# Ultraviolet radiation modulates DNA methylation in melanocytes

**DOI:** 10.1101/2021.10.14.464470

**Authors:** Sarah Preston-Alp, Jaroslav Jelinek, Jean-Pierre Issa, M. Raza Zaidi

## Abstract

Ultraviolet radiation (UVR) is the principal causal factor for melanoma; albeit the underlying mechanisms remain unclear. While the mutagenic properties of UVR are irrefutable, the role of UVR-induced mutations in the initiation of melanoma is controversial which highlights the gap in our knowledge of the initial critical molecular mechanisms of UVR-induced melanomagenesis. To investigate the potential non-mutational mechanisms of UVR-induced melanomagenesis, we studied the role of UVR in modulating DNA methylation changes in melanocytes via next-generation sequencing-based methodologies. Here we show that UVR directly causes stable changes in the DNA methylome and transcriptome, one month after exposure. Genomic features associated with transcription were protected from 5mC alterations whereas CpG sites found in intergenic regions were more likely to be affected. Additionally, the long-term effects of UVR seem to perturb signaling pathways important for melanocyte biology. Interestingly, UVR-sensitive CpG sites were found to be prognostic of overall patient survival and highlighted a subset of CpG sites that may be relevant in melanomagenesis.

**Significance:** We report a novel finding that ultraviolet radiation (UVR) induces DNA methylation changes along with stable alterations in gene expression in cultured melanocytes. Our results provide experimental evidence of UVR-induced epigenetic rewiring, which may be implicated in the susceptibility to melanomagenesis, independently of its mutational effects. These findings offer novel insight into the role of UVR in the initiation and pathogenesis of melanoma via a currently underappreciated mechanism.

## 1 Introduction

Exposure to ultraviolet radiation (UVR) is the greatest risk factor for melanoma development [1]. UVR is considered a complete carcinogen because it is not only mutagenic but also promotes tumor development. It has many complex effects on melanocytes and the skin microenvironment that contribute to melanomagenesis. The most well-studied of these, are the direct and indirect DNA damaging effects [2]. DNA damage, if left unrepaired, is deleterious to the cell by disrupting transcription, replication, and cell cycle progression. The most prevalent mutation induced by UVR-induced DNA damage is the C to T signature mutation, which makes up more than 60% of all UVR-induced mutations. Recently, two genomic studies identified a handful of apparent tumor-suppressor and oncogenes that contain UVR-induced mutations in a low percentage of cases [3, 4]. Paradoxically, two of the most common driver mutations found in human melanoma (BRAF^V600E^ and NRAS^Q61R^) do not carry UVR-signature mutations but another UVR canonical mutation that is induced at a much lower frequency [5]. The mutations found in these common drivers can also be caused by other mutagens and so it remains controversial whether these are induced by UVR since there has been no direct experimental evidence. Some of the other common susceptibility mutations (e.g. in *CDKN2A, TP53*, and *PTEN* genes) are found in non-melanoma cancers of internal tissues that are not exposed to UVR. As such, the role of UVR-induced mutations in melanoma initiation remains debatable. Concurrently, there is a need to investigate whether non-mutational effects of UVR may be involved; specifically, epigenetic modulations, which is a hallmark of melanoma [6].

Epigenetic modifications are heritable chemical changes that do not alter the DNA sequence. This includes changes in the chromatin structure, which are dictated by chemical modifications to the DNA and histone proteins. DNA methylation occurs at cytosines within CpG dinucleotides with the covalent addition of a methyl group to form 5’methyl-cytosine (5mC)[7]. CpG sites can be found grouped in high concentration in areas of the genome called CpG islands (CpGi). Adjacent flanking DNA regions to CpGi make up CpG shores. Many CpGi can be found in the promoter regions of numerous genes where hypomethylation allows for enhanced gene transcription [8]. DNA methylation is an important regulatory mechanism for the genome that can lead to transcriptional activation or repression. DNA methylation can be sensitive to environmental factors so that external stimuli can cause lasting transcriptional effects through alterations of the 5mC. Aberrant 5mC changes are a hallmark of melanoma, showing a global attenuation of 5mC levels and with focal hypermethylation, which suppresses transcription of numerous tumor suppressor genes, including: *PTEN, CDKN2A*, and *RASSF1A* [9-17]. Hypomethylation leading to enhanced transcription of oncogenes is relatively rare in melanoma but instead leads to expression of a number of neoantigens such as, *MAGE-A1* [18]. How these changes arise during melanoma development remains poorly investigated.

Our understanding of UVR in modulating epigenetic features, including DNA methylation, is extremely limited. To date there are no reports on the effect of UVR on the melanocyte epigenome. A recent study identified large hypomethylated blocks in chronically sun-exposed skin that were conserved in skin squamous cell carcinoma [19]. Another study showed an acute methylation and transcriptional response in the epidermis of healthy individuals following UVR exposure that correlated with an individual’s sun sensitivity [20]. However, these changes were likely caused by the recruitment of different cellular populations to the skin after UVR exposure. We hypothesized that UVR directly causes changes in DNA methylation and transcription in melanocytes. Here we report a novel finding that UVR-induces DNA methylation changes in melanocytes along with stable changes in gene expression and that these alterations correlate to methylation changes found in melanoma.

## 2 Material and Methods

### 2.1 Cell culture and UVR exposure

Mouse melan-a cells [21] and human primary epidermal melanocytes (HEMn-DP1 and HEMn-DP2) were used for DNA methylation profiling. Melan-a cells were cultured in RPMI 1640 media supplemented with 10% FBS, 0.1% gentamicin antibiotic, GlutaMAX (Gibco), and phorbol 12-myristate 13-acetate (TPA, 200nM). HEMn cells were obtained from two darkly pigmented neonatal single donors (Gibco #C-202-5C, HEMn-DP1 lot #1216616 & HEMn-DP2 lot #1817883) and cultured in 254 Medium (Gibco), human melanocyte growth supplement (HMGS, Gibco), gentamicin (Gibco) and TPA (200nM). Cells were cultured in a humidified incubator at 37°C and 5% CO_2_. Cells were plated in biological replicates in a 60 mm plate 48 h before UVR exposure. On the day of treatment, adherent cells were washed with DPBS and covered with a thin layer of DPBS. Tissue culture plates were covered with Saran wrap and irradiated with two FS20 bulbs (with the same spectral output as FS40 bulbs used in reference) which emit a broadband UV spectrum containing 35% UVA and 65% UVB (with peak emission at 313nm in the UVB range) [22]. Cells received a total UVR dose of 175 J/m^2^ over 90 s at a rate of 1.94 W/m^2^. Approximately 80% of the melanocytes survive this exposure. Cells were cultured for one month after UVR exposure to detect stable and heritable changes in 5’mC.

### 2.2 DREAM methylation profiling and data analysis

DNA was extracted as follows: cell pellets were lysed in 2% SDS, 25 mM EDTA; proteins were precipitated by the addition of 10 M ammonium acetate and removed by centrifugation. DNA was precipitated by isopropanol, washed with 70% ethanol, and dissolved in TE (Tris 10 mM, EDTA 1 mM). High molecular DNA was recovered by immobilization on 0.4x AMPure XP beads (Beckman Coulter) prior to methylation analysis. Digital Restriction Enzyme Analysis of Methylation (DREAM) was previously described and detects approximately 50,000 unique CpG sites [23, 24]. We detected 15,642 CpG sites in the melan-a cells and 48,927 CpG sites in the HEMn-DP1 cells. Briefly, 2 μg DNA spiked with methylation standards was digested with *SmaI* endonuclease (NEB) at 25°C for 8 h. *SmaI* cuts only the unmethylated recognition site (5’CCCGGG) leaving a 5’GGG signature. Next, followed digestion with *XmaI* (NEB) at 37°C overnight leaving a 5’CCGGG methylation signature. 3’ end filling and dA tailing was performed by Klenow Fragment (3’>5’ exonuclease deficient; NEB). Libraries were prepared by ligation of NEBNext adapters and indexed i7 primers (NEB). Libraries were amplified for 12-14 cycles and cleaned with AMPureXP beads prior to sequencing. DREAM libraries were spiked with 25% phiX Illumina library to ensure sequence diversity and sequenced on Illumina HiSeq 2500 paired-ends for 40 bases. Adapter sequences were removed using TrimGalore! (https://www.bioinformatics.babraham.ac.uk/projects/trim_galore/) [25]. All sequencing was performed at Fox Chase Cancer Center Genomics Facility. Sequences were aligned to the NCBI/mm9 mouse genome or the GRCh37/hg19 human genome using Bowtie2. Signatures corresponding to unmethylated and methylated CpGs were counted, and methylation was calculated as the ratio of methylated read signatures to all of the reads mapping to each CpG site. Methylation was adjusted based on spiked in standards. Coverage thresholds were set at greater than 50 reads per CpG site for DREAM libraries. Data have been deposited in GEO and can be accessed through the following accession numbers: PENDING.

### 2.3 RRBS methylation profiling and data analysis

Reduced Representation Bisulfite Sequencing (RRBS) was previously described [26]. Briefly, genomic DNA was spiked with lambda phage DNA and digested with MspI restriction enzyme. Fragments were ligated with methylated adapters (NEB) to protect adapter sequences from downstream bisulfite treatment. Digested DNA was bisulfite converted using Qiagen’s EpiTect Bisulfite Conversion Kit. All libraries were prepared using NEBNext adapters and indexed with i7 primers. These methylation-based libraries were spiked 25% phiX Illumina library to ensure sequence diversity and sequenced 75 single-end on Illumina HiSeq 2500. Adapter sequences were removed using TrimGalore! (https://www.bioinformatics.babraham.ac.uk/projects/trim_galore/) [25]. RRBS sequence alignment to the human GRCh37/hg19 genome was performed by Bismark [27]. CpG counts were performed using bismark_methylation_extractor function. Coverage thresholds for HEMn-DP2 by RRBS was set at greater than 15 reads per CpG site allowing for detection of 1,018,477 unique CpG sites. Differential methylation analysis was performed using methylKit in R [28, 29]. *P* values were adjusted for multiple testing using SLIM method [30]. Differentially methylated sites were filtered by methylation change > 10% and q < 0.05, unless otherwise stated. Data have been deposited in GEO and can be accessed through the following accession number: GSE169479 (https://www.ncbi.nlm.nih.gov/geo/).

### 2.3 RNA-sequencing

RNA was isolated from HEMn-DP2 cells using Qiagen’s RNeasy Plus Mini kit following manufacturer’s instructions and done in biological triplicates. On column DNase treatment of the samples was included in the protocol. Strand-specific RNA libraries were generated from 1 μg of RNA using NEBNext Ultra II RNA library Prep Kit for Illumina following the manufacturer’s instructions. Libraries were sequenced 75 bases single-end on Illumina HiSeq 2500. Quality monitoring was done using FastQC. Cutadapt was used to remove adapter sequences. Sequence alignment to the human genome GRCh37/hg19 and gene counting was performed by STAR [31]. Genes that had 0 reads across all samples were excluded from the analysis. Differential expression analysis was performed using DESeq2 [32]. Genes were considered upregulated with an FC > 2 and downregulated with an FC < 0.5 and *Q*_*FDR*_ < 0.05, unless otherwise stated. Data have been deposited in GEO and can be accessed through the following accession numbers: GSE169382 (https://www.ncbi.nlm.nih.gov/geo/).

### 2.4 Human epidermal melanocyte datasets from ENCODE

DNase and ChIP-seq data for histone marks H3K9me3, H3K27ac, H3K27me3, H3K4me1, and H3K4me3 were downloaded from the Encyclopedia of DNA Elements (ENCODE)[33, 34] (https://www.encodeproject.org/) for the human epidermal melanocyte cell line using the following identifiers: DNase-seq (ENCF137FNR, ENCFF605NTT), H3K9me3 (ENCFF014FHO, ENCFF849GBJ), H3K4me3 (ENCFF128QSF, ENCFF499KTB, ENCFF317ZWT), H3K4me1 (ENCFF464TQM, ENCFFOVO, ENCFF572TXY), H3K27me3 (ENCFF640WPP), H3K27ac (ENCFF209WNE, ENCFF326ITF). DNase and ChIP-seq profile plots were created for a 10 kb window centered around CpG sites. Profile plots were created by averaging the peak signal using the normalizeToMatrix function in the EnrichedHeatmap package in R with the following parameters: “w0” mean mode and 100 bp bin size [35].

### 2.4 TCGA DNA methylation array data

DNA methylation data for melanoma SKCM cohort (n=462) from the Illumina 450k array platform was downloaded from The Cancer Genome Atlas (TCGA) using TCGAbiolinks R package[36-38]. Publicly available data from the TCGA were sourced from deidentified patients. CpG sites with NA values were excluded leaving a total of 373,814 sites. CpG sites were then filtered to include those that were detectable in either the HEMn-DP1 DREAM dataset or the HEMn-DP2 RRBS dataset leaving a total of 29,666 CpG sites. 896 CpG sites were considered UVR sensitive if they changed (differential methylation > 10% & q < 0.05) in either HEMn-DP1 or HEMn-DP2 after UVR exposure. 25,711 sites were considered UVR insensitive if they did not change (differential methylation < 10%) in both HEMn-DP1 and HEMn-DP2 cells.

### 2.5 Statistical analyses

All analyses were performed using R[28, 29]. Statistical tests used are noted in the figures where they appear. Student’s *t* test was used to compare continuous variables. Comparison between multiple groups of continuous variables was performed by one-way ANOVA followed by post-hoc Tukey test.

Odds ratio (OR) calculations for changes in methylation at specific locations (e.g. CpG islands, promoters, histone marks, etc.) were calculated as:

|  |  | In location |  |
| --- | --- | --- | --- |
|  |  | Yes | No |
| Significantly changed | Yes | a | b |
|  | No | c | d |

OR statistical significance was computed using Fisher’s exact test. Unsupervised hierarchical clustering was performed by Ward’s method using the hclust function in R. Heatmaps were generated using the pheatmap package[39]. Overrepresentation of known TF binding sites were identified using “findMotifGenome.pl” from HOMER[40]. Analysis was conducted on regions ±50 bp surrounding differentially methylated CpG sites within gene promoter regions (100bp upstream of the TSS). Motif enrichment analysis was conducted against background regions containing equivalent GC content. Gene set enrichment analysis (GSEA) was performed on differentially regulated gene transcripts using the hallmark gene set and legacy transcription factor targets using version 7.2 [41, 42]. Kaplan-Meir survival curves and log-rank test statistic was computed using R survival package [43].

## 3 Results

### 3.1 UVR induces DNA methylation changes in melanocytes

We hypothesized that UVR could directly induce methylation of cytosine to 5mC in exposed melanocytes. We irradiated primary human epidermal melanocytes, HEMn-DP1 and HEMn-DP2, with a broadband containing terrestrially relevant UVA and UVB wavebands. Cells were treated in biological replicates and cultured for one month after irradiation to determine stable and heritable changes in methylation. 5mC levels were detected by digital restriction enzyme analysis of methylation (DREAM) and reduced representation bisulfite sequencing (RRBS) methodologies in HEMn-DP1 and HEMn-DP2 cells, respectively. We found that both cell populations were sensitive to UVR-induced gain or loss of DNA methylation at numerous CpG sites (Figure 1a). In the HEMn-DP1 cells 3% of the CpG sites lost methylation and 4% gained methylation (FDR < 0.05 & methylation change > 10%). A similar effect size was seen in the HEMn-DP2 cells, where 2% of the CpG sites lost methylation and 4% gained methylation. From the 9333 CpG sites that we could detect in both HEMn-DP1 and HEMn-DP2 cells, we found that the changes in the two cell populations were distinct and did not correlate with one another (Figure 1b). To expand our study, we tested a mouse melanocyte cell line, melan-a. We found that the melan-a cells were also sensitive to UVR with 10% of the CpG sites losing methylation and 12% gaining methylation (Figure 1c). These also showed a more pronounced trend in hypermethylation. In order to compare between mouse and human samples, we looked at changes in methylation at gene promoters (3000 bp upstream of the transcription start site). These should be similarly regulated in a cell-type specific manner. We found small overlaps in two-way comparisons, however, there were no common changes in promoter methylation between all three cell populations (Figure 1d). It should be noted, that the greater number of differentially methylated promoters in the HEMn-DP2 population is due to the analysis by RRBS which detects more CpG total sites and subsequently more CpG sites within promoter regions. The non-UVR exposed baseline methylation for UVR-insensitive sites were at the extremes of low or highly methylated (Figure 1e). In contrast, CpG sites that were sensitive to UVR were partially methylated non-exposed cells. CpG sites were divided into three groups based on the non-exposed baseline methylation: low (< 20%), medium (20-80%), and high (> 80%). In all three cell populations, a higher percentage of CpG sites that had a medium baseline methylation level were either hyper- or hypomethylated after UVR exposure (Figure 1f).

**Figure 1.**
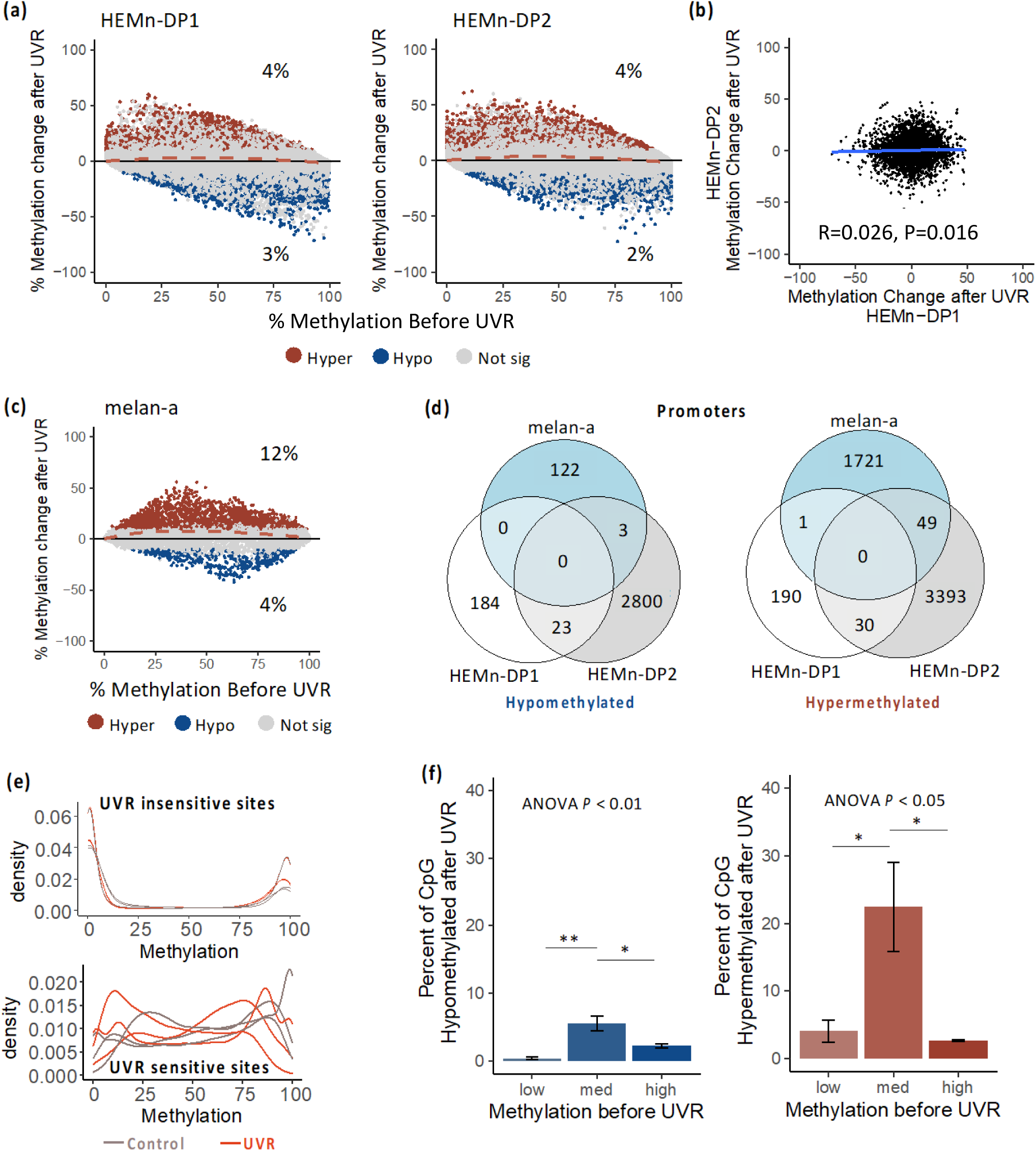
UVR induces unique DNA methylation changes in mouse and human melanocytes at partially methylated CpG sites. (a) Mean-average plot showing the baseline methylation before UVR exposure compared to the change in methylation after UVR exposure for human HEMn-DP1 and HEMn-DP2 cells. Sites that are hypermethylated after UVR (methylation gain > 10% & FDR < 0.05) are highlighted in red and sites that are hypomethylated after UVR (methylation loss > 10% & FDR < 0.05) are highlighted in blue. (b) Scatter-plot comparing differences in 5’mC after UVR HEMn-DP1 cells to HEMn-DP2 cells. (c) MA plot showing the baseline 5’mC levels before UVR to changes in 5’mC after UVR for mouse melan-a cells. (d) Venn diagram showing the lack of overlap between differentially hyper or hypo methylated promoters (1000bp upstream of the TSS) in all three cell lines. (e) Density plot showing distribution of 5’mC levels of UVR insensitive sites and UVR sensitive sites. (f) Bar plot of the percent of CpG sites that are hypomethylated (blue) or hyper methylated (red) after UVR. CpG sites were divided into three groups based on 5’mC levels before UVR: low (< 20%), medium (20-80%), and high (> 80%). Significance calculated by one-way ANOVA followed by Tukey post-hoc (*P* < 0.05 = *, *P* < 0.01 = **).

### 3.2 UVR modulates 5mC levels at different genomic and CpG regions

Next, we analyzed the UVR-induced changes in 5mC with respect to genomic location. We found both hypo and hypermethylated changes occurring at all regions, with no statistically significant difference between the hypo vs. hypermethylated sites (Figure 2a). We compared the ratio of sites that were hypermethylated in promoter, intron, exon, and intergenic regions to the expected ratio (the ratio of all detectable CpG sites in those regions). We found a greater proportion of intergenic CpG sites and a smaller proportion of promoter CpG’s were hypermethylated than what was expected, in all three cell populations (Figure 2b). The same was true for sites that were hypomethylated. Odds ratio analysis identified promoters, and to a lesser extent introns and exons, with an odds ratio less than one as being protected from 5mC changes after UVR (see methods; Figure 2c). Conversely, 5mC changes were more likely to occur at intergenic regions with an odds ratio greater than one (Figure 2c). The same analysis for CpG islands, shores, and open sea regions (sites outside of islands or shores) showed that a greater proportion of open sea sites and a smaller proportion of CpG islands sites were hypermethylated and hypomethylated than expected (Figure 2d). CpG island sites were protected from 5mC changes with an odds ratio less than one, while open sea sites were susceptible with an odds ratio greater than one (Figure 2e). The conclusions remained true for methylation threshold of 15% (Supplemental Figure 1).

**Figure 2.**
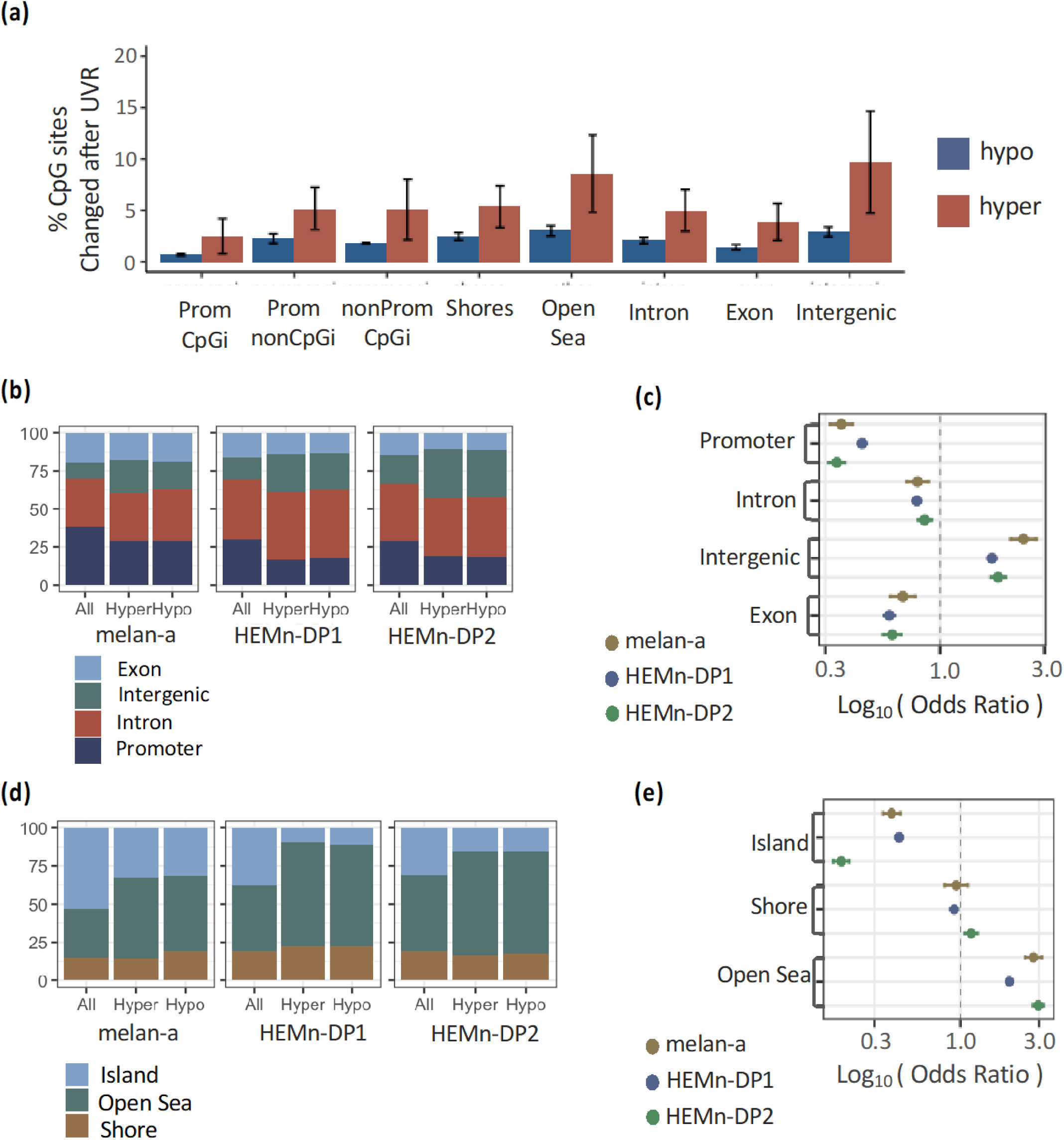
UVR-induced DNA methylation changes occur at distinct genomic regions. (a) Percent of hyper and hypo methylated sites out of the total detectable sites occurring at different regions. (b) The proportion of CpG changes occurring in each cell line for defined genic regions compared to the proportions of regions detected. (c) The odds ratio of changes occurring at defined genic regions. (d) The proportion of CpG changes occurring at regions defined relative to CpG islands compared to proportion of all CpG sites detected. (e) The odds ratio of changes occurring at regions defined relative to CpG islands.

### 3.3 UVR modulates 5mC levels at different chromatin regions

We next explored how these 5mC changes occurred with respect to the chromatin landscape. We obtained DNase and histone ChIP-seq data for human epidermal melanocytes from the ENCODE database. The CpG sites that did not change after UVR exposure were located in areas wither greater DNase sensitivity, H3K27ac, and H3K4me3 as seen in both HEMn-DP1 and HEMn-DP2 cell populations (Figure 3a). The CpG sites that gained methylation were located in regions with lower levels of H3K4me1 than sites that were hypomethylated or did not change after UVR exposure. The CpG sites that gained methylation, particularly the hypermethylated CpG sites in HEMn-DP1 cells, were located in regions with higher levels of H3K9me3 than the CpG sites that did not change. We found no conserved pattern to changes in 5mC with respect to H3K27me3 repressive mark. Odds ratio analysis showed that only the CpG sites in heterochromatic regions marked with H3K9me3 were more likely to have changes in 5mC after UVR exposure (Figure 3b). The CpG sites located in open chromatin (DNase hypersensitivity), actively transcribed (H3K27ac), promoters (H3K4me3), enhancers (H3K4me1), or bivalent promoters (H3K27me3 & H3K4me3) were protected from changes in 5mC after UVR exposure.

**Figure 3.**
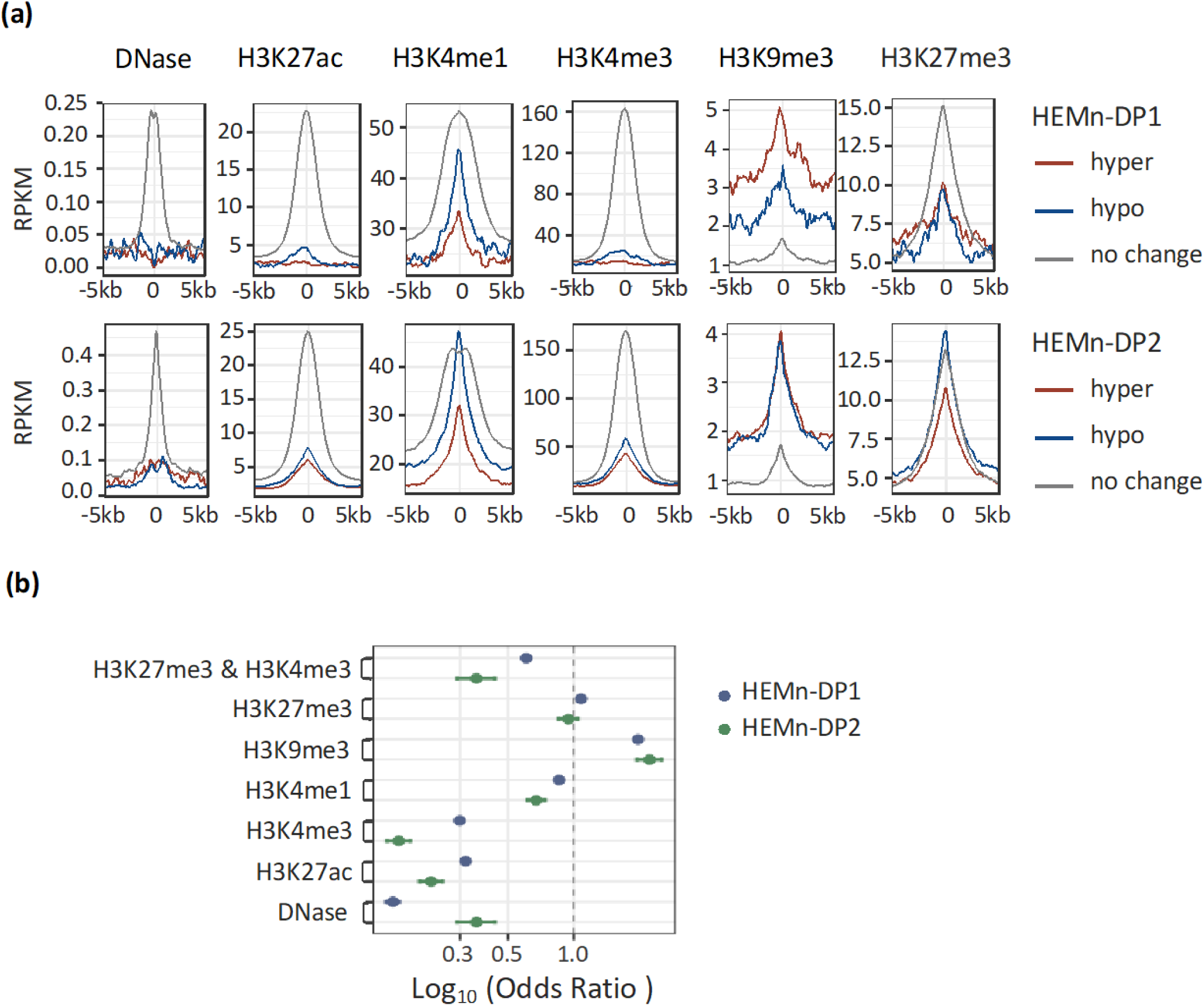
UVR-induced DNA methylation changes occur at distinct chromatin regions. (a) Profile plots of ENCODE sequencing signal from DNAse and histone ChIP-seq experiments from human epidermal melanocytes. Red depicts 10kb regions centered on hypermethylated CpG sites, blue is centered on hypomethylated CpG sites and gray is centered on CpG sites that do not change after UVR exposure. The top panel is relative to changes in HEMn-DP1 cells and the bottom is for HEMn-DP2 cells. (b) The odds ratio of 5’mC changes occurring at regions defined by the chromatin landscape.

### 3.4 Changes in promoter 5mC correlates with UVR-induced transcriptional changes

Hypermethylation of gene promoters is important for silencing gene transcription and vice-versa [8]. In order to understand if UVR changes in 5mC are having a functional effect on transcription we performed RNA-seq analysis in HEMn-DP2 cells one month after UVR exposure (Supplementary File 1). Principal component analysis of the transcriptome showed separation between control and UVR exposed samples (Figure 4a). There were 348 genes up-regulated and 105 genes down-regulated (Q < 0.05 & fold change > 2) after UVR exposure (Figure 4b,c). To determine if gains in promoter methylation would shut down gene transcription or if loss of methylation would aid in transcriptional activation, we compared UVR-induced changes in promoter methylation (500 bp upstream of the transcriptional start site) to changes in gene transcription (Figure 4d) and found a negative correlation between the two. There were 22 genes that increased methylation and were down regulated after UVR exposure and 31 genes in which promoter methylation decreased and were upregulated after UVR exposures (Figure 4e). While a proportion of genes were explained by this relationship in promoter methylation, a greater proportion showed changes in gene expression without any changes in promoter methylation.

**Figure 4.**
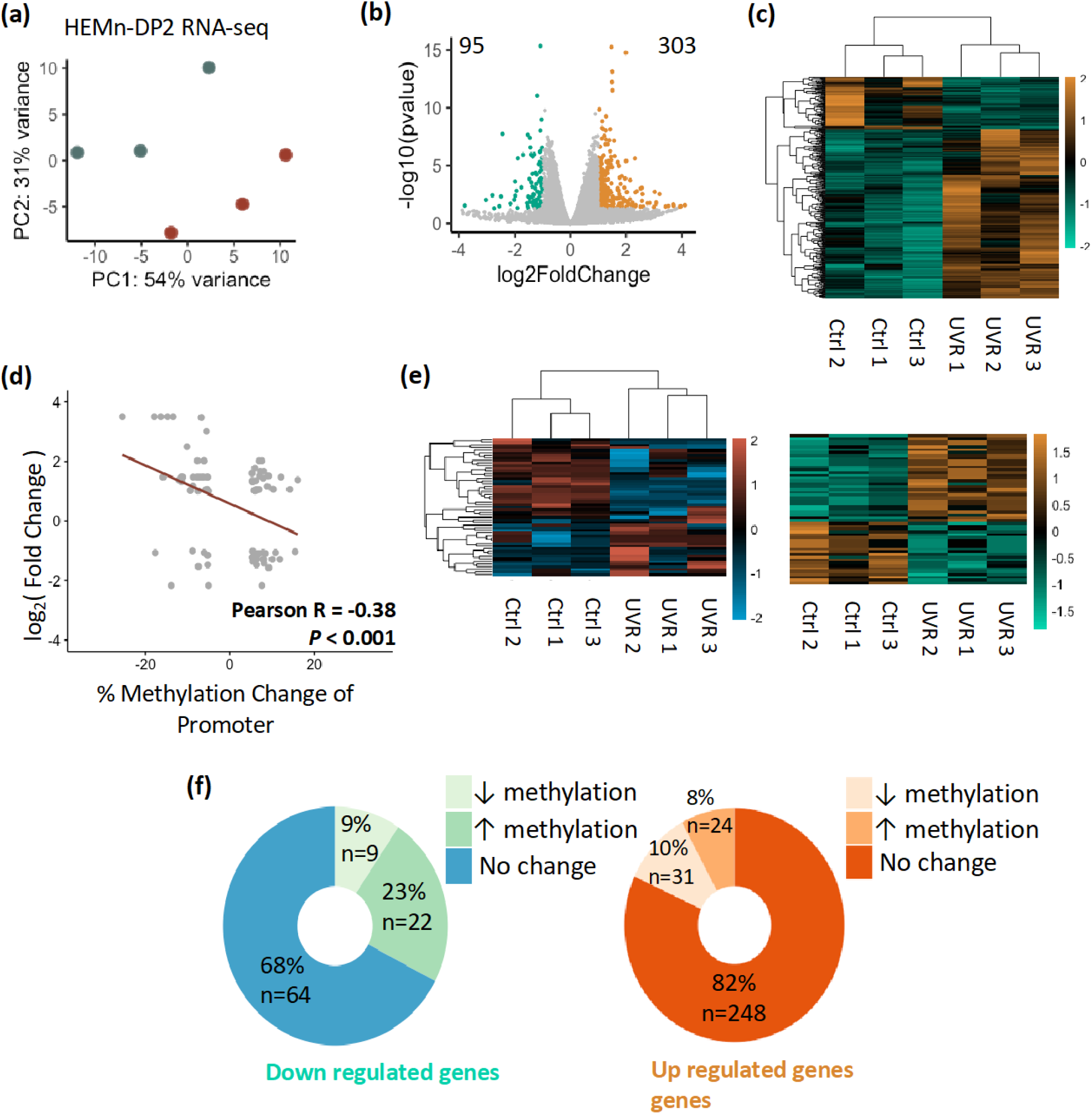
UVR-induced changes in promoter DNA methylation partially explain transcriptional changes. (a) PCA of the transcriptional changes that occur in HEMnDP2 cells after UVR exposure. (b) Volcano plot showing the fold change in transcriptional changes and significance; up regulated genes (n=348) are in green and down regulated genes (n= 105) are in orange. Highlighted genes have a log2(Fold Change) cutoff at ±1 and *P* < 0.05. These points are plotted in a heatmap (c). (d) Scatter plot showing changes in methylation (CpG’s with greater than 10% by RRBS) at the gene promoter (−500 bp of the TSS) and the change in gene expression. Changes in promoter methylation negatively correlate with changes in transcription for 53 genes. (e) A heatmap of promoter methylation of these genes. (f) A heatmap of corresponding gene transcription. Sample and gene clustering are the same as E. (f) A donut plot showing percent of down or up regulated genes with increasing, decreasing, or no methylation changes in the promoter after UVR exposure.

### 3.5 DNA methylation changes occur at transcription factor binding regions that are down-regulated after UVR exposure

To understand the biologically consequences of the UVR-induced methylation changes, we performed gene set enrichment analysis using GSEA. We did not find any significantly enriched pathways for genes that were upregulated after UVR exposure in this analysis. However, we found numerous pathways that were enriched for down-regulated genes (Figure 5a). Pathways related to inflammation were identified, with interferon-alpha response being the top hit. Additionally, a number of different signaling pathways were also identified. Analysis of transcription factor targets revealed a number of potential factors that may be controlling these changes in gene expression (Figure 5b). To determine if methylation could be playing a role in the negative regulation of these transcription factors, we performed motif enrichment analysis on regions containing hyper methylated CpG sites in either HEMnDP1 or HEMnDP2 cells. We identified 45 significantly enriched motifs (Q_FDR_ < 0.0001), including STAT3 and AP2 which were identified by GSEA analysis in Figure 5b (Figure 5C). As expected, many of the transcription factor binding motifs were CG rich. We looked further to see if the AP2 target genes that were down-regulated also contained hypermethylated CpG sites within the gene promoter (3000 bp upstream of the TSS). We found a significant overlap of 23 genes that were down regulated and hypermethylated (Figure 5d). We also identified 18 STAT3 target genes that were down regulated and hypermethylated, but this was not a statistically significant overlap (Figure 5e).

**Figure 5.**
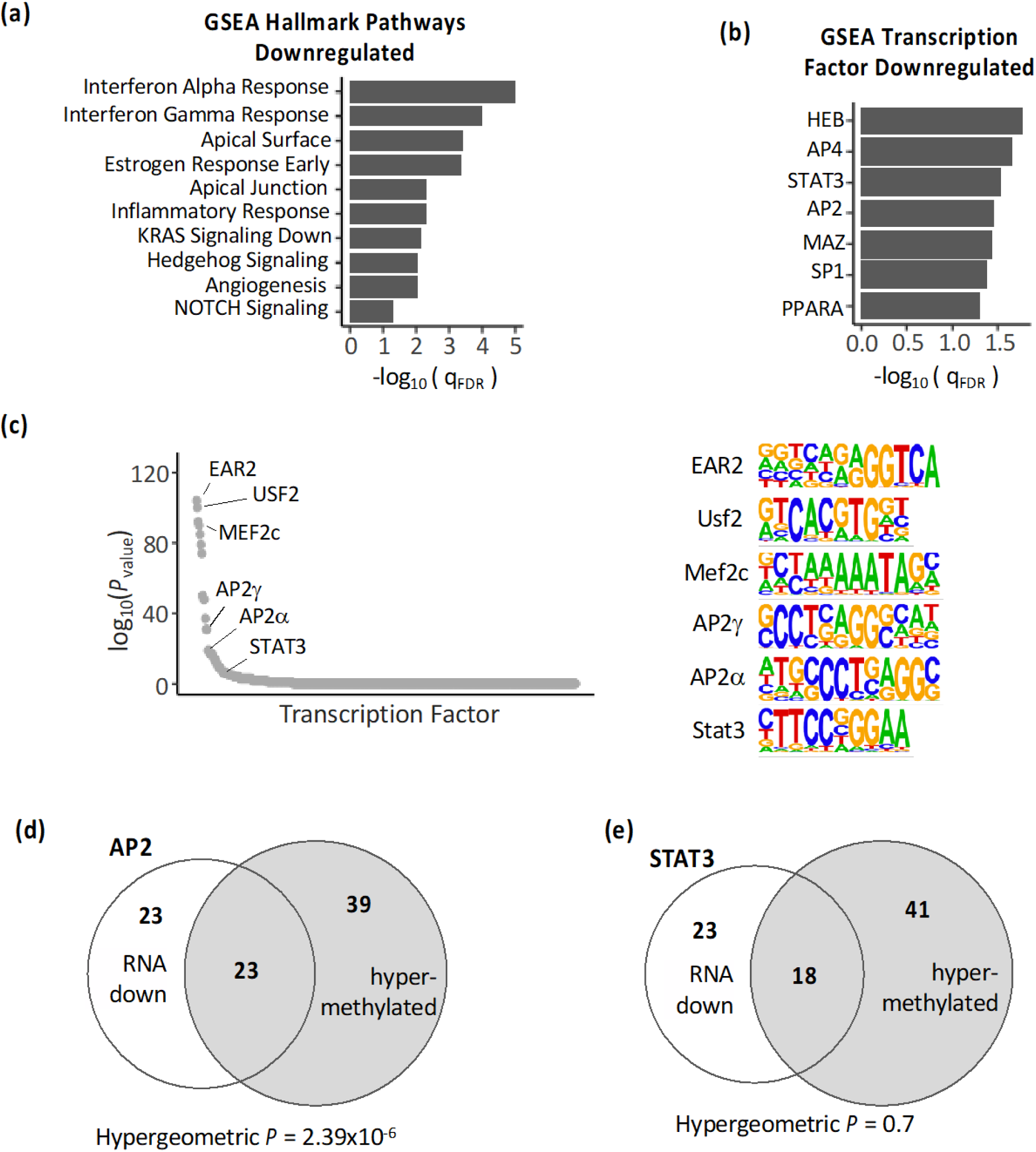
Changes in DNA methylation occur at binding motifs for transcription factors that are down-regulated after UVR. (a) Pathway enrichment analysis by GSEA for HEMn-DP2 cells after UVR. Minus log_10_ of the FDR value of the enrichment scores for each hallmark pathway is shown. These pathways were downregulated after UVR. (b) Minus log_10_ of the FDR value of the enrichment scores of target genes for each transcription factor. Transcription factor target genes were downregulated after UVR. (c) Motif analysis of DNA sequences of the 50bp region flanking significantly differentially methylated CpG sites after UVR identifies enrichment of multiple transcription factor binding sites. Enrichment p_values_ are plotted for all transcription factors. Logo image of transcription factor binding motifs are shown. (d) Venn diagram showing AP2 target genes that are down-regulated after UVR or contain CpG sites which gain methylation in the promoter (3000bp upstream of TSS) after UVR. (e) Venn diagram showing STAT3 target genes.

### 3.6 Methylation of UVR sensitive CpG sites as a prognostic marker for patient survival

To study the disease relevance of the UVR-sensitive sites found in the HEMn-DP cells, we analyzed Illumina 450k arrays from 462 melanoma samples from the TCGA-SKCM cohort[44]. Of the 29,666 CpG sites that were detectable in the either the HEMn-DP1 or the HEMn-DP2 data sets and the 450k array, 896 CpG sites were UVR-sensitive and significantly changed in either HEMn-DP1 or HEMn-DP2 cells after UVR exposure. 27,511 CpG sites were UVR insensitive and did not change in either cell population. Hierarchical clustering of the patient samples based on the UVR-sensitive sites revealed two groups with distinct methylation profiles (Figure 6a). PCA analysis based on these UVR sensitive sites shows separation of these patient samples by cluster (Figure 6b). There were no clinical characteristics that were different between the two group, however cluster 1 had a larger proportion of tumors bearing UVR-signature mutations compared to cluster 2 (Supplementary Table 1). To find if the same changes occurring in melanocytes after UVR exposure also occurred in the melanoma samples, we compared UVR-induced changes to those found in melanoma samples. Patients in Cluster 1 showed the strongest correlation with UVR induced changes (R = 0.47) (Supplementary Figure 1). Survival analysis showed that compared to cluster 1, cluster 2 was associated with worse overall survival (median OS: cluster1 = 104.7 months, cluster 2 = 63.7 months; log-rank *P* = 0.0042; Figure 6d). Hierarchical clustering of the patient samples on the UVR insensitive CpG sites, disrupted the UVR sensitive clusters and the resulting groups no longer showed a difference in overall patient survival (Supplemental Figure 2). To determine if there was any phenotype associated with the two clusters, RNA-seq associated with the patients was analyzed for differential gene expression (FC > 1.5 & Q < 0.05). There were 797 genes upregulated in cluster 1 and 6 genes upregulated in cluster 2 melanomas (Figure 6d,e). Cluster 1 genes are enriched for a number of pathways involved in immunological processes and signaling (Figure 6f).

**Figure 6.**
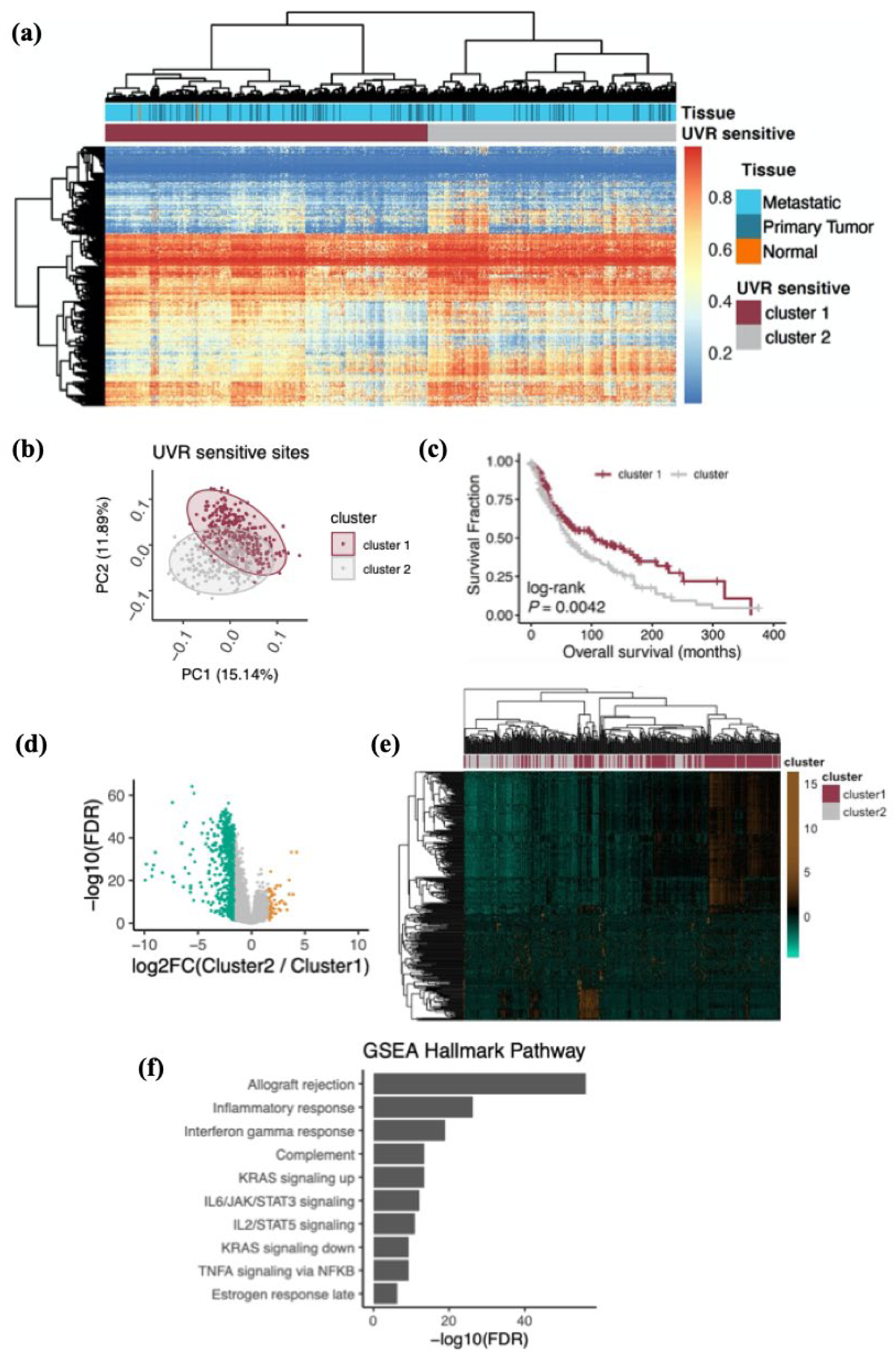
Methylation of UVR sensitive CpG sites as a prognostic marker for patient survival (a) Heatmap of methylation of Illumina 450k array β-values for the TCGA-SKCM cohort. Only the 1510 CpG sites that are sensitive to UVR in the human melanocytes are included. Based on these sites, patients cluster into three distinct groups. (b) Principal component analysis of patients colored by cluster based on UVR sensitive CpG sites. (c) Survival plot based on patient clustering in A. (d) Volcano plot for RNA-seq expression differences between patient clusters. Genes up-regulated in cluster 1 shown in green and genes up-regulated in cluster 2 shown in orange. (e) Heatmap showing gene expression of genes with a fold change greater than 1.5 and Q < 0.05. (f) Gene Set Enrichment Analysis for hallmark pathways for genes upregulated in cluster 1 patients.

## 4 Discussion

Although numerous studies have investigated the roles of UVR-induced genetic mutations in melanomagenesis, there is a significant gap in our knowledge regarding the direct or indirect causal role of the non-mutational events modulated by UVR. Even more profound is the lack of investigation into UVR-induced modulations of epigenetic events. Emerging evidence suggests a role for UVR-modulated epigenetic changes in skin carcinogenesis [20, 45]. These changes can occur at the level of histone modifications, chromatin conformation, or DNA methylation. In this study we focused on the UVR-induced changes occurring in 5mC and found distinct changes in both mouse and human melanocytes. While there were a few concordant changes in 5mC, globally the differences appear to be random and distinct to each melanocyte population, much like the occurrence of UVR-induced DNA mutations. This may provide an additional source of genetic diversity through which selection of biologically advantageous changes in 5mC may be selected for within the melanocyte. Interestingly, the genomic location (i.e. promoter, CpGi, etc.) of the CpG site defined its susceptibility to changes in 5mC. A higher percentage of sites that underwent changes in methylation after UVR exposure were partially methylated (20-80% methylation). These sites likely reflect the plastic regions of the melanocyte genome. All three cell populations showed a propensity for DNA methylation changes to occur within intergenic regions and regions outside of CpGi or shores. These areas are likely exhibit more plasticity with a lower density of regulatory elements located within them; therefore, many of the changes that occur may have no effect on cellular gene expression allowing them to be tolerated by the cell. In line with this notion, regions that are associated with transcriptional regulation (i.e. promoters & CpGi) are protected from these UVR modulated changes. Indeed, when considering where these changes occur relative to the chromatin structure, we found the same pattern. Changes were more likely to occur in heterochromatin marked by H3K9me3 which is associated with silenced gene transcription. Open chromatin (DNAse hypersensitivity), active enhancers (H3K27ac), promoters (H3K4me3), and bivalent promoters (H3K27me3 & H3K4me3) were protected from changes in 5mC.

Even though areas involved in transcriptional control were protected from UVR induced 5mC changes, there were some changes that still occurred. We found stable changes in gene expression in UVR-exposed human melanocytes, which suggests that UVR-induced 5mC changes has functional consequences. A number of these transcriptional changes could be explained by changes in promoter methylation. We found a negative correlation in promoter methylation and gene transcription, which was modest although not unexpected. DNA methylation is one piece in transcriptional control which will also depends on binding of transcription factors and appropriate chromatin conformational changes, and more. Further studies investigating the functional consequences of UVR-induced DNA methylation in transcription factor binding along with concurrent changes in chromatin may better explain the observed transcriptional changes.

We found an enrichment of transcription factor binding motifs in regions where hypermethylation occurred. Of the identified transcription factors, AP2 and STAT3 target genes were downregulated after UVR. The AP2 transcription factor family is made up of five different proteins. The binding motif of two AP2 transcription factors, AP2α and AP2γ, were enriched at regions of hypermethylation. Evidence supports the role of AP2α and AP2γ as a pioneering transcription factors that are essential for normal development of the neural crest cell lineages, which include melanocytes[46, 47]. AP2α is known to bind to both promoter and enhancer sequences up to 150kb from the TSS in melanocytes and works with the master transcriptional regulator for melanocytes, MITF, to coordinate differentiation[48]. The long-term effects of UVR on melanocytes seem to affect gene networks important to melanocyte biology.

The comparison between UVR-induced DNA methylation changes in primary melanocytes with the patient samples from the TCGA-SKCM cohort revealed a positive correlation between UVR-induced 5mC changes and changes found in patient samples. These clusters appear related to UVR exposure since patients in cluster 1 were significantly enriched in the UVR mutational signature and likely greater exposure. Interestingly, UVR-sensitive sites seem to be prognostic for patient survival. Patients that clustered with normal tissue had better overall survival. Additionally, while many of the changes in cluster 1 may be inconsequential, it is likely that some are selected for because they confer increased fitness that is advantageous to the cell. To date there have been several efforts to identify a minimum number of DNA methylation and these data further support the role of locus-specific methylation patterns as a prognostic marker to identify high-risk patients [9, 15, 17, 49, 50]. It is unknown what causal role UVR-induced methylation changes play in melanomagenesis and how they might interact with UVR-induced mutations or other driver mutations of melanoma. Additionally, deeper investigation of the mechanisms of UVR-induced methylation changes is needed.

While UVR is known to have systemic effects, a recent epidemiological study profiled methylation of blood leukocytes in groups of people and found no correlation between 5mC and sun exposure[51]. Leukocyte progenitors do not reside in the skin and taken together with our data, shows that UVR is directly interacting with melanocytes either through direct manipulation of the epigenetic machinery or through selection pressure forcing an outgrowth of clones resistant to UVR stress. These two mechanisms are not mutually exclusive and further studies are needed to understand how these UVR induced methylation changes manifest. It may depend on a number of factors such as cell divisions, time of exposure, pigmentation level, amount of DNA damage, or UVR waveband-specific effects. Understanding how UVR-induced methylation changes interact with age-induced methylation drift would be an interesting area of investigation, since childhood UVR exposure is the greatest causal factor for melanoma development that occurs much later in life. Additionally, these findings have implications for other cellular residents of the skin and may be relevant to other dermatological diseases.

## Acknowledgments

This research was funded by US Department of Defense (#W81XWH-16-1-0177), WW Smith Charitable Trust (#C1508) and American Cancer Society (#IRG-92-027-19). We would like to thank Kelsey Keith for proofreading of the manuscript.

The authors declare no conflict of interest related to these experiments. All sequencing data have been made publicly available on the Gene Expression Omnibus (GEO) database (www.ncbi.nlm.nih.gov). All experiments were approved by the Temple University Institutional Biosafety Committee. Data were sourced from publicly available databases: ENCODE and TCGA. Patient participants gave their informed consent prior to their inclusion in the study and data were obtained from deidentified samples. All experimentation was conducted in accordance with the Declaration of Helsinki; US Federal Policy for the Protection of Human Subjects; and the European Medicines Agency Guidelines for Good Clinical Practice.

---

**Supplementary Figure 1.**
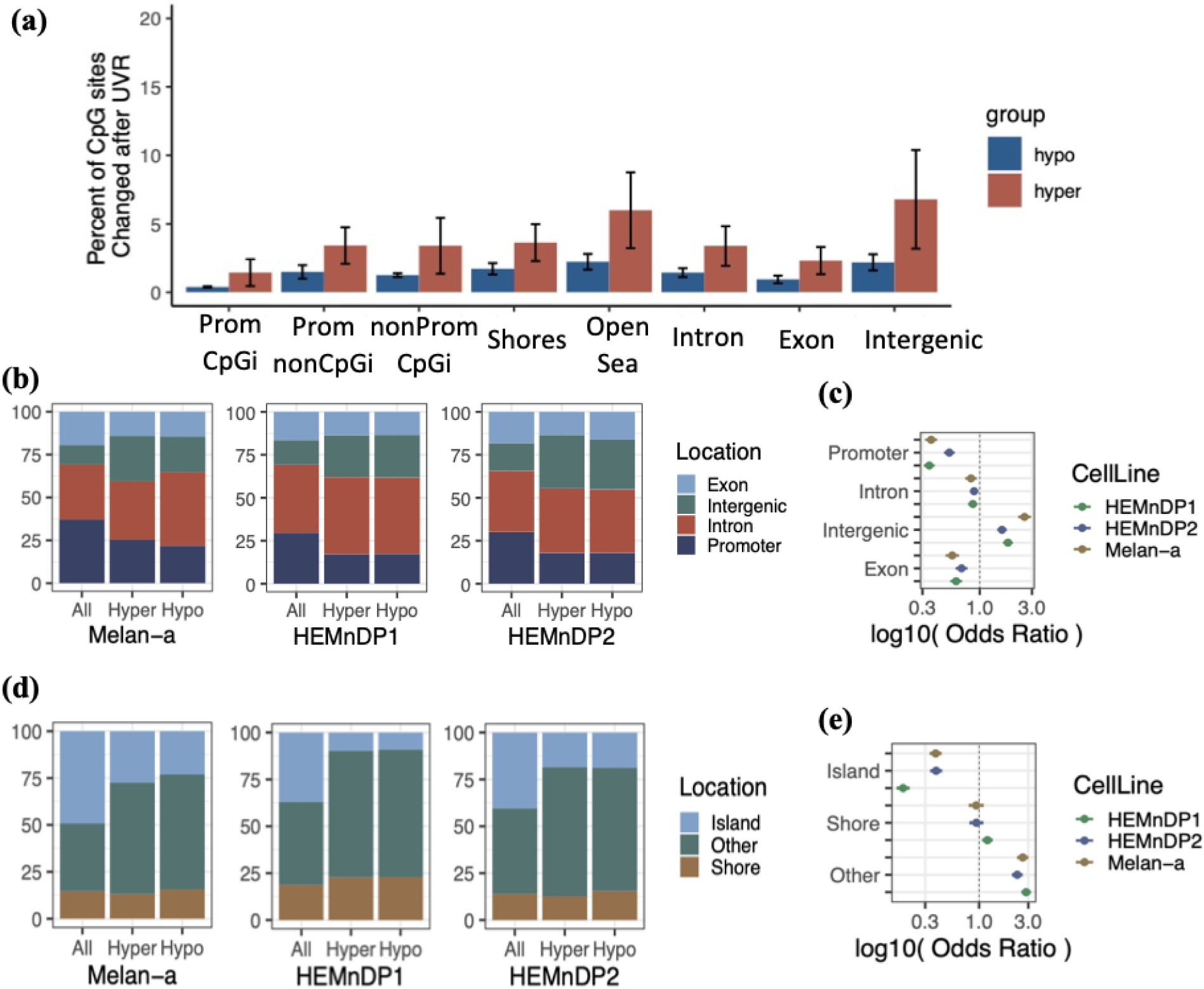
(a) Percent of hyper and hypo methylated sites (methylation difference > 15%) out of the total detectable sites occurring at different regions. (b) The proportion of CpG changes occurring in each cell line for defined genic regions compared to the proportions of regions detected. (c) The odds ratio of changes occurring at defined genic regions. (d) The proportion of CpG changes occurring at regions defined relative to CpG islands compared to proportion of all CpG sites detected. (e) The odds ratio of changes occurring at regions defined relative to CpG islands.

**Supplementary Figure 2.**
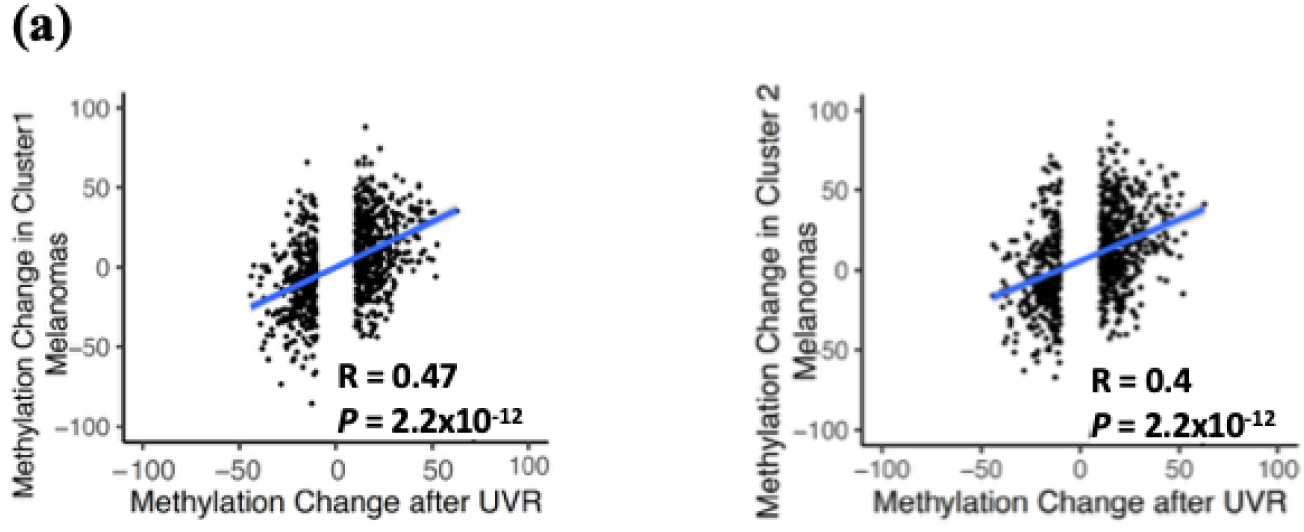
UVR-induced methylation changes correlate with changes found in TCGA-SKCM patients. (a) Plot showing the changes in methylation after UVR exposure in HEMn-DP melanocytes and changes in methylation in melanoma compared to melanocytes for patients in each of the three clusters.

**Supplementary Figure 3.**
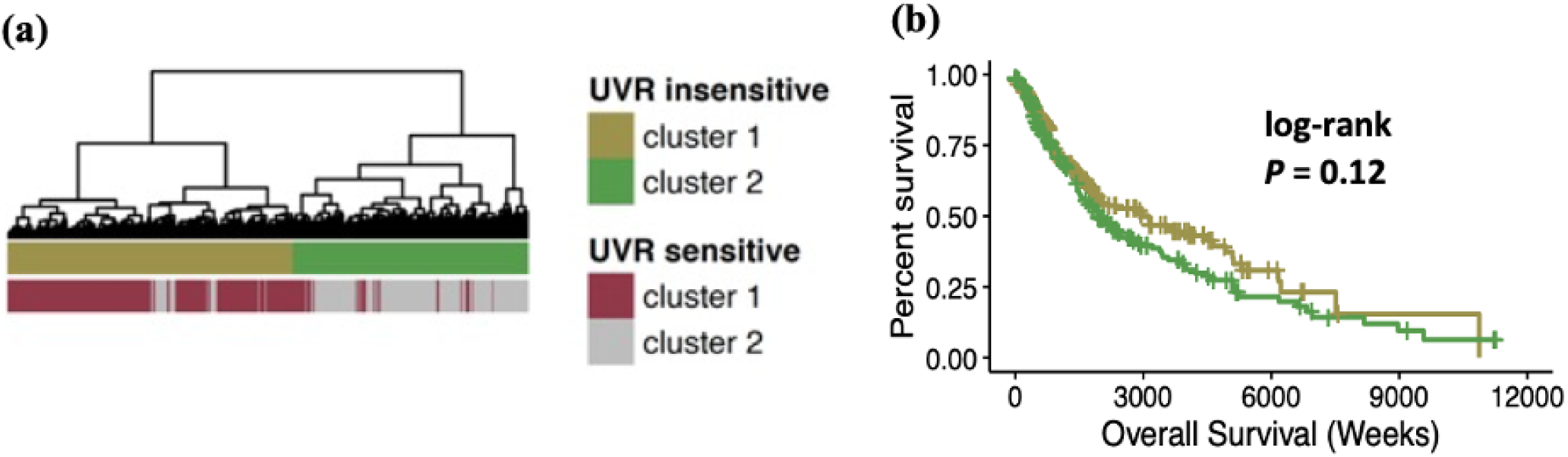
UVR-insensitive CpG sites are not prognostic for patient survival. (a) Dendrogram showing clustering of patient samples based on UVR insensitive CpG sites. Included are annotations of the same patients clustered on UVR sensitive CpG sites from Figure 6A. (b) Overall survival of patients based on UVR insensitive clusters.

**Supplementary Table 1.**
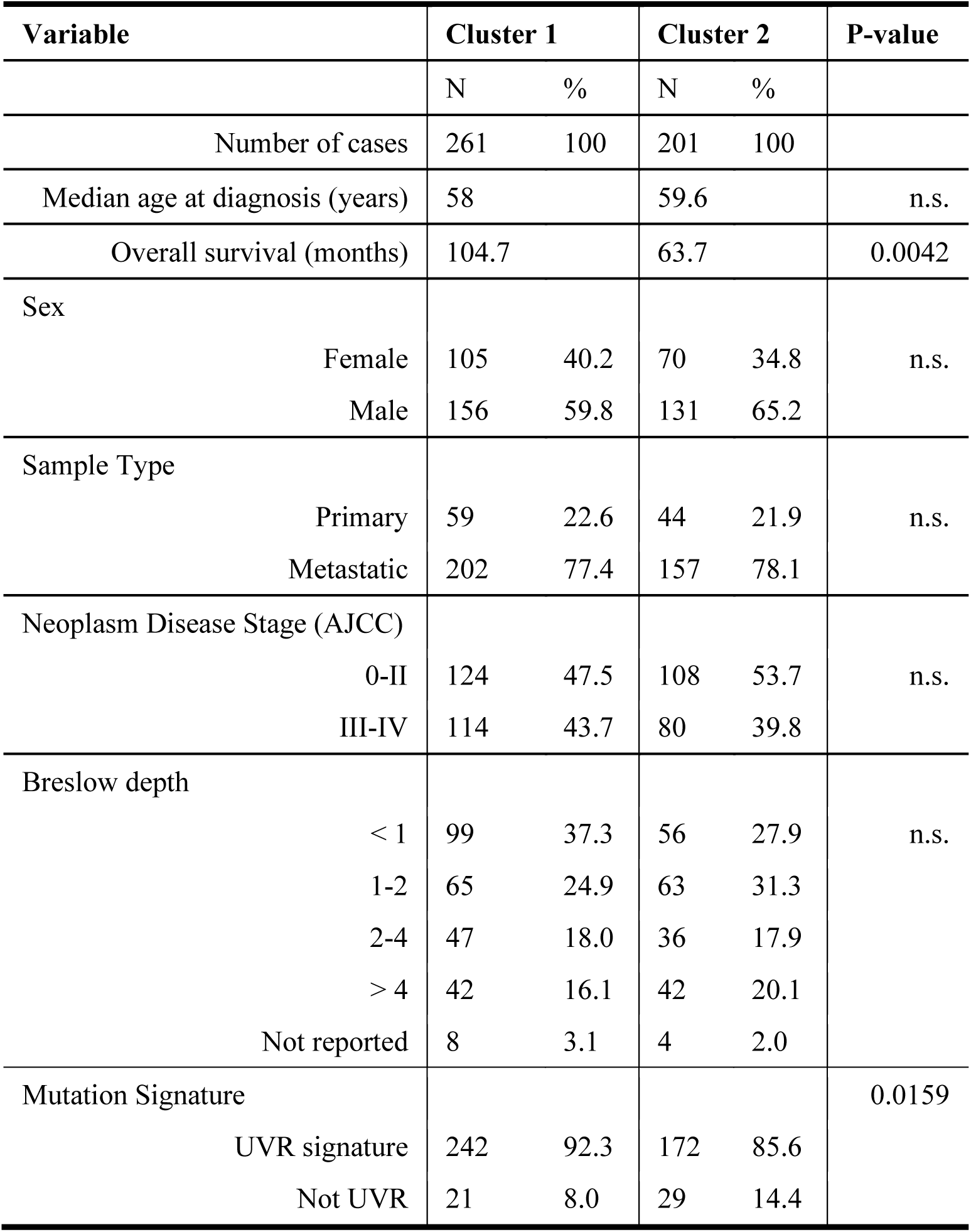
Clinical characteristics of TCGA-SKCM patients clustered by UVR-sensitive CpG sites.

