## Supplemental File 1 for "Ultraviolet radiation modulates DNA methylation in melanocytes"

| hgnc_symb | baseMean | log2FoldCh | lfcSE | stat | pvalue | padj |
|---|---|---|---|---|---|---|
| SCYL3 | 224.1094 | 0.296571 | 0.208426 | 1.422911 | 0.154762 | 0.300109 |
| C1orf112 | 340.09 | -0.05267 | 0.2687 | -0.19603 | 0.844589 | 0.907349 |
| FGR | 0.192403 | 0.975392 | 4.060716 | 0.240202 | 0.810174 | NA |
| CFH | 54.40414 | -0.31781 | 0.475938 | -0.66776 | 0.50429 | 0.653944 |
| STPG1 | 662.8721 | -0.28641 | 0.163665 | -1.74996 | 0.080125 | 0.191663 |
| NIPAL3 | 3455.799 | -0.17816 | 0.163925 | -1.08684 | 0.277106 | 0.443033 |
| AK2 | 4613.099 | 0.026088 | 0.129094 | 0.202086 | 0.83985 | 0.9048 |
| KDM1A | 1777.252 | 0.040052 | 0.151809 | 0.263832 | 0.791909 | 0.87396 |
| ZNF195 | 186.6874 | 0.728132 | 0.22437 | 3.245229 | 0.001174 | 0.011498 |
| ABCC8 | 0.268266 | -2.15396 | 4.038351 | -0.53338 | 0.593773 | NA |
| TMEM132A | 1697.149 | -0.30891 | 0.211333 | -1.4617 | 0.143824 | 0.285467 |
| ST7L | 554.4035 | -0.10197 | 0.163579 | -0.62335 | 0.533058 | 0.677164 |
| PAX6 | 13.66367 | 0.773468 | 0.56264 | 1.374713 | 0.169221 | NA |
| DNAJC11 | 1467.713 | -0.37808 | 0.143247 | -2.63933 | 0.008307 | 0.04157 |
| FMO3 | 0.326536 | 0.037056 | 3.842554 | 0.009644 | 0.992306 | NA |
| E2F2 | 77.92322 | -0.4278 | 0.417172 | -1.02547 | 0.305139 | 0.472372 |
| CAMK1G | 4.321587 | -0.93415 | 1.004304 | -0.93015 | 0.352295 | NA |
| CDK11A | 47.94832 | 0.269555 | 0.312111 | 0.863652 | 0.387779 | 0.553708 |
| NADK | 2883.235 | -0.1923 | 0.132937 | -1.44658 | 0.148016 | 0.291096 |
| CSDE1 | 12047.16 | 0.378952 | 0.170456 | 2.223158 | 0.026205 | 0.090873 |
| MASP2 | 14.17974 | 0.63773 | 0.591921 | 1.077391 | 0.281306 | NA |
| FAM76A | 215.0253 | -0.15573 | 0.204802 | -0.76041 | 0.44701 | 0.606718 |
| TRAF3IP3 | 39.61909 | -0.03943 | 0.358046 | -0.11013 | 0.912307 | NA |
| SPRTN | 400.3169 | 0.450794 | 0.162236 | 2.778633 | 0.005459 | 0.031461 |
| METTL13 | 1851.873 | -0.21536 | 0.142064 | -1.51593 | 0.129538 | 0.266009 |
| SCMH1 | 1245.091 | -0.46551 | 0.1894 | -2.45782 | 0.013978 | 0.05912 |
| FMO1 | 1.557745 | -0.52689 | 1.57873 | -0.33375 | 0.738572 | NA |
| TCEB3 | 2088.541 | -0.02498 | 0.127722 | -0.19558 | 0.844942 | 0.907647 |
| LYPLA2 | 2793.234 | -0.09877 | 0.14562 | -0.67827 | 0.497602 | 0.648606 |
| CLCN6 | 654.699 | -0.16359 | 0.172828 | -0.94653 | 0.343878 | 0.512081 |
| SYT7 | 1.965572 | -1.24964 | 1.632783 | -0.76534 | 0.444067 | NA |
| PIK3C2A | 1168.437 | 0.995101 | 0.271979 | 3.65874 | 2.53E-04 | 0.004485 |
| CD6 | 4.779577 | -0.62493 | 1.242658 | -0.5029 | 0.615037 | NA |
| MTMR11 | 183.345 | -0.23962 | 0.321637 | -0.745 | 0.456274 | 0.614817 |
| COX15 | 1441.797 | -0.38662 | 0.136744 | -2.82734 | 0.004694 | 0.028303 |
| ZMYND11 | 814.2462 | 0.584353 | 0.221495 | 2.638218 | 0.008334 | 0.041654 |
| CLCA1 | 0.516164 | -1.70268 | 2.844782 | -0.59853 | 0.549488 | NA |
| ATP1A2 | 0.593018 | -0.10389 | 3.151334 | -0.03297 | 0.973702 | NA |
| PRDM11 | 100.1727 | -0.6384 | 0.260548 | -2.45022 | 0.014277 | 0.060039 |
| SYT13 | 0.163957 | 0.424913 | 4.079006 | 0.104171 | 0.917034 | NA |
| ZRANB1 | 878.1605 | 0.203755 | 0.168528 | 1.209027 | 0.226653 | 0.388178 |
| NCDN | 2430.419 | -0.34376 | 0.162625 | -2.11382 | 0.034531 | 0.108908 |
| RUNX3 | 3496.193 | -0.20914 | 0.196328 | -1.06527 | 0.286755 | 0.453083 |
| OSBPL5 | 908.2823 | -0.49119 | 0.156566 | -3.13729 | 0.001705 | 0.014606 |
| ZDHHC6 | 701.0937 | 0.484317 | 0.152595 | 3.173874 | 0.001504 | 0.013592 |
| RNH1 | 7172.302 | -0.25439 | 0.12876 | -1.97567 | 0.048193 | 0.135481 |

| | | | | | | |
|---|---|---|---|---|---|---|
| GLRX2 | 477.1498 | 0.615823 | 0.176029 | 3.498412 | 4.68E-04 | 0.006535 |
| ABCC2 | 2839.359 | -0.1978 | 0.196831 | -1.00493 | 0.314933 | 0.482368 |
| PLEKHO1 | 959.474 | 0.338251 | 0.222719 | 1.518737 | 0.128829 | 0.265134 |
| GCLM | 688.7395 | 0.648968 | 0.217749 | 2.980351 | 0.002879 | 0.020562 |
| DEPDC1 | 327.5054 | 0.806586 | 0.173237 | 4.65597 | 3.22E-06 | 3.27E-04 |
| NR1H3 | 357.005 | 0.034052 | 0.190786 | 0.178481 | 0.858345 | 0.91615 |
| KPNA6 | 2222.17 | -0.3888 | 0.15266 | -2.54686 | 0.01087 | 0.049677 |
| VIM | 93043.95 | -0.49073 | 0.140081 | -3.50321 | 4.60E-04 | 0.006449 |
| FAS | 255.552 | 0.394722 | 0.222529 | 1.773802 | 0.076096 | 0.184822 |
| CD44 | 25786.22 | -0.3061 | 0.1156 | -2.64795 | 0.008098 | 0.040922 |
| SLAMF7 | 0.489058 | -0.58569 | 3.427543 | -0.17088 | 0.864321 | NA |
| INSRR | 0.163957 | 0.424913 | 4.079006 | 0.104171 | 0.917034 | NA |
| SH2D2A | 2.000058 | 1.08437 | 1.455747 | 0.744889 | 0.456339 | NA |
| TNFRSF1B | 2.573727 | 1.598062 | 1.375703 | 1.161633 | 0.245385 | NA |
| NUP160 | 1136.715 | -0.00832 | 0.143872 | -0.05786 | 0.953862 | 0.974786 |
| SARS | 4565.998 | -0.09837 | 0.131979 | -0.74533 | 0.456075 | 0.614797 |
| LRRC7 | 48.25972 | 1.114472 | 0.456518 | 2.441245 | 0.014637 | 0.060862 |
| MYOC | 0.3441 | -0.0166 | 3.769682 | -0.0044 | 0.996486 | NA |
| VCL | 4816.629 | -0.23763 | 0.164616 | -1.44355 | 0.148866 | 0.292187 |
| ADSS | 1134.43 | 0.491205 | 0.165861 | 2.961541 | 0.003061 | 0.021524 |
| ZZZ3 | 800.4392 | 0.593776 | 0.236126 | 2.514664 | 0.011915 | 0.053101 |
| FBXO42 | 713.7234 | -0.23557 | 0.140342 | -1.67856 | 0.093237 | 0.213137 |
| PQLC2 | 916.7455 | -0.19091 | 0.144782 | -1.31862 | 0.187296 | 0.341474 |
| THAP3 | 495.2832 | -0.15492 | 0.184352 | -0.84036 | 0.400705 | 0.565475 |
| AIFM2 | 1841.104 | -0.12457 | 0.159004 | -0.78345 | 0.433363 | 0.594551 |
| USH2A | 457.3924 | -0.35188 | 0.175952 | -1.99987 | 0.045515 | 0.13062 |
| TRIT1 | 274.5782 | 0.09941 | 0.206823 | 0.480651 | 0.630765 | 0.753088 |
| ADRB1 | 0.163957 | 0.424913 | 4.079006 | 0.104171 | 0.917034 | NA |
| WDR37 | 1064.755 | -0.17216 | 0.128344 | -1.34141 | 0.179788 | 0.332292 |
| TPR | 947.0658 | 0.888011 | 1.011703 | 0.877739 | 0.380086 | 0.546414 |
| VPS13D | 1662.131 | -0.23389 | 0.146326 | -1.59839 | 0.109957 | 0.237569 |
| CELF2 | 4895.198 | -0.24309 | 0.21502 | -1.13056 | 0.258241 | 0.422501 |
| COL9A2 | 19.46962 | 0.242193 | 0.479963 | 0.504607 | 0.613835 | NA |
| H6PD | 1164.825 | -0.88906 | 0.175159 | -5.07575 | 3.86E-07 | 8.53E-05 |
| VAMP3 | 5593.588 | -0.03889 | 0.136659 | -0.28458 | 0.77597 | 0.861664 |
| PER3 | 326.8472 | -0.20288 | 0.194285 | -1.04424 | 0.296376 | 0.463649 |
| UTS2 | 2.736105 | 1.342371 | 1.508106 | 0.890103 | 0.37341 | NA |
| TNFRSF9 | 0.260158 | 1.240347 | 4.047124 | 0.306476 | 0.759242 | NA |
| RCN1 | 1341.067 | 0.307054 | 0.140398 | 2.187024 | 0.028741 | 0.096331 |
| DKK3 | 16464.73 | 0.021394 | 0.188654 | 0.113403 | 0.909711 | 0.947766 |
| PTGER3 | 1.103647 | 3.314505 | 2.124646 | 1.560027 | 0.118754 | NA |
| FAM160A2 | 473.0779 | -0.20485 | 0.162332 | -1.2619 | 0.206985 | 0.36412 |
| SIKE1 | 348.5181 | 0.577042 | 0.2429 | 2.375635 | 0.017519 | 0.069159 |
| RRP12 | 819.0614 | -0.46504 | 0.17657 | -2.63373 | 0.008445 | 0.042031 |
| TTC17 | 1060.527 | 0.259267 | 0.155488 | 1.667447 | 0.095426 | 0.216181 |
| ALX4 | 0.096202 | 0.408546 | 4.079639 | 0.100143 | 0.920231 | NA |
| AKR7A2 | 1941.63 | -0.4203 | 0.1579 | -2.66181 | 0.007772 | 0.039835 |

| | | | | | | |
|---|---|---|---|---|---|---|
| MRTO4 | 1313.935 | -0.00755 | 0.206454 | -0.03655 | 0.970842 | 0.983955 |
| KCNQ1 | 0.134133 | -1.48909 | 4.077584 | -0.36519 | 0.71497 | NA |
| TRAPPC3 | 2304.898 | 0.008508 | 0.145913 | 0.058311 | 0.953501 | 0.974629 |
| THRAP3 | 3476.593 | 0.022346 | 0.166376 | 0.134311 | 0.893157 | 0.937439 |
| ARID4B | 314.0267 | 0.924091 | 0.807418 | 1.144501 | 0.252416 | 0.416477 |
| OPN3 | 270.9776 | 0.303414 | 0.210233 | 1.443226 | 0.148957 | 0.292317 |
| SDCCAG8 | 408.9498 | 0.695122 | 0.191736 | 3.625419 | 2.88E-04 | 0.00481 |
| HHAT | 470.8397 | 0.004093 | 0.175854 | 0.023276 | 0.98143 | 0.990173 |
| KIF1B | 1527.158 | -0.1194 | 0.186147 | -0.6414 | 0.521261 | 0.668146 |
| SZRD1 | 6369.397 | -0.27431 | 0.171194 | -1.60236 | 0.109077 | 0.236227 |
| MCOLN3 | 1058.49 | 0.645846 | 0.209111 | 3.088538 | 0.002011 | 0.016255 |
| MRPL43 | 1829.402 | -0.07785 | 0.123007 | -0.63286 | 0.526826 | 0.672284 |
| SOAT1 | 4455.221 | 0.034626 | 0.171734 | 0.201624 | 0.840211 | 0.905024 |
| MSH4 | 1.063143 | 1.12382 | 1.855095 | 0.605802 | 0.544646 | NA |
| GDI2 | 7412.943 | 0.066329 | 0.141354 | 0.469242 | 0.638897 | 0.759385 |
| PITHD1 | 1632.523 | -0.18812 | 0.147275 | -1.27737 | 0.201472 | 0.357821 |
| LAMC2 | 4.333408 | 0.145177 | 1.059986 | 0.136962 | 0.891061 | NA |
| CROCC | 586.656 | -0.32207 | 0.262729 | -1.22586 | 0.220251 | 0.379959 |
| ATP2B4 | 3513.382 | -0.29085 | 0.165205 | -1.76054 | 0.078316 | 0.188553 |
| ZC3H11A | 1870.844 | 0.542153 | 0.20725 | 2.615945 | 0.008898 | 0.043752 |
| YIPF1 | 1203.779 | -0.13526 | 0.148804 | -0.90898 | 0.363362 | 0.53062 |
| NDC1 | 720.7316 | 0.11093 | 0.178956 | 0.619875 | 0.53534 | 0.678475 |
| ALDH18A1 | 2538.639 | -0.48984 | 0.124173 | -3.9448 | 7.99E-05 | 0.002212 |
| TARBP1 | 696.1703 | 0.096819 | 0.193465 | 0.500448 | 0.61676 | 0.742224 |
| PSD | 53.09778 | -0.33775 | 0.367055 | -0.92017 | 0.357484 | 0.525021 |
| CCAR1 | 656.3063 | 0.963097 | 0.836269 | 1.151659 | 0.249461 | 0.413328 |
| PIGV | 546.0879 | -0.38697 | 0.156097 | -2.47904 | 0.013174 | 0.056978 |
| PTPRU | 1997.059 | -0.36618 | 0.199006 | -1.84003 | 0.065764 | 0.167212 |
| SNRNP40 | 974.2023 | -0.10176 | 0.184352 | -0.55198 | 0.580965 | 0.714677 |
| COL11A1 | 2257.632 | 0.596693 | 0.542748 | 1.099392 | 0.271597 | 0.436818 |
| QSER1 | 598.0205 | -0.0475 | 0.331235 | -0.1434 | 0.885978 | 0.932547 |
| WAPAL | 1012.368 | 0.599436 | 0.207549 | 2.888161 | 0.003875 | 0.024953 |
| TSPAN32 | 2.976098 | -1.89976 | 1.351807 | -1.40535 | 0.159918 | NA |
| DDX20 | 707.1869 | 0.127934 | 0.166936 | 0.766366 | 0.443458 | 0.603336 |
| CHI3L2 | 3.319069 | -1.01124 | 1.210886 | -0.83513 | 0.403646 | NA |
| GNAI3 | 1712.356 | 0.349094 | 0.171195 | 2.039155 | 0.041435 | 0.123384 |
| OAT | 6278.789 | 0.364821 | 0.161073 | 2.264938 | 0.023516 | 0.084459 |
| WDR3 | 1423.294 | 0.110149 | 0.124192 | 0.886928 | 0.375118 | 0.541491 |
| PKN2 | 747.526 | 0.788338 | 0.31041 | 2.539664 | 0.011096 | 0.050462 |
| MCM10 | 152.2167 | 0.105206 | 0.298927 | 0.351946 | 0.724879 | 0.82527 |
| SPEN | 1106.019 | -0.18268 | 0.254605 | -0.71752 | 0.473053 | 0.628776 |
| TMEM206 | 855.7561 | 0.146665 | 0.15306 | 0.958224 | 0.33795 | 0.506818 |
| SLK | 943.6148 | 0.741753 | 0.89564 | 0.828183 | 0.407567 | 0.571292 |
| COL17A1 | 0.151697 | -1.546 | 4.075469 | -0.37934 | 0.704433 | NA |
| GSTO2 | 96.34809 | -0.11078 | 0.262788 | -0.42156 | 0.673345 | 0.786221 |
| SEC61A2 | 220.2919 | 0.05751 | 0.219541 | 0.261956 | 0.793355 | 0.874742 |
| PRKCQ | 0.83597 | 1.639551 | 2.453777 | 0.668175 | 0.504022 | NA |

| | | | | | | |
|---|---|---|---|---|---|---|
| FAM107B | 368.8103 | 0.814626 | 0.197164 | 4.131714 | 3.60E-05 | 0.001362 |
| YBX1 | 25103.54 | 0.112837 | 0.149106 | 0.756757 | 0.449196 | 0.60872 |
| PPP2R5A | 1397.329 | 0.137218 | 0.152141 | 0.901917 | 0.367101 | 0.534248 |
| TIE1 | 0.326333 | 1.533889 | 3.826878 | 0.40082 | 0.688553 | NA |
| KDM4A | 1436.943 | -0.43176 | 0.15399 | -2.80382 | 0.00505 | 0.029845 |
| NFYC | 1139.06 | -0.2872 | 0.133708 | -2.14795 | 0.031718 | 0.103187 |
| ZMYND12 | 102.8834 | -0.19662 | 0.229375 | -0.85721 | 0.391331 | 0.556629 |
| ASPM | 453.4251 | 0.93395 | 0.695307 | 1.343219 | 0.179201 | 0.331683 |
| CD84 | 0.747876 | -0.62091 | 2.562446 | -0.24231 | 0.80854 | NA |
| ELOVL1 | 3658.18 | -0.18013 | 0.157585 | -1.14304 | 0.253023 | 0.417247 |
| SPI1 | 1.943774 | -0.14746 | 1.446255 | -0.10196 | 0.918789 | NA |
| MPPED2 | 223.4521 | -0.38492 | 0.218127 | -1.76464 | 0.077625 | 0.187485 |
| FGFR2 | 0.654449 | 1.296029 | 2.63083 | 0.492631 | 0.622273 | NA |
| LRRC40 | 334.3283 | 0.848453 | 0.255741 | 3.317629 | 9.08E-04 | 0.009758 |
| PFKP | 8621.362 | -0.22981 | 0.150517 | -1.52679 | 0.126812 | 0.26232 |
| IDI1 | 3672.963 | 0.9749 | 0.226845 | 4.297644 | 1.73E-05 | 8.54E-04 |
| KLF6 | 5334.111 | 0.220934 | 0.152049 | 1.453052 | 0.146209 | 0.288741 |
| EVI5 | 649.1135 | 0.855278 | 0.332242 | 2.574266 | 0.010045 | 0.047332 |
| DNTTIP2 | 676.3698 | 1.118141 | 0.309529 | 3.612393 | 3.03E-04 | 0.004988 |
| RRP15 | 393.6701 | 0.639324 | 0.244946 | 2.610064 | 0.009053 | 0.044218 |
| PRKCZ | 160.6338 | 0.347401 | 0.298681 | 1.163116 | 0.244782 | 0.40757 |
| IARS2 | 3602.44 | 0.217362 | 0.135393 | 1.605413 | 0.108403 | 0.235264 |
| INPP5A | 809.6296 | 0.009972 | 0.172802 | 0.057705 | 0.953983 | 0.974786 |
| NUP133 | 1157.12 | 0.036235 | 0.154234 | 0.234936 | 0.814258 | 0.889353 |
| NUCKS1 | 3620.936 | 0.25145 | 0.173951 | 1.445519 | 0.148312 | 0.291534 |
| KCNAB2 | 19150.65 | -0.60118 | 0.189136 | -3.17854 | 0.00148 | 0.013453 |
| DRD4 | 1.208159 | 0.798263 | 2.146876 | 0.371826 | 0.710023 | NA |
| TGFBR3 | 664.1667 | 0.054336 | 0.224951 | 0.241548 | 0.809131 | 0.885851 |
| KIAA1107 | 25.91016 | 0.49203 | 0.504162 | 0.975936 | 0.329096 | NA |
| HES2 | 8.939077 | -0.10394 | 0.683563 | -0.15205 | 0.879148 | NA |
| SCT | 0.796689 | -1.1728 | 2.541471 | -0.46147 | 0.644464 | NA |
| PHRF1 | 1605.969 | -0.48244 | 0.153683 | -3.13921 | 0.001694 | 0.014563 |
| NUCB2 | 501.5252 | 1.130565 | 0.29258 | 3.864122 | 1.11E-04 | 0.002754 |
| TESK2 | 800.7611 | -0.23612 | 0.254203 | -0.92885 | 0.352968 | 0.521035 |
| EIF2B3 | 790.5521 | 0.033199 | 0.14014 | 0.236898 | 0.812736 | 0.888591 |
| CDC42 | 4393.853 | -0.08167 | 0.145281 | -0.56218 | 0.573997 | 0.709491 |
| EPHA8 | 0.314073 | -0.57308 | 3.898934 | -0.14698 | 0.883146 | NA |
| TRPM5 | 0.315653 | -0.57305 | 3.892639 | -0.14721 | 0.882963 | NA |
| MS4A12 | 0.247898 | -0.5901 | 4.074633 | -0.14482 | 0.884851 | NA |
| UBE2D1 | 393.2383 | 0.468807 | 0.207081 | 2.263888 | 0.023581 | 0.084611 |
| RHOBTB1 | 906.8142 | -0.20296 | 0.207583 | -0.97771 | 0.328216 | 0.496427 |
| ASAH2C | 0.575812 | 0.936922 | 2.776634 | 0.337431 | 0.735792 | NA |
| FCGR2B | 55.72714 | 0.043171 | 0.394984 | 0.109298 | 0.912966 | 0.94937 |
| MRVI1 | 1.961428 | -1.1534 | 1.481335 | -0.77862 | 0.436202 | NA |
| CD5L | 0.652666 | 2.537288 | 2.975981 | 0.852589 | 0.393887 | NA |
| PTGS2 | 292.9033 | 1.823113 | 0.766808 | 2.377535 | 0.017429 | 0.069026 |
| TSG101 | 2135.459 | 0.064754 | 0.14507 | 0.446363 | 0.655335 | 0.772419 |

| | | | | | | |
|---|---|---|---|---|---|---|
| ENO1 | 35010.51 | -0.22869 | 0.124924 | -1.83065 | 0.067153 | 0.169494 |
| ARHGEF10I | 1662.029 | -0.27138 | 0.15941 | -1.70239 | 0.088683 | 0.205499 |
| TACR2 | 41.02514 | -0.28491 | 0.385221 | -0.7396 | 0.459543 | NA |
| EIF4G3 | 2068.28 | 0.30423 | 0.205524 | 1.480264 | 0.138803 | 0.278663 |
| WNT8B | 0.424115 | 1.932531 | 3.45462 | 0.559405 | 0.575885 | NA |
| RASAL2 | 484.6036 | -0.09067 | 0.205175 | -0.44191 | 0.658553 | 0.77483 |
| ZNF37A | 334.1565 | 0.847096 | 0.403166 | 2.10111 | 0.035631 | 0.111248 |
| SEC31B | 42.4341 | 0.459703 | 0.364126 | 1.262482 | 0.206775 | NA |
| PAX2 | 0.096202 | 0.408546 | 4.079639 | 0.100143 | 0.920231 | NA |
| KIFAP3 | 978.538 | 0.410856 | 0.165894 | 2.476619 | 0.013263 | 0.057239 |
| FMO4 | 105.1657 | -0.19638 | 0.261835 | -0.75 | 0.453255 | 0.61221 |
| KLHL20 | 625.8116 | 0.397718 | 0.191218 | 2.079922 | 0.037533 | 0.115123 |
| PLXNA2 | 128.4201 | -0.3063 | 0.259306 | -1.18123 | 0.23751 | 0.400358 |
| NT5C2 | 1116.564 | 0.039242 | 0.130041 | 0.301767 | 0.76283 | 0.852664 |
| RAP1GAP | 2225.06 | -0.09315 | 0.146621 | -0.63533 | 0.525215 | 0.670944 |
| TM9SF3 | 3436.769 | 0.318528 | 0.17283 | 1.843011 | 0.065327 | 0.166729 |
| NFKB2 | 1642.585 | -0.45035 | 0.15318 | -2.93998 | 0.003282 | 0.022409 |
| UBE2T | 533.606 | 0.158202 | 0.179063 | 0.883501 | 0.376966 | 0.543599 |
| PPP1R12B | 494.5571 | -0.09806 | 0.156366 | -0.62714 | 0.530571 | 0.675246 |
| USP33 | 1156.483 | 0.725096 | 0.238686 | 3.037866 | 0.002383 | 0.018106 |
| SPAG6 | 1.306087 | 0.163842 | 1.952563 | 0.083911 | 0.933127 | NA |
| APBB1IP | 0.260158 | 1.240347 | 4.047124 | 0.306476 | 0.759242 | NA |
| ACTN2 | 6.70751 | 0.621525 | 0.844327 | 0.736118 | 0.461659 | NA |
| CAPZB | 7371.348 | -0.15647 | 0.116962 | -1.33778 | 0.180968 | 0.333581 |
| GPR137B | 5863.595 | -0.12399 | 0.13931 | -0.89001 | 0.373461 | 0.539792 |
| ITGA8 | 0.096202 | 0.408546 | 4.079639 | 0.100143 | 0.920231 | NA |
| NEBL | 0.654449 | 1.296029 | 2.63083 | 0.492631 | 0.622273 | NA |
| GNB1 | 17847.98 | -0.19198 | 0.118325 | -1.62251 | 0.104695 | 0.2301 |
| MLLT10 | 847.0715 | 0.442388 | 0.152976 | 2.891875 | 0.00383 | 0.024734 |
| NRD1 | 2582.849 | 0.338769 | 0.143102 | 2.367328 | 0.017917 | 0.070101 |
| SDF4 | 8580.918 | -0.16442 | 0.135798 | -1.21078 | 0.225982 | 0.387196 |
| TP73 | 4.356263 | 1.077093 | 0.988108 | 1.090056 | 0.275688 | NA |
| TOLLIP | 5312.586 | -0.43938 | 0.131314 | -3.34602 | 8.20E-04 | 0.009177 |
| MKNK1 | 632.0065 | -0.03725 | 0.144279 | -0.25818 | 0.796265 | 0.87685 |
| SAR1A | 3237.646 | 0.025863 | 0.156179 | 0.165601 | 0.868471 | 0.921978 |
| CDC14A | 130.9005 | 0.559694 | 0.251355 | 2.226707 | 0.025967 | 0.090324 |
| PGM1 | 3202.355 | -0.22494 | 0.221807 | -1.01414 | 0.310515 | 0.47801 |
| CROCCP3 | 35.38335 | 0.26145 | 0.506027 | 0.516672 | 0.605385 | NA |
| RSBN1 | 320.9092 | 0.287107 | 0.193223 | 1.48588 | 0.137311 | 0.276526 |
| MAGI3 | 184.5694 | 0.463285 | 0.27581 | 1.67973 | 0.09301 | 0.212781 |
| PKP1 | 48.9226 | -1.90187 | 0.403051 | -4.71868 | 2.37E-06 | 2.68E-04 |
| JMJD4 | 224.7279 | -0.16271 | 0.180026 | -0.9038 | 0.366103 | 0.533342 |
| DUSP12 | 377.5518 | 0.034161 | 0.212807 | 0.160526 | 0.872467 | 0.924068 |
| HSPB11 | 450.922 | 0.603552 | 0.166302 | 3.629246 | 2.84E-04 | 0.004766 |
| IL12RB2 | 1068.395 | -0.20833 | 0.244297 | -0.85277 | 0.393786 | 0.558547 |
| KCNK2 | 18.05907 | -1.56756 | 0.660287 | -2.37405 | 0.017594 | NA |
| SERTAD4 | 455.6898 | 1.183559 | 0.382655 | 3.09302 | 0.001981 | 0.016077 |

| | | | | | | |
|---|---|---|---|---|---|---|
| TRAF5 | 250.606 | 0.361221 | 0.211697 | 1.70631 | 0.08795 | 0.204161 |
| PLOD1 | 7737.079 | -0.3603 | 0.148458 | -2.42694 | 0.015227 | 0.062744 |
| SMAP2 | 1358.956 | -0.41913 | 0.13393 | -3.12946 | 0.001751 | 0.014833 |
| PPIE | 1044.924 | -0.02229 | 0.157114 | -0.14187 | 0.887186 | 0.933443 |
| ZMPSTE24 | 1848.631 | 0.270949 | 0.15604 | 1.736413 | 0.082491 | 0.195273 |
| EIF3I | 6674.858 | -0.09626 | 0.133935 | -0.71868 | 0.472337 | 0.628176 |
| NKAIN1 | 3.935479 | -0.24649 | 1.124792 | -0.21914 | 0.826537 | NA |
| COL16A1 | 565.8076 | 0.448116 | 0.254587 | 1.760168 | 0.078379 | 0.188591 |
| TXLNA | 3271.87 | -0.52811 | 0.145704 | -3.62454 | 2.89E-04 | 0.004811 |
| CD59 | 28677.82 | -0.02194 | 0.134494 | -0.16312 | 0.870424 | 0.923107 |
| CD82 | 1756.8 | -0.73499 | 0.210811 | -3.48649 | 4.89E-04 | 0.0067 |
| WDR47 | 924.7529 | 0.291531 | 0.146632 | 1.988177 | 0.046792 | 0.133122 |
| OVGP1 | 13.7793 | 0.381369 | 0.57305 | 0.665508 | 0.505726 | NA |
| SLC25A24 | 1008.879 | 0.630747 | 0.207321 | 3.042364 | 0.002347 | 0.017941 |
| IGSF9 | 0.476449 | 0.524262 | 3.402674 | 0.154073 | 0.877552 | NA |
| TTC39A | 9.225306 | -0.96156 | 0.677468 | -1.41935 | 0.155798 | NA |
| EPS15 | 3582.223 | -0.25509 | 0.19748 | -1.29173 | 0.19645 | 0.351906 |
| ORC1 | 172.6997 | -0.19423 | 0.230349 | -0.8432 | 0.399116 | 0.563814 |
| POMGNT1 | 6815.678 | -0.26643 | 0.1236 | -2.15559 | 0.031116 | 0.101631 |
| RAD54L | 152.7992 | -0.17797 | 0.276955 | -0.64258 | 0.520494 | 0.667581 |
| MAST2 | 2273.184 | -0.3698 | 0.158971 | -2.32624 | 0.020006 | 0.075797 |
| FOLH1 | 0.163957 | 0.424913 | 4.079006 | 0.104171 | 0.917034 | NA |
| SEPHS1 | 1099.379 | -0.0651 | 0.155978 | -0.41734 | 0.676427 | 0.788478 |
| ERO1LB | 198.2328 | 0.974725 | 0.230449 | 4.229671 | 2.34E-05 | 0.001011 |
| ALG6 | 427.1553 | -0.07563 | 0.214248 | -0.35301 | 0.72408 | 0.824799 |
| ASAP3 | 319.2639 | -0.2766 | 0.172741 | -1.60124 | 0.109325 | 0.236504 |
| GANAB | 11806.65 | -0.58978 | 0.144425 | -4.08364 | 4.43E-05 | 0.001546 |
| DHX32 | 1021.155 | -0.01856 | 0.158191 | -0.11733 | 0.906601 | 0.945876 |
| SLC9A1 | 1186.074 | -0.79356 | 0.159772 | -4.96681 | 6.81E-07 | 1.26E-04 |
| RGS1 | 39.77479 | 1.253734 | 0.36076 | 3.475255 | 5.10E-04 | NA |
| NUDC | 3187.382 | -0.18533 | 0.153999 | -1.20348 | 0.228791 | 0.390543 |
| MUL1 | 1487.693 | -0.30461 | 0.132337 | -2.30181 | 0.021346 | 0.079153 |
| PABPC4 | 3521.088 | -0.39356 | 0.139263 | -2.82603 | 0.004713 | 0.028375 |
| USP48 | 1962.807 | 0.217472 | 0.220651 | 0.985593 | 0.324333 | 0.492439 |
| FH | 2570.868 | 0.062607 | 0.130739 | 0.478868 | 0.632033 | 0.754253 |
| SLC17A6 | 7.515663 | 1.277133 | 0.806202 | 1.584135 | 0.113163 | NA |
| TBX15 | 0.096202 | 0.408546 | 4.079639 | 0.100143 | 0.920231 | NA |
| PHGDH | 1767.614 | -0.60728 | 0.145051 | -4.18669 | 2.83E-05 | 0.001145 |
| AGO1 | 1531.536 | -0.38434 | 0.194135 | -1.97974 | 0.047733 | 0.134702 |
| TEKT2 | 0.49029 | 2.123092 | 3.304282 | 0.642527 | 0.520531 | NA |
| CLSPN | 131.4361 | 0.615424 | 0.313644 | 1.962173 | 0.049742 | 0.138006 |
| TGFB2 | 88.82036 | 0.151971 | 0.25755 | 0.590064 | 0.555147 | 0.695073 |
| GPATCH2 | 215.7629 | 0.800599 | 0.229451 | 3.489189 | 4.84E-04 | 0.006671 |
| FMO2 | 0.134133 | -1.48909 | 4.077584 | -0.36519 | 0.71497 | NA |
| SUCO | 535.6015 | 0.577003 | 0.241898 | 2.385312 | 0.017065 | 0.067932 |
| PDE6C | 1.148462 | 1.182044 | 1.944788 | 0.607801 | 0.54332 | NA |
| CWF19L1 | 525.8632 | -0.20481 | 0.173133 | -1.18294 | 0.236833 | 0.399729 |

| | | | | | | |
|---|---|---|---|---|---|---|
| SEMA4G | 181.645 | -0.24722 | 0.203518 | -1.21474 | 0.224467 | 0.385493 |
| BTAF1 | 823.0324 | 0.456537 | 0.175244 | 2.605156 | 0.009183 | 0.044545 |
| IKZF5 | 358.7583 | 0.380481 | 0.178413 | 2.132584 | 0.032959 | 0.105731 |
| BLNK | 0.883146 | 1.112834 | 2.554178 | 0.435692 | 0.663061 | NA |
| TLL2 | 3.30337 | 0.168173 | 1.070454 | 0.157105 | 0.875162 | NA |
| CYP26A1 | 2.243002 | 0.506383 | 1.305121 | 0.387997 | 0.698018 | NA |
| SORBS1 | 3317.642 | -0.01534 | 0.152706 | -0.10047 | 0.919971 | 0.953678 |
| CRTAC1 | 17.30241 | -0.50909 | 0.513878 | -0.99069 | 0.321837 | NA |
| BAMBI | 3560.307 | -0.0449 | 0.143384 | -0.31314 | 0.754175 | 0.846661 |
| WAC | 2483.109 | 0.329821 | 0.143192 | 2.303354 | 0.021259 | 0.078903 |
| CREM | 1315.477 | 0.499129 | 0.182886 | 2.729177 | 0.006349 | 0.034646 |
| SIRT1 | 382.1952 | 0.467826 | 0.197125 | 2.373248 | 0.017632 | 0.069469 |
| HNRNPH3 | 1739.952 | 0.289981 | 0.143321 | 2.0233 | 0.043042 | 0.126367 |
| ACOT7 | 6830.851 | -0.20819 | 0.125845 | -1.65434 | 0.098059 | 0.220378 |
| SH3GLB1 | 2595.973 | 0.454865 | 0.158424 | 2.87119 | 0.004089 | 0.025773 |
| CDC7 | 229.1692 | 0.737148 | 0.213396 | 3.454373 | 5.52E-04 | 0.007178 |
| SYDE2 | 124.8628 | 1.044066 | 0.247771 | 4.213833 | 2.51E-05 | 0.001062 |
| SCD | 27200.89 | -0.44697 | 0.223126 | -2.0032 | 0.045156 | 0.129997 |
| ABLIM1 | 978.2745 | -0.17161 | 0.220875 | -0.77695 | 0.437186 | 0.597569 |
| RAB18 | 1829.451 | 0.799337 | 0.209547 | 3.814596 | 1.36E-04 | 0.003063 |
| NRP1 | 51.71731 | 0.290298 | 0.429641 | 0.675676 | 0.499246 | 0.649678 |
| HSD17B7P2 | 52.8471 | 0.886924 | 0.314877 | 2.816733 | 0.004851 | 0.028931 |
| PRTFDC1 | 1.44374 | -0.31701 | 1.683753 | -0.18828 | 0.85066 | NA |
| PALMD | 156.7238 | -0.05318 | 0.393395 | -0.13519 | 0.892462 | 0.936854 |
| TSPAN15 | 841.4023 | 0.032497 | 0.196648 | 0.165255 | 0.868743 | 0.922011 |
| H2AFY2 | 188.5543 | -0.75843 | 0.27893 | -2.71908 | 0.006546 | 0.035454 |
| FAM21A | 207.7782 | 0.104592 | 0.194505 | 0.537738 | 0.590758 | 0.722451 |
| CDHR5 | 0.162376 | 0.424913 | 4.079006 | 0.104171 | 0.917034 | NA |
| RASSF7 | 316.8127 | 0.397604 | 0.260043 | 1.528992 | 0.126266 | 0.261419 |
| NHP2P1 | 2.164161 | 0.754144 | 1.383147 | 0.545238 | 0.58559 | NA |
| DVL1 | 3109.219 | -0.16786 | 0.177664 | -0.94479 | 0.344764 | 0.512821 |
| PDLIM1 | 2204.781 | 0.177175 | 0.212649 | 0.833179 | 0.404744 | 0.569118 |
| CCNJ | 169.8347 | 0.52343 | 0.234467 | 2.232425 | 0.025587 | 0.089457 |
| GATA3 | 0.096202 | 0.408546 | 4.079639 | 0.100143 | 0.920231 | NA |
| ATRNL1 | 373.7921 | 0.049801 | 0.21419 | 0.232509 | 0.816143 | 0.889858 |
| HPS1 | 7484.871 | -0.34497 | 0.124317 | -2.77493 | 0.005521 | 0.031724 |
| PHYH | 822.9825 | 0.009274 | 0.151209 | 0.061334 | 0.951094 | 0.973008 |
| RASSF4 | 351.0376 | 0.255179 | 0.236037 | 1.081096 | 0.279654 | 0.445605 |
| DNMBP | 476.2675 | -0.47626 | 0.17704 | -2.6901 | 0.007143 | 0.037636 |
| RAB11FIP2 | 707.3659 | 0.972479 | 0.211975 | 4.58771 | 4.48E-06 | 3.99E-04 |
| CXCL12 | 179.0207 | 0.919531 | 0.306232 | 3.002728 | 0.002676 | 0.019678 |
| ERLIN1 | 1467.251 | -0.14601 | 0.13999 | -1.04297 | 0.296962 | 0.464076 |
| EIF3A | 6174.16 | 0.460851 | 0.291465 | 1.581153 | 0.113843 | 0.242954 |
| PKD2L1 | 0.410275 | 0.244283 | 3.100889 | 0.078778 | 0.937209 | NA |
| CUBN | 674.72 | -0.3787 | 0.214078 | -1.76897 | 0.076899 | 0.186314 |
| TRDMT1 | 175.8205 | 0.552311 | 0.281381 | 1.962857 | 0.049663 | 0.137946 |
| GDF10 | 0.163957 | 0.424913 | 4.079006 | 0.104171 | 0.917034 | NA |

| | | | | | | |
|---|---|---|---|---|---|---|
| DDX50 | 807.9426 | 0.5712 | 0.19429 | 2.939932 | 0.003283 | 0.022409 |
| MAPK8 | 347.1366 | 0.56709 | 0.200743 | 2.824955 | 0.004729 | 0.028456 |
| SEC23IP | 1173.948 | 0.204573 | 0.171899 | 1.190079 | 0.234015 | 0.396549 |
| ATE1 | 906.0239 | 0.511026 | 0.196304 | 2.603241 | 0.009235 | 0.044695 |
| NSMCE4A | 428.1949 | 0.3227 | 0.200452 | 1.609864 | 0.107427 | 0.233913 |
| PLEKHA1 | 806.0882 | 0.677838 | 0.183908 | 3.685742 | 2.28E-04 | 0.004196 |
| PALD1 | 0.096202 | 0.408546 | 4.079639 | 0.100143 | 0.920231 | NA |
| UNC5B | 5.034543 | 0.913357 | 0.958711 | 0.952692 | 0.340746 | NA |
| CDH23 | 1.06329 | 0.285492 | 1.942399 | 0.146979 | 0.883149 | NA |
| C10orf54 | 65.82138 | 0.013441 | 0.609006 | 0.02207 | 0.982392 | 0.990412 |
| SPOCK2 | 8.398473 | 0.469623 | 0.724367 | 0.648323 | 0.516776 | NA |
| MICU1 | 1865.723 | -0.19468 | 0.135595 | -1.43574 | 0.151076 | 0.295011 |
| PPP3CB | 1286.488 | 0.05713 | 0.138256 | 0.413217 | 0.679448 | 0.790858 |
| CCSER2 | 1284.089 | 0.614252 | 0.161355 | 3.806846 | 1.41E-04 | 0.003125 |
| BMPR1A | 440.4059 | 0.389566 | 0.192961 | 2.018887 | 0.043499 | 0.127074 |
| MINPP1 | 644.2536 | 0.050085 | 0.153325 | 0.326659 | 0.743926 | 0.83881 |
| ACTA2 | 414.6836 | -0.56257 | 0.208636 | -2.6964 | 0.007009 | 0.037173 |
| LIPA | 13133.16 | 0.162244 | 0.200125 | 0.810711 | 0.417532 | 0.580424 |
| C10orf2 | 760.5592 | -0.08998 | 0.180572 | -0.49828 | 0.618284 | 0.743157 |
| LZTS2 | 3597.062 | -0.27071 | 0.141838 | -1.90859 | 0.056315 | 0.150161 |
| SFXN3 | 4405.225 | -0.32502 | 0.145381 | -2.23567 | 0.025374 | 0.088919 |
| KAZALD1 | 29.28855 | 0.31259 | 0.425432 | 0.73476 | 0.462486 | NA |
| FBXW4 | 947.2473 | -0.23383 | 0.154387 | -1.51455 | 0.129887 | 0.266173 |
| FGF8 | 0.419962 | -2.64111 | 3.429364 | -0.77014 | 0.441214 | NA |
| NPM3 | 608.6067 | -0.11059 | 0.172492 | -0.64113 | 0.521441 | 0.668146 |
| TNKS2 | 2518.445 | 0.359836 | 0.149588 | 2.405515 | 0.01615 | 0.06541 |
| GBF1 | 2955.541 | -0.61194 | 0.154906 | -3.95041 | 7.80E-05 | 0.002183 |
| ARHGAP21 | 1471.884 | 0.319351 | 0.198762 | 1.606702 | 0.10812 | 0.234927 |
| CPEB3 | 47.72626 | 0.317118 | 0.38484 | 0.824026 | 0.409925 | 0.573435 |
| FBXL15 | 701.0305 | -0.0354 | 0.161563 | -0.21912 | 0.826553 | 0.895602 |
| CUEDC2 | 3069.116 | -0.14953 | 0.141101 | -1.05974 | 0.289264 | 0.45559 |
| SUFU | 1130.692 | -0.62405 | 0.146995 | -4.24538 | 2.18E-05 | 9.62E-04 |
| ANKRD26 | 121.8596 | 0.859332 | 1.040276 | 0.826062 | 0.408769 | 0.572402 |
| ACBD5 | 1180.831 | 0.68013 | 0.194595 | 3.495115 | 4.74E-04 | 0.006586 |
| LHPP | 346.7926 | -0.42867 | 0.18394 | -2.3305 | 0.01978 | 0.075109 |
| LARP4B | 1177.14 | -0.03263 | 0.141035 | -0.23138 | 0.817018 | 0.890316 |
| GTPBP4 | 1375.891 | 0.374527 | 0.150927 | 2.481515 | 0.013083 | 0.056665 |
| EDRF1 | 267.5266 | 0.569364 | 0.21186 | 2.687455 | 0.0072 | 0.037853 |
| BCCIP | 668.9777 | 0.822198 | 0.23963 | 3.431112 | 6.01E-04 | 0.007597 |
| MTPAP | 383.953 | 0.247926 | 0.193413 | 1.28185 | 0.199895 | 0.355943 |
| NEURL1 | 13.42221 | 0.348106 | 0.624819 | 0.557131 | 0.577438 | NA |
| SH3PXD2A | 8256.065 | -0.08512 | 0.167527 | -0.50809 | 0.611393 | 0.738281 |
| PITRM1 | 3808.509 | -0.31761 | 0.127786 | -2.48551 | 0.012937 | 0.05628 |
| OBFC1 | 350.4743 | 0.033544 | 0.163051 | 0.205726 | 0.837005 | 0.902391 |
| MAP3K8 | 34.09989 | 0.223912 | 0.388563 | 0.576256 | 0.564442 | NA |
| DKK1 | 1400.923 | 0.60809 | 0.262383 | 2.317564 | 0.020473 | 0.076899 |
| EBF3 | 0.533728 | -1.74213 | 2.820432 | -0.61768 | 0.536785 | NA |

| | | | | | | |
|---|---|---|---|---|---|---|
| GLRX3 | 1433.535 | 0.37059 | 0.164864 | 2.247851 | 0.024586 | 0.086816 |
| SORCS1 | 673.1237 | -0.14342 | 0.168437 | -0.8515 | 0.39449 | 0.559233 |
| FAM208B | 1347.316 | 0.176542 | 0.221166 | 0.798233 | 0.424735 | 0.58672 |
| XPNPEP1 | 3365.562 | -0.29575 | 0.12975 | -2.2794 | 0.022643 | 0.082217 |
| SMC3 | 595.2298 | 0.877652 | 0.796439 | 1.10197 | 0.270475 | 0.435443 |
| SHOC2 | 951.815 | 0.690924 | 0.205125 | 3.36831 | 7.56E-04 | 0.008794 |
| TFAM | 525.7189 | 0.628269 | 0.213776 | 2.938914 | 0.003294 | 0.02243 |
| CCDC6 | 931.3244 | -0.11413 | 0.14941 | -0.76384 | 0.444965 | 0.60481 |
| CUL2 | 969.1345 | 0.665963 | 0.234398 | 2.841165 | 0.004495 | 0.027439 |
| CCNY | 1424.963 | -0.2196 | 0.152824 | -1.43694 | 0.150734 | 0.294487 |
| ZMIZ1 | 2218.713 | -0.77725 | 0.256109 | -3.03482 | 0.002407 | 0.018243 |
| DNAJC12 | 44.57201 | 0.1372 | 0.379698 | 0.361338 | 0.717847 | NA |
| PPIF | 2469.933 | -0.39856 | 0.156612 | -2.54488 | 0.010931 | 0.049887 |
| PBLD | 109.7464 | 0.129801 | 0.277892 | 0.467091 | 0.640435 | 0.760529 |
| TSPAN14 | 3792.086 | -0.34808 | 0.126879 | -2.74341 | 0.006081 | 0.033748 |
| TBC1D12 | 374.7939 | 0.406609 | 0.168622 | 2.411367 | 0.015893 | 0.064533 |
| HTATIP2 | 2462.177 | 0.164757 | 0.169842 | 0.970062 | 0.332016 | 0.500515 |
| CCDC34 | 189.7136 | 0.822355 | 0.361257 | 2.27637 | 0.022824 | 0.08267 |
| ELP4 | 260.3942 | 0.105867 | 0.173076 | 0.611679 | 0.54075 | 0.682777 |
| MTCH2 | 4012.604 | -0.18117 | 0.125696 | -1.44133 | 0.149491 | 0.293079 |
| FNBP4 | 752.2973 | 0.76816 | 0.156459 | 4.909654 | 9.12E-07 | 1.53E-04 |
| P2RX3 | 0.151697 | -1.546 | 4.075469 | -0.37934 | 0.704433 | NA |
| LPXN | 595.4468 | -0.22619 | 0.178756 | -1.26535 | 0.205748 | 0.362805 |
| DTX4 | 742.402 | -0.95424 | 0.214044 | -4.45817 | 8.27E-06 | 5.54E-04 |
| OSBP | 2082.682 | -0.28276 | 0.135327 | -2.08946 | 0.036666 | 0.113473 |
| MS4A6A | 0.163957 | 0.424913 | 4.079006 | 0.104171 | 0.917034 | NA |
| CCDC86 | 1135.239 | -0.14183 | 0.208588 | -0.67996 | 0.49653 | 0.647636 |
| PRPF19 | 2795.056 | -0.25108 | 0.130541 | -1.92335 | 0.054436 | 0.146868 |
| TMEM109 | 4520.339 | -0.28344 | 0.158871 | -1.78409 | 0.074409 | 0.182182 |
| HPX | 2.849643 | 0.101521 | 1.201689 | 0.084482 | 0.932674 | NA |
| TRIM3 | 969.3124 | -0.23964 | 0.15598 | -1.53638 | 0.124446 | 0.259008 |
| RNF141 | 946.641 | 0.40992 | 0.183937 | 2.228596 | 0.025841 | 0.09006 |
| EIF4G2 | 19739.75 | 0.089828 | 0.147924 | 0.607262 | 0.543677 | 0.685451 |
| GALNT18 | 862.0482 | -0.63846 | 0.168193 | -3.79599 | 1.47E-04 | 0.003187 |
| HIPK3 | 1252.228 | 0.462612 | 0.200151 | 2.311312 | 0.020816 | 0.077845 |
| KIAA1549L | 1003.856 | -0.67918 | 0.199684 | -3.40126 | 6.71E-04 | 0.008081 |
| FBXO3 | 762.7446 | 0.49778 | 0.166725 | 2.985634 | 0.00283 | 0.020359 |
| PDHX | 503.5474 | 0.292474 | 0.160621 | 1.820892 | 0.068623 | 0.172248 |
| SLC1A2 | 0.452561 | 2.061505 | 3.349456 | 0.615474 | 0.538241 | NA |
| COMMD9 | 1448.644 | -0.23036 | 0.128869 | -1.78754 | 0.073851 | 0.181188 |
| SLC15A3 | 76.30038 | -0.51531 | 0.436147 | -1.1815 | 0.237402 | 0.400285 |
| CD5 | 0.846649 | 2.927909 | 2.364097 | 1.23849 | 0.215535 | NA |
| ACCS | 167.2434 | 0.245633 | 0.272881 | 0.900148 | 0.368042 | 0.535223 |
| MDK | 123.9906 | -0.69394 | 0.361064 | -1.92194 | 0.054614 | 0.147199 |
| AMBRA1 | 2171.104 | -0.43477 | 0.208727 | -2.08295 | 0.037256 | 0.11457 |
| MADD | 2451.508 | -0.41313 | 0.191238 | -2.1603 | 0.030749 | 0.100783 |
| PTPMT1 | 246.5957 | -0.1533 | 0.197554 | -0.77599 | 0.437754 | 0.597921 |

| | | | | | | |
|---|---|---|---|---|---|---|
| CARS | 1111.167 | 0.052927 | 0.135023 | 0.391986 | 0.695068 | 0.803279 |
| SLC22A18 | 1385.581 | -0.5344 | 0.190759 | -2.80146 | 0.005087 | 0.029945 |
| CD81 | 29879.99 | -0.16079 | 0.172071 | -0.93445 | 0.35007 | 0.518113 |
| CALCA | 0.288605 | 1.443619 | 4.029058 | 0.358302 | 0.720117 | NA |
| SOX6 | 860.2326 | 0.242132 | 0.207486 | 1.166979 | 0.243219 | 0.406164 |
| C11orf58 | 3255.94 | 0.499392 | 0.155785 | 3.205645 | 0.001348 | 0.012589 |
| RPS13 | 8421.119 | 0.171631 | 0.170525 | 1.006489 | 0.31418 | 0.481589 |
| NUP98 | 2596.056 | -0.28251 | 0.142413 | -1.98373 | 0.047286 | 0.133854 |
| HPS5 | 2543.002 | -4.85E-04 | 0.164381 | -0.00295 | 0.997645 | 0.999133 |
| GTF2H1 | 1844.317 | 0.443785 | 0.154859 | 2.865736 | 0.00416 | 0.026068 |
| PTPN5 | 0.096202 | 0.408546 | 4.079639 | 0.100143 | 0.920231 | NA |
| BCL9 | 758.7995 | -0.52102 | 0.175177 | -2.97425 | 0.002937 | 0.02086 |
| PRRX1 | 22.39433 | 1.569443 | 0.583378 | 2.690267 | 0.007139 | NA |
| DHCR24 | 12225.2 | -0.44553 | 0.184196 | -2.41879 | 0.015572 | 0.063581 |
| DNAJC16 | 446.0884 | -0.2288 | 0.166482 | -1.37432 | 0.169342 | 0.319725 |
| MARK1 | 327.6334 | 0.339988 | 0.187205 | 1.81613 | 0.06935 | 0.173413 |
| TNR | 0.601483 | -1.7332 | 2.729494 | -0.63499 | 0.525436 | NA |
| MORN1 | 32.76862 | -0.71208 | 0.373055 | -1.90878 | 0.056291 | NA |
| GPX7 | 177.0585 | -0.10351 | 0.226748 | -0.4565 | 0.648033 | 0.766251 |
| CACYBP | 1499.018 | 0.48465 | 0.152341 | 3.181357 | 0.001466 | 0.013388 |
| SCP2 | 2142.499 | 0.684125 | 0.164019 | 4.171012 | 3.03E-05 | 0.001198 |
| PAPPA2 | 0.722204 | 1.447218 | 2.611582 | 0.554154 | 0.579474 | NA |
| RALGPS2 | 677.5527 | 0.790849 | 0.221587 | 3.569025 | 3.58E-04 | 0.005531 |
| ANGPTL1 | 0.3441 | -0.0166 | 3.769682 | -0.0044 | 0.996486 | NA |
| CEP104 | 1053.116 | -0.04638 | 0.157312 | -0.2948 | 0.768146 | 0.856103 |
| FAM20B | 1490.682 | 0.09515 | 0.178195 | 0.533969 | 0.593363 | 0.723919 |
| TCEANC2 | 421.3588 | 0.090342 | 0.160156 | 0.564084 | 0.572697 | 0.708624 |
| TMEM59 | 8649.736 | 0.21588 | 0.145474 | 1.48398 | 0.137814 | 0.277148 |
| LRRC42 | 788.674 | -0.12703 | 0.153184 | -0.82924 | 0.406968 | 0.570689 |
| WRAP73 | 429.1407 | 0.056879 | 0.156352 | 0.363786 | 0.716018 | 0.818789 |
| MRPL37 | 2904.841 | -0.26536 | 0.148915 | -1.78196 | 0.074755 | 0.182462 |
| ICMT | 6087.934 | -0.23083 | 0.176578 | -1.30724 | 0.191131 | 0.345564 |
| RPL22 | 3833.035 | 0.205844 | 0.181519 | 1.134006 | 0.256792 | 0.42077 |
| CHD5 | 2.037197 | -3.52786 | 2.070223 | -1.7041 | 0.088363 | NA |
| QSOX1 | 5001.716 | -0.44881 | 0.129669 | -3.46117 | 5.38E-04 | 0.007049 |
| STXBP3 | 676.9927 | 0.975911 | 0.275989 | 3.536046 | 4.06E-04 | 0.005981 |
| PHF13 | 720.57 | -0.37239 | 0.190558 | -1.9542 | 0.050677 | 0.139911 |
| ERRFI1 | 151.4258 | 0.296137 | 0.213447 | 1.3874 | 0.16532 | 0.314232 |
| PARK7 | 8646.256 | 0.09645 | 0.141466 | 0.681793 | 0.49537 | 0.646549 |
| KIAA1324 | 7.425292 | 0.206909 | 0.731256 | 0.282951 | 0.777215 | NA |
| OPRD1 | 8.165897 | -0.63668 | 0.767849 | -0.82917 | 0.407008 | NA |
| AMPD2 | 1663.882 | -0.12223 | 0.188273 | -0.64921 | 0.516199 | 0.663725 |
| SRSF4 | 2278.617 | 0.00164 | 0.146764 | 0.011175 | 0.991084 | 0.996125 |
| MECR | 737.5864 | -0.27236 | 0.172001 | -1.58346 | 0.113317 | 0.242093 |
| KCNC4 | 125.224 | -0.29909 | 0.301402 | -0.99233 | 0.321035 | 0.489249 |
| EDEM3 | 947.0091 | 0.748728 | 0.291171 | 2.571435 | 0.010128 | 0.047569 |
| WDR77 | 731.5349 | -0.08403 | 0.143013 | -0.58756 | 0.556825 | 0.696179 |

| | | | | | | |
|---|---|---|---|---|---|---|
| ATP5F1 | 4085.097 | 0.245777 | 0.153661 | 1.599479 | 0.109714 | 0.237131 |
| RAP1A | 1235.035 | 0.529677 | 0.168859 | 3.1368 | 0.001708 | 0.014611 |
| HDAC1 | 4629.776 | 0.033192 | 0.143247 | 0.231709 | 0.816764 | 0.890203 |
| CAPZA1 | 4401.722 | 0.385906 | 0.186619 | 2.067882 | 0.038651 | 0.117459 |
| S100PBP | 446.6626 | -1.60E-04 | 0.165352 | -9.68E-04 | 0.999227 | 0.999472 |
| RNF19B | 1312.647 | -0.57963 | 0.179823 | -3.22332 | 0.001267 | 0.012066 |
| SCAMP3 | 3524.47 | -0.17388 | 0.139284 | -1.24838 | 0.211891 | 0.370332 |
| TRIM62 | 355.9909 | -0.65163 | 0.179203 | -3.63628 | 2.77E-04 | 0.004684 |
| ASH1L | 1080.229 | 0.168565 | 0.195553 | 0.86199 | 0.388693 | 0.554241 |
| DLGAP3 | 1.054044 | 0.320362 | 1.998516 | 0.1603 | 0.872645 | NA |
| SFPQ | 3975.694 | 0.084056 | 0.141271 | 0.595 | 0.551844 | 0.6928 |
| RHOU | 8.364656 | 1.088681 | 1.138959 | 0.955857 | 0.339145 | NA |
| GON4L | 819.6799 | -0.13946 | 0.135736 | -1.02747 | 0.3042 | 0.47141 |
| ARHGEF2 | 3433.122 | -0.45161 | 0.151578 | -2.97935 | 0.002889 | 0.02058 |
| LAMTOR2 | 1546.178 | -0.08166 | 0.13484 | -0.60563 | 0.544759 | 0.686428 |
| MEF2D | 1844.139 | -0.4144 | 0.162768 | -2.54596 | 0.010898 | 0.049765 |
| DOCK7 | 3115.794 | 0.00751 | 0.1923 | 0.039053 | 0.968848 | 0.983028 |
| SRM | 3729.348 | -0.27352 | 0.128657 | -2.12599 | 0.033504 | 0.106886 |
| DLEU2L | 2.164218 | 0.833174 | 1.362062 | 0.611701 | 0.540736 | NA |
| FBXO2 | 1106.706 | -0.04084 | 0.968948 | -0.04214 | 0.966383 | 0.981451 |
| FBXO6 | 320.5197 | -0.49519 | 0.178452 | -2.77492 | 0.005521 | 0.031724 |
| C1orf21 | 2143.516 | 0.047617 | 0.157911 | 0.301542 | 0.763002 | 0.852775 |
| SWT1 | 130.4791 | 0.747131 | 0.332576 | 2.246496 | 0.024672 | 0.086963 |
| MAD2L2 | 958.1752 | -0.09603 | 0.16196 | -0.59292 | 0.553236 | 0.693776 |
| DNAJC6 | 609.7249 | 0.351121 | 0.186066 | 1.887076 | 0.05915 | 0.155655 |
| LEPR | 42.14526 | 1.08978 | 0.374404 | 2.910704 | 0.003606 | NA |
| IVNS1ABP | 3726.974 | 0.278862 | 0.132119 | 2.110689 | 0.034799 | 0.109544 |
| KIAA2013 | 1985.07 | -0.2891 | 0.131755 | -2.1942 | 0.028221 | 0.095208 |
| MFN2 | 4810.798 | -0.40237 | 0.135167 | -2.97681 | 0.002913 | 0.020726 |
| PRG4 | 4.992252 | 0.515492 | 1.19796 | 0.430308 | 0.666972 | NA |
| MIIP | 784.0923 | -0.17084 | 0.144868 | -1.17929 | 0.238284 | 0.401144 |
| SMG7 | 1970.946 | -0.37082 | 0.134872 | -2.74944 | 0.00597 | 0.033288 |
| NCF2 | 0.315653 | -0.57305 | 3.892639 | -0.14721 | 0.882963 | NA |
| PDC | 0.096202 | 0.408546 | 4.079639 | 0.100143 | 0.920231 | NA |
| SLC35D1 | 547.3641 | -0.18906 | 0.197756 | -0.95603 | 0.339058 | 0.507709 |
| PLA2G4A | 1859.441 | 1.129664 | 0.238498 | 4.736575 | 2.17E-06 | 2.63E-04 |
| GADD45A | 856.0797 | 0.10407 | 0.257056 | 0.404852 | 0.685586 | 0.79533 |
| WLS | 3251.465 | -0.14435 | 0.14541 | -0.99271 | 0.320851 | 0.489095 |
| PRDM2 | 974.6858 | -0.03986 | 0.162671 | -0.24503 | 0.806433 | 0.88439 |
| RGS2 | 236.4328 | 1.156681 | 0.334458 | 3.45838 | 5.43E-04 | 0.007103 |
| RPE65 | 6.974697 | 0.539064 | 0.780197 | 0.690932 | 0.489608 | NA |
| TROVE2 | 831.7542 | 0.900859 | 0.294036 | 3.063772 | 0.002186 | 0.017131 |
| UCHL5 | 557.2541 | 0.713695 | 0.270074 | 2.642586 | 0.008228 | 0.041382 |
| BCAS2 | 645.1575 | 0.719251 | 0.196803 | 3.654669 | 2.58E-04 | 0.004536 |
| SRSF11 | 1464.704 | 1.003743 | 0.292233 | 3.434738 | 5.93E-04 | 0.007528 |
| CTH | 166.2951 | -0.47296 | 0.21315 | -2.2189 | 0.026493 | 0.091302 |
| AGMAT | 35.45064 | -0.39581 | 0.35936 | -1.10142 | 0.270713 | NA |

| | | | | | | |
|---|---|---|---|---|---|---|
| OLFML3 | 57.94518 | -0.2397 | 0.326416 | -0.73433 | 0.462746 | 0.620712 |
| CFHR3 | 0.47961 | 0.535796 | 3.39605 | 0.15777 | 0.874638 | NA |
| PLEKHM2 | 7978.005 | -0.28325 | 0.169086 | -1.67521 | 0.093893 | 0.214014 |
| CRYZ | 822.1582 | 0.708512 | 0.205757 | 3.443442 | 5.74E-04 | 0.007385 |
| PHTF1 | 719.9477 | 0.467908 | 0.182094 | 2.569598 | 0.010182 | 0.047747 |
| ZBTB17 | 844.8217 | -0.3321 | 0.180253 | -1.84243 | 0.065412 | 0.166897 |
| CD58 | 358.1293 | 0.646246 | 0.16884 | 3.827568 | 1.29E-04 | 0.002993 |
| TFAP2E | 101.3506 | 0.265906 | 0.255962 | 1.03885 | 0.298875 | 0.465742 |
| CD2 | 0.163957 | 0.424913 | 4.079006 | 0.104171 | 0.917034 | NA |
| TTF2 | 773.4421 | -0.09456 | 0.150515 | -0.62822 | 0.529861 | 0.674874 |
| NR5A2 | 22.1047 | -0.2516 | 0.570447 | -0.44105 | 0.659176 | NA |
| KIF21B | 8.462004 | -1.0269 | 0.883312 | -1.16256 | 0.245008 | NA |
| TMEM9 | 2304.323 | -0.10994 | 0.138046 | -0.79638 | 0.425813 | 0.587731 |
| ADPRHL2 | 968.4038 | -0.12798 | 0.161262 | -0.79364 | 0.427407 | 0.589313 |
| MAP7D1 | 6981.875 | -0.43379 | 0.14543 | -2.98282 | 0.002856 | 0.020475 |
| WARS2 | 318.0667 | -0.01645 | 0.167958 | -0.09795 | 0.921974 | 0.954836 |
| HAO2 | 1.02675 | -3.84073 | 2.498363 | -1.5373 | 0.12422 | NA |
| | 7.070697 | -0.73007 | 0.806922 | -0.90476 | 0.365593 | NA |
| OSCP1 | 217.2568 | -0.21605 | 0.19977 | -1.08152 | 0.279466 | 0.445601 |
| MRPS15 | 2117.967 | 0.24189 | 0.139825 | 1.729947 | 0.08364 | 0.197007 |
| EXOC8 | 432.125 | 0.383626 | 0.219189 | 1.750207 | 0.080083 | 0.1916 |
| GNPAT | 1392.326 | 0.146126 | 0.176518 | 0.827821 | 0.407772 | 0.571343 |
| TSNAX | 1298.338 | 1.008859 | 0.226603 | 4.452106 | 8.50E-06 | 5.66E-04 |
| C1orf109 | 404.0012 | 0.265004 | 0.158542 | 1.671507 | 0.094621 | 0.21514 |
| RRAGC | 1940.592 | 0.068543 | 0.13369 | 0.512702 | 0.60816 | 0.735387 |
| TBCE | 727.0404 | 0.300558 | 0.155362 | 1.934566 | 0.053044 | 0.144021 |
| NID1 | 1657.659 | -0.88932 | 0.188155 | -4.72652 | 2.28E-06 | 2.67E-04 |
| LGALS8 | 1019.753 | 0.339149 | 0.163796 | 2.07056 | 0.0384 | 0.116966 |
| HPCAL4 | 0.163957 | 0.424913 | 4.079006 | 0.104171 | 0.917034 | NA |
| MTR | 1231.126 | 0.389182 | 0.258934 | 1.503018 | 0.132834 | 0.270093 |
| BMP8B | 6.522309 | 0.220497 | 0.924086 | 0.238611 | 0.811407 | NA |
| MYCL | 1.325977 | -1.6864 | 2.032952 | -0.82953 | 0.406804 | NA |
| SIPA1L2 | 1895.038 | -0.02144 | 0.133447 | -0.16067 | 0.872351 | 0.924028 |
| RLF | 286.206 | 0.818439 | 0.252098 | 3.246514 | 0.001168 | 0.011465 |
| KMO | 19.27903 | 0.455864 | 0.582913 | 0.782045 | 0.434188 | NA |
| ZNF684 | 88.54893 | 0.704461 | 0.2558 | 2.753954 | 0.005888 | 0.032988 |
| KCNQ4 | 3.983783 | -0.6257 | 0.993123 | -0.63004 | 0.52867 | NA |
| RIMS3 | 76.08058 | -0.3098 | 0.298353 | -1.03838 | 0.299092 | 0.465931 |
| AKT3 | 1515.688 | 0.49805 | 0.243135 | 2.048452 | 0.040516 | 0.121376 |
| ETV3 | 1565.945 | 0.263417 | 0.204148 | 1.290328 | 0.196937 | 0.352352 |
| ACADM | 1227.878 | 0.67386 | 0.18413 | 3.659696 | 2.53E-04 | 0.004475 |
| ST6GALNA( | 85.02774 | 0.34942 | 0.337906 | 1.034075 | 0.301101 | 0.467954 |
| SLAMF1 | 0.518939 | 0.925178 | 3.271397 | 0.282808 | 0.777324 | NA |
| LPHN2 | 438.1967 | 0.363111 | 0.179303 | 2.025128 | 0.042854 | 0.126001 |
| PADI2 | 20.06008 | -1.00558 | 0.459094 | -2.19037 | 0.028498 | NA |
| SDHB | 2886.259 | 0.134479 | 0.149985 | 0.896619 | 0.369922 | 0.536838 |
| MFAP2 | 294.7351 | -0.02268 | 0.193526 | -0.1172 | 0.906701 | 0.945876 |

| | | | | | | |
|---|---|---|---|---|---|---|
| RPF1 | 988.6311 | 0.529692 | 0.182601 | 2.900818 | 0.003722 | 0.02435 |
| KDM5B | 1510.019 | -0.24974 | 0.148228 | -1.68482 | 0.092024 | 0.210973 |
| UAP1 | 637.7613 | 0.176967 | 0.170341 | 1.038893 | 0.298854 | 0.465742 |
| CTBS | 541.4332 | 0.144008 | 0.196581 | 0.732563 | 0.463825 | 0.621386 |
| RGS4 | 35.5753 | 1.021058 | 0.466866 | 2.187045 | 0.028739 | NA |
| KLHL12 | 1476.144 | -0.03019 | 0.153797 | -0.19629 | 0.844385 | 0.907349 |
| SSX2IP | 328.4171 | 0.704093 | 0.315402 | 2.23237 | 0.025591 | 0.089457 |
| ZNHIT6 | 457.8831 | 0.366016 | 0.23124 | 1.582838 | 0.113458 | 0.242307 |
| RBBP5 | 713.7574 | -0.01675 | 0.17095 | -0.09799 | 0.921943 | 0.954836 |
| GBP3 | 403.8692 | 0.872124 | 0.295819 | 2.948167 | 0.003197 | 0.022084 |
| GBP1 | 186.6728 | -0.14648 | 0.239523 | -0.61154 | 0.540845 | 0.682824 |
| PINK1-AS | 88.61496 | -0.55151 | 0.297065 | -1.85651 | 0.06338 | 0.163381 |
| KIF17 | 1.457231 | 1.67388 | 1.698239 | 0.985656 | 0.324302 | NA |
| GPR89A | 85.81416 | -0.08024 | 0.258078 | -0.31091 | 0.755867 | 0.847421 |
| CDK18 | 432.5186 | -0.45054 | 0.157816 | -2.85486 | 0.004306 | 0.026682 |
| RAB7L1 | 1573.326 | 0.019024 | 0.16643 | 0.114309 | 0.908993 | 0.947483 |
| CD160 | 0.921134 | 0.131863 | 2.11947 | 0.062215 | 0.950391 | NA |
| TXNIP | 2017.913 | -0.8782 | 0.205479 | -4.2739 | 1.92E-05 | 9.04E-04 |
| ECE1 | 1060.177 | -0.1659 | 0.21339 | -0.77746 | 0.436886 | 0.597357 |
| HMGCL | 1051.425 | -0.24093 | 0.160601 | -1.50018 | 0.133569 | 0.271237 |
| GALE | 946.9416 | -0.42944 | 0.153067 | -2.80558 | 0.005023 | 0.029727 |
| ID3 | 520.4595 | 1.02204 | 0.677107 | 1.509421 | 0.131191 | 0.267735 |
| CD46 | 2215.994 | 0.481859 | 0.153527 | 3.138603 | 0.001698 | 0.014564 |
| PRPF3 | 760.1483 | 0.285081 | 0.136538 | 2.087927 | 0.036804 | 0.113713 |
| APH1A | 4232.537 | -0.2505 | 0.123671 | -2.02554 | 0.042812 | 0.125941 |
| LEPRE1 | 1510.974 | -0.32301 | 0.147348 | -2.19216 | 0.028368 | 0.09539 |
| SLC2A1 | 1658.658 | -0.34474 | 0.168444 | -2.04663 | 0.040695 | 0.121674 |
| EBNA1BP2 | 1354.798 | 0.410705 | 0.17401 | 2.360237 | 0.018263 | 0.071011 |
| CDC20 | 1523.597 | -0.53812 | 0.253495 | -2.12278 | 0.033772 | 0.107317 |
| MPL | 0.448206 | -1.2532 | 3.007315 | -0.41672 | 0.676885 | NA |
| ARTN | 7.853323 | -0.08065 | 0.773469 | -0.10427 | 0.916954 | NA |
| IPO13 | 2474.491 | -0.57667 | 0.147213 | -3.91721 | 8.96E-05 | 0.002386 |
| ATP6V0B | 5832.13 | 0.046796 | 0.190675 | 0.245421 | 0.806131 | 0.88422 |
| B4GALT2 | 1767.763 | -0.24194 | 0.147034 | -1.64548 | 0.09987 | 0.223198 |
| ERI3 | 1674.954 | -0.18795 | 0.146626 | -1.28187 | 0.199889 | 0.355943 |
| PTCH2 | 5.84713 | -0.34857 | 0.890473 | -0.39144 | 0.695472 | NA |
| AKR1A1 | 2035.12 | -0.20295 | 0.147898 | -1.37223 | 0.169992 | 0.320363 |
| PRDX1 | 13484.58 | 0.122153 | 0.150245 | 0.813024 | 0.416204 | 0.579324 |
| PIK3R3 | 506.6227 | -0.3538 | 0.296694 | -1.19247 | 0.233078 | 0.395583 |
| TSPAN1 | 9.959758 | -0.56719 | 0.729667 | -0.77732 | 0.43697 | NA |
| BLZF1 | 557.9861 | 0.759469 | 0.244965 | 3.100314 | 0.001933 | 0.015805 |
| CCDC181 | 22.34229 | 0.802168 | 0.684846 | 1.17131 | 0.241474 | NA |
| SLC19A2 | 1297.813 | 0.155453 | 0.17232 | 0.902117 | 0.366995 | 0.534224 |
| FAAH | 24.23153 | 0.622241 | 0.419254 | 1.484162 | 0.137766 | NA |
| NSUN4 | 779.2122 | -0.08262 | 0.171352 | -0.48215 | 0.6297 | 0.752151 |
| TMED5 | 1017.518 | 0.740803 | 0.190859 | 3.881423 | 1.04E-04 | 0.002626 |
| MROH9 | 6.235232 | 0.356648 | 0.871765 | 0.40911 | 0.682459 | NA |

| | | | | | | |
|---|---|---|---|---|---|---|
| DR1 | 1518.519 | 0.25659 | 0.138846 | 1.848016 | 0.0646 | 0.165674 |
| CNN3 | 9536.557 | 0.281396 | 0.165859 | 1.696599 | 0.089772 | 0.207416 |
| PRRC2C | 2639.297 | -0.0371 | 0.333572 | -0.11121 | 0.911453 | 0.948657 |
| F3 | 10.56311 | -0.28438 | 0.660941 | -0.43027 | 0.666998 | NA |
| ABCD3 | 1801.339 | 0.302167 | 0.188677 | 1.601507 | 0.109265 | 0.236504 |
| VAMP4 | 426.9538 | 0.766964 | 0.189408 | 4.049266 | 5.14E-05 | 0.001673 |
| DPH5 | 468.13 | -0.01671 | 0.161048 | -0.10373 | 0.917382 | 0.952076 |
| PTBP2 | 163.9922 | 0.72268 | 0.248523 | 2.907899 | 0.003639 | 0.024058 |
| TNFSF4 | 33.94066 | 0.261133 | 0.374709 | 0.696895 | 0.485869 | NA |
| PRDX6 | 4207.127 | -0.10894 | 0.141657 | -0.76902 | 0.441882 | 0.601869 |
| DARS2 | 780.8194 | -0.18276 | 0.138432 | -1.32024 | 0.186753 | 0.340881 |
| IRF6 | 232.7272 | -0.35068 | 0.183755 | -1.90843 | 0.056336 | 0.150183 |
| DIEXF | 688.9647 | 0.210431 | 0.16019 | 1.313631 | 0.18897 | 0.343263 |
| | 75.56938 | -0.57645 | 0.356171 | -1.61847 | 0.105562 | 0.231319 |
| | 7.158911 | -1.74781 | 1.065998 | -1.6396 | 0.101088 | NA |
| SERPINC1 | 2.760746 | 0.010583 | 1.168747 | 0.009055 | 0.992776 | NA |
| RCAN3 | 809.8339 | -0.01207 | 0.142858 | -0.08452 | 0.932644 | 0.961853 |
| SYF2 | 766.4757 | 0.599586 | 0.164502 | 3.644857 | 2.68E-04 | 0.004599 |
| C1orf63 | 433.7099 | 1.501794 | 0.208515 | 7.202341 | 5.92E-13 | 1.40E-09 |
| SLC35A3 | 295.4181 | 0.852427 | 0.318458 | 2.676732 | 0.007434 | 0.038826 |
| RCOR3 | 646.3948 | 0.124503 | 0.17826 | 0.698435 | 0.484905 | 0.638373 |
| STMN1 | 7210.339 | -0.14898 | 0.188412 | -0.79069 | 0.429125 | 0.590656 |
| MTFR1L | 1824.457 | -0.11163 | 0.141949 | -0.78639 | 0.431636 | 0.592892 |
| MAN1C1 | 283.7036 | -0.407 | 0.173164 | -2.35036 | 0.018755 | 0.072322 |
| NEK2 | 413.3044 | -0.01364 | 0.282028 | -0.04836 | 0.961433 | 0.979108 |
| RPS6KA1 | 585.2275 | -0.01836 | 0.200151 | -0.09174 | 0.926903 | 0.958156 |
| DHDDS | 1243.201 | -0.31501 | 0.140146 | -2.2477 | 0.024595 | 0.086816 |
| NENF | 2273.706 | 0.115703 | 0.163315 | 0.708467 | 0.478655 | 0.633256 |
| NSL1 | 586.8607 | 0.434556 | 0.205836 | 2.111172 | 0.034758 | 0.109442 |
| PROX1 | 12.17747 | -0.64419 | 0.622622 | -1.03464 | 0.300837 | NA |
| ARID1A | 1794.941 | -0.71607 | 0.243458 | -2.94124 | 0.003269 | 0.022362 |
| CENPF | 1257.095 | 0.893761 | 0.860109 | 1.039125 | 0.298747 | 0.465699 |
| RPA2 | 1055.17 | 0.279642 | 0.165853 | 1.686088 | 0.091779 | 0.210711 |
| PPP1R8 | 1108.251 | 0.143128 | 0.142438 | 1.004838 | 0.314975 | 0.48237 |
| STX12 | 1524.224 | 0.206481 | 0.13526 | 1.526556 | 0.126871 | 0.262351 |
| 2-Mar | 237.2537 | 0.02871 | 0.17966 | 0.159803 | 0.873037 | 0.924424 |
| SLC5A9 | 10.10982 | 0.485295 | 0.682166 | 0.711403 | 0.476835 | NA |
| OSBPL9 | 3852.304 | 0.084636 | 0.161826 | 0.523005 | 0.600971 | 0.730247 |
| TXNDC12 | 2022.874 | 0.15059 | 0.140202 | 1.074095 | 0.28278 | 0.449137 |
| MUC5B | 7.56184 | 0.422626 | 0.803924 | 0.525704 | 0.599094 | NA |
| CTSD | 36813.96 | -0.12577 | 0.164121 | -0.76633 | 0.443481 | 0.603336 |
| KIF14 | 165.2351 | 0.732214 | 0.375718 | 1.94884 | 0.051315 | 0.140999 |
| TNNT2 | 0.298089 | -0.51711 | 3.975232 | -0.13008 | 0.896501 | NA |
| DDX59 | 424.8899 | 0.62427 | 0.187056 | 3.337348 | 8.46E-04 | 0.009353 |
| CAMSAP2 | 1235.854 | 0.637782 | 0.209116 | 3.049902 | 0.002289 | 0.017639 |
| ATF6 | 1818.978 | 0.107643 | 0.172351 | 0.624556 | 0.532263 | 0.676528 |
| C1orf54 | 60.08771 | -0.1992 | 0.289404 | -0.6883 | 0.491262 | 0.64317 |

| | | | | | | |
|---|---|---|---|---|---|---|
| CA14 | 2713.078 | -0.68732 | 0.248914 | -2.76128 | 0.005757 | 0.032518 |
| ANKRD13C | 674.2072 | 0.499234 | 0.196558 | 2.539882 | 0.011089 | 0.050462 |
| SGIP1 | 83.82387 | 0.246203 | 0.357542 | 0.688599 | 0.491075 | 0.642998 |
| DCLRE1B | 330.9988 | 0.030098 | 0.208342 | 0.144464 | 0.885134 | 0.932072 |
| CASQ2 | 24.75354 | -1.4498 | 0.524665 | -2.76329 | 0.005722 | NA |
| OLFM3 | 0.422534 | 1.924583 | 3.001758 | 0.641152 | 0.521424 | NA |
| RAB3GAP2 | 1223.566 | 0.644212 | 0.204885 | 3.144254 | 0.001665 | 0.014409 |
| C1orf198 | 4687.301 | -0.05606 | 0.327901 | -0.17096 | 0.864259 | 0.92003 |
| TRIM67 | 21.51096 | -0.67103 | 0.587345 | -1.14248 | 0.253256 | NA |
| HEATR1 | 1198.298 | 0.038741 | 0.139112 | 0.278487 | 0.780639 | 0.86547 |
| FAM178A | 747.9346 | 0.406203 | 0.19609 | 2.071518 | 0.03831 | 0.116813 |
| IDE | 1025.385 | 0.340036 | 0.177454 | 1.916199 | 0.05534 | 0.148596 |
| TECTB | 1.140586 | -0.14889 | 1.823527 | -0.08165 | 0.934926 | NA |
| ELOVL3 | 1.568656 | -2.16843 | 1.872601 | -1.15798 | 0.246874 | NA |
| IFIT3 | 3458.822 | -0.32368 | 0.24567 | -1.31754 | 0.187659 | 0.341836 |
| IFIT2 | 909.9531 | 0.122034 | 0.332686 | 0.366815 | 0.713757 | 0.817308 |
| GPAM | 446.2833 | 0.363543 | 0.206569 | 1.759908 | 0.078423 | 0.188658 |
| CUTC | 310.8974 | 0.224631 | 0.179307 | 1.252773 | 0.210288 | 0.368021 |
| PPP1R3C | 1562.003 | -0.2773 | 0.161392 | -1.71815 | 0.085769 | 0.200354 |
| PYROXD2 | 154.5833 | -0.42051 | 0.256718 | -1.63802 | 0.101419 | 0.225027 |
| CNNM1 | 293.5815 | -0.35893 | 0.296319 | -1.2113 | 0.225782 | 0.386966 |
| MXI1 | 3572.671 | 0.443477 | 0.192106 | 2.308499 | 0.020971 | 0.078206 |
| SMNDC1 | 714.1141 | 0.07772 | 0.153909 | 0.504969 | 0.61358 | 0.740094 |
| C10orf88 | 316.6782 | 0.402732 | 0.193397 | 2.082411 | 0.037305 | 0.114667 |
| HELLS | 173.206 | 0.496472 | 0.25534 | 1.94436 | 0.051852 | 0.141801 |
| TCTN3 | 1580.43 | -0.2668 | 0.149302 | -1.78698 | 0.073941 | 0.181373 |
| FAM45A | 605.1975 | 0.165275 | 0.173054 | 0.955048 | 0.339553 | 0.508003 |
| AVPI1 | 3076.944 | -0.14393 | 0.163119 | -0.88234 | 0.377594 | 0.544108 |
| WDR11 | 1998.94 | 0.281646 | 0.154939 | 1.817786 | 0.069097 | 0.172997 |
| C10orf76 | 1095.772 | -0.26995 | 0.141382 | -1.90937 | 0.056215 | 0.150029 |
| KCNIP2 | 24.95027 | -0.31522 | 0.493477 | -0.63878 | 0.522967 | NA |
| CCDC147 | 3.111381 | 0.316474 | 1.146239 | 0.276098 | 0.782473 | NA |
| GOT1 | 1366.826 | -0.23682 | 0.138139 | -1.71433 | 0.086467 | 0.201508 |
| CPN1 | 220.0631 | 0.472879 | 0.350099 | 1.350698 | 0.176792 | 0.329058 |
| C10orf95 | 0.589223 | -3.04304 | 3.047314 | -0.9986 | 0.317989 | NA |
| SFRP5 | 3.436623 | -0.22799 | 1.523268 | -0.14967 | 0.881026 | NA |
| TNN | 5.18199 | 0.041792 | 0.945093 | 0.04422 | 0.964729 | NA |
| MRPS14 | 1005.215 | 0.262069 | 0.128128 | 2.04536 | 0.040819 | 0.121985 |
| CENPL | 301.8094 | 0.242358 | 0.236108 | 1.026473 | 0.304669 | 0.472013 |
| TNFSF18 | 0.162376 | 0.424913 | 4.079006 | 0.104171 | 0.917034 | NA |
| SEC16B | 1.728229 | -1.16052 | 1.555065 | -0.74628 | 0.455497 | NA |
| GORAB | 148.281 | 1.344287 | 0.255374 | 5.263985 | 1.41E-07 | 4.29E-05 |
| MASTL | 560.2347 | 0.517634 | 0.161803 | 3.199164 | 0.001378 | 0.012785 |
| KIAA1217 | 205.6812 | -0.04788 | 0.210527 | -0.22744 | 0.820078 | 0.892334 |
| SEPT7P9 | 0.49029 | 2.123092 | 3.304282 | 0.642527 | 0.520531 | NA |
| PLXDC2 | 163.3123 | 0.346371 | 0.248645 | 1.393037 | 0.163608 | 0.312181 |
| EPC1 | 341.561 | 0.289132 | 0.202193 | 1.429982 | 0.152722 | 0.297294 |

| | | | | | | |
|---|---|---|---|---|---|---|
| TAF12 | 456.6411 | 0.248984 | 0.174294 | 1.428524 | 0.153141 | 0.297865 |
| NPPB | 0.430642 | -1.1891 | 3.485782 | -0.34113 | 0.733008 | NA |
| UBIAD1 | 610.2707 | -0.27571 | 0.159749 | -1.72588 | 0.084368 | 0.197978 |
| TARDBP | 2300.599 | 0.108241 | 0.140788 | 0.76882 | 0.442 | 0.601943 |
| TNFRSF8 | 0.096202 | 0.408546 | 4.079639 | 0.100143 | 0.920231 | NA |
| TRIM6 | 100.829 | -0.09549 | 0.327146 | -0.29188 | 0.770375 | 0.858023 |
| ECHDC2 | 321.7561 | 0.259044 | 0.18591 | 1.393381 | 0.163505 | 0.312042 |
| LHX4 | 20.37447 | 0.680262 | 0.505713 | 1.345152 | 0.178576 | NA |
| RNF2 | 434.8897 | 0.11716 | 0.194268 | 0.603082 | 0.546454 | 0.688001 |
| TRMT1L | 454.1882 | 0.398671 | 0.225703 | 1.766349 | 0.077337 | 0.186995 |
| KIF18A | 137.272 | 1.406962 | 0.260351 | 5.404106 | 6.51E-08 | 2.57E-05 |
| DESI2 | 1797.262 | 0.183008 | 0.169365 | 1.080556 | 0.279895 | 0.445868 |
| MAPK8IP1 | 343.6062 | -0.17877 | 0.183087 | -0.97641 | 0.328862 | 0.497151 |
| CRY2 | 439.3277 | -0.35835 | 0.161182 | -2.22325 | 0.026199 | 0.090873 |
| PEX16 | 683.9545 | -0.1686 | 0.164754 | -1.02332 | 0.306157 | 0.473267 |
| DEPDC7 | 252.1233 | 0.417844 | 0.199336 | 2.096185 | 0.036066 | 0.112073 |
| CAT | 2110.154 | 0.169185 | 0.157993 | 1.070838 | 0.284242 | 0.450855 |
| BAI2 | 141.4766 | -0.23557 | 0.250933 | -0.93877 | 0.347848 | 0.515984 |
| HCRTR1 | 0.971297 | 3.154786 | 2.249666 | 1.402335 | 0.160815 | NA |
| ZCCHC17 | 1148.342 | 0.212608 | 0.159937 | 1.329325 | 0.183741 | 0.336981 |
| FABP3 | 179.4808 | 0.58221 | 0.393552 | 1.479373 | 0.139041 | 0.279046 |
| KHDRBS1 | 2949.242 | -0.02975 | 0.183676 | -0.16199 | 0.871313 | 0.923353 |
| TMEM39B | 324.2527 | -0.39325 | 0.206449 | -1.90485 | 0.0568 | 0.151046 |
| RNF115 | 1140.241 | -0.13871 | 0.141247 | -0.98202 | 0.326088 | 0.494155 |
| POLR3GL | 786.7416 | 0.18465 | 0.147328 | 1.253328 | 0.210086 | 0.367775 |
| CPXM2 | 3210.285 | 0.011624 | 0.223966 | 0.051899 | 0.958609 | 0.977155 |
| TMEM54 | 936.1956 | -0.07292 | 0.137749 | -0.52934 | 0.596572 | 0.726525 |
| ZSCAN20 | 150.8307 | -0.13403 | 0.202945 | -0.66043 | 0.50898 | 0.657437 |
| CSMD2 | 0.162376 | 0.424913 | 4.079006 | 0.104171 | 0.917034 | NA |
| HPCA | 5.000189 | -0.80439 | 1.018967 | -0.78942 | 0.429866 | NA |
| LRIF1 | 206.7313 | 0.834455 | 0.275997 | 3.023416 | 0.002499 | 0.018801 |
| ADORA3 | 3.309249 | 0.682598 | 1.132778 | 0.602587 | 0.546783 | NA |
| CLCC1 | 696.5738 | 0.243326 | 0.171473 | 1.419038 | 0.155888 | 0.301723 |
| GPSM2 | 498.7101 | 0.407239 | 0.228829 | 1.779662 | 0.075131 | 0.183032 |
| FMOD | 38.06608 | 0.15388 | 0.456213 | 0.337298 | 0.735892 | NA |
| MYOG | 0.163957 | 0.424913 | 4.079006 | 0.104171 | 0.917034 | NA |
| COPA | 8658.801 | -0.29096 | 0.133896 | -2.17307 | 0.029775 | 0.098517 |
| CD244 | 0.915556 | -1.07826 | 2.191337 | -0.49206 | 0.62268 | NA |
| ANXA11 | 8331.034 | -0.21798 | 0.142982 | -1.52452 | 0.12738 | 0.263265 |
| LDB3 | 62.14506 | -1.04672 | 0.428381 | -2.44344 | 0.014548 | 0.06067 |
| FAM35A | 378.1288 | 1.011101 | 0.246603 | 4.100107 | 4.13E-05 | 0.001488 |
| FAM213A | 3026.791 | 0.070449 | 0.146354 | 0.481364 | 0.630258 | 0.752665 |
| RPL5 | 15906.25 | 0.475607 | 0.183355 | 2.593908 | 0.009489 | 0.045544 |
| ODF2L | 223.1283 | 1.079143 | 0.358004 | 3.014333 | 0.002575 | 0.019243 |
| SPATA1 | 5.272178 | 0.624747 | 0.914548 | 0.683121 | 0.49453 | NA |
| TRMT13 | 103.5258 | 1.028722 | 0.354141 | 2.904837 | 0.003674 | 0.024173 |
| LRRC39 | 168.9592 | 1.034133 | 0.268026 | 3.858331 | 1.14E-04 | 0.002779 |

| | | | | | | |
|---|---|---|---|---|---|---|
| RWDD3 | 203.5709 | 0.648293 | 0.219818 | 2.949223 | 0.003186 | 0.022047 |
| ZNF644 | 455.836 | 0.936092 | 0.347884 | 2.690813 | 0.007128 | 0.037572 |
| CCDC18 | 46.23504 | 1.066126 | 1.068949 | 0.997359 | 0.31859 | NA |
| RPAP2 | 163.3506 | 0.610409 | 0.274227 | 2.225926 | 0.026019 | 0.09043 |
| NBPF14 | 59.90566 | 0.495068 | 0.334013 | 1.48218 | 0.138292 | 0.277874 |
| PLAU | 1.095699 | -1.13063 | 2.016328 | -0.56074 | 0.574978 | NA |
| SRGN | 571.2963 | 0.089871 | 0.201263 | 0.446534 | 0.655211 | 0.77237 |
| CHST3 | 2643.51 | -0.46873 | 0.218844 | -2.14184 | 0.032207 | 0.104129 |
| BICC1 | 208.86 | 0.654263 | 0.260725 | 2.509398 | 0.012094 | 0.053697 |
| CISD1 | 663.3329 | 0.332066 | 0.164681 | 2.016421 | 0.043756 | 0.127483 |
| EGR2 | 10.88194 | 0.06974 | 0.665466 | 0.104799 | 0.916535 | NA |
| ECD | 1205.686 | 0.144919 | 0.140861 | 1.028811 | 0.303569 | 0.47084 |
| P4HA1 | 1254.219 | 0.369631 | 0.154496 | 2.392491 | 0.016734 | 0.067227 |
| SLC25A16 | 393.1384 | 0.203948 | 0.193962 | 1.051479 | 0.293039 | 0.460068 |
| ZWINT | 1241.066 | -0.01155 | 0.195563 | -0.05906 | 0.952904 | 0.974355 |
| VPS26A | 1680.266 | 0.49039 | 0.191068 | 2.566579 | 0.010271 | 0.048031 |
| CDKN2C | 1182.849 | -0.53789 | 0.255738 | -2.10329 | 0.03544 | 0.110885 |
| RNF11 | 3613.571 | 0.402819 | 0.149018 | 2.703164 | 0.006868 | 0.036661 |
| OPTN | 5185.925 | 0.469101 | 0.17811 | 2.633775 | 0.008444 | 0.042031 |
| ITIH5 | 60.13219 | 0.694552 | 1.54035 | 0.450905 | 0.652058 | 0.769935 |
| KBTBD4 | 217.6765 | -0.33969 | 0.193943 | -1.75151 | 0.079857 | 0.191148 |
| TYRL | 14.5993 | -0.45067 | 0.572665 | -0.78697 | 0.431298 | NA |
| ATPAF1 | 1442.804 | -0.1373 | 0.143614 | -0.95604 | 0.339051 | 0.507709 |
| STIL | 350.2548 | 0.197254 | 0.204883 | 0.962764 | 0.335666 | 0.504477 |
| LPGAT1 | 1689.123 | 0.401647 | 0.250158 | 1.605572 | 0.108368 | 0.23525 |
| BATF3 | 31.84067 | 0.572776 | 0.429578 | 1.333347 | 0.182418 | NA |
| G0S2 | 3.098036 | -0.9555 | 1.190182 | -0.80282 | 0.422078 | NA |
| PFKFB2 | 758.5855 | -0.05136 | 0.144567 | -0.35524 | 0.72241 | 0.823619 |
| C4BPB | 0.096202 | 0.408546 | 4.079639 | 0.100143 | 0.920231 | NA |
| TRIM51 | 55.1216 | 0.435477 | 0.419207 | 1.038812 | 0.298892 | 0.465742 |
| MYRF | 13.41555 | -0.16899 | 0.549201 | -0.30771 | 0.758303 | NA |
| AHNAK | 48066.94 | -0.20557 | 0.139325 | -1.47545 | 0.140091 | 0.280348 |
| MSTO1 | 287.3216 | -0.13725 | 0.198857 | -0.69018 | 0.490083 | 0.642357 |
| ATG4C | 366.4366 | 0.780265 | 0.239698 | 3.255201 | 0.001133 | 0.011192 |
| HNRNPR | 2833.431 | 0.401952 | 0.142737 | 2.816043 | 0.004862 | 0.028972 |
| ZNF436 | 370.4173 | -0.10681 | 0.227053 | -0.47044 | 0.638041 | 0.758814 |
| PSMB2 | 2849.554 | -0.11904 | 0.130587 | -0.9116 | 0.361977 | 0.529064 |
| AGO3 | 107.4897 | 0.304165 | 0.274424 | 1.108374 | 0.2677 | 0.432663 |
| UROD | 3326.526 | -0.28598 | 0.123911 | -2.30794 | 0.021003 | 0.078273 |
| ST3GAL3 | 755.7712 | -0.44113 | 0.139482 | -3.16261 | 0.001564 | 0.013896 |
| TMEM53 | 347.875 | -0.3372 | 0.180044 | -1.87286 | 0.061087 | 0.159302 |
| HECTD3 | 2996.485 | -0.44842 | 0.1313 | -3.41523 | 6.37E-04 | 0.007829 |
| DNAJC8 | 2939.646 | 0.357445 | 0.171061 | 2.089576 | 0.036656 | 0.113473 |
| AHDC1 | 661.7997 | -0.75355 | 0.22306 | -3.37825 | 7.29E-04 | 0.008554 |
| IFI6 | 26191.13 | -1.10825 | 0.22265 | -4.97753 | 6.44E-07 | 1.21E-04 |
| CPSF3L | 2109.141 | -0.24375 | 0.12074 | -2.0188 | 0.043508 | 0.127074 |
| HIVEP3 | 231.5576 | -0.57163 | 0.219499 | -2.60425 | 0.009207 | 0.044608 |

| | | | | | | |
|---|---|---|---|---|---|---|
| PPCS | 1271.719 | 0.262086 | 0.138911 | 1.886725 | 0.059197 | 0.155744 |
| AUNIP | 117.8429 | 0.079889 | 0.322762 | 0.247518 | 0.804507 | 0.883177 |
| EMC1 | 1660.93 | -0.56802 | 0.149501 | -3.79943 | 1.45E-04 | 0.003166 |
| UBR4 | 4467.114 | -0.56277 | 0.177761 | -3.16587 | 0.001546 | 0.013793 |
| HP1BP3 | 2864.861 | -0.13318 | 0.151952 | -0.87649 | 0.380765 | 0.546883 |
| MACF1 | 3463.991 | 0.251398 | 0.170747 | 1.472343 | 0.140928 | 0.281423 |
| ECHS1 | 4481.88 | -0.19616 | 0.149946 | -1.30823 | 0.190795 | 0.345061 |
| ARHGAP22 | 5.976682 | 0.713856 | 0.892319 | 0.800002 | 0.42371 | NA |
| COPB1 | 2903.384 | 0.270162 | 0.164284 | 1.644485 | 0.100076 | 0.223218 |
| PSMA1 | 512.218 | 0.295902 | 0.152415 | 1.94142 | 0.052207 | 0.14251 |
| BBOX1 | 0.134133 | -1.48909 | 4.077584 | -0.36519 | 0.71497 | NA |
| MYOD1 | 0.260158 | 1.240347 | 4.047124 | 0.306476 | 0.759242 | NA |
| SERGEF | 427.2164 | -0.03472 | 0.181064 | -0.19174 | 0.847944 | 0.909306 |
| KCNC1 | 3.068778 | 2.003469 | 1.500976 | 1.334777 | 0.181949 | NA |
| TPH1 | 1.137783 | 0.40046 | 1.978632 | 0.202392 | 0.83961 | NA |
| CSRP3 | 0.489058 | -0.58569 | 3.427543 | -0.17088 | 0.864321 | NA |
| E2F8 | 12.81442 | 0.354669 | 0.618451 | 0.57348 | 0.56632 | NA |
| CHRNA10 | 9.732896 | -0.36006 | 0.671957 | -0.53584 | 0.59207 | NA |
| CDKN1C | 381.0598 | -0.15657 | 0.187749 | -0.83396 | 0.404306 | 0.569031 |
| STK33 | 2.941917 | 0.754544 | 1.184601 | 0.63696 | 0.524151 | NA |
| LSP1 | 0.326536 | 0.037056 | 3.842554 | 0.009644 | 0.992306 | NA |
| TNNT3 | 0.785714 | -3.2458 | 3.343263 | -0.97085 | 0.331623 | NA |
| TNNI2 | 0.392857 | -1.546 | 4.075469 | -0.37934 | 0.704433 | NA |
| H19 | 2.820211 | 1.069965 | 1.248846 | 0.856763 | 0.391576 | NA |
| TUBGCP2 | 2508.545 | -0.30588 | 0.131222 | -2.33099 | 0.019754 | 0.075058 |
| CALY | 0.192403 | 0.975392 | 4.060716 | 0.240202 | 0.810174 | NA |
| CYP2E1 | 3.617697 | 2.601175 | 1.246578 | 2.086651 | 0.03692 | NA |
| CEP85 | 472.4615 | -0.09944 | 0.152983 | -0.64998 | 0.515705 | 0.663161 |
| ARHGEF16 | 12.87285 | 0.094739 | 0.578241 | 0.163839 | 0.869858 | NA |
| LRRC47 | 1506.425 | -0.42316 | 0.128341 | -3.29717 | 9.77E-04 | 0.010183 |
| SESN2 | 710.3648 | -0.43004 | 0.161575 | -2.66157 | 0.007778 | 0.039847 |
| SMPDL3B | 0.968668 | -0.08001 | 2.195869 | -0.03644 | 0.970935 | NA |
| ATPIF1 | 1885.493 | 0.051897 | 0.13554 | 0.382894 | 0.701799 | 0.808378 |
| MED18 | 647.3899 | -0.14649 | 0.148731 | -0.98494 | 0.324654 | 0.492617 |
| THEMIS2 | 904.2662 | -0.83393 | 0.227812 | -3.6606 | 2.52E-04 | 0.004472 |
| UBE4B | 2611.483 | -0.12472 | 0.142924 | -0.87265 | 0.382856 | 0.548882 |
| CASZ1 | 6.348526 | 0.159266 | 0.924264 | 0.172316 | 0.863189 | NA |
| GJA9 | 0.382031 | -1.26484 | 3.151989 | -0.40128 | 0.688212 | NA |
| CAP1 | 17019.41 | 0.005313 | 0.163251 | 0.032544 | 0.974038 | 0.985529 |
| PPT1 | 5090.419 | -0.17202 | 0.178921 | -0.96144 | 0.33633 | 0.505284 |
| ACAP3 | 2197.274 | -0.0523 | 0.171456 | -0.30502 | 0.760354 | 0.850778 |
| C1orf159 | 472.5651 | -0.05152 | 0.179748 | -0.28663 | 0.774395 | 0.860453 |
| NPHP4 | 526.3485 | -0.28597 | 0.167958 | -1.70265 | 0.088634 | 0.205459 |
| CHD1L | 2103.288 | 0.104563 | 0.14561 | 0.718098 | 0.472697 | 0.628442 |
| PEX11B | 912.4982 | -0.19981 | 0.136693 | -1.46175 | 0.143809 | 0.285467 |
| FMO5 | 94.68091 | 0.291953 | 0.259184 | 1.126434 | 0.259982 | 0.424355 |
| PIAS3 | 517.3753 | -0.34297 | 0.177828 | -1.92867 | 0.053771 | 0.145548 |

| | | | | | | |
|---|---|---|---|---|---|---|
| PRKAB2 | 814.7744 | 0.256235 | 0.161289 | 1.588666 | 0.112136 | 0.240523 |
| RBM8A | 724.1979 | 0.057068 | 0.137712 | 0.414405 | 0.678578 | 0.790385 |
| LIN28A | 0.096202 | 0.408546 | 4.079639 | 0.100143 | 0.920231 | NA |
| TRIM21 | 926.4127 | -0.44456 | 0.158793 | -2.79962 | 0.005116 | 0.030041 |
| SPATA6 | 444.9128 | 0.327432 | 0.182673 | 1.79245 | 0.073061 | 0.180107 |
| LRRC41 | 2593.193 | -0.54873 | 0.12442 | -4.41029 | 1.03E-05 | 6.34E-04 |
| FCRLA | 7180.426 | -0.21928 | 0.167249 | -1.31108 | 0.18983 | 0.34409 |
| HSD17B7 | 483.8921 | 0.33025 | 0.157317 | 2.099266 | 0.035794 | 0.111578 |
| ARFIP2 | 1815.862 | 0.033279 | 0.175308 | 0.189829 | 0.849443 | 0.910419 |
| TRIM5 | 592.8935 | -0.20168 | 0.204032 | -0.98846 | 0.322928 | 0.490809 |
| CNGA4 | 1.061949 | -0.45116 | 2.030098 | -0.22224 | 0.82413 | NA |
| TRIM22 | 356.7487 | -0.40639 | 0.250141 | -1.62463 | 0.104241 | 0.229312 |
| RRP8 | 565.7299 | -0.17479 | 0.14639 | -1.19402 | 0.232469 | 0.394718 |
| TIMM10B | 1494.436 | -0.20554 | 0.137518 | -1.49464 | 0.135008 | 0.273593 |
| PTPRE | 14.53178 | 0.759209 | 0.926839 | 0.819138 | 0.412708 | NA |
| ZRANB2 | 783.7367 | 1.049539 | 0.290835 | 3.608712 | 3.08E-04 | 0.005032 |
| DAP3 | 2465.44 | 0.071987 | 0.131272 | 0.548378 | 0.583433 | 0.716894 |
| RHBG | 1.440017 | 0.403371 | 1.70198 | 0.237001 | 0.812656 | NA |
| KIAA0907 | 763.1213 | 0.377452 | 0.159661 | 2.364081 | 0.018075 | 0.070509 |
| ATP1A4 | 0.58526 | 0.001197 | 3.235889 | 3.70E-04 | 0.999705 | NA |
| NES | 4415.034 | 0.015357 | 0.321511 | 0.047767 | 0.961902 | 0.979417 |
| BCAN | 12442.6 | -0.87595 | 0.201485 | -4.34749 | 1.38E-05 | 7.56E-04 |
| ARHGEF11 | 4258.71 | -0.34541 | 0.157625 | -2.19132 | 0.028429 | 0.095503 |
| RAB25 | 0.162376 | 0.424913 | 4.079006 | 0.104171 | 0.917034 | NA |
| HAPLN2 | 0.949524 | -0.003 | 2.040023 | -0.00147 | 0.998826 | NA |
| FCRL2 | 0.518939 | 0.925178 | 3.271397 | 0.282808 | 0.777324 | NA |
| DCAF8 | 1260.608 | -0.01555 | 0.152746 | -0.1018 | 0.918912 | 0.95308 |
| SYT11 | 1885.524 | -0.14761 | 0.186361 | -0.79204 | 0.428335 | 0.590112 |
| MMACHC | 775.2374 | -0.71799 | 0.194294 | -3.6954 | 2.20E-04 | 0.00412 |
| DPH2 | 960.2618 | -0.11511 | 0.155969 | -0.73804 | 0.460491 | 0.618738 |
| TOE1 | 453.6833 | -0.37095 | 0.152437 | -2.4335 | 0.014954 | 0.061855 |
| NASP | 958.315 | 0.458863 | 0.14499 | 3.164787 | 0.001552 | 0.013833 |
| MUTYH | 322.9853 | -0.34455 | 0.157838 | -2.18296 | 0.029039 | 0.096974 |
| INADL | 138.5263 | 0.187523 | 0.257425 | 0.728457 | 0.466334 | 0.623479 |
| KANK4 | 0.162376 | 0.424913 | 4.079006 | 0.104171 | 0.917034 | NA |
| ANGPTL3 | 0.096202 | 0.408546 | 4.079639 | 0.100143 | 0.920231 | NA |
| FBXO44 | 1222.354 | -0.09199 | 0.157747 | -0.58315 | 0.559794 | 0.69845 |
| RSG1 | 141.4002 | -0.2339 | 0.222987 | -1.04895 | 0.2942 | 0.461219 |
| CASP9 | 336.2478 | -0.17053 | 0.197496 | -0.86345 | 0.387892 | 0.553708 |
| CHRM3 | 0.096202 | 0.408546 | 4.079639 | 0.100143 | 0.920231 | NA |
| MYBPH | 0.096202 | 0.408546 | 4.079639 | 0.100143 | 0.920231 | NA |
| PIK3C2B | 1104.216 | -0.81547 | 0.180973 | -4.50601 | 6.61E-06 | 4.97E-04 |
| DSTYK | 5959.642 | 0.039492 | 0.141889 | 0.278333 | 0.780757 | 0.8655 |
| SLC41A1 | 1153.1 | -0.28408 | 0.161528 | -1.75869 | 0.07863 | 0.18904 |
| LGR6 | 0.163957 | 0.424913 | 4.079006 | 0.104171 | 0.917034 | NA |
| TMCC2 | 3494.838 | -0.22123 | 0.230464 | -0.95995 | 0.337079 | 0.505912 |
| EPHB2 | 0.616518 | 2.49285 | 2.980788 | 0.836306 | 0.402983 | NA |

| | | | | | | |
|---|---|---|---|---|---|---|
| SRRM1 | 1247.63 | -0.18003 | 0.163371 | -1.10195 | 0.270485 | 0.435443 |
| WDR74 | 840.2889 | -0.04303 | 0.192888 | -0.22307 | 0.82348 | 0.893657 |
| SFTPD | 0.230334 | -0.53318 | 4.076749 | -0.13079 | 0.895945 | NA |
| DYDC2 | 1.188177 | -0.03068 | 1.878929 | -0.01633 | 0.986971 | NA |
| TMEM254 | 192.2091 | -0.41441 | 0.234344 | -1.7684 | 0.076995 | 0.186394 |
| SWAP70 | 2260.266 | 0.481458 | 0.145628 | 3.306074 | 9.46E-04 | 0.00998 |
| ARNTL | 301.9651 | 0.410965 | 0.179025 | 2.295581 | 0.0217 | 0.080091 |
| AMPD3 | 62.93996 | -0.18431 | 0.430088 | -0.42854 | 0.668258 | 0.782242 |
| MICALCL | 0.440301 | 0.468383 | 3.449808 | 0.135771 | 0.892003 | NA |
| SBF2 | 1078.404 | -0.05272 | 0.154173 | -0.34196 | 0.732377 | 0.830527 |
| MICAL2 | 53.8623 | -0.83327 | 0.322309 | -2.58532 | 0.009729 | 0.046338 |
| RRAS2 | 296.1896 | 0.064006 | 0.174344 | 0.367123 | 0.713528 | 0.817124 |
| GNAT2 | 4.164569 | 0.455944 | 0.986074 | 0.462383 | 0.643806 | NA |
| GSTM1 | 286.1896 | -0.26724 | 0.200716 | -1.33144 | 0.183045 | 0.336174 |
| PRPF38B | 679.776 | 1.009372 | 0.569716 | 1.771709 | 0.076443 | 0.185473 |
| TSPAN2 | 0.192403 | 0.975392 | 4.060716 | 0.240202 | 0.810174 | NA |
| GSTM5 | 0.422737 | 0.518872 | 3.501356 | 0.148192 | 0.882191 | NA |
| GSTM3 | 1957.581 | 0.208714 | 0.149994 | 1.391482 | 0.164079 | 0.312777 |
| SYT6 | 15.00252 | -0.21773 | 0.575354 | -0.37843 | 0.70511 | NA |
| VAV3 | 138.3962 | 0.20032 | 0.27543 | 0.727298 | 0.467043 | 0.623713 |
| PSRC1 | 974.6654 | -0.30891 | 0.182464 | -1.693 | 0.090456 | 0.208467 |
| PTPN22 | 0.520316 | 2.239969 | 3.173769 | 0.705776 | 0.480328 | NA |
| SORT1 | 11155.05 | -0.27185 | 0.149534 | -1.818 | 0.069064 | 0.172997 |
| WNT2B | 15.1156 | 1.042018 | 0.567392 | 1.836506 | 0.066283 | NA |
| PTGFRN | 1811.119 | -0.70318 | 0.184511 | -3.81103 | 1.38E-04 | 0.00309 |
| LAMTOR5 | 2384.365 | 0.172452 | 0.164367 | 1.049185 | 0.294093 | 0.461117 |
| NOTCH2 | 2841.294 | -0.67777 | 0.192237 | -3.52568 | 4.22E-04 | 0.006131 |
| TRIM45 | 54.69377 | -0.54308 | 0.299409 | -1.81383 | 0.069704 | 0.174038 |
| CEPT1 | 802.6344 | 0.31816 | 0.172796 | 1.841248 | 0.065585 | 0.167055 |
| CD101 | 5.545709 | 0.413153 | 0.890165 | 0.464131 | 0.642554 | NA |
| VTCN1 | 0.399595 | -1.31595 | 3.579466 | -0.36764 | 0.713143 | NA |
| NGF | 24.20227 | 0.841378 | 0.723719 | 1.162576 | 0.245002 | NA |
| AP4B1 | 259.6388 | -0.16852 | 0.18982 | -0.88778 | 0.374658 | 0.54093 |
| LDHA | 30115.96 | 0.122205 | 0.152526 | 0.801209 | 0.423011 | 0.585361 |
| ANO3 | 2.830285 | -0.18544 | 1.311018 | -0.14145 | 0.887518 | NA |
| NAV1 | 2017.724 | -0.7892 | 0.192416 | -4.10155 | 4.10E-05 | 0.001483 |
| CDC73 | 976.4314 | 0.82815 | 0.163487 | 5.065558 | 4.07E-07 | 8.78E-05 |
| TIMM17A | 2437.438 | 0.355829 | 0.159009 | 2.237794 | 0.025234 | 0.088507 |
| CRB1 | 1.195988 | -1.61404 | 2.13645 | -0.75548 | 0.449963 | NA |
| FBXO18 | 2771.9 | -0.07928 | 0.135962 | -0.58311 | 0.559821 | 0.69845 |
| RBM17 | 1090.78 | 0.441649 | 0.154636 | 2.85605 | 0.004289 | 0.026624 |
| ANKRD16 | 154.9643 | -0.06816 | 0.202876 | -0.33596 | 0.736901 | 0.833267 |
| ECHDC3 | 185.4207 | -0.48572 | 0.252634 | -1.92261 | 0.054529 | 0.147055 |
| IL15RA | 2.453932 | 0.61104 | 1.502475 | 0.406689 | 0.684236 | NA |
| LRP4 | 404.0086 | -0.68696 | 0.179239 | -3.83266 | 1.27E-04 | 0.002972 |
| DDB2 | 2001.265 | -0.11108 | 0.171977 | -0.64591 | 0.51834 | 0.665395 |
| ACP2 | 1191.078 | -0.33665 | 0.17401 | -1.93467 | 0.05303 | 0.144021 |

| | | | | | | |
|---|---|---|---|---|---|---|
| PUM1 | 1890.176 | -0.11004 | 0.128972 | -0.85321 | 0.393545 | 0.55842 |
| SPOCD1 | 248.188 | 0.311733 | 0.374886 | 0.831541 | 0.405668 | 0.56959 |
| YARS | 2369.589 | -0.14359 | 0.163967 | -0.87572 | 0.381181 | 0.547413 |
| PHC2 | 4003.727 | -0.13314 | 0.136445 | -0.9758 | 0.329163 | 0.497366 |
| CDCA8 | 487.5601 | -0.41764 | 0.240874 | -1.73386 | 0.082944 | 0.195915 |
| GNL2 | 947.1777 | 0.753649 | 0.202197 | 3.727301 | 1.94E-04 | 0.003807 |
| AGO4 | 583.7519 | 0.198686 | 0.170714 | 1.163849 | 0.244485 | 0.407246 |
| HOOK1 | 27.79773 | 1.321844 | 0.46735 | 2.828381 | 0.004678 | NA |
| CYP2J2 | 138.0198 | -0.11962 | 0.235148 | -0.5087 | 0.610966 | 0.737952 |
| BTF3L4 | 609.7631 | 0.139581 | 0.147416 | 0.946848 | 0.343716 | 0.511904 |
| ZCCHC11 | 378.1266 | 0.409598 | 0.317435 | 1.290336 | 0.196934 | 0.352352 |
| PRPF38A | 530.9575 | 0.327863 | 0.15149 | 2.164251 | 0.030445 | 0.100035 |
| DAGLA | 101.0016 | -0.63087 | 0.2904 | -2.17241 | 0.029825 | 0.098598 |
| SLC43A3 | 3049.359 | -0.15768 | 0.240205 | -0.65644 | 0.51154 | 0.659379 |
| TIMM10 | 698.2364 | 0.11792 | 0.184272 | 0.639924 | 0.522222 | 0.66864 |
| GIF | 0.258578 | 1.233151 | 4.047809 | 0.304646 | 0.760635 | NA |
| APLNR | 0.575812 | 0.936922 | 2.776634 | 0.337431 | 0.735792 | NA |
| FADS2 | 8004.207 | -0.48641 | 0.149067 | -3.26306 | 0.001102 | 0.011045 |
| TMEM258 | 137.4693 | -0.14343 | 0.208093 | -0.68927 | 0.490651 | 0.642797 |
| TCN1 | 44.2246 | 0.047957 | 0.438655 | 0.109328 | 0.912943 | NA |
| PRR5L | 64.5143 | -0.36049 | 0.289559 | -1.24497 | 0.213141 | 0.37186 |
| LMO2 | 153.9101 | 1.323748 | 0.425085 | 3.114078 | 0.001845 | 0.015297 |
| PHF21A | 725.2254 | -0.12785 | 0.164384 | -0.77775 | 0.436715 | 0.597284 |
| NAT10 | 1473.43 | -0.21946 | 0.128368 | -1.70961 | 0.087338 | 0.203019 |
| EHF | 0.096202 | 0.408546 | 4.079639 | 0.100143 | 0.920231 | NA |
| PRRG4 | 209.4455 | 0.915818 | 0.216181 | 4.236351 | 2.27E-05 | 9.97E-04 |
| CAPRIN1 | 7202.886 | -0.14391 | 0.118495 | -1.21447 | 0.224569 | 0.385613 |
| AGT | 4.504847 | 0.4133 | 1.091602 | 0.378618 | 0.704972 | NA |
| ZNF670 | 49.96237 | 0.501046 | 0.317297 | 1.579108 | 0.114311 | 0.243515 |
| PCNXL2 | 289.6966 | -0.48131 | 0.17716 | -2.71681 | 0.006591 | 0.035632 |
| KCNK1 | 0.192403 | 0.975392 | 4.060716 | 0.240202 | 0.810174 | NA |
| URB2 | 479.6845 | -0.14982 | 0.175499 | -0.85371 | 0.393267 | 0.558333 |
| EGLN1 | 2817.243 | 0.040194 | 0.170108 | 0.236285 | 0.813211 | 0.8887 |
| CAPN9 | 4.399315 | 0.886612 | 0.965877 | 0.917935 | 0.358653 | NA |
| COG2 | 1217.409 | 0.011601 | 0.175029 | 0.06628 | 0.947155 | 0.97057 |
| ABCB10 | 344.238 | 0.594443 | 0.241324 | 2.463255 | 0.013768 | 0.058503 |
| NTPCR | 986.0055 | -0.05851 | 0.142683 | -0.41007 | 0.681754 | 0.792683 |
| TAF5L | 699.2931 | 0.078885 | 0.147279 | 0.53562 | 0.592221 | 0.7233 |
| GLUL | 8104.867 | -0.20114 | 0.321272 | -0.62609 | 0.531258 | 0.675613 |
| STX6 | 1365.233 | 0.13312 | 0.181376 | 0.733942 | 0.462984 | 0.620891 |
| RNASEL | 313.9046 | 0.105417 | 0.196434 | 0.536653 | 0.591507 | 0.723 |
| DHX9 | 4593.728 | 0.058422 | 0.128445 | 0.454842 | 0.649223 | 0.767275 |
| KIAA1614 | 319.5987 | -0.55661 | 0.188953 | -2.94577 | 0.003222 | 0.022179 |
| CEP350 | 850.9723 | 0.731829 | 0.269253 | 2.718002 | 0.006568 | 0.035537 |
| NPL | 618.7039 | 0.120932 | 0.242169 | 0.499372 | 0.617518 | 0.742537 |
| FAM129A | 3539.379 | 0.382065 | 0.266175 | 1.435394 | 0.151175 | 0.295106 |
| PIGC | 790.537 | 0.022473 | 0.132597 | 0.169486 | 0.865415 | 0.920317 |

| | | | | | | |
|---|---|---|---|---|---|---|
| ACBD6 | 1166.432 | -0.04278 | 0.132852 | -0.32203 | 0.74743 | 0.84124 |
| LAMC1 | 7137.256 | -0.3659 | 0.127991 | -2.85881 | 0.004252 | 0.026491 |
| RC3H1 | 341.3071 | 0.480104 | 0.20223 | 2.374055 | 0.017594 | 0.069387 |
| EPRS | 2839.071 | 0.624337 | 0.175195 | 3.563677 | 3.66E-04 | 0.005558 |
| HLX | 12.22168 | -0.34024 | 0.634006 | -0.53665 | 0.591513 | NA |
| VPS45 | 1084.518 | 0.26054 | 0.162303 | 1.605272 | 0.108434 | 0.235264 |
| IL10 | 0.151697 | -1.546 | 4.075469 | -0.37934 | 0.704433 | NA |
| KCTD3 | 965.8211 | 0.51577 | 0.241402 | 2.136556 | 0.032634 | 0.10513 |
| RPS6KC1 | 803.2709 | 0.441112 | 0.193957 | 2.274281 | 0.022949 | 0.082916 |
| RASSF5 | 2.707634 | -0.2818 | 1.224144 | -0.2302 | 0.817936 | NA |
| STAM | 2648.585 | -0.02179 | 0.163947 | -0.1329 | 0.89427 | 0.938268 |
| ABI1 | 2475.4 | 0.262771 | 0.142256 | 1.847169 | 0.064723 | 0.165792 |
| YME1L1 | 4575.654 | 0.31811 | 0.162519 | 1.957373 | 0.050304 | 0.139139 |
| DNAJC1 | 769.1839 | 0.675908 | 0.135225 | 4.998395 | 5.78E-07 | 1.12E-04 |
| BCAR3 | 1135.019 | -0.19776 | 0.148816 | -1.32891 | 0.183877 | 0.337126 |
| TTLL7 | 476.0311 | 1.083131 | 0.255003 | 4.247526 | 2.16E-05 | 9.60E-04 |
| FNBP1L | 817.6976 | 0.176737 | 0.326521 | 0.541273 | 0.58832 | 0.720882 |
| CCBL2 | 959.7174 | 0.477861 | 0.173167 | 2.759531 | 0.005788 | 0.032619 |
| GTF2B | 903.8516 | 0.44124 | 0.165661 | 2.663511 | 0.007733 | 0.039779 |
| RABGGTB | 1252.78 | 0.581061 | 0.16956 | 3.426875 | 6.11E-04 | 0.007678 |
| IFI44L | 566.9837 | -0.94571 | 0.811085 | -1.16598 | 0.243624 | 0.406464 |
| GIPC2 | 0.258578 | 1.233151 | 4.047809 | 0.304646 | 0.760635 | NA |
| ARHGAP29 | 27.30991 | 0.358687 | 0.468078 | 0.766297 | 0.443499 | NA |
| IFI44 | 251.6889 | 0.081904 | 0.378453 | 0.216418 | 0.828662 | 0.896738 |
| SLC44A5 | 1.717004 | 3.960539 | 1.870373 | 2.117513 | 0.034216 | NA |
| RPL7P9 | 44.0867 | 0.188184 | 0.362445 | 0.519205 | 0.603618 | NA |
| CLCA2 | 7.246772 | 0.503663 | 1.041565 | 0.483563 | 0.628696 | NA |
| DNASE2B | 17.13681 | -0.03879 | 0.636595 | -0.06094 | 0.951407 | NA |
| DBT | 300.0821 | 0.448716 | 0.279826 | 1.603554 | 0.108812 | 0.235864 |
| RTCA | 623.3005 | 0.339717 | 0.211934 | 1.602941 | 0.108948 | 0.236033 |
| ACTR1A | 8580.893 | -0.36635 | 0.138718 | -2.641 | 0.008266 | 0.041524 |
| TMEM180 | 181.9876 | -0.24983 | 0.214305 | -1.16577 | 0.243709 | 0.406467 |
| MYOF | 12739.05 | 0.5172 | 0.285177 | 1.813609 | 0.069738 | 0.174038 |
| LOXL4 | 2.390983 | 1.906279 | 1.448951 | 1.315627 | 0.188299 | NA |
| STAMBPL1 | 262.6008 | 0.419294 | 0.184322 | 2.274793 | 0.022918 | 0.082895 |
| CH25H | 5.784974 | -1.06173 | 0.905629 | -1.17237 | 0.241049 | NA |
| ATAD1 | 2076.038 | 0.525309 | 0.213869 | 2.456219 | 0.014041 | 0.059342 |
| BTBD16 | 5.044115 | -0.60608 | 0.911673 | -0.6648 | 0.506175 | NA |
| KIF11 | 603.5858 | 0.561575 | 0.231793 | 2.422743 | 0.015404 | 0.063089 |
| CUZD1 | 7.732755 | 1.383471 | 0.798669 | 1.73222 | 0.083234 | NA |
| TACC2 | 405.3469 | -0.46337 | 0.213297 | -2.17242 | 0.029824 | 0.098598 |
| DUSP5 | 141.8687 | 0.06247 | 0.206531 | 0.302473 | 0.762292 | 0.852303 |
| CALHM2 | 639.1693 | -0.05832 | 0.190121 | -0.30677 | 0.759019 | 0.849766 |
| ARL3 | 728.2726 | 0.197683 | 0.182668 | 1.082195 | 0.279166 | 0.445426 |
| CEP55 | 572.0468 | 0.628345 | 0.182421 | 3.444472 | 5.72E-04 | 0.007385 |
| KIF20B | 217.996 | 1.209067 | 1.161031 | 1.041374 | 0.297702 | 0.464828 |
| ENTPD1 | 309.7073 | 0.538445 | 0.196341 | 2.742395 | 0.006099 | 0.033788 |

| | | | | | | |
|---|---|---|---|---|---|---|
| EXOC6 | 176.8589 | 0.843921 | 0.241155 | 3.4995 | 4.66E-04 | 0.006516 |
| PLCE1 | 10.85352 | 1.760769 | 0.752677 | 2.339343 | 0.019318 | NA |
| RBP4 | 0.192403 | 0.975392 | 4.060716 | 0.240202 | 0.810174 | NA |
| ANXA7 | 1953.935 | 0.105095 | 0.139009 | 0.756031 | 0.449631 | 0.609046 |
| FAM149B1 | 571.2285 | 0.227456 | 0.164041 | 1.386584 | 0.165569 | 0.314604 |
| NCOA4 | 7724.073 | -0.03381 | 0.145029 | -0.23312 | 0.815668 | 0.889858 |
| TIMM23 | 1669.6 | 0.155481 | 0.168891 | 0.920601 | 0.357259 | 0.524821 |
| ASCC1 | 836.4917 | -0.00594 | 0.15486 | -0.03838 | 0.969386 | 0.983447 |
| PLA2G12B | 0.163957 | 0.424913 | 4.079006 | 0.104171 | 0.917034 | NA |
| ZNF365 | 36.46303 | 0.081812 | 0.353865 | 0.231194 | 0.817164 | NA |
| OIT3 | 0.326333 | 1.533889 | 3.826878 | 0.40082 | 0.688553 | NA |
| ADAMTS14 | 0.606787 | -3.06078 | 3.567758 | -0.8579 | 0.390947 | NA |
| RPS24 | 10941.58 | 0.567569 | 0.161859 | 3.506564 | 4.54E-04 | 0.00643 |
| TET1 | 41.6392 | 0.37741 | 0.42532 | 0.887354 | 0.374888 | NA |
| DNA2 | 56.29388 | 0.5224 | 0.298727 | 1.748757 | 0.080333 | 0.191928 |
| MYPN | 38.29594 | 0.572018 | 0.389843 | 1.467303 | 0.142294 | NA |
| SIRT3 | 490.5855 | -0.34602 | 0.161125 | -2.1475 | 0.031754 | 0.103214 |
| IFITM3 | 6220.616 | -0.76466 | 0.223499 | -3.42131 | 6.23E-04 | 0.007763 |
| ATHL1 | 156.0527 | 0.292172 | 0.224168 | 1.303364 | 0.19245 | 0.346996 |
| IL19 | 0.134133 | -1.48909 | 4.077584 | -0.36519 | 0.71497 | NA |
| SLC2A5 | 2.305381 | -0.28469 | 1.260946 | -0.22577 | 0.821377 | NA |
| RERE | 2539.008 | -0.80224 | 0.222232 | -3.60992 | 3.06E-04 | 0.005015 |
| MMEL1 | 3.432513 | -1.06801 | 1.106015 | -0.96564 | 0.334226 | NA |
| PRDM16 | 17.15368 | 0.264765 | 0.506579 | 0.522653 | 0.601216 | NA |
| CELA2A | 0.163957 | 0.424913 | 4.079006 | 0.104171 | 0.917034 | NA |
| FHAD1 | 1.410339 | -1.33003 | 1.92711 | -0.69017 | 0.490089 | NA |
| PADI1 | 1.121596 | 1.119295 | 2.311568 | 0.484215 | 0.628233 | NA |
| EPHA2 | 472.5465 | -0.2776 | 0.241299 | -1.15045 | 0.249958 | 0.41369 |
| ARHGEF19 | 141.583 | -0.26052 | 0.300427 | -0.86716 | 0.385853 | 0.551916 |
| EFHD2 | 2836.096 | -0.06497 | 0.165617 | -0.3923 | 0.69484 | 0.80325 |
| PEX14 | 796.6774 | -0.40581 | 0.137454 | -2.95233 | 0.003154 | 0.021929 |
| PGD | 4250.269 | -0.41468 | 0.172675 | -2.40152 | 0.016327 | 0.065972 |
| MYOM3 | 0.314073 | -0.57308 | 3.898934 | -0.14698 | 0.883146 | NA |
| SH3BGRL3 | 4037.813 | -0.18875 | 0.227004 | -0.83148 | 0.405704 | 0.56959 |
| CNKSR1 | 1.441394 | 1.117352 | 1.830657 | 0.610356 | 0.541626 | NA |
| RPL11 | 15869.22 | 0.225719 | 0.129732 | 1.739888 | 0.081879 | 0.194406 |
| IL22RA1 | 0.314073 | -0.57308 | 3.898934 | -0.14698 | 0.883146 | NA |
| ZNF593 | 1.132682 | 0.571077 | 1.952886 | 0.292427 | 0.76996 | NA |
| C1orf216 | 2441.703 | -0.02005 | 0.186588 | -0.10746 | 0.914425 | 0.950505 |
| KIAA0319L | 2653.289 | -0.40467 | 0.145051 | -2.78983 | 0.005274 | 0.030661 |
| EVA1B | 835.0941 | -0.06122 | 0.198903 | -0.30778 | 0.758246 | 0.849302 |
| DMRTA2 | 0.622842 | 1.158846 | 2.735323 | 0.42366 | 0.671814 | NA |
| MAP3K6 | 1059.616 | -0.56573 | 0.173256 | -3.26529 | 0.001094 | 0.010986 |
| FCN3 | 0.841063 | -2.08438 | 2.306687 | -0.90362 | 0.366195 | NA |
| GPN2 | 592.3928 | -0.19709 | 0.145498 | -1.35456 | 0.175558 | 0.327548 |
| SYTL1 | 3.651037 | -0.19113 | 1.102581 | -0.17335 | 0.862378 | NA |
| WDTC1 | 2199.431 | -0.51795 | 0.138643 | -3.73585 | 1.87E-04 | 0.003727 |

| | | | | | | |
|---|---|---|---|---|---|---|
| NBPF3 | 439.887 | -0.11174 | 0.167488 | -0.66712 | 0.504694 | 0.654325 |
| HSPG2 | 9626.487 | -0.51214 | 0.268877 | -1.90472 | 0.056816 | 0.151056 |
| ITGB3BP | 245.52 | 0.646044 | 0.224523 | 2.877407 | 0.00401 | 0.025436 |
| SERBP1 | 4708.802 | 0.277615 | 0.140004 | 1.982913 | 0.047377 | 0.134015 |
| BCL10 | 403.2355 | 0.117416 | 0.165086 | 0.711245 | 0.476933 | 0.631807 |
| CYR61 | 677.1788 | 0.53432 | 0.788129 | 0.677961 | 0.497797 | 0.648717 |
| PRKACB | 1818.437 | 0.62028 | 0.209683 | 2.958181 | 0.003095 | 0.021663 |
| PIGK | 1504.344 | 0.752813 | 0.206521 | 3.645212 | 2.67E-04 | 0.004599 |
| TINAGL1 | 3.052272 | 0.552538 | 1.248387 | 0.442601 | 0.658054 | NA |
| ADC | 306.3739 | 0.092381 | 0.184262 | 0.501358 | 0.616119 | 0.742059 |
| RPS8 | 21285.64 | 0.337768 | 0.142079 | 2.377317 | 0.017439 | 0.069029 |
| KIF2C | 793.0517 | -0.34481 | 0.248462 | -1.38779 | 0.165202 | 0.314059 |
| PTPRF | 324.8093 | -0.37111 | 0.306697 | -1.21003 | 0.226267 | 0.38763 |
| BEST4 | 0.561971 | -0.49218 | 2.810061 | -0.17515 | 0.860962 | NA |
| MOB3C | 253.1684 | -0.11665 | 0.185024 | -0.63046 | 0.528393 | 0.673776 |
| TMEM61 | 0.163957 | 0.424913 | 4.079006 | 0.104171 | 0.917034 | NA |
| LMO4 | 1946.358 | 0.032227 | 0.14953 | 0.215525 | 0.829358 | 0.897165 |
| SYPL2 | 42.82258 | -0.19583 | 0.343699 | -0.56976 | 0.568842 | NA |
| BARHL2 | 0.192403 | 0.975392 | 4.060716 | 0.240202 | 0.810174 | NA |
| MTF2 | 241.3031 | 0.602598 | 0.229096 | 2.630333 | 0.00853 | 0.042312 |
| SLC44A3 | 256.6622 | -0.25992 | 0.229701 | -1.13157 | 0.257814 | 0.421919 |
| IGSF3 | 3616.001 | -0.55056 | 0.154973 | -3.5526 | 3.81E-04 | 0.005731 |
| ZNF697 | 1175.825 | -0.15709 | 0.189143 | -0.83054 | 0.406236 | 0.569932 |
| CTTNBP2NL | 683.7142 | 0.112991 | 0.177902 | 0.635129 | 0.525344 | 0.670971 |
| STRIP1 | 247.3274 | -0.2284 | 0.214709 | -1.06378 | 0.287427 | 0.453587 |
| KCNA10 | 0.163957 | 0.424913 | 4.079006 | 0.104171 | 0.917034 | NA |
| PSMA5 | 3016.11 | 0.109938 | 0.172805 | 0.636195 | 0.524649 | 0.670408 |
| FNDC7 | 0.096202 | 0.408546 | 4.079639 | 0.100143 | 0.920231 | NA |
| C1orf162 | 1.60277 | 1.310065 | 2.178035 | 0.60149 | 0.547514 | NA |
| CD53 | 0.545988 | -0.43913 | 2.541145 | -0.17281 | 0.862803 | NA |
| CELSR2 | 2982.381 | -0.74801 | 0.175481 | -4.26264 | 2.02E-05 | 9.36E-04 |
| ITGA10 | 3.892366 | 0.75777 | 1.301938 | 0.582032 | 0.560545 | NA |
| GJA5 | 15.13021 | 0.019889 | 0.60783 | 0.032721 | 0.973897 | NA |
| GPR161 | 481.5864 | -0.12216 | 0.159396 | -0.76637 | 0.443456 | 0.603336 |
| ALDH9A1 | 2507.681 | -0.29413 | 0.141619 | -2.07691 | 0.03781 | 0.115644 |
| ATP1B1 | 741.847 | 0.328374 | 0.357096 | 0.919567 | 0.357799 | 0.525131 |
| TIPRL | 876.0109 | 0.401279 | 0.142221 | 2.821527 | 0.00478 | 0.028636 |
| NME7 | 233.9264 | 0.360101 | 0.193157 | 1.864293 | 0.062281 | 0.161562 |
| POGK | 1627.712 | -0.1388 | 0.15465 | -0.89753 | 0.369435 | 0.536481 |
| MPC2 | 767.444 | 0.176393 | 0.155521 | 1.134209 | 0.256707 | 0.420707 |
| CREG1 | 1618.867 | 0.177111 | 0.212521 | 0.833382 | 0.404629 | 0.569092 |
| DCAF6 | 1991.808 | 0.373671 | 0.136398 | 2.739568 | 0.006152 | 0.03389 |
| RXRG | 1196.586 | -0.45577 | 0.163822 | -2.78208 | 0.005401 | 0.031234 |
| TBX19 | 21.9291 | 0.412348 | 0.568316 | 0.72556 | 0.468108 | NA |
| UCK2 | 1253.893 | -0.30693 | 0.163044 | -1.88252 | 0.059765 | 0.156926 |
| TMCO1 | 2374.941 | 0.505453 | 0.164537 | 3.071984 | 0.002126 | 0.016816 |
| XCL1 | 0.096202 | 0.408546 | 4.079639 | 0.100143 | 0.920231 | NA |

| | | | | | | |
|---|---|---|---|---|---|---|
| POU2F1 | 372.8568 | 0.069529 | 0.22411 | 0.310245 | 0.756374 | 0.847846 |
| ILDR2 | 1.427407 | 0.984049 | 1.659042 | 0.593143 | 0.553086 | NA |
| MGST3 | 4077.502 | -0.00133 | 0.152977 | -0.00866 | 0.993088 | 0.996912 |
| ADCY10 | 0.619047 | -1.7574 | 3.107232 | -0.56558 | 0.571677 | NA |
| RFWD2 | 949.8406 | 0.129178 | 0.164136 | 0.787021 | 0.43127 | 0.592665 |
| PVRL4 | 3.747367 | -0.81319 | 1.12328 | -0.72394 | 0.469102 | NA |
| UFC1 | 2340.57 | 0.356959 | 0.124221 | 2.873593 | 0.004058 | 0.02562 |
| PPOX | 425.3966 | -0.30184 | 0.151329 | -1.99459 | 0.046087 | 0.131845 |
| FCGR2A | 34.73607 | -0.29482 | 0.385328 | -0.76512 | 0.444197 | NA |
| NUF2 | 212.0165 | 1.148873 | 0.190025 | 6.045917 | 1.49E-09 | 1.35E-06 |
| RGS5 | 8.927165 | 0.650965 | 0.679617 | 0.957841 | 0.338143 | NA |
| SDHC | 2102.271 | -0.08083 | 0.142553 | -0.56702 | 0.570704 | 0.7066 |
| PFDN2 | 1653.695 | 0.260776 | 0.178765 | 1.458767 | 0.144629 | 0.286575 |
| NR1I3 | 12.76885 | 0.134064 | 0.655094 | 0.204648 | 0.837847 | NA |
| USP21 | 631.5313 | -0.35191 | 0.192322 | -1.82979 | 0.067281 | 0.169633 |
| F13B | 0.163957 | 0.424913 | 4.079006 | 0.104171 | 0.917034 | NA |
| PRCC | 1765.395 | -0.28591 | 0.154293 | -1.85302 | 0.063879 | 0.164341 |
| FCRL5 | 0.163957 | 0.424913 | 4.079006 | 0.104171 | 0.917034 | NA |
| RRNAD1 | 418.952 | -0.22042 | 0.177823 | -1.23956 | 0.21514 | 0.374251 |
| MRPL24 | 1919.722 | -0.15617 | 0.170556 | -0.91567 | 0.359841 | 0.526982 |
| PIGM | 356.5289 | -0.15074 | 0.201275 | -0.74894 | 0.453892 | 0.612718 |
| CASQ1 | 0.49187 | 2.136088 | 3.895953 | 0.548284 | 0.583497 | NA |
| ISG20L2 | 838.5181 | -0.19655 | 0.145674 | -1.34921 | 0.17727 | 0.329394 |
| CRABP2 | 8.243275 | 0.609877 | 0.735293 | 0.829434 | 0.406859 | NA |
| HDGF | 8623.122 | -0.14807 | 0.203583 | -0.72733 | 0.467023 | 0.623713 |
| ABL2 | 7440.286 | 0.093261 | 0.17656 | 0.528213 | 0.597352 | 0.727064 |
| XPR1 | 851.5278 | 0.210968 | 0.160627 | 1.313404 | 0.189047 | 0.343314 |
| RGS16 | 511.2001 | 1.700257 | 0.881239 | 1.929393 | 0.053682 | 0.145364 |
| TOR1AIP1 | 1929.525 | 0.065835 | 0.196485 | 0.335062 | 0.737579 | 0.833837 |
| FAM163A | 8.792415 | 0.704529 | 0.717027 | 0.98257 | 0.325819 | NA |
| HMCN1 | 1861.961 | 0.090638 | 0.141355 | 0.641207 | 0.521388 | 0.668146 |
| RGL1 | 602.2438 | -0.32911 | 0.166672 | -1.9746 | 0.048314 | 0.135641 |
| LYPLAL1 | 292.4568 | 0.213716 | 0.312013 | 0.684957 | 0.493371 | 0.64458 |
| LHX9 | 0.489058 | -0.58569 | 3.427543 | -0.17088 | 0.864321 | NA |
| PRUNE | 654.8569 | -0.17314 | 0.17478 | -0.99061 | 0.321875 | 0.490088 |
| RORC | 0.392857 | -1.546 | 4.075469 | -0.37934 | 0.704433 | NA |
| TUFT1 | 382.3039 | -0.62159 | 0.16224 | -3.83128 | 1.27E-04 | 0.002981 |
| SF3B4 | 1648.369 | -0.43498 | 0.152169 | -2.85855 | 0.004256 | 0.026493 |
| ECM1 | 9656.536 | -0.59037 | 0.176699 | -3.3411 | 8.34E-04 | 0.009297 |
| ZNF687 | 1224.466 | -0.67403 | 0.149773 | -4.50032 | 6.79E-06 | 5.06E-04 |
| TARS2 | 1085.351 | -0.37628 | 0.139896 | -2.6897 | 0.007152 | 0.037648 |
| CGN | 255.4162 | -0.19742 | 0.228156 | -0.86527 | 0.386893 | 0.552871 |
| SNX27 | 1427.861 | -0.44354 | 0.226495 | -1.95827 | 0.050198 | 0.138943 |
| SETDB1 | 605.6723 | -0.33309 | 0.180603 | -1.84433 | 0.065135 | 0.166416 |
| ADAMTSL4 | 30.21972 | -0.33206 | 0.377668 | -0.87924 | 0.37927 | NA |
| MCL1 | 4125.283 | 0.00768 | 0.127062 | 0.060445 | 0.951801 | 0.97348 |
| CTSK | 44351.08 | -0.09258 | 0.175472 | -0.52763 | 0.597756 | 0.727481 |

| | | | | | | |
|---|---|---|---|---|---|---|
| RFX5 | 2385.628 | -0.13439 | 0.182844 | -0.73498 | 0.462352 | 0.620395 |
| PI4KB | 3380.785 | -0.40204 | 0.130133 | -3.08945 | 0.002005 | 0.016238 |
| PIP5K1A | 1852.527 | -0.19281 | 0.148772 | -1.29604 | 0.194962 | 0.350068 |
| ANP32E | 636.7749 | 0.476829 | 0.266213 | 1.791152 | 0.073269 | 0.180395 |
| FAM63A | 1225.41 | -0.47231 | 0.283036 | -1.66872 | 0.095172 | 0.215853 |
| ANXA9 | 97.41616 | -0.56725 | 0.372023 | -1.52476 | 0.127319 | 0.263216 |
| SELENBP1 | 2471.14 | -0.61719 | 0.196345 | -3.14337 | 0.00167 | 0.014431 |
| CERS2 | 3263.07 | -0.21289 | 0.165981 | -1.28265 | 0.199615 | 0.355802 |
| ENSA | 4150.586 | -0.02732 | 0.141478 | -0.19308 | 0.846895 | 0.908839 |
| SEMA6C | 112.4745 | -0.26288 | 0.250535 | -1.04926 | 0.294056 | 0.461117 |
| MRPL9 | 643.6555 | 0.1526 | 0.138057 | 1.105344 | 0.269011 | 0.433776 |
| ARNT | 1389.654 | -0.29239 | 0.143366 | -2.03943 | 0.041407 | 0.123369 |
| POGZ | 1155.297 | -0.48484 | 0.135286 | -3.58384 | 3.39E-04 | 0.005324 |
| C1orf56 | 147.3659 | -0.42631 | 0.228793 | -1.86329 | 0.062421 | 0.161813 |
| OAZ3 | 20.28709 | -0.32109 | 0.458668 | -0.70004 | 0.483899 | NA |
| HORMAD1 | 0.268266 | -2.15396 | 4.038351 | -0.53338 | 0.593773 | NA |
| GOLPH3L | 482.4849 | -0.05794 | 0.183164 | -0.31631 | 0.751767 | 0.844731 |
| GABPB2 | 108.0292 | 0.037966 | 0.279672 | 0.135751 | 0.892018 | 0.936771 |
| IKBKE | 434.3926 | -0.09509 | 0.161643 | -0.5883 | 0.556332 | 0.695782 |
| SYT14 | 483.008 | 0.549073 | 0.265207 | 2.070352 | 0.038419 | 0.116995 |
| KCNH1 | 142.7504 | -0.31918 | 0.220868 | -1.44512 | 0.148425 | 0.291611 |
| DTL | 517.6619 | -0.00957 | 0.221572 | -0.04319 | 0.965546 | 0.981198 |
| DYRK3 | 373.363 | 0.143812 | 0.187534 | 0.76686 | 0.443165 | 0.603321 |
| EIF2D | 1184.947 | -0.04896 | 0.133698 | -0.36621 | 0.714211 | 0.81767 |
| INTS7 | 989.4679 | -0.08849 | 0.176885 | -0.50028 | 0.616879 | 0.742224 |
| VASH2 | 9.316558 | 0.030778 | 0.717061 | 0.042923 | 0.965763 | NA |
| TAF1A | 71.55051 | 0.301931 | 0.283015 | 1.066835 | 0.286046 | 0.452446 |
| SMYD2 | 1857.831 | 0.107953 | 0.164306 | 0.657025 | 0.511165 | 0.659182 |
| DUSP10 | 1050.177 | 0.060661 | 0.222183 | 0.273021 | 0.784837 | 0.869149 |
| HHIPL2 | 0.392857 | -1.546 | 4.075469 | -0.37934 | 0.704433 | NA |
| TP53BP2 | 1377.537 | 0.054235 | 0.154572 | 0.350871 | 0.725685 | 0.825703 |
| ATP8B2 | 4693.41 | -0.11409 | 0.219651 | -0.51944 | 0.603456 | 0.732334 |
| FLG2 | 0.82213 | 0.50433 | 2.197682 | 0.229483 | 0.818494 | NA |
| ADAM15 | 5141.859 | -0.36716 | 0.128001 | -2.86842 | 0.004125 | 0.025944 |
| JTB | 3229.917 | -0.09603 | 0.181548 | -0.52898 | 0.596821 | 0.726717 |
| RAB13 | 3371.391 | 0.474659 | 0.189386 | 2.506307 | 0.0122 | 0.054027 |
| TPM3 | 10866.93 | -0.05285 | 0.120642 | -0.43803 | 0.661363 | 0.776975 |
| NUP210L | 1.148665 | 0.530005 | 1.802635 | 0.294017 | 0.768745 | NA |
| SNAPIN | 1207.977 | -0.15315 | 0.152584 | -1.00372 | 0.315515 | 0.482886 |
| SLC27A3 | 896.5635 | -0.67452 | 0.290482 | -2.32207 | 0.020229 | 0.076302 |
| UBAP2L | 5682.438 | -0.48308 | 0.141113 | -3.42337 | 6.18E-04 | 0.007734 |
| SLC39A1 | 5723.467 | -0.12032 | 0.155086 | -0.77584 | 0.437846 | 0.597933 |
| HAX1 | 2558.827 | 0.058186 | 0.13839 | 0.420452 | 0.674155 | 0.786547 |
| CREB3L4 | 144.3961 | -0.31811 | 0.229031 | -1.38892 | 0.164856 | 0.313702 |
| EFNA3 | 14.63853 | 0.183366 | 0.591896 | 0.309794 | 0.756718 | NA |
| AQP10 | 0.163957 | 0.424913 | 4.079006 | 0.104171 | 0.917034 | NA |
| KCNN3 | 14.18208 | 0.01159 | 0.550065 | 0.021071 | 0.983189 | NA |

| | | | | | | |
|---|---|---|---|---|---|---|
| C1orf43 | 9653.78 | -0.04778 | 0.149977 | -0.31859 | 0.750036 | 0.843533 |
| GATAD2B | 976.1301 | -0.51771 | 0.14131 | -3.66367 | 2.49E-04 | 0.004426 |
| ILF2 | 4091.995 | -0.02498 | 0.14143 | -0.17661 | 0.859812 | 0.916725 |
| RIT1 | 756.8501 | 0.25431 | 0.186994 | 1.359994 | 0.173832 | 0.325815 |
| INTS3 | 1492.577 | -0.37027 | 0.152395 | -2.42971 | 0.015111 | 0.062396 |
| PKLR | 4.514655 | -0.22274 | 1.008203 | -0.22093 | 0.825149 | NA |
| HCN3 | 89.8586 | -0.0378 | 0.250841 | -0.15069 | 0.880218 | 0.928462 |
| FLG | 271.6411 | -0.87979 | 0.34079 | -2.58162 | 0.009834 | 0.046688 |
| ACTA1 | 1.794833 | 0.911042 | 1.612736 | 0.564905 | 0.572139 | NA |
| C1orf131 | 152.0711 | 0.806474 | 0.229583 | 3.512783 | 4.43E-04 | 0.006334 |
| GALNT2 | 10280.59 | -0.33532 | 0.183874 | -1.82362 | 0.068209 | 0.171408 |
| TTC13 | 303.2533 | 0.380864 | 0.219945 | 1.731632 | 0.083339 | 0.196653 |
| SCCPDH | 2420.308 | 0.106604 | 0.183647 | 0.580484 | 0.561589 | 0.699721 |
| LYST | 6296.071 | 0.519756 | 0.243541 | 2.134167 | 0.032829 | 0.105579 |
| | 12.8184 | 0.438103 | 0.643793 | 0.680503 | 0.496186 | NA |
| CEP170 | 759.2163 | 0.421662 | 0.279417 | 1.509077 | 0.131279 | 0.267868 |
| SNAP47 | 860.8847 | -0.10058 | 0.179039 | -0.56176 | 0.574276 | 0.709634 |
| SRP9 | 1513.349 | 0.600948 | 0.177447 | 3.386638 | 7.08E-04 | 0.008346 |
| NVL | 356.8128 | 0.38178 | 0.206169 | 1.851783 | 0.064057 | 0.164621 |
| SDE2 | 380.2543 | 0.304463 | 0.220475 | 1.380941 | 0.167297 | 0.316783 |
| DEGS1 | 4082.09 | 0.029586 | 0.158272 | 0.186931 | 0.851715 | 0.911865 |
| FBXO28 | 1090.409 | 0.246073 | 0.167134 | 1.47231 | 0.140937 | 0.281423 |
| ARF1 | 13642.1 | -0.2666 | 0.119965 | -2.22234 | 0.026261 | 0.090994 |
| CNIH4 | 1735.23 | 0.192971 | 0.147266 | 1.310353 | 0.190076 | 0.344286 |
| ITPKB | 10703.62 | -0.56387 | 0.138376 | -4.0749 | 4.60E-05 | 0.001568 |
| GUK1 | 7470.929 | -0.0335 | 0.131592 | -0.25461 | 0.799028 | 0.878627 |
| CDC42BPA | 1375.488 | 0.733694 | 0.307537 | 2.38571 | 0.017046 | 0.067932 |
| CNIH3 | 156.3662 | -0.36306 | 0.203709 | -1.78224 | 0.07471 | 0.182427 |
| C1orf35 | 385.5671 | -0.12304 | 0.160753 | -0.76541 | 0.44403 | 0.603875 |
| PARP1 | 4292.075 | -0.21806 | 0.122682 | -1.77741 | 0.075501 | 0.183754 |
| PSEN2 | 1792.723 | -0.29114 | 0.164158 | -1.77351 | 0.076144 | 0.184863 |
| PYCR2 | 1071.808 | -0.1699 | 0.136084 | -1.24852 | 0.211839 | 0.370324 |
| LBR | 436.3215 | 0.406537 | 0.191716 | 2.120515 | 0.033963 | 0.107625 |
| WNT9A | 0.785567 | -0.70568 | 2.358063 | -0.29926 | 0.764741 | NA |
| EPHX1 | 4384.758 | -0.70582 | 0.184301 | -3.8297 | 1.28E-04 | 0.002993 |
| REN | 0.096202 | 0.408546 | 4.079639 | 0.100143 | 0.920231 | NA |
| SOX13 | 1581.394 | -0.45659 | 0.192478 | -2.37216 | 0.017684 | 0.069628 |
| ETNK2 | 1081.344 | -0.377 | 0.180972 | -2.08319 | 0.037234 | 0.114532 |
| PPFIA4 | 26.49784 | -0.47598 | 0.430829 | -1.1048 | 0.269248 | NA |
| PLEKHA6 | 303.4822 | -1.54252 | 0.275604 | -5.59686 | 2.18E-08 | 1.29E-05 |
| PTPN7 | 0.326333 | 1.533889 | 3.826878 | 0.40082 | 0.688553 | NA |
| SYT2 | 1.716012 | 1.369682 | 1.544299 | 0.886928 | 0.375118 | NA |
| ARL8A | 2184.348 | -0.22481 | 0.133024 | -1.69001 | 0.091026 | 0.209456 |
| ZMYM4 | 883.1561 | 0.268361 | 0.250265 | 1.072307 | 0.283582 | 0.450055 |
| IDI2 | 0.602678 | 1.083611 | 2.70938 | 0.399948 | 0.689195 | NA |
| PROSER2 | 84.01825 | 0.092206 | 0.362847 | 0.254119 | 0.799404 | 0.878959 |
| USP6NL | 626.2258 | 0.07649 | 0.177914 | 0.429926 | 0.66725 | 0.781814 |

| | | | | | | |
|---|---|---|---|---|---|---|
| COMMD3 | 1001.259 | 0.440921 | 0.165793 | 2.659468 | 0.007826 | 0.040061 |
| MSRB2 | 1025.398 | -0.18937 | 0.148493 | -1.27525 | 0.202222 | 0.358669 |
| PDSS1 | 231.8665 | 0.386087 | 0.234048 | 1.649604 | 0.099024 | 0.221873 |
| FAM171A1 | 11.26362 | 0.596917 | 0.672315 | 0.887852 | 0.37462 | NA |
| FAM188A | 1318.732 | 0.116094 | 0.159271 | 0.728911 | 0.466056 | 0.623178 |
| SLC39A12 | 24.86941 | 0.189224 | 0.427403 | 0.442728 | 0.657962 | NA |
| TMEM236 | 0.162376 | 0.424913 | 4.079006 | 0.104171 | 0.917034 | NA |
| RSU1 | 6290.543 | 0.228833 | 0.134463 | 1.701828 | 0.088788 | 0.205541 |
| ST8SIA6 | 878.105 | 0.263049 | 0.265968 | 0.989026 | 0.32265 | 0.490721 |
| PARD3 | 834.1324 | 0.104864 | 0.148442 | 0.706433 | 0.479919 | 0.634563 |
| ANKRD30A | 0.163957 | 0.424913 | 4.079006 | 0.104171 | 0.917034 | NA |
| ZEB1 | 0.096202 | 0.408546 | 4.079639 | 0.100143 | 0.920231 | NA |
| FAM13C | 12.27581 | 0.054146 | 0.632979 | 0.085542 | 0.93183 | NA |
| NRBF2 | 574.5386 | 0.534016 | 0.170645 | 3.129389 | 0.001752 | 0.014833 |
| CDHR1 | 22.93818 | 0.148339 | 0.760409 | 0.195078 | 0.845332 | NA |
| RGR | 2.195937 | -0.78637 | 1.675285 | -0.46939 | 0.638789 | NA |
| POLR3A | 1107.681 | -0.31565 | 0.133293 | -2.36811 | 0.017879 | 0.07 |
| HERC4 | 953.6574 | 0.796025 | 0.216371 | 3.678987 | 2.34E-04 | 0.004291 |
| C10orf11 | 843.0091 | -0.01337 | 0.139434 | -0.09592 | 0.923582 | 0.956 |
| CAMK2G | 577.5129 | -0.31216 | 0.195039 | -1.60051 | 0.109486 | 0.236809 |
| ADIRF | 1.275711 | 1.512025 | 1.883195 | 0.802904 | 0.42203 | NA |
| GLUD1 | 3289.977 | 0.100754 | 0.127364 | 0.791066 | 0.428905 | 0.59062 |
| ANKRD1 | 0.864416 | 1.729451 | 2.409716 | 0.717699 | 0.472943 | NA |
| HTR7 | 37.48487 | 0.941902 | 0.500548 | 1.88174 | 0.059871 | NA |
| RPP30 | 714.3754 | 0.376741 | 0.152827 | 2.465152 | 0.013696 | 0.058277 |
| FRA10AC1 | 232.3804 | 0.494899 | 0.31104 | 1.591112 | 0.111584 | 0.239791 |
| ADD3 | 4957.585 | 0.452758 | 0.247312 | 1.830717 | 0.067143 | 0.169494 |
| VAX1 | 0.096202 | 0.408546 | 4.079639 | 0.100143 | 0.920231 | NA |
| DNAJB12 | 1987.465 | -0.33991 | 0.152148 | -2.23405 | 0.02548 | 0.089236 |
| EIF4EBP2 | 3686.923 | -0.28258 | 0.122257 | -2.31139 | 0.020812 | 0.077845 |
| PLEKHS1 | 0.162376 | 0.424913 | 4.079006 | 0.104171 | 0.917034 | NA |
| TCF7L2 | 374.8013 | -0.35333 | 0.197583 | -1.78825 | 0.073735 | 0.181055 |
| MKI67 | 1611.442 | -0.33303 | 0.259055 | -1.28557 | 0.198592 | 0.354457 |
| CYP17A1 | 0.642189 | 0.046766 | 2.378271 | 0.019664 | 0.984311 | NA |
| INA | 8.833155 | 0.105611 | 0.825886 | 0.127876 | 0.898247 | NA |
| FUOM | 739.2279 | -0.08316 | 0.231659 | -0.35898 | 0.71961 | 0.82133 |
| LRRC27 | 341.7712 | 0.002936 | 0.187732 | 0.015642 | 0.98752 | 0.99347 |
| MTG1 | 80.95469 | -0.07858 | 0.252664 | -0.311 | 0.755798 | 0.847421 |
| PAOX | 127.4225 | -0.02421 | 0.248076 | -0.09758 | 0.922263 | 0.955032 |
| GSTO1 | 14116.17 | 0.181592 | 0.191124 | 0.950124 | 0.342049 | 0.51019 |
| TAF5 | 118.7878 | -0.00923 | 0.269282 | -0.03428 | 0.972653 | 0.985034 |
| PPRC1 | 892.1439 | -0.44401 | 0.18008 | -2.46562 | 0.013678 | 0.058239 |
| ITPRIP | 836.4972 | -0.37618 | 0.176408 | -2.13246 | 0.032969 | 0.105731 |
| CNNM2 | 431.7595 | -0.36313 | 0.163344 | -2.22312 | 0.026208 | 0.090873 |
| PDCD11 | 1974 | -0.41522 | 0.153382 | -2.70707 | 0.006788 | 0.036363 |
| ADAM12 | 144.6709 | -0.76483 | 0.345115 | -2.21616 | 0.026681 | 0.091707 |
| RGS10 | 2581.112 | 0.379162 | 0.178851 | 2.119989 | 0.034007 | 0.107737 |

| | | | | | | |
|---|---|---|---|---|---|---|
| BTBD10 | 626.5265 | 0.380649 | 0.185451 | 2.052554 | 0.040116 | 0.120643 |
| ADM | 120.3731 | -0.44692 | 0.239259 | -1.86795 | 0.061769 | 0.160479 |
| GAS2 | 118.4881 | 0.141953 | 0.302682 | 0.468986 | 0.63908 | 0.759385 |
| SLC5A12 | 0.70851 | -1.62753 | 2.686091 | -0.60591 | 0.544574 | NA |
| LIN7C | 311.1776 | 0.734234 | 0.296425 | 2.476968 | 0.01325 | 0.057233 |
| IMMP1L | 112.8472 | 0.189244 | 0.275655 | 0.686523 | 0.492383 | 0.643732 |
| PGAP2 | 347.993 | -0.25167 | 0.171089 | -1.47101 | 0.14129 | 0.28189 |
| TUT1 | 203.9728 | -0.56804 | 0.188192 | -3.01838 | 0.002541 | 0.019043 |
| SCGB1A1 | 0.47961 | 0.535796 | 3.39605 | 0.15777 | 0.874638 | NA |
| SYT8 | 0.162376 | 0.424913 | 4.079006 | 0.104171 | 0.917034 | NA |
| ZNF214 | 25.68135 | 0.879615 | 0.450301 | 1.953392 | 0.050773 | NA |
| ZNF215 | 78.10623 | 0.641505 | 0.295415 | 2.17154 | 0.02989 | 0.098788 |
| HSD17B12 | 4140.305 | 0.430394 | 0.154207 | 2.791018 | 0.005254 | 0.030564 |
| APIP | 422.7527 | 0.693304 | 0.196619 | 3.526124 | 4.22E-04 | 0.006131 |
| PAMR1 | 27.01302 | -1.17087 | 0.435558 | -2.68821 | 0.007184 | NA |
| DGKZ | 1772.319 | -0.2688 | 0.172615 | -1.55723 | 0.119415 | 0.251264 |
| EIF3M | 2483.562 | 0.264954 | 0.166979 | 1.586754 | 0.112568 | 0.241189 |
| TNKS1BP1 | 2612.334 | -0.52663 | 0.179541 | -2.9332 | 0.003355 | 0.022701 |
| GLYAT | 0.392711 | 0.292538 | 3.145221 | 0.09301 | 0.925895 | NA |
| SERPING1 | 254.7203 | -0.45217 | 0.294296 | -1.53644 | 0.124431 | 0.259008 |
| SSRP1 | 3248.961 | -0.08187 | 0.120959 | -0.67687 | 0.498485 | 0.649308 |
| SLC43A1 | 332.5359 | -0.19528 | 0.176523 | -1.10623 | 0.268627 | 0.433599 |
| PTPRJ | 1832.463 | -0.31148 | 0.153695 | -2.0266 | 0.042703 | 0.125806 |
| C11orf49 | 853.065 | -0.32688 | 0.139473 | -2.34368 | 0.019095 | 0.073307 |
| ARFGAP2 | 2139.23 | -0.38761 | 0.129298 | -2.99782 | 0.002719 | 0.019923 |
| CELF1 | 1217.804 | 0.075589 | 0.163288 | 0.46292 | 0.643422 | 0.762832 |
| DAK | 526.3821 | -0.33266 | 0.20911 | -1.59083 | 0.111647 | 0.239791 |
| MTA2 | 2546.714 | -0.41039 | 0.153013 | -2.68208 | 0.007317 | 0.038363 |
| TMEM138 | 930.7724 | -0.03807 | 0.163313 | -0.23312 | 0.815667 | 0.889858 |
| FADS1 | 3556.222 | -0.26678 | 0.122474 | -2.17823 | 0.029389 | 0.097646 |
| ROM1 | 70.17626 | -0.00173 | 0.285779 | -0.00607 | 0.995157 | 0.997908 |
| EML3 | 1497 | -0.31224 | 0.152325 | -2.04982 | 0.040382 | 0.121106 |
| INCENP | 2138.128 | -0.80751 | 0.17648 | -4.57564 | 4.75E-06 | 4.09E-04 |
| OOSP2 | 0.392857 | -1.546 | 4.075469 | -0.37934 | 0.704433 | NA |
| MS4A3 | 0.134133 | -1.48909 | 4.077584 | -0.36519 | 0.71497 | NA |
| PLCH2 | 1.195639 | 0.515221 | 2.098932 | 0.245468 | 0.806094 | NA |
| CPSF7 | 1431.149 | -0.33567 | 0.151499 | -2.21567 | 0.026714 | 0.091765 |
| MS4A2 | 0.364467 | -1.20166 | 3.685431 | -0.32606 | 0.744381 | NA |
| B3GAT3 | 2057.796 | -0.09469 | 0.151893 | -0.62338 | 0.533037 | 0.677164 |
| MPP7 | 1.909793 | 2.161609 | 1.550034 | 1.394556 | 0.16315 | NA |
| | 1.107959 | -0.3351 | 1.823092 | -0.18381 | 0.854166 | NA |
| ITGB1 | 10751.94 | 0.625094 | 0.17001 | 3.676805 | 2.36E-04 | 0.004314 |
| TRIM48 | 3.425227 | 1.779718 | 1.573138 | 1.131317 | 0.257922 | NA |
| ARID5B | 1106.609 | -0.18257 | 0.174908 | -1.04379 | 0.296585 | 0.463665 |
| PDCD4 | 1305.238 | 0.266623 | 0.195412 | 1.364414 | 0.172437 | 0.323815 |
| ADRA2A | 7.767025 | 2.005369 | 0.978958 | 2.048472 | 0.040514 | NA |
| RGS18 | 0.411855 | 0.250456 | 3.097618 | 0.080854 | 0.935558 | NA |

| | | | | | | |
|---|---|---|---|---|---|---|
| DOCK1 | 1945.047 | 0.238917 | 0.165037 | 1.447661 | 0.147712 | 0.290884 |
| PIP4K2A | 4124.115 | 0.082035 | 0.144227 | 0.56879 | 0.569498 | 0.705805 |
| ENKUR | 1.208102 | 0.723565 | 1.850162 | 0.391082 | 0.695737 | NA |
| GPR158 | 100.3647 | -1.23854 | 0.474157 | -2.61208 | 0.008999 | 0.044103 |
| UEVLD | 555.1056 | 0.681702 | 0.199295 | 3.420573 | 6.25E-04 | 0.007763 |
| TMEM86A | 360.5968 | -0.50307 | 0.182683 | -2.75377 | 0.005891 | 0.032991 |
| ANK3 | 109.2335 | 0.312979 | 0.426287 | 0.734197 | 0.462829 | 0.620753 |
| IPMK | 138.6969 | 0.738875 | 0.334094 | 2.21158 | 0.026996 | 0.092445 |
| DLG5 | 1370.677 | -0.29574 | 0.170348 | -1.73612 | 0.082543 | 0.195357 |
| MAT1A | 3.171021 | 0.089038 | 1.081071 | 0.082361 | 0.93436 | NA |
| DIP2C | 3331.53 | -0.47321 | 0.175677 | -2.69363 | 0.007068 | 0.037373 |
| AGAP11 | 0.47803 | 0.520083 | 2.97881 | 0.174594 | 0.861399 | NA |
| EXT2 | 3369.847 | -0.3169 | 0.144549 | -2.19231 | 0.028357 | 0.09539 |
| NEK7 | 1116.627 | 1.065306 | 0.225507 | 4.724042 | 2.31E-06 | 2.67E-04 |
| UPF2 | 537.6028 | 0.920549 | 0.616741 | 1.492602 | 0.135541 | 0.274393 |
| CDC123 | 1464.365 | 0.253013 | 0.176045 | 1.437203 | 0.15066 | 0.294428 |
| CCDC3 | 23.27401 | 0.764929 | 0.55145 | 1.387122 | 0.165405 | NA |
| FRMD4A | 1113.985 | -0.02489 | 0.200484 | -0.12417 | 0.901177 | 0.942187 |
| VTI1A | 744.7635 | 0.041743 | 0.15085 | 0.276721 | 0.781995 | 0.866568 |
| FAM160B1 | 1000.768 | 0.234759 | 0.177687 | 1.321193 | 0.186437 | 0.340557 |
| AKR1C2 | 48.39518 | 0.741064 | 0.479735 | 1.544736 | 0.12241 | 0.255848 |
| DPYSL4 | 395.1204 | -0.31919 | 0.345838 | -0.92293 | 0.356043 | 0.523878 |
| ADAM8 | 3.193832 | -0.37271 | 1.160792 | -0.32108 | 0.748146 | NA |
| ITIH2 | 1.083511 | 0.500984 | 1.995536 | 0.251053 | 0.801774 | NA |
| KIN | 151.662 | 0.595134 | 0.256338 | 2.321672 | 0.020251 | 0.076358 |
| GFRA1 | 206.9549 | -0.4285 | 0.298972 | -1.43325 | 0.151786 | 0.295909 |
| CACUL1 | 1125.951 | 0.215075 | 0.18727 | 1.148473 | 0.250773 | 0.414408 |
| TIAL1 | 1449.431 | 0.280903 | 0.141148 | 1.99013 | 0.046577 | 0.1327 |
| BAG3 | 1202.6 | -0.42627 | 0.153842 | -2.77087 | 0.005591 | 0.031988 |
| LIX1L | 799.9079 | -0.3923 | 0.145772 | -2.69118 | 0.00712 | 0.037572 |
| NBPF11 | 2.320295 | -0.86825 | 1.786949 | -0.48589 | 0.627048 | NA |
| RABGAP1L | 521.578 | 0.292365 | 0.255987 | 1.142111 | 0.253408 | 0.417475 |
| TMEM56 | 82.24594 | 0.977757 | 0.280832 | 3.481637 | 4.98E-04 | 0.006768 |
| ASTN1 | 1.274772 | -2.84703 | 2.61544 | -1.08855 | 0.276354 | NA |
| PTPN14 | 863.5432 | -0.41232 | 0.152138 | -2.71017 | 0.006725 | 0.036139 |
| ARL14EP | 516.6436 | 0.669745 | 0.193058 | 3.469137 | 5.22E-04 | 0.006967 |
| SPON1 | 1.74341 | -0.25121 | 1.432949 | -0.17531 | 0.860838 | NA |
| PDE3B | 1057.907 | 0.33334 | 0.209101 | 1.59416 | 0.1109 | 0.23892 |
| UHMK1 | 3305.419 | 0.372181 | 0.223255 | 1.667064 | 0.095502 | 0.216311 |
| TADA1 | 216.0206 | 0.180334 | 0.195778 | 0.921117 | 0.356989 | 0.524555 |
| SUV39H2 | 311.012 | 0.436293 | 0.189808 | 2.298598 | 0.021528 | 0.079679 |
| DCLRE1C | 289.6347 | 0.762525 | 0.20918 | 3.645302 | 2.67E-04 | 0.004599 |
| OLAH | 0.096202 | 0.408546 | 4.079639 | 0.100143 | 0.920231 | NA |
| RPP38 | 224.3942 | 0.442141 | 0.177938 | 2.484804 | 0.012962 | 0.05635 |
| NMT2 | 962.2322 | 0.35455 | 0.236073 | 1.501864 | 0.133132 | 0.270625 |
| ARL5B-AS1 | 1.150043 | 1.185274 | 1.979154 | 0.598879 | 0.549253 | NA |
| FAM21B | 122.2669 | -0.05257 | 0.226026 | -0.23261 | 0.816068 | 0.889858 |

| | | | | | | |
|---|---|---|---|---|---|---|
| TCTEX1D1 | 8.941415 | -0.17468 | 0.774991 | -0.22539 | 0.821673 | NA |
| WDR78 | 47.53739 | 0.823304 | 0.34785 | 2.366836 | 0.017941 | 0.070142 |
| ANKRD22 | 0.134133 | -1.48909 | 4.077584 | -0.36519 | 0.71497 | NA |
| IFIT5 | 917.6121 | 0.271194 | 0.235445 | 1.151836 | 0.249389 | 0.41332 |
| PANK1 | 223.6207 | 0.468663 | 0.212935 | 2.200965 | 0.027738 | 0.09425 |
| HHEX | 247.3817 | 0.040495 | 0.193737 | 0.20902 | 0.834432 | 0.900929 |
| GGPS1 | 995.5567 | 0.505933 | 0.177012 | 2.858192 | 0.004261 | 0.0265 |
| MR1 | 1237.384 | -0.47111 | 0.289916 | -1.62497 | 0.104168 | 0.229312 |
| HNRNPU | 5953.878 | -0.0339 | 0.17261 | -0.19637 | 0.844317 | 0.907349 |
| AHCTF1 | 534.7244 | 0.546532 | 0.198097 | 2.75892 | 0.005799 | 0.032657 |
| LINC00467 | 113.0489 | 0.604799 | 0.241451 | 2.504846 | 0.01225 | 0.05417 |
| MCOLN2 | 146.4482 | 0.742166 | 0.289763 | 2.561286 | 0.010429 | 0.048407 |
| DDAH1 | 54.1634 | 0.153331 | 0.408467 | 0.375381 | 0.707377 | 0.812751 |
| HS2ST1 | 971.8609 | 0.211322 | 0.211234 | 1.000414 | 0.31711 | 0.484826 |
| AK5 | 4221.074 | 0.64647 | 0.786116 | 0.82236 | 0.410872 | 0.57424 |
| CC2D1B | 2904.369 | -0.24041 | 0.154662 | -1.55441 | 0.120086 | 0.25228 |
| MIA3 | 1058.597 | 0.452296 | 0.275824 | 1.639797 | 0.101047 | 0.224455 |
| DISP1 | 310.7638 | -0.99881 | 0.209117 | -4.77634 | 1.79E-06 | 2.28E-04 |
| OBSCN | 238.5012 | -0.36982 | 0.334443 | -1.10578 | 0.268823 | 0.433709 |
| TRIM11 | 942.0556 | -0.33059 | 0.153697 | -2.15092 | 0.031483 | 0.102534 |
| ENAH | 1750.294 | 0.425704 | 0.239193 | 1.779752 | 0.075117 | 0.183032 |
| CCSAP | 179.5123 | 0.219813 | 0.198983 | 1.104682 | 0.269297 | 0.433993 |
| GBP5 | 0.096202 | 0.408546 | 4.079639 | 0.100143 | 0.920231 | NA |
| BUB3 | 2402.044 | 0.124234 | 0.158223 | 0.785183 | 0.432347 | 0.5935 |
| GPR26 | 1.162303 | 2.195571 | 2.038153 | 1.077235 | 0.281375 | NA |
| MMP21 | 0.640406 | 1.11402 | 2.710977 | 0.410929 | 0.681124 | NA |
| C10orf90 | 3652.602 | -0.57634 | 0.178732 | -3.22461 | 0.001261 | 0.012031 |
| FAM69A | 534.9017 | 0.018034 | 0.195892 | 0.092059 | 0.926651 | 0.958156 |
| MMS19 | 2747.087 | -0.41515 | 0.127404 | -3.25856 | 0.00112 | 0.01112 |
| PI4K2A | 1828.833 | -0.30485 | 0.151142 | -2.01701 | 0.043695 | 0.127367 |
| MARVELD1 | 6622.643 | -0.50623 | 0.124089 | -4.07959 | 4.51E-05 | 0.001553 |
| ZFYVE27 | 1905.517 | -0.09871 | 0.157168 | -0.62807 | 0.529956 | 0.674874 |
| GOLGA7B | 2199.19 | -0.51006 | 0.125537 | -4.06299 | 4.84E-05 | 0.001613 |
| SLC25A28 | 580.344 | -0.07087 | 0.163195 | -0.43429 | 0.664078 | 0.779393 |
| MOV10 | 1269.951 | -0.65605 | 0.15577 | -4.21168 | 2.53E-05 | 0.001062 |
| RHOC | 4403.059 | -0.04605 | 0.13233 | -0.34801 | 0.727834 | 0.827433 |
| PPM1J | 9.420576 | -0.30549 | 0.701348 | -0.43558 | 0.66314 | NA |
| SLC16A1 | 1330.03 | -0.07968 | 0.164181 | -0.48533 | 0.627446 | 0.750113 |
| PIK3AP1 | 123.7874 | -0.18427 | 0.259003 | -0.71145 | 0.476805 | 0.631755 |
| C10orf12 | 84.78011 | 0.085538 | 0.254102 | 0.336628 | 0.736398 | 0.833121 |
| SPAG17 | 2.197166 | 1.403219 | 1.431805 | 0.980035 | 0.327069 | NA |
| FMN2 | 31.465 | 0.110791 | 0.538074 | 0.205902 | 0.836867 | NA |
| MCU | 466.3226 | 0.033302 | 0.180325 | 0.184678 | 0.853481 | 0.912964 |
| TTC18 | 13.95256 | 0.892382 | 0.575329 | 1.55108 | 0.120883 | NA |
| GPR61 | 1.252771 | 0.053586 | 1.700195 | 0.031518 | 0.974857 | NA |
| ADK | 1199.133 | 0.491524 | 0.200878 | 2.446879 | 0.01441 | 0.060428 |
| KCNMA1 | 12.5928 | -0.78724 | 0.684523 | -1.15005 | 0.250122 | NA |

| | | | | | | |
|---|---|---|---|---|---|---|
| DRAM2 | 1201.248 | 0.195375 | 0.172242 | 1.134308 | 0.256666 | 0.420707 |
| PCGF6 | 162.7525 | 0.456581 | 0.255679 | 1.785758 | 0.074138 | 0.181706 |
| SFR1 | 82.32754 | 0.460981 | 0.311509 | 1.479828 | 0.138919 | 0.27885 |
| SFXN2 | 73.57091 | -0.63969 | 0.401843 | -1.59188 | 0.111412 | 0.239657 |
| SUPV3L1 | 691.7085 | -0.06078 | 0.149545 | -0.40641 | 0.684442 | 0.794639 |
| HKDC1 | 74.97562 | -1.34524 | 0.452839 | -2.97067 | 0.002971 | 0.021006 |
| HK1 | 9082.729 | -0.23364 | 0.16763 | -1.39376 | 0.163391 | 0.311966 |
| TYSND1 | 1004.76 | -0.33136 | 0.14537 | -2.27943 | 0.022642 | 0.082217 |
| NODAL | 0.28583 | -2.21358 | 3.97376 | -0.55705 | 0.577494 | NA |
| UBE2L6 | 1525.838 | -0.89124 | 0.272257 | -3.27354 | 0.001062 | 0.010789 |
| ZDHHC5 | 3611.645 | -0.33001 | 0.135015 | -2.44428 | 0.014514 | 0.060596 |
| MED19 | 555.4246 | -0.08117 | 0.158206 | -0.51309 | 0.607889 | 0.735325 |
| KAT6B | 388.9949 | 0.233055 | 0.204011 | 1.142365 | 0.253302 | 0.417475 |
| SAMD8 | 869.8812 | 0.691972 | 0.220385 | 3.139836 | 0.00169 | 0.014563 |
| GLYATL2 | 14.50518 | 0.809009 | 0.572495 | 1.41313 | 0.157618 | NA |
| MS4A1 | 0.162376 | 0.424913 | 4.079006 | 0.104171 | 0.917034 | NA |
| FRRS1 | 117.4945 | -0.25899 | 0.25626 | -1.01065 | 0.312184 | 0.479394 |
| HIAT1 | 1445.52 | 0.221007 | 0.165028 | 1.339214 | 0.180501 | 0.333033 |
| SASS6 | 77.62934 | 0.887503 | 0.357746 | 2.480823 | 0.013108 | 0.056734 |
| SHCBP1L | 0.437526 | -2.68242 | 3.384147 | -0.79264 | 0.427986 | NA |
| NMNAT2 | 0.875299 | 1.848392 | 2.453783 | 0.753283 | 0.45128 | NA |
| ZFYVE9 | 848.628 | 0.327178 | 0.166888 | 1.960462 | 0.049942 | 0.138396 |
| C1orf27 | 264.0953 | 0.904036 | 0.315256 | 2.867626 | 0.004136 | 0.025982 |
| CPT2 | 572.1598 | -0.36229 | 0.166552 | -2.17524 | 0.029612 | 0.098146 |
| NECAP2 | 3245.992 | -0.11684 | 0.12384 | -0.94348 | 0.345435 | 0.513404 |
| LRP8 | 873.2783 | 0.080848 | 0.143774 | 0.562331 | 0.57389 | 0.709434 |
| CDCP2 | 0.134133 | -1.48909 | 4.077584 | -0.36519 | 0.71497 | NA |
| SSBP3 | 964.2562 | -0.33435 | 0.151959 | -2.20025 | 0.027789 | 0.094335 |
| CREB3L1 | 65.60934 | 0.285911 | 0.976999 | 0.292642 | 0.769796 | 0.85762 |
| FAM213B | 3651.18 | -0.28046 | 0.145177 | -1.93182 | 0.053382 | 0.144782 |
| TNFRSF14 | 9866.44 | -0.17484 | 0.151954 | -1.1506 | 0.249898 | 0.41369 |
| PANK4 | 1121.629 | -0.34092 | 0.152917 | -2.22944 | 0.025784 | 0.089951 |
| PEX10 | 2150.943 | -0.29096 | 0.144069 | -2.01961 | 0.043424 | 0.126951 |
| RER1 | 3099.273 | -0.1606 | 0.129829 | -1.23701 | 0.216084 | 0.37531 |
| SKI | 2057.296 | -0.51499 | 0.178443 | -2.88601 | 0.003902 | 0.025065 |
| LDLRAP1 | 1249.184 | -0.31196 | 0.134992 | -2.31094 | 0.020836 | 0.077896 |
| PAFAH2 | 666.3978 | -0.15239 | 0.152906 | -0.99665 | 0.318933 | 0.486797 |
| EXTL1 | 824.4662 | -0.48067 | 0.202868 | -2.36937 | 0.017818 | 0.06981 |
| SLC30A2 | 12.27355 | -0.69502 | 0.749514 | -0.92729 | 0.353774 | NA |
| TRIM63 | 7846.85 | -0.0594 | 0.144442 | -0.41124 | 0.680899 | 0.792077 |
| MRPL17 | 2576.71 | 0.085158 | 0.175148 | 0.486208 | 0.62682 | 0.749465 |
| GRHL3 | 0.558248 | 0.982076 | 2.804738 | 0.350149 | 0.726227 | NA |
| UBXN11 | 370.9293 | 0.057814 | 0.166385 | 0.34747 | 0.728238 | 0.827654 |
| NLRP14 | 0.314073 | -0.57308 | 3.898934 | -0.14698 | 0.883146 | NA |
| TPRG1L | 3321.51 | -0.10001 | 0.14945 | -0.66915 | 0.503398 | 0.653359 |
| XKR8 | 415.4776 | -0.23235 | 0.17489 | -1.32855 | 0.183995 | 0.337239 |
| EYA3 | 557.4497 | -0.17587 | 0.141743 | -1.24078 | 0.214686 | 0.373586 |

| | | | | | | |
|---|---|---|---|---|---|---|
| WASF2 | 3254.7 | -0.62649 | 0.152881 | -4.09788 | 4.17E-05 | 0.001498 |
| FAM46B | 67.04167 | -0.62976 | 0.317664 | -1.98246 | 0.047427 | 0.134086 |
| RNF207 | 150.3415 | -0.04774 | 0.233169 | -0.20476 | 0.837756 | 0.902976 |
| GPR153 | 2107.469 | -0.33793 | 0.143372 | -2.35698 | 0.018424 | 0.07152 |
| RHBDL2 | 1.670591 | -0.12805 | 1.762369 | -0.07266 | 0.942078 | NA |
| CD1D | 1.288102 | -0.49186 | 1.904342 | -0.25828 | 0.796187 | NA |
| CD1A | 0.163957 | 0.424913 | 4.079006 | 0.104171 | 0.917034 | NA |
| CD1E | 0.162376 | 0.424913 | 4.079006 | 0.104171 | 0.917034 | NA |
| PPP1R15B | 898.7313 | 0.195726 | 0.187421 | 1.044308 | 0.296343 | 0.463649 |
| TAGLN2 | 12668.26 | -0.14344 | 0.137686 | -1.04177 | 0.297517 | 0.464637 |
| ELK4 | 740.1297 | 0.462523 | 0.244025 | 1.895392 | 0.058041 | 0.153519 |
| SLAMF8 | 1.400891 | -0.43398 | 1.760974 | -0.24644 | 0.805341 | NA |
| SLC45A3 | 160.9144 | -0.12547 | 0.300142 | -0.41802 | 0.675931 | 0.788232 |
| DUSP23 | 996.8007 | 0.14551 | 0.16332 | 0.890951 | 0.372956 | 0.539654 |
| NBL1 | 31878.02 | -0.53314 | 0.151316 | -3.52339 | 4.26E-04 | 0.00616 |
| ITLN2 | 1.558526 | -0.08772 | 1.723497 | -0.0509 | 0.959408 | NA |
| F11R | 1998.539 | -0.32359 | 0.207891 | -1.55653 | 0.119582 | 0.251535 |
| USF1 | 1347.902 | -0.2383 | 0.172369 | -1.38249 | 0.166823 | 0.316277 |
| NIT1 | 637.8217 | -0.21169 | 0.149952 | -1.41173 | 0.158029 | 0.304545 |
| DEDD | 569.7809 | -0.20099 | 0.17935 | -1.12063 | 0.262444 | 0.426852 |
| CDA | 3.062502 | -0.46625 | 1.115883 | -0.41783 | 0.67607 | NA |
| PINK1 | 1534.106 | -0.11062 | 0.134038 | -0.82529 | 0.409207 | 0.572795 |
| B4GALT3 | 1206.857 | -0.29969 | 0.151461 | -1.97863 | 0.047858 | 0.134892 |
| ADAMTS4 | 90.49349 | -1.08811 | 0.434484 | -2.50436 | 0.012267 | 0.054213 |
| NDUFS2 | 4035.087 | -0.27048 | 0.11803 | -2.29161 | 0.021928 | 0.080682 |
| FCER1G | 5.064141 | 0.16476 | 0.883912 | 0.186399 | 0.852132 | NA |
| APOA2 | 0.162376 | 0.424913 | 4.079006 | 0.104171 | 0.917034 | NA |
| TOMM40L | 744.0745 | -0.17334 | 0.162506 | -1.06664 | 0.286132 | 0.452521 |
| MPZ | 1266.166 | 0.110268 | 0.803194 | 0.137287 | 0.890804 | 0.936099 |
| CACHD1 | 262.5668 | -0.15742 | 0.237975 | -0.66151 | 0.508287 | 0.657089 |
| EPB41 | 776.7209 | 0.208754 | 0.165102 | 1.264396 | 0.206088 | 0.363081 |
| SV2A | 1271.591 | -0.48673 | 0.199069 | -2.44503 | 0.014484 | 0.060538 |
| TNNI1 | 0.163957 | 0.424913 | 4.079006 | 0.104171 | 0.917034 | NA |
| CSRP1 | 12187.18 | -0.4955 | 0.136911 | -3.61912 | 2.96E-04 | 0.004874 |
| C1QC | 0.163957 | 0.424913 | 4.079006 | 0.104171 | 0.917034 | NA |
| C1orf51 | 209.7135 | -0.11136 | 0.202069 | -0.55109 | 0.58157 | 0.715198 |
| CCDC24 | 69.98769 | 0.052474 | 0.263934 | 0.198814 | 0.842408 | 0.906403 |
| PADI4 | 0.162376 | 0.424913 | 4.079006 | 0.104171 | 0.917034 | NA |
| ADIPOR1 | 5675.109 | -0.13915 | 0.129579 | -1.07388 | 0.282876 | 0.44923 |
| CYB5R1 | 4386.065 | -0.37653 | 0.128418 | -2.93206 | 0.003367 | 0.022712 |
| PSMD4 | 3068.332 | -0.05765 | 0.131414 | -0.4387 | 0.660881 | 0.776641 |
| ATP13A2 | 2634.8 | -0.34949 | 0.158367 | -2.20684 | 0.027325 | 0.093221 |
| PSMB4 | 7930.779 | -0.14098 | 0.141559 | -0.99588 | 0.319308 | 0.487181 |
| BTG2 | 287.0408 | -0.03899 | 0.184112 | -0.21178 | 0.832279 | 0.899259 |
| CELF3 | 0.258578 | 1.233151 | 4.047809 | 0.304646 | 0.760635 | NA |
| ALDH4A1 | 550.8879 | -0.38339 | 0.157577 | -2.43301 | 0.014974 | 0.061916 |
| THEM4 | 1206.754 | 0.447359 | 0.185534 | 2.411188 | 0.015901 | 0.064533 |

| | | | | | | |
|---|---|---|---|---|---|---|
| TCHH | 14.58885 | 1.053477 | 0.751851 | 1.401178 | 0.161161 | NA |
| MED8 | 623.0291 | -0.08973 | 0.144037 | -0.62297 | 0.533307 | 0.677202 |
| SPRR2G | 0.785714 | -3.2458 | 3.343263 | -0.97085 | 0.331623 | NA |
| CCDC17 | 3.333746 | -0.3349 | 1.086799 | -0.30816 | 0.757963 | NA |
| GPBP1L1 | 1593.557 | 0.142541 | 0.162386 | 0.87779 | 0.380058 | 0.546414 |
| TMEM69 | 504.8597 | 0.073103 | 0.157288 | 0.464773 | 0.642094 | 0.761812 |
| EFCAB14 | 3146.463 | -0.00888 | 0.124229 | -0.07151 | 0.942988 | 0.968558 |
| DFFA | 1362.997 | -0.2752 | 0.189271 | -1.45402 | 0.145941 | 0.28834 |
| CCDC28B | 275.4596 | -0.27239 | 0.207843 | -1.31057 | 0.190004 | 0.344236 |
| IQCC | 56.9599 | -0.04557 | 0.333905 | -0.13647 | 0.891449 | 0.936392 |
| TMEM234 | 250.9576 | -0.07138 | 0.186598 | -0.38252 | 0.702074 | 0.808431 |
| BSDC1 | 1986.62 | -0.07593 | 0.126672 | -0.59938 | 0.548917 | 0.690147 |
| ZBTB8A | 14.63497 | 0.643585 | 0.579271 | 1.111025 | 0.266558 | NA |
| ATAD3B | 719.1085 | 0.054832 | 0.205546 | 0.266762 | 0.789653 | 0.872526 |
| SSU72 | 6096.89 | 0.091338 | 0.14856 | 0.614826 | 0.538669 | 0.681093 |
| UBE2J2 | 1623.157 | -0.05628 | 0.135556 | -0.41519 | 0.678005 | 0.789873 |
| ZNF362 | 702.0208 | -0.32379 | 0.141247 | -2.29237 | 0.021884 | 0.080546 |
| FNDC5 | 0.296509 | -0.51709 | 3.982155 | -0.12985 | 0.896684 | NA |
| S100A1 | 277.678 | 0.155123 | 0.204408 | 0.758891 | 0.447918 | 0.60742 |
| CHTOP | 1277.297 | -0.09295 | 0.158106 | -0.58792 | 0.556583 | 0.695949 |
| ZBTB7B | 2103.497 | -0.21229 | 0.184448 | -1.15097 | 0.249747 | 0.41357 |
| FLAD1 | 1192.253 | -0.1761 | 0.133686 | -1.31728 | 0.187744 | 0.341836 |
| SHC1 | 8169.135 | -0.18202 | 0.158946 | -1.14514 | 0.252152 | 0.4161 |
| ADAR | 8806.446 | -0.75826 | 0.156209 | -4.85416 | 1.21E-06 | 1.79E-04 |
| IL6R | 1120.612 | -0.08055 | 0.150378 | -0.53566 | 0.592194 | 0.7233 |
| UBE2Q1 | 3194.589 | -0.0406 | 0.148459 | -0.27351 | 0.784464 | 0.868817 |
| CHRNB2 | 9.422762 | -0.60303 | 0.71663 | -0.84148 | 0.400077 | NA |
| CRTC2 | 1008.621 | -0.17913 | 0.148303 | -1.20784 | 0.227107 | 0.388563 |
| FDPS | 1575.074 | -0.01212 | 0.141871 | -0.0854 | 0.931946 | 0.961401 |
| RUSC1 | 760.3851 | -0.26231 | 0.156743 | -1.67352 | 0.094225 | 0.214616 |
| GBAP1 | 123.5708 | -0.20267 | 0.213592 | -0.94887 | 0.342688 | 0.510822 |
| FAM189B | 1763.604 | -0.3713 | 0.198973 | -1.86611 | 0.062026 | 0.161078 |
| PAQR6 | 15.82584 | -0.32927 | 0.508997 | -0.6469 | 0.517697 | NA |
| PMF1 | 402.841 | -0.21783 | 0.195554 | -1.11389 | 0.265326 | 0.429881 |
| SLC25A44 | 1270.532 | -0.29878 | 0.155034 | -1.92718 | 0.053957 | 0.145874 |
| LMNA | 36654.76 | -0.47746 | 0.153086 | -3.11887 | 0.001815 | 0.015146 |
| UBQLN4 | 1380.794 | -0.38762 | 0.13379 | -2.89723 | 0.003765 | 0.024485 |
| GPATCH4 | 889.8956 | 0.466872 | 0.134593 | 3.468775 | 5.23E-04 | 0.006967 |
| LRRC71 | 0.803131 | -0.76693 | 2.349822 | -0.32638 | 0.744138 | NA |
| FCRL3 | 0.134133 | -1.48909 | 4.077584 | -0.36519 | 0.71497 | NA |
| LRRC56 | 49.51286 | -0.07391 | 0.325541 | -0.22704 | 0.820394 | 0.892334 |
| CYB561A3 | 6378.276 | -0.12465 | 0.139064 | -0.89636 | 0.370061 | 0.536887 |
| PPP1R32 | 3.433897 | 0.80273 | 1.104518 | 0.72677 | 0.467367 | NA |
| ASRGL1 | 122.7195 | 0.002002 | 0.24859 | 0.008052 | 0.993576 | 0.997229 |
| GNG3 | 1.52677 | 1.133152 | 1.652118 | 0.685878 | 0.49279 | NA |
| UBXN1 | 3509.002 | -0.14083 | 0.145496 | -0.96796 | 0.333066 | 0.501397 |
| C11orf48 | 623.4995 | 0.216868 | 0.183864 | 1.179502 | 0.238198 | 0.401059 |

| | | | | | | |
|---|---|---|---|---|---|---|
| TTC9C | 617.5598 | -0.10822 | 0.185755 | -0.58261 | 0.560155 | 0.698472 |
| TAF6L | 588.4864 | -0.28249 | 0.179418 | -1.57446 | 0.115382 | 0.245355 |
| NXF1 | 1329.881 | 0.042207 | 0.153156 | 0.275579 | 0.782872 | 0.867305 |
| STX5 | 1200.36 | -0.20746 | 0.1691 | -1.22682 | 0.21989 | 0.379574 |
| CMPK1 | 2737.263 | 0.356188 | 0.144334 | 2.467803 | 0.013595 | 0.057993 |
| BEND5 | 46.90606 | -0.10048 | 0.324646 | -0.3095 | 0.756942 | NA |
| COA7 | 490.0465 | 0.179919 | 0.167917 | 1.071478 | 0.283954 | 0.450519 |
| ZYG11B | 1104.517 | 0.343214 | 0.185114 | 1.854066 | 0.06373 | 0.1641 |
| SLC1A7 | 0.460669 | -0.62593 | 3.398671 | -0.18417 | 0.853882 | NA |
| C1orf123 | 870.7182 | -0.04556 | 0.133455 | -0.34137 | 0.732821 | 0.830894 |
| MAGOH | 426.9034 | 0.585083 | 0.160493 | 3.64553 | 2.67E-04 | 0.004599 |
| ACOT11 | 29.04059 | -0.10977 | 0.397165 | -0.27638 | 0.782257 | NA |
| FAM151A | 0.585114 | 1.12675 | 2.73598 | 0.411827 | 0.680466 | NA |
| PARS2 | 225.796 | -0.47592 | 0.217348 | -2.18965 | 0.02855 | 0.095853 |
| C1orf177 | 0.096202 | 0.408546 | 4.079639 | 0.100143 | 0.920231 | NA |
| USP24 | 1619.465 | 0.258919 | 0.159956 | 1.61869 | 0.105514 | 0.2313 |
| PPAP2B | 110.1483 | 0.01109 | 0.245676 | 0.045141 | 0.963995 | 0.980386 |
| NOL9 | 449.2494 | -0.62873 | 0.167675 | -3.74972 | 1.77E-04 | 0.0036 |
| PRKAA2 | 2.629185 | 1.279429 | 1.260852 | 1.014734 | 0.310233 | NA |
| KLHL21 | 4619.354 | -0.29931 | 0.161823 | -1.84958 | 0.064374 | 0.16522 |
| ZSWIM5 | 1.791029 | -1.75516 | 1.594408 | -1.10082 | 0.270974 | NA |
| GMEB1 | 469.1677 | -0.42569 | 0.168308 | -2.52923 | 0.011431 | 0.051527 |
| SLC45A1 | 11.71396 | 0.763097 | 0.689115 | 1.107358 | 0.268139 | NA |
| SEPN1 | 9618.045 | -0.55726 | 0.126216 | -4.41512 | 1.01E-05 | 6.32E-04 |
| AK4 | 329.9011 | -0.51336 | 0.296051 | -1.73402 | 0.082915 | 0.195915 |
| JAK1 | 3068.57 | -0.14259 | 0.129586 | -1.10038 | 0.271168 | 0.436364 |
| RAVER2 | 453.7733 | 0.076926 | 0.196742 | 0.390997 | 0.695799 | 0.80381 |
| CTRC | 0.462046 | 0.589069 | 3.442439 | 0.17112 | 0.86413 | NA |
| LZIC | 771.6075 | 0.350174 | 0.15588 | 2.246427 | 0.024677 | 0.086963 |
| RBP7 | 33.34612 | 0.37164 | 0.451722 | 0.822717 | 0.410669 | NA |
| FBLIM1 | 442.8711 | -0.53558 | 0.201889 | -2.65284 | 0.007982 | 0.040559 |
| SLC25A34 | 20.85167 | -0.03599 | 0.506549 | -0.07104 | 0.943365 | NA |
| AKR7A3 | 4.001018 | -0.03053 | 1.035836 | -0.02948 | 0.976485 | NA |
| DRAXIN | 1.393789 | -0.8924 | 1.663459 | -0.53647 | 0.591633 | NA |
| PDPN | 0.517155 | 2.232561 | 3.179631 | 0.702145 | 0.482589 | NA |
| DHRS3 | 772.7476 | -0.36268 | 0.315432 | -1.14979 | 0.250231 | 0.413909 |
| MATN1 | 2.606035 | -0.2512 | 1.263501 | -0.19882 | 0.842407 | NA |
| LAPTM5 | 0.902348 | 0.812045 | 2.125951 | 0.381968 | 0.702485 | NA |
| SDC3 | 8686.333 | -0.4865 | 0.148554 | -3.27488 | 0.001057 | 0.010747 |
| PEF1 | 1714.206 | -0.34037 | 0.12443 | -2.73543 | 0.00623 | 0.034188 |
| SYNC | 1913.76 | 0.300752 | 0.154426 | 1.947544 | 0.05147 | 0.141276 |
| RBBP4 | 2216.453 | -0.26374 | 0.131464 | -2.00618 | 0.044837 | 0.129393 |
| KIAA1522 | 1350.657 | -0.64413 | 0.160721 | -4.00777 | 6.13E-05 | 0.001863 |
| TSSK3 | 2.572664 | -0.5296 | 1.491849 | -0.355 | 0.722592 | NA |
| TMCO4 | 619.7007 | -0.30529 | 0.211042 | -1.44658 | 0.148015 | 0.291096 |
| UBXN10 | 0.46577 | -1.30191 | 3.402879 | -0.38259 | 0.702024 | NA |
| CAMK2N1 | 368.8922 | -0.42482 | 0.257601 | -1.64913 | 0.099121 | 0.222006 |

| | | | | | | |
|---|---|---|---|---|---|---|
| ALPL | 0.700488 | -1.9689 | 2.983028 | -0.66003 | 0.509233 | NA |
| WNT4 | 42.0672 | 1.227662 | 0.5129 | 2.393571 | 0.016685 | NA |
| SCNN1D | 18.78261 | 0.257454 | 0.503224 | 0.51161 | 0.608924 | NA |
| MXRA8 | 24.38134 | 0.018187 | 0.546046 | 0.033306 | 0.973431 | NA |
| C1orf86 | 2521.326 | -0.2187 | 0.142855 | -1.53093 | 0.125788 | 0.260839 |
| MEGF6 | 29.22879 | -0.91081 | 0.519272 | -1.75401 | 0.079429 | NA |
| CCDC27 | 0.163957 | 0.424913 | 4.079006 | 0.104171 | 0.917034 | NA |
| IL23R | 0.162376 | 0.424913 | 4.079006 | 0.104171 | 0.917034 | NA |
| DIRAS3 | 9.092803 | 0.292714 | 0.677059 | 0.432332 | 0.6655 | NA |
| C1orf87 | 0.151697 | -1.546 | 4.075469 | -0.37934 | 0.704433 | NA |
| NFIA | 119.6879 | -0.45166 | 0.370854 | -1.21788 | 0.22327 | 0.384105 |
| OMA1 | 161.7502 | 0.605012 | 0.251446 | 2.406128 | 0.016123 | 0.065355 |
| MYSM1 | 314.9481 | 1.268713 | 0.339446 | 3.737606 | 1.86E-04 | 0.003708 |
| TM2D1 | 682.5092 | 0.368795 | 0.16398 | 2.24902 | 0.024511 | 0.086714 |
| USP1 | 462.834 | 0.916997 | 0.208348 | 4.401284 | 1.08E-05 | 6.41E-04 |
| FUBP1 | 902.1753 | 0.053091 | 0.192965 | 0.275131 | 0.783215 | 0.867596 |
| NEXN | 17.82072 | 0.97946 | 0.809997 | 1.209214 | 0.226581 | NA |
| DNAJB4 | 287.7249 | 1.051385 | 0.240825 | 4.365759 | 1.27E-05 | 7.08E-04 |
| ELTD1 | 244.6992 | 0.900856 | 0.19042 | 4.730896 | 2.24E-06 | 2.66E-04 |
| LRRIQ3 | 33.01778 | 0.301223 | 0.439144 | 0.685932 | 0.492756 | NA |
| TYW3 | 610.7059 | 0.750577 | 0.210374 | 3.567825 | 3.60E-04 | 0.005535 |
| SNX7 | 1344.259 | 0.560758 | 0.153521 | 3.652645 | 2.60E-04 | 0.004545 |
| B3GALT2 | 8.006064 | -0.52421 | 0.739137 | -0.70922 | 0.478187 | NA |
| NTNG1 | 6.345064 | -1.32101 | 1.082531 | -1.2203 | 0.22235 | NA |
| FAM102B | 727.3311 | 0.622023 | 0.22349 | 2.783225 | 0.005382 | 0.031155 |
| HENMT1 | 99.96136 | 0.433294 | 0.334029 | 1.297173 | 0.194572 | 0.349673 |
| AKNAD1 | 14.12094 | 0.16606 | 0.550287 | 0.30177 | 0.762828 | NA |
| C1orf52 | 393.5923 | 0.462303 | 0.199907 | 2.312591 | 0.020745 | 0.07768 |
| WDR63 | 177.5838 | 0.827742 | 0.270103 | 3.064543 | 0.00218 | 0.017115 |
| GBP2 | 220.1442 | 0.125877 | 0.664602 | 0.189402 | 0.849778 | 0.910531 |
| ATXN7L2 | 140.5869 | -0.27106 | 0.261318 | -1.03729 | 0.299599 | 0.466435 |
| GBP4 | 28.55005 | -0.53485 | 0.485715 | -1.10115 | 0.27083 | NA |
| ZNF326 | 307.4404 | 0.74714 | 0.337072 | 2.216557 | 0.026653 | 0.091691 |
| HFM1 | 2.845442 | -0.00832 | 1.367645 | -0.00608 | 0.995148 | NA |
| GFI1 | 4.969337 | 0.333659 | 0.883308 | 0.377738 | 0.705625 | NA |
| KCNT2 | 1.63207 | 1.952588 | 1.715903 | 1.137936 | 0.255147 | NA |
| AGL | 558.1128 | 0.782765 | 0.3417 | 2.290795 | 0.021975 | 0.08078 |
| EXTL2 | 1338.507 | 0.35255 | 0.185151 | 1.904117 | 0.056895 | 0.151187 |
| SLC30A7 | 622.7807 | 0.078992 | 0.19288 | 0.409541 | 0.682143 | 0.792948 |
| ZNF281 | 342.6238 | 0.278839 | 0.248866 | 1.12044 | 0.262526 | 0.426921 |
| ARPC5 | 4574.642 | 0.059859 | 0.135894 | 0.440485 | 0.659586 | 0.775579 |
| CADM3 | 1962.773 | -0.48453 | 0.250601 | -1.93347 | 0.053178 | 0.144294 |
| NLRP3 | 0.452561 | 2.061505 | 3.349456 | 0.615474 | 0.538241 | NA |
| ZNF496 | 907.6615 | -0.53053 | 0.136923 | -3.87469 | 1.07E-04 | 0.002681 |
| TRIM58 | 4.718578 | 1.80318 | 1.166672 | 1.545576 | 0.122207 | NA |
| SLAMF9 | 154.3857 | 0.164394 | 0.32568 | 0.504771 | 0.61372 | 0.740094 |
| KCNJ9 | 0.419962 | -2.64111 | 3.429364 | -0.77014 | 0.441214 | NA |

| | | | | | | |
|---|---|---|---|---|---|---|
| IGSF8 | 4574.616 | -0.66598 | 0.153305 | -4.34415 | 1.40E-05 | 7.64E-04 |
| DDR2 | 1102.097 | 0.276192 | 0.18284 | 1.510564 | 0.1309 | 0.267383 |
| PEA15 | 20132.94 | 0.064581 | 0.199267 | 0.324091 | 0.745869 | 0.839962 |
| PEX19 | 1785.992 | -0.30324 | 0.130281 | -2.32756 | 0.019936 | 0.075556 |
| NCSTN | 3512.365 | -0.39088 | 0.12244 | -3.19238 | 0.001411 | 0.013038 |
| VANGL2 | 938.6525 | -0.46851 | 0.157111 | -2.98201 | 0.002864 | 0.020475 |
| OLFML2B | 116.028 | -0.95663 | 0.332324 | -2.8786 | 0.003994 | 0.025436 |
| FCRLB | 65.20968 | -0.09651 | 0.281547 | -0.34278 | 0.731764 | 0.829991 |
| SLC9C2 | 0.162376 | 0.424913 | 4.079006 | 0.104171 | 0.917034 | NA |
| KLHDC9 | 4.027679 | 0.417834 | 1.077701 | 0.387709 | 0.698232 | NA |
| C1orf74 | 112.8532 | -0.13023 | 0.220984 | -0.58932 | 0.555647 | 0.695357 |
| LRRC52 | 0.35636 | 1.709252 | 3.651833 | 0.468053 | 0.639746 | NA |
| FLVCR1 | 203.8403 | -0.14183 | 0.198117 | -0.71589 | 0.474057 | 0.629544 |
| ATF3 | 848.0191 | 0.004451 | 0.304993 | 0.014592 | 0.988358 | 0.994219 |
| RBM15 | 233.934 | 0.163917 | 0.185639 | 0.882985 | 0.377245 | 0.543869 |
| DENND2D | 84.01853 | -0.34143 | 0.350717 | -0.97353 | 0.33029 | 0.498547 |
| AXDND1 | 2.823169 | 1.434326 | 1.261875 | 1.136663 | 0.255679 | NA |
| TDRD5 | 0.712719 | 2.707521 | 2.836859 | 0.954408 | 0.339877 | NA |
| IER5 | 1858.56 | -0.55192 | 0.220736 | -2.50038 | 0.012406 | 0.054553 |
| BPNT1 | 1503.67 | 0.080791 | 0.143447 | 0.563212 | 0.573291 | 0.70921 |
| SPATA17 | 8.420374 | 1.258089 | 0.743779 | 1.691482 | 0.090745 | NA |
| C1orf115 | 66.892 | -0.05156 | 0.285346 | -0.18069 | 0.856612 | 0.914811 |
| BROX | 876.1069 | 0.369806 | 0.206079 | 1.794491 | 0.072735 | 0.179416 |
| ACP6 | 438.0736 | -0.12399 | 0.179051 | -0.69247 | 0.488639 | 0.641437 |
| WDR64 | 0.298089 | -0.51711 | 3.975232 | -0.13008 | 0.896501 | NA |
| KIF26B | 557.1586 | -1.22346 | 0.304563 | -4.01709 | 5.89E-05 | 0.0018 |
| TFB2M | 457.8926 | 0.310069 | 0.190751 | 1.625515 | 0.104053 | 0.229156 |
| CNST | 832.0883 | 0.454849 | 0.192903 | 2.357918 | 0.018378 | 0.071385 |
| KLHDC8A | 2.495138 | 0.554089 | 1.487489 | 0.372499 | 0.709521 | NA |
| PM20D1 | 2.066076 | 0.869491 | 1.629363 | 0.533639 | 0.593591 | NA |
| B3GALNT2 | 284.0165 | 0.339215 | 0.178069 | 1.904967 | 0.056784 | 0.151046 |
| C1orf147 | 3.113685 | 0.506547 | 1.169058 | 0.433295 | 0.664801 | NA |
| MAPKAPK2 | 3307.13 | -0.20431 | 0.182531 | -1.11931 | 0.263009 | 0.42747 |
| IL20 | 0.75318 | -2.10401 | 2.55086 | -0.82482 | 0.409472 | NA |
| IL24 | 5951.555 | 0.043823 | 0.246351 | 0.177889 | 0.85881 | 0.916352 |
| FAIM3 | 3956.192 | 0.036045 | 0.16874 | 0.213612 | 0.83085 | 0.898124 |
| PIGR | 0.303394 | -2.27006 | 4.032451 | -0.56295 | 0.57347 | NA |
| CAPN2 | 9571.569 | -0.07949 | 0.155023 | -0.51276 | 0.608122 | 0.735387 |
| MRPL55 | 521.3778 | -0.04602 | 0.158796 | -0.28981 | 0.771963 | 0.858857 |
| C1orf145 | 2.105562 | 1.288309 | 1.646982 | 0.782225 | 0.434083 | NA |
| WDR26 | 3655.481 | -0.06754 | 0.139407 | -0.48451 | 0.628023 | 0.75045 |
| TRIM17 | 18.44928 | -0.11527 | 0.49848 | -0.23124 | 0.81713 | NA |
| DISC1 | 992.5471 | -0.41662 | 0.151204 | -2.75534 | 0.005863 | 0.032879 |
| H3F3A | 341.6773 | 0.081368 | 0.158131 | 0.514559 | 0.606861 | 0.734751 |
| ADCK3 | 1020.353 | -0.40689 | 0.167182 | -2.43382 | 0.01494 | 0.061822 |
| OTUD7B | 3199.513 | -0.31414 | 0.136286 | -2.30504 | 0.021164 | 0.07868 |
| RPRD2 | 718.3124 | -0.4163 | 0.166122 | -2.50596 | 0.012212 | 0.05404 |

| | | | | | | |
|---|---|---|---|---|---|---|
| CTSS | 543.6593 | 0.329944 | 0.185359 | 1.78003 | 0.075071 | 0.182994 |
| BNIPL | 3.678218 | 0.629948 | 1.213706 | 0.519028 | 0.603741 | NA |
| TNFAIP8L2 | 0.399595 | -1.31595 | 3.579466 | -0.36764 | 0.713143 | NA |
| LYSMD1 | 275.4599 | -0.23702 | 0.183421 | -1.29221 | 0.196286 | 0.351718 |
| SCNM1 | 1147.499 | 0.121043 | 0.14225 | 0.850919 | 0.394814 | 0.559494 |
| VPS72 | 1344.11 | -0.19039 | 0.15807 | -1.20446 | 0.22841 | 0.390061 |
| S100A11 | 10841.12 | 0.017121 | 0.136053 | 0.125837 | 0.899861 | 0.941392 |
| SPRR2D | 0.192403 | 0.975392 | 4.060716 | 0.240202 | 0.810174 | NA |
| S100A9 | 0.151697 | -1.546 | 4.075469 | -0.37934 | 0.704433 | NA |
| TDRD10 | 0.260158 | 1.240347 | 4.047124 | 0.306476 | 0.759242 | NA |
| C1orf189 | 0.748867 | 2.740193 | 2.468928 | 1.109871 | 0.267054 | NA |
| PMVK | 1311.15 | -0.18945 | 0.154528 | -1.226 | 0.220197 | 0.379921 |
| PBXIP1 | 5047.385 | -0.39434 | 0.161618 | -2.43997 | 0.014688 | 0.061041 |
| PYGO2 | 1303.145 | -0.38891 | 0.165638 | -2.34795 | 0.018877 | 0.072659 |
| HIPK1 | 1532.81 | -0.08608 | 0.138788 | -0.62025 | 0.53509 | 0.678307 |
| LENEP | 0.315653 | -0.57305 | 3.892639 | -0.14721 | 0.882963 | NA |
| DCST2 | 1.335911 | 1.000206 | 1.953834 | 0.51192 | 0.608707 | NA |
| DCST1 | 2.518306 | 0.259631 | 1.443686 | 0.179839 | 0.857279 | NA |
| C1orf106 | 9.40638 | -0.34769 | 0.691602 | -0.50273 | 0.615154 | NA |
| YY1AP1 | 601.2106 | -0.2457 | 0.166353 | -1.47699 | 0.139677 | 0.279803 |
| APOA1BP | 3356.626 | -0.21263 | 0.122879 | -1.73041 | 0.083557 | 0.196933 |
| NBPF10 | 189.8489 | 0.393081 | 0.353934 | 1.110604 | 0.266739 | 0.431639 |
| SLC22A15 | 178.8708 | 0.524823 | 0.20124 | 2.607943 | 0.009109 | 0.044401 |
| ATP1A1 | 55460.57 | -0.42401 | 0.254477 | -1.66622 | 0.09567 | 0.216537 |
| LMOD1 | 48.33969 | -0.22568 | 0.453292 | -0.49787 | 0.618573 | 0.743278 |
| ELF3 | 1.238368 | 0.11839 | 1.910502 | 0.061968 | 0.950588 | NA |
| TMEM183A | 742.7804 | 0.085332 | 0.16954 | 0.503316 | 0.614742 | 0.740853 |
| TRIM46 | 118.5208 | 0.120193 | 0.224528 | 0.535315 | 0.592432 | 0.7233 |
| KRTCAP2 | 1581.679 | -0.12098 | 0.174206 | -0.69446 | 0.487391 | 0.64041 |
| TSACC | 10.45512 | 0.662306 | 0.624378 | 1.060746 | 0.288805 | NA |
| CCT3 | 10475.11 | -0.07121 | 0.128176 | -0.55556 | 0.578514 | 0.71305 |
| TMEM79 | 129.7886 | -0.42062 | 0.223415 | -1.88266 | 0.059746 | 0.156909 |
| SSR2 | 4060.987 | -0.1227 | 0.131538 | -0.93281 | 0.35092 | 0.518852 |
| ADORA1 | 15.28882 | -0.464 | 0.511661 | -0.90684 | 0.364491 | NA |
| SRGAP2 | 1875.177 | -0.36982 | 0.144991 | -2.55065 | 0.010752 | 0.049294 |
| NFASC | 41.61412 | -0.49691 | 0.481627 | -1.03173 | 0.3022 | NA |
| NUAK2 | 8.548466 | -0.86026 | 0.717139 | -1.19958 | 0.230303 | NA |
| SPTA1 | 19.64572 | 1.062922 | 0.535179 | 1.986105 | 0.047022 | NA |
| MNDA | 1.915931 | 3.081113 | 1.977775 | 1.557868 | 0.119265 | NA |
| PYHIN1 | 72.4785 | -0.69617 | 0.432548 | -1.60946 | 0.107515 | 0.233955 |
| IFI16 | 4664.848 | 0.256086 | 0.258166 | 0.991947 | 0.321224 | 0.489411 |
| AIM2 | 1.431763 | 2.610478 | 1.858973 | 1.404258 | 0.160242 | NA |
| SMIM12 | 1392.949 | -0.15919 | 0.144197 | -1.104 | 0.269593 | 0.434302 |
| ZMYM6 | 330.9785 | 0.785818 | 0.228804 | 3.434463 | 5.94E-04 | 0.007528 |
| GRIK3 | 30.11261 | -0.49726 | 0.514482 | -0.96652 | 0.333784 | NA |
| ZC3H12A | 77.8403 | -0.08066 | 0.25663 | -0.31432 | 0.75328 | 0.845894 |
| MEAF6 | 1074.325 | 0.270848 | 0.131691 | 2.056697 | 0.039715 | 0.119833 |

| | | | | | | |
|---|---|---|---|---|---|---|
| SNIP1 | 355.9428 | -0.1176 | 0.184713 | -0.63666 | 0.524347 | 0.670347 |
| DNALI1 | 162.4773 | -0.28147 | 0.194462 | -1.44745 | 0.14777 | 0.290937 |
| HEYL | 12.70978 | -0.20295 | 0.615955 | -0.32949 | 0.741782 | NA |
| EXO5 | 148.1009 | -0.01471 | 0.215 | -0.06842 | 0.945451 | 0.969593 |
| CLDN19 | 0.163957 | 0.424913 | 4.079006 | 0.104171 | 0.917034 | NA |
| C1orf50 | 199.2476 | -0.03182 | 0.195265 | -0.16298 | 0.870535 | 0.923107 |
| ERMAP | 235.2514 | -0.30309 | 0.229481 | -1.32076 | 0.186582 | 0.340716 |
| ZNF691 | 175.7128 | -0.13103 | 0.199901 | -0.65548 | 0.512158 | 0.65973 |
| ARMC3 | 0.162376 | 0.424913 | 4.079006 | 0.104171 | 0.917034 | NA |
| OTUD1 | 388.309 | -0.01047 | 0.183937 | -0.0569 | 0.954622 | 0.974998 |
| ARHGAP12 | 831.752 | 0.94642 | 0.222867 | 4.246574 | 2.17E-05 | 9.60E-04 |
| HECTD2 | 122.4176 | 1.024526 | 0.290376 | 3.528281 | 4.18E-04 | 0.006122 |
| ZNF488 | 0.824933 | -0.18668 | 2.233587 | -0.08358 | 0.933393 | NA |
| 8-Mar | 632.0569 | -0.24965 | 0.154182 | -1.61918 | 0.105409 | 0.231198 |
| ZCCHC24 | 2753.739 | -0.49188 | 0.155496 | -3.16331 | 0.00156 | 0.013873 |
| PHYHIPL | 0.861641 | -0.2592 | 2.180811 | -0.11885 | 0.90539 | NA |
| SLC16A9 | 0.47803 | 0.520083 | 2.97881 | 0.174594 | 0.861399 | NA |
| MBL2 | 1.599821 | -1.04115 | 1.538529 | -0.67672 | 0.498586 | NA |
| REEP3 | 2178.098 | 0.828874 | 0.203252 | 4.078068 | 4.54E-05 | 0.001556 |
| C10orf10 | 59.48175 | 0.002702 | 0.494929 | 0.005459 | 0.995644 | 0.997917 |
| C10orf25 | 12.97946 | 0.44744 | 0.596205 | 0.75048 | 0.452966 | NA |
| ZNF22 | 157.0752 | 0.592916 | 0.301873 | 1.964125 | 0.049516 | 0.137692 |
| AKR1E2 | 35.53611 | -0.21385 | 0.423143 | -0.50539 | 0.613282 | NA |
| NUDT5 | 1207.648 | 0.064223 | 0.153153 | 0.419342 | 0.674967 | 0.787316 |
| BEND7 | 76.60939 | 0.267148 | 0.282317 | 0.946273 | 0.344009 | 0.512167 |
| ATP5C1 | 5567.335 | 0.274959 | 0.131717 | 2.0875 | 0.036843 | 0.113773 |
| PRPF18 | 95.56668 | 0.786353 | 0.26254 | 2.995171 | 0.002743 | 0.019974 |
| TAF3 | 339.3267 | 0.304596 | 0.269794 | 1.128992 | 0.258901 | 0.423177 |
| VDAC2 | 1905.306 | -0.00294 | 0.165285 | -0.01777 | 0.985819 | 0.992517 |
| COMTD1 | 465.5924 | -0.16474 | 0.165883 | -0.99308 | 0.32067 | 0.488882 |
| SLC18A2 | 63.00721 | 0.481896 | 0.337535 | 1.427693 | 0.15338 | 0.298135 |
| PDZD8 | 859.2601 | 0.541373 | 0.216189 | 2.504161 | 0.012274 | 0.054213 |
| ZNF503 | 1198.408 | -0.23471 | 0.155227 | -1.51205 | 0.130522 | 0.26704 |
| FAM175B | 466.4923 | 0.60138 | 0.177849 | 3.381413 | 7.21E-04 | 0.008464 |
| FAM204A | 596.6111 | 0.866117 | 0.183455 | 4.721128 | 2.35E-06 | 2.67E-04 |
| PRDX3 | 4419.357 | 0.186077 | 0.14217 | 1.308836 | 0.19059 | 0.344953 |
| GHITM | 6673.449 | 0.260582 | 0.131296 | 1.984686 | 0.047179 | 0.133679 |
| STOX1 | 1.973773 | 1.114095 | 1.6015 | 0.695657 | 0.486644 | NA |
| RET | 4.609926 | -0.54331 | 1.084162 | -0.50114 | 0.616274 | NA |
| DDX21 | 1983.273 | 0.710176 | 0.185191 | 3.834832 | 1.26E-04 | 0.002961 |
| BMS1 | 930.4146 | 0.595191 | 0.215689 | 2.759489 | 0.005789 | 0.032619 |
| STK32C | 662.208 | -0.18277 | 0.17779 | -1.028 | 0.303951 | 0.471148 |
| KIAA1462 | 7.05429 | 1.469205 | 0.902171 | 1.628521 | 0.103414 | NA |
| CASP7 | 415.4698 | 0.229408 | 0.186567 | 1.229632 | 0.218835 | 0.378502 |
| C10orf118 | 136.7667 | 1.44672 | 0.926898 | 1.56082 | 0.118566 | 0.249961 |
| VWA2 | 0.86916 | 0.686589 | 2.213986 | 0.310114 | 0.756474 | NA |
| PRAP1 | 0.419962 | -2.64111 | 3.429364 | -0.77014 | 0.441214 | NA |

| | | | | | | |
|---|---|---|---|---|---|---|
| TRUB1 | 655.1851 | 0.466496 | 0.192475 | 2.42367 | 0.015365 | 0.062996 |
| C10orf82 | 2.600456 | -0.74513 | 1.242329 | -0.59978 | 0.548652 | NA |
| HSPA12A | 3166.108 | -0.31855 | 0.200223 | -1.59096 | 0.111619 | 0.239791 |
| FAM35BP | 0.096202 | 0.408546 | 4.079639 | 0.100143 | 0.920231 | NA |
| FRAT1 | 191.5294 | -0.30043 | 0.23915 | -1.25623 | 0.209034 | 0.366643 |
| UBTD1 | 2158.068 | -0.07621 | 0.164349 | -0.46371 | 0.642857 | 0.762412 |
| ANKRD2 | 1.159142 | 2.191644 | 1.992053 | 1.100194 | 0.271248 | NA |
| PACSIN3 | 1330.873 | -0.1179 | 0.155932 | -0.75608 | 0.449599 | 0.609046 |
| SLC39A13 | 3622.396 | -0.21828 | 0.135514 | -1.61074 | 0.107237 | 0.233736 |
| PSMC3 | 3705.681 | -0.06229 | 0.144976 | -0.42967 | 0.667437 | 0.781864 |
| RAPSN | 1.448132 | 0.963259 | 1.699702 | 0.566722 | 0.570903 | NA |
| AGBL2 | 6.664764 | -0.35887 | 0.810746 | -0.44264 | 0.658027 | NA |
| NELL1 | 19.90572 | -0.96585 | 0.662691 | -1.45747 | 0.144986 | NA |
| PTER | 326.6116 | 0.674033 | 0.181241 | 3.718985 | 2.00E-04 | 0.003868 |
| C1QL3 | 11.12619 | 1.243588 | 0.634183 | 1.96093 | 0.049887 | NA |
| CACNB2 | 11.20144 | 0.31172 | 0.813154 | 0.383347 | 0.701462 | NA |
| PTPLA | 1406.749 | -0.11779 | 0.20444 | -0.57614 | 0.56452 | 0.702244 |
| ARL5B | 270.2812 | 0.882059 | 0.275089 | 3.206443 | 0.001344 | 0.012564 |
| TRIM51HP | 0.35636 | 1.709252 | 3.651833 | 0.468053 | 0.639746 | NA |
| ABTB2 | 625.2338 | -0.69786 | 0.194167 | -3.59411 | 3.25E-04 | 0.005188 |
| R3HCC1L | 589.5609 | 0.404358 | 0.171601 | 2.356377 | 0.018454 | 0.071588 |
| HTRA1 | 4362.129 | -0.24474 | 0.164416 | -1.48852 | 0.136613 | 0.275639 |
| HIF1AN | 1626.23 | -0.29542 | 0.137192 | -2.1533 | 0.031295 | 0.102008 |
| NDUFB8 | 463.9592 | 0.117609 | 0.149442 | 0.786989 | 0.431288 | 0.592665 |
| BTRC | 739.1406 | -0.19564 | 0.141392 | -1.38371 | 0.166449 | 0.31572 |
| POLL | 462.1721 | -0.30248 | 0.153463 | -1.97104 | 0.048719 | 0.136411 |
| DPCD | 452.0288 | 0.062258 | 0.221905 | 0.280562 | 0.779047 | 0.864433 |
| API5 | 1979.397 | 0.408289 | 0.158414 | 2.577359 | 0.009956 | 0.047051 |
| HPS6 | 1012.121 | -0.37364 | 0.134993 | -2.76788 | 0.005642 | 0.032127 |
| NOLC1 | 2715.721 | 0.011328 | 0.1493 | 0.075871 | 0.939522 | 0.966423 |
| ALKBH3 | 550.2086 | -0.10311 | 0.182558 | -0.56479 | 0.572219 | 0.708254 |
| SGPL1 | 1256.522 | -0.63586 | 0.160692 | -3.95699 | 7.59E-05 | 0.002147 |
| PCBD1 | 832.4119 | -0.25209 | 0.162849 | -1.54803 | 0.121615 | 0.254591 |
| WBP1L | 1699.566 | -0.29429 | 0.165742 | -1.77558 | 0.075802 | 0.184259 |
| C10orf32 | 629.6144 | 0.62369 | 0.161842 | 3.853704 | 1.16E-04 | 0.002815 |
| ANAPC16 | 1271.145 | 0.063464 | 0.134347 | 0.472391 | 0.636648 | 0.757473 |
| SMPD1 | 3797.05 | -0.44018 | 0.124868 | -3.52517 | 4.23E-04 | 0.006131 |
| APBB1 | 3138.534 | -0.41362 | 0.134362 | -3.07839 | 0.002081 | 0.016597 |
| SYNPO2L | 1.265234 | 0.20438 | 1.735412 | 0.11777 | 0.90625 | NA |
| NUDT13 | 69.73631 | 0.561685 | 0.303668 | 1.849667 | 0.064362 | 0.16522 |
| TRIM44 | 1930.053 | 0.148072 | 0.126072 | 1.174504 | 0.240193 | 0.403355 |
| ILK | 21.56117 | -0.4553 | 0.45558 | -0.99939 | 0.317604 | NA |
| TAF10 | 2381.181 | -0.10489 | 0.145722 | -0.71979 | 0.471657 | 0.627552 |
| TPP1 | 7996.695 | -0.641 | 0.150874 | -4.24858 | 2.15E-05 | 9.60E-04 |
| DCHS1 | 382.7907 | -0.74583 | 0.190494 | -3.91523 | 9.03E-05 | 0.002389 |
| MSS51 | 12.73064 | 0.252389 | 0.64739 | 0.389856 | 0.696643 | NA |
| USP54 | 1220.96 | -0.68196 | 0.136739 | -4.9873 | 6.12E-07 | 1.17E-04 |

| | | | | | | |
|---|---|---|---|---|---|---|
| RAG1 | 93.85154 | 0.203443 | 0.253046 | 0.803975 | 0.421411 | 0.584035 |
| C11orf74 | 285.4709 | 0.577823 | 0.18526 | 3.118976 | 0.001815 | 0.015146 |
| PPFIBP2 | 1166.401 | -0.19991 | 0.167372 | -1.19443 | 0.23231 | 0.394566 |
| CYB5R2 | 313.0965 | -0.12652 | 0.183928 | -0.68788 | 0.491528 | 0.643306 |
| TUB | 279.2415 | -0.10146 | 0.192989 | -0.52572 | 0.599084 | 0.728649 |
| RIC3 | 0.834592 | 0.684312 | 2.202411 | 0.31071 | 0.756021 | NA |
| LMO1 | 4.535118 | 0.466124 | 0.962343 | 0.484363 | 0.628128 | NA |
| TRIM66 | 93.94327 | -0.26643 | 0.283932 | -0.93836 | 0.348058 | 0.51605 |
| RPL27A | 18993.16 | 0.216282 | 0.138496 | 1.561648 | 0.118371 | 0.24974 |
| ST5 | 3155.891 | -0.55863 | 0.144699 | -3.86063 | 1.13E-04 | 0.002776 |
| AKIP1 | 755.9424 | -0.06347 | 0.149823 | -0.42362 | 0.671844 | 0.785165 |
| TMEM41B | 400.7442 | 0.454441 | 0.183686 | 2.474013 | 0.01336 | 0.05747 |
| ZNF143 | 348.341 | 0.588656 | 0.185934 | 3.165942 | 0.001546 | 0.013793 |
| WEE1 | 420.6048 | -0.13114 | 0.203119 | -0.64564 | 0.518514 | 0.665474 |
| FAM86GP | 0.516164 | -1.70268 | 2.844782 | -0.59853 | 0.549488 | NA |
| NDST2 | 542.6405 | -0.32789 | 0.15761 | -2.08036 | 0.037492 | 0.115058 |
| PLEKHA7 | 638.2381 | 0.768414 | 0.312415 | 2.459592 | 0.01391 | 0.058918 |
| SAAL1 | 578.9368 | 0.229071 | 0.154186 | 1.485676 | 0.137365 | 0.276573 |
| YPEL4 | 84.3818 | -0.57193 | 0.259621 | -2.20295 | 0.027598 | 0.093918 |
| LDHC | 75.71829 | -0.113 | 0.333764 | -0.33856 | 0.734943 | 0.832209 |
| LDHAL6A | 2.297063 | -0.38665 | 1.314651 | -0.29411 | 0.768675 | NA |
| FAM111A | 793.4508 | 0.494334 | 0.192719 | 2.565049 | 0.010316 | 0.048042 |
| NAV2 | 1933.051 | -0.61222 | 0.220568 | -2.77564 | 0.005509 | 0.031705 |
| GLYATL1 | 2.088502 | 1.209743 | 1.495322 | 0.809018 | 0.418505 | NA |
| PATL1 | 1570.589 | -0.35304 | 0.136626 | -2.58395 | 0.009768 | 0.046467 |
| STX3 | 3742.797 | -0.03458 | 0.168917 | -0.20474 | 0.837776 | 0.902976 |
| MRPL16 | 1149.343 | 0.157246 | 0.17146 | 0.917103 | 0.359089 | 0.526345 |
| MS4A7 | 101.547 | 0.778952 | 0.281351 | 2.768614 | 0.00563 | 0.032101 |
| MS4A14 | 6.21497 | 2.250883 | 1.202412 | 1.871973 | 0.06121 | NA |
| IGF2 | 0.873901 | 0.695147 | 2.212014 | 0.31426 | 0.753324 | NA |
| ART5 | 5.59545 | 0.260272 | 0.885373 | 0.293969 | 0.768781 | NA |
| STIM1 | 2170.782 | -0.33356 | 0.152886 | -2.18178 | 0.029126 | 0.097017 |
| RRM1 | 1403.49 | 0.128613 | 0.16266 | 0.790683 | 0.429129 | 0.590656 |
| OR51E2 | 0.162376 | 0.424913 | 4.079006 | 0.104171 | 0.917034 | NA |
| TRIM68 | 498.9736 | -0.26104 | 0.152478 | -1.71199 | 0.086898 | 0.202353 |
| SDHAF2 | 303.7072 | -0.18887 | 0.185142 | -1.02011 | 0.307675 | 0.474933 |
| DDB1 | 14146.92 | -0.33765 | 0.132543 | -2.54747 | 0.010851 | 0.049628 |
| VPS37C | 913.0536 | -0.51057 | 0.147861 | -3.45303 | 5.54E-04 | 0.007206 |
| VWCE | 13.74387 | 0.486053 | 0.584975 | 0.830895 | 0.406033 | NA |
| RAB3IL1 | 656.7248 | -0.28761 | 0.163051 | -1.7639 | 0.077749 | 0.187683 |
| BEST1 | 1407.728 | -0.07618 | 0.177831 | -0.42839 | 0.668369 | 0.782261 |
| FTH1 | 34941.9 | -0.18025 | 0.143966 | -1.252 | 0.21057 | 0.368427 |
| BSCL2 | 137.0091 | -0.34868 | 0.260102 | -1.34055 | 0.180068 | 0.332612 |
| POLR2G | 1354.882 | -0.10498 | 0.1468 | -0.71514 | 0.474524 | 0.629784 |
| RAB4A | 900.452 | 0.31948 | 0.153748 | 2.077951 | 0.037714 | 0.11547 |
| RNF187 | 5675.672 | -0.15698 | 0.127399 | -1.23219 | 0.217878 | 0.377341 |
| DDIT4 | 3278.144 | -0.29677 | 0.159086 | -1.86547 | 0.062115 | 0.161274 |

| | | | | | | |
|---|---|---|---|---|---|---|
| GNG4 | 6.533293 | 0.731516 | 0.81988 | 0.892224 | 0.372273 | NA |
| IRF2BP2 | 1349.35 | -0.30111 | 0.206885 | -1.45543 | 0.145551 | 0.287692 |
| COA6 | 747.2282 | 0.52746 | 0.170226 | 3.098593 | 0.001944 | 0.015854 |
| BMI1 | 837.7029 | 0.577314 | 0.174286 | 3.312449 | 9.25E-04 | 0.009851 |
| MFSD2A | 89.74612 | -0.06429 | 0.265593 | -0.24207 | 0.808726 | 0.88584 |
| FEN1 | 1566.73 | -0.11998 | 0.192842 | -0.62216 | 0.533839 | 0.677587 |
| SERINC2 | 379.1607 | -0.0293 | 0.184577 | -0.15873 | 0.873879 | 0.924986 |
| TMEM223 | 486.0765 | -0.0558 | 0.146625 | -0.38058 | 0.703513 | 0.809566 |
| NBPF9 | 120.8814 | -0.09108 | 0.259951 | -0.35038 | 0.726056 | 0.826046 |
| NDUFS5 | 4630.381 | 0.332798 | 0.131909 | 2.522927 | 0.011638 | 0.052222 |
| SLC16A4 | 405.42 | 0.177551 | 0.253512 | 0.700366 | 0.483699 | 0.637354 |
| AHCYL1 | 4928.83 | -0.1731 | 0.135455 | -1.27796 | 0.201265 | 0.357599 |
| GSTM4 | 945.9095 | -0.41979 | 0.161797 | -2.59457 | 0.009471 | 0.045531 |
| ARMC4 | 1.28379 | 2.393551 | 1.949457 | 1.227804 | 0.219521 | NA |
| AFAP1L2 | 537.5447 | -0.10889 | 0.203311 | -0.53558 | 0.592247 | 0.7233 |
| PCSK9 | 38.17485 | -0.00337 | 0.418102 | -0.00805 | 0.993575 | NA |
| RAB3B | 28.08211 | -0.93245 | 0.51135 | -1.82351 | 0.068226 | NA |
| RSPO1 | 0.134133 | -1.48909 | 4.077584 | -0.36519 | 0.71497 | NA |
| GCSAML | 0.096202 | 0.408546 | 4.079639 | 0.100143 | 0.920231 | NA |
| THBS3 | 206.6414 | -0.34471 | 0.181715 | -1.897 | 0.057827 | 0.153057 |
| SLC50A1 | 950.3255 | -0.20223 | 0.181827 | -1.11223 | 0.266038 | 0.4308 |
| EFNA1 | 61.31644 | -0.04387 | 0.351419 | -0.12484 | 0.900652 | 0.941822 |
| SHE | 12.94622 | 1.908918 | 0.712713 | 2.678383 | 0.007398 | NA |
| PTAFR | 1.256924 | 2.336092 | 1.930733 | 1.209951 | 0.226298 | NA |
| NPR1 | 4.160617 | -0.14911 | 1.118834 | -0.13327 | 0.893977 | NA |
| CLIC4 | 7410.339 | 0.393897 | 0.189207 | 2.081833 | 0.037358 | 0.114765 |
| METTL15 | 190.279 | 0.533444 | 0.256559 | 2.079224 | 0.037597 | 0.11523 |
| DFFB | 127.1413 | -0.11201 | 0.21622 | -0.51805 | 0.604423 | 0.732942 |
| LUZP1 | 1633.35 | 0.253077 | 0.134924 | 1.875704 | 0.060696 | 0.158526 |
| ZNF32 | 667.0735 | 0.12359 | 0.178148 | 0.693745 | 0.487842 | 0.640674 |
| HNRNPF | 4802.006 | -0.03123 | 0.116161 | -0.26889 | 0.788018 | 0.871362 |
| CSGALNAC | 381.1939 | 0.779893 | 0.20443 | 3.814967 | 1.36E-04 | 0.003063 |
| CALML6 | 1.369137 | 0.32666 | 1.894165 | 0.172456 | 0.863079 | NA |
| TOR1AIP2 | 1180.616 | 0.175981 | 0.172917 | 1.017721 | 0.30881 | 0.476128 |
| OTUD3 | 163.6318 | 0.63491 | 0.269099 | 2.359396 | 0.018305 | 0.071125 |
| TAS1R3 | 11.92187 | -0.40638 | 0.621185 | -0.6542 | 0.512984 | NA |
| PUSL1 | 326.3913 | -0.08813 | 0.206503 | -0.42676 | 0.669556 | 0.78334 |
| IFFO2 | 442.194 | -0.11234 | 0.203481 | -0.5521 | 0.580879 | 0.714667 |
| GPR37L1 | 0.303394 | -2.27006 | 4.032451 | -0.56295 | 0.57347 | NA |
| USP47 | 946.5989 | 0.62447 | 0.328597 | 1.900414 | 0.057379 | 0.152074 |
| CDK1 | 1081.003 | 0.461313 | 0.21182 | 2.177855 | 0.029417 | 0.097713 |
| EMX2 | 0.586491 | 2.39072 | 2.689057 | 0.889055 | 0.373973 | NA |
| SLC30A1 | 433.3919 | 0.485294 | 0.177115 | 2.739986 | 0.006144 | 0.03389 |
| MGMT | 814.5007 | -0.1842 | 0.16825 | -1.09478 | 0.273612 | 0.439461 |
| PFKFB3 | 1925.168 | -0.13604 | 0.207508 | -0.65557 | 0.512103 | 0.65973 |
| SYT9 | 0.288605 | 1.443619 | 4.029058 | 0.358302 | 0.720117 | NA |
| KIF5B | 2931.471 | 0.886829 | 0.612837 | 1.447088 | 0.147872 | 0.291007 |

| | | | | | | |
|---|---|---|---|---|---|---|
| OR10A4 | 0.162376 | 0.424913 | 4.079006 | 0.104171 | 0.917034 | NA |
| DNAJC24 | 210.2739 | 0.681967 | 0.297913 | 2.289153 | 0.02207 | 0.081 |
| PRKCDBP | 5145.796 | -0.14653 | 0.167939 | -0.8725 | 0.382936 | 0.548882 |
| DCDC1 | 0.915759 | -1.79003 | 2.342046 | -0.7643 | 0.444687 | NA |
| S1PR1 | 23.16456 | -0.07114 | 0.479351 | -0.14841 | 0.882016 | NA |
| MORN4 | 212.0405 | -0.43107 | 0.200424 | -2.15077 | 0.031494 | 0.102545 |
| ZNF672 | 1029.132 | -0.37119 | 0.146494 | -2.53382 | 0.011283 | 0.051071 |
| ZNF692 | 291.0407 | 0.477103 | 0.195428 | 2.441323 | 0.014634 | 0.060862 |
| TRIM8 | 1896.572 | -0.18275 | 0.165203 | -1.10619 | 0.268645 | 0.433599 |
| C10orf35 | 2.310757 | 0.560693 | 1.288184 | 0.435259 | 0.663375 | NA |
| ZDHHC16 | 1976.094 | -0.27082 | 0.17451 | -1.5519 | 0.120687 | 0.25305 |
| EXOSC1 | 799.8538 | 0.114045 | 0.165975 | 0.687117 | 0.492009 | 0.643509 |
| PGAM1 | 2889.518 | -0.08503 | 0.132949 | -0.63959 | 0.522441 | 0.668705 |
| LURAP1 | 176.6141 | -0.3044 | 0.225938 | -1.34729 | 0.177888 | 0.330231 |
| KCND3 | 15.02461 | 1.600934 | 0.813639 | 1.967624 | 0.049111 | NA |
| LRRC8C | 542.1801 | 0.098277 | 0.215877 | 0.455244 | 0.648934 | 0.76701 |
| LRRC8D | 919.9994 | -0.25938 | 0.191897 | -1.35168 | 0.176477 | 0.328797 |
| COL24A1 | 19.68969 | -0.24571 | 0.842707 | -0.29157 | 0.770612 | NA |
| LPAR3 | 32.90902 | -0.18459 | 0.41517 | -0.4446 | 0.656605 | NA |
| CLSTN1 | 9502.388 | -0.46035 | 0.158953 | -2.89611 | 0.003778 | 0.024529 |
| PIK3CD | 3996.051 | -0.50618 | 0.154009 | -3.28672 | 0.001014 | 0.010449 |
| SLC25A33 | 331.0315 | -0.13468 | 0.217335 | -0.61968 | 0.535467 | 0.678563 |
| SPSB1 | 2516.847 | -0.49165 | 0.155804 | -3.15554 | 0.001602 | 0.014125 |
| PLEKHG5 | 183.4814 | -0.06659 | 0.270348 | -0.24631 | 0.80544 | 0.883628 |
| ANO5 | 306.9938 | 0.772483 | 0.192024 | 4.022843 | 5.75E-05 | 0.001791 |
| TMEM51 | 607.8277 | -0.67995 | 0.225819 | -3.01104 | 0.002604 | 0.019343 |
| CAMTA1 | 1599.527 | 0.155298 | 0.16401 | 0.946881 | 0.3437 | 0.511904 |
| NHLH1 | 3.366467 | -0.09369 | 1.212209 | -0.07729 | 0.938395 | NA |
| SLFNL1 | 4.269613 | -0.81605 | 0.956627 | -0.85305 | 0.393632 | NA |
| CTPS1 | 1114.116 | -0.1185 | 0.15869 | -0.74677 | 0.455203 | 0.614138 |
| KNDC1 | 8.031159 | 0.656648 | 0.714575 | 0.918934 | 0.35813 | NA |
| METTL18 | 105.3133 | 0.792392 | 0.278944 | 2.840688 | 0.004502 | 0.027466 |
| TTC40 | 35.65064 | 0.288741 | 0.385386 | 0.749226 | 0.453721 | NA |
| COL8A2 | 6.4213 | -0.61136 | 0.863691 | -0.70784 | 0.479042 | NA |
| PWWP2B | 892.5777 | -0.25665 | 0.134251 | -1.91175 | 0.055908 | 0.149547 |
| ANGPTL7 | 352.4162 | 1.120277 | 0.761269 | 1.471591 | 0.141131 | 0.281685 |
| EXOSC10 | 1909.378 | -0.2009 | 0.131806 | -1.52424 | 0.12745 | 0.263318 |
| PTEN | 1746.123 | 0.20644 | 0.256212 | 0.805738 | 0.420394 | 0.583153 |
| KLF17 | 0.804566 | 0.545894 | 2.033578 | 0.26844 | 0.78836 | NA |
| SRGAP2C | 65.83458 | 0.42069 | 0.306016 | 1.374735 | 0.169214 | 0.319584 |
| PPIH | 664.1955 | 0.150861 | 0.156147 | 0.966145 | 0.333972 | 0.502504 |
| JMJD1C | 654.8471 | 0.918631 | 0.32442 | 2.831611 | 0.004631 | 0.027999 |
| EPHX4 | 1.618376 | 0.649083 | 1.559215 | 0.416288 | 0.677199 | NA |
| C1QTNF4 | 1.459963 | -0.77351 | 1.658727 | -0.46633 | 0.64098 | NA |
| NEGR1 | 38.44071 | 0.840616 | 0.376182 | 2.234599 | 0.025444 | NA |
| ALG14 | 355.559 | 0.214784 | 0.182453 | 1.177203 | 0.239115 | 0.402358 |
| OR5B12 | 0.163957 | 0.424913 | 4.079006 | 0.104171 | 0.917034 | NA |

| | | | | | | |
|---|---|---|---|---|---|---|
| GNG12 | 7317.86 | 0.168718 | 0.148565 | 1.135649 | 0.256103 | 0.42028 |
| CLP1 | 575.081 | 0.05675 | 0.172241 | 0.329483 | 0.74179 | 0.837188 |
| INSL5 | 0.163957 | 0.424913 | 4.079006 | 0.104171 | 0.917034 | NA |
| FGGY | 212.4056 | -0.10595 | 0.189555 | -0.55892 | 0.576217 | 0.711346 |
| CHCHD1 | 929.4783 | 0.071708 | 0.146544 | 0.489327 | 0.62461 | 0.748409 |
| AGAP5 | 0.638825 | 1.117948 | 3.106359 | 0.35989 | 0.718929 | NA |
| FAM21C | 677.5293 | 0.268212 | 0.145259 | 1.846441 | 0.064828 | 0.165956 |
| ZFAND4 | 81.45562 | 0.506843 | 0.268681 | 1.886411 | 0.05924 | 0.155821 |
| LRRC20 | 137.0414 | -0.69483 | 0.236061 | -2.94345 | 0.003246 | 0.022307 |
| | 1.037858 | 1.046985 | 2.132048 | 0.49107 | 0.623377 | NA |
| HPSE2 | 40.82952 | -0.38746 | 0.374235 | -1.03533 | 0.300517 | NA |
| C10orf131 | 2.626439 | 1.61026 | 1.352553 | 1.190533 | 0.233837 | NA |
| HSPA6 | 51.17383 | -1.37474 | 0.339093 | -4.05416 | 5.03E-05 | 0.001652 |
| NOC3L | 362.6109 | 1.003713 | 0.321111 | 3.125754 | 0.001773 | 0.014986 |
| MTX1 | 653.9545 | -0.19242 | 0.175996 | -1.09333 | 0.274247 | 0.439886 |
| CKS1B | 745.7955 | -0.17793 | 0.25527 | -0.69703 | 0.485782 | 0.639244 |
| MAB21L3 | 0.518736 | 2.237423 | 2.7713 | 0.807355 | 0.419462 | NA |
| VANGL1 | 1443.817 | -0.43542 | 0.141692 | -3.07299 | 0.002119 | 0.016804 |
| SNCG | 0.585114 | 1.12675 | 2.73598 | 0.411827 | 0.680466 | NA |
| MMRN2 | 2.415212 | -0.54696 | 1.283125 | -0.42627 | 0.669912 | NA |
| DAB1 | 5.226698 | 0.599426 | 1.07565 | 0.557269 | 0.577344 | NA |
| ARV1 | 681.6651 | 0.133759 | 0.156962 | 0.852178 | 0.394115 | 0.55888 |
| SAA1 | 2.827416 | -1.13972 | 1.314006 | -0.86736 | 0.385742 | NA |
| MINOS1 | 2904.374 | -0.08584 | 0.168199 | -0.51037 | 0.609795 | 0.736838 |
| NMNAT1 | 486.2059 | 0.044587 | 0.191332 | 0.233035 | 0.815734 | 0.889858 |
| APOBEC4 | 0.162376 | 0.424913 | 4.079006 | 0.104171 | 0.917034 | NA |
| HSPB7 | 0.640755 | -1.64826 | 2.754778 | -0.59833 | 0.549621 | NA |
| UQCRH | 4316.803 | 0.364577 | 0.14278 | 2.553421 | 0.010667 | 0.049123 |
| TAS1R1 | 1.493931 | -0.17651 | 1.55812 | -0.11328 | 0.909808 | NA |
| HES3 | 0.956409 | -1.29092 | 2.212928 | -0.58335 | 0.559655 | NA |
| TOMM20 | 4870.448 | 0.181128 | 0.1592 | 1.13774 | 0.255229 | 0.419495 |
| C1orf100 | 1.005865 | 3.174325 | 2.260631 | 1.404176 | 0.160266 | NA |
| PLK3 | 437.0649 | 0.043684 | 0.192605 | 0.226805 | 0.820576 | 0.892334 |
| NET1 | 678.8474 | 0.571556 | 0.205573 | 2.780303 | 0.005431 | 0.031364 |
| TUBB8 | 1.017482 | -0.71858 | 2.105405 | -0.3413 | 0.732874 | NA |
| USMG5 | 3000.39 | 0.39565 | 0.168282 | 2.351117 | 0.018717 | 0.072265 |
| PIFO | 44.72523 | 0.058316 | 0.374578 | 0.155685 | 0.876281 | NA |
| GNG5 | 2633.938 | 0.071181 | 0.141914 | 0.501578 | 0.615964 | 0.742024 |
| CD34 | 10.86108 | 0.19727 | 0.65014 | 0.303428 | 0.761564 | NA |
| CYB561D1 | 711.4824 | -0.36066 | 0.168254 | -2.14356 | 0.032068 | 0.103843 |
| AGAP8 | 2.562176 | 0.329363 | 1.283314 | 0.25665 | 0.797449 | NA |
| FAM21D | 0.163957 | 0.424913 | 4.079006 | 0.104171 | 0.917034 | NA |
| PHLDA3 | 5790.5 | -0.01974 | 0.205236 | -0.09617 | 0.923384 | 0.955878 |
| GLIS1 | 4.575678 | 0.974457 | 0.958561 | 1.016582 | 0.309352 | NA |
| PODN | 7.790982 | 0.726191 | 0.737564 | 0.98458 | 0.32483 | NA |
| EXO1 | 297.0712 | 0.304619 | 0.238376 | 1.277893 | 0.201287 | 0.357599 |
| MFSD4 | 18.92454 | 0.073155 | 0.486295 | 0.150433 | 0.880423 | NA |

| | | | | | | |
|---|---|---|---|---|---|---|
| TMEM81 | 36.1864 | 0.297961 | 0.368241 | 0.809147 | 0.418431 | NA |
| GOLT1A | 0.260158 | 1.240347 | 4.047124 | 0.306476 | 0.759242 | NA |
| AKIRIN1 | 2158.296 | 0.153632 | 0.149943 | 1.0246 | 0.305552 | 0.472703 |
| ANGEL2 | 540.3953 | 0.252359 | 0.179936 | 1.402498 | 0.160767 | 0.308346 |
| BRSK2 | 86.62832 | 0.373105 | 0.413634 | 0.902017 | 0.367048 | 0.534236 |
| FGFBP3 | 22.79718 | 0.095428 | 0.514632 | 0.18543 | 0.852892 | NA |
| HRAS | 1585.371 | 0.059303 | 0.175544 | 0.337824 | 0.735496 | 0.832559 |
| PDZK1 | 1.591567 | 0.691343 | 1.560782 | 0.442947 | 0.657804 | NA |
| GLMN | 115.7059 | 1.03728 | 0.292505 | 3.546191 | 3.91E-04 | 0.005836 |
| NLRP6 | 0.163957 | 0.424913 | 4.079006 | 0.104171 | 0.917034 | NA |
| OR9G1 | 4.976811 | -0.37693 | 0.883267 | -0.42674 | 0.669569 | NA |
| PTDSS2 | 1901.397 | -0.16225 | 0.130703 | -1.24139 | 0.214463 | 0.373452 |
| CTBP2 | 1669.555 | 0.041064 | 0.131703 | 0.31179 | 0.7552 | 0.84717 |
| PDIK1L | 93.87062 | 0.72922 | 0.282185 | 2.584193 | 0.009761 | 0.046452 |
| RAG2 | 0.520316 | 2.239969 | 3.173769 | 0.705776 | 0.480328 | NA |
| TRAF6 | 320.7892 | 0.078492 | 0.169622 | 0.462744 | 0.643548 | 0.762832 |
| MARCKSL1 | 15473.39 | -0.43862 | 0.137686 | -3.18565 | 0.001444 | 0.013252 |
| SH3BP5L | 1459.743 | -0.31009 | 0.148141 | -2.0932 | 0.036332 | 0.112722 |
| TMEM51-A | 1.082076 | -0.5518 | 2.085766 | -0.26455 | 0.791353 | NA |
| NPPA | 1.525336 | 0.472832 | 1.602087 | 0.295135 | 0.767891 | NA |
| ZNF408 | 388.7429 | -0.21823 | 0.208934 | -1.04447 | 0.296267 | 0.463602 |
| CKAP5 | 3703.429 | -0.16067 | 0.13378 | -1.20102 | 0.229744 | 0.39183 |
| ARHGAP1 | 2467.312 | -0.6076 | 0.127395 | -4.76941 | 1.85E-06 | 2.33E-04 |
| ATG13 | 2947.191 | -0.21935 | 0.146218 | -1.50017 | 0.133571 | 0.271237 |
| C1orf127 | 3.962703 | -0.05378 | 1.122624 | -0.04791 | 0.961789 | NA |
| CHST1 | 2.255639 | 1.459338 | 1.534785 | 0.950842 | 0.341685 | NA |
| TP53I11 | 242.4185 | -0.58733 | 0.27115 | -2.16609 | 0.030305 | 0.099795 |
| APITD1 | 212.6455 | 0.182288 | 0.198589 | 0.917916 | 0.358663 | 0.525906 |
| TMEM9B | 1794.585 | 0.097208 | 0.151929 | 0.639826 | 0.522286 | 0.66865 |
| NRIP3 | 1029.548 | -0.53357 | 0.167027 | -3.19449 | 0.001401 | 0.012963 |
| SCUBE2 | 2777.263 | -0.14302 | 0.178008 | -0.80345 | 0.421714 | 0.584317 |
| EIF3F | 3526.436 | -0.11409 | 0.14287 | -0.79859 | 0.424528 | 0.586587 |
| ZNF25 | 273.7047 | 1.06597 | 0.277202 | 3.845464 | 1.20E-04 | 0.002893 |
| PPP2R2D | 942.5589 | 0.181753 | 0.14818 | 1.226574 | 0.219983 | 0.379661 |
| UBQLNL | 0.918331 | 0.764804 | 2.118268 | 0.361052 | 0.718061 | NA |
| UBQLN3 | 0.163957 | 0.424913 | 4.079006 | 0.104171 | 0.917034 | NA |
| PNLIP | 0.163957 | 0.424913 | 4.079006 | 0.104171 | 0.917034 | NA |
| AURKAIP1 | 3620.48 | -0.02053 | 0.159452 | -0.12874 | 0.897561 | 0.939815 |
| SFN | 1.6974 | -0.35191 | 1.528515 | -0.23023 | 0.817912 | NA |
| CALCB | 0.354779 | 1.703954 | 3.657232 | 0.465914 | 0.641277 | NA |
| DENND2C | 62.71436 | 0.171816 | 0.312442 | 0.549912 | 0.58238 | 0.715905 |
| B3GALT6 | 1201.233 | -0.24334 | 0.135912 | -1.79042 | 0.073387 | 0.18061 |
| C11orf16 | 1.284176 | 0.715001 | 1.74187 | 0.410479 | 0.681455 | NA |
| CSTF3 | 480.7937 | 0.118293 | 0.196208 | 0.602897 | 0.546577 | 0.688082 |
| TCP11L1 | 426.2896 | -0.11074 | 0.158867 | -0.69708 | 0.485756 | 0.639244 |
| BNIP3 | 986.2079 | 0.373931 | 0.153927 | 2.429278 | 0.015129 | 0.062427 |
| C10orf111 | 2.327386 | -0.35638 | 1.354737 | -0.26306 | 0.792501 | NA |

| | | | | | | |
|---|---|---|---|---|---|---|
| ACBD7 | 7.871062 | -0.56971 | 0.801656 | -0.71067 | 0.477288 | NA |
| ZBTB8OS | 444.5395 | 0.624578 | 0.166411 | 3.75322 | 1.75E-04 | 0.003558 |
| SLC35G1 | 695.8105 | 0.310801 | 0.199565 | 1.557393 | 0.119377 | 0.251237 |
| | 0.162376 | 0.424913 | 4.079006 | 0.104171 | 0.917034 | NA |
| RNPEP | 2584.224 | -0.3801 | 0.142359 | -2.67001 | 0.007585 | 0.039355 |
| CLK2 | 489.3756 | -0.02741 | 0.163113 | -0.16805 | 0.866543 | 0.920772 |
| DMBT1P1 | 0.163957 | 0.424913 | 4.079006 | 0.104171 | 0.917034 | NA |
| BDNF | 166.5819 | 0.412919 | 0.74442 | 0.554685 | 0.57911 | 0.713515 |
| OR10AB1P | 0.096202 | 0.408546 | 4.079639 | 0.100143 | 0.920231 | NA |
| LINC00303 | 1.373421 | -0.39644 | 1.899148 | -0.20875 | 0.834645 | NA |
| TCERG1L | 72.35578 | 1.038451 | 0.372953 | 2.784402 | 0.005363 | 0.031088 |
| FIBIN | 0.548763 | 2.343695 | 3.135478 | 0.747476 | 0.454776 | NA |
| SEC24C | 4123.043 | -0.51759 | 0.12852 | -4.0273 | 5.64E-05 | 0.001779 |
| MTHFR | 903.7463 | -0.57696 | 0.165576 | -3.48454 | 4.93E-04 | 0.006739 |
| DEAF1 | 2200.071 | -0.17423 | 0.140159 | -1.24305 | 0.213848 | 0.372655 |
| TMEM80 | 371.1999 | -0.2873 | 0.173727 | -1.65373 | 0.098183 | 0.220574 |
| ZDHHC13 | 387.6857 | 0.550821 | 0.189464 | 2.907253 | 0.003646 | 0.024068 |
| RHOG | 1255.19 | -0.24602 | 0.15669 | -1.57012 | 0.116388 | 0.246693 |
| EPS8L2 | 1333.256 | -0.37055 | 0.200974 | -1.84376 | 0.065219 | 0.166489 |
| MRVI1-AS1 | 0.298089 | -0.51711 | 3.975232 | -0.13008 | 0.896501 | NA |
| NUDT4P1 | 1.331206 | 0.831477 | 1.740564 | 0.477705 | 0.63286 | NA |
| TALDO1 | 9046.257 | -0.13468 | 0.167429 | -0.8044 | 0.421168 | 0.583902 |
| NAP1L4P1 | 10.84016 | 0.532429 | 0.63469 | 0.83888 | 0.401537 | NA |
| RIMKLA | 1.970021 | 0.196592 | 1.679275 | 0.11707 | 0.906805 | NA |
| PDDC1 | 2226.119 | -0.42148 | 0.169098 | -2.49251 | 0.012684 | 0.055427 |
| C10orf85 | 8.789895 | 0.890326 | 0.767104 | 1.160634 | 0.245791 | NA |
| | 8.535338 | -0.03179 | 0.705405 | -0.04507 | 0.964055 | NA |
| FZD8 | 101.1853 | -0.72494 | 0.268624 | -2.69872 | 0.006961 | 0.037018 |
| GJD4 | 0.862836 | 1.736629 | 2.194323 | 0.791419 | 0.4287 | NA |
| KCNA2 | 116.3617 | -0.32193 | 0.316916 | -1.01582 | 0.309715 | 0.476976 |
| LRRN4CL | 482.6181 | -0.77614 | 0.217895 | -3.56199 | 3.68E-04 | 0.005587 |
| CBX3P1 | 0.682693 | 2.623213 | 2.518338 | 1.041645 | 0.297576 | NA |
| | 0.162376 | 0.424913 | 4.079006 | 0.104171 | 0.917034 | NA |
| SLC25A22 | 564.377 | -0.0874 | 0.181077 | -0.48268 | 0.629324 | 0.751777 |
| NHLH2 | 0.604258 | 1.087325 | 2.707696 | 0.401568 | 0.688002 | NA |
| PIDD | 408.1951 | -0.28261 | 0.172983 | -1.63372 | 0.102318 | 0.226383 |
| RPLP2 | 14492.18 | -0.09442 | 0.155278 | -0.60804 | 0.543161 | 0.685164 |
| JUN | 3930.54 | 0.239225 | 0.337388 | 0.70905 | 0.478294 | 0.633185 |
| CSTF2T | 853.7687 | -0.17275 | 0.184652 | -0.93556 | 0.349502 | 0.517595 |
| PGBD5 | 19.77403 | -0.47637 | 0.586282 | -0.81253 | 0.41649 | NA |
| GBA | 4090.418 | -0.37789 | 0.133587 | -2.82882 | 0.004672 | 0.028187 |
| CASC2 | 32.24267 | 0.269708 | 0.399964 | 0.674332 | 0.5001 | NA |
| PNPLA2 | 1874.341 | -0.11979 | 0.163574 | -0.73236 | 0.463952 | 0.621415 |
| AGTRAP | 1272.309 | -0.29936 | 0.144644 | -2.06965 | 0.038485 | 0.117153 |
| EFCAB4A | 6.088278 | 1.203205 | 0.882252 | 1.363789 | 0.172634 | NA |
| CD151 | 9580.304 | -0.1663 | 0.138798 | -1.19815 | 0.23086 | 0.393112 |
| POLR2L | 6038.2 | -0.05383 | 0.147421 | -0.36513 | 0.715012 | 0.81835 |

| | | | | | | |
|---|---|---|---|---|---|---|
| FAM87B | 1.129667 | -0.3654 | 1.881429 | -0.19421 | 0.846008 | NA |
| CDKN2AIPN | 0.296509 | -0.51709 | 3.982155 | -0.12985 | 0.896684 | NA |
| | 2.05912 | -0.02414 | 1.404518 | -0.01719 | 0.986288 | NA |
| MYOZ1 | 12.19396 | 0.230668 | 0.59308 | 0.388932 | 0.697326 | NA |
| KCNJ10 | 7.697011 | -0.22693 | 0.761108 | -0.29816 | 0.765584 | NA |
| CHID1 | 2247.301 | -0.43931 | 0.131909 | -3.33038 | 8.67E-04 | 0.009476 |
| ZNF518A | 188.3284 | 0.884132 | 0.443645 | 1.992885 | 0.046274 | 0.132188 |
| CCDC23 | 261.3956 | 0.261181 | 0.174314 | 1.498332 | 0.134047 | 0.272017 |
| ZBTB41 | 150.9904 | 0.87639 | 0.342558 | 2.558366 | 0.010517 | 0.048662 |
| BET1L | 950.7381 | -0.29316 | 0.145498 | -2.0149 | 0.043915 | 0.127733 |
| RPS27 | 11833.99 | 0.296316 | 0.161269 | 1.837401 | 0.066151 | 0.16784 |
| RIC8A | 3206.03 | -0.165 | 0.145205 | -1.13636 | 0.255808 | 0.41997 |
| DMAP1 | 858.1471 | -0.223 | 0.168476 | -1.32361 | 0.185632 | 0.339451 |
| BOLA1 | 295.56 | -0.14038 | 0.167086 | -0.84016 | 0.400817 | 0.565475 |
| PDE4DIP | 2893.702 | 0.289899 | 0.131421 | 2.205877 | 0.027393 | 0.093396 |
| | 0.163957 | 0.424913 | 4.079006 | 0.104171 | 0.917034 | NA |
| SH2D4B | 0.303394 | -2.27006 | 4.032451 | -0.56295 | 0.57347 | NA |
| RPS3AP5 | 13.10821 | -0.07038 | 0.602785 | -0.11676 | 0.907047 | NA |
| LINC00843 | 1.440017 | 0.403371 | 1.70198 | 0.237001 | 0.812656 | NA |
| TUBAL3 | 0.399595 | -1.31595 | 3.579466 | -0.36764 | 0.713143 | NA |
| CTNNBIP1 | 744.1739 | -0.14803 | 0.138292 | -1.0704 | 0.284437 | 0.450947 |
| | 11.58567 | 0.145905 | 0.595292 | 0.245099 | 0.80638 | NA |
| | 2.856359 | 1.072283 | 1.222022 | 0.877466 | 0.380233 | NA |
| RIIAD1 | 2.68605 | 0.174248 | 1.321602 | 0.131846 | 0.895106 | NA |
| TMEM52 | 3.808652 | -0.18334 | 1.007611 | -0.18196 | 0.855615 | NA |
| HYI | 210.5502 | -0.29382 | 0.22379 | -1.31295 | 0.189201 | 0.343409 |
| RCC2 | 2892.957 | -0.51299 | 0.15392 | -3.33286 | 8.60E-04 | 0.009421 |
| IGSF22 | 10.78072 | 0.096615 | 0.715059 | 0.135115 | 0.892521 | NA |
| DPM3 | 650.094 | -0.1959 | 0.236759 | -0.82742 | 0.408001 | 0.571572 |
| SPTY2D1 | 465.6971 | 0.426145 | 0.207287 | 2.055818 | 0.0398 | 0.119967 |
| C10orf67 | 5.966277 | -0.01921 | 0.820153 | -0.02342 | 0.981317 | NA |
| FUCA1 | 693.0016 | -0.15629 | 0.157595 | -0.99174 | 0.321324 | 0.489482 |
| HNRNPCL1 | 0.518736 | 2.237423 | 2.7713 | 0.807355 | 0.419462 | NA |
| LDLRAD3 | 1765.038 | -0.54378 | 0.169037 | -3.21694 | 0.001296 | 0.012249 |
| CTGLF12P | 1.86508 | -0.62073 | 1.426117 | -0.43526 | 0.663374 | NA |
| C1orf101 | 11.73064 | 0.92102 | 0.670196 | 1.374255 | 0.169363 | NA |
| VWA1 | 1292.124 | -0.205 | 0.325922 | -0.62898 | 0.529365 | 0.674507 |
| HNRNPCP5 | 0.096202 | 0.408546 | 4.079639 | 0.100143 | 0.920231 | NA |
| FJX1 | 320.0984 | -0.13648 | 0.17625 | -0.77434 | 0.438732 | 0.598798 |
| ZBTB18 | 319.7876 | 0.467316 | 0.254664 | 1.835026 | 0.066502 | 0.168462 |
| DNHD1 | 77.9942 | 0.252756 | 0.290746 | 0.869337 | 0.384663 | 0.550705 |
| HTR1D | 0.163957 | 0.424913 | 4.079006 | 0.104171 | 0.917034 | NA |
| NBPF23 | 1.330186 | 0.778826 | 1.870599 | 0.416351 | 0.677153 | NA |
| | 65.59283 | 0.243847 | 0.300515 | 0.811431 | 0.417118 | 0.580188 |
| ATOH7 | 1.215189 | -0.88793 | 1.977332 | -0.44905 | 0.653393 | NA |
| MRGPRX4 | 0.096202 | 0.408546 | 4.079639 | 0.100143 | 0.920231 | NA |
| C1orf200 | 0.163957 | 0.424913 | 4.079006 | 0.104171 | 0.917034 | NA |

| | | | | | | |
|---|---|---|---|---|---|---|
| CITED4 | 11.8032 | -0.34071 | 0.602673 | -0.56534 | 0.571845 | NA |
| C1orf194 | 0.604258 | 1.087325 | 2.707696 | 0.401568 | 0.688002 | NA |
| ZNF648 | 0.192403 | 0.975392 | 4.060716 | 0.240202 | 0.810174 | NA |
| PSTK | 93.91758 | 0.300453 | 0.238674 | 1.258846 | 0.208086 | 0.365406 |
| C10orf91 | 1.420452 | -0.26596 | 1.837831 | -0.14471 | 0.884936 | NA |
| TRNAU1AP | 467.0033 | 0.124194 | 0.178421 | 0.696074 | 0.486383 | 0.639483 |
| ACTA2-AS1 | 0.899187 | 0.806933 | 2.127165 | 0.379346 | 0.704431 | NA |
| RCC1 | 1067.799 | -0.17181 | 0.159273 | -1.07874 | 0.280706 | 0.4468 |
| TPT1P10 | 0.422534 | 1.924583 | 3.001758 | 0.641152 | 0.521424 | NA |
| NACAP2 | 0.163957 | 0.424913 | 4.079006 | 0.104171 | 0.917034 | NA |
| PLD5 | 0.35636 | 1.709252 | 3.651833 | 0.468053 | 0.639746 | NA |
| HARBI1 | 160.5925 | 0.072827 | 0.22259 | 0.327178 | 0.743533 | 0.838446 |
| FAM73A | 396.9626 | 0.786635 | 0.261203 | 3.011592 | 0.002599 | 0.019343 |
| PRR26 | 3.567128 | -0.72889 | 1.300039 | -0.56067 | 0.575025 | NA |
| SRP9P1 | 2.631997 | 2.105551 | 1.384532 | 1.520767 | 0.128318 | NA |
| SKIDA1 | 9.597225 | 0.377363 | 0.705042 | 0.535234 | 0.592488 | NA |
| PCGF5 | 1184.851 | 0.811153 | 0.197146 | 4.114474 | 3.88E-05 | 0.001429 |
| YOD1 | 264.7967 | 1.068193 | 0.206406 | 5.175197 | 2.28E-07 | 5.51E-05 |
| CHRM4 | 3.016252 | -0.28968 | 1.202452 | -0.24091 | 0.809626 | NA |
| GPR157 | 152.229 | -0.62637 | 0.312911 | -2.00175 | 0.045312 | 0.130349 |
| PIPSL | 12.79081 | -0.02292 | 0.628704 | -0.03646 | 0.970919 | NA |
| PPA1 | 1456.737 | 0.428758 | 0.164751 | 2.602451 | 0.009256 | 0.044768 |
| PDIA3P1 | 49.03917 | -0.05107 | 0.332476 | -0.15359 | 0.877933 | 0.926876 |
| GREM2 | 1.564315 | 1.893687 | 1.744001 | 1.085829 | 0.277555 | NA |
| C11orf42 | 0.422534 | 1.924583 | 3.001758 | 0.641152 | 0.521424 | NA |
| OR56B4 | 0.544554 | -2.65036 | 3.220086 | -0.82307 | 0.410468 | NA |
| FCRL6 | 0.382031 | -1.26484 | 3.151989 | -0.40128 | 0.688212 | NA |
| ANKRD34A | 120.1166 | -0.07326 | 0.296923 | -0.24673 | 0.805116 | 0.883508 |
| SDCCAG3P | 0.392857 | -1.546 | 4.075469 | -0.37934 | 0.704433 | NA |
| DHTKD1 | 693.7992 | -0.20774 | 0.146714 | -1.41596 | 0.156788 | 0.303069 |
| HIST3H2A | 397.0864 | -0.08562 | 0.241768 | -0.35413 | 0.723243 | 0.824269 |
| | 3.676062 | -0.07103 | 1.024706 | -0.06932 | 0.944734 | NA |
| FRAT2 | 419.8273 | -0.19284 | 0.205719 | -0.9374 | 0.348554 | 0.516513 |
| OR9G3P | 0.260158 | 1.240347 | 4.047124 | 0.306476 | 0.759242 | NA |
| ZNF678 | 28.1078 | 0.834381 | 0.594537 | 1.403412 | 0.160494 | NA |
| PHLDA2 | 3011.22 | -0.09805 | 0.174309 | -0.56252 | 0.573765 | 0.709406 |
| AMIGO1 | 221.4 | -0.04351 | 0.244921 | -0.17767 | 0.858983 | 0.916352 |
| GPR3 | 38.55963 | -0.15861 | 0.417502 | -0.37991 | 0.704013 | NA |
| CELF2-AS1 | 13.62277 | 0.448232 | 0.602862 | 0.743507 | 0.457175 | NA |
| LSM10 | 1506.951 | 0.053176 | 0.158728 | 0.33501 | 0.737617 | 0.833837 |
| SLC35C1 | 1793.375 | -0.41796 | 0.131955 | -3.16747 | 0.001538 | 0.013769 |
| IBA57 | 216.6195 | -0.1333 | 0.224306 | -0.59426 | 0.552341 | 0.693167 |
| ADO | 1202.252 | -0.02856 | 0.137427 | -0.20779 | 0.835396 | 0.901477 |
| SNRPE | 662.9335 | 0.508701 | 0.14698 | 3.461027 | 5.38E-04 | 0.007049 |
| RTKN2 | 78.70715 | 0.466254 | 0.295663 | 1.576978 | 0.114801 | 0.244338 |
| CHST15 | 1.075752 | 0.450053 | 2.098769 | 0.214437 | 0.830206 | NA |
| PTCHD3 | 2.071461 | -2.08094 | 1.736734 | -1.19819 | 0.230842 | NA |

| | | | | | | |
|---|---|---|---|---|---|---|
| | 11.31238 | 0.230073 | 0.721412 | 0.31892 | 0.749787 | NA |
| FAM89A | 2567.933 | 0.185433 | 0.163369 | 1.135052 | 0.256353 | 0.420516 |
| TDRKH | 110.2814 | -0.12075 | 0.291703 | -0.41394 | 0.678916 | 0.790468 |
| MRPS16 | 2574.692 | -0.29011 | 0.141505 | -2.05017 | 0.040347 | 0.121032 |
| MOB2 | 762.6699 | -0.04394 | 0.188447 | -0.23317 | 0.815633 | 0.889858 |
| KCNA4 | 0.807523 | 1.550758 | 2.909197 | 0.533053 | 0.593997 | NA |
| NLRP10 | 0.833012 | 0.681553 | 2.203157 | 0.309353 | 0.757053 | NA |
| B4GALNT4 | 639.8767 | -0.39657 | 0.168447 | -2.35426 | 0.018559 | 0.071832 |
| CCNYL2 | 2.631917 | -0.18165 | 1.184684 | -0.15333 | 0.878138 | NA |
| PAQR7 | 340.2849 | -0.44358 | 0.164019 | -2.70443 | 0.006842 | 0.036554 |
| GRID1 | 0.163957 | 0.424913 | 4.079006 | 0.104171 | 0.917034 | NA |
| C1orf116 | 12.70671 | 0.340412 | 0.868005 | 0.392178 | 0.694927 | NA |
| ACBD3 | 1394.373 | 0.41351 | 0.206683 | 2.000703 | 0.045424 | 0.130579 |
| LCK | 0.327913 | 1.543154 | 4.021063 | 0.383768 | 0.701151 | NA |
| | 13.86216 | -0.24804 | 0.588272 | -0.42163 | 0.673292 | NA |
| RGS7 | 0.977621 | 1.971661 | 2.116061 | 0.93176 | 0.351461 | NA |
| AP2A2 | 3051.482 | -0.38237 | 0.175204 | -2.18242 | 0.029078 | 0.096999 |
| CAMK1D | 9.720807 | 0.462975 | 0.768364 | 0.602546 | 0.546811 | NA |
| FAM133CP | 0.192403 | 0.975392 | 4.060716 | 0.240202 | 0.810174 | NA |
| FAM43B | 9.308978 | 0.86244 | 0.913304 | 0.944307 | 0.345013 | NA |
| PTGDR2 | 2.743928 | -0.75182 | 1.236464 | -0.60804 | 0.543161 | NA |
| RABIF | 1001.154 | -0.005 | 0.139289 | -0.03587 | 0.971388 | 0.984257 |
| FANCF | 332.2775 | -0.00973 | 0.175993 | -0.05528 | 0.955916 | 0.975764 |
| CTNNA3 | 0.843488 | 2.923514 | 2.365851 | 1.235713 | 0.216565 | NA |
| | 5046.05 | 0.385652 | 0.191483 | 2.014031 | 0.044006 | 0.127866 |
| | 3.29664 | 0.956323 | 1.130499 | 0.84593 | 0.397592 | NA |
| EPHA10 | 0.096202 | 0.408546 | 4.079639 | 0.100143 | 0.920231 | NA |
| C10orf107 | 3.50681 | 0.817067 | 1.066851 | 0.765868 | 0.443755 | NA |
| OVCH2 | 0.460466 | 0.568655 | 3.014642 | 0.188631 | 0.850382 | NA |
| FHL3 | 619.2228 | -0.03435 | 0.149762 | -0.22937 | 0.818578 | 0.891361 |
| SF3A3 | 2228.946 | 0.002996 | 0.129797 | 0.02308 | 0.981586 | 0.990189 |
| FAM46C | 6.076508 | -0.32626 | 0.809947 | -0.40282 | 0.687079 | NA |
| UTP11L | 702.4078 | 0.467931 | 0.179679 | 2.604267 | 0.009207 | 0.044608 |
| | 0.574434 | -0.17781 | 2.756096 | -0.06452 | 0.948559 | NA |
| HMGN3P1 | 0.314073 | -0.57308 | 3.898934 | -0.14698 | 0.883146 | NA |
| HIST2H3D | 0.162376 | 0.424913 | 4.079006 | 0.104171 | 0.917034 | NA |
| SFXN4 | 425.4375 | -0.04273 | 0.16235 | -0.26322 | 0.792379 | 0.874234 |
| FAM167B | 2429.772 | 0.072473 | 0.299992 | 0.241584 | 0.809103 | 0.885851 |
| ZNF438 | 58.45835 | -0.21497 | 0.317819 | -0.67638 | 0.498799 | 0.649348 |
| PTPN20B | 0.151697 | -1.546 | 4.075469 | -0.37934 | 0.704433 | NA |
| BMP8A | 0.508056 | 0.731032 | 2.861912 | 0.255435 | 0.798387 | NA |
| TMEM50A | 3895.284 | 0.151417 | 0.139047 | 1.088958 | 0.276173 | 0.441779 |
| | 0.096202 | 0.408546 | 4.079639 | 0.100143 | 0.920231 | NA |
| SLC35F3 | 0.096202 | 0.408546 | 4.079639 | 0.100143 | 0.920231 | NA |
| OLFML1 | 1.869078 | 0.043799 | 1.547192 | 0.028309 | 0.977416 | NA |
| LIN9 | 115.8869 | 1.026679 | 0.233773 | 4.391771 | 1.12E-05 | 6.60E-04 |
| ANKRD45 | 7.449626 | -1.0654 | 0.781994 | -1.36241 | 0.173068 | NA |

| | | | | | | |
|---|---|---|---|---|---|---|
| KIRREL | 4277.288 | -0.43387 | 0.149255 | -2.9069 | 0.00365 | 0.024081 |
| IQGAP3 | 929.0558 | -0.95646 | 0.23508 | -4.06867 | 4.73E-05 | 0.001588 |
| LRRC55 | 0.096202 | 0.408546 | 4.079639 | 0.100143 | 0.920231 | NA |
| DUSP5P1 | 1.185162 | -1.08022 | 1.836234 | -0.58828 | 0.556345 | NA |
| ST6GALNA( | 1234.103 | -0.25863 | 0.227324 | -1.13772 | 0.255236 | 0.419495 |
| PTP4A2 | 2810.736 | 0.200095 | 0.145515 | 1.375081 | 0.169106 | 0.319451 |
| DENND5A | 3111.15 | -0.25651 | 0.159302 | -1.61019 | 0.107357 | 0.233913 |
| | 0.651288 | 1.289307 | 2.633717 | 0.489539 | 0.62446 | NA |
| CNTN2 | 0.816825 | 1.673341 | 2.21346 | 0.755984 | 0.449659 | NA |
| FAM132A | 3.461362 | 0.492956 | 1.117179 | 0.441251 | 0.659031 | NA |
| HIST2H2AC | 8.604045 | -1.1528 | 0.690431 | -1.66968 | 0.094983 | NA |
| HIST2H2AB | 0.394291 | 0.298745 | 3.141711 | 0.09509 | 0.924243 | NA |
| TSSC4 | 1047.528 | -0.15182 | 0.154872 | -0.98026 | 0.326957 | 0.495092 |
| TACSTD2 | 1.230224 | 1.989951 | 2.317289 | 0.858741 | 0.390483 | NA |
| MROH7 | 0.192403 | 0.975392 | 4.060716 | 0.240202 | 0.810174 | NA |
| MRGPRE | 0.134133 | -1.48909 | 4.077584 | -0.36519 | 0.71497 | NA |
| PKP3 | 17.41608 | -0.65473 | 0.496007 | -1.32 | 0.186835 | NA |
| CSF1 | 368.9962 | -0.37373 | 0.24719 | -1.51191 | 0.130558 | 0.26704 |
| A3GALT2 | 0.813811 | 0.928591 | 2.591569 | 0.358312 | 0.72011 | NA |
| CEND1 | 6.063121 | 0.803699 | 0.853329 | 0.94184 | 0.346274 | NA |
| DUSP8 | 186.8038 | -0.11462 | 0.236252 | -0.48518 | 0.627551 | 0.750113 |
| PDE4B | 365.0699 | -0.73491 | 0.274062 | -2.68154 | 0.007328 | 0.038391 |
| FAM19A3 | 4.062689 | 0.057964 | 0.996734 | 0.058154 | 0.953626 | NA |
| OR7E14P | 0.296509 | -0.51709 | 3.982155 | -0.12985 | 0.896684 | NA |
| ZBTB40 | 944.8933 | -0.14554 | 0.183453 | -0.79331 | 0.427596 | 0.589436 |
| HIST2H2BE | 153.1103 | -0.17828 | 0.329133 | -0.54167 | 0.588047 | 0.720622 |
| RNLS | 160.8149 | 0.025257 | 0.213705 | 0.118185 | 0.905921 | 0.945645 |
| | 0.392857 | -1.546 | 4.075469 | -0.37934 | 0.704433 | NA |
| CLCNKB | 2.665739 | 0.587723 | 1.236007 | 0.475502 | 0.634429 | NA |
| NUTM2A | 5.340717 | -0.45351 | 0.852268 | -0.53212 | 0.594641 | NA |
| MUC6 | 0.315653 | -0.57305 | 3.892639 | -0.14721 | 0.882963 | NA |
| AP3M1 | 2061.112 | -0.01979 | 0.175597 | -0.1127 | 0.91027 | 0.947811 |
| | 0.52699 | -2.57878 | 3.248478 | -0.79384 | 0.427288 | NA |
| INTS5 | 968.7635 | -0.45879 | 0.135224 | -3.39285 | 6.92E-04 | 0.00825 |
| MANEAL | 22.98747 | -0.40485 | 0.433665 | -0.93355 | 0.350538 | NA |
| ANO9 | 12.20615 | -0.43437 | 0.630867 | -0.68853 | 0.491117 | NA |
| FAF1 | 1131.558 | -0.23244 | 0.134899 | -1.72308 | 0.084873 | 0.198652 |
| MIXL1 | 0.87074 | 0.675988 | 2.012127 | 0.335957 | 0.736903 | NA |
| SIGIRR | 852.5663 | -0.21365 | 0.14548 | -1.46857 | 0.14195 | 0.282921 |
| IFITM2 | 2530.449 | -0.3623 | 0.17406 | -2.08146 | 0.037392 | 0.114841 |
| PGBD2 | 74.90062 | 0.003941 | 0.276065 | 0.014276 | 0.98861 | 0.994313 |
| GAPDHP24 | 0.134133 | -1.48909 | 4.077584 | -0.36519 | 0.71497 | NA |
| PRMT3 | 443.895 | 0.542971 | 0.204931 | 2.649527 | 0.00806 | 0.040801 |
| CDNF | 12.3171 | 0.96735 | 0.634674 | 1.524168 | 0.127467 | NA |
| ZBTB37 | 79.00604 | 0.230326 | 0.297029 | 0.775432 | 0.438085 | 0.598191 |
| SMYD3 | 783.6756 | -0.03482 | 0.147276 | -0.23641 | 0.813115 | 0.888676 |
| IFNLR1 | 76.61821 | 0.230643 | 0.434997 | 0.530217 | 0.595961 | 0.726065 |

| | | | | | | |
|---|---|---|---|---|---|---|
| TMEM179I | 687.7797 | -0.13624 | 0.164671 | -0.82736 | 0.408032 | 0.571572 |
| ROR1 | 238.3469 | 0.163278 | 0.199661 | 0.817779 | 0.413484 | 0.576688 |
| | 0.640406 | 1.11402 | 2.710977 | 0.410929 | 0.681124 | NA |
| MUC1 | 22.13705 | -0.5495 | 0.458429 | -1.19866 | 0.230662 | NA |
| IRF7 | 817.3531 | -0.92142 | 0.292655 | -3.14849 | 0.001641 | 0.014318 |
| FAM131C | 0.162376 | 0.424913 | 4.079006 | 0.104171 | 0.917034 | NA |
| C11orf35 | 43.58819 | -0.42189 | 0.333587 | -1.26472 | 0.205971 | NA |
| C1orf227 | 1.225666 | 0.672437 | 1.847219 | 0.364026 | 0.715838 | NA |
| PRKG1 | 333.2455 | 0.612414 | 0.259148 | 2.363182 | 0.018119 | 0.070588 |
| PSMD13 | 3869.872 | -0.03407 | 0.147022 | -0.23176 | 0.816726 | 0.890203 |
| PBX1 | 274.2478 | -0.34357 | 0.224645 | -1.5294 | 0.126164 | 0.261259 |
| POU3F1 | 3.740693 | -0.0298 | 1.044676 | -0.02853 | 0.977242 | NA |
| ZBTB3 | 107.8589 | -0.42854 | 0.23813 | -1.79962 | 0.071921 | 0.178 |
| ADARB2 | 21.80817 | 0.356049 | 0.584753 | 0.608887 | 0.542599 | NA |
| NRG3 | 141.0937 | -0.26293 | 0.274273 | -0.95866 | 0.33773 | 0.506619 |
| IFIT1 | 1058.112 | -0.02347 | 0.562417 | -0.04173 | 0.966714 | 0.981618 |
| | 11.83765 | -0.37686 | 0.598478 | -0.6297 | 0.528888 | NA |
| DNAH14 | 52.21041 | 1.107787 | 0.332893 | 3.327758 | 8.75E-04 | 0.009546 |
| C1orf110 | 0.134133 | -1.48909 | 4.077584 | -0.36519 | 0.71497 | NA |
| THNSL1 | 151.0785 | -0.08724 | 0.248254 | -0.35143 | 0.725263 | 0.825454 |
| IFITM1 | 549.1323 | -1.34696 | 0.606668 | -2.22026 | 0.026401 | 0.091242 |
| LINC00839 | 588.3663 | -0.04294 | 0.144571 | -0.29699 | 0.766476 | 0.854885 |
| CALHM1 | 1.169179 | -1.04993 | 1.839034 | -0.57091 | 0.568059 | NA |
| RNPC3 | 108.0238 | 0.524757 | 0.276942 | 1.894826 | 0.058115 | 0.153614 |
| LEMD1 | 0.163957 | 0.424913 | 4.079006 | 0.104171 | 0.917034 | NA |
| MATN1-AS | 5.926132 | -0.21669 | 0.811669 | -0.26696 | 0.789497 | NA |
| AIDA | 500.8969 | 0.255739 | 0.180648 | 1.415677 | 0.15687 | 0.303119 |
| NBPF6 | 0.096202 | 0.408546 | 4.079639 | 0.100143 | 0.920231 | NA |
| AGBL4 | 0.825079 | -2.06702 | 2.533819 | -0.81577 | 0.41463 | NA |
| CYP2R1 | 207.972 | 0.447149 | 0.20866 | 2.142956 | 0.032117 | 0.103971 |
| TEX38 | 0.424115 | 1.932531 | 3.45462 | 0.559405 | 0.575885 | NA |
| POLR3C | 759.1248 | -0.05138 | 0.141136 | -0.36404 | 0.715826 | 0.81867 |
| UBL4B | 0.162376 | 0.424913 | 4.079006 | 0.104171 | 0.917034 | NA |
| EDARADD | 26.03697 | -0.42288 | 0.432804 | -0.97707 | 0.328536 | NA |
| 1-Mar | 13.68004 | -0.19485 | 0.557502 | -0.34951 | 0.726707 | NA |
| LCE5A | 0.247898 | -0.5901 | 4.074633 | -0.14482 | 0.884851 | NA |
| NBPF12 | 17.56812 | 0.283009 | 0.505443 | 0.559923 | 0.575532 | NA |
| TOR3A | 675.749 | -0.26559 | 0.142384 | -1.8653 | 0.06214 | 0.161303 |
| MST1P2 | 5.945589 | 0.912862 | 0.85287 | 1.070341 | 0.284466 | NA |
| NUDT17 | 93.74844 | -0.35956 | 0.284318 | -1.26464 | 0.206002 | 0.363038 |
| CYP4X1 | 1.370827 | -0.71211 | 1.970974 | -0.3613 | 0.717877 | NA |
| CCDC30 | 24.44397 | 0.869945 | 0.455516 | 1.909799 | 0.056159 | NA |
| TMEM222 | 1539.269 | -0.1805 | 0.133301 | -1.35406 | 0.175717 | 0.327792 |
| OR9Q1 | 0.096202 | 0.408546 | 4.079639 | 0.100143 | 0.920231 | NA |
| CLCNKA | 1.532066 | -1.16575 | 1.666556 | -0.6995 | 0.484243 | NA |
| ARHGAP30 | 1.853869 | 0.497017 | 1.627184 | 0.305446 | 0.760026 | NA |
| CROCCP5 | 0.134133 | -1.48909 | 4.077584 | -0.36519 | 0.71497 | NA |

| | | | | | | |
|---|---|---|---|---|---|---|
| FOXD2 | 64.6832 | -0.54655 | 0.291334 | -1.87602 | 0.060653 | 0.158483 |
| HPDL | 385.9434 | -0.89031 | 0.308036 | -2.89026 | 0.003849 | 0.024827 |
| PRG2 | 0.134133 | -1.48909 | 4.077584 | -0.36519 | 0.71497 | NA |
| ZFP91 | 2790.032 | 0.161084 | 0.219276 | 0.734616 | 0.462574 | 0.620551 |
| CCDC73 | 0.881566 | 1.109202 | 2.554987 | 0.434132 | 0.664192 | NA |
| MST1L | 2.950775 | 1.963936 | 1.371623 | 1.431834 | 0.152191 | NA |
| FOXE3 | 0.640755 | -1.64826 | 2.754778 | -0.59833 | 0.549621 | NA |
| TNFRSF4 | 0.096202 | 0.408546 | 4.079639 | 0.100143 | 0.920231 | NA |
| PDZD7 | 110.8239 | -0.09654 | 0.256237 | -0.37675 | 0.706362 | 0.81182 |
| TNFRSF18 | 0.708364 | 0.219301 | 2.144998 | 0.102238 | 0.918568 | NA |
| RTN4RL2 | 134.8946 | 0.068389 | 0.265854 | 0.257243 | 0.796991 | 0.87712 |
| FAM183A | 2.422003 | -1.52324 | 1.697968 | -0.89709 | 0.369668 | NA |
| RHD | 5.69752 | -0.53409 | 0.99828 | -0.53501 | 0.592646 | NA |
| ESPN | 16.8923 | -0.10225 | 0.539314 | -0.1896 | 0.849622 | NA |
| PNLIPRP1 | 0.518736 | 2.237423 | 2.7713 | 0.807355 | 0.419462 | NA |
| TMEM216 | 147.6831 | 0.174911 | 0.245112 | 0.713598 | 0.475476 | 0.630722 |
| TEAD1 | 4283.258 | 0.261935 | 0.237054 | 1.104958 | 0.269178 | 0.433868 |
| AKR1C1 | 94.22899 | 0.314905 | 0.319902 | 0.98438 | 0.324929 | 0.492847 |
| FOXD3 | 1748.983 | 0.237657 | 0.243464 | 0.976148 | 0.328991 | 0.49728 |
| MRPS21 | 2297.232 | 0.149919 | 0.151092 | 0.992239 | 0.321081 | 0.489256 |
| RNF220 | 1693.536 | -0.14513 | 0.144364 | -1.00533 | 0.314738 | 0.482131 |
| KIAA1598 | 1131.43 | 0.331743 | 0.548338 | 0.604997 | 0.545181 | 0.686764 |
| LUZP2 | 1.500895 | 3.745333 | 2.013071 | 1.860508 | 0.062814 | NA |
| C11orf96 | 1995.081 | 0.078108 | 0.334273 | 0.233665 | 0.815245 | 0.889858 |
| HSPA14 | 736.4108 | 0.546974 | 0.144621 | 3.782114 | 1.56E-04 | 0.003343 |
| CYP26C1 | 5.061295 | -0.78307 | 0.971042 | -0.80642 | 0.420001 | NA |
| TLR5 | 2.167096 | -1.3286 | 1.41398 | -0.93962 | 0.347415 | NA |
| PLEKHN1 | 20.30243 | 0.106034 | 0.49641 | 0.213602 | 0.830857 | NA |
| ISG15 | 2144.524 | -0.9995 | 0.345755 | -2.89077 | 0.003843 | 0.024801 |
| SAMD11 | 15.17558 | 1.183493 | 0.608231 | 1.945796 | 0.051679 | NA |
| C1orf170 | 5.524773 | 0.097068 | 0.861316 | 0.112698 | 0.91027 | NA |
| SLC18A3 | 0.558248 | 0.982076 | 2.804738 | 0.350149 | 0.726227 | NA |
| PEAR1 | 0.315653 | -0.57305 | 3.892639 | -0.14721 | 0.882963 | NA |
| ZFP69B | 20.05969 | -0.0871 | 0.545082 | -0.15979 | 0.873044 | NA |
| ZFP69 | 86.44575 | 0.800236 | 0.305005 | 2.623679 | 0.008699 | 0.042987 |
| DMBT1 | 0.392857 | -1.546 | 4.075469 | -0.37934 | 0.704433 | NA |
| LDLRAD2 | 4.841472 | 0.16512 | 0.982704 | 0.168026 | 0.866563 | NA |
| HS6ST1P1 | 0.260158 | 1.240347 | 4.047124 | 0.306476 | 0.759242 | NA |
| KLHL17 | 416.4864 | -0.08484 | 0.159738 | -0.53113 | 0.595325 | 0.725499 |
| PLA2G2C | 0.162376 | 0.424913 | 4.079006 | 0.104171 | 0.917034 | NA |
| C1orf204 | 5.288889 | -0.59816 | 0.880006 | -0.67972 | 0.49668 | NA |
| S100A3 | 24.24942 | 0.259546 | 0.44981 | 0.577011 | 0.563932 | NA |
| RAB42 | 25.52126 | -0.99558 | 0.460593 | -2.16152 | 0.030655 | NA |
| GPR89B | 31.95485 | -0.13857 | 0.381041 | -0.36367 | 0.716106 | NA |
| OR2AG2 | 0.735027 | 1.3952 | 2.594798 | 0.537691 | 0.59079 | NA |
| AGRN | 6985.642 | -1.08555 | 0.133624 | -8.12386 | 4.52E-16 | 3.30E-12 |
| NUTM2B | 0.802985 | 0.54709 | 2.208262 | 0.247747 | 0.80433 | NA |

| | | | | | | |
|---|---|---|---|---|---|---|
| HNRNPU-A | 12.57288 | 1.443954 | 0.685539 | 2.106304 | 0.035178 | NA |
| NCR3LG1 | 93.03316 | 0.098165 | 0.291373 | 0.336906 | 0.736188 | 0.833095 |
| AGAP4 | 5.466016 | 0.908925 | 0.927385 | 0.980095 | 0.327039 | NA |
| HES4 | 287.3743 | 0.01604 | 0.281952 | 0.056891 | 0.954632 | 0.974998 |
| ZNF669 | 108.1767 | -0.45486 | 0.236339 | -1.9246 | 0.054279 | 0.146612 |
| ENO4 | 17.97889 | 0.131987 | 0.527394 | 0.250262 | 0.802384 | NA |
| JAKMIP3 | 8.180557 | 1.028849 | 0.820094 | 1.254551 | 0.209642 | NA |
| TCTEX1D4 | 5.478937 | -0.63941 | 0.902463 | -0.70852 | 0.478624 | NA |
| SELL | 4.898297 | -0.2348 | 0.989586 | -0.23727 | 0.812444 | NA |
| ACTBP11 | 0.296509 | -0.51709 | 3.982155 | -0.12985 | 0.896684 | NA |
| INSC | 286.5942 | 0.413482 | 0.292594 | 1.413156 | 0.15761 | 0.304013 |
| SRSF10 | 1842.919 | 0.568442 | 0.157391 | 3.611643 | 3.04E-04 | 0.004996 |
| FAM72B | 35.00636 | 0.326363 | 0.412805 | 0.790599 | 0.429178 | NA |
| ASAH2 | 3.563849 | 0.576979 | 1.126554 | 0.512163 | 0.608537 | NA |
| NANOS1 | 72.67502 | -0.1663 | 0.430067 | -0.38669 | 0.698989 | 0.806395 |
| DPYD | 599.6254 | 0.633531 | 0.205754 | 3.079067 | 0.002076 | 0.016597 |
| S100A16 | 534.5994 | -0.07006 | 0.250567 | -0.2796 | 0.779788 | 0.864932 |
| CC2D2B | 0.654246 | 2.541085 | 3.009764 | 0.84428 | 0.398513 | NA |
| RHCE | 3.8416 | 0.051907 | 0.990999 | 0.052379 | 0.958227 | NA |
| UROS | 2439.476 | -0.17944 | 0.132252 | -1.35682 | 0.174837 | 0.326835 |
| RBM34 | 7.712775 | 0.795235 | 0.838194 | 0.948747 | 0.342749 | NA |
| BCL2L15 | 1.637521 | 0.608302 | 1.617589 | 0.376055 | 0.706876 | NA |
| PRELP | 3467.309 | -0.71236 | 0.226292 | -3.14798 | 0.001644 | 0.014318 |
| MTF1 | 199.2363 | 0.046386 | 0.208307 | 0.222682 | 0.823783 | 0.893663 |
| TMCO2 | 0.096202 | 0.408546 | 4.079639 | 0.100143 | 0.920231 | NA |
| TMEM201 | 1643.491 | -0.49819 | 0.165448 | -3.01114 | 0.002603 | 0.019343 |
| HMX2 | 0.096202 | 0.408546 | 4.079639 | 0.100143 | 0.920231 | NA |
| FAM78B | 203.6792 | -0.28466 | 0.254462 | -1.11866 | 0.263287 | 0.427592 |
| FAM196A | 1.156773 | -0.72344 | 1.907588 | -0.37924 | 0.704509 | NA |
| C1orf192 | 92.50238 | 0.333023 | 0.256628 | 1.297689 | 0.194394 | 0.349475 |
| NOC2L | 2610.527 | -0.30191 | 0.160607 | -1.87981 | 0.060134 | 0.157509 |
| FAM35DP | 0.651288 | 1.289307 | 2.633717 | 0.489539 | 0.62446 | NA |
| VHLL | 0.288605 | 1.443619 | 4.029058 | 0.358302 | 0.720117 | NA |
| FAM111B | 88.80184 | 1.233948 | 0.514226 | 2.399621 | 0.016412 | 0.066156 |
| PLAC9 | 114.8961 | 0.16307 | 0.261138 | 0.624459 | 0.532326 | 0.676529 |
| S100A13 | 2743.532 | 0.26054 | 0.139908 | 1.862223 | 0.062572 | 0.161938 |
| ZNF33A | 208.3111 | 0.847443 | 0.319952 | 2.648654 | 0.008081 | 0.040855 |
| BTBD8 | 6.009777 | 1.246908 | 0.936632 | 1.331269 | 0.183101 | NA |
| PNRC2 | 784.8673 | 0.716385 | 0.210335 | 3.405932 | 6.59E-04 | 0.007993 |
| | 3.181643 | 0.283798 | 1.11508 | 0.254509 | 0.799102 | NA |
| GJB5 | 0.260158 | 1.240347 | 4.047124 | 0.306476 | 0.759242 | NA |
| FAM53B | 5077.418 | -0.65059 | 0.146938 | -4.42766 | 9.53E-06 | 6.14E-04 |
| | 0.230334 | -0.53318 | 4.076749 | -0.13079 | 0.895945 | NA |
| S100A14 | 0.834389 | 1.638864 | 2.19944 | 0.745128 | 0.456195 | NA |
| KAZN | 1503.208 | -0.26202 | 0.146944 | -1.78314 | 0.074564 | 0.182375 |
| SLC35E2B | 1316.074 | -0.53647 | 0.161779 | -3.31604 | 9.13E-04 | 0.009778 |
| MMP23B | 1.033116 | 1.025241 | 1.933005 | 0.530387 | 0.595844 | NA |

| | | | | | | |
|---|---|---|---|---|---|---|
| SH2D5 | 9.278536 | 0.934169 | 0.855354 | 1.092142 | 0.27477 | NA |
| RASSF10 | 0.298089 | -0.51711 | 3.975232 | -0.13008 | 0.896501 | NA |
| RNU1-75P | 0.151697 | -1.546 | 4.075469 | -0.37934 | 0.704433 | NA |
| BLOC1S2 | 1092.86 | 0.690433 | 0.198358 | 3.480745 | 5.00E-04 | 0.006782 |
| AKR1C3 | 16.89467 | 0.161093 | 0.629596 | 0.255868 | 0.798053 | NA |
| S100A4 | 189.3203 | 0.362125 | 0.301038 | 1.202923 | 0.229006 | 0.390822 |
| ACADSB | 820.2983 | 0.802358 | 0.263263 | 3.047743 | 0.002306 | 0.017726 |
| STK40 | 1939.807 | -0.26901 | 0.133296 | -2.01818 | 0.043573 | 0.127168 |
| TMEM63A | 974.4377 | -0.2516 | 0.177801 | -1.41505 | 0.157055 | 0.303288 |
| CTSE | 3.338287 | -0.41406 | 1.142991 | -0.36226 | 0.717158 | NA |
| SEMA4A | 12.22226 | -0.06062 | 0.606676 | -0.09991 | 0.920412 | NA |
| LCOR | 176.9531 | 0.473723 | 0.310319 | 1.526571 | 0.126868 | 0.262351 |
| OR2C3 | 1.024017 | 0.177191 | 1.894522 | 0.093528 | 0.925484 | NA |
| CD55 | 520.8294 | -0.16887 | 0.18291 | -0.92323 | 0.355885 | 0.523878 |
| SRGAP2B | 180.7979 | -0.14132 | 0.196097 | -0.72064 | 0.471128 | 0.627342 |
| ASB13 | 1642.708 | -0.34164 | 0.155724 | -2.19391 | 0.028242 | 0.095224 |
| THEM5 | 6.502437 | -0.21271 | 0.985755 | -0.21578 | 0.829157 | NA |
| ZNF124 | 83.32791 | 0.284246 | 0.267683 | 1.061873 | 0.288293 | 0.454443 |
| S100A5 | 0.47961 | 0.535796 | 3.39605 | 0.15777 | 0.874638 | NA |
| NBPF4 | 0.230334 | -0.53318 | 4.076749 | -0.13079 | 0.895945 | NA |
| YRDC | 574.1877 | 0.004393 | 0.193316 | 0.022726 | 0.981869 | 0.990311 |
| ESRRG | 1.743207 | 0.180904 | 1.513109 | 0.119558 | 0.904833 | NA |
| GDAP2 | 307.9553 | 0.193517 | 0.197746 | 0.978615 | 0.32777 | 0.495937 |
| SLC6A9 | 396.3005 | 0.009549 | 0.67856 | 0.014072 | 0.988773 | 0.994393 |
| FAM72A | 18.51041 | 0.385635 | 0.563811 | 0.683978 | 0.493989 | NA |
| LINC00610 | 39.51917 | 2.76E-04 | 0.336437 | 8.19E-04 | 0.999347 | NA |
| HBG2 | 0.163957 | 0.424913 | 4.079006 | 0.104171 | 0.917034 | NA |
| AJAP1 | 29.04019 | 1.110261 | 0.576522 | 1.925789 | 0.054131 | NA |
| SLC30A10 | 33.42983 | 0.478244 | 0.373369 | 1.28089 | 0.200232 | NA |
| FAM180B | 1.772088 | -2.18236 | 1.792446 | -1.21753 | 0.223402 | NA |
| ZNF33B | 180.709 | 0.774549 | 0.375111 | 2.064854 | 0.038937 | 0.118085 |
| LCE1B | 0.096202 | 0.408546 | 4.079639 | 0.100143 | 0.920231 | NA |
| S100A2 | 32.11113 | 0.186747 | 0.40435 | 0.461844 | 0.644193 | NA |
| ZNF239 | 17.61174 | -0.47304 | 0.510971 | -0.92577 | 0.354567 | NA |
| NHLRC2 | 413.5637 | 0.554351 | 0.212162 | 2.612869 | 0.008979 | 0.04402 |
| LAMB3 | 326.9499 | -0.05782 | 0.213763 | -0.27048 | 0.786793 | 0.870828 |
| HIST3H2BB | 1.435718 | -0.5457 | 1.807394 | -0.30193 | 0.762708 | NA |
| TMEM26 | 78.56975 | -0.53354 | 0.322966 | -1.65201 | 0.098532 | 0.221104 |
| | 0.583919 | -1.6942 | 2.749224 | -0.61625 | 0.537732 | NA |
| FUT11 | 437.4358 | -0.55067 | 0.206143 | -2.67129 | 0.007556 | 0.039253 |
| ZMYM1 | 121.374 | 0.730231 | 0.335033 | 2.179579 | 0.029289 | 0.097369 |
| SLC6A17 | 187.4238 | -0.31711 | 0.199418 | -1.59016 | 0.1118 | 0.240019 |
| ACSL5 | 9.974034 | -0.00545 | 0.638746 | -0.00854 | 0.993187 | NA |
| LRRC8B | 330.2201 | 0.015748 | 0.349116 | 0.045109 | 0.964021 | 0.980386 |
| | 0.995388 | 1.024939 | 2.067489 | 0.495741 | 0.620077 | NA |
| FAM110D | 0.622842 | 1.158846 | 2.735323 | 0.42366 | 0.671814 | NA |
| | 0.927816 | -0.09492 | 2.155471 | -0.04404 | 0.964876 | NA |

| | | | | | | |
|---|---|---|---|---|---|---|
| DDI2 | 958.138 | 0.064658 | 0.174148 | 0.37128 | 0.710429 | 0.814835 |
| SVIL | 3720.584 | 0.10221 | 0.231721 | 0.44109 | 0.659148 | 0.775241 |
| TRIM33 | 753.9546 | 0.522541 | 0.246398 | 2.120718 | 0.033946 | 0.107602 |
| IPP | 280.7205 | -0.17168 | 0.193595 | -0.8868 | 0.375184 | 0.541491 |
| OGDHL | 0.92901 | 1.841773 | 2.123319 | 0.867403 | 0.385721 | NA |
| COL13A1 | 0.303394 | -2.27006 | 4.032451 | -0.56295 | 0.57347 | NA |
| ZNF695 | 8.741731 | 0.241675 | 0.711244 | 0.339793 | 0.734013 | NA |
| FAM177B | 0.134133 | -1.48909 | 4.077584 | -0.36519 | 0.71497 | NA |
| MIB2 | 1223.92 | -0.19203 | 0.162626 | -1.18083 | 0.237671 | 0.400568 |
| HHLA3 | 585.6224 | -0.28589 | 0.165253 | -1.73 | 0.08363 | 0.197007 |
| DMBX1 | 0.134133 | -1.48909 | 4.077584 | -0.36519 | 0.71497 | NA |
| FAR1 | 754.7552 | 0.80547 | 0.36374 | 2.214412 | 0.0268 | 0.091986 |
| | 0.096202 | 0.408546 | 4.079639 | 0.100143 | 0.920231 | NA |
| CDC42SE1 | 3356.519 | -0.29708 | 0.154531 | -1.92244 | 0.05455 | 0.147075 |
| MPEG1 | 0.268266 | -2.15396 | 4.038351 | -0.53338 | 0.593773 | NA |
| PARVA | 11346.77 | -0.19588 | 0.129176 | -1.51635 | 0.12943 | 0.26588 |
| CR1L | 0.151697 | -1.546 | 4.075469 | -0.37934 | 0.704433 | NA |
| PSAP | 113827.3 | -0.40716 | 0.130314 | -3.12448 | 0.001781 | 0.015029 |
| S100A10 | 14245.38 | 0.045604 | 0.151048 | 0.301915 | 0.762717 | 0.852618 |
| WDR96 | 6.828636 | 0.016374 | 0.828137 | 0.019772 | 0.984226 | NA |
| MAP1LC3C | 10.65751 | 0.592827 | 0.814116 | 0.728185 | 0.4665 | NA |
| MCMBP | 1695.552 | 0.211367 | 0.169299 | 1.248481 | 0.211855 | 0.370324 |
| TAF13 | 597.5163 | 0.421937 | 0.254369 | 1.658764 | 0.097163 | 0.218901 |
| ATAD3A | 1565.738 | -0.13317 | 0.194264 | -0.68554 | 0.493005 | 0.644243 |
| FAM212B | 188.4273 | -0.18088 | 0.25239 | -0.71667 | 0.473579 | 0.629168 |
| MEIG1 | 9.154862 | 0.01582 | 0.65349 | 0.024208 | 0.980687 | NA |
| NRAP | 1.924573 | -0.37826 | 1.424337 | -0.26557 | 0.79057 | NA |
| HRNR | 15.11495 | -0.61163 | 0.59513 | -1.02772 | 0.304079 | NA |
| HES5 | 6.754154 | -0.63643 | 0.807409 | -0.78823 | 0.43056 | NA |
| S100A6 | 17987.38 | 0.273747 | 0.20433 | 1.339734 | 0.180332 | 0.332892 |
| DNM3 | 113.9389 | -0.17122 | 0.312055 | -0.5487 | 0.583214 | 0.7167 |
| MPZL1 | 1212.602 | -0.17174 | 0.159114 | -1.07935 | 0.280431 | 0.446542 |
| C1orf122 | 1870.87 | -0.21259 | 0.130238 | -1.63232 | 0.102612 | 0.226809 |
| SNHG12 | 168.4463 | 0.790462 | 0.210179 | 3.760894 | 1.69E-04 | 0.003497 |
| ENTPD7 | 243.3808 | 0.021614 | 0.189182 | 0.114249 | 0.909041 | 0.947483 |
| 5-Mar | 1078.04 | 0.267195 | 0.14865 | 1.797478 | 0.07226 | 0.178615 |
| ZNF248 | 182.3997 | 0.671753 | 0.200469 | 3.350917 | 8.05E-04 | 0.009085 |
| MIER1 | 565.6617 | 1.01169 | 0.245868 | 4.114765 | 3.88E-05 | 0.001429 |
| MAN1A2 | 859.1901 | 0.637069 | 0.226041 | 2.818374 | 0.004827 | 0.028841 |
| SVIP | 663.0259 | 0.760581 | 0.177201 | 4.292187 | 1.77E-05 | 8.56E-04 |
| SZT2 | 827.1508 | -0.53871 | 0.180261 | -2.9885 | 0.002803 | 0.020297 |
| CACNA1E | 0.558248 | 0.982076 | 2.804738 | 0.350149 | 0.726227 | NA |
| SLC29A3 | 37.77396 | -0.07537 | 0.360212 | -0.20925 | 0.834256 | NA |
| ANTXRL | 0.326333 | 1.533889 | 3.826878 | 0.40082 | 0.688553 | NA |
| ZNF485 | 50.52204 | 0.673833 | 0.31766 | 2.12124 | 0.033902 | 0.1076 |
| NTRK1 | 0.862864 | -0.92591 | 2.497199 | -0.37078 | 0.710801 | NA |
| MGEA5 | 1603.943 | 0.332621 | 0.22376 | 1.486506 | 0.137145 | 0.276319 |

| | | | | | | |
|---|---|---|---|---|---|---|
| FLVCR1-AS | 81.15914 | 0.082288 | 0.267484 | 0.307636 | 0.758359 | 0.849319 |
| ANKRD35 | 0.478233 | -0.67301 | 2.937473 | -0.22911 | 0.818782 | NA |
| YTHDF2 | 2419.241 | -0.10554 | 0.124071 | -0.85065 | 0.394962 | 0.559494 |
| C1orf228 | 8.17769 | -0.48609 | 0.78587 | -0.61854 | 0.536219 | NA |
| ZNF511 | 399.0523 | -0.03335 | 0.176035 | -0.18948 | 0.84972 | 0.910531 |
| CTNND1 | 7234.203 | -0.37963 | 0.119693 | -3.17171 | 0.001515 | 0.013631 |
| SH2D1B | 0.35636 | 1.709252 | 3.651833 | 0.468053 | 0.639746 | NA |
| MDM4 | 284.7149 | 0.035263 | 0.198535 | 0.177617 | 0.859024 | 0.916352 |
| RYR2 | 4.586351 | -0.05723 | 0.955659 | -0.05989 | 0.952244 | NA |
| PAPSS2 | 18.16792 | 0.424948 | 0.513236 | 0.827977 | 0.407683 | NA |
| ABCA4 | 0.314073 | -0.57308 | 3.898934 | -0.14698 | 0.883146 | NA |
| IPO9 | 3687.242 | -0.23915 | 0.130252 | -1.83602 | 0.066354 | 0.168227 |
| SSBP3-AS1 | 1.953231 | 0.600653 | 1.420042 | 0.422983 | 0.672308 | NA |
| C1orf85 | 6692.045 | -0.18089 | 0.14768 | -1.22489 | 0.220617 | 0.38052 |
| LDB1 | 2694.506 | -0.40742 | 0.128226 | -3.17733 | 0.001486 | 0.013472 |
| CTR9 | 1097.281 | 0.304158 | 0.256863 | 1.184128 | 0.236362 | 0.399195 |
| | 5184.049 | -0.38998 | 0.134841 | -2.89212 | 0.003827 | 0.024734 |
| GPATCH3 | 436.6463 | -0.07454 | 0.172415 | -0.43232 | 0.665507 | 0.780503 |
| COLGALT2 | 113.5171 | -1.00031 | 0.274205 | -3.64802 | 2.64E-04 | 0.004594 |
| RCSD1 | 1.645628 | -0.4699 | 1.475911 | -0.31838 | 0.750199 | NA |
| MUC2 | 0.586491 | 2.39072 | 2.689057 | 0.889055 | 0.373973 | NA |
| MTOR | 3385.884 | -0.47244 | 0.147218 | -3.20908 | 0.001332 | 0.012499 |
| BRINP2 | 0.959212 | -2.37936 | 2.418243 | -0.98392 | 0.325155 | NA |
| LRIG2 | 161.5023 | -0.09117 | 0.198602 | -0.45906 | 0.646189 | 0.76491 |
| FOXJ3 | 1138.642 | -0.08615 | 0.140393 | -0.61367 | 0.539436 | 0.681722 |
| CD247 | 0.555087 | 0.973802 | 2.808575 | 0.346725 | 0.728798 | NA |
| INPP5F | 1964.06 | 0.232034 | 0.169108 | 1.372103 | 0.170031 | 0.320363 |
| HMGN2 | 1642.805 | -0.56766 | 0.139039 | -4.08275 | 4.45E-05 | 0.001547 |
| GJC2 | 25.28319 | -0.16846 | 0.46355 | -0.36342 | 0.71629 | NA |
| DENND4B | 1354.553 | -0.59206 | 0.159256 | -3.71764 | 2.01E-04 | 0.003869 |
| KTI12 | 446.3019 | -0.189 | 0.180808 | -1.04528 | 0.295893 | 0.463138 |
| DUSP27 | 0.298089 | -0.51711 | 3.975232 | -0.13008 | 0.896501 | NA |
| HSD3BP5 | 20.02961 | 0.392399 | 0.491692 | 0.798059 | 0.424836 | NA |
| TSEN15 | 812.602 | 0.210845 | 0.13914 | 1.515342 | 0.129686 | 0.266024 |
| GRK5 | 867.8451 | 0.108062 | 0.146466 | 0.737795 | 0.460639 | 0.618796 |
| SFMBT2 | 317.7427 | -0.28689 | 0.224126 | -1.28004 | 0.200532 | 0.356579 |
| PRMT6 | 362.7417 | -0.32015 | 0.172047 | -1.86084 | 0.062767 | 0.162216 |
| SHISA4 | 1665.603 | -0.08884 | 0.163481 | -0.54343 | 0.586836 | 0.719553 |
| C1orf174 | 396.7914 | 0.306076 | 0.202005 | 1.515186 | 0.129725 | 0.266024 |
| RASGEF1A | 8.800993 | 0.640948 | 0.703287 | 0.91136 | 0.362106 | NA |
| DCLRE1A | 238.5383 | 0.435373 | 0.196196 | 2.219069 | 0.026482 | 0.091289 |
| NOS1AP | 2.290629 | 0.525815 | 1.348424 | 0.389948 | 0.696575 | NA |
| SMG5 | 5497.958 | -0.29404 | 0.210447 | -1.39721 | 0.16235 | 0.310342 |
| KIAA1279 | 1434.606 | 0.187091 | 0.13475 | 1.38843 | 0.165006 | 0.313807 |
| SGMS1 | 334.7267 | 0.698577 | 0.207618 | 3.364719 | 7.66E-04 | 0.00881 |
| RNU5E-1 | 0.382031 | -1.26484 | 3.151989 | -0.40128 | 0.688212 | NA |
| | 0.151697 | -1.546 | 4.075469 | -0.37934 | 0.704433 | NA |

| | | | | | | |
|---|---|---|---|---|---|---|
| | 0.096202 | 0.408546 | 4.079639 | 0.100143 | 0.920231 | NA |
| | 0.937264 | -1.23834 | 2.063944 | -0.59999 | 0.548514 | NA |
| RNU6-37P | 0.163957 | 0.424913 | 4.079006 | 0.104171 | 0.917034 | NA |
| RNU6-1266 | 0.096202 | 0.408546 | 4.079639 | 0.100143 | 0.920231 | NA |
| RNU4-21P | 0.096202 | 0.408546 | 4.079639 | 0.100143 | 0.920231 | NA |
| RNU6-490F | 0.163957 | 0.424913 | 4.079006 | 0.104171 | 0.917034 | NA |
| | 0.096202 | 0.408546 | 4.079639 | 0.100143 | 0.920231 | NA |
| | 0.162376 | 0.424913 | 4.079006 | 0.104171 | 0.917034 | NA |
| RNU6-403F | 0.096202 | 0.408546 | 4.079639 | 0.100143 | 0.920231 | NA |
| SNORA73B | 5.313327 | 0.535587 | 0.922014 | 0.580888 | 0.561316 | NA |
| | 0.602678 | 1.083611 | 2.70938 | 0.399948 | 0.689195 | NA |
| RNU5D-1 | 0.134133 | -1.48909 | 4.077584 | -0.36519 | 0.71497 | NA |
| | 1.044145 | 0.31036 | 2.099918 | 0.147796 | 0.882504 | NA |
| RNU6-529F | 1.677089 | 0.300749 | 1.519436 | 0.197934 | 0.843096 | NA |
| | 0.096202 | 0.408546 | 4.079639 | 0.100143 | 0.920231 | NA |
| RNU6-833F | 0.151697 | -1.546 | 4.075469 | -0.37934 | 0.704433 | NA |
| RNU5E-10F | 0.192403 | 0.975392 | 4.060716 | 0.240202 | 0.810174 | NA |
| RNU6-1099 | 0.883146 | 1.112834 | 2.554178 | 0.435692 | 0.663061 | NA |
| RNU6-1205 | 0.162376 | 0.424913 | 4.079006 | 0.104171 | 0.917034 | NA |
| | 0.162376 | 0.424913 | 4.079006 | 0.104171 | 0.917034 | NA |
| | 0.151697 | -1.546 | 4.075469 | -0.37934 | 0.704433 | NA |
| RN7SKP160 | 0.586491 | 2.39072 | 2.689057 | 0.889055 | 0.373973 | NA |
| | 0.399595 | -1.31595 | 3.579466 | -0.36764 | 0.713143 | NA |
| RNU6-884F | 0.151697 | -1.546 | 4.075469 | -0.37934 | 0.704433 | NA |
| SNORD46 | 2.031812 | 0.640288 | 1.376527 | 0.465147 | 0.641826 | NA |
| RNU5A-8P | 0.315653 | -0.57305 | 3.892639 | -0.14721 | 0.882963 | NA |
| | 0.096202 | 0.408546 | 4.079639 | 0.100143 | 0.920231 | NA |
| RNU6-151F | 0.162376 | 0.424913 | 4.079006 | 0.104171 | 0.917034 | NA |
| | 0.987658 | -1.76927 | 2.378195 | -0.74395 | 0.456905 | NA |
| RNU4-59P | 0.314073 | -0.57308 | 3.898934 | -0.14698 | 0.883146 | NA |
| RN7SKP37 | 0.096202 | 0.408546 | 4.079639 | 0.100143 | 0.920231 | NA |
| | 0.096202 | 0.408546 | 4.079639 | 0.100143 | 0.920231 | NA |
| | 0.654246 | 2.541085 | 3.009764 | 0.84428 | 0.398513 | NA |
| SNORA55 | 2.341133 | -0.23198 | 1.448554 | -0.16015 | 0.872765 | NA |
| RNA5SP78 | 0.162376 | 0.424913 | 4.079006 | 0.104171 | 0.917034 | NA |
| SNORA16B | 0.3441 | -0.0166 | 3.769682 | -0.0044 | 0.996486 | NA |
| RNVU1-6 | 0.162376 | 0.424913 | 4.079006 | 0.104171 | 0.917034 | NA |
| RN7SKP50 | 0.096202 | 0.408546 | 4.079639 | 0.100143 | 0.920231 | NA |
| RN7SKP78 | 0.326333 | 1.533889 | 3.826878 | 0.40082 | 0.688553 | NA |
| RNU6-740F | 0.096202 | 0.408546 | 4.079639 | 0.100143 | 0.920231 | NA |
| RNU1-91P | 0.151697 | -1.546 | 4.075469 | -0.37934 | 0.704433 | NA |
| RNU6-828F | 0.478233 | -0.67301 | 2.937473 | -0.22911 | 0.818782 | NA |
| RNU5E-4P | 0.151697 | -1.546 | 4.075469 | -0.37934 | 0.704433 | NA |
| RNA5SP298 | 0.640755 | -1.64826 | 2.754778 | -0.59833 | 0.549621 | NA |
| | 0.296509 | -0.51709 | 3.982155 | -0.12985 | 0.896684 | NA |
| | 0.163957 | 0.424913 | 4.079006 | 0.104171 | 0.917034 | NA |
| SNORA23 | 1.0349 | 0.343148 | 1.852366 | 0.185249 | 0.853034 | NA |

| | | | | | | |
|---|---|---|---|---|---|---|
| RNU6-124( | 0.162376 | 0.424913 | 4.079006 | 0.104171 | 0.917034 | NA |
| SNORD38A | 1.885622 | -0.0039 | 1.460299 | -0.00267 | 0.997868 | NA |
| | 0.651435 | 0.249629 | 2.788679 | 0.089515 | 0.928672 | NA |
| | 0.162376 | 0.424913 | 4.079006 | 0.104171 | 0.917034 | NA |
| RN7SKP26( | 1.341068 | 0.883398 | 1.741964 | 0.507128 | 0.612065 | NA |
| | 0.813811 | 0.928591 | 2.591569 | 0.358312 | 0.72011 | NA |
| | 0.192403 | 0.975392 | 4.060716 | 0.240202 | 0.810174 | NA |
| RNVU1-20 | 0.096202 | 0.408546 | 4.079639 | 0.100143 | 0.920231 | NA |
| | 0.247898 | -0.5901 | 4.074633 | -0.14482 | 0.884851 | NA |
| | 0.298089 | -0.51711 | 3.975232 | -0.13008 | 0.896501 | NA |
| | 0.163957 | 0.424913 | 4.079006 | 0.104171 | 0.917034 | NA |
| | 3.74639 | 0.711058 | 1.042145 | 0.682302 | 0.495048 | NA |
| | 3.303167 | 0.400132 | 1.10982 | 0.360538 | 0.718445 | NA |
| | 5.708114 | 0.037733 | 0.815975 | 0.046243 | 0.963116 | NA |
| | 1.611703 | 2.750352 | 1.792131 | 1.534683 | 0.124862 | NA |
| | 0.260158 | 1.240347 | 4.047124 | 0.306476 | 0.759242 | NA |
| BTBD7P1 | 0.096202 | 0.408546 | 4.079639 | 0.100143 | 0.920231 | NA |
| | 0.134133 | -1.48909 | 4.077584 | -0.36519 | 0.71497 | NA |
| | 0.134133 | -1.48909 | 4.077584 | -0.36519 | 0.71497 | NA |
| PDCD4-AS1 | 69.84078 | -0.26756 | 0.309347 | -0.86492 | 0.387085 | 0.552959 |
| | 0.163957 | 0.424913 | 4.079006 | 0.104171 | 0.917034 | NA |
| | 0.452561 | 2.061505 | 3.349456 | 0.615474 | 0.538241 | NA |
| | 0.816825 | 1.673341 | 2.21346 | 0.755984 | 0.449659 | NA |
| EFCAB2 | 104.8894 | -0.07405 | 0.278352 | -0.26601 | 0.790228 | 0.872781 |
| COX20 | 47.77291 | 0.381414 | 0.339652 | 1.122957 | 0.261456 | 0.425939 |
| CHML | 313.0612 | 0.858656 | 0.396964 | 2.163055 | 0.030537 | 0.100281 |
| IBA57-AS1 | 0.6908 | 0.277604 | 2.346083 | 0.118327 | 0.905809 | NA |
| C1orf95 | 5.236644 | -0.84457 | 0.987899 | -0.85492 | 0.392596 | NA |
| | 1.42088 | 2.532189 | 1.880145 | 1.346806 | 0.178043 | NA |
| CAPN8 | 0.705792 | -3.31105 | 2.811825 | -1.17754 | 0.238978 | NA |
| TATDN3 | 307.1796 | 0.25506 | 0.186773 | 1.365615 | 0.17206 | 0.32326 |
| SERTAD4-A | 27.4574 | 1.229009 | 0.492727 | 2.494299 | 0.012621 | NA |
| C1orf132 | 0.777517 | 1.644684 | 2.527228 | 0.650785 | 0.515185 | NA |
| LINC00862 | 0.735616 | -2.07189 | 2.528723 | -0.81934 | 0.412591 | NA |
| C1orf53 | 36.78561 | 0.441912 | 0.350991 | 1.259041 | 0.208016 | NA |
| GPR52 | 0.096202 | 0.408546 | 4.079639 | 0.100143 | 0.920231 | NA |
| | 25.79425 | 0.602791 | 0.448047 | 1.345374 | 0.178504 | NA |
| METTL11B | 0.450981 | 2.057325 | 3.353149 | 0.61355 | 0.539513 | NA |
| MSTO2P | 6.313944 | -0.22484 | 0.794194 | -0.28311 | 0.777094 | NA |
| SPRN | 50.40274 | -0.30465 | 0.365631 | -0.8332 | 0.40473 | 0.569118 |
| FANK1 | 64.10464 | 0.17912 | 0.300022 | 0.597021 | 0.550493 | 0.691764 |
| KPRP | 0.192403 | 0.975392 | 4.060716 | 0.240202 | 0.810174 | NA |
| METTL10 | 79.7017 | 0.257028 | 0.254334 | 1.010593 | 0.312211 | 0.479394 |
| ADAMTSL4 | 0.853323 | -0.46916 | 2.317774 | -0.20242 | 0.839588 | NA |
| PPAPDC1A | 650.1462 | 0.351197 | 0.255691 | 1.37352 | 0.169591 | 0.320041 |
| HIST2H2BF | 1.190749 | 2.261198 | 1.987481 | 1.137721 | 0.255237 | NA |
| FAM231D | 7.24034 | -1.05564 | 0.753874 | -1.40029 | 0.161426 | NA |

| | | | | | | |
|---|---|---|---|---|---|---|
| FAM72C | 1.594488 | 1.432245 | 1.712183 | 0.836503 | 0.402872 | NA |
| HIST2H3PS | 27.11344 | -0.05086 | 0.423517 | -0.12009 | 0.904409 | NA |
| HIST2H2BC | 2.077262 | 0.716114 | 1.425336 | 0.502417 | 0.615374 | NA |
| | 0.096202 | 0.408546 | 4.079639 | 0.100143 | 0.920231 | NA |
| NBPF16 | 5.589522 | 0.386123 | 0.901281 | 0.428416 | 0.668348 | NA |
| NBPF20 | 17.79996 | 0.173737 | 0.553692 | 0.313779 | 0.753689 | NA |
| NBPF24 | 1.658237 | -1.16816 | 1.652219 | -0.70702 | 0.479551 | NA |
| PNLIPRP3 | 69.05303 | -0.13425 | 0.386499 | -0.34735 | 0.728332 | 0.827682 |
| | 5.270793 | 4.66E-04 | 0.967957 | 4.81E-04 | 0.999616 | NA |
| HSD3BP4 | 0.748867 | 2.740193 | 2.468928 | 1.109871 | 0.267054 | NA |
| ATP1A1OS | 36.53134 | -0.17366 | 0.364548 | -0.47638 | 0.633805 | NA |
| RBM20 | 0.544554 | -2.65036 | 3.220086 | -0.82307 | 0.410468 | NA |
| | 7.301399 | -0.31349 | 0.744979 | -0.42081 | 0.673894 | NA |
| SPATA42 | 0.506476 | 0.726418 | 2.864186 | 0.253621 | 0.799788 | NA |
| C1orf146 | 0.420954 | 1.923376 | 3.462803 | 0.555439 | 0.578594 | NA |
| HSP90B3P | 1.62392 | -0.42828 | 1.578049 | -0.2714 | 0.786082 | NA |
| C10orf62 | 0.296509 | -0.51709 | 3.982155 | -0.12985 | 0.896684 | NA |
| SAMD13 | 57.20607 | 0.185511 | 0.46965 | 0.394998 | 0.692844 | 0.801725 |
| C1orf141 | 0.162376 | 0.424913 | 4.079006 | 0.104171 | 0.917034 | NA |
| EFCAB7 | 109.6533 | 0.824086 | 0.36495 | 2.258076 | 0.023941 | 0.085415 |
| ZYG11A | 3.783386 | -0.11777 | 1.013353 | -0.11622 | 0.907479 | NA |
| IFIT1B | 1.173331 | 1.84241 | 2.290367 | 0.804417 | 0.421156 | NA |
| LIPJ | 0.364467 | -1.20166 | 3.685431 | -0.32606 | 0.744381 | NA |
| FOXO6 | 0.815594 | -0.45539 | 2.531219 | -0.17991 | 0.857224 | NA |
| INPP5B | 718.4331 | -0.40848 | 0.160121 | -2.55111 | 0.010738 | 0.049294 |
| RUFY2 | 172.6832 | 0.773778 | 0.321826 | 2.404335 | 0.016202 | 0.065554 |
| PHACTR4 | 994.7948 | -0.49204 | 0.186995 | -2.63131 | 0.008506 | 0.042257 |
| ASAH2B | 9.391628 | 0.79531 | 0.667931 | 1.190706 | 0.233769 | NA |
| AGAP6 | 40.67781 | 1.078156 | 0.362172 | 2.976915 | 0.002912 | NA |
| CTGLF11P | 0.162376 | 0.424913 | 4.079006 | 0.104171 | 0.917034 | NA |
| TIMM23B | 1.673669 | 0.581916 | 1.860244 | 0.312817 | 0.75442 | NA |
| ZDHHC18 | 689.6945 | -0.26169 | 0.16947 | -1.54419 | 0.122543 | 0.256035 |
| BMS1P5 | 2.022327 | 1.148615 | 1.525757 | 0.752817 | 0.45156 | NA |
| SYT15 | 19.30988 | -0.73642 | 0.524853 | -1.4031 | 0.160586 | NA |
| BMS1P1 | 2.500559 | 0.72947 | 1.300262 | 0.561018 | 0.574785 | NA |
| TMEM57 | 656.8915 | 0.224959 | 0.154386 | 1.457126 | 0.145082 | 0.28704 |
| PTPN20A | 1.159491 | 0.557655 | 1.885778 | 0.295716 | 0.767447 | NA |
| TCEA3 | 14.52536 | -0.08983 | 0.542946 | -0.16545 | 0.868589 | NA |
| | 0.983514 | -1.5981 | 2.093157 | -0.76349 | 0.445173 | NA |
| C10orf126 | 0.95764 | 0.792425 | 2.400829 | 0.330063 | 0.741352 | NA |
| C1orf134 | 9.126964 | -0.58554 | 0.742928 | -0.78815 | 0.430608 | NA |
| C1orf195 | 0.327913 | 1.543154 | 4.021063 | 0.383768 | 0.701151 | NA |
| PTCHD2 | 0.814253 | -1.23545 | 2.246777 | -0.54988 | 0.582405 | NA |
| CASC10 | 77.19139 | -0.1362 | 0.264447 | -0.51503 | 0.606529 | 0.734539 |
| MALRD1 | 0.545988 | -0.43913 | 2.541145 | -0.17281 | 0.862803 | NA |
| ST8SIA6-AS | 31.75686 | 0.474274 | 0.431472 | 1.099201 | 0.27168 | NA |
| ZBTB48 | 300.8907 | -0.28671 | 0.168082 | -1.70579 | 0.088046 | 0.204344 |

| | | | | | | |
|---|---|---|---|---|---|---|
| C11orf83 | 358.8059 | -0.19083 | 0.221311 | -0.86225 | 0.388549 | 0.554176 |
| LRRC10B | 0.69625 | -3.13723 | 2.93845 | -1.06765 | 0.285679 | NA |
| TMEM240 | 64.77576 | -0.04363 | 0.303062 | -0.14395 | 0.885537 | 0.932306 |
| | 4.084104 | 0.579678 | 1.07436 | 0.539556 | 0.589503 | NA |
| TMEM88B | 0.583919 | -1.6942 | 2.749224 | -0.61625 | 0.537732 | NA |
| ACCSL | 28.54554 | -1.07032 | 0.476204 | -2.2476 | 0.024602 | NA |
| C11orf91 | 3.373201 | -0.74424 | 1.121021 | -0.6639 | 0.506755 | NA |
| LGR4 | 475.6448 | 0.188549 | 0.250197 | 0.7536 | 0.45109 | 0.610395 |
| IPO7 | 3748.727 | 0.443273 | 0.199515 | 2.221756 | 0.0263 | 0.091046 |
| CALML3-AS | 0.096202 | 0.408546 | 4.079639 | 0.100143 | 0.920231 | NA |
| NAP1L4 | 2226.802 | 0.258212 | 0.162517 | 1.588831 | 0.112099 | 0.2405 |
| | 0.247898 | -0.5901 | 4.074633 | -0.14482 | 0.884851 | NA |
| KRTAP5-1 | 0.921281 | -1.21902 | 2.226885 | -0.54741 | 0.584096 | NA |
| | 0.420954 | 1.923376 | 3.462803 | 0.555439 | 0.578594 | NA |
| RNVU1-18 | 0.258578 | 1.233151 | 4.047809 | 0.304646 | 0.760635 | NA |
| RNU6-418F | 0.298089 | -0.51711 | 3.975232 | -0.13008 | 0.896501 | NA |
| RNU6-949F | 0.096202 | 0.408546 | 4.079639 | 0.100143 | 0.920231 | NA |
| | 0.247898 | -0.5901 | 4.074633 | -0.14482 | 0.884851 | NA |
| RNU6-48P | 0.574231 | 0.93227 | 2.49998 | 0.372911 | 0.709215 | NA |
| RNU6-481F | 0.163957 | 0.424913 | 4.079006 | 0.104171 | 0.917034 | NA |
| RNU6-514F | 0.096202 | 0.408546 | 4.079639 | 0.100143 | 0.920231 | NA |
| | 0.151697 | -1.546 | 4.075469 | -0.37934 | 0.704433 | NA |
| RNU6-43P | 0.230334 | -0.53318 | 4.076749 | -0.13079 | 0.895945 | NA |
| SNORA14B | 2.39569 | 1.551076 | 1.377709 | 1.125837 | 0.260234 | NA |
| RNVU1-15 | 0.260158 | 1.240347 | 4.047124 | 0.306476 | 0.759242 | NA |
| RNU6-125F | 0.487129 | 2.123445 | 3.910539 | 0.543006 | 0.587126 | NA |
| RNU6-15P | 0.327913 | 1.543154 | 4.021063 | 0.383768 | 0.701151 | NA |
| | 0.163957 | 0.424913 | 4.079006 | 0.104171 | 0.917034 | NA |
| SNORA42 | 0.258578 | 1.233151 | 4.047809 | 0.304646 | 0.760635 | NA |
| RNVU1-14 | 0.556813 | -0.57007 | 3.294615 | -0.17303 | 0.862627 | NA |
| SNORA66 | 1.624354 | -3.47806 | 2.019127 | -1.72255 | 0.084969 | NA |
| MIR606 | 0.324752 | 1.53082 | 4.022025 | 0.380609 | 0.703493 | NA |
| MIR197 | 0.49029 | 2.123092 | 3.304282 | 0.642527 | 0.520531 | NA |
| MIR186 | 1.03648 | 0.354657 | 2.004615 | 0.17692 | 0.859571 | NA |
| MIR34A | 2.448816 | -0.35943 | 1.340867 | -0.26806 | 0.788654 | NA |
| MIR181B1 | 0.192403 | 0.975392 | 4.060716 | 0.240202 | 0.810174 | NA |
| MIR130A | 0.392857 | -1.546 | 4.075469 | -0.37934 | 0.704433 | NA |
| SNORD78 | 7.241389 | 1.234784 | 0.806855 | 1.530367 | 0.125926 | NA |
| C11orf31 | 1854.974 | 0.042533 | 0.135536 | 0.31381 | 0.753665 | 0.846169 |
| GNRHR2 | 10.49914 | -0.277 | 0.663916 | -0.41722 | 0.676517 | NA |
| DIO1 | 1.159491 | 0.557655 | 1.885778 | 0.295716 | 0.767447 | NA |
| AKR7L | 2.664102 | 0.494969 | 1.255467 | 0.394251 | 0.693396 | NA |
| MIR760 | 0.544554 | -2.65036 | 3.220086 | -0.82307 | 0.410468 | NA |
| MIR765 | 0.163957 | 0.424913 | 4.079006 | 0.104171 | 0.917034 | NA |
| SNORD67 | 1.607697 | 0.014776 | 1.512674 | 0.009768 | 0.992206 | NA |
| | 0.096202 | 0.408546 | 4.079639 | 0.100143 | 0.920231 | NA |
| RNA5SP339 | 0.163957 | 0.424913 | 4.079006 | 0.104171 | 0.917034 | NA |

| | | | | | | |
|---|---|---|---|---|---|---|
| RNU6-817F | 2.822334 | 0.578323 | 1.217756 | 0.474909 | 0.634852 | NA |
| RNA5SP323 | 0.326333 | 1.533889 | 3.826878 | 0.40082 | 0.688553 | NA |
| RNU11-3P | 0.096202 | 0.408546 | 4.079639 | 0.100143 | 0.920231 | NA |
| RNVU1-13 | 0.230334 | -0.53318 | 4.076749 | -0.13079 | 0.895945 | NA |
| SNORA12 | 0.134133 | -1.48909 | 4.077584 | -0.36519 | 0.71497 | NA |
| RNU6-510F | 0.163957 | 0.424913 | 4.079006 | 0.104171 | 0.917034 | NA |
| SNORA45 | 4.947779 | 0.237085 | 1.218864 | 0.194513 | 0.845774 | NA |
| | 23.86382 | 0.342779 | 0.480464 | 0.713433 | 0.475578 | NA |
| | 1.981677 | 0.663067 | 1.469751 | 0.451143 | 0.651887 | NA |
| | 0.314073 | -0.57308 | 3.898934 | -0.14698 | 0.883146 | NA |
| ST13P5 | 2.212195 | 0.082105 | 1.371686 | 0.059857 | 0.952269 | NA |
| MAP10 | 19.73681 | 0.525998 | 0.63009 | 0.834799 | 0.403831 | NA |
| COX20P1 | 0.134133 | -1.48909 | 4.077584 | -0.36519 | 0.71497 | NA |
| CFL1P4 | 1.46011 | -1.91326 | 1.810838 | -1.05656 | 0.290713 | NA |
| | 0.449786 | -1.25325 | 3.00474 | -0.41709 | 0.676613 | NA |
| AURKAPS1 | 0.192403 | 0.975392 | 4.060716 | 0.240202 | 0.810174 | NA |
| | 0.843488 | 2.923514 | 2.365851 | 1.235713 | 0.216565 | NA |
| | 0.162376 | 0.424913 | 4.079006 | 0.104171 | 0.917034 | NA |
| DENND1B | 178.8826 | 0.746639 | 0.292101 | 2.556094 | 0.010585 | 0.048867 |
| RPL5P5 | 0.715249 | -0.95037 | 2.624937 | -0.36205 | 0.717313 | NA |
| C1orf220 | 2.153967 | 2.417005 | 1.616103 | 1.495576 | 0.134764 | NA |
| | 866.2683 | -0.0076 | 0.162578 | -0.04672 | 0.962733 | 0.979674 |
| | 0.134133 | -1.48909 | 4.077584 | -0.36519 | 0.71497 | NA |
| PFN1P11 | 1.101221 | -0.557 | 1.814706 | -0.30694 | 0.758892 | NA |
| | 24.26739 | -0.24834 | 0.59989 | -0.41397 | 0.678897 | NA |
| RPL29P7 | 1.288531 | 2.391399 | 2.161427 | 1.106399 | 0.268554 | NA |
| SFT2D2 | 158.9627 | -0.01446 | 0.215231 | -0.06719 | 0.946428 | 0.970016 |
| RPL31P11 | 0.544407 | -0.44425 | 2.542878 | -0.1747 | 0.861313 | NA |
| | 16.56424 | 0.434212 | 0.515791 | 0.841836 | 0.39988 | NA |
| CCDC19 | 4.362361 | -0.1453 | 0.981709 | -0.14801 | 0.882334 | NA |
| DARC | 13.85062 | -0.53572 | 0.661575 | -0.80977 | 0.418072 | NA |
| FAM24B | 2.644003 | 1.59733 | 1.479017 | 1.079994 | 0.280145 | NA |
| MLLT11 | 328.252 | -0.23584 | 0.186868 | -1.26207 | 0.206925 | 0.36412 |
| | 0.151697 | -1.546 | 4.075469 | -0.37934 | 0.704433 | NA |
| NOTCH2NL | 94.78146 | 0.148626 | 0.325374 | 0.456786 | 0.647825 | 0.766234 |
| HIST2H3DF | 5.385629 | -0.08047 | 0.929245 | -0.0866 | 0.930991 | NA |
| YWHAZP5 | 0.69625 | -3.13723 | 2.93845 | -1.06765 | 0.285679 | NA |
| NRAS | 1169.386 | 0.318454 | 0.161712 | 1.96926 | 0.048923 | 0.136821 |
| CHUK | 986.4978 | 0.658088 | 0.157742 | 4.171914 | 3.02E-05 | 0.001198 |
| GSTM2 | 456.6602 | -0.23766 | 0.185264 | -1.28279 | 0.199564 | 0.355802 |
| ARHGAP19 | 300.5711 | -0.13271 | 0.20411 | -0.65017 | 0.515585 | 0.663079 |
| HNRNPA1P | 0.392857 | -1.546 | 4.075469 | -0.37934 | 0.704433 | NA |
| | 0.326333 | 1.533889 | 3.826878 | 0.40082 | 0.688553 | NA |
| CFL1P2 | 0.937264 | -1.23834 | 2.063944 | -0.59999 | 0.548514 | NA |
| GBP7 | 6.498148 | -0.38041 | 0.843949 | -0.45075 | 0.652171 | NA |
| | 2.134979 | -2.67184 | 1.875654 | -1.42448 | 0.154307 | NA |
| | 0.516164 | -1.70268 | 2.844782 | -0.59853 | 0.549488 | NA |

| | | | | | | |
|---|---|---|---|---|---|---|
| RBMXL1 | 666.5532 | -0.08153 | 0.174596 | -0.46698 | 0.640517 | 0.76055 |
| KRT8P41 | 0.096202 | 0.408546 | 4.079639 | 0.100143 | 0.920231 | NA |
| DNAJC9 | 153.5355 | 0.424096 | 0.251485 | 1.686366 | 0.091725 | 0.210696 |
| LDHAP5 | 4.42133 | -0.79336 | 1.09671 | -0.7234 | 0.469433 | NA |
| | 0.690946 | -1.56103 | 2.699004 | -0.57837 | 0.563013 | NA |
| | 0.315653 | -0.57305 | 3.892639 | -0.14721 | 0.882963 | NA |
| TMX2 | 3355.11 | -0.19914 | 0.156151 | -1.27528 | 0.20221 | 0.358669 |
| | 10.77556 | 0.17306 | 0.658001 | 0.263009 | 0.792544 | NA |
| | 0.545988 | -0.43913 | 2.541145 | -0.17281 | 0.862803 | NA |
| NDUFS3 | 2060.214 | -0.10832 | 0.168808 | -0.64168 | 0.521084 | 0.668146 |
| LEPROT | 3173.844 | 0.268148 | 0.159126 | 1.685124 | 0.091965 | 0.210878 |
| PGGT1BP1 | 0.151697 | -1.546 | 4.075469 | -0.37934 | 0.704433 | NA |
| SEC14L1P1 | 24.37473 | 0.386137 | 0.451018 | 0.856146 | 0.391917 | NA |
| RPL17P50 | 4.288867 | 0.999767 | 0.997774 | 1.001997 | 0.316345 | NA |
| PIGCP1 | 90.79973 | -0.05839 | 0.251157 | -0.2325 | 0.816148 | 0.889858 |
| PRDX2P2 | 0.163957 | 0.424913 | 4.079006 | 0.104171 | 0.917034 | NA |
| ANAPC10P | 0.162376 | 0.424913 | 4.079006 | 0.104171 | 0.917034 | NA |
| FAM210CP | 0.326333 | 1.533889 | 3.826878 | 0.40082 | 0.688553 | NA |
| | 0.422534 | 1.924583 | 3.001758 | 0.641152 | 0.521424 | NA |
| | 3.082776 | -0.99474 | 1.252559 | -0.79416 | 0.427101 | NA |
| MRPL23 | 1954.46 | -0.03273 | 0.138704 | -0.236 | 0.813434 | 0.888705 |
| TSPAN4 | 3169.915 | -0.3999 | 0.16 | -2.49937 | 0.012441 | 0.054606 |
| | 0.624422 | 1.162588 | 2.733761 | 0.425271 | 0.670639 | NA |
| | 0.162376 | 0.424913 | 4.079006 | 0.104171 | 0.917034 | NA |
| MYCBP | 367.9157 | 0.097617 | 0.196795 | 0.496033 | 0.619871 | 0.74446 |
| | 0.526843 | -0.38719 | 2.855892 | -0.13558 | 0.892156 | NA |
| SH3D21 | 90.77221 | -0.57851 | 0.309436 | -1.86956 | 0.061544 | 0.160095 |
| HNRNPA1P | 0.354779 | 1.703954 | 3.657232 | 0.465914 | 0.641277 | NA |
| ANKRD36B | 87.24764 | 0.036333 | 0.267196 | 0.135978 | 0.891839 | 0.936719 |
| | 10.57975 | 0.668855 | 0.62278 | 1.073984 | 0.28283 | NA |
| ALDOAP2 | 1.621188 | 1.859647 | 1.660407 | 1.119994 | 0.262716 | NA |
| | 0.314073 | -0.57308 | 3.898934 | -0.14698 | 0.883146 | NA |
| BBIP1 | 252.5157 | 0.619676 | 0.182773 | 3.390416 | 6.98E-04 | 0.008275 |
| AS3MT | 191.7705 | -0.50249 | 0.264043 | -1.90308 | 0.05703 | 0.151387 |
| NUTM2D | 7.643819 | -1.10894 | 0.721525 | -1.53694 | 0.124309 | NA |
| PGGT1BP2 | 0.392857 | -1.546 | 4.075469 | -0.37934 | 0.704433 | NA |
| RPSAP6 | 0.163957 | 0.424913 | 4.079006 | 0.104171 | 0.917034 | NA |
| ZSWIM8 | 2746.651 | -0.58959 | 0.146738 | -4.01798 | 5.87E-05 | 0.001798 |
| HNRNPUL2 | 1013.981 | -0.26356 | 0.157966 | -1.66845 | 0.095227 | 0.215867 |
| METTL12 | 125.6215 | -0.28829 | 0.231713 | -1.24418 | 0.213434 | 0.372096 |
| MS4A4E | 0.399595 | -1.31595 | 3.579466 | -0.36764 | 0.713143 | NA |
| OOSP1 | 0.766634 | 1.516849 | 2.262991 | 0.670285 | 0.502676 | NA |
| | 60.98826 | -0.40709 | 0.418751 | -0.97216 | 0.33097 | 0.499446 |
| NPM1P31 | 0.326536 | 0.037056 | 3.842554 | 0.009644 | 0.992306 | NA |
| | 0.411855 | 0.250456 | 3.097618 | 0.080854 | 0.935558 | NA |
| | 15.48958 | -0.17975 | 0.596012 | -0.30159 | 0.762961 | NA |
| SMTNL1 | 0.8427 | -0.93379 | 2.415151 | -0.38664 | 0.699024 | NA |

| | | | | | | |
|---|---|---|---|---|---|---|
| MED28P1 | 0.3441 | -0.0166 | 3.769682 | -0.0044 | 0.996486 | NA |
| TMEM14D | 1.321867 | 0.888794 | 1.796137 | 0.494836 | 0.620716 | NA |
| CHCHD3P3 | 0.862836 | 1.736629 | 2.194323 | 0.791419 | 0.4287 | NA |
| PARGP1 | 26.37699 | 0.247967 | 0.401467 | 0.617651 | 0.536805 | NA |
| | 109.7921 | 0.679772 | 0.482206 | 1.409712 | 0.158625 | 0.305275 |
| NPM1P24 | 0.162376 | 0.424913 | 4.079006 | 0.104171 | 0.917034 | NA |
| | 4.451223 | 1.037885 | 1.176996 | 0.881809 | 0.37788 | NA |
| MUC5AC | 0.134133 | -1.48909 | 4.077584 | -0.36519 | 0.71497 | NA |
| | 0.096202 | 0.408546 | 4.079639 | 0.100143 | 0.920231 | NA |
| | 1.0491 | 1.000662 | 2.336632 | 0.42825 | 0.668469 | NA |
| | 0.163957 | 0.424913 | 4.079006 | 0.104171 | 0.917034 | NA |
| CELA2B | 0.392711 | 0.292538 | 3.145221 | 0.09301 | 0.925895 | NA |
| TMEM167I | 1152.64 | 0.230351 | 0.162647 | 1.416259 | 0.1567 | 0.302948 |
| | 0.268266 | -2.15396 | 4.038351 | -0.53338 | 0.593773 | NA |
| FAM72D | 5.395043 | 0.850198 | 0.941656 | 0.902875 | 0.366592 | NA |
| CFL1P6 | 0.151697 | -1.546 | 4.075469 | -0.37934 | 0.704433 | NA |
| TNFRSF25 | 3.256529 | 1.793141 | 1.342954 | 1.335222 | 0.181804 | NA |
| SLC35E2 | 77.50763 | -0.3794 | 0.27862 | -1.3617 | 0.173294 | 0.325011 |
| | 0.450981 | 2.057325 | 3.353149 | 0.61355 | 0.539513 | NA |
| | 0.162376 | 0.424913 | 4.079006 | 0.104171 | 0.917034 | NA |
| BTNL10 | 0.555087 | 0.973802 | 2.808575 | 0.346725 | 0.728798 | NA |
| ZC3H11B | 15.60696 | 0.290563 | 0.54722 | 0.53098 | 0.595433 | NA |
| QRSL1P1 | 0.151697 | -1.546 | 4.075469 | -0.37934 | 0.704433 | NA |
| | 4.788277 | -1.02052 | 0.938194 | -1.08775 | 0.276705 | NA |
| | 0.844077 | -0.22325 | 2.189502 | -0.10196 | 0.918786 | NA |
| TSTD1 | 26.49565 | -0.51701 | 0.516145 | -1.00168 | 0.3165 | NA |
| | 10.56221 | 0.358211 | 0.7039 | 0.508895 | 0.610826 | NA |
| LINC01138 | 1.932835 | -0.51746 | 1.539434 | -0.33614 | 0.736767 | NA |
| NBPF7 | 0.460466 | 0.568655 | 3.014642 | 0.188631 | 0.850382 | NA |
| | 0.870886 | -0.53073 | 2.310896 | -0.22966 | 0.818353 | NA |
| KRT18P57 | 0.555233 | -0.57009 | 3.296947 | -0.17291 | 0.862719 | NA |
| CYB5RL | 126.4802 | -0.15194 | 0.226398 | -0.67114 | 0.502132 | 0.652287 |
| | 0.163957 | 0.424913 | 4.079006 | 0.104171 | 0.917034 | NA |
| | 2.029037 | -0.33495 | 1.388259 | -0.24128 | 0.809341 | NA |
| | 0.392711 | 0.292538 | 3.145221 | 0.09301 | 0.925895 | NA |
| SEPW1P | 0.247898 | -0.5901 | 4.074633 | -0.14482 | 0.884851 | NA |
| | 4.072088 | 0.36099 | 1.092158 | 0.330529 | 0.741 | NA |
| CROCCP2 | 380.783 | 0.228493 | 0.190562 | 1.199046 | 0.23051 | 0.392685 |
| C1orf167 | 0.162376 | 0.424913 | 4.079006 | 0.104171 | 0.917034 | NA |
| TTC34 | 1.902276 | 0.990844 | 1.560755 | 0.634849 | 0.525527 | NA |
| MMP23A | 0.556813 | -0.57007 | 3.294615 | -0.17303 | 0.862627 | NA |
| ATAD3C | 0.667861 | -2.0782 | 2.593993 | -0.80116 | 0.42304 | NA |
| | 0.720827 | 0.448488 | 2.540859 | 0.17651 | 0.859893 | NA |
| ANXA2P3 | 0.926585 | -2.43791 | 2.203722 | -1.10627 | 0.268609 | NA |
| CCDC7 | 25.52158 | 0.991193 | 0.445721 | 2.223794 | 0.026162 | NA |
| PPIAP31 | 1.39931 | -0.43358 | 1.630816 | -0.26587 | 0.790343 | NA |
| | 0.847947 | -3.45038 | 2.752525 | -1.25353 | 0.210012 | NA |

| | | | | | | |
|---|---|---|---|---|---|---|
| | 13.8515 | 0.28418 | 0.592117 | 0.479939 | 0.631271 | NA |
| MORF4L1P | 265.3204 | 0.289107 | 0.179755 | 1.608339 | 0.107761 | 0.234362 |
| LINC00339 | 325.4586 | -0.29595 | 0.190433 | -1.55412 | 0.120157 | 0.25234 |
| CELA3B | 0.162376 | 0.424913 | 4.079006 | 0.104171 | 0.917034 | NA |
| HNRNPA3P | 3.627277 | -0.64646 | 1.032318 | -0.62623 | 0.531167 | NA |
| | 0.476449 | 0.524262 | 3.402674 | 0.154073 | 0.877552 | NA |
| | 2.087307 | 0.208675 | 1.354452 | 0.154066 | 0.877558 | NA |
| MBL3P | 0.163957 | 0.424913 | 4.079006 | 0.104171 | 0.917034 | NA |
| NBPF1 | 612.5897 | 0.028789 | 0.204837 | 0.140547 | 0.888228 | 0.934251 |
| RPL12P8 | 0.096202 | 0.408546 | 4.079639 | 0.100143 | 0.920231 | NA |
| HIST2H2BD | 1.9781 | 0.647328 | 1.543328 | 0.419436 | 0.674897 | NA |
| DDX18P6 | 0.162376 | 0.424913 | 4.079006 | 0.104171 | 0.917034 | NA |
| RPL21P28 | 4.235679 | -0.32806 | 0.990553 | -0.33119 | 0.740499 | NA |
| MTMR9LP | 1848.017 | 0.593803 | 0.293174 | 2.025429 | 0.042823 | 0.125942 |
| | 18.57263 | 1.072852 | 0.537152 | 1.997297 | 0.045793 | NA |
| | 0.162376 | 0.424913 | 4.079006 | 0.104171 | 0.917034 | NA |
| | 4.173511 | 0.553341 | 1.024973 | 0.539859 | 0.589294 | NA |
| MIR1254-1 | 0.488709 | 2.119056 | 3.307564 | 0.64067 | 0.521737 | NA |
| RNU6ATAC | 0.247898 | -0.5901 | 4.074633 | -0.14482 | 0.884851 | NA |
| | 0.162376 | 0.424913 | 4.079006 | 0.104171 | 0.917034 | NA |
| RNU6ATAC | 0.268266 | -2.15396 | 4.038351 | -0.53338 | 0.593773 | NA |
| | 0.162376 | 0.424913 | 4.079006 | 0.104171 | 0.917034 | NA |
| MIR320B2 | 0.151697 | -1.546 | 4.075469 | -0.37934 | 0.704433 | NA |
| SNORD99 | 4.43424 | 1.064895 | 0.983699 | 1.082541 | 0.279012 | NA |
| SNORA77 | 0.382031 | -1.26484 | 3.151989 | -0.40128 | 0.688212 | NA |
| MIR1290 | 0.134133 | -1.48909 | 4.077584 | -0.36519 | 0.71497 | NA |
| | 16.03305 | 0.380778 | 0.564789 | 0.674197 | 0.500186 | NA |
| C1orf229 | 1.637374 | 1.223851 | 1.56921 | 0.779915 | 0.435441 | NA |
| FADS3 | 1349.321 | -0.44744 | 0.181576 | -2.46422 | 0.013731 | 0.058409 |
| CCNL2 | 1255.425 | 0.635851 | 0.155958 | 4.077072 | 4.56E-05 | 0.001558 |
| MYBPHL | 0.668852 | 1.246964 | 2.610519 | 0.477669 | 0.632886 | NA |
| BTBD19 | 183.3659 | 0.352466 | 0.223434 | 1.577493 | 0.114682 | 0.244173 |
| DCDC2B | 0.880631 | -1.73238 | 2.414456 | -0.7175 | 0.473064 | NA |
| RNU6-1165 | 0.162376 | 0.424913 | 4.079006 | 0.104171 | 0.917034 | NA |
| MIR1915 | 0.28583 | -2.21358 | 3.97376 | -0.55705 | 0.577494 | NA |
| RNA5SP317 | 0.420954 | 1.923376 | 3.462803 | 0.555439 | 0.578594 | NA |
| RN7SKP16 | 1.752566 | 0.483454 | 1.698619 | 0.284616 | 0.775939 | NA |
| SNORA36B | 0.134133 | -1.48909 | 4.077584 | -0.36519 | 0.71497 | NA |
| RNU2-59P | 1.66072 | 1.41188 | 1.770174 | 0.797594 | 0.425106 | NA |
| RNA5SP338 | 0.151697 | -1.546 | 4.075469 | -0.37934 | 0.704433 | NA |
| | 0.162376 | 0.424913 | 4.079006 | 0.104171 | 0.917034 | NA |
| RNU2-42P | 0.162376 | 0.424913 | 4.079006 | 0.104171 | 0.917034 | NA |
| RNU2-70P | 0.096202 | 0.408546 | 4.079639 | 0.100143 | 0.920231 | NA |
| RNU2-38P | 0.096202 | 0.408546 | 4.079639 | 0.100143 | 0.920231 | NA |
| MIR1537 | 0.163957 | 0.424913 | 4.079006 | 0.104171 | 0.917034 | NA |
| RNU6-1245 | 0.163957 | 0.424913 | 4.079006 | 0.104171 | 0.917034 | NA |
| RNA5SP304 | 0.78443 | -2.34592 | 2.507874 | -0.93542 | 0.34957 | NA |

| | | | | | | |
|---|---|---|---|---|---|---|
| | 0.230334 | -0.53318 | 4.076749 | -0.13079 | 0.895945 | NA |
| RNU6-387F | 0.440301 | 0.468383 | 3.449808 | 0.135771 | 0.892003 | NA |
| HIST2H2BA | 1.160722 | 2.193377 | 1.991761 | 1.101225 | 0.270799 | NA |
| | 1.87189 | 1.070262 | 1.669165 | 0.641196 | 0.521395 | NA |
| SEC22B | 2927.984 | 0.08331 | 0.147656 | 0.564217 | 0.572607 | 0.708586 |
| | 22.75773 | 1.130747 | 0.47293 | 2.390942 | 0.016805 | NA |
| | 0.394291 | 0.298745 | 3.141711 | 0.09509 | 0.924243 | NA |
| | 4.35542 | -0.02653 | 1.002167 | -0.02647 | 0.97888 | NA |
| RPS10P7 | 20.00212 | -0.83251 | 0.462331 | -1.80067 | 0.071755 | NA |
| RPS26P15 | 1.510061 | -1.22497 | 1.912065 | -0.64065 | 0.521749 | NA |
| | 0.163957 | 0.424913 | 4.079006 | 0.104171 | 0.917034 | NA |
| | 0.327913 | 1.543154 | 4.021063 | 0.383768 | 0.701151 | NA |
| | 0.315653 | -0.57305 | 3.892639 | -0.14721 | 0.882963 | NA |
| | 11.71504 | 0.027107 | 0.619534 | 0.043754 | 0.965101 | NA |
| NUTM2A-A | 267.149 | 0.169905 | 0.179667 | 0.945668 | 0.344318 | 0.512394 |
| | 0.638825 | 1.117948 | 3.106359 | 0.35989 | 0.718929 | NA |
| | 0.162376 | 0.424913 | 4.079006 | 0.104171 | 0.917034 | NA |
| KIF28P | 1.616593 | 1.302318 | 1.799016 | 0.723906 | 0.469124 | NA |
| | 0.096202 | 0.408546 | 4.079639 | 0.100143 | 0.920231 | NA |
| RPS15AP5 | 0.288605 | 1.443619 | 4.029058 | 0.358302 | 0.720117 | NA |
| RBMX2P3 | 0.163957 | 0.424913 | 4.079006 | 0.104171 | 0.917034 | NA |
| | 1.418308 | 0.350852 | 1.643553 | 0.213472 | 0.830959 | NA |
| | 1.005222 | -1.76557 | 2.135341 | -0.82683 | 0.408331 | NA |
| HBD | 0.392857 | -1.546 | 4.075469 | -0.37934 | 0.704433 | NA |
| | 2.88101 | 0.391059 | 1.222954 | 0.319766 | 0.749146 | NA |
| | 0.163957 | 0.424913 | 4.079006 | 0.104171 | 0.917034 | NA |
| | 0.151697 | -1.546 | 4.075469 | -0.37934 | 0.704433 | NA |
| | 0.327913 | 1.543154 | 4.021063 | 0.383768 | 0.701151 | NA |
| | 0.151697 | -1.546 | 4.075469 | -0.37934 | 0.704433 | NA |
| ADH5P3 | 0.163957 | 0.424913 | 4.079006 | 0.104171 | 0.917034 | NA |
| | 33.30528 | 0.375278 | 0.389871 | 0.962569 | 0.335764 | NA |
| TSSC2 | 0.258578 | 1.233151 | 4.047809 | 0.304646 | 0.760635 | NA |
| | 13.3444 | -0.3589 | 0.559822 | -0.6411 | 0.521455 | NA |
| | 1.143158 | 2.223222 | 2.00111 | 1.110994 | 0.266571 | NA |
| | 0.162376 | 0.424913 | 4.079006 | 0.104171 | 0.917034 | NA |
| LGALS8-AS | 2.056195 | -1.8206 | 1.65918 | -1.09729 | 0.272516 | NA |
| | 3.498731 | 0.553286 | 1.057234 | 0.523334 | 0.600742 | NA |
| | 30.71966 | 0.078972 | 0.374107 | 0.211096 | 0.832813 | NA |
| | 0.452561 | 2.061505 | 3.349456 | 0.615474 | 0.538241 | NA |
| CFL1P1 | 24.71598 | 0.689798 | 0.486117 | 1.418994 | 0.155901 | NA |
| | 0.410275 | 0.244283 | 3.100889 | 0.078778 | 0.937209 | NA |
| | 0.151697 | -1.546 | 4.075469 | -0.37934 | 0.704433 | NA |
| CCNJP2 | 0.096202 | 0.408546 | 4.079639 | 0.100143 | 0.920231 | NA |
| | 2.682109 | -0.11615 | 1.162354 | -0.09992 | 0.920406 | NA |
| | 12.77887 | 0.746213 | 0.638105 | 1.169421 | 0.242234 | NA |
| SNRPD2P2 | 0.162376 | 0.424913 | 4.079006 | 0.104171 | 0.917034 | NA |
| HNRNPRP1 | 0.437526 | -2.68242 | 3.384147 | -0.79264 | 0.427986 | NA |

| | | | | | | |
|---|---|---|---|---|---|---|
| | 1.010809 | 2.054089 | 2.306032 | 0.890746 | 0.373065 | NA |
| | 0.303394 | -2.27006 | 4.032451 | -0.56295 | 0.57347 | NA |
| | 3.407031 | 0.756828 | 1.145738 | 0.660559 | 0.508895 | NA |
| | 0.247898 | -0.5901 | 4.074633 | -0.14482 | 0.884851 | NA |
| HMGN1P4 | 0.163957 | 0.424913 | 4.079006 | 0.104171 | 0.917034 | NA |
| GLTPD1 | 977.8179 | -0.33107 | 0.165614 | -1.99905 | 0.045603 | 0.130744 |
| | 0.616518 | 2.49285 | 2.980788 | 0.836306 | 0.402983 | NA |
| LINC01057 | 33.98436 | -0.46675 | 0.387044 | -1.20594 | 0.22784 | NA |
| | 0.938495 | 0.819106 | 2.155823 | 0.379951 | 0.703982 | NA |
| | 663.3293 | 0.096117 | 0.195926 | 0.49058 | 0.623724 | 0.7478 |
| | 0.096202 | 0.408546 | 4.079639 | 0.100143 | 0.920231 | NA |
| | 0.151697 | -1.546 | 4.075469 | -0.37934 | 0.704433 | NA |
| SDHDP6 | 1.890309 | -0.88832 | 2.005201 | -0.44301 | 0.657761 | NA |
| | 1.501019 | -1.35042 | 1.798237 | -0.75097 | 0.45267 | NA |
| LINC00466 | 7.660573 | -0.03391 | 0.743278 | -0.04562 | 0.963611 | NA |
| | 11.37874 | -0.28913 | 0.61269 | -0.47191 | 0.636993 | NA |
| WARS2-IT1 | 0.162376 | 0.424913 | 4.079006 | 0.104171 | 0.917034 | NA |
| | 0.260158 | 1.240347 | 4.047124 | 0.306476 | 0.759242 | NA |
| PDCL3P6 | 0.134133 | -1.48909 | 4.077584 | -0.36519 | 0.71497 | NA |
| | 37.58193 | 1.003878 | 0.466064 | 2.153949 | 0.031244 | NA |
| | 1.290906 | -0.9649 | 1.844052 | -0.52325 | 0.600799 | NA |
| IFIT6P | 0.096202 | 0.408546 | 4.079639 | 0.100143 | 0.920231 | NA |
| | 0.654246 | 2.541085 | 3.009764 | 0.84428 | 0.398513 | NA |
| | 0.247898 | -0.5901 | 4.074633 | -0.14482 | 0.884851 | NA |
| RPL7P7 | 0.327913 | 1.543154 | 4.021063 | 0.383768 | 0.701151 | NA |
| | 0.324752 | 1.53082 | 4.022025 | 0.380609 | 0.703493 | NA |
| | 2.659023 | -0.34535 | 1.242982 | -0.27784 | 0.781136 | NA |
| | 0.296509 | -0.51709 | 3.982155 | -0.12985 | 0.896684 | NA |
| LINC00703 | 1.505106 | -2.35873 | 2.214667 | -1.06505 | 0.286854 | NA |
| | 0.162376 | 0.424913 | 4.079006 | 0.104171 | 0.917034 | NA |
| | 6.851224 | 0.72748 | 0.858963 | 0.846928 | 0.397035 | NA |
| | 0.151697 | -1.546 | 4.075469 | -0.37934 | 0.704433 | NA |
| | 3.484291 | 0.586232 | 1.126666 | 0.520325 | 0.602837 | NA |
| | 1.39478 | 0.860979 | 1.847866 | 0.465931 | 0.641265 | NA |
| | 0.296509 | -0.51709 | 3.982155 | -0.12985 | 0.896684 | NA |
| HNRNPA3P | 0.162376 | 0.424913 | 4.079006 | 0.104171 | 0.917034 | NA |
| KRT8P45 | 0.75318 | -2.10401 | 2.55086 | -0.82482 | 0.409472 | NA |
| | 0.163957 | 0.424913 | 4.079006 | 0.104171 | 0.917034 | NA |
| | 3.544459 | -0.77103 | 1.091738 | -0.70624 | 0.480037 | NA |
| | 1.804695 | 0.942614 | 1.666796 | 0.565525 | 0.571717 | NA |
| FAM96AP2 | 1.044588 | -1.04753 | 2.032811 | -0.51531 | 0.606337 | NA |
| | 0.326536 | 0.037056 | 3.842554 | 0.009644 | 0.992306 | NA |
| ZMIZ1-AS1 | 2.893024 | -0.88112 | 1.188539 | -0.74134 | 0.458485 | NA |
| PTCHD3P1 | 1465.686 | 0.112515 | 0.162178 | 0.693773 | 0.487825 | 0.640674 |
| | 0.258578 | 1.233151 | 4.047809 | 0.304646 | 0.760635 | NA |
| | 13.18016 | 0.408459 | 0.589154 | 0.693298 | 0.488122 | NA |
| | 0.586491 | 2.39072 | 2.689057 | 0.889055 | 0.373973 | NA |

| | | | | | | |
|---|---|---|---|---|---|---|
| IPO8P1 | 3.812015 | -0.35929 | 1.014609 | -0.35411 | 0.723253 | NA |
| PLA2G12AI | 3.001557 | 0.520879 | 1.164306 | 0.447373 | 0.654606 | NA |
| RASAL2-AS | 15.41322 | 0.027377 | 0.586325 | 0.046693 | 0.962758 | NA |
| UBE2D3P3 | 4.274222 | 1.573184 | 1.139664 | 1.380393 | 0.167466 | NA |
| | 0.382031 | -1.26484 | 3.151989 | -0.40128 | 0.688212 | NA |
| LAMTOR5-/ | 2.096406 | 0.696389 | 1.379677 | 0.504748 | 0.613736 | NA |
| CEP57L1P1 | 0.163957 | 0.424913 | 4.079006 | 0.104171 | 0.917034 | NA |
| FCF1P7 | 0.488709 | 2.119056 | 3.307564 | 0.64067 | 0.521737 | NA |
| | 0.096202 | 0.408546 | 4.079639 | 0.100143 | 0.920231 | NA |
| SHMT1P1 | 0.163957 | 0.424913 | 4.079006 | 0.104171 | 0.917034 | NA |
| FDPSP8 | 0.644964 | 2.579089 | 2.975624 | 0.866739 | 0.386085 | NA |
| | 0.392857 | -1.546 | 4.075469 | -0.37934 | 0.704433 | NA |
| LINC00853 | 0.940076 | 0.824756 | 2.188343 | 0.376886 | 0.706258 | NA |
| | 0.437526 | -2.68242 | 3.384147 | -0.79264 | 0.427986 | NA |
| TMEM72-A | 0.967145 | 0.230519 | 2.075373 | 0.111074 | 0.911558 | NA |
| | 255.4012 | -0.14614 | 0.240197 | -0.60841 | 0.542916 | 0.684928 |
| GCSHP5 | 0.35636 | 1.709252 | 3.651833 | 0.468053 | 0.639746 | NA |
| LINC00502 | 0.096202 | 0.408546 | 4.079639 | 0.100143 | 0.920231 | NA |
| | 457.499 | -0.0647 | 0.15377 | -0.42078 | 0.673918 | 0.786547 |
| BEND3P3 | 5.434059 | -0.35757 | 0.857096 | -0.41719 | 0.67654 | NA |
| | 0.46577 | -1.30191 | 3.402879 | -0.38259 | 0.702024 | NA |
| LINC00863 | 33.87422 | -0.05045 | 0.38866 | -0.12982 | 0.896712 | NA |
| | 3.79911 | -0.05293 | 1.020515 | -0.05187 | 0.958635 | NA |
| LINC00184 | 0.327913 | 1.543154 | 4.021063 | 0.383768 | 0.701151 | NA |
| | 7.10184 | 0.904764 | 1.014074 | 0.892207 | 0.372282 | NA |
| | 1.790798 | 0.241408 | 1.584876 | 0.15232 | 0.878935 | NA |
| KCNC4-AS1 | 0.653015 | 0.2552 | 2.787103 | 0.091565 | 0.927044 | NA |
| | 0.422534 | 1.924583 | 3.001758 | 0.641152 | 0.521424 | NA |
| | 0.528424 | -0.38142 | 2.853651 | -0.13366 | 0.893671 | NA |
| | 0.315653 | -0.57305 | 3.892639 | -0.14721 | 0.882963 | NA |
| PPP1R8P1 | 0.096202 | 0.408546 | 4.079639 | 0.100143 | 0.920231 | NA |
| | 0.258578 | 1.233151 | 4.047809 | 0.304646 | 0.760635 | NA |
| SNX18P15 | 0.298089 | -0.51711 | 3.975232 | -0.13008 | 0.896501 | NA |
| | 0.134133 | -1.48909 | 4.077584 | -0.36519 | 0.71497 | NA |
| | 0.586491 | 2.39072 | 2.689057 | 0.889055 | 0.373973 | NA |
| | 0.654449 | 1.296029 | 2.63083 | 0.492631 | 0.622273 | NA |
| LINC00337 | 9.954373 | -0.62895 | 0.639753 | -0.98312 | 0.325549 | NA |
| DAP3P1 | 0.392857 | -1.546 | 4.075469 | -0.37934 | 0.704433 | NA |
| ATP6V1E1I | 1.460955 | 0.943975 | 1.856661 | 0.508426 | 0.611155 | NA |
| | 3.011174 | -0.98124 | 1.122724 | -0.87398 | 0.382128 | NA |
| LINC00999 | 22.80794 | -0.30621 | 0.463868 | -0.66012 | 0.509178 | NA |
| | 0.096202 | 0.408546 | 4.079639 | 0.100143 | 0.920231 | NA |
| DYNC1I2P1 | 7.241836 | 0.424388 | 0.837642 | 0.506645 | 0.612404 | NA |
| | 7.871297 | -0.71989 | 0.725942 | -0.99166 | 0.321363 | NA |
| NPM1P39 | 11.89634 | 0.548333 | 0.619927 | 0.884513 | 0.376419 | NA |
| BRI3P1 | 0.288605 | 1.443619 | 4.029058 | 0.358302 | 0.720117 | NA |
| DUTP6 | 0.288605 | 1.443619 | 4.029058 | 0.358302 | 0.720117 | NA |

| | | | | | | |
|---|---|---|---|---|---|---|
| | 0.260158 | 1.240347 | 4.047124 | 0.306476 | 0.759242 | NA |
| ZNF33BP1 | 0.096202 | 0.408546 | 4.079639 | 0.100143 | 0.920231 | NA |
| MIR137HG | 16.56734 | 0.141242 | 0.560323 | 0.252073 | 0.800985 | NA |
| | 0.364467 | -1.20166 | 3.685431 | -0.32606 | 0.744381 | NA |
| HSPA7 | 0.151697 | -1.546 | 4.075469 | -0.37934 | 0.704433 | NA |
| | 181.5037 | 0.10629 | 0.217356 | 0.489012 | 0.624833 | 0.74847 |
| | 36.41408 | 0.304915 | 0.364493 | 0.836546 | 0.402848 | NA |
| LINC00705 | 0.096202 | 0.408546 | 4.079639 | 0.100143 | 0.920231 | NA |
| | 0.419962 | -2.64111 | 3.429364 | -0.77014 | 0.441214 | NA |
| | 0.46735 | -1.30193 | 3.399671 | -0.38296 | 0.701751 | NA |
| | 0.354779 | 1.703954 | 3.657232 | 0.465914 | 0.641277 | NA |
| OSTCP2 | 0.162376 | 0.424913 | 4.079006 | 0.104171 | 0.917034 | NA |
| | 5.949922 | -0.52171 | 0.863257 | -0.60435 | 0.545613 | NA |
| | 13.13028 | 0.08415 | 0.642721 | 0.130928 | 0.895832 | NA |
| | 0.324752 | 1.53082 | 4.022025 | 0.380609 | 0.703493 | NA |
| | 2.126672 | 0.355758 | 1.367755 | 0.260104 | 0.794784 | NA |
| ARL6IP1P2 | 0.792452 | -2.15617 | 2.562821 | -0.84133 | 0.400166 | NA |
| SFTA1P | 0.163957 | 0.424913 | 4.079006 | 0.104171 | 0.917034 | NA |
| TGIF2P1 | 0.192403 | 0.975392 | 4.060716 | 0.240202 | 0.810174 | NA |
| TNPO1P1 | 0.900767 | 0.808531 | 1.963257 | 0.411831 | 0.680463 | NA |
| | 0.096202 | 0.408546 | 4.079639 | 0.100143 | 0.920231 | NA |
| RPS15AP1( | 0.765054 | 1.506026 | 2.496832 | 0.603175 | 0.546392 | NA |
| FDPSP1 | 0.382031 | -1.26484 | 3.151989 | -0.40128 | 0.688212 | NA |
| | 1.393935 | -2.09637 | 1.832113 | -1.14424 | 0.252526 | NA |
| | 77.19094 | -0.05395 | 0.292881 | -0.1842 | 0.853853 | 0.913247 |
| | 0.247898 | -0.5901 | 4.074633 | -0.14482 | 0.884851 | NA |
| GBP1P1 | 0.975407 | 0.039022 | 2.314857 | 0.016857 | 0.98655 | NA |
| | 0.260158 | 1.240347 | 4.047124 | 0.306476 | 0.759242 | NA |
| CYP4A22-A | 0.151697 | -1.546 | 4.075469 | -0.37934 | 0.704433 | NA |
| | 3.294766 | -1.1324 | 1.200938 | -0.94293 | 0.345719 | NA |
| | 0.258578 | 1.233151 | 4.047809 | 0.304646 | 0.760635 | NA |
| | 0.717609 | 0.516981 | 3.063469 | 0.168757 | 0.865988 | NA |
| | 0.555233 | -0.57009 | 3.296947 | -0.17291 | 0.862719 | NA |
| C2CD4D | 12.98362 | -0.29053 | 0.7108 | -0.40873 | 0.682735 | NA |
| | 0.58526 | 0.001197 | 3.235889 | 3.70E-04 | 0.999705 | NA |
| | 0.651288 | 1.289307 | 2.633717 | 0.489539 | 0.62446 | NA |
| MTOR-AS1 | 0.134133 | -1.48909 | 4.077584 | -0.36519 | 0.71497 | NA |
| | 0.45509 | -2.71227 | 3.963744 | -0.68427 | 0.493805 | NA |
| | 45.44657 | 0.483869 | 0.358459 | 1.349857 | 0.177062 | NA |
| AGBL4-IT1 | 0.162376 | 0.424913 | 4.079006 | 0.104171 | 0.917034 | NA |
| MTND2P28 | 9.535166 | -0.87784 | 0.6702 | -1.30981 | 0.19026 | NA |
| | 5.095411 | 0.128713 | 0.999332 | 0.128799 | 0.897517 | NA |
| | 0.230334 | -0.53318 | 4.076749 | -0.13079 | 0.895945 | NA |
| | 2.626467 | 0.572296 | 1.252999 | 0.456741 | 0.647857 | NA |
| | 0.163957 | 0.424913 | 4.079006 | 0.104171 | 0.917034 | NA |
| | 0.096202 | 0.408546 | 4.079639 | 0.100143 | 0.920231 | NA |
| | 0.392857 | -1.546 | 4.075469 | -0.37934 | 0.704433 | NA |

| | | | | | | |
|---|---|---|---|---|---|---|
| ATP5G2P1 | 0.819558 | -2.40918 | 2.432038 | -0.9906 | 0.321881 | NA |
| | 1.553782 | 0.573128 | 1.572738 | 0.364414 | 0.715549 | NA |
| NRG3-AS1 | 0.315653 | -0.57305 | 3.892639 | -0.14721 | 0.882963 | NA |
| | 0.096202 | 0.408546 | 4.079639 | 0.100143 | 0.920231 | NA |
| | 4.89692 | -0.39774 | 0.986816 | -0.40306 | 0.686905 | NA |
| | 0.192403 | 0.975392 | 4.060716 | 0.240202 | 0.810174 | NA |
| | 31.9586 | -0.42954 | 0.603782 | -0.71142 | 0.476825 | NA |
| PROSER2-A | 2.115985 | -1.28611 | 1.367822 | -0.94026 | 0.347083 | NA |
| | 0.134133 | -1.48909 | 4.077584 | -0.36519 | 0.71497 | NA |
| | 16.5199 | 0.520484 | 0.509312 | 1.021934 | 0.306812 | NA |
| LINC00626 | 0.163957 | 0.424913 | 4.079006 | 0.104171 | 0.917034 | NA |
| FAM229A | 35.87037 | 0.46041 | 0.415352 | 1.10848 | 0.267654 | NA |
| ERCC6 | 508.6122 | 0.799403 | 0.167017 | 4.786367 | 1.70E-06 | 2.24E-04 |
| | 1.040001 | 0.161297 | 2.078712 | 0.077595 | 0.938151 | NA |
| RUSC1-AS1 | 43.53812 | 0.068076 | 0.326965 | 0.208206 | 0.835068 | NA |
| | 0.163957 | 0.424913 | 4.079006 | 0.104171 | 0.917034 | NA |
| LINC00115 | 5.794556 | -0.11904 | 0.893969 | -0.13316 | 0.89407 | NA |
| | 7.527701 | -0.86029 | 0.779226 | -1.10403 | 0.269582 | NA |
| | 0.508056 | 0.731032 | 2.861912 | 0.255435 | 0.798387 | NA |
| | 0.324752 | 1.53082 | 4.022025 | 0.380609 | 0.703493 | NA |
| RIMKLBP2 | 1.625501 | -0.42008 | 1.608465 | -0.26117 | 0.793964 | NA |
| | 4.211125 | -1.55004 | 1.004879 | -1.54251 | 0.122949 | NA |
| MTND1P23 | 4001.098 | -0.32659 | 0.163256 | -2.00051 | 0.045445 | 0.130606 |
| HMGB1P26 | 0.192403 | 0.975392 | 4.060716 | 0.240202 | 0.810174 | NA |
| | 0.288605 | 1.443619 | 4.029058 | 0.358302 | 0.720117 | NA |
| KCNIP2-AS1 | 0.45509 | -2.71227 | 3.963744 | -0.68427 | 0.493805 | NA |
| CCT8P1 | 18.61826 | 0.679978 | 0.501842 | 1.354964 | 0.175429 | NA |
| | 3.134613 | 0.224642 | 1.137984 | 0.197403 | 0.843512 | NA |
| | 11.03865 | 0.960513 | 0.682043 | 1.408288 | 0.159046 | NA |
| ZNF503-AS | 1.811425 | -0.28142 | 1.691423 | -0.16638 | 0.867858 | NA |
| LINC00869 | 20.85342 | -0.15593 | 0.462084 | -0.33744 | 0.735782 | NA |
| SLC39A12-A | 0.162376 | 0.424913 | 4.079006 | 0.104171 | 0.917034 | NA |
| | 15.06922 | 0.834198 | 0.574591 | 1.451811 | 0.146554 | NA |
| EIF3LP3 | 0.555087 | 0.973802 | 2.808575 | 0.346725 | 0.728798 | NA |
| | 1.957487 | -1.47079 | 1.568454 | -0.93773 | 0.348383 | NA |
| PSMC1P12 | 0.162376 | 0.424913 | 4.079006 | 0.104171 | 0.917034 | NA |
| SLC25A39P | 0.163957 | 0.424913 | 4.079006 | 0.104171 | 0.917034 | NA |
| AP4B1-AS1 | 4.758293 | 0.134486 | 0.948652 | 0.141765 | 0.887266 | NA |
| | 7.089677 | 0.114626 | 0.748586 | 0.153123 | 0.878301 | NA |
| | 0.096202 | 0.408546 | 4.079639 | 0.100143 | 0.920231 | NA |
| | 0.268266 | -2.15396 | 4.038351 | -0.53338 | 0.593773 | NA |
| LEMD1-AS1 | 0.163957 | 0.424913 | 4.079006 | 0.104171 | 0.917034 | NA |
| ZNF32-AS1 | 1.728418 | 1.57973 | 1.632683 | 0.967567 | 0.333261 | NA |
| | 11.59615 | 0.337155 | 0.786766 | 0.428532 | 0.668264 | NA |
| | 0.392857 | -1.546 | 4.075469 | -0.37934 | 0.704433 | NA |
| | 1.255896 | 0.001954 | 2.021155 | 9.67E-04 | 0.999229 | NA |
| | 0.672013 | 1.253691 | 2.607776 | 0.480751 | 0.630693 | NA |

| | | | | | | |
|---|---|---|---|---|---|---|
| | 14.1121 | -1.25852 | 0.564433 | -2.2297 | 0.025767 | NA |
| | 0.163957 | 0.424913 | 4.079006 | 0.104171 | 0.917034 | NA |
| | 48.31458 | -0.06023 | 0.314783 | -0.19133 | 0.84827 | 0.909573 |
| PARD3-AS1 | 1.556514 | -1.92657 | 1.793358 | -1.07428 | 0.282698 | NA |
| RPS14P3 | 13.77757 | 0.712902 | 0.650113 | 1.096582 | 0.272824 | NA |
| RPL23P11 | 0.327913 | 1.543154 | 4.021063 | 0.383768 | 0.701151 | NA |
| TPI1P1 | 70.78455 | -0.13723 | 0.284748 | -0.48195 | 0.629841 | 0.752244 |
| SLC16A1-A | 30.72499 | 0.531216 | 0.41044 | 1.294258 | 0.195576 | NA |
| | 0.815245 | 1.670886 | 2.214146 | 0.754641 | 0.450464 | NA |
| | 0.432222 | -1.18916 | 3.482045 | -0.34151 | 0.732718 | NA |
| | 0.096202 | 0.408546 | 4.079639 | 0.100143 | 0.920231 | NA |
| GAPDHP32 | 0.303394 | -2.27006 | 4.032451 | -0.56295 | 0.57347 | NA |
| NOTCH2P1 | 0.163957 | 0.424913 | 4.079006 | 0.104171 | 0.917034 | NA |
| | 0.392857 | -1.546 | 4.075469 | -0.37934 | 0.704433 | NA |
| | 1.139355 | -2.01403 | 2.05367 | -0.9807 | 0.326743 | NA |
| RPA2P2 | 0.096202 | 0.408546 | 4.079639 | 0.100143 | 0.920231 | NA |
| | 0.288605 | 1.443619 | 4.029058 | 0.358302 | 0.720117 | NA |
| | 0.298089 | -0.51711 | 3.975232 | -0.13008 | 0.896501 | NA |
| | 1.207956 | 1.431596 | 1.927393 | 0.742763 | 0.457625 | NA |
| KIRREL-IT1 | 0.556667 | 0.972496 | 2.522454 | 0.385536 | 0.699841 | NA |
| | 0.096202 | 0.408546 | 4.079639 | 0.100143 | 0.920231 | NA |
| | 0.192403 | 0.975392 | 4.060716 | 0.240202 | 0.810174 | NA |
| TRAF6P1 | 0.392857 | -1.546 | 4.075469 | -0.37934 | 0.704433 | NA |
| | 0.872321 | 0.678609 | 2.011629 | 0.337343 | 0.735858 | NA |
| | 1.476193 | 2.630491 | 1.892307 | 1.390097 | 0.164499 | NA |
| RPL7P37 | 0.47803 | 0.520083 | 2.97881 | 0.174594 | 0.861399 | NA |
| | 1.415497 | -0.96742 | 1.679672 | -0.57596 | 0.564643 | NA |
| | 0.247898 | -0.5901 | 4.074633 | -0.14482 | 0.884851 | NA |
| ENTPD1-AS | 35.88345 | 0.03833 | 0.38358 | 0.099926 | 0.920403 | NA |
| | 37.34339 | -0.48176 | 0.372349 | -1.29384 | 0.19572 | NA |
| | 0.458885 | 0.577525 | 3.449278 | 0.167433 | 0.867029 | NA |
| | 1.18812 | -0.45987 | 2.292512 | -0.2006 | 0.841014 | NA |
| | 0.163957 | 0.424913 | 4.079006 | 0.104171 | 0.917034 | NA |
| | 1.313991 | -0.53321 | 1.770244 | -0.30121 | 0.763258 | NA |
| | 0.957843 | 0.061519 | 2.068681 | 0.029738 | 0.976276 | NA |
| | 20.01975 | -0.21168 | 0.487088 | -0.43457 | 0.663872 | NA |
| DISC1-IT1 | 1.121588 | -1.32591 | 2.108778 | -0.62876 | 0.529507 | NA |
| FABP7P1 | 0.096202 | 0.408546 | 4.079639 | 0.100143 | 0.920231 | NA |
| HNRNPA3P | 0.410275 | 0.244283 | 3.100889 | 0.078778 | 0.937209 | NA |
| LINC00840 | 0.162376 | 0.424913 | 4.079006 | 0.104171 | 0.917034 | NA |
| | 0.163957 | 0.424913 | 4.079006 | 0.104171 | 0.917034 | NA |
| | 0.134133 | -1.48909 | 4.077584 | -0.36519 | 0.71497 | NA |
| | 0.163957 | 0.424913 | 4.079006 | 0.104171 | 0.917034 | NA |
| | 1.045872 | -1.31674 | 2.456371 | -0.53605 | 0.591923 | NA |
| | 5.278252 | 0.290087 | 0.867834 | 0.334266 | 0.738179 | NA |
| | 2.019755 | -0.21719 | 1.385839 | -0.15672 | 0.875466 | NA |
| | 0.163957 | 0.424913 | 4.079006 | 0.104171 | 0.917034 | NA |

| | | | | | | |
|---|---|---|---|---|---|---|
| ATE1-AS1 | 2.669646 | -0.19974 | 1.175537 | -0.16991 | 0.865081 | NA |
| | 2.988009 | -0.62454 | 1.180146 | -0.52921 | 0.596662 | NA |
| | 0.162376 | 0.424913 | 4.079006 | 0.104171 | 0.917034 | NA |
| | 0.096202 | 0.408546 | 4.079639 | 0.100143 | 0.920231 | NA |
| | 0.260158 | 1.240347 | 4.047124 | 0.306476 | 0.759242 | NA |
| | 11.50838 | -0.08492 | 0.631291 | -0.13452 | 0.892995 | NA |
| CEP164P1 | 0.622842 | 1.158846 | 2.735323 | 0.42366 | 0.671814 | NA |
| | 25.92463 | 0.024575 | 0.500031 | 0.049148 | 0.960802 | NA |
| | 82.70647 | -0.07966 | 0.329886 | -0.24149 | 0.809178 | 0.885851 |
| | 0.260158 | 1.240347 | 4.047124 | 0.306476 | 0.759242 | NA |
| | 0.268266 | -2.15396 | 4.038351 | -0.53338 | 0.593773 | NA |
| RHEBP2 | 0.843488 | 2.923514 | 2.365851 | 1.235713 | 0.216565 | NA |
| HAUS4P1 | 0.526843 | -0.38719 | 2.855892 | -0.13558 | 0.892156 | NA |
| | 2.384076 | 0.044259 | 1.355317 | 0.032656 | 0.973949 | NA |
| | 0.947944 | -0.02704 | 2.20277 | -0.01228 | 0.990206 | NA |
| NBPF2P | 1.465688 | -0.81787 | 1.578746 | -0.51805 | 0.604423 | NA |
| | 5.522295 | -0.27325 | 0.860357 | -0.3176 | 0.750789 | NA |
| | 0.49029 | 2.123092 | 3.304282 | 0.642527 | 0.520531 | NA |
| | 0.422534 | 1.924583 | 3.001758 | 0.641152 | 0.521424 | NA |
| | 0.162376 | 0.424913 | 4.079006 | 0.104171 | 0.917034 | NA |
| | 0.738391 | 0.402625 | 2.276191 | 0.176885 | 0.859599 | NA |
| | 2.189144 | 1.004458 | 1.585016 | 0.633721 | 0.526263 | NA |
| HMGB3P8 | 0.572651 | 0.928649 | 2.780261 | 0.334015 | 0.738368 | NA |
| | 0.134133 | -1.48909 | 4.077584 | -0.36519 | 0.71497 | NA |
| LINC00595 | 1.946346 | 1.109615 | 1.463021 | 0.75844 | 0.448187 | NA |
| | 0.748867 | 2.740193 | 2.468928 | 1.109871 | 0.267054 | NA |
| | 0.247898 | -0.5901 | 4.074633 | -0.14482 | 0.884851 | NA |
| WDR11-AS | 1.551007 | -0.72774 | 1.546985 | -0.47043 | 0.638051 | NA |
| MYL6P3 | 0.096202 | 0.408546 | 4.079639 | 0.100143 | 0.920231 | NA |
| | 1.859683 | -0.49983 | 1.84375 | -0.27109 | 0.78632 | NA |
| GLUD1P7 | 0.738188 | 1.39874 | 2.317793 | 0.603479 | 0.54619 | NA |
| PFN1P9 | 0.260158 | 1.240347 | 4.047124 | 0.306476 | 0.759242 | NA |
| RPL31P12 | 0.247898 | -0.5901 | 4.074633 | -0.14482 | 0.884851 | NA |
| PFN1P7 | 0.420954 | 1.923376 | 3.462803 | 0.555439 | 0.578594 | NA |
| | 0.411855 | 0.250456 | 3.097618 | 0.080854 | 0.935558 | NA |
| WASH7P | 35.24847 | 0.059552 | 0.368636 | 0.161547 | 0.871663 | NA |
| | 0.260158 | 1.240347 | 4.047124 | 0.306476 | 0.759242 | NA |
| | 0.303394 | -2.27006 | 4.032451 | -0.56295 | 0.57347 | NA |
| NBPF13P | 4.86153 | 0.164023 | 0.975024 | 0.168225 | 0.866406 | NA |
| LINC00845 | 0.45509 | -2.71227 | 3.963744 | -0.68427 | 0.493805 | NA |
| | 0.3441 | -0.0166 | 3.769682 | -0.0044 | 0.996486 | NA |
| KLLN | 42.81724 | -0.77581 | 0.349018 | -2.22283 | 0.026227 | NA |
| | 0.096202 | 0.408546 | 4.079639 | 0.100143 | 0.920231 | NA |
| ELL2P1 | 0.947944 | -0.02704 | 2.20277 | -0.01228 | 0.990206 | NA |
| | 0.518736 | 2.237423 | 2.7713 | 0.807355 | 0.419462 | NA |
| PARG | 237.4782 | 0.507428 | 0.218193 | 2.32559 | 0.02004 | 0.07588 |
| NANOGP6 | 0.70046 | 1.37107 | 2.594868 | 0.528378 | 0.597237 | NA |

| | | | | | | |
|---|---|---|---|---|---|---|
| LINC00866 | 0.984737 | -2.5904 | 2.366084 | -1.0948 | 0.273603 | NA |
| TP73-AS1 | 1002.105 | -0.16608 | 0.159521 | -1.04109 | 0.297835 | 0.464828 |
| | 0.47803 | 0.520083 | 2.97881 | 0.174594 | 0.861399 | NA |
| | 0.919351 | 0.885831 | 2.421371 | 0.365839 | 0.714485 | NA |
| EIF4A2P2 | 2.587248 | 0.050618 | 1.233937 | 0.041021 | 0.967279 | NA |
| ZMYM4-AS | 0.096202 | 0.408546 | 4.079639 | 0.100143 | 0.920231 | NA |
| | 0.432222 | -1.18916 | 3.482045 | -0.34151 | 0.732718 | NA |
| | 0.411855 | 0.250456 | 3.097618 | 0.080854 | 0.935558 | NA |
| | 0.258578 | 1.233151 | 4.047809 | 0.304646 | 0.760635 | NA |
| | 0.326536 | 0.037056 | 3.842554 | 0.009644 | 0.992306 | NA |
| | 0.096202 | 0.408546 | 4.079639 | 0.100143 | 0.920231 | NA |
| | 1.430735 | 0.40353 | 2.158823 | 0.186922 | 0.851722 | NA |
| | 12.25362 | 0.486641 | 0.790183 | 0.615859 | 0.537988 | NA |
| | 2.792114 | 0.320313 | 1.289509 | 0.248399 | 0.803826 | NA |
| SLC2A1-AS | 23.85525 | -0.36751 | 0.424817 | -0.8651 | 0.386984 | NA |
| | 8.863515 | 0.770768 | 0.72968 | 1.05631 | 0.290827 | NA |
| | 0.392711 | 0.292538 | 3.145221 | 0.09301 | 0.925895 | NA |
| | 0.49029 | 2.123092 | 3.304282 | 0.642527 | 0.520531 | NA |
| | 0.489058 | -0.58569 | 3.427543 | -0.17088 | 0.864321 | NA |
| PHBP11 | 0.554095 | -2.96528 | 3.108533 | -0.95392 | 0.340127 | NA |
| LINC01132 | 0.162376 | 0.424913 | 4.079006 | 0.104171 | 0.917034 | NA |
| | 0.247898 | -0.5901 | 4.074633 | -0.14482 | 0.884851 | NA |
| MIR3916 | 99.3258 | 1.358538 | 0.316157 | 4.297034 | 1.73E-05 | 8.54E-04 |
| DNMBP-AS | 0.096202 | 0.408546 | 4.079639 | 0.100143 | 0.920231 | NA |
| | 2.192608 | 0.816572 | 1.350765 | 0.604526 | 0.545494 | NA |
| | 0.096202 | 0.408546 | 4.079639 | 0.100143 | 0.920231 | NA |
| | 0.151697 | -1.546 | 4.075469 | -0.37934 | 0.704433 | NA |
| | 1.452059 | -0.1661 | 1.670582 | -0.09943 | 0.920799 | NA |
| | 7.998798 | 0.58217 | 0.786164 | 0.74052 | 0.458984 | NA |
| CALR4P | 0.572651 | 0.928649 | 2.780261 | 0.334015 | 0.738368 | NA |
| | 0.096202 | 0.408546 | 4.079639 | 0.100143 | 0.920231 | NA |
| ASH1L-IT1 | 0.096202 | 0.408546 | 4.079639 | 0.100143 | 0.920231 | NA |
| | 119.6894 | -0.382 | 0.228693 | -1.67037 | 0.094845 | 0.215442 |
| GLUD1P2 | 0.392857 | -1.546 | 4.075469 | -0.37934 | 0.704433 | NA |
| FAM212B-/ | 2.51582 | -1.54961 | 1.478323 | -1.04822 | 0.294538 | NA |
| | 1.144885 | 1.758273 | 2.389324 | 0.735887 | 0.461799 | NA |
| SEPT2P1 | 2.787702 | -0.54395 | 1.227666 | -0.44308 | 0.657711 | NA |
| | 0.788379 | 1.583268 | 2.566172 | 0.616976 | 0.53725 | NA |
| RPS26P13 | 0.621261 | 1.168072 | 3.136787 | 0.372379 | 0.709611 | NA |
| | 6.087027 | 0.844595 | 0.81341 | 1.038338 | 0.299113 | NA |
| | 0.258578 | 1.233151 | 4.047809 | 0.304646 | 0.760635 | NA |
| | 0.096202 | 0.408546 | 4.079639 | 0.100143 | 0.920231 | NA |
| | 2.504255 | -0.92795 | 1.27986 | -0.72504 | 0.468429 | NA |
| | 0.35636 | 1.709252 | 3.651833 | 0.468053 | 0.639746 | NA |
| | 3.546119 | 0.816607 | 1.090543 | 0.748808 | 0.453973 | NA |
| | 1.351894 | 0.976798 | 1.815732 | 0.537964 | 0.590602 | NA |
| | 1.142167 | -0.11411 | 2.006648 | -0.05686 | 0.954653 | NA |

| | | | | | | |
|---|---|---|---|---|---|---|
| | 2.806488 | -0.84723 | 1.267362 | -0.6685 | 0.503814 | NA |
| PRKRIRP3 | 0.163957 | 0.424913 | 4.079006 | 0.104171 | 0.917034 | NA |
| | 0.392857 | -1.546 | 4.075469 | -0.37934 | 0.704433 | NA |
| | 14.00651 | 0.279196 | 0.608414 | 0.458892 | 0.646312 | NA |
| HNRNPA1P | 0.163957 | 0.424913 | 4.079006 | 0.104171 | 0.917034 | NA |
| | 0.410275 | 0.244283 | 3.100889 | 0.078778 | 0.937209 | NA |
| | 0.585114 | 1.12675 | 2.73598 | 0.411827 | 0.680466 | NA |
| | 1.281364 | -0.71591 | 1.774413 | -0.40346 | 0.686607 | NA |
| | 0.151697 | -1.546 | 4.075469 | -0.37934 | 0.704433 | NA |
| RANP5 | 0.162376 | 0.424913 | 4.079006 | 0.104171 | 0.917034 | NA |
| CFL1P3 | 0.422737 | 0.518872 | 3.501356 | 0.148192 | 0.882191 | NA |
| SEC63P1 | 4.254832 | 0.286917 | 0.979867 | 0.292812 | 0.769666 | NA |
| | 0.716471 | -2.03238 | 2.582811 | -0.78689 | 0.431348 | NA |
| | 3.009819 | 0.446123 | 1.160579 | 0.384397 | 0.700684 | NA |
| | 11.97778 | -0.05696 | 0.615424 | -0.09256 | 0.926256 | NA |
| | 0.394291 | 0.298745 | 3.141711 | 0.09509 | 0.924243 | NA |
| | 0.162376 | 0.424913 | 4.079006 | 0.104171 | 0.917034 | NA |
| | 5.117625 | -0.50486 | 0.898196 | -0.56208 | 0.57406 | NA |
| | 119.4596 | 0.21868 | 0.242733 | 0.900907 | 0.367638 | 0.534761 |
| ST13P19 | 0.247898 | -0.5901 | 4.074633 | -0.14482 | 0.884851 | NA |
| PHBP3 | 0.288605 | 1.443619 | 4.029058 | 0.358302 | 0.720117 | NA |
| LINC00568 | 1.598452 | 0.120209 | 1.543205 | 0.077896 | 0.937911 | NA |
| | 1.652366 | -0.32069 | 1.531078 | -0.20945 | 0.834095 | NA |
| | 8.209933 | 0.162492 | 0.701023 | 0.231793 | 0.816699 | NA |
| PPIAP19 | 0.134133 | -1.48909 | 4.077584 | -0.36519 | 0.71497 | NA |
| | 10.31474 | -0.11917 | 0.620814 | -0.19196 | 0.847775 | NA |
| GEMIN8P4 | 17.41399 | -0.32736 | 0.548248 | -0.59711 | 0.550437 | NA |
| | 0.392857 | -1.546 | 4.075469 | -0.37934 | 0.704433 | NA |
| | 0.151697 | -1.546 | 4.075469 | -0.37934 | 0.704433 | NA |
| | 1.259656 | -0.65019 | 1.760499 | -0.36932 | 0.71189 | NA |
| | 0.327913 | 1.543154 | 4.021063 | 0.383768 | 0.701151 | NA |
| NPM1P30 | 0.258578 | 1.233151 | 4.047809 | 0.304646 | 0.760635 | NA |
| EFCAB14-A | 0.192403 | 0.975392 | 4.060716 | 0.240202 | 0.810174 | NA |
| UBBP2 | 0.394291 | 0.298745 | 3.141711 | 0.09509 | 0.924243 | NA |
| | 0.719393 | -0.9644 | 2.652838 | -0.36354 | 0.716204 | NA |
| | 1.163601 | -2.7311 | 2.310237 | -1.18217 | 0.237138 | NA |
| | 1.210913 | 2.305897 | 2.03591 | 1.132612 | 0.257377 | NA |
| PCAT6 | 45.84956 | -0.14024 | 0.34336 | -0.40844 | 0.682953 | NA |
| | 3.426838 | 2.869292 | 1.367437 | 2.098299 | 0.035879 | NA |
| | 12.23388 | -0.28294 | 0.635702 | -0.44508 | 0.656265 | NA |
| | 0.899573 | -1.00757 | 2.384851 | -0.42249 | 0.67267 | NA |
| AMD1P1 | 0.192403 | 0.975392 | 4.060716 | 0.240202 | 0.810174 | NA |
| EFCAB14P1 | 0.719043 | 1.433043 | 2.335048 | 0.61371 | 0.539407 | NA |
| ITPKB-IT1 | 0.260158 | 1.240347 | 4.047124 | 0.306476 | 0.759242 | NA |
| | 4.878348 | -0.49565 | 0.992872 | -0.49921 | 0.617631 | NA |
| | 0.956059 | 0.787678 | 2.36595 | 0.332923 | 0.739193 | NA |
| | 3.140462 | 0.966115 | 1.2711 | 0.760062 | 0.447217 | NA |

| | | | | | | |
|---|---|---|---|---|---|---|
| | 0.392711 | 0.292538 | 3.145221 | 0.09301 | 0.925895 | NA |
| | 0.556813 | -0.57007 | 3.294615 | -0.17303 | 0.862627 | NA |
| | 14.94737 | 0.058903 | 0.535977 | 0.109899 | 0.912489 | NA |
| | 3.925127 | -0.41739 | 1.021011 | -0.4088 | 0.682687 | NA |
| | 5.078387 | 0.059412 | 0.865596 | 0.068637 | 0.945279 | NA |
| | 0.424115 | 1.932531 | 3.45462 | 0.559405 | 0.575885 | NA |
| | 8.312605 | -0.04664 | 0.690388 | -0.06756 | 0.946135 | NA |
| | 0.774539 | 0.39797 | 2.296349 | 0.173305 | 0.862411 | NA |
| | 0.411855 | 0.250456 | 3.097618 | 0.080854 | 0.935558 | NA |
| GRK5-IT1 | 0.134133 | -1.48909 | 4.077584 | -0.36519 | 0.71497 | NA |
| EEF1A1P11 | 27.81032 | -0.1723 | 0.421954 | -0.40834 | 0.683022 | NA |
| | 0.096202 | 0.408546 | 4.079639 | 0.100143 | 0.920231 | NA |
| | 6.854165 | 0.061623 | 0.808101 | 0.076256 | 0.939215 | NA |
| | 5.19545 | 1.247508 | 1.03 | 1.211174 | 0.225829 | NA |
| ITPKB-AS1 | 0.247898 | -0.5901 | 4.074633 | -0.14482 | 0.884851 | NA |
| | 1.782739 | -0.05743 | 1.614198 | -0.03558 | 0.971618 | NA |
| SPCS2P4 | 4.579869 | -0.70011 | 1.002728 | -0.69821 | 0.485048 | NA |
| C1orf233 | 1477.519 | -0.23369 | 0.14166 | -1.64967 | 0.099011 | 0.221873 |
| | 5.101233 | 0.368884 | 0.946307 | 0.389814 | 0.696674 | NA |
| | 0.163957 | 0.424913 | 4.079006 | 0.104171 | 0.917034 | NA |
| | 1.496357 | 2.668981 | 1.845797 | 1.445977 | 0.148184 | NA |
| | 13.89481 | -0.18294 | 0.610999 | -0.29941 | 0.764626 | NA |
| | 1.389825 | 0.28982 | 1.646534 | 0.176018 | 0.860279 | NA |
| | 1.170171 | 1.843131 | 2.071651 | 0.889692 | 0.373631 | NA |
| | 0.151697 | -1.546 | 4.075469 | -0.37934 | 0.704433 | NA |
| TNKS2-AS1 | 16.27852 | 0.269276 | 0.569138 | 0.47313 | 0.63612 | NA |
| | 0.772958 | 0.415176 | 2.577252 | 0.161092 | 0.872021 | NA |
| | 0.260158 | 1.240347 | 4.047124 | 0.306476 | 0.759242 | NA |
| | 1.514445 | -1.85382 | 1.951264 | -0.95006 | 0.34208 | NA |
| | 0.70046 | 1.37107 | 2.594868 | 0.528378 | 0.597237 | NA |
| FAF2P1 | 0.163957 | 0.424913 | 4.079006 | 0.104171 | 0.917034 | NA |
| LINC01128 | 187.1272 | -3.32E-04 | 0.191344 | -0.00173 | 0.998617 | 0.999371 |
| AK3P5 | 0.247898 | -0.5901 | 4.074633 | -0.14482 | 0.884851 | NA |
| | 3.399651 | 2.128942 | 1.34967 | 1.577379 | 0.114708 | NA |
| | 0.747636 | 0.735436 | 2.634453 | 0.279161 | 0.780122 | NA |
| | 0.303394 | -2.27006 | 4.032451 | -0.56295 | 0.57347 | NA |
| | 0.506476 | 0.726418 | 2.864186 | 0.253621 | 0.799788 | NA |
| | 1.120219 | 0.417842 | 1.804388 | 0.23157 | 0.816872 | NA |
| | 0.096202 | 0.408546 | 4.079639 | 0.100143 | 0.920231 | NA |
| | 0.247898 | -0.5901 | 4.074633 | -0.14482 | 0.884851 | NA |
| | 43.58277 | 0.088355 | 0.374556 | 0.235893 | 0.813516 | NA |
| SNRPD2P1 | 0.69238 | 0.295083 | 2.603008 | 0.113362 | 0.909743 | NA |
| AKT3-IT1 | 0.151697 | -1.546 | 4.075469 | -0.37934 | 0.704433 | NA |
| | 0.192403 | 0.975392 | 4.060716 | 0.240202 | 0.810174 | NA |
| ACTBP14 | 0.151697 | -1.546 | 4.075469 | -0.37934 | 0.704433 | NA |
| AKR7A2P1 | 1.208102 | 0.723565 | 1.850162 | 0.391082 | 0.695737 | NA |
| | 0.151697 | -1.546 | 4.075469 | -0.37934 | 0.704433 | NA |

| | | | | | | |
|---|---|---|---|---|---|---|
| | 0.096202 | 0.408546 | 4.079639 | 0.100143 | 0.920231 | NA |
| LINC01165 | 1.426176 | -0.29008 | 1.685773 | -0.17207 | 0.86338 | NA |
| CALM2P2 | 1.846496 | -0.60587 | 1.5496 | -0.39099 | 0.695807 | NA |
| | 0.096202 | 0.408546 | 4.079639 | 0.100143 | 0.920231 | NA |
| | 0.162376 | 0.424913 | 4.079006 | 0.104171 | 0.917034 | NA |
| | 0.151697 | -1.546 | 4.075469 | -0.37934 | 0.704433 | NA |
| CYCSP4 | 0.392857 | -1.546 | 4.075469 | -0.37934 | 0.704433 | NA |
| VIM-AS1 | 13.87362 | 0.42329 | 0.558589 | 0.757784 | 0.448581 | NA |
| RPS7P4 | 0.392857 | -1.546 | 4.075469 | -0.37934 | 0.704433 | NA |
| | 3.096913 | -0.62368 | 1.158402 | -0.5384 | 0.590302 | NA |
| | 1.239506 | 1.943067 | 2.060745 | 0.942896 | 0.345734 | NA |
| | 0.230334 | -0.53318 | 4.076749 | -0.13079 | 0.895945 | NA |
| | 0.556813 | -0.57007 | 3.294615 | -0.17303 | 0.862627 | NA |
| | 0.162376 | 0.424913 | 4.079006 | 0.104171 | 0.917034 | NA |
| | 0.392857 | -1.546 | 4.075469 | -0.37934 | 0.704433 | NA |
| | 0.151697 | -1.546 | 4.075469 | -0.37934 | 0.704433 | NA |
| ST13P13 | 0.192403 | 0.975392 | 4.060716 | 0.240202 | 0.810174 | NA |
| | 1.45782 | -0.09859 | 1.838232 | -0.05364 | 0.957226 | NA |
| | 6.71997 | 1.880122 | 0.937054 | 2.006419 | 0.044812 | NA |
| | 0.163957 | 0.424913 | 4.079006 | 0.104171 | 0.917034 | NA |
| | 23.7869 | 0.032541 | 0.425025 | 0.076562 | 0.938972 | NA |
| HYI-AS1 | 2.049607 | -1.04999 | 1.364257 | -0.76964 | 0.441511 | NA |
| HMGN2P1 | 0.162376 | 0.424913 | 4.079006 | 0.104171 | 0.917034 | NA |
| | 13.97455 | -0.76342 | 0.593599 | -1.28609 | 0.198413 | NA |
| | 9.653704 | 0.174483 | 0.743141 | 0.234791 | 0.814371 | NA |
| | 0.162376 | 0.424913 | 4.079006 | 0.104171 | 0.917034 | NA |
| | 61.78128 | 1.531877 | 0.402925 | 3.80189 | 1.44E-04 | 0.003158 |
| | 1.413713 | -0.46869 | 1.759142 | -0.26643 | 0.789905 | NA |
| NPM1P25 | 1.118842 | -0.14871 | 1.94468 | -0.07647 | 0.939045 | NA |
| | 2.423677 | -0.99396 | 1.3533 | -0.73447 | 0.462661 | NA |
| | 0.614937 | 2.493639 | 2.617677 | 0.952615 | 0.340785 | NA |
| LYST-AS1 | 0.327913 | 1.543154 | 4.021063 | 0.383768 | 0.701151 | NA |
| | 1.266612 | 0.746194 | 1.745721 | 0.427442 | 0.669058 | NA |
| | 21.61164 | 0.344614 | 0.523266 | 0.658584 | 0.510163 | NA |
| | 0.719043 | 1.433043 | 2.335048 | 0.61371 | 0.539407 | NA |
| | 0.151697 | -1.546 | 4.075469 | -0.37934 | 0.704433 | NA |
| HNRNPA1P | 1.05103 | -0.7935 | 2.094237 | -0.3789 | 0.704763 | NA |
| | 12.18101 | -0.03736 | 0.571771 | -0.06534 | 0.947903 | NA |
| SSU72P7 | 0.163957 | 0.424913 | 4.079006 | 0.104171 | 0.917034 | NA |
| | 1.488207 | -0.17791 | 1.698666 | -0.10474 | 0.916584 | NA |
| NKX1-2 | 0.545988 | -0.43913 | 2.541145 | -0.17281 | 0.862803 | NA |
| | 1.664626 | 0.191034 | 1.497529 | 0.127566 | 0.898493 | NA |
| | 0.096202 | 0.408546 | 4.079639 | 0.100143 | 0.920231 | NA |
| | 0.586491 | 2.39072 | 2.689057 | 0.889055 | 0.373973 | NA |
| | 2.334257 | 1.49891 | 1.427213 | 1.050236 | 0.29361 | NA |
| | 0.096202 | 0.408546 | 4.079639 | 0.100143 | 0.920231 | NA |
| NAMPTL | 49.15548 | 0.631224 | 0.424006 | 1.488715 | 0.136563 | 0.275613 |

| | | | | | | |
|---|---|---|---|---|---|---|
| | 2.49477 | -0.52702 | 1.401337 | -0.37609 | 0.706853 | NA |
| | 10.1302 | 0.582943 | 0.684877 | 0.851164 | 0.394678 | NA |
| | 0.151697 | -1.546 | 4.075469 | -0.37934 | 0.704433 | NA |
| | 0.151697 | -1.546 | 4.075469 | -0.37934 | 0.704433 | NA |
| ASNSP5 | 0.096202 | 0.408546 | 4.079639 | 0.100143 | 0.920231 | NA |
| | 0.096202 | 0.408546 | 4.079639 | 0.100143 | 0.920231 | NA |
| | 0.324752 | 1.53082 | 4.022025 | 0.380609 | 0.703493 | NA |
| | 0.151697 | -1.546 | 4.075469 | -0.37934 | 0.704433 | NA |
| | 1.364154 | 2.182193 | 1.9921 | 1.095424 | 0.273331 | NA |
| RPL7P10 | 0.326333 | 1.533889 | 3.826878 | 0.40082 | 0.688553 | NA |
| UBE2Q1-AS | 0.963147 | -0.83257 | 2.358641 | -0.35299 | 0.724097 | NA |
| SLC25A38P | 0.163957 | 0.424913 | 4.079006 | 0.104171 | 0.917034 | NA |
| SNRPFP2 | 0.163957 | 0.424913 | 4.079006 | 0.104171 | 0.917034 | NA |
| | 3.010867 | 1.461946 | 1.204451 | 1.213786 | 0.224829 | NA |
| | 0.192403 | 0.975392 | 4.060716 | 0.240202 | 0.810174 | NA |
| ISCA1P2 | 0.163957 | 0.424913 | 4.079006 | 0.104171 | 0.917034 | NA |
| | 4.075388 | 1.659961 | 1.076222 | 1.542397 | 0.122977 | NA |
| | 2.054954 | 1.273794 | 1.497066 | 0.85086 | 0.394847 | NA |
| EMX2OS | 1.326462 | 1.54459 | 1.923183 | 0.803143 | 0.421892 | NA |
| | 0.151697 | -1.546 | 4.075469 | -0.37934 | 0.704433 | NA |
| | 0.572651 | 0.928649 | 2.780261 | 0.334015 | 0.738368 | NA |
| | 19.46373 | -0.46187 | 0.464356 | -0.99464 | 0.319912 | NA |
| HNRNPA1P | 0.314073 | -0.57308 | 3.898934 | -0.14698 | 0.883146 | NA |
| RHEBP1 | 2.137555 | 0.433445 | 1.355399 | 0.319792 | 0.749126 | NA |
| YWHAZP3 | 20.90081 | 0.155686 | 0.493304 | 0.315598 | 0.752308 | NA |
| | 19.72124 | -0.69015 | 0.466397 | -1.47975 | 0.138939 | NA |
| ZRANB2-AS | 10.72946 | -0.26205 | 0.651441 | -0.40226 | 0.687494 | NA |
| | 0.162376 | 0.424913 | 4.079006 | 0.104171 | 0.917034 | NA |
| | 1.307318 | 1.582229 | 1.879481 | 0.841843 | 0.399876 | NA |
| | 0.548763 | 2.343695 | 3.135478 | 0.747476 | 0.454776 | NA |
| MIR181A1I | 0.774742 | -0.40577 | 2.223483 | -0.18249 | 0.855195 | NA |
| AKR1B1P1 | 0.260158 | 1.240347 | 4.047124 | 0.306476 | 0.759242 | NA |
| RPL5P4 | 2.550694 | 0.695799 | 1.394547 | 0.498942 | 0.61782 | NA |
| | 1.494374 | -1.03816 | 1.752549 | -0.59237 | 0.553604 | NA |
| | 0.096202 | 0.408546 | 4.079639 | 0.100143 | 0.920231 | NA |
| | 0.420954 | 1.923376 | 3.462803 | 0.555439 | 0.578594 | NA |
| TMEM254- | 15.50138 | 0.618186 | 0.562973 | 1.098074 | 0.272172 | NA |
| | 1.525336 | 0.472832 | 1.602087 | 0.295135 | 0.767891 | NA |
| | 0.192403 | 0.975392 | 4.060716 | 0.240202 | 0.810174 | NA |
| | 0.096202 | 0.408546 | 4.079639 | 0.100143 | 0.920231 | NA |
| | 15.19014 | 0.685568 | 0.531675 | 1.289449 | 0.197242 | NA |
| | 0.392857 | -1.546 | 4.075469 | -0.37934 | 0.704433 | NA |
| | 22.44107 | 0.182523 | 0.478152 | 0.381726 | 0.702665 | NA |
| RPL22P18 | 0.661122 | -3.06315 | 2.98049 | -1.02773 | 0.304076 | NA |
| | 0.096202 | 0.408546 | 4.079639 | 0.100143 | 0.920231 | NA |
| SMYD3-IT1 | 0.326333 | 1.533889 | 3.826878 | 0.40082 | 0.688553 | NA |
| | 0.162376 | 0.424913 | 4.079006 | 0.104171 | 0.917034 | NA |

| | | | | | | |
|---|---|---|---|---|---|---|
| PHBP9 | 3.59893 | 0.975603 | 1.18979 | 0.819979 | 0.412228 | NA |
| | 0.803131 | -0.76693 | 2.349822 | -0.32638 | 0.744138 | NA |
| HNRNPA1P | 2.179413 | -0.57541 | 1.515979 | -0.37956 | 0.704272 | NA |
| | 0.096202 | 0.408546 | 4.079639 | 0.100143 | 0.920231 | NA |
| | 0.392857 | -1.546 | 4.075469 | -0.37934 | 0.704433 | NA |
| BANF1P2 | 1.110174 | 1.280534 | 2.2388 | 0.571973 | 0.56734 | NA |
| | 0.192403 | 0.975392 | 4.060716 | 0.240202 | 0.810174 | NA |
| | 2.013017 | -0.32834 | 1.438204 | -0.2283 | 0.819412 | NA |
| | 5.613144 | 0.312056 | 0.856669 | 0.364266 | 0.715659 | NA |
| RPL4P3 | 1.389476 | 1.604276 | 1.701484 | 0.942869 | 0.345748 | NA |
| | 3.309869 | 0.431586 | 1.139166 | 0.378861 | 0.704791 | NA |
| | 0.134133 | -1.48909 | 4.077584 | -0.36519 | 0.71497 | NA |
| LINC00856 | 0.260158 | 1.240347 | 4.047124 | 0.306476 | 0.759242 | NA |
| | 3.323157 | 1.60396 | 1.206803 | 1.329098 | 0.183816 | NA |
| RSU1P1 | 0.410275 | 0.244283 | 3.100889 | 0.078778 | 0.937209 | NA |
| ERVMER61 | 2.094183 | -1.68969 | 1.465687 | -1.15283 | 0.24898 | NA |
| | 3.556719 | -0.49511 | 1.050763 | -0.47119 | 0.637505 | NA |
| LRRC37A6F | 1.15555 | -0.20065 | 2.223695 | -0.09023 | 0.928102 | NA |
| PROX1-AS1 | 0.163957 | 0.424913 | 4.079006 | 0.104171 | 0.917034 | NA |
| | 0.681112 | 2.620532 | 2.519661 | 1.040033 | 0.298324 | NA |
| ANKRD30B | 0.163957 | 0.424913 | 4.079006 | 0.104171 | 0.917034 | NA |
| | 2.612424 | 0.50933 | 1.256884 | 0.405233 | 0.685307 | NA |
| | 0.69625 | -3.13723 | 2.93845 | -1.06765 | 0.285679 | NA |
| HMGB1P11 | 0.096202 | 0.408546 | 4.079639 | 0.100143 | 0.920231 | NA |
| | 1.560455 | -1.53619 | 1.708948 | -0.89891 | 0.368702 | NA |
| | 19.73409 | -0.44549 | 0.456286 | -0.97634 | 0.328898 | NA |
| ZNF32-AS2 | 9.013796 | 1.776326 | 0.777176 | 2.285615 | 0.022277 | NA |
| | 0.163957 | 0.424913 | 4.079006 | 0.104171 | 0.917034 | NA |
| PHBP12 | 0.230334 | -0.53318 | 4.076749 | -0.13079 | 0.895945 | NA |
| | 0.629726 | -0.22725 | 2.732935 | -0.08315 | 0.93373 | NA |
| | 2.493575 | -1.37122 | 1.292426 | -1.06096 | 0.288707 | NA |
| | 0.631307 | -0.20384 | 3.097925 | -0.0658 | 0.947537 | NA |
| | 0.419962 | -2.64111 | 3.429364 | -0.77014 | 0.441214 | NA |
| SETSIP | 0.247898 | -0.5901 | 4.074633 | -0.14482 | 0.884851 | NA |
| ENO1-AS1 | 0.777314 | 2.821035 | 2.404647 | 1.17316 | 0.240732 | NA |
| | 1.693073 | 0.276824 | 1.517715 | 0.182395 | 0.855272 | NA |
| LINC01001 | 0.411855 | 0.250456 | 3.097618 | 0.080854 | 0.935558 | NA |
| | 0.151697 | -1.546 | 4.075469 | -0.37934 | 0.704433 | NA |
| | 3.051888 | 0.221958 | 1.236967 | 0.179437 | 0.857594 | NA |
| | 0.162376 | 0.424913 | 4.079006 | 0.104171 | 0.917034 | NA |
| | 3.290033 | -0.26187 | 1.142388 | -0.22923 | 0.818688 | NA |
| PSAT1P3 | 0.440301 | 0.468383 | 3.449808 | 0.135771 | 0.892003 | NA |
| | 343.7728 | 0.237042 | 0.203535 | 1.164627 | 0.24417 | 0.406893 |
| | 0.096202 | 0.408546 | 4.079639 | 0.100143 | 0.920231 | NA |
| | 1.430568 | 0.993923 | 1.70292 | 0.583658 | 0.55945 | NA |
| | 0.247898 | -0.5901 | 4.074633 | -0.14482 | 0.884851 | NA |
| RPEP3 | 0.326333 | 1.533889 | 3.826878 | 0.40082 | 0.688553 | NA |

| | | | | | | |
|---|---|---|---|---|---|---|
| | 0.410275 | 0.244283 | 3.100889 | 0.078778 | 0.937209 | NA |
| GOT2P2 | 0.432222 | -1.18916 | 3.482045 | -0.34151 | 0.732718 | NA |
| | 4.103229 | 0.507794 | 1.077966 | 0.471066 | 0.637593 | NA |
| | 0.162376 | 0.424913 | 4.079006 | 0.104171 | 0.917034 | NA |
| CTGLF10P | 0.736607 | 1.395844 | 2.318642 | 0.602009 | 0.547168 | NA |
| | 0.555087 | 0.973802 | 2.808575 | 0.346725 | 0.728798 | NA |
| | 18.84406 | -0.83674 | 0.469427 | -1.78247 | 0.074673 | NA |
| | 0.49029 | 2.123092 | 3.304282 | 0.642527 | 0.520531 | NA |
| | 0.392857 | -1.546 | 4.075469 | -0.37934 | 0.704433 | NA |
| | 4.537932 | 0.675692 | 0.997928 | 0.677095 | 0.498346 | NA |
| | 0.422534 | 1.924583 | 3.001758 | 0.641152 | 0.521424 | NA |
| | 44.17169 | -0.08102 | 0.316731 | -0.25581 | 0.798095 | NA |
| | 5.61525 | 0.138147 | 0.866339 | 0.159461 | 0.873306 | NA |
| | 0.949727 | -0.95431 | 2.383855 | -0.40032 | 0.688919 | NA |
| | 0.941473 | 1.959123 | 2.152875 | 0.910003 | 0.362821 | NA |
| | 0.845418 | 1.095819 | 2.535106 | 0.432258 | 0.665554 | NA |
| | 7.993088 | 0.183231 | 0.80705 | 0.227039 | 0.820394 | NA |
| | 0.392857 | -1.546 | 4.075469 | -0.37934 | 0.704433 | NA |
| GYG1P3 | 4.295865 | -0.26096 | 1.141591 | -0.2286 | 0.819183 | NA |
| | 1.53222 | -0.24624 | 1.840826 | -0.13376 | 0.893589 | NA |
| | 0.950755 | 1.904765 | 2.40482 | 0.792061 | 0.428325 | NA |
| | 0.392857 | -1.546 | 4.075469 | -0.37934 | 0.704433 | NA |
| | 0.588072 | 2.39955 | 3.098804 | 0.774347 | 0.438726 | NA |
| | 1.103996 | 1.740043 | 2.131488 | 0.816351 | 0.414299 | NA |
| | 0.162376 | 0.424913 | 4.079006 | 0.104171 | 0.917034 | NA |
| | 2.106443 | -1.1289 | 1.368162 | -0.82512 | 0.409302 | NA |
| | 9.894646 | -0.44021 | 0.726392 | -0.60602 | 0.544499 | NA |
| | 0.315653 | -0.57305 | 3.892639 | -0.14721 | 0.882963 | NA |
| | 0.151697 | -1.546 | 4.075469 | -0.37934 | 0.704433 | NA |
| GLUD1P6 | 1.379 | 0.36119 | 1.668785 | 0.216439 | 0.828646 | NA |
| CCDC147-A | 2.724352 | -1.78843 | 1.303057 | -1.37249 | 0.169911 | NA |
| C1DP1 | 20.42699 | 0.840425 | 0.487553 | 1.723763 | 0.084751 | NA |
| | 1.755873 | -0.13678 | 1.568634 | -0.0872 | 0.930513 | NA |
| | 0.163957 | 0.424913 | 4.079006 | 0.104171 | 0.917034 | NA |
| LINC00704 | 0.298089 | -0.51711 | 3.975232 | -0.13008 | 0.896501 | NA |
| RPL34P6 | 0.815245 | 1.670886 | 2.214146 | 0.754641 | 0.450464 | NA |
| | 2.006225 | 0.394572 | 1.60469 | 0.245887 | 0.80577 | NA |
| LINC01160 | 1.331766 | 0.790302 | 2.090457 | 0.378052 | 0.705392 | NA |
| | 2.091894 | -0.46434 | 1.479118 | -0.31393 | 0.753575 | NA |
| | 88.63616 | -0.01137 | 0.274756 | -0.04137 | 0.966997 | 0.981674 |
| | 7.371 | 0.94281 | 0.744899 | 1.265688 | 0.205625 | NA |
| | 1.396003 | 0.40896 | 1.86007 | 0.219863 | 0.825978 | NA |
| | 0.134133 | -1.48909 | 4.077584 | -0.36519 | 0.71497 | NA |
| | 0.28583 | -2.21358 | 3.97376 | -0.55705 | 0.577494 | NA |
| | 1.264828 | 1.410059 | 1.995098 | 0.706762 | 0.479715 | NA |
| | 0.296509 | -0.51709 | 3.982155 | -0.12985 | 0.896684 | NA |
| PRDX3P2 | 0.314073 | -0.57308 | 3.898934 | -0.14698 | 0.883146 | NA |

| | | | | | | |
|---|---|---|---|---|---|---|
| | 0.490493 | 0.776209 | 2.895261 | 0.268096 | 0.788625 | NA |
| | 0.314073 | -0.57308 | 3.898934 | -0.14698 | 0.883146 | NA |
| | 0.096202 | 0.408546 | 4.079639 | 0.100143 | 0.920231 | NA |
| | 1.35356 | -0.61965 | 2.121437 | -0.29209 | 0.770217 | NA |
| | 1.342677 | -0.80138 | 1.862188 | -0.43034 | 0.666947 | NA |
| LINC01037 | 0.162376 | 0.424913 | 4.079006 | 0.104171 | 0.917034 | NA |
| | 0.162376 | 0.424913 | 4.079006 | 0.104171 | 0.917034 | NA |
| | 0.096202 | 0.408546 | 4.079639 | 0.100143 | 0.920231 | NA |
| | 15.0359 | 0.736926 | 0.524274 | 1.405615 | 0.159839 | NA |
| | 1.366683 | -0.61531 | 1.76266 | -0.34908 | 0.727028 | NA |
| | 0.134133 | -1.48909 | 4.077584 | -0.36519 | 0.71497 | NA |
| | 0.757324 | -2.08271 | 2.587512 | -0.80491 | 0.420873 | NA |
| | 0.162376 | 0.424913 | 4.079006 | 0.104171 | 0.917034 | NA |
| | 0.489058 | -0.58569 | 3.427543 | -0.17088 | 0.864321 | NA |
| | 0.096202 | 0.408546 | 4.079639 | 0.100143 | 0.920231 | NA |
| | 1.965637 | 0.575394 | 1.419817 | 0.405259 | 0.685287 | NA |
| EMBP1 | 0.918331 | 0.764804 | 2.118268 | 0.361052 | 0.718061 | NA |
| MIR4426 | 12.75471 | 0.364362 | 0.697669 | 0.522256 | 0.601492 | NA |
| | 282.2498 | -0.0594 | 0.202713 | -0.29304 | 0.769488 | 0.857357 |
| | 2.231916 | 0.810284 | 1.423104 | 0.569378 | 0.5691 | NA |
| | 0.69238 | 0.295083 | 2.603008 | 0.113362 | 0.909743 | NA |
| | 0.096202 | 0.408546 | 4.079639 | 0.100143 | 0.920231 | NA |
| | 0.096202 | 0.408546 | 4.079639 | 0.100143 | 0.920231 | NA |
| IPO9-AS1 | 2.332916 | 0.99215 | 1.398314 | 0.709533 | 0.477994 | NA |
| RPS7P3 | 3.567681 | 1.170429 | 1.072849 | 1.090954 | 0.275293 | NA |
| | 0.258578 | 1.233151 | 4.047809 | 0.304646 | 0.760635 | NA |
| | 0.577209 | 2.423681 | 3.646109 | 0.664731 | 0.506223 | NA |
| | 2.718662 | -0.63292 | 1.203011 | -0.52611 | 0.598809 | NA |
| | 0.096202 | 0.408546 | 4.079639 | 0.100143 | 0.920231 | NA |
| | 49.4659 | -0.12477 | 0.334673 | -0.3728 | 0.709294 | 0.814006 |
| CAP1P2 | 13.13696 | 0.040131 | 0.615219 | 0.065231 | 0.94799 | NA |
| HSPE1P25 | 0.163957 | 0.424913 | 4.079006 | 0.104171 | 0.917034 | NA |
| | 2.998865 | -0.57147 | 1.465583 | -0.38992 | 0.696592 | NA |
| | 1.330121 | -1.83415 | 1.905355 | -0.96263 | 0.335733 | NA |
| | 0.298089 | -0.51711 | 3.975232 | -0.13008 | 0.896501 | NA |
| | 0.544554 | -2.65036 | 3.220086 | -0.82307 | 0.410468 | NA |
| | 0.424115 | 1.932531 | 3.45462 | 0.559405 | 0.575885 | NA |
| MRPL35P2 | 1.784319 | -0.05578 | 1.614116 | -0.03456 | 0.97243 | NA |
| LINC01031 | 0.096202 | 0.408546 | 4.079639 | 0.100143 | 0.920231 | NA |
| | 6.705625 | 0.010557 | 0.808291 | 0.013061 | 0.989579 | NA |
| | 0.096202 | 0.408546 | 4.079639 | 0.100143 | 0.920231 | NA |
| | 0.327913 | 1.543154 | 4.021063 | 0.383768 | 0.701151 | NA |
| RPS15AP12 | 2.217141 | -0.60446 | 1.362968 | -0.44349 | 0.657412 | NA |
| LINC00867 | 0.672013 | 1.253691 | 2.607776 | 0.480751 | 0.630693 | NA |
| | 3.303277 | 0.037364 | 1.188021 | 0.031451 | 0.97491 | NA |
| | 0.163957 | 0.424913 | 4.079006 | 0.104171 | 0.917034 | NA |
| | 0.163957 | 0.424913 | 4.079006 | 0.104171 | 0.917034 | NA |

| | | | | | | |
|---|---|---|---|---|---|---|
| CNOT7P2 | 0.162376 | 0.424913 | 4.079006 | 0.104171 | 0.917034 | NA |
| | 0.163957 | 0.424913 | 4.079006 | 0.104171 | 0.917034 | NA |
| | 0.394291 | 0.298745 | 3.141711 | 0.09509 | 0.924243 | NA |
| LINC00202 | 0.35636 | 1.709252 | 3.651833 | 0.468053 | 0.639746 | NA |
| LINC00865 | 34.07387 | -0.10117 | 0.42399 | -0.23861 | 0.811406 | NA |
| ASCL5 | 0.422737 | 0.518872 | 3.501356 | 0.148192 | 0.882191 | NA |
| FTH1P1 | 0.489058 | -0.58569 | 3.427543 | -0.17088 | 0.864321 | NA |
| | 6.192874 | -0.27407 | 0.786338 | -0.34854 | 0.727438 | NA |
| GNG12-AS1 | 34.80291 | -0.03922 | 0.39103 | -0.1003 | 0.920104 | NA |
| | 0.151697 | -1.546 | 4.075469 | -0.37934 | 0.704433 | NA |
| | 0.544554 | -2.65036 | 3.220086 | -0.82307 | 0.410468 | NA |
| RPL4P2 | 1.015699 | 0.142451 | 2.181052 | 0.065313 | 0.947925 | NA |
| | 0.192403 | 0.975392 | 4.060716 | 0.240202 | 0.810174 | NA |
| | 7.280297 | 0.297124 | 0.739609 | 0.401731 | 0.687882 | NA |
| | 1.281015 | 0.71131 | 1.742215 | 0.408279 | 0.683069 | NA |
| RPL5P6 | 0.314073 | -0.57308 | 3.898934 | -0.14698 | 0.883146 | NA |
| | 6.06069 | 1.705329 | 1.079252 | 1.580102 | 0.114083 | NA |
| | 14.99332 | -0.2464 | 0.584062 | -0.42187 | 0.673118 | NA |
| | 4.149349 | 0.256032 | 1.006476 | 0.254385 | 0.799198 | NA |
| | 0.891311 | -0.69809 | 2.403604 | -0.29043 | 0.771484 | NA |
| | 1.284314 | -2.91715 | 2.227383 | -1.30968 | 0.190306 | NA |
| | 7.623211 | 0.384473 | 0.728031 | 0.5281 | 0.59743 | NA |
| | 0.952539 | 0.987729 | 2.327171 | 0.424433 | 0.67125 | NA |
| | 22.01654 | 0.547966 | 0.47699 | 1.148799 | 0.250639 | NA |
| | 0.4986 | -1.64638 | 3.317584 | -0.49626 | 0.619712 | NA |
| | 0.096202 | 0.408546 | 4.079639 | 0.100143 | 0.920231 | NA |
| RSU1P2 | 0.151697 | -1.546 | 4.075469 | -0.37934 | 0.704433 | NA |
| | 0.833158 | -0.52049 | 2.521552 | -0.20642 | 0.836464 | NA |
| | 0.134133 | -1.48909 | 4.077584 | -0.36519 | 0.71497 | NA |
| | 0.096202 | 0.408546 | 4.079639 | 0.100143 | 0.920231 | NA |
| | 0.247898 | -0.5901 | 4.074633 | -0.14482 | 0.884851 | NA |
| | 12.62988 | 0.140485 | 0.64454 | 0.217962 | 0.827458 | NA |
| | 0.402398 | -2.58771 | 4.018369 | -0.64397 | 0.519595 | NA |
| | 0.697685 | -0.92563 | 2.353044 | -0.39338 | 0.694041 | NA |
| IDI2-AS1 | 17.89073 | -0.32509 | 0.514801 | -0.63149 | 0.52772 | NA |
| | 4.651997 | -0.70875 | 1.047718 | -0.67647 | 0.498743 | NA |
| | 0.151697 | -1.546 | 4.075469 | -0.37934 | 0.704433 | NA |
| | 5.34214 | 0.540786 | 1.000101 | 0.540732 | 0.588692 | NA |
| | 0.653015 | 0.2552 | 2.787103 | 0.091565 | 0.927044 | NA |
| ARID4B-IT1 | 0.163957 | 0.424913 | 4.079006 | 0.104171 | 0.917034 | NA |
| MARK2P9 | 0.230334 | -0.53318 | 4.076749 | -0.13079 | 0.895945 | NA |
| | 1.910345 | 0.57799 | 1.57782 | 0.366322 | 0.714125 | NA |
| ANKRD20A | 0.563552 | -0.48645 | 2.808029 | -0.17324 | 0.862465 | NA |
| | 0.096202 | 0.408546 | 4.079639 | 0.100143 | 0.920231 | NA |
| | 0.096202 | 0.408546 | 4.079639 | 0.100143 | 0.920231 | NA |
| | 0.654246 | 2.541085 | 3.009764 | 0.84428 | 0.398513 | NA |
| | 0.163957 | 0.424913 | 4.079006 | 0.104171 | 0.917034 | NA |

| | | | | | | |
|---|---|---|---|---|---|---|
| | 0.134133 | -1.48909 | 4.077584 | -0.36519 | 0.71497 | NA |
| | 3.891202 | 0.823048 | 1.029009 | 0.799846 | 0.4238 | NA |
| TXNP1 | 0.410275 | 0.244283 | 3.100889 | 0.078778 | 0.937209 | NA |
| | 0.096202 | 0.408546 | 4.079639 | 0.100143 | 0.920231 | NA |
| | 0.394291 | 0.298745 | 3.141711 | 0.09509 | 0.924243 | NA |
| KATNBL1P2 | 0.163957 | 0.424913 | 4.079006 | 0.104171 | 0.917034 | NA |
| SMG7-AS1 | 17.36649 | 0.168162 | 0.484502 | 0.347082 | 0.72853 | NA |
| | 0.747636 | 0.735436 | 2.634453 | 0.279161 | 0.780122 | NA |
| PHKA1P1 | 0.747636 | 0.735436 | 2.634453 | 0.279161 | 0.780122 | NA |
| | 0.230334 | -0.53318 | 4.076749 | -0.13079 | 0.895945 | NA |
| | 0.258578 | 1.233151 | 4.047809 | 0.304646 | 0.760635 | NA |
| HSPE1P6 | 0.151697 | -1.546 | 4.075469 | -0.37934 | 0.704433 | NA |
| | 0.096202 | 0.408546 | 4.079639 | 0.100143 | 0.920231 | NA |
| | 0.268266 | -2.15396 | 4.038351 | -0.53338 | 0.593773 | NA |
| | 17.67456 | 0.739848 | 0.506156 | 1.4617 | 0.143823 | NA |
| | 26.8151 | 0.34558 | 0.397774 | 0.868784 | 0.384965 | NA |
| HDAC1P2 | 0.869306 | -0.5527 | 2.538621 | -0.21772 | 0.82765 | NA |
| | 0.162376 | 0.424913 | 4.079006 | 0.104171 | 0.917034 | NA |
| | 0.260158 | 1.240347 | 4.047124 | 0.306476 | 0.759242 | NA |
| | 1.50496 | -0.97023 | 1.870122 | -0.51881 | 0.603895 | NA |
| EEF1A1P14 | 1.770313 | -0.12937 | 1.519242 | -0.08515 | 0.932138 | NA |
| TTLL7-IT1 | 0.326333 | 1.533889 | 3.826878 | 0.40082 | 0.688553 | NA |
| | 0.46577 | -1.30191 | 3.402879 | -0.38259 | 0.702024 | NA |
| | 0.3441 | -0.0166 | 3.769682 | -0.0044 | 0.996486 | NA |
| LINC00702 | 30.86581 | 1.519735 | 0.54416 | 2.792808 | 0.005225 | NA |
| CTAGE7P | 2.892718 | 1.745396 | 1.287082 | 1.356088 | 0.175071 | NA |
| ACTBP12 | 0.458885 | 0.577525 | 3.449278 | 0.167433 | 0.867029 | NA |
| | 0.247898 | -0.5901 | 4.074633 | -0.14482 | 0.884851 | NA |
| | 0.792306 | -0.44145 | 2.21354 | -0.19943 | 0.841926 | NA |
| | 13.15243 | 0.543958 | 0.600785 | 0.905411 | 0.365248 | NA |
| | 0.260158 | 1.240347 | 4.047124 | 0.306476 | 0.759242 | NA |
| | 75.25933 | 0.387424 | 0.26771 | 1.447179 | 0.147847 | 0.291005 |
| RPL12P29 | 0.420954 | 1.923376 | 3.462803 | 0.555439 | 0.578594 | NA |
| | 0.096202 | 0.408546 | 4.079639 | 0.100143 | 0.920231 | NA |
| | 0.786799 | 1.578877 | 2.532268 | 0.623503 | 0.532954 | NA |
| SNX18P16 | 6.922233 | 1.115741 | 0.803858 | 1.387982 | 0.165143 | NA |
| | 0.258578 | 1.233151 | 4.047809 | 0.304646 | 0.760635 | NA |
| | 0.096202 | 0.408546 | 4.079639 | 0.100143 | 0.920231 | NA |
| PPIAP32 | 0.258578 | 1.233151 | 4.047809 | 0.304646 | 0.760635 | NA |
| PFN1P1 | 9.324748 | -0.11608 | 0.742901 | -0.15625 | 0.875835 | NA |
| FAM53B-AS | 0.596179 | -0.09275 | 3.146759 | -0.02948 | 0.976485 | NA |
| UBE2FP3 | 3.035104 | 0.488613 | 1.118677 | 0.436778 | 0.662273 | NA |
| | 0.151697 | -1.546 | 4.075469 | -0.37934 | 0.704433 | NA |
| | 1.900265 | -0.35855 | 1.466414 | -0.24451 | 0.806839 | NA |
| | 11.73947 | 1.078507 | 0.625533 | 1.72414 | 0.084683 | NA |
| | 13.56446 | -0.58651 | 0.609738 | -0.96191 | 0.336095 | NA |
| PRKAR1AP | 0.803131 | -0.76693 | 2.349822 | -0.32638 | 0.744138 | NA |

| | | | | | | |
|---|---|---|---|---|---|---|
| | 5.26224 | 0.830911 | 0.923374 | 0.899864 | 0.368193 | NA |
| | 3.0665 | -0.05995 | 1.163215 | -0.05154 | 0.958894 | NA |
| | 0.134133 | -1.48909 | 4.077584 | -0.36519 | 0.71497 | NA |
| | 0.69625 | -3.13723 | 2.93845 | -1.06765 | 0.285679 | NA |
| | 0.70693 | -1.62766 | 2.687154 | -0.60572 | 0.544701 | NA |
| AGGF1P2 | 1.308267 | -0.51088 | 1.710881 | -0.2986 | 0.765242 | NA |
| BTBD18 | 2.634075 | 0.430464 | 1.288106 | 0.334183 | 0.738241 | NA |
| | 128.346 | 0.453506 | 0.263615 | 1.720331 | 0.085372 | 0.199625 |
| | 0.096202 | 0.408546 | 4.079639 | 0.100143 | 0.920231 | NA |
| | 0.725928 | 0.181819 | 2.321433 | 0.078322 | 0.937572 | NA |
| DNM3-IT1 | 0.364467 | -1.20166 | 3.685431 | -0.32606 | 0.744381 | NA |
| | 0.296509 | -0.51709 | 3.982155 | -0.12985 | 0.896684 | NA |
| | 0.813462 | 2.857892 | 2.821395 | 1.012936 | 0.311091 | NA |
| | 30.71108 | 0.34151 | 0.433696 | 0.78744 | 0.431024 | NA |
| ERI3-IT1 | 0.87074 | 0.675988 | 2.012127 | 0.335957 | 0.736903 | NA |
| | 0.162376 | 0.424913 | 4.079006 | 0.104171 | 0.917034 | NA |
| LINC01137 | 157.4508 | 0.114893 | 0.202264 | 0.568034 | 0.570012 | 0.706055 |
| PGAM1P11 | 0.258578 | 1.233151 | 4.047809 | 0.304646 | 0.760635 | NA |
| | 0.846852 | 1.76834 | 2.426477 | 0.728769 | 0.466143 | NA |
| GPR158-AS | 0.134133 | -1.48909 | 4.077584 | -0.36519 | 0.71497 | NA |
| | 0.096202 | 0.408546 | 4.079639 | 0.100143 | 0.920231 | NA |
| | 0.162376 | 0.424913 | 4.079006 | 0.104171 | 0.917034 | NA |
| | 0.096202 | 0.408546 | 4.079639 | 0.100143 | 0.920231 | NA |
| CICP27 | 0.247898 | -0.5901 | 4.074633 | -0.14482 | 0.884851 | NA |
| | 0.163957 | 0.424913 | 4.079006 | 0.104171 | 0.917034 | NA |
| LINC01136 | 0.951161 | 0.273761 | 2.339218 | 0.117031 | 0.906835 | NA |
| | 0.58526 | 0.001197 | 3.235889 | 3.70E-04 | 0.999705 | NA |
| EIF3LP2 | 0.668852 | 1.246964 | 2.610519 | 0.477669 | 0.632886 | NA |
| | 0.151697 | -1.546 | 4.075469 | -0.37934 | 0.704433 | NA |
| DLG5-AS1 | 11.5214 | -0.10657 | 0.63534 | -0.16773 | 0.866795 | NA |
| MT1XP1 | 0.517155 | 2.232561 | 3.179631 | 0.702145 | 0.482589 | NA |
| KRTAP5-AS | 2.440991 | 2.626753 | 1.534577 | 1.711712 | 0.08695 | NA |
| | 146.6825 | 0.450109 | 0.227369 | 1.979643 | 0.047744 | 0.134702 |
| | 0.3441 | -0.0166 | 3.769682 | -0.0044 | 0.996486 | NA |
| | 0.883203 | 0.854847 | 2.135496 | 0.400304 | 0.688933 | NA |
| GDI2P2 | 2.731879 | -0.37661 | 1.258909 | -0.29916 | 0.764821 | NA |
| | 0.162376 | 0.424913 | 4.079006 | 0.104171 | 0.917034 | NA |
| CHIAP3 | 0.448206 | -1.2532 | 3.007315 | -0.41672 | 0.676885 | NA |
| | 0.162376 | 0.424913 | 4.079006 | 0.104171 | 0.917034 | NA |
| | 0.260158 | 1.240347 | 4.047124 | 0.306476 | 0.759242 | NA |
| | 6.045033 | 0.290783 | 0.834556 | 0.348428 | 0.727518 | NA |
| | 1.380377 | 0.882103 | 1.689164 | 0.522213 | 0.601522 | NA |
| | 0.324752 | 1.53082 | 4.022025 | 0.380609 | 0.703493 | NA |
| RPS15AP11 | 0.758758 | -0.35643 | 2.514365 | -0.14176 | 0.887273 | NA |
| | 0.134133 | -1.48909 | 4.077584 | -0.36519 | 0.71497 | NA |
| | 0.134133 | -1.48909 | 4.077584 | -0.36519 | 0.71497 | NA |
| RPL13AP6 | 3.247176 | -1.21004 | 1.14015 | -1.0613 | 0.288554 | NA |

| | | | | | | |
|---|---|---|---|---|---|---|
| | 0.998346 | 1.943196 | 2.103833 | 0.923646 | 0.355671 | NA |
| TCEB2P4 | 0.096202 | 0.408546 | 4.079639 | 0.100143 | 0.920231 | NA |
| | 4.08628 | -0.10899 | 1.018444 | -0.10702 | 0.914773 | NA |
| | 0.096202 | 0.408546 | 4.079639 | 0.100143 | 0.920231 | NA |
| | 0.230334 | -0.53318 | 4.076749 | -0.13079 | 0.895945 | NA |
| | 5.210091 | 0.578227 | 0.930141 | 0.621655 | 0.534168 | NA |
| | 3.361612 | 1.321071 | 1.254799 | 1.052815 | 0.292426 | NA |
| KCNH1-IT1 | 10.65541 | 0.512447 | 0.686039 | 0.746965 | 0.455085 | NA |
| | 0.489058 | -0.58569 | 3.427543 | -0.17088 | 0.864321 | NA |
| SOD2P1 | 0.096202 | 0.408546 | 4.079639 | 0.100143 | 0.920231 | NA |
| | 1.747563 | 1.527456 | 1.672994 | 0.913007 | 0.361239 | NA |
| | 0.162376 | 0.424913 | 4.079006 | 0.104171 | 0.917034 | NA |
| | 3.871359 | 1.936975 | 1.206446 | 1.605522 | 0.108379 | NA |
| | 2.916153 | 1.50767 | 1.372294 | 1.098649 | 0.271921 | NA |
| BCAS2P2 | 0.842643 | -2.08432 | 2.306153 | -0.90381 | 0.366097 | NA |
| RPS4XP11 | 0.610582 | -0.18034 | 2.753139 | -0.0655 | 0.947774 | NA |
| PFN1P3 | 0.72077 | 0.527966 | 3.060238 | 0.172524 | 0.863025 | NA |
| TATDN1P1 | 0.452561 | 2.061505 | 3.349456 | 0.615474 | 0.538241 | NA |
| | 0.096202 | 0.408546 | 4.079639 | 0.100143 | 0.920231 | NA |
| CTBP2P8 | 3.299212 | 2.109404 | 1.322635 | 1.594849 | 0.110746 | NA |
| | 2.350232 | 0.186277 | 1.454679 | 0.128054 | 0.898106 | NA |
| | 0.652869 | 1.292592 | 2.374873 | 0.544278 | 0.58625 | NA |
| ZNF37BP | 75.1592 | 1.076277 | 0.30119 | 3.573418 | 3.52E-04 | 0.005475 |
| | 0.151697 | -1.546 | 4.075469 | -0.37934 | 0.704433 | NA |
| ZNF101P2 | 0.462046 | 0.589069 | 3.442439 | 0.17112 | 0.86413 | NA |
| | 0.326333 | 1.533889 | 3.826878 | 0.40082 | 0.688553 | NA |
| | 0.163957 | 0.424913 | 4.079006 | 0.104171 | 0.917034 | NA |
| | 0.268266 | -2.15396 | 4.038351 | -0.53338 | 0.593773 | NA |
| PTGES3P1 | 63.16873 | -0.07325 | 0.276941 | -0.26449 | 0.791403 | 0.873483 |
| NDUFB1P2 | 0.163957 | 0.424913 | 4.079006 | 0.104171 | 0.917034 | NA |
| | 5.306589 | 0.510604 | 0.873259 | 0.584711 | 0.558742 | NA |
| | 2.140476 | 0.841474 | 1.391203 | 0.604854 | 0.545276 | NA |
| LINC00624 | 0.134133 | -1.48909 | 4.077584 | -0.36519 | 0.71497 | NA |
| | 7.337811 | -0.30534 | 0.776059 | -0.39345 | 0.693984 | NA |
| RPL7P11 | 0.151697 | -1.546 | 4.075469 | -0.37934 | 0.704433 | NA |
| | 2.84726 | 0.705717 | 1.16239 | 0.607126 | 0.543767 | NA |
| | 1.801534 | 0.918548 | 1.480142 | 0.620581 | 0.534875 | NA |
| | 5.280436 | -1.16621 | 0.882733 | -1.32113 | 0.186457 | NA |
| | 0.983514 | -1.5981 | 2.093157 | -0.76349 | 0.445173 | NA |
| GAS5 | 4621.383 | 0.163829 | 0.176183 | 0.929881 | 0.352433 | 0.520437 |
| EIF5AP4 | 0.28583 | -2.21358 | 3.97376 | -0.55705 | 0.577494 | NA |
| C11orf94 | 1.525189 | 1.125604 | 1.607983 | 0.70001 | 0.483921 | NA |
| LINC01135 | 3.894501 | -0.26163 | 1.201292 | -0.21779 | 0.827594 | NA |
| | 2.168733 | -0.97571 | 1.562198 | -0.62457 | 0.532252 | NA |
| SVILP1 | 26.49082 | -0.06968 | 0.418647 | -0.16644 | 0.867809 | NA |
| HNRNPUL2 | 13.49995 | -0.33604 | 0.605937 | -0.55458 | 0.579182 | NA |
| | 6.440547 | 0.20414 | 0.817942 | 0.249578 | 0.802914 | NA |

| | | | | | | |
|---|---|---|---|---|---|---|
| | 0.162376 | 0.424913 | 4.079006 | 0.104171 | 0.917034 | NA |
| LINC00700 | 0.096202 | 0.408546 | 4.079639 | 0.100143 | 0.920231 | NA |
| FABP5P7 | 0.163957 | 0.424913 | 4.079006 | 0.104171 | 0.917034 | NA |
| FTH1P2 | 2.966504 | -0.84042 | 1.122743 | -0.74854 | 0.454133 | NA |
| | 16.73214 | 0.454921 | 0.57266 | 0.794399 | 0.426963 | NA |
| | 0.268266 | -2.15396 | 4.038351 | -0.53338 | 0.593773 | NA |
| CUBNP2 | 0.134133 | -1.48909 | 4.077584 | -0.36519 | 0.71497 | NA |
| | 16.84842 | -0.48483 | 0.510143 | -0.95039 | 0.341915 | NA |
| | 1.665821 | 1.285941 | 1.600278 | 0.803574 | 0.421643 | NA |
| | 0.096202 | 0.408546 | 4.079639 | 0.100143 | 0.920231 | NA |
| | 0.641986 | 1.117767 | 2.709494 | 0.412537 | 0.679946 | NA |
| | 48.20781 | 0.338683 | 0.376813 | 0.89881 | 0.368754 | 0.535798 |
| VDAC1P4 | 0.162376 | 0.424913 | 4.079006 | 0.104171 | 0.917034 | NA |
| ZRANB2-AS | 0.096202 | 0.408546 | 4.079639 | 0.100143 | 0.920231 | NA |
| SUMO1P3 | 0.889885 | 0.644651 | 2.1706 | 0.296992 | 0.766472 | NA |
| CHCHD2P6 | 5.092616 | -0.30208 | 0.906899 | -0.33309 | 0.739064 | NA |
| | 0.324752 | 1.53082 | 4.022025 | 0.380609 | 0.703493 | NA |
| ANKRD65 | 1.729367 | -0.39907 | 1.526801 | -0.26137 | 0.793804 | NA |
| | 5.293712 | 1.404263 | 0.995045 | 1.411256 | 0.158169 | NA |
| | 6.299698 | -0.14756 | 0.776483 | -0.19003 | 0.849282 | NA |
| | 0.298089 | -0.51711 | 3.975232 | -0.13008 | 0.896501 | NA |
| | 0.096202 | 0.408546 | 4.079639 | 0.100143 | 0.920231 | NA |
| | 0.260158 | 1.240347 | 4.047124 | 0.306476 | 0.759242 | NA |
| | 0.815391 | 0.931626 | 2.360431 | 0.394685 | 0.693076 | NA |
| | 0.151697 | -1.546 | 4.075469 | -0.37934 | 0.704433 | NA |
| | 0.134133 | -1.48909 | 4.077584 | -0.36519 | 0.71497 | NA |
| SMIM1 | 85.78757 | -0.61354 | 0.294178 | -2.08562 | 0.037013 | 0.11412 |
| LINC00601 | 7.865005 | 0.591988 | 0.717958 | 0.824544 | 0.40963 | NA |
| | 13.50226 | -0.42874 | 0.618151 | -0.69358 | 0.487945 | NA |
| | 10.35266 | 0.309526 | 0.662433 | 0.467256 | 0.640317 | NA |
| RPL5P28 | 0.134133 | -1.48909 | 4.077584 | -0.36519 | 0.71497 | NA |
| | 0.096202 | 0.408546 | 4.079639 | 0.100143 | 0.920231 | NA |
| | 0.392857 | -1.546 | 4.075469 | -0.37934 | 0.704433 | NA |
| MRPL53P1 | 0.192403 | 0.975392 | 4.060716 | 0.240202 | 0.810174 | NA |
| | 0.134133 | -1.48909 | 4.077584 | -0.36519 | 0.71497 | NA |
| | 0.584911 | 2.38756 | 2.690807 | 0.887303 | 0.374916 | NA |
| DUSP8P5 | 18.28592 | 0.359866 | 0.522026 | 0.689364 | 0.490594 | NA |
| | 1.276206 | -1.07269 | 2.030043 | -0.52841 | 0.597216 | NA |
| SNRPGP10 | 29.22525 | 0.646732 | 0.423578 | 1.526831 | 0.126803 | NA |
| | 1.128326 | -0.93784 | 1.813709 | -0.51708 | 0.605099 | NA |
| | 0.458885 | 0.577525 | 3.449278 | 0.167433 | 0.867029 | NA |
| RPEL1 | 1.879969 | 1.528624 | 1.494589 | 1.022772 | 0.306416 | NA |
| MRPS10P1 | 0.735616 | -2.07189 | 2.528723 | -0.81934 | 0.412591 | NA |
| LINC00623 | 0.134133 | -1.48909 | 4.077584 | -0.36519 | 0.71497 | NA |
| | 1.071608 | 0.294623 | 2.235884 | 0.13177 | 0.895166 | NA |
| | 0.610582 | -0.18034 | 2.753139 | -0.0655 | 0.947774 | NA |
| | 0.315653 | -0.57305 | 3.892639 | -0.14721 | 0.882963 | NA |

| | | | | | | |
|---|---|---|---|---|---|---|
| | 1.452059 | -0.1661 | 1.670582 | -0.09943 | 0.920799 | NA |
| | 64.46126 | 0.405952 | 0.28456 | 1.426594 | 0.153697 | 0.298554 |
| DBF4P1 | 1.909939 | 1.719596 | 1.59897 | 1.07544 | 0.282178 | NA |
| | 224.2242 | 0.017146 | 0.200371 | 0.085569 | 0.931809 | 0.961401 |
| | 1.312853 | -1.50193 | 1.923163 | -0.78097 | 0.434822 | NA |
| | 1.63242 | 0.749319 | 1.641395 | 0.456513 | 0.648021 | NA |
| | 3.257331 | 0.957301 | 1.104529 | 0.866705 | 0.386104 | NA |
| | 7.991692 | 0.227289 | 0.813705 | 0.279326 | 0.779994 | NA |
| | 1.827295 | -0.97528 | 1.644179 | -0.59317 | 0.553068 | NA |
| GAPDHP23 | 0.151697 | -1.546 | 4.075469 | -0.37934 | 0.704433 | NA |
| | 0.640755 | -1.64826 | 2.754778 | -0.59833 | 0.549621 | NA |
| NSRP1P1 | 0.163957 | 0.424913 | 4.079006 | 0.104171 | 0.917034 | NA |
| | 7.204324 | -0.54959 | 0.80203 | -0.68525 | 0.493186 | NA |
| | 0.163957 | 0.424913 | 4.079006 | 0.104171 | 0.917034 | NA |
| LDHAP2 | 3.256393 | 2.986394 | 1.451737 | 2.057118 | 0.039675 | NA |
| NPM1P26 | 1.713595 | 0.710076 | 1.748569 | 0.40609 | 0.684677 | NA |
| CYCSP52 | 1.113131 | 2.168349 | 2.301419 | 0.942179 | 0.346101 | NA |
| | 0.134133 | -1.48909 | 4.077584 | -0.36519 | 0.71497 | NA |
| | 0.260158 | 1.240347 | 4.047124 | 0.306476 | 0.759242 | NA |
| KIAA0040 | 1.929221 | -0.83152 | 1.789462 | -0.46468 | 0.642163 | NA |
| | 0.594599 | -0.12152 | 2.771051 | -0.04385 | 0.965021 | NA |
| | 2.760157 | 0.991918 | 1.232547 | 0.804771 | 0.420952 | NA |
| | 0.946806 | -1.54284 | 2.156294 | -0.7155 | 0.474297 | NA |
| PRELID1P3 | 0.096202 | 0.408546 | 4.079639 | 0.100143 | 0.920231 | NA |
| LINC00263 | 196.3694 | -0.13146 | 0.209137 | -0.6286 | 0.529608 | 0.674744 |
| HSP90AB7I | 0.163957 | 0.424913 | 4.079006 | 0.104171 | 0.917034 | NA |
| | 0.392857 | -1.546 | 4.075469 | -0.37934 | 0.704433 | NA |
| | 0.392857 | -1.546 | 4.075469 | -0.37934 | 0.704433 | NA |
| | 12.57796 | 0.025669 | 0.571878 | 0.044886 | 0.964198 | NA |
| ASH1L-AS1 | 18.40878 | -0.21232 | 0.498446 | -0.42596 | 0.670138 | NA |
| NEXN-AS1 | 0.681461 | 0.491544 | 3.066278 | 0.160307 | 0.87264 | NA |
| | 0.096202 | 0.408546 | 4.079639 | 0.100143 | 0.920231 | NA |
| | 0.096202 | 0.408546 | 4.079639 | 0.100143 | 0.920231 | NA |
| | 63.52469 | -0.30729 | 0.294935 | -1.04189 | 0.297462 | 0.464613 |
| | 2.011233 | 0.070694 | 1.456915 | 0.048523 | 0.9613 | NA |
| LINC01036 | 0.70693 | -1.62766 | 2.687154 | -0.60572 | 0.544701 | NA |
| | 1.698391 | 1.510988 | 1.665448 | 0.907257 | 0.364271 | NA |
| | 0.610582 | -0.18034 | 2.753139 | -0.0655 | 0.947774 | NA |
| CHIAP1 | 0.67854 | -0.89371 | 2.366391 | -0.37767 | 0.705677 | NA |
| | 3.638371 | -0.06284 | 1.010725 | -0.06217 | 0.950425 | NA |
| | 0.163957 | 0.424913 | 4.079006 | 0.104171 | 0.917034 | NA |
| | 0.458885 | 0.577525 | 3.449278 | 0.167433 | 0.867029 | NA |
| | 0.780475 | 2.824774 | 2.754691 | 1.025441 | 0.305155 | NA |
| | 0.392857 | -1.546 | 4.075469 | -0.37934 | 0.704433 | NA |
| | 1.024164 | -1.06815 | 2.390105 | -0.4469 | 0.654944 | NA |
| | 2.225438 | 0.52186 | 1.418589 | 0.367873 | 0.712968 | NA |
| | 0.134133 | -1.48909 | 4.077584 | -0.36519 | 0.71497 | NA |

| | | | | | | |
|---|---|---|---|---|---|---|
| | 9.667345 | 0.579356 | 0.687472 | 0.842734 | 0.399377 | NA |
| KDM4A-AS | 7.83113 | 0.515024 | 0.802203 | 0.642012 | 0.520865 | NA |
| | 0.134133 | -1.48909 | 4.077584 | -0.36519 | 0.71497 | NA |
| PPP1R11P1 | 0.096202 | 0.408546 | 4.079639 | 0.100143 | 0.920231 | NA |
| | 3.075177 | 0.213863 | 1.18206 | 0.180924 | 0.856428 | NA |
| EI24P2 | 39.34571 | -0.1914 | 0.374353 | -0.51129 | 0.609145 | NA |
| RPL26P30 | 0.327913 | 1.543154 | 4.021063 | 0.383768 | 0.701151 | NA |
| | 3.090011 | 1.146059 | 1.237554 | 0.926068 | 0.354411 | NA |
| | 2.795911 | 0.019446 | 1.293245 | 0.015036 | 0.988003 | NA |
| ENO1-IT1 | 1.209942 | 0.263859 | 2.515142 | 0.104908 | 0.916449 | NA |
| ZBED5 | 532.787 | 0.71464 | 0.227043 | 3.147593 | 0.001646 | 0.014318 |
| | 0.258578 | 1.233151 | 4.047809 | 0.304646 | 0.760635 | NA |
| | 3.106232 | 1.129267 | 1.302564 | 0.866957 | 0.385966 | NA |
| | 0.382031 | -1.26484 | 3.151989 | -0.40128 | 0.688212 | NA |
| PPIAL4G | 0.151697 | -1.546 | 4.075469 | -0.37934 | 0.704433 | NA |
| | 0.315653 | -0.57305 | 3.892639 | -0.14721 | 0.882963 | NA |
| | 0.151697 | -1.546 | 4.075469 | -0.37934 | 0.704433 | NA |
| | 1.744255 | 2.014588 | 1.598588 | 1.260229 | 0.207587 | NA |
| | 0.096202 | 0.408546 | 4.079639 | 0.100143 | 0.920231 | NA |
| LINC01134 | 30.57846 | 0.994124 | 0.479549 | 2.07304 | 0.038169 | NA |
| | 0.096202 | 0.408546 | 4.079639 | 0.100143 | 0.920231 | NA |
| | 0.887313 | -2.40227 | 2.38425 | -1.00756 | 0.313667 | NA |
| KRT8P29 | 0.35636 | 1.709252 | 3.651833 | 0.468053 | 0.639746 | NA |
| | 0.163957 | 0.424913 | 4.079006 | 0.104171 | 0.917034 | NA |
| | 1.373639 | 1.032867 | 1.793152 | 0.576006 | 0.564611 | NA |
| | 0.506476 | 0.726418 | 2.864186 | 0.253621 | 0.799788 | NA |
| PKMP1 | 4.377828 | -0.2907 | 1.0808 | -0.26897 | 0.787953 | NA |
| RRM2P2 | 0.134133 | -1.48909 | 4.077584 | -0.36519 | 0.71497 | NA |
| EIF2S2P3 | 0.915556 | -1.07826 | 2.191337 | -0.49206 | 0.62268 | NA |
| | 0.260158 | 1.240347 | 4.047124 | 0.306476 | 0.759242 | NA |
| NPM1P40 | 0.35636 | 1.709252 | 3.651833 | 0.468053 | 0.639746 | NA |
| | 0.526843 | -0.38719 | 2.855892 | -0.13558 | 0.892156 | NA |
| RPL13AP5 | 86.22855 | -0.25059 | 0.319135 | -0.78522 | 0.432328 | 0.5935 |
| PRPF38AP2 | 0.887313 | -2.40227 | 2.38425 | -1.00756 | 0.313667 | NA |
| CCDC163P | 51.39035 | -0.55835 | 0.383044 | -1.45766 | 0.144934 | 0.286891 |
| LAMTOR5P | 0.163957 | 0.424913 | 4.079006 | 0.104171 | 0.917034 | NA |
| | 0.096202 | 0.408546 | 4.079639 | 0.100143 | 0.920231 | NA |
| PRKG1-AS1 | 7.067086 | 0.907633 | 0.849166 | 1.068852 | 0.285137 | NA |
| MTX1P1 | 46.25037 | 0.073849 | 0.343173 | 0.215196 | 0.829615 | NA |
| EIF1AXP1 | 1.905786 | 0.007761 | 1.367774 | 0.005674 | 0.995472 | NA |
| | 1.101367 | -2.48182 | 2.397488 | -1.03518 | 0.300587 | NA |
| | 1.182873 | 0.724533 | 2.160891 | 0.335294 | 0.737403 | NA |
| | 0.392857 | -1.546 | 4.075469 | -0.37934 | 0.704433 | NA |
| | 2.028302 | 1.607251 | 1.523631 | 1.054882 | 0.291479 | NA |
| | 0.230334 | -0.53318 | 4.076749 | -0.13079 | 0.895945 | NA |
| | 5.674955 | -0.4284 | 0.880964 | -0.48629 | 0.626764 | NA |
| | 1.464116 | 0.942734 | 1.806734 | 0.521789 | 0.601817 | NA |

| | | | | | | |
|---|---|---|---|---|---|---|
| SNX25P1 | 3.900861 | 1.080774 | 1.080445 | 1.000304 | 0.317163 | NA |
| | 11.58574 | 0.534508 | 0.603844 | 0.885176 | 0.376062 | NA |
| ARL5AP2 | 0.162376 | 0.424913 | 4.079006 | 0.104171 | 0.917034 | NA |
| | 0.585114 | 1.12675 | 2.73598 | 0.411827 | 0.680466 | NA |
| | 0.653015 | 0.2552 | 2.787103 | 0.091565 | 0.927044 | NA |
| TMSB4XP1 | 0.096202 | 0.408546 | 4.079639 | 0.100143 | 0.920231 | NA |
| | 0.162376 | 0.424913 | 4.079006 | 0.104171 | 0.917034 | NA |
| TIMM9P1 | 0.392857 | -1.546 | 4.075469 | -0.37934 | 0.704433 | NA |
| | 0.163957 | 0.424913 | 4.079006 | 0.104171 | 0.917034 | NA |
| | 0.163957 | 0.424913 | 4.079006 | 0.104171 | 0.917034 | NA |
| PTGES3P4 | 0.897403 | 1.74906 | 2.471224 | 0.707771 | 0.479088 | NA |
| | 0.258578 | 1.233151 | 4.047809 | 0.304646 | 0.760635 | NA |
| | 26.62742 | 0.244933 | 0.481711 | 0.508464 | 0.611128 | NA |
| | 1.419881 | -1.51816 | 1.910563 | -0.79461 | 0.42684 | NA |
| | 0.134133 | -1.48909 | 4.077584 | -0.36519 | 0.71497 | NA |
| LINC01141 | 0.392857 | -1.546 | 4.075469 | -0.37934 | 0.704433 | NA |
| | 0.151697 | -1.546 | 4.075469 | -0.37934 | 0.704433 | NA |
| LINC00614 | 0.096202 | 0.408546 | 4.079639 | 0.100143 | 0.920231 | NA |
| | 4.057385 | 0.289849 | 1.022207 | 0.283552 | 0.776754 | NA |
| EDRF1-AS1 | 0.795081 | 1.604357 | 2.508662 | 0.639527 | 0.52248 | NA |
| GAPDHP21 | 0.69625 | -3.13723 | 2.93845 | -1.06765 | 0.285679 | NA |
| | 0.151697 | -1.546 | 4.075469 | -0.37934 | 0.704433 | NA |
| | 0.303394 | -2.27006 | 4.032451 | -0.56295 | 0.57347 | NA |
| RPL34P1 | 0.162376 | 0.424913 | 4.079006 | 0.104171 | 0.917034 | NA |
| | 4.991882 | 0.179316 | 0.92717 | 0.193401 | 0.846645 | NA |
| | 0.440301 | 0.468383 | 3.449808 | 0.135771 | 0.892003 | NA |
| | 0.162376 | 0.424913 | 4.079006 | 0.104171 | 0.917034 | NA |
| | 0.823499 | -2.06708 | 2.534576 | -0.81555 | 0.414756 | NA |
| | 0.134133 | -1.48909 | 4.077584 | -0.36519 | 0.71497 | NA |
| | 0.47803 | 0.520083 | 2.97881 | 0.174594 | 0.861399 | NA |
| ZNF503-AS | 253.725 | 0.179111 | 0.250771 | 0.714242 | 0.475078 | 0.630406 |
| | 0.324752 | 1.53082 | 4.022025 | 0.380609 | 0.703493 | NA |
| | 0.410275 | 0.244283 | 3.100889 | 0.078778 | 0.937209 | NA |
| | 35.07259 | 0.524935 | 0.386044 | 1.359779 | 0.1739 | NA |
| ZBTB40-IT1 | 0.834592 | 0.684312 | 2.202411 | 0.31071 | 0.756021 | NA |
| | 0.096202 | 0.408546 | 4.079639 | 0.100143 | 0.920231 | NA |
| | 0.873515 | 2.9967 | 2.318337 | 1.292607 | 0.196147 | NA |
| ANO7P1 | 63.03336 | -0.60883 | 0.295503 | -2.06033 | 0.039367 | 0.119055 |
| | 0.326333 | 1.533889 | 3.826878 | 0.40082 | 0.688553 | NA |
| | 0.162376 | 0.424913 | 4.079006 | 0.104171 | 0.917034 | NA |
| | 0.544407 | -0.44425 | 2.542878 | -0.1747 | 0.861313 | NA |
| RNF223 | 0.163957 | 0.424913 | 4.079006 | 0.104171 | 0.917034 | NA |
| | 0.096202 | 0.408546 | 4.079639 | 0.100143 | 0.920231 | NA |
| PITRM1-AS | 19.71086 | 1.183778 | 0.505382 | 2.342346 | 0.019163 | NA |
| CAMTA1-IT | 0.488912 | 0.771583 | 2.897678 | 0.266276 | 0.790026 | NA |
| NRBF2P3 | 0.096202 | 0.408546 | 4.079639 | 0.100143 | 0.920231 | NA |
| | 0.288605 | 1.443619 | 4.029058 | 0.358302 | 0.720117 | NA |

| | | | | | | |
|---|---|---|---|---|---|---|
| XRCC6P1 | 0.858627 | -2.14058 | 2.513451 | -0.85165 | 0.394408 | NA |
| FOXD2-AS1 | 120.7544 | -0.37459 | 0.267545 | -1.4001 | 0.161484 | 0.309069 |
| | 0.151697 | -1.546 | 4.075469 | -0.37934 | 0.704433 | NA |
| | 7.117247 | 0.136736 | 0.823013 | 0.166141 | 0.868046 | NA |
| | 0.163957 | 0.424913 | 4.079006 | 0.104171 | 0.917034 | NA |
| | 0.563552 | -0.48645 | 2.808029 | -0.17324 | 0.862465 | NA |
| DCLRE1CP1 | 0.326536 | 0.037056 | 3.842554 | 0.009644 | 0.992306 | NA |
| | 0.716471 | -2.03238 | 2.582811 | -0.78689 | 0.431348 | NA |
| LINC00959 | 15.24585 | 0.231162 | 0.553156 | 0.417898 | 0.676022 | NA |
| | 1.697428 | 1.343932 | 1.617987 | 0.83062 | 0.406188 | NA |
| | 0.230334 | -0.53318 | 4.076749 | -0.13079 | 0.895945 | NA |
| UNC5B-AS1 | 0.096202 | 0.408546 | 4.079639 | 0.100143 | 0.920231 | NA |
| PTP4A1P7 | 0.163957 | 0.424913 | 4.079006 | 0.104171 | 0.917034 | NA |
| LINC00857 | 60.59772 | 0.233287 | 0.402442 | 0.579679 | 0.562131 | 0.700081 |
| | 0.411855 | 0.250456 | 3.097618 | 0.080854 | 0.935558 | NA |
| | 0.364467 | -1.20166 | 3.685431 | -0.32606 | 0.744381 | NA |
| | 0.163957 | 0.424913 | 4.079006 | 0.104171 | 0.917034 | NA |
| | 2.43421 | -0.0563 | 1.224433 | -0.04598 | 0.963328 | NA |
| | 15.55481 | 0.629336 | 0.581723 | 1.081847 | 0.27932 | NA |
| OXCT2P1 | 1.263597 | -0.09324 | 1.722854 | -0.05412 | 0.956842 | NA |
| | 0.096202 | 0.408546 | 4.079639 | 0.100143 | 0.920231 | NA |
| | 1.579453 | -0.56402 | 1.503609 | -0.37511 | 0.707578 | NA |
| | 1.374791 | -2.07862 | 1.792619 | -1.15955 | 0.246234 | NA |
| | 1.610712 | 0.666451 | 1.557285 | 0.427957 | 0.668683 | NA |
| FAM21FP | 0.640609 | 0.042838 | 2.379511 | 0.018003 | 0.985637 | NA |
| | 0.956262 | 0.048146 | 1.9097 | 0.025211 | 0.979886 | NA |
| | 1.005076 | -0.3957 | 2.04069 | -0.1939 | 0.846251 | NA |
| | 0.462046 | 0.589069 | 3.442439 | 0.17112 | 0.86413 | NA |
| | 0.872467 | -0.54278 | 2.536752 | -0.21397 | 0.830573 | NA |
| PPIAP13 | 1.043796 | 2.066306 | 2.144878 | 0.963368 | 0.335363 | NA |
| | 0.162376 | 0.424913 | 4.079006 | 0.104171 | 0.917034 | NA |
| | 3.69128 | 0.404532 | 1.040341 | 0.388846 | 0.69739 | NA |
| | 0.162376 | 0.424913 | 4.079006 | 0.104171 | 0.917034 | NA |
| | 0.163957 | 0.424913 | 4.079006 | 0.104171 | 0.917034 | NA |
| | 0.47803 | 0.520083 | 2.97881 | 0.174594 | 0.861399 | NA |
| | 0.55687 | -0.14239 | 3.170029 | -0.04492 | 0.964172 | NA |
| | 0.854903 | -0.47663 | 2.545657 | -0.18723 | 0.851479 | NA |
| | 9.702686 | -0.25405 | 0.663815 | -0.38271 | 0.701935 | NA |
| PRKCQ-AS1 | 5.603886 | -0.50157 | 0.884566 | -0.56702 | 0.5707 | NA |
| LINC00844 | 1.73954 | 0.950876 | 1.722233 | 0.552118 | 0.580867 | NA |
| | 3.134024 | 1.206366 | 1.28827 | 0.936423 | 0.349055 | NA |
| | 16325.55 | -0.56459 | 0.131519 | -4.29282 | 1.76E-05 | 8.56E-04 |
| FLG-AS1 | 96.77347 | -0.81072 | 0.302422 | -2.68077 | 0.007345 | 0.038445 |
| | 21.792 | 0.735837 | 0.448116 | 1.642069 | 0.100576 | NA |
| | 0.399595 | -1.31595 | 3.579466 | -0.36764 | 0.713143 | NA |
| RPL35P1 | 0.71919 | 0.510978 | 2.728734 | 0.187258 | 0.851458 | NA |
| | 0.163957 | 0.424913 | 4.079006 | 0.104171 | 0.917034 | NA |

| | | | | | | |
|---|---|---|---|---|---|---|
| | 0.411855 | 0.250456 | 3.097618 | 0.080854 | 0.935558 | NA |
| | 0.258578 | 1.233151 | 4.047809 | 0.304646 | 0.760635 | NA |
| | 1.265234 | 0.20438 | 1.735412 | 0.11777 | 0.90625 | NA |
| | 1.631999 | 0.092865 | 1.767507 | 0.05254 | 0.958098 | NA |
| | 0.162376 | 0.424913 | 4.079006 | 0.104171 | 0.917034 | NA |
| ARMC4P1 | 0.163957 | 0.424913 | 4.079006 | 0.104171 | 0.917034 | NA |
| | 0.364467 | -1.20166 | 3.685431 | -0.32606 | 0.744381 | NA |
| | 0.230334 | -0.53318 | 4.076749 | -0.13079 | 0.895945 | NA |
| | 0.422534 | 1.924583 | 3.001758 | 0.641152 | 0.521424 | NA |
| | 0.804769 | -0.21293 | 2.457871 | -0.08663 | 0.930964 | NA |
| | 0.516164 | -1.70268 | 2.844782 | -0.59853 | 0.549488 | NA |
| | 1.14431 | -0.94385 | 1.953694 | -0.48311 | 0.629019 | NA |
| | 0.096202 | 0.408546 | 4.079639 | 0.100143 | 0.920231 | NA |
| SLC25A24P | 0.298089 | -0.51711 | 3.975232 | -0.13008 | 0.896501 | NA |
| ARPC3P2 | 1.248845 | 1.474697 | 2.199721 | 0.670402 | 0.502602 | NA |
| | 1.030544 | -1.32239 | 2.273679 | -0.58161 | 0.560832 | NA |
| | 12.04419 | -0.61971 | 0.642898 | -0.96394 | 0.335078 | NA |
| | 146.7285 | -0.29425 | 0.259257 | -1.13496 | 0.25639 | 0.420517 |
| CD81-AS1 | 4.689626 | 0.752162 | 0.954728 | 0.787828 | 0.430797 | NA |
| | 3.396419 | 1.006368 | 1.164465 | 0.864232 | 0.38746 | NA |
| | 2.610238 | -1.08952 | 1.342129 | -0.81179 | 0.416913 | NA |
| UBE2V2P3 | 0.258578 | 1.233151 | 4.047809 | 0.304646 | 0.760635 | NA |
| | 0.670636 | 0.219794 | 2.632532 | 0.083491 | 0.933461 | NA |
| | 0.326333 | 1.533889 | 3.826878 | 0.40082 | 0.688553 | NA |
| OR2W3 | 0.450981 | 2.057325 | 3.353149 | 0.61355 | 0.539513 | NA |
| HMGN2P17 | 0.410275 | 0.244283 | 3.100889 | 0.078778 | 0.937209 | NA |
| | 0.314073 | -0.57308 | 3.898934 | -0.14698 | 0.883146 | NA |
| | 0.780475 | 2.824774 | 2.754691 | 1.025441 | 0.305155 | NA |
| | 5.897639 | 0.937283 | 0.839566 | 1.11639 | 0.264255 | NA |
| LINC00707 | 1.439814 | 1.120216 | 1.789219 | 0.626092 | 0.531255 | NA |
| | 0.134133 | -1.48909 | 4.077584 | -0.36519 | 0.71497 | NA |
| | 11.15474 | 0.91395 | 0.658719 | 1.387465 | 0.1653 | NA |
| | 1.751075 | -0.45025 | 1.655132 | -0.27203 | 0.785598 | NA |
| | 3.243731 | 0.392625 | 1.132892 | 0.346569 | 0.728915 | NA |
| | 0.096202 | 0.408546 | 4.079639 | 0.100143 | 0.920231 | NA |
| RNU7-57P | 0.544407 | -0.44425 | 2.542878 | -0.1747 | 0.861313 | NA |
| | 2.65893 | -0.58926 | 1.341551 | -0.43924 | 0.660488 | NA |
| | 0.151697 | -1.546 | 4.075469 | -0.37934 | 0.704433 | NA |
| RNY4P25 | 0.151697 | -1.546 | 4.075469 | -0.37934 | 0.704433 | NA |
| | 0.096202 | 0.408546 | 4.079639 | 0.100143 | 0.920231 | NA |
| | 0.151697 | -1.546 | 4.075469 | -0.37934 | 0.704433 | NA |
| | 0.151697 | -1.546 | 4.075469 | -0.37934 | 0.704433 | NA |
| RNU7-163P | 1.313817 | 2.449753 | 1.95775 | 1.25131 | 0.210821 | NA |
| | 1.723043 | 0.159776 | 1.599185 | 0.099911 | 0.920415 | NA |
| | 0.096202 | 0.408546 | 4.079639 | 0.100143 | 0.920231 | NA |
| RN7SL854P | 0.520316 | 2.239969 | 3.173769 | 0.705776 | 0.480328 | NA |
| RN7SL425P | 0.260158 | 1.240347 | 4.047124 | 0.306476 | 0.759242 | NA |

| | | | | | | |
|---|---|---|---|---|---|---|
| | 1.427407 | 0.984049 | 1.659042 | 0.593143 | 0.553086 | NA |
| RN7SL333F | 1.869436 | 0.435321 | 1.4412 | 0.302055 | 0.76261 | NA |
| | 5.80372 | -0.36206 | 0.898322 | -0.40304 | 0.686922 | NA |
| | 55.82682 | 0.843809 | 0.308491 | 2.735278 | 0.006233 | 0.034188 |
| | 2.176072 | 2.355961 | 1.626038 | 1.448897 | 0.147366 | NA |
| RN7SL825F | 0.162376 | 0.424913 | 4.079006 | 0.104171 | 0.917034 | NA |
| | 0.162376 | 0.424913 | 4.079006 | 0.104171 | 0.917034 | NA |
| RN7SL754F | 0.096202 | 0.408546 | 4.079639 | 0.100143 | 0.920231 | NA |
| GAPDHP27 | 0.162376 | 0.424913 | 4.079006 | 0.104171 | 0.917034 | NA |
| | 0.488709 | 2.119056 | 3.307564 | 0.64067 | 0.521737 | NA |
| C1orf226 | 3.668581 | -0.30557 | 1.109632 | -0.27538 | 0.783026 | NA |
| | 0.134133 | -1.48909 | 4.077584 | -0.36519 | 0.71497 | NA |
| | 0.402398 | -2.58771 | 4.018369 | -0.64397 | 0.519595 | NA |
| TEX35 | 0.162376 | 0.424913 | 4.079006 | 0.104171 | 0.917034 | NA |
| | 3.935815 | 0.530981 | 1.031337 | 0.514847 | 0.60666 | NA |
| AMY2B | 6.606616 | 2.214117 | 0.920752 | 2.404684 | 0.016186 | NA |
| GAPDHP33 | 2.066793 | 1.103409 | 1.573842 | 0.701093 | 0.483245 | NA |
| | 1.572394 | 2.735269 | 1.870564 | 1.46227 | 0.143667 | NA |
| RPL36AP4C | 0.163957 | 0.424913 | 4.079006 | 0.104171 | 0.917034 | NA |
| RPS4XP13 | 1.252918 | -0.9216 | 1.717077 | -0.53673 | 0.591455 | NA |
| MTATP8P1 | 2.034487 | -1.68121 | 1.764336 | -0.95288 | 0.340649 | NA |
| UOX | 1.32207 | 0.156255 | 1.806852 | 0.086479 | 0.931085 | NA |
| L1TD1 | 3.58885 | 0.417884 | 1.122952 | 0.37213 | 0.709796 | NA |
| | 10.01969 | 0.280217 | 0.636158 | 0.440484 | 0.659587 | NA |
| RN7SL559F | 0.314073 | -0.57308 | 3.898934 | -0.14698 | 0.883146 | NA |
| | 0.162376 | 0.424913 | 4.079006 | 0.104171 | 0.917034 | NA |
| RN7SL23P | 0.736957 | -1.0303 | 2.640699 | -0.39016 | 0.696418 | NA |
| HIST2H2BB | 41.2129 | 0.096029 | 0.436136 | 0.220182 | 0.82573 | NA |
| | 0.151697 | -1.546 | 4.075469 | -0.37934 | 0.704433 | NA |
| | 2.023144 | 1.869689 | 1.609954 | 1.16133 | 0.245508 | NA |
| | 8.281653 | -1.25368 | 0.7019 | -1.78612 | 0.074079 | NA |
| NSUN6 | 67.71386 | 0.444423 | 0.297224 | 1.495243 | 0.134851 | 0.273368 |
| | 1.206375 | 1.44275 | 2.16406 | 0.666687 | 0.504972 | NA |
| | 0.392857 | -1.546 | 4.075469 | -0.37934 | 0.704433 | NA |
| OR5G3 | 0.096202 | 0.408546 | 4.079639 | 0.100143 | 0.920231 | NA |
| | 6.925979 | -0.70841 | 0.771267 | -0.9185 | 0.358357 | NA |
| CORT | 2.674176 | -0.96082 | 1.297644 | -0.74043 | 0.459039 | NA |
| | 0.096202 | 0.408546 | 4.079639 | 0.100143 | 0.920231 | NA |
| HOGA1 | 25.96961 | -0.51414 | 0.521605 | -0.98569 | 0.324287 | NA |
| TCEB1P19 | 2.537939 | 1.503433 | 1.342756 | 1.119662 | 0.262858 | NA |
| SNHG3 | 424.594 | 0.191963 | 0.182037 | 1.054525 | 0.291643 | 0.458435 |
| | 246.7736 | -0.36291 | 0.231033 | -1.57083 | 0.116222 | 0.246565 |
| RPL5P29 | 0.134133 | -1.48909 | 4.077584 | -0.36519 | 0.71497 | NA |
| BGLAP | 1.158297 | -1.38149 | 2.066675 | -0.66846 | 0.503841 | NA |
| SKINTL | 0.096202 | 0.408546 | 4.079639 | 0.100143 | 0.920231 | NA |
| | 8.486641 | 0.897468 | 0.729375 | 1.230461 | 0.218524 | NA |
| BMS1P4 | 3.703176 | 0.447254 | 1.094534 | 0.408625 | 0.682815 | NA |

| | | | | | | |
|---|---|---|---|---|---|---|
| NPPA-AS1 | 10.18157 | 2.314221 | 0.75099 | 3.081562 | 0.002059 | NA |
| | 0.151697 | -1.546 | 4.075469 | -0.37934 | 0.704433 | NA |
| | 0.268266 | -2.15396 | 4.038351 | -0.53338 | 0.593773 | NA |
| | 1.511146 | 1.03585 | 1.697098 | 0.610365 | 0.54162 | NA |
| | 0.134133 | -1.48909 | 4.077584 | -0.36519 | 0.71497 | NA |
| MRPL20 | 1866.089 | 0.146595 | 0.1488 | 0.985183 | 0.324534 | 0.492617 |
| | 0.260158 | 1.240347 | 4.047124 | 0.306476 | 0.759242 | NA |
| MBL1P | 0.162376 | 0.424913 | 4.079006 | 0.104171 | 0.917034 | NA |
| CNTF | 18.6534 | 0.929782 | 0.507143 | 1.833373 | 0.066747 | NA |
| RN7SL846F | 0.096202 | 0.408546 | 4.079639 | 0.100143 | 0.920231 | NA |
| RN7SL749F | 0.134133 | -1.48909 | 4.077584 | -0.36519 | 0.71497 | NA |
| | 5.797937 | 0.019979 | 0.84726 | 0.02358 | 0.981187 | NA |
| | 6.656703 | -0.28981 | 0.782421 | -0.3704 | 0.711087 | NA |
| PGBD3 | 0.652666 | 2.537288 | 2.975981 | 0.852589 | 0.393887 | NA |
| VSIG8 | 1.064927 | 0.46521 | 2.027265 | 0.229477 | 0.818498 | NA |
| PTPRVP | 0.621261 | 1.168072 | 3.136787 | 0.372379 | 0.709611 | NA |
| EFNA4 | 146.2829 | 0.054359 | 0.235897 | 0.230436 | 0.817753 | 0.890746 |
| NBPF15 | 246.8168 | -0.04331 | 0.178161 | -0.24307 | 0.807953 | 0.885317 |
| ANTXRLP1 | 0.134133 | -1.48909 | 4.077584 | -0.36519 | 0.71497 | NA |
| ZNF487 | 37.61624 | 0.729283 | 0.358655 | 2.033382 | 0.042014 | NA |
| LEFTY1 | 0.995185 | 1.939099 | 2.104716 | 0.921311 | 0.356888 | NA |
| WDR65 | 2.428731 | 1.993731 | 1.397524 | 1.426616 | 0.153691 | NA |
| | 0.508056 | 0.731032 | 2.861912 | 0.255435 | 0.798387 | NA |
| TTC4 | 46.99375 | -0.33878 | 0.314127 | -1.07848 | 0.280819 | NA |
| RPLP0P2 | 6.208323 | -0.51988 | 0.837024 | -0.6211 | 0.534533 | NA |
| ZMYM6NB | 0.162376 | 0.424913 | 4.079006 | 0.104171 | 0.917034 | NA |
| | 3.061539 | -0.30991 | 1.187414 | -0.26099 | 0.794098 | NA |
| | 1.755524 | 0.886375 | 1.510448 | 0.586829 | 0.557319 | NA |
| | 2.012907 | 0.257818 | 1.430096 | 0.18028 | 0.856933 | NA |
| PPIEL | 8.877394 | 0.771699 | 0.67676 | 1.140285 | 0.254168 | NA |
| DDOST | 7676.666 | -0.23008 | 0.121748 | -1.88978 | 0.058788 | 0.154976 |
| | 0.614937 | 2.493639 | 2.617677 | 0.952615 | 0.340785 | NA |
| IFITM10 | 34.73536 | 0.374629 | 0.359194 | 1.042971 | 0.296962 | NA |
| RN7SL130F | 0.74907 | 1.552259 | 2.278648 | 0.681219 | 0.495733 | NA |
| RPL34P22 | 0.941656 | 0.847273 | 2.457626 | 0.344752 | 0.730281 | NA |
| | 52.08748 | 0.059243 | 0.302125 | 0.196088 | 0.844542 | 0.907349 |
| PFN1P8 | 0.424115 | 1.932531 | 3.45462 | 0.559405 | 0.575885 | NA |
| | 1.661894 | 2.829199 | 1.787306 | 1.582941 | 0.113435 | NA |
| | 42.83268 | 0.397836 | 0.343489 | 1.15822 | 0.246774 | NA |
| CFHR1 | 1.043154 | -2.66287 | 2.122813 | -1.25441 | 0.209695 | NA |
| PGBD4P7 | 0.354779 | 1.703954 | 3.657232 | 0.465914 | 0.641277 | NA |
| | 0.163957 | 0.424913 | 4.079006 | 0.104171 | 0.917034 | NA |
| RN7SL532F | 0.354779 | 1.703954 | 3.657232 | 0.465914 | 0.641277 | NA |
| FCGR2C | 0.516164 | -1.70268 | 2.844782 | -0.59853 | 0.549488 | NA |
| | 3.086839 | 0.569446 | 1.177751 | 0.483503 | 0.628739 | NA |
| | 0.410275 | 0.244283 | 3.100889 | 0.078778 | 0.937209 | NA |
| HBB | 0.460669 | -0.62593 | 3.398671 | -0.18417 | 0.853882 | NA |

| | | | | | | |
|---|---|---|---|---|---|---|
| ALKBH3-AS | 9.95667 | 0.017687 | 0.657606 | 0.026895 | 0.978543 | NA |
| | 6.303344 | 0.484933 | 0.839024 | 0.577973 | 0.563282 | NA |
| | 2.837915 | -1.32569 | 1.21841 | -1.08805 | 0.276575 | NA |
| | 192.7352 | -0.41761 | 0.231191 | -1.80635 | 0.070863 | 0.176045 |
| BDNF-AS | 17.39982 | 0.580899 | 0.58154 | 0.998897 | 0.317845 | NA |
| | 2.424965 | 0.039885 | 1.322746 | 0.030153 | 0.975945 | NA |
| | 2.702098 | 1.238568 | 1.248859 | 0.99176 | 0.321315 | NA |
| | 2.552973 | 3.470942 | 1.601323 | 2.167547 | 0.030193 | NA |
| SBF2-AS1 | 47.43875 | 0.00329 | 0.340238 | 0.00967 | 0.992285 | 0.996738 |
| | 1.387896 | 1.605513 | 1.744576 | 0.920288 | 0.357422 | NA |
| MIR210HG | 21.67178 | 0.138277 | 0.434866 | 0.317975 | 0.750504 | NA |
| CSTF3-AS1 | 0.298089 | -0.51711 | 3.975232 | -0.13008 | 0.896501 | NA |
| ZBED5-AS1 | 295.609 | -0.2458 | 0.221668 | -1.10886 | 0.267489 | 0.432558 |
| CARS-AS1 | 0.134133 | -1.48909 | 4.077584 | -0.36519 | 0.71497 | NA |
| SPTY2D1-A | 9.658977 | -0.00533 | 0.637782 | -0.00835 | 0.993334 | NA |
| LRP4-AS1 | 6.426459 | 0.80519 | 0.87946 | 0.915551 | 0.359903 | NA |
| CDK11B | 598.3685 | 0.1966 | 0.150558 | 1.305808 | 0.191618 | 0.346232 |
| | 0.555233 | -0.57009 | 3.296947 | -0.17291 | 0.862719 | NA |
| PCP4L1 | 0.850576 | 0.626886 | 2.42682 | 0.258316 | 0.796163 | NA |
| MTATP6P1 | 3345.227 | -0.51045 | 0.157923 | -3.23223 | 0.001228 | 0.011864 |
| | 139.0862 | -0.31338 | 0.226739 | -1.38212 | 0.166935 | 0.316389 |
| C1orf213 | 13.17539 | 0.199968 | 0.588471 | 0.339809 | 0.734 | NA |
| | 0.948386 | -1.53855 | 2.155471 | -0.71379 | 0.475358 | NA |
| | 17.41645 | 1.043815 | 0.526696 | 1.981816 | 0.0475 | NA |
| | 4.536858 | -0.96879 | 0.966452 | -1.00242 | 0.316141 | NA |
| TRIM51CP | 0.756975 | 0.468225 | 2.554602 | 0.183287 | 0.854573 | NA |
| | 15.51108 | -0.78535 | 0.654795 | -1.19939 | 0.230376 | NA |
| | 15.23696 | -1.09941 | 0.616675 | -1.78281 | 0.074617 | NA |
| | 4.105425 | -0.12674 | 1.057789 | -0.11981 | 0.904633 | NA |
| ABHD17AP | 0.585114 | 1.12675 | 2.73598 | 0.411827 | 0.680466 | NA |
| | 2.069743 | 0.229564 | 1.330067 | 0.172596 | 0.862969 | NA |
| | 0.315653 | -0.57305 | 3.892639 | -0.14721 | 0.882963 | NA |
| | 8.739529 | -0.20174 | 0.725959 | -0.27789 | 0.781094 | NA |
| GLUD1P3 | 19.13469 | 0.012759 | 0.483278 | 0.026402 | 0.978937 | NA |
| | 4.793161 | 1.542891 | 1.070693 | 1.441021 | 0.149579 | NA |
| | 86.58684 | -0.1617 | 0.289656 | -0.55825 | 0.576672 | 0.711611 |
| | 0.096202 | 0.408546 | 4.079639 | 0.100143 | 0.920231 | NA |
| APITD1-CO | 0.394291 | 0.298745 | 3.141711 | 0.09509 | 0.924243 | NA |
| | 0.672013 | 1.253691 | 2.607776 | 0.480751 | 0.630693 | NA |
| HNRNPKP3 | 1.347189 | 0.804826 | 1.780891 | 0.451923 | 0.651324 | NA |
| | 0.954682 | 0.055803 | 2.069758 | 0.026961 | 0.978491 | NA |
| RNU7-123F | 0.258578 | 1.233151 | 4.047809 | 0.304646 | 0.760635 | NA |
| RNU7-8P | 0.096202 | 0.408546 | 4.079639 | 0.100143 | 0.920231 | NA |
| RNU6-117( | 0.392857 | -1.546 | 4.075469 | -0.37934 | 0.704433 | NA |
| RNA5SP20 | 0.392857 | -1.546 | 4.075469 | -0.37934 | 0.704433 | NA |
| | 2.757796 | 0.623063 | 1.21821 | 0.511458 | 0.60903 | NA |
| | 2.050599 | 0.247963 | 1.354617 | 0.183051 | 0.854758 | NA |

| | | | | | | |
|---|---|---|---|---|---|---|
| RMRPP4 | 0.162376 | 0.424913 | 4.079006 | 0.104171 | 0.917034 | NA |
| RNU7-49P | 0.260158 | 1.240347 | 4.047124 | 0.306476 | 0.759242 | NA |
| RNU2-30P | 0.315653 | -0.57305 | 3.892639 | -0.14721 | 0.882963 | NA |
| | 0.096202 | 0.408546 | 4.079639 | 0.100143 | 0.920231 | NA |
| RNA5SP67 | 0.162376 | 0.424913 | 4.079006 | 0.104171 | 0.917034 | NA |
| | 0.162376 | 0.424913 | 4.079006 | 0.104171 | 0.917034 | NA |
| RN7SKP287 | 0.230334 | -0.53318 | 4.076749 | -0.13079 | 0.895945 | NA |
| | 0.096202 | 0.408546 | 4.079639 | 0.100143 | 0.920231 | NA |
| RNU6-118P | 0.422534 | 1.924583 | 3.001758 | 0.641152 | 0.521424 | NA |
| RNU6-885P | 0.765054 | 1.506026 | 2.496832 | 0.603175 | 0.546392 | NA |
| | 0.260158 | 1.240347 | 4.047124 | 0.306476 | 0.759242 | NA |
| | 3.769441 | 0.083809 | 1.242005 | 0.067479 | 0.946201 | NA |
| | 0.392711 | 0.292538 | 3.145221 | 0.09301 | 0.925895 | NA |
| RNU6-1171 | 0.163957 | 0.424913 | 4.079006 | 0.104171 | 0.917034 | NA |
| RNA5SP82 | 0.260158 | 1.240347 | 4.047124 | 0.306476 | 0.759242 | NA |
| RNU6-307P | 0.163957 | 0.424913 | 4.079006 | 0.104171 | 0.917034 | NA |
| RNU7-28P | 0.096202 | 0.408546 | 4.079639 | 0.100143 | 0.920231 | NA |
| | 0.096202 | 0.408546 | 4.079639 | 0.100143 | 0.920231 | NA |
| RN7SKP88 | 0.315653 | -0.57305 | 3.892639 | -0.14721 | 0.882963 | NA |
| SCARNA4 | 0.584911 | 2.38756 | 2.690807 | 0.887303 | 0.374916 | NA |
| | 0.954682 | 0.055803 | 2.069758 | 0.026961 | 0.978491 | NA |
| | 0.151697 | -1.546 | 4.075469 | -0.37934 | 0.704433 | NA |
| RNU1-68P | 1.769082 | -1.26494 | 1.448173 | -0.87347 | 0.382407 | NA |
| | 0.096202 | 0.408546 | 4.079639 | 0.100143 | 0.920231 | NA |
| RNA5SP40 | 0.162376 | 0.424913 | 4.079006 | 0.104171 | 0.917034 | NA |
| | 0.192403 | 0.975392 | 4.060716 | 0.240202 | 0.810174 | NA |
| | 0.28583 | -2.21358 | 3.97376 | -0.55705 | 0.577494 | NA |
| TMEM200B | 208.3179 | -0.11298 | 0.188686 | -0.59879 | 0.549312 | 0.690498 |
| | 0.303394 | -2.27006 | 4.032451 | -0.56295 | 0.57347 | NA |
| TRNP1 | 1235.711 | -0.20817 | 0.228445 | -0.91126 | 0.362158 | 0.529133 |
| EIF5AL1 | 180.2202 | -0.18858 | 0.225651 | -0.8357 | 0.403324 | 0.56793 |
| | 3.384894 | -0.35715 | 1.083062 | -0.32976 | 0.741583 | NA |
| | 1.189372 | 1.382636 | 1.842322 | 0.750486 | 0.452962 | NA |
| | 0.623045 | 0.091811 | 2.666142 | 0.034436 | 0.97253 | NA |
| | 0.096202 | 0.408546 | 4.079639 | 0.100143 | 0.920231 | NA |
| OR10Y1P | 1.954637 | -0.19007 | 1.66658 | -0.11405 | 0.909198 | NA |
| | 5.138679 | -0.66145 | 0.955285 | -0.69241 | 0.488679 | NA |
| | 0.643384 | 2.576138 | 2.977667 | 0.865153 | 0.386955 | NA |
| | 0.230334 | -0.53318 | 4.076749 | -0.13079 | 0.895945 | NA |
| RCC2P6 | 1.094548 | 2.111186 | 2.036733 | 1.036555 | 0.299943 | NA |
| | 0.735027 | 1.3952 | 2.594798 | 0.537691 | 0.59079 | NA |
| PRKRIRP4 | 0.326333 | 1.533889 | 3.826878 | 0.40082 | 0.688553 | NA |
| | 11.94156 | -0.52583 | 0.58099 | -0.90505 | 0.365439 | NA |
| | 0.134133 | -1.48909 | 4.077584 | -0.36519 | 0.71497 | NA |
| | 0.192403 | 0.975392 | 4.060716 | 0.240202 | 0.810174 | NA |
| CBX3P8 | 0.162376 | 0.424913 | 4.079006 | 0.104171 | 0.917034 | NA |
| | 0.163957 | 0.424913 | 4.079006 | 0.104171 | 0.917034 | NA |

| | | | | | | |
|---|---|---|---|---|---|---|
| | 0.298089 | -0.51711 | 3.975232 | -0.13008 | 0.896501 | NA |
| | 0.096202 | 0.408546 | 4.079639 | 0.100143 | 0.920231 | NA |
| | 0.747636 | 0.735436 | 2.634453 | 0.279161 | 0.780122 | NA |
| | 0.324752 | 1.53082 | 4.022025 | 0.380609 | 0.703493 | NA |
| | 0.134133 | -1.48909 | 4.077584 | -0.36519 | 0.71497 | NA |
| | 0.87074 | 0.675988 | 2.012127 | 0.335957 | 0.736903 | NA |
| NAV2-AS3 | 0.096202 | 0.408546 | 4.079639 | 0.100143 | 0.920231 | NA |
| | 2.210403 | -0.75719 | 1.326814 | -0.57068 | 0.568217 | NA |
| | 5.577274 | -0.51081 | 0.957693 | -0.53337 | 0.593776 | NA |
| | 0.394291 | 0.298745 | 3.141711 | 0.09509 | 0.924243 | NA |
| | 0.490493 | 0.776209 | 2.895261 | 0.268096 | 0.788625 | NA |
| | 1.335358 | 3.577115 | 2.0361 | 1.756847 | 0.078944 | NA |
| EIF4A2P3 | 3.972572 | -0.06743 | 0.990119 | -0.06811 | 0.945701 | NA |
| | 10.14264 | 0.852772 | 0.657093 | 1.297796 | 0.194357 | NA |
| CSNK2A3 | 11.33662 | -0.15205 | 0.664656 | -0.22876 | 0.819055 | NA |
| | 0.604258 | 1.087325 | 2.707696 | 0.401568 | 0.688002 | NA |
| WAC-AS1 | 214.3735 | 0.594701 | 0.3165 | 1.878992 | 0.060246 | 0.15768 |
| ARMS2 | 0.46735 | -1.30193 | 3.399671 | -0.38296 | 0.701751 | NA |
| | 0.096202 | 0.408546 | 4.079639 | 0.100143 | 0.920231 | NA |
| | 0.327913 | 1.543154 | 4.021063 | 0.383768 | 0.701151 | NA |
| FPGT | 210.9812 | 0.546256 | 0.266094 | 2.052867 | 0.040085 | 0.120637 |
| | 0.163957 | 0.424913 | 4.079006 | 0.104171 | 0.917034 | NA |
| | 0.134133 | -1.48909 | 4.077584 | -0.36519 | 0.71497 | NA |
| | 0.163957 | 0.424913 | 4.079006 | 0.104171 | 0.917034 | NA |
| | 0.460466 | 0.568655 | 3.014642 | 0.188631 | 0.850382 | NA |
| GLYATL1P2 | 1.551852 | 1.796216 | 1.661946 | 1.080791 | 0.27979 | NA |
| | 0.604258 | 1.087325 | 2.707696 | 0.401568 | 0.688002 | NA |
| MEX3A | 81.2606 | -0.07021 | 0.327424 | -0.21443 | 0.830214 | 0.89752 |
| | 0.87074 | 0.675988 | 2.012127 | 0.335957 | 0.736903 | NA |
| | 0.659753 | -0.00581 | 2.629089 | -0.00221 | 0.998237 | NA |
| OR10V3P | 0.324752 | 1.53082 | 4.022025 | 0.380609 | 0.703493 | NA |
| | 0.134133 | -1.48909 | 4.077584 | -0.36519 | 0.71497 | NA |
| EEF1G | 1.343815 | 0.18908 | 1.751639 | 0.107945 | 0.91404 | NA |
| | 0.785567 | -0.70568 | 2.358063 | -0.29926 | 0.764741 | NA |
| GVINP2 | 0.096202 | 0.408546 | 4.079639 | 0.100143 | 0.920231 | NA |
| FAR1-IT1 | 0.52699 | -2.57878 | 3.248478 | -0.79384 | 0.427288 | NA |
| | 1.968652 | 1.1111 | 1.635176 | 0.679499 | 0.496822 | NA |
| SLC22A18A | 503.5504 | 0.351819 | 0.258997 | 1.358393 | 0.174339 | 0.326301 |
| GVINP1 | 10.49265 | 1.207659 | 0.709153 | 1.702958 | 0.088576 | NA |
| | 1.407272 | -0.69944 | 1.719223 | -0.40683 | 0.684131 | NA |
| TMEM9B-A | 19.23342 | -0.10437 | 0.495479 | -0.21065 | 0.833162 | NA |
| | 0.622842 | 1.158846 | 2.735323 | 0.42366 | 0.671814 | NA |
| | 0.096202 | 0.408546 | 4.079639 | 0.100143 | 0.920231 | NA |
| | 0.163957 | 0.424913 | 4.079006 | 0.104171 | 0.917034 | NA |
| | 0.545988 | -0.43913 | 2.541145 | -0.17281 | 0.862803 | NA |
| | 5.198911 | -0.81674 | 0.894175 | -0.9134 | 0.361031 | NA |
| | 0.324752 | 1.53082 | 4.022025 | 0.380609 | 0.703493 | NA |

| | | | | | | |
|---|---|---|---|---|---|---|
| | 0.247898 | -0.5901 | 4.074633 | -0.14482 | 0.884851 | NA |
| | 2.85952 | 1.075323 | 1.201346 | 0.895099 | 0.370734 | NA |
| | 0.163957 | 0.424913 | 4.079006 | 0.104171 | 0.917034 | NA |
| | 0.151697 | -1.546 | 4.075469 | -0.37934 | 0.704433 | NA |
| | 0.787148 | -0.70094 | 2.357021 | -0.29738 | 0.766174 | NA |
| | 1.410193 | -0.26412 | 1.718689 | -0.15368 | 0.877864 | NA |
| SDHCP4 | 0.258578 | 1.233151 | 4.047809 | 0.304646 | 0.760635 | NA |
| HNRNPRP2 | 0.096202 | 0.408546 | 4.079639 | 0.100143 | 0.920231 | NA |
| | 0.260158 | 1.240347 | 4.047124 | 0.306476 | 0.759242 | NA |
| | 0.488709 | 2.119056 | 3.307564 | 0.64067 | 0.521737 | NA |
| | 0.230334 | -0.53318 | 4.076749 | -0.13079 | 0.895945 | NA |
| | 0.163957 | 0.424913 | 4.079006 | 0.104171 | 0.917034 | NA |
| KIAA0754 | 663.4958 | 0.351373 | 0.18908 | 1.858333 | 0.063122 | 0.162888 |
| | 0.354779 | 1.703954 | 3.657232 | 0.465914 | 0.641277 | NA |
| | 0.162376 | 0.424913 | 4.079006 | 0.104171 | 0.917034 | NA |
| | 2.823112 | 1.407056 | 1.422761 | 0.988962 | 0.322682 | NA |
| HNRNPA1P | 0.096202 | 0.408546 | 4.079639 | 0.100143 | 0.920231 | NA |
| | 1.968166 | -0.93029 | 1.436258 | -0.64772 | 0.517169 | NA |
| | 1.463088 | -0.85499 | 1.867569 | -0.45781 | 0.64709 | NA |
| TOLLIP-AS1 | 54.81778 | 0.067967 | 0.363231 | 0.187119 | 0.851568 | 0.91179 |
| | 1.85064 | -0.65041 | 1.59592 | -0.40755 | 0.683605 | NA |
| | 0.46735 | -1.30193 | 3.399671 | -0.38296 | 0.701751 | NA |
| | 4.514633 | 0.283992 | 0.929091 | 0.305667 | 0.759858 | NA |
| | 2.676951 | -0.19021 | 1.260829 | -0.15086 | 0.880084 | NA |
| NPM1P35 | 0.46577 | -1.30191 | 3.402879 | -0.38259 | 0.702024 | NA |
| | 3.223566 | 0.384848 | 1.152556 | 0.333909 | 0.738449 | NA |
| | 0.268266 | -2.15396 | 4.038351 | -0.53338 | 0.593773 | NA |
| RRM1-AS1 | 0.151697 | -1.546 | 4.075469 | -0.37934 | 0.704433 | NA |
| | 0.096202 | 0.408546 | 4.079639 | 0.100143 | 0.920231 | NA |
| | 37.65602 | 0.01182 | 0.383015 | 0.03086 | 0.975381 | NA |
| | 0.899333 | -0.26387 | 2.222941 | -0.1187 | 0.905512 | NA |
| OR5BA1P | 1.866072 | 0.900686 | 1.586103 | 0.567861 | 0.570129 | NA |
| | 0.399595 | -1.31595 | 3.579466 | -0.36764 | 0.713143 | NA |
| | 0.260158 | 1.240347 | 4.047124 | 0.306476 | 0.759242 | NA |
| | 0.096202 | 0.408546 | 4.079639 | 0.100143 | 0.920231 | NA |
| | 1.0349 | 0.343148 | 1.852366 | 0.185249 | 0.853034 | NA |
| | 0.35636 | 1.709252 | 3.651833 | 0.468053 | 0.639746 | NA |
| | 0.392857 | -1.546 | 4.075469 | -0.37934 | 0.704433 | NA |
| | 0.849381 | -1.30173 | 2.226174 | -0.58474 | 0.558724 | NA |
| | 1.83031 | -0.19201 | 1.590164 | -0.12075 | 0.903888 | NA |
| CCDC179 | 0.163957 | 0.424913 | 4.079006 | 0.104171 | 0.917034 | NA |
| | 16.81538 | 0.097006 | 0.538237 | 0.18023 | 0.856972 | NA |
| | 0.298089 | -0.51711 | 3.975232 | -0.13008 | 0.896501 | NA |
| | 0.815391 | 0.931626 | 2.360431 | 0.394685 | 0.693076 | NA |
| | 0.096202 | 0.408546 | 4.079639 | 0.100143 | 0.920231 | NA |
| | 0.326333 | 1.533889 | 3.826878 | 0.40082 | 0.688553 | NA |
| | 0.354779 | 1.703954 | 3.657232 | 0.465914 | 0.641277 | NA |

| | | | | | | |
|---|---|---|---|---|---|---|
| | 0.163957 | 0.424913 | 4.079006 | 0.104171 | 0.917034 | NA |
| | 137.8484 | 2.310287 | 1.154473 | 2.001162 | 0.045375 | 0.130468 |
| | 3.0782 | 2.704018 | 1.498394 | 1.80461 | 0.071136 | NA |
| | 14.41628 | -0.08755 | 0.577538 | -0.1516 | 0.879506 | NA |
| | 0.151697 | -1.546 | 4.075469 | -0.37934 | 0.704433 | NA |
| | 1.713733 | -1.73465 | 1.625178 | -1.06736 | 0.285811 | NA |
| | 0.422534 | 1.924583 | 3.001758 | 0.641152 | 0.521424 | NA |
| EIF4A2P5 | 0.364467 | -1.20166 | 3.685431 | -0.32606 | 0.744381 | NA |
| | 0.163957 | 0.424913 | 4.079006 | 0.104171 | 0.917034 | NA |
| MIR3654 | 30.97962 | -0.10535 | 0.445176 | -0.23664 | 0.812934 | NA |
| | 0.163957 | 0.424913 | 4.079006 | 0.104171 | 0.917034 | NA |
| | 2.702033 | 0.171052 | 1.232121 | 0.138828 | 0.889586 | NA |
| OR10V2P | 0.192403 | 0.975392 | 4.060716 | 0.240202 | 0.810174 | NA |
| | 3.764974 | 0.717869 | 1.06388 | 0.674765 | 0.499825 | NA |
| | 1.879969 | 1.528624 | 1.494589 | 1.022772 | 0.306416 | NA |
| MTRNR2L8 | 0.624422 | 1.162588 | 2.733761 | 0.425271 | 0.670639 | NA |
| | 0.392857 | -1.546 | 4.075469 | -0.37934 | 0.704433 | NA |
| | 3.555412 | 0.121994 | 1.109578 | 0.109946 | 0.912452 | NA |
| PADI6 | 0.151697 | -1.546 | 4.075469 | -0.37934 | 0.704433 | NA |
| | 1.645482 | 0.21678 | 1.529826 | 0.141702 | 0.887315 | NA |
| | 1.673366 | 0.912533 | 1.950485 | 0.467849 | 0.639893 | NA |
| | 0.230334 | -0.53318 | 4.076749 | -0.13079 | 0.895945 | NA |
| | 0.163957 | 0.424913 | 4.079006 | 0.104171 | 0.917034 | NA |
| | 1.083714 | -0.06832 | 2.255294 | -0.03029 | 0.975832 | NA |
| | 1.179509 | 1.360465 | 1.979386 | 0.687317 | 0.491883 | NA |
| | 1.813822 | 0.133354 | 1.692505 | 0.078791 | 0.937199 | NA |
| | 0.640755 | -1.64826 | 2.754778 | -0.59833 | 0.549621 | NA |
| PGA5 | 0.163957 | 0.424913 | 4.079006 | 0.104171 | 0.917034 | NA |
| | 0.392857 | -1.546 | 4.075469 | -0.37934 | 0.704433 | NA |
| | 0.71919 | 0.510978 | 2.728734 | 0.187258 | 0.851458 | NA |
| | 3.674947 | 0.943153 | 1.083698 | 0.87031 | 0.384131 | NA |
| | 1.416874 | -0.45964 | 1.62883 | -0.28219 | 0.777799 | NA |
| | 4.188357 | 0.174664 | 0.98068 | 0.178105 | 0.85864 | NA |
| | 4.021115 | -0.63212 | 1.239669 | -0.50991 | 0.610113 | NA |
| ZBED6 | 202.8584 | 0.628781 | 0.302748 | 2.076911 | 0.03781 | 0.115644 |
| HLA-AS1 | 0.544554 | -2.65036 | 3.220086 | -0.82307 | 0.410468 | NA |
| | 1.22552 | 1.38393 | 1.789452 | 0.773382 | 0.439296 | NA |
| | 0.3441 | -0.0166 | 3.769682 | -0.0044 | 0.996486 | NA |
| | 1.720122 | -0.23797 | 1.501053 | -0.15853 | 0.874036 | NA |
| | 100.9985 | -0.59591 | 0.235039 | -2.53536 | 0.011233 | 0.050945 |
| TRIM34 | 0.563552 | -0.48645 | 2.808029 | -0.17324 | 0.862465 | NA |
| ERCC6-PGE | 1.30252 | 2.031918 | 2.097119 | 0.96891 | 0.33259 | NA |
| FPGT-TNNI | 0.151697 | -1.546 | 4.075469 | -0.37934 | 0.704433 | NA |
| | 0.163957 | 0.424913 | 4.079006 | 0.104171 | 0.917034 | NA |
| | 24.87144 | 1.528758 | 0.592288 | 2.581107 | 0.009848 | NA |
| | 0.163957 | 0.424913 | 4.079006 | 0.104171 | 0.917034 | NA |
| | 0.69625 | -3.13723 | 2.93845 | -1.06765 | 0.285679 | NA |

| | | | | | | |
|---|---|---|---|---|---|---|
| | 78.05754 | 0.573396 | 0.262547 | 2.183977 | 0.028964 | 0.09686 |
| | 0.518736 | 2.237423 | 2.7713 | 0.807355 | 0.419462 | NA |
| | 6.262366 | -0.17368 | 0.837672 | -0.20733 | 0.83575 | NA |
| | 0.52699 | -2.57878 | 3.248478 | -0.79384 | 0.427288 | NA |
| | 5.171371 | 0.043789 | 0.86521 | 0.050611 | 0.959636 | NA |
| | 0.260158 | 1.240347 | 4.047124 | 0.306476 | 0.759242 | NA |
| | 0.776322 | -0.40209 | 2.222574 | -0.18091 | 0.856435 | NA |
| | 10.17106 | 0.438459 | 0.661456 | 0.66287 | 0.507414 | NA |
| | 0.134133 | -1.48909 | 4.077584 | -0.36519 | 0.71497 | NA |
| | 2.422233 | 1.568442 | 1.522498 | 1.030177 | 0.302927 | NA |
| | 12.05644 | 0.207525 | 0.647128 | 0.320687 | 0.748448 | NA |
| | 0.151697 | -1.546 | 4.075469 | -0.37934 | 0.704433 | NA |
| | 27.09635 | -0.37237 | 0.427358 | -0.87132 | 0.383579 | NA |
| | 13.38374 | -0.12854 | 0.550591 | -0.23345 | 0.81541 | NA |
| | 1.793455 | 0.460616 | 1.498402 | 0.307405 | 0.758535 | NA |
| | 9.999846 | 0.490943 | 0.663688 | 0.73972 | 0.45947 | NA |
| | 2.451774 | -0.07167 | 1.21207 | -0.05913 | 0.952848 | NA |
| | 33.28892 | 0.778953 | 0.404185 | 1.927217 | 0.053953 | NA |
| | 37.75773 | -0.54092 | 0.503069 | -1.07523 | 0.282271 | NA |
| | 14.02046 | 0.855206 | 0.578168 | 1.479164 | 0.139097 | NA |
| | 6.365027 | 0.141653 | 0.864378 | 0.163879 | 0.869827 | NA |
| | 126.7711 | -0.3427 | 0.222615 | -1.53941 | 0.123703 | 0.258051 |
| | 12.40308 | -0.49362 | 0.661573 | -0.74614 | 0.455585 | NA |
| | 1.283964 | -0.71772 | 1.990944 | -0.36049 | 0.718481 | NA |
| | 0.096202 | 0.408546 | 4.079639 | 0.100143 | 0.920231 | NA |
| LINC00622 | 18.64979 | 0.419036 | 0.494281 | 0.847768 | 0.396567 | NA |
| | 7.619397 | 0.700574 | 0.805933 | 0.869271 | 0.384699 | NA |
| | 0.298089 | -0.51711 | 3.975232 | -0.13008 | 0.896501 | NA |
| | 0.096202 | 0.408546 | 4.079639 | 0.100143 | 0.920231 | NA |
| | 0.652869 | 1.292592 | 2.374873 | 0.544278 | 0.58625 | NA |
| | 3.822463 | 0.732383 | 1.03687 | 0.70634 | 0.479977 | NA |
| | 0.736607 | 1.395844 | 2.318642 | 0.602009 | 0.547168 | NA |
| | 2.387757 | 0.565833 | 1.449857 | 0.390268 | 0.696338 | NA |
| | 0.52699 | -2.57878 | 3.248478 | -0.79384 | 0.427288 | NA |
| | 0.586694 | 1.130467 | 2.734203 | 0.413454 | 0.679274 | NA |
| | 1.27255 | 1.529745 | 2.170945 | 0.704645 | 0.481031 | NA |
| | 2.475209 | -0.65254 | 1.450663 | -0.44982 | 0.65284 | NA |
| | 3.907433 | 1.263887 | 1.172934 | 1.077542 | 0.281238 | NA |
| | 0.163957 | 0.424913 | 4.079006 | 0.104171 | 0.917034 | NA |
| | 9.357305 | 1.900038 | 0.797924 | 2.381228 | 0.017255 | NA |
| | 5.989909 | -0.31868 | 0.832194 | -0.38294 | 0.701761 | NA |
| | 13.2196 | 0.637478 | 0.562286 | 1.133724 | 0.25691 | NA |
| | 4.024999 | -0.11911 | 1.087544 | -0.10952 | 0.91279 | NA |
| | 1.446958 | -0.00969 | 1.799806 | -0.00538 | 0.995704 | NA |
| | 0.096202 | 0.408546 | 4.079639 | 0.100143 | 0.920231 | NA |
| | 0.658173 | -0.01016 | 2.630582 | -0.00386 | 0.996919 | NA |
| | 7.872371 | 0.17632 | 0.806355 | 0.218663 | 0.826912 | NA |

| | | | | | | |
|---|---|---|---|---|---|---|
| | 40.05031 | 0.511647 | 0.356713 | 1.43434 | 0.151475 | NA |
| | 20.36629 | -0.13687 | 0.516479 | -0.265 | 0.79101 | NA |
| | 0.260158 | 1.240347 | 4.047124 | 0.306476 | 0.759242 | NA |
| OCLM | 1.294855 | 1.463581 | 1.753443 | 0.83469 | 0.403892 | NA |
| | 0.624422 | 1.162588 | 2.733761 | 0.425271 | 0.670639 | NA |
| MIR3973 | 0.134133 | -1.48909 | 4.077584 | -0.36519 | 0.71497 | NA |
| MIR5187 | 0.544554 | -2.65036 | 3.220086 | -0.82307 | 0.410468 | NA |
| MIR3916 | 3.176967 | 1.190721 | 1.198486 | 0.993521 | 0.320456 | NA |
| RN7SL768F | 0.544554 | -2.65036 | 3.220086 | -0.82307 | 0.410468 | NA |
| MIR5581 | 1.091387 | 2.10753 | 2.037446 | 1.034398 | 0.30095 | NA |
| RN7SL276F | 0.134133 | -1.48909 | 4.077584 | -0.36519 | 0.71497 | NA |
| | 0.785567 | -0.70568 | 2.358063 | -0.29926 | 0.764741 | NA |
| MIR4689 | 0.134133 | -1.48909 | 4.077584 | -0.36519 | 0.71497 | NA |
| MIR4258 | 1.090541 | -1.58559 | 1.951054 | -0.81268 | 0.4164 | NA |
| RN7SL431F | 0.392857 | -1.546 | 4.075469 | -0.37934 | 0.704433 | NA |
| MIR548AC | 0.298089 | -0.51711 | 3.975232 | -0.13008 | 0.896501 | NA |
| RN7SL119F | 0.298089 | -0.51711 | 3.975232 | -0.13008 | 0.896501 | NA |
| | 144.3599 | -0.15023 | 0.214218 | -0.70128 | 0.483126 | 0.637021 |
| RN7SL473F | 0.096202 | 0.408546 | 4.079639 | 0.100143 | 0.920231 | NA |
| RN7SL600F | 0.096202 | 0.408546 | 4.079639 | 0.100143 | 0.920231 | NA |
| RN7SL131F | 0.162376 | 0.424913 | 4.079006 | 0.104171 | 0.917034 | NA |
| MIR4685 | 0.192403 | 0.975392 | 4.060716 | 0.240202 | 0.810174 | NA |
| | 0.134133 | -1.48909 | 4.077584 | -0.36519 | 0.71497 | NA |
| MIR4677 | 0.392857 | -1.546 | 4.075469 | -0.37934 | 0.704433 | NA |
| | 0.163957 | 0.424913 | 4.079006 | 0.104171 | 0.917034 | NA |
| MIR3118-2 | 0.192403 | 0.975392 | 4.060716 | 0.240202 | 0.810174 | NA |
| RN7SL721F | 0.162376 | 0.424913 | 4.079006 | 0.104171 | 0.917034 | NA |
| RN7SL574F | 1.102655 | 0.449766 | 1.809137 | 0.248608 | 0.803664 | NA |
| | 0.419962 | -2.64111 | 3.429364 | -0.77014 | 0.441214 | NA |
| MIR4423 | 0.422534 | 1.924583 | 3.001758 | 0.641152 | 0.521424 | NA |
| RN7SL480F | 0.163957 | 0.424913 | 4.079006 | 0.104171 | 0.917034 | NA |
| MIR3605 | 0.586694 | 1.130467 | 2.734203 | 0.413454 | 0.679274 | NA |
| RN7SL235F | 0.151697 | -1.546 | 4.075469 | -0.37934 | 0.704433 | NA |
| | 0.162376 | 0.424913 | 4.079006 | 0.104171 | 0.917034 | NA |
| MIR4482-1 | 27.39177 | -0.10547 | 0.417263 | -0.25276 | 0.80045 | NA |
| | 5.345973 | -0.23745 | 0.923849 | -0.25702 | 0.797164 | NA |
| | 3.578157 | 1.859715 | 1.183007 | 1.572024 | 0.115945 | NA |
| LINC01140 | 1.561833 | -1.02461 | 1.776553 | -0.57674 | 0.564114 | NA |
| | 0.096202 | 0.408546 | 4.079639 | 0.100143 | 0.920231 | NA |
| | 1.255729 | 0.65706 | 2.007478 | 0.327306 | 0.743436 | NA |
| | 0.430642 | -1.1891 | 3.485782 | -0.34113 | 0.733008 | NA |
| | 2.08364 | 0.821011 | 1.46384 | 0.560861 | 0.574892 | NA |
| | 34.4678 | -0.13268 | 0.399731 | -0.33192 | 0.739947 | NA |
| | 0.151697 | -1.546 | 4.075469 | -0.37934 | 0.704433 | NA |
| | 0.566355 | -1.63598 | 3.185967 | -0.5135 | 0.607605 | NA |
| | 0.134133 | -1.48909 | 4.077584 | -0.36519 | 0.71497 | NA |
| | 2.844594 | -0.44661 | 1.369317 | -0.32616 | 0.744307 | NA |

| | | | | | | |
|---|---|---|---|---|---|---|
| | 19.71062 | 0.065907 | 0.562846 | 0.117096 | 0.906784 | NA |
| | 31.82264 | -0.548 | 0.411479 | -1.33179 | 0.182929 | NA |
| | 5.933764 | 0.083353 | 0.86444 | 0.096424 | 0.923184 | NA |
| ESPNP | 0.354779 | 1.703954 | 3.657232 | 0.465914 | 0.641277 | NA |
| PLCE1-AS1 | 13.16357 | -0.4019 | 0.660348 | -0.60862 | 0.542775 | NA |
| | 9.640674 | 0.184037 | 0.690673 | 0.266461 | 0.789885 | NA |
| TRABD2B | 0.296509 | -0.51709 | 3.982155 | -0.12985 | 0.896684 | NA |
| | 6.595764 | 0.082887 | 0.802434 | 0.103295 | 0.917729 | NA |
| | 1.798759 | -0.05195 | 1.440467 | -0.03606 | 0.971233 | NA |
| | 7.015182 | 0.179962 | 0.918137 | 0.196007 | 0.844604 | NA |
| | 0.326333 | 1.533889 | 3.826878 | 0.40082 | 0.688553 | NA |
| | 0.268266 | -2.15396 | 4.038351 | -0.53338 | 0.593773 | NA |
| | 0.555233 | -0.57009 | 3.296947 | -0.17291 | 0.862719 | NA |
| | 91.41374 | -0.12835 | 0.279103 | -0.45988 | 0.645604 | 0.764599 |
| | 0.774539 | 0.39797 | 2.296349 | 0.173305 | 0.862411 | NA |
| | 28.8134 | -0.23739 | 0.394635 | -0.60153 | 0.547487 | NA |
| | 0.314073 | -0.57308 | 3.898934 | -0.14698 | 0.883146 | NA |
| | 0.151697 | -1.546 | 4.075469 | -0.37934 | 0.704433 | NA |
| | 68.31061 | -0.48955 | 0.298904 | -1.63781 | 0.101462 | 0.22508 |
| | 7.710037 | -0.32477 | 0.718265 | -0.45216 | 0.651154 | NA |
| | 0.315653 | -0.57305 | 3.892639 | -0.14721 | 0.882963 | NA |
| | 29.33753 | 0.428667 | 0.397616 | 1.078095 | 0.280991 | NA |
| | 4.42675 | 0.243623 | 0.941044 | 0.258886 | 0.795723 | NA |
| KCNQ1OT1 | 26.89193 | 1.203826 | 0.542413 | 2.21939 | 0.02646 | NA |
| | 0.163957 | 0.424913 | 4.079006 | 0.104171 | 0.917034 | NA |
| RAD52 | 165.5453 | 0.170529 | 0.214924 | 0.793441 | 0.427521 | 0.589401 |
| BAD | 1058.352 | -0.11685 | 0.157622 | -0.74132 | 0.458501 | 0.616973 |
| M6PR | 4448.222 | -0.0146 | 0.12956 | -0.11266 | 0.910299 | 0.947811 |
| FKBP4 | 3750.871 | -0.24212 | 0.162255 | -1.4922 | 0.135647 | 0.27456 |
| RECQL | 930.1333 | 1.055521 | 0.315762 | 3.342772 | 8.29E-04 | 0.009268 |
| RPAP3 | 282.6718 | 1.0854 | 0.371071 | 2.925049 | 0.003444 | 0.023028 |
| MYCBP2 | 2238.119 | 0.173032 | 0.173185 | 0.999116 | 0.317739 | 0.485536 |
| FBXL3 | 827.5723 | 0.76585 | 0.173744 | 4.407925 | 1.04E-05 | 6.38E-04 |
| MAP3K9 | 50.91429 | -0.31377 | 0.349025 | -0.89899 | 0.368657 | 0.535723 |
| ALDH3B1 | 449.0647 | -0.60175 | 0.186401 | -3.22827 | 0.001245 | 0.011897 |
| ADIPOR2 | 4238.475 | -0.17446 | 0.185502 | -0.9405 | 0.346962 | 0.514991 |
| PLEKHG6 | 3.191351 | -0.16798 | 1.320657 | -0.12719 | 0.898787 | NA |
| MGST1 | 350.4568 | -0.09958 | 0.208957 | -0.47653 | 0.633694 | 0.755323 |
| CRY1 | 283.8806 | 0.644791 | 0.17282 | 3.730984 | 1.91E-04 | 0.003775 |
| POMT2 | 929.0411 | -0.33776 | 0.147809 | -2.28511 | 0.022306 | 0.081613 |
| DYRK4 | 366.2518 | -4.73E-04 | 0.183884 | -0.00257 | 0.997946 | 0.999295 |
| CD9 | 11533.72 | -0.04234 | 0.158019 | -0.26797 | 0.788721 | 0.87174 |
| NCAPD2 | 2727.365 | -0.68252 | 0.194206 | -3.51443 | 4.41E-04 | 0.006302 |
| IFFO1 | 1496.097 | -0.2177 | 0.135241 | -1.60972 | 0.10746 | 0.233921 |
| SLC6A13 | 2.229519 | 0.376185 | 1.445484 | 0.260249 | 0.794672 | NA |
| CD4 | 96.73289 | -0.56582 | 0.345549 | -1.63745 | 0.101536 | 0.225161 |
| LRRC23 | 222.6892 | 0.090744 | 0.185994 | 0.487889 | 0.625628 | 0.748948 |

| | | | | | | |
|---|---|---|---|---|---|---|
| TSPAN9 | 850.2774 | -0.68143 | 0.180861 | -3.76769 | 1.65E-04 | 0.003469 |
| BTBD7 | 934.1856 | 0.079717 | 0.159683 | 0.499223 | 0.617622 | 0.742546 |
| DCN | 2.863629 | 0.031091 | 1.141318 | 0.027241 | 0.978268 | NA |
| NR1H4 | 0.612365 | -1.10341 | 2.696105 | -0.40926 | 0.682349 | NA |
| CALCOCO1 | 1864.032 | -0.07453 | 0.144112 | -0.51716 | 0.605043 | 0.733563 |
| UBR7 | 1099.714 | -0.15108 | 0.1759 | -0.8589 | 0.390398 | 0.555738 |
| MAP4K5 | 728.6928 | 0.933737 | 0.233947 | 3.991239 | 6.57E-05 | 0.001943 |
| POLR3B | 350.7104 | 0.067132 | 0.17892 | 0.375206 | 0.707507 | 0.812765 |
| ANGEL1 | 910.1589 | -0.37741 | 0.134853 | -2.79869 | 0.005131 | 0.030098 |
| DDX11 | 181.1412 | -0.20397 | 0.226002 | -0.90253 | 0.366777 | 0.534116 |
| HEBP1 | 2432.436 | -0.37066 | 0.126146 | -2.93838 | 0.003299 | 0.022439 |
| GPRC5A | 403.2845 | -1.44976 | 0.281776 | -5.14506 | 2.67E-07 | 6.34E-05 |
| POLA2 | 550.3817 | -0.28629 | 0.194744 | -1.47009 | 0.141538 | 0.28229 |
| CAPN1 | 8513.172 | -0.41124 | 0.140059 | -2.93618 | 0.003323 | 0.022535 |
| CCDC88C | 2.691208 | 0.226118 | 1.181666 | 0.191355 | 0.848247 | NA |
| YAF2 | 301.9518 | 0.7924 | 0.201114 | 3.940048 | 8.15E-05 | 0.002244 |
| IGF1 | 14.62989 | 0.535079 | 0.614721 | 0.870443 | 0.384059 | NA |
| CNTN1 | 231.7394 | 0.025413 | 0.291776 | 0.087096 | 0.930595 | 0.960575 |
| PHLDB1 | 1634.454 | -0.24885 | 0.145153 | -1.71442 | 0.086452 | 0.201508 |
| MNAT1 | 192.5385 | 0.577905 | 0.228053 | 2.53408 | 0.011274 | 0.051071 |
| SAMD4A | 1860.614 | 0.615693 | 0.148816 | 4.137271 | 3.51E-05 | 0.001342 |
| MRE11A | 321.4797 | 0.838749 | 0.225438 | 3.720536 | 1.99E-04 | 0.003853 |
| PLEKHB1 | 145.5545 | -0.93577 | 0.392851 | -2.38201 | 0.017219 | 0.068384 |
| NRXN3 | 9.724941 | -0.17719 | 0.639222 | -0.27719 | 0.781631 | NA |
| AQR | 904.5138 | 0.061758 | 0.138546 | 0.445758 | 0.655772 | 0.772781 |
| RNF10 | 3342.116 | -0.43444 | 0.11785 | -3.68641 | 2.27E-04 | 0.004193 |
| ZNF839 | 264.9565 | -0.32756 | 0.185641 | -1.76448 | 0.077651 | 0.187485 |
| GRAMD1B | 20.71113 | -0.51044 | 0.449132 | -1.13651 | 0.255742 | NA |
| BIRC3 | 35.11883 | 0.198982 | 0.499548 | 0.398324 | 0.690392 | NA |
| AKAP11 | 1559.167 | 0.349507 | 0.26272 | 1.330338 | 0.183407 | 0.33663 |
| SNAPC1 | 83.48992 | 0.94116 | 0.28468 | 3.306033 | 9.46E-04 | 0.00998 |
| DERA | 1335.333 | 0.200358 | 0.132208 | 1.515477 | 0.129652 | 0.266013 |
| STRAP | 5155.431 | 0.089574 | 0.149448 | 0.599364 | 0.54893 | 0.690147 |
| HSD17B6 | 39.70661 | 0.369028 | 0.377677 | 0.977098 | 0.328521 | NA |
| MIPEP | 970.6926 | -0.11786 | 0.137955 | -0.85432 | 0.392927 | 0.557996 |
| PRKCH | 560.907 | -0.25619 | 0.173584 | -1.47587 | 0.13998 | 0.280266 |
| VEZT | 662.4892 | 0.721908 | 0.203059 | 3.555165 | 3.78E-04 | 0.00569 |
| SNX1 | 3083.105 | 0.082867 | 0.121853 | 0.680054 | 0.49647 | 0.647636 |
| ARNTL2 | 3.673387 | 1.015712 | 1.106828 | 0.917678 | 0.358787 | NA |
| SLC39A9 | 1994.117 | -0.3019 | 0.154482 | -1.95426 | 0.05067 | 0.139911 |
| NR2E3 | 0.911593 | 1.257757 | 2.250933 | 0.558771 | 0.576318 | NA |
| ARID4A | 258.865 | 0.96067 | 0.87332 | 1.100021 | 0.271323 | 0.436436 |
| IFT88 | 229.9365 | 0.89419 | 0.305855 | 2.923579 | 0.00346 | 0.02312 |
| ZCCHC8 | 249.4172 | 0.167881 | 0.186742 | 0.899002 | 0.368652 | 0.535723 |
| FUT8 | 491.2032 | -0.04207 | 0.190896 | -0.22039 | 0.825566 | 0.895106 |
| GAB2 | 4025.912 | -0.63046 | 0.179059 | -3.52097 | 4.30E-04 | 0.006186 |
| PIAS1 | 488.1163 | 0.268755 | 0.149154 | 1.801861 | 0.071567 | 0.177335 |

| | | | | | | |
|---|---|---|---|---|---|---|
| APBA2 | 1633.907 | -0.20273 | 0.175384 | -1.15591 | 0.247717 | 0.411243 |
| DAPK2 | 17.29517 | 0.205317 | 0.621752 | 0.330223 | 0.741232 | NA |
| CYP46A1 | 0.696104 | -0.93034 | 2.354268 | -0.39517 | 0.692715 | NA |
| USP2 | 34.98202 | 0.157059 | 0.411354 | 0.38181 | 0.702603 | NA |
| METTL1 | 340.0709 | -0.05016 | 0.167197 | -0.29999 | 0.764186 | 0.853696 |
| HOXC8 | 111.7502 | 0.123013 | 0.254189 | 0.483942 | 0.628427 | 0.750857 |
| PSMA4 | 2366.597 | 0.583152 | 0.197092 | 2.958775 | 0.003089 | 0.021653 |
| MYO16 | 136.7234 | -0.20273 | 0.282084 | -0.71869 | 0.472335 | 0.628176 |
| TDP1 | 431.7842 | -0.12328 | 0.186907 | -0.65956 | 0.509538 | 0.657799 |
| SPATA7 | 158.9819 | 0.320906 | 0.23359 | 1.373803 | 0.169503 | 0.319926 |
| MED17 | 680.565 | 0.204189 | 0.167774 | 1.217043 | 0.223588 | 0.384373 |
| FAM214A | 410.3958 | 0.56345 | 0.262276 | 2.148311 | 0.031689 | 0.103151 |
| ANO2 | 12.51369 | 0.805687 | 0.790379 | 1.019367 | 0.308029 | NA |
| C12orf4 | 436.9148 | 0.408196 | 0.205159 | 1.989658 | 0.046629 | 0.132793 |
| USP28 | 390.0964 | 0.118304 | 0.168097 | 0.703785 | 0.481567 | 0.635881 |
| LMO3 | 9.077713 | -1.18502 | 0.760212 | -1.5588 | 0.119044 | NA |
| RSF1 | 922.7683 | 0.734759 | 0.607898 | 1.208688 | 0.226783 | 0.388289 |
| KITLG | 250.507 | 1.057865 | 0.248865 | 4.250753 | 2.13E-05 | 9.59E-04 |
| JKAMP | 1429.529 | 0.25149 | 0.165211 | 1.522237 | 0.12795 | 0.26412 |
| LIMA1 | 5500.38 | 0.675171 | 0.165815 | 4.071825 | 4.66E-05 | 0.00158 |
| LETMD1 | 1176.854 | -0.24567 | 0.146835 | -1.67308 | 0.094311 | 0.214722 |
| SLC4A8 | 6.222722 | -0.98979 | 0.863072 | -1.14682 | 0.251458 | NA |
| RAD51 | 352.3287 | -0.14742 | 0.283541 | -0.51991 | 0.603128 | 0.73214 |
| MPHOSPH | 318.0316 | 0.59731 | 0.247233 | 2.41598 | 0.015693 | 0.063953 |
| PLEKHA5 | 2628.717 | 0.742432 | 0.279621 | 2.655132 | 0.007928 | 0.04044 |
| FOXN3 | 1474.828 | -0.11975 | 0.151402 | -0.79094 | 0.42898 | 0.59062 |
| NRIP2 | 81.70877 | 0.251724 | 0.320834 | 0.784591 | 0.432693 | 0.593839 |
| AP5M1 | 1054.506 | 0.518223 | 0.166571 | 3.111125 | 0.001864 | 0.015428 |
| SYNE2 | 1220.049 | 0.715699 | 0.736634 | 0.971581 | 0.331259 | 0.499755 |
| PLEKHH1 | 543.7269 | -0.78137 | 0.278818 | -2.80246 | 0.005071 | 0.029923 |
| CHRDL2 | 0.544554 | -2.65036 | 3.220086 | -0.82307 | 0.410468 | NA |
| FAM168A | 1684.802 | -0.4437 | 0.144705 | -3.06624 | 0.002168 | 0.017049 |
| RELT | 251.682 | 0.148898 | 0.30374 | 0.490213 | 0.623983 | 0.747838 |
| GALC | 681.1967 | -0.03188 | 0.193071 | -0.16511 | 0.868857 | 0.922011 |
| PKP2 | 0.555233 | -0.57009 | 3.296947 | -0.17291 | 0.862719 | NA |
| F7 | 1.429191 | 0.470595 | 1.645258 | 0.286031 | 0.774855 | NA |
| TMCC3 | 274.2567 | -0.79819 | 0.260408 | -3.06515 | 0.002176 | 0.017092 |
| PPP1R12A | 713.9067 | 0.953646 | 0.602057 | 1.583979 | 0.113198 | 0.241987 |
| RASGRF1 | 25.16172 | -0.22855 | 0.440572 | -0.51875 | 0.603933 | NA |
| CDK17 | 740.5452 | 0.43275 | 0.22292 | 1.941282 | 0.052224 | 0.142522 |
| SLC2A3 | 415.5249 | -0.16592 | 0.192525 | -0.86182 | 0.388788 | 0.554243 |
| YBX3 | 4168.013 | -0.18288 | 0.138949 | -1.31615 | 0.188124 | 0.342213 |
| STYK1 | 0.593018 | -0.10389 | 3.151334 | -0.03297 | 0.973702 | NA |
| WNK1 | 4441.569 | -0.27676 | 0.193531 | -1.43007 | 0.152697 | 0.297294 |
| RIMBP2 | 0.847947 | -3.45038 | 2.752525 | -1.25353 | 0.210012 | NA |
| BCAT1 | 5.140881 | -0.08864 | 1.000696 | -0.08858 | 0.929414 | NA |
| HDAC7 | 1117.887 | -0.30391 | 0.145525 | -2.08834 | 0.036767 | 0.113713 |

| | | | | | | |
|---|---|---|---|---|---|---|
| MRPS35 | 1000.816 | 0.305085 | 0.161084 | 1.893952 | 0.058231 | 0.153818 |
| SFSWAP | 1394.573 | 0.153789 | 0.144111 | 1.067151 | 0.285904 | 0.452344 |
| MON2 | 1049.501 | 0.279431 | 0.159284 | 1.754298 | 0.07938 | 0.190341 |
| DGAT2 | 21.94956 | -0.39025 | 0.46185 | -0.84497 | 0.39813 | NA |
| CS | 3802.855 | -0.4437 | 0.124474 | -3.56462 | 3.64E-04 | 0.005555 |
| LTK | 14.17817 | 0.982511 | 0.599546 | 1.638759 | 0.101263 | NA |
| EIF4B | 8415.224 | 0.029917 | 0.153923 | 0.194361 | 0.845894 | 0.908422 |
| ADCK1 | 420.957 | -0.33399 | 0.155743 | -2.14447 | 0.031995 | 0.103731 |
| ASUN | 676.8278 | 0.441763 | 0.165695 | 2.666124 | 0.007673 | 0.03955 |
| TM7SF3 | 5381.475 | -0.30769 | 0.176903 | -1.73929 | 0.081983 | 0.194498 |
| SPA17 | 88.17011 | 0.207021 | 0.290015 | 0.713827 | 0.475334 | 0.630605 |
| CDON | 646.4127 | -0.64523 | 0.162345 | -3.97441 | 7.06E-05 | 0.002041 |
| BTBD1 | 2363.282 | 0.620757 | 0.178756 | 3.472642 | 5.15E-04 | 0.006915 |
| FAR2 | 377.3031 | -0.17288 | 0.170954 | -1.0113 | 0.311874 | 0.479045 |
| IPO5 | 3986.179 | 0.061649 | 0.122108 | 0.504874 | 0.613648 | 0.740094 |
| DGKA | 363.594 | 0.158757 | 0.221056 | 0.718174 | 0.47265 | 0.628442 |
| ERBB3 | 5380.827 | -0.31386 | 0.143679 | -2.18448 | 0.028927 | 0.096764 |
| FOXJ2 | 480.3768 | -0.58793 | 0.172014 | -3.41794 | 6.31E-04 | 0.007785 |
| DIP2B | 3854.583 | -0.10659 | 0.128431 | -0.82992 | 0.406587 | 0.570289 |
| SMARCD1 | 2248.656 | -0.42968 | 0.125731 | -3.41748 | 6.32E-04 | 0.007789 |
| ATXN3 | 340.1969 | 0.517964 | 0.270631 | 1.913912 | 0.055631 | 0.149143 |
| GOLGA5 | 1187.487 | 0.855773 | 0.165049 | 5.184973 | 2.16E-07 | 5.34E-05 |
| EML1 | 226.0873 | 0.056602 | 0.190211 | 0.297576 | 0.766027 | 0.854786 |
| KIF26A | 1.589987 | 0.694691 | 1.591628 | 0.436466 | 0.662499 | NA |
| ATG2B | 1502.044 | 0.269114 | 0.143572 | 1.874424 | 0.060872 | 0.158881 |
| MYO9A | 599.346 | 0.481188 | 0.24668 | 1.95066 | 0.051098 | 0.140646 |
| NEO1 | 3077.729 | -0.45981 | 0.188287 | -2.44208 | 0.014603 | 0.060786 |
| TNFRSF1A | 1959.864 | -0.30722 | 0.142542 | -2.15533 | 0.031136 | 0.101656 |
| STOML1 | 599.0809 | -0.27777 | 0.147581 | -1.88215 | 0.059816 | 0.157023 |
| PKM | 65874.09 | -0.22006 | 0.153395 | -1.43462 | 0.151395 | 0.295445 |
| TP53BP1 | 1318.768 | -0.05305 | 0.143939 | -0.36856 | 0.712456 | 0.8164 |
| SYT1 | 87.21715 | 0.862344 | 0.336773 | 2.560609 | 0.010449 | 0.048444 |
| NAV3 | 294.3512 | -0.07449 | 0.251114 | -0.29662 | 0.766753 | 0.855114 |
| ATP11A | 868.9975 | -0.43166 | 0.189928 | -2.27276 | 0.023041 | 0.083125 |
| CYFIP1 | 6188.736 | -0.3094 | 0.123764 | -2.49992 | 0.012422 | 0.054583 |
| RASGRP2 | 0.163957 | 0.424913 | 4.079006 | 0.104171 | 0.917034 | NA |
| PPP2R5B | 1689.61 | -0.11484 | 0.136807 | -0.83944 | 0.401223 | 0.565913 |
| PYGM | 0.881566 | 1.109202 | 2.554987 | 0.434132 | 0.664192 | NA |
| GAL | 8.366671 | -1.22017 | 0.727823 | -1.67646 | 0.093647 | NA |
| CLEC2D | 35.49743 | 0.723171 | 0.394334 | 1.833904 | 0.066668 | NA |
| RORA | 54.00936 | 0.690695 | 0.349974 | 1.973564 | 0.048431 | 0.135798 |
| NEDD4 | 341.9331 | 0.642485 | 0.249143 | 2.57878 | 0.009915 | 0.046923 |
| PIGB | 598.6197 | 0.108751 | 0.172243 | 0.631381 | 0.527791 | 0.673299 |
| MAPK6 | 820.509 | 0.71042 | 0.214121 | 3.317841 | 9.07E-04 | 0.009758 |
| GNB5 | 810.1381 | 0.06655 | 0.15781 | 0.421714 | 0.673234 | 0.786169 |
| RAB27A | 5457.574 | 0.206614 | 0.200833 | 1.028788 | 0.303579 | 0.47084 |
| LRP6 | 822.1983 | 0.391398 | 0.184802 | 2.117939 | 0.03418 | 0.10817 |

| | | | | | | |
|---|---|---|---|---|---|---|
| SPTB | 52.94142 | -1.08367 | 0.354221 | -3.05929 | 0.002219 | 0.017292 |
| TMEM260 | 392.8758 | 0.12953 | 0.19151 | 0.676364 | 0.49881 | 0.649348 |
| EXOC5 | 1488.555 | 0.753193 | 0.255996 | 2.942211 | 0.003259 | 0.022344 |
| PTPN21 | 703.3797 | 0.336368 | 0.180611 | 1.862388 | 0.062548 | 0.161937 |
| ATP2B1 | 558.5492 | 0.668418 | 0.261317 | 2.557881 | 0.010531 | 0.048693 |
| VASH1 | 342.6263 | -0.40637 | 0.186198 | -2.18248 | 0.029074 | 0.096999 |
| SEL1L | 3888.833 | 0.056527 | 0.164335 | 0.343975 | 0.730865 | 0.829368 |
| SLC6A15 | 1978.319 | 0.307415 | 0.151569 | 2.028216 | 0.042538 | 0.125476 |
| RDH11 | 2691.827 | 0.073968 | 0.167688 | 0.441108 | 0.659135 | 0.775241 |
| ACTN1 | 7140.691 | -0.02762 | 0.198611 | -0.13906 | 0.889405 | 0.935323 |
| ZFYVE26 | 2602.622 | -0.46414 | 0.148616 | -3.1231 | 0.00179 | 0.015078 |
| MPP5 | 791.6655 | 0.530136 | 0.215803 | 2.456577 | 0.014027 | 0.059304 |
| MARK2 | 1019.445 | -0.49788 | 0.163565 | -3.0439 | 0.002335 | 0.017885 |
| CHFR | 756.7129 | -0.20669 | 0.159806 | -1.29341 | 0.195869 | 0.351223 |
| TRHDE | 0.260158 | 1.240347 | 4.047124 | 0.306476 | 0.759242 | NA |
| SCARB1 | 12247.38 | -0.62923 | 0.136484 | -4.61028 | 4.02E-06 | 3.70E-04 |
| PDE8A | 1159.204 | 0.38624 | 0.15118 | 2.554843 | 0.010624 | 0.048986 |
| KDM5A | 1537.258 | 0.278842 | 0.171489 | 1.626002 | 0.103949 | 0.229013 |
| FERMT2 | 1252.701 | 0.406026 | 0.182012 | 2.230761 | 0.025697 | 0.089705 |
| FRY | 317.2328 | -0.46867 | 0.259105 | -1.80882 | 0.07048 | 0.175377 |
| PICALM | 3672.248 | 0.351118 | 0.181465 | 1.934911 | 0.053001 | 0.144013 |
| CLNS1A | 2091.071 | 0.238895 | 0.144124 | 1.657564 | 0.097405 | 0.219324 |
| EED | 690.72 | 0.046782 | 0.158844 | 0.294515 | 0.768364 | 0.856266 |
| CA12 | 49.43441 | -0.58394 | 0.389692 | -1.49845 | 0.134015 | 0.272 |
| NTN4 | 45.19265 | 1.668366 | 0.618686 | 2.696628 | 0.007005 | NA |
| NUAK1 | 227.0234 | 0.639529 | 0.37947 | 1.685321 | 0.091927 | 0.210878 |
| DPP8 | 1371.533 | 0.471016 | 0.216595 | 2.174638 | 0.029657 | 0.098236 |
| SLC24A1 | 462.5707 | -0.30239 | 0.175364 | -1.72433 | 0.084649 | 0.198309 |
| PTPLAD1 | 5387.761 | -0.01859 | 0.152368 | -0.12199 | 0.90291 | 0.943218 |
| SLC12A1 | 3.013754 | 0.735973 | 1.434827 | 0.512935 | 0.607997 | NA |
| ACTR6 | 446.2838 | 0.918785 | 0.215682 | 4.259914 | 2.05E-05 | 9.40E-04 |
| TIPIN | 166.1403 | 0.32069 | 0.222804 | 1.439334 | 0.150056 | 0.2936 |
| NUP37 | 397.3137 | 0.149439 | 0.163812 | 0.912263 | 0.36163 | 0.528818 |
| ACAT1 | 1838.861 | 0.299433 | 0.183497 | 1.631815 | 0.102719 | 0.226934 |
| MARK3 | 1356.604 | 0.359127 | 0.154847 | 2.319243 | 0.020382 | 0.076683 |
| SLC25A3 | 17818.58 | -0.11721 | 0.143794 | -0.81511 | 0.41501 | 0.578272 |
| ATP12A | 3.950447 | -0.11069 | 1.052486 | -0.10517 | 0.916238 | NA |
| SART3 | 1455.349 | -0.13484 | 0.142825 | -0.94409 | 0.345125 | 0.51312 |
| REXO2 | 1397.47 | 0.392764 | 0.156056 | 2.516822 | 0.011842 | 0.052866 |
| RBM7 | 265.8964 | 0.500131 | 0.212336 | 2.355373 | 0.018504 | 0.071688 |
| RBMS2 | 564.2855 | -0.09482 | 0.170289 | -0.55679 | 0.577671 | 0.712388 |
| BAZ2A | 2900.927 | -0.53831 | 0.149281 | -3.60603 | 3.11E-04 | 0.00505 |
| UNG | 1719.221 | -0.36152 | 0.135516 | -2.66773 | 0.007637 | 0.039439 |
| ANKRD13A | 1321.073 | 0.171925 | 0.175022 | 0.982302 | 0.325951 | 0.494011 |
| ACACB | 868.7104 | -0.47559 | 0.193014 | -2.46404 | 0.013738 | 0.058417 |
| MCAM | 13499.6 | -0.02306 | 0.290823 | -0.0793 | 0.936791 | 0.964637 |
| FAM76B | 244.1951 | 1.004873 | 0.311262 | 3.228383 | 0.001245 | 0.011897 |

| | | | | | | |
|---|---|---|---|---|---|---|
| TYR | 59184.77 | -0.32663 | 0.145743 | -2.24117 | 0.025015 | 0.087841 |
| POLD3 | 323.1764 | 0.332493 | 0.173636 | 1.914886 | 0.055507 | 0.148877 |
| NAALAD2 | 4.82481 | 1.058497 | 0.993451 | 1.065474 | 0.286661 | NA |
| ACER3 | 802.2844 | 0.219811 | 0.194039 | 1.13282 | 0.25729 | 0.421353 |
| C12orf5 | 1118.917 | 0.249952 | 0.152884 | 1.634917 | 0.102066 | 0.226041 |
| TULP3 | 954.4464 | -0.26221 | 0.13451 | -1.94936 | 0.051253 | 0.140974 |
| PPP2R5C | 2582.665 | 0.158307 | 0.148966 | 1.062704 | 0.287916 | 0.454191 |
| RAPGEF3 | 6.455707 | -0.77873 | 0.774893 | -1.00495 | 0.314923 | NA |
| SENP1 | 405.8196 | -0.07614 | 0.18884 | -0.40321 | 0.686795 | 0.796123 |
| DCT | 34472.32 | -0.39424 | 0.574297 | -0.68648 | 0.492411 | 0.643732 |
| RAB21 | 978.9374 | 0.341946 | 0.19674 | 1.738064 | 0.0822 | 0.194856 |
| CPB2 | 7.343857 | 1.096432 | 0.812138 | 1.350057 | 0.176998 | NA |
| CHRNA3 | 5.637092 | 0.388074 | 1.019385 | 0.380695 | 0.70343 | NA |
| PSEN1 | 1903.648 | -0.10378 | 0.140651 | -0.73788 | 0.460587 | 0.618796 |
| MOK | 1133.897 | -0.28581 | 0.23035 | -1.24075 | 0.214698 | 0.373586 |
| HSP90AA1 | 16597.47 | 1.069966 | 0.266146 | 4.020227 | 5.81E-05 | 0.001793 |
| IGSF9B | 1.850145 | 1.001833 | 1.48382 | 0.675172 | 0.499567 | NA |
| AP4E1 | 466.6764 | 0.602978 | 0.193537 | 3.115562 | 0.001836 | 0.015242 |
| EXD2 | 987.704 | -0.25979 | 0.133532 | -1.9455 | 0.051714 | 0.141653 |
| ARG2 | 240.5897 | 0.247607 | 0.402377 | 0.61536 | 0.538317 | 0.681093 |
| AACS | 2386.255 | -0.27954 | 0.122106 | -2.28934 | 0.022059 | 0.081 |
| ERC1 | 1888.241 | -0.43856 | 0.177224 | -2.4746 | 0.013339 | 0.057401 |
| DIS3 | 1090.13 | 0.68031 | 0.242955 | 2.800151 | 0.005108 | 0.030022 |
| PIBF1 | 118.432 | 1.265146 | 0.770358 | 1.642283 | 0.100531 | 0.223686 |
| TDRD3 | 2196.479 | 0.904691 | 0.277584 | 3.259155 | 0.001117 | 0.01112 |
| NUFIP1 | 216.2747 | 0.219344 | 0.221474 | 0.990381 | 0.321988 | 0.490197 |
| PDS5B | 642.7492 | 0.568212 | 0.195669 | 2.903942 | 0.003685 | 0.024216 |
| EPYC | 1.213849 | -1.34398 | 1.887808 | -0.71193 | 0.47651 | NA |
| HAL | 1.681242 | 1.965083 | 1.654241 | 1.187906 | 0.23487 | NA |
| SSH1 | 5651.607 | -0.28688 | 0.125107 | -2.29309 | 0.021843 | 0.080418 |
| GSTP1 | 20584.64 | -0.18829 | 0.201398 | -0.93494 | 0.349821 | 0.517938 |
| APLP2 | 11983.17 | -0.13834 | 0.122106 | -1.13296 | 0.257231 | 0.421315 |
| KIAA1467 | 274.4189 | -0.71669 | 0.173479 | -4.13128 | 3.61E-05 | 0.001362 |
| SLCO1A2 | 0.410275 | 0.244283 | 3.100889 | 0.078778 | 0.937209 | NA |
| WBP11 | 2957.639 | -0.2735 | 0.130365 | -2.09798 | 0.035907 | 0.111755 |
| CTTN | 10315.8 | -0.07987 | 0.159577 | -0.5005 | 0.61672 | 0.742224 |
| WNT11 | 1.866864 | -0.99112 | 1.481314 | -0.66908 | 0.503445 | NA |
| AQP6 | 2.778603 | -0.95543 | 1.330682 | -0.718 | 0.472759 | NA |
| TMED2 | 5017.145 | 0.080725 | 0.155443 | 0.519324 | 0.603535 | 0.732334 |
| ZFAND6 | 1630.025 | 0.392629 | 0.168828 | 2.325614 | 0.020039 | 0.07588 |
| ZW10 | 486.1826 | 0.240157 | 0.151688 | 1.583236 | 0.113368 | 0.242158 |
| ALG9 | 157.3288 | -0.27149 | 0.220341 | -1.23212 | 0.217906 | 0.377341 |
| NOX4 | 16.57423 | 0.699624 | 0.574773 | 1.217217 | 0.223522 | NA |
| MTMR2 | 2181.507 | 0.186316 | 0.161676 | 1.152408 | 0.249154 | 0.412992 |
| L2HGDH | 171.7243 | 0.454361 | 0.274216 | 1.656947 | 0.09753 | 0.219422 |
| TXNDC16 | 261.6903 | 0.745889 | 0.237864 | 3.135784 | 0.001714 | 0.01465 |
| C14orf166 | 2135.586 | 0.426215 | 0.127961 | 3.330813 | 8.66E-04 | 0.00947 |

| | | | | | | |
|---|---|---|---|---|---|---|
| NID2 | 23.95602 | 0.960737 | 0.439406 | 2.186448 | 0.028783 | NA |
| SF3B2 | 6969.833 | -0.22964 | 0.120738 | -1.90199 | 0.057172 | 0.151627 |
| KLHL42 | 1368.704 | 0.039324 | 0.180063 | 0.218389 | 0.827126 | 0.895814 |
| DNM1L | 1917.991 | 0.371563 | 0.174039 | 2.134941 | 0.032766 | 0.105468 |
| PTHLH | 5.929091 | 1.679868 | 0.977748 | 1.718098 | 0.085779 | NA |
| ERGIC2 | 632.8509 | 1.053464 | 0.260379 | 4.045894 | 5.21E-05 | 0.001693 |
| AAMDC | 387.7597 | 0.127066 | 0.184693 | 0.687985 | 0.491462 | 0.643291 |
| DOCK9 | 424.261 | 0.021865 | 0.20454 | 0.106898 | 0.91487 | 0.950884 |
| ANKRD10 | 1011.551 | 0.655011 | 0.155593 | 4.209765 | 2.56E-05 | 0.001067 |
| TGDS | 248.1501 | 0.493837 | 0.226183 | 2.183349 | 0.02901 | 0.096932 |
| PPP1R13B | 257.7981 | -0.07895 | 0.224756 | -0.35126 | 0.72539 | 0.825454 |
| DYNLL1 | 6509.897 | 0.265235 | 0.145836 | 1.818716 | 0.068955 | 0.172824 |
| TESC | 1.178278 | -0.21536 | 1.987735 | -0.10835 | 0.913721 | NA |
| RPL6 | 9474.37 | 0.370247 | 0.139672 | 2.650831 | 0.008029 | 0.040681 |
| MAPKAPK5 | 635.367 | 0.093993 | 0.157696 | 0.596036 | 0.551151 | 0.69215 |
| P2RX7 | 6154.387 | -0.16178 | 0.152533 | -1.06062 | 0.288862 | 0.455138 |
| ANAPC5 | 4817.895 | 0.071216 | 0.142551 | 0.499584 | 0.617368 | 0.742508 |
| SLC8B1 | 458.7326 | -0.20973 | 0.176042 | -1.19137 | 0.233507 | 0.396083 |
| KDM2B | 712.9184 | -0.41998 | 0.157376 | -2.66865 | 0.007616 | 0.039391 |
| LHX5 | 0.327913 | 1.543154 | 4.021063 | 0.383768 | 0.701151 | NA |
| OAS1 | 1570.286 | -1.11311 | 0.294326 | -3.7819 | 1.56E-04 | 0.003343 |
| GCN1L1 | 4789.143 | -0.67554 | 0.167406 | -4.03537 | 5.45E-05 | 0.001728 |
| RPLP0 | 34337.92 | -0.17614 | 0.119376 | -1.47551 | 0.140074 | 0.280348 |
| PXN | 3396.116 | -0.47394 | 0.125989 | -3.76177 | 1.69E-04 | 0.003497 |
| SIRT4 | 52.2776 | 0.018334 | 0.312074 | 0.058749 | 0.953152 | 0.974441 |
| RPH3A | 17.23813 | -0.6431 | 0.614677 | -1.04623 | 0.295453 | NA |
| PEBP1 | 18140.62 | 0.022661 | 0.121343 | 0.186755 | 0.851853 | 0.911895 |
| TBX5 | 0.230334 | -0.53318 | 4.076749 | -0.13079 | 0.895945 | NA |
| BRAP | 493.6574 | 0.241558 | 0.152545 | 1.583523 | 0.113302 | 0.242093 |
| ERP29 | 4016.789 | -0.06834 | 0.150504 | -0.45406 | 0.649786 | 0.767864 |
| LAG3 | 0.919846 | -3.47044 | 2.735469 | -1.26868 | 0.204555 | NA |
| MLF2 | 5958.743 | -0.13546 | 0.143248 | -0.94561 | 0.344348 | 0.512394 |
| OTUB2 | 126.6978 | -0.28503 | 0.226913 | -1.25613 | 0.209069 | 0.366643 |
| DDX24 | 5266.18 | 0.01207 | 0.119823 | 0.100728 | 0.919766 | 0.953549 |
| ZBTB25 | 79.6361 | 0.126522 | 0.314337 | 0.402506 | 0.687312 | 0.796567 |
| NECAP1 | 931.9913 | -0.04293 | 0.137898 | -0.31134 | 0.755544 | 0.847421 |
| RCOR1 | 656.5372 | -0.09509 | 0.162843 | -0.58394 | 0.559263 | 0.698048 |
| GPATCH2L | 895.697 | 0.371536 | 0.267063 | 1.391192 | 0.164167 | 0.312894 |
| PAPOLA | 3818.226 | 0.547276 | 0.143193 | 3.821941 | 1.32E-04 | 0.003029 |
| CCNK | 1408.396 | -0.41118 | 0.162777 | -2.52601 | 0.011537 | 0.051904 |
| IRAK3 | 386.8579 | 0.383437 | 0.234247 | 1.636894 | 0.101653 | 0.225335 |
| LYZ | 1.438436 | 0.401503 | 1.702105 | 0.235886 | 0.813521 | NA |
| PDCD7 | 664.6685 | -0.09814 | 0.146739 | -0.66881 | 0.503616 | 0.653539 |
| SPG21 | 3251.344 | -0.22229 | 0.144347 | -1.53995 | 0.123572 | 0.257867 |
| ZNF268 | 127.304 | 0.556318 | 0.438657 | 1.268232 | 0.204715 | 0.361395 |
| GOLGA3 | 2148.632 | -0.32649 | 0.195452 | -1.67045 | 0.09483 | 0.215442 |
| PITPNM2 | 627.7704 | -0.08836 | 0.271609 | -0.32533 | 0.744934 | 0.839379 |

| | | | | | | |
|---|---|---|---|---|---|---|
| OSBPL8 | 1490.173 | 0.861406 | 0.561642 | 1.533729 | 0.125096 | 0.260016 |
| CMA1 | 7.191762 | 1.214558 | 1.332728 | 0.911332 | 0.36212 | NA |
| PSME1 | 3611.351 | -0.25715 | 0.183122 | -1.40424 | 0.160249 | 0.307602 |
| PPP2R3C | 610.9385 | 0.471432 | 0.186214 | 2.53167 | 0.011352 | 0.051249 |
| HAUS4 | 314.5932 | -0.15577 | 0.16796 | -0.92744 | 0.353697 | 0.521656 |
| JPH4 | 10.53174 | -1.00951 | 0.61878 | -1.63145 | 0.102796 | NA |
| MYH7 | 0.823499 | -2.06708 | 2.534576 | -0.81555 | 0.414756 | NA |
| SLC7A8 | 8024.589 | -0.07543 | 0.199951 | -0.37724 | 0.705993 | 0.811553 |
| OSGEP | 527.2385 | -0.05007 | 0.1638 | -0.30569 | 0.759844 | 0.850495 |
| SLC22A17 | 476.991 | 0.169454 | 0.25115 | 0.674713 | 0.499858 | 0.65026 |
| RNF31 | 718.1545 | -0.48017 | 0.137895 | -3.48215 | 4.97E-04 | 0.006762 |
| SCFD1 | 711.561 | 0.955594 | 0.233437 | 4.093593 | 4.25E-05 | 0.001514 |
| G2E3 | 286.5439 | 0.865888 | 0.267604 | 3.235706 | 0.001213 | 0.01178 |
| HECTD1 | 2442.863 | 0.548694 | 0.204592 | 2.681894 | 0.007321 | 0.038367 |
| HNRNPC | 8509.434 | -0.03592 | 0.163469 | -0.21974 | 0.826077 | 0.895364 |
| RPGRIP1 | 2.593629 | -0.1577 | 1.339341 | -0.11775 | 0.906269 | NA |
| SUPT16H | 2671.368 | 0.412058 | 0.18694 | 2.204231 | 0.027508 | 0.093709 |
| TOX4 | 1734.84 | -0.15137 | 0.125524 | -1.20588 | 0.227862 | 0.389574 |
| GEMIN2 | 120.8416 | 0.303811 | 0.230999 | 1.315207 | 0.18844 | 0.342737 |
| TGM1 | 8.256349 | 0.006869 | 0.718669 | 0.009558 | 0.992374 | NA |
| TINF2 | 1379.183 | -0.22109 | 0.134537 | -1.6433 | 0.10032 | 0.223425 |
| TRPM7 | 1781.92 | 0.959862 | 0.247711 | 3.874919 | 1.07E-04 | 0.002681 |
| TYRO3 | 3332.335 | -0.11913 | 0.158701 | -0.75066 | 0.452857 | 0.611879 |
| WDR76 | 232.7271 | 0.053769 | 0.205545 | 0.261594 | 0.793635 | 0.874943 |
| CAPN3 | 1083.233 | -0.24142 | 0.162154 | -1.48884 | 0.136528 | 0.27559 |
| SNAP23 | 1123.89 | 0.192311 | 0.137193 | 1.401755 | 0.160988 | 0.308572 |
| MYL6 | 18367.12 | 0.143738 | 0.142949 | 1.005522 | 0.314646 | 0.482053 |
| AAAS | 1150.829 | -0.40796 | 0.134231 | -3.03919 | 0.002372 | 0.01805 |
| CBX5 | 1569.268 | -0.26046 | 0.165737 | -1.57151 | 0.116065 | 0.246398 |
| NXPE1 | 0.134133 | -1.48909 | 4.077584 | -0.36519 | 0.71497 | NA |
| ARCN1 | 3697.199 | -0.05003 | 0.136062 | -0.3677 | 0.713098 | 0.816711 |
| CEP170B | 2590.289 | -0.3991 | 0.151146 | -2.64051 | 0.008278 | 0.041548 |
| KCNK10 | 0.556813 | -0.57007 | 3.294615 | -0.17303 | 0.862627 | NA |
| ABHD4 | 1774.805 | 0.021974 | 0.159808 | 0.1375 | 0.890636 | 0.936036 |
| KHNYN | 1444.296 | -0.30018 | 0.140712 | -2.13326 | 0.032904 | 0.105627 |
| FKBP3 | 674.5733 | 0.539166 | 0.209297 | 2.576086 | 0.009993 | 0.047157 |
| SDR39U1 | 210.7141 | 0.25139 | 0.200377 | 1.254586 | 0.209629 | 0.367191 |
| CTSG | 1.826346 | 0.764873 | 2.012993 | 0.379968 | 0.703969 | NA |
| GZMH | 0.192403 | 0.975392 | 4.060716 | 0.240202 | 0.810174 | NA |
| GZMB | 12.41494 | 0.621932 | 0.585658 | 1.061938 | 0.288264 | NA |
| RBM23 | 2228.992 | -0.17202 | 0.169412 | -1.01537 | 0.309931 | 0.477235 |
| PRMT5 | 1408.072 | -0.28153 | 0.159406 | -1.7661 | 0.077379 | 0.187058 |
| COCH | 3.74797 | 0.711688 | 1.042266 | 0.682828 | 0.494716 | NA |
| AP4S1 | 78.1861 | 0.270157 | 0.264559 | 1.021161 | 0.307178 | 0.474475 |
| POLE2 | 121.3341 | 0.255498 | 0.293644 | 0.870096 | 0.384248 | 0.550351 |
| VCPKMT | 90.75812 | 0.652796 | 0.251198 | 2.598728 | 0.009357 | 0.045147 |
| SOS2 | 1371.788 | 0.578177 | 0.221141 | 2.614515 | 0.008935 | 0.043845 |

| | | | | | | |
|---|---|---|---|---|---|---|
| CDKL1 | 26.49563 | 0.179364 | 0.540778 | 0.331678 | 0.740132 | NA |
| NIN | 1403.294 | 0.429042 | 0.646043 | 0.664107 | 0.506622 | 0.655875 |
| PYGL | 2835.397 | -0.15239 | 0.25735 | -0.59215 | 0.553748 | 0.69406 |
| TRIM9 | 75.30382 | -0.77189 | 0.34957 | -2.20811 | 0.027237 | 0.092946 |
| PSMC6 | 914.3287 | 0.889844 | 0.212901 | 4.179609 | 2.92E-05 | 0.00117 |
| GNPNAT1 | 437.2931 | 0.119378 | 0.17062 | 0.699672 | 0.484132 | 0.63777 |
| DDHD1 | 355.405 | 0.365357 | 0.17734 | 2.060206 | 0.039379 | 0.119061 |
| CDKN3 | 706.7297 | 0.269581 | 0.179184 | 1.504493 | 0.132455 | 0.269673 |
| CNIH1 | 2485.261 | 0.222051 | 0.164495 | 1.349898 | 0.177049 | 0.329241 |
| CGRRF1 | 344.584 | 0.513853 | 0.214828 | 2.391922 | 0.01676 | 0.067263 |
| ATP6V1D | 3538.268 | 0.379994 | 0.164293 | 2.312913 | 0.020727 | 0.077653 |
| C14orf105 | 0.163957 | 0.424913 | 4.079006 | 0.104171 | 0.917034 | NA |
| PLEK2 | 261.8968 | -0.01742 | 0.178986 | -0.09733 | 0.922464 | 0.95505 |
| PIGH | 629.4128 | -0.03838 | 0.175268 | -0.219 | 0.82665 | 0.895625 |
| C14orf166I | 0.260158 | 1.240347 | 4.047124 | 0.306476 | 0.759242 | NA |
| PSMA3 | 1326.467 | 0.584763 | 0.170291 | 3.43391 | 5.95E-04 | 0.007535 |
| VTI1B | 2016.545 | 0.136723 | 0.133041 | 1.027677 | 0.304102 | 0.47132 |
| TIMM9 | 377.39 | 0.021983 | 0.168498 | 0.130467 | 0.896197 | 0.939326 |
| GSTZ1 | 525.9814 | -0.36783 | 0.19522 | -1.88416 | 0.059544 | 0.156482 |
| KIAA0586 | 236.7911 | 0.41552 | 0.335443 | 1.238718 | 0.21545 | 0.374583 |
| TMED8 | 1185.01 | -0.03111 | 0.188259 | -0.16523 | 0.868765 | 0.922011 |
| SAMD15 | 16.09067 | -0.87805 | 0.56259 | -1.56072 | 0.118589 | NA |
| AHSA1 | 2993.866 | 0.003039 | 0.135724 | 0.022392 | 0.982135 | 0.990311 |
| DAAM1 | 561.3025 | 0.816865 | 0.24465 | 3.338917 | 8.41E-04 | 0.00931 |
| ISM2 | 1.274922 | -0.93633 | 1.907377 | -0.4909 | 0.623496 | NA |
| SPTLC2 | 1908.816 | -0.24482 | 0.152098 | -1.60962 | 0.107481 | 0.233925 |
| RIN3 | 2565.485 | -0.67024 | 0.144368 | -4.64259 | 3.44E-06 | 3.37E-04 |
| LGMN | 1707.839 | -0.29587 | 0.137779 | -2.14741 | 0.031761 | 0.103214 |
| ALKBH1 | 303.8824 | -0.10018 | 0.174169 | -0.57517 | 0.565177 | 0.702545 |
| SNW1 | 1493.307 | 0.260521 | 0.190569 | 1.367064 | 0.171605 | 0.322661 |
| CHGA | 1.061949 | -0.45116 | 2.030098 | -0.22224 | 0.82413 | NA |
| ITPK1 | 10423.15 | -0.39744 | 0.148959 | -2.66808 | 0.007629 | 0.039423 |
| DHRS7 | 3131.392 | -0.06356 | 0.123802 | -0.51337 | 0.607693 | 0.735198 |
| PPM1A | 1152.158 | 0.265383 | 0.15972 | 1.661551 | 0.096603 | 0.218014 |
| SIX4 | 2.405524 | 0.20377 | 1.29768 | 0.157026 | 0.875224 | NA |
| GALNT16 | 0.247898 | -0.5901 | 4.074633 | -0.14482 | 0.884851 | NA |
| ASB2 | 290.8645 | 0.225952 | 0.303305 | 0.744965 | 0.456293 | 0.614817 |
| CEP128 | 277.9316 | 0.662743 | 0.306092 | 2.165173 | 0.030374 | 0.099941 |
| ERH | 2691.904 | 0.227217 | 0.177313 | 1.281447 | 0.200037 | 0.356067 |
| HIF1A | 2152.921 | 0.436166 | 0.168976 | 2.581233 | 0.009845 | 0.046721 |
| KIAA0247 | 699.6575 | -0.44559 | 0.16186 | -2.75294 | 0.005906 | 0.033043 |
| SRSF5 | 2796.025 | 0.504174 | 0.156964 | 3.212033 | 0.001318 | 0.012423 |
| SLC10A1 | 0.80576 | 2.889259 | 2.725368 | 1.060136 | 0.289083 | NA |
| EIF5 | 3599.943 | 0.369866 | 0.141505 | 2.613801 | 0.008954 | 0.043919 |
| DICER1 | 861.4729 | 0.527832 | 0.308891 | 1.708795 | 0.087489 | 0.203249 |
| ZFYVE21 | 1289.35 | -0.06775 | 0.141789 | -0.4778 | 0.632793 | 0.754704 |
| MTHFD1 | 3353.417 | -0.1944 | 0.124739 | -1.55846 | 0.119123 | 0.250792 |

| | | | | | | |
|---|---|---|---|---|---|---|
| ZC3H14 | 741.426 | 0.521925 | 0.196353 | 2.658099 | 0.007858 | 0.04019 |
| PCNX | 1422.617 | 0.058617 | 0.162715 | 0.360241 | 0.718667 | 0.820552 |
| BDKRB1 | 0.134133 | -1.48909 | 4.077584 | -0.36519 | 0.71497 | NA |
| GSKIP | 664.8355 | 0.62862 | 0.21971 | 2.861139 | 0.004221 | 0.02638 |
| VRK1 | 408.5704 | 0.714737 | 0.200535 | 3.564159 | 3.65E-04 | 0.005555 |
| PSMC1 | 848.9312 | 0.284344 | 0.182419 | 1.558742 | 0.119057 | 0.250759 |
| PAPLN | 113.99 | -0.55356 | 0.349238 | -1.58504 | 0.112957 | 0.241759 |
| RPS6KA5 | 290.7414 | 0.175165 | 0.224136 | 0.781513 | 0.434501 | 0.595561 |
| SMEK1 | 941.6758 | 0.590509 | 0.203996 | 2.894705 | 0.003795 | 0.024612 |
| C14orf93 | 171.5665 | -0.47077 | 0.200305 | -2.35028 | 0.018759 | 0.072322 |
| PSMB5 | 4137.572 | -0.04952 | 0.143652 | -0.34474 | 0.730287 | 0.82887 |
| YY1 | 2593.04 | 0.035358 | 0.141774 | 0.249397 | 0.803054 | 0.882059 |
| ACIN1 | 3966.516 | -0.22657 | 0.13285 | -1.70548 | 0.088105 | 0.204439 |
| CCNB1IP1 | 1304.597 | 0.099119 | 0.180248 | 0.549903 | 0.582386 | 0.715905 |
| TRIP11 | 382.0279 | 0.802418 | 1.020413 | 0.786366 | 0.431653 | 0.592892 |
| APEX1 | 4804.728 | -0.03627 | 0.133961 | -0.27078 | 0.78656 | 0.870699 |
| PABPN1 | 800.2641 | 0.009339 | 0.154375 | 0.060494 | 0.951762 | 0.97348 |
| EFS | 605.5214 | -0.23563 | 0.161253 | -1.46124 | 0.14395 | 0.285539 |
| ARHGAP5 | 1650.933 | 0.920852 | 0.31725 | 2.902602 | 0.003701 | 0.024281 |
| CINP | 1031.086 | -0.28516 | 0.129036 | -2.20993 | 0.02711 | 0.092647 |
| DHRS2 | 3.661405 | -2.57027 | 1.207498 | -2.12859 | 0.033288 | NA |
| SRP54 | 1233.987 | 0.512183 | 0.210332 | 2.435117 | 0.014887 | 0.061701 |
| CPNE6 | 0.449786 | -1.25325 | 3.00474 | -0.41709 | 0.676613 | NA |
| CHD8 | 2517.478 | -0.31306 | 0.123023 | -2.54471 | 0.010937 | 0.049887 |
| PCK2 | 860.4103 | -0.59026 | 0.15938 | -3.70349 | 2.13E-04 | 0.00404 |
| KIAA0391 | 21.77152 | -0.1278 | 0.515354 | -0.24798 | 0.804154 | NA |
| DCAF11 | 2469.573 | -0.30581 | 0.152994 | -1.99884 | 0.045625 | 0.130776 |
| PSMA6 | 595.5763 | 0.180089 | 0.206013 | 0.874167 | 0.382027 | 0.548164 |
| NFKBIA | 1040.967 | -0.0471 | 0.142791 | -0.32985 | 0.741516 | 0.837127 |
| EMC9 | 343.0361 | -0.1968 | 0.206952 | -0.95096 | 0.341623 | 0.509812 |
| PSME2 | 3261.514 | -0.06575 | 0.150496 | -0.43689 | 0.662191 | 0.777717 |
| BRMS1L | 274.4264 | 0.547579 | 0.226672 | 2.415735 | 0.015703 | 0.063974 |
| REC8 | 71.7793 | -0.47463 | 0.342754 | -1.38477 | 0.166124 | 0.315242 |
| TM9SF1 | 246.8525 | -0.28843 | 0.204198 | -1.41248 | 0.157809 | 0.304248 |
| SEC23A | 2243.533 | 0.455936 | 0.162941 | 2.798158 | 0.005139 | 0.030133 |
| GMPR2 | 1716.314 | 1.79E-04 | 0.125853 | 0.001424 | 0.998864 | 0.999384 |
| PNN | 562.4092 | 1.005855 | 0.685226 | 1.467918 | 0.142126 | 0.282988 |
| RABGGTA | 1181.756 | -0.22797 | 0.137291 | -1.6605 | 0.096815 | 0.218284 |
| NFATC4 | 1882.938 | -0.42114 | 0.127416 | -3.30524 | 9.49E-04 | 0.009991 |
| EEA1 | 611.8255 | 1.167205 | 0.946782 | 1.232813 | 0.217646 | 0.377018 |
| KIAA0226L | 23.34107 | -0.9391 | 0.468627 | -2.00393 | 0.045078 | NA |
| NALCN | 0.46735 | -1.30193 | 3.399671 | -0.38296 | 0.701751 | NA |
| FGF14 | 12.9214 | -1.17085 | 0.680331 | -1.72099 | 0.085252 | NA |
| HTR2A | 0.247898 | -0.5901 | 4.074633 | -0.14482 | 0.884851 | NA |
| NDFIP2 | 1395.704 | 0.569593 | 0.163705 | 3.47939 | 5.03E-04 | 0.006793 |
| TNFSF13B | 21.24719 | 0.333122 | 0.533162 | 0.624804 | 0.5321 | NA |
| FNDC3A | 1584.619 | 0.614323 | 0.215309 | 2.853223 | 0.004328 | 0.02675 |

| | | | | | | |
|---|---|---|---|---|---|---|
| MLNR | 5.586379 | -0.38745 | 0.852778 | -0.45434 | 0.649585 | NA |
| CDADC1 | 239.7018 | 0.951316 | 0.250495 | 3.79775 | 1.46E-04 | 0.003182 |
| CAB39L | 1021.369 | 0.709314 | 0.268288 | 2.64385 | 0.008197 | 0.041263 |
| KLF5 | 454.3616 | 0.855664 | 0.242172 | 3.533285 | 4.10E-04 | 0.006022 |
| STK24 | 3489.834 | -0.32023 | 0.154964 | -2.06648 | 0.038784 | 0.117741 |
| DNAJC3 | 668.2013 | 0.577312 | 0.178749 | 3.229741 | 0.001239 | 0.011874 |
| UGGT2 | 400.5675 | 0.813677 | 0.294294 | 2.764845 | 0.005695 | 0.03235 |
| ARHGEF7 | 1746.114 | -0.18323 | 0.122979 | -1.48991 | 0.136248 | 0.275434 |
| FGF9 | 0.096202 | 0.408546 | 4.079639 | 0.100143 | 0.920231 | NA |
| SGCG | 0.268266 | -2.15396 | 4.038351 | -0.53338 | 0.593773 | NA |
| PARP4 | 2288.406 | 0.053741 | 0.166851 | 0.322091 | 0.747384 | 0.84124 |
| SUPT20H | 721.859 | 0.29566 | 0.204899 | 1.442957 | 0.149033 | 0.292417 |
| MRPS31 | 238.7009 | 0.687621 | 0.202098 | 3.402412 | 6.68E-04 | 0.008064 |
| SLC25A15 | 357.15 | -0.47444 | 0.166984 | -2.8412 | 0.004494 | 0.027439 |
| KPNA3 | 1287.454 | 0.42674 | 0.150314 | 2.83899 | 0.004526 | 0.027525 |
| FLT1 | 79.78403 | 1.260187 | 0.481834 | 2.615396 | 0.008912 | 0.043786 |
| RGCC | 45.36337 | 0.616053 | 0.346077 | 1.780105 | 0.075059 | NA |
| VWA8 | 602.9804 | 0.017367 | 0.157622 | 0.110184 | 0.912263 | 0.949227 |
| DGKH | 193.2589 | 0.824826 | 0.218862 | 3.76871 | 1.64E-04 | 0.003461 |
| KATNAL1 | 720.8895 | 0.542755 | 0.232813 | 2.331296 | 0.019738 | 0.075032 |
| INTS6 | 532.8147 | 0.583432 | 0.202222 | 2.885106 | 0.003913 | 0.025101 |
| DHRS12 | 351.767 | -0.0357 | 0.191424 | -0.1865 | 0.85205 | 0.911895 |
| MEDAG | 2.072664 | 0.637648 | 1.573195 | 0.40532 | 0.685242 | NA |
| TSC22D1 | 2449.103 | -0.00853 | 0.13675 | -0.0624 | 0.950244 | 0.972391 |
| CLN5 | 1471.518 | 0.144059 | 0.157296 | 0.915845 | 0.359748 | 0.526911 |
| OLFM4 | 0.134133 | -1.48909 | 4.077584 | -0.36519 | 0.71497 | NA |
| AAGAB | 1130.616 | 0.159527 | 0.154399 | 1.033213 | 0.301504 | 0.468442 |
| IQCH | 19.02849 | 0.291363 | 0.517062 | 0.563496 | 0.573097 | NA |
| LACTB | 839.0588 | 0.69777 | 0.236787 | 2.946819 | 0.003211 | 0.022116 |
| CORO2B | 2089.895 | -1.31012 | 0.268442 | -4.88047 | 1.06E-06 | 1.68E-04 |
| CSK | 2673.798 | -0.44742 | 0.144913 | -3.08751 | 0.002018 | 0.016256 |
| HERC1 | 2039.791 | -0.24174 | 0.16262 | -1.48653 | 0.137138 | 0.276319 |
| TRIP4 | 554.5652 | 0.107287 | 0.174184 | 0.615943 | 0.537932 | 0.681032 |
| MTFMT | 481.2982 | 0.087602 | 0.173652 | 0.504469 | 0.613932 | 0.740239 |
| RASL12 | 7.579471 | 0.665094 | 0.754147 | 0.881915 | 0.377823 | NA |
| AP3B2 | 5.579503 | 0.272157 | 0.849264 | 0.320462 | 0.748618 | NA |
| ACSBG1 | 73.37614 | 0.736622 | 0.462461 | 1.592831 | 0.111198 | 0.239334 |
| IGDCC4 | 2.71165 | -1.70307 | 1.310914 | -1.29915 | 0.193894 | NA |
| RAB11A | 4079.449 | 0.26764 | 0.119715 | 2.235654 | 0.025374 | 0.088919 |
| CTSH | 1790.858 | -0.32832 | 0.139126 | -2.35987 | 0.018282 | 0.071058 |
| GOLGA8UF | 0.315653 | -0.57305 | 3.892639 | -0.14721 | 0.882963 | NA |
| CD276 | 2821.07 | -0.4947 | 0.123884 | -3.99325 | 6.52E-05 | 0.001941 |
| FAH | 1785.766 | -0.30601 | 0.151685 | -2.01737 | 0.043657 | 0.127299 |
| KIAA1199 | 21.35959 | 0.204006 | 0.560682 | 0.363853 | 0.715968 | NA |
| RPAP1 | 1000.04 | -0.52618 | 0.150115 | -3.50519 | 4.56E-04 | 0.00643 |
| HOMER2 | 202.203 | -0.20177 | 0.263294 | -0.76634 | 0.443471 | 0.603336 |
| EHD4 | 6244.73 | -0.49269 | 0.174843 | -2.8179 | 0.004834 | 0.028855 |

| | | | | | | |
|---|---|---|---|---|---|---|
| TMEM87A | 1588.44 | 0.275235 | 0.150931 | 1.823576 | 0.068216 | 0.171408 |
| ZNF106 | 17771.56 | 0.352039 | 0.177091 | 1.987902 | 0.046823 | 0.133177 |
| CEP152 | 107.5289 | 1.212594 | 0.392578 | 3.088799 | 0.00201 | 0.016251 |
| ATP8B4 | 0.747636 | 0.735436 | 2.634453 | 0.279161 | 0.780122 | NA |
| OCA2 | 13170.55 | -0.72188 | 0.183739 | -3.92883 | 8.54E-05 | 0.002317 |
| DTWD1 | 208.6138 | 0.612304 | 0.214964 | 2.848399 | 0.004394 | 0.027074 |
| FAM189A1 | 0.303394 | -2.27006 | 4.032451 | -0.56295 | 0.57347 | NA |
| GABPB1 | 410.1335 | 0.318126 | 0.181571 | 1.752073 | 0.079761 | 0.191024 |
| TJP1 | 1302.366 | -0.13518 | 0.196292 | -0.68868 | 0.491022 | 0.642998 |
| BMF | 810.4359 | 0.494066 | 0.192971 | 2.560311 | 0.010458 | 0.048467 |
| DMXL2 | 1409.055 | 0.411953 | 0.289388 | 1.42353 | 0.154583 | 0.299882 |
| SCG3 | 3.815762 | 0.718638 | 1.100791 | 0.652838 | 0.513861 | NA |
| DNAJC17 | 236.631 | -0.11732 | 0.203778 | -0.57574 | 0.564788 | 0.702356 |
| EIF3J | 979.1652 | 0.616199 | 0.211265 | 2.916718 | 0.003537 | 0.023559 |
| SPG11 | 1222.69 | -0.10373 | 0.208522 | -0.49746 | 0.618868 | 0.743481 |
| RHOV | 1.79044 | -0.15215 | 1.460398 | -0.10418 | 0.917026 | NA |
| VPS18 | 3578.993 | -0.57877 | 0.128455 | -4.50565 | 6.62E-06 | 4.97E-04 |
| OIP5 | 97.03407 | -0.30573 | 0.297183 | -1.02876 | 0.303594 | 0.47084 |
| SLC30A4 | 339.6214 | 0.259695 | 0.201771 | 1.287077 | 0.198067 | 0.353894 |
| BLOC1S6 | 1878.87 | 0.654302 | 0.182444 | 3.58632 | 3.35E-04 | 0.005288 |
| MYEF2 | 964.1347 | 0.371611 | 0.225244 | 1.649818 | 0.09898 | 0.221858 |
| DDX25 | 12.77159 | 0.511044 | 0.609981 | 0.837802 | 0.402142 | NA |
| CRYAB | 13035.24 | 1.042702 | 0.535858 | 1.945856 | 0.051672 | 0.141603 |
| CTSC | 1885.281 | -0.09325 | 0.158046 | -0.59002 | 0.555179 | 0.695073 |
| ZBTB16 | 0.258578 | 1.233151 | 4.047809 | 0.304646 | 0.760635 | NA |
| ZNF259 | 1486.494 | -0.04822 | 0.142547 | -0.33828 | 0.735151 | 0.83232 |
| TECTA | 22.4882 | -0.21121 | 0.447671 | -0.4718 | 0.637067 | NA |
| SC5D | 1510.871 | 0.925844 | 0.233645 | 3.962606 | 7.41E-05 | 0.002116 |
| C11orf63 | 245.0143 | 0.303293 | 0.200583 | 1.512056 | 0.13052 | 0.26704 |
| B3GAT1 | 5.51306 | -0.20413 | 0.951978 | -0.21442 | 0.830217 | NA |
| HSPA8 | 45672.15 | -0.0016 | 0.172116 | -0.00929 | 0.992587 | 0.996738 |
| VWA5A | 105.8564 | -0.26588 | 0.298966 | -0.88935 | 0.373816 | 0.540043 |
| DNAJC4 | 669.6114 | -0.31 | 0.150881 | -2.05459 | 0.039919 | 0.120287 |
| SIAE | 681.93 | 0.176001 | 0.149824 | 1.174713 | 0.24011 | 0.403355 |
| SNX15 | 573.6783 | -0.2336 | 0.196692 | -1.18763 | 0.23498 | 0.397785 |
| ATG2A | 3269.396 | -0.66531 | 0.124586 | -5.34018 | 9.29E-08 | 3.24E-05 |
| EHD1 | 1934.748 | -0.33277 | 0.148758 | -2.23701 | 0.025286 | 0.088662 |
| UNC93B1 | 1049.781 | -0.68365 | 0.154869 | -4.41435 | 1.01E-05 | 6.32E-04 |
| PUS3 | 220.889 | 0.280971 | 0.21374 | 1.314544 | 0.188663 | 0.342879 |
| DCPS | 987.9958 | -0.23866 | 0.141451 | -1.68719 | 0.091566 | 0.210412 |
| SUV420H1 | 516.884 | 0.508375 | 0.225358 | 2.255858 | 0.02408 | 0.085828 |
| FOXRED1 | 627.9511 | -0.2046 | 0.20691 | -0.98882 | 0.322753 | 0.490733 |
| PPP6R3 | 1773.081 | 0.501277 | 0.200356 | 2.501927 | 0.012352 | 0.054456 |
| NRXN2 | 6.7854 | 1.117226 | 0.848078 | 1.317363 | 0.187717 | NA |
| ST3GAL4 | 16740.54 | -0.01291 | 0.128877 | -0.10014 | 0.920231 | 0.953864 |
| CPT1A | 2935.198 | -0.03283 | 0.22581 | -0.14537 | 0.884415 | 0.931563 |
| CCND1 | 42521.56 | 0.199689 | 0.285591 | 0.699213 | 0.484419 | 0.638017 |

| | | | | | | |
|---|---|---|---|---|---|---|
| CHORDC1 | 585.7887 | 1.003884 | 0.226647 | 4.429286 | 9.45E-06 | 6.12E-04 |
| FOLR1 | 0.392857 | -1.546 | 4.075469 | -0.37934 | 0.704433 | NA |
| ANAPC15 | 186.5754 | 0.118437 | 0.21453 | 0.552076 | 0.580897 | 0.714667 |
| FOLR3 | 12.61514 | 0.569088 | 0.755876 | 0.752885 | 0.451519 | NA |
| PANX1 | 1787.137 | -0.02765 | 0.158872 | -0.17405 | 0.861824 | 0.91821 |
| ARHGEF17 | 1744.083 | -0.55912 | 0.175305 | -3.18942 | 0.001426 | 0.013132 |
| CEP164 | 681.0927 | -0.36565 | 0.153424 | -2.38323 | 0.017162 | 0.06825 |
| KIAA1377 | 45.49467 | 0.442785 | 0.526951 | 0.840276 | 0.400754 | NA |
| IL10RA | 0.192403 | 0.975392 | 4.060716 | 0.240202 | 0.810174 | NA |
| BIRC2 | 954.3643 | 0.875951 | 0.255213 | 3.432236 | 5.99E-04 | 0.007574 |
| UBE4A | 1155.352 | -0.14127 | 0.172124 | -0.82073 | 0.411802 | 0.575044 |
| DDX6 | 1842.526 | 0.045009 | 0.186889 | 0.240831 | 0.809686 | 0.886237 |
| UPK2 | 1.967817 | 0.07878 | 1.540974 | 0.051123 | 0.959227 | NA |
| CBL | 1196.918 | -0.21118 | 0.161215 | -1.30992 | 0.190224 | 0.344448 |
| PVRL1 | 649.9437 | -0.62276 | 0.182832 | -3.40618 | 6.59E-04 | 0.007993 |
| NAA40 | 411.0902 | -0.03974 | 0.187077 | -0.21243 | 0.83177 | 0.898873 |
| SLC35F2 | 966.3919 | 0.26177 | 0.194685 | 1.344584 | 0.17876 | 0.33126 |
| ELMOD1 | 0.260158 | 1.240347 | 4.047124 | 0.306476 | 0.759242 | NA |
| PITPNM1 | 3081.317 | -0.25528 | 0.196519 | -1.29899 | 0.193948 | 0.348953 |
| AIP | 2517.344 | -0.12837 | 0.143113 | -0.89699 | 0.369722 | 0.536613 |
| NDUFS8 | 2379.283 | -0.14473 | 0.141741 | -1.02106 | 0.307226 | 0.474487 |
| TCIRG1 | 922.8365 | -0.14919 | 0.172389 | -0.86542 | 0.386809 | 0.552871 |
| CHKA | 661.5905 | -0.13853 | 0.175779 | -0.78809 | 0.430647 | 0.592195 |
| EXPH5 | 3.379497 | -0.22606 | 1.14372 | -0.19765 | 0.84332 | NA |
| POU2AF1 | 0.715249 | -0.95037 | 2.624937 | -0.36205 | 0.717313 | NA |
| VWF | 13.70362 | -0.98159 | 0.589374 | -1.66547 | 0.095818 | NA |
| PSMD9 | 410.5209 | -0.07697 | 0.161783 | -0.47577 | 0.634235 | 0.755664 |
| LEPREL2 | 2025.186 | -0.3142 | 0.156878 | -2.00284 | 0.045195 | 0.130076 |
| PPFIBP1 | 2498.418 | 0.382241 | 0.199069 | 1.920142 | 0.05484 | 0.147555 |
| PRPF40B | 203.5412 | -0.47261 | 0.203933 | -2.31747 | 0.020478 | 0.076899 |
| CD69 | 0.296509 | -0.51709 | 3.982155 | -0.12985 | 0.896684 | NA |
| PRDM4 | 1069.808 | 0.02709 | 0.147587 | 0.18355 | 0.854366 | 0.913506 |
| CLEC2B | 198.5363 | 1.150685 | 0.307097 | 3.746972 | 1.79E-04 | 0.003621 |
| COQ5 | 740.1567 | -0.09462 | 0.143622 | -0.65881 | 0.51002 | 0.65835 |
| SELPLG | 15.68907 | -0.1 | 0.546814 | -0.18288 | 0.854896 | NA |
| CORO1C | 9514.469 | -0.02283 | 0.148493 | -0.15371 | 0.877836 | 0.926876 |
| ASIC1 | 138.1824 | -0.25453 | 0.308914 | -0.82396 | 0.409961 | 0.573435 |
| DAO | 1.692658 | -0.35328 | 1.666493 | -0.21199 | 0.832116 | NA |
| CAPRIN2 | 701.7692 | 0.495291 | 0.170921 | 2.897784 | 0.003758 | 0.024485 |
| TSPAN11 | 269.1288 | 0.104884 | 0.26487 | 0.395984 | 0.692117 | 0.800961 |
| KCTD10 | 3368.363 | -0.07345 | 0.134066 | -0.54787 | 0.583779 | 0.717088 |
| SLC11A2 | 2475.514 | -0.03467 | 0.128682 | -0.26941 | 0.787618 | 0.871252 |
| MLEC | 16526.23 | -0.59412 | 0.13428 | -4.42452 | 9.67E-06 | 6.19E-04 |
| MVK | 1181.438 | -0.35593 | 0.186432 | -1.90916 | 0.056242 | 0.150066 |
| CSRNP2 | 994.3119 | -0.09652 | 0.186196 | -0.51836 | 0.604209 | 0.732942 |
| CAMKK2 | 1993.752 | -0.22569 | 0.129879 | -1.73769 | 0.082265 | 0.194934 |
| IL23A | 6.925069 | -0.41734 | 0.777885 | -0.53651 | 0.591607 | NA |

| | | | | | | |
|---|---|---|---|---|---|---|
| ATP5B | 26943.49 | -0.15373 | 0.119479 | -1.28671 | 0.198197 | 0.35399 |
| PTGES3 | 4714.949 | 0.023568 | 0.149262 | 0.1579 | 0.874535 | 0.925145 |
| BCL7A | 421.0166 | -0.29914 | 0.225589 | -1.32604 | 0.184826 | 0.338239 |
| RSRC2 | 651.4674 | 0.783218 | 0.267005 | 2.933342 | 0.003353 | 0.022701 |
| CYP27B1 | 5.953686 | 0.351317 | 0.834136 | 0.421175 | 0.673627 | NA |
| LIN7A | 0.555087 | 0.973802 | 2.808575 | 0.346725 | 0.728798 | NA |
| KRT18 | 1.088049 | 1.239202 | 2.655459 | 0.466662 | 0.640742 | NA |
| ACSS3 | 0.258578 | 1.233151 | 4.047809 | 0.304646 | 0.760635 | NA |
| TENC1 | 708.2511 | -0.69139 | 0.156936 | -4.40557 | 1.06E-05 | 6.38E-04 |
| GLI1 | 51.87452 | 0.40356 | 0.341345 | 1.182267 | 0.2371 | 0.399946 |
| PPM1H | 1405.645 | -0.5261 | 0.18532 | -2.83886 | 0.004527 | 0.027525 |
| METAP2 | 1333.015 | 0.870489 | 0.249942 | 3.482764 | 4.96E-04 | 0.006762 |
| LTA4H | 2704.446 | -0.02496 | 0.148597 | -0.16798 | 0.866598 | 0.920772 |
| ELK3 | 1037.758 | 0.555584 | 0.182408 | 3.045834 | 0.00232 | 0.017804 |
| SLC6A12 | 3.737079 | -0.07962 | 1.04607 | -0.07611 | 0.939328 | NA |
| WNT5B | 5.313012 | -0.19573 | 0.934487 | -0.20945 | 0.834096 | NA |
| MAGOHB | 166.3203 | 0.470492 | 0.25641 | 1.834918 | 0.066518 | 0.168462 |
| TRPV4 | 38.41131 | 0.386029 | 0.406126 | 0.950517 | 0.34185 | NA |
| ITFG2 | 598.706 | -0.24806 | 0.155416 | -1.59611 | 0.110464 | 0.238361 |
| FOXM1 | 2479.11 | -0.64997 | 0.304798 | -2.13245 | 0.03297 | 0.105731 |
| PRR4 | 38.98555 | 0.562048 | 0.399389 | 1.40727 | 0.159347 | NA |
| PRMT8 | 0.47961 | 0.535796 | 3.39605 | 0.15777 | 0.874638 | NA |
| PARP11 | 154.688 | 0.097135 | 0.329103 | 0.295152 | 0.767878 | 0.855885 |
| ARPC3 | 3472.963 | 0.503159 | 0.149321 | 3.369639 | 7.53E-04 | 0.008782 |
| GPN3 | 469.3864 | 0.55621 | 0.17626 | 3.155623 | 0.001602 | 0.014125 |
| VPS29 | 1850.586 | 0.503482 | 0.154073 | 3.267822 | 0.001084 | 0.010916 |
| RAD51AP1 | 154.191 | 0.920548 | 0.22979 | 4.006046 | 6.17E-05 | 0.001867 |
| CUX2 | 0.096202 | 0.408546 | 4.079639 | 0.100143 | 0.920231 | NA |
| SH2B3 | 1828.671 | -0.21687 | 0.191513 | -1.1324 | 0.257466 | 0.421466 |
| AKAP3 | 48.83099 | 0.080051 | 0.335509 | 0.238594 | 0.81142 | 0.887561 |
| MANSC1 | 121.539 | -0.03451 | 0.266481 | -0.1295 | 0.896965 | 0.939771 |
| KCNA1 | 0.392857 | -1.546 | 4.075469 | -0.37934 | 0.704433 | NA |
| DUSP16 | 936.7709 | -0.14931 | 0.210251 | -0.71014 | 0.477615 | 0.632358 |
| CREBL2 | 3644.465 | -0.01692 | 0.180739 | -0.0936 | 0.925427 | 0.957492 |
| ACAD10 | 666.1145 | -0.32932 | 0.167289 | -1.96859 | 0.049 | 0.136932 |
| ALDH2 | 340.9054 | -0.25469 | 0.226643 | -1.12374 | 0.261123 | 0.425573 |
| CDKN1B | 1658.959 | 0.099577 | 0.187211 | 0.531895 | 0.594798 | 0.725073 |
| GPRC5D | 0.354779 | 1.703954 | 3.657232 | 0.465914 | 0.641277 | NA |
| NAA25 | 560.8217 | 0.457039 | 0.185822 | 2.459559 | 0.013911 | 0.058918 |
| GSG1 | 0.640755 | -1.64826 | 2.754778 | -0.59833 | 0.549621 | NA |
| SCNN1A | 0.163957 | 0.424913 | 4.079006 | 0.104171 | 0.917034 | NA |
| LTBR | 5472.731 | -0.14946 | 0.14773 | -1.0117 | 0.31168 | 0.478933 |
| OGFOD2 | 12.78002 | 0.140989 | 0.55723 | 0.253017 | 0.800255 | NA |
| CDK2AP1 | 4617.949 | 0.189499 | 0.218279 | 0.86815 | 0.385312 | 0.55121 |
| OAS3 | 3931.976 | -1.46735 | 0.267045 | -5.49477 | 3.91E-08 | 1.85E-05 |
| OAS2 | 3347.99 | -1.42022 | 0.291712 | -4.86856 | 1.12E-06 | 1.75E-04 |
| ART4 | 0.506476 | 0.726418 | 2.864186 | 0.253621 | 0.799788 | NA |

| | | | | | | |
|---|---|---|---|---|---|---|
| MGP | 66.64449 | 3.063377 | 1.317757 | 2.32469 | 0.020089 | 0.075965 |
| ARHGDIB | 19.65994 | -0.56751 | 0.45314 | -1.25239 | 0.210428 | NA |
| GTF2H3 | 635.6028 | 0.271558 | 0.22747 | 1.193821 | 0.232548 | 0.394795 |
| EIF2B1 | 1556.349 | 0.02356 | 0.142516 | 0.165313 | 0.868698 | 0.922011 |
| DDX55 | 591.0547 | 0.766741 | 0.171255 | 4.477187 | 7.56E-06 | 5.34E-04 |
| SLC38A1 | 0.324752 | 1.53082 | 4.022025 | 0.380609 | 0.703493 | NA |
| ENDOU | 0.230334 | -0.53318 | 4.076749 | -0.13079 | 0.895945 | NA |
| C12orf49 | 2200.853 | -0.24142 | 0.132248 | -1.82553 | 0.06792 | 0.17081 |
| VDR | 295.9677 | -0.1352 | 0.231403 | -0.58427 | 0.55904 | 0.697891 |
| FZD10 | 3.324243 | -0.48167 | 1.361003 | -0.35391 | 0.723409 | NA |
| RFC5 | 451.2476 | -0.04346 | 0.195028 | -0.22287 | 0.82364 | 0.893657 |
| STX2 | 877.954 | 0.446487 | 0.133087 | 3.354847 | 7.94E-04 | 0.008997 |
| GPR133 | 5.385308 | -0.20423 | 0.936315 | -0.21813 | 0.827331 | NA |
| COPZ1 | 4045.44 | -0.27774 | 0.134355 | -2.06718 | 0.038717 | 0.1176 |
| TBC1D30 | 8.292965 | 0.326313 | 0.754606 | 0.432429 | 0.66543 | NA |
| CAND1 | 3735.698 | 0.375561 | 0.157638 | 2.38243 | 0.017199 | 0.068328 |
| IL26 | 0.163957 | 0.424913 | 4.079006 | 0.104171 | 0.917034 | NA |
| IFNG | 0.163957 | 0.424913 | 4.079006 | 0.104171 | 0.917034 | NA |
| RAB5B | 1996.113 | -0.30826 | 0.126611 | -2.43472 | 0.014903 | 0.061732 |
| MDM1 | 124.9859 | 0.726619 | 0.273577 | 2.65599 | 0.007908 | 0.040371 |
| NUP107 | 563.9707 | 0.547416 | 0.229439 | 2.385889 | 0.017038 | 0.067932 |
| CNOT2 | 1122.474 | 0.393181 | 0.160771 | 2.445606 | 0.014461 | 0.060524 |
| TIMELESS | 1226.272 | -0.34765 | 0.222715 | -1.56095 | 0.118536 | 0.249961 |
| CPSF6 | 1555.734 | 0.13582 | 0.12885 | 1.054099 | 0.291838 | 0.458668 |
| KRR1 | 286.6831 | 1.167391 | 0.332023 | 3.515991 | 4.38E-04 | 0.006274 |
| MRPL51 | 2531.92 | 0.044982 | 0.14461 | 0.311054 | 0.75576 | 0.847421 |
| GAPDH | 55018.16 | -0.23777 | 0.135601 | -1.75343 | 0.079529 | 0.190609 |
| NOP2 | 1746.512 | -0.15667 | 0.216978 | -0.72208 | 0.470248 | 0.626592 |
| CHD4 | 8497.996 | -0.24468 | 0.119197 | -2.05278 | 0.040093 | 0.120637 |
| ACRBP | 1.404614 | -1.20865 | 1.657173 | -0.72935 | 0.465789 | NA |
| UHRF1BP1I | 428.9719 | 0.902517 | 0.201058 | 4.488836 | 7.16E-06 | 5.25E-04 |
| COPS7A | 3863.44 | -0.20555 | 0.148436 | -1.38474 | 0.166132 | 0.315242 |
| ING4 | 727.3382 | -0.03648 | 0.166743 | -0.21876 | 0.826836 | 0.895634 |
| GNB3 | 9.841368 | -0.04706 | 0.667342 | -0.07052 | 0.943778 | NA |
| CDCA3 | 114.0269 | -0.45626 | 0.301661 | -1.5125 | 0.130407 | 0.267007 |
| CHPT1 | 2973.825 | 0.021332 | 0.176827 | 0.120637 | 0.903978 | 0.944033 |
| USP5 | 4048.961 | -0.46621 | 0.124073 | -3.75755 | 1.72E-04 | 0.003525 |
| TPI1 | 14285.65 | -0.11634 | 0.128495 | -0.90543 | 0.365238 | 0.532491 |
| GNPTAB | 5735.272 | 0.293909 | 0.180398 | 1.629222 | 0.103266 | 0.227762 |
| SPSB2 | 353.646 | 0.014814 | 0.157924 | 0.093802 | 0.925266 | 0.957409 |
| ENO2 | 3223.971 | -0.12281 | 0.211454 | -0.58078 | 0.561392 | 0.699601 |
| ATN1 | 3690.911 | -0.54253 | 0.176883 | -3.06718 | 0.002161 | 0.017033 |
| C12orf57 | 2144.137 | 0.023607 | 0.144411 | 0.163472 | 0.870147 | 0.923095 |
| PTPN6 | 2.52498 | -0.94193 | 1.325075 | -0.71085 | 0.477178 | NA |
| LPCAT3 | 954.265 | -0.29376 | 0.212101 | -1.385 | 0.166051 | 0.315242 |
| NT5DC3 | 693.1947 | -0.54961 | 0.239213 | -2.29756 | 0.021587 | 0.079801 |
| SUDS3 | 838.9296 | 0.363556 | 0.169382 | 2.146366 | 0.031844 | 0.103399 |

| | | | | | | |
|---|---|---|---|---|---|---|
| GOLT1B | 1036.993 | 0.550824 | 0.169277 | 3.253988 | 0.001138 | 0.011214 |
| GYS2 | 0.971297 | 3.154786 | 2.249666 | 1.402335 | 0.160815 | NA |
| LDHB | 18503.33 | 0.233339 | 0.153208 | 1.523022 | 0.127753 | 0.263761 |
| PRKAB1 | 484.2883 | -0.35464 | 0.163397 | -2.17045 | 0.029973 | 0.098922 |
| CMAS | 977.5543 | 0.396194 | 0.143693 | 2.757231 | 0.005829 | 0.032752 |
| HCFC2 | 366.7014 | 0.629934 | 0.202227 | 3.114991 | 0.00184 | 0.015261 |
| ST8SIA1 | 479.6627 | 0.35929 | 0.225725 | 1.591714 | 0.111449 | 0.239657 |
| CLEC4A | 1.518479 | 3.806204 | 1.96339 | 1.938588 | 0.052551 | NA |
| C2CD5 | 493.7433 | 0.671555 | 0.229016 | 2.932354 | 0.003364 | 0.022712 |
| AICDA | 0.151697 | -1.546 | 4.075469 | -0.37934 | 0.704433 | NA |
| RAB35 | 1902.175 | -0.22176 | 0.142503 | -1.55616 | 0.119669 | 0.251583 |
| PHC1 | 296.4557 | -0.33451 | 0.22724 | -1.47208 | 0.141001 | 0.281502 |
| COX6A1 | 1082.366 | 0.014761 | 0.147787 | 0.09988 | 0.92044 | 0.953884 |
| RFX4 | 0.820549 | 0.502564 | 2.433834 | 0.206491 | 0.836408 | NA |
| RIC8B | 392.374 | 0.203891 | 0.169382 | 1.203735 | 0.228692 | 0.390429 |
| SRSF9 | 3568.686 | -0.12666 | 0.118419 | -1.06958 | 0.284808 | 0.45116 |
| | 53.7037 | 0.172018 | 0.302513 | 0.56863 | 0.569607 | 0.705813 |
| FGFR1OP2 | 821.6326 | 0.777676 | 0.185225 | 4.198536 | 2.69E-05 | 0.001098 |
| KLRB1 | 2.718671 | 0.259743 | 1.235096 | 0.210302 | 0.833432 | NA |
| FBRSL1 | 1188.909 | -0.46863 | 0.160166 | -2.92589 | 0.003435 | 0.022986 |
| UBE3A | 1405.691 | 0.493479 | 0.227978 | 2.16459 | 0.030419 | 0.099989 |
| MESDC2 | 2027.132 | -0.0265 | 0.122612 | -0.21614 | 0.828875 | 0.896738 |
| RCN2 | 1970.643 | 0.710143 | 0.166751 | 4.258704 | 2.06E-05 | 9.41E-04 |
| CHRNB4 | 7.792552 | -0.22113 | 0.733556 | -0.30145 | 0.763072 | NA |
| KMT2A | 1101.214 | -0.26006 | 0.187069 | -1.39019 | 0.164472 | 0.313274 |
| TREH | 0.424115 | 1.932531 | 3.45462 | 0.559405 | 0.575885 | NA |
| IFT46 | 678.4693 | -0.05124 | 0.161808 | -0.31667 | 0.751491 | 0.844608 |
| MMP8 | 1142.962 | 0.402039 | 0.188014 | 2.138352 | 0.032488 | 0.104859 |
| RPS25 | 12779.69 | 0.524303 | 0.13585 | 3.859432 | 1.14E-04 | 0.002777 |
| CASC1 | 2.394146 | 1.418732 | 1.411215 | 1.005326 | 0.31474 | NA |
| SPCS2 | 1119.406 | -0.1244 | 0.189185 | -0.65757 | 0.510816 | 0.659019 |
| USP35 | 721.1787 | -0.44568 | 0.156945 | -2.8397 | 0.004516 | 0.027525 |
| SLC16A7 | 122.2521 | 0.330877 | 0.309913 | 1.067644 | 0.285681 | 0.452291 |
| TMEM5 | 449.3651 | 0.131314 | 0.201068 | 0.653082 | 0.513704 | 0.660803 |
| KLF12 | 412.8669 | 0.317385 | 0.216841 | 1.463681 | 0.143281 | 0.284682 |
| UCHL3 | 72.77435 | 0.271219 | 0.263428 | 1.029577 | 0.303208 | 0.47046 |
| PCDH17 | 1.920675 | 2.241316 | 1.635771 | 1.370189 | 0.170628 | NA |
| CCND2 | 32.13658 | 0.367371 | 0.559254 | 0.656895 | 0.511248 | NA |
| | 6.706811 | -0.09877 | 0.782859 | -0.12616 | 0.899604 | NA |
| CCDC92 | 1454.005 | -0.16278 | 0.156743 | -1.03851 | 0.299031 | 0.465897 |
| YLPM1 | 1707.582 | -0.45389 | 0.238001 | -1.90708 | 0.056511 | 0.150513 |
| DCAF4 | 727.2682 | -0.10962 | 0.174951 | -0.62657 | 0.530941 | 0.675555 |
| PROX2 | 5.968126 | 0.339973 | 0.818242 | 0.415492 | 0.677782 | NA |
| VSX2 | 0.096202 | 0.408546 | 4.079639 | 0.100143 | 0.920231 | NA |
| FCF1 | 715.3089 | 0.007538 | 0.178344 | 0.042267 | 0.966286 | 0.981451 |
| PGF | 159.8991 | 0.138289 | 0.357951 | 0.386335 | 0.699248 | 0.806443 |
| IFI27L2 | 2814.873 | -0.05518 | 0.137914 | -0.40014 | 0.689056 | 0.797886 |

| | | | | | | |
|---|---|---|---|---|---|---|
| CCDC176 | 18.28887 | 1.268522 | 0.53012 | 2.392895 | 0.016716 | NA |
| NEK9 | 3441.565 | -0.35452 | 0.127747 | -2.77519 | 0.005517 | 0.031724 |
| ACYP1 | 139.8056 | 0.094238 | 0.217277 | 0.433723 | 0.66449 | 0.779567 |
| IFT43 | 377.6751 | -0.14346 | 0.159834 | -0.89758 | 0.369409 | 0.536481 |
| NPC2 | 9035.039 | -0.24411 | 0.12604 | -1.93674 | 0.052778 | 0.143471 |
| DNAL1 | 258.775 | 0.161603 | 0.186657 | 0.865776 | 0.386613 | 0.552737 |
| IRF2BPL | 2350.892 | -0.28322 | 0.149467 | -1.89488 | 0.058108 | 0.153614 |
| ACOT2 | 1121.327 | -0.3308 | 0.15038 | -2.1998 | 0.027821 | 0.094368 |
| LTBP2 | 38.11508 | -0.16167 | 0.362321 | -0.44619 | 0.655458 | NA |
| AREL1 | 2357.521 | -0.27412 | 0.133837 | -2.04818 | 0.040543 | 0.121382 |
| MLH3 | 434.0897 | 0.582335 | 0.210094 | 2.771787 | 0.005575 | 0.031939 |
| TTLL5 | 836.8987 | 0.004926 | 0.174484 | 0.028233 | 0.977476 | 0.987689 |
| FLVCR2 | 39.90603 | -0.03235 | 0.351784 | -0.09196 | 0.926731 | NA |
| ABCD4 | 898.0905 | -0.11255 | 0.137187 | -0.82044 | 0.411966 | 0.575044 |
| DLST | 3108.114 | -0.27941 | 0.118108 | -2.36568 | 0.017997 | 0.070292 |
| PPP4R4 | 1.103647 | 3.314505 | 2.124646 | 1.560027 | 0.118754 | NA |
| TGFB3 | 21.28501 | -0.01333 | 0.491253 | -0.02714 | 0.978347 | NA |
| ZC2HC1C | 42.7938 | 0.223532 | 0.410277 | 0.544833 | 0.585869 | NA |
| SLIRP | 1186.704 | 0.43952 | 0.129439 | 3.395571 | 6.85E-04 | 0.008201 |
| RBM25 | 969.3411 | 0.878908 | 0.607824 | 1.445991 | 0.14818 | 0.291322 |
| ALDH6A1 | 626.8966 | -0.38095 | 0.218374 | -1.74447 | 0.081078 | 0.193008 |
| GPR68 | 1.032125 | -1.35749 | 2.070496 | -0.65563 | 0.51206 | NA |
| ESRRB | 0.957485 | -0.51149 | 2.371634 | -0.21567 | 0.829247 | NA |
| EIF2B2 | 1220.079 | -0.13185 | 0.186685 | -0.70628 | 0.480012 | 0.634563 |
| NRDE2 | 470.0873 | -0.02274 | 0.156146 | -0.14563 | 0.884214 | 0.93146 |
| COQ6 | 232.482 | -0.08439 | 0.2003 | -0.4213 | 0.673532 | 0.786323 |
| ZNF410 | 9.910471 | 0.921759 | 0.682916 | 1.34974 | 0.177099 | NA |
| SNX19 | 2303.959 | -0.36184 | 0.122653 | -2.95012 | 0.003176 | 0.022009 |
| KCNJ5 | 0.653015 | 0.2552 | 2.787103 | 0.091565 | 0.927044 | NA |
| MSANTD2 | 112.5168 | 0.497474 | 0.221121 | 2.249778 | 0.024463 | 0.08666 |
| TP53AIP1 | 0.163957 | 0.424913 | 4.079006 | 0.104171 | 0.917034 | NA |
| IQSEC3 | 0.774888 | -2.09879 | 2.346327 | -0.8945 | 0.371054 | NA |
| CCDC77 | 230.0313 | 0.72113 | 0.188682 | 3.821923 | 1.32E-04 | 0.003029 |
| ENOX1 | 22.32377 | 1.412801 | 0.551441 | 2.562017 | 0.010407 | NA |
| MTRF1 | 143.5894 | 0.917867 | 0.245897 | 3.732724 | 1.89E-04 | 0.003755 |
| SPG20OS | 18.50189 | 0.543062 | 0.615745 | 0.88196 | 0.377798 | NA |
| SOHLH2 | 14.31798 | 0.346377 | 0.632021 | 0.548046 | 0.58366 | NA |
| DNAJC15 | 1664.683 | 0.366331 | 0.147344 | 2.486221 | 0.012911 | 0.056209 |
| PROSER1 | 975.3528 | -0.28795 | 0.187361 | -1.53687 | 0.124326 | 0.258985 |
| UFM1 | 1021.554 | 0.940483 | 0.254914 | 3.689416 | 2.25E-04 | 0.004177 |
| WBP4 | 254.5923 | 0.9292 | 0.267618 | 3.472118 | 5.16E-04 | 0.006917 |
| ELF1 | 2014.56 | 0.331603 | 0.120438 | 2.753298 | 0.0059 | 0.033023 |
| SMAD9 | 26.25068 | -0.32121 | 0.488699 | -0.65727 | 0.511006 | NA |
| HSPH1 | 3366.76 | 0.720227 | 0.192585 | 3.739788 | 1.84E-04 | 0.003682 |
| KBTBD7 | 252.8609 | -0.06175 | 0.213614 | -0.28909 | 0.772511 | 0.85903 |
| ALG5 | 1055.507 | 0.225453 | 0.166204 | 1.356477 | 0.174947 | 0.326872 |
| EXOSC8 | 438.9351 | 0.440774 | 0.19122 | 2.305058 | 0.021163 | 0.07868 |

| | | | | | | |
|---|---|---|---|---|---|---|
| NR2C1 | 341.0419 | 0.407711 | 0.221357 | 1.841872 | 0.065494 | 0.166907 |
| UTP20 | 874.4906 | 0.108422 | 0.202757 | 0.534738 | 0.592831 | 0.723441 |
| TMPO | 1116.625 | 0.051624 | 0.170342 | 0.303063 | 0.761842 | 0.851908 |
| ARL1 | 1756.156 | 0.570571 | 0.175845 | 3.244739 | 0.001176 | 0.011508 |
| GLT8D2 | 5.754573 | 0.742224 | 0.882232 | 0.841303 | 0.400178 | NA |
| MTERFD3 | 154.0211 | 0.563738 | 0.211267 | 2.668365 | 0.007622 | 0.039407 |
| SOCS2 | 1137.735 | 0.662661 | 0.268453 | 2.468441 | 0.01357 | 0.057932 |
| NFYB | 312.0856 | 0.709501 | 0.179286 | 3.957363 | 7.58E-05 | 0.002147 |
| CCDC53 | 730.568 | 0.279356 | 0.155909 | 1.791789 | 0.073167 | 0.180256 |
| APAF1 | 296.0914 | 0.484217 | 0.244444 | 1.980891 | 0.047604 | 0.13441 |
| PLBD1 | 10.1395 | 0.368819 | 0.700858 | 0.52624 | 0.598722 | NA |
| TAS2R10 | 1.094548 | 2.111186 | 2.036733 | 1.036555 | 0.299943 | NA |
| PRB2 | 1.105576 | 1.732871 | 2.324315 | 0.745541 | 0.455945 | NA |
| PYROXD1 | 83.2296 | 1.037447 | 0.395294 | 2.624493 | 0.008678 | 0.042936 |
| BCL2L14 | 0.162376 | 0.424913 | 4.079006 | 0.104171 | 0.917034 | NA |
| PSPC1 | 959.5836 | 0.254879 | 0.146404 | 1.740932 | 0.081695 | 0.194127 |
| ZMYM2 | 1551.915 | 0.530296 | 0.198809 | 2.667359 | 0.007645 | 0.039439 |
| GJB6 | 0.640755 | -1.64826 | 2.754778 | -0.59833 | 0.549621 | NA |
| GJA3 | 658.1006 | 0.06697 | 0.158612 | 0.422225 | 0.672861 | 0.785888 |
| TBC1D15 | 1053.816 | 0.779231 | 0.206492 | 3.773662 | 1.61E-04 | 0.003424 |
| FLT3 | 0.096202 | 0.408546 | 4.079639 | 0.100143 | 0.920231 | NA |
| RPL21 | 5055.703 | 0.386242 | 0.152662 | 2.53004 | 0.011405 | 0.051429 |
| MTIF3 | 519.3202 | 0.709388 | 0.194738 | 3.64277 | 2.70E-04 | 0.004608 |
| GTF3A | 2862.146 | 0.033368 | 0.160835 | 0.207464 | 0.835647 | 0.901643 |
| RASL11A | 12.54502 | 0.07366 | 0.637751 | 0.1155 | 0.908049 | NA |
| UBL3 | 6924.32 | 0.271558 | 0.187466 | 1.448573 | 0.147457 | 0.290626 |
| RBM19 | 1118.662 | -0.0665 | 0.143152 | -0.46457 | 0.642238 | 0.761906 |
| CIT | 2503.75 | -0.6791 | 0.199031 | -3.41202 | 6.45E-04 | 0.007889 |
| IFT81 | 889.14 | 0.996293 | 0.318701 | 3.126101 | 0.001771 | 0.014979 |
| ACADS | 1184.458 | -0.35609 | 0.139386 | -2.5547 | 0.010628 | 0.048986 |
| HVCN1 | 20.79363 | 0.25112 | 0.528757 | 0.474924 | 0.634841 | NA |
| NME2P1 | 37.95209 | 0.348616 | 0.403719 | 0.863511 | 0.387857 | NA |
| DDX54 | 2064.256 | -0.32424 | 0.132379 | -2.44931 | 0.014313 | 0.060106 |
| MED13L | 825.1915 | 0.188606 | 0.208048 | 0.906549 | 0.364646 | 0.531934 |
| RASSF8 | 1147.131 | 1.139413 | 0.2807 | 4.059181 | 4.92E-05 | 0.001622 |
| BHLHE41 | 2004.916 | -1.06788 | 0.291406 | -3.66457 | 2.48E-04 | 0.004422 |
| SSPN | 38.34516 | -0.07295 | 0.416962 | -0.17496 | 0.861107 | NA |
| ITPR2 | 488.4067 | 0.567682 | 0.20502 | 2.768908 | 0.005624 | 0.032088 |
| CCDC91 | 288.442 | 0.972983 | 0.365309 | 2.66345 | 0.007734 | 0.039779 |
| CCDC70 | 0.832809 | 1.636407 | 2.200093 | 0.74379 | 0.457004 | NA |
| SPRYD7 | 466.5101 | 0.248656 | 0.163462 | 1.52118 | 0.128215 | 0.264483 |
| EBPL | 480.9179 | 0.11446 | 0.155628 | 0.735469 | 0.462054 | 0.620275 |
| ATP7B | 931.5927 | -0.41964 | 0.182739 | -2.2964 | 0.021653 | 0.079992 |
| ZC3H13 | 2461.294 | 0.894492 | 0.357867 | 2.499507 | 0.012437 | 0.054606 |
| ATF1 | 294.6867 | 0.668704 | 0.20114 | 3.324561 | 8.86E-04 | 0.009614 |
| TSFM | 622.8788 | 0.009992 | 0.141538 | 0.070596 | 0.94372 | 0.968771 |
| ARHGAP9 | 6.890512 | -0.16069 | 0.83944 | -0.19142 | 0.848194 | NA |

| | | | | | | |
|---|---|---|---|---|---|---|
| NCKAP1L | 0.651085 | 2.534961 | 3.013697 | 0.841147 | 0.400266 | NA |
| MMP19 | 38.87154 | 0.209147 | 0.507301 | 0.412274 | 0.680138 | NA |
| PFDN5 | 6574.106 | 0.315598 | 0.153789 | 2.052149 | 0.040155 | 0.12073 |
| SPATS2 | 709.0126 | -0.07282 | 0.165635 | -0.43965 | 0.660194 | 0.776182 |
| ORMDL2 | 477.3711 | -0.13533 | 0.177169 | -0.76382 | 0.444973 | 0.60481 |
| NR4A1 | 1818.956 | 0.447281 | 0.284905 | 1.56993 | 0.116431 | 0.246701 |
| PDE1B | 27.54376 | 0.712553 | 0.651948 | 1.09296 | 0.274411 | NA |
| HOXC13 | 22.50726 | 0.70973 | 0.480438 | 1.477256 | 0.139607 | NA |
| CDK2 | 9552.682 | -0.3898 | 0.1277 | -3.05248 | 0.00227 | 0.017517 |
| LRP1 | 10307.94 | -0.65155 | 0.209769 | -3.10605 | 0.001896 | 0.015566 |
| HOXC11 | 117.1542 | -0.21623 | 0.230456 | -0.93826 | 0.348111 | 0.51605 |
| C12orf44 | 893.4062 | -0.11543 | 0.200313 | -0.57625 | 0.564443 | 0.702222 |
| NFE2 | 0.641986 | 1.117767 | 2.709494 | 0.412537 | 0.679946 | NA |
| HOXC12 | 0.392711 | 0.292538 | 3.145221 | 0.09301 | 0.925895 | NA |
| IKZF4 | 230.2225 | -0.41827 | 0.20264 | -2.0641 | 0.039008 | 0.118181 |
| SMUG1 | 964.3079 | -0.20255 | 0.139428 | -1.45269 | 0.146309 | 0.28883 |
| TUBA1B | 4670.102 | -0.44157 | 0.190325 | -2.32006 | 0.020337 | 0.076589 |
| METTL21B | 173.1587 | -0.19365 | 0.200964 | -0.96359 | 0.33525 | 0.503981 |
| RAB38 | 9193.989 | 0.005703 | 0.114279 | 0.049907 | 0.960197 | 0.9781 |
| GPR83 | 1.039255 | 2.120498 | 2.300766 | 0.921649 | 0.356712 | NA |
| WNT1 | 0.163957 | 0.424913 | 4.079006 | 0.104171 | 0.917034 | NA |
| PIWIL1 | 0.314073 | -0.57308 | 3.898934 | -0.14698 | 0.883146 | NA |
| GPR18 | 0.998549 | 1.029429 | 2.066501 | 0.498151 | 0.618378 | NA |
| CLYBL | 237.6909 | -0.05903 | 0.201861 | -0.29242 | 0.769966 | 0.857729 |
| TMTC4 | 354.9948 | 0.228937 | 0.188779 | 1.212729 | 0.225234 | 0.386306 |
| RAP2A | 2008.684 | 0.705495 | 0.206335 | 3.419174 | 6.28E-04 | 0.007783 |
| ABCC4 | 921.5012 | 0.185194 | 0.155597 | 1.190216 | 0.233961 | 0.396514 |
| EFNB2 | 48.99859 | -0.60516 | 0.347118 | -1.74338 | 0.081267 | 0.193289 |
| TM9SF2 | 6119.743 | 0.071601 | 0.121429 | 0.589654 | 0.555423 | 0.695305 |
| ATP5S | 156.9884 | 0.43379 | 0.213637 | 2.030501 | 0.042306 | 0.125119 |
| BMP4 | 188.0324 | 0.354488 | 0.371994 | 0.95294 | 0.340621 | 0.5087 |
| MAX | 2289.537 | -0.29111 | 0.130416 | -2.2322 | 0.025602 | 0.089457 |
| KLC1 | 973.9864 | 0.025152 | 0.137141 | 0.183404 | 0.854481 | 0.913506 |
| XRCC3 | 535.1114 | -0.31413 | 0.194287 | -1.61686 | 0.105909 | 0.23178 |
| TUBGCP3 | 595.188 | 0.320396 | 0.158607 | 2.020071 | 0.043376 | 0.126906 |
| MCF2L | 4.777174 | 0.063604 | 0.909166 | 0.069959 | 0.944227 | NA |
| F10 | 3.833521 | -0.18634 | 1.003402 | -0.18571 | 0.852672 | NA |
| PCID2 | 1113.339 | 0.166898 | 0.177803 | 0.938668 | 0.347901 | 0.515998 |
| PROZ | 1.284021 | -0.2497 | 2.112256 | -0.11821 | 0.905899 | NA |
| FRMD8 | 2090.898 | -0.18737 | 0.214417 | -0.87385 | 0.382199 | 0.548277 |
| PRDX5 | 6947.159 | -0.11149 | 0.144825 | -0.76984 | 0.441395 | 0.601534 |
| FLRT1 | 107.365 | 0.042866 | 0.433809 | 0.098813 | 0.921287 | 0.954374 |
| ZNF384 | 891.8545 | -0.20067 | 0.171494 | -1.17013 | 0.24195 | 0.404958 |
| EMG1 | 160.3098 | 0.036755 | 0.226941 | 0.16196 | 0.871338 | 0.923353 |
| PCNXL4 | 1959.815 | 0.707926 | 0.202839 | 3.490082 | 4.83E-04 | 0.006662 |
| ATG14 | 739.3228 | 0.410551 | 0.162803 | 2.521766 | 0.011677 | 0.0523 |
| KTN1 | 1827.958 | 1.12286 | 1.028703 | 1.091531 | 0.275039 | 0.440659 |

| | | | | | | |
|---|---|---|---|---|---|---|
| SIX1 | 2.092829 | 0.67355 | 1.571041 | 0.428729 | 0.668121 | NA |
| RHOJ | 1706.825 | 0.109034 | 0.172434 | 0.632325 | 0.527175 | 0.672657 |
| DLGAP5 | 343.8484 | 0.893819 | 0.204445 | 4.371921 | 1.23E-05 | 7.02E-04 |
| L3HYPDH | 175.5684 | 0.255684 | 0.23604 | 1.083223 | 0.278709 | 0.444877 |
| HSPA2 | 4175.329 | 0.137813 | 0.195357 | 0.705439 | 0.480537 | 0.63495 |
| ZBTB1 | 647.88 | 0.897593 | 0.230791 | 3.889194 | 1.01E-04 | 0.002559 |
| TRMT5 | 423.1803 | 0.448472 | 0.203982 | 2.198586 | 0.027907 | 0.094605 |
| SGPP1 | 563.8257 | 0.167701 | 0.192211 | 0.872484 | 0.382944 | 0.548882 |
| PLEKHG3 | 2079.635 | -0.68204 | 0.166033 | -4.10789 | 3.99E-05 | 0.001466 |
| PZP | 31.00667 | 2.301233 | 0.488131 | 4.714372 | 2.42E-06 | NA |
| HELB | 43.52338 | 0.56424 | 0.327033 | 1.72533 | 0.084468 | NA |
| RAP1B | 1613.019 | 0.2731 | 0.192941 | 1.415459 | 0.156934 | 0.303153 |
| TSPAN8 | 0.162376 | 0.424913 | 4.079006 | 0.104171 | 0.917034 | NA |
| BEST3 | 0.260158 | 1.240347 | 4.047124 | 0.306476 | 0.759242 | NA |
| RAB3IP | 40.07007 | 0.750729 | 0.460364 | 1.630729 | 0.102947 | NA |
| PTPRB | 2.483235 | 0.42635 | 1.317155 | 0.32369 | 0.746173 | NA |
| DYRK2 | 2067.21 | 0.881959 | 0.207155 | 4.257491 | 2.07E-05 | 9.43E-04 |
| YEATS4 | 313.4778 | 0.828953 | 0.198319 | 4.179901 | 2.92E-05 | 0.00117 |
| METTL25 | 98.91054 | 0.472142 | 0.26217 | 1.800899 | 0.071719 | 0.177575 |
| TNFRSF19 | 1070.7 | -0.13369 | 0.160319 | -0.83388 | 0.40435 | 0.569031 |
| RNF6 | 865.5691 | 1.002951 | 0.252363 | 3.97424 | 7.06E-05 | 0.002041 |
| EMC4 | 2694.519 | 0.294242 | 0.130613 | 2.252786 | 0.024273 | 0.086227 |
| HERC2 | 14294.43 | -0.26547 | 0.15842 | -1.6757 | 0.093797 | 0.213921 |
| SNRPN | 22.93679 | -0.10109 | 0.450248 | -0.22451 | 0.822359 | NA |
| EIF2AK4 | 3535.274 | 0.343165 | 0.139861 | 2.453618 | 0.014143 | 0.059645 |
| MYO5C | 14.46411 | 0.348432 | 0.566691 | 0.614853 | 0.538652 | NA |
| CGNL1 | 0.763859 | -0.6945 | 2.307767 | -0.30094 | 0.76346 | NA |
| TMOD2 | 1381.963 | 0.306635 | 0.229593 | 1.335563 | 0.181692 | 0.334622 |
| TTBK2 | 367.4413 | 0.28216 | 0.206583 | 1.365839 | 0.171989 | 0.323179 |
| ELL3 | 6.529656 | -0.11223 | 0.972903 | -0.11536 | 0.908162 | NA |
| C15orf57 | 290.4182 | 0.070078 | 0.178373 | 0.392872 | 0.694414 | 0.802993 |
| INO80 | 1100.44 | 0.185521 | 0.152203 | 1.218905 | 0.22288 | 0.383712 |
| NARG2 | 501.5732 | 0.396188 | 0.222243 | 1.782679 | 0.074639 | 0.182427 |
| DLL4 | 0.096202 | 0.408546 | 4.079639 | 0.100143 | 0.920231 | NA |
| ALDH1A2 | 6.667354 | 0.20773 | 0.780429 | 0.266174 | 0.790106 | NA |
| FAM63B | 568.5663 | 0.647049 | 0.372156 | 1.738651 | 0.082096 | 0.194689 |
| IVD | 2569.461 | -0.243 | 0.1443 | -1.68401 | 0.092181 | 0.21121 |
| KNSTRN | 722.2062 | 0.140871 | 0.208451 | 0.6758 | 0.499168 | 0.649647 |
| DUT | 1580.397 | 0.075765 | 0.144322 | 0.52497 | 0.599604 | 0.729132 |
| CHAC1 | 46.63209 | 0.067794 | 0.314168 | 0.215788 | 0.829153 | NA |
| CLN6 | 1951.572 | -0.44314 | 0.149103 | -2.97202 | 0.002959 | 0.020939 |
| ARPP19 | 4873.827 | 0.30298 | 0.157915 | 1.918622 | 0.055032 | 0.147937 |
| VPS13C | 1142.014 | 0.834471 | 0.327373 | 2.548992 | 0.010803 | 0.04945 |
| CALML4 | 81.43483 | 0.16074 | 0.271199 | 0.5927 | 0.553382 | 0.693776 |
| ISLR | 0.096202 | 0.408546 | 4.079639 | 0.100143 | 0.920231 | NA |
| THAP10 | 60.90156 | 0.36646 | 0.283784 | 1.291333 | 0.196588 | 0.352003 |
| LOXL1 | 22.98339 | -0.94756 | 0.441311 | -2.14716 | 0.031781 | NA |

| | | | | | | |
|---|---|---|---|---|---|---|
| CCNT1 | 864.7781 | 0.16733 | 0.172551 | 0.969742 | 0.332175 | 0.500589 |
| PUS7L | 128.2858 | 1.138676 | 0.311717 | 3.652921 | 2.59E-04 | 0.004545 |
| NGDN | 531.5596 | 0.416874 | 0.169404 | 2.460831 | 0.013862 | 0.058773 |
| RIPK3 | 218.1693 | -0.09313 | 0.225357 | -0.41325 | 0.679424 | 0.790858 |
| ADCY4 | 3.604243 | 1.423059 | 1.129811 | 1.259555 | 0.20783 | NA |
| RAB2B | 893.1573 | -0.27592 | 0.143286 | -1.92568 | 0.054145 | 0.146282 |
| BCL2L2 | 1784.548 | 0.171742 | 0.165151 | 1.03991 | 0.298382 | 0.465314 |
| AJUBA | 463.9365 | 0.041701 | 0.267768 | 0.155735 | 0.876242 | 0.926081 |
| DTD2 | 398.3015 | 0.344484 | 0.16425 | 2.097307 | 0.035966 | 0.111853 |
| PARP2 | 484.8253 | 0.306561 | 0.160499 | 1.910044 | 0.056128 | 0.149964 |
| HEATR5A | 807.3293 | 0.495984 | 0.188055 | 2.637434 | 0.008354 | 0.041733 |
| FOXA1 | 0.555233 | -0.57009 | 3.296947 | -0.17291 | 0.862719 | NA |
| SNX6 | 1515.942 | 0.406697 | 0.208336 | 1.952121 | 0.050924 | 0.140406 |
| EAPP | 945.9148 | 0.412864 | 0.208769 | 1.977614 | 0.047972 | 0.135086 |
| EGLN3 | 380.6492 | -0.0864 | 0.167918 | -0.51451 | 0.606897 | 0.734751 |
| MIS18BP1 | 223.1883 | 1.291388 | 0.439022 | 2.941511 | 0.003266 | 0.022362 |
| NRL | 37.97357 | -0.58208 | 0.342441 | -1.69981 | 0.089167 | NA |
| RNASE1 | 1.375219 | 1.009999 | 1.970616 | 0.512529 | 0.608281 | NA |
| NEDD8 | 1532.549 | 0.216808 | 0.147758 | 1.467317 | 0.14229 | 0.283134 |
| DAD1 | 5412.164 | -0.01493 | 0.149566 | -0.09981 | 0.920495 | 0.953884 |
| TEP1 | 821.6361 | -0.70037 | 0.257332 | -2.72166 | 0.006495 | 0.035242 |
| GALNT8 | 0.912824 | 3.024111 | 2.331252 | 1.297204 | 0.194561 | NA |
| KCNA5 | 6.625106 | 0.038043 | 1.035456 | 0.03674 | 0.970692 | NA |
| EFCAB4B | 28.19624 | 0.386354 | 0.479165 | 0.806306 | 0.420066 | NA |
| CDC16 | 3303.049 | -0.04481 | 0.125605 | -0.35678 | 0.72126 | 0.822641 |
| CLIP1 | 1927.637 | 0.84365 | 0.567234 | 1.487306 | 0.136934 | 0.276127 |
| CCDC62 | 2.57088 | -0.27441 | 1.421289 | -0.19307 | 0.846906 | NA |
| HIP1R | 2685.249 | -0.55467 | 0.240633 | -2.30503 | 0.021165 | 0.07868 |
| C12orf65 | 399.6599 | 0.28453 | 0.17558 | 1.620518 | 0.105121 | 0.230737 |
| TRAF3 | 1969.228 | -0.15347 | 0.148165 | -1.03583 | 0.300282 | 0.467235 |
| PPFIA1 | 2079.466 | 0.685733 | 0.14092 | 4.866119 | 1.14E-06 | 1.75E-04 |
| LRRC9 | 0.518736 | 2.237423 | 2.7713 | 0.807355 | 0.419462 | NA |
| ACTR10 | 2356.993 | 0.38272 | 0.146648 | 2.609788 | 0.00906 | 0.044236 |
| ABHD12B | 5.046634 | 0.241624 | 0.88373 | 0.273414 | 0.784535 | NA |
| GCH1 | 78.56282 | 0.173842 | 0.327619 | 0.530624 | 0.595679 | 0.725848 |
| LGALS3 | 33314.91 | -0.31967 | 0.141521 | -2.25882 | 0.023895 | 0.085385 |
| RAN | 9389.734 | 0.090372 | 0.146308 | 0.617681 | 0.536786 | 0.679944 |
| IGHMBP2 | 730.4038 | -0.34818 | 0.149221 | -2.33335 | 0.01963 | 0.07478 |
| ACY3 | 0.326333 | 1.533889 | 3.826878 | 0.40082 | 0.688553 | NA |
| ALDH3B2 | 0.774742 | -0.40577 | 2.223483 | -0.18249 | 0.855195 | NA |
| MTL5 | 68.1712 | -0.61679 | 0.277128 | -2.22567 | 0.026036 | 0.090437 |
| ATP8A2 | 21.66572 | 0.063959 | 0.466115 | 0.137216 | 0.89086 | NA |
| MTUS2 | 0.192403 | 0.975392 | 4.060716 | 0.240202 | 0.810174 | NA |
| ZMYM5 | 231.1996 | 0.856729 | 0.280347 | 3.055959 | 0.002243 | 0.017406 |
| USPL1 | 269.389 | 0.654744 | 0.205656 | 3.183679 | 0.001454 | 0.013322 |
| XPO4 | 958.5078 | 0.100166 | 0.187158 | 0.535194 | 0.592516 | 0.7233 |
| TPTE2 | 0.520316 | 2.239969 | 3.173769 | 0.705776 | 0.480328 | NA |

| | | | | | | |
|---|---|---|---|---|---|---|
| POMP | 3433.699 | 0.332532 | 0.192897 | 1.723882 | 0.084729 | 0.198393 |
| CDK8 | 468.5461 | 0.226892 | 0.160448 | 1.414113 | 0.157329 | 0.30352 |
| ALOX5AP | 1.927292 | 0.450759 | 1.598627 | 0.281967 | 0.777969 | NA |
| WASF3 | 960.2482 | -0.05442 | 0.20567 | -0.26458 | 0.791329 | 0.873483 |
| DCLK1 | 696.234 | -0.79974 | 0.342875 | -2.33247 | 0.019676 | 0.074931 |
| CCNA1 | 8.886889 | -0.1472 | 0.68824 | -0.21388 | 0.830641 | NA |
| COG6 | 726.3751 | 0.974367 | 0.209261 | 4.656235 | 3.22E-06 | 3.27E-04 |
| SPG20 | 2134.516 | 0.739512 | 0.206572 | 3.579918 | 3.44E-04 | 0.00538 |
| RXFP2 | 2.118136 | -0.47216 | 1.43056 | -0.33005 | 0.741363 | NA |
| EPSTI1 | 85.1687 | -0.98305 | 0.440477 | -2.23178 | 0.025629 | 0.089521 |
| TRPC4 | 67.41424 | 0.435706 | 0.31455 | 1.385175 | 0.165999 | 0.31522 |
| POSTN | 0.46735 | -1.30193 | 3.399671 | -0.38296 | 0.701751 | NA |
| RFXAP | 44.49943 | 0.395433 | 0.371501 | 1.064421 | 0.287138 | NA |
| TPT1 | 60712.41 | 0.510092 | 0.168134 | 3.033838 | 0.002415 | 0.018279 |
| GPALPP1 | 409.5268 | 0.438383 | 0.199552 | 2.196831 | 0.028033 | 0.094788 |
| KL | 0.096202 | 0.408546 | 4.079639 | 0.100143 | 0.920231 | NA |
| RFC3 | 525.4917 | 0.288246 | 0.236459 | 1.21901 | 0.22284 | 0.3837 |
| STARD13 | 1576.516 | 0.498452 | 0.245555 | 2.029899 | 0.042367 | 0.125189 |
| MACROD1 | 834.0294 | -0.29602 | 0.146115 | -2.02596 | 0.042769 | 0.125906 |
| RTN3 | 2019.833 | -0.10923 | 0.140584 | -0.77696 | 0.437183 | 0.597569 |
| RARRES3 | 28.00872 | -0.75445 | 0.457237 | -1.65002 | 0.098938 | NA |
| HRASLS2 | 0.70046 | 1.37107 | 2.594868 | 0.528378 | 0.597237 | NA |
| NTS | 1.877157 | 0.513842 | 1.558459 | 0.329711 | 0.741618 | NA |
| BTG1 | 2028.604 | -0.26281 | 0.152592 | -1.7223 | 0.085015 | 0.198945 |
| LRRIQ1 | 8.17534 | 0.803291 | 1.04736 | 0.766968 | 0.443101 | NA |
| C12orf29 | 298.0917 | 1.003613 | 0.222765 | 4.505257 | 6.63E-06 | 4.97E-04 |
| TMTC1 | 1937.269 | 0.121604 | 0.171649 | 0.708447 | 0.478668 | 0.633256 |
| KRAS | 565.816 | 0.707634 | 0.244705 | 2.891781 | 0.003831 | 0.024734 |
| IPO8 | 1115.527 | 0.171654 | 0.185006 | 0.927829 | 0.353496 | 0.521597 |
| CCDC59 | 272.2005 | 0.792623 | 0.243195 | 3.259215 | 0.001117 | 0.01112 |
| ZFC3H1 | 350.6037 | 0.944685 | 0.616689 | 1.531867 | 0.125555 | 0.26063 |
| DPF2 | 1124.405 | -0.50353 | 0.136247 | -3.69574 | 2.19E-04 | 0.00412 |
| MEN1 | 1086.416 | -0.4183 | 0.160561 | -2.60523 | 0.009181 | 0.044545 |
| C14orf1 | 2306.197 | -0.12677 | 0.192927 | -0.65708 | 0.511128 | 0.659182 |
| C14orf159 | 88.00948 | -0.4969 | 0.286048 | -1.73713 | 0.082363 | 0.195089 |
| UNC79 | 12.51734 | 0.590779 | 0.637208 | 0.927137 | 0.353855 | NA |
| NUMB | 1405.407 | 0.137707 | 0.133757 | 1.029533 | 0.303229 | 0.47046 |
| VRTN | 1.902479 | 0.587734 | 1.628375 | 0.360933 | 0.71815 | NA |
| COX16 | 597.7516 | 0.445438 | 0.192133 | 2.318387 | 0.020428 | 0.076808 |
| TTC9 | 6.499048 | -0.15545 | 0.860001 | -0.18076 | 0.856558 | NA |
| MED6 | 353.486 | 0.366607 | 0.173907 | 2.10807 | 0.035025 | 0.10994 |
| EIF2S1 | 1903.807 | 0.391956 | 0.174385 | 2.247649 | 0.024599 | 0.086816 |
| ADAM20 | 0.584911 | 2.38756 | 2.690807 | 0.887303 | 0.374916 | NA |
| MEIS2 | 0.382031 | -1.26484 | 3.151989 | -0.40128 | 0.688212 | NA |
| DPH6 | 183.2704 | -0.072 | 0.219379 | -0.32821 | 0.742755 | 0.837887 |
| KATNBL1 | 440.2876 | 1.183388 | 0.204298 | 5.79247 | 6.94E-09 | 5.29E-06 |
| EMC7 | 3659.249 | 0.130521 | 0.150649 | 0.866391 | 0.386276 | 0.552388 |

| | | | | | | |
|---|---|---|---|---|---|---|
| TRPM1 | 4255.386 | 0.132404 | 0.177496 | 0.745952 | 0.455697 | 0.614594 |
| PPHLN1 | 1528.874 | 0.379463 | 0.157815 | 2.40448 | 0.016195 | 0.065554 |
| FKBP11 | 890.2408 | -0.01083 | 0.142078 | -0.07624 | 0.939227 | 0.966288 |
| ARF3 | 5137.391 | -0.40211 | 0.139351 | -2.8856 | 0.003907 | 0.025075 |
| TMEM106( | 2914.758 | -0.0442 | 0.182923 | -0.24164 | 0.809061 | 0.885851 |
| SLC38A2 | 4607.806 | 0.584838 | 0.221892 | 2.635687 | 0.008397 | 0.041895 |
| PLEKHA8P1 | 69.81466 | 0.484594 | 0.282078 | 1.717942 | 0.085807 | 0.200403 |
| EMP1 | 11507.65 | -0.16466 | 0.145105 | -1.13478 | 0.256468 | 0.42053 |
| SOX5 | 648.4172 | 0.460245 | 0.261275 | 1.761534 | 0.078148 | 0.188276 |
| RERG | 93.46748 | 1.486096 | 0.975601 | 1.523262 | 0.127693 | 0.263683 |
| KLRC1 | 0.134133 | -1.48909 | 4.077584 | -0.36519 | 0.71497 | NA |
| C12orf39 | 49.71973 | 0.250388 | 0.330802 | 0.756911 | 0.449103 | 0.608679 |
| PRH2 | 0.392711 | 0.292538 | 3.145221 | 0.09301 | 0.925895 | NA |
| PIWIL4 | 20.65009 | -0.12717 | 0.481524 | -0.2641 | 0.791701 | NA |
| GGACT | 78.707 | -0.47064 | 0.341235 | -1.37922 | 0.167827 | 0.317471 |
| COL4A2 | 9017.263 | -0.3705 | 0.214769 | -1.72513 | 0.084504 | 0.198068 |
| CLDN10 | 0.411855 | 0.250456 | 3.097618 | 0.080854 | 0.935558 | NA |
| DZIP1 | 957.8627 | 0.59741 | 0.260971 | 2.289179 | 0.022069 | 0.081 |
| UBAC2 | 1543.546 | -0.13471 | 0.177476 | -0.75904 | 0.44783 | 0.60737 |
| ARGLU1 | 854.2439 | 0.910813 | 0.21882 | 4.162393 | 3.15E-05 | 0.001232 |
| BIVM | 718.707 | 0.589394 | 0.190862 | 3.088068 | 0.002015 | 0.016256 |
| ERCC5 | 1117.084 | 0.490358 | 0.171462 | 2.859872 | 0.004238 | 0.026417 |
| TPP2 | 943.7543 | 0.637166 | 0.260326 | 2.447571 | 0.014382 | 0.060355 |
| KDELC1 | 325.3053 | 0.036605 | 0.184067 | 0.198867 | 0.842367 | 0.906403 |
| CARS2 | 2458.385 | -0.0209 | 0.145286 | -0.14388 | 0.885592 | 0.932306 |
| ARHGAP32 | 770.03 | -0.23055 | 0.181357 | -1.27127 | 0.203631 | 0.360091 |
| STT3A | 5872.566 | -0.30441 | 0.162605 | -1.87208 | 0.061196 | 0.159516 |
| ADAMTS8 | 0.258578 | 1.233151 | 4.047809 | 0.304646 | 0.760635 | NA |
| ACRV1 | 1.498287 | 1.287568 | 1.734062 | 0.742515 | 0.457775 | NA |
| ETS1 | 9420.749 | -0.0387 | 0.225752 | -0.17143 | 0.863885 | 0.919827 |
| SLC37A2 | 152.852 | 0.033079 | 0.273389 | 0.120995 | 0.903695 | 0.94382 |
| TAOK3 | 1453.94 | 0.830438 | 0.179749 | 4.61998 | 3.84E-06 | 3.67E-04 |
| USP30 | 369.589 | -0.45086 | 0.169169 | -2.66514 | 0.007696 | 0.039632 |
| SDS | 2.714496 | 0.228167 | 1.324092 | 0.172319 | 0.863186 | NA |
| MSI1 | 1.62392 | -0.42828 | 1.578049 | -0.2714 | 0.786082 | NA |
| HNF1A | 26.63049 | -0.34342 | 0.420909 | -0.8159 | 0.41456 | NA |
| FBXO21 | 880.3211 | -0.10702 | 0.162962 | -0.65672 | 0.511363 | 0.659366 |
| TBX3 | 10.52912 | 0.667194 | 0.902214 | 0.739507 | 0.459599 | NA |
| OASL | 59.06015 | -1.31719 | 0.44749 | -2.94351 | 0.003245 | 0.022307 |
| HRK | 534.9049 | -0.35415 | 0.175768 | -2.01488 | 0.043917 | 0.127733 |
| RNFT2 | 57.58372 | -0.48511 | 0.361005 | -1.34378 | 0.179021 | 0.331557 |
| P2RX4 | 1587.506 | -0.18923 | 0.150632 | -1.25621 | 0.20904 | 0.366643 |
| CCDC64 | 29.73125 | -0.0111 | 0.428286 | -0.02591 | 0.979326 | NA |
| DTX1 | 1.016048 | -2.46876 | 2.437554 | -1.0128 | 0.311154 | NA |
| TRAFD1 | 823.4018 | -0.31515 | 0.138486 | -2.27571 | 0.022864 | 0.082762 |
| ATP5G2 | 5390.02 | -0.14658 | 0.157573 | -0.93025 | 0.352241 | 0.520351 |
| DNAJC14 | 635.8639 | -0.26299 | 0.164651 | -1.59727 | 0.110205 | 0.237974 |

| | | | | | | |
|---|---|---|---|---|---|---|
| CD63 | 68888.68 | -0.15973 | 0.1336 | -1.19556 | 0.23187 | 0.394209 |
| PRPH | 0.163957 | 0.424913 | 4.079006 | 0.104171 | 0.917034 | NA |
| AVIL | 29.34866 | 0.23232 | 0.384587 | 0.604077 | 0.545792 | NA |
| AMHR2 | 0.096202 | 0.408546 | 4.079639 | 0.100143 | 0.920231 | NA |
| GDF11 | 2180.133 | -0.08138 | 0.141443 | -0.57534 | 0.565061 | 0.702545 |
| GLS2 | 19.96814 | 0.164175 | 0.477563 | 0.343776 | 0.731015 | NA |
| ITGA7 | 92.06731 | -0.72457 | 0.367303 | -1.97267 | 0.048533 | 0.135986 |
| FAM186B | 6.978356 | -0.02285 | 0.769153 | -0.02971 | 0.976302 | NA |
| RDH5 | 1.302723 | 0.907293 | 1.965565 | 0.461594 | 0.644372 | NA |
| AGAP2 | 22.07064 | -0.10258 | 0.474459 | -0.21621 | 0.828828 | NA |
| BLOC1S1 | 216.0712 | 0.085473 | 0.195954 | 0.43619 | 0.662699 | 0.778159 |
| CDK4 | 3403.632 | -0.25181 | 0.126511 | -1.99039 | 0.046548 | 0.132668 |
| PPP1R1A | 0.392857 | -1.546 | 4.075469 | -0.37934 | 0.704433 | NA |
| TROAP | 519.6896 | -0.33563 | 0.249572 | -1.34484 | 0.178677 | 0.33126 |
| TSPAN31 | 1414.379 | -0.20922 | 0.149279 | -1.40151 | 0.161062 | 0.308595 |
| B4GALNT1 | 149.0279 | -0.00255 | 0.273158 | -0.00932 | 0.99256 | 0.996738 |
| TFCP2 | 684.1292 | -0.0561 | 0.172238 | -0.32571 | 0.744643 | 0.839219 |
| COQ10A | 203.9142 | -0.06409 | 0.254205 | -0.2521 | 0.800964 | 0.880141 |
| FAIM2 | 0.134133 | -1.48909 | 4.077584 | -0.36519 | 0.71497 | NA |
| PAN2 | 398.1181 | -0.09142 | 0.192763 | -0.47426 | 0.635311 | 0.756414 |
| ESPL1 | 515.9445 | -0.7192 | 0.22714 | -3.16634 | 0.001544 | 0.013793 |
| KRT7 | 0.815594 | -0.45539 | 2.531219 | -0.17991 | 0.857224 | NA |
| ZC3H10 | 226.0873 | -0.01835 | 0.19539 | -0.0939 | 0.925188 | 0.957409 |
| HNRNPA1 | 8574.722 | 0.028683 | 0.119229 | 0.240569 | 0.80989 | 0.886296 |
| SLC26A10 | 3.245254 | 0.297076 | 1.173785 | 0.253092 | 0.800197 | NA |
| ACVR1B | 693.4776 | -0.24995 | 0.139753 | -1.78853 | 0.073691 | 0.181021 |
| OS9 | 7509.303 | -0.27388 | 0.151091 | -1.81271 | 0.069877 | 0.174179 |
| MIP | 1.252771 | 0.053586 | 1.700195 | 0.031518 | 0.974857 | NA |
| KCNH3 | 0.654449 | 1.296029 | 2.63083 | 0.492631 | 0.622273 | NA |
| KCNMB4 | 20.6996 | 0.099527 | 0.490357 | 0.202968 | 0.83916 | NA |
| USP15 | 523.7807 | 1.139143 | 0.294093 | 3.873403 | 1.07E-04 | 0.002688 |
| GNS | 10325.13 | -0.47615 | 0.116237 | -4.09642 | 4.20E-05 | 0.001503 |
| CPM | 265.7179 | -0.24956 | 0.209453 | -1.1915 | 0.233458 | 0.396057 |
| MDM2 | 1591.508 | 0.341341 | 0.292119 | 1.168499 | 0.242605 | 0.40554 |
| ISCU | 2116.016 | 0.245979 | 0.136323 | 1.804388 | 0.071171 | 0.176549 |
| ALDH1L2 | 1920.318 | -0.44468 | 0.330165 | -1.34684 | 0.178032 | 0.330397 |
| STAB2 | 3.861671 | -0.08078 | 1.099581 | -0.07346 | 0.941437 | NA |
| USP44 | 2.590206 | 0.353836 | 1.320578 | 0.26794 | 0.788745 | NA |
| SCYL2 | 1251.667 | 0.521266 | 0.183983 | 2.833231 | 0.004608 | 0.027914 |
| CKAP4 | 2370.612 | -0.23086 | 0.122204 | -1.88913 | 0.058874 | 0.155139 |
| PLXNC1 | 11277.09 | -0.35543 | 0.237285 | -1.49792 | 0.134153 | 0.272187 |
| APPL2 | 511.7687 | 0.056849 | 0.172562 | 0.329442 | 0.741822 | 0.837188 |
| PWP1 | 2193.594 | 0.281636 | 0.131692 | 2.138593 | 0.032469 | 0.104859 |
| DRAM1 | 1538.825 | 0.693256 | 0.218001 | 3.180059 | 0.001472 | 0.013425 |
| KIAA1033 | 2247.736 | 0.912545 | 0.293887 | 3.105086 | 0.001902 | 0.015585 |
| SLC41A2 | 95.84726 | 0.393216 | 0.263232 | 1.493801 | 0.135228 | 0.273898 |
| NEK3 | 131.9165 | 0.254731 | 0.237984 | 1.070372 | 0.284452 | 0.450947 |

| | | | | | | |
|---|---|---|---|---|---|---|
| PCDH8 | 0.757324 | -2.08271 | 2.587512 | -0.80491 | 0.420873 | NA |
| VPS36 | 1027.287 | 0.426205 | 0.199791 | 2.133249 | 0.032904 | 0.105627 |
| RNASEH2B | 464.147 | 0.18717 | 0.199325 | 0.939017 | 0.347722 | 0.515942 |
| CKAP2 | 1131.882 | 0.538659 | 0.160212 | 3.36216 | 7.73E-04 | 0.008875 |
| LECT1 | 0.096202 | 0.408546 | 4.079639 | 0.100143 | 0.920231 | NA |
| TBC1D4 | 781.3584 | 0.22429 | 0.146259 | 1.533511 | 0.12515 | 0.260016 |
| THSD1 | 18.69169 | -0.25152 | 0.528216 | -0.47616 | 0.633957 | NA |
| BORA | 148.9783 | 0.626575 | 0.210108 | 2.982154 | 0.002862 | 0.020475 |
| LRCH1 | 750.319 | 0.226905 | 0.147939 | 1.533771 | 0.125086 | 0.260016 |
| SUCLA2 | 737.6778 | 0.742823 | 0.20858 | 3.561341 | 3.69E-04 | 0.005593 |
| RCBTB1 | 901.956 | 0.44677 | 0.219656 | 2.033959 | 0.041956 | 0.124471 |
| MED4 | 433.8775 | 0.590899 | 0.174178 | 3.392489 | 6.93E-04 | 0.00825 |
| PHF11 | 171.3548 | -0.23465 | 0.284361 | -0.82517 | 0.409275 | 0.572795 |
| RPL13AP25 | 50.74544 | -0.0944 | 0.315567 | -0.29915 | 0.764825 | 0.853847 |
| COG3 | 538.2223 | 0.490271 | 0.176174 | 2.782885 | 0.005388 | 0.031172 |
| LMO7 | 1194.266 | 0.596292 | 0.261793 | 2.277728 | 0.022743 | 0.082451 |
| SCEL | 66.46984 | 1.705768 | 0.527978 | 3.230753 | 0.001235 | 0.011864 |
| ITM2B | 12872 | 0.128611 | 0.115562 | 1.112924 | 0.265741 | 0.430423 |
| SPRY2 | 1488.049 | 0.512359 | 0.213379 | 2.401169 | 0.016343 | 0.066012 |
| NUDT15 | 729.7206 | 0.298441 | 0.218838 | 1.363756 | 0.172644 | 0.324032 |
| EDNRB | 16625.15 | 0.008332 | 0.201497 | 0.041349 | 0.967017 | 0.981674 |
| RCBTB2 | 903.0472 | 0.046486 | 0.193127 | 0.240701 | 0.809787 | 0.886265 |
| LCP1 | 84.5323 | 0.008022 | 0.308454 | 0.026006 | 0.979253 | 0.988929 |
| SETDB2 | 776.6633 | 0.63768 | 0.217481 | 2.932115 | 0.003367 | 0.022712 |
| | 0.134133 | -1.48909 | 4.077584 | -0.36519 | 0.71497 | NA |
| TTC5 | 583.9749 | -0.15066 | 0.176516 | -0.85349 | 0.393386 | 0.55838 |
| NKX2-8 | 0.163957 | 0.424913 | 4.079006 | 0.104171 | 0.917034 | NA |
| ZFHX2 | 28.18132 | -0.10963 | 0.409868 | -0.26749 | 0.789095 | NA |
| MTHFS | 99.72658 | -0.11885 | 0.25758 | -0.46142 | 0.644497 | 0.763593 |
| ADAMTS7 | 448.062 | -0.01639 | 0.21384 | -0.07663 | 0.938918 | 0.966165 |
| ABHD17C | 298.3971 | -0.02398 | 0.180643 | -0.13276 | 0.894385 | 0.938268 |
| IREB2 | 976.1946 | 0.315172 | 0.236113 | 1.334836 | 0.18193 | 0.334748 |
| ALPK3 | 6.203315 | -0.43997 | 0.871287 | -0.50496 | 0.613587 | NA |
| TM6SF1 | 20.55063 | 0.621657 | 0.682837 | 0.910403 | 0.36261 | NA |
| CIB2 | 356.1107 | -0.27027 | 0.226185 | -1.19493 | 0.232115 | 0.394343 |
| MYO7A | 8.712409 | 0.419245 | 0.69477 | 0.60343 | 0.546223 | NA |
| FCHSD2 | 187.3852 | 0.234979 | 0.24081 | 0.975784 | 0.329172 | 0.497366 |
| ARRB1 | 2286.348 | -0.45511 | 0.127482 | -3.57004 | 3.57E-04 | 0.005524 |
| SLCO2B1 | 32.52346 | -0.39678 | 0.650607 | -0.60986 | 0.541953 | NA |
| PRKRIR | 369.0572 | 0.635542 | 0.173002 | 3.673619 | 2.39E-04 | 0.004335 |
| ANKRD42 | 252.0437 | 0.36816 | 0.211135 | 1.743723 | 0.081207 | 0.193238 |
| IL18BP | 31.11298 | 0.161352 | 0.402454 | 0.40092 | 0.688479 | NA |
| NUMA1 | 3022.111 | -0.55321 | 0.176479 | -3.13468 | 0.00172 | 0.014673 |
| CCDC90B | 710.5313 | 0.550582 | 0.167164 | 3.29366 | 9.89E-04 | 0.010266 |
| SYTL2 | 1108.571 | 0.759735 | 0.352368 | 2.156083 | 0.031077 | 0.101597 |
| RAB30 | 489.1966 | 0.05097 | 0.248336 | 0.205248 | 0.837379 | 0.902712 |
| CREBZF | 716.7625 | 0.786547 | 0.199732 | 3.938008 | 8.22E-05 | 0.002255 |

| | | | | | | |
|---|---|---|---|---|---|---|
| LRRC32 | 0.881566 | 1.109202 | 2.554987 | 0.434132 | 0.664192 | NA |
| PRCP | 4402.435 | -0.09921 | 0.16129 | -0.61512 | 0.538474 | 0.681093 |
| NARS2 | 1013.155 | -0.19145 | 0.1893 | -1.01136 | 0.311846 | 0.479045 |
| RNF121 | 673.2095 | -0.19371 | 0.177403 | -1.09195 | 0.274856 | 0.440564 |
| SORL1 | 11.16238 | 0.969911 | 0.704034 | 1.377648 | 0.168312 | NA |
| BUD13 | 433.1091 | 0.032874 | 0.163536 | 0.20102 | 0.840683 | 0.905184 |
| TRPC6 | 0.392711 | 0.292538 | 3.145221 | 0.09301 | 0.925895 | NA |
| MMP7 | 1.127884 | 0.383009 | 2.112806 | 0.18128 | 0.856148 | NA |
| C11orf70 | 14.13991 | -0.30416 | 0.627233 | -0.48493 | 0.627725 | NA |
| DCUN1D5 | 888.9903 | 0.171861 | 0.167671 | 1.024987 | 0.305369 | 0.472543 |
| YAP1 | 2442.988 | 0.290577 | 0.153104 | 1.897907 | 0.057708 | 0.152776 |
| TRIM29 | 0.327913 | 1.543154 | 4.021063 | 0.383768 | 0.701151 | NA |
| SLC37A4 | 782.8762 | -0.35825 | 0.187624 | -1.9094 | 0.056211 | 0.150029 |
| BTG4 | 1.090395 | -0.30963 | 1.797091 | -0.17229 | 0.863206 | NA |
| POU2F3 | 0.574434 | -0.17781 | 2.756096 | -0.06452 | 0.948559 | NA |
| RDX | 3110.121 | 0.705239 | 0.310146 | 2.273893 | 0.022972 | 0.082954 |
| PPP2R1B | 743.8416 | 0.104182 | 0.167962 | 0.620268 | 0.535082 | 0.678307 |
| FDX1 | 594.0364 | 0.246932 | 0.146107 | 1.690083 | 0.091012 | 0.209456 |
| C11orf1 | 402.6347 | 0.392027 | 0.179235 | 2.187219 | 0.028727 | 0.09631 |
| FXYD6 | 14.84317 | -0.85829 | 0.717933 | -1.1955 | 0.231893 | NA |
| ARHGAP20 | 2.768679 | 0.6833 | 1.247632 | 0.547677 | 0.583914 | NA |
| FXYD2 | 0.134133 | -1.48909 | 4.077584 | -0.36519 | 0.71497 | NA |
| TMPRSS13 | 7.366888 | -0.71575 | 1.202947 | -0.595 | 0.551843 | NA |
| CASP1 | 925.5995 | 0.098557 | 0.20853 | 0.472626 | 0.63648 | 0.757389 |
| ALKBH8 | 183.1117 | 0.744579 | 0.244108 | 3.0502 | 0.002287 | 0.017639 |
| MAP2K5 | 475.8449 | -0.39374 | 0.14945 | -2.63461 | 0.008423 | 0.041958 |
| UNC13C | 0.424115 | 1.932531 | 3.45462 | 0.559405 | 0.575885 | NA |
| SQRDL | 470.4442 | 0.433811 | 0.335297 | 1.293811 | 0.195731 | 0.351042 |
| CTDSPL2 | 376.0091 | 0.650343 | 0.190106 | 3.42095 | 6.24E-04 | 0.007763 |
| SLTM | 777.6877 | 1.014015 | 0.641708 | 1.580183 | 0.114065 | 0.243201 |
| THBS1 | 2284.969 | 0.474383 | 0.844837 | 0.561508 | 0.574451 | 0.709683 |
| MAPKBP1 | 1244.689 | -0.43541 | 0.193995 | -2.24446 | 0.024803 | 0.087277 |
| NUSAP1 | 1253.35 | 0.238094 | 0.178985 | 1.330251 | 0.183436 | 0.33663 |
| NDUFAF1 | 673.1062 | 0.034258 | 0.153771 | 0.222786 | 0.823702 | 0.893657 |
| KIF23 | 541.5085 | 0.658276 | 0.167988 | 3.9186 | 8.91E-05 | 0.002378 |
| ITGA11 | 8.02285 | 1.615897 | 0.90646 | 1.782645 | 0.074644 | NA |
| CASC5 | 259.0868 | 0.644527 | 0.406105 | 1.587095 | 0.112491 | 0.241067 |
| HAUS2 | 519.3357 | 0.237062 | 0.188458 | 1.257907 | 0.208425 | 0.365894 |
| RTF1 | 1662.854 | 0.395942 | 0.195814 | 2.022031 | 0.043173 | 0.126625 |
| PARP6 | 898.547 | -0.13629 | 0.268594 | -0.5074 | 0.611871 | 0.738594 |
| RPLP1 | 32703.37 | 0.001211 | 0.130906 | 0.009252 | 0.992618 | 0.996738 |
| PAQR5 | 11.04298 | 0.622657 | 0.627087 | 0.992935 | 0.320742 | NA |
| LRRC49 | 67.41339 | 0.849721 | 0.327142 | 2.597405 | 0.009393 | 0.045248 |
| TUBGCP4 | 849.0166 | -0.14924 | 0.137029 | -1.08914 | 0.276092 | 0.44171 |
| RMDN3 | 1111.929 | -0.06094 | 0.136455 | -0.44658 | 0.655177 | 0.77237 |
| ITPKA | 4.126278 | 0.720924 | 1.012227 | 0.712216 | 0.476331 | NA |
| UACA | 504.3362 | 0.950039 | 1.143446 | 0.830856 | 0.406055 | 0.569881 |

| | | | | | | |
|---|---|---|---|---|---|---|
| SMAD6 | 155.7391 | 0.428992 | 0.274742 | 1.561437 | 0.118421 | 0.2498 |
| PLCB2 | 856.3529 | -0.36381 | 0.15302 | -2.37757 | 0.017427 | 0.069026 |
| TMEM62 | 489.9838 | -0.04106 | 0.184537 | -0.22249 | 0.823934 | 0.893745 |
| PAK6 | 0.998549 | 1.029429 | 2.066501 | 0.498151 | 0.618378 | NA |
| ADAM10 | 3583.945 | 0.344248 | 0.224901 | 1.530668 | 0.125852 | 0.260925 |
| DUOX1 | 24.43191 | -0.35685 | 0.469141 | -0.76065 | 0.446867 | NA |
| SLC28A2 | 0.298089 | -0.51711 | 3.975232 | -0.13008 | 0.896501 | NA |
| CYP19A1 | 4.99666 | 0.582714 | 0.963819 | 0.604589 | 0.545452 | NA |
| ZNF280D | 137.7263 | 0.545951 | 0.296768 | 1.839657 | 0.065819 | 0.167298 |
| SEMA6D | 1021.836 | -0.14788 | 0.2023 | -0.73099 | 0.464787 | 0.622183 |
| RSL24D1 | 1679.476 | 0.992078 | 0.235739 | 4.208374 | 2.57E-05 | 0.00107 |
| SPTBN5 | 70.68785 | 0.256418 | 0.431524 | 0.594214 | 0.552369 | 0.693167 |
| GCHFR | 109.3459 | 0.061833 | 0.275667 | 0.224304 | 0.822521 | 0.893074 |
| MNS1 | 31.73439 | 0.978665 | 0.696311 | 1.405499 | 0.159873 | NA |
| USP8 | 1000.378 | 0.830986 | 0.302323 | 2.748674 | 0.005984 | 0.033351 |
| SECISBP2L | 2148.128 | 0.769666 | 0.27276 | 2.821776 | 0.004776 | 0.028636 |
| TMOD3 | 701.2771 | 0.655722 | 0.22781 | 2.878374 | 0.003997 | 0.025436 |
| SPPL2A | 843.6936 | 0.173137 | 0.164438 | 1.052901 | 0.292386 | 0.459165 |
| GLCE | 640.2105 | 0.259594 | 0.201528 | 1.288128 | 0.197701 | 0.353347 |
| SHF | 1010.085 | -0.24385 | 0.182308 | -1.33757 | 0.181038 | 0.333581 |
| APH1B | 2257.938 | -0.31244 | 0.195075 | -1.60162 | 0.109239 | 0.236492 |
| VWA9 | 1403.196 | -0.15525 | 0.147186 | -1.0548 | 0.291517 | 0.458407 |
| CILP | 0.326536 | 0.037056 | 3.842554 | 0.009644 | 0.992306 | NA |
| PARP16 | 127.642 | -0.19959 | 0.230668 | -0.86527 | 0.386892 | 0.552871 |
| PPCDC | 278.9236 | -0.14318 | 0.180531 | -0.79311 | 0.427714 | 0.589531 |
| SEMA7A | 148.1448 | -0.72975 | 0.30712 | -2.37611 | 0.017496 | 0.069122 |
| UBL7 | 1501.914 | -0.28523 | 0.154581 | -1.84517 | 0.065013 | 0.166214 |
| B4GALNT3 | 15.21636 | 0.392737 | 0.564461 | 0.695775 | 0.48657 | NA |
| ERP27 | 0.326536 | 0.037056 | 3.842554 | 0.009644 | 0.992306 | NA |
| ETV6 | 254.1318 | -0.3059 | 0.20536 | -1.48958 | 0.136334 | 0.275434 |
| GABARAPL | 2350.674 | 0.199724 | 0.148667 | 1.343433 | 0.179132 | 0.331658 |
| KIF21A | 200.87 | 0.672541 | 0.729221 | 0.922273 | 0.356386 | 0.524101 |
| CPNE8 | 796.1365 | 0.804354 | 0.195941 | 4.105093 | 4.04E-05 | 0.001474 |
| YARS2 | 633.1619 | 0.028101 | 0.140431 | 0.200108 | 0.841396 | 0.905561 |
| FGD4 | 332.6321 | 0.843363 | 0.23092 | 3.65218 | 2.60E-04 | 0.004547 |
| ALG10 | 40.11783 | 0.214147 | 0.337868 | 0.633819 | 0.526199 | NA |
| FAM60A | 552.5103 | 0.233193 | 0.176219 | 1.323312 | 0.185732 | 0.339582 |
| AEBP2 | 456.6002 | 0.157432 | 0.195199 | 0.80652 | 0.419943 | 0.58289 |
| SLCO1C1 | 0.659753 | -0.00581 | 2.629089 | -0.00221 | 0.998237 | NA |
| METTL20 | 66.84537 | -0.07066 | 0.309503 | -0.22831 | 0.819407 | 0.891854 |
| ETNK1 | 552.3863 | 0.61681 | 0.159736 | 3.861432 | 1.13E-04 | 0.002775 |
| ZCRB1 | 822.4974 | 0.647838 | 0.191225 | 3.387835 | 7.04E-04 | 0.008326 |
| TMEM117 | 703.4798 | 0.110349 | 0.175896 | 0.627353 | 0.530428 | 0.675137 |
| PRICKLE1 | 14.50111 | -0.16067 | 0.573708 | -0.28006 | 0.779432 | NA |
| C1RL | 222.1881 | -0.29226 | 0.181049 | -1.61425 | 0.106474 | 0.232543 |
| NDUFA9 | 729.0356 | -0.17035 | 0.158326 | -1.07595 | 0.281951 | 0.448104 |
| CLSTN3 | 173.1225 | 0.036122 | 0.26419 | 0.136726 | 0.891248 | 0.936356 |

| | | | | | | |
|---|---|---|---|---|---|---|
| KLRG1 | 24.92763 | 0.39613 | 0.521674 | 0.759345 | 0.447646 | NA |
| VAMP1 | 52.89476 | 0.214559 | 0.297893 | 0.720256 | 0.471367 | 0.627431 |
| TAPBPL | 233.4904 | -0.30996 | 0.192543 | -1.60983 | 0.107434 | 0.233913 |
| CD27 | 0.813665 | 1.671689 | 2.474058 | 0.675687 | 0.499239 | NA |
| RBP5 | 5.039398 | 0.997438 | 1.012796 | 0.984836 | 0.324705 | NA |
| PEX5 | 1573.515 | -0.34779 | 0.125114 | -2.7798 | 0.005439 | 0.031378 |
| PIANP | 2.719894 | 0.0302 | 1.21839 | 0.024787 | 0.980225 | NA |
| SLC38A4 | 0.162376 | 0.424913 | 4.079006 | 0.104171 | 0.917034 | NA |
| AMIGO2 | 1.392454 | 2.595684 | 1.940577 | 1.337584 | 0.181032 | NA |
| SCAF11 | 729.4203 | 0.818836 | 0.362228 | 2.260553 | 0.023787 | 0.085143 |
| COL2A1 | 5.65939 | -0.1603 | 0.959709 | -0.16703 | 0.867349 | NA |
| PPFIA2 | 33.12995 | 0.930774 | 0.395768 | 2.351818 | 0.018682 | NA |
| LLPH | 362.606 | 0.503477 | 0.191085 | 2.634835 | 0.008418 | 0.041958 |
| RPL14P1 | 114.8333 | 0.204787 | 0.262001 | 0.781626 | 0.434434 | 0.595539 |
| LRIG3 | 174.8885 | 0.123178 | 0.235548 | 0.522943 | 0.601014 | 0.730247 |
| 9-Mar | 1075.357 | -0.16806 | 0.17488 | -0.961 | 0.336551 | 0.505541 |
| INHBE | 5.132156 | 0.118856 | 0.869925 | 0.136628 | 0.891325 | NA |
| GLIPR1 | 28.55839 | 0.727355 | 0.578733 | 1.256806 | 0.208824 | NA |
| TPH2 | 0.096202 | 0.408546 | 4.079639 | 0.100143 | 0.920231 | NA |
| PHLDA1 | 6197.274 | -0.06152 | 0.136178 | -0.45179 | 0.65142 | 0.769259 |
| TMEM19 | 818.4766 | 0.01157 | 0.187465 | 0.061716 | 0.950789 | 0.97278 |
| PTPRQ | 0.162376 | 0.424913 | 4.079006 | 0.104171 | 0.917034 | NA |
| DUSP6 | 3432.412 | 0.019248 | 0.162123 | 0.118724 | 0.905494 | 0.945366 |
| POC1B | 464.8048 | 0.578903 | 0.184391 | 3.139551 | 0.001692 | 0.014563 |
| TMTC3 | 348.9043 | 1.020334 | 0.266047 | 3.835162 | 1.25E-04 | 0.002961 |
| LUM | 26.2727 | 0.283748 | 0.550775 | 0.51518 | 0.606427 | NA |
| SNRPF | 580.0261 | 0.208303 | 0.154834 | 1.345327 | 0.17852 | 0.331094 |
| AMDHD1 | 6.040659 | 1.730504 | 0.911422 | 1.898686 | 0.057606 | NA |
| NEDD1 | 441.2635 | 0.859 | 0.20158 | 4.261339 | 2.03E-05 | 9.37E-04 |
| SYCP3 | 5.139522 | 0.339164 | 0.973167 | 0.348516 | 0.727453 | NA |
| ASCL1 | 0.47961 | 0.535796 | 3.39605 | 0.15777 | 0.874638 | NA |
| GAS2L3 | 1178.747 | 0.166428 | 0.178789 | 0.930864 | 0.351924 | 0.520012 |
| TMEM132I | 6.365156 | -0.25932 | 0.933485 | -0.27779 | 0.781171 | NA |
| SLC15A4 | 1797.156 | -0.34124 | 0.13234 | -2.57849 | 0.009923 | 0.046925 |
| TDG | 214.5096 | 0.896806 | 0.225574 | 3.975662 | 7.02E-05 | 0.002039 |
| RITA1 | 789.0026 | -0.21833 | 0.171323 | -1.27435 | 0.202539 | 0.358825 |
| SDSL | 270.5032 | -0.3233 | 0.272937 | -1.1845 | 0.236214 | 0.399143 |
| MMAB | 1209.736 | -0.16469 | 0.141255 | -1.16591 | 0.243649 | 0.406464 |
| GLTP | 1062.294 | -0.22921 | 0.168944 | -1.35671 | 0.174873 | 0.326835 |
| GIT2 | 580.1113 | -0.23099 | 0.14544 | -1.58824 | 0.112233 | 0.240687 |
| TCHP | 575.1009 | -0.05038 | 0.144265 | -0.34925 | 0.726901 | 0.82669 |
| FAM222A | 217.8988 | -0.72166 | 0.265472 | -2.7184 | 0.00656 | 0.03551 |
| FOXN4 | 0.151697 | -1.546 | 4.075469 | -0.37934 | 0.704433 | NA |
| NUPL1 | 1130.062 | 0.415326 | 0.176828 | 2.348762 | 0.018836 | 0.072547 |
| MTMR6 | 1171.849 | 0.619952 | 0.206133 | 3.007537 | 0.002634 | 0.019441 |
| SLC46A3 | 159.4396 | 0.612376 | 0.254734 | 2.403986 | 0.016217 | 0.065572 |
| SLC7A1 | 1829.635 | -0.52995 | 0.178188 | -2.9741 | 0.002939 | 0.02086 |

| | | | | | | |
|---|---|---|---|---|---|---|
| PDX1 | 0.392857 | -1.546 | 4.075469 | -0.37934 | 0.704433 | NA |
| LNX2 | 374.332 | -0.09059 | 0.211168 | -0.42899 | 0.667931 | 0.782205 |
| SUOX | 543.8106 | -0.44032 | 0.147061 | -2.99415 | 0.002752 | 0.020028 |
| CCDC65 | 19.98623 | 0.346612 | 0.460757 | 0.752268 | 0.45189 | NA |
| SLC39A5 | 0.614937 | 2.493639 | 2.617677 | 0.952615 | 0.340785 | NA |
| TARBP2 | 1325.02 | -0.22824 | 0.145788 | -1.56555 | 0.117454 | 0.248216 |
| RDH16 | 1.562036 | -1.53353 | 1.708649 | -0.89751 | 0.369445 | NA |
| DHH | 11.7182 | 0.071261 | 0.60173 | 0.118427 | 0.90573 | NA |
| ACVRL1 | 135.288 | 0.103668 | 0.254669 | 0.407069 | 0.683957 | 0.794388 |
| NPFF | 8.90935 | 0.570504 | 0.697193 | 0.818287 | 0.413193 | NA |
| NABP2 | 1903.782 | -0.13801 | 0.174951 | -0.78885 | 0.430198 | 0.591716 |
| N4BP2L1 | 38.94385 | 0.123005 | 0.406769 | 0.302396 | 0.76235 | NA |
| SMARCC2 | 2693.37 | -0.45066 | 0.137843 | -3.26937 | 0.001078 | 0.010884 |
| BRCA2 | 107.2159 | 0.952661 | 0.497328 | 1.915559 | 0.055421 | 0.148748 |
| KANSL2 | 478.902 | 0.157165 | 0.14813 | 1.060997 | 0.288691 | 0.45493 |
| CERS5 | 1570.58 | -0.21573 | 0.171128 | -1.26063 | 0.207443 | 0.364547 |
| MAP3K12 | 1170.331 | -0.27968 | 0.132241 | -2.11493 | 0.034436 | 0.108718 |
| ITGB7 | 31.97284 | -0.60251 | 0.41293 | -1.45912 | 0.144533 | NA |
| GALNT6 | 12.52327 | -0.2771 | 0.714127 | -0.38802 | 0.697998 | NA |
| CSAD | 90.65591 | 0.214827 | 0.36725 | 0.584961 | 0.558574 | 0.697556 |
| LMBR1L | 542.3029 | -0.19466 | 0.143264 | -1.35877 | 0.174221 | 0.326183 |
| C12orf10 | 1047.034 | -0.15364 | 0.142655 | -1.07697 | 0.281492 | 0.447691 |
| ESYT1 | 5301.806 | -0.48094 | 0.14611 | -3.29166 | 9.96E-04 | 0.010321 |
| TMBIM6 | 17379.26 | -0.08891 | 0.131466 | -0.6763 | 0.498849 | 0.649348 |
| ANKRD52 | 3616.926 | -0.71911 | 0.150442 | -4.78001 | 1.75E-06 | 2.26E-04 |
| ZNF740 | 603.5094 | -0.15992 | 0.190959 | -0.83746 | 0.402332 | 0.566803 |
| SMIM2 | 0.845069 | 2.925226 | 2.326646 | 1.257272 | 0.208655 | NA |
| WDFY2 | 217.1214 | 0.023048 | 0.218417 | 0.105525 | 0.915959 | 0.951316 |
| HNRNPA1L | 25.43172 | 0.753386 | 0.423915 | 1.777208 | 0.075534 | NA |
| LPAR6 | 430.3259 | 1.063537 | 0.244024 | 4.358327 | 1.31E-05 | 7.23E-04 |
| ESD | 2365.313 | 0.086029 | 0.13754 | 0.625483 | 0.531654 | 0.675972 |
| RB1 | 1963.157 | 0.626344 | 0.206385 | 3.034826 | 0.002407 | 0.018243 |
| SBNO1 | 1105.546 | 0.529098 | 0.215793 | 2.451881 | 0.014211 | 0.05987 |
| MORN3 | 18.02204 | -0.52985 | 0.516093 | -1.02666 | 0.304583 | NA |
| SETD1B | 548.9981 | -0.60277 | 0.20495 | -2.94107 | 0.003271 | 0.022362 |
| VPS33A | 360.1308 | -0.06518 | 0.183674 | -0.35489 | 0.722672 | 0.823698 |
| VPS37B | 3561.347 | -0.53238 | 0.126699 | -4.20193 | 2.65E-05 | 0.001093 |
| RHOF | 7.604379 | -0.59641 | 0.739458 | -0.80655 | 0.419928 | NA |
| DENR | 1088.612 | 0.337206 | 0.17866 | 1.88742 | 0.059104 | 0.155568 |
| DIAPH3 | 356.6716 | 0.592179 | 0.242114 | 2.44587 | 0.01445 | 0.060524 |
| SLAIN1 | 82.17517 | -0.71609 | 0.45014 | -1.59081 | 0.111653 | 0.239791 |
| RBM26 | 475.542 | 0.767338 | 0.340173 | 2.255728 | 0.024088 | 0.085831 |
| SRRM4 | 0.247898 | -0.5901 | 4.074633 | -0.14482 | 0.884851 | NA |
| METTL21C | 0.162376 | 0.424913 | 4.079006 | 0.104171 | 0.917034 | NA |
| MBNL2 | 1309.07 | 0.562878 | 0.322076 | 1.747656 | 0.080524 | 0.19219 |
| ABHD13 | 419.0732 | 0.501771 | 0.174348 | 2.877985 | 0.004002 | 0.025436 |
| RAB20 | 133.9659 | -0.29181 | 0.216926 | -1.34519 | 0.178565 | 0.331125 |

| | | | | | | |
|---|---|---|---|---|---|---|
| GRTP1 | 6.783178 | -0.4024 | 0.775989 | -0.51856 | 0.604067 | NA |
| CUL4A | 1970.138 | 0.399291 | 0.126865 | 3.147363 | 0.001648 | 0.014319 |
| TTC6 | 0.096202 | 0.408546 | 4.079639 | 0.100143 | 0.920231 | NA |
| CDH24 | 126.1189 | -0.13692 | 0.330547 | -0.41423 | 0.678705 | 0.790407 |
| REM2 | 2.625236 | -0.09254 | 1.248331 | -0.07413 | 0.940908 | NA |
| CBLN3 | 35.26339 | -0.41586 | 0.366227 | -1.13554 | 0.256151 | NA |
| TSSK4 | 26.09612 | 0.322014 | 0.416036 | 0.774004 | 0.438928 | NA |
| NOVA1 | 1.717389 | 1.966895 | 1.611749 | 1.220349 | 0.222333 | NA |
| FITM1 | 2.267479 | -1.12802 | 1.320914 | -0.85397 | 0.393123 | NA |
| MDGA2 | 257.8158 | -0.04621 | 0.272669 | -0.16946 | 0.865431 | 0.920317 |
| TMX1 | 954.7275 | 0.738424 | 0.189681 | 3.892976 | 9.90E-05 | 0.00253 |
| FRMD6 | 3032.45 | 0.566422 | 0.178124 | 3.17993 | 0.001473 | 0.013425 |
| PELI2 | 373.0767 | 0.022635 | 0.199358 | 0.113538 | 0.909604 | 0.947766 |
| RTN1 | 3.936649 | 1.141119 | 1.087397 | 1.049405 | 0.293992 | NA |
| C14orf37 | 362.46 | -0.05647 | 0.216953 | -0.26028 | 0.794648 | 0.875678 |
| SLC38A6 | 400.0694 | 0.0503 | 0.198841 | 0.252967 | 0.800294 | 0.879693 |
| NAA30 | 693.2 | 0.338155 | 0.175462 | 1.927232 | 0.053951 | 0.145874 |
| ADAM21 | 12.87973 | 0.049686 | 0.614751 | 0.080822 | 0.935583 | NA |
| RDH12 | 3.546147 | 0.091757 | 1.176519 | 0.077991 | 0.937835 | NA |
| DCAF5 | 1446.09 | -0.15959 | 0.152043 | -1.04961 | 0.293898 | 0.461117 |
| RAB15 | 361.5397 | -0.17191 | 0.197958 | -0.8684 | 0.385177 | 0.551083 |
| WDR89 | 114.8938 | 0.836949 | 0.315275 | 2.654664 | 0.007939 | 0.040458 |
| ESR2 | 78.92564 | -0.39859 | 0.277859 | -1.43451 | 0.151428 | 0.295445 |
| STON2 | 9.748985 | 0.357206 | 0.69387 | 0.514803 | 0.606691 | NA |
| EFCAB11 | 102.407 | 0.436348 | 0.251484 | 1.735091 | 0.082725 | 0.195632 |
| GPR65 | 6.003384 | 0.735223 | 0.969449 | 0.758393 | 0.448216 | NA |
| PTGR2 | 50.23745 | 0.770137 | 0.322009 | 2.391661 | 0.016772 | 0.067265 |
| JDP2 | 363.3441 | 0.146297 | 0.194253 | 0.753126 | 0.451374 | 0.610554 |
| AK7 | 39.11674 | 0.230014 | 0.340352 | 0.675811 | 0.49916 | NA |
| SLC24A4 | 17.51226 | -1.52906 | 0.54188 | -2.82177 | 0.004776 | NA |
| FBLN5 | 273.8083 | -0.03416 | 0.21642 | -0.15786 | 0.874569 | 0.925145 |
| C14orf79 | 285.8626 | -0.2279 | 0.18294 | -1.24574 | 0.21286 | 0.371589 |
| WARS | 13245.22 | -0.48343 | 0.16605 | -2.91137 | 0.003599 | 0.023873 |
| SLC25A47 | 0.883203 | 0.854847 | 2.135496 | 0.400304 | 0.688933 | NA |
| WDR20 | 560.9875 | 0.063105 | 0.14815 | 0.425953 | 0.670142 | 0.783871 |
| NIPA2 | 3026.603 | 0.050705 | 0.135255 | 0.374882 | 0.707748 | 0.812783 |
| HERC2P2 | 169.7848 | 1.168183 | 0.335017 | 3.486934 | 4.89E-04 | 0.006696 |
| SLC12A6 | 206.5567 | -0.42298 | 0.181018 | -2.33667 | 0.019456 | 0.074285 |
| DUOXA1 | 1.246033 | -0.06945 | 1.833478 | -0.03788 | 0.969784 | NA |
| MFAP1 | 1076.111 | 0.210149 | 0.144586 | 1.453452 | 0.146098 | 0.288569 |
| TCF12 | 1191.364 | 0.044403 | 0.195776 | 0.226806 | 0.820574 | 0.892334 |
| SORD | 1845.623 | -0.28636 | 0.130681 | -2.19131 | 0.028429 | 0.095503 |
| SERF2 | 12102.11 | -0.10779 | 0.146899 | -0.73375 | 0.463103 | 0.62097 |
| ZSCAN29 | 605.1661 | 0.145388 | 0.174715 | 0.832146 | 0.405327 | 0.569465 |
| DUOX2 | 0.921281 | -1.21902 | 2.226885 | -0.54741 | 0.584096 | NA |
| LYSMD2 | 407.333 | 0.077865 | 0.16445 | 0.473489 | 0.635864 | 0.756921 |
| FGF7 | 0.585114 | 1.12675 | 2.73598 | 0.411827 | 0.680466 | NA |

| | | | | | | |
|---|---|---|---|---|---|---|
| GCNT3 | 4.502387 | -1.00442 | 0.960317 | -1.04592 | 0.295596 | NA |
| BNIP2 | 1509.203 | 0.717077 | 0.188098 | 3.812244 | 1.38E-04 | 0.003086 |
| GTF2A2 | 1640.662 | 0.319203 | 0.161244 | 1.979632 | 0.047745 | 0.134702 |
| SRP14 | 7286.688 | 0.406093 | 0.148144 | 2.741206 | 0.006121 | 0.033867 |
| BAHD1 | 1229.085 | -0.52462 | 0.167325 | -3.13531 | 0.001717 | 0.014663 |
| DISP2 | 5.020519 | -1.0029 | 1.039216 | -0.96505 | 0.33452 | NA |
| CDAN1 | 399.3268 | -0.46754 | 0.17527 | -2.66756 | 0.00764 | 0.039439 |
| TLE3 | 968.2962 | -0.32319 | 0.183894 | -1.75751 | 0.078832 | 0.189295 |
| ANP32A | 2186.669 | -0.07386 | 0.151048 | -0.48899 | 0.624851 | 0.74847 |
| COMMD4 | 2589.719 | -0.18718 | 0.134615 | -1.39045 | 0.164392 | 0.313223 |
| UBE2Q2 | 1143.721 | 0.422449 | 0.175583 | 2.405985 | 0.016129 | 0.065355 |
| PSTPIP1 | 1.082076 | -0.5518 | 2.085766 | -0.26455 | 0.791353 | NA |
| ETFA | 3888.727 | 0.29293 | 0.148268 | 1.975679 | 0.048191 | 0.135481 |
| BCL2A1 | 2369.376 | 0.803336 | 0.174126 | 4.613539 | 3.96E-06 | 3.70E-04 |
| HMG20A | 1285.916 | -0.12948 | 0.158872 | -0.81499 | 0.415075 | 0.578296 |
| SCAPER | 243.6292 | 0.481481 | 0.331803 | 1.451106 | 0.14675 | 0.28952 |
| TSPAN3 | 7531.332 | -8.82E-05 | 0.14359 | -6.14E-04 | 0.99951 | 0.999594 |
| WDR61 | 2160.725 | 0.280143 | 0.171764 | 1.630976 | 0.102895 | 0.227198 |
| NEIL1 | 93.64644 | 0.487083 | 0.25089 | 1.941418 | 0.052208 | 0.14251 |
| MAN2C1 | 1353.857 | -0.23414 | 0.139572 | -1.6776 | 0.093426 | 0.213322 |
| DNAJA4 | 5511.513 | 0.027917 | 0.140684 | 0.198437 | 0.842703 | 0.906638 |
| MESDC1 | 1400.979 | -0.60236 | 0.180922 | -3.32941 | 8.70E-04 | 0.0095 |
| TPM1 | 1985.74 | 0.654801 | 0.33085 | 1.979144 | 0.0478 | 0.134794 |
| PIF1 | 142.8503 | -0.09755 | 0.274658 | -0.35517 | 0.722464 | 0.823619 |
| USP3 | 325.8248 | 3.41E-04 | 0.177539 | 0.00192 | 0.998468 | 0.999371 |
| BBS4 | 943.6187 | 0.244487 | 0.177304 | 1.378915 | 0.167921 | 0.317598 |
| PML | 2489.413 | -0.81199 | 0.200034 | -4.05927 | 4.92E-05 | 0.001622 |
| CYP1A1 | 46.98859 | -0.14544 | 0.343848 | -0.42297 | 0.672314 | NA |
| ULK3 | 1467.013 | -0.1848 | 0.138521 | -1.33411 | 0.182169 | 0.335032 |
| CELF6 | 0.134133 | -1.48909 | 4.077584 | -0.36519 | 0.71497 | NA |
| SCAMP2 | 2325.234 | -0.19248 | 0.154778 | -1.2436 | 0.213646 | 0.372356 |
| HAPLN3 | 175.613 | 0.679814 | 0.313301 | 2.16984 | 0.030019 | 0.099019 |
| RHCG | 0.987658 | -1.76927 | 2.378195 | -0.74395 | 0.456905 | NA |
| POLG | 1945.541 | -0.31233 | 0.143744 | -2.1728 | 0.029796 | 0.098557 |
| RLBP1 | 2.192993 | -0.09431 | 1.365441 | -0.06907 | 0.944934 | NA |
| FANCI | 1369.515 | 0.041576 | 0.185839 | 0.223722 | 0.822973 | 0.893194 |
| ABHD2 | 3673.382 | 0.018579 | 0.202506 | 0.091748 | 0.926899 | 0.958156 |
| WDR93 | 7.663941 | -0.06801 | 0.775917 | -0.08765 | 0.930158 | NA |
| TICRR | 173.2976 | -0.57157 | 0.278726 | -2.05065 | 0.040301 | 0.120955 |
| NTRK3 | 1.006214 | 1.50962 | 2.386044 | 0.632687 | 0.526938 | NA |
| DET1 | 227.1798 | -0.17941 | 0.194323 | -0.92327 | 0.355864 | 0.523878 |
| MFGE8 | 17780.96 | 0.043317 | 0.192794 | 0.224681 | 0.822227 | 0.893037 |
| ZNF710 | 392.4607 | -0.64979 | 0.18991 | -3.42155 | 6.23E-04 | 0.007763 |
| EFTUD1 | 559.9045 | 0.353018 | 0.160364 | 2.201349 | 0.027711 | 0.094239 |
| SEC11A | 4104.081 | 0.220763 | 0.141538 | 1.559743 | 0.118821 | 0.250377 |
| SCYL1 | 3242.519 | -0.2714 | 0.1353 | -2.00589 | 0.044868 | 0.129419 |
| AKT1 | 8601.395 | -0.19538 | 0.145217 | -1.34541 | 0.178492 | 0.331093 |

| | | | | | | |
|---|---|---|---|---|---|---|
| CDCA5 | 736.9223 | -0.3352 | 0.277054 | -1.20986 | 0.226331 | 0.387683 |
| C11orf73 | 822.3069 | 0.138168 | 0.170429 | 0.810708 | 0.417533 | 0.580424 |
| CCDC81 | 4.956451 | 0.277358 | 1.069143 | 0.25942 | 0.795311 | NA |
| SESN3 | 2621.571 | 0.030477 | 0.279238 | 0.109144 | 0.913089 | 0.94937 |
| ENDOD1 | 4692.867 | -0.08664 | 0.140776 | -0.61543 | 0.53827 | 0.681093 |
| CCDC82 | 305.8255 | 1.282026 | 0.741617 | 1.72869 | 0.083865 | 0.197265 |
| KLHL35 | 43.54699 | -0.45205 | 0.35853 | -1.26086 | 0.207361 | NA |
| TENM4 | 0.296509 | -0.51709 | 3.982155 | -0.12985 | 0.896684 | NA |
| SERPINH1 | 1829.361 | -0.54146 | 0.183111 | -2.95701 | 0.003106 | 0.021663 |
| CAPN5 | 1038.86 | -0.53638 | 0.131312 | -4.08482 | 4.41E-05 | 0.001543 |
| INTS4 | 591.192 | -0.18848 | 0.152459 | -1.23627 | 0.216359 | 0.375491 |
| PAK1 | 1860.219 | -0.31916 | 0.125763 | -2.53778 | 0.011156 | 0.050652 |
| RPS3 | 31633.84 | 0.058557 | 0.144698 | 0.404686 | 0.685708 | 0.79533 |
| ZC3H12C | 1925.151 | 0.603419 | 0.204053 | 2.957171 | 0.003105 | 0.021663 |
| TTC12 | 268.3676 | -0.0697 | 0.202459 | -0.34427 | 0.730646 | 0.829198 |
| NCAM1 | 2062.936 | -0.40821 | 0.171302 | -2.38295 | 0.017174 | 0.068277 |
| DRD2 | 1.243672 | -0.61289 | 1.854147 | -0.33055 | 0.740984 | NA |
| C11orf52 | 0.490493 | 0.776209 | 2.895261 | 0.268096 | 0.788625 | NA |
| NPAT | 925.8281 | 0.495567 | 0.294174 | 1.684608 | 0.092064 | 0.211025 |
| ATM | 3093.208 | 0.664453 | 0.295608 | 2.247749 | 0.024592 | 0.086816 |
| AASDHPPT | 721.5283 | 0.462056 | 0.246262 | 1.876276 | 0.060617 | 0.158426 |
| GLB1L2 | 52.48834 | -0.05045 | 0.322798 | -0.1563 | 0.8758 | 0.925779 |
| LAMTOR1 | 3540.594 | -0.18157 | 0.137832 | -1.3173 | 0.187737 | 0.341836 |
| P4HA3 | 5.798526 | -0.43487 | 0.813514 | -0.53456 | 0.592953 | NA |
| GRIK4 | 1.445321 | -0.30112 | 1.586081 | -0.18985 | 0.849425 | NA |
| HYOU1 | 3957.229 | -0.48285 | 0.138133 | -3.49556 | 4.73E-04 | 0.006586 |
| SLC22A8 | 0.163957 | 0.424913 | 4.079006 | 0.104171 | 0.917034 | NA |
| EI24 | 5398.64 | -0.24134 | 0.144459 | -1.67062 | 0.094796 | 0.215442 |
| CCDC15 | 58.94248 | 0.637817 | 0.329115 | 1.937977 | 0.052626 | 0.143256 |
| CHEK1 | 341.1536 | 0.256068 | 0.16073 | 1.593161 | 0.111124 | 0.239218 |
| FEZ1 | 4008.165 | -0.17044 | 0.127506 | -1.3367 | 0.181321 | 0.333991 |
| ESAM | 0.616518 | 2.49285 | 2.980788 | 0.836306 | 0.402983 | NA |
| KIRREL3 | 33.18748 | 0.347725 | 0.395063 | 0.880174 | 0.378765 | NA |
| MPZL2 | 4.807253 | -0.24307 | 0.940505 | -0.25845 | 0.796059 | NA |
| SCN2B | 0.544407 | -0.44425 | 2.542878 | -0.1747 | 0.861313 | NA |
| SIDT2 | 912.475 | -0.72037 | 0.164184 | -4.38759 | 1.15E-05 | 6.61E-04 |
| TMEM25 | 30.23383 | -0.63913 | 0.420029 | -1.52163 | 0.128103 | NA |
| TAGLN | 774.5028 | 0.192847 | 0.709324 | 0.271874 | 0.785719 | 0.869964 |
| ORAOV1 | 303.0951 | 0.419599 | 0.167598 | 2.503606 | 0.012293 | 0.054239 |
| GPHA2 | 0.555233 | -0.57009 | 3.296947 | -0.17291 | 0.862719 | NA |
| TRPT1 | 510.6318 | -0.05521 | 0.156451 | -0.35287 | 0.724189 | 0.824799 |
| NUDT22 | 808.0105 | -0.19368 | 0.148528 | -1.30403 | 0.192225 | 0.3468 |
| FERMT3 | 141.9899 | -0.53052 | 0.361295 | -1.46838 | 0.142002 | 0.282978 |
| PLCB3 | 1843.667 | -0.27612 | 0.165963 | -1.66376 | 0.09616 | 0.21722 |
| MRPL49 | 1846.297 | -0.11903 | 0.175452 | -0.67841 | 0.497511 | 0.648559 |
| CDC42EP2 | 74.98032 | -0.28669 | 0.268612 | -1.06731 | 0.285834 | 0.452344 |
| FAU | 6437.406 | 0.016733 | 0.126962 | 0.131797 | 0.895145 | 0.938944 |

| | | | | | | |
|---|---|---|---|---|---|---|
| TM7SF2 | 230.2138 | 0.07868 | 0.226239 | 0.347775 | 0.728009 | 0.827474 |
| VPS51 | 2317.677 | -0.40644 | 0.153007 | -2.65633 | 0.0079 | 0.040349 |
| HMGA2 | 4.021676 | 1.675267 | 1.088257 | 1.539404 | 0.123706 | NA |
| MMP3 | 1.97459 | 1.626118 | 1.48773 | 1.093019 | 0.274385 | NA |
| KLRF1 | 2.233342 | 0.108512 | 1.464909 | 0.074074 | 0.940951 | NA |
| CLEC1A | 0.230334 | -0.53318 | 4.076749 | -0.13079 | 0.895945 | NA |
| CWC15 | 1113.607 | 0.656944 | 0.18583 | 3.535196 | 4.07E-04 | 0.005986 |
| KLHL1 | 0.096202 | 0.408546 | 4.079639 | 0.100143 | 0.920231 | NA |
| DCUN1D2 | 228.2241 | 0.201212 | 0.193084 | 1.042091 | 0.297369 | 0.464559 |
| TMCO3 | 1520.747 | 0.144866 | 0.14873 | 0.97402 | 0.330047 | 0.498243 |
| TMEM218 | 559.8087 | -0.16268 | 0.166864 | -0.97492 | 0.3296 | 0.497782 |
| TIRAP | 143.4911 | -0.25327 | 0.221914 | -1.1413 | 0.253745 | 0.417799 |
| N6AMT2 | 165.1476 | 0.134705 | 0.218651 | 0.616071 | 0.537848 | 0.680998 |
| LATS2 | 2103.527 | 0.207436 | 0.205116 | 1.011313 | 0.311867 | 0.479045 |
| SAP18 | 4389.136 | 0.253087 | 0.161195 | 1.570062 | 0.116401 | 0.246693 |
| FAM124A | 342.7341 | -1.06452 | 0.2774 | -3.83751 | 1.24E-04 | 0.002947 |
| CTAGE5 | 19.12986 | 0.170573 | 0.472105 | 0.361303 | 0.717873 | NA |
| FSIP1 | 9.428021 | 0.381464 | 0.715952 | 0.532806 | 0.594168 | NA |
| DLG2 | 10.32307 | 0.690359 | 0.646557 | 1.067746 | 0.285635 | NA |
| CCDC83 | 0.162376 | 0.424913 | 4.079006 | 0.104171 | 0.917034 | NA |
| PRSS23 | 4031.78 | 0.118781 | 0.195593 | 0.607289 | 0.543659 | 0.685451 |
| DIXDC1 | 695.2382 | 0.189515 | 0.190286 | 0.995947 | 0.319276 | 0.487181 |
| DLAT | 1535.526 | 0.173328 | 0.145114 | 1.194422 | 0.232313 | 0.394566 |
| PIH1D2 | 26.5364 | 0.899771 | 0.413563 | 2.175659 | 0.029581 | NA |
| C11orf57 | 728.3127 | 0.510743 | 0.173939 | 2.936333 | 0.003321 | 0.022535 |
| TIMM8B | 1396.392 | 0.236827 | 0.182444 | 1.298081 | 0.19426 | 0.349445 |
| IL18 | 1.132682 | 0.571077 | 1.952886 | 0.292427 | 0.76996 | NA |
| TEX12 | 0.558248 | 0.982076 | 2.804738 | 0.350149 | 0.726227 | NA |
| PTS | 793.6565 | 0.669477 | 0.204057 | 3.280828 | 0.001035 | 0.010582 |
| FREM2 | 67.61936 | -0.17556 | 0.268174 | -0.65464 | 0.5127 | 0.659941 |
| FOXO1 | 352.4641 | 0.08232 | 0.176943 | 0.465233 | 0.641764 | 0.76165 |
| ABCB9 | 513.2968 | -0.66028 | 0.155221 | -4.25382 | 2.10E-05 | 9.51E-04 |
| RILPL2 | 412.5211 | 0.090245 | 0.195966 | 0.460512 | 0.645149 | 0.764212 |
| DHX37 | 1476.969 | -0.30819 | 0.176151 | -1.74958 | 0.080192 | 0.191745 |
| UBC | 22084.98 | -0.21835 | 0.154134 | -1.41666 | 0.156582 | 0.30277 |
| CACNA2D4 | 4.503699 | -0.3195 | 1.019213 | -0.31347 | 0.753921 | NA |
| DCP1B | 275.766 | -0.2619 | 0.187085 | -1.39988 | 0.161549 | 0.309097 |
| CACNA1C | 1.592559 | 2.773497 | 1.813178 | 1.529633 | 0.126108 | NA |
| KCNA6 | 0.260158 | 1.240347 | 4.047124 | 0.306476 | 0.759242 | NA |
| C12orf45 | 147.1416 | 0.445267 | 0.218032 | 2.042209 | 0.041131 | 0.122803 |
| C12orf23 | 1200.125 | 0.603178 | 0.190607 | 3.164511 | 0.001553 | 0.013836 |
| BTBD11 | 0.258578 | 1.233151 | 4.047809 | 0.304646 | 0.760635 | NA |
| UBE3B | 1527.311 | -0.39263 | 0.127479 | -3.07996 | 0.00207 | 0.016597 |
| RAD9B | 11.18987 | 0.611863 | 0.631681 | 0.968627 | 0.332731 | NA |
| PLBD2 | 4181.383 | -0.40831 | 0.138387 | -2.95046 | 0.003173 | 0.022004 |
| SLC2A13 | 688.3862 | 0.41948 | 0.316857 | 1.32388 | 0.185543 | 0.339342 |
| GXYLT1 | 341.641 | 0.669242 | 0.361245 | 1.852599 | 0.06394 | 0.164427 |

| | | | | | | |
|---|---|---|---|---|---|---|
| TWF1 | 1144.214 | 1.041164 | 0.225665 | 4.613756 | 3.95E-06 | 3.70E-04 |
| TEX30 | 479.4794 | 0.598856 | 0.176534 | 3.392296 | 6.93E-04 | 0.00825 |
| AKAP6 | 349.8824 | 0.18226 | 0.212631 | 0.857164 | 0.391354 | 0.556629 |
| NPAS3 | 6.383668 | 0.20968 | 0.818243 | 0.256256 | 0.797753 | NA |
| FAM177A1 | 1760.134 | 0.279797 | 0.210797 | 1.327331 | 0.184399 | 0.337667 |
| MBIP | 268.5165 | 0.82753 | 0.251983 | 3.284076 | 0.001023 | 0.010511 |
| MIPOL1 | 75.69758 | 0.906983 | 0.439093 | 2.065583 | 0.038868 | 0.117937 |
| KCTD14 | 23.43343 | -0.61091 | 0.534866 | -1.14217 | 0.253385 | NA |
| NDUFC2 | 651.7318 | -0.13237 | 0.15164 | -0.87291 | 0.382714 | 0.548751 |
| ME3 | 32.15267 | -0.06096 | 0.386344 | -0.15779 | 0.874623 | NA |
| NUBPL | 404.5518 | -0.00204 | 0.185523 | -0.01102 | 0.99121 | 0.996168 |
| VIPAS39 | 655.9611 | -0.34026 | 0.143565 | -2.37004 | 0.017786 | 0.069751 |
| PTPRO | 108.4237 | 0.942228 | 0.480274 | 1.961858 | 0.049779 | 0.138075 |
| EPS8 | 2650.104 | 0.595009 | 0.180489 | 3.296653 | 9.78E-04 | 0.010189 |
| ACAD8 | 747.9221 | -0.05401 | 0.162372 | -0.33262 | 0.739423 | 0.835401 |
| THYN1 | 514.6569 | 0.237721 | 0.147929 | 1.606997 | 0.108055 | 0.234829 |
| VPS26B | 2543.004 | -0.33592 | 0.138261 | -2.42957 | 0.015117 | 0.062398 |
| NCAPD3 | 1192.536 | 0.020321 | 0.177229 | 0.114657 | 0.908717 | 0.947313 |
| ANO4 | 566.569 | 0.406406 | 0.145426 | 2.79459 | 0.005197 | 0.030347 |
| TEX9 | 35.57315 | 1.046538 | 0.428502 | 2.442316 | 0.014593 | NA |
| FLI1 | 77.18097 | 0.680197 | 0.412436 | 1.649219 | 0.099103 | 0.222006 |
| KCNJ1 | 0.096202 | 0.408546 | 4.079639 | 0.100143 | 0.920231 | NA |
| TMEM45B | 0.804769 | -0.21293 | 2.457871 | -0.08663 | 0.930964 | NA |
| AMN1 | 88.82719 | 0.208584 | 0.251879 | 0.82811 | 0.407608 | 0.571292 |
| BICD1 | 425.362 | 0.100772 | 0.18418 | 0.547137 | 0.584284 | 0.717551 |
| SAV1 | 559.7799 | 0.543454 | 0.17146 | 3.169559 | 0.001527 | 0.013701 |
| CCDC122 | 7.932522 | 0.332414 | 0.781041 | 0.425604 | 0.670397 | NA |
| SERP2 | 258.7758 | -0.08812 | 0.249827 | -0.35272 | 0.724302 | 0.824841 |
| SACS | 778.2974 | 0.878748 | 0.306394 | 2.868032 | 0.00413 | 0.025962 |
| CCDC175 | 7.387064 | 0.377913 | 0.782287 | 0.483088 | 0.629033 | NA |
| PABPC3 | 8.795746 | -0.88842 | 0.752289 | -1.18096 | 0.237619 | NA |
| CENPJ | 153.7816 | 0.95886 | 0.369371 | 2.595925 | 0.009434 | 0.045406 |
| GLT1D1 | 5.439952 | -1.04952 | 0.877183 | -1.19646 | 0.231515 | NA |
| TMEM132I | 910.0127 | -0.37531 | 0.173867 | -2.1586 | 0.030881 | 0.101132 |
| HSPB8 | 85.74673 | -0.82278 | 0.302507 | -2.71988 | 0.006531 | 0.035384 |
| POU4F1 | 1.237926 | 1.940943 | 2.028448 | 0.956861 | 0.338637 | NA |
| RNF219 | 145.9995 | 0.684991 | 0.238155 | 2.876239 | 0.004024 | 0.025476 |
| CYSLTR2 | 208.9792 | 0.279776 | 0.238377 | 1.17367 | 0.240527 | 0.403783 |
| ARL11 | 0.544407 | -0.44425 | 2.542878 | -0.1747 | 0.861313 | NA |
| KCNK13 | 12.43513 | -0.26008 | 0.70838 | -0.36715 | 0.713505 | NA |
| GUCY1A2 | 5.923302 | 0.214784 | 0.855262 | 0.251133 | 0.801712 | NA |
| CWF19L2 | 154.9653 | 0.766542 | 0.341451 | 2.244956 | 0.024771 | 0.08722 |
| USP12 | 826.7121 | 0.790496 | 0.183433 | 4.309456 | 1.64E-05 | 8.29E-04 |
| PAN3 | 554.1984 | 0.280798 | 0.188547 | 1.489273 | 0.136415 | 0.275453 |
| PFKM | 6400.908 | -0.35363 | 0.122058 | -2.89719 | 0.003765 | 0.024485 |
| TMEM123 | 3628.237 | 0.783486 | 0.181572 | 4.315016 | 1.60E-05 | 8.22E-04 |
| GRIA4 | 1.207956 | 1.431596 | 1.927393 | 0.742763 | 0.457625 | NA |

| | | | | | | |
|---|---|---|---|---|---|---|
| GPR180 | 756.919 | 0.470636 | 0.235747 | 1.996359 | 0.045895 | 0.131389 |
| FARP1 | 4512.837 | -0.55738 | 0.155157 | -3.59234 | 3.28E-04 | 0.005201 |
| IFLTD1 | 0.258578 | 1.233151 | 4.047809 | 0.304646 | 0.760635 | NA |
| MED21 | 614.6026 | 0.399185 | 0.192257 | 2.076309 | 0.037865 | 0.115728 |
| RASSF3 | 6658.293 | 0.240758 | 0.190392 | 1.264536 | 0.206038 | 0.363046 |
| PTPRR | 0.488709 | 2.119056 | 3.307564 | 0.64067 | 0.521737 | NA |
| TMEM251 | 487.7416 | 0.176696 | 0.170632 | 1.035539 | 0.300417 | 0.467283 |
| ING1 | 379.8398 | 0.357492 | 0.16333 | 2.188769 | 0.028614 | 0.096013 |
| TEX29 | 1.392783 | 0.953702 | 1.968261 | 0.484541 | 0.628002 | NA |
| ADPRHL1 | 15.14204 | 0.17186 | 0.578984 | 0.29683 | 0.766596 | NA |
| TUBGCP5 | 518.7563 | 0.324569 | 0.153365 | 2.116322 | 0.034317 | 0.10846 |
| GOLGA8I | 1.301528 | -0.75253 | 1.814473 | -0.41474 | 0.678336 | NA |
| PPP2R5E | 1008.507 | 0.269054 | 0.198178 | 1.357637 | 0.174579 | 0.326595 |
| THY1 | 513.3824 | -0.0276 | 0.25547 | -0.10805 | 0.913956 | 0.9501 |
| TBCEL | 616.3452 | 0.636865 | 0.195655 | 3.255034 | 0.001134 | 0.011192 |
| UBASH3B | 946.3995 | 0.088216 | 0.176229 | 0.500577 | 0.616669 | 0.742224 |
| ROBO4 | 0.394291 | 0.298745 | 3.141711 | 0.09509 | 0.924243 | NA |
| ROBO3 | 152.2832 | 0.250801 | 0.201118 | 1.247036 | 0.212384 | 0.370976 |
| TBRG1 | 789.3244 | 0.408384 | 0.152328 | 2.680959 | 0.007341 | 0.038441 |
| NRGN | 139.0806 | -0.04983 | 0.312409 | -0.15951 | 0.873267 | 0.924503 |
| OXA1L | 3464.356 | -0.30433 | 0.142396 | -2.13719 | 0.032583 | 0.105021 |
| SLC7A7 | 34.71597 | -0.81232 | 0.368398 | -2.20501 | 0.027454 | NA |
| TMBIM4 | 512.382 | 0.399819 | 0.25078 | 1.594298 | 0.110869 | 0.23892 |
| GRIP1 | 80.91727 | -0.37039 | 0.45675 | -0.81093 | 0.417406 | 0.580397 |
| KIF5A | 24.38951 | 0.374391 | 0.421419 | 0.888407 | 0.374322 | NA |
| ELMSAN1 | 1727.863 | -0.63678 | 0.24115 | -2.64058 | 0.008276 | 0.041548 |
| FAM161B | 120.7761 | -0.08876 | 0.214816 | -0.41321 | 0.679451 | 0.790858 |
| BATF | 28.4719 | -0.65015 | 0.396486 | -1.63978 | 0.101051 | NA |
| C15orf26 | 0.288605 | 1.443619 | 4.029058 | 0.358302 | 0.720117 | NA |
| ADAMTSL3 | 5.832313 | -0.55584 | 0.848369 | -0.65519 | 0.512349 | NA |
| SLC28A1 | 0.163957 | 0.424913 | 4.079006 | 0.104171 | 0.917034 | NA |
| WHAMM | 301.0686 | 0.309252 | 0.170781 | 1.810816 | 0.070169 | 0.174687 |
| ANKRD9 | 1500.916 | -0.07091 | 0.158423 | -0.44759 | 0.654446 | 0.771756 |
| C14orf2 | 2228.673 | 0.362057 | 0.156376 | 2.315297 | 0.020597 | 0.077246 |
| NPTN | 2999.624 | 0.112262 | 0.133896 | 0.838431 | 0.401789 | 0.566424 |
| GALK2 | 489.6589 | 0.244199 | 0.167123 | 1.461195 | 0.143962 | 0.285539 |
| BUB1B | 478.4703 | 0.24041 | 0.206358 | 1.165013 | 0.244014 | 0.406747 |
| FAM92A1P | 39.16135 | 0.216967 | 0.361205 | 0.600675 | 0.548057 | NA |
| MMP14 | 21841.87 | -0.43552 | 0.166033 | -2.62312 | 0.008713 | 0.04302 |
| | 3.946754 | 0.63627 | 1.006195 | 0.632353 | 0.527156 | NA |
| DHRS4 | 177.437 | -0.18797 | 0.207718 | -0.90493 | 0.365502 | 0.532706 |
| DHRS1 | 861.6824 | -0.30909 | 0.140926 | -2.19326 | 0.028289 | 0.095301 |
| RNF111 | 910.4952 | -0.20564 | 0.148506 | -1.3847 | 0.166144 | 0.315242 |
| CCNB2 | 838.6909 | -0.15934 | 0.173637 | -0.91767 | 0.358792 | 0.52603 |
| FAM81A | 84.64037 | -0.10236 | 0.278375 | -0.36771 | 0.713089 | 0.816711 |
| MYO1E | 5004.951 | -0.05283 | 0.1292 | -0.4089 | 0.682611 | 0.793368 |
| SNX22 | 20.31866 | -0.17297 | 0.491006 | -0.35228 | 0.724626 | NA |

| | | | | | | |
|---|---|---|---|---|---|---|
| ACAN | 198.2298 | 0.959497 | 0.855449 | 1.12163 | 0.26202 | 0.426448 |
| CABP1 | 5.852879 | -1.36285 | 0.930906 | -1.46401 | 0.143192 | NA |
| AP3S2 | 482.9354 | -0.409 | 0.159413 | -2.56565 | 0.010298 | 0.048042 |
| SPPL3 | 1370.844 | -0.39184 | 0.158081 | -2.47871 | 0.013186 | 0.057008 |
| MEGF11 | 1.878681 | 0.344883 | 1.596577 | 0.216014 | 0.828977 | NA |
| C12orf43 | 114.9468 | -0.47335 | 0.230122 | -2.05694 | 0.039692 | 0.119794 |
| WDR66 | 49.68398 | 0.284696 | 0.309614 | 0.919521 | 0.357823 | 0.525131 |
| HPD | 2.582978 | -1.55512 | 1.400481 | -1.11042 | 0.266819 | NA |
| LRRC43 | 21.98637 | -0.51263 | 0.451199 | -1.13615 | 0.255895 | NA |
| FAM86C1 | 126.3565 | -0.29732 | 0.225161 | -1.32047 | 0.186677 | 0.340837 |
| GDPD5 | 533.7829 | -0.86926 | 0.241952 | -3.5927 | 3.27E-04 | 0.005201 |
| C11orf30 | 359.2611 | 0.037696 | 0.163625 | 0.230381 | 0.817796 | 0.890746 |
| ALG8 | 1687.586 | -0.14458 | 0.145669 | -0.99254 | 0.320936 | 0.489161 |
| GJD2 | 1.105576 | 1.732871 | 2.324315 | 0.745541 | 0.455945 | NA |
| ACTC1 | 24.65306 | 2.021568 | 0.653517 | 3.093368 | 0.001979 | NA |
| ADPGK | 1712.862 | 0.101668 | 0.158988 | 0.639467 | 0.522519 | 0.668732 |
| PTMS | 9391.889 | -0.23775 | 0.19011 | -1.25059 | 0.211086 | 0.36923 |
| PLA2G4D | 0.151697 | -1.546 | 4.075469 | -0.37934 | 0.704433 | NA |
| C1R | 144.1058 | -0.48336 | 0.237049 | -2.03908 | 0.041442 | 0.123384 |
| STARD9 | 416.2847 | 0.166145 | 0.169359 | 0.981022 | 0.326582 | 0.494777 |
| UBR1 | 1096.537 | 0.336268 | 0.188826 | 1.780831 | 0.07494 | 0.182764 |
| ISL2 | 0.354779 | 1.703954 | 3.657232 | 0.465914 | 0.641277 | NA |
| FAM86C2P | 32.85406 | -0.47561 | 0.367704 | -1.29346 | 0.195853 | NA |
| SIK3 | 894.2035 | -0.23718 | 0.175738 | -1.34964 | 0.177133 | 0.329294 |
| MPZL3 | 11.5335 | -0.40388 | 0.596266 | -0.67734 | 0.498189 | NA |
| PCSK7 | 2182.718 | -0.25149 | 0.161506 | -1.55716 | 0.119432 | 0.251264 |
| CXCR5 | 0.134133 | -1.48909 | 4.077584 | -0.36519 | 0.71497 | NA |
| VPS11 | 2315.633 | -0.37134 | 0.129036 | -2.8778 | 0.004005 | 0.025436 |
| NLRX1 | 703.5817 | -0.59882 | 0.163763 | -3.65662 | 2.56E-04 | 0.004508 |
| ITGA5 | 3205.96 | -0.4178 | 0.162359 | -2.57332 | 0.010073 | 0.047424 |
| ZNF385A | 490.214 | -0.21444 | 0.174579 | -1.22832 | 0.219328 | 0.379029 |
| FMNL3 | 1238.637 | -0.42925 | 0.141221 | -3.03956 | 0.002369 | 0.018039 |
| AQP5 | 2.046804 | -0.69781 | 1.34901 | -0.51728 | 0.604962 | NA |
| RACGAP1 | 1848.387 | -0.21444 | 0.219074 | -0.97884 | 0.327657 | 0.495835 |
| LARP4 | 1220.184 | 0.712843 | 0.252556 | 2.822513 | 0.004765 | 0.028601 |
| GRASP | 1445.759 | -0.47058 | 0.155556 | -3.02514 | 0.002485 | 0.018718 |
| SHANK2 | 4.128953 | -0.30277 | 1.018811 | -0.29718 | 0.766329 | NA |
| CLPB | 832.4749 | -0.38675 | 0.160376 | -2.41154 | 0.015885 | 0.064533 |
| NEU3 | 446.1017 | -0.72298 | 0.192488 | -3.75595 | 1.73E-04 | 0.003542 |
| SLC25A45 | 157.6597 | -0.31258 | 0.214733 | -1.45566 | 0.145487 | 0.287692 |
| SYVN1 | 1226.661 | -0.49245 | 0.170445 | -2.88918 | 0.003862 | 0.024886 |
| ZFPL1 | 198.9435 | -0.50452 | 0.201689 | -2.50147 | 0.012368 | 0.054506 |
| RPS6KA4 | 2567.275 | -0.35886 | 0.137205 | -2.61553 | 0.008909 | 0.043786 |
| LRP5 | 765.0245 | -0.32095 | 0.161558 | -1.9866 | 0.046967 | 0.133364 |
| TPCN2 | 6285.233 | -0.6623 | 0.17349 | -3.81753 | 1.35E-04 | 0.00305 |
| FAM86FP | 29.46254 | -0.78548 | 0.397973 | -1.97369 | 0.048417 | NA |
| SLITRK5 | 9.271597 | -0.73279 | 0.742952 | -0.98632 | 0.323974 | NA |

| | | | | | | |
|---|---|---|---|---|---|---|
| FAT3 | 4.624772 | -0.81996 | 1.064429 | -0.77033 | 0.441104 | NA |
| FBXO33 | 775.3425 | 0.699544 | 0.149337 | 4.684336 | 2.81E-06 | 2.97E-04 |
| LRFN5 | 11.76148 | -0.2463 | 0.588555 | -0.41849 | 0.675589 | NA |
| SPTSSA | 1951.653 | 0.508655 | 0.196062 | 2.594356 | 0.009477 | 0.045537 |
| TSHR | 1.436653 | 1.111781 | 1.831046 | 0.607184 | 0.543729 | NA |
| CFL2 | 1158.843 | 0.328147 | 0.176522 | 1.858959 | 0.063033 | 0.16273 |
| SUGT1 | 680.3198 | 0.59921 | 0.173979 | 3.444147 | 5.73E-04 | 0.007385 |
| GTF2A1 | 593.6175 | 0.501258 | 0.262149 | 1.912115 | 0.055861 | 0.149468 |
| PGM2L1 | 481.9271 | 1.483415 | 0.183173 | 8.09844 | 5.57E-16 | 3.30E-12 |
| INPPL1 | 4694.166 | -0.24448 | 0.132523 | -1.84485 | 0.065059 | 0.166294 |
| GJB2 | 51.32571 | -0.405 | 0.594456 | -0.68129 | 0.495689 | 0.646895 |
| CRYL1 | 9651.377 | -0.29526 | 0.187167 | -1.57751 | 0.114678 | 0.244173 |
| HEPACAM | 1.258707 | 1.465113 | 1.941661 | 0.754567 | 0.450509 | NA |
| SKA3 | 209.0984 | 0.270021 | 0.209114 | 1.291261 | 0.196613 | 0.352003 |
| MICU2 | 669.8127 | 0.776331 | 0.226736 | 3.423947 | 6.17E-04 | 0.007733 |
| C11orf82 | 145.1754 | 0.466947 | 0.288433 | 1.618911 | 0.105466 | 0.231238 |
| PCF11 | 586.5385 | 0.620231 | 0.272204 | 2.278548 | 0.022694 | 0.082325 |
| PKNOX2 | 1620.178 | -0.66134 | 0.140334 | -4.71265 | 2.45E-06 | 2.71E-04 |
| LRR1 | 377.2548 | -0.06511 | 0.224042 | -0.29063 | 0.771337 | 0.858691 |
| RPL36AL | 3599.164 | 0.196535 | 0.154498 | 1.272084 | 0.203343 | 0.359796 |
| DNAAF2 | 215.1606 | 0.760292 | 0.189136 | 4.019822 | 5.82E-05 | 0.001793 |
| KLHDC2 | 1017.505 | 0.141809 | 0.157649 | 0.899527 | 0.368372 | 0.535506 |
| EML5 | 40.84712 | 0.786677 | 0.437965 | 1.796211 | 0.072461 | NA |
| NEMF | 651.2814 | 0.932923 | 0.686106 | 1.359736 | 0.173914 | 0.325916 |
| RPUSD4 | 823.5779 | 0.020162 | 0.143427 | 0.140575 | 0.888206 | 0.934251 |
| ARF6 | 2277.49 | -0.09965 | 0.128419 | -0.77597 | 0.437764 | 0.597921 |
| TTC8 | 670.4249 | 0.427959 | 0.177353 | 2.413037 | 0.01582 | 0.064361 |
| TMEM63C | 0.192403 | 0.975392 | 4.060716 | 0.240202 | 0.810174 | NA |
| NOXRED1 | 2.912882 | 1.771888 | 1.294643 | 1.368631 | 0.171115 | NA |
| AMER2 | 0.399595 | -1.31595 | 3.579466 | -0.36764 | 0.713143 | NA |
| KBTBD6 | 404.5194 | 0.100594 | 0.217797 | 0.46187 | 0.644175 | 0.763364 |
| DACT1 | 1152.853 | 0.604766 | 0.349802 | 1.728881 | 0.08383 | 0.197223 |
| DACH1 | 0.544554 | -2.65036 | 3.220086 | -0.82307 | 0.410468 | NA |
| LOH12CR1 | 285.9705 | -0.32377 | 0.186112 | -1.73968 | 0.081915 | 0.194416 |
| TMEM55B | 1609.368 | -0.1139 | 0.166747 | -0.68309 | 0.494551 | 0.645623 |
| METTL17 | 485.847 | 0.099574 | 0.158554 | 0.628016 | 0.529994 | 0.674874 |
| SLC39A2 | 1.353328 | 1.603733 | 1.750302 | 0.916261 | 0.35953 | NA |
| NDRG2 | 345.4519 | -0.154 | 0.199938 | -0.77026 | 0.441146 | 0.601277 |
| ARHGEF40 | 2928.155 | -0.86496 | 0.173 | -4.99981 | 5.74E-07 | 1.12E-04 |
| ZNF219 | 1774.627 | -0.06999 | 0.309375 | -0.22625 | 0.82101 | 0.892334 |
| C12orf50 | 0.410275 | 0.244283 | 3.100889 | 0.078778 | 0.937209 | NA |
| PPP1R36 | 53.52765 | 0.119573 | 0.348308 | 0.343296 | 0.731376 | 0.829789 |
| METTL3 | 965.6576 | 0.091654 | 0.156074 | 0.587248 | 0.557037 | 0.696198 |
| SALL2 | 231.4188 | -0.55991 | 0.187706 | -2.98292 | 0.002855 | 0.020475 |
| FAM194B | 0.574434 | -0.17781 | 2.756096 | -0.06452 | 0.948559 | NA |
| ZFYVE1 | 2113.32 | -0.45176 | 0.143327 | -3.15198 | 0.001622 | 0.014248 |
| E2F7 | 1256.089 | -0.04755 | 0.212193 | -0.22409 | 0.822687 | 0.893127 |

| | | | | | | |
|---|---|---|---|---|---|---|
| ARHGAP42 | 610.2853 | 0.649071 | 0.247416 | 2.623401 | 0.008706 | 0.043002 |
| ISCA2 | 452.4445 | -0.05531 | 0.204969 | -0.26987 | 0.787263 | 0.871205 |
| OTOGL | 17.64367 | 1.17725 | 0.742392 | 1.585753 | 0.112795 | NA |
| TTC7B | 1248.373 | -0.32081 | 0.152674 | -2.10125 | 0.035619 | 0.111239 |
| TC2N | 0.096202 | 0.408546 | 4.079639 | 0.100143 | 0.920231 | NA |
| CPSF2 | 1702.267 | 0.493963 | 0.207006 | 2.386228 | 0.017022 | 0.0679 |
| MOAP1 | 1187.807 | 0.110808 | 0.165812 | 0.668272 | 0.50396 | 0.653751 |
| IFI27L1 | 699.5119 | -0.16321 | 0.160504 | -1.01688 | 0.30921 | 0.47662 |
| IFI27 | 1851.045 | -1.33487 | 0.533796 | -2.50072 | 0.012394 | 0.054541 |
| CLMN | 123.5224 | -0.73095 | 0.349996 | -2.08846 | 0.036756 | 0.113713 |
| CCDC38 | 0.913027 | 1.87659 | 2.13519 | 0.878887 | 0.379463 | NA |
| SMCO4 | 90.59454 | 0.172232 | 0.27767 | 0.620278 | 0.535075 | 0.678307 |
| KIAA1731 | 182.3507 | 1.080603 | 0.362318 | 2.982468 | 0.002859 | 0.020475 |
| TAF1D | 530.836 | 0.917357 | 0.267471 | 3.429746 | 6.04E-04 | 0.007625 |
| AMOTL1 | 2128.034 | -0.09077 | 0.15246 | -0.59539 | 0.55158 | 0.692542 |
| LIPC | 0.422534 | 1.924583 | 3.001758 | 0.641152 | 0.521424 | NA |
| CEP57 | 658.1982 | 0.720248 | 0.208049 | 3.46191 | 5.36E-04 | 0.007049 |
| TCP11L2 | 149.1128 | -0.03906 | 0.293041 | -0.13331 | 0.89395 | 0.938105 |
| SPRED1 | 2327.715 | 0.25286 | 0.199095 | 1.270043 | 0.204069 | 0.360758 |
| GPR176 | 901.2023 | 0.221579 | 0.252011 | 0.879242 | 0.37927 | 0.545595 |
| JAM3 | 2486.643 | 0.03896 | 0.146474 | 0.265984 | 0.790251 | 0.872781 |
| CMTM5 | 90.15076 | -0.37818 | 0.268485 | -1.40857 | 0.158964 | 0.305729 |
| | 0.096202 | 0.408546 | 4.079639 | 0.100143 | 0.920231 | NA |
| GLB1L3 | 0.151697 | -1.546 | 4.075469 | -0.37934 | 0.704433 | NA |
| ADAMTS15 | 2.184788 | 0.203834 | 1.444961 | 0.141065 | 0.887819 | NA |
| SVOP | 0.653015 | 0.2552 | 2.787103 | 0.091565 | 0.927044 | NA |
| AMN | 10.59589 | 0.84723 | 0.679202 | 1.247391 | 0.212254 | NA |
| RAB8B | 1246.398 | 0.552671 | 0.187448 | 2.948401 | 0.003194 | 0.02208 |
| IKBIP | 754.3122 | 0.988307 | 0.247655 | 3.990661 | 6.59E-05 | 0.001943 |
| RPUSD2 | 359.8305 | -0.13823 | 0.184412 | -0.7496 | 0.453497 | 0.612394 |
| ZFYVE19 | 863.0248 | -0.1792 | 0.170632 | -1.05023 | 0.293611 | 0.460845 |
| PPP1R14D | 0.163957 | 0.424913 | 4.079006 | 0.104171 | 0.917034 | NA |
| SPINT1 | 403.2246 | -0.0406 | 0.317393 | -0.12793 | 0.898205 | 0.940157 |
| FBN1 | 1330.229 | -0.30586 | 0.143334 | -2.13392 | 0.032849 | 0.105592 |
| AVPR1A | 0.247898 | -0.5901 | 4.074633 | -0.14482 | 0.884851 | NA |
| DEPDC4 | 19.97163 | 0.340882 | 0.461099 | 0.739282 | 0.459736 | NA |
| CKB | 618.1477 | -0.14636 | 0.288088 | -0.50804 | 0.611425 | 0.738281 |
| TRMT61A | 929.9448 | -0.01923 | 0.192863 | -0.09971 | 0.920573 | 0.953884 |
| BAG5 | 887.98 | 0.313648 | 0.19065 | 1.645151 | 0.099939 | 0.223198 |
| LARP6 | 678.3859 | 0.049325 | 0.226355 | 0.21791 | 0.827499 | 0.895973 |
| SENP8 | 56.25996 | 0.621061 | 0.326318 | 1.903241 | 0.057009 | 0.151364 |
| COPS2 | 1429.298 | 1.119833 | 0.304697 | 3.675239 | 2.38E-04 | 0.004321 |
| GABRB3 | 64.28581 | 0.909499 | 0.332653 | 2.73408 | 0.006255 | 0.034285 |
| FRS2 | 567.7467 | 0.589693 | 0.223388 | 2.639776 | 0.008296 | 0.041553 |
| CCT2 | 4294.781 | 0.416441 | 0.141723 | 2.938415 | 0.003299 | 0.022439 |
| ARIH1 | 1634.428 | -0.0175 | 0.133177 | -0.13143 | 0.895432 | 0.939039 |
| CLMP | 4.412962 | -0.35292 | 1.007869 | -0.35016 | 0.726217 | NA |

| | | | | | | |
|---|---|---|---|---|---|---|
| SCN3B | 0.586491 | 2.39072 | 2.689057 | 0.889055 | 0.373973 | NA |
| ZNF202 | 176.6027 | -0.06285 | 0.208767 | -0.30103 | 0.76339 | 0.853129 |
| FAM227B | 54.36149 | 0.541639 | 0.406399 | 1.332776 | 0.182605 | 0.335522 |
| CUL5 | 526.3915 | 0.935092 | 0.326462 | 2.864325 | 0.004179 | 0.026171 |
| MYRFL | 0.288605 | 1.443619 | 4.029058 | 0.358302 | 0.720117 | NA |
| C11orf65 | 6.524359 | 1.085018 | 0.845125 | 1.283855 | 0.199193 | NA |
| IDH3A | 1075.338 | 0.279186 | 0.148887 | 1.875154 | 0.060772 | 0.158689 |
| WDR72 | 43.12786 | 0.996171 | 0.420148 | 2.370998 | 0.01774 | NA |
| CRABP1 | 3.689846 | 0.080366 | 1.07659 | 0.074649 | 0.940494 | NA |
| PLD4 | 2.599116 | -1.00346 | 1.255664 | -0.79915 | 0.424205 | NA |
| XRRA1 | 186.0689 | -0.33012 | 0.201116 | -1.64145 | 0.100704 | 0.223943 |
| RNF169 | 418.351 | 0.108569 | 0.199805 | 0.543376 | 0.586871 | 0.719553 |
| PRTG | 99.70714 | 0.255516 | 0.286848 | 0.890772 | 0.373052 | 0.539727 |
| LEO1 | 346.8165 | 0.831482 | 0.348572 | 2.385395 | 0.017061 | 0.067932 |
| | 739.313 | 0.596424 | 0.245926 | 2.42522 | 0.015299 | 0.062857 |
| RIMKLB | 444.9774 | 0.164027 | 0.244857 | 0.669892 | 0.502927 | 0.652962 |
| A2ML1 | 0.572651 | 0.928649 | 2.780261 | 0.334015 | 0.738368 | NA |
| TMED3 | 6090.906 | -0.02114 | 0.166881 | -0.12666 | 0.89921 | 0.94096 |
| TMEM135 | 787.2609 | 0.186036 | 0.212016 | 0.877463 | 0.380235 | 0.546453 |
| IQCD | 28.51368 | 0.226494 | 0.414596 | 0.5463 | 0.58486 | NA |
| HSP90B1 | 9111.195 | 0.595181 | 0.31508 | 1.888988 | 0.058893 | 0.155152 |
| MMP10 | 0.410275 | 0.244283 | 3.100889 | 0.078778 | 0.937209 | NA |
| TMPRSS5 | 56.87489 | -0.54485 | 0.296254 | -1.83913 | 0.065896 | 0.167388 |
| B2M | 49905.64 | 0.03968 | 0.174305 | 0.227647 | 0.81992 | 0.892331 |
| ZNF592 | 1471.352 | -0.43629 | 0.186856 | -2.33492 | 0.019548 | 0.074586 |
| CASC4 | 687.9601 | 0.96142 | 0.244359 | 3.934453 | 8.34E-05 | 0.002283 |
| NNMT | 9.732849 | 1.592661 | 0.892692 | 1.784111 | 0.074406 | NA |
| AGBL1 | 0.984855 | -1.02676 | 2.366773 | -0.43382 | 0.664418 | NA |
| CATSPER2 | 1.296436 | 1.468688 | 1.796302 | 0.817618 | 0.413576 | NA |
| PPIB | 4154.628 | -0.09068 | 0.18035 | -0.50281 | 0.615096 | 0.741129 |
| FAM96A | 1401.919 | 0.498369 | 0.184052 | 2.707765 | 0.006774 | 0.036304 |
| KIAA0101 | 211.9514 | 0.014703 | 0.220365 | 0.066722 | 0.946803 | 0.970293 |
| KIF7 | 271.8565 | -0.00866 | 0.226452 | -0.03822 | 0.96951 | 0.983447 |
| PLIN1 | 2.436593 | -0.69042 | 1.311432 | -0.52646 | 0.598569 | NA |
| PEX11A | 209.9236 | 0.031247 | 0.186654 | 0.167405 | 0.867052 | 0.921131 |
| MESP1 | 13.03264 | -0.17198 | 0.57806 | -0.29751 | 0.766078 | NA |
| ANPEP | 111.6459 | -0.24954 | 0.351353 | -0.71023 | 0.477562 | 0.632358 |
| RBPMS2 | 42.26399 | -0.44428 | 0.479498 | -0.92654 | 0.354163 | NA |
| ANKDD1A | 85.5578 | 0.635856 | 0.255022 | 2.493339 | 0.012655 | 0.055359 |
| CLPX | 841.3446 | 0.409274 | 0.188156 | 2.175184 | 0.029616 | 0.098146 |
| ZBTB39 | 300.0261 | -0.31908 | 0.187493 | -1.70184 | 0.088785 | 0.205541 |
| TAC3 | 0.096202 | 0.408546 | 4.079639 | 0.100143 | 0.920231 | NA |
| MYO1A | 0.096202 | 0.408546 | 4.079639 | 0.100143 | 0.920231 | NA |
| TMEM194/ | 744.2651 | 0.201857 | 0.197315 | 1.02302 | 0.306298 | 0.473301 |
| NAB2 | 2445.787 | -6.70E-04 | 0.19367 | -0.00346 | 0.997241 | 0.999011 |
| VPS39 | 3000.671 | -0.32911 | 0.142097 | -2.31605 | 0.020555 | 0.07714 |
| STAT6 | 7546.61 | -0.30934 | 0.130412 | -2.37201 | 0.017692 | 0.069634 |

| | | | | | | |
|---|---|---|---|---|---|---|
| XRCC6BP1 | 198.9297 | 0.165868 | 0.227408 | 0.729385 | 0.465766 | 0.622931 |
| PIP4K2C | 1752.765 | -0.31857 | 0.146093 | -2.18058 | 0.029214 | 0.097148 |
| MTMR10 | 1965.396 | 0.317873 | 0.158427 | 2.006431 | 0.04481 | 0.129347 |
| C15orf48 | 1.801563 | -0.37483 | 1.531969 | -0.24467 | 0.806711 | NA |
| SCG5 | 4.50015 | -0.95214 | 0.94131 | -1.01151 | 0.311774 | NA |
| GREM1 | 9.90554 | 1.727078 | 0.813536 | 2.122927 | 0.03376 | NA |
| DIS3L | 534.7849 | -0.05926 | 0.185919 | -0.31873 | 0.749929 | 0.843492 |
| CCNDBP1 | 1988.336 | -0.03967 | 0.136752 | -0.29008 | 0.771753 | 0.858857 |
| EPB42 | 42.88933 | -0.20195 | 0.363193 | -0.55604 | 0.578187 | NA |
| SMAD3 | 593.13 | -0.24942 | 0.217157 | -1.14858 | 0.250729 | 0.414408 |
| MAP1A | 445.4072 | -0.27994 | 0.178762 | -1.566 | 0.117349 | 0.24807 |
| MARS | 3305.382 | -0.21575 | 0.125364 | -1.72095 | 0.085259 | 0.199473 |
| MBD6 | 1329.002 | -0.49875 | 0.163918 | -3.04268 | 0.002345 | 0.017934 |
| PDIA3 | 6758.852 | -0.07547 | 0.124169 | -0.60777 | 0.543342 | 0.68532 |
| C15orf39 | 679.5613 | -0.56645 | 0.178616 | -3.17135 | 0.001517 | 0.013637 |
| ISLR2 | 0.556813 | -0.57007 | 3.294615 | -0.17303 | 0.862627 | NA |
| FBXO22 | 964.2306 | 0.15662 | 0.140318 | 1.116176 | 0.264347 | 0.428902 |
| TBC1D2B | 1817.969 | -0.15164 | 0.139639 | -1.08591 | 0.277519 | 0.443456 |
| RNF214 | 444.1051 | -0.14828 | 0.152074 | -0.97509 | 0.329518 | 0.497761 |
| POP5 | 904.5649 | 0.0538 | 0.141718 | 0.379628 | 0.704221 | 0.810224 |
| ATP5L | 3083.807 | 0.276957 | 0.154548 | 1.792049 | 0.073125 | 0.180191 |
| ZNF641 | 437.2794 | -0.016 | 0.217373 | -0.07363 | 0.941307 | 0.967169 |
| CACNB3 | 1110.218 | -0.09183 | 0.19248 | -0.47707 | 0.633315 | 0.755023 |
| KMT2D | 2305.546 | -0.7251 | 0.275654 | -2.63047 | 0.008527 | 0.042312 |
| RHEBL1 | 87.21659 | 0.617842 | 0.306573 | 2.01532 | 0.043871 | 0.127724 |
| TUBA1A | 9614.322 | -0.38661 | 0.185012 | -2.08966 | 0.036649 | 0.113473 |
| TUBA1C | 1558.577 | -0.29386 | 0.179834 | -1.63408 | 0.102242 | 0.226262 |
| NCKAP5L | 1391.323 | -0.53804 | 0.142993 | -3.76267 | 1.68E-04 | 0.003497 |
| GPD1 | 0.594599 | -0.12152 | 2.771051 | -0.04385 | 0.965021 | NA |
| KRT80 | 9.400658 | 0.538967 | 0.850856 | 0.633442 | 0.526445 | NA |
| OTUB1 | 3636.959 | -0.13877 | 0.129868 | -1.06856 | 0.285269 | 0.451752 |
| RCOR2 | 8.021085 | -1.17148 | 0.803984 | -1.45709 | 0.145092 | NA |
| SPRYD3 | 1614.83 | -0.36393 | 0.200758 | -1.8128 | 0.069863 | 0.174179 |
| IGFBP6 | 92.50253 | -0.2887 | 0.409025 | -0.70583 | 0.480293 | 0.634828 |
| SOAT2 | 4.77459 | 0.402857 | 1.049301 | 0.383929 | 0.701031 | NA |
| NDUFV1 | 5007.267 | -0.21445 | 0.117963 | -1.81791 | 0.069079 | 0.172997 |
| CDK2AP2 | 1545.87 | -0.31551 | 0.156252 | -2.01924 | 0.043462 | 0.127 |
| NUDT8 | 110.7407 | -0.36729 | 0.245657 | -1.49513 | 0.134879 | 0.273379 |
| SLC3A2 | 13408.78 | -0.32906 | 0.145073 | -2.26821 | 0.023316 | 0.083915 |
| HRASLS5 | 0.315653 | -0.57305 | 3.892639 | -0.14721 | 0.882963 | NA |
| C11orf84 | 1370.868 | -0.1265 | 0.15863 | -0.79746 | 0.425183 | 0.587202 |
| ATG16L2 | 144.3751 | 0.101639 | 0.230002 | 0.441907 | 0.658556 | 0.77483 |
| C2CD3 | 640.4646 | -0.2092 | 0.163208 | -1.28182 | 0.199904 | 0.355943 |
| FADD | 758.1709 | -0.29503 | 0.191682 | -1.53916 | 0.123765 | 0.258089 |
| LTBP3 | 3325.129 | -0.64549 | 0.15086 | -4.27873 | 1.88E-05 | 8.88E-04 |
| NAALADL1 | 290.3657 | -0.25846 | 0.166422 | -1.55303 | 0.120415 | 0.252704 |
| SAC3D1 | 491.6702 | 0.012902 | 0.156454 | 0.082463 | 0.934279 | 0.96312 |

| | | | | | | |
|---|---|---|---|---|---|---|
| BATF2 | 135.6082 | -0.89301 | 0.263124 | -3.39387 | 6.89E-04 | 0.008244 |
| SF1 | 4808.933 | -0.44461 | 0.138066 | -3.22026 | 0.001281 | 0.012146 |
| MAP4K2 | 791.1561 | -0.3542 | 0.15987 | -2.21558 | 0.026721 | 0.091765 |
| C11orf85 | 1.20553 | -1.82229 | 1.887291 | -0.96556 | 0.334266 | NA |
| CCDC88B | 69.25158 | -0.25845 | 0.274537 | -0.94141 | 0.346497 | 0.514366 |
| PAFAH1B2 | 2594.666 | -0.01968 | 0.152729 | -0.12884 | 0.897487 | 0.939815 |
| MAPK1IP1l | 1434.488 | -0.16453 | 0.146595 | -1.12234 | 0.26172 | 0.426135 |
| C14orf183 | 0.163957 | 0.424913 | 4.079006 | 0.104171 | 0.917034 | NA |
| MGAT2 | 102.7242 | -0.39064 | 0.245111 | -1.59373 | 0.110997 | 0.238989 |
| DEGS2 | 0.162376 | 0.424913 | 4.079006 | 0.104171 | 0.917034 | NA |
| STIP1 | 6885 | -0.15221 | 0.130441 | -1.16689 | 0.243255 | 0.406166 |
| CHRM1 | 50.04418 | -0.91763 | 0.346201 | -2.65057 | 0.008036 | 0.040692 |
| TCTN2 | 1475.476 | -0.22363 | 0.140995 | -1.58611 | 0.112715 | 0.241329 |
| PPIP5K1 | 357.3848 | -0.43817 | 0.183165 | -2.39223 | 0.016746 | 0.067229 |
| ADAL | 290.3102 | 0.322351 | 0.199636 | 1.614696 | 0.106377 | 0.232417 |
| LCMT2 | 555.2585 | -0.21258 | 0.169795 | -1.25196 | 0.210583 | 0.368427 |
| TPTE2P5 | 0.909021 | -2.40928 | 2.210188 | -1.09008 | 0.275678 | NA |
| ANKRD49 | 195.5015 | 1.001321 | 0.23324 | 4.293102 | 1.76E-05 | 8.56E-04 |
| STXBP6 | 2079.437 | 0.336035 | 0.238966 | 1.406202 | 0.159664 | 0.306827 |
| GRM5 | 41.36026 | 0.642251 | 0.343357 | 1.870506 | 0.061414 | NA |
| JMJD7-PLA | 19.28902 | -0.36129 | 0.475756 | -0.7594 | 0.44761 | NA |
| FEM1B | 2993.408 | 0.553448 | 0.176318 | 3.13892 | 0.001696 | 0.014564 |
| MAP2K1 | 1559.49 | 0.054147 | 0.156005 | 0.347085 | 0.728527 | 0.827745 |
| UPF3A | 349.1706 | 0.717274 | 0.250243 | 2.866315 | 0.004153 | 0.026037 |
| CHST14 | 1114.09 | -0.3888 | 0.177804 | -2.18671 | 0.028764 | 0.096381 |
| CSNK1G1 | 708.6331 | -0.16201 | 0.173541 | -0.93355 | 0.350536 | 0.518479 |
| OR5AU1 | 0.096202 | 0.408546 | 4.079639 | 0.100143 | 0.920231 | NA |
| KIAA1024 | 52.3695 | 0.622611 | 0.34555 | 1.801798 | 0.071577 | 0.177335 |
| SNUPN | 487.1772 | 0.047937 | 0.16884 | 0.283918 | 0.776473 | 0.862136 |
| CRADD | 542.7487 | 0.14079 | 0.227097 | 0.619957 | 0.535286 | 0.678475 |
| SIN3A | 975.2847 | -0.34343 | 0.1385 | -2.47965 | 0.013151 | 0.056901 |
| PTPN9 | 2436.357 | -0.09401 | 0.13321 | -0.70576 | 0.480338 | 0.634828 |
| RNASE6 | 2.775239 | -0.66259 | 1.280711 | -0.51736 | 0.604904 | NA |
| BNC1 | 1.045726 | 0.338916 | 1.94625 | 0.174138 | 0.861757 | NA |
| C15orf40 | 507.9325 | 0.231848 | 0.18164 | 1.276418 | 0.201808 | 0.358267 |
| FAM103A1 | 96.90417 | 0.495872 | 0.25744 | 1.926164 | 0.054084 | 0.146184 |
| CHRNA5 | 43.59983 | 0.475993 | 0.350429 | 1.358315 | 0.174364 | NA |
| NRG4 | 8.304742 | 1.273919 | 0.872156 | 1.460655 | 0.14411 | NA |
| C15orf27 | 9.624717 | 0.936294 | 0.677602 | 1.381776 | 0.167041 | NA |
| LINGO1 | 371.5477 | -1.51145 | 1.028256 | -1.46992 | 0.141584 | 0.282335 |
| ONECUT1 | 8.255362 | -0.25712 | 0.775144 | -0.3317 | 0.740113 | NA |
| AVEN | 574.3662 | 0.187312 | 0.17373 | 1.078178 | 0.280954 | 0.447135 |
| WNT10B | 2.921985 | -0.2809 | 1.218233 | -0.23058 | 0.81764 | NA |
| OTUD7A | 18.17566 | 0.506866 | 0.479558 | 1.056944 | 0.290537 | NA |
| KLF13 | 2402.498 | -0.58985 | 0.16643 | -3.54412 | 3.94E-04 | 0.005852 |
| SERPINA6 | 0.162376 | 0.424913 | 4.079006 | 0.104171 | 0.917034 | NA |
| NIPA1 | 1409.216 | 0.397389 | 0.152533 | 2.605263 | 0.00918 | 0.044545 |

| | | | | | | |
|---|---|---|---|---|---|---|
| SIK2 | 1781.317 | -0.33678 | 0.150258 | -2.24137 | 0.025002 | 0.087822 |
| ANKK1 | 0.411855 | 0.250456 | 3.097618 | 0.080854 | 0.935558 | NA |
| USP50 | 0.394291 | 0.298745 | 3.141711 | 0.09509 | 0.924243 | NA |
| C14orf142 | 351.4246 | 0.431345 | 0.21233 | 2.031485 | 0.042206 | 0.124994 |
| | 19.87159 | -0.06309 | 0.600806 | -0.10501 | 0.91637 | NA |
| SLN | 0.424115 | 1.932531 | 3.45462 | 0.559405 | 0.575885 | NA |
| NFRKB | 952.5823 | -0.36085 | 0.136746 | -2.63881 | 0.00832 | 0.041599 |
| PRDM10 | 369.3027 | 0.150422 | 0.183226 | 0.820968 | 0.411665 | 0.574963 |
| FOS | 180.6565 | 0.436055 | 0.308004 | 1.415747 | 0.15685 | 0.303119 |
| TMED10 | 8083.463 | -0.15148 | 0.135871 | -1.1149 | 0.264892 | 0.429314 |
| KRT8 | 1.459015 | 1.061255 | 1.654615 | 0.641391 | 0.521269 | NA |
| SDR9C7 | 1.087795 | -0.34293 | 2.583446 | -0.13274 | 0.894399 | NA |
| DENND5B | 868.2944 | 0.406528 | 0.182547 | 2.226981 | 0.025949 | 0.090291 |
| WIBG | 628.9102 | 0.001048 | 0.139471 | 0.007517 | 0.994003 | 0.997382 |
| PA2G4 | 2610.95 | -0.09836 | 0.13448 | -0.73138 | 0.464545 | 0.621928 |
| SMAGP | 78.09451 | 0.009044 | 0.286431 | 0.031574 | 0.974812 | 0.98618 |
| STAT2 | 4157.957 | -0.60768 | 0.149982 | -4.05172 | 5.08E-05 | 0.00166 |
| GTSF1 | 0.163957 | 0.424913 | 4.079006 | 0.104171 | 0.917034 | NA |
| RNF34 | 999.2021 | -0.06828 | 0.172291 | -0.39632 | 0.691872 | 0.800756 |
| TMEM133 | 121.2692 | 0.492064 | 0.308436 | 1.595351 | 0.110634 | 0.238553 |
| ATF7 | 433.2475 | -0.38952 | 0.170236 | -2.28811 | 0.022131 | 0.081127 |
| AKAP13 | 2082.048 | -0.09367 | 0.141611 | -0.66145 | 0.508322 | 0.657089 |
| CDCA4 | 536.1916 | -0.33202 | 0.213846 | -1.55263 | 0.120511 | 0.252771 |
| TRIAP1 | 794.2133 | 0.225209 | 0.161841 | 1.391539 | 0.164062 | 0.312777 |
| PLA2G1B | 2.750847 | -0.07032 | 1.360375 | -0.05169 | 0.958777 | NA |
| MSANTD4 | 592.3357 | 0.642586 | 0.193758 | 3.316434 | 9.12E-04 | 0.009773 |
| TPT1-AS1 | 216.8041 | -0.26142 | 0.353986 | -0.73852 | 0.460201 | 0.618489 |
| PDGFD | 78.98 | -0.64692 | 0.398195 | -1.62464 | 0.10424 | 0.229312 |
| PYGO1 | 252.032 | 0.26237 | 0.243543 | 1.077305 | 0.281344 | 0.447516 |
| C11orf24 | 3469.968 | -0.36001 | 0.118475 | -3.03871 | 0.002376 | 0.018067 |
| TMEM126A | 689.5815 | 0.421138 | 0.166253 | 2.533111 | 0.011306 | 0.051105 |
| TMEM126B | 1094.979 | 0.767202 | 0.21911 | 3.50144 | 4.63E-04 | 0.006477 |
| CDC42BPG | 70.0077 | -0.26742 | 0.338555 | -0.78988 | 0.4296 | 0.591099 |
| FAM98B | 446.3947 | 0.325017 | 0.21716 | 1.496672 | 0.134479 | 0.272754 |
| CHST11 | 2098.627 | -0.23067 | 0.164504 | -1.40221 | 0.160851 | 0.308409 |
| KSR2 | 0.28583 | -2.21358 | 3.97376 | -0.55705 | 0.577494 | NA |
| MAP1LC3B | 31.24605 | -0.28466 | 0.393766 | -0.72291 | 0.469732 | NA |
| MAP6 | 66.23122 | 0.349602 | 0.319041 | 1.095789 | 0.273171 | 0.439051 |
| OR2AT4 | 0.47803 | 0.520083 | 2.97881 | 0.174594 | 0.861399 | NA |
| P2RY6 | 0.629929 | -1.1413 | 2.677641 | -0.42623 | 0.669937 | NA |
| ATF7IP | 868.0622 | -0.04114 | 0.230179 | -0.17875 | 0.858136 | 0.916009 |
| GPHN | 682.3999 | -0.29144 | 0.185245 | -1.57328 | 0.115654 | 0.245718 |
| PAH | 0.134133 | -1.48909 | 4.077584 | -0.36519 | 0.71497 | NA |
| SPATA5L1 | 232.4399 | -0.36607 | 0.185985 | -1.96829 | 0.049034 | 0.136938 |
| GATM | 105.508 | -0.26792 | 0.354076 | -0.75667 | 0.44925 | 0.60872 |
| RHNO1 | 531.8931 | -0.25321 | 0.166399 | -1.5217 | 0.128085 | 0.264307 |
| FBXL14 | 303.1763 | -0.27436 | 0.189366 | -1.44881 | 0.147389 | 0.290626 |

| | | | | | | |
|---|---|---|---|---|---|---|
| NINJ2 | 4.14663 | -0.08568 | 0.9959 | -0.08604 | 0.931438 | NA |
| FAM90A1 | 8.486397 | -0.86966 | 0.767061 | -1.13375 | 0.256898 | NA |
| C3AR1 | 26.05207 | -0.62535 | 0.418872 | -1.49294 | 0.135453 | NA |
| FRMD5 | 784.6393 | 0.146878 | 0.190201 | 0.772226 | 0.439981 | 0.59995 |
| TLN2 | 569.3202 | -0.60576 | 0.197526 | -3.06673 | 0.002164 | 0.017036 |
| LDHAL6B | 2.927421 | 0.739947 | 1.224727 | 0.604173 | 0.545728 | NA |
| ISG20 | 81.93496 | 0.140122 | 0.293234 | 0.47785 | 0.632757 | 0.754704 |
| CLEC7A | 0.096202 | 0.408546 | 4.079639 | 0.100143 | 0.920231 | NA |
| DPAGT1 | 1305.223 | -0.42323 | 0.15547 | -2.72227 | 0.006483 | 0.035193 |
| HINFP | 557.5168 | -0.06441 | 0.219172 | -0.29388 | 0.768853 | 0.85673 |
| STARD5 | 102.7 | -0.20101 | 0.238434 | -0.84303 | 0.399209 | 0.563878 |
| IL16 | 717.4785 | -0.85092 | 0.299249 | -2.84352 | 0.004462 | 0.02732 |
| ABCG4 | 43.68638 | -0.27444 | 0.352307 | -0.77897 | 0.435995 | NA |
| PDZD3 | 0.315653 | -0.57305 | 3.892639 | -0.14721 | 0.882963 | NA |
| C2CD2L | 552.1065 | -0.07709 | 0.187595 | -0.41092 | 0.681131 | 0.792212 |
| ARNT2 | 1482.595 | -0.79583 | 0.173848 | -4.57774 | 4.70E-06 | 4.09E-04 |
| TTC36 | 0.965361 | 0.923875 | 2.105329 | 0.438827 | 0.660787 | NA |
| IL17D | 3237.286 | -0.44392 | 0.230708 | -1.92418 | 0.054332 | 0.146721 |
| FIBP | 3287.467 | -0.16679 | 0.156988 | -1.06245 | 0.288029 | 0.454295 |
| CARNS1 | 2.65471 | 0.900497 | 1.246302 | 0.722536 | 0.469965 | NA |
| PPP1CA | 5347.71 | -0.23446 | 0.155119 | -1.51146 | 0.130672 | 0.267092 |
| CTSW | 0.230334 | -0.53318 | 4.076749 | -0.13079 | 0.895945 | NA |
| PDE3A | 232.9854 | -0.66205 | 0.400941 | -1.65123 | 0.098691 | 0.221377 |
| RASGRP1 | 1.48623 | -0.09738 | 1.890909 | -0.0515 | 0.958928 | NA |
| MRPL52 | 1527.09 | 0.045 | 0.154372 | 0.291503 | 0.770667 | 0.858187 |
| RND1 | 9.865322 | -0.82615 | 0.68072 | -1.21364 | 0.224885 | NA |
| RAD9A | 166.7352 | -0.26896 | 0.208094 | -1.29247 | 0.196194 | 0.351608 |
| EFEMP2 | 579.4248 | -0.16993 | 0.180345 | -0.94224 | 0.346072 | 0.513927 |
| OR10AD1 | 0.314073 | -0.57308 | 3.898934 | -0.14698 | 0.883146 | NA |
| TMEM134 | 767.739 | -0.16311 | 0.14334 | -1.13792 | 0.255155 | 0.419478 |
| FAM71D | 2.8894 | 1.458038 | 1.283254 | 1.136203 | 0.255871 | NA |
| CORO1B | 5984.267 | -0.26372 | 0.135101 | -1.95205 | 0.050932 | 0.140406 |
| MUS81 | 805.6708 | -0.13592 | 0.168124 | -0.80844 | 0.41884 | 0.581831 |
| CFL1 | 25438.61 | -0.22972 | 0.139457 | -1.64726 | 0.099505 | 0.222698 |
| NAA16 | 144.1932 | 1.144011 | 0.346062 | 3.305799 | 9.47E-04 | 0.00998 |
| HOXC5 | 112.3323 | -0.20906 | 0.285447 | -0.73239 | 0.463933 | 0.621415 |
| SNX32 | 86.66929 | -0.222 | 0.262147 | -0.84686 | 0.397074 | 0.56175 |
| OVOL1 | 24.90752 | 0.019784 | 0.482056 | 0.041041 | 0.967263 | NA |
| RARG | 1548.533 | -0.40397 | 0.154144 | -2.62072 | 0.008774 | 0.043288 |
| SSH3 | 1014.254 | -0.17809 | 0.157933 | -1.12762 | 0.259479 | 0.423872 |
| NADSYN1 | 1737.727 | -0.25704 | 0.188167 | -1.36605 | 0.171925 | 0.323108 |
| DHCR7 | 2816.452 | -0.34968 | 0.247251 | -1.41428 | 0.157281 | 0.303477 |
| NBEA | 231.8461 | 0.148636 | 0.220257 | 0.674829 | 0.499784 | 0.650235 |
| RNASEH2C | 2055.256 | -0.04719 | 0.165115 | -0.28577 | 0.775053 | 0.860888 |
| MYEOV | 157.6716 | 0.680011 | 0.440002 | 1.545473 | 0.122232 | 0.255566 |
| ANKRD13D | 1577.035 | -0.06442 | 0.164313 | -0.39205 | 0.695022 | 0.803279 |
| MRGPRF | 14.41378 | 0.257799 | 0.542362 | 0.475328 | 0.634553 | NA |

| | | | | | | |
|---|---|---|---|---|---|---|
| KAT5 | 870.0092 | -0.23476 | 0.144162 | -1.62847 | 0.103425 | 0.227984 |
| ADRBK1 | 2081.17 | -0.39896 | 0.130576 | -3.05537 | 0.002248 | 0.017429 |
| RELA | 3180.719 | -0.33394 | 0.139621 | -2.39177 | 0.016767 | 0.067265 |
| HECTD4 | 1240.692 | -0.55187 | 0.209066 | -2.63972 | 0.008297 | 0.041553 |
| TRMT112 | 3539.853 | 0.004087 | 0.145074 | 0.028172 | 0.977525 | 0.987689 |
| KDM2A | 1786.425 | -0.47892 | 0.161943 | -2.95731 | 0.003103 | 0.021663 |
| MRP63 | 1370.795 | 0.04058 | 0.129143 | 0.314227 | 0.753349 | 0.845894 |
| ESRRA | 1798.924 | -0.09358 | 0.140012 | -0.66839 | 0.503883 | 0.653751 |
| RHOD | 134.93 | 0.149268 | 0.236843 | 0.630241 | 0.528537 | 0.673809 |
| ABCD2 | 7.619279 | -0.63938 | 0.785479 | -0.814 | 0.415647 | NA |
| SYT12 | 0.096202 | 0.408546 | 4.079639 | 0.100143 | 0.920231 | NA |
| SLC2A14 | 0.268266 | -2.15396 | 4.038351 | -0.53338 | 0.593773 | NA |
| GPR137 | 1645.778 | -0.24239 | 0.203878 | -1.18889 | 0.234485 | 0.397117 |
| MAP3K11 | 10656.12 | -0.53385 | 0.136898 | -3.89962 | 9.63E-05 | 0.002488 |
| KCNK7 | 1.342381 | -0.86875 | 1.876521 | -0.46296 | 0.643393 | NA |
| OLR1 | 0.096202 | 0.408546 | 4.079639 | 0.100143 | 0.920231 | NA |
| GLIPR1L1 | 1.506338 | -0.39416 | 1.788825 | -0.22035 | 0.8256 | NA |
| EHBP1L1 | 1474.999 | -0.25199 | 0.182482 | -1.38092 | 0.167303 | 0.316783 |
| THAP2 | 51.52727 | 0.879762 | 0.363136 | 2.42268 | 0.015406 | 0.063089 |
| RNF26 | 872.3329 | -0.45249 | 0.147019 | -3.07778 | 0.002085 | 0.016597 |
| PPP1R14B | 2564.756 | 0.1692 | 0.124772 | 1.356077 | 0.175075 | 0.327007 |
| SSSCA1 | 693.2021 | -0.07366 | 0.15586 | -0.47258 | 0.636514 | 0.757389 |
| FKBP2 | 1990.12 | -0.17153 | 0.167181 | -1.02603 | 0.304879 | 0.472185 |
| VEGFB | 4757.089 | -0.31871 | 0.1743 | -1.82851 | 0.067473 | 0.169976 |
| PEAK1 | 1997.636 | 0.028331 | 0.177751 | 0.159383 | 0.873367 | 0.924526 |
| CSPG4 | 4556.297 | -0.92395 | 0.144666 | -6.38678 | 1.69E-10 | 2.23E-07 |
| SNX33 | 1941.522 | -0.49192 | 0.125486 | -3.92014 | 8.85E-05 | 0.002374 |
| CCDC41 | 70.95705 | 0.900812 | 0.457298 | 1.96986 | 0.048854 | 0.136693 |
| NUDT4 | 900.475 | 0.451299 | 0.182501 | 2.472855 | 0.013404 | 0.057636 |
| PC | 1071.893 | -0.81603 | 0.193214 | -4.22344 | 2.41E-05 | 0.001026 |
| LRFN4 | 651.9097 | -0.14621 | 0.155018 | -0.9432 | 0.345581 | 0.513496 |
| RCE1 | 409.6002 | -0.0029 | 0.155407 | -0.01866 | 0.985111 | 0.992141 |
| C11orf80 | 148.8501 | 0.017136 | 0.216456 | 0.079168 | 0.936899 | 0.964646 |
| | 45.81893 | 0.592553 | 0.343899 | 1.723043 | 0.084881 | NA |
| TIGD3 | 5.53682 | 0.086628 | 0.956586 | 0.090559 | 0.927843 | NA |
| | 1.62174 | 0.128717 | 1.625655 | 0.079179 | 0.936891 | NA |
| SPTBN2 | 841.89 | -0.74305 | 0.189511 | -3.92091 | 8.82E-05 | 0.002371 |
| RBM4B | 394.5849 | -0.01927 | 0.154519 | -0.12473 | 0.900736 | 0.941822 |
| RBM4 | 94.30335 | 0.014054 | 0.253981 | 0.055333 | 0.955873 | 0.975764 |
| LRRC63 | 9.369396 | 0.467242 | 0.662631 | 0.705132 | 0.480728 | NA |
| CCS | 703.3496 | -0.10907 | 0.155639 | -0.70079 | 0.483436 | 0.637289 |
| SLC25A30 | 826.9483 | 0.312777 | 0.161685 | 1.934491 | 0.053053 | 0.144021 |
| CTSF | 6023.383 | -0.29874 | 0.12749 | -2.34323 | 0.019118 | 0.073347 |
| MSRB3 | 1112.702 | 0.479489 | 0.184434 | 2.599788 | 0.009328 | 0.045045 |
| LEMD3 | 680.9963 | 0.496045 | 0.179434 | 2.764491 | 0.005701 | 0.032354 |
| ZDHHC24 | 734.9905 | -0.17261 | 0.170218 | -1.01403 | 0.310567 | 0.478027 |
| | 68.56374 | -0.2337 | 0.29972 | -0.77973 | 0.43555 | 0.596586 |

| | | | | | | |
|---|---|---|---|---|---|---|
| MGA | 747.6071 | 0.109742 | 0.191339 | 0.573546 | 0.566275 | 0.703394 |
| C12orf66 | 125.1555 | -0.06834 | 0.22102 | -0.30918 | 0.757183 | 0.848416 |
| ADCY6 | 1813.958 | -0.19003 | 0.163191 | -1.16449 | 0.244226 | 0.406914 |
| REP15 | 2.786186 | 0.582628 | 1.208666 | 0.482042 | 0.629776 | NA |
| DDX23 | 2640.368 | -0.10845 | 0.154109 | -0.7037 | 0.481618 | 0.635881 |
| ZNHIT2 | 266.9018 | -0.21381 | 0.18935 | -1.12919 | 0.258818 | 0.423177 |
| C11orf45 | 58.10464 | -0.24662 | 0.308358 | -0.79977 | 0.423845 | 0.585969 |
| RALGAPA1 | 184.5044 | 0.313107 | 0.296462 | 1.056147 | 0.290901 | 0.457864 |
| LIG4 | 287.6093 | 1.516024 | 0.684224 | 2.215682 | 0.026713 | 0.091765 |
| RPL35P9 | 0.816972 | 0.936457 | 2.589651 | 0.361615 | 0.71764 | NA |
| ATP2A2 | 10284.32 | -0.40654 | 0.158435 | -2.56596 | 0.010289 | 0.048042 |
| ZWILCH | 357.0353 | 0.664402 | 0.19175 | 3.464949 | 5.30E-04 | 0.007022 |
| RPL4 | 30736.17 | -3.85E-04 | 0.133543 | -0.00288 | 0.997701 | 0.999133 |
| SNAPC5 | 584.7289 | 0.441917 | 0.168005 | 2.630381 | 0.008529 | 0.042312 |
| C12orf76 | 351.1855 | 0.116514 | 0.204927 | 0.568565 | 0.569651 | 0.705813 |
| BBS1 | 8.070571 | -0.15001 | 0.707438 | -0.21204 | 0.832076 | NA |
| DENND4A | 521.1064 | 0.572848 | 0.212847 | 2.691357 | 0.007116 | 0.037572 |
| IGDCC3 | 0.162376 | 0.424913 | 4.079006 | 0.104171 | 0.917034 | NA |
| PELI3 | 779.9869 | -0.33493 | 0.157214 | -2.1304 | 0.033139 | 0.106017 |
| MYO1H | 2.261098 | -0.97306 | 1.490013 | -0.65305 | 0.513722 | NA |
| MRPL11 | 776.0155 | -0.12653 | 0.162496 | -0.7787 | 0.436159 | 0.597005 |
| CMKLR1 | 0.151697 | -1.546 | 4.075469 | -0.37934 | 0.704433 | NA |
| SLC29A2 | 99.93426 | 0.094134 | 0.334451 | 0.281459 | 0.778359 | 0.863912 |
| B3GNT1 | 1561.862 | -0.26431 | 0.14979 | -1.76451 | 0.077646 | 0.187485 |
| KIAA1551 | 172.7779 | 0.927989 | 0.71878 | 1.291061 | 0.196683 | 0.352003 |
| BRMS1 | 1942.51 | -0.09081 | 0.164672 | -0.55148 | 0.581305 | 0.714966 |
| RIN1 | 267.7993 | -0.31183 | 0.18548 | -1.6812 | 0.092723 | 0.212249 |
| FZD4 | 176.5211 | -0.33402 | 0.216797 | -1.54073 | 0.123384 | 0.25752 |
| YIF1A | 2364.968 | -0.19822 | 0.186124 | -1.06499 | 0.286881 | 0.453221 |
| CNIH2 | 42.39657 | -0.3957 | 0.351442 | -1.12595 | 0.260188 | NA |
| RAB1B | 5702.142 | -0.27882 | 0.123105 | -2.26492 | 0.023517 | 0.084459 |
| FBXW8 | 657.0832 | -0.27246 | 0.170736 | -1.59578 | 0.110537 | 0.238431 |
| KLC2 | 1743.573 | -0.26145 | 0.184763 | -1.41506 | 0.15705 | 0.303288 |
| PACS1 | 1745.337 | -0.46394 | 0.160892 | -2.88353 | 0.003932 | 0.025186 |
| CSRP2 | 1307.392 | 0.466488 | 0.26468 | 1.762456 | 0.077992 | 0.188118 |
| INHBC | 0.315653 | -0.57305 | 3.892639 | -0.14721 | 0.882963 | NA |
| DDIT3 | 366.2918 | 0.161462 | 0.203022 | 0.795291 | 0.426445 | 0.588328 |
| PCCA | 390.5679 | -0.00275 | 0.168093 | -0.01638 | 0.986928 | 0.993212 |
| HIGD2B | 0.151697 | -1.546 | 4.075469 | -0.37934 | 0.704433 | NA |
| DCTN2 | 5015.695 | -0.15634 | 0.136315 | -1.14687 | 0.251434 | 0.415377 |
| CTDSP2 | 4407.97 | -0.41128 | 0.124535 | -3.3025 | 9.58E-04 | 0.010071 |
| GOLGA8A | 47.44897 | 1.70662 | 0.515181 | 3.31266 | 9.24E-04 | 0.009851 |
| CATSPER1 | 9.199294 | -0.01706 | 0.863373 | -0.01975 | 0.984239 | NA |
| CST6 | 0.670433 | 1.255281 | 2.356401 | 0.532711 | 0.594234 | NA |
| GRAMD2 | 1.681242 | 1.965083 | 1.654241 | 1.187906 | 0.23487 | NA |
| BANF1 | 2258.9 | -0.30756 | 0.139051 | -2.21188 | 0.026975 | 0.092425 |
| APOF | 0.247898 | -0.5901 | 4.074633 | -0.14482 | 0.884851 | NA |

| | | | | | | |
|---|---|---|---|---|---|---|
| CHRNA7 | 0.419962 | -2.64111 | 3.429364 | -0.77014 | 0.441214 | NA |
| EIF1AD | 571.388 | -0.09424 | 0.185071 | -0.5092 | 0.610614 | 0.737603 |
| TBC1D10C | 0.558248 | 0.982076 | 2.804738 | 0.350149 | 0.726227 | NA |
| SART1 | 2803.418 | -0.53102 | 0.14582 | -3.6416 | 2.71E-04 | 0.004615 |
| POLD4 | 349.6261 | -0.07135 | 0.184737 | -0.38624 | 0.69932 | 0.806443 |
| CLCF1 | 17.02367 | 0.056835 | 0.596726 | 0.095246 | 0.92412 | NA |
| TSGA10IP | 1.909793 | 2.161609 | 1.550034 | 1.394556 | 0.16315 | NA |
| LIPT2 | 19.18733 | -0.21817 | 0.472471 | -0.46176 | 0.644257 | NA |
| KCNE3 | 13.26789 | -0.11316 | 0.56945 | -0.19871 | 0.842489 | NA |
| ALG10B | 50.33524 | 0.262615 | 0.327999 | 0.800657 | 0.42333 | 0.585598 |
| DRAP1 | 2829.318 | 0.136525 | 0.130478 | 1.046345 | 0.295402 | 0.462675 |
| UCP3 | 6.572201 | 0.598467 | 0.812179 | 0.736866 | 0.461204 | NA |
| UCP2 | 21.86427 | -0.67546 | 0.581048 | -1.16248 | 0.245039 | NA |
| C11orf68 | 1697.544 | -0.03964 | 0.173418 | -0.22858 | 0.819192 | 0.891783 |
| PAAF1 | 733.5905 | -0.3007 | 0.14906 | -2.01733 | 0.043661 | 0.127299 |
| MRPL48 | 762.0708 | 0.054602 | 0.17049 | 0.320267 | 0.748766 | 0.842272 |
| RAB6A | 2453.282 | 0.297333 | 0.173813 | 1.710652 | 0.087145 | 0.202702 |
| FOSL1 | 1154.724 | 0.25413 | 0.341157 | 0.744906 | 0.456329 | 0.614817 |
| CCDC85B | 2189.989 | -0.02996 | 0.184819 | -0.1621 | 0.871231 | 0.923353 |
| RPS6KB2 | 1139.644 | -0.14842 | 0.166703 | -0.89033 | 0.373289 | 0.539792 |
| GOLGA8EP | 0.67854 | -0.89371 | 2.366391 | -0.37767 | 0.705677 | NA |
| MLXIP | 1910.287 | -0.75452 | 0.202904 | -3.71859 | 2.00E-04 | 0.003868 |
| C15orf54 | 0.623045 | 0.091811 | 2.666142 | 0.034436 | 0.97253 | NA |
| | 1.306087 | 0.163842 | 1.952563 | 0.083911 | 0.933127 | NA |
| SLC35E3 | 485.3845 | -0.18231 | 0.166586 | -1.09442 | 0.273772 | 0.439659 |
| PRIMA1 | 712.1819 | 0.638982 | 0.661517 | 0.965934 | 0.334077 | 0.502536 |
| A2M | 18010.98 | 0.278278 | 0.260925 | 1.066503 | 0.286196 | 0.452562 |
| UNC119B | 1734.132 | -0.27679 | 0.140623 | -1.96831 | 0.049033 | 0.136938 |
| PLEKHD1 | 0.679532 | 2.619206 | 2.901822 | 0.902607 | 0.366734 | NA |
| | 0.392857 | -1.546 | 4.075469 | -0.37934 | 0.704433 | NA |
| DLEU1 | 356.3478 | 0.234532 | 0.181803 | 1.290037 | 0.197038 | 0.35248 |
| GPX2 | 0.324752 | 1.53082 | 4.022025 | 0.380609 | 0.703493 | NA |
| FOXR1 | 0.134133 | -1.48909 | 4.077584 | -0.36519 | 0.71497 | NA |
| COX8A | 2884.875 | -0.1439 | 0.174409 | -0.82506 | 0.40934 | 0.572795 |
| RPL37AP8 | 1.088612 | 0.323874 | 2.06822 | 0.156596 | 0.875564 | NA |
| ZSCAN2 | 265.4577 | -0.3922 | 0.186698 | -2.10073 | 0.035664 | 0.111306 |
| B3GNT4 | 5.328646 | 0.174498 | 0.869271 | 0.20074 | 0.840902 | NA |
| SPRYD4 | 426.217 | -0.31365 | 0.166929 | -1.87894 | 0.060252 | 0.15768 |
| SYNE3 | 249.9261 | -0.37078 | 0.206216 | -1.79804 | 0.072171 | 0.178471 |
| LPCAT4 | 479.1611 | -0.18289 | 0.17589 | -1.03982 | 0.298425 | 0.46532 |
| WDR25 | 354.5356 | -0.17603 | 0.184086 | -0.95624 | 0.338952 | 0.507678 |
| PLA2G16 | 1375.074 | 0.120473 | 0.153546 | 0.784609 | 0.432683 | 0.593839 |
| NANOGP1 | 0.162376 | 0.424913 | 4.079006 | 0.104171 | 0.917034 | NA |
| SCAND2P | 192.5637 | 0.153348 | 0.188842 | 0.812045 | 0.416766 | 0.579969 |
| C15orf56 | 1.150035 | -1.04642 | 2.019749 | -0.51809 | 0.604393 | NA |
| VSIG10 | 390.8508 | 0.062697 | 0.209395 | 0.299418 | 0.764621 | 0.853834 |
| WSB2 | 4677.624 | -0.17439 | 0.126372 | -1.38001 | 0.167583 | 0.317106 |

| | | | | | | |
|---|---|---|---|---|---|---|
| | 0.303394 | -2.27006 | 4.032451 | -0.56295 | 0.57347 | NA |
| PXMP2 | 49.68881 | -0.37659 | 0.320993 | -1.17319 | 0.240718 | 0.403979 |
| PNMA1 | 1797.851 | -0.16484 | 0.139755 | -1.1795 | 0.2382 | 0.401059 |
| ANKLE2 | 1926.562 | 0.200255 | 0.155416 | 1.288507 | 0.197569 | 0.353177 |
| FAM89B | 228.6077 | -0.07738 | 0.217516 | -0.35576 | 0.722021 | 0.823351 |
| WDR73 | 420.0794 | 0.162059 | 0.170041 | 0.953058 | 0.340561 | 0.5087 |
| POLE | 911.9116 | -0.51779 | 0.226814 | -2.28289 | 0.022437 | 0.081935 |
| SCN4B | 4.046348 | -0.08523 | 1.042717 | -0.08174 | 0.934854 | NA |
| ZDHHC22 | 22.89108 | -0.18879 | 0.798409 | -0.23646 | 0.813073 | NA |
| ANO6 | 2843.637 | 0.174207 | 0.195184 | 0.892531 | 0.372108 | 0.53876 |
| ULK1 | 878.464 | -0.43161 | 0.162655 | -2.65351 | 0.007966 | 0.040531 |
| PUS1 | 427.9386 | -0.04602 | 0.188193 | -0.24456 | 0.806797 | 0.884627 |
| PCNPP5 | 5.127063 | 0.642706 | 0.937001 | 0.685917 | 0.492765 | NA |
| | 2.34924 | -0.95648 | 1.467294 | -0.65186 | 0.514488 | NA |
| RPL13AP3 | 1.291897 | 0.841043 | 1.879006 | 0.4476 | 0.654442 | NA |
| | 5.181767 | 1.15753 | 0.949156 | 1.219537 | 0.222641 | NA |
| | 19.45938 | -0.49198 | 0.524631 | -0.93776 | 0.348366 | NA |
| PAWR | 297.7452 | 0.707597 | 0.235036 | 3.010595 | 0.002607 | 0.019343 |
| ACOT4 | 104.4477 | -0.11348 | 0.228388 | -0.49687 | 0.61928 | 0.743902 |
| CD163L1 | 0.096202 | 0.408546 | 4.079639 | 0.100143 | 0.920231 | NA |
| | 1.302374 | 2.434983 | 1.999113 | 1.218032 | 0.223212 | NA |
| C12orf68 | 160.3014 | -0.13514 | 0.303814 | -0.44481 | 0.656456 | 0.77328 |
| UBE2N | 3330.703 | 0.160098 | 0.153839 | 1.040688 | 0.29802 | 0.465056 |
| IMP3 | 1924.697 | -0.0968 | 0.145866 | -0.66365 | 0.506918 | 0.65606 |
| ASB8 | 866.2516 | 0.014547 | 0.160272 | 0.090766 | 0.927679 | 0.958566 |
| DPY19L2 | 35.46564 | 0.364342 | 0.394691 | 0.923108 | 0.355951 | NA |
| DDX10 | 411.2182 | 0.624602 | 0.212096 | 2.944904 | 0.003231 | 0.022228 |
| KDELC2 | 978.2139 | 0.464603 | 0.190082 | 2.444224 | 0.014516 | 0.060596 |
| AQP11 | 97.45504 | -0.3564 | 0.245938 | -1.44916 | 0.147293 | 0.290493 |
| DNAJC22 | 16.7366 | 0.045615 | 0.548902 | 0.083102 | 0.93377 | NA |
| COX14 | 993.0334 | 0.260222 | 0.16092 | 1.617083 | 0.10586 | 0.231729 |
| DTX3 | 549.2364 | -0.48081 | 0.229224 | -2.09755 | 0.035945 | 0.111821 |
| KCTD12 | 615.9444 | -0.33139 | 0.279254 | -1.18669 | 0.235348 | 0.398124 |
| RPP25 | 2390.415 | -0.17602 | 0.154169 | -1.14177 | 0.253551 | 0.417595 |
| C13orf45 | 0.411855 | 0.250456 | 3.097618 | 0.080854 | 0.935558 | NA |
| COX5A | 3515.702 | -0.03603 | 0.146848 | -0.24533 | 0.806203 | 0.88422 |
| FAM219B | 1932.777 | -0.02498 | 0.171329 | -0.14582 | 0.884063 | 0.931441 |
| MPI | 831.0838 | -0.47383 | 0.16531 | -2.86629 | 0.004153 | 0.026037 |
| APOLD1 | 6231.917 | 0.173971 | 0.160461 | 1.084195 | 0.278279 | 0.444453 |
| FAM101A | 11.38461 | -1.29598 | 0.673663 | -1.92379 | 0.054381 | NA |
| FBXO34 | 736.9588 | 0.216316 | 0.197741 | 1.093936 | 0.273983 | 0.439786 |
| EXD1 | 1.92343 | 4.105171 | 1.852697 | 2.215781 | 0.026706 | NA |
| | 0.192403 | 0.975392 | 4.060716 | 0.240202 | 0.810174 | NA |
| CCDC89 | 0.163957 | 0.424913 | 4.079006 | 0.104171 | 0.917034 | NA |
| C12orf42 | 1.295204 | 0.045195 | 1.784804 | 0.025322 | 0.979798 | NA |
| TMTC2 | 2713.43 | -0.04293 | 0.126707 | -0.3388 | 0.734764 | 0.832199 |
| EDC3 | 1079.855 | -0.3596 | 0.155943 | -2.30596 | 0.021113 | 0.07861 |

| | | | | | | |
|---|---|---|---|---|---|---|
| ZNF664 | 3087.873 | 0.308437 | 0.16327 | 1.88912 | 0.058876 | 0.155139 |
| | 23.23297 | 0.153329 | 0.462953 | 0.331197 | 0.740495 | NA |
| SMCO3 | 22.39009 | -0.29608 | 0.523665 | -0.56541 | 0.571795 | NA |
| TMEM151/ | 4.090811 | 1.695162 | 1.145185 | 1.480252 | 0.138806 | NA |
| PTPN11 | 4446.572 | 0.663147 | 0.229232 | 2.892908 | 0.003817 | 0.024686 |
| RAB39A | 4.649602 | 1.733492 | 1.083202 | 1.600341 | 0.109523 | NA |
| CLK3 | 1188.602 | -0.15576 | 0.15053 | -1.03473 | 0.300797 | 0.467542 |
| ARID3B | 22.43161 | 0.523022 | 0.448325 | 1.166613 | 0.243367 | NA |
| HMGN2P4( | 7.146764 | 0.227674 | 0.771111 | 0.295254 | 0.7678 | NA |
| PACS2 | 2437.82 | -0.44575 | 0.149597 | -2.9797 | 0.002885 | 0.020568 |
| GPC5 | 1.149832 | -0.38667 | 2.069202 | -0.18687 | 0.851763 | NA |
| KLHL28 | 136.3217 | 0.427047 | 0.230023 | 1.856536 | 0.063377 | 0.163381 |
| C14orf28 | 90.49628 | 0.240018 | 0.2585 | 0.928503 | 0.353147 | 0.521234 |
| EIF3J-AS1 | 138.0795 | 0.231455 | 0.223892 | 1.033779 | 0.30124 | 0.468108 |
| DGKZP1 | 5.215421 | 0.035742 | 0.939476 | 0.038045 | 0.969652 | NA |
| ZBTB42 | 341.7561 | -0.09384 | 0.189658 | -0.4948 | 0.620745 | 0.745208 |
| LACC1 | 253.5673 | 1.012087 | 0.367456 | 2.754312 | 0.005882 | 0.032967 |
| AKAP5 | 1.70513 | 1.285072 | 1.554544 | 0.826656 | 0.408432 | NA |
| PHC1P1 | 17.68505 | -0.39482 | 0.478314 | -0.82545 | 0.409115 | NA |
| R3HDM2 | 546.8218 | -0.25385 | 0.211498 | -1.20026 | 0.230039 | 0.392221 |
| C14orf119 | 1689.793 | 0.032075 | 0.136725 | 0.234598 | 0.814521 | 0.889545 |
| BBS10 | 258.5471 | 0.360143 | 0.247041 | 1.457827 | 0.144888 | 0.286849 |
| SOCS4 | 773.6293 | 1.024451 | 0.248357 | 4.124916 | 3.71E-05 | 0.001391 |
| C12orf40 | 0.298089 | -0.51711 | 3.975232 | -0.13008 | 0.896501 | NA |
| CSNK1A1L | 0.134133 | -1.48909 | 4.077584 | -0.36519 | 0.71497 | NA |
| RPS12P23 | 1.617111 | -3.3093 | 2.151623 | -1.53805 | 0.124037 | NA |
| HMGB1P14 | 1.539942 | -0.09296 | 1.64952 | -0.05636 | 0.955058 | NA |
| HERC2P3 | 39.14978 | 1.307622 | 0.398804 | 3.278856 | 0.001042 | NA |
| FGD6 | 85.45423 | 0.459068 | 0.320991 | 1.430156 | 0.152672 | 0.297294 |
| OAZ2 | 2138.09 | -0.08329 | 0.150715 | -0.55261 | 0.580534 | 0.714368 |
| ALX1 | 617.2539 | 0.163649 | 0.253641 | 0.645197 | 0.518799 | 0.665768 |
| KCTD4 | 0.430642 | -1.1891 | 3.485782 | -0.34113 | 0.733008 | NA |
| ZNF609 | 1262.079 | -0.51486 | 0.160759 | -3.2027 | 0.001361 | 0.012689 |
| ATP5EP2 | 7.347821 | 0.793834 | 0.762098 | 1.041643 | 0.297577 | NA |
| C11orf71 | 78.76077 | -0.21123 | 0.280873 | -0.75203 | 0.452032 | 0.610972 |
| GLIPR1L2 | 11.25722 | 0.33281 | 0.615085 | 0.54108 | 0.588453 | NA |
| | 3.089165 | -0.15942 | 1.157552 | -0.13772 | 0.890461 | NA |
| MAB21L1 | 1.176909 | 1.262874 | 1.942452 | 0.650144 | 0.515599 | NA |
| SHISA2 | 17.53011 | -2.5095 | 0.887634 | -2.82717 | 0.004696 | NA |
| SRSF8 | 2122.756 | -0.02439 | 0.143847 | -0.16957 | 0.865349 | 0.920317 |
| SLC36A4 | 371.9805 | 0.840343 | 0.226763 | 3.705816 | 2.11E-04 | 0.004029 |
| ZDHHC20 | 509.8659 | 0.761394 | 0.236611 | 3.21791 | 0.001291 | 0.012237 |
| HOXC9 | 545.8576 | 0.189931 | 0.148869 | 1.275826 | 0.202017 | 0.35852 |
| HOXC10 | 684.085 | -0.1902 | 0.137258 | -1.38574 | 0.165827 | 0.314993 |
| C12orf36 | 0.758555 | 0.471868 | 2.553489 | 0.184794 | 0.853391 | NA |
| CAPS2 | 21.96885 | 0.217083 | 0.498881 | 0.43514 | 0.663461 | NA |
| ST20 | 51.20304 | 0.107894 | 0.318551 | 0.338701 | 0.734835 | 0.8322 |

| | | | | | | |
|---|---|---|---|---|---|---|
| LRRC57 | 376.4561 | -0.02885 | 0.178516 | -0.16162 | 0.871602 | 0.923482 |
| GPR137C | 24.69111 | 0.737013 | 0.481003 | 1.532244 | 0.125462 | NA |
| AEN | 1679.322 | -0.08143 | 0.182369 | -0.44651 | 0.655228 | 0.77237 |
| TMEM132( | 0.096202 | 0.408546 | 4.079639 | 0.100143 | 0.920231 | NA |
| TMEM136 | 221.9117 | 0.167326 | 0.17686 | 0.946089 | 0.344103 | 0.512195 |
| HEPHL1 | 6.961029 | 0.008073 | 0.994165 | 0.008121 | 0.993521 | NA |
| DDN | 4.022025 | 1.17277 | 1.0424 | 1.125067 | 0.26056 | NA |
| GPR135 | 16.16807 | 0.712886 | 0.540661 | 1.318543 | 0.187322 | NA |
| RFX7 | 678.2467 | 0.11056 | 0.176226 | 0.627378 | 0.530411 | 0.675137 |
| RNF41 | 1365.12 | -0.18391 | 0.136883 | -1.34358 | 0.179083 | 0.33162 |
| | 0.096202 | 0.408546 | 4.079639 | 0.100143 | 0.920231 | NA |
| COA4 | 1279.963 | -0.09515 | 0.161391 | -0.58954 | 0.555501 | 0.695329 |
| PRKAG1 | 2621.419 | -0.13253 | 0.132033 | -1.00377 | 0.31549 | 0.482886 |
| MRPS11 | 1490.58 | -0.03037 | 0.146587 | -0.20718 | 0.835871 | 0.901643 |
| MGAT4C | 0.096202 | 0.408546 | 4.079639 | 0.100143 | 0.920231 | NA |
| IDH2 | 1803.536 | -0.30633 | 0.166468 | -1.84015 | 0.065747 | 0.167212 |
| FAM181B | 33.62164 | -0.05964 | 0.37893 | -0.15739 | 0.874939 | NA |
| TMEM30B | 1.810576 | 1.595827 | 1.631515 | 0.978126 | 0.328012 | NA |
| NOP10 | 3109.515 | 0.232789 | 0.181793 | 1.280514 | 0.200365 | 0.356417 |
| RAD51B | 200.6724 | -0.05427 | 0.183283 | -0.29608 | 0.767168 | 0.855496 |
| ARL6IP4 | 8.375433 | 0.212134 | 0.688736 | 0.308005 | 0.758079 | NA |
| SHMT2 | 4338.366 | -0.27557 | 0.137816 | -1.99957 | 0.045547 | 0.130677 |
| HHIPL1 | 83.20556 | -0.23943 | 0.268251 | -0.89257 | 0.372086 | 0.53876 |
| GABRG3 | 13.99254 | 0.393414 | 0.574566 | 0.684714 | 0.493524 | NA |
| C1S | 31.88444 | -0.3878 | 0.390225 | -0.99379 | 0.320326 | NA |
| DAOA | 0.163957 | 0.424913 | 4.079006 | 0.104171 | 0.917034 | NA |
| KBTBD3 | 94.76337 | 0.414826 | 0.242158 | 1.713038 | 0.086706 | 0.201984 |
| NXPH4 | 19.29992 | -0.17579 | 0.498437 | -0.35269 | 0.724319 | NA |
| TRAPPC6B | 483.1919 | 0.793317 | 0.265697 | 2.985799 | 0.002828 | 0.020359 |
| PGBD4 | 64.35332 | 0.224485 | 0.322809 | 0.695413 | 0.486797 | 0.639869 |
| KCNK4 | 0.327913 | 1.543154 | 4.021063 | 0.383768 | 0.701151 | NA |
| ORAI1 | 1680.647 | -0.16732 | 0.153456 | -1.09032 | 0.275573 | 0.441265 |
| GLRX5 | 1793.309 | -0.1338 | 0.141428 | -0.94604 | 0.344128 | 0.512195 |
| MFSD5 | 1566.087 | -0.34241 | 0.15147 | -2.26061 | 0.023784 | 0.085143 |
| RNASE10 | 0.096202 | 0.408546 | 4.079639 | 0.100143 | 0.920231 | NA |
| | 0.151697 | -1.546 | 4.075469 | -0.37934 | 0.704433 | NA |
| NDN | 1317.273 | -0.21061 | 0.142833 | -1.47451 | 0.140345 | 0.280568 |
| NTM | 1423.104 | 0.725674 | 0.240874 | 3.012666 | 0.00259 | 0.01932 |
| TSKU | 782.952 | -0.40197 | 0.162294 | -2.47679 | 0.013257 | 0.057233 |
| ANXA2 | 44878.57 | 0.091257 | 0.199568 | 0.457274 | 0.647474 | 0.766201 |
| HCAR2 | 1.564701 | 0.631505 | 1.640308 | 0.384991 | 0.700244 | NA |
| CCDC87 | 6.931283 | -0.85729 | 0.794195 | -1.07945 | 0.280389 | NA |
| TMEM198I | 355.8365 | -0.34522 | 0.167873 | -2.05645 | 0.039739 | 0.119837 |
| CRIP2 | 3802.612 | -0.06621 | 0.128915 | -0.51356 | 0.607559 | 0.73511 |
| GALNT9 | 0.452561 | 2.061505 | 3.349456 | 0.615474 | 0.538241 | NA |
| C11orf54 | 554.6508 | 0.784412 | 0.233406 | 3.360724 | 7.77E-04 | 0.008896 |
| SRPR | 4818.619 | -0.29747 | 0.124593 | -2.38755 | 0.016961 | 0.067758 |

| | | | | | | |
|---|---|---|---|---|---|---|
| ODF3L1 | 7.173345 | -1.09971 | 0.781974 | -1.40633 | 0.159626 | NA |
| SPATA13 | 413.3062 | -0.12646 | 0.224319 | -0.56377 | 0.572911 | 0.708814 |
| SOX1 | 1.114915 | 1.271012 | 1.991514 | 0.638214 | 0.523334 | NA |
| MTA1 | 1422.615 | -0.27942 | 0.140735 | -1.98542 | 0.047098 | 0.133662 |
| CADM1 | 2243.479 | 0.409639 | 0.234284 | 1.748472 | 0.080382 | 0.192008 |
| C12orf60 | 78.26084 | 0.34023 | 0.267009 | 1.274229 | 0.202582 | 0.358825 |
| SLC25A21 | 1.859833 | 0.149049 | 1.479696 | 0.100729 | 0.919765 | NA |
| GAS6 | 329.9803 | -0.28854 | 0.203194 | -1.42004 | 0.155596 | 0.301452 |
| BEGAIN | 17.4554 | -0.09803 | 0.497719 | -0.19695 | 0.843865 | NA |
| GPC6 | 1.353328 | 1.603733 | 1.750302 | 0.916261 | 0.35953 | NA |
| GPR19 | 117.1226 | -0.30629 | 0.301376 | -1.01629 | 0.309489 | 0.476864 |
| TMEM119 | 0.230334 | -0.53318 | 4.076749 | -0.13079 | 0.895945 | NA |
| DAZAP2 | 4788.714 | -0.15354 | 0.124713 | -1.23115 | 0.218267 | 0.377746 |
| JRKL | 145.4363 | 0.673633 | 0.345993 | 1.946957 | 0.05154 | 0.141393 |
| SYNDIG1L | 0.746498 | -1.433 | 2.535747 | -0.56512 | 0.571994 | NA |
| PMCH | 1.843415 | 2.949948 | 1.749784 | 1.685893 | 0.091816 | NA |
| SH2D7 | 5.210826 | -0.20642 | 0.931077 | -0.2217 | 0.824544 | NA |
| GPR132 | 0.163957 | 0.424913 | 4.079006 | 0.104171 | 0.917034 | NA |
| EP400 | 1246.024 | -0.72954 | 0.230992 | -3.15831 | 0.001587 | 0.01406 |
| MEX3B | 166.465 | 0.012306 | 0.222675 | 0.055264 | 0.955928 | 0.975764 |
| FOLR4 | 0.134133 | -1.48909 | 4.077584 | -0.36519 | 0.71497 | NA |
| SETD3 | 1646.613 | 0.3646 | 0.155187 | 2.34942 | 0.018803 | 0.072466 |
| TNFAIP8L3 | 6.89169 | 2.00217 | 0.970789 | 2.062416 | 0.039168 | NA |
| C11orf88 | 0.260158 | 1.240347 | 4.047124 | 0.306476 | 0.759242 | NA |
| NDUFB1 | 2490.556 | 0.213951 | 0.184777 | 1.157888 | 0.24691 | 0.410366 |
| KLHL25 | 228.9239 | -0.42 | 0.239553 | -1.75327 | 0.079556 | 0.190609 |
| OPCML | 2.241072 | 1.35339 | 1.434445 | 0.943494 | 0.345428 | NA |
| LHFP | 397.4525 | 0.282792 | 0.231433 | 1.221917 | 0.221739 | 0.382136 |
| TBK1 | 690.1675 | 1.115736 | 0.246395 | 4.528239 | 5.95E-06 | 4.70E-04 |
| NUDT14 | 486.6608 | -0.18666 | 0.185824 | -1.00451 | 0.315133 | 0.482549 |
| HTR7P1 | 107.3002 | -0.29965 | 0.250022 | -1.19847 | 0.230733 | 0.392952 |
| SETD8 | 503.0984 | 0.085906 | 0.164829 | 0.521181 | 0.602241 | 0.731363 |
| DIABLO | 165.247 | -0.20578 | 0.215522 | -0.9548 | 0.33968 | 0.508003 |
| OR7E87P | 0.163957 | 0.424913 | 4.079006 | 0.104171 | 0.917034 | NA |
| LRTOMT | 99.60515 | -0.21361 | 0.242304 | -0.88158 | 0.378006 | 0.544304 |
| GOLGA6L4 | 0.260158 | 1.240347 | 4.047124 | 0.306476 | 0.759242 | NA |
| SNRNP35 | 738.0974 | 0.098888 | 0.176659 | 0.559769 | 0.575637 | 0.710703 |
| C11orf72 | 0.458885 | 0.577525 | 3.449278 | 0.167433 | 0.867029 | NA |
| PCDH9 | 82.82406 | 0.203311 | 0.297242 | 0.683991 | 0.493981 | 0.645092 |
| ACOT1 | 244.8069 | -0.23866 | 0.249033 | -0.95834 | 0.33789 | 0.506793 |
| OAF | 2457.641 | -0.21469 | 0.154981 | -1.38524 | 0.165979 | 0.31522 |
| POU6F1 | 282.603 | -0.49039 | 0.234622 | -2.09012 | 0.036607 | 0.113399 |
| PRKD1 | 1254.77 | -0.23355 | 0.13287 | -1.75773 | 0.078794 | 0.189295 |
| MAML2 | 276.8442 | -0.04649 | 0.205224 | -0.22655 | 0.820772 | 0.892334 |
| KNTC1 | 463.1678 | 0.418183 | 0.18085 | 2.312317 | 0.02076 | 0.077712 |
| TMEM255B | 27.49865 | 0.168748 | 0.441796 | 0.38196 | 0.702491 | NA |
| SLITRK6 | 98.16466 | 1.732462 | 0.906993 | 1.910116 | 0.056118 | 0.149964 |

| | | | | | | |
|---|---|---|---|---|---|---|
| XPOT | 2262.842 | 0.387158 | 0.174674 | 2.216457 | 0.02666 | 0.091691 |
| NELL2 | 5.554469 | 0.668316 | 0.969881 | 0.68907 | 0.490779 | NA |
| ATL3 | 1436.074 | 0.363593 | 0.209674 | 1.734088 | 0.082902 | 0.195915 |
| NDUFA12 | 1227.729 | 0.199365 | 0.152191 | 1.309964 | 0.190208 | 0.344448 |
| RPS17 | 0.134133 | -1.48909 | 4.077584 | -0.36519 | 0.71497 | NA |
| BTBD6 | 1899.848 | -0.08865 | 0.215562 | -0.41124 | 0.680899 | 0.792077 |
| JAG2 | 37.64613 | 0.104875 | 0.352581 | 0.29745 | 0.766123 | NA |
| NOC4L | 711.228 | -0.31885 | 0.154086 | -2.0693 | 0.038518 | 0.117204 |
| CHRM5 | 0.096202 | 0.408546 | 4.079639 | 0.100143 | 0.920231 | NA |
| TMEM121 | 306.8858 | -0.01216 | 0.188117 | -0.06465 | 0.948455 | 0.971147 |
| SIVA1 | 1450.062 | 0.138814 | 0.179211 | 0.774588 | 0.438583 | 0.598665 |
| BRI3BP | 843.6848 | -0.1667 | 0.248337 | -0.67126 | 0.502053 | 0.65228 |
| BRF1 | 1085.184 | -0.34863 | 0.152708 | -2.28298 | 0.022431 | 0.081935 |
| SEMA4B | 532.3591 | -0.67876 | 0.171832 | -3.95012 | 7.81E-05 | 0.002183 |
| ANKS1B | 51.51004 | -1.25774 | 1.234445 | -1.01887 | 0.308266 | 0.475536 |
| FLRT2 | 184.8707 | 0.654911 | 0.286384 | 2.286833 | 0.022206 | 0.08135 |
| RPS27L | 6649.652 | 0.547375 | 0.158133 | 3.461477 | 5.37E-04 | 0.007049 |
| ADSSL1 | 471.9925 | -0.21642 | 0.186016 | -1.16345 | 0.244645 | 0.407398 |
| NDNL2 | 783.1924 | 0.001043 | 0.139021 | 0.007501 | 0.994015 | 0.997382 |
| DDX51 | 662.9677 | -0.24244 | 0.155174 | -1.5624 | 0.118194 | 0.249499 |
| TNFAIP2 | 101.0297 | 0.29698 | 0.331983 | 0.894564 | 0.37102 | 0.53772 |
| PRPF39 | 341.3734 | 1.436388 | 0.287246 | 5.000557 | 5.72E-07 | 1.12E-04 |
| ATP6V0A2 | 751.906 | 0.404647 | 0.165512 | 2.44482 | 0.014492 | 0.060538 |
| C14orf80 | 321.5422 | -0.46959 | 0.282976 | -1.65948 | 0.097019 | 0.218619 |
| HS6ST3 | 0.096202 | 0.408546 | 4.079639 | 0.100143 | 0.920231 | NA |
| METTL7A | 382.7004 | -0.78962 | 0.292279 | -2.70158 | 0.006901 | 0.036755 |
| PARPBP | 283.4155 | 0.91809 | 0.210101 | 4.369759 | 1.24E-05 | 7.06E-04 |
| STAC3 | 3.742926 | 0.814627 | 1.111397 | 0.732976 | 0.463573 | NA |
| AHNAK2 | 81.49012 | -0.32642 | 0.437355 | -0.74634 | 0.455461 | 0.614347 |
| SP1 | 1648.894 | -0.29202 | 0.140594 | -2.07708 | 0.037794 | 0.115644 |
| ACTBP7 | 3.763486 | 0.769247 | 1.126925 | 0.682606 | 0.494856 | NA |
| NDUFA4L2 | 3.50523 | 0.816397 | 1.066676 | 0.765365 | 0.444054 | NA |
| SHC4 | 2758.883 | 0.341249 | 0.179782 | 1.898127 | 0.057679 | 0.152734 |
| KRT79 | 0.536531 | -2.91136 | 3.721629 | -0.78228 | 0.434049 | NA |
| ZFP36L1 | 1822.463 | -0.14562 | 0.161146 | -0.90368 | 0.366164 | 0.533342 |
| NTF3 | 378.3107 | 0.64431 | 0.341171 | 1.888523 | 0.058956 | 0.155268 |
| PMEL | 251738 | -0.69197 | 0.208454 | -3.31953 | 9.02E-04 | 0.009727 |
| EP400NL | 100.1439 | -0.00162 | 0.250732 | -0.00646 | 0.994842 | 0.997872 |
| C11orf87 | 0.298089 | -0.51711 | 3.975232 | -0.13008 | 0.896501 | NA |
| MORF4L1 | 6352.231 | 0.252722 | 0.148878 | 1.697509 | 0.089601 | 0.20714 |
| TRIM69 | 928.8567 | -0.66304 | 0.19255 | -3.44347 | 5.74E-04 | 0.007385 |
| LAMP1 | 9771.273 | -0.33884 | 0.148431 | -2.28281 | 0.022441 | 0.081935 |
| IRS2 | 3417.029 | -0.40717 | 0.289683 | -1.40555 | 0.159857 | 0.306994 |
| FAM186A | 0.916751 | 0.760304 | 2.340291 | 0.324876 | 0.745275 | NA |
| GRK1 | 0.096202 | 0.408546 | 4.079639 | 0.100143 | 0.920231 | NA |
| RASA3 | 2347.123 | 0.04949 | 0.188451 | 0.262613 | 0.792849 | 0.874397 |
| ATP4B | 0.134133 | -1.48909 | 4.077584 | -0.36519 | 0.71497 | NA |

| | | | | | | |
|---|---|---|---|---|---|---|
| DLEU7 | 93.49168 | 0.716044 | 0.276268 | 2.591843 | 0.009546 | 0.045744 |
| C15orf41 | 227.8306 | 0.050265 | 0.209473 | 0.239959 | 0.810362 | 0.886567 |
| | 0.574231 | 0.93227 | 2.49998 | 0.372911 | 0.709215 | NA |
| TAS2R42 | 0.151697 | -1.546 | 4.075469 | -0.37934 | 0.704433 | NA |
| CCDC84 | 118.7787 | 0.676716 | 0.219826 | 3.078411 | 0.002081 | 0.016597 |
| BCL9L | 1234.95 | -0.5036 | 0.190598 | -2.64222 | 0.008237 | 0.041392 |
| POLR1D | 2589.001 | 0.02922 | 0.160902 | 0.181599 | 0.855898 | 0.914361 |
| SLC51B | 0.096202 | 0.408546 | 4.079639 | 0.100143 | 0.920231 | NA |
| KDM4D | 62.43489 | -0.45424 | 0.321965 | -1.41085 | 0.158289 | 0.304828 |
| GABRA5 | 7.050363 | 1.325147 | 0.815911 | 1.624131 | 0.104348 | NA |
| PPP1CC | 3081.958 | 0.198171 | 0.138402 | 1.431853 | 0.152186 | 0.296494 |
| BACE1 | 1023.9 | -0.3224 | 0.140166 | -2.30011 | 0.021442 | 0.07946 |
| GOLGA6L1 | 0.70693 | -1.62766 | 2.687154 | -0.60572 | 0.544701 | NA |
| GOLGA8R | 0.75318 | -2.10401 | 2.55086 | -0.82482 | 0.409472 | NA |
| GNG2 | 1929.597 | 0.086236 | 0.195519 | 0.441062 | 0.659168 | 0.775241 |
| KTN1-AS1 | 60.12202 | 0.026819 | 0.293001 | 0.091531 | 0.92707 | 0.958156 |
| FSD2 | 9.319227 | 1.054989 | 0.718792 | 1.467725 | 0.142179 | NA |
| ARAP1 | 3025.542 | -0.4669 | 0.139066 | -3.35738 | 7.87E-04 | 0.008969 |
| PDE2A | 4.09818 | 0.18726 | 1.089342 | 0.171902 | 0.863515 | NA |
| LRRC16B | 59.28639 | -0.29936 | 0.308881 | -0.96917 | 0.332463 | 0.500806 |
| BCDIN3D | 62.80136 | -0.04655 | 0.287189 | -0.16208 | 0.871241 | 0.923353 |
| CCDC42B | 0.247898 | -0.5901 | 4.074633 | -0.14482 | 0.884851 | NA |
| TPCN1 | 1525.19 | -0.52313 | 0.131065 | -3.99137 | 6.57E-05 | 0.001943 |
| C1QL4 | 4.730481 | 0.543931 | 0.982782 | 0.553461 | 0.579948 | NA |
| ZDHHC17 | 417.9057 | 0.900061 | 0.250364 | 3.595012 | 3.24E-04 | 0.005188 |
| C14orf23 | 0.260158 | 1.240347 | 4.047124 | 0.306476 | 0.759242 | NA |
| TMEM262 | 9.509042 | 0.232588 | 0.666436 | 0.349003 | 0.727087 | NA |
| ENTPD5 | 609.9072 | -0.21533 | 0.172968 | -1.24489 | 0.213173 | 0.37186 |
| HEATR4 | 6.678365 | -0.12493 | 0.783399 | -0.15947 | 0.873301 | NA |
| NAP1L1 | 6061.075 | 0.581354 | 0.218248 | 2.663736 | 0.007728 | 0.039779 |
| H1FNT | 0.162376 | 0.424913 | 4.079006 | 0.104171 | 0.917034 | NA |
| DYNC2H1 | 754.2679 | 0.718492 | 0.564859 | 1.271984 | 0.203379 | 0.359805 |
| CHP1 | 8655.889 | -0.2268 | 0.142964 | -1.58641 | 0.112646 | 0.241224 |
| COL4A1 | 2482.307 | 0.155688 | 0.258537 | 0.602189 | 0.547049 | 0.688457 |
| COX8C | 0.555233 | -0.57009 | 3.296947 | -0.17291 | 0.862719 | NA |
| TCL6 | 0.096202 | 0.408546 | 4.079639 | 0.100143 | 0.920231 | NA |
| DHRS4L2 | 166.424 | -0.15746 | 0.207325 | -0.7595 | 0.447553 | 0.607134 |
| WHAMMP | 16.92909 | 0.806827 | 0.531104 | 1.519153 | 0.128724 | NA |
| B3GALTL | 1233.402 | 0.368136 | 0.167074 | 2.203432 | 0.027564 | 0.093874 |
| KRT18P59 | 1.993937 | 1.135122 | 1.450207 | 0.782731 | 0.433785 | NA |
| THSD4 | 7.245904 | -0.20652 | 0.76072 | -0.27147 | 0.786027 | NA |
| MCRS1 | 1901.099 | -0.19384 | 0.123695 | -1.56707 | 0.117099 | 0.247716 |
| FANCM | 124.0972 | 1.212565 | 0.418335 | 2.89855 | 0.003749 | 0.02446 |
| | 0.096202 | 0.408546 | 4.079639 | 0.100143 | 0.920231 | NA |
| OVCH1 | 1.21484 | 0.621271 | 1.748506 | 0.355315 | 0.722353 | NA |
| ARHGAP11 | 59.13938 | 0.241854 | 0.312144 | 0.774814 | 0.43845 | 0.59858 |
| RILPL1 | 433.2861 | -0.5009 | 0.224877 | -2.22744 | 0.025918 | 0.09021 |

| | | | | | | |
|---|---|---|---|---|---|---|
| C11orf95 | 1400.096 | -0.03831 | 0.212399 | -0.18037 | 0.856861 | 0.914942 |
| PLA2G4E | 0.163957 | 0.424913 | 4.079006 | 0.104171 | 0.917034 | NA |
| MESP2 | 0.787148 | -0.70094 | 2.357021 | -0.29738 | 0.766174 | NA |
| COMMD6 | 1085.856 | 0.407017 | 0.172282 | 2.3625 | 0.018152 | 0.070694 |
| HYKK | 95.44828 | 0.217316 | 0.316145 | 0.687393 | 0.491835 | 0.643353 |
| C15orf62 | 23.32191 | 0.184968 | 0.481543 | 0.384115 | 0.700894 | NA |
| OR6C6 | 0.989036 | -0.99378 | 2.406119 | -0.41302 | 0.67959 | NA |
| GTF2F2 | 1091.987 | 0.472807 | 0.195499 | 2.41846 | 0.015586 | 0.063606 |
| H3F3C | 1.954397 | 0.097524 | 1.395751 | 0.069872 | 0.944295 | NA |
| CSPG4P8 | 0.162376 | 0.424913 | 4.079006 | 0.104171 | 0.917034 | NA |
| SLC24A5 | 5331.681 | 0.107654 | 0.197692 | 0.544556 | 0.586059 | 0.71894 |
| H2AFX | 1956.179 | -0.29107 | 0.20197 | -1.44114 | 0.149544 | 0.293079 |
| SERPINA5 | 0.096202 | 0.408546 | 4.079639 | 0.100143 | 0.920231 | NA |
| LCTL | 36.76497 | 0.282067 | 0.415268 | 0.679241 | 0.496985 | NA |
| C15orf52 | 3131.555 | -0.0053 | 0.224974 | -0.02356 | 0.981203 | 0.990173 |
| C12orf55 | 2.066793 | 1.103409 | 1.573842 | 0.701093 | 0.483245 | NA |
| FAM154B | 11.14673 | 0.379304 | 0.707141 | 0.53639 | 0.591689 | NA |
| TMEM120B | 404.7434 | -0.46854 | 0.212877 | -2.201 | 0.027736 | 0.09425 |
| PLET1 | 0.096202 | 0.408546 | 4.079639 | 0.100143 | 0.920231 | NA |
| SKOR1 | 5.520903 | 0.476744 | 0.863515 | 0.552097 | 0.580882 | NA |
| NHLRC3 | 809.6816 | 0.314937 | 0.155863 | 2.0206 | 0.043321 | 0.126879 |
| TMC3 | 2.564593 | 0.807492 | 1.44533 | 0.558691 | 0.576373 | NA |
| LRRK2 | 563.931 | 0.413577 | 0.323142 | 1.279862 | 0.200594 | 0.356605 |
| KCTD21 | 347.4668 | -0.23591 | 0.171613 | -1.37463 | 0.169245 | 0.319593 |
| ALKBH2 | 287.1438 | -0.05959 | 0.200382 | -0.29739 | 0.766172 | 0.854855 |
| ARID2 | 624.7348 | 0.187434 | 0.202047 | 0.927679 | 0.353574 | 0.521606 |
| UBE2Q2P1 | 14.55667 | 1.021659 | 0.59087 | 1.729076 | 0.083795 | NA |
| ZAR1L | 0.162376 | 0.424913 | 4.079006 | 0.104171 | 0.917034 | NA |
| C15orf61 | 98.92931 | -0.33158 | 0.23868 | -1.38923 | 0.164762 | 0.313625 |
| HMGB1 | 2417.557 | 0.648593 | 0.168902 | 3.84005 | 1.23E-04 | 0.002922 |
| | 0.163957 | 0.424913 | 4.079006 | 0.104171 | 0.917034 | NA |
| MYBPC1 | 0.518736 | 2.237423 | 2.7713 | 0.807355 | 0.419462 | NA |
| OR6M1 | 0.096202 | 0.408546 | 4.079639 | 0.100143 | 0.920231 | NA |
| SERPINA3 | 0.260158 | 1.240347 | 4.047124 | 0.306476 | 0.759242 | NA |
| RPS2P4 | 0.163957 | 0.424913 | 4.079006 | 0.104171 | 0.917034 | NA |
| MPHOSPH8 | 783.4712 | 0.782712 | 0.353504 | 2.214151 | 0.026818 | 0.091994 |
| ZBTB44 | 573.126 | 0.30126 | 0.184357 | 1.634115 | 0.102235 | 0.226262 |
| FUT4 | 129.4718 | -0.26455 | 0.244649 | -1.08133 | 0.279549 | 0.445601 |
| ZNF140 | 265.4556 | 0.459649 | 0.176358 | 2.606346 | 0.009151 | 0.044481 |
| EVL | 588.8199 | -0.58875 | 0.221064 | -2.66327 | 0.007739 | 0.039783 |
| ZNF605 | 140.7782 | 0.940984 | 0.854658 | 1.101006 | 0.270894 | 0.436042 |
| MYL6B | 1343.483 | -0.11915 | 0.171895 | -0.69313 | 0.488226 | 0.641037 |
| IPO4 | 14.48218 | -0.47862 | 0.539085 | -0.88784 | 0.374626 | NA |
| NCOR2 | 4584.271 | -0.55772 | 0.178482 | -3.12479 | 0.001779 | 0.015024 |
| ANAPC7 | 1154.269 | -0.02381 | 0.147556 | -0.16136 | 0.871806 | 0.923616 |
| NACA | 8307.338 | 0.484547 | 0.18881 | 2.566323 | 0.010278 | 0.048042 |
| SLC22A25 | 0.28583 | -2.21358 | 3.97376 | -0.55705 | 0.577494 | NA |

| | | | | | | |
|---|---|---|---|---|---|---|
| MMP1 | 293.9648 | 2.292503 | 1.511451 | 1.516756 | 0.129328 | 0.265763 |
| GOLGA6L2( | 0.727151 | -0.55687 | 2.954176 | -0.1885 | 0.850483 | NA |
| TRAPPC4 | 1345.677 | -0.11681 | 0.168769 | -0.69211 | 0.488867 | 0.641594 |
| TECPR2 | 2378.586 | -0.58526 | 0.167702 | -3.48987 | 4.83E-04 | 0.006662 |
| LINC00173 | 4.644877 | -0.35725 | 0.961209 | -0.37166 | 0.710144 | NA |
| STRN3 | 606.6009 | 0.538018 | 0.226647 | 2.373812 | 0.017606 | 0.069409 |
| PPTC7 | 935.8432 | 0.426499 | 0.143438 | 2.973406 | 0.002945 | 0.020882 |
| TOMM20L | 2.534778 | 1.509919 | 1.390916 | 1.085558 | 0.277675 | NA |
| SCN8A | 44.98724 | 0.401741 | 0.389642 | 1.031053 | 0.302516 | NA |
| ARHGEF12 | 5082.856 | 0.354682 | 0.149934 | 2.365589 | 0.018001 | 0.070292 |
| HCAR1 | 1.939565 | -1.92319 | 1.540988 | -1.24802 | 0.212022 | NA |
| SRGAP1 | 383.0869 | -0.59551 | 0.233454 | -2.55085 | 0.010746 | 0.049294 |
| NOP9 | 1199.437 | -0.37417 | 0.173178 | -2.16062 | 0.030725 | 0.100759 |
| ZNF705A | 0.096202 | 0.408546 | 4.079639 | 0.100143 | 0.920231 | NA |
| CASP4 | 1575.373 | 0.461086 | 0.160225 | 2.877737 | 0.004005 | 0.025436 |
| GMFB | 2214.178 | 0.67373 | 0.19292 | 3.492269 | 4.79E-04 | 0.006641 |
| DYNC1H1 | 16930.59 | -0.50353 | 0.171885 | -2.92945 | 0.003396 | 0.022819 |
| PCBP2 | 3040.867 | -0.42397 | 0.145114 | -2.92165 | 0.003482 | 0.023241 |
| SLC25A29 | 676.9305 | -0.23964 | 0.154286 | -1.55321 | 0.120373 | 0.25266 |
| PCNXL3 | 1966.686 | -0.69346 | 0.15515 | -4.4696 | 7.84E-06 | 5.40E-04 |
| NEK5 | 12.15423 | 0.879906 | 0.612211 | 1.437259 | 0.150644 | NA |
| SERPINA1 | 87.42776 | -0.17732 | 1.054699 | -0.16812 | 0.866486 | 0.920772 |
| LRP10 | 9537.304 | -0.35223 | 0.180922 | -1.94686 | 0.051552 | 0.141393 |
| MRPL21 | 1034.869 | 0.085867 | 0.134814 | 0.63693 | 0.52417 | 0.670307 |
| BNIP3P1 | 4.365928 | -0.44958 | 1.00189 | -0.44874 | 0.653622 | NA |
| FBXL22 | 2.201304 | -1.21337 | 1.317832 | -0.92073 | 0.35719 | NA |
| MYO5A | 20529.35 | 0.325443 | 0.251143 | 1.295844 | 0.195029 | 0.350068 |
| SIPA1L1 | 618.5515 | 0.0376 | 0.172479 | 0.217996 | 0.827432 | 0.895973 |
| BCO2 | 4.373266 | 1.095796 | 1.047106 | 1.046499 | 0.295331 | NA |
| MFAP5 | 0.326333 | 1.533889 | 3.826878 | 0.40082 | 0.688553 | NA |
| MYH6 | 0.162376 | 0.424913 | 4.079006 | 0.104171 | 0.917034 | NA |
| DNAH10 | 17.12546 | 0.075668 | 0.519811 | 0.145569 | 0.884262 | NA |
| CBX3P7 | 46.97023 | 0.564041 | 0.5317 | 1.060824 | 0.28877 | NA |
| NMB | 218.6908 | -0.13963 | 0.201876 | -0.69165 | 0.48916 | 0.641907 |
| RPS26 | 4364.121 | 0.06439 | 0.151599 | 0.424743 | 0.671024 | 0.784504 |
| C14orf178 | 0.163957 | 0.424913 | 4.079006 | 0.104171 | 0.917034 | NA |
| HOXC6 | 379.1764 | -0.03451 | 0.171723 | -0.20097 | 0.840725 | 0.905184 |
| KLHDC1 | 55.25219 | 0.595445 | 0.328028 | 1.815227 | 0.069489 | 0.173679 |
| FAM118B | 961.0234 | 0.176031 | 0.165112 | 1.066131 | 0.286364 | 0.4527 |
| HIST4H4 | 12.58821 | 0.316697 | 0.591251 | 0.535639 | 0.592208 | NA |
| SLC22A20 | 0.628146 | -0.21495 | 3.102079 | -0.06929 | 0.944756 | NA |
| PRB3 | 0.785567 | -0.70568 | 2.358063 | -0.29926 | 0.764741 | NA |
| TEAD4 | 455.166 | -0.25003 | 0.191785 | -1.30371 | 0.192333 | 0.346889 |
| ERO1L | 1097.577 | 0.274579 | 0.193572 | 1.418487 | 0.156049 | 0.301904 |
| PCDH20 | 249.2389 | 1.223993 | 0.29357 | 4.169345 | 3.05E-05 | 0.001199 |
| IRAK4 | 527.7625 | 0.641376 | 0.192335 | 3.334691 | 8.54E-04 | 0.0094 |
| MRPL42 | 1147.283 | 0.704572 | 0.18384 | 3.832531 | 1.27E-04 | 0.002972 |

| | | | | | | |
|---|---|---|---|---|---|---|
| TUBA3C | 0.151697 | -1.546 | 4.075469 | -0.37934 | 0.704433 | NA |
| ZNF84 | 224.7956 | 0.974169 | 0.32495 | 2.997908 | 0.002718 | 0.019923 |
| PRIM1 | 155.6022 | 0.332376 | 0.218928 | 1.518197 | 0.128965 | 0.265292 |
| TMEM229I | 1634.042 | -0.6084 | 0.154697 | -3.93282 | 8.40E-05 | 0.002288 |
| | 22.50343 | 0.044595 | 0.463525 | 0.096208 | 0.923355 | NA |
| ZNF770 | 355.0509 | 0.890627 | 0.243982 | 3.650376 | 2.62E-04 | 0.004572 |
| TFDP1 | 2404.005 | 0.004815 | 0.144351 | 0.033355 | 0.973391 | 0.985192 |
| RPS6KL1 | 130.3467 | -0.00478 | 0.279977 | -0.01708 | 0.98637 | 0.992734 |
| STYX | 423.1448 | 0.680844 | 0.21684 | 3.139852 | 0.00169 | 0.014563 |
| TMEM116 | 89.61963 | 0.073919 | 0.273932 | 0.269843 | 0.787281 | 0.871205 |
| FAM109A | 592.2171 | -0.08216 | 0.178839 | -0.45943 | 0.645929 | 0.764681 |
| HYLS1 | 353.6419 | 0.038498 | 0.215888 | 0.178326 | 0.858467 | 0.916198 |
| HOXC4 | 171.0157 | -0.54755 | 0.218929 | -2.50104 | 0.012383 | 0.054536 |
| UVRAG | 751.4198 | 0.008466 | 0.148977 | 0.056825 | 0.954684 | 0.974998 |
| TPTE2P3 | 0.151697 | -1.546 | 4.075469 | -0.37934 | 0.704433 | NA |
| ZNF26 | 109.577 | 0.490483 | 0.235339 | 2.084155 | 0.037146 | 0.114411 |
| TXNRD1 | 7550.004 | 0.353832 | 0.197738 | 1.789401 | 0.07355 | 0.180809 |
| B3GNT6 | 0.324752 | 1.53082 | 4.022025 | 0.380609 | 0.703493 | NA |
| ATL1 | 279.3248 | 0.221937 | 0.196559 | 1.12911 | 0.258851 | 0.423177 |
| ITGBL1 | 1.139997 | 2.221426 | 2.048614 | 1.084356 | 0.278207 | NA |
| KCNRG | 1.15657 | -0.18697 | 1.786779 | -0.10464 | 0.916661 | NA |
| WDHD1 | 233.3563 | 0.732398 | 0.288163 | 2.541606 | 0.011034 | 0.050236 |
| MMP17 | 11520.93 | -0.17037 | 0.135433 | -1.25799 | 0.208397 | 0.365894 |
| BAZ1A | 565.283 | 1.000728 | 0.5901 | 1.69586 | 0.089912 | 0.207626 |
| CALM1 | 9252.841 | 0.11586 | 0.149027 | 0.777442 | 0.436898 | 0.597357 |
| FAM19A2 | 2.682954 | 1.258151 | 1.258455 | 0.999758 | 0.317428 | NA |
| FAN1 | 443.1137 | 0.074549 | 0.168268 | 0.443037 | 0.657739 | 0.774252 |
| CEP290 | 102.2863 | 0.730706 | 1.283534 | 0.569292 | 0.569158 | 0.705795 |
| FAM179B | 420.2129 | 0.645371 | 0.23465 | 2.750352 | 0.005953 | 0.033243 |
| SMOC1 | 3.421976 | 2.168003 | 1.359694 | 1.594478 | 0.110829 | NA |
| CDC42BPB | 4364.88 | -0.40734 | 0.155528 | -2.61907 | 0.008817 | 0.043426 |
| RASSF9 | 0.096202 | 0.408546 | 4.079639 | 0.100143 | 0.920231 | NA |
| SCAMP5 | 336.3794 | -0.1717 | 0.196259 | -0.87485 | 0.381655 | 0.547763 |
| PNP | 717.0787 | 0.121509 | 0.182985 | 0.664041 | 0.506664 | 0.655875 |
| CHAMP1 | 587.5388 | -0.79889 | 0.227136 | -3.51721 | 4.36E-04 | 0.006267 |
| ARHGAP11 | 507.0831 | 0.619664 | 0.232445 | 2.665857 | 0.007679 | 0.039564 |
| RYR3 | 22.93417 | 0.515837 | 0.444885 | 1.159483 | 0.246259 | NA |
| FICD | 207.1002 | -0.2564 | 0.199284 | -1.28658 | 0.19824 | 0.35399 |
| CIPC | 551.119 | -0.00577 | 0.160361 | -0.03598 | 0.971296 | 0.984247 |
| MIR412 | 0.096202 | 0.408546 | 4.079639 | 0.100143 | 0.920231 | NA |
| MIR148B | 0.162376 | 0.424913 | 4.079006 | 0.104171 | 0.917034 | NA |
| MIRLET7I | 2.753587 | -0.41429 | 1.161068 | -0.35682 | 0.721226 | NA |
| RNU6-834I | 0.163957 | 0.424913 | 4.079006 | 0.104171 | 0.917034 | NA |
| RNU4-39P | 0.230334 | -0.53318 | 4.076749 | -0.13079 | 0.895945 | NA |
| SNORD9 | 0.315653 | -0.57305 | 3.892639 | -0.14721 | 0.882963 | NA |
| RNU5A-1 | 0.258578 | 1.233151 | 4.047809 | 0.304646 | 0.760635 | NA |
| RNU4-23P | 1.08386 | -1.26078 | 2.279484 | -0.5531 | 0.580197 | NA |

| | | | | | | |
|---|---|---|---|---|---|---|
| SNORD115 | 0.096202 | 0.408546 | 4.079639 | 0.100143 | 0.920231 | NA |
| RNA5SP38: | 1.867644 | -0.67834 | 1.721248 | -0.3941 | 0.693507 | NA |
| RNU1-47P | 0.151697 | -1.546 | 4.075469 | -0.37934 | 0.704433 | NA |
| | 0.230334 | -0.53318 | 4.076749 | -0.13079 | 0.895945 | NA |
| | 0.162376 | 0.424913 | 4.079006 | 0.104171 | 0.917034 | NA |
| | 0.420954 | 1.923376 | 3.462803 | 0.555439 | 0.578594 | NA |
| RN7SKP24: | 0.096202 | 0.408546 | 4.079639 | 0.100143 | 0.920231 | NA |
| | 0.162376 | 0.424913 | 4.079006 | 0.104171 | 0.917034 | NA |
| RNU4-80P | 0.096202 | 0.408546 | 4.079639 | 0.100143 | 0.920231 | NA |
| | 0.096202 | 0.408546 | 4.079639 | 0.100143 | 0.920231 | NA |
| RNU6-216I | 0.327913 | 1.543154 | 4.021063 | 0.383768 | 0.701151 | NA |
| RNU5B-1 | 0.096202 | 0.408546 | 4.079639 | 0.100143 | 0.920231 | NA |
| RNU4-32P | 0.682693 | 2.623213 | 2.518338 | 1.041645 | 0.297576 | NA |
| RNU6-485I | 0.314073 | -0.57308 | 3.898934 | -0.14698 | 0.883146 | NA |
| | 0.575812 | 0.936922 | 2.776634 | 0.337431 | 0.735792 | NA |
| RNU6-1017 | 0.162376 | 0.424913 | 4.079006 | 0.104171 | 0.917034 | NA |
| RNA5SP34! | 0.151697 | -1.546 | 4.075469 | -0.37934 | 0.704433 | NA |
| | 9.767547 | 0.42182 | 0.686862 | 0.614126 | 0.539132 | NA |
| SNORD8 | 0.096202 | 0.408546 | 4.079639 | 0.100143 | 0.920231 | NA |
| RNU4-1 | 1.957172 | 1.15993 | 1.516365 | 0.764941 | 0.444307 | NA |
| | 0.296509 | -0.51709 | 3.982155 | -0.12985 | 0.896684 | NA |
| SNORD14E | 1.446755 | 0.439331 | 1.610042 | 0.272869 | 0.784954 | NA |
| | 0.134133 | -1.48909 | 4.077584 | -0.36519 | 0.71497 | NA |
| | 0.096202 | 0.408546 | 4.079639 | 0.100143 | 0.920231 | NA |
| RNU6-188I | 0.096202 | 0.408546 | 4.079639 | 0.100143 | 0.920231 | NA |
| RNU6-879I | 0.163957 | 0.424913 | 4.079006 | 0.104171 | 0.917034 | NA |
| | 0.392857 | -1.546 | 4.075469 | -0.37934 | 0.704433 | NA |
| RN7SKP71 | 0.162376 | 0.424913 | 4.079006 | 0.104171 | 0.917034 | NA |
| RNU4-24P | 0.096202 | 0.408546 | 4.079639 | 0.100143 | 0.920231 | NA |
| | 0.162376 | 0.424913 | 4.079006 | 0.104171 | 0.917034 | NA |
| | 0.298089 | -0.51711 | 3.975232 | -0.13008 | 0.896501 | NA |
| RNU6-343I | 0.296509 | -0.51709 | 3.982155 | -0.12985 | 0.896684 | NA |
| | 0.940279 | 0.09645 | 2.075894 | 0.046462 | 0.962942 | NA |
| RN7SKP23! | 0.314073 | -0.57308 | 3.898934 | -0.14698 | 0.883146 | NA |
| RN7SKP9 | 0.163957 | 0.424913 | 4.079006 | 0.104171 | 0.917034 | NA |
| | 0.096202 | 0.408546 | 4.079639 | 0.100143 | 0.920231 | NA |
| RNU6-113I | 0.162376 | 0.424913 | 4.079006 | 0.104171 | 0.917034 | NA |
| RNA5SP37 | 0.35636 | 1.709252 | 3.651833 | 0.468053 | 0.639746 | NA |
| SNORD6 | 1.092967 | 2.10982 | 2.003899 | 1.052857 | 0.292406 | NA |
| | 0.653072 | 0.258115 | 3.011926 | 0.085698 | 0.931707 | NA |
| RNU6-8 | 3.223784 | 0.989208 | 1.13607 | 0.870728 | 0.383902 | NA |
| RNA5SP30 | 0.096202 | 0.408546 | 4.079639 | 0.100143 | 0.920231 | NA |
| RNU4-2 | 1.668917 | -0.52368 | 1.657435 | -0.31596 | 0.752033 | NA |
| | 80.51089 | 0.107507 | 0.347801 | 0.309107 | 0.75724 | 0.848416 |
| | 11.12161 | 0.558783 | 0.701847 | 0.796159 | 0.425939 | NA |
| | 0.162376 | 0.424913 | 4.079006 | 0.104171 | 0.917034 | NA |
| | 0.096202 | 0.408546 | 4.079639 | 0.100143 | 0.920231 | NA |

| | | | | | | |
|---|---|---|---|---|---|---|
| | 0.134133 | -1.48909 | 4.077584 | -0.36519 | 0.71497 | NA |
| LINC00449 | 6.596207 | -0.14137 | 0.764298 | -0.18496 | 0.853257 | NA |
| INF2 | 6473.493 | -0.4089 | 0.175024 | -2.33623 | 0.019479 | 0.074349 |
| | 0.151697 | -1.546 | 4.075469 | -0.37934 | 0.704433 | NA |
| | 4.220616 | -0.39002 | 0.977645 | -0.39894 | 0.689935 | NA |
| | 0.689366 | -1.56115 | 2.700123 | -0.57818 | 0.563145 | NA |
| | 0.096202 | 0.408546 | 4.079639 | 0.100143 | 0.920231 | NA |
| | 0.247898 | -0.5901 | 4.074633 | -0.14482 | 0.884851 | NA |
| | 0.392857 | -1.546 | 4.075469 | -0.37934 | 0.704433 | NA |
| | 2.656648 | 1.201711 | 1.390789 | 0.86405 | 0.38756 | NA |
| | 13.90016 | 0.539555 | 0.595396 | 0.906212 | 0.364824 | NA |
| | 1.1119 | 0.46633 | 1.933926 | 0.241131 | 0.809453 | NA |
| | 3.464515 | -0.24952 | 1.089822 | -0.22896 | 0.818902 | NA |
| NXPE2 | 0.096202 | 0.408546 | 4.079639 | 0.100143 | 0.920231 | NA |
| SDHD | 908.6419 | 0.110113 | 0.192479 | 0.572078 | 0.567269 | 0.704113 |
| LAYN | 447.0075 | 0.710214 | 0.353596 | 2.008548 | 0.044585 | 0.129054 |
| CARD16 | 29.90437 | 0.497359 | 0.471979 | 1.053773 | 0.291987 | NA |
| CASP12 | 0.616518 | 2.49285 | 2.980788 | 0.836306 | 0.402983 | NA |
| LRCOL1 | 1.771507 | 0.854675 | 1.477668 | 0.578395 | 0.562998 | NA |
| CNN2P12 | 0.392857 | -1.546 | 4.075469 | -0.37934 | 0.704433 | NA |
| ATXN2 | 895.0035 | -0.27242 | 0.186174 | -1.46324 | 0.143403 | 0.284812 |
| TCTN1 | 673.1315 | -0.26912 | 0.182193 | -1.47713 | 0.13964 | 0.279776 |
| FAM216A | 149.3689 | 0.271517 | 0.214491 | 1.265866 | 0.205561 | 0.362584 |
| MZT1 | 436.1566 | 0.441724 | 0.15919 | 2.774821 | 0.005523 | 0.031724 |
| C12orf73 | 151.196 | 0.395649 | 0.211144 | 1.873831 | 0.060954 | 0.158996 |
| TRIM13 | 624.4405 | 0.219342 | 0.182136 | 1.204277 | 0.228482 | 0.390128 |
| COX17P1 | 4.959446 | 0.86577 | 0.944757 | 0.916395 | 0.35946 | NA |
| GOLGA6L1( | 1.033263 | 0.082415 | 2.175992 | 0.037875 | 0.969788 | NA |
| SARNP | 92.00666 | 0.229418 | 0.238746 | 0.960932 | 0.336587 | 0.505541 |
| OR6C72P | 0.162376 | 0.424913 | 4.079006 | 0.104171 | 0.917034 | NA |
| PRR13 | 1069.384 | -0.10151 | 0.144346 | -0.70323 | 0.48191 | 0.636073 |
| C15orf59 | 832.4065 | -1.11864 | 0.24544 | -4.55768 | 5.17E-06 | 4.24E-04 |
| CCDC85C | 232.765 | -0.66229 | 0.18581 | -3.56431 | 3.65E-04 | 0.005555 |
| | 1.725846 | -0.23167 | 1.459515 | -0.15873 | 0.873881 | NA |
| DYNLL1P1 | 1.181439 | -0.20977 | 1.987054 | -0.10557 | 0.915925 | NA |
| MUC19 | 0.813811 | 0.928591 | 2.591569 | 0.358312 | 0.72011 | NA |
| LIN52 | 343.4439 | 0.104634 | 0.197004 | 0.531126 | 0.595331 | 0.725499 |
| DPF3 | 16.5688 | -0.82846 | 0.536853 | -1.54318 | 0.122786 | NA |
| LYRM5 | 210.602 | 0.499279 | 0.194703 | 2.564317 | 0.010338 | 0.048099 |
| CATSPER2F | 15.17374 | -0.2393 | 0.558316 | -0.42862 | 0.668201 | NA |
| LOH12CR2 | 41.8635 | -0.09783 | 0.381973 | -0.25611 | 0.797863 | NA |
| TPTE2P6 | 0.908083 | 3.025337 | 2.728641 | 1.108734 | 0.267545 | NA |
| C1QTNF9B- | 1.197073 | 1.305651 | 1.836533 | 0.710932 | 0.477126 | NA |
| C1QTNF9B | 0.298089 | -0.51711 | 3.975232 | -0.13008 | 0.896501 | NA |
| RPS3AP47 | 7.956367 | 0.253152 | 0.794826 | 0.3185 | 0.750106 | NA |
| C1RL-AS1 | 19.90167 | 0.367579 | 0.482267 | 0.762191 | 0.445946 | NA |
| NYNRIN | 532.2352 | -0.91977 | 0.206048 | -4.46387 | 8.05E-06 | 5.46E-04 |

| | | | | | | |
|---|---|---|---|---|---|---|
| | 0.813811 | 0.928591 | 2.591569 | 0.358312 | 0.72011 | NA |
| GOLGA8O | 0.096202 | 0.408546 | 4.079639 | 0.100143 | 0.920231 | NA |
| HERC2P9 | 47.27451 | 0.816001 | 0.384056 | 2.124692 | 0.033612 | 0.107059 |
| | 1.099787 | -2.48189 | 2.397845 | -1.03505 | 0.300646 | NA |
| ATP10A | 2496.005 | -0.45755 | 0.207966 | -2.2001 | 0.0278 | 0.094335 |
| RNU6-1 | 0.096202 | 0.408546 | 4.079639 | 0.100143 | 0.920231 | NA |
| SNORA70G | 0.096202 | 0.408546 | 4.079639 | 0.100143 | 0.920231 | NA |
| SNORD116 | 0.162376 | 0.424913 | 4.079006 | 0.104171 | 0.917034 | NA |
| | 0.096202 | 0.408546 | 4.079639 | 0.100143 | 0.920231 | NA |
| SNORD116 | 0.163957 | 0.424913 | 4.079006 | 0.104171 | 0.917034 | NA |
| | 0.163957 | 0.424913 | 4.079006 | 0.104171 | 0.917034 | NA |
| | 0.163957 | 0.424913 | 4.079006 | 0.104171 | 0.917034 | NA |
| | 0.572651 | 0.928649 | 2.780261 | 0.334015 | 0.738368 | NA |
| | 0.163957 | 0.424913 | 4.079006 | 0.104171 | 0.917034 | NA |
| | 0.851596 | 0.74818 | 2.455927 | 0.304643 | 0.760638 | NA |
| RNU6-36P | 0.402398 | -2.58771 | 4.018369 | -0.64397 | 0.519595 | NA |
| RNY1P6 | 1.38705 | -1.18656 | 1.632795 | -0.7267 | 0.467408 | NA |
| RNU6-80P | 0.162376 | 0.424913 | 4.079006 | 0.104171 | 0.917034 | NA |
| RNU6-689P | 0.163957 | 0.424913 | 4.079006 | 0.104171 | 0.917034 | NA |
| RNU6-574P | 0.134133 | -1.48909 | 4.077584 | -0.36519 | 0.71497 | NA |
| SNORD59A | 1.770073 | 0.275437 | 1.470334 | 0.187329 | 0.851402 | NA |
| SNORD116 | 0.35636 | 1.709252 | 3.651833 | 0.468053 | 0.639746 | NA |
| SNORD116 | 0.192403 | 0.975392 | 4.060716 | 0.240202 | 0.810174 | NA |
| RNU6-16P | 0.163957 | 0.424913 | 4.079006 | 0.104171 | 0.917034 | NA |
| | 0.708364 | 0.219301 | 2.144998 | 0.102238 | 0.918568 | NA |
| SNORD116 | 0.192403 | 0.975392 | 4.060716 | 0.240202 | 0.810174 | NA |
| RNU6-1157 | 0.3441 | -0.0166 | 3.769682 | -0.0044 | 0.996486 | NA |
| RNU6-45P | 0.162376 | 0.424913 | 4.079006 | 0.104171 | 0.917034 | NA |
| SNORD116 | 0.230334 | -0.53318 | 4.076749 | -0.13079 | 0.895945 | NA |
| SNORD116 | 1.531863 | -0.70736 | 1.574423 | -0.44928 | 0.653231 | NA |
| RNU6-83P | 0.163957 | 0.424913 | 4.079006 | 0.104171 | 0.917034 | NA |
| SNORA2B | 0.640755 | -1.64826 | 2.754778 | -0.59833 | 0.549621 | NA |
| RNY1P4 | 0.258578 | 1.233151 | 4.047809 | 0.304646 | 0.760635 | NA |
| SNORD116 | 0.151697 | -1.546 | 4.075469 | -0.37934 | 0.704433 | NA |
| SNORD116 | 0.804712 | -0.7622 | 2.348804 | -0.3245 | 0.745556 | NA |
| SNORD15B | 1.036241 | 1.610618 | 2.405425 | 0.669577 | 0.503127 | NA |
| RNU6-19P | 0.134133 | -1.48909 | 4.077584 | -0.36519 | 0.71497 | NA |
| SNORD116 | 0.258578 | 1.233151 | 4.047809 | 0.304646 | 0.760635 | NA |
| MIR615 | 0.3441 | -0.0166 | 3.769682 | -0.0044 | 0.996486 | NA |
| MIR619 | 0.230334 | -0.53318 | 4.076749 | -0.13079 | 0.895945 | NA |
| MIR621 | 20.2572 | -0.71416 | 0.504451 | -1.41572 | 0.156859 | NA |
| MIR211 | 9.558167 | 0.385079 | 0.667123 | 0.577224 | 0.563788 | NA |
| MIR26A2 | 0.28583 | -2.21358 | 3.97376 | -0.55705 | 0.577494 | NA |
| MIR34B | 0.096202 | 0.408546 | 4.079639 | 0.100143 | 0.920231 | NA |
| MIR613 | 0.230334 | -0.53318 | 4.076749 | -0.13079 | 0.895945 | NA |
| MIR616 | 4.258344 | -0.40106 | 0.965309 | -0.41547 | 0.677796 | NA |
| DIO2 | 3.034513 | 1.968463 | 1.374074 | 1.432574 | 0.151979 | NA |

| | | | | | | |
|---|---|---|---|---|---|---|
| STK38L | 350.986 | 0.841751 | 0.251857 | 3.342173 | 8.31E-04 | 0.009276 |
| SLC48A1 | 504.053 | -0.35203 | 0.218074 | -1.61425 | 0.106472 | 0.232543 |
| TRAV18 | 0.706784 | 0.225544 | 2.334352 | 0.096619 | 0.923029 | NA |
| TRAV19 | 0.556667 | 0.972496 | 2.522454 | 0.385536 | 0.699841 | NA |
| TRAV20 | 0.162376 | 0.424913 | 4.079006 | 0.104171 | 0.917034 | NA |
| TRAV38-2D | 0.096202 | 0.408546 | 4.079639 | 0.100143 | 0.920231 | NA |
| TRDC | 0.384806 | 1.846216 | 3.999752 | 0.461582 | 0.644381 | NA |
| TRAJ13 | 1.05261 | -0.79 | 2.093679 | -0.37733 | 0.705932 | NA |
| TRAJ10 | 0.096202 | 0.408546 | 4.079639 | 0.100143 | 0.920231 | NA |
| TRAJ6 | 1.099787 | -2.48189 | 2.397845 | -1.03505 | 0.300646 | NA |
| TRAJ5 | 0.163957 | 0.424913 | 4.079006 | 0.104171 | 0.917034 | NA |
| TRAJ3 | 0.162376 | 0.424913 | 4.079006 | 0.104171 | 0.917034 | NA |
| IGHV3-53 | 0.162376 | 0.424913 | 4.079006 | 0.104171 | 0.917034 | NA |
| | 0.096202 | 0.408546 | 4.079639 | 0.100143 | 0.920231 | NA |
| | 21.89628 | -0.69866 | 0.500479 | -1.39597 | 0.162722 | NA |
| TAS2R19 | 1.335561 | 2.492706 | 1.894034 | 1.316083 | 0.188146 | NA |
| TAS2R15 | 2.02629 | 0.206442 | 1.452317 | 0.142147 | 0.886964 | NA |
| TAS2R50 | 0.162376 | 0.424913 | 4.079006 | 0.104171 | 0.917034 | NA |
| TAS2R14 | 16.46604 | 0.692878 | 0.593931 | 1.166597 | 0.243373 | NA |
| TAS2R13 | 0.624422 | 1.162588 | 2.733761 | 0.425271 | 0.670639 | NA |
| RNA5SP37 | 0.096202 | 0.408546 | 4.079639 | 0.100143 | 0.920231 | NA |
| RNU1-149P | 3.223152 | 0.068747 | 1.110916 | 0.061883 | 0.950656 | NA |
| RNY3P6 | 0.163957 | 0.424913 | 4.079006 | 0.104171 | 0.917034 | NA |
| | 0.260158 | 1.240347 | 4.047124 | 0.306476 | 0.759242 | NA |
| | 0.151697 | -1.546 | 4.075469 | -0.37934 | 0.704433 | NA |
| | 1.238275 | -0.45084 | 2.273975 | -0.19826 | 0.842842 | NA |
| | 0.096202 | 0.408546 | 4.079639 | 0.100143 | 0.920231 | NA |
| SNORA53 | 1.323098 | 2.407088 | 1.923019 | 1.251723 | 0.210671 | NA |
| RNU6-466P | 0.422534 | 1.924583 | 3.001758 | 0.641152 | 0.521424 | NA |
| RNU6-808P | 0.096202 | 0.408546 | 4.079639 | 0.100143 | 0.920231 | NA |
| | 3.168197 | 0.207413 | 1.302884 | 0.159196 | 0.873515 | NA |
| | 56.59085 | 0.255853 | 0.364003 | 0.702886 | 0.482127 | 0.636186 |
| LINC00277 | 0.28583 | -2.21358 | 3.97376 | -0.55705 | 0.577494 | NA |
| HNRNPA1P | 0.151697 | -1.546 | 4.075469 | -0.37934 | 0.704433 | NA |
| | 1.951447 | 1.007412 | 1.471129 | 0.684788 | 0.493477 | NA |
| CRIP1 | 4.347302 | -0.03171 | 1.054136 | -0.03008 | 0.976002 | NA |
| RPL7AP60 | 0.930591 | 1.843923 | 2.122799 | 0.868628 | 0.385051 | NA |
| RPL13P6 | 0.394291 | 0.298745 | 3.141711 | 0.09509 | 0.924243 | NA |
| | 0.151697 | -1.546 | 4.075469 | -0.37934 | 0.704433 | NA |
| ST13P10 | 0.72077 | 0.527966 | 3.060238 | 0.172524 | 0.863025 | NA |
| EEF1A1P16 | 2.535716 | -0.30891 | 1.198026 | -0.25785 | 0.796522 | NA |
| RBMS2P1 | 8.040457 | -0.27504 | 0.783429 | -0.35107 | 0.725532 | NA |
| | 0.476449 | 0.524262 | 3.402674 | 0.154073 | 0.877552 | NA |
| RPL6P25 | 0.326536 | 0.037056 | 3.842554 | 0.009644 | 0.992306 | NA |
| | 0.162376 | 0.424913 | 4.079006 | 0.104171 | 0.917034 | NA |
| IFITM9P | 2.316301 | -0.16482 | 1.342383 | -0.12278 | 0.902282 | NA |
| | 0.096202 | 0.408546 | 4.079639 | 0.100143 | 0.920231 | NA |

| | | | | | | |
|---|---|---|---|---|---|---|
| HNRNPCP6 | 0.324752 | 1.53082 | 4.022025 | 0.380609 | 0.703493 | NA |
| | 25.6154 | 0.142729 | 0.413147 | 0.345468 | 0.729743 | NA |
| NPM1P50 | 1.025415 | 1.11532 | 2.099549 | 0.531219 | 0.595267 | NA |
| PCNPP3 | 0.816972 | 0.936457 | 2.589651 | 0.361615 | 0.71764 | NA |
| GAPDHP44 | 0.488709 | 2.119056 | 3.307564 | 0.64067 | 0.521737 | NA |
| RPS3P6 | 0.622842 | 1.158846 | 2.735323 | 0.42366 | 0.671814 | NA |
| | 0.162376 | 0.424913 | 4.079006 | 0.104171 | 0.917034 | NA |
| ST13P11 | 0.424115 | 1.932531 | 3.45462 | 0.559405 | 0.575885 | NA |
| PTPRCAP | 14.73882 | -0.43931 | 0.603179 | -0.72832 | 0.46642 | NA |
| | 0.70693 | -1.62766 | 2.687154 | -0.60572 | 0.544701 | NA |
| RPL18AP3 | 134.8566 | -0.05328 | 0.236173 | -0.2256 | 0.821513 | 0.892561 |
| | 16.50028 | 0.885645 | 0.624066 | 1.419154 | 0.155854 | NA |
| SIPA1 | 3078.898 | -0.01179 | 0.231514 | -0.05092 | 0.959387 | 0.977677 |
| SYNJ2BP | 359.2711 | 0.328136 | 0.208529 | 1.573574 | 0.115586 | 0.245656 |
| ARL2 | 2700.767 | -0.17234 | 0.13921 | -1.23799 | 0.215719 | 0.374868 |
| | 6.974466 | 0.234262 | 0.801796 | 0.292171 | 0.770156 | NA |
| HEXA | 2535.365 | -0.30293 | 0.14437 | -2.09826 | 0.035882 | 0.111736 |
| RPSAP54 | 3.930524 | -0.51793 | 1.00031 | -0.51777 | 0.604621 | NA |
| RPS29 | 7954.279 | 0.302907 | 0.175176 | 1.729159 | 0.083781 | 0.197146 |
| DDX47 | 12.4221 | -0.25422 | 0.599993 | -0.4237 | 0.671781 | NA |
| | 1.802483 | -0.52384 | 1.643994 | -0.31864 | 0.750001 | NA |
| | 1.405846 | 0.267018 | 1.884093 | 0.141722 | 0.887299 | NA |
| LTB4R | 19.05159 | 0.109581 | 0.504205 | 0.217335 | 0.827947 | NA |
| LTB4R2 | 3.235558 | 0.010078 | 1.112256 | 0.009061 | 0.992771 | NA |
| MDP1 | 3.864504 | -0.2023 | 1.057335 | -0.19133 | 0.848266 | NA |
| IRF9 | 36.83968 | 0.316178 | 0.389466 | 0.811823 | 0.416893 | NA |
| | 1.176909 | 1.262874 | 1.942452 | 0.650144 | 0.515599 | NA |
| | 0.544407 | -0.44425 | 2.542878 | -0.1747 | 0.861313 | NA |
| AP1G2 | 358.0889 | -0.19568 | 0.238954 | -0.8189 | 0.412842 | 0.576064 |
| CARKD | 1096.995 | -0.16173 | 0.138344 | -1.16904 | 0.242389 | 0.40535 |
| GANC | 1160.563 | -0.07668 | 0.164267 | -0.46683 | 0.640625 | 0.760601 |
| ZNF891 | 33.47736 | 0.314831 | 0.38663 | 0.814295 | 0.415476 | NA |
| PTMAP5 | 15.84053 | 0.211538 | 0.603626 | 0.350446 | 0.726004 | NA |
| | 3.437285 | -0.40546 | 1.18878 | -0.34107 | 0.733052 | NA |
| RPS4XP1 | 0.478233 | -0.67301 | 2.937473 | -0.22911 | 0.818782 | NA |
| TUBBP2 | 0.46577 | -1.30191 | 3.402879 | -0.38259 | 0.702024 | NA |
| | 0.096202 | 0.408546 | 4.079639 | 0.100143 | 0.920231 | NA |
| SNURF | 12.71502 | 1.715751 | 0.632838 | 2.711199 | 0.006704 | NA |
| STARP1 | 0.096202 | 0.408546 | 4.079639 | 0.100143 | 0.920231 | NA |
| LGMNP1 | 15.49644 | 0.178889 | 0.535786 | 0.333882 | 0.738469 | NA |
| ANG | 101.0851 | -0.28282 | 0.258147 | -1.09557 | 0.273265 | 0.439144 |
| RPL18P10 | 0.392857 | -1.546 | 4.075469 | -0.37934 | 0.704433 | NA |
| VSTM5 | 0.424115 | 1.932531 | 3.45462 | 0.559405 | 0.575885 | NA |
| | 7.587319 | -0.67706 | 0.757599 | -0.8937 | 0.371485 | NA |
| RCN1P2 | 48.75925 | 0.275458 | 0.330413 | 0.833679 | 0.404462 | 0.569031 |
| PPME1 | 1446.39 | -0.19632 | 0.143597 | -1.36713 | 0.171585 | 0.322661 |
| STARD10 | 814.2569 | 0.149724 | 0.272723 | 0.548995 | 0.583009 | 0.716522 |

| | | | | | | |
|---|---|---|---|---|---|---|
| ZNF705E | 0.706784 | 0.225544 | 2.334352 | 0.096619 | 0.923029 | NA |
| CPEB1 | 188.9966 | -0.13417 | 0.194804 | -0.68873 | 0.490993 | 0.642998 |
| | 0.28583 | -2.21358 | 3.97376 | -0.55705 | 0.577494 | NA |
| | 0.968668 | -0.08001 | 2.195869 | -0.03644 | 0.970935 | NA |
| RPL21P1 | 0.803131 | -0.76693 | 2.349822 | -0.32638 | 0.744138 | NA |
| | 1.670148 | 0.750303 | 1.621208 | 0.462805 | 0.643504 | NA |
| | 31.98526 | -0.21183 | 0.441478 | -0.47982 | 0.631355 | NA |
| DDX12P | 85.487 | -0.05052 | 0.264652 | -0.19091 | 0.848596 | 0.909758 |
| LINC00612 | 3.217323 | -0.63032 | 1.178317 | -0.53493 | 0.592699 | NA |
| RPSAP55 | 0.134133 | -1.48909 | 4.077584 | -0.36519 | 0.71497 | NA |
| C14orf182 | 79.02082 | 0.558636 | 0.328246 | 1.701882 | 0.088777 | 0.205541 |
| PHB2 | 6685.312 | -0.24616 | 0.124038 | -1.98455 | 0.047195 | 0.13369 |
| CD27-AS1 | 163.4341 | 0.058054 | 0.206424 | 0.281239 | 0.778527 | 0.864018 |
| PRDX2P1 | 1.118436 | 1.112087 | 2.263211 | 0.491376 | 0.623161 | NA |
| | 0.298089 | -0.51711 | 3.975232 | -0.13008 | 0.896501 | NA |
| | 3.539394 | -0.66879 | 1.051204 | -0.63621 | 0.52464 | NA |
| GOLGA8B | 33.54609 | 1.947515 | 0.580513 | 3.354818 | 7.94E-04 | NA |
| DHRS4-AS1 | 141.03 | -0.0332 | 0.254874 | -0.13027 | 0.896352 | 0.939326 |
| HOMEZ | 163.0035 | -0.42948 | 0.234625 | -1.83048 | 0.067178 | 0.169519 |
| C14orf164 | 10.7926 | -0.92675 | 0.678416 | -1.36606 | 0.171922 | NA |
| | 2.347465 | 0.521868 | 1.301692 | 0.400915 | 0.688482 | NA |
| MRPL3P1 | 1.101075 | 0.447616 | 1.809417 | 0.247381 | 0.804613 | NA |
| | 0.258578 | 1.233151 | 4.047809 | 0.304646 | 0.760635 | NA |
| MIR17HG | 8.585096 | 1.045771 | 0.726367 | 1.439729 | 0.149944 | NA |
| | 0.163957 | 0.424913 | 4.079006 | 0.104171 | 0.917034 | NA |
| SIAH3 | 1.618965 | -1.05912 | 1.584264 | -0.66853 | 0.503798 | NA |
| OR7E37P | 0.096202 | 0.408546 | 4.079639 | 0.100143 | 0.920231 | NA |
| CALM2P3 | 2.088733 | -0.46612 | 1.368405 | -0.34063 | 0.733384 | NA |
| LINC00598 | 0.448206 | -1.2532 | 3.007315 | -0.41672 | 0.676885 | NA |
| | 0.986086 | 0.883356 | 2.062196 | 0.428357 | 0.668391 | NA |
| GRK6P1 | 2.223077 | 0.160383 | 1.360044 | 0.117925 | 0.906127 | NA |
| ESRRAP1 | 0.672013 | 1.253691 | 2.607776 | 0.480751 | 0.630693 | NA |
| | 1.667584 | 0.663098 | 1.683543 | 0.393871 | 0.693676 | NA |
| | 55.71008 | 0.545087 | 0.315666 | 1.726781 | 0.084207 | 0.197795 |
| ADAMTS7F | 5.536364 | -0.5861 | 0.938607 | -0.62444 | 0.53234 | NA |
| H2AFZP3 | 6.203955 | -0.76943 | 0.890187 | -0.86435 | 0.387398 | NA |
| RPL31P52 | 0.134133 | -1.48909 | 4.077584 | -0.36519 | 0.71497 | NA |
| | 0.490493 | 0.776209 | 2.895261 | 0.268096 | 0.788625 | NA |
| TEX40 | 0.096202 | 0.408546 | 4.079639 | 0.100143 | 0.920231 | NA |
| | 4.179201 | -0.13697 | 0.975509 | -0.14041 | 0.888339 | NA |
| | 1.079942 | 3.273783 | 2.61046 | 1.254102 | 0.209805 | NA |
| | 0.749217 | 0.7397 | 2.633289 | 0.280903 | 0.778784 | NA |
| | 0.324752 | 1.53082 | 4.022025 | 0.380609 | 0.703493 | NA |
| MIR548L | 0.163957 | 0.424913 | 4.079006 | 0.104171 | 0.917034 | NA |
| | 0.162376 | 0.424913 | 4.079006 | 0.104171 | 0.917034 | NA |
| MIR548K | 0.134133 | -1.48909 | 4.077584 | -0.36519 | 0.71497 | NA |
| SNORA34 | 0.651435 | 0.249629 | 2.788679 | 0.089515 | 0.928672 | NA |

| | | | | | | |
|---|---|---|---|---|---|---|
| | 8.452807 | -0.341 | 0.792082 | -0.43051 | 0.666822 | NA |
| C12orf61 | 15.71428 | 0.092675 | 0.518779 | 0.178641 | 0.858219 | NA |
| PSMB11 | 0.163957 | 0.424913 | 4.079006 | 0.104171 | 0.917034 | NA |
| POTEG | 0.134133 | -1.48909 | 4.077584 | -0.36519 | 0.71497 | NA |
| RNU6-262F | 0.134133 | -1.48909 | 4.077584 | -0.36519 | 0.71497 | NA |
| RNU6-101F | 0.151697 | -1.546 | 4.075469 | -0.37934 | 0.704433 | NA |
| RNU2-33P | 0.096202 | 0.408546 | 4.079639 | 0.100143 | 0.920231 | NA |
| RNY5P8 | 0.899333 | -0.26387 | 2.222941 | -0.1187 | 0.905512 | NA |
| RNU6-554F | 0.096202 | 0.408546 | 4.079639 | 0.100143 | 0.920231 | NA |
| | 1.933118 | 2.224537 | 1.741093 | 1.277667 | 0.201367 | NA |
| | 0.268266 | -2.15396 | 4.038351 | -0.53338 | 0.593773 | NA |
| | 0.247898 | -0.5901 | 4.074633 | -0.14482 | 0.884851 | NA |
| RNU2-51P | 0.394291 | 0.298745 | 3.141711 | 0.09509 | 0.924243 | NA |
| RNU2-7P | 1.024164 | -1.06815 | 2.390105 | -0.4469 | 0.654944 | NA |
| RN7SKP8 | 0.163957 | 0.424913 | 4.079006 | 0.104171 | 0.917034 | NA |
| RN7SKP17 | 0.162376 | 0.424913 | 4.079006 | 0.104171 | 0.917034 | NA |
| RNY3P8 | 1.621188 | 1.859647 | 1.660407 | 1.119994 | 0.262716 | NA |
| RNU6-516F | 6.195513 | 0.267439 | 0.977125 | 0.2737 | 0.784315 | NA |
| RNU2-6P | 0.134133 | -1.48909 | 4.077584 | -0.36519 | 0.71497 | NA |
| GAPDHP69 | 0.134133 | -1.48909 | 4.077584 | -0.36519 | 0.71497 | NA |
| | 5.950091 | 1.090689 | 0.87453 | 1.247172 | 0.212335 | NA |
| CSNK1A1P1 | 0.096202 | 0.408546 | 4.079639 | 0.100143 | 0.920231 | NA |
| | 0.296509 | -0.51709 | 3.982155 | -0.12985 | 0.896684 | NA |
| LINC00571 | 1.771654 | 0.279558 | 1.444842 | 0.193487 | 0.846577 | NA |
| DNAJA1P1 | 0.134133 | -1.48909 | 4.077584 | -0.36519 | 0.71497 | NA |
| | 11.92172 | -0.73798 | 0.594995 | -1.2403 | 0.214863 | NA |
| | 0.681112 | 2.620532 | 2.519661 | 1.040033 | 0.298324 | NA |
| ATP5C1P1 | 1.639904 | -0.43485 | 1.659691 | -0.262 | 0.793318 | NA |
| GAPDHP55 | 0.163957 | 0.424913 | 4.079006 | 0.104171 | 0.917034 | NA |
| SNHG14 | 845.937 | 0.37831 | 0.187895 | 2.013408 | 0.044072 | 0.127962 |
| PTPN2P2 | 0.096202 | 0.408546 | 4.079639 | 0.100143 | 0.920231 | NA |
| LINC00403 | 0.162376 | 0.424913 | 4.079006 | 0.104171 | 0.917034 | NA |
| LINC00297 | 0.774742 | -0.40577 | 2.223483 | -0.18249 | 0.855195 | NA |
| | 0.487129 | 2.123445 | 3.910539 | 0.543006 | 0.587126 | NA |
| STK24-AS1 | 3.616451 | -0.55447 | 1.027565 | -0.53959 | 0.589478 | NA |
| | 1.94629 | 1.079575 | 1.452696 | 0.743153 | 0.457389 | NA |
| | 12.63693 | 0.148744 | 0.645216 | 0.230534 | 0.817677 | NA |
| ATP5F1P1 | 0.230334 | -0.53318 | 4.076749 | -0.13079 | 0.895945 | NA |
| LINC00365 | 1.848857 | -0.19382 | 1.495245 | -0.12962 | 0.896865 | NA |
| PDIA3P2 | 0.324752 | 1.53082 | 4.022025 | 0.380609 | 0.703493 | NA |
| | 7.24198 | 0.6665 | 0.796021 | 0.837289 | 0.40243 | NA |
| TEX26-AS1 | 0.785714 | -3.2458 | 3.343263 | -0.97085 | 0.331623 | NA |
| | 12.9738 | 0.424364 | 0.63978 | 0.663297 | 0.507141 | NA |
| COL4A2-AS | 0.556813 | -0.57007 | 3.294615 | -0.17303 | 0.862627 | NA |
| | 0.162376 | 0.424913 | 4.079006 | 0.104171 | 0.917034 | NA |
| YBX1P1 | 14.61979 | -0.2136 | 0.555639 | -0.38443 | 0.700662 | NA |
| RPS4XP16 | 3.574565 | 0.849777 | 1.065626 | 0.797444 | 0.425193 | NA |

| | | | | | | |
|---|---|---|---|---|---|---|
| | 0.911244 | 3.022036 | 2.332005 | 1.295896 | 0.195011 | NA |
| LINC01034 | 1.078361 | 3.265254 | 2.293219 | 1.423874 | 0.154483 | NA |
| TMEM233 | 1.544269 | -1.01379 | 1.908871 | -0.53109 | 0.595354 | NA |
| | 6.692583 | 0.177249 | 0.779117 | 0.2275 | 0.820035 | NA |
| LINC01058 | 0.947944 | -0.02704 | 2.20277 | -0.01228 | 0.990206 | NA |
| PSME2P2 | 62.0673 | 0.084732 | 0.318431 | 0.266092 | 0.790168 | 0.872781 |
| GOLGA2P7 | 30.175 | -0.33331 | 0.37785 | -0.88211 | 0.377715 | NA |
| LINC00618 | 0.354779 | 1.703954 | 3.657232 | 0.465914 | 0.641277 | NA |
| | 0.096202 | 0.408546 | 4.079639 | 0.100143 | 0.920231 | NA |
| RPS12P26 | 0.583919 | -1.6942 | 2.749224 | -0.61625 | 0.537732 | NA |
| | 18.5892 | 0.125417 | 0.555248 | 0.225875 | 0.821299 | NA |
| | 0.28583 | -2.21358 | 3.97376 | -0.55705 | 0.577494 | NA |
| SMARCE1P | 0.096202 | 0.408546 | 4.079639 | 0.100143 | 0.920231 | NA |
| | 0.247898 | -0.5901 | 4.074633 | -0.14482 | 0.884851 | NA |
| | 0.163957 | 0.424913 | 4.079006 | 0.104171 | 0.917034 | NA |
| CT62 | 0.134133 | -1.48909 | 4.077584 | -0.36519 | 0.71497 | NA |
| RBMS1P1 | 11.05148 | 0.943177 | 0.640735 | 1.472024 | 0.141014 | NA |
| EDNRB-AS1 | 0.52699 | -2.57878 | 3.248478 | -0.79384 | 0.427288 | NA |
| IPO7P2 | 0.192403 | 0.975392 | 4.060716 | 0.240202 | 0.810174 | NA |
| | 1.139355 | -2.01403 | 2.05367 | -0.9807 | 0.326743 | NA |
| | 0.69625 | -3.13723 | 2.93845 | -1.06765 | 0.285679 | NA |
| | 0.736957 | -1.0303 | 2.640699 | -0.39016 | 0.696418 | NA |
| | 145.8737 | 0.53197 | 0.209479 | 2.539494 | 0.011101 | 0.050462 |
| LINC00937 | 3.598859 | 0.138804 | 1.022933 | 0.135693 | 0.892064 | NA |
| | 2.530704 | -0.99584 | 1.401627 | -0.71049 | 0.477402 | NA |
| CLCP2 | 0.384806 | 1.846216 | 3.999752 | 0.461582 | 0.644381 | NA |
| EIF4A1P5 | 0.296509 | -0.51709 | 3.982155 | -0.12985 | 0.896684 | NA |
| TEX22 | 10.03624 | 0.178812 | 0.653379 | 0.273673 | 0.784336 | NA |
| | 136.1179 | -0.04253 | 0.212099 | -0.20053 | 0.84107 | 0.905336 |
| | 1.760025 | 1.602275 | 1.820053 | 0.880345 | 0.378672 | NA |
| OR8T1P | 0.260158 | 1.240347 | 4.047124 | 0.306476 | 0.759242 | NA |
| ADAM1B | 1.131248 | -0.36273 | 1.88105 | -0.19283 | 0.847091 | NA |
| | 0.458885 | 0.577525 | 3.449278 | 0.167433 | 0.867029 | NA |
| RPS7P10 | 3.165239 | -0.04759 | 1.141752 | -0.04168 | 0.966751 | NA |
| RPL21P110 | 0.134133 | -1.48909 | 4.077584 | -0.36519 | 0.71497 | NA |
| LINC00343 | 0.163957 | 0.424913 | 4.079006 | 0.104171 | 0.917034 | NA |
| | 2.336437 | 0.902978 | 1.396602 | 0.646553 | 0.517921 | NA |
| IGBP1P1 | 0.672013 | 1.253691 | 2.607776 | 0.480751 | 0.630693 | NA |
| PWWP2AP | 0.163957 | 0.424913 | 4.079006 | 0.104171 | 0.917034 | NA |
| ANKRD20A | 0.260158 | 1.240347 | 4.047124 | 0.306476 | 0.759242 | NA |
| FAM66C | 16.015 | -0.13606 | 0.506307 | -0.26873 | 0.788136 | NA |
| OR7E101P | 1.654727 | 0.113166 | 1.647441 | 0.068692 | 0.945235 | NA |
| C14orf132 | 2745.773 | -0.36345 | 0.13303 | -2.73209 | 0.006293 | 0.034429 |
| | 106.1338 | 0.188543 | 0.284512 | 0.662688 | 0.50753 | 0.656489 |
| | 550.2232 | 0.020524 | 0.20012 | 0.102559 | 0.918313 | 0.952709 |
| PARP1P1 | 0.326333 | 1.533889 | 3.826878 | 0.40082 | 0.688553 | NA |
| SMIM2-AS1 | 9.39097 | 1.006931 | 0.770238 | 1.307298 | 0.191112 | NA |

| | | | | | | |
|---|---|---|---|---|---|---|
| LINC00434 | 0.096202 | 0.408546 | 4.079639 | 0.100143 | 0.920231 | NA |
| RBM26-AS: | 28.46376 | 0.752959 | 0.41055 | 1.834025 | 0.06665 | NA |
| FTH1P16 | 10.29492 | -0.30592 | 0.628367 | -0.48685 | 0.626365 | NA |
| | 5.893531 | -0.24375 | 0.950035 | -0.25657 | 0.79751 | NA |
| NCAM1-AS | 1.1988 | 0.520582 | 2.098288 | 0.248099 | 0.804058 | NA |
| RPL7P6 | 1.140347 | 0.579353 | 2.036091 | 0.284542 | 0.775995 | NA |
| LINC00376 | 0.544554 | -2.65036 | 3.220086 | -0.82307 | 0.410468 | NA |
| ATP5G1P5 | 0.151697 | -1.546 | 4.075469 | -0.37934 | 0.704433 | NA |
| | 119.506 | 0.260156 | 0.222145 | 1.17111 | 0.241554 | 0.404634 |
| SLC9A7P1 | 7.403608 | -0.66507 | 0.856118 | -0.77684 | 0.437251 | NA |
| SUCLA2-AS | 5.864462 | 0.231329 | 0.94249 | 0.245444 | 0.806112 | NA |
| PSPC1P1 | 0.508056 | 0.731032 | 2.861912 | 0.255435 | 0.798387 | NA |
| PRKRIRP6 | 0.151697 | -1.546 | 4.075469 | -0.37934 | 0.704433 | NA |
| | 0.584911 | 2.38756 | 2.690807 | 0.887303 | 0.374916 | NA |
| NPM1P22 | 0.163957 | 0.424913 | 4.079006 | 0.104171 | 0.917034 | NA |
| PIN4P1 | 1.806984 | -0.21108 | 1.611054 | -0.13102 | 0.895759 | NA |
| DHX9P1 | 1.762197 | -0.6745 | 1.464846 | -0.46046 | 0.645189 | NA |
| C14orf144 | 0.478233 | -0.67301 | 2.937473 | -0.22911 | 0.818782 | NA |
| | 0.151697 | -1.546 | 4.075469 | -0.37934 | 0.704433 | NA |
| | 0.938698 | 0.081169 | 1.915817 | 0.042368 | 0.966205 | NA |
| | 1.492911 | -0.25016 | 1.872222 | -0.13362 | 0.893707 | NA |
| RPL15P18 | 2.821489 | -0.82645 | 1.204481 | -0.68615 | 0.492621 | NA |
| | 0.354779 | 1.703954 | 3.657232 | 0.465914 | 0.641277 | NA |
| HOTAIR | 9.982744 | 0.366337 | 0.63859 | 0.573666 | 0.566194 | NA |
| | 0.151697 | -1.546 | 4.075469 | -0.37934 | 0.704433 | NA |
| | 4.789858 | -1.02054 | 0.949192 | -1.07517 | 0.282299 | NA |
| FAM207BP | 0.624422 | 1.162588 | 2.733761 | 0.425271 | 0.670639 | NA |
| HMGB3P4 | 0.314073 | -0.57308 | 3.898934 | -0.14698 | 0.883146 | NA |
| | 0.298089 | -0.51711 | 3.975232 | -0.13008 | 0.896501 | NA |
| UBAC2-AS1 | 26.56922 | -0.75171 | 0.42953 | -1.75008 | 0.080105 | NA |
| OR7E128P | 1.443391 | 0.958388 | 1.70004 | 0.563744 | 0.572928 | NA |
| RPL5P32 | 0.382031 | -1.26484 | 3.151989 | -0.40128 | 0.688212 | NA |
| MED4-AS1 | 0.422534 | 1.924583 | 3.001758 | 0.641152 | 0.521424 | NA |
| RPL41 | 5993.863 | -0.07112 | 0.192291 | -0.36986 | 0.711487 | 0.815575 |
| | 4.938747 | -0.90559 | 1.042728 | -0.86848 | 0.385133 | NA |
| ANKRD10-I | 22.33485 | 1.2129 | 0.496676 | 2.442035 | 0.014605 | NA |
| TRAC | 27.49807 | 0.251473 | 0.474186 | 0.530325 | 0.595887 | NA |
| ADAM1A | 15.90435 | 0.81391 | 0.538568 | 1.511249 | 0.130725 | NA |
| | 20.95357 | -0.32449 | 0.487118 | -0.66613 | 0.505326 | NA |
| LINC00459 | 0.163957 | 0.424913 | 4.079006 | 0.104171 | 0.917034 | NA |
| LINC00452 | 0.163957 | 0.424913 | 4.079006 | 0.104171 | 0.917034 | NA |
| ANKRD26P | 0.965158 | 1.861755 | 2.440023 | 0.763007 | 0.445459 | NA |
| RGS17P1 | 0.47961 | 0.535796 | 3.39605 | 0.15777 | 0.874638 | NA |
| PATL2 | 10.60637 | 0.232215 | 0.630013 | 0.368587 | 0.712435 | NA |
| MYCBP2-A: | 0.424115 | 1.932531 | 3.45462 | 0.559405 | 0.575885 | NA |
| LINC00428 | 0.162376 | 0.424913 | 4.079006 | 0.104171 | 0.917034 | NA |
| SACS-AS1 | 1.179712 | 0.701541 | 1.930555 | 0.363389 | 0.716315 | NA |

| | | | | | | |
|---|---|---|---|---|---|---|
| LINC00341 | 107.9972 | -0.38183 | 0.252783 | -1.5105 | 0.130917 | 0.267383 |
| KARSP1 | 0.096202 | 0.408546 | 4.079639 | 0.100143 | 0.920231 | NA |
| EEF1DP3 | 21.10675 | -0.15646 | 0.443123 | -0.35308 | 0.724027 | NA |
| ATP8A2P2 | 0.766634 | 1.516849 | 2.262991 | 0.670285 | 0.502676 | NA |
| LIPT1P1 | 0.096202 | 0.408546 | 4.079639 | 0.100143 | 0.920231 | NA |
| FNTAP2 | 0.162376 | 0.424913 | 4.079006 | 0.104171 | 0.917034 | NA |
| CNOT6LP1 | 0.402398 | -2.58771 | 4.018369 | -0.64397 | 0.519595 | NA |
| FKBP1AP3 | 0.162376 | 0.424913 | 4.079006 | 0.104171 | 0.917034 | NA |
| | 2.946864 | 0.225162 | 1.229126 | 0.183189 | 0.85465 | NA |
| STARD13-I | 0.096202 | 0.408546 | 4.079639 | 0.100143 | 0.920231 | NA |
| OR6C5P | 0.162376 | 0.424913 | 4.079006 | 0.104171 | 0.917034 | NA |
| GOLGA6L5 | 5.662086 | 0.766998 | 0.946355 | 0.810476 | 0.417667 | NA |
| | 0.096202 | 0.408546 | 4.079639 | 0.100143 | 0.920231 | NA |
| | 67.00966 | 0.048367 | 0.30236 | 0.159965 | 0.872909 | 0.924371 |
| | 0.260158 | 1.240347 | 4.047124 | 0.306476 | 0.759242 | NA |
| CCT5P2 | 0.802985 | 0.54709 | 2.208262 | 0.247747 | 0.80433 | NA |
| | 2.652138 | -0.01633 | 1.243077 | -0.01314 | 0.989516 | NA |
| USP12-AS2 | 1.007648 | 2.051938 | 2.077812 | 0.987548 | 0.323374 | NA |
| SMARCE1P | 0.614937 | 2.493639 | 2.617677 | 0.952615 | 0.340785 | NA |
| RPS6P23 | 0.151697 | -1.546 | 4.075469 | -0.37934 | 0.704433 | NA |
| PPIHP1 | 0.260158 | 1.240347 | 4.047124 | 0.306476 | 0.759242 | NA |
| | 17.64901 | -0.1929 | 0.611569 | -0.31541 | 0.752448 | NA |
| EEF1B2P1 | 1.078282 | -2.71064 | 2.113412 | -1.28259 | 0.199636 | NA |
| FAM58DP | 9.012845 | -0.40247 | 0.703851 | -0.5718 | 0.567454 | NA |
| CLIC1P1 | 1.366537 | 0.235538 | 1.674427 | 0.140668 | 0.888132 | NA |
| | 0.517155 | 2.232561 | 3.179631 | 0.702145 | 0.482589 | NA |
| IRX1P1 | 0.574231 | 0.93227 | 2.49998 | 0.372911 | 0.709215 | NA |
| LINC00441 | 0.999644 | -3.70618 | 2.616772 | -1.41632 | 0.156682 | NA |
| PTMAP4 | 24.84219 | -0.09912 | 0.46694 | -0.21227 | 0.831895 | NA |
| | 0.526843 | -0.38719 | 2.855892 | -0.13558 | 0.892156 | NA |
| DLEU2 | 29.60532 | 0.593646 | 0.390237 | 1.521243 | 0.128199 | NA |
| | 6.743483 | -0.12688 | 0.850949 | -0.14911 | 0.88147 | NA |
| OGFOD1P1 | 3.907185 | 0.80505 | 1.047921 | 0.768236 | 0.442347 | NA |
| | 0.134133 | -1.48909 | 4.077584 | -0.36519 | 0.71497 | NA |
| TSPAN19 | 4.943722 | 0.738953 | 0.914493 | 0.808047 | 0.419063 | NA |
| DOC2GP | 1.055267 | -0.21843 | 2.044568 | -0.10683 | 0.914922 | NA |
| | 18.5375 | 0.112112 | 0.534723 | 0.209664 | 0.83393 | NA |
| | 19.14701 | -0.25647 | 0.510941 | -0.50196 | 0.615693 | NA |
| SIK3-IT1 | 0.913173 | 1.256193 | 2.465498 | 0.509509 | 0.610396 | NA |
| TMEM253 | 3.396914 | 0.128457 | 1.11968 | 0.114727 | 0.908662 | NA |
| HCFC2P1 | 0.46735 | -1.30193 | 3.399671 | -0.38296 | 0.701751 | NA |
| LINC00384 | 0.096202 | 0.408546 | 4.079639 | 0.100143 | 0.920231 | NA |
| ST13P4 | 3.405654 | 0.501711 | 1.101469 | 0.455492 | 0.648755 | NA |
| RPLP1P13 | 0.258578 | 1.233151 | 4.047809 | 0.304646 | 0.760635 | NA |
| FTH1P7 | 4.411179 | -0.19126 | 1.072553 | -0.17832 | 0.858472 | NA |
| FTLP8 | 0.134133 | -1.48909 | 4.077584 | -0.36519 | 0.71497 | NA |
| LINC01047 | 0.873901 | 0.695147 | 2.212014 | 0.31426 | 0.753324 | NA |

| | | | | | | |
|---|---|---|---|---|---|---|
| ZDHHC20P- | 2.764323 | 0.075324 | 1.259021 | 0.059827 | 0.952293 | NA |
| | 0.162376 | 0.424913 | 4.079006 | 0.104171 | 0.917034 | NA |
| RASA3-IT1 | 0.392857 | -1.546 | 4.075469 | -0.37934 | 0.704433 | NA |
| RPL3P4 | 57.16923 | -0.17392 | 0.321592 | -0.5408 | 0.588642 | 0.721053 |
| EEF1A1P3 | 2.149758 | 0.779682 | 1.404031 | 0.555317 | 0.578678 | NA |
| LDHBP1 | 1.022583 | -1.06903 | 2.182302 | -0.48986 | 0.62423 | NA |
| | 4.301862 | 0.328265 | 0.950415 | 0.345391 | 0.7298 | NA |
| COL4A2-AS | 1.425973 | 0.261345 | 1.671539 | 0.15635 | 0.875757 | NA |
| RPL34P27 | 1.071397 | -1.5616 | 1.9555 | -0.79857 | 0.424542 | NA |
| CTAGE3P | 0.458885 | 0.577525 | 3.449278 | 0.167433 | 0.867029 | NA |
| RPS11P5 | 21.85286 | 0.135008 | 0.458002 | 0.294776 | 0.768165 | NA |
| MRPS31P2 | 0.258578 | 1.233151 | 4.047809 | 0.304646 | 0.760635 | NA |
| LINC00327 | 18.87619 | 0.700313 | 0.477408 | 1.466907 | 0.142401 | NA |
| | 1.821901 | 0.527314 | 1.486958 | 0.354626 | 0.72287 | NA |
| LINC00456 | 12.38673 | -0.69583 | 0.588124 | -1.18313 | 0.236757 | NA |
| LINC00167 | 1.947724 | 1.580121 | 1.487596 | 1.062198 | 0.288146 | NA |
| MIPEPP3 | 7.615881 | 0.118808 | 0.823674 | 0.144242 | 0.885309 | NA |
| | 0.298089 | -0.51711 | 3.975232 | -0.13008 | 0.896501 | NA |
| AK4P3 | 0.260158 | 1.240347 | 4.047124 | 0.306476 | 0.759242 | NA |
| HMGA1P6 | 0.162376 | 0.424913 | 4.079006 | 0.104171 | 0.917034 | NA |
| LINC01077 | 12.90717 | -0.06252 | 0.620995 | -0.10067 | 0.91981 | NA |
| ST6GALNA( | 0.816825 | 1.673341 | 2.21346 | 0.755984 | 0.449659 | NA |
| RANP8 | 0.640406 | 1.11402 | 2.710977 | 0.410929 | 0.681124 | NA |
| LINC00460 | 0.247898 | -0.5901 | 4.074633 | -0.14482 | 0.884851 | NA |
| KRT8P39 | 0.913027 | 1.87659 | 2.13519 | 0.878887 | 0.379463 | NA |
| CYP51A1P2 | 1.787491 | 0.838926 | 1.549277 | 0.541495 | 0.588166 | NA |
| LINC00462 | 453.0131 | -0.10942 | 0.223165 | -0.49033 | 0.6239 | 0.747838 |
| CALM2P4 | 0.326536 | 0.037056 | 3.842554 | 0.009644 | 0.992306 | NA |
| RNASEH2B- | 1.385616 | -2.9218 | 2.008356 | -1.45482 | 0.145718 | NA |
| GAS6-AS1 | 0.134133 | -1.48909 | 4.077584 | -0.36519 | 0.71497 | NA |
| GOT2P3 | 1.785145 | 2.02495 | 1.632855 | 1.240129 | 0.214928 | NA |
| LINC00284 | 0.327913 | 1.543154 | 4.021063 | 0.383768 | 0.701151 | NA |
| | 0.392857 | -1.546 | 4.075469 | -0.37934 | 0.704433 | NA |
| ATP8A2P3 | 0.096202 | 0.408546 | 4.079639 | 0.100143 | 0.920231 | NA |
| RPS3AP44 | 0.631307 | -0.20384 | 3.097925 | -0.0658 | 0.947537 | NA |
| | 0.096202 | 0.408546 | 4.079639 | 0.100143 | 0.920231 | NA |
| | 3.631044 | 0.651097 | 1.064004 | 0.611931 | 0.540584 | NA |
| NAP1L4P3 | 1.137783 | 0.40046 | 1.978632 | 0.202392 | 0.83961 | NA |
| TCEB2P1 | 0.151697 | -1.546 | 4.075469 | -0.37934 | 0.704433 | NA |
| FHP1 | 0.392857 | -1.546 | 4.075469 | -0.37934 | 0.704433 | NA |
| | 2.192993 | -0.09431 | 1.365441 | -0.06907 | 0.944934 | NA |
| RPS26P47 | 1.357235 | 0.140439 | 1.935419 | 0.072562 | 0.942154 | NA |
| RNF219-AS | 1.082491 | 0.43074 | 2.021318 | 0.213098 | 0.83125 | NA |
| | 0.162376 | 0.424913 | 4.079006 | 0.104171 | 0.917034 | NA |
| RPL13AP20 | 5.68644 | 0.545043 | 0.848966 | 0.642009 | 0.520868 | NA |
| | 35.39683 | 0.121877 | 0.457429 | 0.266439 | 0.789902 | NA |
| | 1.67288 | -1.61584 | 1.528478 | -1.05716 | 0.290441 | NA |

| | | | | | | |
|---|---|---|---|---|---|---|
| MAPKAPK5 | 96.63708 | 0.094074 | 0.252592 | 0.372436 | 0.709569 | 0.814163 |
| OR2AT2P | 0.488709 | 2.119056 | 3.307564 | 0.64067 | 0.521737 | NA |
| | 4.739025 | -0.10566 | 1.010789 | -0.10453 | 0.916746 | NA |
| PCCA-AS1 | 0.151697 | -1.546 | 4.075469 | -0.37934 | 0.704433 | NA |
| HMGN2P5 | 36.43418 | -0.36278 | 0.408841 | -0.88733 | 0.374901 | NA |
| | 1.867652 | 0.904046 | 1.572196 | 0.575021 | 0.565277 | NA |
| LINC00412 | 0.296509 | -0.51709 | 3.982155 | -0.12985 | 0.896684 | NA |
| RPS3AP6 | 48.89546 | 0.30645 | 0.328154 | 0.933861 | 0.350376 | 0.518371 |
| PHBP13 | 0.314073 | -0.57308 | 3.898934 | -0.14698 | 0.883146 | NA |
| TEX21P | 0.303394 | -2.27006 | 4.032451 | -0.56295 | 0.57347 | NA |
| | 1.068447 | 0.289228 | 2.237018 | 0.129292 | 0.897127 | NA |
| LINC00940 | 0.151697 | -1.546 | 4.075469 | -0.37934 | 0.704433 | NA |
| GAPDHP52 | 1.018642 | -1.62775 | 2.390605 | -0.68089 | 0.495938 | NA |
| CHCHD2P8 | 0.151697 | -1.546 | 4.075469 | -0.37934 | 0.704433 | NA |
| | 0.151697 | -1.546 | 4.075469 | -0.37934 | 0.704433 | NA |
| C12orf75 | 994.3988 | 0.14879 | 0.241929 | 0.615017 | 0.538544 | 0.681093 |
| PPP1R3E | 238.4399 | -0.39543 | 0.214872 | -1.84031 | 0.065723 | 0.167212 |
| KDM4E | 0.628146 | -0.21495 | 3.102079 | -0.06929 | 0.944756 | NA |
| | 0.134133 | -1.48909 | 4.077584 | -0.36519 | 0.71497 | NA |
| MCF2L-AS1 | 0.260158 | 1.240347 | 4.047124 | 0.306476 | 0.759242 | NA |
| SMIM2-IT1 | 0.192403 | 0.975392 | 4.060716 | 0.240202 | 0.810174 | NA |
| | 0.506476 | 0.726418 | 2.864186 | 0.253621 | 0.799788 | NA |
| RPL23AP69 | 0.315653 | -0.57305 | 3.892639 | -0.14721 | 0.882963 | NA |
| DNM1P51 | 5.275331 | 0.136208 | 0.864166 | 0.157618 | 0.874758 | NA |
| | 0.163957 | 0.424913 | 4.079006 | 0.104171 | 0.917034 | NA |
| RPL29P30 | 0.134133 | -1.48909 | 4.077584 | -0.36519 | 0.71497 | NA |
| | 90.83934 | -0.03735 | 0.316819 | -0.1179 | 0.906147 | 0.945762 |
| ESRRAP2 | 0.298089 | -0.51711 | 3.975232 | -0.13008 | 0.896501 | NA |
| EIF4A1P7 | 0.814253 | -1.23545 | 2.246777 | -0.54988 | 0.582405 | NA |
| SNX19P2 | 2.836377 | 0.644165 | 1.159887 | 0.555369 | 0.578642 | NA |
| | 0.544407 | -0.44425 | 2.542878 | -0.1747 | 0.861313 | NA |
| POU5F1P3 | 0.096202 | 0.408546 | 4.079639 | 0.100143 | 0.920231 | NA |
| DICER1-AS1 | 16.29554 | 0.596445 | 0.516638 | 1.154473 | 0.248306 | NA |
| ADAM21P1 | 1.654967 | -0.2547 | 1.735485 | -0.14676 | 0.883319 | NA |
| | 2.542728 | 0.557988 | 1.345426 | 0.41473 | 0.67834 | NA |
| LINC00941 | 4.872573 | 0.487219 | 0.914862 | 0.53256 | 0.594339 | NA |
| CPB2-AS1 | 12.40572 | 0.734894 | 0.654648 | 1.122579 | 0.261616 | NA |
| COX7CP1 | 2.542397 | -0.4133 | 1.282105 | -0.32236 | 0.747178 | NA |
| FABP5P2 | 0.392857 | -1.546 | 4.075469 | -0.37934 | 0.704433 | NA |
| MYCBP2-AS | 0.162376 | 0.424913 | 4.079006 | 0.104171 | 0.917034 | NA |
| LINC00545 | 0.096202 | 0.408546 | 4.079639 | 0.100143 | 0.920231 | NA |
| LINC01069 | 0.151697 | -1.546 | 4.075469 | -0.37934 | 0.704433 | NA |
| MYO16-AS | 0.35636 | 1.709252 | 3.651833 | 0.468053 | 0.639746 | NA |
| SUCLG2P2 | 3.76165 | 0.5145 | 1.119782 | 0.459465 | 0.645901 | NA |
| UBE2L5P | 0.623045 | 0.091811 | 2.666142 | 0.034436 | 0.97253 | NA |
| H3F3BP1 | 0.134133 | -1.48909 | 4.077584 | -0.36519 | 0.71497 | NA |
| HNRNPA3P | 0.47961 | 0.535796 | 3.39605 | 0.15777 | 0.874638 | NA |

| | | | | | | |
|---|---|---|---|---|---|---|
| SNRPGP14 | 1.399513 | -0.94631 | 1.712153 | -0.5527 | 0.580467 | NA |
| STARD13-A | 2.993413 | 1.191992 | 1.335601 | 0.892476 | 0.372138 | NA |
| | 2.262146 | 0.489305 | 1.281353 | 0.381866 | 0.702561 | NA |
| INTS6-AS1 | 6.895391 | 0.074815 | 0.851476 | 0.087865 | 0.929984 | NA |
| | 0.489058 | -0.58569 | 3.427543 | -0.17088 | 0.864321 | NA |
| | 12.51679 | 0.33972 | 0.609607 | 0.557278 | 0.577338 | NA |
| FABP5P1 | 0.096202 | 0.408546 | 4.079639 | 0.100143 | 0.920231 | NA |
| PRMT5-AS: | 1.052464 | 0.310783 | 1.847086 | 0.168256 | 0.866382 | NA |
| COX5BP6 | 3.230276 | -0.39527 | 1.199369 | -0.32957 | 0.741728 | NA |
| LINC01065 | 0.096202 | 0.408546 | 4.079639 | 0.100143 | 0.920231 | NA |
| DLEU7-AS1 | 1.282798 | 0.172952 | 1.69878 | 0.10181 | 0.918908 | NA |
| | 1.160722 | 2.193377 | 1.991761 | 1.101225 | 0.270799 | NA |
| | 0.448206 | -1.2532 | 3.007315 | -0.41672 | 0.676885 | NA |
| LINC00987 | 14.29693 | 0.91093 | 0.612104 | 1.488195 | 0.1367 | NA |
| MAPK6PS3 | 0.623191 | -1.58103 | 2.770593 | -0.57065 | 0.568239 | NA |
| CKMT1B | 0.151697 | -1.546 | 4.075469 | -0.37934 | 0.704433 | NA |
| | 11.34758 | -0.19578 | 0.619549 | -0.316 | 0.752001 | NA |
| LINC01071 | 0.163957 | 0.424913 | 4.079006 | 0.104171 | 0.917034 | NA |
| | 2.597964 | 0.416761 | 1.251379 | 0.333041 | 0.739103 | NA |
| UBE2Q2P6 | 2.731508 | 1.316359 | 1.28786 | 1.022129 | 0.30672 | NA |
| | 0.163957 | 0.424913 | 4.079006 | 0.104171 | 0.917034 | NA |
| FRY-AS1 | 0.151697 | -1.546 | 4.075469 | -0.37934 | 0.704433 | NA |
| KRR1P1 | 0.392711 | 0.292538 | 3.145221 | 0.09301 | 0.925895 | NA |
| GGTA2P | 0.096202 | 0.408546 | 4.079639 | 0.100143 | 0.920231 | NA |
| | 0.640609 | 0.042838 | 2.379511 | 0.018003 | 0.985637 | NA |
| GOLGA2P5 | 8.871856 | 0.688393 | 0.769291 | 0.894841 | 0.370872 | NA |
| | 1.427903 | -1.31794 | 1.795076 | -0.7342 | 0.462829 | NA |
| LINC00426 | 2.951455 | -0.08069 | 1.27144 | -0.06347 | 0.949395 | NA |
| CASC4P1 | 0.096202 | 0.408546 | 4.079639 | 0.100143 | 0.920231 | NA |
| | 0.230334 | -0.53318 | 4.076749 | -0.13079 | 0.895945 | NA |
| TARDBPP2 | 0.411855 | 0.250456 | 3.097618 | 0.080854 | 0.935558 | NA |
| SLC35E1P1 | 1.103647 | 3.314505 | 2.124646 | 1.560027 | 0.118754 | NA |
| | 0.162376 | 0.424913 | 4.079006 | 0.104171 | 0.917034 | NA |
| | 1.140733 | -1.35268 | 2.071057 | -0.65314 | 0.513669 | NA |
| | 1.139152 | -1.36402 | 1.928942 | -0.70714 | 0.479482 | NA |
| RNU7-166F | 0.134133 | -1.48909 | 4.077584 | -0.36519 | 0.71497 | NA |
| | 0.3441 | -0.0166 | 3.769682 | -0.0044 | 0.996486 | NA |
| | 0.096202 | 0.408546 | 4.079639 | 0.100143 | 0.920231 | NA |
| SCARNA12 | 1.85861 | 0.482468 | 1.468988 | 0.328436 | 0.742582 | NA |
| RNU7-59P | 0.162376 | 0.424913 | 4.079006 | 0.104171 | 0.917034 | NA |
| | 0.163957 | 0.424913 | 4.079006 | 0.104171 | 0.917034 | NA |
| SNORD108 | 1.443594 | 0.453431 | 1.768231 | 0.256432 | 0.797617 | NA |
| | 0.650297 | -2.03865 | 2.649751 | -0.76937 | 0.441671 | NA |
| SNORD127 | 0.324752 | 1.53082 | 4.022025 | 0.380609 | 0.703493 | NA |
| | 0.260158 | 1.240347 | 4.047124 | 0.306476 | 0.759242 | NA |
| RPSAP4 | 2.126965 | -0.80875 | 1.567017 | -0.51611 | 0.605777 | NA |
| RPL21P10 | 0.852156 | 0.629888 | 2.425889 | 0.259653 | 0.795132 | NA |

| | | | | | | |
|---|---|---|---|---|---|---|
| RBM14 | 895.3801 | -0.60097 | 0.149426 | -4.02182 | 5.78E-05 | 0.001791 |
| | 7.864145 | -0.2059 | 0.721675 | -0.28531 | 0.775405 | NA |
| | 1.321867 | 0.888794 | 1.796137 | 0.494836 | 0.620716 | NA |
| | 0.326333 | 1.533889 | 3.826878 | 0.40082 | 0.688553 | NA |
| RPL9P5 | 0.151697 | -1.546 | 4.075469 | -0.37934 | 0.704433 | NA |
| | 0.134133 | -1.48909 | 4.077584 | -0.36519 | 0.71497 | NA |
| RPL37P2 | 48.33873 | -0.27015 | 0.346597 | -0.77945 | 0.435715 | 0.59671 |
| | 0.71919 | 0.510978 | 2.728734 | 0.187258 | 0.851458 | NA |
| | 97.34957 | 0.660228 | 0.354404 | 1.862925 | 0.062473 | 0.161813 |
| RN7SL213F | 0.518736 | 2.237423 | 2.7713 | 0.807355 | 0.419462 | NA |
| SUGT1P3 | 5.041982 | -0.06011 | 0.936039 | -0.06422 | 0.948798 | NA |
| RPL23AP1C | 0.458885 | 0.577525 | 3.449278 | 0.167433 | 0.867029 | NA |
| | 2.20918 | -0.44242 | 1.287664 | -0.34358 | 0.73116 | NA |
| RN7SL351F | 0.324752 | 1.53082 | 4.022025 | 0.380609 | 0.703493 | NA |
| | 2.655003 | 0.108486 | 1.212894 | 0.089444 | 0.928729 | NA |
| | 0.450981 | 2.057325 | 3.353149 | 0.61355 | 0.539513 | NA |
| | 0.163957 | 0.424913 | 4.079006 | 0.104171 | 0.917034 | NA |
| RN7SL587F | 0.163957 | 0.424913 | 4.079006 | 0.104171 | 0.917034 | NA |
| RPS2P5 | 3126.885 | -0.26798 | 0.138826 | -1.93031 | 0.053569 | 0.145156 |
| RPL36AP45 | 0.326333 | 1.533889 | 3.826878 | 0.40082 | 0.688553 | NA |
| RPL13P5 | 90.66876 | -0.11663 | 0.272186 | -0.4285 | 0.668288 | 0.782242 |
| RPL9P3 | 0.601483 | -1.7332 | 2.729494 | -0.63499 | 0.525436 | NA |
| | 2.632763 | 1.204797 | 1.291257 | 0.933042 | 0.350798 | NA |
| | 0.326333 | 1.533889 | 3.826878 | 0.40082 | 0.688553 | NA |
| RN7SL472F | 0.096202 | 0.408546 | 4.079639 | 0.100143 | 0.920231 | NA |
| RPS19P3 | 15.04233 | 0.287811 | 0.555866 | 0.517771 | 0.604618 | NA |
| RN7SL331F | 0.258578 | 1.233151 | 4.047809 | 0.304646 | 0.760635 | NA |
| RN7SL38P | 0.151697 | -1.546 | 4.075469 | -0.37934 | 0.704433 | NA |
| | 8.451452 | -1.00783 | 0.717621 | -1.40441 | 0.160198 | NA |
| | 0.296509 | -0.51709 | 3.982155 | -0.12985 | 0.896684 | NA |
| ARHGEF25 | 387.8174 | -0.18014 | 0.161579 | -1.11488 | 0.264901 | 0.429314 |
| RPL15P2 | 0.326536 | 0.037056 | 3.842554 | 0.009644 | 0.992306 | NA |
| RPL4P1 | 0.096202 | 0.408546 | 4.079639 | 0.100143 | 0.920231 | NA |
| RPL23AP64 | 7.838431 | 0.287132 | 0.746616 | 0.384578 | 0.70055 | NA |
| RN7SL459F | 0.096202 | 0.408546 | 4.079639 | 0.100143 | 0.920231 | NA |
| RPL29P24 | 0.968668 | -0.08001 | 2.195869 | -0.03644 | 0.970935 | NA |
| | 0.096202 | 0.408546 | 4.079639 | 0.100143 | 0.920231 | NA |
| RPL5P1 | 2.783208 | 0.178505 | 1.260212 | 0.141646 | 0.887359 | NA |
| | 6.273313 | 0.330079 | 0.891758 | 0.370144 | 0.711275 | NA |
| | 2.883233 | 2.214892 | 1.438507 | 1.539715 | 0.12363 | NA |
| RN7SL809F | 0.586491 | 2.39072 | 2.689057 | 0.889055 | 0.373973 | NA |
| | 2.720828 | 0.886028 | 1.22997 | 0.720366 | 0.4713 | NA |
| RPL36AP43 | 0.652869 | 1.292592 | 2.374873 | 0.544278 | 0.58625 | NA |
| | 8.577369 | 0.37333 | 0.748145 | 0.499008 | 0.617774 | NA |
| | 0.327913 | 1.543154 | 4.021063 | 0.383768 | 0.701151 | NA |
| | 1.101075 | 0.447616 | 1.809417 | 0.247381 | 0.804613 | NA |
| | 0.134133 | -1.48909 | 4.077584 | -0.36519 | 0.71497 | NA |

| | | | | | | |
|---|---|---|---|---|---|---|
| RPL31P49 | 5.052267 | -0.38978 | 0.930686 | -0.41881 | 0.675356 | NA |
| RN7SL10P | 0.163957 | 0.424913 | 4.079006 | 0.104171 | 0.917034 | NA |
| RPSAP52 | 0.247898 | -0.5901 | 4.074633 | -0.14482 | 0.884851 | NA |
| | 0.096202 | 0.408546 | 4.079639 | 0.100143 | 0.920231 | NA |
| | 1.018713 | 1.086497 | 2.092548 | 0.519222 | 0.603606 | NA |
| RN7SL390F | 0.134133 | -1.48909 | 4.077584 | -0.36519 | 0.71497 | NA |
| | 0.452561 | 2.061505 | 3.349456 | 0.615474 | 0.538241 | NA |
| PLEKHO2 | 2848.939 | -0.44905 | 0.130414 | -3.44327 | 5.75E-04 | 0.007385 |
| RPL32P26 | 0.326333 | 1.533889 | 3.826878 | 0.40082 | 0.688553 | NA |
| RPL7AP2 | 0.736607 | 1.395844 | 2.318642 | 0.602009 | 0.547168 | NA |
| HYPK | 14.20291 | 0.302233 | 0.626659 | 0.482293 | 0.629598 | NA |
| | 0.096202 | 0.408546 | 4.079639 | 0.100143 | 0.920231 | NA |
| RPL7AP6 | 18.42248 | 0.085006 | 0.58065 | 0.146398 | 0.883607 | NA |
| RPS20P33 | 2.041297 | 0.149622 | 1.411528 | 0.106 | 0.915582 | NA |
| | 0.326536 | 0.037056 | 3.842554 | 0.009644 | 0.992306 | NA |
| RPL5P3 | 0.586491 | 2.39072 | 2.689057 | 0.889055 | 0.373973 | NA |
| | 0.631307 | -0.20384 | 3.097925 | -0.0658 | 0.947537 | NA |
| | 0.314073 | -0.57308 | 3.898934 | -0.14698 | 0.883146 | NA |
| RN7SL683F | 0.096202 | 0.408546 | 4.079639 | 0.100143 | 0.920231 | NA |
| | 0.659753 | -0.00581 | 2.629089 | -0.00221 | 0.998237 | NA |
| | 0.162376 | 0.424913 | 4.079006 | 0.104171 | 0.917034 | NA |
| | 0.247898 | -0.5901 | 4.074633 | -0.14482 | 0.884851 | NA |
| C15orf38 | 494.7522 | -0.50551 | 0.168927 | -2.99246 | 0.002767 | 0.020111 |
| RPL21P11 | 0.394291 | 0.298745 | 3.141711 | 0.09509 | 0.924243 | NA |
| | 0.831228 | 1.631574 | 2.456512 | 0.664183 | 0.506573 | NA |
| RN7SL529F | 3.986369 | -1.24414 | 0.989977 | -1.25674 | 0.208849 | NA |
| CCDC169 | 30.38053 | 1.351557 | 0.406236 | 3.327023 | 8.78E-04 | NA |
| | 0.394291 | 0.298745 | 3.141711 | 0.09509 | 0.924243 | NA |
| | 3.697751 | -0.20305 | 1.102831 | -0.18412 | 0.853922 | NA |
| STRC | 0.162376 | 0.424913 | 4.079006 | 0.104171 | 0.917034 | NA |
| | 0.163957 | 0.424913 | 4.079006 | 0.104171 | 0.917034 | NA |
| | 0.392857 | -1.546 | 4.075469 | -0.37934 | 0.704433 | NA |
| | 0.392711 | 0.292538 | 3.145221 | 0.09301 | 0.925895 | NA |
| | 0.420954 | 1.923376 | 3.462803 | 0.555439 | 0.578594 | NA |
| RN7SL239F | 0.432222 | -1.18916 | 3.482045 | -0.34151 | 0.732718 | NA |
| | 0.258578 | 1.233151 | 4.047809 | 0.304646 | 0.760635 | NA |
| RN7SL737F | 4.010794 | 1.638452 | 1.088017 | 1.505907 | 0.132091 | NA |
| UBA52P3 | 0.631307 | -0.20384 | 3.097925 | -0.0658 | 0.947537 | NA |
| RPS11P6 | 4.544631 | -0.35603 | 1.05273 | -0.3382 | 0.735215 | NA |
| | 0.3441 | -0.0166 | 3.769682 | -0.0044 | 0.996486 | NA |
| | 0.506476 | 0.726418 | 2.864186 | 0.253621 | 0.799788 | NA |
| | 0.163957 | 0.424913 | 4.079006 | 0.104171 | 0.917034 | NA |
| RPS6P16 | 0.162376 | 0.424913 | 4.079006 | 0.104171 | 0.917034 | NA |
| FGF14-IT1 | 0.096202 | 0.408546 | 4.079639 | 0.100143 | 0.920231 | NA |
| RPL3P3 | 1.099787 | -2.48189 | 2.397845 | -1.03505 | 0.300646 | NA |
| | 1.239709 | 0.808051 | 1.989464 | 0.406165 | 0.684621 | NA |
| RPL35AP32 | 0.640755 | -1.64826 | 2.754778 | -0.59833 | 0.549621 | NA |

| | | | | | | |
|---|---|---|---|---|---|---|
| MRPS31P5 | 0.162376 | 0.424913 | 4.079006 | 0.104171 | 0.917034 | NA |
| RPL5P30 | 1.39773 | -0.43577 | 1.63096 | -0.26719 | 0.789323 | NA |
| | 6.393591 | -0.49725 | 0.839425 | -0.59237 | 0.553606 | NA |
| RPS4XP14 | 1.306686 | -0.48448 | 1.881729 | -0.25747 | 0.796819 | NA |
| PLA2G4B | 2.019792 | -0.18807 | 1.448619 | -0.12983 | 0.896704 | NA |
| RPL21P116 | 0.449786 | -1.25325 | 3.00474 | -0.41709 | 0.676613 | NA |
| JMJD7 | 2.207037 | 0.007309 | 1.447076 | 0.005051 | 0.99597 | NA |
| | 0.410275 | 0.244283 | 3.100889 | 0.078778 | 0.937209 | NA |
| | 0.804566 | 0.545894 | 2.033578 | 0.26844 | 0.78836 | NA |
| RN7SL67P | 6.16097 | 1.17684 | 0.969312 | 1.214099 | 0.22471 | NA |
| | 0.258578 | 1.233151 | 4.047809 | 0.304646 | 0.760635 | NA |
| | 0.258578 | 1.233151 | 4.047809 | 0.304646 | 0.760635 | NA |
| | 0.46577 | -1.30191 | 3.402879 | -0.38259 | 0.702024 | NA |
| RPL13AP2 | 0.824933 | -0.18668 | 2.233587 | -0.08358 | 0.933393 | NA |
| RN7SL417P | 0.163957 | 0.424913 | 4.079006 | 0.104171 | 0.917034 | NA |
| RPS12P20 | 0.151697 | -1.546 | 4.075469 | -0.37934 | 0.704433 | NA |
| | 0.793683 | 0.370036 | 2.527443 | 0.146407 | 0.8836 | NA |
| RPS6P22 | 0.163957 | 0.424913 | 4.079006 | 0.104171 | 0.917034 | NA |
| RPL30P11 | 0.489058 | -0.58569 | 3.427543 | -0.17088 | 0.864321 | NA |
| RPL12P33 | 0.35636 | 1.709252 | 3.651833 | 0.468053 | 0.639746 | NA |
| | 1.060368 | -0.45434 | 2.030513 | -0.22376 | 0.822947 | NA |
| | 1.832453 | -0.75045 | 1.601513 | -0.46859 | 0.639363 | NA |
| N4BP2L2 | 487.1816 | 0.940265 | 0.279405 | 3.365244 | 7.65E-04 | 0.00881 |
| GABPB1-AS | 72.43765 | 0.154469 | 0.305352 | 0.505871 | 0.612947 | 0.739743 |
| | 2.414891 | 1.483519 | 1.358407 | 1.092102 | 0.274788 | NA |
| | 3.654921 | -0.1225 | 1.10877 | -0.11048 | 0.91203 | NA |
| A2M-AS1 | 11.69529 | -0.67408 | 0.665803 | -1.01244 | 0.311329 | NA |
| ARAP1-AS2 | 1.033466 | -0.7302 | 1.946441 | -0.37515 | 0.707552 | NA |
| | 4.84627 | 0.245993 | 0.889018 | 0.276702 | 0.782009 | NA |
| EEF1A1P4 | 5.915188 | -0.38467 | 0.85364 | -0.45062 | 0.652263 | NA |
| USP2-AS1 | 2.280196 | 2.527188 | 1.647676 | 1.533789 | 0.125082 | NA |
| | 6.204949 | -0.77867 | 0.866429 | -0.89871 | 0.368805 | NA |
| NEAT1 | 2431.345 | 1.288125 | 0.207646 | 6.203472 | 5.52E-10 | 5.95E-07 |
| | 1.98685 | 2.287057 | 1.615946 | 1.415305 | 0.156979 | NA |
| | 30.53494 | 0.47002 | 0.390137 | 1.204757 | 0.228297 | NA |
| DDX11-AS1 | 13.79744 | -0.53183 | 0.553149 | -0.96145 | 0.336325 | NA |
| | 3.734383 | -1.03366 | 1.112721 | -0.92895 | 0.352914 | NA |
| | 19.86412 | -0.1936 | 0.45775 | -0.42295 | 0.672333 | NA |
| | 3.008385 | 0.062782 | 1.190885 | 0.052719 | 0.957956 | NA |
| | 0.162376 | 0.424913 | 4.079006 | 0.104171 | 0.917034 | NA |
| | 1.526916 | 0.456861 | 1.739807 | 0.262593 | 0.792864 | NA |
| | 0.555233 | -0.57009 | 3.296947 | -0.17291 | 0.862719 | NA |
| RAD51-AS1 | 39.5717 | 0.138551 | 0.403099 | 0.343714 | 0.731061 | NA |
| | 13.40024 | 0.179171 | 0.601108 | 0.298067 | 0.765652 | NA |
| | 3.752429 | -0.64717 | 1.101257 | -0.58766 | 0.55676 | NA |
| RAB30-AS1 | 47.5124 | 0.421885 | 0.323678 | 1.303411 | 0.192434 | 0.346996 |
| LINC00900 | 82.57411 | -0.48444 | 0.258004 | -1.87764 | 0.060431 | 0.158077 |

| | | | | | | |
|---|---|---|---|---|---|---|
| | 0.326333 | 1.533889 | 3.826878 | 0.40082 | 0.688553 | NA |
| KCTD21-AS | 26.70192 | -0.32951 | 0.400648 | -0.82244 | 0.410826 | NA |
| | 0.247898 | -0.5901 | 4.074633 | -0.14482 | 0.884851 | NA |
| | 1.335764 | 1.644383 | 1.880121 | 0.874615 | 0.381783 | NA |
| | 0.096202 | 0.408546 | 4.079639 | 0.100143 | 0.920231 | NA |
| | 43.41838 | 0.418262 | 0.331751 | 1.260771 | 0.207391 | NA |
| | 8.043721 | -0.19341 | 0.849656 | -0.22764 | 0.819928 | NA |
| | 1.118785 | -0.60441 | 1.92382 | -0.31417 | 0.753389 | NA |
| RASSF8-AS: | 272.2867 | 0.157309 | 0.245696 | 0.640259 | 0.522004 | 0.668506 |
| H2AFJ | 3640.073 | -0.29408 | 0.181986 | -1.61593 | 0.10611 | 0.231919 |
| | 1.285748 | -1.30211 | 1.957858 | -0.66507 | 0.506007 | NA |
| | 0.806146 | 0.553068 | 2.206737 | 0.250627 | 0.802102 | NA |
| DNM1P35 | 4.537878 | -0.97632 | 0.933645 | -1.04571 | 0.295696 | NA |
| | 11.16733 | -0.21274 | 0.707579 | -0.30066 | 0.763674 | NA |
| UBAP1L | 440.4702 | -0.23267 | 0.31279 | -0.74384 | 0.456972 | 0.615377 |
| SOCS2-AS1 | 235.2461 | 0.573432 | 0.287539 | 1.994279 | 0.046122 | 0.131911 |
| PGAM5 | 769.9283 | -0.38783 | 0.177785 | -2.18149 | 0.029148 | 0.097036 |
| SNHG10 | 49.48539 | -0.19786 | 0.3224 | -0.61371 | 0.539408 | 0.681722 |
| | 0.163957 | 0.424913 | 4.079006 | 0.104171 | 0.917034 | NA |
| | 14.28345 | 0.430259 | 0.563529 | 0.763508 | 0.445161 | NA |
| | 3.574768 | 0.615445 | 1.045416 | 0.588708 | 0.556057 | NA |
| UBL7-AS1 | 48.778 | 0.007333 | 0.328571 | 0.022318 | 0.982194 | 0.990311 |
| | 1.401882 | 2.193078 | 1.965211 | 1.11595 | 0.264443 | NA |
| | 0.640406 | 1.11402 | 2.710977 | 0.410929 | 0.681124 | NA |
| | 13.47165 | 0.517551 | 0.608774 | 0.850153 | 0.39524 | NA |
| DNAJC3-AS | 17.26775 | 0.773752 | 0.560197 | 1.381214 | 0.167213 | NA |
| | 16.03473 | 0.113446 | 0.499901 | 0.226937 | 0.820473 | NA |
| | 5.624361 | -0.38084 | 0.901343 | -0.42253 | 0.672639 | NA |
| OIP5-AS1 | 2555.608 | 0.399291 | 0.163321 | 2.444832 | 0.014492 | 0.060538 |
| | 0.853176 | 0.733892 | 2.189324 | 0.335214 | 0.737464 | NA |
| | 3.283206 | 0.164504 | 1.078272 | 0.152563 | 0.878743 | NA |
| | 4.01558 | 0.500928 | 1.319698 | 0.379578 | 0.704259 | NA |
| | 184.6749 | -0.06356 | 0.228034 | -0.27874 | 0.780442 | 0.865414 |
| | 6.862706 | 0.605981 | 0.87752 | 0.690561 | 0.489842 | NA |
| | 2.170891 | -0.41034 | 1.428667 | -0.28722 | 0.773943 | NA |
| LINC00926 | 6.135989 | 0.085949 | 0.854683 | 0.100563 | 0.919897 | NA |
| DYNLL1-AS | 185.9854 | -0.03494 | 0.206434 | -0.16925 | 0.8656 | 0.920414 |
| | 4.167163 | 0.471009 | 1.116196 | 0.421977 | 0.673042 | NA |
| DPH6-AS1 | 4.612581 | 0.165864 | 1.003248 | 0.165327 | 0.868686 | NA |
| | 6.115823 | 0.550027 | 0.838459 | 0.655997 | 0.511826 | NA |
| | 7.440335 | -0.04402 | 0.846462 | -0.05201 | 0.958522 | NA |
| WHAMMP: | 8.785492 | 2.1251 | 0.801806 | 2.650392 | 0.00804 | NA |
| GAPDHP61 | 0.747876 | -0.62091 | 2.562446 | -0.24231 | 0.80854 | NA |
| SRP14-AS1 | 126.7058 | 0.205954 | 0.225526 | 0.913219 | 0.361128 | 0.528409 |
| | 0.850576 | 0.626886 | 2.42682 | 0.258316 | 0.796163 | NA |
| DSTNP2 | 99.07425 | 0.097534 | 0.264835 | 0.368283 | 0.712662 | 0.816474 |
| | 0.096202 | 0.408546 | 4.079639 | 0.100143 | 0.920231 | NA |

| | | | | | | |
|---|---|---|---|---|---|---|
| RBM14-RBI | 5.305933 | 0.676901 | 0.919363 | 0.736271 | 0.461565 | NA |
| | 0.544407 | -0.44425 | 2.542878 | -0.1747 | 0.861313 | NA |
| CCDC153 | 11.08961 | 0.316564 | 0.669713 | 0.472686 | 0.636437 | NA |
| ACTN3 | 1.370624 | -0.08357 | 2.026746 | -0.04123 | 0.967111 | NA |
| FMN1 | 9019.585 | -0.16739 | 0.199497 | -0.83906 | 0.401434 | 0.566143 |
| | 0.163957 | 0.424913 | 4.079006 | 0.104171 | 0.917034 | NA |
| | 0.134133 | -1.48909 | 4.077584 | -0.36519 | 0.71497 | NA |
| | 0.555087 | 0.973802 | 2.808575 | 0.346725 | 0.728798 | NA |
| | 0.163957 | 0.424913 | 4.079006 | 0.104171 | 0.917034 | NA |
| PGAM1P9 | 0.096202 | 0.408546 | 4.079639 | 0.100143 | 0.920231 | NA |
| | 0.258578 | 1.233151 | 4.047809 | 0.304646 | 0.760635 | NA |
| | 0.192403 | 0.975392 | 4.060716 | 0.240202 | 0.810174 | NA |
| GAPDHP70 | 1.296007 | -1.20838 | 1.886673 | -0.64048 | 0.521858 | NA |
| | 78.64024 | -0.11298 | 0.372835 | -0.30304 | 0.761858 | 0.851908 |
| LINC00942 | 1.341885 | 1.595113 | 1.923255 | 0.829382 | 0.406888 | NA |
| HOXC13-AS | 1.465513 | 1.689182 | 1.843746 | 0.916168 | 0.359579 | NA |
| | 0.392857 | -1.546 | 4.075469 | -0.37934 | 0.704433 | NA |
| | 1.901714 | 1.59349 | 1.67874 | 0.949218 | 0.34251 | NA |
| | 159.6858 | -0.56331 | 0.290291 | -1.94052 | 0.052317 | 0.142676 |
| | 0.326536 | 0.037056 | 3.842554 | 0.009644 | 0.992306 | NA |
| C15orf38-A | 1.114915 | 1.271012 | 1.991514 | 0.638214 | 0.523334 | NA |
| | 0.910455 | -0.74751 | 2.116266 | -0.35322 | 0.723924 | NA |
| DNAH10OS | 5.740727 | -0.32516 | 0.853599 | -0.38093 | 0.703254 | NA |
| | 0.555233 | -0.57009 | 3.296947 | -0.17291 | 0.862719 | NA |
| | 0.506476 | 0.726418 | 2.864186 | 0.253621 | 0.799788 | NA |
| | 18.97635 | 0.16757 | 0.489203 | 0.342537 | 0.731947 | NA |
| HOXC-AS2 | 7.700602 | -0.84264 | 0.715364 | -1.17791 | 0.238832 | NA |
| FZD10-AS1 | 1.640738 | 0.753093 | 1.709688 | 0.440486 | 0.659585 | NA |
| | 0.260158 | 1.240347 | 4.047124 | 0.306476 | 0.759242 | NA |
| | 52.94507 | 0.191165 | 0.296827 | 0.644027 | 0.519558 | 0.666597 |
| | 5.983388 | -0.06815 | 0.876239 | -0.07777 | 0.93801 | NA |
| LINC00617 | 0.096202 | 0.408546 | 4.079639 | 0.100143 | 0.920231 | NA |
| | 0.096202 | 0.408546 | 4.079639 | 0.100143 | 0.920231 | NA |
| | 1.024164 | -1.06815 | 2.390105 | -0.4469 | 0.654944 | NA |
| HOXC-AS1 | 22.05389 | -0.07859 | 0.434976 | -0.18068 | 0.856621 | NA |
| | 11.33578 | 0.440564 | 0.891553 | 0.494153 | 0.621198 | NA |
| GPR162 | 17.24243 | 0.157003 | 0.507177 | 0.309563 | 0.756894 | NA |
| | 1.834308 | 0.474853 | 1.555301 | 0.305313 | 0.760128 | NA |
| | 0.58526 | 0.001197 | 3.235889 | 3.70E-04 | 0.999705 | NA |
| | 0.163957 | 0.424913 | 4.079006 | 0.104171 | 0.917034 | NA |
| | 0.6908 | 0.277604 | 2.346083 | 0.118327 | 0.905809 | NA |
| | 626.289 | -0.25269 | 0.140576 | -1.79756 | 0.072247 | 0.178615 |
| | 6.14547 | 0.54166 | 0.972095 | 0.557209 | 0.577385 | NA |
| | 0.162376 | 0.424913 | 4.079006 | 0.104171 | 0.917034 | NA |
| | 1.55407 | -1.15016 | 2.33721 | -0.49211 | 0.622642 | NA |
| | 201.0937 | -0.17619 | 0.23492 | -0.75 | 0.453257 | 0.61221 |
| | 16.95504 | 0.741511 | 0.562345 | 1.318605 | 0.187301 | NA |

| | | | | | | |
|---|---|---|---|---|---|---|
| | 1.227246 | 0.674628 | 1.847001 | 0.365256 | 0.71492 | NA |
| SLC25A30-/ | 1.580685 | 0.588029 | 1.561672 | 0.376538 | 0.706517 | NA |
| | 3.242499 | -0.16236 | 1.077264 | -0.15071 | 0.880201 | NA |
| HOXC-AS3 | 7.916615 | -0.67556 | 0.761367 | -0.8873 | 0.374918 | NA |
| | 1.347981 | -1.54787 | 1.827445 | -0.84701 | 0.396988 | NA |
| | 1.820264 | 0.374755 | 1.447466 | 0.258904 | 0.795709 | NA |
| | 1.214637 | 1.27501 | 1.967912 | 0.6479 | 0.51705 | NA |
| | 3.694193 | -0.19766 | 1.251748 | -0.15791 | 0.87453 | NA |
| MALAT1 | 2179.287 | 1.162575 | 0.292179 | 3.978975 | 6.92E-05 | 0.002018 |
| | 62.20918 | -0.10712 | 0.294003 | -0.36433 | 0.715608 | 0.81867 |
| | 0.163957 | 0.424913 | 4.079006 | 0.104171 | 0.917034 | NA |
| PRB1 | 0.151697 | -1.546 | 4.075469 | -0.37934 | 0.704433 | NA |
| RNA5SP38! | 0.260158 | 1.240347 | 4.047124 | 0.306476 | 0.759242 | NA |
| | 5.066916 | 0.546293 | 0.879206 | 0.621349 | 0.53437 | NA |
| SNORD116 | 0.096202 | 0.408546 | 4.079639 | 0.100143 | 0.920231 | NA |
| SCARNA11 | 0.162376 | 0.424913 | 4.079006 | 0.104171 | 0.917034 | NA |
| RNU6-415F | 1.812737 | -1.90001 | 1.779242 | -1.06788 | 0.285576 | NA |
| RNU6-327F | 0.163957 | 0.424913 | 4.079006 | 0.104171 | 0.917034 | NA |
| | 0.162376 | 0.424913 | 4.079006 | 0.104171 | 0.917034 | NA |
| | 0.422534 | 1.924583 | 3.001758 | 0.641152 | 0.521424 | NA |
| RNU7-40P | 0.096202 | 0.408546 | 4.079639 | 0.100143 | 0.920231 | NA |
| | 0.096202 | 0.408546 | 4.079639 | 0.100143 | 0.920231 | NA |
| RNA5SP39! | 0.162376 | 0.424913 | 4.079006 | 0.104171 | 0.917034 | NA |
| RNU6-539F | 0.260158 | 1.240347 | 4.047124 | 0.306476 | 0.759242 | NA |
| SCARNA13 | 2.712042 | -0.27924 | 1.363112 | -0.20486 | 0.837685 | NA |
| RNU6-126F | 0.49029 | 2.123092 | 3.304282 | 0.642527 | 0.520531 | NA |
| | 0.096202 | 0.408546 | 4.079639 | 0.100143 | 0.920231 | NA |
| RNU7-111F | 14.29487 | -0.5343 | 0.568534 | -0.93979 | 0.347327 | NA |
| SCARNA14 | 0.096202 | 0.408546 | 4.079639 | 0.100143 | 0.920231 | NA |
| RNU6-273F | 0.163957 | 0.424913 | 4.079006 | 0.104171 | 0.917034 | NA |
| RNA5SP36/ | 0.134133 | -1.48909 | 4.077584 | -0.36519 | 0.71497 | NA |
| | 0.151697 | -1.546 | 4.075469 | -0.37934 | 0.704433 | NA |
| RN7SKP11 | 0.096202 | 0.408546 | 4.079639 | 0.100143 | 0.920231 | NA |
| RNU6-64P | 0.096202 | 0.408546 | 4.079639 | 0.100143 | 0.920231 | NA |
| RNU6-131! | 0.162376 | 0.424913 | 4.079006 | 0.104171 | 0.917034 | NA |
| MIR2116 | 0.422534 | 1.924583 | 3.001758 | 0.641152 | 0.521424 | NA |
| | 10.98513 | 0.61326 | 0.614899 | 0.997334 | 0.318603 | NA |
| | 0.778894 | 2.823374 | 2.403613 | 1.174637 | 0.24014 | NA |
| SERPINE3 | 16.96413 | 0.520713 | 0.527817 | 0.986541 | 0.323868 | NA |
| | 4.3544 | -0.03958 | 0.937379 | -0.04223 | 0.966318 | NA |
| IGHV3-52 | 0.096202 | 0.408546 | 4.079639 | 0.100143 | 0.920231 | NA |
| ALG11 | 246.2074 | 0.446854 | 0.184697 | 2.419384 | 0.015547 | 0.063532 |
| ATXN7L3B | 1877.046 | -0.24604 | 0.134177 | -1.83369 | 0.0667 | 0.168708 |
| TPTE2P1 | 6.192955 | -0.08995 | 0.912038 | -0.09863 | 0.921432 | NA |
| UTP14C | 564.7907 | 0.60425 | 0.196547 | 3.074325 | 0.00211 | 0.016753 |
| ALG1L10P | 3.234473 | -1.11855 | 1.212694 | -0.92237 | 0.356336 | NA |
| | 0.432222 | -1.18916 | 3.482045 | -0.34151 | 0.732718 | NA |

| | | | | | | |
|---|---|---|---|---|---|---|
| | 1.750092 | -0.41542 | 1.662801 | -0.24983 | 0.802719 | NA |
| | 468.3954 | -0.69719 | 0.258637 | -2.69564 | 0.007025 | 0.037215 |
| | 0.096202 | 0.408546 | 4.079639 | 0.100143 | 0.920231 | NA |
| | 7.458852 | 0.695051 | 0.844705 | 0.822833 | 0.410603 | NA |
| | 109.458 | -0.69897 | 0.277332 | -2.52034 | 0.011724 | 0.05241 |
| | 2.141707 | 1.764114 | 1.462683 | 1.206081 | 0.227786 | NA |
| OR7E11P | 0.151697 | -1.546 | 4.075469 | -0.37934 | 0.704433 | NA |
| SF3A3P2 | 0.544554 | -2.65036 | 3.220086 | -0.82307 | 0.410468 | NA |
| | 0.162376 | 0.424913 | 4.079006 | 0.104171 | 0.917034 | NA |
| HIGD1AP1( | 0.162376 | 0.424913 | 4.079006 | 0.104171 | 0.917034 | NA |
| | 0.096202 | 0.408546 | 4.079639 | 0.100143 | 0.920231 | NA |
| | 2.648569 | 0.862988 | 1.338033 | 0.644968 | 0.518948 | NA |
| | 20.55458 | 0.426989 | 0.501 | 0.852274 | 0.394062 | NA |
| AP5B1 | 1270.694 | -0.44486 | 0.160891 | -2.76497 | 0.005693 | 0.03235 |
| OR2AT1P | 0.151697 | -1.546 | 4.075469 | -0.37934 | 0.704433 | NA |
| PTP4A2P2 | 0.314073 | -0.57308 | 3.898934 | -0.14698 | 0.883146 | NA |
| | 0.296509 | -0.51709 | 3.982155 | -0.12985 | 0.896684 | NA |
| | 0.163957 | 0.424913 | 4.079006 | 0.104171 | 0.917034 | NA |
| RANP3 | 0.162376 | 0.424913 | 4.079006 | 0.104171 | 0.917034 | NA |
| | 1.120365 | -0.61682 | 2.073914 | -0.29742 | 0.766148 | NA |
| CHMP4A | 12.62081 | 0.461139 | 0.591333 | 0.779831 | 0.435491 | NA |
| | 0.862836 | 1.736629 | 2.194323 | 0.791419 | 0.4287 | NA |
| | 0.268266 | -2.15396 | 4.038351 | -0.53338 | 0.593773 | NA |
| OMP | 1.645774 | -1.39946 | 1.608559 | -0.87001 | 0.384296 | NA |
| MTND2P2( | 0.326333 | 1.533889 | 3.826878 | 0.40082 | 0.688553 | NA |
| MAGEL2 | 1.454862 | -0.58515 | 1.616608 | -0.36196 | 0.717381 | NA |
| | 0.315653 | -0.57305 | 3.892639 | -0.14721 | 0.882963 | NA |
| OR7E126P | 0.793886 | -0.43778 | 2.212663 | -0.19785 | 0.84316 | NA |
| | 4.142692 | 1.533768 | 1.164086 | 1.317573 | 0.187647 | NA |
| | 0.792306 | -0.44145 | 2.21354 | -0.19943 | 0.841926 | NA |
| | 0.151697 | -1.546 | 4.075469 | -0.37934 | 0.704433 | NA |
| | 9.700948 | 0.102745 | 0.690477 | 0.148802 | 0.88171 | NA |
| | 47.04734 | 0.192203 | 0.33435 | 0.574857 | 0.565388 | 0.702734 |
| | 0.725928 | 0.181819 | 2.321433 | 0.078322 | 0.937572 | NA |
| | 0.162376 | 0.424913 | 4.079006 | 0.104171 | 0.917034 | NA |
| | 0.314073 | -0.57308 | 3.898934 | -0.14698 | 0.883146 | NA |
| | 0.420954 | 1.923376 | 3.462803 | 0.555439 | 0.578594 | NA |
| | 0.163957 | 0.424913 | 4.079006 | 0.104171 | 0.917034 | NA |
| | 3.311384 | -0.51928 | 1.250045 | -0.41541 | 0.677841 | NA |
| | 0.717609 | 0.516981 | 3.063469 | 0.168757 | 0.865988 | NA |
| STT3A-AS1 | 0.82213 | 0.50433 | 2.197682 | 0.229483 | 0.818494 | NA |
| | 0.449786 | -1.25325 | 3.00474 | -0.41709 | 0.676613 | NA |
| | 1.910944 | 0.085569 | 1.452676 | 0.058904 | 0.953028 | NA |
| OSBPL9P2 | 0.163957 | 0.424913 | 4.079006 | 0.104171 | 0.917034 | NA |
| | 0.096202 | 0.408546 | 4.079639 | 0.100143 | 0.920231 | NA |
| | 15.30307 | 0.347611 | 0.557908 | 0.623062 | 0.533244 | NA |
| | 0.163957 | 0.424913 | 4.079006 | 0.104171 | 0.917034 | NA |

| | | | | | | |
|---|---|---|---|---|---|---|
| | 3.336611 | 3.058139 | 1.369167 | 2.233577 | 0.025511 | NA |
| COPS8P3 | 0.668852 | 1.246964 | 2.610519 | 0.477669 | 0.632886 | NA |
| | 0.192403 | 0.975392 | 4.060716 | 0.240202 | 0.810174 | NA |
| SENCR | 0.883349 | -0.20011 | 2.228601 | -0.08979 | 0.928451 | NA |
| | 1.694885 | -1.50313 | 1.723959 | -0.87191 | 0.383259 | NA |
| PTP4A1P6 | 0.247898 | -0.5901 | 4.074633 | -0.14482 | 0.884851 | NA |
| | 0.163957 | 0.424913 | 4.079006 | 0.104171 | 0.917034 | NA |
| | 1.102655 | 0.449766 | 1.809137 | 0.248608 | 0.803664 | NA |
| | 0.163957 | 0.424913 | 4.079006 | 0.104171 | 0.917034 | NA |
| NAP1L1P1 | 2.652053 | 0.796077 | 1.344113 | 0.59227 | 0.55367 | NA |
| | 0.545988 | -0.43913 | 2.541145 | -0.17281 | 0.862803 | NA |
| | 0.151697 | -1.546 | 4.075469 | -0.37934 | 0.704433 | NA |
| | 9.637127 | 0.204644 | 0.695706 | 0.294154 | 0.768641 | NA |
| FDPSP4 | 0.096202 | 0.408546 | 4.079639 | 0.100143 | 0.920231 | NA |
| TFAMP2 | 0.163957 | 0.424913 | 4.079006 | 0.104171 | 0.917034 | NA |
| | 0.247898 | -0.5901 | 4.074633 | -0.14482 | 0.884851 | NA |
| | 39.91681 | 0.760697 | 0.406286 | 1.872319 | 0.061163 | NA |
| | 0.624422 | 1.162588 | 2.733761 | 0.425271 | 0.670639 | NA |
| | 0.28583 | -2.21358 | 3.97376 | -0.55705 | 0.577494 | NA |
| | 14.80269 | -0.45989 | 0.551817 | -0.83341 | 0.404611 | NA |
| | 0.68922 | 0.287127 | 2.605675 | 0.110193 | 0.912256 | NA |
| | 2.700895 | -0.37548 | 1.18548 | -0.31673 | 0.751447 | NA |
| | 0.162376 | 0.424913 | 4.079006 | 0.104171 | 0.917034 | NA |
| | 0.315653 | -0.57305 | 3.892639 | -0.14721 | 0.882963 | NA |
| | 3.490623 | 1.092801 | 1.098535 | 0.99478 | 0.319843 | NA |
| SCARNA9 | 4.277702 | 1.118365 | 1.121699 | 0.997028 | 0.318751 | NA |
| | 0.71919 | 0.510978 | 2.728734 | 0.187258 | 0.851458 | NA |
| | 14.49186 | 0.282269 | 0.5659 | 0.498797 | 0.617923 | NA |
| | 0.134133 | -1.48909 | 4.077584 | -0.36519 | 0.71497 | NA |
| | 0.52699 | -2.57878 | 3.248478 | -0.79384 | 0.427288 | NA |
| | 1.263248 | 1.409426 | 2.009206 | 0.701484 | 0.483001 | NA |
| ATP5F1P5 | 12.97561 | 0.505837 | 0.578037 | 0.875096 | 0.381522 | NA |
| ALG1L9P | 4.705116 | -0.76797 | 0.978669 | -0.7847 | 0.432627 | NA |
| RSF1-IT2 | 0.815391 | 0.931626 | 2.360431 | 0.394685 | 0.693076 | NA |
| DPP3 | 2539.475 | -0.43245 | 0.130466 | -3.31468 | 9.17E-04 | 0.009804 |
| | 0.422534 | 1.924583 | 3.001758 | 0.641152 | 0.521424 | NA |
| TCEB1P22 | 0.096202 | 0.408546 | 4.079639 | 0.100143 | 0.920231 | NA |
| ARL6IP1P3 | 0.162376 | 0.424913 | 4.079006 | 0.104171 | 0.917034 | NA |
| | 0.192403 | 0.975392 | 4.060716 | 0.240202 | 0.810174 | NA |
| | 23.52934 | -0.04539 | 0.434524 | -0.10446 | 0.916804 | NA |
| | 2.981043 | 1.120101 | 1.193578 | 0.93844 | 0.348018 | NA |
| | 0.258578 | 1.233151 | 4.047809 | 0.304646 | 0.760635 | NA |
| | 0.296509 | -0.51709 | 3.982155 | -0.12985 | 0.896684 | NA |
| | 0.134133 | -1.48909 | 4.077584 | -0.36519 | 0.71497 | NA |
| | 9.335594 | -0.29312 | 0.64205 | -0.45653 | 0.648007 | NA |
| GRM5-AS1 | 0.394291 | 0.298745 | 3.141711 | 0.09509 | 0.924243 | NA |
| | 0.230334 | -0.53318 | 4.076749 | -0.13079 | 0.895945 | NA |

| | | | | | | |
|---|---|---|---|---|---|---|
| | 4.254992 | -1.20277 | 1.001714 | -1.20071 | 0.229863 | NA |
| | 3.020512 | -0.84213 | 1.218712 | -0.691 | 0.489568 | NA |
| | 0.708364 | 0.219301 | 2.144998 | 0.102238 | 0.918568 | NA |
| OVOL1-AS1 | 0.629726 | -0.22725 | 2.732935 | -0.08315 | 0.93373 | NA |
| | 66.8297 | -0.57532 | 0.287749 | -1.99937 | 0.045568 | 0.130706 |
| | 9.221146 | -0.10059 | 0.68248 | -0.14738 | 0.882828 | NA |
| | 48.70628 | -0.27296 | 0.325474 | -0.83867 | 0.401656 | 0.566389 |
| | 10.60164 | -0.37679 | 0.656795 | -0.57368 | 0.566184 | NA |
| EID3 | 11.21681 | 0.571513 | 0.775332 | 0.737121 | 0.461049 | NA |
| | 1.493785 | 0.546363 | 1.627789 | 0.335647 | 0.737137 | NA |
| | 3.329673 | 0.51225 | 1.128749 | 0.453821 | 0.649958 | NA |
| | 0.420954 | 1.923376 | 3.462803 | 0.555439 | 0.578594 | NA |
| DDX18P5 | 0.94967 | -2.20819 | 2.817377 | -0.78378 | 0.433171 | NA |
| SETP17 | 0.096202 | 0.408546 | 4.079639 | 0.100143 | 0.920231 | NA |
| | 0.47961 | 0.535796 | 3.39605 | 0.15777 | 0.874638 | NA |
| | 1.331352 | 0.018035 | 1.822963 | 0.009893 | 0.992107 | NA |
| | 1.073823 | 2.138376 | 2.047615 | 1.044325 | 0.296335 | NA |
| | 66.46906 | -0.07546 | 0.29352 | -0.25707 | 0.797121 | 0.877181 |
| | 0.230334 | -0.53318 | 4.076749 | -0.13079 | 0.895945 | NA |
| TCEB2P2 | 0.448206 | -1.2532 | 3.007315 | -0.41672 | 0.676885 | NA |
| WTAPP1 | 0.681461 | 0.491544 | 3.066278 | 0.160307 | 0.87264 | NA |
| EID1 | 2058.087 | 0.635073 | 0.226239 | 2.80709 | 0.004999 | 0.029618 |
| | 2.505689 | -0.39071 | 1.208862 | -0.32321 | 0.746537 | NA |
| | 0.096202 | 0.408546 | 4.079639 | 0.100143 | 0.920231 | NA |
| | 0.230334 | -0.53318 | 4.076749 | -0.13079 | 0.895945 | NA |
| ENPP7P8 | 0.162376 | 0.424913 | 4.079006 | 0.104171 | 0.917034 | NA |
| | 16.3695 | -0.48125 | 0.530973 | -0.90635 | 0.364748 | NA |
| | 3.588566 | -0.60795 | 1.08586 | -0.55988 | 0.575562 | NA |
| | 1.130898 | 1.215321 | 1.814636 | 0.669732 | 0.503028 | NA |
| | 2.336428 | -0.1932 | 1.251564 | -0.15437 | 0.877321 | NA |
| NOX5 | 0.892891 | -0.69406 | 2.402721 | -0.28886 | 0.772685 | NA |
| | 0.392857 | -1.546 | 4.075469 | -0.37934 | 0.704433 | NA |
| | 0.096202 | 0.408546 | 4.079639 | 0.100143 | 0.920231 | NA |
| | 0.163957 | 0.424913 | 4.079006 | 0.104171 | 0.917034 | NA |
| | 0.134133 | -1.48909 | 4.077584 | -0.36519 | 0.71497 | NA |
| TAS2R43 | 0.151697 | -1.546 | 4.075469 | -0.37934 | 0.704433 | NA |
| | 0.162376 | 0.424913 | 4.079006 | 0.104171 | 0.917034 | NA |
| | 0.422534 | 1.924583 | 3.001758 | 0.641152 | 0.521424 | NA |
| | 2.33742 | 0.998267 | 1.333492 | 0.748612 | 0.454091 | NA |
| TBX5-AS1 | 0.151697 | -1.546 | 4.075469 | -0.37934 | 0.704433 | NA |
| RSF1-IT1 | 0.536503 | 0.87599 | 3.23231 | 0.27101 | 0.786383 | NA |
| | 0.315653 | -0.57305 | 3.892639 | -0.14721 | 0.882963 | NA |
| | 0.862836 | 1.736629 | 2.194323 | 0.791419 | 0.4287 | NA |
| | 0.749217 | 0.7397 | 2.633289 | 0.280903 | 0.778784 | NA |
| | 0.422534 | 1.924583 | 3.001758 | 0.641152 | 0.521424 | NA |
| | 0.709944 | 0.232855 | 2.332364 | 0.099837 | 0.920474 | NA |
| | 61.63181 | 0.112617 | 0.286721 | 0.392775 | 0.694486 | 0.802997 |

| | | | | | | |
|---|---|---|---|---|---|---|
| | 32.27793 | 0.210788 | 0.474413 | 0.444313 | 0.656816 | NA |
| | 1.434495 | -0.14593 | 1.626958 | -0.08969 | 0.928531 | NA |
| | 0.682896 | 1.412658 | 2.61672 | 0.539858 | 0.589295 | NA |
| | 0.151697 | -1.546 | 4.075469 | -0.37934 | 0.704433 | NA |
| | 0.326536 | 0.037056 | 3.842554 | 0.009644 | 0.992306 | NA |
| | 2.86757 | 0.334717 | 1.197728 | 0.27946 | 0.779892 | NA |
| | 55.24623 | 0.205429 | 0.327238 | 0.627766 | 0.530157 | 0.675004 |
| POLR2M | 367.5482 | 0.517764 | 0.190145 | 2.722995 | 0.006469 | 0.035132 |
| | 2.200512 | 0.353205 | 1.395128 | 0.25317 | 0.800137 | NA |
| | 0.163957 | 0.424913 | 4.079006 | 0.104171 | 0.917034 | NA |
| FDXACB1 | 48.99082 | -0.02005 | 0.31521 | -0.06359 | 0.949293 | 0.9717 |
| | 0.247898 | -0.5901 | 4.074633 | -0.14482 | 0.884851 | NA |
| | 0.162376 | 0.424913 | 4.079006 | 0.104171 | 0.917034 | NA |
| LINC00925 | 1.503831 | -0.03766 | 1.734351 | -0.02172 | 0.982674 | NA |
| | 0.163957 | 0.424913 | 4.079006 | 0.104171 | 0.917034 | NA |
| | 0.258578 | 1.233151 | 4.047809 | 0.304646 | 0.760635 | NA |
| | 0.151697 | -1.546 | 4.075469 | -0.37934 | 0.704433 | NA |
| | 1.033319 | 0.349745 | 2.005511 | 0.174392 | 0.861557 | NA |
| | 23.103 | -0.06937 | 0.461544 | -0.1503 | 0.88053 | NA |
| | 0.162376 | 0.424913 | 4.079006 | 0.104171 | 0.917034 | NA |
| | 0.163957 | 0.424913 | 4.079006 | 0.104171 | 0.917034 | NA |
| PABPC1P4 | 5.191893 | 0.182245 | 0.915436 | 0.199079 | 0.842201 | NA |
| FAM222A-/ | 3.491767 | -0.74146 | 1.069324 | -0.69339 | 0.488065 | NA |
| | 1.180005 | -1.57876 | 2.139972 | -0.73775 | 0.460668 | NA |
| | 0.314073 | -0.57308 | 3.898934 | -0.14698 | 0.883146 | NA |
| | 0.192403 | 0.975392 | 4.060716 | 0.240202 | 0.810174 | NA |
| | 0.134133 | -1.48909 | 4.077584 | -0.36519 | 0.71497 | NA |
| | 0.303394 | -2.27006 | 4.032451 | -0.56295 | 0.57347 | NA |
| | 0.574231 | 0.93227 | 2.49998 | 0.372911 | 0.709215 | NA |
| | 0.392857 | -1.546 | 4.075469 | -0.37934 | 0.704433 | NA |
| | 0.3441 | -0.0166 | 3.769682 | -0.0044 | 0.996486 | NA |
| SNHG1 | 813.6945 | 0.223641 | 0.199888 | 1.11883 | 0.263213 | 0.427592 |
| | 0.096202 | 0.408546 | 4.079639 | 0.100143 | 0.920231 | NA |
| HNRNPABF | 0.420954 | 1.923376 | 3.462803 | 0.555439 | 0.578594 | NA |
| AGAP2-AS1 | 258.8503 | -0.10708 | 0.191707 | -0.55856 | 0.576462 | 0.711426 |
| | 1.269344 | -2.06734 | 1.94024 | -1.06551 | 0.286646 | NA |
| | 0.151697 | -1.546 | 4.075469 | -0.37934 | 0.704433 | NA |
| | 1.590133 | 0.039996 | 1.513994 | 0.026417 | 0.978925 | NA |
| | 0.162376 | 0.424913 | 4.079006 | 0.104171 | 0.917034 | NA |
| RMST | 0.163957 | 0.424913 | 4.079006 | 0.104171 | 0.917034 | NA |
| TAS2R20 | 0.584911 | 2.38756 | 2.690807 | 0.887303 | 0.374916 | NA |
| | 0.853323 | -0.46916 | 2.317774 | -0.20242 | 0.839588 | NA |
| TMEM5-AS | 0.315653 | -0.57305 | 3.892639 | -0.14721 | 0.882963 | NA |
| PXN-AS1 | 42.48497 | 0.002375 | 0.334914 | 0.007093 | 0.994341 | NA |
| | 0.151697 | -1.546 | 4.075469 | -0.37934 | 0.704433 | NA |
| | 0.162376 | 0.424913 | 4.079006 | 0.104171 | 0.917034 | NA |
| | 18.82327 | -0.06498 | 0.474275 | -0.13701 | 0.891023 | NA |

| | | | | | | |
|---|---|---|---|---|---|---|
| | 0.096202 | 0.408546 | 4.079639 | 0.100143 | 0.920231 | NA |
| DENND5B-/ | 2.939049 | 0.014517 | 1.164267 | 0.012469 | 0.990052 | NA |
| LINC00346 | 6.565849 | 0.311811 | 0.785221 | 0.3971 | 0.691294 | NA |
| | 2.337834 | 1.382708 | 1.380207 | 1.001812 | 0.316434 | NA |
| | 0.582339 | -1.69412 | 2.750801 | -0.61586 | 0.537985 | NA |
| | 27.64948 | -0.24131 | 0.439537 | -0.54901 | 0.582999 | NA |
| | 0.804712 | -0.7622 | 2.348804 | -0.3245 | 0.745556 | NA |
| | 0.326333 | 1.533889 | 3.826878 | 0.40082 | 0.688553 | NA |
| | 0.163957 | 0.424913 | 4.079006 | 0.104171 | 0.917034 | NA |
| | 0.49029 | 2.123092 | 3.304282 | 0.642527 | 0.520531 | NA |
| | 5.819104 | -0.23193 | 0.818327 | -0.28342 | 0.776853 | NA |
| | 0.747636 | 0.735436 | 2.634453 | 0.279161 | 0.780122 | NA |
| | 1.362427 | 2.544836 | 1.926007 | 1.321302 | 0.186401 | NA |
| | 0.247898 | -0.5901 | 4.074633 | -0.14482 | 0.884851 | NA |
| | 1.98028 | 0.149202 | 1.410595 | 0.105773 | 0.915763 | NA |
| PSMC1P9 | 0.651085 | 2.534961 | 3.013697 | 0.841147 | 0.400266 | NA |
| | 0.392857 | -1.546 | 4.075469 | -0.37934 | 0.704433 | NA |
| | 1.754301 | 1.32931 | 1.726412 | 0.769984 | 0.441309 | NA |
| TAS2R63P | 0.326333 | 1.533889 | 3.826878 | 0.40082 | 0.688553 | NA |
| TCEB1P31 | 0.192403 | 0.975392 | 4.060716 | 0.240202 | 0.810174 | NA |
| | 1.05261 | -0.79 | 2.093679 | -0.37733 | 0.705932 | NA |
| CBX3P4 | 1.254352 | 0.05565 | 1.700006 | 0.032735 | 0.973886 | NA |
| | 0.151697 | -1.546 | 4.075469 | -0.37934 | 0.704433 | NA |
| MRPL40P1 | 0.326333 | 1.533889 | 3.826878 | 0.40082 | 0.688553 | NA |
| | 43.68378 | 0.114629 | 0.368422 | 0.311136 | 0.755697 | NA |
| APOPT1 | 9.5576 | 0.404069 | 0.693873 | 0.582338 | 0.560339 | NA |
| | 0.70046 | 1.37107 | 2.594868 | 0.528378 | 0.597237 | NA |
| DYX1C1 | 2.955555 | 1.419847 | 1.291632 | 1.099266 | 0.271652 | NA |
| A2MP1 | 3.417796 | 2.100238 | 1.248363 | 1.682394 | 0.092492 | NA |
| | 0.260158 | 1.240347 | 4.047124 | 0.306476 | 0.759242 | NA |
| | 0.424115 | 1.932531 | 3.45462 | 0.559405 | 0.575885 | NA |
| | 0.394291 | 0.298745 | 3.141711 | 0.09509 | 0.924243 | NA |
| | 0.392857 | -1.546 | 4.075469 | -0.37934 | 0.704433 | NA |
| | 0.626003 | 1.180003 | 3.130502 | 0.376937 | 0.70622 | NA |
| | 18.99129 | 0.465008 | 0.465821 | 0.998255 | 0.318156 | NA |
| | 13.80225 | -0.14391 | 0.581585 | -0.24744 | 0.804565 | NA |
| | 0.518736 | 2.237423 | 2.7713 | 0.807355 | 0.419462 | NA |
| | 0.096202 | 0.408546 | 4.079639 | 0.100143 | 0.920231 | NA |
| | 0.163957 | 0.424913 | 4.079006 | 0.104171 | 0.917034 | NA |
| | 1.400505 | 1.12359 | 1.76195 | 0.637697 | 0.523671 | NA |
| | 0.096202 | 0.408546 | 4.079639 | 0.100143 | 0.920231 | NA |
| | 0.192403 | 0.975392 | 4.060716 | 0.240202 | 0.810174 | NA |
| | 25.9307 | -0.6215 | 0.412984 | -1.5049 | 0.132349 | NA |
| | 32.99617 | 0.67054 | 0.812217 | 0.825568 | 0.409049 | NA |
| | 0.260158 | 1.240347 | 4.047124 | 0.306476 | 0.759242 | NA |
| | 0.134133 | -1.48909 | 4.077584 | -0.36519 | 0.71497 | NA |
| | 0.163957 | 0.424913 | 4.079006 | 0.104171 | 0.917034 | NA |

| | | | | | | |
|---|---|---|---|---|---|---|
| CCND2-AS1 | 0.736957 | -1.0303 | 2.640699 | -0.39016 | 0.696418 | NA |
| ATF4P4 | 1.199003 | -0.22526 | 1.838234 | -0.12254 | 0.902472 | NA |
| | 7.299809 | 0.613015 | 0.817199 | 0.750142 | 0.453169 | NA |
| TAS2R30 | 0.134133 | -1.48909 | 4.077584 | -0.36519 | 0.71497 | NA |
| | 0.720624 | 1.435938 | 2.334155 | 0.615185 | 0.538432 | NA |
| | 0.774742 | -0.40577 | 2.223483 | -0.18249 | 0.855195 | NA |
| ZNF10 | 80.36831 | 0.499544 | 0.269133 | 1.856124 | 0.063436 | 0.16345 |
| | 0.392857 | -1.546 | 4.075469 | -0.37934 | 0.704433 | NA |
| | 0.230334 | -0.53318 | 4.076749 | -0.13079 | 0.895945 | NA |
| | 24.86703 | 0.374204 | 0.469188 | 0.797556 | 0.425128 | NA |
| RPL7AP3 | 0.247898 | -0.5901 | 4.074633 | -0.14482 | 0.884851 | NA |
| | 0.93741 | -3.49975 | 2.723608 | -1.28497 | 0.198804 | NA |
| | 0.495797 | -0.73607 | 3.324282 | -0.22142 | 0.824763 | NA |
| | 4.189955 | -0.9329 | 1.039406 | -0.89753 | 0.369436 | NA |
| | 24.83666 | 1.344025 | 0.442596 | 3.036683 | 0.002392 | NA |
| HMBS | 929.7599 | -0.15751 | 0.163273 | -0.96468 | 0.334706 | 0.50329 |
| CACNA1C-/ | 0.134133 | -1.48909 | 4.077584 | -0.36519 | 0.71497 | NA |
| TAS2R64P | 0.47803 | 0.520083 | 2.97881 | 0.174594 | 0.861399 | NA |
| | 1.776037 | -0.1261 | 1.592469 | -0.07918 | 0.936887 | NA |
| | 0.096202 | 0.408546 | 4.079639 | 0.100143 | 0.920231 | NA |
| OSBPL9P5 | 0.096202 | 0.408546 | 4.079639 | 0.100143 | 0.920231 | NA |
| | 33.61992 | 0.968203 | 0.456599 | 2.120465 | 0.033967 | NA |
| | 0.163957 | 0.424913 | 4.079006 | 0.104171 | 0.917034 | NA |
| | 3.024854 | 1.545652 | 1.233907 | 1.252649 | 0.210333 | NA |
| | 1.425973 | 0.261345 | 1.671539 | 0.15635 | 0.875757 | NA |
| | 0.298089 | -0.51711 | 3.975232 | -0.13008 | 0.896501 | NA |
| | 0.163957 | 0.424913 | 4.079006 | 0.104171 | 0.917034 | NA |
| | 1.991105 | 0.057673 | 1.395223 | 0.041336 | 0.967028 | NA |
| | 1.045579 | 1.171193 | 2.006879 | 0.583589 | 0.559497 | NA |
| RPL41P2 | 8.696822 | 0.393644 | 0.673746 | 0.584261 | 0.559045 | NA |
| | 0.727508 | 0.192954 | 2.572892 | 0.074995 | 0.940219 | NA |
| NTAN1P3 | 0.192403 | 0.975392 | 4.060716 | 0.240202 | 0.810174 | NA |
| HSPA8P5 | 2.747897 | 0.613672 | 1.228266 | 0.499624 | 0.61734 | NA |
| | 6.35499 | -0.15884 | 0.803564 | -0.19767 | 0.843302 | NA |
| | 2.373275 | -2.31356 | 1.482837 | -1.56023 | 0.118707 | NA |
| | 1.334367 | 0.842194 | 1.784035 | 0.472073 | 0.636875 | NA |
| | 20.66547 | -0.45205 | 0.547198 | -0.82611 | 0.40874 | NA |
| | 0.354779 | 1.703954 | 3.657232 | 0.465914 | 0.641277 | NA |
| | 1.859776 | -0.06877 | 1.46217 | -0.04703 | 0.962486 | NA |
| | 5.9772 | 2.527435 | 1.164373 | 2.170641 | 0.029958 | NA |
| | 3.430519 | 0.207856 | 1.20887 | 0.171942 | 0.863483 | NA |
| TAS2R31 | 0.778894 | 2.823374 | 2.403613 | 1.174637 | 0.24014 | NA |
| | 0.096202 | 0.408546 | 4.079639 | 0.100143 | 0.920231 | NA |
| TRAV11 | 0.163957 | 0.424913 | 4.079006 | 0.104171 | 0.917034 | NA |
| | 2.170139 | -1.99351 | 1.634851 | -1.21938 | 0.222699 | NA |
| | 0.653015 | 0.2552 | 2.787103 | 0.091565 | 0.927044 | NA |
| | 0.258578 | 1.233151 | 4.047809 | 0.304646 | 0.760635 | NA |

| | | | | | | |
|---|---|---|---|---|---|---|
| | 0.392857 | -1.546 | 4.075469 | -0.37934 | 0.704433 | NA |
| | 0.134133 | -1.48909 | 4.077584 | -0.36519 | 0.71497 | NA |
| | 0.096202 | 0.408546 | 4.079639 | 0.100143 | 0.920231 | NA |
| | 0.92743 | 1.834049 | 2.364191 | 0.775762 | 0.43789 | NA |
| | 0.247898 | -0.5901 | 4.074633 | -0.14482 | 0.884851 | NA |
| | 0.096202 | 0.408546 | 4.079639 | 0.100143 | 0.920231 | NA |
| | 11.24705 | -0.16569 | 0.650411 | -0.25474 | 0.798923 | NA |
| | 0.298089 | -0.51711 | 3.975232 | -0.13008 | 0.896501 | NA |
| | 1.011758 | -0.66314 | 1.918131 | -0.34572 | 0.729551 | NA |
| NANOGNBI | 0.134133 | -1.48909 | 4.077584 | -0.36519 | 0.71497 | NA |
| | 0.151697 | -1.546 | 4.075469 | -0.37934 | 0.704433 | NA |
| | 12.22339 | -0.69134 | 0.582714 | -1.18641 | 0.235461 | NA |
| NLRP9P | 0.134133 | -1.48909 | 4.077584 | -0.36519 | 0.71497 | NA |
| | 172.2912 | 0.09644 | 0.19261 | 0.500701 | 0.616582 | 0.742224 |
| | 0.45509 | -2.71227 | 3.963744 | -0.68427 | 0.493805 | NA |
| | 0.151697 | -1.546 | 4.075469 | -0.37934 | 0.704433 | NA |
| TAS2R67P | 0.162376 | 0.424913 | 4.079006 | 0.104171 | 0.917034 | NA |
| | 2.464091 | 0.438611 | 1.29761 | 0.338014 | 0.735352 | NA |
| | 0.260158 | 1.240347 | 4.047124 | 0.306476 | 0.759242 | NA |
| CLEC12B | 0.488912 | 0.771583 | 2.897678 | 0.266276 | 0.790026 | NA |
| | 11.5745 | 0.07812 | 0.638559 | 0.122338 | 0.902631 | NA |
| KLRAP1 | 6.511756 | 2.552621 | 1.006884 | 2.535169 | 0.011239 | NA |
| | 0.258578 | 1.233151 | 4.047809 | 0.304646 | 0.760635 | NA |
| | 2148.182 | -0.25102 | 0.229815 | -1.09226 | 0.27472 | 0.440406 |
| | 0.134133 | -1.48909 | 4.077584 | -0.36519 | 0.71497 | NA |
| | 0.298089 | -0.51711 | 3.975232 | -0.13008 | 0.896501 | NA |
| | 0.163957 | 0.424913 | 4.079006 | 0.104171 | 0.917034 | NA |
| | 0.392857 | -1.546 | 4.075469 | -0.37934 | 0.704433 | NA |
| | 0.392857 | -1.546 | 4.075469 | -0.37934 | 0.704433 | NA |
| | 0.803131 | -0.76693 | 2.349822 | -0.32638 | 0.744138 | NA |
| | 0.096202 | 0.408546 | 4.079639 | 0.100143 | 0.920231 | NA |
| | 38.95826 | 0.232285 | 0.350783 | 0.66219 | 0.507849 | NA |
| | 0.612162 | -0.19706 | 2.47693 | -0.07956 | 0.936588 | NA |
| | 0.096202 | 0.408546 | 4.079639 | 0.100143 | 0.920231 | NA |
| RBBP4P5 | 0.134133 | -1.48909 | 4.077584 | -0.36519 | 0.71497 | NA |
| | 3.843836 | -0.07511 | 1.060062 | -0.07085 | 0.943515 | NA |
| | 10.90545 | 0.354549 | 0.769914 | 0.460505 | 0.645154 | NA |
| | 0.437526 | -2.68242 | 3.384147 | -0.79264 | 0.427986 | NA |
| | 1.383152 | 2.519898 | 1.91781 | 1.313945 | 0.188865 | NA |
| KLRF2 | 0.096202 | 0.408546 | 4.079639 | 0.100143 | 0.920231 | NA |
| | 0.162376 | 0.424913 | 4.079006 | 0.104171 | 0.917034 | NA |
| | 9.107713 | -0.53307 | 0.776085 | -0.68687 | 0.492167 | NA |
| TPT1P12 | 0.808921 | 2.894015 | 2.722603 | 1.062959 | 0.287801 | NA |
| | 3.945785 | 1.390314 | 1.09909 | 1.264969 | 0.205883 | NA |
| | 1.139209 | -0.69433 | 1.964144 | -0.3535 | 0.723712 | NA |
| SLC5A8 | 3.765879 | 0.018359 | 1.169792 | 0.015694 | 0.987478 | NA |
| | 6.456118 | 0.434152 | 0.885734 | 0.490161 | 0.62402 | NA |

| | | | | | | |
|---|---|---|---|---|---|---|
| | 21.1608 | -0.93177 | 0.464294 | -2.00686 | 0.044764 | NA |
| | 0.870886 | -0.53073 | 2.310896 | -0.22966 | 0.818353 | NA |
| | 0.869306 | -0.5527 | 2.538621 | -0.21772 | 0.82765 | NA |
| | 1.758242 | 2.645987 | 1.831913 | 1.444385 | 0.148631 | NA |
| | 0.096202 | 0.408546 | 4.079639 | 0.100143 | 0.920231 | NA |
| | 0.096202 | 0.408546 | 4.079639 | 0.100143 | 0.920231 | NA |
| | 28.63005 | 0.697428 | 0.399914 | 1.743943 | 0.081169 | NA |
| | 0.354779 | 1.703954 | 3.657232 | 0.465914 | 0.641277 | NA |
| | 1.999698 | 1.226422 | 1.658952 | 0.739275 | 0.45974 | NA |
| | 2.493326 | 2.029023 | 1.384698 | 1.465317 | 0.142834 | NA |
| | 0.768215 | 1.51173 | 2.494681 | 0.605981 | 0.544527 | NA |
| | 0.258578 | 1.233151 | 4.047809 | 0.304646 | 0.760635 | NA |
| TAS2R18 | 1.11032 | 0.442168 | 2.116951 | 0.20887 | 0.83455 | NA |
| | 1.837265 | 0.948736 | 1.814134 | 0.522969 | 0.600996 | NA |
| | 0.230334 | -0.53318 | 4.076749 | -0.13079 | 0.895945 | NA |
| | 0.096202 | 0.408546 | 4.079639 | 0.100143 | 0.920231 | NA |
| | 0.096202 | 0.408546 | 4.079639 | 0.100143 | 0.920231 | NA |
| | 0.586694 | 1.130467 | 2.734203 | 0.413454 | 0.679274 | NA |
| | 38.67527 | -0.28446 | 0.336472 | -0.84541 | 0.397883 | NA |
| | 16.30939 | -0.2695 | 0.515764 | -0.52253 | 0.601303 | NA |
| | 3.42311 | -2.79658 | 1.275524 | -2.19249 | 0.028344 | NA |
| | 0.163957 | 0.424913 | 4.079006 | 0.104171 | 0.917034 | NA |
| | 0.296509 | -0.51709 | 3.982155 | -0.12985 | 0.896684 | NA |
| RPEP6 | 0.151697 | -1.546 | 4.075469 | -0.37934 | 0.704433 | NA |
| | 13.31092 | -0.1186 | 0.547623 | -0.21657 | 0.828545 | NA |
| | 0.163957 | 0.424913 | 4.079006 | 0.104171 | 0.917034 | NA |
| | 0.163957 | 0.424913 | 4.079006 | 0.104171 | 0.917034 | NA |
| KRT127P | 0.096202 | 0.408546 | 4.079639 | 0.100143 | 0.920231 | NA |
| CLLU1 | 0.134133 | -1.48909 | 4.077584 | -0.36519 | 0.71497 | NA |
| PWAR6 | 25.24956 | 0.844875 | 0.542315 | 1.557905 | 0.119256 | NA |
| | 0.192403 | 0.975392 | 4.060716 | 0.240202 | 0.810174 | NA |
| TMPO-AS1 | 46.38746 | -0.48496 | 0.344853 | -1.40629 | 0.159638 | NA |
| | 0.163957 | 0.424913 | 4.079006 | 0.104171 | 0.917034 | NA |
| | 4.47014 | 0.19181 | 1.06581 | 0.179966 | 0.857179 | NA |
| HMGN2P6 | 0.162376 | 0.424913 | 4.079006 | 0.104171 | 0.917034 | NA |
| | 0.163957 | 0.424913 | 4.079006 | 0.104171 | 0.917034 | NA |
| HNRNPA1P | 0.134133 | -1.48909 | 4.077584 | -0.36519 | 0.71497 | NA |
| | 0.422534 | 1.924583 | 3.001758 | 0.641152 | 0.521424 | NA |
| | 0.424115 | 1.932531 | 3.45462 | 0.559405 | 0.575885 | NA |
| GATC | 616.3201 | -0.03749 | 0.186573 | -0.20095 | 0.840741 | 0.905184 |
| | 2.99108 | -0.03517 | 1.142884 | -0.03077 | 0.975453 | NA |
| | 0.411855 | 0.250456 | 3.097618 | 0.080854 | 0.935558 | NA |
| | 2.149721 | 0.810232 | 1.528411 | 0.530114 | 0.596033 | NA |
| | 0.096202 | 0.408546 | 4.079639 | 0.100143 | 0.920231 | NA |
| | 0.163957 | 0.424913 | 4.079006 | 0.104171 | 0.917034 | NA |
| C12orf79 | 3.103119 | 0.388405 | 1.185552 | 0.327615 | 0.743203 | NA |
| | 1.645774 | -1.39946 | 1.608559 | -0.87001 | 0.384296 | NA |

| | | | | | | |
|---|---|---|---|---|---|---|
| | 0.46577 | -1.30191 | 3.402879 | -0.38259 | 0.702024 | NA |
| | 2.89453 | 0.478374 | 1.211725 | 0.394787 | 0.693 | NA |
| | 0.46735 | -1.30193 | 3.399671 | -0.38296 | 0.701751 | NA |
| | 0.096202 | 0.408546 | 4.079639 | 0.100143 | 0.920231 | NA |
| | 0.526843 | -0.38719 | 2.855892 | -0.13558 | 0.892156 | NA |
| | 13.80147 | -0.06144 | 0.604236 | -0.10168 | 0.91901 | NA |
| | 5.610114 | -0.30155 | 0.880564 | -0.34245 | 0.73201 | NA |
| | 0.096202 | 0.408546 | 4.079639 | 0.100143 | 0.920231 | NA |
| | 0.096202 | 0.408546 | 4.079639 | 0.100143 | 0.920231 | NA |
| | 6.140178 | -0.66175 | 0.861774 | -0.76789 | 0.442552 | NA |
| | 3.845395 | 0.569024 | 1.05206 | 0.540866 | 0.5886 | NA |
| | 0.163957 | 0.424913 | 4.079006 | 0.104171 | 0.917034 | NA |
| | 8.93058 | 0.884925 | 0.743157 | 1.190766 | 0.233746 | NA |
| | 4.108285 | -0.17235 | 0.955853 | -0.18031 | 0.856906 | NA |
| | 0.911593 | 1.257757 | 2.250933 | 0.558771 | 0.576318 | NA |
| | 0.830383 | -3.4083 | 2.430438 | -1.40234 | 0.160814 | NA |
| | 2.442224 | -1.07758 | 1.439207 | -0.74873 | 0.454018 | NA |
| PRELID2P1 | 0.422534 | 1.924583 | 3.001758 | 0.641152 | 0.521424 | NA |
| | 82.05029 | -0.07922 | 0.277571 | -0.28542 | 0.775326 | 0.861029 |
| | 1.752712 | -0.14013 | 1.568628 | -0.08933 | 0.928817 | NA |
| | 9.014089 | 0.322867 | 0.835802 | 0.386296 | 0.699278 | NA |
| | 0.45509 | -2.71227 | 3.963744 | -0.68427 | 0.493805 | NA |
| FNTB | 59.55158 | -0.49829 | 0.297462 | -1.67512 | 0.09391 | 0.214014 |
| | 0.163957 | 0.424913 | 4.079006 | 0.104171 | 0.917034 | NA |
| | 0.518736 | 2.237423 | 2.7713 | 0.807355 | 0.419462 | NA |
| | 0.151697 | -1.546 | 4.075469 | -0.37934 | 0.704433 | NA |
| | 0.392857 | -1.546 | 4.075469 | -0.37934 | 0.704433 | NA |
| | 2.420341 | 0.591326 | 1.450258 | 0.407738 | 0.683466 | NA |
| | 0.192403 | 0.975392 | 4.060716 | 0.240202 | 0.810174 | NA |
| | 0.678686 | -3.08037 | 2.621268 | -1.17514 | 0.239937 | NA |
| | 4.25003 | 0.069896 | 1.06142 | 0.065851 | 0.947496 | NA |
| | 0.506476 | 0.726418 | 2.864186 | 0.253621 | 0.799788 | NA |
| | 0.260158 | 1.240347 | 4.047124 | 0.306476 | 0.759242 | NA |
| | 0.151697 | -1.546 | 4.075469 | -0.37934 | 0.704433 | NA |
| | 0.3441 | -0.0166 | 3.769682 | -0.0044 | 0.996486 | NA |
| | 0.488912 | 0.771583 | 2.897678 | 0.266276 | 0.790026 | NA |
| | 0.949727 | -0.95431 | 2.383855 | -0.40032 | 0.688919 | NA |
| MRPL2P1 | 0.392857 | -1.546 | 4.075469 | -0.37934 | 0.704433 | NA |
| | 1.110734 | 1.171607 | 2.014126 | 0.581695 | 0.560772 | NA |
| | 0.392857 | -1.546 | 4.075469 | -0.37934 | 0.704433 | NA |
| | 2.58982 | 1.232038 | 1.429809 | 0.86168 | 0.388864 | NA |
| | 0.440301 | 0.468383 | 3.449808 | 0.135771 | 0.892003 | NA |
| | 0.392857 | -1.546 | 4.075469 | -0.37934 | 0.704433 | NA |
| | 1.436075 | -0.1339 | 1.755147 | -0.07629 | 0.939188 | NA |
| | 23.64998 | 0.746438 | 0.470618 | 1.586079 | 0.112721 | NA |
| | 0.392711 | 0.292538 | 3.145221 | 0.09301 | 0.925895 | NA |
| | 1.123225 | -0.64885 | 2.232612 | -0.29062 | 0.771339 | NA |

| | | | | | | |
|---|---|---|---|---|---|---|
| | 0.394291 | 0.298745 | 3.141711 | 0.09509 | 0.924243 | NA |
| | 2.129382 | -0.16373 | 1.460928 | -0.11207 | 0.910768 | NA |
| HSPA8P14 | 0.460466 | 0.568655 | 3.014642 | 0.188631 | 0.850382 | NA |
| OR7E47P | 0.192403 | 0.975392 | 4.060716 | 0.240202 | 0.810174 | NA |
| | 1.872599 | -0.0909 | 1.613914 | -0.05632 | 0.955084 | NA |
| | 3.807099 | 0.555626 | 1.110579 | 0.500303 | 0.616862 | NA |
| | 4.395388 | 1.361139 | 1.018859 | 1.335945 | 0.181567 | NA |
| | 0.192403 | 0.975392 | 4.060716 | 0.240202 | 0.810174 | NA |
| GSTP1P1 | 0.690946 | -1.56103 | 2.699004 | -0.57837 | 0.563013 | NA |
| | 0.883 | 1.784395 | 2.188528 | 0.81534 | 0.414878 | NA |
| | 0.163957 | 0.424913 | 4.079006 | 0.104171 | 0.917034 | NA |
| GALNT4 | 8.760222 | 0.194988 | 0.684422 | 0.284895 | 0.775725 | NA |
| | 1.6849 | -0.55131 | 1.656497 | -0.33282 | 0.739274 | NA |
| | 0.795081 | 1.604357 | 2.508662 | 0.639527 | 0.52248 | NA |
| | 28.09123 | 0.499132 | 0.410356 | 1.216338 | 0.223856 | NA |
| | 0.096202 | 0.408546 | 4.079639 | 0.100143 | 0.920231 | NA |
| | 0.392857 | -1.546 | 4.075469 | -0.37934 | 0.704433 | NA |
| | 0.162376 | 0.424913 | 4.079006 | 0.104171 | 0.917034 | NA |
| | 269.0018 | 0.9217 | 0.247306 | 3.726954 | 1.94E-04 | 0.003807 |
| | 0.192403 | 0.975392 | 4.060716 | 0.240202 | 0.810174 | NA |
| | 0.640755 | -1.64826 | 2.754778 | -0.59833 | 0.549621 | NA |
| | 0.163957 | 0.424913 | 4.079006 | 0.104171 | 0.917034 | NA |
| | 0.909866 | 1.872373 | 2.379543 | 0.786862 | 0.431362 | NA |
| | 0.096202 | 0.408546 | 4.079639 | 0.100143 | 0.920231 | NA |
| | 0.871089 | -1.46164 | 2.262872 | -0.64592 | 0.518329 | NA |
| | 0.151697 | -1.546 | 4.075469 | -0.37934 | 0.704433 | NA |
| | 0.096202 | 0.408546 | 4.079639 | 0.100143 | 0.920231 | NA |
| | 8.029319 | 0.734772 | 0.721381 | 1.018563 | 0.30841 | NA |
| | 200.6517 | 0.07234 | 0.228639 | 0.316395 | 0.751703 | 0.844731 |
| | 0.508056 | 0.731032 | 2.861912 | 0.255435 | 0.798387 | NA |
| | 6.989616 | -0.64745 | 0.767859 | -0.84319 | 0.399122 | NA |
| CNPY2 | 187.903 | 0.244656 | 0.190956 | 1.281214 | 0.200119 | 0.35611 |
| LSM6P2 | 0.096202 | 0.408546 | 4.079639 | 0.100143 | 0.920231 | NA |
| | 2.026158 | 2.299413 | 1.641253 | 1.40101 | 0.161211 | NA |
| | 0.162376 | 0.424913 | 4.079006 | 0.104171 | 0.917034 | NA |
| | 7.500492 | 0.086219 | 0.748457 | 0.115195 | 0.90829 | NA |
| | 41.06222 | 1.060732 | 0.441645 | 2.401775 | 0.016316 | NA |
| | 0.163957 | 0.424913 | 4.079006 | 0.104171 | 0.917034 | NA |
| | 0.651085 | 2.534961 | 3.013697 | 0.841147 | 0.400266 | NA |
| | 0.392857 | -1.546 | 4.075469 | -0.37934 | 0.704433 | NA |
| | 0.162376 | 0.424913 | 4.079006 | 0.104171 | 0.917034 | NA |
| ST13P3 | 0.678686 | -3.08037 | 2.621268 | -1.17514 | 0.239937 | NA |
| | 0.572651 | 0.928649 | 2.780261 | 0.334015 | 0.738368 | NA |
| | 1.398079 | -2.25327 | 1.966256 | -1.14597 | 0.251807 | NA |
| MRS2P2 | 0.528424 | -0.38142 | 2.853651 | -0.13366 | 0.893671 | NA |
| | 1.567836 | 1.766079 | 1.822638 | 0.968968 | 0.332561 | NA |
| | 0.3441 | -0.0166 | 3.769682 | -0.0044 | 0.996486 | NA |

| | | | | | | |
|---|---|---|---|---|---|---|
| | 0.394291 | 0.298745 | 3.141711 | 0.09509 | 0.924243 | NA |
| | 15.2287 | 0.176048 | 0.528469 | 0.333128 | 0.739038 | NA |
| | 0.163957 | 0.424913 | 4.079006 | 0.104171 | 0.917034 | NA |
| | 8.9783 | 0.808923 | 0.705082 | 1.147274 | 0.251268 | NA |
| | 0.096202 | 0.408546 | 4.079639 | 0.100143 | 0.920231 | NA |
| | 0.096202 | 0.408546 | 4.079639 | 0.100143 | 0.920231 | NA |
| HNRNPA3P | 0.096202 | 0.408546 | 4.079639 | 0.100143 | 0.920231 | NA |
| CASC18 | 1.878884 | -0.09816 | 1.5128 | -0.06488 | 0.948266 | NA |
| | 0.162376 | 0.424913 | 4.079006 | 0.104171 | 0.917034 | NA |
| | 0.134133 | -1.48909 | 4.077584 | -0.36519 | 0.71497 | NA |
| | 0.774742 | -0.40577 | 2.223483 | -0.18249 | 0.855195 | NA |
| | 0.151697 | -1.546 | 4.075469 | -0.37934 | 0.704433 | NA |
| | 0.192403 | 0.975392 | 4.060716 | 0.240202 | 0.810174 | NA |
| | 0.096202 | 0.408546 | 4.079639 | 0.100143 | 0.920231 | NA |
| | 0.725928 | 0.181819 | 2.321433 | 0.078322 | 0.937572 | NA |
| | 0.392857 | -1.546 | 4.075469 | -0.37934 | 0.704433 | NA |
| EEF1A1P17 | 0.815042 | 2.859272 | 2.76927 | 1.0325 | 0.301838 | NA |
| | 0.70046 | 1.37107 | 2.594868 | 0.528378 | 0.597237 | NA |
| | 2.583 | 3.498257 | 1.603106 | 2.182174 | 0.029097 | NA |
| | 1.419889 | 0.381214 | 1.819664 | 0.209497 | 0.83406 | NA |
| | 2.598138 | 1.174145 | 1.379767 | 0.850973 | 0.394784 | NA |
| | 0.324752 | 1.53082 | 4.022025 | 0.380609 | 0.703493 | NA |
| | 0.258578 | 1.233151 | 4.047809 | 0.304646 | 0.760635 | NA |
| | 0.163957 | 0.424913 | 4.079006 | 0.104171 | 0.917034 | NA |
| | 0.260158 | 1.240347 | 4.047124 | 0.306476 | 0.759242 | NA |
| | 1.941991 | 0.19281 | 1.456829 | 0.132349 | 0.894708 | NA |
| | 0.872321 | 0.678609 | 2.011629 | 0.337343 | 0.735858 | NA |
| | 3.502066 | 1.101642 | 1.213332 | 0.907947 | 0.363906 | NA |
| QRSL1P3 | 0.508056 | 0.731032 | 2.861912 | 0.255435 | 0.798387 | NA |
| | 0.558248 | 0.982076 | 2.804738 | 0.350149 | 0.726227 | NA |
| OLA1P3 | 0.134133 | -1.48909 | 4.077584 | -0.36519 | 0.71497 | NA |
| | 1.06329 | 0.285492 | 1.942399 | 0.146979 | 0.883149 | NA |
| | 0.437526 | -2.68242 | 3.384147 | -0.79264 | 0.427986 | NA |
| | 0.151697 | -1.546 | 4.075469 | -0.37934 | 0.704433 | NA |
| | 1.009228 | 2.053856 | 2.077371 | 0.98868 | 0.322819 | NA |
| HMGA1P3 | 1.144942 | 1.358971 | 2.181094 | 0.623068 | 0.53324 | NA |
| | 0.763859 | -0.6945 | 2.307767 | -0.30094 | 0.76346 | NA |
| HIGD1AP1 | 0.696104 | -0.93034 | 2.354268 | -0.39517 | 0.692715 | NA |
| | 1.19906 | 0.118246 | 1.94637 | 0.060752 | 0.951557 | NA |
| | 0.616518 | 2.49285 | 2.980788 | 0.836306 | 0.402983 | NA |
| | 0.392857 | -1.546 | 4.075469 | -0.37934 | 0.704433 | NA |
| | 0.260158 | 1.240347 | 4.047124 | 0.306476 | 0.759242 | NA |
| | 0.151697 | -1.546 | 4.075469 | -0.37934 | 0.704433 | NA |
| | 123.7106 | -0.03993 | 0.214883 | -0.18584 | 0.852568 | 0.912284 |
| BCDIN3D-A | 8.691469 | -0.08522 | 0.74103 | -0.115 | 0.908447 | NA |
| | 0.758758 | -0.35643 | 2.514365 | -0.14176 | 0.887273 | NA |
| | 0.096202 | 0.408546 | 4.079639 | 0.100143 | 0.920231 | NA |

| | | | | | | |
|---|---|---|---|---|---|---|
| | 0.151697 | -1.546 | 4.075469 | -0.37934 | 0.704433 | NA |
| | 13.63292 | 0.304796 | 0.585314 | 0.520739 | 0.602549 | NA |
| | 0.437526 | -2.68242 | 3.384147 | -0.79264 | 0.427986 | NA |
| | 11.78575 | 0.80302 | 0.654567 | 1.226796 | 0.219899 | NA |
| | 0.640755 | -1.64826 | 2.754778 | -0.59833 | 0.549621 | NA |
| | 0.919911 | 0.765837 | 2.338731 | 0.327458 | 0.743321 | NA |
| | 3.124771 | 0.29748 | 1.15812 | 0.256865 | 0.797283 | NA |
| | 0.392857 | -1.546 | 4.075469 | -0.37934 | 0.704433 | NA |
| | 2.918527 | -0.25001 | 1.288043 | -0.1941 | 0.846098 | NA |
| MKRN9P | 0.314073 | -0.57308 | 3.898934 | -0.14698 | 0.883146 | NA |
| | 0.192403 | 0.975392 | 4.060716 | 0.240202 | 0.810174 | NA |
| | 0.162376 | 0.424913 | 4.079006 | 0.104171 | 0.917034 | NA |
| | 0.096202 | 0.408546 | 4.079639 | 0.100143 | 0.920231 | NA |
| | 3.56843 | -1.34096 | 1.178044 | -1.1383 | 0.254997 | NA |
| LINC00485 | 0.134133 | -1.48909 | 4.077584 | -0.36519 | 0.71497 | NA |
| | 0.489058 | -0.58569 | 3.427543 | -0.17088 | 0.864321 | NA |
| | 0.192403 | 0.975392 | 4.060716 | 0.240202 | 0.810174 | NA |
| | 0.750651 | 1.552121 | 2.514127 | 0.61736 | 0.536998 | NA |
| | 32.68951 | -0.64335 | 0.399392 | -1.61082 | 0.107218 | NA |
| | 1.045872 | -1.31674 | 2.456371 | -0.53605 | 0.591923 | NA |
| | 0.162376 | 0.424913 | 4.079006 | 0.104171 | 0.917034 | NA |
| | 1.132479 | 1.219292 | 1.844561 | 0.66102 | 0.508599 | NA |
| | 0.162376 | 0.424913 | 4.079006 | 0.104171 | 0.917034 | NA |
| | 0.151697 | -1.546 | 4.075469 | -0.37934 | 0.704433 | NA |
| | 0.684273 | 2.627785 | 2.896325 | 0.907282 | 0.364257 | NA |
| | 0.327913 | 1.543154 | 4.021063 | 0.383768 | 0.701151 | NA |
| | 5.025427 | 0.768916 | 0.963542 | 0.79801 | 0.424865 | NA |
| | 0.151697 | -1.546 | 4.075469 | -0.37934 | 0.704433 | NA |
| | 0.392857 | -1.546 | 4.075469 | -0.37934 | 0.704433 | NA |
| | 0.247898 | -0.5901 | 4.074633 | -0.14482 | 0.884851 | NA |
| | 1.730947 | -0.39777 | 1.500857 | -0.26503 | 0.790989 | NA |
| | 0.460669 | -0.62593 | 3.398671 | -0.18417 | 0.853882 | NA |
| CHURC1 | 1586.483 | 0.891174 | 0.213355 | 4.176946 | 2.95E-05 | 0.001179 |
| | 47.88276 | 0.570931 | 0.320821 | 1.779592 | 0.075143 | 0.183032 |
| NUTF2P2 | 0.314073 | -0.57308 | 3.898934 | -0.14698 | 0.883146 | NA |
| | 55.60719 | -0.44539 | 0.307577 | -1.44807 | 0.147597 | 0.290713 |
| | 1.055239 | 2.092943 | 2.097897 | 0.997638 | 0.318455 | NA |
| | 4.50713 | -0.46311 | 1.230134 | -0.37647 | 0.706568 | NA |
| | 0.555233 | -0.57009 | 3.296947 | -0.17291 | 0.862719 | NA |
| | 11.56565 | -0.28827 | 0.679701 | -0.42411 | 0.671486 | NA |
| | 1.836451 | -0.07811 | 1.514376 | -0.05158 | 0.958865 | NA |
| | 0.354779 | 1.703954 | 3.657232 | 0.465914 | 0.641277 | NA |
| | 2.019341 | -0.87103 | 1.521974 | -0.5723 | 0.567119 | NA |
| | 0.096202 | 0.408546 | 4.079639 | 0.100143 | 0.920231 | NA |
| | 0.163957 | 0.424913 | 4.079006 | 0.104171 | 0.917034 | NA |
| | 0.422534 | 1.924583 | 3.001758 | 0.641152 | 0.521424 | NA |
| | 0.326536 | 0.037056 | 3.842554 | 0.009644 | 0.992306 | NA |

| | | | | | | |
|---|---|---|---|---|---|---|
| | 0.163957 | 0.424913 | 4.079006 | 0.104171 | 0.917034 | NA |
| | 0.151697 | -1.546 | 4.075469 | -0.37934 | 0.704433 | NA |
| | 0.315653 | -0.57305 | 3.892639 | -0.14721 | 0.882963 | NA |
| PCNPP1 | 1.187588 | 2.257887 | 1.988061 | 1.135723 | 0.256072 | NA |
| | 0.506476 | 0.726418 | 2.864186 | 0.253621 | 0.799788 | NA |
| SLC25A3P2 | 0.725928 | 0.181819 | 2.321433 | 0.078322 | 0.937572 | NA |
| | 113.4921 | -0.48168 | 0.266536 | -1.8072 | 0.070731 | 0.17579 |
| RPA2P1 | 0.260158 | 1.240347 | 4.047124 | 0.306476 | 0.759242 | NA |
| | 0.096202 | 0.408546 | 4.079639 | 0.100143 | 0.920231 | NA |
| | 0.163957 | 0.424913 | 4.079006 | 0.104171 | 0.917034 | NA |
| | 0.58526 | 0.001197 | 3.235889 | 3.70E-04 | 0.999705 | NA |
| | 0.260158 | 1.240347 | 4.047124 | 0.306476 | 0.759242 | NA |
| | 3.844228 | 0.819205 | 1.021673 | 0.801827 | 0.422653 | NA |
| RBM8B | 2.098336 | -0.26092 | 1.394053 | -0.18717 | 0.851529 | NA |
| | 0.230334 | -0.53318 | 4.076749 | -0.13079 | 0.895945 | NA |
| | 0.096202 | 0.408546 | 4.079639 | 0.100143 | 0.920231 | NA |
| | 1.197625 | -0.85962 | 1.981435 | -0.43384 | 0.664407 | NA |
| | 0.151697 | -1.546 | 4.075469 | -0.37934 | 0.704433 | NA |
| LINC00641 | 54.05232 | 0.942007 | 0.414833 | 2.270811 | 0.023158 | 0.083463 |
| | 0.096202 | 0.408546 | 4.079639 | 0.100143 | 0.920231 | NA |
| | 1.034697 | 1.024735 | 1.902596 | 0.538598 | 0.590164 | NA |
| | 0.582339 | -1.69412 | 2.750801 | -0.61586 | 0.537985 | NA |
| | 0.315653 | -0.57305 | 3.892639 | -0.14721 | 0.882963 | NA |
| | 0.47961 | 0.535796 | 3.39605 | 0.15777 | 0.874638 | NA |
| | 0.698879 | 1.36795 | 2.596167 | 0.526912 | 0.598255 | NA |
| | 0.192403 | 0.975392 | 4.060716 | 0.240202 | 0.810174 | NA |
| | 5.347756 | -0.37464 | 0.885326 | -0.42316 | 0.672177 | NA |
| | 0.134133 | -1.48909 | 4.077584 | -0.36519 | 0.71497 | NA |
| | 44.55358 | 0.238562 | 0.377262 | 0.632351 | 0.527157 | NA |
| | 0.192403 | 0.975392 | 4.060716 | 0.240202 | 0.810174 | NA |
| CHMP4BP1 | 0.750448 | 2.743526 | 2.504729 | 1.095338 | 0.273368 | NA |
| | 0.58526 | 0.001197 | 3.235889 | 3.70E-04 | 0.999705 | NA |
| | 0.296509 | -0.51709 | 3.982155 | -0.12985 | 0.896684 | NA |
| PPIAP6 | 0.392857 | -1.546 | 4.075469 | -0.37934 | 0.704433 | NA |
| LINC00640 | 1.045376 | 2.066118 | 2.09889 | 0.984386 | 0.324926 | NA |
| SPESP1 | 16.69185 | 1.094745 | 0.565352 | 1.936396 | 0.052819 | NA |
| | 0.163957 | 0.424913 | 4.079006 | 0.104171 | 0.917034 | NA |
| DIO3OS | 0.71919 | 0.510978 | 2.728734 | 0.187258 | 0.851458 | NA |
| | 0.151697 | -1.546 | 4.075469 | -0.37934 | 0.704433 | NA |
| | 0.163957 | 0.424913 | 4.079006 | 0.104171 | 0.917034 | NA |
| EIF3LP1 | 0.399595 | -1.31595 | 3.579466 | -0.36764 | 0.713143 | NA |
| | 0.162376 | 0.424913 | 4.079006 | 0.104171 | 0.917034 | NA |
| | 0.526843 | -0.38719 | 2.855892 | -0.13558 | 0.892156 | NA |
| | 0.162376 | 0.424913 | 4.079006 | 0.104171 | 0.917034 | NA |
| | 0.28583 | -2.21358 | 3.97376 | -0.55705 | 0.577494 | NA |
| | 0.584911 | 2.38756 | 2.690807 | 0.887303 | 0.374916 | NA |
| | 2.712961 | 1.226314 | 1.347066 | 0.910359 | 0.362633 | NA |

| | | | | | | |
|---|---|---|---|---|---|---|
| NT5CP1 | 1.410404 | 0.968834 | 1.838328 | 0.527019 | 0.59818 | NA |
| | 1.441811 | 0.977266 | 1.883072 | 0.518974 | 0.603779 | NA |
| | 0.163957 | 0.424913 | 4.079006 | 0.104171 | 0.917034 | NA |
| | 0.747636 | 0.735436 | 2.634453 | 0.279161 | 0.780122 | NA |
| | 0.230334 | -0.53318 | 4.076749 | -0.13079 | 0.895945 | NA |
| | 0.163957 | 0.424913 | 4.079006 | 0.104171 | 0.917034 | NA |
| | 6.401733 | -0.32591 | 0.896981 | -0.36334 | 0.716353 | NA |
| | 4.639366 | -0.06815 | 0.999854 | -0.06816 | 0.945656 | NA |
| BLZF2P | 0.298089 | -0.51711 | 3.975232 | -0.13008 | 0.896501 | NA |
| RHOQP1 | 23.79738 | 0.090358 | 0.456957 | 0.19774 | 0.843249 | NA |
| | 0.096202 | 0.408546 | 4.079639 | 0.100143 | 0.920231 | NA |
| PTTG4P | 0.326536 | 0.037056 | 3.842554 | 0.009644 | 0.992306 | NA |
| | 4.332838 | -0.30815 | 0.949418 | -0.32457 | 0.745507 | NA |
| | 44.45893 | 0.501341 | 0.470409 | 1.065756 | 0.286534 | NA |
| | 0.151697 | -1.546 | 4.075469 | -0.37934 | 0.704433 | NA |
| | 0.096202 | 0.408546 | 4.079639 | 0.100143 | 0.920231 | NA |
| NBEAP1 | 1.589987 | 0.694691 | 1.591628 | 0.436466 | 0.662499 | NA |
| | 0.151697 | -1.546 | 4.075469 | -0.37934 | 0.704433 | NA |
| | 0.544554 | -2.65036 | 3.220086 | -0.82307 | 0.410468 | NA |
| | 0.162376 | 0.424913 | 4.079006 | 0.104171 | 0.917034 | NA |
| | 0.162376 | 0.424913 | 4.079006 | 0.104171 | 0.917034 | NA |
| | 0.258578 | 1.233151 | 4.047809 | 0.304646 | 0.760635 | NA |
| | 0.734035 | -2.0718 | 2.529697 | -0.81899 | 0.41279 | NA |
| DNAJC19P9 | 1.179509 | 1.360465 | 1.979386 | 0.687317 | 0.491883 | NA |
| RPL21P7 | 1.663192 | -0.49205 | 1.500975 | -0.32782 | 0.743046 | NA |
| | 8.296205 | -0.25486 | 0.964051 | -0.26437 | 0.791497 | NA |
| | 2.027456 | -0.33651 | 1.388219 | -0.2424 | 0.80847 | NA |
| OR11H5P | 0.151697 | -1.546 | 4.075469 | -0.37934 | 0.704433 | NA |
| COX7A2P1 | 1.525539 | -0.06231 | 1.650827 | -0.03775 | 0.969889 | NA |
| | 0.162376 | 0.424913 | 4.079006 | 0.104171 | 0.917034 | NA |
| | 0.163957 | 0.424913 | 4.079006 | 0.104171 | 0.917034 | NA |
| | 1.956752 | 0.523504 | 1.674627 | 0.312609 | 0.754578 | NA |
| | 0.192403 | 0.975392 | 4.060716 | 0.240202 | 0.810174 | NA |
| BCL2L2-PAI | 6.377438 | 1.884526 | 0.917569 | 2.053826 | 0.039993 | NA |
| HSPE1P2 | 1.0491 | 1.000662 | 2.336632 | 0.42825 | 0.668469 | NA |
| | 1.037472 | 3.237257 | 2.161528 | 1.49767 | 0.134219 | NA |
| UBE2CP1 | 0.151697 | -1.546 | 4.075469 | -0.37934 | 0.704433 | NA |
| | 0.134133 | -1.48909 | 4.077584 | -0.36519 | 0.71497 | NA |
| ARHGAP5-/ | 78.39591 | 0.209861 | 0.316116 | 0.663874 | 0.506771 | 0.655941 |
| | 253.2828 | 0.204481 | 0.201963 | 1.012466 | 0.311315 | 0.478807 |
| | 0.47803 | 0.520083 | 2.97881 | 0.174594 | 0.861399 | NA |
| | 45.99527 | 0.400054 | 0.352437 | 1.135109 | 0.25633 | NA |
| HIF1A-AS2 | 0.83597 | 1.639551 | 2.453777 | 0.668175 | 0.504022 | NA |
| | 0.462046 | 0.589069 | 3.442439 | 0.17112 | 0.86413 | NA |
| | 0.163957 | 0.424913 | 4.079006 | 0.104171 | 0.917034 | NA |
| | 3.952204 | 0.113403 | 1.013837 | 0.111856 | 0.910938 | NA |
| | 0.163957 | 0.424913 | 4.079006 | 0.104171 | 0.917034 | NA |

| | | | | | | |
|---|---|---|---|---|---|---|
| LINC00871 | 0.572651 | 0.928649 | 2.780261 | 0.334015 | 0.738368 | NA |
| LINC00638 | 23.39985 | 0.303632 | 0.448374 | 0.677184 | 0.498289 | NA |
| | 2.573501 | -0.14231 | 1.243414 | -0.11445 | 0.908883 | NA |
| | 18.27639 | 0.390942 | 0.541608 | 0.721817 | 0.470407 | NA |
| SLC25A21-/ | 4.179707 | -0.50494 | 0.999095 | -0.50539 | 0.613282 | NA |
| | 0.842643 | -2.08432 | 2.306153 | -0.90381 | 0.366097 | NA |
| | 7.609678 | 1.564299 | 0.81565 | 1.917857 | 0.055129 | NA |
| ITPK1-AS1 | 0.411855 | 0.250456 | 3.097618 | 0.080854 | 0.935558 | NA |
| | 0.908875 | -0.75101 | 2.116968 | -0.35476 | 0.722771 | NA |
| | 14.93381 | 0.180179 | 0.54011 | 0.333597 | 0.738683 | NA |
| | 17.38958 | 0.213482 | 0.490176 | 0.435522 | 0.663183 | NA |
| LINC00637 | 0.260158 | 1.240347 | 4.047124 | 0.306476 | 0.759242 | NA |
| | 18.06122 | -0.08896 | 0.518257 | -0.17165 | 0.863716 | NA |
| | 0.815391 | 0.931626 | 2.360431 | 0.394685 | 0.693076 | NA |
| | 1.503445 | 1.098331 | 1.654119 | 0.663997 | 0.506692 | NA |
| | 0.422534 | 1.924583 | 3.001758 | 0.641152 | 0.521424 | NA |
| | 0.247898 | -0.5901 | 4.074633 | -0.14482 | 0.884851 | NA |
| DIO2-AS1 | 0.258578 | 1.233151 | 4.047809 | 0.304646 | 0.760635 | NA |
| | 11.7156 | -0.47396 | 0.651464 | -0.72753 | 0.466901 | NA |
| RAP1AP | 2.303947 | -0.85625 | 1.34365 | -0.63726 | 0.523957 | NA |
| HIF1A-AS1 | 0.151697 | -1.546 | 4.075469 | -0.37934 | 0.704433 | NA |
| BMS1P15 | 0.134133 | -1.48909 | 4.077584 | -0.36519 | 0.71497 | NA |
| | 0.096202 | 0.408546 | 4.079639 | 0.100143 | 0.920231 | NA |
| | 0.163957 | 0.424913 | 4.079006 | 0.104171 | 0.917034 | NA |
| | 6.218328 | 0.386852 | 0.803329 | 0.481561 | 0.630118 | NA |
| | 5.527089 | 0.963196 | 0.914233 | 1.053556 | 0.292086 | NA |
| LINC00520 | 1854.438 | 0.007292 | 0.123201 | 0.059186 | 0.952804 | 0.974337 |
| ADIPOR1P2 | 0.134133 | -1.48909 | 4.077584 | -0.36519 | 0.71497 | NA |
| | 0.303394 | -2.27006 | 4.032451 | -0.56295 | 0.57347 | NA |
| | 0.327913 | 1.543154 | 4.021063 | 0.383768 | 0.701151 | NA |
| | 0.151697 | -1.546 | 4.075469 | -0.37934 | 0.704433 | NA |
| RNASE4 | 0.230334 | -0.53318 | 4.076749 | -0.13079 | 0.895945 | NA |
| | 17.31485 | -0.40308 | 0.528128 | -0.76322 | 0.445334 | NA |
| | 0.096202 | 0.408546 | 4.079639 | 0.100143 | 0.920231 | NA |
| | 9.221276 | 0.534909 | 0.678777 | 0.788049 | 0.430668 | NA |
| | 0.134133 | -1.48909 | 4.077584 | -0.36519 | 0.71497 | NA |
| TRAV28 | 0.162376 | 0.424913 | 4.079006 | 0.104171 | 0.917034 | NA |
| | 0.545988 | -0.43913 | 2.541145 | -0.17281 | 0.862803 | NA |
| EEF1A1P33 | 0.314073 | -0.57308 | 3.898934 | -0.14698 | 0.883146 | NA |
| | 0.296509 | -0.51709 | 3.982155 | -0.12985 | 0.896684 | NA |
| | 0.520316 | 2.239969 | 3.173769 | 0.705776 | 0.480328 | NA |
| | 2.087161 | 0.707052 | 1.371442 | 0.515554 | 0.606166 | NA |
| | 1.18109 | 1.362393 | 1.979079 | 0.688397 | 0.491203 | NA |
| | 0.163957 | 0.424913 | 4.079006 | 0.104171 | 0.917034 | NA |
| EIF4EBP1P1 | 0.399595 | -1.31595 | 3.579466 | -0.36764 | 0.713143 | NA |
| | 0.134133 | -1.48909 | 4.077584 | -0.36519 | 0.71497 | NA |
| | 0.162376 | 0.424913 | 4.079006 | 0.104171 | 0.917034 | NA |

| | | | | | | |
|---|---|---|---|---|---|---|
| | 5.356833 | 0.282257 | 0.911454 | 0.309678 | 0.756806 | NA |
| | 0.296509 | -0.51709 | 3.982155 | -0.12985 | 0.896684 | NA |
| SCOCP1 | 0.586491 | 2.39072 | 2.689057 | 0.889055 | 0.373973 | NA |
| EGLN3-AS1 | 0.163957 | 0.424913 | 4.079006 | 0.104171 | 0.917034 | NA |
| HNRNPCP1 | 14.82971 | 1.544073 | 0.618203 | 2.497681 | 0.012501 | NA |
| | 1.605554 | 0.650643 | 1.821017 | 0.357297 | 0.72087 | NA |
| | 1.20711 | -1.81886 | 1.886695 | -0.96404 | 0.335024 | NA |
| | 0.661122 | -3.06315 | 2.98049 | -1.02773 | 0.304076 | NA |
| | 1.044291 | -1.32184 | 2.457199 | -0.53794 | 0.590615 | NA |
| | 0.247898 | -0.5901 | 4.074633 | -0.14482 | 0.884851 | NA |
| ZMYND19F | 0.162376 | 0.424913 | 4.079006 | 0.104171 | 0.917034 | NA |
| | 0.326333 | 1.533889 | 3.826878 | 0.40082 | 0.688553 | NA |
| FOXN3-AS1 | 72.9889 | -0.07341 | 0.279502 | -0.26265 | 0.792817 | 0.874397 |
| | 0.268266 | -2.15396 | 4.038351 | -0.53338 | 0.593773 | NA |
| | 0.162376 | 0.424913 | 4.079006 | 0.104171 | 0.917034 | NA |
| | 19.00403 | 0.34094 | 0.482227 | 0.707012 | 0.479559 | NA |
| | 0.096202 | 0.408546 | 4.079639 | 0.100143 | 0.920231 | NA |
| | 0.697685 | -0.92563 | 2.353044 | -0.39338 | 0.694041 | NA |
| | 7.222713 | -0.07022 | 0.779678 | -0.09006 | 0.928238 | NA |
| | 4.547042 | -0.05974 | 0.915673 | -0.06524 | 0.94798 | NA |
| | 2.478051 | 1.097295 | 1.471185 | 0.745858 | 0.455753 | NA |
| | 1.522764 | -1.40258 | 1.668578 | -0.84059 | 0.40058 | NA |
| | 5.03101 | 0.212717 | 0.937266 | 0.226955 | 0.820459 | NA |
| KRT18P7 | 0.247898 | -0.5901 | 4.074633 | -0.14482 | 0.884851 | NA |
| | 0.096202 | 0.408546 | 4.079639 | 0.100143 | 0.920231 | NA |
| | 1.161072 | 0.527093 | 2.031776 | 0.259425 | 0.795308 | NA |
| | 1.561687 | -0.08099 | 1.556458 | -0.05204 | 0.9585 | NA |
| | 0.392857 | -1.546 | 4.075469 | -0.37934 | 0.704433 | NA |
| HIF1AP1 | 0.230334 | -0.53318 | 4.076749 | -0.13079 | 0.895945 | NA |
| | 0.296509 | -0.51709 | 3.982155 | -0.12985 | 0.896684 | NA |
| COX5AP2 | 0.163957 | 0.424913 | 4.079006 | 0.104171 | 0.917034 | NA |
| | 2.530355 | -0.09898 | 1.275318 | -0.07761 | 0.938135 | NA |
| | 0.489058 | -0.58569 | 3.427543 | -0.17088 | 0.864321 | NA |
| | 3.356907 | 1.27795 | 1.239575 | 1.030958 | 0.302561 | NA |
| TMEM179 | 1.083803 | -2.45142 | 2.403819 | -1.0198 | 0.307822 | NA |
| | 4.713177 | 0.43764 | 0.992699 | 0.440858 | 0.659315 | NA |
| | 0.096202 | 0.408546 | 4.079639 | 0.100143 | 0.920231 | NA |
| | 0.46735 | -1.30193 | 3.399671 | -0.38296 | 0.701751 | NA |
| | 1.828546 | 0.414547 | 1.589727 | 0.260766 | 0.794273 | NA |
| | 0.151697 | -1.546 | 4.075469 | -0.37934 | 0.704433 | NA |
| RPPH1 | 0.134133 | -1.48909 | 4.077584 | -0.36519 | 0.71497 | NA |
| | 0.28583 | -2.21358 | 3.97376 | -0.55705 | 0.577494 | NA |
| | 2.726685 | -0.50315 | 1.221344 | -0.41197 | 0.680363 | NA |
| | 0.096202 | 0.408546 | 4.079639 | 0.100143 | 0.920231 | NA |
| | 1.802629 | -1.30223 | 1.594252 | -0.81683 | 0.414026 | NA |
| | 0.71919 | 0.510978 | 2.728734 | 0.187258 | 0.851458 | NA |
| DNAJC8P1 | 0.298089 | -0.51711 | 3.975232 | -0.13008 | 0.896501 | NA |

| | | | | | | |
|---|---|---|---|---|---|---|
| | 0.163957 | 0.424913 | 4.079006 | 0.104171 | 0.917034 | NA |
| ENSAP2 | 3.608685 | -1.46153 | 1.184653 | -1.23372 | 0.217307 | NA |
| | 3.6153 | 0.417124 | 1.062798 | 0.392477 | 0.694706 | NA |
| | 0.476449 | 0.524262 | 3.402674 | 0.154073 | 0.877552 | NA |
| CHORDC2P | 0.162376 | 0.424913 | 4.079006 | 0.104171 | 0.917034 | NA |
| HNRNPUP1 | 1.050883 | 0.315451 | 1.99938 | 0.157774 | 0.874635 | NA |
| | 0.392857 | -1.546 | 4.075469 | -0.37934 | 0.704433 | NA |
| ACTN1-AS1 | 0.192403 | 0.975392 | 4.060716 | 0.240202 | 0.810174 | NA |
| | 41.7291 | -0.27388 | 0.329234 | -0.83187 | 0.405482 | NA |
| | 2.462271 | 0.753131 | 1.440002 | 0.523007 | 0.60097 | NA |
| LINC00639 | 1.202377 | 0.510208 | 2.000895 | 0.25499 | 0.798731 | NA |
| | 14.60367 | 0.400936 | 0.537221 | 0.746314 | 0.455478 | NA |
| POC1B-GAI | 1.657888 | 0.125143 | 1.487942 | 0.084105 | 0.932973 | NA |
| PTBP1P | 0.134133 | -1.48909 | 4.077584 | -0.36519 | 0.71497 | NA |
| | 0.870886 | -0.53073 | 2.310896 | -0.22966 | 0.818353 | NA |
| | 0.46735 | -1.30193 | 3.399671 | -0.38296 | 0.701751 | NA |
| | 0.774742 | -0.40577 | 2.223483 | -0.18249 | 0.855195 | NA |
| | 0.163957 | 0.424913 | 4.079006 | 0.104171 | 0.917034 | NA |
| | 0.096202 | 0.408546 | 4.079639 | 0.100143 | 0.920231 | NA |
| | 4.185913 | 0.280677 | 1.081071 | 0.259628 | 0.795151 | NA |
| | 0.260158 | 1.240347 | 4.047124 | 0.306476 | 0.759242 | NA |
| TRAV30 | 0.096202 | 0.408546 | 4.079639 | 0.100143 | 0.920231 | NA |
| FAM35CP | 1.609131 | 0.660667 | 1.527112 | 0.432625 | 0.665287 | NA |
| | 0.230334 | -0.53318 | 4.076749 | -0.13079 | 0.895945 | NA |
| | 0.151697 | -1.546 | 4.075469 | -0.37934 | 0.704433 | NA |
| | 0.47961 | 0.535796 | 3.39605 | 0.15777 | 0.874638 | NA |
| | 0.977767 | 1.39453 | 2.230419 | 0.625232 | 0.531819 | NA |
| | 2.416618 | 1.077017 | 1.31645 | 0.818122 | 0.413287 | NA |
| | 0.258578 | 1.233151 | 4.047809 | 0.304646 | 0.760635 | NA |
| | 0.394291 | 0.298745 | 3.141711 | 0.09509 | 0.924243 | NA |
| | 0.162376 | 0.424913 | 4.079006 | 0.104171 | 0.917034 | NA |
| LINC00648 | 2.764884 | 0.056249 | 1.324736 | 0.042461 | 0.966131 | NA |
| | 0.506476 | 0.726418 | 2.864186 | 0.253621 | 0.799788 | NA |
| | 0.28583 | -2.21358 | 3.97376 | -0.55705 | 0.577494 | NA |
| | 1.63787 | -0.73465 | 1.830211 | -0.4014 | 0.688123 | NA |
| RANBP20P | 0.315653 | -0.57305 | 3.892639 | -0.14721 | 0.882963 | NA |
| | 0.875053 | -3.61314 | 2.623168 | -1.3774 | 0.168389 | NA |
| | 3.080515 | 0.898939 | 1.228642 | 0.731652 | 0.464381 | NA |
| | 0.134133 | -1.48909 | 4.077584 | -0.36519 | 0.71497 | NA |
| ADAM20P1 | 0.151697 | -1.546 | 4.075469 | -0.37934 | 0.704433 | NA |
| | 0.324752 | 1.53082 | 4.022025 | 0.380609 | 0.703493 | NA |
| DDX18P1 | 0.682896 | 1.412658 | 2.61672 | 0.539858 | 0.589295 | NA |
| | 0.163957 | 0.424913 | 4.079006 | 0.104171 | 0.917034 | NA |
| | 0.392857 | -1.546 | 4.075469 | -0.37934 | 0.704433 | NA |
| | 0.46577 | -1.30191 | 3.402879 | -0.38259 | 0.702024 | NA |
| | 0.420954 | 1.923376 | 3.462803 | 0.555439 | 0.578594 | NA |
| | 0.134133 | -1.48909 | 4.077584 | -0.36519 | 0.71497 | NA |

| | | | | | | |
|---|---|---|---|---|---|---|
| | 0.364467 | -1.20166 | 3.685431 | -0.32606 | 0.744381 | NA |
| | 0.096202 | 0.408546 | 4.079639 | 0.100143 | 0.920231 | NA |
| | 0.613357 | 2.486614 | 2.985155 | 0.832993 | 0.404848 | NA |
| | 3.01392 | -1.52565 | 1.291801 | -1.18103 | 0.237591 | NA |
| | 0.518736 | 2.237423 | 2.7713 | 0.807355 | 0.419462 | NA |
| | 0.749217 | 0.7397 | 2.633289 | 0.280903 | 0.778784 | NA |
| | 0.757324 | -2.08271 | 2.587512 | -0.80491 | 0.420873 | NA |
| | 0.354779 | 1.703954 | 3.657232 | 0.465914 | 0.641277 | NA |
| | 1.197422 | -0.20967 | 1.730805 | -0.12114 | 0.903582 | NA |
| GOLGA6L1! | 0.652869 | 1.292592 | 2.374873 | 0.544278 | 0.58625 | NA |
| | 0.163957 | 0.424913 | 4.079006 | 0.104171 | 0.917034 | NA |
| USP3-AS1 | 1.781138 | -0.27647 | 1.592047 | -0.17366 | 0.862134 | NA |
| | 13.85264 | 0.649279 | 0.586898 | 1.106288 | 0.268602 | NA |
| | 0.422534 | 1.924583 | 3.001758 | 0.641152 | 0.521424 | NA |
| | 0.296509 | -0.51709 | 3.982155 | -0.12985 | 0.896684 | NA |
| NPM1P42 | 0.162376 | 0.424913 | 4.079006 | 0.104171 | 0.917034 | NA |
| | 0.711139 | 2.710446 | 2.504485 | 1.082237 | 0.279147 | NA |
| | 4.749331 | 1.22067 | 0.968876 | 1.259883 | 0.207712 | NA |
| | 0.134133 | -1.48909 | 4.077584 | -0.36519 | 0.71497 | NA |
| | 3.477374 | -0.23186 | 1.191682 | -0.19457 | 0.845733 | NA |
| CSPG4P12 | 0.410275 | 0.244283 | 3.100889 | 0.078778 | 0.937209 | NA |
| | 0.151697 | -1.546 | 4.075469 | -0.37934 | 0.704433 | NA |
| | 1.255343 | 2.334528 | 1.930962 | 1.208997 | 0.226664 | NA |
| | 1.235354 | -0.89516 | 1.719639 | -0.52055 | 0.60268 | NA |
| PLCB2-AS1 | 3.382649 | -1.1119 | 1.173628 | -0.9474 | 0.343435 | NA |
| | 0.588072 | 2.39955 | 3.098804 | 0.774347 | 0.438726 | NA |
| | 0.984855 | -1.02676 | 2.366773 | -0.43382 | 0.664418 | NA |
| | 1.285748 | -1.30211 | 1.957858 | -0.66507 | 0.506007 | NA |
| | 4.673294 | 1.189217 | 0.987794 | 1.203911 | 0.228624 | NA |
| | 10.06864 | -0.13586 | 0.645414 | -0.2105 | 0.833279 | NA |
| | 4.225239 | 0.558606 | 1.056329 | 0.528819 | 0.596931 | NA |
| | 0.863039 | 0.783382 | 2.464829 | 0.317824 | 0.750618 | NA |
| | 0.314073 | -0.57308 | 3.898934 | -0.14698 | 0.883146 | NA |
| | 2.614996 | 1.595137 | 1.312255 | 1.215569 | 0.224149 | NA |
| | 0.419962 | -2.64111 | 3.429364 | -0.77014 | 0.441214 | NA |
| LINC00984 | 169.5306 | -0.08013 | 0.226837 | -0.35325 | 0.723905 | 0.824785 |
| | 0.096202 | 0.408546 | 4.079639 | 0.100143 | 0.920231 | NA |
| | 17.30094 | -0.23594 | 0.531475 | -0.44393 | 0.657095 | NA |
| | 17.70169 | 0.363114 | 0.532988 | 0.681279 | 0.495695 | NA |
| | 2.903986 | 1.032296 | 1.195705 | 0.863337 | 0.387952 | NA |
| | 0.096202 | 0.408546 | 4.079639 | 0.100143 | 0.920231 | NA |
| | 0.303394 | -2.27006 | 4.032451 | -0.56295 | 0.57347 | NA |
| | 0.162376 | 0.424913 | 4.079006 | 0.104171 | 0.917034 | NA |
| | 0.134133 | -1.48909 | 4.077584 | -0.36519 | 0.71497 | NA |
| | 0.858627 | -2.14058 | 2.513451 | -0.85165 | 0.394408 | NA |
| | 1.205355 | 1.329837 | 2.166752 | 0.613747 | 0.539383 | NA |
| | 0.096202 | 0.408546 | 4.079639 | 0.100143 | 0.920231 | NA |

| | | | | | | |
|---|---|---|---|---|---|---|
| | 0.719043 | 1.433043 | 2.335048 | 0.61371 | 0.539407 | NA |
| | 0.192403 | 0.975392 | 4.060716 | 0.240202 | 0.810174 | NA |
| DCAF13P3 | 0.520316 | 2.239969 | 3.173769 | 0.705776 | 0.480328 | NA |
| | 4.695119 | -0.17001 | 1.03572 | -0.16415 | 0.869612 | NA |
| | 5.112502 | 0.011782 | 0.947005 | 0.012442 | 0.990073 | NA |
| H3F3AP1 | 0.314073 | -0.57308 | 3.898934 | -0.14698 | 0.883146 | NA |
| | 3.325529 | 0.483026 | 1.128751 | 0.42793 | 0.668702 | NA |
| | 1.519814 | -0.07644 | 1.621533 | -0.04714 | 0.962401 | NA |
| | 0.163957 | 0.424913 | 4.079006 | 0.104171 | 0.917034 | NA |
| | 0.230334 | -0.53318 | 4.076749 | -0.13079 | 0.895945 | NA |
| | 0.303394 | -2.27006 | 4.032451 | -0.56295 | 0.57347 | NA |
| EFTUD1P1 | 0.163957 | 0.424913 | 4.079006 | 0.104171 | 0.917034 | NA |
| ISCA1P4 | 1.921624 | 0.541658 | 1.46181 | 0.370539 | 0.710981 | NA |
| | 0.918331 | 0.764804 | 2.118268 | 0.361052 | 0.718061 | NA |
| | 1.93072 | 1.568832 | 1.705316 | 0.919966 | 0.357591 | NA |
| HNRNPA1P | 0.556667 | 0.972496 | 2.522454 | 0.385536 | 0.699841 | NA |
| HERC2P7 | 3.238398 | 1.553647 | 1.217746 | 1.275839 | 0.202012 | NA |
| | 1.099787 | -2.48189 | 2.397845 | -1.03505 | 0.300646 | NA |
| | 661.7948 | -0.31825 | 0.161848 | -1.96637 | 0.049255 | 0.13727 |
| | 0.326333 | 1.533889 | 3.826878 | 0.40082 | 0.688553 | NA |
| HNRNPCP3 | 0.192403 | 0.975392 | 4.060716 | 0.240202 | 0.810174 | NA |
| | 102.4265 | -0.84023 | 0.362255 | -2.31945 | 0.02037 | 0.076664 |
| | 0.583919 | -1.6942 | 2.749224 | -0.61625 | 0.537732 | NA |
| | 2.271391 | 0.449361 | 1.368753 | 0.3283 | 0.742685 | NA |
| | 28.26053 | 0.84794 | 0.534966 | 1.585035 | 0.112958 | NA |
| THTPA | 228.1685 | -0.3274 | 0.185205 | -1.76776 | 0.077101 | 0.186614 |
| | 0.096202 | 0.408546 | 4.079639 | 0.100143 | 0.920231 | NA |
| | 0.490493 | 0.776209 | 2.895261 | 0.268096 | 0.788625 | NA |
| | 0.162376 | 0.424913 | 4.079006 | 0.104171 | 0.917034 | NA |
| | 4.286293 | 0.57239 | 1.007633 | 0.568054 | 0.569998 | NA |
| | 6.40045 | 1.024871 | 0.949747 | 1.079098 | 0.280544 | NA |
| | 0.230334 | -0.53318 | 4.076749 | -0.13079 | 0.895945 | NA |
| | 0.096202 | 0.408546 | 4.079639 | 0.100143 | 0.920231 | NA |
| ANP32BP3 | 0.326333 | 1.533889 | 3.826878 | 0.40082 | 0.688553 | NA |
| | 1.740542 | -2.32937 | 1.935186 | -1.20369 | 0.228708 | NA |
| | 0.452561 | 2.061505 | 3.349456 | 0.615474 | 0.538241 | NA |
| AHCYP7 | 0.163957 | 0.424913 | 4.079006 | 0.104171 | 0.917034 | NA |
| NDUFAF4P | 0.162376 | 0.424913 | 4.079006 | 0.104171 | 0.917034 | NA |
| | 59.6638 | -0.71498 | 0.299245 | -2.38927 | 0.016882 | 0.067589 |
| | 1.144942 | 1.358971 | 2.181094 | 0.623068 | 0.53324 | NA |
| | 0.326536 | 0.037056 | 3.842554 | 0.009644 | 0.992306 | NA |
| MRPL46 | 191.0821 | 0.181443 | 0.22297 | 0.813756 | 0.415785 | 0.578944 |
| | 2.393061 | 0.101261 | 1.341689 | 0.075473 | 0.939839 | NA |
| | 11.43701 | 2.490035 | 0.767299 | 3.245196 | 0.001174 | NA |
| | 0.151697 | -1.546 | 4.075469 | -0.37934 | 0.704433 | NA |
| | 0.460466 | 0.568655 | 3.014642 | 0.188631 | 0.850382 | NA |
| UBE2Q2L | 4.842263 | -0.55125 | 0.894359 | -0.61636 | 0.537654 | NA |

| | | | | | | |
|---|---|---|---|---|---|---|
| HNRNPA1P | 0.151697 | -1.546 | 4.075469 | -0.37934 | 0.704433 | NA |
| | 0.151697 | -1.546 | 4.075469 | -0.37934 | 0.704433 | NA |
| | 2.136267 | -0.64375 | 1.350128 | -0.4768 | 0.633501 | NA |
| ANP32AP1 | 0.192403 | 0.975392 | 4.060716 | 0.240202 | 0.810174 | NA |
| | 1.522175 | 0.469421 | 1.602229 | 0.29298 | 0.769538 | NA |
| | 1.470789 | -1.06553 | 1.748153 | -0.60952 | 0.542183 | NA |
| | 0.260158 | 1.240347 | 4.047124 | 0.306476 | 0.759242 | NA |
| | 5.216244 | -0.21017 | 0.850786 | -0.24703 | 0.804887 | NA |
| LINC00052 | 3.661422 | -0.62858 | 1.149676 | -0.54675 | 0.584552 | NA |
| | 0.35636 | 1.709252 | 3.651833 | 0.468053 | 0.639746 | NA |
| | 6.53117 | 0.064543 | 0.785065 | 0.082213 | 0.934477 | NA |
| | 1.235004 | 0.68061 | 1.965017 | 0.346363 | 0.72907 | NA |
| RPL21P12 | 0.163957 | 0.424913 | 4.079006 | 0.104171 | 0.917034 | NA |
| | 2.929521 | -1.72955 | 1.299834 | -1.3306 | 0.183322 | NA |
| UBE2Q2P1 | 0.556667 | 0.972496 | 2.522454 | 0.385536 | 0.699841 | NA |
| | 0.096202 | 0.408546 | 4.079639 | 0.100143 | 0.920231 | NA |
| | 0.134133 | -1.48909 | 4.077584 | -0.36519 | 0.71497 | NA |
| | 0.096202 | 0.408546 | 4.079639 | 0.100143 | 0.920231 | NA |
| | 2.404212 | 1.045664 | 1.336098 | 0.782625 | 0.433847 | NA |
| HMGN1P2( | 0.151697 | -1.546 | 4.075469 | -0.37934 | 0.704433 | NA |
| | 0.096202 | 0.408546 | 4.079639 | 0.100143 | 0.920231 | NA |
| | 8.928684 | 1.122113 | 0.80144 | 1.400121 | 0.161477 | NA |
| | 0.326536 | 0.037056 | 3.842554 | 0.009644 | 0.992306 | NA |
| | 2.091262 | -2.0689 | 1.502393 | -1.37707 | 0.168491 | NA |
| | 0.258578 | 1.233151 | 4.047809 | 0.304646 | 0.760635 | NA |
| | 0.46735 | -1.30193 | 3.399671 | -0.38296 | 0.701751 | NA |
| | 5.668023 | 0.378774 | 0.969005 | 0.390889 | 0.695879 | NA |
| TYRO3P | 0.968522 | 0.920016 | 1.915592 | 0.480278 | 0.63103 | NA |
| | 5.524224 | -0.60908 | 0.857683 | -0.71015 | 0.477612 | NA |
| | 0.134133 | -1.48909 | 4.077584 | -0.36519 | 0.71497 | NA |
| | 0.151697 | -1.546 | 4.075469 | -0.37934 | 0.704433 | NA |
| PRELID1P4 | 0.547182 | 2.340248 | 3.137689 | 0.745851 | 0.455757 | NA |
| | 0.682693 | 2.623213 | 2.518338 | 1.041645 | 0.297576 | NA |
| | 0.162376 | 0.424913 | 4.079006 | 0.104171 | 0.917034 | NA |
| | 1.178424 | -1.58299 | 2.140626 | -0.7395 | 0.459604 | NA |
| | 5.29916 | 0.055735 | 0.946293 | 0.058898 | 0.953033 | NA |
| EEF1A1P22 | 11.25408 | 0.547455 | 0.725594 | 0.754492 | 0.450554 | NA |
| | 0.151697 | -1.546 | 4.075469 | -0.37934 | 0.704433 | NA |
| | 0.096202 | 0.408546 | 4.079639 | 0.100143 | 0.920231 | NA |
| | 0.230334 | -0.53318 | 4.076749 | -0.13079 | 0.895945 | NA |
| | 0.096202 | 0.408546 | 4.079639 | 0.100143 | 0.920231 | NA |
| | 0.364467 | -1.20166 | 3.685431 | -0.32606 | 0.744381 | NA |
| C15orf37 | 8.612945 | -0.19159 | 0.715505 | -0.26777 | 0.788879 | NA |
| | 0.314073 | -0.57308 | 3.898934 | -0.14698 | 0.883146 | NA |
| | 0.422534 | 1.924583 | 3.001758 | 0.641152 | 0.521424 | NA |
| | 0.28583 | -2.21358 | 3.97376 | -0.55705 | 0.577494 | NA |
| | 1.965223 | -0.02685 | 1.585705 | -0.01693 | 0.986492 | NA |

| | | | | | | |
|---|---|---|---|---|---|---|
| | 1.005019 | -1.03066 | 2.033647 | -0.50681 | 0.612291 | NA |
| | 0.736607 | 1.395844 | 2.318642 | 0.602009 | 0.547168 | NA |
| PIGHP1 | 1.281015 | 0.71131 | 1.742215 | 0.408279 | 0.683069 | NA |
| | 2.681463 | 0.796338 | 1.389329 | 0.573182 | 0.566522 | NA |
| | 0.327913 | 1.543154 | 4.021063 | 0.383768 | 0.701151 | NA |
| | 3.572396 | 1.806576 | 1.24243 | 1.454066 | 0.145928 | NA |
| IQCH-AS1 | 25.84053 | 0.998999 | 0.418227 | 2.388654 | 0.01691 | NA |
| | 6.573904 | 2.844181 | 1.084775 | 2.621908 | 0.008744 | NA |
| | 1.252771 | 0.053586 | 1.700195 | 0.031518 | 0.974857 | NA |
| | 0.392857 | -1.546 | 4.075469 | -0.37934 | 0.704433 | NA |
| | 7.529124 | -0.16489 | 0.755857 | -0.21815 | 0.827314 | NA |
| | 4.960305 | 0.553316 | 1.055331 | 0.524306 | 0.600066 | NA |
| | 7.433428 | 0.367405 | 0.731911 | 0.501981 | 0.615681 | NA |
| | 0.151697 | -1.546 | 4.075469 | -0.37934 | 0.704433 | NA |
| HMGB1P8 | 4.882269 | 0.664693 | 0.964065 | 0.689469 | 0.490528 | NA |
| | 0.163957 | 0.424913 | 4.079006 | 0.104171 | 0.917034 | NA |
| | 5.698826 | -0.83055 | 0.878005 | -0.94595 | 0.344173 | NA |
| | 0.326333 | 1.533889 | 3.826878 | 0.40082 | 0.688553 | NA |
| | 0.74907 | 1.552259 | 2.278648 | 0.681219 | 0.495733 | NA |
| | 0.411855 | 0.250456 | 3.097618 | 0.080854 | 0.935558 | NA |
| | 2.506295 | 1.550002 | 1.638967 | 0.945719 | 0.344292 | NA |
| | 12.29968 | 0.818275 | 0.640288 | 1.27798 | 0.201257 | NA |
| | 0.460466 | 0.568655 | 3.014642 | 0.188631 | 0.850382 | NA |
| CSPG4P11 | 26.80473 | -0.26896 | 0.495116 | -0.54323 | 0.586971 | NA |
| | 0.508056 | 0.731032 | 2.861912 | 0.255435 | 0.798387 | NA |
| LINC00933 | 5.4778 | 0.131157 | 0.879395 | 0.149145 | 0.881439 | NA |
| | 2.621909 | 0.195456 | 1.295417 | 0.150883 | 0.880068 | NA |
| | 0.392857 | -1.546 | 4.075469 | -0.37934 | 0.704433 | NA |
| | 0.258578 | 1.233151 | 4.047809 | 0.304646 | 0.760635 | NA |
| | 0.410275 | 0.244283 | 3.100889 | 0.078778 | 0.937209 | NA |
| | 0.327913 | 1.543154 | 4.021063 | 0.383768 | 0.701151 | NA |
| | 0.326333 | 1.533889 | 3.826878 | 0.40082 | 0.688553 | NA |
| | 0.096202 | 0.408546 | 4.079639 | 0.100143 | 0.920231 | NA |
| | 1.590076 | -0.3384 | 1.915172 | -0.1767 | 0.859748 | NA |
| | 1.292043 | 0.040764 | 1.785235 | 0.022834 | 0.981783 | NA |
| | 3.294178 | -0.25529 | 1.127593 | -0.2264 | 0.820889 | NA |
| | 4.548265 | 0.948465 | 1.048301 | 0.904764 | 0.36559 | NA |
| ANP32BP1 | 1.450276 | 0.320155 | 1.846699 | 0.173366 | 0.862364 | NA |
| | 1.696234 | 0.294515 | 1.666475 | 0.176729 | 0.859721 | NA |
| | 1.538158 | 0.399163 | 1.868829 | 0.21359 | 0.830867 | NA |
| LINC00567 | 0.162376 | 0.424913 | 4.079006 | 0.104171 | 0.917034 | NA |
| TCEB1P2 | 0.967291 | -0.94461 | 2.160526 | -0.43721 | 0.661958 | NA |
| HERC2P10 | 1.149832 | -0.38667 | 2.069202 | -0.18687 | 0.851763 | NA |
| | 1.725846 | -0.23167 | 1.459515 | -0.15873 | 0.873881 | NA |
| | 0.846649 | 2.927909 | 2.364097 | 1.23849 | 0.215535 | NA |
| HNRNPLP2 | 17.01841 | -0.35958 | 0.49859 | -0.72119 | 0.470793 | NA |
| | 0.422534 | 1.924583 | 3.001758 | 0.641152 | 0.521424 | NA |

| | | | | | | |
|---|---|---|---|---|---|---|
| | 0.45509 | -2.71227 | 3.963744 | -0.68427 | 0.493805 | NA |
| | 0.260158 | 1.240347 | 4.047124 | 0.306476 | 0.759242 | NA |
| | 9.081005 | 0.476881 | 0.702355 | 0.678974 | 0.497154 | NA |
| | 1.411499 | -2.12166 | 2.00599 | -1.05766 | 0.29021 | NA |
| | 0.192403 | 0.975392 | 4.060716 | 0.240202 | 0.810174 | NA |
| | 0.489058 | -0.58569 | 3.427543 | -0.17088 | 0.864321 | NA |
| | 0.830237 | -1.27668 | 2.202503 | -0.57965 | 0.562151 | NA |
| | 0.555233 | -0.57009 | 3.296947 | -0.17291 | 0.862719 | NA |
| | 1.120219 | 0.417842 | 1.804388 | 0.23157 | 0.816872 | NA |
| | 0.151697 | -1.546 | 4.075469 | -0.37934 | 0.704433 | NA |
| | 0.746295 | -0.65731 | 2.318283 | -0.28353 | 0.776767 | NA |
| AGBL1-AS1 | 0.303394 | -2.27006 | 4.032451 | -0.56295 | 0.57347 | NA |
| CSPG4P13 | 13.21597 | -0.38344 | 0.591137 | -0.64865 | 0.516562 | NA |
| | 0.518736 | 2.237423 | 2.7713 | 0.807355 | 0.419462 | NA |
| | 0.392711 | 0.292538 | 3.145221 | 0.09301 | 0.925895 | NA |
| | 0.258578 | 1.233151 | 4.047809 | 0.304646 | 0.760635 | NA |
| | 1.587707 | -3.26952 | 1.872881 | -1.74572 | 0.08086 | NA |
| SSSCA1-AS: | 36.0452 | 0.217623 | 0.359136 | 0.605962 | 0.54454 | NA |
| | 0.965508 | -0.08709 | 2.197017 | -0.03964 | 0.968381 | NA |
| | 6.874439 | -0.13721 | 0.85936 | -0.15967 | 0.873143 | NA |
| | 20.99496 | 0.343964 | 0.472453 | 0.728038 | 0.46659 | NA |
| | 17.74357 | -0.07467 | 0.516102 | -0.14469 | 0.884959 | NA |
| | 0.841063 | -2.08438 | 2.306687 | -0.90362 | 0.366195 | NA |
| NTRK3-AS1 | 0.096202 | 0.408546 | 4.079639 | 0.100143 | 0.920231 | NA |
| | 12.89136 | -0.67553 | 0.609147 | -1.10898 | 0.26744 | NA |
| HEXA-AS1 | 1.215189 | -0.88793 | 1.977332 | -0.44905 | 0.653393 | NA |
| LINC01043 | 0.134133 | -1.48909 | 4.077584 | -0.36519 | 0.71497 | NA |
| LINC00562 | 1.534877 | 0.084903 | 1.62645 | 0.052202 | 0.958368 | NA |
| | 0.162376 | 0.424913 | 4.079006 | 0.104171 | 0.917034 | NA |
| | 7.420055 | 0.612123 | 0.805413 | 0.760011 | 0.447248 | NA |
| | 1108.612 | -0.19364 | 0.227158 | -0.85245 | 0.393964 | 0.558732 |
| | 0.994194 | -0.63787 | 1.893391 | -0.33689 | 0.736198 | NA |
| | 2.739001 | 0.003038 | 1.222432 | 0.002485 | 0.998017 | NA |
| | 1.987199 | 1.212507 | 1.528164 | 0.793441 | 0.427521 | NA |
| | 1.820467 | -0.04814 | 1.430357 | -0.03366 | 0.973151 | NA |
| | 21.09833 | 0.267726 | 0.46172 | 0.579846 | 0.562019 | NA |
| | 3.295116 | 0.995266 | 1.133385 | 0.878135 | 0.37987 | NA |
| | 0.096202 | 0.408546 | 4.079639 | 0.100143 | 0.920231 | NA |
| | 5.951279 | 0.427489 | 0.859821 | 0.497184 | 0.619059 | NA |
| TPST2P1 | 0.326333 | 1.533889 | 3.826878 | 0.40082 | 0.688553 | NA |
| | 0.4986 | -1.64638 | 3.317584 | -0.49626 | 0.619712 | NA |
| | 38.68426 | 0.401952 | 0.387305 | 1.037819 | 0.299354 | NA |
| | 0.134133 | -1.48909 | 4.077584 | -0.36519 | 0.71497 | NA |
| | 1.977162 | 4.164432 | 1.778566 | 2.341455 | 0.019209 | NA |
| | 0.192403 | 0.975392 | 4.060716 | 0.240202 | 0.810174 | NA |
| | 1.314202 | 0.813471 | 1.874167 | 0.434044 | 0.664256 | NA |
| | 6.512862 | -0.04364 | 0.818562 | -0.05331 | 0.957484 | NA |

| | | | | | | |
|---|---|---|---|---|---|---|
| | 0.096202 | 0.408546 | 4.079639 | 0.100143 | 0.920231 | NA |
| HNRNPA3P | 1.195639 | 0.515221 | 2.098932 | 0.245468 | 0.806094 | NA |
| | 0.096202 | 0.408546 | 4.079639 | 0.100143 | 0.920231 | NA |
| LINC00543 | 1.181439 | -0.20977 | 1.987054 | -0.10557 | 0.915925 | NA |
| | 3.077559 | -0.28713 | 1.213075 | -0.23669 | 0.812894 | NA |
| | 0.448206 | -1.2532 | 3.007315 | -0.41672 | 0.676885 | NA |
| | 0.134133 | -1.48909 | 4.077584 | -0.36519 | 0.71497 | NA |
| | 7.418965 | 1.401452 | 0.819189 | 1.710779 | 0.087122 | NA |
| | 0.096202 | 0.408546 | 4.079639 | 0.100143 | 0.920231 | NA |
| KRT8P50 | 1.429191 | 0.470595 | 1.645258 | 0.286031 | 0.774855 | NA |
| | 17.26914 | -0.02544 | 0.525533 | -0.04841 | 0.961392 | NA |
| | 0.888304 | 0.641829 | 2.171212 | 0.295609 | 0.767529 | NA |
| | 2.786143 | -1.41 | 1.235015 | -1.14169 | 0.253585 | NA |
| | 0.096202 | 0.408546 | 4.079639 | 0.100143 | 0.920231 | NA |
| | 2.064642 | 0.337739 | 1.492697 | 0.226261 | 0.820998 | NA |
| | 0.192403 | 0.975392 | 4.060716 | 0.240202 | 0.810174 | NA |
| | 1.649984 | 0.73597 | 1.549673 | 0.474919 | 0.634844 | NA |
| | 2.012464 | 1.134837 | 1.467439 | 0.773345 | 0.439318 | NA |
| ADPGK-AS1 | 10.46814 | 1.269757 | 0.746738 | 1.700405 | 0.089055 | NA |
| LINC00565 | 1.657153 | 2.827967 | 1.867826 | 1.514041 | 0.130015 | NA |
| CCPG1 | 210.3128 | 0.991411 | 0.322828 | 3.071023 | 0.002133 | 0.016837 |
| | 0.134133 | -1.48909 | 4.077584 | -0.36519 | 0.71497 | NA |
| SPCS2P1 | 0.151697 | -1.546 | 4.075469 | -0.37934 | 0.704433 | NA |
| | 0.134133 | -1.48909 | 4.077584 | -0.36519 | 0.71497 | NA |
| | 23.412 | 0.482744 | 0.577454 | 0.835986 | 0.403163 | NA |
| | 0.258578 | 1.233151 | 4.047809 | 0.304646 | 0.760635 | NA |
| DOCK9-AS2 | 13.54738 | -0.2184 | 0.564481 | -0.3869 | 0.698831 | NA |
| | 1.007648 | 2.051938 | 2.077812 | 0.987548 | 0.323374 | NA |
| LINC00555 | 0.151697 | -1.546 | 4.075469 | -0.37934 | 0.704433 | NA |
| | 5.887394 | 0.055572 | 0.93027 | 0.059737 | 0.952365 | NA |
| SNORD116 | 25.63856 | 0.687665 | 0.429564 | 1.600843 | 0.109412 | NA |
| SNORD116 | 25.63856 | 0.687665 | 0.429564 | 1.600843 | 0.109412 | NA |
| | 0.422534 | 1.924583 | 3.001758 | 0.641152 | 0.521424 | NA |
| | 0.881915 | -2.2305 | 2.871853 | -0.77668 | 0.437349 | NA |
| | 5.602069 | 0.79565 | 1.209131 | 0.658035 | 0.510516 | NA |
| ATP5J2P6 | 0.258578 | 1.233151 | 4.047809 | 0.304646 | 0.760635 | NA |
| | 12.19533 | 0.433793 | 0.691208 | 0.627586 | 0.530275 | NA |
| | 0.096202 | 0.408546 | 4.079639 | 0.100143 | 0.920231 | NA |
| | 4.243499 | 2.411174 | 1.198977 | 2.011026 | 0.044323 | NA |
| ADAMTS7P | 1.047103 | 1.538196 | 2.447568 | 0.628459 | 0.529703 | NA |
| | 6.863991 | 0.07245 | 0.8159 | 0.088797 | 0.929243 | NA |
| | 0.247898 | -0.5901 | 4.074633 | -0.14482 | 0.884851 | NA |
| | 0.096202 | 0.408546 | 4.079639 | 0.100143 | 0.920231 | NA |
| LINC00561 | 0.163957 | 0.424913 | 4.079006 | 0.104171 | 0.917034 | NA |
| | 1.415736 | -1.38965 | 1.816509 | -0.76501 | 0.444265 | NA |
| | 0.788582 | 0.608095 | 2.456267 | 0.247569 | 0.804468 | NA |
| GOLGA8T | 3.754993 | 1.855738 | 1.163988 | 1.594293 | 0.11087 | NA |

| | | | | | | |
|---|---|---|---|---|---|---|
| | 0.296509 | -0.51709 | 3.982155 | -0.12985 | 0.896684 | NA |
| LARP4P | 0.163957 | 0.424913 | 4.079006 | 0.104171 | 0.917034 | NA |
| | 12.34366 | 1.441162 | 0.626702 | 2.299597 | 0.021471 | NA |
| CD24P2 | 0.163957 | 0.424913 | 4.079006 | 0.104171 | 0.917034 | NA |
| | 12.97556 | -0.59152 | 0.574139 | -1.03028 | 0.302881 | NA |
| | 2.867499 | -0.74884 | 1.22395 | -0.61182 | 0.540657 | NA |
| | 36.00742 | 0.477399 | 0.430804 | 1.108159 | 0.267793 | NA |
| | 0.750651 | 1.552121 | 2.514127 | 0.61736 | 0.536998 | NA |
| | 6.998656 | 0.837063 | 0.764518 | 1.094891 | 0.273564 | NA |
| | 0.506476 | 0.726418 | 2.864186 | 0.253621 | 0.799788 | NA |
| | 2.243205 | 0.161484 | 1.348132 | 0.119784 | 0.904654 | NA |
| LINC00559 | 0.315653 | -0.57305 | 3.892639 | -0.14721 | 0.882963 | NA |
| | 7.172614 | 0.375512 | 0.773976 | 0.485173 | 0.627554 | NA |
| | 0.163957 | 0.424913 | 4.079006 | 0.104171 | 0.917034 | NA |
| PAN3-AS1 | 14.64417 | 0.353407 | 0.529693 | 0.667193 | 0.504649 | NA |
| | 0.163957 | 0.424913 | 4.079006 | 0.104171 | 0.917034 | NA |
| | 0.327913 | 1.543154 | 4.021063 | 0.383768 | 0.701151 | NA |
| | 1.272345 | 3.519403 | 2.400212 | 1.466288 | 0.14257 | NA |
| | 0.460466 | 0.568655 | 3.014642 | 0.188631 | 0.850382 | NA |
| | 26.3913 | -0.11372 | 0.50405 | -0.22562 | 0.8215 | NA |
| TPBGL | 96.05394 | -0.49851 | 0.279148 | -1.78581 | 0.07413 | 0.181706 |
| | 6.033529 | 0.985786 | 0.868393 | 1.135184 | 0.256298 | NA |
| | 0.096202 | 0.408546 | 4.079639 | 0.100143 | 0.920231 | NA |
| | 0.096202 | 0.408546 | 4.079639 | 0.100143 | 0.920231 | NA |
| | 0.298089 | -0.51711 | 3.975232 | -0.13008 | 0.896501 | NA |
| | 0.163957 | 0.424913 | 4.079006 | 0.104171 | 0.917034 | NA |
| DISC1FP1 | 0.326333 | 1.533889 | 3.826878 | 0.40082 | 0.688553 | NA |
| GOLGA6L7 | 0.520316 | 2.239969 | 3.173769 | 0.705776 | 0.480328 | NA |
| | 0.555233 | -0.57009 | 3.296947 | -0.17291 | 0.862719 | NA |
| C15orf65 | 36.88605 | 0.426539 | 0.374101 | 1.140171 | 0.254215 | NA |
| ATP5HP1 | 0.162376 | 0.424913 | 4.079006 | 0.104171 | 0.917034 | NA |
| | 5.165275 | 0.985104 | 0.969506 | 1.016088 | 0.309587 | NA |
| | 0.163957 | 0.424913 | 4.079006 | 0.104171 | 0.917034 | NA |
| GOLGA8S | 0.979551 | 0.16329 | 2.152655 | 0.075855 | 0.939534 | NA |
| | 9.476189 | -0.25275 | 0.69512 | -0.3636 | 0.716155 | NA |
| | 0.402398 | -2.58771 | 4.018369 | -0.64397 | 0.519595 | NA |
| GOLGA8H | 2.252836 | -0.80767 | 1.342103 | -0.6018 | 0.547309 | NA |
| | 69.89156 | 0.040777 | 0.296521 | 0.137519 | 0.890621 | 0.936036 |
| LOXL1-AS1 | 1.494281 | -1.73103 | 1.875863 | -0.92279 | 0.356115 | NA |
| | 0.382031 | -1.26484 | 3.151989 | -0.40128 | 0.688212 | NA |
| DNM1P49 | 0.162376 | 0.424913 | 4.079006 | 0.104171 | 0.917034 | NA |
| | 0.327913 | 1.543154 | 4.021063 | 0.383768 | 0.701151 | NA |
| | 673.5109 | -0.19938 | 0.17869 | -1.11579 | 0.264513 | 0.428974 |
| | 0.134133 | -1.48909 | 4.077584 | -0.36519 | 0.71497 | NA |
| | 1.752655 | -0.41495 | 1.535934 | -0.27016 | 0.787037 | NA |
| RN7SL80P | 1.602596 | 0.149842 | 1.657784 | 0.090387 | 0.92798 | NA |
| RN7SL755P | 0.163957 | 0.424913 | 4.079006 | 0.104171 | 0.917034 | NA |

| | | | | | | |
|---|---|---|---|---|---|---|
| MIR3165 | 0.096202 | 0.408546 | 4.079639 | 0.100143 | 0.920231 | NA |
| | 0.163957 | 0.424913 | 4.079006 | 0.104171 | 0.917034 | NA |
| RN7SL747F | 0.163957 | 0.424913 | 4.079006 | 0.104171 | 0.917034 | NA |
| RN7SL86P | 0.163957 | 0.424913 | 4.079006 | 0.104171 | 0.917034 | NA |
| | 1.071251 | -0.2597 | 1.986911 | -0.1307 | 0.89601 | NA |
| RN7SL744F | 0.52699 | -2.57878 | 3.248478 | -0.79384 | 0.427288 | NA |
| | 26.84876 | -0.35969 | 0.402318 | -0.89403 | 0.371304 | NA |
| RN7SL312F | 0.640609 | 0.042838 | 2.379511 | 0.018003 | 0.985637 | NA |
| | 0.758758 | -0.35643 | 2.514365 | -0.14176 | 0.887273 | NA |
| MIR4712 | 0.506476 | 0.726418 | 2.864186 | 0.253621 | 0.799788 | NA |
| MIR5006 | 0.096202 | 0.408546 | 4.079639 | 0.100143 | 0.920231 | NA |
| RN7SL613F | 0.162376 | 0.424913 | 4.079006 | 0.104171 | 0.917034 | NA |
| MIR4513 | 0.134133 | -1.48909 | 4.077584 | -0.36519 | 0.71497 | NA |
| RN7SL179F | 0.192403 | 0.975392 | 4.060716 | 0.240202 | 0.810174 | NA |
| RN7SL596F | 0.192403 | 0.975392 | 4.060716 | 0.240202 | 0.810174 | NA |
| MIR548AR | 0.561971 | -0.49218 | 2.810061 | -0.17515 | 0.860962 | NA |
| RN7SL595F | 0.134133 | -1.48909 | 4.077584 | -0.36519 | 0.71497 | NA |
| | 0.71919 | 0.510978 | 2.728734 | 0.187258 | 0.851458 | NA |
| RN7SL534F | 0.247898 | -0.5901 | 4.074633 | -0.14482 | 0.884851 | NA |
| MIR3613 | 0.163957 | 0.424913 | 4.079006 | 0.104171 | 0.917034 | NA |
| RN7SL348F | 0.096202 | 0.408546 | 4.079639 | 0.100143 | 0.920231 | NA |
| | 0.422737 | 0.518872 | 3.501356 | 0.148192 | 0.882191 | NA |
| MIR3942 | 0.134133 | -1.48909 | 4.077584 | -0.36519 | 0.71497 | NA |
| RN7SL2 | 104.7469 | -1.04736 | 0.353919 | -2.95932 | 0.003083 | 0.021631 |
| MIR4312 | 0.258578 | 1.233151 | 4.047809 | 0.304646 | 0.760635 | NA |
| | 0.785567 | -0.70568 | 2.358063 | -0.29926 | 0.764741 | NA |
| MIR5188 | 0.096202 | 0.408546 | 4.079639 | 0.100143 | 0.920231 | NA |
| MIR4502 | 0.096202 | 0.408546 | 4.079639 | 0.100143 | 0.920231 | NA |
| MIR3164 | 2.147115 | -2.09721 | 1.547467 | -1.35525 | 0.175336 | NA |
| RN7SL133F | 0.326333 | 1.533889 | 3.826878 | 0.40082 | 0.688553 | NA |
| RN7SL166F | 0.096202 | 0.408546 | 4.079639 | 0.100143 | 0.920231 | NA |
| | 0.690946 | -1.56103 | 2.699004 | -0.57837 | 0.563013 | NA |
| | 32.05604 | -0.42311 | 0.446232 | -0.94818 | 0.343039 | NA |
| RN7SL660F | 0.162376 | 0.424913 | 4.079006 | 0.104171 | 0.917034 | NA |
| RN7SL49P | 0.096202 | 0.408546 | 4.079639 | 0.100143 | 0.920231 | NA |
| MIR4489 | 0.096202 | 0.408546 | 4.079639 | 0.100143 | 0.920231 | NA |
| MIR3685 | 0.547182 | 2.340248 | 3.137689 | 0.745851 | 0.455757 | NA |
| RN7SL3 | 5.203893 | -1.04841 | 0.974329 | -1.07603 | 0.281912 | NA |
| MIR4690 | 0.162376 | 0.424913 | 4.079006 | 0.104171 | 0.917034 | NA |
| | 0.936915 | 0.819377 | 2.189457 | 0.374237 | 0.708228 | NA |
| MIR3657 | 0.134133 | -1.48909 | 4.077584 | -0.36519 | 0.71497 | NA |
| | 0.268266 | -2.15396 | 4.038351 | -0.53338 | 0.593773 | NA |
| | 0.163957 | 0.424913 | 4.079006 | 0.104171 | 0.917034 | NA |
| MIR4512 | 0.670433 | 1.255281 | 2.356401 | 0.532711 | 0.594234 | NA |
| MIR4498 | 0.134133 | -1.48909 | 4.077584 | -0.36519 | 0.71497 | NA |
| | 0.73681 | 0.399395 | 2.277135 | 0.175393 | 0.86077 | NA |
| | 0.268266 | -2.15396 | 4.038351 | -0.53338 | 0.593773 | NA |

| | | | | | | |
|---|---|---|---|---|---|---|
| | 0.163957 | 0.424913 | 4.079006 | 0.104171 | 0.917034 | NA |
| | 0.151697 | -1.546 | 4.075469 | -0.37934 | 0.704433 | NA |
| | 0.230334 | -0.53318 | 4.076749 | -0.13079 | 0.895945 | NA |
| | 0.326333 | 1.533889 | 3.826878 | 0.40082 | 0.688553 | NA |
| | 0.096202 | 0.408546 | 4.079639 | 0.100143 | 0.920231 | NA |
| | 2.788961 | 1.334717 | 1.347333 | 0.990636 | 0.321863 | NA |
| | 0.151697 | -1.546 | 4.075469 | -0.37934 | 0.704433 | NA |
| | 0.725928 | 0.181819 | 2.321433 | 0.078322 | 0.937572 | NA |
| | 1.076781 | 3.261813 | 2.235782 | 1.458914 | 0.144589 | NA |
| | 1.798962 | -0.39956 | 1.497704 | -0.26678 | 0.78964 | NA |
| | 1.966789 | -1.2874 | 1.510993 | -0.85202 | 0.394203 | NA |
| | 0.258578 | 1.233151 | 4.047809 | 0.304646 | 0.760635 | NA |
| | 0.096202 | 0.408546 | 4.079639 | 0.100143 | 0.920231 | NA |
| | 6.149866 | 0.55819 | 0.869189 | 0.642196 | 0.520746 | NA |
| | 2.469192 | 0.347835 | 1.260964 | 0.275849 | 0.782664 | NA |
| | 0.163957 | 0.424913 | 4.079006 | 0.104171 | 0.917034 | NA |
| | 0.258578 | 1.233151 | 4.047809 | 0.304646 | 0.760635 | NA |
| | 0.399595 | -1.31595 | 3.579466 | -0.36764 | 0.713143 | NA |
| LASP1 | 7177.739 | -0.37636 | 0.155181 | -2.42527 | 0.015297 | 0.062857 |
| SNX11 | 945.6622 | -0.16566 | 0.139756 | -1.18538 | 0.235867 | 0.398845 |
| DBNDD1 | 2891.959 | -0.27082 | 0.247311 | -1.09508 | 0.273481 | 0.43932 |
| SARM1 | 311.4514 | -0.51655 | 0.174148 | -2.96613 | 0.003016 | 0.021243 |
| POLDIP2 | 4099.975 | -0.31428 | 0.138841 | -2.26362 | 0.023597 | 0.084618 |
| CAMKK1 | 173.6305 | -0.71762 | 0.220521 | -3.25419 | 0.001137 | 0.011214 |
| HSPB6 | 11.91893 | -1.13541 | 0.619709 | -1.83216 | 0.066928 | NA |
| ARHGAP33 | 118.0783 | -0.48278 | 0.302142 | -1.59787 | 0.110073 | 0.237763 |
| NDUFAB1 | 1845.606 | 0.055508 | 0.13513 | 0.410775 | 0.681237 | 0.792212 |
| CDC27 | 1655.428 | 0.416659 | 0.144786 | 2.877764 | 0.004005 | 0.025436 |
| DVL2 | 2088.588 | -0.49325 | 0.139894 | -3.52589 | 4.22E-04 | 0.006131 |
| UPF1 | 3720.761 | -0.55761 | 0.163456 | -3.41137 | 6.46E-04 | 0.007892 |
| DHX33 | 1460.995 | -0.17256 | 0.149911 | -1.15111 | 0.249685 | 0.41357 |
| LIG3 | 715.6009 | -0.30079 | 0.134873 | -2.2302 | 0.025734 | 0.089808 |
| ACSM3 | 16.38822 | 0.478268 | 0.52119 | 0.917646 | 0.358804 | NA |
| | 89.79565 | -0.63083 | 0.244302 | -2.58217 | 0.009818 | 0.046669 |
| CIAPIN1 | 2396.849 | -0.17646 | 0.13912 | -1.26839 | 0.204658 | 0.361375 |
| SPPL2B | 777.5852 | -0.43875 | 0.171571 | -2.55722 | 0.010551 | 0.048747 |
| COPZ2 | 1015.453 | -0.06641 | 0.134726 | -0.49291 | 0.622079 | 0.746432 |
| CREBBP | 1423.746 | -0.57424 | 0.217749 | -2.63719 | 0.00836 | 0.041745 |
| BZRAP1 | 34.71172 | -0.15544 | 0.514345 | -0.30221 | 0.76249 | NA |
| MPO | 1.213406 | -0.28643 | 1.978822 | -0.14475 | 0.884908 | NA |
| SOX8 | 230.7714 | 0.368237 | 0.340559 | 1.081271 | 0.279577 | 0.445601 |
| ITGAL | 0.260158 | 1.240347 | 4.047124 | 0.306476 | 0.759242 | NA |
| PDK2 | 940.0834 | -0.16818 | 0.13072 | -1.2866 | 0.198233 | 0.35399 |
| ITGA3 | 7562.794 | -0.15329 | 0.329583 | -0.4651 | 0.641857 | 0.761683 |
| GGNBP2 | 1090.392 | 0.509762 | 0.175776 | 2.900065 | 0.003731 | 0.024376 |
| ITGA2B | 2.244436 | 0.956595 | 1.371961 | 0.697247 | 0.485648 | NA |
| GDE1 | 4753.102 | -0.08883 | 0.149861 | -0.59276 | 0.553339 | 0.693776 |

| | | | | | | |
|---|---|---|---|---|---|---|
| C19orf60 | 1002.909 | -0.16824 | 0.159632 | -1.05389 | 0.291931 | 0.458755 |
| CRLF1 | 9.691536 | 0.417674 | 0.703632 | 0.593597 | 0.552782 | NA |
| OSBPL7 | 227.7669 | -0.51777 | 0.179944 | -2.87737 | 0.00401 | 0.025436 |
| TMEM98 | 4149.372 | -0.25198 | 0.122221 | -2.06172 | 0.039235 | 0.118715 |
| YBX2 | 3.088873 | 0.570823 | 1.2046 | 0.47387 | 0.635593 | NA |
| MAP3K14 | 154.4831 | -0.37858 | 0.23288 | -1.62563 | 0.104029 | 0.229146 |
| CCL18 | 690.9575 | 0.555668 | 0.578438 | 0.960635 | 0.336736 | 0.505637 |
| CCL3 | 129.5902 | 1.233022 | 0.845776 | 1.457859 | 0.144879 | 0.286849 |
| SYNRG | 1491.945 | -0.45124 | 0.137959 | -3.27086 | 0.001072 | 0.010848 |
| AP2B1 | 10468.13 | -0.50642 | 0.114603 | -4.41889 | 9.92E-06 | 6.26E-04 |
| ZNF263 | 907.7474 | 0.023254 | 0.148049 | 0.157068 | 0.875191 | 0.925383 |
| CX3CL1 | 72.8889 | -2.45522 | 0.436132 | -5.62955 | 1.81E-08 | 1.13E-05 |
| SPATA20 | 2224.133 | -0.35786 | 0.14409 | -2.48356 | 0.013008 | 0.056459 |
| CACNA1G | 1.469987 | 0.110263 | 2.154214 | 0.051185 | 0.959178 | NA |
| TNFRSF12A | 2092.034 | 0.783633 | 0.718945 | 1.089975 | 0.275724 | 0.4413 |
| TBXA2R | 41.65907 | -0.05022 | 0.346117 | -0.1451 | 0.884628 | NA |
| COX10 | 249.2949 | -0.32463 | 0.182133 | -1.78236 | 0.07469 | 0.182427 |
| PAF1 | 2160.84 | -0.25994 | 0.154462 | -1.68285 | 0.092404 | 0.211681 |
| ARHGAP44 | 11.09565 | -0.26281 | 0.617104 | -0.42587 | 0.670204 | NA |
| ELAC2 | 3135.22 | -0.29121 | 0.12687 | -2.29538 | 0.021712 | 0.080102 |
| PRSS21 | 0.163957 | 0.424913 | 4.079006 | 0.104171 | 0.917034 | NA |
| CCDC124 | 3091.217 | -0.26378 | 0.148553 | -1.7757 | 0.075783 | 0.184251 |
| CEACAM21 | 60.26511 | 1.854226 | 0.470647 | 3.93974 | 8.16E-05 | 0.002244 |
| PAFAH1B1 | 3525.493 | 0.17622 | 0.146933 | 1.199321 | 0.230403 | 0.392623 |
| NOS2 | 2.311106 | -0.28038 | 1.329609 | -0.21087 | 0.832985 | NA |
| DNAH9 | 2.986597 | -2.10126 | 1.271864 | -1.65211 | 0.098512 | NA |
| KIAA0100 | 8590.239 | -0.33305 | 0.171075 | -1.94679 | 0.05156 | 0.141393 |
| GAS7 | 11237.01 | -0.32523 | 0.16928 | -1.92128 | 0.054696 | 0.147336 |
| MATK | 88.03537 | 0.142644 | 0.359561 | 0.396717 | 0.691576 | 0.800491 |
| CD79B | 11.47476 | -0.04336 | 0.841263 | -0.05154 | 0.958897 | NA |
| RPUSD1 | 1491.919 | -0.14133 | 0.176429 | -0.80105 | 0.423104 | 0.585422 |
| RHBDF1 | 1883.363 | -0.40983 | 0.128761 | -3.18285 | 0.001458 | 0.013329 |
| LUC7L | 665.9153 | 0.440031 | 0.181273 | 2.427446 | 0.015206 | 0.0627 |
| BAIAP3 | 6.305248 | 0.45637 | 0.795892 | 0.573408 | 0.566369 | NA |
| TSR3 | 1515.333 | -0.04678 | 0.126759 | -0.36904 | 0.712095 | 0.816193 |
| PIGQ | 2791.551 | -0.42604 | 0.13117 | -3.24801 | 0.001162 | 0.011424 |
| CRAMP1L | 267.9718 | -0.41192 | 0.24291 | -1.69578 | 0.089928 | 0.207626 |
| CYB561 | 1079.08 | -0.25136 | 0.155548 | -1.61598 | 0.1061 | 0.231919 |
| SPAG9 | 2145.605 | 0.395422 | 0.273413 | 1.446243 | 0.148109 | 0.291231 |
| MPND | 538.6723 | -0.30564 | 0.174738 | -1.74916 | 0.080264 | 0.19184 |
| NFIX | 3876.603 | -0.52505 | 0.170142 | -3.08594 | 0.002029 | 0.016331 |
| MMP25 | 3.873581 | 0.767519 | 1.124396 | 0.682606 | 0.494856 | NA |
| IL32 | 8.134361 | -0.70799 | 0.924297 | -0.76597 | 0.443692 | NA |
| PKD1 | 2718.885 | -0.40412 | 0.190542 | -2.12088 | 0.033932 | 0.107602 |
| MED24 | 2097.306 | -0.35976 | 0.122707 | -2.93184 | 0.00337 | 0.022712 |
| ZNF207 | 2406.105 | 0.070433 | 0.122639 | 0.574308 | 0.56576 | 0.702867 |
| ZNF200 | 171.49 | 0.208061 | 0.192285 | 1.082046 | 0.279232 | 0.445471 |

| | | | | | | |
|---|---|---|---|---|---|---|
| MRC2 | 41.49896 | 0.84178 | 0.397885 | 2.115637 | 0.034376 | NA |
| APBA3 | 508.4602 | -0.15846 | 0.167314 | -0.94706 | 0.343611 | 0.511875 |
| MKS1 | 215.4389 | -0.46461 | 0.222532 | -2.08783 | 0.036813 | 0.113713 |
| AKAP8L | 814.142 | -0.17933 | 0.148836 | -1.2049 | 0.228241 | 0.389884 |
| MBTD1 | 190.9717 | 0.439664 | 0.230782 | 1.905105 | 0.056766 | 0.151046 |
| UTP18 | 1246.934 | 0.177209 | 0.14032 | 1.262897 | 0.206626 | 0.363668 |
| TTC19 | 1404.576 | 0.126348 | 0.151955 | 0.831479 | 0.405703 | 0.56959 |
| PTBP1 | 7636.596 | -0.46331 | 0.138166 | -3.35327 | 7.99E-04 | 0.009034 |
| DPF1 | 55.40492 | -0.40724 | 0.29183 | -1.39548 | 0.162871 | 0.311174 |
| PLAUR | 303.8243 | 0.271397 | 0.237626 | 1.142119 | 0.253405 | 0.417475 |
| WIZ | 1994.253 | -0.46809 | 0.150052 | -3.11953 | 0.001811 | 0.015146 |
| ZBTB32 | 1.519814 | -0.07644 | 1.621533 | -0.04714 | 0.962401 | NA |
| TMEM159 | 1027.677 | -0.35787 | 0.162884 | -2.19706 | 0.028016 | 0.09476 |
| BRCA1 | 541.7413 | 0.683684 | 0.28757 | 2.377454 | 0.017433 | 0.069026 |
| CD22 | 7.490461 | 0.583562 | 0.793675 | 0.735267 | 0.462177 | NA |
| PSMC4 | 2652.226 | -0.16166 | 0.132773 | -1.21753 | 0.223403 | 0.384238 |
| SLC25A39 | 3772.438 | -0.1215 | 0.131815 | -0.92173 | 0.356672 | 0.524283 |
| MVP | 7864.005 | -0.41553 | 0.144087 | -2.88386 | 0.003928 | 0.025173 |
| DPEP1 | 0.28583 | -2.21358 | 3.97376 | -0.55705 | 0.577494 | NA |
| XYLT2 | 1301.054 | -0.49783 | 0.150484 | -3.30821 | 9.39E-04 | 0.009948 |
| SRCIN1 | 145.9467 | 0.439456 | 0.329477 | 1.333798 | 0.18227 | 0.335165 |
| RALBP1 | 2148.083 | 0.575372 | 0.177641 | 3.238961 | 0.0012 | 0.011696 |
| SLC7A9 | 1.198617 | 1.916623 | 2.276885 | 0.841774 | 0.399915 | NA |
| POU2F2 | 5.032808 | 1.84834 | 0.986511 | 1.873613 | 0.060984 | NA |
| RABEP1 | 1839.923 | 0.674864 | 0.219421 | 3.075658 | 0.0021 | 0.016689 |
| GRN | 21051.74 | -0.43316 | 0.142992 | -3.02927 | 0.002451 | 0.018505 |
| RANBP3 | 1954.72 | -0.38638 | 0.133817 | -2.88741 | 0.003884 | 0.024986 |
| PNPLA6 | 5321.479 | -0.43073 | 0.202833 | -2.12354 | 0.033709 | 0.107222 |
| ALG1 | 986.756 | -0.32494 | 0.157589 | -2.06197 | 0.03921 | 0.118672 |
| ATP6V0A1 | 5162.058 | -0.55191 | 0.156685 | -3.52241 | 4.28E-04 | 0.006167 |
| UHRF1 | 179.2254 | -0.17356 | 0.294114 | -0.59011 | 0.55512 | 0.695073 |
| MAP2K3 | 2368.934 | -0.23988 | 0.171795 | -1.39629 | 0.162627 | 0.310808 |
| GABARAPL | 3044.97 | 0.163997 | 0.136347 | 1.202788 | 0.229058 | 0.39083 |
| TIMP2 | 51038.81 | -0.44461 | 0.161667 | -2.75017 | 0.005957 | 0.033246 |
| TUBG2 | 582.5384 | -0.09311 | 0.168034 | -0.55414 | 0.579483 | 0.713595 |
| MRI1 | 273.8837 | -0.04575 | 0.198071 | -0.23098 | 0.817331 | 0.890494 |
| EDC4 | 1067.375 | -0.5346 | 0.14607 | -3.65993 | 2.52E-04 | 0.004475 |
| CLEC16A | 1351.469 | -0.51241 | 0.160975 | -3.18314 | 0.001457 | 0.013326 |
| CDH1 | 1426.609 | -1.47544 | 0.540702 | -2.72875 | 0.006358 | 0.034652 |
| FAM65A | 2513.576 | -0.50538 | 0.156159 | -3.23629 | 0.001211 | 0.011777 |
| BEST2 | 0.678686 | -3.08037 | 2.621268 | -1.17514 | 0.239937 | NA |
| PHLPP2 | 199.3803 | 0.050646 | 0.211034 | 0.239988 | 0.810339 | 0.886567 |
| CTNS | 1640.773 | -0.23912 | 0.167362 | -1.42876 | 0.153074 | 0.297783 |
| PHF23 | 2634.79 | -0.21444 | 0.12756 | -1.68107 | 0.09275 | 0.212269 |
| RAB27B | 2.857202 | 3.685639 | 1.624768 | 2.26841 | 0.023304 | NA |
| DSG2 | 650.1259 | 0.269821 | 0.169292 | 1.593818 | 0.110977 | 0.238988 |
| KIAA0556 | 1833.2 | -0.82891 | 0.187571 | -4.4192 | 9.91E-06 | 6.26E-04 |

| | | | | | | |
|---|---|---|---|---|---|---|
| TNFRSF17 | 0.850604 | -2.33906 | 2.457452 | -0.95182 | 0.341186 | NA |
| SNX29 | 888.439 | -0.56869 | 0.149382 | -3.80696 | 1.41E-04 | 0.003125 |
| EPN3 | 0.69238 | 0.295083 | 2.603008 | 0.113362 | 0.909743 | NA |
| NEDD4L | 3504.214 | -0.26417 | 0.170846 | -1.54626 | 0.122042 | 0.255395 |
| BCAR1 | 2158.459 | -0.36401 | 0.153493 | -2.3715 | 0.017716 | 0.069636 |
| HERPUD1 | 1957.411 | -0.12809 | 0.149227 | -0.85838 | 0.390685 | 0.555877 |
| HOMER3 | 1255.163 | -0.16983 | 0.145671 | -1.16586 | 0.243673 | 0.406464 |
| CYBA | 9940.062 | -0.09057 | 0.155776 | -0.58138 | 0.560982 | 0.699164 |
| PRSS8 | 0.151697 | -1.546 | 4.075469 | -0.37934 | 0.704433 | NA |
| USE1 | 759.1509 | -0.02685 | 0.180034 | -0.14915 | 0.881437 | 0.929417 |
| LAMA3 | 18.16285 | 1.348893 | 0.521036 | 2.588869 | 0.009629 | NA |
| USP36 | 1289.243 | -0.26228 | 0.174273 | -1.50496 | 0.132333 | 0.269509 |
| PCGF2 | 1453.781 | -0.31204 | 0.176572 | -1.76722 | 0.077191 | 0.186754 |
| POLR3E | 951.4961 | -0.0234 | 0.16072 | -0.1456 | 0.884239 | 0.93146 |
| FLYWCH1 | 3997.4 | -0.5001 | 0.136415 | -3.666 | 2.46E-04 | 0.004412 |
| UNKL | 411.4327 | -0.1561 | 0.204807 | -0.76219 | 0.445946 | 0.605647 |
| CTDP1 | 1176.299 | -0.29503 | 0.143159 | -2.06082 | 0.03932 | 0.118943 |
| CREB3L3 | 0.424115 | 1.932531 | 3.45462 | 0.559405 | 0.575885 | NA |
| CDH3 | 5039.443 | -0.7866 | 0.234655 | -3.35216 | 8.02E-04 | 0.009053 |
| VMP1 | 3423.992 | 0.072788 | 0.124886 | 0.582838 | 0.560002 | 0.698472 |
| APPBP2 | 1201.19 | 0.266928 | 0.187059 | 1.426971 | 0.153588 | 0.298392 |
| SEZ6 | 0.556813 | -0.57007 | 3.294615 | -0.17303 | 0.862627 | NA |
| MED29 | 1437.777 | -0.13201 | 0.132335 | -0.99757 | 0.318487 | 0.486349 |
| HAGH | 1756.606 | 0.077471 | 0.174853 | 0.443066 | 0.657718 | 0.774252 |
| DLX3 | 0.919554 | 0.143913 | 2.343626 | 0.061406 | 0.951036 | NA |
| ATP2C2 | 1.356286 | 2.46466 | 2.147149 | 1.147875 | 0.25102 | NA |
| NGFR | 12188.7 | 0.314793 | 0.56426 | 0.557886 | 0.576922 | 0.711845 |
| RFXANK | 949.8087 | -0.33478 | 0.17284 | -1.93691 | 0.052757 | 0.143471 |
| TMEM161/ | 672.732 | -0.33292 | 0.152908 | -2.17725 | 0.029462 | 0.097807 |
| LPAR2 | 209.4335 | -0.43239 | 0.217572 | -1.98734 | 0.046884 | 0.133225 |
| SUGP2 | 1768.666 | -0.16195 | 0.169208 | -0.95709 | 0.338524 | 0.507304 |
| CNN2 | 1444.709 | -0.31387 | 0.191099 | -1.64247 | 0.100493 | 0.223642 |
| ABCA7 | 755.2735 | -0.48051 | 0.19027 | -2.52544 | 0.011555 | 0.051969 |
| SBNO2 | 3364.857 | -0.54379 | 0.16283 | -3.33963 | 8.39E-04 | 0.00931 |
| HMG20B | 31122.53 | -0.20884 | 0.121405 | -1.72021 | 0.085394 | 0.199635 |
| AP3D1 | 6732.612 | -0.44006 | 0.131102 | -3.35662 | 7.89E-04 | 0.008985 |
| SLC9A3R2 | 908.9821 | 0.073889 | 0.176228 | 0.419278 | 0.675013 | 0.787316 |
| NTHL1 | 450.361 | -0.12757 | 0.163459 | -0.78046 | 0.435121 | 0.596166 |
| WDR18 | 1524.497 | -0.22548 | 0.123385 | -1.82742 | 0.067636 | 0.170276 |
| NTN1 | 8.314333 | 0.390938 | 0.798685 | 0.489477 | 0.624504 | NA |
| GLP2R | 0.288605 | 1.443619 | 4.029058 | 0.358302 | 0.720117 | NA |
| KARS | 3948.512 | 0.024301 | 0.130275 | 0.186537 | 0.852024 | 0.911895 |
| ADAT1 | 747.7546 | -0.37099 | 0.186285 | -1.99154 | 0.046422 | 0.132419 |
| MAP2K4 | 657.8379 | 0.279281 | 0.170163 | 1.641259 | 0.100744 | 0.22399 |
| TLE2 | 272.2016 | -0.15176 | 0.213194 | -0.71185 | 0.476556 | 0.631541 |
| PDE4A | 1013.398 | -0.27609 | 0.223555 | -1.23499 | 0.216835 | 0.37618 |
| ELAVL1 | 2284.704 | -0.27275 | 0.120742 | -2.25899 | 0.023884 | 0.085385 |

| | | | | | | |
|---|---|---|---|---|---|---|
| THUMPD1 | 585.8597 | 0.648193 | 0.211957 | 3.058136 | 0.002227 | 0.017335 |
| ACSM2B | 0.288605 | 1.443619 | 4.029058 | 0.358302 | 0.720117 | NA |
| FECH | 1404.69 | -0.10327 | 0.14736 | -0.7008 | 0.483429 | 0.637289 |
| CACNB1 | 151.9644 | -0.00976 | 0.213147 | -0.0458 | 0.963466 | 0.980168 |
| METTL22 | 264.5854 | -0.19362 | 0.171695 | -1.12771 | 0.259441 | 0.423872 |
| DHX8 | 2173.742 | -0.37484 | 0.14548 | -2.57656 | 0.009979 | 0.047131 |
| ROGDI | 523.0759 | -0.36539 | 0.190283 | -1.92023 | 0.054829 | 0.147555 |
| ROCK1 | 602.1651 | 0.78342 | 0.989982 | 0.791348 | 0.428741 | 0.590534 |
| CBFB | 1820.828 | 0.344606 | 0.163027 | 2.113798 | 0.034533 | 0.108908 |
| IFI35 | 1058.372 | -0.77206 | 0.197268 | -3.91375 | 9.09E-05 | 0.002389 |
| HEATR6 | 486.0522 | -0.28124 | 0.160253 | -1.75496 | 0.079266 | 0.190145 |
| COASY | 3211.343 | -0.32193 | 0.207346 | -1.55264 | 0.120509 | 0.252771 |
| PLEKHH3 | 669.8565 | -0.07918 | 0.160488 | -0.4934 | 0.621731 | 0.746165 |
| MEF2A | 991.2641 | 0.783871 | 0.159528 | 4.913703 | 8.94E-07 | 1.53E-04 |
| PRR11 | 884.0308 | -0.35433 | 0.21552 | -1.64405 | 0.100165 | 0.223272 |
| SIRT2 | 3801.681 | -0.17224 | 0.130789 | -1.31696 | 0.187851 | 0.341909 |
| SDK2 | 58.55295 | -0.77934 | 0.359657 | -2.16688 | 0.030244 | 0.099677 |
| VPS35 | 4264.146 | 0.189635 | 0.150488 | 1.260132 | 0.207622 | 0.364808 |
| DNAJA2 | 1959.411 | 0.209959 | 0.140981 | 1.489273 | 0.136416 | 0.275453 |
| PLA2G10 | 0.911593 | 1.257757 | 2.250933 | 0.558771 | 0.576318 | NA |
| SMG6 | 1356.071 | -0.45138 | 0.146853 | -3.07369 | 0.002114 | 0.016777 |
| FGF22 | 7.698358 | 0.023891 | 0.770368 | 0.031013 | 0.975259 | NA |
| FSTL3 | 2225.779 | 0.435599 | 0.634534 | 0.686487 | 0.492406 | 0.643732 |
| RNF126 | 1490.017 | -0.16921 | 0.144762 | -1.16886 | 0.24246 | 0.405411 |
| MNT | 493.4098 | -0.74631 | 0.224711 | -3.32121 | 8.96E-04 | 0.009708 |
| JMJD6 | 719.3924 | 0.069998 | 0.208132 | 0.336317 | 0.736632 | 0.833121 |
| WIPI1 | 7251.177 | -0.07281 | 0.138439 | -0.52591 | 0.598954 | 0.728565 |
| ST6GALNA( | 4756.531 | -0.47413 | 0.190104 | -2.49406 | 0.012629 | 0.055268 |
| C16orf80 | 738.6426 | 0.033469 | 0.1496 | 0.223722 | 0.822974 | 0.893194 |
| CSNK2A2 | 1663.413 | -0.19098 | 0.130858 | -1.45947 | 0.144436 | 0.286335 |
| TCF3 | 1648.962 | -0.21085 | 0.147303 | -1.43142 | 0.15231 | 0.296688 |
| DAZAP1 | 2925.844 | -0.16947 | 0.152288 | -1.11286 | 0.265769 | 0.430423 |
| MBD3 | 2539.33 | -0.20416 | 0.170078 | -1.20038 | 0.229992 | 0.392198 |
| CDH19 | 914.9091 | 1.991757 | 0.250353 | 7.95579 | 1.78E-15 | 7.03E-12 |
| PRKACA | 2419.981 | -0.41734 | 0.125976 | -3.31287 | 9.23E-04 | 0.009851 |
| LPHN1 | 684.5287 | -0.53029 | 0.173877 | -3.04981 | 0.00229 | 0.017639 |
| EPN2 | 466.9214 | -0.20803 | 0.162213 | -1.28244 | 0.19969 | 0.355882 |
| ALDH3A2 | 1175.365 | 0.011213 | 0.148864 | 0.075326 | 0.939956 | 0.966534 |
| SREBF1 | 1983.074 | -0.80709 | 0.148742 | -5.42611 | 5.76E-08 | 2.35E-05 |
| NFATC3 | 346.5416 | -0.12649 | 0.155394 | -0.81401 | 0.415639 | 0.578809 |
| ACADVL | 7345.683 | 0.085336 | 0.136673 | 0.62438 | 0.532378 | 0.676529 |
| ACAP1 | 38.20801 | 0.287078 | 0.380989 | 0.753505 | 0.451146 | NA |
| DERL2 | 992.8631 | -0.00228 | 0.165193 | -0.01379 | 0.988998 | 0.994535 |
| NDE1 | 925.0197 | -0.48861 | 0.174906 | -2.79356 | 0.005213 | 0.030399 |
| TMEM38A | 159.0896 | -0.12016 | 0.231916 | -0.51812 | 0.604374 | 0.732942 |
| AP1M1 | 2996.104 | -0.39062 | 0.127138 | -3.0724 | 0.002123 | 0.016805 |
| XRCC1 | 636.7995 | -0.48162 | 0.168886 | -2.85176 | 0.004348 | 0.026845 |

| | | | | | | |
|---|---|---|---|---|---|---|
| LLGL2 | 118.2702 | 0.387433 | 0.311284 | 1.244627 | 0.213269 | 0.371973 |
| NLE1 | 542.1213 | -0.38405 | 0.150544 | -2.55107 | 0.010739 | 0.049294 |
| SMARCE1 | 1200.205 | 0.335132 | 0.165054 | 2.030445 | 0.042311 | 0.125119 |
| FNDC8 | 0.804712 | -0.7622 | 2.348804 | -0.3245 | 0.745556 | NA |
| GSDMB | 19.06567 | 0.356151 | 0.504207 | 0.706359 | 0.479965 | NA |
| ADAM11 | 69.12433 | -0.4466 | 0.335883 | -1.32963 | 0.183641 | 0.336932 |
| TBX21 | 0.52699 | -2.57878 | 3.248478 | -0.79384 | 0.427288 | NA |
| NSF | 1637.097 | 0.013277 | 0.128861 | 0.103036 | 0.917935 | 0.952399 |
| MRPS34 | 3671.548 | -0.0741 | 0.148481 | -0.49903 | 0.617761 | 0.742604 |
| NOTCH3 | 58.84949 | -0.44614 | 0.403924 | -1.10453 | 0.269365 | 0.433993 |
| C17orf85 | 676.7396 | 0.226268 | 0.193126 | 1.171611 | 0.241353 | 0.404426 |
| ATP2A3 | 28.59779 | -0.12161 | 0.398564 | -0.30513 | 0.760267 | NA |
| ZNF532 | 666.2102 | -0.07177 | 0.188351 | -0.38103 | 0.703178 | 0.809337 |
| SCARF1 | 9.803091 | -0.4562 | 0.633562 | -0.72006 | 0.471486 | NA |
| LMAN1 | 1747.017 | 0.42864 | 0.150811 | 2.842226 | 0.00448 | 0.027376 |
| ZZEF1 | 2490.924 | -0.49744 | 0.146684 | -3.39127 | 6.96E-04 | 0.008269 |
| C19orf10 | 4605.456 | -0.09138 | 0.173576 | -0.52644 | 0.598584 | 0.728207 |
| ANO8 | 311.8335 | -0.15254 | 0.224563 | -0.67928 | 0.496962 | 0.648128 |
| TIMM21 | 460.7873 | 0.499748 | 0.182439 | 2.739265 | 0.006158 | 0.033906 |
| VPS9D1 | 325.8217 | -0.26433 | 0.191412 | -1.38096 | 0.167293 | 0.316783 |
| CACNG4 | 0.384806 | 1.846216 | 3.999752 | 0.461582 | 0.644381 | NA |
| MOCOS | 84.02738 | -0.32194 | 0.281061 | -1.14543 | 0.252029 | 0.415977 |
| WDR62 | 447.0978 | -0.53059 | 0.273196 | -1.94216 | 0.052118 | 0.142371 |
| RGS11 | 76.31178 | -0.03161 | 0.457648 | -0.06906 | 0.94494 | 0.969441 |
| SLC46A1 | 99.3018 | -0.00533 | 0.23662 | -0.02254 | 0.982014 | 0.990311 |
| SPAG5 | 348.1858 | -0.54934 | 0.311798 | -1.76184 | 0.078097 | 0.188276 |
| TRAF4 | 1854.11 | -0.05184 | 0.125676 | -0.41251 | 0.679966 | 0.791153 |
| GPATCH1 | 250.5032 | 0.006423 | 0.188547 | 0.034067 | 0.972824 | 0.985082 |
| ICAM3 | 439.2771 | -0.02009 | 0.180975 | -0.11101 | 0.911607 | 0.948657 |
| CAMSAP3 | 2.447462 | 1.62042 | 1.409383 | 1.149737 | 0.250252 | NA |
| XAB2 | 1315.8 | -0.46077 | 0.137353 | -3.35466 | 7.95E-04 | 0.008997 |
| ARHGEF1 | 2652.706 | -0.2838 | 0.14947 | -1.8987 | 0.057604 | 0.152569 |
| STXBP2 | 489.8604 | -0.51191 | 0.18731 | -2.73298 | 0.006276 | 0.034384 |
| MAP2K7 | 1160.973 | -0.33166 | 0.141061 | -2.35116 | 0.018715 | 0.072265 |
| NMRK2 | 1.740418 | -2.59874 | 1.85112 | -1.40387 | 0.160356 | NA |
| GTF3C1 | 3807.315 | -0.70669 | 0.150527 | -4.69477 | 2.67E-06 | 2.93E-04 |
| IL4R | 430.2945 | -0.02831 | 0.23522 | -0.12035 | 0.904204 | 0.944186 |
| SNRPA | 1176.897 | -0.41637 | 0.157704 | -2.64023 | 0.008285 | 0.041548 |
| EXOSC5 | 716.1892 | -0.1561 | 0.195152 | -0.79987 | 0.423788 | 0.585958 |
| SIRT6 | 603.8759 | -0.27814 | 0.181735 | -1.53049 | 0.125896 | 0.260971 |
| PIAS2 | 608.6033 | 0.310637 | 0.167612 | 1.853313 | 0.063837 | 0.164285 |
| PIK3C3 | 1099.14 | 0.653748 | 0.184426 | 3.544765 | 3.93E-04 | 0.005852 |
| TNRC6C | 160.05 | 0.015706 | 0.243954 | 0.064381 | 0.948667 | 0.971279 |
| CLUL1 | 0.258578 | 1.233151 | 4.047809 | 0.304646 | 0.760635 | NA |
| THOC1 | 227.3836 | 0.917271 | 0.245074 | 3.742829 | 1.82E-04 | 0.003659 |
| REXO1 | 890.0126 | -0.44191 | 0.171526 | -2.57633 | 0.009986 | 0.047145 |
| CEACAM1 | 1169.942 | -0.1941 | 0.772455 | -0.25128 | 0.801601 | 0.880721 |

| | | | | | | |
|---|---|---|---|---|---|---|
| CIC | 5116.774 | -0.5254 | 0.185169 | -2.83742 | 0.004548 | 0.027635 |
| LIPE | 953.3048 | -0.34287 | 0.257151 | -1.33333 | 0.182422 | 0.335341 |
| PAFAH1B3 | 634.3358 | -0.25946 | 0.221358 | -1.17214 | 0.241141 | 0.404398 |
| KIF22 | 1103.622 | -0.37753 | 0.173428 | -2.17688 | 0.02949 | 0.097872 |
| DNM2 | 6129.106 | -0.35537 | 0.128123 | -2.77371 | 0.005542 | 0.031817 |
| KEAP1 | 2483.404 | -0.26893 | 0.198318 | -1.35608 | 0.175073 | 0.327007 |
| RDH8 | 1.856284 | 1.630743 | 1.919775 | 0.849445 | 0.395634 | NA |
| COL5A3 | 635.7648 | -0.40005 | 0.285603 | -1.40072 | 0.161297 | 0.308963 |
| SRCAP | 3288.938 | -0.76507 | 0.204886 | -3.73413 | 1.88E-04 | 0.003747 |
| NDC80 | 209.66 | 0.851189 | 0.275042 | 3.094764 | 0.00197 | 0.016038 |
| CDH7 | 6.922962 | 0.515956 | 0.814335 | 0.633593 | 0.526347 | NA |
| ZNF506 | 123.1055 | 0.165048 | 0.252082 | 0.654737 | 0.512637 | 0.659932 |
| PHLPP1 | 1610.303 | -0.78697 | 0.214422 | -3.6702 | 2.42E-04 | 0.004373 |
| ATP8B1 | 205.3258 | 0.675421 | 0.226253 | 2.985246 | 0.002834 | 0.020371 |
| ME2 | 1530.258 | 0.42495 | 0.160434 | 2.648749 | 0.008079 | 0.040855 |
| EPB41L3 | 4256.791 | 0.325076 | 0.14469 | 2.246701 | 0.024659 | 0.086963 |
| NFE2L1 | 11924.88 | -0.29028 | 0.14864 | -1.95291 | 0.05083 | 0.140234 |
| PALB2 | 410.0601 | 0.489737 | 0.193392 | 2.532351 | 0.01133 | 0.051175 |
| ULK2 | 965.2404 | -0.14328 | 0.175417 | -0.81678 | 0.414053 | 0.577278 |
| ITGAE | 782.2597 | 0.300607 | 0.175663 | 1.711269 | 0.087031 | 0.20251 |
| CYLD | 1286.207 | 0.664388 | 0.179967 | 3.691723 | 2.23E-04 | 0.004158 |
| SEH1L | 613.3652 | 0.319565 | 0.148228 | 2.155894 | 0.031092 | 0.101597 |
| ZNF213 | 406.7957 | -0.43149 | 0.180056 | -2.39639 | 0.016557 | 0.066625 |
| RRN3 | 1112.013 | 0.333373 | 0.152499 | 2.186073 | 0.02881 | 0.096427 |
| CHERP | 853.3212 | -0.63129 | 0.205464 | -3.0725 | 0.002123 | 0.016805 |
| MRPL28 | 2036.392 | -0.14509 | 0.138177 | -1.05005 | 0.293697 | 0.460918 |
| HBQ1 | 2.337659 | 0.598536 | 1.358735 | 0.440509 | 0.659568 | NA |
| ITPKC | 596.9116 | -0.2803 | 0.183949 | -1.5238 | 0.127557 | 0.263448 |
| CEACAM6 | 0.163957 | 0.424913 | 4.079006 | 0.104171 | 0.917034 | NA |
| HSD17B2 | 0.163957 | 0.424913 | 4.079006 | 0.104171 | 0.917034 | NA |
| NLK | 804.7145 | 0.210763 | 0.198097 | 1.063942 | 0.287355 | 0.453569 |
| PIGS | 1793.603 | -0.40386 | 0.125265 | -3.22401 | 0.001264 | 0.012046 |
| ATXN7L3 | 2302.023 | -0.18446 | 0.154738 | -1.19206 | 0.233237 | 0.395796 |
| PGS1 | 688.504 | -0.12369 | 0.15532 | -0.79639 | 0.425808 | 0.587731 |
| PSMC5 | 2334.8 | 0.079171 | 0.134003 | 0.590819 | 0.554642 | 0.694914 |
| CETP | 0.163957 | 0.424913 | 4.079006 | 0.104171 | 0.917034 | NA |
| MMP2 | 6837.1 | -1.05196 | 0.231899 | -4.53629 | 5.73E-06 | 4.59E-04 |
| LPCAT2 | 2121.075 | 0.311567 | 0.16722 | 1.863211 | 0.062433 | 0.161813 |
| GNAO1 | 9.048338 | -1.54171 | 0.727946 | -2.11789 | 0.034184 | NA |
| OGFOD1 | 1433.984 | -0.09981 | 0.154362 | -0.64662 | 0.517875 | 0.665014 |
| RFX2 | 85.78793 | -0.02807 | 0.306931 | -0.09147 | 0.92712 | 0.958156 |
| METTL2A | 347.0087 | -0.02557 | 0.18855 | -0.13561 | 0.892131 | 0.936777 |
| KHSRP | 6678.083 | -0.55093 | 0.128283 | -4.29464 | 1.75E-05 | 8.56E-04 |
| GNA11 | 2696.56 | -0.43287 | 0.148444 | -2.91604 | 0.003545 | 0.023597 |
| COQ9 | 1150.304 | -0.16777 | 0.128635 | -1.30422 | 0.192158 | 0.346798 |
| ARHGAP28 | 7.162083 | -0.06835 | 0.810679 | -0.08431 | 0.932812 | NA |
| FUS | 5248.436 | -0.36743 | 0.176448 | -2.08239 | 0.037307 | 0.114667 |

| | | | | | | |
|---|---|---|---|---|---|---|
| FXYD5 | 3181.744 | -0.10914 | 0.211524 | -0.51597 | 0.605876 | 0.734172 |
| ZNF302 | 320.2302 | 0.982367 | 0.265844 | 3.695271 | 2.20E-04 | 0.00412 |
| GRAMD1A | 1528.71 | -0.2744 | 0.148634 | -1.84612 | 0.064875 | 0.166004 |
| FXYD3 | 1320.093 | -0.14744 | 0.162609 | -0.90669 | 0.364573 | 0.531934 |
| CDIP1 | 519.372 | -0.87906 | 0.237525 | -3.70093 | 2.15E-04 | 0.004068 |
| CMTM1 | 15.55637 | 0.842529 | 0.595323 | 1.415246 | 0.156996 | NA |
| KCNH4 | 3.148685 | -0.55529 | 1.348711 | -0.41172 | 0.680548 | NA |
| GMIP | 1222.27 | -0.45625 | 0.166338 | -2.74289 | 0.00609 | 0.033785 |
| BIRC5 | 598.6172 | -0.45869 | 0.267152 | -1.71696 | 0.085986 | 0.200742 |
| ANKRD24 | 3.494367 | -0.73973 | 1.137243 | -0.65046 | 0.515397 | NA |
| LTBP4 | 1601.383 | -0.36173 | 0.129599 | -2.79112 | 0.005253 | 0.030564 |
| BLVRB | 1713.807 | -0.12098 | 0.151825 | -0.79687 | 0.425525 | 0.587538 |
| YPEL3 | 352.2475 | 0.285668 | 0.230088 | 1.241557 | 0.2144 | 0.373397 |
| ICAM1 | 669.3405 | -0.56274 | 0.154959 | -3.63156 | 2.82E-04 | 0.004744 |
| TFAP4 | 320.2602 | -0.4771 | 0.179458 | -2.65855 | 0.007848 | 0.040153 |
| RAB11FIP3 | 1699.219 | -0.26116 | 0.167547 | -1.55871 | 0.119066 | 0.250759 |
| GNPTG | 2615.359 | -0.01151 | 0.124512 | -0.09244 | 0.926347 | 0.958156 |
| CERS4 | 1658.532 | -0.55238 | 0.235947 | -2.34112 | 0.019226 | 0.073619 |
| MCOLN1 | 2330.784 | -0.28095 | 0.169157 | -1.6609 | 0.096733 | 0.21817 |
| PDPR | 1149.541 | -0.52391 | 0.189264 | -2.76814 | 0.005638 | 0.032121 |
| AARS | 5343.845 | -0.42619 | 0.132736 | -3.21083 | 0.001324 | 0.012443 |
| GLG1 | 3393.794 | -0.46033 | 0.141137 | -3.26161 | 0.001108 | 0.01108 |
| TNRC6A | 1144.861 | 0.221867 | 0.195338 | 1.13581 | 0.256036 | 0.420246 |
| FCGBP | 0.668852 | 1.246964 | 2.610519 | 0.477669 | 0.632886 | NA |
| PLEKHG2 | 429.8733 | -0.33028 | 0.167045 | -1.97718 | 0.048021 | 0.135192 |
| DLL3 | 534.9751 | -1.30318 | 0.642658 | -2.0278 | 0.042581 | 0.125539 |
| WDR7 | 887.9932 | -0.02646 | 0.167365 | -0.15808 | 0.874392 | 0.925145 |
| TXNL1 | 1451.969 | 0.082861 | 0.139132 | 0.595556 | 0.551472 | 0.692479 |
| ABCC6 | 3.507438 | -0.09287 | 1.398622 | -0.0664 | 0.947059 | NA |
| MYO15A | 4.869449 | 0.482956 | 0.948644 | 0.509102 | 0.610681 | NA |
| ALKBH5 | 1743.014 | -0.49721 | 0.16516 | -3.01048 | 0.002608 | 0.019343 |
| NLRP1 | 493.9614 | 0.30983 | 0.358473 | 0.864305 | 0.38742 | 0.553358 |
| PITPNM3 | 7.88232 | 0.095783 | 0.736425 | 0.130065 | 0.896515 | NA |
| SPAG7 | 1655.231 | 0.047521 | 0.138472 | 0.343184 | 0.73146 | 0.829805 |
| ORC6 | 446.4876 | 0.572578 | 0.221781 | 2.581721 | 0.009831 | 0.046688 |
| TMEM101 | 1315.38 | -0.23151 | 0.140298 | -1.65012 | 0.098918 | 0.221761 |
| RFFL | 128.854 | -0.17605 | 0.216187 | -0.81434 | 0.415448 | 0.578747 |
| UNC13D | 20.54652 | -0.70641 | 0.480808 | -1.46922 | 0.141773 | NA |
| MFSD11 | 604.7169 | 0.193789 | 0.213602 | 0.907245 | 0.364277 | 0.531639 |
| CDC6 | 555.4706 | 0.408802 | 0.258718 | 1.580108 | 0.114082 | 0.243201 |
| DHPS | 1640.175 | -0.15517 | 0.159286 | -0.97417 | 0.329974 | 0.498197 |
| HOOK2 | 302.0364 | -0.10637 | 0.169774 | -0.62655 | 0.530957 | 0.675555 |
| NUBP2 | 1729.03 | -0.15718 | 0.155238 | -1.0125 | 0.3113 | 0.478807 |
| C19orf77 | 0.938495 | 0.819106 | 2.155823 | 0.379951 | 0.703982 | NA |
| IL12RB1 | 32.65073 | -0.02356 | 0.419904 | -0.05611 | 0.955256 | NA |
| TMED1 | 537.3295 | -0.27609 | 0.19309 | -1.42988 | 0.152753 | 0.297301 |
| MAST3 | 484.7764 | -0.68305 | 0.179021 | -3.81546 | 1.36E-04 | 0.003063 |

| | | | | | | |
|---|---|---|---|---|---|---|
| OCEL1 | 736.6253 | -0.09367 | 0.181427 | -0.5163 | 0.605647 | 0.734104 |
| MYO9B | 12261.14 | -0.36118 | 0.168758 | -2.14022 | 0.032337 | 0.104486 |
| KCNK6 | 94.86872 | 0.534028 | 0.26949 | 1.981624 | 0.047521 | 0.134231 |
| CATSPERG | 4.17502 | -0.14877 | 0.984148 | -0.15116 | 0.879847 | NA |
| PSMD8 | 8977.649 | 0.081437 | 0.167322 | 0.48671 | 0.626464 | 0.749266 |
| FBXL19 | 1197.936 | -0.32429 | 0.161477 | -2.00825 | 0.044616 | 0.129054 |
| STX1B | 23.714 | -0.38019 | 0.422619 | -0.89962 | 0.368325 | NA |
| HSD3B7 | 544.2087 | -0.22012 | 0.204922 | -1.07417 | 0.282748 | 0.449137 |
| SETD1A | 1098.941 | -0.45024 | 0.189505 | -2.37589 | 0.017507 | 0.069134 |
| BCL7C | 734.3155 | -0.29149 | 0.15693 | -1.85742 | 0.063252 | 0.163152 |
| EFNA2 | 5.476309 | 0.47379 | 1.061186 | 0.446472 | 0.655256 | NA |
| CIRBP | 3404.09 | -0.40715 | 0.160448 | -2.53757 | 0.011162 | 0.050662 |
| ATP5D | 2973.63 | -0.17438 | 0.149697 | -1.16489 | 0.244063 | 0.406771 |
| C19orf26 | 262.0672 | -0.01738 | 0.231591 | -0.07503 | 0.940188 | 0.966605 |
| IGFALS | 1.958606 | 1.668876 | 1.598728 | 1.043877 | 0.296542 | NA |
| HNRNPM | 5092.607 | -0.29178 | 0.149143 | -1.9564 | 0.050419 | 0.139392 |
| 2-Mar | 1540.877 | -0.14362 | 0.15958 | -0.9 | 0.368119 | 0.535269 |
| NDUFB7 | 3392.163 | 0.052544 | 0.161733 | 0.324879 | 0.745273 | 0.83953 |
| TECR | 4026.946 | -0.17845 | 0.18583 | -0.96028 | 0.336916 | 0.505844 |
| TIMM13 | 2905.105 | -0.13785 | 0.154094 | -0.8946 | 0.371003 | 0.53772 |
| CDC34 | 2105.232 | -0.02884 | 0.149124 | -0.19338 | 0.846665 | 0.908806 |
| MISP | 0.296509 | -0.51709 | 3.982155 | -0.12985 | 0.896684 | NA |
| POLR2E | 6595.136 | -0.24068 | 0.13227 | -1.81961 | 0.068818 | 0.172582 |
| POLRMT | 1444.851 | -0.47694 | 0.136067 | -3.50519 | 4.56E-04 | 0.00643 |
| HCN2 | 118.3863 | -0.00676 | 0.268416 | -0.02517 | 0.97992 | 0.98935 |
| IZUMO4 | 6.417751 | -0.18881 | 1.026509 | -0.18393 | 0.854067 | NA |
| GADD45B | 450.1923 | 0.470651 | 0.343939 | 1.368417 | 0.171182 | 0.322069 |
| PALM | 1298.253 | -0.30806 | 0.14259 | -2.16048 | 0.030736 | 0.100766 |
| MADCAM1 | 19.13758 | 0.09434 | 0.476145 | 0.198132 | 0.842941 | NA |
| MKNK2 | 2104.828 | -0.55398 | 0.167008 | -3.3171 | 9.10E-04 | 0.009767 |
| TELO2 | 1322.259 | -0.42525 | 0.146361 | -2.90549 | 0.003667 | 0.024136 |
| CELF4 | 0.629726 | -0.22725 | 2.732935 | -0.08315 | 0.93373 | NA |
| ZNF516 | 847.0703 | -0.32861 | 0.202564 | -1.62223 | 0.104754 | 0.230145 |
| CDH20 | 4.440836 | 1.670105 | 1.163784 | 1.435064 | 0.151269 | NA |
| ADNP2 | 526.437 | -0.28535 | 0.180736 | -1.57882 | 0.114377 | 0.243611 |
| RBFA | 402.4224 | -0.03724 | 0.179005 | -0.20806 | 0.835182 | 0.901409 |
| USP14 | 2295.147 | 0.143563 | 0.13802 | 1.040159 | 0.298266 | 0.465256 |
| VAPA | 2361.365 | 0.164485 | 0.133116 | 1.235656 | 0.216586 | 0.375824 |
| METTL4 | 232.3618 | 0.51496 | 0.212657 | 2.421553 | 0.015454 | 0.06322 |
| LPIN2 | 501.2739 | 0.109212 | 0.185343 | 0.589242 | 0.555699 | 0.695357 |
| SMCHD1 | 748.3107 | 0.562762 | 0.343006 | 1.640676 | 0.100865 | 0.22419 |
| MYOM1 | 8.491556 | 0.231801 | 0.765857 | 0.302669 | 0.762142 | NA |
| MYL12A | 8945.158 | 0.334832 | 0.210259 | 1.592472 | 0.111279 | 0.239458 |
| CEP76 | 149.2275 | 0.386401 | 0.234894 | 1.645001 | 0.09997 | 0.223198 |
| ST8SIA5 | 20.69308 | 1.008262 | 0.632652 | 1.593706 | 0.111002 | NA |
| CEP192 | 465.9637 | 0.43662 | 0.237994 | 1.834583 | 0.066567 | 0.168551 |
| RNMT | 891.0538 | 0.586106 | 0.19477 | 3.009216 | 0.002619 | 0.019392 |

| | | | | | | |
|---|---|---|---|---|---|---|
| SMAD7 | 192.1765 | 0.237896 | 0.206976 | 1.149386 | 0.250397 | 0.41401 |
| LIPG | 83.09492 | -0.96367 | 0.380305 | -2.53394 | 0.011279 | 0.051071 |
| LAMA1 | 2763.708 | -0.2976 | 0.147623 | -2.01592 | 0.043809 | 0.127605 |
| RNF125 | 288.6433 | 0.276788 | 0.218848 | 1.264753 | 0.20596 | 0.363018 |
| ANKRD12 | 302.171 | 0.933785 | 1.138047 | 0.820515 | 0.411922 | 0.575044 |
| POLI | 288.4957 | 0.681579 | 0.288694 | 2.360903 | 0.018231 | 0.070953 |
| MIB1 | 1331.444 | 0.464658 | 0.26591 | 1.747429 | 0.080563 | 0.192203 |
| RBBP8 | 364.8359 | 1.134861 | 0.252437 | 4.495619 | 6.94E-06 | 5.14E-04 |
| RIOK3 | 906.0433 | 0.805215 | 0.170234 | 4.730048 | 2.24E-06 | 2.66E-04 |
| ACP5 | 6036.816 | -0.62975 | 0.163408 | -3.85384 | 1.16E-04 | 0.002815 |
| MSLN | 1.724412 | -1.02404 | 1.531319 | -0.66873 | 0.503667 | NA |
| MGRN1 | 2651.753 | -0.73768 | 0.148771 | -4.95851 | 7.10E-07 | 1.26E-04 |
| ZNF629 | 782.8514 | -0.48327 | 0.167419 | -2.88659 | 0.003894 | 0.025037 |
| TRADD | 1151.435 | -0.21207 | 0.153568 | -1.38092 | 0.167302 | 0.316783 |
| HSF4 | 445.0446 | 0.244244 | 0.2319 | 1.053233 | 0.292234 | 0.458987 |
| CORO1A | 18.07488 | -0.40891 | 0.535979 | -0.76292 | 0.445514 | NA |
| MAPK3 | 2333.18 | -0.33732 | 0.166025 | -2.03177 | 0.042177 | 0.124939 |
| GDPD3 | 5.138027 | -0.08834 | 0.889347 | -0.09933 | 0.920874 | NA |
| ELMO3 | 32.547 | 0.062398 | 0.418341 | 0.149155 | 0.881431 | NA |
| PHKB | 837.9457 | 0.19741 | 0.220069 | 0.897039 | 0.369698 | 0.536613 |
| LYRM1 | 308.583 | 0.32629 | 0.177763 | 1.835531 | 0.066427 | 0.168375 |
| NUTF2 | 1311.434 | -0.26968 | 0.195019 | -1.38283 | 0.166716 | 0.316127 |
| NUP93 | 1995.15 | -0.41603 | 0.130358 | -3.19142 | 0.001416 | 0.013072 |
| CENPT | 623.3415 | 0.031248 | 0.155843 | 0.200508 | 0.841083 | 0.905336 |
| TSNAXIP1 | 46.02354 | 0.575503 | 0.333324 | 1.726555 | 0.084248 | NA |
| NFAT5 | 696.0417 | 0.3485 | 0.190566 | 1.828768 | 0.067434 | 0.169948 |
| LONP2 | 3118.571 | -0.15856 | 0.131414 | -1.20654 | 0.22761 | 0.389254 |
| N4BP1 | 1245.961 | -0.19968 | 0.166393 | -1.20006 | 0.230116 | 0.392295 |
| CBLN1 | 0.134133 | -1.48909 | 4.077584 | -0.36519 | 0.71497 | NA |
| ARL2BP | 1333.352 | -0.06966 | 0.143095 | -0.48679 | 0.62641 | 0.749266 |
| PLLP | 28.4561 | 0.354211 | 0.389587 | 0.909196 | 0.363246 | NA |
| ZNF423 | 2.311106 | -0.28038 | 1.329609 | -0.21087 | 0.832985 | NA |
| DHODH | 209.9448 | -0.23769 | 0.192656 | -1.23377 | 0.217287 | 0.37671 |
| CCL17 | 2.307108 | -0.86877 | 1.445999 | -0.60081 | 0.547966 | NA |
| CTCF | 1384.79 | -0.11058 | 0.134018 | -0.8251 | 0.409313 | 0.572795 |
| ACD | 584.9111 | -0.31247 | 0.170645 | -1.8311 | 0.067086 | 0.169467 |
| POLR2C | 1901.445 | -0.05695 | 0.149598 | -0.38071 | 0.70342 | 0.809538 |
| PARD6A | 45.58421 | 0.708188 | 0.39201 | 1.806557 | 0.070831 | NA |
| ZNF821 | 202.9321 | -0.31949 | 0.195441 | -1.63474 | 0.102104 | 0.226082 |
| MMP15 | 905.4114 | -0.52239 | 0.153296 | -3.40768 | 6.55E-04 | 0.007966 |
| USB1 | 1403.933 | -0.06936 | 0.159248 | -0.43557 | 0.663147 | 0.778531 |
| CYB5B | 3094.961 | -0.049 | 0.147656 | -0.33183 | 0.740019 | 0.835857 |
| CCDC113 | 16.16604 | 0.30412 | 0.612156 | 0.496801 | 0.61933 | NA |
| NME3 | 762.3376 | 0.030855 | 0.183709 | 0.167954 | 0.866619 | 0.920772 |
| NDRG4 | 16.80936 | 0.107449 | 0.525977 | 0.204285 | 0.838131 | NA |
| PSMD7 | 1324.385 | 0.371188 | 0.128784 | 2.882239 | 0.003949 | 0.025262 |
| SETD6 | 202.7712 | 0.153506 | 0.188279 | 0.815312 | 0.414894 | 0.578178 |

| | | | | | | |
|---|---|---|---|---|---|---|
| SLC38A7 | 1187.523 | -0.40027 | 0.165041 | -2.42525 | 0.015298 | 0.062857 |
| VAC14 | 2240.531 | -0.48306 | 0.143969 | -3.35527 | 7.93E-04 | 0.008995 |
| HAS3 | 25.50552 | -0.43199 | 0.441962 | -0.97743 | 0.328358 | NA |
| TANGO6 | 1072.553 | -0.22874 | 0.175666 | -1.30215 | 0.192867 | 0.347688 |
| COG4 | 2179.381 | -0.29388 | 0.130908 | -2.24494 | 0.024772 | 0.08722 |
| SMPD3 | 8.395275 | 0.460372 | 0.77451 | 0.594404 | 0.552242 | NA |
| SLC7A6OS | 303.7404 | 0.02561 | 0.1672 | 0.153169 | 0.878265 | 0.926986 |
| SLC7A6 | 926.8276 | -0.18837 | 0.200935 | -0.93748 | 0.348512 | 0.516513 |
| PLA2G15 | 2498.464 | -0.28295 | 0.152166 | -1.85946 | 0.062962 | 0.162583 |
| ESRP2 | 2.422097 | -0.98393 | 1.262327 | -0.77946 | 0.435711 | NA |
| FA2H | 0.096202 | 0.408546 | 4.079639 | 0.100143 | 0.920231 | NA |
| WDR59 | 919.9561 | -0.12837 | 0.13905 | -0.92316 | 0.355923 | 0.523878 |
| MON1B | 1854.507 | -0.23157 | 0.142471 | -1.62541 | 0.104076 | 0.229164 |
| CMC2 | 901.2349 | 0.135952 | 0.149912 | 0.90688 | 0.36447 | 0.531856 |
| AXIN1 | 1258.081 | -0.16232 | 0.179928 | -0.90216 | 0.366974 | 0.534224 |
| HCFC1R1 | 2370.634 | 0.045887 | 0.203458 | 0.225536 | 0.821563 | 0.892561 |
| NPRL3 | 1589.023 | -0.48763 | 0.163121 | -2.9894 | 0.002795 | 0.020255 |
| MLYCD | 589.8599 | -0.03234 | 0.171174 | -0.18896 | 0.850125 | 0.910747 |
| MPG | 1551.149 | -0.04436 | 0.197177 | -0.22496 | 0.822012 | 0.892885 |
| NECAB2 | 12.52999 | -0.62601 | 0.638083 | -0.98108 | 0.326552 | NA |
| HSDL1 | 1022.935 | 0.136952 | 0.1502 | 0.911798 | 0.361875 | 0.52898 |
| TAF1C | 703.4771 | -0.05699 | 0.172199 | -0.33094 | 0.74069 | 0.836509 |
| NAGPA | 765.3604 | -0.36337 | 0.159434 | -2.27912 | 0.02266 | 0.082252 |
| WFDC1 | 496.3632 | -0.08672 | 0.223142 | -0.38862 | 0.697559 | 0.805171 |
| SEC14L5 | 22.32269 | -2.03201 | 0.775529 | -2.62016 | 0.008789 | NA |
| COTL1 | 5413.771 | -0.49376 | 0.201538 | -2.44997 | 0.014287 | 0.060039 |
| USP10 | 1446.485 | -0.3018 | 0.127016 | -2.37608 | 0.017498 | 0.069122 |
| CRISPLD2 | 12.37998 | 0.216115 | 0.690652 | 0.312915 | 0.754345 | NA |
| TSC2 | 2420.701 | -0.53091 | 0.133991 | -3.96226 | 7.42E-05 | 0.002116 |
| ZNF500 | 357.1766 | -0.34651 | 0.186093 | -1.86201 | 0.062602 | 0.161941 |
| NME4 | 5735.867 | -0.18039 | 0.126369 | -1.42747 | 0.153444 | 0.298209 |
| ABCC1 | 967.3039 | -0.62003 | 0.177988 | -3.48353 | 4.95E-04 | 0.006751 |
| NOMO3 | 186.2936 | -0.36515 | 0.238752 | -1.52939 | 0.126167 | 0.261259 |
| LMF1 | 757.7953 | -0.4967 | 0.135406 | -3.66822 | 2.44E-04 | 0.004387 |
| FOXF1 | 0.815245 | 1.670886 | 2.214146 | 0.754641 | 0.450464 | NA |
| NARFL | 828.0422 | -0.17582 | 0.149061 | -1.17952 | 0.23819 | 0.401059 |
| MTHFSD | 282.6403 | -0.28619 | 0.171812 | -1.66574 | 0.095766 | 0.21662 |
| CLCN7 | 12825.84 | -0.53933 | 0.161513 | -3.33922 | 8.40E-04 | 0.00931 |
| HAGHL | 1225.057 | -0.18952 | 0.157055 | -1.20669 | 0.22755 | 0.389209 |
| FAM173A | 810.2287 | -0.02395 | 0.145082 | -0.16507 | 0.868891 | 0.922011 |
| SLC7A5 | 19180.39 | -0.80037 | 0.153819 | -5.20329 | 1.96E-07 | 5.16E-05 |
| METRN | 2601.844 | -0.21475 | 0.149722 | -1.4343 | 0.151487 | 0.295469 |
| FBXO31 | 1421.96 | -0.45668 | 0.142208 | -3.21137 | 0.001321 | 0.012431 |
| STUB1 | 2317.138 | -0.06644 | 0.136505 | -0.48671 | 0.626464 | 0.749266 |
| RHBDL1 | 27.03245 | -0.2898 | 0.420822 | -0.68865 | 0.491044 | NA |
| NUBP1 | 414.2923 | 0.066971 | 0.18495 | 0.362105 | 0.717273 | 0.819751 |
| UBE2I | 2675.499 | -0.25017 | 0.130492 | -1.91711 | 0.055224 | 0.148327 |

| | | | | | | |
|---|---|---|---|---|---|---|
| MEFV | 0.163957 | 0.424913 | 4.079006 | 0.104171 | 0.917034 | NA |
| CRYM | 20.40004 | -0.18259 | 0.544146 | -0.33556 | 0.737203 | NA |
| EEF2K | 724.8643 | -0.71267 | 0.152317 | -4.67888 | 2.88E-06 | 2.97E-04 |
| CAPN15 | 973.6524 | -0.3884 | 0.169046 | -2.29762 | 0.021584 | 0.079801 |
| PIEZO1 | 4195.57 | -0.39366 | 0.187153 | -2.1034 | 0.035431 | 0.110885 |
| GSPT1 | 3659.481 | 0.016482 | 0.124193 | 0.132711 | 0.894422 | 0.938268 |
| ZNF174 | 320.5175 | -0.28926 | 0.180803 | -1.59987 | 0.109627 | 0.237028 |
| CLUAP1 | 306.8252 | 0.465573 | 0.225268 | 2.066757 | 0.038757 | 0.117691 |
| UBFD1 | 2322.246 | -0.38073 | 0.132178 | -2.88042 | 0.003972 | 0.025354 |
| EARS2 | 931.1938 | -0.22888 | 0.158711 | -1.44211 | 0.14927 | 0.292786 |
| TCEB2 | 5818.775 | -0.05249 | 0.181049 | -0.28995 | 0.771857 | 0.858857 |
| GGA2 | 3132.787 | -0.29412 | 0.136483 | -2.15496 | 0.031165 | 0.101725 |
| CPPED1 | 2249.447 | -0.16352 | 0.157997 | -1.03495 | 0.300691 | 0.4675 |
| USP31 | 338.5518 | -0.1043 | 0.195904 | -0.53239 | 0.594453 | 0.724801 |
| HMOX2 | 2331.294 | 0.005477 | 0.127972 | 0.042802 | 0.96586 | 0.981339 |
| DNAJA3 | 1477.161 | -0.03285 | 0.126645 | -0.25938 | 0.795345 | 0.876284 |
| BFAR | 3039.133 | -0.141 | 0.132511 | -1.06409 | 0.287288 | 0.453569 |
| SALL1 | 1.180005 | -1.57876 | 2.139972 | -0.73775 | 0.460668 | NA |
| TOX3 | 69.99333 | -0.47358 | 0.329479 | -1.43737 | 0.150612 | 0.294428 |
| RRN3P2 | 5.762652 | -0.05091 | 0.861995 | -0.05906 | 0.952903 | NA |
| RBL2 | 1353.538 | 0.389025 | 0.158762 | 2.450367 | 0.014271 | 0.060039 |
| QPRT | 10470.01 | -0.28427 | 0.152113 | -1.86882 | 0.061648 | 0.16029 |
| XYLT1 | 1246.726 | 0.227293 | 0.197681 | 1.149798 | 0.250227 | 0.413909 |
| PYCARD | 1073.424 | -0.5624 | 0.261169 | -2.1534 | 0.031287 | 0.102008 |
| RPGRIP1L | 174.2649 | 0.6484 | 0.300651 | 2.156652 | 0.031033 | 0.101516 |
| MAZ | 1127.925 | -0.45872 | 0.208258 | -2.20267 | 0.027618 | 0.093949 |
| STX4 | 1171.004 | -0.01485 | 0.144143 | -0.10304 | 0.917935 | 0.952399 |
| CDIPT | 4210.498 | -0.134 | 0.159359 | -0.84087 | 0.400421 | 0.565252 |
| BCKDK | 1828.102 | -0.29478 | 0.141681 | -2.08059 | 0.037472 | 0.115025 |
| KAT8 | 643.4628 | -0.23385 | 0.143077 | -1.63443 | 0.102168 | 0.226182 |
| NOMO1 | 1313.442 | -0.40635 | 0.202244 | -2.00922 | 0.044514 | 0.128924 |
| SYT17 | 0.853323 | -0.46916 | 2.317774 | -0.20242 | 0.839588 | NA |
| TMC5 | 2.801193 | 0.592885 | 1.253673 | 0.472918 | 0.636271 | NA |
| CCP110 | 275.752 | 0.856578 | 0.342078 | 2.50404 | 0.012278 | 0.054213 |
| C16orf62 | 2270.113 | -0.46787 | 0.139186 | -3.36144 | 7.75E-04 | 0.00889 |
| RNF40 | 3352.551 | -0.4057 | 0.142631 | -2.84437 | 0.00445 | 0.027291 |
| KNOP1 | 225.7628 | -0.03541 | 0.183164 | -0.1933 | 0.84672 | 0.908806 |
| TTC23 | 643.0176 | 0.167808 | 0.182117 | 0.921428 | 0.356827 | 0.524381 |
| KLHDC4 | 323.7416 | -0.30495 | 0.186316 | -1.63675 | 0.101682 | 0.225359 |
| MAN2B1 | 3775.785 | -0.44904 | 0.136411 | -3.29179 | 9.96E-04 | 0.010321 |
| KCNN4 | 27.5129 | 0.345294 | 0.691353 | 0.499448 | 0.617464 | NA |
| MAP4K1 | 9.154904 | -0.7914 | 0.69599 | -1.13709 | 0.255501 | NA |
| ECH1 | 1682.126 | -3.27E-04 | 0.140308 | -0.00233 | 0.998138 | 0.999318 |
| HNRNPL | 4923.5 | -0.46575 | 0.155994 | -2.98568 | 0.002829 | 0.020359 |
| NFKBIB | 1734.56 | 0.121854 | 0.216855 | 0.561917 | 0.574173 | 0.709634 |
| TUBB4A | 1929.507 | -0.83176 | 0.158634 | -5.24329 | 1.58E-07 | 4.45E-05 |
| SARS2 | 19.01493 | -0.26512 | 0.486809 | -0.5446 | 0.586028 | NA |

| | | | | | | |
|---|---|---|---|---|---|---|
| ARHGEF18 | 881.2572 | -0.46774 | 0.155466 | -3.00864 | 0.002624 | 0.019406 |
| PEX11G | 74.95599 | -0.35312 | 0.272375 | -1.29646 | 0.194817 | 0.349933 |
| DOT1L | 578.6836 | -0.63298 | 0.204997 | -3.08775 | 0.002017 | 0.016256 |
| PLEKHJ1 | 1026.876 | -0.26374 | 0.151349 | -1.74257 | 0.081409 | 0.193523 |
| RNASEH2A | 974.0827 | -0.1981 | 0.15578 | -1.27168 | 0.203486 | 0.359941 |
| SF3A2 | 1100.151 | -0.49035 | 0.167442 | -2.92847 | 0.003406 | 0.022866 |
| AMH | 4.809433 | -0.45179 | 0.957788 | -0.4717 | 0.637141 | NA |
| LYL1 | 110.3173 | -0.12685 | 0.25131 | -0.50474 | 0.613738 | 0.740094 |
| OAZ1 | 14803.74 | -0.00881 | 0.136172 | -0.06469 | 0.948422 | 0.971147 |
| TRMT1 | 912.199 | -0.17894 | 0.142579 | -1.25503 | 0.209468 | 0.367072 |
| STX10 | 1549.586 | 0.034655 | 0.160166 | 0.21637 | 0.828699 | 0.896738 |
| TLE6 | 39.51403 | -0.3937 | 0.353879 | -1.11254 | 0.265906 | NA |
| CCDC130 | 681.5034 | 3.11E-04 | 0.152778 | 0.002033 | 0.998378 | 0.999371 |
| AES | 8713.714 | -0.08058 | 0.151421 | -0.53214 | 0.594628 | 0.724939 |
| SGTA | 3057.688 | -0.23469 | 0.127707 | -1.83769 | 0.066108 | 0.167818 |
| SNAPC2 | 1021.692 | -0.12347 | 0.138017 | -0.89464 | 0.370982 | 0.53772 |
| C19orf53 | 3584.488 | 0.081122 | 0.178828 | 0.45363 | 0.650095 | 0.768075 |
| TIMM44 | 1010.056 | 0.009947 | 0.144154 | 0.069 | 0.94499 | 0.969441 |
| IL27RA | 88.8479 | 0.164141 | 0.287075 | 0.571771 | 0.567477 | 0.704296 |
| ASF1B | 533.1976 | -0.32231 | 0.23815 | -1.35339 | 0.175931 | 0.328088 |
| FAM32A | 3307.633 | 0.171977 | 0.132531 | 1.297629 | 0.194415 | 0.349475 |
| C19orf44 | 150.4041 | -0.10065 | 0.21059 | -0.47795 | 0.632685 | 0.754704 |
| MED26 | 174.238 | -0.41138 | 0.214312 | -1.91955 | 0.054915 | 0.147724 |
| OLFM2 | 99.30955 | -0.07375 | 0.358296 | -0.20583 | 0.836925 | 0.902391 |
| RASAL3 | 3.174787 | 1.674832 | 1.208817 | 1.385513 | 0.165896 | NA |
| AKAP8 | 741.7989 | 0.10755 | 0.166925 | 0.644302 | 0.51938 | 0.666441 |
| EPHX3 | 3.435195 | -0.69272 | 1.112681 | -0.62257 | 0.533566 | NA |
| ILVBL | 5781.975 | -0.52164 | 0.162583 | -3.20843 | 0.001335 | 0.012508 |
| SYDE1 | 4389.122 | -0.29251 | 0.141167 | -2.07208 | 0.038258 | 0.116684 |
| POP4 | 1011.428 | 0.154228 | 0.14696 | 1.049458 | 0.293967 | 0.461117 |
| CCNE1 | 343.5553 | 0.102045 | 0.19972 | 0.510941 | 0.609393 | 0.736502 |
| URI1 | 1478.217 | 0.479501 | 0.186322 | 2.573504 | 0.010067 | 0.047417 |
| PDCD5 | 816.048 | 0.614703 | 0.169742 | 3.621397 | 2.93E-04 | 0.004852 |
| ANKRD27 | 783.3068 | 0.125106 | 0.138132 | 0.9057 | 0.365094 | 0.532373 |
| RPS16 | 25739.14 | -0.05529 | 0.158657 | -0.34847 | 0.72749 | 0.827121 |
| TIMM50 | 3182.474 | -0.31967 | 0.123666 | -2.58497 | 0.009739 | 0.046367 |
| FBL | 2709.581 | -0.31433 | 0.167074 | -1.88136 | 0.059922 | 0.157094 |
| DYRK1B | 1339.222 | -0.42698 | 0.195774 | -2.18099 | 0.029184 | 0.097105 |
| CNTD2 | 0.87074 | 0.675988 | 2.012127 | 0.335957 | 0.736903 | NA |
| GPI | 9383.406 | -0.39672 | 0.125304 | -3.16605 | 0.001545 | 0.013793 |
| AKT2 | 4211.479 | -0.46513 | 0.143942 | -3.23135 | 0.001232 | 0.011864 |
| PLD3 | 9500.401 | -0.37299 | 0.113168 | -3.2959 | 9.81E-04 | 0.010195 |
| PRX | 149.1331 | -0.50098 | 0.239953 | -2.08782 | 0.036814 | 0.113713 |
| PIAS4 | 579.0217 | -0.3946 | 0.169334 | -2.33032 | 0.019789 | 0.075121 |
| NUMBL | 485.0037 | -0.18011 | 0.210892 | -0.85404 | 0.393084 | 0.558152 |
| EBI3 | 1.009228 | 2.053856 | 2.077371 | 0.98868 | 0.322819 | NA |
| CCDC94 | 625.3503 | -0.22486 | 0.175307 | -1.28265 | 0.199615 | 0.355802 |

| | | | | | | |
|---|---|---|---|---|---|---|
| SHD | 0.327913 | 1.543154 | 4.021063 | 0.383768 | 0.701151 | NA |
| TBCB | 2536.765 | -0.1964 | 0.129209 | -1.52002 | 0.128505 | 0.264779 |
| FSD1 | 63.40428 | 0.435061 | 0.322437 | 1.349287 | 0.177245 | 0.329394 |
| POLR2I | 221.6747 | 0.0307 | 0.199851 | 0.153612 | 0.877916 | 0.926876 |
| OVOL3 | 0.258578 | 1.233151 | 4.047809 | 0.304646 | 0.760635 | NA |
| CLIP3 | 1106.629 | -0.58162 | 0.167693 | -3.46837 | 5.24E-04 | 0.006967 |
| ZFR2 | 1.656722 | 0.713654 | 1.587909 | 0.44943 | 0.653122 | NA |
| TJP3 | 2.658018 | 0.523981 | 1.240973 | 0.422234 | 0.672854 | NA |
| APLP1 | 70.41939 | 0.363324 | 0.515524 | 0.704766 | 0.480956 | 0.635361 |
| CACTIN | 369.0683 | -0.20541 | 0.192426 | -1.06747 | 0.285758 | 0.452292 |
| HNRNPUL1 | 7026.251 | -0.40777 | 0.136461 | -2.98819 | 0.002806 | 0.020298 |
| FZR1 | 1578.588 | -0.45168 | 0.179935 | -2.51024 | 0.012065 | 0.053649 |
| TGFB1 | 2909.643 | 0.043641 | 0.294682 | 0.148094 | 0.882268 | 0.930128 |
| ATP5SL | 1652.466 | -0.20815 | 0.126471 | -1.64584 | 0.099796 | 0.223181 |
| PLIN3 | 3388.652 | -0.24637 | 0.147063 | -1.67525 | 0.093886 | 0.214014 |
| MRPL4 | 1237.774 | -0.12434 | 0.156422 | -0.79492 | 0.426658 | 0.588485 |
| CD79A | 4.13239 | 7.15E-04 | 1.170226 | 6.11E-04 | 0.999512 | NA |
| RPS19 | 26096.01 | 0.076786 | 0.153212 | 0.501173 | 0.616249 | 0.742141 |
| ICAM5 | 24.11163 | 0.616008 | 0.44055 | 1.39827 | 0.162032 | NA |
| BABAM1 | 656.4684 | -0.07516 | 0.144018 | -0.52189 | 0.601748 | 0.730914 |
| TYK2 | 2070.904 | -0.33508 | 0.138611 | -2.41738 | 0.015633 | 0.063752 |
| CDC37 | 7992.396 | -0.07971 | 0.138411 | -0.57591 | 0.564674 | 0.702288 |
| RABAC1 | 5916.723 | 0.051173 | 0.125609 | 0.407398 | 0.683716 | 0.794262 |
| ATP1A3 | 13.77016 | -0.11748 | 0.601986 | -0.19515 | 0.845273 | NA |
| PTPRS | 3431.341 | -0.65361 | 0.170518 | -3.83307 | 1.27E-04 | 0.002972 |
| CNFN | 35.67279 | 0.237304 | 0.407065 | 0.582963 | 0.559918 | NA |
| MEGF8 | 6294.716 | -0.66155 | 0.165767 | -3.99087 | 6.58E-05 | 0.001943 |
| RAB3D | 575.4899 | -0.38967 | 0.188285 | -2.06959 | 0.038491 | 0.117153 |
| TMEM205 | 698.4835 | -0.12057 | 0.151729 | -0.79461 | 0.426838 | 0.588596 |
| CAPS | 83.53 | 0.286943 | 0.2492 | 1.151456 | 0.249545 | 0.413409 |
| | 2171.285 | -0.19284 | 0.171011 | -1.12765 | 0.259467 | 0.423872 |
| MIER2 | 460.5619 | -0.25816 | 0.150387 | -1.71661 | 0.08605 | 0.200851 |
| TNPO2 | 3800.516 | -0.45915 | 0.144759 | -3.17184 | 0.001515 | 0.013631 |
| WDR83OS | 881.3872 | -0.08335 | 0.168163 | -0.49565 | 0.620143 | 0.744712 |
| GCDH | 875.8716 | -0.30928 | 0.133161 | -2.32261 | 0.0202 | 0.076241 |
| KLF1 | 0.163957 | 0.424913 | 4.079006 | 0.104171 | 0.917034 | NA |
| DNASE2 | 3102.623 | -0.16886 | 0.155753 | -1.08416 | 0.278294 | 0.444453 |
| MAST1 | 13.91642 | -0.5068 | 0.596965 | -0.84896 | 0.395904 | NA |
| JAK3 | 16.01564 | -0.6436 | 0.53323 | -1.20698 | 0.227438 | NA |
| RPL18A | 11504.7 | -0.06423 | 0.155521 | -0.41302 | 0.679591 | 0.790866 |
| SLC5A5 | 0.35636 | 1.709252 | 3.651833 | 0.468053 | 0.639746 | NA |
| KCNN1 | 12.75078 | 0.294865 | 0.693373 | 0.425261 | 0.670646 | NA |
| ARRDC2 | 852.747 | 0.187633 | 0.202664 | 0.925835 | 0.354532 | 0.522758 |
| PIK3R2 | 2.855716 | -0.54733 | 1.242175 | -0.44062 | 0.659486 | NA |
| RAB3A | 352.6529 | -0.47061 | 0.174448 | -2.6977 | 0.006982 | 0.037101 |
| PDE4C | 2.417995 | 1.510691 | 1.442345 | 1.047385 | 0.294922 | NA |
| ISYNA1 | 1056.081 | -0.48849 | 0.151623 | -3.22174 | 0.001274 | 0.01212 |

| | | | | | | |
|---|---|---|---|---|---|---|
| ELL | 412.7169 | -0.32101 | 0.170655 | -1.88107 | 0.059962 | 0.157164 |
| CRTC1 | 762.745 | -0.54172 | 0.18251 | -2.96815 | 0.002996 | 0.02114 |
| KMT2B | 1036.318 | -0.61988 | 0.198055 | -3.12985 | 0.001749 | 0.014831 |
| COMP | 0.315653 | -0.57305 | 3.892639 | -0.14721 | 0.882963 | NA |
| UPK1A | 7.274719 | -0.63835 | 0.830899 | -0.76827 | 0.442328 | NA |
| COPE | 3411.725 | -0.18667 | 0.132516 | -1.40865 | 0.158938 | 0.305729 |
| DDX49 | 1428.128 | -0.14318 | 0.146458 | -0.97761 | 0.328265 | 0.496438 |
| ETV2 | 72.6702 | -0.45587 | 0.275134 | -1.65689 | 0.097542 | 0.219422 |
| ATP4A | 0.303394 | -2.27006 | 4.032451 | -0.56295 | 0.57347 | NA |
| ARMC6 | 1092.209 | -0.27807 | 0.153357 | -1.81324 | 0.069794 | 0.174046 |
| TMEM147 | 3478.68 | -0.15757 | 0.151345 | -1.04111 | 0.297825 | 0.464828 |
| GAPDHS | 55.03542 | -0.44027 | 0.435773 | -1.01032 | 0.312344 | 0.479394 |
| TCEB1P28 | 0.682693 | 2.623213 | 2.518338 | 1.041645 | 0.297576 | NA |
| MAG | 0.508056 | 0.731032 | 2.861912 | 0.255435 | 0.798387 | NA |
| TMEM59L | 6.198377 | 0.575006 | 1.075545 | 0.534619 | 0.592914 | NA |
| HAMP | 0.096202 | 0.408546 | 4.079639 | 0.100143 | 0.920231 | NA |
| USF2 | 8254.556 | -0.20627 | 0.147825 | -1.39538 | 0.162901 | 0.311181 |
| LSR | 53.87344 | 0.090148 | 0.327132 | 0.275569 | 0.782879 | 0.867305 |
| KXD1 | 2013.921 | -0.09565 | 0.142458 | -0.67141 | 0.501957 | 0.65228 |
| FKBP8 | 10634.01 | -0.25663 | 0.13159 | -1.95025 | 0.051146 | 0.140741 |
| SUGP1 | 921.2549 | -0.24887 | 0.154598 | -1.60982 | 0.107436 | 0.233913 |
| HPN | 0.260158 | 1.240347 | 4.047124 | 0.306476 | 0.759242 | NA |
| ZNF14 | 66.6702 | 0.403297 | 0.300472 | 1.342212 | 0.179527 | 0.332131 |
| SCN1B | 103.7922 | -0.53234 | 0.274639 | -1.93832 | 0.052584 | 0.143198 |
| PBX4 | 42.92076 | 0.208188 | 0.325008 | 0.640565 | 0.521806 | NA |
| ERF | 2645.786 | -0.19018 | 0.15366 | -1.23765 | 0.215847 | 0.375035 |
| GSK3A | 1685.981 | -0.20954 | 0.159391 | -1.31462 | 0.188638 | 0.342879 |
| ATP13A1 | 2392.958 | -0.43227 | 0.151816 | -2.84732 | 0.004409 | 0.027138 |
| ZNF574 | 357.5302 | -0.51124 | 0.161829 | -3.15914 | 0.001582 | 0.014031 |
| GRIK5 | 1.313788 | 0.047668 | 1.966741 | 0.024237 | 0.980663 | NA |
| SIPA1L3 | 479.2218 | -0.58135 | 0.219248 | -2.65156 | 0.008012 | 0.040636 |
| ZNF85 | 42.56964 | 0.620602 | 0.50511 | 1.228647 | 0.219204 | NA |
| ETHE1 | 976.406 | -0.27821 | 0.160511 | -1.73328 | 0.083047 | 0.196119 |
| CADM4 | 4027.528 | 0.075032 | 0.196642 | 0.381566 | 0.702784 | 0.808962 |
| SMG9 | 1135.286 | -0.21339 | 0.167397 | -1.27478 | 0.202386 | 0.358744 |
| KRT23 | 0.92901 | 1.841773 | 2.123319 | 0.867403 | 0.385721 | NA |
| CRYBA1 | 0.192403 | 0.975392 | 4.060716 | 0.240202 | 0.810174 | NA |
| NUFIP2 | 1880.705 | 0.48598 | 0.184061 | 2.640329 | 0.008283 | 0.041548 |
| GIT1 | 4047.539 | -0.16791 | 0.176515 | -0.95123 | 0.341486 | 0.509671 |
| TADA2A | 280.905 | -0.18765 | 0.196548 | -0.95471 | 0.339726 | 0.508003 |
| AATF | 1601.802 | -0.09339 | 0.146745 | -0.63642 | 0.524502 | 0.670401 |
| DHRS11 | 153.678 | -0.16429 | 0.280014 | -0.58674 | 0.557379 | 0.696283 |
| ZNHIT3 | 693.7494 | 0.269936 | 0.177602 | 1.519898 | 0.128537 | 0.264779 |
| MLLT6 | 1843.732 | -0.34225 | 0.189587 | -1.80522 | 0.07104 | 0.176299 |
| PSMB3 | 2994.513 | -0.10189 | 0.150613 | -0.67653 | 0.498706 | 0.649348 |
| CWC25 | 388.7616 | 0.46881 | 0.151808 | 3.08818 | 0.002014 | 0.016256 |
| RPL19 | 27517.74 | 0.153107 | 0.157156 | 0.974235 | 0.32994 | 0.498197 |

| | | | | | | |
|---|---|---|---|---|---|---|
| FBXL20 | 211.0173 | -0.06673 | 0.205288 | -0.32506 | 0.745135 | 0.839454 |
| RUNDC3A | 42.04081 | 0.059216 | 0.403412 | 0.146788 | 0.883299 | NA |
| UBTF | 2571.438 | -0.17626 | 0.127607 | -1.3813 | 0.167188 | 0.316767 |
| PSMD3 | 4221.249 | -0.16088 | 0.138943 | -1.15789 | 0.246907 | 0.410366 |
| CASC3 | 2292.061 | -0.38101 | 0.145222 | -2.62367 | 0.008699 | 0.042987 |
| RAPGEFL1 | 37.34926 | 0.215521 | 0.387669 | 0.555942 | 0.578251 | NA |
| RGS9 | 24.26831 | 0.303507 | 0.54659 | 0.555274 | 0.578708 | NA |
| RNF43 | 69.05498 | -1.41364 | 0.389356 | -3.6307 | 2.83E-04 | 0.004753 |
| WNT3 | 22.96015 | 0.188456 | 0.532964 | 0.353601 | 0.723638 | NA |
| ASPA | 3288.741 | 0.154033 | 0.167746 | 0.918253 | 0.358486 | 0.525712 |
| RAD51C | 403.023 | 0.277829 | 0.171989 | 1.615386 | 0.106227 | 0.232133 |
| 4-Sep | 205.3091 | -0.9495 | 0.687123 | -1.38185 | 0.167018 | 0.316495 |
| MTMR4 | 1528.227 | -0.40536 | 0.128393 | -3.15716 | 0.001593 | 0.014094 |
| TRIM37 | 1017.324 | 0.416086 | 0.17487 | 2.379398 | 0.017341 | 0.068778 |
| DHX40 | 1261.706 | 0.411553 | 0.199494 | 2.062985 | 0.039114 | 0.118441 |
| TUBD1 | 86.53642 | 0.538995 | 0.306213 | 1.760196 | 0.078375 | 0.188591 |
| KPNB1 | 6881.057 | 0.077322 | 0.13889 | 0.556715 | 0.577722 | 0.712388 |
| GOSR2 | 399.3746 | -0.14368 | 0.166602 | -0.86239 | 0.388473 | 0.554176 |
| PNPO | 551.6463 | -0.47285 | 0.198719 | -2.37947 | 0.017338 | 0.068778 |
| RPS6KB1 | 377.9187 | 0.615923 | 0.176449 | 3.490646 | 4.82E-04 | 0.006662 |
| TRIM16L | 365.3067 | -0.48531 | 0.296716 | -1.63561 | 0.101922 | 0.225805 |
| ZNF29P | 0.460466 | 0.568655 | 3.014642 | 0.188631 | 0.850382 | NA |
| CDK5RAP3 | 912.6855 | -0.05911 | 0.133787 | -0.4418 | 0.658633 | 0.774844 |
| CBX1 | 1772.961 | 0.083555 | 0.125118 | 0.667808 | 0.504256 | 0.653944 |
| RECQL5 | 540.2168 | -0.27383 | 0.158477 | -1.7279 | 0.084007 | 0.197482 |
| PIGL | 77.36449 | 0.484017 | 0.309212 | 1.565328 | 0.117506 | 0.248269 |
| GALK1 | 1173.427 | -0.18715 | 0.179603 | -1.04205 | 0.297388 | 0.464559 |
| INTS2 | 265.1307 | 0.554748 | 0.233908 | 2.371653 | 0.017709 | 0.069636 |
| CAMTA2 | 1467.778 | -0.44593 | 0.171825 | -2.59523 | 0.009453 | 0.045467 |
| MED13 | 937.1159 | 0.313425 | 0.202273 | 1.549514 | 0.121258 | 0.254023 |
| HOXB6 | 183.7114 | 0.136414 | 0.233987 | 0.582998 | 0.559895 | 0.698469 |
| ENO3 | 77.94722 | 0.1936 | 0.277176 | 0.698473 | 0.484882 | 0.638373 |
| PFN1 | 15809.19 | -0.14015 | 0.118901 | -1.17875 | 0.238499 | 0.401449 |
| RNF167 | 2827.023 | -0.17978 | 0.158411 | -1.13488 | 0.256425 | 0.420517 |
| SLC25A11 | 3641.429 | -0.29071 | 0.132398 | -2.19576 | 0.028109 | 0.094913 |
| RASD1 | 2.953092 | 0.092653 | 1.150143 | 0.080558 | 0.935794 | NA |
| CHRNE | 2.382439 | -0.1301 | 1.252185 | -0.1039 | 0.91725 | NA |
| RAI1 | 3004.522 | -0.6625 | 0.193175 | -3.42953 | 6.05E-04 | 0.007625 |
| NUP88 | 1058.155 | 0.209587 | 0.148181 | 1.414399 | 0.157245 | 0.303457 |
| C1QBP | 4473.162 | -0.07154 | 0.128018 | -0.55881 | 0.57629 | 0.711361 |
| SLC6A4 | 1.44374 | -0.31701 | 1.683753 | -0.18828 | 0.85066 | NA |
| BLMH | 1098.419 | -0.14802 | 0.132765 | -1.11488 | 0.264903 | 0.429314 |
| CPD | 1925.673 | -0.03208 | 0.152357 | -0.21058 | 0.833218 | 0.899945 |
| GOSR1 | 801.7323 | -0.19576 | 0.144386 | -1.35579 | 0.175165 | 0.327064 |
| CCDC47 | 1687.938 | 0.642641 | 0.177124 | 3.628204 | 2.85E-04 | 0.004779 |
| MED31 | 254.4583 | 0.529343 | 0.216952 | 2.439906 | 0.014691 | 0.061041 |
| DRG2 | 999.9402 | -0.19884 | 0.151952 | -1.30855 | 0.190687 | 0.345024 |

| | | | | | | |
|---|---|---|---|---|---|---|
| FTSJ3 | 1688.084 | -0.13672 | 0.133934 | -1.02078 | 0.30736 | 0.474571 |
| AKAP10 | 629.45 | 0.372314 | 0.164247 | 2.266792 | 0.023403 | 0.084142 |
| ALDH3A1 | 4.402805 | 0.392202 | 1.002412 | 0.391258 | 0.695606 | NA |
| SMARCD2 | 3168.879 | -0.52335 | 0.15876 | -3.29652 | 9.79E-04 | 0.010189 |
| SYNGR2 | 1020.634 | 0.152801 | 0.22137 | 0.690253 | 0.490035 | 0.642357 |
| B9D1 | 252.6088 | -0.11438 | 0.228404 | -0.50079 | 0.61652 | 0.742224 |
| UTP6 | 915.3188 | 0.384429 | 0.143207 | 2.68442 | 0.007266 | 0.038129 |
| DDX5 | 9281.009 | 0.565264 | 0.139905 | 4.040346 | 5.34E-05 | 0.001715 |
| C17orf75 | 224.4408 | 0.267852 | 0.215156 | 1.244922 | 0.21316 | 0.37186 |
| CYTH1 | 917.6255 | -0.11347 | 0.140373 | -0.80832 | 0.418907 | 0.581856 |
| PSMD11 | 2186.119 | -0.15672 | 0.130816 | -1.19802 | 0.230909 | 0.393132 |
| LGALS3BP | 26816.55 | -0.8257 | 0.149628 | -5.51831 | 3.42E-08 | 1.69E-05 |
| ASIC2 | 0.392857 | -1.546 | 4.075469 | -0.37934 | 0.704433 | NA |
| CCL7 | 0.392857 | -1.546 | 4.075469 | -0.37934 | 0.704433 | NA |
| CCL2 | 1136.378 | -0.10245 | 0.214317 | -0.47803 | 0.632629 | 0.754704 |
| CCL8 | 0.883203 | 0.854847 | 2.135496 | 0.400304 | 0.688933 | NA |
| CCL1 | 0.134133 | -1.48909 | 4.077584 | -0.36519 | 0.71497 | NA |
| PEX12 | 162.7013 | -0.13819 | 0.224516 | -0.61549 | 0.538229 | 0.681093 |
| HNF1B | 0.430642 | -1.1891 | 3.485782 | -0.34113 | 0.733008 | NA |
| DHX58 | 452.714 | -1.26061 | 0.307163 | -4.10402 | 4.06E-05 | 0.001477 |
| KAT2A | 649.3007 | -0.10407 | 0.173248 | -0.60073 | 0.548023 | 0.689317 |
| RAB5C | 3186.26 | -0.21962 | 0.143583 | -1.52954 | 0.12613 | 0.261259 |
| NAGLU | 3686.948 | -0.38384 | 0.17206 | -2.23082 | 0.025693 | 0.089705 |
| HSD17B1P1 | 20.73462 | 0.517589 | 0.457219 | 1.132039 | 0.257618 | NA |
| HSD17B1 | 81.38566 | -0.67833 | 0.297408 | -2.2808 | 0.022561 | 0.082016 |
| MLX | 2005.752 | -0.08766 | 0.16044 | -0.54639 | 0.584798 | 0.717944 |
| CNTNAP1 | 783.6345 | -0.90547 | 0.185558 | -4.87971 | 1.06E-06 | 1.68E-04 |
| EZH1 | 610.7419 | -0.07228 | 0.146194 | -0.49444 | 0.620994 | 0.745432 |
| DLX4 | 0.716471 | -2.03238 | 2.582811 | -0.78689 | 0.431348 | NA |
| PPP1R9B | 3243.574 | -0.45097 | 0.168962 | -2.66903 | 0.007607 | 0.039383 |
| COL1A1 | 2.58453 | -0.5009 | 1.220575 | -0.41038 | 0.681526 | NA |
| SGCA | 9.818575 | 0.543852 | 0.704473 | 0.771999 | 0.440115 | NA |
| MRPL27 | 1112.977 | 0.221122 | 0.141119 | 1.566915 | 0.117135 | 0.247716 |
| VAT1 | 48045.93 | -0.51016 | 0.125513 | -4.06461 | 4.81E-05 | 0.001611 |
| LRRC59 | 4517.061 | -0.15715 | 0.186082 | -0.84453 | 0.398373 | 0.563233 |
| RND2 | 140.0146 | 0.250715 | 0.374159 | 0.670078 | 0.502808 | 0.65288 |
| ALOX12 | 31.29843 | -0.24438 | 0.398355 | -0.61347 | 0.539567 | NA |
| HDAC5 | 2896.227 | -0.00389 | 0.203088 | -0.01917 | 0.984707 | 0.991987 |
| ABCC3 | 0.845475 | 0.825047 | 2.791858 | 0.295519 | 0.767597 | NA |
| LUC7L3 | 863.7448 | 1.414958 | 0.736236 | 1.921881 | 0.054621 | 0.147199 |
| MPP2 | 99.75846 | -0.46837 | 0.239469 | -1.95586 | 0.050482 | 0.139472 |
| SMURF2 | 414.7978 | 0.456793 | 0.214452 | 2.130046 | 0.033168 | 0.106042 |
| DUSP3 | 6948.139 | -0.13987 | 0.130665 | -1.07044 | 0.284422 | 0.450947 |
| CACNG1 | 0.096202 | 0.408546 | 4.079639 | 0.100143 | 0.920231 | NA |
| EFTUD2 | 4398.447 | -0.26842 | 0.132175 | -2.03076 | 0.042279 | 0.125117 |
| HLF | 0.977767 | 1.39453 | 2.230419 | 0.625232 | 0.531819 | NA |
| SLC16A6 | 11778.88 | 0.281807 | 0.229204 | 1.229505 | 0.218883 | 0.378502 |

| | | | | | | |
|---|---|---|---|---|---|---|
| PRKAR1A | 7420.349 | 0.34476 | 0.170191 | 2.025722 | 0.042793 | 0.125933 |
| EFNB3 | 932.5969 | -0.31052 | 0.154669 | -2.00761 | 0.044685 | 0.129144 |
| FAM20A | 6.777794 | -1.12877 | 0.808393 | -1.39632 | 0.162619 | NA |
| YWHAE | 16685.39 | -0.08187 | 0.144659 | -0.56598 | 0.571409 | 0.707325 |
| | 1.498886 | 0.419584 | 1.712531 | 0.245008 | 0.80645 | NA |
| MMD | 692.3949 | 0.094439 | 0.184263 | 0.512526 | 0.608283 | 0.735461 |
| RANGRF | 104.162 | -0.01355 | 0.248649 | -0.05451 | 0.956533 | 0.976145 |
| DPH1 | 623.822 | -0.07701 | 0.175193 | -0.4396 | 0.66023 | 0.776182 |
| MAP2K6 | 193.8833 | 0.024286 | 0.230277 | 0.105464 | 0.916007 | 0.951316 |
| DHRS7B | 794.0317 | 0.0368 | 0.162347 | 0.226673 | 0.820678 | 0.892334 |
| WSB1 | 1689.311 | 0.536694 | 0.141949 | 3.780887 | 1.56E-04 | 0.00335 |
| RCVRN | 7.459299 | 0.226774 | 0.862872 | 0.262813 | 0.792695 | NA |
| MYH1 | 0.52699 | -2.57878 | 3.248478 | -0.79384 | 0.427288 | NA |
| SLC9A3R1 | 692.5983 | -0.10344 | 0.155283 | -0.66615 | 0.505314 | 0.65473 |
| MYH3 | 4.267151 | 0.369863 | 1.016902 | 0.363715 | 0.716071 | NA |
| NAT9 | 887.4746 | -0.05339 | 0.142868 | -0.37367 | 0.708652 | 0.813584 |
| TMEM104 | 1633.839 | -0.17951 | 0.231922 | -0.774 | 0.438933 | 0.599004 |
| TNFAIP1 | 2278.548 | -0.07251 | 0.161092 | -0.45013 | 0.652619 | 0.770362 |
| IFT20 | 333.754 | 0.65525 | 0.203928 | 3.213149 | 0.001313 | 0.012392 |
| TMEM97 | 1986.939 | -0.34907 | 0.157011 | -2.22321 | 0.026201 | 0.090873 |
| CDR2L | 994.1406 | -0.29987 | 0.220852 | -1.35779 | 0.17453 | 0.326554 |
| PMP22 | 8324.613 | -0.02608 | 0.208987 | -0.12481 | 0.90067 | 0.941822 |
| UNC119 | 933.7266 | -0.14655 | 0.153481 | -0.95484 | 0.339656 | 0.508003 |
| ALDOC | 623.608 | -0.13346 | 0.172509 | -0.77364 | 0.439142 | 0.599011 |
| SUPT6H | 5384.764 | -0.29631 | 0.133822 | -2.21418 | 0.026817 | 0.091994 |
| RAB34 | 3710.038 | -0.17064 | 0.163489 | -1.04377 | 0.296593 | 0.463665 |
| PHF12 | 823.3167 | -0.36383 | 0.159487 | -2.28125 | 0.022533 | 0.081968 |
| REEP6 | 249.607 | -0.12555 | 0.205243 | -0.61171 | 0.540727 | 0.682777 |
| PCSK4 | 76.66566 | -0.27503 | 0.291773 | -0.94263 | 0.34587 | 0.513692 |
| APC2 | 128.7802 | -0.35751 | 0.233869 | -1.52866 | 0.126348 | 0.261544 |
| RPS15 | 10959.77 | -0.06373 | 0.142033 | -0.44871 | 0.653641 | 0.77119 |
| NDUFS7 | 1273.614 | -0.28894 | 0.142381 | -2.02937 | 0.04242 | 0.125269 |
| KISS1R | 0.163957 | 0.424913 | 4.079006 | 0.104171 | 0.917034 | NA |
| ARID3A | 203.1781 | -0.00512 | 0.229707 | -0.0223 | 0.982208 | 0.990311 |
| GRIN3B | 1.252974 | -0.44216 | 1.86133 | -0.23755 | 0.812231 | NA |
| STK11 | 2274.165 | -0.30997 | 0.139675 | -2.2192 | 0.026473 | 0.091289 |
| TTR | 79.7382 | 1.439653 | 0.453037 | 3.177783 | 0.001484 | 0.013461 |
| B4GALT6 | 736.2017 | 0.987445 | 0.187085 | 5.278042 | 1.31E-07 | 4.07E-05 |
| PMFBP1 | 4.898962 | 0.219989 | 0.948762 | 0.23187 | 0.816639 | NA |
| ZNF430 | 78.49146 | 1.025481 | 0.346945 | 2.95575 | 0.003119 | 0.021713 |
| MYL12B | 10051.33 | 0.31368 | 0.18228 | 1.720869 | 0.085275 | 0.199473 |
| FAM86A | 259.5309 | -0.33798 | 0.170547 | -1.98177 | 0.047505 | 0.134217 |
| PPL | 3.455562 | -0.90989 | 1.073091 | -0.84791 | 0.396486 | NA |
| UBN1 | 2037.261 | -0.34104 | 0.154321 | -2.20994 | 0.02711 | 0.092647 |
| KDSR | 3476.624 | 0.283326 | 0.134149 | 2.112024 | 0.034684 | 0.109309 |
| VPS4B | 1478.915 | 0.563824 | 0.187577 | 3.00583 | 0.002649 | 0.019519 |
| ONECUT2 | 2.623599 | -0.25975 | 1.227421 | -0.21163 | 0.832399 | NA |

| | | | | | | |
|---|---|---|---|---|---|---|
| C19orf25 | 1221.322 | -0.12325 | 0.148436 | -0.83033 | 0.406351 | 0.570027 |
| GNA13 | 1875.911 | 0.101761 | 0.173302 | 0.58719 | 0.557076 | 0.696198 |
| HOXB8 | 68.07159 | -0.21481 | 0.308722 | -0.69579 | 0.486561 | 0.63963 |
| KANSL1 | 648.9187 | -0.22839 | 0.155076 | -1.47275 | 0.140818 | 0.28128 |
| HOXB5 | 61.50988 | -0.16106 | 0.28898 | -0.55734 | 0.577293 | 0.712154 |
| HOXB3 | 106.8773 | -0.01755 | 0.23287 | -0.07536 | 0.939929 | 0.966534 |
| ZFP30 | 193.5316 | 0.717821 | 0.249539 | 2.876591 | 0.00402 | 0.025471 |
| EPX | 1.020857 | 0.19325 | 2.085078 | 0.092682 | 0.926156 | NA |
| AKAP1 | 968.9712 | -0.01411 | 0.143238 | -0.09847 | 0.921556 | 0.95457 |
| COIL | 370.1109 | 0.367105 | 0.16615 | 2.209478 | 0.027141 | 0.092684 |
| TRIM25 | 3076.174 | -0.53652 | 0.144894 | -3.70288 | 2.13E-04 | 0.004043 |
| SCPEP1 | 3303.427 | 7.16E-04 | 0.125444 | 0.005706 | 0.995447 | 0.997908 |
| SPOP | 949.104 | 0.2636 | 0.171664 | 1.535554 | 0.124648 | 0.259382 |
| TBX2 | 4793.98 | -0.38543 | 0.137887 | -2.79526 | 0.005186 | 0.030299 |
| SLC35B1 | 1736.804 | 0.011182 | 0.150439 | 0.074331 | 0.940747 | 0.966779 |
| TBX4 | 1.724558 | -2.27467 | 1.823655 | -1.24731 | 0.212282 | NA |
| TEX14 | 2.404598 | 0.3092 | 1.330977 | 0.232311 | 0.816297 | NA |
| FAM117A | 283.2754 | -0.40499 | 0.187038 | -2.1653 | 0.030365 | 0.099938 |
| ABCC11 | 7.078242 | -0.8332 | 0.813598 | -1.02409 | 0.305792 | NA |
| PAPD5 | 339.4679 | 0.596867 | 0.21052 | 2.835204 | 0.00458 | 0.027771 |
| ADCY7 | 18.9641 | -0.45402 | 0.500421 | -0.90728 | 0.364259 | NA |
| CEP89 | 462.7777 | 0.060329 | 0.19761 | 0.305291 | 0.760144 | 0.850634 |
| TSHZ3 | 318.1169 | -0.2056 | 0.195898 | -1.04954 | 0.293932 | 0.461117 |
| HS3ST2 | 241.5603 | 0.386221 | 0.427226 | 0.904021 | 0.365984 | 0.533277 |
| RBBP6 | 643.0145 | 0.528316 | 0.348996 | 1.513816 | 0.130072 | 0.26643 |
| ZC3H7A | 1616.22 | 0.686265 | 0.173605 | 3.953029 | 7.72E-05 | 0.002172 |
| ZNF205 | 467.0619 | -0.18986 | 0.206552 | -0.91918 | 0.358001 | 0.525131 |
| NAA60 | 286.1693 | -0.46864 | 0.172033 | -2.72413 | 0.006447 | 0.035027 |
| PQLC1 | 3193.842 | -0.24618 | 0.138229 | -1.78099 | 0.074915 | 0.182739 |
| DDX39A | 3118.793 | -0.07188 | 0.250469 | -0.28699 | 0.774121 | 0.860327 |
| PKN1 | 7777.436 | -0.20358 | 0.171458 | -1.18736 | 0.235085 | 0.397849 |
| C19orf43 | 5258.953 | 0.012134 | 0.146837 | 0.082635 | 0.934142 | 0.963062 |
| CD97 | 1492.052 | -0.49428 | 0.168295 | -2.93701 | 0.003314 | 0.022501 |
| WDR83 | 174.4787 | -0.06401 | 0.205792 | -0.31103 | 0.755781 | 0.847421 |
| GIPC1 | 10462.62 | -0.24682 | 0.158713 | -1.55511 | 0.119919 | 0.252019 |
| KCNJ2 | 48.13584 | 0.240659 | 0.39809 | 0.604535 | 0.545488 | 0.687003 |
| B9D2 | 109.7711 | -0.08866 | 0.231045 | -0.38373 | 0.701175 | 0.807895 |
| ADCK4 | 1028.281 | -0.40113 | 0.153651 | -2.61064 | 0.009037 | 0.044218 |
| SLC12A4 | 2653.263 | -0.00115 | 0.203669 | -0.00565 | 0.995492 | 0.997908 |
| ENKD1 | 414.1854 | -0.21517 | 0.175953 | -1.22286 | 0.221382 | 0.381687 |
| PEPD | 8846.757 | -0.24662 | 0.130377 | -1.89157 | 0.058548 | 0.154515 |
| CHST8 | 1.080707 | 1.085144 | 1.993395 | 0.54437 | 0.586187 | NA |
| USP22 | 7523.758 | -0.4725 | 0.123837 | -3.81548 | 1.36E-04 | 0.003063 |
| ZNF576 | 371.1103 | 0.195914 | 0.16391 | 1.195254 | 0.231988 | 0.394248 |
| ZNF45 | 326.1218 | 0.2377 | 0.199673 | 1.190448 | 0.23387 | 0.396416 |
| LYPD3 | 78.80432 | -0.47199 | 0.266113 | -1.77365 | 0.07612 | 0.184844 |
| CNOT1 | 5044.187 | -0.20188 | 0.125619 | -1.60708 | 0.108037 | 0.234829 |

| | | | | | | |
|---|---|---|---|---|---|---|
| LRRC29 | 58.20426 | -0.55279 | 0.333087 | -1.65959 | 0.096998 | 0.218613 |
| BBS2 | 1201.313 | 0.216733 | 0.187997 | 1.152858 | 0.248969 | 0.412801 |
| MT1G | 2.79602 | -0.44088 | 1.200066 | -0.36738 | 0.713338 | NA |
| MT2A | 2072.711 | -0.023 | 0.148708 | -0.15465 | 0.877095 | 0.926461 |
| C16orf70 | 426.499 | -0.27037 | 0.171205 | -1.57924 | 0.114281 | 0.243493 |
| GOT2 | 4411.178 | -0.32504 | 0.154872 | -2.09875 | 0.035839 | 0.111632 |
| DOK4 | 724.9933 | -0.00597 | 0.183632 | -0.03252 | 0.974057 | 0.985529 |
| C17orf53 | 236.3933 | -0.36703 | 0.227558 | -1.6129 | 0.106765 | 0.233009 |
| SOX9 | 40.59763 | -1.07033 | 0.442104 | -2.42099 | 0.015478 | NA |
| TEKT3 | 3.590723 | -0.22496 | 1.078005 | -0.20868 | 0.834698 | NA |
| HS3ST3B1 | 1.03648 | 0.354657 | 2.004615 | 0.17692 | 0.859571 | NA |
| SLC25A35 | 170.0091 | -0.29778 | 0.211844 | -1.40565 | 0.159828 | 0.306992 |
| MRPS7 | 2000.694 | 0.054145 | 0.129586 | 0.417833 | 0.67607 | 0.788316 |
| GGA3 | 1247.365 | -0.56617 | 0.16846 | -3.36087 | 7.77E-04 | 0.008896 |
| ARMC7 | 511.5699 | -0.10771 | 0.181625 | -0.59305 | 0.553151 | 0.693776 |
| NUP85 | 918.8248 | -0.01235 | 0.166172 | -0.07431 | 0.940764 | 0.966779 |
| SLC25A19 | 199.3827 | -0.00151 | 0.255539 | -0.00591 | 0.995282 | 0.997908 |
| MIF4GD | 329.9275 | 0.004959 | 0.182965 | 0.027101 | 0.978379 | 0.988384 |
| NT5C | 699.1617 | -0.10911 | 0.146399 | -0.74532 | 0.456078 | 0.614797 |
| SLC25A23 | 2618.824 | -0.50009 | 0.160364 | -3.11845 | 0.001818 | 0.015146 |
| PSPN | 58.85503 | -0.40816 | 0.309937 | -1.31691 | 0.18787 | 0.341909 |
| GTF2F1 | 3235.947 | -0.30789 | 0.147693 | -2.08465 | 0.037101 | 0.114349 |
| ALKBH7 | 1319.542 | -0.25901 | 0.15097 | -1.71564 | 0.086229 | 0.20115 |
| CLPP | 2198.265 | -0.22358 | 0.130836 | -1.70886 | 0.087476 | 0.203249 |
| TNFSF9 | 100.7814 | -0.12919 | 0.274615 | -0.47043 | 0.638048 | 0.758814 |
| MED1 | 1336.326 | 0.043071 | 0.172757 | 0.249316 | 0.803116 | 0.882059 |
| RPL23 | 18791.77 | 0.512252 | 0.137687 | 3.7204 | 1.99E-04 | 0.003853 |
| CD70 | 38.81849 | -0.09035 | 0.58916 | -0.15335 | 0.878124 | NA |
| C3 | 12.42398 | -0.03917 | 0.979147 | -0.04 | 0.968092 | NA |
| SH2D3A | 6.619504 | 0.253643 | 0.773981 | 0.327713 | 0.743129 | NA |
| TRIP10 | 1050.946 | 0.027407 | 0.174375 | 0.157174 | 0.875108 | 0.925378 |
| GPR108 | 1572.539 | -0.16417 | 0.182579 | -0.89917 | 0.368562 | 0.535717 |
| S1PR4 | 0.430642 | -1.1891 | 3.485782 | -0.34113 | 0.733008 | NA |
| NCLN | 3902.015 | -0.3233 | 0.144801 | -2.23269 | 0.025569 | 0.089457 |
| LRFN3 | 785.0617 | -0.35854 | 0.163907 | -2.18745 | 0.02871 | 0.09628 |
| IGFLR1 | 2.502168 | -0.28004 | 1.24218 | -0.22544 | 0.821633 | NA |
| CAPNS1 | 12257.4 | -0.20292 | 0.151473 | -1.33961 | 0.180371 | 0.332913 |
| PDCD2L | 79.43409 | -0.24244 | 0.280776 | -0.86348 | 0.387874 | 0.553708 |
| RBM42 | 2716.373 | -0.30549 | 0.145695 | -2.09676 | 0.036015 | 0.111944 |
| KIRREL2 | 2.317924 | 2.53317 | 1.596966 | 1.58624 | 0.112685 | NA |
| UBA2 | 2537.079 | 0.363507 | 0.162097 | 2.242531 | 0.024927 | 0.087611 |
| FFAR2 | 0.45509 | -2.71227 | 3.963744 | -0.68427 | 0.493805 | NA |
| HCST | 10.57887 | 0.315754 | 0.615096 | 0.513341 | 0.607713 | NA |
| COX6B1 | 5135.16 | 0.030071 | 0.130027 | 0.231266 | 0.817108 | 0.890333 |
| KRT36 | 0.392857 | -1.546 | 4.075469 | -0.37934 | 0.704433 | NA |
| THRA | 1980.788 | -0.46467 | 0.124156 | -3.74265 | 1.82E-04 | 0.003659 |
| CCR7 | 0.422534 | 1.924583 | 3.001758 | 0.641152 | 0.521424 | NA |

| | | | | | | |
|---|---|---|---|---|---|---|
| NR1D1 | 482.5814 | -0.31306 | 0.179454 | -1.74452 | 0.081069 | 0.193008 |
| STAT5A | 2499.787 | -0.49923 | 0.139998 | -3.56599 | 3.62E-04 | 0.005542 |
| WNK4 | 62.2606 | -0.83961 | 0.33379 | -2.51538 | 0.01189 | 0.053032 |
| BECN1 | 2273.474 | 0.164319 | 0.145855 | 1.126591 | 0.259915 | 0.424304 |
| TRAP1 | 944.7845 | -0.34578 | 0.169687 | -2.03777 | 0.041572 | 0.123674 |
| GLIS2 | 120.9259 | -0.29569 | 0.21692 | -1.36313 | 0.172842 | 0.324267 |
| NSRP1 | 450.4733 | 0.723054 | 0.706512 | 1.023414 | 0.306112 | 0.473261 |
| PRDM7 | 1555.692 | -0.26678 | 0.208383 | -1.28025 | 0.200456 | 0.356497 |
| RHOT1 | 722.0437 | 0.445458 | 0.163225 | 2.729107 | 0.006351 | 0.034646 |
| EVI2A | 93.49639 | 1.151549 | 0.419876 | 2.742589 | 0.006096 | 0.033788 |
| OMG | 7.439697 | -0.62791 | 0.90306 | -0.69531 | 0.486859 | NA |
| MAP2K2 | 7821.315 | -0.14282 | 0.132234 | -1.08004 | 0.280125 | 0.446115 |
| ABHD8 | 397.238 | -0.0458 | 0.206839 | -0.22143 | 0.824756 | 0.894421 |
| PIN1 | 3266.291 | -0.13432 | 0.168258 | -0.79828 | 0.424708 | 0.58672 |
| FBXL12 | 480.4575 | -0.16237 | 0.155537 | -1.04392 | 0.296521 | 0.463665 |
| EMR2 | 27.51858 | -0.10037 | 0.418458 | -0.23985 | 0.810443 | NA |
| SIN3B | 1307.655 | -0.24102 | 0.140518 | -1.71521 | 0.086307 | 0.201293 |
| SLC35E1 | 1188.83 | -0.41479 | 0.159533 | -2.60003 | 0.009321 | 0.045031 |
| EPS15L1 | 895.4124 | -0.35029 | 0.140741 | -2.48893 | 0.012813 | 0.055885 |
| KLF2 | 68.46477 | -0.33176 | 0.300094 | -1.10551 | 0.268937 | 0.433776 |
| OR7C1 | 2.838307 | -0.05032 | 1.310603 | -0.03839 | 0.969375 | NA |
| UQCR11 | 11.03257 | -0.38766 | 0.603588 | -0.64226 | 0.520704 | NA |
| GFER | 657.1147 | -0.15659 | 0.175193 | -0.89382 | 0.371419 | 0.538088 |
| SYNGR3 | 21.97473 | 0.541716 | 0.54211 | 0.999273 | 0.317663 | NA |
| PKMYT1 | 572.0261 | -0.5066 | 0.288826 | -1.75398 | 0.079434 | 0.190434 |
| WFIKKN1 | 4.826469 | 0.391654 | 1.019052 | 0.384332 | 0.700733 | NA |
| WDR24 | 394.1893 | -0.26837 | 0.189105 | -1.41914 | 0.155859 | 0.301715 |
| FBXL16 | 7.629539 | 0.139709 | 1.058877 | 0.13194 | 0.895031 | NA |
| CHTF18 | 386.8703 | -0.36764 | 0.211513 | -1.73815 | 0.082184 | 0.194856 |
| SMARCA4 | 4862.285 | -0.65668 | 0.15275 | -4.29905 | 1.72E-05 | 8.54E-04 |
| KDM4B | 1017.084 | -0.51916 | 0.155586 | -3.33683 | 8.47E-04 | 0.009354 |
| TICAM1 | 1078.252 | -0.22837 | 0.168138 | -1.35823 | 0.174389 | 0.326343 |
| EMC6 | 71.80145 | 0.047486 | 0.307379 | 0.154485 | 0.877227 | 0.926461 |
| METTL16 | 578.5339 | -0.08526 | 0.149233 | -0.57135 | 0.56776 | 0.704501 |
| ZNF780B | 129.8063 | 0.612408 | 0.331528 | 1.847229 | 0.064714 | 0.165792 |
| LRFN1 | 78.34324 | -0.29756 | 0.30729 | -0.96835 | 0.33287 | 0.501166 |
| ZFP36 | 262.0629 | -0.02664 | 0.195037 | -0.13661 | 0.891335 | 0.936356 |
| KRT17 | 4.800401 | -0.08021 | 1.051788 | -0.07626 | 0.939208 | NA |
| RNF112 | 0.815391 | 0.931626 | 2.360431 | 0.394685 | 0.693076 | NA |
| SPECC1 | 1073.739 | 0.160692 | 0.138112 | 1.163486 | 0.244632 | 0.407398 |
| MRPS12 | 1367.24 | -0.03318 | 0.157537 | -0.21063 | 0.833176 | 0.899945 |
| PSMG2 | 75.8862 | -0.0347 | 0.309248 | -0.11221 | 0.910658 | 0.948085 |
| TWSG1 | 1418.368 | 0.718831 | 0.18297 | 3.928684 | 8.54E-05 | 0.002317 |
| SOX15 | 16.97119 | -0.23645 | 0.542949 | -0.43549 | 0.663205 | NA |
| FAM64A | 636.2298 | -0.55336 | 0.223615 | -2.47463 | 0.013337 | 0.057401 |
| RPAIN | 588.7682 | 0.459313 | 0.155628 | 2.951358 | 0.003164 | 0.021972 |
| USP6 | 9.769836 | 0.757996 | 0.670404 | 1.130655 | 0.2582 | NA |

| | | | | | | |
|---|---|---|---|---|---|---|
| SHBG | 5.348839 | 0.733494 | 0.876264 | 0.83707 | 0.402553 | NA |
| PLD2 | 684.2291 | -0.24772 | 0.143556 | -1.72562 | 0.084415 | 0.197995 |
| AIPL1 | 0.134133 | -1.48909 | 4.077584 | -0.36519 | 0.71497 | NA |
| CD68 | 2.905851 | -0.6846 | 1.243442 | -0.55057 | 0.581931 | NA |
| TXNDC17 | 866.799 | 0.334002 | 0.174093 | 1.918527 | 0.055044 | 0.147937 |
| ATP1B2 | 33.51146 | 0.350663 | 0.426655 | 0.821889 | 0.41114 | NA |
| FXR2 | 1972.161 | -0.24065 | 0.14457 | -1.66457 | 0.095998 | 0.21698 |
| KIF1C | 15541.66 | -0.2999 | 0.160348 | -1.87029 | 0.061443 | 0.160019 |
| MPDU1 | 2815.126 | -0.2364 | 0.14923 | -1.58412 | 0.113167 | 0.241987 |
| MMP28 | 7.145811 | -0.10329 | 1.036802 | -0.09962 | 0.920645 | NA |
| CCL4 | 0.520316 | 2.239969 | 3.173769 | 0.705776 | 0.480328 | NA |
| MRM1 | 261.7921 | -0.16265 | 0.197679 | -0.82278 | 0.410633 | 0.573995 |
| KRI1 | 771.1398 | -0.1899 | 0.147917 | -1.2838 | 0.199214 | 0.355407 |
| ILF3 | 6684.785 | -0.47685 | 0.127955 | -3.72672 | 1.94E-04 | 0.003807 |
| SLC44A2 | 3129.934 | -0.23977 | 0.168516 | -1.42286 | 0.154776 | 0.300109 |
| AP1M2 | 0.86916 | 0.686589 | 2.213986 | 0.310114 | 0.756474 | NA |
| CDKN2D | 555.7236 | -0.18329 | 0.180829 | -1.01363 | 0.310761 | 0.478203 |
| ITFG1 | 1532.171 | 0.178024 | 0.151683 | 1.173656 | 0.240533 | 0.403783 |
| QRICH2 | 18.29177 | -1.28701 | 0.559331 | -2.30097 | 0.021393 | NA |
| FOXJ1 | 0.258578 | 1.233151 | 4.047809 | 0.304646 | 0.760635 | NA |
| SEC14L1 | 2640.897 | -0.08523 | 0.21741 | -0.39204 | 0.695031 | 0.803279 |
| RHBDF2 | 617.5626 | -0.57979 | 0.161075 | -3.59949 | 3.19E-04 | 0.005122 |
| AANAT | 0.314073 | -0.57308 | 3.898934 | -0.14698 | 0.883146 | NA |
| CDH15 | 2.380537 | -0.29443 | 1.65885 | -0.17749 | 0.859125 | NA |
| KLF16 | 989.3192 | -0.27781 | 0.158941 | -1.74787 | 0.080486 | 0.192178 |
| TMEM8A | 6478.956 | -0.22074 | 0.167345 | -1.31906 | 0.187151 | 0.341386 |
| DOHH | 779.4343 | -0.21156 | 0.165039 | -1.28185 | 0.199894 | 0.355943 |
| MAU2 | 1298.054 | -0.35188 | 0.142438 | -2.47044 | 0.013495 | 0.057755 |
| SHC2 | 718.0499 | 0.093688 | 0.201376 | 0.465239 | 0.641761 | 0.76165 |
| | 1.166229 | 0.502497 | 1.978277 | 0.254007 | 0.79949 | NA |
| ABHD17A | 1701.058 | -0.24333 | 0.146039 | -1.6662 | 0.095674 | 0.216537 |
| GAMT | 1046.942 | -0.02076 | 0.15189 | -0.13665 | 0.891309 | 0.936356 |
| DOCK6 | 995.3216 | -0.50996 | 0.154964 | -3.29085 | 9.99E-04 | 0.010333 |
| ECSIT | 832.7259 | -0.27655 | 0.15504 | -1.78372 | 0.074469 | 0.182252 |
| LDLR | 3830.328 | -0.57998 | 0.206917 | -2.80299 | 0.005063 | 0.029892 |
| ELOF1 | 1355.848 | -0.09189 | 0.152708 | -0.60176 | 0.547331 | 0.688738 |
| C19orf80 | 3.537121 | -0.61686 | 1.181234 | -0.52222 | 0.601517 | NA |
| PRKCSH | 8231.685 | -0.47479 | 0.157283 | -3.01868 | 0.002539 | 0.019039 |
| CNN1 | 59.54041 | 0.185723 | 1.1473 | 0.161878 | 0.871402 | 0.923353 |
| ZSCAN10 | 0.163957 | 0.424913 | 4.079006 | 0.104171 | 0.917034 | NA |
| FAM98C | 494.88 | -0.20546 | 0.149756 | -1.37197 | 0.170073 | 0.32037 |
| SAFB2 | 1421.214 | -0.16604 | 0.19259 | -0.86216 | 0.388601 | 0.554176 |
| RPL36 | 11434.63 | 0.08832 | 0.164612 | 0.536533 | 0.59159 | 0.723 |
| ATP8B3 | 103.8608 | -0.30889 | 0.263242 | -1.17343 | 0.240625 | 0.40388 |
| NCAN | 0.247898 | -0.5901 | 4.074633 | -0.14482 | 0.884851 | NA |
| GTPBP3 | 456.218 | -0.25305 | 0.166803 | -1.51706 | 0.129253 | 0.2657 |
| BST2 | 294.9094 | -1.70982 | 0.351783 | -4.86044 | 1.17E-06 | 1.78E-04 |

| | | | | | | |
|---|---|---|---|---|---|---|
| SLC27A1 | 1133.545 | -0.45175 | 0.13823 | -3.2681 | 0.001083 | 0.010915 |
| USHBP1 | 3.26039 | -0.10407 | 1.385659 | -0.0751 | 0.940134 | NA |
| COLGALT1 | 3339.423 | -0.37719 | 0.143352 | -2.6312 | 0.008508 | 0.042257 |
| DDA1 | 2121.196 | -0.22429 | 0.157294 | -1.42595 | 0.153883 | 0.298867 |
| MRPL34 | 2320.39 | 0.100677 | 0.133829 | 0.752277 | 0.451884 | 0.610912 |
| PGLS | 1541.926 | -0.21163 | 0.138162 | -1.53177 | 0.125578 | 0.260631 |
| LSM7 | 1186.638 | -0.08977 | 0.190854 | -0.47036 | 0.638095 | 0.758814 |
| ACSBG2 | 0.949524 | -0.003 | 2.040023 | -0.00147 | 0.998826 | NA |
| MLLT1 | 998.4875 | -0.49975 | 0.178566 | -2.79869 | 0.005131 | 0.030098 |
| ACTN4 | 11353.06 | -0.09978 | 0.168524 | -0.5921 | 0.553784 | 0.69406 |
| FCHO1 | 20.77847 | 0.017942 | 0.48871 | 0.036713 | 0.970714 | NA |
| UNC13A | 4.148014 | 1.56405 | 1.130456 | 1.383556 | 0.166494 | NA |
| MAP1S | 1141.571 | -0.50338 | 0.140142 | -3.59195 | 3.28E-04 | 0.005201 |
| SSBP4 | 1840.793 | -0.23876 | 0.139593 | -1.71044 | 0.087185 | 0.202702 |
| GDF15 | 2527.985 | -0.18042 | 0.270902 | -0.66598 | 0.505422 | 0.654767 |
| PGPEP1 | 2624.972 | -0.50538 | 0.149263 | -3.38585 | 7.10E-04 | 0.008362 |
| KIAA1683 | 15.2255 | -0.38636 | 0.559268 | -0.69082 | 0.489676 | NA |
| LSM4 | 4918.465 | -0.17624 | 0.130428 | -1.35121 | 0.176628 | 0.328872 |
| JUND | 3970.477 | -0.24123 | 0.17444 | -1.38291 | 0.166693 | 0.316127 |
| ZNF557 | 197.3401 | -0.04945 | 0.203516 | -0.24297 | 0.808025 | 0.885317 |
| CRB3 | 1.033263 | 0.082415 | 2.175992 | 0.037875 | 0.969788 | NA |
| HBZ | 0.134133 | -1.48909 | 4.077584 | -0.36519 | 0.71497 | NA |
| PAK4 | 2105.699 | -0.59061 | 0.13781 | -4.28565 | 1.82E-05 | 8.74E-04 |
| C16orf13 | 1391.684 | -0.10206 | 0.150304 | -0.67899 | 0.497143 | 0.648243 |
| YIPF2 | 1734.731 | -0.13474 | 0.156117 | -0.86308 | 0.388093 | 0.553851 |
| ATG4D | 829.019 | -0.23755 | 0.16987 | -1.39841 | 0.161989 | 0.309788 |
| GMFG | 6.55017 | 0.086949 | 0.893409 | 0.097322 | 0.922471 | NA |
| MAP3K10 | 1270.138 | -0.12004 | 0.182388 | -0.65816 | 0.510437 | 0.658745 |
| ZNF317 | 1138.133 | -0.32494 | 0.15336 | -2.1188 | 0.034107 | 0.107997 |
| PPAN | 117.2522 | -0.15 | 0.244346 | -0.6139 | 0.539284 | 0.681722 |
| EIF3G | 4540.302 | -0.14003 | 0.130834 | -1.07027 | 0.284497 | 0.450958 |
| ANGPTL6 | 11.55107 | -0.18283 | 0.586867 | -0.31154 | 0.755392 | NA |
| C19orf66 | 1152.586 | -0.11331 | 0.164836 | -0.68743 | 0.491812 | 0.643353 |
| DNMT1 | 2770.015 | -0.25739 | 0.15338 | -1.67812 | 0.093324 | 0.213265 |
| ZNF426 | 179.4333 | 0.294333 | 0.250274 | 1.176042 | 0.239578 | 0.402786 |
| ZNF236 | 334.968 | 0.235613 | 0.180465 | 1.305593 | 0.191691 | 0.346259 |
| SLC7A10 | 10.69122 | -0.05699 | 0.660302 | -0.08631 | 0.931222 | NA |
| LRP3 | 3082.094 | -0.23108 | 0.14589 | -1.58395 | 0.113206 | 0.241987 |
| NOL11 | 653.6066 | 0.55465 | 0.186708 | 2.970677 | 0.002971 | 0.021006 |
| C1QL1 | 25.06385 | -0.11778 | 0.495809 | -0.23756 | 0.812224 | NA |
| GFAP | 2.086949 | -0.12983 | 1.399804 | -0.09275 | 0.926106 | NA |
| PYY | 0.382031 | -1.26484 | 3.151989 | -0.40128 | 0.688212 | NA |
| HIGD1B | 0.288605 | 1.443619 | 4.029058 | 0.358302 | 0.720117 | NA |
| ZNF428 | 1057.276 | -0.12274 | 0.132712 | -0.92487 | 0.355036 | 0.523241 |
| CCL25 | 0.327913 | 1.543154 | 4.021063 | 0.383768 | 0.701151 | NA |
| COX4I1 | 10217.15 | -0.01206 | 0.19285 | -0.06254 | 0.950129 | 0.972357 |
| EMC8 | 1003.296 | -0.16553 | 0.133893 | -1.23626 | 0.216363 | 0.375491 |

| | | | | | | |
|---|---|---|---|---|---|---|
| GSE1 | 839.9626 | -0.19785 | 0.235512 | -0.84008 | 0.400864 | 0.565475 |
| GINS2 | 361.584 | -0.15096 | 0.2503 | -0.60311 | 0.546435 | 0.688001 |
| CHMP1A | 3237.327 | -0.30313 | 0.139342 | -2.17546 | 0.029596 | 0.098146 |
| NFATC1 | 401.4153 | -0.40151 | 0.180889 | -2.21966 | 0.026442 | 0.091282 |
| RAB11FIP4 | 1323.602 | -0.31578 | 0.174512 | -1.80951 | 0.070372 | 0.175154 |
| HAUS8 | 260.1027 | 0.179185 | 0.248863 | 0.720016 | 0.471515 | 0.627434 |
| EMR3 | 0.382031 | -1.26484 | 3.151989 | -0.40128 | 0.688212 | NA |
| TUBG1 | 1501.098 | -0.16122 | 0.20766 | -0.77636 | 0.437536 | 0.597921 |
| PSME3 | 5463.732 | -0.27796 | 0.14877 | -1.86837 | 0.06171 | 0.160363 |
| RPL27 | 20569.81 | 0.288375 | 0.167163 | 1.725116 | 0.084507 | 0.198068 |
| PSMC3IP | 195.7163 | 0.147747 | 0.230428 | 0.641184 | 0.521403 | 0.668146 |
| AOC3 | 5.186136 | 0.285546 | 1.145668 | 0.249239 | 0.803176 | NA |
| ACLY | 10712.08 | -0.34335 | 0.125233 | -2.74173 | 0.006112 | 0.033841 |
| VPS25 | 1824.667 | -0.19708 | 0.147354 | -1.33747 | 0.181069 | 0.333581 |
| RAMP2 | 49.7793 | 0.158128 | 0.301848 | 0.523867 | 0.600371 | 0.729915 |
| AOC2 | 13.27409 | 0.470248 | 0.625387 | 0.751931 | 0.452093 | NA |
| | 0.162376 | 0.424913 | 4.079006 | 0.104171 | 0.917034 | NA |
| TMEM204 | 169.6082 | -0.41607 | 0.223587 | -1.86087 | 0.062762 | 0.162216 |
| KREMEN2 | 40.06797 | 0.164299 | 0.402488 | 0.408209 | 0.683121 | NA |
| THOC6 | 664.4705 | -0.02586 | 0.169104 | -0.15295 | 0.878437 | 0.926996 |
| TRAF7 | 3630.957 | -0.36344 | 0.129937 | -2.79706 | 0.005157 | 0.0302 |
| KRT34 | 0.163957 | 0.424913 | 4.079006 | 0.104171 | 0.917034 | NA |
| TOP2A | 1884.445 | 0.405637 | 0.137977 | 2.939886 | 0.003283 | 0.022409 |
| STARD3 | 2013.332 | -0.1267 | 0.135031 | -0.9383 | 0.348088 | 0.51605 |
| RARA | 809.3006 | -0.49727 | 0.142998 | -3.47749 | 5.06E-04 | 0.006834 |
| PPP1R1B | 2.286237 | -0.27061 | 1.271382 | -0.21284 | 0.831449 | NA |
| CLUHP3 | 32.75899 | 0.008416 | 0.418596 | 0.020104 | 0.98396 | NA |
| VIMP | 1498.382 | 0.321719 | 0.138824 | 2.317466 | 0.020478 | 0.076899 |
| CHSY1 | 654.7541 | -0.01472 | 0.19392 | -0.07589 | 0.939505 | 0.966423 |
| SNRPA1 | 450.4307 | 0.131867 | 0.186003 | 0.708947 | 0.478358 | 0.633189 |
| KRT17P1 | 0.162376 | 0.424913 | 4.079006 | 0.104171 | 0.917034 | NA |
| LLGL1 | 1471.921 | -0.59579 | 0.164138 | -3.62982 | 2.84E-04 | 0.004762 |
| RHPN2 | 227.2799 | -0.04688 | 0.196727 | -0.23832 | 0.811633 | 0.887711 |
| C19orf12 | 5177.675 | -0.05539 | 0.137073 | -0.40409 | 0.686143 | 0.795602 |
| C19orf40 | 155.0997 | -0.07903 | 0.267135 | -0.29585 | 0.767342 | 0.85561 |
| PODNL1 | 5.691221 | 0.389678 | 0.898986 | 0.433464 | 0.664678 | NA |
| DNAJB1 | 4713.489 | -0.20272 | 0.158545 | -1.27865 | 0.20102 | 0.357285 |
| ZSWIM4 | 545.62 | -0.18807 | 0.193282 | -0.97303 | 0.330537 | 0.498856 |
| FBXW9 | 210.4158 | -0.40694 | 0.218377 | -1.86348 | 0.062394 | 0.161786 |
| RFX1 | 435.7512 | -0.4905 | 0.190217 | -2.57864 | 0.009919 | 0.046924 |
| ZNF20 | 2.61063 | 0.195781 | 1.272841 | 0.153814 | 0.877756 | NA |
| C19orf57 | 39.21178 | -0.20556 | 0.377592 | -0.54439 | 0.58617 | NA |
| DCAF15 | 701.6523 | -0.39954 | 0.177148 | -2.25541 | 0.024107 | 0.085854 |
| CC2D1A | 1283.416 | -0.3546 | 0.17265 | -2.05386 | 0.039989 | 0.120414 |
| RTBDN | 0.230334 | -0.53318 | 4.076749 | -0.13079 | 0.895945 | NA |
| LHX1 | 3.091445 | 1.483234 | 1.238772 | 1.197342 | 0.231173 | NA |
| CCT6B | 69.98802 | 0.354922 | 0.303085 | 1.171032 | 0.241586 | 0.404634 |

| | | | | | | |
|---|---|---|---|---|---|---|
| ACACA | 3590.086 | -0.52901 | 0.124347 | -4.25431 | 2.10E-05 | 9.51E-04 |
| ENOSF1 | 667.4693 | 0.074202 | 0.14037 | 0.528615 | 0.597073 | 0.726799 |
| LINC00470 | 49.98648 | 0.443655 | 0.380281 | 1.166649 | 0.243352 | 0.406201 |
| EMILIN2 | 981.186 | -0.05279 | 0.232462 | -0.22709 | 0.820352 | 0.892334 |
| RAP1GAP2 | 9.539528 | 0.254813 | 0.772338 | 0.329924 | 0.741457 | NA |
| CLUH | 3627.487 | -0.4707 | 0.154164 | -3.05324 | 0.002264 | 0.017517 |
| INPP5K | 2375.741 | -0.16565 | 0.144233 | -1.14846 | 0.250777 | 0.414408 |
| MYBBP1A | 1677.559 | -0.63235 | 0.161563 | -3.91398 | 9.08E-05 | 0.002389 |
| RPA1 | 3549.526 | -0.2214 | 0.165062 | -1.34134 | 0.179811 | 0.332292 |
| SERPINF1 | 5904.942 | -0.56838 | 0.184229 | -3.08516 | 0.002034 | 0.016362 |
| UBE2G1 | 1156.657 | -0.20255 | 0.133795 | -1.51388 | 0.130055 | 0.26643 |
| ITGB4 | 33.19143 | -0.21752 | 0.40649 | -0.53513 | 0.592563 | NA |
| WBP2 | 4945.744 | -0.3424 | 0.134942 | -2.5374 | 0.011168 | 0.050668 |
| H3F3B | 6774.177 | -0.0799 | 0.142232 | -0.56174 | 0.574292 | 0.709634 |
| UNK | 1120.827 | -0.43329 | 0.140645 | -3.08071 | 0.002065 | 0.016587 |
| TRIM47 | 419.0232 | -0.4462 | 0.37221 | -1.19878 | 0.230614 | 0.392807 |
| EIF5A | 13782.11 | -0.33969 | 0.172466 | -1.96961 | 0.048884 | 0.136742 |
| KDM6B | 633.9689 | -0.47319 | 0.212934 | -2.22226 | 0.026266 | 0.090994 |
| GPS2 | 73.46561 | 0.471513 | 0.38127 | 1.23669 | 0.216202 | 0.375377 |
| XAF1 | 371.2166 | -0.19394 | 0.378087 | -0.51294 | 0.607993 | 0.735336 |
| DLG4 | 751.2345 | -0.26764 | 0.16508 | -1.62127 | 0.104959 | 0.230452 |
| SDF2 | 1618.857 | 0.086452 | 0.148938 | 0.580457 | 0.561607 | 0.699721 |
| FLOT2 | 3063.724 | -0.41218 | 0.127538 | -3.23185 | 0.00123 | 0.011864 |
| ERAL1 | 1117.565 | -0.28129 | 0.131132 | -2.14507 | 0.031947 | 0.103652 |
| PRMT7 | 1371.488 | -0.36279 | 0.146863 | -2.47026 | 0.013501 | 0.057763 |
| NIP7 | 1137.026 | 0.215087 | 0.149663 | 1.437139 | 0.150679 | 0.294428 |
| TERF2 | 578.3494 | -0.32372 | 0.141359 | -2.29006 | 0.022018 | 0.080912 |
| VPS4A | 2876.365 | -0.25135 | 0.139295 | -1.80444 | 0.071162 | 0.176549 |
| MTSS1L | 1400.425 | -0.45647 | 0.148294 | -3.07813 | 0.002083 | 0.016597 |
| SYT4 | 0.162376 | 0.424913 | 4.079006 | 0.104171 | 0.917034 | NA |
| SLC14A2 | 0.163957 | 0.424913 | 4.079006 | 0.104171 | 0.917034 | NA |
| MYH10 | 11952.52 | -0.06909 | 0.171479 | -0.40293 | 0.687001 | 0.796285 |
| PEMT | 461.6371 | -0.29519 | 0.155154 | -1.90258 | 0.057096 | 0.151527 |
| SCO1 | 733.4922 | 0.069841 | 0.142159 | 0.491288 | 0.623223 | 0.747426 |
| MPRIP | 6300.128 | -0.13876 | 0.164488 | -0.84357 | 0.39891 | 0.563724 |
| FAM104A | 605.526 | 0.003105 | 0.15177 | 0.020459 | 0.983677 | 0.991371 |
| SLC39A11 | 715.8626 | -0.50164 | 0.143112 | -3.50523 | 4.56E-04 | 0.00643 |
| BTBD2 | 3045.356 | -0.47194 | 0.140255 | -3.36486 | 7.66E-04 | 0.00881 |
| PRAM1 | 0.488709 | 2.119056 | 3.307564 | 0.64067 | 0.521737 | NA |
| ZNF414 | 326.5935 | -0.34134 | 0.18216 | -1.87387 | 0.060949 | 0.158996 |
| CSNK1G2 | 3292.535 | -0.17231 | 0.158987 | -1.08379 | 0.278459 | 0.444537 |
| CNDP2 | 4612.227 | -0.36518 | 0.131551 | -2.77595 | 0.005504 | 0.031691 |
| MYH11 | 18.08413 | -0.25259 | 0.548596 | -0.46043 | 0.64521 | NA |
| FOPNL | 791.9555 | 0.619999 | 0.2031 | 3.052677 | 0.002268 | 0.017517 |
| CTIF | 1298.802 | -0.41274 | 0.139099 | -2.96724 | 0.003005 | 0.021179 |
| MRO | 3.34123 | -0.90316 | 1.128352 | -0.80043 | 0.423463 | NA |
| MBD2 | 1253.79 | 0.281487 | 0.142503 | 1.975311 | 0.048233 | 0.135498 |

| | | | | | | |
|---|---|---|---|---|---|---|
| IER3IP1 | 736.1215 | 0.500606 | 0.153294 | 3.26566 | 0.001092 | 0.010985 |
| NAPG | 694.8595 | 0.597725 | 0.181056 | 3.301334 | 9.62E-04 | 0.010104 |
| SPIRE1 | 1667.02 | 0.214817 | 0.149639 | 1.435571 | 0.151125 | 0.295056 |
| ERN2 | 0.750448 | 2.743526 | 2.504729 | 1.095338 | 0.273368 | NA |
| RPS15A | 2053.512 | 0.276154 | 0.155145 | 1.779971 | 0.075081 | 0.182994 |
| NARS | 4548.018 | 0.57631 | 0.165058 | 3.491562 | 4.80E-04 | 0.006651 |
| KIAA1468 | 771.9128 | 0.443912 | 0.204256 | 2.173314 | 0.029757 | 0.098483 |
| HRH4 | 4.850408 | 0.256905 | 0.944793 | 0.271916 | 0.785686 | NA |
| TMEM241 | 278.9644 | -0.43353 | 0.202143 | -2.14465 | 0.031981 | 0.10373 |
| KCTD1 | 676.8826 | -0.35959 | 0.181782 | -1.97815 | 0.047912 | 0.134949 |
| CABLES1 | 13885.97 | -0.46238 | 0.194972 | -2.37152 | 0.017715 | 0.069636 |
| RNF138 | 194.621 | 0.451224 | 0.230162 | 1.960464 | 0.049942 | 0.138396 |
| ELP2 | 2092.297 | 0.158553 | 0.156086 | 1.015804 | 0.309723 | 0.476976 |
| DSG1 | 0.163957 | 0.424913 | 4.079006 | 0.104171 | 0.917034 | NA |
| DSC3 | 0.096202 | 0.408546 | 4.079639 | 0.100143 | 0.920231 | NA |
| DTNA | 282.2529 | 0.201958 | 0.187136 | 1.079201 | 0.280498 | 0.446589 |
| FHOD3 | 2146.92 | -0.18656 | 0.225058 | -0.82892 | 0.40715 | 0.570809 |
| TPGS2 | 3727.284 | -0.12327 | 0.137645 | -0.89553 | 0.370504 | 0.537354 |
| KLHL36 | 1502.354 | -0.27711 | 0.165435 | -1.67503 | 0.093928 | 0.214015 |
| BCMO1 | 0.399595 | -1.31595 | 3.579466 | -0.36764 | 0.713143 | NA |
| MPHOSPH( | 455.3712 | 0.496183 | 0.206232 | 2.405944 | 0.016131 | 0.065355 |
| CHST5 | 0.70693 | -1.62766 | 2.687154 | -0.60572 | 0.544701 | NA |
| KIAA0513 | 623.0058 | -0.54893 | 0.143421 | -3.8274 | 1.30E-04 | 0.002993 |
| DYNC1LI2 | 4598.202 | 0.434287 | 0.173291 | 2.506107 | 0.012207 | 0.054038 |
| FBXL8 | 129.4635 | -0.05053 | 0.235592 | -0.21449 | 0.830164 | 0.89752 |
| FHOD1 | 1476.073 | -0.138 | 0.175102 | -0.78814 | 0.430615 | 0.592195 |
| CCDC102A | 370.5208 | -0.10239 | 0.179647 | -0.56995 | 0.568713 | 0.705388 |
| SLC9A5 | 145.3986 | -0.01344 | 0.25749 | -0.05221 | 0.958358 | 0.977053 |
| CALCOCO2 | 1359.452 | 0.075027 | 0.145485 | 0.515706 | 0.60606 | 0.734271 |
| RSAD1 | 1263.353 | -0.13981 | 0.169373 | -0.82546 | 0.409108 | 0.572742 |
| NMT1 | 4736.957 | -0.37534 | 0.129027 | -2.90902 | 0.003626 | 0.024006 |
| MYCBPAP | 0.816825 | 1.673341 | 2.21346 | 0.755984 | 0.449659 | NA |
| SRSF1 | 3074.177 | 0.19188 | 0.146075 | 1.313573 | 0.18899 | 0.343263 |
| VEZF1 | 1538.969 | 0.161508 | 0.13485 | 1.197686 | 0.231039 | 0.393158 |
| CHAD | 18.90266 | -0.18877 | 0.518696 | -0.36392 | 0.715916 | NA |
| TACO1 | 1025.722 | -0.10927 | 0.148247 | -0.73705 | 0.461092 | 0.619194 |
| TEX2 | 2820.022 | -0.06593 | 0.156761 | -0.42055 | 0.674087 | 0.786547 |
| DCAF7 | 5832.639 | -0.50452 | 0.122385 | -4.12238 | 3.75E-05 | 0.0014 |
| LIMD2 | 2254.995 | -0.53689 | 0.196081 | -2.73811 | 0.006179 | 0.033993 |
| BRIP1 | 148.1806 | 0.551474 | 0.300636 | 1.834357 | 0.066601 | 0.1686 |
| KAT7 | 1235.797 | -0.39836 | 0.13996 | -2.84621 | 0.004424 | 0.027205 |
| MAPK8IP3 | 1017.035 | 0.014892 | 0.211118 | 0.070538 | 0.943765 | 0.968771 |
| IGF1R | 1519.059 | -0.66399 | 0.200655 | -3.30909 | 9.36E-04 | 0.009929 |
| ARRDC4 | 1602.149 | -0.40662 | 0.222558 | -1.82705 | 0.067692 | 0.17038 |
| ADAMTS17 | 13.84789 | -1.05391 | 0.54945 | -1.91812 | 0.055096 | NA |
| LINS | 296.0777 | 0.280125 | 0.203628 | 1.375668 | 0.168924 | 0.31919 |
| PCSK6 | 36.80669 | -0.05538 | 0.414617 | -0.13356 | 0.893749 | NA |

| | | | | | | |
|---|---|---|---|---|---|---|
| UNC45A | 4265.146 | -0.38838 | 0.143186 | -2.71239 | 0.00668 | 0.035967 |
| ST8SIA2 | 579.9247 | 0.839281 | 0.882669 | 0.950845 | 0.341683 | 0.509837 |
| MCTP2 | 5.020371 | 1.360092 | 0.951355 | 1.429637 | 0.152821 | NA |
| FURIN | 4429.567 | -0.22764 | 0.219379 | -1.03765 | 0.299432 | 0.466338 |
| IQGAP1 | 7577.798 | 0.149787 | 0.151771 | 0.986928 | 0.323678 | 0.491623 |
| CRTC3 | 1245.096 | -0.37508 | 0.134929 | -2.77984 | 0.005439 | 0.031378 |
| GLYR1 | 1296.648 | -0.18043 | 0.14688 | -1.22845 | 0.21928 | 0.379 |
| PMM2 | 338.666 | -0.16166 | 0.172197 | -0.93884 | 0.347814 | 0.515984 |
| SLC5A2 | 1.304543 | 0.199695 | 1.761391 | 0.113373 | 0.909734 | NA |
| ITGAX | 2.97067 | -1.66397 | 1.199888 | -1.38677 | 0.165512 | NA |
| TGFB1I1 | 2364.867 | -0.33211 | 0.141789 | -2.34225 | 0.019168 | 0.073491 |
| C16orf58 | 1235.507 | -0.30278 | 0.161747 | -1.87193 | 0.061216 | 0.159533 |
| ARMC5 | 427.8794 | -0.32961 | 0.222658 | -1.48035 | 0.13878 | 0.278663 |
| PARN | 1029.261 | 0.036733 | 0.155041 | 0.236925 | 0.812715 | 0.888591 |
| FTO | 1054.462 | -0.50969 | 0.140043 | -3.63951 | 2.73E-04 | 0.004639 |
| UQCRC2 | 4124.909 | 0.01371 | 0.160209 | 0.085578 | 0.931802 | 0.961401 |
| CDR2 | 560.9048 | 0.381132 | 0.199436 | 1.911045 | 0.055999 | 0.149722 |
| IGSF6 | 4.658332 | 0.28294 | 1.008635 | 0.280517 | 0.779081 | NA |
| ARHGAP17 | 1211.856 | -0.36419 | 0.155444 | -2.34289 | 0.019135 | 0.073388 |
| MYLK3 | 1.389622 | 0.986783 | 1.761847 | 0.560085 | 0.575422 | NA |
| NKD1 | 104.8708 | -0.22615 | 0.318283 | -0.71053 | 0.477375 | 0.632252 |
| DHX38 | 1912.096 | -0.34792 | 0.16073 | -2.16463 | 0.030416 | 0.099989 |
| TXNL4B | 263.3329 | 0.092464 | 0.18062 | 0.511927 | 0.608702 | 0.735818 |
| MARVELD3 | 0.326333 | 1.533889 | 3.826878 | 0.40082 | 0.688553 | NA |
| ZFHX3 | 620.4579 | -0.15647 | 0.322493 | -0.48519 | 0.627544 | 0.750113 |
| CLEC18B | 0.134133 | -1.48909 | 4.077584 | -0.36519 | 0.71497 | NA |
| CPNE2 | 718.1206 | -0.11949 | 0.151683 | -0.78778 | 0.430824 | 0.592371 |
| NLRC5 | 302.8194 | -0.92854 | 0.24162 | -3.84298 | 1.22E-04 | 0.0029 |
| KATNB1 | 1544.308 | -0.19883 | 0.152151 | -1.30678 | 0.191288 | 0.34574 |
| KIFC3 | 4060.178 | -0.14024 | 0.154837 | -0.90574 | 0.365073 | 0.532373 |
| NUDT7 | 32.82425 | 0.263048 | 0.393749 | 0.668061 | 0.504095 | NA |
| GCSH | 75.24389 | 0.365257 | 0.289071 | 1.263555 | 0.20639 | 0.363559 |
| CMTM3 | 1709.049 | -0.01784 | 0.162914 | -0.1095 | 0.912806 | 0.94937 |
| CMTM2 | 0.258578 | 1.233151 | 4.047809 | 0.304646 | 0.760635 | NA |
| CDH11 | 171.3404 | 0.693555 | 0.26127 | 2.654553 | 0.007941 | 0.040458 |
| NOL3 | 572.0371 | -0.05858 | 0.16971 | -0.34518 | 0.729957 | 0.828734 |
| MAP1LC3B | 4991.937 | 0.102486 | 0.150031 | 0.683097 | 0.494546 | 0.645623 |
| MBTPS1 | 4430.516 | -0.01896 | 0.134685 | -0.14079 | 0.888032 | 0.934211 |
| CDH13 | 264.0501 | -0.28237 | 0.255317 | -1.10594 | 0.268753 | 0.433655 |
| ZCCHC14 | 1091.464 | -0.03116 | 0.201261 | -0.15482 | 0.876961 | 0.926427 |
| TLDC1 | 485.3341 | -0.42741 | 0.144421 | -2.95949 | 0.003081 | 0.021631 |
| OSGIN1 | 1119.033 | -0.49737 | 0.27232 | -1.82642 | 0.067788 | 0.170548 |
| RHOT2 | 2476.239 | -0.20458 | 0.134798 | -1.51771 | 0.129088 | 0.265408 |
| RPL3L | 4.523207 | -0.22046 | 1.004538 | -0.21946 | 0.826289 | NA |
| ZSCAN32 | 286.4232 | 0.002581 | 0.178023 | 0.014499 | 0.988432 | 0.994219 |
| RPS2 | 27447.23 | -0.33582 | 0.149742 | -2.24264 | 0.02492 | 0.087611 |
| NDUFB10 | 3051.265 | -0.02507 | 0.13523 | -0.18536 | 0.852943 | 0.912603 |

| | | | | | | |
|---|---|---|---|---|---|---|
| PDPK1 | 869.8565 | -0.21581 | 0.140968 | -1.53093 | 0.125786 | 0.260839 |
| TIGD7 | 17.8806 | 0.721079 | 0.485576 | 1.484997 | 0.137545 | NA |
| DEF8 | 2970.351 | -0.08464 | 0.137205 | -0.61692 | 0.537287 | 0.680433 |
| TCF25 | 4894.547 | -0.28592 | 0.11606 | -2.46357 | 0.013756 | 0.058473 |
| GALNS | 1320.377 | -0.45357 | 0.143447 | -3.16193 | 0.001567 | 0.013907 |
| GAS8 | 698.4938 | -0.18283 | 0.223444 | -0.81825 | 0.413216 | 0.576519 |
| MED9 | 396.2707 | -0.32493 | 0.17715 | -1.83424 | 0.066618 | 0.168608 |
| NCOR1 | 1693.158 | 0.111555 | 0.254911 | 0.437623 | 0.66166 | 0.777247 |
| CDRT15P1 | 2.846037 | 0.963741 | 1.337804 | 0.72039 | 0.471285 | NA |
| COPS3 | 1346.632 | 0.157464 | 0.145647 | 1.081134 | 0.279638 | 0.445605 |
| GID4 | 450.2324 | 0.019268 | 0.147948 | 0.130235 | 0.896381 | 0.939326 |
| ZNF287 | 93.19733 | 0.425248 | 0.262833 | 1.617938 | 0.105676 | 0.231481 |
| KSR1 | 336.7891 | -0.37967 | 0.192201 | -1.97538 | 0.048225 | 0.135498 |
| CIRH1A | 2265.734 | -0.15986 | 0.156301 | -1.02275 | 0.306424 | 0.473434 |
| RANBP10 | 1700.802 | -0.61052 | 0.185091 | -3.29848 | 9.72E-04 | 0.010161 |
| CTRL | 0.296509 | -0.51709 | 3.982155 | -0.12985 | 0.896684 | NA |
| DPEP3 | 0.151697 | -1.546 | 4.075469 | -0.37934 | 0.704433 | NA |
| GFOD2 | 445.7454 | -0.18488 | 0.193263 | -0.9566 | 0.338769 | 0.507469 |
| NOB1 | 1072.703 | -0.02422 | 0.151373 | -0.16 | 0.872881 | 0.924371 |
| PRPSAP2 | 611.593 | 0.06099 | 0.164547 | 0.370653 | 0.710896 | 0.815056 |
| MYO19 | 735.0567 | -0.08927 | 0.170582 | -0.52332 | 0.600749 | 0.73015 |
| DDX52 | 613.5185 | 0.594966 | 0.181213 | 3.283243 | 0.001026 | 0.010515 |
| RASL10B | 13.35331 | 0.332287 | 0.569002 | 0.583981 | 0.559233 | NA |
| PCTP | 690.9314 | -0.24574 | 0.157004 | -1.56519 | 0.117538 | 0.248292 |
| TOM1L1 | 810.1978 | 0.41059 | 0.163736 | 2.507628 | 0.012154 | 0.053906 |
| C17orf80 | 430.3665 | 0.588442 | 0.182257 | 3.228643 | 0.001244 | 0.011897 |
| TOB1 | 600.1155 | 0.101926 | 0.139737 | 0.729414 | 0.465748 | 0.622931 |
| VPS53 | 3232.368 | -0.67737 | 0.127821 | -5.29933 | 1.16E-07 | 3.83E-05 |
| SGSM2 | 1460.487 | -0.47352 | 0.144162 | -3.28463 | 0.001021 | 0.010509 |
| NPEPPS | 1674.339 | -0.00823 | 0.141609 | -0.05811 | 0.95366 | 0.974708 |
| SKAP1 | 0.315653 | -0.57305 | 3.892639 | -0.14721 | 0.882963 | NA |
| LRRC46 | 14.68614 | -0.73021 | 0.539634 | -1.35317 | 0.176002 | NA |
| SCRN2 | 685.4606 | -0.35286 | 0.152844 | -2.30865 | 0.020963 | 0.078199 |
| SSH2 | 1164.844 | -0.08164 | 0.151354 | -0.5394 | 0.589614 | 0.721647 |
| RHBDL3 | 10.87254 | 0.127715 | 0.662289 | 0.192839 | 0.847085 | NA |
| SPACA3 | 64.1724 | -0.03293 | 0.469668 | -0.0701 | 0.944111 | 0.969042 |
| ARSG | 742.6262 | -0.23994 | 0.162575 | -1.4759 | 0.13997 | 0.280266 |
| ABCA8 | 37.38088 | -0.23395 | 0.387402 | -0.6039 | 0.545912 | NA |
| G6PC3 | 3534.902 | -0.25531 | 0.155755 | -1.63917 | 0.101178 | 0.224703 |
| CLTC | 12266.23 | -0.08065 | 0.140941 | -0.57221 | 0.567182 | 0.704078 |
| C17orf64 | 0.326333 | 1.533889 | 3.826878 | 0.40082 | 0.688553 | NA |
| BCAS3 | 1318.736 | -0.40922 | 0.135094 | -3.02918 | 0.002452 | 0.018505 |
| PTRH2 | 830.002 | 0.478116 | 0.162012 | 2.951106 | 0.003166 | 0.021977 |
| SS18 | 965.7846 | 0.589176 | 0.187426 | 3.143502 | 0.001669 | 0.014431 |
| TAF4B | 508.4363 | 0.004151 | 0.201989 | 0.020549 | 0.983605 | 0.991371 |
| AFG3L2 | 2320.827 | 0.046712 | 0.135846 | 0.343864 | 0.730949 | 0.829383 |
| SLMO1 | 76.13042 | 0.243008 | 0.2652 | 0.916322 | 0.359498 | 0.526609 |

| | | | | | | |
|---|---|---|---|---|---|---|
| IMPA2 | 1265.171 | 0.215615 | 0.246446 | 0.874898 | 0.381629 | 0.547763 |
| GNAL | 740.2129 | 0.16736 | 0.248152 | 0.674425 | 0.500041 | 0.650427 |
| SLC39A6 | 6239.429 | 0.550904 | 0.184866 | 2.980023 | 0.002882 | 0.020568 |
| RPRD1A | 1009.708 | 0.519067 | 0.193115 | 2.687871 | 0.007191 | 0.037838 |
| C18orf21 | 152.9579 | 0.300554 | 0.223532 | 1.344565 | 0.178766 | 0.33126 |
| GALNT1 | 1233.515 | 0.662598 | 0.197435 | 3.356031 | 7.91E-04 | 0.008987 |
| ASXL3 | 0.096202 | 0.408546 | 4.079639 | 0.100143 | 0.920231 | NA |
| MEP1B | 0.134133 | -1.48909 | 4.077584 | -0.36519 | 0.71497 | NA |
| GAREM | 232.8334 | -0.20848 | 0.226773 | -0.91932 | 0.357926 | 0.525131 |
| ESCO1 | 310.4094 | 1.210118 | 0.81194 | 1.490404 | 0.136118 | 0.275278 |
| OSBPL1A | 3054.2 | 0.209846 | 0.182541 | 1.149582 | 0.250316 | 0.413938 |
| GATA6 | 0.230334 | -0.53318 | 4.076749 | -0.13079 | 0.895945 | NA |
| GREB1L | 63.1502 | -0.08815 | 0.303224 | -0.29071 | 0.771275 | 0.858691 |
| C18orf8 | 727.8338 | -0.01018 | 0.187306 | -0.05435 | 0.956655 | 0.976145 |
| PELP1 | 1544.239 | -0.60031 | 0.144801 | -4.14571 | 3.39E-05 | 0.001308 |
| NPC1 | 1686.913 | -0.44468 | 0.202157 | -2.19969 | 0.027829 | 0.094368 |
| SLC14A1 | 0.247898 | -0.5901 | 4.074633 | -0.14482 | 0.884851 | NA |
| ARRB2 | 626.7817 | -0.18352 | 0.154598 | -1.1871 | 0.235187 | 0.397965 |
| SLC13A5 | 0.258578 | 1.233151 | 4.047809 | 0.304646 | 0.760635 | NA |
| ZMYND15 | 1.655373 | -1.06548 | 1.720195 | -0.6194 | 0.535655 | NA |
| WRAP53 | 445.6729 | -0.27248 | 0.205953 | -1.32301 | 0.185833 | 0.339662 |
| MINK1 | 3592.987 | -0.46505 | 0.140499 | -3.30998 | 9.33E-04 | 0.00992 |
| SAT2 | 1854.819 | -0.03361 | 0.143355 | -0.23442 | 0.814658 | 0.889545 |
| ASGR1 | 1.144942 | 1.358971 | 2.181094 | 0.623068 | 0.53324 | NA |
| PIK3R5 | 2.031454 | 0.266464 | 1.51793 | 0.175544 | 0.860652 | NA |
| TP53 | 2305.397 | -0.5146 | 0.142809 | -3.60342 | 3.14E-04 | 0.005079 |
| CCDC40 | 80.86317 | 0.288775 | 0.289118 | 0.998813 | 0.317885 | 0.485635 |
| ARHGDIA | 13762.85 | -0.38206 | 0.151521 | -2.52148 | 0.011686 | 0.052315 |
| TMC6 | 1051.678 | -0.31415 | 0.174911 | -1.79608 | 0.072482 | 0.178968 |
| SLC16A3 | 1706.143 | 0.128466 | 0.252164 | 0.509456 | 0.610433 | 0.737459 |
| CARD14 | 16.14605 | -1.47402 | 0.599309 | -2.45953 | 0.013912 | NA |
| TTYH2 | 5925.498 | -0.58523 | 0.172665 | -3.38937 | 7.01E-04 | 0.008296 |
| RAB40B | 347.4353 | -0.15605 | 0.182281 | -0.85611 | 0.391935 | 0.557055 |
| EIF4A3 | 2401.748 | -0.00591 | 0.143386 | -0.04124 | 0.967101 | 0.981675 |
| CSNK1D | 4696.472 | -0.35933 | 0.124156 | -2.89415 | 0.003802 | 0.024629 |
| ANAPC11 | 2600.947 | -0.132 | 0.176546 | -0.74771 | 0.454637 | 0.613515 |
| TBCD | 3689.085 | -0.5562 | 0.140087 | -3.97041 | 7.17E-05 | 0.002065 |
| FN3KRP | 1127.121 | -0.26268 | 0.170804 | -1.53789 | 0.124074 | 0.258597 |
| NARF | 3259.872 | 0.088404 | 0.136276 | 0.648707 | 0.516528 | 0.664003 |
| RPTOR | 2780.368 | -0.54086 | 0.194669 | -2.77837 | 0.005463 | 0.031471 |
| FOXK2 | 1823.29 | -0.30925 | 0.158452 | -1.95166 | 0.050978 | 0.140481 |
| TRIM65 | 795.2259 | -0.27505 | 0.157191 | -1.7498 | 0.080153 | 0.191691 |
| CBX8 | 375.1512 | -0.23212 | 0.178849 | -1.29783 | 0.194344 | 0.349475 |
| SECTM1 | 236.8556 | -0.10217 | 0.306158 | -0.3337 | 0.738603 | 0.834633 |
| RNF157 | 888.41 | -0.78947 | 0.157852 | -5.00135 | 5.69E-07 | 1.12E-04 |
| AZI1 | 642.4974 | -0.23145 | 0.194201 | -1.19181 | 0.233336 | 0.395906 |
| WDR45B | 2584.327 | -0.14345 | 0.166285 | -0.86265 | 0.388329 | 0.554122 |

| | | | | | | |
|---|---|---|---|---|---|---|
| CBX4 | 1879.125 | -0.41744 | 0.121237 | -3.44315 | 5.75E-04 | 0.007385 |
| RNF165 | 6.248912 | -0.61233 | 0.890375 | -0.68773 | 0.491625 | NA |
| DYM | 915.4552 | -0.18889 | 0.144448 | -1.30768 | 0.190983 | 0.345349 |
| MAPK4 | 3.024222 | 0.429684 | 1.16008 | 0.370391 | 0.711091 | NA |
| ELAC1 | 129.3437 | -0.11183 | 0.216649 | -0.51617 | 0.605736 | 0.734104 |
| MBD1 | 1606.853 | -0.15805 | 0.132074 | -1.19667 | 0.231434 | 0.393637 |
| SMAD4 | 996.7509 | 0.212053 | 0.139968 | 1.515012 | 0.129769 | 0.266024 |
| TNFRSF11A | 32.57774 | -0.1723 | 0.418139 | -0.41207 | 0.68029 | NA |
| ZCCHC2 | 320.4531 | -0.01945 | 0.188534 | -0.10315 | 0.917844 | 0.952399 |
| FBXO15 | 11.31273 | 0.002756 | 0.604259 | 0.004561 | 0.996361 | NA |
| CBLN2 | 0.298089 | -0.51711 | 3.975232 | -0.13008 | 0.896501 | NA |
| PMAIP1 | 23.39882 | 0.756679 | 0.497166 | 1.521985 | 0.128013 | NA |
| LEPREL4 | 928.5177 | -0.30765 | 0.156338 | -1.96783 | 0.049087 | 0.137021 |
| NT5C3B | 2033.846 | 0.044685 | 0.149933 | 0.298035 | 0.765677 | 0.854637 |
| FAM134C | 1796.249 | -0.54958 | 0.150971 | -3.64029 | 2.72E-04 | 0.004632 |
| PIP4K2B | 4000.223 | -0.67634 | 0.160533 | -4.2131 | 2.52E-05 | 0.001062 |
| ERBB2 | 2836.533 | -0.3837 | 0.132237 | -2.90164 | 0.003712 | 0.02432 |
| GRB7 | 3.007853 | 1.131564 | 1.17366 | 0.964132 | 0.33498 | NA |
| MIEN1 | 1556.895 | 0.00738 | 0.145158 | 0.050842 | 0.959451 | 0.977677 |
| ARL5C | 1.87576 | -0.09267 | 1.436063 | -0.06453 | 0.948549 | NA |
| STAC2 | 1.552348 | -0.27508 | 1.744736 | -0.15766 | 0.874724 | NA |
| IGFBP4 | 4274.288 | -0.23952 | 0.151077 | -1.58539 | 0.112878 | 0.241634 |
| FKBP10 | 7546.929 | -0.32204 | 0.112836 | -2.85408 | 0.004316 | 0.02672 |
| TXNL4A | 2646.945 | -0.0585 | 0.143887 | -0.40655 | 0.68434 | 0.794605 |
| CACNA1A | 270.1059 | -0.14072 | 0.288476 | -0.48782 | 0.625678 | 0.748948 |
| | 46.22141 | 0.205633 | 0.325771 | 0.631221 | 0.527896 | NA |
| SAMD1 | 502.675 | -0.28978 | 0.184367 | -1.57176 | 0.116007 | 0.24633 |
| BRD4 | 1547.158 | -0.59901 | 0.1738 | -3.44654 | 5.68E-04 | 0.007341 |
| SLC39A3 | 711.9997 | -0.35395 | 0.144256 | -2.45363 | 0.014142 | 0.059645 |
| NFIC | 1680.594 | -0.52296 | 0.189472 | -2.7601 | 0.005778 | 0.032602 |
| TPGS1 | 492.5259 | 0.17235 | 0.161194 | 1.069214 | 0.284973 | 0.45135 |
| PPAP2C | 5.796096 | -1.10378 | 0.913688 | -1.20805 | 0.227029 | NA |
| FEM1A | 41.21598 | -0.64149 | 0.341224 | -1.87996 | 0.060114 | NA |
| VAV1 | 0.096202 | 0.408546 | 4.079639 | 0.100143 | 0.920231 | NA |
| MVB12A | 1545.982 | -0.19202 | 0.136571 | -1.406 | 0.159724 | 0.306892 |
| CIB3 | 0.151697 | -1.546 | 4.075469 | -0.37934 | 0.704433 | NA |
| SH3GL1 | 2474.389 | -0.21624 | 0.133117 | -1.62441 | 0.104288 | 0.229312 |
| DUS3L | 544.2647 | -0.09827 | 0.165374 | -0.59421 | 0.55237 | 0.693167 |
| DPP9 | 2296.823 | -0.30781 | 0.142141 | -2.1655 | 0.030349 | 0.099913 |
| CCDC97 | 798.0274 | -0.42355 | 0.134524 | -3.14852 | 0.001641 | 0.014318 |
| TMEM91 | 9.87501 | 0.469964 | 0.678226 | 0.692931 | 0.488353 | NA |
| ZFP14 | 85.589 | 0.802215 | 0.328475 | 2.442239 | 0.014596 | 0.060786 |
| WTIP | 765.0034 | 0.187617 | 0.186288 | 1.007136 | 0.31387 | 0.481299 |
| ADAMTS10 | 18.38734 | -0.3411 | 0.588885 | -0.57922 | 0.562438 | NA |
| MYO1F | 37.02773 | -0.71073 | 0.367916 | -1.93176 | 0.053389 | NA |
| C19orf52 | 557.2308 | -0.28323 | 0.157018 | -1.80381 | 0.07126 | 0.176735 |
| FBN3 | 0.326333 | 1.533889 | 3.826878 | 0.40082 | 0.688553 | NA |

| | | | | | | |
|---|---|---|---|---|---|---|
| CARM1 | 3019.041 | -0.50869 | 0.119312 | -4.26354 | 2.01E-05 | 9.36E-04 |
| EVI5L | 1891.228 | -0.47609 | 0.137372 | -3.46569 | 5.29E-04 | 0.007021 |
| SLC47A1 | 275.7699 | -0.25194 | 0.193906 | -1.29926 | 0.193854 | 0.348951 |
| PSMB6 | 2429.677 | -0.16648 | 0.153606 | -1.08384 | 0.278435 | 0.444537 |
| TLK2 | 399.038 | 0.63155 | 0.258131 | 2.44663 | 0.01442 | 0.060448 |
| TBX6 | 36.03574 | -0.17555 | 0.400463 | -0.43838 | 0.661112 | NA |
| PPP4C | 3447.41 | -0.30925 | 0.141556 | -2.18462 | 0.028917 | 0.096756 |
| ALDOA | 33410.79 | -0.26216 | 0.124708 | -2.10218 | 0.035538 | 0.111053 |
| FAM57B | 0.739585 | 2.778432 | 2.852922 | 0.97389 | 0.330111 | NA |
| DOC2A | 0.750448 | 2.743526 | 2.504729 | 1.095338 | 0.273368 | NA |
| HIRIP3 | 464.5635 | -0.24205 | 0.17138 | -1.41237 | 0.157842 | 0.304264 |
| TAOK2 | 2753.616 | -0.4721 | 0.147343 | -3.20411 | 0.001355 | 0.012647 |
| TMEM219 | 2472.238 | -0.16643 | 0.142075 | -1.17139 | 0.241442 | 0.404507 |
| CTF1 | 32.43662 | -0.78233 | 0.38803 | -2.01617 | 0.043783 | NA |
| CDH8 | 0.890088 | -0.08229 | 2.128047 | -0.03867 | 0.969154 | NA |
| KIAA1328 | 108.7215 | 0.13583 | 0.263281 | 0.515911 | 0.605916 | 0.734172 |
| CCDC102B | 9.589508 | 0.453936 | 0.678214 | 0.669311 | 0.503297 | NA |
| CD226 | 2.682568 | 2.1703 | 1.448071 | 1.498753 | 0.133938 | NA |
| CNDP1 | 0.230334 | -0.53318 | 4.076749 | -0.13079 | 0.895945 | NA |
| PRSS53 | 5.910213 | 0.28957 | 0.8639 | 0.335189 | 0.737483 | NA |
| SETBP1 | 66.73856 | -0.21867 | 0.280563 | -0.7794 | 0.435742 | 0.59671 |
| EPG5 | 2477.114 | -0.32433 | 0.159409 | -2.03458 | 0.041893 | 0.124348 |
| PSTPIP2 | 37.30212 | -0.47538 | 0.441632 | -1.07641 | 0.281745 | NA |
| ATP5A1 | 16234.39 | -0.16161 | 0.122634 | -1.31778 | 0.187577 | 0.341795 |
| HAUS1 | 561.0262 | 0.601629 | 0.19992 | 3.009345 | 0.002618 | 0.019392 |
| C18orf25 | 599.1315 | 0.084789 | 0.196514 | 0.431466 | 0.66613 | 0.780873 |
| CNTNAP4 | 4.247695 | -0.49087 | 1.201298 | -0.40862 | 0.682822 | NA |
| CARHSP1 | 1686.229 | 0.024309 | 0.182828 | 0.13296 | 0.894225 | 0.938268 |
| TEKT5 | 0.096202 | 0.408546 | 4.079639 | 0.100143 | 0.920231 | NA |
| TXNDC11 | 1180.991 | -0.10815 | 0.170635 | -0.63381 | 0.526207 | 0.671711 |
| TRAPPC8 | 1025.477 | 0.596934 | 0.203527 | 2.932942 | 0.003358 | 0.022707 |
| INO80C | 250.9359 | -0.02181 | 0.192257 | -0.11347 | 0.90966 | 0.947766 |
| NMRAL1 | 1256.555 | -0.27035 | 0.150129 | -1.80077 | 0.071739 | 0.177587 |
| UBALD1 | 1114.991 | -0.13769 | 0.174856 | -0.78746 | 0.431011 | 0.592439 |
| C16orf89 | 7.910083 | -3.05234 | 1.171084 | -2.60643 | 0.009149 | NA |
| CFDP1 | 569.9778 | 0.354293 | 0.24787 | 1.429349 | 0.152904 | 0.297502 |
| ZDHHC7 | 2712.516 | -0.28378 | 0.128989 | -2.20005 | 0.027803 | 0.094335 |
| CMIP | 683.5238 | -0.31266 | 0.209841 | -1.48998 | 0.13623 | 0.275434 |
| KCNJ16 | 0.583919 | -1.6942 | 2.749224 | -0.61625 | 0.537732 | NA |
| CEBPG | 353.1868 | 0.686656 | 0.165383 | 4.151918 | 3.30E-05 | 0.001286 |
| KCTD15 | 2221.741 | -0.1287 | 0.148161 | -0.86868 | 0.385022 | 0.550927 |
| ZNF599 | 105.583 | -0.00042 | 0.232884 | -0.00182 | 0.99855 | 0.999371 |
| LGI4 | 41.67974 | 0.182882 | 0.39281 | 0.465574 | 0.64152 | NA |
| ANKFN1 | 0.134133 | -1.48909 | 4.077584 | -0.36519 | 0.71497 | NA |
| DGKE | 72.54098 | 1.474836 | 0.317074 | 4.651399 | 3.30E-06 | 3.28E-04 |
| MSI2 | 2088.079 | 0.397505 | 0.208317 | 1.908171 | 0.056369 | 0.150237 |
| HS3ST3A1 | 8.596956 | 0.929521 | 0.999664 | 0.929833 | 0.352457 | NA |

| | | | | | | |
|---|---|---|---|---|---|---|
| GDPD1 | 63.94401 | 0.050005 | 0.281369 | 0.177721 | 0.858942 | 0.916352 |
| GRAP | 0.163957 | 0.424913 | 4.079006 | 0.104171 | 0.917034 | NA |
| SLC5A10 | 201.7954 | -0.53 | 0.295097 | -1.79601 | 0.072493 | 0.178968 |
| C17orf103 | 786.04 | -0.43378 | 0.146706 | -2.95683 | 0.003108 | 0.021663 |
| CABYR | 64.15415 | -0.09181 | 0.295348 | -0.31084 | 0.755924 | 0.847421 |
| IMPACT | 949.9679 | 0.410436 | 0.201076 | 2.0412 | 0.041231 | 0.123029 |
| ANKRD29 | 0.980782 | 1.975754 | 2.115136 | 0.934103 | 0.350251 | NA |
| CHST9 | 73.45371 | -0.02552 | 0.317351 | -0.08041 | 0.935912 | 0.964049 |
| DNAAF1 | 12.31717 | 0.137425 | 0.681533 | 0.201641 | 0.840197 | NA |
| C16orf74 | 141.4134 | -0.31722 | 0.259895 | -1.22057 | 0.222247 | 0.382845 |
| JPH3 | 0.908875 | -0.75101 | 2.116968 | -0.35476 | 0.722771 | NA |
| PITPNC1 | 407.6551 | 0.167406 | 0.219602 | 0.762316 | 0.445871 | 0.605614 |
| CERS3 | 0.585114 | 1.12675 | 2.73598 | 0.411827 | 0.680466 | NA |
| PRKCA | 1278.877 | -0.10863 | 0.182876 | -0.59401 | 0.552502 | 0.69323 |
| LRRK1 | 1028.604 | -0.38676 | 0.169493 | -2.28186 | 0.022498 | 0.081968 |
| CEP112 | 162.214 | 1.104485 | 0.729617 | 1.513788 | 0.13008 | 0.26643 |
| ABCA9 | 11.09955 | 0.655656 | 0.639917 | 1.024594 | 0.305555 | NA |
| ABCA6 | 10.70214 | 0.014938 | 0.623922 | 0.023942 | 0.980899 | NA |
| ABCA10 | 17.27802 | 1.740975 | 0.546798 | 3.183946 | 0.001453 | NA |
| ABCA5 | 593.2477 | 0.705319 | 0.285302 | 2.472183 | 0.013429 | 0.05764 |
| PSMA8 | 3.732761 | 0.911123 | 1.202203 | 0.757878 | 0.448524 | NA |
| L3MBTL4 | 2.962017 | 1.937686 | 1.350031 | 1.43529 | 0.151204 | NA |
| SLFN13 | 9.690987 | -0.37073 | 0.702686 | -0.52759 | 0.597782 | NA |
| C17orf50 | 0.134133 | -1.48909 | 4.077584 | -0.36519 | 0.71497 | NA |
| FLCN | 740.3012 | -0.35703 | 0.137967 | -2.58777 | 0.00966 | 0.046071 |
| CXXC1 | 1538.478 | -0.10701 | 0.144073 | -0.74271 | 0.457655 | 0.616114 |
| SKA1 | 117.3551 | -0.01908 | 0.274787 | -0.06943 | 0.944651 | 0.969345 |
| PPP4R1 | 1705.667 | 0.047474 | 0.14975 | 0.317019 | 0.751229 | 0.844474 |
| APCDD1 | 290.0879 | -0.58639 | 0.228393 | -2.56744 | 0.010245 | 0.047932 |
| PIEZO2 | 0.151697 | -1.546 | 4.075469 | -0.37934 | 0.704433 | NA |
| CCDC144B | 0.847801 | -1.30616 | 2.227118 | -0.58648 | 0.557554 | NA |
| MPPE1 | 471.6716 | -0.0145 | 0.158314 | -0.09156 | 0.927048 | 0.958156 |
| CCDC144CI | 0.623191 | -1.58103 | 2.770593 | -0.57065 | 0.568239 | NA |
| USP43 | 15.77106 | 0.243552 | 0.551384 | 0.44171 | 0.658699 | NA |
| EME1 | 127.0934 | 0.014343 | 0.237186 | 0.060473 | 0.951779 | 0.97348 |
| ANKRD40 | 2374.649 | 0.050266 | 0.158464 | 0.317206 | 0.751087 | 0.844394 |
| ZNF18 | 186.7157 | -0.04651 | 0.204516 | -0.22744 | 0.820085 | 0.892334 |
| C16orf87 | 153.3101 | 0.803037 | 0.229063 | 3.505752 | 4.55E-04 | 0.00643 |
| HEATR3 | 595.8561 | -0.06815 | 0.186769 | -0.3649 | 0.715185 | 0.818469 |
| ZKSCAN2 | 158.4575 | -0.14476 | 0.228364 | -0.63391 | 0.526137 | 0.671695 |
| KDM8 | 57.44527 | -0.33293 | 0.324204 | -1.02692 | 0.30446 | 0.471751 |
| PDZD9 | 0.288605 | 1.443619 | 4.029058 | 0.358302 | 0.720117 | NA |
| OTOA | 0.326536 | 0.037056 | 3.842554 | 0.009644 | 0.992306 | NA |
| ZNF689 | 502.4719 | -0.37118 | 0.173974 | -2.13353 | 0.032881 | 0.105609 |
| PRR14 | 954.88 | -0.40319 | 0.171689 | -2.34836 | 0.018857 | 0.072603 |
| FBRS | 2273.157 | -0.59184 | 0.151717 | -3.90095 | 9.58E-05 | 0.00248 |
| PHKG2 | 565.5938 | -0.01388 | 0.194615 | -0.07131 | 0.943149 | 0.968558 |

| | | | | | | |
|---|---|---|---|---|---|---|
| COX6A2 | 0.52699 | -2.57878 | 3.248478 | -0.79384 | 0.427288 | NA |
| ITGAD | 0.134133 | -1.48909 | 4.077584 | -0.36519 | 0.71497 | NA |
| MPV17L | 0.096202 | 0.408546 | 4.079639 | 0.100143 | 0.920231 | NA |
| NTAN1 | 2406.693 | 0.099671 | 0.184964 | 0.538866 | 0.58998 | 0.721887 |
| SMG1 | 1513.908 | 0.497348 | 0.206201 | 2.411954 | 0.015867 | 0.064508 |
| TMED6 | 8.891183 | -0.04045 | 0.671642 | -0.06022 | 0.951977 | NA |
| CLEC18A | 0.151697 | -1.546 | 4.075469 | -0.37934 | 0.704433 | NA |
| DDX19B | 205.837 | -0.15931 | 0.184537 | -0.86332 | 0.387961 | 0.55373 |
| ST3GAL2 | 1555.29 | -0.42553 | 0.143328 | -2.96891 | 0.002989 | 0.021102 |
| FUK | 740.9296 | -0.36752 | 0.135726 | -2.70783 | 0.006773 | 0.036304 |
| IL34 | 29.77196 | -1.95104 | 0.61492 | -3.17284 | 0.00151 | NA |
| HYDIN | 10.6705 | -0.01181 | 0.628458 | -0.0188 | 0.985002 | NA |
| ZNF19 | 46.41791 | -0.06588 | 0.33595 | -0.1961 | 0.844534 | NA |
| SLC38A10 | 5200.114 | -0.43654 | 0.160961 | -2.71206 | 0.006687 | 0.035967 |
| ABHD3 | 335.5981 | 0.535641 | 0.188382 | 2.843378 | 0.004464 | 0.02732 |
| COLEC12 | 0.793886 | -0.43778 | 2.212663 | -0.19785 | 0.84316 | NA |
| DNAH3 | 1.116067 | -1.84515 | 2.007497 | -0.91913 | 0.358027 | NA |
| ZC3H18 | 1379.692 | -0.34763 | 0.152003 | -2.28701 | 0.022195 | 0.081338 |
| RNF166 | 386.0581 | -0.20414 | 0.158774 | -1.2857 | 0.198547 | 0.354431 |
| SPATA2L | 469.2372 | -0.17111 | 0.172543 | -0.99168 | 0.321353 | 0.489482 |
| ZNF276 | 272.6181 | -0.03744 | 0.173828 | -0.21539 | 0.82946 | 0.897194 |
| SLC5A11 | 0.563552 | -0.48645 | 2.808029 | -0.17324 | 0.862465 | NA |
| WNT9B | 0.151697 | -1.546 | 4.075469 | -0.37934 | 0.704433 | NA |
| MRPL10 | 1109.769 | -0.26386 | 0.158427 | -1.6655 | 0.095813 | 0.216643 |
| PRAC1 | 0.783004 | -0.66714 | 2.898037 | -0.2302 | 0.817934 | NA |
| HOXB13 | 501.0679 | -0.10755 | 0.201919 | -0.53266 | 0.594272 | 0.724655 |
| ATP5G1 | 1977.416 | -0.10108 | 0.133439 | -0.75749 | 0.448756 | 0.608279 |
| UBE2Z | 5140.808 | -0.34963 | 0.123395 | -2.83346 | 0.004605 | 0.027908 |
| SNF8 | 1712.052 | -0.08802 | 0.140137 | -0.62811 | 0.52993 | 0.674874 |
| IGF2BP1 | 315.4141 | -0.07524 | 0.202984 | -0.37069 | 0.710866 | 0.815056 |
| ARHGAP27 | 14.67201 | -0.43871 | 0.539244 | -0.81357 | 0.415892 | NA |
| IRX6 | 358.634 | -1.05839 | 0.297301 | -3.56002 | 3.71E-04 | 0.005601 |
| AMFR | 2745.014 | -0.24664 | 0.165569 | -1.48966 | 0.136313 | 0.275434 |
| RSPRY1 | 504.1672 | 0.185829 | 0.177225 | 1.048548 | 0.294386 | 0.461451 |
| NAE1 | 964.864 | 0.835889 | 0.178611 | 4.679932 | 2.87E-06 | 2.97E-04 |
| GPR114 | 1.224528 | -0.60718 | 1.721431 | -0.35272 | 0.724301 | NA |
| CCDC135 | 0.314073 | -0.57308 | 3.898934 | -0.14698 | 0.883146 | NA |
| ACE | 7.787013 | 0.250265 | 0.711289 | 0.351847 | 0.724953 | NA |
| TEPP | 3.785341 | 0.493848 | 1.098804 | 0.449441 | 0.653114 | NA |
| LRRC36 | 1.506403 | 1.704481 | 1.818534 | 0.937283 | 0.348613 | NA |
| TPPP3 | 14.82505 | 0.210092 | 0.575996 | 0.364745 | 0.715302 | NA |
| ZDHHC1 | 620.1331 | 0.008898 | 0.167237 | 0.053206 | 0.957568 | 0.976514 |
| ATP6V0D1 | 6241.274 | -0.36652 | 0.140421 | -2.61017 | 0.00905 | 0.044218 |
| AGRP | 1.418511 | -0.10897 | 1.758181 | -0.06198 | 0.950581 | NA |
| RLTPR | 6.447728 | -0.00332 | 0.779743 | -0.00425 | 0.996606 | NA |
| C16orf86 | 271.9525 | -0.31065 | 0.192517 | -1.6136 | 0.106614 | 0.232765 |
| PSKH1 | 2618.592 | -0.27279 | 0.15246 | -1.78924 | 0.073577 | 0.180809 |

| | | | | | | |
|---|---|---|---|---|---|---|
| ABR | 11969.1 | -0.58318 | 0.166324 | -3.50631 | 4.54E-04 | 0.00643 |
| LYPD5 | 4.855515 | 0.215408 | 0.921023 | 0.23388 | 0.815078 | NA |
| ZNF230 | 95.7004 | 0.770881 | 0.252409 | 3.054097 | 0.002257 | 0.01748 |
| ZNF221 | 30.67488 | 0.697306 | 0.488713 | 1.42682 | 0.153632 | NA |
| CPAMD8 | 3.595881 | -0.16574 | 1.059027 | -0.1565 | 0.875637 | NA |
| NR2F6 | 1738.247 | -0.24994 | 0.139408 | -1.79286 | 0.072996 | 0.179985 |
| ANKLE1 | 19.71551 | 0.477488 | 0.536415 | 0.890147 | 0.373387 | NA |
| CILP2 | 27.04253 | -0.72216 | 0.488017 | -1.47978 | 0.138931 | NA |
| ZNF66 | 6.059757 | 0.95456 | 0.915162 | 1.043051 | 0.296925 | NA |
| ZNF714 | 91.50285 | 0.521969 | 0.34566 | 1.510064 | 0.131027 | 0.267538 |
| C19orf47 | 555.7052 | -0.19854 | 0.178405 | -1.11289 | 0.265757 | 0.430423 |
| HIPK4 | 0.696104 | -0.93034 | 2.354268 | -0.39517 | 0.692715 | NA |
| SHKBP1 | 3050.317 | -0.37564 | 0.128375 | -2.92609 | 0.003433 | 0.022986 |
| SPTBN4 | 28.83067 | -0.19727 | 0.404502 | -0.48769 | 0.62577 | NA |
| TAOK1 | 2136.149 | 0.508417 | 0.253572 | 2.005024 | 0.044961 | 0.129622 |
| DEDD2 | 635.604 | 0.109075 | 0.232438 | 0.469263 | 0.638881 | 0.759385 |
| NEK8 | 81.22614 | -0.45124 | 0.268765 | -1.67892 | 0.093167 | 0.213018 |
| TLCD1 | 90.62936 | -0.40281 | 0.265778 | -1.51559 | 0.129624 | 0.266013 |
| SAFB | 2246.043 | -0.28703 | 0.156637 | -1.83245 | 0.066885 | 0.169066 |
| NACC1 | 2124.184 | -0.5443 | 0.160019 | -3.40146 | 6.70E-04 | 0.008081 |
| IER2 | 531.212 | -0.02753 | 0.198416 | -0.13877 | 0.889632 | 0.935478 |
| PTGER1 | 9.838178 | 0.243782 | 0.692766 | 0.351897 | 0.724916 | NA |
| MUM1 | 1687.858 | -0.29481 | 0.137436 | -2.14506 | 0.031948 | 0.103652 |
| ZNF333 | 234.6453 | -0.01544 | 0.222821 | -0.06931 | 0.944746 | 0.969358 |
| CELF5 | 4.381652 | -0.42709 | 0.926823 | -0.46081 | 0.644934 | NA |
| MFSD12 | 69360.31 | -0.15409 | 0.135472 | -1.1374 | 0.255371 | 0.419543 |
| FBXO27 | 121.0576 | -0.4193 | 0.247691 | -1.69284 | 0.090486 | 0.208495 |
| DMKN | 154.2912 | -0.35179 | 0.218628 | -1.60908 | 0.107599 | 0.234081 |
| U2AF1L4 | 45.45618 | 0.260919 | 0.375211 | 0.695391 | 0.48681 | NA |
| NPHS1 | 0.162376 | 0.424913 | 4.079006 | 0.104171 | 0.917034 | NA |
| THAP8 | 286.7964 | -0.30536 | 0.17625 | -1.73255 | 0.083176 | 0.196347 |
| COX7A1 | 133.3179 | 0.443323 | 0.325948 | 1.360101 | 0.173798 | 0.325802 |
| ZNF382 | 32.7445 | 0.25464 | 0.405704 | 0.62765 | 0.530233 | NA |
| DUSP14 | 1510.04 | 0.349457 | 0.212291 | 1.646123 | 0.099738 | 0.223094 |
| PLXDC1 | 3.184229 | -0.49504 | 1.088395 | -0.45483 | 0.64923 | NA |
| PGAP3 | 408.8978 | -0.30715 | 0.166929 | -1.83999 | 0.06577 | 0.167212 |
| GRIN2C | 4.272038 | 0.110166 | 0.940685 | 0.117113 | 0.90677 | NA |
| FDXR | 867.5706 | -0.3461 | 0.169002 | -2.04793 | 0.040567 | 0.121415 |
| SAP30BP | 1709.739 | 5.26E-05 | 0.160163 | 3.28E-04 | 0.999738 | 0.999738 |
| ACOX1 | 1519.618 | -0.26945 | 0.147706 | -1.82424 | 0.068116 | 0.17123 |
| PRPSAP1 | 998.2354 | -0.03504 | 0.147788 | -0.23708 | 0.812594 | 0.888591 |
| CYGB | 25347.38 | -0.28292 | 0.191317 | -1.47878 | 0.1392 | 0.279177 |
| SRSF2 | 2031.313 | -3.64E-04 | 0.153221 | -0.00237 | 0.998106 | 0.999318 |
| CCL5 | 60.62018 | -0.09349 | 0.34812 | -0.26857 | 0.788263 | 0.871434 |
| CCL16 | 0.163957 | 0.424913 | 4.079006 | 0.104171 | 0.917034 | NA |
| KLHL10 | 0.720624 | 1.435938 | 2.334155 | 0.615185 | 0.538432 | NA |
| HCRT | 0.096202 | 0.408546 | 4.079639 | 0.100143 | 0.920231 | NA |

| | | | | | | |
|---|---|---|---|---|---|---|
| MPP3 | 72.33945 | 0.143155 | 0.270038 | 0.530129 | 0.596023 | 0.726065 |
| NAGS | 68.84413 | -0.04659 | 0.322503 | -0.14447 | 0.885128 | 0.932072 |
| LSM12 | 123.2811 | -0.14766 | 0.231075 | -0.63903 | 0.522806 | 0.669027 |
| ASB16 | 5.704136 | -1.11004 | 0.852704 | -1.30178 | 0.192991 | NA |
| FAM171A2 | 241.5603 | -0.11742 | 0.17438 | -0.67335 | 0.500725 | 0.651245 |
| DBF4B | 408.5861 | -0.42503 | 0.212518 | -1.99995 | 0.045506 | 0.13062 |
| PLCD3 | 1756.951 | -0.16779 | 0.14306 | -1.17288 | 0.240842 | 0.404073 |
| OR7G1 | 0.260158 | 1.240347 | 4.047124 | 0.306476 | 0.759242 | NA |
| RAVER1 | 738.578 | -0.63534 | 0.178044 | -3.56844 | 3.59E-04 | 0.005535 |
| SYCE2 | 3.612285 | 0.108506 | 1.055075 | 0.102842 | 0.918088 | NA |
| SPC24 | 382.0926 | -0.66429 | 0.292797 | -2.26876 | 0.023283 | 0.08387 |
| ALOX15 | 0.163957 | 0.424913 | 4.079006 | 0.104171 | 0.917034 | NA |
| ZNF653 | 139.5539 | 0.222741 | 0.23497 | 0.947953 | 0.343154 | 0.511307 |
| MED11 | 291.7345 | 0.080658 | 0.173989 | 0.463581 | 0.642948 | 0.762444 |
| CXCL16 | 63.7131 | 0.252424 | 0.299487 | 0.842855 | 0.39931 | 0.563892 |
| SCIMP | 0.774539 | 0.39797 | 2.296349 | 0.173305 | 0.862411 | NA |
| RNASEK-C1 | 17.8458 | 0.551052 | 0.489391 | 1.125996 | 0.260167 | NA |
| BCL6B | 1.393935 | -2.09637 | 1.832113 | -1.14424 | 0.252526 | NA |
| TNFSF13 | 63.55021 | -0.2785 | 0.292227 | -0.95301 | 0.340583 | 0.5087 |
| SENP3 | 779.9757 | -0.19062 | 0.170809 | -1.11598 | 0.264432 | 0.428902 |
| FGF11 | 10.42709 | 0.231059 | 0.657958 | 0.351176 | 0.725456 | NA |
| EIF4A1 | 51.55799 | 0.061687 | 0.331669 | 0.18599 | 0.852452 | 0.912243 |
| RPL26 | 3023.107 | 0.593228 | 0.132172 | 4.488318 | 7.18E-06 | 5.25E-04 |
| POLR3K | 573.2507 | 0.067634 | 0.204734 | 0.330352 | 0.741134 | 0.836776 |
| SNRNP25 | 1158.607 | 0.002721 | 0.156084 | 0.017433 | 0.986091 | 0.992622 |
| WDR90 | 289.5006 | -0.41451 | 0.191754 | -2.16165 | 0.030645 | 0.100548 |
| JMJD8 | 959.8958 | -0.27879 | 0.227089 | -1.22769 | 0.219565 | 0.379257 |
| CCDC78 | 48.25427 | 0.176942 | 0.310991 | 0.568961 | 0.569382 | 0.705805 |
| SPSB3 | 4.32768 | -0.36301 | 0.966051 | -0.37577 | 0.707089 | NA |
| MEIOB | 1.653947 | -0.31892 | 1.531024 | -0.20831 | 0.834989 | NA |
| HS3ST6 | 0.260158 | 1.240347 | 4.047124 | 0.306476 | 0.759242 | NA |
| C16orf59 | 199.1988 | -0.3638 | 0.245248 | -1.48338 | 0.137973 | 0.277326 |
| CCNF | 497.957 | -0.57832 | 0.204319 | -2.83046 | 0.004648 | 0.028075 |
| TBC1D24 | 401.0178 | -0.34548 | 0.198791 | -1.73792 | 0.082224 | 0.194875 |
| AMDHD2 | 1173.379 | -0.16207 | 0.163886 | -0.98895 | 0.32269 | 0.490721 |
| NTN3 | 2.015828 | 0.661606 | 1.414901 | 0.467599 | 0.640072 | NA |
| CCDC64B | 0.151697 | -1.546 | 4.075469 | -0.37934 | 0.704433 | NA |
| PAQR4 | 527.9352 | -0.03697 | 0.249509 | -0.14819 | 0.882194 | 0.930128 |
| FLYWCH2 | 696.4659 | -0.1994 | 0.143806 | -1.3866 | 0.165565 | 0.314604 |
| ZG16B | 0.151697 | -1.546 | 4.075469 | -0.37934 | 0.704433 | NA |
| ZNF75A | 187.2889 | 0.452641 | 0.206423 | 2.192781 | 0.028323 | 0.095362 |
| ADCY9 | 3770.049 | -0.6226 | 0.195557 | -3.18374 | 0.001454 | 0.013322 |
| SNHG16 | 1503.142 | -0.15524 | 0.190213 | -0.81613 | 0.414426 | 0.57773 |
| SLC35G3 | 0.151697 | -1.546 | 4.075469 | -0.37934 | 0.704433 | NA |
| GPT2 | 273.8453 | -0.05512 | 0.183619 | -0.30021 | 0.764016 | 0.853667 |
| BRD7 | 704.6696 | 0.458344 | 0.18925 | 2.421896 | 0.01544 | 0.063199 |
| ZNF319 | 342.8351 | -0.41463 | 0.17097 | -2.42515 | 0.015302 | 0.062857 |

| | | | | | | |
|---|---|---|---|---|---|---|
| C16orf71 | 15.77033 | 0.211595 | 0.560921 | 0.377227 | 0.706005 | NA |
| COX11 | 856.442 | 0.225642 | 0.14768 | 1.527909 | 0.126535 | 0.261839 |
| STXBP4 | 184.5479 | 0.431585 | 0.211384 | 2.041708 | 0.041181 | 0.122909 |
| PLEKHF1 | 476.6063 | -0.06655 | 0.172446 | -0.38593 | 0.699551 | 0.806494 |
| TMEM100 | 0.516164 | -1.70268 | 2.844782 | -0.59853 | 0.549488 | NA |
| CYB5A | 995.2898 | 0.274795 | 0.13905 | 1.976229 | 0.048129 | 0.135397 |
| WDR88 | 3.589549 | -0.62427 | 1.024673 | -0.60924 | 0.542366 | NA |
| ATP9B | 502.9259 | -0.28282 | 0.16184 | -1.74756 | 0.080541 | 0.192192 |
| KIAA0355 | 617.7343 | -0.07919 | 0.173442 | -0.45661 | 0.647952 | 0.766251 |
| SERPINB8 | 443.1379 | 0.40572 | 0.206312 | 1.966533 | 0.049237 | 0.13727 |
| CDYL2 | 64.84865 | -0.17627 | 0.300711 | -0.58617 | 0.557764 | 0.69668 |
| CENPN | 732.6256 | -0.06174 | 0.177825 | -0.3472 | 0.728442 | 0.827728 |
| ATMIN | 1298.556 | 0.038283 | 0.171759 | 0.22289 | 0.823621 | 0.893657 |
| C16orf46 | 22.46713 | -0.78451 | 0.446403 | -1.7574 | 0.07885 | NA |
| PKD1L2 | 6.489683 | -0.23053 | 0.7899 | -0.29185 | 0.770401 | NA |
| TMX3 | 947.5539 | 1.001996 | 0.215766 | 4.643902 | 3.42E-06 | 3.37E-04 |
| MFAP4 | 3.260346 | 1.263551 | 1.150494 | 1.098269 | 0.272087 | NA |
| MAPK7 | 797.9642 | -0.44271 | 0.225749 | -1.96109 | 0.049869 | 0.138259 |
| PRKCB | 2447.499 | 0.007161 | 0.148719 | 0.048152 | 0.961595 | 0.979188 |
| CCDC68 | 2.698724 | 0.94992 | 1.28577 | 0.738794 | 0.460032 | NA |
| BEAN1 | 92.56035 | -0.48752 | 0.319875 | -1.52411 | 0.127481 | 0.263336 |
| TK2 | 1078.554 | -0.39803 | 0.141458 | -2.81379 | 0.004896 | 0.029107 |
| SLC38A8 | 89.53987 | -1.15837 | 1.09831 | -1.05468 | 0.291571 | 0.458432 |
| SEC11C | 1626.158 | 0.485557 | 0.168643 | 2.879192 | 0.003987 | 0.025425 |
| CPLX4 | 0.096202 | 0.408546 | 4.079639 | 0.100143 | 0.920231 | NA |
| NDEL1 | 1204.915 | 0.274606 | 0.164249 | 1.671885 | 0.094547 | 0.215011 |
| CENPV | 33.50242 | 0.116065 | 0.417681 | 0.27788 | 0.781104 | NA |
| CDH16 | 0.163957 | 0.424913 | 4.079006 | 0.104171 | 0.917034 | NA |
| RRAD | 0.392711 | 0.292538 | 3.145221 | 0.09301 | 0.925895 | NA |
| FAM96B | 2088.053 | -0.13247 | 0.170223 | -0.77821 | 0.436444 | 0.597188 |
| WDR16 | 1.51462 | -0.4102 | 2.147409 | -0.19102 | 0.848508 | NA |
| MC4R | 13.10437 | -0.41273 | 0.576099 | -0.71643 | 0.473726 | NA |
| ATF7IP2 | 183.0002 | 0.369995 | 0.253145 | 1.461592 | 0.143853 | 0.285467 |
| TVP23A | 4.456448 | 0.747356 | 1.02381 | 0.729975 | 0.465405 | NA |
| COG1 | 793.4136 | -0.43271 | 0.143553 | -3.01426 | 0.002576 | 0.019243 |
| AP1G1 | 2311.428 | -0.08615 | 0.134486 | -0.64062 | 0.521772 | 0.668425 |
| SLFN5 | 3015.301 | 0.469644 | 0.285566 | 1.644605 | 0.100051 | 0.223218 |
| C16orf45 | 313.7764 | -0.04212 | 0.212741 | -0.19801 | 0.84304 | 0.906755 |
| KIAA0430 | 2219.943 | -0.27565 | 0.156383 | -1.76269 | 0.077953 | 0.188086 |
| LDHD | 172.4653 | -0.58882 | 0.210451 | -2.7979 | 0.005144 | 0.030142 |
| TMEM170/ | 212.8614 | 0.136179 | 0.181854 | 0.748839 | 0.453955 | 0.612733 |
| C18orf54 | 181.6925 | 0.469137 | 0.260471 | 1.801108 | 0.071686 | 0.17753 |
| DCTN5 | 2737.858 | -0.15716 | 0.170274 | -0.92296 | 0.356026 | 0.523878 |
| TERF2IP | 2005.351 | 0.097995 | 0.155508 | 0.630161 | 0.528589 | 0.673809 |
| PLK1 | 1447.79 | -0.6326 | 0.279446 | -2.26377 | 0.023588 | 0.084611 |
| CHP2 | 0.394291 | 0.298745 | 3.141711 | 0.09509 | 0.924243 | NA |
| CCDC178 | 0.682693 | 2.623213 | 2.518338 | 1.041645 | 0.297576 | NA |

| | | | | | | |
|---|---|---|---|---|---|---|
| RCCD1 | 842.024 | -0.24396 | 0.175471 | -1.39029 | 0.164441 | 0.313265 |
| AKTIP | 159.209 | 0.233241 | 0.210005 | 1.110649 | 0.266719 | 0.431639 |
| MAPRE2 | 1510.817 | -0.0522 | 0.14561 | -0.35852 | 0.719955 | 0.821628 |
| NUDT21 | 1951.839 | 0.272618 | 0.171335 | 1.59114 | 0.111578 | 0.239791 |
| B4GALNT2 | 0.096202 | 0.408546 | 4.079639 | 0.100143 | 0.920231 | NA |
| PHB | 4581.261 | -0.14044 | 0.163361 | -0.85967 | 0.389968 | 0.555391 |
| SNRPD1 | 1135.088 | 0.343497 | 0.16735 | 2.052566 | 0.040115 | 0.120643 |
| SAMD14 | 93.98675 | -0.16078 | 0.241536 | -0.66567 | 0.505621 | 0.654811 |
| TMEM92 | 0.096202 | 0.408546 | 4.079639 | 0.100143 | 0.920231 | NA |
| ACSF2 | 2704.897 | -0.19198 | 0.191654 | -1.00172 | 0.316478 | 0.483985 |
| LINC00483 | 0.096202 | 0.408546 | 4.079639 | 0.100143 | 0.920231 | NA |
| SP2 | 725.8871 | -0.36482 | 0.169024 | -2.15838 | 0.030898 | 0.10116 |
| PRR15L | 2.064156 | -1.93642 | 1.516065 | -1.27727 | 0.201509 | NA |
| COQ7 | 415.3415 | 0.262263 | 0.169681 | 1.545624 | 0.122195 | 0.255544 |
| GPRC5B | 268.4399 | -0.06005 | 0.291824 | -0.20578 | 0.836965 | 0.902391 |
| CRK | 2277.504 | 0.197798 | 0.165488 | 1.195242 | 0.231992 | 0.394248 |
| NOD2 | 11.32747 | -0.97757 | 0.658117 | -1.48541 | 0.137436 | NA |
| SNX20 | 0.452561 | 2.061505 | 3.349456 | 0.615474 | 0.538241 | NA |
| KATNAL2 | 50.90407 | 0.54661 | 0.307193 | 1.779369 | 0.075179 | 0.183083 |
| HDHD2 | 518.4354 | 0.157047 | 0.198134 | 0.792633 | 0.427992 | 0.589776 |
| ZNF91 | 73.82684 | 1.17967 | 0.510497 | 2.310827 | 0.020842 | 0.077896 |
| CCL23 | 1.078282 | -2.71064 | 2.113412 | -1.28259 | 0.199636 | NA |
| CDK12 | 1403.609 | -0.08002 | 0.149563 | -0.53505 | 0.592612 | 0.7233 |
| DPEP2 | 0.163957 | 0.424913 | 4.079006 | 0.104171 | 0.917034 | NA |
| DUS2 | 275.4532 | -0.39142 | 0.18267 | -2.14276 | 0.032132 | 0.103994 |
| ENGASE | 542.5059 | 0.001092 | 0.195786 | 0.005576 | 0.995551 | 0.997908 |
| RBFOX3 | 0.584911 | 2.38756 | 2.690807 | 0.887303 | 0.374916 | NA |
| TBC1D16 | 71296.3 | -0.52462 | 0.16305 | -3.21757 | 0.001293 | 0.012237 |
| ENTHD2 | 195.9362 | 0.118062 | 0.22855 | 0.516567 | 0.605459 | 0.733992 |
| MYO5B | 1.996495 | -1.46063 | 1.495385 | -0.97676 | 0.328688 | NA |
| ACAA2 | 1551.361 | 0.154729 | 0.122614 | 1.261916 | 0.206979 | 0.36412 |
| FN3K | 173.8269 | -0.52703 | 0.310774 | -1.69587 | 0.089911 | 0.207626 |
| PRRT2 | 26.70055 | -0.333 | 0.427678 | -0.77861 | 0.436207 | NA |
| ZNF23 | 8.413513 | 0.51893 | 0.768666 | 0.675105 | 0.499609 | NA |
| IRGQ | 692.1798 | -0.54382 | 0.154066 | -3.52981 | 4.16E-04 | 0.006094 |
| ZNF668 | 335.519 | -0.37004 | 0.18062 | -2.04873 | 0.040488 | 0.121331 |
| ZNF646 | 831.8007 | -0.51252 | 0.162376 | -3.1564 | 0.001597 | 0.014121 |
| VKORC1 | 571.4335 | -0.0221 | 0.171435 | -0.12891 | 0.897426 | 0.939815 |
| LPO | 0.613743 | -0.17004 | 2.749758 | -0.06184 | 0.950691 | NA |
| SMG8 | 759.0731 | -0.2016 | 0.155164 | -1.29925 | 0.193859 | 0.348951 |
| TPM4 | 19572.47 | 0.038694 | 0.172548 | 0.224251 | 0.822562 | 0.893074 |
| RAB8A | 1079.55 | -0.34683 | 0.171419 | -2.0233 | 0.043043 | 0.126367 |
| GPX4 | 6461.865 | -0.07222 | 0.177935 | -0.4059 | 0.684813 | 0.794992 |
| MIDN | 612.864 | -0.64455 | 0.207023 | -3.11343 | 0.001849 | 0.01532 |
| JSRP1 | 0.690946 | -1.56103 | 2.699004 | -0.57837 | 0.563013 | NA |
| FAM129C | 2.107674 | -0.14896 | 1.323619 | -0.11254 | 0.910393 | NA |
| KLHL26 | 140.9337 | -0.65776 | 0.266103 | -2.47184 | 0.013442 | 0.057654 |

| | | | | | | |
|---|---|---|---|---|---|---|
| GATAD2A | 4695.401 | -0.33962 | 0.161191 | -2.10692 | 0.035125 | 0.110159 |
| MVD | 1985.003 | -0.24245 | 0.157336 | -1.54098 | 0.123322 | 0.257437 |
| CDT1 | 360.912 | -0.41261 | 0.239327 | -1.72404 | 0.0847 | 0.198364 |
| TRAPPC2L | 1674.027 | 0.109873 | 0.135446 | 0.811195 | 0.417254 | 0.580308 |
| ANKRD11 | 3076.819 | 0.117214 | 0.198652 | 0.590046 | 0.55516 | 0.695073 |
| SPATA33 | 176.4841 | 0.167462 | 0.228516 | 0.732826 | 0.463665 | 0.621241 |
| | 2.828658 | 0.246865 | 1.306914 | 0.188892 | 0.850178 | NA |
| PROCA1 | 24.21066 | 0.482502 | 0.508028 | 0.949753 | 0.342238 | NA |
| RPL13 | 29574.98 | -0.0052 | 0.144205 | -0.03609 | 0.971212 | 0.984247 |
| DHRS13 | 116.1193 | -0.71643 | 0.259414 | -2.76174 | 0.005749 | 0.032518 |
| TP53I13 | 1391.227 | -0.1088 | 0.126474 | -0.86024 | 0.389655 | 0.555087 |
| CORO6 | 41.43218 | 0.19611 | 0.360471 | 0.544037 | 0.586416 | NA |
| SERTAD3 | 602.9621 | -0.14491 | 0.158279 | -0.91556 | 0.3599 | 0.527003 |
| RAB4B | 238.086 | -0.16988 | 0.183877 | -0.92389 | 0.355545 | 0.523732 |
| C19orf55 | 161.4982 | -0.82263 | 0.244438 | -3.36539 | 7.64E-04 | 0.00881 |
| AXL | 750.1144 | -0.4494 | 0.199508 | -2.25252 | 0.02429 | 0.086239 |
| NFKBID | 63.77454 | -0.18641 | 0.291038 | -0.64048 | 0.521857 | 0.668462 |
| TMEM145 | 21.24412 | -0.83146 | 0.468617 | -1.77429 | 0.076015 | NA |
| ZNF526 | 534.3336 | -0.43474 | 0.165007 | -2.63468 | 0.008422 | 0.041958 |
| ZNF146 | 707.8174 | 0.724613 | 0.257073 | 2.818709 | 0.004822 | 0.028826 |
| ZNF283 | 90.47926 | 0.571671 | 0.270952 | 2.109859 | 0.03487 | 0.109623 |
| PPP1R14A | 12.04344 | -0.20984 | 0.613326 | -0.34214 | 0.732244 | NA |
| SPINT2 | 0.997115 | 0.104049 | 2.1472 | 0.048458 | 0.961351 | NA |
| C19orf33 | 1.099787 | -2.48189 | 2.397845 | -1.03505 | 0.300646 | NA |
| YIF1B | 2749.909 | -0.14237 | 0.124737 | -1.14137 | 0.253716 | 0.417799 |
| ATCAY | 0.260158 | 1.240347 | 4.047124 | 0.306476 | 0.759242 | NA |
| DAPK3 | 3025.812 | -0.05144 | 0.191368 | -0.2688 | 0.788084 | 0.871362 |
| EEF2 | 84952.05 | -0.41014 | 0.134132 | -3.05776 | 0.00223 | 0.017335 |
| TMIGD2 | 0.823352 | -0.17011 | 2.472341 | -0.06881 | 0.945144 | NA |
| CHAF1A | 747.369 | -0.20368 | 0.198745 | -1.02481 | 0.305451 | 0.472607 |
| UBXN6 | 4961.52 | -0.17043 | 0.118482 | -1.43842 | 0.150315 | 0.294008 |
| | 1815.737 | -0.40762 | 0.130659 | -3.11972 | 0.00181 | 0.015146 |
| PLIN4 | 2.633238 | -0.0914 | 1.465737 | -0.06236 | 0.950275 | NA |
| SEMA6B | 22.91647 | -0.03441 | 0.609294 | -0.05647 | 0.954964 | NA |
| NXN | 490.0246 | -0.36518 | 0.172595 | -2.1158 | 0.034362 | 0.108572 |
| FAM57A | 830.4154 | -0.22501 | 0.137445 | -1.6371 | 0.101609 | 0.225279 |
| GLOD4 | 1552.325 | 0.110569 | 0.125203 | 0.883115 | 0.377174 | 0.543833 |
| SLC43A2 | 36.42524 | -0.09877 | 0.370037 | -0.26693 | 0.789523 | NA |
| RILP | 332.6132 | -0.0227 | 0.16003 | -0.14182 | 0.887224 | 0.933443 |
| SERPINF2 | 0.957843 | 0.061519 | 2.068681 | 0.029738 | 0.976276 | NA |
| WDR81 | 4507.488 | -0.6095 | 0.174313 | -3.4966 | 4.71E-04 | 0.006572 |
| SRR | 243.8408 | -0.11299 | 0.205069 | -0.55098 | 0.581646 | 0.715217 |
| TSR1 | 1160.589 | 0.119663 | 0.135831 | 0.88097 | 0.378334 | 0.544645 |
| TRPV3 | 111.8742 | 0.048311 | 0.302563 | 0.159672 | 0.87314 | 0.92445 |
| HSD11B1L | 285.183 | 0.135696 | 0.168159 | 0.806949 | 0.419696 | 0.582747 |
| CYB5D2 | 892.2802 | -0.15444 | 0.163814 | -0.94278 | 0.345791 | 0.51364 |
| ACER1 | 0.163957 | 0.424913 | 4.079006 | 0.104171 | 0.917034 | NA |

| | | | | | | |
|---|---|---|---|---|---|---|
| ANGPTL4 | 247.9567 | 1.560416 | 0.675828 | 2.308895 | 0.020949 | 0.078197 |
| | 0.46735 | -1.30193 | 3.399671 | -0.38296 | 0.701751 | NA |
| CD320 | 2546.266 | -0.2663 | 0.140589 | -1.89417 | 0.058202 | 0.153798 |
| ZNF558 | 171.1278 | 0.277964 | 0.245276 | 1.133269 | 0.257101 | 0.42116 |
| C3P1 | 0.163957 | 0.424913 | 4.079006 | 0.104171 | 0.917034 | NA |
| PRDX2 | 4811.311 | -0.04571 | 0.136055 | -0.33596 | 0.736902 | 0.833267 |
| ZNF232 | 149.6398 | 0.25306 | 0.208817 | 1.211878 | 0.225559 | 0.386778 |
| MIS12 | 392.5459 | 0.399034 | 0.168135 | 2.373289 | 0.01763 | 0.069469 |
| CD300C | 0.096202 | 0.408546 | 4.079639 | 0.100143 | 0.920231 | NA |
| HID1 | 9.794912 | 0.785744 | 0.80308 | 0.978413 | 0.32787 | NA |
| ICT1 | 741.5716 | 0.23169 | 0.165476 | 1.400143 | 0.16147 | 0.309069 |
| ATP5H | 3004.755 | 0.36666 | 0.170416 | 2.15156 | 0.031432 | 0.102411 |
| TMEM88 | 3.926707 | -0.41365 | 1.00175 | -0.41293 | 0.679658 | NA |
| EVPL | 0.163957 | 0.424913 | 4.079006 | 0.104171 | 0.917034 | NA |
| SRP68 | 2104.946 | -0.16326 | 0.141103 | -1.15702 | 0.247263 | 0.410746 |
| MGAT5B | 1108.957 | -1.02805 | 0.216962 | -4.7384 | 2.15E-06 | 2.63E-04 |
| TMC8 | 8.350422 | -1.15187 | 0.734767 | -1.56767 | 0.116959 | NA |
| TK1 | 1424.216 | -0.43318 | 0.192548 | -2.24974 | 0.024465 | 0.08666 |
| GSDMA | 0.134133 | -1.48909 | 4.077584 | -0.36519 | 0.71497 | NA |
| TMEM99 | 342.0168 | -0.23143 | 0.172479 | -1.34176 | 0.179673 | 0.332225 |
| GHDC | 1582.97 | -0.36898 | 0.159873 | -2.30795 | 0.021002 | 0.078273 |
| ITFG3 | 1640.5 | -0.4334 | 0.141731 | -3.05792 | 0.002229 | 0.017335 |
| PRR25 | 1.743613 | -0.66006 | 1.560343 | -0.42302 | 0.67228 | NA |
| ZNF598 | 1374.817 | -0.31833 | 0.170276 | -1.8695 | 0.061553 | 0.160095 |
| RAB26 | 22.8114 | 0.20335 | 0.464993 | 0.437319 | 0.66188 | NA |
| MLST8 | 2430.283 | -0.13235 | 0.15384 | -0.8603 | 0.389623 | 0.555087 |
| E4F1 | 621.9934 | -0.1432 | 0.164192 | -0.87215 | 0.383129 | 0.549014 |
| DNASE1L2 | 6.533516 | -0.21659 | 0.842943 | -0.25694 | 0.797224 | NA |
| ECI1 | 683.5386 | -0.24159 | 0.180061 | -1.34171 | 0.17969 | 0.332225 |
| CASKIN1 | 13.31733 | -0.17511 | 0.648778 | -0.26991 | 0.78723 | NA |
| ABCA3 | 32.96932 | 0.385645 | 0.558165 | 0.690916 | 0.489618 | NA |
| KCTD5 | 1050.532 | -0.11493 | 0.164107 | -0.70036 | 0.483701 | 0.637354 |
| SRRM2 | 5478.045 | -0.57561 | 0.220607 | -2.6092 | 0.009075 | 0.044294 |
| ZNF597 | 41.01763 | 0.103047 | 0.355851 | 0.28958 | 0.772138 | NA |
| NLRC3 | 9.338266 | 0.603765 | 0.671017 | 0.899775 | 0.36824 | NA |
| ANKS3 | 409.6063 | 0.011861 | 0.206342 | 0.057483 | 0.95416 | 0.974799 |
| NUDT16L1 | 956.9019 | -0.14342 | 0.164032 | -0.87435 | 0.381925 | 0.548084 |
| OR1F1 | 1.212268 | -1.34711 | 1.888238 | -0.71342 | 0.475586 | NA |
| VASN | 82.61998 | -0.70016 | 0.300339 | -2.33124 | 0.019741 | 0.075032 |
| OR2C1 | 0.392857 | -1.546 | 4.075469 | -0.37934 | 0.704433 | NA |
| TTC39C | 254.6647 | 0.059071 | 0.277538 | 0.212839 | 0.831452 | 0.89868 |
| NKIRAS2 | 834.1 | -0.28031 | 0.137785 | -2.03438 | 0.041914 | 0.124377 |
| DNAJC7 | 941.4004 | 0.5546 | 0.18231 | 3.042076 | 0.00235 | 0.017947 |
| THAP11 | 880.1578 | -0.06669 | 0.173757 | -0.3838 | 0.701129 | 0.807895 |
| MLKL | 171.3255 | -0.11907 | 0.264658 | -0.4499 | 0.652786 | 0.770411 |
| RFWD3 | 1058.348 | -0.26118 | 0.191528 | -1.36369 | 0.172666 | 0.324032 |
| COG7 | 369.2623 | -0.29985 | 0.172683 | -1.73644 | 0.082485 | 0.195273 |

| | | | | | | |
|---|---|---|---|---|---|---|
| TXNDC2 | 0.641986 | 1.117767 | 2.709494 | 0.412537 | 0.679946 | NA |
| RAB31 | 1816.99 | 0.450453 | 0.238264 | 1.890559 | 0.058683 | 0.154802 |
| ATXN2L | 2653.659 | -0.50238 | 0.157237 | -3.19505 | 0.001398 | 0.012949 |
| SOGA2 | 562.4007 | -0.46153 | 0.193323 | -2.38737 | 0.01697 | 0.067758 |
| HEXIM2 | 75.41388 | -0.21307 | 0.273555 | -0.7789 | 0.436036 | 0.596949 |
| DYNLRB2 | 1.670148 | 0.750303 | 1.621208 | 0.462805 | 0.643504 | NA |
| TMUB2 | 1269.399 | -0.22745 | 0.175347 | -1.29714 | 0.194584 | 0.349673 |
| STAT3 | 3377.884 | -0.4354 | 0.119169 | -3.65362 | 2.59E-04 | 0.004545 |
| AXIN2 | 160.3611 | -0.13921 | 0.197682 | -0.70422 | 0.481295 | 0.635739 |
| ZNF30 | 63.73729 | 0.327709 | 0.307404 | 1.066056 | 0.286398 | 0.4527 |
| LDLRAD4 | 1.720268 | -0.9865 | 1.532984 | -0.64351 | 0.51989 | NA |
| KCTD19 | 1.590133 | 0.039996 | 1.513994 | 0.026417 | 0.978925 | NA |
| TMEM208 | 1297.894 | 0.042332 | 0.149146 | 0.28383 | 0.77654 | 0.862136 |
| CA7 | 0.586491 | 2.39072 | 2.689057 | 0.889055 | 0.373973 | NA |
| ABHD15 | 432.877 | -0.25799 | 0.181001 | -1.42537 | 0.15405 | 0.299094 |
| CHTF8 | 1019.33 | -0.46956 | 0.156979 | -2.99124 | 0.002778 | 0.020158 |
| SNTB2 | 1044.049 | -0.05513 | 0.154236 | -0.35742 | 0.72078 | 0.82241 |
| ZNF507 | 372.1255 | 0.46339 | 0.23487 | 1.972968 | 0.048499 | 0.135924 |
| DDX19A | 1245.365 | -0.03514 | 0.132635 | -0.26494 | 0.791052 | 0.873258 |
| LRRC28 | 491.2435 | -0.08014 | 0.164412 | -0.48746 | 0.625933 | 0.749009 |
| CTRB2 | 0.556813 | -0.57007 | 3.294615 | -0.17303 | 0.862627 | NA |
| LGALS9 | 8.08693 | -2.81941 | 0.972111 | -2.90029 | 0.003728 | NA |
| UQCRFS1 | 1946.712 | -0.16573 | 0.14425 | -1.14891 | 0.250594 | 0.414278 |
| XPO6 | 3909.113 | -0.37981 | 0.122303 | -3.10543 | 0.0019 | 0.015577 |
| GSG1L | 95.08646 | -0.45371 | 0.295271 | -1.53659 | 0.124394 | 0.258991 |
| NSMCE1 | 1170.963 | -0.1552 | 0.133563 | -1.16201 | 0.245231 | 0.408145 |
| | 0.230334 | -0.53318 | 4.076749 | -0.13079 | 0.895945 | NA |
| CD2BP2 | 2929.779 | -0.19394 | 0.156519 | -1.2391 | 0.215309 | 0.374429 |
| TBC1D10B | 2471.594 | -0.33768 | 0.167162 | -2.02007 | 0.043376 | 0.126906 |
| NPIPB3 | 76.59327 | 0.476049 | 0.381606 | 1.247487 | 0.212219 | 0.370797 |
| PDILT | 0.424115 | 1.932531 | 3.45462 | 0.559405 | 0.575885 | NA |
| UMOD | 2.601125 | 0.424042 | 1.33029 | 0.318759 | 0.74991 | NA |
| GP2 | 27.1222 | -0.4636 | 0.723558 | -0.64072 | 0.521703 | NA |
| INO80E | 1302.782 | 0.004777 | 0.151624 | 0.031505 | 0.974867 | 0.98618 |
| HEXDC | 270.5198 | -0.18508 | 0.199064 | -0.92975 | 0.352499 | 0.520437 |
| SPNS1 | 19.3737 | -0.13602 | 0.494712 | -0.27495 | 0.783353 | NA |
| LRRC45 | 823.3918 | -0.21849 | 0.160432 | -1.3619 | 0.17323 | 0.324944 |
| STRA13 | 1397.221 | 0.034527 | 0.179398 | 0.192461 | 0.847381 | 0.909114 |
| ASPSCR1 | 824.2025 | -0.38203 | 0.173619 | -2.20039 | 0.027779 | 0.094335 |
| FASN | 12257.73 | -0.82336 | 0.180808 | -4.55378 | 5.27E-06 | 4.28E-04 |
| MT1E | 119.1389 | -0.4644 | 0.349603 | -1.32838 | 0.184054 | 0.337295 |
| DUS1L | 2398.84 | -0.03729 | 0.139274 | -0.26774 | 0.788903 | 0.87186 |
| GPS1 | 3799.033 | -0.24692 | 0.121567 | -2.03117 | 0.042238 | 0.125025 |
| RFNG | 1022.089 | -0.20918 | 0.176863 | -1.18274 | 0.236911 | 0.399741 |
| DCXR | 1632.254 | -0.16615 | 0.156148 | -1.06405 | 0.287305 | 0.453569 |
| RAC3 | 563.8368 | 0.166993 | 0.202767 | 0.82357 | 0.410184 | 0.573638 |
| AHSP | 0.162376 | 0.424913 | 4.079006 | 0.104171 | 0.917034 | NA |

| | | | | | | |
|---|---|---|---|---|---|---|
| ITGAM | 1.740846 | -0.21584 | 1.754359 | -0.12303 | 0.902081 | NA |
| PYDC1 | 0.613743 | -0.17004 | 2.749758 | -0.06184 | 0.950691 | NA |
| ZNF764 | 159.1341 | -0.3122 | 0.213041 | -1.46546 | 0.142795 | 0.283795 |
| ZNF747 | 175.2574 | -0.47951 | 0.219758 | -2.18201 | 0.029109 | 0.09701 |
| ZNF768 | 1130.433 | -0.3443 | 0.162334 | -2.12094 | 0.033927 | 0.107602 |
| NLGN2 | 882.2065 | -0.66708 | 0.177372 | -3.76094 | 1.69E-04 | 0.003497 |
| CHD3 | 11222.32 | -0.24741 | 0.181236 | -1.36515 | 0.172207 | 0.323484 |
| CNTROB | 1103.09 | -0.35756 | 0.169638 | -2.10778 | 0.03505 | 0.109983 |
| TRAPPC1 | 3697.399 | -0.1575 | 0.136136 | -1.1569 | 0.247314 | 0.410746 |
| KCNAB3 | 22.52702 | -0.12323 | 0.461594 | -0.26696 | 0.789503 | NA |
| ZNF778 | 69.20478 | 0.301931 | 0.284297 | 1.062028 | 0.288223 | 0.454433 |
| CHRNB1 | 316.2424 | 0.007352 | 0.211131 | 0.034822 | 0.972221 | 0.984681 |
| SLC16A5 | 408.035 | -0.52689 | 0.197862 | -2.66294 | 0.007746 | 0.039802 |
| ADPRM | 298.5605 | 0.455911 | 0.19394 | 2.350785 | 0.018734 | 0.072294 |
| ELP5 | 912.7446 | 0.002572 | 0.159682 | 0.016108 | 0.987148 | 0.993273 |
| GABARAP | 1184.355 | -0.20854 | 0.187748 | -1.11074 | 0.266682 | 0.431639 |
| STX8 | 851.0859 | 0.174672 | 0.180115 | 0.969781 | 0.332156 | 0.500589 |
| UBB | 16926.98 | -0.27699 | 0.186603 | -1.48438 | 0.137709 | 0.277124 |
| GPRC5C | 1.307261 | 2.037817 | 2.096389 | 0.972061 | 0.33102 | NA |
| ADORA2B | 234.3144 | -0.08032 | 0.24439 | -0.32868 | 0.742401 | 0.837721 |
| TMC7 | 4.532546 | -0.14519 | 0.930678 | -0.15601 | 0.876026 | NA |
| ARL6IP1 | 4610.237 | -0.16351 | 0.205751 | -0.79471 | 0.426784 | 0.58859 |
| CDH2 | 714.393 | 0.050677 | 0.198687 | 0.255059 | 0.798677 | 0.878323 |
| DLGAP1 | 485.3312 | 0.116169 | 0.187001 | 0.621221 | 0.534454 | 0.678129 |
| SOCS6 | 1119.4 | 0.136146 | 0.252787 | 0.538582 | 0.590175 | 0.721887 |
| HOXB9 | 33.45683 | 0.104313 | 0.39268 | 0.265644 | 0.790513 | NA |
| TTLL6 | 19.77029 | 0.556594 | 0.543922 | 1.023299 | 0.306167 | NA |
| USP32 | 1773.663 | 0.344889 | 0.142434 | 2.421391 | 0.015461 | 0.063226 |
| PPM1D | 436.3557 | 0.086689 | 0.219693 | 0.394592 | 0.693144 | 0.801993 |
| PSG6 | 0.192403 | 0.975392 | 4.060716 | 0.240202 | 0.810174 | NA |
| TANC2 | 1166.102 | -0.37216 | 0.150368 | -2.475 | 0.013324 | 0.057401 |
| MBD3L1 | 0.151697 | -1.546 | 4.075469 | -0.37934 | 0.704433 | NA |
| LRRC8E | 16.35059 | -0.37606 | 0.506599 | -0.74233 | 0.457887 | NA |
| INSR | 269.9216 | -0.27671 | 0.217142 | -1.27431 | 0.202554 | 0.358825 |
| NRTN | 88.52868 | -0.18989 | 0.277295 | -0.68479 | 0.493476 | 0.644607 |
| NETO2 | 143.2646 | 0.022089 | 0.217506 | 0.101556 | 0.919109 | 0.953118 |
| JUNB | 872.0325 | -0.10123 | 0.174455 | -0.58026 | 0.561741 | 0.699815 |
| SHCBP1 | 707.222 | 0.031226 | 0.25507 | 0.122421 | 0.902566 | 0.943056 |
| NPTX1 | 2612.11 | 0.208965 | 0.859151 | 0.243223 | 0.807833 | 0.885317 |
| | 1601.083 | -0.80176 | 0.228752 | -3.50493 | 4.57E-04 | 0.00643 |
| ZNF439 | 170.5131 | 0.130316 | 0.269931 | 0.482775 | 0.629255 | 0.751771 |
| ZNF440 | 283.5138 | -0.15712 | 0.222094 | -0.70747 | 0.479274 | 0.633846 |
| GAA | 3443.984 | -0.52469 | 0.123964 | -4.2326 | 2.31E-05 | 0.001003 |
| CANT1 | 2327.138 | -0.46195 | 0.140163 | -3.29585 | 9.81E-04 | 0.010195 |
| KRT19 | 0.460669 | -0.62593 | 3.398671 | -0.18417 | 0.853882 | NA |
| KRT9 | 26.25622 | 0.539687 | 0.672053 | 0.803042 | 0.421951 | NA |
| KRT27 | 23.69956 | -0.54474 | 0.51662 | -1.05444 | 0.291682 | NA |

| | | | | | | |
|---|---|---|---|---|---|---|
| DSEL | 244.6531 | 0.953872 | 0.238794 | 3.994537 | 6.48E-05 | 0.001936 |
| ZNF562 | 479.9404 | -0.22657 | 0.187525 | -1.20824 | 0.226957 | 0.388458 |
| ZNF561 | 759.6662 | 0.145087 | 0.1807 | 0.802918 | 0.422022 | 0.584608 |
| WIPF2 | 1616.195 | -0.5131 | 0.151446 | -3.38799 | 7.04E-04 | 0.008326 |
| RSL1D1 | 2714.739 | 0.405825 | 0.18352 | 2.211337 | 0.027012 | 0.092446 |
| NEUROD2 | 0.151697 | -1.546 | 4.075469 | -0.37934 | 0.704433 | NA |
| DNAI2 | 0.163957 | 0.424913 | 4.079006 | 0.104171 | 0.917034 | NA |
| BPTF | 1179.845 | 0.39942 | 0.660204 | 0.604995 | 0.545182 | 0.686764 |
| LGALS4 | 0.134133 | -1.48909 | 4.077584 | -0.36519 | 0.71497 | NA |
| RASGRP4 | 0.968319 | 1.86533 | 2.390384 | 0.780348 | 0.435186 | NA |
| BCL2 | 474.7605 | -1.53E-04 | 0.174938 | -8.74E-04 | 0.999303 | 0.999472 |
| ZNF540 | 25.47874 | 0.84929 | 0.489024 | 1.736705 | 0.082439 | NA |
| ZNF570 | 54.71832 | 0.099885 | 0.304685 | 0.32783 | 0.74304 | 0.838049 |
| RNMTL1 | 570.6666 | -0.0924 | 0.168674 | -0.54782 | 0.583813 | 0.717088 |
| AQP4 | 1.656519 | 1.208323 | 1.708219 | 0.707358 | 0.479344 | NA |
| CYP4F11 | 1.630839 | 0.747778 | 1.64147 | 0.455554 | 0.648711 | NA |
| LGALS9C | 0.315653 | -0.57305 | 3.892639 | -0.14721 | 0.882963 | NA |
| TVP23B | 361.3052 | 0.845224 | 0.209929 | 4.026232 | 5.67E-05 | 0.001782 |
| FBXW10 | 0.545988 | -0.43913 | 2.541145 | -0.17281 | 0.862803 | NA |
| ATPAF2 | 386.9738 | -0.27937 | 0.159735 | -1.74897 | 0.080297 | 0.19188 |
| LRRC48 | 39.10222 | -0.28337 | 0.360595 | -0.78583 | 0.431967 | NA |
| ZNF57 | 66.09312 | 0.137389 | 0.279017 | 0.492404 | 0.622434 | 0.746665 |
| ZNF554 | 95.21832 | -0.23327 | 0.252968 | -0.92213 | 0.356459 | 0.524101 |
| THOP1 | 1691.855 | -0.25366 | 0.185438 | -1.36788 | 0.17135 | 0.322285 |
| ORMDL3 | 1789.021 | -5.75E-04 | 0.145353 | -0.00396 | 0.996842 | 0.998779 |
| MOB3A | 798.9604 | -0.49018 | 0.164352 | -2.98253 | 0.002859 | 0.020475 |
| SLFN12 | 236.1803 | 0.73371 | 0.231206 | 3.173403 | 0.001507 | 0.013595 |
| CALB2 | 0.230334 | -0.53318 | 4.076749 | -0.13079 | 0.895945 | NA |
| TEFM | 256.6068 | 0.3999 | 0.194469 | 2.056371 | 0.039747 | 0.119837 |
| MALT1 | 398.4485 | 0.847946 | 0.1671 | 5.074466 | 3.89E-07 | 8.53E-05 |
| AZU1 | 0.163957 | 0.424913 | 4.079006 | 0.104171 | 0.917034 | NA |
| BSG | 18902.55 | -0.40741 | 0.154167 | -2.64266 | 0.008226 | 0.041382 |
| COPRS | 1570.896 | -0.03258 | 0.142685 | -0.22832 | 0.819398 | 0.891854 |
| CCDC11 | 10.38969 | 0.446658 | 0.62509 | 0.71455 | 0.474887 | NA |
| FAM195A | 707.8871 | 0.013709 | 0.18232 | 0.075189 | 0.940064 | 0.966562 |
| PRSS27 | 21.493 | -0.81254 | 0.467475 | -1.73815 | 0.082184 | NA |
| PRSS30P | 0.617467 | -1.75734 | 3.109154 | -0.56522 | 0.571927 | NA |
| ZNF24 | 2499.423 | 0.638457 | 0.217763 | 2.931886 | 0.003369 | 0.022712 |
| BANP | 639.3526 | 0.058632 | 0.181774 | 0.322557 | 0.747031 | 0.84103 |
| C17orf66 | 0.134133 | -1.48909 | 4.077584 | -0.36519 | 0.71497 | NA |
| TAF15 | 2710.38 | -0.10932 | 0.150693 | -0.72542 | 0.468195 | 0.624629 |
| ZNF738 | 50.38079 | 0.696408 | 0.378067 | 1.842021 | 0.065472 | 0.166907 |
| SLFN11 | 7.68328 | -0.54074 | 0.772199 | -0.70026 | 0.483764 | NA |
| FAM192A | 1578.582 | -0.00634 | 0.149017 | -0.04253 | 0.966079 | 0.981394 |
| RAB37 | 2.867381 | -1.23554 | 1.213096 | -1.0185 | 0.30844 | NA |
| RPL38 | 10105.03 | 0.256417 | 0.178741 | 1.434576 | 0.151408 | 0.295445 |
| CES4A | 15.35837 | 0.096542 | 0.520153 | 0.185604 | 0.852755 | NA |

| | | | | | | |
|---|---|---|---|---|---|---|
| CES3 | 3.668398 | -0.01606 | 1.03245 | -0.01555 | 0.987591 | NA |
| CES2 | 1164.76 | -0.22974 | 0.158586 | -1.44868 | 0.147427 | 0.290626 |
| PDP2 | 381.1962 | -0.0451 | 0.17552 | -0.25695 | 0.797214 | 0.877202 |
| DCAKD | 640.1833 | -0.75687 | 0.194197 | -3.89745 | 9.72E-05 | 0.0025 |
| FAM222B | 649.2125 | -0.50531 | 0.171177 | -2.95198 | 0.003157 | 0.021941 |
| | 1.893177 | 0.476377 | 1.467625 | 0.324591 | 0.745491 | NA |
| PTPRM | 6728.138 | 0.034014 | 0.14767 | 0.230336 | 0.817831 | 0.890746 |
| CHD2 | 1585.821 | 0.31137 | 0.209863 | 1.483681 | 0.137893 | 0.277213 |
| WFIKKN2 | 0.096202 | 0.408546 | 4.079639 | 0.100143 | 0.920231 | NA |
| STAT5B | 1687.67 | -0.03696 | 0.12746 | -0.28995 | 0.771853 | 0.858857 |
| CD7 | 0.709944 | 0.232855 | 2.332364 | 0.099837 | 0.920474 | NA |
| CNP | 9998.743 | -0.54834 | 0.121038 | -4.53031 | 5.89E-06 | 4.69E-04 |
| JUP | 161.4812 | -0.60504 | 0.218318 | -2.77138 | 0.005582 | 0.031952 |
| HAP1 | 3.23156 | -0.3842 | 1.247451 | -0.30799 | 0.75809 | NA |
| TDRD12 | 1.32425 | -0.50336 | 1.927261 | -0.26118 | 0.793956 | NA |
| EIF1 | 12134.45 | -0.16452 | 0.150391 | -1.09397 | 0.273969 | 0.439786 |
| ENDOV | 216.0586 | -0.35848 | 0.192504 | -1.86219 | 0.062576 | 0.161938 |
| RNF213 | 4747.641 | -0.90088 | 0.181459 | -4.96464 | 6.88E-07 | 1.26E-04 |
| 10-Mar | 2.365232 | 0.168011 | 1.321467 | 0.12714 | 0.89883 | NA |
| PHOSPHO1 | 6.011485 | 0.304738 | 0.84316 | 0.361424 | 0.717783 | NA |
| ZNF791 | 314.3178 | 0.534635 | 0.200849 | 2.661876 | 0.007771 | 0.039835 |
| CBX2 | 349.1034 | -0.37084 | 0.183233 | -2.02385 | 0.042985 | 0.126323 |
| HOXB2 | 239.6841 | 0.085432 | 0.197105 | 0.433434 | 0.664699 | 0.779659 |
| C1QTNF1 | 2133.52 | -0.5181 | 0.206424 | -2.50989 | 0.012077 | 0.053662 |
| SWSAP1 | 94.80944 | -0.20885 | 0.236611 | -0.88268 | 0.377407 | 0.543904 |
| RAX2 | 0.151697 | -1.546 | 4.075469 | -0.37934 | 0.704433 | NA |
| TCAP | 12.23593 | 0.003673 | 0.602455 | 0.006096 | 0.995136 | NA |
| PIK3R6 | 0.162376 | 0.424913 | 4.079006 | 0.104171 | 0.917034 | NA |
| | 47.24161 | 0.109969 | 0.312553 | 0.351842 | 0.724957 | 0.82527 |
| MRPL45 | 1008.928 | 0.211807 | 0.158726 | 1.334418 | 0.182067 | 0.334896 |
| C16orf91 | 302.4021 | 0.083459 | 0.173832 | 0.480117 | 0.631144 | 0.75342 |
| SOCS7 | 276.563 | -0.22898 | 0.169565 | -1.35037 | 0.176898 | 0.329058 |
| CTU2 | 589.0256 | -0.22583 | 0.149226 | -1.51337 | 0.130186 | 0.266601 |
| PRPF8 | 9295.123 | -0.69569 | 0.132969 | -5.23199 | 1.68E-07 | 4.62E-05 |
| PITPNA | 4147.868 | -0.23739 | 0.119751 | -1.98238 | 0.047436 | 0.134086 |
| ZBTB4 | 3758.372 | -0.48108 | 0.133327 | -3.60828 | 3.08E-04 | 0.005033 |
| TNK1 | 51.14625 | 0.339495 | 0.437528 | 0.775939 | 0.437785 | 0.597921 |
| SLC16A11 | 2.318422 | 0.491792 | 1.496177 | 0.328699 | 0.742383 | NA |
| SLC16A13 | 38.18315 | -0.23917 | 0.391174 | -0.61141 | 0.540931 | NA |
| STARD6 | 1.020857 | 0.19325 | 2.085078 | 0.092682 | 0.926156 | NA |
| TTC9B | 7.08884 | -0.0493 | 0.760709 | -0.06481 | 0.948325 | NA |
| IQCK | 61.66951 | 0.039494 | 0.333056 | 0.11858 | 0.905608 | 0.945402 |
| ZNF266 | 301.0176 | -0.11052 | 0.168896 | -0.65437 | 0.512875 | 0.660023 |
| PCP2 | 2.710295 | 0.070169 | 1.266431 | 0.055407 | 0.955815 | NA |
| EMR1 | 0.163957 | 0.424913 | 4.079006 | 0.104171 | 0.917034 | NA |
| NDUFA11 | 551.3599 | 0.087637 | 0.152 | 0.576559 | 0.564238 | 0.702086 |
| C19orf70 | 1737.006 | -0.04546 | 0.15721 | -0.28917 | 0.772449 | 0.85903 |

| | | | | | | |
|---|---|---|---|---|---|---|
| SEZ6L2 | 588.7539 | -0.49841 | 0.201721 | -2.4708 | 0.013481 | 0.05774 |
| ASPHD1 | 278.1291 | 0.22319 | 0.265565 | 0.840434 | 0.400665 | 0.565475 |
| KCTD13 | 549.3549 | -0.13621 | 0.166133 | -0.81986 | 0.412295 | 0.575436 |
| | 8.790834 | 0.704451 | 0.721635 | 0.976188 | 0.328971 | NA |
| FAM211A-/ | 12074.5 | -0.01434 | 0.162724 | -0.08814 | 0.929762 | 0.960189 |
| TVP23C | 30.36174 | -0.07909 | 0.411202 | -0.19233 | 0.847481 | NA |
| YPEL2 | 379.0235 | 0.268758 | 0.207099 | 1.297724 | 0.194382 | 0.349475 |
| PPM1E | 40.11079 | 0.345633 | 0.350065 | 0.987341 | 0.323475 | NA |
| MED16 | 1223.138 | -0.56525 | 0.152996 | -3.69455 | 2.20E-04 | 0.004125 |
| ZNF519 | 29.98527 | 0.927205 | 0.410684 | 2.25771 | 0.023964 | NA |
| PTPN2 | 268.8296 | 0.595679 | 0.247777 | 2.404092 | 0.016213 | 0.065572 |
| SMAD2 | 2063.27 | 0.134567 | 0.147094 | 0.914842 | 0.360275 | 0.527421 |
| ERCC4 | 234.4081 | 0.061377 | 0.225412 | 0.272287 | 0.785401 | 0.869693 |
| | 0.151697 | -1.546 | 4.075469 | -0.37934 | 0.704433 | NA |
| RMI2 | 202.545 | -0.27072 | 0.216094 | -1.25277 | 0.210289 | 0.368021 |
| TOM1L2 | 3800.915 | -0.49076 | 0.198759 | -2.46911 | 0.013545 | 0.057906 |
| ZNF77 | 172.0977 | 0.135406 | 0.208896 | 0.648198 | 0.516857 | 0.664138 |
| B3GNTL1 | 73.6397 | -0.20726 | 0.2871 | -0.72191 | 0.470348 | 0.626655 |
| CTDNEP1 | 1828.557 | -0.3262 | 0.162428 | -2.00827 | 0.044615 | 0.129054 |
| ETV4 | 224.0926 | -0.51798 | 0.263203 | -1.968 | 0.049068 | 0.137 |
| BAIAP2 | 1810.832 | -0.58305 | 0.156524 | -3.725 | 1.95E-04 | 0.003827 |
| RPL7AP66 | 1.560106 | -0.0832 | 1.582614 | -0.05257 | 0.958072 | NA |
| | 245.0524 | -0.3867 | 0.207555 | -1.86311 | 0.062447 | 0.161813 |
| ARL4D | 292.6961 | -0.06636 | 0.222769 | -0.29788 | 0.765794 | 0.854687 |
| | 17.8928 | -0.02088 | 0.51703 | -0.04039 | 0.967779 | NA |
| UBE2O | 1528.285 | -0.62822 | 0.154611 | -4.0632 | 4.84E-05 | 0.001613 |
| ORAI3 | 416.5844 | -0.6619 | 0.180017 | -3.6769 | 2.36E-04 | 0.004314 |
| TUBB6 | 4623.209 | -0.08621 | 0.198585 | -0.43412 | 0.664203 | 0.779426 |
| NUPR1 | 161.1582 | 0.46683 | 0.250427 | 1.864131 | 0.062303 | 0.161585 |
| YES1 | 700.5738 | 0.71191 | 0.250339 | 2.843785 | 0.004458 | 0.027313 |
| CHMP6 | 795.4578 | -0.2655 | 0.144796 | -1.8336 | 0.066713 | 0.168708 |
| CCDC57 | 259.4563 | 0.184938 | 0.208304 | 0.887824 | 0.374636 | 0.54093 |
| HSF5 | 0.326333 | 1.533889 | 3.826878 | 0.40082 | 0.688553 | NA |
| SPHK1 | 1216.342 | 0.289888 | 0.291578 | 0.994202 | 0.320125 | 0.488238 |
| CIDEA | 1.257127 | 1.462356 | 1.777172 | 0.822855 | 0.41059 | NA |
| ATAD5 | 51.35305 | 1.391614 | 0.539614 | 2.578909 | 0.009911 | 0.046923 |
| ZNF404 | 19.86067 | 0.654308 | 0.479852 | 1.363561 | 0.172706 | NA |
| RTTN | 467.9378 | -0.25138 | 0.163508 | -1.53743 | 0.124188 | 0.258742 |
| TAC4 | 0.891108 | 0.008384 | 2.435479 | 0.003442 | 0.997253 | NA |
| HSD11B2 | 1.075752 | 0.450053 | 2.098769 | 0.214437 | 0.830206 | NA |
| CRLF3 | 239.02 | 0.453994 | 0.197818 | 2.295011 | 0.021732 | 0.080136 |
| EID2 | 344.9173 | 0.386576 | 0.183872 | 2.102423 | 0.035516 | 0.111035 |
| EID2B | 34.1741 | 0.186383 | 0.366948 | 0.507928 | 0.611504 | NA |
| SLCO3A1 | 573.729 | 0.145669 | 0.179955 | 0.809478 | 0.41824 | 0.581245 |
| ZNF575 | 75.59794 | -0.45432 | 0.308731 | -1.47156 | 0.141139 | 0.281685 |
| CCDC101 | 167.7733 | -0.29496 | 0.268697 | -1.09776 | 0.272311 | 0.437848 |
| DIRAS1 | 1267.983 | -0.18152 | 0.198048 | -0.91656 | 0.359373 | 0.526491 |

| | | | | | | |
|---|---|---|---|---|---|---|
| PHLDB3 | 199.8717 | -0.30733 | 0.216662 | -1.41846 | 0.156057 | 0.301904 |
| GNG7 | 1500.153 | -0.46231 | 0.141638 | -3.26401 | 0.001098 | 0.011027 |
| CNTD1 | 12.39559 | 0.675336 | 0.650691 | 1.037875 | 0.299328 | NA |
| LMNB2 | 2700.639 | -0.48175 | 0.219129 | -2.19848 | 0.027915 | 0.094605 |
| MEX3C | 1098.514 | 0.418072 | 0.149784 | 2.791168 | 0.005252 | 0.030564 |
| RNF152 | 64.71525 | 0.629809 | 0.312152 | 2.017638 | 0.043629 | 0.127269 |
| MYO1D | 10082.82 | -0.5256 | 0.181446 | -2.89671 | 0.003771 | 0.024496 |
| FOXL1 | 1.431763 | 2.610478 | 1.858973 | 1.404258 | 0.160242 | NA |
| LRRC37A | 5.491862 | 0.917401 | 0.978245 | 0.937803 | 0.348346 | NA |
| FOXC2 | 1.044145 | 0.31036 | 2.099918 | 0.147796 | 0.882504 | NA |
| ACSF3 | 670.1388 | -0.24766 | 0.141348 | -1.75215 | 0.079749 | 0.191024 |
| ZNF843 | 21.26856 | -0.25242 | 0.478045 | -0.52802 | 0.597488 | NA |
| CDK5R1 | 1475.631 | -0.01736 | 0.166058 | -0.10452 | 0.91676 | 0.951845 |
| LRRC37A3 | 17.09575 | 0.097072 | 0.500143 | 0.194089 | 0.846106 | NA |
| IRX5 | 153.0566 | -0.30509 | 0.268198 | -1.13754 | 0.255311 | 0.419503 |
| METRNL | 513.3266 | -0.11413 | 0.219472 | -0.52 | 0.603064 | 0.732138 |
| TYMS | 2078.727 | -0.18229 | 0.204786 | -0.89015 | 0.373385 | 0.539792 |
| C18orf56 | 54.22354 | -0.52493 | 0.355809 | -1.47531 | 0.140129 | 0.280366 |
| EFCAB5 | 16.84879 | 0.896843 | 0.556775 | 1.61078 | 0.107228 | NA |
| NFATC2IP | 1006.451 | -0.12063 | 0.134584 | -0.8963 | 0.370092 | 0.536887 |
| SHMT1 | 680.6237 | -0.29742 | 0.203204 | -1.46365 | 0.143289 | 0.284682 |
| SMCR8 | 608.7687 | -0.41445 | 0.16089 | -2.57596 | 0.009996 | 0.047157 |
| FAM210A | 445.8608 | 0.389068 | 0.188932 | 2.059303 | 0.039465 | 0.119231 |
| B3GNT8 | 30.14725 | -0.53701 | 0.428604 | -1.25292 | 0.210235 | NA |
| CHD9 | 698.269 | 0.882803 | 0.326191 | 2.706403 | 0.006802 | 0.03642 |
| TRIM72 | 0.247898 | -0.5901 | 4.074633 | -0.14482 | 0.884851 | NA |
| FBXO39 | 0.315653 | -0.57305 | 3.892639 | -0.14721 | 0.882963 | NA |
| TOP3A | 791.0101 | -0.39531 | 0.160714 | -2.45969 | 0.013906 | 0.058918 |
| CASKIN2 | 673.0642 | -0.50514 | 0.17494 | -2.88749 | 0.003883 | 0.024986 |
| DLGAP1-AS | 84.0007 | 0.190093 | 0.263966 | 0.720142 | 0.471438 | 0.627431 |
| TIMM22 | 948.5439 | -0.30891 | 0.161754 | -1.90973 | 0.056167 | 0.150029 |
| HIC1 | 6.000843 | 0.137366 | 0.840059 | 0.16352 | 0.870109 | NA |
| TGIF1 | 1227.662 | 0.328592 | 0.178764 | 1.838135 | 0.066043 | 0.167724 |
| MIEF2 | 324.4099 | -0.05023 | 0.185193 | -0.27125 | 0.786198 | 0.870413 |
| CD19 | 0.706784 | 0.225544 | 2.334352 | 0.096619 | 0.923029 | NA |
| PTRF | 14286.51 | -0.48203 | 0.1613 | -2.9884 | 0.002804 | 0.020297 |
| IRX3 | 392.3131 | -0.12154 | 0.207058 | -0.58697 | 0.557225 | 0.696283 |
| ST8SIA3 | 0.382031 | -1.26484 | 3.151989 | -0.40128 | 0.688212 | NA |
| RABEP2 | 1293.534 | -0.26247 | 0.180084 | -1.45751 | 0.144976 | 0.286927 |
| C18orf32 | 85.03774 | 0.14637 | 0.273795 | 0.534597 | 0.592928 | 0.723463 |
| ZNF491 | 58.71431 | -0.13184 | 0.325745 | -0.40474 | 0.685671 | 0.79533 |
| GSG2 | 123.0864 | -0.47377 | 0.292207 | -1.62137 | 0.104939 | 0.230452 |
| KIAA0195 | 2336.486 | -0.64584 | 0.148854 | -4.3387 | 1.43E-05 | 7.69E-04 |
| FLII | 5913.701 | -0.35999 | 0.145892 | -2.46749 | 0.013606 | 0.058023 |
| GRB2 | 2835.594 | -0.14696 | 0.134628 | -1.09158 | 0.275016 | 0.440659 |
| CENPBD1 | 254.6339 | -0.15257 | 0.221048 | -0.69023 | 0.490048 | 0.642357 |
| STAP2 | 601.0946 | -0.08743 | 0.174762 | -0.50027 | 0.616882 | 0.742224 |

| | | | | | | |
|---|---|---|---|---|---|---|
| TWF1P1 | 12.44896 | 0.709534 | 0.628121 | 1.129615 | 0.258639 | NA |
| TSSK6 | 60.82931 | -0.34436 | 0.289239 | -1.19057 | 0.233822 | 0.396391 |
| NDUFV2 | 6.875594 | 0.434384 | 0.800025 | 0.542963 | 0.587155 | NA |
| PARD6G | 1276.635 | -0.28007 | 0.156442 | -1.79025 | 0.073413 | 0.180638 |
| SH2B1 | 1424.598 | -0.24965 | 0.130076 | -1.91925 | 0.054952 | 0.14779 |
| PRSS36 | 13.66487 | -0.26704 | 0.574163 | -0.46509 | 0.641864 | NA |
| TMPRSS9 | 2.276549 | 0.46223 | 1.343335 | 0.344092 | 0.730777 | NA |
| TMEM11 | 846.3345 | -0.03214 | 0.15108 | -0.21276 | 0.831516 | 0.89868 |
| KCNG2 | 3.102859 | 0.529001 | 1.117184 | 0.473513 | 0.635847 | NA |
| | 9.006416 | -0.11015 | 0.70382 | -0.1565 | 0.87564 | NA |
| | 0.900564 | 1.750062 | 2.175753 | 0.804348 | 0.421196 | NA |
| | 1.082223 | -2.45149 | 2.404193 | -1.01967 | 0.307883 | NA |
| | 10.53045 | -0.52133 | 0.671283 | -0.77662 | 0.437384 | NA |
| KLHL11 | 70.57645 | 0.08844 | 0.31576 | 0.280084 | 0.779413 | 0.864677 |
| CTXN1 | 290.2326 | 0.144473 | 0.205293 | 0.703741 | 0.481594 | 0.635881 |
| MAF | 1119.831 | -0.32201 | 0.184737 | -1.74308 | 0.081321 | 0.193352 |
| ERN1 | 115.4873 | -0.5544 | 0.2392 | -2.31774 | 0.020463 | 0.076899 |
| SUZ12 | 643.8392 | 0.733125 | 0.306576 | 2.391336 | 0.016787 | 0.067285 |
| CPNE7 | 798.572 | 0.695326 | 0.583804 | 1.191026 | 0.233643 | 0.396258 |
| EFCAB13 | 12.02747 | 1.113729 | 0.607768 | 1.83249 | 0.066878 | NA |
| DPY19L3 | 1836.165 | 0.496268 | 0.15667 | 3.167609 | 0.001537 | 0.013769 |
| PFAS | 1075.65 | -0.76674 | 0.220713 | -3.47391 | 5.13E-04 | 0.00691 |
| C17orf62 | 1355.256 | -0.30067 | 0.129655 | -2.319 | 0.020395 | 0.076709 |
| ZBTB7A | 1923.174 | -0.22338 | 0.165029 | -1.3536 | 0.175865 | 0.328018 |
| TUFM | 8489.081 | -0.13293 | 0.120346 | -1.10454 | 0.269359 | 0.433993 |
| CTC1 | 374.2137 | -0.47193 | 0.204047 | -2.31284 | 0.020731 | 0.077653 |
| LINC00324 | 68.24171 | 0.473732 | 0.583586 | 0.81176 | 0.41693 | 0.580061 |
| EIF3K | 4530.468 | -0.17872 | 0.136608 | -1.30824 | 0.190792 | 0.345061 |
| AURKB | 626.1516 | -0.47536 | 0.252986 | -1.87901 | 0.060243 | 0.15768 |
| TMEM107 | 401.3422 | -0.33225 | 0.165116 | -2.01222 | 0.044197 | 0.128169 |
| EXOC3L1 | 20.3057 | -0.11245 | 0.481504 | -0.23354 | 0.815338 | NA |
| | 1.705682 | -0.24761 | 1.597667 | -0.15498 | 0.876835 | NA |
| PER1 | 802.447 | -0.53566 | 0.248503 | -2.15554 | 0.031119 | 0.101631 |
| HES7 | 83.60612 | 0.369486 | 0.40363 | 0.915409 | 0.359977 | 0.527051 |
| FARSA | 3140.087 | -0.28114 | 0.143276 | -1.96224 | 0.049734 | 0.138006 |
| SAMD4B | 2861.762 | -0.29939 | 0.171736 | -1.74332 | 0.081277 | 0.193289 |
| ALOXE3 | 2.025073 | 0.612299 | 1.55562 | 0.393604 | 0.693873 | NA |
| GGN | 9.713374 | 0.123061 | 0.667491 | 0.184363 | 0.853729 | NA |
| CALR | 30755.43 | -0.49643 | 0.151706 | -3.2723 | 0.001067 | 0.010822 |
| RAD23A | 4906.946 | -0.16146 | 0.150064 | -1.07594 | 0.281953 | 0.448104 |
| GADD45GI | 2428.397 | -0.0699 | 0.171007 | -0.40874 | 0.682729 | 0.793428 |
| MEIS3P1 | 101.9522 | 0.393574 | 0.286466 | 1.373893 | 0.169475 | 0.319924 |
| DAND5 | 0.162376 | 0.424913 | 4.079006 | 0.104171 | 0.917034 | NA |
| C17orf96 | 132.2862 | -0.06226 | 0.322239 | -0.19322 | 0.846788 | 0.908806 |
| WSCD1 | 9.974803 | -0.09875 | 0.675321 | -0.14623 | 0.883743 | NA |
| GEMIN4 | 811.3965 | -0.50181 | 0.159587 | -3.14442 | 0.001664 | 0.014409 |
| RNF151 | 0.488709 | 2.119056 | 3.307564 | 0.64067 | 0.521737 | NA |

| | | | | | | |
|---|---|---|---|---|---|---|
| CIITA | 45.64817 | -0.43329 | 0.32757 | -1.32273 | 0.185925 | NA |
| ZFPM1 | 269.0231 | -0.35564 | 0.20186 | -1.76182 | 0.0781 | 0.188276 |
| ALOX15B | 0.495797 | -0.73607 | 3.324282 | -0.22142 | 0.824763 | NA |
| PLD6 | 56.66412 | -0.58756 | 0.373068 | -1.57494 | 0.115269 | 0.245203 |
| CDC42EP4 | 441.2247 | -0.44372 | 0.154413 | -2.87357 | 0.004059 | 0.02562 |
| SYCN | 0.151697 | -1.546 | 4.075469 | -0.37934 | 0.704433 | NA |
| PIPOX | 14.32909 | -0.16471 | 0.65591 | -0.25112 | 0.801719 | NA |
| CDH5 | 0.162376 | 0.424913 | 4.079006 | 0.104171 | 0.917034 | NA |
| GIPC3 | 1005.522 | -0.18104 | 0.243796 | -0.7426 | 0.457722 | 0.616134 |
| | 78.4486 | -0.44608 | 0.274609 | -1.62441 | 0.104288 | 0.229312 |
| PDXDC1 | 5293.597 | -0.11867 | 0.187887 | -0.63162 | 0.527638 | 0.673176 |
| B3GNT3 | 0.520316 | 2.239969 | 3.173769 | 0.705776 | 0.480328 | NA |
| SEPHS2 | 3535.239 | 0.117808 | 0.139595 | 0.843927 | 0.39871 | 0.563508 |
| DCTPP1 | 1371.586 | -0.02313 | 0.171124 | -0.13514 | 0.892497 | 0.936854 |
| ZNF771 | 298.5072 | -0.08036 | 0.171559 | -0.46843 | 0.639474 | 0.75954 |
| TSHZ1 | 995.7915 | -0.22313 | 0.213125 | -1.04694 | 0.295129 | 0.462431 |
| ZADH2 | 1628.694 | 0.02795 | 0.147353 | 0.18968 | 0.84956 | 0.910463 |
| ZNF48 | 220.6419 | -0.27281 | 0.210532 | -1.2958 | 0.195046 | 0.350068 |
| 1-Sep | 50.56 | -0.04354 | 0.305326 | -0.1426 | 0.886606 | 0.933124 |
| FAHD1 | 1307.51 | -0.06127 | 0.142965 | -0.42858 | 0.668227 | 0.782242 |
| MYLPF | 2.190881 | 1.283853 | 1.452526 | 0.883876 | 0.376763 | NA |
| CCDC43 | 608.0858 | 0.406172 | 0.165458 | 2.454831 | 0.014095 | 0.059508 |
| C17orf104 | 9.579516 | 1.383086 | 0.699117 | 1.978332 | 0.047891 | NA |
| FZD2 | 232.423 | -0.22321 | 0.218745 | -1.02039 | 0.307543 | 0.474791 |
| LINC00304 | 2.727412 | 0.363803 | 1.32179 | 0.275236 | 0.783135 | NA |
| HMHA1 | 394.1917 | -0.15469 | 0.182472 | -0.84775 | 0.396579 | 0.561232 |
| | 2.320658 | 0.254809 | 1.5859 | 0.160671 | 0.872352 | NA |
| ZNF571 | 56.38809 | 0.321685 | 0.316602 | 1.016053 | 0.309604 | 0.476958 |
| SSTR2 | 8.300069 | 2.33008 | 1.05057 | 2.21792 | 0.02656 | NA |
| ZNF594 | 46.08915 | 0.586307 | 0.357177 | 1.641501 | 0.100694 | NA |
| SLC47A2 | 2.773577 | 0.898 | 1.292386 | 0.694839 | 0.487156 | NA |
| S1PR5 | 3.844874 | 0.253917 | 1.187509 | 0.213823 | 0.830685 | NA |
| | 56.44728 | 0.053336 | 0.339344 | 0.157173 | 0.875108 | 0.925378 |
| ANKRD30B | 34.31932 | 1.812223 | 0.510829 | 3.547611 | 3.89E-04 | NA |
| ZFP3 | 33.63836 | -0.20826 | 0.517435 | -0.40248 | 0.68733 | NA |
| CSNK1G2-A | 0.976776 | -2.39789 | 2.178705 | -1.1006 | 0.271069 | NA |
| ZNF443 | 24.93723 | 0.827047 | 0.451689 | 1.831011 | 0.067099 | NA |
| ZNF792 | 128.4856 | -0.11604 | 0.238288 | -0.48697 | 0.626281 | 0.749266 |
| CUEDC1 | 1241.857 | -0.31071 | 0.128676 | -2.41466 | 0.01575 | 0.064127 |
| KCTD2 | 1948.775 | -0.48529 | 0.156002 | -3.11077 | 0.001866 | 0.015428 |
| CMTR2 | 579.4446 | 0.497373 | 0.198429 | 2.506561 | 0.012191 | 0.054008 |
| ZFP82 | 72.15901 | 0.081435 | 0.275512 | 0.295577 | 0.767553 | 0.855764 |
| NQO1 | 37221.55 | -0.05776 | 0.19806 | -0.29162 | 0.770581 | 0.858171 |
| TRAPPC5 | 23.37412 | -0.25569 | 0.513747 | -0.49769 | 0.618705 | NA |
| RPH3AL | 3.284234 | 0.952345 | 1.150697 | 0.827624 | 0.407883 | NA |
| SLC25A42 | 214.0947 | -0.37359 | 0.216451 | -1.72599 | 0.084349 | 0.197978 |
| METTL23 | 589.0319 | 0.056539 | 0.196063 | 0.288373 | 0.773061 | 0.8594 |

| | | | | | | |
|---|---|---|---|---|---|---|
| SLC26A11 | 582.6662 | -0.36205 | 0.161303 | -2.24452 | 0.024799 | 0.087277 |
| MUC16 | 0.151697 | -1.546 | 4.075469 | -0.37934 | 0.704433 | NA |
| POLR2A | 5345.916 | -0.86968 | 0.184521 | -4.71316 | 2.44E-06 | 2.71E-04 |
| SLC25A41 | 1.228827 | 0.64505 | 2.019407 | 0.319426 | 0.749404 | NA |
| TMEM102 | 515.8914 | -0.15603 | 0.214493 | -0.72744 | 0.466957 | 0.623713 |
| TMEM132I | 0.327913 | 1.543154 | 4.021063 | 0.383768 | 0.701151 | NA |
| FAM211A | 186.0432 | 0.060632 | 0.324986 | 0.186568 | 0.852 | 0.911895 |
| CCL13 | 0.096202 | 0.408546 | 4.079639 | 0.100143 | 0.920231 | NA |
| SYNE4 | 1.247817 | -0.68254 | 1.954986 | -0.34913 | 0.726992 | NA |
| OGFOD3 | 819.5596 | -0.415 | 0.161966 | -2.56229 | 0.010398 | 0.048343 |
| AATK | 112.9088 | -1.23693 | 0.348718 | -3.54707 | 3.90E-04 | 0.005824 |
| RNF135 | 539.481 | -0.24986 | 0.155595 | -1.60583 | 0.108311 | 0.23517 |
| ACBD4 | 333.3813 | -0.23218 | 0.204501 | -1.13537 | 0.25622 | 0.420414 |
| SGSH | 1575.579 | -0.42463 | 0.163948 | -2.59001 | 0.009597 | 0.045907 |
| MEX3D | 689.853 | -0.22288 | 0.207913 | -1.07199 | 0.283723 | 0.450213 |
| MRPS23 | 1384.07 | 0.436965 | 0.187384 | 2.331917 | 0.019705 | 0.074993 |
| SLX1B | 0.392711 | 0.292538 | 3.145221 | 0.09301 | 0.925895 | NA |
| HKR1 | 472.1401 | -0.21666 | 0.152614 | -1.41969 | 0.155699 | 0.301553 |
| ODF3L2 | 0.612162 | -0.19706 | 2.47693 | -0.07956 | 0.936588 | NA |
| SLC2A4 | 5.847236 | 0.609023 | 0.841038 | 0.724133 | 0.468984 | NA |
| CLDN7 | 13.32979 | -0.01134 | 0.608683 | -0.01862 | 0.985142 | NA |
| ZNF101 | 185.8563 | -0.00118 | 0.200867 | -0.00585 | 0.995329 | 0.997908 |
| GINS3 | 252.2809 | -0.13428 | 0.209291 | -0.64162 | 0.521123 | 0.668146 |
| TMEM259 | 4538.872 | -0.17772 | 0.14271 | -1.24534 | 0.213005 | 0.371732 |
| DEXI | 10.86358 | -0.33549 | 0.624375 | -0.53732 | 0.591044 | NA |
| ZNF708 | 54.18009 | 0.597683 | 0.546573 | 1.093511 | 0.274169 | 0.439821 |
| IST1 | 2396.247 | -0.1621 | 0.13622 | -1.18998 | 0.234056 | 0.39656 |
| TSEN54 | 473.0904 | -0.00309 | 0.172149 | -0.01797 | 0.985659 | 0.99244 |
| RGMA | 13.31775 | -1.99578 | 0.694836 | -2.87231 | 0.004075 | NA |
| CYB5D1 | 256.9034 | 0.049648 | 0.217183 | 0.228598 | 0.819181 | 0.891783 |
| SYNM | 703.4479 | 0.098091 | 0.185547 | 0.528658 | 0.597042 | 0.726799 |
| GLTPD2 | 15.87243 | -0.23149 | 0.529483 | -0.4372 | 0.661969 | NA |
| | 0.883146 | 1.112834 | 2.554178 | 0.435692 | 0.663061 | NA |
| DNM1P46 | 27.81323 | 0.369157 | 0.415866 | 0.887683 | 0.374711 | NA |
| NPLOC4 | 6025.604 | -0.38859 | 0.130304 | -2.98222 | 0.002862 | 0.020475 |
| TEX19 | 0.599903 | -1.73313 | 2.730982 | -0.63462 | 0.525678 | NA |
| CAPN12 | 1.44476 | -0.28802 | 2.162196 | -0.13321 | 0.89403 | NA |
| EXOC7 | 2681.835 | -0.34064 | 0.127585 | -2.66993 | 0.007587 | 0.039355 |
| KPNA2 | 7260.809 | -0.34846 | 0.1711 | -2.0366 | 0.04169 | 0.123969 |
| FES | 850.9716 | -0.42899 | 0.142738 | -3.00545 | 0.002652 | 0.019527 |
| MXRA7 | 3509.267 | -0.21536 | 0.176734 | -1.21857 | 0.223008 | 0.38382 |
| NXPH3 | 2.519101 | -1.49614 | 1.35817 | -1.10158 | 0.270642 | NA |
| HS3ST4 | 0.268266 | -2.15396 | 4.038351 | -0.53338 | 0.593773 | NA |
| TSPAN10 | 5142.975 | -0.69435 | 0.261712 | -2.65311 | 0.007975 | 0.040544 |
| SKA2 | 1063.352 | 0.471368 | 0.15313 | 3.078221 | 0.002082 | 0.016597 |
| PPP1R27 | 1.793601 | -0.1489 | 1.460382 | -0.10196 | 0.918788 | NA |
| BRICD5 | 46.07352 | -0.07398 | 0.383828 | -0.19274 | 0.847165 | NA |

| | | | | | | |
|---|---|---|---|---|---|---|
| GALR2 | 2.241218 | 0.891611 | 1.362709 | 0.654293 | 0.512923 | NA |
| HOXB4 | 262.0135 | -0.19818 | 0.184057 | -1.07675 | 0.281592 | 0.44773 |
| NGRN | 1773.299 | -0.08777 | 0.168221 | -0.52175 | 0.601844 | 0.730956 |
| DDX28 | 360.9948 | -0.1959 | 0.161726 | -1.21132 | 0.225771 | 0.386966 |
| C16orf72 | 944.428 | -0.02428 | 0.138092 | -0.17581 | 0.860446 | 0.917154 |
| VMO1 | 13.5104 | 1.083488 | 0.603268 | 1.79603 | 0.07249 | NA |
| TMEM95 | 0.163957 | 0.424913 | 4.079006 | 0.104171 | 0.917034 | NA |
| GJC1 | 1204.5 | -0.16122 | 0.225374 | -0.71536 | 0.474387 | 0.629745 |
| PYCR1 | 464.1535 | -0.15495 | 0.275202 | -0.56305 | 0.573399 | 0.70927 |
| LSMD1 | 1639.17 | 0.011405 | 0.133291 | 0.085563 | 0.931813 | 0.961401 |
| SPNS2 | 6.097093 | -0.17053 | 0.843966 | -0.20206 | 0.839872 | NA |
| C19orf59 | 0.392857 | -1.546 | 4.075469 | -0.37934 | 0.704433 | NA |
| OTOP2 | 0.268266 | -2.15396 | 4.038351 | -0.53338 | 0.593773 | NA |
| ABAT | 516.1524 | -0.46832 | 0.257619 | -1.81786 | 0.069086 | 0.172997 |
| SLC25A10 | 124.3241 | -0.45866 | 0.250548 | -1.83064 | 0.067154 | 0.169494 |
| LYSMD4 | 178.3773 | -0.0519 | 0.212421 | -0.24434 | 0.80697 | 0.884734 |
| AFMID | 803.3721 | -0.18044 | 0.135212 | -1.33449 | 0.182042 | 0.334896 |
| LYPD4 | 0.327913 | 1.543154 | 4.021063 | 0.383768 | 0.701151 | NA |
| GJD3 | 7.395332 | -0.74531 | 0.748358 | -0.99592 | 0.319289 | NA |
| C2CD4C | 19.39513 | 0.558339 | 0.689188 | 0.810141 | 0.417859 | NA |
| CHST6 | 114.6085 | -1.01441 | 0.230562 | -4.39972 | 1.08E-05 | 6.42E-04 |
| GDPGP1 | 54.22987 | -0.27519 | 0.307554 | -0.89476 | 0.370915 | 0.53772 |
| | 1.614435 | 0.100692 | 1.58733 | 0.063435 | 0.94942 | NA |
| CCBE1 | 0.192403 | 0.975392 | 4.060716 | 0.240202 | 0.810174 | NA |
| C19orf71 | 2485.473 | -0.60039 | 0.219817 | -2.73134 | 0.006308 | 0.034476 |
| CCDC159 | 373.8692 | -0.22691 | 0.202674 | -1.11956 | 0.262902 | 0.427413 |
| RPS7P1 | 41.56131 | 0.36178 | 0.359761 | 1.005612 | 0.314602 | NA |
| NPIPA1 | 269.3121 | -0.12257 | 0.256313 | -0.47821 | 0.6325 | 0.754704 |
| GRIN2A | 0.298089 | -0.51711 | 3.975232 | -0.13008 | 0.896501 | NA |
| | 52.39726 | -0.32883 | 0.43282 | -0.75974 | 0.447412 | 0.607081 |
| ASB7 | 413.2467 | -0.03251 | 0.190213 | -0.17089 | 0.864308 | 0.92003 |
| ACSM5 | 21.7637 | -0.07932 | 0.43957 | -0.18045 | 0.856797 | NA |
| PGPEP1L | 0.544554 | -2.65036 | 3.220086 | -0.82307 | 0.410468 | NA |
| | 15.22963 | 0.33896 | 0.526748 | 0.643496 | 0.519903 | NA |
| MRPL54 | 864.2614 | 0.108814 | 0.160773 | 0.676814 | 0.498524 | 0.649308 |
| C15orf32 | 0.327913 | 1.543154 | 4.021063 | 0.383768 | 0.701151 | NA |
| ALYREF | 1434.608 | -0.22802 | 0.162589 | -1.40243 | 0.160786 | 0.308346 |
| FAM101B | 4113.223 | -0.44208 | 0.135787 | -3.25567 | 0.001131 | 0.011192 |
| CMTM4 | 1019.337 | -0.04984 | 0.141902 | -0.35126 | 0.725397 | 0.825454 |
| TBL3 | 1395.765 | -0.37704 | 0.145085 | -2.59875 | 0.009357 | 0.045147 |
| | 26.76023 | -0.49616 | 0.586471 | -0.84601 | 0.397549 | NA |
| NPIPA5 | 21.59285 | -0.15373 | 0.532144 | -0.2889 | 0.772661 | NA |
| ZNF730 | 6.454116 | 1.281575 | 1.034515 | 1.238817 | 0.215413 | NA |
| | 0.230334 | -0.53318 | 4.076749 | -0.13079 | 0.895945 | NA |
| DNAH2 | 1.344026 | 1.504564 | 2.095704 | 0.717928 | 0.472802 | NA |
| NPW | 12.49568 | -1.05481 | 0.779117 | -1.35385 | 0.175783 | NA |
| COA3 | 1804.352 | -0.05413 | 0.130528 | -0.41471 | 0.678356 | 0.790205 |

| | | | | | | |
|---|---|---|---|---|---|---|
| ACTG1 | 93611.85 | -0.32251 | 0.159674 | -2.01979 | 0.043405 | 0.126951 |
| VPS33B | 774.1455 | -0.34925 | 0.139029 | -2.5121 | 0.012002 | 0.053428 |
| ADAP2 | 0.989093 | -0.34325 | 2.31721 | -0.14813 | 0.882238 | NA |
| EIF3C | 149.4424 | -0.63669 | 0.309317 | -2.05838 | 0.039553 | 0.119436 |
| NR2C2AP | 339.0827 | -0.09535 | 0.203916 | -0.46759 | 0.640077 | 0.76018 |
| KCNJ12 | 4.850742 | -1.91533 | 0.959526 | -1.99612 | 0.04592 | NA |
| PGP | 1370.058 | -0.05633 | 0.160073 | -0.35189 | 0.724919 | 0.82527 |
| ALDH1A3 | 781.5386 | 0.567129 | 0.287263 | 1.974253 | 0.048353 | 0.135674 |
| TM2D3 | 704.4873 | 0.195951 | 0.180245 | 1.087137 | 0.276976 | 0.442933 |
| SPATA32 | 0.938698 | 0.081169 | 1.915817 | 0.042368 | 0.966205 | NA |
| CCR10 | 24.11936 | -0.47472 | 0.425946 | -1.1145 | 0.265066 | NA |
| HDDC3 | 220.9124 | -0.36621 | 0.177231 | -2.06626 | 0.038804 | 0.117772 |
| ZFP1 | 212.6768 | 0.401026 | 0.255718 | 1.568236 | 0.116826 | 0.247228 |
| | 6.244676 | 0.086618 | 0.949012 | 0.091272 | 0.927276 | NA |
| SOCS3 | 21.61883 | 0.418804 | 0.454772 | 0.920908 | 0.357098 | NA |
| SNN | 2637.694 | -0.32909 | 0.154146 | -2.13495 | 0.032765 | 0.105468 |
| KRBA2 | 11.20732 | -0.1704 | 0.628939 | -0.27092 | 0.786449 | NA |
| ZNF93 | 44.5009 | 1.171609 | 0.398211 | 2.942182 | 0.003259 | NA |
| 9-Sep | 27290.34 | -0.6988 | 0.165921 | -4.21162 | 2.54E-05 | 0.001062 |
| CLDN6 | 1.931684 | 1.755502 | 1.70602 | 1.029004 | 0.303478 | NA |
| APOBR | 2.329755 | 0.974366 | 1.316456 | 0.740143 | 0.459213 | NA |
| ZBTB7C | 0.94809 | -2.20833 | 2.818137 | -0.78361 | 0.433267 | NA |
| TMEM186 | 179.2589 | 0.109795 | 0.20857 | 0.526418 | 0.598598 | 0.728207 |
| SDR42E1 | 1.269533 | 2.018767 | 2.236927 | 0.902474 | 0.366805 | NA |
| PIGW | 264.1005 | -0.01467 | 0.196262 | -0.07477 | 0.940397 | 0.966736 |
| FMNL1 | 103.7882 | -0.65962 | 0.331883 | -1.98752 | 0.046865 | 0.133202 |
| ZFP90 | 94.53485 | -0.13215 | 0.261463 | -0.50542 | 0.613262 | 0.740047 |
| TMEM106/ | 169.0361 | -0.63333 | 0.208282 | -3.04072 | 0.00236 | 0.017993 |
| CIB1 | 1751.699 | -0.13262 | 0.150413 | -0.8817 | 0.377939 | 0.544274 |
| MYADML2 | 0.410275 | 0.244283 | 3.100889 | 0.078778 | 0.937209 | NA |
| TBC1D3F | 5.579871 | 0.707105 | 0.972349 | 0.727213 | 0.467095 | NA |
| LRRC37B | 68.70076 | -0.00877 | 0.326995 | -0.02683 | 0.978598 | 0.988521 |
| NOMO2 | 492.456 | -0.50529 | 0.221474 | -2.28147 | 0.022521 | 0.081968 |
| LINC00482 | 16.37015 | 1.109878 | 0.540346 | 2.054013 | 0.039974 | NA |
| PRSS57 | 0.517155 | 2.232561 | 3.179631 | 0.702145 | 0.482589 | NA |
| MC2R | 0.151697 | -1.546 | 4.075469 | -0.37934 | 0.704433 | NA |
| RAB11B | 2997.429 | -0.16243 | 0.133013 | -1.22118 | 0.222016 | 0.382559 |
| GP1BA | 13.49112 | -1.05043 | 0.584999 | -1.7956 | 0.072557 | NA |
| UBALD2 | 1877.462 | -0.08223 | 0.173095 | -0.47505 | 0.634751 | 0.75605 |
| NOTUM | 2.946439 | 1.173852 | 1.279053 | 0.917751 | 0.358749 | NA |
| CCDC137 | 1070.364 | 0.009197 | 0.169444 | 0.054279 | 0.956713 | 0.976145 |
| CDK10 | 1113.623 | 0.034119 | 0.156451 | 0.218078 | 0.827368 | 0.895973 |
| SOCS1 | 24.74643 | -1.14699 | 0.524187 | -2.18812 | 0.028661 | NA |
| HGS | 3174.011 | -0.37852 | 0.134713 | -2.80983 | 0.004957 | 0.029396 |
| TNFAIP8L1 | 517.4517 | -0.46415 | 0.219976 | -2.11 | 0.034858 | 0.109623 |
| RAD51D | 180.7649 | -0.47498 | 0.24469 | -1.94116 | 0.052239 | 0.142531 |
| TARSL2 | 177.8877 | 0.47999 | 0.210377 | 2.28157 | 0.022515 | 0.081968 |

| | | | | | | |
|---|---|---|---|---|---|---|
| FAM174B | 999.0098 | -0.40682 | 0.149325 | -2.72439 | 0.006442 | 0.035027 |
| C17orf70 | 1182.649 | -0.3775 | 0.184086 | -2.05065 | 0.040301 | 0.120955 |
| SV2B | 1614.443 | -0.67713 | 0.21327 | -3.175 | 0.001498 | 0.013563 |
| PDE6G | 16.4432 | -0.47321 | 0.519706 | -0.91054 | 0.362538 | NA |
| NR2F2 | 788.5528 | -0.18492 | 0.241889 | -0.76447 | 0.444586 | 0.604492 |
| TLCD2 | 22.85019 | -0.25061 | 0.450461 | -0.55633 | 0.577982 | NA |
| WASH3P | 99.52332 | 0.091065 | 0.237764 | 0.383005 | 0.701716 | 0.808361 |
| P4HB | 19456.81 | -0.30572 | 0.123014 | -2.48526 | 0.012946 | 0.056298 |
| SNAI3 | 8.869512 | -0.0259 | 0.71271 | -0.03634 | 0.97101 | NA |
| C16orf52 | 291.5498 | 0.635628 | 0.192728 | 3.298056 | 9.74E-04 | 0.010161 |
| ANKFY1 | 3340.676 | -0.50326 | 0.145822 | -3.45119 | 5.58E-04 | 0.007239 |
| SRL | 0.096202 | 0.408546 | 4.079639 | 0.100143 | 0.920231 | NA |
| ADAMTSL5 | 328.0939 | 0.027323 | 0.250352 | 0.109138 | 0.913093 | 0.94937 |
| PCYT2 | 3120.146 | -0.26238 | 0.136469 | -1.9226 | 0.05453 | 0.147055 |
| ARL17A | 3.205172 | 0.035747 | 1.19021 | 0.030034 | 0.97604 | NA |
| EVI2B | 61.88611 | 0.138321 | 0.322642 | 0.428712 | 0.668133 | 0.782242 |
| NPIPB4 | 16.54936 | 0.126004 | 0.613115 | 0.205514 | 0.83717 | NA |
| ZNF829 | 42.45053 | 0.700523 | 0.555261 | 1.261611 | 0.207089 | NA |
| ATP6V0C | 55.55524 | -0.44958 | 0.330863 | -1.3588 | 0.174211 | 0.326183 |
| C16orf54 | 0.163957 | 0.424913 | 4.079006 | 0.104171 | 0.917034 | NA |
| RTN4RL1 | 255.5871 | -0.22908 | 0.186535 | -1.22805 | 0.219427 | 0.379144 |
| PAGR1 | 396.6908 | -0.23424 | 0.219749 | -1.06592 | 0.286458 | 0.452735 |
| ZNF267 | 19.49198 | 0.683468 | 0.544018 | 1.256334 | 0.208995 | NA |
| RPS4XP21 | 0.163957 | 0.424913 | 4.079006 | 0.104171 | 0.917034 | NA |
| NDUFA13 | 111.4519 | -0.06599 | 0.244728 | -0.26966 | 0.787425 | 0.871252 |
| ZNF566 | 121.1583 | 0.530935 | 0.240643 | 2.206315 | 0.027362 | 0.093319 |
| ZNF529 | 136.1057 | 0.478346 | 0.290873 | 1.644518 | 0.100069 | 0.223218 |
| PIP5K1C | 3144.655 | -0.49398 | 0.141774 | -3.48429 | 4.93E-04 | 0.006739 |
| WWOX | 369.3187 | -0.17124 | 0.152627 | -1.12193 | 0.261892 | 0.426298 |
| KIF18B | 255.8995 | -0.7227 | 0.300575 | -2.40439 | 0.0162 | 0.065554 |
| ZNRF1 | 370.949 | -0.38166 | 0.179162 | -2.13028 | 0.033149 | 0.106017 |
| MKL2 | 604.4976 | -0.34801 | 0.198321 | -1.7548 | 0.079294 | 0.190173 |
| ZNF555 | 126.6701 | -0.06406 | 0.221269 | -0.28952 | 0.772184 | 0.858857 |
| RGS9BP | 12.1994 | -0.30449 | 0.634616 | -0.4798 | 0.631371 | NA |
| KRT10 | 1388.204 | 0.052965 | 0.160761 | 0.329465 | 0.741804 | 0.837188 |
| ANKRD20A | 1.159694 | -0.24887 | 2.055096 | -0.1211 | 0.903612 | NA |
| ZNF396 | 41.47874 | -0.20002 | 0.372557 | -0.5369 | 0.591339 | NA |
| SMYD4 | 541.344 | -0.0487 | 0.170495 | -0.28565 | 0.775145 | 0.860909 |
| GPATCH8 | 1250.414 | -0.09041 | 0.144891 | -0.62401 | 0.532622 | 0.676767 |
| MIR22HG | 498.0497 | 0.519273 | 0.191408 | 2.712914 | 0.006669 | 0.035967 |
| C17orf58 | 198.2276 | -0.09325 | 0.243804 | -0.38248 | 0.702106 | 0.808431 |
| FSCN2 | 3.82523 | 0.560588 | 1.038966 | 0.539563 | 0.589498 | NA |
| ZNF397 | 233.4431 | 0.23637 | 0.259087 | 0.912322 | 0.361599 | 0.528818 |
| ZSCAN30 | 263.1181 | 0.170117 | 0.223595 | 0.760828 | 0.44676 | 0.606544 |
| HEXIM1 | 2389.079 | 0.036049 | 0.119959 | 0.300513 | 0.763786 | 0.85349 |
| KRT14 | 14.5453 | 0.30899 | 0.680406 | 0.454126 | 0.649738 | NA |
| MAPT | 110.5376 | -1.48496 | 0.390519 | -3.80254 | 1.43E-04 | 0.003156 |

| | | | | | | |
|---|---|---|---|---|---|---|
| ZACN | 1.937385 | -1.41272 | 1.911357 | -0.73912 | 0.459835 | NA |
| KANK3 | 22.97608 | -0.38973 | 0.50364 | -0.77383 | 0.439033 | NA |
| PKD1L3 | 5.12799 | 0.614834 | 0.880377 | 0.698376 | 0.484942 | NA |
| C17orf82 | 61.46491 | -0.03856 | 0.335699 | -0.11486 | 0.908554 | 0.947226 |
| VSTM2B | 13.29234 | 0.467825 | 0.630199 | 0.742345 | 0.457879 | NA |
| | 0.556813 | -0.57007 | 3.294615 | -0.17303 | 0.862627 | NA |
| ZNF546 | 47.12841 | 0.506643 | 0.342504 | 1.479231 | 0.139078 | 0.279075 |
| MT1X | 31.02081 | -0.61382 | 0.372212 | -1.64911 | 0.099125 | NA |
| KRT12 | 0.296509 | -0.51709 | 3.982155 | -0.12985 | 0.896684 | NA |
| EPOR | 612.108 | -0.38099 | 0.176816 | -2.15475 | 0.031182 | 0.10175 |
| RDM1 | 23.4307 | -0.28482 | 0.514491 | -0.5536 | 0.579853 | NA |
| RPL7P48 | 0.096202 | 0.408546 | 4.079639 | 0.100143 | 0.920231 | NA |
| SIRT7 | 448.929 | 0.040225 | 0.183927 | 0.218699 | 0.826884 | 0.895634 |
| | 31.18565 | -0.05114 | 0.389801 | -0.1312 | 0.895615 | NA |
| IFT140 | 716.7107 | -0.45211 | 0.149423 | -3.02574 | 0.00248 | 0.018705 |
| USP7 | 1843.016 | 0.122775 | 0.15258 | 0.80466 | 0.421016 | 0.58376 |
| NANOS3 | 6.887129 | -0.70056 | 0.793309 | -0.88309 | 0.377188 | NA |
| ZNF385C | 1.591713 | 0.041659 | 1.513932 | 0.027517 | 0.978047 | NA |
| | 94.57298 | 0.31422 | 0.287907 | 1.091392 | 0.275101 | 0.440659 |
| C17orf97 | 34.18922 | -0.4251 | 0.367656 | -1.15625 | 0.247581 | NA |
| VMAC | 109.1017 | -0.14622 | 0.229054 | -0.63838 | 0.523225 | 0.669419 |
| HAPLN4 | 0.432222 | -1.18916 | 3.482045 | -0.34151 | 0.732718 | NA |
| TRPV2 | 6095.618 | -0.32661 | 0.146307 | -2.23238 | 0.02559 | 0.089457 |
| FANCA | 278.4184 | -0.42465 | 0.244209 | -1.73889 | 0.082054 | 0.194628 |
| FAM69C | 1487.218 | -0.3983 | 0.226417 | -1.75913 | 0.078556 | 0.188901 |
| DNAH17 | 16.0936 | 0.115907 | 0.530692 | 0.218407 | 0.827112 | NA |
| TMEM220 | 121.6827 | 0.161479 | 0.301772 | 0.535101 | 0.59258 | 0.7233 |
| TMEM256- | 5.989862 | -0.06492 | 0.834623 | -0.07779 | 0.937999 | NA |
| PALM3 | 2.471756 | 0.38649 | 1.294845 | 0.298484 | 0.765334 | NA |
| CLEC17A | 0.938845 | -1.23445 | 2.063177 | -0.59832 | 0.549624 | NA |
| RINL | 493.3368 | 0.140877 | 0.168275 | 0.837184 | 0.402489 | 0.566889 |
| OR7D2 | 9.857115 | -0.67816 | 0.657082 | -1.03207 | 0.302039 | NA |
| MEIS3P2 | 127.4285 | 0.409334 | 0.27922 | 1.465992 | 0.142651 | 0.283603 |
| C19orf67 | 0.921281 | -1.21902 | 2.226885 | -0.54741 | 0.584096 | NA |
| ZNF490 | 38.38958 | -0.61968 | 0.345361 | -1.79429 | 0.072766 | NA |
| TMEM221 | 4.232021 | 0.604413 | 1.013648 | 0.596275 | 0.550991 | NA |
| ZNF626 | 35.79495 | 0.662542 | 0.474669 | 1.395798 | 0.162775 | NA |
| SMTNL2 | 0.382031 | -1.26484 | 3.151989 | -0.40128 | 0.688212 | NA |
| DCUN1D3 | 181.6858 | -0.0989 | 0.251545 | -0.39315 | 0.694207 | 0.802988 |
| ZNF793 | 1.79002 | -0.62742 | 1.802477 | -0.34809 | 0.727773 | NA |
| OR7A5 | 6.370754 | -0.27517 | 0.89021 | -0.30911 | 0.757239 | NA |
| ZNF383 | 99.18667 | 0.418707 | 0.242384 | 1.727449 | 0.084087 | 0.197553 |
| C19orf35 | 0.411855 | 0.250456 | 3.097618 | 0.080854 | 0.935558 | NA |
| ZNF559 | 160.9035 | 0.349602 | 0.249278 | 1.402458 | 0.160779 | 0.308346 |
| SBK1 | 7.714043 | 0.176137 | 0.716814 | 0.245722 | 0.805897 | NA |
| PRR19 | 46.3873 | 0.068415 | 0.360072 | 0.190005 | 0.849305 | NA |
| C19orf54 | 304.9823 | -0.57225 | 0.171589 | -3.33503 | 8.53E-04 | 0.009397 |

| | | | | | | |
|---|---|---|---|---|---|---|
| NCCRP1 | 1056.992 | -1.2198 | 0.610494 | -1.99806 | 0.04571 | 0.130925 |
| FAM83G | 214.729 | -0.37554 | 0.200573 | -1.87231 | 0.061163 | 0.159466 |
| NBR1 | 4996.027 | 0.14297 | 0.144572 | 0.988916 | 0.322704 | 0.490721 |
| NPIPP1 | 64.68168 | 0.015569 | 0.332848 | 0.046775 | 0.962692 | 0.979674 |
| CLN3 | 110.1478 | -0.18074 | 0.237962 | -0.75954 | 0.44753 | 0.607134 |
| SUMO2 | 2039.075 | 0.222569 | 0.146793 | 1.516214 | 0.129465 | 0.265906 |
| ZNF177 | 20.14523 | 0.080289 | 0.569485 | 0.140985 | 0.887882 | NA |
| HILS1 | 66.21171 | 0.275492 | 0.287778 | 0.957308 | 0.338412 | 0.507254 |
| SPRED3 | 7.635982 | -0.57193 | 0.823844 | -0.69422 | 0.487543 | NA |
| SHISA6 | 0.163957 | 0.424913 | 4.079006 | 0.104171 | 0.917034 | NA |
| LINC00910 | 166.8081 | -0.06436 | 0.255325 | -0.25207 | 0.800987 | 0.880141 |
| SLX4 | 360.8485 | -0.75884 | 0.201899 | -3.7585 | 1.71E-04 | 0.003521 |
| ZNF563 | 41.26696 | 0.317871 | 0.385927 | 0.823655 | 0.410136 | NA |
| FBF1 | 89.1926 | -0.34564 | 0.311066 | -1.11114 | 0.266509 | 0.431444 |
| GPR179 | 0.151697 | -1.546 | 4.075469 | -0.37934 | 0.704433 | NA |
| MSL1 | 2312.477 | -0.00498 | 0.134068 | -0.03713 | 0.970382 | 0.983911 |
| | 4.347647 | -0.64483 | 1.092113 | -0.59045 | 0.554891 | NA |
| DHFRP1 | 4.750133 | 0.788689 | 0.983111 | 0.802238 | 0.422415 | NA |
| SBSN | 1.646997 | -2.04733 | 2.063777 | -0.99203 | 0.321182 | NA |
| ZNF567 | 80.60938 | 0.594871 | 0.296429 | 2.006794 | 0.044772 | 0.129302 |
| RNFT1 | 205.2386 | 0.523722 | 0.219443 | 2.386594 | 0.017005 | 0.067878 |
| LITAF | 5692.885 | -0.28392 | 0.172191 | -1.64887 | 0.099174 | 0.222084 |
| SF3B3 | 4141.477 | -0.52339 | 0.13012 | -4.02239 | 5.76E-05 | 0.001791 |
| SP6 | 0.571659 | -3.00664 | 3.038724 | -0.98944 | 0.322447 | NA |
| ZNF573 | 40.5552 | -0.15013 | 0.388944 | -0.386 | 0.699499 | NA |
| CRYM-AS1 | 11.06214 | 0.554588 | 0.709713 | 0.781425 | 0.434552 | NA |
| HN1 | 2120.603 | -0.34784 | 0.143111 | -2.43058 | 0.015075 | 0.062268 |
| ZNF527 | 63.36681 | 0.341442 | 0.295446 | 1.155683 | 0.247811 | 0.411341 |
| RPS2P46 | 80.78632 | -0.5918 | 0.301348 | -1.96384 | 0.049548 | 0.137692 |
| SPATA41 | 3.125213 | -0.16906 | 1.131669 | -0.14939 | 0.881246 | NA |
| USP32P3 | 0.399595 | -1.31595 | 3.579466 | -0.36764 | 0.713143 | NA |
| ZNF724P | 17.9268 | 0.93616 | 0.661092 | 1.416081 | 0.156752 | NA |
| ZNF699 | 43.20356 | -0.05447 | 0.370756 | -0.14691 | 0.8832 | NA |
| C16orf93 | 22.6253 | 0.126428 | 0.456288 | 0.277079 | 0.781719 | NA |
| KIAA0895L | 397.6373 | 0.048019 | 0.197598 | 0.243016 | 0.807993 | 0.885317 |
| PLEKHG4 | 306.7026 | -0.50549 | 0.23256 | -2.1736 | 0.029735 | 0.098467 |
| KIF19 | 0.096202 | 0.408546 | 4.079639 | 0.100143 | 0.920231 | NA |
| ZNF681 | 11.50453 | 1.427618 | 0.868075 | 1.64458 | 0.100056 | NA |
| RYR1 | 35.58175 | 0.105225 | 0.405221 | 0.259674 | 0.795115 | NA |
| SUPT5H | 4757.338 | -0.37852 | 0.134421 | -2.81596 | 0.004863 | 0.028972 |
| ZNF493 | 41.40127 | 0.352928 | 0.334235 | 1.055929 | 0.291 | NA |
| ATP2A1 | 6.65476 | 0.740658 | 0.785532 | 0.942874 | 0.345745 | NA |
| ZNF565 | 37.41542 | 0.698956 | 0.373767 | 1.870032 | 0.061479 | NA |
| ELAVL3 | 0.476449 | 0.524262 | 3.402674 | 0.154073 | 0.877552 | NA |
| LONP1 | 5187.17 | -0.31934 | 0.139955 | -2.28177 | 0.022503 | 0.081968 |
| ZNF781 | 6.273071 | 0.571724 | 0.862172 | 0.663121 | 0.507253 | NA |
| INCA1 | 34.77037 | -0.13344 | 0.375298 | -0.35556 | 0.72217 | NA |

| | | | | | | |
|---|---|---|---|---|---|---|
| ZNF774 | 20.60349 | -0.14978 | 0.501281 | -0.29879 | 0.765103 | NA |
| NOXO1 | 6.6445 | -0.34253 | 0.837808 | -0.40884 | 0.682655 | NA |
| PRTN3 | 0.327913 | 1.543154 | 4.021063 | 0.383768 | 0.701151 | NA |
| NPIPB15 | 17.6568 | 1.006111 | 0.575431 | 1.748447 | 0.080387 | NA |
| ZNF569 | 56.01445 | 0.857819 | 0.372204 | 2.304701 | 0.021183 | 0.078699 |
| ZNF799 | 33.9852 | 1.113346 | 0.389269 | 2.860092 | 0.004235 | NA |
| SIAH1 | 192.7673 | 0.292619 | 0.19147 | 1.528281 | 0.126443 | 0.261693 |
| SULT1A1 | 3.103422 | 0.213398 | 1.342045 | 0.159009 | 0.873661 | NA |
| MYO18A | 3144.007 | -0.6842 | 0.170997 | -4.00123 | 6.30E-05 | 0.001896 |
| C17orf59 | 737.3658 | -0.1646 | 0.186467 | -0.88275 | 0.377369 | 0.543904 |
| MAN2A2 | 4347.887 | -0.37344 | 0.279683 | -1.33524 | 0.181799 | 0.334668 |
| CACNA1H | 0.661122 | -3.06315 | 2.98049 | -1.02773 | 0.304076 | NA |
| ZNF846 | 89.91484 | 0.183619 | 0.250273 | 0.733674 | 0.463147 | 0.62097 |
| TCF4 | 276.8897 | 0.192679 | 0.269448 | 0.715086 | 0.474556 | 0.629784 |
| ZNF136 | 113.4602 | 0.494716 | 0.237893 | 2.079574 | 0.037565 | 0.115179 |
| | 0.134133 | -1.48909 | 4.077584 | -0.36519 | 0.71497 | NA |
| ERI2 | 203.8134 | 0.64847 | 0.272841 | 2.376732 | 0.017467 | 0.069115 |
| HSH2D | 0.392857 | -1.546 | 4.075469 | -0.37934 | 0.704433 | NA |
| TRPV1 | 1.222745 | -0.03123 | 1.954581 | -0.01598 | 0.98725 | NA |
| PDXDC2P | 9.455456 | 0.670656 | 0.734431 | 0.913165 | 0.361156 | NA |
| AMZ2 | 1898.179 | -0.05956 | 0.133671 | -0.44555 | 0.655919 | 0.772809 |
| ZNF431 | 63.94692 | 0.902099 | 0.319152 | 2.82655 | 0.004705 | 0.028358 |
| NF1 | 1770.906 | 0.159217 | 0.204128 | 0.779989 | 0.435398 | 0.596445 |
| ZNF700 | 141.902 | 0.793842 | 0.25848 | 3.071197 | 0.002132 | 0.016837 |
| | 12.28946 | -0.82779 | 0.644226 | -1.28493 | 0.198816 | NA |
| | 0.574231 | 0.93227 | 2.49998 | 0.372911 | 0.709215 | NA |
| ZNF585A | 76.53701 | 0.516886 | 0.33458 | 1.544882 | 0.122375 | 0.25582 |
| METTL9 | 6123.14 | 0.066136 | 0.155484 | 0.425356 | 0.670577 | 0.784225 |
| ZNF429 | 70.70707 | -0.20196 | 0.325201 | -0.62104 | 0.53457 | 0.678152 |
| SERTAD1 | 577.415 | 0.16858 | 0.213497 | 0.789614 | 0.429753 | 0.591241 |
| ZNF100 | 50.61591 | 0.75054 | 0.486377 | 1.543125 | 0.1228 | 0.256483 |
| ZNF441 | 70.94198 | 1.474345 | 0.480729 | 3.066891 | 0.002163 | 0.017036 |
| SIGLEC15 | 1.39126 | 0.989136 | 1.681347 | 0.5883 | 0.556331 | NA |
| ZNF420 | 118.0556 | 0.456285 | 0.240569 | 1.896692 | 0.057869 | 0.153117 |
| ZNF763 | 10.72775 | 0.514603 | 0.710017 | 0.724776 | 0.468589 | NA |
| MAFG | 2271.287 | -0.02502 | 0.145511 | -0.17195 | 0.863479 | 0.91956 |
| ZNF682 | 24.49149 | 0.751948 | 0.478166 | 1.572567 | 0.115819 | NA |
| ZNF257 | 23.35526 | 0.530851 | 0.448118 | 1.184623 | 0.236167 | NA |
| ZNF785 | 126.3297 | -0.57469 | 0.263252 | -2.18304 | 0.029033 | 0.096974 |
| PSMD12 | 950.1281 | 0.492789 | 0.211163 | 2.333688 | 0.019612 | 0.074735 |
| KANK2 | 8054.248 | -0.57524 | 0.1833 | -3.13824 | 0.0017 | 0.014569 |
| IL27 | 0.151697 | -1.546 | 4.075469 | -0.37934 | 0.704433 | NA |
| RAMP2-AS: | 2.345178 | 1.027271 | 1.462557 | 0.70238 | 0.482442 | NA |
| BLM | 163.153 | 0.399927 | 0.263087 | 1.520133 | 0.128477 | 0.264779 |
| ZNF720 | 164.9514 | 0.743079 | 0.327722 | 2.267406 | 0.023365 | 0.084066 |
| ZNF833P | 4.811198 | 0.33303 | 0.927731 | 0.358973 | 0.719615 | NA |
| ZNF98 | 0.096202 | 0.408546 | 4.079639 | 0.100143 | 0.920231 | NA |

| | | | | | | |
|---|---|---|---|---|---|---|
| ZNF675 | 47.63168 | 0.990966 | 0.460012 | 2.154218 | 0.031223 | 0.101857 |
| SHPK | 149.1059 | -0.16777 | 0.21237 | -0.78998 | 0.429541 | 0.591086 |
| C16orf47 | 8.278543 | -0.21431 | 0.689525 | -0.31081 | 0.755948 | NA |
| SPN | 0.533728 | -1.74213 | 2.820432 | -0.61768 | 0.536785 | NA |
| RAB40C | 457.9568 | -0.31222 | 0.153702 | -2.03133 | 0.042222 | 0.12501 |
| PIGN | 904.8822 | 0.17495 | 0.171158 | 1.02216 | 0.306705 | 0.473806 |
| ZNF624 | 55.69186 | 0.467915 | 0.493059 | 0.949004 | 0.342619 | 0.510783 |
| CCDC154 | 38.04706 | 0.145322 | 0.362123 | 0.401306 | 0.688195 | NA |
| SERPINB2 | 26.38998 | 1.476851 | 0.583387 | 2.531512 | 0.011357 | NA |
| ZNF433 | 57.45935 | 0.330082 | 0.294664 | 1.120199 | 0.262629 | 0.427029 |
| TBC1D3 | 0.327913 | 1.543154 | 4.021063 | 0.383768 | 0.701151 | NA |
| KLHL14 | 6.7713 | 1.075662 | 0.784998 | 1.370274 | 0.170601 | NA |
| HSPB9 | 2.576503 | -1.97666 | 1.500193 | -1.3176 | 0.187637 | NA |
| | 0.258578 | 1.233151 | 4.047809 | 0.304646 | 0.760635 | NA |
| CFD | 6.306961 | 0.389523 | 0.921592 | 0.422663 | 0.672541 | NA |
| EME2 | 238.0642 | 0.014036 | 0.21333 | 0.065797 | 0.94754 | 0.97088 |
| ZNF780A | 189.8376 | 0.587052 | 0.279713 | 2.098766 | 0.035837 | 0.111632 |
| ZNF461 | 105.8763 | 0.32305 | 0.235436 | 1.372135 | 0.170021 | 0.320363 |
| | 11.99152 | 0.477138 | 0.623298 | 0.765506 | 0.443971 | NA |
| ZNF181 | 107.6467 | 1.047534 | 0.315376 | 3.321541 | 8.95E-04 | 0.009708 |
| ZNF44 | 84.27876 | 0.547727 | 0.279714 | 1.958171 | 0.05021 | 0.138945 |
| ZNF790 | 75.82487 | 0.540969 | 0.281494 | 1.921775 | 0.054634 | 0.147202 |
| MYO1C | 13501.68 | -0.42581 | 0.127764 | -3.33278 | 8.60E-04 | 0.009421 |
| SPG7 | 1659.187 | -0.25096 | 0.146717 | -1.71047 | 0.087179 | 0.202702 |
| ZNF823 | 77.19918 | 0.692573 | 0.32915 | 2.104124 | 0.035368 | 0.110746 |
| PLCG2 | 180.0878 | -0.2038 | 0.20329 | -1.00249 | 0.316105 | 0.483539 |
| ZNF121 | 202.6146 | 0.654799 | 0.275752 | 2.374595 | 0.017568 | 0.069308 |
| MBP | 14604.2 | -0.35235 | 0.171748 | -2.05157 | 0.040211 | 0.120837 |
| CCDC151 | 13.46421 | -0.31382 | 0.564176 | -0.55624 | 0.578049 | NA |
| ZNF560 | 2.084378 | -1.4482 | 1.518173 | -0.95391 | 0.34013 | NA |
| | 0.489058 | -0.58569 | 3.427543 | -0.17088 | 0.864321 | NA |
| ZBTB14 | 330.0921 | 0.436524 | 0.193776 | 2.25273 | 0.024276 | 0.086227 |
| SNX29P2 | 2.137658 | -3.78876 | 1.733952 | -2.18504 | 0.028886 | NA |
| | 6.376974 | 0.590149 | 0.844253 | 0.699018 | 0.48454 | NA |
| ZNF607 | 78.28973 | 0.021638 | 0.30393 | 0.071194 | 0.943243 | 0.968571 |
| DDX42 | 2337.121 | 0.125834 | 0.158091 | 0.795961 | 0.426055 | 0.587965 |
| RPL23A | 4312.515 | 0.265396 | 0.196534 | 1.350382 | 0.176893 | 0.329058 |
| UBL5 | 4559.732 | 0.139728 | 0.162711 | 0.858749 | 0.390479 | 0.555738 |
| HELZ | 1224.893 | 0.034218 | 0.163143 | 0.209739 | 0.833871 | 0.900487 |
| MYL4 | 0.803131 | -0.76693 | 2.349822 | -0.32638 | 0.744138 | NA |
| ZNF442 | 9.863829 | 0.602043 | 0.643304 | 0.935861 | 0.349345 | NA |
| ASNA1 | 4473.361 | -0.14871 | 0.160891 | -0.92428 | 0.355342 | 0.523627 |
| WWP2 | 1495.328 | -0.57234 | 0.141615 | -4.04153 | 5.31E-05 | 0.001711 |
| MT1F | 58.11559 | 0.022386 | 0.344787 | 0.064926 | 0.948233 | 0.971147 |
| ZNF69 | 161.187 | 0.378636 | 0.205362 | 1.843752 | 0.065219 | 0.166489 |
| ZNF568 | 137.9762 | 0.426246 | 0.252372 | 1.688958 | 0.091227 | 0.209796 |
| NBR2 | 50.4681 | -0.18197 | 0.333074 | -0.54635 | 0.584826 | 0.717944 |

| | | | | | | |
|---|---|---|---|---|---|---|
| ZNF43 | 94.95523 | 0.384732 | 0.400791 | 0.959932 | 0.33709 | 0.505912 |
| ZNF627 | 323.5522 | 0.108321 | 0.181512 | 0.596769 | 0.550662 | 0.691902 |
| ZNF536 | 417.4583 | -0.23749 | 0.201173 | -1.18052 | 0.237794 | 0.400717 |
| TAT | 0.35636 | 1.709252 | 3.651833 | 0.468053 | 0.639746 | NA |
| ANKRD13B | 360.107 | -0.3698 | 0.168251 | -2.1979 | 0.027957 | 0.094666 |
| C19orf45 | 0.327913 | 1.543154 | 4.021063 | 0.383768 | 0.701151 | NA |
| MSRB1 | 799.4208 | -0.13919 | 0.163556 | -0.851 | 0.394767 | 0.559494 |
| ZNF652 | 347.7235 | -0.00673 | 0.234177 | -0.02872 | 0.977087 | 0.987584 |
| ZNF830 | 305.1799 | 0.522783 | 0.170313 | 3.06954 | 0.002144 | 0.01691 |
| ZNF521 | 168.8817 | -0.8116 | 0.275371 | -2.94728 | 0.003206 | 0.0221 |
| ZNF358 | 1640.089 | -0.24635 | 0.226973 | -1.08536 | 0.277764 | 0.443726 |
| ARHGEF15 | 0.134133 | -1.48909 | 4.077584 | -0.36519 | 0.71497 | NA |
| CES1 | 55.76336 | 0.635536 | 1.143245 | 0.555906 | 0.578275 | 0.712922 |
| R3HDM4 | 1284.322 | -0.20618 | 0.128141 | -1.60902 | 0.107612 | 0.234081 |
| RUNDC1 | 665.0092 | -0.28341 | 0.167145 | -1.69558 | 0.089966 | 0.207661 |
| PRC1 | 2284.94 | 0.025186 | 0.245827 | 0.102453 | 0.918397 | 0.952713 |
| MAP3K3 | 3757.623 | -0.53363 | 0.137192 | -3.88964 | 1.00E-04 | 0.002559 |
| KIAA0753 | 307.1641 | 0.162351 | 0.176119 | 0.921824 | 0.35662 | 0.524272 |
| APRT | 2618.268 | -0.14403 | 0.150958 | -0.95414 | 0.340015 | 0.508371 |
| TBKBP1 | 442.0744 | -0.27285 | 0.154893 | -1.76153 | 0.078148 | 0.188276 |
| MIR324 | 1.373899 | 0.534052 | 2.128914 | 0.250857 | 0.801925 | NA |
| MIR365B | 0.163957 | 0.424913 | 4.079006 | 0.104171 | 0.917034 | NA |
| SNORD104 | 33.39325 | 0.234187 | 0.396415 | 0.590762 | 0.55468 | NA |
| | 0.096202 | 0.408546 | 4.079639 | 0.100143 | 0.920231 | NA |
| RNA5SP452 | 0.096202 | 0.408546 | 4.079639 | 0.100143 | 0.920231 | NA |
| SNORD1B | 4.013075 | 0.404453 | 0.991036 | 0.408111 | 0.683192 | NA |
| RNU6-623P | 0.422534 | 1.924583 | 3.001758 | 0.641152 | 0.521424 | NA |
| | 0.096202 | 0.408546 | 4.079639 | 0.100143 | 0.920231 | NA |
| RNU4-58P | 0.192403 | 0.975392 | 4.060716 | 0.240202 | 0.810174 | NA |
| RNU6-181P | 0.260158 | 1.240347 | 4.047124 | 0.306476 | 0.759242 | NA |
| RNA5SP440 | 0.163957 | 0.424913 | 4.079006 | 0.104171 | 0.917034 | NA |
| | 0.192403 | 0.975392 | 4.060716 | 0.240202 | 0.810174 | NA |
| | 0.134133 | -1.48909 | 4.077584 | -0.36519 | 0.71497 | NA |
| RNU6-97P | 0.096202 | 0.408546 | 4.079639 | 0.100143 | 0.920231 | NA |
| SNORA38B | 0.134133 | -1.48909 | 4.077584 | -0.36519 | 0.71497 | NA |
| RNA5-8SP2 | 63.30495 | 0.012479 | 0.329776 | 0.03784 | 0.969815 | 0.983611 |
| RN7SKP44 | 0.163957 | 0.424913 | 4.079006 | 0.104171 | 0.917034 | NA |
| RNU6-288P | 0.162376 | 0.424913 | 4.079006 | 0.104171 | 0.917034 | NA |
| | 1.214074 | 2.321363 | 2.323069 | 0.999266 | 0.317666 | NA |
| | 0.096202 | 0.408546 | 4.079639 | 0.100143 | 0.920231 | NA |
| | 0.096202 | 0.408546 | 4.079639 | 0.100143 | 0.920231 | NA |
| | 0.163957 | 0.424913 | 4.079006 | 0.104171 | 0.917034 | NA |
| RNU1-60P | 1.237926 | 1.940943 | 2.028448 | 0.956861 | 0.338637 | NA |
| | 0.163957 | 0.424913 | 4.079006 | 0.104171 | 0.917034 | NA |
| | 0.558248 | 0.982076 | 2.804738 | 0.350149 | 0.726227 | NA |
| | 9.327467 | -0.15949 | 0.69308 | -0.23012 | 0.817996 | NA |
| | 0.937264 | -1.23834 | 2.063944 | -0.59999 | 0.548514 | NA |

| | | | | | | |
|---|---|---|---|---|---|---|
| OR1F2P | 0.28583 | -2.21358 | 3.97376 | -0.55705 | 0.577494 | NA |
| OXLD1 | 569.6384 | -0.1129 | 0.150036 | -0.75251 | 0.451744 | 0.610791 |
| TNRC6C-AS | 15.77342 | 0.064306 | 0.560718 | 0.114684 | 0.908695 | NA |
| MRPL38 | 79.86152 | -0.12259 | 0.300346 | -0.40816 | 0.683159 | 0.79385 |
| SMIM5 | 8.346302 | 0.080316 | 0.794352 | 0.101109 | 0.919464 | NA |
| | 0.420954 | 1.923376 | 3.462803 | 0.555439 | 0.578594 | NA |
| CRHR1-IT1 | 119.7695 | -0.38814 | 0.236668 | -1.64004 | 0.100997 | 0.224425 |
| TTC25 | 22.0921 | 0.306416 | 0.466608 | 0.656688 | 0.511381 | NA |
| LRRC3C | 0.326333 | 1.533889 | 3.826878 | 0.40082 | 0.688553 | NA |
| ZNF155 | 85.85491 | 0.222992 | 0.285445 | 0.781206 | 0.434681 | 0.595739 |
| CD177 | 0.151697 | -1.546 | 4.075469 | -0.37934 | 0.704433 | NA |
| FBXO47 | 0.555087 | 0.973802 | 2.808575 | 0.346725 | 0.728798 | NA |
| | 9.28157 | -0.07425 | 0.687651 | -0.10798 | 0.914015 | NA |
| SPIRE2 | 378.7447 | -0.11627 | 0.19496 | -0.59638 | 0.550922 | 0.692009 |
| | 0.394291 | 0.298745 | 3.141711 | 0.09509 | 0.924243 | NA |
| | 3.689279 | 0.108067 | 1.126196 | 0.095958 | 0.923554 | NA |
| SLFN12L | 1.268901 | -0.96607 | 1.991761 | -0.48503 | 0.627654 | NA |
| | 31.8265 | -0.17993 | 0.408045 | -0.44097 | 0.659237 | NA |
| SYCE1L | 19.11847 | 0.334084 | 0.46221 | 0.722797 | 0.469805 | NA |
| TMEM231 | 227.7298 | -0.50259 | 0.188157 | -2.6711 | 0.00756 | 0.039258 |
| SDHAF1 | 401.2434 | -0.08872 | 0.178336 | -0.49749 | 0.618844 | 0.743481 |
| | 1.406049 | -0.26601 | 1.627656 | -0.16343 | 0.870178 | NA |
| PSENEN | 376.1298 | -0.07447 | 0.196058 | -0.37986 | 0.704048 | 0.810103 |
| SCGB2B2 | 2.414083 | -0.03119 | 1.350453 | -0.02309 | 0.981576 | NA |
| PSMB10 | 518.6682 | -0.28811 | 0.164493 | -1.75149 | 0.079861 | 0.191148 |
| RPSAP58 | 14.77933 | -0.08879 | 0.530329 | -0.16743 | 0.867033 | NA |
| E2F4 | 1594.289 | -0.05561 | 0.142859 | -0.38927 | 0.697077 | 0.804815 |
| NT5M | 108.5885 | -0.90043 | 0.257803 | -3.49269 | 4.78E-04 | 0.006638 |
| GPR56 | 16137.23 | -1.0276 | 0.200263 | -5.13124 | 2.88E-07 | 6.69E-05 |
| MT1DP | 0.096202 | 0.408546 | 4.079639 | 0.100143 | 0.920231 | NA |
| MT1A | 9.773541 | -0.32501 | 0.65462 | -0.49649 | 0.619552 | NA |
| MT1M | 1.312853 | -1.50193 | 1.923163 | -0.78097 | 0.434822 | NA |
| | 3.307514 | 0.181502 | 1.246266 | 0.145636 | 0.884208 | NA |
| CNEP1R1 | 440.9966 | 0.27292 | 0.169397 | 1.611126 | 0.107152 | 0.233595 |
| RGL3 | 105.2815 | 0.13614 | 0.245877 | 0.553693 | 0.579789 | 0.713749 |
| | 30.62381 | -0.24461 | 0.433548 | -0.56421 | 0.572609 | NA |
| TMEM256 | 253.3787 | 0.007451 | 0.197597 | 0.037707 | 0.969921 | 0.983611 |
| EIF3CL | 0.545988 | -0.43913 | 2.541145 | -0.17281 | 0.862803 | NA |
| LCMT1 | 943.2055 | -0.15079 | 0.159665 | -0.94442 | 0.344954 | 0.512974 |
| C17orf107 | 9.837432 | 0.443386 | 0.823385 | 0.538492 | 0.590238 | NA |
| ITPRIPL2 | 1768.938 | -0.02335 | 0.180508 | -0.12933 | 0.897094 | 0.939771 |
| DENND1C | 12.17063 | -0.66178 | 0.804321 | -0.82278 | 0.410634 | NA |
| | 16.88463 | -0.295 | 0.494175 | -0.59696 | 0.550537 | NA |
| | 0.744715 | -0.63075 | 2.564852 | -0.24592 | 0.805743 | NA |
| | 2.024984 | 1.25763 | 1.59508 | 0.788443 | 0.430438 | NA |
| | 3.664471 | 0.443989 | 1.062389 | 0.417916 | 0.676009 | NA |
| C16orf96 | 2.505857 | -1.86969 | 1.465604 | -1.27571 | 0.202058 | NA |

| | | | | | | |
|---|---|---|---|---|---|---|
| SRRM2-AS: | 2.907285 | -0.22405 | 1.221114 | -0.18348 | 0.85442 | NA |
| | 1.693073 | 0.276824 | 1.517715 | 0.182395 | 0.855272 | NA |
| RNPS1 | 1723.826 | -0.37973 | 0.139314 | -2.7257 | 0.006416 | 0.034941 |
| DOK6 | 0.488912 | 0.771583 | 2.897678 | 0.266276 | 0.790026 | NA |
| HN1L | 3197.249 | -0.15682 | 0.178001 | -0.88098 | 0.378329 | 0.544645 |
| | 1.880172 | 1.059817 | 1.495568 | 0.708638 | 0.478549 | NA |
| | 2.128816 | -0.06384 | 1.426692 | -0.04475 | 0.964311 | NA |
| HBA1 | 0.422534 | 1.924583 | 3.001758 | 0.641152 | 0.521424 | NA |
| HBM | 0.134133 | -1.48909 | 4.077584 | -0.36519 | 0.71497 | NA |
| TCEB3B | 0.489058 | -0.58569 | 3.427543 | -0.17088 | 0.864321 | NA |
| RAB12 | 753.8383 | 0.221033 | 0.140147 | 1.577146 | 0.114762 | 0.244299 |
| TMEM200( | 0.854757 | 0.736707 | 2.188665 | 0.336601 | 0.736418 | NA |
| SNORD60 | 2.173877 | 1.254109 | 1.428055 | 0.878194 | 0.379839 | NA |
| RNU6-945I | 0.162376 | 0.424913 | 4.079006 | 0.104171 | 0.917034 | NA |
| SNORA30 | 0.230334 | -0.53318 | 4.076749 | -0.13079 | 0.895945 | NA |
| SNORA50 | 0.975407 | 0.039022 | 2.314857 | 0.016857 | 0.98655 | NA |
| | 0.163957 | 0.424913 | 4.079006 | 0.104171 | 0.917034 | NA |
| | 0.096202 | 0.408546 | 4.079639 | 0.100143 | 0.920231 | NA |
| RNU6-196I | 0.162376 | 0.424913 | 4.079006 | 0.104171 | 0.917034 | NA |
| RNU6-39P | 0.258578 | 1.233151 | 4.047809 | 0.304646 | 0.760635 | NA |
| | 0.151697 | -1.546 | 4.075469 | -0.37934 | 0.704433 | NA |
| RNU6-128‹ | 0.151697 | -1.546 | 4.075469 | -0.37934 | 0.704433 | NA |
| RNU6-106! | 0.163957 | 0.424913 | 4.079006 | 0.104171 | 0.917034 | NA |
| RNU6-100! | 0.613357 | 2.486614 | 2.985155 | 0.832993 | 0.404848 | NA |
| SNORD7 | 0.448206 | -1.2532 | 3.007315 | -0.41672 | 0.676885 | NA |
| | 0.134133 | -1.48909 | 4.077584 | -0.36519 | 0.71497 | NA |
| | 9.946209 | 0.058155 | 0.719013 | 0.080882 | 0.935536 | NA |
| SNORA46 | 0.508056 | 0.731032 | 2.861912 | 0.255435 | 0.798387 | NA |
| MIR635 | 120.6537 | -0.41613 | 0.244924 | -1.69902 | 0.089316 | 0.206563 |
| MIR193A | 1.645774 | -1.39946 | 1.608559 | -0.87001 | 0.384296 | NA |
| MIR641 | 1.966803 | -0.02513 | 1.585714 | -0.01585 | 0.987354 | NA |
| | 0.163957 | 0.424913 | 4.079006 | 0.104171 | 0.917034 | NA |
| MIR199A1 | 0.096202 | 0.408546 | 4.079639 | 0.100143 | 0.920231 | NA |
| MIR27A | 1.717795 | 0.894528 | 1.576519 | 0.567407 | 0.570438 | NA |
| MIR640 | 0.163957 | 0.424913 | 4.079006 | 0.104171 | 0.917034 | NA |
| MIR33B | 0.134133 | -1.48909 | 4.077584 | -0.36519 | 0.71497 | NA |
| MIR152 | 0.314073 | -0.57308 | 3.898934 | -0.14698 | 0.883146 | NA |
| MIR328 | 0.162376 | 0.424913 | 4.079006 | 0.104171 | 0.917034 | NA |
| MIR23A | 2.164713 | -0.53863 | 1.366844 | -0.39407 | 0.69353 | NA |
| SNORD105 | 0.096202 | 0.408546 | 4.079639 | 0.100143 | 0.920231 | NA |
| SNORD41 | 0.556813 | -0.57007 | 3.294615 | -0.17303 | 0.862627 | NA |
| RNU6ATAC | 0.162376 | 0.424913 | 4.079006 | 0.104171 | 0.917034 | NA |
| MIR320C2 | 0.096202 | 0.408546 | 4.079639 | 0.100143 | 0.920231 | NA |
| PRR22 | 32.40959 | -0.4405 | 0.426484 | -1.03285 | 0.301672 | NA |
| | 0.163957 | 0.424913 | 4.079006 | 0.104171 | 0.917034 | NA |
| | 0.151697 | -1.546 | 4.075469 | -0.37934 | 0.704433 | NA |
| RNA5SP46! | 0.28583 | -2.21358 | 3.97376 | -0.55705 | 0.577494 | NA |

| | | | | | | |
|---|---|---|---|---|---|---|
| SNORD91B | 0.096202 | 0.408546 | 4.079639 | 0.100143 | 0.920231 | NA |
| C17orf51 | 702.3763 | -0.31079 | 0.164938 | -1.88427 | 0.059528 | 0.156476 |
| KRTAP2-3 | 1.763631 | -4.79E-04 | 1.512262 | -3.17E-04 | 0.999747 | NA |
| C17orf100 | 0.162376 | 0.424913 | 4.079006 | 0.104171 | 0.917034 | NA |
| KRTAP3-2 | 0.134133 | -1.48909 | 4.077584 | -0.36519 | 0.71497 | NA |
| FAM106A | 0.315653 | -0.57305 | 3.892639 | -0.14721 | 0.882963 | NA |
| ZNF254 | 79.42482 | 1.006538 | 0.505892 | 1.989631 | 0.046632 | 0.132793 |
| RPL32P31 | 0.327913 | 1.543154 | 4.021063 | 0.383768 | 0.701151 | NA |
| | 0.526843 | -0.38719 | 2.855892 | -0.13558 | 0.892156 | NA |
| RPL36AP48 | 0.324752 | 1.53082 | 4.022025 | 0.380609 | 0.703493 | NA |
| RPL12P38 | 1.724412 | -1.02404 | 1.531319 | -0.66873 | 0.503667 | NA |
| SUPT4H1 | 1629.91 | 0.004408 | 0.14174 | 0.031102 | 0.975188 | 0.986377 |
| | 1.157911 | 0.522058 | 2.032481 | 0.256858 | 0.797289 | NA |
| PGK1P2 | 1.943129 | 1.068393 | 1.509387 | 0.707832 | 0.479049 | NA |
| | 1.529502 | -1.07961 | 1.722295 | -0.62685 | 0.53076 | NA |
| | 0.709944 | 0.232855 | 2.332364 | 0.099837 | 0.920474 | NA |
| | 1.270335 | 0.030019 | 1.726924 | 0.017383 | 0.986131 | NA |
| | 0.268266 | -2.15396 | 4.038351 | -0.53338 | 0.593773 | NA |
| RPS7P11 | 4.164605 | 0.457383 | 1.07088 | 0.427109 | 0.6693 | NA |
| QTRT1 | 658.0981 | -0.17031 | 0.153644 | -1.10845 | 0.267667 | 0.432663 |
| COG8 | 511.4913 | -0.47339 | 0.151792 | -3.11864 | 0.001817 | 0.015146 |
| LCAT | 75.07554 | 0.014904 | 0.258339 | 0.05769 | 0.953995 | 0.974786 |
| KRT222 | 2.064018 | 0.206892 | 1.466235 | 0.141104 | 0.887788 | NA |
| TTLL13 | 0.162376 | 0.424913 | 4.079006 | 0.104171 | 0.917034 | NA |
| ADAT3 | 231.0565 | -0.20552 | 0.232977 | -0.88217 | 0.377686 | 0.544108 |
| LAT | 0.163957 | 0.424913 | 4.079006 | 0.104171 | 0.917034 | NA |
| EMP2 | 921.9207 | -0.14749 | 0.208176 | -0.70848 | 0.478649 | 0.633256 |
| KCTD11 | 279.2475 | -0.24202 | 0.184607 | -1.31102 | 0.189852 | 0.34409 |
| LIPE-AS1 | 13.17637 | -0.37639 | 0.582986 | -0.64563 | 0.518519 | NA |
| DNASE1 | 258.7354 | -0.02141 | 0.220098 | -0.09728 | 0.922503 | 0.95505 |
| | 0.690946 | -1.56103 | 2.699004 | -0.57837 | 0.563013 | NA |
| | 4.331672 | -0.04699 | 0.989526 | -0.04749 | 0.962122 | NA |
| NUDT19 | 420.8049 | -0.19875 | 0.171769 | -1.1571 | 0.247232 | 0.410746 |
| ZNF726 | 29.67853 | -0.07699 | 0.499644 | -0.15409 | 0.877542 | NA |
| | 31.39552 | -0.18663 | 0.3926 | -0.47536 | 0.634532 | NA |
| | 5.389001 | 2.126793 | 0.99418 | 2.139244 | 0.032416 | NA |
| TAX1BP3 | 606.1178 | -0.13283 | 0.205458 | -0.6465 | 0.517957 | 0.665047 |
| | 7.093517 | 0.741193 | 0.781445 | 0.948491 | 0.34288 | NA |
| ZNF90 | 24.76349 | 0.52561 | 0.49313 | 1.065866 | 0.286484 | NA |
| TM6SF2 | 7.17186 | -0.57849 | 0.828351 | -0.69837 | 0.484947 | NA |
| MEF2B | 1.424835 | -0.73179 | 1.663339 | -0.43995 | 0.659972 | NA |
| SMIM7 | 1030.44 | -0.09057 | 0.153061 | -0.59175 | 0.554019 | 0.69428 |
| ARL16 | 819.5969 | 0.212463 | 0.172115 | 1.234427 | 0.217044 | 0.376453 |
| | 0.134133 | -1.48909 | 4.077584 | -0.36519 | 0.71497 | NA |
| | 0.247898 | -0.5901 | 4.074633 | -0.14482 | 0.884851 | NA |
| PRCD | 196.5916 | -0.28343 | 0.230369 | -1.23034 | 0.218569 | 0.378116 |
| AMZ2P1 | 251.5978 | 0.299919 | 0.195658 | 1.532876 | 0.125306 | 0.260204 |

| | | | | | | |
|---|---|---|---|---|---|---|
| PLEKHM1P | 120.6409 | 0.362395 | 0.284136 | 1.275427 | 0.202158 | 0.358669 |
| ZNF788 | 1.372261 | 0.301516 | 1.751077 | 0.172189 | 0.863289 | NA |
| C19orf38 | 4.788056 | 0.102852 | 0.900699 | 0.114192 | 0.909086 | NA |
| HNRNPA1P | 0.151697 | -1.546 | 4.075469 | -0.37934 | 0.704433 | NA |
| C17orf67 | 71.61464 | 0.282603 | 0.285493 | 0.989877 | 0.322234 | 0.490321 |
| | 3.802894 | -0.01117 | 1.257688 | -0.00888 | 0.992913 | NA |
| | 3.354296 | -0.93218 | 1.156844 | -0.8058 | 0.42036 | NA |
| | 28.19209 | -0.05207 | 0.408269 | -0.12753 | 0.898524 | NA |
| OR4F13P | 0.162376 | 0.424913 | 4.079006 | 0.104171 | 0.917034 | NA |
| | 0.151697 | -1.546 | 4.075469 | -0.37934 | 0.704433 | NA |
| VAC14-AS1 | 3.640004 | -0.59395 | 1.192152 | -0.49822 | 0.618332 | NA |
| KANSL1-AS | 11.60049 | -0.08246 | 0.630613 | -0.13075 | 0.895969 | NA |
| FAM149B1 | 0.392857 | -1.546 | 4.075469 | -0.37934 | 0.704433 | NA |
| LRRC37A4F | 43.64164 | 0.247313 | 0.33464 | 0.739042 | 0.459881 | NA |
| | 2.913288 | 1.087029 | 1.211981 | 0.896902 | 0.369771 | NA |
| | 0.315653 | -0.57305 | 3.892639 | -0.14721 | 0.882963 | NA |
| PLIN5 | 0.268266 | -2.15396 | 4.038351 | -0.53338 | 0.593773 | NA |
| KRT42P | 0.134133 | -1.48909 | 4.077584 | -0.36519 | 0.71497 | NA |
| | 0.134133 | -1.48909 | 4.077584 | -0.36519 | 0.71497 | NA |
| C17orf98 | 1.926154 | -0.37773 | 1.398683 | -0.27006 | 0.787112 | NA |
| HMGN2P1! | 0.749217 | 0.7397 | 2.633289 | 0.280903 | 0.778784 | NA |
| | 5.470759 | -0.8813 | 0.8618 | -1.02263 | 0.306484 | NA |
| | 164.5331 | -0.12647 | 0.218159 | -0.57969 | 0.562121 | 0.700081 |
| CDIPT-AS1 | 19.39623 | 0.199519 | 0.493724 | 0.40411 | 0.686132 | NA |
| CDRT15L2 | 0.258578 | 1.233151 | 4.047809 | 0.304646 | 0.760635 | NA |
| UPF3AP2 | 1.077932 | -0.48016 | 2.057314 | -0.23339 | 0.815457 | NA |
| | 0.163957 | 0.424913 | 4.079006 | 0.104171 | 0.917034 | NA |
| ZSWIM7 | 760.6361 | 0.014029 | 0.148943 | 0.094192 | 0.924957 | 0.957339 |
| RPL15P20 | 0.096202 | 0.408546 | 4.079639 | 0.100143 | 0.920231 | NA |
| RPL13P12 | 1814.293 | 0.108864 | 0.147586 | 0.737632 | 0.460738 | 0.618859 |
| NEURL4 | 316.5737 | -0.42119 | 0.189633 | -2.22105 | 0.026347 | 0.091117 |
| | 61.57278 | 0.06 | 0.28733 | 0.208819 | 0.834589 | 0.901016 |
| C16orf90 | 0.151697 | -1.546 | 4.075469 | -0.37934 | 0.704433 | NA |
| PRSS41 | 0.134133 | -1.48909 | 4.077584 | -0.36519 | 0.71497 | NA |
| | 67.37649 | -0.01658 | 0.292856 | -0.05661 | 0.954857 | 0.975091 |
| ZNF407 | 318.9104 | 0.064236 | 0.216197 | 0.297117 | 0.766377 | 0.854855 |
| HNRNPA1P | 1.387253 | -1.71455 | 1.84915 | -0.92721 | 0.353817 | NA |
| | 31.5921 | -0.3006 | 0.387067 | -0.7766 | 0.437395 | NA |
| | 4.059361 | 0.254981 | 1.06889 | 0.238548 | 0.811456 | NA |
| IFI30 | 0.192403 | 0.975392 | 4.060716 | 0.240202 | 0.810174 | NA |
| CKLF | 165.8811 | 0.391681 | 0.215787 | 1.815128 | 0.069504 | 0.173679 |
| PAM16 | 27.03644 | -0.47488 | 0.424746 | -1.11804 | 0.26355 | NA |
| RPS3AP2 | 0.096202 | 0.408546 | 4.079639 | 0.100143 | 0.920231 | NA |
| RNASEK | 21.91499 | -0.50737 | 0.470185 | -1.07908 | 0.280552 | NA |
| RPS20P2 | 2.201586 | 0.98381 | 1.629419 | 0.603779 | 0.54599 | NA |
| | 148.8911 | 0.798841 | 0.204352 | 3.909139 | 9.26E-05 | 0.002414 |
| | 17.92191 | -0.65283 | 0.570746 | -1.14382 | 0.2527 | NA |

| | | | | | | |
|---|---|---|---|---|---|---|
| | 0.612162 | -0.19706 | 2.47693 | -0.07956 | 0.936588 | NA |
| ZGLP1 | 4.674555 | 0.118404 | 0.931054 | 0.127172 | 0.898805 | NA |
| VAMP2 | 961.3332 | -0.36839 | 0.141383 | -2.60566 | 0.00917 | 0.044534 |
| | 0.556813 | -0.57007 | 3.294615 | -0.17303 | 0.862627 | NA |
| RPL21P119 | 0.804363 | 1.537223 | 2.514087 | 0.611444 | 0.540906 | NA |
| | 1.020857 | 0.19325 | 2.085078 | 0.092682 | 0.926156 | NA |
| | 1.053841 | 0.997099 | 2.078418 | 0.479739 | 0.631413 | NA |
| | 1.777406 | 2.716315 | 1.959292 | 1.386376 | 0.165632 | NA |
| | 0.392857 | -1.546 | 4.075469 | -0.37934 | 0.704433 | NA |
| MIR1227 | 0.392857 | -1.546 | 4.075469 | -0.37934 | 0.704433 | NA |
| MIR320C1 | 0.096202 | 0.408546 | 4.079639 | 0.100143 | 0.920231 | NA |
| MIR1180 | 0.315653 | -0.57305 | 3.892639 | -0.14721 | 0.882963 | NA |
| | 0.162376 | 0.424913 | 4.079006 | 0.104171 | 0.917034 | NA |
| C16orf3 | 3.528961 | 0.37363 | 1.062702 | 0.351585 | 0.72515 | NA |
| PSG3 | 0.162376 | 0.424913 | 4.079006 | 0.104171 | 0.917034 | NA |
| KRTAP1-5 | 0.741166 | 2.78101 | 2.851312 | 0.975344 | 0.32939 | NA |
| HMSD | 13.97156 | 0.548128 | 0.575418 | 0.952574 | 0.340806 | NA |
| TRIM16 | 295.4213 | -0.44972 | 0.199494 | -2.2543 | 0.024178 | 0.085996 |
| FXYD7 | 3.627691 | -0.3067 | 1.030789 | -0.29754 | 0.766052 | NA |
| UBA52 | 15308.7 | -0.06 | 0.149173 | -0.40221 | 0.687532 | 0.796745 |
| TIAF1 | 42.51629 | -0.77886 | 0.337831 | -2.30547 | 0.02114 | NA |
| URAHP | 25.06378 | -0.52271 | 0.485182 | -1.07734 | 0.281327 | NA |
| RNU4-30P | 0.163957 | 0.424913 | 4.079006 | 0.104171 | 0.917034 | NA |
| MIR1910 | 0.151697 | -1.546 | 4.075469 | -0.37934 | 0.704433 | NA |
| | 0.162376 | 0.424913 | 4.079006 | 0.104171 | 0.917034 | NA |
| RNU4-47P | 1.045726 | 0.338916 | 1.94625 | 0.174138 | 0.861757 | NA |
| | 0.354779 | 1.703954 | 3.657232 | 0.465914 | 0.641277 | NA |
| RNU6-981F | 0.163957 | 0.424913 | 4.079006 | 0.104171 | 0.917034 | NA |
| RNA5SP45( | 0.162376 | 0.424913 | 4.079006 | 0.104171 | 0.917034 | NA |
| SNORD71 | 0.096202 | 0.408546 | 4.079639 | 0.100143 | 0.920231 | NA |
| RNU6-195F | 0.610582 | -0.18034 | 2.753139 | -0.0655 | 0.947774 | NA |
| EXOSC6 | 345.5734 | -0.46676 | 0.173463 | -2.69085 | 0.007127 | 0.037572 |
| CDRT15 | 0.163957 | 0.424913 | 4.079006 | 0.104171 | 0.917034 | NA |
| ZNF844 | 88.3196 | -0.3568 | 0.308869 | -1.15518 | 0.248018 | 0.411627 |
| TINCR | 115.0135 | 0.438408 | 0.435737 | 1.006129 | 0.314354 | 0.48173 |
| CERS1 | 475.4029 | -0.27804 | 0.254698 | -1.09163 | 0.274997 | 0.440659 |
| | 1.221725 | -0.08735 | 1.955093 | -0.04468 | 0.964363 | NA |
| AFG3L1P | 228.3841 | 0.336158 | 0.202913 | 1.656659 | 0.097588 | 0.219455 |
| | 0.162376 | 0.424913 | 4.079006 | 0.104171 | 0.917034 | NA |
| | 22.23102 | -0.56895 | 0.455801 | -1.24823 | 0.211945 | NA |
| UBE2SP2 | 1.079513 | -0.47694 | 2.056893 | -0.23187 | 0.816636 | NA |
| | 2.613035 | 1.637854 | 1.511179 | 1.083825 | 0.278442 | NA |
| OACYLP | 3.495928 | 0.768956 | 1.122683 | 0.684927 | 0.49339 | NA |
| C17orf72 | 0.640755 | -1.64826 | 2.754778 | -0.59833 | 0.549621 | NA |
| | 1.568425 | 0.022422 | 1.54585 | 0.014505 | 0.988427 | NA |
| ATXN1L | 1535.807 | -0.36052 | 0.159051 | -2.26669 | 0.023409 | 0.084142 |
| | 0.918331 | 0.764804 | 2.118268 | 0.361052 | 0.718061 | NA |

| | | | | | | |
|---|---|---|---|---|---|---|
| SNRPGP15 | 33.6911 | -0.1902 | 0.480006 | -0.39624 | 0.691928 | NA |
| HNRNPA1P | 42.94844 | -0.01211 | 0.345923 | -0.03501 | 0.972071 | NA |
| | 17.11241 | 0.345689 | 0.633595 | 0.5456 | 0.585341 | NA |
| | 25.78285 | -0.13562 | 0.440209 | -0.30808 | 0.758018 | NA |
| C17orf89 | 1259.42 | 0.03503 | 0.161201 | 0.217304 | 0.827972 | 0.896239 |
| | 4.454002 | -0.12176 | 0.95338 | -0.12771 | 0.898376 | NA |
| | 0.163957 | 0.424913 | 4.079006 | 0.104171 | 0.917034 | NA |
| RPL23AP2 | 2.455191 | 3.407096 | 1.621984 | 2.100573 | 0.035678 | NA |
| RPSAP56 | 1.407243 | 0.957292 | 1.678143 | 0.570447 | 0.568374 | NA |
| AATK-AS1 | 0.392857 | -1.546 | 4.075469 | -0.37934 | 0.704433 | NA |
| PLEKHM1 | 1134.755 | -0.4177 | 0.161416 | -2.58773 | 0.009661 | 0.046071 |
| WASH5P | 1.167745 | -2.70405 | 2.12796 | -1.27072 | 0.203828 | NA |
| MPRIP-AS1 | 0.260158 | 1.240347 | 4.047124 | 0.306476 | 0.759242 | NA |
| ARHGAP23 | 696.3463 | -0.30173 | 0.245873 | -1.22718 | 0.219756 | 0.379436 |
| ZNF469 | 3.128677 | -0.4151 | 1.135073 | -0.3657 | 0.714586 | NA |
| FAM195B | 487.905 | -0.24853 | 0.186695 | -1.33119 | 0.183127 | 0.336255 |
| | 0.096202 | 0.408546 | 4.079639 | 0.100143 | 0.920231 | NA |
| | 0.162376 | 0.424913 | 4.079006 | 0.104171 | 0.917034 | NA |
| | 3.457019 | 1.088867 | 1.230579 | 0.884842 | 0.376242 | NA |
| | 0.151697 | -1.546 | 4.075469 | -0.37934 | 0.704433 | NA |
| | 14.4027 | 0.231327 | 0.553327 | 0.418065 | 0.675899 | NA |
| TLK2P1 | 4.784329 | -0.14117 | 1.031973 | -0.1368 | 0.891188 | NA |
| | 0.151697 | -1.546 | 4.075469 | -0.37934 | 0.704433 | NA |
| | 0.162376 | 0.424913 | 4.079006 | 0.104171 | 0.917034 | NA |
| BAIAP2-AS1 | 1328.274 | -0.41951 | 0.16519 | -2.53956 | 0.011099 | 0.050462 |
| | 0.192403 | 0.975392 | 4.060716 | 0.240202 | 0.810174 | NA |
| RPL23AP76 | 0.096202 | 0.408546 | 4.079639 | 0.100143 | 0.920231 | NA |
| | 3.09529 | -3.29418 | 1.435157 | -2.29535 | 0.021713 | NA |
| | 0.314073 | -0.57308 | 3.898934 | -0.14698 | 0.883146 | NA |
| | 1.598249 | 0.567223 | 1.588614 | 0.357055 | 0.72105 | NA |
| UPF3AP1 | 1.282595 | 0.709442 | 1.711365 | 0.414547 | 0.678473 | NA |
| UPK1A-AS1 | 12.33722 | 0.071377 | 0.60198 | 0.11857 | 0.905616 | NA |
| SCDP1 | 4.589687 | 0.322806 | 0.987378 | 0.326932 | 0.743719 | NA |
| | 2.520955 | -0.55493 | 1.273981 | -0.43559 | 0.663137 | NA |
| HSBP1L1 | 372.8444 | 0.216373 | 0.197988 | 1.092861 | 0.274455 | 0.4401 |
| SMCR5 | 0.450981 | 2.057325 | 3.353149 | 0.61355 | 0.539513 | NA |
| SRRM5 | 4.78832 | 0.16687 | 0.951875 | 0.175307 | 0.860839 | NA |
| CACTIN-AS1 | 2.728147 | -0.99709 | 1.187266 | -0.83982 | 0.401011 | NA |
| ABHD17AP | 0.448206 | -1.2532 | 3.007315 | -0.41672 | 0.676885 | NA |
| IL6STP1 | 0.096202 | 0.408546 | 4.079639 | 0.100143 | 0.920231 | NA |
| LINC00511 | 1599.368 | -0.32898 | 0.146132 | -2.25129 | 0.024367 | 0.086453 |
| | 0.776119 | 0.401378 | 2.295521 | 0.174853 | 0.861196 | NA |
| | 0.410275 | 0.244283 | 3.100889 | 0.078778 | 0.937209 | NA |
| HPN-AS1 | 0.134133 | -1.48909 | 4.077584 | -0.36519 | 0.71497 | NA |
| | 2.54546 | -0.69669 | 1.360464 | -0.51209 | 0.608585 | NA |
| | 0.288605 | 1.443619 | 4.029058 | 0.358302 | 0.720117 | NA |
| SCAMP4 | 1503.304 | -0.6307 | 0.133592 | -4.72107 | 2.35E-06 | 2.67E-04 |

| | | | | | | |
|---|---|---|---|---|---|---|
| | 0.163957 | 0.424913 | 4.079006 | 0.104171 | 0.917034 | NA |
| SPAG5-AS1 | 3.075097 | -1.675 | 1.4379 | -1.16489 | 0.244064 | NA |
| | 0.506476 | 0.726418 | 2.864186 | 0.253621 | 0.799788 | NA |
| | 0.247898 | -0.5901 | 4.074633 | -0.14482 | 0.884851 | NA |
| | 7.451333 | -0.22594 | 0.787597 | -0.28687 | 0.774212 | NA |
| | 0.260158 | 1.240347 | 4.047124 | 0.306476 | 0.759242 | NA |
| RPL7P47 | 0.326536 | 0.037056 | 3.842554 | 0.009644 | 0.992306 | NA |
| | 2.363835 | -0.30712 | 1.425613 | -0.21543 | 0.829432 | NA |
| | 0.588072 | 2.39955 | 3.098804 | 0.774347 | 0.438726 | NA |
| CASP16 | 3.242371 | -1.28545 | 1.371672 | -0.93714 | 0.348686 | NA |
| | 4.488975 | -0.43695 | 0.928976 | -0.47035 | 0.638102 | NA |
| | 2.827075 | 0.581194 | 1.244514 | 0.467004 | 0.640497 | NA |
| ATF4P3 | 0.846852 | 1.76834 | 2.426477 | 0.728769 | 0.466143 | NA |
| C19orf24 | 333.6383 | 0.066372 | 0.184674 | 0.359401 | 0.719295 | 0.821112 |
| | 0.314073 | -0.57308 | 3.898934 | -0.14698 | 0.883146 | NA |
| CES1P1 | 5.986285 | 0.82383 | 0.860169 | 0.957754 | 0.338187 | NA |
| ARL17B | 4.906548 | -0.28261 | 1.036403 | -0.27268 | 0.785099 | NA |
| MTND2P12 | 0.096202 | 0.408546 | 4.079639 | 0.100143 | 0.920231 | NA |
| | 0.555233 | -0.57009 | 3.296947 | -0.17291 | 0.862719 | NA |
| | 0.452561 | 2.061505 | 3.349456 | 0.615474 | 0.538241 | NA |
| MRPL45P2 | 71.42061 | 0.177408 | 0.276818 | 0.640883 | 0.521599 | 0.668276 |
| | 0.900913 | -0.26014 | 2.222223 | -0.11706 | 0.90681 | NA |
| EEF1DP1 | 1.921421 | 0.960699 | 1.530318 | 0.627777 | 0.53015 | NA |
| | 0.47961 | 0.535796 | 3.39605 | 0.15777 | 0.874638 | NA |
| | 138.6033 | -0.21681 | 0.233654 | -0.92791 | 0.353454 | 0.521597 |
| THEMIS3P | 0.096202 | 0.408546 | 4.079639 | 0.100143 | 0.920231 | NA |
| | 0.096202 | 0.408546 | 4.079639 | 0.100143 | 0.920231 | NA |
| | 0.162376 | 0.424913 | 4.079006 | 0.104171 | 0.917034 | NA |
| PRAC2 | 20.30031 | 0.014766 | 0.477743 | 0.030907 | 0.975343 | NA |
| ZNF492 | 6.136394 | 0.805709 | 0.97565 | 0.825817 | 0.408908 | NA |
| ZNF688 | 663.9068 | -0.33572 | 0.165518 | -2.02829 | 0.04253 | 0.125476 |
| PET100 | 157.6711 | 0.436046 | 0.219435 | 1.98713 | 0.046908 | 0.133248 |
| | 1.50225 | -0.05913 | 1.577722 | -0.03748 | 0.970105 | NA |
| RPL7P46 | 0.163957 | 0.424913 | 4.079006 | 0.104171 | 0.917034 | NA |
| | 6.201264 | 0.205707 | 0.832268 | 0.247164 | 0.804781 | NA |
| | 1.075752 | 0.450053 | 2.098769 | 0.214437 | 0.830206 | NA |
| TOB1-AS1 | 15.18391 | 0.182191 | 0.548627 | 0.332085 | 0.739825 | NA |
| CISD3 | 1601.492 | -0.05818 | 0.166571 | -0.34927 | 0.72689 | 0.82669 |
| HOXB-AS1 | 3.241311 | 1.016377 | 1.202619 | 0.845136 | 0.398035 | NA |
| ATP6V0CP1 | 0.096202 | 0.408546 | 4.079639 | 0.100143 | 0.920231 | NA |
| | 0.134133 | -1.48909 | 4.077584 | -0.36519 | 0.71497 | NA |
| HERC2P4 | 1.620759 | -0.39583 | 1.782798 | -0.22203 | 0.824291 | NA |
| | 2.02629 | 0.206442 | 1.452317 | 0.142147 | 0.886964 | NA |
| HMGN2P3 | 0.792452 | -2.15617 | 2.562821 | -0.84133 | 0.400166 | NA |
| | 0.096202 | 0.408546 | 4.079639 | 0.100143 | 0.920231 | NA |
| TNPO1P2 | 0.151697 | -1.546 | 4.075469 | -0.37934 | 0.704433 | NA |
| | 1.514445 | -1.85382 | 1.951264 | -0.95006 | 0.34208 | NA |

| | | | | | | |
|---|---|---|---|---|---|---|
| | 0.832809 | 1.636407 | 2.200093 | 0.74379 | 0.457004 | NA |
| RPS18P12 | 5.951147 | 1.080792 | 0.858301 | 1.259222 | 0.20795 | NA |
| | 0.296509 | -0.51709 | 3.982155 | -0.12985 | 0.896684 | NA |
| HSBP1 | 5700.967 | 0.286798 | 0.174452 | 1.644001 | 0.100176 | 0.223272 |
| C17orf105 | 1.789506 | -1.22275 | 1.690248 | -0.72342 | 0.469423 | NA |
| WASIR2 | 0.134133 | -1.48909 | 4.077584 | -0.36519 | 0.71497 | NA |
| RNASEH1P: | 0.247898 | -0.5901 | 4.074633 | -0.14482 | 0.884851 | NA |
| CEACAMP5 | 0.599903 | -1.73313 | 2.730982 | -0.63462 | 0.525678 | NA |
| | 4.956957 | 0.055715 | 0.950028 | 0.058646 | 0.953234 | NA |
| | 0.151697 | -1.546 | 4.075469 | -0.37934 | 0.704433 | NA |
| DBIL5P | 5.445936 | -1.0604 | 0.922993 | -1.14887 | 0.250609 | NA |
| | 0.151697 | -1.546 | 4.075469 | -0.37934 | 0.704433 | NA |
| MTRNR2L4 | 0.46735 | -1.30193 | 3.399671 | -0.38296 | 0.701751 | NA |
| FAM215B | 0.260158 | 1.240347 | 4.047124 | 0.306476 | 0.759242 | NA |
| | 5.752771 | -0.97316 | 0.892867 | -1.08993 | 0.275744 | NA |
| | 19.70551 | -0.36327 | 0.516162 | -0.70379 | 0.481563 | NA |
| RPL21P131 | 0.163957 | 0.424913 | 4.079006 | 0.104171 | 0.917034 | NA |
| SLC16A6P1 | 0.652869 | 1.292592 | 2.374873 | 0.544278 | 0.58625 | NA |
| | 0.134133 | -1.48909 | 4.077584 | -0.36519 | 0.71497 | NA |
| LINC00665 | 229.6013 | -0.3057 | 0.192537 | -1.58775 | 0.112343 | 0.24088 |
| | 0.28583 | -2.21358 | 3.97376 | -0.55705 | 0.577494 | NA |
| LYRM9 | 112.5823 | -0.20119 | 0.229535 | -0.87653 | 0.380743 | 0.546883 |
| | 0.162376 | 0.424913 | 4.079006 | 0.104171 | 0.917034 | NA |
| | 28.97437 | 0.610332 | 0.430811 | 1.416704 | 0.15657 | NA |
| | 14.66444 | 1.735916 | 0.700383 | 2.478523 | 0.013193 | NA |
| HOXB-AS3 | 25.94034 | 1.261716 | 0.429897 | 2.934924 | 0.003336 | NA |
| | 0.640609 | 0.042838 | 2.379511 | 0.018003 | 0.985637 | NA |
| | 102.2461 | -0.11865 | 0.27311 | -0.43444 | 0.663971 | 0.779344 |
| NPIPB7 | 0.096202 | 0.408546 | 4.079639 | 0.100143 | 0.920231 | NA |
| SNRPEP4 | 2.611201 | 0.774644 | 1.313868 | 0.58959 | 0.555465 | NA |
| USP32P2 | 1.977505 | -0.69819 | 1.634598 | -0.42714 | 0.669281 | NA |
| RPS2P48 | 0.825079 | -2.06702 | 2.533819 | -0.81577 | 0.41463 | NA |
| | 2.934545 | 0.091001 | 1.323007 | 0.068784 | 0.945162 | NA |
| | 39.06095 | -0.16635 | 0.349274 | -0.47627 | 0.633884 | NA |
| DDX11L10 | 0.324752 | 1.53082 | 4.022025 | 0.380609 | 0.703493 | NA |
| CYP2T2P | 16.42522 | 0.012457 | 0.49639 | 0.025095 | 0.979979 | NA |
| | 0.163957 | 0.424913 | 4.079006 | 0.104171 | 0.917034 | NA |
| RPS28 | 8297.119 | 0.006865 | 0.154758 | 0.04436 | 0.964617 | 0.980626 |
| UBE2SP1 | 14.93936 | -0.45626 | 0.530999 | -0.85924 | 0.390206 | NA |
| | 0.324752 | 1.53082 | 4.022025 | 0.380609 | 0.703493 | NA |
| RPL9P2 | 0.151697 | -1.546 | 4.075469 | -0.37934 | 0.704433 | NA |
| | 39.42814 | 0.268033 | 0.361249 | 0.741961 | 0.458111 | NA |
| | 3.880348 | 0.058672 | 1.088936 | 0.05388 | 0.95703 | NA |
| PINLYP | 349.1832 | 0.065096 | 0.171587 | 0.379377 | 0.704408 | 0.81036 |
| | 40.91765 | -0.12744 | 0.340593 | -0.37416 | 0.708283 | NA |
| | 8.504856 | -0.22528 | 0.840167 | -0.26814 | 0.788591 | NA |
| | 1.007794 | 1.520243 | 2.181859 | 0.696765 | 0.48595 | NA |

| | | | | | | |
|---|---|---|---|---|---|---|
| WASH4P | 8.298218 | -0.26121 | 0.746407 | -0.34995 | 0.726375 | NA |
| | 3.577379 | 1.130377 | 1.209498 | 0.934583 | 0.350003 | NA |
| | 0.163957 | 0.424913 | 4.079006 | 0.104171 | 0.917034 | NA |
| | 0.163957 | 0.424913 | 4.079006 | 0.104171 | 0.917034 | NA |
| SOX9-AS1 | 3.968168 | 0.004427 | 1.145687 | 0.003864 | 0.996917 | NA |
| LINC00338 | 50.53457 | 0.265847 | 0.335305 | 0.792851 | 0.427865 | 0.589669 |
| TBC1D3C | 0.449786 | -1.25325 | 3.00474 | -0.41709 | 0.676613 | NA |
| HMGN1P3( | 0.163957 | 0.424913 | 4.079006 | 0.104171 | 0.917034 | NA |
| | 0.326333 | 1.533889 | 3.826878 | 0.40082 | 0.688553 | NA |
| FAUP1 | 5.010108 | 0.061041 | 0.956789 | 0.063798 | 0.949131 | NA |
| | 1.185016 | -0.03538 | 1.879555 | -0.01882 | 0.984982 | NA |
| | 0.247898 | -0.5901 | 4.074633 | -0.14482 | 0.884851 | NA |
| RPL6P27 | 9.155305 | 0.439546 | 0.691931 | 0.635246 | 0.525268 | NA |
| | 0.134133 | -1.48909 | 4.077584 | -0.36519 | 0.71497 | NA |
| | 18.05252 | 0.369964 | 0.500168 | 0.73968 | 0.459494 | NA |
| | 0.555233 | -0.57009 | 3.296947 | -0.17291 | 0.862719 | NA |
| | 1.082288 | 1.087249 | 1.993006 | 0.545532 | 0.585387 | NA |
| COX10-AS1 | 0.3441 | -0.0166 | 3.769682 | -0.0044 | 0.996486 | NA |
| | 53.75441 | 0.304944 | 0.355891 | 0.856847 | 0.391529 | 0.556781 |
| | 0.640755 | -1.64826 | 2.754778 | -0.59833 | 0.549621 | NA |
| | 0.247898 | -0.5901 | 4.074633 | -0.14482 | 0.884851 | NA |
| | 0.658173 | -0.01016 | 2.630582 | -0.00386 | 0.996919 | NA |
| | 1.143508 | 0.584377 | 2.035365 | 0.287112 | 0.774027 | NA |
| LINC00854 | 4.372138 | -0.9656 | 0.995469 | -0.97 | 0.332049 | NA |
| | 0.134133 | -1.48909 | 4.077584 | -0.36519 | 0.71497 | NA |
| PITPNA-AS: | 33.7982 | 0.193527 | 0.384318 | 0.50356 | 0.614571 | NA |
| | 15.20595 | 0.143856 | 0.572519 | 0.251268 | 0.801607 | NA |
| | 46.46249 | -0.03809 | 0.31959 | -0.11917 | 0.90514 | NA |
| CYP2F2P | 8.752894 | -0.57446 | 0.731509 | -0.7853 | 0.432275 | NA |
| B3GNT9 | 123.6051 | -0.08665 | 0.25539 | -0.33928 | 0.734399 | 0.832129 |
| RSL24D1P8 | 0.326536 | 0.037056 | 3.842554 | 0.009644 | 0.992306 | NA |
| SMG1P1 | 0.914404 | 3.02753 | 2.381561 | 1.271238 | 0.203644 | NA |
| RAI1-AS1 | 0.096202 | 0.408546 | 4.079639 | 0.100143 | 0.920231 | NA |
| RPL21P121 | 0.096202 | 0.408546 | 4.079639 | 0.100143 | 0.920231 | NA |
| ZNF737 | 44.4063 | 0.311296 | 0.430064 | 0.723836 | 0.469167 | NA |
| SHISA9 | 1.526587 | 1.88588 | 1.829401 | 1.030873 | 0.3026 | NA |
| PYY2 | 0.422534 | 1.924583 | 3.001758 | 0.641152 | 0.521424 | NA |
| | 1.726838 | 1.578405 | 1.632716 | 0.966736 | 0.333676 | NA |
| LINC00674 | 120.4796 | -0.10845 | 0.242158 | -0.44784 | 0.654271 | 0.771702 |
| | 7.310894 | -1.62588 | 0.788438 | -2.06215 | 0.039193 | NA |
| LRRC37A2 | 14.09357 | 0.186254 | 0.541418 | 0.344012 | 0.730837 | NA |
| ABCA17P | 1.237982 | 1.502941 | 1.913566 | 0.785414 | 0.432211 | NA |
| GABARAPL | 0.422534 | 1.924583 | 3.001758 | 0.641152 | 0.521424 | NA |
| | 1.245013 | -0.14306 | 1.979798 | -0.07226 | 0.942396 | NA |
| RNU7-140F | 0.096202 | 0.408546 | 4.079639 | 0.100143 | 0.920231 | NA |
| | 0.162376 | 0.424913 | 4.079006 | 0.104171 | 0.917034 | NA |
| RNU7-186F | 0.586491 | 2.39072 | 2.689057 | 0.889055 | 0.373973 | NA |

| | | | | | | |
|---|---|---|---|---|---|---|
| SNORD42B | 0.163957 | 0.424913 | 4.079006 | 0.104171 | 0.917034 | NA |
| RNU7-134F | 0.096202 | 0.408546 | 4.079639 | 0.100143 | 0.920231 | NA |
| | 1.519603 | -1.40785 | 1.668985 | -0.84353 | 0.39893 | NA |
| MIR1972-1 | 0.516164 | -1.70268 | 2.844782 | -0.59853 | 0.549488 | NA |
| | 0.096202 | 0.408546 | 4.079639 | 0.100143 | 0.920231 | NA |
| | 0.096202 | 0.408546 | 4.079639 | 0.100143 | 0.920231 | NA |
| SNORD124 | 0.151697 | -1.546 | 4.075469 | -0.37934 | 0.704433 | NA |
| | 0.853379 | -0.00837 | 2.414748 | -0.00347 | 0.997234 | NA |
| | 0.096202 | 0.408546 | 4.079639 | 0.100143 | 0.920231 | NA |
| | 0.096202 | 0.408546 | 4.079639 | 0.100143 | 0.920231 | NA |
| RNU7-125F | 0.162376 | 0.424913 | 4.079006 | 0.104171 | 0.917034 | NA |
| | 0.162376 | 0.424913 | 4.079006 | 0.104171 | 0.917034 | NA |
| | 0.163957 | 0.424913 | 4.079006 | 0.104171 | 0.917034 | NA |
| | 0.394291 | 0.298745 | 3.141711 | 0.09509 | 0.924243 | NA |
| | 1.686391 | 0.414168 | 1.73616 | 0.238554 | 0.811451 | NA |
| RPL34P31 | 0.943799 | -0.10552 | 2.406177 | -0.04385 | 0.965022 | NA |
| | 1.86508 | -0.62073 | 1.426117 | -0.43526 | 0.663374 | NA |
| | 1.278872 | 1.519834 | 1.931715 | 0.78678 | 0.431411 | NA |
| ALKBH6 | 40.39009 | 0.039153 | 0.329966 | 0.118658 | 0.905547 | NA |
| RPL17P42 | 0.518736 | 2.237423 | 2.7713 | 0.807355 | 0.419462 | NA |
| RPL32P34 | 0.163957 | 0.424913 | 4.079006 | 0.104171 | 0.917034 | NA |
| HOXB-AS2 | 2.594269 | 2.102981 | 1.401351 | 1.500682 | 0.133438 | NA |
| | 0.314073 | -0.57308 | 3.898934 | -0.14698 | 0.883146 | NA |
| NME1 | 1412.303 | 0.077074 | 0.172432 | 0.446983 | 0.654888 | 0.772199 |
| TNFSF12 | 1260.804 | -0.17912 | 0.199931 | -0.89589 | 0.370311 | 0.537139 |
| CDRT4 | 1.073823 | 2.138376 | 2.047615 | 1.044325 | 0.296335 | NA |
| | 0.963578 | 1.864506 | 2.778761 | 0.670985 | 0.50223 | NA |
| | 1.104199 | 0.511516 | 2.339905 | 0.218605 | 0.826957 | NA |
| | 0.643384 | 2.576138 | 2.977667 | 0.865153 | 0.386955 | NA |
| RPS4XP22 | 1.684697 | -0.06999 | 1.739411 | -0.04024 | 0.967905 | NA |
| RPL23AP72 | 0.096202 | 0.408546 | 4.079639 | 0.100143 | 0.920231 | NA |
| RN7SL608F | 0.230334 | -0.53318 | 4.076749 | -0.13079 | 0.895945 | NA |
| RN7SL792F | 0.163957 | 0.424913 | 4.079006 | 0.104171 | 0.917034 | NA |
| RN7SL606F | 0.163957 | 0.424913 | 4.079006 | 0.104171 | 0.917034 | NA |
| RN7SL368F | 0.163957 | 0.424913 | 4.079006 | 0.104171 | 0.917034 | NA |
| RPL9P30 | 0.134133 | -1.48909 | 4.077584 | -0.36519 | 0.71497 | NA |
| | 0.544554 | -2.65036 | 3.220086 | -0.82307 | 0.410468 | NA |
| RN7SL146F | 0.544554 | -2.65036 | 3.220086 | -0.82307 | 0.410468 | NA |
| | 0.296509 | -0.51709 | 3.982155 | -0.12985 | 0.896684 | NA |
| RPS27P29 | 0.556813 | -0.57007 | 3.294615 | -0.17303 | 0.862627 | NA |
| RPL12P37 | 0.452561 | 2.061505 | 3.349456 | 0.615474 | 0.538241 | NA |
| TCAM1P | 1.530429 | -1.87841 | 1.918298 | -0.97921 | 0.327478 | NA |
| | 0.533728 | -1.74213 | 2.820432 | -0.61768 | 0.536785 | NA |
| | 1.25178 | -2.04774 | 1.94381 | -1.05346 | 0.292128 | NA |
| | 0.544407 | -0.44425 | 2.542878 | -0.1747 | 0.861313 | NA |
| RPS12P27 | 0.151697 | -1.546 | 4.075469 | -0.37934 | 0.704433 | NA |
| FOXO3B | 32.04818 | -0.07709 | 0.37261 | -0.20689 | 0.836097 | NA |

| | | | | | | |
|---|---|---|---|---|---|---|
| RPL29P2 | 1.401868 | -0.28042 | 2.112032 | -0.13277 | 0.894375 | NA |
| RPL7AP10 | 0.315653 | -0.57305 | 3.892639 | -0.14721 | 0.882963 | NA |
| RPL21P123 | 0.35636 | 1.709252 | 3.651833 | 0.468053 | 0.639746 | NA |
| | 6.247731 | 0.356774 | 0.862949 | 0.413436 | 0.679287 | NA |
| | 0.638825 | 1.117948 | 3.106359 | 0.35989 | 0.718929 | NA |
| | 0.624422 | 1.162588 | 2.733761 | 0.425271 | 0.670639 | NA |
| UBA52P8 | 2.011501 | 1.112925 | 1.476593 | 0.753711 | 0.451023 | NA |
| | 0.096202 | 0.408546 | 4.079639 | 0.100143 | 0.920231 | NA |
| RPL21P4 | 0.230334 | -0.53318 | 4.076749 | -0.13079 | 0.895945 | NA |
| | 3.054655 | -0.00752 | 1.133328 | -0.00664 | 0.994703 | NA |
| | 0.326333 | 1.533889 | 3.826878 | 0.40082 | 0.688553 | NA |
| RPL21P122 | 0.303394 | -2.27006 | 4.032451 | -0.56295 | 0.57347 | NA |
| RN7SL39P | 0.163957 | 0.424913 | 4.079006 | 0.104171 | 0.917034 | NA |
| CDRT1 | 0.326333 | 1.533889 | 3.826878 | 0.40082 | 0.688553 | NA |
| RPL35AP33 | 0.134133 | -1.48909 | 4.077584 | -0.36519 | 0.71497 | NA |
| RPL39P38 | 0.629726 | -0.22725 | 2.732935 | -0.08315 | 0.93373 | NA |
| RPL7P5 | 0.163957 | 0.424913 | 4.079006 | 0.104171 | 0.917034 | NA |
| RN7SL316F | 0.392857 | -1.546 | 4.075469 | -0.37934 | 0.704433 | NA |
| RN7SL129F | 0.151697 | -1.546 | 4.075469 | -0.37934 | 0.704433 | NA |
| RPS4XP19 | 0.096202 | 0.408546 | 4.079639 | 0.100143 | 0.920231 | NA |
| RPS3AP49 | 2.578645 | 2.040025 | 1.364441 | 1.495136 | 0.134879 | NA |
| RPL9P28 | 0.315653 | -0.57305 | 3.892639 | -0.14721 | 0.882963 | NA |
| RPL9P32 | 0.556667 | 0.972496 | 2.522454 | 0.385536 | 0.699841 | NA |
| | 0.392857 | -1.546 | 4.075469 | -0.37934 | 0.704433 | NA |
| ARHGDIG | 1.945904 | 2.844225 | 1.832967 | 1.551705 | 0.120733 | NA |
| | 0.096202 | 0.408546 | 4.079639 | 0.100143 | 0.920231 | NA |
| | 0.096202 | 0.408546 | 4.079639 | 0.100143 | 0.920231 | NA |
| RPS12P31 | 0.775733 | 2.817284 | 2.75909 | 1.021092 | 0.307211 | NA |
| RN7SL800F | 0.327913 | 1.543154 | 4.021063 | 0.383768 | 0.701151 | NA |
| DECR2 | 351.2577 | -0.14683 | 0.180336 | -0.81419 | 0.415536 | 0.578766 |
| | 9.134901 | -0.06429 | 0.65846 | -0.09764 | 0.922218 | NA |
| RPL21P129 | 0.327913 | 1.543154 | 4.021063 | 0.383768 | 0.701151 | NA |
| RN7SL862F | 0.151697 | -1.546 | 4.075469 | -0.37934 | 0.704433 | NA |
| RPS27AP1 | 0.49187 | 2.136088 | 3.895953 | 0.548284 | 0.583497 | NA |
| ZNF709 | 0.74749 | 1.546424 | 2.516383 | 0.614543 | 0.538857 | NA |
| RN7SL449F | 0.410275 | 0.244283 | 3.100889 | 0.078778 | 0.937209 | NA |
| RPL31P58 | 0.096202 | 0.408546 | 4.079639 | 0.100143 | 0.920231 | NA |
| RPS3P7 | 0.28583 | -2.21358 | 3.97376 | -0.55705 | 0.577494 | NA |
| RPS10P28 | 1.129464 | 0.412965 | 1.930323 | 0.213936 | 0.830597 | NA |
| RPL31P61 | 0.151697 | -1.546 | 4.075469 | -0.37934 | 0.704433 | NA |
| RN7SL775F | 0.776119 | 0.401378 | 2.295521 | 0.174853 | 0.861196 | NA |
| | 0.487129 | 2.123445 | 3.910539 | 0.543006 | 0.587126 | NA |
| RPS18P13 | 0.314073 | -0.57308 | 3.898934 | -0.14698 | 0.883146 | NA |
| RPL32P33 | 0.163957 | 0.424913 | 4.079006 | 0.104171 | 0.917034 | NA |
| NME1-NMI | 36.10042 | -0.00968 | 0.381731 | -0.02537 | 0.979759 | NA |
| | 0.628146 | -0.21495 | 3.102079 | -0.06929 | 0.944756 | NA |
| | 0.163957 | 0.424913 | 4.079006 | 0.104171 | 0.917034 | NA |

| | | | | | | |
|---|---|---|---|---|---|---|
| NPIPB5 | 39.21969 | 0.472103 | 0.395423 | 1.19392 | 0.232509 | NA |
| RN7SL844P | 0.976776 | -2.39789 | 2.178705 | -1.1006 | 0.271069 | NA |
| | 0.804712 | -0.7622 | 2.348804 | -0.3245 | 0.745556 | NA |
| RPS24P17 | 0.410275 | 0.244283 | 3.100889 | 0.078778 | 0.937209 | NA |
| RPS27P27 | 0.096202 | 0.408546 | 4.079639 | 0.100143 | 0.920231 | NA |
| TMEM199 | 372.8348 | -0.07904 | 0.177753 | -0.44464 | 0.656578 | 0.773346 |
| RPS20P35 | 0.162376 | 0.424913 | 4.079006 | 0.104171 | 0.917034 | NA |
| RPS4XP17 | 1.007794 | 1.520243 | 2.181859 | 0.696765 | 0.48595 | NA |
| P2RY11 | 50.61718 | -0.51844 | 0.330572 | -1.56832 | 0.116808 | 0.247228 |
| | 2.038753 | 0.289574 | 1.442866 | 0.200694 | 0.840938 | NA |
| RPL26P35 | 0.163957 | 0.424913 | 4.079006 | 0.104171 | 0.917034 | NA |
| RPS2P45 | 0.871089 | -1.46164 | 2.262872 | -0.64592 | 0.518329 | NA |
| RPL18P13 | 2.69007 | -0.26589 | 1.24433 | -0.21368 | 0.830797 | NA |
| RPL21P120 | 1.935464 | 1.025732 | 1.42996 | 0.717315 | 0.47318 | NA |
| | 0.394291 | 0.298745 | 3.141711 | 0.09509 | 0.924243 | NA |
| | 1.2271 | 1.384124 | 1.759418 | 0.786694 | 0.431461 | NA |
| | 1.091976 | -0.2898 | 1.942755 | -0.14917 | 0.881418 | NA |
| | 3.864539 | 0.56212 | 1.014242 | 0.554227 | 0.579424 | NA |
| ZNF585B | 81.23346 | 0.152337 | 0.324527 | 0.469414 | 0.638774 | 0.759385 |
| CRNDE | 275.9735 | 0.396798 | 0.20889 | 1.899556 | 0.057491 | 0.152304 |
| | 0.327913 | 1.543154 | 4.021063 | 0.383768 | 0.701151 | NA |
| CEBPA | 0.561971 | -0.49218 | 2.810061 | -0.17515 | 0.860962 | NA |
| | 2.247322 | 0.060802 | 1.454546 | 0.041802 | 0.966657 | NA |
| | 0.096202 | 0.408546 | 4.079639 | 0.100143 | 0.920231 | NA |
| | 0.392857 | -1.546 | 4.075469 | -0.37934 | 0.704433 | NA |
| | 3.015411 | -0.74847 | 1.207536 | -0.61983 | 0.535368 | NA |
| | 110.5236 | -0.83295 | 0.320212 | -2.60125 | 0.009289 | 0.04489 |
| | 5.589993 | 0.617629 | 0.916919 | 0.673592 | 0.500571 | NA |
| LINC00920 | 203.6178 | 0.465778 | 0.218551 | 2.131203 | 0.033072 | 0.105994 |
| | 0.748867 | 2.740193 | 2.468928 | 1.109871 | 0.267054 | NA |
| | 136.7735 | -0.416 | 0.20799 | -2.00012 | 0.045487 | 0.13062 |
| | 0.489058 | -0.58569 | 3.427543 | -0.17088 | 0.864321 | NA |
| | 0.712719 | 2.707521 | 2.836859 | 0.954408 | 0.339877 | NA |
| | 7.597451 | -0.43649 | 0.754706 | -0.57836 | 0.563023 | NA |
| NR2F2-AS1 | 54.11087 | 0.27186 | 0.327899 | 0.829095 | 0.407051 | 0.570737 |
| | 1.579453 | -0.56402 | 1.503609 | -0.37511 | 0.707578 | NA |
| BCKDHA | 66.83338 | -0.54582 | 0.332287 | -1.64261 | 0.100464 | 0.223642 |
| SMURF2P1 | 35.65904 | -0.10079 | 0.368575 | -0.27344 | 0.784511 | NA |
| RRN3P1 | 25.15024 | 0.453101 | 0.428543 | 1.057307 | 0.290371 | NA |
| SUMO2P17 | 1.693073 | 0.276824 | 1.517715 | 0.182395 | 0.855272 | NA |
| | 3.755377 | 1.537176 | 1.195151 | 1.286177 | 0.198381 | NA |
| | 3.035693 | -0.36101 | 1.109794 | -0.32529 | 0.744961 | NA |
| FAM138E | 0.134133 | -1.48909 | 4.077584 | -0.36519 | 0.71497 | NA |
| HAUS5 | 333.3714 | -0.58025 | 0.204374 | -2.83917 | 0.004523 | 0.027525 |
| | 0.260158 | 1.240347 | 4.047124 | 0.306476 | 0.759242 | NA |
| CASC16 | 0.247898 | -0.5901 | 4.074633 | -0.14482 | 0.884851 | NA |
| | 0.163957 | 0.424913 | 4.079006 | 0.104171 | 0.917034 | NA |

| | | | | | | |
|---|---|---|---|---|---|---|
| ZNF286B | 29.75467 | 0.370624 | 0.406667 | 0.911368 | 0.362102 | NA |
| ZNF564 | 1.778392 | 0.325778 | 1.507564 | 0.216096 | 0.828913 | NA |
| | 8.859647 | 0.906469 | 0.712261 | 1.272665 | 0.203137 | NA |
| CCDC79 | 0.883 | 1.784395 | 2.188528 | 0.81534 | 0.414878 | NA |
| RPS4XP20 | 0.151697 | -1.546 | 4.075469 | -0.37934 | 0.704433 | NA |
| YJEFN3 | 5.951756 | 0.651991 | 0.827487 | 0.787917 | 0.430745 | NA |
| CACNA1G-/ | 0.490493 | 0.776209 | 2.895261 | 0.268096 | 0.788625 | NA |
| PKD1P6 | 110.5913 | -0.27086 | 0.33626 | -0.80551 | 0.420527 | 0.583218 |
| | 0.096202 | 0.408546 | 4.079639 | 0.100143 | 0.920231 | NA |
| | 0.873901 | 0.695147 | 2.212014 | 0.31426 | 0.753324 | NA |
| LRRC37BP1 | 2.038339 | -0.21984 | 1.344596 | -0.1635 | 0.870123 | NA |
| CDK3 | 0.659753 | -0.00581 | 2.629089 | -0.00221 | 0.998237 | NA |
| | 11.31807 | 0.488799 | 0.641446 | 0.762027 | 0.446044 | NA |
| | 0.134133 | -1.48909 | 4.077584 | -0.36519 | 0.71497 | NA |
| PGAM1P12 | 0.096202 | 0.408546 | 4.079639 | 0.100143 | 0.920231 | NA |
| GAPDHP66 | 1.266372 | 2.02221 | 2.026982 | 0.997646 | 0.318451 | NA |
| | 1.51427 | 1.23776 | 1.732335 | 0.714504 | 0.474915 | NA |
| | 0.877625 | -0.2954 | 2.206963 | -0.13385 | 0.893522 | NA |
| LINC00923 | 2.008726 | 0.218933 | 1.399242 | 0.156465 | 0.875666 | NA |
| ZNF345 | 48.74145 | 0.842809 | 0.32809 | 2.568833 | 0.010204 | 0.047814 |
| | 4.184733 | 0.463208 | 1.036206 | 0.447023 | 0.654858 | NA |
| | 0.162376 | 0.424913 | 4.079006 | 0.104171 | 0.917034 | NA |
| | 1.916815 | -0.52082 | 1.457929 | -0.35723 | 0.720917 | NA |
| PTX4 | 0.260158 | 1.240347 | 4.047124 | 0.306476 | 0.759242 | NA |
| RNU6-322F | 1.042804 | -0.41495 | 2.072413 | -0.20022 | 0.841305 | NA |
| RNU7-71P | 0.392857 | -1.546 | 4.075469 | -0.37934 | 0.704433 | NA |
| RNU2-69P | 0.134133 | -1.48909 | 4.077584 | -0.36519 | 0.71497 | NA |
| SCARNA17 | 2.713302 | -0.48302 | 1.228329 | -0.39323 | 0.69415 | NA |
| RNU2-27P | 2.21112 | -0.5549 | 1.752187 | -0.31669 | 0.751479 | NA |
| RNU6-1262 | 0.151697 | -1.546 | 4.075469 | -0.37934 | 0.704433 | NA |
| | 0.096202 | 0.408546 | 4.079639 | 0.100143 | 0.920231 | NA |
| | 0.614937 | 2.493639 | 2.617677 | 0.952615 | 0.340785 | NA |
| | 0.490493 | 0.776209 | 2.895261 | 0.268096 | 0.788625 | NA |
| RNU1-103F | 0.74749 | 1.546424 | 2.516383 | 0.614543 | 0.538857 | NA |
| | 0.591998 | -0.2499 | 3.117006 | -0.08017 | 0.936101 | NA |
| RNU6-1002 | 0.134133 | -1.48909 | 4.077584 | -0.36519 | 0.71497 | NA |
| RNU6-638F | 0.162376 | 0.424913 | 4.079006 | 0.104171 | 0.917034 | NA |
| RNA5SP434 | 0.096202 | 0.408546 | 4.079639 | 0.100143 | 0.920231 | NA |
| RNU6-1153 | 0.230334 | -0.53318 | 4.076749 | -0.13079 | 0.895945 | NA |
| SCARNA20 | 0.392857 | -1.546 | 4.075469 | -0.37934 | 0.704433 | NA |
| RNU7-117F | 0.392711 | 0.292538 | 3.145221 | 0.09301 | 0.925895 | NA |
| RNU6-807F | 0.134133 | -1.48909 | 4.077584 | -0.36519 | 0.71497 | NA |
| | 0.506476 | 0.726418 | 2.864186 | 0.253621 | 0.799788 | NA |
| | 0.096202 | 0.408546 | 4.079639 | 0.100143 | 0.920231 | NA |
| SCARNA21 | 0.815391 | 0.931626 | 2.360431 | 0.394685 | 0.693076 | NA |
| RN7SKP23 | 0.296509 | -0.51709 | 3.982155 | -0.12985 | 0.896684 | NA |
| | 0.163957 | 0.424913 | 4.079006 | 0.104171 | 0.917034 | NA |

| | | | | | | |
|---|---|---|---|---|---|---|
| RNU6-1223 | 0.163957 | 0.424913 | 4.079006 | 0.104171 | 0.917034 | NA |
| RNU6-993F | 0.096202 | 0.408546 | 4.079639 | 0.100143 | 0.920231 | NA |
| RNU6-1057 | 0.151697 | -1.546 | 4.075469 | -0.37934 | 0.704433 | NA |
| RNU6-840F | 1.003293 | 0.228546 | 2.091683 | 0.109264 | 0.912993 | NA |
| | 1.104032 | 1.152394 | 2.115117 | 0.544837 | 0.585866 | NA |
| | 0.815245 | 1.670886 | 2.214146 | 0.754641 | 0.450464 | NA |
| | 0.967088 | -0.05827 | 2.035183 | -0.02863 | 0.977158 | NA |
| | 0.898984 | 1.748354 | 2.20932 | 0.791354 | 0.428737 | NA |
| HMGN1P38 | 3.093907 | -0.15646 | 1.157911 | -0.13512 | 0.892518 | NA |
| ZNF260 | 218.2159 | 0.870179 | 0.395867 | 2.19816 | 0.027938 | 0.094656 |
| | 0.163957 | 0.424913 | 4.079006 | 0.104171 | 0.917034 | NA |
| NPIPB11 | 4.98643 | 0.008113 | 1.017479 | 0.007974 | 0.993638 | NA |
| | 1.347707 | -4.22129 | 2.39527 | -1.76234 | 0.078011 | NA |
| | 0.420954 | 1.923376 | 3.462803 | 0.555439 | 0.578594 | NA |
| PKD1P5 | 0.162376 | 0.424913 | 4.079006 | 0.104171 | 0.917034 | NA |
| | 0.556813 | -0.57007 | 3.294615 | -0.17303 | 0.862627 | NA |
| NPIPA2 | 0.096202 | 0.408546 | 4.079639 | 0.100143 | 0.920231 | NA |
| MPV17L2 | 767.1049 | -0.11748 | 0.210679 | -0.55762 | 0.577102 | 0.711993 |
| | 0.749273 | 0.569822 | 2.889797 | 0.197184 | 0.843683 | NA |
| MEF2BNB | 329.0013 | 0.017127 | 0.164838 | 0.103899 | 0.91725 | 0.952022 |
| HMGB1P24 | 0.640755 | -1.64826 | 2.754778 | -0.59833 | 0.549621 | NA |
| CHMP1B | 3055.972 | 0.236398 | 0.149012 | 1.586431 | 0.112642 | 0.241224 |
| | 1.228827 | 0.64505 | 2.019407 | 0.319426 | 0.749404 | NA |
| SNHG9 | 28.03361 | 0.113932 | 0.405035 | 0.281288 | 0.77849 | NA |
| ABCC6P2 | 0.384806 | 1.846216 | 3.999752 | 0.461582 | 0.644381 | NA |
| | 0.096202 | 0.408546 | 4.079639 | 0.100143 | 0.920231 | NA |
| CYP2A6 | 0.192403 | 0.975392 | 4.060716 | 0.240202 | 0.810174 | NA |
| | 0.606787 | -3.06078 | 3.567758 | -0.8579 | 0.390947 | NA |
| LINC01152 | 8.518145 | -0.52735 | 0.704477 | -0.74856 | 0.45412 | NA |
| | 0.715249 | -0.95037 | 2.624937 | -0.36205 | 0.717313 | NA |
| ZNF486 | 77.91851 | 0.448998 | 0.375408 | 1.196027 | 0.231686 | 0.394009 |
| | 2.967758 | 1.893378 | 1.575191 | 1.201999 | 0.229364 | NA |
| | 13.76889 | -0.36894 | 0.563247 | -0.65502 | 0.512452 | NA |
| SALL3 | 147.0157 | -0.40038 | 0.303949 | -1.31726 | 0.18775 | 0.341836 |
| POLG2 | 150.885 | 0.214708 | 0.197511 | 1.08707 | 0.277006 | 0.442933 |
| CYP2B7P | 1.064724 | 1.125776 | 1.854789 | 0.606957 | 0.54388 | NA |
| | 2.664902 | 0.147767 | 1.34749 | 0.109661 | 0.912678 | NA |
| MTRNR2L1 | 0.258578 | 1.233151 | 4.047809 | 0.304646 | 0.760635 | NA |
| LINC00273 | 3.500661 | 0.016287 | 1.103296 | 0.014762 | 0.988222 | NA |
| SDCCAG3P | 0.162376 | 0.424913 | 4.079006 | 0.104171 | 0.917034 | NA |
| ZNF253 | 62.93087 | 0.303998 | 0.389531 | 0.78042 | 0.435143 | 0.596166 |
| CAPNS2 | 4.231781 | 0.816898 | 1.02846 | 0.794292 | 0.427025 | NA |
| | 0.163957 | 0.424913 | 4.079006 | 0.104171 | 0.917034 | NA |
| HP | 0.163957 | 0.424913 | 4.079006 | 0.104171 | 0.917034 | NA |
| RPL29P33 | 0.327913 | 1.543154 | 4.021063 | 0.383768 | 0.701151 | NA |
| LSM14A | 1573.966 | 0.408094 | 0.165316 | 2.468575 | 0.013565 | 0.057932 |
| NHLRC4 | 68.40061 | -0.5543 | 0.293934 | -1.8858 | 0.059321 | 0.155967 |

| | | | | | | |
|---|---|---|---|---|---|---|
| RRN3P3 | 105.8982 | 0.405681 | 0.318859 | 1.272287 | 0.203271 | 0.359796 |
| MIR10A | 1.495512 | -0.17463 | 1.558026 | -0.11208 | 0.910756 | NA |
| | 0.151697 | -1.546 | 4.075469 | -0.37934 | 0.704433 | NA |
| MIR3179-1 | 0.881915 | -2.2305 | 2.871853 | -0.77668 | 0.437349 | NA |
| ZNF271 | 870.1736 | 0.584303 | 0.193128 | 3.025466 | 0.002483 | 0.018709 |
| MIR3180-4 | 0.096202 | 0.408546 | 4.079639 | 0.100143 | 0.920231 | NA |
| | 0.489058 | -0.58569 | 3.427543 | -0.17088 | 0.864321 | NA |
| ZNF878 | 4.474129 | -0.12922 | 0.917483 | -0.14085 | 0.887992 | NA |
| NPIPB1P | 0.163957 | 0.424913 | 4.079006 | 0.104171 | 0.917034 | NA |
| ZNF625 | 3.727383 | -0.45558 | 1.415775 | -0.32179 | 0.747614 | NA |
| | 0.096202 | 0.408546 | 4.079639 | 0.100143 | 0.920231 | NA |
| | 0.096202 | 0.408546 | 4.079639 | 0.100143 | 0.920231 | NA |
| | 0.296509 | -0.51709 | 3.982155 | -0.12985 | 0.896684 | NA |
| TEN1 | 0.449786 | -1.25325 | 3.00474 | -0.41709 | 0.676613 | NA |
| P2RX5-TAX | 20.45085 | -0.21454 | 0.482757 | -0.44441 | 0.656745 | NA |
| | 0.247898 | -0.5901 | 4.074633 | -0.14482 | 0.884851 | NA |
| | 0.449786 | -1.25325 | 3.00474 | -0.41709 | 0.676613 | NA |
| | 0.096202 | 0.408546 | 4.079639 | 0.100143 | 0.920231 | NA |
| C17orf49 | 2.29519 | 0.624482 | 1.540198 | 0.405456 | 0.685142 | NA |
| MIR3180-1 | 0.574231 | 0.93227 | 2.49998 | 0.372911 | 0.709215 | NA |
| | 8.008484 | -0.22621 | 0.724357 | -0.31229 | 0.754822 | NA |
| PDF | 281.4919 | -0.33342 | 0.187492 | -1.77833 | 0.07535 | 0.183424 |
| | 9.603648 | -0.57616 | 0.654052 | -0.88091 | 0.378367 | NA |
| | 0.460669 | -0.62593 | 3.398671 | -0.18417 | 0.853882 | NA |
| ASB9P1 | 0.326333 | 1.533889 | 3.826878 | 0.40082 | 0.688553 | NA |
| | 0.163957 | 0.424913 | 4.079006 | 0.104171 | 0.917034 | NA |
| | 0.151697 | -1.546 | 4.075469 | -0.37934 | 0.704433 | NA |
| PRC1-AS1 | 2.233846 | -0.11844 | 1.400399 | -0.08457 | 0.932599 | NA |
| ENO1P2 | 1.173388 | 1.426393 | 2.178451 | 0.654774 | 0.512613 | NA |
| | 2.046804 | -0.69781 | 1.34901 | -0.51728 | 0.604962 | NA |
| | 0.651435 | 0.249629 | 2.788679 | 0.089515 | 0.928672 | NA |
| | 0.162376 | 0.424913 | 4.079006 | 0.104171 | 0.917034 | NA |
| MC1R | 46.86515 | -0.2172 | 0.365173 | -0.59479 | 0.551983 | NA |
| CEP95 | 187.0985 | 0.927965 | 0.296289 | 3.131964 | 0.001736 | 0.01475 |
| | 0.162376 | 0.424913 | 4.079006 | 0.104171 | 0.917034 | NA |
| | 0.354779 | 1.703954 | 3.657232 | 0.465914 | 0.641277 | NA |
| NPM1P5 | 1.274691 | 1.421878 | 1.902542 | 0.747357 | 0.454848 | NA |
| | 210.8488 | -0.4397 | 0.258649 | -1.7 | 0.089131 | 0.206216 |
| | 0.720624 | 1.435938 | 2.334155 | 0.615185 | 0.538432 | NA |
| | 1.679661 | 1.962788 | 1.612764 | 1.217034 | 0.223591 | NA |
| SMIM6 | 0.69625 | -3.13723 | 2.93845 | -1.06765 | 0.285679 | NA |
| | 0.28583 | -2.21358 | 3.97376 | -0.55705 | 0.577494 | NA |
| | 2.125295 | 0.031146 | 1.347263 | 0.023118 | 0.981556 | NA |
| | 0.777314 | 2.821035 | 2.404647 | 1.17316 | 0.240732 | NA |
| PRKXP1 | 9.061775 | 0.124644 | 0.74158 | 0.168079 | 0.866521 | NA |
| ITGB3 | 2759.433 | -0.22479 | 0.211283 | -1.06391 | 0.287369 | 0.453569 |
| | 0.792452 | -2.15617 | 2.562821 | -0.84133 | 0.400166 | NA |

| | | | | | | |
|---|---|---|---|---|---|---|
| | 5.257995 | 0.134226 | 1.073174 | 0.125074 | 0.900465 | NA |
| SLC35G6 | 1.208102 | 0.723565 | 1.850162 | 0.391082 | 0.695737 | NA |
| HNRNPA1P | 0.327913 | 1.543154 | 4.021063 | 0.383768 | 0.701151 | NA |
| | 0.508056 | 0.731032 | 2.861912 | 0.255435 | 0.798387 | NA |
| | 0.392711 | 0.292538 | 3.145221 | 0.09301 | 0.925895 | NA |
| | 0.399595 | -1.31595 | 3.579466 | -0.36764 | 0.713143 | NA |
| NDUFA3P4 | 0.096202 | 0.408546 | 4.079639 | 0.100143 | 0.920231 | NA |
| | 0.399595 | -1.31595 | 3.579466 | -0.36764 | 0.713143 | NA |
| | 0.681315 | 1.40954 | 2.618083 | 0.538386 | 0.59031 | NA |
| | 0.326536 | 0.037056 | 3.842554 | 0.009644 | 0.992306 | NA |
| | 0.815391 | 0.931626 | 2.360431 | 0.394685 | 0.693076 | NA |
| | 0.096202 | 0.408546 | 4.079639 | 0.100143 | 0.920231 | NA |
| | 2.305024 | -0.57525 | 1.402311 | -0.41022 | 0.681646 | NA |
| | 0.382031 | -1.26484 | 3.151989 | -0.40128 | 0.688212 | NA |
| | 14.28919 | 0.250671 | 0.55148 | 0.454542 | 0.649439 | NA |
| | 0.682693 | 2.623213 | 2.518338 | 1.041645 | 0.297576 | NA |
| GH1 | 0.327913 | 1.543154 | 4.021063 | 0.383768 | 0.701151 | NA |
| | 0.151697 | -1.546 | 4.075469 | -0.37934 | 0.704433 | NA |
| | 0.883146 | 1.112834 | 2.554178 | 0.435692 | 0.663061 | NA |
| | 0.651288 | 1.289307 | 2.633717 | 0.489539 | 0.62446 | NA |
| | 54.21734 | 0.189066 | 0.31064 | 0.608633 | 0.542768 | 0.684848 |
| | 5.679485 | 0.203882 | 0.822385 | 0.247916 | 0.804199 | NA |
| | 106.9952 | 0.099765 | 0.239455 | 0.416633 | 0.676947 | 0.788873 |
| | 13.71705 | 0.828682 | 0.596122 | 1.390121 | 0.164492 | NA |
| DNM1P47 | 1.962127 | 1.55416 | 1.508101 | 1.030541 | 0.302756 | NA |
| | 0.163957 | 0.424913 | 4.079006 | 0.104171 | 0.917034 | NA |
| HSP90B2P | 2.779498 | -1.2191 | 1.32034 | -0.92333 | 0.355838 | NA |
| | 0.900564 | 1.750062 | 2.175753 | 0.804348 | 0.421196 | NA |
| | 0.163957 | 0.424913 | 4.079006 | 0.104171 | 0.917034 | NA |
| | 2.829879 | 0.325508 | 1.207529 | 0.269565 | 0.787495 | NA |
| HSPE1P3 | 0.842643 | -2.08432 | 2.306153 | -0.90381 | 0.366097 | NA |
| | 0.303394 | -2.27006 | 4.032451 | -0.56295 | 0.57347 | NA |
| | 0.69238 | 0.295083 | 2.603008 | 0.113362 | 0.909743 | NA |
| | 0.410275 | 0.244283 | 3.100889 | 0.078778 | 0.937209 | NA |
| | 12.2512 | 0.207726 | 0.616588 | 0.336896 | 0.736196 | NA |
| | 0.382031 | -1.26484 | 3.151989 | -0.40128 | 0.688212 | NA |
| | 0.260158 | 1.240347 | 4.047124 | 0.306476 | 0.759242 | NA |
| | 0.096202 | 0.408546 | 4.079639 | 0.100143 | 0.920231 | NA |
| SLC22A31 | 2.027659 | -0.68335 | 1.335538 | -0.51167 | 0.608883 | NA |
| | 1.075752 | 0.450053 | 2.098769 | 0.214437 | 0.830206 | NA |
| | 39.13946 | -0.17838 | 0.347881 | -0.51275 | 0.608128 | NA |
| | 0.151697 | -1.546 | 4.075469 | -0.37934 | 0.704433 | NA |
| | 0.851742 | -0.48655 | 2.547586 | -0.19099 | 0.848537 | NA |
| RAB43P1 | 0.402398 | -2.58771 | 4.018369 | -0.64397 | 0.519595 | NA |
| | 0.382031 | -1.26484 | 3.151989 | -0.40128 | 0.688212 | NA |
| | 0.162376 | 0.424913 | 4.079006 | 0.104171 | 0.917034 | NA |
| | 0.327913 | 1.543154 | 4.021063 | 0.383768 | 0.701151 | NA |

| | | | | | | |
|---|---|---|---|---|---|---|
| | 20.41429 | 0.283728 | 0.464254 | 0.611149 | 0.541101 | NA |
| | 0.777314 | 2.821035 | 2.404647 | 1.17316 | 0.240732 | NA |
| | 2.066525 | -0.02235 | 1.74566 | -0.0128 | 0.989787 | NA |
| | 61.69004 | -0.70488 | 0.297769 | -2.3672 | 0.017923 | 0.070101 |
| | 0.096202 | 0.408546 | 4.079639 | 0.100143 | 0.920231 | NA |
| NDUFA5P1 | 0.303394 | -2.27006 | 4.032451 | -0.56295 | 0.57347 | NA |
| | 0.833158 | -0.52049 | 2.521552 | -0.20642 | 0.836464 | NA |
| | 0.163957 | 0.424913 | 4.079006 | 0.104171 | 0.917034 | NA |
| | 0.951104 | -0.01996 | 2.201588 | -0.00907 | 0.992765 | NA |
| | 0.162376 | 0.424913 | 4.079006 | 0.104171 | 0.917034 | NA |
| | 0.601483 | -1.7332 | 2.729494 | -0.63499 | 0.525436 | NA |
| | 3.779762 | 0.161804 | 1.094867 | 0.147784 | 0.882513 | NA |
| VN1R68P | 0.151697 | -1.546 | 4.075469 | -0.37934 | 0.704433 | NA |
| CDC37P1 | 0.151697 | -1.546 | 4.075469 | -0.37934 | 0.704433 | NA |
| | 81.67283 | 0.108865 | 0.261237 | 0.41673 | 0.676876 | 0.788868 |
| | 0.260158 | 1.240347 | 4.047124 | 0.306476 | 0.759242 | NA |
| TGFBR3L | 39.55638 | 0.204956 | 0.34453 | 0.594886 | 0.55192 | NA |
| | 11.00529 | 0.465625 | 0.657869 | 0.707777 | 0.479084 | NA |
| ZNF720P1 | 0.47961 | 0.535796 | 3.39605 | 0.15777 | 0.874638 | NA |
| LSM3P5 | 0.163957 | 0.424913 | 4.079006 | 0.104171 | 0.917034 | NA |
| | 38.00235 | -0.27808 | 0.390767 | -0.71163 | 0.476695 | NA |
| | 0.163957 | 0.424913 | 4.079006 | 0.104171 | 0.917034 | NA |
| | 2.13478 | -0.36016 | 1.376212 | -0.2617 | 0.793551 | NA |
| | 4.486682 | -0.47681 | 1.001574 | -0.47606 | 0.63403 | NA |
| | 2.86268 | 1.072281 | 1.272156 | 0.842885 | 0.399293 | NA |
| | 1.73954 | 0.950876 | 1.722233 | 0.552118 | 0.580867 | NA |
| | 0.163957 | 0.424913 | 4.079006 | 0.104171 | 0.917034 | NA |
| | 25.99999 | -0.37582 | 0.475558 | -0.79027 | 0.429373 | NA |
| | 4.016447 | 0.853425 | 1.038836 | 0.82152 | 0.41135 | NA |
| | 0.096202 | 0.408546 | 4.079639 | 0.100143 | 0.920231 | NA |
| | 1.752655 | -0.41495 | 1.535934 | -0.27016 | 0.787037 | NA |
| NSFP1 | 2.173632 | -0.80894 | 1.454172 | -0.55629 | 0.578014 | NA |
| | 0.612162 | -0.19706 | 2.47693 | -0.07956 | 0.936588 | NA |
| MIR4519 | 8.701434 | 1.435529 | 0.730666 | 1.964686 | 0.049451 | NA |
| | 64.67535 | 0.162657 | 0.277528 | 0.586092 | 0.557814 | 0.69668 |
| | 1.949022 | -0.91689 | 1.390122 | -0.65957 | 0.509528 | NA |
| AOC4P | 0.163957 | 0.424913 | 4.079006 | 0.104171 | 0.917034 | NA |
| | 0.162376 | 0.424913 | 4.079006 | 0.104171 | 0.917034 | NA |
| | 34.55028 | 0.260745 | 0.358456 | 0.727413 | 0.466973 | NA |
| | 13.50829 | -0.57028 | 0.588475 | -0.96908 | 0.332503 | NA |
| | 0.424115 | 1.932531 | 3.45462 | 0.559405 | 0.575885 | NA |
| CA5AP1 | 0.096202 | 0.408546 | 4.079639 | 0.100143 | 0.920231 | NA |
| | 18.33886 | 0.494061 | 0.489936 | 1.008421 | 0.313253 | NA |
| | 0.258578 | 1.233151 | 4.047809 | 0.304646 | 0.760635 | NA |
| | 0.392857 | -1.546 | 4.075469 | -0.37934 | 0.704433 | NA |
| | 1.490136 | -1.58638 | 1.719755 | -0.92245 | 0.356296 | NA |
| | 0.738391 | 0.402625 | 2.276191 | 0.176885 | 0.859599 | NA |

| | | | | | | |
|---|---|---|---|---|---|---|
| | 0.162376 | 0.424913 | 4.079006 | 0.104171 | 0.917034 | NA |
| | 0.296509 | -0.51709 | 3.982155 | -0.12985 | 0.896684 | NA |
| | 0.162376 | 0.424913 | 4.079006 | 0.104171 | 0.917034 | NA |
| | 0.096202 | 0.408546 | 4.079639 | 0.100143 | 0.920231 | NA |
| | 2.525225 | 1.149076 | 1.501647 | 0.765211 | 0.444146 | NA |
| | 17.198 | 0.095788 | 0.510366 | 0.187686 | 0.851123 | NA |
| | 0.260158 | 1.240347 | 4.047124 | 0.306476 | 0.759242 | NA |
| | 0.678686 | -3.08037 | 2.621268 | -1.17514 | 0.239937 | NA |
| SUB1P4 | 0.314073 | -0.57308 | 3.898934 | -0.14698 | 0.883146 | NA |
| | 1.075752 | 0.450053 | 2.098769 | 0.214437 | 0.830206 | NA |
| | 0.651288 | 1.289307 | 2.633717 | 0.489539 | 0.62446 | NA |
| | 0.844077 | -0.22325 | 2.189502 | -0.10196 | 0.918786 | NA |
| | 3.813117 | -0.00516 | 1.065024 | -0.00484 | 0.996136 | NA |
| | 108.4308 | 0.124319 | 0.254135 | 0.489186 | 0.62471 | 0.748452 |
| | 73.97407 | -0.44869 | 0.275051 | -1.6313 | 0.102827 | 0.227098 |
| LINC00919 | 35.00198 | -0.45973 | 0.509505 | -0.9023 | 0.366898 | NA |
| | 10.3787 | -0.29082 | 0.632873 | -0.45953 | 0.645857 | NA |
| | 0.260158 | 1.240347 | 4.047124 | 0.306476 | 0.759242 | NA |
| | 2.78706 | 0.981564 | 1.245966 | 0.787793 | 0.430818 | NA |
| | 1.858399 | -0.4589 | 1.422607 | -0.32258 | 0.747017 | NA |
| | 18.70002 | 0.190779 | 0.570458 | 0.334431 | 0.738055 | NA |
| | 0.555087 | 0.973802 | 2.808575 | 0.346725 | 0.728798 | NA |
| | 0.52699 | -2.57878 | 3.248478 | -0.79384 | 0.427288 | NA |
| | 0.746498 | -1.433 | 2.535747 | -0.56512 | 0.571994 | NA |
| | 1.539593 | 1.098873 | 1.649451 | 0.666205 | 0.50528 | NA |
| | 0.518736 | 2.237423 | 2.7713 | 0.807355 | 0.419462 | NA |
| COX6CP1 | 26.15081 | 0.596757 | 0.474748 | 1.256999 | 0.208754 | NA |
| | 0.410275 | 0.244283 | 3.100889 | 0.078778 | 0.937209 | NA |
| | 0.766634 | 1.516849 | 2.262991 | 0.670285 | 0.502676 | NA |
| | 26.81401 | -0.18074 | 0.429986 | -0.42035 | 0.674233 | NA |
| | 4.227775 | -0.10388 | 1.0071 | -0.10315 | 0.917846 | NA |
| | 1.045523 | 1.534279 | 2.413816 | 0.635624 | 0.525022 | NA |
| | 1.582111 | -0.20376 | 1.782133 | -0.11433 | 0.908974 | NA |
| | 1.713586 | -0.7846 | 1.451865 | -0.54041 | 0.588917 | NA |
| | 1.007648 | 2.051938 | 2.077812 | 0.987548 | 0.323374 | NA |
| AQP4-AS1 | 2.347457 | -0.62172 | 1.316207 | -0.47236 | 0.636671 | NA |
| | 0.986086 | 0.883356 | 2.062196 | 0.428357 | 0.668391 | NA |
| WBP11P1 | 0.096202 | 0.408546 | 4.079639 | 0.100143 | 0.920231 | NA |
| | 6.19662 | 0.374285 | 0.810192 | 0.461971 | 0.644102 | NA |
| | 0.096202 | 0.408546 | 4.079639 | 0.100143 | 0.920231 | NA |
| | 0.410275 | 0.244283 | 3.100889 | 0.078778 | 0.937209 | NA |
| | 0.096202 | 0.408546 | 4.079639 | 0.100143 | 0.920231 | NA |
| | 1.132625 | 0.391474 | 2.111679 | 0.185385 | 0.852927 | NA |
| | 2.03799 | 0.682895 | 1.447426 | 0.4718 | 0.63707 | NA |
| | 4.323635 | 1.026699 | 0.970174 | 1.058263 | 0.289936 | NA |
| | 1.659229 | 0.61197 | 1.860398 | 0.328946 | 0.742197 | NA |
| | 1.373842 | 0.279944 | 1.886333 | 0.148406 | 0.882022 | NA |

| | | | | | | |
|---|---|---|---|---|---|---|
| | 14.13077 | -0.50564 | 0.581619 | -0.86936 | 0.384651 | NA |
| | 0.247898 | -0.5901 | 4.074633 | -0.14482 | 0.884851 | NA |
| | 0.555233 | -0.57009 | 3.296947 | -0.17291 | 0.862719 | NA |
| | 16.75055 | 0.436839 | 0.517437 | 0.844235 | 0.398538 | NA |
| TPRKBP2 | 0.163957 | 0.424913 | 4.079006 | 0.104171 | 0.917034 | NA |
| C16orf95 | 77.40956 | 0.068696 | 0.316106 | 0.217319 | 0.827959 | 0.896239 |
| FTLP14 | 5.875241 | -1.20562 | 0.926409 | -1.30139 | 0.193125 | NA |
| | 1.380377 | 0.882103 | 1.689164 | 0.522213 | 0.601522 | NA |
| | 0.452561 | 2.061505 | 3.349456 | 0.615474 | 0.538241 | NA |
| | 0.151697 | -1.546 | 4.075469 | -0.37934 | 0.704433 | NA |
| | 0.844077 | -0.22325 | 2.189502 | -0.10196 | 0.918786 | NA |
| | 224.8437 | -0.10289 | 0.198592 | -0.51808 | 0.6044 | 0.732942 |
| | 2.227201 | 0.035543 | 1.594631 | 0.022289 | 0.982218 | NA |
| | 0.163957 | 0.424913 | 4.079006 | 0.104171 | 0.917034 | NA |
| | 0.629929 | -1.1413 | 2.677641 | -0.42623 | 0.669937 | NA |
| | 0.163957 | 0.424913 | 4.079006 | 0.104171 | 0.917034 | NA |
| | 0.268266 | -2.15396 | 4.038351 | -0.53338 | 0.593773 | NA |
| | 0.898195 | -1.76199 | 2.351517 | -0.7493 | 0.453676 | NA |
| BMS1P8 | 0.980782 | 1.975754 | 2.115136 | 0.934103 | 0.350251 | NA |
| | 2.725141 | -0.4901 | 1.19359 | -0.41061 | 0.681357 | NA |
| | 0.162376 | 0.424913 | 4.079006 | 0.104171 | 0.917034 | NA |
| FAM157C | 0.303394 | -2.27006 | 4.032451 | -0.56295 | 0.57347 | NA |
| | 0.430642 | -1.1891 | 3.485782 | -0.34113 | 0.733008 | NA |
| | 201.5808 | -0.62385 | 0.191453 | -3.25851 | 0.00112 | 0.01112 |
| | 1.223736 | 2.27097 | 2.079939 | 1.091845 | 0.274901 | NA |
| | 0.47961 | 0.535796 | 3.39605 | 0.15777 | 0.874638 | NA |
| MT1L | 16.96543 | -0.06403 | 0.550246 | -0.11636 | 0.907369 | NA |
| | 17.36563 | 0.011153 | 0.563509 | 0.019793 | 0.984209 | NA |
| | 0.392857 | -1.546 | 4.075469 | -0.37934 | 0.704433 | NA |
| | 18.48208 | 0.350709 | 0.598601 | 0.585881 | 0.557955 | NA |
| ERVK13-1 | 92.4274 | 0.485922 | 0.248937 | 1.95199 | 0.050939 | 0.140406 |
| | 96.74727 | -1.12023 | 0.306297 | -3.65735 | 2.55E-04 | 0.004502 |
| | 0.151697 | -1.546 | 4.075469 | -0.37934 | 0.704433 | NA |
| | 0.162376 | 0.424913 | 4.079006 | 0.104171 | 0.917034 | NA |
| | 0.163957 | 0.424913 | 4.079006 | 0.104171 | 0.917034 | NA |
| | 2.672807 | -0.19295 | 1.233288 | -0.15645 | 0.875676 | NA |
| | 0.163957 | 0.424913 | 4.079006 | 0.104171 | 0.917034 | NA |
| | 1.798317 | 0.827623 | 1.614273 | 0.512691 | 0.608168 | NA |
| SNAI3-AS1 | 129.9189 | -0.35554 | 0.223068 | -1.59384 | 0.110971 | 0.238988 |
| | 0.162376 | 0.424913 | 4.079006 | 0.104171 | 0.917034 | NA |
| | 1.295204 | 0.045195 | 1.784804 | 0.025322 | 0.979798 | NA |
| | 0.247898 | -0.5901 | 4.074633 | -0.14482 | 0.884851 | NA |
| | 2.308039 | -0.06851 | 1.331323 | -0.05146 | 0.958956 | NA |
| | 4.745231 | 0.288732 | 1.042426 | 0.276981 | 0.781795 | NA |
| | 1.039255 | 2.120498 | 2.300766 | 0.921649 | 0.356712 | NA |
| | 0.096202 | 0.408546 | 4.079639 | 0.100143 | 0.920231 | NA |
| | 1.045726 | 0.338916 | 1.94625 | 0.174138 | 0.861757 | NA |

| | | | | | | |
|---|---|---|---|---|---|---|
| | 0.424115 | 1.932531 | 3.45462 | 0.559405 | 0.575885 | NA |
| | 2.450047 | 0.365243 | 1.275853 | 0.286274 | 0.774668 | NA |
| CYCSP39 | 1.090395 | -0.30963 | 1.797091 | -0.17229 | 0.863206 | NA |
| | 0.422737 | 0.518872 | 3.501356 | 0.148192 | 0.882191 | NA |
| SLC7A5P1 | 73.49839 | -0.47935 | 0.287912 | -1.66492 | 0.095929 | 0.216865 |
| | 2.000639 | -1.54444 | 1.550255 | -0.99625 | 0.319131 | NA |
| | 0.399595 | -1.31595 | 3.579466 | -0.36764 | 0.713143 | NA |
| | 0.327913 | 1.543154 | 4.021063 | 0.383768 | 0.701151 | NA |
| | 4.357609 | -0.51648 | 1.033074 | -0.49995 | 0.617111 | NA |
| | 10.53938 | 0.143377 | 0.624157 | 0.229713 | 0.818315 | NA |
| | 3.854313 | 0.332139 | 1.07691 | 0.308419 | 0.757764 | NA |
| MIR940 | 8.721525 | 0.491284 | 0.706035 | 0.695835 | 0.486532 | NA |
| | 0.296509 | -0.51709 | 3.982155 | -0.12985 | 0.896684 | NA |
| CASC17 | 0.151697 | -1.546 | 4.075469 | -0.37934 | 0.704433 | NA |
| | 59.16305 | 2.134162 | 0.991937 | 2.151509 | 0.031436 | 0.102411 |
| | 0.476449 | 0.524262 | 3.402674 | 0.154073 | 0.877552 | NA |
| | 9.701216 | 0.331995 | 0.685212 | 0.484514 | 0.628021 | NA |
| | 0.096202 | 0.408546 | 4.079639 | 0.100143 | 0.920231 | NA |
| | 1.181236 | 0.579619 | 2.102215 | 0.275718 | 0.782765 | NA |
| | 86.79399 | 0.179113 | 0.306562 | 0.584265 | 0.559042 | 0.697891 |
| | 0.392857 | -1.546 | 4.075469 | -0.37934 | 0.704433 | NA |
| | 0.163957 | 0.424913 | 4.079006 | 0.104171 | 0.917034 | NA |
| | 0.096202 | 0.408546 | 4.079639 | 0.100143 | 0.920231 | NA |
| FBXL19-AS: | 52.22095 | 0.789922 | 0.362518 | 2.178988 | 0.029333 | 0.097487 |
| | 9.191802 | 0.897862 | 0.837476 | 1.072105 | 0.283673 | NA |
| | 0.911832 | -0.02323 | 2.42372 | -0.00958 | 0.992354 | NA |
| | 8.343255 | -0.03514 | 0.754539 | -0.04658 | 0.962851 | NA |
| | 2.196 | -0.73478 | 1.378535 | -0.53302 | 0.594022 | NA |
| | 1.129813 | -1.87082 | 2.359654 | -0.79284 | 0.427873 | NA |
| HCCAT5 | 4.595314 | -1.24777 | 1.050555 | -1.18773 | 0.234942 | NA |
| | 0.163957 | 0.424913 | 4.079006 | 0.104171 | 0.917034 | NA |
| | 1.393586 | -0.37783 | 1.885612 | -0.20038 | 0.841185 | NA |
| | 1.013338 | -0.63846 | 2.07512 | -0.30768 | 0.75833 | NA |
| | 0.354779 | 1.703954 | 3.657232 | 0.465914 | 0.641277 | NA |
| | 0.151697 | -1.546 | 4.075469 | -0.37934 | 0.704433 | NA |
| | 0.766634 | 1.516849 | 2.262991 | 0.670285 | 0.502676 | NA |
| | 4.73862 | -0.61014 | 1.029084 | -0.5929 | 0.553247 | NA |
| | 1.235354 | -0.89516 | 1.719639 | -0.52055 | 0.60268 | NA |
| | 0.489058 | -0.58569 | 3.427543 | -0.17088 | 0.864321 | NA |
| | 0.134133 | -1.48909 | 4.077584 | -0.36519 | 0.71497 | NA |
| FTO-IT1 | 1.3408 | -0.91042 | 2.016493 | -0.45149 | 0.65164 | NA |
| | 0.096202 | 0.408546 | 4.079639 | 0.100143 | 0.920231 | NA |
| | 0.162376 | 0.424913 | 4.079006 | 0.104171 | 0.917034 | NA |
| | 1.090395 | -0.30963 | 1.797091 | -0.17229 | 0.863206 | NA |
| | 5.16304 | -0.99379 | 1.028853 | -0.96592 | 0.334085 | NA |
| | 1.3507 | -0.58284 | 1.734422 | -0.33604 | 0.736839 | NA |
| | 35.65302 | 0.18171 | 0.35147 | 0.517001 | 0.605156 | NA |

| | | | | | | |
|---|---|---|---|---|---|---|
| | 5.778957 | 0.615569 | 0.949138 | 0.648555 | 0.516626 | NA |
| | 2.9895 | -0.03609 | 1.142782 | -0.03158 | 0.974807 | NA |
| | 0.420954 | 1.923376 | 3.462803 | 0.555439 | 0.578594 | NA |
| SULT1A3 | 0.938845 | -1.23445 | 2.063177 | -0.59832 | 0.549624 | NA |
| | 0.911593 | 1.257757 | 2.250933 | 0.558771 | 0.576318 | NA |
| | 1.456146 | -0.54961 | 1.792133 | -0.30668 | 0.759086 | NA |
| | 130.1536 | 0.009688 | 0.242464 | 0.039958 | 0.968127 | 0.982548 |
| | 4.291093 | 0.433256 | 0.999594 | 0.433432 | 0.664701 | NA |
| | 0.230334 | -0.53318 | 4.076749 | -0.13079 | 0.895945 | NA |
| | 5.121477 | -0.05316 | 0.870952 | -0.06103 | 0.951333 | NA |
| | 131.0698 | 0.033821 | 0.208128 | 0.162503 | 0.87091 | 0.923244 |
| | 0.247898 | -0.5901 | 4.074633 | -0.14482 | 0.884851 | NA |
| | 8.39255 | 0.306771 | 0.759101 | 0.404124 | 0.686122 | NA |
| | 7.956511 | -0.28685 | 0.786515 | -0.36471 | 0.715325 | NA |
| | 2.150763 | -0.37881 | 1.366729 | -0.27717 | 0.781653 | NA |
| | 3.794163 | 0.407584 | 1.106588 | 0.368325 | 0.712631 | NA |
| | 0.785714 | -3.2458 | 3.343263 | -0.97085 | 0.331623 | NA |
| | 0.861641 | -0.2592 | 2.180811 | -0.11885 | 0.90539 | NA |
| RBFADN | 0.602678 | 1.083611 | 2.70938 | 0.399948 | 0.689195 | NA |
| | 1.239709 | 0.808051 | 1.989464 | 0.406165 | 0.684621 | NA |
| | 1.284322 | -0.14931 | 2.000728 | -0.07463 | 0.940512 | NA |
| | 0.460669 | -0.62593 | 3.398671 | -0.18417 | 0.853882 | NA |
| | 0.268266 | -2.15396 | 4.038351 | -0.53338 | 0.593773 | NA |
| | 0.806146 | 0.553068 | 2.206737 | 0.250627 | 0.802102 | NA |
| C16orf97 | 2.365832 | -0.19289 | 1.442079 | -0.13376 | 0.893592 | NA |
| | 0.394291 | 0.298745 | 3.141711 | 0.09509 | 0.924243 | NA |
| | 10.24388 | -0.00122 | 0.646886 | -0.00189 | 0.998495 | NA |
| UBE2FP2 | 0.096202 | 0.408546 | 4.079639 | 0.100143 | 0.920231 | NA |
| | 0.575812 | 0.936922 | 2.776634 | 0.337431 | 0.735792 | NA |
| | 3.454027 | -0.08071 | 1.230601 | -0.06559 | 0.947705 | NA |
| | 0.556667 | 0.972496 | 2.522454 | 0.385536 | 0.699841 | NA |
| | 0.163957 | 0.424913 | 4.079006 | 0.104171 | 0.917034 | NA |
| | 1.110909 | -2.65317 | 2.097332 | -1.26502 | 0.205864 | NA |
| | 0.816972 | 0.936457 | 2.589651 | 0.361615 | 0.71764 | NA |
| | 19.53367 | 0.651127 | 0.507401 | 1.28326 | 0.199401 | NA |
| | 0.192403 | 0.975392 | 4.060716 | 0.240202 | 0.810174 | NA |
| | 0.58526 | 0.001197 | 3.235889 | 3.70E-04 | 0.999705 | NA |
| | 3.932562 | 1.095479 | 1.079882 | 1.014443 | 0.310371 | NA |
| | 2.603434 | -0.2949 | 1.366905 | -0.21574 | 0.829187 | NA |
| | 0.67854 | -0.89371 | 2.366391 | -0.37767 | 0.705677 | NA |
| | 0.162376 | 0.424913 | 4.079006 | 0.104171 | 0.917034 | NA |
| | 0.163957 | 0.424913 | 4.079006 | 0.104171 | 0.917034 | NA |
| | 1.389972 | -0.74064 | 1.967559 | -0.37643 | 0.7066 | NA |
| | 0.163957 | 0.424913 | 4.079006 | 0.104171 | 0.917034 | NA |
| | 73.44925 | -0.63245 | 0.31542 | -2.00511 | 0.044952 | 0.129622 |
| | 1.923204 | 0.535803 | 1.417119 | 0.378093 | 0.705361 | NA |
| EIF4BP5 | 0.629726 | -0.22725 | 2.732935 | -0.08315 | 0.93373 | NA |

| | | | | | | |
|---|---|---|---|---|---|---|
| | 1.434292 | 0.3217 | 1.639728 | 0.196191 | 0.844461 | NA |
| | 7.110337 | -0.31902 | 0.771063 | -0.41374 | 0.679062 | NA |
| C16orf98 | 1.167249 | 0.55518 | 1.831153 | 0.303186 | 0.761748 | NA |
| | 0.556813 | -0.57007 | 3.294615 | -0.17303 | 0.862627 | NA |
| VPS9D1-AS | 35.98301 | -0.38496 | 0.503176 | -0.76507 | 0.444231 | NA |
| | 2.368036 | -0.11252 | 1.265119 | -0.08894 | 0.929129 | NA |
| HSPE1P5 | 0.163957 | 0.424913 | 4.079006 | 0.104171 | 0.917034 | NA |
| | 0.258578 | 1.233151 | 4.047809 | 0.304646 | 0.760635 | NA |
| | 0.526843 | -0.38719 | 2.855892 | -0.13558 | 0.892156 | NA |
| | 1.06329 | 0.285492 | 1.942399 | 0.146979 | 0.883149 | NA |
| TEN1-CDK3 | 4.453819 | 0.115849 | 0.936593 | 0.123692 | 0.901559 | NA |
| ATP5F1P7 | 0.163957 | 0.424913 | 4.079006 | 0.104171 | 0.917034 | NA |
| | 0.28583 | -2.21358 | 3.97376 | -0.55705 | 0.577494 | NA |
| | 3.977673 | -0.13348 | 1.018003 | -0.13112 | 0.89568 | NA |
| | 0.258578 | 1.233151 | 4.047809 | 0.304646 | 0.760635 | NA |
| UBE2MP1 | 1.845944 | 0.850627 | 1.744226 | 0.487682 | 0.625775 | NA |
| | 0.296509 | -0.51709 | 3.982155 | -0.12985 | 0.896684 | NA |
| | 1.171751 | 1.845228 | 2.071346 | 0.890835 | 0.373018 | NA |
| | 4.127519 | -0.64195 | 1.067764 | -0.60121 | 0.547699 | NA |
| | 0.528424 | -0.38142 | 2.853651 | -0.13366 | 0.893671 | NA |
| | 0.258578 | 1.233151 | 4.047809 | 0.304646 | 0.760635 | NA |
| | 0.096202 | 0.408546 | 4.079639 | 0.100143 | 0.920231 | NA |
| | 0.623045 | 0.091811 | 2.666142 | 0.034436 | 0.97253 | NA |
| | 39.51607 | 0.390432 | 0.381136 | 1.024391 | 0.30565 | NA |
| | 42.55407 | -0.15032 | 0.375273 | -0.40056 | 0.688748 | NA |
| | 1.446755 | 0.439331 | 1.610042 | 0.272869 | 0.784954 | NA |
| | 0.268266 | -2.15396 | 4.038351 | -0.53338 | 0.593773 | NA |
| DLGAP1-AS | 0.163957 | 0.424913 | 4.079006 | 0.104171 | 0.917034 | NA |
| | 24.96469 | 0.27925 | 0.411374 | 0.678822 | 0.497251 | NA |
| | 0.162376 | 0.424913 | 4.079006 | 0.104171 | 0.917034 | NA |
| | 0.326333 | 1.533889 | 3.826878 | 0.40082 | 0.688553 | NA |
| | 0.746295 | -0.65731 | 2.318283 | -0.28353 | 0.776767 | NA |
| | 0.163957 | 0.424913 | 4.079006 | 0.104171 | 0.917034 | NA |
| | 3.325822 | -0.188 | 1.138868 | -0.16508 | 0.868884 | NA |
| | 4.792369 | -0.60317 | 0.90617 | -0.66563 | 0.505648 | NA |
| | 11.33232 | 0.846793 | 0.630195 | 1.343702 | 0.179045 | NA |
| EEF1A1P38 | 1.2271 | 1.384124 | 1.759418 | 0.786694 | 0.431461 | NA |
| | 1.437067 | 1.633888 | 1.799299 | 0.908069 | 0.363842 | NA |
| | 0.151697 | -1.546 | 4.075469 | -0.37934 | 0.704433 | NA |
| | 2.468778 | -0.06491 | 1.319921 | -0.04918 | 0.960778 | NA |
| | 0.258578 | 1.233151 | 4.047809 | 0.304646 | 0.760635 | NA |
| | 15.96635 | 0.98271 | 0.5426 | 1.811112 | 0.070124 | NA |
| | 0.192403 | 0.975392 | 4.060716 | 0.240202 | 0.810174 | NA |
| | 3.891551 | 0.300811 | 1.048584 | 0.286874 | 0.774209 | NA |
| HERC2P8 | 0.134133 | -1.48909 | 4.077584 | -0.36519 | 0.71497 | NA |
| GAN | 87.28824 | 0.675117 | 0.34367 | 1.964431 | 0.04948 | 0.137692 |
| | 9.017983 | 0.182591 | 0.662426 | 0.27564 | 0.782825 | NA |

| | | | | | | |
|---|---|---|---|---|---|---|
| YBX3P1 | 0.815042 | 2.859272 | 2.76927 | 1.0325 | 0.301838 | NA |
| | 7.381608 | 0.652192 | 0.771448 | 0.845414 | 0.39788 | NA |
| HMGN2P4: | 0.096202 | 0.408546 | 4.079639 | 0.100143 | 0.920231 | NA |
| | 0.819558 | -2.40918 | 2.432038 | -0.9906 | 0.321881 | NA |
| | 1.928929 | 0.614588 | 1.501461 | 0.409327 | 0.6823 | NA |
| IMPDH1P1 | 0.230334 | -0.53318 | 4.076749 | -0.13079 | 0.895945 | NA |
| | 0.314073 | -0.57308 | 3.898934 | -0.14698 | 0.883146 | NA |
| | 23.61359 | -0.29934 | 0.419742 | -0.71316 | 0.475747 | NA |
| | 8.413393 | -0.16995 | 0.706622 | -0.24051 | 0.809932 | NA |
| | 0.440301 | 0.468383 | 3.449808 | 0.135771 | 0.892003 | NA |
| | 41.34939 | -0.12416 | 0.392969 | -0.31596 | 0.752031 | NA |
| | 32.96979 | 0.164305 | 0.394433 | 0.416559 | 0.677001 | NA |
| | 0.575812 | 0.936922 | 2.776634 | 0.337431 | 0.735792 | NA |
| HPR | 0.230334 | -0.53318 | 4.076749 | -0.13079 | 0.895945 | NA |
| | 74.39508 | 0.005475 | 0.25769 | 0.021248 | 0.983048 | 0.990989 |
| | 0.230334 | -0.53318 | 4.076749 | -0.13079 | 0.895945 | NA |
| | 55.00802 | -0.8722 | 0.602326 | -1.44806 | 0.147601 | 0.290713 |
| | 0.315653 | -0.57305 | 3.892639 | -0.14721 | 0.882963 | NA |
| | 0.392711 | 0.292538 | 3.145221 | 0.09301 | 0.925895 | NA |
| | 2.020426 | 0.714606 | 1.492083 | 0.478932 | 0.631987 | NA |
| | 2.152939 | 0.882335 | 1.661577 | 0.531022 | 0.595403 | NA |
| | 0.163957 | 0.424913 | 4.079006 | 0.104171 | 0.917034 | NA |
| | 0.163957 | 0.424913 | 4.079006 | 0.104171 | 0.917034 | NA |
| | 1.602539 | -0.12674 | 1.714409 | -0.07393 | 0.941069 | NA |
| | 0.230334 | -0.53318 | 4.076749 | -0.13079 | 0.895945 | NA |
| | 2.098189 | 0.306117 | 1.31934 | 0.232022 | 0.816521 | NA |
| | 4.925421 | 1.793918 | 1.033635 | 1.735542 | 0.082645 | NA |
| | 0.162376 | 0.424913 | 4.079006 | 0.104171 | 0.917034 | NA |
| | 9.771284 | -0.15735 | 0.702446 | -0.22401 | 0.822751 | NA |
| | 5.450605 | -0.45901 | 0.920693 | -0.49854 | 0.6181 | NA |
| | 0.162376 | 0.424913 | 4.079006 | 0.104171 | 0.917034 | NA |
| | 0.151697 | -1.546 | 4.075469 | -0.37934 | 0.704433 | NA |
| TUBB8P7 | 19.67994 | -0.57399 | 0.509684 | -1.12616 | 0.260097 | NA |
| | 3.543803 | -0.70909 | 1.19969 | -0.59106 | 0.554481 | NA |
| LINC00662 | 204.7099 | 0.137138 | 0.179813 | 0.762669 | 0.445661 | 0.605468 |
| | 0.096202 | 0.408546 | 4.079639 | 0.100143 | 0.920231 | NA |
| | 0.163957 | 0.424913 | 4.079006 | 0.104171 | 0.917034 | NA |
| | 18.13016 | -0.67182 | 0.590585 | -1.13756 | 0.255305 | NA |
| MIA | 20.59 | 0.040668 | 0.560538 | 0.072552 | 0.942163 | NA |
| | 0.831228 | 1.631574 | 2.456512 | 0.664183 | 0.506573 | NA |
| | 0.096202 | 0.408546 | 4.079639 | 0.100143 | 0.920231 | NA |
| | 6.876318 | -0.29719 | 0.842423 | -0.35278 | 0.724255 | NA |
| | 19.96563 | 0.062895 | 0.488147 | 0.128845 | 0.897481 | NA |
| | 0.556813 | -0.57007 | 3.294615 | -0.17303 | 0.862627 | NA |
| | 16.08288 | -0.34078 | 0.583316 | -0.58421 | 0.559076 | NA |
| | 1.938064 | 1.141618 | 1.716815 | 0.664963 | 0.506074 | NA |
| | 0.134133 | -1.48909 | 4.077584 | -0.36519 | 0.71497 | NA |

| | | | | | | |
|---|---|---|---|---|---|---|
| | 0.768215 | 1.51173 | 2.494681 | 0.605981 | 0.544527 | NA |
| | 6.030238 | -0.31978 | 0.812846 | -0.39341 | 0.694017 | NA |
| | 4.571048 | 0.034242 | 0.975375 | 0.035106 | 0.971995 | NA |
| | 6.90539 | 0.15417 | 0.745965 | 0.206672 | 0.836266 | NA |
| | 0.861641 | -0.2592 | 2.180811 | -0.11885 | 0.90539 | NA |
| | 4.070159 | 0.841953 | 1.069278 | 0.787404 | 0.431045 | NA |
| DLGAP1-AS | 23.24446 | 1.084718 | 0.489848 | 2.214396 | 0.026802 | NA |
| | 0.163957 | 0.424913 | 4.079006 | 0.104171 | 0.917034 | NA |
| | 0.096202 | 0.408546 | 4.079639 | 0.100143 | 0.920231 | NA |
| | 0.096202 | 0.408546 | 4.079639 | 0.100143 | 0.920231 | NA |
| | 1.979049 | -0.78005 | 1.386138 | -0.56275 | 0.573606 | NA |
| | 37.37857 | -0.33472 | 0.341569 | -0.97993 | 0.327119 | NA |
| | 4.024692 | -0.50352 | 1.008377 | -0.49934 | 0.617541 | NA |
| | 0.654449 | 1.296029 | 2.63083 | 0.492631 | 0.622273 | NA |
| | 11.91493 | -0.59675 | 0.584288 | -1.02132 | 0.307101 | NA |
| | 1.793804 | -0.54764 | 1.551304 | -0.35302 | 0.724074 | NA |
| | 0.163957 | 0.424913 | 4.079006 | 0.104171 | 0.917034 | NA |
| BCAR4 | 0.45509 | -2.71227 | 3.963744 | -0.68427 | 0.493805 | NA |
| | 0.555233 | -0.57009 | 3.296947 | -0.17291 | 0.862719 | NA |
| | 7.505329 | 0.735183 | 0.863755 | 0.851148 | 0.394687 | NA |
| | 2.719894 | 0.0302 | 1.21839 | 0.024787 | 0.980225 | NA |
| LINC00514 | 1.336939 | 3.579225 | 2.06038 | 1.737167 | 0.082358 | NA |
| | 2.717236 | -0.14276 | 1.185372 | -0.12044 | 0.904136 | NA |
| | 2.081857 | 1.209762 | 1.428786 | 0.846706 | 0.397159 | NA |
| | 0.192403 | 0.975392 | 4.060716 | 0.240202 | 0.810174 | NA |
| | 6.780705 | 0.09184 | 0.752543 | 0.12204 | 0.902868 | NA |
| | 3.086805 | -0.39256 | 1.145211 | -0.34278 | 0.731761 | NA |
| | 2.0698 | 0.379819 | 1.382616 | 0.27471 | 0.783539 | NA |
| SNORD3D | 0.162376 | 0.424913 | 4.079006 | 0.104171 | 0.917034 | NA |
| | 135.893 | -0.71734 | 0.295976 | -2.42364 | 0.015366 | 0.062996 |
| | 593.5812 | -0.3482 | 0.207845 | -1.67529 | 0.093878 | 0.214014 |
| | 7.720404 | 0.059284 | 0.772879 | 0.076706 | 0.938858 | NA |
| CORO7 | 354.5373 | -0.36799 | 0.217339 | -1.69315 | 0.090428 | 0.208467 |
| | 1.072389 | 1.632428 | 2.363252 | 0.690755 | 0.48972 | NA |
| | 4.050508 | 1.165508 | 1.087108 | 1.072118 | 0.283667 | NA |
| | 0.315653 | -0.57305 | 3.892639 | -0.14721 | 0.882963 | NA |
| | 0.298089 | -0.51711 | 3.975232 | -0.13008 | 0.896501 | NA |
| | 0.096202 | 0.408546 | 4.079639 | 0.100143 | 0.920231 | NA |
| | 0.151697 | -1.546 | 4.075469 | -0.37934 | 0.704433 | NA |
| HNRNPA1P | 3.864651 | -0.52494 | 1.103454 | -0.47573 | 0.63427 | NA |
| | 0.47803 | 0.520083 | 2.97881 | 0.174594 | 0.861399 | NA |
| | 1.335708 | 2.111616 | 2.003911 | 1.053747 | 0.291999 | NA |
| | 0.528424 | -0.38142 | 2.853651 | -0.13366 | 0.893671 | NA |
| | 0.518939 | 0.925178 | 3.271397 | 0.282808 | 0.777324 | NA |
| | 1.962505 | -0.77218 | 1.464239 | -0.52736 | 0.597943 | NA |
| | 2.39365 | -1.12277 | 1.328429 | -0.84518 | 0.398009 | NA |
| | 0.432222 | -1.18916 | 3.482045 | -0.34151 | 0.732718 | NA |

| | | | | | | |
|---|---|---|---|---|---|---|
| | 8.709521 | 1.162048 | 0.725652 | 1.601384 | 0.109292 | NA |
| | 9.995378 | 0.503282 | 0.69403 | 0.725159 | 0.468355 | NA |
| | 1.755727 | 0.469734 | 1.546775 | 0.303686 | 0.761367 | NA |
| CCER2 | 0.096202 | 0.408546 | 4.079639 | 0.100143 | 0.920231 | NA |
| | 0.151697 | -1.546 | 4.075469 | -0.37934 | 0.704433 | NA |
| | 0.096202 | 0.408546 | 4.079639 | 0.100143 | 0.920231 | NA |
| | 0.258578 | 1.233151 | 4.047809 | 0.304646 | 0.760635 | NA |
| TXNP4 | 0.681112 | 2.620532 | 2.519661 | 1.040033 | 0.298324 | NA |
| | 0.364467 | -1.20166 | 3.685431 | -0.32606 | 0.744381 | NA |
| | 0.096202 | 0.408546 | 4.079639 | 0.100143 | 0.920231 | NA |
| | 18.51758 | -0.41438 | 0.471335 | -0.87916 | 0.379312 | NA |
| | 0.55687 | -0.14239 | 3.170029 | -0.04492 | 0.964172 | NA |
| | 0.72077 | 0.527966 | 3.060238 | 0.172524 | 0.863025 | NA |
| | 12.79215 | -0.4605 | 0.592757 | -0.77687 | 0.437234 | NA |
| BTF3P14 | 0.151697 | -1.546 | 4.075469 | -0.37934 | 0.704433 | NA |
| | 0.518736 | 2.237423 | 2.7713 | 0.807355 | 0.419462 | NA |
| | 1.001078 | -1.62003 | 2.135717 | -0.75854 | 0.448126 | NA |
| | 0.192403 | 0.975392 | 4.060716 | 0.240202 | 0.810174 | NA |
| | 1.034843 | 0.085767 | 2.175521 | 0.039424 | 0.968553 | NA |
| | 4.869652 | 0.322708 | 0.921036 | 0.350375 | 0.726057 | NA |
| | 3.745223 | -0.53295 | 1.117835 | -0.47677 | 0.633529 | NA |
| | 3.914245 | -0.48465 | 1.029466 | -0.47078 | 0.637797 | NA |
| | 8.632703 | -0.01516 | 0.717709 | -0.02113 | 0.983145 | NA |
| | 2.789192 | 0.067163 | 1.187257 | 0.05657 | 0.954888 | NA |
| | 0.69625 | -3.13723 | 2.93845 | -1.06765 | 0.285679 | NA |
| | 5.630985 | -0.52176 | 0.904187 | -0.57705 | 0.563909 | NA |
| | 0.096202 | 0.408546 | 4.079639 | 0.100143 | 0.920231 | NA |
| | 5.895241 | -0.1152 | 0.851715 | -0.13526 | 0.89241 | NA |
| | 1.130898 | 1.215321 | 1.814636 | 0.669732 | 0.503028 | NA |
| | 1.053979 | -3.69847 | 2.572645 | -1.43761 | 0.150544 | NA |
| | 111.5942 | -0.19527 | 0.271047 | -0.72044 | 0.471253 | 0.627368 |
| MRPS21P9 | 0.296509 | -0.51709 | 3.982155 | -0.12985 | 0.896684 | NA |
| | 3.065168 | 0.533651 | 1.202032 | 0.443958 | 0.657073 | NA |
| | 0.096202 | 0.408546 | 4.079639 | 0.100143 | 0.920231 | NA |
| | 2.736598 | 0.533229 | 1.309967 | 0.407055 | 0.683968 | NA |
| | 0.163957 | 0.424913 | 4.079006 | 0.104171 | 0.917034 | NA |
| MRPL12 | 550.3136 | -0.19978 | 0.150488 | -1.32752 | 0.184336 | 0.337654 |
| | 0.517155 | 2.232561 | 3.179631 | 0.702145 | 0.482589 | NA |
| | 1.619957 | 0.655051 | 1.533765 | 0.427087 | 0.669316 | NA |
| | 14.43836 | 0.686968 | 0.549223 | 1.2508 | 0.211007 | NA |
| | 30.62901 | -0.26119 | 0.47096 | -0.55459 | 0.579178 | NA |
| | 212.2235 | -0.14399 | 0.204902 | -0.70273 | 0.482224 | 0.636186 |
| | 2.87841 | -1.69901 | 1.199019 | -1.417 | 0.156483 | NA |
| | 8.793356 | -0.58249 | 0.720195 | -0.8088 | 0.418631 | NA |
| | 8.92833 | -0.46897 | 0.711101 | -0.6595 | 0.509578 | NA |
| | 1.160925 | 1.304491 | 1.810948 | 0.720336 | 0.471318 | NA |
| | 1.242238 | -1.86491 | 1.881316 | -0.99128 | 0.321549 | NA |

| | | | | | | |
|---|---|---|---|---|---|---|
| ALOX12P2 | 0.449786 | -1.25325 | 3.00474 | -0.41709 | 0.676613 | NA |
| | 175.2114 | -0.28543 | 0.198585 | -1.43731 | 0.150631 | 0.294428 |
| | 0.258578 | 1.233151 | 4.047809 | 0.304646 | 0.760635 | NA |
| EIF4A1P9 | 0.151697 | -1.546 | 4.075469 | -0.37934 | 0.704433 | NA |
| | 1.862697 | 0.366375 | 1.626114 | 0.225307 | 0.82174 | NA |
| | 0.163957 | 0.424913 | 4.079006 | 0.104171 | 0.917034 | NA |
| | 29.81954 | -0.35763 | 0.391922 | -0.91249 | 0.361509 | NA |
| ROCK1P1 | 2.339203 | 0.618811 | 1.28931 | 0.479955 | 0.631259 | NA |
| | 0.162376 | 0.424913 | 4.079006 | 0.104171 | 0.917034 | NA |
| | 43.90737 | -0.03547 | 0.329981 | -0.1075 | 0.914394 | NA |
| | 0.096202 | 0.408546 | 4.079639 | 0.100143 | 0.920231 | NA |
| | 4.562926 | -1.43068 | 1.149626 | -1.24448 | 0.213323 | NA |
| | 7.557873 | -0.18468 | 0.78601 | -0.23496 | 0.814243 | NA |
| | 48.12244 | -0.4049 | 0.330991 | -1.22331 | 0.221214 | 0.381453 |
| | 2.294288 | -1.37299 | 1.467774 | -0.93542 | 0.349571 | NA |
| | 0.392857 | -1.546 | 4.075469 | -0.37934 | 0.704433 | NA |
| | 6.172539 | -0.54425 | 0.894896 | -0.60817 | 0.543077 | NA |
| | 63.71483 | 0.215982 | 0.308369 | 0.7004 | 0.483677 | 0.637354 |
| | 4.350633 | -1.10369 | 0.959624 | -1.15013 | 0.250092 | NA |
| | 0.875299 | 1.848392 | 2.453783 | 0.753283 | 0.45128 | NA |
| | 2.103725 | -1.81171 | 1.466282 | -1.23558 | 0.216614 | NA |
| | 1.893955 | 1.755314 | 1.648659 | 1.064692 | 0.287015 | NA |
| | 1.877194 | 0.496221 | 1.45427 | 0.341217 | 0.73294 | NA |
| LRRC37A17 | 123.5567 | -0.1917 | 0.26889 | -0.71292 | 0.475896 | 0.630941 |
| | 44.74483 | -0.01336 | 0.346922 | -0.0385 | 0.969292 | NA |
| | 4.934635 | -1.51572 | 1.049029 | -1.44488 | 0.148492 | NA |
| | 0.151697 | -1.546 | 4.075469 | -0.37934 | 0.704433 | NA |
| | 0.601483 | -1.7332 | 2.729494 | -0.63499 | 0.525436 | NA |
| HNRNPCP4 | 0.192403 | 0.975392 | 4.060716 | 0.240202 | 0.810174 | NA |
| | 0.727508 | 0.192954 | 2.572892 | 0.074995 | 0.940219 | NA |
| ZNF205-AS | 5.195998 | -0.41305 | 0.963049 | -0.4289 | 0.667995 | NA |
| RYKP1 | 0.545988 | -0.43913 | 2.541145 | -0.17281 | 0.862803 | NA |
| | 0.162376 | 0.424913 | 4.079006 | 0.104171 | 0.917034 | NA |
| ATP5A1P3 | 1.613001 | -0.63725 | 1.794857 | -0.35504 | 0.722558 | NA |
| | 2.229454 | -1.17346 | 1.513553 | -0.7753 | 0.438162 | NA |
| | 0.658173 | -0.01016 | 2.630582 | -0.00386 | 0.996919 | NA |
| | 360.3159 | -0.1083 | 0.222108 | -0.48761 | 0.625826 | 0.748957 |
| | 0.162376 | 0.424913 | 4.079006 | 0.104171 | 0.917034 | NA |
| | 0.096202 | 0.408546 | 4.079639 | 0.100143 | 0.920231 | NA |
| | 0.258578 | 1.233151 | 4.047809 | 0.304646 | 0.760635 | NA |
| | 0.151697 | -1.546 | 4.075469 | -0.37934 | 0.704433 | NA |
| | 7.613917 | 0.244391 | 0.81363 | 0.300372 | 0.763894 | NA |
| | 0.296509 | -0.51709 | 3.982155 | -0.12985 | 0.896684 | NA |
| | 0.163957 | 0.424913 | 4.079006 | 0.104171 | 0.917034 | NA |
| | 11.38957 | 0.71091 | 0.673605 | 1.055381 | 0.291251 | NA |
| | 1.237926 | 1.940943 | 2.028448 | 0.956861 | 0.338637 | NA |
| | 46.72576 | -0.55669 | 0.344385 | -1.61649 | 0.105989 | NA |

| | | | | | | |
|---|---|---|---|---|---|---|
| | 10.41667 | -0.78746 | 0.663777 | -1.18633 | 0.23549 | NA |
| | 5.526565 | -0.58592 | 0.902359 | -0.64932 | 0.516134 | NA |
| | 0.162376 | 0.424913 | 4.079006 | 0.104171 | 0.917034 | NA |
| | 0.640406 | 1.11402 | 2.710977 | 0.410929 | 0.681124 | NA |
| | 0.296509 | -0.51709 | 3.982155 | -0.12985 | 0.896684 | NA |
| | 0.327913 | 1.543154 | 4.021063 | 0.383768 | 0.701151 | NA |
| | 0.288605 | 1.443619 | 4.029058 | 0.358302 | 0.720117 | NA |
| | 14.0023 | 0.139588 | 0.677163 | 0.206137 | 0.836684 | NA |
| | 9.316283 | 0.365258 | 0.671903 | 0.543616 | 0.586705 | NA |
| | 2.749086 | -0.60639 | 1.322411 | -0.45855 | 0.646558 | NA |
| | 0.096202 | 0.408546 | 4.079639 | 0.100143 | 0.920231 | NA |
| | 0.392857 | -1.546 | 4.075469 | -0.37934 | 0.704433 | NA |
| | 1.168976 | -0.41512 | 1.914592 | -0.21682 | 0.82835 | NA |
| | 35.06982 | 0.424305 | 0.423409 | 1.002115 | 0.316288 | NA |
| | 0.260158 | 1.240347 | 4.047124 | 0.306476 | 0.759242 | NA |
| RN7SL155F | 0.151697 | -1.546 | 4.075469 | -0.37934 | 0.704433 | NA |
| RN7SL94P | 0.134133 | -1.48909 | 4.077584 | -0.36519 | 0.71497 | NA |
| | 0.096202 | 0.408546 | 4.079639 | 0.100143 | 0.920231 | NA |
| | 0.096202 | 0.408546 | 4.079639 | 0.100143 | 0.920231 | NA |
| | 0.303394 | -2.27006 | 4.032451 | -0.56295 | 0.57347 | NA |
| | 0.518736 | 2.237423 | 2.7713 | 0.807355 | 0.419462 | NA |
| AK4P1 | 0.680121 | -0.88898 | 2.365123 | -0.37587 | 0.707013 | NA |
| RN7SL716F | 0.151697 | -1.546 | 4.075469 | -0.37934 | 0.704433 | NA |
| UBBP4 | 15.02865 | -0.01345 | 0.543328 | -0.02476 | 0.980246 | NA |
| | 0.392711 | 0.292538 | 3.145221 | 0.09301 | 0.925895 | NA |
| | 0.258578 | 1.233151 | 4.047809 | 0.304646 | 0.760635 | NA |
| | 0.364467 | -1.20166 | 3.685431 | -0.32606 | 0.744381 | NA |
| | 0.979202 | 1.972754 | 2.08204 | 0.94751 | 0.343379 | NA |
| | 1.03648 | 0.354657 | 2.004615 | 0.17692 | 0.859571 | NA |
| | 1.446515 | 1.109807 | 1.923134 | 0.577082 | 0.563884 | NA |
| | 0.134133 | -1.48909 | 4.077584 | -0.36519 | 0.71497 | NA |
| | 0.384806 | 1.846216 | 3.999752 | 0.461582 | 0.644381 | NA |
| | 0.096202 | 0.408546 | 4.079639 | 0.100143 | 0.920231 | NA |
| | 0.815391 | 0.931626 | 2.360431 | 0.394685 | 0.693076 | NA |
| | 1.530697 | 0.042995 | 1.600009 | 0.026872 | 0.978562 | NA |
| | 0.096202 | 0.408546 | 4.079639 | 0.100143 | 0.920231 | NA |
| | 23.70808 | -0.29975 | 0.5017 | -0.59747 | 0.550193 | NA |
| SLC25A51P | 0.134133 | -1.48909 | 4.077584 | -0.36519 | 0.71497 | NA |
| | 5.617836 | -0.63907 | 1.019855 | -0.62663 | 0.530902 | NA |
| | 0.163957 | 0.424913 | 4.079006 | 0.104171 | 0.917034 | NA |
| DLGAP1-AS | 0.163957 | 0.424913 | 4.079006 | 0.104171 | 0.917034 | NA |
| FTLP13 | 0.096202 | 0.408546 | 4.079639 | 0.100143 | 0.920231 | NA |
| | 0.430642 | -1.1891 | 3.485782 | -0.34113 | 0.733008 | NA |
| | 10.67697 | -0.19909 | 0.623422 | -0.31935 | 0.749461 | NA |
| OOSP1P2 | 0.561971 | -0.49218 | 2.810061 | -0.17515 | 0.860962 | NA |
| LINC00667 | 671.2974 | 0.190827 | 0.163116 | 1.169891 | 0.242045 | 0.405002 |
| | 6.184352 | -0.19522 | 0.797281 | -0.24486 | 0.806564 | NA |

| | | | | | | |
|---|---|---|---|---|---|---|
| | 1.185162 | -1.08022 | 1.836234 | -0.58828 | 0.556345 | NA |
| | 0.816825 | 1.673341 | 2.21346 | 0.755984 | 0.449659 | NA |
| | 2.151755 | 0.765568 | 1.39866 | 0.547358 | 0.584133 | NA |
| | 1.569997 | -1.71637 | 1.688941 | -1.01624 | 0.309515 | NA |
| | 0.599903 | -1.73313 | 2.730982 | -0.63462 | 0.525678 | NA |
| | 5.471427 | -0.31212 | 0.894981 | -0.34875 | 0.727278 | NA |
| | 0.151697 | -1.546 | 4.075469 | -0.37934 | 0.704433 | NA |
| FAM60BP | 1.360148 | -1.53382 | 1.974839 | -0.77668 | 0.437346 | NA |
| | 15.41925 | -0.33055 | 0.573206 | -0.57666 | 0.564166 | NA |
| CIAPIN1P | 0.58526 | 0.001197 | 3.235889 | 3.70E-04 | 0.999705 | NA |
| | 0.163957 | 0.424913 | 4.079006 | 0.104171 | 0.917034 | NA |
| | 0.230334 | -0.53318 | 4.076749 | -0.13079 | 0.895945 | NA |
| | 0.163957 | 0.424913 | 4.079006 | 0.104171 | 0.917034 | NA |
| LINC00672 | 1.892791 | 1.520634 | 1.724951 | 0.881552 | 0.378019 | NA |
| | 0.392711 | 0.292538 | 3.145221 | 0.09301 | 0.925895 | NA |
| | 1.517285 | 1.795409 | 1.693559 | 1.060139 | 0.289081 | NA |
| | 0.096202 | 0.408546 | 4.079639 | 0.100143 | 0.920231 | NA |
| | 2.309491 | -1.65158 | 1.395625 | -1.1834 | 0.236651 | NA |
| | 0.517155 | 2.232561 | 3.179631 | 0.702145 | 0.482589 | NA |
| | 11.3439 | -0.67215 | 0.672748 | -0.9991 | 0.317744 | NA |
| | 0.096202 | 0.408546 | 4.079639 | 0.100143 | 0.920231 | NA |
| RN7SL381F | 0.979202 | 1.972754 | 2.08204 | 0.94751 | 0.343379 | NA |
| | 1.361233 | 0.907194 | 1.662201 | 0.545779 | 0.585218 | NA |
| | 0.528424 | -0.38142 | 2.853651 | -0.13366 | 0.893671 | NA |
| | 1.689495 | 0.214366 | 1.601324 | 0.133868 | 0.893507 | NA |
| | 0.163957 | 0.424913 | 4.079006 | 0.104171 | 0.917034 | NA |
| | 0.151697 | -1.546 | 4.075469 | -0.37934 | 0.704433 | NA |
| | 2.996531 | -0.75577 | 1.148112 | -0.65827 | 0.510366 | NA |
| | 0.736957 | -1.0303 | 2.640699 | -0.39016 | 0.696418 | NA |
| MIR4737 | 5.083498 | -1.61825 | 0.992569 | -1.63036 | 0.103024 | NA |
| | 0.326333 | 1.533889 | 3.826878 | 0.40082 | 0.688553 | NA |
| | 65.58055 | -0.32721 | 0.310141 | -1.05503 | 0.291413 | 0.458333 |
| | 0.096202 | 0.408546 | 4.079639 | 0.100143 | 0.920231 | NA |
| DND1P1 | 12.81869 | -0.75853 | 0.602383 | -1.25922 | 0.20795 | NA |
| | 7.996246 | -0.63278 | 0.746834 | -0.84729 | 0.396836 | NA |
| | 4.032761 | 1.472242 | 1.148916 | 1.281418 | 0.200047 | NA |
| | 39.22442 | 1.364574 | 0.437685 | 3.117708 | 0.001823 | NA |
| | 0.134133 | -1.48909 | 4.077584 | -0.36519 | 0.71497 | NA |
| | 6.771606 | -0.04538 | 0.758112 | -0.05987 | 0.952263 | NA |
| | 0.096202 | 0.408546 | 4.079639 | 0.100143 | 0.920231 | NA |
| | 0.162376 | 0.424913 | 4.079006 | 0.104171 | 0.917034 | NA |
| MIR3175 | 0.162376 | 0.424913 | 4.079006 | 0.104171 | 0.917034 | NA |
| MIR3189 | 0.966942 | 0.917791 | 1.915974 | 0.47902 | 0.631924 | NA |
| MAGOH2 | 1.668982 | 1.28852 | 1.600275 | 0.805187 | 0.420712 | NA |
| | 0.162376 | 0.424913 | 4.079006 | 0.104171 | 0.917034 | NA |
| | 0.92936 | -0.12162 | 2.442792 | -0.04979 | 0.960292 | NA |
| | 0.488709 | 2.119056 | 3.307564 | 0.64067 | 0.521737 | NA |

| | | | | | | |
|---|---|---|---|---|---|---|
| | 0.326333 | 1.533889 | 3.826878 | 0.40082 | 0.688553 | NA |
| | 0.192403 | 0.975392 | 4.060716 | 0.240202 | 0.810174 | NA |
| | 0.162376 | 0.424913 | 4.079006 | 0.104171 | 0.917034 | NA |
| | 1.147231 | -0.4269 | 2.00528 | -0.21289 | 0.831415 | NA |
| | 4.923887 | 0.300131 | 0.89385 | 0.335774 | 0.737042 | NA |
| LINC00909 | 218.1805 | 0.420393 | 0.217553 | 1.932374 | 0.053313 | 0.144629 |
| MIR4322 | 0.163957 | 0.424913 | 4.079006 | 0.104171 | 0.917034 | NA |
| | 0.288605 | 1.443619 | 4.029058 | 0.358302 | 0.720117 | NA |
| | 2.634985 | -0.49847 | 1.581439 | -0.3152 | 0.752608 | NA |
| | 52.73325 | -0.47557 | 0.331541 | -1.43444 | 0.151448 | 0.295445 |
| | 2.483438 | 0.165792 | 1.476545 | 0.112284 | 0.910598 | NA |
| | 0.517155 | 2.232561 | 3.179631 | 0.702145 | 0.482589 | NA |
| | 1.873819 | 0.045943 | 1.713684 | 0.02681 | 0.978611 | NA |
| | 1.781601 | -0.72609 | 1.91181 | -0.37979 | 0.7041 | NA |
| CCDC58P1 | 0.096202 | 0.408546 | 4.079639 | 0.100143 | 0.920231 | NA |
| SNRPGP2 | 75.95222 | 0.341466 | 0.265013 | 1.288487 | 0.197577 | 0.353177 |
| DYNLL2 | 329.712 | -0.14062 | 0.209082 | -0.67258 | 0.501217 | 0.651742 |
| | 3.787218 | 0.392155 | 1.137444 | 0.344769 | 0.730268 | NA |
| MYH4 | 0.134133 | -1.48909 | 4.077584 | -0.36519 | 0.71497 | NA |
| MIR3183 | 0.326333 | 1.533889 | 3.826878 | 0.40082 | 0.688553 | NA |
| | 0.096202 | 0.408546 | 4.079639 | 0.100143 | 0.920231 | NA |
| | 3.050874 | 0.188739 | 1.178886 | 0.160099 | 0.872803 | NA |
| | 22.75799 | 0.369254 | 0.485211 | 0.761017 | 0.446647 | NA |
| | 0.326536 | 0.037056 | 3.842554 | 0.009644 | 0.992306 | NA |
| | 0.488912 | 0.771583 | 2.897678 | 0.266276 | 0.790026 | NA |
| | 0.151697 | -1.546 | 4.075469 | -0.37934 | 0.704433 | NA |
| | 87.12941 | 0.37322 | 0.262073 | 1.424109 | 0.154415 | 0.299654 |
| RN7SL405F | 0.192403 | 0.975392 | 4.060716 | 0.240202 | 0.810174 | NA |
| | 0.162376 | 0.424913 | 4.079006 | 0.104171 | 0.917034 | NA |
| | 0.46735 | -1.30193 | 3.399671 | -0.38296 | 0.701751 | NA |
| | 3.637233 | -0.39882 | 1.104536 | -0.36108 | 0.718041 | NA |
| SNX19P3 | 6.589737 | 0.167552 | 0.761926 | 0.219906 | 0.825944 | NA |
| LINC00526 | 28.51667 | 0.235657 | 0.415072 | 0.56775 | 0.570204 | NA |
| | 0.478233 | -0.67301 | 2.937473 | -0.22911 | 0.818782 | NA |
| | 1.289268 | -1.64884 | 1.960876 | -0.84087 | 0.400421 | NA |
| | 0.163957 | 0.424913 | 4.079006 | 0.104171 | 0.917034 | NA |
| | 1.673872 | 0.096871 | 1.512454 | 0.064049 | 0.948931 | NA |
| | 0.096202 | 0.408546 | 4.079639 | 0.100143 | 0.920231 | NA |
| | 0.508056 | 0.731032 | 2.861912 | 0.255435 | 0.798387 | NA |
| KRT18P8 | 0.163957 | 0.424913 | 4.079006 | 0.104171 | 0.917034 | NA |
| | 0.151697 | -1.546 | 4.075469 | -0.37934 | 0.704433 | NA |
| | 0.151697 | -1.546 | 4.075469 | -0.37934 | 0.704433 | NA |
| | 0.556813 | -0.57007 | 3.294615 | -0.17303 | 0.862627 | NA |
| | 0.096202 | 0.408546 | 4.079639 | 0.100143 | 0.920231 | NA |
| | 1.332989 | 0.293283 | 1.682898 | 0.174273 | 0.861651 | NA |
| DPRXP4 | 0.096202 | 0.408546 | 4.079639 | 0.100143 | 0.920231 | NA |
| | 0.717609 | 0.516981 | 3.063469 | 0.168757 | 0.865988 | NA |

| | | | | | | |
|---|---|---|---|---|---|---|
| RN7SL805F | 2.04344 | -0.35152 | 1.464824 | -0.23998 | 0.810349 | NA |
| | 2.941781 | -1.32267 | 1.198684 | -1.10343 | 0.269839 | NA |
| SNORA67 | 0.163957 | 0.424913 | 4.079006 | 0.104171 | 0.917034 | NA |
| PPIAP14 | 0.327913 | 1.543154 | 4.021063 | 0.383768 | 0.701151 | NA |
| | 0.134133 | -1.48909 | 4.077584 | -0.36519 | 0.71497 | NA |
| MIR4523 | 1.14589 | -0.96272 | 1.809626 | -0.532 | 0.594725 | NA |
| | 0.247898 | -0.5901 | 4.074633 | -0.14482 | 0.884851 | NA |
| | 0.151697 | -1.546 | 4.075469 | -0.37934 | 0.704433 | NA |
| | 0.327913 | 1.543154 | 4.021063 | 0.383768 | 0.701151 | NA |
| | 0.327913 | 1.543154 | 4.021063 | 0.383768 | 0.701151 | NA |
| RN7SL45P | 0.192403 | 0.975392 | 4.060716 | 0.240202 | 0.810174 | NA |
| | 0.163957 | 0.424913 | 4.079006 | 0.104171 | 0.917034 | NA |
| | 36.96743 | 0.056549 | 0.358875 | 0.157572 | 0.874794 | NA |
| | 0.884581 | 1.788183 | 2.221027 | 0.805115 | 0.420753 | NA |
| | 113.2055 | 0.050649 | 0.285968 | 0.177116 | 0.859418 | 0.916432 |
| | 0.49187 | 2.136088 | 3.895953 | 0.548284 | 0.583497 | NA |
| | 0.162376 | 0.424913 | 4.079006 | 0.104171 | 0.917034 | NA |
| SH3GL1P2 | 1.156919 | -2.03273 | 2.016184 | -1.0082 | 0.313356 | NA |
| | 0.6882 | 0.151421 | 2.969628 | 0.05099 | 0.959334 | NA |
| | 0.163957 | 0.424913 | 4.079006 | 0.104171 | 0.917034 | NA |
| | 14.89469 | 0.420071 | 0.552207 | 0.760713 | 0.446828 | NA |
| | 3.015426 | 1.909797 | 1.291355 | 1.478909 | 0.139165 | NA |
| | 0.163957 | 0.424913 | 4.079006 | 0.104171 | 0.917034 | NA |
| | 2.87001 | -0.12341 | 1.199607 | -0.10287 | 0.918065 | NA |
| | 0.999498 | -1.62429 | 2.136491 | -0.76026 | 0.4471 | NA |
| | 1.687712 | 0.72863 | 1.534051 | 0.474971 | 0.634807 | NA |
| | 7.072639 | -0.27436 | 0.806032 | -0.34038 | 0.733568 | NA |
| | 0.230334 | -0.53318 | 4.076749 | -0.13079 | 0.895945 | NA |
| | 0.162376 | 0.424913 | 4.079006 | 0.104171 | 0.917034 | NA |
| | 0.096202 | 0.408546 | 4.079639 | 0.100143 | 0.920231 | NA |
| FTLP12 | 8.072267 | -0.3475 | 0.780736 | -0.44509 | 0.656253 | NA |
| C1QTNF1-A | 0.247898 | -0.5901 | 4.074633 | -0.14482 | 0.884851 | NA |
| | 1.869078 | 0.043799 | 1.547192 | 0.028309 | 0.977416 | NA |
| MIR4731 | 0.134133 | -1.48909 | 4.077584 | -0.36519 | 0.71497 | NA |
| | 0.096202 | 0.408546 | 4.079639 | 0.100143 | 0.920231 | NA |
| | 0.134133 | -1.48909 | 4.077584 | -0.36519 | 0.71497 | NA |
| | 0.162376 | 0.424913 | 4.079006 | 0.104171 | 0.917034 | NA |
| | 0.572651 | 0.928649 | 2.780261 | 0.334015 | 0.738368 | NA |
| | 2.383816 | 0.17615 | 1.248176 | 0.141126 | 0.88777 | NA |
| MIR133A1 | 0.561971 | -0.49218 | 2.810061 | -0.17515 | 0.860962 | NA |
| MIR1-2 | 0.561971 | -0.49218 | 2.810061 | -0.17515 | 0.860962 | NA |
| BZRAP1-AS | 2.896242 | -0.65652 | 1.266018 | -0.51857 | 0.60406 | NA |
| LRRC37A7F | 17.38973 | 0.020879 | 0.504447 | 0.041391 | 0.966984 | NA |
| MIR4728 | 0.247898 | -0.5901 | 4.074633 | -0.14482 | 0.884851 | NA |
| | 0.327913 | 1.543154 | 4.021063 | 0.383768 | 0.701151 | NA |
| SRP72P1 | 0.096202 | 0.408546 | 4.079639 | 0.100143 | 0.920231 | NA |
| SNORD3B-1 | 0.58333 | 2.389479 | 3.105916 | 0.769331 | 0.441697 | NA |

| | | | | | | |
|---|---|---|---|---|---|---|
| | 2.282513 | 0.155166 | 1.292723 | 0.12003 | 0.904459 | NA |
| | 0.151697 | -1.546 | 4.075469 | -0.37934 | 0.704433 | NA |
| | 32.78001 | 0.569241 | 0.458321 | 1.242015 | 0.214231 | NA |
| | 13.03046 | -0.65593 | 0.576857 | -1.13708 | 0.255507 | NA |
| | 0.163957 | 0.424913 | 4.079006 | 0.104171 | 0.917034 | NA |
| | 0.49187 | 2.136088 | 3.895953 | 0.548284 | 0.583497 | NA |
| PGDP1 | 2.783208 | 0.178505 | 1.260212 | 0.141646 | 0.887359 | NA |
| | 0.134133 | -1.48909 | 4.077584 | -0.36519 | 0.71497 | NA |
| ARGFXP2 | 1.27606 | 0.077176 | 1.788447 | 0.043153 | 0.96558 | NA |
| FEM1AP2 | 0.151697 | -1.546 | 4.075469 | -0.37934 | 0.704433 | NA |
| | 3.912526 | 0.508559 | 1.06795 | 0.476201 | 0.633931 | NA |
| | 0.162376 | 0.424913 | 4.079006 | 0.104171 | 0.917034 | NA |
| | 0.162376 | 0.424913 | 4.079006 | 0.104171 | 0.917034 | NA |
| | 0.296509 | -0.51709 | 3.982155 | -0.12985 | 0.896684 | NA |
| | 2.137169 | 1.307794 | 1.475997 | 0.886041 | 0.375595 | NA |
| | 0.151697 | -1.546 | 4.075469 | -0.37934 | 0.704433 | NA |
| RN7SL219F | 0.162376 | 0.424913 | 4.079006 | 0.104171 | 0.917034 | NA |
| MIR4999 | 0.163957 | 0.424913 | 4.079006 | 0.104171 | 0.917034 | NA |
| | 0.508056 | 0.731032 | 2.861912 | 0.255435 | 0.798387 | NA |
| | 22.80864 | 0.505959 | 0.435648 | 1.161394 | 0.245482 | NA |
| | 0.096202 | 0.408546 | 4.079639 | 0.100143 | 0.920231 | NA |
| | 11.54896 | -0.67551 | 0.716898 | -0.94226 | 0.346058 | NA |
| | 72.33402 | -0.24428 | 0.321356 | -0.76017 | 0.447156 | 0.606803 |
| | 10.63373 | 0.576007 | 0.676529 | 0.851414 | 0.394539 | NA |
| | 0.392857 | -1.546 | 4.075469 | -0.37934 | 0.704433 | NA |
| | 0.247898 | -0.5901 | 4.074633 | -0.14482 | 0.884851 | NA |
| | 1.312622 | 0.801445 | 1.708401 | 0.46912 | 0.638984 | NA |
| | 5.736617 | 0.132558 | 0.851515 | 0.155673 | 0.876291 | NA |
| | 0.151697 | -1.546 | 4.075469 | -0.37934 | 0.704433 | NA |
| | 2.697101 | -1.0829 | 1.195349 | -0.90593 | 0.364973 | NA |
| | 0.392857 | -1.546 | 4.075469 | -0.37934 | 0.704433 | NA |
| | 2.048669 | 1.176779 | 1.517324 | 0.775562 | 0.438008 | NA |
| | 0.544554 | -2.65036 | 3.220086 | -0.82307 | 0.410468 | NA |
| | 0.604258 | 1.087325 | 2.707696 | 0.401568 | 0.688002 | NA |
| MIR1539 | 0.655827 | 2.55247 | 3.531744 | 0.722722 | 0.46985 | NA |
| SENP3-EIF4 | 0.594599 | -0.12152 | 2.771051 | -0.04385 | 0.965021 | NA |
| | 0.162376 | 0.424913 | 4.079006 | 0.104171 | 0.917034 | NA |
| | 2.955475 | -0.49386 | 1.124699 | -0.43911 | 0.660584 | NA |
| | 88.05794 | -0.0818 | 0.261417 | -0.31291 | 0.754348 | 0.846776 |
| | 0.162376 | 0.424913 | 4.079006 | 0.104171 | 0.917034 | NA |
| | 0.411855 | 0.250456 | 3.097618 | 0.080854 | 0.935558 | NA |
| | 0.326333 | 1.533889 | 3.826878 | 0.40082 | 0.688553 | NA |
| MIR5010 | 0.785567 | -0.70568 | 2.358063 | -0.29926 | 0.764741 | NA |
| | 0.863039 | 0.783382 | 2.464829 | 0.317824 | 0.750618 | NA |
| | 3.913281 | -0.35004 | 1.008996 | -0.34692 | 0.728653 | NA |
| TUFMP1 | 0.247898 | -0.5901 | 4.074633 | -0.14482 | 0.884851 | NA |
| | 0.623191 | -1.58103 | 2.770593 | -0.57065 | 0.568239 | NA |

| | | | | | | |
|---|---|---|---|---|---|---|
| | 46.33191 | -0.43456 | 0.352052 | -1.23435 | 0.217072 | NA |
| | 3.077919 | -0.33604 | 1.409086 | -0.23848 | 0.811511 | NA |
| RPL17 | 235.9615 | 0.304933 | 0.246174 | 1.23869 | 0.21546 | 0.374583 |
| | 2.767428 | -0.07853 | 1.161193 | -0.06763 | 0.946082 | NA |
| MAFG-AS1 | 43.56932 | -0.22943 | 0.357048 | -0.64258 | 0.520496 | NA |
| | 0.478233 | -0.67301 | 2.937473 | -0.22911 | 0.818782 | NA |
| | 0.327913 | 1.543154 | 4.021063 | 0.383768 | 0.701151 | NA |
| | 0.682693 | 2.623213 | 2.518338 | 1.041645 | 0.297576 | NA |
| | 0.096202 | 0.408546 | 4.079639 | 0.100143 | 0.920231 | NA |
| | 0.096202 | 0.408546 | 4.079639 | 0.100143 | 0.920231 | NA |
| | 0.151697 | -1.546 | 4.075469 | -0.37934 | 0.704433 | NA |
| | 5.260597 | 0.734904 | 0.978597 | 0.750978 | 0.452666 | NA |
| | 6.242527 | 1.55332 | 0.842705 | 1.843256 | 0.065292 | NA |
| CXADRP3 | 0.162376 | 0.424913 | 4.079006 | 0.104171 | 0.917034 | NA |
| | 23.66599 | 0.224549 | 0.487108 | 0.460985 | 0.644809 | NA |
| | 1.033116 | 1.025241 | 1.933005 | 0.530387 | 0.595844 | NA |
| | 0.682693 | 2.623213 | 2.518338 | 1.041645 | 0.297576 | NA |
| | 0.354779 | 1.703954 | 3.657232 | 0.465914 | 0.641277 | NA |
| | 1.056848 | -0.19221 | 2.254508 | -0.08525 | 0.932059 | NA |
| | 0.508056 | 0.731032 | 2.861912 | 0.255435 | 0.798387 | NA |
| MIR3177 | 0.392857 | -1.546 | 4.075469 | -0.37934 | 0.704433 | NA |
| | 0.096202 | 0.408546 | 4.079639 | 0.100143 | 0.920231 | NA |
| RN7SL226F | 0.151697 | -1.546 | 4.075469 | -0.37934 | 0.704433 | NA |
| | 0.654449 | 1.296029 | 2.63083 | 0.492631 | 0.622273 | NA |
| MIR4734 | 0.096202 | 0.408546 | 4.079639 | 0.100143 | 0.920231 | NA |
| | 0.774539 | 0.39797 | 2.296349 | 0.173305 | 0.862411 | NA |
| UBE2CP2 | 0.163957 | 0.424913 | 4.079006 | 0.104171 | 0.917034 | NA |
| | 0.748867 | 2.740193 | 2.468928 | 1.109871 | 0.267054 | NA |
| | 1.349265 | -1.88672 | 2.030501 | -0.92919 | 0.352791 | NA |
| | 0.163957 | 0.424913 | 4.079006 | 0.104171 | 0.917034 | NA |
| | 0.260158 | 1.240347 | 4.047124 | 0.306476 | 0.759242 | NA |
| MIR4725 | 0.163957 | 0.424913 | 4.079006 | 0.104171 | 0.917034 | NA |
| | 2.718228 | 0.842344 | 1.32905 | 0.633794 | 0.526216 | NA |
| MIR3615 | 0.134133 | -1.48909 | 4.077584 | -0.36519 | 0.71497 | NA |
| | 4.033828 | -0.25627 | 0.992674 | -0.25817 | 0.796279 | NA |
| | 10.27448 | -0.0644 | 0.765414 | -0.08414 | 0.932948 | NA |
| POLRMTP1 | 10.36413 | -0.6804 | 0.641997 | -1.05982 | 0.289227 | NA |
| | 1.02264 | -0.42936 | 1.996699 | -0.21504 | 0.82974 | NA |
| | 0.956262 | 0.048146 | 1.9097 | 0.025211 | 0.979886 | NA |
| | 0.520316 | 2.239969 | 3.173769 | 0.705776 | 0.480328 | NA |
| | 0.151697 | -1.546 | 4.075469 | -0.37934 | 0.704433 | NA |
| RHOT1P1 | 0.392857 | -1.546 | 4.075469 | -0.37934 | 0.704433 | NA |
| | 0.192403 | 0.975392 | 4.060716 | 0.240202 | 0.810174 | NA |
| | 0.096202 | 0.408546 | 4.079639 | 0.100143 | 0.920231 | NA |
| | 0.327913 | 1.543154 | 4.021063 | 0.383768 | 0.701151 | NA |
| RN7SL758F | 0.151697 | -1.546 | 4.075469 | -0.37934 | 0.704433 | NA |
| | 0.296509 | -0.51709 | 3.982155 | -0.12985 | 0.896684 | NA |

| | | | | | | |
|---|---|---|---|---|---|---|
| STRADA | 101.3146 | -0.09327 | 0.277168 | -0.33652 | 0.736481 | 0.833121 |
| | 0.653072 | 0.258115 | 3.011926 | 0.085698 | 0.931707 | NA |
| | 0.394291 | 0.298745 | 3.141711 | 0.09509 | 0.924243 | NA |
| MIR3186 | 0.45509 | -2.71227 | 3.963744 | -0.68427 | 0.493805 | NA |
| | 122.0852 | -0.33082 | 0.229114 | -1.4439 | 0.148768 | 0.292187 |
| MIR3178 | 0.134133 | -1.48909 | 4.077584 | -0.36519 | 0.71497 | NA |
| MIR3176 | 0.422737 | 0.518872 | 3.501356 | 0.148192 | 0.882191 | NA |
| NARF-IT1 | 3.748311 | -0.61749 | 1.190982 | -0.51847 | 0.604132 | NA |
| GRAMD4P7 | 0.811881 | 2.862678 | 3.297846 | 0.868045 | 0.38537 | NA |
| | 1.671942 | 1.245692 | 1.722473 | 0.7232 | 0.469557 | NA |
| | 5.912277 | 0.517229 | 0.897838 | 0.576083 | 0.564559 | NA |
| | 0.460466 | 0.568655 | 3.014642 | 0.188631 | 0.850382 | NA |
| | 0.192403 | 0.975392 | 4.060716 | 0.240202 | 0.810174 | NA |
| UBL5P2 | 2.712881 | -0.84497 | 1.209025 | -0.69888 | 0.484625 | NA |
| | 0.804712 | -0.7622 | 2.348804 | -0.3245 | 0.745556 | NA |
| | 0.326333 | 1.533889 | 3.826878 | 0.40082 | 0.688553 | NA |
| ARIH2P1 | 1.06329 | 0.285492 | 1.942399 | 0.146979 | 0.883149 | NA |
| | 0.651435 | 0.249629 | 2.788679 | 0.089515 | 0.928672 | NA |
| | 1.868058 | 0.021519 | 1.434582 | 0.015 | 0.988032 | NA |
| RN7SL192P | 0.678686 | -3.08037 | 2.621268 | -1.17514 | 0.239937 | NA |
| | 0.151697 | -1.546 | 4.075469 | -0.37934 | 0.704433 | NA |
| | 0.61514 | 1.240942 | 3.104385 | 0.399738 | 0.689349 | NA |
| | 0.096202 | 0.408546 | 4.079639 | 0.100143 | 0.920231 | NA |
| | 0.134133 | -1.48909 | 4.077584 | -0.36519 | 0.71497 | NA |
| MIR4740 | 3.204954 | -0.5026 | 1.095676 | -0.45872 | 0.646438 | NA |
| | 0.192403 | 0.975392 | 4.060716 | 0.240202 | 0.810174 | NA |
| | 0.488709 | 2.119056 | 3.307564 | 0.64067 | 0.521737 | NA |
| SNORA76 | 23.51996 | 0.159622 | 0.460971 | 0.346273 | 0.729138 | NA |
| CBX3P2 | 5.708392 | -0.80233 | 0.898388 | -0.89308 | 0.371813 | NA |
| | 0.49029 | 2.123092 | 3.304282 | 0.642527 | 0.520531 | NA |
| RN7SL663P | 0.134133 | -1.48909 | 4.077584 | -0.36519 | 0.71497 | NA |
| | 0.839896 | -0.29864 | 2.789385 | -0.10706 | 0.914738 | NA |
| | 0.787148 | -0.70094 | 2.357021 | -0.29738 | 0.766174 | NA |
| | 2.165392 | 1.715894 | 1.561644 | 1.098774 | 0.271867 | NA |
| | 183.925 | -0.23878 | 0.208469 | -1.14541 | 0.252039 | 0.415977 |
| | 5.60953 | 0.298923 | 0.854079 | 0.349995 | 0.726343 | NA |
| | 5.263079 | 0.442544 | 0.9051 | 0.488945 | 0.624881 | NA |
| | 1.0066 | -1.02879 | 2.187904 | -0.47022 | 0.638201 | NA |
| | 0.296509 | -0.51709 | 3.982155 | -0.12985 | 0.896684 | NA |
| | 0.678686 | -3.08037 | 2.621268 | -1.17514 | 0.239937 | NA |
| | 0.556813 | -0.57007 | 3.294615 | -0.17303 | 0.862627 | NA |
| | 0.230334 | -0.53318 | 4.076749 | -0.13079 | 0.895945 | NA |
| | 0.424115 | 1.932531 | 3.45462 | 0.559405 | 0.575885 | NA |
| | 0.260158 | 1.240347 | 4.047124 | 0.306476 | 0.759242 | NA |
| | 70.92569 | 0.001973 | 0.529416 | 0.003727 | 0.997027 | 0.99888 |
| MIR4530 | 0.230334 | -0.53318 | 4.076749 | -0.13079 | 0.895945 | NA |
| | 0.163957 | 0.424913 | 4.079006 | 0.104171 | 0.917034 | NA |

| | | | | | | |
|---|---|---|---|---|---|---|
| | 0.162376 | 0.424913 | 4.079006 | 0.104171 | 0.917034 | NA |
| | 2.082352 | -0.26786 | 1.502059 | -0.17833 | 0.858466 | NA |
| | 1.459015 | 1.061255 | 1.654615 | 0.641391 | 0.521269 | NA |
| | 4.794084 | -0.02453 | 0.976326 | -0.02513 | 0.979953 | NA |
| RFWD2P1 | 0.151697 | -1.546 | 4.075469 | -0.37934 | 0.704433 | NA |
| | 0.096202 | 0.408546 | 4.079639 | 0.100143 | 0.920231 | NA |
| | 0.162376 | 0.424913 | 4.079006 | 0.104171 | 0.917034 | NA |
| MIR3677 | 0.488709 | 2.119056 | 3.307564 | 0.64067 | 0.521737 | NA |
| | 0.096202 | 0.408546 | 4.079639 | 0.100143 | 0.920231 | NA |
| | 5.660072 | -0.19348 | 0.893068 | -0.21665 | 0.828485 | NA |
| | 9.571418 | -0.22341 | 0.758462 | -0.29456 | 0.768331 | NA |
| | 0.163957 | 0.424913 | 4.079006 | 0.104171 | 0.917034 | NA |
| | 0.163957 | 0.424913 | 4.079006 | 0.104171 | 0.917034 | NA |
| | 0.298089 | -0.51711 | 3.975232 | -0.13008 | 0.896501 | NA |
| | 1.60814 | -0.79697 | 1.672191 | -0.4766 | 0.633645 | NA |
| PSMD7P1 | 0.584911 | 2.38756 | 2.690807 | 0.887303 | 0.374916 | NA |
| | 1.63242 | 0.749319 | 1.641395 | 0.456513 | 0.648021 | NA |
| | 0.134133 | -1.48909 | 4.077584 | -0.36519 | 0.71497 | NA |
| MYO15B | 1.989671 | -0.61782 | 1.573501 | -0.39264 | 0.694586 | NA |
| | 0.561971 | -0.49218 | 2.810061 | -0.17515 | 0.860962 | NA |
| | 4.14127 | 0.045396 | 0.981659 | 0.046244 | 0.963116 | NA |
| MIR5695 | 0.326333 | 1.533889 | 3.826878 | 0.40082 | 0.688553 | NA |
| | 46.83153 | -0.15692 | 0.360073 | -0.4358 | 0.66298 | NA |
| MIR3680-2 | 0.163957 | 0.424913 | 4.079006 | 0.104171 | 0.917034 | NA |
| MIR3680-1 | 0.163957 | 0.424913 | 4.079006 | 0.104171 | 0.917034 | NA |
| SH3GL1P1 | 4.424121 | 0.802111 | 1.188266 | 0.675026 | 0.499659 | NA |
| | 1.63594 | 0.601505 | 1.647219 | 0.365164 | 0.714989 | NA |
| | 2.527405 | 0.655359 | 1.371876 | 0.47771 | 0.632857 | NA |
| | 0.260158 | 1.240347 | 4.047124 | 0.306476 | 0.759242 | NA |
| | 0.610582 | -0.18034 | 2.753139 | -0.0655 | 0.947774 | NA |
| | 1.006454 | 0.21027 | 1.900041 | 0.110666 | 0.911881 | NA |
| | 0.884727 | 1.125991 | 2.88877 | 0.389782 | 0.696698 | NA |
| | 106.6638 | -0.22423 | 0.297272 | -0.75431 | 0.450664 | 0.609958 |
| | 0.35636 | 1.709252 | 3.651833 | 0.468053 | 0.639746 | NA |
| | 0.192403 | 0.975392 | 4.060716 | 0.240202 | 0.810174 | NA |
| | 0.162376 | 0.424913 | 4.079006 | 0.104171 | 0.917034 | NA |
| ITM2BP1 | 0.518736 | 2.237423 | 2.7713 | 0.807355 | 0.419462 | NA |
| | 64.39698 | 0.107137 | 0.275166 | 0.389356 | 0.697013 | 0.804815 |
| | 0.815391 | 0.931626 | 2.360431 | 0.394685 | 0.693076 | NA |
| | 0.258578 | 1.233151 | 4.047809 | 0.304646 | 0.760635 | NA |
| | 0.574231 | 0.93227 | 2.49998 | 0.372911 | 0.709215 | NA |
| | 1.397876 | -1.6426 | 1.882481 | -0.87257 | 0.382897 | NA |
| AKR1B1P7 | 0.35636 | 1.709252 | 3.651833 | 0.468053 | 0.639746 | NA |
| | 0.392711 | 0.292538 | 3.145221 | 0.09301 | 0.925895 | NA |
| LINC00663 | 21.54501 | -0.18733 | 0.511518 | -0.36622 | 0.714202 | NA |
| SLC25A6P4 | 0.096202 | 0.408546 | 4.079639 | 0.100143 | 0.920231 | NA |
| | 0.28583 | -2.21358 | 3.97376 | -0.55705 | 0.577494 | NA |

| | | | | | | |
|---|---|---|---|---|---|---|
| | 1.522175 | 0.469421 | 1.602229 | 0.29298 | 0.769538 | NA |
| | 0.885732 | -2.40221 | 2.384877 | -1.00727 | 0.313806 | NA |
| | 0.096202 | 0.408546 | 4.079639 | 0.100143 | 0.920231 | NA |
| ACTBP9 | 0.410275 | 0.244283 | 3.100889 | 0.078778 | 0.937209 | NA |
| | 15.97993 | 1.076306 | 0.573613 | 1.876363 | 0.060606 | NA |
| | 2.73364 | 0.23429 | 1.220763 | 0.191921 | 0.847804 | NA |
| | 0.134133 | -1.48909 | 4.077584 | -0.36519 | 0.71497 | NA |
| | 23.02616 | 0.32922 | 0.579377 | 0.568231 | 0.569878 | NA |
| | 2.775818 | 1.021775 | 1.330991 | 0.76768 | 0.442678 | NA |
| | 3.379379 | -0.64657 | 1.097621 | -0.58907 | 0.555818 | NA |
| | 1.961282 | -0.41505 | 1.36029 | -0.30512 | 0.760277 | NA |
| | 0.298089 | -0.51711 | 3.975232 | -0.13008 | 0.896501 | NA |
| | 0.096202 | 0.408546 | 4.079639 | 0.100143 | 0.920231 | NA |
| | 0.298089 | -0.51711 | 3.975232 | -0.13008 | 0.896501 | NA |
| | 3.669206 | -0.50547 | 1.1305 | -0.44712 | 0.654786 | NA |
| MRPL37P1 | 0.162376 | 0.424913 | 4.079006 | 0.104171 | 0.917034 | NA |
| | 22.89693 | -0.68564 | 0.457473 | -1.49876 | 0.133936 | NA |
| | 0.163957 | 0.424913 | 4.079006 | 0.104171 | 0.917034 | NA |
| | 0.134133 | -1.48909 | 4.077584 | -0.36519 | 0.71497 | NA |
| | 0.506476 | 0.726418 | 2.864186 | 0.253621 | 0.799788 | NA |
| | 48.19835 | -0.43516 | 0.33149 | -1.31273 | 0.189273 | 0.343409 |
| | 0.326333 | 1.533889 | 3.826878 | 0.40082 | 0.688553 | NA |
| FXYD1 | 1.025561 | 0.190158 | 2.384522 | 0.079747 | 0.936439 | NA |
| AARSD1 | 8.404492 | 0.53358 | 0.75005 | 0.711393 | 0.476841 | NA |
| | 0.298089 | -0.51711 | 3.975232 | -0.13008 | 0.896501 | NA |
| | 2.038339 | -0.21984 | 1.344596 | -0.1635 | 0.870123 | NA |
| | 0.288605 | 1.443619 | 4.029058 | 0.358302 | 0.720117 | NA |
| LRRC37A9P | 1.268755 | 0.049606 | 1.899823 | 0.026111 | 0.979169 | NA |
| | 0.544554 | -2.65036 | 3.220086 | -0.82307 | 0.410468 | NA |
| | 0.3441 | -0.0166 | 3.769682 | -0.0044 | 0.996486 | NA |
| | 0.392711 | 0.292538 | 3.145221 | 0.09301 | 0.925895 | NA |
| | 1.687494 | -0.33602 | 1.768527 | -0.19 | 0.849309 | NA |
| | 0.163957 | 0.424913 | 4.079006 | 0.104171 | 0.917034 | NA |
| | 2.193103 | -0.79153 | 1.563251 | -0.50634 | 0.612621 | NA |
| | 0.163957 | 0.424913 | 4.079006 | 0.104171 | 0.917034 | NA |
| | 0.163957 | 0.424913 | 4.079006 | 0.104171 | 0.917034 | NA |
| | 0.268266 | -2.15396 | 4.038351 | -0.53338 | 0.593773 | NA |
| DHX40P1 | 0.151697 | -1.546 | 4.075469 | -0.37934 | 0.704433 | NA |
| | 0.258578 | 1.233151 | 4.047809 | 0.304646 | 0.760635 | NA |
| | 0.440301 | 0.468383 | 3.449808 | 0.135771 | 0.892003 | NA |
| | 0.52699 | -2.57878 | 3.248478 | -0.79384 | 0.427288 | NA |
| | 1.354277 | -0.37751 | 1.844618 | -0.20465 | 0.837843 | NA |
| | 24.87575 | 0.517603 | 0.455578 | 1.136146 | 0.255896 | NA |
| | 1.402122 | 0.958574 | 1.966468 | 0.48746 | 0.625933 | NA |
| | 0.698879 | 1.36795 | 2.596167 | 0.526912 | 0.598255 | NA |
| | 10.04629 | 0.273448 | 0.683214 | 0.400238 | 0.688981 | NA |
| | 9.266682 | 2.049499 | 0.781477 | 2.622598 | 0.008726 | NA |

| | | | | | | |
|---|---|---|---|---|---|---|
| | 5.757131 | -0.17545 | 0.833236 | -0.21056 | 0.833231 | NA |
| | 0.327913 | 1.543154 | 4.021063 | 0.383768 | 0.701151 | NA |
| | 0.314073 | -0.57308 | 3.898934 | -0.14698 | 0.883146 | NA |
| | 0.163957 | 0.424913 | 4.079006 | 0.104171 | 0.917034 | NA |
| | 2.861 | -1.12209 | 1.256704 | -0.89289 | 0.371918 | NA |
| LRRC37A16 | 665.6814 | -0.14642 | 0.15782 | -0.92777 | 0.353525 | 0.521597 |
| | 1.147085 | 0.53291 | 1.948376 | 0.273515 | 0.784457 | NA |
| | 1.721113 | 1.271654 | 1.706047 | 0.74538 | 0.456042 | NA |
| | 2.413211 | -0.9507 | 1.424441 | -0.66742 | 0.504506 | NA |
| | 1.169179 | -1.04993 | 1.839034 | -0.57091 | 0.568059 | NA |
| | 69.64668 | 0.122049 | 0.27013 | 0.451816 | 0.651401 | 0.769259 |
| ZNF850 | 53.66576 | 0.291369 | 0.360008 | 0.809339 | 0.41832 | 0.581245 |
| | 1.70671 | 1.288284 | 1.568841 | 0.82117 | 0.41155 | NA |
| | 1.465485 | -0.31895 | 1.726907 | -0.18469 | 0.85347 | NA |
| | 0.70851 | -1.62753 | 2.686091 | -0.60591 | 0.544574 | NA |
| | 0.92901 | 1.841773 | 2.123319 | 0.867403 | 0.385721 | NA |
| | 0.162376 | 0.424913 | 4.079006 | 0.104171 | 0.917034 | NA |
| | 13.61595 | 0.173826 | 0.587252 | 0.295998 | 0.767231 | NA |
| | 4.813983 | -0.772 | 0.969028 | -0.79667 | 0.42564 | NA |
| PTGES3L | 2.118878 | 0.16818 | 1.744666 | 0.096397 | 0.923205 | NA |
| | 0.641986 | 1.117767 | 2.709494 | 0.412537 | 0.679946 | NA |
| | 5.17595 | -0.77842 | 1.028855 | -0.75659 | 0.449295 | NA |
| | 0.096202 | 0.408546 | 4.079639 | 0.100143 | 0.920231 | NA |
| | 0.163957 | 0.424913 | 4.079006 | 0.104171 | 0.917034 | NA |
| CCDC58P3 | 0.096202 | 0.408546 | 4.079639 | 0.100143 | 0.920231 | NA |
| | 0.260158 | 1.240347 | 4.047124 | 0.306476 | 0.759242 | NA |
| | 3.491718 | -0.1222 | 1.175017 | -0.104 | 0.917171 | NA |
| | 5.554397 | -0.80556 | 0.914931 | -0.88046 | 0.378611 | NA |
| KRT18P61 | 0.134133 | -1.48909 | 4.077584 | -0.36519 | 0.71497 | NA |
| | 0.725928 | 0.181819 | 2.321433 | 0.078322 | 0.937572 | NA |
| CTBP2P7 | 0.134133 | -1.48909 | 4.077584 | -0.36519 | 0.71497 | NA |
| | 0.151697 | -1.546 | 4.075469 | -0.37934 | 0.704433 | NA |
| | 0.151697 | -1.546 | 4.075469 | -0.37934 | 0.704433 | NA |
| ILF3-AS1 | 251.095 | 0.114159 | 0.209282 | 0.545478 | 0.585425 | 0.718394 |
| | 12.90917 | 1.431228 | 0.760156 | 1.882808 | 0.059726 | NA |
| C19orf82 | 263.0963 | 0.004251 | 0.217091 | 0.019583 | 0.984376 | 0.991773 |
| | 0.163957 | 0.424913 | 4.079006 | 0.104171 | 0.917034 | NA |
| | 1.70513 | 1.285072 | 1.554544 | 0.826656 | 0.408432 | NA |
| | 0.727508 | 0.192954 | 2.572892 | 0.074995 | 0.940219 | NA |
| | 0.096202 | 0.408546 | 4.079639 | 0.100143 | 0.920231 | NA |
| | 14.36088 | -0.7354 | 0.530059 | -1.38739 | 0.165323 | NA |
| RNF157-AS | 1.598249 | 0.567223 | 1.588614 | 0.357055 | 0.72105 | NA |
| | 0.096202 | 0.408546 | 4.079639 | 0.100143 | 0.920231 | NA |
| | 4.863125 | 0.67049 | 0.940336 | 0.713032 | 0.475826 | NA |
| | 0.151697 | -1.546 | 4.075469 | -0.37934 | 0.704433 | NA |
| | 0.096202 | 0.408546 | 4.079639 | 0.100143 | 0.920231 | NA |
| | 1.458812 | 1.676572 | 1.711787 | 0.979428 | 0.327369 | NA |

| | | | | | | |
|---|---|---|---|---|---|---|
| | 2.014191 | 0.480495 | 1.639894 | 0.293004 | 0.769519 | NA |
| | 0.354779 | 1.703954 | 3.657232 | 0.465914 | 0.641277 | NA |
| | 7.694489 | 0.265617 | 0.826278 | 0.321462 | 0.74786 | NA |
| TPMTP1 | 1.083685 | 2.141036 | 2.332782 | 0.917804 | 0.358721 | NA |
| | 0.151697 | -1.546 | 4.075469 | -0.37934 | 0.704433 | NA |
| | 2.251117 | 0.861744 | 1.341313 | 0.642463 | 0.520573 | NA |
| | 35.61229 | -0.41448 | 0.44215 | -0.93743 | 0.348539 | NA |
| | 0.528424 | -0.38142 | 2.853651 | -0.13366 | 0.893671 | NA |
| | 49.36093 | 0.011551 | 0.331467 | 0.03485 | 0.9722 | 0.984681 |
| CALM2P1 | 0.096202 | 0.408546 | 4.079639 | 0.100143 | 0.920231 | NA |
| | 0.672013 | 1.253691 | 2.607776 | 0.480751 | 0.630693 | NA |
| | 0.74749 | 1.546424 | 2.516383 | 0.614543 | 0.538857 | NA |
| PHF5CP | 0.096202 | 0.408546 | 4.079639 | 0.100143 | 0.920231 | NA |
| | 0.682896 | 1.412658 | 2.61672 | 0.539858 | 0.589295 | NA |
| PTP4A2P1 | 1.945261 | -0.45366 | 1.579784 | -0.28717 | 0.773983 | NA |
| ATP5LP7 | 0.134133 | -1.48909 | 4.077584 | -0.36519 | 0.71497 | NA |
| | 0.872321 | 0.678609 | 2.011629 | 0.337343 | 0.735858 | NA |
| | 0.162376 | 0.424913 | 4.079006 | 0.104171 | 0.917034 | NA |
| | 22.68814 | 0.407791 | 0.522114 | 0.781038 | 0.43478 | NA |
| | 2.047649 | 1.111208 | 1.445131 | 0.768932 | 0.441934 | NA |
| | 1.593507 | 0.596556 | 1.813973 | 0.328867 | 0.742256 | NA |
| | 5.775418 | -0.13384 | 0.846633 | -0.15808 | 0.874392 | NA |
| SNRPGP4 | 2.101361 | 1.122709 | 1.659381 | 0.676583 | 0.498671 | NA |
| | 0.230334 | -0.53318 | 4.076749 | -0.13079 | 0.895945 | NA |
| | 0.489058 | -0.58569 | 3.427543 | -0.17088 | 0.864321 | NA |
| | 6.278755 | -0.40218 | 0.855057 | -0.47035 | 0.638102 | NA |
| | 5.360882 | -0.44905 | 0.898553 | -0.49974 | 0.617255 | NA |
| | 1.210485 | -0.74676 | 2.171272 | -0.34393 | 0.7309 | NA |
| | 0.364467 | -1.20166 | 3.685431 | -0.32606 | 0.744381 | NA |
| | 0.384806 | 1.846216 | 3.999752 | 0.461582 | 0.644381 | NA |
| | 2.837986 | -0.25817 | 1.219204 | -0.21176 | 0.832298 | NA |
| | 0.134133 | -1.48909 | 4.077584 | -0.36519 | 0.71497 | NA |
| | 74.64515 | -0.54021 | 0.344768 | -1.56689 | 0.11714 | 0.247716 |
| | 0.162376 | 0.424913 | 4.079006 | 0.104171 | 0.917034 | NA |
| | 17.15483 | -0.72353 | 0.515454 | -1.40368 | 0.160415 | NA |
| | 14.70337 | -0.60305 | 0.625688 | -0.96382 | 0.335135 | NA |
| | 0.719043 | 1.433043 | 2.335048 | 0.61371 | 0.539407 | NA |
| | 0.134133 | -1.48909 | 4.077584 | -0.36519 | 0.71497 | NA |
| | 24.65357 | 0.164544 | 0.457524 | 0.359641 | 0.719116 | NA |
| | 7.695947 | 0.57135 | 0.747093 | 0.764763 | 0.444412 | NA |
| | 5.544789 | 0.470174 | 0.881767 | 0.533218 | 0.593883 | NA |
| | 0.49187 | 2.136088 | 3.895953 | 0.548284 | 0.583497 | NA |
| | 44.35985 | 0.321185 | 0.357295 | 0.898936 | 0.368687 | NA |
| | 3.485094 | 0.068354 | 1.03024 | 0.066348 | 0.947101 | NA |
| | 0.162376 | 0.424913 | 4.079006 | 0.104171 | 0.917034 | NA |
| | 1.216035 | 2.309201 | 2.192558 | 1.053199 | 0.29225 | NA |
| | 39.1865 | 0.423732 | 0.343036 | 1.235238 | 0.216742 | NA |

| | | | | | | |
|---|---|---|---|---|---|---|
| | 0.163957 | 0.424913 | 4.079006 | 0.104171 | 0.917034 | NA |
| | 2.838978 | 0.674599 | 1.221319 | 0.552353 | 0.580706 | NA |
| | 5.759886 | 0.070468 | 0.836531 | 0.084238 | 0.932867 | NA |
| MAP3K14-/ | 9.792401 | 0.165158 | 0.666267 | 0.247886 | 0.804223 | NA |
| | 129.1104 | 0.219518 | 0.286317 | 0.766698 | 0.443261 | 0.603336 |
| | 98.92001 | -0.49962 | 0.240368 | -2.07858 | 0.037656 | 0.115375 |
| | 2.812004 | -0.45922 | 1.221993 | -0.3758 | 0.707067 | NA |
| | 0.151697 | -1.546 | 4.075469 | -0.37934 | 0.704433 | NA |
| | 7.252257 | 0.897188 | 0.940553 | 0.953894 | 0.340137 | NA |
| | 0.872321 | 0.678609 | 2.011629 | 0.337343 | 0.735858 | NA |
| | 0.192403 | 0.975392 | 4.060716 | 0.240202 | 0.810174 | NA |
| | 0.192403 | 0.975392 | 4.060716 | 0.240202 | 0.810174 | NA |
| RPL23AP77 | 0.394291 | 0.298745 | 3.141711 | 0.09509 | 0.924243 | NA |
| | 1.851522 | 1.481591 | 1.522604 | 0.973064 | 0.330521 | NA |
| | 1.04716 | 1.17329 | 2.006462 | 0.584756 | 0.558712 | NA |
| | 0.151697 | -1.546 | 4.075469 | -0.37934 | 0.704433 | NA |
| | 22.13307 | -0.38281 | 0.448427 | -0.85367 | 0.393285 | NA |
| | 0.314073 | -0.57308 | 3.898934 | -0.14698 | 0.883146 | NA |
| | 5.447125 | 0.7082 | 0.947773 | 0.747225 | 0.454927 | NA |
| | 121.2241 | -0.60043 | 0.237873 | -2.52415 | 0.011598 | 0.0521 |
| | 1.401939 | 1.712015 | 1.808509 | 0.946644 | 0.34382 | NA |
| | 144.548 | 0.151496 | 0.227433 | 0.666115 | 0.505338 | 0.65473 |
| | 0.69238 | 0.295083 | 2.603008 | 0.113362 | 0.909743 | NA |
| SLC25A1P5 | 0.681461 | 0.491544 | 3.066278 | 0.160307 | 0.87264 | NA |
| | 0.134133 | -1.48909 | 4.077584 | -0.36519 | 0.71497 | NA |
| | 0.162376 | 0.424913 | 4.079006 | 0.104171 | 0.917034 | NA |
| | 0.134133 | -1.48909 | 4.077584 | -0.36519 | 0.71497 | NA |
| | 0.162376 | 0.424913 | 4.079006 | 0.104171 | 0.917034 | NA |
| | 0.834389 | 1.638864 | 2.19944 | 0.745128 | 0.456195 | NA |
| LINC00906 | 38.40108 | -0.49335 | 0.484277 | -1.01874 | 0.308325 | NA |
| | 1.1119 | 0.46633 | 1.933926 | 0.241131 | 0.809453 | NA |
| | 4.374021 | 0.272612 | 1.024158 | 0.266181 | 0.7901 | NA |
| | 0.096202 | 0.408546 | 4.079639 | 0.100143 | 0.920231 | NA |
| | 0.230334 | -0.53318 | 4.076749 | -0.13079 | 0.895945 | NA |
| | 0.47803 | 0.520083 | 2.97881 | 0.174594 | 0.861399 | NA |
| | 677.1642 | -0.40304 | 0.150269 | -2.68215 | 0.007315 | 0.038363 |
| SH3GL1P3 | 15.49571 | 0.897078 | 0.567186 | 1.581629 | 0.113734 | NA |
| RPL9P29 | 2.475063 | -0.0703 | 1.293957 | -0.05433 | 0.956675 | NA |
| | 0.096202 | 0.408546 | 4.079639 | 0.100143 | 0.920231 | NA |
| | 0.489058 | -0.58569 | 3.427543 | -0.17088 | 0.864321 | NA |
| | 0.162376 | 0.424913 | 4.079006 | 0.104171 | 0.917034 | NA |
| KCNJ2-AS1 | 2.471967 | 1.085049 | 1.29937 | 0.835057 | 0.403685 | NA |
| | 0.551292 | -1.76878 | 3.219819 | -0.54934 | 0.582772 | NA |
| | 2.483959 | 0.688835 | 1.405878 | 0.489968 | 0.624157 | NA |
| | 0.460669 | -0.62593 | 3.398671 | -0.18417 | 0.853882 | NA |
| | 0.162376 | 0.424913 | 4.079006 | 0.104171 | 0.917034 | NA |
| LINC00669 | 9.381958 | 0.530982 | 0.715187 | 0.742438 | 0.457822 | NA |

| | | | | | | | |
|---|---|---|---|---|---|---|---|
| | 0.994194 | -0.63787 | 1.893391 | -0.33689 | 0.736198 | NA | |
| | 10.22396 | 0.695279 | 0.893195 | 0.778418 | 0.436323 | NA | |
| | 2.966358 | -0.41178 | 1.111022 | -0.37063 | 0.710909 | NA | |
| | 2.609049 | 0.194659 | 1.234738 | 0.157652 | 0.874731 | NA | |
| | 0.424115 | 1.932531 | 3.45462 | 0.559405 | 0.575885 | NA | |
| | 16.39913 | 0.105211 | 0.534987 | 0.196661 | 0.844093 | NA | |
| | 11.89516 | -0.3051 | 0.610057 | -0.50012 | 0.616988 | NA | |
| | 1.911373 | 2.164016 | 1.571809 | 1.376768 | 0.168584 | NA | |
| | 0.096202 | 0.408546 | 4.079639 | 0.100143 | 0.920231 | NA | |
| | 1.36261 | 1.545046 | 1.790867 | 0.862736 | 0.388282 | NA | |
| | 4.000069 | 0.692654 | 1.072448 | 0.645862 | 0.518369 | NA | |
| | 0.49029 | 2.123092 | 3.304282 | 0.642527 | 0.520531 | NA | |
| | 0.163957 | 0.424913 | 4.079006 | 0.104171 | 0.917034 | NA | |
| | 3.577572 | 0.419799 | 1.057528 | 0.396962 | 0.691395 | NA | |
| | 5.363563 | -0.20328 | 0.904002 | -0.22486 | 0.822086 | NA | |
| | 0.738188 | 1.39874 | 2.317793 | 0.603479 | 0.54619 | NA | |
| | 12.14299 | 0.225915 | 0.603546 | 0.374313 | 0.708171 | NA | |
| | 0.134133 | -1.48909 | 4.077584 | -0.36519 | 0.71497 | NA | |
| | 1.921973 | -0.39508 | 1.48345 | -0.26632 | 0.789989 | NA | |
| | 1.102655 | 0.449766 | 1.809137 | 0.248608 | 0.803664 | NA | |
| | 1.878884 | -0.09816 | 1.5128 | -0.06488 | 0.948266 | NA | |
| | 0.096202 | 0.408546 | 4.079639 | 0.100143 | 0.920231 | NA | |
| | 1031.162 | -0.51485 | 0.156109 | -3.29802 | 9.74E-04 | | 0.010161 |
| | 0.586491 | 2.39072 | 2.689057 | 0.889055 | 0.373973 | NA | |
| | 23.87511 | -0.23393 | 0.461365 | -0.50704 | 0.612124 | NA | |
| | 0.151697 | -1.546 | 4.075469 | -0.37934 | 0.704433 | NA | |
| | 0.247898 | -0.5901 | 4.074633 | -0.14482 | 0.884851 | NA | |
| | 0.163957 | 0.424913 | 4.079006 | 0.104171 | 0.917034 | NA | |
| ZNF571-AS | 25.81548 | -0.15685 | 0.51323 | -0.30562 | 0.759893 | NA | |
| | 0.247898 | -0.5901 | 4.074633 | -0.14482 | 0.884851 | NA | |
| | 7.200528 | 0.033883 | 0.888804 | 0.038122 | 0.96959 | NA | |
| | 1.312622 | 0.801445 | 1.708401 | 0.46912 | 0.638984 | NA | |
| | 22.99658 | 0.166786 | 0.449851 | 0.370759 | 0.710817 | NA | |
| | 0.303394 | -2.27006 | 4.032451 | -0.56295 | 0.57347 | NA | |
| | 0.163957 | 0.424913 | 4.079006 | 0.104171 | 0.917034 | NA | |
| | 12.18294 | -0.46438 | 0.704324 | -0.65932 | 0.509688 | NA | |
| | 8.703154 | 0.583345 | 0.704372 | 0.828178 | 0.40757 | NA | |
| | 1.754236 | -0.4131 | 1.535881 | -0.26897 | 0.787956 | NA | |
| CIRBP-AS1 | 16.13526 | 0.049529 | 0.595367 | 0.08319 | 0.9337 | NA | |
| | 2.922863 | 0.302936 | 1.270293 | 0.238478 | 0.811511 | NA | |
| | 6.575728 | 0.590007 | 0.850216 | 0.693949 | 0.487714 | NA | |
| | 0.096202 | 0.408546 | 4.079639 | 0.100143 | 0.920231 | NA | |
| | 0.516164 | -1.70268 | 2.844782 | -0.59853 | 0.549488 | NA | |
| | 4.147151 | 0.187879 | 1.008974 | 0.186208 | 0.852282 | NA | |
| | 4.917602 | 0.294334 | 0.911693 | 0.322843 | 0.746814 | NA | |
| | 0.448206 | -1.2532 | 3.007315 | -0.41672 | 0.676885 | NA | |
| MIR24-2 | 123.5918 | 0.527495 | 0.267658 | 1.970782 | 0.048749 | | 0.136462 |

| | | | | | | |
|---|---|---|---|---|---|---|
| | 0.096202 | 0.408546 | 4.079639 | 0.100143 | 0.920231 | NA |
| | 106.0453 | 4.53E-04 | 0.321814 | 0.001406 | 0.998878 | 0.999384 |
| | 0.856337 | 0.757304 | 2.453242 | 0.308695 | 0.757553 | NA |
| MIR497HG | 0.841063 | -2.08438 | 2.306687 | -0.90362 | 0.366195 | NA |
| | 1.75409 | 0.304005 | 1.445805 | 0.210267 | 0.83346 | NA |
| S1PR2 | 133.3907 | -0.44506 | 0.226364 | -1.96614 | 0.049282 | 0.137306 |
| | 4.699603 | -0.31381 | 1.057638 | -0.29671 | 0.76669 | NA |
| | 0.696104 | -0.93034 | 2.354268 | -0.39517 | 0.692715 | NA |
| | 2.469446 | 1.232821 | 1.849878 | 0.666433 | 0.505134 | NA |
| | 6.892755 | 0.530413 | 0.884009 | 0.600009 | 0.5485 | NA |
| | 0.247898 | -0.5901 | 4.074633 | -0.14482 | 0.884851 | NA |
| | 0.314073 | -0.57308 | 3.898934 | -0.14698 | 0.883146 | NA |
| | 3.000933 | 0.439498 | 1.288042 | 0.341214 | 0.732942 | NA |
| | 0.558248 | 0.982076 | 2.804738 | 0.350149 | 0.726227 | NA |
| | 0.314073 | -0.57308 | 3.898934 | -0.14698 | 0.883146 | NA |
| | 0.70851 | -1.62753 | 2.686091 | -0.60591 | 0.544574 | NA |
| | 0.984506 | 0.880958 | 2.062676 | 0.427094 | 0.669311 | NA |
| | 95.09136 | -0.26611 | 0.25063 | -1.06176 | 0.288344 | 0.454443 |
| | 0.489058 | -0.58569 | 3.427543 | -0.17088 | 0.864321 | NA |
| | 0.096202 | 0.408546 | 4.079639 | 0.100143 | 0.920231 | NA |
| | 0.162376 | 0.424913 | 4.079006 | 0.104171 | 0.917034 | NA |
| LINC00907 | 1.249194 | -0.07766 | 2.009285 | -0.03865 | 0.969167 | NA |
| | 0.258578 | 1.233151 | 4.047809 | 0.304646 | 0.760635 | NA |
| | 0.489058 | -0.58569 | 3.427543 | -0.17088 | 0.864321 | NA |
| CYP4F24P | 0.096202 | 0.408546 | 4.079639 | 0.100143 | 0.920231 | NA |
| | 0.420954 | 1.923376 | 3.462803 | 0.555439 | 0.578594 | NA |
| CCL15 | 0.134133 | -1.48909 | 4.077584 | -0.36519 | 0.71497 | NA |
| | 0.879205 | -0.27615 | 2.437194 | -0.11331 | 0.909788 | NA |
| | 0.151697 | -1.546 | 4.075469 | -0.37934 | 0.704433 | NA |
| | 0.489058 | -0.58569 | 3.427543 | -0.17088 | 0.864321 | NA |
| | 0.163957 | 0.424913 | 4.079006 | 0.104171 | 0.917034 | NA |
| | 0.192403 | 0.975392 | 4.060716 | 0.240202 | 0.810174 | NA |
| | 0.096202 | 0.408546 | 4.079639 | 0.100143 | 0.920231 | NA |
| | 8.98481 | -0.18923 | 0.77819 | -0.24316 | 0.80788 | NA |
| | 0.28583 | -2.21358 | 3.97376 | -0.55705 | 0.577494 | NA |
| | 0.162376 | 0.424913 | 4.079006 | 0.104171 | 0.917034 | NA |
| | 0.162376 | 0.424913 | 4.079006 | 0.104171 | 0.917034 | NA |
| | 0.163957 | 0.424913 | 4.079006 | 0.104171 | 0.917034 | NA |
| | 0.488709 | 2.119056 | 3.307564 | 0.64067 | 0.521737 | NA |
| | 2.69718 | 0.917905 | 1.306194 | 0.702733 | 0.482222 | NA |
| | 4.92034 | 0.681943 | 0.972015 | 0.701576 | 0.482943 | NA |
| | 3.04257 | -1.64089 | 1.17621 | -1.39506 | 0.162997 | NA |
| | 1.668917 | -0.52368 | 1.657435 | -0.31596 | 0.752033 | NA |
| | 4.752471 | -0.07868 | 0.996345 | -0.07897 | 0.937056 | NA |
| | 0.162376 | 0.424913 | 4.079006 | 0.104171 | 0.917034 | NA |
| | 0.096202 | 0.408546 | 4.079639 | 0.100143 | 0.920231 | NA |
| FDX1L | 5.525269 | -0.50035 | 0.92654 | -0.54002 | 0.58918 | NA |

| | | | | | | |
|---|---|---|---|---|---|---|
| THA1P | 0.520316 | 2.239969 | 3.173769 | 0.705776 | 0.480328 | NA |
| EIF5AP2 | 2.239849 | 1.762932 | 1.548313 | 1.138615 | 0.254864 | NA |
| | 2.608215 | -0.61314 | 1.305052 | -0.46982 | 0.638481 | NA |
| | 2.896309 | -0.57467 | 1.356537 | -0.42363 | 0.671834 | NA |
| | 80.0589 | 0.332255 | 0.293576 | 1.13175 | 0.257739 | 0.421856 |
| SHC1P2 | 2.982677 | 0.677647 | 1.383657 | 0.489751 | 0.62431 | NA |
| TAF9P3 | 0.640406 | 1.11402 | 2.710977 | 0.410929 | 0.681124 | NA |
| | 1.656308 | 0.122105 | 1.513683 | 0.080667 | 0.935707 | NA |
| | 1.460595 | 1.079353 | 1.826252 | 0.591021 | 0.554506 | NA |
| | 0.260158 | 1.240347 | 4.047124 | 0.306476 | 0.759242 | NA |
| | 1.007445 | 3.175243 | 2.206758 | 1.438872 | 0.150187 | NA |
| | 0.096202 | 0.408546 | 4.079639 | 0.100143 | 0.920231 | NA |
| | 0.314073 | -0.57308 | 3.898934 | -0.14698 | 0.883146 | NA |
| | 2.060404 | 0.103335 | 1.531879 | 0.067456 | 0.946219 | NA |
| | 0.096202 | 0.408546 | 4.079639 | 0.100143 | 0.920231 | NA |
| | 0.163957 | 0.424913 | 4.079006 | 0.104171 | 0.917034 | NA |
| | 0.298089 | -0.51711 | 3.975232 | -0.13008 | 0.896501 | NA |
| | 0.489058 | -0.58569 | 3.427543 | -0.17088 | 0.864321 | NA |
| UBE2L4 | 0.558248 | 0.982076 | 2.804738 | 0.350149 | 0.726227 | NA |
| FAM60CP | 0.833012 | 0.681553 | 2.203157 | 0.309353 | 0.757053 | NA |
| | 10.96746 | 0.144534 | 0.750727 | 0.192525 | 0.847331 | NA |
| | 0.151697 | -1.546 | 4.075469 | -0.37934 | 0.704433 | NA |
| | 1.667722 | -2.06177 | 1.8579 | -1.10973 | 0.267114 | NA |
| | 10.5736 | 0.214699 | 0.705771 | 0.304205 | 0.760972 | NA |
| RPS10P27 | 0.096202 | 0.408546 | 4.079639 | 0.100143 | 0.920231 | NA |
| | 1.425973 | 0.261345 | 1.671539 | 0.15635 | 0.875757 | NA |
| | 0.327913 | 1.543154 | 4.021063 | 0.383768 | 0.701151 | NA |
| | 2.579033 | 0.807412 | 1.498533 | 0.538801 | 0.590024 | NA |
| | 0.924655 | -0.07191 | 2.416297 | -0.02976 | 0.976257 | NA |
| | 1.675509 | 0.299267 | 1.519489 | 0.196952 | 0.843865 | NA |
| | 0.096202 | 0.408546 | 4.079639 | 0.100143 | 0.920231 | NA |
| | 1.702704 | -0.96991 | 1.494987 | -0.64877 | 0.516485 | NA |
| | 0.151697 | -1.546 | 4.075469 | -0.37934 | 0.704433 | NA |
| | 0.392857 | -1.546 | 4.075469 | -0.37934 | 0.704433 | NA |
| | 4.730268 | 0.32851 | 1.029675 | 0.319043 | 0.749694 | NA |
| | 0.163957 | 0.424913 | 4.079006 | 0.104171 | 0.917034 | NA |
| | 0.163957 | 0.424913 | 4.079006 | 0.104171 | 0.917034 | NA |
| SMIM22 | 1.045523 | 1.534279 | 2.413816 | 0.635624 | 0.525022 | NA |
| LIN37 | 55.41202 | -0.01664 | 0.309556 | -0.05375 | 0.957137 | 0.976409 |
| | 0.151697 | -1.546 | 4.075469 | -0.37934 | 0.704433 | NA |
| | 0.096202 | 0.408546 | 4.079639 | 0.100143 | 0.920231 | NA |
| | 0.096202 | 0.408546 | 4.079639 | 0.100143 | 0.920231 | NA |
| | 4.439708 | -0.40936 | 1.093232 | -0.37445 | 0.708072 | NA |
| NDUFA7 | 18.88465 | -0.16808 | 0.488377 | -0.34415 | 0.730731 | NA |
| | 0.134133 | -1.48909 | 4.077584 | -0.36519 | 0.71497 | NA |
| | 6.163083 | 0.672026 | 0.903656 | 0.743675 | 0.457073 | NA |
| | 0.268266 | -2.15396 | 4.038351 | -0.53338 | 0.593773 | NA |

| | | | | | | |
|---|---|---|---|---|---|---|
| | 1.404468 | -0.26803 | 1.627787 | -0.16466 | 0.869211 | NA |
| | 36.93261 | -0.40937 | 0.36562 | -1.11967 | 0.262855 | NA |
| SNX6P1 | 0.192403 | 0.975392 | 4.060716 | 0.240202 | 0.810174 | NA |
| | 0.326333 | 1.533889 | 3.826878 | 0.40082 | 0.688553 | NA |
| | 0.448206 | -1.2532 | 3.007315 | -0.41672 | 0.676885 | NA |
| | 5.524828 | -0.77281 | 0.965633 | -0.80032 | 0.423526 | NA |
| MIR3188 | 3.718735 | -0.27607 | 1.052401 | -0.26233 | 0.79307 | NA |
| | 3.533501 | 0.327098 | 1.060446 | 0.308453 | 0.757738 | NA |
| | 21.01519 | -0.11726 | 0.471104 | -0.24891 | 0.803432 | NA |
| | 0.97663 | -0.60197 | 1.931737 | -0.31162 | 0.755327 | NA |
| | 6.52162 | -0.30015 | 0.869449 | -0.34522 | 0.72993 | NA |
| | 0.424115 | 1.932531 | 3.45462 | 0.559405 | 0.575885 | NA |
| | 0.35636 | 1.709252 | 3.651833 | 0.468053 | 0.639746 | NA |
| | 0.870886 | -0.53073 | 2.310896 | -0.22966 | 0.818353 | NA |
| | 0.163957 | 0.424913 | 4.079006 | 0.104171 | 0.917034 | NA |
| | 24.08065 | -0.15607 | 0.492331 | -0.317 | 0.751243 | NA |
| PCGF7P | 0.47803 | 0.520083 | 2.97881 | 0.174594 | 0.861399 | NA |
| | 0.697685 | -0.92563 | 2.353044 | -0.39338 | 0.694041 | NA |
| | 4.556933 | -0.54654 | 0.937162 | -0.58319 | 0.559765 | NA |
| | 0.230334 | -0.53318 | 4.076749 | -0.13079 | 0.895945 | NA |
| | 0.87074 | 0.675988 | 2.012127 | 0.335957 | 0.736903 | NA |
| | 8.40724 | 0.988444 | 0.704051 | 1.403938 | 0.160337 | NA |
| EEF1A1P7 | 1.327411 | -0.52458 | 1.752811 | -0.29928 | 0.764727 | NA |
| | 5.353123 | -0.51005 | 0.852881 | -0.59803 | 0.549822 | NA |
| | 0.230334 | -0.53318 | 4.076749 | -0.13079 | 0.895945 | NA |
| | 5.476242 | 1.138662 | 0.874968 | 1.301376 | 0.19313 | NA |
| VN1R81P | 3.086141 | 1.915777 | 1.285352 | 1.490469 | 0.136101 | NA |
| | 34.68971 | 0.255394 | 0.3681 | 0.693815 | 0.487798 | NA |
| | 1.935667 | 0.621898 | 1.421215 | 0.437582 | 0.661689 | NA |
| FENDRR | 0.134133 | -1.48909 | 4.077584 | -0.36519 | 0.71497 | NA |
| | 0.163957 | 0.424913 | 4.079006 | 0.104171 | 0.917034 | NA |
| HAVCR1P1 | 0.596179 | -0.09275 | 3.146759 | -0.02948 | 0.976485 | NA |
| | 0.303394 | -2.27006 | 4.032451 | -0.56295 | 0.57347 | NA |
| | 0.268266 | -2.15396 | 4.038351 | -0.53338 | 0.593773 | NA |
| VN1R83P | 1.06136 | 2.032712 | 2.134262 | 0.952419 | 0.340884 | NA |
| | 1.757959 | -1.11865 | 1.765356 | -0.63367 | 0.526298 | NA |
| | 0.151697 | -1.546 | 4.075469 | -0.37934 | 0.704433 | NA |
| | 4.365372 | -0.38823 | 0.998997 | -0.38862 | 0.697554 | NA |
| | 34.41337 | 0.357834 | 0.367152 | 0.974621 | 0.329748 | NA |
| | 1.340865 | 1.498962 | 1.835156 | 0.816804 | 0.41404 | NA |
| LINC00664 | 0.096202 | 0.408546 | 4.079639 | 0.100143 | 0.920231 | NA |
| | 1.710076 | 0.325008 | 1.719316 | 0.189033 | 0.850067 | NA |
| | 0.162376 | 0.424913 | 4.079006 | 0.104171 | 0.917034 | NA |
| | 0.192403 | 0.975392 | 4.060716 | 0.240202 | 0.810174 | NA |
| | 0.151697 | -1.546 | 4.075469 | -0.37934 | 0.704433 | NA |
| | 0.489058 | -0.58569 | 3.427543 | -0.17088 | 0.864321 | NA |
| | 0.802985 | 0.54709 | 2.208262 | 0.247747 | 0.80433 | NA |

| | | | | | | |
|---|---|---|---|---|---|---|
| SCGB1B2P | 2.640025 | -0.78976 | 1.201548 | -0.65728 | 0.510998 | NA |
| | 0.980928 | 1.403548 | 2.471832 | 0.567817 | 0.570159 | NA |
| | 0.247898 | -0.5901 | 4.074633 | -0.14482 | 0.884851 | NA |
| | 17.36913 | -0.60087 | 0.519539 | -1.15655 | 0.247457 | NA |
| | 1.12045 | -2.80351 | 2.29454 | -1.22182 | 0.221777 | NA |
| | 2.916778 | 0.365893 | 1.286424 | 0.284426 | 0.776084 | NA |
| | 0.163957 | 0.424913 | 4.079006 | 0.104171 | 0.917034 | NA |
| | 6.455038 | 1.650753 | 0.869854 | 1.897735 | 0.057731 | NA |
| | 2.604609 | 0.322508 | 1.355816 | 0.23787 | 0.811982 | NA |
| | 0.872467 | -0.54278 | 2.536752 | -0.21397 | 0.830573 | NA |
| | 2.13647 | -1.01226 | 1.463792 | -0.69153 | 0.48923 | NA |
| | 1.024017 | 0.177191 | 1.894522 | 0.093528 | 0.925484 | NA |
| | 3.847304 | -0.05962 | 1.155266 | -0.05161 | 0.958843 | NA |
| | 0.096202 | 0.408546 | 4.079639 | 0.100143 | 0.920231 | NA |
| | 0.096202 | 0.408546 | 4.079639 | 0.100143 | 0.920231 | NA |
| VN1R82P | 1.880917 | 0.013191 | 1.501705 | 0.008784 | 0.992991 | NA |
| KRT18P40 | 0.162376 | 0.424913 | 4.079006 | 0.104171 | 0.917034 | NA |
| | 4.556986 | -0.9973 | 0.972209 | -1.02581 | 0.30498 | NA |
| | 0.096202 | 0.408546 | 4.079639 | 0.100143 | 0.920231 | NA |
| | 0.260158 | 1.240347 | 4.047124 | 0.306476 | 0.759242 | NA |
| | 65.35623 | -0.70487 | 0.36181 | -1.94819 | 0.051392 | 0.141162 |
| | 0.096202 | 0.408546 | 4.079639 | 0.100143 | 0.920231 | NA |
| CALR3 | 0.096202 | 0.408546 | 4.079639 | 0.100143 | 0.920231 | NA |
| | 1.433526 | 1.662906 | 2.25556 | 0.737247 | 0.460972 | NA |
| | 1.946529 | 0.580056 | 1.523353 | 0.380776 | 0.70337 | NA |
| HNRNPA1P | 0.096202 | 0.408546 | 4.079639 | 0.100143 | 0.920231 | NA |
| | 148.8384 | -0.15707 | 0.250777 | -0.62633 | 0.531098 | 0.675555 |
| | 0.163957 | 0.424913 | 4.079006 | 0.104171 | 0.917034 | NA |
| | 0.096202 | 0.408546 | 4.079639 | 0.100143 | 0.920231 | NA |
| | 2.035535 | 0.108936 | 1.514714 | 0.071918 | 0.942667 | NA |
| | 1.292283 | -0.48427 | 1.713883 | -0.28256 | 0.777517 | NA |
| | 0.134133 | -1.48909 | 4.077584 | -0.36519 | 0.71497 | NA |
| FBXO17 | 366.0585 | -0.47234 | 0.183008 | -2.581 | 0.009851 | 0.046728 |
| | 0.163957 | 0.424913 | 4.079006 | 0.104171 | 0.917034 | NA |
| | 0.247898 | -0.5901 | 4.074633 | -0.14482 | 0.884851 | NA |
| | 5.767615 | 0.534273 | 0.839593 | 0.636348 | 0.52455 | NA |
| | 3.09931 | 1.591108 | 1.214998 | 1.309556 | 0.190346 | NA |
| | 0.394291 | 0.298745 | 3.141711 | 0.09509 | 0.924243 | NA |
| | 0.134133 | -1.48909 | 4.077584 | -0.36519 | 0.71497 | NA |
| | 0.151697 | -1.546 | 4.075469 | -0.37934 | 0.704433 | NA |
| | 2.472105 | -0.40473 | 1.415547 | -0.28592 | 0.77494 | NA |
| | 0.392857 | -1.546 | 4.075469 | -0.37934 | 0.704433 | NA |
| | 0.192403 | 0.975392 | 4.060716 | 0.240202 | 0.810174 | NA |
| ITGB1P1 | 21.13367 | 1.391622 | 0.472599 | 2.944616 | 0.003234 | NA |
| RAB11B-AS | 66.95044 | 0.313139 | 0.326852 | 0.958044 | 0.33804 | 0.506825 |
| | 2.631357 | -0.16789 | 1.252809 | -0.13401 | 0.893397 | NA |
| | 11.76312 | 0.426432 | 0.597901 | 0.713215 | 0.475713 | NA |

| | | | | | | |
|---|---|---|---|---|---|---|
| | 5.357264 | -1.17127 | 1.116132 | -1.0494 | 0.293993 | NA |
| | 0.151697 | -1.546 | 4.075469 | -0.37934 | 0.704433 | NA |
| | 0.303394 | -2.27006 | 4.032451 | -0.56295 | 0.57347 | NA |
| | 1.684994 | -0.19875 | 1.503821 | -0.13217 | 0.894853 | NA |
| | 19.41254 | 0.475435 | 0.532904 | 0.892158 | 0.372308 | NA |
| | 17.05399 | 0.272332 | 0.549136 | 0.495928 | 0.619945 | NA |
| | 0.556667 | 0.972496 | 2.522454 | 0.385536 | 0.699841 | NA |
| | 1.204307 | -1.1284 | 2.006461 | -0.56238 | 0.573855 | NA |
| | 12.91087 | -0.09067 | 0.592552 | -0.15302 | 0.878383 | NA |
| | 3.172629 | -0.68317 | 1.163065 | -0.58739 | 0.556942 | NA |
| | 0.096202 | 0.408546 | 4.079639 | 0.100143 | 0.920231 | NA |
| | 0.650297 | -2.03865 | 2.649751 | -0.76937 | 0.441671 | NA |
| TDGF1P7 | 0.162376 | 0.424913 | 4.079006 | 0.104171 | 0.917034 | NA |
| | 13.79184 | -0.78766 | 0.725959 | -1.085 | 0.277923 | NA |
| | 0.268266 | -2.15396 | 4.038351 | -0.53338 | 0.593773 | NA |
| | 0.814253 | -1.23545 | 2.246777 | -0.54988 | 0.582405 | NA |
| | 3.508187 | 1.077206 | 1.132163 | 0.951458 | 0.341372 | NA |
| | 0.575812 | 0.936922 | 2.776634 | 0.337431 | 0.735792 | NA |
| | 4.990142 | 1.55494 | 0.960961 | 1.618109 | 0.105639 | NA |
| | 8.767932 | -0.76135 | 0.796575 | -0.95578 | 0.339184 | NA |
| | 0.87074 | 0.675988 | 2.012127 | 0.335957 | 0.736903 | NA |
| | 0.956409 | -1.29092 | 2.212928 | -0.58335 | 0.559655 | NA |
| | 0.163957 | 0.424913 | 4.079006 | 0.104171 | 0.917034 | NA |
| | 0.096202 | 0.408546 | 4.079639 | 0.100143 | 0.920231 | NA |
| PLEKHA3P1 | 2.912646 | -0.38294 | 1.188788 | -0.32212 | 0.747358 | NA |
| | 0.392857 | -1.546 | 4.075469 | -0.37934 | 0.704433 | NA |
| | 1.931458 | -0.90132 | 1.377069 | -0.65452 | 0.512777 | NA |
| | 1.768879 | -0.85345 | 1.581407 | -0.53968 | 0.58942 | NA |
| | 64.90373 | -0.42551 | 0.337934 | -1.25916 | 0.207972 | 0.365314 |
| | 7.153534 | 0.025206 | 0.818849 | 0.030782 | 0.975444 | NA |
| | 0.824933 | -0.18668 | 2.233587 | -0.08358 | 0.933393 | NA |
| EGLN2 | 293.6081 | -0.28271 | 0.183353 | -1.5419 | 0.123097 | 0.257058 |
| DPM1 | 683.5853 | 0.662703 | 0.165784 | 3.997397 | 6.40E-05 | 0.001922 |
| SEMA3F | 6.839683 | 0.531363 | 0.818812 | 0.648944 | 0.516375 | NA |
| CYP26B1 | 3.317413 | 0.228639 | 1.083213 | 0.211075 | 0.832829 | NA |
| ALS2 | 1036.35 | 0.124799 | 0.140209 | 0.890095 | 0.373415 | 0.539792 |
| CASP10 | 44.02085 | -0.42194 | 0.37375 | -1.12895 | 0.25892 | NA |
| CFLAR | 532.2376 | -0.00122 | 0.179168 | -0.00682 | 0.99456 | 0.997674 |
| TFPI | 125.8824 | 0.414499 | 0.270042 | 1.534945 | 0.124797 | 0.259602 |
| NDUFAF7 | 391.6278 | 1.00315 | 0.211165 | 4.750542 | 2.03E-06 | 2.51E-04 |
| RBM5 | 1451.444 | 0.280171 | 0.138568 | 2.021901 | 0.043187 | 0.126625 |
| RBM6 | 893.6013 | 0.50127 | 0.157906 | 3.174484 | 0.001501 | 0.013573 |
| ZMYND10 | 32.33838 | -0.30597 | 0.445227 | -0.68722 | 0.491947 | NA |
| GCFC2 | 279.4384 | 0.623558 | 0.213735 | 2.917443 | 0.003529 | 0.023518 |
| WDR54 | 603.9808 | -0.21116 | 0.160133 | -1.31865 | 0.187286 | 0.341474 |
| FARP2 | 1959.656 | -0.54617 | 0.163214 | -3.34636 | 8.19E-04 | 0.009177 |
| UPP2 | 0.314073 | -0.57308 | 3.898934 | -0.14698 | 0.883146 | NA |

| | | | | | | |
|---|---|---|---|---|---|---|
| MARK4 | 1918.521 | -0.29085 | 0.145897 | -1.99352 | 0.046205 | 0.132085 |
| TRAPPC6A | 591.7147 | -0.10252 | 0.193494 | -0.52986 | 0.596208 | 0.726194 |
| CACNA2D2 | 5.701587 | 0.329671 | 0.848434 | 0.388564 | 0.697599 | NA |
| DLEC1 | 2.192461 | 1.275037 | 1.395526 | 0.91366 | 0.360895 | NA |
| CELSR3 | 87.72729 | -0.43028 | 0.285236 | -1.50851 | 0.131424 | 0.268072 |
| SS18L2 | 371.6285 | 0.436748 | 0.210152 | 2.078247 | 0.037687 | 0.115416 |
| PGLYRP1 | 0.298089 | -0.51711 | 3.975232 | -0.13008 | 0.896501 | NA |
| MAPK8IP2 | 349.6312 | -0.74755 | 0.1668 | -4.4817 | 7.41E-06 | 5.34E-04 |
| HEATR5B | 798.0647 | 0.11064 | 0.151579 | 0.729914 | 0.465443 | 0.622857 |
| UQCRC1 | 7034.218 | -0.12806 | 0.145994 | -0.87714 | 0.380413 | 0.546642 |
| HHATL | 0.163957 | 0.424913 | 4.079006 | 0.104171 | 0.917034 | NA |
| GIPR | 59.7092 | 0.561842 | 0.308783 | 1.81954 | 0.068829 | 0.172582 |
| PHF7 | 86.78278 | -0.16151 | 0.296106 | -0.54545 | 0.585441 | 0.718394 |
| SEMA3G | 1.716508 | -0.4344 | 1.679058 | -0.25872 | 0.795853 | NA |
| NISCH | 2021.707 | -0.46639 | 0.168898 | -2.76136 | 0.005756 | 0.032518 |
| STAB1 | 0.610582 | -0.18034 | 2.753139 | -0.0655 | 0.947774 | NA |
| FUZ | 424.5154 | -0.3127 | 0.17176 | -1.82058 | 0.068671 | 0.172333 |
| ABHD5 | 219.1596 | 0.15874 | 0.185585 | 0.855353 | 0.392356 | 0.557385 |
| LARS2 | 991.5442 | -0.32466 | 0.136396 | -2.3803 | 0.017298 | 0.068678 |
| QPCTL | 1791.206 | -0.2303 | 0.14177 | -1.62447 | 0.104276 | 0.229312 |
| PPP5C | 2406.048 | -0.08571 | 0.134728 | -0.63615 | 0.524677 | 0.670408 |
| CEP68 | 464.978 | -0.35871 | 0.184454 | -1.94471 | 0.05181 | 0.141718 |
| MAP4K3 | 1157.186 | 0.936163 | 0.176251 | 5.311542 | 1.09E-07 | 3.68E-05 |
| ERCC1 | 1887.326 | -0.16693 | 0.142743 | -1.16943 | 0.242231 | 0.405185 |
| SEMA3B | 1237.641 | 0.312578 | 0.627707 | 0.497967 | 0.618507 | 0.743274 |
| LTF | 2.373805 | 1.048392 | 1.648031 | 0.636148 | 0.52468 | NA |
| EHD3 | 916.2879 | -0.41356 | 0.157051 | -2.63328 | 0.008456 | 0.042069 |
| CLK1 | 873.9734 | 1.518504 | 0.217776 | 6.972791 | 3.11E-12 | 6.14E-09 |
| MDH1 | 5490.535 | 0.105928 | 0.13885 | 0.762898 | 0.445524 | 0.605351 |
| BID | 733.9275 | 0.458049 | 0.18809 | 2.435264 | 0.014881 | 0.061701 |
| CHDH | 12.00986 | -0.07489 | 0.744959 | -0.10053 | 0.919923 | NA |
| GLT8D1 | 1269.346 | 0.098432 | 0.13448 | 0.731949 | 0.464199 | 0.621537 |
| SLC11A1 | 8.106394 | 0.922808 | 0.774783 | 1.191054 | 0.233632 | NA |
| AGPS | 1431.498 | 0.425557 | 0.248788 | 1.710521 | 0.08717 | 0.202702 |
| ZNF806 | 0.996766 | 1.94108 | 2.070839 | 0.93734 | 0.348584 | NA |
| TTC27 | 437.1012 | -0.11552 | 0.175086 | -0.65978 | 0.509392 | 0.657683 |
| ZNF582 | 63.39683 | 0.602391 | 0.33814 | 1.781482 | 0.074834 | 0.18258 |
| CYP24A1 | 7.431959 | 3.191505 | 1.072623 | 2.97542 | 0.002926 | NA |
| ZFP64 | 577.8389 | -0.35447 | 0.165488 | -2.14199 | 0.032194 | 0.104129 |
| SPAST | 317.8265 | 0.403332 | 0.238019 | 1.694535 | 0.090164 | 0.207996 |
| CPS1 | 1033.705 | -0.17252 | 0.166321 | -1.03727 | 0.299612 | 0.466435 |
| RTFDC1 | 3400.582 | -0.01459 | 0.139946 | -0.10424 | 0.916979 | 0.951908 |
| NLRP2 | 3.646653 | -0.02289 | 1.09334 | -0.02094 | 0.983295 | NA |
| NDUFS1 | 3072.425 | 0.109372 | 0.133707 | 0.817999 | 0.413358 | 0.576581 |
| ALAS1 | 2291.129 | -0.17784 | 0.152074 | -1.16943 | 0.242229 | 0.405185 |
| EHD2 | 3589.114 | -0.53624 | 0.145455 | -3.68663 | 2.27E-04 | 0.004193 |
| PHF20 | 986.1271 | 0.131537 | 0.231526 | 0.568129 | 0.569947 | 0.706055 |

| | | | | | | |
|---|---|---|---|---|---|---|
| TYMP | 381.6914 | -0.06098 | 0.179758 | -0.33925 | 0.734421 | 0.832129 |
| NCAPH2 | 1093.159 | -0.26407 | 0.161818 | -1.6319 | 0.1027 | 0.226934 |
| TOMM34 | 1188.933 | -0.16864 | 0.176466 | -0.95566 | 0.339244 | 0.507923 |
| RTEL1-TNFI | 25.26852 | 0.083656 | 0.419445 | 0.199444 | 0.841916 | NA |
| KCNG1 | 39.50324 | 0.384395 | 0.432962 | 0.887828 | 0.374633 | NA |
| VRK2 | 306.6764 | 0.793897 | 0.251956 | 3.15094 | 0.001627 | 0.014249 |
| IKZF2 | 32.19331 | 0.404545 | 0.50812 | 0.79616 | 0.425939 | NA |
| TSSC1 | 1040.127 | -0.16627 | 0.135237 | -1.22947 | 0.218896 | 0.378502 |
| SLC4A7 | 730.5294 | 0.382645 | 0.309073 | 1.23804 | 0.215701 | 0.374868 |
| TMSB10 | 28503.75 | 0.061711 | 0.1763 | 0.350036 | 0.726312 | 0.826178 |
| SH3YL1 | 227.2554 | 0.709014 | 0.287541 | 2.465781 | 0.013671 | 0.058239 |
| FAM136A | 770.9124 | 0.226572 | 0.156949 | 1.4436 | 0.148852 | 0.292187 |
| CUL3 | 1324.824 | 0.573121 | 0.225491 | 2.541658 | 0.011033 | 0.050236 |
| PNKP | 1292.734 | -0.16217 | 0.157501 | -1.02967 | 0.303166 | 0.47046 |
| RTN4R | 52.30136 | -0.05396 | 0.317794 | -0.16981 | 0.865161 | 0.920317 |
| INPP4A | 788.0935 | -0.07449 | 0.178952 | -0.41628 | 0.677209 | 0.789101 |
| PARP3 | 395.9921 | -0.00499 | 0.189267 | -0.02639 | 0.978948 | 0.988706 |
| FAM65C | 8.378283 | -0.04796 | 0.730026 | -0.0657 | 0.947616 | NA |
| RETSAT | 6267.83 | -0.42355 | 0.140525 | -3.01407 | 0.002578 | 0.019243 |
| CAPG | 5052.373 | -0.14443 | 0.194181 | -0.74379 | 0.457004 | 0.615377 |
| AP2S1 | 5007.673 | -0.21208 | 0.153884 | -1.37817 | 0.168151 | 0.317983 |
| MAP4 | 16914.03 | -0.41347 | 0.16021 | -2.58077 | 0.009858 | 0.046728 |
| R3HDM1 | 604.0996 | 0.096211 | 0.221875 | 0.433628 | 0.664559 | 0.779571 |
| LTBP1 | 10.41132 | -0.24257 | 0.671983 | -0.36098 | 0.718113 | NA |
| NNAT | 5.739124 | 0.075109 | 0.841998 | 0.089204 | 0.92892 | NA |
| LY75 | 1.699972 | 1.5106 | 1.678814 | 0.899802 | 0.368225 | NA |
| PTPRN | 1.791001 | -0.18165 | 1.567091 | -0.11592 | 0.907718 | NA |
| TBC1D22A | 1156.895 | -0.3738 | 0.137657 | -2.71546 | 0.006618 | 0.035745 |
| ATP9A | 2554.915 | -0.57388 | 0.158275 | -3.62586 | 2.88E-04 | 0.004809 |
| CBLN4 | 0.260158 | 1.240347 | 4.047124 | 0.306476 | 0.759242 | NA |
| NOP58 | 507.419 | 0.944882 | 0.334703 | 2.823042 | 0.004757 | 0.028569 |
| EIF2AK2 | 1847.073 | -0.03031 | 0.257693 | -0.1176 | 0.906382 | 0.945876 |
| CCDC85A | 109.032 | 0.707025 | 0.258426 | 2.73589 | 0.006221 | 0.034176 |
| PUM2 | 1863.963 | 0.483651 | 0.195647 | 2.472056 | 0.013434 | 0.05764 |
| ITIH4 | 1.349265 | -1.88672 | 2.030501 | -0.92919 | 0.352791 | NA |
| ITIH1 | 0.151697 | -1.546 | 4.075469 | -0.37934 | 0.704433 | NA |
| PHF21B | 1.133876 | 2.273059 | 2.072112 | 1.096977 | 0.272651 | NA |
| IL17RB | 60.35348 | -0.06495 | 0.28688 | -0.22639 | 0.820898 | 0.892334 |
| MTA3 | 993.6143 | -0.05878 | 0.162161 | -0.36247 | 0.717002 | 0.81952 |
| MXD1 | 234.6403 | 0.193014 | 0.195954 | 0.984998 | 0.324625 | 0.492617 |
| OGFR | 2226.6 | -0.30254 | 0.147715 | -2.04814 | 0.040546 | 0.121382 |
| ACAA1 | 1742.2 | -0.05878 | 0.191265 | -0.30734 | 0.758583 | 0.849439 |
| SPAG4 | 22.4326 | -0.4794 | 0.459497 | -1.04331 | 0.296804 | NA |
| NCKAP1 | 4183.563 | 0.493038 | 0.18078 | 2.727278 | 0.006386 | 0.034791 |
| ZNF112 | 78.87234 | 1.029944 | 0.347018 | 2.967982 | 0.002998 | 0.02114 |
| ELMO2 | 1572.566 | -0.46705 | 0.127316 | -3.66843 | 2.44E-04 | 0.004387 |
| POLD1 | 1034.988 | -0.43239 | 0.172572 | -2.50557 | 0.012225 | 0.054079 |

| | | | | | | |
|---|---|---|---|---|---|---|
| SLC6A16 | 5.531737 | 0.954414 | 0.857282 | 1.113302 | 0.265579 | NA |
| GLTSCR1 | 468.1374 | -0.56488 | 0.200661 | -2.81509 | 0.004876 | 0.029006 |
| SPHK2 | 1167.863 | -0.36482 | 0.170054 | -2.1453 | 0.031929 | 0.103646 |
| RPL18 | 16886.79 | -0.06071 | 0.137721 | -0.4408 | 0.659359 | 0.77539 |
| CA11 | 565.5366 | -0.17847 | 0.183858 | -0.97068 | 0.331706 | 0.500238 |
| ISOC2 | 1811.368 | -0.27741 | 0.137093 | -2.02356 | 0.043016 | 0.126367 |
| U2AF2 | 6038.568 | -0.35235 | 0.153929 | -2.28906 | 0.022076 | 0.081 |
| EPN1 | 7181.13 | -0.33721 | 0.178375 | -1.89047 | 0.058695 | 0.154802 |
| GSC2 | 0.134133 | -1.48909 | 4.077584 | -0.36519 | 0.71497 | NA |
| GPC1 | 2646.143 | -0.22758 | 0.165824 | -1.37239 | 0.169941 | 0.320363 |
| CASP8 | 303.2704 | -0.14285 | 0.185757 | -0.769 | 0.441895 | 0.601869 |
| WISP2 | 563.1183 | -0.05461 | 0.765165 | -0.07138 | 0.943099 | 0.968558 |
| CTSA | 13225.6 | -0.2925 | 0.154591 | -1.89211 | 0.058477 | 0.154396 |
| EYA2 | 0.594802 | -1.06492 | 3.125073 | -0.34077 | 0.73328 | NA |
| BCAS1 | 50.91668 | 0.745199 | 0.599675 | 1.242671 | 0.213989 | 0.37274 |
| PMS1 | 228.0468 | 0.921688 | 0.301604 | 3.055951 | 0.002243 | 0.017406 |
| CALCRL | 30.36395 | 0.65725 | 0.391132 | 1.680379 | 0.092884 | NA |
| ANKRD44 | 482.9763 | 0.461308 | 0.177866 | 2.593574 | 0.009498 | 0.04557 |
| ZC3H15 | 890.2989 | 1.041185 | 0.252379 | 4.125473 | 3.70E-05 | 0.001391 |
| ASB1 | 1034.96 | -0.18477 | 0.152473 | -1.21184 | 0.225574 | 0.386778 |
| MTHFD2 | 587.5073 | 0.274867 | 0.153849 | 1.786599 | 0.074002 | 0.181485 |
| CTNNA2 | 62.4348 | 0.198523 | 0.338314 | 0.586802 | 0.557337 | 0.696283 |
| NGEF | 0.230334 | -0.53318 | 4.076749 | -0.13079 | 0.895945 | NA |
| SP100 | 2023.702 | 0.576229 | 0.229886 | 2.506579 | 0.012191 | 0.054008 |
| RHOA | 19397.94 | -0.20116 | 0.145983 | -1.37797 | 0.168212 | 0.318046 |
| HYAL2 | 1096.616 | -0.22559 | 0.139998 | -1.61141 | 0.10709 | 0.233566 |
| HDAC4 | 1034.34 | -0.31346 | 0.154415 | -2.03 | 0.042356 | 0.125189 |
| RASSF1 | 2146.86 | 0.042501 | 0.204431 | 0.207898 | 0.835309 | 0.901464 |
| REEP1 | 1.6765 | 1.961523 | 1.654439 | 1.185612 | 0.235775 | NA |
| POLR1A | 2988.509 | -0.61956 | 0.16811 | -3.68544 | 2.28E-04 | 0.004196 |
| LAPTM4A | 9609.698 | -0.05224 | 0.135162 | -0.38652 | 0.699112 | 0.806443 |
| TTC7A | 305.5758 | -0.10057 | 0.190236 | -0.52868 | 0.597025 | 0.726799 |
| IP6K2 | 1262.312 | 0.162353 | 0.138187 | 1.174878 | 0.240044 | 0.403304 |
| STON1-GTI | 0.296509 | -0.51709 | 3.982155 | -0.12985 | 0.896684 | NA |
| SRBD1 | 477.898 | 0.301636 | 0.1575 | 1.915155 | 0.055473 | 0.148819 |
| PSME4 | 1528.075 | 0.142446 | 0.151947 | 0.937469 | 0.348517 | 0.516513 |
| ERLEC1 | 1474.623 | 0.487238 | 0.147247 | 3.308971 | 9.36E-04 | 0.009929 |
| BCL3 | 120.2924 | -0.85179 | 0.27066 | -3.14708 | 0.001649 | 0.014322 |
| CECR5 | 1315.993 | -0.17393 | 0.168348 | -1.03315 | 0.301533 | 0.468442 |
| UFD1L | 2204.537 | 0.208834 | 0.163812 | 1.274843 | 0.202365 | 0.358744 |
| CLTCL1 | 256.728 | -0.37969 | 0.180435 | -2.10431 | 0.035351 | 0.110724 |
| DGCR2 | 2466.303 | -0.40228 | 0.132606 | -3.03365 | 0.002416 | 0.018279 |
| RAD18 | 443.3956 | 0.691333 | 0.201774 | 3.426265 | 6.12E-04 | 0.007685 |
| NCK2 | 3396.681 | 0.002089 | 0.156893 | 0.013317 | 0.989375 | 0.994829 |
| MAP4K4 | 4201.226 | 0.110464 | 0.13098 | 0.843372 | 0.39902 | 0.563807 |
| MGAT4A | 108.4141 | 0.457094 | 0.309998 | 1.474507 | 0.140345 | 0.280568 |
| RPL31 | 17300.95 | 0.448776 | 0.16114 | 2.785015 | 0.005353 | 0.031044 |

| | | | | | | |
|---|---|---|---|---|---|---|
| LMCD1 | 569.1795 | -0.12878 | 0.154957 | -0.83108 | 0.405928 | 0.56977 |
| TRIB2 | 5631.448 | -0.35692 | 0.171279 | -2.08387 | 0.037172 | 0.11443 |
| MYO3B | 20.11441 | 0.14271 | 0.490772 | 0.290787 | 0.771214 | NA |
| CYBRD1 | 8159.668 | 0.442613 | 0.190493 | 2.323514 | 0.020152 | 0.07613 |
| PTPN18 | 1586.827 | -0.28478 | 0.13836 | -2.05825 | 0.039566 | 0.119444 |
| LIMS2 | 1360.444 | -0.25208 | 0.158637 | -1.58902 | 0.112056 | 0.240482 |
| ASIC4 | 1.189169 | 2.260443 | 1.954925 | 1.156281 | 0.247566 | NA |
| SPEG | 34.48522 | 0.414857 | 0.414693 | 1.000396 | 0.317119 | NA |
| TRNT1 | 285.2607 | 0.70532 | 0.189158 | 3.728731 | 1.92E-04 | 0.003796 |
| PVR | 1026.964 | -0.20616 | 0.136779 | -1.50726 | 0.131743 | 0.268537 |
| PANX2 | 182.3974 | 0.152792 | 0.264357 | 0.577976 | 0.56328 | 0.701302 |
| | 875.1048 | -0.28932 | 0.158034 | -1.83072 | 0.067142 | 0.169494 |
| GLI2 | 6.286122 | -0.29733 | 0.900236 | -0.33028 | 0.741188 | NA |
| CLASP1 | 3043.796 | -0.18553 | 0.137103 | -1.35322 | 0.175984 | 0.328121 |
| TEAD2 | 1253.621 | -0.18171 | 0.16249 | -1.11826 | 0.263455 | 0.427785 |
| BCS1L | 813.2662 | -0.23722 | 0.135091 | -1.75599 | 0.07909 | 0.1898 |
| GTSE1 | 754.5004 | -0.50089 | 0.266622 | -1.87866 | 0.060291 | 0.157746 |
| TTC38 | 1065.602 | -0.31965 | 0.157892 | -2.0245 | 0.042919 | 0.12616 |
| GRAMD4 | 1384.207 | -0.33308 | 0.20669 | -1.61147 | 0.107077 | 0.233566 |
| CELSR1 | 409.08 | -0.831 | 0.266661 | -3.11631 | 0.001831 | 0.015235 |
| ZNF638 | 987.2873 | 0.74611 | 0.322783 | 2.311488 | 0.020806 | 0.077845 |
| ADD2 | 0.662557 | -0.83483 | 2.671964 | -0.31244 | 0.754706 | NA |
| FOSL2 | 1415.498 | -0.3206 | 0.194865 | -1.64525 | 0.099919 | 0.223198 |
| TMEM131 | 1157.589 | -0.1234 | 0.188061 | -0.65614 | 0.511731 | 0.659436 |
| ARHGAP15 | 146.4261 | 0.490722 | 0.279879 | 1.753339 | 0.079544 | 0.190609 |
| TUBA3D | 2.2421 | -1.81358 | 1.684218 | -1.07681 | 0.281565 | NA |
| EXOSC7 | 737.4966 | -0.01357 | 0.139143 | -0.09751 | 0.922325 | 0.955032 |
| MKRN2 | 1164.639 | 0.010645 | 0.139246 | 0.076448 | 0.939062 | 0.966202 |
| MCM6 | 785.9761 | 0.105952 | 0.229971 | 0.46072 | 0.645 | 0.764112 |
| PTPN23 | 2466.046 | -0.68439 | 0.152781 | -4.47952 | 7.48E-06 | 5.34E-04 |
| MLH1 | 1190.152 | -0.1049 | 0.137324 | -0.76391 | 0.444922 | 0.60481 |
| DGKD | 534.431 | -0.32556 | 0.203032 | -1.60347 | 0.10883 | 0.235864 |
| RARB | 384.7089 | 0.314562 | 0.271063 | 1.160475 | 0.245855 | 0.409012 |
| TOP2B | 1746.349 | 0.56309 | 0.575718 | 0.978065 | 0.328042 | 0.496228 |
| DNAJC10 | 1696.429 | 0.730184 | 0.23866 | 3.059514 | 0.002217 | 0.017291 |
| DYNC1I2 | 2099.082 | 0.390409 | 0.163661 | 2.385482 | 0.017057 | 0.067932 |
| SMC1B | 0.533728 | -1.74213 | 2.820432 | -0.61768 | 0.536785 | NA |
| FBLN1 | 146.0093 | 0.082512 | 0.306908 | 0.268849 | 0.788046 | 0.871362 |
| CST7 | 0.163957 | 0.424913 | 4.079006 | 0.104171 | 0.917034 | NA |
| MAP2 | 1059.652 | 0.428453 | 0.254251 | 1.685158 | 0.091958 | 0.210878 |
| FAP | 912.2307 | 0.999529 | 0.192437 | 5.194059 | 2.06E-07 | 5.27E-05 |
| CBFA2T2 | 534.9899 | -0.55623 | 0.170549 | -3.26144 | 0.001108 | 0.01108 |
| ITCH | 1330.246 | 0.43658 | 0.170164 | 2.565638 | 0.010299 | 0.048042 |
| TP53INP2 | 2556.13 | -0.24164 | 0.183453 | -1.3172 | 0.187772 | 0.341836 |
| MYH7B | 6.412846 | -0.16944 | 0.858777 | -0.19731 | 0.843587 | NA |
| FKBP7 | 1020.106 | 0.428459 | 0.167031 | 2.565145 | 0.010313 | 0.048042 |
| OSBPL6 | 254.5149 | -0.11563 | 0.211923 | -0.54561 | 0.585336 | 0.718394 |

| | | | | | | |
|---|---|---|---|---|---|---|
| XRCC5 | 9544.016 | 0.108512 | 0.13881 | 0.78173 | 0.434373 | 0.595539 |
| SP140 | 0.739585 | 2.778432 | 2.852922 | 0.97389 | 0.330111 | NA |
| TNS1 | 6748.315 | -1.14915 | 0.256519 | -4.47977 | 7.47E-06 | 5.34E-04 |
| DDX1 | 2779.503 | 0.464391 | 0.157569 | 2.947228 | 0.003206 | 0.0221 |
| RABL2B | 137.9196 | 0.041032 | 0.214738 | 0.191078 | 0.848464 | 0.909699 |
| PTPRH | 338.3053 | -0.28267 | 0.201695 | -1.40145 | 0.161078 | 0.308595 |
| SLC35C2 | 1569.704 | -0.19592 | 0.138121 | -1.41846 | 0.156058 | 0.301904 |
| RIF1 | 493.3022 | 0.944425 | 0.771068 | 1.224826 | 0.220641 | 0.38052 |
| RBL1 | 275.6138 | 0.595341 | 0.226676 | 2.626395 | 0.008629 | 0.042733 |
| DLGAP4 | 2165.55 | -0.16526 | 0.206098 | -0.80184 | 0.422644 | 0.58499 |
| COL4A4 | 3.672979 | 1.928605 | 1.26448 | 1.525215 | 0.127205 | NA |
| STK17B | 597.8744 | 0.620858 | 0.197239 | 3.147744 | 0.001645 | 0.014318 |
| LRP2 | 21.96771 | -0.95508 | 0.595465 | -1.60392 | 0.108732 | NA |
| MPP4 | 21.94812 | -0.25916 | 0.489652 | -0.52928 | 0.596613 | NA |
| STRADB | 1494.868 | -0.18343 | 0.129842 | -1.41274 | 0.157734 | 0.304153 |
| BZW1 | 1420.73 | 0.399004 | 0.204006 | 1.955849 | 0.050483 | 0.139472 |
| CCNT2 | 484.9817 | 0.973775 | 0.208047 | 4.680555 | 2.86E-06 | 2.97E-04 |
| COBLL1 | 529.1731 | 0.418524 | 0.276913 | 1.511389 | 0.130689 | 0.267092 |
| XPO1 | 3908.567 | 0.514444 | 0.186676 | 2.755811 | 0.005855 | 0.032848 |
| SLC27A5 | 167.3652 | -0.36415 | 0.244531 | -1.48919 | 0.136437 | 0.275453 |
| ZNF324 | 308.0152 | -0.3294 | 0.190609 | -1.72812 | 0.083966 | 0.197425 |
| ZNF671 | 238.6439 | -0.12741 | 0.182735 | -0.69724 | 0.485651 | 0.639244 |
| ZNF416 | 150.7913 | -0.23211 | 0.212161 | -1.09402 | 0.273946 | 0.439786 |
| ZNF586 | 148.8713 | 0.043151 | 0.208543 | 0.206916 | 0.836075 | 0.901643 |
| ZNF446 | 136.1291 | -0.21779 | 0.226995 | -0.95943 | 0.337342 | 0.506162 |
| | 10.60205 | -0.5146 | 0.638857 | -0.80549 | 0.420534 | NA |
| ZNF264 | 131.2006 | -0.51159 | 0.268639 | -1.90438 | 0.05686 | 0.151138 |
| RPS5 | 17049.43 | -0.00637 | 0.147921 | -0.04308 | 0.965638 | 0.981198 |
| STARD7 | 5312.181 | -0.40166 | 0.142654 | -2.81564 | 0.004868 | 0.028977 |
| NCOA1 | 732.3359 | -0.03216 | 0.177237 | -0.18146 | 0.856006 | 0.914395 |
| AGBL5 | 1343.415 | -0.20027 | 0.171131 | -1.17026 | 0.241895 | 0.404924 |
| EFR3B | 603.684 | -1.08932 | 0.217797 | -5.00155 | 5.69E-07 | 1.12E-04 |
| KIF3C | 3721.713 | -0.39054 | 0.175312 | -2.22769 | 0.025901 | 0.090178 |
| RAB10 | 3318.361 | 0.142243 | 0.181436 | 0.783984 | 0.43305 | 0.59425 |
| HADHA | 12600.94 | -0.16082 | 0.129741 | -1.23953 | 0.21515 | 0.374251 |
| MAPRE3 | 766.2214 | -0.26311 | 0.162444 | -1.61971 | 0.105294 | 0.231073 |
| CAD | 2265.767 | -0.67549 | 0.172953 | -3.9056 | 9.40E-05 | 0.002438 |
| WDFY1 | 5490.399 | 0.101033 | 0.241936 | 0.417604 | 0.676237 | 0.788433 |
| MTIF2 | 614.5464 | 0.598059 | 0.199864 | 2.992325 | 0.002769 | 0.020111 |
| ATG16L1 | 888.2918 | -0.03339 | 0.155339 | -0.21495 | 0.829805 | 0.897322 |
| USP40 | 777.8736 | -0.10272 | 0.190225 | -0.53999 | 0.589207 | 0.721298 |
| PPP1R15A | 5915.513 | 0.032916 | 0.181103 | 0.181754 | 0.855776 | 0.914347 |
| HSD17B14 | 532.9394 | -0.10635 | 0.158268 | -0.67197 | 0.501603 | 0.652172 |
| FTL | 50435.1 | -0.00278 | 0.150982 | -0.01839 | 0.985326 | 0.992273 |
| BAX | 4182.086 | -0.23158 | 0.165179 | -1.40197 | 0.160924 | 0.308498 |
| GMCL1 | 602.7353 | 0.289083 | 0.151389 | 1.909539 | 0.056193 | 0.150029 |
| GNAS | 30789.78 | 0.04693 | 0.146539 | 0.320257 | 0.748773 | 0.842272 |

| | | | | | | |
|---|---|---|---|---|---|---|
| PHACTR3 | 20.49229 | 0.809677 | 0.592772 | 1.365918 | 0.171965 | NA |
| TFAP2C | 766.5722 | 0.331396 | 0.212932 | 1.556348 | 0.119625 | 0.251537 |
| AURKA | 1035.993 | -0.0187 | 0.232146 | -0.08056 | 0.935793 | 0.964049 |
| CASS4 | 0.096202 | 0.408546 | 4.079639 | 0.100143 | 0.920231 | NA |
| SULT2B1 | 9.14016 | -0.26618 | 0.760316 | -0.3501 | 0.726266 | NA |
| CNOT3 | 1104.895 | -0.39823 | 0.158361 | -2.5147 | 0.011914 | 0.053101 |
| GP6 | 0.303394 | -2.27006 | 4.032451 | -0.56295 | 0.57347 | NA |
| PTPN4 | 296.0567 | 0.773673 | 0.199856 | 3.871159 | 1.08E-04 | 0.002703 |
| DDX18 | 1216.103 | 0.752654 | 0.191691 | 3.926391 | 8.62E-05 | 0.002334 |
| EDEM2 | 1247.996 | -0.26084 | 0.154175 | -1.69185 | 0.090674 | 0.208808 |
| DNMT3B | 55.90225 | -0.17943 | 0.343746 | -0.52198 | 0.601682 | 0.730909 |
| REM1 | 2.275383 | 0.887859 | 1.613783 | 0.550173 | 0.582201 | NA |
| TPX2 | 2369.1 | -0.29209 | 0.18701 | -1.56191 | 0.11831 | 0.249699 |
| FER1L4 | 2.403594 | 1.045512 | 1.350445 | 0.774198 | 0.438814 | NA |
| PDRG1 | 549.8635 | 0.135543 | 0.195192 | 0.69441 | 0.487425 | 0.64041 |
| EPB41L1 | 330.7488 | 0.005997 | 0.177878 | 0.033712 | 0.973107 | 0.985082 |
| DOCK3 | 9.731736 | 0.678875 | 0.6911 | 0.982311 | 0.325947 | NA |
| C3orf18 | 656.4549 | -0.33768 | 0.152305 | -2.21715 | 0.026613 | 0.091632 |
| TMEM40 | 0.661122 | -3.06315 | 2.98049 | -1.02773 | 0.304076 | NA |
| KIF9 | 97.4837 | -0.17496 | 0.241649 | -0.72401 | 0.469062 | 0.625504 |
| CRLS1 | 647.8311 | 0.162017 | 0.146188 | 1.108277 | 0.267742 | 0.432672 |
| ATRN | 2699.093 | -0.21643 | 0.13797 | -1.56865 | 0.11673 | 0.247157 |
| SMOX | 395.1699 | -0.33291 | 0.156331 | -2.1295 | 0.033213 | 0.106157 |
| SIGLEC1 | 0.134133 | -1.48909 | 4.077584 | -0.36519 | 0.71497 | NA |
| FKBP1A | 16466.71 | -0.07433 | 0.179504 | -0.41409 | 0.678808 | 0.790421 |
| NSFL1C | 1396.15 | -0.19558 | 0.139363 | -1.40337 | 0.160507 | 0.308029 |
| SLC4A11 | 379.0373 | -0.05217 | 0.276269 | -0.18885 | 0.85021 | 0.910747 |
| C20orf194 | 769.1359 | -0.19956 | 0.187434 | -1.06468 | 0.287019 | 0.453378 |
| ZNF343 | 177.5546 | 0.19466 | 0.191311 | 1.017504 | 0.308914 | 0.476225 |
| EBF4 | 49.74195 | 0.379456 | 0.380703 | 0.996724 | 0.318898 | 0.486797 |
| CPXM1 | 1.42761 | 0.468916 | 1.645385 | 0.284989 | 0.775653 | NA |
| MAVS | 2204.538 | -0.56846 | 0.164082 | -3.4645 | 5.31E-04 | 0.007022 |
| | 374.7149 | -0.1982 | 0.177096 | -1.11918 | 0.263063 | 0.4275 |
| XRN2 | 2865.68 | 0.269096 | 0.143015 | 1.881592 | 0.059891 | 0.157094 |
| PLK1S1 | 204.8344 | 0.624907 | 0.243629 | 2.564999 | 0.010318 | 0.048042 |
| SNX5 | 1759.181 | 0.296348 | 0.140868 | 2.103722 | 0.035403 | 0.110826 |
| ESF1 | 239.0991 | 1.01507 | 0.985353 | 1.030158 | 0.302936 | 0.470374 |
| RBBP9 | 657.4135 | -0.13572 | 0.150821 | -0.89985 | 0.368198 | 0.535318 |
| SLC23A2 | 7032.093 | -0.17253 | 0.178118 | -0.96864 | 0.332725 | 0.50106 |
| TMEM230 | 4067.859 | -0.09498 | 0.131742 | -0.72095 | 0.47094 | 0.627233 |
| DZANK1 | 53.80778 | 0.040034 | 0.316408 | 0.126526 | 0.899315 | 0.940983 |
| C20orf26 | 673.2339 | -0.79105 | 0.263109 | -3.00655 | 0.002642 | 0.019492 |
| TASP1 | 329.4156 | 0.515435 | 0.187706 | 2.745967 | 0.006033 | 0.03358 |
| KIF16B | 647.6182 | 0.396533 | 0.214177 | 1.851425 | 0.064108 | 0.164717 |
| TRMT6 | 390.5727 | 0.540823 | 0.185643 | 2.913237 | 0.003577 | 0.02377 |
| CHGB | 0.652666 | 2.537288 | 2.975981 | 0.852589 | 0.393887 | NA |
| PCBP4 | 1996.344 | -0.15124 | 0.235413 | -0.64246 | 0.520576 | 0.667614 |

| | | | | | | |
|---|---|---|---|---|---|---|
| STRN4 | 3232.433 | -0.4152 | 0.137038 | -3.02985 | 0.002447 | 0.018487 |
| FLT3LG | 97.34813 | -0.00128 | 0.280394 | -0.00455 | 0.996366 | 0.998387 |
| NAT14 | 2071.426 | -0.1566 | 0.186848 | -0.83809 | 0.401979 | 0.566508 |
| NLRC4 | 4.34678 | 1.079787 | 1.068213 | 1.010835 | 0.312095 | NA |
| CMTM6 | 1294.651 | 0.212322 | 0.181968 | 1.166807 | 0.243288 | 0.406166 |
| ITGA6 | 2570.712 | 0.315964 | 0.184639 | 1.711254 | 0.087034 | 0.20251 |
| RAPGEF4 | 51.11787 | 0.591355 | 0.327205 | 1.807291 | 0.070717 | 0.17579 |
| | 777.0765 | 0.445395 | 0.160691 | 2.771754 | 0.005576 | 0.031939 |
| DAZL | 0.096202 | 0.408546 | 4.079639 | 0.100143 | 0.920231 | NA |
| COL9A3 | 1483.783 | 0.88294 | 0.353822 | 2.495434 | 0.01258 | 0.055074 |
| NUP50 | 936.5829 | 0.471853 | 0.16601 | 2.842325 | 0.004479 | 0.027376 |
| CDC45 | 282.0384 | -0.15125 | 0.242969 | -0.62251 | 0.533607 | 0.677477 |
| COMT | 4876.74 | -0.17694 | 0.14031 | -1.26107 | 0.207284 | 0.364376 |
| CECR1 | 1059.876 | -0.75896 | 0.171688 | -4.42059 | 9.84E-06 | 6.26E-04 |
| | 5.364803 | 0.666836 | 0.881766 | 0.75625 | 0.449499 | NA |
| LRRFIP2 | 1906.72 | 0.696352 | 0.161498 | 4.311824 | 1.62E-05 | 8.27E-04 |
| SEC22C | 2056.947 | 0.028241 | 0.143981 | 0.196142 | 0.844499 | 0.907349 |
| XYLB | 266.1104 | -0.10003 | 0.254076 | -0.3937 | 0.693805 | 0.802601 |
| MSH2 | 411.7459 | 0.411921 | 0.271888 | 1.515038 | 0.129763 | 0.266024 |
| IL11 | 7.048518 | 1.175 | 0.80953 | 1.45146 | 0.146652 | NA |
| MZF1 | 165.4147 | -0.18895 | 0.277786 | -0.68021 | 0.496374 | 0.647636 |
| ARVCF | 183.0082 | -0.18551 | 0.254495 | -0.72892 | 0.46605 | 0.623178 |
| TRMT2A | 1121.899 | -0.07598 | 0.137489 | -0.55265 | 0.580501 | 0.714368 |
| RANBP1 | 2378.094 | -0.06821 | 0.187379 | -0.36402 | 0.715844 | 0.81867 |
| ZDHHC8 | 1373.656 | -0.38694 | 0.13129 | -2.94723 | 0.003206 | 0.0221 |
| KLHL22 | 699.2547 | -0.45404 | 0.158128 | -2.87134 | 0.004087 | 0.025773 |
| MED15 | 5692.999 | -0.41281 | 0.18356 | -2.24889 | 0.02452 | 0.086719 |
| SERPIND1 | 0.816972 | 0.936457 | 2.589651 | 0.361615 | 0.71764 | NA |
| SNAP29 | 1951.791 | -0.09782 | 0.132924 | -0.73592 | 0.46178 | 0.620048 |
| CRKL | 1814.286 | -0.30224 | 0.171361 | -1.76375 | 0.077774 | 0.187707 |
| LZTR1 | 2811.641 | -0.23262 | 0.125877 | -1.84797 | 0.064607 | 0.165674 |
| MMP11 | 55.17287 | 0.140821 | 0.353814 | 0.398008 | 0.690625 | 0.799546 |
| CECR2 | 57.99951 | -0.64021 | 0.322546 | -1.98486 | 0.04716 | 0.133679 |
| SMARCB1 | 2626.646 | -0.28381 | 0.120488 | -2.35553 | 0.018496 | 0.071682 |
| P2RX6 | 132.1389 | -0.13793 | 0.224674 | -0.6139 | 0.53928 | 0.681722 |
| DERL3 | 22.4545 | -0.09899 | 0.432088 | -0.22911 | 0.818787 | NA |
| SLC7A4 | 177.6395 | -0.20388 | 0.218417 | -0.93345 | 0.350589 | 0.518492 |
| BCL2L13 | 3053.455 | -0.15504 | 0.149437 | -1.03747 | 0.299518 | 0.466411 |
| DDTL | 23.63038 | 0.318821 | 0.446014 | 0.714823 | 0.474719 | NA |
| DDT | 1.147231 | -0.4269 | 2.00528 | -0.21289 | 0.831415 | NA |
| GSTT2 | 42.44455 | 0.322327 | 0.338696 | 0.95167 | 0.341264 | NA |
| CABIN1 | 2090.309 | -0.72463 | 0.166209 | -4.35971 | 1.30E-05 | 7.21E-04 |
| TBC1D10A | 635.7927 | -0.26492 | 0.149775 | -1.76878 | 0.07693 | 0.186347 |
| SUSD2 | 45.42746 | -0.64775 | 0.39129 | -1.65541 | 0.09784 | NA |
| SF3A1 | 3452.411 | -0.57786 | 0.144203 | -4.00723 | 6.14E-05 | 0.001863 |
| RNF215 | 154.5786 | -0.07161 | 0.211968 | -0.33781 | 0.735503 | 0.832559 |
| SEC14L2 | 273.4851 | -0.01965 | 0.247805 | -0.07928 | 0.936809 | 0.964637 |

| | | | | | | |
|---|---|---|---|---|---|---|
| SPECC1L | 786.5571 | 0.100944 | 0.147053 | 0.686441 | 0.492435 | 0.643732 |
| PPIL2 | 1040.232 | -0.43394 | 0.144073 | -3.01192 | 0.002596 | 0.019343 |
| UPB1 | 0.096202 | 0.408546 | 4.079639 | 0.100143 | 0.920231 | NA |
| YPEL1 | 58.80584 | -0.14041 | 0.362659 | -0.38718 | 0.698626 | 0.806212 |
| SNRPD3 | 1482.703 | 0.01907 | 0.152905 | 0.124715 | 0.900749 | 0.941822 |
| PES1 | 2554.931 | -0.2916 | 0.147749 | -1.97362 | 0.048425 | 0.135798 |
| MAPK1 | 6941.204 | 0.040515 | 0.113553 | 0.356795 | 0.721245 | 0.822641 |
| GGT1 | 83.49724 | -0.5492 | 0.271637 | -2.02182 | 0.043195 | 0.126625 |
| PRODH | 11.1522 | 0.062772 | 0.740137 | 0.084811 | 0.932412 | NA |
| PPM1F | 2069.246 | -0.49402 | 0.143647 | -3.4391 | 5.84E-04 | 0.007456 |
| SLC35E4 | 319.7166 | -0.07021 | 0.224561 | -0.31267 | 0.754528 | 0.846873 |
| TOP3B | 462.016 | -0.24344 | 0.185583 | -1.31176 | 0.1896 | 0.343844 |
| CYTH4 | 0.134133 | -1.48909 | 4.077584 | -0.36519 | 0.71497 | NA |
| DGCR14 | 755.8093 | -0.31937 | 0.152683 | -2.09173 | 0.036463 | 0.113084 |
| CRYBB2P1 | 550.7038 | -0.01636 | 0.14606 | -0.11204 | 0.910789 | 0.948139 |
| CARD10 | 37.04387 | -0.58163 | 0.473617 | -1.22807 | 0.219421 | NA |
| LRP5L | 23.29146 | -0.11624 | 0.435361 | -0.26701 | 0.789463 | NA |
| SLC25A1 | 4576.228 | -0.32259 | 0.132668 | -2.43154 | 0.015035 | 0.062124 |
| ADRBK2 | 219.6128 | -0.13677 | 0.23977 | -0.57043 | 0.568389 | 0.705133 |
| LGALS2 | 7.476712 | 0.61834 | 0.835061 | 0.740473 | 0.459013 | NA |
| GGA1 | 2007.339 | -0.44288 | 0.153894 | -2.87786 | 0.004004 | 0.025436 |
| HIRA | 295.9834 | -0.16346 | 0.183549 | -0.89056 | 0.373168 | 0.539764 |
| SH3BP1 | 383.4107 | -0.22836 | 0.163199 | -1.39924 | 0.161741 | 0.309415 |
| SEZ6L | 0.134133 | -1.48909 | 4.077584 | -0.36519 | 0.71497 | NA |
| LGALS1 | 42924.75 | -0.13236 | 0.154699 | -0.85561 | 0.392214 | 0.557321 |
| HPS4 | 5023.625 | -0.32242 | 0.254005 | -1.26933 | 0.204323 | 0.360991 |
| PIK3IP1 | 1019.414 | -0.1394 | 0.160408 | -0.86904 | 0.384827 | 0.550782 |
| SRRD | 517.2346 | 0.027158 | 0.187015 | 0.14522 | 0.884537 | 0.931609 |
| PATZ1 | 1088.681 | -0.69897 | 0.14528 | -4.81116 | 1.50E-06 | 2.07E-04 |
| TRIOBP | 552.7021 | -0.27997 | 0.175607 | -1.59429 | 0.110871 | 0.23892 |
| TFIP11 | 1939.047 | -0.11787 | 0.194649 | -0.60557 | 0.544799 | 0.686428 |
| GCAT | 307.4934 | -0.48409 | 0.212488 | -2.27818 | 0.022716 | 0.082378 |
| CRYBB1 | 0.437526 | -2.68242 | 3.384147 | -0.79264 | 0.427986 | NA |
| ANKRD54 | 1069.124 | -0.39758 | 0.18137 | -2.19211 | 0.028371 | 0.09539 |
| EIF3L | 8518.491 | -0.17518 | 0.140085 | -1.25054 | 0.211104 | 0.36923 |
| NHP2L1 | 4552.117 | -0.17338 | 0.146942 | -1.17993 | 0.238029 | 0.401057 |
| MICALL1 | 2805.547 | -0.3783 | 0.211077 | -1.79226 | 0.073091 | 0.180143 |
| POLR2F | 1990.899 | 0.105178 | 0.158755 | 0.662521 | 0.507637 | 0.656489 |
| SOX10 | 23924.6 | -0.41861 | 0.142254 | -2.9427 | 0.003254 | 0.022326 |
| CCDC134 | 164.126 | -0.64331 | 0.262395 | -2.45168 | 0.014219 | 0.059882 |
| DEPDC5 | 435.9137 | -0.26974 | 0.160378 | -1.68193 | 0.092582 | 0.212008 |
| PICK1 | 743.9582 | -0.1733 | 0.14114 | -1.22789 | 0.219488 | 0.379195 |
| TTC28 | 408.2836 | -0.60662 | 0.247605 | -2.44997 | 0.014287 | 0.060039 |
| SLC16A8 | 10.57772 | 0.580784 | 0.653007 | 0.889399 | 0.373789 | NA |
| CENPM | 406.6 | -0.39266 | 0.23702 | -1.65664 | 0.097593 | 0.219455 |
| 3-Sep | 386.3211 | -0.9504 | 0.221626 | -4.28829 | 1.80E-05 | 8.68E-04 |
| SLC5A1 | 0.192403 | 0.975392 | 4.060716 | 0.240202 | 0.810174 | NA |

| | | | | | | |
|---|---|---|---|---|---|---|
| TPTEP1 | 1.870833 | 1.056985 | 1.563681 | 0.67596 | 0.499066 | NA |
| SLC5A4 | 80.8712 | -0.3588 | 0.431409 | -0.8317 | 0.405577 | 0.56959 |
| KDELR3 | 1088.705 | 0.138859 | 0.195011 | 0.712055 | 0.476431 | 0.631494 |
| CYP2D6 | 2.606992 | -0.351 | 1.300043 | -0.26999 | 0.787167 | NA |
| DDX17 | 6647.668 | 0.15126 | 0.165946 | 0.9115 | 0.362032 | 0.529079 |
| DMC1 | 6.845703 | 0.017809 | 0.893584 | 0.01993 | 0.9841 | NA |
| TCF20 | 997.5049 | -0.23492 | 0.175135 | -1.34135 | 0.179808 | 0.332292 |
| HSCB | 156.3769 | 0.432222 | 0.222449 | 1.943021 | 0.052014 | 0.142177 |
| CBY1 | 345.523 | -0.1208 | 0.163303 | -0.73972 | 0.459471 | 0.617857 |
| TOMM22 | 1981.394 | -0.19736 | 0.197876 | -0.99738 | 0.318581 | 0.486385 |
| RTDR1 | 0.526843 | -0.38719 | 2.855892 | -0.13558 | 0.892156 | NA |
| XBP1 | 1505.61 | -0.0016 | 0.185069 | -0.00864 | 0.993109 | 0.996912 |
| RTCB | 3138.821 | -0.04945 | 0.140124 | -0.35289 | 0.724169 | 0.824799 |
| JOSD1 | 1826.28 | -0.20897 | 0.13993 | -1.49336 | 0.135344 | 0.27404 |
| FBXO7 | 7874.327 | -0.24755 | 0.153014 | -1.61785 | 0.105695 | 0.231481 |
| GTPBP1 | 1442.525 | -0.54986 | 0.149054 | -3.68898 | 2.25E-04 | 0.004177 |
| POLDIP3 | 3122.043 | -0.43244 | 0.142481 | -3.03509 | 0.002405 | 0.018243 |
| RAB36 | 146.4474 | -0.50268 | 0.288998 | -1.7394 | 0.081964 | 0.194493 |
| TIMP3 | 18395.72 | 0.350819 | 0.333074 | 1.053276 | 0.292215 | 0.458987 |
| PPP6R2 | 2314.875 | -0.3075 | 0.144201 | -2.13242 | 0.032972 | 0.105731 |
| SBF1 | 6769.986 | -0.43575 | 0.176509 | -2.46871 | 0.01356 | 0.057932 |
| SUN2 | 2516.347 | -0.49182 | 0.254876 | -1.92965 | 0.05365 | 0.145343 |
| CYB5R3 | 17994.41 | -0.3411 | 0.131831 | -2.58744 | 0.009669 | 0.046091 |
| DNAL4 | 398.9228 | -0.36426 | 0.162419 | -2.24273 | 0.024914 | 0.087611 |
| C22orf31 | 0.488912 | 0.771583 | 2.897678 | 0.266276 | 0.790026 | NA |
| LMF2 | 2845.257 | -0.37466 | 0.176498 | -2.12275 | 0.033775 | 0.107317 |
| RHBDD3 | 676.9771 | -0.26507 | 0.155962 | -1.69958 | 0.089209 | 0.206357 |
| PACSIN2 | 3385.958 | -0.18175 | 0.134344 | -1.35289 | 0.176091 | 0.32818 |
| TTLL1 | 405.2306 | -0.36022 | 0.154457 | -2.33218 | 0.019691 | 0.074965 |
| RASL10A | 0.364467 | -1.20166 | 3.685431 | -0.32606 | 0.744381 | NA |
| AP1B1 | 4073.333 | -0.49383 | 0.121227 | -4.07364 | 4.63E-05 | 0.001572 |
| HMGXB4 | 698.7769 | 0.152969 | 0.143 | 1.069714 | 0.284748 | 0.45116 |
| TOM1 | 3929.305 | -0.17921 | 0.135563 | -1.32198 | 0.186176 | 0.340184 |
| NEFH | 47.81888 | -0.0558 | 0.525103 | -0.10627 | 0.91537 | 0.951141 |
| CHKB | 10.96108 | 0.399894 | 0.685686 | 0.583202 | 0.559757 | NA |
| BIK | 90.31456 | 0.830699 | 0.380465 | 2.183376 | 0.029008 | 0.096932 |
| HMOX1 | 1146.328 | 0.182242 | 0.28723 | 0.634481 | 0.525767 | 0.671366 |
| MCAT | 729.7086 | -0.23296 | 0.176107 | -1.32284 | 0.185888 | 0.339711 |
| THOC5 | 1056.776 | -0.17234 | 0.147051 | -1.172 | 0.241197 | 0.404398 |
| MCM5 | 1649.75 | -0.3157 | 0.197759 | -1.59637 | 0.110405 | 0.238287 |
| APOBEC3H | 3.3084 | -0.75637 | 1.272253 | -0.59451 | 0.55217 | NA |
| ARSA | 1655.334 | -0.44462 | 0.140112 | -3.17336 | 0.001507 | 0.013595 |
| TSPO | 11527.7 | -0.14656 | 0.133819 | -1.09522 | 0.273419 | 0.43932 |
| RASD2 | 1.129907 | -0.91109 | 1.957893 | -0.46534 | 0.641688 | NA |
| TTLL12 | 3073.854 | -0.17614 | 0.152628 | -1.15404 | 0.248483 | 0.412284 |
| CBX7 | 384.6874 | -0.07484 | 0.177024 | -0.42275 | 0.672476 | 0.785573 |
| PDGFB | 18.03449 | 0.522798 | 0.68239 | 0.766127 | 0.443601 | NA |

| | | | | | | |
|---|---|---|---|---|---|---|
| ACR | 0.928019 | -0.78227 | 2.07134 | -0.37767 | 0.705679 | NA |
| CABP7 | 3.736352 | -0.97232 | 1.458408 | -0.6667 | 0.504965 | NA |
| RPL3 | 41634.03 | -0.08373 | 0.122503 | -0.68353 | 0.494271 | 0.6454 |
| ZMAT5 | 305.5286 | -0.19216 | 0.216674 | -0.88685 | 0.375158 | 0.541491 |
| RBFOX2 | 3102.244 | -0.11895 | 0.142923 | -0.83225 | 0.405269 | 0.569451 |
| SYNGR1 | 4598.232 | -0.41526 | 0.180267 | -2.3036 | 0.021245 | 0.078903 |
| TAB1 | 1073.849 | -0.39601 | 0.138018 | -2.86926 | 0.004114 | 0.025903 |
| ASCC2 | 1600.598 | -0.27959 | 0.132472 | -2.11057 | 0.034809 | 0.109548 |
| MTMR3 | 966.2169 | -0.31759 | 0.173065 | -1.83508 | 0.066494 | 0.168462 |
| MIEF1 | 2086.783 | -0.29519 | 0.150782 | -1.95773 | 0.050261 | 0.139054 |
| APOL1 | 12.79325 | -1.15537 | 0.664389 | -1.739 | 0.082035 | NA |
| PNPLA3 | 393.0694 | -0.38847 | 0.212141 | -1.83116 | 0.067076 | 0.169467 |
| MYH9 | 25779.47 | -0.66254 | 0.184216 | -3.59654 | 3.22E-04 | 0.005173 |
| CACNA1I | 0.134133 | -1.48909 | 4.077584 | -0.36519 | 0.71497 | NA |
| SAMM50 | 2741.675 | -0.17754 | 0.138382 | -1.28298 | 0.199497 | 0.355802 |
| TXN2 | 2370.882 | -0.12382 | 0.124266 | -0.99644 | 0.319035 | 0.48689 |
| FOXRED2 | 1341.308 | -0.66587 | 0.165488 | -4.02369 | 5.73E-05 | 0.001791 |
| GRAP2 | 0.888653 | -1.5355 | 2.47141 | -0.62131 | 0.534398 | NA |
| EIF3D | 5442.974 | -0.10658 | 0.135672 | -0.78558 | 0.432116 | 0.59339 |
| TNRC6B | 990.9371 | 0.04448 | 0.173917 | 0.255756 | 0.79814 | 0.877894 |
| SGSM3 | 3703.889 | -0.26995 | 0.134952 | -2.00034 | 0.045464 | 0.130611 |
| IFT27 | 752.4851 | -0.08958 | 0.141394 | -0.63356 | 0.526369 | 0.671846 |
| PVALB | 0.151697 | -1.546 | 4.075469 | -0.37934 | 0.704433 | NA |
| KIAA0930 | 24623.6 | -0.56163 | 0.142323 | -3.94615 | 7.94E-05 | 0.002205 |
| SLC25A17 | 608.8494 | -0.07126 | 0.155103 | -0.45942 | 0.645931 | 0.764681 |
| FAM118A | 243.5809 | -0.09396 | 0.183138 | -0.51304 | 0.607923 | 0.735325 |
| KCTD17 | 572.6413 | -0.07625 | 0.177 | -0.43077 | 0.666639 | 0.781315 |
| ST13 | 2955.036 | 0.292667 | 0.139761 | 2.094051 | 0.036255 | 0.112574 |
| IL2RB | 0.601483 | -1.7332 | 2.729494 | -0.63499 | 0.525436 | NA |
| RBX1 | 166.2238 | -0.11095 | 0.260942 | -0.42518 | 0.670706 | 0.784298 |
| EP300 | 1746.994 | -0.48889 | 0.24022 | -2.03517 | 0.041833 | 0.124232 |
| L3MBTL2 | 748.6317 | -0.34725 | 0.144853 | -2.39724 | 0.016519 | 0.066542 |
| CHADL | 14.11892 | -0.38842 | 0.610143 | -0.6366 | 0.524383 | NA |
| RANGAP1 | 4271.831 | -0.4066 | 0.182609 | -2.22661 | 0.025973 | 0.090324 |
| ZC3H7B | 5548.466 | -0.63288 | 0.154863 | -4.08675 | 4.37E-05 | 0.001538 |
| PHF5A | 706.3562 | 0.00589 | 0.15773 | 0.037339 | 0.970214 | 0.983824 |
| ACO2 | 3164.058 | -0.49182 | 0.120804 | -4.07119 | 4.68E-05 | 0.00158 |
| POLR3H | 2035.81 | -0.25308 | 0.182255 | -1.3886 | 0.164954 | 0.313788 |
| TRMU | 827.86 | 0.005306 | 0.185649 | 0.028583 | 0.977197 | 0.98761 |
| PMM1 | 1497.88 | -0.1503 | 0.132394 | -1.13524 | 0.256273 | 0.420442 |
| DESI1 | 2179.798 | -0.30579 | 0.155336 | -1.96857 | 0.049002 | 0.136932 |
| CERK | 3655.946 | 0.135687 | 0.177969 | 0.76242 | 0.445809 | 0.6056 |
| BRD1 | 952.1621 | -0.27506 | 0.173845 | -1.58222 | 0.1136 | 0.242522 |
| ZBED4 | 842.2449 | -0.18768 | 0.139754 | -1.34296 | 0.179287 | 0.331737 |
| HDAC10 | 36.86294 | -0.34532 | 0.368949 | -0.93596 | 0.349292 | NA |
| PLTP | 4898.444 | -0.48731 | 0.129447 | -3.76453 | 1.67E-04 | 0.003495 |
| PCIF1 | 1209.24 | -0.46075 | 0.197881 | -2.32841 | 0.01989 | 0.075456 |

| | | | | | | |
|---|---|---|---|---|---|---|
| GSS | 1838.481 | -0.36271 | 0.12325 | -2.94292 | 0.003251 | 0.022326 |
| TRPC4AP | 3674.05 | -0.28262 | 0.144549 | -1.95516 | 0.050564 | 0.139665 |
| PYGB | 11339.78 | -0.1259 | 0.192037 | -0.65561 | 0.512074 | 0.65973 |
| ABHD12 | 3371.53 | -0.22413 | 0.148996 | -1.5043 | 0.132503 | 0.269681 |
| PROCR | 569.9329 | 0.186569 | 0.226668 | 0.823094 | 0.410455 | 0.573881 |
| GINS1 | 290.3214 | -0.41808 | 0.2296 | -1.82092 | 0.06862 | 0.172248 |
| NINL | 266.8598 | -0.1184 | 0.203649 | -0.58138 | 0.560981 | 0.699164 |
| CD40 | 5.43439 | 0.12428 | 0.873347 | 0.142304 | 0.88684 | NA |
| UQCC1 | 1315.861 | -0.08007 | 0.131947 | -0.60683 | 0.543961 | 0.685662 |
| ZMYND8 | 2477.381 | -0.02594 | 0.153802 | -0.16867 | 0.866059 | 0.920654 |
| SGK2 | 2.887517 | -0.30445 | 1.174222 | -0.25928 | 0.795421 | NA |
| IFT52 | 891.6143 | 0.013738 | 0.152065 | 0.090344 | 0.928014 | 0.958745 |
| MYBL2 | 425.9409 | -0.25991 | 0.277712 | -0.9359 | 0.349324 | 0.51746 |
| NDRG3 | 1835.122 | -0.26032 | 0.147094 | -1.76975 | 0.076768 | 0.186035 |
| SLA2 | 0.230334 | -0.53318 | 4.076749 | -0.13079 | 0.895945 | NA |
| C20orf24 | 600.9271 | -0.06507 | 0.17479 | -0.3723 | 0.709667 | 0.814198 |
| NFATC2 | 4677.129 | -0.34752 | 0.220434 | -1.57653 | 0.114903 | 0.244486 |
| RIMS4 | 5.085703 | 0.392794 | 0.881004 | 0.445848 | 0.655707 | NA |
| PABPC1L | 91.43365 | 1.004483 | 0.306292 | 3.279496 | 0.00104 | 0.01061 |
| STK4 | 1094.012 | 0.323594 | 0.166745 | 1.94065 | 0.052301 | 0.142666 |
| SALL4 | 40.76832 | -0.30492 | 0.34696 | -0.87882 | 0.379498 | NA |
| ADNP | 2510.763 | -0.32957 | 0.146444 | -2.25045 | 0.02442 | 0.086599 |
| PFDN4 | 341.4056 | 1.29801 | 0.283107 | 4.584868 | 4.54E-06 | 3.99E-04 |
| DOK5 | 364.465 | -0.63916 | 0.23047 | -2.77327 | 0.00555 | 0.031844 |
| CSTF1 | 1107.592 | -0.11651 | 0.154902 | -0.75214 | 0.451967 | 0.610954 |
| BMP7 | 5.03502 | -0.30254 | 0.978505 | -0.30918 | 0.757183 | NA |
| RAE1 | 897.2946 | -0.13179 | 0.165196 | -0.7978 | 0.424987 | 0.587001 |
| TPD52L2 | 3736.358 | -0.17251 | 0.154784 | -1.11454 | 0.265048 | 0.42949 |
| DNAJC5 | 3657.789 | -0.39045 | 0.142891 | -2.73248 | 0.006286 | 0.034405 |
| NELFCD | 2535.76 | -0.17607 | 0.137508 | -1.28047 | 0.200381 | 0.356417 |
| CTSZ | 8596.657 | -0.09462 | 0.147178 | -0.64292 | 0.520279 | 0.667378 |
| PRPF6 | 3018.853 | -0.31354 | 0.126814 | -2.47246 | 0.013419 | 0.057638 |
| TUBB1 | 3.95361 | -0.35544 | 1.003844 | -0.35408 | 0.723277 | NA |
| SLMO2 | 1401.821 | 0.660403 | 0.186015 | 3.550268 | 3.85E-04 | 0.005768 |
| HRH3 | 0.151697 | -1.546 | 4.075469 | -0.37934 | 0.704433 | NA |
| MTG2 | 892.0526 | -0.21934 | 0.167994 | -1.30561 | 0.191684 | 0.346259 |
| PSMA7 | 6504.316 | 0.084912 | 0.129343 | 0.656486 | 0.511511 | 0.659379 |
| SLCO4A1 | 818.1837 | -1.35932 | 0.283191 | -4.80001 | 1.59E-06 | 2.16E-04 |
| NTSR1 | 2.484548 | -0.48062 | 1.325183 | -0.36268 | 0.716844 | NA |
| MRGBP | 629.9737 | 0.036189 | 0.182594 | 0.198196 | 0.842892 | 0.906755 |
| TCFL5 | 1344.337 | -0.1423 | 0.137373 | -1.03584 | 0.300275 | 0.467235 |
| DIDO1 | 1828.515 | -0.39975 | 0.145276 | -2.75166 | 0.005929 | 0.033157 |
| GID8 | 649.0397 | -0.27233 | 0.158026 | -1.72329 | 0.084836 | 0.198603 |
| SLC17A9 | 61.60077 | 0.075817 | 0.346617 | 0.218735 | 0.826856 | 0.895634 |
| BIRC7 | 46.26161 | -0.51052 | 0.487853 | -1.04646 | 0.29535 | NA |
| NKAIN4 | 2.133346 | -0.97961 | 1.345693 | -0.72796 | 0.46664 | NA |
| ARFGAP1 | 2498.063 | -0.32113 | 0.142556 | -2.25269 | 0.024279 | 0.086227 |

| | | | | | | |
|---|---|---|---|---|---|---|
| COL20A1 | 0.163957 | 0.424913 | 4.079006 | 0.104171 | 0.917034 | NA |
| CHRNA4 | 0.163957 | 0.424913 | 4.079006 | 0.104171 | 0.917034 | NA |
| EEF1A2 | 36.0274 | 0.389927 | 0.492876 | 0.791126 | 0.428871 | NA |
| PTK6 | 6.380828 | -0.55255 | 0.823559 | -0.67093 | 0.502264 | NA |
| GMEB2 | 436.6872 | -0.39054 | 0.163318 | -2.39131 | 0.016789 | 0.067285 |
| C20orf27 | 1777.656 | -0.30291 | 0.130276 | -2.32511 | 0.020066 | 0.075931 |
| SPEF1 | 9.996749 | -0.7114 | 0.690326 | -1.03053 | 0.30276 | NA |
| CDC25B | 3762.966 | -0.45234 | 0.150964 | -2.99633 | 0.002732 | 0.019947 |
| ISM1 | 103.222 | 0.002347 | 0.32132 | 0.007305 | 0.994171 | 0.997453 |
| RNF24 | 752.1348 | -0.37602 | 0.203892 | -1.84423 | 0.06515 | 0.166418 |
| ARFRP1 | 887.5121 | -0.04775 | 0.141674 | -0.33705 | 0.736082 | 0.833056 |
| NDUFAF5 | 536.6163 | 0.316059 | 0.177939 | 1.776222 | 0.075696 | 0.184078 |
| SEL1L2 | 0.247898 | -0.5901 | 4.074633 | -0.14482 | 0.884851 | NA |
| TRIB3 | 897.1714 | -0.12438 | 0.183565 | -0.67755 | 0.498054 | 0.648931 |
| RASSF2 | 143.392 | -0.4052 | 0.360488 | -1.12404 | 0.260995 | 0.425422 |
| CSNK2A1 | 1949.72 | -0.20593 | 0.131357 | -1.56771 | 0.116949 | 0.247445 |
| SLC52A3 | 7.34869 | -1.21834 | 0.868748 | -1.40241 | 0.160793 | NA |
| | 0.97663 | -0.60197 | 1.931737 | -0.31162 | 0.755327 | NA |
| RSPO4 | 2.430604 | 0.986118 | 1.668526 | 0.591011 | 0.554513 | NA |
| CDS2 | 3841.914 | -0.30524 | 0.153896 | -1.98342 | 0.04732 | 0.133917 |
| HM13 | 6309.771 | -0.14414 | 0.159896 | -0.90145 | 0.367349 | 0.534412 |
| SNPH | 771.0008 | -0.59001 | 0.14922 | -3.95399 | 7.69E-05 | 0.002169 |
| MYLK2 | 1.273342 | -0.93915 | 1.907721 | -0.49229 | 0.622516 | NA |
| SIRPB1 | 134.3374 | -0.62671 | 0.245105 | -2.55691 | 0.010561 | 0.048772 |
| SEC23B | 1694.108 | -0.07899 | 0.147536 | -0.53541 | 0.592369 | 0.7233 |
| FERMT1 | 0.410275 | 0.244283 | 3.100889 | 0.078778 | 0.937209 | NA |
| PDYN | 0.354779 | 1.703954 | 3.657232 | 0.465914 | 0.641277 | NA |
| CCM2L | 0.134133 | -1.48909 | 4.077584 | -0.36519 | 0.71497 | NA |
| PLCB4 | 3.332028 | 0.727512 | 1.127894 | 0.645018 | 0.518915 | NA |
| MYL9 | 2096.092 | -0.02721 | 0.258474 | -0.10527 | 0.916159 | 0.95139 |
| TM9SF4 | 1950.425 | -0.49113 | 0.13156 | -3.73311 | 1.89E-04 | 0.003755 |
| TLDC2 | 9.726134 | 0.989254 | 0.729076 | 1.35686 | 0.174826 | NA |
| CRNKL1 | 698.1459 | 0.483057 | 0.219907 | 2.196641 | 0.028046 | 0.094807 |
| POFUT1 | 1773.331 | -0.39144 | 0.151147 | -2.58979 | 0.009603 | 0.045907 |
| SAMHD1 | 2352.37 | 0.15556 | 0.174291 | 0.892527 | 0.37211 | 0.53876 |
| PAK7 | 0.096202 | 0.408546 | 4.079639 | 0.100143 | 0.920231 | NA |
| KIF3B | 1990.495 | 0.083291 | 0.149592 | 0.556786 | 0.577674 | 0.712388 |
| MROH8 | 35.75787 | -0.24301 | 0.365614 | -0.66467 | 0.506262 | NA |
| NOP56 | 2347.637 | -0.0249 | 0.141251 | -0.17628 | 0.860071 | 0.916919 |
| MANBAL | 2062.68 | -0.24546 | 0.17908 | -1.37066 | 0.17048 | 0.32085 |
| IDH3B | 2809.778 | -0.06745 | 0.133231 | -0.50628 | 0.61266 | 0.739472 |
| MAPRE1 | 4381.774 | 0.103132 | 0.127996 | 0.805741 | 0.420392 | 0.583153 |
| JAG1 | 575.0155 | -0.49788 | 0.201349 | -2.47273 | 0.013409 | 0.057636 |
| CDK5RAP1 | 551.1745 | -0.13259 | 0.146075 | -0.90769 | 0.36404 | 0.531423 |
| SNTA1 | 950.6274 | -0.19795 | 0.133275 | -1.48524 | 0.137481 | 0.276759 |
| OXT | 0.681112 | 2.620532 | 2.519661 | 1.040033 | 0.298324 | NA |
| TTI1 | 502.8602 | -0.47772 | 0.172992 | -2.7615 | 0.005754 | 0.032518 |

| | | | | | | |
|---|---|---|---|---|---|---|
| E2F1 | 1026.108 | -0.15021 | 0.236828 | -0.63425 | 0.525917 | 0.671486 |
| RPRD1B | 539.0863 | -0.23513 | 0.146495 | -1.60505 | 0.108482 | 0.235327 |
| PXMP4 | 353.6835 | -0.50194 | 0.22846 | -2.19707 | 0.028016 | 0.09476 |
| CHMP4B | 3304.752 | -0.09867 | 0.13609 | -0.72502 | 0.468439 | 0.624884 |
| CST3 | 13325.67 | -0.2744 | 0.162815 | -1.68535 | 0.091921 | 0.210878 |
| ASIP | 0.162376 | 0.424913 | 4.079006 | 0.104171 | 0.917034 | NA |
| ACTR5 | 196.8592 | -0.03767 | 0.1853 | -0.2033 | 0.8389 | 0.903953 |
| WFDC2 | 8.969902 | 0.141471 | 0.728564 | 0.194179 | 0.846036 | NA |
| AHCY | 6135.551 | -0.21729 | 0.138872 | -1.56469 | 0.117655 | 0.248495 |
| PPP1R16B | 2.908162 | -2.75694 | 1.399282 | -1.97025 | 0.048809 | NA |
| FAM83D | 481.5493 | -0.53321 | 0.241113 | -2.21146 | 0.027004 | 0.092445 |
| DHX35 | 305.3501 | -0.00845 | 0.187878 | -0.04496 | 0.964138 | 0.980386 |
| DNTTIP1 | 615.8772 | -0.02399 | 0.154306 | -0.15545 | 0.876465 | 0.926153 |
| MAP1LC3A | 1076.306 | -0.05662 | 0.191712 | -0.29535 | 0.767726 | 0.855796 |
| PIGU | 611.0415 | -0.31717 | 0.150897 | -2.10192 | 0.03556 | 0.111086 |
| TNNC2 | 1.711657 | 0.295749 | 1.545121 | 0.191408 | 0.848206 | NA |
| ACOT8 | 731.6287 | -0.19688 | 0.140299 | -1.40331 | 0.160523 | 0.308029 |
| APMAP | 3893.675 | -0.06337 | 0.136521 | -0.46418 | 0.642521 | 0.762089 |
| | 13.76456 | -0.04221 | 0.585402 | -0.0721 | 0.94252 | NA |
| TULP2 | 1.712152 | -1.73709 | 1.625446 | -1.06869 | 0.285211 | NA |
| NUCB1 | 8763.438 | -0.50156 | 0.151642 | -3.30754 | 9.41E-04 | 0.009957 |
| DHDH | 19.62454 | -0.37117 | 0.647898 | -0.57289 | 0.566722 | NA |
| GYS1 | 3255.031 | -0.41038 | 0.127901 | -3.20857 | 0.001334 | 0.012508 |
| LHB | 7.181079 | 0.230916 | 0.780062 | 0.296023 | 0.767213 | NA |
| KCNA7 | 1.311188 | 0.009168 | 1.781747 | 0.005146 | 0.995894 | NA |
| SNRNP70 | 4672.503 | -0.03638 | 0.145238 | -0.25049 | 0.802206 | 0.881304 |
| CLPTM1 | 4724.131 | -0.34099 | 0.181692 | -1.87678 | 0.060548 | 0.15835 |
| RELB | 527.9995 | -0.44807 | 0.195271 | -2.2946 | 0.021756 | 0.080199 |
| CLASRP | 591.1288 | -0.13709 | 0.153129 | -0.89524 | 0.370657 | 0.537509 |
| LIN7B | 127.8375 | 0.114619 | 0.222784 | 0.514483 | 0.606914 | 0.734751 |
| PPP1R37 | 3372.484 | -0.42879 | 0.139201 | -3.08037 | 0.002067 | 0.016594 |
| FCGRT | 1980.674 | -0.44038 | 0.121884 | -3.61311 | 3.03E-04 | 0.004981 |
| PIH1D1 | 1795.019 | -0.06825 | 0.168905 | -0.40409 | 0.686144 | 0.795602 |
| CKM | 1.391203 | 0.983095 | 1.790872 | 0.548947 | 0.583042 | NA |
| PPP1R13L | 66.07878 | -0.18596 | 0.32238 | -0.57683 | 0.564051 | 0.702029 |
| ERCC2 | 1254.982 | -0.39056 | 0.129457 | -3.01689 | 0.002554 | 0.019126 |
| SLC17A7 | 100.3878 | 0.611024 | 0.45634 | 1.338967 | 0.180581 | 0.333093 |
| KLC3 | 1.713586 | -0.7846 | 1.451865 | -0.54041 | 0.588917 | NA |
| CD37 | 6.611053 | -0.35411 | 0.774172 | -0.4574 | 0.647385 | NA |
| DKKL1 | 3.812964 | -1.11982 | 1.031899 | -1.0852 | 0.277833 | NA |
| DMPK | 1005.76 | -0.08963 | 0.241994 | -0.37038 | 0.711097 | 0.815207 |
| RSPH6A | 0.192403 | 0.975392 | 4.060716 | 0.240202 | 0.810174 | NA |
| TBC1D17 | 1786.189 | -0.05665 | 0.257306 | -0.22018 | 0.825728 | 0.895199 |
| IL4I1 | 6.958559 | -1.47821 | 0.850549 | -1.73795 | 0.08222 | NA |
| PTOV1 | 1334.053 | -0.15043 | 0.155045 | -0.97022 | 0.331936 | 0.500458 |
| NOVA2 | 0.883203 | 0.854847 | 2.135496 | 0.400304 | 0.688933 | NA |
| MED25 | 1221.826 | -0.45897 | 0.148531 | -3.09003 | 0.002001 | 0.016217 |

| | | | | | | |
|---|---|---|---|---|---|---|
| CCDC61 | 265.9475 | -0.36041 | 0.226842 | -1.5888 | 0.112105 | 0.2405 |
| TNNT1 | 513.1281 | 0.315248 | 0.288839 | 1.091429 | 0.275084 | 0.440659 |
| VRK3 | 843.3536 | -0.3379 | 0.13942 | -2.4236 | 0.015368 | 0.062996 |
| PPP6R1 | 4854.377 | -0.38594 | 0.138036 | -2.79597 | 0.005174 | 0.030263 |
| ZNF419 | 198.0023 | 0.104631 | 0.19476 | 0.537229 | 0.591109 | 0.722706 |
| AURKC | 3.966365 | -0.99333 | 1.056926 | -0.93983 | 0.347307 | NA |
| SLC1A5 | 1105.119 | -0.42523 | 0.189571 | -2.2431 | 0.02489 | 0.087559 |
| PRKD2 | 574.0944 | -0.49835 | 0.182905 | -2.72465 | 0.006437 | 0.03502 |
| CCDC9 | 460.5058 | -0.31948 | 0.17214 | -1.85592 | 0.063465 | 0.16349 |
| BBC3 | 620.4452 | -0.52063 | 0.202634 | -2.5693 | 0.01019 | 0.047769 |
| MYH14 | 65.46752 | -0.63876 | 0.355103 | -1.79881 | 0.072049 | 0.178242 |
| GLTSCR2 | 6460.877 | -0.38475 | 0.140146 | -2.74536 | 0.006044 | 0.03362 |
| NKG7 | 0.46577 | -1.30191 | 3.402879 | -0.38259 | 0.702024 | NA |
| ETFB | 900.7617 | -0.20918 | 0.157848 | -1.32519 | 0.18511 | 0.338653 |
| NAPA | 7178.754 | -0.27154 | 0.129689 | -2.09382 | 0.036276 | 0.112606 |
| MEIS3 | 1352.249 | -0.03291 | 0.174507 | -0.1886 | 0.850406 | 0.910875 |
| KDELR1 | 5360.142 | -0.31045 | 0.146609 | -2.11756 | 0.034213 | 0.108216 |
| CYTH2 | 1395.272 | -0.03871 | 0.147161 | -0.26306 | 0.792507 | 0.874295 |
| GRWD1 | 1746.836 | -0.26297 | 0.162622 | -1.61706 | 0.105866 | 0.231729 |
| GRIN2D | 98.99575 | -0.72346 | 0.252728 | -2.86261 | 0.004202 | 0.026285 |
| SYNGR4 | 0.72077 | 0.527966 | 3.060238 | 0.172524 | 0.863025 | NA |
| CLEC11A | 5413.052 | -0.3155 | 0.137248 | -2.29874 | 0.02152 | 0.079674 |
| CCDC114 | 0.936915 | 0.819377 | 2.189457 | 0.374237 | 0.708228 | NA |
| CARD8 | 440.1655 | 0.383673 | 0.169838 | 2.259046 | 0.023881 | 0.085385 |
| LIG1 | 780.2278 | -0.42205 | 0.152231 | -2.77241 | 0.005564 | 0.031913 |
| ZNF175 | 197.1032 | 0.052138 | 0.199648 | 0.26115 | 0.793977 | 0.875021 |
| PLA2G4C | 538.8327 | -0.27177 | 0.226657 | -1.19905 | 0.230507 | 0.392685 |
| DBP | 260.8371 | -0.52234 | 0.260544 | -2.00481 | 0.044983 | 0.129624 |
| FAM83E | 0.314073 | -0.57308 | 3.898934 | -0.14698 | 0.883146 | NA |
| RASIP1 | 25.4228 | -0.72845 | 0.415327 | -1.75393 | 0.079442 | NA |
| BCAT2 | 729.2478 | -0.33208 | 0.19816 | -1.67583 | 0.093772 | 0.213905 |
| PLEKHA4 | 5692.249 | -0.30893 | 0.152315 | -2.02825 | 0.042534 | 0.125476 |
| PPP2R1A | 8877.494 | -0.34712 | 0.12876 | -2.69584 | 0.007021 | 0.037208 |
| LENG1 | 350.0915 | -0.06377 | 0.195525 | -0.32616 | 0.744307 | 0.838959 |
| PRPF31 | 1187.869 | -0.16251 | 0.154109 | -1.05451 | 0.291651 | 0.458435 |
| TFPT | 1456.802 | -0.10261 | 0.167236 | -0.61357 | 0.539497 | 0.681722 |
| UBE2S | 3771.294 | -0.3392 | 0.184995 | -1.83359 | 0.066715 | 0.168708 |
| RPL28 | 18523.47 | -0.11766 | 0.158796 | -0.74093 | 0.458735 | 0.617112 |
| | 1385.844 | 0.457747 | 0.167746 | 2.728804 | 0.006356 | 0.034652 |
| ACTR8 | 933.8306 | 0.023442 | 0.144231 | 0.162534 | 0.870886 | 0.923244 |
| CRBN | 713.0074 | 0.79567 | 0.223052 | 3.567202 | 3.61E-04 | 0.005541 |
| OGG1 | 379.3035 | -0.20874 | 0.161969 | -1.28879 | 0.197472 | 0.353175 |
| KAT2B | 12688.78 | 0.390257 | 0.160125 | 2.437202 | 0.014801 | 0.061418 |
| WNT5A | 8.151007 | 0.788983 | 0.768358 | 1.026843 | 0.304494 | NA |
| PFKFB4 | 742.0347 | 0.062948 | 0.225922 | 0.278625 | 0.780532 | 0.865433 |
| COL7A1 | 51.96083 | 0.885121 | 0.429862 | 2.059082 | 0.039486 | 0.119264 |
| PRKAR2A | 1799.137 | -0.21444 | 0.145211 | -1.47676 | 0.139739 | 0.27988 |

| | | | | | | |
|---|---|---|---|---|---|---|
| USP4 | 1712.114 | -0.18928 | 0.141788 | -1.33492 | 0.181902 | 0.334748 |
| GNAI2 | 14524.85 | -0.34031 | 0.125476 | -2.71217 | 0.006684 | 0.035967 |
| HYAL1 | 39.70691 | -0.66476 | 0.376871 | -1.76389 | 0.07775 | NA |
| TUSC2 | 1258.981 | 0.020731 | 0.163131 | 0.127079 | 0.898878 | 0.940695 |
| NPRL2 | 499.5752 | -0.02368 | 0.154681 | -0.15306 | 0.87835 | 0.926986 |
| CYB561D2 | 727.4607 | -0.11138 | 0.161056 | -0.69154 | 0.489224 | 0.641921 |
| C3orf14 | 474.1 | 0.488343 | 0.171104 | 2.854072 | 0.004316 | 0.02672 |
| CSPG5 | 13.85786 | 0.351533 | 0.732512 | 0.479901 | 0.631298 | NA |
| KLHL18 | 826.3958 | -0.03175 | 0.144305 | -0.22005 | 0.825832 | 0.89523 |
| SCAP | 2556.149 | -0.43948 | 0.164754 | -2.66751 | 0.007642 | 0.039439 |
| HEMK1 | 459.1978 | -0.22575 | 0.200922 | -1.12357 | 0.261195 | 0.425632 |
| CISH | 22.79921 | -0.69418 | 0.453361 | -1.53118 | 0.125726 | NA |
| MAPKAPK3 | 2281.081 | -0.2301 | 0.159616 | -1.4416 | 0.149416 | 0.293024 |
| ACVR2B | 54.50921 | 0.10982 | 0.324095 | 0.33885 | 0.734722 | 0.832199 |
| WDR48 | 1071.645 | 0.379001 | 0.186709 | 2.029902 | 0.042367 | 0.125189 |
| GORASP1 | 1933.957 | -0.26944 | 0.137514 | -1.95939 | 0.050067 | 0.138646 |
| RRP9 | 946.0295 | -0.18025 | 0.188266 | -0.95743 | 0.33835 | 0.507226 |
| ABHD14B | 1565.392 | -0.43748 | 0.161802 | -2.70378 | 0.006856 | 0.03661 |
| EIF1B | 1195.806 | 0.080551 | 0.159585 | 0.504753 | 0.613732 | 0.740094 |
| ABHD14A-/ | 0.151697 | -1.546 | 4.075469 | -0.37934 | 0.704433 | NA |
| VIPR1 | 0.424115 | 1.932531 | 3.45462 | 0.559405 | 0.575885 | NA |
| DNAH1 | 63.74217 | -0.1332 | 0.303432 | -0.43899 | 0.660667 | 0.776536 |
| ZBTB47 | 2335.878 | -0.16782 | 0.203138 | -0.82612 | 0.408737 | 0.572402 |
| TNNC1 | 1.534288 | 1.842014 | 1.773824 | 1.038442 | 0.299064 | NA |
| NKTR | 470.6986 | 1.094232 | 0.300955 | 3.635868 | 2.77E-04 | 0.004685 |
| SPCS1 | 3088.169 | -0.15363 | 0.135455 | -1.13418 | 0.256718 | 0.420707 |
| NEK4 | 412.8484 | 0.371234 | 0.190226 | 1.95154 | 0.050993 | 0.140488 |
| SLC4A3 | 683.4809 | -0.15113 | 0.157652 | -0.95866 | 0.337732 | 0.506619 |
| INO80D | 224.0708 | 0.482221 | 0.276537 | 1.743786 | 0.081196 | 0.193238 |
| EEF1B2 | 7430.233 | 0.403479 | 0.134007 | 3.010871 | 0.002605 | 0.019343 |
| ADAM23 | 4842.968 | 0.253725 | 0.149099 | 1.701726 | 0.088807 | 0.205546 |
| DGUOK | 1294.433 | -0.02428 | 0.143808 | -0.16886 | 0.865903 | 0.920653 |
| MOB1A | 1466.063 | 0.421842 | 0.156086 | 2.702621 | 0.00688 | 0.036671 |
| KANSL3 | 2101.654 | -0.31734 | 0.188201 | -1.68617 | 0.091763 | 0.210711 |
| LMAN2L | 572.7032 | -0.2347 | 0.146259 | -1.60471 | 0.108557 | 0.235403 |
| RTKN | 1687.293 | -0.39232 | 0.148888 | -2.63502 | 0.008413 | 0.041958 |
| TTL | 3074.124 | 0.131763 | 0.164197 | 0.80247 | 0.422281 | 0.584768 |
| IL1A | 0.326536 | 0.037056 | 3.842554 | 0.009644 | 0.992306 | NA |
| CCL20 | 0.863039 | 0.783382 | 2.464829 | 0.317824 | 0.750618 | NA |
| PIKFYVE | 1076.603 | 0.497535 | 0.219351 | 2.26821 | 0.023316 | 0.083915 |
| KCNIP3 | 1692.248 | -0.82037 | 0.225229 | -3.64239 | 2.70E-04 | 0.004608 |
| FAHD2A | 635.5765 | -0.19756 | 0.142271 | -1.38861 | 0.164951 | 0.313788 |
| NCL | 7808.996 | 0.426665 | 0.29749 | 1.434217 | 0.15151 | 0.295469 |
| ACTR1B | 2730.681 | -0.36885 | 0.185829 | -1.98487 | 0.047159 | 0.133679 |
| SLC35F5 | 3559.443 | 0.737426 | 0.178018 | 4.14243 | 3.44E-05 | 0.001323 |
| ZAP70 | 0.971254 | -2.67708 | 2.319442 | -1.15419 | 0.248421 | NA |
| ACTR3 | 7405.046 | 0.703974 | 0.155044 | 4.540475 | 5.61E-06 | 4.53E-04 |

| | | | | | | |
|---|---|---|---|---|---|---|
| STEAP3 | 100.096 | 0.158853 | 0.269111 | 0.59029 | 0.554996 | 0.695073 |
| EPB41L5 | 705.3235 | 0.254379 | 0.170233 | 1.494303 | 0.135096 | 0.273679 |
| TFCP2L1 | 0.258578 | 1.233151 | 4.047809 | 0.304646 | 0.760635 | NA |
| | 1495.452 | 0.582657 | 0.142789 | 4.080556 | 4.49E-05 | 0.001553 |
| TP53I3 | 918.7123 | -0.16382 | 0.178158 | -0.91952 | 0.357826 | 0.525131 |
| DNAJC27 | 116.0712 | 0.560543 | 0.252849 | 2.216906 | 0.026629 | 0.091638 |
| POMC | 62.18208 | -0.53738 | 0.293907 | -1.82841 | 0.067489 | 0.169976 |
| STAM2 | 647.2515 | 0.772703 | 0.219771 | 3.515944 | 4.38E-04 | 0.006274 |
| OTOF | 1.982494 | 1.115242 | 1.486995 | 0.749997 | 0.453257 | NA |
| GPD2 | 1435.806 | 0.20327 | 0.177466 | 1.145402 | 0.252043 | 0.415977 |
| CENPA | 351.2506 | -0.05813 | 0.273809 | -0.21229 | 0.83188 | 0.89891 |
| CYTIP | 0.586694 | 1.130467 | 2.734203 | 0.413454 | 0.679274 | NA |
| ACVR1 | 909.9887 | -0.39239 | 0.153075 | -2.56336 | 0.010366 | 0.048212 |
| TANC1 | 4235.004 | 0.113662 | 0.157336 | 0.722413 | 0.470041 | 0.626386 |
| SLC30A3 | 3.96192 | -0.96838 | 1.007122 | -0.96153 | 0.336285 | NA |
| MPV17 | 3055.433 | -0.2157 | 0.15072 | -1.4311 | 0.152402 | 0.296817 |
| GTF3C2 | 1652.078 | -0.28299 | 0.169169 | -1.67281 | 0.094364 | 0.214796 |
| EIF2B4 | 1089.897 | -0.05881 | 0.161974 | -0.3631 | 0.716532 | 0.81922 |
| NRBP1 | 2781.049 | -0.09107 | 0.127197 | -0.71599 | 0.474 | 0.629544 |
| ITGB6 | 0.713668 | -0.95576 | 2.626349 | -0.36391 | 0.715923 | NA |
| FNDC4 | 245.2875 | 0.219537 | 0.353431 | 0.62116 | 0.534495 | 0.678129 |
| ITGA4 | 492.8631 | 0.873352 | 0.28422 | 3.072806 | 0.002121 | 0.016804 |
| PSMD14 | 1805.159 | 0.350141 | 0.159918 | 2.189503 | 0.02856 | 0.095861 |
| SNX17 | 7079.757 | -0.22215 | 0.153889 | -1.44359 | 0.148854 | 0.292187 |
| GPR75-ASB | 429.0868 | 0.188221 | 0.167149 | 1.126067 | 0.260137 | 0.424389 |
| PPM1G | 3484.652 | -0.15558 | 0.1668 | -0.93272 | 0.350966 | 0.518856 |
| PDE1A | 0.602678 | 1.083611 | 2.70938 | 0.399948 | 0.689195 | NA |
| IFIH1 | 499.6444 | -0.1545 | 0.621624 | -0.24854 | 0.803716 | 0.882554 |
| GCA | 150.2786 | 0.363749 | 0.20051 | 1.814117 | 0.06966 | 0.173966 |
| INO80B | 893.5039 | -0.20635 | 0.210774 | -0.97902 | 0.327568 | 0.495828 |
| MOGS | 1959.651 | -0.26851 | 0.153841 | -1.74535 | 0.080924 | 0.192758 |
| TTC31 | 661.0813 | -0.3474 | 0.145578 | -2.38638 | 0.017015 | 0.067895 |
| PCGF1 | 444.6946 | 0.112074 | 0.179051 | 0.625934 | 0.531358 | 0.675668 |
| GRB14 | 82.97314 | 0.588096 | 0.258546 | 2.274629 | 0.022928 | 0.082895 |
| CLIP4 | 988.5909 | 0.45205 | 0.181726 | 2.487539 | 0.012863 | 0.056084 |
| TLX2 | 7.011286 | 0.749953 | 0.804765 | 0.931891 | 0.351393 | NA |
| SPTBN1 | 36356.93 | -0.00944 | 0.145789 | -0.06474 | 0.948383 | 0.971147 |
| AUP1 | 3917.327 | 0.009774 | 0.144951 | 0.067432 | 0.946238 | 0.970016 |
| RTN4 | 9997.013 | 0.022358 | 0.126022 | 0.177417 | 0.859181 | 0.916352 |
| HTRA2 | 895.4553 | -0.0278 | 0.201376 | -0.13807 | 0.890182 | 0.935891 |
| LOXL3 | 1423.222 | 0.476679 | 0.604036 | 0.789156 | 0.430021 | 0.591541 |
| DOK1 | 434.5311 | 0.331979 | 0.270233 | 1.22849 | 0.219263 | 0.379 |
| GALNT3 | 1290.591 | 0.203793 | 0.240418 | 0.847659 | 0.396628 | 0.561234 |
| POLE4 | 1302.312 | 0.106718 | 0.162826 | 0.655413 | 0.512202 | 0.65973 |
| TACR1 | 0.151697 | -1.546 | 4.075469 | -0.37934 | 0.704433 | NA |
| CCDC88A | 311.8917 | 0.73098 | 1.11908 | 0.653198 | 0.513629 | 0.660803 |
| ACADL | 1.866864 | -0.99112 | 1.481314 | -0.66908 | 0.503445 | NA |

| | | | | | | |
|---|---|---|---|---|---|---|
| EVA1A | 1879.899 | -0.05101 | 0.14421 | -0.35371 | 0.723553 | 0.824463 |
| MRPL19 | 856.8151 | 0.460441 | 0.158376 | 2.907265 | 0.003646 | 0.024068 |
| LANCL1 | 1856.891 | 0.119517 | 0.18473 | 0.646984 | 0.517642 | 0.664972 |
| WDR75 | 572.01 | 0.646896 | 0.165286 | 3.913798 | 9.09E-05 | 0.002389 |
| EFEMP1 | 82.40891 | 0.025399 | 0.341357 | 0.074407 | 0.940687 | 0.966779 |
| FANCL | 219.5216 | 0.560409 | 0.215747 | 2.597525 | 0.00939 | 0.045248 |
| FN1 | 51501.47 | -0.34058 | 0.259698 | -1.31146 | 0.189701 | 0.343974 |
| STAT1 | 6172.155 | -0.48968 | 0.244252 | -2.00483 | 0.044981 | 0.129624 |
| GLS | 1615.387 | 0.662007 | 0.201788 | 3.280711 | 0.001035 | 0.010582 |
| PAPOLG | 317.1263 | 0.204211 | 0.182882 | 1.116628 | 0.264153 | 0.428684 |
| DNAH6 | 2.214612 | 0.568451 | 1.3908 | 0.408722 | 0.682744 | NA |
| PECR | 561.288 | 0.103254 | 0.158095 | 0.653115 | 0.513682 | 0.660803 |
| UNC50 | 953.2846 | 0.336041 | 0.137411 | 2.445522 | 0.014464 | 0.060524 |
| IGFBP2 | 324.4969 | 0.717161 | 0.401803 | 1.784854 | 0.074285 | 0.18199 |
| ELMOD3 | 616.8155 | -0.02731 | 0.152268 | -0.17934 | 0.857673 | 0.91568 |
| IGFBP5 | 283.5202 | 0.410451 | 0.407725 | 1.006686 | 0.314085 | 0.481568 |
| USP34 | 2479.193 | 0.67014 | 0.202639 | 3.307068 | 9.43E-04 | 0.009961 |
| EFHD1 | 256.1099 | -0.56157 | 0.254088 | -2.21015 | 0.027095 | 0.092647 |
| KCNJ13 | 145.753 | -1.3198 | 0.854579 | -1.54439 | 0.122495 | 0.25598 |
| CCT4 | 4932.488 | 0.086384 | 0.147865 | 0.584211 | 0.559078 | 0.697891 |
| GGCX | 1713.979 | -0.33332 | 0.156008 | -2.13659 | 0.032631 | 0.10513 |
| EHBP1 | 1182.822 | 0.472224 | 0.206958 | 2.281741 | 0.022505 | 0.081968 |
| OTX1 | 34.30333 | -0.08084 | 0.452046 | -0.17883 | 0.858069 | NA |
| TXNDC9 | 743.1653 | 0.574503 | 0.212344 | 2.705531 | 0.00682 | 0.03645 |
| COQ10B | 1113.398 | 0.783804 | 0.160108 | 4.895477 | 9.81E-07 | 1.61E-04 |
| GNLY | 0.230334 | -0.53318 | 4.076749 | -0.13079 | 0.895945 | NA |
| SF3B1 | 4456.63 | 0.685646 | 0.188764 | 3.632285 | 2.81E-04 | 0.004737 |
| ST3GAL5 | 2649.554 | -0.2713 | 0.173714 | -1.56175 | 0.118346 | 0.249732 |
| CHST10 | 594.0245 | -0.23054 | 0.146403 | -1.57466 | 0.115335 | 0.2453 |
| PDCL3 | 312.233 | 0.360272 | 0.203506 | 1.770327 | 0.076673 | 0.18588 |
| MOB4 | 757.1963 | 0.738208 | 0.193145 | 3.822045 | 1.32E-04 | 0.003029 |
| HSPE1 | 2042.658 | 0.501991 | 0.172481 | 2.910419 | 0.003609 | 0.023918 |
| KDM3A | 1024.644 | 0.308013 | 0.147256 | 2.091679 | 0.036467 | 0.113084 |
| PLCD4 | 365.1179 | -0.56344 | 0.311851 | -1.80675 | 0.070802 | 0.175929 |
| CHMP3 | 3143.368 | 0.152696 | 0.151745 | 1.006272 | 0.314285 | 0.481686 |
| ZNF142 | 505.681 | -0.52172 | 0.204627 | -2.54959 | 0.010785 | 0.049404 |
| PRKAG3 | 1.086672 | 0.527563 | 2.235195 | 0.236026 | 0.813413 | NA |
| SMYD1 | 0.944149 | -2.4805 | 2.433399 | -1.01936 | 0.308033 | NA |
| IL1R1 | 146.9931 | -0.17724 | 0.248623 | -0.71291 | 0.475904 | 0.630941 |
| WNT6 | 14.17338 | 0.22344 | 0.619793 | 0.360507 | 0.718468 | NA |
| IL1RL2 | 3.317036 | 1.622005 | 1.169564 | 1.386845 | 0.165489 | NA |
| IL1RL1 | 0.096202 | 0.408546 | 4.079639 | 0.100143 | 0.920231 | NA |
| IL18R1 | 0.326333 | 1.533889 | 3.826878 | 0.40082 | 0.688553 | NA |
| FHL2 | 1558.596 | -0.09014 | 0.21461 | -0.42003 | 0.674462 | 0.786828 |
| MLPH | 8990.975 | -0.4642 | 0.150826 | -3.07773 | 0.002086 | 0.016597 |
| CNPPD1 | 3742.825 | -0.5908 | 0.129231 | -4.57168 | 4.84E-06 | 4.10E-04 |
| UXS1 | 1150.806 | 0.137707 | 0.151906 | 0.906524 | 0.364658 | 0.531934 |

| | | | | | | |
|---|---|---|---|---|---|---|
| ABCB6 | 48.63816 | -0.21639 | 0.316165 | -0.68442 | 0.493711 | 0.644811 |
| STK16 | 1322.219 | -0.15015 | 0.155695 | -0.9644 | 0.334844 | 0.503433 |
| SLC5A7 | 11.18354 | 2.502891 | 0.758619 | 3.299271 | 9.69E-04 | NA |
| HDLBP | 22866.81 | -0.20443 | 0.134791 | -1.51666 | 0.129353 | 0.265768 |
| PPP1R7 | 1882.235 | -0.05885 | 0.137689 | -0.42741 | 0.669083 | 0.782942 |
| PASK | 229.5102 | -0.29786 | 0.258972 | -1.15016 | 0.250077 | 0.41377 |
| STK25 | 3637.923 | -0.1102 | 0.12677 | -0.86928 | 0.384695 | 0.550705 |
| PROC | 0.968522 | 0.920016 | 1.915592 | 0.480278 | 0.63103 | NA |
| ID2 | 929.4601 | 0.85454 | 0.19236 | 4.442392 | 8.90E-06 | 5.86E-04 |
| TAF1B | 401.8292 | 0.6193 | 0.20665 | 2.99686 | 0.002728 | 0.019937 |
| HPCAL1 | 7457.682 | 0.11736 | 0.164349 | 0.714092 | 0.47517 | 0.630458 |
| ODC1 | 412.3638 | 0.09693 | 0.18657 | 0.519534 | 0.603388 | 0.732334 |
| BIRC6 | 2249.199 | 0.170546 | 0.156426 | 1.090267 | 0.275596 | 0.441265 |
| NOL10 | 489.3343 | 0.719441 | 0.218957 | 3.285757 | 0.001017 | 0.010476 |
| PLEKHB2 | 7768.963 | -0.13605 | 0.164327 | -0.82791 | 0.407721 | 0.571339 |
| GORASP2 | 2893.317 | -0.14263 | 0.152009 | -0.93828 | 0.348099 | 0.51605 |
| STRN | 728.2571 | 0.420534 | 0.171358 | 2.454123 | 0.014123 | 0.059604 |
| CEBPZ | 646.9744 | 1.260279 | 0.316757 | 3.978698 | 6.93E-05 | 0.002018 |
| PRKD3 | 4304.729 | 0.08429 | 0.200089 | 0.421261 | 0.673565 | 0.786323 |
| DCAF17 | 339.5589 | 0.49792 | 0.178031 | 2.796819 | 0.005161 | 0.0302 |
| QPCT | 17786.34 | -0.20699 | 0.150831 | -1.37236 | 0.169953 | 0.320363 |
| RAB3GAP1 | 2176.71 | 0.182156 | 0.140618 | 1.29539 | 0.195186 | 0.350192 |
| SLC25A12 | 766.2323 | -0.07281 | 0.1632 | -0.44614 | 0.655495 | 0.772532 |
| RMDN2 | 189.5985 | 1.263153 | 0.240271 | 5.257195 | 1.46E-07 | 4.34E-05 |
| DLX2 | 44.18598 | 0.301016 | 0.478218 | 0.629453 | 0.529053 | NA |
| DARS | 1027.616 | 0.696133 | 0.199169 | 3.495193 | 4.74E-04 | 0.006586 |
| SRSF7 | 1852.315 | 0.354397 | 0.157839 | 2.245299 | 0.024749 | 0.087191 |
| SDC1 | 212.7046 | -0.63273 | 0.326195 | -1.93974 | 0.052411 | 0.142855 |
| PLCL1 | 115.0384 | 0.22733 | 0.336749 | 0.675072 | 0.49963 | 0.650106 |
| SLC1A4 | 24314.26 | -0.765 | 0.150603 | -5.07957 | 3.78E-07 | 8.53E-05 |
| SOS1 | 626.2041 | 0.490113 | 0.300567 | 1.630627 | 0.102969 | 0.227301 |
| KYNU | 320.1748 | 0.332948 | 0.23383 | 1.423888 | 0.154479 | 0.299729 |
| WIPF1 | 1861.331 | -0.11292 | 0.168209 | -0.67133 | 0.502013 | 0.65228 |
| ORC2 | 353.4229 | 0.191843 | 0.188176 | 1.019488 | 0.307971 | 0.475143 |
| COX7A2L | 2523.708 | 0.05073 | 0.169095 | 0.300008 | 0.764171 | 0.853696 |
| PNO1 | 778.3033 | 0.205698 | 0.161007 | 1.277568 | 0.201402 | 0.357749 |
| ORC4 | 480.1365 | 0.420072 | 0.198441 | 2.116858 | 0.034272 | 0.108345 |
| PLEK | 0.230334 | -0.53318 | 4.076749 | -0.13079 | 0.895945 | NA |
| RND3 | 1258.763 | 1.149705 | 0.270648 | 4.247968 | 2.16E-05 | 9.60E-04 |
| ATF2 | 731.4554 | 0.626888 | 0.202674 | 3.093082 | 0.001981 | 0.016077 |
| THADA | 699.5274 | -0.21317 | 0.149529 | -1.42559 | 0.153985 | 0.299017 |
| AAK1 | 226.8353 | -0.21654 | 0.203228 | -1.06552 | 0.28664 | 0.452961 |
| TRAK2 | 5361.301 | 0.338464 | 0.141621 | 2.389925 | 0.016852 | 0.067493 |
| C2orf42 | 349.1221 | -0.04641 | 0.179696 | -0.25828 | 0.796194 | 0.87685 |
| TIA1 | 774.3301 | 0.812807 | 0.24136 | 3.36762 | 7.58E-04 | 0.008794 |
| PCYOX1 | 2693.969 | 0.035618 | 0.132473 | 0.268867 | 0.788032 | 0.871362 |
| EPAS1 | 3518.761 | -0.55448 | 0.17399 | -3.18686 | 0.001438 | 0.013218 |

| | | | | | | |
|---|---|---|---|---|---|---|
| SUMO1 | 1742.798 | 0.538617 | 0.155904 | 3.454793 | 5.51E-04 | 0.007174 |
| CD207 | 1.286979 | 0.266221 | 1.956478 | 0.136072 | 0.891765 | NA |
| VAX2 | 73.64879 | -0.55784 | 0.313341 | -1.78031 | 0.075025 | 0.182934 |
| ATP6V1B1 | 17.14997 | -0.59021 | 0.540751 | -1.09147 | 0.275068 | NA |
| NFE2L2 | 2335.729 | 0.534091 | 0.187163 | 2.85362 | 0.004322 | 0.02673 |
| MSH6 | 1054.629 | 0.199179 | 0.148418 | 1.342014 | 0.179591 | 0.332198 |
| PLEKHA3 | 533.8067 | 0.368494 | 0.156735 | 2.351058 | 0.01872 | 0.072265 |
| SPR | 1829.393 | -0.00749 | 0.163971 | -0.04568 | 0.963563 | 0.980182 |
| EPHA4 | 28.31947 | -0.27836 | 0.431839 | -0.64459 | 0.519194 | NA |
| PARD3B | 535.2771 | 0.143726 | 0.163179 | 0.880792 | 0.37843 | 0.544717 |
| FARSB | 1200.853 | -0.06897 | 0.156019 | -0.44209 | 0.658423 | 0.774827 |
| ALMS1 | 256.0589 | 0.054121 | 0.241828 | 0.223797 | 0.822915 | 0.893194 |
| CD3EAP | 72.84467 | -0.28921 | 0.314576 | -0.91938 | 0.357897 | 0.525131 |
| COLEC11 | 0.640609 | 0.042838 | 2.379511 | 0.018003 | 0.985637 | NA |
| ZNF541 | 2.002937 | -1.34767 | 1.541055 | -0.87451 | 0.381838 | NA |
| SLC8A2 | 4.827714 | -0.28483 | 0.89285 | -0.31901 | 0.749716 | NA |
| KPTN | 264.3234 | -0.24497 | 0.205714 | -1.19083 | 0.233722 | 0.396277 |
| CRYGD | 0.298089 | -0.51711 | 3.975232 | -0.13008 | 0.896501 | NA |
| MREG | 1645.396 | -0.6237 | 0.208844 | -2.98646 | 0.002822 | 0.020352 |
| FASTKD2 | 734.6776 | 0.067275 | 0.170569 | 0.394417 | 0.693273 | 0.802064 |
| NRP2 | 4953.05 | -0.95717 | 0.189577 | -5.04896 | 4.44E-07 | 9.40E-05 |
| CREB1 | 661.8865 | 0.697805 | 0.185896 | 3.753736 | 1.74E-04 | 0.003558 |
| KLF7 | 257.65 | 0.186072 | 0.185445 | 1.003383 | 0.315676 | 0.483007 |
| VAMP8 | 1002.175 | 0.119289 | 0.145298 | 0.820996 | 0.411649 | 0.574963 |
| RPN2 | 6430.104 | -0.23353 | 0.126484 | -1.84634 | 0.064843 | 0.165956 |
| TGIF2 | 506.414 | -0.72333 | 0.159832 | -4.52557 | 6.02E-06 | 4.73E-04 |
| HS1BP3 | 2549.338 | -0.29137 | 0.132874 | -2.19282 | 0.028321 | 0.095362 |
| C2orf43 | 662.0059 | 0.047876 | 0.184912 | 0.258912 | 0.795703 | 0.876597 |
| WDR35 | 584.9084 | 0.38898 | 0.21408 | 1.816984 | 0.06922 | 0.173232 |
| DNAH7 | 4.468811 | -0.43226 | 0.951733 | -0.45418 | 0.649696 | NA |
| CYP20A1 | 308.6465 | 0.138456 | 0.161706 | 0.856225 | 0.391874 | 0.557055 |
| NDUFB3 | 1469.741 | 0.470007 | 0.151189 | 3.108738 | 0.001879 | 0.015479 |
| GTF3C3 | 953.008 | 0.345455 | 0.162629 | 2.124195 | 0.033654 | 0.107125 |
| SATB2 | 124.464 | -0.04455 | 0.262085 | -0.17 | 0.865014 | 0.920317 |
| C2orf40 | 3.147287 | 0.859007 | 1.363056 | 0.630206 | 0.528559 | NA |
| ITGB1BP1 | 4174.538 | 0.084076 | 0.145507 | 0.577818 | 0.563387 | 0.70135 |
| CPSF3 | 1287.932 | 0.16913 | 0.14811 | 1.141922 | 0.253487 | 0.417547 |
| ZBTB45 | 685.504 | -0.49474 | 0.154501 | -3.20219 | 0.001364 | 0.012701 |
| RHOQ | 4326.212 | 0.443896 | 0.175694 | 2.526535 | 0.011519 | 0.051866 |
| GPR75 | 6.319708 | 0.487556 | 0.841988 | 0.579053 | 0.562553 | NA |
| SUPT7L | 1049.916 | 0.163473 | 0.145079 | 1.126784 | 0.259834 | 0.42423 |
| KLHL29 | 990.3582 | -0.4145 | 0.176474 | -2.34879 | 0.018835 | 0.072547 |
| DNMT3A | 595.2588 | -0.33726 | 0.172937 | -1.95017 | 0.051156 | 0.140741 |
| TMEM214 | 2834.301 | -0.32628 | 0.145031 | -2.24972 | 0.024466 | 0.08666 |
| ATAD2B | 127.4984 | 0.657174 | 0.389865 | 1.685647 | 0.091864 | 0.210851 |
| FKBP1B | 253.1351 | -0.06596 | 0.177025 | -0.37262 | 0.709433 | 0.814086 |
| ATL2 | 668.1251 | 0.597578 | 0.15329 | 3.898346 | 9.69E-05 | 0.002496 |

| | | | | | | |
|---|---|---|---|---|---|---|
| YPEL5 | 3163.705 | 0.352959 | 0.157421 | 2.242137 | 0.024953 | 0.087674 |
| FAM98A | 2377.823 | 0.175205 | 0.173013 | 1.01267 | 0.311218 | 0.478781 |
| YIPF4 | 1634.731 | 0.406096 | 0.146516 | 2.771677 | 0.005577 | 0.031939 |
| AFTPH | 587.8703 | 0.716521 | 0.190357 | 3.764079 | 1.67E-04 | 0.003495 |
| LGALSL | 88.47161 | -0.02945 | 0.256245 | -0.11493 | 0.908504 | 0.947226 |
| CNRIP1 | 1152.277 | -0.10353 | 0.167442 | -0.61832 | 0.536362 | 0.679625 |
| BCL11A | 28.15894 | -0.09247 | 0.454559 | -0.20344 | 0.838793 | NA |
| CRIPT | 464.0212 | 0.129489 | 0.173746 | 0.745273 | 0.456106 | 0.614797 |
| NCAPH | 628.0487 | -0.2672 | 0.225936 | -1.18265 | 0.236946 | 0.399744 |
| ZNF549 | 262.1863 | 0.049787 | 0.204258 | 0.243747 | 0.807427 | 0.885096 |
| A1BG | 19.59517 | -0.0462 | 0.461118 | -0.10019 | 0.920195 | NA |
| ZSCAN18 | 280.5955 | -0.47403 | 0.284162 | -1.66817 | 0.095283 | 0.215939 |
| ZNF211 | 256.7933 | 0.083052 | 0.184542 | 0.450043 | 0.652679 | 0.770362 |
| GTDC1 | 555.8702 | 0.199336 | 0.173184 | 1.151004 | 0.249731 | 0.41357 |
| CXCR4 | 0.596179 | -0.09275 | 3.146759 | -0.02948 | 0.976485 | NA |
| ZRANB3 | 87.68746 | 0.408453 | 0.293142 | 1.393364 | 0.16351 | 0.312042 |
| ACVR2A | 121.4134 | -0.1162 | 0.238943 | -0.48633 | 0.626735 | 0.74944 |
| MTERFD2 | 550.2448 | 0.037212 | 0.144373 | 0.257752 | 0.796598 | 0.87685 |
| HJURP | 479.9533 | -0.04244 | 0.247501 | -0.17149 | 0.863842 | 0.919827 |
| METTL8 | 260.0703 | 0.229636 | 0.170792 | 1.344531 | 0.178777 | 0.33126 |
| TTC21B | 409.4641 | 0.825961 | 0.19145 | 4.314249 | 1.60E-05 | 8.22E-04 |
| NMI | 53.68312 | 0.102508 | 0.395019 | 0.259501 | 0.795249 | 0.876259 |
| TNFAIP6 | 146.5106 | 0.812506 | 0.277578 | 2.927126 | 0.003421 | 0.022949 |
| ACVR1C | 138.0865 | 0.301557 | 0.312407 | 0.965271 | 0.334409 | 0.502972 |
| BAZ2B | 251.9643 | 0.705438 | 0.653027 | 1.080259 | 0.280027 | 0.446019 |
| ZNF137P | 16.21175 | 0.759253 | 0.611152 | 1.242331 | 0.214115 | NA |
| ACSL3 | 4131.076 | 0.199021 | 0.246806 | 0.806387 | 0.42002 | 0.582924 |
| CHPF | 15943.62 | -0.2118 | 0.186305 | -1.13685 | 0.255601 | 0.419747 |
| DNPEP | 1561.135 | -0.22533 | 0.1302 | -1.73066 | 0.083512 | 0.196865 |
| INHA | 3.050336 | -0.54935 | 1.21517 | -0.45208 | 0.651213 | NA |
| MOGAT1 | 0.448206 | -1.2532 | 3.007315 | -0.41672 | 0.676885 | NA |
| OBSL1 | 3618.795 | -0.24455 | 0.128128 | -1.90864 | 0.056309 | 0.150161 |
| FAM124B | 1.391057 | 1.605737 | 1.701366 | 0.943793 | 0.345276 | NA |
| GCNT7 | 2.574785 | -0.04064 | 1.25544 | -0.03237 | 0.974175 | NA |
| CTCFL | 0.392857 | -1.546 | 4.075469 | -0.37934 | 0.704433 | NA |
| HMGB1P1 | 28.36367 | 0.969995 | 0.487768 | 1.988639 | 0.046741 | NA |
| FAM210B | 7344.757 | 0.666374 | 0.215437 | 3.093127 | 0.001981 | 0.016077 |
| PI3 | 0.84823 | 2.93367 | 2.713279 | 1.081227 | 0.279596 | NA |
| FAM209A | 0.247898 | -0.5901 | 4.074633 | -0.14482 | 0.884851 | NA |
| SNX21 | 477.1087 | -0.24448 | 0.170229 | -1.43616 | 0.150957 | 0.294826 |
| SLPI | 0.163957 | 0.424913 | 4.079006 | 0.104171 | 0.917034 | NA |
| WFDC3 | 2.029094 | -0.10546 | 1.369715 | -0.07699 | 0.938628 | NA |
| TTPAL | 689.9136 | -0.26322 | 0.151241 | -1.7404 | 0.081788 | 0.194269 |
| PREX1 | 316.6543 | -2.1689 | 0.930592 | -2.33067 | 0.019771 | 0.075098 |
| KCNS1 | 20.28448 | 0.23584 | 0.539444 | 0.43719 | 0.661973 | NA |
| SLC12A5 | 0.163957 | 0.424913 | 4.079006 | 0.104171 | 0.917034 | NA |
| ARHGAP40 | 0.851742 | -0.48655 | 2.547586 | -0.19099 | 0.848537 | NA |

| | | | | | | |
|---|---|---|---|---|---|---|
| SDC4 | 312.8745 | -0.62898 | 0.231885 | -2.71246 | 0.006679 | 0.035967 |
| NCOA3 | 1021.484 | 0.178413 | 0.18118 | 0.984727 | 0.324758 | 0.492707 |
| PIGT | 4454.531 | -0.4219 | 0.142685 | -2.95688 | 0.003108 | 0.021663 |
| MATN4 | 1.401533 | 3.641611 | 2.075973 | 1.754171 | 0.079401 | NA |
| NCOA5 | 881.1832 | -0.31406 | 0.19265 | -1.6302 | 0.103059 | 0.227389 |
| VAPB | 2477.509 | -0.11794 | 0.151085 | -0.7806 | 0.435038 | 0.596159 |
| PARD6B | 8.815948 | 0.343273 | 0.69224 | 0.495888 | 0.619974 | NA |
| ATP5E | 4136.76 | 0.500399 | 0.129948 | 3.850769 | 1.18E-04 | 0.002843 |
| CHD6 | 1481.659 | 0.056481 | 0.167762 | 0.336671 | 0.736365 | 0.833121 |
| PLCG1 | 3535.471 | -0.39638 | 0.143154 | -2.76894 | 0.005624 | 0.032088 |
| TOX2 | 1.774311 | 0.514284 | 1.66994 | 0.307966 | 0.758108 | NA |
| SRSF6 | 2520.225 | 0.11334 | 0.156697 | 0.723304 | 0.469493 | 0.625866 |
| GDAP1L1 | 0.411855 | 0.250456 | 3.097618 | 0.080854 | 0.935558 | NA |
| ARFGEF2 | 2026.75 | 0.285271 | 0.186011 | 1.533624 | 0.125122 | 0.260016 |
| ZNFX1 | 3474.016 | -0.48641 | 0.12706 | -3.82819 | 1.29E-04 | 0.002993 |
| ZNF831 | 0.096202 | 0.408546 | 4.079639 | 0.100143 | 0.920231 | NA |
| CSE1L | 2599.122 | 0.154396 | 0.181246 | 0.85186 | 0.394292 | 0.559064 |
| RAB22A | 1362.907 | 0.475564 | 0.15627 | 3.043212 | 0.002341 | 0.017914 |
| PTGIS | 0.134133 | -1.48909 | 4.077584 | -0.36519 | 0.71497 | NA |
| STAU1 | 2854.497 | -0.14049 | 0.123127 | -1.14101 | 0.253867 | 0.417883 |
| CDH26 | 0.860207 | -2.14053 | 2.512764 | -0.85186 | 0.394291 | NA |
| SNAI1 | 3.575599 | -0.49632 | 1.132444 | -0.43827 | 0.661191 | NA |
| MOCS3 | 369.1506 | -0.31003 | 0.177565 | -1.74602 | 0.080808 | 0.192597 |
| STX16 | 1333.21 | 0.157298 | 0.152914 | 1.028673 | 0.303634 | 0.47084 |
| PPP4R1L | 455.5192 | 0.06529 | 0.219254 | 0.297782 | 0.765869 | 0.854691 |
| PMEPA1 | 2230.44 | 0.825874 | 0.585023 | 1.411695 | 0.15804 | 0.304545 |
| RNF114 | 2374.545 | -0.26859 | 0.127715 | -2.10302 | 0.035464 | 0.11093 |
| DDX27 | 823.5408 | 0.366669 | 0.175128 | 2.093722 | 0.036285 | 0.112606 |
| RBPJL | 0.448206 | -1.2532 | 3.007315 | -0.41672 | 0.676885 | NA |
| BCAS4 | 151.811 | -0.44393 | 0.232999 | -1.90528 | 0.056744 | 0.151033 |
| KCNK15 | 11.60576 | -0.15568 | 0.657849 | -0.23665 | 0.81293 | NA |
| TP53TG5 | 1.674314 | -0.71508 | 1.5215 | -0.46998 | 0.638367 | NA |
| NEURL2 | 61.97705 | -0.20884 | 0.291352 | -0.71678 | 0.473509 | 0.629168 |
| STAMBP | 1180.612 | 0.011164 | 0.14441 | 0.077304 | 0.938381 | 0.965761 |
| NAGK | 3442.588 | -0.20075 | 0.154835 | -1.29656 | 0.194784 | 0.349926 |
| MCEE | 146.5227 | 0.421146 | 0.236988 | 1.777079 | 0.075555 | 0.18381 |
| PAIP2B | 79.48499 | 0.096557 | 0.250096 | 0.386078 | 0.699439 | 0.806443 |
| SNRNP27 | 389.1796 | 0.701964 | 0.172082 | 4.079247 | 4.52E-05 | 0.001553 |
| MPHOSPH: | 258.808 | 1.084749 | 0.698591 | 1.552768 | 0.120479 | 0.252771 |
| LRRFIP1 | 1799.582 | 0.636988 | 0.261646 | 2.43454 | 0.014911 | 0.061742 |
| | 0.096202 | 0.408546 | 4.079639 | 0.100143 | 0.920231 | NA |
| RAB17 | 711.812 | -0.01864 | 0.348913 | -0.05342 | 0.957399 | 0.976509 |
| EMC3 | 2433.155 | 0.105546 | 0.155451 | 0.678966 | 0.497159 | 0.648243 |
| SSUH2 | 159.3668 | -0.13651 | 0.250998 | -0.54388 | 0.586521 | 0.719339 |
| PPP1R12C | 4226.521 | -0.13241 | 0.150148 | -0.8819 | 0.377833 | 0.544236 |
| MBOAT7 | 2465.678 | -0.48808 | 0.139232 | -3.50554 | 4.56E-04 | 0.00643 |
| SRMS | 1.052261 | 0.993047 | 2.045208 | 0.485548 | 0.627288 | NA |

| | | | | | | |
|---|---|---|---|---|---|---|
| OPRL1 | 87.2988 | -0.52179 | 0.292628 | -1.78313 | 0.074565 | 0.182375 |
| SLC2A4RG | 955.5834 | -0.2749 | 0.143769 | -1.91208 | 0.055866 | 0.149468 |
| C20orf195 | 3.259058 | 0.624146 | 1.148946 | 0.543234 | 0.586969 | NA |
| PPDPF | 3270.445 | -0.20933 | 0.135101 | -1.54942 | 0.121281 | 0.254026 |
| IL1B | 0.326333 | 1.533889 | 3.826878 | 0.40082 | 0.688553 | NA |
| PLGLB2 | 0.096202 | 0.408546 | 4.079639 | 0.100143 | 0.920231 | NA |
| CHCHD5 | 604.985 | -0.11169 | 0.14983 | -0.74547 | 0.455986 | 0.614797 |
| PAX8 | 74.26128 | -0.41045 | 0.286336 | -1.43345 | 0.15173 | 0.295848 |
| INSIG2 | 287.8759 | 0.566884 | 0.204643 | 2.770117 | 0.005604 | 0.03203 |
| POLR1B | 926.0584 | -0.14587 | 0.14883 | -0.98009 | 0.32704 | 0.495154 |
| HTR5BP | 0.614937 | 2.493639 | 2.617677 | 0.952615 | 0.340785 | NA |
| CCDC93 | 692.5219 | 0.434156 | 0.174556 | 2.487207 | 0.012875 | 0.056115 |
| PSD4 | 50.08175 | 0.205402 | 0.557912 | 0.368161 | 0.712753 | 0.816474 |
| FOSB | 32.2889 | -0.32548 | 0.408288 | -0.79719 | 0.425343 | NA |
| OPA3 | 1384.347 | -0.49031 | 0.150607 | -3.25555 | 0.001132 | 0.011192 |
| SNRPD2 | 6178.091 | 0.147606 | 0.160276 | 0.920947 | 0.357078 | 0.52462 |
| RTN2 | 1217.664 | -0.04486 | 0.156367 | -0.2869 | 0.774185 | 0.860327 |
| EML2 | 2786.805 | -0.2843 | 0.145886 | -1.94879 | 0.051321 | 0.140999 |
| VASP | 2336.39 | -0.18276 | 0.155971 | -1.17177 | 0.24129 | 0.404426 |
| SYMPK | 2334.116 | -0.55521 | 0.138178 | -4.0181 | 5.87E-05 | 0.001798 |
| GPCPD1 | 172.9738 | 0.252536 | 0.273855 | 0.922155 | 0.356447 | 0.524101 |
| SDCBP2 | 74.38096 | 0.390605 | 0.299519 | 1.304107 | 0.192197 | 0.3468 |
| PANK2 | 1199.708 | -0.07075 | 0.130788 | -0.54097 | 0.58853 | 0.721053 |
| GNRH2 | 0.914607 | 1.879338 | 2.377229 | 0.790558 | 0.429202 | NA |
| DEFB126 | 0.88142 | 1.783485 | 2.222213 | 0.802571 | 0.422222 | NA |
| FAM182A | 0.3441 | -0.0166 | 3.769682 | -0.0044 | 0.996486 | NA |
| CD93 | 0.096202 | 0.408546 | 4.079639 | 0.100143 | 0.920231 | NA |
| GZF1 | 691.4207 | 0.121189 | 0.141167 | 0.858476 | 0.39063 | 0.555865 |
| NAPB | 399.8773 | 0.311147 | 0.194734 | 1.597802 | 0.110087 | 0.237763 |
| NKX2-4 | 0.653015 | 0.2552 | 2.787103 | 0.091565 | 0.927044 | NA |
| CENPB | 6370.629 | -0.27877 | 0.144486 | -1.92942 | 0.053679 | 0.145364 |
| PSMF1 | 3114.738 | -0.30771 | 0.120513 | -2.55333 | 0.01067 | 0.049123 |
| NKX2-2 | 0.394291 | 0.298745 | 3.141711 | 0.09509 | 0.924243 | NA |
| DTD1 | 1440.68 | -0.06278 | 0.154418 | -0.40654 | 0.684346 | 0.794605 |
| RBCK1 | 2770.723 | -0.49074 | 0.136251 | -3.60173 | 3.16E-04 | 0.005092 |
| TMX4 | 1323.537 | -0.08213 | 0.204467 | -0.40166 | 0.687935 | 0.796977 |
| STK35 | 661.8003 | -0.65681 | 0.167769 | -3.91498 | 9.04E-05 | 0.002389 |
| SNRPB | 3836.999 | -0.13063 | 0.15726 | -0.83063 | 0.406183 | 0.569932 |
| NRSN2 | 2273.859 | -0.38648 | 0.145632 | -2.65384 | 0.007958 | 0.040509 |
| AP5S1 | 888.3541 | -0.35253 | 0.15282 | -2.30686 | 0.021063 | 0.078448 |
| RRBP1 | 3997.816 | -0.25513 | 0.199025 | -1.28191 | 0.199873 | 0.355943 |
| BMP2 | 13.64711 | 1.318993 | 0.614272 | 2.147246 | 0.031774 | NA |
| ZNF133 | 293.1409 | 0.189057 | 0.175568 | 1.076834 | 0.281554 | 0.44773 |
| FLRT3 | 244.2264 | -0.33587 | 0.335438 | -1.00127 | 0.316694 | 0.484252 |
| PCSK2 | 389.8081 | 2.057779 | 1.071583 | 1.920317 | 0.054818 | 0.147555 |
| MKKS | 1285.353 | 0.327201 | 0.145086 | 2.255222 | 0.024119 | 0.085854 |
| BFSP1 | 152.0946 | 0.02259 | 0.727466 | 0.031052 | 0.975228 | 0.986377 |

| | | | | | | |
|---|---|---|---|---|---|---|
| DSTN | 6843.808 | 0.423727 | 0.165194 | 2.565032 | 0.010317 | 0.048042 |
| SNRPB2 | 1204.163 | 0.59132 | 0.183036 | 3.230627 | 0.001235 | 0.011864 |
| MGME1 | 1072.073 | 0.269345 | 0.157785 | 1.707044 | 0.087814 | 0.203885 |
| LRRN4 | 0.719246 | 0.444943 | 2.542118 | 0.175028 | 0.861057 | NA |
| TBC1D20 | 2814.139 | -0.32908 | 0.16379 | -2.00914 | 0.044523 | 0.128924 |
| ITPA | 2233.668 | 0.014037 | 0.160533 | 0.08744 | 0.930322 | 0.960504 |
| TCF15 | 0.151697 | -1.546 | 4.075469 | -0.37934 | 0.704433 | NA |
| MCM8 | 278.2001 | 0.47582 | 0.228293 | 2.084255 | 0.037137 | 0.114411 |
| TMEM74B | 8.080034 | 0.040001 | 0.727406 | 0.054991 | 0.956146 | NA |
| FAM110A | 98.24489 | -0.31881 | 0.253496 | -1.25767 | 0.208512 | 0.365923 |
| MRPS26 | 1332.83 | -0.05727 | 0.14046 | -0.40774 | 0.683463 | 0.794046 |
| MMP24 | 15.8973 | -0.23362 | 0.518886 | -0.45024 | 0.65254 | NA |
| NECAB3 | 432.0465 | -0.04794 | 0.161307 | -0.29721 | 0.766304 | 0.854855 |
| ID1 | 19.66448 | 0.823406 | 0.851978 | 0.966464 | 0.333812 | NA |
| RALY | 4618.692 | -0.36815 | 0.138847 | -2.65148 | 0.008014 | 0.040636 |
| DYNLRB1 | 4948.772 | -0.04006 | 0.16364 | -0.24479 | 0.806618 | 0.884511 |
| EIF2S2 | 1175.317 | 0.740648 | 0.210997 | 3.510228 | 4.48E-04 | 0.006372 |
| ERGIC3 | 6401.777 | -0.14436 | 0.139761 | -1.03292 | 0.301643 | 0.468492 |
| ROMO1 | 1981.401 | -0.02672 | 0.168668 | -0.15843 | 0.874114 | 0.925069 |
| BPIFB9P | 0.163957 | 0.424913 | 4.079006 | 0.104171 | 0.917034 | NA |
| FAM83C | 0.134133 | -1.48909 | 4.077584 | -0.36519 | 0.71497 | NA |
| CEP250 | 802.1949 | -0.32029 | 0.214825 | -1.49092 | 0.135984 | 0.275053 |
| PLAGL2 | 379.2146 | -0.44486 | 0.178171 | -2.4968 | 0.012532 | 0.054882 |
| MMP24-AS | 1913.642 | -0.00838 | 0.121745 | -0.06881 | 0.94514 | 0.969511 |
| TMEM115 | 1764.123 | -0.25681 | 0.158881 | -1.61638 | 0.106012 | 0.231871 |
| BCL2L12 | 545.4893 | -0.21339 | 0.189979 | -1.12325 | 0.261329 | 0.425792 |
| IRF3 | 942.2917 | -0.06238 | 0.164695 | -0.37876 | 0.704866 | 0.810651 |
| PRMT1 | 4614.531 | -0.17768 | 0.13721 | -1.29493 | 0.195343 | 0.3504 |
| RRAS | 2195.752 | -0.08622 | 0.237322 | -0.36332 | 0.716366 | 0.819109 |
| PRRG2 | 21.6072 | 0.082233 | 0.499843 | 0.164518 | 0.869323 | NA |
| SCAF1 | 3173.142 | -0.57246 | 0.144938 | -3.94967 | 7.83E-05 | 0.002183 |
| PRR12 | 1247.899 | -0.76191 | 0.168453 | -4.52301 | 6.10E-06 | 4.76E-04 |
| TUBA4A | 89.00479 | 0.282326 | 0.270507 | 1.04369 | 0.296629 | 0.463665 |
| AAMP | 2643.087 | -0.12049 | 0.131436 | -0.91671 | 0.359296 | 0.526452 |
| PNKD | 1928.01 | -0.12351 | 0.129354 | -0.95482 | 0.339667 | 0.508003 |
| ZNF835 | 0.815391 | 0.931626 | 2.360431 | 0.394685 | 0.693076 | NA |
| TUBGCP6 | 1110.177 | -0.39245 | 0.187942 | -2.08815 | 0.036784 | 0.113713 |
| ADM2 | 2.621466 | 0.792797 | 1.312902 | 0.603851 | 0.545943 | NA |
| DGCR6L | 1554.558 | -0.29783 | 0.14756 | -2.01837 | 0.043552 | 0.127152 |
| DGCR8 | 769.3867 | -0.11675 | 0.168046 | -0.69477 | 0.487199 | 0.640255 |
| ASPHD2 | 127.5882 | -0.09768 | 0.240841 | -0.40557 | 0.685058 | 0.795121 |
| VPREB3 | 1.79623 | 1.522559 | 1.657218 | 0.918744 | 0.358229 | NA |
| SDF2L1 | 906.9186 | -0.32099 | 0.214563 | -1.49601 | 0.134651 | 0.273009 |
| GAL3ST1 | 0.591998 | -0.2499 | 3.117006 | -0.08017 | 0.936101 | NA |
| YWHAH | 4119.228 | -0.15992 | 0.149466 | -1.06995 | 0.284643 | 0.451128 |
| RFPL2 | 18.7435 | -0.28996 | 0.787741 | -0.36809 | 0.712805 | NA |
| C22orf24 | 1.973339 | 0.491936 | 1.602661 | 0.30695 | 0.758882 | NA |

| | | | | | | |
|---|---|---|---|---|---|---|
| POM121L9 | 0.162376 | 0.424913 | 4.079006 | 0.104171 | 0.917034 | NA |
| GNAZ | 7.529601 | 0.023681 | 0.730046 | 0.032438 | 0.974123 | NA |
| MGAT3 | 1.550621 | 0.545407 | 1.695251 | 0.321726 | 0.74766 | NA |
| ADORA2A | 0.736957 | -1.0303 | 2.640699 | -0.39016 | 0.696418 | NA |
| ATF4 | 5908.372 | 0.287447 | 0.208239 | 1.380374 | 0.167472 | 0.317051 |
| A4GALT | 0.526843 | -0.38719 | 2.855892 | -0.13558 | 0.892156 | NA |
| CDC42EP1 | 3516.198 | -0.08765 | 0.193121 | -0.45388 | 0.649913 | 0.767937 |
| APOL3 | 8.24849 | -0.18141 | 0.823991 | -0.22015 | 0.825751 | NA |
| MCHR1 | 64.46676 | -0.54457 | 0.401241 | -1.35722 | 0.174711 | 0.326741 |
| TPST2 | 1666.216 | 0.149363 | 0.214585 | 0.696054 | 0.486395 | 0.639483 |
| BAIAP2L2 | 470.809 | 0.11456 | 0.206536 | 0.554674 | 0.579118 | 0.713515 |
| MPST | 1548.812 | -0.17145 | 0.16479 | -1.04042 | 0.298146 | 0.465146 |
| TST | 529.0721 | -0.17591 | 0.150087 | -1.17204 | 0.241181 | 0.404398 |
| APOL5 | 0.162376 | 0.424913 | 4.079006 | 0.104171 | 0.917034 | NA |
| APOL2 | 1242.566 | -0.43188 | 0.146931 | -2.93933 | 0.003289 | 0.022423 |
| RAC2 | 0.949524 | -0.003 | 2.040023 | -0.00147 | 0.998826 | NA |
| LIF | 2.72356 | -0.48968 | 1.206281 | -0.40594 | 0.684784 | NA |
| C22orf23 | 155.8129 | -0.70366 | 0.281532 | -2.49939 | 0.012441 | 0.054606 |
| APOBEC3A | 0.134133 | -1.48909 | 4.077584 | -0.36519 | 0.71497 | NA |
| APOBEC3F | 87.04915 | -0.80169 | 0.254574 | -3.14916 | 0.001637 | 0.014318 |
| RIBC2 | 8.212371 | -0.28561 | 0.819916 | -0.34834 | 0.727583 | NA |
| MYO1B | 2838.53 | 0.40605 | 0.187039 | 2.17094 | 0.029936 | 0.098867 |
| HOXD1 | 51.80741 | 0.973136 | 0.567018 | 1.716234 | 0.086119 | 0.200973 |
| HOXD3 | 33.36177 | 0.089523 | 0.455889 | 0.196371 | 0.84432 | NA |
| MTX2 | 793.7332 | 0.335783 | 0.13581 | 2.472454 | 0.013419 | 0.057638 |
| PDE11A | 12.10583 | -0.2722 | 0.615685 | -0.4421 | 0.658413 | NA |
| CHN1 | 155.1153 | -0.43325 | 0.325992 | -1.32901 | 0.183844 | 0.337118 |
| GAD1 | 158.425 | 0.954572 | 0.243777 | 3.915764 | 9.01E-05 | 0.002389 |
| EIF2S2P4 | 4.344419 | 0.840459 | 1.040496 | 0.807748 | 0.419236 | NA |
| OSGEPL1 | 238.6638 | 0.289546 | 0.200987 | 1.440618 | 0.149693 | 0.293211 |
| ORMDL1 | 813.7099 | 0.746909 | 0.152165 | 4.908543 | 9.18E-07 | 1.53E-04 |
| HAT1 | 944.8817 | 0.879114 | 0.227295 | 3.867715 | 1.10E-04 | 0.00273 |
| HOXD9 | 18.79977 | 0.354487 | 0.526594 | 0.67317 | 0.500839 | NA |
| HOXD13 | 0.162376 | 0.424913 | 4.079006 | 0.104171 | 0.917034 | NA |
| KLK14 | 0.314073 | -0.57308 | 3.898934 | -0.14698 | 0.883146 | NA |
| SIGLEC9 | 0.268266 | -2.15396 | 4.038351 | -0.53338 | 0.593773 | NA |
| KLK10 | 0.544554 | -2.65036 | 3.220086 | -0.82307 | 0.410468 | NA |
| SGOL1 | 85.04633 | 0.467604 | 0.280587 | 1.666521 | 0.09561 | 0.216515 |
| LBP | 0.134133 | -1.48909 | 4.077584 | -0.36519 | 0.71497 | NA |
| SYT5 | 23.60212 | -0.46068 | 0.445961 | -1.033 | 0.301604 | NA |
| TNNI3 | 69.87148 | -0.06241 | 0.32481 | -0.19215 | 0.847624 | 0.909194 |
| SH3BP4 | 880.6295 | -0.09203 | 0.160804 | -0.57233 | 0.567096 | 0.704045 |
| EXOC3L2 | 0.555087 | 0.973802 | 2.808575 | 0.346725 | 0.728798 | NA |
| PVRL2 | 2235.686 | -0.32129 | 0.154725 | -2.0765 | 0.037848 | 0.115728 |
| APOE | 19395.05 | -0.22137 | 0.193169 | -1.14601 | 0.25179 | 0.415791 |
| TOMM40 | 2132.527 | -0.01327 | 0.17329 | -0.0766 | 0.938944 | 0.966165 |
| APOC1 | 87.63815 | -0.24858 | 0.340659 | -0.72971 | 0.46557 | 0.622879 |

| | | | | | | |
|---|---|---|---|---|---|---|
| KIF1A | 6.036618 | -0.39027 | 0.82189 | -0.47485 | 0.634894 | NA |
| NDUFA10 | 2852.793 | -0.02586 | 0.128666 | -0.20102 | 0.840686 | 0.905184 |
| SCO2 | 1032.01 | -0.05741 | 0.154846 | -0.37074 | 0.710828 | 0.815056 |
| PXDN | 3197.738 | -0.36828 | 0.185637 | -1.98386 | 0.047272 | 0.133854 |
| HRC | 0.315653 | -0.57305 | 3.892639 | -0.14721 | 0.882963 | NA |
| TRPM4 | 1580.545 | -0.34194 | 0.171911 | -1.98906 | 0.046695 | 0.132894 |
| ZBTB46 | 81.85059 | -0.25124 | 0.261519 | -0.96071 | 0.336698 | 0.505637 |
| HELZ2 | 2219.235 | -1.20131 | 0.17622 | -6.8171 | 9.29E-12 | 1.57E-08 |
| SAMD10 | 96.1628 | -0.0553 | 0.256955 | -0.2152 | 0.829612 | 0.897277 |
| ATXN10 | 2379.19 | -0.11922 | 0.1352 | -0.88183 | 0.377866 | 0.544236 |
| ZNF337 | 23.42223 | 0.166182 | 0.444439 | 0.373914 | 0.708468 | NA |
| TAF4 | 367.1414 | -0.33584 | 0.243356 | -1.38005 | 0.16757 | 0.317106 |
| GATA5 | 0.096202 | 0.408546 | 4.079639 | 0.100143 | 0.920231 | NA |
| LAMA5 | 176.6948 | -0.64587 | 0.259459 | -2.48929 | 0.0128 | 0.055849 |
| OSBPL2 | 991.8041 | -0.09008 | 0.137512 | -0.65505 | 0.512435 | 0.659815 |
| ADRM1 | 2155.332 | -0.0924 | 0.125747 | -0.73478 | 0.462471 | 0.620483 |
| CHMP2A | 3161.297 | -0.08779 | 0.127161 | -0.6904 | 0.48994 | 0.642357 |
| UBE2M | 3039.643 | -0.17019 | 0.157257 | -1.08225 | 0.279142 | 0.445426 |
| TRIM28 | 6775.226 | -0.51691 | 0.126404 | -4.08932 | 4.33E-05 | 0.001531 |
| TMEM160 | 1121.314 | -0.03634 | 0.160128 | -0.22694 | 0.820471 | 0.892334 |
| ZC3H4 | 2356.099 | -0.62658 | 0.183655 | -3.4117 | 6.46E-04 | 0.00789 |
| NPAS1 | 38.34955 | 0.232445 | 0.466576 | 0.498194 | 0.618347 | NA |
| ZNF331 | 143.3607 | 0.343434 | 0.224466 | 1.530001 | 0.126017 | 0.26113 |
| PKDREJ | 6.06565 | 0.311836 | 0.877886 | 0.355212 | 0.722431 | NA |
| EPS8L1 | 357.3822 | -0.18643 | 0.195722 | -0.95251 | 0.340837 | 0.508949 |
| AAR2 | 1474.554 | -0.44361 | 0.132978 | -3.33598 | 8.50E-04 | 0.009373 |
| TTLL9 | 5.013204 | 0.558837 | 0.881791 | 0.633751 | 0.526243 | NA |
| RBM39 | 2113.333 | 0.868056 | 0.250909 | 3.459649 | 5.41E-04 | 0.007077 |
| ZNF341 | 116.7698 | -0.45206 | 0.248105 | -1.82204 | 0.068449 | 0.171957 |
| GGT7 | 941.5841 | -0.22488 | 0.159307 | -1.4116 | 0.158066 | 0.304547 |
| ACSS2 | 1494.589 | -0.29581 | 0.207286 | -1.42704 | 0.153569 | 0.298392 |
| ATP6V1E1 | 6025.494 | 0.310999 | 0.145936 | 2.13107 | 0.033083 | 0.106001 |
| ZNF227 | 193.2399 | 0.129323 | 0.21902 | 0.59046 | 0.554882 | 0.695073 |
| MRPS25 | 1267.446 | 0.063803 | 0.143906 | 0.443367 | 0.6575 | 0.774125 |
| SH3BP5 | 574.1772 | 0.336454 | 0.186968 | 1.799525 | 0.071936 | 0.178 |
| HACL1 | 974.2132 | 0.177115 | 0.173971 | 1.01807 | 0.308645 | 0.476058 |
| TBC1D5 | 1174.276 | -0.21055 | 0.164126 | -1.28285 | 0.199544 | 0.355802 |
| CAPN7 | 651.1625 | 0.510553 | 0.216275 | 2.360664 | 0.018242 | 0.070975 |
| RFTN1 | 299.0881 | -0.28866 | 0.344462 | -0.838 | 0.402029 | 0.56651 |
| C3orf20 | 5.161872 | -1.43807 | 0.91778 | -1.5669 | 0.117138 | NA |
| ZFYVE20 | 924.5142 | -0.40451 | 0.166749 | -2.42586 | 0.015272 | 0.062844 |
| GALNT15 | 73.50999 | -0.14508 | 0.259658 | -0.55872 | 0.576355 | 0.711368 |
| SLC6A6 | 3084.534 | -0.45724 | 0.209152 | -2.18618 | 0.028802 | 0.096427 |
| KCNC3 | 4.375813 | 0.676853 | 1.011737 | 0.669001 | 0.503495 | NA |
| NAPSA | 6.938498 | -0.58553 | 0.788086 | -0.74297 | 0.457498 | NA |
| NAPSB | 0.163957 | 0.424913 | 4.079006 | 0.104171 | 0.917034 | NA |
| NR1H2 | 3432.299 | -0.32624 | 0.149419 | -2.1834 | 0.029006 | 0.096932 |

| | | | | | | |
|---|---|---|---|---|---|---|
| LRRC4B | 9.392518 | 0.357472 | 0.921127 | 0.388081 | 0.697956 | NA |
| ZNF304 | 304.9947 | 0.005542 | 0.164968 | 0.033596 | 0.973199 | 0.985082 |
| ZSCAN5A | 60.19506 | -0.07124 | 0.307828 | -0.23142 | 0.816985 | 0.890316 |
| ZNF132 | 113.4318 | -0.37186 | 0.247844 | -1.50038 | 0.133516 | 0.271218 |
| USP29 | 0.392857 | -1.546 | 4.075469 | -0.37934 | 0.704433 | NA |
| MATN3 | 87.24314 | -0.01147 | 0.375243 | -0.03057 | 0.975612 | 0.986598 |
| DHX30 | 3838.201 | -0.41839 | 0.143601 | -2.91352 | 0.003574 | 0.023762 |
| RAF1 | 3204.275 | -0.07667 | 0.204074 | -0.37571 | 0.707136 | 0.812552 |
| PPARG | 273.4437 | 0.154834 | 0.22435 | 0.690142 | 0.490105 | 0.642357 |
| NUP210 | 1.119861 | -0.05288 | 2.270917 | -0.02329 | 0.981422 | NA |
| PTCD3 | 2360.131 | 0.182284 | 0.139996 | 1.302069 | 0.192893 | 0.347688 |
| IMMT | 3337.792 | -0.07091 | 0.118124 | -0.60033 | 0.548284 | 0.689499 |
| MRPL35 | 1213.651 | 0.431838 | 0.153244 | 2.817978 | 0.004833 | 0.028855 |
| IQCA1 | 0.192403 | 0.975392 | 4.060716 | 0.240202 | 0.810174 | NA |
| ILKAP | 1558.324 | -0.15111 | 0.129664 | -1.16542 | 0.243848 | 0.406585 |
| PER2 | 18.10775 | -0.63598 | 0.564156 | -1.12732 | 0.259609 | NA |
| RAMP1 | 46.23514 | -1.01696 | 0.591622 | -1.71894 | 0.085625 | NA |
| SCLY | 74.1414 | -0.25742 | 0.285522 | -0.90157 | 0.367286 | 0.534385 |
| HSPA12B | 61.92697 | -0.07075 | 0.292831 | -0.2416 | 0.809093 | 0.885851 |
| ANKEF1 | 51.25883 | -0.30436 | 0.365465 | -0.83281 | 0.404952 | 0.569272 |
| PCED1A | 960.2338 | -0.15421 | 0.166912 | -0.92389 | 0.355546 | 0.523732 |
| SNAP25 | 78.17632 | -0.04947 | 0.302977 | -0.16329 | 0.87029 | 0.923107 |
| BTBD3 | 1702.476 | 0.479918 | 0.16883 | 2.842613 | 0.004475 | 0.027371 |
| PCNA | 3200.544 | 0.246976 | 0.148438 | 1.663833 | 0.096146 | 0.21722 |
| NXT1 | 464.8407 | -0.08254 | 0.220629 | -0.37412 | 0.708318 | 0.813279 |
| POLR3F | 234.2487 | 0.4036 | 0.17747 | 2.274185 | 0.022955 | 0.082916 |
| RIN2 | 35.95076 | 0.189973 | 0.364582 | 0.521072 | 0.602317 | NA |
| PTPRA | 3013.167 | -0.08027 | 0.139525 | -0.57533 | 0.565067 | 0.702545 |
| SSTR4 | 9.005877 | 0.269818 | 0.740381 | 0.364432 | 0.715536 | NA |
| CTNNBL1 | 1395.045 | -0.18123 | 0.134511 | -1.34735 | 0.177868 | 0.330231 |
| LPIN3 | 496.3892 | -0.21045 | 0.180576 | -1.16546 | 0.243833 | 0.406585 |
| ZSWIM3 | 117.9249 | -0.25028 | 0.251567 | -0.9949 | 0.319785 | 0.487783 |
| RBM38 | 230.6607 | -0.28699 | 0.204993 | -1.40002 | 0.161509 | 0.309069 |
| VSTM2L | 11.88974 | -0.68267 | 0.719733 | -0.9485 | 0.342876 | NA |
| OSER1 | 1563.722 | 0.253485 | 0.168637 | 1.503141 | 0.132803 | 0.270093 |
| SERINC3 | 3869.839 | 0.260153 | 0.136024 | 1.912552 | 0.055805 | 0.149437 |
| PPP1R3D | 402.6861 | -0.12129 | 0.15588 | -0.77811 | 0.436504 | 0.597202 |
| | 13.96696 | 0.003386 | 0.692771 | 0.004888 | 0.9961 | NA |
| HMGB1P5 | 6.237208 | 0.326263 | 0.855511 | 0.381366 | 0.702932 | NA |
| SUV420H2 | 121.7774 | -0.09251 | 0.252581 | -0.36627 | 0.714163 | 0.81767 |
| HSPBP1 | 1290.769 | -0.28737 | 0.126268 | -2.2759 | 0.022852 | 0.082745 |
| MORC2 | 947.8296 | -0.34172 | 0.168949 | -2.02263 | 0.043111 | 0.126537 |
| LARGE | 2016.539 | -0.63967 | 0.132637 | -4.82269 | 1.42E-06 | 2.02E-04 |
| GSTT2B | 8.327107 | 0.641343 | 0.706236 | 0.908113 | 0.363818 | NA |
| SLC2A11 | 137.8539 | 0.249265 | 0.236045 | 1.056007 | 0.290965 | 0.457904 |
| C1QTNF6 | 303.7121 | -0.44818 | 0.191568 | -2.33954 | 0.019308 | 0.073836 |
| FAM83F | 0.247898 | -0.5901 | 4.074633 | -0.14482 | 0.884851 | NA |

| | | | | | | |
|---|---|---|---|---|---|---|
| SEC14L4 | 0.6908 | 0.277604 | 2.346083 | 0.118327 | 0.905809 | NA |
| IRAK2 | 45.16764 | -0.52108 | 0.351255 | -1.48348 | 0.137947 | NA |
| CAMK1 | 219.7807 | -0.32821 | 0.253784 | -1.29326 | 0.195921 | 0.351223 |
| THUMPD3 | 563.7368 | 0.517299 | 0.188463 | 2.744822 | 0.006054 | 0.033634 |
| VHL | 679.7119 | -0.27352 | 0.184629 | -1.48145 | 0.138488 | 0.278172 |
| BHLHE40 | 1435.409 | -0.40542 | 0.207665 | -1.9523 | 0.050903 | 0.140403 |
| ARL8B | 3054.357 | 0.124876 | 0.140479 | 0.888931 | 0.37404 | 0.540301 |
| EDEM1 | 1875.488 | 0.041837 | 0.19511 | 0.214426 | 0.830215 | 0.89752 |
| CNTN6 | 1.390854 | 2.478543 | 1.961142 | 1.263826 | 0.206292 | NA |
| CHL1 | 722.4564 | 0.302944 | 0.263231 | 1.150867 | 0.249787 | 0.413579 |
| YWHAQ | 9947.318 | 1.34E-04 | 0.150713 | 8.88E-04 | 0.999292 | 0.999472 |
| KIDINS220 | 2345.461 | 0.377653 | 0.213962 | 1.765043 | 0.077557 | 0.18741 |
| GRHL1 | 7.931914 | 0.125478 | 0.704805 | 0.178033 | 0.858697 | NA |
| ROCK2 | 1320.643 | 1.07122 | 0.843401 | 1.27012 | 0.204042 | 0.360758 |
| RSAD2 | 69.37704 | -1.3044 | 0.506112 | -2.57729 | 0.009958 | 0.047051 |
| LPIN1 | 2551.854 | 0.085467 | 0.155606 | 0.549251 | 0.582833 | 0.71638 |
| CMPK2 | 402.5147 | -1.0051 | 0.310668 | -3.23529 | 0.001215 | 0.01178 |
| IAH1 | 1736.494 | 0.141595 | 0.129597 | 1.092585 | 0.274576 | 0.440235 |
| DHX34 | 415.5965 | -0.43065 | 0.168581 | -2.55453 | 0.010633 | 0.048992 |
| PRADC1 | 809.9243 | -0.20189 | 0.166 | -1.21618 | 0.223917 | 0.384772 |
| SEMA4F | 1208.675 | -0.39355 | 0.178796 | -2.20111 | 0.027728 | 0.09425 |
| CCT7 | 10015.37 | -0.15311 | 0.129169 | -1.18536 | 0.235875 | 0.398845 |
| EGR4 | 0.163957 | 0.424913 | 4.079006 | 0.104171 | 0.917034 | NA |
| RAB11FIP5 | 2670.655 | -0.35978 | 0.18785 | -1.91527 | 0.055457 | 0.148811 |
| SMYD5 | 868.4293 | -0.36543 | 0.156143 | -2.34036 | 0.019265 | 0.073746 |
| DYSF | 12.67371 | -0.25859 | 0.594958 | -0.43463 | 0.663833 | NA |
| CCDC142 | 186.1621 | -0.1227 | 0.198965 | -0.61667 | 0.537455 | 0.680573 |
| GPR55 | 3.388535 | -0.19891 | 1.161905 | -0.17119 | 0.864075 | NA |
| SP110 | 856.0045 | -0.46123 | 0.237788 | -1.93968 | 0.052418 | 0.142855 |
| MRPL44 | 3397.733 | -0.31293 | 0.137232 | -2.28029 | 0.02259 | 0.082075 |
| CHRND | 0.134133 | -1.48909 | 4.077584 | -0.36519 | 0.71497 | NA |
| PAX3 | 2041.899 | -0.20055 | 0.148527 | -1.35029 | 0.176922 | 0.329058 |
| DOCK10 | 892.5146 | 0.366402 | 0.251412 | 1.457373 | 0.145013 | 0.286953 |
| TTLL4 | 927.0064 | -0.55368 | 0.15621 | -3.54446 | 3.93E-04 | 0.005852 |
| USP37 | 156.3211 | 0.545017 | 0.237608 | 2.293765 | 0.021804 | 0.08035 |
| HTR2B | 4308.474 | 0.897067 | 0.277161 | 3.236623 | 0.00121 | 0.011773 |
| ITM2C | 20544.98 | -0.10456 | 0.135164 | -0.77356 | 0.43919 | 0.599011 |
| SLC19A3 | 22.11972 | -0.58365 | 0.507577 | -1.14987 | 0.250199 | NA |
| SERPINE2 | 21122.89 | 0.281572 | 0.221085 | 1.273595 | 0.202807 | 0.359115 |
| DNAJB2 | 3124.265 | -0.07313 | 0.150838 | -0.48483 | 0.627795 | 0.750253 |
| TMBIM1 | 8253.349 | -0.16028 | 0.154458 | -1.03768 | 0.299418 | 0.466338 |
| CYP27A1 | 8048.371 | -0.55986 | 0.17305 | -3.23527 | 0.001215 | 0.01178 |
| EIF4E2 | 1580.371 | -0.02378 | 0.130073 | -0.18286 | 0.85491 | 0.913749 |
| ARMC9 | 7052.404 | 0.19425 | 0.136348 | 1.424672 | 0.154252 | 0.299387 |
| CAB39 | 1384.498 | 0.405031 | 0.179308 | 2.258856 | 0.023892 | 0.085385 |
| COX5B | 6241.924 | 0.00557 | 0.159587 | 0.034899 | 0.97216 | 0.984681 |
| REV1 | 528.9749 | 0.620101 | 0.168917 | 3.671036 | 2.42E-04 | 0.004366 |

| | | | | | | |
|---|---|---|---|---|---|---|
| TSGA10 | 49.56112 | 1.115019 | 0.435188 | 2.562154 | 0.010403 | 0.048343 |
| MFSD9 | 377.771 | 0.189586 | 0.158259 | 1.197946 | 0.230938 | 0.393132 |
| TMEM127 | 2593.182 | -0.3741 | 0.16707 | -2.23917 | 0.025145 | 0.088218 |
| EDAR | 0.326333 | 1.533889 | 3.826878 | 0.40082 | 0.688553 | NA |
| TGFBRAP1 | 2817.758 | -0.22998 | 0.146399 | -1.57094 | 0.116197 | 0.246557 |
| GCC2 | 355.4854 | 1.027308 | 1.125916 | 0.91242 | 0.361548 | 0.528818 |
| MRPS9 | 361.4625 | 0.30133 | 0.224878 | 1.339969 | 0.180255 | 0.332803 |
| GPR45 | 0.258578 | 1.233151 | 4.047809 | 0.304646 | 0.760635 | NA |
| C2orf49 | 428.2546 | 0.674373 | 0.19548 | 3.449832 | 5.61E-04 | 0.007268 |
| ANKRD36 | 38.04553 | 0.772081 | 0.561727 | 1.374477 | 0.169294 | NA |
| EPC2 | 413.0524 | 0.704286 | 0.164515 | 4.280986 | 1.86E-05 | 8.82E-04 |
| ARHGEF4 | 2.975254 | 0.137013 | 1.1669 | 0.117416 | 0.90653 | NA |
| VILL | 181.4889 | -0.12807 | 0.199153 | -0.64305 | 0.520191 | 0.667338 |
| FLNB | 618.0205 | -0.53227 | 0.240938 | -2.20918 | 0.027162 | 0.092718 |
| SCN2A | 12.24352 | 0.0934 | 0.589273 | 0.158501 | 0.874062 | NA |
| TBR1 | 0.653072 | 0.258115 | 3.011926 | 0.085698 | 0.931707 | NA |
| 7-Mar | 1137.078 | 0.543975 | 0.193722 | 2.808021 | 0.004985 | 0.029547 |
| ERMN | 0.582339 | -1.69412 | 2.750801 | -0.61586 | 0.537985 | NA |
| GALNT5 | 57.69749 | 0.856932 | 0.460094 | 1.862513 | 0.062531 | 0.161927 |
| SCN7A | 1.490236 | 2.70501 | 1.918689 | 1.409822 | 0.158592 | NA |
| TANK | 877.7139 | 1.006434 | 0.219502 | 4.585068 | 4.54E-06 | 3.99E-04 |
| CBWD2 | 139.2516 | 0.520206 | 0.266916 | 1.948947 | 0.051302 | 0.140999 |
| IL1RN | 0.84823 | 2.93367 | 2.713279 | 1.081227 | 0.279596 | NA |
| SMPD4 | 1359.466 | -0.37026 | 0.166818 | -2.21956 | 0.026449 | 0.091282 |
| WDR33 | 1021.147 | 0.098731 | 0.135894 | 0.726526 | 0.467516 | 0.624004 |
| CCDC115 | 1324.537 | -0.29875 | 0.146376 | -2.041 | 0.041251 | 0.123057 |
| SAP130 | 799.5784 | -0.46084 | 0.147624 | -3.12173 | 0.001798 | 0.015096 |
| BIN1 | 1169.227 | -0.43147 | 0.145802 | -2.95927 | 0.003084 | 0.021631 |
| IMP4 | 2244.646 | -0.08918 | 0.160054 | -0.55722 | 0.577378 | 0.712186 |
| HS6ST1 | 335.0599 | -0.25964 | 0.207144 | -1.25343 | 0.210048 | 0.367763 |
| UGGT1 | 2569.48 | -0.28006 | 0.133226 | -2.10214 | 0.035541 | 0.111053 |
| GYPC | 6096.033 | -0.00539 | 0.147401 | -0.03656 | 0.970834 | 0.983955 |
| IFT172 | 679.8241 | -0.3817 | 0.161008 | -2.37068 | 0.017755 | 0.069688 |
| EPT1 | 888.3038 | 0.452154 | 0.238716 | 1.894106 | 0.058211 | 0.153798 |
| CGREF1 | 88.71383 | 0.001482 | 0.257086 | 0.005766 | 0.9954 | 0.997908 |
| HADHB | 5740.807 | -0.15901 | 0.129243 | -1.23032 | 0.218577 | 0.378116 |
| KHK | 137.0024 | -0.00475 | 0.27253 | -0.01744 | 0.986084 | 0.992622 |
| ADCY3 | 1449.075 | -0.22941 | 0.155522 | -1.47511 | 0.140184 | 0.280391 |
| PPM1B | 1089.065 | 0.495862 | 0.16544 | 2.99722 | 0.002725 | 0.019937 |
| PNPT1 | 889.1343 | 0.572798 | 0.245402 | 2.334116 | 0.01959 | 0.074698 |
| DYNC2LI1 | 239.8853 | 0.714389 | 0.229973 | 3.106404 | 0.001894 | 0.015558 |
| LHCGR | 0.192403 | 0.975392 | 4.060716 | 0.240202 | 0.810174 | NA |
| SMEK2 | 1110.908 | 0.68955 | 0.198136 | 3.48019 | 5.01E-04 | 0.006784 |
| THUMPD2 | 98.87816 | 1.027521 | 0.310903 | 3.304956 | 9.50E-04 | 0.009992 |
| CYP1B1 | 3269.513 | 0.496396 | 0.334466 | 1.484146 | 0.13777 | 0.277148 |
| SULT6B1 | 0.326333 | 1.533889 | 3.826878 | 0.40082 | 0.688553 | NA |
| RAB1A | 5300.02 | 0.114058 | 0.138051 | 0.826199 | 0.408691 | 0.572402 |

| | | | | | | |
|---|---|---|---|---|---|---|
| ACTR2 | 7326.49 | 0.470103 | 0.172565 | 2.724207 | 0.006446 | 0.035027 |
| PREB | 2269.8 | -0.14693 | 0.131788 | -1.11494 | 0.264878 | 0.429314 |
| SLC5A6 | 1123.003 | -0.36001 | 0.176271 | -2.04238 | 0.041114 | 0.122803 |
| ABCG5 | 1.143305 | 1.751855 | 2.151234 | 0.814349 | 0.415445 | NA |
| PREPL | 1650.525 | 0.328019 | 0.186633 | 1.757567 | 0.078821 | 0.189295 |
| SLC3A1 | 0.594599 | -0.12152 | 2.771051 | -0.04385 | 0.965021 | NA |
| EMILIN1 | 286.6661 | -0.72072 | 0.185688 | -3.88137 | 1.04E-04 | 0.002626 |
| FBXO11 | 1102.891 | 0.280729 | 0.225372 | 1.245629 | 0.212901 | 0.371604 |
| SIX3 | 0.326536 | 0.037056 | 3.842554 | 0.009644 | 0.992306 | NA |
| ATRAID | 2488 | -0.25164 | 0.148395 | -1.69575 | 0.089934 | 0.207626 |
| CENPO | 429.9939 | -0.21861 | 0.202851 | -1.07769 | 0.28117 | 0.4473 |
| LRPPRC | 3475.989 | 0.329512 | 0.201082 | 1.638691 | 0.101278 | 0.22484 |
| TRIM54 | 0.162376 | 0.424913 | 4.079006 | 0.104171 | 0.917034 | NA |
| DTNB | 412.7463 | -0.30963 | 0.178576 | -1.73387 | 0.082942 | 0.195915 |
| AOX1 | 22.60601 | 1.15677 | 0.524661 | 2.204796 | 0.027468 | NA |
| ATIC | 3728.931 | -0.39938 | 0.16332 | -2.44541 | 0.014469 | 0.060524 |
| SMARCAL1 | 937.8664 | -0.45965 | 0.165307 | -2.78057 | 0.005426 | 0.031364 |
| BARD1 | 163.9245 | 0.169045 | 0.239345 | 0.706283 | 0.480012 | 0.634563 |
| STAT4 | 5.861493 | 0.780095 | 0.879749 | 0.886725 | 0.375227 | NA |
| MSTN | 3.395551 | 0.720309 | 1.271361 | 0.566565 | 0.57101 | NA |
| CARF | 99.46749 | 0.620222 | 0.2857 | 2.170889 | 0.02994 | 0.098867 |
| ASNSD1 | 951.5385 | 0.790612 | 0.211987 | 3.729537 | 1.92E-04 | 0.00379 |
| METTL5 | 576.3409 | 0.553953 | 0.174705 | 3.170796 | 0.00152 | 0.013653 |
| SSB | 894.0389 | 1.141769 | 0.655892 | 1.740787 | 0.081721 | 0.194149 |
| NAB1 | 507.7519 | 0.591372 | 0.269722 | 2.19252 | 0.028342 | 0.09539 |
| CDK15 | 113.8589 | -0.03423 | 0.266535 | -0.12844 | 0.897804 | 0.939904 |
| PPIG | 363.177 | 0.880889 | 0.95728 | 0.920199 | 0.357469 | 0.525021 |
| FASTKD1 | 455.7435 | 0.708937 | 0.2128 | 3.33148 | 8.64E-04 | 0.009456 |
| MDH1B | 28.55908 | 0.10885 | 0.401575 | 0.271059 | 0.786346 | NA |
| HECW2 | 0.296509 | -0.51709 | 3.982155 | -0.12985 | 0.896684 | NA |
| IDH1 | 3868.425 | 0.272833 | 0.146973 | 1.856346 | 0.063404 | 0.163404 |
| OLA1 | 1398.83 | 0.328906 | 0.161773 | 2.033135 | 0.042039 | 0.124655 |
| CIR1 | 398.3278 | 0.943429 | 0.334309 | 2.822028 | 0.004772 | 0.02863 |
| SSFA2 | 3953.71 | 0.71322 | 0.19379 | 3.68038 | 2.33E-04 | 0.004274 |
| CHRNA1 | 16.84002 | -0.0637 | 0.582574 | -0.10934 | 0.912934 | NA |
| FAM117B | 203.9826 | -0.22586 | 0.228091 | -0.99022 | 0.322069 | 0.490258 |
| WDR12 | 1125.99 | 0.343102 | 0.128885 | 2.662071 | 0.007766 | 0.039835 |
| ABI2 | 809.7636 | 0.011842 | 0.182808 | 0.064779 | 0.94835 | 0.971147 |
| ITGAV | 2714.173 | 0.736196 | 0.28107 | 2.619263 | 0.008812 | 0.043419 |
| SLC40A1 | 3.31461 | 0.433739 | 1.069846 | 0.405422 | 0.685167 | NA |
| GUCD1 | 3303.35 | -0.54531 | 0.126312 | -4.31719 | 1.58E-05 | 8.22E-04 |
| RNF185 | 1242.618 | -0.34518 | 0.13534 | -2.5505 | 0.010757 | 0.049294 |
| KIAA1644 | 2.137295 | 0.691348 | 1.581597 | 0.43712 | 0.662024 | NA |
| PARVG | 0.574231 | 0.93227 | 2.49998 | 0.372911 | 0.709215 | NA |
| PRDM15 | 225.7343 | -0.09281 | 0.20657 | -0.44927 | 0.653233 | 0.770862 |
| PFKL | 6783.42 | -0.39772 | 0.116261 | -3.42091 | 6.24E-04 | 0.007763 |
| HUNK | 123.2616 | 0.195687 | 0.285391 | 0.685679 | 0.492915 | 0.644197 |

| | | | | | | |
|---|---|---|---|---|---|---|
| COL6A1 | 2258.483 | -0.47982 | 0.156658 | -3.06288 | 0.002192 | 0.017165 |
| IFNAR1 | 1429.635 | 0.212946 | 0.159006 | 1.339236 | 0.180494 | 0.333033 |
| SOD1 | 7096.327 | 0.148853 | 0.144739 | 1.028423 | 0.303751 | 0.470961 |
| COL6A2 | 1339.079 | -0.01852 | 0.200165 | -0.09252 | 0.926285 | 0.958156 |
| SIK1 | 121.3059 | -0.53322 | 0.34551 | -1.5433 | 0.122759 | 0.256442 |
| TRPM2 | 0.298089 | -0.51711 | 3.975232 | -0.13008 | 0.896501 | NA |
| TMEM50B | 920.1705 | 0.172639 | 0.172638 | 1.000005 | 0.317308 | 0.485004 |
| APP | 14735.79 | -0.12324 | 0.118499 | -1.04 | 0.29834 | 0.46531 |
| DOPEY2 | 495.1476 | -0.72287 | 0.193051 | -3.74446 | 1.81E-04 | 0.003645 |
| URB1 | 2144.019 | -0.54669 | 0.176683 | -3.0942 | 0.001973 | 0.016057 |
| EMP3 | 7166.195 | -0.33715 | 0.174347 | -1.93379 | 0.053139 | 0.144222 |
| SAE1 | 5013.211 | -0.16073 | 0.136801 | -1.17491 | 0.240032 | 0.403304 |
| NTN5 | 1.465099 | 1.176849 | 1.91655 | 0.614046 | 0.539185 | NA |
| LMTK3 | 36.94602 | -0.40812 | 0.347629 | -1.17401 | 0.240389 | NA |
| GEMIN7 | 240.8261 | -0.20668 | 0.18805 | -1.09904 | 0.271749 | 0.437003 |
| RNPEPL1 | 4583.408 | -0.41802 | 0.152427 | -2.74244 | 0.006098 | 0.033788 |
| CAPN10 | 643.2009 | -0.10603 | 0.174594 | -0.60732 | 0.543638 | 0.685451 |
| ERVK3-1 | 158.1958 | 0.555328 | 0.209787 | 2.647103 | 0.008118 | 0.04099 |
| ZNF787 | 1227.594 | -0.33926 | 0.150066 | -2.26075 | 0.023774 | 0.085143 |
| ACPT | 1.035046 | -0.72706 | 1.945928 | -0.37363 | 0.708677 | NA |
| ZNF473 | 245.7824 | -0.06386 | 0.211408 | -0.30209 | 0.762584 | 0.85255 |
| FAM71E1 | 3.096941 | 0.328808 | 1.177613 | 0.279216 | 0.780079 | NA |
| RPS11 | 33531.3 | 0.260528 | 0.138449 | 1.881759 | 0.059869 | 0.157093 |
| RPL13A | 49570.3 | -0.06065 | 0.121388 | -0.49967 | 0.617307 | 0.742508 |
| CTU1 | 271.1721 | -0.25949 | 0.200226 | -1.29598 | 0.194981 | 0.350068 |
| NOSIP | 1748.064 | -0.03657 | 0.150047 | -0.24372 | 0.80745 | 0.885096 |
| IGLON5 | 26.52116 | 1.181085 | 0.661378 | 1.785795 | 0.074132 | NA |
| RCN3 | 182.9088 | -0.13765 | 0.298208 | -0.46158 | 0.644385 | 0.763536 |
| ZNF614 | 126.5015 | 0.516691 | 0.235245 | 2.196399 | 0.028063 | 0.094838 |
| | 109.9269 | 0.203824 | 0.228993 | 0.890088 | 0.373419 | 0.539792 |
| ACP1 | 1553.693 | 0.64819 | 0.163993 | 3.952541 | 7.73E-05 | 0.002172 |
| MBOAT2 | 658.357 | 0.306776 | 0.17692 | 1.733981 | 0.082921 | 0.195915 |
| OSR1 | 0.652869 | 1.292592 | 2.374873 | 0.544278 | 0.58625 | NA |
| GDF7 | 0.420954 | 1.923376 | 3.462803 | 0.555439 | 0.578594 | NA |
| PDIA6 | 5800.36 | -0.13959 | 0.123622 | -1.12915 | 0.258834 | 0.423177 |
| RHOB | 4990.295 | -0.0832 | 0.205999 | -0.40389 | 0.686297 | 0.795623 |
| ATP6V1C2 | 15.46788 | 0.805779 | 0.537941 | 1.497896 | 0.13416 | NA |
| HNRNPLL | 1535.197 | 0.133585 | 0.147573 | 0.905213 | 0.365353 | 0.532553 |
| GALM | 2159.889 | -0.06879 | 0.133828 | -0.514 | 0.607251 | 0.735038 |
| CAMKMT | 273.805 | -0.0075 | 0.175945 | -0.04263 | 0.965996 | 0.981393 |
| EML4 | 661.0573 | 0.119871 | 0.188185 | 0.636987 | 0.524133 | 0.670307 |
| CALM2 | 11686.83 | 0.232642 | 0.148603 | 1.565523 | 0.11746 | 0.248216 |
| CHAC2 | 73.01953 | 0.461211 | 0.303445 | 1.519919 | 0.128531 | 0.264779 |
| RPS27A | 12772.86 | 0.443473 | 0.158745 | 2.793613 | 0.005212 | 0.030399 |
| WDPCP | 123.2946 | 0.004223 | 0.262334 | 0.016098 | 0.987156 | 0.993273 |
| VPS54 | 498.2694 | 0.426191 | 0.219829 | 1.938735 | 0.052534 | 0.143104 |
| ASXL2 | 1188.29 | -0.12343 | 0.153318 | -0.80503 | 0.420805 | 0.583536 |

| | | | | | | |
|---|---|---|---|---|---|---|
| ETAA1 | 173.5946 | 1.009792 | 0.378627 | 2.666986 | 0.007653 | 0.039466 |
| SNRPG | 1003.545 | 0.42504 | 0.16305 | 2.606811 | 0.009139 | 0.044457 |
| ABHD1 | 9.92242 | 0.212346 | 0.66439 | 0.31961 | 0.749264 | NA |
| MEIS1 | 129.8157 | -0.17499 | 0.262618 | -0.66632 | 0.505206 | 0.654702 |
| TRIM43B | 0.134133 | -1.48909 | 4.077584 | -0.36519 | 0.71497 | NA |
| TRIM43 | 0.151697 | -1.546 | 4.075469 | -0.37934 | 0.704433 | NA |
| CIAO1 | 2506.709 | -0.11912 | 0.11683 | -1.01959 | 0.307925 | 0.475133 |
| ZNF514 | 147.3799 | 0.215525 | 0.240953 | 0.89447 | 0.37107 | 0.53772 |
| SNRNP200 | 6753.174 | -0.46447 | 0.131293 | -3.53766 | 4.04E-04 | 0.005952 |
| MRPS5 | 1663.035 | 0.163434 | 0.128469 | 1.272163 | 0.203315 | 0.359796 |
| ANKRD53 | 46.85149 | -0.69422 | 0.337495 | -2.05696 | 0.03969 | NA |
| TPRKB | 362.9247 | 0.550445 | 0.176318 | 3.121897 | 0.001797 | 0.015096 |
| NAT8 | 0.526843 | -0.38719 | 2.855892 | -0.13558 | 0.892156 | NA |
| EXOC6B | 1583.561 | -0.14041 | 0.143414 | -0.97907 | 0.327544 | 0.495828 |
| SFXN5 | 313.6226 | -0.09287 | 0.167811 | -0.55342 | 0.579977 | 0.713906 |
| TEX261 | 3303.278 | -0.27138 | 0.166875 | -1.62623 | 0.1039 | 0.228948 |
| DQX1 | 0.987484 | 1.983755 | 2.381011 | 0.833157 | 0.404756 | NA |
| DUSP11 | 453.9281 | 0.316026 | 0.182077 | 1.735672 | 0.082622 | 0.195427 |
| ST6GAL2 | 0.783004 | -0.66714 | 2.898037 | -0.2302 | 0.817934 | NA |
| NPHP1 | 100.5893 | 0.693026 | 0.270975 | 2.557529 | 0.010542 | 0.048723 |
| MALL | 5.428448 | -0.15699 | 0.963128 | -0.163 | 0.870516 | NA |
| THNSL2 | 438.7062 | -0.41737 | 0.19639 | -2.12524 | 0.033567 | 0.107001 |
| RALB | 1620.275 | 0.144003 | 0.143457 | 1.003804 | 0.315473 | 0.482886 |
| TMEM177 | 462.8887 | -0.21598 | 0.182788 | -1.18159 | 0.237368 | 0.400283 |
| NT5DC4 | 0.163957 | 0.424913 | 4.079006 | 0.104171 | 0.917034 | NA |
| RABL2A | 138.478 | 0.042255 | 0.225189 | 0.18764 | 0.851158 | 0.911516 |
| SLC20A1 | 1404.946 | -0.05671 | 0.159764 | -0.35496 | 0.722622 | 0.823698 |
| FBLN7 | 73.60003 | -0.79312 | 0.270967 | -2.927 | 0.003423 | 0.022949 |
| | 0.151697 | -1.546 | 4.075469 | -0.37934 | 0.704433 | NA |
| ZC3H8 | 59.36606 | 0.818428 | 0.325739 | 2.512527 | 0.011987 | 0.053383 |
| LIPT1 | 82.91151 | 0.688199 | 0.254317 | 2.706072 | 0.006808 | 0.036429 |
| CNGA3 | 0.096202 | 0.408546 | 4.079639 | 0.100143 | 0.920231 | NA |
| FAHD2B | 301.0608 | -0.38271 | 0.215339 | -1.77724 | 0.075529 | 0.183785 |
| LYG1 | 4.607726 | -0.05412 | 1.088989 | -0.0497 | 0.96036 | NA |
| AFF3 | 89.15721 | -0.74814 | 0.341276 | -2.19218 | 0.028367 | 0.09539 |
| UBXN4 | 1214.584 | 0.935021 | 0.317089 | 2.948766 | 0.00319 | 0.022067 |
| SPOPL | 233.9118 | 0.696601 | 0.191104 | 3.645129 | 2.67E-04 | 0.004599 |
| THSD7B | 0.440301 | 0.468383 | 3.449808 | 0.135771 | 0.892003 | NA |
| GPR17 | 6.142073 | 0.044894 | 0.834691 | 0.053785 | 0.957107 | NA |
| POLR2D | 539.1971 | -0.1489 | 0.184602 | -0.80658 | 0.419911 | 0.58289 |
| AMMECR1 | 1870.37 | -0.30404 | 0.132984 | -2.28628 | 0.022238 | 0.081443 |
| PKP4 | 1281.068 | 0.066097 | 0.140966 | 0.468886 | 0.639151 | 0.759385 |
| SCN1A | 0.163957 | 0.424913 | 4.079006 | 0.104171 | 0.917034 | NA |
| SLC4A10 | 1.376428 | -1.32951 | 2.007901 | -0.66214 | 0.507882 | NA |
| SCRN3 | 324.899 | 0.668953 | 0.254344 | 2.630108 | 0.008536 | 0.042322 |
| KIAA1715 | 830.4767 | 0.574806 | 0.2253 | 2.551296 | 0.010732 | 0.049294 |
| ZNF385B | 0.258578 | 1.233151 | 4.047809 | 0.304646 | 0.760635 | NA |

| | | | | | | |
|---|---|---|---|---|---|---|
| TMEFF2 | 0.163957 | 0.424913 | 4.079006 | 0.104171 | 0.917034 | NA |
| CDCA7 | 123.5023 | 0.096178 | 0.317387 | 0.30303 | 0.761867 | 0.851908 |
| DLX1 | 128.3298 | 0.112541 | 0.258199 | 0.435868 | 0.662932 | 0.778356 |
| UBR3 | 1036.702 | 0.500795 | 0.219774 | 2.278676 | 0.022686 | 0.082322 |
| PHOSPHO2 | 25.09976 | 0.777427 | 0.445123 | 1.746546 | 0.080716 | NA |
| GULP1 | 1616.456 | 1.028954 | 0.232213 | 4.431078 | 9.38E-06 | 6.11E-04 |
| FAM171B | 300.5383 | 0.686002 | 0.272468 | 2.517729 | 0.011811 | 0.05276 |
| HSPD1 | 6494.428 | 0.293683 | 0.164226 | 1.788288 | 0.07373 | 0.181055 |
| CCDC150 | 26.87282 | 1.777278 | 0.458657 | 3.874964 | 1.07E-04 | NA |
| METTL21A | 214.8516 | 0.127716 | 0.17915 | 0.712902 | 0.475907 | 0.630941 |
| UNC80 | 7.999178 | -0.34415 | 0.77978 | -0.44134 | 0.658966 | NA |
| PTH2R | 8.446619 | -0.64792 | 0.762368 | -0.84988 | 0.395394 | NA |
| CPO | 2.642062 | 1.686433 | 1.542721 | 1.093155 | 0.274326 | NA |
| NBEAL1 | 415.7627 | 0.9859 | 0.612404 | 1.609885 | 0.107423 | 0.233913 |
| KANSL1L | 294.2237 | 0.551053 | 0.227318 | 2.424152 | 0.015344 | 0.062965 |
| SPAG16 | 241.3742 | 0.187328 | 0.213283 | 0.878307 | 0.379777 | 0.546126 |
| ABCA12 | 43.66635 | -0.37914 | 0.44245 | -0.85692 | 0.391491 | NA |
| SUMF1 | 1623.076 | -0.124 | 0.167615 | -0.73977 | 0.45944 | 0.617857 |
| RHBDD1 | 414.6351 | -0.03914 | 0.177496 | -0.22049 | 0.825493 | 0.895106 |
| ACKR3 | 0.327913 | 1.543154 | 4.021063 | 0.383768 | 0.701151 | NA |
| TRPM8 | 45.65487 | 1.479759 | 0.413261 | 3.580688 | 3.43E-04 | NA |
| HES6 | 5688.674 | -0.41099 | 0.191478 | -2.14641 | 0.03184 | 0.103399 |
| ESPNL | 1.586192 | -0.40493 | 1.57816 | -0.25658 | 0.797501 | NA |
| ANKMY1 | 161.1998 | -0.14259 | 0.247236 | -0.57672 | 0.56413 | 0.702054 |
| COPS7B | 780.0015 | -0.32247 | 0.142489 | -2.26316 | 0.023626 | 0.084695 |
| DIS3L2 | 1096.043 | -0.3524 | 0.156476 | -2.2521 | 0.024316 | 0.086308 |
| CPNE9 | 0.096202 | 0.408546 | 4.079639 | 0.100143 | 0.920231 | NA |
| FANCD2 | 628.6819 | -0.42544 | 0.214511 | -1.98331 | 0.047332 | 0.13392 |
| TAMM41 | 215.0552 | -0.27954 | 0.206255 | -1.35532 | 0.175315 | 0.327146 |
| VGLL4 | 2084.108 | 0.090743 | 0.197872 | 0.458593 | 0.646527 | 0.765233 |
| RAB5A | 1789.652 | 0.554262 | 0.180193 | 3.075937 | 0.002098 | 0.016685 |
| FAM134A | 5797.673 | -0.35429 | 0.144894 | -2.4452 | 0.014477 | 0.060538 |
| CTDSP1 | 2378.295 | -0.26344 | 0.146636 | -1.79653 | 0.07241 | 0.178874 |
| RQCD1 | 1706.692 | -0.22841 | 0.140323 | -1.62773 | 0.103581 | 0.228287 |
| 4-Mar | 12.95796 | 0.577097 | 1.005255 | 0.57408 | 0.565913 | NA |
| STK11IP | 946.2693 | -0.43447 | 0.157553 | -2.75762 | 0.005822 | 0.032734 |
| GMPPA | 1231.057 | -0.25146 | 0.143594 | -1.75121 | 0.07991 | 0.191226 |
| GRIP2 | 0.719393 | -0.9644 | 2.652838 | -0.36354 | 0.716204 | NA |
| EAF1 | 455.8167 | 0.102612 | 0.166871 | 0.614917 | 0.538609 | 0.681093 |
| CNTN4 | 59.02051 | 0.977862 | 0.333535 | 2.93181 | 0.00337 | 0.022712 |
| DYNC1LI1 | 1274.873 | 0.732995 | 0.189253 | 3.873092 | 1.07E-04 | 0.002688 |
| RBMS3 | 79.67314 | -0.12672 | 0.365517 | -0.34669 | 0.728827 | 0.827926 |
| GADL1 | 2.94063 | 0.01549 | 1.164381 | 0.013303 | 0.989386 | NA |
| OSBPL10 | 1416.897 | 0.06267 | 0.161942 | 0.386993 | 0.698762 | 0.806239 |
| POMGNT2 | 949.8415 | -0.33811 | 0.16892 | -2.00162 | 0.045326 | 0.130359 |
| ACKR2 | 4.28352 | -0.06196 | 1.002882 | -0.06178 | 0.950737 | NA |
| FAM198A | 6.279148 | -0.76742 | 0.861271 | -0.89103 | 0.372912 | NA |

| | | | | | | |
|---|---|---|---|---|---|---|
| CSRNP1 | 215.7663 | 0.1896 | 0.321243 | 0.590209 | 0.55505 | 0.695073 |
| SLC25A38 | 688.3896 | -0.35263 | 0.15378 | -2.29309 | 0.021843 | 0.080418 |
| ITGA9 | 80.31749 | -0.81812 | 0.347556 | -2.35392 | 0.018576 | 0.071874 |
| SLC22A14 | 1.945123 | 1.531652 | 1.629791 | 0.939784 | 0.347328 | NA |
| GOLGA4 | 873.1381 | 1.012422 | 1.300792 | 0.778312 | 0.436385 | 0.597177 |
| CTDSPL | 1448.221 | -0.13349 | 0.130119 | -1.02587 | 0.304951 | 0.472185 |
| STAC | 22.25644 | -0.37709 | 0.504536 | -0.74739 | 0.454825 | NA |
| IQSEC1 | 1082.086 | -0.50901 | 0.2152 | -2.36527 | 0.018017 | 0.07033 |
| CAND2 | 156.1207 | -0.48037 | 0.255248 | -1.88199 | 0.059837 | 0.157045 |
| RPL32 | 14828.65 | 0.255243 | 0.156569 | 1.630228 | 0.103053 | 0.227389 |
| PTPRG | 697.4879 | -0.15064 | 0.19967 | -0.75447 | 0.450569 | 0.609899 |
| IL17RD | 109.6929 | -0.67432 | 0.252649 | -2.66901 | 0.007608 | 0.039383 |
| SLC25A26 | 832.689 | 0.030777 | 0.160039 | 0.192308 | 0.847501 | 0.909159 |
| TMF1 | 390.8645 | 1.186449 | 0.738194 | 1.607233 | 0.108003 | 0.234803 |
| LRIG1 | 242.3113 | 0.091341 | 0.187547 | 0.487031 | 0.626236 | 0.749266 |
| LIMD1 | 2834.499 | -0.97241 | 0.218693 | -4.44649 | 8.73E-06 | 5.78E-04 |
| ZNF660 | 16.78369 | 0.327867 | 0.567174 | 0.578071 | 0.563216 | NA |
| AMT | 37.60942 | -0.44288 | 0.381246 | -1.16167 | 0.245369 | NA |
| TCTA | 1789.28 | -0.24115 | 0.119177 | -2.02346 | 0.043026 | 0.126367 |
| NICN1 | 549.154 | -0.52802 | 0.154719 | -3.41276 | 6.43E-04 | 0.007876 |
| UCN2 | 480.0695 | 0.918368 | 0.633983 | 1.448568 | 0.147458 | 0.290626 |
| VPRBP | 742.7472 | -0.22364 | 0.16973 | -1.31762 | 0.187629 | 0.341836 |
| MANF | 2027.703 | 0.141224 | 0.160933 | 0.877535 | 0.380196 | 0.546453 |
| | 1.710434 | 0.70104 | 1.69845 | 0.412753 | 0.679788 | NA |
| ANO7 | 102.0865 | -0.22043 | 0.302984 | -0.72752 | 0.466907 | 0.623713 |
| WASH2P | 111.2058 | 0.040005 | 0.236059 | 0.169468 | 0.865428 | 0.920317 |
| SLX4IP | 26.86509 | -0.11441 | 0.437581 | -0.26147 | 0.79373 | NA |
| ADAM33 | 1.493931 | -0.17651 | 1.55812 | -0.11328 | 0.909808 | NA |
| CSRP2BP | 682.4753 | -0.23957 | 0.144223 | -1.66107 | 0.096699 | 0.21817 |
| TMC2 | 0.610582 | -0.18034 | 2.753139 | -0.0655 | 0.947774 | NA |
| FRG1B | 271.8091 | 0.05564 | 0.223379 | 0.249083 | 0.803296 | 0.882175 |
| JPH2 | 249.8865 | -0.01073 | 0.303184 | -0.0354 | 0.971758 | 0.984547 |
| DUSP15 | 276.2334 | -0.19065 | 0.188076 | -1.01369 | 0.310729 | 0.478203 |
| COMMD7 | 1759.808 | -0.10881 | 0.16652 | -0.65345 | 0.513468 | 0.660715 |
| C20orf144 | 0.326333 | 1.533889 | 3.826878 | 0.40082 | 0.688553 | NA |
| KIAA1755 | 78.02091 | -0.05388 | 0.417802 | -0.12895 | 0.897395 | 0.939815 |
| SPATA25 | 6.974181 | -0.0644 | 0.83478 | -0.07715 | 0.938504 | NA |
| DSN1 | 412.15 | 0.324652 | 0.200111 | 1.622355 | 0.104727 | 0.230128 |
| SOGA1 | 2293.368 | -0.73284 | 0.250177 | -2.9293 | 0.003397 | 0.022819 |
| CNBD2 | 3.338084 | -0.18775 | 1.206716 | -0.15558 | 0.876361 | NA |
| LSM14B | 990.6095 | -0.14989 | 0.149928 | -0.99974 | 0.317436 | 0.485136 |
| YTHDF1 | 1244.582 | -0.19895 | 0.133222 | -1.49341 | 0.135331 | 0.27404 |
| CABLES2 | 247.0948 | -0.48541 | 0.195291 | -2.48558 | 0.012934 | 0.05628 |
| HNMT | 1080.519 | 0.512842 | 0.260146 | 1.971361 | 0.048683 | 0.136341 |
| LYPD1 | 804.2559 | 0.739407 | 0.574523 | 1.286992 | 0.198097 | 0.353894 |
| LYPD6B | 1.209536 | 1.431081 | 1.892393 | 0.756228 | 0.449512 | NA |
| PPP1R1C | 5.890479 | -0.81139 | 1.012334 | -0.80151 | 0.422838 | NA |

| | | | | | | |
|---|---|---|---|---|---|---|
| C2orf50 | 0.528424 | -0.38142 | 2.853651 | -0.13366 | 0.893671 | NA |
| CRIM1 | 1020.903 | 0.370416 | 0.185935 | 1.992186 | 0.046351 | 0.132315 |
| ITPR1 | 419.6001 | 0.026471 | 0.158911 | 0.166575 | 0.867705 | 0.921425 |
| NGLY1 | 403.435 | 0.458116 | 0.196304 | 2.333706 | 0.019611 | 0.074735 |
| OXSM | 306.0267 | 0.099457 | 0.167791 | 0.592745 | 0.553352 | 0.693776 |
| MAGI1 | 23.56808 | -0.02893 | 0.695055 | -0.04163 | 0.966798 | NA |
| TMEM18 | 639.5 | 0.018445 | 0.143806 | 0.12826 | 0.897943 | 0.939966 |
| PIGF | 246.9053 | 0.198742 | 0.199586 | 0.995772 | 0.319361 | 0.487199 |
| ANKAR | 11.12163 | 0.887991 | 0.617652 | 1.437689 | 0.150522 | NA |
| INPP1 | 726.2152 | 0.077044 | 0.216748 | 0.355456 | 0.722248 | 0.823531 |
| MFSD6 | 634.4537 | 0.298535 | 0.151798 | 1.966665 | 0.049222 | 0.137268 |
| RNF144A | 1395.187 | -0.56285 | 0.210902 | -2.66877 | 0.007613 | 0.039391 |
| ASAP2 | 1535.443 | 0.250463 | 0.178159 | 1.40584 | 0.159772 | 0.306935 |
| ADAM17 | 431.7217 | 0.456834 | 0.176324 | 2.590882 | 0.009573 | 0.045835 |
| NBAS | 1506.793 | -0.2822 | 0.131013 | -2.15396 | 0.031243 | 0.101895 |
| KCNE4 | 2353.98 | -0.56835 | 0.207547 | -2.73841 | 0.006174 | 0.033979 |
| AP1S3 | 124.1859 | 0.225647 | 0.22562 | 1.000118 | 0.317253 | 0.484982 |
| CCDC74B | 62.34053 | 0.165031 | 0.345624 | 0.477486 | 0.633016 | 0.754818 |
| MZT2B | 1289.863 | -0.10798 | 0.149289 | -0.7233 | 0.469498 | 0.625866 |
| FAM168B | 3055.543 | -0.07891 | 0.12528 | -0.62985 | 0.528794 | 0.673961 |
| | 138.2613 | -0.01683 | 0.22592 | -0.07449 | 0.940617 | 0.966779 |
| MGAT5 | 3814.072 | -0.14101 | 0.193887 | -0.72729 | 0.467047 | 0.623713 |
| TMEM163 | 1.216567 | -0.28085 | 1.97818 | -0.14197 | 0.887102 | NA |
| GPATCH11 | 254.3596 | 1.226258 | 0.220458 | 5.562317 | 2.66E-08 | 1.50E-05 |
| GEMIN6 | 505.8092 | 0.373828 | 0.212438 | 1.759708 | 0.078457 | 0.188702 |
| TMEM178A | 2.374817 | 2.591957 | 1.575311 | 1.645362 | 0.099895 | NA |
| SPC25 | 149.691 | -0.09101 | 0.28778 | -0.31625 | 0.751815 | 0.844731 |
| PDK1 | 141.6999 | 0.162299 | 0.242685 | 0.668763 | 0.503647 | 0.653539 |
| TCF7L1 | 458.3126 | -0.45069 | 0.194761 | -2.31405 | 0.020665 | 0.077477 |
| TGOLN2 | 6705.606 | -0.30778 | 0.157162 | -1.95835 | 0.050189 | 0.138943 |
| SH2D6 | 2.789432 | -0.1464 | 1.272725 | -0.11503 | 0.908423 | NA |
| ZNF547 | 54.01912 | 0.548958 | 0.321366 | 1.708204 | 0.087598 | 0.203424 |
| ZNF773 | 145.4273 | -0.13721 | 0.249285 | -0.5504 | 0.582043 | 0.715632 |
| ZNF776 | 375.2133 | 0.307421 | 0.194516 | 1.58044 | 0.114006 | 0.243127 |
| ZNF256 | 78.31338 | 0.07667 | 0.31994 | 0.239638 | 0.810611 | 0.886758 |
| ZNF837 | 105.5965 | -0.44616 | 0.232305 | -1.92057 | 0.054786 | 0.14751 |
| ZFP36L2 | 1363.389 | -0.50185 | 0.20313 | -2.47059 | 0.013489 | 0.057751 |
| PLEKHH2 | 131.2494 | 0.95846 | 0.278745 | 3.438486 | 5.85E-04 | 0.007457 |
| GPD1L | 775.8945 | -0.04994 | 0.154988 | -0.32221 | 0.747296 | 0.84124 |
| CLEC4F | 0.096202 | 0.408546 | 4.079639 | 0.100143 | 0.920231 | NA |
| SLC30A6 | 694.4137 | 0.277919 | 0.196325 | 1.415607 | 0.156891 | 0.303119 |
| RASGRP3 | 696.9078 | 0.616534 | 0.217947 | 2.828822 | 0.004672 | 0.028187 |
| ACMSD | 0.298089 | -0.51711 | 3.975232 | -0.13008 | 0.896501 | NA |
| ACOXL | 0.585114 | 1.12675 | 2.73598 | 0.411827 | 0.680466 | NA |
| BCL2L11 | 117.3665 | -0.39968 | 0.29537 | -1.35316 | 0.176003 | 0.328121 |
| ANAPC1 | 858.3907 | -0.17447 | 0.159326 | -1.09507 | 0.273486 | 0.43932 |
| RGPD3 | 0.748867 | 2.740193 | 2.468928 | 1.109871 | 0.267054 | NA |

| | | | | | | |
|---|---|---|---|---|---|---|
| RANBP2 | 1386.806 | 0.541949 | 0.287781 | 1.883196 | 0.059674 | 0.156754 |
| MERTK | 123.8198 | 0.362015 | 0.236062 | 1.533557 | 0.125139 | 0.260016 |
| TMEM87B | 602.1014 | 0.428662 | 0.150394 | 2.85026 | 0.004368 | 0.026944 |
| NR4A2 | 342.6674 | 0.17997 | 0.209399 | 0.859459 | 0.390087 | 0.555494 |
| CCDC148 | 7.922402 | 0.990735 | 0.816615 | 1.213221 | 0.225045 | NA |
| PLA2R1 | 402.134 | 0.486355 | 0.17423 | 2.791461 | 0.005247 | 0.030564 |
| RBMS1 | 1137.103 | 0.465712 | 0.251873 | 1.848991 | 0.064459 | 0.165367 |
| SCN3A | 5.081602 | 1.592675 | 0.936965 | 1.699824 | 0.089164 | NA |
| CMTM7 | 460.4764 | -0.18601 | 0.165256 | -1.12558 | 0.260343 | 0.424593 |
| FBXL2 | 512.7023 | 0.133894 | 0.160627 | 0.833568 | 0.404525 | 0.569031 |
| UBP1 | 2032.962 | 0.033182 | 0.129979 | 0.255284 | 0.798503 | 0.878213 |
| RMND5A | 862.9789 | 0.15476 | 0.212808 | 0.727226 | 0.467088 | 0.623713 |
| CD8A | 0.364467 | -1.20166 | 3.685431 | -0.32606 | 0.744381 | NA |
| RPIA | 318.9731 | 0.109894 | 0.165274 | 0.664923 | 0.5061 | 0.655288 |
| SPHKAP | 1.24767 | 0.253795 | 1.932703 | 0.131316 | 0.895525 | NA |
| PID1 | 0.613743 | -0.17004 | 2.749758 | -0.06184 | 0.950691 | NA |
| TRIP12 | 4827.188 | 0.201036 | 0.145685 | 1.379942 | 0.167604 | 0.317106 |
| FBXO36 | 68.26428 | -0.1163 | 0.289257 | -0.40208 | 0.687628 | 0.796777 |
| GAL3ST2 | 0.689366 | -1.56115 | 2.700123 | -0.57818 | 0.563145 | NA |
| CCDC173 | 1.71604 | -0.01617 | 1.776198 | -0.00911 | 0.992735 | NA |
| ATP5G3 | 6538.456 | -0.01894 | 0.146499 | -0.1293 | 0.897122 | 0.939771 |
| CXADR | 32.3714 | 0.24154 | 0.412122 | 0.586088 | 0.557816 | NA |
| BTG3 | 1168.041 | 0.770304 | 0.147906 | 5.208062 | 1.91E-07 | 5.14E-05 |
| C21orf91 | 881.1706 | 0.576796 | 0.268591 | 2.147486 | 0.031755 | 0.103214 |
| CHODL | 6.120558 | -0.3267 | 0.855491 | -0.38189 | 0.702542 | NA |
| NCAM2 | 694.2676 | 0.525127 | 0.293199 | 1.791024 | 0.073289 | 0.180408 |
| MRPL39 | 588.9505 | 0.365877 | 0.1597 | 2.291024 | 0.021962 | 0.080757 |
| JAM2 | 4.435655 | 1.633151 | 1.204899 | 1.355425 | 0.175282 | NA |
| ATP5J | 2192.482 | 0.273965 | 0.145608 | 1.881529 | 0.0599 | 0.157094 |
| GABPA | 328.629 | 0.813092 | 0.316633 | 2.567934 | 0.010231 | 0.047882 |
| ADAMTS1 | 236.2828 | 0.150378 | 0.182754 | 0.822842 | 0.410598 | 0.573995 |
| ADAMTS5 | 4.980102 | 1.156546 | 1.06022 | 1.090855 | 0.275337 | NA |
| TSEN2 | 329.6294 | 0.265325 | 0.18058 | 1.469296 | 0.141753 | 0.282575 |
| XPC | 1308.098 | 0.167189 | 0.179829 | 0.929714 | 0.352519 | 0.520437 |
| CCDC174 | 335.1057 | 0.753997 | 0.208041 | 3.62427 | 2.90E-04 | 0.004811 |
| FGD5 | 0.288605 | 1.443619 | 4.029058 | 0.358302 | 0.720117 | NA |
| DPH3 | 1221.275 | 0.443665 | 0.181664 | 2.442234 | 0.014597 | 0.060786 |
| OXNAD1 | 377.9143 | 0.1457 | 0.161454 | 0.902427 | 0.36683 | 0.534116 |
| PLCL2 | 173.121 | 0.129002 | 0.242719 | 0.531487 | 0.595081 | 0.725343 |
| ACSS1 | 894.3565 | -0.67904 | 0.215025 | -3.15798 | 0.001589 | 0.014065 |
| CNTNAP5 | 0.096202 | 0.408546 | 4.079639 | 0.100143 | 0.920231 | NA |
| PROM2 | 2.614297 | -0.35047 | 1.216186 | -0.28817 | 0.773216 | NA |
| HSPA13 | 1216.391 | 0.72489 | 0.191187 | 3.791517 | 1.50E-04 | 0.003239 |
| USP25 | 712.9634 | 0.552953 | 0.234948 | 2.353511 | 0.018597 | 0.071931 |
| DBI | 4476.611 | 0.63816 | 0.162562 | 3.925633 | 8.65E-05 | 0.002336 |
| NIFK | 538.8157 | 0.887016 | 0.235602 | 3.764886 | 1.67E-04 | 0.003495 |
| RBM45 | 250.0677 | 0.416442 | 0.226683 | 1.837114 | 0.066193 | 0.16789 |

| | | | | | | |
|---|---|---|---|---|---|---|
| TTN | 148.0957 | 0.932997 | 0.231189 | 4.035654 | 5.45E-05 | 0.001728 |
| KCTD18 | 322.9648 | 0.148612 | 0.173802 | 0.855068 | 0.392514 | 0.557543 |
| FAM126B | 118.0701 | 0.243893 | 0.287453 | 0.848462 | 0.396181 | 0.56087 |
| ALS2CR12 | 12.74255 | 1.045826 | 0.639823 | 1.634554 | 0.102143 | NA |
| ALS2CR11 | 15.5973 | 0.745515 | 0.552708 | 1.348842 | 0.177388 | NA |
| TMEM237 | 428.9862 | 0.219608 | 0.170747 | 1.286165 | 0.198386 | 0.354196 |
| FZD7 | 427.065 | -0.185 | 0.388423 | -0.4763 | 0.633863 | 0.755448 |
| N6AMT1 | 264.9358 | 0.500791 | 0.206557 | 2.424471 | 0.015331 | 0.062931 |
| RWDD2B | 713.9112 | 0.119082 | 0.160729 | 0.740889 | 0.458761 | 0.617112 |
| USP16 | 567.493 | 1.214178 | 0.303876 | 3.995642 | 6.45E-05 | 0.001931 |
| CCT8 | 4676.723 | 0.30025 | 0.143796 | 2.088032 | 0.036795 | 0.113713 |
| MAP3K7CL | 15.49427 | 0.618208 | 0.663146 | 0.932235 | 0.351215 | NA |
| BACH1 | 668.9186 | 0.895318 | 0.184397 | 4.855393 | 1.20E-06 | 1.79E-04 |
| TIAM1 | 394.1118 | -0.29211 | 0.180728 | -1.6163 | 0.106029 | 0.231871 |
| SCAF4 | 855.1391 | -0.54844 | 0.166675 | -3.29048 | 0.001 | 0.010337 |
| LHFPL4 | 0.096202 | 0.408546 | 4.079639 | 0.100143 | 0.920231 | NA |
| B3GNT7 | 15.91213 | -0.77474 | 0.507387 | -1.52693 | 0.126779 | NA |
| PDE6D | 846.3557 | -0.02822 | 0.182566 | -0.15457 | 0.87716 | 0.926461 |
| BRPF1 | 643.9631 | -0.30019 | 0.152665 | -1.96636 | 0.049257 | 0.13727 |
| RPUSD3 | 1215.339 | -0.21706 | 0.139526 | -1.55572 | 0.119774 | 0.251758 |
| TATDN2 | 2042.227 | -0.28967 | 0.201136 | -1.44015 | 0.149825 | 0.293243 |
| GHRL | 1.154989 | -0.15951 | 1.966205 | -0.08112 | 0.935343 | NA |
| SEC13 | 2750.207 | -0.06808 | 0.14285 | -0.47661 | 0.633642 | 0.755323 |
| EXOG | 342.9944 | -0.03117 | 0.177498 | -0.17562 | 0.860592 | 0.917227 |
| TIMP4 | 63.90203 | 0.387465 | 0.47514 | 0.815477 | 0.414799 | 0.578115 |
| SYN2 | 5.495967 | 0.110851 | 0.888614 | 0.124746 | 0.900725 | NA |
| CACNA1D | 34.70031 | 0.067817 | 0.378472 | 0.179187 | 0.857791 | NA |
| CACNA2D3 | 0.151697 | -1.546 | 4.075469 | -0.37934 | 0.704433 | NA |
| APPL1 | 559.4901 | 0.646914 | 0.286507 | 2.257937 | 0.02395 | 0.085416 |
| DSCR3 | 3298.258 | -0.01077 | 0.131311 | -0.08203 | 0.934624 | 0.963308 |
| DYRK1A | 780.3181 | 0.133757 | 0.152775 | 0.875517 | 0.381293 | 0.547441 |
| ERG | 0.682693 | 2.623213 | 2.518338 | 1.041645 | 0.297576 | NA |
| ETS2 | 346.5483 | 0.048972 | 0.189916 | 0.25786 | 0.796515 | 0.87685 |
| LCA5L | 7.595756 | 0.722768 | 0.747439 | 0.966993 | 0.333548 | NA |
| MX1 | 6898.78 | -1.14641 | 0.310741 | -3.68928 | 2.25E-04 | 0.004177 |
| C2CD2 | 270.7961 | -0.30013 | 0.195854 | -1.53243 | 0.125417 | 0.260388 |
| FMNL2 | 254.2926 | 0.018195 | 0.270371 | 0.067295 | 0.946347 | 0.970016 |
| GAREML | 196.269 | -0.22483 | 0.23615 | -0.95208 | 0.341056 | 0.509157 |
| DRC1 | 1.439667 | 1.722755 | 1.816404 | 0.948443 | 0.342904 | NA |
| AGAP1 | 939.5245 | 0.150188 | 0.160399 | 0.936343 | 0.349097 | 0.517189 |
| KRTCAP3 | 2.08364 | 0.821011 | 1.46384 | 0.560861 | 0.574892 | NA |
| BRE | 1045.184 | -0.39922 | 0.163472 | -2.44214 | 0.0146 | 0.060786 |
| DUSP2 | 9.280623 | 0.12476 | 0.698272 | 0.17867 | 0.858197 | NA |
| GALNT14 | 156.2274 | -0.41589 | 0.315933 | -1.31638 | 0.188048 | 0.342128 |
| CNNM4 | 411.3362 | -0.59414 | 0.176355 | -3.36899 | 7.54E-04 | 0.008786 |
| SLC13A3 | 125.9975 | -0.51098 | 0.250215 | -2.04217 | 0.041135 | 0.122803 |
| MITD1 | 213.8396 | 1.049256 | 0.25474 | 4.118936 | 3.81E-05 | 0.001415 |

| | | | | | | |
|---|---|---|---|---|---|---|
| EIF5B | 1540.712 | 0.761038 | 1.015885 | 0.749138 | 0.453774 | 0.612629 |
| C2orf62 | 0.58526 | 0.001197 | 3.235889 | 3.70E-04 | 0.999705 | NA |
| CNOT11 | 1294.379 | -0.31356 | 0.141915 | -2.20953 | 0.027138 | 0.092684 |
| KCNB1 | 0.163957 | 0.424913 | 4.079006 | 0.104171 | 0.917034 | NA |
| B4GALT5 | 3095.862 | -0.53878 | 0.143779 | -3.74729 | 1.79E-04 | 0.003621 |
| SPATA2 | 347.6494 | -0.45723 | 0.192409 | -2.37634 | 0.017485 | 0.069119 |
| ZFAND2B | 854.5722 | -0.10069 | 0.176167 | -0.57154 | 0.567636 | 0.70442 |
| MIS18A | 225.8312 | 0.039118 | 0.198202 | 0.197365 | 0.843542 | 0.906882 |
| C21orf59 | 32.07704 | 0.386745 | 0.406501 | 0.951401 | 0.341401 | NA |
| SYNJ1 | 479.2123 | -0.06069 | 0.180462 | -0.33632 | 0.736628 | 0.833121 |
| PAXBP1 | 297.7144 | 1.159581 | 0.324899 | 3.569051 | 3.58E-04 | 0.005531 |
| IFNAR2 | 259.8209 | 0.073952 | 0.171493 | 0.431227 | 0.666303 | 0.780999 |
| IFNGR2 | 1382.111 | 0.124496 | 0.133728 | 0.930965 | 0.351872 | 0.52 |
| GART | 2495.654 | -0.01036 | 0.140563 | -0.07368 | 0.941266 | 0.967169 |
| SON | 3990.456 | -0.07194 | 0.197518 | -0.36424 | 0.715679 | 0.81867 |
| DONSON | 319.4789 | 0.346845 | 0.194978 | 1.778889 | 0.075258 | 0.183237 |
| KCNE2 | 3.878343 | 0.808628 | 1.20882 | 0.66894 | 0.503534 | NA |
| RCAN1 | 10373.7 | 0.187936 | 0.205571 | 0.914218 | 0.360602 | 0.52777 |
| CLIC6 | 43.55567 | 0.735031 | 0.613231 | 1.19862 | 0.230676 | NA |
| RUNX1 | 958.9044 | -0.23196 | 0.215543 | -1.07615 | 0.281861 | 0.448098 |
| CBR1 | 2984.334 | -0.18047 | 0.145045 | -1.24421 | 0.21342 | 0.372096 |
| CBR3 | 777.7217 | -0.2267 | 0.156282 | -1.4506 | 0.14689 | 0.289747 |
| C2orf81 | 134.616 | -0.44721 | 0.26024 | -1.71844 | 0.085717 | 0.200271 |
| MORC3 | 634.3709 | 0.819104 | 0.329716 | 2.484271 | 0.012982 | 0.05641 |
| CHAF1B | 309.0343 | -0.18305 | 0.21306 | -0.85914 | 0.390266 | 0.555681 |
| CLDN14 | 20.62101 | 0.81463 | 0.462055 | 1.763059 | 0.07789 | NA |
| SIM2 | 3.4531 | 0.55852 | 1.251527 | 0.446271 | 0.655402 | NA |
| HLCS | 752.5962 | 0.002508 | 0.14554 | 0.017235 | 0.986249 | 0.992697 |
| SCUBE1 | 0.488912 | 0.771583 | 2.897678 | 0.266276 | 0.790026 | NA |
| M1AP | 11.13466 | 0.120114 | 0.66524 | 0.180557 | 0.856715 | NA |
| HK2 | 2227.67 | -0.38732 | 0.145798 | -2.65655 | 0.007895 | 0.040341 |
| RGL4 | 15.55322 | -0.16684 | 0.620974 | -0.26867 | 0.788181 | NA |
| CCDC117 | 442.1044 | 0.250374 | 0.186752 | 1.340679 | 0.180025 | 0.332584 |
| ZNF222 | 68.15122 | 0.84373 | 0.287977 | 2.929858 | 0.003391 | 0.022819 |
| ZNF233 | 48.06442 | 0.074332 | 0.339018 | 0.219258 | 0.826449 | 0.895572 |
| ZNF235 | 89.08839 | 0.381667 | 0.251309 | 1.518716 | 0.128834 | 0.265134 |
| TNFRSF13C | 1.467268 | -0.8203 | 1.674139 | -0.48998 | 0.624146 | NA |
| ARHGAP35 | 3447.224 | -0.28846 | 0.15644 | -1.84389 | 0.065199 | 0.166489 |
| PTGIR | 2.276578 | -0.67486 | 1.369131 | -0.49291 | 0.622078 | NA |
| CALM3 | 7998.055 | -0.50313 | 0.155745 | -3.23047 | 0.001236 | 0.011864 |
| ABCG1 | 4.072117 | -0.27203 | 1.015369 | -0.26792 | 0.788764 | NA |
| TFF3 | 263.5033 | -0.36762 | 0.214048 | -1.71748 | 0.085891 | 0.200558 |
| TFF2 | 1.946346 | 1.109615 | 1.463021 | 0.75844 | 0.448187 | NA |
| RSPH1 | 5.267224 | 0.461082 | 0.864758 | 0.533192 | 0.593901 | NA |
| SLC37A1 | 499.6478 | -0.12941 | 0.161341 | -0.80206 | 0.422518 | 0.584968 |
| PDE9A | 1.333192 | -0.17938 | 1.88139 | -0.09534 | 0.924042 | NA |
| WDR4 | 337.118 | -0.31323 | 0.199575 | -1.56948 | 0.116537 | 0.246836 |

| | | | | | | |
|---|---|---|---|---|---|---|
| NDUFV3 | 1119.239 | -0.11617 | 0.145477 | -0.79857 | 0.42454 | 0.586587 |
| PKNOX1 | 223.0911 | 0.131698 | 0.275953 | 0.477248 | 0.633185 | 0.754944 |
| CBS | 341.3515 | 0.178417 | 0.190996 | 0.934141 | 0.350231 | 0.518221 |
| U2AF1 | 1692.757 | 0.086549 | 0.131112 | 0.660118 | 0.509178 | 0.657545 |
| HSF2BP | 16.49601 | 0.471613 | 0.533286 | 0.884353 | 0.376506 | NA |
| RRP1B | 931.5942 | -0.20461 | 0.164879 | -1.24095 | 0.214626 | 0.373586 |
| PDXK | 1737.25 | -0.33445 | 0.193207 | -1.73106 | 0.083442 | 0.196777 |
| CSTB | 6374.015 | -0.04528 | 0.194558 | -0.23272 | 0.815979 | 0.889858 |
| RRP1 | 1183.85 | -0.19502 | 0.167105 | -1.16703 | 0.243198 | 0.406164 |
| AGPAT3 | 3402.644 | -0.4563 | 0.133482 | -3.4184 | 6.30E-04 | 0.007785 |
| TRAPPC10 | 1531.066 | 0.08865 | 0.144522 | 0.613401 | 0.539611 | 0.681774 |
| C21orf33 | 1303.842 | -0.29805 | 0.150665 | -1.97822 | 0.047904 | 0.134949 |
| ICOSLG | 9.427146 | 0.079445 | 0.756742 | 0.104983 | 0.91639 | NA |
| AIRE | 0.392857 | -1.546 | 4.075469 | -0.37934 | 0.704433 | NA |
| C21orf2 | 386.7072 | -0.20085 | 0.183656 | -1.09361 | 0.274126 | 0.439821 |
| LRRC3 | 109.1021 | -0.49535 | 0.413716 | -1.19731 | 0.231185 | 0.393327 |
| ITGB2 | 2.912992 | 1.48646 | 1.407855 | 1.055833 | 0.291044 | NA |
| FAM207A | 582.9244 | -0.20985 | 0.180679 | -1.16143 | 0.245467 | 0.408479 |
| FTCD | 1.187588 | 2.257887 | 1.988061 | 1.135723 | 0.256072 | NA |
| SPATC1L | 421.9755 | -0.34548 | 0.162998 | -2.11952 | 0.034047 | 0.107834 |
| LSS | 7433.024 | -0.56393 | 0.139148 | -4.05273 | 5.06E-05 | 0.001658 |
| MCM3AP | 2242.646 | -0.35026 | 0.148742 | -2.3548 | 0.018533 | 0.071752 |
| C21orf58 | 148.7237 | -0.14338 | 0.246127 | -0.58256 | 0.560191 | 0.698472 |
| PCNT | 785.1696 | 0.004057 | 0.207636 | 0.019539 | 0.984411 | 0.991773 |
| DIP2A | 930.1956 | -0.0425 | 0.172986 | -0.24566 | 0.805942 | 0.884098 |
| S100B | 57275.32 | 0.20371 | 0.241319 | 0.844153 | 0.398584 | 0.563397 |
| PRMT2 | 4375.279 | -0.04235 | 0.141518 | -0.29927 | 0.764737 | 0.853834 |
| CLDND2 | 8.354036 | 1.118433 | 0.806799 | 1.38626 | 0.165668 | NA |
| ZNF761 | 107.3522 | 0.686537 | 0.345166 | 1.989006 | 0.046701 | 0.132894 |
| RDH13 | 263.5359 | -0.34407 | 0.184906 | -1.86075 | 0.062779 | 0.162216 |
| BRSK1 | 381.5925 | 0.073915 | 0.18596 | 0.397476 | 0.691016 | 0.799922 |
| COX6B2 | 0.134133 | -1.48909 | 4.077584 | -0.36519 | 0.71497 | NA |
| ANO10 | 1672.846 | -0.15783 | 0.147559 | -1.06961 | 0.284794 | 0.45116 |
| NBEAL2 | 547.1419 | -0.31113 | 0.263029 | -1.18287 | 0.236859 | 0.399729 |
| CCDC12 | 653.4129 | 0.069014 | 0.155505 | 0.443806 | 0.657183 | 0.773905 |
| PTH1R | 3.523716 | 0.540587 | 1.457378 | 0.370931 | 0.710689 | NA |
| MYL3 | 1.006656 | -0.39276 | 2.040151 | -0.19251 | 0.847339 | NA |
| | 0.096202 | 0.408546 | 4.079639 | 0.100143 | 0.920231 | NA |
| | 3.710345 | -1.18879 | 1.095121 | -1.08553 | 0.277687 | NA |
| TUBA3FP | 14.2814 | 0.122055 | 0.566431 | 0.21548 | 0.829393 | NA |
| YDJC | 610.5633 | -0.36674 | 0.156644 | -2.34125 | 0.019219 | 0.073617 |
| CCDC116 | 2.564951 | 1.131087 | 1.385234 | 0.816531 | 0.414196 | NA |
| ZNF577 | 76.74944 | 0.804511 | 0.287019 | 2.802993 | 0.005063 | 0.029892 |
| TMEM143 | 219.5014 | -0.43089 | 0.196769 | -2.18984 | 0.028536 | 0.095833 |
| ALDH16A1 | 1422.867 | -0.51166 | 0.136029 | -3.76141 | 1.69E-04 | 0.003497 |
| SIGLEC16 | 1.276206 | -1.07269 | 2.030043 | -0.52841 | 0.597216 | NA |
| IZUMO2 | 0.35636 | 1.709252 | 3.651833 | 0.468053 | 0.639746 | NA |

| | | | | | | |
|---|---|---|---|---|---|---|
| EMC10 | 4970.984 | -0.31039 | 0.12057 | -2.57437 | 0.010042 | 0.047332 |
| JOSD2 | 1196.966 | -0.29986 | 0.157277 | -1.90659 | 0.056574 | 0.150648 |
| SHANK1 | 355.2375 | -0.25662 | 0.199701 | -1.28501 | 0.198787 | 0.354753 |
| RPL29 | 14027 | 0.017902 | 0.169475 | 0.105633 | 0.915873 | 0.951316 |
| ITIH3 | 32.91392 | -0.39307 | 0.583289 | -0.67389 | 0.50038 | NA |
| DCP1A | 688.5955 | -0.2588 | 0.15865 | -1.63128 | 0.102831 | 0.227098 |
| SNED1 | 11.04745 | -0.17573 | 0.648494 | -0.27099 | 0.7864 | NA |
| PPP1R21 | 634.0437 | 0.396351 | 0.1846 | 2.147083 | 0.031787 | 0.10327 |
| PKDCC | 59.83067 | -0.16229 | 0.298283 | -0.54409 | 0.586381 | 0.71925 |
| OXER1 | 1.024017 | 0.177191 | 1.894522 | 0.093528 | 0.925484 | NA |
| HAAO | 7.272616 | -0.67743 | 0.85204 | -0.79507 | 0.426575 | NA |
| REL | 57.4676 | 0.831576 | 0.318531 | 2.610656 | 0.009037 | 0.044218 |
| PUS10 | 63.36788 | 0.382487 | 0.294132 | 1.300393 | 0.193466 | 0.348531 |
| PEX13 | 281.0334 | 0.372494 | 0.181795 | 2.048982 | 0.040464 | 0.121289 |
| KIAA1841 | 247.5737 | 0.627641 | 0.22819 | 2.750515 | 0.00595 | 0.033242 |
| RFTN2 | 381.4681 | 0.659684 | 0.266268 | 2.47752 | 0.01323 | 0.057179 |
| LRRTM1 | 0.678686 | -3.08037 | 2.621268 | -1.17514 | 0.239937 | NA |
| MEMO1 | 68.73646 | -0.06996 | 0.29843 | -0.23442 | 0.81466 | 0.889545 |
| DPY30 | 892.9102 | 0.26803 | 0.190496 | 1.407011 | 0.159424 | 0.306416 |
| TYW5 | 167.9014 | 0.734499 | 0.207761 | 3.535317 | 4.07E-04 | 0.005986 |
| C2orf47 | 284.2944 | 0.045706 | 0.184925 | 0.247158 | 0.804786 | 0.88332 |
| KCNF1 | 0.845069 | 2.925226 | 2.326646 | 1.257272 | 0.208655 | NA |
| PQLC3 | 1098.813 | -0.06485 | 0.153742 | -0.42182 | 0.673159 | 0.786159 |
| ARL5A | 1206.873 | 0.734631 | 0.175052 | 4.196637 | 2.71E-05 | 0.001104 |
| FAM84A | 0.489058 | -0.58569 | 3.427543 | -0.17088 | 0.864321 | NA |
| KCNJ3 | 1.107756 | 0.313909 | 2.268235 | 0.138394 | 0.889929 | NA |
| CLHC1 | 25.51217 | 0.807933 | 0.422638 | 1.911642 | 0.055922 | NA |
| PRORSD1P | 11.30348 | 0.964854 | 0.668671 | 1.442943 | 0.149037 | NA |
| FRZB | 6.603366 | 0.87179 | 0.816226 | 1.068074 | 0.285487 | NA |
| DUSP19 | 59.07804 | 0.223009 | 0.340801 | 0.654368 | 0.512875 | 0.660023 |
| CCDC104 | 571.1926 | 0.990375 | 0.22115 | 4.478287 | 7.52E-06 | 5.34E-04 |
| NUP35 | 339.3791 | 0.191999 | 0.179512 | 1.069566 | 0.284815 | 0.45116 |
| CCDC138 | 43.47795 | 0.993172 | 0.35812 | 2.773294 | 0.005549 | NA |
| C2orf48 | 1.296153 | -2.92934 | 2.02948 | -1.44339 | 0.14891 | NA |
| FBXO41 | 586.5436 | -0.41217 | 0.182777 | -2.25505 | 0.02413 | 0.085854 |
| ALMS1P | 2.554511 | 0.386807 | 1.281927 | 0.301739 | 0.762851 | NA |
| ACTG2 | 14.97735 | 0.460028 | 0.613102 | 0.750329 | 0.453056 | NA |
| C2orf44 | 349.5284 | -0.10075 | 0.163831 | -0.61494 | 0.538596 | 0.681093 |
| SMC6 | 393.0805 | 0.857221 | 1.035222 | 0.828055 | 0.407639 | 0.571292 |
| VSNL1 | 2.863483 | 0.401522 | 1.182071 | 0.339677 | 0.7341 | NA |
| CCDC74A | 140.2035 | 0.011898 | 0.242307 | 0.049104 | 0.960836 | 0.978668 |
| ANKRD30B | 56.44146 | 0.213277 | 0.301983 | 0.706253 | 0.480031 | 0.634563 |
| SLC16A14 | 0.315653 | -0.57305 | 3.892639 | -0.14721 | 0.882963 | NA |
| TEKT4 | 0.392857 | -1.546 | 4.075469 | -0.37934 | 0.704433 | NA |
| EN1 | 0.096202 | 0.408546 | 4.079639 | 0.100143 | 0.920231 | NA |
| ZNF2 | 241.1452 | -0.09548 | 0.188003 | -0.50785 | 0.611558 | 0.738367 |
| NOSTRIN | 6.257068 | 0.90057 | 0.849699 | 1.059869 | 0.289204 | NA |

| | | | | | | |
|---|---|---|---|---|---|---|
| | 0.163957 | 0.424913 | 4.079006 | 0.104171 | 0.917034 | NA |
| CCDC140 | 53.00839 | -0.01125 | 0.304702 | -0.03692 | 0.970547 | 0.983948 |
| SGPP2 | 2.574988 | -0.33668 | 1.250299 | -0.26928 | 0.787717 | NA |
| INHBB | 0.134133 | -1.48909 | 4.077584 | -0.36519 | 0.71497 | NA |
| XIRP2 | 0.911244 | 3.022036 | 2.332005 | 1.295896 | 0.195011 | NA |
| BBS5 | 733.9145 | -0.05362 | 0.343347 | -0.15617 | 0.875895 | 0.925797 |
| NEURL3 | 0.719246 | 0.444943 | 2.542118 | 0.175028 | 0.861057 | NA |
| ANKRD23 | 5.109057 | 0.972273 | 0.886705 | 1.096501 | 0.27286 | NA |
| ERCC3 | 1817.782 | 0.080439 | 0.145058 | 0.554528 | 0.579217 | 0.713563 |
| RNF149 | 646.3434 | 0.155609 | 0.157471 | 0.988178 | 0.323066 | 0.490955 |
| IWS1 | 895.9011 | 0.670687 | 0.24785 | 2.706021 | 0.006809 | 0.036429 |
| BOLA3 | 703.3241 | 0.334894 | 0.168694 | 1.985213 | 0.047121 | 0.133662 |
| CDC42EP3 | 1493.782 | -0.26959 | 0.214726 | -1.25553 | 0.209287 | 0.366863 |
| DHX57 | 565.6817 | 0.14146 | 0.148125 | 0.955001 | 0.339577 | 0.508003 |
| ARHGAP25 | 0.35636 | 1.709252 | 3.651833 | 0.468053 | 0.639746 | NA |
| TGFA | 4.375545 | 0.188785 | 1.027869 | 0.183666 | 0.854275 | NA |
| CCNYL1 | 371.4216 | 0.120643 | 0.16775 | 0.719183 | 0.472028 | 0.627976 |
| FZD5 | 83.14643 | -0.2962 | 0.290508 | -1.01961 | 0.307915 | 0.475133 |
| NPPC | 0.774539 | 0.39797 | 2.296349 | 0.173305 | 0.862411 | NA |
| ALPP | 0.424115 | 1.932531 | 3.45462 | 0.559405 | 0.575885 | NA |
| GPR155 | 2509.454 | 0.157145 | 0.191749 | 0.819534 | 0.412482 | 0.57563 |
| DAPL1 | 16.61454 | -0.92842 | 0.588548 | -1.57748 | 0.114686 | NA |
| COL6A3 | 13.45162 | 0.131437 | 0.551636 | 0.238267 | 0.811674 | NA |
| LINC01116 | 89.91242 | 0.292322 | 0.261323 | 1.118626 | 0.2633 | 0.427592 |
| KBTBD8 | 20.67915 | 0.916518 | 0.502145 | 1.825205 | 0.06797 | NA |
| FAM19A4 | 0.392857 | -1.546 | 4.075469 | -0.37934 | 0.704433 | NA |
| EOGT | 427.4002 | 0.168954 | 0.237907 | 0.710167 | 0.477601 | 0.632358 |
| TMEM169 | 228.468 | 0.002868 | 0.2365 | 0.012125 | 0.990326 | 0.995532 |
| ARPC2 | 11364.38 | 0.126772 | 0.136302 | 0.930089 | 0.352325 | 0.52041 |
| RNF25 | 879.5026 | -0.14636 | 0.156678 | -0.93414 | 0.35023 | 0.518221 |
| STK36 | 493.9085 | -0.24725 | 0.188861 | -1.30917 | 0.190476 | 0.344799 |
| NEK10 | 8.818382 | 1.066783 | 0.786438 | 1.356475 | 0.174948 | NA |
| FEV | 0.392857 | -1.546 | 4.075469 | -0.37934 | 0.704433 | NA |
| CRYBA2 | 0.478233 | -0.67301 | 2.937473 | -0.22911 | 0.818782 | NA |
| IHH | 0.640609 | 0.042838 | 2.379511 | 0.018003 | 0.985637 | NA |
| EOMES | 232.9978 | -0.54356 | 0.30259 | -1.79635 | 0.072439 | 0.17891 |
| CWC22 | 196.8974 | 1.155235 | 0.730678 | 1.581044 | 0.113868 | 0.242963 |
| AZI2 | 670.5785 | 0.440906 | 0.174603 | 2.525195 | 0.011563 | 0.051985 |
| TGFBR2 | 5415.017 | -0.04819 | 0.213882 | -0.2253 | 0.821748 | 0.89268 |
| ANKZF1 | 530.1908 | -0.01042 | 0.152955 | -0.0681 | 0.945708 | 0.969758 |
| HDAC11 | 1035.494 | -0.24816 | 0.14853 | -1.67079 | 0.094763 | 0.21542 |
| FBLN2 | 191.8235 | 0.098969 | 1.118802 | 0.08846 | 0.929511 | 0.960041 |
| GLB1L | 155.5703 | -0.6883 | 0.209842 | -3.28009 | 0.001038 | 0.010596 |
| STT3B | 3609.341 | 0.300875 | 0.179812 | 1.673274 | 0.094273 | 0.214677 |
| CHCHD4 | 402.6123 | 0.118325 | 0.165278 | 0.715914 | 0.474045 | 0.629544 |
| SGOL2 | 171.9029 | 1.218354 | 0.729482 | 1.670163 | 0.094887 | 0.215459 |
| CLASP2 | 1882.085 | 0.238403 | 0.173257 | 1.376003 | 0.168821 | 0.319045 |

| | | | | | | |
|---|---|---|---|---|---|---|
| SUCLG1 | 2081.618 | 0.016017 | 0.132302 | 0.121063 | 0.903641 | 0.94382 |
| EFHB | 4.815426 | 0.047293 | 0.900985 | 0.05249 | 0.958138 | NA |
| FABP1 | 0.096202 | 0.408546 | 4.079639 | 0.100143 | 0.920231 | NA |
| ICA1L | 36.83056 | 0.732334 | 0.35697 | 2.051529 | 0.040215 | NA |
| CTLA4 | 8.746229 | 0.759947 | 0.810605 | 0.937506 | 0.348498 | NA |
| ICOS | 66.11862 | 0.005048 | 0.5228 | 0.009655 | 0.992296 | 0.996738 |
| CADPS | 69.73955 | -0.30658 | 0.346959 | -0.88362 | 0.376902 | 0.543599 |
| SYNPR | 12.8939 | 2.368043 | 0.753647 | 3.142112 | 0.001677 | NA |
| C3orf49 | 1.588764 | 1.141653 | 1.736673 | 0.657379 | 0.510937 | NA |
| THOC7 | 823.667 | 0.491395 | 0.163337 | 3.008473 | 0.002626 | 0.019406 |
| ATXN7 | 374.3643 | 0.2761 | 0.209029 | 1.320872 | 0.186544 | 0.340699 |
| PSMD6 | 1961.333 | 0.402626 | 0.142775 | 2.820014 | 0.004802 | 0.028752 |
| PRICKLE2 | 351.4364 | -0.3779 | 0.169202 | -2.23341 | 0.025522 | 0.089331 |
| ADAMTS9 | 129.8399 | -0.48527 | 0.316042 | -1.53545 | 0.124674 | 0.259392 |
| HESX1 | 2.711583 | 0.884043 | 1.242322 | 0.711605 | 0.476709 | NA |
| SLMAP | 889.447 | 0.61417 | 0.209503 | 2.931558 | 0.003373 | 0.022718 |
| RPP14 | 13.70661 | 0.314424 | 0.594835 | 0.52859 | 0.59709 | NA |
| ABHD6 | 2057.975 | -0.0778 | 0.140509 | -0.55371 | 0.579776 | 0.713749 |
| C3orf67 | 117.316 | 0.323542 | 0.297947 | 1.085907 | 0.27752 | 0.443456 |
| IL17RE | 66.35432 | -0.13361 | 0.271175 | -0.49271 | 0.622219 | 0.746524 |
| IL17RC | 1227.206 | -0.35003 | 0.131886 | -2.65401 | 0.007954 | 0.040505 |
| CRELD1 | 516.1599 | -0.29652 | 0.166041 | -1.78579 | 0.074133 | 0.181706 |
| PRRT3 | 187.0235 | -0.76307 | 0.212335 | -3.5937 | 3.26E-04 | 0.005188 |
| MTMR14 | 1551.678 | -0.27698 | 0.130609 | -2.12071 | 0.033946 | 0.107602 |
| SNRK | 1445.446 | 0.215292 | 0.184298 | 1.168176 | 0.242736 | 0.4057 |
| DNAJC5G | 0.230334 | -0.53318 | 4.076749 | -0.13079 | 0.895945 | NA |
| UCN | 14.16027 | -0.1941 | 0.528988 | -0.36693 | 0.713669 | NA |
| ZNF513 | 729.2306 | 0.023319 | 0.184439 | 0.126431 | 0.899391 | 0.940983 |
| SLC4A1AP | 639.4647 | 0.591668 | 0.204026 | 2.899968 | 0.003732 | 0.024376 |
| PLB1 | 4.8131 | 0.479285 | 0.935797 | 0.512167 | 0.608534 | NA |
| SPDYA | 2.451774 | -0.07167 | 1.21207 | -0.05913 | 0.952848 | NA |
| KIAA1143 | 271.4593 | 0.758443 | 0.334517 | 2.267279 | 0.023373 | 0.084068 |
| KIF15 | 100.3504 | 0.897727 | 0.305165 | 2.941779 | 0.003263 | 0.022362 |
| TGM4 | 0.651435 | 0.249629 | 2.788679 | 0.089515 | 0.928672 | NA |
| WDR43 | 1010.809 | 0.723119 | 0.220562 | 3.278532 | 0.001043 | 0.010637 |
| ZDHHC3 | 1835.797 | -0.20733 | 0.13284 | -1.56076 | 0.118581 | 0.249961 |
| CDCP1 | 3.375736 | 0.021243 | 1.285112 | 0.01653 | 0.986812 | NA |
| CLEC3B | 0.382031 | -1.26484 | 3.151989 | -0.40128 | 0.688212 | NA |
| SLC6A20 | 0.629726 | -0.22725 | 2.732935 | -0.08315 | 0.93373 | NA |
| LZTFL1 | 313.0507 | 0.876946 | 0.247644 | 3.541156 | 3.98E-04 | 0.00591 |
| FYCO1 | 5327.033 | -0.31811 | 0.157393 | -2.02114 | 0.043266 | 0.126802 |
| CCR1 | 27.85061 | 0.042977 | 0.544671 | 0.078905 | 0.937108 | NA |
| RTP3 | 0.096202 | 0.408546 | 4.079639 | 0.100143 | 0.920231 | NA |
| LRRC2 | 263.0431 | 0.604625 | 0.239748 | 2.521919 | 0.011672 | 0.0523 |
| ELP6 | 1179.289 | -0.02038 | 0.131986 | -0.15438 | 0.877311 | 0.926467 |
| BAP1 | 4398.791 | -0.20905 | 0.154503 | -1.35305 | 0.176041 | 0.328139 |
| TKT | 15077.54 | -0.34689 | 0.158648 | -2.18655 | 0.028775 | 0.096393 |

| | | | | | | |
|---|---|---|---|---|---|---|
| PRKCD | 2117.003 | -0.41356 | 0.137364 | -3.01066 | 0.002607 | 0.019343 |
| RFT1 | 719.0743 | -0.20002 | 0.17495 | -1.14329 | 0.252918 | 0.417132 |
| SFMBT1 | 385.2341 | -0.21882 | 0.163442 | -1.33884 | 0.180624 | 0.33312 |
| GNL3 | 1690.487 | 0.899143 | 0.143286 | 6.275177 | 3.49E-10 | 4.14E-07 |
| PBRM1 | 1976.109 | 0.362961 | 0.248844 | 1.458586 | 0.144679 | 0.286626 |
| FAM208A | 715.1896 | 0.531006 | 0.31166 | 1.703796 | 0.088419 | 0.205049 |
| ARHGEF3 | 966.072 | -0.26305 | 0.1925 | -1.3665 | 0.171781 | 0.32289 |
| CDC25A | 255.8763 | -0.12239 | 0.260855 | -0.4692 | 0.638928 | 0.759385 |
| ZNF589 | 746.3019 | 0.099857 | 0.195982 | 0.509518 | 0.610389 | 0.737459 |
| FBXW12 | 0.752028 | 2.752519 | 2.876634 | 0.956854 | 0.338641 | NA |
| PLXNB1 | 2787.194 | -0.32037 | 0.19662 | -1.62936 | 0.103236 | 0.227739 |
| CCDC51 | 598.2098 | -0.10747 | 0.154323 | -0.69639 | 0.486183 | 0.639469 |
| ATRIP | 254.4735 | -0.24253 | 0.185961 | -1.3042 | 0.192165 | 0.346798 |
| SHISA5 | 6401.039 | -0.545 | 0.163832 | -3.32656 | 8.79E-04 | 0.009556 |
| BSN | 15.31282 | -0.40116 | 0.670692 | -0.59813 | 0.549751 | NA |
| APEH | 6283.659 | -0.27928 | 0.121254 | -2.30325 | 0.021265 | 0.078903 |
| RNF123 | 1591.703 | -0.49077 | 0.135465 | -3.62286 | 2.91E-04 | 0.004831 |
| MON1A | 740.0358 | -0.11234 | 0.182661 | -0.615 | 0.538553 | 0.681093 |
| MST1R | 4.932527 | -0.67028 | 0.959106 | -0.69885 | 0.484643 | NA |
| RAD54L2 | 529.7106 | -0.34184 | 0.179032 | -1.90937 | 0.056214 | 0.150029 |
| TEX264 | 2499.984 | -0.14299 | 0.141243 | -1.01236 | 0.311365 | 0.478811 |
| GRM2 | 0.662557 | -0.83483 | 2.671964 | -0.31244 | 0.754706 | NA |
| DUSP7 | 703.3788 | -0.16985 | 0.228212 | -0.74427 | 0.456711 | 0.615154 |
| POC1A | 709.0716 | -0.27203 | 0.222609 | -1.22203 | 0.221698 | 0.382121 |
| PPM1M | 524.9594 | -0.10577 | 0.222174 | -0.47606 | 0.63403 | 0.755518 |
| WDR82 | 3187.098 | -0.1854 | 0.138555 | -1.33811 | 0.180861 | 0.333453 |
| BLCAP | 1413.944 | -0.38512 | 0.14559 | -2.64522 | 0.008164 | 0.041148 |
| ZNF606 | 90.08153 | 0.471119 | 0.313929 | 1.500717 | 0.133429 | 0.271088 |
| ZNF667-AS | 343.9922 | -0.1088 | 0.265697 | -0.40949 | 0.682183 | 0.792948 |
| CACNG2 | 0.392857 | -1.546 | 4.075469 | -0.37934 | 0.704433 | NA |
| YWHAB | 11393.65 | -0.07308 | 0.145328 | -0.50285 | 0.615068 | 0.741129 |
| EVA1C | 371.7354 | 0.015119 | 0.220867 | 0.068453 | 0.945425 | 0.969593 |
| SGSM1 | 0.392711 | 0.292538 | 3.145221 | 0.09301 | 0.925895 | NA |
| | 3.71852 | -1.98365 | 1.403315 | -1.41355 | 0.157495 | NA |
| DUSP18 | 60.06827 | -0.26337 | 0.295261 | -0.89198 | 0.372404 | 0.539054 |
| TEF | 398.9894 | -0.57366 | 0.155089 | -3.69892 | 2.17E-04 | 0.004081 |
| MEI1 | 1.093402 | -1.45329 | 2.572346 | -0.56497 | 0.572096 | NA |
| ZNF226 | 337.7948 | 0.657877 | 0.198909 | 3.307428 | 0.000942 | 0.009957 |
| ZNF229 | 127.5627 | 0.193887 | 0.264507 | 0.733014 | 0.46355 | 0.621213 |
| ZNF180 | 94.61734 | 0.640642 | 0.286102 | 2.239209 | 0.025142 | 0.088218 |
| ZNF610 | 41.30666 | -0.0751 | 0.572873 | -0.13109 | 0.895701 | NA |
| ZNF528 | 190.3686 | 0.191081 | 0.341257 | 0.559931 | 0.575526 | 0.710649 |
| ZNF701 | 82.71295 | 0.092207 | 0.258994 | 0.35602 | 0.721826 | 0.823208 |
| TMC4 | 8.945608 | 0.437286 | 0.730579 | 0.598548 | 0.549475 | NA |
| TTYH1 | 3.713791 | -0.08076 | 1.011309 | -0.07986 | 0.936351 | NA |
| LENG8 | 966.4044 | -0.06919 | 0.224947 | -0.30758 | 0.758405 | 0.849319 |
| CDC42EP5 | 1035.913 | -0.08254 | 0.196295 | -0.42051 | 0.674112 | 0.786547 |

| | | | | | | |
|---|---|---|---|---|---|---|
| NLRP7 | 0.163957 | 0.424913 | 4.079006 | 0.104171 | 0.917034 | NA |
| DNAAF3 | 51.90718 | 0.090763 | 0.39118 | 0.232023 | 0.81652 | 0.890173 |
| ZNF444 | 834.1602 | -0.35548 | 0.151674 | -2.34373 | 0.019092 | 0.073307 |
| C19orf48 | 880.5168 | -0.2309 | 0.20479 | -1.12749 | 0.259534 | 0.423872 |
| KLK1 | 3.138611 | 0.613958 | 1.243352 | 0.493792 | 0.621453 | NA |
| KLK4 | 0.556813 | -0.57007 | 3.294615 | -0.17303 | 0.862627 | NA |
| KLK2 | 0.163957 | 0.424913 | 4.079006 | 0.104171 | 0.917034 | NA |
| KLK13 | 0.298089 | -0.51711 | 3.975232 | -0.13008 | 0.896501 | NA |
| | 18.27553 | -0.21502 | 0.471686 | -0.45586 | 0.648493 | NA |
| ZNF83 | 165.4837 | 0.83762 | 0.265816 | 3.151131 | 0.001626 | 0.014249 |
| TRANK1 | 74.52227 | -0.38441 | 0.276159 | -1.39199 | 0.163925 | 0.312633 |
| TTC21A | 56.73661 | -0.14421 | 0.287438 | -0.50171 | 0.615873 | 0.741989 |
| RPSA | 12218.75 | -0.15049 | 0.132853 | -1.13272 | 0.257334 | 0.421366 |
| ENTPD3 | 0.968725 | 0.233151 | 2.074813 | 0.112372 | 0.910528 | NA |
| CTNNB1 | 25727.76 | -0.01662 | 0.142129 | -0.11696 | 0.906889 | 0.945907 |
| ULK4 | 149.7108 | -0.0391 | 0.231826 | -0.16867 | 0.866056 | 0.920654 |
| KCNJ4 | 0.134133 | -1.48909 | 4.077584 | -0.36519 | 0.71497 | NA |
| SETD5 | 1610.943 | -0.01517 | 0.134391 | -0.11287 | 0.910131 | 0.947811 |
| GLYCTK | 192.1249 | -0.2456 | 0.194165 | -1.26492 | 0.2059 | 0.362965 |
| NT5DC2 | 2754.014 | -0.05519 | 0.172185 | -0.3205 | 0.748589 | 0.842225 |
| SMIM4 | 428.2587 | 0.095641 | 0.192788 | 0.496091 | 0.61983 | 0.74446 |
| KIF5C | 260.7511 | 1.11255 | 0.35006 | 3.178171 | 0.001482 | 0.013453 |
| MMADHC | 1907.058 | 0.096695 | 0.161004 | 0.600574 | 0.548124 | 0.689371 |
| PDHB | 1858.141 | 0.211504 | 0.156518 | 1.351311 | 0.176596 | 0.328863 |
| PXK | 621.0969 | 0.004702 | 0.20263 | 0.023205 | 0.981487 | 0.990173 |
| KCTD6 | 124.4637 | 0.290253 | 0.228914 | 1.267956 | 0.204814 | 0.361427 |
| ACOX2 | 0.86916 | 0.686589 | 2.213986 | 0.310114 | 0.756474 | NA |
| CX3CR1 | 0.151697 | -1.546 | 4.075469 | -0.37934 | 0.704433 | NA |
| XIRP1 | 0.394291 | 0.298745 | 3.141711 | 0.09509 | 0.924243 | NA |
| SCN11A | 0.930794 | 0.917844 | 2.145018 | 0.427896 | 0.668727 | NA |
| ARF4 | 5961.017 | -0.02436 | 0.127035 | -0.19175 | 0.847937 | 0.909306 |
| 2-Sep | 10892.43 | 0.252928 | 0.138505 | 1.826129 | 0.067831 | 0.170621 |
| DTYMK | 1965.139 | -0.23898 | 0.164041 | -1.45683 | 0.145165 | 0.287108 |
| ING5 | 300.8406 | -0.12955 | 0.170606 | -0.75936 | 0.447636 | 0.607177 |
| ATG4B | 1776.185 | -0.1186 | 0.129027 | -0.91922 | 0.357982 | 0.525131 |
| KLHL30 | 0.247898 | -0.5901 | 4.074633 | -0.14482 | 0.884851 | NA |
| GBX2 | 0.315653 | -0.57305 | 3.892639 | -0.14721 | 0.882963 | NA |
| MYL1 | 0.326333 | 1.533889 | 3.826878 | 0.40082 | 0.688553 | NA |
| COL3A1 | 17.22689 | 0.682579 | 0.542999 | 1.257054 | 0.208734 | NA |
| ZSWIM1 | 276.4524 | -0.53325 | 0.21322 | -2.50095 | 0.012386 | 0.054536 |
| LRP1B | 0.096202 | 0.408546 | 4.079639 | 0.100143 | 0.920231 | NA |
| PKIG | 1440.594 | -0.08657 | 0.177281 | -0.4883 | 0.625334 | 0.748746 |
| FAM178B | 36.46315 | -0.94775 | 0.440138 | -2.1533 | 0.031295 | NA |
| SEMA4C | 2602.137 | -0.37487 | 0.186449 | -2.01057 | 0.044371 | 0.128578 |
| CNNM3 | 652.9964 | -0.29618 | 0.210473 | -1.40723 | 0.159359 | 0.30634 |
| ATOH8 | 439.2495 | -0.15125 | 0.219642 | -0.6886 | 0.491072 | 0.642998 |
| SFTPB | 0.230334 | -0.53318 | 4.076749 | -0.13079 | 0.895945 | NA |

| | | | | | | |
|---|---|---|---|---|---|---|
| USP39 | 2069.075 | -0.18566 | 0.155594 | -1.19326 | 0.232768 | 0.395112 |
| C2orf68 | 403.86 | -0.31742 | 0.191567 | -1.65697 | 0.097525 | 0.219422 |
| TMEM150/ | 581.455 | -0.23875 | 0.191858 | -1.24442 | 0.213346 | 0.372053 |
| RNF181 | 1913.58 | 0.162687 | 0.134012 | 1.213974 | 0.224758 | 0.385824 |
| VAMP5 | 3340.58 | -0.11963 | 0.178361 | -0.6707 | 0.502413 | 0.652458 |
| MAT2A | 3948.29 | -0.01075 | 0.121262 | -0.08866 | 0.929352 | 0.959961 |
| INPP5D | 1.06329 | 0.285492 | 1.942399 | 0.146979 | 0.883149 | NA |
| TM4SF20 | 0.411855 | 0.250456 | 3.097618 | 0.080854 | 0.935558 | NA |
| MFF | 2083.69 | 0.061049 | 0.131963 | 0.462623 | 0.643634 | 0.762832 |
| OR7E102P | 0.096202 | 0.408546 | 4.079639 | 0.100143 | 0.920231 | NA |
| E2F6 | 325.8062 | 0.364235 | 0.18792 | 1.938252 | 0.052592 | 0.143198 |
| COL4A3 | 4.333681 | 0.779565 | 1.057861 | 0.736926 | 0.461168 | NA |
| IRS1 | 1075.544 | -0.28386 | 0.18167 | -1.56251 | 0.118167 | 0.249487 |
| ATF5 | 796.37 | -0.0802 | 0.170975 | -0.46907 | 0.639021 | 0.759385 |
| CPT1C | 187.7302 | -0.04229 | 0.311635 | -0.13572 | 0.892046 | 0.936771 |
| MN1 | 7.909093 | 0.055987 | 0.76027 | 0.07364 | 0.941297 | NA |
| C22orf15 | 8.054734 | -0.29232 | 0.731766 | -0.39947 | 0.689548 | NA |
| ELSPBP1 | 30.34978 | 0.201531 | 1.763885 | 0.114254 | 0.909036 | NA |
| SCN9A | 131.8035 | 1.328777 | 0.318248 | 4.175282 | 2.98E-05 | 0.001184 |
| SLC38A11 | 4.42275 | 0.666304 | 1.040149 | 0.640585 | 0.521792 | NA |
| CCDC8 | 61.59727 | -0.23802 | 0.280039 | -0.84997 | 0.395341 | 0.559815 |
| ZEB2 | 7349.055 | 0.356518 | 0.212729 | 1.675922 | 0.093753 | 0.213904 |
| PCBP1 | 4965.561 | -0.30854 | 0.128558 | -2.4 | 0.016395 | 0.066132 |
| NFU1 | 977.6186 | 0.295662 | 0.18707 | 1.580484 | 0.113996 | 0.243127 |
| ANTXR1 | 6147.132 | 0.003162 | 0.130323 | 0.024266 | 0.98064 | 0.989974 |
| CKAP2L | 209.8178 | 0.629542 | 0.218826 | 2.876911 | 0.004016 | 0.025459 |
| PROKR1 | 4.210329 | 0.214424 | 1.123542 | 0.190847 | 0.848646 | NA |
| APLF | 105.3124 | 1.155655 | 0.317161 | 3.643744 | 2.69E-04 | 0.004608 |
| RGPD8 | 6.687921 | 1.033438 | 0.791343 | 1.305929 | 0.191577 | NA |
| HIC2 | 163.1498 | -0.47115 | 0.217715 | -2.16409 | 0.030458 | 0.100049 |
| BUB1 | 812.9069 | 0.054686 | 0.206125 | 0.265307 | 0.790773 | 0.873094 |
| LIMS1 | 577.5442 | 0.566842 | 0.255954 | 2.214628 | 0.026786 | 0.091962 |
| UGP2 | 835.8474 | 0.372175 | 0.157385 | 2.36475 | 0.018042 | 0.070405 |
| BTD | 1536.516 | -0.25264 | 0.227566 | -1.11017 | 0.266925 | 0.431764 |
| TMEM42 | 135.1208 | -0.38399 | 0.23579 | -1.62852 | 0.103415 | 0.227984 |
| MAP3K2 | 674.107 | 0.801701 | 0.293386 | 2.732586 | 0.006284 | 0.034405 |
| ZNF35 | 216.9235 | 0.236283 | 0.219634 | 1.075801 | 0.282016 | 0.448104 |
| MYO7B | 45.31717 | -0.7594 | 0.484123 | -1.5686 | 0.116741 | NA |
| MYRIP | 1.033466 | -0.7302 | 1.946441 | -0.37515 | 0.707552 | NA |
| UBE2E3 | 1277.009 | -0.17969 | 0.156702 | -1.14671 | 0.251501 | 0.415431 |
| UBE2E1 | 2327.284 | 0.366977 | 0.158464 | 2.315844 | 0.020567 | 0.077158 |
| HNRNPA3 | 2171.733 | 0.135016 | 0.195656 | 0.690066 | 0.490152 | 0.642357 |
| HOXD4 | 34.53 | 0.71622 | 0.428585 | 1.671129 | 0.094696 | NA |
| NANP | 280.8198 | 0.517169 | 0.177616 | 2.911732 | 0.003594 | 0.023861 |
| PDCD6IP | 2823.026 | 0.15003 | 0.155306 | 0.966028 | 0.33403 | 0.502529 |
| MRAP | 0.382031 | -1.26484 | 3.151989 | -0.40128 | 0.688212 | NA |
| FAM161A | 109.1494 | 0.578983 | 0.441268 | 1.31209 | 0.18949 | 0.343696 |

| | | | | | | |
|---|---|---|---|---|---|---|
| GLB1 | 2952.286 | -0.38834 | 0.145879 | -2.6621 | 0.007766 | 0.039835 |
| CRTAP | 14239.8 | -0.20086 | 0.148157 | -1.35572 | 0.175188 | 0.327064 |
| CMTM8 | 38.1648 | 0.527209 | 0.371688 | 1.418418 | 0.156069 | NA |
| B3GNT2 | 837.4815 | 0.348761 | 0.167973 | 2.076296 | 0.037867 | 0.115728 |
| SETMAR | 443.4167 | 0.186371 | 0.154672 | 1.204943 | 0.228225 | 0.389884 |
| CST2 | 4.291343 | -1.40232 | 0.972263 | -1.44233 | 0.14921 | NA |
| CST1 | 125.6431 | -1.19526 | 0.383271 | -3.11858 | 0.001817 | 0.015146 |
| ZNF804A | 3.782003 | 1.971981 | 1.149193 | 1.715969 | 0.086168 | NA |
| TMEM182 | 190.6183 | 0.22008 | 0.190868 | 1.153049 | 0.24889 | 0.412786 |
| RALGAPB | 1758.288 | -0.0141 | 0.132815 | -0.10618 | 0.915439 | 0.951141 |
| NPAS2 | 420.3456 | -0.07154 | 0.176396 | -0.40558 | 0.685053 | 0.795121 |
| LONRF2 | 0.326333 | 1.533889 | 3.826878 | 0.40082 | 0.688553 | NA |
| SIX2 | 0.288605 | 1.443619 | 4.029058 | 0.358302 | 0.720117 | NA |
| IRF2BP1 | 1014.829 | -0.40765 | 0.183548 | -2.22094 | 0.026355 | 0.091117 |
| FOXA3 | 0.596179 | -0.09275 | 3.146759 | -0.02948 | 0.976485 | NA |
| ACYP2 | 261.4094 | 0.462818 | 0.234916 | 1.970145 | 0.048822 | 0.136634 |
| TRABD | 1650.093 | -0.13615 | 0.144573 | -0.94173 | 0.346333 | 0.514186 |
| ZNF296 | 5.004451 | -1.33372 | 0.977327 | -1.36466 | 0.172361 | NA |
| KCNS3 | 1993.017 | -0.05287 | 0.229574 | -0.23029 | 0.817864 | 0.890746 |
| FOXN2 | 598.5715 | 0.727895 | 0.250257 | 2.908586 | 0.003631 | 0.024019 |
| LSM3 | 1653.069 | 0.552609 | 0.162418 | 3.402395 | 6.68E-04 | 0.008064 |
| TMEM43 | 2711.291 | -0.16939 | 0.132122 | -1.28204 | 0.199827 | 0.355943 |
| RPS9 | 27613.33 | 0.026041 | 0.15292 | 0.170293 | 0.86478 | 0.920317 |
| TSEN34 | 1459.979 | -0.29582 | 0.136253 | -2.17111 | 0.029923 | 0.098867 |
| NDUFA3 | 1198.384 | -0.00744 | 0.182017 | -0.04087 | 0.967398 | 0.981893 |
| OSCAR | 4.103164 | 1.153916 | 1.090408 | 1.058243 | 0.289945 | NA |
| ZNF160 | 256.8172 | 0.379307 | 0.24541 | 1.545607 | 0.1222 | 0.255544 |
| ZNF415 | 8.440509 | 0.588019 | 0.798981 | 0.735961 | 0.461755 | NA |
| FEZ2 | 1048.353 | 0.205576 | 0.1834 | 1.120916 | 0.262324 | 0.426852 |
| ALK | 44.12645 | -0.8407 | 0.432969 | -1.94171 | 0.052173 | NA |
| SIGLEC17P | 0.680121 | -0.88898 | 2.365123 | -0.37587 | 0.707013 | NA |
| TRMT61B | 300.8175 | 0.321042 | 0.178134 | 1.80225 | 0.071506 | 0.17727 |
| KCNG3 | 0.774888 | -2.09879 | 2.346327 | -0.8945 | 0.371054 | NA |
| PRKCE | 763.7574 | -0.49115 | 0.175103 | -2.80493 | 0.005033 | 0.029758 |
| JAGN1 | 1518.158 | -0.1827 | 0.173404 | -1.05359 | 0.292071 | 0.458913 |
| TADA3 | 3494.36 | -0.22156 | 0.16139 | -1.3728 | 0.169814 | 0.320309 |
| SOCS5 | 461.9634 | 0.474924 | 0.194183 | 2.445755 | 0.014455 | 0.060524 |
| RBKS | 40.90197 | -0.01688 | 0.372818 | -0.04528 | 0.963881 | NA |
| GRIK1 | 1.428602 | 2.607905 | 1.905474 | 1.368639 | 0.171112 | NA |
| SCAND1 | 2034.65 | 0.015629 | 0.148776 | 0.105049 | 0.916337 | 0.951491 |
| TMEM37 | 0.298089 | -0.51711 | 3.975232 | -0.13008 | 0.896501 | NA |
| ZNF581 | 878.1987 | -0.21366 | 0.220532 | -0.96883 | 0.332632 | 0.500997 |
| ZNF524 | 313.6953 | -0.28794 | 0.193483 | -1.48819 | 0.136701 | 0.275751 |
| CDK5R2 | 0.825079 | -2.06702 | 2.533819 | -0.81577 | 0.41463 | NA |
| ASXL1 | 2493.226 | -0.37173 | 0.18005 | -2.06459 | 0.038961 | 0.11813 |
| BCL2L1 | 3274.831 | -0.27664 | 0.163194 | -1.69519 | 0.090039 | 0.207788 |
| ZNF584 | 361.8621 | -0.0318 | 0.190936 | -0.16656 | 0.867717 | 0.921425 |

| | | | | | | |
|---|---|---|---|---|---|---|
| NMUR1 | 0.230334 | -0.53318 | 4.076749 | -0.13079 | 0.895945 | NA |
| ZNF274 | 639.6578 | -0.04364 | 0.194454 | -0.2244 | 0.822448 | 0.893074 |
| ZIK1 | 204.3594 | -0.05046 | 0.243856 | -0.20691 | 0.836082 | 0.901643 |
| C20orf201 | 4.405822 | 0.424092 | 0.964325 | 0.439781 | 0.660096 | NA |
| RGS19 | 512.0451 | 0.105559 | 0.185422 | 0.569288 | 0.569161 | 0.705795 |
| TCEA2 | 800.59 | -0.3779 | 0.152686 | -2.47502 | 0.013323 | 0.057401 |
| RRM2 | 1142.117 | -0.22229 | 0.315744 | -0.70402 | 0.481423 | 0.635836 |
| TRAPPC12 | 1691.296 | -0.31515 | 0.128649 | -2.4497 | 0.014297 | 0.060062 |
| RPS21 | 6725.856 | 0.249148 | 0.165407 | 1.506271 | 0.131998 | 0.268871 |
| RPS7 | 10212.99 | 0.315416 | 0.16568 | 1.903767 | 0.056941 | 0.151216 |
| RNASEH1 | 644.0765 | 0.169998 | 0.163274 | 1.041182 | 0.297791 | 0.464828 |
| PRNP | 10669.33 | 0.511171 | 0.25063 | 2.039541 | 0.041396 | 0.123366 |
| ADRA1D | 0.163957 | 0.424913 | 4.079006 | 0.104171 | 0.917034 | NA |
| ZNF217 | 810.9761 | 0.222479 | 0.167655 | 1.327004 | 0.184507 | 0.337769 |
| SCG2 | 139.301 | 0.541949 | 0.263124 | 2.059672 | 0.03943 | 0.119184 |
| C20orf196 | 119.9913 | -0.15737 | 0.243368 | -0.64663 | 0.517868 | 0.665014 |
| MAL | 9025.522 | -0.25325 | 0.16102 | -1.57281 | 0.115762 | 0.245898 |
| LAMB2 | 10836.96 | -0.48717 | 0.150853 | -3.22943 | 0.00124 | 0.011878 |
| USP19 | 2213.728 | -0.44059 | 0.133136 | -3.30932 | 9.35E-04 | 0.009929 |
| QARS | 6177.177 | -0.27993 | 0.126143 | -2.21918 | 0.026474 | 0.091289 |
| KLF11 | 498.9344 | -0.09466 | 0.162587 | -0.58224 | 0.560406 | 0.698666 |
| EIF2AK3 | 666.7419 | 0.23779 | 0.168493 | 1.411274 | 0.158164 | 0.304685 |
| TEX37 | 0.096202 | 0.408546 | 4.079639 | 0.100143 | 0.920231 | NA |
| KRCC1 | 522.539 | 1.040672 | 0.273876 | 3.79979 | 1.45E-04 | 0.003166 |
| NME6 | 545.4581 | -0.1945 | 0.145893 | -1.33317 | 0.182475 | 0.335386 |
| CD8B | 1.351894 | 0.976798 | 1.815732 | 0.537964 | 0.590602 | NA |
| CXCR6 | 5.881259 | 0.008707 | 0.866379 | 0.01005 | 0.991981 | NA |
| CEBPB | 1425.457 | 0.133164 | 0.185212 | 0.718983 | 0.472151 | 0.628069 |
| SERHL | 7.72197 | -0.25801 | 0.73218 | -0.35239 | 0.724546 | NA |
| MACROD2 | 41.58728 | 0.516428 | 0.366868 | 1.407669 | 0.159229 | NA |
| CERS6 | 524.1019 | 0.067582 | 0.192347 | 0.351353 | 0.725324 | 0.825454 |
| SPTLC3 | 8.976657 | 0.130955 | 0.677852 | 0.19319 | 0.84681 | NA |
| TP53RK | 1184.718 | 0.077282 | 0.136921 | 0.564426 | 0.572464 | 0.708483 |
| B3GALT1 | 32.60779 | -0.26637 | 0.4031 | -0.6608 | 0.50874 | NA |
| SUCLG2 | 2407.622 | 0.412421 | 0.165034 | 2.499002 | 0.012454 | 0.054643 |
| CSDC2 | 0.096202 | 0.408546 | 4.079639 | 0.100143 | 0.920231 | NA |
| DNAJB7 | 0.681112 | 2.620532 | 2.519661 | 1.040033 | 0.298324 | NA |
| MYEOV2 | 2059.83 | 0.189876 | 0.205567 | 0.92367 | 0.355658 | 0.523768 |
| AGXT | 0.554095 | -2.96528 | 3.108533 | -0.95392 | 0.340127 | NA |
| ZBTB8OSP2 | 23.92628 | 0.555587 | 0.453977 | 1.223822 | 0.221019 | NA |
| SP3 | 983.4823 | 0.694142 | 0.206828 | 3.356137 | 7.90E-04 | 0.008987 |
| METAP1D | 207.9098 | 0.036238 | 0.231696 | 0.156404 | 0.875715 | 0.925771 |
| ZNF621 | 429.2732 | 0.474924 | 0.231234 | 2.053871 | 0.039988 | 0.120414 |
| MYD88 | 1935.189 | -0.3821 | 0.125477 | -3.04514 | 0.002326 | 0.017834 |
| OXSR1 | 1778.328 | 0.344704 | 0.174573 | 1.97456 | 0.048318 | 0.135641 |
| SLC22A13 | 0.508056 | 0.731032 | 2.861912 | 0.255435 | 0.798387 | NA |
| LCLAT1 | 240.29 | -0.01923 | 0.236645 | -0.08128 | 0.935223 | 0.963758 |

| | | | | | | |
|---|---|---|---|---|---|---|
| MIR4435-1 | 681.4258 | 0.215015 | 0.208771 | 1.029908 | 0.303053 | 0.47046 |
| | 42.52463 | -0.04445 | 0.35891 | -0.12386 | 0.901429 | NA |
| SH3RF3 | 259.2319 | 0.338757 | 0.362404 | 0.934749 | 0.349917 | 0.517951 |
| ARPP21 | 3.747263 | 1.018348 | 1.10352 | 0.922818 | 0.356102 | NA |
| COMMD1 | 788.2733 | 0.041703 | 0.175219 | 0.238003 | 0.811879 | 0.887899 |
| RAPH1 | 679.6847 | -0.32969 | 0.258576 | -1.27501 | 0.202305 | 0.358723 |
| AHSA2 | 186.6247 | 1.19426 | 0.246671 | 4.841503 | 1.29E-06 | 1.86E-04 |
| TERF1P1 | 0.162376 | 0.424913 | 4.079006 | 0.104171 | 0.917034 | NA |
| MZT2A | 1092.157 | -0.09304 | 0.140427 | -0.66254 | 0.507627 | 0.656489 |
| ZBTB21 | 474.0456 | 0.514133 | 0.168038 | 3.059628 | 0.002216 | 0.017291 |
| DAG1 | 9546.375 | -0.48421 | 0.203049 | -2.38471 | 0.017092 | 0.06802 |
| INSM1 | 12.81815 | -0.14031 | 0.589165 | -0.23815 | 0.811765 | NA |
| NAA20 | 1428.122 | 0.275215 | 0.161104 | 1.70831 | 0.087579 | 0.203418 |
| CCDC36 | 29.57723 | -0.42961 | 0.413575 | -1.03877 | 0.298913 | NA |
| SMARCC1 | 2381.185 | -0.0435 | 0.221806 | -0.19612 | 0.844519 | 0.907349 |
| ZNF417 | 114.8811 | 0.263992 | 0.262848 | 1.00435 | 0.31521 | 0.482605 |
| MST1 | 62.26767 | 0.155671 | 0.328046 | 0.474541 | 0.635114 | 0.756331 |
| GMPPB | 35.44036 | -0.47583 | 0.384347 | -1.23802 | 0.215709 | NA |
| C2orf70 | 0.869749 | -2.37356 | 2.394068 | -0.99144 | 0.321473 | NA |
| NABP1 | 252.9657 | 0.6541 | 0.232753 | 2.810276 | 0.00495 | 0.02937 |
| GPR113 | 5.355384 | -0.45412 | 0.934486 | -0.48596 | 0.626995 | NA |
| XCR1 | 0.288605 | 1.443619 | 4.029058 | 0.358302 | 0.720117 | NA |
| CCDC106 | 1295.565 | -0.18975 | 0.165842 | -1.14417 | 0.252552 | 0.416644 |
| CCR9 | 0.382031 | -1.26484 | 3.151989 | -0.40128 | 0.688212 | NA |
| SLC19A1 | 614.8502 | -0.48422 | 0.145 | -3.33943 | 8.40E-04 | 0.00931 |
| PSMD1 | 2778.353 | 0.025775 | 0.181485 | 0.142025 | 0.88706 | 0.933443 |
| SUSD5 | 1051.089 | -0.23507 | 0.215702 | -1.08979 | 0.275807 | 0.441374 |
| AGFG1 | 1629.127 | -0.02565 | 0.133126 | -0.19267 | 0.847214 | 0.909017 |
| CCDC13-AS | 0.681315 | 1.40954 | 2.618083 | 0.538386 | 0.59031 | NA |
| UBXN2A | 1034.938 | 0.448724 | 0.169372 | 2.649334 | 0.008065 | 0.040807 |
| EVX2 | 0.151697 | -1.546 | 4.075469 | -0.37934 | 0.704433 | NA |
| ZHX3 | 1061.026 | -0.21847 | 0.16016 | -1.36405 | 0.172552 | 0.323928 |
| SNHG11 | 253.6346 | -0.01278 | 0.183467 | -0.06967 | 0.944457 | 0.96923 |
| ANKRD36C | 18.1232 | 0.597633 | 0.536488 | 1.113973 | 0.265291 | NA |
| ZNF497 | 29.5162 | -0.6658 | 0.398465 | -1.67091 | 0.094739 | NA |
| GRIK1-AS1 | 0.230334 | -0.53318 | 4.076749 | -0.13079 | 0.895945 | NA |
| NR1D2 | 530.2592 | 0.708282 | 0.224928 | 3.148928 | 0.001639 | 0.014318 |
| RPL15 | 17874.53 | 0.063435 | 0.138857 | 0.456836 | 0.647789 | 0.766234 |
| DENND6A | 521.6502 | -0.00631 | 0.210457 | -0.03 | 0.976065 | 0.986887 |
| PDE12 | 678.5482 | 0.238784 | 0.197818 | 1.207089 | 0.227398 | 0.389004 |
| FUT1 | 52.23434 | -0.2792 | 0.364651 | -0.76567 | 0.443874 | 0.603732 |
| UBE2C | 927.073 | -0.45923 | 0.215958 | -2.1265 | 0.033461 | 0.106779 |
| DES | 0.258578 | 1.233151 | 4.047809 | 0.304646 | 0.760635 | NA |
| FAM182B | 4.924129 | -0.02667 | 0.940521 | -0.02835 | 0.977382 | NA |
| ANKRD30B | 1.345046 | 1.598466 | 1.922802 | 0.831321 | 0.405792 | NA |
| DPP10 | 19.77021 | 0.126597 | 0.506585 | 0.249903 | 0.802662 | NA |
| | 11.91946 | -0.06855 | 0.682601 | -0.10043 | 0.920006 | NA |

| | | | | | | |
|---|---|---|---|---|---|---|
| LINC00116 | 553.509 | 0.059227 | 0.144158 | 0.410851 | 0.681182 | 0.792212 |
| BAK1P1 | 18.50893 | 0.24051 | 0.537137 | 0.447762 | 0.654325 | NA |
| CREG2 | 11.43609 | -0.80508 | 0.617792 | -1.30316 | 0.192519 | NA |
| HOXD8 | 224.5424 | 0.196445 | 0.177191 | 1.108664 | 0.267575 | 0.432638 |
| TSPEAR | 0.640406 | 1.11402 | 2.710977 | 0.410929 | 0.681124 | NA |
| ZNF613 | 110.8333 | 0.554192 | 0.258002 | 2.148015 | 0.031713 | 0.103187 |
| RPL23P2 | 2.272754 | -0.39359 | 1.487681 | -0.26456 | 0.791346 | NA |
| IP6K1 | 3156.445 | -0.41326 | 0.166998 | -2.47462 | 0.013338 | 0.057401 |
| ENTHD1 | 392.5939 | 0.202914 | 0.231868 | 0.875128 | 0.381505 | 0.547679 |
| MYPOP | 406.912 | -0.17378 | 0.235226 | -0.73876 | 0.460053 | 0.618366 |
| LRRTM4 | 0.883349 | -0.20011 | 2.228601 | -0.08979 | 0.928451 | NA |
| ZNF135 | 42.44616 | -0.44746 | 0.458512 | -0.9759 | 0.329116 | NA |
| | 125.6308 | -0.2933 | 0.225959 | -1.29802 | 0.19428 | 0.349445 |
| KCMF1 | 1565.417 | 0.215398 | 0.150846 | 1.42794 | 0.153309 | 0.298135 |
| | 45.05158 | 0.166787 | 0.328329 | 0.507988 | 0.611462 | NA |
| MAP3K19 | 0.096202 | 0.408546 | 4.079639 | 0.100143 | 0.920231 | NA |
| CCDC121 | 34.61516 | 0.077541 | 0.453814 | 0.170864 | 0.864331 | NA |
| BOK | 307.8378 | -0.17136 | 0.250132 | -0.68507 | 0.493303 | 0.644561 |
| PFN4 | 6.549752 | 0.211703 | 0.828528 | 0.255517 | 0.798324 | NA |
| ZNF285B | 5.397779 | 0.347577 | 0.883731 | 0.393306 | 0.694093 | NA |
| NCKAP5 | 77.10238 | 0.050668 | 0.343925 | 0.147322 | 0.882878 | 0.930536 |
| | 2.007575 | 2.252112 | 1.550518 | 1.45249 | 0.146365 | NA |
| SOX11 | 83.98917 | -0.36477 | 0.386755 | -0.94314 | 0.345608 | 0.513496 |
| MAMSTR | 39.6716 | -0.35535 | 0.345033 | -1.0299 | 0.303058 | NA |
| FUT2 | 6.366729 | -0.15908 | 0.821185 | -0.19372 | 0.846399 | NA |
| THAP4 | 1505.36 | -0.24461 | 0.158253 | -1.54569 | 0.12218 | 0.255544 |
| C19orf18 | 2.480046 | 1.487004 | 1.533155 | 0.969898 | 0.332098 | NA |
| SIX5 | 146.3052 | -0.39578 | 0.224209 | -1.76522 | 0.077526 | 0.187374 |
| FBXO46 | 280.8479 | -0.40281 | 0.173296 | -2.32439 | 0.020104 | 0.076001 |
| FAM109B | 1028.915 | -0.1212 | 0.15506 | -0.78165 | 0.434423 | 0.595539 |
| SPACA4 | 0.096202 | 0.408546 | 4.079639 | 0.100143 | 0.920231 | NA |
| CCDC71 | 665.7166 | -0.35601 | 0.167825 | -2.12131 | 0.033896 | 0.1076 |
| PPFIA3 | 546.2704 | -0.3229 | 0.196307 | -1.64489 | 0.099992 | 0.223198 |
| UMODL1 | 2.198501 | -0.87571 | 1.324322 | -0.66125 | 0.508449 | NA |
| ZFAS1 | 3139.386 | 0.193437 | 0.167056 | 1.157921 | 0.246896 | 0.410366 |
| NR2C2 | 313.6768 | -0.05964 | 0.16381 | -0.36405 | 0.715819 | 0.81867 |
| GPR4 | 15.55366 | -0.26811 | 0.555954 | -0.48226 | 0.629623 | NA |
| ARIH2 | 2091.453 | 0.113642 | 0.174487 | 0.651294 | 0.514857 | 0.662214 |
| RBM44 | 0.984506 | 0.880958 | 2.062676 | 0.427094 | 0.669311 | NA |
| RPRM | 0.096202 | 0.408546 | 4.079639 | 0.100143 | 0.920231 | NA |
| IL17RA | 1152.15 | -0.51109 | 0.170389 | -2.99955 | 0.002704 | 0.019859 |
| DNAJC28 | 19.20669 | 1.236392 | 0.57736 | 2.141459 | 0.032237 | NA |
| SOX12 | 2220.5 | -0.5262 | 0.183065 | -2.87439 | 0.004048 | 0.025585 |
| ZCCHC3 | 105.3477 | -0.3187 | 0.276361 | -1.15322 | 0.248822 | 0.412786 |
| ZNF620 | 24.18006 | 0.19174 | 0.47867 | 0.400569 | 0.688737 | NA |
| CACYBPP2 | 8.877632 | 0.361713 | 0.756247 | 0.4783 | 0.632437 | NA |
| ZNF619 | 120.5673 | -0.42476 | 0.215269 | -1.97314 | 0.048479 | 0.1359 |

| | | | | | | |
|---|---|---|---|---|---|---|
| ARL6IP6 | 332.6701 | 0.230379 | 0.177343 | 1.299056 | 0.193925 | 0.348953 |
| ODF3B | 50.77129 | -0.47946 | 0.305328 | -1.57031 | 0.116344 | 0.246692 |
| ZNRF3-AS1 | 0.134133 | -1.48909 | 4.077584 | -0.36519 | 0.71497 | NA |
| C2orf73 | 2.37594 | -0.51067 | 1.267993 | -0.40274 | 0.687138 | NA |
| FAM211B | 98.8834 | -0.2732 | 0.268711 | -1.01671 | 0.309289 | 0.476621 |
| IMPDH2 | 4307.679 | -0.26299 | 0.118214 | -2.22473 | 0.026099 | 0.090629 |
| ALS2CL | 63.24305 | 0.25281 | 0.294392 | 0.858752 | 0.390477 | 0.555738 |
| PRSS42 | 0.489058 | -0.58569 | 3.427543 | -0.17088 | 0.864321 | NA |
| NDUFAF3 | 2218.665 | -0.13145 | 0.155341 | -0.84618 | 0.397455 | 0.562203 |
| C2orf69 | 275.3963 | 0.334763 | 0.259124 | 1.291901 | 0.196391 | 0.351854 |
| DALRD3 | 777.8184 | -0.26331 | 0.143277 | -1.83779 | 0.066093 | 0.167815 |
| ZNF114 | 50.20196 | 0.236132 | 0.316264 | 0.746629 | 0.455288 | 0.614182 |
| FAR2P2 | 0.651435 | 0.249629 | 2.788679 | 0.089515 | 0.928672 | NA |
| VN1R1 | 3.171232 | 0.630437 | 1.182542 | 0.53312 | 0.59395 | NA |
| ZNF543 | 90.18313 | -0.38778 | 0.257429 | -1.50636 | 0.131975 | 0.268871 |
| WDR6 | 2879.865 | -0.48135 | 0.217584 | -2.21223 | 0.02695 | 0.092394 |
| GEN1 | 253.8222 | 0.603336 | 0.349909 | 1.724266 | 0.08466 | 0.198309 |
| PLEKHM3 | 146.9604 | 0.084121 | 0.214066 | 0.392966 | 0.694345 | 0.802993 |
| ZNF223 | 20.69348 | 0.364804 | 0.518385 | 0.703732 | 0.481599 | NA |
| P4HTM | 210.6716 | -0.15452 | 0.190011 | -0.81322 | 0.416095 | 0.579239 |
| SLC25A20 | 763.6231 | 0.02362 | 0.155371 | 0.15202 | 0.879171 | 0.927523 |
| CD28 | 0.55687 | -0.14239 | 3.170029 | -0.04492 | 0.964172 | NA |
| EPM2AIP1 | 373.7407 | 0.438264 | 0.27534 | 1.591717 | 0.111448 | 0.239657 |
| ERBB4 | 4.038972 | 1.173534 | 1.072249 | 1.09446 | 0.273753 | NA |
| OTOS | 0.303394 | -2.27006 | 4.032451 | -0.56295 | 0.57347 | NA |
| GPR35 | 1.185016 | -0.03538 | 1.879555 | -0.01882 | 0.984982 | NA |
| CSRNP3 | 154.0095 | 0.061886 | 0.281688 | 0.219698 | 0.826107 | 0.895364 |
| THBD | 3054.337 | 0.618401 | 0.307712 | 2.009675 | 0.044466 | 0.128822 |
| FAM132B | 1.696031 | 0.715654 | 1.584139 | 0.451762 | 0.65144 | NA |
| ADORA2A-/ | 1.928872 | 0.45231 | 1.598625 | 0.282937 | 0.777225 | NA |
| ZNF852 | 34.88722 | 0.254555 | 0.357356 | 0.712329 | 0.476261 | NA |
| ZNF552 | 133.9871 | 0.074845 | 0.225571 | 0.331803 | 0.740038 | 0.835857 |
| SEPW1 | 7176.587 | 0.007427 | 0.140341 | 0.052923 | 0.957793 | 0.976659 |
| | 0.382031 | -1.26484 | 3.151989 | -0.40128 | 0.688212 | NA |
| TCAIM | 567.7971 | 0.608978 | 0.211019 | 2.885894 | 0.003903 | 0.025065 |
| CDH4 | 1.015902 | -0.7221 | 2.105992 | -0.34288 | 0.731689 | NA |
| C2orf71 | 0.298089 | -0.51711 | 3.975232 | -0.13008 | 0.896501 | NA |
| | 0.247898 | -0.5901 | 4.074633 | -0.14482 | 0.884851 | NA |
| LBX2 | 71.65716 | -0.35934 | 0.297428 | -1.20818 | 0.22698 | 0.388458 |
| LSMEM2 | 1.076781 | 3.261813 | 2.235782 | 1.458914 | 0.144589 | NA |
| NLRP8 | 0.315653 | -0.57305 | 3.892639 | -0.14721 | 0.882963 | NA |
| APOBEC3B | 239.5115 | -0.60215 | 0.238633 | -2.52334 | 0.011625 | 0.052182 |
| PCBP1-AS1 | 184.3993 | 0.131957 | 0.224058 | 0.588941 | 0.555901 | 0.695462 |
| MYADM | 1191.465 | -0.46027 | 0.158366 | -2.90636 | 0.003657 | 0.024096 |
| SERTAD2 | 1293.906 | 0.02549 | 0.221738 | 0.114955 | 0.908481 | 0.947226 |
| RBM15B | 2026.368 | -0.46473 | 0.142191 | -3.26833 | 0.001082 | 0.010915 |
| NKPD1 | 2.939695 | -0.70666 | 1.150051 | -0.61446 | 0.538912 | NA |

| | | | | | | |
|---|---|---|---|---|---|---|
| NLRP11 | 0.3441 | -0.0166 | 3.769682 | -0.0044 | 0.996486 | NA |
| ZNF154 | 208.2604 | 0.197878 | 0.193402 | 1.02314 | 0.306241 | 0.473301 |
| NRXN1 | 2.509132 | 1.575381 | 1.457675 | 1.080749 | 0.279809 | NA |
| GPBAR1 | 3.061564 | 1.110828 | 1.252755 | 0.886708 | 0.375236 | NA |
| ZNF784 | 183.6278 | -0.47845 | 0.198199 | -2.414 | 0.015778 | 0.064213 |
| LINC00652 | 2.657632 | 1.318474 | 1.268818 | 1.039136 | 0.298741 | NA |
| FIZ1 | 428.4212 | -0.25877 | 0.171287 | -1.51072 | 0.13086 | 0.267382 |
| SSC5D | 785.5037 | -0.25034 | 0.305029 | -0.82069 | 0.411821 | 0.575044 |
| TMEM150I | 0.326333 | 1.533889 | 3.826878 | 0.40082 | 0.688553 | NA |
| TMEM86B | 11.85718 | -0.1421 | 0.598704 | -0.23735 | 0.812389 | NA |
| | 0.163957 | 0.424913 | 4.079006 | 0.104171 | 0.917034 | NA |
| FAR2P1 | 0.096202 | 0.408546 | 4.079639 | 0.100143 | 0.920231 | NA |
| PRKRA | 988.651 | 0.265986 | 0.154129 | 1.725731 | 0.084396 | 0.197995 |
| SLC9A4 | 0.748867 | 2.740193 | 2.468928 | 1.109871 | 0.267054 | NA |
| ZNF816 | 68.87286 | 0.2328 | 0.290984 | 0.800044 | 0.423685 | 0.585953 |
| | 0.354779 | 1.703954 | 3.657232 | 0.465914 | 0.641277 | NA |
| CCDC66 | 121.4628 | 1.105405 | 0.447846 | 2.468268 | 0.013577 | 0.057939 |
| EMC3-AS1 | 97.25671 | -0.15121 | 0.264113 | -0.57252 | 0.566971 | 0.704045 |
| MCFD2 | 2455.824 | 0.190421 | 0.144794 | 1.315113 | 0.188472 | 0.342742 |
| TPRXL | 2.905861 | -0.06101 | 1.328316 | -0.04593 | 0.963367 | NA |
| KCNE1 | 5.929201 | -0.50177 | 0.928901 | -0.54017 | 0.589079 | NA |
| NRIP1 | 669.9986 | 0.536688 | 0.244467 | 2.195343 | 0.028139 | 0.094961 |
| | 300.5496 | -0.90957 | 0.223915 | -4.06213 | 4.86E-05 | 0.001615 |
| CXCR2 | 0.151697 | -1.546 | 4.075469 | -0.37934 | 0.704433 | NA |
| D2HGDH | 332.8073 | -0.15071 | 0.174776 | -0.8623 | 0.388525 | 0.554176 |
| OXTR | 36.69379 | 1.515148 | 0.703535 | 2.153621 | 0.03127 | NA |
| GPR62 | 0.667861 | -2.0782 | 2.593993 | -0.80116 | 0.42304 | NA |
| PITPNB | 1571.637 | 0.51352 | 0.183287 | 2.801735 | 0.005083 | 0.029934 |
| FKRP | 663.7889 | -0.36356 | 0.164437 | -2.21091 | 0.027042 | 0.092521 |
| HIGD1A | 3114.209 | 0.451512 | 0.180325 | 2.503876 | 0.012284 | 0.054217 |
| CCDC108 | 1.388631 | -1.18407 | 1.727625 | -0.68538 | 0.493107 | NA |
| SETD2 | 853.2048 | 0.54844 | 0.222767 | 2.461945 | 0.013819 | 0.058675 |
| TMIE | 0.640755 | -1.64826 | 2.754778 | -0.59833 | 0.549621 | NA |
| FDX1P1 | 0.163957 | 0.424913 | 4.079006 | 0.104171 | 0.917034 | NA |
| LINC00471 | 3.5525 | 0.61808 | 1.100255 | 0.561761 | 0.574279 | NA |
| ZNF329 | 312.6591 | -0.29032 | 0.183 | -1.58643 | 0.112642 | 0.241224 |
| PNMAL1 | 251.1785 | -0.17969 | 0.182808 | -0.98295 | 0.325634 | 0.493656 |
| | 5.51732 | 0.444081 | 0.882978 | 0.502935 | 0.61501 | NA |
| WRB | 1206.372 | 0.058128 | 0.134622 | 0.431789 | 0.665895 | 0.780675 |
| ASB18 | 0.296509 | -0.51709 | 3.982155 | -0.12985 | 0.896684 | NA |
| UBA7 | 1282.608 | -0.81823 | 0.239206 | -3.4206 | 6.25E-04 | 0.007763 |
| BACE2 | 8685.605 | -0.3591 | 0.116675 | -3.07778 | 0.002085 | 0.016597 |
| UBE2E2 | 518.7137 | -0.05747 | 0.174455 | -0.3294 | 0.741853 | 0.837188 |
| C22orf26 | 7.046159 | -0.30979 | 0.77992 | -0.39721 | 0.691214 | NA |
| FIGN | 308.1399 | 0.342226 | 0.22137 | 1.545947 | 0.122117 | 0.255507 |
| IZUMO1 | 0.622842 | 1.158846 | 2.735323 | 0.42366 | 0.671814 | NA |
| SPACA6P | 48.74787 | 0.184817 | 0.341696 | 0.540881 | 0.58859 | 0.721053 |

| | | | | | | |
|---|---|---|---|---|---|---|
| ZSCAN22 | 175.1494 | -0.40667 | 0.210944 | -1.92785 | 0.053873 | 0.145748 |
| KCNJ14 | 3.687973 | 0.690237 | 1.025257 | 0.673234 | 0.500799 | NA |
| | 0.46577 | -1.30191 | 3.402879 | -0.38259 | 0.702024 | NA |
| YBEY | 382.2695 | -0.03585 | 0.185098 | -0.19367 | 0.846431 | 0.908806 |
| CACNB4 | 21.59296 | 0.498667 | 0.576652 | 0.864762 | 0.38717 | NA |
| TSHZ2 | 6.493762 | 0.129476 | 0.920058 | 0.140726 | 0.888086 | NA |
| ABHD17AP | 0.298089 | -0.51711 | 3.975232 | -0.13008 | 0.896501 | NA |
| LIMK2 | 3983.449 | 0.058831 | 0.173369 | 0.339338 | 0.734355 | 0.832129 |
| ADI1 | 7592.262 | -0.08074 | 0.149411 | -0.54039 | 0.588931 | 0.721298 |
| SATB1 | 421.6979 | 0.187974 | 0.203211 | 0.925019 | 0.354956 | 0.523188 |
| ACTL10 | 74.80303 | -0.40494 | 0.318098 | -1.273 | 0.203016 | 0.359432 |
| LINC00334 | 1.075549 | 1.639348 | 2.132574 | 0.768718 | 0.442061 | NA |
| C2orf82 | 13.07983 | 0.452461 | 0.594038 | 0.76167 | 0.446257 | NA |
| TRAK1 | 2760.147 | -0.27924 | 0.154975 | -1.80186 | 0.071568 | 0.177335 |
| PLCB1 | 40.31307 | 0.319372 | 0.426803 | 0.748289 | 0.454286 | NA |
| TTC3 | 3138.299 | 0.323985 | 0.341779 | 0.947938 | 0.343161 | 0.511307 |
| RRP7B | 136.0335 | -0.13491 | 0.22822 | -0.59112 | 0.554439 | 0.694733 |
| ALG12 | 867.3546 | -0.2578 | 0.168471 | -1.53027 | 0.125951 | 0.26104 |
| COL18A1 | 6354.379 | -0.4517 | 0.213612 | -2.11458 | 0.034465 | 0.108758 |
| SLC25A18 | 20.67379 | 0.248328 | 0.494499 | 0.502181 | 0.61554 | NA |
| LENG9 | 252.0037 | -0.34126 | 0.206417 | -1.65327 | 0.098276 | 0.220698 |
| C21orf90 | 0.660976 | -0.84029 | 2.673548 | -0.3143 | 0.753294 | NA |
| EWSR1 | 4512.695 | -0.3041 | 0.133288 | -2.28155 | 0.022516 | 0.081968 |
| CNOT10 | 542.1924 | 0.188748 | 0.152104 | 1.240915 | 0.214637 | 0.373586 |
| ZNF662 | 19.29686 | 0.048742 | 0.5778 | 0.084358 | 0.932772 | NA |
| ZNF320 | 143.5246 | 0.1699 | 0.364602 | 0.465986 | 0.641225 | 0.761162 |
| TPM3P8 | 1.675801 | -1.18946 | 1.651352 | -0.7203 | 0.471342 | NA |
| SLC8A1 | 61.03477 | -0.08287 | 0.328584 | -0.25221 | 0.800878 | 0.880141 |
| WBP2NL | 3.131666 | 0.505914 | 1.407369 | 0.359475 | 0.71924 | NA |
| IGSF5 | 1.621777 | 0.185689 | 1.804641 | 0.102895 | 0.918046 | NA |
| NEB | 6.305303 | 1.169258 | 1.007067 | 1.161053 | 0.24562 | NA |
| RIPPLY3 | 4.218823 | 1.491436 | 1.05795 | 1.409741 | 0.158616 | NA |
| SMDT1 | 1166.387 | -0.08001 | 0.153647 | -0.52072 | 0.602564 | 0.731681 |
| RUVBL2 | 2917.047 | -0.2973 | 0.130881 | -2.2715 | 0.023117 | 0.083373 |
| C21orf67 | 63.84944 | -0.12163 | 0.300415 | -0.40486 | 0.685582 | 0.79533 |
| PTTG1IP | 20735.79 | -0.49063 | 0.128737 | -3.81114 | 1.38E-04 | 0.00309 |
| ABHD16B | 97.14777 | -0.23446 | 0.267138 | -0.87768 | 0.380116 | 0.546414 |
| | 0.392857 | -1.546 | 4.075469 | -0.37934 | 0.704433 | NA |
| CECR6 | 21.30937 | -0.33402 | 0.449772 | -0.74265 | 0.457693 | NA |
| | 0.162376 | 0.424913 | 4.079006 | 0.104171 | 0.917034 | NA |
| TMEM89 | 0.163957 | 0.424913 | 4.079006 | 0.104171 | 0.917034 | NA |
| RIPK4 | 1050.943 | -0.70251 | 0.210321 | -3.3402 | 8.37E-04 | 0.00931 |
| MX2 | 4114.337 | -0.37246 | 0.140413 | -2.65261 | 0.007987 | 0.040569 |
| PI4KAP2 | 99.33911 | -0.08071 | 0.233432 | -0.34573 | 0.729543 | 0.828343 |
| COA5 | 572.7428 | 0.518494 | 0.18139 | 2.858448 | 0.004257 | 0.026493 |
| PSMG1 | 978.4478 | 0.208476 | 0.149976 | 1.390063 | 0.16451 | 0.313294 |
| PRR14L | 553.5818 | 0.06428 | 0.237438 | 0.270724 | 0.786603 | 0.870699 |

| | | | | | | |
|---|---|---|---|---|---|---|
| COL18A1-A | 0.616518 | 2.49285 | 2.980788 | 0.836306 | 0.402983 | NA |
| SERHL2 | 63.59326 | -0.24747 | 0.29528 | -0.8381 | 0.401976 | 0.566508 |
| PCBP3 | 3.161361 | -0.18095 | 1.119589 | -0.16162 | 0.871602 | NA |
| ZNRF3 | 106.6792 | -0.48644 | 0.335888 | -1.44821 | 0.147559 | 0.290713 |
| TANGO2 | 2375.727 | -0.43781 | 0.126043 | -3.47351 | 5.14E-04 | 0.006912 |
| DGCR6 | 164.88 | -0.08132 | 0.234398 | -0.34694 | 0.728639 | 0.827792 |
| ZNF530 | 113.3433 | -0.01004 | 0.296207 | -0.0339 | 0.972959 | 0.985082 |
| FAM19A1 | 0.913173 | 1.256193 | 2.465498 | 0.509509 | 0.610396 | NA |
| GPR1 | 7.934447 | -1.19238 | 0.834716 | -1.42848 | 0.153153 | NA |
| CBX6 | 3954.906 | -0.6667 | 0.183293 | -3.63735 | 2.75E-04 | 0.004672 |
| KREMEN1 | 1674.981 | -0.49891 | 0.19162 | -2.60364 | 0.009224 | 0.044669 |
| TRAIP | 185.9482 | -0.21042 | 0.289062 | -0.72795 | 0.466645 | 0.623684 |
| CHEK2 | 282.0523 | 0.239479 | 0.210639 | 1.136919 | 0.255572 | 0.419747 |
| AIFM3 | 64.25286 | -0.1987 | 0.274875 | -0.72288 | 0.469755 | 0.626075 |
| TUBA8 | 2.282485 | 1.411313 | 1.467837 | 0.961492 | 0.336305 | NA |
| EMILIN3 | 0.938845 | -1.23445 | 2.063177 | -0.59832 | 0.549624 | NA |
| CCR4 | 0.303394 | -2.27006 | 4.032451 | -0.56295 | 0.57347 | NA |
| GPR39 | 0.488709 | 2.119056 | 3.307564 | 0.64067 | 0.521737 | NA |
| FAM3B | 0.364467 | -1.20166 | 3.685431 | -0.32606 | 0.744381 | NA |
| TOB2 | 985.4482 | -0.39649 | 0.151611 | -2.61518 | 0.008918 | 0.043796 |
| SCN5A | 9.186919 | 1.443008 | 0.791044 | 1.824182 | 0.068125 | NA |
| TTC32 | 18.61747 | 0.534491 | 0.543875 | 0.982746 | 0.325732 | NA |
| KCNH8 | 51.40156 | 0.990635 | 0.346371 | 2.860043 | 0.004236 | 0.026417 |
| SMTN | 2131.414 | -0.38572 | 0.273972 | -1.40789 | 0.159163 | 0.306013 |
| PP2D1 | 7.182898 | 1.009665 | 0.819672 | 1.231791 | 0.218027 | NA |
| TMPRSS2 | 4.212277 | 2.648905 | 1.392932 | 1.901676 | 0.057214 | NA |
| TBX1 | 0.303394 | -2.27006 | 4.032451 | -0.56295 | 0.57347 | NA |
| | 59.33648 | -0.3608 | 0.326095 | -1.10641 | 0.268547 | 0.433559 |
| UQCR10 | 1417.603 | -5.27E-04 | 0.160906 | -0.00328 | 0.997384 | 0.999065 |
| CLDN5 | 2.462397 | 0.159807 | 1.370046 | 0.116643 | 0.907143 | NA |
| NIPSNAP1 | 1331.528 | -0.43907 | 0.197645 | -2.2215 | 0.026317 | 0.091064 |
| CRELD2 | 869.5626 | -0.19503 | 0.162618 | -1.19931 | 0.230407 | 0.392623 |
| UBE2F | 367.0976 | 0.027995 | 0.183706 | 0.15239 | 0.878879 | 0.927317 |
| C22orf46 | 224.4377 | 0.396275 | 0.259036 | 1.529805 | 0.126065 | 0.261185 |
| KCNK12 | 0.151697 | -1.546 | 4.075469 | -0.37934 | 0.704433 | NA |
| LINC00315 | 3.97596 | 0.856131 | 1.111051 | 0.77056 | 0.440968 | NA |
| DRD5P1 | 0.392857 | -1.546 | 4.075469 | -0.37934 | 0.704433 | NA |
| RPL23AP82 | 364.1619 | 0.329024 | 0.178141 | 1.846981 | 0.06475 | 0.165826 |
| KRTAP19-1 | 28.92925 | 0.421074 | 0.511764 | 0.82279 | 0.410627 | NA |
| PLA2G6 | 201.3393 | -0.21881 | 0.220768 | -0.99111 | 0.32163 | 0.489777 |
| SS18L1 | 234.2757 | 0.33997 | 0.213141 | 1.59505 | 0.110701 | 0.238655 |
| | 0.71919 | 0.510978 | 2.728734 | 0.187258 | 0.851458 | NA |
| THAP7 | 840.1729 | -0.06302 | 0.196278 | -0.32107 | 0.748154 | 0.841815 |
| | 32.08593 | -0.2735 | 0.375891 | -0.72761 | 0.466854 | NA |
| BPIFC | 0.258578 | 1.233151 | 4.047809 | 0.304646 | 0.760635 | NA |
| TXNRD2 | 1394.33 | -0.1242 | 0.161491 | -0.76911 | 0.441831 | 0.601869 |
| FOXD4L1 | 1.945444 | -1.14135 | 1.564851 | -0.72937 | 0.465776 | NA |

| | | | | | | |
|---|---|---|---|---|---|---|
| PIWIL3 | 0.247898 | -0.5901 | 4.074633 | -0.14482 | 0.884851 | NA |
| KRTAP7-1 | 0.163957 | 0.424913 | 4.079006 | 0.104171 | 0.917034 | NA |
| KCNH7 | 0.392711 | 0.292538 | 3.145221 | 0.09301 | 0.925895 | NA |
| ZNF840 | 0.81145 | -0.46945 | 2.476203 | -0.18958 | 0.849635 | NA |
| 5-Sep | 131.2108 | -0.11536 | 0.228705 | -0.5044 | 0.613983 | 0.740239 |
| EIF4ENIF1 | 557.5501 | -0.23353 | 0.167455 | -1.39457 | 0.163145 | 0.311548 |
| UBE2G2 | 2992.949 | -0.00972 | 0.152538 | -0.06373 | 0.949184 | 0.9717 |
| OSBP2 | 47.38274 | -0.44034 | 0.335332 | -1.31316 | 0.18913 | 0.343385 |
| RBM43 | 139.009 | -0.4087 | 0.344354 | -1.18685 | 0.235286 | 0.398076 |
| SUMO3 | 5049.94 | -0.20132 | 0.121737 | -1.65372 | 0.098184 | 0.220574 |
| PTRHD1 | 396.2756 | 0.120496 | 0.15926 | 0.756604 | 0.449287 | 0.60872 |
| FAM227A | 18.14799 | 0.540992 | 0.488536 | 1.107373 | 0.268133 | NA |
| USP18 | 414.5118 | -1.05432 | 0.241584 | -4.36419 | 1.28E-05 | 7.10E-04 |
| NDUFA6 | 2131.459 | 0.22736 | 0.15585 | 1.458836 | 0.14461 | 0.286575 |
| NT5C1B | 1.267603 | 3.51044 | 2.032346 | 1.727284 | 0.084117 | NA |
| UBOX5 | 256.0344 | -0.5014 | 0.174731 | -2.86956 | 0.00411 | 0.025893 |
| MAFF | 2591.888 | -0.25541 | 0.155908 | -1.6382 | 0.10138 | 0.225006 |
| SLC24A3 | 0.230334 | -0.53318 | 4.076749 | -0.13079 | 0.895945 | NA |
| | 2.182371 | -0.29308 | 1.361793 | -0.21521 | 0.829601 | NA |
| INPP5J | 14.99298 | -0.0483 | 0.695723 | -0.06942 | 0.944658 | NA |
| ZNF445 | 410.8829 | 0.068164 | 0.188682 | 0.361262 | 0.717903 | 0.820154 |
| ZNF74 | 429.0879 | -0.31184 | 0.194579 | -1.60266 | 0.10901 | 0.236125 |
| RBM11 | 8.714746 | 0.363828 | 0.731798 | 0.497171 | 0.619069 | NA |
| TCN2 | 701.6178 | -0.76974 | 0.229907 | -3.34806 | 8.14E-04 | 0.009153 |
| GAS2L1 | 1505.205 | -0.26735 | 0.177865 | -1.50311 | 0.13281 | 0.270093 |
| MAPK11 | 116.5592 | -0.06702 | 0.310083 | -0.21613 | 0.828887 | 0.896738 |
| SP140L | 293.5001 | -0.07254 | 0.165989 | -0.43704 | 0.662084 | 0.777669 |
| MRPL30 | 1.913748 | -0.23169 | 1.455845 | -0.15915 | 0.873553 | NA |
| LINC00158 | 0.815245 | 1.670886 | 2.214146 | 0.754641 | 0.450464 | NA |
| SH3BGR | 56.31473 | 0.33065 | 0.310645 | 1.064396 | 0.287149 | 0.453464 |
| C19orf68 | 170.9524 | -0.44224 | 0.246392 | -1.79488 | 0.072673 | 0.179299 |
| L3MBTL1 | 150.9431 | 0.051178 | 0.255288 | 0.200473 | 0.841111 | 0.905336 |
| MRPL40 | 862.091 | 0.123988 | 0.145338 | 0.853102 | 0.393603 | 0.55842 |
| FAM212A | 377.7841 | -0.3457 | 0.213907 | -1.61611 | 0.106071 | 0.231919 |
| UBE2L3 | 3405.812 | -0.10566 | 0.131758 | -0.80196 | 0.422578 | 0.584968 |
| BRWD1 | 578.6773 | 0.533128 | 0.299089 | 1.782504 | 0.074667 | 0.182427 |
| SYN3 | 35.94917 | 0.165102 | 0.415426 | 0.397429 | 0.691051 | NA |
| LYG2 | 0.327913 | 1.543154 | 4.021063 | 0.383768 | 0.701151 | NA |
| PRAME | 0.46735 | -1.30193 | 3.399671 | -0.38296 | 0.701751 | NA |
| DRG1 | 1554.018 | 0.177544 | 0.162458 | 1.092861 | 0.274455 | 0.4401 |
| NLRP9 | 0.715249 | -0.95037 | 2.624937 | -0.36205 | 0.717313 | NA |
| DMWD | 1015.988 | -0.20042 | 0.143952 | -1.39224 | 0.16385 | 0.31254 |
| PIGP | 385.7687 | 0.18598 | 0.169462 | 1.097476 | 0.272433 | 0.437971 |
| RPL12P4 | 14.96086 | 0.051615 | 0.551957 | 0.093513 | 0.925496 | NA |
| CECR5-AS1 | 3.130461 | -0.64363 | 1.092136 | -0.58934 | 0.555636 | NA |
| GNB1L | 163.1389 | 0.029483 | 0.237836 | 0.123966 | 0.901343 | 0.942193 |
| KLHDC8B | 1252.445 | -0.1484 | 0.151307 | -0.98081 | 0.326685 | 0.494807 |

| | | | | | | |
|---|---|---|---|---|---|---|
| SETD4 | 467.5009 | 0.145344 | 0.150854 | 0.963472 | 0.335311 | 0.504007 |
| | 14.33859 | 0.505867 | 0.547704 | 0.923614 | 0.355687 | NA |
| ZNF284 | 56.76914 | 0.539353 | 0.302508 | 1.782939 | 0.074596 | 0.182413 |
| C2orf76 | 189.6825 | 0.452324 | 0.216725 | 2.087089 | 0.03688 | 0.113858 |
| CIDECP | 191.946 | -0.06687 | 0.191568 | -0.34904 | 0.727056 | 0.826786 |
| BPIFB4 | 0.096202 | 0.408546 | 4.079639 | 0.100143 | 0.920231 | NA |
| ZNF749 | 398.532 | -0.16215 | 0.239329 | -0.67752 | 0.498078 | 0.648931 |
| | 249.454 | -0.25497 | 0.191222 | -1.33335 | 0.182418 | 0.335341 |
| ZNF17 | 58.84211 | 0.294988 | 0.308973 | 0.954738 | 0.33971 | 0.508003 |
| GPAT2 | 22.98335 | -0.24802 | 0.473394 | -0.52391 | 0.600341 | NA |
| ZNF501 | 32.92905 | 0.255178 | 0.372143 | 0.685699 | 0.492903 | NA |
| ZNF197 | 315.2958 | 0.464789 | 0.278983 | 1.666012 | 0.095711 | 0.216579 |
| SPATA12 | 0.315653 | -0.57305 | 3.892639 | -0.14721 | 0.882963 | NA |
| FAM228A | 2.783062 | 0.573833 | 1.268947 | 0.452212 | 0.651116 | NA |
| MYT1L | 0.326333 | 1.533889 | 3.826878 | 0.40082 | 0.688553 | NA |
| 10-Sep | 1089.652 | 0.439859 | 0.16867 | 2.607808 | 0.009112 | 0.044401 |
| CEACAM19 | 1.114712 | 2.170961 | 2.315032 | 0.937767 | 0.348364 | NA |
| NF2 | 1348.84 | -0.47234 | 0.180329 | -2.61935 | 0.00881 | 0.043419 |
| PRR5 | 565.4272 | -0.04672 | 0.254063 | -0.18389 | 0.8541 | 0.913428 |
| CYP27C1 | 60.51516 | 0.511674 | 0.430721 | 1.187948 | 0.234854 | 0.397628 |
| BCR | 1223.895 | -0.41749 | 0.167189 | -2.49713 | 0.01252 | 0.054851 |
| HYAL3 | 132.8242 | 0.002358 | 0.263662 | 0.008943 | 0.992864 | 0.996901 |
| VSIG10L | 222.844 | -0.3959 | 0.193022 | -2.05104 | 0.040263 | 0.120949 |
| LINC00846 | 0.778894 | 2.823374 | 2.403613 | 1.174637 | 0.24014 | NA |
| TRABD2A | 6.407353 | 0.179155 | 0.828611 | 0.216211 | 0.828823 | NA |
| POFUT2 | 1175.784 | -0.17345 | 0.143948 | -1.20492 | 0.228233 | 0.389884 |
| TMEM17 | 131.4714 | 0.126983 | 0.236701 | 0.536472 | 0.591632 | 0.723 |
| PPARA | 917.194 | -0.21681 | 0.153725 | -1.41039 | 0.158425 | 0.30499 |
| EFCAB6 | 12.93886 | 0.100725 | 0.582799 | 0.17283 | 0.862785 | NA |
| EMID1 | 0.314073 | -0.57308 | 3.898934 | -0.14698 | 0.883146 | NA |
| KRTAP21-2 | 0.476449 | 0.524262 | 3.402674 | 0.154073 | 0.877552 | NA |
| TMPRSS6 | 0.440301 | 0.468383 | 3.449808 | 0.135771 | 0.892003 | NA |
| RPS19BP1 | 2798.318 | 0.115567 | 0.16714 | 0.691437 | 0.489291 | 0.641937 |
| PLCD1 | 841.9477 | -0.22708 | 0.135038 | -1.68161 | 0.092645 | 0.212111 |
| CMC1 | 671.1425 | 0.289196 | 0.162887 | 1.775437 | 0.075826 | 0.184279 |
| LYPD6 | 507.5951 | 0.735723 | 0.308405 | 2.385573 | 0.017053 | 0.067932 |
| SESTD1 | 478.2506 | 0.170233 | 0.233943 | 0.727669 | 0.466816 | 0.623713 |
| BCAM | 139.5751 | -0.24264 | 0.24849 | -0.97647 | 0.328829 | 0.497151 |
| CIDEC | 1.49294 | -2.24184 | 1.750328 | -1.28081 | 0.20026 | NA |
| CDHR4 | 0.151697 | -1.546 | 4.075469 | -0.37934 | 0.704433 | NA |
| PTMA | 11493.19 | 0.008317 | 0.179454 | 0.046345 | 0.963035 | 0.979897 |
| TPM3P7 | 0.247898 | -0.5901 | 4.074633 | -0.14482 | 0.884851 | NA |
| TET3 | 85.25096 | -0.52084 | 0.294663 | -1.76758 | 0.077132 | 0.18665 |
| C2orf88 | 134.8209 | -0.02954 | 0.37208 | -0.07939 | 0.936719 | 0.964637 |
| NHEJ1 | 18.3214 | -0.12865 | 0.486235 | -0.26459 | 0.791328 | NA |
| ZNF70 | 132.4748 | -0.63907 | 0.244844 | -2.61011 | 0.009051 | 0.044218 |
| CCDC157 | 103.0049 | 0.10775 | 0.227979 | 0.472631 | 0.636476 | 0.757389 |

| | | | | | | |
|---|---|---|---|---|---|---|
| SHISA7 | 2.497224 | -1.83E-04 | 1.274863 | -1.43E-04 | 0.999886 | NA |
| | 0.134133 | -1.48909 | 4.077584 | -0.36519 | 0.71497 | NA |
| C2orf66 | 0.763859 | -0.6945 | 2.307767 | -0.30094 | 0.76346 | NA |
| DNER | 0.638825 | 1.117948 | 3.106359 | 0.35989 | 0.718929 | NA |
| MORN2 | 419.0815 | 0.43706 | 0.165623 | 2.638887 | 0.008318 | 0.041599 |
| ARL4C | 3863.786 | 0.329871 | 0.261549 | 1.26122 | 0.20723 | 0.364335 |
| | 0.46577 | -1.30191 | 3.402879 | -0.38259 | 0.702024 | NA |
| MAPK12 | 181.9059 | -0.03225 | 0.219689 | -0.1468 | 0.883288 | 0.930872 |
| TMPPE | 53.06842 | -0.35445 | 0.302438 | -1.17198 | 0.241206 | 0.404398 |
| ZC3H6 | 147.9997 | 0.472317 | 0.358395 | 1.317865 | 0.187549 | 0.341795 |
| POTEE | 0.151697 | -1.546 | 4.075469 | -0.37934 | 0.704433 | NA |
| IL17REL | 0.151697 | -1.546 | 4.075469 | -0.37934 | 0.704433 | NA |
| C3orf62 | 151.238 | -0.11256 | 0.208601 | -0.53957 | 0.589493 | 0.721573 |
| BSPH1 | 0.096202 | 0.408546 | 4.079639 | 0.100143 | 0.920231 | NA |
| SLC38A3 | 0.315653 | -0.57305 | 3.892639 | -0.14721 | 0.882963 | NA |
| | 0.420954 | 1.923376 | 3.462803 | 0.555439 | 0.578594 | NA |
| PDCD1 | 0.163957 | 0.424913 | 4.079006 | 0.104171 | 0.917034 | NA |
| CERKL | 82.97668 | 1.030266 | 0.303043 | 3.399732 | 6.75E-04 | 0.008102 |
| DUSP28 | 78.94267 | -0.12823 | 0.298311 | -0.42985 | 0.667305 | 0.781814 |
| RALGAPA2 | 487.9244 | -0.23515 | 0.155851 | -1.50881 | 0.131346 | 0.267959 |
| LDOC1L | 1615.427 | -0.49132 | 0.156631 | -3.13677 | 0.001708 | 0.014611 |
| LINC00319 | 0.134133 | -1.48909 | 4.077584 | -0.36519 | 0.71497 | NA |
| PARVB | 4648.984 | -0.25346 | 0.122826 | -2.06353 | 0.039062 | 0.118314 |
| TEKT4P2 | 42.10026 | 0.248747 | 0.36414 | 0.683108 | 0.494539 | NA |
| SLC4A5 | 2.940695 | 1.014936 | 1.277125 | 0.794703 | 0.426786 | NA |
| FSIP2 | 1.512544 | 1.793508 | 1.857062 | 0.965777 | 0.334156 | NA |
| TMEM198 | 251.3693 | -0.02789 | 0.189363 | -0.14731 | 0.88289 | 0.930536 |
| TMSB4XP2 | 0.096202 | 0.408546 | 4.079639 | 0.100143 | 0.920231 | NA |
| ZNF548 | 135.1053 | 0.152593 | 0.248705 | 0.613551 | 0.539512 | 0.681722 |
| RPL14 | 14120.66 | 0.225126 | 0.149392 | 1.506947 | 0.131824 | 0.26861 |
| | 237.218 | -0.27166 | 0.230787 | -1.17708 | 0.239163 | 0.402358 |
| H1F0 | 6622.255 | -0.17953 | 0.203502 | -0.88221 | 0.377666 | 0.544108 |
| RIMKLBP1 | 0.803131 | -0.76693 | 2.349822 | -0.32638 | 0.744138 | NA |
| BLOC1S3 | 440.2819 | -0.38655 | 0.18247 | -2.11841 | 0.03414 | 0.108072 |
| ZNF600 | 65.78503 | 0.529784 | 0.305987 | 1.731395 | 0.083381 | 0.196714 |
| PAX8-AS1 | 324.4435 | 0.075443 | 0.196358 | 0.384212 | 0.700821 | 0.807644 |
| | 5.500383 | -0.12256 | 0.928581 | -0.13198 | 0.894999 | NA |
| C22orf43 | 0.162376 | 0.424913 | 4.079006 | 0.104171 | 0.917034 | NA |
| FHIT | 250.2298 | -0.21922 | 0.185145 | -1.18407 | 0.236385 | 0.399195 |
| FAM150B | 3.846479 | 1.353942 | 1.144252 | 1.183256 | 0.236708 | NA |
| RRP7A | 1496.948 | -0.30328 | 0.205315 | -1.47716 | 0.139632 | 0.279776 |
| FAM179A | 4.669382 | -0.47417 | 0.98863 | -0.47963 | 0.631493 | NA |
| TMEM194I | 295.8612 | 0.11324 | 0.238291 | 0.475218 | 0.634631 | 0.755984 |
| SYCP2 | 35.22193 | 1.100294 | 0.588764 | 1.868819 | 0.061648 | NA |
| PTPRT | 0.134133 | -1.48909 | 4.077584 | -0.36519 | 0.71497 | NA |
| | 0.096202 | 0.408546 | 4.079639 | 0.100143 | 0.920231 | NA |
| MYT1 | 3.688615 | -0.50513 | 1.11215 | -0.45419 | 0.64969 | NA |

| | | | | | | |
|---|---|---|---|---|---|---|
| SPATS2L | 1727.446 | 0.111694 | 0.15679 | 0.71238 | 0.476229 | 0.631298 |
| WDSUB1 | 232.4572 | 0.002 | 0.20905 | 0.009569 | 0.992365 | 0.996738 |
| GREB1 | 3282.868 | -0.69212 | 0.18598 | -3.7215 | 1.98E-04 | 0.003853 |
| ZNF766 | 194.5453 | 0.490661 | 0.274222 | 1.789283 | 0.073569 | 0.180809 |
| SRGAP3 | 42.26572 | -0.90174 | 0.363754 | -2.47899 | 0.013176 | NA |
| FAM217B | 459.496 | 0.60434 | 0.22059 | 2.73965 | 0.00615 | 0.03389 |
| XPNPEP3 | 299.0986 | -0.16628 | 0.183661 | -0.90538 | 0.365265 | 0.532491 |
| ZNF471 | 33.25134 | 1.217997 | 0.518156 | 2.350638 | 0.018741 | NA |
| ZNF836 | 68.07101 | -0.13078 | 0.313416 | -0.41728 | 0.676473 | 0.788478 |
| NIF3L1 | 587.6249 | 0.150823 | 0.176327 | 0.855359 | 0.392352 | 0.557385 |
| CGB7 | 6.617679 | 0.263935 | 0.850222 | 0.310431 | 0.756233 | NA |
| ZKSCAN7 | 62.28348 | -0.46679 | 0.30141 | -1.54869 | 0.121457 | 0.254304 |
| PTPN1 | 3756.536 | -0.53357 | 0.124578 | -4.28299 | 1.84E-05 | 8.78E-04 |
| ZNF765 | 101.394 | 0.556988 | 0.317942 | 1.751856 | 0.079799 | 0.191075 |
| XRCC6 | 7586.388 | -0.09598 | 0.143755 | -0.66763 | 0.504367 | 0.653972 |
| LINC00176 | 4.792016 | -0.73524 | 0.981694 | -0.74895 | 0.453889 | NA |
| RFX8 | 0.163957 | 0.424913 | 4.079006 | 0.104171 | 0.917034 | NA |
| C20orf96 | 348.0428 | -0.01725 | 0.187993 | -0.09177 | 0.92688 | 0.958156 |
| PRPF40A | 1178.82 | 0.949548 | 0.653077 | 1.453959 | 0.145958 | 0.28834 |
| SULF2 | 35.94157 | -0.67522 | 0.404602 | -1.66885 | 0.095147 | NA |
| PLXNB2 | 6554.076 | -0.52572 | 0.148529 | -3.53949 | 4.01E-04 | 0.005923 |
| MKL1 | 1208.815 | -0.53864 | 0.139233 | -3.8686 | 1.09E-04 | 0.002726 |
| POTEF | 0.937118 | 0.09073 | 2.07701 | 0.043683 | 0.965157 | NA |
| RIMBP3 | 7.109639 | 0.222472 | 0.751773 | 0.29593 | 0.767284 | NA |
| HRH1 | 0.556667 | 0.972496 | 2.522454 | 0.385536 | 0.699841 | NA |
| ZNF502 | 30.14269 | 0.641987 | 0.438423 | 1.464312 | 0.143109 | NA |
| TTC30B | 95.83457 | -0.00387 | 0.246588 | -0.0157 | 0.98747 | 0.99347 |
| ZNF512B | 964.37 | -0.37496 | 0.160634 | -2.33424 | 0.019583 | 0.074698 |
| ZNF418 | 58.27558 | 0.305722 | 0.368463 | 0.82972 | 0.406697 | 0.570377 |
| SNHG17 | 692.8432 | -0.16054 | 0.159067 | -1.00927 | 0.312846 | 0.48004 |
| CHRNG | 0.853176 | 0.733892 | 2.189324 | 0.335214 | 0.737464 | NA |
| ADA | 202.5835 | -0.37041 | 0.246239 | -1.50428 | 0.132509 | 0.269681 |
| ARID5A | 703.4044 | 0.137293 | 0.260574 | 0.526887 | 0.598272 | 0.728035 |
| RGPD4 | 0.096202 | 0.408546 | 4.079639 | 0.100143 | 0.920231 | NA |
| ZFP28 | 157.6119 | 0.197168 | 0.284173 | 0.69383 | 0.487789 | 0.640674 |
| KIAA1211L | 8.887573 | -0.90556 | 0.755883 | -1.19801 | 0.230913 | NA |
| ANKRD36B | 22.20557 | 0.928071 | 0.522186 | 1.77728 | 0.075522 | NA |
| RIMBP3B | 0.162376 | 0.424913 | 4.079006 | 0.104171 | 0.917034 | NA |
| SLC39A10 | 526.6736 | 0.727549 | 0.256255 | 2.839159 | 0.004523 | 0.027525 |
| AP2A1 | 6008.048 | -0.32916 | 0.129841 | -2.53508 | 0.011242 | 0.050962 |
| ANXA4 | 3345.085 | 0.23242 | 0.131076 | 1.773168 | 0.076201 | 0.184963 |
| ZNF470 | 132.6582 | 0.320899 | 0.378939 | 0.846836 | 0.397087 | 0.56175 |
| | 0.230334 | -0.53318 | 4.076749 | -0.13079 | 0.895945 | NA |
| KIAA1671 | 213.3477 | -0.34481 | 0.237537 | -1.4516 | 0.146613 | 0.289346 |
| ZGPAT | 648.6934 | -0.23208 | 0.15472 | -1.5 | 0.133615 | 0.27128 |
| PGAP1 | 313.4354 | 0.226815 | 0.260849 | 0.869527 | 0.384559 | 0.550664 |
| SRC | 1354.437 | -0.33422 | 0.140991 | -2.37049 | 0.017764 | 0.069688 |

| | | | | | | |
|---|---|---|---|---|---|---|
| ZNF772 | 139.9251 | 0.027232 | 0.2451 | 0.111106 | 0.911532 | 0.948657 |
| | 20.02639 | 0.33854 | 0.481647 | 0.702879 | 0.482131 | NA |
| C20orf112 | 361.3733 | -0.58449 | 0.205349 | -2.84635 | 0.004422 | 0.027205 |
| ZSCAN5B | 1.047306 | 0.316192 | 2.099018 | 0.150638 | 0.880261 | NA |
| C1D | 340.0802 | 0.686675 | 0.188525 | 3.642352 | 2.70E-04 | 0.004608 |
| FITM2 | 1117.485 | -0.50225 | 0.168803 | -2.97537 | 0.002926 | 0.020798 |
| PELI1 | 330.4046 | 0.41954 | 0.253395 | 1.655672 | 0.097788 | 0.219811 |
| DACT3 | 134.4713 | -0.18906 | 0.212887 | -0.88808 | 0.374495 | 0.540843 |
| ADARB1 | 1231.062 | -0.14911 | 0.134726 | -1.10673 | 0.268411 | 0.433457 |
| ZNF860 | 52.88868 | 0.677849 | 0.348412 | 1.945541 | 0.05171 | 0.141653 |
| C5AR1 | 0.654449 | 1.296029 | 2.63083 | 0.492631 | 0.622273 | NA |
| STMN3 | 651.5118 | -0.06667 | 0.281883 | -0.23653 | 0.81302 | 0.888676 |
| ZNF628 | 331.3688 | -0.57105 | 0.190129 | -3.00347 | 0.002669 | 0.019642 |
| SLC2A10 | 407.7405 | -0.1743 | 0.228887 | -0.76152 | 0.446347 | 0.606122 |
| ZNF665 | 17.76221 | -0.07621 | 0.551794 | -0.13811 | 0.890152 | NA |
| ATG7 | 465.4979 | -0.23847 | 0.159304 | -1.49698 | 0.134399 | 0.272639 |
| TTC30A | 101.3398 | 0.030328 | 0.268629 | 0.1129 | 0.91011 | 0.947811 |
| | 0.260158 | 1.240347 | 4.047124 | 0.306476 | 0.759242 | NA |
| ENTPD6 | 2653.655 | -0.05672 | 0.134179 | -0.42269 | 0.672525 | 0.785573 |
| KLKP1 | 0.488709 | 2.119056 | 3.307564 | 0.64067 | 0.521737 | NA |
| ZNF841 | 104.676 | 0.413119 | 0.285831 | 1.445325 | 0.148367 | 0.291544 |
| ZNF615 | 96.57883 | 0.776098 | 0.259769 | 2.987643 | 0.002811 | 0.020314 |
| DPP4 | 3983.416 | -0.15805 | 0.193327 | -0.81755 | 0.413617 | 0.576757 |
| | 3.289142 | 0.728857 | 1.118029 | 0.651912 | 0.514458 | NA |
| RPE | 651.7711 | 0.414033 | 0.166719 | 2.483413 | 0.013013 | 0.056459 |
| ZNF460 | 38.71958 | 0.144244 | 0.39379 | 0.366296 | 0.714144 | NA |
| RPL37A | 27232.45 | 0.211181 | 0.158431 | 1.332948 | 0.182549 | 0.33547 |
| | 7.77511 | -0.47217 | 0.754999 | -0.62539 | 0.531715 | NA |
| SLC9A8 | 865.6522 | -0.57475 | 0.157694 | -3.64473 | 2.68E-04 | 0.004599 |
| FAM49A | 12.00353 | 0.130448 | 0.734287 | 0.177653 | 0.858995 | NA |
| NKIRAS1 | 379.0811 | 0.643674 | 0.192342 | 3.346508 | 8.18E-04 | 0.009177 |
| C2orf27A | 14.06475 | 0.013826 | 0.543078 | 0.025459 | 0.979689 | NA |
| ZNF677 | 162.6369 | 0.202897 | 0.316344 | 0.641382 | 0.521274 | 0.668146 |
| ZNF347 | 80.95212 | 0.211045 | 0.353833 | 0.596454 | 0.550872 | 0.692009 |
| ZNF71 | 491.1627 | -0.17639 | 0.164489 | -1.07238 | 0.283549 | 0.450055 |
| ZNF335 | 678.2095 | -0.32748 | 0.159723 | -2.05033 | 0.040332 | 0.121017 |
| ZNF667 | 104.274 | 0.75943 | 0.386681 | 1.963968 | 0.049534 | 0.137692 |
| SIRPA | 4852.321 | -0.62443 | 0.188021 | -3.32109 | 8.97E-04 | 0.009708 |
| SULT1C4 | 0.394291 | 0.298745 | 3.141711 | 0.09509 | 0.924243 | NA |
| SFI1 | 405.1663 | -0.30915 | 0.174792 | -1.7687 | 0.076944 | 0.186347 |
| ZNF649 | 216.5632 | -0.04737 | 0.209524 | -0.22609 | 0.821128 | 0.892334 |
| HIBCH | 945.7739 | 0.314984 | 0.185628 | 1.69686 | 0.089723 | 0.207343 |
| ZNF544 | 305.2574 | -0.13371 | 0.165793 | -0.80651 | 0.419946 | 0.58289 |
| SOWAHC | 295.5556 | 0.390487 | 0.200341 | 1.949109 | 0.051282 | 0.140999 |
| DDRGK1 | 1749.779 | -0.15358 | 0.174609 | -0.87959 | 0.379082 | 0.545391 |
| ZNF334 | 44.08821 | 0.483253 | 0.509185 | 0.949072 | 0.342584 | NA |
| SULT1C2 | 104.659 | -0.24285 | 0.461167 | -0.5266 | 0.59847 | 0.7282 |

| | | | | | | |
|---|---|---|---|---|---|---|
| QRICH1 | 1820.268 | -0.31154 | 0.125497 | -2.48241 | 0.01305 | 0.056565 |
| UCKL1 | 770.6606 | -0.25041 | 0.161517 | -1.55038 | 0.121051 | 0.253678 |
| PEG3 | 11.50172 | 0.600176 | 0.655749 | 0.915253 | 0.360059 | NA |
| TMEM239 | 0.134133 | -1.48909 | 4.077584 | -0.36519 | 0.71497 | NA |
| ZNF813 | 81.25736 | 0.674816 | 0.47856 | 1.410097 | 0.158511 | 0.305106 |
| PIM3 | 2204.816 | -0.05556 | 0.219543 | -0.25308 | 0.800205 | 0.879677 |
| SPRED2 | 902.483 | -0.1971 | 0.168217 | -1.1717 | 0.241317 | 0.404426 |
| GFPT1 | 1725.985 | 0.359528 | 0.154428 | 2.328125 | 0.019905 | 0.075484 |
| ITSN2 | 558.8557 | 0.799342 | 0.347208 | 2.302201 | 0.021324 | 0.079097 |
| ZNF583 | 88.70664 | 0.538917 | 0.26752 | 2.014492 | 0.043958 | 0.127788 |
| ZNF480 | 225.5226 | 0.945942 | 0.342342 | 2.763149 | 0.005725 | 0.032425 |
| ZNF587 | 298.223 | -0.1066 | 0.206825 | -0.51543 | 0.606254 | 0.734356 |
| ZNF280B | 207.2176 | -0.22502 | 0.26453 | -0.85063 | 0.394973 | 0.559494 |
| ZNF808 | 67.77139 | 0.075143 | 0.318166 | 0.236177 | 0.813295 | 0.888705 |
| GPN1 | 851.0018 | 0.08532 | 0.142005 | 0.600821 | 0.547959 | 0.689317 |
| PABPC1P2 | 0.70851 | -1.62753 | 2.686091 | -0.60591 | 0.544574 | NA |
| ZNF28 | 118.7888 | 0.522451 | 0.3314 | 1.576495 | 0.114912 | 0.244486 |
| TLK1 | 456.9685 | 0.532809 | 0.287485 | 1.853346 | 0.063833 | 0.164285 |
| C3orf35 | 1.854793 | 0.825848 | 2.000198 | 0.412883 | 0.679692 | NA |
| COPS8 | 3384.627 | 0.205904 | 0.135045 | 1.524699 | 0.127334 | 0.263216 |
| PPIAP22 | 101.6692 | 0.281253 | 0.259125 | 1.085393 | 0.277748 | 0.443726 |
| ZNF534 | 3.532335 | 0.602567 | 1.085964 | 0.554869 | 0.578984 | NA |
| FAM3D | 0.298089 | -0.51711 | 3.975232 | -0.13008 | 0.896501 | NA |
| NCOA6 | 990.9433 | -0.475 | 0.213067 | -2.22936 | 0.02579 | 0.089951 |
| STK39 | 750.5775 | 0.300133 | 0.145554 | 2.062008 | 0.039207 | 0.118672 |
| SLC5A3 | 2731.627 | 0.84337 | 0.927328 | 0.909462 | 0.363106 | 0.530322 |
| TMEM184I | 2273.344 | -0.44262 | 0.144616 | -3.06066 | 0.002209 | 0.017259 |
| | 424.033 | 0.769377 | 0.31514 | 2.44138 | 0.014631 | 0.060862 |
| LTN1 | 752.279 | 0.795534 | 0.266627 | 2.983697 | 0.002848 | 0.020461 |
| ITPRIPL1 | 221.0795 | -0.16608 | 0.219827 | -0.75553 | 0.449933 | 0.609317 |
| TOP1 | 948.8962 | 0.574893 | 0.602911 | 0.953529 | 0.340322 | 0.508571 |
| SREBF2 | 10863.44 | -0.49886 | 0.121025 | -4.12198 | 3.76E-05 | 0.0014 |
| POU3F3 | 0.586491 | 2.39072 | 2.689057 | 0.889055 | 0.373973 | NA |
| ATG9A | 3161.901 | -0.5329 | 0.167968 | -3.17262 | 0.001511 | 0.013619 |
| NAGA | 1681.45 | -0.47437 | 0.143704 | -3.30103 | 9.63E-04 | 0.010106 |
| TGM2 | 12.70821 | -0.5593 | 0.5946 | -0.94063 | 0.346897 | NA |
| MIR103A2 | 4.493666 | 0.6875 | 1.067422 | 0.644075 | 0.519527 | NA |
| MIR425 | 0.096202 | 0.408546 | 4.079639 | 0.100143 | 0.920231 | NA |
| MIR26A1 | 0.162376 | 0.424913 | 4.079006 | 0.104171 | 0.917034 | NA |
| MIR26B | 0.096202 | 0.408546 | 4.079639 | 0.100143 | 0.920231 | NA |
| MIRLET7G | 0.151697 | -1.546 | 4.075469 | -0.37934 | 0.704433 | NA |
| SNORA60 | 1.118988 | -1.27548 | 2.033024 | -0.62738 | 0.530411 | NA |
| | 1.292043 | 0.040764 | 1.785235 | 0.022834 | 0.981783 | NA |
| | 0.151697 | -1.546 | 4.075469 | -0.37934 | 0.704433 | NA |
| RNA5SP47 | 0.163957 | 0.424913 | 4.079006 | 0.104171 | 0.917034 | NA |
| RNU6-228I | 0.151697 | -1.546 | 4.075469 | -0.37934 | 0.704433 | NA |
| RNU6-1301 | 0.151697 | -1.546 | 4.075469 | -0.37934 | 0.704433 | NA |

| | | | | | | |
|---|---|---|---|---|---|---|
| | 0.151697 | -1.546 | 4.075469 | -0.37934 | 0.704433 | NA |
| RN7SKP185 | 0.096202 | 0.408546 | 4.079639 | 0.100143 | 0.920231 | NA |
| RN7SKP173 | 0.192403 | 0.975392 | 4.060716 | 0.240202 | 0.810174 | NA |
| RN7SKP74 | 0.738188 | 1.39874 | 2.317793 | 0.603479 | 0.54619 | NA |
| RNU1-134P | 0.134133 | -1.48909 | 4.077584 | -0.36519 | 0.71497 | NA |
| | 0.096202 | 0.408546 | 4.079639 | 0.100143 | 0.920231 | NA |
| RNU6-1290 | 0.134133 | -1.48909 | 4.077584 | -0.36519 | 0.71497 | NA |
| RNU5B-2P | 0.303394 | -2.27006 | 4.032451 | -0.56295 | 0.57347 | NA |
| RNU6-433P | 0.327913 | 1.543154 | 4.021063 | 0.383768 | 0.701151 | NA |
| RNA5SP123 | 0.163957 | 0.424913 | 4.079006 | 0.104171 | 0.917034 | NA |
| RNU6-1147 | 0.163957 | 0.424913 | 4.079006 | 0.104171 | 0.917034 | NA |
| SNORA71D | 0.260158 | 1.240347 | 4.047124 | 0.306476 | 0.759242 | NA |
| | 0.163957 | 0.424913 | 4.079006 | 0.104171 | 0.917034 | NA |
| RNU6-137P | 0.162376 | 0.424913 | 4.079006 | 0.104171 | 0.917034 | NA |
| | 0.804712 | -0.7622 | 2.348804 | -0.3245 | 0.745556 | NA |
| RNU6-674P | 0.163957 | 0.424913 | 4.079006 | 0.104171 | 0.917034 | NA |
| RNU6-1048 | 0.314073 | -0.57308 | 3.898934 | -0.14698 | 0.883146 | NA |
| | 0.392857 | -1.546 | 4.075469 | -0.37934 | 0.704433 | NA |
| RNU6-454P | 0.260158 | 1.240347 | 4.047124 | 0.306476 | 0.759242 | NA |
| RNU6-1019 | 0.163957 | 0.424913 | 4.079006 | 0.104171 | 0.917034 | NA |
| RNU6-312P | 0.422534 | 1.924583 | 3.001758 | 0.641152 | 0.521424 | NA |
| SNORA71C | 0.842497 | -0.24499 | 2.018471 | -0.12137 | 0.903397 | NA |
| RNA5SP132 | 0.163957 | 0.424913 | 4.079006 | 0.104171 | 0.917034 | NA |
| RNA5SP479 | 0.260158 | 1.240347 | 4.047124 | 0.306476 | 0.759242 | NA |
| RNU6-384P | 0.134133 | -1.48909 | 4.077584 | -0.36519 | 0.71497 | NA |
| RNU4-8P | 0.653015 | 0.2552 | 2.787103 | 0.091565 | 0.927044 | NA |
| RN7SKP80 | 0.526843 | -0.38719 | 2.855892 | -0.13558 | 0.892156 | NA |
| RNU6-407P | 0.162376 | 0.424913 | 4.079006 | 0.104171 | 0.917034 | NA |
| | 0.585114 | 1.12675 | 2.73598 | 0.411827 | 0.680466 | NA |
| SNORD82 | 0.258578 | 1.233151 | 4.047809 | 0.304646 | 0.760635 | NA |
| RNU6-108P | 0.134133 | -1.48909 | 4.077584 | -0.36519 | 0.71497 | NA |
| RNU6-395P | 27.04871 | -0.55569 | 0.401149 | -1.38525 | 0.165976 | NA |
| | 0.096202 | 0.408546 | 4.079639 | 0.100143 | 0.920231 | NA |
| ZNF525 | 51.94492 | 0.366017 | 0.415837 | 0.880195 | 0.378754 | 0.544984 |
| | 0.096202 | 0.408546 | 4.079639 | 0.100143 | 0.920231 | NA |
| | 0.163957 | 0.424913 | 4.079006 | 0.104171 | 0.917034 | NA |
| | 0.296509 | -0.51709 | 3.982155 | -0.12985 | 0.896684 | NA |
| RBMS3-AS2 | 0.163957 | 0.424913 | 4.079006 | 0.104171 | 0.917034 | NA |
| | 0.303394 | -2.27006 | 4.032451 | -0.56295 | 0.57347 | NA |
| | 0.162376 | 0.424913 | 4.079006 | 0.104171 | 0.917034 | NA |
| PCMTD2 | 695.4753 | 0.404308 | 0.204383 | 1.978189 | 0.047907 | 0.134949 |
| SOX18 | 9.322111 | -0.32028 | 0.653111 | -0.4904 | 0.623852 | NA |
| | 3.298775 | -0.18716 | 1.253306 | -0.14933 | 0.881291 | NA |
| | 0.296509 | -0.51709 | 3.982155 | -0.12985 | 0.896684 | NA |
| SYS1 | 691.337 | -0.13647 | 0.184595 | -0.73927 | 0.459743 | 0.618083 |
| MAFB | 315.5927 | 0.036974 | 0.318537 | 0.116074 | 0.907594 | 0.946475 |
| TRAF3IP1 | 359.0687 | 0.440305 | 0.219409 | 2.006781 | 0.044773 | 0.129302 |

| | | | | | | |
|---|---|---|---|---|---|---|
| | 0.151697 | -1.546 | 4.075469 | -0.37934 | 0.704433 | NA |
| GIGYF2 | 2110.581 | -0.09118 | 0.132898 | -0.68609 | 0.492657 | 0.64393 |
| C2orf72 | 243.0503 | 0.281985 | 0.22671 | 1.243817 | 0.213567 | 0.372274 |
| ZDBF2 | 137.1154 | 0.687688 | 0.349873 | 1.965534 | 0.049352 | 0.137438 |
| BMPR2 | 2104.434 | 0.188139 | 0.284558 | 0.661163 | 0.508508 | 0.657257 |
| HNRNPCP2 | 10.60422 | -0.48156 | 0.690903 | -0.697 | 0.485803 | NA |
| COL5A2 | 3514.869 | -0.05178 | 0.20079 | -0.25789 | 0.796491 | 0.87685 |
| DFNB59 | 8.77964 | 1.082939 | 0.697865 | 1.551788 | 0.120713 | NA |
| ERICH2 | 2.19147 | 0.087988 | 1.326957 | 0.066308 | 0.947133 | NA |
| SP5 | 2.251467 | 0.062178 | 1.297425 | 0.047924 | 0.961777 | NA |
| | 4.873253 | 1.869503 | 1.224237 | 1.527076 | 0.126742 | NA |
| MBD5 | 303.8509 | 0.083519 | 0.187453 | 0.445545 | 0.655926 | 0.772809 |
| POTEKP | 0.854757 | 0.736707 | 2.188665 | 0.336601 | 0.736418 | NA |
| ZNF814 | 93.45528 | 0.166684 | 0.251663 | 0.662329 | 0.507761 | 0.656577 |
| ZNF551 | 17.63827 | -0.10876 | 0.559217 | -0.19449 | 0.845794 | NA |
| ZNF805 | 56.65299 | 0.155329 | 0.308539 | 0.503434 | 0.614659 | 0.740828 |
| ZSCAN5C | 0.680121 | -0.88898 | 2.365123 | -0.37587 | 0.707013 | NA |
| | 4.109902 | -0.161 | 1.007253 | -0.15985 | 0.873003 | NA |
| | 0.163957 | 0.424913 | 4.079006 | 0.104171 | 0.917034 | NA |
| | 14.4191 | 0.96592 | 0.562407 | 1.717475 | 0.085892 | NA |
| | 0.096202 | 0.408546 | 4.079639 | 0.100143 | 0.920231 | NA |
| LINC01123 | 0.230334 | -0.53318 | 4.076749 | -0.13079 | 0.895945 | NA |
| ZNF468 | 109.4752 | 0.39157 | 0.254394 | 1.539226 | 0.123749 | 0.258089 |
| ZNF616 | 190.2282 | 0.332496 | 0.246707 | 1.347736 | 0.177743 | 0.330119 |
| TBC1D8 | 529.3584 | 0.120553 | 0.204864 | 0.588451 | 0.55623 | 0.695727 |
| | 0.326333 | 1.533889 | 3.826878 | 0.40082 | 0.688553 | NA |
| ASPDH | 2.887306 | -1.01849 | 1.378645 | -0.73876 | 0.460053 | NA |
| | 0.489058 | -0.58569 | 3.427543 | -0.17088 | 0.864321 | NA |
| AKT1S1 | 2556.712 | -0.20914 | 0.147857 | -1.41448 | 0.157222 | 0.303457 |
| STARD7-AS | 112.0643 | -0.17494 | 0.232347 | -0.7529 | 0.451507 | 0.610554 |
| | 1.482903 | 0.434893 | 1.641474 | 0.26494 | 0.791055 | NA |
| | 14.05327 | 1.073555 | 0.665688 | 1.6127 | 0.10681 | NA |
| MRPL53 | 35.22366 | -0.20656 | 0.374698 | -0.55127 | 0.581449 | NA |
| DCTN1 | 7951.339 | -0.32986 | 0.140055 | -2.35519 | 0.018513 | 0.0717 |
| PNMAL2 | 54.25327 | -0.68456 | 0.321887 | -2.12671 | 0.033444 | 0.10676 |
| IGFL2 | 0.192403 | 0.975392 | 4.060716 | 0.240202 | 0.810174 | NA |
| IGFL4 | 7.931441 | 1.564366 | 1.047382 | 1.493596 | 0.135281 | NA |
| FBXO48 | 52.63254 | 0.232117 | 0.308828 | 0.751607 | 0.452287 | 0.611248 |
| | 2.008661 | -1.42041 | 1.485751 | -0.95602 | 0.339061 | NA |
| | 0.258578 | 1.233151 | 4.047809 | 0.304646 | 0.760635 | NA |
| LINC01121 | 0.653015 | 0.2552 | 2.787103 | 0.091565 | 0.927044 | NA |
| C2orf91 | 0.134133 | -1.48909 | 4.077584 | -0.36519 | 0.71497 | NA |
| CDKL4 | 1.148665 | 0.530005 | 1.802635 | 0.294017 | 0.768745 | NA |
| LINC00654 | 153.1202 | -0.2785 | 0.221764 | -1.25584 | 0.209173 | 0.366717 |
| VIT | 0.162376 | 0.424913 | 4.079006 | 0.104171 | 0.917034 | NA |
| | 0.392857 | -1.546 | 4.075469 | -0.37934 | 0.704433 | NA |
| CHKB-AS1 | 38.84085 | 0.037393 | 0.336578 | 0.111098 | 0.911539 | NA |

| | | | | | | |
|---|---|---|---|---|---|---|
| CPT1B | 10.0238 | -0.24533 | 0.717245 | -0.34205 | 0.732313 | NA |
| HMGN1 | 1527.748 | 0.262532 | 0.144496 | 1.816884 | 0.069235 | 0.173233 |
| DENND6B | 64.89032 | 0.005453 | 0.271349 | 0.020096 | 0.983967 | 0.991494 |
| | 3.712034 | -0.3207 | 1.16677 | -0.27486 | 0.783425 | NA |
| | 0.163957 | 0.424913 | 4.079006 | 0.104171 | 0.917034 | NA |
| MFSD2B | 0.709559 | 2.702128 | 2.840259 | 0.951367 | 0.341418 | NA |
| CDPF1 | 160.5866 | -0.2126 | 0.216319 | -0.98282 | 0.325696 | 0.493687 |
| SMIM11 | 580.4195 | 0.320302 | 0.179688 | 1.78255 | 0.07466 | 0.182427 |
| CYP2D7P | 4.297625 | 0.197216 | 1.143534 | 0.172461 | 0.863075 | NA |
| LINC00634 | 3.230422 | -0.80657 | 1.177974 | -0.68471 | 0.493529 | NA |
| ITSN1 | 714.1793 | 0.394914 | 0.23707 | 1.665813 | 0.095751 | 0.21662 |
| CRYZL1 | 560.0829 | 0.348973 | 0.188367 | 1.852628 | 0.063936 | 0.164427 |
| CYS1 | 17.33995 | -0.59412 | 0.521725 | -1.13876 | 0.254805 | NA |
| RFPL3S | 0.601483 | -1.7332 | 2.729494 | -0.63499 | 0.525436 | NA |
| C21orf62 | 0.354779 | 1.703954 | 3.657232 | 0.465914 | 0.641277 | NA |
| C21orf49 | 14.28863 | 0.639434 | 0.558612 | 1.144684 | 0.25234 | NA |
| | 0.096202 | 0.408546 | 4.079639 | 0.100143 | 0.920231 | NA |
| IGLL3P | 1.733779 | 0.863739 | 1.543172 | 0.559716 | 0.575673 | NA |
| TMEM191( | 9.611356 | -0.03818 | 0.719487 | -0.05307 | 0.957674 | NA |
| P2RX6P | 0.661122 | -3.06315 | 2.98049 | -1.02773 | 0.304076 | NA |
| | 14.84438 | -0.147 | 0.520801 | -0.28225 | 0.77775 | NA |
| | 0.247898 | -0.5901 | 4.074633 | -0.14482 | 0.884851 | NA |
| TRIM71 | 1.015699 | 0.142451 | 2.181052 | 0.065313 | 0.947925 | NA |
| ZCWPW2 | 9.441945 | 0.441029 | 0.803143 | 0.549129 | 0.582917 | NA |
| ANKRD28 | 2803.688 | 0.426795 | 0.235342 | 1.813507 | 0.069754 | 0.174038 |
| COLQ | 27.94037 | 0.216727 | 0.485106 | 0.446762 | 0.655047 | NA |
| METTL6 | 292.7089 | 0.165841 | 0.175563 | 0.944626 | 0.34485 | 0.512884 |
| | 25.16112 | -0.74276 | 0.418118 | -1.77643 | 0.075662 | NA |
| SETD5-AS1 | 135.2905 | 0.85895 | 0.258142 | 3.327432 | 8.77E-04 | 0.009546 |
| | 0.394291 | 0.298745 | 3.141711 | 0.09509 | 0.924243 | NA |
| | 0.151697 | -1.546 | 4.075469 | -0.37934 | 0.704433 | NA |
| | 0.162376 | 0.424913 | 4.079006 | 0.104171 | 0.917034 | NA |
| | 0.825079 | -2.06702 | 2.533819 | -0.81577 | 0.41463 | NA |
| SNORA70F | 0.162376 | 0.424913 | 4.079006 | 0.104171 | 0.917034 | NA |
| SNORA75 | 0.520316 | 2.239969 | 3.173769 | 0.705776 | 0.480328 | NA |
| | 0.162376 | 0.424913 | 4.079006 | 0.104171 | 0.917034 | NA |
| | 0.163957 | 0.424913 | 4.079006 | 0.104171 | 0.917034 | NA |
| RNU6-1024 | 0.487129 | 2.123445 | 3.910539 | 0.543006 | 0.587126 | NA |
| SNORA70B | 0.392857 | -1.546 | 4.075469 | -0.37934 | 0.704433 | NA |
| | 0.437526 | -2.68242 | 3.384147 | -0.79264 | 0.427986 | NA |
| RNU6-5P | 0.623191 | -1.58103 | 2.770593 | -0.57065 | 0.568239 | NA |
| RNU6-611F | 0.641986 | 1.117767 | 2.709494 | 0.412537 | 0.679946 | NA |
| SNORD51 | 1.595437 | -0.5998 | 1.62242 | -0.3697 | 0.711608 | NA |
| | 0.096202 | 0.408546 | 4.079639 | 0.100143 | 0.920231 | NA |
| | 0.163957 | 0.424913 | 4.079006 | 0.104171 | 0.917034 | NA |
| RNU6-31P | 0.163957 | 0.424913 | 4.079006 | 0.104171 | 0.917034 | NA |
| RNU6-900F | 0.163957 | 0.424913 | 4.079006 | 0.104171 | 0.917034 | NA |

| | | | | | | |
|---|---|---|---|---|---|---|
| SNORD20 | 0.392711 | 0.292538 | 3.145221 | 0.09301 | 0.925895 | NA |
| RNU6-851F | 0.096202 | 0.408546 | 4.079639 | 0.100143 | 0.920231 | NA |
| | 3.518438 | 0.256826 | 1.091358 | 0.235327 | 0.813955 | NA |
| | 0.162376 | 0.424913 | 4.079006 | 0.104171 | 0.917034 | NA |
| RNU6-476F | 0.394291 | 0.298745 | 3.141711 | 0.09509 | 0.924243 | NA |
| MIR647 | 0.326333 | 1.533889 | 3.826878 | 0.40082 | 0.688553 | NA |
| MIR191 | 0.162376 | 0.424913 | 4.079006 | 0.104171 | 0.917034 | NA |
| MIR499A | 0.162376 | 0.424913 | 4.079006 | 0.104171 | 0.917034 | NA |
| MIR659 | 2.145512 | -0.8614 | 1.478923 | -0.58245 | 0.560262 | NA |
| MIR642A | 0.315653 | -0.57305 | 3.892639 | -0.14721 | 0.882963 | NA |
| MIR646 | 0.096202 | 0.408546 | 4.079639 | 0.100143 | 0.920231 | NA |
| MIR138-1 | 0.764062 | -1.46884 | 2.483016 | -0.59155 | 0.55415 | NA |
| MIR644A | 0.327913 | 1.543154 | 4.021063 | 0.383768 | 0.701151 | NA |
| MIR645 | 0.096202 | 0.408546 | 4.079639 | 0.100143 | 0.920231 | NA |
| MIR185 | 0.163957 | 0.424913 | 4.079006 | 0.104171 | 0.917034 | NA |
| SNORD94 | 2.608257 | 1.742849 | 1.402152 | 1.242982 | 0.213875 | NA |
| SNORD83A | 3.256689 | -0.45393 | 1.107272 | -0.40996 | 0.681837 | NA |
| SACM1L | 2081.834 | 0.681881 | 0.160988 | 4.235615 | 2.28E-05 | 9.97E-04 |
| TSN | 2648.913 | 0.040193 | 0.131659 | 0.30528 | 0.760153 | 0.850634 |
| IGLV5-52 | 0.327913 | 1.543154 | 4.021063 | 0.383768 | 0.701151 | NA |
| IGLV1-51 | 0.584911 | 2.38756 | 2.690807 | 0.887303 | 0.374916 | NA |
| | 0.163957 | 0.424913 | 4.079006 | 0.104171 | 0.917034 | NA |
| | 0.268266 | -2.15396 | 4.038351 | -0.53338 | 0.593773 | NA |
| SNORD17 | 10.65257 | 0.667961 | 0.625026 | 1.068692 | 0.285208 | NA |
| SNORD89 | 1.865712 | 1.010494 | 1.752093 | 0.576736 | 0.564118 | NA |
| SNORD12 | 1.988227 | 3.102483 | 1.717293 | 1.806612 | 0.070823 | NA |
| RNU6-882F | 1.171751 | 1.845228 | 2.071346 | 0.890835 | 0.373018 | NA |
| SNORD69 | 1.455009 | -1.54356 | 1.678891 | -0.91939 | 0.357891 | NA |
| SNORD19 | 0.921134 | 0.131863 | 2.11947 | 0.062215 | 0.950391 | NA |
| | 0.151697 | -1.546 | 4.075469 | -0.37934 | 0.704433 | NA |
| RNA5SP474 | 0.247898 | -0.5901 | 4.074633 | -0.14482 | 0.884851 | NA |
| | 365.1518 | -0.24891 | 0.157052 | -1.58489 | 0.112991 | 0.241789 |
| RPS15AP36 | 1.278443 | -1.18833 | 1.874683 | -0.63388 | 0.526158 | NA |
| VN2R17P | 0.303394 | -2.27006 | 4.032451 | -0.56295 | 0.57347 | NA |
| ZNF580 | 804.5622 | -0.22649 | 0.140407 | -1.6131 | 0.106722 | 0.232957 |
| ZNF611 | 97.41737 | 0.232765 | 0.289588 | 0.80378 | 0.421524 | 0.584123 |
| SYT3 | 7.368645 | 0.843199 | 0.807038 | 1.044808 | 0.296112 | NA |
| NUP62 | 1268.881 | -0.40634 | 0.134609 | -3.01864 | 0.002539 | 0.019039 |
| RPL23AP80 | 0.151697 | -1.546 | 4.075469 | -0.37934 | 0.704433 | NA |
| | 0.46735 | -1.30193 | 3.399671 | -0.38296 | 0.701751 | NA |
| PPP1R14BP | 30.31373 | -0.29079 | 0.402015 | -0.72333 | 0.469475 | NA |
| | 4.299544 | 1.534052 | 1.034316 | 1.483155 | 0.138033 | NA |
| NPM1P46 | 0.162376 | 0.424913 | 4.079006 | 0.104171 | 0.917034 | NA |
| AHCYP5 | 0.096202 | 0.408546 | 4.079639 | 0.100143 | 0.920231 | NA |
| | 0.134133 | -1.48909 | 4.077584 | -0.36519 | 0.71497 | NA |
| | 0.151697 | -1.546 | 4.075469 | -0.37934 | 0.704433 | NA |
| KLHL23 | 21.9493 | 0.011669 | 0.467752 | 0.024947 | 0.980097 | NA |

| | | | | | | |
|---|---|---|---|---|---|---|
| BTF3L4P2 | 3.577326 | -0.87674 | 1.075871 | -0.81491 | 0.415122 | NA |
| | 0.448206 | -1.2532 | 3.007315 | -0.41672 | 0.676885 | NA |
| | 0.696104 | -0.93034 | 2.354268 | -0.39517 | 0.692715 | NA |
| | 18.80446 | 0.716831 | 0.545538 | 1.313989 | 0.18885 | NA |
| | 5.66767 | -0.31984 | 1.070817 | -0.29869 | 0.765178 | NA |
| YWHAZP2 | 7.72682 | -0.42193 | 0.731239 | -0.57701 | 0.563934 | NA |
| ANKRD39 | 423.8485 | -0.11299 | 0.16067 | -0.70321 | 0.481924 | 0.636073 |
| | 0.392711 | 0.292538 | 3.145221 | 0.09301 | 0.925895 | NA |
| | 1.282944 | -0.71339 | 1.77411 | -0.40211 | 0.687601 | NA |
| | 0.258578 | 1.233151 | 4.047809 | 0.304646 | 0.760635 | NA |
| H2AFZP1 | 0.096202 | 0.408546 | 4.079639 | 0.100143 | 0.920231 | NA |
| FTH1P3 | 2.488511 | -1.15753 | 1.43419 | -0.80709 | 0.419612 | NA |
| PPIAP16 | 0.841063 | -2.08438 | 2.306687 | -0.90362 | 0.366195 | NA |
| TMEM110 | 104.1072 | -0.27218 | 0.256772 | -1.05999 | 0.28915 | 0.455471 |
| RPLP0P6 | 444.5745 | -0.26242 | 0.166935 | -1.572 | 0.11595 | 0.246254 |
| | 0.151697 | -1.546 | 4.075469 | -0.37934 | 0.704433 | NA |
| LBH | 594.5226 | -0.62242 | 0.156967 | -3.96528 | 7.33E-05 | 0.002099 |
| PPP1CB | 2113.337 | 0.393199 | 0.183098 | 2.147482 | 0.031755 | 0.103214 |
| NCKIPSD | 753.1469 | -0.27223 | 0.178885 | -1.5218 | 0.12806 | 0.264301 |
| | 0.926438 | -0.78569 | 2.072011 | -0.37919 | 0.704544 | NA |
| TREX1 | 746.6165 | -0.35593 | 0.148366 | -2.39899 | 0.01644 | 0.066248 |
| SLC35F6 | 1110.466 | -0.57118 | 0.151451 | -3.77136 | 1.62E-04 | 0.003443 |
| HMGB1P1( | 1.937191 | 0.501769 | 1.44322 | 0.347673 | 0.728086 | NA |
| FAM209B | 1.007997 | 0.255096 | 2.389543 | 0.106755 | 0.914983 | NA |
| | 0.162376 | 0.424913 | 4.079006 | 0.104171 | 0.917034 | NA |
| ZNF337-AS | 18.77407 | 0.189628 | 0.626536 | 0.302662 | 0.762148 | NA |
| CENPBD1P | 753.4406 | -0.21269 | 0.144564 | -1.47128 | 0.141216 | 0.281791 |
| ZNF134 | 484.4823 | 0.021221 | 0.200599 | 0.105786 | 0.915752 | 0.9513 |
| | 0.162376 | 0.424913 | 4.079006 | 0.104171 | 0.917034 | NA |
| | 0.163957 | 0.424913 | 4.079006 | 0.104171 | 0.917034 | NA |
| | 0.813811 | 0.928591 | 2.591569 | 0.358312 | 0.72011 | NA |
| OLA1P1 | 1.610509 | 1.167658 | 1.618286 | 0.72154 | 0.470577 | NA |
| ZNF888 | 0.571659 | -3.00664 | 3.038724 | -0.98944 | 0.322447 | NA |
| ZNF845 | 96.17599 | 0.623624 | 0.379672 | 1.642536 | 0.100479 | 0.223642 |
| RPL13P2 | 0.096202 | 0.408546 | 4.079639 | 0.100143 | 0.920231 | NA |
| | 0.163957 | 0.424913 | 4.079006 | 0.104171 | 0.917034 | NA |
| SUGT1P2 | 0.35636 | 1.709252 | 3.651833 | 0.468053 | 0.639746 | NA |
| | 78.68394 | -0.42345 | 0.278957 | -1.51798 | 0.129018 | 0.265356 |
| | 0.192403 | 0.975392 | 4.060716 | 0.240202 | 0.810174 | NA |
| | 0.151697 | -1.546 | 4.075469 | -0.37934 | 0.704433 | NA |
| TAF9BP1 | 0.354779 | 1.703954 | 3.657232 | 0.465914 | 0.641277 | NA |
| | 0.326536 | 0.037056 | 3.842554 | 0.009644 | 0.992306 | NA |
| CFL1P7 | 0.162376 | 0.424913 | 4.079006 | 0.104171 | 0.917034 | NA |
| | 0.134133 | -1.48909 | 4.077584 | -0.36519 | 0.71497 | NA |
| RPL13AP7 | 3.872583 | 0.021082 | 1.11819 | 0.018854 | 0.984958 | NA |
| | 92.47403 | -0.41402 | 0.249286 | -1.66084 | 0.096746 | 0.21817 |
| PPM1N | 51.85221 | 0.651305 | 0.341421 | 1.90763 | 0.056439 | 0.150368 |

| | | | | | | |
|---|---|---|---|---|---|---|
| SLC23A3 | 9.736634 | 0.368699 | 0.765117 | 0.481886 | 0.629887 | NA |
| RPL5P8 | 0.392857 | -1.546 | 4.075469 | -0.37934 | 0.704433 | NA |
| CSNK1E | 5571.971 | -0.09167 | 0.138684 | -0.66097 | 0.508634 | 0.657276 |
| SEPHS1P6 | 0.555233 | -0.57009 | 3.296947 | -0.17291 | 0.862719 | NA |
| DNAJB1P1 | 0.096202 | 0.408546 | 4.079639 | 0.100143 | 0.920231 | NA |
| RPS10P2 | 0.448206 | -1.2532 | 3.007315 | -0.41672 | 0.676885 | NA |
| | 0.163957 | 0.424913 | 4.079006 | 0.104171 | 0.917034 | NA |
| | 0.151697 | -1.546 | 4.075469 | -0.37934 | 0.704433 | NA |
| API5P2 | 0.392857 | -1.546 | 4.075469 | -0.37934 | 0.704433 | NA |
| | 33.84223 | -0.145 | 0.410929 | -0.35286 | 0.724195 | NA |
| RPL7AP14 | 0.315653 | -0.57305 | 3.892639 | -0.14721 | 0.882963 | NA |
| | 1.209682 | 0.725753 | 1.849937 | 0.392312 | 0.694828 | NA |
| TTLL3 | 70.92416 | 0.122261 | 0.301591 | 0.405387 | 0.685193 | 0.795122 |
| PGAM1P4 | 0.134133 | -1.48909 | 4.077584 | -0.36519 | 0.71497 | NA |
| CPSF1P1 | 46.26413 | -1.39885 | 0.503724 | -2.77702 | 0.005486 | NA |
| GNAQP1 | 0.744715 | -0.63075 | 2.564852 | -0.24592 | 0.805743 | NA |
| CPNE1 | 2795.862 | -0.18268 | 0.216163 | -0.84509 | 0.398062 | 0.56286 |
| | 0.315653 | -0.57305 | 3.892639 | -0.14721 | 0.882963 | NA |
| | 10.4878 | -0.70901 | 0.634379 | -1.11765 | 0.263717 | NA |
| | 2.724395 | 0.283408 | 1.215847 | 0.233095 | 0.815688 | NA |
| XPOTP1 | 3.586802 | -0.29828 | 1.095928 | -0.27217 | 0.785491 | NA |
| EEF1A1P12 | 33.18315 | -0.25856 | 0.476063 | -0.54311 | 0.587052 | NA |
| PDCL3P3 | 0.296509 | -0.51709 | 3.982155 | -0.12985 | 0.896684 | NA |
| | 0.151697 | -1.546 | 4.075469 | -0.37934 | 0.704433 | NA |
| CYCSP6 | 0.151697 | -1.546 | 4.075469 | -0.37934 | 0.704433 | NA |
| SMARCE1P | 0.258578 | 1.233151 | 4.047809 | 0.304646 | 0.760635 | NA |
| RPS15AP1 | 6.51075 | 0.396523 | 0.792247 | 0.500504 | 0.61672 | NA |
| RPS4XP3 | 5.471844 | -0.2732 | 0.848492 | -0.32198 | 0.747468 | NA |
| COPS8P2 | 1.424835 | -0.73179 | 1.663339 | -0.43995 | 0.659972 | NA |
| EML6 | 17.88121 | 0.644486 | 0.539233 | 1.195189 | 0.232013 | NA |
| RPS19P1 | 6.244503 | 0.019255 | 0.861823 | 0.022342 | 0.982175 | NA |
| | 0.230334 | -0.53318 | 4.076749 | -0.13079 | 0.895945 | NA |
| ARHGEF33 | 13.02809 | 0.880751 | 0.601521 | 1.464205 | 0.143138 | NA |
| IFRD2 | 1953.058 | -0.21087 | 0.154638 | -1.36361 | 0.17269 | 0.324032 |
| CAPN14 | 64.04248 | -0.22246 | 0.289276 | -0.76902 | 0.441883 | 0.601869 |
| RPL5P35 | 0.096202 | 0.408546 | 4.079639 | 0.100143 | 0.920231 | NA |
| | 0.556667 | 0.972496 | 2.522454 | 0.385536 | 0.699841 | NA |
| MPRIPP1 | 3.822252 | 0.260935 | 0.999985 | 0.260939 | 0.794139 | NA |
| HMGB1P4 | 0.315653 | -0.57305 | 3.892639 | -0.14721 | 0.882963 | NA |
| APOC1P1 | 0.096202 | 0.408546 | 4.079639 | 0.100143 | 0.920231 | NA |
| SRSF9P1 | 0.915556 | -1.07826 | 2.191337 | -0.49206 | 0.62268 | NA |
| RPL23AP3 | 0.260158 | 1.240347 | 4.047124 | 0.306476 | 0.759242 | NA |
| | 8.583584 | 1.147005 | 0.880866 | 1.302135 | 0.19287 | NA |
| C22orf29 | 533.3202 | -0.46569 | 0.187464 | -2.48417 | 0.012985 | 0.05641 |
| DSTNP4 | 0.162376 | 0.424913 | 4.079006 | 0.104171 | 0.917034 | NA |
| OR7E122P | 0.545988 | -0.43913 | 2.541145 | -0.17281 | 0.862803 | NA |
| PEX26 | 2207.466 | -0.2443 | 0.160965 | -1.51771 | 0.129089 | 0.265408 |

| | | | | | | |
|---|---|---|---|---|---|---|
| FASTKD5 | 691.9547 | -0.00597 | 0.162843 | -0.03669 | 0.970735 | 0.983955 |
| | 0.163957 | 0.424913 | 4.079006 | 0.104171 | 0.917034 | NA |
| VPS16 | 1035.472 | -0.29358 | 0.130005 | -2.25823 | 0.023931 | 0.085415 |
| GPX1P2 | 0.392857 | -1.546 | 4.075469 | -0.37934 | 0.704433 | NA |
| SLC25A5P1 | 0.354779 | 1.703954 | 3.657232 | 0.465914 | 0.641277 | NA |
| LINC00478 | 5.751486 | 0.733216 | 0.900538 | 0.814198 | 0.415532 | NA |
| ACTG1P3 | 0.720624 | 1.435938 | 2.334155 | 0.615185 | 0.538432 | NA |
| | 0.163957 | 0.424913 | 4.079006 | 0.104171 | 0.917034 | NA |
| MCM3AP-A | 59.40956 | 0.339872 | 0.299241 | 1.135783 | 0.256047 | 0.420246 |
| NPEPL1 | 209.0547 | -0.27487 | 0.196319 | -1.40012 | 0.161479 | 0.309069 |
| | 0.151697 | -1.546 | 4.075469 | -0.37934 | 0.704433 | NA |
| | 131.2096 | -0.2057 | 0.343242 | -0.5993 | 0.548974 | 0.690147 |
| ZNF663P | 1.266023 | 3.509235 | 2.049126 | 1.712552 | 0.086795 | NA |
| | 3.627127 | -1.12929 | 1.286379 | -0.87788 | 0.380007 | NA |
| | 0.151697 | -1.546 | 4.075469 | -0.37934 | 0.704433 | NA |
| RPL27AP | 0.968668 | -0.08001 | 2.195869 | -0.03644 | 0.970935 | NA |
| | 4.164851 | 1.80197 | 1.077332 | 1.672622 | 0.094402 | NA |
| PI4KAP1 | 73.08972 | 0.365615 | 0.29343 | 1.246005 | 0.212763 | 0.371473 |
| EFCAB8 | 0.163957 | 0.424913 | 4.079006 | 0.104171 | 0.917034 | NA |
| ANKRD20A | 3.603012 | 0.882982 | 1.058219 | 0.834404 | 0.404054 | NA |
| GAB4 | 4.464965 | -1.11824 | 0.993995 | -1.12499 | 0.260592 | NA |
| C20orf202 | 1.506403 | 1.704481 | 1.818534 | 0.937283 | 0.348613 | NA |
| | 0.526843 | -0.38719 | 2.855892 | -0.13558 | 0.892156 | NA |
| MIR941-1 | 0.163957 | 0.424913 | 4.079006 | 0.104171 | 0.917034 | NA |
| MIR933 | 0.096202 | 0.408546 | 4.079639 | 0.100143 | 0.920231 | NA |
| MIR298 | 0.162376 | 0.424913 | 4.079006 | 0.104171 | 0.917034 | NA |
| RPL10P1 | 0.841063 | -2.08438 | 2.306687 | -0.90362 | 0.366195 | NA |
| | 0.258578 | 1.233151 | 4.047809 | 0.304646 | 0.760635 | NA |
| | 0.247898 | -0.5901 | 4.074633 | -0.14482 | 0.884851 | NA |
| SYCE3 | 0.28583 | -2.21358 | 3.97376 | -0.55705 | 0.577494 | NA |
| | 0.758758 | -0.35643 | 2.514365 | -0.14176 | 0.887273 | NA |
| | 1.741627 | 0.179272 | 1.513121 | 0.118478 | 0.905689 | NA |
| | 0.163957 | 0.424913 | 4.079006 | 0.104171 | 0.917034 | NA |
| | 19.01966 | -0.42416 | 0.542066 | -0.78248 | 0.433933 | NA |
| | 0.571659 | -3.00664 | 3.038724 | -0.98944 | 0.322447 | NA |
| | 8.950751 | -0.45148 | 0.664162 | -0.67977 | 0.496648 | NA |
| | 0.151697 | -1.546 | 4.075469 | -0.37934 | 0.704433 | NA |
| CEBPZ-AS1 | 725.7043 | 0.07159 | 0.178681 | 0.400661 | 0.68867 | 0.797595 |
| TDRD15 | 0.163957 | 0.424913 | 4.079006 | 0.104171 | 0.917034 | NA |
| ZNF579 | 545.5455 | -0.41147 | 0.171444 | -2.40005 | 0.016393 | 0.066132 |
| | 0.392857 | -1.546 | 4.075469 | -0.37934 | 0.704433 | NA |
| FAM19A5 | 398.5576 | -0.39019 | 0.291737 | -1.33746 | 0.181071 | 0.333581 |
| FAM228B | 41.77065 | 0.849211 | 0.417976 | 2.031724 | 0.042182 | NA |
| | 0.151697 | -1.546 | 4.075469 | -0.37934 | 0.704433 | NA |
| | 0.298089 | -0.51711 | 3.975232 | -0.13008 | 0.896501 | NA |
| | 17.15323 | 0.40571 | 0.524465 | 0.77357 | 0.439185 | NA |
| MIR1302-3 | 0.096202 | 0.408546 | 4.079639 | 0.100143 | 0.920231 | NA |

| | | | | | | |
|---|---|---|---|---|---|---|
| | 0.28583 | -2.21358 | 3.97376 | -0.55705 | 0.577494 | NA |
| SNORD88A | 0.544407 | -0.44425 | 2.542878 | -0.1747 | 0.861313 | NA |
| MIR663B | 3.417318 | -0.06746 | 1.169148 | -0.0577 | 0.953988 | NA |
| | 0.820549 | 0.502564 | 2.433834 | 0.206491 | 0.836408 | NA |
| MIR1301 | 0.163957 | 0.424913 | 4.079006 | 0.104171 | 0.917034 | NA |
| RNU6ATAC | 0.354779 | 1.703954 | 3.657232 | 0.465914 | 0.641277 | NA |
| MIR1249 | 7.524266 | -1.17654 | 0.811673 | -1.44953 | 0.14719 | NA |
| PPP3R1 | 1917.982 | -0.12638 | 0.127856 | -0.98848 | 0.322918 | 0.490809 |
| C2orf16 | 32.02894 | -0.58856 | 0.38148 | -1.54282 | 0.122874 | NA |
| ARIH2OS | 20.66694 | -0.33273 | 0.4472 | -0.74404 | 0.456853 | NA |
| NPTXR | 52.47306 | -0.64346 | 0.325507 | -1.9768 | 0.048064 | 0.13528 |
| C19orf73 | 15.20893 | 0.114403 | 0.542982 | 0.210694 | 0.833126 | NA |
| ZNF880 | 171.5192 | 0.048825 | 0.303734 | 0.160748 | 0.872292 | 0.924028 |
| TIGD1 | 31.08026 | 0.804543 | 0.38371 | 2.096745 | 0.036016 | NA |
| APOL6 | 279.2797 | -0.5594 | 0.231474 | -2.41671 | 0.015662 | 0.063848 |
| | 0.230334 | -0.53318 | 4.076749 | -0.13079 | 0.895945 | NA |
| LINC01118 | 0.804712 | -0.7622 | 2.348804 | -0.3245 | 0.745556 | NA |
| RAB6C | 0.096202 | 0.408546 | 4.079639 | 0.100143 | 0.920231 | NA |
| C21orf140 | 15.84576 | -0.05611 | 0.542177 | -0.1035 | 0.917569 | NA |
| | 7.103232 | 0.548589 | 0.818807 | 0.669985 | 0.502867 | NA |
| | 0.162376 | 0.424913 | 4.079006 | 0.104171 | 0.917034 | NA |
| | 0.326333 | 1.533889 | 3.826878 | 0.40082 | 0.688553 | NA |
| | 0.096202 | 0.408546 | 4.079639 | 0.100143 | 0.920231 | NA |
| | 0.096202 | 0.408546 | 4.079639 | 0.100143 | 0.920231 | NA |
| LINC01124 | 0.489058 | -0.58569 | 3.427543 | -0.17088 | 0.864321 | NA |
| | 0.163957 | 0.424913 | 4.079006 | 0.104171 | 0.917034 | NA |
| ADRA2B | 2.024856 | -0.35634 | 1.529265 | -0.23301 | 0.815751 | NA |
| LINC00152 | 371.3248 | 0.123204 | 0.207618 | 0.593415 | 0.552904 | 0.693616 |
| | 2.825495 | 0.580163 | 1.217955 | 0.476342 | 0.633831 | NA |
| | 2.803778 | -0.34892 | 1.145644 | -0.30456 | 0.760701 | NA |
| | 2.102342 | 1.932711 | 1.554638 | 1.24319 | 0.213798 | NA |
| SNORD12B | 0.561971 | -0.49218 | 2.810061 | -0.17515 | 0.860962 | NA |
| RN7SKP27 | 0.315653 | -0.57305 | 3.892639 | -0.14721 | 0.882963 | NA |
| RNA5SP49 | 0.192403 | 0.975392 | 4.060716 | 0.240202 | 0.810174 | NA |
| RNU2-52P | 0.449786 | -1.25325 | 3.00474 | -0.41709 | 0.676613 | NA |
| | 0.134133 | -1.48909 | 4.077584 | -0.36519 | 0.71497 | NA |
| RN7SKP38 | 0.096202 | 0.408546 | 4.079639 | 0.100143 | 0.920231 | NA |
| RNU4-78P | 0.555233 | -0.57009 | 3.296947 | -0.17291 | 0.862719 | NA |
| RN7SKP11 | 0.163957 | 0.424913 | 4.079006 | 0.104171 | 0.917034 | NA |
| RNA5SP10 | 0.163957 | 0.424913 | 4.079006 | 0.104171 | 0.917034 | NA |
| | 0.490493 | 0.776209 | 2.895261 | 0.268096 | 0.788625 | NA |
| | 0.151697 | -1.546 | 4.075469 | -0.37934 | 0.704433 | NA |
| | 1.010809 | 2.054089 | 2.306032 | 0.890746 | 0.373065 | NA |
| | 0.134133 | -1.48909 | 4.077584 | -0.36519 | 0.71497 | NA |
| | 3.038265 | 0.490262 | 1.118905 | 0.438163 | 0.661268 | NA |
| ATP6V0E1 | 0.096202 | 0.408546 | 4.079639 | 0.100143 | 0.920231 | NA |
| | 2.416995 | -0.8668 | 1.411984 | -0.61389 | 0.539288 | NA |

| | | | | | | |
|---|---|---|---|---|---|---|
| MTND5P28 | 0.096202 | 0.408546 | 4.079639 | 0.100143 | 0.920231 | NA |
| | 0.268266 | -2.15396 | 4.038351 | -0.53338 | 0.593773 | NA |
| | 0.162376 | 0.424913 | 4.079006 | 0.104171 | 0.917034 | NA |
| | 0.096202 | 0.408546 | 4.079639 | 0.100143 | 0.920231 | NA |
| RFPL4A | 0.134133 | -1.48909 | 4.077584 | -0.36519 | 0.71497 | NA |
| | 9.376012 | 0.131641 | 0.668433 | 0.196939 | 0.843875 | NA |
| | 0.392857 | -1.546 | 4.075469 | -0.37934 | 0.704433 | NA |
| EEF1A1P24 | 2.164713 | -0.53863 | 1.366844 | -0.39407 | 0.69353 | NA |
| DIP2A-IT1 | 0.744715 | -0.63075 | 2.564852 | -0.24592 | 0.805743 | NA |
| | 0.134133 | -1.48909 | 4.077584 | -0.36519 | 0.71497 | NA |
| | 0.900913 | -0.26014 | 2.222223 | -0.11706 | 0.90681 | NA |
| | 0.653015 | 0.2552 | 2.787103 | 0.091565 | 0.927044 | NA |
| | 0.947594 | 1.902131 | 2.454497 | 0.774958 | 0.438365 | NA |
| | 0.315653 | -0.57305 | 3.892639 | -0.14721 | 0.882963 | NA |
| | 0.786799 | 1.578877 | 2.532268 | 0.623503 | 0.532954 | NA |
| LINC00205 | 72.87691 | -0.25071 | 0.373334 | -0.67156 | 0.501867 | 0.65228 |
| UBE2E1-AS | 3.553381 | -0.76523 | 1.132848 | -0.67549 | 0.499363 | NA |
| ENTPD3-AS | 95.52466 | 0.041716 | 0.273786 | 0.152366 | 0.878898 | 0.927317 |
| IL10RB-AS1 | 7.697188 | -0.10631 | 0.741225 | -0.14343 | 0.885952 | NA |
| LINC00114 | 0.3441 | -0.0166 | 3.769682 | -0.0044 | 0.996486 | NA |
| EEF1A1P1 | 0.303394 | -2.27006 | 4.032451 | -0.56295 | 0.57347 | NA |
| DAZAP2P1 | 61.85578 | -0.3127 | 0.326363 | -0.95814 | 0.337993 | 0.506819 |
| | 0.476449 | 0.524262 | 3.402674 | 0.154073 | 0.877552 | NA |
| | 0.883 | 1.784395 | 2.188528 | 0.81534 | 0.414878 | NA |
| | 0.572651 | 0.928649 | 2.780261 | 0.334015 | 0.738368 | NA |
| OSER1-AS1 | 246.434 | -0.10986 | 0.175023 | -0.62769 | 0.53021 | 0.675004 |
| SEC13P1 | 0.230334 | -0.53318 | 4.076749 | -0.13079 | 0.895945 | NA |
| | 0.327913 | 1.543154 | 4.021063 | 0.383768 | 0.701151 | NA |
| | 0.096202 | 0.408546 | 4.079639 | 0.100143 | 0.920231 | NA |
| ASS1P2 | 0.796661 | 1.607053 | 2.50763 | 0.640865 | 0.52161 | NA |
| | 0.151697 | -1.546 | 4.075469 | -0.37934 | 0.704433 | NA |
| | 3.434882 | -0.07673 | 1.183693 | -0.06482 | 0.948317 | NA |
| | 210.4043 | 0.057235 | 0.199852 | 0.286387 | 0.774582 | 0.860453 |
| | 0.134133 | -1.48909 | 4.077584 | -0.36519 | 0.71497 | NA |
| | 0.709944 | 0.232855 | 2.332364 | 0.099837 | 0.920474 | NA |
| | 0.151697 | -1.546 | 4.075469 | -0.37934 | 0.704433 | NA |
| | 0.096202 | 0.408546 | 4.079639 | 0.100143 | 0.920231 | NA |
| CCNT2-AS1 | 20.41633 | 0.575894 | 0.484963 | 1.187502 | 0.23503 | NA |
| | 2.357956 | -0.8582 | 1.388485 | -0.61808 | 0.536521 | NA |
| | 13.4436 | 0.751213 | 0.661464 | 1.135683 | 0.256089 | NA |
| | 0.162376 | 0.424913 | 4.079006 | 0.104171 | 0.917034 | NA |
| | 12.86923 | 0.183569 | 0.577704 | 0.317757 | 0.750669 | NA |
| UBE2FP1 | 16.96237 | 0.839818 | 0.530698 | 1.582478 | 0.113541 | NA |
| | 15.86924 | 0.194101 | 0.542349 | 0.35789 | 0.720426 | NA |
| RPS24P8 | 1.667133 | -0.03825 | 1.741107 | -0.02197 | 0.982471 | NA |
| ATG12P2 | 0.096202 | 0.408546 | 4.079639 | 0.100143 | 0.920231 | NA |
| | 0.727508 | 0.192954 | 2.572892 | 0.074995 | 0.940219 | NA |

| | | | | | | |
|---|---|---|---|---|---|---|
| | 33.28908 | -0.19588 | 0.376931 | -0.51966 | 0.603299 | NA |
| HMGB1P9 | 0.096202 | 0.408546 | 4.079639 | 0.100143 | 0.920231 | NA |
| DNAJC27-A | 4.236693 | -0.27344 | 0.955014 | -0.28632 | 0.774632 | NA |
| | 109.2778 | 0.003299 | 0.305452 | 0.010802 | 0.991382 | 0.996256 |
| HOXD-AS1 | 312.486 | 0.808672 | 0.35261 | 2.293392 | 0.021825 | 0.080404 |
| | 1.481322 | 0.442244 | 1.686201 | 0.262272 | 0.793111 | NA |
| MINOS1P3 | 24.96345 | 0.215786 | 0.427423 | 0.504854 | 0.613661 | NA |
| | 0.163957 | 0.424913 | 4.079006 | 0.104171 | 0.917034 | NA |
| | 0.392857 | -1.546 | 4.075469 | -0.37934 | 0.704433 | NA |
| MSL3P1 | 52.69824 | 0.231217 | 0.317533 | 0.728167 | 0.466511 | 0.623646 |
| | 0.478233 | -0.67301 | 2.937473 | -0.22911 | 0.818782 | NA |
| CHL1-AS2 | 3.335577 | -0.0373 | 1.16813 | -0.03194 | 0.974524 | NA |
| | 0.162376 | 0.424913 | 4.079006 | 0.104171 | 0.917034 | NA |
| | 1.06294 | 2.033685 | 2.088387 | 0.973807 | 0.330152 | NA |
| | 0.915759 | -1.79003 | 2.342046 | -0.7643 | 0.444687 | NA |
| | 16.90646 | -0.4537 | 0.510162 | -0.88932 | 0.37383 | NA |
| BACE2-IT1 | 0.987309 | 0.267164 | 2.098144 | 0.127333 | 0.898677 | NA |
| | 0.134133 | -1.48909 | 4.077584 | -0.36519 | 0.71497 | NA |
| | 1437.954 | -0.3914 | 0.228992 | -1.70925 | 0.087405 | 0.203133 |
| | 0.096202 | 0.408546 | 4.079639 | 0.100143 | 0.920231 | NA |
| ADM5 | 4.51147 | 0.489996 | 1.039548 | 0.471355 | 0.637387 | NA |
| PRKAR2A-A | 13.62684 | -0.66995 | 0.545107 | -1.22902 | 0.219065 | NA |
| | 0.424115 | 1.932531 | 3.45462 | 0.559405 | 0.575885 | NA |
| RSL24D1P6 | 1.233919 | -2.69077 | 2.106908 | -1.27712 | 0.20156 | NA |
| PGAM1P6 | 0.162376 | 0.424913 | 4.079006 | 0.104171 | 0.917034 | NA |
| | 0.599903 | -1.73313 | 2.730982 | -0.63462 | 0.525678 | NA |
| TTC21B-AS | 0.096202 | 0.408546 | 4.079639 | 0.100143 | 0.920231 | NA |
| | 0.432222 | -1.18916 | 3.482045 | -0.34151 | 0.732718 | NA |
| LINC01087 | 0.326333 | 1.533889 | 3.826878 | 0.40082 | 0.688553 | NA |
| | 0.58526 | 0.001197 | 3.235889 | 3.70E-04 | 0.999705 | NA |
| LINC01117 | 8.102691 | -0.33521 | 0.727383 | -0.46085 | 0.644907 | NA |
| RPS5P2 | 0.163957 | 0.424913 | 4.079006 | 0.104171 | 0.917034 | NA |
| | 0.163957 | 0.424913 | 4.079006 | 0.104171 | 0.917034 | NA |
| | 0.52699 | -2.57878 | 3.248478 | -0.79384 | 0.427288 | NA |
| | 5.226267 | 0.070078 | 0.920841 | 0.076102 | 0.939338 | NA |
| | 0.268266 | -2.15396 | 4.038351 | -0.53338 | 0.593773 | NA |
| SH3BP5-AS | 40.5083 | 1.067619 | 0.356643 | 2.993523 | 0.002758 | NA |
| EIF3EP2 | 1.526916 | 0.456861 | 1.739807 | 0.262593 | 0.792864 | NA |
| | 2.525679 | 1.072278 | 1.336133 | 0.802523 | 0.42225 | NA |
| | 2.011684 | 0.595727 | 1.515957 | 0.392971 | 0.694341 | NA |
| | 0.134133 | -1.48909 | 4.077584 | -0.36519 | 0.71497 | NA |
| | 0.430642 | -1.1891 | 3.485782 | -0.34113 | 0.733008 | NA |
| | 1.25731 | 0.686444 | 2.256212 | 0.304246 | 0.76094 | NA |
| RPL29P11 | 2.77536 | 0.611145 | 1.24288 | 0.491717 | 0.622919 | NA |
| | 0.134133 | -1.48909 | 4.077584 | -0.36519 | 0.71497 | NA |
| TSSC1-IT1 | 2.156045 | 0.295738 | 1.51456 | 0.195263 | 0.845187 | NA |
| | 0.162376 | 0.424913 | 4.079006 | 0.104171 | 0.917034 | NA |

| | | | | | | |
|---|---|---|---|---|---|---|
| VPS26BP1 | 0.163957 | 0.424913 | 4.079006 | 0.104171 | 0.917034 | NA |
| | 8.440534 | 0.065924 | 0.702225 | 0.093879 | 0.925205 | NA |
| | 4.077027 | -0.70736 | 1.016579 | -0.69583 | 0.486537 | NA |
| | 0.410275 | 0.244283 | 3.100889 | 0.078778 | 0.937209 | NA |
| | 4.492008 | -1.23723 | 1.020162 | -1.21278 | 0.225215 | NA |
| | 0.327913 | 1.543154 | 4.021063 | 0.383768 | 0.701151 | NA |
| LARGE-AS1 | 0.163957 | 0.424913 | 4.079006 | 0.104171 | 0.917034 | NA |
| UBE2D3P1 | 2.964631 | -0.01655 | 1.235813 | -0.01339 | 0.989317 | NA |
| | 0.298089 | -0.51711 | 3.975232 | -0.13008 | 0.896501 | NA |
| RPL18AP2 | 0.096202 | 0.408546 | 4.079639 | 0.100143 | 0.920231 | NA |
| | 0.260158 | 1.240347 | 4.047124 | 0.306476 | 0.759242 | NA |
| | 0.604258 | 1.087325 | 2.707696 | 0.401568 | 0.688002 | NA |
| | 0.315653 | -0.57305 | 3.892639 | -0.14721 | 0.882963 | NA |
| | 0.430642 | -1.1891 | 3.485782 | -0.34113 | 0.733008 | NA |
| | 0.134133 | -1.48909 | 4.077584 | -0.36519 | 0.71497 | NA |
| | 0.192403 | 0.975392 | 4.060716 | 0.240202 | 0.810174 | NA |
| | 0.324752 | 1.53082 | 4.022025 | 0.380609 | 0.703493 | NA |
| | 3.985335 | 0.039176 | 1.093674 | 0.035821 | 0.971425 | NA |
| | 4.461695 | 0.466163 | 0.947645 | 0.491918 | 0.622777 | NA |
| | 0.162376 | 0.424913 | 4.079006 | 0.104171 | 0.917034 | NA |
| ARL2BPP1( | 2.292212 | 0.216442 | 1.666511 | 0.129877 | 0.896663 | NA |
| TRAPPC12- | 12.77156 | -0.74681 | 0.679062 | -1.09977 | 0.271433 | NA |
| RPS2P1 | 0.151697 | -1.546 | 4.075469 | -0.37934 | 0.704433 | NA |
| RPL8P2 | 0.684273 | 2.627785 | 2.896325 | 0.907282 | 0.364257 | NA |
| | 0.162376 | 0.424913 | 4.079006 | 0.104171 | 0.917034 | NA |
| | 5.189175 | -0.10831 | 0.869303 | -0.1246 | 0.900842 | NA |
| | 0.230334 | -0.53318 | 4.076749 | -0.13079 | 0.895945 | NA |
| | 3.217856 | 1.197227 | 1.206904 | 0.991982 | 0.321206 | NA |
| RPF2P1 | 1.331206 | 0.831477 | 1.740564 | 0.477705 | 0.63286 | NA |
| | 48.8941 | -0.09679 | 0.337803 | -0.28654 | 0.774468 | 0.860453 |
| | 0.326333 | 1.533889 | 3.826878 | 0.40082 | 0.688553 | NA |
| | 5.185079 | -0.49664 | 0.877128 | -0.56621 | 0.571252 | NA |
| | 0.151697 | -1.546 | 4.075469 | -0.37934 | 0.704433 | NA |
| | 0.804566 | 0.545894 | 2.033578 | 0.26844 | 0.78836 | NA |
| | 0.315653 | -0.57305 | 3.892639 | -0.14721 | 0.882963 | NA |
| BOLA3-AS1 | 228.9678 | -0.2367 | 0.187426 | -1.26292 | 0.206617 | 0.363668 |
| RAB6C-AS1 | 0.247898 | -0.5901 | 4.074633 | -0.14482 | 0.884851 | NA |
| | 0.260158 | 1.240347 | 4.047124 | 0.306476 | 0.759242 | NA |
| RFPL1S | 4.707867 | -0.46749 | 0.991037 | -0.47172 | 0.637126 | NA |
| | 4.192641 | -0.05789 | 1.025018 | -0.05648 | 0.954961 | NA |
| | 0.803131 | -0.76693 | 2.349822 | -0.32638 | 0.744138 | NA |
| C3orf83 | 4.348746 | 0.615662 | 0.978036 | 0.629488 | 0.529029 | NA |
| | 0.752028 | 2.752519 | 2.876634 | 0.956854 | 0.338641 | NA |
| | 0.163957 | 0.424913 | 4.079006 | 0.104171 | 0.917034 | NA |
| | 0.28583 | -2.21358 | 3.97376 | -0.55705 | 0.577494 | NA |
| UBE2D3P2 | 0.163957 | 0.424913 | 4.079006 | 0.104171 | 0.917034 | NA |
| | 5.742115 | 0.093431 | 0.968974 | 0.096422 | 0.923185 | NA |

| | | | | | | |
|---|---|---|---|---|---|---|
| | 1.027741 | -0.65078 | 2.331852 | -0.27908 | 0.780182 | NA |
| | 0.247898 | -0.5901 | 4.074633 | -0.14482 | 0.884851 | NA |
| | 5.139995 | 0.426608 | 0.939713 | 0.453977 | 0.649846 | NA |
| HNRNPA1P | 0.45509 | -2.71227 | 3.963744 | -0.68427 | 0.493805 | NA |
| SLC26A6 | 501.9946 | 0.137948 | 0.158127 | 0.872386 | 0.382998 | 0.548893 |
| FGD5-AS1 | 3250.513 | 0.123095 | 0.14097 | 0.873203 | 0.382552 | 0.548652 |
| PLAC4 | 4.316897 | 1.060859 | 0.993355 | 1.067956 | 0.28554 | NA |
| SIRPAP1 | 156.2177 | -0.57205 | 0.249144 | -2.29607 | 0.021672 | 0.080037 |
| MIAT | 3.209952 | 1.52726 | 1.226588 | 1.245129 | 0.213084 | NA |
| LSM3P3 | 0.411855 | 0.250456 | 3.097618 | 0.080854 | 0.935558 | NA |
| DNAJC19P5 | 1.225869 | -0.07951 | 1.838357 | -0.04325 | 0.965503 | NA |
| MKRN7P | 1.908212 | 2.160586 | 1.550034 | 1.393896 | 0.163349 | NA |
| | 0.326333 | 1.533889 | 3.826878 | 0.40082 | 0.688553 | NA |
| | 4.191367 | -0.03314 | 1.062858 | -0.03118 | 0.975127 | NA |
| HSPE1P13 | 0.096202 | 0.408546 | 4.079639 | 0.100143 | 0.920231 | NA |
| NTF4 | 1.063143 | 1.12382 | 1.855095 | 0.605802 | 0.544646 | NA |
| SATB2-AS1 | 0.708567 | -0.60605 | 2.553496 | -0.23734 | 0.812392 | NA |
| | 5.652336 | 0.278619 | 1.069802 | 0.260439 | 0.794525 | NA |
| | 0.641986 | 1.117767 | 2.709494 | 0.412537 | 0.679946 | NA |
| | 0.163957 | 0.424913 | 4.079006 | 0.104171 | 0.917034 | NA |
| HAR1A | 1.293421 | 0.777973 | 1.840346 | 0.422732 | 0.672491 | NA |
| PRICKLE2-A | 0.247898 | -0.5901 | 4.074633 | -0.14482 | 0.884851 | NA |
| PPIAP21 | 8.10848 | 0.062185 | 0.716326 | 0.08681 | 0.930822 | NA |
| MEMO1P1 | 8.248039 | 0.282944 | 0.704068 | 0.40187 | 0.687779 | NA |
| PRSS44 | 0.163957 | 0.424913 | 4.079006 | 0.104171 | 0.917034 | NA |
| | 0.298089 | -0.51711 | 3.975232 | -0.13008 | 0.896501 | NA |
| | 1.101221 | -0.557 | 1.814706 | -0.30694 | 0.758892 | NA |
| UQCRFS1P1 | 39.9702 | -0.40685 | 0.351645 | -1.15698 | 0.247281 | NA |
| | 0.162376 | 0.424913 | 4.079006 | 0.104171 | 0.917034 | NA |
| AHCTF1P1 | 6.043371 | 0.949469 | 0.840244 | 1.129992 | 0.25848 | NA |
| FTCDNL1 | 3.942407 | 0.807763 | 1.070238 | 0.754751 | 0.450399 | NA |
| | 0.556813 | -0.57007 | 3.294615 | -0.17303 | 0.862627 | NA |
| RPLP0P1 | 0.163957 | 0.424913 | 4.079006 | 0.104171 | 0.917034 | NA |
| | 0.230334 | -0.53318 | 4.076749 | -0.13079 | 0.895945 | NA |
| RPL37AP1 | 8.006967 | -0.27846 | 0.879441 | -0.31663 | 0.751525 | NA |
| | 0.192403 | 0.975392 | 4.060716 | 0.240202 | 0.810174 | NA |
| GRM7-AS3 | 0.410275 | 0.244283 | 3.100889 | 0.078778 | 0.937209 | NA |
| | 0.364467 | -1.20166 | 3.685431 | -0.32606 | 0.744381 | NA |
| | 12.30302 | 0.170611 | 0.611627 | 0.278946 | 0.780286 | NA |
| | 0.151697 | -1.546 | 4.075469 | -0.37934 | 0.704433 | NA |
| | 1.199003 | -0.22526 | 1.838234 | -0.12254 | 0.902472 | NA |
| ARPC3P1 | 1.224085 | 0.637824 | 2.020305 | 0.315707 | 0.752225 | NA |
| TMEM191A | 30.48928 | 0.61656 | 0.420572 | 1.466004 | 0.142647 | NA |
| | 0.327913 | 1.543154 | 4.021063 | 0.383768 | 0.701151 | NA |
| CFLAR-AS1 | 0.563552 | -0.48645 | 2.808029 | -0.17324 | 0.862465 | NA |
| | 118.3259 | -0.18456 | 0.28171 | -0.65516 | 0.512366 | 0.659798 |
| | 5.5215 | 0.458027 | 0.868923 | 0.52712 | 0.598111 | NA |

| | | | | | | |
|---|---|---|---|---|---|---|
| | 0.545988 | -0.43913 | 2.541145 | -0.17281 | 0.862803 | NA |
| | 0.162376 | 0.424913 | 4.079006 | 0.104171 | 0.917034 | NA |
| RPL10AP6 | 14.37375 | 0.703941 | 0.54286 | 1.296725 | 0.194726 | NA |
| | 8.622137 | -0.02553 | 0.68459 | -0.03729 | 0.970254 | NA |
| | 0.46735 | -1.30193 | 3.399671 | -0.38296 | 0.701751 | NA |
| | 0.296509 | -0.51709 | 3.982155 | -0.12985 | 0.896684 | NA |
| | 0.46577 | -1.30191 | 3.402879 | -0.38259 | 0.702024 | NA |
| HMGN2P7 | 0.096202 | 0.408546 | 4.079639 | 0.100143 | 0.920231 | NA |
| PLCL2-AS1 | 0.096202 | 0.408546 | 4.079639 | 0.100143 | 0.920231 | NA |
| CYP2D8P | 0.776322 | -0.40209 | 2.222574 | -0.18091 | 0.856435 | NA |
| | 8.318872 | 0.457008 | 0.720059 | 0.63468 | 0.525637 | NA |
| | 3.433716 | 0.237336 | 1.194573 | 0.198678 | 0.842514 | NA |
| TMEM185E | 499.0433 | -0.32769 | 0.154742 | -2.11767 | 0.034203 | 0.108214 |
| FTOP1 | 0.35636 | 1.709252 | 3.651833 | 0.468053 | 0.639746 | NA |
| LINC00323 | 2.645372 | 0.890839 | 1.237315 | 0.719977 | 0.471539 | NA |
| USF1P1 | 0.162376 | 0.424913 | 4.079006 | 0.104171 | 0.917034 | NA |
| | 2.364672 | 0.192374 | 1.260595 | 0.152606 | 0.878709 | NA |
| | 0.151697 | -1.546 | 4.075469 | -0.37934 | 0.704433 | NA |
| FAM138B | 0.151697 | -1.546 | 4.075469 | -0.37934 | 0.704433 | NA |
| | 0.81145 | -0.46945 | 2.476203 | -0.18958 | 0.849635 | NA |
| MYL6P1 | 1.053979 | -3.69847 | 2.572645 | -1.43761 | 0.150544 | NA |
| | 0.134133 | -1.48909 | 4.077584 | -0.36519 | 0.71497 | NA |
| FTH1P20 | 8.753558 | -0.29529 | 0.700183 | -0.42174 | 0.673218 | NA |
| FTLP3 | 47.47783 | -0.12304 | 0.318587 | -0.38621 | 0.699341 | 0.806443 |
| | 0.268266 | -2.15396 | 4.038351 | -0.53338 | 0.593773 | NA |
| UBE2V1P1 | 0.096202 | 0.408546 | 4.079639 | 0.100143 | 0.920231 | NA |
| | 0.878848 | -0.97535 | 2.435542 | -0.40046 | 0.688814 | NA |
| | 0.58526 | 0.001197 | 3.235889 | 3.70E-04 | 0.999705 | NA |
| HSPA9P1 | 1.827206 | 0.036111 | 1.461036 | 0.024716 | 0.980282 | NA |
| TEX41 | 935.2692 | 0.510794 | 0.226208 | 2.258077 | 0.023941 | 0.085415 |
| LENG8-AS1 | 10.42249 | -0.71959 | 0.639368 | -1.12547 | 0.260389 | NA |
| | 0.420954 | 1.923376 | 3.462803 | 0.555439 | 0.578594 | NA |
| | 2.4883 | 0.333004 | 1.333482 | 0.249725 | 0.8028 | NA |
| | 0.288605 | 1.443619 | 4.029058 | 0.358302 | 0.720117 | NA |
| | 1.762555 | -0.28036 | 1.575099 | -0.17799 | 0.858728 | NA |
| | 1.437298 | -0.55785 | 1.673581 | -0.33333 | 0.738887 | NA |
| MEIS1-AS3 | 0.489058 | -0.58569 | 3.427543 | -0.17088 | 0.864321 | NA |
| | 3.510176 | 0.321416 | 1.114129 | 0.288491 | 0.772971 | NA |
| EEF1GP3 | 0.260158 | 1.240347 | 4.047124 | 0.306476 | 0.759242 | NA |
| | 4.87421 | 0.545155 | 0.95358 | 0.571693 | 0.56753 | NA |
| | 0.296509 | -0.51709 | 3.982155 | -0.12985 | 0.896684 | NA |
| | 0.151697 | -1.546 | 4.075469 | -0.37934 | 0.704433 | NA |
| GSTM3P2 | 0.258578 | 1.233151 | 4.047809 | 0.304646 | 0.760635 | NA |
| | 0.3441 | -0.0166 | 3.769682 | -0.0044 | 0.996486 | NA |
| BSN-AS2 | 0.689366 | -1.56115 | 2.700123 | -0.57818 | 0.563145 | NA |
| | 0.096202 | 0.408546 | 4.079639 | 0.100143 | 0.920231 | NA |
| RPS14P4 | 0.096202 | 0.408546 | 4.079639 | 0.100143 | 0.920231 | NA |

| | | | | | | |
|---|---|---|---|---|---|---|
| GTF2IP2 | 0.151697 | -1.546 | 4.075469 | -0.37934 | 0.704433 | NA |
| NRBF2P2 | 0.151697 | -1.546 | 4.075469 | -0.37934 | 0.704433 | NA |
| LINC00161 | 0.392857 | -1.546 | 4.075469 | -0.37934 | 0.704433 | NA |
| RPS4XP2 | 2.095707 | -2.44478 | 1.780965 | -1.37273 | 0.169836 | NA |
| | 0.315653 | -0.57305 | 3.892639 | -0.14721 | 0.882963 | NA |
| | 0.192403 | 0.975392 | 4.060716 | 0.240202 | 0.810174 | NA |
| | 0.52699 | -2.57878 | 3.248478 | -0.79384 | 0.427288 | NA |
| | 0.163957 | 0.424913 | 4.079006 | 0.104171 | 0.917034 | NA |
| | 0.096202 | 0.408546 | 4.079639 | 0.100143 | 0.920231 | NA |
| SLC8A1-AS1 | 0.163957 | 0.424913 | 4.079006 | 0.104171 | 0.917034 | NA |
| ITGB2-AS1 | 0.096202 | 0.408546 | 4.079639 | 0.100143 | 0.920231 | NA |
| RPL41P1 | 0.936915 | 0.819377 | 2.189457 | 0.374237 | 0.708228 | NA |
| | 1.855835 | -0.4368 | 1.475879 | -0.29596 | 0.767264 | NA |
| LMCD1-AS1 | 11.86244 | -0.60902 | 0.637065 | -0.95597 | 0.339086 | NA |
| RPL12P44 | 0.260158 | 1.240347 | 4.047124 | 0.306476 | 0.759242 | NA |
| | 0.314073 | -0.57308 | 3.898934 | -0.14698 | 0.883146 | NA |
| MIR663A | 311.2959 | 0.169004 | 0.244691 | 0.690682 | 0.489765 | 0.642357 |
| | 2.381447 | -1.21496 | 1.279327 | -0.94968 | 0.342273 | NA |
| TTC39DP | 0.46735 | -1.30193 | 3.399671 | -0.38296 | 0.701751 | NA |
| | 0.151697 | -1.546 | 4.075469 | -0.37934 | 0.704433 | NA |
| | 11.29417 | -0.27977 | 0.638946 | -0.43787 | 0.661482 | NA |
| MIS18A-AS | 1.303954 | 2.434324 | 1.945662 | 1.251154 | 0.210878 | NA |
| | 9.93106 | -0.1217 | 0.688689 | -0.17672 | 0.85973 | NA |
| | 0.096202 | 0.408546 | 4.079639 | 0.100143 | 0.920231 | NA |
| | 0.651288 | 1.289307 | 2.633717 | 0.489539 | 0.62446 | NA |
| THRAP3P1 | 0.730312 | -0.54604 | 2.950954 | -0.18504 | 0.853199 | NA |
| HNRNPKP2 | 9.477839 | -0.48798 | 0.687723 | -0.70956 | 0.477979 | NA |
| | 72.50654 | 0.056783 | 0.273846 | 0.207355 | 0.835733 | 0.901643 |
| | 0.151697 | -1.546 | 4.075469 | -0.37934 | 0.704433 | NA |
| | 0.586491 | 2.39072 | 2.689057 | 0.889055 | 0.373973 | NA |
| PPIAP2 | 0.162376 | 0.424913 | 4.079006 | 0.104171 | 0.917034 | NA |
| KIF9-AS1 | 5.707539 | -0.35604 | 0.866648 | -0.41083 | 0.681199 | NA |
| RNF6P1 | 0.096202 | 0.408546 | 4.079639 | 0.100143 | 0.920231 | NA |
| | 0.151697 | -1.546 | 4.075469 | -0.37934 | 0.704433 | NA |
| | 0.151697 | -1.546 | 4.075469 | -0.37934 | 0.704433 | NA |
| | 0.151697 | -1.546 | 4.075469 | -0.37934 | 0.704433 | NA |
| | 0.096202 | 0.408546 | 4.079639 | 0.100143 | 0.920231 | NA |
| LINC00310 | 47.43115 | 0.763325 | 0.42832 | 1.782138 | 0.074727 | 0.182431 |
| | 0.628146 | -0.21495 | 3.102079 | -0.06929 | 0.944756 | NA |
| STK4-AS1 | 5.495189 | -0.19269 | 0.835002 | -0.23076 | 0.8175 | NA |
| CCDC148-A | 0.324752 | 1.53082 | 4.022025 | 0.380609 | 0.703493 | NA |
| | 0.163957 | 0.424913 | 4.079006 | 0.104171 | 0.917034 | NA |
| PABPC1P6 | 0.163957 | 0.424913 | 4.079006 | 0.104171 | 0.917034 | NA |
| | 10.30609 | -0.18157 | 0.650374 | -0.27918 | 0.78011 | NA |
| CERS6-AS1 | 0.588072 | 2.39955 | 3.098804 | 0.774347 | 0.438726 | NA |
| | 2.596586 | 0.117223 | 1.189184 | 0.098574 | 0.921476 | NA |
| ATP5A1P2 | 0.096202 | 0.408546 | 4.079639 | 0.100143 | 0.920231 | NA |

| | | | | | | |
|---|---|---|---|---|---|---|
| HNRNPA3P | 0.268266 | -2.15396 | 4.038351 | -0.53338 | 0.593773 | NA |
| MED28P3 | 0.998346 | 1.943196 | 2.103833 | 0.923646 | 0.355671 | NA |
| GSTM3P1 | 1.312825 | 0.283311 | 1.873516 | 0.151219 | 0.879803 | NA |
| | 0.876044 | -0.28385 | 2.43892 | -0.11638 | 0.907348 | NA |
| | 0.162376 | 0.424913 | 4.079006 | 0.104171 | 0.917034 | NA |
| | 0.268266 | -2.15396 | 4.038351 | -0.53338 | 0.593773 | NA |
| RPL7P12 | 0.162376 | 0.424913 | 4.079006 | 0.104171 | 0.917034 | NA |
| | 13.03138 | 0.082942 | 0.601853 | 0.13781 | 0.89039 | NA |
| | 0.298089 | -0.51711 | 3.975232 | -0.13008 | 0.896501 | NA |
| PSMA2P3 | 0.258578 | 1.233151 | 4.047809 | 0.304646 | 0.760635 | NA |
| | 0.298089 | -0.51711 | 3.975232 | -0.13008 | 0.896501 | NA |
| SNAP25-AS | 0.69625 | -3.13723 | 2.93845 | -1.06765 | 0.285679 | NA |
| | 47.49639 | -0.24934 | 0.357704 | -0.69705 | 0.485772 | 0.639244 |
| | 9.987987 | 0.167888 | 0.680069 | 0.246868 | 0.80501 | NA |
| | 1.042804 | -0.41495 | 2.072413 | -0.20022 | 0.841305 | NA |
| | 57.27763 | 0.60381 | 0.309837 | 1.948802 | 0.051319 | 0.140999 |
| | 6.850557 | -0.3093 | 0.874937 | -0.35351 | 0.723703 | NA |
| | 2.121931 | 0.362628 | 1.420635 | 0.255258 | 0.798524 | NA |
| | 0.49187 | 2.136088 | 3.895953 | 0.548284 | 0.583497 | NA |
| | 0.134133 | -1.48909 | 4.077584 | -0.36519 | 0.71497 | NA |
| | 1.84895 | 0.046134 | 1.458861 | 0.031623 | 0.974772 | NA |
| RPL3P1 | 0.831817 | -1.27234 | 2.201546 | -0.57793 | 0.563311 | NA |
| RNF144A-A | 105.7804 | -0.14657 | 0.257519 | -0.56916 | 0.569251 | 0.705805 |
| LINC00693 | 10.22333 | 0.773024 | 0.827119 | 0.934599 | 0.349995 | NA |
| | 6.71462 | -0.17143 | 0.79799 | -0.21483 | 0.8299 | NA |
| | 2.032161 | -0.37398 | 1.522723 | -0.2456 | 0.805993 | NA |
| | 0.757324 | -2.08271 | 2.587512 | -0.80491 | 0.420873 | NA |
| | 20.90864 | 0.31253 | 0.465055 | 0.672028 | 0.501566 | NA |
| | 32.12106 | -0.11303 | 0.381032 | -0.29665 | 0.766735 | NA |
| | 0.096202 | 0.408546 | 4.079639 | 0.100143 | 0.920231 | NA |
| | 11.0559 | 0.098936 | 0.597953 | 0.165457 | 0.868584 | NA |
| CYP4F29P | 0.324752 | 1.53082 | 4.022025 | 0.380609 | 0.703493 | NA |
| GUSBP11 | 145.6132 | 0.132563 | 0.210355 | 0.630186 | 0.528573 | 0.673809 |
| | 0.672013 | 1.253691 | 2.607776 | 0.480751 | 0.630693 | NA |
| | 0.392857 | -1.546 | 4.075469 | -0.37934 | 0.704433 | NA |
| | 19.5318 | -0.04042 | 0.467716 | -0.08641 | 0.931139 | NA |
| RPS26P4 | 0.151697 | -1.546 | 4.075469 | -0.37934 | 0.704433 | NA |
| | 1.811165 | -0.16997 | 1.434073 | -0.11852 | 0.905655 | NA |
| | 1.846927 | 0.935885 | 1.55051 | 0.603599 | 0.546111 | NA |
| | 0.623191 | -1.58103 | 2.770593 | -0.57065 | 0.568239 | NA |
| | 0.478233 | -0.67301 | 2.937473 | -0.22911 | 0.818782 | NA |
| | 2.766621 | 0.250459 | 1.548165 | 0.161778 | 0.871481 | NA |
| | 3.251984 | -0.44676 | 1.083476 | -0.41234 | 0.68009 | NA |
| LINC00687 | 0.327913 | 1.543154 | 4.021063 | 0.383768 | 0.701151 | NA |
| | 0.654246 | 2.541085 | 3.009764 | 0.84428 | 0.398513 | NA |
| OST4 | 2938.589 | 0.139734 | 0.1377 | 1.014772 | 0.310215 | 0.47761 |
| | 0.163957 | 0.424913 | 4.079006 | 0.104171 | 0.917034 | NA |

| | | | | | | |
|---|---|---|---|---|---|---|
| LINC01125 | 12.88229 | 1.098667 | 0.66195 | 1.659744 | 0.096966 | NA |
| | 0.096202 | 0.408546 | 4.079639 | 0.100143 | 0.920231 | NA |
| TMSB10P1 | 2.623838 | -0.52493 | 1.185717 | -0.44271 | 0.657977 | NA |
| DAP3P2 | 0.996766 | 1.94108 | 2.070839 | 0.93734 | 0.348584 | NA |
| | 0.151697 | -1.546 | 4.075469 | -0.37934 | 0.704433 | NA |
| SNRPGP9 | 0.151697 | -1.546 | 4.075469 | -0.37934 | 0.704433 | NA |
| | 8.48495 | 1.920008 | 0.823733 | 2.330861 | 0.019761 | NA |
| | 0.096202 | 0.408546 | 4.079639 | 0.100143 | 0.920231 | NA |
| | 0.134133 | -1.48909 | 4.077584 | -0.36519 | 0.71497 | NA |
| FCF1P2 | 57.7607 | -0.01499 | 0.329184 | -0.04555 | 0.963673 | 0.980209 |
| | 0.151697 | -1.546 | 4.075469 | -0.37934 | 0.704433 | NA |
| PSMD10P1 | 0.738188 | 1.39874 | 2.317793 | 0.603479 | 0.54619 | NA |
| | 0.163957 | 0.424913 | 4.079006 | 0.104171 | 0.917034 | NA |
| TTC3-AS1 | 0.364467 | -1.20166 | 3.685431 | -0.32606 | 0.744381 | NA |
| LINC00659 | 0.162376 | 0.424913 | 4.079006 | 0.104171 | 0.917034 | NA |
| | 2.189327 | 0.530062 | 1.457209 | 0.363751 | 0.716044 | NA |
| | 0.151697 | -1.546 | 4.075469 | -0.37934 | 0.704433 | NA |
| | 0.096202 | 0.408546 | 4.079639 | 0.100143 | 0.920231 | NA |
| LINC00954 | 1.216209 | -0.8802 | 2.00349 | -0.43933 | 0.66042 | NA |
| | 0.096202 | 0.408546 | 4.079639 | 0.100143 | 0.920231 | NA |
| | 0.47961 | 0.535796 | 3.39605 | 0.15777 | 0.874638 | NA |
| | 0.163957 | 0.424913 | 4.079006 | 0.104171 | 0.917034 | NA |
| BACH1-IT2 | 0.392711 | 0.292538 | 3.145221 | 0.09301 | 0.925895 | NA |
| | 2.658727 | -0.29798 | 1.26623 | -0.23533 | 0.813956 | NA |
| | 5.707846 | -0.37555 | 0.855721 | -0.43887 | 0.660757 | NA |
| | 0.151697 | -1.546 | 4.075469 | -0.37934 | 0.704433 | NA |
| | 4.32507 | 1.260866 | 1.019325 | 1.236962 | 0.216101 | NA |
| | 1.593148 | 0.697664 | 1.591488 | 0.438372 | 0.661117 | NA |
| | 0.134133 | -1.48909 | 4.077584 | -0.36519 | 0.71497 | NA |
| UBTFL6 | 0.327913 | 1.543154 | 4.021063 | 0.383768 | 0.701151 | NA |
| | 45.4099 | -0.0855 | 0.317983 | -0.2689 | 0.788009 | NA |
| MTND1P9 | 0.314073 | -0.57308 | 3.898934 | -0.14698 | 0.883146 | NA |
| SEPT7P8 | 0.163957 | 0.424913 | 4.079006 | 0.104171 | 0.917034 | NA |
| EXOSC3P1 | 4.524146 | 0.713389 | 0.92907 | 0.767853 | 0.442574 | NA |
| | 0.326333 | 1.533889 | 3.826878 | 0.40082 | 0.688553 | NA |
| ABHD17AP | 0.689366 | -1.56115 | 2.700123 | -0.57818 | 0.563145 | NA |
| AGBL5-IT1 | 1.760763 | -1.59021 | 1.646198 | -0.96599 | 0.33405 | NA |
| | 1.972806 | 2.209214 | 1.577792 | 1.400193 | 0.161455 | NA |
| CRYGEP | 0.134133 | -1.48909 | 4.077584 | -0.36519 | 0.71497 | NA |
| | 0.247898 | -0.5901 | 4.074633 | -0.14482 | 0.884851 | NA |
| | 0.163957 | 0.424913 | 4.079006 | 0.104171 | 0.917034 | NA |
| | 0.268266 | -2.15396 | 4.038351 | -0.53338 | 0.593773 | NA |
| | 75.96791 | 0.2702 | 0.354128 | 0.763001 | 0.445463 | 0.605337 |
| HNRNPA1P | 0.134133 | -1.48909 | 4.077584 | -0.36519 | 0.71497 | NA |
| KRT18P4 | 1.580685 | 0.588029 | 1.561672 | 0.376538 | 0.706517 | NA |
| FEM1AP1 | 0.134133 | -1.48909 | 4.077584 | -0.36519 | 0.71497 | NA |
| | 1.025598 | 0.201544 | 2.083681 | 0.096725 | 0.922945 | NA |

| | | | | | | |
|---|---|---|---|---|---|---|
| | 1.327411 | -0.52458 | 1.752811 | -0.29928 | 0.764727 | NA |
| | 0.640755 | -1.64826 | 2.754778 | -0.59833 | 0.549621 | NA |
| | 0.151697 | -1.546 | 4.075469 | -0.37934 | 0.704433 | NA |
| LRRC3-AS1 | 0.162376 | 0.424913 | 4.079006 | 0.104171 | 0.917034 | NA |
| PPP1R2P5 | 0.162376 | 0.424913 | 4.079006 | 0.104171 | 0.917034 | NA |
| | 0.774539 | 0.39797 | 2.296349 | 0.173305 | 0.862411 | NA |
| SPINK8 | 0.163957 | 0.424913 | 4.079006 | 0.104171 | 0.917034 | NA |
| | 0.326333 | 1.533889 | 3.826878 | 0.40082 | 0.688553 | NA |
| | 0.096202 | 0.408546 | 4.079639 | 0.100143 | 0.920231 | NA |
| | 6.504503 | -0.13836 | 0.85256 | -0.16229 | 0.871075 | NA |
| | 0.162376 | 0.424913 | 4.079006 | 0.104171 | 0.917034 | NA |
| | 0.296509 | -0.51709 | 3.982155 | -0.12985 | 0.896684 | NA |
| | 0.163957 | 0.424913 | 4.079006 | 0.104171 | 0.917034 | NA |
| | 1.787637 | 0.238177 | 1.584929 | 0.150276 | 0.880547 | NA |
| | 0.314073 | -0.57308 | 3.898934 | -0.14698 | 0.883146 | NA |
| | 8.109571 | -0.48913 | 0.71427 | -0.6848 | 0.493469 | NA |
| ACVR2B-AS | 23.84064 | -1.25047 | 0.49544 | -2.52395 | 0.011604 | NA |
| SUCLA2P3 | 0.151697 | -1.546 | 4.075469 | -0.37934 | 0.704433 | NA |
| PRDX3P1 | 1.562036 | -1.53353 | 1.708649 | -0.89751 | 0.369445 | NA |
| | 0.678686 | -3.08037 | 2.621268 | -1.17514 | 0.239937 | NA |
| | 0.096202 | 0.408546 | 4.079639 | 0.100143 | 0.920231 | NA |
| PKP4P1 | 0.134133 | -1.48909 | 4.077584 | -0.36519 | 0.71497 | NA |
| | 87.14688 | 0.524694 | 0.260797 | 2.011889 | 0.044232 | 0.128238 |
| SOS1-IT1 | 5.150291 | 0.429044 | 1.032528 | 0.415527 | 0.677756 | NA |
| | 3.978408 | -1.16153 | 1.01341 | -1.14616 | 0.251731 | NA |
| | 0.298089 | -0.51711 | 3.975232 | -0.13008 | 0.896501 | NA |
| DYNLT3P2 | 0.258578 | 1.233151 | 4.047809 | 0.304646 | 0.760635 | NA |
| MRPS18AP | 0.719043 | 1.433043 | 2.335048 | 0.61371 | 0.539407 | NA |
| | 0.163957 | 0.424913 | 4.079006 | 0.104171 | 0.917034 | NA |
| RPS15P5 | 9.887331 | -0.61215 | 0.673726 | -0.9086 | 0.363559 | NA |
| DDX50P1 | 2.642889 | -0.80496 | 1.30695 | -0.61591 | 0.537953 | NA |
| | 0.096202 | 0.408546 | 4.079639 | 0.100143 | 0.920231 | NA |
| | 1.484097 | 1.734562 | 1.70674 | 1.016301 | 0.309486 | NA |
| | 0.096202 | 0.408546 | 4.079639 | 0.100143 | 0.920231 | NA |
| | 0.162376 | 0.424913 | 4.079006 | 0.104171 | 0.917034 | NA |
| STEAP3-AS | 1.784263 | -0.32213 | 1.625942 | -0.19812 | 0.84295 | NA |
| OGFR-AS1 | 6.277041 | 0.301388 | 0.789029 | 0.381973 | 0.702481 | NA |
| IMMTP1 | 6.680847 | -0.83044 | 0.800471 | -1.03745 | 0.299528 | NA |
| | 31.9107 | -0.59986 | 0.499315 | -1.20137 | 0.229608 | NA |
| | 0.67854 | -0.89371 | 2.366391 | -0.37767 | 0.705677 | NA |
| | 0.096202 | 0.408546 | 4.079639 | 0.100143 | 0.920231 | NA |
| | 0.574231 | 0.93227 | 2.49998 | 0.372911 | 0.709215 | NA |
| | 3.643155 | 1.563256 | 1.224453 | 1.276698 | 0.201709 | NA |
| | 0.151697 | -1.546 | 4.075469 | -0.37934 | 0.704433 | NA |
| | 0.163957 | 0.424913 | 4.079006 | 0.104171 | 0.917034 | NA |
| ALMS1-IT1 | 4.763536 | -0.26761 | 0.914365 | -0.29267 | 0.769775 | NA |
| ANKRD36B | 0.260158 | 1.240347 | 4.047124 | 0.306476 | 0.759242 | NA |

| | | | | | | |
|---|---|---|---|---|---|---|
| | 0.163957 | 0.424913 | 4.079006 | 0.104171 | 0.917034 | NA |
| UBBP1 | 0.884581 | 1.788183 | 2.221027 | 0.805115 | 0.420753 | NA |
| BIRC6-AS1 | 0.096202 | 0.408546 | 4.079639 | 0.100143 | 0.920231 | NA |
| | 0.192403 | 0.975392 | 4.060716 | 0.240202 | 0.810174 | NA |
| HSPD1P6 | 0.610582 | -0.18034 | 2.753139 | -0.0655 | 0.947774 | NA |
| RPL4P6 | 4.622694 | -0.78594 | 0.981179 | -0.80102 | 0.423122 | NA |
| | 11.91069 | -0.41809 | 0.616044 | -0.67867 | 0.49735 | NA |
| PRRT3-AS1 | 29.30106 | 8.14E-04 | 0.417927 | 0.001948 | 0.998445 | NA |
| | 6.361296 | 1.255933 | 0.883915 | 1.420875 | 0.155353 | NA |
| | 0.151697 | -1.546 | 4.075469 | -0.37934 | 0.704433 | NA |
| | 1.551852 | 1.796216 | 1.661946 | 1.080791 | 0.27979 | NA |
| | 0.096202 | 0.408546 | 4.079639 | 0.100143 | 0.920231 | NA |
| | 0.162376 | 0.424913 | 4.079006 | 0.104171 | 0.917034 | NA |
| | 0.096202 | 0.408546 | 4.079639 | 0.100143 | 0.920231 | NA |
| | 0.151697 | -1.546 | 4.075469 | -0.37934 | 0.704433 | NA |
| | 3.327044 | -0.39903 | 1.120445 | -0.35614 | 0.721737 | NA |
| PHBP4 | 0.392857 | -1.546 | 4.075469 | -0.37934 | 0.704433 | NA |
| | 0.163957 | 0.424913 | 4.079006 | 0.104171 | 0.917034 | NA |
| | 3.444184 | -0.03491 | 1.072795 | -0.03255 | 0.974037 | NA |
| FANCD2P2 | 4.491468 | -1.09294 | 1.056511 | -1.03449 | 0.300909 | NA |
| DSCR9 | 2.274552 | 0.43835 | 1.406043 | 0.311761 | 0.755222 | NA |
| | 1.022583 | -1.06903 | 2.182302 | -0.48986 | 0.62423 | NA |
| | 0.096202 | 0.408546 | 4.079639 | 0.100143 | 0.920231 | NA |
| TCEA1P2 | 3.258847 | 0.008462 | 1.165955 | 0.007258 | 0.994209 | NA |
| | 3.302568 | 0.735889 | 1.224447 | 0.600997 | 0.547842 | NA |
| | 0.314073 | -0.57308 | 3.898934 | -0.14698 | 0.883146 | NA |
| | 0.6908 | 0.277604 | 2.346083 | 0.118327 | 0.905809 | NA |
| | 7.449344 | 0.098624 | 0.762663 | 0.129315 | 0.897108 | NA |
| PPP5D1 | 5.739033 | -0.51187 | 0.848038 | -0.6036 | 0.546111 | NA |
| THAP7-AS1 | 60.12073 | 0.057232 | 0.293722 | 0.194852 | 0.845509 | 0.908091 |
| COL6A4P1 | 0.162376 | 0.424913 | 4.079006 | 0.104171 | 0.917034 | NA |
| LIMD1-AS1 | 0.288605 | 1.443619 | 4.029058 | 0.358302 | 0.720117 | NA |
| | 0.151697 | -1.546 | 4.075469 | -0.37934 | 0.704433 | NA |
| FAM133DP | 0.488912 | 0.771583 | 2.897678 | 0.266276 | 0.790026 | NA |
| SF3A3P1 | 0.659753 | -0.00581 | 2.629089 | -0.00221 | 0.998237 | NA |
| | 1.849097 | -0.59873 | 1.451994 | -0.41235 | 0.680083 | NA |
| | 1.195493 | 1.307783 | 1.879298 | 0.695889 | 0.486498 | NA |
| HMGB1P27 | 1.216218 | 1.276946 | 1.967625 | 0.648978 | 0.516352 | NA |
| HM13-AS1 | 2.309977 | 0.223757 | 1.504738 | 0.148701 | 0.881789 | NA |
| | 0.163957 | 0.424913 | 4.079006 | 0.104171 | 0.917034 | NA |
| KLF2P2 | 0.162376 | 0.424913 | 4.079006 | 0.104171 | 0.917034 | NA |
| DDTP1 | 0.596179 | -0.09275 | 3.146759 | -0.02948 | 0.976485 | NA |
| | 0.134133 | -1.48909 | 4.077584 | -0.36519 | 0.71497 | NA |
| | 0.096202 | 0.408546 | 4.079639 | 0.100143 | 0.920231 | NA |
| FBXW4P1 | 4.536727 | -0.14121 | 0.993339 | -0.14215 | 0.88696 | NA |
| | 0.612162 | -0.19706 | 2.47693 | -0.07956 | 0.936588 | NA |
| | 0.739585 | 2.778432 | 2.852922 | 0.97389 | 0.330111 | NA |

| | | | | | | |
|---|---|---|---|---|---|---|
| | 0.979202 | 1.972754 | 2.08204 | 0.94751 | 0.343379 | NA |
| | 17.73676 | 0.244374 | 0.571507 | 0.427597 | 0.668945 | NA |
| MEIS1-AS2 | 0.151697 | -1.546 | 4.075469 | -0.37934 | 0.704433 | NA |
| SDAD1P2 | 0.392711 | 0.292538 | 3.145221 | 0.09301 | 0.925895 | NA |
| | 0.192403 | 0.975392 | 4.060716 | 0.240202 | 0.810174 | NA |
| RHOQP3 | 7.15226 | -0.22837 | 0.758403 | -0.30111 | 0.763327 | NA |
| NFU1P1 | 0.192403 | 0.975392 | 4.060716 | 0.240202 | 0.810174 | NA |
| SNAP23P | 0.71919 | 0.510978 | 2.728734 | 0.187258 | 0.851458 | NA |
| | 0.354779 | 1.703954 | 3.657232 | 0.465914 | 0.641277 | NA |
| VN1R108P | 2.095471 | -0.34436 | 1.392102 | -0.24737 | 0.804624 | NA |
| | 0.134133 | -1.48909 | 4.077584 | -0.36519 | 0.71497 | NA |
| | 0.556813 | -0.57007 | 3.294615 | -0.17303 | 0.862627 | NA |
| | 0.151697 | -1.546 | 4.075469 | -0.37934 | 0.704433 | NA |
| | 1.529059 | -0.23872 | 1.67839 | -0.14223 | 0.886895 | NA |
| | 0.134133 | -1.48909 | 4.077584 | -0.36519 | 0.71497 | NA |
| | 0.4986 | -1.64638 | 3.317584 | -0.49626 | 0.619712 | NA |
| | 17.40025 | 0.938215 | 0.647881 | 1.44813 | 0.147581 | NA |
| RPL31P2 | 0.588072 | 2.39955 | 3.098804 | 0.774347 | 0.438726 | NA |
| | 0.163957 | 0.424913 | 4.079006 | 0.104171 | 0.917034 | NA |
| | 0.162376 | 0.424913 | 4.079006 | 0.104171 | 0.917034 | NA |
| FBXW11P1 | 1.061709 | 0.25203 | 2.095453 | 0.120275 | 0.904266 | NA |
| | 0.918331 | 0.764804 | 2.118268 | 0.361052 | 0.718061 | NA |
| | 8.883173 | 0.269725 | 0.719775 | 0.374735 | 0.707857 | NA |
| | 0.096202 | 0.408546 | 4.079639 | 0.100143 | 0.920231 | NA |
| | 0.954682 | 0.055803 | 2.069758 | 0.026961 | 0.978491 | NA |
| | 0.574231 | 0.93227 | 2.49998 | 0.372911 | 0.709215 | NA |
| SNX18P13 | 0.230334 | -0.53318 | 4.076749 | -0.13079 | 0.895945 | NA |
| | 8.903579 | -0.3201 | 0.763521 | -0.41924 | 0.675039 | NA |
| DSTNP1 | 12.12045 | 0.847303 | 0.709864 | 1.193613 | 0.232629 | NA |
| FAM201B | 1.139363 | 0.418105 | 2.214864 | 0.188772 | 0.850271 | NA |
| RPL12P15 | 0.888304 | 0.641829 | 2.171212 | 0.295609 | 0.767529 | NA |
| | 2.586163 | 2.662875 | 1.487078 | 1.790676 | 0.073345 | NA |
| | 2.002833 | 2.249182 | 1.55051 | 1.450608 | 0.146889 | NA |
| | 13.73635 | -0.34064 | 0.543857 | -0.62634 | 0.531091 | NA |
| | 0.230334 | -0.53318 | 4.076749 | -0.13079 | 0.895945 | NA |
| | 0.774742 | -0.40577 | 2.223483 | -0.18249 | 0.855195 | NA |
| TSSK1A | 0.628146 | -0.21495 | 3.102079 | -0.06929 | 0.944756 | NA |
| | 2.348513 | 1.894099 | 1.407134 | 1.346068 | 0.17828 | NA |
| | 2.688201 | 0.798118 | 1.352306 | 0.590191 | 0.555063 | NA |
| HAR1B | 0.314073 | -0.57308 | 3.898934 | -0.14698 | 0.883146 | NA |
| TCF7L1-IT1 | 0.096202 | 0.408546 | 4.079639 | 0.100143 | 0.920231 | NA |
| ARHGAP42 | 1.091976 | -0.2898 | 1.942755 | -0.14917 | 0.881418 | NA |
| PTP4A1P1 | 0.162376 | 0.424913 | 4.079006 | 0.104171 | 0.917034 | NA |
| | 0.35636 | 1.709252 | 3.651833 | 0.468053 | 0.639746 | NA |
| LINC00852 | 2.186312 | 0.02037 | 1.320744 | 0.015423 | 0.987694 | NA |
| | 0.134133 | -1.48909 | 4.077584 | -0.36519 | 0.71497 | NA |
| | 0.508056 | 0.731032 | 2.861912 | 0.255435 | 0.798387 | NA |

| | | | | | | |
|---|---|---|---|---|---|---|
| KRT8P10 | 0.162376 | 0.424913 | 4.079006 | 0.104171 | 0.917034 | NA |
| | 0.735027 | 1.3952 | 2.594798 | 0.537691 | 0.59079 | NA |
| ITPR1-AS1 | 3.906273 | 1.677951 | 1.102371 | 1.522129 | 0.127977 | NA |
| | 118.2131 | 0.141354 | 0.269963 | 0.523604 | 0.600554 | 0.729987 |
| APCDD1L-A | 0.162376 | 0.424913 | 4.079006 | 0.104171 | 0.917034 | NA |
| | 0.296509 | -0.51709 | 3.982155 | -0.12985 | 0.896684 | NA |
| EZH2P1 | 0.746498 | -1.433 | 2.535747 | -0.56512 | 0.571994 | NA |
| RPL36P2 | 0.162376 | 0.424913 | 4.079006 | 0.104171 | 0.917034 | NA |
| RPS3P2 | 0.096202 | 0.408546 | 4.079639 | 0.100143 | 0.920231 | NA |
| | 289.0244 | 0.168332 | 0.214734 | 0.783909 | 0.433093 | 0.59425 |
| | 9.03475 | 0.154378 | 0.793313 | 0.194599 | 0.845707 | NA |
| | 0.555233 | -0.57009 | 3.296947 | -0.17291 | 0.862719 | NA |
| | 0.230334 | -0.53318 | 4.076749 | -0.13079 | 0.895945 | NA |
| | 0.268266 | -2.15396 | 4.038351 | -0.53338 | 0.593773 | NA |
| | 0.314073 | -0.57308 | 3.898934 | -0.14698 | 0.883146 | NA |
| | 3.196314 | 0.906285 | 1.181678 | 0.766947 | 0.443113 | NA |
| | 0.230334 | -0.53318 | 4.076749 | -0.13079 | 0.895945 | NA |
| | 0.096202 | 0.408546 | 4.079639 | 0.100143 | 0.920231 | NA |
| | 0.424115 | 1.932531 | 3.45462 | 0.559405 | 0.575885 | NA |
| TIMM8AP1 | 9.005095 | -0.13812 | 0.662554 | -0.20846 | 0.834868 | NA |
| | 0.70851 | -1.62753 | 2.686091 | -0.60591 | 0.544574 | NA |
| | 1.813308 | -0.73304 | 1.579784 | -0.46401 | 0.642639 | NA |
| | 0.134133 | -1.48909 | 4.077584 | -0.36519 | 0.71497 | NA |
| RPL3P5 | 0.134133 | -1.48909 | 4.077584 | -0.36519 | 0.71497 | NA |
| PTCHD3P2 | 0.314073 | -0.57308 | 3.898934 | -0.14698 | 0.883146 | NA |
| IGBP1P3 | 0.096202 | 0.408546 | 4.079639 | 0.100143 | 0.920231 | NA |
| FAHD2CP | 33.56403 | -0.6317 | 0.42282 | -1.49402 | 0.135171 | NA |
| | 6.187946 | 0.677285 | 0.972292 | 0.696586 | 0.486062 | NA |
| AGBL5-AS1 | 0.258578 | 1.233151 | 4.047809 | 0.304646 | 0.760635 | NA |
| | 0.192403 | 0.975392 | 4.060716 | 0.240202 | 0.810174 | NA |
| DIRC3 | 3.154651 | 0.681991 | 1.175834 | 0.580006 | 0.56191 | NA |
| LINC01090 | 0.096202 | 0.408546 | 4.079639 | 0.100143 | 0.920231 | NA |
| LINC00899 | 4.059617 | -0.33783 | 0.991674 | -0.34067 | 0.733355 | NA |
| COX6CP2 | 0.419962 | -2.64111 | 3.429364 | -0.77014 | 0.441214 | NA |
| | 1.157561 | 2.190278 | 2.039086 | 1.074147 | 0.282757 | NA |
| GAPDHP59 | 0.591998 | -0.2499 | 3.117006 | -0.08017 | 0.936101 | NA |
| | 2.768622 | 0.601503 | 1.17228 | 0.513105 | 0.607878 | NA |
| | 1.962622 | -0.06096 | 1.552879 | -0.03925 | 0.968688 | NA |
| | 1.876605 | 2.119198 | 1.61187 | 1.314744 | 0.188596 | NA |
| | 0.096202 | 0.408546 | 4.079639 | 0.100143 | 0.920231 | NA |
| | 0.907091 | -0.03396 | 2.426081 | -0.014 | 0.988832 | NA |
| | 1.361161 | -1.04569 | 2.138753 | -0.48893 | 0.624893 | NA |
| | 0.162376 | 0.424913 | 4.079006 | 0.104171 | 0.917034 | NA |
| | 0.096202 | 0.408546 | 4.079639 | 0.100143 | 0.920231 | NA |
| NDUFB1P1 | 1.291491 | 2.354367 | 2.238911 | 1.051568 | 0.292998 | NA |
| | 7.333295 | -0.72463 | 0.770579 | -0.94038 | 0.347024 | NA |
| | 3.91202 | 0.832749 | 1.067824 | 0.779856 | 0.435476 | NA |

| | | | | | | |
|---|---|---|---|---|---|---|
| | 0.422534 | 1.924583 | 3.001758 | 0.641152 | 0.521424 | NA |
| GAPDHP37 | 0.642189 | 0.046766 | 2.378271 | 0.019664 | 0.984311 | NA |
| IDH1-AS1 | 15.86006 | -0.49752 | 0.561945 | -0.88536 | 0.375962 | NA |
| HS1BP3-IT1 | 1.284176 | 0.715001 | 1.74187 | 0.410479 | 0.681455 | NA |
| | 40.92999 | 0.094952 | 0.367051 | 0.258688 | 0.795876 | NA |
| | 5.453477 | 0.047336 | 0.851564 | 0.055588 | 0.95567 | NA |
| | 1.091618 | -0.82092 | 2.284043 | -0.35941 | 0.719285 | NA |
| | 2.040212 | -1.79536 | 1.632227 | -1.09995 | 0.271356 | NA |
| | 3.448889 | -0.0326 | 1.060385 | -0.03074 | 0.975473 | NA |
| | 0.888304 | 0.641829 | 2.171212 | 0.295609 | 0.767529 | NA |
| | 1.295261 | 0.315793 | 1.914152 | 0.164978 | 0.868961 | NA |
| | 217.3076 | -0.32311 | 0.18551 | -1.74174 | 0.081553 | 0.193828 |
| TMA7 | 466.5285 | 0.333828 | 0.173528 | 1.923774 | 0.054383 | 0.146816 |
| BACH1-AS1 | 0.324752 | 1.53082 | 4.022025 | 0.380609 | 0.703493 | NA |
| | 0.151697 | -1.546 | 4.075469 | -0.37934 | 0.704433 | NA |
| IMPDH1P1 | 0.937264 | -1.23834 | 2.063944 | -0.59999 | 0.548514 | NA |
| DUXAP7 | 0.230334 | -0.53318 | 4.076749 | -0.13079 | 0.895945 | NA |
| | 0.3441 | -0.0166 | 3.769682 | -0.0044 | 0.996486 | NA |
| | 1.636529 | -1.07836 | 1.58333 | -0.68107 | 0.495829 | NA |
| | 0.638825 | 1.117948 | 3.106359 | 0.35989 | 0.718929 | NA |
| BTF3L4P1 | 0.163957 | 0.424913 | 4.079006 | 0.104171 | 0.917034 | NA |
| | 8.77332 | -0.24681 | 0.701907 | -0.35162 | 0.72512 | NA |
| | 0.327913 | 1.543154 | 4.021063 | 0.383768 | 0.701151 | NA |
| RPS27AP2 | 1.175714 | -0.20823 | 1.812447 | -0.11489 | 0.908534 | NA |
| | 6.692833 | 0.31336 | 0.839039 | 0.373474 | 0.708795 | NA |
| SEMA3B-AS | 0.315653 | -0.57305 | 3.892639 | -0.14721 | 0.882963 | NA |
| VIPR1-AS1 | 0.151697 | -1.546 | 4.075469 | -0.37934 | 0.704433 | NA |
| | 0.151697 | -1.546 | 4.075469 | -0.37934 | 0.704433 | NA |
| | 1.033466 | -0.7302 | 1.946441 | -0.37515 | 0.707552 | NA |
| LINC00493 | 725.6341 | 0.292324 | 0.174337 | 1.676776 | 0.093586 | 0.213567 |
| | 62.86003 | 0.447927 | 0.343226 | 1.305048 | 0.191876 | 0.346488 |
| RPL31P15 | 0.163957 | 0.424913 | 4.079006 | 0.104171 | 0.917034 | NA |
| RPL18AP7 | 3.244522 | -0.53296 | 1.192784 | -0.44682 | 0.655005 | NA |
| | 64.96735 | 0.081611 | 0.290534 | 0.280899 | 0.778788 | 0.864226 |
| | 0.296509 | -0.51709 | 3.982155 | -0.12985 | 0.896684 | NA |
| LARS2-AS1 | 0.162376 | 0.424913 | 4.079006 | 0.104171 | 0.917034 | NA |
| NCKAP5-IT1 | 0.134133 | -1.48909 | 4.077584 | -0.36519 | 0.71497 | NA |
| | 4.514722 | -0.08652 | 1.042883 | -0.08297 | 0.933878 | NA |
| | 5.601593 | -0.22332 | 0.869725 | -0.25677 | 0.797355 | NA |
| RPL12P11 | 0.268266 | -2.15396 | 4.038351 | -0.53338 | 0.593773 | NA |
| | 16.67423 | -0.53355 | 0.499552 | -1.06807 | 0.28549 | NA |
| | 0.526843 | -0.38719 | 2.855892 | -0.13558 | 0.892156 | NA |
| | 1.434698 | -0.57873 | 2.005846 | -0.28852 | 0.772949 | NA |
| | 5.685357 | -0.4785 | 0.892355 | -0.53622 | 0.591807 | NA |
| | 0.163957 | 0.424913 | 4.079006 | 0.104171 | 0.917034 | NA |
| | 0.653015 | 0.2552 | 2.787103 | 0.091565 | 0.927044 | NA |
| | 0.096202 | 0.408546 | 4.079639 | 0.100143 | 0.920231 | NA |

| | | | | | | |
|---|---|---|---|---|---|---|
| | 0.096202 | 0.408546 | 4.079639 | 0.100143 | 0.920231 | NA |
| | 0.28583 | -2.21358 | 3.97376 | -0.55705 | 0.577494 | NA |
| | 0.47803 | 0.520083 | 2.97881 | 0.174594 | 0.861399 | NA |
| | 5.629925 | -0.29338 | 0.932903 | -0.31448 | 0.75316 | NA |
| | 0.096202 | 0.408546 | 4.079639 | 0.100143 | 0.920231 | NA |
| RPL12P9 | 0.134133 | -1.48909 | 4.077584 | -0.36519 | 0.71497 | NA |
| | 0.163957 | 0.424913 | 4.079006 | 0.104171 | 0.917034 | NA |
| | 0.476449 | 0.524262 | 3.402674 | 0.154073 | 0.877552 | NA |
| | 1.206668 | -0.63372 | 2.320936 | -0.27305 | 0.784818 | NA |
| | 1.336859 | -1.35856 | 1.768035 | -0.7684 | 0.442247 | NA |
| TRIM51JP | 0.914607 | 1.879338 | 2.377229 | 0.790558 | 0.429202 | NA |
| | 0.422534 | 1.924583 | 3.001758 | 0.641152 | 0.521424 | NA |
| RHOQP2 | 12.52566 | 0.097749 | 0.609141 | 0.16047 | 0.872511 | NA |
| | 0.162376 | 0.424913 | 4.079006 | 0.104171 | 0.917034 | NA |
| MRPL51P2 | 0.151697 | -1.546 | 4.075469 | -0.37934 | 0.704433 | NA |
| | 0.354779 | 1.703954 | 3.657232 | 0.465914 | 0.641277 | NA |
| | 517.4764 | -1.30641 | 0.297704 | -4.38828 | 1.14E-05 | 6.61E-04 |
| | 0.151697 | -1.546 | 4.075469 | -0.37934 | 0.704433 | NA |
| PET117 | 13.84906 | 0.388333 | 0.589686 | 0.658542 | 0.51019 | NA |
| | 0.134133 | -1.48909 | 4.077584 | -0.36519 | 0.71497 | NA |
| SEC1P | 0.422534 | 1.924583 | 3.001758 | 0.641152 | 0.521424 | NA |
| CYCSP10 | 1.891889 | -0.87458 | 1.795209 | -0.48717 | 0.626136 | NA |
| | 1.522524 | -0.95549 | 1.738521 | -0.5496 | 0.582592 | NA |
| | 3.816111 | 0.206557 | 1.080563 | 0.191157 | 0.848403 | NA |
| LINC00342 | 10.00121 | 1.331456 | 0.982283 | 1.355471 | 0.175267 | NA |
| HSPA8P18 | 0.327913 | 1.543154 | 4.021063 | 0.383768 | 0.701151 | NA |
| | 0.411855 | 0.250456 | 3.097618 | 0.080854 | 0.935558 | NA |
| POLHP1 | 0.258578 | 1.233151 | 4.047809 | 0.304646 | 0.760635 | NA |
| CYP1B1-AS | 13.30331 | -0.30234 | 0.569979 | -0.53045 | 0.595801 | NA |
| | 0.288605 | 1.443619 | 4.029058 | 0.358302 | 0.720117 | NA |
| RPEP4 | 0.260158 | 1.240347 | 4.047124 | 0.306476 | 0.759242 | NA |
| | 2.300317 | -0.13427 | 1.404442 | -0.0956 | 0.923835 | NA |
| | 0.162376 | 0.424913 | 4.079006 | 0.104171 | 0.917034 | NA |
| ERVH48-1 | 0.458885 | 0.577525 | 3.449278 | 0.167433 | 0.867029 | NA |
| | 0.326333 | 1.533889 | 3.826878 | 0.40082 | 0.688553 | NA |
| | 0.163957 | 0.424913 | 4.079006 | 0.104171 | 0.917034 | NA |
| | 0.096202 | 0.408546 | 4.079639 | 0.100143 | 0.920231 | NA |
| | 3.218494 | -0.25215 | 1.171048 | -0.21532 | 0.829521 | NA |
| | 0.247898 | -0.5901 | 4.074633 | -0.14482 | 0.884851 | NA |
| BPIFB5P | 0.326333 | 1.533889 | 3.826878 | 0.40082 | 0.688553 | NA |
| HMGA1P8 | 2.007146 | 0.22136 | 1.429046 | 0.1549 | 0.8769 | NA |
| RPS24P6 | 0.163957 | 0.424913 | 4.079006 | 0.104171 | 0.917034 | NA |
| | 0.449786 | -1.25325 | 3.00474 | -0.41709 | 0.676613 | NA |
| | 34.13883 | 0.37753 | 0.399981 | 0.94387 | 0.345236 | NA |
| | 3.476037 | 1.481057 | 1.146155 | 1.292196 | 0.196289 | NA |
| | 0.424115 | 1.932531 | 3.45462 | 0.559405 | 0.575885 | NA |
| HMGB1P31 | 0.969717 | 3.152914 | 2.250337 | 1.401085 | 0.161189 | NA |

| | | | | | | |
|---|---|---|---|---|---|---|
| | 0.192403 | 0.975392 | 4.060716 | 0.240202 | 0.810174 | NA |
| GPX1 | 14091.8 | -0.16434 | 0.138303 | -1.18828 | 0.234723 | 0.397463 |
| | 0.258578 | 1.233151 | 4.047809 | 0.304646 | 0.760635 | NA |
| EIF4A1P1 | 0.096202 | 0.408546 | 4.079639 | 0.100143 | 0.920231 | NA |
| EEF1A1P34 | 0.096202 | 0.408546 | 4.079639 | 0.100143 | 0.920231 | NA |
| ERP29P1 | 1.117204 | -0.62358 | 2.074777 | -0.30055 | 0.763757 | NA |
| LINC00028 | 1.11032 | 0.442168 | 2.116951 | 0.20887 | 0.83455 | NA |
| | 0.163957 | 0.424913 | 4.079006 | 0.104171 | 0.917034 | NA |
| | 0.610582 | -0.18034 | 2.753139 | -0.0655 | 0.947774 | NA |
| | 0.518939 | 0.925178 | 3.271397 | 0.282808 | 0.777324 | NA |
| | 550.0178 | 0.284332 | 0.21144 | 1.344742 | 0.178709 | 0.33126 |
| | 0.996766 | 1.94108 | 2.070839 | 0.93734 | 0.348584 | NA |
| EIF3FP3 | 19.66005 | -0.13418 | 0.455481 | -0.29458 | 0.768314 | NA |
| | 0.151697 | -1.546 | 4.075469 | -0.37934 | 0.704433 | NA |
| | 0.163957 | 0.424913 | 4.079006 | 0.104171 | 0.917034 | NA |
| | 0.558248 | 0.982076 | 2.804738 | 0.350149 | 0.726227 | NA |
| KRT18P62 | 0.163957 | 0.424913 | 4.079006 | 0.104171 | 0.917034 | NA |
| TMEM238 | 23.63094 | 0.546422 | 0.524391 | 1.042013 | 0.297406 | NA |
| | 0.151697 | -1.546 | 4.075469 | -0.37934 | 0.704433 | NA |
| ZNF197-AS | 1.678227 | 1.465617 | 1.653772 | 0.886227 | 0.375495 | NA |
| | 0.096202 | 0.408546 | 4.079639 | 0.100143 | 0.920231 | NA |
| EIF3KP2 | 0.163957 | 0.424913 | 4.079006 | 0.104171 | 0.917034 | NA |
| | 0.602678 | 1.083611 | 2.70938 | 0.399948 | 0.689195 | NA |
| | 0.35636 | 1.709252 | 3.651833 | 0.468053 | 0.639746 | NA |
| BTF3P5 | 1.782573 | 0.375106 | 1.545899 | 0.242646 | 0.80828 | NA |
| | 0.258578 | 1.233151 | 4.047809 | 0.304646 | 0.760635 | NA |
| TWIST2 | 5.927287 | 0.193859 | 1.090318 | 0.177801 | 0.85888 | NA |
| | 3.548251 | 0.038109 | 1.203409 | 0.031668 | 0.974737 | NA |
| | 0.151697 | -1.546 | 4.075469 | -0.37934 | 0.704433 | NA |
| ANAPC1P1 | 0.096202 | 0.408546 | 4.079639 | 0.100143 | 0.920231 | NA |
| FDPSP6 | 0.162376 | 0.424913 | 4.079006 | 0.104171 | 0.917034 | NA |
| | 1.268009 | 1.569749 | 1.956286 | 0.802413 | 0.422314 | NA |
| EIF4EBP2P1 | 0.162376 | 0.424913 | 4.079006 | 0.104171 | 0.917034 | NA |
| LINC01122 | 0.815391 | 0.931626 | 2.360431 | 0.394685 | 0.693076 | NA |
| | 0.096202 | 0.408546 | 4.079639 | 0.100143 | 0.920231 | NA |
| H2AFZP6 | 0.162376 | 0.424913 | 4.079006 | 0.104171 | 0.917034 | NA |
| | 0.420954 | 1.923376 | 3.462803 | 0.555439 | 0.578594 | NA |
| RPL36AP13 | 0.411855 | 0.250456 | 3.097618 | 0.080854 | 0.935558 | NA |
| | 19.33636 | 0.579799 | 0.493095 | 1.175836 | 0.23966 | NA |
| | 75.30911 | 0.119225 | 0.356678 | 0.334265 | 0.738179 | 0.834314 |
| | 0.151697 | -1.546 | 4.075469 | -0.37934 | 0.704433 | NA |
| | 0.926585 | -2.43791 | 2.203722 | -1.10627 | 0.268609 | NA |
| | 0.303394 | -2.27006 | 4.032451 | -0.56295 | 0.57347 | NA |
| DPH3P1 | 0.303394 | -2.27006 | 4.032451 | -0.56295 | 0.57347 | NA |
| | 0.151697 | -1.546 | 4.075469 | -0.37934 | 0.704433 | NA |
| | 0.651435 | 0.249629 | 2.788679 | 0.089515 | 0.928672 | NA |
| | 18.68268 | 0.088396 | 0.465948 | 0.189713 | 0.849534 | NA |

| | | | | | | |
|---|---|---|---|---|---|---|
| | 0.850576 | 0.626886 | 2.42682 | 0.258316 | 0.796163 | NA |
| | 0.938698 | 0.081169 | 1.915817 | 0.042368 | 0.966205 | NA |
| | 5.185868 | 0.109678 | 0.884501 | 0.124 | 0.901316 | NA |
| | 4.915751 | 0.038728 | 0.933992 | 0.041465 | 0.966925 | NA |
| | 1.392637 | 1.61 | 1.744172 | 0.923074 | 0.355969 | NA |
| | 0.670433 | 1.255281 | 2.356401 | 0.532711 | 0.594234 | NA |
| BTF3P6 | 0.192403 | 0.975392 | 4.060716 | 0.240202 | 0.810174 | NA |
| RPS20P10 | 1.282595 | 0.709442 | 1.711365 | 0.414547 | 0.678473 | NA |
| | 1.982423 | -0.27454 | 1.675555 | -0.16385 | 0.869848 | NA |
| RPL5P34 | 1.359799 | 0.16786 | 1.649101 | 0.101789 | 0.918924 | NA |
| | 0.888653 | -1.5355 | 2.47141 | -0.62131 | 0.534398 | NA |
| | 8.94174 | 0.343128 | 0.694063 | 0.494376 | 0.621041 | NA |
| | 0.326536 | 0.037056 | 3.842554 | 0.009644 | 0.992306 | NA |
| | 0.134133 | -1.48909 | 4.077584 | -0.36519 | 0.71497 | NA |
| MTND4P26 | 0.163957 | 0.424913 | 4.079006 | 0.104171 | 0.917034 | NA |
| | 38.30433 | 0.086831 | 0.373756 | 0.23232 | 0.81629 | NA |
| | 0.151697 | -1.546 | 4.075469 | -0.37934 | 0.704433 | NA |
| RPEP2 | 0.134133 | -1.48909 | 4.077584 | -0.36519 | 0.71497 | NA |
| RNASEH1-A | 163.7588 | 0.096952 | 0.212647 | 0.455928 | 0.648442 | 0.766581 |
| | 0.696104 | -0.93034 | 2.354268 | -0.39517 | 0.692715 | NA |
| | 0.834592 | 0.684312 | 2.202411 | 0.31071 | 0.756021 | NA |
| | 0.260158 | 1.240347 | 4.047124 | 0.306476 | 0.759242 | NA |
| | 14.77633 | 0.058985 | 0.553404 | 0.106585 | 0.915118 | NA |
| | 0.296509 | -0.51709 | 3.982155 | -0.12985 | 0.896684 | NA |
| RPL7P13 | 0.163957 | 0.424913 | 4.079006 | 0.104171 | 0.917034 | NA |
| | 0.162376 | 0.424913 | 4.079006 | 0.104171 | 0.917034 | NA |
| | 0.422534 | 1.924583 | 3.001758 | 0.641152 | 0.521424 | NA |
| RPL30P2 | 0.096202 | 0.408546 | 4.079639 | 0.100143 | 0.920231 | NA |
| | 2.239638 | 0.887786 | 1.412205 | 0.628652 | 0.529577 | NA |
| | 0.392857 | -1.546 | 4.075469 | -0.37934 | 0.704433 | NA |
| | 1.641702 | 0.656084 | 1.865072 | 0.351774 | 0.725008 | NA |
| | 1.625115 | 0.641078 | 1.584788 | 0.40452 | 0.68583 | NA |
| | 0.163957 | 0.424913 | 4.079006 | 0.104171 | 0.917034 | NA |
| | 6.035329 | 0.80957 | 0.908118 | 0.891481 | 0.372671 | NA |
| | 15.05183 | 0.434391 | 0.584899 | 0.742676 | 0.457678 | NA |
| | 19.05216 | 0.038307 | 0.609621 | 0.062838 | 0.949896 | NA |
| | 0.315653 | -0.57305 | 3.892639 | -0.14721 | 0.882963 | NA |
| | 0.151697 | -1.546 | 4.075469 | -0.37934 | 0.704433 | NA |
| | 35.01526 | 0.256331 | 0.364326 | 0.703575 | 0.481698 | NA |
| | 1.954011 | 1.124182 | 1.723765 | 0.652167 | 0.514294 | NA |
| HNRNPA1P | 0.134133 | -1.48909 | 4.077584 | -0.36519 | 0.71497 | NA |
| | 1.041581 | 0.164062 | 2.078273 | 0.078942 | 0.937079 | NA |
| TPMTP2 | 0.151697 | -1.546 | 4.075469 | -0.37934 | 0.704433 | NA |
| | 0.096202 | 0.408546 | 4.079639 | 0.100143 | 0.920231 | NA |
| | 0.151697 | -1.546 | 4.075469 | -0.37934 | 0.704433 | NA |
| SNRK-AS1 | 0.303394 | -2.27006 | 4.032451 | -0.56295 | 0.57347 | NA |
| | 0.736607 | 1.395844 | 2.318642 | 0.602009 | 0.547168 | NA |

| | | | | | | | |
|---|---|---|---|---|---|---|---|
| EIF3FP1 | 0.096202 | 0.408546 | 4.079639 | 0.100143 | 0.920231 | NA | |
| YWHAEP5 | 1.332786 | 0.833291 | 1.740529 | 0.478757 | 0.632111 | NA | |
| | 1.329828 | 0.312177 | 1.851641 | 0.168595 | 0.866115 | NA | |
| CHL1-AS1 | 1.847727 | 0.402595 | 1.539106 | 0.261577 | 0.793647 | NA | |
| | 0.452561 | 2.061505 | 3.349456 | 0.615474 | 0.538241 | NA | |
| SDCBP2-AS | 42.3512 | 0.170994 | 0.410354 | 0.4167 | 0.676898 | NA | |
| | 0.575812 | 0.936922 | 2.776634 | 0.337431 | 0.735792 | NA | |
| | 0.163957 | 0.424913 | 4.079006 | 0.104171 | 0.917034 | NA | |
| | 0.163957 | 0.424913 | 4.079006 | 0.104171 | 0.917034 | NA | |
| | 1.385121 | 0.296115 | 1.904095 | 0.155515 | 0.876416 | NA | |
| | 0.163957 | 0.424913 | 4.079006 | 0.104171 | 0.917034 | NA | |
| | 0.506476 | 0.726418 | 2.864186 | 0.253621 | 0.799788 | NA | |
| | 1.195842 | -0.23035 | 1.838794 | -0.12527 | 0.900307 | NA | |
| | 0.599903 | -1.73313 | 2.730982 | -0.63462 | 0.525678 | NA | |
| | 1.575098 | -2.07544 | 1.826684 | -1.13618 | 0.255882 | NA | |
| LINC00163 | 0.788582 | 0.608095 | 2.456267 | 0.247569 | 0.804468 | NA | |
| MIR155HG | 20.06643 | 0.968682 | 0.514825 | 1.881576 | 0.059894 | NA | |
| | 0.303394 | -2.27006 | 4.032451 | -0.56295 | 0.57347 | NA | |
| | 0.162376 | 0.424913 | 4.079006 | 0.104171 | 0.917034 | NA | |
| | 0.586491 | 2.39072 | 2.689057 | 0.889055 | 0.373973 | NA | |
| CDC42P1 | 0.260158 | 1.240347 | 4.047124 | 0.306476 | 0.759242 | NA | |
| | 1.380369 | -0.94489 | 1.897497 | -0.49796 | 0.618509 | NA | |
| | 0.192403 | 0.975392 | 4.060716 | 0.240202 | 0.810174 | NA | |
| | 0.424115 | 1.932531 | 3.45462 | 0.559405 | 0.575885 | NA | |
| SDHCP3 | 0.315653 | -0.57305 | 3.892639 | -0.14721 | 0.882963 | NA | |
| | 49.91606 | -0.12513 | 0.378217 | -0.33085 | 0.740757 | | 0.836509 |
| | 0.162376 | 0.424913 | 4.079006 | 0.104171 | 0.917034 | NA | |
| COX20P2 | 0.296509 | -0.51709 | 3.982155 | -0.12985 | 0.896684 | NA | |
| | 9.048576 | 0.193613 | 0.684438 | 0.282878 | 0.77727 | NA | |
| DPP10-AS1 | 0.258578 | 1.233151 | 4.047809 | 0.304646 | 0.760635 | NA | |
| C19orf81 | 2.952706 | 0.782588 | 1.219776 | 0.641583 | 0.521144 | NA | |
| | 1.332786 | 0.833291 | 1.740529 | 0.478757 | 0.632111 | NA | |
| PPIAP3 | 0.303394 | -2.27006 | 4.032451 | -0.56295 | 0.57347 | NA | |
| BCRP5 | 0.258578 | 1.233151 | 4.047809 | 0.304646 | 0.760635 | NA | |
| SLC25A5P2 | 1.59186 | -0.82019 | 1.702621 | -0.48172 | 0.630002 | NA | |
| RPL24P2 | 39.55892 | 0.232918 | 0.390705 | 0.596147 | 0.551077 | NA | |
| | 0.096202 | 0.408546 | 4.079639 | 0.100143 | 0.920231 | NA | |
| | 1.30445 | -0.21678 | 2.036318 | -0.10646 | 0.915221 | NA | |
| | 10.68623 | -0.29543 | 0.750086 | -0.39387 | 0.69368 | NA | |
| | 0.096202 | 0.408546 | 4.079639 | 0.100143 | 0.920231 | NA | |
| | 0.163957 | 0.424913 | 4.079006 | 0.104171 | 0.917034 | NA | |
| | 12.00435 | 0.36313 | 0.637981 | 0.569186 | 0.56923 | NA | |
| | 1.061709 | 0.25203 | 2.095453 | 0.120275 | 0.904266 | NA | |
| | 3.114558 | -3.13293 | 1.644033 | -1.90564 | 0.056697 | NA | |
| FABP7P2 | 0.424115 | 1.932531 | 3.45462 | 0.559405 | 0.575885 | NA | |
| | 0.151697 | -1.546 | 4.075469 | -0.37934 | 0.704433 | NA | |
| | 1.186394 | 0.520608 | 1.795836 | 0.289897 | 0.771895 | NA | |

| | | | | | | |
|---|---|---|---|---|---|---|
| NUCB1-AS1 | 0.134133 | -1.48909 | 4.077584 | -0.36519 | 0.71497 | NA |
| | 13.49138 | -0.15158 | 0.617584 | -0.24544 | 0.806114 | NA |
| | 0.392857 | -1.546 | 4.075469 | -0.37934 | 0.704433 | NA |
| FAM83C-AS | 0.35636 | 1.709252 | 3.651833 | 0.468053 | 0.639746 | NA |
| TSPY26P | 62.06467 | -0.50052 | 0.289178 | -1.73082 | 0.083483 | 0.196836 |
| | 3.705537 | 0.678072 | 1.026525 | 0.660551 | 0.5089 | NA |
| | 0.382031 | -1.26484 | 3.151989 | -0.40128 | 0.688212 | NA |
| | 0.911649 | 0.958357 | 2.37748 | 0.403098 | 0.686876 | NA |
| | 0.096202 | 0.408546 | 4.079639 | 0.100143 | 0.920231 | NA |
| | 66.1522 | -0.23422 | 0.272279 | -0.86023 | 0.389661 | 0.555087 |
| HM13-IT1 | 8.356237 | 0.201677 | 0.901843 | 0.223628 | 0.823047 | NA |
| | 2.856113 | -0.61814 | 1.220246 | -0.50657 | 0.612456 | NA |
| | 0.096202 | 0.408546 | 4.079639 | 0.100143 | 0.920231 | NA |
| | 0.230334 | -0.53318 | 4.076749 | -0.13079 | 0.895945 | NA |
| | 0.096202 | 0.408546 | 4.079639 | 0.100143 | 0.920231 | NA |
| | 0.163957 | 0.424913 | 4.079006 | 0.104171 | 0.917034 | NA |
| | 0.662557 | -0.83483 | 2.671964 | -0.31244 | 0.754706 | NA |
| RPL36AP15 | 0.314073 | -0.57308 | 3.898934 | -0.14698 | 0.883146 | NA |
| SSR4P1 | 25.23863 | -0.17256 | 0.531422 | -0.32471 | 0.745401 | NA |
| | 3.119339 | -0.51944 | 1.192618 | -0.43555 | 0.663163 | NA |
| | 1.759044 | 0.775977 | 1.554492 | 0.499184 | 0.61765 | NA |
| | 0.163957 | 0.424913 | 4.079006 | 0.104171 | 0.917034 | NA |
| | 0.260158 | 1.240347 | 4.047124 | 0.306476 | 0.759242 | NA |
| | 0.163957 | 0.424913 | 4.079006 | 0.104171 | 0.917034 | NA |
| | 0.303394 | -2.27006 | 4.032451 | -0.56295 | 0.57347 | NA |
| | 0.162376 | 0.424913 | 4.079006 | 0.104171 | 0.917034 | NA |
| | 0.705792 | -3.31105 | 2.811825 | -1.17754 | 0.238978 | NA |
| | 5.312242 | -0.01676 | 0.842213 | -0.0199 | 0.984122 | NA |
| RPS2P7 | 2.816038 | -0.01639 | 1.153967 | -0.0142 | 0.988669 | NA |
| | 7.636515 | 0.132198 | 0.896914 | 0.147393 | 0.882822 | NA |
| | 0.192403 | 0.975392 | 4.060716 | 0.240202 | 0.810174 | NA |
| FTLP1 | 0.151697 | -1.546 | 4.075469 | -0.37934 | 0.704433 | NA |
| AGAP1-IT1 | 2.761591 | 1.35653 | 1.286842 | 1.054155 | 0.291812 | NA |
| NFAM1 | 1.977119 | 0.154188 | 1.45511 | 0.105963 | 0.915612 | NA |
| | 2.751444 | -0.06133 | 1.204921 | -0.0509 | 0.959408 | NA |
| GNAS-AS1 | 31.39835 | -0.49931 | 0.381159 | -1.30997 | 0.190206 | NA |
| LINC01102 | 5.022415 | -0.06489 | 0.91709 | -0.07075 | 0.943594 | NA |
| | 47.1564 | 0.88404 | 0.33908 | 2.607175 | 0.009129 | 0.044457 |
| | 1.551649 | 2.656399 | 1.922175 | 1.381975 | 0.166979 | NA |
| | 0.296509 | -0.51709 | 3.982155 | -0.12985 | 0.896684 | NA |
| | 0.151697 | -1.546 | 4.075469 | -0.37934 | 0.704433 | NA |
| | 0.392711 | 0.292538 | 3.145221 | 0.09301 | 0.925895 | NA |
| H3F3AP4 | 66.21314 | -0.07702 | 0.292269 | -0.26354 | 0.792134 | 0.874127 |
| RPL21P40 | 0.162376 | 0.424913 | 4.079006 | 0.104171 | 0.917034 | NA |
| | 1.379817 | 1.008459 | 2.131447 | 0.473134 | 0.636118 | NA |
| PA2G4P2 | 0.680121 | -0.88898 | 2.365123 | -0.37587 | 0.707013 | NA |
| PCBP2P1 | 1.329975 | -0.56057 | 1.737247 | -0.32268 | 0.746938 | NA |

| | | | | | | |
|---|---|---|---|---|---|---|
| | 0.298089 | -0.51711 | 3.975232 | -0.13008 | 0.896501 | NA |
| | 0.574231 | 0.93227 | 2.49998 | 0.372911 | 0.709215 | NA |
| | 0.163957 | 0.424913 | 4.079006 | 0.104171 | 0.917034 | NA |
| HMGN1P3( | 0.528424 | -0.38142 | 2.853651 | -0.13366 | 0.893671 | NA |
| LINC00607 | 27.22122 | 1.34623 | 0.526608 | 2.556416 | 0.010576 | NA |
| | 2.623046 | 0.793201 | 1.284228 | 0.617648 | 0.536808 | NA |
| | 3.325233 | 0.66382 | 1.141487 | 0.58154 | 0.560877 | NA |
| | 0.151697 | -1.546 | 4.075469 | -0.37934 | 0.704433 | NA |
| MYL6P2 | 1.120365 | -0.61682 | 2.073914 | -0.29742 | 0.766148 | NA |
| | 0.096202 | 0.408546 | 4.079639 | 0.100143 | 0.920231 | NA |
| BHLHE40-A | 1.923204 | 0.535803 | 1.417119 | 0.378093 | 0.705361 | NA |
| | 0.162376 | 0.424913 | 4.079006 | 0.104171 | 0.917034 | NA |
| LDHAP7 | 0.647493 | -1.18339 | 3.051782 | -0.38777 | 0.698186 | NA |
| RMDN2-AS | 0.326333 | 1.533889 | 3.826878 | 0.40082 | 0.688553 | NA |
| | 1.101278 | -0.11563 | 1.987513 | -0.05818 | 0.953605 | NA |
| | 12.58292 | 0.145458 | 0.615421 | 0.236355 | 0.813157 | NA |
| | 0.162376 | 0.424913 | 4.079006 | 0.104171 | 0.917034 | NA |
| TSPEAR-AS | 1.21937 | -0.87362 | 2.002767 | -0.43621 | 0.662686 | NA |
| RBMS3-AS3 | 0.987309 | 0.267164 | 2.098144 | 0.127333 | 0.898677 | NA |
| RHOA-IT1 | 0.45509 | -2.71227 | 3.963744 | -0.68427 | 0.493805 | NA |
| MACROD2- | 0.670433 | 1.255281 | 2.356401 | 0.532711 | 0.594234 | NA |
| | 1.971815 | 0.617709 | 1.421036 | 0.43469 | 0.663788 | NA |
| TTC28-AS1 | 78.09734 | 0.240511 | 0.268003 | 0.897419 | 0.369495 | 0.536481 |
| UBOX5-AS1 | 1.462885 | -0.34575 | 1.712353 | -0.20192 | 0.839981 | NA |
| | 0.422534 | 1.924583 | 3.001758 | 0.641152 | 0.521424 | NA |
| MORC2-AS | 63.65945 | -0.32092 | 0.329189 | -0.97489 | 0.329615 | 0.497782 |
| | 0.162376 | 0.424913 | 4.079006 | 0.104171 | 0.917034 | NA |
| | 0.134133 | -1.48909 | 4.077584 | -0.36519 | 0.71497 | NA |
| | 0.986086 | 0.883356 | 2.062196 | 0.428357 | 0.668391 | NA |
| | 0.298089 | -0.51711 | 3.975232 | -0.13008 | 0.896501 | NA |
| | 2.020921 | -0.85308 | 1.414228 | -0.60321 | 0.546366 | NA |
| LDHAP3 | 1.017279 | 0.1458 | 2.180567 | 0.066863 | 0.946691 | NA |
| HAUS6P2 | 0.096202 | 0.408546 | 4.079639 | 0.100143 | 0.920231 | NA |
| | 0.392857 | -1.546 | 4.075469 | -0.37934 | 0.704433 | NA |
| MRPS36P1 | 0.162376 | 0.424913 | 4.079006 | 0.104171 | 0.917034 | NA |
| | 0.096202 | 0.408546 | 4.079639 | 0.100143 | 0.920231 | NA |
| PRADC1P1 | 15.7421 | 0.245259 | 0.598237 | 0.40997 | 0.681828 | NA |
| | 0.162376 | 0.424913 | 4.079006 | 0.104171 | 0.917034 | NA |
| | 4.899599 | -0.73086 | 0.89438 | -0.81717 | 0.413829 | NA |
| | 0.163957 | 0.424913 | 4.079006 | 0.104171 | 0.917034 | NA |
| | 19.31368 | 0.179235 | 0.479511 | 0.373787 | 0.708563 | NA |
| | 0.314073 | -0.57308 | 3.898934 | -0.14698 | 0.883146 | NA |
| | 0.096202 | 0.408546 | 4.079639 | 0.100143 | 0.920231 | NA |
| | 0.096202 | 0.408546 | 4.079639 | 0.100143 | 0.920231 | NA |
| | 1.253121 | -1.51159 | 1.920912 | -0.78691 | 0.431334 | NA |
| RPL5P9 | 1.647006 | -0.01044 | 1.67675 | -0.00623 | 0.995031 | NA |
| DDX11L2 | 56.22829 | 0.188551 | 0.410277 | 0.45957 | 0.645825 | 0.764681 |

| | | | | | | |
|---|---|---|---|---|---|---|
| NRADDP | 0.891457 | -2.39215 | 2.460214 | -0.97233 | 0.330884 | NA |
| NDUFAF4P | 1.913516 | 1.485373 | 1.634512 | 0.908756 | 0.363479 | NA |
| | 8.42455 | 0.071999 | 0.686549 | 0.104871 | 0.916478 | NA |
| | 45.20494 | 0.286456 | 0.331672 | 0.863674 | 0.387767 | NA |
| | 0.544407 | -0.44425 | 2.542878 | -0.1747 | 0.861313 | NA |
| | 0.452561 | 2.061505 | 3.349456 | 0.615474 | 0.538241 | NA |
| | 6.643885 | 1.128681 | 0.844801 | 1.336032 | 0.181539 | NA |
| | 0.096202 | 0.408546 | 4.079639 | 0.100143 | 0.920231 | NA |
| | 0.651435 | 0.249629 | 2.788679 | 0.089515 | 0.928672 | NA |
| LINC00896 | 0.755955 | 0.348448 | 2.512583 | 0.138681 | 0.889702 | NA |
| | 0.134133 | -1.48909 | 4.077584 | -0.36519 | 0.71497 | NA |
| | 2.719451 | 0.566644 | 1.195923 | 0.473813 | 0.635633 | NA |
| | 4.257221 | 1.278572 | 1.052541 | 1.214748 | 0.224462 | NA |
| ARF4P2 | 0.247898 | -0.5901 | 4.074633 | -0.14482 | 0.884851 | NA |
| | 0.978999 | 3.124447 | 2.610028 | 1.197093 | 0.23127 | NA |
| | 0.640755 | -1.64826 | 2.754778 | -0.59833 | 0.549621 | NA |
| | 107.2012 | -0.51817 | 0.267245 | -1.93894 | 0.052509 | 0.143068 |
| MED28P7 | 0.163957 | 0.424913 | 4.079006 | 0.104171 | 0.917034 | NA |
| | 0.096202 | 0.408546 | 4.079639 | 0.100143 | 0.920231 | NA |
| | 0.28583 | -2.21358 | 3.97376 | -0.55705 | 0.577494 | NA |
| | 0.260158 | 1.240347 | 4.047124 | 0.306476 | 0.759242 | NA |
| YWHAQP5 | 0.134133 | -1.48909 | 4.077584 | -0.36519 | 0.71497 | NA |
| RAD23BP1 | 1.16883 | 0.552533 | 1.800196 | 0.306929 | 0.758897 | NA |
| | 0.134133 | -1.48909 | 4.077584 | -0.36519 | 0.71497 | NA |
| MED15P5 | 0.995185 | 1.939099 | 2.104716 | 0.921311 | 0.356888 | NA |
| | 1.975967 | 2.208389 | 1.54315 | 1.431091 | 0.152404 | NA |
| | 0.588072 | 2.39955 | 3.098804 | 0.774347 | 0.438726 | NA |
| KRT18P5 | 2.788933 | 0.211304 | 1.204028 | 0.175497 | 0.860689 | NA |
| | 0.602678 | 1.083611 | 2.70938 | 0.399948 | 0.689195 | NA |
| | 6.764147 | -0.24744 | 0.796185 | -0.31078 | 0.755968 | NA |
| | 0.096202 | 0.408546 | 4.079639 | 0.100143 | 0.920231 | NA |
| | 1.14213 | -0.25238 | 2.265158 | -0.11142 | 0.911283 | NA |
| | 0.382031 | -1.26484 | 3.151989 | -0.40128 | 0.688212 | NA |
| | 2.639582 | -0.29606 | 1.334442 | -0.22186 | 0.82442 | NA |
| | 0.096202 | 0.408546 | 4.079639 | 0.100143 | 0.920231 | NA |
| BCRP8 | 0.096202 | 0.408546 | 4.079639 | 0.100143 | 0.920231 | NA |
| GAPDHP2 | 1.024164 | -1.06815 | 2.390105 | -0.4469 | 0.654944 | NA |
| BCYRN1 | 24.23991 | -0.38606 | 0.75405 | -0.51199 | 0.60866 | NA |
| CBR3-AS1 | 10.0504 | -0.20271 | 0.624806 | -0.32443 | 0.745611 | NA |
| | 0.096202 | 0.408546 | 4.079639 | 0.100143 | 0.920231 | NA |
| | 6.333745 | 2.083279 | 0.937185 | 2.222911 | 0.026222 | NA |
| BMS1P20 | 253.8178 | 0.589193 | 0.198195 | 2.972802 | 0.002951 | 0.020911 |
| NIFK-AS1 | 155.0911 | 0.171319 | 0.236584 | 0.724133 | 0.468984 | 0.62547 |
| | 2.128013 | 0.763944 | 1.355313 | 0.563666 | 0.572981 | NA |
| | 0.151697 | -1.546 | 4.075469 | -0.37934 | 0.704433 | NA |
| | 0.489058 | -0.58569 | 3.427543 | -0.17088 | 0.864321 | NA |
| | 0.422534 | 1.924583 | 3.001758 | 0.641152 | 0.521424 | NA |

| | | | | | | |
|---|---|---|---|---|---|---|
| | 0.49029 | 2.123092 | 3.304282 | 0.642527 | 0.520531 | NA |
| | 0.247898 | -0.5901 | 4.074633 | -0.14482 | 0.884851 | NA |
| GGT8P | 0.151697 | -1.546 | 4.075469 | -0.37934 | 0.704433 | NA |
| ANKRD44-I | 0.71919 | 0.510978 | 2.728734 | 0.187258 | 0.851458 | NA |
| | 0.134133 | -1.48909 | 4.077584 | -0.36519 | 0.71497 | NA |
| | 1.473564 | 0.334948 | 1.696825 | 0.197397 | 0.843517 | NA |
| | 6.170349 | 0.863743 | 0.854953 | 1.010281 | 0.31236 | NA |
| | 8.917465 | 0.330692 | 0.791232 | 0.417946 | 0.675986 | NA |
| | 2.206302 | 1.802392 | 1.42796 | 1.262215 | 0.206872 | NA |
| NDUFA6-A! | 17.21984 | -0.22584 | 0.507273 | -0.44519 | 0.65618 | NA |
| FUCA1P1 | 0.151697 | -1.546 | 4.075469 | -0.37934 | 0.704433 | NA |
| | 0.315653 | -0.57305 | 3.892639 | -0.14721 | 0.882963 | NA |
| | 0.096202 | 0.408546 | 4.079639 | 0.100143 | 0.920231 | NA |
| | 6.443839 | -0.3711 | 0.793805 | -0.46749 | 0.64015 | NA |
| | 0.624422 | 1.162588 | 2.733761 | 0.425271 | 0.670639 | NA |
| | 0.392857 | -1.546 | 4.075469 | -0.37934 | 0.704433 | NA |
| SNAI1P1 | 0.096202 | 0.408546 | 4.079639 | 0.100143 | 0.920231 | NA |
| | 11.49105 | -0.20443 | 0.629132 | -0.32494 | 0.745228 | NA |
| SULT1C2P1 | 0.45509 | -2.71227 | 3.963744 | -0.68427 | 0.493805 | NA |
| ZNF295-AS | 19.76518 | 0.323167 | 0.632723 | 0.510756 | 0.609522 | NA |
| | 0.134133 | -1.48909 | 4.077584 | -0.36519 | 0.71497 | NA |
| LINC00851 | 0.163957 | 0.424913 | 4.079006 | 0.104171 | 0.917034 | NA |
| | 0.134133 | -1.48909 | 4.077584 | -0.36519 | 0.71497 | NA |
| TTN-AS1 | 30.90531 | 0.515164 | 0.389331 | 1.323204 | 0.185768 | NA |
| | 0.163957 | 0.424913 | 4.079006 | 0.104171 | 0.917034 | NA |
| | 0.134133 | -1.48909 | 4.077584 | -0.36519 | 0.71497 | NA |
| HOXD-AS2 | 65.51509 | 0.068405 | 0.276494 | 0.247401 | 0.804598 | 0.883196 |
| | 0.163957 | 0.424913 | 4.079006 | 0.104171 | 0.917034 | NA |
| NDUFA9P1 | 0.35636 | 1.709252 | 3.651833 | 0.468053 | 0.639746 | NA |
| | 0.458885 | 0.577525 | 3.449278 | 0.167433 | 0.867029 | NA |
| CECR7 | 61.96524 | -0.05035 | 0.296073 | -0.17006 | 0.864963 | 0.920317 |
| | 0.151697 | -1.546 | 4.075469 | -0.37934 | 0.704433 | NA |
| | 76.03828 | -0.32302 | 0.264314 | -1.22212 | 0.221663 | 0.382116 |
| SGOL1-AS1 | 1.754301 | 1.32931 | 1.726412 | 0.769984 | 0.441309 | NA |
| DGCR5 | 16.12519 | -0.69462 | 0.609473 | -1.13971 | 0.254409 | NA |
| NONOP2 | 2.346687 | -0.18054 | 1.432587 | -0.12603 | 0.899712 | NA |
| | 0.192403 | 0.975392 | 4.060716 | 0.240202 | 0.810174 | NA |
| | 0.162376 | 0.424913 | 4.079006 | 0.104171 | 0.917034 | NA |
| | 5.057403 | 0.13977 | 0.90946 | 0.153684 | 0.877859 | NA |
| | 1.765212 | 0.00666 | 1.543016 | 0.004316 | 0.996556 | NA |
| | 0.314073 | -0.57308 | 3.898934 | -0.14698 | 0.883146 | NA |
| | 0.422534 | 1.924583 | 3.001758 | 0.641152 | 0.521424 | NA |
| RPS15AP38 | 0.520316 | 2.239969 | 3.173769 | 0.705776 | 0.480328 | NA |
| RPL30P4 | 2.407316 | 0.987739 | 1.458328 | 0.677309 | 0.49821 | NA |
| LINC00312 | 4.483339 | 1.908502 | 1.168247 | 1.633646 | 0.102333 | NA |
| | 23.97804 | -0.6964 | 0.476114 | -1.46267 | 0.143558 | NA |
| DCTN1-AS1 | 0.364467 | -1.20166 | 3.685431 | -0.32606 | 0.744381 | NA |

| | | | | | | |
|---|---|---|---|---|---|---|
| | 0.151697 | -1.546 | 4.075469 | -0.37934 | 0.704433 | NA |
| LINC01143 | 0.296509 | -0.51709 | 3.982155 | -0.12985 | 0.896684 | NA |
| | 27.73922 | -0.15547 | 0.42151 | -0.36885 | 0.712243 | NA |
| BANF1P3 | 0.792452 | -2.15617 | 2.562821 | -0.84133 | 0.400166 | NA |
| | 1.212268 | -1.34711 | 1.888238 | -0.71342 | 0.475586 | NA |
| | 0.096202 | 0.408546 | 4.079639 | 0.100143 | 0.920231 | NA |
| LINC00211 | 0.28583 | -2.21358 | 3.97376 | -0.55705 | 0.577494 | NA |
| | 0.096202 | 0.408546 | 4.079639 | 0.100143 | 0.920231 | NA |
| | 1.226015 | -1.35692 | 2.261032 | -0.60013 | 0.548416 | NA |
| | 0.163957 | 0.424913 | 4.079006 | 0.104171 | 0.917034 | NA |
| | 0.163957 | 0.424913 | 4.079006 | 0.104171 | 0.917034 | NA |
| LINC00322 | 0.134133 | -1.48909 | 4.077584 | -0.36519 | 0.71497 | NA |
| | 5.915641 | 0.373422 | 0.884913 | 0.421987 | 0.673035 | NA |
| DGUOK-AS | 13.70977 | 1.764828 | 0.593022 | 2.97599 | 0.00292 | NA |
| KLF7-IT1 | 1.369312 | 1.561252 | 1.870566 | 0.834642 | 0.403919 | NA |
| | 0.424115 | 1.932531 | 3.45462 | 0.559405 | 0.575885 | NA |
| | 0.410275 | 0.244283 | 3.100889 | 0.078778 | 0.937209 | NA |
| | 0.624422 | 1.162588 | 2.733761 | 0.425271 | 0.670639 | NA |
| LINC00649 | 11.71583 | 0.055946 | 0.842916 | 0.066372 | 0.947081 | NA |
| | 0.163957 | 0.424913 | 4.079006 | 0.104171 | 0.917034 | NA |
| EIF4HP2 | 0.430642 | -1.1891 | 3.485782 | -0.34113 | 0.733008 | NA |
| | 0.589223 | -3.04304 | 3.047314 | -0.9986 | 0.317989 | NA |
| | 133.5602 | -0.49577 | 0.243009 | -2.04014 | 0.041337 | 0.123231 |
| CNTN4-AS1 | 0.163957 | 0.424913 | 4.079006 | 0.104171 | 0.917034 | NA |
| | 20.31379 | 0.914537 | 0.476074 | 1.920997 | 0.054732 | NA |
| ZEB2-AS1 | 3.815745 | 0.808612 | 1.096738 | 0.737288 | 0.460947 | NA |
| SPATA3-AS | 0.324752 | 1.53082 | 4.022025 | 0.380609 | 0.703493 | NA |
| | 5.688496 | -0.06809 | 0.950477 | -0.07164 | 0.942887 | NA |
| | 0.162376 | 0.424913 | 4.079006 | 0.104171 | 0.917034 | NA |
| MLK7-AS1 | 0.877625 | -0.2954 | 2.206963 | -0.13385 | 0.893522 | NA |
| | 0.888304 | 0.641829 | 2.171212 | 0.295609 | 0.767529 | NA |
| MLLT10P1 | 3.352338 | 0.427864 | 1.068489 | 0.400438 | 0.688834 | NA |
| | 0.841063 | -2.08438 | 2.306687 | -0.90362 | 0.366195 | NA |
| | 2.091516 | 1.692039 | 1.468596 | 1.152147 | 0.249261 | NA |
| | 0.096202 | 0.408546 | 4.079639 | 0.100143 | 0.920231 | NA |
| | 0.163957 | 0.424913 | 4.079006 | 0.104171 | 0.917034 | NA |
| PAXBP1-AS | 16.63012 | -0.2834 | 0.523687 | -0.54115 | 0.588401 | NA |
| | 0.096202 | 0.408546 | 4.079639 | 0.100143 | 0.920231 | NA |
| | 3.761957 | 0.66749 | 1.166195 | 0.572366 | 0.567074 | NA |
| SNORD11 | 0.422534 | 1.924583 | 3.001758 | 0.641152 | 0.521424 | NA |
| | 0.392857 | -1.546 | 4.075469 | -0.37934 | 0.704433 | NA |
| | 0.162376 | 0.424913 | 4.079006 | 0.104171 | 0.917034 | NA |
| | 0.162376 | 0.424913 | 4.079006 | 0.104171 | 0.917034 | NA |
| | 1.778181 | -0.69451 | 1.536305 | -0.45206 | 0.651223 | NA |
| | 0.774742 | -0.40577 | 2.223483 | -0.18249 | 0.855195 | NA |
| | 0.096202 | 0.408546 | 4.079639 | 0.100143 | 0.920231 | NA |
| | 0.096202 | 0.408546 | 4.079639 | 0.100143 | 0.920231 | NA |

| | | | | | | |
|---|---|---|---|---|---|---|
| | 0.296509 | -0.51709 | 3.982155 | -0.12985 | 0.896684 | NA |
| | 1.380369 | -0.94489 | 1.897497 | -0.49796 | 0.618509 | NA |
| SNORD125 | 4.265649 | -0.40988 | 1.012666 | -0.40475 | 0.685658 | NA |
| | 0.163957 | 0.424913 | 4.079006 | 0.104171 | 0.917034 | NA |
| | 0.151697 | -1.546 | 4.075469 | -0.37934 | 0.704433 | NA |
| | 0.134133 | -1.48909 | 4.077584 | -0.36519 | 0.71497 | NA |
| GATSL3 | 7.149136 | -0.22349 | 0.739466 | -0.30224 | 0.762471 | NA |
| | 1.543159 | 0.16312 | 1.777685 | 0.09176 | 0.926889 | NA |
| RNF103 | 388.4479 | 0.170036 | 0.165117 | 1.029789 | 0.303109 | 0.47046 |
| | 25.69351 | -0.2142 | 0.493277 | -0.43424 | 0.664117 | NA |
| LINC01119 | 2.502739 | 0.330592 | 1.359654 | 0.243144 | 0.807894 | NA |
| ASB14 | 2.927432 | 1.379921 | 1.384344 | 0.996805 | 0.318859 | NA |
| | 22.74158 | -0.22067 | 0.441949 | -0.49931 | 0.617558 | NA |
| | 13.21969 | 0.466233 | 0.585889 | 0.795771 | 0.426165 | NA |
| KLHL41 | 15.6393 | -0.17917 | 0.628251 | -0.28519 | 0.7755 | NA |
| | 0.747876 | -0.62091 | 2.562446 | -0.24231 | 0.80854 | NA |
| COX6CP14 | 0.151697 | -1.546 | 4.075469 | -0.37934 | 0.704433 | NA |
| C2orf61 | 0.162376 | 0.424913 | 4.079006 | 0.104171 | 0.917034 | NA |
| PSMD6-AS: | 6.800283 | 0.882823 | 1.040615 | 0.848367 | 0.396234 | NA |
| APOBEC3G | 143.3544 | -0.30136 | 0.225034 | -1.33915 | 0.180521 | 0.333033 |
| TLR9 | 0.411855 | 0.250456 | 3.097618 | 0.080854 | 0.935558 | NA |
| WBP1 | 181.0821 | -0.6535 | 0.208498 | -3.13431 | 0.001723 | 0.014681 |
| ITIH4-AS1 | 0.192403 | 0.975392 | 4.060716 | 0.240202 | 0.810174 | NA |
| RNF7P1 | 0.134133 | -1.48909 | 4.077584 | -0.36519 | 0.71497 | NA |
| RN7SL674F | 0.399595 | -1.31595 | 3.579466 | -0.36764 | 0.713143 | NA |
| ADSL | 1678.077 | -0.17097 | 0.127282 | -1.34325 | 0.17919 | 0.331683 |
| RPL39P36 | 0.192403 | 0.975392 | 4.060716 | 0.240202 | 0.810174 | NA |
| PRDX3P4 | 0.354779 | 1.703954 | 3.657232 | 0.465914 | 0.641277 | NA |
| | 0.354779 | 1.703954 | 3.657232 | 0.465914 | 0.641277 | NA |
| LILRA2 | 0.134133 | -1.48909 | 4.077584 | -0.36519 | 0.71497 | NA |
| | 0.096202 | 0.408546 | 4.079639 | 0.100143 | 0.920231 | NA |
| RN7SL263F | 0.490493 | 0.776209 | 2.895261 | 0.268096 | 0.788625 | NA |
| ZNF542 | 182.0128 | -0.1811 | 0.236867 | -0.76458 | 0.444519 | 0.604471 |
| GHRLOS | 0.881623 | 0.857232 | 2.360567 | 0.363147 | 0.716495 | NA |
| PPIL3 | 206.0792 | 0.54738 | 0.24751 | 2.21155 | 0.026998 | 0.092445 |
| | 0.096202 | 0.408546 | 4.079639 | 0.100143 | 0.920231 | NA |
| RPL23AP7 | 765.6842 | 0.393054 | 0.186375 | 2.108935 | 0.03495 | 0.109816 |
| | 0.968522 | 0.920016 | 1.915592 | 0.480278 | 0.63103 | NA |
| THOC7-AS1 | 0.258578 | 1.233151 | 4.047809 | 0.304646 | 0.760635 | NA |
| | 5.29181 | -0.4684 | 0.883326 | -0.53027 | 0.595927 | NA |
| KRBOX1 | 41.32919 | -0.11173 | 0.333632 | -0.33489 | 0.737707 | NA |
| | 40.01202 | 0.092699 | 0.352443 | 0.26302 | 0.792535 | NA |
| C21orf91-C | 0.392857 | -1.546 | 4.075469 | -0.37934 | 0.704433 | NA |
| TMEM189 | 728.7637 | -0.2303 | 0.142366 | -1.6177 | 0.105728 | 0.231513 |
| RDH14 | 371.1914 | 0.461355 | 0.168306 | 2.741168 | 0.006122 | 0.033867 |
| MIF | 10.4822 | -0.61372 | 0.656739 | -0.93449 | 0.350049 | NA |
| TPM3P9 | 180.1897 | 0.230769 | 0.199464 | 1.156941 | 0.247297 | 0.410746 |

| | | | | | | |
|---|---|---|---|---|---|---|
| | 0.162376 | 0.424913 | 4.079006 | 0.104171 | 0.917034 | NA |
| ADAMTS9-/ | 0.096202 | 0.408546 | 4.079639 | 0.100143 | 0.920231 | NA |
| | 52.74827 | 0.608639 | 0.341058 | 1.784559 | 0.074333 | 0.182069 |
| PDXP | 176.7585 | 0.093895 | 0.208328 | 0.450707 | 0.6522 | 0.76995 |
| CD302 | 255.7826 | 0.445204 | 0.242038 | 1.839397 | 0.065857 | 0.16736 |
| | 0.096202 | 0.408546 | 4.079639 | 0.100143 | 0.920231 | NA |
| PTPRG-AS1 | 42.29009 | -0.21002 | 0.379912 | -0.55282 | 0.580384 | NA |
| ARHGAP8 | 0.096202 | 0.408546 | 4.079639 | 0.100143 | 0.920231 | NA |
| PSMC1P1 | 7.807809 | 0.6223 | 0.757252 | 0.821787 | 0.411198 | NA |
| | 0.902145 | 1.753058 | 2.208187 | 0.79389 | 0.427259 | NA |
| | 5.404363 | 0.095891 | 0.877528 | 0.109274 | 0.912985 | NA |
| ARPC4 | 3329.372 | -0.2575 | 0.207501 | -1.24096 | 0.214622 | 0.373586 |
| ADAMTS9-/ | 8.530407 | 1.874196 | 0.788463 | 2.377025 | 0.017453 | NA |
| RN7SL704F | 0.163957 | 0.424913 | 4.079006 | 0.104171 | 0.917034 | NA |
| VWFP1 | 0.230334 | -0.53318 | 4.076749 | -0.13079 | 0.895945 | NA |
| ATP5O | 1079.08 | 0.272592 | 0.167814 | 1.624366 | 0.104298 | 0.229312 |
| PISD | 772.8765 | -0.10583 | 0.16505 | -0.64119 | 0.521396 | 0.668146 |
| | 0.230334 | -0.53318 | 4.076749 | -0.13079 | 0.895945 | NA |
| RN7SL517F | 0.096202 | 0.408546 | 4.079639 | 0.100143 | 0.920231 | NA |
| PWP2 | 878.89 | -0.35145 | 0.154508 | -2.27466 | 0.022926 | 0.082895 |
| | 0.394291 | 0.298745 | 3.141711 | 0.09509 | 0.924243 | NA |
| C2orf15 | 12.3317 | 0.578652 | 0.629378 | 0.919404 | 0.357884 | NA |
| PI4KA | 4307.392 | -0.5292 | 0.163364 | -3.23938 | 0.001198 | 0.011696 |
| | 57.78529 | 0.236683 | 0.352583 | 0.671283 | 0.50204 | 0.65228 |
| | 7.826727 | -1.66151 | 0.994502 | -1.67069 | 0.094783 | NA |
| RN7SL113F | 1.061709 | 0.25203 | 2.095453 | 0.120275 | 0.904266 | NA |
| MTFP1 | 103.0687 | -0.04732 | 0.290497 | -0.16289 | 0.870604 | 0.923107 |
| TCP10L | 0.247898 | -0.5901 | 4.074633 | -0.14482 | 0.884851 | NA |
| ARFGAP3 | 656.0561 | 0.413425 | 0.174935 | 2.363309 | 0.018113 | 0.070587 |
| C22orf39 | 350.5664 | -0.01593 | 0.197904 | -0.08047 | 0.935861 | 0.964049 |
| | 22.72505 | -0.27239 | 0.490181 | -0.55568 | 0.578427 | NA |
| | 0.151697 | -1.546 | 4.075469 | -0.37934 | 0.704433 | NA |
| EIF6 | 2014.101 | -0.17979 | 0.169089 | -1.06331 | 0.287642 | 0.453819 |
| | 1.195988 | -1.61404 | 2.13645 | -0.75548 | 0.449963 | NA |
| DBIL5P2 | 0.440301 | 0.468383 | 3.449808 | 0.135771 | 0.892003 | NA |
| RN7SL313F | 0.096202 | 0.408546 | 4.079639 | 0.100143 | 0.920231 | NA |
| GTF2A1L | 0.134133 | -1.48909 | 4.077584 | -0.36519 | 0.71497 | NA |
| NDUFB4P1 | 0.096202 | 0.408546 | 4.079639 | 0.100143 | 0.920231 | NA |
| UGT1A10 | 0.096202 | 0.408546 | 4.079639 | 0.100143 | 0.920231 | NA |
| | 0.392711 | 0.292538 | 3.145221 | 0.09301 | 0.925895 | NA |
| | 7.009004 | 1.467877 | 0.8424 | 1.742495 | 0.081422 | NA |
| | 0.096202 | 0.408546 | 4.079639 | 0.100143 | 0.920231 | NA |
| RN7SL40P | 0.096202 | 0.408546 | 4.079639 | 0.100143 | 0.920231 | NA |
| RPL23AP81 | 0.247898 | -0.5901 | 4.074633 | -0.14482 | 0.884851 | NA |
| ZNF702P | 4.064526 | 1.677979 | 1.072401 | 1.564693 | 0.117655 | NA |
| GLYCTK-AS | 1.636529 | -1.07836 | 1.58333 | -0.68107 | 0.495829 | NA |
| ALDOAP1 | 1.178075 | 0.574265 | 2.102875 | 0.273086 | 0.784787 | NA |

| | | | | | | |
|---|---|---|---|---|---|---|
| RN7SL812F | 0.462046 | 0.589069 | 3.442439 | 0.17112 | 0.86413 | NA |
| MRPL33 | 1218.943 | 0.38112 | 0.168592 | 2.260608 | 0.023784 | 0.085143 |
| MICAL3 | 1689.506 | -0.64216 | 0.189613 | -3.3867 | 7.07E-04 | 0.008346 |
| | 4.674197 | -0.03658 | 0.932897 | -0.03922 | 0.968719 | NA |
| STON1 | 274.2874 | 0.301837 | 0.271582 | 1.111399 | 0.266396 | 0.431321 |
| | 1.3816 | 0.438123 | 1.829949 | 0.239418 | 0.810781 | NA |
| NAT6 | 184.6419 | -0.15618 | 0.217721 | -0.71733 | 0.473171 | 0.628861 |
| | 0.096202 | 0.408546 | 4.079639 | 0.100143 | 0.920231 | NA |
| | 1.180005 | -1.57876 | 2.139972 | -0.73775 | 0.460668 | NA |
| IL10RB | 600.7657 | -0.17012 | 0.149548 | -1.13757 | 0.255299 | 0.419503 |
| RN7SL834F | 1.6482 | 1.396964 | 1.664639 | 0.839199 | 0.401358 | NA |
| WDR92 | 179.4606 | 0.167332 | 0.238089 | 0.702812 | 0.482173 | 0.636186 |
| RPL37P23 | 5.371825 | -0.24857 | 0.876211 | -0.28369 | 0.776646 | NA |
| | 16.70474 | -0.30186 | 0.602613 | -0.50092 | 0.616426 | NA |
| | 0.134133 | -1.48909 | 4.077584 | -0.36519 | 0.71497 | NA |
| APOBEC3D | 121.0794 | -0.69868 | 0.217852 | -3.20712 | 0.001341 | 0.012544 |
| RN7SL832F | 1.034697 | 1.024735 | 1.902596 | 0.538598 | 0.590164 | NA |
| | 6.297002 | 0.435294 | 0.843849 | 0.515843 | 0.605964 | NA |
| RN7SL610F | 0.46577 | -1.30191 | 3.402879 | -0.38259 | 0.702024 | NA |
| TUBA4B | 0.134133 | -1.48909 | 4.077584 | -0.36519 | 0.71497 | NA |
| MRPS6 | 2349.492 | 0.530851 | 0.849858 | 0.624635 | 0.532211 | 0.676528 |
| ZNF512 | 521.2398 | 0.01786 | 0.165222 | 0.108096 | 0.913919 | 0.9501 |
| ACY1 | 21.26765 | -0.46289 | 0.494946 | -0.93523 | 0.34967 | NA |
| NFS1 | 604.7382 | -0.21414 | 0.148603 | -1.44104 | 0.149573 | 0.293079 |
| | 0.897992 | -1.0518 | 2.16635 | -0.48552 | 0.627309 | NA |
| FLNB-AS1 | 0.556813 | -0.57007 | 3.294615 | -0.17303 | 0.862627 | NA |
| | 0.096202 | 0.408546 | 4.079639 | 0.100143 | 0.920231 | NA |
| ILF2P1 | 0.163957 | 0.424913 | 4.079006 | 0.104171 | 0.917034 | NA |
| DBNDD2 | 30.2084 | 0.229197 | 0.428478 | 0.534909 | 0.592712 | NA |
| ECEL1P2 | 9.458062 | 0.111951 | 1.160171 | 0.096495 | 0.923127 | NA |
| | 0.162376 | 0.424913 | 4.079006 | 0.104171 | 0.917034 | NA |
| LINC00698 | 0.163957 | 0.424913 | 4.079006 | 0.104171 | 0.917034 | NA |
| RN7SL145F | 0.411855 | 0.250456 | 3.097618 | 0.080854 | 0.935558 | NA |
| RN7SL636F | 0.162376 | 0.424913 | 4.079006 | 0.104171 | 0.917034 | NA |
| | 0.192403 | 0.975392 | 4.060716 | 0.240202 | 0.810174 | NA |
| RN7SL268F | 1.399513 | -0.94631 | 1.712153 | -0.5527 | 0.580467 | NA |
| RBM12 | 1270.692 | -0.17832 | 0.246549 | -0.72325 | 0.469528 | 0.625866 |
| SCARF2 | 2421.743 | -0.03707 | 0.239337 | -0.15488 | 0.876914 | 0.926427 |
| | 0.394291 | 0.298745 | 3.141711 | 0.09509 | 0.924243 | NA |
| APOBEC3C | 1872.343 | -0.54804 | 0.124242 | -4.41103 | 1.03E-05 | 6.34E-04 |
| | 9.055222 | 1.149442 | 0.700157 | 1.64169 | 0.100654 | NA |
| | 19.02703 | 0.388897 | 0.662083 | 0.587383 | 0.556946 | NA |
| | 0.35636 | 1.709252 | 3.651833 | 0.468053 | 0.639746 | NA |
| | 58.85195 | -0.10339 | 0.288992 | -0.35775 | 0.720532 | 0.822207 |
| WNT5A-AS | 1.367914 | 0.791109 | 1.898548 | 0.416692 | 0.676904 | NA |
| RAD21L1 | 0.258578 | 1.233151 | 4.047809 | 0.304646 | 0.760635 | NA |
| CCDC13 | 7.033615 | 0.237522 | 0.789917 | 0.300692 | 0.763649 | NA |

| | | | | | | |
|---|---|---|---|---|---|---|
| ASPRV1 | 14.18485 | -0.04303 | 0.578247 | -0.07442 | 0.940674 | NA |
| | 61.14853 | -0.1125 | 0.311197 | -0.36151 | 0.717721 | 0.82009 |
| | 19.01683 | -0.09684 | 0.500772 | -0.19338 | 0.846663 | NA |
| RN7SL280F | 0.247898 | -0.5901 | 4.074633 | -0.14482 | 0.884851 | NA |
| | 0.815391 | 0.931626 | 2.360431 | 0.394685 | 0.693076 | NA |
| UBE2V1 | 43.1443 | 0.008559 | 0.406988 | 0.021031 | 0.983221 | NA |
| ASLP1 | 0.82213 | 0.50433 | 2.197682 | 0.229483 | 0.818494 | NA |
| CRYBB2 | 7.027669 | -0.23208 | 0.776669 | -0.29881 | 0.765082 | NA |
| DACT3-AS1 | 1.625501 | -0.42008 | 1.608465 | -0.26117 | 0.793964 | NA |
| | 3.024571 | -0.20468 | 1.130771 | -0.18101 | 0.856357 | NA |
| TWF2 | 856.7871 | -0.16384 | 0.173145 | -0.94625 | 0.344022 | 0.512167 |
| MARS2 | 424.0348 | -0.14525 | 0.168471 | -0.86219 | 0.388584 | 0.554176 |
| BACH1-IT1 | 1.794942 | 0.281406 | 1.639616 | 0.171629 | 0.863729 | NA |
| ABHD14A | 291.2581 | -0.26908 | 0.184251 | -1.46039 | 0.144182 | 0.285928 |
| GAPDHP38 | 0.096202 | 0.408546 | 4.079639 | 0.100143 | 0.920231 | NA |
| ATP5L2 | 0.640406 | 1.11402 | 2.710977 | 0.410929 | 0.681124 | NA |
| ZNF324B | 124.0953 | -0.16395 | 0.211885 | -0.77376 | 0.439075 | 0.599011 |
| ANKRD20A | 0.392711 | 0.292538 | 3.145221 | 0.09301 | 0.925895 | NA |
| | 5.446869 | 1.093521 | 0.941897 | 1.160977 | 0.245651 | NA |
| FER1L5 | 2.036758 | -0.22167 | 1.464317 | -0.15138 | 0.879675 | NA |
| EAF1-AS1 | 0.55687 | -0.14239 | 3.170029 | -0.04492 | 0.964172 | NA |
| | 0.163957 | 0.424913 | 4.079006 | 0.104171 | 0.917034 | NA |
| TMEM158 | 17.57114 | -0.48865 | 0.526277 | -0.9285 | 0.353147 | NA |
| HMGB1P3 | 0.288605 | 1.443619 | 4.029058 | 0.358302 | 0.720117 | NA |
| | 5.617353 | 0.83049 | 0.90767 | 0.914969 | 0.360208 | NA |
| | 6.700535 | 0.305515 | 0.890686 | 0.343011 | 0.73159 | NA |
| FGD5P1 | 2.329755 | 0.974366 | 1.316456 | 0.740143 | 0.459213 | NA |
| CHCHD10 | 1531.066 | -0.13952 | 0.223177 | -0.62515 | 0.53187 | 0.676174 |
| ATP6V1E2 | 91.20387 | -0.07208 | 0.271596 | -0.26539 | 0.790712 | 0.873094 |
| | 0.162376 | 0.424913 | 4.079006 | 0.104171 | 0.917034 | NA |
| SHANK3 | 1047.594 | -0.343 | 0.180306 | -1.90235 | 0.057126 | 0.151573 |
| ZNF550 | 159.0276 | 0.597776 | 0.225303 | 2.653214 | 0.007973 | 0.040544 |
| | 4.081208 | -0.70658 | 1.023659 | -0.69024 | 0.49004 | NA |
| RNU6-377F | 0.151697 | -1.546 | 4.075469 | -0.37934 | 0.704433 | NA |
| | 0.192403 | 0.975392 | 4.060716 | 0.240202 | 0.810174 | NA |
| SNORD119 | 0.134133 | -1.48909 | 4.077584 | -0.36519 | 0.71497 | NA |
| RNU6ATAC | 0.162376 | 0.424913 | 4.079006 | 0.104171 | 0.917034 | NA |
| | 0.640755 | -1.64826 | 2.754778 | -0.59833 | 0.549621 | NA |
| SCARNA5 | 0.096202 | 0.408546 | 4.079639 | 0.100143 | 0.920231 | NA |
| | 0.516164 | -1.70268 | 2.844782 | -0.59853 | 0.549488 | NA |
| | 0.268266 | -2.15396 | 4.038351 | -0.53338 | 0.593773 | NA |
| RNU6-1206 | 0.392857 | -1.546 | 4.075469 | -0.37934 | 0.704433 | NA |
| RNU7-64P | 0.162376 | 0.424913 | 4.079006 | 0.104171 | 0.917034 | NA |
| RN7SKP83 | 0.096202 | 0.408546 | 4.079639 | 0.100143 | 0.920231 | NA |
| | 1.008034 | 0.23684 | 2.090235 | 0.113308 | 0.909786 | NA |
| RNU6-100F | 0.288605 | 1.443619 | 4.029058 | 0.358302 | 0.720117 | NA |
| RNU6-107F | 0.096202 | 0.408546 | 4.079639 | 0.100143 | 0.920231 | NA |

| | | | | | | |
|---|---|---|---|---|---|---|
| RMRPP3 | 0.096202 | 0.408546 | 4.079639 | 0.100143 | 0.920231 | NA |
| | 0.584911 | 2.38756 | 2.690807 | 0.887303 | 0.374916 | NA |
| RNU7-162F | 0.163957 | 0.424913 | 4.079006 | 0.104171 | 0.917034 | NA |
| RNU6-759F | 0.151697 | -1.546 | 4.075469 | -0.37934 | 0.704433 | NA |
| | 0.420954 | 1.923376 | 3.462803 | 0.555439 | 0.578594 | NA |
| RNU6-856F | 0.3441 | -0.0166 | 3.769682 | -0.0044 | 0.996486 | NA |
| | 0.247898 | -0.5901 | 4.074633 | -0.14482 | 0.884851 | NA |
| RNA5SP101 | 1.262459 | -1.18346 | 1.757563 | -0.67335 | 0.500723 | NA |
| RNU6-548F | 1.484446 | 0.457722 | 1.720573 | 0.266028 | 0.790217 | NA |
| | 0.614937 | 2.493639 | 2.617677 | 0.952615 | 0.340785 | NA |
| RNU6-762F | 0.555087 | 0.973802 | 2.808575 | 0.346725 | 0.728798 | NA |
| RNA5SP49( | 0.354779 | 1.703954 | 3.657232 | 0.465914 | 0.641277 | NA |
| | 0.520316 | 2.239969 | 3.173769 | 0.705776 | 0.480328 | NA |
| RNU6-902F | 0.163957 | 0.424913 | 4.079006 | 0.104171 | 0.917034 | NA |
| TUG1 | 1779.43 | 0.01746 | 0.191412 | 0.091216 | 0.927321 | 0.95828 |
| IGKV1OR22 | 0.247898 | -0.5901 | 4.074633 | -0.14482 | 0.884851 | NA |
| | 0.35636 | 1.709252 | 3.651833 | 0.468053 | 0.639746 | NA |
| OSGEPL1-A | 1.198654 | 1.307481 | 1.836319 | 0.712012 | 0.476457 | NA |
| SIGLEC14 | 0.151697 | -1.546 | 4.075469 | -0.37934 | 0.704433 | NA |
| | 1.044145 | 0.31036 | 2.099918 | 0.147796 | 0.882504 | NA |
| | 0.448206 | -1.2532 | 3.007315 | -0.41672 | 0.676885 | NA |
| XIRP2-AS1 | 0.134133 | -1.48909 | 4.077584 | -0.36519 | 0.71497 | NA |
| STX16-NPE | 1.399164 | 0.376665 | 1.647821 | 0.228584 | 0.819192 | NA |
| BRK1 | 4593.777 | -0.02752 | 0.177039 | -0.15545 | 0.876467 | 0.926153 |
| | 0.392857 | -1.546 | 4.075469 | -0.37934 | 0.704433 | NA |
| | 2.548924 | -1.08944 | 1.264576 | -0.86151 | 0.388958 | NA |
| | 0.151697 | -1.546 | 4.075469 | -0.37934 | 0.704433 | NA |
| BRWD1-IT2 | 5.523001 | -0.49122 | 0.878058 | -0.55943 | 0.575865 | NA |
| | 0.134133 | -1.48909 | 4.077584 | -0.36519 | 0.71497 | NA |
| | 4.599698 | -0.09121 | 0.979239 | -0.09315 | 0.925787 | NA |
| TRAPPC2P1 | 91.04261 | 0.480653 | 0.260947 | 1.841955 | 0.065482 | 0.166907 |
| C21orf119 | 53.56228 | 0.157796 | 0.305673 | 0.516226 | 0.605696 | 0.734104 |
| ZNF432 | 105.8417 | 0.593941 | 0.295389 | 2.010704 | 0.044357 | 0.128569 |
| MTRNR2L3 | 0.437526 | -2.68242 | 3.384147 | -0.79264 | 0.427986 | NA |
| ZNF225 | 103.2632 | 0.441661 | 0.241304 | 1.830308 | 0.067204 | 0.169547 |
| | 0.891457 | -2.39215 | 2.460214 | -0.97233 | 0.330884 | NA |
| ZNF350 | 126.7497 | 0.46619 | 0.232651 | 2.003813 | 0.04509 | 0.129901 |
| | 0.096202 | 0.408546 | 4.079639 | 0.100143 | 0.920231 | NA |
| | 0.766634 | 1.516849 | 2.262991 | 0.670285 | 0.502676 | NA |
| | 2.420167 | -0.1478 | 1.257336 | -0.11755 | 0.906425 | NA |
| LBX2-AS1 | 14.87409 | 0.72108 | 0.563412 | 1.279847 | 0.200599 | NA |
| PRR24 | 140.644 | 0.017352 | 0.229337 | 0.075662 | 0.939688 | 0.96651 |
| FNBP1P1 | 6.34969 | 0.322625 | 1.050891 | 0.307001 | 0.758843 | NA |
| | 0.162376 | 0.424913 | 4.079006 | 0.104171 | 0.917034 | NA |
| RTEL1 | 26.00738 | -0.50507 | 0.472769 | -1.06832 | 0.285378 | NA |
| | 0.162376 | 0.424913 | 4.079006 | 0.104171 | 0.917034 | NA |
| ZNF578 | 16.69825 | -0.43594 | 0.550109 | -0.79247 | 0.428087 | NA |

| | | | | | | |
|---|---|---|---|---|---|---|
| | 5.103827 | 0.370771 | 0.894675 | 0.41442 | 0.678567 | NA |
| UBE2F-SCL' | 0.716471 | -2.03238 | 2.582811 | -0.78689 | 0.431348 | NA |
| | 0.163957 | 0.424913 | 4.079006 | 0.104171 | 0.917034 | NA |
| | 4.258489 | 1.720456 | 1.091448 | 1.576306 | 0.114955 | NA |
| | 27.02723 | -0.49274 | 0.414856 | -1.18774 | 0.234936 | NA |
| | 0.163957 | 0.424913 | 4.079006 | 0.104171 | 0.917034 | NA |
| | 0.326333 | 1.533889 | 3.826878 | 0.40082 | 0.688553 | NA |
| SH3RF3-AS | 26.57976 | 0.323607 | 0.532901 | 0.607256 | 0.543681 | NA |
| | 0.574231 | 0.93227 | 2.49998 | 0.372911 | 0.709215 | NA |
| | 13.11849 | 0.581479 | 0.642055 | 0.905653 | 0.365119 | NA |
| | 18.61663 | 0.51258 | 0.502839 | 1.019372 | 0.308026 | NA |
| LINC00657 | 3797.991 | 0.223641 | 0.242398 | 0.922616 | 0.356208 | 0.523926 |
| | 150.069 | 0.20262 | 0.26738 | 0.757796 | 0.448573 | 0.608203 |
| | 42.0055 | -0.63593 | 0.372653 | -1.7065 | 0.087914 | NA |
| | 8.565859 | 1.040058 | 0.800475 | 1.299301 | 0.193841 | NA |
| | 4.428276 | -0.04009 | 1.091329 | -0.03674 | 0.970694 | NA |
| | 0.658173 | -0.01016 | 2.630582 | -0.00386 | 0.996919 | NA |
| | 11.10437 | -0.06513 | 0.681613 | -0.09556 | 0.923873 | NA |
| | 13.84505 | 0.986569 | 0.598751 | 1.647712 | 0.099412 | NA |
| | 1.208248 | -0.56723 | 2.124937 | -0.26694 | 0.789515 | NA |
| | 0.696104 | -0.93034 | 2.354268 | -0.39517 | 0.692715 | NA |
| | 1.292043 | 0.040764 | 1.785235 | 0.022834 | 0.981783 | NA |
| | 0.711139 | 2.710446 | 2.504485 | 1.082237 | 0.279147 | NA |
| LINC00515 | 0.422534 | 1.924583 | 3.001758 | 0.641152 | 0.521424 | NA |
| | 0.247898 | -0.5901 | 4.074633 | -0.14482 | 0.884851 | NA |
| | 0.419962 | -2.64111 | 3.429364 | -0.77014 | 0.441214 | NA |
| | 35.83004 | 0.525714 | 0.375081 | 1.401601 | 0.161034 | NA |
| | 4.444074 | 0.41613 | 0.938019 | 0.443627 | 0.657313 | NA |
| | 189.0091 | 0.121546 | 0.3148 | 0.386107 | 0.699418 | 0.806443 |
| | 17.10346 | -0.11523 | 0.608357 | -0.18941 | 0.849771 | NA |
| XKR7 | 0.392857 | -1.546 | 4.075469 | -0.37934 | 0.704433 | NA |
| | 17.36517 | 0.537086 | 0.491297 | 1.0932 | 0.274306 | NA |
| CAPN10-AS | 24.22567 | 0.027873 | 0.470239 | 0.059274 | 0.952734 | NA |
| | 0.911447 | 1.874441 | 2.135629 | 0.8777 | 0.380107 | NA |
| | 2.196774 | -0.30951 | 1.370947 | -0.22576 | 0.821385 | NA |
| | 4.385241 | 0.402622 | 1.008534 | 0.399216 | 0.689734 | NA |
| | 0.460466 | 0.568655 | 3.014642 | 0.188631 | 0.850382 | NA |
| | 28.96621 | 0.267886 | 0.404075 | 0.662963 | 0.507354 | NA |
| | 2.094826 | 0.691584 | 1.424787 | 0.485394 | 0.627397 | NA |
| | 0.096202 | 0.408546 | 4.079639 | 0.100143 | 0.920231 | NA |
| ZNF865 | 768.9855 | -0.38638 | 0.175957 | -2.19591 | 0.028099 | 0.094903 |
| | 2.84171 | -0.61273 | 1.162219 | -0.52721 | 0.59805 | NA |
| | 58.41643 | 0.038237 | 0.309405 | 0.123583 | 0.901645 | 0.942427 |
| | 0.314073 | -0.57308 | 3.898934 | -0.14698 | 0.883146 | NA |
| | 0.315653 | -0.57305 | 3.892639 | -0.14721 | 0.882963 | NA |
| | 9.493688 | 0.224798 | 0.851692 | 0.263943 | 0.791824 | NA |
| | 1.770073 | 0.275437 | 1.470334 | 0.187329 | 0.851402 | NA |

| | | | | | | |
|---|---|---|---|---|---|---|
| | 300.8944 | -1.03158 | 0.226001 | -4.56449 | 5.01E-06 | 4.21E-04 |
| | 14.66828 | 0.597312 | 0.553678 | 1.078808 | 0.280673 | NA |
| | 0.327913 | 1.543154 | 4.021063 | 0.383768 | 0.701151 | NA |
| | 2.37081 | 0.670197 | 1.331783 | 0.503233 | 0.614801 | NA |
| | 1.137783 | 0.40046 | 1.978632 | 0.202392 | 0.83961 | NA |
| | 0.35636 | 1.709252 | 3.651833 | 0.468053 | 0.639746 | NA |
| | 0.572651 | 0.928649 | 2.780261 | 0.334015 | 0.738368 | NA |
| | 100.645 | 0.466335 | 0.323773 | 1.440316 | 0.149778 | 0.293243 |
| | 2.611914 | 0.304918 | 1.49116 | 0.204484 | 0.837975 | NA |
| GFY | 0.163957 | 0.424913 | 4.079006 | 0.104171 | 0.917034 | NA |
| ZNF234 | 150.5379 | 0.425813 | 0.217668 | 1.956256 | 0.050435 | 0.139405 |
| RN7SL821F | 0.163957 | 0.424913 | 4.079006 | 0.104171 | 0.917034 | NA |
| MIR378I | 0.163957 | 0.424913 | 4.079006 | 0.104171 | 0.917034 | NA |
| RN7SL533F | 0.151697 | -1.546 | 4.075469 | -0.37934 | 0.704433 | NA |
| MIR4777 | 1.312411 | -0.54592 | 1.909738 | -0.28586 | 0.774985 | NA |
| | 0.526843 | -0.38719 | 2.855892 | -0.13558 | 0.892156 | NA |
| MIR3197 | 1.847719 | -0.97377 | 1.415589 | -0.68789 | 0.49152 | NA |
| RN7SL4P | 0.96595 | -1.57133 | 2.099557 | -0.74841 | 0.454214 | NA |
| | 0.163957 | 0.424913 | 4.079006 | 0.104171 | 0.917034 | NA |
| MIR3679 | 0.134133 | -1.48909 | 4.077584 | -0.36519 | 0.71497 | NA |
| RN7SL678F | 0.151697 | -1.546 | 4.075469 | -0.37934 | 0.704433 | NA |
| MIR3687 | 23.74419 | 0.374012 | 0.498759 | 0.749885 | 0.453324 | NA |
| | 0.134133 | -1.48909 | 4.077584 | -0.36519 | 0.71497 | NA |
| MIR4442 | 1.775617 | -0.64845 | 1.543375 | -0.42015 | 0.674373 | NA |
| MIR4786 | 0.544554 | -2.65036 | 3.220086 | -0.82307 | 0.410468 | NA |
| MIR2467 | 0.163957 | 0.424913 | 4.079006 | 0.104171 | 0.917034 | NA |
| | 0.448206 | -1.2532 | 3.007315 | -0.41672 | 0.676885 | NA |
| RN7SL602F | 0.096202 | 0.408546 | 4.079639 | 0.100143 | 0.920231 | NA |
| MIR3125 | 4.912997 | -0.26298 | 0.882144 | -0.29811 | 0.765618 | NA |
| MIR3648 | 45.74526 | 0.845743 | 0.555112 | 1.523554 | 0.12762 | NA |
| | 0.151697 | -1.546 | 4.075469 | -0.37934 | 0.704433 | NA |
| MIR4534 | 1.265031 | 0.740117 | 1.715192 | 0.431507 | 0.6661 | NA |
| | 0.096202 | 0.408546 | 4.079639 | 0.100143 | 0.920231 | NA |
| | 0.151697 | -1.546 | 4.075469 | -0.37934 | 0.704433 | NA |
| RN7SL690F | 0.151697 | -1.546 | 4.075469 | -0.37934 | 0.704433 | NA |
| RN7SL525F | 0.268266 | -2.15396 | 4.038351 | -0.53338 | 0.593773 | NA |
| RN7SL701F | 0.162376 | 0.424913 | 4.079006 | 0.104171 | 0.917034 | NA |
| MIR4431 | 0.096202 | 0.408546 | 4.079639 | 0.100143 | 0.920231 | NA |
| MIR4765 | 0.163957 | 0.424913 | 4.079006 | 0.104171 | 0.917034 | NA |
| MIR4761 | 0.163957 | 0.424913 | 4.079006 | 0.104171 | 0.917034 | NA |
| MIR3192 | 0.096202 | 0.408546 | 4.079639 | 0.100143 | 0.920231 | NA |
| | 0.096202 | 0.408546 | 4.079639 | 0.100143 | 0.920231 | NA |
| MIR4263 | 1.576679 | -2.08983 | 1.715854 | -1.21795 | 0.223242 | NA |
| MIR3128 | 0.134133 | -1.48909 | 4.077584 | -0.36519 | 0.71497 | NA |
| MIR4782 | 0.162376 | 0.424913 | 4.079006 | 0.104171 | 0.917034 | NA |
| | 0.096202 | 0.408546 | 4.079639 | 0.100143 | 0.920231 | NA |
| | 13.03768 | 0.925137 | 0.736818 | 1.255585 | 0.209266 | NA |

| | | | | | | |
|---|---|---|---|---|---|---|
| MIR3682 | 0.364467 | -1.20166 | 3.685431 | -0.32606 | 0.744381 | NA |
| RN7SL243F | 0.268266 | -2.15396 | 4.038351 | -0.53338 | 0.593773 | NA |
| | 0.495797 | -0.73607 | 3.324282 | -0.22142 | 0.824763 | NA |
| | 0.096202 | 0.408546 | 4.079639 | 0.100143 | 0.920231 | NA |
| | 0.192403 | 0.975392 | 4.060716 | 0.240202 | 0.810174 | NA |
| RN7SL53P | 0.163957 | 0.424913 | 4.079006 | 0.104171 | 0.917034 | NA |
| ESRG | 0.919846 | -3.47044 | 2.735469 | -1.26868 | 0.204555 | NA |
| MIR4326 | 0.69625 | -3.13723 | 2.93845 | -1.06765 | 0.285679 | NA |
| MIR4440 | 0.151697 | -1.546 | 4.075469 | -0.37934 | 0.704433 | NA |
| RN7SL514F | 0.163957 | 0.424913 | 4.079006 | 0.104171 | 0.917034 | NA |
| RN7SL349F | 0.096202 | 0.408546 | 4.079639 | 0.100143 | 0.920231 | NA |
| RN7SL389F | 0.628146 | -0.21495 | 3.102079 | -0.06929 | 0.944756 | NA |
| | 0.163957 | 0.424913 | 4.079006 | 0.104171 | 0.917034 | NA |
| MIR4766 | 0.46735 | -1.30193 | 3.399671 | -0.38296 | 0.701751 | NA |
| | 0.192403 | 0.975392 | 4.060716 | 0.240202 | 0.810174 | NA |
| MIR4754 | 3.549271 | 0.069978 | 1.08178 | 0.064688 | 0.948423 | NA |
| MIR3135A | 0.151697 | -1.546 | 4.075469 | -0.37934 | 0.704433 | NA |
| MIR3194 | 1.116213 | -3.87733 | 2.172197 | -1.78498 | 0.074264 | NA |
| | 0.163957 | 0.424913 | 4.079006 | 0.104171 | 0.917034 | NA |
| | 0.623191 | -1.58103 | 2.770593 | -0.57065 | 0.568239 | NA |
| | 0.640406 | 1.11402 | 2.710977 | 0.410929 | 0.681124 | NA |
| | 0.134133 | -1.48909 | 4.077584 | -0.36519 | 0.71497 | NA |
| | 1.457581 | 0.387035 | 1.603931 | 0.241304 | 0.80932 | NA |
| | 33.64823 | -0.64858 | 0.453328 | -1.43072 | 0.152511 | NA |
| | 0.411855 | 0.250456 | 3.097618 | 0.080854 | 0.935558 | NA |
| | 0.258578 | 1.233151 | 4.047809 | 0.304646 | 0.760635 | NA |
| | 4.927625 | 0.666712 | 0.933969 | 0.713848 | 0.475321 | NA |
| | 0.298089 | -0.51711 | 3.975232 | -0.13008 | 0.896501 | NA |
| | 1.170556 | -0.4175 | 2.096325 | -0.19916 | 0.84214 | NA |
| | 0.556813 | -0.57007 | 3.294615 | -0.17303 | 0.862627 | NA |
| | 0.394291 | 0.298745 | 3.141711 | 0.09509 | 0.924243 | NA |
| | 2.131774 | 0.283052 | 1.512425 | 0.187151 | 0.851542 | NA |
| | 1.10954 | -0.31713 | 1.971981 | -0.16082 | 0.872235 | NA |
| | 0.392711 | 0.292538 | 3.145221 | 0.09301 | 0.925895 | NA |
| | 9.667163 | 0.084391 | 0.715011 | 0.118027 | 0.906046 | NA |
| | 4.818697 | 1.679054 | 1.022421 | 1.642234 | 0.100542 | NA |
| | 11.57075 | -1.13334 | 0.595206 | -1.90412 | 0.056894 | NA |
| | 0.624422 | 1.162588 | 2.733761 | 0.425271 | 0.670639 | NA |
| | 0.096202 | 0.408546 | 4.079639 | 0.100143 | 0.920231 | NA |
| EIF5AP3 | 0.288605 | 1.443619 | 4.029058 | 0.358302 | 0.720117 | NA |
| | 0.163957 | 0.424913 | 4.079006 | 0.104171 | 0.917034 | NA |
| | 4.490259 | -0.40596 | 0.922618 | -0.44001 | 0.659931 | NA |
| | 5.664818 | 0.135163 | 0.955107 | 0.141516 | 0.887463 | NA |
| | 0.545988 | -0.43913 | 2.541145 | -0.17281 | 0.862803 | NA |
| ZNF582-AS | 58.43549 | -0.0184 | 0.295772 | -0.0622 | 0.9504 | 0.972466 |
| | 0.163957 | 0.424913 | 4.079006 | 0.104171 | 0.917034 | NA |
| ZNF285 | 63.18521 | -0.00946 | 0.306815 | -0.03084 | 0.975398 | 0.986466 |

| | | | | | | |
|---|---|---|---|---|---|---|
| | 36.00373 | 0.633398 | 0.48735 | 1.299678 | 0.193711 | NA |
| | 4.808541 | 0.251617 | 0.885675 | 0.284096 | 0.776337 | NA |
| | 0.938845 | -1.23445 | 2.063177 | -0.59832 | 0.549624 | NA |
| | 2.485799 | 0.454375 | 1.396144 | 0.32545 | 0.744841 | NA |
| | 0.096202 | 0.408546 | 4.079639 | 0.100143 | 0.920231 | NA |
| | 4.08872 | -0.39114 | 1.046468 | -0.37377 | 0.708575 | NA |
| ZNF224 | 26.89289 | 1.242227 | 0.421058 | 2.95025 | 0.003175 | NA |
| | 1.878738 | 0.532302 | 1.445272 | 0.368305 | 0.712646 | NA |
| | 2.392712 | 0.970407 | 1.417933 | 0.684382 | 0.493734 | NA |
| | 0.640406 | 1.11402 | 2.710977 | 0.410929 | 0.681124 | NA |
| SLC25A36P | 0.162376 | 0.424913 | 4.079006 | 0.104171 | 0.917034 | NA |
| | 0.411855 | 0.250456 | 3.097618 | 0.080854 | 0.935558 | NA |
| NDUFV2P1 | 9.453555 | 0.601295 | 0.661797 | 0.90858 | 0.363572 | NA |
| | 0.411855 | 0.250456 | 3.097618 | 0.080854 | 0.935558 | NA |
| | 0.411855 | 0.250456 | 3.097618 | 0.080854 | 0.935558 | NA |
| | 0.163957 | 0.424913 | 4.079006 | 0.104171 | 0.917034 | NA |
| | 35.33172 | 0.113074 | 0.389669 | 0.290179 | 0.771679 | NA |
| | 1.993937 | 1.135122 | 1.450207 | 0.782731 | 0.433785 | NA |
| | 1.130695 | 2.130887 | 2.289427 | 0.930751 | 0.351982 | NA |
| | 2.110246 | 1.254374 | 1.426464 | 0.879359 | 0.379207 | NA |
| | 2.035818 | 3.11688 | 1.697119 | 1.836572 | 0.066273 | NA |
| ZSCAN5D | 0.151697 | -1.546 | 4.075469 | -0.37934 | 0.704433 | NA |
| | 0.402398 | -2.58771 | 4.018369 | -0.64397 | 0.519595 | NA |
| | 2.388374 | 0.65264 | 1.307415 | 0.499184 | 0.61765 | NA |
| | 0.315653 | -0.57305 | 3.892639 | -0.14721 | 0.882963 | NA |
| | 0.096202 | 0.408546 | 4.079639 | 0.100143 | 0.920231 | NA |
| | 54.6728 | 0.666515 | 0.328568 | 2.028546 | 0.042505 | 0.125471 |
| PTOV1-AS1 | 177.76 | -0.28908 | 0.213067 | -1.35677 | 0.174855 | 0.326835 |
| | 3.891951 | -1.563 | 1.038649 | -1.50484 | 0.132366 | NA |
| | 0.258578 | 1.233151 | 4.047809 | 0.304646 | 0.760635 | NA |
| | 1.044145 | 0.31036 | 2.099918 | 0.147796 | 0.882504 | NA |
| | 10.64079 | -0.48724 | 0.618431 | -0.78787 | 0.430772 | NA |
| NAPA-AS1 | 16.88403 | -0.22951 | 0.51654 | -0.44432 | 0.656808 | NA |
| | 2.544398 | -0.03055 | 1.41344 | -0.02161 | 0.982756 | NA |
| | 0.162376 | 0.424913 | 4.079006 | 0.104171 | 0.917034 | NA |
| | 0.298089 | -0.51711 | 3.975232 | -0.13008 | 0.896501 | NA |
| | 0.392857 | -1.546 | 4.075469 | -0.37934 | 0.704433 | NA |
| | 0.518736 | 2.237423 | 2.7713 | 0.807355 | 0.419462 | NA |
| | 0.717609 | 0.516981 | 3.063469 | 0.168757 | 0.865988 | NA |
| SMIM17 | 0.572651 | 0.928649 | 2.780261 | 0.334015 | 0.738368 | NA |
| | 0.192403 | 0.975392 | 4.060716 | 0.240202 | 0.810174 | NA |
| | 1.866157 | -0.27478 | 1.606737 | -0.17101 | 0.864212 | NA |
| | 88.20155 | 0.113382 | 0.309826 | 0.365956 | 0.714398 | 0.817726 |
| | 2.341586 | -0.0989 | 1.335614 | -0.07405 | 0.940973 | NA |
| | 32.49863 | 0.427797 | 0.431581 | 0.991232 | 0.321572 | NA |
| | 1.142073 | -0.81184 | 2.330623 | -0.34833 | 0.727589 | NA |
| | 3.877961 | 0.281994 | 1.159108 | 0.243286 | 0.807784 | NA |

| | | | | | | |
|---|---|---|---|---|---|---|
| | 1.87245 | 1.005198 | 1.659443 | 0.605744 | 0.544685 | NA |
| | 0.162376 | 0.424913 | 4.079006 | 0.104171 | 0.917034 | NA |
| | 0.392857 | -1.546 | 4.075469 | -0.37934 | 0.704433 | NA |
| | 0.296509 | -0.51709 | 3.982155 | -0.12985 | 0.896684 | NA |
| | 0.162376 | 0.424913 | 4.079006 | 0.104171 | 0.917034 | NA |
| | 0.489058 | -0.58569 | 3.427543 | -0.17088 | 0.864321 | NA |
| | 1.609066 | -1.39607 | 1.853511 | -0.7532 | 0.451329 | NA |
| | 2.010242 | -1.40707 | 1.41265 | -0.99605 | 0.319225 | NA |
| | 0.314073 | -0.57308 | 3.898934 | -0.14698 | 0.883146 | NA |
| | 0.162376 | 0.424913 | 4.079006 | 0.104171 | 0.917034 | NA |
| | 0.163957 | 0.424913 | 4.079006 | 0.104171 | 0.917034 | NA |
| | 1.092967 | 2.10982 | 2.003899 | 1.052857 | 0.292406 | NA |
| | 1.716398 | 0.300458 | 1.545048 | 0.194465 | 0.845812 | NA |
| | 0.545988 | -0.43913 | 2.541145 | -0.17281 | 0.862803 | NA |
| RPL7AP69 | 0.096202 | 0.408546 | 4.079639 | 0.100143 | 0.920231 | NA |
| | 0.247898 | -0.5901 | 4.074633 | -0.14482 | 0.884851 | NA |
| | 86.78446 | -0.17586 | 0.302265 | -0.58182 | 0.560689 | 0.698945 |
| | 0.599903 | -1.73313 | 2.730982 | -0.63462 | 0.525678 | NA |
| | 0.793886 | -0.43778 | 2.212663 | -0.19785 | 0.84316 | NA |
| | 0.28583 | -2.21358 | 3.97376 | -0.55705 | 0.577494 | NA |
| | 0.151697 | -1.546 | 4.075469 | -0.37934 | 0.704433 | NA |
| | 8.580253 | 0.450086 | 0.681441 | 0.660491 | 0.508939 | NA |
| VN1R107P | 0.315653 | -0.57305 | 3.892639 | -0.14721 | 0.882963 | NA |
| | 16.62088 | -0.56572 | 0.500714 | -1.12983 | 0.258548 | NA |
| | 0.738391 | 0.402625 | 2.276191 | 0.176885 | 0.859599 | NA |
| | 0.162376 | 0.424913 | 4.079006 | 0.104171 | 0.917034 | NA |
| | 1.774778 | 0.157681 | 1.859216 | 0.08481 | 0.932412 | NA |
| | 1.943515 | -0.00541 | 1.550248 | -0.00349 | 0.997218 | NA |
| | 0.873864 | 1.205392 | 2.855153 | 0.422181 | 0.672893 | NA |
| | 0.163957 | 0.424913 | 4.079006 | 0.104171 | 0.917034 | NA |
| MIR296 | 0.845418 | 1.095819 | 2.535106 | 0.432258 | 0.665554 | NA |
| | 6.823343 | -0.42044 | 0.825844 | -0.5091 | 0.610683 | NA |
| | 0.326333 | 1.533889 | 3.826878 | 0.40082 | 0.688553 | NA |
| | 1.866275 | 0.432622 | 1.441196 | 0.300183 | 0.764038 | NA |
| | 0.163957 | 0.424913 | 4.079006 | 0.104171 | 0.917034 | NA |
| | 1.239563 | 1.504644 | 1.913282 | 0.786421 | 0.431621 | NA |
| | 12.41695 | 0.115433 | 0.628342 | 0.18371 | 0.854241 | NA |
| | 0.134133 | -1.48909 | 4.077584 | -0.36519 | 0.71497 | NA |
| | 0.354779 | 1.703954 | 3.657232 | 0.465914 | 0.641277 | NA |
| | 0.260158 | 1.240347 | 4.047124 | 0.306476 | 0.759242 | NA |
| | 0.422534 | 1.924583 | 3.001758 | 0.641152 | 0.521424 | NA |
| | 14.04323 | 0.067879 | 0.570255 | 0.119033 | 0.905249 | NA |
| | 0.582339 | -1.69412 | 2.750801 | -0.61586 | 0.537985 | NA |
| | 0.096202 | 0.408546 | 4.079639 | 0.100143 | 0.920231 | NA |
| SIGLEC22P | 0.260158 | 1.240347 | 4.047124 | 0.306476 | 0.759242 | NA |
| | 6.346297 | -0.53098 | 0.778523 | -0.68204 | 0.495213 | NA |
| IGFL1P1 | 0.096202 | 0.408546 | 4.079639 | 0.100143 | 0.920231 | NA |

| | | | | | | |
|---|---|---|---|---|---|---|
| A1BG-AS1 | 119.8421 | -0.14459 | 0.238577 | -0.60606 | 0.544475 | 0.686238 |
| | 28.94988 | -0.25517 | 0.405209 | -0.62972 | 0.528875 | NA |
| | 2.075104 | -0.04243 | 1.404124 | -0.03022 | 0.975893 | NA |
| | 5.454979 | 0.617273 | 0.944818 | 0.653325 | 0.513547 | NA |
| ZNF818P | 2.093688 | -0.03423 | 1.403269 | -0.02439 | 0.980541 | NA |
| | 0.134133 | -1.48909 | 4.077584 | -0.36519 | 0.71497 | NA |
| | 4.283963 | -0.34721 | 1.04811 | -0.33127 | 0.740438 | NA |
| | 4.622537 | -1.01428 | 0.999887 | -1.01439 | 0.310396 | NA |
| | 1.60219 | 1.363436 | 1.739646 | 0.783743 | 0.433191 | NA |
| | 0.490493 | 0.776209 | 2.895261 | 0.268096 | 0.788625 | NA |
| | 0.096202 | 0.408546 | 4.079639 | 0.100143 | 0.920231 | NA |
| | 0.584911 | 2.38756 | 2.690807 | 0.887303 | 0.374916 | NA |
| | 0.327913 | 1.543154 | 4.021063 | 0.383768 | 0.701151 | NA |
| | 1.49965 | -0.07978 | 2.07973 | -0.03836 | 0.969401 | NA |
| | 4.587117 | -0.41298 | 0.967476 | -0.42686 | 0.669479 | NA |
| | 2.070791 | 1.716585 | 1.470558 | 1.167302 | 0.243088 | NA |
| | 7.734557 | -0.07463 | 0.739878 | -0.10087 | 0.919654 | NA |
| LINC00528 | 0.324752 | 1.53082 | 4.022025 | 0.380609 | 0.703493 | NA |
| HCCAT3 | 0.853176 | 0.733892 | 2.189324 | 0.335214 | 0.737464 | NA |
| | 1.475145 | 0.336818 | 1.696708 | 0.198512 | 0.842644 | NA |
| ZNF587B | 146.0861 | -0.18047 | 0.214274 | -0.84223 | 0.399661 | 0.564314 |
| | 1.784716 | -0.14025 | 1.577434 | -0.08891 | 0.929155 | NA |
| | 10.77733 | 0.319277 | 0.638566 | 0.49999 | 0.617082 | NA |
| | 1.107959 | -0.3351 | 1.823092 | -0.18381 | 0.854166 | NA |
| | 58.43228 | -0.47478 | 0.357097 | -1.32957 | 0.183661 | 0.336932 |
| | 0.966942 | 0.917791 | 1.915974 | 0.47902 | 0.631924 | NA |
| | 0.326333 | 1.533889 | 3.826878 | 0.40082 | 0.688553 | NA |
| SPIB | 1.192047 | -2.11452 | 2.271965 | -0.9307 | 0.352008 | NA |
| | 0.3441 | -0.0166 | 3.769682 | -0.0044 | 0.996486 | NA |
| NPCDR1 | 0.163957 | 0.424913 | 4.079006 | 0.104171 | 0.917034 | NA |
| | 1.1119 | 0.46633 | 1.933926 | 0.241131 | 0.809453 | NA |
| | 0.151697 | -1.546 | 4.075469 | -0.37934 | 0.704433 | NA |
| LRRC3DN | 76.30696 | -0.56726 | 0.529017 | -1.0723 | 0.283586 | 0.450055 |
| | 5.000827 | 0.082522 | 0.903795 | 0.091306 | 0.927249 | NA |
| | 0.35636 | 1.709252 | 3.651833 | 0.468053 | 0.639746 | NA |
| | 2.431407 | 0.225417 | 1.273728 | 0.176975 | 0.859528 | NA |
| | 0.97663 | -0.60197 | 1.931737 | -0.31162 | 0.755327 | NA |
| | 0.545988 | -0.43913 | 2.541145 | -0.17281 | 0.862803 | NA |
| | 0.151697 | -1.546 | 4.075469 | -0.37934 | 0.704433 | NA |
| | 0.230334 | -0.53318 | 4.076749 | -0.13079 | 0.895945 | NA |
| | 0.151697 | -1.546 | 4.075469 | -0.37934 | 0.704433 | NA |
| RPL12P41 | 0.260158 | 1.240347 | 4.047124 | 0.306476 | 0.759242 | NA |
| | 0.489058 | -0.58569 | 3.427543 | -0.17088 | 0.864321 | NA |
| | 7.635363 | -0.60015 | 0.719059 | -0.83463 | 0.403924 | NA |
| | 12.07104 | 0.103175 | 0.579153 | 0.178147 | 0.858607 | NA |
| | 0.134133 | -1.48909 | 4.077584 | -0.36519 | 0.71497 | NA |
| | 2.880779 | -0.3917 | 1.222068 | -0.32052 | 0.748572 | NA |

| | | | | | | |
|---|---|---|---|---|---|---|
| | 2.92619 | -5.69E-05 | 1.276199 | -4.46E-05 | 0.999964 | NA |
| | 33.49778 | 0.819219 | 0.44472 | 1.842101 | 0.06546 | NA |
| | 37.79193 | 0.163843 | 0.400018 | 0.409589 | 0.682107 | NA |
| MIR519A2 | 0.134133 | -1.48909 | 4.077584 | -0.36519 | 0.71497 | NA |
| | 0.392711 | 0.292538 | 3.145221 | 0.09301 | 0.925895 | NA |
| | 0.230334 | -0.53318 | 4.076749 | -0.13079 | 0.895945 | NA |
| | 34.61105 | -0.14691 | 0.403599 | -0.36401 | 0.71585 | NA |
| | 0.134133 | -1.48909 | 4.077584 | -0.36519 | 0.71497 | NA |
| FUCA2 | 1586.446 | -0.13482 | 0.142019 | -0.94929 | 0.342475 | 0.510698 |
| GCLC | 919.2599 | 0.329031 | 0.15258 | 2.156454 | 0.031048 | 0.101538 |
| NFYA | 767.7125 | -0.02343 | 0.136885 | -0.17113 | 0.864122 | 0.919997 |
| ENPP4 | 1.249048 | 0.800669 | 1.919369 | 0.417152 | 0.676567 | NA |
| LAP3 | 3115.041 | -0.17221 | 0.163869 | -1.05093 | 0.293291 | 0.460403 |
| HS3ST1 | 1.937191 | 0.501769 | 1.44322 | 0.347673 | 0.728086 | NA |
| PLXND1 | 2906.194 | -0.33705 | 0.195101 | -1.72755 | 0.08407 | 0.197553 |
| CD38 | 0.860207 | -2.14053 | 2.512764 | -0.85186 | 0.394291 | NA |
| SLC22A16 | 0.151697 | -1.546 | 4.075469 | -0.37934 | 0.704433 | NA |
| CCDC109B | 142.1583 | 0.067873 | 0.223787 | 0.303291 | 0.761668 | 0.851908 |
| IBTK | 900.8505 | 0.55804 | 0.29781 | 1.873812 | 0.060956 | 0.158996 |
| CDKL3 | 37.98882 | 0.711924 | 0.389774 | 1.826506 | 0.067774 | NA |
| PROM1 | 26.61304 | 0.096155 | 0.450577 | 0.213403 | 0.831012 | NA |
| TEAD3 | 588.3462 | -0.31426 | 0.153541 | -2.04674 | 0.040683 | 0.121674 |
| MYLIP | 575.2741 | -0.05042 | 0.148796 | -0.33887 | 0.734706 | 0.832199 |
| JARID2 | 1110.263 | -0.3082 | 0.140052 | -2.20062 | 0.027763 | 0.094307 |
| TFAP2B | 3.18775 | -0.56873 | 1.217754 | -0.46704 | 0.640474 | NA |
| TFAP2D | 0.096202 | 0.408546 | 4.079639 | 0.100143 | 0.920231 | NA |
| SEC62 | 1630.358 | 0.93749 | 0.580965 | 1.613678 | 0.106597 | 0.232765 |
| REV3L | 804.2154 | 0.274545 | 0.59646 | 0.460291 | 0.645308 | 0.764324 |
| VTA1 | 998.2584 | 0.580346 | 0.200247 | 2.89815 | 0.003754 | 0.024478 |
| RANBP9 | 1322.655 | 0.217577 | 0.135444 | 1.606398 | 0.108187 | 0.234942 |
| ETV7 | 61.0803 | -0.8523 | 0.357728 | -2.38252 | 0.017194 | 0.068328 |
| HFE | 407.0791 | -0.20884 | 0.160055 | -1.30481 | 0.191959 | 0.346567 |
| FYN | 3473.809 | -0.10692 | 0.144269 | -0.7411 | 0.458635 | 0.617083 |
| HIVEP2 | 732.4387 | 0.177223 | 0.153716 | 1.152925 | 0.248941 | 0.412801 |
| ELOVL5 | 2780.551 | 0.016648 | 0.120982 | 0.137611 | 0.890548 | 0.936036 |
| MAN2B2 | 2173.387 | -0.45628 | 0.133816 | -3.40974 | 6.50E-04 | 0.007923 |
| SLC7A14 | 124.1301 | 2.153344 | 1.102251 | 1.953589 | 0.05075 | 0.140047 |
| CLDN11 | 282.0927 | 0.656305 | 1.166863 | 0.562452 | 0.573808 | 0.709406 |
| PGM3 | 730.9543 | 0.561533 | 0.185694 | 3.02397 | 0.002495 | 0.018778 |
| RWDD2A | 208.6721 | 0.28603 | 0.193468 | 1.478431 | 0.139292 | 0.279268 |
| RNF14 | 2611.175 | 0.089508 | 0.13346 | 0.670673 | 0.502429 | 0.652458 |
| TACC3 | 972.4209 | -0.56682 | 0.234996 | -2.41205 | 0.015863 | 0.064508 |
| UFL1 | 460.5 | 0.768706 | 0.610343 | 1.259464 | 0.207863 | 0.365177 |
| ACPP | 5.998947 | -0.29473 | 0.857545 | -0.34369 | 0.731082 | NA |
| SLC30A9 | 1844.291 | 0.506549 | 0.209163 | 2.421796 | 0.015444 | 0.063199 |
| MATR3 | 4158.843 | 0.622988 | 0.283948 | 2.194024 | 0.028234 | 0.095223 |
| ATP2C1 | 4092.518 | 0.161097 | 0.134733 | 1.195679 | 0.231822 | 0.394183 |

| | | | | | | |
|---|---|---|---|---|---|---|
| RUFY3 | 652.4729 | 0.322477 | 0.288645 | 1.117212 | 0.263904 | 0.428396 |
| WWTR1 | 2620.833 | 0.167393 | 0.229172 | 0.730426 | 0.46513 | 0.622571 |
| CD74 | 26.07483 | -0.16296 | 0.454817 | -0.3583 | 0.72012 | NA |
| SERPINB1 | 627.1426 | 0.037533 | 0.154153 | 0.243481 | 0.807633 | 0.885215 |
| DEF6 | 49.3895 | 0.188144 | 0.383385 | 0.490743 | 0.623608 | 0.747737 |
| UBR2 | 1019.969 | 0.207569 | 0.187406 | 1.107592 | 0.268038 | 0.433032 |
| CCDC28A | 497.749 | 0.106571 | 0.162371 | 0.656343 | 0.511604 | 0.659389 |
| RRAGD | 3556.979 | -0.00515 | 0.119021 | -0.04324 | 0.96551 | 0.981198 |
| HSF2 | 527.9416 | 0.501297 | 0.174061 | 2.880008 | 0.003977 | 0.025373 |
| SEC63 | 1395.195 | 0.393645 | 0.229602 | 1.714469 | 0.086443 | 0.201508 |
| BTN3A1 | 505 | -0.50302 | 0.230097 | -2.1861 | 0.028808 | 0.096427 |
| IFNGR1 | 1262.981 | 0.619108 | 0.149709 | 4.135404 | 3.54E-05 | 0.001346 |
| B4GALT7 | 750.5233 | -0.21338 | 0.140348 | -1.52033 | 0.128427 | 0.264779 |
| BRD9 | 875.4039 | 0.050048 | 0.132064 | 0.378965 | 0.704714 | 0.810633 |
| TBPL1 | 337.0992 | 0.309851 | 0.172436 | 1.796908 | 0.07235 | 0.178764 |
| BCLAF1 | 1922.449 | 0.585274 | 0.327078 | 1.789401 | 0.07355 | 0.180809 |
| IBSP | 0.518939 | 0.925178 | 3.271397 | 0.282808 | 0.777324 | NA |
| BAK1 | 536.5866 | -0.34539 | 0.173438 | -1.99143 | 0.046434 | 0.132422 |
| FAM13B | 1209.314 | 0.48757 | 0.187803 | 2.596186 | 0.009427 | 0.04539 |
| ARHGAP31 | 2785.989 | -0.57088 | 0.149484 | -3.81903 | 1.34E-04 | 0.003043 |
| CENPQ | 179.3642 | 1.28651 | 0.236972 | 5.42895 | 5.67E-08 | 2.35E-05 |
| UBA6 | 693.8851 | 0.782891 | 0.269823 | 2.901502 | 0.003714 | 0.02432 |
| ASTE1 | 117.8967 | 0.175947 | 0.24546 | 0.716806 | 0.473494 | 0.629168 |
| PEX3 | 286.151 | 0.246729 | 0.166002 | 1.4863 | 0.1372 | 0.276381 |
| DEPDC1B | 304.0393 | 0.369141 | 0.229107 | 1.61122 | 0.107132 | 0.233594 |
| RFC1 | 798.5903 | 0.765115 | 0.315292 | 2.426688 | 0.015237 | 0.062765 |
| TBC1D23 | 860.6802 | 0.979007 | 0.22692 | 4.314332 | 1.60E-05 | 8.22E-04 |
| RPL26L1 | 529.9437 | 0.417423 | 0.201232 | 2.074332 | 0.038048 | 0.116164 |
| FLT4 | 0.324752 | 1.53082 | 4.022025 | 0.380609 | 0.703493 | NA |
| NSUN2 | 1891.971 | 0.197661 | 0.123801 | 1.596606 | 0.110354 | 0.238252 |
| MFAP3 | 878.6442 | 0.100415 | 0.186278 | 0.539058 | 0.589847 | 0.721858 |
| AGA | 724.3791 | 0.197031 | 0.196617 | 1.002106 | 0.316292 | 0.483763 |
| PI4K2B | 501.2562 | 0.680378 | 0.208349 | 3.265566 | 0.001092 | 0.010985 |
| BOD1L1 | 790.195 | 0.777378 | 1.038538 | 0.748531 | 0.45414 | 0.612913 |
| MAT2B | 1433.671 | 0.518385 | 0.187962 | 2.757927 | 0.005817 | 0.032729 |
| TRIO | 3855.6 | -0.10108 | 0.173468 | -0.58269 | 0.560101 | 0.698472 |
| VCAN | 365.9386 | -0.60586 | 0.401111 | -1.51045 | 0.130928 | 0.267383 |
| SKIV2L2 | 1025.476 | 0.629876 | 0.263711 | 2.388513 | 0.016917 | 0.067684 |
| DNAH5 | 1.401736 | 2.557161 | 1.873156 | 1.365162 | 0.172202 | NA |
| ZFYVE16 | 4678.843 | 0.932369 | 0.334758 | 2.785202 | 0.005349 | 0.031041 |
| RAI14 | 2651.829 | 0.571682 | 0.230415 | 2.481098 | 0.013098 | 0.056711 |
| SOX30 | 5.325266 | 0.010376 | 0.914028 | 0.011352 | 0.990943 | NA |
| SPDL1 | 171.5345 | 1.205413 | 0.391624 | 3.077988 | 0.002084 | 0.016597 |
| CDH10 | 0.163957 | 0.424913 | 4.079006 | 0.104171 | 0.917034 | NA |
| LSG1 | 1192.253 | 0.428195 | 0.133933 | 3.197079 | 0.001388 | 0.012868 |
| DCUN1D1 | 432.8935 | 0.840725 | 0.289767 | 2.901381 | 0.003715 | 0.02432 |
| JADE2 | 3010.682 | -0.31125 | 0.305828 | -1.01774 | 0.3088 | 0.476128 |

| | | | | | | |
|---|---|---|---|---|---|---|
| LCP2 | 1.600812 | 0.66716 | 1.651524 | 0.403966 | 0.686238 | NA |
| CUL7 | 1275.296 | -0.59285 | 0.149718 | -3.95978 | 7.50E-05 | 0.002133 |
| CTNNA1 | 18178.74 | 0.079017 | 0.129837 | 0.608592 | 0.542795 | 0.684848 |
| EPHA3 | 4.586454 | -0.27843 | 1.046999 | -0.26594 | 0.790289 | NA |
| YTHDC2 | 543.6999 | 0.781823 | 0.236198 | 3.310036 | 9.33E-04 | 0.00992 |
| POLR2B | 3587.335 | 0.371125 | 0.154888 | 2.396095 | 0.016571 | 0.066637 |
| ARAP2 | 88.58401 | 0.798905 | 0.423535 | 1.886278 | 0.059258 | 0.155834 |
| CP | 0.35636 | 1.709252 | 3.651833 | 0.468053 | 0.639746 | NA |
| DTNBP1 | 595.139 | -0.16697 | 0.148721 | -1.12268 | 0.261575 | 0.426022 |
| FAM184B | 142.3985 | 0.374252 | 0.212891 | 1.75795 | 0.078756 | 0.189267 |
| GOPC | 744.592 | 0.630692 | 0.202425 | 3.115687 | 0.001835 | 0.015242 |
| TSPAN17 | 2650.354 | -0.36387 | 0.127645 | -2.85063 | 0.004363 | 0.026927 |
| NOP16 | 472.4544 | 0.461252 | 0.180071 | 2.561503 | 0.010422 | 0.048395 |
| CC2D2A | 1224.217 | 0.12106 | 0.153936 | 0.786434 | 0.431613 | 0.592892 |
| MRPS10 | 1113.009 | 0.155652 | 0.151166 | 1.029673 | 0.303164 | 0.47046 |
| GUCA1A | 4.858375 | 1.300778 | 0.994027 | 1.308594 | 0.190672 | NA |
| ERCC8 | 235.7273 | 0.274506 | 0.209446 | 1.310628 | 0.189984 | 0.344236 |
| ADAMTS6 | 4.434857 | 1.079029 | 1.095704 | 0.984781 | 0.324731 | NA |
| CLPTM1L | 3936.355 | -0.11019 | 0.151388 | -0.72784 | 0.466712 | 0.623704 |
| HEXB | 6568.79 | 0.089985 | 0.145079 | 0.620251 | 0.535093 | 0.678307 |
| PTCD2 | 260.7526 | -0.08641 | 0.187981 | -0.45968 | 0.645743 | 0.764681 |
| MCUR1 | 712.1722 | 0.357064 | 0.145726 | 2.450239 | 0.014276 | 0.060039 |
| TNIP3 | 0.162376 | 0.424913 | 4.079006 | 0.104171 | 0.917034 | NA |
| MAPK9 | 1078.419 | 0.303497 | 0.153354 | 1.979067 | 0.047808 | 0.134794 |
| POLQ | 162.277 | 0.989493 | 0.286158 | 3.457863 | 5.44E-04 | 0.007109 |
| PIK3CB | 1476.907 | -0.10547 | 0.191252 | -0.55146 | 0.581321 | 0.714966 |
| THOC3 | 304.5649 | 0.004144 | 0.185107 | 0.02239 | 0.982137 | 0.990311 |
| HEBP2 | 612.4116 | 0.113345 | 0.170258 | 0.665727 | 0.505586 | 0.654811 |
| FNIP2 | 2355.183 | 0.382191 | 0.153405 | 2.49138 | 0.012725 | 0.055583 |
| MSMO1 | 3213.454 | 0.251487 | 0.183459 | 1.370803 | 0.170436 | 0.32082 |
| FSTL4 | 4.422597 | -0.32034 | 0.934232 | -0.3429 | 0.731677 | NA |
| METTL24 | 0.151697 | -1.546 | 4.075469 | -0.37934 | 0.704433 | NA |
| MCF2L2 | 25.81497 | 0.160874 | 0.419172 | 0.383789 | 0.701135 | NA |
| ANAPC4 | 496.8055 | 0.526259 | 0.151789 | 3.46703 | 5.26E-04 | 0.006994 |
| FOXC1 | 64.61663 | 0.046338 | 0.296175 | 0.156454 | 0.875675 | 0.925771 |
| FAM114A2 | 615.3801 | 0.29402 | 0.181537 | 1.619612 | 0.105316 | 0.231078 |
| CYFIP2 | 302.9481 | 0.00176 | 0.23466 | 0.007499 | 0.994017 | 0.997382 |
| TAB2 | 1849.174 | 0.013333 | 0.131678 | 0.101253 | 0.919349 | 0.953283 |
| GINM1 | 734.0359 | 0.171179 | 0.149235 | 1.147044 | 0.251363 | 0.415319 |
| C4orf27 | 432.1287 | 0.630868 | 0.188553 | 3.345842 | 8.20E-04 | 0.009177 |
| ZFR | 3144.131 | 0.324227 | 0.159075 | 2.038206 | 0.041529 | 0.123608 |
| NPFFR2 | 157.8919 | -0.71576 | 0.288225 | -2.48333 | 0.013016 | 0.056459 |
| TRAF3IP2 | 1312.369 | -0.38141 | 0.18783 | -2.03063 | 0.042292 | 0.125119 |
| DCBLD2 | 2130.939 | 0.242274 | 0.160725 | 1.507377 | 0.131714 | 0.268537 |
| PRDM1 | 9.050699 | 0.041624 | 0.667472 | 0.062361 | 0.950276 | NA |
| ATG5 | 631.4566 | 0.737735 | 0.22506 | 3.277955 | 0.001046 | 0.010649 |
| USP13 | 824.8244 | -0.08215 | 0.204902 | -0.40092 | 0.688481 | 0.797454 |

| | | | | | | |
|---|---|---|---|---|---|---|
| ATP11B | 1362.286 | 0.703575 | 0.219092 | 3.211321 | 0.001321 | 0.012431 |
| SEC61A1 | 8820.335 | -0.38385 | 0.155446 | -2.46934 | 0.013536 | 0.05789 |
| RIOK2 | 331.0769 | 0.858534 | 0.224661 | 3.821461 | 1.33E-04 | 0.003029 |
| DGKG | 13.75978 | -0.65837 | 0.636324 | -1.03465 | 0.300831 | NA |
| PET112 | 885.3892 | -0.10174 | 0.132066 | -0.77034 | 0.441096 | 0.601277 |
| RPS17P5 | 0.327913 | 1.543154 | 4.021063 | 0.383768 | 0.701151 | NA |
| PRDM6 | 1.855741 | -0.82082 | 1.601184 | -0.51263 | 0.608209 | NA |
| GUCY1B3 | 115.8765 | 0.731447 | 0.305936 | 2.390851 | 0.016809 | 0.067346 |
| TNK2 | 1416.162 | -0.27119 | 0.197222 | -1.37505 | 0.169116 | 0.319451 |
| GPBP1 | 843.1175 | 0.711527 | 0.212539 | 3.347755 | 8.15E-04 | 0.009155 |
| AHRR | 2323.793 | -0.3056 | 0.168527 | -1.81337 | 0.069775 | 0.174038 |
| RNF4 | 2115.731 | -0.21687 | 0.142536 | -1.52153 | 0.128126 | 0.264346 |
| LIMCH1 | 65.50675 | 1.360056 | 0.360285 | 3.774942 | 1.60E-04 | 0.003412 |
| ST3GAL6 | 1538.312 | 0.423541 | 0.158656 | 2.669553 | 0.007595 | 0.03937 |
| SLC12A2 | 659.4911 | 0.259239 | 0.250979 | 1.032912 | 0.301645 | 0.468492 |
| SNX24 | 395.3558 | 0.510738 | 0.207375 | 2.462871 | 0.013783 | 0.058544 |
| SNCAIP | 0.096202 | 0.408546 | 4.079639 | 0.100143 | 0.920231 | NA |
| POU1F1 | 0.162376 | 0.424913 | 4.079006 | 0.104171 | 0.917034 | NA |
| TAF11 | 565.1711 | 0.212784 | 0.157812 | 1.348338 | 0.17755 | 0.329811 |
| ANKS1A | 3451.803 | -0.2753 | 0.187601 | -1.46747 | 0.142248 | 0.283134 |
| ZNF76 | 771.8313 | -0.18398 | 0.144373 | -1.27431 | 0.202553 | 0.358825 |
| UHRF1BP1 | 1000.525 | -0.32711 | 0.153554 | -2.13025 | 0.033151 | 0.106017 |
| TRAM2 | 3459.809 | -0.45104 | 0.148329 | -3.04081 | 0.002359 | 0.017993 |
| ROPN1 | 445.8781 | -0.18138 | 0.195641 | -0.9271 | 0.353877 | 0.521857 |
| PDIA5 | 623.7357 | -0.08305 | 0.161181 | -0.51525 | 0.606382 | 0.734435 |
| TBC1D22B | 467.561 | -0.0406 | 0.168386 | -0.24111 | 0.80947 | 0.886082 |
| NDUFB4 | 3219.098 | -0.02028 | 0.1662 | -0.12203 | 0.902871 | 0.943218 |
| MYLK | 594.8607 | -0.35473 | 0.226429 | -1.56662 | 0.117204 | 0.247806 |
| SNAP91 | 0.163957 | 0.424913 | 4.079006 | 0.104171 | 0.917034 | NA |
| CYB5R4 | 425.7091 | 0.571257 | 0.202593 | 2.819723 | 0.004807 | 0.028764 |
| ME1 | 1064.441 | 0.450467 | 0.145623 | 3.093381 | 0.001979 | 0.016077 |
| TBC1D1 | 2182.341 | 0.143672 | 0.222478 | 0.64578 | 0.518422 | 0.665428 |
| SLC9A3 | 35.01134 | -0.06123 | 0.451977 | -0.13548 | 0.892232 | NA |
| ZNRD1 | 194.2801 | -0.14486 | 0.188071 | -0.77024 | 0.441157 | 0.601277 |
| CLDN18 | 7.444634 | 1.245235 | 0.81121 | 1.535034 | 0.124775 | NA |
| ZBTB11 | 396.4308 | 0.663941 | 0.204485 | 3.246899 | 0.001167 | 0.011459 |
| ISOC1 | 185.7115 | 0.111176 | 0.20581 | 0.540188 | 0.589068 | 0.721298 |
| TRMT11 | 182.738 | 0.948006 | 0.21709 | 4.366881 | 1.26E-05 | 7.08E-04 |
| PPAP2A | 989.7099 | 0.459527 | 0.265714 | 1.729403 | 0.083737 | 0.197146 |
| DHX29 | 549.3833 | 0.797955 | 0.298306 | 2.674955 | 0.007474 | 0.038996 |
| FGFR3 | 4.081179 | -0.03727 | 0.990527 | -0.03763 | 0.969983 | NA |
| KIF2A | 562.622 | 0.635773 | 0.229546 | 2.76969 | 0.005611 | 0.032054 |
| IFT80 | 161.7394 | 1.200579 | 0.37089 | 3.237022 | 0.001208 | 0.011766 |
| PITX1 | 0.787148 | -0.70094 | 2.357021 | -0.29738 | 0.766174 | NA |
| MAST4 | 140.8988 | -0.67258 | 0.25455 | -2.64222 | 0.008236 | 0.041392 |
| GPR116 | 0.314073 | -0.57308 | 3.898934 | -0.14698 | 0.883146 | NA |
| ATP1B3 | 7753.813 | 0.082825 | 0.202138 | 0.409746 | 0.681992 | 0.792882 |

| | | | | | | |
|---|---|---|---|---|---|---|
| PFN2 | 4963.938 | -0.36529 | 0.307136 | -1.18933 | 0.234309 | 0.396876 |
| DAPP1 | 0.096202 | 0.408546 | 4.079639 | 0.100143 | 0.920231 | NA |
| ZXDC | 424.255 | -0.16031 | 0.189853 | -0.84439 | 0.398454 | 0.563281 |
| NDST1 | 6105.354 | -0.51185 | 0.143539 | -3.56595 | 3.63E-04 | 0.005542 |
| CAMK2A | 12.71722 | -0.34066 | 0.638378 | -0.53363 | 0.593595 | NA |
| TCOF1 | 1656.269 | -0.27185 | 0.171841 | -1.582 | 0.11365 | 0.242586 |
| WDR1 | 17266.34 | -0.17671 | 0.141811 | -1.24609 | 0.212731 | 0.371473 |
| ARHGAP10 | 472.4795 | 0.289343 | 0.15102 | 1.915922 | 0.055375 | 0.148657 |
| TRIP13 | 644.7418 | -0.13495 | 0.238044 | -0.56692 | 0.570765 | 0.706602 |
| HLTF | 873.5427 | 0.781927 | 0.318054 | 2.45847 | 0.013953 | 0.059055 |
| LNX1 | 6.916809 | 0.04757 | 0.931729 | 0.051055 | 0.959282 | NA |
| TFRC | 3227.072 | 0.175066 | 0.210237 | 0.832709 | 0.405009 | 0.569272 |
| AFF4 | 3430.738 | 0.244635 | 0.166735 | 1.467209 | 0.142319 | 0.283134 |
| HMMR | 234.1803 | 1.117189 | 0.296387 | 3.769359 | 1.64E-04 | 0.003461 |
| P4HA2 | 1537.829 | -0.1343 | 0.195368 | -0.6874 | 0.491832 | 0.643353 |
| STK10 | 4756.262 | -0.40548 | 0.130484 | -3.10753 | 0.001887 | 0.015523 |
| FBXW11 | 2458.402 | -0.0327 | 0.166295 | -0.19663 | 0.844117 | 0.907349 |
| CRMP1 | 207.8506 | -0.42722 | 0.212043 | -2.01476 | 0.04393 | 0.127739 |
| EVC | 2902.791 | -0.62674 | 0.152787 | -4.10205 | 4.10E-05 | 0.001483 |
| SIDT1 | 29.99277 | -0.71784 | 0.462907 | -1.55072 | 0.120968 | NA |
| MCM2 | 981.4633 | -0.4871 | 0.227864 | -2.13769 | 0.032542 | 0.105005 |
| TP63 | 0.754374 | 0.344906 | 2.513743 | 0.137208 | 0.890866 | NA |
| ALPK1 | 477.1858 | -0.12765 | 0.17367 | -0.735 | 0.462339 | 0.620395 |
| SDHA | 4295.652 | -0.26868 | 0.13815 | -1.94486 | 0.051791 | 0.141718 |
| PPP2R3A | 796.3116 | 0.60062 | 0.186466 | 3.221075 | 0.001277 | 0.012122 |
| IGF2BP2 | 1241.116 | -0.0672 | 0.171554 | -0.39173 | 0.695257 | 0.803408 |
| MAP3K13 | 776.6438 | 0.265641 | 0.141689 | 1.874817 | 0.060818 | 0.158775 |
| ST6GAL1 | 3101.85 | -0.29705 | 0.147291 | -2.01674 | 0.043723 | 0.127417 |
| PPP2R2C | 0.651288 | 1.289307 | 2.633717 | 0.489539 | 0.62446 | NA |
| CDHR2 | 12.86336 | -0.50989 | 0.58696 | -0.86869 | 0.385016 | NA |
| SNCB | 45.87398 | -1.23936 | 0.571702 | -2.16783 | 0.030171 | NA |
| MGLL | 213.6793 | 0.08809 | 0.886399 | 0.099379 | 0.920837 | 0.953991 |
| TUBE1 | 180.4881 | 0.648099 | 0.202446 | 3.201344 | 0.001368 | 0.012729 |
| TXK | 3.71531 | -0.28034 | 1.138387 | -0.24626 | 0.805483 | NA |
| FNDC3B | 1310.134 | 0.13187 | 0.192581 | 0.684751 | 0.493501 | 0.644607 |
| FRYL | 1290.441 | 0.295687 | 0.142027 | 2.081904 | 0.037351 | 0.114765 |
| PLD1 | 848.9125 | -0.13918 | 0.184865 | -0.75289 | 0.451516 | 0.610554 |
| DLG1 | 1362.154 | 0.673058 | 0.229418 | 2.933763 | 0.003349 | 0.022686 |
| RAB7A | 20682.44 | -0.06388 | 0.127975 | -0.49918 | 0.61765 | 0.742546 |
| JADE1 | 284.04 | 0.280432 | 0.2215 | 1.266058 | 0.205492 | 0.362516 |
| MCCC1 | 708.7849 | 0.126332 | 0.18378 | 0.687407 | 0.491826 | 0.643353 |
| LAMP3 | 0.884727 | 1.125991 | 2.88877 | 0.389782 | 0.696698 | NA |
| UBE2K | 1939.78 | 0.286878 | 0.144373 | 1.987066 | 0.046915 | 0.133248 |
| N4BP2 | 50.46297 | 0.634905 | 0.476967 | 1.331131 | 0.183146 | 0.336255 |
| ADCY2 | 562.7294 | -0.49234 | 0.189883 | -2.59284 | 0.009519 | 0.045649 |
| PKD2L2 | 2.077676 | 1.160533 | 1.441605 | 0.805029 | 0.420803 | NA |
| SLC1A3 | 71.90752 | 0.588829 | 0.33996 | 1.732053 | 0.083264 | 0.196515 |

| | | | | | | |
|---|---|---|---|---|---|---|
| LXN | 392.2295 | 0.470523 | 0.191859 | 2.452437 | 0.014189 | 0.059806 |
| SCGN | 1.38705 | -1.18656 | 1.632795 | -0.7267 | 0.467408 | NA |
| LRRC16A | 868.9875 | -0.27615 | 0.179994 | -1.5342 | 0.12498 | 0.259891 |
| EPB41L2 | 1536.676 | 0.071807 | 0.220088 | 0.326263 | 0.744226 | 0.838959 |
| MOXD1 | 381.1448 | 0.120203 | 0.229563 | 0.523617 | 0.600545 | 0.729987 |
| STX7 | 6363.56 | 0.397133 | 0.186738 | 2.126685 | 0.033446 | 0.10676 |
| DDX43 | 19.00863 | 0.777574 | 0.509449 | 1.526303 | 0.126934 | NA |
| CRYBG3 | 282.4831 | 0.589698 | 0.316823 | 1.861285 | 0.062704 | 0.162128 |
| EPHA6 | 0.392857 | -1.546 | 4.075469 | -0.37934 | 0.704433 | NA |
| SLC4A4 | 9.339025 | -0.1431 | 0.674163 | -0.21227 | 0.831898 | NA |
| SESN1 | 478.7142 | -0.04021 | 0.203052 | -0.198 | 0.843041 | 0.906755 |
| KCNN2 | 82.75127 | -1.27506 | 0.384261 | -3.31822 | 9.06E-04 | 0.009758 |
| CPOX | 1109.565 | 0.081143 | 0.150956 | 0.537525 | 0.590905 | 0.722556 |
| CLDND1 | 1139.999 | 0.528219 | 0.168846 | 3.128398 | 0.001758 | 0.014873 |
| CXCL2 | 18.99524 | -1.52623 | 0.777631 | -1.96267 | 0.049685 | NA |
| TCF7 | 95.98481 | -0.14345 | 0.264593 | -0.54216 | 0.587705 | 0.720278 |
| OSTM1 | 12193.6 | 0.50684 | 0.182299 | 2.780267 | 0.005431 | 0.031364 |
| IMPG2 | 2.502197 | 0.811632 | 1.388482 | 0.584547 | 0.558853 | NA |
| PCNP | 2164.527 | 0.581444 | 0.19562 | 2.972308 | 0.002956 | 0.020932 |
| MEF2C | 1851.459 | 0.661699 | 0.205179 | 3.224979 | 0.00126 | 0.012025 |
| UBA5 | 479.1081 | 0.493592 | 0.238323 | 2.071106 | 0.038349 | 0.11687 |
| KIAA0141 | 1822.571 | -0.24959 | 0.120082 | -2.0785 | 0.037663 | 0.115375 |
| PCDHB4 | 0.814253 | -1.23545 | 2.246777 | -0.54988 | 0.582405 | NA |
| PCDHA6 | 0.622842 | 1.158846 | 2.735323 | 0.42366 | 0.671814 | NA |
| PCDHGA2 | 44.62724 | -0.49215 | 0.337274 | -1.4592 | 0.144511 | NA |
| WDR70 | 616.687 | 0.466438 | 0.167585 | 2.783295 | 0.005381 | 0.031155 |
| FYB | 10.68047 | 0.084523 | 0.794305 | 0.106411 | 0.915256 | NA |
| C1QTNF3 | 7.41121 | 0.31041 | 0.756851 | 0.410134 | 0.681708 | NA |
| C5orf22 | 1221.326 | -0.03605 | 0.197174 | -0.18284 | 0.854926 | 0.913749 |
| FAM135A | 188.9079 | 0.802218 | 0.267645 | 2.997325 | 0.002724 | 0.019937 |
| COL19A1 | 29.54896 | 0.335381 | 0.459156 | 0.73043 | 0.465127 | NA |
| MRPL22 | 1009.57 | 0.26149 | 0.178234 | 1.467118 | 0.142344 | 0.283135 |
| GEMIN5 | 662.0322 | -0.30136 | 0.15191 | -1.9838 | 0.047278 | 0.133854 |
| GSK3B | 1612.148 | -0.0671 | 0.171323 | -0.39165 | 0.695316 | 0.803408 |
| ITGB5 | 5819.378 | -0.03401 | 0.167495 | -0.20307 | 0.839077 | 0.904049 |
| C4orf6 | 0.555087 | 0.973802 | 2.808575 | 0.346725 | 0.728798 | NA |
| RNF13 | 1965.699 | 0.45593 | 0.165264 | 2.7588 | 0.005801 | 0.032657 |
| DOPEY1 | 343.6667 | 0.202663 | 0.219918 | 0.921536 | 0.356771 | 0.524363 |
| LYRM2 | 1049.118 | 0.630747 | 0.178133 | 3.54088 | 3.99E-04 | 0.00591 |
| BCKDHB | 663.5624 | 0.103667 | 0.248436 | 0.417278 | 0.676475 | 0.788478 |
| TNPO1 | 5670.671 | 0.719139 | 0.189404 | 3.796861 | 1.47E-04 | 0.003187 |
| OXCT1 | 2484.232 | 0.263664 | 0.176999 | 1.489631 | 0.136321 | 0.275434 |
| FAT1 | 3489.787 | -0.38226 | 0.167238 | -2.28572 | 0.022271 | 0.081537 |
| YTHDC1 | 911.955 | 0.60053 | 0.258474 | 2.323365 | 0.02016 | 0.076136 |
| CHMP2B | 771.9123 | 0.926589 | 0.2026 | 4.573494 | 4.80E-06 | 4.09E-04 |
| NOA1 | 797.4221 | -0.2634 | 0.147218 | -1.78916 | 0.073589 | 0.180809 |
| REST | 466.056 | 0.58799 | 0.294127 | 1.999104 | 0.045597 | 0.130744 |

| | | | | | | |
|---|---|---|---|---|---|---|
| TAF9 | 1484.533 | 0.528567 | 0.18191 | 2.90566 | 0.003665 | 0.024136 |
| MYNN | 398.9494 | 0.666078 | 0.226879 | 2.935827 | 0.003327 | 0.022548 |
| MECOM | 0.640755 | -1.64826 | 2.754778 | -0.59833 | 0.549621 | NA |
| SCAMP1 | 924.8011 | 0.640458 | 0.189399 | 3.381535 | 7.21E-04 | 0.008464 |
| PREP | 1963.301 | -0.49644 | 0.145654 | -3.40833 | 6.54E-04 | 0.007955 |
| HACE1 | 318.7201 | 0.44563 | 0.223388 | 1.994873 | 0.046057 | 0.131789 |
| MGST2 | 307.6902 | -0.11135 | 0.189049 | -0.58901 | 0.555854 | 0.695462 |
| DIMT1 | 766.2639 | 0.228552 | 0.149941 | 1.524282 | 0.127438 | 0.263318 |
| IPO11 | 601.6733 | 0.194348 | 0.183979 | 1.056362 | 0.290803 | 0.45777 |
| FAT2 | 15.64352 | 0.016576 | 0.54634 | 0.03034 | 0.975796 | NA |
| RBM22 | 998.2796 | -0.14441 | 0.139443 | -1.0356 | 0.300389 | 0.467283 |
| ACOX3 | 486.637 | -0.34788 | 0.173206 | -2.00848 | 0.044592 | 0.129054 |
| ADAMTS2 | 134.7966 | -0.77896 | 0.396987 | -1.96217 | 0.049743 | 0.138006 |
| TMPRSS11I | 0.899187 | 0.806933 | 2.127165 | 0.379346 | 0.704431 | NA |
| UIMC1 | 580.7494 | 0.314352 | 0.200381 | 1.568775 | 0.1167 | 0.247139 |
| SH3BP2 | 1370.292 | -0.37719 | 0.14857 | -2.53883 | 0.011122 | 0.050539 |
| NOP14 | 1309.497 | 0.071148 | 0.145655 | 0.488473 | 0.625215 | 0.748721 |
| ADD1 | 8420.74 | -0.51488 | 0.115043 | -4.47552 | 7.62E-06 | 5.35E-04 |
| F11 | 0.394291 | 0.298745 | 3.141711 | 0.09509 | 0.924243 | NA |
| MAEA | 2852.721 | -0.26121 | 0.138879 | -1.88087 | 0.059989 | 0.157199 |
| DNAJB11 | 1639.331 | 0.468752 | 0.157114 | 2.983523 | 0.00285 | 0.020461 |
| LEPREL1 | 1399.954 | 0.980227 | 0.668055 | 1.467285 | 0.142299 | 0.283134 |
| CHRD | 1.780992 | 0.370243 | 1.531458 | 0.241758 | 0.808968 | NA |
| EXOC1 | 895.5533 | 0.744495 | 0.264891 | 2.810569 | 0.004945 | 0.029358 |
| RBM27 | 497.3019 | 0.628442 | 0.245031 | 2.564749 | 0.010325 | 0.048058 |
| POU4F3 | 0.410275 | 0.244283 | 3.100889 | 0.078778 | 0.937209 | NA |
| SEL1L3 | 186.8964 | 0.408921 | 0.301712 | 1.355336 | 0.175311 | 0.327146 |
| TF | 1.643845 | -0.01408 | 1.719549 | -0.00819 | 0.993465 | NA |
| CDV3 | 3113.925 | 0.220049 | 0.146263 | 1.504475 | 0.132459 | 0.269673 |
| CD200 | 328.2241 | -0.88664 | 0.283091 | -3.13199 | 0.001736 | 0.01475 |
| CCDC80 | 1884.535 | -0.18811 | 0.236411 | -0.7957 | 0.426204 | 0.588064 |
| SEMA6A | 2784.455 | -0.23431 | 0.181482 | -1.29109 | 0.196673 | 0.352003 |
| ECHDC1 | 1917.713 | 0.636058 | 0.173025 | 3.676116 | 2.37E-04 | 0.004317 |
| GABRP | 0.545988 | -0.43913 | 2.541145 | -0.17281 | 0.862803 | NA |
| CDC23 | 778.7543 | -0.07805 | 0.158641 | -0.49197 | 0.622742 | 0.746925 |
| MAP3K1 | 615.2008 | -0.13353 | 0.244083 | -0.54707 | 0.58433 | 0.717551 |
| HIVEP1 | 310.3068 | -0.05575 | 0.23087 | -0.24148 | 0.809184 | 0.885851 |
| FKBP5 | 709.6094 | -0.72921 | 0.250886 | -2.90652 | 0.003655 | 0.024096 |
| SRPK1 | 1145.414 | 0.383355 | 0.174623 | 2.195329 | 0.02814 | 0.094961 |
| BRPF3 | 1210.901 | -0.34247 | 0.169493 | -2.02057 | 0.043324 | 0.126879 |
| MRPS18A | 789.3449 | -0.03944 | 0.154214 | -0.25578 | 0.798121 | 0.877894 |
| TMEM14A | 927.9692 | 0.178669 | 0.152306 | 1.173095 | 0.240758 | 0.403989 |
| EFHC1 | 335.9232 | 0.384903 | 0.184215 | 2.089425 | 0.036669 | 0.113473 |
| HSP90AB1 | 24931.76 | 0.023424 | 0.140139 | 0.167146 | 0.867256 | 0.921265 |
| MLN | 0.162376 | 0.424913 | 4.079006 | 0.104171 | 0.917034 | NA |
| CDC5L | 1027.016 | 0.74819 | 0.233848 | 3.199474 | 0.001377 | 0.012785 |
| ITPR3 | 2960.593 | -0.42807 | 0.177597 | -2.41033 | 0.015938 | 0.064664 |

| | | | | | | |
|---|---|---|---|---|---|---|
| ZNF184 | 185.5905 | 0.523051 | 0.221 | 2.366744 | 0.017945 | 0.070142 |
| DSP | 2.215953 | 0.955495 | 1.484968 | 0.643445 | 0.519936 | NA |
| TMEM33 | 2172.495 | 0.161202 | 0.177358 | 0.908906 | 0.3634 | 0.53062 |
| GNRHR | 1.033116 | 1.025241 | 1.933005 | 0.530387 | 0.595844 | NA |
| SLAIN2 | 1105.098 | 0.217555 | 0.164786 | 1.320229 | 0.186759 | 0.340881 |
| OCIAD1 | 2201.038 | 0.377777 | 0.154114 | 2.451275 | 0.014235 | 0.059928 |
| DCUN1D4 | 815.5187 | 0.655181 | 0.235346 | 2.783908 | 0.005371 | 0.03112 |
| USP46 | 560.251 | 0.104125 | 0.21408 | 0.486381 | 0.626697 | 0.74944 |
| SULT1E1 | 0.151697 | -1.546 | 4.075469 | -0.37934 | 0.704433 | NA |
| CHIC2 | 283.8303 | 0.120838 | 0.182159 | 0.663368 | 0.507095 | 0.656218 |
| NMU | 6.368434 | 0.025781 | 0.841692 | 0.03063 | 0.975565 | NA |
| KIAA1211 | 1040.316 | -0.42877 | 0.158646 | -2.70268 | 0.006878 | 0.036671 |
| LAMTOR3 | 814.4725 | 0.672565 | 0.177164 | 3.796278 | 1.47E-04 | 0.003187 |
| PF4V1 | 0.151697 | -1.546 | 4.075469 | -0.37934 | 0.704433 | NA |
| NFKB1 | 1741.683 | -0.27042 | 0.136418 | -1.98227 | 0.047449 | 0.134091 |
| MANBA | 1209.435 | -0.10388 | 0.137927 | -0.75315 | 0.451362 | 0.610554 |
| UBE2D3 | 6765.19 | 0.063848 | 0.134125 | 0.476035 | 0.634049 | 0.755518 |
| MAPK10 | 13.12067 | -0.0447 | 0.575461 | -0.07768 | 0.938086 | NA |
| ELF2 | 315.7446 | 0.782953 | 0.196284 | 3.988886 | 6.64E-05 | 0.001953 |
| NDUFC1 | 1374.852 | 0.268517 | 0.170986 | 1.570403 | 0.116321 | 0.246689 |
| UCP1 | 0.151697 | -1.546 | 4.075469 | -0.37934 | 0.704433 | NA |
| TBC1D9 | 795.9136 | 0.252722 | 0.16336 | 1.547022 | 0.121858 | 0.255054 |
| ZNF330 | 1198.207 | 0.26978 | 0.178463 | 1.511684 | 0.130614 | 0.267063 |
| INPP4B | 6108.326 | 0.773494 | 0.17155 | 4.508845 | 6.52E-06 | 4.97E-04 |
| GAB1 | 548.9982 | 0.223443 | 0.226159 | 0.987988 | 0.323158 | 0.491033 |
| KLHL2 | 459.6415 | 0.638222 | 0.178982 | 3.565847 | 3.63E-04 | 0.005542 |
| CPE | 4.193981 | 0.087923 | 1.069757 | 0.08219 | 0.934496 | NA |
| RPL34 | 7729.884 | 0.526801 | 0.197094 | 2.672837 | 0.007521 | 0.039107 |
| WFS1 | 3194.472 | -0.46241 | 0.241915 | -1.91148 | 0.055943 | 0.149607 |
| ANXA10 | 0.096202 | 0.408546 | 4.079639 | 0.100143 | 0.920231 | NA |
| GRPEL1 | 1302.683 | 0.009323 | 0.155973 | 0.059773 | 0.952337 | 0.973943 |
| GAR1 | 435.0446 | -0.15839 | 0.16299 | -0.9718 | 0.331149 | 0.499652 |
| FRG1 | 535.7073 | 0.646705 | 0.230934 | 2.800391 | 0.005104 | 0.030014 |
| CLCN3 | 4513.428 | 0.289122 | 0.191194 | 1.512193 | 0.130485 | 0.26704 |
| AADAT | 222.3631 | -0.00129 | 0.215931 | -0.00597 | 0.995238 | 0.997908 |
| GALNT7 | 800.76 | 0.424441 | 0.179391 | 2.366008 | 0.017981 | 0.070259 |
| DHX15 | 4063.63 | 0.31702 | 0.13479 | 2.35196 | 0.018675 | 0.072184 |
| SOD3 | 18.93969 | -0.1695 | 0.53964 | -0.3141 | 0.753446 | NA |
| SEPSECS | 213.191 | 0.628568 | 0.236553 | 2.657198 | 0.007879 | 0.04028 |
| TRIM2 | 4039.217 | 0.14414 | 0.260073 | 0.554229 | 0.579422 | 0.713593 |
| SLC2A9 | 35.33256 | -0.15246 | 0.372564 | -0.40923 | 0.682371 | NA |
| FBXW7 | 313.4663 | 0.42454 | 0.195181 | 2.17511 | 0.029622 | 0.098146 |
| NEIL3 | 76.0719 | 0.13037 | 0.302855 | 0.430469 | 0.666855 | 0.781491 |
| TBC1D19 | 301.5843 | 0.138019 | 0.209526 | 0.65872 | 0.510076 | 0.65835 |
| WHSC1 | 1916.34 | -0.32657 | 0.216001 | -1.51192 | 0.130555 | 0.26704 |
| SH3D19 | 1348.446 | 0.091623 | 0.210694 | 0.43486 | 0.663664 | 0.779061 |
| STIM2 | 462.585 | 0.220842 | 0.173004 | 1.276513 | 0.201774 | 0.358267 |

| | | | | | | |
|---|---|---|---|---|---|---|
| NKX3-2 | 3.989361 | -0.30952 | 1.060279 | -0.29192 | 0.770346 | NA |
| MFSD10 | 1983.649 | -0.1482 | 0.16926 | -0.87556 | 0.381272 | 0.547441 |
| GLRB | 74.48803 | 0.943828 | 0.315029 | 2.996008 | 0.002735 | 0.019954 |
| BST1 | 818.8125 | -0.1184 | 0.152013 | -0.77885 | 0.436068 | 0.596949 |
| RAPGEF2 | 1146.355 | 0.074208 | 0.133849 | 0.554414 | 0.579295 | 0.713563 |
| HGFAC | 0.912824 | 3.024111 | 2.331252 | 1.297204 | 0.194561 | NA |
| SNX25 | 1650.008 | 0.792144 | 0.163406 | 4.84772 | 1.25E-06 | 1.83E-04 |
| LRP2BP | 53.00272 | 0.837335 | 0.371947 | 2.251221 | 0.024372 | 0.086453 |
| UFSP2 | 550.1789 | 0.391826 | 0.179538 | 2.182411 | 0.029079 | 0.096999 |
| KLF3 | 656.4804 | 0.343112 | 0.201499 | 1.702796 | 0.088606 | 0.205443 |
| KLHL5 | 671.0018 | 0.362352 | 0.176515 | 2.052813 | 0.040091 | 0.120637 |
| FAM149A | 738.0111 | -0.34721 | 0.195872 | -1.77266 | 0.076286 | 0.185131 |
| NCAPG | 319.1286 | 0.732178 | 0.214345 | 3.415884 | 6.36E-04 | 0.007818 |
| UGDH | 661.8757 | 0.023067 | 0.189195 | 0.121924 | 0.902959 | 0.943218 |
| PPARGC1A | 75.00745 | -0.67315 | 0.514153 | -1.30924 | 0.190452 | 0.344799 |
| COL12A1 | 585.467 | -0.56292 | 0.20426 | -2.7559 | 0.005853 | 0.032848 |
| BTN3A3 | 475.6047 | -0.45499 | 0.214163 | -2.12451 | 0.033628 | 0.107079 |
| TDP2 | 735.4877 | 0.681293 | 0.154628 | 4.406001 | 1.05E-05 | 6.38E-04 |
| FRK | 31.93434 | 0.06483 | 0.398047 | 0.16287 | 0.870621 | NA |
| DSE | 134.4042 | 0.476774 | 0.258951 | 1.841175 | 0.065596 | 0.167055 |
| RWDD1 | 1576.463 | 0.846307 | 0.218927 | 3.865707 | 1.11E-04 | 0.002747 |
| RSPH4A | 4.788028 | 0.623077 | 0.914052 | 0.681665 | 0.495451 | NA |
| MAK | 1.649634 | 1.909873 | 1.612286 | 1.184575 | 0.236186 | NA |
| TMEM14C | 3889.007 | 0.155139 | 0.134133 | 1.156607 | 0.247433 | 0.410886 |
| PAK1IP1 | 486.0193 | 0.601439 | 0.153954 | 3.906618 | 9.36E-05 | 0.002433 |
| GCNT2 | 3137.737 | 0.04193 | 0.208616 | 0.200994 | 0.840704 | 0.905184 |
| SMIM8 | 101.1787 | 0.37249 | 0.247823 | 1.503046 | 0.132827 | 0.270093 |
| NEDD9 | 469.7843 | -0.18321 | 0.165983 | -1.10376 | 0.269697 | 0.434351 |
| CEP85L | 97.56302 | 0.743597 | 0.451632 | 1.646465 | 0.099668 | 0.222979 |
| ADTRP | 11.54799 | -0.27716 | 0.596794 | -0.46441 | 0.642353 | NA |
| ASF1A | 411.0904 | 0.715209 | 0.16756 | 4.268374 | 1.97E-05 | 9.23E-04 |
| MCM9 | 219.5001 | 0.254874 | 0.181462 | 1.404557 | 0.160153 | 0.307468 |
| FAM184A | 2.865049 | 3.506201 | 1.637196 | 2.141589 | 0.032227 | NA |
| RNGTT | 571.8088 | 0.227472 | 0.168673 | 1.3486 | 0.177465 | 0.329706 |
| MAN1A1 | 509.6807 | 0.594912 | 0.173065 | 3.437504 | 5.87E-04 | 0.007468 |
| GABRR2 | 2.262349 | 0.147942 | 1.309881 | 0.112943 | 0.910076 | NA |
| SERINC1 | 8046.415 | 0.669763 | 0.186757 | 3.586287 | 3.35E-04 | 0.005288 |
| HDDC2 | 2164.811 | 0.263414 | 0.137904 | 1.910134 | 0.056116 | 0.149964 |
| TPD52L1 | 70.28062 | 0.602816 | 1.108514 | 0.543806 | 0.586575 | 0.719339 |
| HINT3 | 359.191 | 0.37515 | 0.25919 | 1.447391 | 0.147788 | 0.290937 |
| NCOA7 | 481.5984 | 0.661631 | 0.209934 | 3.151614 | 0.001624 | 0.014248 |
| FAM65B | 268.6309 | -1.12526 | 0.365607 | -3.07778 | 0.002085 | 0.016597 |
| SASH1 | 5265.923 | 0.223689 | 0.21462 | 1.042255 | 0.297293 | 0.464533 |
| UST | 827.6468 | -0.00696 | 0.15579 | -0.04469 | 0.964353 | 0.980481 |
| ULBP1 | 7.109806 | 1.360974 | 0.915105 | 1.487233 | 0.136953 | NA |
| PPARD | 1335.763 | -0.2699 | 0.151354 | -1.78326 | 0.074544 | 0.182375 |
| FANCE | 224.913 | -0.40916 | 0.203919 | -2.00649 | 0.044805 | 0.129347 |

| | | | | | | |
|---|---|---|---|---|---|---|
| TULP1 | 0.623045 | 0.091811 | 2.666142 | 0.034436 | 0.97253 | NA |
| SLC26A8 | 1.56189 | -0.54495 | 1.531142 | -0.35591 | 0.721905 | NA |
| MAPK14 | 1834.232 | -0.09953 | 0.13718 | -0.72557 | 0.468103 | 0.624576 |
| KCTD20 | 5616.1 | -0.01986 | 0.157999 | -0.12569 | 0.899978 | 0.941432 |
| STK38 | 1189.251 | 0.001309 | 0.134942 | 0.009701 | 0.99226 | 0.996738 |
| SRSF3 | 4327.71 | 0.099688 | 0.14319 | 0.696194 | 0.486307 | 0.639483 |
| MCM3 | 3072.387 | -0.14539 | 0.181378 | -0.80156 | 0.422807 | 0.585148 |
| RNF8 | 528.5092 | 0.298127 | 0.149853 | 1.989468 | 0.04665 | 0.132812 |
| PHACTR1 | 5743.062 | 0.575294 | 0.266656 | 2.157439 | 0.030971 | 0.101371 |
| MDGA1 | 103.0302 | -0.6046 | 0.299553 | -2.01833 | 0.043557 | 0.127152 |
| ICK | 509.1572 | 0.328301 | 0.24 | 1.367922 | 0.171337 | 0.322285 |
| FBXO9 | 1369.022 | 0.095086 | 0.128127 | 0.742122 | 0.458013 | 0.616456 |
| CD83 | 427.7479 | -0.08713 | 0.159672 | -0.54568 | 0.585283 | 0.718394 |
| MDN1 | 1331.547 | -0.02341 | 0.163151 | -0.14348 | 0.885915 | 0.932547 |
| GLP1R | 0.151697 | -1.546 | 4.075469 | -0.37934 | 0.704433 | NA |
| SAYSD1 | 226.2792 | -0.1974 | 0.206293 | -0.9569 | 0.338616 | 0.507304 |
| BACH2 | 29.14915 | -0.28827 | 0.440528 | -0.65437 | 0.512876 | NA |
| RBM24 | 2.880633 | 0.035057 | 1.181172 | 0.02968 | 0.976322 | NA |
| CAP2 | 674.9739 | 0.146756 | 0.175049 | 0.838368 | 0.401824 | 0.566424 |
| TREML2 | 0.163957 | 0.424913 | 4.079006 | 0.104171 | 0.917034 | NA |
| ZNF451 | 476.5638 | 0.667629 | 0.215456 | 3.098686 | 0.001944 | 0.015854 |
| BAG2 | 75.405 | 0.332697 | 0.271009 | 1.227623 | 0.219588 | 0.379257 |
| RAB23 | 249.5899 | 0.729149 | 0.232022 | 3.142589 | 0.001675 | 0.014459 |
| TSPO2 | 0.452561 | 2.061505 | 3.349456 | 0.615474 | 0.538241 | NA |
| GPR63 | 42.39654 | 0.243122 | 0.38197 | 0.636494 | 0.524454 | NA |
| KHDRBS2 | 0.247898 | -0.5901 | 4.074633 | -0.14482 | 0.884851 | NA |
| FBXL4 | 600.26 | 0.178584 | 0.183905 | 0.971068 | 0.331514 | 0.500013 |
| CCNC | 809.6509 | 0.565558 | 0.209575 | 2.698591 | 0.006963 | 0.037018 |
| PRDM13 | 0.298089 | -0.51711 | 3.975232 | -0.13008 | 0.896501 | NA |
| E2F3 | 616.6046 | 0.103973 | 0.169538 | 0.613271 | 0.539697 | 0.681811 |
| PTP4A1 | 2435.444 | 0.221484 | 0.169752 | 1.304748 | 0.191979 | 0.346567 |
| ASCC3 | 1285.657 | 0.460601 | 0.227348 | 2.025972 | 0.042768 | 0.125906 |
| HDGFL1 | 0.151697 | -1.546 | 4.075469 | -0.37934 | 0.704433 | NA |
| BVES | 1102.178 | -0.11336 | 0.161263 | -0.70294 | 0.482093 | 0.636186 |
| COL9A1 | 0.096202 | 0.408546 | 4.079639 | 0.100143 | 0.920231 | NA |
| MED23 | 669.3518 | 0.137555 | 0.180723 | 0.761136 | 0.446576 | 0.606363 |
| WASF1 | 216.2147 | -0.04872 | 0.260397 | -0.18712 | 0.851568 | 0.91179 |
| GPLD1 | 3.058247 | 0.878725 | 1.162511 | 0.755885 | 0.449718 | NA |
| ALDH5A1 | 76.30187 | -0.42003 | 0.332532 | -1.26312 | 0.206546 | 0.363668 |
| AIM1 | 4.18377 | 0.507874 | 0.962285 | 0.527779 | 0.597653 | NA |
| VNN2 | 2.621815 | 0.026494 | 1.280344 | 0.020693 | 0.983491 | NA |
| ACOT13 | 604.3494 | 0.287416 | 0.163191 | 1.761223 | 0.078201 | 0.188314 |
| SMAP1 | 249.7082 | 0.080011 | 0.190234 | 0.420593 | 0.674052 | 0.786547 |
| RPS12 | 31861.85 | 0.173489 | 0.159162 | 1.090015 | 0.275706 | 0.4413 |
| C6orf62 | 3218.022 | -0.40039 | 0.15381 | -2.60316 | 0.009237 | 0.044695 |
| B3GAT2 | 3.146306 | 0.541236 | 1.25772 | 0.430331 | 0.666955 | NA |
| GMNN | 495.4953 | 0.469429 | 0.157862 | 2.97366 | 0.002943 | 0.020877 |

| | | | | | | |
|---|---|---|---|---|---|---|
| EYA4 | 11.07236 | 0.537756 | 0.741061 | 0.725657 | 0.468049 | NA |
| SNX3 | 6588.153 | 0.193003 | 0.135724 | 1.422026 | 0.155019 | 0.300482 |
| HBS1L | 2096.197 | 0.456027 | 0.172441 | 2.644533 | 0.00818 | 0.041197 |
| TRIM38 | 746.5932 | 0.197426 | 0.157378 | 1.254474 | 0.20967 | 0.367209 |
| PEX7 | 232.4915 | 9.62E-04 | 0.184788 | 0.005207 | 0.995845 | 0.998034 |
| ZBTB24 | 1040.731 | 0.592824 | 0.157148 | 3.772397 | 1.62E-04 | 0.003435 |
| FIG4 | 1078.138 | 0.023675 | 0.171445 | 0.13809 | 0.890169 | 0.935891 |
| PERP | 3423.552 | -0.03669 | 0.145562 | -0.25205 | 0.800999 | 0.880141 |
| KIAA1244 | 71.3004 | -0.1093 | 0.313524 | -0.34861 | 0.727383 | 0.827079 |
| SLC16A10 | 187.2043 | -0.3393 | 0.292767 | -1.15894 | 0.246482 | 0.409996 |
| HECA | 490.5128 | 0.563524 | 0.215869 | 2.610487 | 0.009041 | 0.044218 |
| GPR126 | 908.4889 | 0.137072 | 0.190831 | 0.718294 | 0.472576 | 0.628423 |
| PHACTR2 | 442.5623 | 0.715731 | 0.268081 | 2.669836 | 0.007589 | 0.039355 |
| EPM2A | 537.4827 | 0.075758 | 0.180094 | 0.420657 | 0.674006 | 0.786547 |
| SLC39A7 | 4380.597 | -0.07413 | 0.144104 | -0.51443 | 0.606952 | 0.734751 |
| PHF1 | 1543.656 | 0.035252 | 0.185343 | 0.190197 | 0.849155 | 0.910193 |
| CUTA | 4955.286 | -0.15476 | 0.138336 | -1.11875 | 0.263246 | 0.427592 |
| MDFI | 1.99927 | -0.50401 | 1.413276 | -0.35663 | 0.72137 | NA |
| TFEB | 162.6172 | -0.26737 | 0.219515 | -1.21801 | 0.223222 | 0.384077 |
| CCND3 | 3491.901 | -0.22995 | 0.143533 | -1.6021 | 0.109134 | 0.236308 |
| BYSL | 1030.241 | -0.30068 | 0.180812 | -1.66296 | 0.096321 | 0.217461 |
| GUCA1B | 8.743757 | 0.495457 | 0.704708 | 0.703068 | 0.482013 | NA |
| PRPH2 | 0.785567 | -0.70568 | 2.358063 | -0.29926 | 0.764741 | NA |
| GLTSCR1L | 332.8359 | -0.27014 | 0.17623 | -1.5329 | 0.1253 | 0.260204 |
| PPP2R5D | 2221.304 | -0.33006 | 0.120452 | -2.74018 | 0.006141 | 0.03389 |
| MRPL2 | 871.9336 | -0.13546 | 0.189977 | -0.71302 | 0.475833 | 0.630941 |
| PTK7 | 1984.741 | -0.36476 | 0.134636 | -2.70921 | 0.006744 | 0.036212 |
| SRF | 1436.198 | -0.23683 | 0.144016 | -1.64449 | 0.100075 | 0.223218 |
| CUL9 | 1003.835 | -0.59389 | 0.165527 | -3.58786 | 3.33E-04 | 0.00527 |
| DNPH1 | 1583.149 | -0.29975 | 0.214332 | -1.39851 | 0.161959 | 0.309782 |
| DUSP22 | 1855.2 | -0.12289 | 0.136956 | -0.89727 | 0.369576 | 0.536532 |
| EXOC2 | 1410.846 | 0.116814 | 0.148324 | 0.787557 | 0.430956 | 0.592439 |
| COX7A2 | 5884.554 | 0.195681 | 0.173324 | 1.128987 | 0.258903 | 0.423177 |
| TMEM30A | 3050.591 | 0.640012 | 0.172769 | 3.704436 | 2.12E-04 | 0.00404 |
| GMDS | 627.4728 | -0.11988 | 0.161629 | -0.74171 | 0.458262 | 0.616722 |
| SENP6 | 638.4791 | 0.82405 | 0.620793 | 1.327414 | 0.184372 | 0.337667 |
| IMPG1 | 0.3441 | -0.0166 | 3.769682 | -0.0044 | 0.996486 | NA |
| VEGFA | 192.4268 | 0.494518 | 0.248307 | 1.99156 | 0.046419 | 0.132419 |
| PRPF4B | 1014.266 | 1.108726 | 0.320048 | 3.464252 | 5.32E-04 | 0.007022 |
| TTK | 282.6046 | 0.814772 | 0.217585 | 3.744613 | 1.81E-04 | 0.003645 |
| SLC29A1 | 1505.577 | -0.39105 | 0.169609 | -2.30558 | 0.021134 | 0.078663 |
| WISP3 | 0.886724 | 0.648622 | 2.434305 | 0.26645 | 0.789892 | NA |
| BTN2A1 | 809.8648 | 0.308749 | 0.17835 | 1.73114 | 0.083427 | 0.196777 |
| LAMA4 | 1879.587 | -0.36336 | 0.179831 | -2.02057 | 0.043324 | 0.126879 |
| FAM46A | 568.3668 | -0.28038 | 0.210693 | -1.33073 | 0.183277 | 0.336442 |
| ENPP5 | 23.59907 | 0.016717 | 0.453367 | 0.036874 | 0.970586 | NA |
| PRSS16 | 0.564775 | -1.6359 | 3.188247 | -0.5131 | 0.607879 | NA |

| | | | | | | |
|---|---|---|---|---|---|---|
| TBX18 | 8.0246 | 0.397117 | 0.778292 | 0.510241 | 0.609882 | NA |
| ERBB2IP | 1451.611 | 0.816544 | 0.302383 | 2.700363 | 0.006926 | 0.036855 |
| PCDHB2 | 2.085966 | -0.1142 | 1.452328 | -0.07863 | 0.937325 | NA |
| HARS2 | 824.0106 | 0.026183 | 0.147876 | 0.177062 | 0.859459 | 0.916432 |
| NUDT12 | 353.0599 | 0.852915 | 0.32163 | 2.651851 | 0.008005 | 0.040626 |
| CEP72 | 143.737 | -0.54095 | 0.249899 | -2.16467 | 0.030413 | 0.099989 |
| MAN2A1 | 713.9005 | 0.15668 | 0.180893 | 0.866146 | 0.38641 | 0.552513 |
| SEMA5A | 2544.619 | -0.55016 | 0.211992 | -2.59519 | 0.009454 | 0.045467 |
| C7 | 1.63049 | 1.95266 | 1.731691 | 1.127603 | 0.259488 | NA |
| PAPD7 | 518.3228 | -0.07489 | 0.176938 | -0.42325 | 0.672109 | 0.78532 |
| GHR | 167.6349 | 0.095729 | 0.287321 | 0.333179 | 0.738999 | 0.835002 |
| HMGCS1 | 2061.046 | 0.456373 | 0.255035 | 1.789452 | 0.073542 | 0.180809 |
| DAP | 4403.658 | -0.2255 | 0.198055 | -1.13859 | 0.254875 | 0.419192 |
| NME5 | 129.0512 | 0.643172 | 0.255164 | 2.520616 | 0.011715 | 0.052388 |
| BRD8 | 1385.031 | -0.05405 | 0.186715 | -0.28949 | 0.772208 | 0.858857 |
| KIF20A | 1357.157 | -0.52731 | 0.243856 | -2.16237 | 0.030589 | 0.100398 |
| NNT | 2825.297 | 0.068499 | 0.148035 | 0.462725 | 0.643562 | 0.762832 |
| MRPS30 | 918.5643 | -0.00771 | 0.17158 | -0.04491 | 0.964177 | 0.980386 |
| HSPA9 | 8042.223 | 0.278553 | 0.125471 | 2.220066 | 0.026414 | 0.091242 |
| MRPS27 | 1742.709 | -0.27744 | 0.14605 | -1.89962 | 0.057483 | 0.152304 |
| PFDN1 | 1408.998 | 0.255484 | 0.146053 | 1.749251 | 0.080248 | 0.19184 |
| HBEGF | 265.2514 | 0.15406 | 0.293183 | 0.525474 | 0.599254 | 0.728781 |
| SLC4A9 | 0.651435 | 0.249629 | 2.788679 | 0.089515 | 0.928672 | NA |
| LOX | 361.9088 | 0.240081 | 0.184692 | 1.299897 | 0.193636 | 0.348763 |
| APBB3 | 173.6388 | 0.14598 | 0.291777 | 0.500312 | 0.616856 | 0.742224 |
| TMCO6 | 189.4526 | -0.21101 | 0.184727 | -1.1423 | 0.25333 | 0.417475 |
| SPARC | 31938.83 | -0.09374 | 0.245127 | -0.3824 | 0.702167 | 0.808431 |
| IK | 2111.23 | 0.306836 | 0.160736 | 1.908946 | 0.056269 | 0.150106 |
| HMGCR | 3976.825 | 0.084399 | 0.17278 | 0.488474 | 0.625214 | 0.748721 |
| COL4A3BP | 5822.515 | 0.239986 | 0.293369 | 0.818036 | 0.413337 | 0.576581 |
| FAF2 | 1624.617 | -0.17415 | 0.134267 | -1.29702 | 0.194624 | 0.349693 |
| PCDHB5 | 19.05335 | 0.055941 | 0.519493 | 0.107684 | 0.914246 | NA |
| PCDHB6 | 3.532713 | -0.58767 | 1.04039 | -0.56485 | 0.572173 | NA |
| PCDHB7 | 5.136339 | -0.00468 | 1.048828 | -0.00446 | 0.996441 | NA |
| PDE8B | 37.78456 | 0.988172 | 0.367791 | 2.686779 | 0.007214 | NA |
| CLK4 | 217.8155 | 1.437867 | 0.248444 | 5.787493 | 7.14E-09 | 5.29E-06 |
| PCDHB15 | 9.338158 | -0.61497 | 0.917023 | -0.67062 | 0.502465 | NA |
| GRM6 | 0.134133 | -1.48909 | 4.077584 | -0.36519 | 0.71497 | NA |
| ITK | 2.522361 | 0.824874 | 1.302023 | 0.633533 | 0.526386 | NA |
| RNF130 | 2621.637 | 0.015045 | 0.132553 | 0.113501 | 0.909633 | 0.947766 |
| THG1L | 364.7671 | 0.193381 | 0.178909 | 1.080889 | 0.279746 | 0.445692 |
| ARSB | 1846.906 | -0.37999 | 0.151529 | -2.50773 | 0.012151 | 0.053906 |
| CLINT1 | 1580.746 | 0.456994 | 0.186104 | 2.45558 | 0.014066 | 0.059426 |
| THBS4 | 28.89142 | 0.103269 | 0.386546 | 0.267157 | 0.789348 | NA |
| CNOT6 | 804.214 | -0.1168 | 0.222824 | -0.52416 | 0.600166 | 0.72974 |
| TTC1 | 1268.707 | 0.693105 | 0.178017 | 3.893467 | 9.88E-05 | 0.00253 |
| MSH3 | 444.3784 | 0.236836 | 0.225938 | 1.048236 | 0.29453 | 0.461615 |

| | | | | | | |
|---|---|---|---|---|---|---|
| RASGRF2 | 25.54178 | 0.586261 | 0.480494 | 1.220122 | 0.222419 | NA |
| CCNG1 | 5038.848 | 0.126513 | 0.199155 | 0.635249 | 0.525266 | 0.670944 |
| POLR3G | 487.8133 | 0.840447 | 0.344834 | 2.43725 | 0.014799 | 0.061418 |
| DROSHA | 1663.725 | -0.2362 | 0.136028 | -1.73642 | 0.08249 | 0.195273 |
| CDH6 | 14.77112 | 1.926026 | 0.730202 | 2.637664 | 0.008348 | NA |
| LMNB1 | 301.2663 | -0.17808 | 0.222695 | -0.79964 | 0.423917 | 0.586 |
| ARRDC3 | 315.5646 | 0.049076 | 0.291675 | 0.168256 | 0.866382 | 0.920772 |
| GOLPH3 | 2190.231 | 0.23692 | 0.150591 | 1.573271 | 0.115656 | 0.245718 |
| SUB1 | 1577.081 | 0.491466 | 0.189683 | 2.590986 | 0.00957 | 0.045835 |
| NPR3 | 0.151697 | -1.546 | 4.075469 | -0.37934 | 0.704433 | NA |
| FAM172A | 492.6574 | 0.30211 | 0.214367 | 1.40931 | 0.158743 | 0.305454 |
| SLC27A6 | 0.163957 | 0.424913 | 4.079006 | 0.104171 | 0.917034 | NA |
| TARS | 3022.409 | 0.433876 | 0.140021 | 3.09865 | 0.001944 | 0.015854 |
| IRX4 | 0.288605 | 1.443619 | 4.029058 | 0.358302 | 0.720117 | NA |
| LNPEP | 676.1742 | 0.370066 | 0.233517 | 1.584752 | 0.113023 | 0.241813 |
| PDE4D | 786.6491 | 0.179553 | 0.255579 | 0.702533 | 0.482347 | 0.636247 |
| RAD1 | 664.5784 | 0.21959 | 0.149745 | 1.466428 | 0.142532 | 0.283414 |
| BRIX1 | 408.0614 | 0.733293 | 0.211274 | 3.470818 | 5.19E-04 | 0.006943 |
| PRLR | 0.326536 | 0.037056 | 3.842554 | 0.009644 | 0.992306 | NA |
| SLC12A7 | 671.438 | -0.51391 | 0.186882 | -2.74991 | 0.005961 | 0.033256 |
| IL4 | 0.163957 | 0.424913 | 4.079006 | 0.104171 | 0.917034 | NA |
| RAD50 | 390.3272 | 0.937786 | 1.099674 | 0.852785 | 0.393778 | 0.558547 |
| ST8SIA4 | 89.74367 | 0.6336 | 0.29684 | 2.134486 | 0.032803 | 0.10553 |
| GNPDA1 | 2952.931 | -0.1197 | 0.140693 | -0.85078 | 0.394894 | 0.559494 |
| PCDH12 | 24.56414 | -0.6732 | 0.439052 | -1.53331 | 0.1252 | NA |
| SKP1 | 6926.746 | 0.001275 | 0.13046 | 0.009774 | 0.992201 | 0.996738 |
| NUP155 | 995.4175 | -0.0923 | 0.160104 | -0.5765 | 0.564275 | 0.702086 |
| PPP2CA | 2795.275 | 0.267034 | 0.129667 | 2.059374 | 0.039458 | 0.119231 |
| FGF1 | 86.242 | -0.06934 | 0.279855 | -0.24776 | 0.804323 | 0.883057 |
| NR3C1 | 906.2927 | 0.269946 | 0.169522 | 1.592395 | 0.111296 | 0.239458 |
| C5orf15 | 1464.869 | 0.340721 | 0.16504 | 2.064469 | 0.038973 | 0.118135 |
| PPWD1 | 220.3716 | 0.777754 | 0.263015 | 2.957069 | 0.003106 | 0.021663 |
| LIFR | 0.875096 | 2.998778 | 2.317512 | 1.293965 | 0.195678 | NA |
| TRIM23 | 287.4772 | 0.984953 | 0.297141 | 3.314763 | 9.17E-04 | 0.009804 |
| TRAPPC13 | 328.4924 | 0.765999 | 0.206803 | 3.704006 | 0.000212 | 0.00404 |
| C9 | 0.640755 | -1.64826 | 2.754778 | -0.59833 | 0.549621 | NA |
| SEC24A | 898.4982 | 0.12082 | 0.169836 | 0.711394 | 0.47684 | 0.631755 |
| TXNDC15 | 976.2661 | 0.00281 | 0.150371 | 0.018685 | 0.985093 | 0.992141 |
| TTC33 | 309.3254 | 1.026577 | 0.238409 | 4.30595 | 1.66E-05 | 8.32E-04 |
| RARS | 1935.113 | 0.426917 | 0.175573 | 2.431569 | 0.015034 | 0.062124 |
| WWC1 | 438.8702 | 0.293182 | 0.280285 | 1.046013 | 0.295555 | 0.462854 |
| H2AFY | 2736.596 | -0.11799 | 0.126216 | -0.9348 | 0.34989 | 0.517951 |
| TCERG1 | 670.3373 | 0.607444 | 0.264111 | 2.299956 | 0.021451 | 0.079468 |
| DPYSL3 | 177.0354 | -0.61622 | 0.236415 | -2.60651 | 0.009147 | 0.044479 |
| SMAD5 | 1196.076 | 0.374886 | 0.228596 | 1.639952 | 0.101015 | 0.224425 |
| CSNK1A1 | 2164.193 | 0.26908 | 0.154079 | 1.746379 | 0.080745 | 0.192525 |
| HMGXB3 | 3520.074 | -0.27283 | 0.155174 | -1.75822 | 0.078709 | 0.189193 |

| | | | | | | |
|---|---|---|---|---|---|---|
| ERGIC1 | 3866.936 | -0.34433 | 0.127154 | -2.70798 | 0.006769 | 0.036304 |
| PDGFRB | 1.930363 | 1.160669 | 1.615427 | 0.718491 | 0.472455 | NA |
| ATP6V0E1 | 6950.13 | 0.001025 | 0.149912 | 0.00684 | 0.994543 | 0.997674 |
| BNIP1 | 248.4661 | 0.074313 | 0.175533 | 0.423355 | 0.672036 | 0.785312 |
| STC2 | 2.49803 | 2.344218 | 1.523164 | 1.539045 | 0.123793 | NA |
| CPEB4 | 1188.502 | 0.68831 | 0.183146 | 3.75826 | 1.71E-04 | 0.003521 |
| HRH2 | 607.7421 | 0.362003 | 0.328872 | 1.100741 | 0.271009 | 0.436168 |
| DBN1 | 3260.75 | -0.40809 | 0.192321 | -2.1219 | 0.033846 | 0.107486 |
| ZNF346 | 281.501 | -0.34985 | 0.164576 | -2.12575 | 0.033524 | 0.106893 |
| EHHADH | 386.341 | 0.093653 | 0.20723 | 0.451927 | 0.651322 | 0.769259 |
| CNTN3 | 3.359711 | 1.034846 | 1.163471 | 0.889448 | 0.373763 | NA |
| SMC4 | 509.104 | 1.17002 | 0.886647 | 1.3196 | 0.186969 | 0.341107 |
| TBCCD1 | 245.1701 | -0.12101 | 0.205293 | -0.58944 | 0.555567 | 0.695339 |
| TIMMDC1 | 2109.21 | 0.164575 | 0.144801 | 1.136557 | 0.255723 | 0.419889 |
| BCL6 | 645.444 | 0.35531 | 0.167226 | 2.124733 | 0.033609 | 0.107059 |
| HGD | 0.163957 | 0.424913 | 4.079006 | 0.104171 | 0.917034 | NA |
| CLDN16 | 0.458885 | 0.577525 | 3.449278 | 0.167433 | 0.867029 | NA |
| ARL6 | 120.3031 | 0.906339 | 0.296884 | 3.052837 | 0.002267 | 0.017517 |
| NPHP3 | 326.2726 | 0.575585 | 0.283634 | 2.029322 | 0.042426 | 0.125269 |
| AMOTL2 | 1362.206 | -0.08081 | 0.242136 | -0.33376 | 0.738563 | 0.834633 |
| NIT2 | 1582.111 | 0.162632 | 0.187011 | 0.86964 | 0.384497 | 0.550642 |
| FAM162A | 1140.52 | 0.325166 | 0.14059 | 2.312867 | 0.02073 | 0.077653 |
| KPNA1 | 2536.951 | -0.14142 | 0.148814 | -0.9503 | 0.341957 | 0.510118 |
| PCCB | 1323.156 | -0.20135 | 0.159411 | -1.26308 | 0.206562 | 0.363668 |
| ARMC8 | 562.1976 | 0.211467 | 0.154237 | 1.371049 | 0.17036 | 0.320797 |
| CEP70 | 247.2168 | 0.800688 | 0.245968 | 3.255255 | 0.001133 | 0.011192 |
| RBP2 | 0.260158 | 1.240347 | 4.047124 | 0.306476 | 0.759242 | NA |
| RBP1 | 0.604258 | 1.087325 | 2.707696 | 0.401568 | 0.688002 | NA |
| SLC25A36 | 1337.308 | 0.748862 | 0.205333 | 3.64707 | 2.65E-04 | 0.004599 |
| GRK7 | 0.787148 | -0.70094 | 2.357021 | -0.29738 | 0.766174 | NA |
| RNF7 | 1823.974 | 0.165435 | 0.13588 | 1.217504 | 0.223413 | 0.384238 |
| TFDP2 | 960.7408 | -0.08605 | 0.141088 | -0.60988 | 0.541942 | 0.684136 |
| XRN1 | 529.2029 | 0.473555 | 0.265309 | 1.784919 | 0.074274 | 0.18199 |
| BCHE | 1775.794 | 0.647004 | 0.191748 | 3.37425 | 7.40E-04 | 0.008671 |
| PDCD10 | 966.1968 | 1.062894 | 0.242086 | 4.39056 | 1.13E-05 | 6.60E-04 |
| FGF12 | 3.317616 | 0.016454 | 1.118979 | 0.014705 | 0.988268 | NA |
| HES1 | 1423.866 | 0.117975 | 0.136292 | 0.865604 | 0.386708 | 0.552806 |
| ACAP2 | 1072.992 | 0.527477 | 0.322118 | 1.637527 | 0.10152 | 0.225161 |
| ECT2 | 849.5672 | 0.881395 | 0.164518 | 5.357437 | 8.44E-08 | 3.13E-05 |
| TFG | 2329.298 | 0.011997 | 0.139064 | 0.08627 | 0.931251 | 0.961169 |
| RPL24 | 13170.78 | 0.403771 | 0.178847 | 2.257627 | 0.023969 | 0.085459 |
| FXR1 | 2732.392 | 0.726337 | 0.217934 | 3.332831 | 8.60E-04 | 0.009421 |
| CBLB | 481.4404 | 0.044081 | 0.171356 | 0.257249 | 0.796987 | 0.87712 |
| BBX | 1255.121 | 0.505495 | 0.281254 | 1.797292 | 0.072289 | 0.178651 |
| IFT57 | 499.6705 | 0.602581 | 0.184165 | 3.271967 | 0.001068 | 0.010822 |
| GNB4 | 810.926 | 0.409024 | 0.240311 | 1.702063 | 0.088744 | 0.205541 |
| IQCG | 87.99696 | 0.411355 | 0.267405 | 1.538321 | 0.12397 | 0.258425 |

| | | | | | | |
|---|---|---|---|---|---|---|
| GBE1 | 1504.472 | 0.54199 | 0.199213 | 2.720651 | 0.006515 | 0.035318 |
| MORC1 | 1.347189 | 0.804826 | 1.780891 | 0.451923 | 0.651324 | NA |
| UMPS | 652.7886 | 0.221454 | 0.157996 | 1.401646 | 0.161021 | 0.308584 |
| NCBP2 | 1742.685 | 0.338149 | 0.163737 | 2.065188 | 0.038905 | 0.11802 |
| SNX4 | 838.0933 | 0.892569 | 0.271855 | 3.283255 | 0.001026 | 0.010515 |
| C3orf52 | 101.3017 | 0.085171 | 0.272859 | 0.312142 | 0.754933 | 0.847111 |
| FRMD4B | 481.0811 | 0.104805 | 0.290583 | 0.360673 | 0.718344 | 0.820342 |
| SLC41A3 | 1638.879 | -0.36319 | 0.130319 | -2.78693 | 0.005321 | 0.030907 |
| ROPN1B | 221.1035 | -0.13314 | 0.190266 | -0.69977 | 0.484068 | 0.637767 |
| PLXNA1 | 6801.174 | -0.92027 | 0.196621 | -4.68042 | 2.86E-06 | 2.97E-04 |
| ATP6V1A | 5794.842 | 0.131752 | 0.167527 | 0.786452 | 0.431603 | 0.592892 |
| ABTB1 | 1030.419 | 0.094627 | 0.159745 | 0.592361 | 0.553609 | 0.693987 |
| PODXL2 | 228.895 | -0.72789 | 0.344156 | -2.115 | 0.03443 | 0.108718 |
| EFCC1 | 6.547771 | -0.76776 | 0.795793 | -0.96478 | 0.334657 | NA |
| KIAA1257 | 1.380174 | 1.51503 | 1.913079 | 0.791933 | 0.4284 | NA |
| NEK11 | 73.24137 | 0.152003 | 0.310023 | 0.490297 | 0.623924 | 0.747838 |
| MRPL3 | 1629.439 | 0.400231 | 0.132494 | 3.020738 | 0.002522 | 0.018956 |
| PLSCR4 | 381.7787 | 0.593322 | 0.207652 | 2.857287 | 0.004273 | 0.026562 |
| COMMD2 | 597.4061 | 0.556431 | 0.207196 | 2.685528 | 0.007242 | 0.038036 |
| PEX5L | 17.10063 | -1.12803 | 0.772529 | -1.46018 | 0.144241 | NA |
| ABCC5 | 1229.841 | -0.00638 | 0.181268 | -0.0352 | 0.971921 | 0.984629 |
| ARHGEF26 | 284.0418 | 0.082295 | 0.215476 | 0.381923 | 0.702518 | 0.808735 |
| KLHL24 | 896.6482 | 0.507585 | 0.201313 | 2.521373 | 0.01169 | 0.052315 |
| PLCH1 | 1.766406 | 0.99678 | 1.673542 | 0.595611 | 0.551435 | NA |
| SSR3 | 4314.252 | 0.059853 | 0.199827 | 0.299524 | 0.76454 | 0.853834 |
| CLCN2 | 104.4638 | -0.23785 | 0.227629 | -1.04489 | 0.296072 | 0.463358 |
| FOXP1 | 154.7562 | -0.07262 | 0.219815 | -0.33036 | 0.741129 | 0.836776 |
| EIF4G1 | 14363.2 | -0.49427 | 0.15493 | -3.19026 | 0.001421 | 0.013114 |
| STAG1 | 513.9278 | 0.637794 | 0.184078 | 3.464814 | 5.31E-04 | 0.007022 |
| A4GNT | 0.572651 | 0.928649 | 2.780261 | 0.334015 | 0.738368 | NA |
| ATP10B | 165.1397 | -0.67226 | 0.267804 | -2.51026 | 0.012064 | 0.053649 |
| ELOVL4 | 215.6376 | 0.355428 | 0.205978 | 1.725566 | 0.084425 | 0.197995 |
| FILIP1 | 1.120365 | -0.61682 | 2.073914 | -0.29742 | 0.766148 | NA |
| CASP8AP2 | 180.171 | 1.080988 | 0.848739 | 1.273639 | 0.202791 | 0.359115 |
| HMGN3 | 1041.622 | 0.434866 | 0.15732 | 2.764202 | 0.005706 | 0.032367 |
| UBE3D | 128.3165 | -0.05728 | 0.255144 | -0.22449 | 0.822379 | 0.893074 |
| CNR1 | 0.096202 | 0.408546 | 4.079639 | 0.100143 | 0.920231 | NA |
| PHF3 | 627.187 | 0.87178 | 0.707671 | 1.231899 | 0.217987 | 0.377425 |
| ZC2HC1B | 0.096202 | 0.408546 | 4.079639 | 0.100143 | 0.920231 | NA |
| ADGB | 0.258578 | 1.233151 | 4.047809 | 0.304646 | 0.760635 | NA |
| PLAGL1 | 0.67854 | -0.89371 | 2.366391 | -0.37767 | 0.705677 | NA |
| FBXO30 | 544.3912 | 0.512186 | 0.185032 | 2.768101 | 0.005638 | 0.032121 |
| TNFAIP3 | 710.9264 | -0.38126 | 0.173059 | -2.20307 | 0.02759 | 0.093918 |
| AKAP7 | 172.0071 | 0.23842 | 0.306529 | 0.777805 | 0.436684 | 0.597284 |
| RAB32 | 17364.11 | 0.127881 | 0.166854 | 0.766423 | 0.443425 | 0.603336 |
| MYB | 13.28514 | 0.019364 | 0.621652 | 0.03115 | 0.97515 | NA |
| ALDH8A1 | 2.220006 | -0.5859 | 1.353543 | -0.43287 | 0.665111 | NA |

| | | | | | | |
|---|---|---|---|---|---|---|
| SGK1 | 12949.9 | -0.01426 | 0.261725 | -0.05449 | 0.956546 | 0.976145 |
| RNF146 | 871.4667 | 0.534321 | 0.161976 | 3.298758 | 9.71E-04 | 0.010161 |
| ARG1 | 0.640609 | 0.042838 | 2.379511 | 0.018003 | 0.985637 | NA |
| CTGF | 56.9022 | 0.767964 | 0.388765 | 1.975394 | 0.048223 | 0.135498 |
| FBXL5 | 2155.548 | 0.050153 | 0.191068 | 0.262487 | 0.792946 | 0.874397 |
| MED28 | 732.7065 | 0.003743 | 0.160623 | 0.023303 | 0.981409 | 0.990173 |
| FOXO3 | 610.5767 | -0.48613 | 0.19412 | -2.50426 | 0.012271 | 0.054213 |
| ARMC2 | 35.59993 | 0.143034 | 0.377423 | 0.378974 | 0.704707 | NA |
| PKD2 | 3508.71 | 0.456574 | 0.143581 | 3.179898 | 0.001473 | 0.013425 |
| ABCG2 | 62.61801 | 0.739359 | 0.344832 | 2.144113 | 0.032024 | 0.103756 |
| SPP1 | 53.80006 | -1.04145 | 0.527012 | -1.97613 | 0.04814 | 0.135397 |
| FAM47E-ST | 74.57004 | 0.014547 | 0.270498 | 0.053777 | 0.957113 | 0.976409 |
| CCNI | 5131.77 | -0.40534 | 0.120385 | -3.36704 | 7.60E-04 | 0.008794 |
| RARRES1 | 22.31499 | -0.09095 | 0.473638 | -0.19203 | 0.847717 | NA |
| MFSD1 | 2703.668 | 0.197559 | 0.13468 | 1.466877 | 0.14241 | 0.283219 |
| ELL2 | 397.4689 | 0.458311 | 0.165686 | 2.766143 | 0.005672 | 0.032268 |
| GLRXP3 | 0.315653 | -0.57305 | 3.892639 | -0.14721 | 0.882963 | NA |
| UBE2B | 1289.84 | 0.505406 | 0.143955 | 3.510857 | 4.47E-04 | 0.006364 |
| PIGZ | 197.7822 | -0.32271 | 0.244774 | -1.3184 | 0.187369 | 0.341521 |
| SENP5 | 854.4017 | 0.19811 | 0.181104 | 1.093899 | 0.273999 | 0.439786 |
| SLC17A5 | 1314.145 | 0.218053 | 0.181384 | 1.202163 | 0.2293 | 0.39113 |
| OGFRL1 | 151.4809 | 0.096937 | 0.284191 | 0.341098 | 0.73303 | 0.831029 |
| DUSP1 | 279.2147 | 0.025589 | 0.309398 | 0.082706 | 0.934085 | 0.963062 |
| PANK3 | 1955.527 | 0.630152 | 0.247897 | 2.541994 | 0.011022 | 0.050218 |
| MSX2 | 55.4149 | -0.30943 | 0.348431 | -0.88806 | 0.374507 | 0.540843 |
| GRIA2 | 0.151697 | -1.546 | 4.075469 | -0.37934 | 0.704433 | NA |
| NUP43 | 639.672 | 0.131674 | 0.224138 | 0.587468 | 0.556889 | 0.696185 |
| MTHFD1L | 2162.374 | -0.11076 | 0.203885 | -0.54326 | 0.586951 | 0.719576 |
| LRP11 | 445.9335 | -0.05918 | 0.172634 | -0.34278 | 0.731763 | 0.829991 |
| PCMT1 | 5260.66 | 0.134513 | 0.159741 | 0.842066 | 0.399751 | 0.564374 |
| PLEKHG1 | 323.2185 | -0.01168 | 0.224078 | -0.05213 | 0.958427 | 0.977053 |
| CYSTM1 | 2673.982 | -0.05407 | 0.156097 | -0.34639 | 0.729052 | 0.827945 |
| WDR55 | 1381.79 | -0.16967 | 0.158409 | -1.07106 | 0.284142 | 0.450757 |
| ARAP3 | 1746.564 | -0.51164 | 0.139372 | -3.67103 | 2.42E-04 | 0.004366 |
| PCDHB8 | 2.127664 | 1.692754 | 1.452671 | 1.165269 | 0.24391 | NA |
| PCDHB10 | 1.932835 | -0.51746 | 1.539434 | -0.33614 | 0.736767 | NA |
| PCDHB14 | 10.93435 | 0.508748 | 0.708758 | 0.717803 | 0.472879 | NA |
| PCDHB12 | 2.010535 | 2.225923 | 1.666585 | 1.33562 | 0.181674 | NA |
| SLC25A2 | 0.096202 | 0.408546 | 4.079639 | 0.100143 | 0.920231 | NA |
| SLC10A7 | 104.9285 | 0.486945 | 0.249588 | 1.950993 | 0.051058 | 0.140634 |
| ETF1 | 4994.126 | 0.483927 | 0.177819 | 2.721453 | 0.0065 | 0.035248 |
| TGFBI | 237.6707 | 1.751346 | 0.397909 | 4.401372 | 1.08E-05 | 6.41E-04 |
| FAM53C | 2151.781 | -0.29798 | 0.149664 | -1.99102 | 0.046479 | 0.132518 |
| SIL1 | 2006.209 | -0.37568 | 0.143911 | -2.61049 | 0.009041 | 0.044218 |
| PAIP2 | 1444.539 | 0.483729 | 0.168228 | 2.87543 | 0.004035 | 0.025524 |
| MYOT | 70.0903 | 1.971143 | 0.514966 | 3.827714 | 1.29E-04 | 0.002993 |
| KDM3B | 2351.822 | -0.42054 | 0.126021 | -3.3371 | 8.47E-04 | 0.009353 |

| | | | | | | |
|---|---|---|---|---|---|---|
| EGR1 | 130.4803 | 0.701052 | 0.455347 | 1.539599 | 0.123658 | 0.258002 |
| SERP1 | 2411.566 | 0.048102 | 0.179862 | 0.267438 | 0.789132 | 0.872032 |
| PLS1 | 79.89346 | 0.748968 | 0.350237 | 2.138463 | 0.032479 | 0.104859 |
| NACA3P | 3.204965 | 0.061803 | 1.388805 | 0.044501 | 0.964505 | NA |
| LRAT | 24.49904 | 0.424196 | 0.493009 | 0.860422 | 0.389557 | NA |
| KIAA0922 | 325.7859 | -0.37653 | 0.207641 | -1.81339 | 0.069772 | 0.174038 |
| MND1 | 50.15761 | 0.828344 | 0.316441 | 2.617686 | 0.008853 | 0.043566 |
| PDZRN3 | 2535.48 | 0.473966 | 0.21071 | 2.249378 | 0.024488 | 0.086686 |
| SEC22A | 263.3508 | 0.281679 | 0.192948 | 1.459869 | 0.144326 | 0.286166 |
| DPPA4 | 0.47803 | 0.520083 | 2.97881 | 0.174594 | 0.861399 | NA |
| POPDC2 | 12.83261 | -0.30108 | 0.622656 | -0.48354 | 0.628714 | NA |
| B4GALT4 | 672.6729 | 0.361725 | 0.165714 | 2.18282 | 0.029049 | 0.09698 |
| NAA50 | 2454.838 | 0.384463 | 0.164309 | 2.339881 | 0.01929 | 0.073814 |
| CD80 | 0.230334 | -0.53318 | 4.076749 | -0.13079 | 0.895945 | NA |
| TNFSF10 | 10.51905 | -0.1957 | 0.673689 | -0.29049 | 0.771438 | NA |
| ZNF639 | 348.4077 | 0.298839 | 0.186906 | 1.598869 | 0.10985 | 0.23738 |
| SLITRK3 | 1.113684 | -0.35671 | 2.036264 | -0.17518 | 0.86094 | NA |
| PIK3CA | 418.2185 | 0.778789 | 0.279594 | 2.785422 | 0.005346 | 0.031036 |
| PDS5A | 2328.757 | 0.548988 | 0.176205 | 3.115626 | 0.001836 | 0.015242 |
| LIAS | 150.4034 | 0.082384 | 0.241348 | 0.341349 | 0.732841 | 0.830894 |
| POLK | 446.9837 | 1.340858 | 0.328137 | 4.086271 | 4.38E-05 | 0.001538 |
| SV2C | 0.644964 | 2.579089 | 2.975624 | 0.866739 | 0.386085 | NA |
| FYTTD1 | 1581.692 | 0.609792 | 0.187493 | 3.25234 | 0.001145 | 0.01127 |
| KIAA1191 | 4785.317 | -0.0869 | 0.146117 | -0.59473 | 0.552021 | 0.692949 |
| NLN | 820.2666 | 0.197971 | 0.207637 | 0.953449 | 0.340363 | 0.508571 |
| CENPK | 85.2852 | 1.468199 | 0.306845 | 4.784823 | 1.71E-06 | 2.24E-04 |
| COL10A1 | 0.314073 | -0.57308 | 3.898934 | -0.14698 | 0.883146 | NA |
| AMD1 | 2821.01 | 0.092501 | 0.204361 | 0.452633 | 0.650813 | 0.768772 |
| NDUFAF4 | 435.3489 | 0.689288 | 0.153705 | 4.484474 | 7.31E-06 | 5.32E-04 |
| USP45 | 192.6273 | 0.921924 | 0.277877 | 3.317738 | 9.07E-04 | 0.009758 |
| SLC36A1 | 630.4396 | -0.01513 | 0.181931 | -0.08317 | 0.933715 | 0.962789 |
| EXOSC9 | 516.335 | 0.511504 | 0.175106 | 2.921115 | 0.003488 | 0.023268 |
| PLA2G12A | 760.6644 | 0.205496 | 0.205889 | 0.998093 | 0.318234 | 0.486043 |
| MXD4 | 6714.316 | -0.2528 | 0.125588 | -2.01291 | 0.044125 | 0.128084 |
| MTRR | 1219.047 | 0.291966 | 0.16459 | 1.773904 | 0.076079 | 0.184819 |
| FASTKD3 | 256.1745 | 0.534121 | 0.189449 | 2.819337 | 0.004812 | 0.028784 |
| | 0.601483 | -1.7332 | 2.729494 | -0.63499 | 0.525436 | NA |
| ATP8A1 | 34.68855 | 0.818601 | 0.379341 | 2.157956 | 0.030931 | NA |
| GRM4 | 0.163957 | 0.424913 | 4.079006 | 0.104171 | 0.917034 | NA |
| TRERF1 | 8.056973 | -0.4465 | 0.765069 | -0.5836 | 0.559487 | NA |
| BTN2A2 | 865.0319 | -0.10595 | 0.175808 | -0.60267 | 0.546727 | 0.688197 |
| SIRT5 | 267.6906 | 0.032575 | 0.167967 | 0.193936 | 0.846226 | 0.908697 |
| HIST1H4B | 0.163957 | 0.424913 | 4.079006 | 0.104171 | 0.917034 | NA |
| MRS2 | 916.3212 | 0.192593 | 0.159034 | 1.211017 | 0.225889 | 0.387093 |
| WRNIP1 | 2011.893 | -0.04366 | 0.177118 | -0.2465 | 0.805294 | 0.88355 |
| RRP36 | 1680.749 | 0.097722 | 0.132854 | 0.735558 | 0.462 | 0.620272 |
| BTN2A3P | 98.40804 | 0.611738 | 0.268007 | 2.282548 | 0.022457 | 0.081967 |

| | | | | | | |
|---|---|---|---|---|---|---|
| BTN1A1 | 1.714724 | 0.059918 | 1.668681 | 0.035907 | 0.971356 | NA |
| SNRPC | 1914.574 | 0.003471 | 0.130583 | 0.026579 | 0.978796 | 0.988636 |
| SLC17A3 | 0.151697 | -1.546 | 4.075469 | -0.37934 | 0.704433 | NA |
| SERPINB6 | 4614.464 | -0.16077 | 0.126782 | -1.26804 | 0.204783 | 0.361426 |
| XPO5 | 1650.564 | -0.37275 | 0.145302 | -2.56535 | 0.010307 | 0.048042 |
| ABCC10 | 1121.666 | -0.34615 | 0.1566 | -2.21038 | 0.027079 | 0.09262 |
| HIST1H1D | 2.28652 | 1.927295 | 1.528093 | 1.261242 | 0.207222 | NA |
| PEX6 | 1305.149 | -0.41125 | 0.161513 | -2.54623 | 0.010889 | 0.049746 |
| NQO2 | 1675.352 | 0.134761 | 0.159803 | 0.843294 | 0.399064 | 0.563807 |
| PRICKLE4 | 154.402 | 0.141066 | 0.207319 | 0.68043 | 0.496232 | 0.647533 |
| OARD1 | 497.4302 | 0.285159 | 0.171015 | 1.667445 | 0.095426 | 0.216181 |
| UNC5CL | 9.572125 | 0.028355 | 0.739349 | 0.038352 | 0.969407 | NA |
| AARS2 | 718.5941 | -0.38279 | 0.169046 | -2.2644 | 0.02355 | 0.084549 |
| HIST1H1A | 0.314073 | -0.57308 | 3.898934 | -0.14698 | 0.883146 | NA |
| ZNF391 | 45.69331 | 0.490918 | 0.432142 | 1.136011 | 0.255952 | NA |
| RPS10 | 8842.5 | 0.070822 | 0.155204 | 0.456314 | 0.648164 | 0.76633 |
| MOCS1 | 863.4787 | -0.34971 | 0.138584 | -2.52347 | 0.01162 | 0.052181 |
| HIST1H2BJ | 30.31982 | 0.450669 | 0.629311 | 0.716131 | 0.47391 | NA |
| MED20 | 485.6729 | -0.33676 | 0.165243 | -2.03796 | 0.041554 | 0.123649 |
| OR2B6 | 0.392857 | -1.546 | 4.075469 | -0.37934 | 0.704433 | NA |
| TBCC | 842.4761 | -0.22525 | 0.136999 | -1.64417 | 0.100142 | 0.223272 |
| SPDEF | 0.911593 | 1.257757 | 2.250933 | 0.558771 | 0.576318 | NA |
| TCP11 | 0.303394 | -2.27006 | 4.032451 | -0.56295 | 0.57347 | NA |
| MAD2L1BP | 1426.042 | 0.062616 | 0.143494 | 0.436366 | 0.662571 | 0.778086 |
| HIST1H3B | 1.449704 | -0.80325 | 1.854551 | -0.43313 | 0.664924 | NA |
| APOBEC2 | 0.881769 | -0.20384 | 2.229339 | -0.09144 | 0.927147 | NA |
| KLHDC3 | 2394.409 | -0.19923 | 0.132103 | -1.5081 | 0.131528 | 0.268237 |
| GNMT | 3.137402 | -0.76134 | 1.141897 | -0.66673 | 0.504945 | NA |
| DNAH8 | 0.268266 | -2.15396 | 4.038351 | -0.53338 | 0.593773 | NA |
| MEA1 | 1568.58 | -0.01533 | 0.163 | -0.09407 | 0.925053 | 0.957356 |
| KLHL31 | 0.614937 | 2.493639 | 2.617677 | 0.952615 | 0.340785 | NA |
| COL21A1 | 0.096202 | 0.408546 | 4.079639 | 0.100143 | 0.920231 | NA |
| CDKN1A | 7407.1 | -0.05651 | 0.156118 | -0.36197 | 0.717371 | 0.819783 |
| SOX4 | 2407.972 | -0.41176 | 0.169815 | -2.42475 | 0.015319 | 0.062905 |
| GLO1 | 3758.032 | 0.111958 | 0.156921 | 0.713467 | 0.475557 | 0.63076 |
| CPNE5 | 0.495797 | -0.73607 | 3.324282 | -0.22142 | 0.824763 | NA |
| RREB1 | 1028.051 | -0.65387 | 0.186608 | -3.50399 | 4.58E-04 | 0.006438 |
| SSR1 | 3043.082 | 0.149304 | 0.147758 | 1.010461 | 0.312274 | 0.479394 |
| RIOK1 | 365.4446 | 0.576777 | 0.224222 | 2.572353 | 0.010101 | 0.047462 |
| NRN1 | 13.76436 | 2.549635 | 0.763249 | 3.340504 | 8.36E-04 | NA |
| SLC35B3 | 890.373 | 0.523099 | 0.155264 | 3.369085 | 7.54E-04 | 0.008786 |
| RPP40 | 140.7268 | 0.078383 | 0.226354 | 0.346286 | 0.729127 | 0.827951 |
| ATXN1 | 816.1245 | -0.07032 | 0.202974 | -0.34645 | 0.729008 | 0.827945 |
| NUP153 | 1386.471 | -0.1878 | 0.144489 | -1.29975 | 0.193687 | 0.348801 |
| DEK | 1148.275 | 1.048258 | 0.277064 | 3.783446 | 1.55E-04 | 0.003334 |
| EEF1E1 | 139.0423 | 0.940437 | 0.222446 | 4.227712 | 2.36E-05 | 0.001014 |
| RUNX2 | 65.68573 | -0.22285 | 0.333491 | -0.66825 | 0.503975 | 0.653751 |

| | | | | | | |
|---|---|---|---|---|---|---|
| OPN5 | 0.35636 | 1.709252 | 3.651833 | 0.468053 | 0.639746 | NA |
| CXCL6 | 8.628908 | -1.09391 | 0.755043 | -1.44881 | 0.147392 | NA |
| EREG | 0.192403 | 0.975392 | 4.060716 | 0.240202 | 0.810174 | NA |
| SH3TC1 | 18.05295 | -0.86785 | 0.533265 | -1.62743 | 0.103646 | NA |
| IRF1 | 162.9769 | -0.30376 | 0.26631 | -1.14064 | 0.254021 | 0.418046 |
| FAM193A | 715.0561 | -0.51475 | 0.165547 | -3.10941 | 0.001875 | 0.015465 |
| GRK4 | 299.3721 | -0.13361 | 0.165216 | -0.80872 | 0.418676 | 0.581671 |
| STATH | 0.096202 | 0.408546 | 4.079639 | 0.100143 | 0.920231 | NA |
| CANX | 14632.42 | 0.13347 | 0.122047 | 1.093594 | 0.274133 | 0.439821 |
| COX7C | 5372.61 | 0.349345 | 0.13781 | 2.534975 | 0.011246 | 0.050962 |
| MASP1 | 1.605125 | -1.74683 | 1.620538 | -1.07793 | 0.281064 | NA |
| HRASLS | 18.8955 | -0.30084 | 0.482925 | -0.62296 | 0.53331 | NA |
| IDUA | 244.5049 | -0.06765 | 0.187988 | -0.35986 | 0.718953 | 0.8208 |
| FGFRL1 | 336.2286 | 0.09361 | 0.223355 | 0.41911 | 0.675136 | 0.787382 |
| TMEM175 | 713.2383 | -0.41232 | 0.154261 | -2.67288 | 0.00752 | 0.039107 |
| SRD5A3 | 1083.781 | -0.0721 | 0.139194 | -0.51799 | 0.604468 | 0.732942 |
| SPINK2 | 10.93856 | 1.083652 | 0.802678 | 1.350046 | 0.177001 | NA |
| RASL11B | 4.157926 | 0.323471 | 1.279083 | 0.252893 | 0.800351 | NA |
| PAICS | 4041.41 | 0.209859 | 0.142215 | 1.475648 | 0.140038 | 0.280337 |
| KDR | 264.1325 | -0.18731 | 0.350986 | -0.53366 | 0.593579 | 0.724107 |
| PPAT | 389.8005 | 0.561393 | 0.218229 | 2.572494 | 0.010097 | 0.047462 |
| ACKR4 | 2.853527 | 0.255871 | 1.229786 | 0.208062 | 0.835181 | NA |
| ANAPC13 | 1586.94 | 0.213954 | 0.141489 | 1.512157 | 0.130494 | 0.26704 |
| MBD4 | 413.3845 | 0.633082 | 0.178872 | 3.539299 | 4.01E-04 | 0.005923 |
| PALLD | 687.7807 | 0.507184 | 0.220541 | 2.299729 | 0.021464 | 0.079491 |
| SPCS3 | 2603.816 | 0.51398 | 0.189477 | 2.712629 | 0.006675 | 0.035967 |
| DCTD | 2478.858 | -0.20642 | 0.126391 | -1.63321 | 0.102425 | 0.226539 |
| EPB41L4A | 262.1974 | -0.03994 | 0.176627 | -0.2261 | 0.821124 | 0.892334 |
| CDO1 | 12.21506 | -0.04637 | 0.625303 | -0.07416 | 0.940883 | NA |
| REEP5 | 3247.369 | 0.256004 | 0.142575 | 1.795575 | 0.072562 | 0.179101 |
| RTN4IP1 | 204.0697 | -0.26933 | 0.198485 | -1.35695 | 0.174798 | 0.326835 |
| QRSL1 | 574.5099 | 0.342338 | 0.149009 | 2.297426 | 0.021594 | 0.079801 |
| C6orf203 | 305.1116 | 0.382318 | 0.18385 | 2.079511 | 0.03757 | 0.115179 |
| ZSWIM6 | 550.9141 | 0.118291 | 0.188841 | 0.626408 | 0.531047 | 0.675555 |
| POLN | 42.80983 | 0.430584 | 0.344502 | 1.249874 | 0.211346 | NA |
| PPIL4 | 375.4117 | 1.109032 | 0.322608 | 3.43771 | 5.87E-04 | 0.007468 |
| ULBP2 | 74.68838 | 0.213466 | 0.305531 | 0.698672 | 0.484757 | 0.63832 |
| ULBP3 | 19.4946 | 0.200218 | 0.466633 | 0.429071 | 0.667872 | NA |
| LATS1 | 465.791 | 0.081967 | 0.251932 | 0.325352 | 0.744914 | 0.839379 |
| ZNF141 | 46.73472 | 0.666205 | 0.417338 | 1.596319 | 0.110418 | NA |
| SLC34A1 | 1.153998 | -2.85566 | 2.193533 | -1.30185 | 0.192966 | NA |
| F12 | 10.66307 | -0.20899 | 0.700732 | -0.29824 | 0.765519 | NA |
| PRR7 | 142.0493 | -0.79685 | 0.349137 | -2.28235 | 0.022469 | 0.081968 |
| PDLIM4 | 201.8072 | -0.37831 | 0.195454 | -1.93554 | 0.052924 | 0.143837 |
| KIF3A | 508.6539 | 0.645408 | 0.341424 | 1.890342 | 0.058712 | 0.154812 |
| MGAT1 | 8228.958 | -0.38988 | 0.121714 | -3.20325 | 0.001359 | 0.012675 |
| GFPT2 | 56.40127 | -0.60621 | 0.308688 | -1.96383 | 0.04955 | 0.137692 |

| | | | | | | |
|---|---|---|---|---|---|---|
| NDUFA2 | 1920.867 | 0.131566 | 0.15929 | 0.825956 | 0.408829 | 0.572419 |
| ANKHD1 | 16.1308 | 0.27792 | 0.528995 | 0.525373 | 0.599324 | NA |
| DIAPH1 | 4561.391 | -0.29966 | 0.159699 | -1.87638 | 0.060604 | 0.158426 |
| NDFIP1 | 3838.538 | 0.011493 | 0.148695 | 0.077294 | 0.938389 | 0.965761 |
| UBE2D2 | 2931.053 | -0.12259 | 0.143282 | -0.8556 | 0.392217 | 0.557321 |
| MAP1B | 1708.15 | 0.042941 | 0.27318 | 0.157191 | 0.875094 | 0.925378 |
| CKMT2 | 13.46129 | 0.318266 | 0.55988 | 0.568453 | 0.569727 | NA |
| ZCCHC9 | 735.5231 | 0.919083 | 0.166189 | 5.530334 | 3.20E-08 | 1.65E-05 |
| MCCC2 | 1393.351 | -0.32709 | 0.133036 | -2.45869 | 0.013944 | 0.05904 |
| PRKAA1 | 1139.186 | 0.638934 | 0.213088 | 2.998452 | 0.002714 | 0.019907 |
| CARD6 | 331.2424 | 0.281554 | 0.245635 | 1.146229 | 0.251701 | 0.415702 |
| EEFSEC | 412.6329 | -0.52632 | 0.200603 | -2.62369 | 0.008698 | 0.042987 |
| TBC1D14 | 1566.596 | -0.27232 | 0.16497 | -1.6507 | 0.0988 | 0.221581 |
| TMEM128 | 575.9747 | 0.175775 | 0.15858 | 1.10843 | 0.267676 | 0.432663 |
| COQ3 | 198.5121 | 0.251594 | 0.212482 | 1.184074 | 0.236384 | 0.399195 |
| PNISR | 487.7329 | 1.257176 | 0.640083 | 1.964082 | 0.049521 | 0.137692 |
| POPDC3 | 1015.02 | 0.346703 | 0.13498 | 2.568543 | 0.010213 | 0.047836 |
| GRSF1 | 3384.141 | 0.023477 | 0.148611 | 0.157974 | 0.874477 | 0.925145 |
| ENAM | 0.096202 | 0.408546 | 4.079639 | 0.100143 | 0.920231 | NA |
| IGJ | 0.315653 | -0.57305 | 3.892639 | -0.14721 | 0.882963 | NA |
| ANKRD17 | 2393.033 | 0.055701 | 0.192208 | 0.289795 | 0.771973 | 0.858857 |
| UTP3 | 672.9604 | 0.265031 | 0.231788 | 1.143421 | 0.252864 | 0.4171 |
| REEP2 | 363.0245 | -0.14709 | 0.259051 | -0.56781 | 0.570161 | 0.706149 |
| PCBD2 | 87.42219 | 0.159303 | 0.256667 | 0.620661 | 0.534823 | 0.678307 |
| DMGDH | 6.088804 | -0.16832 | 0.825402 | -0.20393 | 0.83841 | NA |
| BHMT2 | 2.924557 | 0.520008 | 1.207106 | 0.430789 | 0.666621 | NA |
| AP3B1 | 1829.162 | 0.191014 | 0.134463 | 1.420566 | 0.155443 | 0.301206 |
| ZBED3 | 526.5973 | -0.36682 | 0.202484 | -1.81159 | 0.070049 | 0.174461 |
| DCTN4 | 2152.327 | -0.155 | 0.140607 | -1.10239 | 0.270291 | 0.435249 |
| PDE6A | 0.638825 | 1.117948 | 3.106359 | 0.35989 | 0.718929 | NA |
| PDE6B | 2.946864 | 0.225162 | 1.229126 | 0.183189 | 0.85465 | NA |
| ANKRD32 | 148.863 | 1.187084 | 0.289098 | 4.106165 | 4.02E-05 | 0.001472 |
| MED10 | 630.1268 | 0.421991 | 0.152694 | 2.763638 | 0.005716 | 0.032408 |
| PDZD2 | 57.892 | -1.7729 | 0.526817 | -3.3653 | 7.65E-04 | 0.00881 |
| ATP13A3 | 1478.616 | 0.764985 | 0.250567 | 3.053022 | 0.002265 | 0.017517 |
| LARS | 3145.771 | 0.397986 | 0.142373 | 2.79537 | 0.005184 | 0.030299 |
| SPINK5 | 3.113539 | 0.860327 | 1.14635 | 0.750492 | 0.452958 | NA |
| HSD17B4 | 3638.061 | 0.143709 | 0.154156 | 0.932236 | 0.351214 | 0.519158 |
| MRPS36 | 480.4661 | 0.547465 | 0.152331 | 3.593916 | 3.26E-04 | 0.005188 |
| CCNB1 | 2632.018 | -0.21879 | 0.216204 | -1.01196 | 0.311557 | 0.478921 |
| CDK7 | 432.6552 | 0.509761 | 0.176807 | 2.883145 | 0.003937 | 0.025203 |
| CD180 | 1.15657 | -0.18697 | 1.786779 | -0.10464 | 0.916661 | NA |
| IL6ST | 1843.407 | 0.945216 | 0.241673 | 3.911128 | 9.19E-05 | 0.002404 |
| FST | 74.336 | 0.937396 | 0.444438 | 2.10917 | 0.03493 | 0.109781 |
| CCNH | 1003.347 | 0.423679 | 0.165572 | 2.558887 | 0.010501 | 0.048647 |
| DOCK2 | 47.82316 | 0.353442 | 0.481926 | 0.733394 | 0.463318 | 0.621129 |
| TMEM165 | 2852.339 | 0.411861 | 0.137866 | 2.987404 | 0.002814 | 0.020314 |

| | | | | | | |
|---|---|---|---|---|---|---|
| CLOCK | 911.9717 | 0.649938 | 0.223429 | 2.908928 | 0.003627 | 0.024006 |
| PDGFRA | 2.502168 | -0.28004 | 1.24218 | -0.22544 | 0.821633 | NA |
| KLB | 1.708254 | 1.515646 | 1.692807 | 0.895345 | 0.370603 | NA |
| TMED7 | 1489.959 | 0.295877 | 0.199339 | 1.48429 | 0.137732 | 0.277124 |
| APC | 1045.86 | 0.75822 | 0.339678 | 2.232176 | 0.025603 | 0.089457 |
| NREP | 1.06329 | 0.285492 | 1.942399 | 0.146979 | 0.883149 | NA |
| WDR36 | 554.3421 | 0.437026 | 0.20822 | 2.098864 | 0.035829 | 0.111632 |
| ADAM19 | 480.5836 | 0.02303 | 0.251434 | 0.091596 | 0.927019 | 0.958156 |
| HAVCR2 | 0.556667 | 0.972496 | 2.522454 | 0.385536 | 0.699841 | NA |
| CCNJL | 548.5122 | -0.33709 | 0.16256 | -2.07361 | 0.038116 | 0.116339 |
| MTO1 | 449.6229 | 0.242475 | 0.180695 | 1.341907 | 0.179626 | 0.332211 |
| BAI3 | 19.78432 | -0.30506 | 0.548346 | -0.55633 | 0.577987 | NA |
| ANKRD6 | 276.8034 | 0.583473 | 0.169005 | 3.452398 | 5.56E-04 | 0.007215 |
| HTR1B | 0.411855 | 0.250456 | 3.097618 | 0.080854 | 0.935558 | NA |
| KHDC1 | 5.685096 | 0.068273 | 0.910546 | 0.07498 | 0.940231 | NA |
| KIAA1009 | 66.14402 | 1.026699 | 1.043032 | 0.984341 | 0.324948 | 0.492847 |
| SYNCRIP | 2621.505 | 0.3773 | 0.153031 | 2.465517 | 0.013682 | 0.058239 |
| SNX14 | 1200.826 | 0.45146 | 0.16631 | 2.714566 | 0.006636 | 0.035809 |
| NT5E | 825.096 | 0.371044 | 0.257035 | 1.443553 | 0.148865 | 0.292187 |
| MRAP2 | 118.7027 | 0.044262 | 0.255099 | 0.173511 | 0.86225 | 0.918473 |
| AKIRIN2 | 562.2163 | -0.23324 | 0.159577 | -1.46159 | 0.143853 | 0.285467 |
| ORC3 | 477.0586 | 0.516068 | 0.234826 | 2.197661 | 0.027973 | 0.094695 |
| LCA5 | 90.00538 | 0.990387 | 0.454101 | 2.180982 | 0.029185 | 0.097105 |
| MAP3K7 | 907.0234 | 0.63327 | 0.167731 | 3.775512 | 1.60E-04 | 0.003412 |
| LTV1 | 458.8138 | 0.933933 | 0.250994 | 3.72093 | 1.98E-04 | 0.003853 |
| MAP7 | 74.83942 | 0.311893 | 0.315208 | 0.989485 | 0.322426 | 0.490549 |
| CD164 | 6422.352 | 0.301013 | 0.143475 | 2.098016 | 0.035904 | 0.111755 |
| LACE1 | 148.5571 | 0.465874 | 0.244568 | 1.904885 | 0.056795 | 0.151046 |
| NHSL1 | 2787.732 | -0.6422 | 0.2002 | -3.20778 | 0.001338 | 0.012526 |
| AHI1 | 496.6457 | 1.141756 | 0.240054 | 4.756245 | 1.97E-06 | 2.46E-04 |
| HEY2 | 1.359945 | -0.90023 | 1.873741 | -0.48045 | 0.63091 | NA |
| PKIB | 343.2161 | 0.52552 | 0.247435 | 2.123866 | 0.033681 | 0.107164 |
| SMPD2 | 977.9126 | -0.19877 | 0.176607 | -1.12549 | 0.260379 | 0.424594 |
| MICAL1 | 8511.348 | -0.47893 | 0.189844 | -2.52276 | 0.011644 | 0.052224 |
| REPS1 | 907.4195 | -0.24439 | 0.13712 | -1.78231 | 0.074699 | 0.182427 |
| STX11 | 73.81687 | 0.526668 | 0.282219 | 1.866167 | 0.062018 | 0.161078 |
| TEC | 45.44948 | 0.675798 | 0.333781 | 2.024675 | 0.042901 | NA |
| RTP4 | 311.8111 | -0.76817 | 0.269167 | -2.85387 | 0.004319 | 0.026723 |
| ACTL6A | 1461.403 | 0.131234 | 0.169383 | 0.774778 | 0.438471 | 0.59858 |
| NDUFB5 | 2469.982 | 0.130415 | 0.163152 | 0.799345 | 0.42409 | 0.586153 |
| MRPL47 | 842.256 | 0.606058 | 0.174409 | 3.474931 | 5.11E-04 | 0.006892 |
| TRA2B | 2512.733 | 0.063978 | 0.149166 | 0.428905 | 0.667992 | 0.782205 |
| SKIL | 272.4891 | 1.509825 | 0.347654 | 4.342903 | 1.41E-05 | 7.65E-04 |
| CNPY3 | 2537.365 | -0.28201 | 0.189459 | -1.48849 | 0.136622 | 0.275639 |
| FOXP4 | 762.1052 | -0.38475 | 0.149108 | -2.58033 | 0.00987 | 0.046768 |
| PPIL1 | 1091.079 | 7.69E-04 | 0.159048 | 0.004832 | 0.996145 | 0.99825 |
| KLC4 | 517.5287 | -0.25359 | 0.17258 | -1.46943 | 0.141717 | 0.282551 |

| | | | | | | |
|---|---|---|---|---|---|---|
| KIF13A | 2450.262 | -0.02817 | 0.152391 | -0.18487 | 0.853333 | 0.912938 |
| ZSCAN9 | 183.0523 | 0.022057 | 0.21694 | 0.101674 | 0.919015 | 0.953103 |
| PIM1 | 100.5421 | -0.19255 | 0.314236 | -0.61274 | 0.540046 | 0.682173 |
| GMPR | 8289.199 | -0.62448 | 0.198142 | -3.15165 | 0.001623 | 0.014248 |
| CMTR1 | 2052.513 | -0.3716 | 0.128045 | -2.90212 | 0.003706 | 0.024303 |
| TFAP2A | 10099.18 | -0.3289 | 0.163447 | -2.01229 | 0.04419 | 0.128169 |
| YIPF3 | 3936.3 | -0.20628 | 0.138911 | -1.485 | 0.137543 | 0.276837 |
| TMEM14B | 1002.877 | 0.152333 | 0.13593 | 1.120674 | 0.262427 | 0.426852 |
| TMEM63B | 2412.598 | -0.27244 | 0.146317 | -1.86199 | 0.062604 | 0.161941 |
| FRS3 | 134.2992 | -0.41729 | 0.254011 | -1.6428 | 0.100424 | 0.223616 |
| TJAP1 | 763.1979 | -0.34568 | 0.142676 | -2.4228 | 0.015401 | 0.063089 |
| CAPN11 | 2.004414 | 2.249048 | 1.52881 | 1.47111 | 0.141261 | NA |
| HIST1H2AB | 0.833158 | -0.52049 | 2.521552 | -0.20642 | 0.836464 | NA |
| KIAA0319 | 1.169033 | -0.01411 | 1.776406 | -0.00794 | 0.993662 | NA |
| IRF4 | 7289.164 | -0.45974 | 0.150859 | -3.04751 | 0.002307 | 0.017728 |
| SLC22A23 | 342.513 | -0.11527 | 0.230719 | -0.49962 | 0.617344 | 0.742508 |
| TUBB2A | 1144.688 | -0.26589 | 0.188849 | -1.40795 | 0.159147 | 0.306013 |
| LRRC1 | 153.4008 | 0.416153 | 0.299747 | 1.388348 | 0.165031 | 0.313807 |
| GCM1 | 0.163957 | 0.424913 | 4.079006 | 0.104171 | 0.917034 | NA |
| FOXF2 | 391.5908 | 0.20332 | 0.183622 | 1.107272 | 0.268176 | 0.433138 |
| BPHL | 752.15 | -0.27382 | 0.148302 | -1.84638 | 0.064837 | 0.165956 |
| RIPK1 | 1352.289 | 0.037761 | 0.147451 | 0.256091 | 0.797881 | 0.877773 |
| TUBB2B | 616.831 | -0.35523 | 0.203214 | -1.74808 | 0.08045 | 0.192131 |
| UQCC2 | 1856.089 | -0.0185 | 0.14278 | -0.1296 | 0.89688 | 0.939766 |
| HMGA1 | 5722.691 | -0.29585 | 0.126465 | -2.33935 | 0.019317 | 0.07385 |
| TCF19 | 530.269 | -0.24695 | 0.294961 | -0.83722 | 0.402467 | 0.566889 |
| FLOT1 | 7655.127 | -0.13606 | 0.136207 | -0.99895 | 0.317818 | 0.485595 |
| IER3 | 1740.162 | 0.258519 | 0.255822 | 1.010543 | 0.312235 | 0.479394 |
| MDC1 | 763.7894 | -0.59991 | 0.1921 | -3.12291 | 0.001791 | 0.015078 |
| PGBD1 | 254.5109 | 0.337229 | 0.193134 | 1.746086 | 0.080796 | 0.192597 |
| ATAT1 | 268.0798 | -0.21926 | 0.169218 | -1.29573 | 0.195069 | 0.350068 |
| TPMT | 658.5219 | 0.667899 | 0.17789 | 3.754566 | 1.74E-04 | 0.003555 |
| RNF144B | 326.2578 | -0.58007 | 0.243803 | -2.37927 | 0.017347 | 0.068779 |
| NRM | 386.7236 | -0.49471 | 0.169053 | -2.92638 | 0.003429 | 0.022982 |
| MTCH1 | 9469.644 | -0.21505 | 0.135212 | -1.59046 | 0.111731 | 0.239914 |
| VARS2 | 1222.048 | -0.37671 | 0.148717 | -2.53306 | 0.011307 | 0.051105 |
| TAF8 | 666.9725 | -0.14559 | 0.159754 | -0.91131 | 0.362131 | 0.529133 |
| FAM8A1 | 897.8388 | 0.305416 | 0.182501 | 1.673503 | 0.094228 | 0.214616 |
| C6orf52 | 23.67649 | 0.320889 | 0.442615 | 0.724985 | 0.468461 | NA |
| FGFBP2 | 32.34869 | -1.0762 | 0.488783 | -2.2018 | 0.02768 | NA |
| CPEB2 | 1824.007 | 0.041549 | 0.227628 | 0.18253 | 0.855166 | 0.91391 |
| FHDC1 | 1582.337 | -0.08703 | 0.204249 | -0.4261 | 0.670033 | 0.783821 |
| TLR2 | 15.62294 | -0.56405 | 0.578563 | -0.97492 | 0.329602 | NA |
| MGARP | 7.20559 | 0.1137 | 0.860036 | 0.132203 | 0.894824 | NA |
| NEK1 | 271.9725 | 0.505011 | 0.399236 | 1.264943 | 0.205892 | 0.362965 |
| DDX60 | 497.2371 | -0.47485 | 0.645767 | -0.73533 | 0.462136 | 0.620315 |
| DBR1 | 614.4211 | 0.245972 | 0.179291 | 1.371916 | 0.17009 | 0.32037 |

| | | | | | | |
|---|---|---|---|---|---|---|
| DNAJC13 | 4150.377 | 0.540752 | 0.230042 | 2.350666 | 0.01874 | 0.072294 |
| SLC35A5 | 1411.092 | 0.287826 | 0.17533 | 1.641623 | 0.100668 | 0.223907 |
| DIRC2 | 995.6766 | 0.05429 | 0.167839 | 0.323467 | 0.746341 | 0.840414 |
| SENP7 | 193.7841 | 0.910518 | 0.329354 | 2.764559 | 0.0057 | 0.032354 |
| GUCA1C | 0.162376 | 0.424913 | 4.079006 | 0.104171 | 0.917034 | NA |
| CCDC54 | 2.183362 | 0.811377 | 1.346294 | 0.602674 | 0.546726 | NA |
| COX17 | 1004.95 | 0.608943 | 0.20786 | 2.929575 | 0.003394 | 0.022819 |
| PARP9 | 1099.566 | -0.47923 | 0.289449 | -1.65567 | 0.097788 | 0.219811 |
| ARHGAP24 | 1476.83 | 0.213816 | 0.196762 | 1.086676 | 0.27718 | 0.44307 |
| FAM13A | 700.4105 | 0.222352 | 0.182719 | 1.216902 | 0.223642 | 0.384409 |
| HERC3 | 676.3502 | 0.112187 | 0.154508 | 0.726096 | 0.46778 | 0.624246 |
| HERC6 | 972.1377 | -0.63022 | 0.267542 | -2.35558 | 0.018494 | 0.071682 |
| HERC5 | 469.7823 | -0.52902 | 0.311705 | -1.69719 | 0.089661 | 0.207239 |
| PCDH10 | 16.74887 | -0.38463 | 0.540628 | -0.71144 | 0.476809 | NA |
| NDST4 | 0.613743 | -0.17004 | 2.749758 | -0.06184 | 0.950691 | NA |
| C4orf21 | 92.20552 | 1.113245 | 0.426797 | 2.608369 | 0.009097 | 0.044383 |
| AP1AR | 601.1019 | 0.858848 | 0.241712 | 3.553185 | 3.81E-04 | 0.005726 |
| COPS4 | 1246.787 | 0.297468 | 0.175701 | 1.693038 | 0.090448 | 0.208467 |
| HNRNPD | 3660.597 | -0.22257 | 0.163119 | -1.36446 | 0.172422 | 0.323815 |
| RASGEF1B | 41.14487 | 0.005266 | 0.397034 | 0.013264 | 0.989417 | NA |
| SEC31A | 4775.311 | -0.19517 | 0.122688 | -1.59082 | 0.11165 | 0.239791 |
| FGF5 | 0.837367 | 2.9578 | 2.751439 | 1.075001 | 0.282374 | NA |
| AGPAT9 | 64.69772 | 0.442304 | 0.28884 | 1.531313 | 0.125692 | 0.260777 |
| FGF2 | 82.22987 | 0.597112 | 0.299567 | 1.993249 | 0.046234 | 0.132137 |
| BBS7 | 439.0394 | 0.785773 | 0.243178 | 3.231263 | 0.001232 | 0.011864 |
| KIAA1109 | 1230.49 | 0.527785 | 0.250367 | 2.108047 | 0.035027 | 0.10994 |
| BMPR1B | 196.9617 | 0.79662 | 0.224047 | 3.555591 | 3.77E-04 | 0.005688 |
| RAP1GDS1 | 1545.197 | 0.196197 | 0.145178 | 1.351421 | 0.176561 | 0.328856 |
| LARP1B | 194.7263 | 0.844455 | 0.21947 | 3.847699 | 1.19E-04 | 0.002873 |
| MMRN1 | 0.151697 | -1.546 | 4.075469 | -0.37934 | 0.704433 | NA |
| PDE5A | 171.1016 | 0.597676 | 0.210078 | 2.845016 | 0.004441 | 0.02725 |
| PRDM5 | 150.243 | 0.81508 | 0.225206 | 3.619263 | 2.95E-04 | 0.004874 |
| NAAA | 313.4224 | 0.276951 | 0.214136 | 1.293344 | 0.195892 | 0.351223 |
| NUP54 | 413.2013 | 0.559278 | 0.195501 | 2.860749 | 0.004226 | 0.026398 |
| CXCL9 | 3.232475 | 2.761465 | 1.472997 | 1.874725 | 0.060831 | NA |
| BMP2K | 524.905 | 0.379505 | 0.148793 | 2.550564 | 0.010755 | 0.049294 |
| G3BP2 | 5611.448 | 0.175323 | 0.160929 | 1.089441 | 0.275959 | 0.441557 |
| 11-Sep | 2090.726 | 0.283095 | 0.173437 | 1.632266 | 0.102624 | 0.226809 |
| FRAS1 | 432.5173 | -0.32108 | 0.290404 | -1.10562 | 0.268892 | 0.433761 |
| SCARB2 | 9125.7 | 0.300671 | 0.161396 | 1.86294 | 0.062471 | 0.161813 |
| CCNG2 | 762.6128 | 0.119353 | 0.180687 | 0.660553 | 0.508899 | 0.657414 |
| CNOT6L | 365.828 | 0.338442 | 0.205876 | 1.643911 | 0.100195 | 0.223272 |
| USO1 | 1324.22 | 0.727949 | 0.282959 | 2.572632 | 0.010093 | 0.047462 |
| CDKL2 | 109.4119 | 0.444407 | 0.295937 | 1.501695 | 0.133176 | 0.270643 |
| SHROOM3 | 5.052651 | -0.2078 | 0.934223 | -0.22243 | 0.823977 | NA |
| ANXA3 | 117.3924 | 0.31964 | 0.480457 | 0.665283 | 0.50587 | 0.655061 |
| PPA2 | 1495.312 | 0.040276 | 0.138799 | 0.290172 | 0.771685 | 0.858857 |

| | | | | | | |
|---|---|---|---|---|---|---|
| CENPE | 124.8624 | 0.940991 | 1.494099 | 0.629805 | 0.528822 | 0.673961 |
| GSTCD | 357.7451 | 0.456403 | 0.220336 | 2.071392 | 0.038322 | 0.116819 |
| INTS12 | 591.5285 | 0.297415 | 0.166685 | 1.784299 | 0.074375 | 0.182135 |
| CASP6 | 325.2076 | 0.129911 | 0.178926 | 0.726059 | 0.467803 | 0.624246 |
| LEF1 | 736.6444 | -0.41307 | 0.179192 | -2.30516 | 0.021158 | 0.07868 |
| HADH | 1373.896 | -0.12676 | 0.139002 | -0.91194 | 0.361802 | 0.52898 |
| EGF | 13.53347 | 0.855701 | 0.672352 | 1.272697 | 0.203126 | NA |
| PAPSS1 | 2900.012 | 0.048491 | 0.153621 | 0.315655 | 0.752264 | 0.844956 |
| SEC24B | 1148.827 | -0.04091 | 0.184798 | -0.2214 | 0.824784 | 0.894421 |
| PPP3CA | 861.7073 | 0.372328 | 0.174729 | 2.130881 | 0.033099 | 0.106006 |
| SLC39A8 | 713.8584 | 0.245114 | 0.168577 | 1.454018 | 0.145941 | 0.28834 |
| MTTP | 5.245513 | 0.635819 | 0.984898 | 0.645569 | 0.518559 | NA |
| FBN2 | 89.40881 | -0.68373 | 0.320071 | -2.13619 | 0.032664 | 0.105198 |
| SLC6A3 | 0.926585 | -2.43791 | 2.203722 | -1.10627 | 0.268609 | NA |
| PLK4 | 218.9313 | 0.8069 | 0.228885 | 3.525357 | 4.23E-04 | 0.006131 |
| SHQ1 | 573.0568 | -0.10246 | 0.15591 | -0.65715 | 0.511083 | 0.659182 |
| UBA3 | 1401.725 | 0.605238 | 0.165856 | 3.649172 | 2.63E-04 | 0.004587 |
| ARL6IP5 | 5228.486 | 0.355969 | 0.153343 | 2.321398 | 0.020265 | 0.07639 |
| NFKBIZ | 452.5361 | 0.619548 | 0.242169 | 2.558332 | 0.010518 | 0.048662 |
| COL8A1 | 73.70996 | 2.128631 | 0.568881 | 3.741787 | 1.83E-04 | 0.003662 |
| NXPE3 | 982.9138 | 0.204813 | 0.15533 | 1.318568 | 0.187314 | 0.341474 |
| MYH15 | 3.144872 | 0.186687 | 1.300233 | 0.14358 | 0.885832 | NA |
| PHLDB2 | 165.4758 | 0.071967 | 0.352538 | 0.204141 | 0.838244 | 0.903398 |
| ABHD10 | 875.3908 | 0.304715 | 0.178841 | 1.703833 | 0.088412 | 0.205049 |
| TAGLN3 | 1.159694 | -0.24887 | 2.055096 | -0.1211 | 0.903612 | NA |
| PLA1A | 2739.916 | -0.51939 | 0.223842 | -2.32033 | 0.020323 | 0.076558 |
| RABL3 | 421.9663 | 0.274409 | 0.203347 | 1.34946 | 0.177189 | 0.329348 |
| ADPRH | 450.4901 | -0.27704 | 0.254116 | -1.09019 | 0.275628 | 0.441265 |
| IGSF11 | 1569.744 | -0.8596 | 0.253487 | -3.39111 | 6.96E-04 | 0.008269 |
| ATG3 | 1012.799 | 0.454049 | 0.149375 | 3.039653 | 0.002369 | 0.018039 |
| NR1I2 | 6.971389 | -0.57426 | 0.88723 | -0.64725 | 0.517468 | NA |
| BOC | 1610.166 | -0.22999 | 0.250098 | -0.91959 | 0.357789 | 0.525131 |
| SRPRB | 1290.948 | -0.1461 | 0.147973 | -0.98737 | 0.323462 | 0.491431 |
| TMEM108 | 170.9257 | 0.153294 | 0.214305 | 0.715306 | 0.47442 | 0.629745 |
| AGTR1 | 52.92918 | 0.467851 | 0.56128 | 0.833544 | 0.404538 | 0.569031 |
| MED12L | 28.05565 | 0.055906 | 0.44773 | 0.124866 | 0.90063 | NA |
| EIF2A | 1549.283 | 0.595023 | 0.215857 | 2.756566 | 0.005841 | 0.032803 |
| ALDH1L1 | 14.78697 | 0.302515 | 0.616919 | 0.490365 | 0.623876 | NA |
| OSBPL11 | 431.2547 | 0.278183 | 0.163224 | 1.704301 | 0.088325 | 0.20491 |
| TRPC1 | 180.7214 | 0.631771 | 0.27497 | 2.297599 | 0.021585 | 0.079801 |
| NCEH1 | 476.8608 | 0.395208 | 0.184906 | 2.137349 | 0.03257 | 0.105021 |
| LPP | 1542.181 | 0.114554 | 0.283247 | 0.40443 | 0.685897 | 0.795471 |
| TMEM44 | 343.0971 | -0.20342 | 0.232763 | -0.87396 | 0.382142 | 0.548263 |
| KIAA0226 | 1383.432 | -0.3838 | 0.222365 | -1.72601 | 0.084346 | 0.197978 |
| CCDC39 | 12.76142 | 1.39568 | 0.669058 | 2.086039 | 0.036975 | NA |
| EAF2 | 24.85065 | 0.76581 | 0.463416 | 1.652532 | 0.098426 | NA |
| TM4SF19 | 0.921281 | -1.21902 | 2.226885 | -0.54741 | 0.584096 | NA |

| | | | | | | |
|---|---|---|---|---|---|---|
| MUC4 | 0.247898 | -0.5901 | 4.074633 | -0.14482 | 0.884851 | NA |
| SLIT2 | 16.36964 | -0.14963 | 0.527046 | -0.2839 | 0.776487 | NA |
| EIF2B5 | 2083.933 | 0.009289 | 0.146161 | 0.063556 | 0.949324 | 0.9717 |
| AHSG | 0.162376 | 0.424913 | 4.079006 | 0.104171 | 0.917034 | NA |
| ECE2 | 263.9804 | 0.11486 | 0.210236 | 0.54634 | 0.584832 | 0.717944 |
| VWA5B2 | 2.488483 | -0.10059 | 1.309775 | -0.0768 | 0.938783 | NA |
| DGKQ | 1251.686 | -0.29036 | 0.15444 | -1.88012 | 0.060092 | 0.157435 |
| FIP1L1 | 711.0661 | 0.226565 | 0.163195 | 1.388309 | 0.165043 | 0.313807 |
| SLC26A1 | 79.0613 | -0.32613 | 0.278893 | -1.16937 | 0.242256 | 0.405185 |
| LYAR | 421.7037 | 0.607684 | 0.203538 | 2.985606 | 0.00283 | 0.020359 |
| CENPC | 145.4515 | 0.859994 | 0.423248 | 2.031894 | 0.042164 | 0.124934 |
| EPHA5 | 443.7734 | 0.40767 | 0.230529 | 1.768408 | 0.076993 | 0.186394 |
| CORIN | 2.376115 | 0.248207 | 1.301898 | 0.19065 | 0.8488 | NA |
| ATP10D | 563.9026 | 0.269121 | 0.162923 | 1.651834 | 0.098568 | 0.221145 |
| OCIAD2 | 81.66211 | -0.01915 | 0.325076 | -0.05892 | 0.953012 | 0.974382 |
| SLC10A4 | 0.911593 | 1.257757 | 2.250933 | 0.558771 | 0.576318 | NA |
| SCD5 | 662.7733 | -0.31505 | 0.229513 | -1.37267 | 0.169855 | 0.320335 |
| PLAC8 | 1.723043 | 0.159776 | 1.599185 | 0.099911 | 0.920415 | NA |
| ENOPH1 | 1216.125 | 0.500055 | 0.145326 | 3.440916 | 5.80E-04 | 0.007426 |
| TRMT10A | 52.89247 | 1.265092 | 0.3766 | 3.359247 | 7.82E-04 | 0.008918 |
| KLHL8 | 314.3254 | 0.530185 | 0.188209 | 2.817006 | 0.004847 | 0.028921 |
| SNCA | 5587.783 | -0.01638 | 0.13267 | -0.12346 | 0.901746 | 0.942449 |
| PYURF | 0.230334 | -0.53318 | 4.076749 | -0.13079 | 0.895945 | NA |
| TBCK | 568.9151 | 0.02442 | 0.194829 | 0.125341 | 0.900254 | 0.941637 |
| CAMK2D | 1137.57 | 0.076542 | 0.180888 | 0.423145 | 0.672189 | 0.785336 |
| CISD2 | 652.8381 | 0.329265 | 0.15384 | 2.140305 | 0.03233 | 0.104486 |
| DDIT4L | 181.4765 | 0.263586 | 0.338248 | 0.779267 | 0.435822 | 0.596751 |
| ANK2 | 1892.49 | 0.09389 | 0.220983 | 0.424872 | 0.67093 | 0.784483 |
| TIFA | 266.5176 | 0.357752 | 0.205062 | 1.744605 | 0.081054 | 0.193008 |
| SPATA5 | 141.7756 | 0.349359 | 0.236447 | 1.477538 | 0.139531 | 0.279653 |
| FABP2 | 0.163957 | 0.424913 | 4.079006 | 0.104171 | 0.917034 | NA |
| CCNA2 | 584.8611 | -0.05685 | 0.236842 | -0.24004 | 0.810302 | 0.886567 |
| METTL14 | 308.8746 | 0.602359 | 0.233935 | 2.574897 | 0.010027 | 0.047283 |
| USP53 | 779.8244 | 1.509975 | 0.201817 | 7.481911 | 7.32E-14 | 2.17E-10 |
| SETD7 | 2391.91 | 0.345053 | 0.189684 | 1.819098 | 0.068896 | 0.172715 |
| NAF1 | 276.4191 | -0.1319 | 0.168591 | -0.78236 | 0.434002 | 0.595222 |
| 1-Mar | 10.03912 | 1.43329 | 0.702975 | 2.038892 | 0.041461 | NA |
| SFRP2 | 0.134133 | -1.48909 | 4.077584 | -0.36519 | 0.71497 | NA |
| RPS3A | 8587.193 | 0.359559 | 0.18691 | 1.923703 | 0.054392 | 0.146816 |
| RNF175 | 0.658173 | -0.01016 | 2.630582 | -0.00386 | 0.996919 | NA |
| PDGFC | 33.94324 | 1.055077 | 0.399479 | 2.641132 | 0.008263 | NA |
| CBR4 | 416.41 | 0.560636 | 0.227748 | 2.461647 | 0.01383 | 0.058681 |
| GLRA3 | 0.324752 | 1.53082 | 4.022025 | 0.380609 | 0.703493 | NA |
| CYP4V2 | 440.6761 | 0.131891 | 0.234987 | 0.561267 | 0.574615 | 0.709812 |
| ROPN1L | 3.903518 | 1.131075 | 1.067727 | 1.05933 | 0.289449 | NA |
| NDUFS6 | 3049.777 | -0.09315 | 0.141005 | -0.66063 | 0.508849 | 0.657414 |
| 6-Mar | 1990.495 | -0.01063 | 0.181411 | -0.05858 | 0.953288 | 0.974496 |

| | | | | | | |
|---|---|---|---|---|---|---|
| NKD2 | 0.823352 | -0.17011 | 2.472341 | -0.06881 | 0.945144 | NA |
| CDH18 | 262.8343 | 0.409099 | 0.702223 | 0.582577 | 0.560178 | 0.698472 |
| ADAMTS16 | 7.451567 | 0.723867 | 0.746061 | 0.970251 | 0.331921 | NA |
| SRD5A1 | 573.0615 | -0.07067 | 0.149434 | -0.47295 | 0.63625 | 0.757303 |
| MYO10 | 20587.83 | -0.21191 | 0.156222 | -1.35648 | 0.174947 | 0.326872 |
| FAM105A | 0.134133 | -1.48909 | 4.077584 | -0.36519 | 0.71497 | NA |
| RPL37 | 11168.87 | 0.304686 | 0.180438 | 1.688591 | 0.091298 | 0.209918 |
| SKP2 | 347.0685 | 0.058067 | 0.248637 | 0.233541 | 0.815341 | 0.889858 |
| OSMR | 202.6961 | 0.059802 | 0.256903 | 0.232782 | 0.815931 | 0.889858 |
| PLK2 | 1019.804 | -0.21185 | 0.239227 | -0.88557 | 0.37585 | 0.542371 |
| PIK3R1 | 1823.666 | 0.488612 | 0.287085 | 1.701978 | 0.08876 | 0.205541 |
| LHFPL2 | 3299.803 | 0.002683 | 0.14161 | 0.018946 | 0.984884 | 0.992081 |
| SSBP2 | 391.7045 | -0.25882 | 0.177401 | -1.45898 | 0.144569 | 0.286552 |
| ANKRD31 | 2.11319 | 0.276435 | 1.648585 | 0.16768 | 0.866835 | NA |
| IQGAP2 | 94.9362 | 0.126429 | 0.425382 | 0.297212 | 0.766305 | 0.854855 |
| CRHBP | 5.381714 | 1.095163 | 0.945603 | 1.158163 | 0.246797 | NA |
| RASA1 | 654.8692 | 0.319844 | 0.240219 | 1.331467 | 0.183036 | 0.336174 |
| LIX1 | 13.27377 | 0.626861 | 2.279637 | 0.274983 | 0.783329 | NA |
| GIN1 | 88.89661 | 0.977456 | 0.318794 | 3.066102 | 0.002169 | 0.017049 |
| PPIP5K2 | 1068.617 | 1.003378 | 0.291612 | 3.440792 | 5.80E-04 | 0.007426 |
| PAM | 5458.667 | 0.096655 | 0.131085 | 0.737345 | 0.460913 | 0.619023 |
| BDP1 | 444.0439 | 1.037772 | 0.88803 | 1.168622 | 0.242556 | 0.405514 |
| GTF2H2 | 28.93185 | 0.703049 | 0.43842 | 1.603597 | 0.108803 | NA |
| SLC30A5 | 1072.503 | 0.339399 | 0.137365 | 2.470786 | 0.013482 | 0.05774 |
| BTF3 | 9190.895 | 0.184812 | 0.169 | 1.093562 | 0.274147 | 0.439821 |
| FBXL17 | 503.7685 | 0.140435 | 0.178793 | 0.785458 | 0.432185 | 0.593416 |
| SPATA9 | 10.32549 | 0.172038 | 0.654792 | 0.262738 | 0.792753 | NA |
| TSLP | 0.614937 | 2.493639 | 2.617677 | 0.952615 | 0.340785 | NA |
| TNFAIP8 | 49.02622 | 0.613792 | 0.340753 | 1.801285 | 0.071658 | 0.177498 |
| FEM1C | 561.2375 | 0.158031 | 0.178464 | 0.885504 | 0.375885 | 0.542371 |
| COMMD10 | 707.2644 | 0.543399 | 0.204681 | 2.654857 | 0.007934 | 0.040456 |
| ATG12 | 883.4816 | 0.535282 | 0.181009 | 2.957209 | 0.003104 | 0.021663 |
| MEGF10 | 2.070895 | -1.60252 | 1.459124 | -1.09827 | 0.272085 | NA |
| ADAMTS19 | 1.843132 | -0.21958 | 1.589334 | -0.13816 | 0.890117 | NA |
| YIPF5 | 1522.942 | 0.222139 | 0.159757 | 1.390483 | 0.164382 | 0.313223 |
| ARHGAP26 | 230.689 | -0.13274 | 0.202004 | -0.65711 | 0.511108 | 0.659182 |
| CXCL14 | 0.623045 | 0.091811 | 2.666142 | 0.034436 | 0.97253 | NA |
| SLC25A48 | 0.684273 | 2.627785 | 2.896325 | 0.907282 | 0.364257 | NA |
| DDX46 | 927.8878 | 0.804233 | 0.339234 | 2.370731 | 0.017753 | 0.069688 |
| RNF145 | 5585.615 | -0.01833 | 0.147743 | -0.12405 | 0.901278 | 0.942193 |
| C1QTNF2 | 78.481 | -0.30023 | 0.339806 | -0.88354 | 0.376944 | 0.543599 |
| GABRB2 | 0.47961 | 0.535796 | 3.39605 | 0.15777 | 0.874638 | NA |
| FBXO38 | 997.7644 | 0.256116 | 0.1454 | 1.761458 | 0.078161 | 0.188276 |
| PCYOX1L | 361.4359 | -0.60661 | 0.220903 | -2.74606 | 0.006032 | 0.03358 |
| GLRA1 | 0.163957 | 0.424913 | 4.079006 | 0.104171 | 0.917034 | NA |
| TNIP1 | 3164.345 | -0.29695 | 0.117808 | -2.52066 | 0.011714 | 0.052388 |
| G3BP1 | 4871.163 | 0.226728 | 0.150854 | 1.502965 | 0.132848 | 0.270093 |

| | | | | | | |
|---|---|---|---|---|---|---|
| ZNF300 | 140.6535 | 1.105695 | 0.270467 | 4.088096 | 4.35E-05 | 0.001535 |
| N4BP3 | 86.40043 | -0.65248 | 0.305114 | -2.13849 | 0.032477 | 0.104859 |
| NHP2 | 1731.277 | -0.0178 | 0.176138 | -0.10104 | 0.919517 | 0.953373 |
| RMND5B | 1064.106 | -0.30539 | 0.134637 | -2.26824 | 0.023314 | 0.083915 |
| BOD1 | 1160.123 | -0.07772 | 0.167198 | -0.46485 | 0.642041 | 0.761812 |
| TENM2 | 2.891858 | -0.44544 | 1.348405 | -0.33034 | 0.741141 | NA |
| KCNMB1 | 0.151697 | -1.546 | 4.075469 | -0.37934 | 0.704433 | NA |
| FAM50B | 320.789 | -0.2177 | 0.208183 | -1.04573 | 0.295687 | 0.462999 |
| MYLK4 | 27.15463 | -0.02953 | 0.450423 | -0.06557 | 0.947719 | NA |
| | 0.520316 | 2.239969 | 3.173769 | 0.705776 | 0.480328 | NA |
| FAM217A | 0.745706 | 2.741883 | 2.882857 | 0.951099 | 0.341554 | NA |
| TBC1D7 | 7229.29 | -0.10974 | 0.132115 | -0.83062 | 0.406188 | 0.569932 |
| FARS2 | 373.116 | -0.1593 | 0.169644 | -0.939 | 0.347733 | 0.515942 |
| GFOD1 | 215.0347 | 0.033133 | 0.18963 | 0.174722 | 0.861298 | 0.917815 |
| CDKAL1 | 306.3339 | -0.33936 | 0.165852 | -2.04617 | 0.040739 | 0.121776 |
| PCDHB18 | 2.964225 | 0.463071 | 1.2684 | 0.365083 | 0.71505 | NA |
| PSD2 | 0.919554 | 0.143913 | 2.343626 | 0.061406 | 0.951036 | NA |
| LRRTM2 | 3.189542 | 0.027668 | 1.286588 | 0.021505 | 0.982843 | NA |
| ZMAT2 | 1803.107 | 0.105024 | 0.14426 | 0.728022 | 0.4666 | 0.623684 |
| GFRA3 | 161.5847 | 0.688918 | 0.410236 | 1.679319 | 0.09309 | 0.212923 |
| KLHL3 | 65.37756 | -0.41229 | 0.372999 | -1.10535 | 0.269008 | 0.433776 |
| DCDC2 | 4.198742 | -0.1703 | 1.121114 | -0.15191 | 0.879261 | NA |
| SLC17A4 | 0.163957 | 0.424913 | 4.079006 | 0.104171 | 0.917034 | NA |
| KAAG1 | 0.555233 | -0.57009 | 3.296947 | -0.17291 | 0.862719 | NA |
| TRIM7 | 203.6431 | 0.12692 | 0.190471 | 0.666349 | 0.505188 | 0.654702 |
| TRIM41 | 1137.115 | -0.32555 | 0.143804 | -2.26381 | 0.023586 | 0.084611 |
| HIGD2A | 1174.593 | -0.1791 | 0.145273 | -1.23285 | 0.217631 | 0.377018 |
| FAM193B | 411.7343 | -0.03704 | 0.157253 | -0.23552 | 0.813806 | 0.888941 |
| PLA2G7 | 0.326536 | 0.037056 | 3.842554 | 0.009644 | 0.992306 | NA |
| TNFRSF21 | 47.84048 | -2.00159 | 0.542907 | -3.6868 | 2.27E-04 | 0.004193 |
| RNF44 | 984.778 | -0.38487 | 0.18525 | -2.07755 | 0.03775 | 0.115552 |
| MUT | 1136.415 | 0.262055 | 0.157556 | 1.663251 | 0.096262 | 0.217411 |
| RASGEF1C | 76.24601 | -0.02831 | 0.405712 | -0.06978 | 0.944372 | 0.969226 |
| DOK3 | 29.20677 | -0.29762 | 0.515517 | -0.57732 | 0.563723 | NA |
| ABT1 | 857.4315 | 0.07283 | 0.15241 | 0.477856 | 0.632753 | 0.754704 |
| PPP1R18 | 2890.875 | -0.26359 | 0.149202 | -1.76669 | 0.077281 | 0.186896 |
| DAAM2 | 4306.607 | -0.63718 | 0.129764 | -4.91028 | 9.09E-07 | 1.53E-04 |
| PRIM2 | 345.1993 | -0.07216 | 0.191388 | -0.37705 | 0.706138 | 0.811641 |
| MLIP | 768.3848 | 0.064324 | 0.174545 | 0.368525 | 0.712482 | 0.8164 |
| FGD2 | 0.488709 | 2.119056 | 3.307564 | 0.64067 | 0.521737 | NA |
| SCUBE3 | 37.16674 | 0.033668 | 0.440854 | 0.076369 | 0.939125 | NA |
| CRIP3 | 0.163957 | 0.424913 | 4.079006 | 0.104171 | 0.917034 | NA |
| TTBK1 | 0.151697 | -1.546 | 4.075469 | -0.37934 | 0.704433 | NA |
| TCTE1 | 0.399595 | -1.31595 | 3.579466 | -0.36764 | 0.713143 | NA |
| RPL7L1 | 2283.883 | 0.073203 | 0.139956 | 0.523046 | 0.600943 | 0.730247 |
| NFKBIE | 563.3411 | -0.31642 | 0.147612 | -2.1436 | 0.032065 | 0.103843 |
| CYP39A1 | 51.56842 | 0.218428 | 0.381601 | 0.572399 | 0.567052 | 0.704045 |

| | | | | | | |
|---|---|---|---|---|---|---|
| TPBG | 467.2463 | 0.222764 | 0.217066 | 1.02625 | 0.304774 | 0.472114 |
| IRAK1BP1 | 56.24664 | 0.356173 | 0.311717 | 1.142618 | 0.253197 | 0.417418 |
| PHIP | 468.7759 | 0.828597 | 0.327738 | 2.528227 | 0.011464 | 0.051636 |
| PRSS35 | 22.94127 | 3.015514 | 0.969289 | 3.111057 | 0.001864 | NA |
| MMS22L | 173.8513 | 0.538959 | 0.277509 | 1.942135 | 0.052121 | 0.142371 |
| FAXC | 290.352 | 0.13115 | 0.176145 | 0.744556 | 0.45654 | 0.615032 |
| GABRR1 | 0.162376 | 0.424913 | 4.079006 | 0.104171 | 0.917034 | NA |
| PNRC1 | 1919.405 | -0.17031 | 0.163127 | -1.04406 | 0.296456 | 0.463653 |
| PM20D2 | 110.1867 | 0.217707 | 0.423859 | 0.513631 | 0.60751 | 0.73511 |
| RARS2 | 950.6124 | 0.484751 | 0.168287 | 2.880495 | 0.003971 | 0.025354 |
| SCML4 | 0.488709 | 2.119056 | 3.307564 | 0.64067 | 0.521737 | NA |
| TBC1D32 | 109.775 | 0.835717 | 0.39803 | 2.099631 | 0.035761 | 0.111536 |
| RNF217 | 265.3884 | 0.453821 | 0.198607 | 2.285015 | 0.022312 | 0.081613 |
| RSPO3 | 2.84792 | 3.695545 | 1.702753 | 2.170336 | 0.029981 | NA |
| ARHGAP18 | 810.6299 | 0.605959 | 0.174004 | 3.482452 | 4.97E-04 | 0.006762 |
| ABRACL | 1823.772 | 0.49918 | 0.176253 | 2.832184 | 0.004623 | 0.027963 |
| SLC18B1 | 5252.647 | 0.018863 | 0.166901 | 0.113022 | 0.910013 | 0.947811 |
| MTFR2 | 199.7819 | 0.305228 | 0.320082 | 0.953592 | 0.34029 | 0.508571 |
| SLC2A12 | 7.035025 | 0.89405 | 0.827917 | 1.07988 | 0.280196 | NA |
| SHPRH | 298.5076 | 0.672029 | 0.321372 | 2.091125 | 0.036517 | 0.113179 |
| AIG1 | 1061.021 | -0.00111 | 0.138495 | -0.00801 | 0.993611 | 0.997229 |
| LPHN3 | 1.883489 | 1.441511 | 1.673794 | 0.861224 | 0.389115 | NA |
| GPM6A | 6.506648 | -0.09909 | 0.859511 | -0.11528 | 0.90822 | NA |
| WDR17 | 185.7493 | 0.759378 | 0.289326 | 2.624645 | 0.008674 | 0.042935 |
| SPATA4 | 0.642189 | 0.046766 | 2.378271 | 0.019664 | 0.984311 | NA |
| VEGFC | 21.71695 | 1.061272 | 0.648579 | 1.636303 | 0.101776 | NA |
| MTMR12 | 1100.552 | -0.08825 | 0.159682 | -0.55268 | 0.58048 | 0.714368 |
| CCT5 | 6107.063 | 0.051458 | 0.143352 | 0.35896 | 0.719625 | 0.82133 |
| FAM173B | 328.5823 | 0.297174 | 0.182251 | 1.630573 | 0.10298 | 0.227301 |
| SEC24D | 1240.302 | -0.02563 | 0.138811 | -0.18464 | 0.853512 | 0.912964 |
| SLC7A11 | 1559.872 | 0.9599 | 0.303544 | 3.162305 | 0.001565 | 0.0139 |
| CCRN4L | 107.3758 | -0.62613 | 0.271021 | -2.31026 | 0.020874 | 0.077988 |
| EIF4E | 491.8587 | 0.429137 | 0.198078 | 2.166505 | 0.030273 | 0.099745 |
| CSNK1G3 | 542.9633 | 0.744333 | 0.259643 | 2.866756 | 0.004147 | 0.026026 |
| SRFBP1 | 242.9587 | 0.942384 | 0.277725 | 3.393227 | 6.91E-04 | 0.00825 |
| ADAMTS12 | 7.096302 | 0.026427 | 0.75083 | 0.035197 | 0.971923 | NA |
| FER | 674.3186 | 0.744321 | 0.572941 | 1.299123 | 0.193902 | 0.348953 |
| ANKRD50 | 1829.84 | 0.713319 | 0.22142 | 3.221573 | 0.001275 | 0.01212 |
| SCLT1 | 142.8032 | 0.580376 | 0.327614 | 1.771526 | 0.076473 | 0.185473 |
| C4orf33 | 282.5549 | 0.683616 | 0.226642 | 3.016278 | 0.002559 | 0.019152 |
| QDPR | 5749.907 | -0.31168 | 0.128465 | -2.42616 | 0.01526 | 0.062814 |
| QTRTD1 | 761.5902 | 0.347581 | 0.187349 | 1.855266 | 0.063558 | 0.163694 |
| DRD3 | 0.162376 | 0.424913 | 4.079006 | 0.104171 | 0.917034 | NA |
| MMAA | 107.4261 | 0.417409 | 0.26162 | 1.59548 | 0.110605 | 0.238534 |
| ZNF827 | 506.2869 | -0.05505 | 0.196712 | -0.27983 | 0.77961 | 0.864814 |
| EDNRA | 1.009228 | 2.053856 | 2.077371 | 0.98868 | 0.322819 | NA |
| NR3C2 | 29.03244 | -0.14743 | 0.425377 | -0.34659 | 0.728903 | NA |

| | | | | | | |
|---|---|---|---|---|---|---|
| WWC2 | 889.3041 | 0.678168 | 0.253209 | 2.678291 | 0.0074 | 0.03868 |
| CENPU | 246.9855 | 1.061019 | 0.23281 | 4.557444 | 5.18E-06 | 4.24E-04 |
| ACSL1 | 5069.824 | -0.05382 | 0.131046 | -0.41071 | 0.681282 | 0.792212 |
| SLC25A4 | 2917.222 | 0.076804 | 0.123476 | 0.622019 | 0.533929 | 0.677629 |
| TDO2 | 1.266226 | 2.422108 | 1.957907 | 1.237091 | 0.216053 | NA |
| GUF1 | 532.2683 | 0.431601 | 0.239058 | 1.805421 | 0.071009 | 0.176295 |
| FBXO4 | 145.6156 | 0.108046 | 0.250117 | 0.431981 | 0.665755 | 0.780588 |
| C5orf28 | 264.3541 | 0.742636 | 0.203862 | 3.642837 | 2.70E-04 | 0.004608 |
| CCL28 | 722.8642 | 0.095649 | 0.205168 | 0.466198 | 0.641074 | 0.761059 |
| PARP8 | 826.434 | 0.408714 | 0.152854 | 2.673887 | 0.007498 | 0.039071 |
| DST | 2635.288 | 1.12501 | 0.756727 | 1.486679 | 0.1371 | 0.276319 |
| BEND6 | 66.6934 | 1.707211 | 0.374627 | 4.5571 | 5.19E-06 | 4.24E-04 |
| RBM46 | 0.096202 | 0.408546 | 4.079639 | 0.100143 | 0.920231 | NA |
| SCHIP1 | 12.00245 | 0.450707 | 0.605758 | 0.744037 | 0.456854 | NA |
| GRID2 | 3.163758 | 1.939479 | 1.228603 | 1.578606 | 0.114427 | NA |
| ATG10 | 164.1611 | 0.261442 | 0.221817 | 1.178636 | 0.238543 | 0.401466 |
| POC5 | 215.1608 | 0.334934 | 0.20352 | 1.645701 | 0.099825 | 0.223192 |
| SPOCK1 | 172.5714 | -0.03212 | 0.219397 | -0.14639 | 0.883612 | 0.931048 |
| FAM151B | 47.49691 | 0.732887 | 0.325793 | 2.24955 | 0.024478 | 0.086673 |
| JMY | 394.2831 | 0.663458 | 0.2596 | 2.555694 | 0.010598 | 0.048904 |
| HOMER1 | 237.9869 | 0.647577 | 0.263774 | 2.455042 | 0.014087 | 0.059494 |
| XRCC4 | 119.0719 | 0.93819 | 0.26959 | 3.480063 | 5.01E-04 | 0.006784 |
| CCDC50 | 3073.596 | 0.240066 | 0.149427 | 1.606579 | 0.108147 | 0.234942 |
| CAMK4 | 0.563552 | -0.48645 | 2.808029 | -0.17324 | 0.862465 | NA |
| TRIM36 | 91.59458 | 0.344205 | 0.282046 | 1.220388 | 0.222318 | 0.382911 |
| IGSF10 | 39.39773 | -0.74756 | 0.416618 | -1.79435 | 0.072758 | NA |
| SPEF2 | 24.48949 | 1.014083 | 0.53766 | 1.886104 | 0.059281 | NA |
| SPARCL1 | 41.59974 | -0.43226 | 0.579121 | -0.7464 | 0.455425 | NA |
| MBNL1 | 1572.814 | 0.707156 | 0.264507 | 2.673482 | 0.007507 | 0.039084 |
| NADK2 | 211.7854 | 0.526692 | 0.307669 | 1.711882 | 0.086918 | 0.20236 |
| GJA1 | 275.4418 | -1.09152 | 0.380159 | -2.8712 | 0.004089 | 0.025773 |
| CCNO | 20.73795 | 0.759941 | 0.578924 | 1.312679 | 0.189291 | NA |
| DDX4 | 0.151697 | -1.546 | 4.075469 | -0.37934 | 0.704433 | NA |
| PELO | 695.4569 | 0.03544 | 0.15485 | 0.228864 | 0.818974 | 0.89171 |
| SAR1B | 1631.064 | 0.388506 | 0.156856 | 2.476838 | 0.013255 | 0.057233 |
| CATSPER3 | 0.774742 | -0.40577 | 2.223483 | -0.18249 | 0.855195 | NA |
| HNRNPDL | 3112.954 | 0.465876 | 0.152379 | 3.057362 | 0.002233 | 0.017347 |
| UTRN | 2839.337 | 0.533488 | 0.252849 | 2.109907 | 0.034866 | 0.109623 |
| GRM1 | 70.11977 | 0.32482 | 0.402454 | 0.807099 | 0.419609 | 0.582695 |
| PTPRK | 1056.031 | -0.02646 | 0.135781 | -0.19487 | 0.845497 | 0.908091 |
| PART1 | 16.94883 | -0.46589 | 0.563425 | -0.82689 | 0.408299 | NA |
| RAB3C | 14.37188 | 0.483996 | 0.626721 | 0.772268 | 0.439956 | NA |
| MARVELD2 | 19.12094 | -0.45405 | 0.475549 | -0.9548 | 0.33968 | NA |
| RAD17 | 617.2103 | 0.664052 | 0.220684 | 3.009064 | 0.002621 | 0.019392 |
| PLOD2 | 1142.913 | 0.83262 | 0.238716 | 3.487909 | 4.87E-04 | 0.00668 |
| STK32B | 664.2673 | 0.118103 | 0.242129 | 0.487769 | 0.625713 | 0.748948 |
| ZIC1 | 0.096202 | 0.408546 | 4.079639 | 0.100143 | 0.920231 | NA |

| | | | | | | |
|---|---|---|---|---|---|---|
| GPR125 | 1535.655 | -0.19413 | 0.249519 | -0.77801 | 0.436564 | 0.597215 |
| CPB1 | 0.555233 | -0.57009 | 3.296947 | -0.17291 | 0.862719 | NA |
| SREK1IP1 | 251.953 | 0.970251 | 0.290575 | 3.339067 | 8.41E-04 | 0.00931 |
| LGI2 | 8.395325 | 1.036955 | 1.047288 | 0.990133 | 0.322109 | NA |
| CWC27 | 241.5609 | 0.796896 | 0.373328 | 2.134576 | 0.032796 | 0.10553 |
| SRP19 | 229.0299 | 0.650085 | 0.198671 | 3.272165 | 0.001067 | 0.010822 |
| CENPH | 183.1761 | 0.504174 | 0.209877 | 2.402234 | 0.016295 | 0.065864 |
| CDYL | 1279.85 | -0.00662 | 0.131144 | -0.05046 | 0.959758 | 0.977906 |
| BANK1 | 4.241892 | -0.00716 | 1.133692 | -0.00631 | 0.994964 | NA |
| DAB2 | 7242.497 | -0.26849 | 0.117482 | -2.28533 | 0.022293 | 0.081595 |
| CAST | 4096.956 | 0.39877 | 0.138022 | 2.889172 | 0.003863 | 0.024886 |
| SCOC | 1219.261 | 1.093085 | 0.244494 | 4.470803 | 7.79E-06 | 5.40E-04 |
| CLGN | 0.230334 | -0.53318 | 4.076749 | -0.13079 | 0.895945 | NA |
| CETN3 | 203.1394 | 0.687815 | 0.242284 | 2.838877 | 0.004527 | 0.027525 |
| SMARCA5 | 1769.683 | 0.753773 | 0.303179 | 2.48623 | 0.01291 | 0.056209 |
| SYCP2L | 0.815245 | 1.670886 | 2.214146 | 0.754641 | 0.450464 | NA |
| BMP6 | 0.584911 | 2.38756 | 2.690807 | 0.887303 | 0.374916 | NA |
| SLC25A27 | 33.87473 | 0.751727 | 0.484318 | 1.552135 | 0.12063 | NA |
| FAM81B | 0.096202 | 0.408546 | 4.079639 | 0.100143 | 0.920231 | NA |
| LPCAT1 | 3159.837 | -0.6141 | 0.17293 | -3.55116 | 3.84E-04 | 0.005756 |
| PLEKHG4B | 6.93878 | 0.940799 | 0.83421 | 1.127773 | 0.259416 | NA |
| GTF2E1 | 338.2598 | 0.040625 | 0.196001 | 0.207267 | 0.835801 | 0.901643 |
| TMPRSS11I | 0.162376 | 0.424913 | 4.079006 | 0.104171 | 0.917034 | NA |
| SREK1 | 506.83 | 1.054619 | 0.678801 | 1.553649 | 0.120268 | 0.252484 |
| CHD1 | 795.4362 | 1.116642 | 0.34969 | 3.193234 | 0.001407 | 0.01301 |
| ZUFSP | 181.9238 | 0.656741 | 0.24166 | 2.717628 | 0.006575 | 0.035561 |
| NUS1 | 507.8634 | 0.313908 | 0.179811 | 1.745764 | 0.080852 | 0.192625 |
| C6orf57 | 343.3996 | -0.12015 | 0.222435 | -0.54014 | 0.589103 | 0.721298 |
| ANKH | 614.7304 | 0.0215 | 0.164121 | 0.130999 | 0.895776 | 0.939107 |
| FAM105B | 470.5899 | 0.187163 | 0.159117 | 1.176264 | 0.239489 | 0.402773 |
| FAM134B | 141.5294 | 0.161199 | 0.233425 | 0.690581 | 0.489829 | 0.642357 |
| CDH12 | 0.612365 | -1.10341 | 2.696105 | -0.40926 | 0.682349 | NA |
| TOMM70A | 1427.18 | 0.399811 | 0.21708 | 1.841764 | 0.06551 | 0.166907 |
| ABI3BP | 2.592056 | 0.786877 | 1.28561 | 0.612065 | 0.540495 | NA |
| ENPP3 | 6.365945 | -0.65205 | 1.028194 | -0.63417 | 0.525968 | NA |
| C4orf19 | 23.97933 | -1.36613 | 0.562823 | -2.42728 | 0.015213 | NA |
| UCHL1 | 1808.745 | 0.166538 | 0.18351 | 0.907516 | 0.364134 | 0.531495 |
| TNIK | 970.7036 | 0.210834 | 0.152251 | 1.38478 | 0.16612 | 0.315242 |
| SH3RF1 | 447.5064 | 0.19713 | 0.186608 | 1.056386 | 0.290792 | 0.45777 |
| SRSF12 | 17.2774 | 0.246109 | 0.544029 | 0.452382 | 0.650994 | NA |
| PDLIM3 | 696.2998 | 0.326499 | 0.24294 | 1.343951 | 0.178964 | 0.331503 |
| SORBS2 | 2184.821 | 0.096181 | 0.17954 | 0.535711 | 0.592158 | 0.7233 |
| CEP170P1 | 81.98693 | 0.321106 | 0.427282 | 0.751508 | 0.452347 | 0.611259 |
| RAB6B | 609.5973 | -0.31183 | 0.166786 | -1.86967 | 0.061529 | 0.160095 |
| EPHB1 | 27.50757 | -0.63658 | 0.458191 | -1.38934 | 0.164729 | NA |
| DKK2 | 3.746069 | 0.490952 | 1.093864 | 0.448824 | 0.653559 | NA |
| CYP2U1 | 4652.762 | 0.342659 | 0.178568 | 1.918928 | 0.054993 | 0.147867 |

| | | | | | | |
|---|---|---|---|---|---|---|
| AK9 | 129.5272 | 0.502674 | 0.25491 | 1.971969 | 0.048613 | 0.136179 |
| CDK19 | 839.0945 | 0.087101 | 0.188177 | 0.462865 | 0.643461 | 0.762832 |
| GTF3C6 | 1268.39 | 0.061629 | 0.184075 | 0.334806 | 0.737772 | 0.833932 |
| MARCKS | 10174.86 | -0.04299 | 0.204367 | -0.21037 | 0.833377 | 0.900035 |
| TRMT44 | 298.6034 | -0.27094 | 0.18257 | -1.48405 | 0.137796 | 0.277148 |
| GRAMD3 | 588.5789 | 0.307252 | 0.231443 | 1.327552 | 0.184326 | 0.337654 |
| ZCCHC10 | 218.2068 | 0.988704 | 0.235158 | 4.204418 | 2.62E-05 | 0.001085 |
| LARP1 | 7194.05 | -0.51355 | 0.148269 | -3.46366 | 5.33E-04 | 0.007027 |
| CNOT8 | 1139.427 | 0.155598 | 0.173369 | 0.897494 | 0.369455 | 0.536481 |
| GRIA1 | 0.163957 | 0.424913 | 4.079006 | 0.104171 | 0.917034 | NA |
| SETD9 | 307.6763 | 0.061431 | 0.189251 | 0.324602 | 0.745482 | 0.839686 |
| MIER3 | 461.3175 | 0.712608 | 0.263374 | 2.70569 | 0.006816 | 0.036449 |
| PPARGC1B | 493.1843 | -0.4358 | 0.159231 | -2.73691 | 0.006202 | 0.034086 |
| SLC26A2 | 1164.325 | 0.362841 | 0.177945 | 2.039062 | 0.041444 | 0.123384 |
| LSM11 | 405.726 | 0.052891 | 0.166961 | 0.316788 | 0.751404 | 0.844591 |
| MED7 | 181.8759 | 0.320184 | 0.197874 | 1.618121 | 0.105637 | 0.23144 |
| ACPL2 | 156.8359 | 0.153118 | 0.209123 | 0.732188 | 0.464054 | 0.621482 |
| RASA2 | 255.3939 | 0.821909 | 0.244613 | 3.36003 | 7.79E-04 | 0.008909 |
| RAET1L | 0.774888 | -2.09879 | 2.346327 | -0.8945 | 0.371054 | NA |
| DCK | 572.6862 | 0.382188 | 0.184749 | 2.068692 | 0.038575 | 0.117311 |
| ADAMTS3 | 19.23739 | -0.36101 | 0.558232 | -0.6467 | 0.517825 | NA |
| PPEF2 | 0.151697 | -1.546 | 4.075469 | -0.37934 | 0.704433 | NA |
| ART3 | 5.636294 | 0.974853 | 0.938581 | 1.038645 | 0.29897 | NA |
| CXCL13 | 2.435167 | -0.1181 | 1.474488 | -0.08009 | 0.936164 | NA |
| FGF18 | 2.049579 | 0.21725 | 1.39331 | 0.155924 | 0.876093 | NA |
| PCDH1 | 964.1774 | 0.281128 | 0.297786 | 0.94406 | 0.345139 | 0.51312 |
| SH3RF2 | 2.486545 | -0.37573 | 1.244864 | -0.30183 | 0.762784 | NA |
| PPP2R2B | 49.17611 | -0.01139 | 0.347522 | -0.03277 | 0.973857 | 0.985496 |
| EEF1A1 | 108801.4 | 0.133883 | 0.156838 | 0.853637 | 0.393306 | 0.558333 |
| CD109 | 5961.479 | 0.240334 | 0.193402 | 1.242665 | 0.213991 | 0.37274 |
| ZFAND3 | 2622.845 | -0.28855 | 0.1185 | -2.43503 | 0.01489 | 0.061701 |
| MAPK13 | 1.680813 | -0.20962 | 1.673961 | -0.12522 | 0.900347 | NA |
| VPS8 | 1399.483 | 0.148535 | 0.181029 | 0.820503 | 0.41193 | 0.575044 |
| EIF4A2 | 6824.409 | 0.259475 | 0.171643 | 1.511714 | 0.130607 | 0.267063 |
| FCHO2 | 440.7108 | 1.010114 | 0.317314 | 3.183324 | 0.001456 | 0.013326 |
| TMEM171 | 1777.579 | 0.148867 | 0.23888 | 0.623185 | 0.533163 | 0.677164 |
| ARMC12 | 2.192644 | 0.820157 | 1.628928 | 0.503495 | 0.614616 | NA |
| KIT | 5435.243 | 0.21034 | 0.154769 | 1.359051 | 0.17413 | 0.326179 |
| AASDH | 284.7203 | 0.361098 | 0.219522 | 1.644932 | 0.099984 | 0.223198 |
| AFAP1L1 | 1958.331 | 0.050753 | 0.161267 | 0.314711 | 0.752981 | 0.845641 |
| SLC35B2 | 5069.767 | -0.37642 | 0.13081 | -2.87764 | 0.004007 | 0.025436 |
| SLC34A2 | 0.52699 | -2.57878 | 3.248478 | -0.79384 | 0.427288 | NA |
| WDR19 | 736.9108 | 0.392443 | 0.191744 | 2.046699 | 0.040688 | 0.121674 |
| RAB28 | 343.5707 | 0.247536 | 0.176732 | 1.400633 | 0.161324 | 0.308965 |
| NCK1 | 698.2924 | 0.61733 | 0.152888 | 4.037792 | 5.40E-05 | 0.001726 |
| DZIP1L | 139.1463 | 0.188938 | 0.229952 | 0.82164 | 0.411282 | 0.574631 |
| MRAS | 2747.39 | -0.14416 | 0.16727 | -0.86186 | 0.388767 | 0.554243 |

| | | | | | | |
|---|---|---|---|---|---|---|
| ESYT3 | 13.38499 | -0.09851 | 0.676531 | -0.1456 | 0.884234 | NA |
| FAIM | 201.5815 | 0.491094 | 0.251357 | 1.953772 | 0.050728 | 0.140019 |
| CLSTN2 | 3.463713 | 0.232996 | 1.229333 | 0.189531 | 0.849677 | NA |
| HIST1H2BD | 104.2642 | 0.424816 | 0.820041 | 0.518042 | 0.604429 | 0.732942 |
| CDC25C | 168.9608 | -0.32333 | 0.289546 | -1.11667 | 0.264137 | 0.428684 |
| HIST1H4H | 29.22609 | 0.684803 | 0.571391 | 1.198484 | 0.230729 | NA |
| NRG2 | 49.82936 | -0.64463 | 0.313474 | -2.05641 | 0.039743 | 0.119837 |
| ZSCAN12 | 200.5056 | 0.628426 | 0.224025 | 2.805163 | 0.005029 | 0.029751 |
| CDC42SE2 | 682.7305 | 0.189958 | 0.182018 | 1.043624 | 0.29666 | 0.463665 |
| RAPGEF6 | 110.427 | 0.641858 | 0.24473 | 2.622721 | 0.008723 | 0.043052 |
| SPON2 | 332.5022 | -0.28516 | 0.323023 | -0.8828 | 0.377346 | 0.543904 |
| CHCHD6 | 1610.213 | -0.67761 | 0.262561 | -2.58079 | 0.009858 | 0.046728 |
| CTBP1 | 3638.445 | -0.35405 | 0.151318 | -2.33977 | 0.019296 | 0.073814 |
| ZFYVE28 | 574.348 | -0.20775 | 0.202956 | -1.02362 | 0.306013 | 0.473169 |
| RGS12 | 3432.92 | -0.33655 | 0.152619 | -2.20514 | 0.027444 | 0.093545 |
| CCDC58 | 218.8027 | 0.495006 | 0.220083 | 2.249178 | 0.024501 | 0.086705 |
| KALRN | 70.76813 | -0.65568 | 0.373321 | -1.75635 | 0.079028 | 0.189689 |
| FGFR4 | 40.85482 | -0.32583 | 0.33639 | -0.9686 | 0.332742 | NA |
| HK3 | 0.268266 | -2.15396 | 4.038351 | -0.53338 | 0.593773 | NA |
| C5orf45 | 211.8835 | 0.234321 | 0.201907 | 1.160537 | 0.24583 | 0.409012 |
| SQSTM1 | 11889.11 | -0.37621 | 0.134254 | -2.80225 | 0.005075 | 0.029923 |
| MGAT4B | 7138.442 | -0.2177 | 0.118165 | -1.84237 | 0.065421 | 0.166897 |
| MAML1 | 854.4968 | -0.33755 | 0.148639 | -2.27097 | 0.023149 | 0.083463 |
| SCGB3A1 | 1.275914 | 0.876396 | 1.883164 | 0.465385 | 0.641656 | NA |
| DVL3 | 4378.442 | -0.40322 | 0.148445 | -2.71628 | 0.006602 | 0.035673 |
| AP2M1 | 16903.29 | -0.2695 | 0.136177 | -1.97904 | 0.047812 | 0.134794 |
| ABCF3 | 1812.736 | -0.27038 | 0.145894 | -1.85327 | 0.063843 | 0.164285 |
| PCYT1A | 2693.224 | -0.11496 | 0.141672 | -0.81147 | 0.417098 | 0.580188 |
| BDH1 | 1092.31 | -0.29505 | 0.135218 | -2.18204 | 0.029106 | 0.09701 |
| IP6K3 | 361.8185 | -0.70079 | 0.264676 | -2.64771 | 0.008104 | 0.040934 |
| LEMD2 | 1661.479 | -0.1658 | 0.164108 | -1.01032 | 0.312344 | 0.479394 |
| TREML1 | 0.260158 | 1.240347 | 4.047124 | 0.306476 | 0.759242 | NA |
| ADCY10P1 | 34.59204 | 1.059848 | 0.39452 | 2.686423 | 0.007222 | NA |
| MT2P1 | 0.614937 | 2.493639 | 2.617677 | 0.952615 | 0.340785 | NA |
| SGCB | 2682.596 | 0.312424 | 0.148807 | 2.099519 | 0.035771 | 0.111538 |
| SPATA18 | 702.3294 | 0.060447 | 0.233978 | 0.258345 | 0.796141 | 0.87685 |
| SMARCAD1 | 607.5536 | 0.911518 | 0.319498 | 2.852967 | 0.004331 | 0.026758 |
| PDLIM5 | 1152.811 | 0.406054 | 0.146769 | 2.766614 | 0.005664 | 0.032237 |
| STPG2 | 0.91323 | 0.960961 | 2.376661 | 0.404332 | 0.685968 | NA |
| MSX1 | 20.83135 | 0.298283 | 0.455936 | 0.654222 | 0.512969 | NA |
| PACRGL | 297.1313 | 0.372231 | 0.177462 | 2.097528 | 0.035947 | 0.111821 |
| C1QTNF7 | 0.260158 | 1.240347 | 4.047124 | 0.306476 | 0.759242 | NA |
| DCAF16 | 623.8862 | -0.04631 | 0.185312 | -0.24989 | 0.802676 | 0.881739 |
| GNPDA2 | 321.743 | 0.620745 | 0.202751 | 3.061607 | 0.002202 | 0.01722 |
| GABRB1 | 5.906067 | 0.777578 | 0.843832 | 0.921484 | 0.356798 | NA |
| PAQR3 | 455.38 | 0.485984 | 0.178344 | 2.724988 | 0.00643 | 0.035001 |
| NIPAL1 | 25.34972 | 0.621742 | 0.499736 | 1.24414 | 0.213448 | NA |

| | | | | | | |
|---|---|---|---|---|---|---|
| ANTXR2 | 402.7761 | -0.11654 | 0.173184 | -0.67292 | 0.500999 | 0.65153 |
| HELQ | 247.5924 | 0.628947 | 0.235413 | 2.671678 | 0.007547 | 0.039226 |
| MRPS18C | 424.63 | 0.425725 | 0.150317 | 2.832181 | 0.004623 | 0.027963 |
| CGGBP1 | 3239.922 | 0.30968 | 0.156843 | 1.974452 | 0.04833 | 0.135643 |
| FAM175A | 162.499 | 0.50658 | 0.297876 | 1.700641 | 0.08901 | 0.205977 |
| CLDN1 | 222.4245 | 0.449786 | 0.265539 | 1.693862 | 0.090292 | 0.20825 |
| LMOD3 | 1.035891 | 3.235292 | 2.200744 | 1.47009 | 0.141537 | NA |
| POGLUT1 | 486.01 | -0.00653 | 0.194008 | -0.03367 | 0.973138 | 0.985082 |
| SLC15A2 | 5.060003 | 1.367331 | 0.976149 | 1.40074 | 0.161292 | NA |
| EIF4E3 | 109.8803 | -0.25669 | 0.264734 | -0.96962 | 0.332234 | 0.500589 |
| LRRC58 | 1216.64 | 0.689872 | 0.280285 | 2.461324 | 0.013843 | 0.058713 |
| FSTL1 | 10886.33 | -0.6072 | 0.22157 | -2.74045 | 0.006136 | 0.03389 |
| IGFBP7 | 10727.36 | 0.156532 | 0.138849 | 1.127358 | 0.259591 | 0.423892 |
| KIAA1524 | 189.6839 | 1.065579 | 0.297972 | 3.576099 | 3.49E-04 | 0.005433 |
| SERPINI1 | 593.1174 | 1.599938 | 0.332393 | 4.813399 | 1.48E-06 | 2.07E-04 |
| PRKCI | 402.7725 | 0.666113 | 0.206198 | 3.230448 | 0.001236 | 0.011864 |
| EIF5A2 | 605.5823 | 0.652999 | 0.160066 | 4.079571 | 4.51E-05 | 0.001553 |
| RPL22L1 | 2149.042 | 0.57666 | 0.188357 | 3.061536 | 0.002202 | 0.01722 |
| PPM1L | 288.9294 | 0.076742 | 0.205557 | 0.373336 | 0.708899 | 0.813788 |
| RYBP | 964.3564 | 0.310009 | 0.20107 | 1.541802 | 0.123122 | 0.257064 |
| PPP4R2 | 846.9277 | 0.687087 | 0.218046 | 3.151115 | 0.001626 | 0.014249 |
| GTPBP8 | 325.19 | 0.337275 | 0.175351 | 1.923425 | 0.054427 | 0.146868 |
| C3orf17 | 1119.188 | 0.429715 | 0.157308 | 2.731682 | 0.006301 | 0.034456 |
| SPICE1 | 282.6119 | 0.521067 | 0.200593 | 2.597633 | 0.009387 | 0.045248 |
| FAM86KP | 0.804712 | -0.7622 | 2.348804 | -0.3245 | 0.745556 | NA |
| KIAA1407 | 42.08893 | 0.66245 | 0.491986 | 1.346481 | 0.178147 | NA |
| NKX6-1 | 0.327913 | 1.543154 | 4.021063 | 0.383768 | 0.701151 | NA |
| CDS1 | 14.83017 | 0.30337 | 0.729897 | 0.415633 | 0.677678 | NA |
| WDFY3 | 1624.758 | 0.173916 | 0.147921 | 1.175735 | 0.239701 | 0.4029 |
| COX18 | 534.8009 | -0.25075 | 0.149522 | -1.67704 | 0.093534 | 0.213528 |
| PTPN13 | 1158.875 | 0.274635 | 0.252461 | 1.087832 | 0.276669 | 0.442514 |
| C4orf36 | 94.33649 | 0.406655 | 0.264226 | 1.539041 | 0.123794 | 0.258104 |
| PPM1K | 318.9855 | 0.238899 | 0.238037 | 1.00362 | 0.315562 | 0.482895 |
| FAM194A | 1.486027 | 0.48257 | 1.621445 | 0.297617 | 0.765995 | NA |
| GMPS | 1513.283 | 0.113576 | 0.150733 | 0.753493 | 0.451154 | 0.610412 |
| TIPARP | 1517.172 | 0.388422 | 0.15649 | 2.482095 | 0.013061 | 0.056594 |
| CCNL1 | 685.4511 | 0.984731 | 0.177373 | 5.551737 | 2.83E-08 | 1.52E-05 |
| PTX3 | 39.08182 | 0.126675 | 0.35358 | 0.358265 | 0.720145 | NA |
| RPL9 | 9714.425 | 0.30183 | 0.166186 | 1.816223 | 0.069336 | 0.173413 |
| SMIM14 | 235.5512 | -0.0027 | 0.214731 | -0.0126 | 0.98995 | 0.995239 |
| RBM47 | 2.740661 | 2.242175 | 1.490258 | 1.504554 | 0.132439 | NA |
| APBB2 | 837.3169 | -0.20127 | 0.192758 | -1.04415 | 0.296415 | 0.463649 |
| PCOLCE2 | 112.9084 | 0.126779 | 0.286317 | 0.442792 | 0.657916 | 0.774307 |
| U2SURP | 657.4868 | 0.967063 | 0.606636 | 1.59414 | 0.110905 | 0.23892 |
| TTC14 | 279.7145 | 1.268258 | 0.337147 | 3.76174 | 1.69E-04 | 0.003497 |
| CXCL3 | 27.77847 | -0.83034 | 0.436548 | -1.90206 | 0.057163 | NA |
| CXCL5 | 63.65826 | -1.28692 | 0.600167 | -2.14427 | 0.032011 | 0.103742 |

| | | | | | | |
|---|---|---|---|---|---|---|
| PF4 | 8.866015 | 0.100663 | 0.692627 | 0.145336 | 0.884446 | NA |
| MTHFD2L | 102.9254 | 0.659549 | 0.243223 | 2.711705 | 0.006694 | 0.035989 |
| CXCL1 | 162.2694 | -0.76792 | 0.775766 | -0.98989 | 0.32223 | 0.490321 |
| RCHY1 | 439.0293 | 0.399664 | 0.172547 | 2.316261 | 0.020544 | 0.077121 |
| PLSCR2 | 0.326333 | 1.533889 | 3.826878 | 0.40082 | 0.688553 | NA |
| CCDC158 | 2.068374 | 1.106444 | 1.590177 | 0.695799 | 0.486554 | NA |
| GYG1 | 1873.187 | -0.13505 | 0.162247 | -0.83236 | 0.405207 | 0.569433 |
| HPS3 | 1367.697 | 0.013923 | 0.163558 | 0.085128 | 0.93216 | 0.96152 |
| TM4SF18 | 7.859517 | 0.595821 | 0.77924 | 0.764618 | 0.444499 | NA |
| TOPBP1 | 1059 | 0.63414 | 0.17283 | 3.669156 | 2.43E-04 | 0.004384 |
| RYK | 1008.878 | 0.361428 | 0.168749 | 2.141804 | 0.032209 | 0.104129 |
| DTX3L | 1050.591 | -0.69146 | 0.231234 | -2.99031 | 0.002787 | 0.020207 |
| ZNF148 | 610.3889 | 0.728113 | 0.242515 | 3.002339 | 0.002679 | 0.019691 |
| NMNAT3 | 1.140347 | 0.579353 | 2.036091 | 0.284542 | 0.775995 | NA |
| TPRA1 | 1078.531 | -0.19903 | 0.15264 | -1.30391 | 0.192265 | 0.34682 |
| YEATS2 | 1703.776 | -0.0753 | 0.132397 | -0.56875 | 0.569526 | 0.705805 |
| POLR2H | 1237.031 | 0.065964 | 0.164484 | 0.401035 | 0.688395 | 0.797432 |
| KLF15 | 341.7243 | -0.12219 | 0.214331 | -0.57009 | 0.568616 | 0.70534 |
| CCDC37 | 0.296509 | -0.51709 | 3.982155 | -0.12985 | 0.896684 | NA |
| CAMK2N2 | 22.61937 | 0.242599 | 0.490572 | 0.494523 | 0.620937 | NA |
| LIPH | 1.381958 | 0.900194 | 1.866056 | 0.482405 | 0.629519 | NA |
| TMEM41A | 405.8939 | -0.12628 | 0.180497 | -0.6996 | 0.484178 | 0.63777 |
| RPN1 | 7239.93 | -0.22817 | 0.143515 | -1.58984 | 0.11187 | 0.240128 |
| SENP2 | 1217.346 | 0.171044 | 0.168062 | 1.017743 | 0.3088 | 0.476128 |
| IFT122 | 546.7114 | -0.44309 | 0.142899 | -3.10073 | 0.00193 | 0.015794 |
| C3orf65 | 0.394291 | 0.298745 | 3.141711 | 0.09509 | 0.924243 | NA |
| RFC4 | 355.5224 | 0.498434 | 0.18452 | 2.701245 | 0.006908 | 0.036774 |
| RPL39L | 674.1131 | -0.08813 | 0.244306 | -0.36074 | 0.718292 | 0.820342 |
| UVSSA | 202.2431 | -0.07135 | 0.181771 | -0.39252 | 0.694677 | 0.80314 |
| SLBP | 1248.667 | 0.231359 | 0.13063 | 1.7711 | 0.076544 | 0.185607 |
| LRPAP1 | 5739.542 | -0.3353 | 0.154274 | -2.17343 | 0.029748 | 0.098482 |
| ZDHHC19 | 0.448206 | -1.2532 | 3.007315 | -0.41672 | 0.676885 | NA |
| SLC51A | 52.2442 | -0.18801 | 0.30573 | -0.61494 | 0.538592 | 0.681093 |
| UBXN7 | 851.5121 | 0.157056 | 0.181896 | 0.863434 | 0.387899 | 0.553708 |
| RNF168 | 297.9749 | 0.245636 | 0.200342 | 1.226088 | 0.220166 | 0.379921 |
| PIGX | 649.4408 | 0.334321 | 0.186433 | 1.793255 | 0.072932 | 0.179865 |
| MFI2 | 3565.531 | -1.03036 | 0.241686 | -4.26323 | 2.01E-05 | 9.36E-04 |
| S100P | 0.928019 | -0.78227 | 2.07134 | -0.37767 | 0.705679 | NA |
| ABLIM2 | 1.706702 | -0.21718 | 1.592383 | -0.13639 | 0.891516 | NA |
| AIMP1 | 575.2155 | 0.812461 | 0.210536 | 3.859011 | 1.14E-04 | 0.002777 |
| SGMS2 | 268.9839 | 0.608785 | 0.178054 | 3.419099 | 6.28E-04 | 0.007783 |
| METAP1 | 1153.192 | 0.134209 | 0.148859 | 0.901586 | 0.367277 | 0.534385 |
| DNAJB14 | 744.8993 | 1.058441 | 0.338895 | 3.123212 | 0.001789 | 0.015078 |
| H2AFZ | 12500.49 | -0.03078 | 0.135912 | -0.22646 | 0.820845 | 0.892334 |
| EMCN | 2.334113 | 1.474346 | 1.633851 | 0.902375 | 0.366858 | NA |
| SLC9B1 | 20.436 | 0.147126 | 0.462818 | 0.317893 | 0.750566 | NA |
| SLC9B2 | 723.7026 | 0.011718 | 0.146778 | 0.079833 | 0.93637 | 0.964437 |

| | | | | | | |
|---|---|---|---|---|---|---|
| BDH2 | 510.1394 | 0.513514 | 0.154929 | 3.314523 | 9.18E-04 | 0.009804 |
| PGRMC2 | 2354.341 | -0.02339 | 0.199426 | -0.11727 | 0.906643 | 0.945876 |
| SPRY1 | 79.10515 | -0.31104 | 0.266954 | -1.16515 | 0.243959 | 0.406712 |
| INTU | 59.00657 | 0.531117 | 0.302567 | 1.755371 | 0.079196 | 0.190015 |
| HSPA4L | 66.70433 | 0.722915 | 0.371118 | 1.947936 | 0.051423 | 0.141213 |
| MFSD8 | 203.4554 | 0.585983 | 0.206588 | 2.836487 | 0.004561 | 0.027674 |
| C4orf29 | 130.1138 | 0.543605 | 0.225131 | 2.414617 | 0.015752 | 0.064127 |
| ETNPPL | 1.51293 | 0.51091 | 1.562353 | 0.327013 | 0.743658 | NA |
| PITX2 | 1.668624 | 0.880224 | 1.734128 | 0.507589 | 0.611742 | NA |
| C4orf3 | 4981.146 | 0.250485 | 0.140091 | 1.788019 | 0.073773 | 0.181081 |
| PRSS12 | 508.2437 | 0.136215 | 0.151974 | 0.896306 | 0.370089 | 0.536887 |
| NDST3 | 7.875792 | 1.997371 | 0.792217 | 2.521244 | 0.011694 | NA |
| HMGB2 | 768.6242 | 0.489864 | 0.185208 | 2.644948 | 0.00817 | 0.041164 |
| SAP30 | 503.3826 | 0.349114 | 0.192092 | 1.81743 | 0.069151 | 0.173097 |
| SCRG1 | 156.7591 | 1.462319 | 0.326842 | 4.474084 | 7.67E-06 | 5.35E-04 |
| HAND2 | 0.392857 | -1.546 | 4.075469 | -0.37934 | 0.704433 | NA |
| MAD2L1 | 795.0097 | 0.263854 | 0.218974 | 1.204957 | 0.22822 | 0.389884 |
| ANXA5 | 42900.68 | 0.119441 | 0.136735 | 0.873521 | 0.382379 | 0.54847 |
| TMEM155 | 0.724348 | 0.185001 | 2.57534 | 0.071836 | 0.942733 | NA |
| MAP9 | 81.08385 | 1.132944 | 0.962596 | 1.176967 | 0.239209 | 0.402358 |
| FBXO8 | 553.4821 | 0.527154 | 0.181965 | 2.897002 | 0.003767 | 0.024487 |
| CEP44 | 123.0098 | 0.996653 | 0.291906 | 3.414301 | 6.39E-04 | 0.007848 |
| HPGD | 4442.402 | 0.671996 | 0.304148 | 2.209433 | 0.027145 | 0.092684 |
| C4orf45 | 0.315653 | -0.57305 | 3.892639 | -0.14721 | 0.882963 | NA |
| TMEM144 | 77.57131 | -0.43181 | 0.317735 | -1.35903 | 0.174136 | 0.326179 |
| FAM198B | 84.47556 | -1.27014 | 0.922498 | -1.37684 | 0.16856 | 0.318655 |
| NPY1R | 0.163957 | 0.424913 | 4.079006 | 0.104171 | 0.917034 | NA |
| NAA15 | 802.7047 | 0.668599 | 0.299085 | 2.235482 | 0.025386 | 0.088933 |
| IL15 | 22.25475 | -0.13625 | 0.588141 | -0.23166 | 0.816802 | NA |
| FAM160A1 | 5.283286 | -0.20458 | 0.990655 | -0.20651 | 0.836396 | NA |
| ARFIP1 | 996.4539 | 0.675829 | 0.196539 | 3.438659 | 5.85E-04 | 0.007457 |
| KIAA0947 | 920.0421 | 0.664138 | 0.232203 | 2.860158 | 0.004234 | 0.026417 |
| HHIP | 0.670433 | 1.255281 | 2.356401 | 0.532711 | 0.594234 | NA |
| ANAPC10 | 138.5543 | 0.701014 | 0.244971 | 2.861614 | 0.004215 | 0.026354 |
| ABCE1 | 1326.682 | 0.731129 | 0.170688 | 4.283426 | 1.84E-05 | 8.78E-04 |
| OTUD4 | 740.7418 | 0.308702 | 0.168479 | 1.832281 | 0.06691 | 0.169092 |
| LSM6 | 243.2718 | 0.350409 | 0.208979 | 1.67677 | 0.093588 | 0.213567 |
| TMEM184( | 976.9107 | 0.15715 | 0.155275 | 1.012074 | 0.311503 | 0.478908 |
| PRMT10 | 239.0836 | 0.368338 | 0.233208 | 1.579443 | 0.114235 | 0.243483 |
| ITGA2 | 511.0964 | 0.978187 | 0.28996 | 3.373526 | 7.42E-04 | 0.008681 |
| MOCS2 | 955.9294 | 0.703344 | 0.205395 | 3.424352 | 6.16E-04 | 0.007731 |
| SLC45A2 | 721.591 | -1.10617 | 0.266774 | -4.14647 | 3.38E-05 | 0.001308 |
| EDIL3 | 0.288605 | 1.443619 | 4.029058 | 0.358302 | 0.720117 | NA |
| TMEM161I | 355.5636 | 0.752802 | 0.19797 | 3.802614 | 1.43E-04 | 0.003156 |
| ELOVL7 | 5.583458 | 0.466374 | 0.891437 | 0.523171 | 0.600855 | NA |
| NDUFAF2 | 411.3333 | 0.680467 | 0.239875 | 2.836755 | 0.004557 | 0.027664 |
| ZNF474 | 1.043999 | 1.178957 | 2.24578 | 0.524966 | 0.599607 | NA |

| | | | | | | |
|---|---|---|---|---|---|---|
| LMBRD2 | 258.0998 | 0.792297 | 0.23324 | 3.396919 | 6.81E-04 | 0.008177 |
| RANBP3L | 0.9715 | 2.024222 | 2.39445 | 0.845381 | 0.397898 | NA |
| NIPBL | 661.1087 | 0.644866 | 0.659009 | 0.978539 | 0.327808 | 0.495937 |
| RNF180 | 128.1117 | 0.909486 | 0.286121 | 3.178672 | 0.00148 | 0.013453 |
| GPR98 | 3.984775 | 0.074762 | 1.017175 | 0.0735 | 0.941408 | NA |
| SLC25A46 | 1397.25 | 0.555635 | 0.206455 | 2.691318 | 0.007117 | 0.037572 |
| STARD4 | 557.2878 | 0.818741 | 0.255866 | 3.199881 | 0.001375 | 0.012783 |
| PGGT1B | 263.3706 | 0.421581 | 0.252382 | 1.670408 | 0.094839 | 0.215442 |
| F2RL2 | 0.787148 | -0.70094 | 2.357021 | -0.29738 | 0.766174 | NA |
| CCDC112 | 110.9962 | 0.669377 | 1.014149 | 0.660038 | 0.50923 | 0.657545 |
| ANKRD33B | 0.72077 | 0.527966 | 3.060238 | 0.172524 | 0.863025 | NA |
| CMBL | 928.3009 | -0.25884 | 0.145711 | -1.77637 | 0.075673 | 0.184058 |
| C5orf63 | 22.55941 | 0.174111 | 0.440381 | 0.395364 | 0.692574 | NA |
| PRRC1 | 1413.824 | 0.074937 | 0.183998 | 0.40727 | 0.68381 | 0.794294 |
| F2RL1 | 0.719393 | -0.9644 | 2.652838 | -0.36354 | 0.716204 | NA |
| AGGF1 | 746.7095 | 0.950763 | 0.236431 | 4.021304 | 5.79E-05 | 0.001791 |
| WDR41 | 1347.965 | 0.259234 | 0.159901 | 1.621214 | 0.104972 | 0.230452 |
| NDUFS4 | 942.7826 | 0.403883 | 0.207656 | 1.944962 | 0.05178 | 0.141718 |
| SPINK1 | 8.479087 | 1.15163 | 0.747713 | 1.540202 | 0.123511 | NA |
| HTR4 | 0.096202 | 0.408546 | 4.079639 | 0.100143 | 0.920231 | NA |
| GRPEL2 | 245.5394 | 0.473802 | 0.215561 | 2.197994 | 0.02795 | 0.094666 |
| CDC20B | 0.315653 | -0.57305 | 3.892639 | -0.14721 | 0.882963 | NA |
| ARSK | 340.0583 | 0.917756 | 0.226076 | 4.059495 | 4.92E-05 | 0.001622 |
| RHOBTB3 | 1657.501 | 0.45527 | 0.212304 | 2.144429 | 0.031999 | 0.103731 |
| GPX8 | 1725.397 | 0.416888 | 0.152166 | 2.739698 | 0.00615 | 0.03389 |
| TIGD6 | 174.2259 | 0.219091 | 0.223487 | 0.980333 | 0.326922 | 0.495092 |
| SERINC5 | 914.3312 | -0.31436 | 0.147638 | -2.12924 | 0.033234 | 0.106198 |
| CAGE1 | 0.258578 | 1.233151 | 4.047809 | 0.304646 | 0.760635 | NA |
| CASP3 | 638.055 | 0.984598 | 0.228642 | 4.306282 | 1.66E-05 | 8.32E-04 |
| PRIMPOL | 202.1393 | 0.536625 | 0.230829 | 2.324773 | 0.020084 | 0.075965 |
| ERAP1 | 1258.953 | -0.20235 | 0.131842 | -1.53477 | 0.12484 | 0.259646 |
| ERAP2 | 90.71468 | 0.267815 | 0.325763 | 0.822117 | 0.411011 | 0.57432 |
| CMYA5 | 6.008879 | -0.63929 | 0.824285 | -0.77557 | 0.438004 | NA |
| EGFLAM | 0.151697 | -1.546 | 4.075469 | -0.37934 | 0.704433 | NA |
| KIAA1430 | 1652.832 | 0.849893 | 0.190368 | 4.464488 | 8.03E-06 | 5.46E-04 |
| RICTOR | 713.9576 | 0.956707 | 0.248913 | 3.843537 | 1.21E-04 | 0.0029 |
| PAPD4 | 1083.992 | 0.696216 | 0.177986 | 3.911637 | 9.17E-05 | 0.002404 |
| EBF1 | 7.847192 | 0.800597 | 0.815991 | 0.981135 | 0.326526 | NA |
| ANKRA2 | 212.8552 | 0.489357 | 0.203867 | 2.400378 | 0.016378 | 0.066132 |
| UBLCP1 | 680.0792 | 0.71395 | 0.279079 | 2.558238 | 0.01052 | 0.048662 |
| UTP15 | 407.4495 | 0.486005 | 0.16489 | 2.947453 | 0.003204 | 0.0221 |
| TLR3 | 141.5768 | 0.310434 | 0.313789 | 0.989307 | 0.322513 | 0.490619 |
| KLKB1 | 4.810782 | 1.525338 | 1.048407 | 1.45491 | 0.145694 | NA |
| NSA2 | 892.7863 | 0.894248 | 0.235282 | 3.80075 | 1.44E-04 | 0.003162 |
| GFM2 | 879.1991 | 0.350499 | 0.187295 | 1.871376 | 0.061293 | 0.159664 |
| TERT | 0.162376 | 0.424913 | 4.079006 | 0.104171 | 0.917034 | NA |
| CCDC127 | 575.28 | -0.0109 | 0.159335 | -0.0684 | 0.945465 | 0.969593 |

| | | | | | | |
|---|---|---|---|---|---|---|
| FOXQ1 | 163.7724 | 0.818631 | 0.707157 | 1.157637 | 0.247012 | 0.410475 |
| C6orf195 | 0.326333 | 1.533889 | 3.826878 | 0.40082 | 0.688553 | NA |
| GPR111 | 0.327913 | 1.543154 | 4.021063 | 0.383768 | 0.701151 | NA |
| ACSL6 | 11.98768 | 0.26444 | 0.756967 | 0.349342 | 0.726833 | NA |
| 8-Sep | 2028.766 | -0.11595 | 0.204129 | -0.56801 | 0.570025 | 0.706055 |
| SHROOM1 | 98.48175 | -0.44429 | 0.329703 | -1.34754 | 0.177805 | 0.330181 |
| GDF9 | 1.974736 | 1.142697 | 1.441282 | 0.792834 | 0.427875 | NA |
| UQCRQ | 3299.581 | 0.194118 | 0.144866 | 1.339982 | 0.180251 | 0.332803 |
| LEAP2 | 8.710063 | 0.631627 | 0.716497 | 0.881549 | 0.378021 | NA |
| GJB7 | 0.778894 | 2.823374 | 2.403613 | 1.174637 | 0.24014 | NA |
| SLC35A1 | 150.2309 | 0.574328 | 0.240553 | 2.38753 | 0.016962 | 0.067758 |
| GRIK2 | 867.2174 | 0.641381 | 0.205458 | 3.121718 | 0.001798 | 0.015096 |
| MB21D1 | 78.05171 | 0.394248 | 0.300239 | 1.313115 | 0.189144 | 0.343385 |
| FABP7 | 19.54905 | -0.45966 | 0.568321 | -0.8088 | 0.418628 | NA |
| TLX3 | 0.327913 | 1.543154 | 4.021063 | 0.383768 | 0.701151 | NA |
| TXLNB | 1.512858 | -1.25993 | 2.088308 | -0.60332 | 0.546293 | NA |
| CITED2 | 450.1961 | -0.47803 | 0.179542 | -2.66251 | 0.007756 | 0.039822 |
| FAM26D | 0.134133 | -1.48909 | 4.077584 | -0.36519 | 0.71497 | NA |
| CREBRF | 632.2266 | 0.9251 | 0.24979 | 3.703511 | 2.13E-04 | 0.00404 |
| DCBLD1 | 246.2143 | 0.138659 | 0.178361 | 0.777405 | 0.43692 | 0.597357 |
| SFXN1 | 801.1539 | 0.148831 | 0.171318 | 0.868743 | 0.384987 | 0.550927 |
| SAMD3 | 0.324752 | 1.53082 | 4.022025 | 0.380609 | 0.703493 | NA |
| TMEM200A | 0.162376 | 0.424913 | 4.079006 | 0.104171 | 0.917034 | NA |
| PDSS2 | 439.3225 | 0.095521 | 0.156756 | 0.60936 | 0.542286 | 0.684424 |
| STXBP5 | 1455.813 | 0.4124 | 0.185887 | 2.218559 | 0.026517 | 0.091356 |
| IL31RA | 1.943275 | 0.401937 | 1.639001 | 0.245233 | 0.806276 | NA |
| ANKRD55 | 1.047103 | 1.538196 | 2.447568 | 0.628459 | 0.529703 | NA |
| RAET1E | 0.548763 | 2.343695 | 3.135478 | 0.747476 | 0.454776 | NA |
| PI16 | 0.151697 | -1.546 | 4.075469 | -0.37934 | 0.704433 | NA |
| GALNT10 | 847.0717 | -0.29906 | 0.136678 | -2.18808 | 0.028664 | 0.096154 |
| SAP30L | 661.2821 | 0.117641 | 0.164163 | 0.71661 | 0.473614 | 0.629168 |
| RPS14 | 18931.74 | 0.020976 | 0.191069 | 0.10978 | 0.912584 | 0.949346 |
| MYOZ3 | 30.25733 | -0.0598 | 0.412618 | -0.14492 | 0.884775 | NA |
| SLU7 | 1010.174 | 0.873159 | 0.235127 | 3.713557 | 2.04E-04 | 0.00392 |
| PTTG1 | 2559.003 | -0.12214 | 0.215321 | -0.56724 | 0.570548 | 0.706481 |
| CAMLG | 1259.789 | 0.290211 | 0.151335 | 1.917678 | 0.055152 | 0.148193 |
| RELL2 | 461.4291 | 0.029514 | 0.197304 | 0.149588 | 0.881089 | 0.929216 |
| SMAD5-AS1 | 1.330966 | 2.102973 | 2.223531 | 0.945781 | 0.344261 | NA |
| KCNK5 | 1.143361 | 1.348976 | 1.954071 | 0.690341 | 0.48998 | NA |
| KIF6 | 356.9631 | 0.053879 | 0.186666 | 0.28864 | 0.772857 | 0.859277 |
| USP49 | 232.3672 | 0.123578 | 0.244672 | 0.505075 | 0.613506 | 0.740094 |
| PHAX | 797.7291 | 0.459072 | 0.246195 | 1.864669 | 0.062228 | 0.161495 |
| ALDH7A1 | 1525.777 | 0.017174 | 0.143395 | 0.119769 | 0.904666 | 0.944585 |
| KDM1B | 469.1932 | 0.166567 | 0.217502 | 0.765819 | 0.443784 | 0.603679 |
| NSD1 | 1314.168 | -0.17592 | 0.165608 | -1.06228 | 0.288108 | 0.454312 |
| C2 | 2.088887 | 0.208509 | 1.392705 | 0.149715 | 0.880989 | NA |
| HSBP1P2 | 0.624422 | 1.162588 | 2.733761 | 0.425271 | 0.670639 | NA |

| | | | | | | |
|---|---|---|---|---|---|---|
| KIAA1586 | 96.31121 | 1.280904 | 0.471845 | 2.71467 | 0.006634 | 0.035809 |
| OR2W6P | 0.314073 | -0.57308 | 3.898934 | -0.14698 | 0.883146 | NA |
| FAM83B | 0.411855 | 0.250456 | 3.097618 | 0.080854 | 0.935558 | NA |
| THAP9 | 65.72214 | 1.055818 | 0.332549 | 3.174921 | 0.001499 | 0.013563 |
| RBPJ | 1319.32 | 0.207437 | 0.178511 | 1.162042 | 0.245219 | 0.408145 |
| LMBRD1 | 1715.872 | 0.491683 | 0.172119 | 2.856641 | 0.004281 | 0.026588 |
| ZCCHC4 | 205.4997 | 0.163483 | 0.206624 | 0.791209 | 0.428822 | 0.590577 |
| HIST1H2BI | 0.941473 | 1.959123 | 2.152875 | 0.910003 | 0.362821 | NA |
| UBTD2 | 638.412 | 0.016984 | 0.152489 | 0.111379 | 0.911316 | 0.948604 |
| HIST1H2AE | 8.069769 | 0.056421 | 0.741445 | 0.076096 | 0.939342 | NA |
| HIST1H1E | 2.221294 | 0.463708 | 1.319619 | 0.351396 | 0.725292 | NA |
| IRF2 | 526.9501 | 0.039259 | 0.150237 | 0.261316 | 0.793849 | 0.874961 |
| FILIP1L | 3.955504 | 1.385986 | 1.132884 | 1.223414 | 0.221174 | NA |
| TAP1 | 3032.891 | -0.65137 | 0.182309 | -3.57287 | 3.53E-04 | 0.005479 |
| CMAHP | 5.605651 | 1.230508 | 0.953153 | 1.290987 | 0.196708 | NA |
| CDC40 | 382.8811 | 0.797259 | 0.226074 | 3.526542 | 4.21E-04 | 0.006131 |
| TNXB | 10.80352 | -0.418 | 0.651186 | -0.64191 | 0.520934 | NA |
| CCDC110 | 22.62453 | 1.07755 | 1.521553 | 0.708191 | 0.478827 | NA |
| TRAPPC11 | 758.8255 | 0.367119 | 0.233672 | 1.571088 | 0.116162 | 0.246527 |
| ING2 | 143.2928 | 0.293833 | 0.215963 | 1.360574 | 0.173648 | 0.325574 |
| CDKN2AIP | 319.8899 | 0.61289 | 0.211765 | 2.894198 | 0.003801 | 0.024629 |
| SNRNP48 | 224.8266 | 1.028221 | 0.272829 | 3.768732 | 1.64E-04 | 0.003461 |
| ADAM29 | 0.326333 | 1.533889 | 3.826878 | 0.40082 | 0.688553 | NA |
| GDNF | 363.5341 | 0.365917 | 0.323912 | 1.129678 | 0.258612 | 0.422992 |
| UGT3A2 | 6.457834 | -0.01963 | 0.908879 | -0.02159 | 0.982772 | NA |
| IL7R | 10.31216 | 0.670491 | 0.793347 | 0.845142 | 0.398032 | NA |
| DNAJC21 | 708.1097 | 0.790176 | 0.293595 | 2.69138 | 0.007116 | 0.037572 |
| NPNT | 29.72297 | 1.2034 | 0.630756 | 1.907868 | 0.056408 | NA |
| TET2 | 267.154 | 0.59404 | 0.243176 | 2.442834 | 0.014572 | 0.060744 |
| CXXC4 | 10.54029 | 0.271173 | 0.679358 | 0.39916 | 0.689775 | NA |
| SHOX2 | 5.880093 | 0.200048 | 0.858352 | 0.233061 | 0.815714 | NA |
| TSPAN5 | 456.0588 | 0.009049 | 0.228968 | 0.039523 | 0.968474 | 0.982732 |
| IL12A | 6.53687 | -0.06444 | 0.86917 | -0.07414 | 0.940902 | NA |
| STX18 | 928.3671 | 0.103033 | 0.129256 | 0.797124 | 0.425379 | 0.587405 |
| | 6266.093 | -0.32844 | 0.178758 | -1.83737 | 0.066155 | 0.16784 |
| ZBTB49 | 86.07233 | 0.156907 | 0.28023 | 0.559922 | 0.575533 | 0.710649 |
| GFM1 | 1765.377 | 0.304685 | 0.12601 | 2.417942 | 0.015609 | 0.063675 |
| FSTL5 | 9.625695 | 0.370985 | 0.722446 | 0.513513 | 0.607593 | NA |
| TNIP2 | 1648.945 | 0.048195 | 0.179483 | 0.268523 | 0.788297 | 0.871434 |
| ZNF608 | 717.8003 | -0.05916 | 0.15949 | -0.37095 | 0.710677 | 0.81504 |
| SLC35G2 | 77.32865 | 0.775875 | 0.359306 | 2.159374 | 0.030821 | 0.100963 |
| LETM1 | 2035.144 | -0.54113 | 0.142376 | -3.80071 | 1.44E-04 | 0.003162 |
| TMEM129 | 1715.637 | -0.35414 | 0.132664 | -2.66941 | 0.007598 | 0.03937 |
| PPIC | 255.3315 | -0.00727 | 0.19726 | -0.03687 | 0.970585 | 0.983948 |
| CEP120 | 285.2775 | 0.880687 | 0.28135 | 3.130223 | 0.001747 | 0.014823 |
| CPLX1 | 4.156839 | -0.24362 | 0.990595 | -0.24593 | 0.805734 | NA |
| PXDC1 | 1259.731 | 0.086095 | 0.160741 | 0.535615 | 0.592225 | 0.7233 |

| | | | | | | |
|---|---|---|---|---|---|---|
| COMMD8 | 444.5555 | 1.125192 | 0.292786 | 3.843049 | 1.22E-04 | 0.0029 |
| ATP5I | 973.8608 | 0.278027 | 0.164492 | 1.690222 | 0.090986 | 0.209443 |
| MFSD7 | 149.8768 | 0.099801 | 0.272611 | 0.366092 | 0.714296 | 0.817689 |
| HNRNPH1 | 3038.96 | 0.377202 | 0.145695 | 2.588984 | 0.009626 | 0.045996 |
| ZBBX | 0.47803 | 0.520083 | 2.97881 | 0.174594 | 0.861399 | NA |
| ACTBL2 | 1.617239 | -0.2809 | 1.534249 | -0.18308 | 0.854732 | NA |
| BRD7P3 | 0.588072 | 2.39955 | 3.098804 | 0.774347 | 0.438726 | NA |
| HSPBAP1 | 337.0782 | 0.381446 | 0.190375 | 2.00366 | 0.045106 | 0.129917 |
| PARM1 | 0.096202 | 0.408546 | 4.079639 | 0.100143 | 0.920231 | NA |
| ZNF354A | 112.5654 | 0.985676 | 0.276857 | 3.560237 | 3.71E-04 | 0.005601 |
| RGS14 | 544.2698 | 0.25772 | 0.186454 | 1.382217 | 0.166905 | 0.316383 |
| LMAN2 | 5039.625 | -0.30903 | 0.12666 | -2.43981 | 0.014695 | 0.061041 |
| RAB24 | 239.1474 | -0.14294 | 0.176267 | -0.81091 | 0.417416 | 0.580397 |
| PRELID1 | 3082.415 | -0.10191 | 0.160198 | -0.63616 | 0.524673 | 0.670408 |
| CXCL10 | 8.004064 | 0.739983 | 0.962935 | 0.768466 | 0.44221 | NA |
| SH3TC2 | 1034.56 | -0.14405 | 0.159908 | -0.90083 | 0.367679 | 0.534761 |
| CXCL11 | 2.344747 | -0.10449 | 1.291193 | -0.08092 | 0.935504 | NA |
| NMD3 | 868.5617 | 0.765046 | 0.246852 | 3.099212 | 0.00194 | 0.015853 |
| ADRB2 | 428.0396 | 0.470298 | 0.340812 | 1.379932 | 0.167608 | 0.317106 |
| B3GALNT1 | 401.1412 | 0.252407 | 0.160765 | 1.570037 | 0.116406 | 0.246693 |
| GPRIN1 | 720.3848 | -0.49708 | 0.158042 | -3.14523 | 0.00166 | 0.014396 |
| HSPB3 | 7.658997 | 2.18399 | 1.026729 | 2.127134 | 0.033409 | NA |
| KCNAB1 | 47.32775 | 0.33764 | 0.328411 | 1.028102 | 0.303902 | 0.471133 |
| MRPL1 | 365.5676 | 0.728439 | 0.255951 | 2.846011 | 0.004427 | 0.027207 |
| PGM2 | 781.0389 | 0.594979 | 0.169768 | 3.504649 | 4.57E-04 | 0.00643 |
| STK32A | 1199.253 | -0.81183 | 0.287731 | -2.82148 | 0.00478 | 0.028636 |
| P2RY12 | 23.15588 | 0.981692 | 0.58728 | 1.671592 | 0.094605 | NA |
| SLC33A1 | 870.6462 | 0.276264 | 0.161089 | 1.714974 | 0.08635 | 0.201353 |
| ARL13B | 177.2987 | 1.110646 | 0.29008 | 3.828751 | 1.29E-04 | 0.002993 |
| IL8 | 86.94404 | 0.589311 | 0.449677 | 1.310522 | 0.190019 | 0.344236 |
| RASSF6 | 0.392857 | -1.546 | 4.075469 | -0.37934 | 0.704433 | NA |
| HINT1 | 6630.828 | 0.107818 | 0.116536 | 0.925191 | 0.354866 | 0.523121 |
| DTWD2 | 279.7765 | 0.587272 | 0.216873 | 2.707907 | 0.006771 | 0.036304 |
| CNBP | 5827.801 | 0.357893 | 0.142584 | 2.510045 | 0.012072 | 0.053659 |
| LDB2 | 0.392857 | -1.546 | 4.075469 | -0.37934 | 0.704433 | NA |
| NLGN1 | 665.4053 | 0.479376 | 0.237881 | 2.015195 | 0.043884 | 0.127729 |
| TAPT1 | 581.8088 | 0.222655 | 0.195611 | 1.138254 | 0.255014 | 0.419306 |
| PCDH7 | 1212.557 | -0.21166 | 0.241363 | -0.87696 | 0.38051 | 0.546716 |
| ROBO1 | 1076.836 | -0.04182 | 0.197577 | -0.21165 | 0.832383 | 0.899289 |
| P2RY1 | 53.29295 | 0.488707 | 0.435366 | 1.12252 | 0.261641 | 0.426066 |
| CTNND2 | 0.52699 | -2.57878 | 3.248478 | -0.79384 | 0.427288 | NA |
| TM4SF1 | 20.06853 | -1.21206 | 0.529337 | -2.28977 | 0.022034 | NA |
| SF3B5 | 2387.38 | -0.09033 | 0.126211 | -0.71569 | 0.47418 | 0.629587 |
| TIGD4 | 14.54307 | 1.80872 | 0.654738 | 2.762509 | 0.005736 | NA |
| TMEM154 | 11.51156 | -0.03905 | 0.622538 | -0.06273 | 0.94998 | NA |
| ALCAM | 3636.123 | 1.138472 | 0.19523 | 5.831426 | 5.50E-09 | 4.65E-06 |
| ZPLD1 | 0.163957 | 0.424913 | 4.079006 | 0.104171 | 0.917034 | NA |

| | | | | | | |
|---|---|---|---|---|---|---|
| SIMC1 | 364.122 | -0.21845 | 0.172085 | -1.26941 | 0.204296 | 0.360991 |
| TMEM192 | 1223.941 | 0.043634 | 0.127485 | 0.342266 | 0.732151 | 0.83035 |
| | 124.5653 | 0.472278 | 0.240954 | 1.960037 | 0.049991 | 0.138502 |
| | 0.52699 | -2.57878 | 3.248478 | -0.79384 | 0.427288 | NA |
| RNF150 | 395.877 | -0.17091 | 0.160106 | -1.06748 | 0.285755 | 0.452292 |
| USP38 | 601.5796 | 0.082315 | 0.166205 | 0.495261 | 0.620416 | 0.744964 |
| ADRA1B | 19.75906 | -0.5473 | 0.523168 | -1.04612 | 0.295506 | NA |
| PWWP2A | 251.2146 | 0.39683 | 0.209892 | 1.890637 | 0.058673 | 0.154802 |
| FAXDC2 | 937.1583 | -0.18139 | 0.169974 | -1.06715 | 0.285906 | 0.452344 |
| SMAD1 | 138.5073 | -0.13601 | 0.23734 | -0.57307 | 0.566598 | 0.703721 |
| DCLK2 | 191.9975 | -0.35092 | 0.211058 | -1.66265 | 0.096382 | 0.217558 |
| HARS | 3105.454 | -0.0143 | 0.143598 | -0.09958 | 0.920677 | 0.953909 |
| NFXL1 | 154.244 | 0.579343 | 0.251783 | 2.30096 | 0.021394 | 0.079307 |
| CD14 | 48.68346 | -0.22061 | 0.371419 | -0.59396 | 0.552537 | 0.69323 |
| DNAJC18 | 681.8624 | -0.18613 | 0.159483 | -1.16706 | 0.243188 | 0.406164 |
| SPATA24 | 58.22904 | 0.363862 | 0.309506 | 1.175623 | 0.239745 | 0.402918 |
| MZB1 | 0.230334 | -0.53318 | 4.076749 | -0.13079 | 0.895945 | NA |
| SLC23A1 | 3.311945 | -0.51366 | 1.173421 | -0.43774 | 0.661572 | NA |
| NUDT9 | 1217.204 | 0.259066 | 0.140206 | 1.847753 | 0.064638 | 0.165719 |
| HSD17B13 | 0.738391 | 0.402625 | 2.276191 | 0.176885 | 0.859599 | NA |
| ELOVL6 | 1413.863 | 0.293931 | 0.202345 | 1.452626 | 0.146328 | 0.28883 |
| SERPINB9 | 14.64308 | -0.64028 | 0.736315 | -0.86957 | 0.384535 | NA |
| IRX1 | 85.76595 | -0.41914 | 0.275842 | -1.51948 | 0.128643 | 0.264951 |
| IRX2 | 0.382031 | -1.26484 | 3.151989 | -0.40128 | 0.688212 | NA |
| EMB | 272.45 | 0.774257 | 0.201535 | 3.841809 | 1.22E-04 | 0.002907 |
| NUDCD2 | 639.9587 | 0.358103 | 0.173167 | 2.067968 | 0.038643 | 0.117459 |
| HSPA4 | 4713.508 | 0.312364 | 0.192996 | 1.618501 | 0.105555 | 0.231319 |
| FAM71B | 0.392857 | -1.546 | 4.075469 | -0.37934 | 0.704433 | NA |
| SGCD | 738.6885 | -0.21843 | 0.231652 | -0.94291 | 0.345725 | 0.513605 |
| POLH | 561.1929 | -0.35935 | 0.186039 | -1.93156 | 0.053414 | 0.144835 |
| HTRA3 | 0.487129 | 2.123445 | 3.910539 | 0.543006 | 0.587126 | NA |
| BFSP2 | 8.316907 | -0.74082 | 0.799546 | -0.92655 | 0.35416 | NA |
| GPR27 | 1.817839 | -2.25663 | 1.669281 | -1.35186 | 0.176421 | NA |
| | 353.3146 | 0.017698 | 0.2032 | 0.087097 | 0.930594 | 0.960575 |
| MINA | 559.6198 | 0.281772 | 0.170505 | 1.65257 | 0.098418 | 0.220892 |
| KIAA0232 | 812.9187 | 0.55705 | 0.214059 | 2.602323 | 0.009259 | 0.044768 |
| CYTL1 | 949.2256 | 0.00664 | 0.140471 | 0.047268 | 0.9623 | 0.979485 |
| TRH | 0.096202 | 0.408546 | 4.079639 | 0.100143 | 0.920231 | NA |
| GSTA4 | 362.2559 | 0.132504 | 0.181575 | 0.729749 | 0.465544 | 0.622879 |
| PAQR8 | 3574.474 | -0.52201 | 0.190064 | -2.74648 | 0.006024 | 0.033559 |
| NUDT6 | 80.87613 | 0.280094 | 0.282552 | 0.9913 | 0.321539 | 0.489703 |
| PKHD1 | 0.956409 | -1.29092 | 2.212928 | -0.58335 | 0.559655 | NA |
| PGK2 | 0.162376 | 0.424913 | 4.079006 | 0.104171 | 0.917034 | NA |
| FAM86JP | 28.12583 | -0.46809 | 0.407062 | -1.14991 | 0.25018 | NA |
| MFN1 | 1113.927 | 0.720248 | 0.214659 | 3.355314 | 7.93E-04 | 0.008995 |
| KCNMB3 | 5.295926 | 1.923806 | 1.011083 | 1.902718 | 0.057077 | NA |
| MUC7 | 0.327913 | 1.543154 | 4.021063 | 0.383768 | 0.701151 | NA |

| | | | | | | |
|---|---|---|---|---|---|---|
| PROL1 | 0.163957 | 0.424913 | 4.079006 | 0.104171 | 0.917034 | NA |
| UGT2B7 | 38.36616 | -0.12327 | 0.374991 | -0.32873 | 0.742362 | NA |
| TPPP | 44.81987 | -0.58514 | 0.426025 | -1.37349 | 0.169599 | NA |
| PDE7B | 82.98728 | 0.472169 | 0.403396 | 1.170486 | 0.241805 | 0.404887 |
| MRPL36 | 832.9092 | 0.235093 | 0.163464 | 1.438198 | 0.150378 | 0.294052 |
| MCC | 2447.726 | -0.12084 | 0.197245 | -0.61266 | 0.540099 | 0.682173 |
| POLR1C | 850.2402 | -0.0911 | 0.180787 | -0.50393 | 0.614308 | 0.74048 |
| DLK2 | 9.928371 | -0.75382 | 0.725013 | -1.03974 | 0.298461 | NA |
| ZNF318 | 990.5901 | -0.26264 | 0.162492 | -1.61634 | 0.106021 | 0.231871 |
| HOPX | 1.732381 | 0.267512 | 1.498441 | 0.178527 | 0.858309 | NA |
| MROH2B | 0.163957 | 0.424913 | 4.079006 | 0.104171 | 0.917034 | NA |
| PPID | 595.0502 | 0.54022 | 0.247617 | 2.181673 | 0.029134 | 0.097017 |
| ETFDH | 680.7712 | 0.440046 | 0.146827 | 2.99703 | 0.002726 | 0.019937 |
| PTGER4 | 21.54678 | 0.382792 | 0.451689 | 0.847468 | 0.396734 | NA |
| TBCA | 2120.005 | 0.662162 | 0.212229 | 3.120029 | 0.001808 | 0.015146 |
| PLRG1 | 1690.563 | 0.337101 | 0.135346 | 2.49065 | 0.012751 | 0.055677 |
| CXXC5 | 686.2213 | 0.029113 | 0.2561 | 0.113678 | 0.909493 | 0.947766 |
| PTCRA | 14.30962 | 0.444002 | 0.633038 | 0.701382 | 0.483065 | NA |
| ENC1 | 1758.726 | -0.39107 | 0.138166 | -2.83041 | 0.004649 | 0.028075 |
| | 74.24455 | 0.338433 | 0.358208 | 0.944794 | 0.344764 | 0.512821 |
| HDAC3 | 1829.378 | -0.12778 | 0.168048 | -0.76036 | 0.447042 | 0.606718 |
| LRRC34 | 47.0021 | 0.418949 | 0.333435 | 1.256463 | 0.208948 | 0.366595 |
| SYNPO | 250.0851 | -0.78212 | 0.247865 | -3.15544 | 0.001603 | 0.014125 |
| RAB33B | 106.4331 | 0.866777 | 0.29352 | 2.953036 | 0.003147 | 0.021892 |
| GAP43 | 0.70693 | -1.62766 | 2.687154 | -0.60572 | 0.544701 | NA |
| LRRC15 | 7.439618 | 0.890303 | 0.833246 | 1.068476 | 0.285306 | NA |
| SMN1 | 57.07163 | 0.343949 | 0.290248 | 1.185018 | 0.23601 | 0.399016 |
| SLC9C1 | 1.417077 | -0.96506 | 1.679421 | -0.57464 | 0.565534 | NA |
| PRL | 5.446058 | 0.975211 | 0.98651 | 0.988547 | 0.322885 | NA |
| MBOAT1 | 302.7069 | -0.19511 | 0.172611 | -1.13034 | 0.258334 | 0.422595 |
| ID4 | 591.2141 | 0.595834 | 0.229815 | 2.592673 | 0.009523 | 0.045653 |
| PAIP1 | 1021.681 | 0.569198 | 0.153122 | 3.717283 | 2.01E-04 | 0.003869 |
| C5orf34 | 89.58584 | 0.610532 | 0.266731 | 2.288942 | 0.022083 | 0.081 |
| ZNF131 | 428.4777 | 0.772662 | 0.183987 | 4.199539 | 2.67E-05 | 0.001097 |
| RCAN2 | 92.43243 | -0.31032 | 0.33241 | -0.93355 | 0.350534 | 0.518479 |
| MYOZ2 | 12.51772 | 1.197529 | 0.716665 | 1.670974 | 0.094727 | NA |
| SYNPO2 | 23.77767 | 0.330318 | 0.462491 | 0.714215 | 0.475094 | NA |
| RSPH9 | 56.74306 | -0.17842 | 0.298446 | -0.59783 | 0.549951 | 0.691229 |
| GTPBP2 | 1884.008 | -0.09517 | 0.129816 | -0.73315 | 0.463467 | 0.621188 |
| MANEA | 314.155 | 0.927939 | 0.307735 | 3.015386 | 0.002567 | 0.019196 |
| AFF1 | 1182.395 | -0.11955 | 0.164435 | -0.72705 | 0.467196 | 0.623787 |
| ACOT12 | 11.26481 | 1.058247 | 0.731497 | 1.446686 | 0.147985 | NA |
| NIPAL4 | 13.69793 | -1.27341 | 0.604009 | -2.10826 | 0.035008 | NA |
| FNDC9 | 0.134133 | -1.48909 | 4.077584 | -0.36519 | 0.71497 | NA |
| KLHL6 | 35.54598 | -0.06757 | 0.492676 | -0.13715 | 0.89091 | NA |
| SMPDL3A | 567.825 | 0.337751 | 0.170365 | 1.982509 | 0.047422 | 0.134086 |
| ZMAT3 | 4020.203 | 0.275055 | 0.19193 | 1.433106 | 0.151828 | 0.295942 |

| | | | | | | |
|---|---|---|---|---|---|---|
| THEMIS | 0.096202 | 0.408546 | 4.079639 | 0.100143 | 0.920231 | NA |
| TMEM217 | 15.70089 | 0.516542 | 0.5317 | 0.97149 | 0.331304 | NA |
| COL6A5 | 0.556667 | 0.972496 | 2.522454 | 0.385536 | 0.699841 | NA |
| TMCC1 | 457.0529 | -0.03423 | 0.176811 | -0.19359 | 0.846499 | 0.908806 |
| EFCAB12 | 2.176827 | 0.450669 | 1.328484 | 0.339235 | 0.734433 | NA |
| RAB43 | 211.5916 | -0.30768 | 0.213487 | -1.44119 | 0.14953 | 0.293079 |
| DCP2 | 275.8784 | 0.51579 | 0.262219 | 1.967024 | 0.04918 | 0.137198 |
| DMXL1 | 1611.562 | 0.465945 | 0.265116 | 1.757512 | 0.078831 | 0.189295 |
| | 1.327657 | 3.605863 | 2.059194 | 1.751104 | 0.079928 | NA |
| ADH6 | 1.063143 | 1.12382 | 1.855095 | 0.605802 | 0.544646 | NA |
| FRG2C | 0.247898 | -0.5901 | 4.074633 | -0.14482 | 0.884851 | NA |
| UNC93B3 | 0.192403 | 0.975392 | 4.060716 | 0.240202 | 0.810174 | NA |
| GXYLT2 | 82.27133 | -1.3775 | 0.404652 | -3.40417 | 6.64E-04 | 0.008036 |
| TADA2B | 668.1402 | -0.27949 | 0.167085 | -1.67273 | 0.09438 | 0.214796 |
| CCDC96 | 50.54604 | -0.30943 | 0.309837 | -0.99867 | 0.317954 | 0.485678 |
| EVC2 | 282.3381 | -0.19602 | 0.199016 | -0.98493 | 0.324658 | 0.492617 |
| HPSE | 116.5236 | 0.36354 | 0.309087 | 1.176174 | 0.239525 | 0.402776 |
| COQ2 | 439.3954 | -0.00634 | 0.209242 | -0.0303 | 0.975827 | 0.986731 |
| ADCY5 | 72.29781 | -0.23799 | 0.267538 | -0.88954 | 0.373712 | 0.539959 |
| PARP14 | 1518.827 | 0.067323 | 0.290264 | 0.231937 | 0.816587 | 0.890173 |
| PARP15 | 18.90867 | 0.49496 | 0.552422 | 0.895982 | 0.370262 | NA |
| ABLIM3 | 2700.169 | -0.27427 | 0.129915 | -2.11119 | 0.034756 | 0.109442 |
| KIAA1919 | 131.8479 | 0.280942 | 0.217603 | 1.291074 | 0.196678 | 0.352003 |
| GLRX | 705.8561 | 0.500812 | 0.206496 | 2.425285 | 0.015296 | 0.062857 |
| IQCB1 | 273.1377 | 0.375254 | 0.182897 | 2.051719 | 0.040197 | 0.120825 |
| GOLGB1 | 1166.883 | 0.903318 | 0.934902 | 0.966216 | 0.333936 | 0.502504 |
| GPR151 | 0.134133 | -1.48909 | 4.077584 | -0.36519 | 0.71497 | NA |
| PLAC8L1 | 1.513921 | 2.643365 | 1.83893 | 1.437448 | 0.150591 | NA |
| STOX2 | 267.9452 | 0.227069 | 0.267736 | 0.848108 | 0.396378 | 0.561082 |
| NDNF | 72.46457 | 1.308121 | 0.484949 | 2.697442 | 0.006987 | 0.037109 |
| MOB1B | 591.7481 | 0.7621 | 0.604184 | 1.261372 | 0.207175 | 0.364292 |
| ZNF622 | 1586.16 | 0.064082 | 0.155349 | 0.412503 | 0.679971 | 0.791153 |
| HEG1 | 559.6745 | -0.16675 | 0.245228 | -0.67997 | 0.496525 | 0.647636 |
| PHC3 | 395.2294 | 0.439773 | 0.234382 | 1.876306 | 0.060613 | 0.158426 |
| GPR160 | 22.83276 | 0.759992 | 0.464115 | 1.637509 | 0.101524 | NA |
| GOLIM4 | 840.4152 | 0.754625 | 0.32223 | 2.341887 | 0.019187 | 0.073519 |
| 3-Mar | 127.2889 | -0.24286 | 0.229083 | -1.06015 | 0.289076 | 0.455414 |
| SLCO4C1 | 4.89561 | 1.218572 | 1.199455 | 1.015939 | 0.309659 | NA |
| XXYLT1 | 559.2929 | -0.37111 | 0.192422 | -1.92865 | 0.053775 | 0.145548 |
| NRROS | 226.346 | -0.34569 | 0.191179 | -1.80822 | 0.070572 | 0.175542 |
| CEP19 | 224.5981 | 0.841436 | 0.252386 | 3.333925 | 8.56E-04 | 0.009417 |
| FBXO45 | 343.0743 | 0.348049 | 0.213195 | 1.632537 | 0.102566 | 0.226809 |
| TLR10 | 0.327913 | 1.543154 | 4.021063 | 0.383768 | 0.701151 | NA |
| TLR1 | 46.50266 | -0.42086 | 0.352498 | -1.19394 | 0.232503 | NA |
| TLR6 | 83.50799 | -0.18292 | 0.352205 | -0.51937 | 0.603504 | 0.732334 |
| FAM174A | 408.2055 | 0.057261 | 0.202387 | 0.282926 | 0.777233 | 0.862824 |
| RGMB | 584.18 | -0.39319 | 0.183526 | -2.14241 | 0.03216 | 0.104055 |

| | | | | | | |
|---|---|---|---|---|---|---|
| FAM53A | 30.96579 | -0.15378 | 0.382106 | -0.40246 | 0.687346 | NA |
| TRMT10C | 642.7592 | 0.604961 | 0.194497 | 3.11039 | 0.001868 | 0.015436 |
| PIGG | 1521.948 | -0.17472 | 0.138767 | -1.25907 | 0.208004 | 0.365316 |
| CHRNA9 | 0.162376 | 0.424913 | 4.079006 | 0.104171 | 0.917034 | NA |
| GALNTL6 | 58.76875 | -0.05974 | 0.316195 | -0.18892 | 0.850154 | 0.910747 |
| GCSAM | 7.167498 | 0.044162 | 0.805532 | 0.054823 | 0.956279 | NA |
| IL20RB | 3.42026 | 0.249042 | 1.092565 | 0.227942 | 0.819691 | NA |
| | 0.163957 | 0.424913 | 4.079006 | 0.104171 | 0.917034 | NA |
| MSL2 | 478.8849 | -0.08999 | 0.153904 | -0.5847 | 0.558749 | 0.6977 |
| TRAM1L1 | 83.63983 | 0.227836 | 0.264497 | 0.861393 | 0.389022 | 0.554509 |
| UGT8 | 605.9908 | -0.00695 | 0.175875 | -0.03954 | 0.968459 | 0.982732 |
| KY | 1.610669 | -3.3877 | 2.011403 | -1.68424 | 0.092134 | NA |
| SLCO2A1 | 0.134133 | -1.48909 | 4.077584 | -0.36519 | 0.71497 | NA |
| TMEM167A | 1919.647 | 0.596308 | 0.241292 | 2.471317 | 0.013462 | 0.057696 |
| SH3PXD2B | 3385.849 | -0.77216 | 0.182412 | -4.23304 | 2.31E-05 | 0.001003 |
| | 0.449786 | -1.25325 | 3.00474 | -0.41709 | 0.676613 | NA |
| LARP7 | 409.7095 | 1.189306 | 0.613896 | 1.937308 | 0.052708 | 0.14338 |
| C4orf32 | 80.19097 | 0.436745 | 0.262095 | 1.666365 | 0.095641 | 0.216537 |
| WDR49 | 0.163957 | 0.424913 | 4.079006 | 0.104171 | 0.917034 | NA |
| SRP72 | 2431.711 | 0.428239 | 0.162712 | 2.631885 | 0.008491 | 0.042207 |
| THAP6 | 241.5801 | 0.814321 | 0.253527 | 3.211973 | 0.001318 | 0.012423 |
| CEP135 | 82.1746 | 1.423117 | 0.913761 | 1.557429 | 0.119369 | 0.251237 |
| BTC | 0.911244 | 3.022036 | 2.332005 | 1.295896 | 0.195011 | NA |
| RSRC1 | 329.6154 | 0.93444 | 0.228286 | 4.093295 | 4.25E-05 | 0.001514 |
| C3orf55 | 37.72509 | -0.22635 | 0.467808 | -0.48386 | 0.628489 | NA |
| METTL15P1 | 3.617055 | 0.929025 | 1.086095 | 0.855381 | 0.39234 | NA |
| C3orf33 | 56.16612 | 0.8288 | 0.363044 | 2.282919 | 0.022435 | 0.081935 |
| P2RY14 | 0.096202 | 0.408546 | 4.079639 | 0.100143 | 0.920231 | NA |
| GPR171 | 0.260158 | 1.240347 | 4.047124 | 0.306476 | 0.759242 | NA |
| DHX36 | 1173.746 | 0.627465 | 0.243035 | 2.581784 | 0.009829 | 0.046688 |
| ZIC4 | 0.326333 | 1.533889 | 3.826878 | 0.40082 | 0.688553 | NA |
| CHST2 | 858.416 | -0.42844 | 0.160897 | -2.66282 | 0.007749 | 0.039802 |
| ATR | 660.6155 | 0.530653 | 0.23239 | 2.283458 | 0.022403 | 0.081897 |
| GK5 | 112.7273 | 0.587116 | 0.295814 | 1.98475 | 0.047172 | 0.133679 |
| SPSB4 | 1.537313 | -3.12242 | 1.946201 | -1.60437 | 0.108634 | NA |
| ZNF654 | 304.8091 | 0.94645 | 0.219546 | 4.310948 | 1.63E-05 | 8.27E-04 |
| MRPS22 | 835.2885 | 0.476938 | 0.155675 | 3.063678 | 0.002186 | 0.017131 |
| CADM2 | 0.314073 | -0.57308 | 3.898934 | -0.14698 | 0.883146 | NA |
| PSMD2 | 10783.71 | -0.16409 | 0.130768 | -1.25479 | 0.209557 | 0.367119 |
| FAM131A | 688.8032 | -0.31388 | 0.147309 | -2.13072 | 0.033112 | 0.106006 |
| PARL | 1778.27 | -0.055 | 0.12805 | -0.42951 | 0.667553 | 0.781923 |
| PHYKPL | 436.443 | -0.11053 | 0.151006 | -0.73196 | 0.464191 | 0.621537 |
| ARL10 | 786.9937 | -0.29527 | 0.242056 | -1.21983 | 0.222527 | 0.383217 |
| CLTB | 4028.283 | 0.120661 | 0.151755 | 0.795103 | 0.426553 | 0.588409 |
| PCSK1 | 2.807545 | 1.376467 | 1.242047 | 1.108225 | 0.267765 | NA |
| RFESD | 10.64805 | -1.04472 | 0.626201 | -1.66834 | 0.095248 | NA |
| CCDC14 | 484.7353 | 1.040267 | 0.383872 | 2.709933 | 0.00673 | 0.036149 |

| | | | | | | |
|---|---|---|---|---|---|---|
| MCTP1 | 289.0416 | 0.436109 | 0.234733 | 1.857895 | 0.063184 | 0.163013 |
| GPR156 | 3.393351 | -1.49503 | 1.100765 | -1.35817 | 0.174409 | NA |
| NR2F1 | 63.58586 | 0.231763 | 0.328476 | 0.70557 | 0.480456 | 0.634913 |
| EIF3KP1 | 0.709944 | 0.232855 | 2.332364 | 0.099837 | 0.920474 | NA |
| RUVBL1 | 1420.188 | -0.15546 | 0.130546 | -1.19088 | 0.233702 | 0.396277 |
| | 0.3441 | -0.0166 | 3.769682 | -0.0044 | 0.996486 | NA |
| LYSMD3 | 0.354779 | 1.703954 | 3.657232 | 0.465914 | 0.641277 | NA |
| JAKMIP2 | 177.8567 | 0.57595 | 0.403396 | 1.427756 | 0.153362 | 0.298135 |
| MBLAC2 | 4.636571 | 0.582373 | 0.969873 | 0.600463 | 0.548198 | NA |
| SLC35A4 | 4001.789 | -0.31629 | 0.142366 | -2.22169 | 0.026304 | 0.091046 |
| TMEM39A | 496.3491 | 0.286545 | 0.172516 | 1.660979 | 0.096718 | 0.21817 |
| | 0.162376 | 0.424913 | 4.079006 | 0.104171 | 0.917034 | NA |
| CDV3P1 | 0.096202 | 0.408546 | 4.079639 | 0.100143 | 0.920231 | NA |
| | 3.983747 | 1.664671 | 1.14941 | 1.448283 | 0.147538 | NA |
| KIAA2018 | 462.5801 | -0.0937 | 0.259488 | -0.36108 | 0.71804 | 0.820231 |
| B3GNT5 | 108.3926 | 0.646837 | 0.23649 | 2.735162 | 0.006235 | 0.034188 |
| RUFY1 | 1018.214 | 0.32634 | 0.152954 | 2.133581 | 0.032877 | 0.105609 |
| BASP1 | 3007.282 | -0.01673 | 0.16018 | -0.10442 | 0.916838 | 0.951845 |
| GJA1P1 | 0.096202 | 0.408546 | 4.079639 | 0.100143 | 0.920231 | NA |
| GCNT4 | 22.27971 | -0.09581 | 0.475129 | -0.20165 | 0.84019 | NA |
| TOB2P1 | 3.408122 | -0.57029 | 1.145049 | -0.49805 | 0.618451 | NA |
| MUC20 | 5.008496 | 0.102312 | 1.158946 | 0.088281 | 0.929654 | NA |
| HCG4 | 114.1356 | -0.38836 | 0.275334 | -1.41051 | 0.15839 | 0.304971 |
| MTX3 | 101.3228 | 0.548254 | 0.428382 | 1.279823 | 0.200607 | 0.356605 |
| SLC38A9 | 264.4534 | 0.474937 | 0.184594 | 2.57287 | 0.010086 | 0.047448 |
| ZBTB38 | 1349.189 | 0.467807 | 0.183396 | 2.550798 | 0.010748 | 0.049294 |
| MAGEF1 | 817.9651 | 0.142006 | 0.154489 | 0.919199 | 0.357991 | 0.525131 |
| NAP1L5 | 277.4845 | 0.288161 | 0.270057 | 1.067038 | 0.285955 | 0.452361 |
| NIM1K | 74.96334 | 0.252793 | 0.273898 | 0.922944 | 0.356037 | 0.523878 |
| ZBED2 | 0.192403 | 0.975392 | 4.060716 | 0.240202 | 0.810174 | NA |
| ATOX1 | 5860.073 | 0.077753 | 0.168084 | 0.462586 | 0.643661 | 0.762832 |
| TBL1XR1 | 1234.522 | 0.479218 | 0.208061 | 2.303256 | 0.021264 | 0.078903 |
| ACAD9 | 1143.578 | -0.161 | 0.141102 | -1.14105 | 0.253849 | 0.417883 |
| SUMO4 | 0.163957 | 0.424913 | 4.079006 | 0.104171 | 0.917034 | NA |
| NAALADL2 | 79.17504 | 0.368355 | 0.268379 | 1.372517 | 0.169903 | 0.320363 |
| PVRL3 | 292.184 | 0.551134 | 0.248037 | 2.221988 | 0.026284 | 0.091031 |
| ANXA2R | 9.942282 | 0.243251 | 0.71667 | 0.339418 | 0.734295 | NA |
| HNRNPA0 | 3973.11 | -0.37202 | 0.128391 | -2.89752 | 0.003761 | 0.024485 |
| | 62.66926 | 0.189645 | 0.297164 | 0.638182 | 0.523355 | 0.669513 |
| YIPF7 | 6.232227 | -1.06113 | 0.865949 | -1.22539 | 0.220428 | NA |
| | 0.82213 | 0.50433 | 2.197682 | 0.229483 | 0.818494 | NA |
| | 183.1582 | -0.24217 | 0.199236 | -1.21548 | 0.224185 | 0.385176 |
| PCDHB9 | 2.036409 | 0.693963 | 1.504799 | 0.461166 | 0.644679 | NA |
| AP3S1 | 1284.273 | 0.228292 | 0.166747 | 1.36909 | 0.170971 | 0.321724 |
| ZNF354C | 104.1267 | 0.732408 | 0.321655 | 2.276998 | 0.022786 | 0.082561 |
| GPR150 | 38.33838 | -0.12481 | 0.415277 | -0.30054 | 0.763766 | NA |
| FAM26E | 91.76608 | 0.507546 | 0.380958 | 1.332288 | 0.182766 | 0.335764 |

| | | | | | | |
|---|---|---|---|---|---|---|
| MLF1 | 193.094 | 0.376138 | 0.24133 | 1.558607 | 0.119089 | 0.250765 |
| GRAMD1C | 52.44807 | 0.652216 | 0.326374 | 1.998373 | 0.045676 | 0.130858 |
| HTR3C | 0.392857 | -1.546 | 4.075469 | -0.37934 | 0.704433 | NA |
| ZNF518B | 735.8877 | 0.189939 | 0.158593 | 1.19765 | 0.231053 | 0.393158 |
| ZNF366 | 18.71662 | 0.03633 | 0.49511 | 0.073377 | 0.941506 | NA |
| LCORL | 147.0577 | 0.520486 | 0.254614 | 2.044213 | 0.040932 | 0.122292 |
| ZNF454 | 6.651945 | -1.49251 | 0.866741 | -1.72198 | 0.085074 | NA |
| ZC3H12D | 0.096202 | 0.408546 | 4.079639 | 0.100143 | 0.920231 | NA |
| RNF212 | 0.151697 | -1.546 | 4.075469 | -0.37934 | 0.704433 | NA |
| TMEM151E | 1.796976 | 0.349301 | 1.544305 | 0.226187 | 0.821056 | NA |
| ZNF354B | 60.70105 | 0.842199 | 0.312598 | 2.694188 | 0.007056 | 0.037327 |
| NEUROG2 | 1.372408 | -0.70922 | 1.970671 | -0.35989 | 0.718932 | NA |
| BEND3 | 141.7003 | -0.67411 | 0.235011 | -2.86843 | 0.004125 | 0.025944 |
| NT5DC1 | 682.5435 | -0.04428 | 0.14826 | -0.29867 | 0.765188 | 0.854172 |
| H3F3AP6 | 0.163957 | 0.424913 | 4.079006 | 0.104171 | 0.917034 | NA |
| AMBN | 0.326536 | 0.037056 | 3.842554 | 0.009644 | 0.992306 | NA |
| PSAPL1 | 0.162376 | 0.424913 | 4.079006 | 0.104171 | 0.917034 | NA |
| ACTG1P1 | 1.675655 | -0.33821 | 1.472486 | -0.22968 | 0.818338 | NA |
| | 0.258578 | 1.233151 | 4.047809 | 0.304646 | 0.760635 | NA |
| ARMC10P1 | 1.311188 | 0.009168 | 1.781747 | 0.005146 | 0.995894 | NA |
| NSUN3 | 235.4067 | 0.591539 | 0.182628 | 3.239039 | 0.001199 | 0.011696 |
| DHFRL1 | 268.5014 | 0.303019 | 0.310448 | 0.976067 | 0.329031 | 0.49728 |
| C5orf64 | 0.314073 | -0.57308 | 3.898934 | -0.14698 | 0.883146 | NA |
| GP5 | 0.490493 | 0.776209 | 2.895261 | 0.268096 | 0.788625 | NA |
| C5orf46 | 0.230334 | -0.53318 | 4.076749 | -0.13079 | 0.895945 | NA |
| TAF7 | 1390.302 | 0.442174 | 0.198601 | 2.226448 | 0.025984 | 0.090335 |
| GAK | 2690.199 | -0.34159 | 0.13242 | -2.57962 | 0.009891 | 0.046847 |
| | 10.48487 | -0.05067 | 0.633098 | -0.08004 | 0.936209 | NA |
| MRFAP1L1 | 1226.801 | 0.278287 | 0.17379 | 1.601288 | 0.109313 | 0.236504 |
| SNX18 | 666.1268 | 0.048866 | 0.215025 | 0.227259 | 0.820222 | 0.892334 |
| MRFAP1 | 9485.275 | -0.05927 | 0.138656 | -0.42746 | 0.669048 | 0.782942 |
| C3orf38 | 798.1165 | 0.359607 | 0.178051 | 2.019688 | 0.043416 | 0.126951 |
| TRIML2 | 0.548763 | 2.343695 | 3.135478 | 0.747476 | 0.454776 | NA |
| ZFP42 | 0.489058 | -0.58569 | 3.427543 | -0.17088 | 0.864321 | NA |
| TAAR3 | 0.096202 | 0.408546 | 4.079639 | 0.100143 | 0.920231 | NA |
| HTR1F | 0.162376 | 0.424913 | 4.079006 | 0.104171 | 0.917034 | NA |
| RPS2P28 | 0.314073 | -0.57308 | 3.898934 | -0.14698 | 0.883146 | NA |
| HLA-DQB1 | 0.941473 | 1.959123 | 2.152875 | 0.910003 | 0.362821 | NA |
| GATA2 | 0.151697 | -1.546 | 4.075469 | -0.37934 | 0.704433 | NA |
| ELMOD2 | 411.1678 | 0.545361 | 0.229296 | 2.378414 | 0.017387 | 0.068915 |
| DNAJB8 | 0.162376 | 0.424913 | 4.079006 | 0.104171 | 0.917034 | NA |
| PPP1R14BP | 84.82351 | 0.119644 | 0.255367 | 0.468515 | 0.639416 | 0.75954 |
| | 1.404468 | -0.26803 | 1.627787 | -0.16466 | 0.869211 | NA |
| CRIPAK | 81.05084 | -0.16141 | 0.299693 | -0.53859 | 0.59017 | 0.721887 |
| | 34.65176 | 0.747613 | 0.389116 | 1.921309 | 0.054693 | NA |
| C3orf80 | 1.04716 | 1.17329 | 2.006462 | 0.584756 | 0.558712 | NA |
| EXOC3 | 1572.602 | -0.21831 | 0.149708 | -1.45825 | 0.144772 | 0.286715 |

| | | | | | | |
|---|---|---|---|---|---|---|
| TDRD6 | 4.778622 | 2.244701 | 1.051376 | 2.135012 | 0.03276 | NA |
| | 3.407042 | 1.301714 | 1.30407 | 0.998193 | 0.318186 | NA |
| RRH | 1.736737 | 1.348229 | 1.585181 | 0.850521 | 0.395036 | NA |
| PNPLA1 | 1.464254 | -2.05134 | 1.820308 | -1.12692 | 0.259777 | NA |
| TIGD2 | 110.1171 | -0.07446 | 0.266633 | -0.27925 | 0.780056 | 0.865117 |
| HCLS1 | 0.296509 | -0.51709 | 3.982155 | -0.12985 | 0.896684 | NA |
| PAK2 | 2158.745 | 0.132693 | 0.151063 | 0.878393 | 0.37973 | 0.546124 |
| RNF182 | 599.8303 | 0.54155 | 0.239754 | 2.258772 | 0.023898 | 0.085385 |
| HIST1H2AC | 526.7255 | 0.309915 | 0.36993 | 0.837767 | 0.402162 | 0.56663 |
| HIST1H2BC | 5.930152 | 0.922258 | 0.917382 | 1.005315 | 0.314745 | NA |
| ZBTB12P1 | 0.952336 | 1.909203 | 2.452323 | 0.778528 | 0.436258 | NA |
| MB21D2 | 142.2631 | -0.24019 | 0.257427 | -0.93305 | 0.350792 | 0.518727 |
| EXOC5P1 | 1.35035 | 0.802108 | 1.736953 | 0.46179 | 0.644232 | NA |
| | 0.324752 | 1.53082 | 4.022025 | 0.380609 | 0.703493 | NA |
| WDFY3-AS2 | 72.1216 | 0.187844 | 0.277285 | 0.677442 | 0.498125 | 0.648931 |
| ARSJ | 128.4746 | 0.59848 | 0.240057 | 2.493078 | 0.012664 | 0.05536 |
| PSMG4 | 297.1881 | -0.08526 | 0.18914 | -0.45076 | 0.65216 | 0.76995 |
| MAP6D1 | 89.45627 | -0.18973 | 0.293529 | -0.64637 | 0.518037 | 0.665078 |
| MRPL14 | 1254.148 | 0.135489 | 0.16884 | 0.802472 | 0.42228 | 0.584768 |
| BBS12 | 360.1157 | 0.3379 | 0.221181 | 1.527709 | 0.126585 | 0.261896 |
| F2R | 2067.044 | 0.409759 | 0.146797 | 2.791324 | 0.005249 | 0.030564 |
| HLA-V | 23.98273 | -0.3121 | 0.460623 | -0.67756 | 0.49805 | NA |
| NPM1 | 11323.89 | 0.505317 | 0.187411 | 2.696303 | 0.007011 | 0.037173 |
| MTHFD2P7 | 0.162376 | 0.424913 | 4.079006 | 0.104171 | 0.917034 | NA |
| ZNF322 | 83.38615 | 0.255246 | 0.310775 | 0.821321 | 0.411463 | 0.574817 |
| NME9 | 3.860212 | 0.806016 | 1.030839 | 0.781903 | 0.434272 | NA |
| OFCC1 | 8.273207 | 0.145169 | 0.739872 | 0.196208 | 0.844447 | NA |
| HSP90AA6I | 0.750448 | 2.743526 | 2.504729 | 1.095338 | 0.273368 | NA |
| DDX60L | 639.1686 | -0.34412 | 0.628189 | -0.54779 | 0.583833 | 0.717088 |
| SOX2 | 395.8812 | -0.14779 | 0.235148 | -0.62852 | 0.529664 | 0.674744 |
| TMEM45A | 360.2201 | 0.199277 | 0.18427 | 1.081439 | 0.279502 | 0.445601 |
| RAP2B | 2689.285 | -0.23773 | 0.206291 | -1.15242 | 0.24915 | 0.412992 |
| RPL24P4 | 16.12341 | 0.322679 | 0.579321 | 0.556996 | 0.57753 | NA |
| MAB21L2 | 0.450981 | 2.057325 | 3.353149 | 0.61355 | 0.539513 | NA |
| C6orf223 | 1.371709 | 2.506436 | 1.969102 | 1.272883 | 0.203059 | NA |
| FDCSP | 0.712719 | 2.707521 | 2.836859 | 0.954408 | 0.339877 | NA |
| ZBTB20 | 22.51447 | -0.18332 | 0.541269 | -0.33868 | 0.734848 | NA |
| C3orf58 | 196.5539 | 0.19161 | 0.248746 | 0.770303 | 0.44112 | 0.601277 |
| C5orf30 | 532.6673 | 0.328184 | 0.178299 | 1.840635 | 0.065675 | 0.167149 |
| SIAH2 | 368.9086 | -0.16484 | 0.205018 | -0.804 | 0.421395 | 0.584035 |
| COPG1 | 7971.698 | -0.46666 | 0.117191 | -3.9821 | 6.83E-05 | 0.002 |
| SLC9A9 | 209.0825 | -0.14294 | 0.248745 | -0.57466 | 0.565522 | 0.702826 |
| RELL1 | 84.49668 | -0.46516 | 0.292271 | -1.59154 | 0.111489 | 0.2397 |
| TIGIT | 0.163957 | 0.424913 | 4.079006 | 0.104171 | 0.917034 | NA |
| C5orf24 | 1579.472 | 0.684718 | 0.209352 | 3.270656 | 0.001073 | 0.010848 |
| CCDC149 | 1245.537 | 0.165834 | 0.196167 | 0.845373 | 0.397903 | 0.562703 |
| | 0.58526 | 0.001197 | 3.235889 | 3.70E-04 | 0.999705 | NA |

| | | | | | | |
|---|---|---|---|---|---|---|
| UNC5C | 30.31922 | 0.018123 | 0.42934 | 0.042211 | 0.966331 | NA |
| ZAR1 | 0.518736 | 2.237423 | 2.7713 | 0.807355 | 0.419462 | NA |
| DCAF4L1 | 2.341198 | 1.022627 | 1.545554 | 0.661658 | 0.508191 | NA |
| | 0.162376 | 0.424913 | 4.079006 | 0.104171 | 0.917034 | NA |
| CEP97 | 358.2008 | 0.588889 | 0.247481 | 2.379536 | 0.017334 | 0.068778 |
| RWDD4 | 220.2118 | 0.718837 | 0.206986 | 3.472873 | 5.15E-04 | 0.006915 |
| HIST1H3I | 1.20553 | -1.82229 | 1.887291 | -0.96556 | 0.334266 | NA |
| CSF1R | 2.365167 | -1.28602 | 1.366733 | -0.94095 | 0.346732 | NA |
| EPHB3 | 26.95316 | -0.64515 | 0.41818 | -1.54275 | 0.122891 | NA |
| HIST1H2AJ | 2.029451 | 0.228988 | 1.643419 | 0.139336 | 0.889184 | NA |
| IGIP | 126.0178 | 0.801156 | 0.376005 | 2.130703 | 0.033114 | 0.106006 |
| SLC35D3 | 0.586491 | 2.39072 | 2.689057 | 0.889055 | 0.373973 | NA |
| PLCXD3 | 0.096202 | 0.408546 | 4.079639 | 0.100143 | 0.920231 | NA |
| RPL35A | 13168.24 | 0.420024 | 0.17736 | 2.368204 | 0.017875 | 0.07 |
| ZNF721 | 91.76296 | 1.165698 | 0.460549 | 2.531103 | 0.01137 | 0.051312 |
| CEP63 | 350.995 | 0.726221 | 0.283075 | 2.565472 | 0.010304 | 0.048042 |
| HMGN4 | 1511.235 | 0.106609 | 0.171284 | 0.622413 | 0.53367 | 0.677477 |
| FREM3 | 0.163957 | 0.424913 | 4.079006 | 0.104171 | 0.917034 | NA |
| ARHGEF37 | 332.8212 | -0.82058 | 0.290549 | -2.82425 | 0.004739 | 0.02849 |
| CEP57L1 | 127.6445 | 0.850857 | 0.228505 | 3.72358 | 1.96E-04 | 0.00384 |
| HSP90AB3I | 11.65337 | -0.19993 | 0.667897 | -0.29934 | 0.764681 | NA |
| | 1.44669 | -2.05828 | 1.971394 | -1.04407 | 0.296452 | NA |
| DDX41 | 2859.201 | -0.1031 | 0.166822 | -0.61803 | 0.536555 | 0.679797 |
| CCDC125 | 41.46991 | -0.31838 | 0.351568 | -0.90559 | 0.365151 | NA |
| LRIT3 | 0.84823 | 2.93367 | 2.713279 | 1.081227 | 0.279596 | NA |
| TRIM61 | 61.21953 | 0.657824 | 0.312845 | 2.102713 | 0.035491 | 0.110985 |
| GTF2H2C | 68.85321 | 0.486862 | 0.294523 | 1.653051 | 0.098321 | 0.220722 |
| FBXL7 | 1376.933 | -0.51467 | 0.166619 | -3.0889 | 0.002009 | 0.016251 |
| HMCES | 1090.031 | -0.37585 | 0.150448 | -2.49823 | 0.012482 | 0.054702 |
| 11-Mar | 0.785567 | -0.70568 | 2.358063 | -0.29926 | 0.764741 | NA |
| | 2.251467 | 0.062178 | 1.297425 | 0.047924 | 0.961777 | NA |
| GUSBP1 | 129.8058 | 0.24016 | 0.237082 | 1.012982 | 0.311069 | 0.478614 |
| LINC00518 | 4396.465 | 0.419976 | 0.211489 | 1.985807 | 0.047055 | 0.133581 |
| TRIM52 | 75.66585 | 0.31791 | 0.288013 | 1.103804 | 0.269678 | 0.434351 |
| FOXL2 | 34.71527 | 0.391448 | 0.471328 | 0.830523 | 0.406243 | NA |
| KCTD16 | 3.720383 | 0.262403 | 1.077384 | 0.243556 | 0.807575 | NA |
| FAM162B | 0.134133 | -1.48909 | 4.077584 | -0.36519 | 0.71497 | NA |
| BTBD9 | 670.8326 | -0.56704 | 0.143634 | -3.94784 | 7.89E-05 | 0.002195 |
| MAATS1 | 3.141839 | 1.22336 | 1.197961 | 1.021202 | 0.307159 | NA |
| ARSI | 880.1302 | -0.78305 | 0.318093 | -2.4617 | 0.013828 | 0.058681 |
| BRD7P2 | 1.871229 | 0.897865 | 1.717095 | 0.522898 | 0.601046 | NA |
| TREML3P | 0.162376 | 0.424913 | 4.079006 | 0.104171 | 0.917034 | NA |
| RPL7AP28 | 3.276971 | -0.0773 | 1.323546 | -0.0584 | 0.953428 | NA |
| ADRA2C | 2.687449 | -0.40843 | 1.355159 | -0.30139 | 0.763118 | NA |
| SCFD2 | 1320.441 | -0.32699 | 0.140339 | -2.32998 | 0.019807 | 0.075165 |
| | 0.382031 | -1.26484 | 3.151989 | -0.40128 | 0.688212 | NA |
| PPP1R2 | 790.5371 | 0.374087 | 0.203213 | 1.840866 | 0.065641 | 0.167108 |

| | | | | | | |
|---|---|---|---|---|---|---|
| CMSS1 | 651.2068 | 0.402501 | 0.189756 | 2.121151 | 0.033909 | 0.1076 |
| CCSER1 | 16.35122 | 0.990771 | 0.56969 | 1.73914 | 0.08201 | NA |
| ZDHHC23 | 61.58376 | 0.465368 | 0.377397 | 1.233098 | 0.217539 | 0.376981 |
| HIST1H2AK | 1.426388 | 0.934706 | 1.835245 | 0.509308 | 0.610536 | NA |
| EFNA5 | 122.8777 | -0.46022 | 0.260806 | -1.76461 | 0.077629 | 0.187485 |
| HIST1H1B | 0.640406 | 1.11402 | 2.710977 | 0.410929 | 0.681124 | NA |
| ACTRT3 | 97.81976 | 0.246951 | 0.259601 | 0.95127 | 0.341467 | 0.509671 |
| COPB2 | 4320.424 | 0.383698 | 0.151521 | 2.532304 | 0.011332 | 0.051175 |
| POU3F2 | 482.2217 | 0.312624 | 0.190209 | 1.643583 | 0.100262 | 0.223381 |
| PROS1 | 7033.57 | 0.846228 | 0.161343 | 5.24489 | 1.56E-07 | 4.45E-05 |
| C6orf58 | 10.89398 | -1.06693 | 0.703424 | -1.51677 | 0.129325 | NA |
| TMEM173 | 1879.425 | -0.09426 | 0.147647 | -0.63841 | 0.523209 | 0.669419 |
| RPL7P26 | 1.36261 | 1.545046 | 1.790867 | 0.862736 | 0.388282 | NA |
| HIST1H2AH | 1.282741 | -0.13551 | 1.82613 | -0.07421 | 0.940846 | NA |
| PRR16 | 198.6169 | 0.154981 | 0.211381 | 0.733181 | 0.463448 | 0.621188 |
| TMED9 | 6381.583 | 0.024483 | 0.149533 | 0.163727 | 0.869946 | 0.922965 |
| H1FX | 2184.884 | -0.39191 | 0.150281 | -2.60788 | 0.00911 | 0.044401 |
| SORCS2 | 342.9099 | -0.72908 | 0.700183 | -1.04127 | 0.297751 | 0.464828 |
| ROBO2 | 465.8702 | -0.07877 | 0.269834 | -0.29192 | 0.770349 | 0.858023 |
| LRRC14B | 0.163957 | 0.424913 | 4.079006 | 0.104171 | 0.917034 | NA |
| NELFA | 690.4299 | -0.40417 | 0.162116 | -2.49307 | 0.012664 | 0.05536 |
| C5orf47 | 0.354779 | 1.703954 | 3.657232 | 0.465914 | 0.641277 | NA |
| FAM43A | 2259.407 | -0.48554 | 0.274078 | -1.77155 | 0.07647 | 0.185473 |
| PURA | 532.2991 | 0.298747 | 0.204446 | 1.461248 | 0.143947 | 0.285539 |
| HIST1H2BL | 6.481694 | -0.67326 | 0.945975 | -0.71171 | 0.476644 | NA |
| PPIL6 | 40.07056 | -0.13442 | 0.377681 | -0.35591 | 0.721906 | NA |
| KIAA0825 | 36.53858 | 1.122742 | 0.390261 | 2.876904 | 0.004016 | NA |
| ARL15 | 342.2465 | 0.421541 | 0.194263 | 2.169951 | 0.030011 | 0.099019 |
| GPRIN3 | 276.5441 | -0.08054 | 0.325992 | -0.24706 | 0.80486 | 0.88332 |
| SDHAP1 | 65.57517 | 0.447257 | 0.296247 | 1.509742 | 0.131109 | 0.26766 |
| LSAMP | 676.4005 | -0.23284 | 0.183688 | -1.26756 | 0.204955 | 0.361622 |
| PCGF3 | 1394.369 | 0.128171 | 0.143294 | 0.894463 | 0.371074 | 0.53772 |
| LMLN | 216.3499 | 0.128464 | 0.228259 | 0.562799 | 0.573572 | 0.709406 |
| | 13.04996 | 0.486672 | 0.60098 | 0.809797 | 0.418057 | NA |
| C6orf201 | 1.045376 | 2.066118 | 2.09889 | 0.984386 | 0.324926 | NA |
| CLDN24 | 0.749217 | 0.7397 | 2.633289 | 0.280903 | 0.778784 | NA |
| KCNQ5 | 123.034 | 0.361866 | 0.219508 | 1.648533 | 0.099243 | 0.222196 |
| KCNIP4 | 13.43802 | 1.064083 | 0.661774 | 1.607925 | 0.107852 | NA |
| WDR53 | 158.4213 | 0.383312 | 0.200432 | 1.912429 | 0.055821 | 0.149437 |
| NAT8L | 743.0926 | -0.51901 | 0.164031 | -3.16411 | 0.001556 | 0.013845 |
| SDHAP3 | 64.24474 | 0.369752 | 0.328296 | 1.126276 | 0.260049 | 0.424389 |
| LRCH3 | 1390.738 | -0.38106 | 0.171576 | -2.22097 | 0.026353 | 0.091117 |
| HTR3E | 0.151697 | -1.546 | 4.075469 | -0.37934 | 0.704433 | NA |
| LRRC70 | 0.096202 | 0.408546 | 4.079639 | 0.100143 | 0.920231 | NA |
| SOWAHB | 0.134133 | -1.48909 | 4.077584 | -0.36519 | 0.71497 | NA |
| BLOC1S4 | 343.5145 | 0.035568 | 0.162132 | 0.219377 | 0.826357 | 0.895553 |
| KLHL32 | 10.4235 | -1.07439 | 0.742231 | -1.44751 | 0.147754 | NA |

| | | | | | | |
|---|---|---|---|---|---|---|
| PRELID2 | 85.75247 | 0.072806 | 0.243287 | 0.299261 | 0.764741 | 0.853834 |
| SLC36A3 | 0.544554 | -2.65036 | 3.220086 | -0.82307 | 0.410468 | NA |
| SLC36A2 | 0.70693 | -1.62766 | 2.687154 | -0.60572 | 0.544701 | NA |
| ANKRD37 | 180.4575 | 0.174734 | 0.187768 | 0.930581 | 0.35207 | 0.520164 |
| KPNA4 | 2078.799 | 0.215863 | 0.164258 | 1.314175 | 0.188787 | 0.343053 |
| RPS23 | 12211.98 | 0.209749 | 0.160278 | 1.308659 | 0.19065 | 0.345009 |
| BTN3A2 | 1415.85 | -0.43487 | 0.230288 | -1.88835 | 0.058979 | 0.155272 |
| C5orf38 | 0.911649 | 0.958357 | 2.37748 | 0.403098 | 0.686876 | NA |
| C6orf1 | 1938.928 | -0.29775 | 0.147958 | -2.01241 | 0.044177 | 0.128169 |
| KATNA1 | 380.766 | 0.302922 | 0.184076 | 1.645636 | 0.099839 | 0.223192 |
| LYRM7 | 378.7569 | 0.664882 | 0.33642 | 1.976344 | 0.048116 | 0.135394 |
| TPI1P3 | 0.134133 | -1.48909 | 4.077584 | -0.36519 | 0.71497 | NA |
| ZNF732 | 0.69625 | -3.13723 | 2.93845 | -1.06765 | 0.285679 | NA |
| QRFPR | 0.134133 | -1.48909 | 4.077584 | -0.36519 | 0.71497 | NA |
| TMEM232 | 18.22937 | 0.54765 | 0.51176 | 1.070131 | 0.28456 | NA |
| C3orf70 | 385.4329 | -0.40846 | 0.238458 | -1.71292 | 0.086727 | 0.201995 |
| MITF | 11034.11 | 0.185973 | 0.182916 | 1.016711 | 0.309291 | 0.476621 |
| TSPYL4 | 423.5362 | -0.06864 | 0.225654 | -0.30419 | 0.760981 | 0.851319 |
| PCDHB13 | 29.09597 | -0.84524 | 0.43188 | -1.95711 | 0.050335 | NA |
| | 0.230334 | -0.53318 | 4.076749 | -0.13079 | 0.895945 | NA |
| ATP13A5 | 0.151697 | -1.546 | 4.075469 | -0.37934 | 0.704433 | NA |
| NHLRC1 | 132.9089 | -0.44063 | 0.211825 | -2.08015 | 0.037511 | 0.115087 |
| ZKSCAN4 | 173.516 | -0.016 | 0.212354 | -0.07535 | 0.939936 | 0.966534 |
| TMSB4XP8 | 83.94182 | 0.039008 | 0.302952 | 0.128759 | 0.897549 | 0.939815 |
| C5orf52 | 0.303394 | -2.27006 | 4.032451 | -0.56295 | 0.57347 | NA |
| SPRY4 | 880.3387 | 0.056838 | 0.156699 | 0.36272 | 0.716814 | 0.819463 |
| | 0.890088 | -0.08229 | 2.128047 | -0.03867 | 0.969154 | NA |
| KBTBD12 | 0.096202 | 0.408546 | 4.079639 | 0.100143 | 0.920231 | NA |
| HSPE1P11 | 0.258578 | 1.233151 | 4.047809 | 0.304646 | 0.760635 | NA |
| HIST1H1C | 272.3368 | 0.364339 | 0.29902 | 1.218446 | 0.223055 | 0.383846 |
| | 0.192403 | 0.975392 | 4.060716 | 0.240202 | 0.810174 | NA |
| ZSCAN23 | 17.87488 | 0.277884 | 0.582958 | 0.47668 | 0.63359 | NA |
| HIST1H2BG | 6.288645 | 0.566892 | 0.922908 | 0.614245 | 0.539053 | NA |
| TPRG1 | 1.216624 | 0.085515 | 1.905007 | 0.04489 | 0.964195 | NA |
| | 130.617 | 0.259583 | 0.27361 | 0.948733 | 0.342756 | 0.51086 |
| EYS | 3.897095 | -0.15637 | 0.980399 | -0.15949 | 0.873279 | NA |
| | 89.4468 | -0.03071 | 0.272617 | -0.11264 | 0.910315 | 0.947811 |
| LRRIQ4 | 0.460669 | -0.62593 | 3.398671 | -0.18417 | 0.853882 | NA |
| PLSCR1 | 1825.222 | -0.1452 | 0.194867 | -0.74514 | 0.456188 | 0.614817 |
| BLOC1S5 | 262.703 | 0.239266 | 0.233433 | 1.024988 | 0.305369 | 0.472543 |
| COL25A1 | 7.509845 | -0.50426 | 0.783906 | -0.64327 | 0.52005 | NA |
| FBLL1 | 139.748 | 0.200383 | 0.233874 | 0.856801 | 0.391555 | 0.556781 |
| NKAIN2 | 2.597724 | 0.799084 | 1.309333 | 0.610298 | 0.541664 | NA |
| SMIM15 | 762.0997 | 0.745973 | 0.159432 | 4.678957 | 2.88E-06 | 2.97E-04 |
| OSTN | 0.354779 | 1.703954 | 3.657232 | 0.465914 | 0.641277 | NA |
| ZDHHC11 | 2.308185 | -0.57946 | 1.263128 | -0.45875 | 0.646414 | NA |
| FAM26F | 2.653508 | -0.68815 | 1.287749 | -0.53438 | 0.593076 | NA |

| | | | | | | |
|---|---|---|---|---|---|---|
| | 0.151697 | -1.546 | 4.075469 | -0.37934 | 0.704433 | NA |
| UTS2B | 16.35766 | -0.02297 | 0.594699 | -0.03863 | 0.969184 | NA |
| MSANTD1 | 5.426721 | -0.23109 | 0.926272 | -0.24948 | 0.802986 | NA |
| HIST1H4D | 1.772885 | 1.348764 | 1.569085 | 0.859586 | 0.390017 | NA |
| LRRC66 | 6.906402 | 0.575173 | 0.805573 | 0.713992 | 0.475232 | NA |
| ZNF292 | 219.2973 | 0.764781 | 0.68148 | 1.122236 | 0.261762 | 0.426146 |
| HUS1B | 1.685985 | 1.314117 | 1.571458 | 0.836241 | 0.403019 | NA |
| PROSP | 0.260158 | 1.240347 | 4.047124 | 0.306476 | 0.759242 | NA |
| ADAT2 | 114.9229 | 0.623172 | 0.233137 | 2.672985 | 0.007518 | 0.039107 |
| ANKDD1B | 17.32053 | -0.3098 | 0.54601 | -0.56739 | 0.570447 | NA |
| APOD | 745.7263 | -0.67522 | 0.333902 | -2.0222 | 0.043156 | 0.126606 |
| PRSS48 | 0.296509 | -0.51709 | 3.982155 | -0.12985 | 0.896684 | NA |
| ANKRD34B | 0.788582 | 0.608095 | 2.456267 | 0.247569 | 0.804468 | NA |
| NKAPL | 20.36508 | 0.471986 | 0.50357 | 0.937281 | 0.348614 | NA |
| FAM47E | 13.07636 | 0.413216 | 0.555865 | 0.743375 | 0.457255 | NA |
| PCDH18 | 24.09204 | 0.144737 | 0.516933 | 0.279992 | 0.779483 | NA |
| TSPYL1 | 2022.518 | -0.27994 | 0.127608 | -2.19377 | 0.028252 | 0.095227 |
| ZKSCAN3 | 95.78449 | -0.23497 | 0.238721 | -0.98428 | 0.324976 | 0.492847 |
| LIN54 | 310.1351 | 0.851902 | 0.183042 | 4.654146 | 3.25E-06 | 3.27E-04 |
| C6orf222 | 0.556813 | -0.57007 | 3.294615 | -0.17303 | 0.862627 | NA |
| ALG1L | 3.788633 | -0.49455 | 0.996657 | -0.49621 | 0.619746 | NA |
| KIAA0408 | 1.455651 | 1.673699 | 1.711992 | 0.977633 | 0.328256 | NA |
| IL1RAP | 1235.104 | 0.60359 | 0.194891 | 3.097065 | 0.001954 | 0.015925 |
| SPOCK3 | 0.616518 | 2.49285 | 2.980788 | 0.836306 | 0.402983 | NA |
| | 1.685286 | -2.08097 | 1.824913 | -1.14031 | 0.254157 | NA |
| HLA-DRB1 | 100.323 | 0.16927 | 0.272883 | 0.6203 | 0.53506 | 0.678307 |
| FAT4 | 11.87504 | -0.13289 | 0.688026 | -0.19315 | 0.846845 | NA |
| HIST1H4A | 0.230334 | -0.53318 | 4.076749 | -0.13079 | 0.895945 | NA |
| HIST1H2BB | 0.096202 | 0.408546 | 4.079639 | 0.100143 | 0.920231 | NA |
| TUBB | 22784.91 | -0.49061 | 0.174249 | -2.81557 | 0.004869 | 0.028977 |
| SUPT3H | 87.74266 | 0.131135 | 0.308962 | 0.424439 | 0.671246 | 0.78462 |
| HIST1H2BC | 1.561743 | 0.170449 | 1.59364 | 0.106956 | 0.914824 | NA |
| CPNE4 | 0.134133 | -1.48909 | 4.077584 | -0.36519 | 0.71497 | NA |
| | 0.82213 | 0.50433 | 2.197682 | 0.229483 | 0.818494 | NA |
| HIST1H2BN | 0.3441 | -0.0166 | 3.769682 | -0.0044 | 0.996486 | NA |
| SLC35F1 | 375.3082 | 0.308382 | 0.287475 | 1.072727 | 0.283394 | 0.44993 |
| TSC22D2 | 607.3449 | -0.09399 | 0.231929 | -0.40525 | 0.685293 | 0.79516 |
| PIK3R4 | 1319.123 | -0.01265 | 0.139868 | -0.09043 | 0.927946 | 0.958745 |
| ARL9 | 23.36279 | -0.37467 | 0.494523 | -0.75765 | 0.448663 | NA |
| AFAP1 | 979.6266 | -0.2316 | 0.172324 | -1.34397 | 0.178959 | 0.331503 |
| HIST1H3C | 2.421508 | 0.172314 | 1.33497 | 0.129077 | 0.897297 | NA |
| MME | 2535.43 | 0.82357 | 0.150057 | 5.488388 | 4.06E-08 | 1.85E-05 |
| LAMA2 | 101.3153 | 0.060988 | 0.305512 | 0.199625 | 0.841774 | 0.905885 |
| MYO6 | 547.8252 | 0.414912 | 0.280616 | 1.478576 | 0.139254 | 0.279238 |
| HDAC2 | 2018.128 | 0.66613 | 0.193664 | 3.439613 | 5.83E-04 | 0.00745 |
| | 1.113131 | 2.168349 | 2.301419 | 0.942179 | 0.346101 | NA |
| | 0.586694 | 1.130467 | 2.734203 | 0.413454 | 0.679274 | NA |

| | | | | | | |
|---|---|---|---|---|---|---|
| ZFP62 | 173.9947 | 0.556533 | 0.201405 | 2.76325 | 0.005723 | 0.032425 |
| GM2A | 6335.925 | -0.60725 | 0.138728 | -4.3773 | 1.20E-05 | 6.88E-04 |
| HIST1H2AI | 4.159782 | -0.65853 | 0.973205 | -0.67666 | 0.498619 | NA |
| CLPSL2 | 0.411855 | 0.250456 | 3.097618 | 0.080854 | 0.935558 | NA |
| CD47 | 3077.28 | 0.2122 | 0.132668 | 1.59948 | 0.109714 | 0.237131 |
| MAML3 | 340.1539 | 0.189671 | 0.162114 | 1.169987 | 0.242006 | 0.404995 |
| HIST1H2AG | 51.90682 | 0.283114 | 0.401591 | 0.704982 | 0.480822 | 0.635255 |
| CTBP1-AS2 | 328.6785 | -0.27782 | 0.178075 | -1.56012 | 0.118732 | 0.250234 |
| ZSCAN16 | 32.1333 | 1.259964 | 0.402573 | 3.129779 | 0.001749 | NA |
| C6orf106 | 5808.632 | -0.22128 | 0.204556 | -1.08174 | 0.279366 | 0.445506 |
| KPNA5 | 104.6809 | 1.158502 | 0.281426 | 4.116537 | 3.85E-05 | 0.001425 |
| PDLIM7 | 904.8199 | -0.20805 | 0.154903 | -1.34308 | 0.179247 | 0.331717 |
| | 15.27971 | 0.713342 | 0.534159 | 1.335449 | 0.181729 | NA |
| PCDHB16 | 9.145477 | 0.125206 | 0.743845 | 0.168323 | 0.866329 | NA |
| HIST1H3E | 23.64039 | 0.510213 | 0.598678 | 0.852233 | 0.394085 | NA |
| WDR5B | 163.5686 | 0.251058 | 0.24594 | 1.020812 | 0.307343 | 0.474571 |
| ANXA6 | 21808.68 | -0.20788 | 0.130981 | -1.58714 | 0.112481 | 0.241067 |
| DTHD1 | 0.506476 | 0.726418 | 2.864186 | 0.253621 | 0.799788 | NA |
| HIST1H4C | 3.523439 | 0.163673 | 1.12748 | 0.145167 | 0.884579 | NA |
| | 319.728 | 0.368126 | 0.19829 | 1.856505 | 0.063382 | 0.163381 |
| ZNF300P1 | 10.43737 | 1.1802 | 0.704284 | 1.675744 | 0.093788 | NA |
| | 0.258578 | 1.233151 | 4.047809 | 0.304646 | 0.760635 | NA |
| HIST1H3J | 2.182371 | -0.29308 | 1.361793 | -0.21521 | 0.829601 | NA |
| SLC22A4 | 3.533836 | -0.85602 | 1.198539 | -0.71422 | 0.475094 | NA |
| TBC1D9B | 7244.141 | -0.33385 | 0.165459 | -2.01771 | 0.043621 | 0.127269 |
| HIST1H4J | 1.110762 | -0.90437 | 1.863208 | -0.48538 | 0.627405 | NA |
| ZNF165 | 10.53161 | 0.379405 | 0.728727 | 0.520641 | 0.602617 | NA |
| SYNGAP1 | 568.5487 | -0.27652 | 0.151965 | -1.81966 | 0.06881 | 0.172582 |
| | 0.162376 | 0.424913 | 4.079006 | 0.104171 | 0.917034 | NA |
| SLC22A5 | 217.999 | -0.14841 | 0.203333 | -0.7299 | 0.465449 | 0.622857 |
| HTT | 8736.762 | -0.54071 | 0.168221 | -3.21431 | 0.001308 | 0.012352 |
| HIST1H3D | 4.00347 | 0.490809 | 1.109391 | 0.442413 | 0.65819 | NA |
| DCHS2 | 0.298089 | -0.51711 | 3.975232 | -0.13008 | 0.896501 | NA |
| VEPH1 | 2769.761 | 0.416293 | 0.273881 | 1.519981 | 0.128516 | 0.264779 |
| MAP3K5 | 36.77371 | 0.02458 | 0.382258 | 0.064303 | 0.948729 | NA |
| HNRNPAB | 5009.033 | -0.13313 | 0.155029 | -0.85872 | 0.390493 | 0.555738 |
| HIST1H2BH | 8.079288 | -0.21198 | 0.882703 | -0.24015 | 0.810214 | NA |
| GYPE | 4.587592 | 0.087024 | 1.168872 | 0.074451 | 0.940652 | NA |
| PCDHB11 | 2.300923 | 1.882141 | 1.41558 | 1.32959 | 0.183653 | NA |
| RPF2 | 377.196 | 0.679095 | 0.186124 | 3.64861 | 2.64E-04 | 0.00459 |
| C5orf56 | 34.24861 | -0.16553 | 0.388829 | -0.42572 | 0.670314 | NA |
| RPS17P2 | 0.392857 | -1.546 | 4.075469 | -0.37934 | 0.704433 | NA |
| KCNMB2 | 0.162376 | 0.424913 | 4.079006 | 0.104171 | 0.917034 | NA |
| ENPP1 | 115.6499 | -0.16115 | 0.274604 | -0.58686 | 0.557299 | 0.696283 |
| C5orf42 | 314.95 | 0.669275 | 0.322439 | 2.075665 | 0.037925 | 0.115846 |
| HIST1H2BE | 1.407592 | -0.33704 | 1.911227 | -0.17635 | 0.86002 | NA |
| ZNF595 | 14.12287 | 0.835383 | 0.614605 | 1.359219 | 0.174077 | NA |

| | | | | | | |
|---|---|---|---|---|---|---|
| FAM114A1 | 1273.792 | 0.18745 | 0.196212 | 0.955347 | 0.339402 | 0.508003 |
| PTMAP2 | 3.888967 | 0.1813 | 1.243853 | 0.145757 | 0.884113 | NA |
| TXNRD3 | 406.6684 | 0.078061 | 0.152343 | 0.512402 | 0.608369 | 0.73549 |
| OCLN | 2.20928 | 2.480595 | 1.590007 | 1.560116 | 0.118732 | NA |
| | 1.618579 | 0.125281 | 1.62584 | 0.077056 | 0.938579 | NA |
| HIST1H2BF | 1.855449 | 0.479808 | 1.468999 | 0.326623 | 0.743953 | NA |
| SGTB | 672.6956 | 0.626971 | 0.274824 | 2.281358 | 0.022527 | 0.081968 |
| ADH5 | 3997.731 | 0.1964 | 0.14302 | 1.373239 | 0.169678 | 0.320155 |
| HIST1H2BK | 390.0756 | 0.390905 | 0.352047 | 1.110379 | 0.266836 | 0.431705 |
| HIST1H4K | 3.277056 | -0.67446 | 1.1063 | -0.60966 | 0.542088 | NA |
| ZNF311 | 42.80347 | 0.144007 | 0.329731 | 0.43674 | 0.6623 | NA |
| FCHSD1 | 840.7391 | -0.08468 | 0.151123 | -0.56034 | 0.575248 | 0.710519 |
| ELOVL2 | 14.28874 | -0.41512 | 0.639921 | -0.64871 | 0.516524 | NA |
| LEKR1 | 7.734089 | 0.694564 | 0.720954 | 0.963396 | 0.335349 | NA |
| GRK6 | 1565.618 | -0.22814 | 0.15863 | -1.43817 | 0.150387 | 0.294052 |
| CD2AP | 499.4171 | 0.943335 | 0.381663 | 2.471643 | 0.013449 | 0.057665 |
| ADH4 | 0.448206 | -1.2532 | 3.007315 | -0.41672 | 0.676885 | NA |
| CHSY3 | 395.4941 | -0.29128 | 0.173604 | -1.67782 | 0.093383 | 0.213265 |
| ZNF876P | 1.441957 | 0.19835 | 1.953344 | 0.101544 | 0.919119 | NA |
| HSD17B11 | 1004.361 | 0.328132 | 0.204401 | 1.60534 | 0.108419 | 0.235264 |
| FKBP1C | 37.75436 | -0.31663 | 0.400893 | -0.7898 | 0.429643 | NA |
| SDAD1 | 861.9987 | 0.447191 | 0.172639 | 2.590317 | 0.009589 | 0.045892 |
| ZKSCAN8 | 393.0863 | 0.034014 | 0.208377 | 0.163235 | 0.870333 | 0.923107 |
| HIST1H4F | 0.921281 | -1.21902 | 2.226885 | -0.54741 | 0.584096 | NA |
| HIST1H4I | 10.0394 | 0.006993 | 0.687738 | 0.010168 | 0.991887 | NA |
| HIST1H3A | 2.497667 | -0.51615 | 1.29423 | -0.39881 | 0.690033 | NA |
| HIST1H2AL | 2.765461 | 0.596538 | 1.229063 | 0.48536 | 0.627421 | NA |
| TMEM207 | 0.327913 | 1.543154 | 4.021063 | 0.383768 | 0.701151 | NA |
| BZW1P2 | 2.698984 | 0.745853 | 1.560075 | 0.478088 | 0.632588 | NA |
| SH3BGRL2 | 111.7631 | 0.400318 | 0.246849 | 1.621713 | 0.104865 | 0.230345 |
| TMA16 | 239.4179 | 0.623082 | 0.24074 | 2.588193 | 0.009648 | 0.046046 |
| HLA-DRB5 | 4.488516 | 1.35536 | 1.235613 | 1.096912 | 0.27268 | NA |
| CNGA1 | 0.862864 | -0.92591 | 2.497199 | -0.37078 | 0.710801 | NA |
| HIST1H4E | 3.222147 | 0.950603 | 1.186084 | 0.801464 | 0.422863 | NA |
| PLN | 1.469897 | 1.171853 | 1.816097 | 0.645259 | 0.518759 | NA |
| DDX39B | 843.0799 | 0.470179 | 0.170207 | 2.762391 | 0.005738 | 0.03247 |
| NUDT16 | 1217.062 | 0.031332 | 0.175041 | 0.178997 | 0.85794 | 0.915883 |
| LRBA | 3259.651 | 0.144797 | 0.156842 | 0.923199 | 0.355903 | 0.523878 |
| CCDC69 | 77.26935 | 0.110181 | 0.42145 | 0.261433 | 0.793759 | 0.874943 |
| C6orf89 | 1381.73 | -0.36291 | 0.131037 | -2.76957 | 0.005613 | 0.032054 |
| TTC37 | 2032.035 | 0.971053 | 0.288303 | 3.368165 | 7.57E-04 | 0.008794 |
| ECI2 | 991.4436 | 0.117346 | 0.139681 | 0.840101 | 0.400852 | 0.565475 |
| PPP1R14C | 0.134133 | -1.48909 | 4.077584 | -0.36519 | 0.71497 | NA |
| SMIM11P1 | 0.162376 | 0.424913 | 4.079006 | 0.104171 | 0.917034 | NA |
| RPL10A | 14464.73 | 0.147029 | 0.154164 | 0.95372 | 0.340225 | 0.508558 |
| FAM169A | 18.85146 | 0.63942 | 0.480724 | 1.330119 | 0.183479 | NA |
| SUCNR1 | 0.134133 | -1.48909 | 4.077584 | -0.36519 | 0.71497 | NA |

| | | | | | | |
|---|---|---|---|---|---|---|
| UBE2J1 | 2444.186 | 0.359909 | 0.142314 | 2.528977 | 0.01144 | 0.051545 |
| OPA1 | 1484.843 | 0.733165 | 0.243754 | 3.007803 | 0.002631 | 0.019437 |
| | 2158.098 | 0.25245 | 0.171125 | 1.475242 | 0.140147 | 0.280366 |
| OSTC | 1625.614 | 0.305402 | 0.156897 | 1.946511 | 0.051593 | 0.141452 |
| CCDC152 | 1.454862 | -0.58515 | 1.616608 | -0.36196 | 0.717381 | NA |
| DZIP3 | 329.3937 | 0.696267 | 0.396387 | 1.756535 | 0.078997 | 0.189654 |
| CCDC167 | 578.7086 | -0.13412 | 0.212471 | -0.63123 | 0.52789 | 0.673353 |
| ZFP2 | 14.4836 | -0.0468 | 0.745855 | -0.06275 | 0.949964 | NA |
| SOWAHA | 1.72739 | -0.40933 | 2.025112 | -0.20212 | 0.839819 | NA |
| L3MBTL3 | 135.7847 | 0.40565 | 0.285256 | 1.422055 | 0.15501 | 0.300482 |
| MFAP3L | 943.1538 | 0.275098 | 0.159323 | 1.726666 | 0.084228 | 0.197805 |
| PJA2 | 2633.62 | 0.477042 | 0.190941 | 2.498368 | 0.012477 | 0.054701 |
| MIR219-1 | 0.682896 | 1.412658 | 2.61672 | 0.539858 | 0.589295 | NA |
| MIR378A | 1.266612 | 0.746194 | 1.745721 | 0.427442 | 0.669058 | NA |
| MIR30C2 | 0.163957 | 0.424913 | 4.079006 | 0.104171 | 0.917034 | NA |
| | 0.096202 | 0.408546 | 4.079639 | 0.100143 | 0.920231 | NA |
| RNU6-204F | 0.096202 | 0.408546 | 4.079639 | 0.100143 | 0.920231 | NA |
| | 0.260158 | 1.240347 | 4.047124 | 0.306476 | 0.759242 | NA |
| RNA5SP174 | 3.862514 | 1.282392 | 1.377812 | 0.930745 | 0.351985 | NA |
| | 0.163957 | 0.424913 | 4.079006 | 0.104171 | 0.917034 | NA |
| RNU6-1079 | 0.096202 | 0.408546 | 4.079639 | 0.100143 | 0.920231 | NA |
| RNU6-888F | 0.134133 | -1.48909 | 4.077584 | -0.36519 | 0.71497 | NA |
| VTRNA1-1 | 0.268266 | -2.15396 | 4.038351 | -0.53338 | 0.593773 | NA |
| RNA5SP214 | 0.163957 | 0.424913 | 4.079006 | 0.104171 | 0.917034 | NA |
| RNU6-1285 | 0.163957 | 0.424913 | 4.079006 | 0.104171 | 0.917034 | NA |
| SNORA33 | 10.87054 | 1.30781 | 0.653151 | 2.00231 | 0.045251 | NA |
| RNU6-226F | 0.58526 | 0.001197 | 3.235889 | 3.70E-04 | 0.999705 | NA |
| RNU6-681F | 0.096202 | 0.408546 | 4.079639 | 0.100143 | 0.920231 | NA |
| | 0.163957 | 0.424913 | 4.079006 | 0.104171 | 0.917034 | NA |
| RNU6-553F | 0.096202 | 0.408546 | 4.079639 | 0.100143 | 0.920231 | NA |
| SNORA74A | 0.296509 | -0.51709 | 3.982155 | -0.12985 | 0.896684 | NA |
| RNU1-33P | 0.134133 | -1.48909 | 4.077584 | -0.36519 | 0.71497 | NA |
| RNU6-853F | 0.163957 | 0.424913 | 4.079006 | 0.104171 | 0.917034 | NA |
| | 3.290657 | -0.19437 | 1.200431 | -0.16192 | 0.871368 | NA |
| | 0.736607 | 1.395844 | 2.318642 | 0.602009 | 0.547168 | NA |
| RNU6-1305 | 0.096202 | 0.408546 | 4.079639 | 0.100143 | 0.920231 | NA |
| | 0.230334 | -0.53318 | 4.076749 | -0.13079 | 0.895945 | NA |
| | 0.163957 | 0.424913 | 4.079006 | 0.104171 | 0.917034 | NA |
| RNU6-646F | 0.096202 | 0.408546 | 4.079639 | 0.100143 | 0.920231 | NA |
| RN7SKP275 | 4.243848 | -0.57255 | 1.075902 | -0.53216 | 0.594618 | NA |
| | 1.48623 | -0.09738 | 1.890909 | -0.0515 | 0.958928 | NA |
| RNU6-1187 | 0.134133 | -1.48909 | 4.077584 | -0.36519 | 0.71497 | NA |
| | 0.096202 | 0.408546 | 4.079639 | 0.100143 | 0.920231 | NA |
| | 0.163957 | 0.424913 | 4.079006 | 0.104171 | 0.917034 | NA |
| SNORD117 | 0.163957 | 0.424913 | 4.079006 | 0.104171 | 0.917034 | NA |
| RNY4P17 | 0.327913 | 1.543154 | 4.021063 | 0.383768 | 0.701151 | NA |
| SNORD48 | 0.967291 | -0.94461 | 2.160526 | -0.43721 | 0.661958 | NA |

| | | | | | | |
|---|---|---|---|---|---|---|
| | 0.883146 | 1.112834 | 2.554178 | 0.435692 | 0.663061 | NA |
| RNU1-36P | 0.556813 | -0.57007 | 3.294615 | -0.17303 | 0.862627 | NA |
| VTRNA1-2 | 0.162376 | 0.424913 | 4.079006 | 0.104171 | 0.917034 | NA |
| RNU1-100F | 0.134133 | -1.48909 | 4.077584 | -0.36519 | 0.71497 | NA |
| RN7SK | 17.49874 | 0.872519 | 0.549461 | 1.587953 | 0.112297 | NA |
| | 1.412576 | -1.39568 | 1.81698 | -0.76813 | 0.442409 | NA |
| | 0.3441 | -0.0166 | 3.769682 | -0.0044 | 0.996486 | NA |
| | 0.506476 | 0.726418 | 2.864186 | 0.253621 | 0.799788 | NA |
| RN7SKP292 | 0.096202 | 0.408546 | 4.079639 | 0.100143 | 0.920231 | NA |
| RNY4P10 | 1.697254 | 0.321736 | 1.546936 | 0.207983 | 0.835242 | NA |
| RNA5SP151 | 0.162376 | 0.424913 | 4.079006 | 0.104171 | 0.917034 | NA |
| | 0.162376 | 0.424913 | 4.079006 | 0.104171 | 0.917034 | NA |
| | 0.268266 | -2.15396 | 4.038351 | -0.53338 | 0.593773 | NA |
| | 0.162376 | 0.424913 | 4.079006 | 0.104171 | 0.917034 | NA |
| | 8.173542 | -0.48708 | 0.758782 | -0.64192 | 0.520925 | NA |
| | 0.096202 | 0.408546 | 4.079639 | 0.100143 | 0.920231 | NA |
| | 0.727508 | 0.192954 | 2.572892 | 0.074995 | 0.940219 | NA |
| | 0.096202 | 0.408546 | 4.079639 | 0.100143 | 0.920231 | NA |
| | 3.195026 | 0.215958 | 1.271548 | 0.169839 | 0.865137 | NA |
| | 32.93969 | 0.405149 | 0.400802 | 1.010844 | 0.312091 | NA |
| RAET1G | 14.69223 | 0.383111 | 0.813588 | 0.47089 | 0.637719 | NA |
| SAMD5 | 359.7456 | 0.926169 | 0.20994 | 4.41159 | 1.03E-05 | 6.34E-04 |
| ECT2L | 0.548763 | 2.343695 | 3.135478 | 0.747476 | 0.454776 | NA |
| TMEM244 | 0.163957 | 0.424913 | 4.079006 | 0.104171 | 0.917034 | NA |
| CENPW | 398.8299 | 0.076738 | 0.205011 | 0.374311 | 0.708173 | 0.813192 |
| FAM229B | 881.1281 | 0.013623 | 0.17373 | 0.078417 | 0.937496 | 0.965178 |
| DDO | 0.556813 | -0.57007 | 3.294615 | -0.17303 | 0.862627 | NA |
| CCDC162P | 0.163957 | 0.424913 | 4.079006 | 0.104171 | 0.917034 | NA |
| BVES-AS1 | 31.54095 | 0.451579 | 0.38067 | 1.186277 | 0.235513 | NA |
| HIST1H3H | 11.339 | -0.52371 | 0.621335 | -0.84287 | 0.399299 | NA |
| | 0.411855 | 0.250456 | 3.097618 | 0.080854 | 0.935558 | NA |
| C6orf164 | 2.255611 | 0.057932 | 1.466288 | 0.039509 | 0.968485 | NA |
| C6orf163 | 4.342597 | 1.295587 | 1.01208 | 1.280123 | 0.200502 | NA |
| SNHG5 | 920.5441 | 0.281763 | 0.207243 | 1.359578 | 0.173964 | 0.325958 |
| RIPPLY2 | 0.163957 | 0.424913 | 4.079006 | 0.104171 | 0.917034 | NA |
| OOEP | 0.411855 | 0.250456 | 3.097618 | 0.080854 | 0.935558 | NA |
| LRRC73 | 15.37378 | 0.591832 | 0.659509 | 0.897382 | 0.369515 | NA |
| GGNBP1 | 0.708364 | 0.219301 | 2.144998 | 0.102238 | 0.918568 | NA |
| RPL12P1 | 0.096202 | 0.408546 | 4.079639 | 0.100143 | 0.920231 | NA |
| DAXX | 1902.229 | -0.09839 | 0.146574 | -0.67123 | 0.502071 | 0.65228 |
| PFDN6 | 1002.494 | 0.194388 | 0.172621 | 1.126095 | 0.260125 | 0.424389 |
| RING1 | 1829.873 | -0.18266 | 0.156584 | -1.16652 | 0.243403 | 0.406201 |
| HSD17B8 | 236.7387 | -0.45692 | 0.26433 | -1.72859 | 0.083882 | 0.197268 |
| RXRB | 1393.261 | -0.31421 | 0.15808 | -1.98767 | 0.046848 | 0.133187 |
| COL11A2 | 24.79772 | 0.161663 | 0.474513 | 0.340693 | 0.733335 | NA |
| HLA-DOA | 2.272972 | 0.45053 | 1.368805 | 0.329141 | 0.742049 | NA |
| BRD2 | 4585.879 | -0.18926 | 0.153441 | -1.23343 | 0.217417 | 0.376879 |

| | | | | | | |
|---|---|---|---|---|---|---|
| HLA-DMA | 608.7171 | -0.22704 | 0.157081 | -1.44539 | 0.148349 | 0.291544 |
| TAPSAR1 | 20.42414 | -0.48077 | 0.512858 | -0.93744 | 0.348533 | NA |
| PSMB8 | 1961.375 | -0.44306 | 0.187448 | -2.36366 | 0.018096 | 0.070544 |
| TAP2 | 400.8281 | -0.58192 | 0.178402 | -3.26186 | 0.001107 | 0.01108 |
| HLA-DRA | 4.526614 | 0.773459 | 1.439306 | 0.537383 | 0.591003 | NA |
| BTNL2 | 0.47961 | 0.535796 | 3.39605 | 0.15777 | 0.874638 | NA |
| C6orf10 | 0.589223 | -3.04304 | 3.047314 | -0.9986 | 0.317989 | NA |
| NOTCH4 | 78.80127 | -0.14844 | 0.272371 | -0.54498 | 0.58577 | 0.718723 |
| PBX2 | 870.9265 | -0.29925 | 0.138084 | -2.16717 | 0.030222 | 0.099633 |
| AGER | 11.10172 | -0.02947 | 0.651795 | -0.04522 | 0.963933 | NA |
| RNF5 | 1890.548 | -0.06548 | 0.211514 | -0.30956 | 0.756895 | 0.848189 |
| AGPAT1 | 2703.579 | -0.38834 | 0.12504 | -3.10574 | 0.001898 | 0.015572 |
| PRRT1 | 86.84734 | -0.10122 | 0.328423 | -0.30821 | 0.757925 | 0.849022 |
| FKBPL | 273.551 | -0.08965 | 0.221097 | -0.40547 | 0.685134 | 0.795122 |
| CYP21A1P | 0.28583 | -2.21358 | 3.97376 | -0.55705 | 0.577494 | NA |
| STK19 | 48.57111 | 0.146064 | 0.332658 | 0.439081 | 0.660603 | 0.776536 |
| DXO | 365.788 | -0.01764 | 0.200275 | -0.08808 | 0.929816 | 0.960189 |
| SKIV2L | 1861.377 | -0.4 | 0.127178 | -3.14516 | 0.00166 | 0.014396 |
| NELFE | 1848.131 | -0.08639 | 0.149543 | -0.57771 | 0.56346 | 0.701367 |
| ZBTB12 | 113.2881 | -0.38615 | 0.258085 | -1.4962 | 0.134602 | 0.272956 |
| EHMT2 | 1905.137 | -0.42026 | 0.133494 | -3.14818 | 0.001643 | 0.014318 |
| SLC44A4 | 0.192403 | 0.975392 | 4.060716 | 0.240202 | 0.810174 | NA |
| NEU1 | 2954.043 | -0.15573 | 0.128352 | -1.2133 | 0.225015 | 0.386043 |
| C6orf48 | 4638.386 | -0.12867 | 0.144608 | -0.88981 | 0.373565 | 0.539878 |
| HSPA1B | 708.9694 | -0.44761 | 0.179768 | -2.48996 | 0.012776 | 0.055765 |
| HSPA1A | 730.6663 | -0.33037 | 0.182394 | -1.8113 | 0.070095 | 0.174538 |
| HSPA1L | 34.84706 | 0.125856 | 0.379349 | 0.331769 | 0.740063 | NA |
| LSM2 | 928.1304 | -0.17436 | 0.143903 | -1.21163 | 0.225655 | 0.38686 |
| VARS | 4859.73 | -0.54251 | 0.144995 | -3.74159 | 1.83E-04 | 0.003662 |
| VWA7 | 4.003647 | 0.695978 | 1.093243 | 0.636618 | 0.524374 | NA |
| MSH5 | 10.21191 | 0.58341 | 0.653755 | 0.892399 | 0.372179 | NA |
| C6orf25 | 0.296509 | -0.51709 | 3.982155 | -0.12985 | 0.896684 | NA |
| ABHD16A | 73.19949 | -0.26492 | 0.264078 | -1.0032 | 0.315765 | 0.483082 |
| LY6G5C | 56.11626 | 0.016342 | 0.349185 | 0.046801 | 0.962672 | 0.979674 |
| CSNK2B | 469.7469 | -0.13993 | 0.172202 | -0.8126 | 0.416449 | 0.579597 |
| GPANK1 | 595.1797 | -0.13329 | 0.175298 | -0.76037 | 0.447032 | 0.606718 |
| C6orf47 | 614.6189 | -0.05043 | 0.153486 | -0.32859 | 0.742466 | 0.837721 |
| APOM | 15.07329 | 0.116739 | 0.572081 | 0.204059 | 0.838307 | NA |
| BAG6 | 7975.404 | -0.49798 | 0.14545 | -3.42371 | 6.18E-04 | 0.007733 |
| PRRC2A | 8400.823 | -0.77669 | 0.145356 | -5.34338 | 9.12E-08 | 3.24E-05 |
| NCR3 | 0.303394 | -2.27006 | 4.032451 | -0.56295 | 0.57347 | NA |
| LST1 | 0.151697 | -1.546 | 4.075469 | -0.37934 | 0.704433 | NA |
| NFKBIL1 | 541.4423 | -0.20002 | 0.178163 | -1.12267 | 0.261579 | 0.426022 |
| MICB | 446.5374 | -0.34155 | 0.202023 | -1.69065 | 0.090903 | 0.209295 |
| MICA | 422.6807 | -0.00451 | 0.191918 | -0.02351 | 0.98124 | 0.990173 |
| HLA-C | 17051.6 | -0.57443 | 0.184881 | -3.10701 | 0.00189 | 0.015537 |
| POU5F1 | 2.901256 | -1.27773 | 1.398493 | -0.91365 | 0.360903 | NA |

| | | | | | | |
|---|---|---|---|---|---|---|
| CCHCR1 | 373.6535 | -0.22631 | 0.183573 | -1.23278 | 0.217656 | 0.377018 |
| PSORS1C2 | 0.354779 | 1.703954 | 3.657232 | 0.465914 | 0.641277 | NA |
| PSORS1C1 | 5.725448 | 0.492322 | 0.982305 | 0.50119 | 0.616237 | NA |
| DHX16 | 1560.002 | -0.23377 | 0.15721 | -1.48697 | 0.137022 | 0.276211 |
| C6orf136 | 292.3209 | -0.27608 | 0.189694 | -1.45541 | 0.145557 | 0.287692 |
| MRPS18B | 1873.71 | -0.33389 | 0.158943 | -2.10066 | 0.035671 | 0.111306 |
| PPP1R10 | 1229.375 | -0.49976 | 0.17493 | -2.85693 | 0.004278 | 0.026578 |
| ABCF1 | 3122.521 | -0.02605 | 0.128129 | -0.20329 | 0.838911 | 0.903953 |
| PRR3 | 204.3973 | -0.19986 | 0.183315 | -1.09026 | 0.2756 | 0.441265 |
| DDR1 | 2648.911 | -0.12282 | 0.189852 | -0.64693 | 0.517674 | 0.664972 |
| GNL1 | 2750.141 | -0.1104 | 0.136989 | -0.80588 | 0.420315 | 0.583153 |
| HLA-E | 17823.57 | -0.45521 | 0.126979 | -3.58495 | 3.37E-04 | 0.005308 |
| TRIM39 | 256.6078 | 0.005135 | 0.216758 | 0.023691 | 0.981099 | 0.990173 |
| TRIM15 | 0.134133 | -1.48909 | 4.077584 | -0.36519 | 0.71497 | NA |
| TRIM31 | 0.258578 | 1.233151 | 4.047809 | 0.304646 | 0.760635 | NA |
| RNF39 | 21.80129 | 0.223771 | 0.449042 | 0.498329 | 0.618252 | NA |
| PPP1R11 | 1611.98 | -0.19295 | 0.153426 | -1.25761 | 0.208534 | 0.365923 |
| HLA-J | 5.399039 | 1.273498 | 0.883948 | 1.440694 | 0.149671 | NA |
| ZNRD1-AS1 | 71.25524 | 0.05517 | 0.311069 | 0.177355 | 0.859229 | 0.916352 |
| HCG9 | 0.134133 | -1.48909 | 4.077584 | -0.36519 | 0.71497 | NA |
| GNB2L1 | 30450.22 | -0.17217 | 0.139415 | -1.23494 | 0.216855 | 0.37618 |
| HLA-G | 266.42 | -0.24528 | 0.325721 | -0.75304 | 0.451427 | 0.610554 |
| HLA-F | 730.6566 | -0.45083 | 0.192381 | -2.34343 | 0.019107 | 0.073331 |
| CBY3 | 1.380523 | 0.101537 | 1.91231 | 0.053097 | 0.957655 | NA |
| C5orf60 | 0.192403 | 0.975392 | 4.060716 | 0.240202 | 0.810174 | NA |
| GABBR1 | 303.9217 | 0.26215 | 0.187627 | 1.397188 | 0.162357 | 0.310342 |
| C6orf100 | 1.864934 | 0.0222 | 1.442892 | 0.015386 | 0.987725 | NA |
| TRIM27 | 1966.323 | -0.23666 | 0.119892 | -1.97392 | 0.04839 | 0.135747 |
| | 3.655724 | -0.58082 | 1.01832 | -0.57037 | 0.568424 | NA |
| RANBP17 | 87.16078 | 0.370665 | 0.256163 | 1.446987 | 0.147901 | 0.291014 |
| FAM196B | 1.534806 | -1.77475 | 1.804126 | -0.98372 | 0.325255 | NA |
| ZNF204P | 29.14356 | 1.530879 | 0.543776 | 2.815273 | 0.004874 | NA |
| SPINK9 | 0.315653 | -0.57305 | 3.892639 | -0.14721 | 0.882963 | NA |
| PCDHGA1 | 6.119528 | 0.165538 | 0.916747 | 0.180571 | 0.856704 | NA |
| PCDHA7 | 0.574231 | 0.93227 | 2.49998 | 0.372911 | 0.709215 | NA |
| PCDHA4 | 3.328656 | 0.220561 | 1.247445 | 0.17681 | 0.859658 | NA |
| PCDHA2 | 0.162376 | 0.424913 | 4.079006 | 0.104171 | 0.917034 | NA |
| PCDHA1 | 1.40704 | 1.571999 | 1.812999 | 0.867071 | 0.385903 | NA |
| CCNI2 | 1.930655 | -0.04837 | 1.699957 | -0.02845 | 0.9773 | NA |
| FRG2 | 0.288605 | 1.443619 | 4.029058 | 0.358302 | 0.720117 | NA |
| HSP90AA4I | 0.260158 | 1.240347 | 4.047124 | 0.306476 | 0.759242 | NA |
| C4orf47 | 3.116685 | 0.03368 | 1.179472 | 0.028555 | 0.977219 | NA |
| C4orf46 | 320.9953 | 0.034677 | 0.182112 | 0.190416 | 0.848983 | 0.910091 |
| TMEM170I | 142.5163 | 0.590261 | 0.252354 | 2.339019 | 0.019334 | 0.073891 |
| SNX2 | 1755.97 | 0.622037 | 0.200021 | 3.109849 | 0.001872 | 0.015453 |
| GCNT6 | 3.051823 | -0.79466 | 1.126139 | -0.70565 | 0.480405 | NA |
| CFI | 0.096202 | 0.408546 | 4.079639 | 0.100143 | 0.920231 | NA |

| | | | | | | |
|---|---|---|---|---|---|---|
| HNRNPA1P | 0.151697 | -1.546 | 4.075469 | -0.37934 | 0.704433 | NA |
| | 312.2247 | -0.82116 | 0.245706 | -3.34202 | 8.32E-04 | 0.009276 |
| ATP6AP1L | 12.02485 | 0.394337 | 0.595042 | 0.662704 | 0.50752 | NA |
| SMN2 | 12.59166 | 0.614178 | 0.576728 | 1.064936 | 0.286905 | NA |
| SERF1B | 7.241468 | 0.104997 | 0.767478 | 0.136808 | 0.891182 | NA |
| C5orf51 | 1006.819 | 0.376497 | 0.228855 | 1.645131 | 0.099943 | 0.223198 |
| | 0.506476 | 0.726418 | 2.864186 | 0.253621 | 0.799788 | NA |
| GMNC | 1.113481 | 0.468899 | 1.933629 | 0.242497 | 0.808395 | NA |
| TTC23L | 21.52355 | -0.49649 | 0.461445 | -1.07594 | 0.281954 | NA |
| HSP90AB2I | 1.990077 | -1.4068 | 1.647196 | -0.85406 | 0.393073 | NA |
| | 0.411855 | 0.250456 | 3.097618 | 0.080854 | 0.935558 | NA |
| DNAJC19 | 431.0719 | 0.488902 | 0.184085 | 2.655857 | 0.007911 | 0.040371 |
| ZDHHC11B | 0.660976 | -0.84029 | 2.673548 | -0.3143 | 0.753294 | NA |
| | 1.094751 | 1.216857 | 2.01996 | 0.602417 | 0.546897 | NA |
| EGFEM1P | 8.55123 | 1.281917 | 0.779823 | 1.643856 | 0.100206 | NA |
| ANKUB1 | 0.354779 | 1.703954 | 3.657232 | 0.465914 | 0.641277 | NA |
| C3orf72 | 1.966482 | 3.071493 | 1.706538 | 1.799839 | 0.071886 | NA |
| HCP5 | 7.343005 | -1.25261 | 0.814687 | -1.53753 | 0.124163 | NA |
| HLA-H | 1432.002 | -0.57746 | 0.160321 | -3.60188 | 3.16E-04 | 0.005092 |
| HCG27 | 2.605832 | 0.048378 | 1.259015 | 0.038425 | 0.969349 | NA |
| COL6A6 | 3.62086 | 2.155508 | 1.215216 | 1.773766 | 0.076102 | NA |
| H1FX-AS1 | 36.04964 | -0.0413 | 0.385589 | -0.1071 | 0.914706 | NA |
| OR10C1 | 0.151697 | -1.546 | 4.075469 | -0.37934 | 0.704433 | NA |
| HLA-A | 19754.57 | -0.52199 | 0.16473 | -3.16874 | 0.001531 | 0.01373 |
| PTPLB | 2454.69 | 0.15195 | 0.166296 | 0.913735 | 0.360856 | 0.528077 |
| WDR52 | 59.24532 | 1.997901 | 0.433239 | 4.611541 | 4.00E-06 | 3.70E-04 |
| | 0.937264 | -1.23834 | 2.063944 | -0.59999 | 0.548514 | NA |
| LNP1 | 399.6037 | 0.139046 | 0.174561 | 0.796543 | 0.425716 | 0.587731 |
| VGLL3 | 62.38432 | 0.808741 | 0.456876 | 1.770155 | 0.076701 | 0.185912 |
| RNU6-431I | 0.314073 | -0.57308 | 3.898934 | -0.14698 | 0.883146 | NA |
| RNU1-73P | 0.162376 | 0.424913 | 4.079006 | 0.104171 | 0.917034 | NA |
| RNU1-11P | 4.747072 | 0.922713 | 1.145363 | 0.805607 | 0.420469 | NA |
| RNU6-26P | 0.448206 | -1.2532 | 3.007315 | -0.41672 | 0.676885 | NA |
| RNU6-484I | 0.096202 | 0.408546 | 4.079639 | 0.100143 | 0.920231 | NA |
| SNORD101 | 7.707138 | 1.025635 | 0.771796 | 1.328895 | 0.183883 | NA |
| RNU6-701I | 0.162376 | 0.424913 | 4.079006 | 0.104171 | 0.917034 | NA |
| | 0.458885 | 0.577525 | 3.449278 | 0.167433 | 0.867029 | NA |
| RNU6-190I | 0.163957 | 0.424913 | 4.079006 | 0.104171 | 0.917034 | NA |
| RNU6-42P | 0.555087 | 0.973802 | 2.808575 | 0.346725 | 0.728798 | NA |
| RNU6-4P | 0.151697 | -1.546 | 4.075469 | -0.37934 | 0.704433 | NA |
| SNORD63 | 1.406049 | -0.26601 | 1.627656 | -0.16343 | 0.870178 | NA |
| | 0.163957 | 0.424913 | 4.079006 | 0.104171 | 0.917034 | NA |
| SNORA7B | 0.288605 | 1.443619 | 4.029058 | 0.358302 | 0.720117 | NA |
| RNA5SP187 | 0.162376 | 0.424913 | 4.079006 | 0.104171 | 0.917034 | NA |
| | 0.258578 | 1.233151 | 4.047809 | 0.304646 | 0.760635 | NA |
| | 0.162376 | 0.424913 | 4.079006 | 0.104171 | 0.917034 | NA |
| | 0.096202 | 0.408546 | 4.079639 | 0.100143 | 0.920231 | NA |

| | | | | | | |
|---|---|---|---|---|---|---|
| RNU6-35P | 0.192403 | 0.975392 | 4.060716 | 0.240202 | 0.810174 | NA |
| | 0.46735 | -1.30193 | 3.399671 | -0.38296 | 0.701751 | NA |
| RNU6-658F | 0.298089 | -0.51711 | 3.975232 | -0.13008 | 0.896501 | NA |
| | 0.163957 | 0.424913 | 4.079006 | 0.104171 | 0.917034 | NA |
| | 0.096202 | 0.408546 | 4.079639 | 0.100143 | 0.920231 | NA |
| RNU6-33P | 0.640755 | -1.64826 | 2.754778 | -0.59833 | 0.549621 | NA |
| MIR570 | 1.430532 | 1.196549 | 1.793968 | 0.666985 | 0.504782 | NA |
| MIR573 | 1.116855 | 1.112287 | 2.31317 | 0.48085 | 0.630623 | NA |
| MIR584 | 0.230334 | -0.53318 | 4.076749 | -0.13079 | 0.895945 | NA |
| MIR572 | 0.392857 | -1.546 | 4.075469 | -0.37934 | 0.704433 | NA |
| MIR580 | 0.151697 | -1.546 | 4.075469 | -0.37934 | 0.704433 | NA |
| MIR568 | 1.232404 | 0.600643 | 1.916349 | 0.313431 | 0.753953 | NA |
| MIR302A | 0.096202 | 0.408546 | 4.079639 | 0.100143 | 0.920231 | NA |
| MIR567 | 0.096202 | 0.408546 | 4.079639 | 0.100143 | 0.920231 | NA |
| MIR574 | 0.315653 | -0.57305 | 3.892639 | -0.14721 | 0.882963 | NA |
| MIR579 | 0.247898 | -0.5901 | 4.074633 | -0.14482 | 0.884851 | NA |
| MIR576 | 0.163957 | 0.424913 | 4.079006 | 0.104171 | 0.917034 | NA |
| GPX3 | 77.27391 | 0.228085 | 0.477607 | 0.477557 | 0.632965 | 0.754818 |
| | 0.096202 | 0.408546 | 4.079639 | 0.100143 | 0.920231 | NA |
| | 0.134133 | -1.48909 | 4.077584 | -0.36519 | 0.71497 | NA |
| | 0.420954 | 1.923376 | 3.462803 | 0.555439 | 0.578594 | NA |
| RNU6-930F | 0.506476 | 0.726418 | 2.864186 | 0.253621 | 0.799788 | NA |
| SNORD72 | 0.315653 | -0.57305 | 3.892639 | -0.14721 | 0.882963 | NA |
| SNORA74B | 0.422534 | 1.924583 | 3.001758 | 0.641152 | 0.521424 | NA |
| RNU6-460F | 0.326333 | 1.533889 | 3.826878 | 0.40082 | 0.688553 | NA |
| SNORA26 | 3.306277 | -0.34359 | 1.263574 | -0.27192 | 0.785683 | NA |
| RPL15P3 | 32.47328 | 0.045508 | 0.370685 | 0.122767 | 0.902292 | NA |
| PDCL3P5 | 2.747906 | 1.639263 | 1.425566 | 1.149903 | 0.250184 | NA |
| COX5BP2 | 0.192403 | 0.975392 | 4.060716 | 0.240202 | 0.810174 | NA |
| | 0.162376 | 0.424913 | 4.079006 | 0.104171 | 0.917034 | NA |
| TCTEX1D2 | 36.35875 | 0.497085 | 0.413701 | 1.201558 | 0.229535 | NA |
| EEF1DP5 | 9.883981 | -0.66187 | 0.682693 | -0.9695 | 0.332293 | NA |
| YWHAZP4 | 0.506476 | 0.726418 | 2.864186 | 0.253621 | 0.799788 | NA |
| | 0.315653 | -0.57305 | 3.892639 | -0.14721 | 0.882963 | NA |
| CRYGS | 12.41449 | 0.836218 | 0.646014 | 1.294427 | 0.195518 | NA |
| CNN2P9 | 3.444294 | -0.40636 | 1.134588 | -0.35816 | 0.720224 | NA |
| | 0.399595 | -1.31595 | 3.579466 | -0.36764 | 0.713143 | NA |
| | 1.135211 | -1.86222 | 2.017292 | -0.92313 | 0.355939 | NA |
| PRDX2P4 | 0.28583 | -2.21358 | 3.97376 | -0.55705 | 0.577494 | NA |
| | 7.678566 | 0.432225 | 0.748775 | 0.577243 | 0.563775 | NA |
| RPS7P8 | 0.096202 | 0.408546 | 4.079639 | 0.100143 | 0.920231 | NA |
| TRIM59 | 99.58416 | 0.34214 | 0.253984 | 1.34709 | 0.177951 | 0.330298 |
| C6orf165 | 10.68988 | 0.55847 | 0.680914 | 0.820177 | 0.412115 | NA |
| HNRNPA3P | 11.88749 | 0.93451 | 0.6788 | 1.376709 | 0.168602 | NA |
| RPL9P18 | 0.82371 | 0.509057 | 2.431911 | 0.209324 | 0.834195 | NA |
| RPL7AP34 | 0.392711 | 0.292538 | 3.145221 | 0.09301 | 0.925895 | NA |
| | 0.314073 | -0.57308 | 3.898934 | -0.14698 | 0.883146 | NA |

| | | | | | | |
|---|---|---|---|---|---|---|
| MXD3 | 247.6203 | -0.32797 | 0.236911 | -1.38437 | 0.166246 | 0.315386 |
| RANP6 | 0.994194 | -0.63787 | 1.893391 | -0.33689 | 0.736198 | NA |
| NAP1L1P3 | 1.216218 | 1.276946 | 1.967625 | 0.648978 | 0.516352 | NA |
| GAPDHP71 | 0.247898 | -0.5901 | 4.074633 | -0.14482 | 0.884851 | NA |
| | 6.143556 | -0.10362 | 0.844142 | -0.12275 | 0.902305 | NA |
| | 0.096202 | 0.408546 | 4.079639 | 0.100143 | 0.920231 | NA |
| ANXA2P1 | 7.404549 | -0.43696 | 0.814993 | -0.53615 | 0.591854 | NA |
| | 0.151697 | -1.546 | 4.075469 | -0.37934 | 0.704433 | NA |
| HSPD1P1 | 1.986013 | 1.549979 | 1.697517 | 0.913086 | 0.361197 | NA |
| | 5.256711 | 0.126269 | 0.917946 | 0.137556 | 0.890592 | NA |
| VTI1BP2 | 0.411855 | 0.250456 | 3.097618 | 0.080854 | 0.935558 | NA |
| ATP6V0CP3 | 1.141927 | 0.609996 | 1.889109 | 0.322901 | 0.74677 | NA |
| RPS23P2 | 0.326333 | 1.533889 | 3.826878 | 0.40082 | 0.688553 | NA |
| VDAC1P7 | 0.46577 | -1.30191 | 3.402879 | -0.38259 | 0.702024 | NA |
| AKR1B1P2 | 0.652869 | 1.292592 | 2.374873 | 0.544278 | 0.58625 | NA |
| | 0.45509 | -2.71227 | 3.963744 | -0.68427 | 0.493805 | NA |
| NT5C3AP1 | 6.019508 | 0.383078 | 0.916551 | 0.417956 | 0.67598 | NA |
| ACTN4P1 | 1.760763 | -1.59021 | 1.646198 | -0.96599 | 0.33405 | NA |
| LAP3P2 | 0.915759 | -1.79003 | 2.342046 | -0.7643 | 0.444687 | NA |
| | 0.096202 | 0.408546 | 4.079639 | 0.100143 | 0.920231 | NA |
| RAC1P5 | 0.163957 | 0.424913 | 4.079006 | 0.104171 | 0.917034 | NA |
| SRA1 | 1932.946 | 0.058116 | 0.162875 | 0.356812 | 0.721233 | 0.822641 |
| HNRNPA1P | 0.163957 | 0.424913 | 4.079006 | 0.104171 | 0.917034 | NA |
| VDAC1 | 7553.44 | -0.25349 | 0.130513 | -1.94228 | 0.052103 | 0.142371 |
| ZBTB9 | 394.8666 | -0.52851 | 0.156671 | -3.37338 | 7.43E-04 | 0.008681 |
| GPSM3 | 1571.769 | -0.35997 | 0.195718 | -1.83924 | 0.065879 | 0.167381 |
| | 0.315653 | -0.57305 | 3.892639 | -0.14721 | 0.882963 | NA |
| ATF6B | 2521.404 | -0.35057 | 0.146838 | -2.38744 | 0.016966 | 0.067758 |
| SETP22 | 0.938845 | -1.23445 | 2.063177 | -0.59832 | 0.549624 | NA |
| CLIC1 | 11800.46 | 2.67E-04 | 0.163235 | 0.001633 | 0.998697 | 0.999371 |
| DDAH2 | 834.1257 | -0.0671 | 0.212302 | -0.31605 | 0.751963 | 0.844818 |
| POLD2P1 | 0.967088 | -0.05827 | 2.035183 | -0.02863 | 0.977158 | NA |
| | 0.260158 | 1.240347 | 4.047124 | 0.306476 | 0.759242 | NA |
| | 0.785421 | 0.60161 | 2.458349 | 0.244721 | 0.806672 | NA |
| ATP6V1G2 | 26.81712 | -0.28299 | 0.401237 | -0.70529 | 0.480632 | NA |
| ACTBP2 | 7.82054 | 0.201149 | 0.746562 | 0.269434 | 0.787596 | NA |
| GTF2H4 | 13.58965 | 0.222273 | 0.574458 | 0.386926 | 0.698811 | NA |
| CFL1P5 | 0.624422 | 1.162588 | 2.733761 | 0.425271 | 0.670639 | NA |
| | 1.118639 | 0.415692 | 1.804658 | 0.230344 | 0.817825 | NA |
| EEF1B2P2 | 0.900767 | 0.808531 | 1.963257 | 0.411831 | 0.680463 | NA |
| RPL7AP7 | 1.04035 | -1.85633 | 2.355797 | -0.78798 | 0.430706 | NA |
| UBD | 1.390425 | -0.34271 | 2.156269 | -0.15894 | 0.873718 | NA |
| RPL3P6 | 1.500816 | -0.86149 | 1.576095 | -0.5466 | 0.584655 | NA |
| | 0.162376 | 0.424913 | 4.079006 | 0.104171 | 0.917034 | NA |
| RPS17P1 | 0.640609 | 0.042838 | 2.379511 | 0.018003 | 0.985637 | NA |
| ITGA1 | 1380.995 | 0.880328 | 0.230585 | 3.817799 | 1.35E-04 | 0.00305 |
| | 0.326536 | 0.037056 | 3.842554 | 0.009644 | 0.992306 | NA |

| | | | | | | |
|---|---|---|---|---|---|---|
| RPL7P17 | 0.096202 | 0.408546 | 4.079639 | 0.100143 | 0.920231 | NA |
| TPT1P5 | 1.05103 | -0.7935 | 2.094237 | -0.3789 | 0.704763 | NA |
| LYRM4 | 931.9421 | 0.043586 | 0.142224 | 0.306463 | 0.759252 | 0.849946 |
| | 154.5409 | 0.372949 | 0.205668 | 1.813358 | 0.069777 | 0.174038 |
| LINC00887 | 0.94127 | 3.089692 | 2.23227 | 1.384103 | 0.166327 | NA |
| | 6.103354 | -0.3713 | 0.798426 | -0.46504 | 0.641902 | NA |
| ALG3 | 1522.12 | -0.12808 | 0.140468 | -0.91184 | 0.361852 | 0.52898 |
| UBE2V1P2 | 0.877625 | -0.2954 | 2.206963 | -0.13385 | 0.893522 | NA |
| SETP21 | 0.192403 | 0.975392 | 4.060716 | 0.240202 | 0.810174 | NA |
| AGGF1P1 | 0.326536 | 0.037056 | 3.842554 | 0.009644 | 0.992306 | NA |
| | 0.672013 | 1.253691 | 2.607776 | 0.480751 | 0.630693 | NA |
| | 0.096202 | 0.408546 | 4.079639 | 0.100143 | 0.920231 | NA |
| RPL39P5 | 0.258578 | 1.233151 | 4.047809 | 0.304646 | 0.760635 | NA |
| SOGA3 | 3.651376 | -0.37341 | 1.088889 | -0.34293 | 0.731652 | NA |
| NEURL1B | 203.667 | -1.00424 | 0.296814 | -3.3834 | 7.16E-04 | 0.00842 |
| EFCAB9 | 1.523755 | 0.474747 | 1.576468 | 0.301146 | 0.763303 | NA |
| HAUS3 | 195.224 | 0.887163 | 0.288199 | 3.078302 | 0.002082 | 0.016597 |
| LINC00488 | 4.247744 | 0.736631 | 1.008318 | 0.730555 | 0.465051 | NA |
| | 3.620255 | 0.6565 | 1.113929 | 0.589355 | 0.555623 | NA |
| TPT1P6 | 2.395727 | 1.418403 | 1.40559 | 1.009116 | 0.312919 | NA |
| RPL7P1 | 68.10987 | 0.129895 | 0.36006 | 0.360758 | 0.718281 | 0.820342 |
| SPINK13 | 0.151697 | -1.546 | 4.075469 | -0.37934 | 0.704433 | NA |
| | 4.731484 | 0.182957 | 0.904669 | 0.202236 | 0.839732 | NA |
| | 0.410275 | 0.244283 | 3.100889 | 0.078778 | 0.937209 | NA |
| RBBP4P4 | 0.134133 | -1.48909 | 4.077584 | -0.36519 | 0.71497 | NA |
| RPL6P12 | 0.096202 | 0.408546 | 4.079639 | 0.100143 | 0.920231 | NA |
| RPL9P16 | 0.163957 | 0.424913 | 4.079006 | 0.104171 | 0.917034 | NA |
| | 1.987595 | 1.031131 | 1.587234 | 0.64964 | 0.515925 | NA |
| CYCSP55 | 0.230334 | -0.53318 | 4.076749 | -0.13079 | 0.895945 | NA |
| | 6.287306 | 0.291528 | 0.8445 | 0.345208 | 0.729938 | NA |
| SHFM1P1 | 0.230334 | -0.53318 | 4.076749 | -0.13079 | 0.895945 | NA |
| FAM177A1 | 0.392857 | -1.546 | 4.075469 | -0.37934 | 0.704433 | NA |
| | 0.096202 | 0.408546 | 4.079639 | 0.100143 | 0.920231 | NA |
| USP8P1 | 0.593018 | -0.10389 | 3.151334 | -0.03297 | 0.973702 | NA |
| LINC00243 | 0.547182 | 2.340248 | 3.137689 | 0.745851 | 0.455757 | NA |
| HLA-F-AS1 | 150.3441 | 0.027993 | 0.269239 | 0.103972 | 0.917192 | 0.952022 |
| ARHGEF28 | 513.7442 | 0.195977 | 0.161475 | 1.213663 | 0.224876 | 0.385916 |
| PPIAP29 | 39.67779 | -0.27771 | 0.358294 | -0.7751 | 0.438282 | NA |
| CHCHD2P2 | 57.8522 | -0.13665 | 0.351813 | -0.38842 | 0.697706 | 0.805229 |
| | 0.833158 | -0.52049 | 2.521552 | -0.20642 | 0.836464 | NA |
| GNL3LP1 | 0.35636 | 1.709252 | 3.651833 | 0.468053 | 0.639746 | NA |
| PKMP5 | 2.096995 | -0.60359 | 1.338602 | -0.45091 | 0.652055 | NA |
| | 18.51274 | 0.755743 | 0.477914 | 1.581336 | 0.113801 | NA |
| | 0.967088 | -0.05827 | 2.035183 | -0.02863 | 0.977158 | NA |
| | 8.63544 | 1.406924 | 0.735982 | 1.911628 | 0.055924 | NA |
| LINC00680 | 171.2142 | 0.252481 | 0.216443 | 1.1665 | 0.243413 | 0.406201 |
| | 20.29516 | -0.71047 | 0.477447 | -1.48805 | 0.136737 | NA |

| | | | | | | |
|---|---|---|---|---|---|---|
| C5orf49 | 18.22402 | -0.07747 | 0.526352 | -0.14718 | 0.882986 | NA |
| UBE2QL1 | 0.258578 | 1.233151 | 4.047809 | 0.304646 | 0.760635 | NA |
| CYP51A1P3 | 0.462046 | 0.589069 | 3.442439 | 0.17112 | 0.86413 | NA |
| | 2.493511 | 1.151572 | 1.712904 | 0.672292 | 0.501398 | NA |
| ASS1P10 | 0.134133 | -1.48909 | 4.077584 | -0.36519 | 0.71497 | NA |
| MYL5 | 126.3032 | -0.17953 | 0.309577 | -0.57991 | 0.561974 | 0.700032 |
| HMX1 | 0.327913 | 1.543154 | 4.021063 | 0.383768 | 0.701151 | NA |
| GUSBP9 | 0.947797 | 0.979959 | 2.329332 | 0.420704 | 0.673971 | NA |
| MARCKSL1 | 0.096202 | 0.408546 | 4.079639 | 0.100143 | 0.920231 | NA |
| | 0.392857 | -1.546 | 4.075469 | -0.37934 | 0.704433 | NA |
| | 0.096202 | 0.408546 | 4.079639 | 0.100143 | 0.920231 | NA |
| | 0.096202 | 0.408546 | 4.079639 | 0.100143 | 0.920231 | NA |
| HIST1H1PS | 1.115061 | 0.450621 | 2.115791 | 0.21298 | 0.831343 | NA |
| | 0.151697 | -1.546 | 4.075469 | -0.37934 | 0.704433 | NA |
| | 0.629929 | -1.1413 | 2.677641 | -0.42623 | 0.669937 | NA |
| | 0.392857 | -1.546 | 4.075469 | -0.37934 | 0.704433 | NA |
| RPL12P2 | 0.736957 | -1.0303 | 2.640699 | -0.39016 | 0.696418 | NA |
| | 0.162376 | 0.424913 | 4.079006 | 0.104171 | 0.917034 | NA |
| | 0.520316 | 2.239969 | 3.173769 | 0.705776 | 0.480328 | NA |
| | 0.631307 | -0.20384 | 3.097925 | -0.0658 | 0.947537 | NA |
| | 0.258578 | 1.233151 | 4.047809 | 0.304646 | 0.760635 | NA |
| | 0.134133 | -1.48909 | 4.077584 | -0.36519 | 0.71497 | NA |
| | 1.208102 | 0.723565 | 1.850162 | 0.391082 | 0.695737 | NA |
| | 55.42466 | 0.435309 | 0.322303 | 1.350621 | 0.176817 | 0.329058 |
| | 0.28583 | -2.21358 | 3.97376 | -0.55705 | 0.577494 | NA |
| | 0.766634 | 1.516849 | 2.262991 | 0.670285 | 0.502676 | NA |
| TUBB2BP1 | 0.268266 | -2.15396 | 4.038351 | -0.53338 | 0.593773 | NA |
| RBMXP1 | 0.163957 | 0.424913 | 4.079006 | 0.104171 | 0.917034 | NA |
| RPL17P26 | 1.294855 | 1.463581 | 1.753443 | 0.83469 | 0.403892 | NA |
| LY86-AS1 | 0.163957 | 0.424913 | 4.079006 | 0.104171 | 0.917034 | NA |
| | 1.015755 | 0.398771 | 2.259962 | 0.17645 | 0.85994 | NA |
| | 7.229254 | -1.00565 | 0.747073 | -1.34611 | 0.178266 | NA |
| | 0.096202 | 0.408546 | 4.079639 | 0.100143 | 0.920231 | NA |
| RPL21P65 | 0.096202 | 0.408546 | 4.079639 | 0.100143 | 0.920231 | NA |
| | 0.28583 | -2.21358 | 3.97376 | -0.55705 | 0.577494 | NA |
| TPT1P4 | 4.731216 | -0.3231 | 0.960087 | -0.33653 | 0.736469 | NA |
| | 0.74749 | 1.546424 | 2.516383 | 0.614543 | 0.538857 | NA |
| | 0.192403 | 0.975392 | 4.060716 | 0.240202 | 0.810174 | NA |
| FNIP1 | 1149.812 | 0.809502 | 0.186642 | 4.337186 | 1.44E-05 | 7.71E-04 |
| | 2.428632 | -0.55253 | 1.269524 | -0.43523 | 0.663399 | NA |
| LARP1P1 | 0.411855 | 0.250456 | 3.097618 | 0.080854 | 0.935558 | NA |
| ANKRD18E | 19.36366 | 0.524888 | 0.477182 | 1.099976 | 0.271343 | NA |
| | 0.326536 | 0.037056 | 3.842554 | 0.009644 | 0.992306 | NA |
| | 0.382031 | -1.26484 | 3.151989 | -0.40128 | 0.688212 | NA |
| CBX3P9 | 13.48165 | 0.850087 | 0.591678 | 1.436738 | 0.150792 | NA |
| | 0.6908 | 0.277604 | 2.346083 | 0.118327 | 0.905809 | NA |
| PRELID1P1 | 2.866644 | 0.419087 | 1.265513 | 0.33116 | 0.740524 | NA |

| | | | | | | |
|---|---|---|---|---|---|---|
| RPS4XP9 | 0.163957 | 0.424913 | 4.079006 | 0.104171 | 0.917034 | NA |
| ISCA1P1 | 0.260158 | 1.240347 | 4.047124 | 0.306476 | 0.759242 | NA |
| | 0.096202 | 0.408546 | 4.079639 | 0.100143 | 0.920231 | NA |
| | 60.30377 | -0.00909 | 0.304776 | -0.02984 | 0.976194 | 0.986933 |
| CCT7P1 | 0.258578 | 1.233151 | 4.047809 | 0.304646 | 0.760635 | NA |
| HIST1H4PS | 0.489058 | -0.58569 | 3.427543 | -0.17088 | 0.864321 | NA |
| | 0.247898 | -0.5901 | 4.074633 | -0.14482 | 0.884851 | NA |
| RSL24D1P1 | 0.942851 | 3.091781 | 2.270302 | 1.361837 | 0.173249 | NA |
| | 1.536161 | 0.286569 | 1.969673 | 0.145491 | 0.884324 | NA |
| | 2.192405 | 1.262367 | 1.52066 | 0.830144 | 0.406457 | NA |
| | 22.33139 | 0.276189 | 0.490831 | 0.562696 | 0.573642 | NA |
| RPS4XP7 | 0.247898 | -0.5901 | 4.074633 | -0.14482 | 0.884851 | NA |
| HIST1H2AP | 0.163957 | 0.424913 | 4.079006 | 0.104171 | 0.917034 | NA |
| | 0.327913 | 1.543154 | 4.021063 | 0.383768 | 0.701151 | NA |
| TENM3 | 1193.746 | -0.04633 | 0.156794 | -0.29548 | 0.767625 | 0.855764 |
| LYPLA1P3 | 0.258578 | 1.233151 | 4.047809 | 0.304646 | 0.760635 | NA |
| RAET1K | 0.096202 | 0.408546 | 4.079639 | 0.100143 | 0.920231 | NA |
| | 1.21326 | 0.631805 | 1.921077 | 0.32888 | 0.742246 | NA |
| | 5.324291 | -0.18659 | 0.868957 | -0.21473 | 0.829978 | NA |
| RPS27P15 | 0.163957 | 0.424913 | 4.079006 | 0.104171 | 0.917034 | NA |
| | 0.096202 | 0.408546 | 4.079639 | 0.100143 | 0.920231 | NA |
| | 0.192403 | 0.975392 | 4.060716 | 0.240202 | 0.810174 | NA |
| HNRNPA1P | 0.151697 | -1.546 | 4.075469 | -0.37934 | 0.704433 | NA |
| GAPDHP63 | 3.933096 | 0.120985 | 0.992682 | 0.121877 | 0.902996 | NA |
| RPL7P28 | 0.096202 | 0.408546 | 4.079639 | 0.100143 | 0.920231 | NA |
| BRD7P4 | 0.545988 | -0.43913 | 2.541145 | -0.17281 | 0.862803 | NA |
| HIST1H2AP | 0.314073 | -0.57308 | 3.898934 | -0.14698 | 0.883146 | NA |
| ST13P16 | 0.096202 | 0.408546 | 4.079639 | 0.100143 | 0.920231 | NA |
| | 0.163957 | 0.424913 | 4.079006 | 0.104171 | 0.917034 | NA |
| | 2.951366 | 0.463022 | 1.129732 | 0.409852 | 0.681915 | NA |
| DBIP1 | 0.163957 | 0.424913 | 4.079006 | 0.104171 | 0.917034 | NA |
| RPL18AP8 | 0.163957 | 0.424913 | 4.079006 | 0.104171 | 0.917034 | NA |
| | 0.298089 | -0.51711 | 3.975232 | -0.13008 | 0.896501 | NA |
| GSTM2P1 | 0.802985 | 0.54709 | 2.208262 | 0.247747 | 0.80433 | NA |
| | 0.258578 | 1.233151 | 4.047809 | 0.304646 | 0.760635 | NA |
| CNN3P1 | 0.151697 | -1.546 | 4.075469 | -0.37934 | 0.704433 | NA |
| NUFIP1P | 0.93741 | -3.49975 | 2.723608 | -1.28497 | 0.198804 | NA |
| TUBB8P2 | 0.748867 | 2.740193 | 2.468928 | 1.109871 | 0.267054 | NA |
| FTH1P15 | 20.81038 | -1.01423 | 0.580795 | -1.74628 | 0.080763 | NA |
| | 0.640755 | -1.64826 | 2.754778 | -0.59833 | 0.549621 | NA |
| | 1.75567 | 0.303256 | 1.471306 | 0.206114 | 0.836702 | NA |
| | 3.197424 | 0.52441 | 1.222737 | 0.428882 | 0.668009 | NA |
| NPM1P37 | 0.900913 | -0.26014 | 2.222223 | -0.11706 | 0.90681 | NA |
| HMGB1P2( | 1.170556 | -0.4175 | 2.096325 | -0.19916 | 0.84214 | NA |
| | 8.184532 | 0.406414 | 0.818845 | 0.496326 | 0.619664 | NA |
| NPM1P38 | 0.327913 | 1.543154 | 4.021063 | 0.383768 | 0.701151 | NA |
| PIP5K1P1 | 0.424115 | 1.932531 | 3.45462 | 0.559405 | 0.575885 | NA |

| | | | | | | |
|---|---|---|---|---|---|---|
| EEF1A1P42 | 0.151697 | -1.546 | 4.075469 | -0.37934 | 0.704433 | NA |
| | 5.951992 | 0.239761 | 0.848209 | 0.282667 | 0.777432 | NA |
| | 0.883 | 1.784395 | 2.188528 | 0.81534 | 0.414878 | NA |
| | 2.654857 | 0.523648 | 1.183497 | 0.442459 | 0.658157 | NA |
| | 0.594599 | -0.12152 | 2.771051 | -0.04385 | 0.965021 | NA |
| | 0.151697 | -1.546 | 4.075469 | -0.37934 | 0.704433 | NA |
| RPL17P25 | 0.163957 | 0.424913 | 4.079006 | 0.104171 | 0.917034 | NA |
| | 0.162376 | 0.424913 | 4.079006 | 0.104171 | 0.917034 | NA |
| ZNF259P1 | 0.804566 | 0.545894 | 2.033578 | 0.26844 | 0.78836 | NA |
| PPP1R3G | 0.134133 | -1.48909 | 4.077584 | -0.36519 | 0.71497 | NA |
| BMPR1APS | 11.40474 | 0.2734 | 0.635751 | 0.430043 | 0.667164 | NA |
| | 0.842643 | -2.08432 | 2.306153 | -0.90381 | 0.366097 | NA |
| | 0.151697 | -1.546 | 4.075469 | -0.37934 | 0.704433 | NA |
| | 0.151697 | -1.546 | 4.075469 | -0.37934 | 0.704433 | NA |
| | 8.015478 | -0.54992 | 0.809874 | -0.67902 | 0.497125 | NA |
| | 0.715249 | -0.95037 | 2.624937 | -0.36205 | 0.717313 | NA |
| PPIAP9 | 0.613743 | -0.17004 | 2.749758 | -0.06184 | 0.950691 | NA |
| ATP5F1P6 | 0.35636 | 1.709252 | 3.651833 | 0.468053 | 0.639746 | NA |
| GAPDHP42 | 0.643567 | 1.129905 | 3.100382 | 0.364441 | 0.715529 | NA |
| ZSCAN12P1 | 69.74991 | -0.04059 | 0.276965 | -0.14654 | 0.883495 | 0.931007 |
| RPL35P3 | 0.613743 | -0.17004 | 2.749758 | -0.06184 | 0.950691 | NA |
| | 0.382031 | -1.26484 | 3.151989 | -0.40128 | 0.688212 | NA |
| BTF3P7 | 0.392857 | -1.546 | 4.075469 | -0.37934 | 0.704433 | NA |
| | 0.815391 | 0.931626 | 2.360431 | 0.394685 | 0.693076 | NA |
| LAP3P1 | 0.096202 | 0.408546 | 4.079639 | 0.100143 | 0.920231 | NA |
| | 1.626695 | 0.638819 | 1.554182 | 0.411032 | 0.681049 | NA |
| HNRNPA1P | 0.658173 | -0.01016 | 2.630582 | -0.00386 | 0.996919 | NA |
| RPS24P12 | 0.096202 | 0.408546 | 4.079639 | 0.100143 | 0.920231 | NA |
| ACTBP8 | 0.448206 | -1.2532 | 3.007315 | -0.41672 | 0.676885 | NA |
| | 0.096202 | 0.408546 | 4.079639 | 0.100143 | 0.920231 | NA |
| | 0.258578 | 1.233151 | 4.047809 | 0.304646 | 0.760635 | NA |
| | 0.134133 | -1.48909 | 4.077584 | -0.36519 | 0.71497 | NA |
| KRT8P42 | 3.904118 | 0.836012 | 1.094141 | 0.764081 | 0.444819 | NA |
| | 4.414001 | 2.169623 | 1.109848 | 1.954883 | 0.050597 | NA |
| | 6.969485 | 0.346012 | 0.806392 | 0.429086 | 0.66786 | NA |
| YAP1P1 | 2.289455 | -0.07193 | 1.364419 | -0.05272 | 0.957956 | NA |
| | 0.619047 | -1.7574 | 3.107232 | -0.56558 | 0.571677 | NA |
| | 2.284939 | 1.924848 | 1.518308 | 1.267758 | 0.204884 | NA |
| PKMP3 | 0.411855 | 0.250456 | 3.097618 | 0.080854 | 0.935558 | NA |
| RPL35P2 | 18.31067 | 0.158573 | 0.490578 | 0.323237 | 0.746516 | NA |
| | 0.096202 | 0.408546 | 4.079639 | 0.100143 | 0.920231 | NA |
| RCC2P7 | 0.096202 | 0.408546 | 4.079639 | 0.100143 | 0.920231 | NA |
| | 2.410682 | 0.240607 | 1.27308 | 0.188996 | 0.850096 | NA |
| | 0.324752 | 1.53082 | 4.022025 | 0.380609 | 0.703493 | NA |
| OR1F12 | 0.162376 | 0.424913 | 4.079006 | 0.104171 | 0.917034 | NA |
| RPL5P18 | 1.294855 | 1.463581 | 1.753443 | 0.83469 | 0.403892 | NA |
| | 0.258578 | 1.233151 | 4.047809 | 0.304646 | 0.760635 | NA |

| | | | | | | |
|---|---|---|---|---|---|---|
| RPS18P9 | 14.70275 | 0.101451 | 0.533612 | 0.19012 | 0.849215 | NA |
| HIST1H3PS | 0.327913 | 1.543154 | 4.021063 | 0.383768 | 0.701151 | NA |
| | 0.899187 | 0.806933 | 2.127165 | 0.379346 | 0.704431 | NA |
| OSTCP4 | 8.063276 | -1.00088 | 0.760046 | -1.31686 | 0.187885 | NA |
| | 0.134133 | -1.48909 | 4.077584 | -0.36519 | 0.71497 | NA |
| MIR1271 | 0.096202 | 0.408546 | 4.079639 | 0.100143 | 0.920231 | NA |
| SNORD100 | 6.238898 | 0.139359 | 0.800708 | 0.174045 | 0.86183 | NA |
| | 0.151697 | -1.546 | 4.075469 | -0.37934 | 0.704433 | NA |
| MIR1303 | 0.678686 | -3.08037 | 2.621268 | -1.17514 | 0.239937 | NA |
| RNU6ATAC | 0.163957 | 0.424913 | 4.079006 | 0.104171 | 0.917034 | NA |
| | 0.520316 | 2.239969 | 3.173769 | 0.705776 | 0.480328 | NA |
| | 0.096202 | 0.408546 | 4.079639 | 0.100143 | 0.920231 | NA |
| MIR548I1 | 0.298089 | -0.51711 | 3.975232 | -0.13008 | 0.896501 | NA |
| | 1.304653 | -0.93789 | 2.220522 | -0.42237 | 0.672753 | NA |
| MIR1324 | 0.260158 | 1.240347 | 4.047124 | 0.306476 | 0.759242 | NA |
| C6orf226 | 140.8616 | -0.23621 | 0.219173 | -1.07774 | 0.281149 | 0.4473 |
| C5orf54 | 162.9435 | 0.443821 | 0.225994 | 1.963862 | 0.049546 | 0.137692 |
| SLC12A8 | 40.0451 | 1.068237 | 0.466205 | 2.291343 | 0.021944 | NA |
| PPT2 | 1130.669 | -0.12864 | 0.170429 | -0.75481 | 0.450365 | 0.609693 |
| C5orf55 | 106.9727 | 0.138227 | 0.227714 | 0.607018 | 0.543839 | 0.685582 |
| RNU6-757F | 0.260158 | 1.240347 | 4.047124 | 0.306476 | 0.759242 | NA |
| | 0.392857 | -1.546 | 4.075469 | -0.37934 | 0.704433 | NA |
| | 0.162376 | 0.424913 | 4.079006 | 0.104171 | 0.917034 | NA |
| RNA5SP203 | 0.410275 | 0.244283 | 3.100889 | 0.078778 | 0.937209 | NA |
| RNU2-28P | 0.163957 | 0.424913 | 4.079006 | 0.104171 | 0.917034 | NA |
| RN7SKP150 | 1.2638 | -0.68861 | 1.777184 | -0.38747 | 0.698407 | NA |
| RNU4-72P | 0.096202 | 0.408546 | 4.079639 | 0.100143 | 0.920231 | NA |
| RNU4-14P | 0.260158 | 1.240347 | 4.047124 | 0.306476 | 0.759242 | NA |
| | 1.674075 | -0.35038 | 1.560479 | -0.22453 | 0.822342 | NA |
| | 0.845272 | 1.770638 | 2.207891 | 0.801959 | 0.422577 | NA |
| RNU2-61P | 0.096202 | 0.408546 | 4.079639 | 0.100143 | 0.920231 | NA |
| RNA5SP155 | 0.555087 | 0.973802 | 2.808575 | 0.346725 | 0.728798 | NA |
| RN7SKP296 | 0.966942 | 0.917791 | 1.915974 | 0.47902 | 0.631924 | NA |
| RNA5SP221 | 0.096202 | 0.408546 | 4.079639 | 0.100143 | 0.920231 | NA |
| RNU6-1054 | 0.096202 | 0.408546 | 4.079639 | 0.100143 | 0.920231 | NA |
| RNU2-64P | 0.163957 | 0.424913 | 4.079006 | 0.104171 | 0.917034 | NA |
| RN7SKP30 | 0.35636 | 1.709252 | 3.651833 | 0.468053 | 0.639746 | NA |
| | 0.134133 | -1.48909 | 4.077584 | -0.36519 | 0.71497 | NA |
| EHHADH-A | 1.414302 | -3.09041 | 1.920811 | -1.60891 | 0.107636 | NA |
| FTH1P10 | 0.151697 | -1.546 | 4.075469 | -0.37934 | 0.704433 | NA |
| | 1.569416 | 1.765492 | 1.765492 | 1 | 0.31731 | NA |
| | 0.162376 | 0.424913 | 4.079006 | 0.104171 | 0.917034 | NA |
| VPS52 | 1128.948 | -0.33597 | 0.128925 | -2.60591 | 0.009163 | 0.04452 |
| | 1.809439 | 0.428428 | 1.452082 | 0.295044 | 0.76796 | NA |
| EEF1A1P8 | 5.896336 | 0.128022 | 0.840339 | 0.152346 | 0.878914 | NA |
| | 0.230334 | -0.53318 | 4.076749 | -0.13079 | 0.895945 | NA |
| | 0.162376 | 0.424913 | 4.079006 | 0.104171 | 0.917034 | NA |

| | | | | | | |
|---|---|---|---|---|---|---|
| RAET1E-AS | 2.170475 | -0.56273 | 1.654061 | -0.34021 | 0.733697 | NA |
| | 0.921281 | -1.21902 | 2.226885 | -0.54741 | 0.584096 | NA |
| | 0.163957 | 0.424913 | 4.079006 | 0.104171 | 0.917034 | NA |
| RPS20P14 | 26.45355 | 0.529631 | 0.450529 | 1.175577 | 0.239764 | NA |
| | 3.660981 | 1.022934 | 1.170908 | 0.873625 | 0.382323 | NA |
| | 2.331096 | 1.510116 | 1.371203 | 1.101307 | 0.270763 | NA |
| HLA-DPB1 | 589.3493 | -0.02394 | 0.156362 | -0.15311 | 0.878311 | 0.926986 |
| ABCC5-AS1 | 0.230334 | -0.53318 | 4.076749 | -0.13079 | 0.895945 | NA |
| | 0.28583 | -2.21358 | 3.97376 | -0.55705 | 0.577494 | NA |
| | 0.162376 | 0.424913 | 4.079006 | 0.104171 | 0.917034 | NA |
| | 0.134133 | -1.48909 | 4.077584 | -0.36519 | 0.71497 | NA |
| EPB41L4A-. | 646.9651 | 0.104151 | 0.181333 | 0.574361 | 0.565723 | 0.702867 |
| | 1.05103 | -0.7935 | 2.094237 | -0.3789 | 0.704763 | NA |
| HCG14 | 1.131248 | -0.36273 | 1.88105 | -0.19283 | 0.847091 | NA |
| C5orf66 | 11.15196 | 0.708857 | 0.646181 | 1.096995 | 0.272644 | NA |
| | 0.392857 | -1.546 | 4.075469 | -0.37934 | 0.704433 | NA |
| | 0.303394 | -2.27006 | 4.032451 | -0.56295 | 0.57347 | NA |
| MCCD1P2 | 0.151697 | -1.546 | 4.075469 | -0.37934 | 0.704433 | NA |
| | 6.234458 | 0.191948 | 0.831528 | 0.230838 | 0.81744 | NA |
| | 0.134133 | -1.48909 | 4.077584 | -0.36519 | 0.71497 | NA |
| SLC31A1P1 | 0.096202 | 0.408546 | 4.079639 | 0.100143 | 0.920231 | NA |
| | 5.93898 | 0.412252 | 0.980538 | 0.420435 | 0.674168 | NA |
| | 4.258844 | -0.58597 | 0.964177 | -0.60774 | 0.543358 | NA |
| SMIM13 | 503.4579 | 0.366421 | 0.173081 | 2.117053 | 0.034255 | 0.108321 |
| HLA-DPB2 | 0.460669 | -0.62593 | 3.398671 | -0.18417 | 0.853882 | NA |
| LPP-AS1 | 0.518736 | 2.237423 | 2.7713 | 0.807355 | 0.419462 | NA |
| | 0.314073 | -0.57308 | 3.898934 | -0.14698 | 0.883146 | NA |
| TNK2-AS1 | 0.327913 | 1.543154 | 4.021063 | 0.383768 | 0.701151 | NA |
| | 0.258578 | 1.233151 | 4.047809 | 0.304646 | 0.760635 | NA |
| | 0.628146 | -0.21495 | 3.102079 | -0.06929 | 0.944756 | NA |
| E2F3-IT1 | 0.163957 | 0.424913 | 4.079006 | 0.104171 | 0.917034 | NA |
| CETN4P | 10.23449 | 0.042476 | 0.647685 | 0.065581 | 0.947711 | NA |
| RPL32P1 | 0.162376 | 0.424913 | 4.079006 | 0.104171 | 0.917034 | NA |
| DUX4L9 | 0.653015 | 0.2552 | 2.787103 | 0.091565 | 0.927044 | NA |
| | 7.100219 | 0.747005 | 0.762272 | 0.979972 | 0.3271 | NA |
| | 0.162376 | 0.424913 | 4.079006 | 0.104171 | 0.917034 | NA |
| LINC00240 | 2.51156 | -0.96251 | 1.329763 | -0.72382 | 0.469178 | NA |
| | 0.096202 | 0.408546 | 4.079639 | 0.100143 | 0.920231 | NA |
| OPA1-AS1 | 0.629726 | -0.22725 | 2.732935 | -0.08315 | 0.93373 | NA |
| SUCLA2P1 | 3.010731 | -0.5376 | 1.177488 | -0.45657 | 0.647982 | NA |
| PSAT1P4 | 0.134133 | -1.48909 | 4.077584 | -0.36519 | 0.71497 | NA |
| RPL29P12 | 1.466477 | 2.270865 | 2.005054 | 1.13257 | 0.257395 | NA |
| HMGB3P2 | 1.582025 | 1.324554 | 1.679402 | 0.788706 | 0.430284 | NA |
| | 0.448206 | -1.2532 | 3.007315 | -0.41672 | 0.676885 | NA |
| RPL3P7 | 9.170146 | 0.065917 | 0.728337 | 0.090503 | 0.927887 | NA |
| SETP20 | 0.462046 | 0.589069 | 3.442439 | 0.17112 | 0.86413 | NA |
| | 8.151096 | -0.10541 | 0.80381 | -0.13113 | 0.895671 | NA |

| | | | | | | |
|---|---|---|---|---|---|---|
| | 45.74805 | 0.832225 | 0.339771 | 2.44937 | 0.014311 | NA |
| | 38.67062 | -0.243 | 0.42619 | -0.57017 | 0.56856 | NA |
| | 1.244103 | 1.47325 | 2.257332 | 0.652651 | 0.513981 | NA |
| OSTM1-AS: | 0.134133 | -1.48909 | 4.077584 | -0.36519 | 0.71497 | NA |
| | 666.5014 | 0.130334 | 0.172572 | 0.755245 | 0.450102 | 0.609421 |
| | 1.186597 | -0.04644 | 1.739087 | -0.0267 | 0.978695 | NA |
| | 1922.372 | -0.36082 | 0.175942 | -2.05081 | 0.040285 | 0.120955 |
| | 16.91443 | 0.031665 | 0.500317 | 0.063291 | 0.949535 | NA |
| | 0.288605 | 1.443619 | 4.029058 | 0.358302 | 0.720117 | NA |
| NCBP2-AS1 | 1.679458 | 2.805265 | 1.781235 | 1.574899 | 0.11528 | NA |
| | 2.514439 | 0.732437 | 1.372022 | 0.533838 | 0.593454 | NA |
| | 0.392857 | -1.546 | 4.075469 | -0.37934 | 0.704433 | NA |
| LEPREL1-AS | 6.004566 | 0.916195 | 1.055627 | 0.867915 | 0.385441 | NA |
| | 0.28583 | -2.21358 | 3.97376 | -0.55705 | 0.577494 | NA |
| | 4.962207 | -0.41815 | 0.929998 | -0.44963 | 0.652977 | NA |
| TRAM2-AS: | 36.6623 | -0.24888 | 0.374192 | -0.66511 | 0.505979 | NA |
| | 4.568476 | -0.53421 | 0.942642 | -0.56672 | 0.570906 | NA |
| UBXN7-AS1 | 3.673085 | -0.39705 | 1.039182 | -0.38207 | 0.702406 | NA |
| | 0.134133 | -1.48909 | 4.077584 | -0.36519 | 0.71497 | NA |
| | 13.76607 | -0.22975 | 0.596223 | -0.38534 | 0.699987 | NA |
| | 0.296509 | -0.51709 | 3.982155 | -0.12985 | 0.896684 | NA |
| NOL7 | 230.3944 | 0.064715 | 0.198975 | 0.325242 | 0.744998 | 0.839379 |
| | 1.809788 | -0.55855 | 1.421691 | -0.39288 | 0.69441 | NA |
| | 0.585114 | 1.12675 | 2.73598 | 0.411827 | 0.680466 | NA |
| | 0.162376 | 0.424913 | 4.079006 | 0.104171 | 0.917034 | NA |
| | 0.192403 | 0.975392 | 4.060716 | 0.240202 | 0.810174 | NA |
| | 0.315653 | -0.57305 | 3.892639 | -0.14721 | 0.882963 | NA |
| | 0.151697 | -1.546 | 4.075469 | -0.37934 | 0.704433 | NA |
| | 5.560854 | 1.355211 | 0.91678 | 1.47823 | 0.139346 | NA |
| GTF2H2B | 4.063506 | 0.24849 | 0.998654 | 0.248825 | 0.803496 | NA |
| ARHGAP26 | 0.151697 | -1.546 | 4.075469 | -0.37934 | 0.704433 | NA |
| NPY6R | 0.162376 | 0.424913 | 4.079006 | 0.104171 | 0.917034 | NA |
| ZNF192P1 | 12.43983 | 0.329711 | 0.607982 | 0.542304 | 0.587609 | NA |
| | 0.096202 | 0.408546 | 4.079639 | 0.100143 | 0.920231 | NA |
| | 2.562864 | 1.620294 | 1.339764 | 1.209387 | 0.226514 | NA |
| | 0.314073 | -0.57308 | 3.898934 | -0.14698 | 0.883146 | NA |
| | 0.9197 | -1.22346 | 2.227737 | -0.54919 | 0.582872 | NA |
| | 0.096202 | 0.408546 | 4.079639 | 0.100143 | 0.920231 | NA |
| | 2.756391 | -0.6597 | 1.168654 | -0.56449 | 0.572419 | NA |
| GAPDHP73 | 0.709944 | 0.232855 | 2.332364 | 0.099837 | 0.920474 | NA |
| | 0.151697 | -1.546 | 4.075469 | -0.37934 | 0.704433 | NA |
| KIF4B | 0.788379 | 1.583268 | 2.566172 | 0.616976 | 0.53725 | NA |
| PSMD10P2 | 0.163957 | 0.424913 | 4.079006 | 0.104171 | 0.917034 | NA |
| | 0.823499 | -2.06708 | 2.534576 | -0.81555 | 0.414756 | NA |
| FGF12-AS3 | 0.096202 | 0.408546 | 4.079639 | 0.100143 | 0.920231 | NA |
| | 0.670433 | 1.255281 | 2.356401 | 0.532711 | 0.594234 | NA |
| | 0.134133 | -1.48909 | 4.077584 | -0.36519 | 0.71497 | NA |

| | | | | | | |
|---|---|---|---|---|---|---|
| | 2.812849 | 0.924313 | 1.337776 | 0.690933 | 0.489608 | NA |
| | 0.489058 | -0.58569 | 3.427543 | -0.17088 | 0.864321 | NA |
| | 0.489058 | -0.58569 | 3.427543 | -0.17088 | 0.864321 | NA |
| ERVMER34 | 2.947222 | 0.430338 | 1.182873 | 0.363807 | 0.716002 | NA |
| | 2.692439 | 0.903518 | 1.256763 | 0.718925 | 0.472187 | NA |
| DANCR | 1156.027 | 0.146781 | 0.148473 | 0.988602 | 0.322858 | 0.490809 |
| COX6A1P2 | 40.4327 | 0.294363 | 0.332174 | 0.886173 | 0.375525 | NA |
| | 0.134133 | -1.48909 | 4.077584 | -0.36519 | 0.71497 | NA |
| WDR46 | 1817.193 | -0.11356 | 0.133753 | -0.849 | 0.395881 | 0.560513 |
| RBMX2P5 | 0.151697 | -1.546 | 4.075469 | -0.37934 | 0.704433 | NA |
| BTF3P13 | 0.382031 | -1.26484 | 3.151989 | -0.40128 | 0.688212 | NA |
| ZNF717 | 94.47693 | -0.21562 | 0.277886 | -0.77594 | 0.437786 | 0.597921 |
| | 0.977824 | 1.072535 | 2.286794 | 0.469013 | 0.639061 | NA |
| HCG15 | 7.504297 | -0.6424 | 0.766654 | -0.83792 | 0.402073 | NA |
| | 0.420954 | 1.923376 | 3.462803 | 0.555439 | 0.578594 | NA |
| HCG4B | 0.612162 | -0.19706 | 2.47693 | -0.07956 | 0.936588 | NA |
| | 1.198654 | 1.307481 | 1.836319 | 0.712012 | 0.476457 | NA |
| DLG1-AS1 | 0.517155 | 2.232561 | 3.179631 | 0.702145 | 0.482589 | NA |
| ATP5G1P4 | 2.907054 | 1.029835 | 1.344847 | 0.765764 | 0.443817 | NA |
| | 1.631867 | 2.786079 | 1.838189 | 1.515665 | 0.129604 | NA |
| LTB | 2.109486 | -1.84844 | 1.748283 | -1.05729 | 0.290379 | NA |
| | 0.162376 | 0.424913 | 4.079006 | 0.104171 | 0.917034 | NA |
| | 0.096202 | 0.408546 | 4.079639 | 0.100143 | 0.920231 | NA |
| | 1.108105 | -1.62856 | 2.120834 | -0.76789 | 0.442554 | NA |
| | 0.162376 | 0.424913 | 4.079006 | 0.104171 | 0.917034 | NA |
| | 0.258578 | 1.233151 | 4.047809 | 0.304646 | 0.760635 | NA |
| | 0.163957 | 0.424913 | 4.079006 | 0.104171 | 0.917034 | NA |
| | 0.096202 | 0.408546 | 4.079639 | 0.100143 | 0.920231 | NA |
| HCG9P5 | 0.796661 | 1.607053 | 2.50763 | 0.640865 | 0.52161 | NA |
| | 0.315653 | -0.57305 | 3.892639 | -0.14721 | 0.882963 | NA |
| HCG4P5 | 0.554095 | -2.96528 | 3.108533 | -0.95392 | 0.340127 | NA |
| | 0.192403 | 0.975392 | 4.060716 | 0.240202 | 0.810174 | NA |
| | 0.845418 | 1.095819 | 2.535106 | 0.432258 | 0.665554 | NA |
| | 0.230334 | -0.53318 | 4.076749 | -0.13079 | 0.895945 | NA |
| RPL3P2 | 16.58096 | -0.81404 | 0.512456 | -1.58851 | 0.112171 | NA |
| HCG20 | 5.958541 | -0.45983 | 0.817146 | -0.56272 | 0.573624 | NA |
| | 0.392857 | -1.546 | 4.075469 | -0.37934 | 0.704433 | NA |
| HLA-U | 90.66031 | -0.7175 | 0.268045 | -2.67681 | 0.007433 | 0.038826 |
| MFI2-AS1 | 42.65202 | -0.79831 | 0.365814 | -2.18229 | 0.029088 | NA |
| | 2.809357 | 0.073441 | 1.197082 | 0.06135 | 0.95108 | NA |
| | 0.134133 | -1.48909 | 4.077584 | -0.36519 | 0.71497 | NA |
| | 9.980403 | 0.169921 | 0.687924 | 0.247006 | 0.804904 | NA |
| NLGN1-AS1 | 0.247898 | -0.5901 | 4.074633 | -0.14482 | 0.884851 | NA |
| HCG11 | 287.2634 | 0.218684 | 0.194205 | 1.126046 | 0.260146 | 0.424389 |
| COL6A4P2 | 0.631307 | -0.20384 | 3.097925 | -0.0658 | 0.947537 | NA |
| | 0.134133 | -1.48909 | 4.077584 | -0.36519 | 0.71497 | NA |
| KATNBL1P6 | 0.096202 | 0.408546 | 4.079639 | 0.100143 | 0.920231 | NA |

| | | | | | | |
|---|---|---|---|---|---|---|
| LYPLA2P1 | 1.697051 | 0.81389 | 1.790776 | 0.45449 | 0.649476 | NA |
| | 0.151697 | -1.546 | 4.075469 | -0.37934 | 0.704433 | NA |
| | 0.134133 | -1.48909 | 4.077584 | -0.36519 | 0.71497 | NA |
| | 0.910252 | -0.05889 | 2.163226 | -0.02722 | 0.978283 | NA |
| | 0.913027 | 1.87659 | 2.13519 | 0.878887 | 0.379463 | NA |
| | 0.096202 | 0.408546 | 4.079639 | 0.100143 | 0.920231 | NA |
| PTMAP1 | 0.612162 | -0.19706 | 2.47693 | -0.07956 | 0.936588 | NA |
| TSTD3 | 23.84965 | 0.024294 | 0.458278 | 0.053013 | 0.957722 | NA |
| | 5.405245 | 0.800359 | 0.994406 | 0.804861 | 0.4209 | NA |
| LINC01013 | 21.86111 | 0.176224 | 0.486204 | 0.362448 | 0.717017 | NA |
| | 37.65015 | 0.35545 | 0.363112 | 0.9789 | 0.327629 | NA |
| | 0.394291 | 0.298745 | 3.141711 | 0.09509 | 0.924243 | NA |
| | 9.411475 | 0.057859 | 0.765753 | 0.075559 | 0.93977 | NA |
| PROB1 | 92.51709 | -0.11281 | 0.361674 | -0.31191 | 0.755107 | 0.847146 |
| DHFR | 249.3756 | 0.447936 | 0.188608 | 2.374962 | 0.017551 | 0.069262 |
| | 0.162376 | 0.424913 | 4.079006 | 0.104171 | 0.917034 | NA |
| SAPCD1 | 1.286951 | 2.396584 | 1.948979 | 1.229661 | 0.218824 | NA |
| | 0.495797 | -0.73607 | 3.324282 | -0.22142 | 0.824763 | NA |
| RPL7L1P8 | 2.07627 | -0.5899 | 1.351578 | -0.43646 | 0.662506 | NA |
| HCG22 | 0.651435 | 0.249629 | 2.788679 | 0.089515 | 0.928672 | NA |
| | 0.134133 | -1.48909 | 4.077584 | -0.36519 | 0.71497 | NA |
| | 0.623191 | -1.58103 | 2.770593 | -0.57065 | 0.568239 | NA |
| | 38.52021 | 0.349481 | 0.380253 | 0.919074 | 0.358057 | NA |
| | 2.59384 | 0.49647 | 1.262873 | 0.393128 | 0.694225 | NA |
| | 0.575812 | 0.936922 | 2.776634 | 0.337431 | 0.735792 | NA |
| | 0.096202 | 0.408546 | 4.079639 | 0.100143 | 0.920231 | NA |
| SUMO2P8 | 0.437526 | -2.68242 | 3.384147 | -0.79264 | 0.427986 | NA |
| | 8.514852 | 0.527132 | 0.751598 | 0.701348 | 0.483086 | NA |
| | 0.574231 | 0.93227 | 2.49998 | 0.372911 | 0.709215 | NA |
| | 9.016758 | 1.632965 | 0.771379 | 2.116941 | 0.034265 | NA |
| TMPOP1 | 4.654925 | -0.82433 | 1.027659 | -0.80215 | 0.422469 | NA |
| | 3.128088 | 0.477643 | 1.233967 | 0.387079 | 0.698697 | NA |
| HCG4P8 | 1.102801 | -0.56687 | 1.927854 | -0.29404 | 0.768725 | NA |
| | 0.801202 | 1.538543 | 2.914591 | 0.527876 | 0.597585 | NA |
| PTGES3P3 | 6.450147 | -0.19477 | 0.912985 | -0.21333 | 0.831068 | NA |
| REV3L-IT1 | 0.326333 | 1.533889 | 3.826878 | 0.40082 | 0.688553 | NA |
| | 0.162376 | 0.424913 | 4.079006 | 0.104171 | 0.917034 | NA |
| KRT8P12 | 103.0449 | 0.39639 | 0.254173 | 1.55953 | 0.118871 | 0.250438 |
| ACAP2-IT1 | 1.850291 | 0.46146 | 1.416703 | 0.325728 | 0.74463 | NA |
| | 10.96236 | 0.518315 | 0.649437 | 0.798099 | 0.424813 | NA |
| ACTG1P9 | 1.113481 | 0.468899 | 1.933629 | 0.242497 | 0.808395 | NA |
| MICD | 0.556813 | -0.57007 | 3.294615 | -0.17303 | 0.862627 | NA |
| HLA-DRB6 | 0.49029 | 2.123092 | 3.304282 | 0.642527 | 0.520531 | NA |
| | 16.04323 | 0.397894 | 0.510379 | 0.779605 | 0.435624 | NA |
| RPL21P44 | 0.192403 | 0.975392 | 4.060716 | 0.240202 | 0.810174 | NA |
| RPL4P4 | 18.49011 | -0.54553 | 0.539858 | -1.0105 | 0.312254 | NA |
| | 0.303394 | -2.27006 | 4.032451 | -0.56295 | 0.57347 | NA |

| | | | | | | |
|---|---|---|---|---|---|---|
| MAST4-AS1 | 1.225869 | -0.07951 | 1.838357 | -0.04325 | 0.965503 | NA |
| | 0.392711 | 0.292538 | 3.145221 | 0.09301 | 0.925895 | NA |
| EIF2S2P2 | 0.258578 | 1.233151 | 4.047809 | 0.304646 | 0.760635 | NA |
| | 0.134133 | -1.48909 | 4.077584 | -0.36519 | 0.71497 | NA |
| | 133.5213 | 0.697628 | 0.225689 | 3.091105 | 0.001994 | 0.01617 |
| | 3.707 | 0.307065 | 1.123666 | 0.273271 | 0.784645 | NA |
| | 0.151697 | -1.546 | 4.075469 | -0.37934 | 0.704433 | NA |
| | 0.134133 | -1.48909 | 4.077584 | -0.36519 | 0.71497 | NA |
| | 3.562151 | 0.161277 | 1.054983 | 0.152871 | 0.8785 | NA |
| TFAP2A-AS | 191.3245 | -0.03021 | 0.218977 | -0.13795 | 0.890278 | 0.935909 |
| LRRC37A15 | 1.378943 | 0.121795 | 1.745607 | 0.069772 | 0.944375 | NA |
| MTAPP2 | 1.900491 | 2.094705 | 1.703433 | 1.229696 | 0.218811 | NA |
| | 2.663322 | 0.149457 | 1.22865 | 0.121644 | 0.903181 | NA |
| | 4.624886 | 0.954021 | 0.979745 | 0.973745 | 0.330183 | NA |
| | 672.0349 | 0.137348 | 0.170477 | 0.805674 | 0.420431 | 0.583153 |
| FTH1P5 | 1.11225 | -1.84067 | 2.365829 | -0.77802 | 0.436554 | NA |
| XXYLT1-AS2 | 1.774668 | 0.864468 | 1.522063 | 0.567958 | 0.570063 | NA |
| | 94.89486 | -0.48561 | 0.259436 | -1.87179 | 0.061236 | 0.15955 |
| ELOVL2-AS | 0.314073 | -0.57308 | 3.898934 | -0.14698 | 0.883146 | NA |
| | 1.33457 | 0.31828 | 1.850972 | 0.171953 | 0.863475 | NA |
| | 0.163957 | 0.424913 | 4.079006 | 0.104171 | 0.917034 | NA |
| | 0.134133 | -1.48909 | 4.077584 | -0.36519 | 0.71497 | NA |
| | 0.708364 | 0.219301 | 2.144998 | 0.102238 | 0.918568 | NA |
| | 0.574231 | 0.93227 | 2.49998 | 0.372911 | 0.709215 | NA |
| PA2G4P4 | 2.485596 | 0.786393 | 1.355943 | 0.57996 | 0.561942 | NA |
| HCG4P7 | 1.003439 | -1.03737 | 2.189171 | -0.47387 | 0.635596 | NA |
| | 122.2749 | 1.169037 | 0.316463 | 3.694076 | 2.21E-04 | 0.004127 |
| | 1.100083 | -1.81673 | 2.026823 | -0.89634 | 0.37007 | NA |
| OR2H4P | 0.163957 | 0.424913 | 4.079006 | 0.104171 | 0.917034 | NA |
| | 1.018713 | 1.086497 | 2.092548 | 0.519222 | 0.603606 | NA |
| UQCRHP1 | 0.303394 | -2.27006 | 4.032451 | -0.56295 | 0.57347 | NA |
| | 2.135532 | 1.299057 | 1.458087 | 0.890932 | 0.372966 | NA |
| | 0.640406 | 1.11402 | 2.710977 | 0.410929 | 0.681124 | NA |
| HLA-K | 5.744518 | -0.40101 | 0.873098 | -0.4593 | 0.646019 | NA |
| STMND1 | 0.134133 | -1.48909 | 4.077584 | -0.36519 | 0.71497 | NA |
| | 0.399595 | -1.31595 | 3.579466 | -0.36764 | 0.713143 | NA |
| | 1.04874 | 1.185767 | 2.243995 | 0.528418 | 0.597209 | NA |
| | 10.1389 | 0.794476 | 0.665603 | 1.193618 | 0.232628 | NA |
| | 2.089668 | 0.670778 | 1.571007 | 0.426973 | 0.669399 | NA |
| | 0.260158 | 1.240347 | 4.047124 | 0.306476 | 0.759242 | NA |
| LINC00271 | 1.691436 | 0.062501 | 1.630675 | 0.038328 | 0.969426 | NA |
| | 2.333865 | -0.17768 | 1.351318 | -0.13149 | 0.895389 | NA |
| GCNT1P3 | 0.258578 | 1.233151 | 4.047809 | 0.304646 | 0.760635 | NA |
| HCG18 | 330.4346 | 0.048221 | 0.160861 | 0.29977 | 0.764353 | 0.853784 |
| | 0.260158 | 1.240347 | 4.047124 | 0.306476 | 0.759242 | NA |
| | 19.85086 | -0.22042 | 0.47254 | -0.46647 | 0.640883 | NA |
| BTF3P10 | 1.088815 | -0.29525 | 1.943541 | -0.15191 | 0.879257 | NA |

| | | | | | | |
|---|---|---|---|---|---|---|
| HLA-T | 0.489058 | -0.58569 | 3.427543 | -0.17088 | 0.864321 | NA |
| | 3.688404 | -1.17069 | 1.210494 | -0.96712 | 0.333486 | NA |
| COX11P1 | 0.096202 | 0.408546 | 4.079639 | 0.100143 | 0.920231 | NA |
| | 29.26188 | 0.195067 | 0.436508 | 0.446881 | 0.654961 | NA |
| RPL13AP | 0.096202 | 0.408546 | 4.079639 | 0.100143 | 0.920231 | NA |
| | 2.668697 | 0.909012 | 1.246004 | 0.729542 | 0.46567 | NA |
| TBL1XR1-A | 0.315653 | -0.57305 | 3.892639 | -0.14721 | 0.882963 | NA |
| | 0.47961 | 0.535796 | 3.39605 | 0.15777 | 0.874638 | NA |
| | 0.163957 | 0.424913 | 4.079006 | 0.104171 | 0.917034 | NA |
| HLA-DPA1 | 728.6409 | -0.42618 | 0.177603 | -2.39962 | 0.016412 | 0.066156 |
| | 1.196082 | -0.75368 | 1.944802 | -0.38754 | 0.698359 | NA |
| | 0.151697 | -1.546 | 4.075469 | -0.37934 | 0.704433 | NA |
| | 0.134133 | -1.48909 | 4.077584 | -0.36519 | 0.71497 | NA |
| RPS18 | 34397.79 | 0.178395 | 0.175587 | 1.015997 | 0.309631 | 0.476958 |
| | 0.162376 | 0.424913 | 4.079006 | 0.104171 | 0.917034 | NA |
| RPL7P21 | 6.88189 | -0.10157 | 0.759313 | -0.13377 | 0.893585 | NA |
| | 5.409101 | 1.532357 | 0.987967 | 1.55102 | 0.120897 | NA |
| | 2.233168 | 1.843355 | 1.445539 | 1.275202 | 0.202238 | NA |
| | 4.861749 | 0.327265 | 0.962958 | 0.339853 | 0.733967 | NA |
| | 0.260158 | 1.240347 | 4.047124 | 0.306476 | 0.759242 | NA |
| TMEM44-A | 66.54687 | 0.088811 | 0.272059 | 0.326441 | 0.744091 | 0.838915 |
| | 93.826 | -0.72656 | 0.854371 | -0.8504 | 0.395103 | 0.559611 |
| | 5.507944 | -0.59407 | 0.862485 | -0.68879 | 0.490958 | NA |
| | 0.452561 | 2.061505 | 3.349456 | 0.615474 | 0.538241 | NA |
| HMGB3P14 | 0.096202 | 0.408546 | 4.079639 | 0.100143 | 0.920231 | NA |
| MTND5P15 | 0.260158 | 1.240347 | 4.047124 | 0.306476 | 0.759242 | NA |
| TRAF3IP2-A | 63.99283 | 0.164003 | 0.288358 | 0.56875 | 0.569526 | 0.705805 |
| TAPBP | 3209.821 | -0.76564 | 0.163305 | -4.68843 | 2.75E-06 | 2.97E-04 |
| | 15.78969 | 0.904608 | 0.526836 | 1.717057 | 0.085969 | NA |
| | 10.71928 | 0.554838 | 0.664106 | 0.835466 | 0.403455 | NA |
| PPP1R2P3 | 2.889343 | 1.474422 | 1.231237 | 1.197512 | 0.231107 | NA |
| | 0.572651 | 0.928649 | 2.780261 | 0.334015 | 0.738368 | NA |
| LEF1-AS1 | 42.06996 | -0.05622 | 0.337398 | -0.16663 | 0.867663 | NA |
| SCAND3 | 0.163957 | 0.424913 | 4.079006 | 0.104171 | 0.917034 | NA |
| LINC01063 | 1.320673 | -0.77722 | 1.811537 | -0.42904 | 0.667896 | NA |
| | 0.288605 | 1.443619 | 4.029058 | 0.358302 | 0.720117 | NA |
| NCOA7-AS1 | 0.134133 | -1.48909 | 4.077584 | -0.36519 | 0.71497 | NA |
| OR2H5P | 0.096202 | 0.408546 | 4.079639 | 0.100143 | 0.920231 | NA |
| | 0.3441 | -0.0166 | 3.769682 | -0.0044 | 0.996486 | NA |
| | 1.101221 | -0.557 | 1.814706 | -0.30694 | 0.758892 | NA |
| | 0.326333 | 1.533889 | 3.826878 | 0.40082 | 0.688553 | NA |
| | 20.47729 | 0.444254 | 0.468402 | 0.948445 | 0.342903 | NA |
| | 109.3485 | 0.03267 | 0.265466 | 0.123066 | 0.902055 | 0.942689 |
| | 0.399595 | -1.31595 | 3.579466 | -0.36764 | 0.713143 | NA |
| | 3.7347 | 0.898481 | 1.318204 | 0.681595 | 0.495495 | NA |
| ATP5LP2 | 0.739768 | 1.404572 | 2.591328 | 0.542028 | 0.587799 | NA |
| GPR79 | 0.096202 | 0.408546 | 4.079639 | 0.100143 | 0.920231 | NA |

| | | | | | | |
|---|---|---|---|---|---|---|
| | 0.315653 | -0.57305 | 3.892639 | -0.14721 | 0.882963 | NA |
| KNOP1P4 | 0.096202 | 0.408546 | 4.079639 | 0.100143 | 0.920231 | NA |
| | 0.096202 | 0.408546 | 4.079639 | 0.100143 | 0.920231 | NA |
| | 0.163957 | 0.424913 | 4.079006 | 0.104171 | 0.917034 | NA |
| CCT6P4 | 0.096202 | 0.408546 | 4.079639 | 0.100143 | 0.920231 | NA |
| | 1.393832 | 3.667627 | 1.97519 | 1.856848 | 0.063333 | NA |
| | 1.454659 | -0.14314 | 1.627714 | -0.08794 | 0.929924 | NA |
| | 9.862314 | 0.332864 | 0.661776 | 0.502985 | 0.614975 | NA |
| BDH2P1 | 0.488709 | 2.119056 | 3.307564 | 0.64067 | 0.521737 | NA |
| | 0.978999 | 3.124447 | 2.610028 | 1.197093 | 0.23127 | NA |
| ETF1P1 | 0.824933 | -0.18668 | 2.233587 | -0.08358 | 0.933393 | NA |
| TNF | 1.922661 | 1.226147 | 1.926956 | 0.636313 | 0.524572 | NA |
| GSTO3P | 1.150043 | 1.185274 | 1.979154 | 0.598879 | 0.549253 | NA |
| | 0.096202 | 0.408546 | 4.079639 | 0.100143 | 0.920231 | NA |
| | 0.314073 | -0.57308 | 3.898934 | -0.14698 | 0.883146 | NA |
| | 7.669688 | -0.38914 | 0.742775 | -0.52391 | 0.600343 | NA |
| | 0.096202 | 0.408546 | 4.079639 | 0.100143 | 0.920231 | NA |
| | 0.392857 | -1.546 | 4.075469 | -0.37934 | 0.704433 | NA |
| HCG25 | 3.631486 | 0.195864 | 1.019652 | 0.192089 | 0.847673 | NA |
| | 19.85474 | 0.104139 | 0.528648 | 0.196991 | 0.843834 | NA |
| LINC00884 | 6.537965 | 0.141867 | 0.7758 | 0.182866 | 0.854903 | NA |
| | 0.134133 | -1.48909 | 4.077584 | -0.36519 | 0.71497 | NA |
| | 5.233741 | 1.018761 | 0.928801 | 1.096856 | 0.272704 | NA |
| HIST1H2AN | 0.192403 | 0.975392 | 4.060716 | 0.240202 | 0.810174 | NA |
| LINC00472 | 40.53082 | 0.308371 | 0.41606 | 0.741171 | 0.45859 | NA |
| MICF | 0.258578 | 1.233151 | 4.047809 | 0.304646 | 0.760635 | NA |
| | 154.856 | 0.605859 | 0.349648 | 1.732767 | 0.083137 | 0.196293 |
| XXYLT1-AS: | 0.096202 | 0.408546 | 4.079639 | 0.100143 | 0.920231 | NA |
| OSTN-AS1 | 0.897992 | -1.0518 | 2.16635 | -0.48552 | 0.627309 | NA |
| | 7.946065 | -0.38447 | 0.760611 | -0.50547 | 0.613228 | NA |
| | 0.792452 | -2.15617 | 2.562821 | -0.84133 | 0.400166 | NA |
| STXBP5-AS | 13.65322 | 0.662022 | 0.559392 | 1.183468 | 0.236624 | NA |
| | 0.096202 | 0.408546 | 4.079639 | 0.100143 | 0.920231 | NA |
| HNRNPLP1 | 0.93741 | -3.49975 | 2.723608 | -1.28497 | 0.198804 | NA |
| | 0.298089 | -0.51711 | 3.975232 | -0.13008 | 0.896501 | NA |
| | 2.39751 | 1.022432 | 1.321087 | 0.773932 | 0.438971 | NA |
| | 0.660976 | -0.84029 | 2.673548 | -0.3143 | 0.753294 | NA |
| UFL1-AS1 | 0.151697 | -1.546 | 4.075469 | -0.37934 | 0.704433 | NA |
| | 1.367914 | 0.791109 | 1.898548 | 0.416692 | 0.676904 | NA |
| HIST1H2BN | 7.655654 | -0.6177 | 0.862747 | -0.71597 | 0.474011 | NA |
| KCNQ5-IT1 | 1.045376 | 2.066118 | 2.09889 | 0.984386 | 0.324926 | NA |
| | 0.326333 | 1.533889 | 3.826878 | 0.40082 | 0.688553 | NA |
| ADH5P4 | 1.477948 | -0.15579 | 1.683756 | -0.09253 | 0.92628 | NA |
| YEATS2-AS: | 3.591044 | -0.07257 | 1.239442 | -0.05855 | 0.953311 | NA |
| PAIP1P1 | 2.052179 | 0.249241 | 1.35466 | 0.183988 | 0.854023 | NA |
| | 6.530629 | -0.16029 | 0.807109 | -0.1986 | 0.842573 | NA |
| ZDHHC20P | 2.084981 | 1.313786 | 1.543526 | 0.851159 | 0.394681 | NA |

| | | | | | | |
|---|---|---|---|---|---|---|
| | 44.01998 | 0.166537 | 0.350721 | 0.474843 | 0.634899 | NA |
| | 2.781524 | 3.594603 | 1.581572 | 2.272804 | 0.023038 | NA |
| | 0.768215 | 1.51173 | 2.494681 | 0.605981 | 0.544527 | NA |
| DDX39B-AS | 0.258578 | 1.233151 | 4.047809 | 0.304646 | 0.760635 | NA |
| | 0.096202 | 0.408546 | 4.079639 | 0.100143 | 0.920231 | NA |
| TXNP6 | 1.205384 | -0.56038 | 1.915155 | -0.2926 | 0.769824 | NA |
| TPRG1-AS1 | 0.315653 | -0.57305 | 3.892639 | -0.14721 | 0.882963 | NA |
| | 1.756901 | 1.38236 | 1.593708 | 0.867386 | 0.385731 | NA |
| | 0.544554 | -2.65036 | 3.220086 | -0.82307 | 0.410468 | NA |
| TRIM26 | 1664.026 | -0.31735 | 0.139588 | -2.27347 | 0.022998 | 0.08302 |
| | 2.19418 | -0.47604 | 1.606374 | -0.29634 | 0.766968 | NA |
| ETV5-AS1 | 5.298093 | 1.447839 | 0.94588 | 1.53068 | 0.125849 | NA |
| | 1.949471 | 1.180269 | 1.64521 | 0.717397 | 0.473129 | NA |
| | 0.134133 | -1.48909 | 4.077584 | -0.36519 | 0.71497 | NA |
| ZNF879 | 52.37654 | 0.311434 | 0.358292 | 0.869219 | 0.384727 | 0.550705 |
| | 4.115381 | 0.031086 | 1.033234 | 0.030086 | 0.975998 | NA |
| | 1.213646 | -0.78014 | 1.940269 | -0.40208 | 0.687627 | NA |
| | 0.314073 | -0.57308 | 3.898934 | -0.14698 | 0.883146 | NA |
| | 0.247898 | -0.5901 | 4.074633 | -0.14482 | 0.884851 | NA |
| RPL34-AS1 | 4.276196 | -1.06195 | 0.997418 | -1.0647 | 0.287011 | NA |
| C5orf58 | 0.230334 | -0.53318 | 4.076749 | -0.13079 | 0.895945 | NA |
| PPP1R2P1 | 0.151697 | -1.546 | 4.075469 | -0.37934 | 0.704433 | NA |
| | 0.258578 | 1.233151 | 4.047809 | 0.304646 | 0.760635 | NA |
| | 0.834389 | 1.638864 | 2.19944 | 0.745128 | 0.456195 | NA |
| MCIDAS | 0.47961 | 0.535796 | 3.39605 | 0.15777 | 0.874638 | NA |
| WDR82P1 | 0.411855 | 0.250456 | 3.097618 | 0.080854 | 0.935558 | NA |
| | 5.780923 | 1.219494 | 0.890504 | 1.369443 | 0.170861 | NA |
| HLA-B | 7455.81 | -0.71859 | 0.138422 | -5.1913 | 2.09E-07 | 5.27E-05 |
| | 1.661851 | -0.83498 | 1.718581 | -0.48585 | 0.62707 | NA |
| | 0.163957 | 0.424913 | 4.079006 | 0.104171 | 0.917034 | NA |
| MORF4 | 0.70693 | -1.62766 | 2.687154 | -0.60572 | 0.544701 | NA |
| | 0.096202 | 0.408546 | 4.079639 | 0.100143 | 0.920231 | NA |
| | 3.65106 | 1.249513 | 1.158914 | 1.078176 | 0.280955 | NA |
| | 0.967291 | -0.94461 | 2.160526 | -0.43721 | 0.661958 | NA |
| | 48.39735 | 0.505767 | 0.351117 | 1.440453 | 0.149739 | 0.293221 |
| EIF3EP1 | 3.403172 | 2.039051 | 1.256529 | 1.622764 | 0.10464 | NA |
| | 0.151697 | -1.546 | 4.075469 | -0.37934 | 0.704433 | NA |
| | 1.84713 | 0.442645 | 1.569134 | 0.282095 | 0.777871 | NA |
| TECRP1 | 17.74614 | -0.85265 | 0.576704 | -1.47848 | 0.139279 | NA |
| ZSCAN31 | 57.48412 | 0.05223 | 0.336848 | 0.155056 | 0.876777 | 0.926399 |
| | 0.303394 | -2.27006 | 4.032451 | -0.56295 | 0.57347 | NA |
| | 0.096202 | 0.408546 | 4.079639 | 0.100143 | 0.920231 | NA |
| | 19.80667 | 0.28715 | 0.483128 | 0.594355 | 0.552275 | NA |
| RPL39P3 | 3.498863 | -0.46199 | 1.086705 | -0.42513 | 0.670745 | NA |
| SUMO2P1 | 1.033466 | -0.7302 | 1.946441 | -0.37515 | 0.707552 | NA |
| | 0.460466 | 0.568655 | 3.014642 | 0.188631 | 0.850382 | NA |
| HLA-W | 13.50611 | -0.41201 | 0.627174 | -0.65693 | 0.511226 | NA |

| | | | | | | |
|---|---|---|---|---|---|---|
| HLA-Z | 0.556813 | -0.57007 | 3.294615 | -0.17303 | 0.862627 | NA |
| KRT18P63 | 1.607843 | -0.84781 | 1.701599 | -0.49824 | 0.618313 | NA |
| JARID2-AS1 | 0.892688 | 0.011962 | 2.434661 | 0.004913 | 0.99608 | NA |
| | 12.13133 | 0.304461 | 0.592021 | 0.514274 | 0.60706 | NA |
| | 0.163957 | 0.424913 | 4.079006 | 0.104171 | 0.917034 | NA |
| | 9.647339 | -0.60443 | 0.698154 | -0.86576 | 0.386624 | NA |
| SAPCD1-AS | 0.260158 | 1.240347 | 4.047124 | 0.306476 | 0.759242 | NA |
| | 0.268266 | -2.15396 | 4.038351 | -0.53338 | 0.593773 | NA |
| | 0.192403 | 0.975392 | 4.060716 | 0.240202 | 0.810174 | NA |
| | 2.789225 | -0.88523 | 1.536804 | -0.57602 | 0.564604 | NA |
| IFITM4P | 2.886568 | 0.701713 | 1.159402 | 0.605237 | 0.545021 | NA |
| B3GALT4 | 270.2376 | -0.34665 | 0.253655 | -1.3666 | 0.17175 | 0.322883 |
| TM4SF19-A | 0.754374 | 0.344906 | 2.513743 | 0.137208 | 0.890866 | NA |
| | 6.444567 | -0.0035 | 0.787085 | -0.00444 | 0.996456 | NA |
| | 5.392143 | -1.41197 | 0.92897 | -1.51993 | 0.128529 | NA |
| | 0.096202 | 0.408546 | 4.079639 | 0.100143 | 0.920231 | NA |
| HMGN2P28 | 0.192403 | 0.975392 | 4.060716 | 0.240202 | 0.810174 | NA |
| ZBTB22 | 693.3715 | -0.46332 | 0.159622 | -2.90259 | 0.003701 | 0.024281 |
| | 0.727508 | 0.192954 | 2.572892 | 0.074995 | 0.940219 | NA |
| | 0.961566 | -0.83657 | 2.359415 | -0.35457 | 0.722915 | NA |
| VEZF1P1 | 0.28583 | -2.21358 | 3.97376 | -0.55705 | 0.577494 | NA |
| GUSBP5 | 5.679196 | 0.736868 | 1.028749 | 0.716276 | 0.473821 | NA |
| | 4.003647 | 0.695978 | 1.093243 | 0.636618 | 0.524374 | NA |
| | 0.823352 | -0.17011 | 2.472341 | -0.06881 | 0.945144 | NA |
| | 5.717556 | -1.08812 | 0.884465 | -1.23025 | 0.218602 | NA |
| | 0.460466 | 0.568655 | 3.014642 | 0.188631 | 0.850382 | NA |
| | 2.856448 | 0.530055 | 1.408254 | 0.376392 | 0.706626 | NA |
| | 0.52699 | -2.57878 | 3.248478 | -0.79384 | 0.427288 | NA |
| | 0.230334 | -0.53318 | 4.076749 | -0.13079 | 0.895945 | NA |
| RANP1 | 3.606764 | -0.11106 | 1.023458 | -0.10852 | 0.913584 | NA |
| BZW1P1 | 0.601483 | -1.7332 | 2.729494 | -0.63499 | 0.525436 | NA |
| ARHGEF38 | 1.965694 | 0.686335 | 1.470951 | 0.466592 | 0.640792 | NA |
| LINC01010 | 25.45437 | 0.561012 | 0.563974 | 0.994748 | 0.319859 | NA |
| | 0.420954 | 1.923376 | 3.462803 | 0.555439 | 0.578594 | NA |
| | 0.489058 | -0.58569 | 3.427543 | -0.17088 | 0.864321 | NA |
| CTAGE9 | 0.151697 | -1.546 | 4.075469 | -0.37934 | 0.704433 | NA |
| COX7A2P2 | 1.168976 | -0.41512 | 1.914592 | -0.21682 | 0.82835 | NA |
| OR2AI1P | 0.096202 | 0.408546 | 4.079639 | 0.100143 | 0.920231 | NA |
| | 0.096202 | 0.408546 | 4.079639 | 0.100143 | 0.920231 | NA |
| | 0.956262 | 0.048146 | 1.9097 | 0.025211 | 0.979886 | NA |
| C5orf27 | 3.787767 | 1.642894 | 1.209595 | 1.358219 | 0.174394 | NA |
| SULT1D1P | 0.151697 | -1.546 | 4.075469 | -0.37934 | 0.704433 | NA |
| | 1.347189 | 0.804826 | 1.780891 | 0.451923 | 0.651324 | NA |
| MICG | 0.881769 | -0.20384 | 2.229339 | -0.09144 | 0.927147 | NA |
| EHMT2-AS1 | 0.162376 | 0.424913 | 4.079006 | 0.104171 | 0.917034 | NA |
| TAAR9 | 0.162376 | 0.424913 | 4.079006 | 0.104171 | 0.917034 | NA |
| | 0.727508 | 0.192954 | 2.572892 | 0.074995 | 0.940219 | NA |

| | | | | | | |
|---|---|---|---|---|---|---|
| MCFD2P1 | 0.28583 | -2.21358 | 3.97376 | -0.55705 | 0.577494 | NA |
| RPS7P15 | 0.151697 | -1.546 | 4.075469 | -0.37934 | 0.704433 | NA |
| NR2F1-AS1 | 68.20927 | 0.376588 | 0.331255 | 1.136852 | 0.2556 | 0.419747 |
| CDKN2AIPI | 820.8977 | -0.09897 | 0.201903 | -0.49018 | 0.624008 | 0.747838 |
| | 0.653015 | 0.2552 | 2.787103 | 0.091565 | 0.927044 | NA |
| | 0.488709 | 2.119056 | 3.307564 | 0.64067 | 0.521737 | NA |
| HNRNPA1P | 0.163957 | 0.424913 | 4.079006 | 0.104171 | 0.917034 | NA |
| CDC42P6 | 11.48739 | 0.377958 | 0.588433 | 0.642313 | 0.52067 | NA |
| HLA-DPA3 | 0.192403 | 0.975392 | 4.060716 | 0.240202 | 0.810174 | NA |
| | 0.163957 | 0.424913 | 4.079006 | 0.104171 | 0.917034 | NA |
| | 0.151697 | -1.546 | 4.075469 | -0.37934 | 0.704433 | NA |
| RPSAP2 | 0.134133 | -1.48909 | 4.077584 | -0.36519 | 0.71497 | NA |
| RGL2 | 2284.953 | -0.27695 | 0.136238 | -2.03285 | 0.042068 | 0.124711 |
| | 7.880549 | -0.0868 | 0.718233 | -0.12085 | 0.903809 | NA |
| | 0.296509 | -0.51709 | 3.982155 | -0.12985 | 0.896684 | NA |
| | 0.096202 | 0.408546 | 4.079639 | 0.100143 | 0.920231 | NA |
| | 0.488912 | 0.771583 | 2.897678 | 0.266276 | 0.790026 | NA |
| KIFC1 | 809.6308 | -0.45306 | 0.226166 | -2.00322 | 0.045154 | 0.129997 |
| HCG4P3 | 15.11254 | -0.26415 | 0.606221 | -0.43573 | 0.663035 | NA |
| GSTA7P | 0.258578 | 1.233151 | 4.047809 | 0.304646 | 0.760635 | NA |
| | 1.826011 | -0.91092 | 1.478551 | -0.61609 | 0.537833 | NA |
| | 0.324752 | 1.53082 | 4.022025 | 0.380609 | 0.703493 | NA |
| IRGM | 0.163957 | 0.424913 | 4.079006 | 0.104171 | 0.917034 | NA |
| | 0.260158 | 1.240347 | 4.047124 | 0.306476 | 0.759242 | NA |
| P4HA2-AS1 | 0.162376 | 0.424913 | 4.079006 | 0.104171 | 0.917034 | NA |
| | 0.096202 | 0.408546 | 4.079639 | 0.100143 | 0.920231 | NA |
| | 0.670433 | 1.255281 | 2.356401 | 0.532711 | 0.594234 | NA |
| FAM200B | 515.5418 | 0.684468 | 0.169526 | 4.037532 | 5.40E-05 | 0.001726 |
| GFOD1-AS1 | 0.162376 | 0.424913 | 4.079006 | 0.104171 | 0.917034 | NA |
| C3orf79 | 29.82975 | 0.825345 | 0.689953 | 1.196234 | 0.231605 | NA |
| | 0.584911 | 2.38756 | 2.690807 | 0.887303 | 0.374916 | NA |
| | 0.92743 | 1.834049 | 2.364191 | 0.775762 | 0.43789 | NA |
| | 0.151697 | -1.546 | 4.075469 | -0.37934 | 0.704433 | NA |
| OR2I1P | 12.8956 | -1.13386 | 0.621243 | -1.82515 | 0.067978 | NA |
| | 21.29106 | 0.451478 | 0.467037 | 0.966687 | 0.333701 | NA |
| DDX39BP2 | 0.392857 | -1.546 | 4.075469 | -0.37934 | 0.704433 | NA |
| | 0.327913 | 1.543154 | 4.021063 | 0.383768 | 0.701151 | NA |
| | 1.380515 | -2.27951 | 2.164938 | -1.05292 | 0.292376 | NA |
| | 0.260158 | 1.240347 | 4.047124 | 0.306476 | 0.759242 | NA |
| MRPL48P1 | 1.245013 | -0.14306 | 1.979798 | -0.07226 | 0.942396 | NA |
| NMNAT1P3 | 0.162376 | 0.424913 | 4.079006 | 0.104171 | 0.917034 | NA |
| | 0.096202 | 0.408546 | 4.079639 | 0.100143 | 0.920231 | NA |
| | 0.303394 | -2.27006 | 4.032451 | -0.56295 | 0.57347 | NA |
| | 0.151697 | -1.546 | 4.075469 | -0.37934 | 0.704433 | NA |
| ALG1L6P | 14.59141 | -0.65619 | 0.559116 | -1.17363 | 0.240544 | NA |
| | 1.641905 | 0.146254 | 1.634613 | 0.089473 | 0.928706 | NA |
| SNORA13 | 0.35636 | 1.709252 | 3.651833 | 0.468053 | 0.639746 | NA |

| | | | | | | |
|---|---|---|---|---|---|---|
| | 3.339839 | 0.381464 | 1.146708 | 0.33266 | 0.739391 | NA |
| | 0.151697 | -1.546 | 4.075469 | -0.37934 | 0.704433 | NA |
| | 0.392711 | 0.292538 | 3.145221 | 0.09301 | 0.925895 | NA |
| | 0.852156 | 0.629888 | 2.425889 | 0.259653 | 0.795132 | NA |
| TRNAI2 | 0.422534 | 1.924583 | 3.001758 | 0.641152 | 0.521424 | NA |
| TRNAI6 | 0.162376 | 0.424913 | 4.079006 | 0.104171 | 0.917034 | NA |
| | 0.096202 | 0.408546 | 4.079639 | 0.100143 | 0.920231 | NA |
| SCARNA7 | 1.878388 | 1.529417 | 1.509029 | 1.01351 | 0.310816 | NA |
| | 0.163957 | 0.424913 | 4.079006 | 0.104171 | 0.917034 | NA |
| SCARNA18 | 0.096202 | 0.408546 | 4.079639 | 0.100143 | 0.920231 | NA |
| | 0.096202 | 0.408546 | 4.079639 | 0.100143 | 0.920231 | NA |
| SNORA47 | 0.979202 | 1.972754 | 2.08204 | 0.94751 | 0.343379 | NA |
| | 0.556813 | -0.57007 | 3.294615 | -0.17303 | 0.862627 | NA |
| | 0.314073 | -0.57308 | 3.898934 | -0.14698 | 0.883146 | NA |
| | 1.471218 | 2.277312 | 2.055633 | 1.10784 | 0.267931 | NA |
| | 0.561971 | -0.49218 | 2.810061 | -0.17515 | 0.860962 | NA |
| | 0.735027 | 1.3952 | 2.594798 | 0.537691 | 0.59079 | NA |
| SNORD123 | 0.449786 | -1.25325 | 3.00474 | -0.41709 | 0.676613 | NA |
| RNU2-11P | 0.151697 | -1.546 | 4.075469 | -0.37934 | 0.704433 | NA |
| | 0.162376 | 0.424913 | 4.079006 | 0.104171 | 0.917034 | NA |
| GAPDHP39 | 0.096202 | 0.408546 | 4.079639 | 0.100143 | 0.920231 | NA |
| | 26.47223 | 0.4434 | 0.446298 | 0.993508 | 0.320463 | NA |
| | 0.162376 | 0.424913 | 4.079006 | 0.104171 | 0.917034 | NA |
| RN7SL589F | 0.460466 | 0.568655 | 3.014642 | 0.188631 | 0.850382 | NA |
| RN7SL271F | 0.096202 | 0.408546 | 4.079639 | 0.100143 | 0.920231 | NA |
| TXNDC5 | 208.0429 | -0.30542 | 0.228335 | -1.33759 | 0.18103 | 0.333581 |
| CLRN1-AS1 | 0.392857 | -1.546 | 4.075469 | -0.37934 | 0.704433 | NA |
| | 1.052261 | 0.993047 | 2.045208 | 0.485548 | 0.627288 | NA |
| | 0.163957 | 0.424913 | 4.079006 | 0.104171 | 0.917034 | NA |
| | 0.163957 | 0.424913 | 4.079006 | 0.104171 | 0.917034 | NA |
| RN7SL752F | 0.260158 | 1.240347 | 4.047124 | 0.306476 | 0.759242 | NA |
| PABPC1P1( | 0.392857 | -1.546 | 4.075469 | -0.37934 | 0.704433 | NA |
| ST3GAL6-A | 82.88213 | 0.110453 | 0.292012 | 0.378247 | 0.705247 | 0.810853 |
| | 0.700488 | -1.9689 | 2.983028 | -0.66003 | 0.509233 | NA |
| | 0.288605 | 1.443619 | 4.029058 | 0.358302 | 0.720117 | NA |
| RN7SL552F | 0.134133 | -1.48909 | 4.077584 | -0.36519 | 0.71497 | NA |
| | 1.23121 | -0.79704 | 1.984723 | -0.40159 | 0.687987 | NA |
| RPS15AP16 | 0.83597 | 1.639551 | 2.453777 | 0.668175 | 0.504022 | NA |
| MARK2P8 | 0.584911 | 2.38756 | 2.690807 | 0.887303 | 0.374916 | NA |
| PLCH1-AS1 | 1.7399 | 0.805252 | 1.580326 | 0.509548 | 0.610368 | NA |
| FLYWCH1P | 0.296509 | -0.51709 | 3.982155 | -0.12985 | 0.896684 | NA |
| MYLK-AS1 | 19.43096 | 0.326219 | 0.462791 | 0.704895 | 0.480876 | NA |
| | 0.582339 | -1.69412 | 2.750801 | -0.61586 | 0.537985 | NA |
| | 0.448206 | -1.2532 | 3.007315 | -0.41672 | 0.676885 | NA |
| | 0.555233 | -0.57009 | 3.296947 | -0.17291 | 0.862719 | NA |
| RPSAP41 | 0.134133 | -1.48909 | 4.077584 | -0.36519 | 0.71497 | NA |
| GUSBP4 | 0.682693 | 2.623213 | 2.518338 | 1.041645 | 0.297576 | NA |

| | | | | | | |
|---|---|---|---|---|---|---|
| PDZRN3-AS | 0.52699 | -2.57878 | 3.248478 | -0.79384 | 0.427288 | NA |
| | 0.326536 | 0.037056 | 3.842554 | 0.009644 | 0.992306 | NA |
| | 0.247898 | -0.5901 | 4.074633 | -0.14482 | 0.884851 | NA |
| RPL21P39 | 0.544554 | -2.65036 | 3.220086 | -0.82307 | 0.410468 | NA |
| | 1.683616 | -0.56017 | 1.665479 | -0.33634 | 0.736614 | NA |
| | 0.163957 | 0.424913 | 4.079006 | 0.104171 | 0.917034 | NA |
| RN7SL273F | 0.096202 | 0.408546 | 4.079639 | 0.100143 | 0.920231 | NA |
| ENPP7P2 | 6.807348 | 0.169578 | 0.790775 | 0.214446 | 0.8302 | NA |
| | 0.296509 | -0.51709 | 3.982155 | -0.12985 | 0.896684 | NA |
| RPL7P24 | 1.378797 | 0.877327 | 1.658782 | 0.528898 | 0.596876 | NA |
| | 3.312184 | -0.69391 | 1.094709 | -0.63387 | 0.526163 | NA |
| SLC9A9-AS | 0.096202 | 0.408546 | 4.079639 | 0.100143 | 0.920231 | NA |
| LINC00888 | 190.1276 | 0.16182 | 0.298409 | 0.542278 | 0.587627 | 0.720256 |
| | 3.994731 | 0.199059 | 0.979733 | 0.203177 | 0.838997 | NA |
| DDX50P2 | 1.893876 | -1.70301 | 1.672434 | -1.01828 | 0.308543 | NA |
| | 2.189889 | 0.094883 | 1.371711 | 0.069171 | 0.944853 | NA |
| | 0.162376 | 0.424913 | 4.079006 | 0.104171 | 0.917034 | NA |
| LY6G5B | 12.98889 | -0.08024 | 0.56716 | -0.14147 | 0.887499 | NA |
| | 8.198811 | 0.228459 | 0.687325 | 0.332389 | 0.739595 | NA |
| | 0.163957 | 0.424913 | 4.079006 | 0.104171 | 0.917034 | NA |
| PSMB9 | 592.0713 | -0.63923 | 0.184303 | -3.46838 | 5.24E-04 | 0.006967 |
| RPL21P42 | 0.151697 | -1.546 | 4.075469 | -0.37934 | 0.704433 | NA |
| | 16.82849 | 0.402908 | 0.527172 | 0.764282 | 0.444699 | NA |
| PCDHGC3 | 7363.149 | -0.34375 | 0.169361 | -2.0297 | 0.042387 | 0.125217 |
| | 3.03962 | -0.89323 | 1.149327 | -0.77718 | 0.437054 | NA |
| | 0.162376 | 0.424913 | 4.079006 | 0.104171 | 0.917034 | NA |
| | 0.392711 | 0.292538 | 3.145221 | 0.09301 | 0.925895 | NA |
| ACAD11 | 0.163957 | 0.424913 | 4.079006 | 0.104171 | 0.917034 | NA |
| SEC62-AS1 | 1.629295 | 0.702424 | 1.625006 | 0.432259 | 0.665553 | NA |
| | 12.70012 | -0.17972 | 0.640719 | -0.2805 | 0.779096 | NA |
| | 0.163957 | 0.424913 | 4.079006 | 0.104171 | 0.917034 | NA |
| | 0.096202 | 0.408546 | 4.079639 | 0.100143 | 0.920231 | NA |
| | 2.619601 | -0.75992 | 1.264 | -0.60121 | 0.547703 | NA |
| LINC00636 | 0.623191 | -1.58103 | 2.770593 | -0.57065 | 0.568239 | NA |
| LRRFIP1P1 | 6.567426 | 1.491078 | 0.832243 | 1.791638 | 0.073191 | NA |
| | 0.163957 | 0.424913 | 4.079006 | 0.104171 | 0.917034 | NA |
| LINC00973 | 6.341329 | 1.608617 | 0.871056 | 1.846743 | 0.064784 | NA |
| | 0.326333 | 1.533889 | 3.826878 | 0.40082 | 0.688553 | NA |
| SETP14 | 5.666565 | -1.02345 | 0.919999 | -1.11245 | 0.265946 | NA |
| | 4.800515 | -0.30897 | 0.989605 | -0.31221 | 0.754878 | NA |
| | 0.162376 | 0.424913 | 4.079006 | 0.104171 | 0.917034 | NA |
| TM4SF1-AS | 0.163957 | 0.424913 | 4.079006 | 0.104171 | 0.917034 | NA |
| RPL7P15 | 0.162376 | 0.424913 | 4.079006 | 0.104171 | 0.917034 | NA |
| RN7SL403F | 0.624422 | 1.162588 | 2.733761 | 0.425271 | 0.670639 | NA |
| | 0.230334 | -0.53318 | 4.076749 | -0.13079 | 0.895945 | NA |
| LSP1P2 | 0.162376 | 0.424913 | 4.079006 | 0.104171 | 0.917034 | NA |
| MME-AS1 | 0.096202 | 0.408546 | 4.079639 | 0.100143 | 0.920231 | NA |

| | | | | | | |
|---|---|---|---|---|---|---|
| KRT8P36 | 1.250628 | 0.827774 | 2.160628 | 0.383117 | 0.701633 | NA |
| | 0.792306 | -0.44145 | 2.21354 | -0.19943 | 0.841926 | NA |
| ISY1 | 242.0279 | 0.178096 | 0.187335 | 0.95068 | 0.341767 | 0.509897 |
| | 1.481525 | -0.01879 | 1.737485 | -0.01081 | 0.991373 | NA |
| RN7SL382F | 0.162376 | 0.424913 | 4.079006 | 0.104171 | 0.917034 | NA |
| | 0.49029 | 2.123092 | 3.304282 | 0.642527 | 0.520531 | NA |
| RN7SL502F | 0.134133 | -1.48909 | 4.077584 | -0.36519 | 0.71497 | NA |
| | 0.47961 | 0.535796 | 3.39605 | 0.15777 | 0.874638 | NA |
| PCDHGC5 | 9.795327 | 0.061802 | 0.743121 | 0.083165 | 0.93372 | NA |
| | 4.73021 | -1.45756 | 0.949298 | -1.5354 | 0.124685 | NA |
| RPL36AP21 | 0.324752 | 1.53082 | 4.022025 | 0.380609 | 0.703493 | NA |
| | 17.49293 | 0.295485 | 0.501819 | 0.588828 | 0.555977 | NA |
| | 1.204161 | -0.07212 | 1.765777 | -0.04084 | 0.967423 | NA |
| RN7SL128F | 0.584911 | 2.38756 | 2.690807 | 0.887303 | 0.374916 | NA |
| LINC00886 | 18.1732 | -0.06837 | 0.52489 | -0.13026 | 0.89636 | NA |
| | 1.194704 | -1.32372 | 1.908222 | -0.69369 | 0.487874 | NA |
| | 0.3441 | -0.0166 | 3.769682 | -0.0044 | 0.996486 | NA |
| PLCXD2 | 1.533443 | -0.70521 | 1.574317 | -0.44795 | 0.654191 | NA |
| LSAMP-AS1 | 0.52699 | -2.57878 | 3.248478 | -0.79384 | 0.427288 | NA |
| | 4.962913 | -0.68426 | 0.906595 | -0.75476 | 0.450394 | NA |
| RN7SL751F | 0.151697 | -1.546 | 4.075469 | -0.37934 | 0.704433 | NA |
| CYP51A1P1 | 0.151697 | -1.546 | 4.075469 | -0.37934 | 0.704433 | NA |
| | 0.432222 | -1.18916 | 3.482045 | -0.34151 | 0.732718 | NA |
| | 0.096202 | 0.408546 | 4.079639 | 0.100143 | 0.920231 | NA |
| RPL29P14 | 0.3441 | -0.0166 | 3.769682 | -0.0044 | 0.996486 | NA |
| HMGN1P8 | 0.162376 | 0.424913 | 4.079006 | 0.104171 | 0.917034 | NA |
| | 0.555233 | -0.57009 | 3.296947 | -0.17291 | 0.862719 | NA |
| ARHGAP31 | 2.52382 | -0.55776 | 1.44051 | -0.38719 | 0.698612 | NA |
| LINC00877 | 1.894758 | 0.474687 | 1.452938 | 0.326708 | 0.743888 | NA |
| | 0.151697 | -1.546 | 4.075469 | -0.37934 | 0.704433 | NA |
| | 0.162376 | 0.424913 | 4.079006 | 0.104171 | 0.917034 | NA |
| IQCJ-SCHIP | 0.134133 | -1.48909 | 4.077584 | -0.36519 | 0.71497 | NA |
| | 0.096202 | 0.408546 | 4.079639 | 0.100143 | 0.920231 | NA |
| | 0.314073 | -0.57308 | 3.898934 | -0.14698 | 0.883146 | NA |
| | 0.303394 | -2.27006 | 4.032451 | -0.56295 | 0.57347 | NA |
| ENPP7P4 | 5.528533 | -0.07842 | 0.849676 | -0.0923 | 0.926461 | NA |
| | 3.278019 | 0.952522 | 1.274432 | 0.747409 | 0.454817 | NA |
| | 0.134133 | -1.48909 | 4.077584 | -0.36519 | 0.71497 | NA |
| | 5.449222 | 0.794537 | 0.93127 | 0.853176 | 0.393562 | NA |
| PPATP1 | 0.382031 | -1.26484 | 3.151989 | -0.40128 | 0.688212 | NA |
| | 0.162376 | 0.424913 | 4.079006 | 0.104171 | 0.917034 | NA |
| WWTR1-AS | 16.51195 | 0.610113 | 0.521812 | 1.16922 | 0.242315 | NA |
| SETP6 | 0.096202 | 0.408546 | 4.079639 | 0.100143 | 0.920231 | NA |
| | 0.163957 | 0.424913 | 4.079006 | 0.104171 | 0.917034 | NA |
| CT64 | 0.163957 | 0.424913 | 4.079006 | 0.104171 | 0.917034 | NA |
| RPP21 | 23.6141 | 0.049336 | 0.422522 | 0.116765 | 0.907046 | NA |
| EGFL8 | 17.37276 | 0.060279 | 0.498328 | 0.120963 | 0.90372 | NA |

| | | | | | | |
|---|---|---|---|---|---|---|
| | 3.840515 | -0.83162 | 1.113759 | -0.74668 | 0.455257 | NA |
| KRT8P13 | 0.163957 | 0.424913 | 4.079006 | 0.104171 | 0.917034 | NA |
| EEF1A1P25 | 1.631225 | -0.44746 | 1.622828 | -0.27573 | 0.782756 | NA |
| | 4.497572 | 0.194235 | 1.005335 | 0.193204 | 0.846799 | NA |
| RPL7P19 | 0.697685 | -0.92563 | 2.353044 | -0.39338 | 0.694041 | NA |
| HSPA8P9 | 49.41385 | -0.01626 | 0.303864 | -0.0535 | 0.95733 | 0.976509 |
| | 0.134133 | -1.48909 | 4.077584 | -0.36519 | 0.71497 | NA |
| CBX5P1 | 0.315653 | -0.57305 | 3.892639 | -0.14721 | 0.882963 | NA |
| | 0.298089 | -0.51711 | 3.975232 | -0.13008 | 0.896501 | NA |
| GUSBP2 | 3.005106 | 1.434534 | 1.295379 | 1.107424 | 0.268111 | NA |
| | 0.46735 | -1.30193 | 3.399671 | -0.38296 | 0.701751 | NA |
| | 0.096202 | 0.408546 | 4.079639 | 0.100143 | 0.920231 | NA |
| | 0.163957 | 0.424913 | 4.079006 | 0.104171 | 0.917034 | NA |
| | 1.840989 | 0.350993 | 1.484648 | 0.236415 | 0.813111 | NA |
| RARRES2P1 | 0.134133 | -1.48909 | 4.077584 | -0.36519 | 0.71497 | NA |
| UBQLN4P1 | 3.233132 | -1.339 | 1.175417 | -1.13917 | 0.254632 | NA |
| | 3.514658 | 0.457525 | 1.182819 | 0.386809 | 0.698898 | NA |
| | 0.419962 | -2.64111 | 3.429364 | -0.77014 | 0.441214 | NA |
| | 2.942267 | 0.145592 | 1.125565 | 0.12935 | 0.897081 | NA |
| | 0.803131 | -0.76693 | 2.349822 | -0.32638 | 0.744138 | NA |
| | 18.55973 | 0.47244 | 0.718837 | 0.657228 | 0.511034 | NA |
| RPL7AP30 | 37.79676 | 0.138955 | 0.372617 | 0.372918 | 0.70921 | NA |
| | 0.327913 | 1.543154 | 4.021063 | 0.383768 | 0.701151 | NA |
| | 0.163957 | 0.424913 | 4.079006 | 0.104171 | 0.917034 | NA |
| | 3.415766 | 0.751898 | 1.147708 | 0.65513 | 0.512384 | NA |
| RPS20P4 | 0.488912 | 0.771583 | 2.897678 | 0.266276 | 0.790026 | NA |
| | 3.922809 | 0.856527 | 1.026477 | 0.834434 | 0.404037 | NA |
| WWTR1-IT | 0.476449 | 0.524262 | 3.402674 | 0.154073 | 0.877552 | NA |
| ALG1L15P | 1.140347 | 0.579353 | 2.036091 | 0.284542 | 0.775995 | NA |
| RN7SL68P | 1.695819 | -0.35492 | 1.502388 | -0.23624 | 0.813248 | NA |
| | 0.460466 | 0.568655 | 3.014642 | 0.188631 | 0.850382 | NA |
| | 2.2076 | -0.44346 | 1.312082 | -0.33799 | 0.735374 | NA |
| RPL7AP31 | 0.602678 | 1.083611 | 2.70938 | 0.399948 | 0.689195 | NA |
| LINC00969 | 47.81045 | 0.928873 | 0.339022 | 2.739863 | 0.006146 | 0.03389 |
| FOXP1-IT1 | 0.134133 | -1.48909 | 4.077584 | -0.36519 | 0.71497 | NA |
| | 0.392857 | -1.546 | 4.075469 | -0.37934 | 0.704433 | NA |
| AMACR | 58.05572 | -0.04798 | 0.289731 | -0.16561 | 0.868465 | 0.921978 |
| RPL5P13 | 1.073969 | 1.636961 | 2.132951 | 0.767463 | 0.442806 | NA |
| | 0.151697 | -1.546 | 4.075469 | -0.37934 | 0.704433 | NA |
| ABCF2P1 | 0.163957 | 0.424913 | 4.079006 | 0.104171 | 0.917034 | NA |
| | 0.162376 | 0.424913 | 4.079006 | 0.104171 | 0.917034 | NA |
| RN7SL127F | 0.163957 | 0.424913 | 4.079006 | 0.104171 | 0.917034 | NA |
| | 8.372494 | 0.616584 | 0.749537 | 0.82262 | 0.410724 | NA |
| | 0.296509 | -0.51709 | 3.982155 | -0.12985 | 0.896684 | NA |
| PVRL3-AS1 | 1.339488 | 0.884591 | 1.876659 | 0.471365 | 0.63738 | NA |
| | 3.254759 | 0.116353 | 1.062243 | 0.109535 | 0.912778 | NA |
| | 0.162376 | 0.424913 | 4.079006 | 0.104171 | 0.917034 | NA |

| | | | | | | |
|---|---|---|---|---|---|---|
| | 48.10772 | -0.0682 | 0.325809 | -0.20933 | 0.83419 | 0.900749 |
| | 0.096202 | 0.408546 | 4.079639 | 0.100143 | 0.920231 | NA |
| TFP1 | 1.378797 | 0.877327 | 1.658782 | 0.528898 | 0.596876 | NA |
| | 0.151697 | -1.546 | 4.075469 | -0.37934 | 0.704433 | NA |
| PCDHGC4 | 6.404464 | 0.661434 | 0.795048 | 0.831942 | 0.405442 | NA |
| RPL7AP11 | 0.288605 | 1.443619 | 4.029058 | 0.358302 | 0.720117 | NA |
| RBBP4P2 | 2.943404 | 0.725767 | 1.205259 | 0.602167 | 0.547063 | NA |
| | 0.99434 | -2.42645 | 2.17316 | -1.11655 | 0.264186 | NA |
| | 0.096202 | 0.408546 | 4.079639 | 0.100143 | 0.920231 | NA |
| LINC00960 | 0.688228 | -3.2761 | 2.887102 | -1.13474 | 0.256486 | NA |
| | 0.765054 | 1.506026 | 2.496832 | 0.603175 | 0.546392 | NA |
| | 0.096202 | 0.408546 | 4.079639 | 0.100143 | 0.920231 | NA |
| | 136.9521 | 0.231972 | 0.333827 | 0.694887 | 0.487126 | 0.640231 |
| POU5F1P6 | 0.3441 | -0.0166 | 3.769682 | -0.0044 | 0.996486 | NA |
| HLA-DMB | 66.55424 | 0.298524 | 0.327016 | 0.912872 | 0.36131 | 0.52861 |
| | 0.096202 | 0.408546 | 4.079639 | 0.100143 | 0.920231 | NA |
| | 0.326333 | 1.533889 | 3.826878 | 0.40082 | 0.688553 | NA |
| | 13.46575 | 0.128368 | 0.640331 | 0.200471 | 0.841113 | NA |
| LINC00971 | 0.151697 | -1.546 | 4.075469 | -0.37934 | 0.704433 | NA |
| | 3.556847 | 2.153553 | 1.308506 | 1.645811 | 0.099803 | NA |
| | 0.327913 | 1.543154 | 4.021063 | 0.383768 | 0.701151 | NA |
| | 0.163957 | 0.424913 | 4.079006 | 0.104171 | 0.917034 | NA |
| | 0.45509 | -2.71227 | 3.963744 | -0.68427 | 0.493805 | NA |
| RPL5P12 | 15.23883 | 0.599794 | 0.556628 | 1.077549 | 0.281235 | NA |
| | 0.096202 | 0.408546 | 4.079639 | 0.100143 | 0.920231 | NA |
| LINC00882 | 11.51119 | 0.910326 | 0.776053 | 1.17302 | 0.240788 | NA |
| ZBTB20-AS | 0.487129 | 2.123445 | 3.910539 | 0.543006 | 0.587126 | NA |
| SOX2-OT | 1.079716 | -1.10401 | 1.970373 | -0.56031 | 0.57527 | NA |
| | 4.620459 | -0.07983 | 0.99678 | -0.08009 | 0.936169 | NA |
| RPS26P21 | 1.284139 | 0.830248 | 2.021758 | 0.410657 | 0.681324 | NA |
| | 3.306677 | -0.17432 | 1.089825 | -0.15995 | 0.872918 | NA |
| RPL5P11 | 0.574231 | 0.93227 | 2.49998 | 0.372911 | 0.709215 | NA |
| | 0.096202 | 0.408546 | 4.079639 | 0.100143 | 0.920231 | NA |
| PLCH1-AS2 | 0.096202 | 0.408546 | 4.079639 | 0.100143 | 0.920231 | NA |
| | 0.70693 | -1.62766 | 2.687154 | -0.60572 | 0.544701 | NA |
| FTH1P23 | 34.36026 | 0.153932 | 0.423869 | 0.363158 | 0.716487 | NA |
| | 0.096202 | 0.408546 | 4.079639 | 0.100143 | 0.920231 | NA |
| RPS15AP18 | 0.163957 | 0.424913 | 4.079006 | 0.104171 | 0.917034 | NA |
| FTH1P4 | 3.011174 | -0.98124 | 1.122724 | -0.87398 | 0.382128 | NA |
| PTMAP8 | 0.326333 | 1.533889 | 3.826878 | 0.40082 | 0.688553 | NA |
| EIF4EBP3 | 61.37437 | -0.35967 | 0.356765 | -1.00813 | 0.313393 | 0.480754 |
| ARHGEF26- | 6.118693 | 0.748305 | 0.938242 | 0.797561 | 0.425125 | NA |
| LINC00870 | 0.872321 | 0.678609 | 2.011629 | 0.337343 | 0.735858 | NA |
| | 0.151697 | -1.546 | 4.075469 | -0.37934 | 0.704433 | NA |
| | 1.201429 | 3.443164 | 2.06292 | 1.669073 | 0.095103 | NA |
| | 5.441449 | 0.266912 | 0.884482 | 0.301772 | 0.762826 | NA |
| | 14.99569 | -0.17831 | 0.607237 | -0.29364 | 0.769031 | NA |

| | | | | | | |
|---|---|---|---|---|---|---|
| RN7SL558F | 0.163957 | 0.424913 | 4.079006 | 0.104171 | 0.917034 | NA |
| | 0.134133 | -1.48909 | 4.077584 | -0.36519 | 0.71497 | NA |
| | 3.054306 | 0.562517 | 1.156758 | 0.486287 | 0.626763 | NA |
| | 9.665501 | 0.550886 | 0.76821 | 0.717104 | 0.47331 | NA |
| | 0.629726 | -0.22725 | 2.732935 | -0.08315 | 0.93373 | NA |
| RN7SL174F | 0.163957 | 0.424913 | 4.079006 | 0.104171 | 0.917034 | NA |
| RN7SL738F | 2.259225 | 0.145935 | 1.31686 | 0.110821 | 0.911759 | NA |
| RN7SL815F | 0.28583 | -2.21358 | 3.97376 | -0.55705 | 0.577494 | NA |
| MCCC1-AS1 | 0.296509 | -0.51709 | 3.982155 | -0.12985 | 0.896684 | NA |
| RN7SL141F | 0.096202 | 0.408546 | 4.079639 | 0.100143 | 0.920231 | NA |
| TICAM2 | 0.163957 | 0.424913 | 4.079006 | 0.104171 | 0.917034 | NA |
| | 0.450981 | 2.057325 | 3.353149 | 0.61355 | 0.539513 | NA |
| | 0.151697 | -1.546 | 4.075469 | -0.37934 | 0.704433 | NA |
| | 21.43248 | 0.563249 | 0.573506 | 0.982116 | 0.326043 | NA |
| C4orf48 | 1795.165 | 0.018175 | 0.178322 | 0.101924 | 0.918817 | 0.953065 |
| GMFBP1 | 0.096202 | 0.408546 | 4.079639 | 0.100143 | 0.920231 | NA |
| | 1.101075 | 0.447616 | 1.809417 | 0.247381 | 0.804613 | NA |
| | 0.816825 | 1.673341 | 2.21346 | 0.755984 | 0.449659 | NA |
| HNRNPKP4 | 6.470015 | 0.625752 | 0.832768 | 0.751412 | 0.452405 | NA |
| C6orf183 | 1.146882 | 1.181239 | 1.979819 | 0.59664 | 0.550748 | NA |
| RPL17P22 | 0.247898 | -0.5901 | 4.074633 | -0.14482 | 0.884851 | NA |
| | 0.392857 | -1.546 | 4.075469 | -0.37934 | 0.704433 | NA |
| | 0.450981 | 2.057325 | 3.353149 | 0.61355 | 0.539513 | NA |
| CFB | 5.145903 | 0.238868 | 0.875845 | 0.272729 | 0.785062 | NA |
| LINC00883 | 286.3293 | 0.349236 | 0.20121 | 1.735686 | 0.082619 | 0.195427 |
| OR5V1 | 0.258578 | 1.233151 | 4.047809 | 0.304646 | 0.760635 | NA |
| HLA-L | 162.1612 | -0.20276 | 0.238361 | -0.85065 | 0.394964 | 0.559494 |
| ST13P15 | 0.476449 | 0.524262 | 3.402674 | 0.154073 | 0.877552 | NA |
| | 6.834663 | 0.804808 | 0.782207 | 1.028894 | 0.30353 | NA |
| | 0.151697 | -1.546 | 4.075469 | -0.37934 | 0.704433 | NA |
| PSMC1P7 | 0.163957 | 0.424913 | 4.079006 | 0.104171 | 0.917034 | NA |
| RPL5P17 | 0.972246 | 0.032713 | 2.31623 | 0.014124 | 0.988731 | NA |
| | 5.047223 | -0.24262 | 0.891093 | -0.27228 | 0.785409 | NA |
| TIPARP-AS1 | 7.400589 | 0.102476 | 0.755517 | 0.135637 | 0.892108 | NA |
| | 0.326333 | 1.533889 | 3.826878 | 0.40082 | 0.688553 | NA |
| RN7SL684F | 0.162376 | 0.424913 | 4.079006 | 0.104171 | 0.917034 | NA |
| VTI1BP1 | 0.708567 | -0.60605 | 2.553496 | -0.23734 | 0.812392 | NA |
| | 0.326333 | 1.533889 | 3.826878 | 0.40082 | 0.688553 | NA |
| ENO1P3 | 0.096202 | 0.408546 | 4.079639 | 0.100143 | 0.920231 | NA |
| | 0.314073 | -0.57308 | 3.898934 | -0.14698 | 0.883146 | NA |
| | 1.506338 | -0.39416 | 1.788825 | -0.22035 | 0.8256 | NA |
| RPS2P24 | 0.151697 | -1.546 | 4.075469 | -0.37934 | 0.704433 | NA |
| | 0.134133 | -1.48909 | 4.077584 | -0.36519 | 0.71497 | NA |
| FAM86DP | 339.6342 | -0.32083 | 0.158384 | -2.02567 | 0.042799 | 0.125933 |
| IL12A-AS1 | 0.134133 | -1.48909 | 4.077584 | -0.36519 | 0.71497 | NA |
| LINC01011 | 21.11607 | 0.350225 | 0.44918 | 0.779699 | 0.435568 | NA |
| | 0.151697 | -1.546 | 4.075469 | -0.37934 | 0.704433 | NA |

| | | | | | | |
|---|---|---|---|---|---|---|
| RPS4XP6 | 0.802782 | 1.535222 | 2.550022 | 0.602043 | 0.547146 | NA |
| HMGN1P1( | 0.163957 | 0.424913 | 4.079006 | 0.104171 | 0.917034 | NA |
| PDCL3P4 | 1.254352 | 0.05565 | 1.700006 | 0.032735 | 0.973886 | NA |
| ATP1B3-AS | 3.01274 | 0.761226 | 1.23108 | 0.61834 | 0.536352 | NA |
| | 0.134133 | -1.48909 | 4.077584 | -0.36519 | 0.71497 | NA |
| WWP1P1 | 0.640406 | 1.11402 | 2.710977 | 0.410929 | 0.681124 | NA |
| PBX2P1 | 2.145366 | -0.16838 | 1.383752 | -0.12169 | 0.903147 | NA |
| | 2.194145 | -1.96166 | 1.513853 | -1.29581 | 0.195042 | NA |
| PPIAP15 | 0.314073 | -0.57308 | 3.898934 | -0.14698 | 0.883146 | NA |
| | 1.235207 | 0.087213 | 1.736669 | 0.050218 | 0.959948 | NA |
| SIAH2-AS1 | 1.54826 | 0.022494 | 1.643784 | 0.013684 | 0.989082 | NA |
| | 5.086548 | -0.63702 | 0.940312 | -0.67745 | 0.498118 | NA |
| ITGB5-AS1 | 0.326333 | 1.533889 | 3.826878 | 0.40082 | 0.688553 | NA |
| | 2.069357 | 1.246945 | 1.461082 | 0.85344 | 0.393415 | NA |
| RN7SL762F | 0.096202 | 0.408546 | 4.079639 | 0.100143 | 0.920231 | NA |
| RN7SL36P | 0.247898 | -0.5901 | 4.074633 | -0.14482 | 0.884851 | NA |
| | 0.192403 | 0.975392 | 4.060716 | 0.240202 | 0.810174 | NA |
| | 0.849381 | -1.30173 | 2.226174 | -0.58474 | 0.558724 | NA |
| | 0.327913 | 1.543154 | 4.021063 | 0.383768 | 0.701151 | NA |
| | 0.192403 | 0.975392 | 4.060716 | 0.240202 | 0.810174 | NA |
| HCG16 | 4.303363 | -0.99787 | 0.962434 | -1.03682 | 0.299819 | NA |
| RPL7P23 | 2.531789 | 0.365928 | 1.246092 | 0.293661 | 0.769017 | NA |
| | 0.298089 | -0.51711 | 3.975232 | -0.13008 | 0.896501 | NA |
| ETV5 | 4625.634 | -0.76071 | 0.165824 | -4.58743 | 4.49E-06 | 3.99E-04 |
| RPL38P3 | 0.326333 | 1.533889 | 3.826878 | 0.40082 | 0.688553 | NA |
| | 10.77882 | 0.212768 | 0.629606 | 0.337937 | 0.73541 | NA |
| ERVFRD-1 | 0.326333 | 1.533889 | 3.826878 | 0.40082 | 0.688553 | NA |
| SLC9A9-AS: | 0.162376 | 0.424913 | 4.079006 | 0.104171 | 0.917034 | NA |
| KRT18P34 | 0.964516 | -3.64949 | 2.277655 | -1.6023 | 0.109089 | NA |
| | 0.452561 | 2.061505 | 3.349456 | 0.615474 | 0.538241 | NA |
| | 0.096202 | 0.408546 | 4.079639 | 0.100143 | 0.920231 | NA |
| | 0.594599 | -0.12152 | 2.771051 | -0.04385 | 0.965021 | NA |
| | 0.163957 | 0.424913 | 4.079006 | 0.104171 | 0.917034 | NA |
| | 0.545988 | -0.43913 | 2.541145 | -0.17281 | 0.862803 | NA |
| | 0.162376 | 0.424913 | 4.079006 | 0.104171 | 0.917034 | NA |
| PTCHD4 | 314.7944 | 0.358359 | 0.237396 | 1.50954 | 0.131161 | 0.267719 |
| | 0.638825 | 1.117948 | 3.106359 | 0.35989 | 0.718929 | NA |
| RN7SL47P | 0.096202 | 0.408546 | 4.079639 | 0.100143 | 0.920231 | NA |
| | 0.324752 | 1.53082 | 4.022025 | 0.380609 | 0.703493 | NA |
| C4A | 0.898195 | -1.76199 | 2.351517 | -0.7493 | 0.453676 | NA |
| | 0.260158 | 1.240347 | 4.047124 | 0.306476 | 0.759242 | NA |
| | 6.874738 | 0.518226 | 0.798065 | 0.649353 | 0.51611 | NA |
| | 3.227893 | 0.068535 | 1.084808 | 0.063177 | 0.949626 | NA |
| LIFR-AS1 | 0.392857 | -1.546 | 4.075469 | -0.37934 | 0.704433 | NA |
| LINC00847 | 705.9594 | -0.28335 | 0.169781 | -1.66893 | 0.095131 | 0.215801 |
| IGFBP7-AS: | 14.2844 | 0.025271 | 0.53954 | 0.046839 | 0.962642 | NA |
| SMARCA5-. | 1.014533 | 0.983298 | 2.05986 | 0.477361 | 0.633105 | NA |

| | | | | | | |
|---|---|---|---|---|---|---|
| LINC01024 | 16.38376 | 0.675208 | 0.528285 | 1.278111 | 0.20121 | NA |
| | 27.77862 | 0.2734 | 0.419229 | 0.65215 | 0.514305 | NA |
| SAP30L-AS1 | 9.32563 | -0.06139 | 0.720622 | -0.0852 | 0.932106 | NA |
| | 4.53809 | -0.51703 | 0.940875 | -0.54952 | 0.582649 | NA |
| | 7.155781 | -0.26424 | 0.760553 | -0.34743 | 0.728267 | NA |
| | 3.904439 | -0.38671 | 1.129753 | -0.34229 | 0.73213 | NA |
| | 0.247898 | -0.5901 | 4.074633 | -0.14482 | 0.884851 | NA |
| LINC00461 | 0.84823 | 2.93367 | 2.713279 | 1.081227 | 0.279596 | NA |
| | 255.6774 | -0.0314 | 0.23239 | -0.13512 | 0.892521 | 0.936854 |
| | 0.268266 | -2.15396 | 4.038351 | -0.53338 | 0.593773 | NA |
| | 1.241087 | 1.94541 | 2.287778 | 0.850349 | 0.395131 | NA |
| NADK2-AS1 | 2.915161 | 0.375982 | 1.176759 | 0.319506 | 0.749343 | NA |
| | 0.163957 | 0.424913 | 4.079006 | 0.104171 | 0.917034 | NA |
| | 1.91957 | 0.26705 | 1.881816 | 0.141911 | 0.887151 | NA |
| | 0.394291 | 0.298745 | 3.141711 | 0.09509 | 0.924243 | NA |
| | 0.134133 | -1.48909 | 4.077584 | -0.36519 | 0.71497 | NA |
| | 346.4864 | 0.027239 | 0.163243 | 0.166859 | 0.867481 | 0.921339 |
| | 0.891457 | -2.39215 | 2.460214 | -0.97233 | 0.330884 | NA |
| | 89.70795 | 0.172588 | 0.261461 | 0.66009 | 0.509196 | 0.657545 |
| ALDH1L1-A | 0.192403 | 0.975392 | 4.060716 | 0.240202 | 0.810174 | NA |
| NUDT16P1 | 1.754033 | 8.20E-04 | 1.86519 | 4.40E-04 | 0.999649 | NA |
| | 8.754495 | 0.524851 | 0.741322 | 0.707993 | 0.478949 | NA |
| | 1.005019 | -1.03066 | 2.033647 | -0.50681 | 0.612291 | NA |
| | 0.163957 | 0.424913 | 4.079006 | 0.104171 | 0.917034 | NA |
| PRR7-AS1 | 2.759174 | 0.908186 | 1.240891 | 0.731883 | 0.46424 | NA |
| | 9.706349 | 0.448246 | 0.737298 | 0.607958 | 0.543215 | NA |
| | 0.096202 | 0.408546 | 4.079639 | 0.100143 | 0.920231 | NA |
| | 2.035767 | -1.57535 | 1.459165 | -1.07963 | 0.280309 | NA |
| | 0.163957 | 0.424913 | 4.079006 | 0.104171 | 0.917034 | NA |
| | 56.10076 | -0.3025 | 0.292336 | -1.03477 | 0.300778 | 0.467542 |
| | 9.901308 | 0.041081 | 0.735193 | 0.055878 | 0.955439 | NA |
| RGMB-AS1 | 12.32237 | -0.50456 | 0.587921 | -0.85821 | 0.390779 | NA |
| | 0.096202 | 0.408546 | 4.079639 | 0.100143 | 0.920231 | NA |
| STARD4-AS | 1.22868 | 1.387311 | 1.789047 | 0.775447 | 0.438075 | NA |
| | 0.783638 | 1.586366 | 2.937049 | 0.540122 | 0.589113 | NA |
| | 15.24669 | -0.40519 | 0.570769 | -0.7099 | 0.477768 | NA |
| | 0.392857 | -1.546 | 4.075469 | -0.37934 | 0.704433 | NA |
| | 3.088659 | 0.295417 | 1.130612 | 0.26129 | 0.793869 | NA |
| | 0.392711 | 0.292538 | 3.145221 | 0.09301 | 0.925895 | NA |
| | 0.326536 | 0.037056 | 3.842554 | 0.009644 | 0.992306 | NA |
| MIR4458H( | 68.08379 | 0.345682 | 0.312784 | 1.105177 | 0.269083 | 0.433778 |
| CKMT2-AS1 | 347.3615 | 0.413607 | 0.198256 | 2.08623 | 0.036958 | 0.114008 |
| CPEB2-AS1 | 2.381673 | 0.616161 | 1.364308 | 0.451629 | 0.651536 | NA |
| MTND4P12 | 1953.569 | -0.66779 | 0.166844 | -4.00247 | 6.27E-05 | 0.001891 |
| | 24.05611 | -0.10328 | 0.437021 | -0.23633 | 0.813178 | NA |
| STX18-AS1 | 10.70323 | -0.08643 | 0.681943 | -0.12674 | 0.899147 | NA |
| | 0.556667 | 0.972496 | 2.522454 | 0.385536 | 0.699841 | NA |

| | | | | | | |
|---|---|---|---|---|---|---|
| | 2.096552 | 0.10239 | 1.514549 | 0.067604 | 0.946101 | NA |
| | 81.8317 | 0.021654 | 0.253634 | 0.085375 | 0.931963 | 0.961401 |
| TMEM161E | 57.7499 | 0.99056 | 0.316117 | 3.133523 | 0.001727 | 0.01471 |
| | 0.382031 | -1.26484 | 3.151989 | -0.40128 | 0.688212 | NA |
| HMGN1P1: | 0.163957 | 0.424913 | 4.079006 | 0.104171 | 0.917034 | NA |
| | 0.460466 | 0.568655 | 3.014642 | 0.188631 | 0.850382 | NA |
| SEC24B-AS | 7.916697 | -0.09054 | 0.714825 | -0.12666 | 0.899206 | NA |
| FAM13A-A: | 5.172096 | 1.828486 | 1.05034 | 1.740852 | 0.08171 | NA |
| UBA6-AS1 | 432.958 | -0.13239 | 0.182207 | -0.72662 | 0.46746 | 0.624 |
| NNT-AS1 | 107.7293 | 0.359408 | 0.258587 | 1.389891 | 0.164562 | 0.313294 |
| | 0.658173 | -0.01016 | 2.630582 | -0.00386 | 0.996919 | NA |
| | 0.258578 | 1.233151 | 4.047809 | 0.304646 | 0.760635 | NA |
| | 0.968725 | 0.233151 | 2.074813 | 0.112372 | 0.910528 | NA |
| | 0.162376 | 0.424913 | 4.079006 | 0.104171 | 0.917034 | NA |
| ADH1C | 0.296509 | -0.51709 | 3.982155 | -0.12985 | 0.896684 | NA |
| | 0.096202 | 0.408546 | 4.079639 | 0.100143 | 0.920231 | NA |
| | 0.489058 | -0.58569 | 3.427543 | -0.17088 | 0.864321 | NA |
| | 0.583919 | -1.6942 | 2.749224 | -0.61625 | 0.537732 | NA |
| | 10.15731 | -0.23632 | 0.661819 | -0.35708 | 0.721033 | NA |
| | 29.04465 | 0.916802 | 0.569734 | 1.609176 | 0.107578 | NA |
| GAPDHP60 | 1.998193 | -0.8045 | 1.519753 | -0.52936 | 0.596556 | NA |
| | 0.871935 | 2.99121 | 2.649305 | 1.129055 | 0.258875 | NA |
| PES1P1 | 0.392857 | -1.546 | 4.075469 | -0.37934 | 0.704433 | NA |
| | 0.653015 | 0.2552 | 2.787103 | 0.091565 | 0.927044 | NA |
| | 0.096202 | 0.408546 | 4.079639 | 0.100143 | 0.920231 | NA |
| | 0.28583 | -2.21358 | 3.97376 | -0.55705 | 0.577494 | NA |
| | 2.576422 | 0.114583 | 1.20406 | 0.095164 | 0.924185 | NA |
| CICP16 | 0.992613 | -0.6177 | 2.048917 | -0.30148 | 0.763051 | NA |
| | 0.881769 | -0.20384 | 2.229339 | -0.09144 | 0.927147 | NA |
| | 5.602348 | -2.16847 | 0.968295 | -2.23948 | 0.025125 | NA |
| | 0.47803 | 0.520083 | 2.97881 | 0.174594 | 0.861399 | NA |
| | 0.394291 | 0.298745 | 3.141711 | 0.09509 | 0.924243 | NA |
| OCIAD1-AS | 0.748867 | 2.740193 | 2.468928 | 1.109871 | 0.267054 | NA |
| | 0.258578 | 1.233151 | 4.047809 | 0.304646 | 0.760635 | NA |
| TRIM52-AS | 145.1184 | -0.20358 | 0.202113 | -1.00726 | 0.31381 | 0.48127 |
| | 4.366842 | -0.08054 | 1.017927 | -0.07912 | 0.936935 | NA |
| | 0.230334 | -0.53318 | 4.076749 | -0.13079 | 0.895945 | NA |
| | 0.096202 | 0.408546 | 4.079639 | 0.100143 | 0.920231 | NA |
| LINC00616 | 0.134133 | -1.48909 | 4.077584 | -0.36519 | 0.71497 | NA |
| MEF2C-AS1 | 14.10237 | -0.19507 | 0.552835 | -0.35285 | 0.7242 | NA |
| LUCAT1 | 1.982108 | 2.303502 | 1.902751 | 1.210616 | 0.226042 | NA |
| | 0.151697 | -1.546 | 4.075469 | -0.37934 | 0.704433 | NA |
| | 0.49029 | 2.123092 | 3.304282 | 0.642527 | 0.520531 | NA |
| ARL2BPP6 | 0.096202 | 0.408546 | 4.079639 | 0.100143 | 0.920231 | NA |
| | 0.303394 | -2.27006 | 4.032451 | -0.56295 | 0.57347 | NA |
| LINC00504 | 26.39613 | 0.393636 | 0.523953 | 0.751281 | 0.452484 | NA |
| | 1.66204 | 2.53215 | 1.869432 | 1.354502 | 0.175576 | NA |

| | | | | | | |
|---|---|---|---|---|---|---|
| | 0.967291 | -0.94461 | 2.160526 | -0.43721 | 0.661958 | NA |
| PCDHAC1 | 0.162376 | 0.424913 | 4.079006 | 0.104171 | 0.917034 | NA |
| FOSL1P1 | 0.162376 | 0.424913 | 4.079006 | 0.104171 | 0.917034 | NA |
| | 0.096202 | 0.408546 | 4.079639 | 0.100143 | 0.920231 | NA |
| | 0.713668 | -0.95576 | 2.626349 | -0.36391 | 0.715923 | NA |
| | 28.46628 | 0.490202 | 0.46569 | 1.052636 | 0.292508 | NA |
| PCDHGB8P | 2.716151 | -1.47835 | 1.429716 | -1.03402 | 0.301128 | NA |
| | 0.303394 | -2.27006 | 4.032451 | -0.56295 | 0.57347 | NA |
| | 19.5271 | 0.477876 | 0.506062 | 0.944304 | 0.345014 | NA |
| | 0.096202 | 0.408546 | 4.079639 | 0.100143 | 0.920231 | NA |
| | 1.564315 | 1.893687 | 1.744001 | 1.085829 | 0.277555 | NA |
| | 1.856964 | -1.24048 | 1.565636 | -0.79232 | 0.428174 | NA |
| | 4.444545 | 0.695965 | 1.059433 | 0.656921 | 0.511231 | NA |
| POU5F2 | 0.163957 | 0.424913 | 4.079006 | 0.104171 | 0.917034 | NA |
| | 5.86673 | 0.284285 | 0.830021 | 0.342504 | 0.731972 | NA |
| | 0.134133 | -1.48909 | 4.077584 | -0.36519 | 0.71497 | NA |
| | 0.58526 | 0.001197 | 3.235889 | 3.70E-04 | 0.999705 | NA |
| ANP32C | 0.410275 | 0.244283 | 3.100889 | 0.078778 | 0.937209 | NA |
| | 0.720624 | 1.435938 | 2.334155 | 0.615185 | 0.538432 | NA |
| | 0.937118 | 0.09073 | 2.07701 | 0.043683 | 0.965157 | NA |
| | 5.868853 | 0.046458 | 0.829239 | 0.056025 | 0.955322 | NA |
| | 1.576532 | -0.9628 | 1.600171 | -0.60169 | 0.547384 | NA |
| | 0.623045 | 0.091811 | 2.666142 | 0.034436 | 0.97253 | NA |
| GDNF-AS1 | 28.46136 | 0.786005 | 0.423605 | 1.855512 | 0.063523 | NA |
| | 0.096202 | 0.408546 | 4.079639 | 0.100143 | 0.920231 | NA |
| | 0.900767 | 0.808531 | 1.963257 | 0.411831 | 0.680463 | NA |
| | 2.259154 | -1.0427 | 1.434319 | -0.72696 | 0.467248 | NA |
| HSPA8P4 | 12.6107 | -0.61776 | 0.581404 | -1.06253 | 0.287993 | NA |
| | 0.096202 | 0.408546 | 4.079639 | 0.100143 | 0.920231 | NA |
| | 0.096202 | 0.408546 | 4.079639 | 0.100143 | 0.920231 | NA |
| | 0.151697 | -1.546 | 4.075469 | -0.37934 | 0.704433 | NA |
| GAPDHP40 | 1.197276 | 0.652468 | 1.752695 | 0.372265 | 0.709695 | NA |
| | 55.9349 | 0.435449 | 0.324704 | 1.341066 | 0.179899 | 0.332404 |
| HMGA1P2 | 1.021003 | -1.07763 | 2.391516 | -0.45061 | 0.652273 | NA |
| | 0.096202 | 0.408546 | 4.079639 | 0.100143 | 0.920231 | NA |
| | 0.163957 | 0.424913 | 4.079006 | 0.104171 | 0.917034 | NA |
| | 0.82213 | 0.50433 | 2.197682 | 0.229483 | 0.818494 | NA |
| | 0.260158 | 1.240347 | 4.047124 | 0.306476 | 0.759242 | NA |
| LINC00992 | 36.86071 | 1.26502 | 0.707409 | 1.788245 | 0.073736 | NA |
| | 1.639101 | 0.604773 | 1.647095 | 0.367176 | 0.713488 | NA |
| OXCT1-AS1 | 2.915025 | -2.22698 | 1.443804 | -1.54244 | 0.122967 | NA |
| | 0.749217 | 0.7397 | 2.633289 | 0.280903 | 0.778784 | NA |
| | 0.411855 | 0.250456 | 3.097618 | 0.080854 | 0.935558 | NA |
| TOX4P1 | 0.298089 | -0.51711 | 3.975232 | -0.13008 | 0.896501 | NA |
| LINC01085 | 1.710986 | -0.83266 | 1.793773 | -0.46419 | 0.642509 | NA |
| | 0.134133 | -1.48909 | 4.077584 | -0.36519 | 0.71497 | NA |
| | 12.03058 | -0.42061 | 0.634219 | -0.66319 | 0.507207 | NA |

| | | | | | | |
|---|---|---|---|---|---|---|
| NPHP3-AS1 | 1.67044 | -1.14501 | 2.427098 | -0.47176 | 0.637097 | NA |
| | 1.074172 | 0.44736 | 2.099228 | 0.213107 | 0.831244 | NA |
| EGFLAM-AS | 0.392857 | -1.546 | 4.075469 | -0.37934 | 0.704433 | NA |
| | 3.154391 | 0.913608 | 1.180311 | 0.774041 | 0.438907 | NA |
| | 0.134133 | -1.48909 | 4.077584 | -0.36519 | 0.71497 | NA |
| | 6.998969 | 0.572561 | 0.789295 | 0.725408 | 0.468202 | NA |
| | 1.310839 | 1.430611 | 1.872879 | 0.763857 | 0.444953 | NA |
| | 0.785218 | 1.577495 | 2.568228 | 0.614235 | 0.53906 | NA |
| | 0.260158 | 1.240347 | 4.047124 | 0.306476 | 0.759242 | NA |
| HIGD1AP14 | 0.162376 | 0.424913 | 4.079006 | 0.104171 | 0.917034 | NA |
| | 4.287167 | 0.839222 | 1.036486 | 0.80968 | 0.418124 | NA |
| | 3.74075 | 0.087807 | 1.078755 | 0.081397 | 0.935127 | NA |
| | 0.652869 | 1.292592 | 2.374873 | 0.544278 | 0.58625 | NA |
| | 0.440301 | 0.468383 | 3.449808 | 0.135771 | 0.892003 | NA |
| | 0.096202 | 0.408546 | 4.079639 | 0.100143 | 0.920231 | NA |
| | 0.162376 | 0.424913 | 4.079006 | 0.104171 | 0.917034 | NA |
| PCBP2P3 | 0.162376 | 0.424913 | 4.079006 | 0.104171 | 0.917034 | NA |
| | 0.326536 | 0.037056 | 3.842554 | 0.009644 | 0.992306 | NA |
| | 0.162376 | 0.424913 | 4.079006 | 0.104171 | 0.917034 | NA |
| | 0.096202 | 0.408546 | 4.079639 | 0.100143 | 0.920231 | NA |
| | 1.2271 | 1.384124 | 1.759418 | 0.786694 | 0.431461 | NA |
| USP46-AS1 | 57.32345 | -0.04902 | 0.328298 | -0.14933 | 0.881296 | 0.929351 |
| | 0.096202 | 0.408546 | 4.079639 | 0.100143 | 0.920231 | NA |
| SERBP1P6 | 0.766634 | 1.516849 | 2.262991 | 0.670285 | 0.502676 | NA |
| C5orf17 | 19.43644 | 0.160608 | 0.555771 | 0.288982 | 0.772595 | NA |
| | 0.192403 | 0.975392 | 4.060716 | 0.240202 | 0.810174 | NA |
| HHIP-AS1 | 0.593018 | -0.10389 | 3.151334 | -0.03297 | 0.973702 | NA |
| | 0.268266 | -2.15396 | 4.038351 | -0.53338 | 0.593773 | NA |
| HMGB1P21 | 0.911593 | 1.257757 | 2.250933 | 0.558771 | 0.576318 | NA |
| NUP210P3 | 0.448206 | -1.2532 | 3.007315 | -0.41672 | 0.676885 | NA |
| MTND5P11 | 5.232345 | -1.11846 | 0.932379 | -1.19958 | 0.230302 | NA |
| | 12.7377 | -0.13311 | 0.646498 | -0.20589 | 0.836878 | NA |
| | 5.012229 | 1.637029 | 1.027239 | 1.593621 | 0.111021 | NA |
| | 0.681461 | 0.491544 | 3.066278 | 0.160307 | 0.87264 | NA |
| | 28.29816 | -0.06456 | 0.417571 | -0.1546 | 0.877138 | NA |
| | 0.134133 | -1.48909 | 4.077584 | -0.36519 | 0.71497 | NA |
| | 0.488912 | 0.771583 | 2.897678 | 0.266276 | 0.790026 | NA |
| | 0.574231 | 0.93227 | 2.49998 | 0.372911 | 0.709215 | NA |
| | 0.162376 | 0.424913 | 4.079006 | 0.104171 | 0.917034 | NA |
| | 0.823352 | -0.17011 | 2.472341 | -0.06881 | 0.945144 | NA |
| KRT8P46 | 1.802729 | 2.106935 | 1.642366 | 1.282866 | 0.199539 | NA |
| LAMTOR3P | 0.134133 | -1.48909 | 4.077584 | -0.36519 | 0.71497 | NA |
| | 2.006518 | -0.86451 | 1.680207 | -0.51453 | 0.606882 | NA |
| | 0.096202 | 0.408546 | 4.079639 | 0.100143 | 0.920231 | NA |
| CLUHP4 | 0.420954 | 1.923376 | 3.462803 | 0.555439 | 0.578594 | NA |
| | 0.163957 | 0.424913 | 4.079006 | 0.104171 | 0.917034 | NA |
| | 0.489058 | -0.58569 | 3.427543 | -0.17088 | 0.864321 | NA |

| | | | | | | |
|---|---|---|---|---|---|---|
| FTH1P21 | 0.230334 | -0.53318 | 4.076749 | -0.13079 | 0.895945 | NA |
| HMGN2P4 | 0.354779 | 1.703954 | 3.657232 | 0.465914 | 0.641277 | NA |
| | 3.239621 | 1.280837 | 1.16409 | 1.10029 | 0.271206 | NA |
| | 3.153603 | -0.25614 | 1.108678 | -0.23103 | 0.817289 | NA |
| SUMO2P6 | 0.659753 | -0.00581 | 2.629089 | -0.00221 | 0.998237 | NA |
| | 1.483309 | -0.45049 | 1.77171 | -0.25427 | 0.799286 | NA |
| | 49.77682 | 0.146642 | 0.327088 | 0.448324 | 0.653919 | 0.771441 |
| COX6B1P5 | 0.354779 | 1.703954 | 3.657232 | 0.465914 | 0.641277 | NA |
| TBCAP3 | 0.298089 | -0.51711 | 3.975232 | -0.13008 | 0.896501 | NA |
| | 0.162376 | 0.424913 | 4.079006 | 0.104171 | 0.917034 | NA |
| | 0.623191 | -1.58103 | 2.770593 | -0.57065 | 0.568239 | NA |
| | 0.162376 | 0.424913 | 4.079006 | 0.104171 | 0.917034 | NA |
| | 0.134133 | -1.48909 | 4.077584 | -0.36519 | 0.71497 | NA |
| | 0.163957 | 0.424913 | 4.079006 | 0.104171 | 0.917034 | NA |
| | 0.096202 | 0.408546 | 4.079639 | 0.100143 | 0.920231 | NA |
| MTND6P4 | 1619.758 | -0.49666 | 0.190183 | -2.61146 | 0.009016 | 0.044166 |
| | 0.392857 | -1.546 | 4.075469 | -0.37934 | 0.704433 | NA |
| | 0.392711 | 0.292538 | 3.145221 | 0.09301 | 0.925895 | NA |
| SUDS3P1 | 5.046017 | 1.334355 | 0.971955 | 1.372857 | 0.169797 | NA |
| | 0.268266 | -2.15396 | 4.038351 | -0.53338 | 0.593773 | NA |
| | 0.151697 | -1.546 | 4.075469 | -0.37934 | 0.704433 | NA |
| PCDHA11 | 0.247898 | -0.5901 | 4.074633 | -0.14482 | 0.884851 | NA |
| | 4.865057 | 0.093651 | 0.900732 | 0.103972 | 0.917192 | NA |
| | 0.163957 | 0.424913 | 4.079006 | 0.104171 | 0.917034 | NA |
| | 0.162376 | 0.424913 | 4.079006 | 0.104171 | 0.917034 | NA |
| | 0.258578 | 1.233151 | 4.047809 | 0.304646 | 0.760635 | NA |
| | 0.162376 | 0.424913 | 4.079006 | 0.104171 | 0.917034 | NA |
| HSPD1P5 | 0.563552 | -0.48645 | 2.808029 | -0.17324 | 0.862465 | NA |
| | 0.883146 | 1.112834 | 2.554178 | 0.435692 | 0.663061 | NA |
| ATP1B1P1 | 0.3441 | -0.0166 | 3.769682 | -0.0044 | 0.996486 | NA |
| | 0.911593 | 1.257757 | 2.250933 | 0.558771 | 0.576318 | NA |
| | 0.162376 | 0.424913 | 4.079006 | 0.104171 | 0.917034 | NA |
| | 0.448206 | -1.2532 | 3.007315 | -0.41672 | 0.676885 | NA |
| | 0.422737 | 0.518872 | 3.501356 | 0.148192 | 0.882191 | NA |
| | 0.134133 | -1.48909 | 4.077584 | -0.36519 | 0.71497 | NA |
| | 0.629929 | -1.1413 | 2.677641 | -0.42623 | 0.669937 | NA |
| TMEM150( | 20.6002 | -0.62364 | 0.490349 | -1.27182 | 0.203436 | NA |
| | 2.744139 | -0.05867 | 1.249087 | -0.04697 | 0.962535 | NA |
| | 0.35636 | 1.709252 | 3.651833 | 0.468053 | 0.639746 | NA |
| | 4.451739 | 0.381338 | 1.000333 | 0.381211 | 0.703047 | NA |
| | 1.848564 | 1.007892 | 1.528455 | 0.659419 | 0.509627 | NA |
| ATP5LP3 | 0.163957 | 0.424913 | 4.079006 | 0.104171 | 0.917034 | NA |
| EEF1A1P9 | 19.97686 | 0.025286 | 0.536699 | 0.047114 | 0.962422 | NA |
| PDLIM1P4 | 10.0401 | -0.19342 | 0.678527 | -0.28505 | 0.775603 | NA |
| | 0.258578 | 1.233151 | 4.047809 | 0.304646 | 0.760635 | NA |
| | 5.070142 | 1.115229 | 0.992928 | 1.123172 | 0.261364 | NA |
| | 0.096202 | 0.408546 | 4.079639 | 0.100143 | 0.920231 | NA |

| | | | | | | |
|---|---|---|---|---|---|---|
| FTH1P24 | 0.192403 | 0.975392 | 4.060716 | 0.240202 | 0.810174 | NA |
| | 0.487129 | 2.123445 | 3.910539 | 0.543006 | 0.587126 | NA |
| | 2.61566 | -0.99899 | 1.32438 | -0.75431 | 0.450664 | NA |
| LINC01088 | 3.366367 | -0.20305 | 1.22708 | -0.16548 | 0.868568 | NA |
| | 0.163957 | 0.424913 | 4.079006 | 0.104171 | 0.917034 | NA |
| | 0.392857 | -1.546 | 4.075469 | -0.37934 | 0.704433 | NA |
| UGDH-AS1 | 11.61041 | -0.49234 | 0.620219 | -0.79382 | 0.4273 | NA |
| | 0.258578 | 1.233151 | 4.047809 | 0.304646 | 0.760635 | NA |
| NPM1P27 | 48.56559 | 0.738941 | 0.412799 | 1.790075 | 0.073442 | 0.180671 |
| | 5.455529 | 0.307004 | 0.926017 | 0.331532 | 0.740243 | NA |
| | 0.288605 | 1.443619 | 4.029058 | 0.358302 | 0.720117 | NA |
| | 6.515389 | -0.04509 | 0.781394 | -0.05771 | 0.953981 | NA |
| | 0.315653 | -0.57305 | 3.892639 | -0.14721 | 0.882963 | NA |
| | 1.255778 | -0.9021 | 2.16945 | -0.41582 | 0.677543 | NA |
| | 0.516164 | -1.70268 | 2.844782 | -0.59853 | 0.549488 | NA |
| LINC01060 | 0.260158 | 1.240347 | 4.047124 | 0.306476 | 0.759242 | NA |
| | 0.35636 | 1.709252 | 3.651833 | 0.468053 | 0.639746 | NA |
| LINC00500 | 0.326536 | 0.037056 | 3.842554 | 0.009644 | 0.992306 | NA |
| | 0.151697 | -1.546 | 4.075469 | -0.37934 | 0.704433 | NA |
| | 1.338237 | -0.80071 | 1.980184 | -0.40436 | 0.685947 | NA |
| | 1.311042 | 0.799739 | 1.708573 | 0.468074 | 0.639731 | NA |
| NAIP | 7.840282 | 0.441871 | 0.729643 | 0.605599 | 0.544781 | NA |
| HMGN1P14 | 0.163957 | 0.424913 | 4.079006 | 0.104171 | 0.917034 | NA |
| | 0.151697 | -1.546 | 4.075469 | -0.37934 | 0.704433 | NA |
| MLLT10P2 | 0.288605 | 1.443619 | 4.029058 | 0.358302 | 0.720117 | NA |
| LINC01091 | 0.296509 | -0.51709 | 3.982155 | -0.12985 | 0.896684 | NA |
| RBMXP4 | 1.189169 | 2.260443 | 1.954925 | 1.156281 | 0.247566 | NA |
| | 2.572552 | 0.892075 | 1.33224 | 0.669605 | 0.50311 | NA |
| | 0.47961 | 0.535796 | 3.39605 | 0.15777 | 0.874638 | NA |
| RBBP4P1 | 2.953692 | -0.24328 | 1.224822 | -0.19862 | 0.842559 | NA |
| | 6.909105 | -0.34483 | 0.777142 | -0.44372 | 0.657246 | NA |
| | 0.258578 | 1.233151 | 4.047809 | 0.304646 | 0.760635 | NA |
| | 3.198565 | -1.29594 | 1.216539 | -1.06527 | 0.286753 | NA |
| | 0.260158 | 1.240347 | 4.047124 | 0.306476 | 0.759242 | NA |
| MIR302B | 0.260158 | 1.240347 | 4.047124 | 0.306476 | 0.759242 | NA |
| | 0.601483 | -1.7332 | 2.729494 | -0.63499 | 0.525436 | NA |
| RCC2P4 | 0.151697 | -1.546 | 4.075469 | -0.37934 | 0.704433 | NA |
| SERBP1P5 | 4.062902 | 0.276069 | 1.081682 | 0.255222 | 0.798551 | NA |
| | 0.787148 | -0.70094 | 2.357021 | -0.29738 | 0.766174 | NA |
| | 17.54684 | 0.644772 | 0.517901 | 1.244971 | 0.213143 | NA |
| | 0.096202 | 0.408546 | 4.079639 | 0.100143 | 0.920231 | NA |
| | 4.686145 | 1.887386 | 1.078233 | 1.750443 | 0.080042 | NA |
| | 0.612365 | -1.10341 | 2.696105 | -0.40926 | 0.682349 | NA |
| | 0.260158 | 1.240347 | 4.047124 | 0.306476 | 0.759242 | NA |
| | 0.163957 | 0.424913 | 4.079006 | 0.104171 | 0.917034 | NA |
| | 7.866782 | -0.19265 | 0.72669 | -0.26511 | 0.790924 | NA |
| | 0.258578 | 1.233151 | 4.047809 | 0.304646 | 0.760635 | NA |

| | | | | | | |
|---|---|---|---|---|---|---|
| MRPS33P2 | 0.392857 | -1.546 | 4.075469 | -0.37934 | 0.704433 | NA |
| | 0.268266 | -2.15396 | 4.038351 | -0.53338 | 0.593773 | NA |
| | 0.35636 | 1.709252 | 3.651833 | 0.468053 | 0.639746 | NA |
| TNRC18P1 | 0.096202 | 0.408546 | 4.079639 | 0.100143 | 0.920231 | NA |
| NOP14-AS1 | 354.0448 | -0.26105 | 0.17707 | -1.47428 | 0.140406 | 0.280568 |
| | 8.945823 | 0.206303 | 0.710873 | 0.290211 | 0.771655 | NA |
| | 2.920398 | 2.848391 | 1.413805 | 2.014699 | 0.043936 | NA |
| | 0.134133 | -1.48909 | 4.077584 | -0.36519 | 0.71497 | NA |
| | 0.163957 | 0.424913 | 4.079006 | 0.104171 | 0.917034 | NA |
| | 0.134133 | -1.48909 | 4.077584 | -0.36519 | 0.71497 | NA |
| THEGL | 9.27551 | -0.45226 | 0.837802 | -0.53982 | 0.589322 | NA |
| SRD5A3-AS | 13.68893 | 0.571346 | 0.569321 | 1.003556 | 0.315593 | NA |
| | 0.162376 | 0.424913 | 4.079006 | 0.104171 | 0.917034 | NA |
| | 0.151697 | -1.546 | 4.075469 | -0.37934 | 0.704433 | NA |
| | 0.096202 | 0.408546 | 4.079639 | 0.100143 | 0.920231 | NA |
| | 3.533851 | -0.20236 | 1.030113 | -0.19645 | 0.84426 | NA |
| | 0.162376 | 0.424913 | 4.079006 | 0.104171 | 0.917034 | NA |
| | 0.558248 | 0.982076 | 2.804738 | 0.350149 | 0.726227 | NA |
| | 0.662557 | -0.83483 | 2.671964 | -0.31244 | 0.754706 | NA |
| | 4.757731 | 0.351212 | 0.994487 | 0.353159 | 0.723969 | NA |
| | 0.354779 | 1.703954 | 3.657232 | 0.465914 | 0.641277 | NA |
| ENPP7P10 | 0.518736 | 2.237423 | 2.7713 | 0.807355 | 0.419462 | NA |
| HSPE1P10 | 0.162376 | 0.424913 | 4.079006 | 0.104171 | 0.917034 | NA |
| | 0.230334 | -0.53318 | 4.076749 | -0.13079 | 0.895945 | NA |
| | 0.979348 | 1.397336 | 2.229938 | 0.626625 | 0.530905 | NA |
| | 0.558248 | 0.982076 | 2.804738 | 0.350149 | 0.726227 | NA |
| | 0.260158 | 1.240347 | 4.047124 | 0.306476 | 0.759242 | NA |
| | 0.096202 | 0.408546 | 4.079639 | 0.100143 | 0.920231 | NA |
| | 0.163957 | 0.424913 | 4.079006 | 0.104171 | 0.917034 | NA |
| | 7.657888 | 0.240229 | 0.748511 | 0.320942 | 0.748254 | NA |
| | 0.151697 | -1.546 | 4.075469 | -0.37934 | 0.704433 | NA |
| | 0.096202 | 0.408546 | 4.079639 | 0.100143 | 0.920231 | NA |
| | 67.31888 | -0.08634 | 0.278609 | -0.3099 | 0.75664 | 0.848063 |
| PGBD4P3 | 0.508056 | 0.731032 | 2.861912 | 0.255435 | 0.798387 | NA |
| VCAN-AS1 | 0.096202 | 0.408546 | 4.079639 | 0.100143 | 0.920231 | NA |
| | 0.586491 | 2.39072 | 2.689057 | 0.889055 | 0.373973 | NA |
| | 9.589087 | 0.343634 | 0.680132 | 0.505247 | 0.613386 | NA |
| KRT18P31 | 0.450981 | 2.057325 | 3.353149 | 0.61355 | 0.539513 | NA |
| HS3ST5 | 3.486617 | -0.04339 | 1.059191 | -0.04097 | 0.967324 | NA |
| EEF1A1P19 | 17.16125 | 0.335673 | 0.57098 | 0.587889 | 0.556607 | NA |
| SNX5P1 | 1.009228 | 2.053856 | 2.077371 | 0.98868 | 0.322819 | NA |
| | 6.713467 | 0.100053 | 1.108258 | 0.090279 | 0.928065 | NA |
| | 0.163957 | 0.424913 | 4.079006 | 0.104171 | 0.917034 | NA |
| | 0.096202 | 0.408546 | 4.079639 | 0.100143 | 0.920231 | NA |
| | 0.432222 | -1.18916 | 3.482045 | -0.34151 | 0.732718 | NA |
| PDCD6 | 3064.535 | 0.246853 | 0.147108 | 1.678044 | 0.093339 | 0.213265 |
| HNRNPA1P | 0.28583 | -2.21358 | 3.97376 | -0.55705 | 0.577494 | NA |

| | | | | | | |
|---|---|---|---|---|---|---|
| | 1.036277 | 1.02959 | 1.932342 | 0.53282 | 0.594158 | NA |
| | 0.134133 | -1.48909 | 4.077584 | -0.36519 | 0.71497 | NA |
| RAC1P2 | 39.2415 | 0.161742 | 0.332238 | 0.486827 | 0.626381 | NA |
| | 0.49029 | 2.123092 | 3.304282 | 0.642527 | 0.520531 | NA |
| XBP1P1 | 1.537449 | 1.844769 | 1.773573 | 1.040142 | 0.298274 | NA |
| GBA3 | 0.163957 | 0.424913 | 4.079006 | 0.104171 | 0.917034 | NA |
| | 0.315653 | -0.57305 | 3.892639 | -0.14721 | 0.882963 | NA |
| | 0.303394 | -2.27006 | 4.032451 | -0.56295 | 0.57347 | NA |
| CDC42P4 | 0.586694 | 1.130467 | 2.734203 | 0.413454 | 0.679274 | NA |
| | 0.69625 | -3.13723 | 2.93845 | -1.06765 | 0.285679 | NA |
| | 0.260158 | 1.240347 | 4.047124 | 0.306476 | 0.759242 | NA |
| | 3.321364 | -2.28275 | 1.350286 | -1.69057 | 0.090919 | NA |
| | 0.096202 | 0.408546 | 4.079639 | 0.100143 | 0.920231 | NA |
| | 3.552792 | -0.00282 | 1.066512 | -0.00264 | 0.997892 | NA |
| | 0.698879 | 1.36795 | 2.596167 | 0.526912 | 0.598255 | NA |
| SNX18P23 | 0.134133 | -1.48909 | 4.077584 | -0.36519 | 0.71497 | NA |
| | 0.163957 | 0.424913 | 4.079006 | 0.104171 | 0.917034 | NA |
| | 0.653015 | 0.2552 | 2.787103 | 0.091565 | 0.927044 | NA |
| SLC7A11-A | 19.84753 | 0.587333 | 0.481804 | 1.219028 | 0.222834 | NA |
| | 0.727508 | 0.192954 | 2.572892 | 0.074995 | 0.940219 | NA |
| | 4.588738 | -2.13792 | 1.202899 | -1.7773 | 0.075518 | NA |
| CBX3P3 | 0.260158 | 1.240347 | 4.047124 | 0.306476 | 0.759242 | NA |
| | 0.326536 | 0.037056 | 3.842554 | 0.009644 | 0.992306 | NA |
| LINC01018 | 0.096202 | 0.408546 | 4.079639 | 0.100143 | 0.920231 | NA |
| | 0.134133 | -1.48909 | 4.077584 | -0.36519 | 0.71497 | NA |
| | 1.36103 | 1.54208 | 1.777738 | 0.867439 | 0.385701 | NA |
| | 0.134133 | -1.48909 | 4.077584 | -0.36519 | 0.71497 | NA |
| | 0.394291 | 0.298745 | 3.141711 | 0.09509 | 0.924243 | NA |
| | 35.9847 | 0.562047 | 0.42462 | 1.323648 | 0.18562 | NA |
| | 1.111754 | 1.266994 | 1.992225 | 0.635969 | 0.524797 | NA |
| | 0.162376 | 0.424913 | 4.079006 | 0.104171 | 0.917034 | NA |
| | 0.151697 | -1.546 | 4.075469 | -0.37934 | 0.704433 | NA |
| | 0.392711 | 0.292538 | 3.145221 | 0.09301 | 0.925895 | NA |
| PCDHA10 | 0.392711 | 0.292538 | 3.145221 | 0.09301 | 0.925895 | NA |
| | 0.314073 | -0.57308 | 3.898934 | -0.14698 | 0.883146 | NA |
| | 1.331409 | 0.291422 | 1.683059 | 0.17315 | 0.862533 | NA |
| | 2.896228 | 1.036532 | 1.223345 | 0.847293 | 0.396832 | NA |
| | 0.151697 | -1.546 | 4.075469 | -0.37934 | 0.704433 | NA |
| | 2.636447 | -0.93085 | 1.316035 | -0.70732 | 0.479371 | NA |
| | 0.326333 | 1.533889 | 3.826878 | 0.40082 | 0.688553 | NA |
| | 0.327913 | 1.543154 | 4.021063 | 0.383768 | 0.701151 | NA |
| | 3.336614 | 0.327401 | 1.168255 | 0.280248 | 0.779287 | NA |
| MYLK-AS2 | 0.096202 | 0.408546 | 4.079639 | 0.100143 | 0.920231 | NA |
| EEF1A1P13 | 5386.05 | 0.087772 | 0.163397 | 0.537171 | 0.59115 | 0.722706 |
| | 0.134133 | -1.48909 | 4.077584 | -0.36519 | 0.71497 | NA |
| | 0.247898 | -0.5901 | 4.074633 | -0.14482 | 0.884851 | NA |
| HMGN1P1! | 0.495797 | -0.73607 | 3.324282 | -0.22142 | 0.824763 | NA |

| | | | | | | |
|---|---|---|---|---|---|---|
| | 7.569306 | -0.44638 | 0.768714 | -0.58068 | 0.561455 | NA |
| | 0.392857 | -1.546 | 4.075469 | -0.37934 | 0.704433 | NA |
| | 0.268266 | -2.15396 | 4.038351 | -0.53338 | 0.593773 | NA |
| CIR1P2 | 0.163957 | 0.424913 | 4.079006 | 0.104171 | 0.917034 | NA |
| LTV1P1 | 0.096202 | 0.408546 | 4.079639 | 0.100143 | 0.920231 | NA |
| | 4.86571 | 0.181157 | 0.974447 | 0.185908 | 0.852517 | NA |
| | 6.052527 | 0.274691 | 0.89477 | 0.306996 | 0.758846 | NA |
| TRIM60P14 | 1.373493 | 1.652235 | 1.822255 | 0.906698 | 0.364567 | NA |
| | 0.134133 | -1.48909 | 4.077584 | -0.36519 | 0.71497 | NA |
| | 1.239506 | 1.943067 | 2.060745 | 0.942896 | 0.345734 | NA |
| | 0.096202 | 0.408546 | 4.079639 | 0.100143 | 0.920231 | NA |
| | 6.926749 | 0.422972 | 0.751619 | 0.562747 | 0.573607 | NA |
| | 2.461076 | 0.011697 | 1.250833 | 0.009351 | 0.992539 | NA |
| PTTG2 | 0.096202 | 0.408546 | 4.079639 | 0.100143 | 0.920231 | NA |
| | 0.776119 | 0.401378 | 2.295521 | 0.174853 | 0.861196 | NA |
| ZBED1P1 | 0.151697 | -1.546 | 4.075469 | -0.37934 | 0.704433 | NA |
| ZNF718 | 37.12175 | 0.728183 | 0.450697 | 1.615681 | 0.106163 | NA |
| SMIM20 | 623.9003 | 0.375211 | 0.152093 | 2.466982 | 0.013626 | 0.058084 |
| | 7.779649 | 0.319224 | 0.763634 | 0.418033 | 0.675923 | NA |
| IGBP1P4 | 0.096202 | 0.408546 | 4.079639 | 0.100143 | 0.920231 | NA |
| | 9.002948 | -0.12068 | 0.668747 | -0.18046 | 0.85679 | NA |
| | 43.87041 | -0.07525 | 0.363277 | -0.20715 | 0.835893 | NA |
| KDELC1P1 | 0.151697 | -1.546 | 4.075469 | -0.37934 | 0.704433 | NA |
| LINC01021 | 40.27848 | 1.12577 | 0.383458 | 2.935837 | 0.003326 | NA |
| | 0.162376 | 0.424913 | 4.079006 | 0.104171 | 0.917034 | NA |
| | 12.60104 | -0.66478 | 0.600678 | -1.10671 | 0.268418 | NA |
| EEF1GP2 | 0.096202 | 0.408546 | 4.079639 | 0.100143 | 0.920231 | NA |
| | 11.63557 | 0.350364 | 0.594659 | 0.589184 | 0.555738 | NA |
| UNC93B4 | 0.162376 | 0.424913 | 4.079006 | 0.104171 | 0.917034 | NA |
| UBE2CP3 | 0.420954 | 1.923376 | 3.462803 | 0.555439 | 0.578594 | NA |
| | 0.096202 | 0.408546 | 4.079639 | 0.100143 | 0.920231 | NA |
| MTND6P2 | 0.151697 | -1.546 | 4.075469 | -0.37934 | 0.704433 | NA |
| | 0.756975 | 0.468225 | 2.554602 | 0.183287 | 0.854573 | NA |
| | 2.77262 | 1.045132 | 1.334725 | 0.783031 | 0.433609 | NA |
| | 0.096202 | 0.408546 | 4.079639 | 0.100143 | 0.920231 | NA |
| KLHL2P1 | 0.719246 | 0.444943 | 2.542118 | 0.175028 | 0.861057 | NA |
| | 0.354779 | 1.703954 | 3.657232 | 0.465914 | 0.641277 | NA |
| CLSTN2-AS | 0.544554 | -2.65036 | 3.220086 | -0.82307 | 0.410468 | NA |
| | 0.324752 | 1.53082 | 4.022025 | 0.380609 | 0.703493 | NA |
| EIF3KP3 | 0.163957 | 0.424913 | 4.079006 | 0.104171 | 0.917034 | NA |
| | 7.092863 | -0.59847 | 0.834255 | -0.71737 | 0.473146 | NA |
| CCT5P1 | 1.198857 | 0.654362 | 1.752472 | 0.373393 | 0.708856 | NA |
| | 0.192403 | 0.975392 | 4.060716 | 0.240202 | 0.810174 | NA |
| GMPSP1 | 1.174337 | -0.74927 | 1.902857 | -0.39376 | 0.693757 | NA |
| DUTP7 | 0.247898 | -0.5901 | 4.074633 | -0.14482 | 0.884851 | NA |
| FAM218A | 15.29816 | 0.613108 | 0.637789 | 0.961303 | 0.3364 | NA |
| INTS6P1 | 0.774888 | -2.09879 | 2.346327 | -0.8945 | 0.371054 | NA |

| | | | | | | |
|---|---|---|---|---|---|---|
| | 63.31501 | 0.346133 | 0.279546 | 1.238196 | 0.215643 | 0.374847 |
| | 0.47803 | 0.520083 | 2.97881 | 0.174594 | 0.861399 | NA |
| | 0.096202 | 0.408546 | 4.079639 | 0.100143 | 0.920231 | NA |
| | 0.526843 | -0.38719 | 2.855892 | -0.13558 | 0.892156 | NA |
| CCT6P2 | 0.354779 | 1.703954 | 3.657232 | 0.465914 | 0.641277 | NA |
| KRT8P33 | 8.881109 | 0.357502 | 0.689798 | 0.51827 | 0.60427 | NA |
| | 2.803393 | 0.311363 | 1.198606 | 0.259771 | 0.79504 | NA |
| | 0.640406 | 1.11402 | 2.710977 | 0.410929 | 0.681124 | NA |
| | 0.624422 | 1.162588 | 2.733761 | 0.425271 | 0.670639 | NA |
| IQCJ-SCHIP | 91.42227 | 0.903258 | 0.393465 | 2.29565 | 0.021696 | 0.080091 |
| | 1.708291 | 1.296247 | 1.620313 | 0.799998 | 0.423712 | NA |
| | 1.809439 | 0.428428 | 1.452082 | 0.295044 | 0.76796 | NA |
| | 0.163957 | 0.424913 | 4.079006 | 0.104171 | 0.917034 | NA |
| | 0.163957 | 0.424913 | 4.079006 | 0.104171 | 0.917034 | NA |
| | 4.377696 | 0.572006 | 0.972224 | 0.588348 | 0.556299 | NA |
| | 0.163957 | 0.424913 | 4.079006 | 0.104171 | 0.917034 | NA |
| | 0.449786 | -1.25325 | 3.00474 | -0.41709 | 0.676613 | NA |
| | 5.270819 | -0.29879 | 0.878263 | -0.3402 | 0.733702 | NA |
| | 0.258578 | 1.233151 | 4.047809 | 0.304646 | 0.760635 | NA |
| PABPC1P7 | 0.392711 | 0.292538 | 3.145221 | 0.09301 | 0.925895 | NA |
| ST3GAL1P1 | 0.738188 | 1.39874 | 2.317793 | 0.603479 | 0.54619 | NA |
| | 0.162376 | 0.424913 | 4.079006 | 0.104171 | 0.917034 | NA |
| | 0.162376 | 0.424913 | 4.079006 | 0.104171 | 0.917034 | NA |
| SEPP1 | 68.06511 | 0.358362 | 0.320327 | 1.118736 | 0.263253 | 0.427592 |
| | 0.303394 | -2.27006 | 4.032451 | -0.56295 | 0.57347 | NA |
| | 0.298089 | -0.51711 | 3.975232 | -0.13008 | 0.896501 | NA |
| | 1.005865 | 3.174325 | 2.260631 | 1.404176 | 0.160266 | NA |
| | 0.134133 | -1.48909 | 4.077584 | -0.36519 | 0.71497 | NA |
| SNHG18 | 272.7895 | -0.0052 | 0.214864 | -0.02418 | 0.980705 | 0.989974 |
| | 0.163957 | 0.424913 | 4.079006 | 0.104171 | 0.917034 | NA |
| | 1.483106 | -0.03797 | 1.580038 | -0.02403 | 0.98083 | NA |
| ZBED3-AS1 | 44.33995 | 0.035771 | 0.352086 | 0.101597 | 0.919077 | NA |
| | 0.134133 | -1.48909 | 4.077584 | -0.36519 | 0.71497 | NA |
| DCAF13P2 | 1.016113 | 0.982108 | 2.268429 | 0.432946 | 0.665054 | NA |
| | 0.623191 | -1.58103 | 2.770593 | -0.57065 | 0.568239 | NA |
| | 0.163957 | 0.424913 | 4.079006 | 0.104171 | 0.917034 | NA |
| | 0.506476 | 0.726418 | 2.864186 | 0.253621 | 0.799788 | NA |
| | 15.65864 | 0.01456 | 0.562414 | 0.025888 | 0.979347 | NA |
| | 0.987455 | -0.99643 | 2.195011 | -0.45395 | 0.649863 | NA |
| RNF138P1 | 0.096202 | 0.408546 | 4.079639 | 0.100143 | 0.920231 | NA |
| HNRNPKP1 | 2.6145 | -0.63073 | 1.219449 | -0.51722 | 0.605001 | NA |
| | 0.489058 | -0.58569 | 3.427543 | -0.17088 | 0.864321 | NA |
| | 3.180231 | -0.87995 | 1.157109 | -0.76047 | 0.446972 | NA |
| | 1.285719 | 0.832387 | 2.021451 | 0.411777 | 0.680503 | NA |
| RNPS1P1 | 7.619247 | -1.06877 | 0.749662 | -1.42567 | 0.153962 | NA |
| | 1.211116 | 1.435192 | 1.926921 | 0.744811 | 0.456386 | NA |
| GMDS-AS1 | 35.19599 | 0.231493 | 0.379747 | 0.609599 | 0.542128 | NA |

| | | | | | | | |
|---|---|---|---|---|---|---|---|
| | 0.134133 | -1.48909 | 4.077584 | -0.36519 | 0.71497 | NA | |
| | 1.293624 | 0.063882 | 1.680651 | 0.038011 | 0.969679 | NA | |
| | 0.392711 | 0.292538 | 3.145221 | 0.09301 | 0.925895 | NA | |
| | 0.134133 | -1.48909 | 4.077584 | -0.36519 | 0.71497 | NA | |
| | 1.568628 | -0.38598 | 1.564893 | -0.24665 | 0.80518 | NA | |
| | 1.800543 | -0.38901 | 1.612386 | -0.24126 | 0.80935 | NA | |
| | 0.298089 | -0.51711 | 3.975232 | -0.13008 | 0.896501 | NA | |
| | 3.073539 | 0.074083 | 1.305487 | 0.056747 | 0.954747 | NA | |
| | 0.151697 | -1.546 | 4.075469 | -0.37934 | 0.704433 | NA | |
| | 2.140622 | 0.264634 | 1.544978 | 0.171287 | 0.863998 | NA | |
| | 1.353125 | 2.465848 | 1.88624 | 1.307282 | 0.191117 | NA | |
| ORAOV1P1 | 0.422534 | 1.924583 | 3.001758 | 0.641152 | 0.521424 | NA | |
| | 0.230334 | -0.53318 | 4.076749 | -0.13079 | 0.895945 | NA | |
| | 0.096202 | 0.408546 | 4.079639 | 0.100143 | 0.920231 | NA | |
| | 2.185089 | 0.343291 | 1.364346 | 0.251616 | 0.801338 | NA | |
| | 0.258578 | 1.233151 | 4.047809 | 0.304646 | 0.760635 | NA | |
| THAP9-AS1 | 110.2629 | 1.356176 | 0.300056 | 4.519747 | 6.19E-06 | | 4.77E-04 |
| | 2.084991 | 2.265563 | 1.69856 | 1.333814 | 0.182265 | NA | |
| | 0.163957 | 0.424913 | 4.079006 | 0.104171 | 0.917034 | NA | |
| | 4.896424 | 0.290666 | 0.910783 | 0.319139 | 0.749621 | NA | |
| | 4.206855 | 0.741075 | 0.975815 | 0.759442 | 0.447588 | NA | |
| | 0.399595 | -1.31595 | 3.579466 | -0.36764 | 0.713143 | NA | |
| | 0.096202 | 0.408546 | 4.079639 | 0.100143 | 0.920231 | NA | |
| HTT-AS | 1.361379 | 0.169816 | 1.648967 | 0.102983 | 0.917976 | NA | |
| ALG1L3P | 0.134133 | -1.48909 | 4.077584 | -0.36519 | 0.71497 | NA | |
| | 0.354779 | 1.703954 | 3.657232 | 0.465914 | 0.641277 | NA | |
| CTBP2P4 | 0.449786 | -1.25325 | 3.00474 | -0.41709 | 0.676613 | NA | |
| | 0.162376 | 0.424913 | 4.079006 | 0.104171 | 0.917034 | NA | |
| FCF1P8 | 0.260158 | 1.240347 | 4.047124 | 0.306476 | 0.759242 | NA | |
| | 0.192403 | 0.975392 | 4.060716 | 0.240202 | 0.810174 | NA | |
| | 0.163957 | 0.424913 | 4.079006 | 0.104171 | 0.917034 | NA | |
| | 0.162376 | 0.424913 | 4.079006 | 0.104171 | 0.917034 | NA | |
| | 0.327913 | 1.543154 | 4.021063 | 0.383768 | 0.701151 | NA | |
| | 17.78236 | 1.161027 | 0.524867 | 2.212042 | 0.026964 | NA | |
| | 0.096202 | 0.408546 | 4.079639 | 0.100143 | 0.920231 | NA | |
| | 0.096202 | 0.408546 | 4.079639 | 0.100143 | 0.920231 | NA | |
| HULC | 0.979202 | 1.972754 | 2.08204 | 0.94751 | 0.343379 | NA | |
| | 0.163957 | 0.424913 | 4.079006 | 0.104171 | 0.917034 | NA | |
| | 0.258578 | 1.233151 | 4.047809 | 0.304646 | 0.760635 | NA | |
| | 0.422534 | 1.924583 | 3.001758 | 0.641152 | 0.521424 | NA | |
| TMED7-TIC | 0.096202 | 0.408546 | 4.079639 | 0.100143 | 0.920231 | NA | |
| | 0.162376 | 0.424913 | 4.079006 | 0.104171 | 0.917034 | NA | |
| GOLGA5P1 | 0.392857 | -1.546 | 4.075469 | -0.37934 | 0.704433 | NA | |
| | 0.163957 | 0.424913 | 4.079006 | 0.104171 | 0.917034 | NA | |
| | 1.033116 | 1.025241 | 1.933005 | 0.530387 | 0.595844 | NA | |
| | 0.134133 | -1.48909 | 4.077584 | -0.36519 | 0.71497 | NA | |
| | 0.394291 | 0.298745 | 3.141711 | 0.09509 | 0.924243 | NA | |

| | | | | | | |
|---|---|---|---|---|---|---|
| | 0.163957 | 0.424913 | 4.079006 | 0.104171 | 0.917034 | NA |
| | 0.163957 | 0.424913 | 4.079006 | 0.104171 | 0.917034 | NA |
| GTF2F2P1 | 5.473331 | -0.37392 | 0.874241 | -0.42771 | 0.66886 | NA |
| RFPL4B | 0.096202 | 0.408546 | 4.079639 | 0.100143 | 0.920231 | NA |
| | 0.163957 | 0.424913 | 4.079006 | 0.104171 | 0.917034 | NA |
| | 0.314073 | -0.57308 | 3.898934 | -0.14698 | 0.883146 | NA |
| ALG1L7P | 0.151697 | -1.546 | 4.075469 | -0.37934 | 0.704433 | NA |
| | 0.163957 | 0.424913 | 4.079006 | 0.104171 | 0.917034 | NA |
| | 0.298089 | -0.51711 | 3.975232 | -0.13008 | 0.896501 | NA |
| ALG1L2 | 22.2078 | -0.28888 | 0.447044 | -0.6462 | 0.518151 | NA |
| | 0.162376 | 0.424913 | 4.079006 | 0.104171 | 0.917034 | NA |
| | 0.258578 | 1.233151 | 4.047809 | 0.304646 | 0.760635 | NA |
| TUBB7P | 0.288605 | 1.443619 | 4.029058 | 0.358302 | 0.720117 | NA |
| | 1.101075 | 0.447616 | 1.809417 | 0.247381 | 0.804613 | NA |
| | 0.556813 | -0.57007 | 3.294615 | -0.17303 | 0.862627 | NA |
| | 0.134133 | -1.48909 | 4.077584 | -0.36519 | 0.71497 | NA |
| | 2.772563 | 0.997863 | 1.334056 | 0.747992 | 0.454465 | NA |
| | 0.315653 | -0.57305 | 3.892639 | -0.14721 | 0.882963 | NA |
| | 1.919773 | -0.14214 | 1.626033 | -0.08742 | 0.93034 | NA |
| | 0.258578 | 1.233151 | 4.047809 | 0.304646 | 0.760635 | NA |
| RTN3P1 | 0.28583 | -2.21358 | 3.97376 | -0.55705 | 0.577494 | NA |
| HSPD1P11 | 3.925032 | 2.497054 | 1.195761 | 2.088256 | 0.036775 | NA |
| WWC2-AS2 | 23.68843 | 0.012812 | 0.442278 | 0.028968 | 0.97689 | NA |
| | 0.163957 | 0.424913 | 4.079006 | 0.104171 | 0.917034 | NA |
| DBIP2 | 0.096202 | 0.408546 | 4.079639 | 0.100143 | 0.920231 | NA |
| | 0.55687 | -0.14239 | 3.170029 | -0.04492 | 0.964172 | NA |
| LINC00499 | 0.596179 | -0.09275 | 3.146759 | -0.02948 | 0.976485 | NA |
| | 0.151697 | -1.546 | 4.075469 | -0.37934 | 0.704433 | NA |
| | 0.517155 | 2.232561 | 3.179631 | 0.702145 | 0.482589 | NA |
| ALDH7A1P1 | 0.968522 | 0.920016 | 1.915592 | 0.480278 | 0.63103 | NA |
| | 0.392711 | 0.292538 | 3.145221 | 0.09301 | 0.925895 | NA |
| | 0.151697 | -1.546 | 4.075469 | -0.37934 | 0.704433 | NA |
| | 2.339443 | 0.284033 | 1.324263 | 0.214484 | 0.83017 | NA |
| | 1.803318 | 0.520739 | 1.666337 | 0.312505 | 0.754657 | NA |
| | 0.096202 | 0.408546 | 4.079639 | 0.100143 | 0.920231 | NA |
| | 40.91463 | 0.770586 | 0.347412 | 2.218073 | 0.02655 | NA |
| RTEL1P1 | 0.870886 | -0.53073 | 2.310896 | -0.22966 | 0.818353 | NA |
| LINC01094 | 0.718052 | -2.03248 | 2.581778 | -0.78724 | 0.431142 | NA |
| | 0.096202 | 0.408546 | 4.079639 | 0.100143 | 0.920231 | NA |
| HAUS1P1 | 0.462046 | 0.589069 | 3.442439 | 0.17112 | 0.86413 | NA |
| | 0.870886 | -0.53073 | 2.310896 | -0.22966 | 0.818353 | NA |
| | 0.163957 | 0.424913 | 4.079006 | 0.104171 | 0.917034 | NA |
| | 0.392857 | -1.546 | 4.075469 | -0.37934 | 0.704433 | NA |
| FKBP4P1 | 0.096202 | 0.408546 | 4.079639 | 0.100143 | 0.920231 | NA |
| | 0.860207 | -2.14053 | 2.512764 | -0.85186 | 0.394291 | NA |
| RPL32P3 | 120.2086 | 0.53018 | 0.230039 | 2.30474 | 0.021181 | 0.078699 |
| | 0.151697 | -1.546 | 4.075469 | -0.37934 | 0.704433 | NA |

| | | | | | | |
|---|---|---|---|---|---|---|
| LYPLA1P2 | 0.162376 | 0.424913 | 4.079006 | 0.104171 | 0.917034 | NA |
| | 0.096202 | 0.408546 | 4.079639 | 0.100143 | 0.920231 | NA |
| FOXD1 | 0.096202 | 0.408546 | 4.079639 | 0.100143 | 0.920231 | NA |
| PPIAP11 | 2.790424 | 0.696829 | 1.22324 | 0.569658 | 0.568909 | NA |
| LINC01099 | 0.296509 | -0.51709 | 3.982155 | -0.12985 | 0.896684 | NA |
| | 0.162376 | 0.424913 | 4.079006 | 0.104171 | 0.917034 | NA |
| | 0.163957 | 0.424913 | 4.079006 | 0.104171 | 0.917034 | NA |
| | 0.151697 | -1.546 | 4.075469 | -0.37934 | 0.704433 | NA |
| | 0.260158 | 1.240347 | 4.047124 | 0.306476 | 0.759242 | NA |
| | 0.998346 | 1.943196 | 2.103833 | 0.923646 | 0.355671 | NA |
| | 0.3441 | -0.0166 | 3.769682 | -0.0044 | 0.996486 | NA |
| | 0.162376 | 0.424913 | 4.079006 | 0.104171 | 0.917034 | NA |
| | 0.668852 | 1.246964 | 2.610519 | 0.477669 | 0.632886 | NA |
| DDX3YP3 | 0.46735 | -1.30193 | 3.399671 | -0.38296 | 0.701751 | NA |
| | 0.162376 | 0.424913 | 4.079006 | 0.104171 | 0.917034 | NA |
| | 0.151697 | -1.546 | 4.075469 | -0.37934 | 0.704433 | NA |
| | 19.14379 | 0.108988 | 0.482948 | 0.225673 | 0.821456 | NA |
| | 0.096202 | 0.408546 | 4.079639 | 0.100143 | 0.920231 | NA |
| | 6.1007 | 0.499387 | 1.311484 | 0.38078 | 0.703366 | NA |
| TET2-AS1 | 0.260158 | 1.240347 | 4.047124 | 0.306476 | 0.759242 | NA |
| LDHAP1 | 0.689366 | -1.56115 | 2.700123 | -0.57818 | 0.563145 | NA |
| MSNP1 | 3.783295 | -0.98583 | 1.117732 | -0.88199 | 0.37778 | NA |
| ABCA11P | 5.070142 | 1.115229 | 0.992928 | 1.123172 | 0.261364 | NA |
| HADHAP1 | 0.879205 | -0.27615 | 2.437194 | -0.11331 | 0.909788 | NA |
| | 1.307521 | 0.965064 | 2.104457 | 0.458581 | 0.646535 | NA |
| | 1.595437 | -0.5998 | 1.62242 | -0.3697 | 0.711608 | NA |
| | 0.134133 | -1.48909 | 4.077584 | -0.36519 | 0.71497 | NA |
| | 0.096202 | 0.408546 | 4.079639 | 0.100143 | 0.920231 | NA |
| | 0.192403 | 0.975392 | 4.060716 | 0.240202 | 0.810174 | NA |
| | 4.936326 | 0.132181 | 1.110924 | 0.118983 | 0.905289 | NA |
| | 0.490493 | 0.776209 | 2.895261 | 0.268096 | 0.788625 | NA |
| ZNF346-IT1 | 0.924655 | -0.07191 | 2.416297 | -0.02976 | 0.976257 | NA |
| | 0.556667 | 0.972496 | 2.522454 | 0.385536 | 0.699841 | NA |
| FAM86EP | 13.59733 | -0.6405 | 0.576352 | -1.1113 | 0.266441 | NA |
| | 38.50134 | -0.08549 | 0.400281 | -0.21356 | 0.830888 | NA |
| | 0.096202 | 0.408546 | 4.079639 | 0.100143 | 0.920231 | NA |
| | 0.163957 | 0.424913 | 4.079006 | 0.104171 | 0.917034 | NA |
| | 0.296509 | -0.51709 | 3.982155 | -0.12985 | 0.896684 | NA |
| | 1.154989 | -0.15951 | 1.966205 | -0.08112 | 0.935343 | NA |
| | 0.163957 | 0.424913 | 4.079006 | 0.104171 | 0.917034 | NA |
| RNU7-175F | 0.151697 | -1.546 | 4.075469 | -0.37934 | 0.704433 | NA |
| RNU6-1053 | 0.315653 | -0.57305 | 3.892639 | -0.14721 | 0.882963 | NA |
| RNA5SP217 | 0.520316 | 2.239969 | 3.173769 | 0.705776 | 0.480328 | NA |
| RNU7-47P | 0.096202 | 0.408546 | 4.079639 | 0.100143 | 0.920231 | NA |
| RNU6-583F | 0.134133 | -1.48909 | 4.077584 | -0.36519 | 0.71497 | NA |
| RNA5SP162 | 0.162376 | 0.424913 | 4.079006 | 0.104171 | 0.917034 | NA |
| | 0.487129 | 2.123445 | 3.910539 | 0.543006 | 0.587126 | NA |

| | | | | | | |
|---|---|---|---|---|---|---|
| | 0.315653 | -0.57305 | 3.892639 | -0.14721 | 0.882963 | NA |
| | 3.175037 | -1.03161 | 1.175742 | -0.87741 | 0.380262 | NA |
| RNU7-18P | 0.422534 | 1.924583 | 3.001758 | 0.641152 | 0.521424 | NA |
| | 1.071251 | -0.2597 | 1.986911 | -0.1307 | 0.89601 | NA |
| | 0.410275 | 0.244283 | 3.100889 | 0.078778 | 0.937209 | NA |
| RNU6-736F | 0.410275 | 0.244283 | 3.100889 | 0.078778 | 0.937209 | NA |
| RMRPP2 | 12.87014 | -1.09949 | 0.594682 | -1.84886 | 0.064478 | NA |
| RN7SKP70 | 0.755955 | 0.348448 | 2.512583 | 0.138681 | 0.889702 | NA |
| RNU6-1016 | 0.641986 | 1.117767 | 2.709494 | 0.412537 | 0.679946 | NA |
| RNU6-531F | 1.602044 | 1.906086 | 1.769986 | 1.076893 | 0.281528 | NA |
| RNU6-1098 | 0.28583 | -2.21358 | 3.97376 | -0.55705 | 0.577494 | NA |
| RNU6ATAC | 0.258578 | 1.233151 | 4.047809 | 0.304646 | 0.760635 | NA |
| RNU6-678F | 0.3441 | -0.0166 | 3.769682 | -0.0044 | 0.996486 | NA |
| RN7SKP209 | 0.151697 | -1.546 | 4.075469 | -0.37934 | 0.704433 | NA |
| RNU6-850F | 0.151697 | -1.546 | 4.075469 | -0.37934 | 0.704433 | NA |
| RNU6-1142 | 0.163957 | 0.424913 | 4.079006 | 0.104171 | 0.917034 | NA |
| RNU6-1096 | 0.096202 | 0.408546 | 4.079639 | 0.100143 | 0.920231 | NA |
| | 1.189518 | 0.629177 | 2.053952 | 0.306325 | 0.759357 | NA |
| RN7SKP234 | 0.096202 | 0.408546 | 4.079639 | 0.100143 | 0.920231 | NA |
| RN7SKP64 | 0.096202 | 0.408546 | 4.079639 | 0.100143 | 0.920231 | NA |
| | 0.995534 | 0.10093 | 2.147755 | 0.046993 | 0.962518 | NA |
| HNRNPA3P | 0.096202 | 0.408546 | 4.079639 | 0.100143 | 0.920231 | NA |
| CIR1P1 | 0.247898 | -0.5901 | 4.074633 | -0.14482 | 0.884851 | NA |
| PCDHGA12 | 1553.47 | -0.4157 | 0.167359 | -2.48385 | 0.012997 | 0.05644 |
| | 0.411855 | 0.250456 | 3.097618 | 0.080854 | 0.935558 | NA |
| GUSBP3 | 1.408759 | -1.33322 | 1.927461 | -0.6917 | 0.489126 | NA |
| | 0.324752 | 1.53082 | 4.022025 | 0.380609 | 0.703493 | NA |
| | 24.16838 | 0.786246 | 0.490343 | 1.60346 | 0.108833 | NA |
| | 0.392857 | -1.546 | 4.075469 | -0.37934 | 0.704433 | NA |
| | 0.326333 | 1.533889 | 3.826878 | 0.40082 | 0.688553 | NA |
| PCDHGB6 | 95.87025 | -0.48018 | 0.271743 | -1.76704 | 0.077222 | 0.186792 |
| | 4.331357 | 1.063246 | 0.977113 | 1.08815 | 0.276529 | NA |
| MIR4454 | 0.134133 | -1.48909 | 4.077584 | -0.36519 | 0.71497 | NA |
| | 0.298089 | -0.51711 | 3.975232 | -0.13008 | 0.896501 | NA |
| | 13.94872 | -0.17359 | 0.563419 | -0.30811 | 0.758001 | NA |
| | 1.365791 | 1.704791 | 2.102832 | 0.810712 | 0.417531 | NA |
| | 0.392857 | -1.546 | 4.075469 | -0.37934 | 0.704433 | NA |
| CDC42P5 | 0.163957 | 0.424913 | 4.079006 | 0.104171 | 0.917034 | NA |
| PCDHGA5 | 5.453656 | -0.28122 | 0.983789 | -0.28586 | 0.774987 | NA |
| | 0.326536 | 0.037056 | 3.842554 | 0.009644 | 0.992306 | NA |
| | 0.449786 | -1.25325 | 3.00474 | -0.41709 | 0.676613 | NA |
| MIR146A | 1035.138 | 0.826514 | 0.171575 | 4.817216 | 1.46E-06 | 2.05E-04 |
| PCDHGA7 | 11.73314 | -0.87334 | 0.593468 | -1.47158 | 0.141134 | NA |
| FAM86HP | 20.28031 | -0.59473 | 0.460747 | -1.29079 | 0.196775 | NA |
| | 0.162376 | 0.424913 | 4.079006 | 0.104171 | 0.917034 | NA |
| GRPEL2-AS | 0.258578 | 1.233151 | 4.047809 | 0.304646 | 0.760635 | NA |
| | 0.422737 | 0.518872 | 3.501356 | 0.148192 | 0.882191 | NA |

| | | | | | | |
|---|---|---|---|---|---|---|
| | 0.163957 | 0.424913 | 4.079006 | 0.104171 | 0.917034 | NA |
| | 3.820526 | 0.539865 | 1.024331 | 0.527041 | 0.598165 | NA |
| | 1.134622 | 0.410451 | 2.216341 | 0.185193 | 0.853078 | NA |
| | 0.843838 | 1.092325 | 2.536021 | 0.430724 | 0.666669 | NA |
| PCDHGA6 | 59.70356 | -0.68163 | 0.338434 | -2.01409 | 0.044001 | 0.127866 |
| | 1.181293 | 0.703609 | 1.930238 | 0.364519 | 0.71547 | NA |
| PCDHGA8 | 11.35845 | 0.033545 | 0.725938 | 0.046209 | 0.963144 | NA |
| | 0.096202 | 0.408546 | 4.079639 | 0.100143 | 0.920231 | NA |
| | 0.096202 | 0.408546 | 4.079639 | 0.100143 | 0.920231 | NA |
| | 1.268192 | 0.770271 | 1.929297 | 0.39925 | 0.689709 | NA |
| LINC01170 | 0.260158 | 1.240347 | 4.047124 | 0.306476 | 0.759242 | NA |
| PCDHGA10 | 377.8245 | -0.5718 | 0.191021 | -2.99339 | 0.002759 | 0.020066 |
| | 0.151697 | -1.546 | 4.075469 | -0.37934 | 0.704433 | NA |
| | 0.247898 | -0.5901 | 4.074633 | -0.14482 | 0.884851 | NA |
| PIGFP1 | 0.296509 | -0.51709 | 3.982155 | -0.12985 | 0.896684 | NA |
| PCDHGA11 | 411.8221 | -0.34471 | 0.186902 | -1.84434 | 0.065133 | 0.166416 |
| | 0.392857 | -1.546 | 4.075469 | -0.37934 | 0.704433 | NA |
| | 0.151697 | -1.546 | 4.075469 | -0.37934 | 0.704433 | NA |
| PCDHGB2 | 21.27038 | -0.14368 | 0.662048 | -0.21703 | 0.828187 | NA |
| | 2.568343 | -0.22511 | 1.202074 | -0.18727 | 0.851451 | NA |
| PCDHGB4 | 71.08965 | -0.2557 | 0.489393 | -0.52249 | 0.601332 | 0.730558 |
| | 6.460291 | 0.040528 | 0.802343 | 0.050512 | 0.959714 | NA |
| | 0.134133 | -1.48909 | 4.077584 | -0.36519 | 0.71497 | NA |
| | 1.952239 | -0.58283 | 1.566042 | -0.37217 | 0.709767 | NA |
| | 7.009717 | -0.68929 | 0.776844 | -0.8873 | 0.374919 | NA |
| | 0.556813 | -0.57007 | 3.294615 | -0.17303 | 0.862627 | NA |
| | 0.589223 | -3.04304 | 3.047314 | -0.9986 | 0.317989 | NA |
| | 0.654246 | 2.541085 | 3.009764 | 0.84428 | 0.398513 | NA |
| | 0.872524 | -0.04714 | 2.136288 | -0.02207 | 0.982395 | NA |
| PCDHGB7 | 97.65051 | -0.82905 | 0.313395 | -2.6454 | 0.008159 | 0.041144 |
| MTND6P3 | 0.151697 | -1.546 | 4.075469 | -0.37934 | 0.704433 | NA |
| | 0.163957 | 0.424913 | 4.079006 | 0.104171 | 0.917034 | NA |
| | 0.652869 | 1.292592 | 2.374873 | 0.544278 | 0.58625 | NA |
| PCDHGB1 | 6.949805 | -0.95166 | 0.782005 | -1.21695 | 0.223621 | NA |
| | 4.215523 | 0.291166 | 0.973814 | 0.298995 | 0.764944 | NA |
| | 0.163957 | 0.424913 | 4.079006 | 0.104171 | 0.917034 | NA |
| PCDHGA3 | 10.05404 | -0.48789 | 0.778098 | -0.62703 | 0.530641 | NA |
| | 0.134133 | -1.48909 | 4.077584 | -0.36519 | 0.71497 | NA |
| | 0.488912 | 0.771583 | 2.897678 | 0.266276 | 0.790026 | NA |
| | 0.392857 | -1.546 | 4.075469 | -0.37934 | 0.704433 | NA |
| | 0.324752 | 1.53082 | 4.022025 | 0.380609 | 0.703493 | NA |
| | 1.003439 | -1.03737 | 2.189171 | -0.47387 | 0.635596 | NA |
| | 0.422534 | 1.924583 | 3.001758 | 0.641152 | 0.521424 | NA |
| | 0.163957 | 0.424913 | 4.079006 | 0.104171 | 0.917034 | NA |
| | 0.303394 | -2.27006 | 4.032451 | -0.56295 | 0.57347 | NA |
| | 5.734873 | 0.158238 | 1.016471 | 0.155674 | 0.87629 | NA |
| | 1.104053 | 1.332031 | 2.269145 | 0.587019 | 0.557191 | NA |

| | | | | | | |
|---|---|---|---|---|---|---|
| | 0.392857 | -1.546 | 4.075469 | -0.37934 | 0.704433 | NA |
| | 1.607697 | 0.014776 | 1.512674 | 0.009768 | 0.992206 | NA |
| | 1.580648 | 0.625838 | 1.698867 | 0.368386 | 0.712586 | NA |
| PBOV1 | 0.134133 | -1.48909 | 4.077584 | -0.36519 | 0.71497 | NA |
| | 61.39397 | -0.2293 | 0.295151 | -0.77689 | 0.437226 | 0.597569 |
| PABPC4L | 2.353401 | 1.422982 | 1.657097 | 0.85872 | 0.390495 | NA |
| | 2.302649 | 1.463891 | 1.374986 | 1.064659 | 0.28703 | NA |
| | 5.38793 | 0.370864 | 0.869996 | 0.426282 | 0.669902 | NA |
| ANKHD1-EI | 5.959216 | 0.954077 | 0.864964 | 1.103025 | 0.270016 | NA |
| | 18.17214 | 0.461071 | 0.503484 | 0.915762 | 0.359792 | NA |
| PCDHA3 | 0.392711 | 0.292538 | 3.145221 | 0.09301 | 0.925895 | NA |
| EBLN2 | 8.042895 | 1.512583 | 0.803256 | 1.883065 | 0.059692 | NA |
| | 9.760148 | 0.372906 | 0.650032 | 0.573672 | 0.566189 | NA |
| MTRNR2L9 | 0.303394 | -2.27006 | 4.032451 | -0.56295 | 0.57347 | NA |
| | 0.747876 | -0.62091 | 2.562446 | -0.24231 | 0.80854 | NA |
| | 1.225869 | -0.07951 | 1.838357 | -0.04325 | 0.965503 | NA |
| HIST1H3G | 4.925403 | -0.3214 | 0.917835 | -0.35017 | 0.726213 | NA |
| CTSO | 694.6304 | -0.07323 | 0.190994 | -0.38341 | 0.701418 | 0.808096 |
| | 0.916751 | 0.760304 | 2.340291 | 0.324876 | 0.745275 | NA |
| SMLR1 | 0.49029 | 2.123092 | 3.304282 | 0.642527 | 0.520531 | NA |
| SMIM3 | 0.163957 | 0.424913 | 4.079006 | 0.104171 | 0.917034 | NA |
| HIST1H3F | 1.168976 | -0.41512 | 1.914592 | -0.21682 | 0.82835 | NA |
| ASIC5 | 0.163957 | 0.424913 | 4.079006 | 0.104171 | 0.917034 | NA |
| ZBTB11-AS | 10.61125 | 0.410626 | 0.644292 | 0.637329 | 0.52391 | NA |
| | 0.096202 | 0.408546 | 4.079639 | 0.100143 | 0.920231 | NA |
| PPT2-EGFL | 8.607057 | -0.65551 | 0.7139 | -0.91821 | 0.358509 | NA |
| | 28.78026 | -0.01828 | 0.394005 | -0.0464 | 0.962988 | NA |
| | 9.658823 | -0.32896 | 0.851872 | -0.38616 | 0.699379 | NA |
| | 11.78962 | -0.10605 | 0.671805 | -0.15786 | 0.87457 | NA |
| | 13.51637 | -0.31689 | 0.696481 | -0.45498 | 0.649122 | NA |
| | 11.64108 | 0.2022 | 0.591799 | 0.341671 | 0.732599 | NA |
| | 2.230336 | 0.808908 | 1.430626 | 0.565423 | 0.571786 | NA |
| | 0.260158 | 1.240347 | 4.047124 | 0.306476 | 0.759242 | NA |
| | 0.392857 | -1.546 | 4.075469 | -0.37934 | 0.704433 | NA |
| | 4.945474 | 1.596145 | 1.039831 | 1.535005 | 0.124783 | NA |
| | 9.100059 | -0.29641 | 0.695792 | -0.426 | 0.670106 | NA |
| | 4.929446 | 0.476532 | 0.938333 | 0.507849 | 0.611559 | NA |
| | 1.282392 | 1.378416 | 2.174636 | 0.633861 | 0.526172 | NA |
| | 0.610582 | -0.18034 | 2.753139 | -0.0655 | 0.947774 | NA |
| | 27.89983 | 0.061513 | 0.403858 | 0.152314 | 0.87894 | NA |
| | 0.096202 | 0.408546 | 4.079639 | 0.100143 | 0.920231 | NA |
| CASC14 | 14.51396 | 0.614411 | 0.624783 | 0.983398 | 0.325411 | NA |
| | 0.298089 | -0.51711 | 3.975232 | -0.13008 | 0.896501 | NA |
| | 0.134133 | -1.48909 | 4.077584 | -0.36519 | 0.71497 | NA |
| | 9.599923 | 0.500169 | 0.700893 | 0.713617 | 0.475464 | NA |
| | 5.842596 | -0.35685 | 0.823273 | -0.43345 | 0.664686 | NA |
| | 0.162376 | 0.424913 | 4.079006 | 0.104171 | 0.917034 | NA |

| | | | | | | |
|---|---|---|---|---|---|---|
| | 0.247898 | -0.5901 | 4.074633 | -0.14482 | 0.884851 | NA |
| | 1.094751 | 1.216857 | 2.01996 | 0.602417 | 0.546897 | NA |
| | 0.891465 | 0.657772 | 2.431814 | 0.270486 | 0.786786 | NA |
| | 1.467911 | 2.620575 | 1.882259 | 1.39225 | 0.163847 | NA |
| | 17.95018 | 0.145094 | 0.545652 | 0.26591 | 0.790309 | NA |
| | 0.46735 | -1.30193 | 3.399671 | -0.38296 | 0.701751 | NA |
| | 9.045203 | 1.070395 | 0.839629 | 1.274842 | 0.202365 | NA |
| | 1.539593 | 1.098873 | 1.649451 | 0.666205 | 0.50528 | NA |
| | 5.307608 | -1.24694 | 0.93795 | -1.32943 | 0.183706 | NA |
| | 0.096202 | 0.408546 | 4.079639 | 0.100143 | 0.920231 | NA |
| | 115.6025 | -0.48148 | 0.261648 | -1.84019 | 0.06574 | 0.167212 |
| | 0.315653 | -0.57305 | 3.892639 | -0.14721 | 0.882963 | NA |
| | 0.450981 | 2.057325 | 3.353149 | 0.61355 | 0.539513 | NA |
| | 16.07569 | 0.584645 | 0.511677 | 1.142606 | 0.253202 | NA |
| | 33.58044 | 0.729831 | 0.444271 | 1.642762 | 0.100432 | NA |
| | 2.085341 | 1.166815 | 1.698518 | 0.68696 | 0.492108 | NA |
| | 0.854903 | -0.47663 | 2.545657 | -0.18723 | 0.851479 | NA |
| | 36.26005 | -0.20714 | 0.354349 | -0.58456 | 0.558841 | NA |
| | 0.192403 | 0.975392 | 4.060716 | 0.240202 | 0.810174 | NA |
| | 0.315653 | -0.57305 | 3.892639 | -0.14721 | 0.882963 | NA |
| | 0.151697 | -1.546 | 4.075469 | -0.37934 | 0.704433 | NA |
| | 0.516164 | -1.70268 | 2.844782 | -0.59853 | 0.549488 | NA |
| | 2.516368 | -0.01785 | 1.217462 | -0.01466 | 0.9883 | NA |
| | 18.43099 | 0.836958 | 0.529877 | 1.579534 | 0.114214 | NA |
| | 0.956059 | 0.787678 | 2.36595 | 0.332923 | 0.739193 | NA |
| | 18.91754 | -0.60904 | 0.536832 | -1.13451 | 0.256579 | NA |
| | 6.606234 | 1.186104 | 0.876505 | 1.353221 | 0.175985 | NA |
| | 1.689495 | 0.214366 | 1.601324 | 0.133868 | 0.893507 | NA |
| | 3.481561 | 0.812005 | 1.305668 | 0.621908 | 0.534002 | NA |
| MANEA-AS | 3.678285 | 1.214851 | 1.071437 | 1.133852 | 0.256857 | NA |
| | 2.580428 | 1.594005 | 1.311578 | 1.215333 | 0.224239 | NA |
| | 18.89766 | 1.442852 | 0.522077 | 2.763676 | 0.005715 | NA |
| | 0.738188 | 1.39874 | 2.317793 | 0.603479 | 0.54619 | NA |
| | 0.134133 | -1.48909 | 4.077584 | -0.36519 | 0.71497 | NA |
| | 4.287715 | -0.3054 | 1.150611 | -0.26542 | 0.790683 | NA |
| HLA-P | 0.872321 | 0.678609 | 2.011629 | 0.337343 | 0.735858 | NA |
| | 0.801202 | 1.538543 | 2.914591 | 0.527876 | 0.597585 | NA |
| | 0.096202 | 0.408546 | 4.079639 | 0.100143 | 0.920231 | NA |
| | 2.172683 | 0.396959 | 1.354498 | 0.293067 | 0.769471 | NA |
| | 6.389541 | -0.38733 | 0.829666 | -0.46685 | 0.64061 | NA |
| | 0.684273 | 2.627785 | 2.896325 | 0.907282 | 0.364257 | NA |
| | 0.354779 | 1.703954 | 3.657232 | 0.465914 | 0.641277 | NA |
| | 38.50746 | 0.713851 | 0.440574 | 1.620276 | 0.105173 | NA |
| | 0.096202 | 0.408546 | 4.079639 | 0.100143 | 0.920231 | NA |
| | 0.804712 | -0.7622 | 2.348804 | -0.3245 | 0.745556 | NA |
| | 0.327913 | 1.543154 | 4.021063 | 0.383768 | 0.701151 | NA |
| PCDHGA9 | 44.77516 | -0.59923 | 0.388461 | -1.54257 | 0.122936 | NA |

| | | | | | | |
|---|---|---|---|---|---|---|
| | 0.260158 | 1.240347 | 4.047124 | 0.306476 | 0.759242 | NA |
| PCDHB19P | 0.629726 | -0.22725 | 2.732935 | -0.08315 | 0.93373 | NA |
| PCDHGB3 | 24.31768 | -0.16629 | 0.442959 | -0.3754 | 0.707365 | NA |
| | 0.096202 | 0.408546 | 4.079639 | 0.100143 | 0.920231 | NA |
| PCDHGA4 | 82.11342 | -0.53463 | 0.320417 | -1.66853 | 0.095211 | 0.215867 |
| | 0.162376 | 0.424913 | 4.079006 | 0.104171 | 0.917034 | NA |
| TAPT1-AS1 | 24.03619 | 0.108044 | 0.423384 | 0.255191 | 0.798576 | NA |
| MIR378H | 0.247898 | -0.5901 | 4.074633 | -0.14482 | 0.884851 | NA |
| | 1.950602 | -0.91518 | 1.390108 | -0.65836 | 0.51031 | NA |
| RN7SL471F | 1.666353 | -0.49253 | 1.620555 | -0.30393 | 0.761183 | NA |
| RN7SL689F | 4.061117 | 0.664279 | 1.083133 | 0.613294 | 0.539682 | NA |
| MIR4804 | 0.151697 | -1.546 | 4.075469 | -0.37934 | 0.704433 | NA |
| MIR3936 | 1.552202 | 0.57156 | 1.57281 | 0.363401 | 0.716306 | NA |
| MIR4639 | 0.28583 | -2.21358 | 3.97376 | -0.55705 | 0.577494 | NA |
| | 8.343067 | 1.127719 | 0.827427 | 1.362924 | 0.172907 | NA |
| RN7SL275F | 0.151697 | -1.546 | 4.075469 | -0.37934 | 0.704433 | NA |
| RN7SL336F | 1.101075 | 0.447616 | 1.809417 | 0.247381 | 0.804613 | NA |
| MIR5685 | 0.096202 | 0.408546 | 4.079639 | 0.100143 | 0.920231 | NA |
| | 26.9792 | 0.426681 | 0.423473 | 1.007576 | 0.313658 | NA |
| | 0.410275 | 0.244283 | 3.100889 | 0.078778 | 0.937209 | NA |
| MIR4449 | 1.551007 | -0.72774 | 1.546985 | -0.47043 | 0.638051 | NA |
| | 0.096202 | 0.408546 | 4.079639 | 0.100143 | 0.920231 | NA |
| MIR4798 | 0.327913 | 1.543154 | 4.021063 | 0.383768 | 0.701151 | NA |
| MIR3138 | 0.258578 | 1.233151 | 4.047809 | 0.304646 | 0.760635 | NA |
| RN7SL200F | 0.163957 | 0.424913 | 4.079006 | 0.104171 | 0.917034 | NA |
| MIR3136 | 0.326333 | 1.533889 | 3.826878 | 0.40082 | 0.688553 | NA |
| | 0.49029 | 2.123092 | 3.304282 | 0.642527 | 0.520531 | NA |
| | 0.46735 | -1.30193 | 3.399671 | -0.38296 | 0.701751 | NA |
| RN7SL300F | 0.258578 | 1.233151 | 4.047809 | 0.304646 | 0.760635 | NA |
| MIR544B | 0.163957 | 0.424913 | 4.079006 | 0.104171 | 0.917034 | NA |
| | 2.784809 | 0.299865 | 1.227652 | 0.244259 | 0.80703 | NA |
| | 0.163957 | 0.424913 | 4.079006 | 0.104171 | 0.917034 | NA |
| MIR5091 | 0.134133 | -1.48909 | 4.077584 | -0.36519 | 0.71497 | NA |
| | 0.163957 | 0.424913 | 4.079006 | 0.104171 | 0.917034 | NA |
| RN7SL682F | 0.247898 | -0.5901 | 4.074633 | -0.14482 | 0.884851 | NA |
| MIR4451 | 0.192403 | 0.975392 | 4.060716 | 0.240202 | 0.810174 | NA |
| | 2.174052 | -0.41142 | 1.286989 | -0.31968 | 0.749212 | NA |
| MIR3934 | 1.723246 | -0.27497 | 1.627026 | -0.169 | 0.865793 | NA |
| MIR3145 | 0.460669 | -0.62593 | 3.398671 | -0.18417 | 0.853882 | NA |
| MIR4642 | 0.134133 | -1.48909 | 4.077584 | -0.36519 | 0.71497 | NA |
| RN7SL637F | 0.526843 | -0.38719 | 2.855892 | -0.13558 | 0.892156 | NA |
| | 16.68747 | -0.18518 | 0.559222 | -0.33113 | 0.740545 | NA |
| RN7SL773F | 0.163957 | 0.424913 | 4.079006 | 0.104171 | 0.917034 | NA |
| MIR4444-2 | 0.134133 | -1.48909 | 4.077584 | -0.36519 | 0.71497 | NA |
| MIR4444-1 | 0.134133 | -1.48909 | 4.077584 | -0.36519 | 0.71497 | NA |
| | 0.326333 | 1.533889 | 3.826878 | 0.40082 | 0.688553 | NA |
| | 0.151697 | -1.546 | 4.075469 | -0.37934 | 0.704433 | NA |

| | | | | | | |
|---|---|---|---|---|---|---|
| | 0.973826 | 0.026039 | 2.097476 | 0.012415 | 0.990095 | NA |
| | 0.151697 | -1.546 | 4.075469 | -0.37934 | 0.704433 | NA |
| | 30.45755 | -0.12652 | 0.410196 | -0.30843 | 0.757755 | NA |
| | 76.69961 | 0.656791 | 0.422583 | 1.554229 | 0.12013 | 0.252328 |
| | 0.096202 | 0.408546 | 4.079639 | 0.100143 | 0.920231 | NA |
| | 0.096202 | 0.408546 | 4.079639 | 0.100143 | 0.920231 | NA |
| EPB41L4A-, | 43.17021 | -0.05765 | 0.365711 | -0.15764 | 0.874737 | NA |
| | 35.63555 | -0.02616 | 0.396739 | -0.06593 | 0.94743 | NA |
| | 2.214352 | 0.834755 | 1.423656 | 0.586346 | 0.557643 | NA |
| MIR4453 | 24.85527 | 0.733315 | 0.515368 | 1.422896 | 0.154766 | NA |
| | 28.10256 | 0.179434 | 0.517602 | 0.346664 | 0.728844 | NA |
| | 0.151697 | -1.546 | 4.075469 | -0.37934 | 0.704433 | NA |
| | 0.327913 | 1.543154 | 4.021063 | 0.383768 | 0.701151 | NA |
| | 19.74762 | -1.64345 | 0.538484 | -3.052 | 0.002273 | NA |
| ZSCAN16-A | 239.8899 | 0.1556 | 0.246583 | 0.631027 | 0.528023 | 0.673449 |
| | 10.80906 | -0.26236 | 0.622712 | -0.42132 | 0.673519 | NA |
| | 0.613743 | -0.17004 | 2.749758 | -0.06184 | 0.950691 | NA |
| | 11.10862 | 0.96807 | 0.658313 | 1.470533 | 0.141417 | NA |
| | 0.163957 | 0.424913 | 4.079006 | 0.104171 | 0.917034 | NA |
| | 0.563552 | -0.48645 | 2.808029 | -0.17324 | 0.862465 | NA |
| | 161.3085 | 0.28908 | 0.277856 | 1.040394 | 0.298157 | 0.465146 |
| TSPAN6 | 837.751 | 0.050341 | 0.191806 | 0.262458 | 0.792968 | 0.874397 |
| TNMD | 1.365734 | 2.175162 | 2.194216 | 0.991316 | 0.321531 | NA |
| LAS1L | 923.4749 | -0.15174 | 0.165486 | -0.91693 | 0.359178 | 0.5264 |
| CFTR | 0.816825 | 1.673341 | 2.21346 | 0.755984 | 0.449659 | NA |
| ANKIB1 | 1577.583 | 0.302061 | 0.163678 | 1.845464 | 0.06497 | 0.166175 |
| CYP51A1 | 397.5706 | 0.243799 | 0.192244 | 1.268178 | 0.204734 | 0.361395 |
| KRIT1 | 402.2434 | 0.799668 | 0.308951 | 2.588337 | 0.009644 | 0.046045 |
| MYH16 | 0.326333 | 1.533889 | 3.826878 | 0.40082 | 0.688553 | NA |
| HECW1 | 0.937264 | -1.23834 | 2.063944 | -0.59999 | 0.548514 | NA |
| MAD1L1 | 2405.938 | -0.37882 | 0.143128 | -2.64673 | 0.008127 | 0.041007 |
| TMEM176/ | 0.134133 | -1.48909 | 4.077584 | -0.36519 | 0.71497 | NA |
| KLHL13 | 223.2867 | -0.01132 | 0.238025 | -0.04757 | 0.962059 | 0.97942 |
| ICA1 | 342.609 | 0.218033 | 0.174442 | 1.249891 | 0.211339 | 0.369587 |
| MTMR7 | 11.51808 | 1.273177 | 0.679386 | 1.874011 | 0.060929 | NA |
| SLC7A2 | 189.6212 | 0.272864 | 0.261033 | 1.045324 | 0.295873 | 0.463138 |
| ARF5 | 2824.615 | -0.18297 | 0.124625 | -1.46819 | 0.142052 | 0.282988 |
| CCDC132 | 397.6046 | 0.775465 | 0.304394 | 2.547567 | 0.010848 | 0.049628 |
| PDK4 | 1733.487 | 0.923598 | 0.193081 | 4.783469 | 1.72E-06 | 2.24E-04 |
| ABCB5 | 5680.624 | 0.040721 | 0.151274 | 0.269188 | 0.787785 | 0.871356 |
| ARX | 7.359414 | 0.695478 | 1.083728 | 0.641746 | 0.521038 | NA |
| SLC25A13 | 1548.724 | -0.09905 | 0.157013 | -0.63082 | 0.528156 | 0.673547 |
| ST7 | 1102.834 | -0.53789 | 0.176463 | -3.0482 | 0.002302 | 0.01771 |
| HCCS | 1099.576 | -0.01483 | 0.16173 | -0.09171 | 0.92693 | 0.958156 |
| SKAP2 | 696.173 | 0.467275 | 0.210048 | 2.224608 | 0.026108 | 0.090631 |
| SLC25A5 | 12676.18 | -0.19462 | 0.128417 | -1.51556 | 0.129632 | 0.266013 |
| HOXA11 | 8.62723 | -0.34912 | 0.667281 | -0.5232 | 0.600834 | NA |

| | | | | | | |
|---|---|---|---|---|---|---|
| POLR2J | 923.0884 | -0.02622 | 0.143429 | -0.18282 | 0.854939 | 0.913749 |
| THSD7A | 6.457292 | -0.41165 | 1.036076 | -0.39732 | 0.691133 | NA |
| FAM214B | 1828.202 | -0.21315 | 0.162361 | -1.31284 | 0.189238 | 0.343409 |
| PRKAR2B | 317.4093 | 0.612979 | 0.238945 | 2.565352 | 0.010307 | 0.048042 |
| MSL3 | 644.3999 | 0.442422 | 0.167783 | 2.636864 | 0.008368 | 0.041768 |
| CROT | 463.8183 | 0.224458 | 0.181471 | 1.236879 | 0.216132 | 0.37531 |
| ABCB4 | 565.7136 | 0.294199 | 0.204123 | 1.44128 | 0.149505 | 0.293079 |
| KMT2E | 850.2961 | 0.472356 | 0.319561 | 1.478138 | 0.139371 | 0.279378 |
| RHBDD2 | 2727.731 | -0.15947 | 0.165305 | -0.96468 | 0.334704 | 0.50329 |
| ZFX | 548.1399 | 0.374527 | 0.192671 | 1.943864 | 0.051912 | 0.141932 |
| LAMP2 | 9335.705 | 0.289801 | 0.156483 | 1.85196 | 0.064032 | 0.164591 |
| ASB4 | 4.272855 | -0.65514 | 1.422684 | -0.4605 | 0.645161 | NA |
| TAC1 | 737.1808 | 2.021636 | 0.808371 | 2.500878 | 0.012389 | 0.054536 |
| RALA | 1163.275 | 0.556363 | 0.186227 | 2.987552 | 0.002812 | 0.020314 |
| BAIAP2L1 | 665.2864 | -0.29759 | 0.19945 | -1.49206 | 0.135684 | 0.274564 |
| KDM7A | 151.8716 | 0.879817 | 0.29372 | 2.995433 | 0.002741 | 0.019969 |
| ETV1 | 718.9206 | 0.37027 | 0.200034 | 1.85103 | 0.064165 | 0.164827 |
| AGK | 770.2767 | -0.0509 | 0.139699 | -0.36437 | 0.715582 | 0.81867 |
| PHTF2 | 823.1341 | 0.687688 | 0.182718 | 3.763659 | 1.67E-04 | 0.003495 |
| GGCT | 683.6302 | 0.594717 | 0.190726 | 3.118169 | 0.00182 | 0.01515 |
| DBF4 | 237.3153 | 0.966764 | 0.210845 | 4.585196 | 4.54E-06 | 3.99E-04 |
| IFRD1 | 514.2342 | 1.136228 | 0.207796 | 5.467997 | 4.55E-08 | 2.00E-05 |
| GTF2IRD1 | 700.0179 | -0.31156 | 0.161777 | -1.92586 | 0.054122 | 0.146255 |
| VPS41 | 3727.713 | 0.279619 | 0.141598 | 1.974738 | 0.048298 | 0.135641 |
| SCIN | 111.972 | 1.332164 | 0.30827 | 4.321423 | 1.55E-05 | 8.10E-04 |
| ARSD | 695.5398 | -0.53498 | 0.15172 | -3.52609 | 4.22E-04 | 0.006131 |
| PNPLA4 | 525.206 | -0.03946 | 0.15997 | -0.24665 | 0.805181 | 0.883508 |
| TKTL1 | 1.743199 | -1.5863 | 1.808093 | -0.87733 | 0.380306 | NA |
| NOX1 | 4.252441 | 0.023067 | 1.18631 | 0.019444 | 0.984487 | NA |
| PSMB1 | 3347.651 | 0.172674 | 0.14355 | 1.202885 | 0.229021 | 0.390822 |
| SYN1 | 2.649378 | 2.720186 | 1.435542 | 1.894885 | 0.058108 | NA |
| CDKL5 | 112.0184 | 0.052509 | 0.24419 | 0.215032 | 0.829743 | 0.897322 |
| CYTH3 | 7887.838 | 0.035731 | 0.165254 | 0.216218 | 0.828818 | 0.896738 |
| ADAM22 | 113.0013 | 0.508817 | 0.270118 | 1.883685 | 0.059608 | 0.156615 |
| SYPL1 | 2479.956 | -0.08699 | 0.123839 | -0.70248 | 0.482377 | 0.636247 |
| AASS | 306.6684 | 0.498245 | 0.227128 | 2.193675 | 0.028259 | 0.095227 |
| ST3GAL1 | 1235.021 | -0.18575 | 0.20582 | -0.90249 | 0.366795 | 0.534116 |
| RHOBTB2 | 654.1446 | -0.46798 | 0.157818 | -2.96533 | 0.003024 | 0.021273 |
| RPS20 | 16797.1 | 0.269118 | 0.15276 | 1.761707 | 0.078119 | 0.188276 |
| UBE3C | 3437.638 | 0.048321 | 0.145409 | 0.332311 | 0.739655 | 0.835583 |
| MLXIPL | 0.392857 | -1.546 | 4.075469 | -0.37934 | 0.704433 | NA |
| BAZ1B | 2548.605 | 0.399313 | 0.216135 | 1.847513 | 0.064673 | 0.165736 |
| STARD3NL | 1916.439 | 0.368138 | 0.137399 | 2.679332 | 0.007377 | 0.038577 |
| IDS | 932.3283 | 0.154965 | 0.19388 | 0.799283 | 0.424126 | 0.586153 |
| PRSS3 | 4.917271 | 0.90985 | 0.982399 | 0.92615 | 0.354368 | NA |
| BTK | 8.801728 | -1.17863 | 0.787553 | -1.49658 | 0.134504 | NA |
| KAL1 | 190.7025 | 0.507644 | 0.209215 | 2.426427 | 0.015248 | 0.062789 |

| | | | | | | |
|---|---|---|---|---|---|---|
| RNF216 | 1210.123 | -0.31212 | 0.135251 | -2.30771 | 0.021015 | 0.078296 |
| ANLN | 1588.702 | 0.600503 | 0.193512 | 3.103183 | 0.001915 | 0.015674 |
| RABGAP1 | 1238.408 | 0.317706 | 0.168862 | 1.88145 | 0.059911 | 0.157094 |
| MBTPS2 | 966.9871 | 0.073657 | 0.18866 | 0.390419 | 0.696227 | 0.804148 |
| PRICKLE3 | 445.5319 | -0.44308 | 0.158426 | -2.7968 | 0.005161 | 0.0302 |
| EXTL3 | 5111.699 | -0.43852 | 0.15413 | -2.84514 | 0.004439 | 0.02725 |
| KDM5D | 759.0214 | -0.07417 | 0.174279 | -0.42558 | 0.670411 | 0.784109 |
| NUB1 | 1533.506 | 0.149918 | 0.135286 | 1.108154 | 0.267795 | 0.432699 |
| DNASE1L1 | 795.5681 | -0.10996 | 0.159298 | -0.69029 | 0.490011 | 0.642357 |
| MAMLD1 | 584.5745 | -0.48882 | 0.15338 | -3.18701 | 0.001438 | 0.013218 |
| ZC3H3 | 479.6751 | -0.46636 | 0.173725 | -2.68447 | 0.007264 | 0.038129 |
| WAS | 1.223094 | -1.84374 | 1.855462 | -0.99368 | 0.320379 | NA |
| NUDCD3 | 3002.209 | -0.4807 | 0.128225 | -3.74888 | 1.78E-04 | 0.003606 |
| SLC38A5 | 27.05048 | 0.490853 | 0.545544 | 0.89975 | 0.368253 | NA |
| MAGIX | 39.84572 | 0.117651 | 0.458538 | 0.256579 | 0.797504 | NA |
| CXorf56 | 392.6665 | -0.10688 | 0.159604 | -0.66968 | 0.503062 | 0.653066 |
| SNAI2 | 4523.163 | 0.028527 | 0.164568 | 0.173345 | 0.862381 | 0.918473 |
| GPR124 | 2168.999 | -0.7923 | 0.207976 | -3.80959 | 1.39E-04 | 0.003102 |
| FHL1 | 1302.566 | 0.079441 | 0.159511 | 0.498028 | 0.618464 | 0.743274 |
| SLC45A4 | 199.6811 | -0.3768 | 0.206295 | -1.8265 | 0.067775 | 0.170548 |
| RB1CC1 | 634.7771 | 1.074208 | 0.833585 | 1.288661 | 0.197516 | 0.353175 |
| ERP44 | 1433.964 | 0.167801 | 0.160493 | 1.045537 | 0.295775 | 0.463076 |
| RNASET2 | 152.8699 | -0.09713 | 0.335527 | -0.28948 | 0.772211 | 0.858857 |
| AGPAT4 | 230.5297 | 0.027205 | 0.189061 | 0.143894 | 0.885584 | 0.932306 |
| ANK1 | 1.180005 | -1.57876 | 2.139972 | -0.73775 | 0.460668 | NA |
| TFB1M | 334.5535 | 0.17254 | 0.168198 | 1.025815 | 0.304979 | 0.472185 |
| HMGB3 | 663.254 | 0.068341 | 0.152238 | 0.448907 | 0.653499 | 0.771099 |
| ABCF2 | 2478.413 | -0.17328 | 0.144666 | -1.19782 | 0.230987 | 0.393158 |
| CHPF2 | 2650.149 | -0.25159 | 0.157648 | -1.59588 | 0.110516 | 0.238429 |
| EFCAB1 | 27.64383 | -0.00527 | 0.472276 | -0.01115 | 0.991102 | NA |
| RNF19A | 1071.867 | 0.649848 | 0.223243 | 2.910952 | 0.003603 | 0.023891 |
| NSMAF | 1265.555 | 0.181309 | 0.161788 | 1.120653 | 0.262436 | 0.426852 |
| MYOM2 | 17.08325 | 0.082456 | 0.487793 | 0.169039 | 0.865766 | NA |
| OTC | 1.079513 | -0.47694 | 2.056893 | -0.23187 | 0.816636 | NA |
| STAU2 | 730.7829 | 0.526566 | 0.194148 | 2.712181 | 0.006684 | 0.035967 |
| TNC | 1230.386 | -0.63575 | 1.167486 | -0.54454 | 0.586068 | 0.71894 |
| TG | 1.938554 | -0.35547 | 1.901301 | -0.18696 | 0.851691 | NA |
| ADAM28 | 0.096202 | 0.408546 | 4.079639 | 0.100143 | 0.920231 | NA |
| PHKA2 | 482.6871 | -0.26136 | 0.157155 | -1.66307 | 0.096298 | 0.21745 |
| CNTLN | 114.6977 | 0.901002 | 0.783768 | 1.149577 | 0.250318 | 0.413938 |
| HSPA5 | 10895.01 | -0.21186 | 0.160665 | -1.31863 | 0.187292 | 0.341474 |
| GEMIN8 | 190.9857 | -0.14594 | 0.228052 | -0.63995 | 0.522208 | 0.66864 |
| OFD1 | 99.97476 | 1.053819 | 0.503873 | 2.091439 | 0.036489 | 0.113121 |
| GPM6B | 6614.288 | 0.418298 | 0.185896 | 2.250178 | 0.024438 | 0.086635 |
| PREX2 | 10.5654 | -3.04049 | 3.465027 | -0.87748 | 0.380227 | NA |
| CTPS2 | 695.9956 | -0.11397 | 0.163087 | -0.69884 | 0.484655 | 0.638256 |
| ATP6V1H | 3911.204 | -0.12903 | 0.116207 | -1.11034 | 0.266852 | 0.431705 |

| | | | | | | |
|---|---|---|---|---|---|---|
| XK | 1.127884 | 0.383009 | 2.112806 | 0.18128 | 0.856148 | NA |
| SCML1 | 158.5804 | 0.817265 | 0.242776 | 3.366329 | 7.62E-04 | 0.008802 |
| WWC3 | 2206.722 | -0.20742 | 0.152643 | -1.35888 | 0.174185 | 0.326183 |
| ARHGAP6 | 27.38406 | 0.327407 | 0.543459 | 0.602451 | 0.546874 | NA |
| HDAC9 | 871.7665 | 1.115701 | 0.311433 | 3.58248 | 3.40E-04 | 0.005344 |
| RRM2B | 1769.219 | 0.735293 | 0.185094 | 3.97254 | 7.11E-05 | 0.002051 |
| ZNF800 | 326.1145 | 0.517184 | 0.232362 | 2.22577 | 0.02603 | 0.090437 |
| FAM120A | 3738.72 | -0.05191 | 0.169844 | -0.30563 | 0.759886 | 0.850495 |
| ELN | 0.851742 | -0.48655 | 2.547586 | -0.19099 | 0.848537 | NA |
| RFC2 | 929.4984 | -0.04429 | 0.223948 | -0.19775 | 0.843239 | 0.906757 |
| ARID1B | 2498.452 | -0.22911 | 0.164479 | -1.39292 | 0.163644 | 0.312198 |
| FOXP3 | 0.640406 | 1.11402 | 2.710977 | 0.410929 | 0.681124 | NA |
| PPP1R3F | 227.96 | 0.16526 | 0.260294 | 0.6349 | 0.525494 | 0.67109 |
| ARHGEF5 | 87.74116 | -0.60907 | 0.286917 | -2.12281 | 0.03377 | 0.107317 |
| NFE2L3 | 186.2549 | -0.51404 | 0.363807 | -1.41294 | 0.157674 | 0.304087 |
| LAMC3 | 21.44975 | -0.74016 | 0.600069 | -1.23346 | 0.217404 | NA |
| PHPT1 | 2992.352 | -0.04469 | 0.13888 | -0.32179 | 0.747614 | 0.841287 |
| ENTPD2 | 1.599821 | -1.04115 | 1.538529 | -0.67672 | 0.498586 | NA |
| CUL1 | 1473.834 | 0.501461 | 0.173821 | 2.88492 | 0.003915 | 0.025102 |
| KMT2C | 1075.976 | 0.247006 | 0.249476 | 0.990099 | 0.322126 | 0.490281 |
| ZNF280C | 47.01575 | 0.411566 | 0.371624 | 1.10748 | 0.268087 | 0.433052 |
| TRAF1 | 155.1239 | 0.403962 | 0.255235 | 1.582707 | 0.113488 | 0.242328 |
| RC3H2 | 1810.922 | 0.320764 | 0.174159 | 1.84179 | 0.065506 | 0.166907 |
| GYG2 | 2651.854 | -0.55889 | 0.172899 | -3.23248 | 0.001227 | 0.011864 |
| CDK14 | 130.8672 | 0.751576 | 0.294807 | 2.549384 | 0.010791 | 0.049413 |
| CAMK2B | 0.624422 | 1.162588 | 2.733761 | 0.425271 | 0.670639 | NA |
| TBXAS1 | 36.13718 | -0.63427 | 0.430418 | -1.47363 | 0.140583 | NA |
| PARP12 | 377.6839 | -1.05069 | 0.307849 | -3.413 | 6.43E-04 | 0.007876 |
| DNAJC25 | 200.1163 | 0.545031 | 0.202807 | 2.687438 | 0.0072 | 0.037853 |
| MPC1 | 1091.673 | 0.325019 | 0.173812 | 1.869947 | 0.061491 | 0.160074 |
| LZTS1 | 9288.532 | -1.04555 | 0.171292 | -6.10389 | 1.04E-09 | 1.02E-06 |
| ARSF | 0.489058 | -0.58569 | 3.427543 | -0.17088 | 0.864321 | NA |
| MRPS24 | 3303.851 | 0.187204 | 0.181392 | 1.032039 | 0.302054 | 0.469066 |
| ZNF275 | 327.6209 | -0.21417 | 0.193792 | -1.10517 | 0.269085 | 0.433778 |
| MTMR1 | 745.9618 | -0.13288 | 0.155213 | -0.85613 | 0.391927 | 0.557055 |
| TAF2 | 708.2287 | 0.522206 | 0.175819 | 2.970133 | 0.002977 | 0.02103 |
| HIPK2 | 2550.008 | -0.66523 | 0.277176 | -2.40002 | 0.016394 | 0.066132 |
| TNPO3 | 2084.732 | -0.11073 | 0.127176 | -0.87065 | 0.383946 | 0.550052 |
| CDK13 | 960.2716 | -0.02284 | 0.174054 | -0.13123 | 0.895595 | 0.939084 |
| SLC9A7 | 3.282809 | 0.182155 | 1.367638 | 0.13319 | 0.894043 | NA |
| MSANTD3 | 1460.673 | 0.261363 | 0.171922 | 1.520242 | 0.12845 | 0.264779 |
| ARFGEF1 | 884.6795 | 0.673162 | 0.239363 | 2.812303 | 0.004919 | 0.029215 |
| ZFAT | 510.0653 | -0.33741 | 0.226815 | -1.48759 | 0.136859 | 0.276023 |
| MTFR1 | 974.454 | 0.433452 | 0.150397 | 2.882057 | 0.003951 | 0.025263 |
| STAG3 | 27.05752 | -0.31985 | 0.409521 | -0.78103 | 0.434783 | NA |
| DDX3Y | 1255.678 | 0.923104 | 0.192704 | 4.790269 | 1.67E-06 | 2.24E-04 |
| TRAM1 | 3675.455 | 0.554693 | 0.153853 | 3.605343 | 3.12E-04 | 0.005056 |

| | | | | | | |
|---|---|---|---|---|---|---|
| PHKA1 | 204.5377 | -0.01896 | 0.399129 | -0.0475 | 0.962112 | 0.97942 |
| TRO | 839.0634 | -0.24784 | 0.189907 | -1.30507 | 0.19187 | 0.346488 |
| PMS2P4 | 68.33531 | -0.19411 | 0.286272 | -0.67808 | 0.497722 | 0.648692 |
| ZFY | 298.3509 | 0.992598 | 0.252972 | 3.923742 | 8.72E-05 | 0.002349 |
| IDH3G | 2133.246 | -0.13082 | 0.143819 | -0.9096 | 0.363036 | 0.530284 |
| PDZD4 | 267.8704 | -0.38316 | 0.267459 | -1.43258 | 0.151979 | 0.296188 |
| PDK3 | 298.9677 | -0.21416 | 0.181307 | -1.18123 | 0.237513 | 0.400358 |
| OTUD5 | 1989.82 | -0.2647 | 0.125733 | -2.10529 | 0.035266 | 0.110487 |
| TFE3 | 2451.088 | -0.19811 | 0.16436 | -1.20531 | 0.228084 | 0.389884 |
| TBC1D25 | 819.4172 | -0.37842 | 0.192216 | -1.96873 | 0.048984 | 0.136932 |
| ACSL4 | 1397.665 | 0.784919 | 0.224889 | 3.490246 | 4.83E-04 | 0.006662 |
| GPKOW | 600.5011 | -0.27797 | 0.149556 | -1.85861 | 0.063083 | 0.162824 |
| GRIPAP1 | 956.3238 | -0.22216 | 0.16205 | -1.37093 | 0.170397 | 0.320797 |
| FTSJ1 | 1025.983 | -0.1383 | 0.157404 | -0.87862 | 0.379608 | 0.546015 |
| FUNDC1 | 454.9033 | 0.617663 | 0.191229 | 3.229966 | 0.001238 | 0.011874 |
| MAOB | 13.97488 | -0.53307 | 0.601587 | -0.8861 | 0.375563 | NA |
| IKBKAP | 1378.594 | -0.07455 | 0.155435 | -0.47965 | 0.631478 | 0.753743 |
| PTPN3 | 584.213 | 0.201689 | 0.240519 | 0.838556 | 0.401718 | 0.566409 |
| SLC44A1 | 1440.626 | 0.106202 | 0.20754 | 0.511718 | 0.608848 | 0.735919 |
| POLB | 1337.102 | 0.564711 | 0.225587 | 2.503292 | 0.012304 | 0.054267 |
| FRMPD1 | 0.488912 | 0.771583 | 2.897678 | 0.266276 | 0.790026 | NA |
| GBA2 | 1268.856 | -0.12369 | 0.148732 | -0.83165 | 0.405608 | 0.56959 |
| ASNS | 171.8694 | 0.309745 | 0.248344 | 1.247242 | 0.212309 | 0.370899 |
| AP3M2 | 1431.342 | -0.2661 | 0.159768 | -1.66553 | 0.095807 | 0.216643 |
| PABPC1 | 21967.07 | -0.20972 | 0.187911 | -1.11605 | 0.264399 | 0.428902 |
| OSBPL3 | 683.8737 | 0.272685 | 0.16386 | 1.664131 | 0.096086 | 0.217137 |
| SNX13 | 752.2533 | 0.76201 | 0.224107 | 3.400209 | 6.73E-04 | 0.008102 |
| RPS6KA2 | 8048.108 | -0.0601 | 0.254849 | -0.23581 | 0.813577 | 0.888773 |
| ING3 | 191.6004 | 0.596794 | 0.223132 | 2.674622 | 0.007481 | 0.039002 |
| WBSCR22 | 2394.351 | 0.024635 | 0.135568 | 0.181714 | 0.855807 | 0.914347 |
| ATP6AP1 | 7434.317 | -0.42279 | 0.145414 | -2.90753 | 0.003643 | 0.024068 |
| FAM50A | 2579.332 | 0.086063 | 0.138131 | 0.623052 | 0.53325 | 0.677202 |
| FAM3A | 1839.667 | -0.15194 | 0.128586 | -1.18161 | 0.237361 | 0.400283 |
| CPSF1 | 2232.723 | -0.45352 | 0.134689 | -3.36714 | 7.60E-04 | 0.008794 |
| PDCD2 | 1511.248 | 0.097098 | 0.14788 | 0.656599 | 0.511439 | 0.659379 |
| SMC1A | 2121.27 | -0.1217 | 0.143172 | -0.85003 | 0.395308 | 0.559815 |
| HSD17B10 | 2510.95 | -0.09098 | 0.150654 | -0.60393 | 0.545892 | 0.687439 |
| IKBKG | 513.0939 | -0.21382 | 0.15344 | -1.39352 | 0.163462 | 0.312042 |
| CYP2W1 | 0.545988 | -0.43913 | 2.541145 | -0.17281 | 0.862803 | NA |
| CLCN4 | 31.78505 | -1.06504 | 0.474683 | -2.24369 | 0.024852 | NA |
| VDAC1P1 | 4.267201 | 0.20827 | 1.11369 | 0.187009 | 0.851653 | NA |
| IPCEF1 | 0.661122 | -3.06315 | 2.98049 | -1.02773 | 0.304076 | NA |
| SRI | 2428.943 | 0.205021 | 0.171525 | 1.195286 | 0.231975 | 0.394248 |
| SEMA3A | 31.01026 | 0.418631 | 0.462436 | 0.905273 | 0.365321 | NA |
| SEMA3C | 2387.122 | 0.455109 | 0.291258 | 1.562564 | 0.118155 | 0.249487 |
| SLC25A40 | 413.0223 | 0.433906 | 0.172423 | 2.516521 | 0.011852 | 0.052881 |
| FSCN1 | 570.5439 | -0.6114 | 0.579235 | -1.05552 | 0.291185 | 0.458082 |

| | | | | | | |
|---|---|---|---|---|---|---|
| ACTB | 81234.85 | -0.48208 | 0.175606 | -2.74526 | 0.006046 | 0.03362 |
| BCAP29 | 960.4632 | 0.953174 | 0.227619 | 4.187585 | 2.82E-05 | 0.001145 |
| TPD52 | 219.787 | 0.06839 | 0.193316 | 0.353775 | 0.723508 | 0.824463 |
| PAG1 | 2925.308 | 0.162474 | 0.195028 | 0.833082 | 0.404799 | 0.569128 |
| GPC4 | 269.756 | -0.02173 | 0.307502 | -0.07065 | 0.943675 | 0.968771 |
| MBNL3 | 185.1699 | 0.371096 | 0.250421 | 1.481885 | 0.138371 | 0.277985 |
| CTTNBP2 | 524.6493 | 0.21246 | 0.166689 | 1.274587 | 0.202456 | 0.358815 |
| LRCH4 | 1907.643 | -0.4141 | 0.139131 | -2.97634 | 0.002917 | 0.020745 |
| SLC25A43 | 1206.05 | 0.025735 | 0.147858 | 0.174054 | 0.861823 | 0.91821 |
| UBE2A | 2002.7 | 0.046945 | 0.124046 | 0.378451 | 0.705095 | 0.810836 |
| FGFR1 | 2635.891 | -0.28081 | 0.150263 | -1.86877 | 0.061655 | 0.16029 |
| FKBP6 | 0.651435 | 0.249629 | 2.788679 | 0.089515 | 0.928672 | NA |
| GTF2I | 1445.917 | -0.25515 | 0.173216 | -1.47299 | 0.140752 | 0.281197 |
| AMPH | 32.9514 | -0.67021 | 0.519902 | -1.28911 | 0.197359 | NA |
| ARAF | 1523.425 | -0.32433 | 0.143595 | -2.25862 | 0.023907 | 0.085393 |
| SYNJ2 | 119.8293 | 0.157036 | 0.354103 | 0.443476 | 0.657421 | 0.774109 |
| PMS2P1 | 160.4469 | -0.10106 | 0.217594 | -0.46444 | 0.642329 | 0.761939 |
| HOXA9 | 8.357303 | 0.228355 | 0.720828 | 0.316795 | 0.751399 | NA |
| ZCWPW1 | 59.53623 | 0.48693 | 0.312867 | 1.556345 | 0.119626 | 0.251537 |
| FGF20 | 0.679532 | 2.619206 | 2.901822 | 0.902607 | 0.366734 | NA |
| ITM2A | 1611.142 | -0.06723 | 0.21506 | -0.31261 | 0.754578 | 0.846873 |
| VDAC3 | 3090.876 | 0.078208 | 0.135775 | 0.57601 | 0.564608 | 0.70228 |
| PCM1 | 987.7899 | 0.850299 | 0.608249 | 1.397946 | 0.162129 | 0.310007 |
| BRINP1 | 0.460466 | 0.568655 | 3.014642 | 0.188631 | 0.850382 | NA |
| UBE2D4 | 160.3738 | -0.07287 | 0.210359 | -0.34643 | 0.729022 | 0.827945 |
| RUNX1T1 | 6.961074 | 0.689183 | 0.765769 | 0.899989 | 0.368126 | NA |
| CDH17 | 0.326333 | 1.533889 | 3.826878 | 0.40082 | 0.688553 | NA |
| FDFT1 | 11215.62 | -0.18199 | 0.177524 | -1.02516 | 0.305287 | 0.472538 |
| OPHN1 | 298.1804 | -0.3914 | 0.217003 | -1.80365 | 0.071286 | 0.176762 |
| RFX3 | 92.93182 | 0.658736 | 0.316003 | 2.084587 | 0.037107 | 0.114349 |
| SMARCA2 | 2120.336 | 0.088374 | 0.187265 | 0.471916 | 0.636987 | 0.7578 |
| MID2 | 168.5838 | -0.19641 | 0.285002 | -0.68916 | 0.490722 | 0.642818 |
| KIAA0020 | 434.6099 | 0.978329 | 0.328314 | 2.979856 | 0.002884 | 0.020568 |
| CNOT4 | 469.2061 | 0.125549 | 0.205895 | 0.609771 | 0.542014 | 0.684154 |
| CDC14B | 691.1007 | 0.121694 | 0.149907 | 0.811799 | 0.416907 | 0.580061 |
| ZNF510 | 189.12 | 0.861614 | 0.259499 | 3.320302 | 8.99E-04 | 0.009725 |
| CADPS2 | 217.2217 | 0.012009 | 0.238563 | 0.050339 | 0.959852 | 0.977917 |
| SMARCD3 | 766.4616 | -0.11736 | 0.178438 | -0.65773 | 0.510711 | 0.659019 |
| DLG3 | 207.6504 | 0.178008 | 0.201415 | 0.883787 | 0.376811 | 0.543599 |
| TRPM3 | 68.14391 | 0.688724 | 0.330156 | 2.086058 | 0.036973 | 0.114027 |
| KAT6A | 1259.211 | -0.2563 | 0.187895 | -1.36405 | 0.172551 | 0.323928 |
| ZCCHC6 | 414.6003 | 0.82016 | 0.262646 | 3.122679 | 0.001792 | 0.015079 |
| GRHL2 | 0.096202 | 0.408546 | 4.079639 | 0.100143 | 0.920231 | NA |
| RRAGB | 325.5917 | -0.05926 | 0.186526 | -0.31769 | 0.750722 | 0.844063 |
| BCORL1 | 486.9329 | -0.67647 | 0.198551 | -3.40701 | 6.57E-04 | 0.007978 |
| ATRX | 830.1101 | 0.721121 | 0.997097 | 0.723221 | 0.469544 | 0.625866 |
| MAP3K4 | 346.4718 | 0.288516 | 0.172865 | 1.669024 | 0.095113 | 0.215801 |

| | | | | | | |
|---|---|---|---|---|---|---|
| PILRA | 0.47961 | 0.535796 | 3.39605 | 0.15777 | 0.874638 | NA |
| ABCB1 | 32.42389 | 0.702091 | 0.485283 | 1.446767 | 0.147962 | NA |
| AKR1B1 | 10074.63 | 0.290496 | 0.345782 | 0.840114 | 0.400844 | 0.565475 |
| CPNE3 | 2604.498 | 0.120348 | 0.171461 | 0.701899 | 0.482742 | 0.636586 |
| DDHD2 | 1894.847 | 0.270891 | 0.156071 | 1.735689 | 0.082619 | 0.195427 |
| DNAJA1 | 5599.169 | 0.645947 | 0.178403 | 3.620727 | 2.94E-04 | 0.004857 |
| B4GALT1 | 2322.775 | -0.45443 | 0.144362 | -3.14787 | 0.001645 | 0.014318 |
| CHMP5 | 2190.814 | 0.679639 | 0.175834 | 3.865226 | 1.11E-04 | 0.002747 |
| NFX1 | 1101.677 | 0.088316 | 0.143067 | 0.617306 | 0.537033 | 0.680184 |
| EIF2AK1 | 3619.151 | -0.04522 | 0.137877 | -0.32799 | 0.742922 | 0.837997 |
| EPDR1 | 2164.381 | 0.022733 | 0.144122 | 0.157736 | 0.874665 | 0.925156 |
| SNX10 | 1107.197 | 0.643178 | 0.254622 | 2.526007 | 0.011537 | 0.051904 |
| TXLNG | 288.8186 | 0.574749 | 0.204807 | 2.806293 | 0.005012 | 0.029676 |
| PPEF1 | 22.26925 | 0.094135 | 0.500439 | 0.188104 | 0.850795 | NA |
| LAT2 | 6.487656 | -0.07935 | 0.822632 | -0.09646 | 0.923157 | NA |
| HUWE1 | 8298.023 | -0.60242 | 0.167219 | -3.60258 | 3.15E-04 | 0.005089 |
| TRIP6 | 4817.494 | -0.19359 | 0.136841 | -1.41474 | 0.157146 | 0.303367 |
| ACHE | 98.77913 | -0.25754 | 0.239885 | -1.0736 | 0.283002 | 0.449369 |
| SRRT | 3084.151 | -0.20898 | 0.163092 | -1.28137 | 0.200065 | 0.356067 |
| PIR | 7921.261 | 0.063021 | 0.154957 | 0.406701 | 0.684228 | 0.794605 |
| IGBP1 | 1695.9 | 0.040253 | 0.129024 | 0.311977 | 0.755058 | 0.847146 |
| HEPH | 963.0458 | -0.52803 | 0.131257 | -4.0229 | 5.75E-05 | 0.001791 |
| RBM41 | 164.3729 | 0.786876 | 0.34486 | 2.281726 | 0.022506 | 0.081968 |
| ARHGAP4 | 34.86343 | -0.76464 | 0.389259 | -1.96436 | 0.049489 | NA |
| SPTLC1 | 1293.619 | 0.398018 | 0.147329 | 2.701559 | 0.006902 | 0.036755 |
| MRPS33 | 1032.533 | 0.314143 | 0.196308 | 1.600254 | 0.109542 | 0.236889 |
| NDUFB2 | 2189.934 | 0.078379 | 0.131222 | 0.597301 | 0.550306 | 0.691602 |
| EFNB1 | 203.0256 | -0.59104 | 0.202008 | -2.9258 | 0.003436 | 0.022986 |
| KIF4A | 945.9934 | -0.07504 | 0.169451 | -0.44283 | 0.657892 | 0.774307 |
| DTX2 | 257.4404 | -0.18066 | 0.216721 | -0.83362 | 0.404495 | 0.569031 |
| PUS7 | 297.2731 | 0.601254 | 0.182757 | 3.289903 | 0.001002 | 0.01035 |
| LAMB4 | 2.080569 | 0.151284 | 1.434694 | 0.105447 | 0.916021 | NA |
| NRCAM | 3079.502 | 0.732132 | 0.20451 | 3.579941 | 3.44E-04 | 0.00538 |
| LAMB1 | 3039.141 | -0.05253 | 0.188809 | -0.27821 | 0.780854 | 0.8655 |
| SLC26A4 | 16.74205 | 1.088323 | 0.574227 | 1.895284 | 0.058055 | NA |
| DLD | 1822.144 | 0.508596 | 0.185923 | 2.735519 | 0.006228 | 0.034188 |
| SMPX | 0.624625 | 0.096176 | 2.664509 | 0.036095 | 0.971207 | NA |
| ZFHX4 | 20.27027 | 0.125778 | 0.515454 | 0.244014 | 0.80722 | NA |
| ZC3HC1 | 603.7643 | -0.2991 | 0.152615 | -1.95985 | 0.050013 | 0.138529 |
| ESR1 | 68.48403 | -0.22081 | 0.292373 | -0.75523 | 0.450113 | 0.609421 |
| RGS17 | 29.32743 | 0.918365 | 0.436694 | 2.102994 | 0.035466 | NA |
| ANGPT2 | 88.54695 | 0.107968 | 0.461723 | 0.233837 | 0.815112 | 0.889858 |
| TBL1Y | 0.162376 | 0.424913 | 4.079006 | 0.104171 | 0.917034 | NA |
| EZR | 6700.798 | -0.30778 | 0.118908 | -2.5884 | 0.009642 | 0.046045 |
| DPYSL2 | 3728.017 | -0.39873 | 0.197347 | -2.02045 | 0.043336 | 0.126884 |
| HDAC6 | 1433.264 | -0.46149 | 0.131049 | -3.52148 | 4.29E-04 | 0.006182 |
| UPRT | 389.7973 | 0.212791 | 0.181791 | 1.170526 | 0.241789 | 0.404887 |

| | | | | | | |
|---|---|---|---|---|---|---|
| EPB41L4B | 2.298883 | -0.68755 | 1.348458 | -0.50988 | 0.610137 | NA |
| TMEM38B | 366.7962 | 0.582678 | 0.191588 | 3.041298 | 0.002356 | 0.017982 |
| PSMD5 | 1186.532 | 0.080536 | 0.16489 | 0.488424 | 0.62525 | 0.748721 |
| PTGS1 | 6.26539 | -0.18249 | 0.844213 | -0.21616 | 0.82886 | NA |
| NUP188 | 1677.722 | -0.5003 | 0.140946 | -3.54961 | 3.86E-04 | 0.005775 |
| CRAT | 1134.478 | -0.30616 | 0.145078 | -2.11032 | 0.034831 | 0.109586 |
| SH2D3C | 17.25636 | -0.34544 | 0.540258 | -0.6394 | 0.522562 | NA |
| NANS | 1175.012 | -0.01019 | 0.142607 | -0.07148 | 0.943012 | 0.968558 |
| TBC1D2 | 207.8852 | -0.18341 | 0.206036 | -0.89021 | 0.373355 | 0.539792 |
| DFNB31 | 1221.794 | 0.048167 | 0.186812 | 0.257835 | 0.796534 | 0.87685 |
| IFT74 | 103.6431 | 1.10025 | 0.891775 | 1.233776 | 0.217286 | 0.37671 |
| JAK2 | 226.5514 | 0.599564 | 0.309393 | 1.937872 | 0.052639 | 0.143258 |
| ABL1 | 3851.503 | -0.48547 | 0.135009 | -3.59582 | 3.23E-04 | 0.005181 |
| PCSK5 | 17.0311 | -0.3516 | 0.526473 | -0.66784 | 0.504238 | NA |
| ERMP1 | 771.6512 | -0.25282 | 0.157018 | -1.61012 | 0.107372 | 0.233913 |
| AMELY | 0.364467 | -1.20166 | 3.685431 | -0.32606 | 0.744381 | NA |
| PRKY | 213.8389 | 0.576412 | 0.200408 | 2.876191 | 0.004025 | 0.025476 |
| MTAP | 843.8772 | 0.202187 | 0.18758 | 1.077873 | 0.28109 | 0.447292 |
| CSTF2 | 506.218 | -0.09342 | 0.151949 | -0.61484 | 0.53866 | 0.681093 |
| MXRA5 | 0.230334 | -0.53318 | 4.076749 | -0.13079 | 0.895945 | NA |
| VSIG1 | 0.949378 | 0.969416 | 2.078125 | 0.466486 | 0.640868 | NA |
| PSMD10 | 1717.985 | 0.239115 | 0.161666 | 1.479067 | 0.139122 | 0.279116 |
| ATG4A | 574.5234 | 0.40859 | 0.189421 | 2.157051 | 0.031002 | 0.101442 |
| STS | 524.1822 | 0.100841 | 0.194499 | 0.518465 | 0.604134 | 0.732942 |
| TBL1X | 941.4216 | -0.41373 | 0.164392 | -2.51676 | 0.011844 | 0.052866 |
| GPR143 | 7159.618 | -0.4716 | 0.206082 | -2.28841 | 0.022114 | 0.081088 |
| PGRMC1 | 2671.335 | 0.102689 | 0.142523 | 0.720506 | 0.471213 | 0.627368 |
| POLA1 | 522.2301 | 0.238753 | 0.144651 | 1.65055 | 0.098831 | 0.221607 |
| MID1 | 1973.509 | 0.116532 | 0.168236 | 0.69267 | 0.488516 | 0.641347 |
| NKAP | 305.0306 | 0.614859 | 0.244181 | 2.518046 | 0.011801 | 0.052732 |
| NXT2 | 213.0721 | 0.537348 | 0.225035 | 2.387842 | 0.016948 | 0.067758 |
| ALG13 | 334.3298 | 0.201269 | 0.189556 | 1.061797 | 0.288328 | 0.454443 |
| PRPS2 | 967.7819 | -0.03361 | 0.185668 | -0.18104 | 0.856339 | 0.914668 |
| MOSPD1 | 505.5896 | 0.503567 | 0.188262 | 2.674825 | 0.007477 | 0.038996 |
| AMMECR1 | 366.6302 | 0.255784 | 0.213226 | 1.199591 | 0.230298 | 0.39255 |
| WDR13 | 3075.758 | -0.15471 | 0.11968 | -1.29274 | 0.1961 | 0.351492 |
| SUV39H1 | 369.2349 | -0.5496 | 0.218606 | -2.5141 | 0.011934 | 0.053166 |
| SRPX | 4375.946 | -0.50333 | 0.143155 | -3.51602 | 4.38E-04 | 0.006274 |
| GLRA2 | 0.452561 | 2.061505 | 3.349456 | 0.615474 | 0.538241 | NA |
| XIAP | 543.7149 | 0.417184 | 0.263379 | 1.583968 | 0.113201 | 0.241987 |
| STAG2 | 1149.449 | 0.845526 | 0.253086 | 3.34087 | 8.35E-04 | 0.009297 |
| ATP11C | 440.3061 | 0.58697 | 0.250642 | 2.341869 | 0.019187 | 0.073519 |
| MCF2 | 68.24651 | 0.845958 | 0.370873 | 2.280994 | 0.022549 | 0.081999 |
| ABCD1 | 2078.922 | -0.62644 | 0.24556 | -2.55108 | 0.010739 | 0.049294 |
| CCDC22 | 510.0384 | -0.4441 | 0.174568 | -2.54399 | 0.010959 | 0.049971 |
| CACNA1F | 2.53155 | 0.749721 | 1.312559 | 0.57119 | 0.567871 | NA |
| SYP | 127.3598 | -0.05248 | 0.248719 | -0.21101 | 0.832878 | 0.899742 |

| | | | | | | |
|---|---|---|---|---|---|---|
| PLP2 | 1769.878 | -0.07912 | 0.235905 | -0.33538 | 0.737337 | 0.83368 |
| BMX | 0.766634 | 1.516849 | 2.262991 | 0.670285 | 0.502676 | NA |
| LUZP4 | 1.571281 | -2.11612 | 2.025059 | -1.04497 | 0.296038 | NA |
| PLS3 | 5265.623 | 0.322516 | 0.178382 | 1.808005 | 0.070606 | 0.175588 |
| NAA10 | 1322.812 | 0.086956 | 0.137347 | 0.633108 | 0.526663 | 0.672149 |
| RENBP | 2288.427 | -0.18538 | 0.181792 | -1.01976 | 0.307841 | 0.475128 |
| ELF4 | 671.3589 | -0.42344 | 0.173017 | -2.44738 | 0.01439 | 0.060365 |
| SMARCA1 | 1755.392 | 0.595701 | 0.255502 | 2.331493 | 0.019727 | 0.07503 |
| MTMR8 | 44.51572 | 0.180937 | 0.333416 | 0.542677 | 0.587352 | NA |
| ASB9 | 420.0977 | -0.18394 | 0.193015 | -0.95301 | 0.340587 | 0.5087 |
| ZC3H12B | 4.101372 | 0.708971 | 1.008181 | 0.703218 | 0.48192 | NA |
| RBBP7 | 3269.424 | -0.00788 | 0.125649 | -0.06271 | 0.949994 | 0.972303 |
| KCND1 | 64.68888 | -0.69458 | 0.335769 | -2.06861 | 0.038582 | 0.117311 |
| UBE2NL | 2.026079 | -0.68562 | 1.438825 | -0.47651 | 0.63371 | NA |
| SLC25A14 | 276.2973 | 0.243204 | 0.193933 | 1.254064 | 0.209819 | 0.367415 |
| FMR1 | 729.1718 | 0.318892 | 0.237969 | 1.340057 | 0.180227 | 0.332803 |
| PIM2 | 334.2989 | -0.23072 | 0.194735 | -1.18478 | 0.236104 | 0.399117 |
| SCML2 | 34.32495 | -0.10634 | 0.366771 | -0.28994 | 0.771862 | NA |
| SLC35A2 | 906.9189 | -0.11882 | 0.174021 | -0.68278 | 0.494746 | 0.645807 |
| PQBP1 | 1615 | 0.005381 | 0.158645 | 0.033916 | 0.972944 | 0.985082 |
| PCSK1N | 547.7257 | 0.330752 | 0.205262 | 1.611369 | 0.107099 | 0.233566 |
| EMD | 1768.637 | -0.23971 | 0.149418 | -1.60428 | 0.108653 | 0.235566 |
| TAZ | 521.2926 | -0.12668 | 0.168925 | -0.7499 | 0.453312 | 0.612214 |
| RAB40AL | 0.163957 | 0.424913 | 4.079006 | 0.104171 | 0.917034 | NA |
| PGK1 | 17042.21 | -0.05477 | 0.139891 | -0.39155 | 0.695393 | 0.803419 |
| MAGT1 | 2648.255 | -0.15123 | 0.143226 | -1.05589 | 0.291019 | 0.457928 |
| SMS | 2563.124 | 0.146001 | 0.136643 | 1.068482 | 0.285303 | 0.451752 |
| PHEX | 5.096126 | 0.39375 | 0.944519 | 0.416879 | 0.676767 | NA |
| UBL4A | 2263.818 | -0.24839 | 0.148876 | -1.66842 | 0.095233 | 0.215867 |
| CD99L2 | 4768.554 | -0.49399 | 0.12801 | -3.85903 | 1.14E-04 | 0.002777 |
| RP2 | 696.251 | 0.768488 | 0.185692 | 4.13851 | 3.50E-05 | 0.001341 |
| JADE3 | 180.422 | -0.11378 | 0.212082 | -0.53647 | 0.591634 | 0.723 |
| CDK16 | 3863.04 | -0.37732 | 0.129066 | -2.92346 | 0.003462 | 0.02312 |
| USP11 | 2549.607 | -0.42203 | 0.198271 | -2.12857 | 0.03329 | 0.106347 |
| PCYT1B | 32.72735 | -0.84735 | 0.387352 | -2.18755 | 0.028702 | NA |
| BRS3 | 0.096202 | 0.408546 | 4.079639 | 0.100143 | 0.920231 | NA |
| HTATSF1 | 826.857 | 0.666199 | 0.331074 | 2.012234 | 0.044195 | 0.128169 |
| CD40LG | 0.096202 | 0.408546 | 4.079639 | 0.100143 | 0.920231 | NA |
| TIMP1 | 5981.372 | -0.05856 | 0.251834 | -0.23253 | 0.816125 | 0.889858 |
| PCDH11X | 1.203958 | 0.509109 | 1.966676 | 0.258868 | 0.795737 | NA |
| FGD1 | 1242.565 | -0.59 | 0.140485 | -4.19971 | 2.67E-05 | 0.001097 |
| PIN4 | 425.747 | 0.523542 | 0.156328 | 3.348999 | 8.11E-04 | 0.009131 |
| PORCN | 302.8689 | -0.1949 | 0.181399 | -1.07444 | 0.282628 | 0.449015 |
| ITIH6 | 40.01613 | 0.866926 | 0.500034 | 1.733734 | 0.082965 | NA |
| MAGED2 | 10752.08 | -0.08813 | 0.138662 | -0.63558 | 0.525048 | 0.67081 |
| RBM3 | 7213.895 | -0.05984 | 0.227229 | -0.26335 | 0.792281 | 0.874208 |
| KLF8 | 185.1535 | -0.052 | 0.214517 | -0.24239 | 0.808478 | 0.88565 |

| | | | | | | |
|---|---|---|---|---|---|---|
| SRPX2 | 175.6535 | 0.189628 | 0.351935 | 0.538816 | 0.590014 | 0.721887 |
| SYTL4 | 254.5292 | 0.272459 | 0.233454 | 1.167077 | 0.243179 | 0.406164 |
| ZDHHC15 | 0.192403 | 0.975392 | 4.060716 | 0.240202 | 0.810174 | NA |
| CENPI | 265.1846 | 0.245164 | 0.22789 | 1.0758 | 0.282016 | 0.448104 |
| DRP2 | 3.917637 | 0.114344 | 1.045479 | 0.10937 | 0.912909 | NA |
| TAF7L | 66.41931 | 0.340828 | 0.28957 | 1.177014 | 0.23919 | 0.402358 |
| PBDC1 | 432.593 | 0.570882 | 0.249956 | 2.283927 | 0.022376 | 0.081821 |
| GLA | 3651.04 | 0.150314 | 0.184304 | 0.815573 | 0.414744 | 0.578106 |
| ARMCX3 | 2021.99 | 0.440823 | 0.268981 | 1.638865 | 0.101241 | 0.224802 |
| BEX4 | 2185.715 | 0.133571 | 0.191822 | 0.696329 | 0.486223 | 0.639469 |
| SGK3 | 390.6725 | 0.417165 | 0.241782 | 1.725377 | 0.08446 | 0.198036 |
| PDGFRL | 5.904064 | -0.15757 | 0.936619 | -0.16823 | 0.8664 | NA |
| CSPP1 | 150.4404 | 0.668976 | 0.293799 | 2.276987 | 0.022787 | 0.082561 |
| ZDHHC2 | 412.0563 | 0.368122 | 0.200858 | 1.832745 | 0.066841 | 0.16899 |
| BRF2 | 230.2297 | -0.25968 | 0.17747 | -1.46325 | 0.143398 | 0.284812 |
| TRIM35 | 1168.378 | -0.29399 | 0.151399 | -1.94185 | 0.052155 | 0.142432 |
| ZFAND1 | 388.9245 | 0.612984 | 0.200077 | 3.063744 | 0.002186 | 0.017131 |
| RP1 | 10.17903 | 0.560651 | 0.711944 | 0.787494 | 0.430993 | NA |
| FZD3 | 677.9309 | 0.236275 | 0.199444 | 1.184671 | 0.236148 | 0.399134 |
| INTS9 | 843.4445 | -0.30596 | 0.156834 | -1.95086 | 0.051074 | 0.140645 |
| RIPK2 | 496.5182 | 0.108145 | 0.178048 | 0.607389 | 0.543593 | 0.685451 |
| EYA1 | 0.793683 | 0.370036 | 2.527443 | 0.146407 | 0.8836 | NA |
| NBN | 817.6449 | 0.840732 | 0.257061 | 3.270559 | 0.001073 | 0.010848 |
| TRPA1 | 4.988372 | 0.546363 | 0.962155 | 0.567853 | 0.570135 | NA |
| CPQ | 5486.006 | -0.16668 | 0.213125 | -0.78208 | 0.434168 | 0.59538 |
| DECR1 | 2206.404 | 0.056086 | 0.135408 | 0.414197 | 0.67873 | 0.790407 |
| CALB1 | 0.629726 | -0.22725 | 2.732935 | -0.08315 | 0.93373 | NA |
| IMPAD1 | 2368.81 | 0.338103 | 0.163783 | 2.06433 | 0.038986 | 0.118145 |
| SFRP1 | 118.6302 | 0.344736 | 0.311709 | 1.105955 | 0.268746 | 0.433655 |
| LAPTM4B | 5294.544 | -0.27382 | 0.125493 | -2.18192 | 0.029115 | 0.09701 |
| UBE2W | 426.1053 | 0.572556 | 0.233847 | 2.448425 | 0.014348 | 0.060233 |
| POP1 | 651.819 | 0.050104 | 0.188906 | 0.265234 | 0.790829 | 0.873094 |
| NIPAL2 | 600.0351 | 0.200656 | 0.173296 | 1.157883 | 0.246912 | 0.410366 |
| IKBKB | 1759.787 | -0.06746 | 0.142289 | -0.47413 | 0.635411 | 0.756456 |
| PLAT | 2960.322 | -0.32233 | 0.139569 | -2.30948 | 0.020917 | 0.078113 |
| JPH1 | 844.4943 | 0.584649 | 0.30741 | 1.901856 | 0.05719 | 0.151641 |
| DKK4 | 0.247898 | -0.5901 | 4.074633 | -0.14482 | 0.884851 | NA |
| STK3 | 405.3035 | 0.719765 | 0.221038 | 3.2563 | 0.001129 | 0.011188 |
| GDAP1 | 357.7112 | 0.553676 | 0.205424 | 2.695288 | 0.007033 | 0.037237 |
| RAB2A | 3561.18 | 0.317839 | 0.133968 | 2.372503 | 0.017668 | 0.069586 |
| EIF3E | 4398.405 | 0.691135 | 0.230484 | 2.998621 | 0.002712 | 0.019907 |
| EMC2 | 526.3137 | 0.741201 | 0.305342 | 2.427445 | 0.015206 | 0.0627 |
| ESRP1 | 1161.531 | 0.535153 | 0.209164 | 2.558536 | 0.010511 | 0.048662 |
| WISP1 | 7.10926 | -0.41189 | 0.814471 | -0.50572 | 0.613056 | NA |
| NDRG1 | 1846.995 | -0.47024 | 0.143275 | -3.28206 | 0.001031 | 0.01055 |
| ZC2HC1A | 116.032 | 0.659871 | 0.234525 | 2.813647 | 0.004898 | 0.029107 |
| IL7 | 19.35184 | 1.168669 | 0.523135 | 2.233974 | 0.025485 | NA |

| | | | | | | |
|---|---|---|---|---|---|---|
| STMN2 | 5.722143 | 1.046796 | 0.927234 | 1.128944 | 0.258921 | NA |
| ARMC1 | 691.851 | 0.65124 | 0.176024 | 3.699724 | 2.16E-04 | 0.004081 |
| TRPS1 | 253.4777 | 0.355623 | 0.235299 | 1.511364 | 0.130696 | 0.267092 |
| SPAG1 | 190.7482 | 0.713235 | 0.27018 | 2.639852 | 0.008294 | 0.041553 |
| CHRAC1 | 402.2631 | 0.123566 | 0.168091 | 0.735113 | 0.462271 | 0.620395 |
| NCALD | 488.0154 | -0.57245 | 0.21223 | -2.69733 | 0.00699 | 0.037109 |
| SNX16 | 188.7524 | 0.719151 | 0.204144 | 3.522758 | 4.27E-04 | 0.006167 |
| UBR5 | 2041.138 | 0.022422 | 0.170661 | 0.131382 | 0.895473 | 0.939039 |
| GSDMD | 1327.53 | -0.19977 | 0.151935 | -1.31487 | 0.188555 | 0.342841 |
| TSTA3 | 1909.438 | -0.12608 | 0.125265 | -1.00653 | 0.314163 | 0.481589 |
| PYCRL | 421.71 | -0.23429 | 0.250466 | -0.93543 | 0.349567 | 0.517627 |
| EEF1D | 5414.103 | -0.11342 | 0.127015 | -0.893 | 0.371859 | 0.538594 |
| ANXA13 | 4.583956 | -0.39228 | 1.138868 | -0.34445 | 0.730507 | NA |
| SQLE | 3591.17 | 0.407879 | 0.175424 | 2.325098 | 0.020067 | 0.075931 |
| SH2D4A | 37.67944 | -0.18164 | 0.466572 | -0.38931 | 0.697048 | NA |
| INTS10 | 1347.971 | 0.355943 | 0.166545 | 2.137213 | 0.032581 | 0.105021 |
| ERI1 | 411.9537 | 0.484238 | 0.16477 | 2.938882 | 0.003294 | 0.02243 |
| SLC39A14 | 2340.897 | -0.11617 | 0.254148 | -0.45709 | 0.647603 | 0.766201 |
| MTMR9 | 317.7453 | 0.250787 | 0.211443 | 1.186072 | 0.235594 | 0.398483 |
| LEPROTL1 | 1746.814 | 0.137807 | 0.131617 | 1.047027 | 0.295087 | 0.462427 |
| DCTN6 | 417.9535 | 0.747533 | 0.180266 | 4.146826 | 3.37E-05 | 0.001308 |
| R3HCC1 | 856.0179 | -0.17341 | 0.150395 | -1.15306 | 0.248886 | 0.412786 |
| GSR | 2413.757 | -0.24356 | 0.141951 | -1.71577 | 0.086205 | 0.201133 |
| TNFRSF10A | 67.55718 | 0.061474 | 0.340921 | 0.180317 | 0.856904 | 0.914942 |
| UBXN8 | 535.8622 | 0.448618 | 0.187773 | 2.389143 | 0.016888 | 0.067591 |
| PPP2CB | 2718.169 | 0.09653 | 0.144535 | 0.667865 | 0.50422 | 0.653944 |
| ERICH1 | 629.7139 | 0.384371 | 0.148181 | 2.593929 | 0.009489 | 0.045544 |
| NEFM | 34.9088 | 0.990931 | 0.442351 | 2.240148 | 0.025081 | NA |
| TUSC3 | 1177.98 | 0.082824 | 0.215333 | 0.384633 | 0.700509 | 0.807363 |
| NEFL | 32.58922 | 0.978859 | 0.568028 | 1.723258 | 0.084842 | NA |
| ARHGEF10 | 1008.629 | 0.13459 | 0.191339 | 0.703411 | 0.4818 | 0.636051 |
| MCM4 | 1621.402 | -0.00254 | 0.217446 | -0.01168 | 0.990682 | 0.995806 |
| KCTD9 | 820.0101 | 0.631655 | 0.171303 | 3.68735 | 2.27E-04 | 0.004193 |
| FGL1 | 0.327913 | 1.543154 | 4.021063 | 0.383768 | 0.701151 | NA |
| ASAH1 | 25293.74 | 0.238319 | 0.145999 | 1.632332 | 0.10261 | 0.226809 |
| BNIP3L | 3201.356 | 0.308717 | 0.158514 | 1.94757 | 0.051466 | 0.141276 |
| DENND3 | 116.8594 | -0.01602 | 0.238448 | -0.06718 | 0.946436 | 0.970016 |
| AVL9 | 950.5005 | 0.083339 | 0.154317 | 0.540055 | 0.589159 | 0.721298 |
| RUNDC3B | 45.54433 | 0.150776 | 0.357412 | 0.421854 | 0.673131 | NA |
| C7orf63 | 28.2176 | 1.291742 | 0.480304 | 2.689426 | 0.007157 | NA |
| GTPBP10 | 268.6668 | 0.699364 | 0.246924 | 2.832305 | 0.004621 | 0.027963 |
| RASA4 | 0.508056 | 0.731032 | 2.861912 | 0.255435 | 0.798387 | NA |
| CDK6 | 588.0753 | 0.325682 | 0.291281 | 1.1181 | 0.263524 | 0.427839 |
| PMPCB | 1564.754 | 0.046275 | 0.17933 | 0.258041 | 0.796375 | 0.87685 |
| DNAJC2 | 176.9481 | 1.276316 | 0.914872 | 1.395077 | 0.162993 | 0.311306 |
| TFPI2 | 5.333902 | 1.537235 | 1.068743 | 1.438358 | 0.150332 | NA |
| BET1 | 330.4191 | 0.683231 | 0.238904 | 2.859856 | 0.004238 | 0.026417 |

| | | | | | | |
|---|---|---|---|---|---|---|
| NAMPT | 1656.275 | 0.479164 | 0.24769 | 1.934529 | 0.053048 | 0.144021 |
| TWISTNB | 370.597 | 0.788778 | 0.221063 | 3.568107 | 3.60E-04 | 0.005535 |
| PIK3CG | 0.545988 | -0.43913 | 2.541145 | -0.17281 | 0.862803 | NA |
| PON3 | 5.745362 | 2.754252 | 1.162132 | 2.37 | 0.017788 | NA |
| PON2 | 2661.42 | 0.446483 | 0.14827 | 3.011276 | 0.002602 | 0.019343 |
| ITGB8 | 1561.098 | 0.921768 | 0.220355 | 4.183098 | 2.88E-05 | 0.00116 |
| HBP1 | 1689.002 | 0.246902 | 0.175208 | 1.409194 | 0.158778 | 0.305471 |
| DUS4L | 63.14323 | 0.030807 | 0.280728 | 0.109741 | 0.912615 | 0.949346 |
| SP4 | 85.59471 | 0.628838 | 0.276494 | 2.274325 | 0.022946 | 0.082916 |
| WDR91 | 1323.742 | -0.31847 | 0.195543 | -1.62866 | 0.103385 | 0.227981 |
| DNAH11 | 17.23253 | 0.247266 | 0.546692 | 0.452294 | 0.651057 | NA |
| CBLL1 | 380.4551 | -0.02898 | 0.167158 | -0.17338 | 0.862354 | 0.918473 |
| DLX5 | 4.72409 | 2.836474 | 1.189271 | 2.385053 | 0.017077 | NA |
| MTPN | 3420.114 | 0.165655 | 0.191511 | 0.864993 | 0.387043 | 0.552959 |
| STEAP1B | 202.9789 | 0.03334 | 0.239224 | 0.139368 | 0.889159 | 0.935148 |
| PTN | 10.11324 | 0.676791 | 0.711409 | 0.951339 | 0.341433 | NA |
| MPP6 | 96.25681 | 0.576998 | 0.260807 | 2.212356 | 0.026942 | 0.092392 |
| DFNA5 | 1025.122 | 0.028298 | 0.140268 | 0.201744 | 0.840117 | 0.905006 |
| ATP6V0A4 | 212.0279 | -0.61394 | 0.258336 | -2.3765 | 0.017478 | 0.069119 |
| ZC3HAV1 | 1181.266 | -0.02459 | 0.193364 | -0.12718 | 0.898797 | 0.940693 |
| TTC26 | 232.1562 | 0.145783 | 0.182017 | 0.800931 | 0.423172 | 0.585448 |
| OGDH | 4756.67 | -0.52457 | 0.178442 | -2.9397 | 0.003285 | 0.022409 |
| ADAP1 | 18.91072 | -0.26363 | 0.596854 | -0.44169 | 0.658711 | NA |
| TFEC | 1.688557 | 3.924679 | 1.871699 | 2.096854 | 0.036006 | NA |
| H2AFV | 2319.453 | 0.087305 | 0.14265 | 0.612021 | 0.540524 | 0.682636 |
| CAV2 | 906.2315 | 0.240044 | 0.202918 | 1.182959 | 0.236826 | 0.399729 |
| CAV1 | 3497.379 | -0.66752 | 0.165102 | -4.04306 | 5.28E-05 | 0.001709 |
| MET | 2986.017 | 0.22663 | 0.191598 | 1.182846 | 0.23687 | 0.399729 |
| RNF32 | 16.19118 | 0.531305 | 0.509853 | 1.042074 | 0.297377 | NA |
| LMBR1 | 936.6967 | 0.043799 | 0.157456 | 0.278167 | 0.780884 | 0.8655 |
| WNT2 | 0.258578 | 1.233151 | 4.047809 | 0.304646 | 0.760635 | NA |
| HOXA1 | 49.64864 | -0.04451 | 0.32874 | -0.13539 | 0.892306 | 0.936854 |
| DNAJB6 | 3121.92 | 0.197525 | 0.149654 | 1.319878 | 0.186876 | 0.34099 |
| HOXA2 | 0.326333 | 1.533889 | 3.826878 | 0.40082 | 0.688553 | NA |
| HOXA3 | 4.370826 | -0.3618 | 0.942007 | -0.38407 | 0.700926 | NA |
| LFNG | 42.16189 | 0.479196 | 0.403835 | 1.186613 | 0.23538 | NA |
| HOXA5 | 26.3311 | 0.01109 | 0.41634 | 0.026637 | 0.97875 | NA |
| HOXA6 | 4.418067 | 0.086859 | 1.001761 | 0.086706 | 0.930905 | NA |
| BRAT1 | 1544.893 | -0.17407 | 0.183394 | -0.94917 | 0.342536 | 0.510724 |
| IQCE | 2478.506 | -0.21656 | 0.219867 | -0.98495 | 0.324648 | 0.492617 |
| ANKRD7 | 16.81892 | 0.18499 | 0.489322 | 0.378053 | 0.705391 | NA |
| TSPAN12 | 79.22331 | 0.907717 | 0.297937 | 3.046673 | 0.002314 | 0.017766 |
| SSBP1 | 1691.208 | 0.198459 | 0.146968 | 1.350356 | 0.176902 | 0.329058 |
| HOXA13 | 1.062152 | -1.07787 | 1.975873 | -0.54551 | 0.5854 | NA |
| CPED1 | 70.15448 | 0.42543 | 0.324497 | 1.311045 | 0.189842 | 0.34409 |
| HIBADH | 2472.245 | 0.003968 | 0.15644 | 0.025366 | 0.979763 | 0.989276 |
| TAX1BP1 | 1747.357 | 1.172905 | 0.574925 | 2.040101 | 0.04134 | 0.123231 |

| | | | | | | |
|---|---|---|---|---|---|---|
| CPVL | 40.09095 | 0.565652 | 0.372679 | 1.517799 | 0.129065 | NA |
| CHN2 | 51.28155 | 1.006516 | 0.6643 | 1.515154 | 0.129733 | 0.266024 |
| GRB10 | 509.5719 | -0.0651 | 0.247606 | -0.26292 | 0.792614 | 0.874331 |
| ABHD11 | 301.2971 | -0.22272 | 0.221494 | -1.00554 | 0.314635 | 0.482053 |
| COBL | 679.8308 | -0.0196 | 0.166832 | -0.11746 | 0.906496 | 0.945876 |
| FKBP14 | 842.9117 | 0.541164 | 0.173505 | 3.119007 | 0.001815 | 0.015146 |
| PLEKHA8 | 292.7634 | 0.353959 | 0.182665 | 1.937744 | 0.052654 | 0.143268 |
| STX1A | 113.1526 | -0.25192 | 0.26383 | -0.95484 | 0.339657 | 0.508003 |
| NOD1 | 261.2957 | -0.13957 | 0.223947 | -0.62322 | 0.53314 | 0.677164 |
| GARS | 3259.396 | 0.117762 | 0.13019 | 0.904538 | 0.36571 | 0.532944 |
| CRHR2 | 1.582375 | -0.21967 | 1.900405 | -0.11559 | 0.907979 | NA |
| EPHB6 | 3.054691 | 0.010752 | 1.178416 | 0.009125 | 0.99272 | NA |
| FAM188B | 6.260454 | 0.31391 | 0.812609 | 0.386299 | 0.699275 | NA |
| GHRHR | 0.785714 | -3.2458 | 3.343263 | -0.97085 | 0.331623 | NA |
| NSUN5P2 | 0.096202 | 0.408546 | 4.079639 | 0.100143 | 0.920231 | NA |
| CASP2 | 633.3292 | -0.52501 | 0.168312 | -3.11924 | 0.001813 | 0.015146 |
| CHCHD2 | 6591.337 | 0.013631 | 0.162534 | 0.083868 | 0.933162 | 0.962303 |
| HSPB1 | 11930.93 | -0.1681 | 0.143126 | -1.17446 | 0.24021 | 0.403355 |
| NPTX2 | 2.072867 | 0.185838 | 1.57462 | 0.118021 | 0.906051 | NA |
| PDAP1 | 3081.021 | 0.064873 | 0.131092 | 0.49487 | 0.620692 | 0.745208 |
| BUD31 | 1023.83 | 0.093976 | 0.169628 | 0.55401 | 0.579572 | 0.71363 |
| PTCD1 | 83.43697 | -0.41976 | 0.265586 | -1.58049 | 0.113995 | 0.243127 |
| CYP3A5 | 7.781544 | 1.067068 | 0.83054 | 1.284787 | 0.198867 | NA |
| ZKSCAN1 | 2313.418 | -0.1511 | 0.264524 | -0.57122 | 0.567849 | 0.704537 |
| EIF3B | 6437.955 | -0.20116 | 0.125474 | -1.60318 | 0.108895 | 0.235962 |
| SNX8 | 4447.728 | -0.33355 | 0.147912 | -2.25505 | 0.02413 | 0.085854 |
| NUDT1 | 922.5279 | -0.217 | 0.143989 | -1.50707 | 0.131792 | 0.268592 |
| PTPRZ1 | 1958.738 | 0.744253 | 0.213347 | 3.488466 | 4.86E-04 | 0.00668 |
| TAF6 | 1236.081 | -0.4265 | 0.173913 | -2.45239 | 0.014191 | 0.059806 |
| WASL | 1124.883 | 0.403599 | 0.237679 | 1.698081 | 0.089492 | 0.20693 |
| AIMP2 | 515.1346 | -0.16524 | 0.161427 | -1.02365 | 0.306 | 0.473169 |
| TFR2 | 25.94772 | 0.068893 | 0.429888 | 0.160258 | 0.872678 | NA |
| FSCN3 | 0.49187 | 2.136088 | 3.895953 | 0.548284 | 0.583497 | NA |
| MOSPD3 | 753.2798 | -0.20051 | 0.147882 | -1.35587 | 0.175142 | 0.327064 |
| PCOLCE | 152.8804 | -0.11092 | 0.277036 | -0.40039 | 0.688872 | 0.797751 |
| FBXO24 | 2.039773 | 0.314136 | 1.379409 | 0.227732 | 0.819854 | NA |
| RBM28 | 877.6614 | 0.554624 | 0.143073 | 3.876523 | 1.06E-04 | 0.002673 |
| USP42 | 278.7488 | -0.18686 | 0.200651 | -0.93127 | 0.351715 | 0.519833 |
| IMPDH1 | 2976.581 | -0.1313 | 0.150886 | -0.87022 | 0.384183 | 0.550325 |
| AGFG2 | 708.3941 | -0.58176 | 0.152145 | -3.82372 | 1.31E-04 | 0.003029 |
| LSM5 | 566.1271 | 0.66473 | 0.1564 | 4.250194 | 2.14E-05 | 9.59E-04 |
| SERPINE1 | 22.87454 | 1.120079 | 0.689879 | 1.623587 | 0.104464 | NA |
| AP1S1 | 5204.489 | -0.04121 | 0.15761 | -0.26149 | 0.793718 | 0.874943 |
| C1GALT1 | 742.72 | 1.27952 | 0.222589 | 5.748359 | 9.01E-09 | 6.20E-06 |
| PLOD3 | 17370.01 | -0.31193 | 0.148142 | -2.10563 | 0.035236 | 0.110422 |
| RPA3 | 544.1305 | 0.266495 | 0.142504 | 1.870082 | 0.061472 | 0.16006 |
| ZNHIT1 | 4523.405 | -0.12892 | 0.193324 | -0.66688 | 0.504851 | 0.654386 |

| | | | | | | |
|---|---|---|---|---|---|---|
| CLDN15 | 198.7864 | -0.01444 | 0.265209 | -0.05443 | 0.956593 | 0.976145 |
| GLCCI1 | 137.9928 | -0.20522 | 0.245962 | -0.83436 | 0.404077 | 0.568856 |
| PHF14 | 325.004 | 0.740641 | 0.410268 | 1.805263 | 0.071034 | 0.176299 |
| NRF1 | 309.5659 | -0.34091 | 0.193544 | -1.76141 | 0.078169 | 0.188276 |
| TMEM106I | 1148.478 | 0.487714 | 0.241418 | 2.020204 | 0.043362 | 0.126906 |
| EZH2 | 333.1205 | 0.229039 | 0.201029 | 1.139334 | 0.254564 | 0.418798 |
| CEP41 | 443.1613 | -0.18677 | 0.166852 | -1.11936 | 0.262988 | 0.42747 |
| ZNF862 | 313.9641 | -0.00858 | 0.197686 | -0.04339 | 0.96539 | 0.981198 |
| SFRP4 | 1.076984 | 2.142029 | 2.046972 | 1.046438 | 0.295359 | NA |
| MEST | 253.6092 | -0.06595 | 0.194831 | -0.3385 | 0.734983 | 0.832209 |
| MEOX2 | 6.134303 | 0.085455 | 1.151236 | 0.074229 | 0.940828 | NA |
| ANKMY2 | 495.4576 | 0.314486 | 0.207084 | 1.518643 | 0.128852 | 0.265134 |
| ACTR3C | 6.951136 | 0.789503 | 0.939586 | 0.840267 | 0.400759 | NA |
| POU6F2 | 0.495797 | -0.73607 | 3.324282 | -0.22142 | 0.824763 | NA |
| TSPAN13 | 101.4905 | 0.70372 | 0.317518 | 2.216319 | 0.02667 | 0.091696 |
| RARRES2 | 42.93069 | 1.755422 | 0.643648 | 2.7273 | 0.006385 | NA |
| AHR | 1134.504 | -0.16121 | 0.231331 | -0.69687 | 0.485887 | 0.639256 |
| CHCHD3 | 2208.764 | 0.285584 | 0.155138 | 1.84084 | 0.065645 | 0.167108 |
| GIMAP2 | 32.64213 | 0.239761 | 0.403412 | 0.594332 | 0.55229 | NA |
| TMEM176I | 1.411424 | 1.039904 | 1.817075 | 0.572296 | 0.567122 | NA |
| GLI3 | 758.6588 | -0.56005 | 0.176864 | -3.16657 | 0.001542 | 0.013793 |
| PSMA2 | 158.5886 | 0.603021 | 0.230409 | 2.617174 | 0.008866 | 0.043613 |
| MRPL32 | 1172.537 | 0.612004 | 0.160814 | 3.805677 | 1.41E-04 | 0.003128 |
| COA1 | 1055.757 | 0.14233 | 0.159487 | 0.892419 | 0.372168 | 0.538778 |
| BLVRA | 2777.802 | -0.02544 | 0.129765 | -0.19606 | 0.844563 | 0.907349 |
| URGCP | 2300.19 | -0.10482 | 0.174058 | -0.60222 | 0.547031 | 0.688457 |
| TMEM248 | 2888.943 | -0.18222 | 0.117494 | -1.5509 | 0.120927 | 0.253463 |
| STAG3L4 | 280.1228 | 0.170362 | 0.170787 | 0.99751 | 0.318517 | 0.486349 |
| RHEB | 2183.481 | 0.268289 | 0.13431 | 1.997531 | 0.045768 | 0.131057 |
| PRKAG2 | 260.3824 | 0.001683 | 0.195373 | 0.008613 | 0.993128 | 0.996912 |
| AEBP1 | 19452.45 | -0.02501 | 0.203807 | -0.12271 | 0.902336 | 0.9429 |
| POLD2 | 2993.774 | -0.31981 | 0.13498 | -2.36936 | 0.017819 | 0.06981 |
| GCK | 15.82947 | 0.017425 | 0.519672 | 0.03353 | 0.973252 | NA |
| BCL7B | 1450.153 | 0.015877 | 0.181692 | 0.087387 | 0.930364 | 0.960504 |
| YKT6 | 6303.627 | -0.04127 | 0.155288 | -0.26575 | 0.79043 | 0.872896 |
| TBL2 | 1137.709 | -0.20714 | 0.147701 | -1.40241 | 0.160793 | 0.308346 |
| GALNTL5 | 0.163957 | 0.424913 | 4.079006 | 0.104171 | 0.917034 | NA |
| CLIP2 | 2375.551 | -0.34345 | 0.147775 | -2.32414 | 0.020118 | 0.076028 |
| EIF4H | 4261.595 | -0.19784 | 0.136178 | -1.4528 | 0.146278 | 0.288829 |
| LIMK1 | 1956.165 | -0.25536 | 0.120285 | -2.12295 | 0.033758 | 0.107317 |
| SPATA6L | 16.19462 | 0.514259 | 0.532312 | 0.966087 | 0.334001 | NA |
| SLC1A1 | 98.79307 | 0.659822 | 0.246792 | 2.673591 | 0.007504 | 0.039084 |
| LHX2 | 0.883146 | 1.112834 | 2.554178 | 0.435692 | 0.663061 | NA |
| FKTN | 614.5809 | 0.388344 | 0.233964 | 1.659848 | 0.096945 | 0.218537 |
| FSD1L | 100.7137 | 1.579459 | 0.293173 | 5.387467 | 7.15E-08 | 2.73E-05 |
| CNTNAP3 | 123.1237 | -0.10527 | 0.213081 | -0.49403 | 0.621282 | 0.745702 |
| SPIN1 | 1856.741 | 0.073145 | 0.164754 | 0.443968 | 0.657066 | 0.773844 |

| | | | | | | |
|---|---|---|---|---|---|---|
| NMRK1 | 522.2689 | 0.376644 | 0.167519 | 2.248366 | 0.024553 | 0.086758 |
| TMEM245 | 4661.877 | 0.083924 | 0.12791 | 0.656113 | 0.511752 | 0.659436 |
| PRUNE2 | 9704.193 | -0.03353 | 0.199668 | -0.16793 | 0.866636 | 0.920772 |
| MEGF9 | 3165.056 | 0.433913 | 0.2163 | 2.006064 | 0.044849 | 0.129396 |
| TRIM14 | 1450.324 | -1.08866 | 0.189626 | -5.74109 | 9.41E-09 | 6.20E-06 |
| CORO2A | 463.2818 | -0.19299 | 0.160125 | -1.20522 | 0.22812 | 0.389884 |
| TGFBR1 | 1705.056 | 0.22714 | 0.224472 | 1.011888 | 0.311592 | 0.478921 |
| SEC61B | 2025.89 | 0.16447 | 0.152037 | 1.081779 | 0.279351 | 0.445506 |
| C5 | 69.96525 | 0.848072 | 0.277232 | 3.059067 | 0.00222 | 0.017294 |
| OGN | 0.82371 | 0.509057 | 2.431911 | 0.209324 | 0.834195 | NA |
| ASPN | 0.326333 | 1.533889 | 3.826878 | 0.40082 | 0.688553 | NA |
| ECM2 | 0.518736 | 2.237423 | 2.7713 | 0.807355 | 0.419462 | NA |
| TLE4 | 157.3337 | 0.216885 | 0.21482 | 1.009614 | 0.31268 | 0.479848 |
| LHX6 | 1.236585 | 0.682769 | 1.964737 | 0.347512 | 0.728207 | NA |
| PTGR1 | 2017.455 | 0.079344 | 0.137283 | 0.577962 | 0.56329 | 0.701302 |
| SUSD1 | 267.9917 | -0.09331 | 0.194151 | -0.4806 | 0.630802 | 0.753088 |
| AMBP | 0.134133 | -1.48909 | 4.077584 | -0.36519 | 0.71497 | NA |
| AKNA | 580.7329 | -0.28853 | 0.159812 | -1.80546 | 0.071002 | 0.176295 |
| DNM1 | 19.62089 | -0.04047 | 0.504537 | -0.08021 | 0.936069 | NA |
| ENG | 2481.785 | -0.43119 | 0.167897 | -2.56816 | 0.010224 | 0.047869 |
| AK1 | 411.1352 | -0.39944 | 0.188817 | -2.11546 | 0.03439 | 0.108632 |
| CDC37L1 | 287.0656 | 0.789974 | 0.220869 | 3.576665 | 3.48E-04 | 0.005433 |
| RLN2 | 31.85084 | 0.485575 | 0.381775 | 1.271886 | 0.203414 | NA |
| RLN1 | 7.244414 | 0.790598 | 0.783787 | 1.00869 | 0.313124 | NA |
| PLGRKT | 522.8944 | 0.068148 | 0.197659 | 0.344776 | 0.730263 | 0.82887 |
| TBC1D13 | 1240.254 | -0.41847 | 0.135656 | -3.08483 | 0.002037 | 0.01637 |
| KIAA1432 | 838.6011 | 0.335894 | 0.161221 | 2.083434 | 0.037212 | 0.114494 |
| KDM4C | 507.0808 | 0.02974 | 0.175011 | 0.16993 | 0.865065 | 0.920317 |
| DOCK8 | 824.9548 | -1.44E-04 | 0.158414 | -9.09E-04 | 0.999274 | 0.999472 |
| KANK1 | 2497.866 | -0.16544 | 0.142082 | -1.16443 | 0.244251 | 0.406914 |
| ELAVL2 | 0.134133 | -1.48909 | 4.077584 | -0.36519 | 0.71497 | NA |
| NCS1 | 4401.755 | -0.33577 | 0.215954 | -1.55484 | 0.119985 | 0.252113 |
| TESK1 | 865.4722 | -0.05093 | 0.215098 | -0.23675 | 0.812847 | 0.88863 |
| KCNT1 | 0.881566 | 1.109202 | 2.554987 | 0.434132 | 0.664192 | NA |
| CA9 | 0.926235 | -0.09821 | 2.156052 | -0.04555 | 0.963667 | NA |
| FUBP3 | 1345.745 | 0.207268 | 0.170379 | 1.216512 | 0.22379 | 0.384608 |
| TYRP1 | 422913.1 | -0.05737 | 0.135099 | -0.42467 | 0.67108 | 0.784504 |
| CREB3 | 2104.054 | 0.028664 | 0.157307 | 0.182215 | 0.855414 | 0.914092 |
| RGP1 | 1224.742 | -0.50283 | 0.130432 | -3.85511 | 1.16E-04 | 0.00281 |
| MPDZ | 2004.829 | 0.314915 | 0.23527 | 1.338526 | 0.180725 | 0.333255 |
| LHX3 | 0.314073 | -0.57308 | 3.898934 | -0.14698 | 0.883146 | NA |
| DDX58 | 420.5681 | -0.1996 | 0.301445 | -0.66214 | 0.507882 | 0.656663 |
| EDF1 | 5231.736 | -0.03008 | 0.156615 | -0.19207 | 0.847687 | 0.909194 |
| GLIS3 | 15.388 | 0.12226 | 0.517853 | 0.236091 | 0.813362 | NA |
| BAG1 | 3434.041 | 0.240303 | 0.180852 | 1.328731 | 0.183937 | 0.337184 |
| RAPGEF1 | 3436.245 | -0.66665 | 0.195459 | -3.41071 | 6.48E-04 | 0.007902 |
| NPDC1 | 347.8122 | -0.02178 | 0.204388 | -0.10657 | 0.915128 | 0.950986 |

| | | | | | | |
|---|---|---|---|---|---|---|
| APBA1 | 46.90994 | -0.74156 | 0.363475 | -2.04019 | 0.041331 | NA |
| SETX | 1662.275 | 0.591347 | 0.306807 | 1.927425 | 0.053927 | 0.145859 |
| SH3GL2 | 177.1438 | 0.143615 | 0.268069 | 0.535738 | 0.592139 | 0.7233 |
| PTGDS | 10.82253 | -1.23244 | 0.75177 | -1.63938 | 0.101134 | NA |
| ABCA2 | 2303.631 | -0.55646 | 0.145576 | -3.82247 | 1.32E-04 | 0.003029 |
| SHB | 75.87793 | -0.38339 | 0.276522 | -1.38647 | 0.165602 | 0.314617 |
| UBE2R2 | 2368.58 | -0.3469 | 0.126083 | -2.75132 | 0.005936 | 0.033176 |
| ABHD17B | 428.3188 | 0.22322 | 0.19488 | 1.145419 | 0.252036 | 0.415977 |
| EXOSC3 | 274.0022 | 0.211985 | 0.179213 | 1.182863 | 0.236863 | 0.399729 |
| ZFAND5 | 4925.586 | 0.227924 | 0.13854 | 1.645182 | 0.099932 | 0.223198 |
| FBXO5 | 217.1158 | 0.756558 | 0.208724 | 3.624688 | 2.89E-04 | 0.004811 |
| MTRF1L | 426.1018 | 0.675258 | 0.189124 | 3.570452 | 3.56E-04 | 0.005522 |
| SOD2 | 3597.515 | 0.206576 | 0.177678 | 1.162641 | 0.244975 | 0.407833 |
| MRPL18 | 1618.102 | 0.248671 | 0.151463 | 1.641787 | 0.100634 | 0.223873 |
| CCR6 | 0.763859 | -0.6945 | 2.307767 | -0.30094 | 0.76346 | NA |
| SLC22A2 | 0.163957 | 0.424913 | 4.079006 | 0.104171 | 0.917034 | NA |
| PACRG | 1.862957 | 0.101849 | 1.772401 | 0.057464 | 0.954176 | NA |
| QKI | 3735.866 | 0.57058 | 0.223607 | 2.551714 | 0.010719 | 0.049294 |
| PDE10A | 120.3493 | 0.484727 | 0.316176 | 1.533092 | 0.125253 | 0.260185 |
| FAM120B | 1317.841 | -0.13345 | 0.125452 | -1.06372 | 0.287457 | 0.453587 |
| TBP | 333.7528 | -0.05107 | 0.185208 | -0.27575 | 0.782744 | 0.867305 |
| USP9Y | 414.3873 | 0.7278 | 0.299856 | 2.427168 | 0.015217 | 0.062726 |
| ESYT2 | 3641.208 | 0.332533 | 0.131248 | 2.533624 | 0.011289 | 0.05108 |
| TRPM6 | 21.3649 | -0.68696 | 0.503297 | -1.36492 | 0.172277 | NA |
| KLF9 | 2027.475 | 0.718509 | 0.306075 | 2.34749 | 0.0189 | 0.072701 |
| TJP2 | 737.1588 | -0.50025 | 0.153548 | -3.25795 | 0.001122 | 0.011133 |
| PTBP3 | 729.4769 | 0.281658 | 0.231395 | 1.217215 | 0.223523 | 0.384316 |
| RAD23B | 3791.401 | 0.122732 | 0.126362 | 0.971272 | 0.331413 | 0.499924 |
| FKBP15 | 1725.938 | -0.10144 | 0.129543 | -0.78309 | 0.433574 | 0.594703 |
| CTNNAL1 | 1140.281 | 0.752908 | 0.197091 | 3.820111 | 1.33E-04 | 0.003035 |
| FAM206A | 190.2221 | 0.44962 | 0.217502 | 2.067201 | 0.038715 | 0.1176 |
| WDR34 | 1905.358 | -0.167 | 0.180974 | -0.92281 | 0.356107 | 0.523903 |
| SET | 4362.751 | 0.199558 | 0.164466 | 1.213373 | 0.224987 | 0.386043 |
| PPP2R4 | 5734.222 | -0.32581 | 0.161817 | -2.01344 | 0.044068 | 0.127962 |
| GLE1 | 1326.183 | -0.27893 | 0.123774 | -2.25356 | 0.024224 | 0.086136 |
| RAB14 | 3254.082 | 0.020784 | 0.122001 | 0.170361 | 0.864726 | 0.920317 |
| CNTRL | 174.8136 | 0.73382 | 0.458255 | 1.601336 | 0.109303 | 0.236504 |
| TRIM32 | 2096.647 | -0.29549 | 0.121118 | -2.43965 | 0.014702 | 0.061046 |
| FBXW2 | 961.5871 | 0.058538 | 0.156931 | 0.373015 | 0.709137 | 0.813905 |
| PHF19 | 1894.959 | -0.16735 | 0.228321 | -0.73295 | 0.463591 | 0.621213 |
| NEK6 | 2842.342 | -0.3078 | 0.129525 | -2.37634 | 0.017485 | 0.069119 |
| BSPRY | 3.964917 | -2.17056 | 1.193877 | -1.81807 | 0.069053 | NA |
| PPP6C | 1471.547 | 0.039842 | 0.145499 | 0.27383 | 0.784215 | 0.868623 |
| NDUFA8 | 2227.646 | -0.00876 | 0.16439 | -0.05327 | 0.957515 | 0.976514 |
| HDHD3 | 446.2398 | -0.18061 | 0.169658 | -1.06455 | 0.287079 | 0.453413 |
| RBM18 | 620.4072 | 0.469163 | 0.189775 | 2.472211 | 0.013428 | 0.05764 |
| SLC46A2 | 3.207814 | -0.51709 | 1.174608 | -0.44022 | 0.659776 | NA |

| | | | | | | |
|---|---|---|---|---|---|---|
| HSDL2 | 914.0981 | 0.651029 | 0.195709 | 3.326512 | 8.79E-04 | 0.009556 |
| MAPKAP1 | 3509.034 | 0.045749 | 0.126911 | 0.360483 | 0.718486 | 0.820425 |
| NR4A3 | 836.4377 | 0.716324 | 0.328988 | 2.177357 | 0.029454 | 0.097807 |
| INVS | 608.672 | 0.112622 | 0.155526 | 0.724136 | 0.468982 | 0.62547 |
| GALNT12 | 126.514 | 0.077784 | 0.244814 | 0.317728 | 0.750691 | 0.844063 |
| DENND1A | 640.8786 | -0.57459 | 0.180328 | -3.18633 | 0.001441 | 0.013231 |
| ALG2 | 824.2926 | 0.06926 | 0.159559 | 0.43407 | 0.664238 | 0.779426 |
| TEK | 6.199297 | 0.491256 | 0.853123 | 0.575832 | 0.564729 | NA |
| RCL1 | 241.7694 | -0.26463 | 0.179917 | -1.47087 | 0.141326 | 0.281915 |
| CAAP1 | 372.7148 | 0.557453 | 0.192254 | 2.899567 | 0.003737 | 0.024394 |
| EQTN | 0.096202 | 0.408546 | 4.079639 | 0.100143 | 0.920231 | NA |
| MOB3B | 594.6344 | -0.57915 | 0.259938 | -2.22802 | 0.025879 | 0.090129 |
| MLANA | 60192.4 | -0.09033 | 0.137353 | -0.65763 | 0.510774 | 0.659019 |
| CD274 | 36.35392 | -0.67554 | 0.416512 | -1.6219 | 0.104825 | NA |
| CCDC170 | 3.502105 | 0.802226 | 1.148776 | 0.698331 | 0.48497 | NA |
| CXorf21 | 0.151697 | -1.546 | 4.075469 | -0.37934 | 0.704433 | NA |
| ACAT2 | 2184.562 | 0.111714 | 0.161995 | 0.689614 | 0.490437 | 0.642588 |
| TCP1 | 5606.951 | 0.190236 | 0.149054 | 1.276289 | 0.201853 | 0.358283 |
| TEX11 | 0.134133 | -1.48909 | 4.077584 | -0.36519 | 0.71497 | NA |
| ARR3 | 2.104871 | 0.18276 | 1.473042 | 0.12407 | 0.90126 | NA |
| PDZD11 | 1688.359 | -0.08813 | 0.140686 | -0.62642 | 0.53104 | 0.675555 |
| NUDCD1 | 367.5374 | 0.945607 | 0.218371 | 4.33027 | 1.49E-05 | 7.85E-04 |
| ENY2 | 1248.24 | 0.666915 | 0.155226 | 4.296416 | 1.74E-05 | 8.54E-04 |
| DUSP4 | 9863.455 | -0.27203 | 0.171354 | -1.58755 | 0.112388 | 0.240934 |
| CLU | 17.81468 | 0.345669 | 0.929968 | 0.3717 | 0.710116 | NA |
| TNFRSF10B | 4966.312 | -0.08227 | 0.194149 | -0.42376 | 0.671738 | 0.785118 |
| SORBS3 | 3448.453 | -0.48613 | 0.156438 | -3.10748 | 0.001887 | 0.015523 |
| PTK2B | 329.5127 | -0.49103 | 0.162591 | -3.02001 | 0.002528 | 0.01898 |
| PPP3CC | 607.7594 | 0.04586 | 0.171016 | 0.268165 | 0.788573 | 0.871658 |
| PDLIM2 | 538.5144 | 0.053038 | 0.182369 | 0.290827 | 0.771184 | 0.858681 |
| EPHX2 | 175.6334 | -0.58353 | 0.236391 | -2.4685 | 0.013568 | 0.057932 |
| RNF170 | 526.6581 | 0.485285 | 0.169984 | 2.854879 | 0.004305 | 0.026682 |
| ZNF706 | 2277.751 | 0.008522 | 0.147945 | 0.057602 | 0.954065 | 0.974786 |
| LYPLA1 | 1426.273 | 0.403127 | 0.186397 | 2.162732 | 0.030562 | 0.100335 |
| CRISPLD1 | 216.3282 | -0.16273 | 0.289273 | -0.56256 | 0.573735 | 0.709406 |
| COPS5 | 1115.512 | 0.321908 | 0.14256 | 2.258057 | 0.023942 | 0.085415 |
| RDH10 | 179.7138 | 0.032625 | 0.191939 | 0.169976 | 0.865029 | 0.920317 |
| PILRB | 206.9757 | 0.608313 | 0.232352 | 2.618066 | 0.008843 | 0.043536 |
| OCRL | 1192.082 | -0.39511 | 0.13605 | -2.90418 | 0.003682 | 0.024211 |
| PAEP | 4.933614 | -0.19649 | 0.980152 | -0.20047 | 0.841114 | NA |
| MRPS2 | 1085.152 | -0.09049 | 0.145389 | -0.62237 | 0.533696 | 0.677477 |
| TBX22 | 0.820549 | 0.502564 | 2.433834 | 0.206491 | 0.836408 | NA |
| SERAC1 | 474.6533 | 0.343761 | 0.168934 | 2.034881 | 0.041863 | 0.124289 |
| BBS9 | 457.1551 | -0.07007 | 0.188597 | -0.37154 | 0.710234 | 0.814697 |
| PMS2 | 409.522 | -0.09767 | 0.160486 | -0.60861 | 0.542786 | 0.684848 |
| ZMIZ2 | 1725.054 | -0.4297 | 0.161494 | -2.6608 | 0.007795 | 0.03992 |
| OCM | 0.487129 | 2.123445 | 3.910539 | 0.543006 | 0.587126 | NA |

| | | | | | | |
|---|---|---|---|---|---|---|
| 7-Sep | 3126.798 | 0.849095 | 0.254707 | 3.333616 | 8.57E-04 | 0.009419 |
| EEPD1 | 29.06146 | -0.22478 | 0.400732 | -0.56093 | 0.574843 | NA |
| KLHL7 | 1028.178 | 0.294621 | 0.178231 | 1.653032 | 0.098324 | 0.220722 |
| HERPUD2 | 696.6272 | -0.01745 | 0.149032 | -0.1171 | 0.906781 | 0.945876 |
| CBX3 | 4193.925 | 0.899117 | 0.23286 | 3.86119 | 1.13E-04 | 0.002775 |
| HNRNPA2E | 12250.88 | 0.378937 | 0.147932 | 2.561565 | 0.01042 | 0.048395 |
| WIPF3 | 854.3991 | -0.55825 | 0.233774 | -2.38801 | 0.01694 | 0.067755 |
| NXPH1 | 0.096202 | 0.408546 | 4.079639 | 0.100143 | 0.920231 | NA |
| FAM126A | 2774.425 | 0.619072 | 0.20333 | 3.044664 | 0.002329 | 0.017851 |
| HOXA7 | 4.7236 | -0.71487 | 0.944544 | -0.75684 | 0.449145 | NA |
| INHBA | 0.655827 | 2.55247 | 3.531744 | 0.722722 | 0.46985 | NA |
| FKBP9 | 3881.497 | -0.47933 | 0.120284 | -3.985 | 6.75E-05 | 0.00198 |
| NT5C3A | 475.7187 | 0.583023 | 0.221473 | 2.632476 | 0.008477 | 0.042151 |
| ARL4A | 352.8488 | 0.654723 | 0.326255 | 2.006786 | 0.044772 | 0.129302 |
| CCZ1 | 119.3793 | 0.563487 | 0.260883 | 2.159924 | 0.030779 | 0.100851 |
| POLM | 427.6746 | -0.16441 | 0.158139 | -1.03968 | 0.29849 | 0.46536 |
| RAMP3 | 2.865063 | 0.407234 | 1.206676 | 0.337484 | 0.735752 | NA |
| FTSJ2 | 869.0696 | -0.03739 | 0.147317 | -0.25384 | 0.79962 | 0.879115 |
| TWIST1 | 4.234253 | 0.024601 | 0.976711 | 0.025188 | 0.979905 | NA |
| SMU1 | 1596.223 | -0.03602 | 0.152331 | -0.23648 | 0.813059 | 0.888676 |
| GLIPR2 | 219.9136 | -0.47926 | 0.264063 | -1.81496 | 0.06953 | 0.173679 |
| SLC25A51 | 266.6642 | -0.00983 | 0.190954 | -0.05146 | 0.958956 | 0.977425 |
| CLTA | 6818.333 | -0.02145 | 0.131717 | -0.16286 | 0.870625 | 0.923107 |
| RECK | 358.8488 | 0.202699 | 0.195847 | 1.034983 | 0.300677 | 0.4675 |
| SPINK4 | 0.162376 | 0.424913 | 4.079006 | 0.104171 | 0.917034 | NA |
| ACO1 | 1768.823 | -0.25408 | 0.130546 | -1.94627 | 0.051623 | 0.1415 |
| KIAA1045 | 141.4327 | -0.59958 | 0.204079 | -2.93795 | 0.003304 | 0.022446 |
| DNAI1 | 0.672013 | 1.253691 | 2.607776 | 0.480751 | 0.630693 | NA |
| DCAF10 | 770.2619 | 0.209277 | 0.181235 | 1.154727 | 0.248202 | 0.411875 |
| CNTFR | 0.640406 | 1.11402 | 2.710977 | 0.410929 | 0.681124 | NA |
| KIAA1549 | 223.8603 | -0.67987 | 0.198883 | -3.41846 | 6.30E-04 | 0.007785 |
| TRIM24 | 346.1213 | 0.221001 | 0.265909 | 0.831116 | 0.405908 | 0.56977 |
| C7orf49 | 2282.555 | -0.17406 | 0.144485 | -1.20467 | 0.228332 | 0.389984 |
| CALD1 | 5906.149 | 0.704224 | 0.304935 | 2.309421 | 0.02092 | 0.078113 |
| AKR1D1 | 0.258578 | 1.233151 | 4.047809 | 0.304646 | 0.760635 | NA |
| NUDT10 | 122.4145 | -0.15881 | 0.226757 | -0.70036 | 0.483701 | 0.637354 |
| NECAB1 | 56.52685 | 0.54234 | 0.312142 | 1.73748 | 0.082303 | 0.194983 |
| WWP1 | 1120.024 | 0.547711 | 0.18568 | 2.949762 | 0.00318 | 0.022022 |
| ACOT9 | 1977.625 | 0.076756 | 0.14337 | 0.53537 | 0.592394 | 0.7233 |
| PRDX4 | 3978.117 | -0.18552 | 0.12308 | -1.50728 | 0.13174 | 0.268537 |
| SARDH | 34.95628 | -1.32926 | 0.457192 | -2.90745 | 0.003644 | NA |
| DBH | 0.928019 | -0.78227 | 2.07134 | -0.37767 | 0.705679 | NA |
| IL13RA2 | 5.34595 | 3.067102 | 1.257722 | 2.438617 | 0.014744 | NA |
| PLP1 | 16701.31 | -0.00404 | 0.199038 | -0.0203 | 0.983801 | 0.991411 |
| SERPINA7 | 0.134133 | -1.48909 | 4.077584 | -0.36519 | 0.71497 | NA |
| MORF4L2 | 5683.275 | 0.276139 | 0.148455 | 1.860091 | 0.062873 | 0.162387 |
| RAB9B | 63.7054 | 0.437322 | 0.321898 | 1.358571 | 0.174283 | 0.326247 |

| | | | | | | |
|---|---|---|---|---|---|---|
| FAM199X | 923.1419 | 0.544362 | 0.219285 | 2.482443 | 0.013048 | 0.056565 |
| ESX1 | 0.151697 | -1.546 | 4.075469 | -0.37934 | 0.704433 | NA |
| RAB9A | 1248.258 | 0.392383 | 0.181999 | 2.155966 | 0.031086 | 0.101597 |
| RAP2C | 430.9537 | 0.340119 | 0.202173 | 1.682315 | 0.092508 | 0.211878 |
| AGO2 | 429.6863 | 0.073045 | 0.222447 | 0.32837 | 0.742632 | 0.837829 |
| PMS2P5 | 48.23909 | -0.58603 | 0.335479 | -1.74685 | 0.080664 | 0.19237 |
| CKS2 | 2678.108 | 0.270424 | 0.202551 | 1.335092 | 0.181846 | 0.334698 |
| IQSEC2 | 456.8486 | -0.36711 | 0.156105 | -2.3517 | 0.018688 | 0.072212 |
| XG | 18.84873 | -0.11322 | 0.625784 | -0.18093 | 0.856422 | NA |
| USP9X | 3241.55 | 0.136955 | 0.20248 | 0.676388 | 0.498794 | 0.649348 |
| KIF25 | 1.465688 | -0.81787 | 1.578746 | -0.51805 | 0.604423 | NA |
| UPF3B | 124.4254 | 1.178405 | 0.830185 | 1.419448 | 0.155768 | 0.301639 |
| RNF113A | 500.296 | 0.006067 | 0.160737 | 0.037742 | 0.969893 | 0.983611 |
| 6-Sep | 1345.425 | -0.01014 | 0.155416 | -0.06523 | 0.94799 | 0.971147 |
| TMEM255A | 41.64733 | -0.27183 | 0.347183 | -0.78297 | 0.433643 | NA |
| NDUFA1 | 2882.298 | 0.259204 | 0.156756 | 1.653555 | 0.098218 | 0.220609 |
| TTF1 | 242.4595 | 0.663949 | 0.280082 | 2.370553 | 0.017761 | 0.069688 |
| GTF3C4 | 1193.366 | 0.05371 | 0.145869 | 0.368204 | 0.712721 | 0.816474 |
| DDX31 | 498.5547 | -0.03552 | 0.170543 | -0.20825 | 0.835034 | 0.901331 |
| GRIA3 | 104.3011 | 0.236579 | 0.245998 | 0.961712 | 0.336194 | 0.505208 |
| THOC2 | 665.5612 | 0.938711 | 0.336562 | 2.789116 | 0.005285 | 0.030714 |
| CITED1 | 398.8072 | -0.34074 | 0.212114 | -1.60643 | 0.108181 | 0.234942 |
| ARMCX5 | 380.3317 | 0.245965 | 0.208521 | 1.179568 | 0.238172 | 0.401059 |
| GRPR | 6.636206 | 0.21217 | 0.841059 | 0.252265 | 0.800836 | NA |
| KDM5C | 2491.396 | -0.47034 | 0.146186 | -3.21743 | 0.001293 | 0.012237 |
| AMOT | 607.6972 | -0.25587 | 0.174316 | -1.46784 | 0.142147 | 0.282988 |
| ASL | 368.1631 | -0.17268 | 0.163596 | -1.05551 | 0.291194 | 0.458082 |
| SBDS | 1888.92 | 0.615741 | 0.195232 | 3.153894 | 0.001611 | 0.01418 |
| SSX1 | 2.960997 | -0.24818 | 1.173428 | -0.2115 | 0.832496 | NA |
| UXT | 952.9182 | 0.070151 | 0.142478 | 0.492364 | 0.622462 | 0.746665 |
| CFP | 4.007043 | 0.017945 | 1.132721 | 0.015842 | 0.98736 | NA |
| ELK1 | 1755.833 | -0.27119 | 0.161628 | -1.67785 | 0.093376 | 0.213265 |
| TIMM17B | 578.4515 | -0.20425 | 0.162387 | -1.25779 | 0.208467 | 0.365913 |
| WDR60 | 386.9008 | 0.472604 | 0.226501 | 2.086543 | 0.03693 | 0.113951 |
| AIF1L | 899.9472 | -0.77213 | 0.272144 | -2.8372 | 0.004551 | 0.02764 |
| FAM78A | 550.7038 | -0.40836 | 0.278171 | -1.46801 | 0.1421 | 0.282988 |
| NUP214 | 1174.719 | -0.77795 | 0.170104 | -4.5734 | 4.80E-06 | 4.09E-04 |
| AVPR2 | 1.405663 | 0.963764 | 1.838941 | 0.524086 | 0.600218 | NA |
| SLC10A3 | 415.6931 | -0.03226 | 0.171761 | -0.18784 | 0.851005 | 0.911434 |
| HNRNPH2 | 1815.16 | 0.267542 | 0.205745 | 1.300358 | 0.193478 | 0.348531 |
| ARMCX1 | 1956.117 | 0.231284 | 0.137061 | 1.687454 | 0.091516 | 0.210338 |
| TMEM35 | 2.339518 | -1.47824 | 1.425448 | -1.03704 | 0.299719 | NA |
| TIMM8A | 229.5535 | 0.015038 | 0.208005 | 0.072297 | 0.942366 | 0.968089 |
| ZC4H2 | 366.4509 | -0.30908 | 0.188678 | -1.63815 | 0.101391 | 0.225006 |
| IPPK | 152.0111 | 0.184863 | 0.204556 | 0.903726 | 0.366141 | 0.533342 |
| ZNF484 | 75.856 | 1.011267 | 0.298277 | 3.39036 | 6.98E-04 | 0.008275 |
| OMD | 0.722204 | 1.447218 | 2.611582 | 0.554154 | 0.579474 | NA |

| | | | | | | |
|---|---|---|---|---|---|---|
| FGD3 | 1.289683 | -0.48937 | 1.904039 | -0.25702 | 0.797167 | NA |
| TRAF2 | 925.2191 | -0.37862 | 0.157022 | -2.41128 | 0.015897 | 0.064533 |
| TAS2R3 | 0.411855 | 0.250456 | 3.097618 | 0.080854 | 0.935558 | NA |
| TAS2R4 | 0.448206 | -1.2532 | 3.007315 | -0.41672 | 0.676885 | NA |
| TAS2R5 | 1.621391 | 1.253556 | 1.571839 | 0.79751 | 0.425155 | NA |
| CRYGN | 0.452561 | 2.061505 | 3.349456 | 0.615474 | 0.538241 | NA |
| LRRC61 | 51.90936 | 0.01311 | 0.506962 | 0.02586 | 0.979369 | 0.988962 |
| TUBBP1 | 10.02414 | -0.24029 | 0.686395 | -0.35008 | 0.726281 | NA |
| AKAP9 | 392.616 | 0.593739 | 1.25031 | 0.474874 | 0.634877 | 0.756125 |
| GNG11 | 2245.782 | 0.602989 | 0.156907 | 3.842956 | 1.22E-04 | 0.0029 |
| SHFM1 | 1962.789 | 0.546187 | 0.149531 | 3.652676 | 2.60E-04 | 0.004545 |
| HIP1 | 1244.151 | -0.55376 | 0.166958 | -3.31675 | 9.11E-04 | 0.00977 |
| PTPN12 | 1527.914 | 0.742898 | 0.162954 | 4.558941 | 5.14E-06 | 4.24E-04 |
| POR | 3082.888 | -0.3101 | 0.149597 | -2.07287 | 0.038184 | 0.116488 |
| FGL2 | 87.2575 | 0.981947 | 1.028435 | 0.954797 | 0.339681 | 0.508003 |
| STYXL1 | 495.9546 | 0.067014 | 0.214508 | 0.312409 | 0.754729 | 0.846963 |
| STEAP4 | 0.134133 | -1.48909 | 4.077584 | -0.36519 | 0.71497 | NA |
| GNAI1 | 702.3573 | 0.57196 | 0.201113 | 2.843976 | 0.004455 | 0.027311 |
| PMS2P3 | 49.11595 | 0.436094 | 0.306059 | 1.424868 | 0.154195 | 0.299326 |
| PEX1 | 377.1504 | 0.372053 | 0.237217 | 1.568412 | 0.116785 | 0.247228 |
| MTERF | 212.2806 | 1.004062 | 0.307953 | 3.260444 | 0.001112 | 0.011104 |
| SGCE | 1184.668 | 0.548012 | 0.176168 | 3.110734 | 0.001866 | 0.015428 |
| RBM48 | 222.254 | 0.510229 | 0.206141 | 2.475147 | 0.013318 | 0.057401 |
| CASD1 | 705.087 | 0.555686 | 0.213147 | 2.607057 | 0.009132 | 0.044457 |
| CPA4 | 2.955538 | 1.566172 | 1.410237 | 1.110573 | 0.266752 | NA |
| DOCK4 | 617.9404 | 0.244808 | 0.179859 | 1.361114 | 0.173478 | 0.325305 |
| POT1 | 383.3862 | 0.386296 | 0.202504 | 1.907596 | 0.056443 | 0.150368 |
| ATP6V1F | 4914.538 | 0.053559 | 0.12224 | 0.438151 | 0.661277 | 0.776951 |
| NAA38 | 561.3307 | 0.564685 | 0.170082 | 3.32008 | 9.00E-04 | 0.009725 |
| CDHR3 | 122.6678 | 0.097202 | 0.250148 | 0.388577 | 0.697589 | 0.805171 |
| PRKRIP1 | 480.7112 | 0.335136 | 0.148139 | 2.262317 | 0.023678 | 0.084855 |
| VGF | 92.5812 | -0.17914 | 0.41442 | -0.43226 | 0.665551 | 0.780503 |
| PODXL | 2.056563 | 3.167197 | 1.798235 | 1.761281 | 0.078191 | NA |
| FOXP2 | 18.69816 | 0.010778 | 0.557661 | 0.019327 | 0.984581 | NA |
| STRIP2 | 486.172 | 0.624739 | 0.23314 | 2.67967 | 0.007369 | 0.038555 |
| RABL5 | 570.2442 | 0.057748 | 0.141589 | 0.407856 | 0.68338 | 0.794028 |
| MKLN1 | 1290.993 | 0.307044 | 0.175721 | 1.747343 | 0.080578 | 0.192203 |
| DNAJB9 | 994.4074 | 0.360809 | 0.177283 | 2.035216 | 0.041829 | 0.124232 |
| FLNC | 104.3006 | -0.40298 | 0.50365 | -0.80012 | 0.423639 | 0.585953 |
| LRRC4 | 78.23343 | -0.00842 | 0.296739 | -0.02836 | 0.977374 | 0.987689 |
| CALU | 16556.27 | 0.252002 | 0.149462 | 1.686055 | 0.091785 | 0.210711 |
| CCDC136 | 138.8421 | 0.301996 | 0.232421 | 1.299349 | 0.193824 | 0.348951 |
| SMO | 1284.041 | -0.37328 | 0.160932 | -2.31951 | 0.020368 | 0.076664 |
| IRF5 | 88.20561 | 0.078626 | 0.280472 | 0.280335 | 0.77922 | 0.864544 |
| LRRC17 | 1.293478 | 0.830419 | 1.712299 | 0.484973 | 0.627696 | NA |
| KLHDC10 | 1025.344 | -0.09617 | 0.143232 | -0.67143 | 0.501944 | 0.65228 |
| NDUFA5 | 809.8517 | 0.984005 | 0.29372 | 3.350152 | 8.08E-04 | 0.009102 |

| | | | | | | |
|---|---|---|---|---|---|---|
| OPN1SW | 52.98644 | 0.712087 | 0.372309 | 1.912626 | 0.055796 | 0.149437 |
| SUMF2 | 7631.555 | -0.37905 | 0.130584 | -2.90272 | 0.003699 | 0.024281 |
| PHF20L1 | 871.9361 | 0.897197 | 0.320228 | 2.801746 | 0.005083 | 0.029934 |
| LRRC6 | 44.62113 | 0.845352 | 0.380026 | 2.224461 | 0.026117 | NA |
| MTUS1 | 448.9423 | 0.276199 | 0.293805 | 0.940079 | 0.347177 | 0.515246 |
| ARHGEF6 | 709.0499 | 0.499722 | 0.174278 | 2.867382 | 0.004139 | 0.025988 |
| MAP7D3 | 424.1131 | 0.561871 | 0.290108 | 1.936763 | 0.052774 | 0.143471 |
| FGF13 | 11.64606 | -0.44024 | 0.814209 | -0.5407 | 0.588715 | NA |
| ASH2L | 1845.044 | -0.25413 | 0.13067 | -1.9448 | 0.051799 | 0.141718 |
| TTI2 | 391.9107 | -0.18654 | 0.189003 | -0.98694 | 0.323671 | 0.491623 |
| RPS4Y1 | 6837.209 | 0.111075 | 0.167585 | 0.662799 | 0.50746 | 0.656489 |
| HDHD1 | 549.6433 | -0.15215 | 0.150912 | -1.00823 | 0.313346 | 0.480746 |
| ERMARD | 366.67 | 0.111454 | 0.166516 | 0.669325 | 0.503288 | 0.653289 |
| PHF10 | 520.2713 | 0.482711 | 0.24132 | 2.000292 | 0.045469 | 0.130611 |
| PRRG3 | 33.56589 | 0.471374 | 0.511436 | 0.921668 | 0.356702 | NA |
| NXNL2 | 2.19576 | -0.45144 | 1.466513 | -0.30783 | 0.758208 | NA |
| STARD8 | 637.9173 | -0.66607 | 0.188183 | -3.53948 | 4.01E-04 | 0.005923 |
| FAM155B | 2.060055 | 1.158055 | 1.493704 | 0.775291 | 0.438168 | NA |
| GDPD2 | 4.841276 | 2.319018 | 1.087817 | 2.131808 | 0.033023 | NA |
| SAT1 | 14428.47 | 0.206655 | 0.279161 | 0.740274 | 0.459134 | 0.617544 |
| GNL3L | 484.3875 | -0.56818 | 0.165696 | -3.42906 | 6.06E-04 | 0.007631 |
| MOSPD2 | 737.1359 | 0.812808 | 0.277437 | 2.929702 | 0.003393 | 0.022819 |
| THEM6 | 554.3449 | -0.24337 | 0.163395 | -1.48946 | 0.136367 | 0.275452 |
| GADD45G | 6.814529 | -0.00965 | 0.809195 | -0.01193 | 0.990481 | NA |
| LRCH2 | 121.7224 | 0.611774 | 0.353655 | 1.729859 | 0.083655 | 0.197007 |
| DPP6 | 2.189379 | -0.32354 | 1.71792 | -0.18833 | 0.850617 | NA |
| XPO7 | 3529.925 | -0.22484 | 0.12619 | -1.78175 | 0.07479 | 0.18251 |
| ACE2 | 0.314073 | -0.57308 | 3.898934 | -0.14698 | 0.883146 | NA |
| NSUN5 | 636.2933 | -0.29963 | 0.157366 | -1.90405 | 0.056904 | 0.151187 |
| TULP4 | 1523.007 | -0.20744 | 0.197721 | -1.04918 | 0.294096 | 0.461117 |
| SNX9 | 5724.381 | 0.568903 | 0.146444 | 3.884793 | 1.02E-04 | 0.0026 |
| RSPH3 | 248.6511 | 0.271626 | 0.201045 | 1.351066 | 0.176674 | 0.328906 |
| MLLT4 | 1133.227 | 0.264755 | 0.161373 | 1.640643 | 0.100871 | 0.22419 |
| EPO | 0.163957 | 0.424913 | 4.079006 | 0.104171 | 0.917034 | NA |
| ARPC1B | 14155.46 | -0.22554 | 0.111923 | -2.0151 | 0.043894 | 0.127729 |
| OLFM1 | 1267.149 | -0.0305 | 0.166151 | -0.18357 | 0.854354 | 0.913506 |
| CAMSAP1 | 1267.442 | -0.3032 | 0.14224 | -2.13162 | 0.033038 | 0.105914 |
| UBAC1 | 1275.079 | 0.021021 | 0.153186 | 0.137224 | 0.890853 | 0.936099 |
| COL5A1 | 120.654 | 0.298403 | 0.49672 | 0.600747 | 0.548008 | 0.689317 |
| PNPLA7 | 77.65465 | -0.35656 | 0.284101 | -1.25503 | 0.209467 | 0.367072 |
| MNX1 | 0.162376 | 0.424913 | 4.079006 | 0.104171 | 0.917034 | NA |
| ASS1 | 1.884281 | -0.34254 | 1.443049 | -0.23737 | 0.81237 | NA |
| PRDM12 | 0.096202 | 0.408546 | 4.079639 | 0.100143 | 0.920231 | NA |
| EXOSC2 | 602.9185 | -0.079 | 0.184046 | -0.42926 | 0.667735 | 0.782059 |
| POMT1 | 1008.609 | -0.28266 | 0.145156 | -1.94729 | 0.0515 | 0.141327 |
| UCK1 | 2352.88 | -0.23525 | 0.148502 | -1.58413 | 0.113163 | 0.241987 |
| FIBCD1 | 1.118988 | -1.27548 | 2.033024 | -0.62738 | 0.530411 | NA |

| | | | | | | |
|---|---|---|---|---|---|---|
| PRRC2B | 5108.655 | -0.7845 | 0.158194 | -4.95909 | 7.08E-07 | 1.26E-04 |
| EIF2S3 | 2269.472 | -0.00956 | 0.196403 | -0.04869 | 0.961165 | 0.978918 |
| SLC6A8 | 6660.238 | -0.48201 | 0.140658 | -3.42681 | 6.11E-04 | 0.007678 |
| PNCK | 32.46489 | -0.03142 | 0.441299 | -0.07121 | 0.943231 | NA |
| DKC1 | 1377.289 | 0.296784 | 0.125975 | 2.355902 | 0.018478 | 0.071657 |
| PLXNA3 | 512.1517 | -0.32895 | 0.197094 | -1.66898 | 0.095121 | 0.215801 |
| DUSP9 | 134.5108 | -0.62327 | 0.360799 | -1.72748 | 0.084082 | 0.197553 |
| MPP1 | 777.9126 | -0.19977 | 0.149611 | -1.33523 | 0.181802 | 0.334668 |
| HSD17B3 | 0.247898 | -0.5901 | 4.074633 | -0.14482 | 0.884851 | NA |
| NUTM2F | 1.839612 | -0.06541 | 1.404043 | -0.04659 | 0.962841 | NA |
| HABP4 | 1906.697 | 0.423939 | 0.167901 | 2.52494 | 0.011572 | 0.052003 |
| SLC35D2 | 1158.813 | -0.13875 | 0.144517 | -0.96011 | 0.336998 | 0.505902 |
| PRRG1 | 412.7056 | 0.498188 | 0.176272 | 2.826237 | 0.00471 | 0.028372 |
| UBA1 | 10318.91 | -0.53685 | 0.146131 | -3.67374 | 2.39E-04 | 0.004335 |
| RGN | 0.623045 | 0.091811 | 2.666142 | 0.034436 | 0.97253 | NA |
| TXLNG2P | 338.3129 | 1.024799 | 0.297195 | 3.448244 | 5.64E-04 | 0.007303 |
| AKAP12 | 4328.424 | -0.11119 | 0.294516 | -0.37755 | 0.705765 | 0.811369 |
| SYNE1 | 737.8212 | 0.270423 | 0.213204 | 1.268379 | 0.204663 | 0.361375 |
| EDA2R | 422.2196 | 0.173998 | 0.193653 | 0.898508 | 0.368915 | 0.535966 |
| ARHGEF9 | 500.196 | -0.23304 | 0.165151 | -1.41105 | 0.158229 | 0.30476 |
| SH3BGRL | 3214.519 | 0.487188 | 0.190232 | 2.561016 | 0.010437 | 0.048415 |
| COX7B | 3488.404 | 0.279073 | 0.18808 | 1.483797 | 0.137863 | 0.277199 |
| IDO1 | 0.134133 | -1.48909 | 4.077584 | -0.36519 | 0.71497 | NA |
| RLIM | 1182.168 | 0.359901 | 0.237268 | 1.516853 | 0.129304 | 0.265759 |
| ABCB7 | 422.7849 | 0.112605 | 0.176807 | 0.636882 | 0.524202 | 0.670307 |
| EXOC4 | 3218.024 | -0.06656 | 0.132044 | -0.5041 | 0.614191 | 0.740414 |
| BARX1 | 11.95718 | 1.612837 | 0.845227 | 1.90817 | 0.056369 | NA |
| NINJ1 | 3818.217 | -0.24572 | 0.1318 | -1.86436 | 0.062271 | 0.161562 |
| IL13RA1 | 1486.749 | -0.04403 | 0.14218 | -0.30965 | 0.756824 | 0.848189 |
| WDR44 | 412.6275 | 0.494857 | 0.197913 | 2.500376 | 0.012406 | 0.054553 |
| KHDRBS3 | 753.3465 | 0.541705 | 0.209415 | 2.586752 | 0.009689 | 0.046165 |
| PDHA1 | 1473.441 | 0.158084 | 0.128455 | 1.230657 | 0.218451 | 0.378009 |
| RAI2 | 12.96343 | -0.1839 | 0.604714 | -0.30411 | 0.761042 | NA |
| THAP1 | 181.1685 | 0.806077 | 0.230132 | 3.502674 | 4.61E-04 | 0.006455 |
| EFR3A | 1755.283 | 0.423704 | 0.180642 | 2.34554 | 0.019 | 0.073035 |
| SEC61G | 1061.962 | 0.236398 | 0.150209 | 1.573797 | 0.115534 | 0.245635 |
| LANCL2 | 699.3848 | -0.31136 | 0.139699 | -2.2288 | 0.025827 | 0.090054 |
| FIGNL1 | 249.5643 | 0.647246 | 0.22337 | 2.897636 | 0.00376 | 0.024485 |
| DDC | 0.162376 | 0.424913 | 4.079006 | 0.104171 | 0.917034 | NA |
| HRSP12 | 199.8142 | 0.825831 | 0.235465 | 3.50723 | 4.53E-04 | 0.00643 |
| VPS13B | 902.5556 | 0.179244 | 0.175199 | 1.023091 | 0.306265 | 0.473301 |
| MATN2 | 775.9522 | -0.06018 | 0.288847 | -0.20834 | 0.834963 | 0.901331 |
| IRS4 | 25.51753 | -0.60219 | 0.446274 | -1.34938 | 0.177214 | NA |
| MORC4 | 748.6096 | -0.07079 | 0.149241 | -0.47434 | 0.635257 | 0.756414 |
| BEX2 | 58.25924 | 0.734555 | 0.347378 | 2.114569 | 0.034467 | 0.108758 |
| RNF128 | 269.314 | 0.78143 | 0.188891 | 4.136927 | 3.52E-05 | 0.001342 |
| GNG5P2 | 1.017482 | -0.71858 | 2.105405 | -0.3413 | 0.732874 | NA |

| | | | | | | |
|---|---|---|---|---|---|---|
| TBC1D8B | 372.0021 | 0.886871 | 0.209926 | 4.224688 | 2.39E-05 | 0.001024 |
| TCEAL4 | 1722.533 | 0.594096 | 0.23273 | 2.552722 | 0.010688 | 0.049189 |
| BEX1 | 154.9105 | 0.736878 | 0.450311 | 1.636378 | 0.101761 | 0.22549 |
| ADCK2 | 575.2929 | -0.10655 | 0.153463 | -0.69427 | 0.487511 | 0.640453 |
| MKRN1 | 2863.468 | -0.14459 | 0.123895 | -1.16704 | 0.243195 | 0.406164 |
| AGAP3 | 4114.462 | -0.24748 | 0.162968 | -1.51859 | 0.128866 | 0.265134 |
| KRBA1 | 406.5291 | -0.33517 | 0.171814 | -1.95076 | 0.051085 | 0.140645 |
| ZNF767 | 127.7627 | 0.812017 | 0.21809 | 3.723315 | 1.97E-04 | 0.00384 |
| ACTR3B | 107.0018 | 0.298337 | 0.24119 | 1.236937 | 0.216111 | 0.37531 |
| IMPA1 | 609.4157 | 0.696543 | 0.212614 | 3.276092 | 0.001053 | 0.010711 |
| LRRCC1 | 70.74331 | 1.200903 | 0.977102 | 1.229046 | 0.219055 | 0.378722 |
| E2F5 | 102.6152 | 0.416802 | 0.279461 | 1.491448 | 0.135844 | 0.274817 |
| TEX15 | 17.96326 | 0.670052 | 1.660018 | 0.403642 | 0.686476 | NA |
| TMEM66 | 4500.289 | 0.247266 | 0.124086 | 1.992697 | 0.046295 | 0.132215 |
| RNF122 | 186.7884 | -0.53758 | 0.195839 | -2.74498 | 0.006051 | 0.033633 |
| LOXL2 | 256.2341 | 0.352993 | 0.287576 | 1.22748 | 0.219642 | 0.379295 |
| ELP3 | 1226.322 | -0.08996 | 0.124739 | -0.7212 | 0.470786 | 0.627098 |
| PEBP4 | 0.755955 | 0.348448 | 2.512583 | 0.138681 | 0.889702 | NA |
| FAM127A | 4719.17 | -0.23548 | 0.128686 | -1.82991 | 0.067264 | 0.169633 |
| RAB33A | 1484.459 | 0.534167 | 0.297577 | 1.795051 | 0.072646 | 0.17927 |
| RBMX2 | 459.5457 | 0.210703 | 0.156512 | 1.34624 | 0.178225 | 0.330702 |
| | 463.4738 | 0.781435 | 0.159897 | 4.887115 | 1.02E-06 | 1.66E-04 |
| OSTF1 | 1540.136 | 0.174905 | 0.136832 | 1.278246 | 0.201163 | 0.357485 |
| RFK | 779.9195 | 0.235099 | 0.176834 | 1.329493 | 0.183685 | 0.336932 |
| UBQLN1 | 3902.449 | 0.120637 | 0.135605 | 0.889622 | 0.373669 | 0.539959 |
| NAA35 | 359.8866 | 0.299694 | 0.203267 | 1.474383 | 0.140379 | 0.280568 |
| C9orf40 | 303.3454 | -0.12255 | 0.202305 | -0.60578 | 0.54466 | 0.686398 |
| ANXA1 | 5003.638 | 0.830445 | 0.221272 | 3.753049 | 1.75E-04 | 0.003558 |
| CTSL | 5322.333 | -0.01704 | 0.130789 | -0.13032 | 0.896314 | 0.939326 |
| TMEM2 | 534.1031 | 0.013485 | 0.164267 | 0.082093 | 0.934573 | 0.963308 |
| AGTPBP1 | 397.0666 | 0.792038 | 0.20099 | 3.940688 | 8.12E-05 | 0.002244 |
| GOLM1 | 3508.616 | 0.215295 | 0.192897 | 1.116118 | 0.264371 | 0.428902 |
| FAM189A2 | 223.7735 | -0.94699 | 0.268654 | -3.52496 | 4.24E-04 | 0.006131 |
| PSAT1 | 3332.795 | -0.33432 | 0.180951 | -1.84756 | 0.064667 | 0.165736 |
| ISCA1 | 762.7951 | 0.406538 | 0.169939 | 2.39225 | 0.016745 | 0.067229 |
| DMTF1 | 377.5485 | 1.02516 | 0.243333 | 4.21299 | 2.52E-05 | 0.001062 |
| TMEM243 | 457.9562 | 0.159239 | 0.187015 | 0.851474 | 0.394506 | 0.559233 |
| CCDC146 | 13.10471 | 0.99229 | 0.728649 | 1.361822 | 0.173254 | NA |
| TMEM60 | 795.766 | 0.170651 | 0.173947 | 0.981049 | 0.326568 | 0.494777 |
| POM121C | 600.1661 | -0.6487 | 0.195412 | -3.31965 | 9.01E-04 | 0.009727 |
| CD36 | 13622.65 | 1.428965 | 0.329311 | 4.339262 | 1.43E-05 | 7.69E-04 |
| PNPLA8 | 783.2201 | 1.101644 | 0.2374 | 4.640453 | 3.48E-06 | 3.38E-04 |
| HILPDA | 192.8544 | 0.006482 | 0.195403 | 0.033173 | 0.973537 | 0.985255 |
| FAM71F1 | 0.354779 | 1.703954 | 3.657232 | 0.465914 | 0.641277 | NA |
| RINT1 | 506.1309 | 0.392168 | 0.204158 | 1.920902 | 0.054744 | 0.147431 |
| SRPK2 | 1372.976 | 0.246726 | 0.188817 | 1.306695 | 0.191316 | 0.34574 |
| KCP | 22.72309 | -0.77594 | 0.487118 | -1.59291 | 0.11118 | NA |

| | | | | | | |
|---|---|---|---|---|---|---|
| TES | 941.7306 | 0.083621 | 0.170726 | 0.489794 | 0.62428 | 0.748088 |
| MDFIC | 973.8015 | 0.435446 | 0.205152 | 2.122553 | 0.033791 | 0.107341 |
| SCRN1 | 7581.914 | 0.031974 | 0.165778 | 0.192873 | 0.847058 | 0.908932 |
| C7orf25 | 10.74812 | 0.094956 | 0.671706 | 0.141365 | 0.887581 | NA |
| TNS3 | 6881.898 | -0.31798 | 0.173398 | -1.8338 | 0.066683 | 0.168708 |
| SPDYE1 | 1.493931 | -0.17651 | 1.55812 | -0.11328 | 0.909808 | NA |
| CHST12 | 791.1117 | -0.34408 | 0.176919 | -1.94484 | 0.051794 | 0.141718 |
| IGF2BP3 | 344.0547 | 0.093429 | 0.274016 | 0.340961 | 0.733133 | 0.831067 |
| GPNMB | 61853.12 | -0.27805 | 0.136799 | -2.03255 | 0.042098 | 0.124768 |
| RAPGEF5 | 45.95752 | 0.43692 | 0.51206 | 0.85326 | 0.393515 | NA |
| RAC1 | 6486.353 | -0.23614 | 0.121393 | -1.94521 | 0.05175 | 0.141717 |
| KDELR2 | 4311.562 | -0.14788 | 0.146089 | -1.01229 | 0.311399 | 0.478811 |
| NUPL2 | 447.4644 | 0.544507 | 0.168278 | 3.235764 | 0.001213 | 0.01178 |
| IL6 | 4.84592 | 0.631249 | 0.92718 | 0.680827 | 0.495981 | NA |
| ZDHHC4 | 887.9844 | -0.23783 | 0.165096 | -1.44056 | 0.149709 | 0.293211 |
| AOAH | 17.7255 | -0.39467 | 0.520042 | -0.75893 | 0.447897 | NA |
| BZW2 | 1779.708 | -0.18291 | 0.12888 | -1.41926 | 0.155823 | 0.301695 |
| DGKB | 11.83407 | 1.231619 | 0.672086 | 1.832532 | 0.066872 | NA |
| TBRG4 | 2422.551 | -0.30763 | 0.132456 | -2.32248 | 0.020207 | 0.076242 |
| DDX56 | 1888.575 | -0.25773 | 0.135365 | -1.90393 | 0.056919 | 0.151192 |
| HUS1 | 520.2442 | 0.335039 | 0.164553 | 2.036058 | 0.041745 | 0.124043 |
| NACAD | 271.1411 | -0.27973 | 0.264599 | -1.05719 | 0.290426 | 0.457298 |
| C7orf69 | 0.260158 | 1.240347 | 4.047124 | 0.306476 | 0.759242 | NA |
| DBNL | 5350.335 | -0.34387 | 0.157599 | -2.18192 | 0.029115 | 0.09701 |
| CCM2 | 1498.006 | -0.1756 | 0.165778 | -1.05923 | 0.289497 | 0.455896 |
| MYO1G | 11.05213 | 1.095517 | 0.683447 | 1.602931 | 0.10895 | NA |
| TTYH3 | 24549.91 | -0.38817 | 0.185095 | -2.09715 | 0.03598 | 0.111865 |
| NIPSNAP3A | 575.9876 | 0.065386 | 0.167518 | 0.390321 | 0.696299 | 0.804152 |
| LRRC8A | 2184.655 | -0.3011 | 0.164758 | -1.82752 | 0.067622 | 0.170276 |
| CDK9 | 1551.496 | -0.35609 | 0.14556 | -2.44635 | 0.014431 | 0.060473 |
| TXN | 4214.509 | 0.33695 | 0.141318 | 2.384335 | 0.01711 | 0.068067 |
| ODF2 | 779.8149 | -0.18415 | 0.163663 | -1.1252 | 0.260506 | 0.424718 |
| KIAA0368 | 2324.521 | 0.071854 | 0.160501 | 0.447687 | 0.654379 | 0.771753 |
| TOR1B | 955.8118 | -0.056 | 0.200582 | -0.27919 | 0.780101 | 0.865117 |
| C9orf78 | 1981.319 | 0.318103 | 0.141061 | 2.255079 | 0.024128 | 0.085854 |
| SMC2 | 467.8131 | 1.198422 | 0.377791 | 3.17218 | 0.001513 | 0.01363 |
| KLF4 | 250.9452 | 0.38972 | 0.280396 | 1.389893 | 0.164561 | 0.313294 |
| TOR1A | 1268.488 | 0.018354 | 0.162062 | 0.113255 | 0.909828 | 0.947804 |
| RALGPS1 | 62.94642 | -0.43274 | 0.27555 | -1.57045 | 0.11631 | 0.246689 |
| FAM129B | 11385.2 | -0.32155 | 0.143012 | -2.24839 | 0.024551 | 0.086758 |
| ST6GALNA( | 337.5264 | -0.21577 | 0.16412 | -1.3147 | 0.188612 | 0.342879 |
| TMOD1 | 180.0876 | -0.68273 | 0.347668 | -1.96374 | 0.04956 | 0.137692 |
| DAB2IP | 1218.299 | -0.46437 | 0.13405 | -3.46416 | 5.32E-04 | 0.007022 |
| STXBP1 | 6503.91 | -0.36059 | 0.158798 | -2.27074 | 0.023163 | 0.083463 |
| SLC2A8 | 547.9901 | -0.40642 | 0.155438 | -2.61471 | 0.00893 | 0.043838 |
| ANGPTL2 | 1562.373 | -0.47119 | 0.179379 | -2.62677 | 0.00862 | 0.042703 |
| CDK5RAP2 | 1688.677 | 0.362502 | 0.18415 | 1.968513 | 0.049009 | 0.136932 |

| | | | | | | |
|---|---|---|---|---|---|---|
| ZFP37 | 40.13385 | 0.855826 | 0.404902 | 2.113661 | 0.034544 | NA |
| SLC31A2 | 38.08937 | -0.14343 | 0.352834 | -0.40651 | 0.684367 | NA |
| SLC31A1 | 2056.153 | -0.20848 | 0.139509 | -1.49436 | 0.135081 | 0.273679 |
| TLR4 | 264.456 | -0.15456 | 0.231741 | -0.66697 | 0.504792 | 0.654381 |
| ZNF189 | 310.5 | 0.480657 | 0.213924 | 2.246854 | 0.024649 | 0.086963 |
| ALDOB | 3.27791 | 1.250648 | 1.16492 | 1.073592 | 0.283006 | NA |
| STX17 | 325.2179 | 0.494261 | 0.232309 | 2.127606 | 0.03337 | 0.106573 |
| PRPF4 | 1055.488 | -0.05744 | 0.158449 | -0.36254 | 0.716946 | 0.81952 |
| FPGS | 2200.305 | -0.26567 | 0.148926 | -1.78391 | 0.074438 | 0.182214 |
| USP20 | 1394.594 | -0.61209 | 0.151423 | -4.04229 | 5.29E-05 | 0.00171 |
| BAAT | 2.886861 | -0.14642 | 1.309128 | -0.11184 | 0.910949 | NA |
| KIF12 | 1.724209 | -0.58773 | 1.697585 | -0.34622 | 0.729179 | NA |
| ATP6V1G1 | 6510.296 | 0.732532 | 0.175678 | 4.169748 | 3.05E-05 | 0.001199 |
| TEX10 | 1095.278 | 0.144722 | 0.157593 | 0.918325 | 0.358449 | 0.525712 |
| GARNL3 | 168.6444 | -0.51414 | 0.234 | -2.1972 | 0.028006 | 0.09476 |
| MRPL50 | 539.6536 | 0.654165 | 0.205234 | 3.187413 | 0.001436 | 0.013213 |
| DPM2 | 1397.407 | -0.0728 | 0.182237 | -0.39949 | 0.68953 | 0.798357 |
| WDR38 | 2.012464 | 1.134837 | 1.467439 | 0.773345 | 0.439318 | NA |
| TSTD2 | 452.943 | -0.1057 | 0.168819 | -0.62612 | 0.531235 | 0.675613 |
| GABBR2 | 25.71585 | -0.27614 | 0.565273 | -0.48851 | 0.625185 | NA |
| HEMGN | 0.268266 | -2.15396 | 4.038351 | -0.53338 | 0.593773 | NA |
| PSMB7 | 6685.189 | -0.05319 | 0.124524 | -0.42712 | 0.669294 | 0.783111 |
| NR5A1 | 0.392857 | -1.546 | 4.075469 | -0.37934 | 0.704433 | NA |
| C9orf156 | 346.1423 | 0.141535 | 0.176484 | 0.801969 | 0.422571 | 0.584968 |
| RABEPK | 605.6271 | -0.1939 | 0.184794 | -1.04925 | 0.294061 | 0.461117 |
| GOLGA1 | 642.631 | 0.187183 | 0.155165 | 1.206352 | 0.227682 | 0.389322 |
| XPA | 151.8882 | 0.530787 | 0.26182 | 2.027297 | 0.042632 | 0.12566 |
| NCBP1 | 1283.078 | 0.150271 | 0.187111 | 0.803108 | 0.421912 | 0.584524 |
| ANP32B | 2598.523 | 0.253089 | 0.164594 | 1.537659 | 0.124132 | 0.258672 |
| PDCL | 594.5307 | 0.287649 | 0.160547 | 1.791679 | 0.073184 | 0.180262 |
| RPL35 | 15140.04 | 0.078024 | 0.205964 | 0.378823 | 0.704819 | 0.810651 |
| CTSV | 182.9757 | 0.343788 | 0.204008 | 1.685174 | 0.091955 | 0.210878 |
| LMX1B | 0.258578 | 1.233151 | 4.047809 | 0.304646 | 0.760635 | NA |
| ARPC5L | 1073.394 | 0.02667 | 0.160729 | 0.165934 | 0.868209 | 0.921864 |
| ENPP2 | 858.1499 | -0.95323 | 0.260143 | -3.66424 | 2.48E-04 | 0.004422 |
| DSCC1 | 146.108 | 0.527053 | 0.285001 | 1.849302 | 0.064414 | 0.165288 |
| DERL1 | 1911.989 | -0.24063 | 0.144139 | -1.66943 | 0.095033 | 0.215702 |
| MYC | 4525.719 | -0.41556 | 0.154073 | -2.69714 | 0.006994 | 0.037113 |
| NOV | 22728.5 | 0.064335 | 0.203732 | 0.315782 | 0.752168 | 0.844956 |
| IL33 | 6.832173 | 0.790186 | 0.907603 | 0.870629 | 0.383957 | NA |
| TMEM261 | 840.4223 | -0.14628 | 0.216316 | -0.67625 | 0.498883 | 0.649348 |
| RANBP6 | 406.2544 | 0.557377 | 0.263852 | 2.112459 | 0.034647 | 0.10924 |
| POLR1E | 639.4509 | -0.15662 | 0.164442 | -0.95244 | 0.340873 | 0.508949 |
| PLAA | 874.5074 | 0.186915 | 0.163874 | 1.1406 | 0.254037 | 0.418046 |
| IL11RA | 665.6384 | -0.36092 | 0.153793 | -2.34679 | 0.018936 | 0.072813 |
| UBAP2 | 1577.256 | -0.49408 | 0.152109 | -3.24821 | 0.001161 | 0.011424 |
| APTX | 867.0715 | 0.119896 | 0.158226 | 0.757754 | 0.448598 | 0.608203 |

| | | | | | | |
|---|---|---|---|---|---|---|
| RNF38 | 630.7144 | -0.27029 | 0.18487 | -1.46205 | 0.143728 | 0.285409 |
| TLN1 | 16108.53 | -0.6675 | 0.176235 | -3.78756 | 1.52E-04 | 0.003285 |
| DNAJB5 | 575.7752 | -0.43571 | 0.149228 | -2.91978 | 0.003503 | 0.023355 |
| SPAG8 | 13.66764 | 0.292971 | 0.585618 | 0.500276 | 0.616881 | NA |
| DCTN3 | 2893.136 | -0.04343 | 0.14145 | -0.30707 | 0.758793 | 0.849592 |
| CD72 | 6.232182 | -0.28028 | 0.788761 | -0.35534 | 0.722333 | NA |
| TMEM8B | 421.3995 | -0.44932 | 0.169503 | -2.65081 | 0.00803 | 0.040681 |
| GRHPR | 3439.368 | -0.13473 | 0.143771 | -0.93713 | 0.348694 | 0.516656 |
| ALDH1B1 | 849.6859 | -0.46704 | 0.189424 | -2.46557 | 0.013679 | 0.058239 |
| HINT2 | 505.2616 | 0.053821 | 0.17685 | 0.304332 | 0.760875 | 0.851281 |
| ARHGEF39 | 114.3003 | -0.13249 | 0.301869 | -0.43891 | 0.660727 | 0.776536 |
| IGFBPL1 | 0.260158 | 1.240347 | 4.047124 | 0.306476 | 0.759242 | NA |
| DENND4C | 596.755 | 0.458532 | 0.214908 | 2.133618 | 0.032874 | 0.105609 |
| RPS6 | 36162.39 | 0.336316 | 0.150101 | 2.240602 | 0.025052 | 0.087945 |
| MRPL15 | 1306.154 | 0.076551 | 0.141682 | 0.5403 | 0.58899 | 0.721298 |
| PI15 | 64.7616 | -0.01384 | 0.502199 | -0.02756 | 0.978011 | 0.988096 |
| TTPA | 63.2398 | 0.800243 | 0.314183 | 2.547057 | 0.010864 | 0.049667 |
| GGH | 3163.578 | 0.555379 | 0.176002 | 3.155529 | 0.001602 | 0.014125 |
| SLCO5A1 | 0.096202 | 0.408546 | 4.079639 | 0.100143 | 0.920231 | NA |
| SULF1 | 0.260158 | 1.240347 | 4.047124 | 0.306476 | 0.759242 | NA |
| TGS1 | 512.2482 | 0.724119 | 0.221868 | 3.263732 | 0.0011 | 0.011028 |
| SDCBP | 17170.61 | -0.30163 | 0.171358 | -1.76022 | 0.078371 | 0.188591 |
| RGS3 | 1580.88 | 0.558485 | 0.284208 | 1.965055 | 0.049408 | 0.13756 |
| NCOA2 | 1149.184 | -0.04831 | 0.199002 | -0.24278 | 0.808172 | 0.885397 |
| FAM86B2 | 0.968725 | 0.233151 | 2.074813 | 0.112372 | 0.910528 | NA |
| DYNLT1 | 1622.145 | 0.235741 | 0.168342 | 1.400369 | 0.161403 | 0.309067 |
| TIAM2 | 36.73744 | 0.040118 | 0.349104 | 0.114918 | 0.90851 | NA |
| TMEM181 | 1451.188 | 0.43685 | 0.219283 | 1.992172 | 0.046352 | 0.132315 |
| PNLDC1 | 0.888653 | -1.5355 | 2.47141 | -0.62131 | 0.534398 | NA |
| WTAP | 1290.069 | 0.28013 | 0.162441 | 1.724506 | 0.084617 | 0.198286 |
| C6orf211 | 483.2946 | 0.896885 | 0.260659 | 3.440841 | 5.80E-04 | 0.007426 |
| SLC22A3 | 17.17941 | 0.619292 | 0.588434 | 1.05244 | 0.292598 | NA |
| VWDE | 0.536503 | 0.87599 | 3.23231 | 0.27101 | 0.786383 | NA |
| GNA12 | 2795.413 | -0.28931 | 0.133558 | -2.16617 | 0.030298 | 0.099795 |
| C7orf50 | 1022.792 | -0.23113 | 0.204097 | -1.13243 | 0.257455 | 0.421466 |
| SDK1 | 227.5982 | -0.91862 | 0.235775 | -3.89615 | 9.77E-05 | 0.002508 |
| CCZ1B | 119.8601 | 0.350936 | 0.237306 | 1.478835 | 0.139185 | 0.279177 |
| C7orf26 | 983.6711 | -0.16822 | 0.17494 | -0.96157 | 0.336267 | 0.505253 |
| RBAK | 319.7787 | 0.972395 | 0.335243 | 2.900564 | 0.003725 | 0.024357 |
| CREB5 | 51.31576 | 0.638845 | 0.420573 | 1.518989 | 0.128765 | 0.265134 |
| EGFR | 9.04004 | -0.33363 | 1.000866 | -0.33334 | 0.738879 | NA |
| LINC00525 | 0.096202 | 0.408546 | 4.079639 | 0.100143 | 0.920231 | NA |
| IGFBP3 | 179.6271 | -0.78578 | 0.310543 | -2.53033 | 0.011395 | 0.051405 |
| PURB | 1059.49 | 0.126563 | 0.19876 | 0.636763 | 0.524279 | 0.670333 |
| | 17.69193 | 0.096012 | 0.500217 | 0.19194 | 0.847789 | NA |
| IGFBP1 | 0.134133 | -1.48909 | 4.077584 | -0.36519 | 0.71497 | NA |
| SRCRB4D | 6.520021 | 0.136313 | 0.829493 | 0.164333 | 0.869469 | NA |

| | | | | | | |
|---|---|---|---|---|---|---|
| MDH2 | 7435.62 | -0.2519 | 0.129865 | -1.93974 | 0.052411 | 0.142855 |
| POMZP3 | 40.39498 | -0.20171 | 0.382947 | -0.52674 | 0.598373 | NA |
| GBAS | 1381.621 | 0.274612 | 0.180621 | 1.520379 | 0.128416 | 0.264779 |
| CCT6A | 5401.91 | 0.337783 | 0.176196 | 1.917087 | 0.055227 | 0.148327 |
| PSPH | 715.6428 | -0.43174 | 0.144557 | -2.98661 | 0.002821 | 0.020352 |
| ZNF92 | 52.15364 | 0.957428 | 0.512411 | 1.868476 | 0.061696 | 0.16036 |
| ATXN7L1 | 202.7512 | -0.16546 | 0.198716 | -0.83265 | 0.405045 | 0.569272 |
| TMEM168 | 786.1694 | 0.39179 | 0.156125 | 2.509464 | 0.012091 | 0.053697 |
| C7orf43 | 593.1992 | -0.159 | 0.180588 | -0.88044 | 0.378619 | 0.544856 |
| SLC12A9 | 1271.551 | -0.28793 | 0.167974 | -1.71416 | 0.0865 | 0.201545 |
| GIGYF1 | 1070.102 | -0.34177 | 0.167513 | -2.04028 | 0.041322 | 0.123231 |
| TRIM4 | 1755.186 | -0.20062 | 0.127488 | -1.57362 | 0.115576 | 0.245656 |
| MEPCE | 2401.588 | -0.40863 | 0.176082 | -2.3207 | 0.020303 | 0.076507 |
| TMEM209 | 287.3221 | -0.05675 | 0.228752 | -0.2481 | 0.804054 | 0.882843 |
| AGBL3 | 26.60742 | 0.839062 | 0.428473 | 1.958258 | 0.0502 | NA |
| STRA8 | 0.134133 | -1.48909 | 4.077584 | -0.36519 | 0.71497 | NA |
| ZC3HAV1L | 100.1396 | 0.513082 | 0.309349 | 1.658588 | 0.097199 | 0.218901 |
| TMEM140 | 1792.342 | -0.24932 | 0.160899 | -1.54956 | 0.121247 | 0.254023 |
| EPHA1 | 2.608753 | 0.32184 | 1.449872 | 0.221978 | 0.824331 | NA |
| NOM1 | 542.9514 | 0.042715 | 0.182657 | 0.233851 | 0.815101 | 0.889858 |
| NCAPG2 | 988.4249 | 0.127849 | 0.177542 | 0.720107 | 0.471459 | 0.627431 |
| NLGN4X | 201.1359 | 0.009162 | 0.23935 | 0.038279 | 0.969465 | 0.983447 |
| SHROOM2 | 252.9109 | -0.95968 | 0.280578 | -3.42036 | 6.25E-04 | 0.007763 |
| RAB19 | 0.163957 | 0.424913 | 4.079006 | 0.104171 | 0.917034 | NA |
| C7orf55-LU | 1859.735 | 0.233819 | 0.128572 | 1.81859 | 0.068974 | 0.172836 |
| DENND2A | 784.5963 | -0.32957 | 0.167551 | -1.96698 | 0.049185 | 0.137198 |
| TMEM27 | 2.994995 | 0.153163 | 1.334711 | 0.114753 | 0.908641 | NA |
| SH3KBP1 | 1448.222 | -0.01584 | 0.180869 | -0.08759 | 0.930202 | 0.960503 |
| TMEM47 | 1178.53 | 1.378304 | 0.298936 | 4.610695 | 4.01E-06 | 3.70E-04 |
| SYTL5 | 8.028479 | -0.0265 | 0.794037 | -0.03337 | 0.973376 | NA |
| CASK | 409.4907 | 0.148651 | 0.184424 | 0.806029 | 0.420226 | 0.583142 |
| KDM6A | 329.4683 | 0.089318 | 0.183138 | 0.48771 | 0.625755 | 0.748948 |
| SPIN2A | 2.057629 | -0.887 | 1.414579 | -0.62704 | 0.530634 | NA |
| MSN | 17004.09 | -0.41014 | 0.144145 | -2.84537 | 0.004436 | 0.027248 |
| CCNB3 | 34.45923 | 0.104774 | 0.399753 | 0.262098 | 0.793246 | NA |
| HDAC8 | 486.2682 | 0.339545 | 0.150996 | 2.248701 | 0.024532 | 0.086735 |
| SLC16A2 | 198.2532 | -0.54331 | 0.219558 | -2.47458 | 0.013339 | 0.057401 |
| CXorf36 | 0.488709 | 2.119056 | 3.307564 | 0.64067 | 0.521737 | NA |
| ZNF157 | 4.726529 | -0.0092 | 0.944493 | -0.00974 | 0.992232 | NA |
| ZNF182 | 90.18095 | 0.617496 | 0.310648 | 1.987768 | 0.046837 | 0.133187 |
| CHST7 | 144.4651 | -0.60566 | 0.250352 | -2.41923 | 0.015553 | 0.063537 |
| KRBOX4 | 129.1811 | 0.442954 | 0.233744 | 1.89504 | 0.058087 | 0.153608 |
| NDUFB11 | 2596.123 | -0.07201 | 0.16665 | -0.43213 | 0.665644 | 0.780535 |
| ZNF41 | 195.6038 | 0.262831 | 0.215895 | 1.217402 | 0.223451 | 0.384249 |
| RAB41 | 11.78168 | -0.0542 | 0.650679 | -0.0833 | 0.933611 | NA |
| ZMYM3 | 1102.765 | -0.64403 | 0.148684 | -4.33152 | 1.48E-05 | 7.84E-04 |
| TAF1 | 565.9755 | 0.312275 | 0.21737 | 1.436607 | 0.15083 | 0.294626 |

| | | | | | | |
|---|---|---|---|---|---|---|
| NONO | 8008.202 | -0.27923 | 0.125315 | -2.22825 | 0.025864 | 0.090102 |
| CCDC120 | 157.5089 | -0.55202 | 0.214547 | -2.57297 | 0.010083 | 0.047448 |
| LPAR4 | 0.735027 | 1.3952 | 2.594798 | 0.537691 | 0.59079 | NA |
| EBP | 2010.281 | -0.16482 | 0.144697 | -1.13911 | 0.254659 | 0.418896 |
| OGT | 2813.724 | 0.144866 | 0.208939 | 0.693337 | 0.488098 | 0.640939 |
| SNX12 | 3575.065 | -0.27247 | 0.134726 | -2.02244 | 0.043131 | 0.126565 |
| ITGB1BP2 | 0.897403 | 1.74906 | 2.471224 | 0.707771 | 0.479088 | NA |
| IL2RG | 0.303394 | -2.27006 | 4.032451 | -0.56295 | 0.57347 | NA |
| ACRC | 22.80006 | 0.736727 | 0.526664 | 1.398855 | 0.161856 | NA |
| ZNF711 | 0.47803 | 0.520083 | 2.97881 | 0.174594 | 0.861399 | NA |
| DIAPH2 | 440.2427 | 0.501273 | 0.347094 | 1.444199 | 0.148683 | 0.292069 |
| NXF3 | 4.566664 | 0.239389 | 1.005414 | 0.2381 | 0.811804 | NA |
| RIPPLY1 | 0.096202 | 0.408546 | 4.079639 | 0.100143 | 0.920231 | NA |
| PRPS1 | 1464.986 | 0.038531 | 0.17388 | 0.221594 | 0.82463 | 0.894418 |
| CXorf57 | 215.053 | 0.351536 | 0.208697 | 1.684434 | 0.092098 | 0.211061 |
| FRMPD3 | 0.735616 | -2.07189 | 2.528723 | -0.81934 | 0.412591 | NA |
| DOCK11 | 248.7053 | 0.582767 | 0.296698 | 1.964177 | 0.04951 | 0.137692 |
| IGSF1 | 0.162376 | 0.424913 | 4.079006 | 0.104171 | 0.917034 | NA |
| GPC3 | 69.38272 | -0.23624 | 0.327734 | -0.72082 | 0.471022 | 0.62727 |
| RBMX | 2634.431 | 0.079428 | 0.138314 | 0.574261 | 0.565791 | 0.702867 |
| MCPH1 | 298.3322 | 0.38372 | 0.181687 | 2.111987 | 0.034688 | 0.109309 |
| MFHAS1 | 844.4083 | -0.25493 | 0.168309 | -1.51465 | 0.129862 | 0.266168 |
| FBXO25 | 1255.14 | 0.322431 | 0.161055 | 2.00199 | 0.045286 | 0.130307 |
| FAM58A | 825.642 | 0.103267 | 0.145678 | 0.708873 | 0.478403 | 0.633189 |
| NSDHL | 1476.388 | -0.05382 | 0.148901 | -0.36143 | 0.717778 | 0.82009 |
| ZNF185 | 98.24439 | 0.060712 | 0.279293 | 0.217379 | 0.827913 | 0.896239 |
| CETN2 | 1230.307 | 0.226509 | 0.174774 | 1.296011 | 0.194972 | 0.350068 |
| RPL10 | 20142.25 | -0.15147 | 0.156199 | -0.96969 | 0.332199 | 0.500589 |
| CSGALNAC | 110.5208 | -0.65491 | 0.378578 | -1.72992 | 0.083645 | 0.197007 |
| ATP6V1B2 | 19278.06 | -0.18885 | 0.14852 | -1.27153 | 0.203539 | 0.359981 |
| CCDC25 | 877.7566 | 0.649059 | 0.19768 | 3.283381 | 0.001026 | 0.010515 |
| HMBOX1 | 455.1024 | -0.20487 | 0.184666 | -1.10942 | 0.267249 | 0.432229 |
| CHRNB3 | 0.163957 | 0.424913 | 4.079006 | 0.104171 | 0.917034 | NA |
| CHRNA6 | 46.48819 | -1.11471 | 0.380221 | -2.93173 | 0.003371 | NA |
| GNRH1 | 12.99407 | 1.50141 | 0.671452 | 2.236063 | 0.025348 | NA |
| BIN3 | 1538.418 | -0.09098 | 0.13885 | -0.65523 | 0.512318 | 0.659798 |
| DOK2 | 17.75886 | -0.14898 | 0.565783 | -0.26332 | 0.7923 | NA |
| SLC25A37 | 468.0375 | 0.36843 | 0.187242 | 1.967665 | 0.049107 | 0.137043 |
| CHMP7 | 1614.743 | -0.07629 | 0.154647 | -0.49329 | 0.621807 | 0.746181 |
| DOCK5 | 869.5517 | 0.154785 | 0.175752 | 0.880701 | 0.37848 | 0.544722 |
| STAR | 0.80987 | -0.47359 | 2.47725 | -0.19118 | 0.848387 | NA |
| PROSC | 965.7013 | 0.008144 | 0.18867 | 0.043167 | 0.965569 | 0.981198 |
| ERLIN2 | 926.5309 | -0.09034 | 0.178781 | -0.50529 | 0.613354 | 0.740083 |
| PXDNL | 0.324752 | 1.53082 | 4.022025 | 0.380609 | 0.703493 | NA |
| ST18 | 0.151697 | -1.546 | 4.075469 | -0.37934 | 0.704433 | NA |
| RGS20 | 1191.894 | 0.158985 | 0.173822 | 0.914644 | 0.360378 | 0.527508 |
| TACC1 | 9367.65 | 0.069313 | 0.145783 | 0.475455 | 0.634462 | 0.755859 |

| | | | | | | |
|---|---|---|---|---|---|---|
| GOLGA7 | 1962.882 | 0.265219 | 0.146126 | 1.815007 | 0.069523 | 0.173679 |
| PPAPDC1B | 253.7999 | 0.170819 | 0.19484 | 0.876717 | 0.380641 | 0.546836 |
| GINS4 | 313.6364 | -0.3515 | 0.253063 | -1.38897 | 0.164841 | 0.313702 |
| WHSC1L1 | 1101.499 | 0.349931 | 0.173287 | 2.019369 | 0.043449 | 0.126994 |
| TRIM55 | 0.096202 | 0.408546 | 4.079639 | 0.100143 | 0.920231 | NA |
| ADHFE1 | 47.15133 | 0.145472 | 0.387996 | 0.374931 | 0.707712 | 0.812783 |
| MRPS28 | 547.5722 | 0.35623 | 0.15093 | 2.360233 | 0.018263 | 0.071011 |
| PMP2 | 28.929 | 0.634006 | 0.717849 | 0.883202 | 0.377127 | NA |
| LACTB2 | 253.7849 | 0.525879 | 0.185205 | 2.839443 | 0.004519 | 0.027525 |
| TERF1 | 263.4695 | 0.719036 | 0.326402 | 2.202911 | 0.027601 | 0.093918 |
| RPL7 | 11433.82 | 0.484711 | 0.177072 | 2.737371 | 0.006193 | 0.034054 |
| SLC26A7 | 0.162376 | 0.424913 | 4.079006 | 0.104171 | 0.917034 | NA |
| ATP6V0D2 | 12.23306 | -0.95863 | 0.65045 | -1.47379 | 0.140539 | NA |
| SYBU | 250.2913 | -0.12182 | 0.175967 | -0.69226 | 0.488771 | 0.641539 |
| DPYS | 4.040792 | 0.898349 | 1.086002 | 0.827208 | 0.408119 | NA |
| MTDH | 4054.164 | 0.556142 | 0.168596 | 3.298669 | 9.71E-04 | 0.010161 |
| LRP12 | 2313.819 | 0.642771 | 0.197167 | 3.260034 | 0.001114 | 0.011107 |
| EBAG9 | 245.0259 | 0.422796 | 0.210597 | 2.007602 | 0.044686 | 0.129144 |
| POLR2K | 1694.654 | 0.630815 | 0.14133 | 4.463411 | 8.07E-06 | 5.46E-04 |
| MAL2 | 116.94 | 0.977935 | 0.311521 | 3.139229 | 0.001694 | 0.014563 |
| EIF3H | 4362.469 | 0.421855 | 0.1633 | 2.58332 | 0.009785 | 0.046533 |
| UTP23 | 269.3624 | 0.519759 | 0.208012 | 2.498702 | 0.012465 | 0.054669 |
| NDUFB9 | 2533.761 | 0.101565 | 0.161379 | 0.629355 | 0.529117 | 0.674264 |
| TATDN1 | 138.5356 | 0.588361 | 0.252729 | 2.32803 | 0.019911 | 0.075484 |
| FAM83A | 0.134133 | -1.48909 | 4.077584 | -0.36519 | 0.71497 | NA |
| FAM135B | 8.900899 | 0.101275 | 0.682896 | 0.148303 | 0.882104 | NA |
| ZNF7 | 336.9891 | 0.110872 | 0.178605 | 0.620764 | 0.534755 | 0.678307 |
| ARHGAP39 | 281.3138 | -0.66249 | 0.205047 | -3.23091 | 0.001234 | 0.011864 |
| SLC39A4 | 186.2301 | 0.113313 | 0.203987 | 0.555489 | 0.57856 | 0.71305 |
| NAPRT1 | 1268.874 | -0.1535 | 0.141614 | -1.0839 | 0.27841 | 0.444537 |
| VLDLR | 1614.872 | -0.40734 | 0.151179 | -2.69445 | 0.00705 | 0.037314 |
| AK3 | 3420.577 | 0.394114 | 0.173403 | 2.27282 | 0.023037 | 0.083125 |
| UHRF2 | 1104.345 | 0.451123 | 0.175545 | 2.569844 | 0.010174 | 0.047732 |
| NFIB | 421.266 | -0.20018 | 0.25264 | -0.79235 | 0.428158 | 0.589936 |
| PLIN2 | 1946.445 | -0.00889 | 0.160209 | -0.05546 | 0.95577 | 0.975764 |
| IFNA5 | 0.392857 | -1.546 | 4.075469 | -0.37934 | 0.704433 | NA |
| HAUS6 | 654.7757 | 0.79085 | 0.246343 | 3.210362 | 0.001326 | 0.012453 |
| CDKN2B | 179.0977 | 0.649724 | 0.238039 | 2.729482 | 0.006343 | 0.034639 |
| CDKN2A | 1584.245 | -0.05309 | 0.268507 | -0.19771 | 0.843273 | 0.906757 |
| C9orf72 | 140.9155 | 0.504847 | 0.236564 | 2.134082 | 0.032836 | 0.105579 |
| ZCCHC7 | 238.7528 | 0.571895 | 0.208675 | 2.740601 | 0.006133 | 0.03389 |
| FBXO10 | 106.1018 | -0.40635 | 0.237427 | -1.71148 | 0.086992 | 0.202491 |
| SIGMAR1 | 4563.846 | -0.27467 | 0.162542 | -1.68982 | 0.091063 | 0.2095 |
| CBWD5 | 52.08658 | 0.374346 | 0.308132 | 1.21489 | 0.224408 | 0.385447 |
| CEP78 | 496.8421 | 0.682729 | 0.243766 | 2.800756 | 0.005098 | 0.029995 |
| NTRK2 | 4.483854 | 0.696399 | 1.036839 | 0.671656 | 0.501803 | NA |
| IDNK | 66.91514 | 0.251019 | 0.29144 | 0.861308 | 0.389069 | 0.554509 |

| | | | | | | |
|---|---|---|---|---|---|---|
| SHC3 | 6.816372 | -0.4225 | 0.852761 | -0.49545 | 0.620279 | NA |
| AUH | 348.7908 | 0.335709 | 0.206648 | 1.624548 | 0.104259 | 0.229312 |
| HIATL1 | 8339.971 | 0.157059 | 0.121248 | 1.295356 | 0.195198 | 0.350192 |
| C9orf3 | 713.0692 | 0.015915 | 0.145703 | 0.109231 | 0.913019 | 0.94937 |
| ZNF462 | 586.2103 | -0.43797 | 0.191142 | -2.29134 | 0.021944 | 0.080716 |
| INIP | 344.7503 | 0.25043 | 0.166769 | 1.501652 | 0.133187 | 0.270643 |
| UGCG | 3240.707 | 0.280305 | 0.18303 | 1.531468 | 0.125654 | 0.260743 |
| SNX30 | 1885.527 | -0.01445 | 0.185144 | -0.07804 | 0.937799 | 0.965405 |
| STOM | 4996.245 | -0.14709 | 0.138448 | -1.06239 | 0.288058 | 0.454295 |
| GSN | 25174.99 | -0.278 | 0.125252 | -2.21953 | 0.02645 | 0.091282 |
| MRRF | 519.9273 | 0.154848 | 0.160219 | 0.966476 | 0.333806 | 0.502383 |
| NR6A1 | 73.2951 | -0.53573 | 0.294897 | -1.81665 | 0.06927 | 0.173285 |
| ALAD | 3081.916 | -0.16817 | 0.143539 | -1.1716 | 0.241359 | 0.404426 |
| ASTN2 | 290.499 | -0.37358 | 0.178387 | -2.09419 | 0.036243 | 0.112566 |
| WDR31 | 93.33169 | -0.6083 | 0.305055 | -1.99407 | 0.046144 | 0.131945 |
| POLE3 | 1728.94 | -0.13258 | 0.181737 | -0.72953 | 0.46568 | 0.622931 |
| SURF4 | 9135.579 | -0.30923 | 0.148381 | -2.08403 | 0.037158 | 0.114417 |
| GBGT1 | 172.3392 | -0.29434 | 0.214412 | -1.3728 | 0.169814 | 0.320309 |
| SURF1 | 935.1908 | -0.07554 | 0.157524 | -0.47955 | 0.631544 | 0.753746 |
| SURF2 | 397.3502 | 0.05127 | 0.164872 | 0.310971 | 0.755822 | 0.847421 |
| SURF6 | 958.1576 | 0.001742 | 0.185608 | 0.009384 | 0.992513 | 0.996738 |
| MED22 | 1530.701 | -0.44171 | 0.159713 | -2.76564 | 0.005681 | 0.032302 |
| REXO4 | 1294.671 | 0.006297 | 0.14989 | 0.042014 | 0.966488 | 0.981473 |
| RPL7A | 23746.57 | 0.07396 | 0.144861 | 0.510555 | 0.609663 | 0.736753 |
| GTF3C5 | 1612.283 | -0.30233 | 0.13668 | -2.21195 | 0.02697 | 0.092425 |
| ASB6 | 850.2583 | -0.26651 | 0.149063 | -1.7879 | 0.073792 | 0.181081 |
| PTGES2 | 2524.819 | -0.20441 | 0.140167 | -1.45835 | 0.144745 | 0.286709 |
| NTMT1 | 1347.695 | -0.01564 | 0.184017 | -0.085 | 0.932261 | 0.961541 |
| CIZ1 | 3698.042 | -0.29808 | 0.17655 | -1.68835 | 0.091344 | 0.209983 |
| SLC25A25 | 1303.165 | -0.01664 | 0.203901 | -0.08159 | 0.934972 | 0.963583 |
| SH3GLB2 | 1400.617 | -0.065 | 0.174949 | -0.37153 | 0.71024 | 0.814697 |
| FAM73B | 703.2631 | -0.16128 | 0.15411 | -1.04651 | 0.295328 | 0.46262 |
| PTGES | 30.71952 | -0.12125 | 0.416773 | -0.29092 | 0.771112 | NA |
| LRSAM1 | 789.4704 | -0.68704 | 0.158891 | -4.32397 | 1.53E-05 | 8.04E-04 |
| HMCN2 | 0.392857 | -1.546 | 4.075469 | -0.37934 | 0.704433 | NA |
| GPR107 | 3700.977 | -0.34585 | 0.116618 | -2.96566 | 0.00302 | 0.021262 |
| C9orf142 | 466.2464 | -0.13099 | 0.156616 | -0.83638 | 0.402939 | 0.567456 |
| INPP5E | 547.7886 | -0.34554 | 0.190311 | -1.81564 | 0.069425 | 0.173563 |
| SEC16A | 2421.385 | -0.64733 | 0.206505 | -3.13468 | 0.00172 | 0.014673 |
| DPH7 | 402.0745 | 0.061285 | 0.171617 | 0.357105 | 0.721013 | 0.822598 |
| NOTCH1 | 489.7435 | -0.82185 | 0.291076 | -2.82348 | 0.004751 | 0.028544 |
| CACNA1B | 1.876019 | -0.20672 | 1.620349 | -0.12758 | 0.898484 | NA |
| NACC2 | 2410.884 | -0.41725 | 0.208796 | -1.99837 | 0.045676 | 0.130858 |
| CNKSR2 | 23.05165 | 0.435228 | 0.48296 | 0.901168 | 0.367499 | NA |
| ZNF117 | 31.39135 | 0.15651 | 0.445406 | 0.351386 | 0.725299 | NA |
| FAM49B | 1023.883 | 0.294011 | 0.176054 | 1.670004 | 0.094919 | 0.215484 |
| ASAP1 | 1941.166 | -0.27929 | 0.209158 | -1.3353 | 0.181779 | 0.334668 |

| | | | | | | |
|---|---|---|---|---|---|---|
| PTPRD | 0.258578 | 1.233151 | 4.047809 | 0.304646 | 0.760635 | NA |
| LURAP1L | 134.5582 | 0.457627 | 0.360696 | 1.268734 | 0.204536 | 0.361313 |
| CNKSR3 | 167.1325 | -0.52148 | 0.222507 | -2.34367 | 0.019095 | 0.073307 |
| C7orf31 | 352.4385 | -0.20067 | 0.225194 | -0.89112 | 0.372866 | 0.539649 |
| JAZF1 | 503.1603 | 0.074375 | 0.158679 | 0.468712 | 0.639276 | 0.759456 |
| CACNA2D1 | 212.8356 | 0.268481 | 0.361132 | 0.743442 | 0.457214 | 0.615591 |
| SEMA3D | 37.06952 | 0.952808 | 0.375042 | 2.540534 | 0.011068 | NA |
| TDH | 0.327913 | 1.543154 | 4.021063 | 0.383768 | 0.701151 | NA |
| FAM167A | 2.40811 | -0.82326 | 1.338187 | -0.61521 | 0.538417 | NA |
| NEIL2 | 707.5179 | -0.21105 | 0.166199 | -1.26986 | 0.204136 | 0.360822 |
| PGM5 | 26.04562 | -0.61128 | 0.456298 | -1.33965 | 0.18036 | NA |
| LONRF1 | 3291.903 | 5.47E-04 | 0.172041 | 0.003179 | 0.997464 | 0.999065 |
| CNTNAP3B | 12.20541 | 0.012481 | 0.590858 | 0.021124 | 0.983147 | NA |
| FAM27C | 0.845272 | 1.770638 | 2.207891 | 0.801959 | 0.422577 | NA |
| MAGED4 | 0.163957 | 0.424913 | 4.079006 | 0.104171 | 0.917034 | NA |
| TCEB1 | 2008.265 | 0.48337 | 0.161319 | 2.996359 | 0.002732 | 0.019947 |
| LY96 | 898.6184 | 0.399284 | 0.208723 | 1.912981 | 0.05575 | 0.149428 |
| TMSB4Y | 0.61514 | 1.240942 | 3.104385 | 0.399738 | 0.689349 | NA |
| PDE1C | 8.170795 | 2.078686 | 0.892939 | 2.327914 | 0.019917 | NA |
| RABGEF1 | 2.651062 | -0.19575 | 1.230885 | -0.15903 | 0.873643 | NA |
| VOPP1 | 807.6358 | -0.02117 | 0.161483 | -0.13108 | 0.895709 | 0.939107 |
| 14-Sep | 0.326333 | 1.533889 | 3.826878 | 0.40082 | 0.688553 | NA |
| APOOL | 129.66 | 0.206049 | 0.24151 | 0.853169 | 0.393565 | 0.55842 |
| RSPH10B | 1.131045 | 0.415538 | 1.930034 | 0.215301 | 0.829533 | NA |
| FBXL18 | 410.4726 | -0.67746 | 0.187818 | -3.607 | 3.10E-04 | 0.005044 |
| UNC93B2 | 0.298089 | -0.51711 | 3.975232 | -0.13008 | 0.896501 | NA |
| KLF10 | 410.5272 | 0.151139 | 0.275861 | 0.54788 | 0.583775 | 0.717088 |
| PTPRN2 | 63.0659 | 0.088198 | 0.304408 | 0.289737 | 0.772017 | 0.858857 |
| AZIN1 | 3003.505 | 0.018099 | 0.16503 | 0.109672 | 0.912669 | 0.949346 |
| ATP6V1C1 | 2732.738 | 0.738101 | 0.16634 | 4.437302 | 9.11E-06 | 5.97E-04 |
| TMEM55A | 899.0608 | 0.151325 | 0.163791 | 0.923894 | 0.355542 | 0.523732 |
| OTUD6B | 287.8873 | 0.939731 | 0.270621 | 3.472495 | 5.16E-04 | 0.006915 |
| TTC39B | 41.92527 | 0.509416 | 0.424403 | 1.20031 | 0.230019 | NA |
| AGPAT5 | 891.2082 | 0.34458 | 0.162088 | 2.125884 | 0.033513 | 0.106886 |
| | 0.162376 | 0.424913 | 4.079006 | 0.104171 | 0.917034 | NA |
| TRIM74 | 0.326333 | 1.533889 | 3.826878 | 0.40082 | 0.688553 | NA |
| LRGUK | 6.445109 | -0.20501 | 0.845039 | -0.24261 | 0.80831 | NA |
| NUP205 | 2536.019 | -0.21096 | 0.163828 | -1.28769 | 0.197853 | 0.353565 |
| C9orf85 | 177.908 | 0.296003 | 0.239839 | 1.234174 | 0.217138 | 0.376561 |
| VSIG4 | 0.364467 | -1.20166 | 3.685431 | -0.32606 | 0.744381 | NA |
| PDIA4 | 2519.807 | -0.43541 | 0.138727 | -3.13857 | 0.001698 | 0.014564 |
| DEPTOR | 107.3224 | -0.017 | 0.301446 | -0.05641 | 0.955016 | 0.975169 |
| RNF20 | 1360.628 | 0.40444 | 0.221193 | 1.828448 | 0.067482 | 0.169976 |
| ELMO1 | 72.40613 | 0.431963 | 0.32905 | 1.312755 | 0.189266 | 0.343409 |
| FAM154A | 0.163957 | 0.424913 | 4.079006 | 0.104171 | 0.917034 | NA |
| RRAGA | 3678.754 | -0.20532 | 0.126512 | -1.62296 | 0.104599 | 0.229931 |
| SLC24A2 | 0.151697 | -1.546 | 4.075469 | -0.37934 | 0.704433 | NA |

| | | | | | | |
|---|---|---|---|---|---|---|
| ADCY8 | 1.168976 | -0.41512 | 1.914592 | -0.21682 | 0.82835 | NA |
| RMND1 | 466.2955 | 0.304457 | 0.168052 | 1.811682 | 0.070035 | 0.174461 |
| SLA | 0.260158 | 1.240347 | 4.047124 | 0.306476 | 0.759242 | NA |
| VBP1 | 1050.797 | 0.736566 | 0.174164 | 4.229161 | 2.35E-05 | 0.001011 |
| RAB39B | 41.58535 | 0.218965 | 0.360208 | 0.607885 | 0.543264 | NA |
| CLIC2 | 0.940279 | 0.09645 | 2.075894 | 0.046462 | 0.962942 | NA |
| AFF2 | 52.94357 | 0.14952 | 0.317334 | 0.471174 | 0.637516 | 0.758354 |
| MICU3 | 76.26969 | 0.729156 | 0.403766 | 1.805887 | 0.070936 | 0.176188 |
| VPS37A | 796.6613 | 0.373214 | 0.14411 | 2.589788 | 0.009603 | 0.045907 |
| TMEM185/ | 425.9182 | -0.21328 | 0.188555 | -1.13111 | 0.258008 | 0.422179 |
| NAT2 | 0.392711 | 0.292538 | 3.145221 | 0.09301 | 0.925895 | NA |
| PSD3 | 497.0505 | 0.205384 | 0.182164 | 1.127472 | 0.259543 | 0.423872 |
| C9orf41 | 273.401 | 0.514786 | 0.286254 | 1.798351 | 0.072121 | 0.178385 |
| GNA14 | 1.134023 | 1.814051 | 2.35922 | 0.76892 | 0.441941 | NA |
| GNAQ | 1647.467 | 0.043191 | 0.258647 | 0.166988 | 0.86738 | 0.921314 |
| MMP16 | 132.0035 | 0.431631 | 0.305098 | 1.414733 | 0.157147 | 0.303367 |
| DPY19L4 | 495.2927 | 0.640443 | 0.183607 | 3.488118 | 4.86E-04 | 0.00668 |
| NDUFAF6 | 273.2113 | 0.177552 | 0.171411 | 1.035826 | 0.300284 | 0.467235 |
| C8orf37 | 44.4432 | 0.250503 | 0.33001 | 0.759077 | 0.447807 | NA |
| TSPAN7 | 39.10826 | -0.28786 | 0.384856 | -0.74798 | 0.454472 | NA |
| RPGR | 91.16487 | 1.007897 | 0.436572 | 2.308665 | 0.020962 | 0.078199 |
| CDK20 | 294.4044 | -0.15863 | 0.179539 | -0.88352 | 0.376957 | 0.543599 |
| GDF6 | 14.73241 | 1.373047 | 1.013553 | 1.354686 | 0.175518 | NA |
| UQCRB | 2741.057 | 0.734244 | 0.169494 | 4.331972 | 1.48E-05 | 7.84E-04 |
| MTERFD1 | 314.5418 | 0.701576 | 0.212672 | 3.298868 | 9.71E-04 | 0.010161 |
| PTDSS1 | 4233.975 | -0.17759 | 0.18819 | -0.94368 | 0.345333 | 0.513344 |
| RPL30 | 8818.782 | 0.246042 | 0.166893 | 1.47425 | 0.140414 | 0.280568 |
| FAM122C | 37.10366 | 0.040857 | 0.348444 | 0.117255 | 0.906658 | NA |
| FAM122B | 717.7329 | -0.26566 | 0.210309 | -1.26319 | 0.206522 | 0.363668 |
| FBXO43 | 16.20985 | 0.78959 | 0.518349 | 1.523277 | 0.127689 | NA |
| PHF6 | 525.0452 | 0.832215 | 0.297249 | 2.799723 | 0.005115 | 0.030041 |
| RAB11FIP1 | 1002.396 | -0.32379 | 0.148828 | -2.17558 | 0.029586 | 0.098146 |
| UNC5D | 0.268266 | -2.15396 | 4.038351 | -0.53338 | 0.593773 | NA |
| UTP14A | 543.7804 | 0.174445 | 0.172423 | 1.011726 | 0.311669 | 0.478933 |
| AIFM1 | 2698.957 | 0.078657 | 0.136755 | 0.57517 | 0.565177 | 0.702545 |
| BAG4 | 513.7139 | 0.466155 | 0.161783 | 2.881362 | 0.00396 | 0.025305 |
| TBC1D31 | 203.4855 | 0.376193 | 0.259784 | 1.448101 | 0.147589 | 0.290713 |
| WDYHV1 | 221.8123 | -0.07096 | 0.194876 | -0.36412 | 0.715772 | 0.81867 |
| ATAD2 | 471.0617 | 0.930886 | 0.339692 | 2.740383 | 0.006137 | 0.03389 |
| FBXO32 | 2593.869 | -0.08888 | 0.172379 | -0.51559 | 0.606139 | 0.734291 |
| NSMCE2 | 722.0682 | 0.499944 | 0.199396 | 2.507293 | 0.012166 | 0.053937 |
| GPR112 | 0.134133 | -1.48909 | 4.077584 | -0.36519 | 0.71497 | NA |
| ZIC3 | 0.163957 | 0.424913 | 4.079006 | 0.104171 | 0.917034 | NA |
| MALSU1 | 799.7961 | 0.05358 | 0.178779 | 0.299697 | 0.764408 | 0.853784 |
| RBPMS | 1262.665 | -0.31283 | 0.148134 | -2.11181 | 0.034702 | 0.109327 |
| NRG1 | 0.28583 | -2.21358 | 3.97376 | -0.55705 | 0.577494 | NA |
| PAXIP1 | 278.2146 | 0.102092 | 0.183756 | 0.555584 | 0.578495 | 0.71305 |

| | | | | | | |
|---|---|---|---|---|---|---|
| STEAP2 | 6.00388 | 0.229836 | 0.867643 | 0.264897 | 0.791089 | NA |
| CLDN12 | 459.4722 | 0.436816 | 0.191545 | 2.28049 | 0.022579 | 0.082057 |
| FZD1 | 387.6378 | -0.10422 | 0.188006 | -0.55435 | 0.579337 | 0.713563 |
| GATAD1 | 544.3288 | 0.280785 | 0.160824 | 1.745909 | 0.080827 | 0.192603 |
| SUSD3 | 1.442399 | -0.7557 | 1.646343 | -0.45902 | 0.646222 | NA |
| ARSE | 239.2555 | -0.3061 | 0.215843 | -1.41816 | 0.156143 | 0.30202 |
| TSC22D3 | 2280.14 | -0.39953 | 0.276943 | -1.44264 | 0.149121 | 0.292541 |
| TMEM164 | 764.7051 | -0.57515 | 0.160136 | -3.59165 | 3.29E-04 | 0.005201 |
| TAB3 | 1005.974 | 0.355013 | 0.19486 | 1.821887 | 0.068472 | 0.171978 |
| C9orf43 | 82.8688 | 0.441945 | 0.431476 | 1.024265 | 0.30571 | 0.472886 |
| PALM2-AK/ | 5.773446 | -0.76635 | 0.905879 | -0.84598 | 0.397567 | NA |
| ZNF618 | 589.7589 | -0.52818 | 0.147802 | -3.57357 | 3.52E-04 | 0.005475 |
| DGKI | 15.14691 | 0.638579 | 0.53478 | 1.194096 | 0.232441 | NA |
| C9orf91 | 1411.415 | -0.53464 | 0.175023 | -3.05469 | 0.002253 | 0.017456 |
| UBN2 | 228.8079 | 0.499244 | 0.24964 | 1.999855 | 0.045516 | 0.13062 |
| BRAF | 296.3981 | 0.166995 | 0.208764 | 0.79992 | 0.423757 | 0.585958 |
| PSMG3 | 695.1804 | 0.123469 | 0.181919 | 0.678706 | 0.497324 | 0.648387 |
| SLC37A3 | 1096.706 | -0.0441 | 0.156314 | -0.28211 | 0.777859 | 0.863438 |
| RPS4Y2 | 0.162376 | 0.424913 | 4.079006 | 0.104171 | 0.917034 | NA |
| RADIL | 2.046097 | -0.06666 | 1.528709 | -0.04361 | 0.965218 | NA |
| WIPI2 | 2880.547 | -0.17379 | 0.137009 | -1.26849 | 0.204622 | 0.361375 |
| ANKRD61 | 3.357902 | 0.048848 | 1.155402 | 0.042278 | 0.966277 | NA |
| PTPDC1 | 502.1078 | 0.385887 | 0.183036 | 2.108261 | 0.035008 | 0.10994 |
| RHPN1 | 32.19098 | 0.275514 | 0.380837 | 0.723443 | 0.469408 | NA |
| AAED1 | 535.6454 | 0.211095 | 0.155717 | 1.355631 | 0.175216 | 0.327065 |
| TMSB15A | 81.1359 | 0.216975 | 0.265406 | 0.817521 | 0.413631 | 0.576757 |
| FANCC | 291.8802 | -0.09772 | 0.184621 | -0.52929 | 0.596602 | 0.726525 |
| CUL4B | 1136.218 | 0.662819 | 0.246731 | 2.686407 | 0.007223 | 0.037953 |
| GPRASP2 | 203.8431 | -0.24548 | 0.209378 | -1.17243 | 0.241026 | 0.404325 |
| AUTS2 | 560.6453 | -0.54645 | 0.259182 | -2.10837 | 0.034999 | 0.10994 |
| SHROOM4 | 425.6122 | -0.32113 | 0.186743 | -1.71964 | 0.085499 | 0.19984 |
| RIBC1 | 12.07106 | -0.02321 | 0.608801 | -0.03813 | 0.969586 | NA |
| TMSB15B | 249.8935 | 0.118909 | 0.183856 | 0.646749 | 0.517794 | 0.665014 |
| TSPAN33 | 488.645 | -0.28214 | 0.154607 | -1.82488 | 0.068019 | 0.171022 |
| AHCYL2 | 1530.337 | -0.32696 | 0.167099 | -1.95667 | 0.050386 | 0.139334 |
| CPA2 | 20.82464 | 0.080895 | 0.572883 | 0.141206 | 0.887707 | NA |
| NCF1 | 0.163957 | 0.424913 | 4.079006 | 0.104171 | 0.917034 | NA |
| TSR2 | 1595.024 | -0.10835 | 0.146666 | -0.73875 | 0.460058 | 0.618366 |
| PPP1R9A | 20.4142 | 0.945217 | 0.532306 | 1.775703 | 0.075782 | NA |
| DYNC1I1 | 149.5042 | 0.472629 | 0.255427 | 1.850346 | 0.064264 | 0.165041 |
| PFKFB1 | 1.2036 | 0.044837 | 2.244311 | 0.019978 | 0.984061 | NA |
| ALAS2 | 0.258578 | 1.233151 | 4.047809 | 0.304646 | 0.760635 | NA |
| TMED4 | 2231.712 | -0.24672 | 0.130084 | -1.89664 | 0.057876 | 0.153117 |
| COPG2 | 406.3303 | -0.43908 | 0.215595 | -2.03658 | 0.041692 | 0.123969 |
| AGPAT6 | 10941.99 | -0.30448 | 0.132524 | -2.29752 | 0.021589 | 0.079801 |
| PKD1L1 | 10.69426 | -0.40713 | 0.695033 | -0.58577 | 0.558028 | NA |
| NPM2 | 422.068 | 0.004661 | 0.258303 | 0.018046 | 0.985602 | 0.99244 |

| | | | | | | |
|---|---|---|---|---|---|---|
| EDA | 91.60582 | -0.60377 | 0.278482 | -2.16806 | 0.030154 | 0.099437 |
| FGF17 | 4.891297 | 0.258729 | 0.959161 | 0.269745 | 0.787357 | NA |
| DMTN | 1320.785 | 0.052607 | 0.166683 | 0.315609 | 0.752299 | 0.844956 |
| FAM160B2 | 1152.013 | -0.38688 | 0.129937 | -2.97747 | 0.002906 | 0.020694 |
| CCAR2 | 2243.301 | -0.31805 | 0.142718 | -2.22855 | 0.025844 | 0.09006 |
| FBXW5 | 9174.607 | -0.18782 | 0.158605 | -1.18417 | 0.236344 | 0.399195 |
| STC1 | 33.41624 | 0.593619 | 0.525627 | 1.129352 | 0.258749 | NA |
| TUBBP5 | 1.578084 | 0.558508 | 1.77995 | 0.313778 | 0.75369 | NA |
| ANKRD18C | 0.725928 | 0.181819 | 2.321433 | 0.078322 | 0.937572 | NA |
| FAM131B | 2.211191 | 0.832036 | 1.455796 | 0.571533 | 0.567638 | NA |
| ZYX | 2304.286 | -0.17865 | 0.152089 | -1.17463 | 0.240141 | 0.403355 |
| CCDC107 | 248.7788 | 0.056257 | 0.217541 | 0.258602 | 0.795942 | 0.876779 |
| NPR2 | 853.1323 | -0.45994 | 0.141069 | -3.26035 | 0.001113 | 0.011104 |
| GNE | 979.6099 | -0.32693 | 0.156918 | -2.08346 | 0.037209 | 0.114494 |
| VMA21 | 730.0758 | 0.334265 | 0.261881 | 1.276399 | 0.201814 | 0.358267 |
| G6PD | 4724.485 | -0.27733 | 0.151597 | -1.82941 | 0.067338 | 0.169741 |
| GAB3 | 26.02696 | 0.04893 | 0.432826 | 0.113047 | 0.909993 | NA |
| RALGDS | 1236.236 | -0.33394 | 0.198868 | -1.67922 | 0.09311 | 0.212928 |
| VAV2 | 365.7249 | -0.2781 | 0.22669 | -1.22679 | 0.2199 | 0.379574 |
| ADAMTS13 | 84.74809 | -0.05031 | 0.430955 | -0.11673 | 0.907072 | 0.946014 |
| CACFD1 | 420.8718 | -0.20087 | 0.156459 | -1.28385 | 0.199193 | 0.355407 |
| SLC2A6 | 971.5901 | -0.38423 | 0.153679 | -2.50023 | 0.012411 | 0.054556 |
| C9orf116 | 107.0311 | 0.235276 | 0.24821 | 0.947889 | 0.343186 | 0.511307 |
| GPSM1 | 1614.707 | -0.21814 | 0.148827 | -1.46571 | 0.142729 | 0.28371 |
| C9orf117 | 59.28721 | -0.38054 | 0.317868 | -1.19715 | 0.231249 | 0.393379 |
| TOR2A | 221.8983 | -0.21132 | 0.194476 | -1.08662 | 0.277204 | 0.44307 |
| ST6GALNA( | 1442.528 | -0.12064 | 0.181205 | -0.66575 | 0.505568 | 0.654811 |
| ZER1 | 1834.07 | -0.47181 | 0.139368 | -3.38534 | 7.11E-04 | 0.008369 |
| ZDHHC12 | 1106.345 | -0.21713 | 0.150858 | -1.43933 | 0.150057 | 0.2936 |
| PKN3 | 377.7594 | -0.28058 | 0.226763 | -1.2373 | 0.215974 | 0.375202 |
| PPAPDC3 | 83.63976 | -0.28665 | 0.268817 | -1.06633 | 0.286273 | 0.452623 |
| MED27 | 400.1228 | -0.20945 | 0.158032 | -1.32538 | 0.185047 | 0.338591 |
| PPP1R35 | 269.0754 | 0.123433 | 0.19061 | 0.647571 | 0.517263 | 0.664587 |
| STAG3L2 | 145.3765 | -0.21443 | 0.271117 | -0.7909 | 0.429003 | 0.59062 |
| AZGP1 | 179.0029 | -0.37996 | 0.234694 | -1.61897 | 0.105454 | 0.231238 |
| CYP3A4 | 0.736957 | -1.0303 | 2.640699 | -0.39016 | 0.696418 | NA |
| CYP3A7 | 1.867855 | 0.43726 | 1.415907 | 0.30882 | 0.757459 | NA |
| ZNF394 | 791.4299 | -0.12096 | 0.170154 | -0.71092 | 0.477137 | 0.632006 |
| CPSF4 | 513.0038 | -0.33919 | 0.165507 | -2.04937 | 0.040426 | 0.121205 |
| LY6E | 6227.775 | -0.71843 | 0.135702 | -5.29421 | 1.20E-07 | 3.83E-05 |
| VPS28 | 2573.848 | -0.20976 | 0.143977 | -1.45692 | 0.14514 | 0.287107 |
| TONSL | 616.781 | -0.51402 | 0.194732 | -2.63962 | 0.0083 | 0.041553 |
| RECQL4 | 653.6995 | -0.28346 | 0.217828 | -1.30128 | 0.193162 | 0.348068 |
| LRRC14 | 732.5864 | -0.48502 | 0.159095 | -3.0486 | 0.002299 | 0.017699 |
| COL26A1 | 3.008049 | -0.8782 | 1.432188 | -0.61319 | 0.539751 | NA |
| PPP1R16A | 456.3444 | -0.23247 | 0.166522 | -1.39606 | 0.162695 | 0.310889 |
| FOXH1 | 1.009375 | 1.515121 | 2.384802 | 0.635323 | 0.525218 | NA |

| | | | | | | |
|---|---|---|---|---|---|---|
| ORAI2 | 2220.556 | -0.20345 | 0.175952 | -1.15628 | 0.247567 | 0.411052 |
| ALKBH4 | 373.8126 | -0.09707 | 0.165074 | -0.58803 | 0.556513 | 0.695935 |
| SH2B2 | 119.2262 | -0.46174 | 0.25355 | -1.82111 | 0.06859 | 0.172238 |
| RPL8 | 34494.15 | -0.06686 | 0.125452 | -0.53299 | 0.594041 | 0.724448 |
| LRWD1 | 889.8901 | -0.1936 | 0.171656 | -1.12786 | 0.259378 | 0.423872 |
| FBXL13 | 11.30116 | -0.09297 | 0.681677 | -0.13638 | 0.891523 | NA |
| NAPEPLD | 288.6425 | 0.125142 | 0.230698 | 0.542449 | 0.587509 | 0.720186 |
| PSMC2 | 2978.767 | 0.477616 | 0.153636 | 3.108748 | 0.001879 | 0.015479 |
| TBX20 | 0.162376 | 0.424913 | 4.079006 | 0.104171 | 0.917034 | NA |
| DAGLB | 585.2896 | -0.17442 | 0.191116 | -0.91262 | 0.361442 | 0.528738 |
| KIAA0895 | 106.8402 | 0.606658 | 0.301292 | 2.013523 | 0.04406 | 0.127962 |
| STK17A | 571.8387 | 0.568799 | 0.153768 | 3.699083 | 2.16E-04 | 0.004081 |
| TRA2A | 770.1978 | 0.011911 | 0.163576 | 0.072819 | 0.94195 | 0.967746 |
| COG5 | 927.1356 | 0.230538 | 0.167461 | 1.376667 | 0.168615 | 0.318707 |
| C7orf60 | 372.1753 | 0.733647 | 0.242931 | 3.019977 | 0.002528 | 0.01898 |
| GPR85 | 433.2018 | 0.713777 | 0.191829 | 3.720901 | 1.99E-04 | 0.003853 |
| RP9 | 230.8094 | 0.382892 | 0.200179 | 1.912753 | 0.05578 | 0.149437 |
| BMPER | 3.609392 | 0.680843 | 1.091073 | 0.624013 | 0.532619 | NA |
| ZNF12 | 391.0936 | 0.555569 | 0.265745 | 2.090608 | 0.036563 | 0.113292 |
| SLC29A4 | 30.42056 | -0.13446 | 0.377501 | -0.3562 | 0.721693 | NA |
| STEAP1 | 74.16448 | 0.564376 | 0.296703 | 1.902156 | 0.057151 | 0.151605 |
| CDCA7L | 593.6528 | -0.06123 | 0.18067 | -0.33893 | 0.734661 | 0.832199 |
| SP8 | 0.258578 | 1.233151 | 4.047809 | 0.304646 | 0.760635 | NA |
| MIOS | 507.4303 | 0.611324 | 0.17964 | 3.403041 | 6.66E-04 | 0.008061 |
| KIAA1324L | 26.69733 | -0.19249 | 0.475257 | -0.40503 | 0.685458 | NA |
| INTS4L1 | 0.315653 | -0.57305 | 3.892639 | -0.14721 | 0.882963 | NA |
| SYTL3 | 54.76402 | -0.27012 | 0.319338 | -0.84586 | 0.397629 | 0.562383 |
| IQUB | 4.443311 | 0.618512 | 1.00459 | 0.615685 | 0.538102 | NA |
| HEY1 | 228.4687 | 0.344762 | 0.232799 | 1.480943 | 0.138622 | 0.278394 |
| ZNF704 | 800.3387 | -0.09299 | 0.241623 | -0.38484 | 0.700356 | 0.807265 |
| FABP5 | 506.008 | 0.289094 | 0.165823 | 1.743396 | 0.081264 | 0.193289 |
| SHH | 0.162376 | 0.424913 | 4.079006 | 0.104171 | 0.917034 | NA |
| TAGAP | 2.31254 | 0.221919 | 1.320408 | 0.168069 | 0.866529 | NA |
| COL1A2 | 4121.091 | -0.25296 | 0.200305 | -1.26287 | 0.206636 | 0.363668 |
| FNDC1 | 211.1827 | 1.505462 | 0.805189 | 1.869701 | 0.061525 | 0.160095 |
| CHMP4C | 0.192403 | 0.975392 | 4.060716 | 0.240202 | 0.810174 | NA |
| SLC13A4 | 2.267653 | -0.27177 | 1.342478 | -0.20244 | 0.839576 | NA |
| BRI3 | 5908.96 | -0.11076 | 0.134437 | -0.82391 | 0.40999 | 0.573435 |
| LMTK2 | 507.2847 | -0.117 | 0.180313 | -0.64885 | 0.516434 | 0.663954 |
| CTSB | 52601.45 | -0.2918 | 0.147312 | -1.98085 | 0.047608 | 0.13441 |
| SOX17 | 0.134133 | -1.48909 | 4.077584 | -0.36519 | 0.71497 | NA |
| DLC1 | 1857.747 | 0.1502 | 0.231569 | 0.64862 | 0.516584 | 0.664003 |
| ADCY1 | 1125.395 | -0.83716 | 0.23614 | -3.54519 | 3.92E-04 | 0.00585 |
| C8orf48 | 65.1982 | 0.759161 | 0.298593 | 2.542461 | 0.011007 | 0.050171 |
| SUN3 | 2.14873 | -0.54655 | 1.472639 | -0.37113 | 0.710538 | NA |
| C7orf57 | 27.86556 | -0.01298 | 0.419553 | -0.03094 | 0.975315 | NA |
| PEX2 | 1182.97 | 0.256843 | 0.174981 | 1.46783 | 0.14215 | 0.282988 |

| | | | | | | |
|---|---|---|---|---|---|---|
| RAD21 | 2813.359 | 0.429548 | 0.145591 | 2.950376 | 0.003174 | 0.022004 |
| SLC30A8 | 0.151697 | -1.546 | 4.075469 | -0.37934 | 0.704433 | NA |
| MED30 | 95.16662 | 0.204408 | 0.260972 | 0.783259 | 0.433475 | 0.594636 |
| TNFRSF11B | 82.54817 | 1.319953 | 1.092431 | 1.208271 | 0.226943 | 0.388458 |
| SBSPON | 1.006068 | 2.047881 | 2.307936 | 0.887321 | 0.374906 | NA |
| PHKG1 | 12.3155 | 0.213012 | 0.616833 | 0.345332 | 0.729845 | NA |
| EN2 | 204.1703 | 0.596089 | 0.252378 | 2.361891 | 0.018182 | 0.070787 |
| CSMD3 | 1.215043 | 0.083298 | 1.905339 | 0.043718 | 0.965129 | NA |
| SPIDR | 759.5545 | 0.075904 | 0.147764 | 0.513683 | 0.607473 | 0.73511 |
| ORC5 | 520.8786 | 0.332254 | 0.21621 | 1.536714 | 0.124363 | 0.258991 |
| HEATR2 | 845.6899 | -0.4956 | 0.171307 | -2.89306 | 0.003815 | 0.024686 |
| OSGIN2 | 695.4029 | 0.616132 | 0.196127 | 3.1415 | 0.001681 | 0.014503 |
| SUN1 | 1326.66 | 0.10916 | 0.138627 | 0.787441 | 0.431024 | 0.592439 |
| OXR1 | 733.1621 | 0.877119 | 0.296718 | 2.956066 | 0.003116 | 0.021703 |
| TMEM74 | 0.860264 | -0.96741 | 2.363935 | -0.40924 | 0.682365 | NA |
| GPR146 | 12.78639 | -0.25132 | 0.578413 | -0.4345 | 0.663927 | NA |
| GPER1 | 468.4643 | -0.26202 | 0.212746 | -1.23163 | 0.218089 | 0.377547 |
| UNCX | 0.151697 | -1.546 | 4.075469 | -0.37934 | 0.704433 | NA |
| TMEM184A | 0.163957 | 0.424913 | 4.079006 | 0.104171 | 0.917034 | NA |
| NOS3 | 0.303394 | -2.27006 | 4.032451 | -0.56295 | 0.57347 | NA |
| MICALL2 | 218.9348 | -0.06992 | 0.230364 | -0.30354 | 0.76148 | 0.851797 |
| INTS1 | 5235.025 | -0.54947 | 0.152481 | -3.6035 | 3.14E-04 | 0.005079 |
| CDK5 | 1249.706 | -0.09675 | 0.180939 | -0.53471 | 0.592849 | 0.723441 |
| SLC4A2 | 9550.146 | -0.35678 | 0.158348 | -2.25316 | 0.024249 | 0.0862 |
| SLC7A13 | 0.162376 | 0.424913 | 4.079006 | 0.104171 | 0.917034 | NA |
| FASTK | 2350.617 | -0.24498 | 0.126861 | -1.93112 | 0.053469 | 0.144935 |
| TMUB1 | 999.4128 | -0.24399 | 0.150921 | -1.6167 | 0.105943 | 0.231811 |
| C7orf55 | 57.66995 | -0.11623 | 0.297639 | -0.39052 | 0.696155 | 0.804143 |
| GBX1 | 0.134133 | -1.48909 | 4.077584 | -0.36519 | 0.71497 | NA |
| FOXK1 | 1096.058 | -0.82061 | 0.227453 | -3.60782 | 3.09E-04 | 0.005035 |
| COX6C | 2173.411 | 0.487232 | 0.119705 | 4.070265 | 4.70E-05 | 0.001582 |
| OSR2 | 1.064927 | 0.46521 | 2.027265 | 0.229477 | 0.818498 | NA |
| YWHAZ | 17370.12 | -0.04762 | 0.135145 | -0.35234 | 0.724582 | 0.825081 |
| BAALC | 478.5001 | -0.57494 | 0.175054 | -3.28439 | 0.001022 | 0.010509 |
| FZD6 | 676.8622 | 0.473806 | 0.256709 | 1.845691 | 0.064937 | 0.166126 |
| CTHRC1 | 65.9506 | -0.19641 | 0.37052 | -0.5301 | 0.596041 | 0.726065 |
| SLC25A32 | 666.602 | 0.214034 | 0.20448 | 1.046722 | 0.295228 | 0.462525 |
| DCAF13 | 868.8982 | 0.695816 | 0.158603 | 4.387163 | 1.15E-05 | 6.61E-04 |
| DCSTAMP | 6.275674 | 0.554513 | 1.203738 | 0.460659 | 0.645043 | NA |
| TP53INP1 | 1108.431 | -0.10182 | 0.211851 | -0.48061 | 0.630795 | 0.753088 |
| INTS8 | 645.9493 | 0.408943 | 0.167811 | 2.436932 | 0.014812 | 0.061442 |
| KIAA1429 | 1105.858 | 0.29673 | 0.204461 | 1.451281 | 0.146702 | 0.289472 |
| FREM1 | 2.952918 | 1.449616 | 1.315877 | 1.101635 | 0.27062 | NA |
| GEM | 69.14272 | 0.495617 | 0.313828 | 1.579261 | 0.114276 | 0.243493 |
| PDP1 | 734.6325 | 0.346407 | 0.183433 | 1.888462 | 0.058964 | 0.155268 |
| TMEM67 | 193.2213 | 0.742938 | 0.229877 | 3.231898 | 0.00123 | 0.011864 |
| KIAA0196 | 2410.492 | 0.124133 | 0.150769 | 0.823335 | 0.410318 | 0.573757 |

| | | | | | | |
|---|---|---|---|---|---|---|
| RPP25L | 296.7991 | -0.25252 | 0.192376 | -1.31263 | 0.189306 | 0.343416 |
| FAM219A | 1492.969 | -0.20991 | 0.155521 | -1.34973 | 0.177104 | 0.329292 |
| C9orf24 | 0.647493 | -1.18339 | 3.051782 | -0.38777 | 0.698186 | NA |
| SNAPC3 | 1195.049 | 0.247846 | 0.158785 | 1.560889 | 0.11855 | 0.249961 |
| KIAA1161 | 50.28585 | -0.8055 | 0.392741 | -2.05098 | 0.040269 | 0.120949 |
| NUDT2 | 340.9571 | 0.008825 | 0.172156 | 0.051262 | 0.959117 | 0.977505 |
| TMEM65 | 618.9611 | 0.718224 | 0.199527 | 3.599637 | 3.19E-04 | 0.005122 |
| PSIP1 | 644.37 | 0.771797 | 0.608001 | 1.2694 | 0.204298 | 0.360991 |
| CCDC171 | 1594.523 | 0.741568 | 0.258925 | 2.864023 | 0.004183 | 0.026182 |
| UBAP1 | 1582.448 | -0.11131 | 0.143864 | -0.7737 | 0.43911 | 0.599011 |
| SYK | 3347.77 | -0.06146 | 0.213373 | -0.28803 | 0.773324 | 0.859531 |
| NIPSNAP3E | 11.04504 | 0.846022 | 0.6467 | 1.308216 | 0.1908 | NA |
| ABCA1 | 312.9469 | 0.18446 | 0.203517 | 0.906362 | 0.364744 | 0.531993 |
| NFIL3 | 615.1713 | 0.170376 | 0.238065 | 0.71567 | 0.474195 | 0.629587 |
| LETM2 | 58.71549 | 0.410739 | 0.322058 | 1.275355 | 0.202184 | 0.358669 |
| METTL2B | 570.7108 | -0.23995 | 0.184322 | -1.3018 | 0.192985 | 0.347801 |
| FXN | 351.1209 | -0.20852 | 0.171618 | -1.21503 | 0.224356 | 0.385414 |
| ZMAT4 | 0.096202 | 0.408546 | 4.079639 | 0.100143 | 0.920231 | NA |
| TMEM71 | 57.61666 | -0.45319 | 0.429564 | -1.05499 | 0.291431 | 0.458333 |
| MAMDC2 | 61.77649 | 0.364367 | 0.952914 | 0.382371 | 0.702186 | 0.808431 |
| C8orf34 | 0.411855 | 0.250456 | 3.097618 | 0.080854 | 0.935558 | NA |
| TMC1 | 0.411855 | 0.250456 | 3.097618 | 0.080854 | 0.935558 | NA |
| ALDH1A1 | 91.90445 | 0.03799 | 0.35624 | 0.10664 | 0.915074 | 0.950986 |
| HGSNAT | 1867.678 | -0.27624 | 0.144442 | -1.91243 | 0.05582 | 0.149437 |
| RASEF | 52.19952 | 0.99451 | 0.322108 | 3.087508 | 0.002018 | 0.016256 |
| GKAP1 | 5.5181 | 0.607832 | 0.848387 | 0.716456 | 0.47371 | NA |
| KIF27 | 23.40688 | 0.386258 | 0.479698 | 0.80521 | 0.420699 | NA |
| C9orf64 | 289.2343 | -0.12935 | 0.185618 | -0.69685 | 0.485899 | 0.639256 |
| HNRNPK | 12026.44 | 0.022267 | 0.121364 | 0.183475 | 0.854425 | 0.913506 |
| | 11.6988 | 1.311216 | 0.690501 | 1.898936 | 0.057573 | NA |
| SVEP1 | 1.343961 | -0.90382 | 2.015879 | -0.44835 | 0.653901 | NA |
| ANKS6 | 960.9408 | -0.28619 | 0.175041 | -1.63498 | 0.102054 | 0.226041 |
| FBP1 | 4.278134 | -0.89396 | 0.984971 | -0.9076 | 0.364089 | NA |
| TMEM246 | 909.9613 | -0.55005 | 0.15451 | -3.55998 | 3.71E-04 | 0.005601 |
| ZHX1 | 780.6832 | 0.651173 | 0.289871 | 2.246423 | 0.024677 | 0.086963 |
| DYNLT3 | 578.8628 | 0.844133 | 0.2748 | 3.071807 | 0.002128 | 0.016816 |
| WBSCR27 | 6.548573 | -0.14452 | 0.834433 | -0.17319 | 0.8625 | NA |
| MID1IP1 | 1466.471 | -0.49123 | 0.146519 | -3.35264 | 8.00E-04 | 0.009046 |
| NCF1C | 0.134133 | -1.48909 | 4.077584 | -0.36519 | 0.71497 | NA |
| C9orf84 | 1.359799 | 0.16786 | 1.649101 | 0.101789 | 0.918924 | NA |
| CXorf58 | 0.506476 | 0.726418 | 2.864186 | 0.253621 | 0.799788 | NA |
| KIAA1958 | 85.69705 | 0.142887 | 0.27467 | 0.520211 | 0.602916 | 0.732033 |
| PTCHD1 | 0.508056 | 0.731032 | 2.861912 | 0.255435 | 0.798387 | NA |
| RNF183 | 0.419962 | -2.64111 | 3.429364 | -0.77014 | 0.441214 | NA |
| ASB11 | 6.267335 | -1.19849 | 0.963305 | -1.24414 | 0.213448 | NA |
| PCDH19 | 0.792452 | -2.15617 | 2.562821 | -0.84133 | 0.400166 | NA |
| PIGA | 242.4476 | 0.705051 | 0.193344 | 3.646619 | 2.66E-04 | 0.004599 |

| | | | | | | |
|---|---|---|---|---|---|---|
| FIGF | 11.12887 | 0.77106 | 0.620105 | 1.243434 | 0.213708 | NA |
| STRBP | 168.664 | 0.647039 | 0.27674 | 2.338079 | 0.019383 | 0.074029 |
| CLDN3 | 0.096202 | 0.408546 | 4.079639 | 0.100143 | 0.920231 | NA |
| GAPVD1 | 1524.111 | 0.196689 | 0.149726 | 1.313662 | 0.18896 | 0.343263 |
| C9orf89 | 1263.106 | -2.79E-04 | 0.169152 | -0.00165 | 0.998686 | 0.999371 |
| WNK2 | 11.26933 | -0.40238 | 0.65335 | -0.61587 | 0.537982 | NA |
| ATP7A | 1546.526 | 0.27243 | 0.181216 | 1.503341 | 0.132751 | 0.270093 |
| ZNF367 | 222.2579 | 0.637714 | 0.307225 | 2.075721 | 0.03792 | 0.115846 |
| NLGN4Y | 36.78098 | 0.352988 | 0.421835 | 0.836792 | 0.402709 | NA |
| HDX | 66.70513 | 0.728439 | 0.406668 | 1.791239 | 0.073255 | 0.180395 |
| NDUFB6 | 1346.733 | 0.395865 | 0.147113 | 2.690895 | 0.007126 | 0.037572 |
| AQP7 | 0.327913 | 1.543154 | 4.021063 | 0.383768 | 0.701151 | NA |
| NOL6 | 1724.604 | -0.46771 | 0.155605 | -3.00576 | 0.002649 | 0.019519 |
| AQP3 | 9.004298 | -0.81582 | 0.721392 | -1.13089 | 0.2581 | NA |
| TRMT10B | 164.2004 | 0.238221 | 0.210089 | 1.133905 | 0.256834 | 0.420781 |
| VCP | 13340.84 | -0.30134 | 0.123302 | -2.44396 | 0.014527 | 0.060619 |
| PIGO | 869.9995 | -0.44979 | 0.175635 | -2.56096 | 0.010438 | 0.048415 |
| STOML2 | 3264.353 | -0.20312 | 0.143071 | -1.41972 | 0.155689 | 0.301553 |
| BRWD3 | 338.8392 | 0.734817 | 0.242206 | 3.033848 | 0.002415 | 0.018279 |
| MELK | 840.4454 | 0.311741 | 0.14422 | 2.16156 | 0.030652 | 0.100548 |
| DDX26B | 109.5332 | 0.855262 | 0.25402 | 3.366905 | 7.60E-04 | 0.008794 |
| CLDN2 | 0.631307 | -0.20384 | 3.097925 | -0.0658 | 0.947537 | NA |
| WRN | 136.5729 | 0.636607 | 0.286747 | 2.220101 | 0.026412 | 0.091242 |
| SSX5 | 5.586922 | -0.67566 | 0.93289 | -0.72427 | 0.468902 | NA |
| FAAH2 | 0.192403 | 0.975392 | 4.060716 | 0.240202 | 0.810174 | NA |
| QSOX2 | 712.681 | -0.22982 | 0.202609 | -1.13432 | 0.25666 | 0.420707 |
| ENOX2 | 336.5619 | 0.086719 | 0.231198 | 0.375087 | 0.707596 | 0.812765 |
| SNAPC4 | 472.7772 | -0.44438 | 0.170462 | -2.6069 | 0.009137 | 0.044457 |
| PMPCA | 2108.007 | -0.20929 | 0.141962 | -1.47426 | 0.140411 | 0.280568 |
| SDCCAG3 | 1526.124 | -0.10623 | 0.162234 | -0.65478 | 0.512609 | 0.659932 |
| FRMD7 | 3.383802 | 0.541899 | 1.244303 | 0.435504 | 0.663197 | NA |
| AK8 | 7.974993 | -0.15778 | 0.734398 | -0.21485 | 0.829886 | NA |
| C9orf9 | 156.0569 | -0.45878 | 0.229893 | -1.99561 | 0.045976 | 0.131591 |
| TSC1 | 946.9268 | -0.05676 | 0.140511 | -0.40396 | 0.686241 | 0.795623 |
| GFI1B | 0.192403 | 0.975392 | 4.060716 | 0.240202 | 0.810174 | NA |
| HPRT1 | 497.2162 | 0.111957 | 0.162129 | 0.690544 | 0.489852 | 0.642357 |
| FAM69B | 110.9591 | -0.67848 | 0.405704 | -1.67236 | 0.094454 | 0.21488 |
| ZMYND19 | 602.6532 | 0.133837 | 0.165349 | 0.80942 | 0.418274 | 0.581245 |
| FUNDC2 | 1532.764 | 0.241333 | 0.153574 | 1.571441 | 0.11608 | 0.246398 |
| NSMF | 5992.275 | -0.10961 | 0.146672 | -0.7473 | 0.454884 | 0.613778 |
| ZMAT1 | 0.561971 | -0.49218 | 2.810061 | -0.17515 | 0.860962 | NA |
| TMEM130 | 0.151697 | -1.546 | 4.075469 | -0.37934 | 0.704433 | NA |
| MCM7 | 2938.409 | -0.21886 | 0.170571 | -1.28311 | 0.199455 | 0.355784 |
| ZNF3 | 536.2174 | -0.18455 | 0.146287 | -1.26157 | 0.207104 | 0.364221 |
| ZSCAN21 | 159.117 | 0.022709 | 0.214304 | 0.105968 | 0.915608 | 0.951234 |
| SPDYE6 | 5.838249 | -0.23533 | 0.841662 | -0.2796 | 0.779786 | NA |
| NGFRAP1 | 5461.974 | 0.039913 | 0.123453 | 0.323309 | 0.746461 | 0.840469 |

| | | | | | | |
|---|---|---|---|---|---|---|
| ZCCHC18 | 31.85647 | 0.059529 | 0.36973 | 0.161006 | 0.872088 | NA |
| NYAP1 | 103.1165 | -0.04584 | 0.258187 | -0.17753 | 0.859095 | 0.916352 |
| TSC22D4 | 3410.36 | -0.46397 | 0.127664 | -3.6343 | 2.79E-04 | 0.004707 |
| CNPY4 | 406.8085 | -0.17145 | 0.184369 | -0.92994 | 0.352403 | 0.520437 |
| NAT16 | 2.865266 | 0.172729 | 1.249795 | 0.138206 | 0.890078 | NA |
| NKX3-1 | 8.799898 | 0.127474 | 0.700482 | 0.18198 | 0.855598 | NA |
| PBX3 | 422.5366 | 0.137456 | 0.149947 | 0.916697 | 0.359302 | 0.526452 |
| TTC16 | 0.612365 | -1.10341 | 2.696105 | -0.40926 | 0.682349 | NA |
| PIP5KL1 | 8.967054 | 0.074948 | 0.670898 | 0.111712 | 0.911052 | NA |
| FAM102A | 2084.763 | -0.40101 | 0.180877 | -2.21704 | 0.02662 | 0.091632 |
| GOLGA2 | 2138.045 | -0.195 | 0.161428 | -1.20797 | 0.22706 | 0.388539 |
| TRUB2 | 1547.596 | -0.00662 | 0.126651 | -0.05223 | 0.958342 | 0.977053 |
| COQ4 | 736.2083 | -0.33613 | 0.198632 | -1.69221 | 0.090606 | 0.208729 |
| SLC27A4 | 2256.662 | -0.28274 | 0.142054 | -1.99033 | 0.046554 | 0.132668 |
| URM1 | 3423.993 | -0.25142 | 0.1338 | -1.87906 | 0.060236 | 0.15768 |
| CERCAM | 4949.292 | -0.33416 | 0.119251 | -2.80218 | 0.005076 | 0.029923 |
| DOLPP1 | 423.8988 | -0.38596 | 0.178312 | -2.16454 | 0.030423 | 0.099989 |
| ENDOG | 362.0991 | 0.007941 | 0.172523 | 0.046029 | 0.963287 | 0.980069 |
| PRRX2 | 0.88142 | 1.783485 | 2.222213 | 0.802571 | 0.422222 | NA |
| TRAPPC9 | 1331.415 | -0.5413 | 0.125643 | -4.30825 | 1.65E-05 | 8.30E-04 |
| PSCA | 3.234034 | 0.13129 | 1.139422 | 0.115225 | 0.908267 | NA |
| MFSD3 | 1172.383 | -0.23049 | 0.189145 | -1.21858 | 0.223003 | 0.38382 |
| GPT | 12.0033 | 0.305724 | 0.584556 | 0.523002 | 0.600973 | NA |
| KIFC2 | 489.5135 | 0.20848 | 0.198119 | 1.052299 | 0.292662 | 0.459538 |
| TMEM68 | 304.861 | 0.586131 | 0.210922 | 2.778899 | 0.005454 | 0.03145 |
| CYP7A1 | 1.347539 | -0.58768 | 1.73484 | -0.33875 | 0.734797 | NA |
| | 20.11241 | 0.58915 | 0.467282 | 1.260801 | 0.20738 | NA |
| SCARA3 | 13.84301 | -1.29633 | 0.645392 | -2.00859 | 0.044581 | NA |
| PBK | 878.8646 | 0.34205 | 0.258255 | 1.324465 | 0.185349 | 0.339038 |
| SCARA5 | 0.410275 | 0.244283 | 3.100889 | 0.078778 | 0.937209 | NA |
| COPS6 | 3142.093 | -0.10258 | 0.141121 | -0.72689 | 0.467296 | 0.62385 |
| HOOK3 | 453.6965 | 0.754739 | 0.362712 | 2.080822 | 0.03745 | 0.114989 |
| POLR2J3 | 34.01472 | 0.282588 | 0.376241 | 0.751083 | 0.452602 | NA |
| KCNV2 | 1.22552 | 1.38393 | 1.789452 | 0.773382 | 0.439296 | NA |
| PCMTD1 | 518.5561 | 1.050909 | 0.380401 | 2.762636 | 0.005734 | 0.032461 |
| MPLKIP | 573.1624 | 0.048921 | 0.153869 | 0.317937 | 0.750533 | 0.844011 |
| HR | 1257.296 | -0.3641 | 0.201447 | -1.80744 | 0.070694 | 0.175771 |
| REEP4 | 1447.862 | -0.3559 | 0.167507 | -2.12471 | 0.033611 | 0.107059 |
| LGI3 | 16282.97 | -0.64885 | 0.16494 | -3.93384 | 8.36E-05 | 0.002284 |
| SFTPC | 385.4036 | -0.26 | 0.171178 | -1.51891 | 0.128785 | 0.265134 |
| BMP1 | 2867.384 | -0.56618 | 0.128564 | -4.40388 | 1.06E-05 | 6.40E-04 |
| PHYHIP | 34.88634 | 0.116507 | 0.365244 | 0.318985 | 0.749738 | NA |
| POLR3D | 964.6985 | -0.16699 | 0.133994 | -1.24623 | 0.212679 | 0.371436 |
| FNTA | 850.6909 | 0.34241 | 0.162486 | 2.107328 | 0.035089 | 0.110077 |
| GFRA2 | 228.2379 | 0.240763 | 0.86606 | 0.277998 | 0.781014 | 0.865562 |
| SLC20A2 | 932.4868 | 0.009985 | 0.154211 | 0.064747 | 0.948376 | 0.971147 |
| ADAM9 | 2528.792 | 0.522428 | 0.350704 | 1.489656 | 0.136315 | 0.275434 |

| | | | | | | |
|---|---|---|---|---|---|---|
| FAM84B | 404.3777 | -0.38634 | 0.222008 | -1.74021 | 0.081822 | 0.19431 |
| ZBTB5 | 337.3213 | -0.18576 | 0.195593 | -0.94971 | 0.342258 | 0.510438 |
| ENHO | 13.87887 | -0.51611 | 0.570561 | -0.90456 | 0.3657 | NA |
| SPRY3 | 0.134133 | -1.48909 | 4.077584 | -0.36519 | 0.71497 | NA |
| MECP2 | 3014.369 | -0.28962 | 0.13956 | -2.07526 | 0.037962 | 0.11593 |
| AR | 15.02469 | -0.78134 | 0.635964 | -1.2286 | 0.219222 | NA |
| C8orf46 | 486.9731 | -0.33623 | 0.180757 | -1.86014 | 0.062865 | 0.162387 |
| FAM110B | 220.5098 | -0.27279 | 0.188543 | -1.44684 | 0.147942 | 0.291047 |
| UBE2V2 | 679.3983 | 0.653228 | 0.222099 | 2.941154 | 0.00327 | 0.022362 |
| ZBTB43 | 340.6287 | 0.162562 | 0.181801 | 0.894173 | 0.37123 | 0.537879 |
| APEX2 | 830.3007 | -0.22867 | 0.135135 | -1.69212 | 0.090622 | 0.208729 |
| CCDC126 | 240.4315 | 0.973693 | 0.241211 | 4.036687 | 5.42E-05 | 0.001728 |
| CA5B | 446.1863 | -0.14422 | 0.218332 | -0.66054 | 0.508906 | 0.657414 |
| ZRSR2 | 140.5459 | 0.599108 | 0.250213 | 2.394388 | 0.016648 | 0.066926 |
| | 1.421286 | 1.067504 | 1.85287 | 0.576135 | 0.564524 | NA |
| NR0B1 | 200.3442 | -0.05537 | 0.335994 | -0.16481 | 0.869095 | 0.922145 |
| IL1RAPL1 | 145.1426 | 1.129467 | 0.399893 | 2.824422 | 0.004737 | 0.028489 |
| PTK2 | 2519.401 | 0.178411 | 0.14835 | 1.202635 | 0.229118 | 0.390875 |
| RSPH10B2 | 0.28583 | -2.21358 | 3.97376 | -0.55705 | 0.577494 | NA |
| KCNK9 | 1.757047 | 0.794655 | 1.616013 | 0.491738 | 0.622904 | NA |
| COL22A1 | 84.63595 | -0.50874 | 0.33108 | -1.53659 | 0.124393 | 0.258991 |
| SDC2 | 1886.683 | -0.13456 | 0.195665 | -0.6877 | 0.491639 | 0.643353 |
| MMGT1 | 1110.245 | -0.09957 | 0.161192 | -0.61771 | 0.536767 | 0.679944 |
| TM2D2 | 1456.104 | -0.01956 | 0.148644 | -0.13159 | 0.895311 | 0.939035 |
| HTRA4 | 5.257347 | -0.03752 | 0.916184 | -0.04095 | 0.967338 | NA |
| PLEKHA2 | 1130.445 | -0.42681 | 0.176367 | -2.42001 | 0.01552 | 0.063444 |
| GJB1 | 169.8099 | -0.52836 | 0.40999 | -1.2887 | 0.197502 | 0.353175 |
| CLIC3 | 0.35636 | 1.709252 | 3.651833 | 0.468053 | 0.639746 | NA |
| AGPAT2 | 2933.046 | -0.22416 | 0.157748 | -1.42102 | 0.155311 | 0.300999 |
| PRY | 0.151697 | -1.546 | 4.075469 | -0.37934 | 0.704433 | NA |
| TRIM56 | 551.2011 | -0.25965 | 0.21728 | -1.19499 | 0.232093 | 0.394343 |
| REPS2 | 317.5912 | -0.54213 | 0.208973 | -2.59424 | 0.00948 | 0.045537 |
| SYAP1 | 1302.577 | 0.502191 | 0.17066 | 2.942644 | 0.003254 | 0.022326 |
| TPST1 | 431.0661 | 0.380025 | 0.231161 | 1.643987 | 0.100179 | 0.223272 |
| S100G | 0.163957 | 0.424913 | 4.079006 | 0.104171 | 0.917034 | NA |
| GUSB | 1883.472 | -0.49079 | 0.156707 | -3.13188 | 0.001737 | 0.01475 |
| BRD3 | 1100.844 | -0.06062 | 0.161606 | -0.37514 | 0.707558 | 0.812765 |
| FRMPD4 | 5.051668 | -0.20524 | 0.923802 | -0.22217 | 0.824179 | NA |
| ZFPM2 | 4.879953 | 1.47321 | 0.977888 | 1.506522 | 0.131933 | NA |
| YWHAG | 6515.854 | -0.21941 | 0.139778 | -1.56971 | 0.116483 | 0.246766 |
| | 1.322107 | 0.164454 | 1.725698 | 0.095297 | 0.924079 | NA |
| FOXD4 | 4.226631 | 1.579694 | 1.109266 | 1.424089 | 0.154421 | NA |
| | 15.87858 | -0.06624 | 0.560821 | -0.11811 | 0.905984 | NA |
| | 0.162376 | 0.424913 | 4.079006 | 0.104171 | 0.917034 | NA |
| ZNF212 | 313.2548 | -0.21483 | 0.167622 | -1.28165 | 0.199965 | 0.355997 |
| ZNF282 | 1031.181 | -0.35184 | 0.184607 | -1.90589 | 0.056664 | 0.150854 |
| CNGB3 | 0.392857 | -1.546 | 4.075469 | -0.37934 | 0.704433 | NA |

| | | | | | | |
|---|---|---|---|---|---|---|
| FABP4 | 0.096202 | 0.408546 | 4.079639 | 0.100143 | 0.920231 | NA |
| FAM115C | 34.88258 | 0.18232 | 0.38922 | 0.468425 | 0.639481 | NA |
| SEMA3E | 572.7275 | 1.042941 | 0.162593 | 6.414445 | 1.41E-10 | 2.09E-07 |
| | 0.163957 | 0.424913 | 4.079006 | 0.104171 | 0.917034 | NA |
| SLC26A5 | 1.216567 | -0.28085 | 1.97818 | -0.14197 | 0.887102 | NA |
| COMMD5 | 896.8106 | -0.12683 | 0.148409 | -0.85457 | 0.392789 | 0.557866 |
| DPY19L2P2 | 88.99401 | -0.21772 | 0.287416 | -0.75751 | 0.448743 | 0.608279 |
| ZNF16 | 150.9761 | -0.27423 | 0.202921 | -1.35141 | 0.176564 | 0.328856 |
| ARMC10 | 578.9518 | 0.121635 | 0.157241 | 0.773563 | 0.439189 | 0.599011 |
| RASA4B | 0.762279 | -0.66959 | 2.551343 | -0.26245 | 0.792978 | NA |
| MURC | 26.4307 | 0.545117 | 0.41431 | 1.315722 | 0.188268 | NA |
| BOP1 | 1306.005 | -0.38419 | 0.163032 | -2.35652 | 0.018447 | 0.071585 |
| GPR37 | 419.5388 | 0.251649 | 0.29265 | 0.859897 | 0.389846 | 0.555283 |
| SDR16C5 | 0.163957 | 0.424913 | 4.079006 | 0.104171 | 0.917034 | NA |
| CHCHD7 | 686.4954 | 0.332312 | 0.147029 | 2.260174 | 0.02381 | 0.085202 |
| LMOD2 | 0.163957 | 0.424913 | 4.079006 | 0.104171 | 0.917034 | NA |
| CEL | 2.298351 | 0.590632 | 1.32559 | 0.445562 | 0.655914 | NA |
| KBTBD2 | 1403.808 | 0.378865 | 0.159742 | 2.371726 | 0.017705 | 0.069636 |
| MTSS1 | 669.5695 | -0.27683 | 0.2097 | -1.32013 | 0.186793 | 0.340892 |
| RNF139 | 755.722 | 0.091968 | 0.172385 | 0.533503 | 0.593685 | 0.724163 |
| HAS2 | 390.5446 | -0.24505 | 0.221441 | -1.10659 | 0.268469 | 0.433493 |
| PLAC1 | 1.186394 | 0.520608 | 1.795836 | 0.289897 | 0.771895 | NA |
| HS6ST2 | 0.622842 | 1.158846 | 2.735323 | 0.42366 | 0.671814 | NA |
| PKIA | 17.27081 | 0.181355 | 0.500888 | 0.362067 | 0.717302 | NA |
| XKR6 | 107.3372 | -0.326 | 0.229101 | -1.42294 | 0.154753 | 0.300109 |
| TSNARE1 | 311.24 | -0.59014 | 0.216072 | -2.73119 | 0.006311 | 0.034476 |
| SOX7 | 4.822823 | -0.55162 | 1.034483 | -0.53323 | 0.593874 | NA |
| CCBL1 | 231.6156 | -0.28423 | 0.183172 | -1.55171 | 0.120731 | 0.253097 |
| MTM1 | 516.2365 | 0.688211 | 0.201437 | 3.416509 | 6.34E-04 | 0.007808 |
| GIMAP8 | 0.096202 | 0.408546 | 4.079639 | 0.100143 | 0.920231 | NA |
| ATP6V0E2 | 1503.019 | -0.35503 | 0.140146 | -2.53325 | 0.011301 | 0.051105 |
| C1GALT1C1 | 678.0764 | 0.447498 | 0.186919 | 2.394077 | 0.016662 | 0.06696 |
| C9orf16 | 1067.081 | -0.14187 | 0.14545 | -0.97542 | 0.329351 | 0.497573 |
| NAIF1 | 361.0382 | -0.27186 | 0.183292 | -1.48323 | 0.138013 | 0.27736 |
| CLDN20 | 2.765315 | 0.970657 | 1.30076 | 0.746223 | 0.455532 | NA |
| SOSTDC1 | 0.533728 | -1.74213 | 2.820432 | -0.61768 | 0.536785 | NA |
| CHD7 | 701.7356 | 0.041974 | 0.238563 | 0.175947 | 0.860335 | 0.917119 |
| ESCO2 | 145.3103 | 0.664965 | 0.248478 | 2.676146 | 0.007447 | 0.038877 |
| CLCN5 | 611.5207 | -0.15581 | 0.15023 | -1.03712 | 0.299678 | 0.466476 |
| APLN | 5.327669 | -1.2528 | 0.896185 | -1.39793 | 0.162134 | NA |
| NAT1 | 262.9974 | 0.792553 | 0.244301 | 3.244169 | 0.001178 | 0.011522 |
| ZBTB26 | 76.73882 | 0.48385 | 0.29879 | 1.619364 | 0.105369 | 0.231153 |
| OR1L8 | 0.163957 | 0.424913 | 4.079006 | 0.104171 | 0.917034 | NA |
| GPR82 | 0.162376 | 0.424913 | 4.079006 | 0.104171 | 0.917034 | NA |
| GPR34 | 0.151697 | -1.546 | 4.075469 | -0.37934 | 0.704433 | NA |
| MLLT3 | 468.4411 | 0.600392 | 0.202869 | 2.959513 | 0.003081 | 0.021631 |
| IFNB1 | 0.134133 | -1.48909 | 4.077584 | -0.36519 | 0.71497 | NA |

| | | | | | | |
|---|---|---|---|---|---|---|
| MIR31HG | 1.284176 | 0.715001 | 1.74187 | 0.410479 | 0.681455 | NA |
| ANKRD20A | 1.022583 | -1.06903 | 2.182302 | -0.48986 | 0.62423 | NA |
| CYCS | 3858.149 | 0.408173 | 0.161803 | 2.522651 | 0.011647 | 0.052224 |
| FRMD3 | 7.572116 | 1.454692 | 0.777123 | 1.871892 | 0.061221 | NA |
| SNTB1 | 545.5376 | 0.203174 | 0.207134 | 0.980884 | 0.32665 | 0.494807 |
| MTBP | 65.35559 | 0.703027 | 0.32073 | 2.191958 | 0.028383 | 0.0954 |
| MRPL13 | 718.0955 | 0.594529 | 0.166733 | 3.565763 | 3.63E-04 | 0.005542 |
| HMGN1P3! | 0.46735 | -1.30193 | 3.399671 | -0.38296 | 0.701751 | NA |
| GPR22 | 0.315653 | -0.57305 | 3.892639 | -0.14721 | 0.882963 | NA |
| BPGM | 908.3353 | 0.675884 | 0.202012 | 3.345768 | 8.21E-04 | 0.009177 |
| POP7 | 890.088 | -0.00712 | 0.16868 | -0.04223 | 0.966314 | 0.981451 |
| GNB2 | 8655.671 | -0.25757 | 0.126678 | -2.03325 | 0.042027 | 0.12465 |
| TCEAL1 | 536.7128 | 0.740162 | 0.220329 | 3.359347 | 7.81E-04 | 0.008918 |
| RAB40A | 4.386089 | 1.080769 | 1.040994 | 1.038209 | 0.299173 | NA |
| HCFC1 | 2189.893 | -0.88164 | 0.195072 | -4.51954 | 6.20E-06 | 4.77E-04 |
| FUT10 | 365.0443 | -0.12718 | 0.16536 | -0.76909 | 0.441841 | 0.601869 |
| PURG | 2.810669 | 1.424188 | 1.437512 | 0.990731 | 0.321817 | NA |
| ZNF596 | 37.36833 | 0.597894 | 0.404202 | 1.479195 | 0.139088 | NA |
| CBWD1 | 181.6669 | 0.503724 | 0.212562 | 2.369772 | 0.017799 | 0.069778 |
| CYP7B1 | 163.1127 | 0.627598 | 0.212207 | 2.957473 | 0.003102 | 0.021663 |
| EGFL7 | 341.7629 | 0.196636 | 0.303325 | 0.648269 | 0.516811 | 0.664138 |
| PHF8 | 672.2313 | -0.47228 | 0.162029 | -2.91479 | 0.003559 | 0.023678 |
| ZNF680 | 37.45895 | 0.759586 | 0.400241 | 1.897822 | 0.05772 | NA |
| BNC2 | 834.1406 | -0.35823 | 0.212343 | -1.68703 | 0.091597 | 0.210443 |
| 1-Dec | 3.313232 | 0.209363 | 1.092636 | 0.191613 | 0.848045 | NA |
| LRRN3 | 63.69783 | 0.105186 | 0.452221 | 0.232599 | 0.816073 | 0.889858 |
| ADCK5 | 309.6387 | -0.19211 | 0.170746 | -1.12515 | 0.260527 | 0.424718 |
| CYSLTR1 | 0.162376 | 0.424913 | 4.079006 | 0.104171 | 0.917034 | NA |
| DMRT2 | 0.258578 | 1.233151 | 4.047809 | 0.304646 | 0.760635 | NA |
| ZNF483 | 6.554617 | 0.558894 | 0.791864 | 0.705795 | 0.480316 | NA |
| TNKS | 1600.924 | -0.38517 | 0.137852 | -2.79405 | 0.005205 | 0.030383 |
| ZNF449 | 203.2923 | 0.782692 | 0.248205 | 3.153404 | 0.001614 | 0.014192 |
| PPP1R3B | 304.8905 | 0.138138 | 0.171114 | 0.807287 | 0.419501 | 0.582614 |
| FAM86B3P | 33.71187 | -0.82363 | 0.394039 | -2.09022 | 0.036598 | NA |
| TRIB1 | 513.565 | -0.73908 | 0.219199 | -3.37172 | 7.47E-04 | 0.008725 |
| | 0.134133 | -1.48909 | 4.077584 | -0.36519 | 0.71497 | NA |
| TNFRSF10D | 356.4025 | 0.438538 | 0.253595 | 1.729283 | 0.083758 | 0.197146 |
| TNFRSF10C | 88.93608 | -0.27588 | 0.300024 | -0.91954 | 0.357812 | 0.525131 |
| NUDT18 | 610.0615 | 0.034212 | 0.192867 | 0.177387 | 0.859205 | 0.916352 |
| SCAI | 47.9956 | 0.289175 | 0.317829 | 0.909844 | 0.362905 | 0.530159 |
| EIF1AX | 690.9538 | 0.605096 | 0.177037 | 3.417908 | 6.31E-04 | 0.007785 |
| CXorf23 | 74.02523 | 0.884249 | 0.514001 | 1.720326 | 0.085373 | 0.199625 |
| GPR64 | 4.064483 | 0.270134 | 1.046568 | 0.258114 | 0.796319 | NA |
| DPY19L1 | 1065.918 | 0.505193 | 0.166545 | 3.033361 | 0.002418 | 0.018285 |
| KLHL15 | 278.3069 | 0.275124 | 0.219055 | 1.255958 | 0.209131 | 0.366699 |
| FAM46D | 0.548763 | 2.343695 | 3.135478 | 0.747476 | 0.454776 | NA |
| FAM3C2 | 17.30257 | 0.498847 | 0.562786 | 0.886389 | 0.375408 | NA |

| | | | | | | |
|---|---|---|---|---|---|---|
| C9orf131 | 0.354779 | 1.703954 | 3.657232 | 0.465914 | 0.641277 | NA |
| ARL13A | 0.096202 | 0.408546 | 4.079639 | 0.100143 | 0.920231 | NA |
| STAG3L3 | 9.777769 | 0.399885 | 0.649848 | 0.615351 | 0.538323 | NA |
| WBSCR16 | 1146.704 | -0.17496 | 0.147345 | -1.18741 | 0.235068 | 0.397849 |
| GTF2IRD2B | 45.98815 | 0.21132 | 0.347917 | 0.607386 | 0.543595 | NA |
| ABRA | 0.162376 | 0.424913 | 4.079006 | 0.104171 | 0.917034 | NA |
| ZCCHC12 | 0.685425 | -2.11121 | 2.613539 | -0.8078 | 0.419208 | NA |
| CNTNAP2 | 0.096202 | 0.408546 | 4.079639 | 0.100143 | 0.920231 | NA |
| LINGO2 | 133.1796 | 0.043742 | 0.285002 | 0.153478 | 0.878021 | 0.926887 |
| | 0.162376 | 0.424913 | 4.079006 | 0.104171 | 0.917034 | NA |
| KLF14 | 3.766757 | 0.490819 | 1.024048 | 0.479293 | 0.63173 | NA |
| LEP | 3.271578 | 0.742827 | 1.129443 | 0.657693 | 0.510735 | NA |
| PABPC5 | 132.7428 | 0.210674 | 0.27954 | 0.753646 | 0.451062 | 0.610395 |
| AMZ1 | 13.67145 | 0.181427 | 0.544366 | 0.33328 | 0.738923 | NA |
| SLC22A1 | 5.59921 | 0.079747 | 0.848007 | 0.094041 | 0.925077 | NA |
| ZDHHC14 | 438.5116 | -0.46634 | 0.175852 | -2.65186 | 0.008005 | 0.040626 |
| VCPIP1 | 561.8181 | 0.438107 | 0.237813 | 1.842233 | 0.065441 | 0.166907 |
| DOLK | 824.0749 | -0.25186 | 0.139006 | -1.81186 | 0.070009 | 0.174433 |
| PHYHD1 | 104.4894 | -0.36731 | 0.279016 | -1.31644 | 0.188026 | 0.342128 |
| CCNE2 | 48.85924 | 1.041969 | 0.396468 | 2.628129 | 0.008586 | 0.042551 |
| LSM1 | 664.8311 | 0.663844 | 0.189034 | 3.511773 | 4.45E-04 | 0.00635 |
| LPL | 65.40546 | -0.81073 | 0.441475 | -1.83641 | 0.066297 | 0.168118 |
| LONRF3 | 135.9995 | 0.517507 | 0.296101 | 1.747741 | 0.080509 | 0.19219 |
| SUGCT | 179.0459 | -0.0237 | 0.20113 | -0.11784 | 0.906192 | 0.945762 |
| TMEM70 | 937.3658 | 0.507984 | 0.175413 | 2.895928 | 0.00378 | 0.02453 |
| LINC00476 | 84.14814 | 0.250582 | 0.265606 | 0.943433 | 0.34546 | 0.513404 |
| RWDD4P2 | 2.2965 | 0.031519 | 1.626595 | 0.019377 | 0.98454 | NA |
| TTLL11 | 130.8845 | 0.053022 | 0.227992 | 0.232559 | 0.816104 | 0.889858 |
| TOMM5 | 312.8005 | 0.040069 | 0.265348 | 0.151004 | 0.879973 | 0.928286 |
| ZNF169 | 54.44484 | 0.020193 | 0.300652 | 0.067165 | 0.94645 | 0.970016 |
| MSRA | 725.0138 | -0.25718 | 0.21202 | -1.21299 | 0.225132 | 0.386187 |
| SWI5 | 472.6976 | 0.004644 | 0.197369 | 0.023528 | 0.981229 | 0.990173 |
| | 65.02766 | -0.66191 | 0.279607 | -2.3673 | 0.017919 | 0.070101 |
| ZDHHC21 | 139.9017 | 0.899274 | 0.251215 | 3.579698 | 3.44E-04 | 0.00538 |
| PLEKHF2 | 866.2322 | 0.244737 | 0.214618 | 1.140339 | 0.254145 | 0.418166 |
| KLHL38 | 55.00843 | -0.59623 | 0.327662 | -1.81965 | 0.068812 | 0.172582 |
| TPRN | 581.2024 | -0.44577 | 0.156104 | -2.85562 | 0.004295 | 0.026646 |
| KCNE1L | 1.237779 | 2.36266 | 1.939468 | 1.2182 | 0.223148 | NA |
| SSNA1 | 2350.842 | -0.07329 | 0.146755 | -0.49943 | 0.617475 | 0.742537 |
| UFSP1 | 65.35258 | -0.41578 | 0.277202 | -1.49991 | 0.133638 | 0.27128 |
| SMIM19 | 416.6661 | 0.074725 | 0.20023 | 0.373198 | 0.709001 | 0.813827 |
| ANAPC2 | 930.4751 | -0.3262 | 0.140433 | -2.32282 | 0.020189 | 0.076222 |
| SLC25A53 | 71.44251 | -0.04734 | 0.299894 | -0.15785 | 0.874576 | 0.925145 |
| | 3.790431 | -0.04002 | 1.113123 | -0.03595 | 0.971321 | NA |
| PRR18 | 0.258578 | 1.233151 | 4.047809 | 0.304646 | 0.760635 | NA |
| CDC26 | 106.3457 | 0.335297 | 0.281838 | 1.189681 | 0.234172 | 0.3967 |
| DMRTA1 | 0.585114 | 1.12675 | 2.73598 | 0.411827 | 0.680466 | NA |

| | | | | | | |
|---|---|---|---|---|---|---|
| GJC3 | 14.5004 | -0.23516 | 0.604551 | -0.38899 | 0.697286 | NA |
| RIMS2 | 29.04945 | 0.856067 | 0.406977 | 2.103479 | 0.035424 | NA |
| DNAJC30 | 438.1433 | -0.29461 | 0.154053 | -1.91237 | 0.055829 | 0.149437 |
| VPS37D | 158.7451 | -0.15931 | 0.206252 | -0.7724 | 0.439878 | 0.59988 |
| PRR15 | 0.424115 | 1.932531 | 3.45462 | 0.559405 | 0.575885 | NA |
| CNBD1 | 0.489058 | -0.58569 | 3.427543 | -0.17088 | 0.864321 | NA |
| KBTBD11 | 18.38145 | 0.524127 | 0.604823 | 0.86658 | 0.386172 | NA |
| RMDN1 | 1321.174 | 0.310765 | 0.190229 | 1.633636 | 0.102335 | 0.226383 |
| TTTY14 | 31.95297 | 0.326861 | 0.406177 | 0.804727 | 0.420977 | NA |
| C8orf59 | 266.812 | 0.57949 | 0.263877 | 2.196063 | 0.028087 | 0.094892 |
| FKBP9L | 61.26737 | -0.59936 | 0.28131 | -2.13062 | 0.033121 | 0.106006 |
| FAM91A1 | 2049.146 | 0.535457 | 0.187894 | 2.849782 | 0.004375 | 0.026971 |
| GRIN1 | 3.44057 | -0.08563 | 1.094018 | -0.07827 | 0.937611 | NA |
| TCEANC | 52.98596 | -0.14966 | 0.329982 | -0.45354 | 0.650159 | 0.768075 |
| C8G | 5.766238 | 0.402223 | 0.82988 | 0.484676 | 0.627906 | NA |
| DPP7 | 6183.987 | -0.15627 | 0.128695 | -1.2143 | 0.224634 | 0.385667 |
| FMR1NB | 0.163957 | 0.424913 | 4.079006 | 0.104171 | 0.917034 | NA |
| IFNW1 | 0.392857 | -1.546 | 4.075469 | -0.37934 | 0.704433 | NA |
| ACER2 | 18.44084 | -0.40395 | 0.494593 | -0.81674 | 0.414078 | NA |
| ZBTB34 | 236.1407 | 0.393595 | 0.23879 | 1.648288 | 0.099294 | 0.222267 |
| FAM9B | 0.826513 | -0.16238 | 2.470404 | -0.06573 | 0.947593 | NA |
| CLVS1 | 0.411855 | 0.250456 | 3.097618 | 0.080854 | 0.935558 | NA |
| RPS6KA3 | 1494.059 | 0.438776 | 0.206568 | 2.124119 | 0.03366 | 0.107125 |
| MAN1B1 | 3115.358 | -0.31305 | 0.129807 | -2.41161 | 0.015882 | 0.064533 |
| C8orf31 | 0.296509 | -0.51709 | 3.982155 | -0.12985 | 0.896684 | NA |
| SAMD9L | 93.1963 | -0.00694 | 0.542387 | -0.0128 | 0.989789 | 0.995162 |
| | 0.392857 | -1.546 | 4.075469 | -0.37934 | 0.704433 | NA |
| C8orf47 | 0.490493 | 0.776209 | 2.895261 | 0.268096 | 0.788625 | NA |
| ZBTB33 | 730.5777 | 0.721735 | 0.219465 | 3.288606 | 0.001007 | 0.010388 |
| SAMD12 | 5.098543 | 0.626474 | 0.936822 | 0.668722 | 0.503673 | NA |
| MBOAT4 | 1.005019 | -1.03066 | 2.033647 | -0.50681 | 0.612291 | NA |
| SRRM3 | 493.8098 | 0.160271 | 0.173497 | 0.923769 | 0.355607 | 0.523757 |
| THAP5 | 322.6184 | 1.095358 | 0.348343 | 3.144479 | 0.001664 | 0.014409 |
| FAM20C | 520.3034 | -0.25587 | 0.200684 | -1.27499 | 0.202313 | 0.358723 |
| SLC35G5 | 0.643567 | 1.129905 | 3.100382 | 0.364441 | 0.715529 | NA |
| | 0.489058 | -0.58569 | 3.427543 | -0.17088 | 0.864321 | NA |
| TMEM187 | 199.6446 | -0.43082 | 0.189341 | -2.27537 | 0.022884 | 0.08281 |
| MAMDC4 | 31.54767 | -0.02276 | 0.423531 | -0.05375 | 0.957138 | NA |
| LCN15 | 0.096202 | 0.408546 | 4.079639 | 0.100143 | 0.920231 | NA |
| ADAMTSL1 | 4.37404 | -0.80785 | 1.002487 | -0.80584 | 0.420334 | NA |
| PPP1R42 | 2.38367 | 0.617124 | 1.290909 | 0.478054 | 0.632612 | NA |
| | 4.616701 | -0.08772 | 1.007955 | -0.08702 | 0.930653 | NA |
| PLEC | 13845.48 | -0.61186 | 0.240441 | -2.54473 | 0.010936 | 0.049887 |
| GALNT11 | 1316.16 | 0.331452 | 0.148391 | 2.23363 | 0.025507 | 0.089306 |
| ZFAND2A | 518.1794 | 0.198809 | 0.169056 | 1.17599 | 0.239599 | 0.402786 |
| FAM220A | 585.0578 | -0.10354 | 0.150144 | -0.68964 | 0.490422 | 0.642588 |
| GLDC | 69.81381 | -0.37852 | 0.285251 | -1.32699 | 0.184512 | 0.337769 |

| | | | | | | |
|---|---|---|---|---|---|---|
| MCMDC2 | 20.86119 | 0.265726 | 0.462986 | 0.57394 | 0.566008 | NA |
| CA8 | 3691.21 | 0.300269 | 0.164094 | 1.829858 | 0.067271 | 0.169633 |
| ZNF713 | 38.02492 | 0.120685 | 0.403873 | 0.29882 | 0.765077 | NA |
| PARP10 | 2059.805 | -0.96604 | 0.209622 | -4.60849 | 4.06E-06 | 3.70E-04 |
| GRINA | 6667.16 | -0.50493 | 0.145507 | -3.47012 | 5.20E-04 | 0.006953 |
| GLULP4 | 0.544554 | -2.65036 | 3.220086 | -0.82307 | 0.410468 | NA |
| ZHX2 | 249.498 | -0.35208 | 0.187684 | -1.87591 | 0.060667 | 0.158485 |
| TRIM73 | 0.096202 | 0.408546 | 4.079639 | 0.100143 | 0.920231 | NA |
| OPLAH | 145.4744 | -0.67433 | 0.212171 | -3.17824 | 0.001482 | 0.013453 |
| TMEM139 | 1.349176 | -0.20983 | 1.827014 | -0.11485 | 0.908563 | NA |
| MSC | 2959.509 | -0.18426 | 0.185449 | -0.99356 | 0.320436 | 0.48865 |
| EXOSC4 | 818.0153 | 0.004436 | 0.151769 | 0.029228 | 0.976683 | 0.987343 |
| LINC00086 | 7.803317 | -0.36862 | 0.70683 | -0.52152 | 0.602006 | NA |
| RMI1 | 118.9368 | 0.943281 | 0.295727 | 3.189701 | 0.001424 | 0.013129 |
| | 0.096202 | 0.408546 | 4.079639 | 0.100143 | 0.920231 | NA |
| RRS1 | 538.2436 | -0.22435 | 0.156663 | -1.43205 | 0.152131 | 0.296435 |
| C9orf50 | 4.145233 | -0.34696 | 0.993058 | -0.34938 | 0.726804 | NA |
| C9orf106 | 0.192403 | 0.975392 | 4.060716 | 0.240202 | 0.810174 | NA |
| FAM133A | 0.163957 | 0.424913 | 4.079006 | 0.104171 | 0.917034 | NA |
| CYC1 | 4919.554 | -0.14453 | 0.151013 | -0.95704 | 0.338549 | 0.507304 |
| | 1.06487 | 0.25787 | 2.094579 | 0.123113 | 0.902018 | NA |
| | 0.593018 | -0.10389 | 3.151334 | -0.03297 | 0.973702 | NA |
| MAGED1 | 6151.527 | -0.50028 | 0.139168 | -3.59476 | 3.25E-04 | 0.005188 |
| ZCCHC5 | 0.096202 | 0.408546 | 4.079639 | 0.100143 | 0.920231 | NA |
| FAM156B | 14.89821 | 0.474312 | 0.58996 | 0.803972 | 0.421413 | NA |
| TMEM31 | 0.298089 | -0.51711 | 3.975232 | -0.13008 | 0.896501 | NA |
| EGR3 | 167.681 | -0.02227 | 0.276259 | -0.08062 | 0.935745 | 0.964049 |
| LINC00174 | 10.55709 | 0.501132 | 0.754107 | 0.664537 | 0.506347 | NA |
| | 0.314073 | -0.57308 | 3.898934 | -0.14698 | 0.883146 | NA |
| EEF1A1P27 | 0.856337 | 0.757304 | 2.453242 | 0.308695 | 0.757553 | NA |
| OR9A2 | 0.162376 | 0.424913 | 4.079006 | 0.104171 | 0.917034 | NA |
| SHARPIN | 1674.656 | -0.26507 | 0.148696 | -1.78264 | 0.074645 | 0.182427 |
| SLITRK4 | 1.037858 | 1.046985 | 2.132048 | 0.49107 | 0.623377 | NA |
| GCC1 | 1304.141 | -0.20765 | 0.151455 | -1.37102 | 0.17037 | 0.320797 |
| MAF1 | 3992.445 | -0.14522 | 0.152236 | -0.95394 | 0.340113 | 0.508454 |
| KIAA1875 | 1.484244 | 1.158505 | 1.742637 | 0.6648 | 0.506178 | NA |
| ATP8B5P | 3.517343 | 1.405365 | 1.115964 | 1.259329 | 0.207912 | NA |
| MROH1 | 762.8897 | -0.46147 | 0.176164 | -2.61953 | 0.008805 | 0.043419 |
| ABCA13 | 17.39966 | -0.09244 | 0.538774 | -0.17157 | 0.863777 | NA |
| TIGD5 | 377.4736 | -0.37018 | 0.191695 | -1.93107 | 0.053475 | 0.144935 |
| PUF60 | 4729.616 | -0.36245 | 0.125283 | -2.89308 | 0.003815 | 0.024686 |
| ANKRD18A | 3.151324 | 0.926551 | 1.210612 | 0.765357 | 0.444059 | NA |
| LYNX1 | 203.2761 | -0.18718 | 0.210074 | -0.89104 | 0.372906 | 0.539649 |
| MED14 | 1009.087 | -0.05321 | 0.144377 | -0.36856 | 0.712453 | 0.8164 |
| TDRP | 536.3283 | 0.239896 | 0.158287 | 1.515578 | 0.129626 | 0.266013 |
| ZNRF2 | 450.3747 | 0.174474 | 0.195331 | 0.893224 | 0.371737 | 0.538483 |
| GPR144 | 0.134133 | -1.48909 | 4.077584 | -0.36519 | 0.71497 | NA |

| | | | | | | |
|---|---|---|---|---|---|---|
| | 0.134133 | -1.48909 | 4.077584 | -0.36519 | 0.71497 | NA |
| MTURN | 1704.549 | 0.323845 | 0.148877 | 2.17525 | 0.029611 | 0.098146 |
| GAS1 | 407.7501 | -0.50921 | 0.229078 | -2.22286 | 0.026225 | 0.090906 |
| BHLHA15 | 0.298089 | -0.51711 | 3.975232 | -0.13008 | 0.896501 | NA |
| TSPYL5 | 464.7693 | -0.5004 | 0.154682 | -3.23504 | 0.001216 | 0.01178 |
| TMEM64 | 1079.214 | -0.2392 | 0.217428 | -1.10012 | 0.27128 | 0.436426 |
| MAP3K15 | 2.381527 | 1.059729 | 1.393844 | 0.760292 | 0.44708 | NA |
| BHLHE22 | 1.203009 | 3.444822 | 2.116698 | 1.627451 | 0.103641 | NA |
| SSR4 | 4488.896 | -0.04717 | 0.138807 | -0.33981 | 0.734002 | 0.831892 |
| SCRIB | 2317.008 | -0.49082 | 0.137242 | -3.57629 | 3.48E-04 | 0.005433 |
| FAM83H | 1075.92 | -0.4515 | 0.163565 | -2.76037 | 0.005774 | 0.032593 |
| ZNF572 | 41.38184 | 0.230425 | 0.338436 | 0.680851 | 0.495966 | NA |
| TCEAL8 | 2161.076 | 0.638553 | 0.189659 | 3.36684 | 7.60E-04 | 0.008794 |
| LSMEM1 | 5.161673 | 1.844917 | 0.988272 | 1.866811 | 0.061928 | NA |
| MAPK15 | 1.455651 | 1.673699 | 1.711992 | 0.977633 | 0.328256 | NA |
| EHMT1 | 1078.315 | -0.33955 | 0.138152 | -2.45783 | 0.013978 | 0.05912 |
| | 14.07034 | 0.275522 | 0.553166 | 0.498082 | 0.618426 | NA |
| ZNF707 | 382.2509 | -0.12674 | 0.170299 | -0.74424 | 0.456734 | 0.615154 |
| PJA1 | 627.0309 | -0.34314 | 0.136644 | -2.51118 | 0.012033 | 0.053547 |
| PENK | 1.373842 | 0.279944 | 1.886333 | 0.148406 | 0.882022 | NA |
| ZNF746 | 815.6411 | -0.54966 | 0.149973 | -3.66503 | 2.47E-04 | 0.004422 |
| | 127.4821 | -0.20767 | 0.212587 | -0.97688 | 0.32863 | 0.496926 |
| ZNF467 | 4.07945 | 0.450304 | 1.1741 | 0.383531 | 0.701326 | NA |
| ZBTB2 | 320.8762 | 0.298281 | 0.188329 | 1.58383 | 0.113232 | 0.241999 |
| FANCB | 95.85858 | 1.153935 | 0.300906 | 3.83487 | 1.26E-04 | 0.002961 |
| TNFSF15 | 1.602044 | 1.906086 | 1.769986 | 1.076893 | 0.281528 | NA |
| ZFP41 | 216.0316 | -0.43145 | 0.208093 | -2.07337 | 0.038138 | 0.116376 |
| ATG9B | 0.162376 | 0.424913 | 4.079006 | 0.104171 | 0.917034 | NA |
| PLAG1 | 91.57033 | 0.578067 | 0.287853 | 2.008203 | 0.044622 | 0.129054 |
| YIPF6 | 1256.844 | 0.376692 | 0.191933 | 1.962623 | 0.04969 | 0.137989 |
| BAI1 | 17.81821 | -0.021 | 0.476113 | -0.04411 | 0.964816 | NA |
| KCTD9P2 | 2.73364 | 0.23429 | 1.220763 | 0.191921 | 0.847804 | NA |
| TNRC18 | 1245.958 | -0.74322 | 0.219048 | -3.39296 | 6.91E-04 | 0.00825 |
| ZNF716 | 0.096202 | 0.408546 | 4.079639 | 0.100143 | 0.920231 | NA |
| ERCC6L2 | 411.2752 | 0.684553 | 0.300015 | 2.28173 | 0.022505 | 0.081968 |
| MRPL41 | 2989.487 | -0.11232 | 0.169928 | -0.66101 | 0.508609 | 0.657276 |
| CREB3L2 | 2121.563 | -0.42698 | 0.214341 | -1.99205 | 0.046365 | 0.13232 |
| TP53TG1 | 481.4521 | -0.20634 | 0.156108 | -1.32175 | 0.18625 | 0.340267 |
| LDOC1 | 462.3857 | -0.5547 | 0.177087 | -3.13233 | 0.001734 | 0.01475 |
| EXT1 | 894.3018 | -0.24917 | 0.135035 | -1.84521 | 0.065007 | 0.166214 |
| ATP6AP2 | 4011.753 | 0.317761 | 0.124376 | 2.55485 | 0.010623 | 0.048986 |
| AP1S2 | 5188.173 | 0.655935 | 0.21923 | 2.991991 | 0.002772 | 0.020121 |
| C8orf33 | 2680.211 | -0.05895 | 0.152797 | -0.3858 | 0.699646 | 0.806526 |
| | 394.3334 | -0.12612 | 0.218013 | -0.57852 | 0.562915 | 0.700983 |
| FBXL6 | 260.0368 | -0.22796 | 0.207724 | -1.09741 | 0.272462 | 0.437971 |
| | 0.260158 | 1.240347 | 4.047124 | 0.306476 | 0.759242 | NA |
| FAM87A | 0.163957 | 0.424913 | 4.079006 | 0.104171 | 0.917034 | NA |

| | | | | | | |
|---|---|---|---|---|---|---|
| FAM27A | 13.80312 | 0.02074 | 0.585204 | 0.03544 | 0.971729 | NA |
| CLN8 | 826.1331 | -0.06815 | 0.221046 | -0.30831 | 0.757849 | 0.849017 |
| XKRX | 1.115061 | 0.450621 | 2.115791 | 0.21298 | 0.831343 | NA |
| BGN | 10.18698 | 0.557137 | 0.693591 | 0.803264 | 0.421822 | NA |
| LHFPL1 | 19.57608 | 0.045276 | 0.507043 | 0.089295 | 0.928848 | NA |
| FAM104B | 276.0565 | 0.175569 | 0.185726 | 0.945315 | 0.344498 | 0.512554 |
| FAM156A | 14.57635 | 0.322833 | 0.535077 | 0.603339 | 0.546283 | NA |
| LINC01006 | 14.17554 | 0.28627 | 0.589601 | 0.485532 | 0.627299 | NA |
| SEPHS1P1 | 0.392857 | -1.546 | 4.075469 | -0.37934 | 0.704433 | NA |
| PAPPA | 21.20513 | 1.343563 | 0.614272 | 2.187245 | 0.028725 | NA |
| MAFA | 0.967088 | -0.05827 | 2.035183 | -0.02863 | 0.977158 | NA |
| MAGEB17 | 1.254112 | 0.747071 | 1.987113 | 0.375958 | 0.706948 | NA |
| GPIHBP1 | 0.151697 | -1.546 | 4.075469 | -0.37934 | 0.704433 | NA |
| RBM10 | 1358.998 | -0.53798 | 0.137281 | -3.91882 | 8.90E-05 | 0.002378 |
| | 0.230334 | -0.53318 | 4.076749 | -0.13079 | 0.895945 | NA |
| GLUD2 | 186.6973 | 0.084706 | 0.241885 | 0.350191 | 0.726195 | 0.826125 |
| TCEAL7 | 481.4565 | 0.530137 | 0.20623 | 2.57061 | 0.010152 | 0.047664 |
| CSMD1 | 11.99565 | 0.207913 | 0.769103 | 0.270331 | 0.786905 | NA |
| ANKRD20A | 0.315653 | -0.57305 | 3.892639 | -0.14721 | 0.882963 | NA |
| | 18.03365 | -0.34214 | 0.48195 | -0.7099 | 0.477765 | NA |
| CALN1 | 8.357508 | 0.059645 | 0.813048 | 0.07336 | 0.94152 | NA |
| | 0.681461 | 0.491544 | 3.066278 | 0.160307 | 0.87264 | NA |
| ZNF623 | 519.2821 | -0.23122 | 0.149261 | -1.54912 | 0.121353 | 0.254133 |
| BCOR | 990.5713 | -0.40392 | 0.232954 | -1.73389 | 0.082937 | 0.195915 |
| KIAA2026 | 666.038 | 0.345442 | 0.203794 | 1.695056 | 0.090065 | 0.207808 |
| OR7E38P | 99.16199 | -0.10869 | 0.238052 | -0.45658 | 0.647974 | 0.766251 |
| TREX2 | 9.675869 | -0.50499 | 0.688596 | -0.73337 | 0.463336 | NA |
| RP1L1 | 2.125238 | -0.16649 | 1.323752 | -0.12577 | 0.899914 | NA |
| TRMT12 | 440.309 | 0.116237 | 0.166851 | 0.696648 | 0.486023 | 0.639348 |
| EFHC2 | 14.3324 | 0.9704 | 0.569454 | 1.704089 | 0.088365 | NA |
| UPP1 | 3982.452 | -0.23862 | 0.148677 | -1.60495 | 0.108504 | 0.235331 |
| MACC1 | 0.460466 | 0.568655 | 3.014642 | 0.188631 | 0.850382 | NA |
| ZNF703 | 1273.644 | -0.69959 | 0.185764 | -3.766 | 1.66E-04 | 0.003487 |
| C9orf66 | 20.05975 | -0.09072 | 0.50953 | -0.17806 | 0.858679 | NA |
| RBM12B | 204.4296 | 0.398424 | 0.401139 | 0.993231 | 0.320598 | 0.488834 |
| PNMA3 | 9.566258 | 0.575412 | 0.794266 | 0.724457 | 0.468785 | NA |
| UTY | 284.451 | 0.382809 | 0.206895 | 1.850259 | 0.064276 | 0.165041 |
| SH2D1A | 0.096202 | 0.408546 | 4.079639 | 0.100143 | 0.920231 | NA |
| TRBV21OR9 | 0.096202 | 0.408546 | 4.079639 | 0.100143 | 0.920231 | NA |
| PRKX | 428.972 | -0.06579 | 0.229749 | -0.28638 | 0.774589 | 0.860453 |
| FAM120C | 231.1353 | -0.45156 | 0.192352 | -2.34758 | 0.018896 | 0.072701 |
| GPR173 | 215.3493 | -0.55615 | 0.193487 | -2.87434 | 0.004049 | 0.025585 |
| TSPYL2 | 426.4608 | 0.166997 | 0.185882 | 0.898402 | 0.368971 | 0.535982 |
| IRAK1 | 2589.361 | -0.08608 | 0.206973 | -0.41591 | 0.677476 | 0.789334 |
| CDR1 | 64.62606 | 1.170933 | 0.385533 | 3.037183 | 0.002388 | 0.018136 |
| SRPK3 | 4.917951 | -0.07986 | 0.882145 | -0.09053 | 0.927865 | NA |
| MAP7D2 | 46.67806 | -0.19929 | 0.457884 | -0.43525 | 0.663381 | NA |

| | | | | | | |
|---|---|---|---|---|---|---|
| KCND2 | 0.163957 | 0.424913 | 4.079006 | 0.104171 | 0.917034 | NA |
| | 0.572651 | 0.928649 | 2.780261 | 0.334015 | 0.738368 | NA |
| TOP1MT | 649.39 | -0.09565 | 0.170325 | -0.56155 | 0.574421 | 0.709683 |
| LRRC19 | 5.755195 | 1.592818 | 0.889668 | 1.790351 | 0.073398 | NA |
| WDR27 | 147.7968 | 0.342372 | 0.207919 | 1.64666 | 0.099628 | 0.222931 |
| FOXO4 | 846.237 | -0.15576 | 0.180095 | -0.8649 | 0.387095 | 0.552959 |
| PTP4A3 | 219.3484 | -0.66427 | 0.31831 | -2.08686 | 0.036901 | 0.113893 |
| BEX5 | 0.096202 | 0.408546 | 4.079639 | 0.100143 | 0.920231 | NA |
| | 0.871089 | -1.46164 | 2.262872 | -0.64592 | 0.518329 | NA |
| | 7.486644 | -0.38088 | 0.71703 | -0.5312 | 0.595282 | NA |
| MED12 | 1624.487 | -0.82204 | 0.177561 | -4.62961 | 3.66E-06 | 3.53E-04 |
| CDCA2 | 290.8315 | 0.154391 | 0.18736 | 0.824033 | 0.409921 | 0.573435 |
| AMER1 | 126.2281 | -0.51023 | 0.22511 | -2.2666 | 0.023415 | 0.084142 |
| LRRC26 | 0.268266 | -2.15396 | 4.038351 | -0.53338 | 0.593773 | NA |
| SMIM10 | 134.8853 | 0.660173 | 0.275498 | 2.396289 | 0.016562 | 0.066625 |
| TCTE3 | 4.689443 | 1.014293 | 0.945744 | 1.072481 | 0.283504 | NA |
| SATL1 | 48.05705 | -0.20062 | 0.40083 | -0.50051 | 0.616716 | 0.742224 |
| APOO | 470.7826 | 0.086657 | 0.160859 | 0.538713 | 0.590085 | 0.721887 |
| CYCSP45 | 0.478233 | -0.67301 | 2.937473 | -0.22911 | 0.818782 | NA |
| RBM33 | 799.0399 | -0.19401 | 0.153783 | -1.26158 | 0.207101 | 0.364221 |
| ARMCX2 | 738.3603 | -0.54765 | 0.198598 | -2.75757 | 0.005823 | 0.032734 |
| | 2.002337 | -0.85041 | 1.514479 | -0.56152 | 0.574445 | NA |
| IMMP2L | 339.2982 | 0.462666 | 0.195149 | 2.370834 | 0.017748 | 0.069688 |
| TCEAL2 | 0.258578 | 1.233151 | 4.047809 | 0.304646 | 0.760635 | NA |
| LCN12 | 0.420954 | 1.923376 | 3.462803 | 0.555439 | 0.578594 | NA |
| IFNE | 0.096202 | 0.408546 | 4.079639 | 0.100143 | 0.920231 | NA |
| DGAT1 | 1011.758 | -0.13625 | 0.145602 | -0.93579 | 0.349384 | 0.517484 |
| F8 | 70.78402 | 0.031278 | 0.284036 | 0.11012 | 0.912315 | 0.949227 |
| CA13 | 80.22155 | 0.192932 | 0.257091 | 0.750443 | 0.452988 | 0.611985 |
| | 5.097146 | 0.416648 | 0.874831 | 0.476261 | 0.633889 | NA |
| | 0.392857 | -1.546 | 4.075469 | -0.37934 | 0.704433 | NA |
| EFCAB10 | 5.312631 | 0.481753 | 1.082114 | 0.445196 | 0.656178 | NA |
| HSF1 | 2582.622 | -0.29257 | 0.151718 | -1.9284 | 0.053805 | 0.145597 |
| C6orf120 | 729.702 | 0.124847 | 0.20932 | 0.59644 | 0.550881 | 0.692009 |
| NRBP2 | 1137.985 | -0.11843 | 0.264402 | -0.44792 | 0.654208 | 0.771702 |
| WBP5 | 1217.185 | 0.900315 | 0.205053 | 4.390651 | 1.13E-05 | 6.60E-04 |
| WBSCR17 | 0.247898 | -0.5901 | 4.074633 | -0.14482 | 0.884851 | NA |
| CD24P4 | 0.462046 | 0.589069 | 3.442439 | 0.17112 | 0.86413 | NA |
| PARK2 | 83.1603 | 0.119092 | 0.296339 | 0.401877 | 0.687774 | 0.796869 |
| BRCC3 | 348.6124 | 0.847326 | 0.249225 | 3.399837 | 6.74E-04 | 0.008102 |
| OLFML2A | 1115.511 | 0.786429 | 0.811203 | 0.96946 | 0.332316 | 0.500648 |
| MORN5 | 1.574361 | 1.179246 | 1.789405 | 0.659016 | 0.509886 | NA |
| MYBL1 | 186.422 | 1.079633 | 0.306246 | 3.525382 | 4.23E-04 | 0.006131 |
| YTHDF3 | 2581.812 | 0.177872 | 0.150168 | 1.184488 | 0.23622 | 0.399143 |
| ZNF696 | 221.4413 | -0.20553 | 0.189865 | -1.08253 | 0.279019 | 0.445311 |
| CXorf38 | 434.6817 | -0.06782 | 0.192201 | -0.35286 | 0.724195 | 0.824799 |
| SLC52A2 | 1264.789 | -0.13982 | 0.15246 | -0.91709 | 0.359095 | 0.526345 |

| | | | | | | |
|---|---|---|---|---|---|---|
| IKZF1 | 0.096202 | 0.408546 | 4.079639 | 0.100143 | 0.920231 | NA |
| BCAP31 | 7899.68 | -0.22739 | 0.150022 | -1.51573 | 0.129587 | 0.266013 |
| POMK | 58.27194 | 0.012832 | 0.30836 | 0.041612 | 0.966808 | 0.981629 |
| KLHL34 | 82.55749 | -0.17617 | 0.358885 | -0.49087 | 0.623521 | 0.747708 |
| PTCH1 | 1232.161 | -0.03222 | 0.148485 | -0.21698 | 0.828227 | 0.896434 |
| NKAIN3 | 17.89544 | -0.34713 | 0.52018 | -0.66734 | 0.504558 | NA |
| C7orf61 | 1.503619 | -1.38768 | 1.61906 | -0.85709 | 0.391395 | NA |
| BICD2 | 1779.731 | -0.2113 | 0.16609 | -1.27218 | 0.203308 | 0.359796 |
| TMLHE | 351.5711 | -0.24069 | 0.16185 | -1.48713 | 0.136979 | 0.276172 |
| SLITRK2 | 646.6182 | -0.28933 | 0.14744 | -1.96239 | 0.049717 | 0.138006 |
| TAL2 | 0.326536 | 0.037056 | 3.842554 | 0.009644 | 0.992306 | NA |
| GSAP | 94.65394 | 0.19984 | 0.27841 | 0.717792 | 0.472885 | 0.628623 |
| ANKRD46 | 217.9706 | 0.358543 | 0.213428 | 1.679925 | 0.092972 | 0.212735 |
| ZBTB6 | 285.5697 | 0.574354 | 0.21701 | 2.64667 | 0.008129 | 0.041007 |
| SAPCD2 | 184.5613 | -0.3506 | 0.282118 | -1.24276 | 0.213957 | 0.37274 |
| NAP1L3 | 12.44878 | -0.03119 | 0.60663 | -0.05142 | 0.958988 | NA |
| CA5BP1 | 284.2147 | -0.19142 | 0.195551 | -0.97889 | 0.327632 | 0.495835 |
| THBS2 | 1050.512 | -0.23486 | 0.306233 | -0.76693 | 0.443121 | 0.603321 |
| RXRA | 1892.759 | -0.32212 | 0.153347 | -2.10058 | 0.035678 | 0.111306 |
| C9orf47 | 1.721948 | 2.913894 | 1.832123 | 1.590447 | 0.111734 | NA |
| ZNF75D | 219.873 | 0.481374 | 0.204175 | 2.357654 | 0.018391 | 0.071413 |
| NKRF | 417.769 | 0.003168 | 0.152872 | 0.020726 | 0.983464 | 0.991324 |
| AKAP14 | 0.392857 | -1.546 | 4.075469 | -0.37934 | 0.704433 | NA |
| PCLO | 0.653072 | 0.258115 | 3.011926 | 0.085698 | 0.931707 | NA |
| INSIG1 | 2419.751 | 0.08889 | 0.156676 | 0.567351 | 0.570476 | 0.706465 |
| FAM86B1 | 6.65468 | -0.08699 | 0.760863 | -0.11433 | 0.908974 | NA |
| XKR5 | 9.31477 | -0.95619 | 0.668275 | -1.43084 | 0.152477 | NA |
| UBE2H | 2102.652 | -0.01351 | 0.190767 | -0.0708 | 0.943558 | 0.968771 |
| KIF24 | 279.0157 | -0.37414 | 0.266201 | -1.40547 | 0.15988 | 0.306994 |
| SPDYE9P | 0.134133 | -1.48909 | 4.077584 | -0.36519 | 0.71497 | NA |
| EEF1GP1 | 0.151697 | -1.546 | 4.075469 | -0.37934 | 0.704433 | NA |
| DTX2P1 | 7.454188 | -0.02386 | 0.809774 | -0.02946 | 0.976497 | NA |
| SPIN4 | 32.48193 | -0.50079 | 0.384249 | -1.30329 | 0.192475 | NA |
| SPIN2B | 83.59535 | -0.15497 | 0.261195 | -0.59331 | 0.552971 | 0.693627 |
| CXCR3 | 0.7777 | 0.426209 | 2.574008 | 0.165582 | 0.868486 | NA |
| ERCC6L | 116.3534 | 0.492218 | 0.277268 | 1.775243 | 0.075858 | 0.184319 |
| P2RY4 | 0.162376 | 0.424913 | 4.079006 | 0.104171 | 0.917034 | NA |
| ZNF395 | 705.891 | -0.33776 | 0.165024 | -2.04673 | 0.040685 | 0.121674 |
| CHCHD2P9 | 1.267763 | -2.07106 | 1.940816 | -1.06711 | 0.285923 | NA |
| GPR141 | 0.163957 | 0.424913 | 4.079006 | 0.104171 | 0.917034 | NA |
| | 0.604258 | 1.087325 | 2.707696 | 0.401568 | 0.688002 | NA |
| | 5.707562 | 0.493024 | 0.865218 | 0.569827 | 0.568795 | NA |
| GCNT1 | 7.859686 | 0.375818 | 0.726004 | 0.517653 | 0.604701 | NA |
| | 0.326333 | 1.533889 | 3.826878 | 0.40082 | 0.688553 | NA |
| FNBP1 | 1128.653 | -0.39312 | 0.205329 | -1.91457 | 0.055547 | 0.148951 |
| MAGED4B | 0.717609 | 0.516981 | 3.063469 | 0.168757 | 0.865988 | NA |
| RSBN1L | 313.5441 | 0.321073 | 0.245385 | 1.308443 | 0.190723 | 0.345036 |

| | | | | | | |
|---|---|---|---|---|---|---|
| WDR86 | 1.869436 | 0.435321 | 1.4412 | 0.302055 | 0.76261 | NA |
| TAF9B | 279.7049 | 0.435059 | 0.213682 | 2.036015 | 0.041749 | 0.124043 |
| MAGI2 | 289.8623 | -0.37368 | 0.206177 | -1.81243 | 0.06992 | 0.174248 |
| LHFPL3 | 36.09416 | 0.242089 | 0.352315 | 0.687139 | 0.491995 | NA |
| AGMO | 92.75162 | 0.260052 | 0.392342 | 0.662821 | 0.507445 | 0.656489 |
| MAGEH1 | 602.0836 | 0.081009 | 0.162007 | 0.500033 | 0.617052 | 0.742353 |
| EXD3 | 242.4585 | -0.14907 | 0.175796 | -0.84799 | 0.396444 | 0.561108 |
| CXorf67 | 0.151697 | -1.546 | 4.075469 | -0.37934 | 0.704433 | NA |
| TMEM203 | 1016.336 | -0.17026 | 0.149562 | -1.13838 | 0.254963 | 0.419279 |
| TCEA1 | 585.2704 | 0.560523 | 0.177725 | 3.153869 | 0.001611 | 0.01418 |
| SECISBP2 | 940.8195 | 0.378205 | 0.149978 | 2.521741 | 0.011678 | 0.0523 |
| C9orf153 | 0.46735 | -1.30193 | 3.399671 | -0.38296 | 0.701751 | NA |
| SEMA4D | 31.28257 | 0.010454 | 0.396729 | 0.02635 | 0.978978 | NA |
| FAM205CP | 0.640406 | 1.11402 | 2.710977 | 0.410929 | 0.681124 | NA |
| CARD9 | 11.39093 | -0.19711 | 0.601506 | -0.3277 | 0.74314 | NA |
| SOWAHD | 5.151796 | -0.47062 | 0.892288 | -0.52743 | 0.597892 | NA |
| EIF4EBP1 | 2032.353 | -0.31082 | 0.135781 | -2.28912 | 0.022072 | 0.081 |
| FAM122A | 452.189 | 0.030077 | 0.152345 | 0.197429 | 0.843492 | 0.906882 |
| LCN10 | 0.163957 | 0.424913 | 4.079006 | 0.104171 | 0.917034 | NA |
| PMS2CL | 115.0541 | 0.090681 | 0.219518 | 0.41309 | 0.679541 | 0.790866 |
| CYHR1 | 1310.766 | -0.16337 | 0.132111 | -1.23658 | 0.216243 | 0.375393 |
| COL14A1 | 188.1979 | 0.038905 | 0.452508 | 0.085977 | 0.931485 | 0.961326 |
| ANKRD19P | 2.544894 | 2.669624 | 1.521895 | 1.754144 | 0.079406 | NA |
| UBQLN2 | 992.5378 | -0.3583 | 0.211602 | -1.69325 | 0.090408 | 0.208467 |
| RNF133 | 0.258578 | 1.233151 | 4.047809 | 0.304646 | 0.760635 | NA |
| TMEM215 | 225.3702 | 0.420065 | 0.3818 | 1.10022 | 0.271236 | 0.436415 |
| NUTM2G | 31.34266 | 0.258138 | 0.431688 | 0.597973 | 0.549858 | NA |
| COL4A5 | 23.5433 | -0.8406 | 0.449262 | -1.87107 | 0.061335 | NA |
| NHS | 11.51554 | 1.137973 | 0.715426 | 1.590622 | 0.111695 | NA |
| FAM166A | 0.424115 | 1.932531 | 3.45462 | 0.559405 | 0.575885 | NA |
| HEPACAM2 | 0.192403 | 0.975392 | 4.060716 | 0.240202 | 0.810174 | NA |
| LINC00265 | 86.08479 | -0.55191 | 0.343298 | -1.60766 | 0.10791 | 0.234643 |
| LAMTOR4 | 2486.463 | -0.13669 | 0.164386 | -0.83152 | 0.405678 | 0.56959 |
| PRKAR1B | 409.1166 | -0.12487 | 0.168858 | -0.73951 | 0.459595 | 0.617954 |
| TUBB4B | 7277.078 | -0.40101 | 0.204475 | -1.96116 | 0.04986 | 0.138259 |
| CENPP | 146.976 | 0.13811 | 0.213017 | 0.648351 | 0.516758 | 0.664138 |
| FAM92A1 | 448.6499 | 0.664338 | 0.179974 | 3.691299 | 2.23E-04 | 0.004159 |
| FOCAD | 1560.829 | -0.33588 | 0.14015 | -2.39658 | 0.016549 | 0.066618 |
| | 2.418944 | 0.14189 | 1.407484 | 0.100811 | 0.919701 | NA |
| ZP3 | 5.67285 | -1.0361 | 0.836173 | -1.23909 | 0.215311 | NA |
| CHM | 655.1651 | 0.999471 | 0.249368 | 4.008017 | 6.12E-05 | 0.001863 |
| SRP72P2 | 0.394291 | 0.298745 | 3.141711 | 0.09509 | 0.924243 | NA |
| WASF4P | 4.823265 | -0.83338 | 0.930864 | -0.89527 | 0.370642 | NA |
| IER5L | 307.9922 | 0.199912 | 0.233075 | 0.857716 | 0.391049 | 0.556329 |
| | 2.433936 | -0.96658 | 1.339859 | -0.7214 | 0.470662 | NA |
| C9orf171 | 0.162376 | 0.424913 | 4.079006 | 0.104171 | 0.917034 | NA |
| NDOR1 | 782.9327 | -0.37882 | 0.162469 | -2.33165 | 0.019719 | 0.075022 |

| | | | | | | |
|---|---|---|---|---|---|---|
| PTAR1 | 1096.488 | 0.844587 | 0.283381 | 2.98039 | 0.002879 | 0.020562 |
| | 2.071526 | -0.13194 | 1.440511 | -0.09159 | 0.927022 | NA |
| ZDHHC9 | 1252.339 | -0.31354 | 0.133713 | -2.34487 | 0.019034 | 0.073142 |
| ZBED6CL | 27.13592 | 0.024856 | 0.48959 | 0.050769 | 0.95951 | NA |
| QRFP | 1.823628 | -0.05328 | 1.545351 | -0.03448 | 0.972496 | NA |
| FAM221A | 34.16125 | 1.179904 | 0.390768 | 3.019445 | 0.002532 | NA |
| NOXA1 | 14.52472 | -0.03573 | 0.571162 | -0.06255 | 0.950126 | NA |
| FZD9 | 192.9581 | -0.10308 | 0.273761 | -0.37652 | 0.706532 | 0.811936 |
| ZNF322P1 | 9.137335 | 0.611446 | 0.674634 | 0.906337 | 0.364758 | NA |
| ENTPD8 | 1.083511 | 0.500984 | 1.995536 | 0.251053 | 0.801774 | NA |
| RPSAP47 | 0.298089 | -0.51711 | 3.975232 | -0.13008 | 0.896501 | NA |
| RPL10AP2 | 4.377542 | 0.223368 | 0.950927 | 0.234895 | 0.81429 | NA |
| KLRG2 | 0.096202 | 0.408546 | 4.079639 | 0.100143 | 0.920231 | NA |
| TRMT2B | 552.9102 | -0.15351 | 0.172359 | -0.89065 | 0.373116 | 0.539755 |
| PTPLAD2 | 583.3701 | 0.096281 | 0.167593 | 0.574494 | 0.565634 | 0.702867 |
| NYX | 2.209623 | -0.99081 | 1.504018 | -0.65878 | 0.510038 | NA |
| FAM120AC | 632.1572 | 0.129669 | 0.159269 | 0.814149 | 0.41556 | 0.578766 |
| C9orf152 | 0.806146 | 0.553068 | 2.206737 | 0.250627 | 0.802102 | NA |
| NELFB | 2536.522 | -0.25207 | 0.137195 | -1.83734 | 0.066159 | 0.16784 |
| | 0.247898 | -0.5901 | 4.074633 | -0.14482 | 0.884851 | NA |
| NDUFA4 | 6911.048 | 0.348922 | 0.157282 | 2.21844 | 0.026525 | 0.091357 |
| RELN | 9.554546 | 0.378721 | 0.71556 | 0.529265 | 0.596621 | NA |
| TMEM120/ | 1893.581 | -0.11687 | 0.150579 | -0.77616 | 0.437653 | 0.597921 |
| IL1RAPL2 | 0.260158 | 1.240347 | 4.047124 | 0.306476 | 0.759242 | NA |
| CLDN4 | 0.728731 | -0.58211 | 2.613299 | -0.22275 | 0.823732 | NA |
| DPY19L2P1 | 2.196963 | 2.030191 | 1.525428 | 1.330899 | 0.183222 | NA |
| MAOA | 429.3267 | -0.28853 | 0.173961 | -1.65859 | 0.097198 | 0.218901 |
| NUGGC | 0.163957 | 0.424913 | 4.079006 | 0.104171 | 0.917034 | NA |
| FAM180A | 0.943054 | 1.955489 | 2.35722 | 0.829574 | 0.40678 | NA |
| GSPT2 | 319.0836 | 0.101444 | 0.178342 | 0.568816 | 0.569481 | 0.705805 |
| C8orf76 | 120.6991 | 0.073734 | 0.255804 | 0.288246 | 0.773159 | 0.859428 |
| ZFP92 | 6.045368 | 0.090142 | 0.903763 | 0.099741 | 0.92055 | NA |
| PAX5 | 0.163957 | 0.424913 | 4.079006 | 0.104171 | 0.917034 | NA |
| ADAM5 | 0.096202 | 0.408546 | 4.079639 | 0.100143 | 0.920231 | NA |
| TDRD7 | 3765.56 | 0.165029 | 0.170041 | 0.970528 | 0.331783 | 0.500291 |
| ZNF250 | 171.382 | 0.017971 | 0.194854 | 0.092227 | 0.926518 | 0.958156 |
| ZNF79 | 209.5687 | -0.24038 | 0.232175 | -1.03533 | 0.300516 | 0.467351 |
| C8orf86 | 0.134133 | -1.48909 | 4.077584 | -0.36519 | 0.71497 | NA |
| HRCT1 | 2.475158 | -0.16612 | 1.645936 | -0.10093 | 0.919608 | NA |
| RNF216P1 | 228.5958 | -0.10514 | 0.183694 | -0.57238 | 0.567062 | 0.704045 |
| EEF1A1P5 | 77.78796 | 0.114907 | 0.330395 | 0.347785 | 0.728001 | 0.827474 |
| ZNF107 | 58.07316 | 1.092482 | 0.544054 | 2.00804 | 0.044639 | 0.129072 |
| PPIA | 14806.83 | 0.024815 | 0.14153 | 0.175331 | 0.860819 | 0.917387 |
| GTF2IRD2 | 48.66262 | -0.097 | 0.363772 | -0.26664 | 0.789745 | 0.872547 |
| | 118.749 | 0.81743 | 0.2555 | 3.199334 | 0.001377 | 0.012785 |
| IARS | 5215.1 | 0.14069 | 0.143 | 0.983845 | 0.325192 | 0.493049 |
| HIATL2 | 122.8778 | 0.422597 | 0.293708 | 1.438834 | 0.150198 | 0.293827 |

| | | | | | | |
|---|---|---|---|---|---|---|
| POM121 | 833.7208 | -0.56565 | 0.19427 | -2.91169 | 0.003595 | 0.023861 |
| GIMAP5 | 3.990806 | 0.409521 | 1.061586 | 0.385763 | 0.699672 | NA |
| STK31 | 0.260158 | 1.240347 | 4.047124 | 0.306476 | 0.759242 | NA |
| NLGN3 | 28.41319 | -0.45056 | 0.455921 | -0.98825 | 0.32303 | NA |
| NTNG2 | 31.45137 | -0.35449 | 0.390924 | -0.90681 | 0.364509 | NA |
| WDR5 | 1128.499 | -0.25407 | 0.147762 | -1.71943 | 0.085536 | 0.199887 |
| C9orf163 | 6.112948 | -0.74708 | 0.80296 | -0.93041 | 0.352158 | NA |
| TRRAP | 1719.715 | -0.71125 | 0.179306 | -3.96666 | 7.29E-05 | 0.002092 |
| NUDT11 | 250.5169 | -0.15333 | 0.17822 | -0.86035 | 0.389595 | 0.555087 |
| ZNF34 | 91.45041 | -0.10833 | 0.252032 | -0.42982 | 0.667329 | 0.781814 |
| ZNF658 | 25.48304 | 0.655266 | 0.465631 | 1.407265 | 0.159349 | NA |
| EPHB4 | 1674.534 | -0.52343 | 0.171077 | -3.05961 | 0.002216 | 0.017291 |
| PPP1R26 | 1041.198 | -0.20335 | 0.170301 | -1.19407 | 0.232449 | 0.394718 |
| ARMCX4 | 882.9063 | -0.09114 | 0.240908 | -0.3783 | 0.705207 | 0.810853 |
| ZNF777 | 695.9316 | -0.41148 | 0.152232 | -2.703 | 0.006872 | 0.036662 |
| ZNF775 | 372.6143 | -0.23234 | 0.183613 | -1.26536 | 0.205742 | 0.362805 |
| TRAPPC2 | 128.2672 | 0.045993 | 0.264453 | 0.173918 | 0.86193 | 0.91824 |
| TCEAL3 | 845.96 | 0.344179 | 0.170773 | 2.01542 | 0.043861 | 0.127724 |
| TPK1 | 50.31048 | -0.14093 | 0.364193 | -0.38696 | 0.698786 | 0.806239 |
| XRCC2 | 59.66672 | 0.797108 | 0.445814 | 1.787982 | 0.073779 | 0.181081 |
| ZNF782 | 55.08433 | 0.548035 | 0.32772 | 1.67227 | 0.094471 | 0.21488 |
| WNK3 | 84.76561 | 0.216059 | 0.289786 | 0.74558 | 0.455921 | 0.614797 |
| ACN9 | 414.5155 | 0.50283 | 0.231658 | 2.170568 | 0.029964 | 0.09892 |
| RABL6 | 7472.703 | -0.32231 | 0.133256 | -2.41873 | 0.015575 | 0.063581 |
| ZKSCAN5 | 617.6723 | 0.124464 | 0.14917 | 0.834381 | 0.404066 | 0.568856 |
| TLR7 | 0.419962 | -2.64111 | 3.429364 | -0.77014 | 0.441214 | NA |
| TOMM7 | 6202.385 | 0.117733 | 0.167642 | 0.702286 | 0.482501 | 0.636338 |
| VKORC1L1 | 1165.018 | 0.026697 | 0.180803 | 0.14766 | 0.882611 | 0.930406 |
| DAPK1 | 2579.518 | -0.34801 | 0.129379 | -2.68986 | 0.007148 | 0.037647 |
| COL27A1 | 1101.825 | 0.094412 | 0.242248 | 0.389734 | 0.696733 | 0.804576 |
| CXorf24 | 24.8752 | -0.23258 | 0.421242 | -0.55212 | 0.580868 | NA |
| ANKRD20A | 0.096202 | 0.408546 | 4.079639 | 0.100143 | 0.920231 | NA |
| TLE1 | 2632.931 | -0.43409 | 0.14774 | -2.93822 | 0.003301 | 0.022439 |
| MVB12B | 116.1606 | -0.31856 | 0.243723 | -1.30705 | 0.191196 | 0.345629 |
| CBWD3 | 2.774144 | 0.998761 | 1.334142 | 0.748617 | 0.454088 | NA |
| ZNF252P | 468.9558 | 0.494009 | 0.178333 | 2.770142 | 0.005603 | 0.03203 |
| FLNA | 60315.68 | -0.585 | 0.162046 | -3.61007 | 3.06E-04 | 0.005015 |
| RPS26P11 | 0.260158 | 1.240347 | 4.047124 | 0.306476 | 0.759242 | NA |
| FAM3C | 482.6963 | 0.407002 | 0.196552 | 2.070713 | 0.038386 | 0.116952 |
| LINC00087 | 72.6446 | -0.20661 | 0.265382 | -0.77854 | 0.43625 | 0.597062 |
| LAGE3 | 1147.337 | -0.1972 | 0.148639 | -1.3267 | 0.184607 | 0.337891 |
| | 0.402398 | -2.58771 | 4.018369 | -0.64397 | 0.519595 | NA |
| | 1.691289 | 0.711275 | 1.584267 | 0.448962 | 0.653459 | NA |
| WDR45 | 1021.485 | 0.124789 | 0.149676 | 0.833726 | 0.404436 | 0.569031 |
| ZNF138 | 33.77647 | 0.71345 | 0.380039 | 1.877309 | 0.060476 | NA |
| CXorf40B | 565.988 | -0.09017 | 0.14652 | -0.61542 | 0.538277 | 0.681093 |
| ZNF398 | 589.5246 | -0.33366 | 0.184471 | -1.80874 | 0.070491 | 0.175377 |

| | | | | | | |
|---|---|---|---|---|---|---|
| ZSCAN25 | 528.2087 | -0.02344 | 0.156693 | -0.14959 | 0.881087 | 0.929216 |
| ARRDC1 | 519.3386 | 0.08366 | 0.182622 | 0.458108 | 0.646875 | 0.765569 |
| IGF2R | 9552.918 | -0.57144 | 0.190744 | -2.99585 | 0.002737 | 0.019954 |
| GAL3ST4 | 372.6194 | -0.60971 | 0.21411 | -2.84766 | 0.004404 | 0.027123 |
| ADAM32 | 1.731433 | 2.063348 | 1.755809 | 1.175155 | 0.239933 | NA |
| | 0.558248 | 0.982076 | 2.804738 | 0.350149 | 0.726227 | NA |
| ABCB8 | 1395.01 | -0.42089 | 0.127891 | -3.291 | 9.98E-04 | 0.010333 |
| SND1 | 13014.2 | -0.36988 | 0.142007 | -2.60468 | 0.009196 | 0.044588 |
| | 68.58318 | -0.15215 | 0.313788 | -0.4849 | 0.62775 | 0.750253 |
| PIWIL2 | 0.134133 | -1.48909 | 4.077584 | -0.36519 | 0.71497 | NA |
| C9orf169 | 20.26633 | -0.13558 | 0.484281 | -0.27996 | 0.779507 | NA |
| ENTPD4 | 2014.994 | 0.01536 | 0.16561 | 0.092751 | 0.926101 | 0.958106 |
| EIF4BP6 | 8.815607 | -0.77362 | 0.699586 | -1.10582 | 0.268805 | NA |
| GTF2E2 | 984.083 | 0.278935 | 0.139947 | 1.993149 | 0.046245 | 0.132137 |
| RAD54B | 53.55463 | 0.743834 | 0.318044 | 2.338774 | 0.019347 | 0.073916 |
| | 0.260158 | 1.240347 | 4.047124 | 0.306476 | 0.759242 | NA |
| ZNF655 | 728.3326 | 0.416331 | 0.197672 | 2.106169 | 0.03519 | 0.110334 |
| UAP1L1 | 2367.311 | -0.27827 | 0.157874 | -1.76263 | 0.077963 | 0.188086 |
| ZNF786 | 138.938 | -0.33996 | 0.237771 | -1.4298 | 0.152776 | 0.297301 |
| ZNF517 | 218.9639 | -0.37905 | 0.180006 | -2.10575 | 0.035226 | 0.110418 |
| FABP12 | 0.151697 | -1.546 | 4.075469 | -0.37934 | 0.704433 | NA |
| GSTK1 | 2954.71 | -0.10409 | 0.129712 | -0.80246 | 0.422286 | 0.584768 |
| PDGFA | 120.6919 | 0.209868 | 0.378249 | 0.554841 | 0.579004 | 0.713515 |
| SLC28A3 | 1.579307 | 0.141238 | 1.545188 | 0.091405 | 0.927171 | NA |
| SSPO | 0.364467 | -1.20166 | 3.685431 | -0.32606 | 0.744381 | NA |
| COL4A6 | 2.481598 | 0.296742 | 1.233098 | 0.240647 | 0.809829 | NA |
| HOXA4 | 4.740898 | -0.62462 | 0.968224 | -0.64512 | 0.518847 | NA |
| TOPORS | 455.2293 | 0.826701 | 0.274614 | 3.010406 | 0.002609 | 0.019343 |
| GPX1P1 | 6.399117 | -0.03017 | 0.781148 | -0.03862 | 0.969191 | NA |
| CXorf40A | 257.1238 | -0.16304 | 0.168326 | -0.96857 | 0.332758 | 0.50106 |
| PDCD1LG2 | 92.96481 | 0.303025 | 0.31573 | 0.959758 | 0.337177 | 0.505979 |
| SPTAN1 | 11723.07 | -0.47928 | 0.147644 | -3.24619 | 0.00117 | 0.011469 |
| PHF2 | 1209.214 | -0.31417 | 0.154303 | -2.03604 | 0.041746 | 0.124043 |
| C9orf173 | 1.293864 | -0.45062 | 1.884934 | -0.23906 | 0.811057 | NA |
| ZNF81 | 165.2439 | 0.674322 | 0.277037 | 2.434049 | 0.014931 | 0.061804 |
| CCDC180 | 8.564079 | 1.022293 | 0.798734 | 1.279891 | 0.200583 | NA |
| GPAA1 | 4874.737 | -0.40257 | 0.124972 | -3.22125 | 0.001276 | 0.012122 |
| KIF13B | 2846.921 | -0.28387 | 0.169966 | -1.67015 | 0.094889 | 0.215459 |
| IFNA1 | 0.268266 | -2.15396 | 4.038351 | -0.53338 | 0.593773 | NA |
| F8A1 | 23.18235 | -0.18084 | 0.440968 | -0.4101 | 0.681734 | NA |
| RPL12 | 19935.27 | 0.198764 | 0.139214 | 1.427765 | 0.153359 | 0.298135 |
| VPS13A | 493.721 | 0.90177 | 0.359441 | 2.50881 | 0.012114 | 0.053766 |
| KEL | 1.10954 | -0.31713 | 1.971981 | -0.16082 | 0.872235 | NA |
| NOL8 | 464.9935 | 0.853546 | 0.592654 | 1.440211 | 0.149808 | 0.293243 |
| DLGAP2 | 2.795954 | 2.205551 | 1.374407 | 1.604729 | 0.108554 | NA |
| RPS4X | 20721.19 | 0.083221 | 0.163644 | 0.508551 | 0.611067 | 0.737999 |
| ZNF273 | 28.57082 | 0.50191 | 0.480385 | 1.044809 | 0.296111 | NA |

| | | | | | | |
|---|---|---|---|---|---|---|
| MAK16 | 320.2722 | 0.582517 | 0.17982 | 3.239451 | 0.001198 | 0.011696 |
| AKR1B10 | 11.05702 | 0.72358 | 0.692826 | 1.04439 | 0.296305 | NA |
| NUP62CL | 0.610582 | -0.18034 | 2.753139 | -0.0655 | 0.947774 | NA |
| TOR4A | 933.8831 | -0.21647 | 0.188183 | -1.15032 | 0.250012 | 0.41372 |
| LPAR1 | 38.14314 | 0.181089 | 0.435037 | 0.41626 | 0.677219 | NA |
| HMGN5 | 56.15696 | 0.728946 | 1.59533 | 0.456925 | 0.647725 | 0.766234 |
| ZNF251 | 182.3513 | 0.588936 | 0.230863 | 2.551017 | 0.010741 | 0.049294 |
| ZXDA | 103.0771 | -0.05722 | 0.253038 | -0.22611 | 0.821113 | 0.892334 |
| MLLT4-AS1 | 9.007245 | 0.515732 | 0.784907 | 0.657062 | 0.511141 | NA |
| CARD11 | 0.162376 | 0.424913 | 4.079006 | 0.104171 | 0.917034 | NA |
| DCAF12L2 | 0.192403 | 0.975392 | 4.060716 | 0.240202 | 0.810174 | NA |
| ASPH | 3475.366 | 0.645198 | 0.19183 | 3.363378 | 7.70E-04 | 0.008845 |
| | 0.096202 | 0.408546 | 4.079639 | 0.100143 | 0.920231 | NA |
| FAM115A | 383.7968 | 0.52906 | 0.389812 | 1.357218 | 0.174712 | 0.326741 |
| NRARP | 1.676529 | 0.385402 | 1.772898 | 0.217385 | 0.827908 | NA |
| ZXDB | 168.9773 | -0.47862 | 0.201905 | -2.37054 | 0.017762 | 0.069688 |
| TPM2 | 159.8888 | 0.670704 | 0.217924 | 3.077704 | 0.002086 | 0.016597 |
| MAFK | 599.5634 | -0.02112 | 0.235945 | -0.0895 | 0.928688 | 0.959358 |
| ZNF789 | 207.0207 | -0.20518 | 0.222363 | -0.92273 | 0.356148 | 0.523903 |
| SLC34A3 | 6.42176 | 0.329272 | 0.803253 | 0.409922 | 0.681863 | NA |
| ARC | 1.619811 | 1.252241 | 1.571886 | 0.796649 | 0.425655 | NA |
| | 1.400302 | 2.191364 | 1.96538 | 1.114982 | 0.264858 | NA |
| KLHL9 | 1299.533 | 0.233935 | 0.273217 | 0.856224 | 0.391874 | 0.557055 |
| TUSC1 | 647.6314 | -0.23958 | 0.172151 | -1.39169 | 0.164017 | 0.312758 |
| SLC9A6 | 968.9116 | -0.18727 | 0.153377 | -1.22096 | 0.2221 | 0.382647 |
| EIF1AY | 383.8885 | 0.692598 | 0.250944 | 2.759973 | 0.005781 | 0.032602 |
| DLL1 | 70.98745 | 0.369795 | 0.433445 | 0.853154 | 0.393574 | 0.55842 |
| UNC13B | 2859.244 | -0.26238 | 0.140962 | -1.86134 | 0.062696 | 0.162128 |
| SMURF1 | 1253.524 | -0.17843 | 0.142943 | -1.24826 | 0.211934 | 0.370354 |
| GATSL2 | 1.369486 | -1.16441 | 1.660465 | -0.70125 | 0.483145 | NA |
| PLXNB3 | 1340.474 | -0.64059 | 0.192523 | -3.32731 | 8.77E-04 | 0.009546 |
| EGFL6 | 3.355745 | 2.238261 | 1.346575 | 1.662188 | 0.096475 | NA |
| GRIN3A | 6.735497 | -0.17383 | 0.769727 | -0.22584 | 0.821329 | NA |
| CNOT7 | 1177.587 | 0.257773 | 0.155538 | 1.657298 | 0.097459 | 0.219362 |
| GK | 718.586 | 0.519563 | 0.143292 | 3.625915 | 2.88E-04 | 0.004809 |
| SFT2D1 | 960.7801 | 0.098301 | 0.168392 | 0.583762 | 0.55938 | 0.698121 |
| ZNF277 | 311.8797 | 0.855771 | 0.357018 | 2.396998 | 0.01653 | 0.066564 |
| TOX | 24.94265 | -0.30288 | 0.458596 | -0.66045 | 0.508968 | NA |
| RUSC2 | 1517.955 | -0.42151 | 0.18364 | -2.29529 | 0.021716 | 0.080102 |
| C9orf96 | 76.84536 | 0.035668 | 0.273349 | 0.130486 | 0.896182 | 0.939326 |
| TYW1 | 320.7246 | 0.164885 | 0.16708 | 0.986858 | 0.323712 | 0.491623 |
| DCAF12 | 1574.276 | -0.13508 | 0.160265 | -0.84285 | 0.399314 | 0.563892 |
| SMC5 | 234.4885 | 1.102952 | 0.710629 | 1.552079 | 0.120643 | 0.253003 |
| DCAF12L1 | 8.933959 | -0.966 | 0.789053 | -1.22425 | 0.220857 | NA |
| CAPZA2 | 2283.259 | 0.964373 | 0.246699 | 3.909102 | 9.26E-05 | 0.002414 |
| BHLHB9 | 93.14029 | 0.368968 | 0.28572 | 1.291362 | 0.196578 | 0.352003 |
| L1CAM | 17183.07 | -0.55892 | 0.146272 | -3.82112 | 1.33E-04 | 0.003029 |

| | | | | | | |
|---|---|---|---|---|---|---|
| C9orf114 | 862.7619 | -0.25312 | 0.150849 | -1.67795 | 0.093357 | 0.213265 |
| RPL39 | 241.7984 | 0.19721 | 0.213149 | 0.925222 | 0.35485 | 0.523121 |
| GPRASP1 | 21.64218 | 0.255568 | 0.454338 | 0.562506 | 0.573771 | NA |
| MAGEE1 | 21.75209 | -0.5007 | 0.453978 | -1.10292 | 0.270061 | NA |
| DMD | 389.3193 | 0.204116 | 0.276512 | 0.738183 | 0.460403 | 0.61869 |
| ARMCX6 | 547.8016 | -0.12578 | 0.166466 | -0.75556 | 0.449914 | 0.609317 |
| RORB | 12.24928 | -0.44776 | 0.645195 | -0.69398 | 0.487692 | NA |
| MIR148A | 0.134133 | -1.48909 | 4.077584 | -0.36519 | 0.71497 | NA |
| MIR450A1 | 0.134133 | -1.48909 | 4.077584 | -0.36519 | 0.71497 | NA |
| MIRLET7D | 0.777314 | 2.821035 | 2.404647 | 1.17316 | 0.240732 | NA |
| | 0.392857 | -1.546 | 4.075469 | -0.37934 | 0.704433 | NA |
| RNU4-82P | 0.258578 | 1.233151 | 4.047809 | 0.304646 | 0.760635 | NA |
| | 0.260158 | 1.240347 | 4.047124 | 0.306476 | 0.759242 | NA |
| | 0.134133 | -1.48909 | 4.077584 | -0.36519 | 0.71497 | NA |
| RNA5SP29! | 0.151697 | -1.546 | 4.075469 | -0.37934 | 0.704433 | NA |
| RNU6-223I | 0.162376 | 0.424913 | 4.079006 | 0.104171 | 0.917034 | NA |
| | 0.096202 | 0.408546 | 4.079639 | 0.100143 | 0.920231 | NA |
| RNU1-72P | 0.354779 | 1.703954 | 3.657232 | 0.465914 | 0.641277 | NA |
| | 2.153335 | 0.761958 | 1.375318 | 0.554023 | 0.579563 | NA |
| | 0.151697 | -1.546 | 4.075469 | -0.37934 | 0.704433 | NA |
| RNU6-824I | 0.58526 | 0.001197 | 3.235889 | 3.70E-04 | 0.999705 | NA |
| | 1.255343 | 2.334528 | 1.930962 | 1.208997 | 0.226664 | NA |
| RNA5SP26( | 0.096202 | 0.408546 | 4.079639 | 0.100143 | 0.920231 | NA |
| | 0.392857 | -1.546 | 4.075469 | -0.37934 | 0.704433 | NA |
| | 0.288605 | 1.443619 | 4.029058 | 0.358302 | 0.720117 | NA |
| RNU6-1091 | 0.096202 | 0.408546 | 4.079639 | 0.100143 | 0.920231 | NA |
| | 1.090192 | 0.321964 | 2.034163 | 0.158278 | 0.874237 | NA |
| | 0.096202 | 0.408546 | 4.079639 | 0.100143 | 0.920231 | NA |
| RNA5SP24: | 0.651435 | 0.249629 | 2.788679 | 0.089515 | 0.928672 | NA |
| RNY1 | 1.127738 | 1.224352 | 2.034449 | 0.60181 | 0.5473 | NA |
| | 0.575812 | 0.936922 | 2.776634 | 0.337431 | 0.735792 | NA |
| RNA5-8SP3 | 1.533443 | -0.70521 | 1.574317 | -0.44795 | 0.654191 | NA |
| RNU4-50P | 0.298089 | -0.51711 | 3.975232 | -0.13008 | 0.896501 | NA |
| SNORA65 | 5.53545 | 0.435282 | 0.921217 | 0.472508 | 0.636564 | NA |
| RNU6-108: | 0.096202 | 0.408546 | 4.079639 | 0.100143 | 0.920231 | NA |
| | 0.151697 | -1.546 | 4.075469 | -0.37934 | 0.704433 | NA |
| SNORA14A | 0.949524 | -0.003 | 2.040023 | -0.00147 | 0.998826 | NA |
| SNORA5C | 6.063088 | -0.325 | 0.842472 | -0.38577 | 0.699667 | NA |
| | 0.096202 | 0.408546 | 4.079639 | 0.100143 | 0.920231 | NA |
| | 0.326333 | 1.533889 | 3.826878 | 0.40082 | 0.688553 | NA |
| | 0.162376 | 0.424913 | 4.079006 | 0.104171 | 0.917034 | NA |
| | 0.151697 | -1.546 | 4.075469 | -0.37934 | 0.704433 | NA |
| RNA5SP25! | 0.151697 | -1.546 | 4.075469 | -0.37934 | 0.704433 | NA |
| RNA5SP28: | 5.429561 | 0.724098 | 0.927571 | 0.780638 | 0.435015 | NA |
| | 0.163957 | 0.424913 | 4.079006 | 0.104171 | 0.917034 | NA |
| | 1.590279 | -0.7938 | 1.607826 | -0.49371 | 0.621512 | NA |
| | 1.033263 | 0.082415 | 2.175992 | 0.037875 | 0.969788 | NA |

| | | | | | | |
|---|---|---|---|---|---|---|
| | 1.772096 | -0.48395 | 1.620622 | -0.29862 | 0.765231 | NA |
| | 0.096202 | 0.408546 | 4.079639 | 0.100143 | 0.920231 | NA |
| | 0.653015 | 0.2552 | 2.787103 | 0.091565 | 0.927044 | NA |
| | 195.0215 | -0.09066 | 0.26863 | -0.33749 | 0.735748 | 0.832757 |
| FAM83H-A | 44.7185 | -0.35429 | 0.353826 | -1.0013 | 0.316679 | NA |
| | 0.298089 | -0.51711 | 3.975232 | -0.13008 | 0.896501 | NA |
| | 0.956059 | 0.787678 | 2.36595 | 0.332923 | 0.739193 | NA |
| | 0.163957 | 0.424913 | 4.079006 | 0.104171 | 0.917034 | NA |
| IGBP1-AS1 | 0.162376 | 0.424913 | 4.079006 | 0.104171 | 0.917034 | NA |
| | 5.413185 | 0.042152 | 0.866614 | 0.04864 | 0.961206 | NA |
| | 0.717463 | 1.437863 | 2.615223 | 0.549805 | 0.582453 | NA |
| C6orf99 | 3.776591 | -0.29511 | 1.040166 | -0.28372 | 0.776627 | NA |
| GDI1 | 5595.701 | -0.26768 | 0.14151 | -1.89158 | 0.058547 | 0.154515 |
| LINC00632 | 22.29087 | 1.015484 | 0.514972 | 1.971922 | 0.048618 | NA |
| FAM127B | 1177.163 | -0.23003 | 0.150228 | -1.53118 | 0.125726 | 0.260802 |
| CCDC160 | 7.171478 | 1.825651 | 0.917456 | 1.989907 | 0.046601 | NA |
| C9orf37 | 275.2029 | -0.19674 | 0.181865 | -1.08177 | 0.279356 | 0.445506 |
| GLT6D1 | 0.163957 | 0.424913 | 4.079006 | 0.104171 | 0.917034 | NA |
| COL5A1-AS | 0.162376 | 0.424913 | 4.079006 | 0.104171 | 0.917034 | NA |
| LINC00963 | 1046.096 | -0.1655 | 0.172955 | -0.95691 | 0.338612 | 0.507304 |
| | 6.336146 | -1.30605 | 0.87466 | -1.49321 | 0.135383 | NA |
| TCEAL5 | 0.162376 | 0.424913 | 4.079006 | 0.104171 | 0.917034 | NA |
| TCEAL6 | 0.70693 | -1.62766 | 2.687154 | -0.60572 | 0.544701 | NA |
| RPA4 | 0.939893 | 1.95109 | 2.358696 | 0.82719 | 0.408129 | NA |
| CHIC1 | 156.3753 | 0.532871 | 0.31547 | 1.689134 | 0.091194 | 0.20976 |
| NAP1L6 | 0.096202 | 0.408546 | 4.079639 | 0.100143 | 0.920231 | NA |
| NHSL2 | 0.804712 | -0.7622 | 2.348804 | -0.3245 | 0.745556 | NA |
| GGTA1P | 0.354779 | 1.703954 | 3.657232 | 0.465914 | 0.641277 | NA |
| CXorf65 | 8.616975 | -1.21994 | 0.762171 | -1.60062 | 0.109462 | NA |
| LRRC37A5F | 10.78275 | 0.204102 | 0.687982 | 0.296667 | 0.766721 | NA |
| SPIN3 | 71.12099 | 0.291005 | 0.288835 | 1.007513 | 0.313688 | 0.481146 |
| | 917.6503 | -0.10778 | 0.152244 | -0.70794 | 0.47898 | 0.633527 |
| PAGE3 | 0.096202 | 0.408546 | 4.079639 | 0.100143 | 0.920231 | NA |
| COL15A1 | 872.5397 | 0.307682 | 0.577048 | 0.533199 | 0.593896 | 0.724345 |
| C9orf129 | 0.28583 | -2.21358 | 3.97376 | -0.55705 | 0.577494 | NA |
| | 0.162376 | 0.424913 | 4.079006 | 0.104171 | 0.917034 | NA |
| C9orf170 | 5.836878 | -0.39858 | 0.862041 | -0.46237 | 0.643816 | NA |
| | 8.905883 | -0.07016 | 0.691681 | -0.10143 | 0.919211 | NA |
| SSX9 | 0.151697 | -1.546 | 4.075469 | -0.37934 | 0.704433 | NA |
| MAMDC2-/ | 2.126075 | 0.393395 | 1.65399 | 0.237846 | 0.812001 | NA |
| C9orf135 | 0.720624 | 1.435938 | 2.334155 | 0.615185 | 0.538432 | NA |
| | 0.422737 | 0.518872 | 3.501356 | 0.148192 | 0.882191 | NA |
| | 0.134133 | -1.48909 | 4.077584 | -0.36519 | 0.71497 | NA |
| CBWD6 | 13.11368 | 0.606523 | 0.582577 | 1.041104 | 0.297827 | NA |
| | 0.410275 | 0.244283 | 3.100889 | 0.078778 | 0.937209 | NA |
| PGM5P1 | 3.347063 | 1.650862 | 1.179703 | 1.399389 | 0.161696 | NA |
| FAM27E4 | 1.317366 | 0.190486 | 1.911887 | 0.099632 | 0.920636 | NA |

| | | | | | | |
|---|---|---|---|---|---|---|
| FAM27E2 | 3.708982 | -0.71544 | 1.093841 | -0.65406 | 0.513071 | NA |
| | 4.362761 | 1.310113 | 0.978317 | 1.33915 | 0.180522 | NA |
| MROH6 | 94.92151 | -0.23224 | 0.246512 | -0.94209 | 0.346147 | 0.513975 |
| FAM201A | 4.13263 | 1.21096 | 1.004818 | 1.205153 | 0.228144 | NA |
| | 18.68046 | 0.220236 | 0.506112 | 0.435154 | 0.663451 | NA |
| CXorf31 | 1.262017 | -0.10925 | 1.859842 | -0.05874 | 0.95316 | NA |
| ATP6V0E2- | 12.6377 | -0.11987 | 0.565764 | -0.21188 | 0.832201 | NA |
| ZNF783 | 135.0009 | -0.47316 | 0.274149 | -1.72591 | 0.084363 | 0.197978 |
| ZNF425 | 107.2951 | 0.140032 | 0.2331 | 0.600736 | 0.548016 | 0.689317 |
| ARHGEF34I | 16.9437 | -0.50554 | 0.540934 | -0.93457 | 0.350012 | NA |
| PRSS3P4 | 0.096202 | 0.408546 | 4.079639 | 0.100143 | 0.920231 | NA |
| AARD | 18.57848 | 0.568765 | 0.582832 | 0.975865 | 0.329131 | NA |
| PKHD1L1 | 0.230334 | -0.53318 | 4.076749 | -0.13079 | 0.895945 | NA |
| SLC35B4 | 1586.716 | -0.20291 | 0.160934 | -1.2608 | 0.207381 | 0.364492 |
| FAM71F2 | 6.247363 | -0.01866 | 0.788384 | -0.02367 | 0.981115 | NA |
| FAM205A | 0.151697 | -1.546 | 4.075469 | -0.37934 | 0.704433 | NA |
| TRIQK | 385.8764 | 0.756941 | 0.238155 | 3.178358 | 0.001481 | 0.013453 |
| ARID3C | 2.251467 | 0.062178 | 1.297425 | 0.047924 | 0.961777 | NA |
| ZBTB10 | 486.2807 | 0.346838 | 0.256543 | 1.351967 | 0.176386 | 0.328679 |
| SPDYE2 | 0.420954 | 1.923376 | 3.462803 | 0.555439 | 0.578594 | NA |
| DGAT2L7P | 0.134133 | -1.48909 | 4.077584 | -0.36519 | 0.71497 | NA |
| PDE7A | 198.5925 | 0.054768 | 0.213615 | 0.256387 | 0.797652 | 0.877602 |
| TRBV20OR! | 0.430642 | -1.1891 | 3.485782 | -0.34113 | 0.733008 | NA |
| MUC12 | 3.438787 | 0.143424 | 1.133422 | 0.12654 | 0.899304 | NA |
| | 14.22953 | 0.919389 | 0.644341 | 1.426868 | 0.153618 | NA |
| GOLGA2P1 | 0.096202 | 0.408546 | 4.079639 | 0.100143 | 0.920231 | NA |
| TECPR1 | 1676.248 | -0.57307 | 0.150567 | -3.8061 | 1.41E-04 | 0.003128 |
| SAMD9 | 292.7716 | 0.501788 | 0.784387 | 0.639719 | 0.522355 | 0.668666 |
| | 0.163957 | 0.424913 | 4.079006 | 0.104171 | 0.917034 | NA |
| | 35.89805 | 0.243736 | 0.377114 | 0.646319 | 0.518073 | NA |
| | 0.134133 | -1.48909 | 4.077584 | -0.36519 | 0.71497 | NA |
| TMSB4X | 3094.366 | 0.407276 | 0.198829 | 2.048373 | 0.040523 | 0.121376 |
| C9orf92 | 0.392711 | 0.292538 | 3.145221 | 0.09301 | 0.925895 | NA |
| POM121B | 45.17695 | -0.57381 | 0.354284 | -1.61964 | 0.105309 | NA |
| STAG3L1 | 17.75556 | -0.05076 | 0.502741 | -0.10097 | 0.919572 | NA |
| | 0.163957 | 0.424913 | 4.079006 | 0.104171 | 0.917034 | NA |
| | 5.674431 | 0.539744 | 0.930096 | 0.580309 | 0.561706 | NA |
| | 0.151697 | -1.546 | 4.075469 | -0.37934 | 0.704433 | NA |
| | 220.2556 | -0.20689 | 0.185482 | -1.11541 | 0.264676 | 0.42918 |
| ARSH | 0.303394 | -2.27006 | 4.032451 | -0.56295 | 0.57347 | NA |
| RP9P | 216.5822 | 0.484934 | 0.231465 | 2.095064 | 0.036165 | 0.112353 |
| PPAPDC2 | 234.0406 | -0.29597 | 0.17696 | -1.67253 | 0.094421 | 0.214847 |
| FAM90A2P | 0.163957 | 0.424913 | 4.079006 | 0.104171 | 0.917034 | NA |
| DEFB134 | 4.172499 | 1.180712 | 1.115748 | 1.058224 | 0.289953 | NA |
| DEFB135 | 0.458885 | 0.577525 | 3.449278 | 0.167433 | 0.867029 | NA |
| DEFB136 | 0.488709 | 2.119056 | 3.307564 | 0.64067 | 0.521737 | NA |
| | 0.192403 | 0.975392 | 4.060716 | 0.240202 | 0.810174 | NA |

| | | | | | | |
|---|---|---|---|---|---|---|
| ZNF316 | 779.9616 | -0.36599 | 0.18062 | -2.02627 | 0.042737 | 0.125875 |
| | 0.640406 | 1.11402 | 2.710977 | 0.410929 | 0.681124 | NA |
| OR7E161P | 2.294599 | 2.485977 | 1.485749 | 1.673215 | 0.094285 | NA |
| | 1.009025 | 3.177087 | 2.206167 | 1.440093 | 0.149841 | NA |
| GYG2P1 | 0.708364 | 0.219301 | 2.144998 | 0.102238 | 0.918568 | NA |
| HNRNPA1P | 0.260158 | 1.240347 | 4.047124 | 0.306476 | 0.759242 | NA |
| XKR4 | 11.98745 | 2.569433 | 0.863369 | 2.976056 | 0.00292 | NA |
| | 1.09159 | 1.212842 | 2.020776 | 0.600186 | 0.548382 | NA |
| SNORA69 | 0.134133 | -1.48909 | 4.077584 | -0.36519 | 0.71497 | NA |
| SNORA22 | 0.230334 | -0.53318 | 4.076749 | -0.13079 | 0.895945 | NA |
| RNU6-10P | 0.556813 | -0.57007 | 3.294615 | -0.17303 | 0.862627 | NA |
| | 0.151697 | -1.546 | 4.075469 | -0.37934 | 0.704433 | NA |
| SNORA5A | 8.569254 | -0.18227 | 0.784453 | -0.23236 | 0.816261 | NA |
| SNORA36A | 0.268266 | -2.15396 | 4.038351 | -0.53338 | 0.593773 | NA |
| RNU6-13P | 0.096202 | 0.408546 | 4.079639 | 0.100143 | 0.920231 | NA |
| RNU6-3P | 0.163957 | 0.424913 | 4.079006 | 0.104171 | 0.917034 | NA |
| RNU1-106P | 1.569619 | 1.16325 | 1.623046 | 0.716708 | 0.473555 | NA |
| RNU6-729P | 0.096202 | 0.408546 | 4.079639 | 0.100143 | 0.920231 | NA |
| | 0.151697 | -1.546 | 4.075469 | -0.37934 | 0.704433 | NA |
| RNU1-67P | 0.392857 | -1.546 | 4.075469 | -0.37934 | 0.704433 | NA |
| RNU6-800P | 0.326333 | 1.533889 | 3.826878 | 0.40082 | 0.688553 | NA |
| RNA5SP284 | 0.697685 | -0.92563 | 2.353044 | -0.39338 | 0.694041 | NA |
| RNU6-30P | 0.624422 | 1.162588 | 2.733761 | 0.425271 | 0.670639 | NA |
| RNU6-925P | 0.258578 | 1.233151 | 4.047809 | 0.304646 | 0.760635 | NA |
| RNU6-178P | 0.584911 | 2.38756 | 2.690807 | 0.887303 | 0.374916 | NA |
| SNORA20 | 0.890088 | -0.08229 | 2.128047 | -0.03867 | 0.969154 | NA |
| RNU6-1011 | 0.422534 | 1.924583 | 3.001758 | 0.641152 | 0.521424 | NA |
| | 0.096202 | 0.408546 | 4.079639 | 0.100143 | 0.920231 | NA |
| MIR25 | 5.1054 | -0.12976 | 0.875037 | -0.14829 | 0.882112 | NA |
| MIR23B | 0.678686 | -3.08037 | 2.621268 | -1.17514 | 0.239937 | NA |
| MIR593 | 0.326333 | 1.533889 | 3.826878 | 0.40082 | 0.688553 | NA |
| MIR508 | 0.151697 | -1.546 | 4.075469 | -0.37934 | 0.704433 | NA |
| MIR32 | 0.096202 | 0.408546 | 4.079639 | 0.100143 | 0.920231 | NA |
| MIR548AA1 | 0.162376 | 0.424913 | 4.079006 | 0.104171 | 0.917034 | NA |
| MIR548D1 | 0.162376 | 0.424913 | 4.079006 | 0.104171 | 0.917034 | NA |
| MIR222 | 0.162376 | 0.424913 | 4.079006 | 0.104171 | 0.917034 | NA |
| MIR455 | 0.151697 | -1.546 | 4.075469 | -0.37934 | 0.704433 | NA |
| MIR181B2 | 0.096202 | 0.408546 | 4.079639 | 0.100143 | 0.920231 | NA |
| MIR590 | 1.296253 | 2.478474 | 1.96601 | 1.260662 | 0.207431 | NA |
| MIR93 | 0.950958 | 0.971806 | 2.077609 | 0.467752 | 0.639962 | NA |
| MIR27B | 1.533057 | 0.55892 | 1.825656 | 0.306147 | 0.759492 | NA |
| MIR221 | 1.311188 | 0.009168 | 1.781747 | 0.005146 | 0.995894 | NA |
| MIR219-2 | 0.268266 | -2.15396 | 4.038351 | -0.53338 | 0.593773 | NA |
| MIR589 | 1.860032 | -2.16427 | 2.345446 | -0.92275 | 0.356135 | NA |
| MIR503 | 0.162376 | 0.424913 | 4.079006 | 0.104171 | 0.917034 | NA |
| MIR548A3 | 0.392857 | -1.546 | 4.075469 | -0.37934 | 0.704433 | NA |
| MIR92A2 | 0.163957 | 0.424913 | 4.079006 | 0.104171 | 0.917034 | NA |

| | | | | | | |
|---|---|---|---|---|---|---|
| MIR106B | 0.720624 | 1.435938 | 2.334155 | 0.615185 | 0.538432 | NA |
| MIR718 | 0.162376 | 0.424913 | 4.079006 | 0.104171 | 0.917034 | NA |
| TRGV5 | 0.151697 | -1.546 | 4.075469 | -0.37934 | 0.704433 | NA |
| TRGV4 | 0.392857 | -1.546 | 4.075469 | -0.37934 | 0.704433 | NA |
| TRBV7-3 | 0.392857 | -1.546 | 4.075469 | -0.37934 | 0.704433 | NA |
| MIR374B | 0.260158 | 1.240347 | 4.047124 | 0.306476 | 0.759242 | NA |
| | 0.151697 | -1.546 | 4.075469 | -0.37934 | 0.704433 | NA |
| | 0.642189 | 0.046766 | 2.378271 | 0.019664 | 0.984311 | NA |
| FAM127C | 586.32 | -0.19335 | 0.159937 | -1.20893 | 0.226688 | 0.388183 |
| RPS26P3 | 1.674018 | -0.7301 | 1.698253 | -0.42991 | 0.667257 | NA |
| RNF208 | 89.28344 | -0.21871 | 0.268821 | -0.8136 | 0.415876 | 0.579003 |
| POU5F1B | 0.096202 | 0.408546 | 4.079639 | 0.100143 | 0.920231 | NA |
| RPS26P6 | 0.096202 | 0.408546 | 4.079639 | 0.100143 | 0.920231 | NA |
| RBM12B-A: | 4.205368 | 0.76966 | 1.022802 | 0.752501 | 0.45175 | NA |
| BTF3P12 | 0.134133 | -1.48909 | 4.077584 | -0.36519 | 0.71497 | NA |
| PTTG3P | 0.314073 | -0.57308 | 3.898934 | -0.14698 | 0.883146 | NA |
| | 0.35636 | 1.709252 | 3.651833 | 0.468053 | 0.639746 | NA |
| FGFR1OP | 236.3738 | 1.012052 | 0.233242 | 4.339072 | 1.43E-05 | 7.69E-04 |
| LPAL2 | 4.785702 | 1.077113 | 0.95359 | 1.129534 | 0.258673 | NA |
| | 76.77499 | 0.199067 | 0.290009 | 0.686416 | 0.492451 | 0.643732 |
| | 0.162376 | 0.424913 | 4.079006 | 0.104171 | 0.917034 | NA |
| SCAF8 | 798.5966 | -0.12277 | 0.173375 | -0.70813 | 0.478867 | 0.633449 |
| | 4.144726 | 1.43462 | 1.14871 | 1.248897 | 0.211703 | NA |
| KRT18P68 | 0.162376 | 0.424913 | 4.079006 | 0.104171 | 0.917034 | NA |
| YBX1P4 | 0.327913 | 1.543154 | 4.021063 | 0.383768 | 0.701151 | NA |
| ASIC3 | 16.58078 | 0.272625 | 0.499948 | 0.545308 | 0.585542 | NA |
| NCLP1 | 2.94221 | 0.01418 | 1.184645 | 0.01197 | 0.99045 | NA |
| CCDC183 | 41.81402 | -0.15486 | 0.333783 | -0.46397 | 0.642671 | NA |
| ARHGEF35 | 15.91185 | 0.271307 | 0.563356 | 0.481591 | 0.630097 | NA |
| OR2F1 | 0.392857 | -1.546 | 4.075469 | -0.37934 | 0.704433 | NA |
| | 0.555233 | -0.57009 | 3.296947 | -0.17291 | 0.862719 | NA |
| DNLZ | 6.041723 | -0.31429 | 0.92574 | -0.3395 | 0.734236 | NA |
| | 1.959984 | 2.251507 | 1.641087 | 1.371961 | 0.170076 | NA |
| | 7.411785 | 2.083851 | 0.846083 | 2.462939 | 0.01378 | NA |
| | 0.847801 | -1.30616 | 2.227118 | -0.58648 | 0.557554 | NA |
| | 3.31765 | 2.476355 | 1.363578 | 1.816071 | 0.069359 | NA |
| | 0.298089 | -0.51711 | 3.975232 | -0.13008 | 0.896501 | NA |
| | 0.163957 | 0.424913 | 4.079006 | 0.104171 | 0.917034 | NA |
| FTH1P12 | 5.726165 | -1.01596 | 0.886593 | -1.14591 | 0.251833 | NA |
| | 1.703752 | 0.798229 | 1.53059 | 0.521517 | 0.602007 | NA |
| | 0.831817 | -1.27234 | 2.201546 | -0.57793 | 0.563311 | NA |
| HAUS7 | 5.16276 | 0.453789 | 0.90233 | 0.502908 | 0.615029 | NA |
| PVRIG | 2.95265 | 0.634121 | 1.3868 | 0.457255 | 0.647488 | NA |
| GPC2 | 41.82946 | 0.139 | 0.347821 | 0.399631 | 0.689428 | NA |
| ERV3-1 | 183.1598 | -0.23268 | 0.227309 | -1.02364 | 0.306006 | 0.473169 |
| HMGB1P37 | 0.410275 | 0.244283 | 3.100889 | 0.078778 | 0.937209 | NA |
| | 9.452649 | 0.054672 | 0.689515 | 0.079291 | 0.936801 | NA |

| | | | | | | |
|---|---|---|---|---|---|---|
| | 0.096202 | 0.408546 | 4.079639 | 0.100143 | 0.920231 | NA |
| TPT1P13 | 0.230334 | -0.53318 | 4.076749 | -0.13079 | 0.895945 | NA |
| SETP8 | 0.296509 | -0.51709 | 3.982155 | -0.12985 | 0.896684 | NA |
| YBX1P6 | 0.315653 | -0.57305 | 3.892639 | -0.14721 | 0.882963 | NA |
| | 2.190182 | -1.05595 | 1.345622 | -0.78473 | 0.432611 | NA |
| | 1.300939 | 2.033521 | 2.329095 | 0.873095 | 0.382611 | NA |
| | 0.162376 | 0.424913 | 4.079006 | 0.104171 | 0.917034 | NA |
| C8orf82 | 1173.648 | -0.13116 | 0.195538 | -0.67077 | 0.502366 | 0.652458 |
| | 0.46577 | -1.30191 | 3.402879 | -0.38259 | 0.702024 | NA |
| | 0.326333 | 1.533889 | 3.826878 | 0.40082 | 0.688553 | NA |
| | 0.52699 | -2.57878 | 3.248478 | -0.79384 | 0.427288 | NA |
| | 0.490493 | 0.776209 | 2.895261 | 0.268096 | 0.788625 | NA |
| SLC25A5P3 | 0.420954 | 1.923376 | 3.462803 | 0.555439 | 0.578594 | NA |
| LDHBP2 | 1.863914 | 0.005204 | 1.528388 | 0.003405 | 0.997283 | NA |
| S1PR3 | 162.6844 | 0.928535 | 0.392812 | 2.363813 | 0.018088 | 0.070537 |
| RPS7P14 | 1.334367 | 0.842194 | 1.784035 | 0.472073 | 0.636875 | NA |
| HMGN2P3( | 0.096202 | 0.408546 | 4.079639 | 0.100143 | 0.920231 | NA |
| SERBP1P1 | 1.136405 | -0.17844 | 1.973701 | -0.09041 | 0.927961 | NA |
| RPS10P14 | 0.162376 | 0.424913 | 4.079006 | 0.104171 | 0.917034 | NA |
| | 0.422534 | 1.924583 | 3.001758 | 0.641152 | 0.521424 | NA |
| | 2.883464 | 0.777594 | 1.235375 | 0.629439 | 0.529061 | NA |
| RPL35P4 | 0.163957 | 0.424913 | 4.079006 | 0.104171 | 0.917034 | NA |
| NHP2P2 | 0.134133 | -1.48909 | 4.077584 | -0.36519 | 0.71497 | NA |
| | 0.623191 | -1.58103 | 2.770593 | -0.57065 | 0.568239 | NA |
| | 0.230334 | -0.53318 | 4.076749 | -0.13079 | 0.895945 | NA |
| CNN2P4 | 12.60682 | -0.03503 | 0.583261 | -0.06006 | 0.952107 | NA |
| | 57.54291 | 0.366778 | 0.318451 | 1.151755 | 0.249422 | 0.41332 |
| VDAC1P2 | 1.415294 | -0.46183 | 1.62897 | -0.28351 | 0.776784 | NA |
| RPL21P75 | 1.754504 | 0.853428 | 1.719701 | 0.496265 | 0.619707 | NA |
| C8orf44 | 27.12826 | 0.351271 | 0.399481 | 0.879318 | 0.379229 | NA |
| YBX1P10 | 7.821239 | -0.28459 | 0.715961 | -0.3975 | 0.691 | NA |
| | 0.096202 | 0.408546 | 4.079639 | 0.100143 | 0.920231 | NA |
| NPM1P6 | 6.236561 | 0.866281 | 0.869107 | 0.996747 | 0.318887 | NA |
| GALT | 606.8119 | -0.14605 | 0.14806 | -0.98646 | 0.32391 | 0.49186 |
| | 0.490493 | 0.776209 | 2.895261 | 0.268096 | 0.788625 | NA |
| | 0.230334 | -0.53318 | 4.076749 | -0.13079 | 0.895945 | NA |
| | 0.803131 | -0.76693 | 2.349822 | -0.32638 | 0.744138 | NA |
| | 1.244664 | 1.349438 | 1.881361 | 0.717267 | 0.473209 | NA |
| RRM2P3 | 2.384864 | 1.444326 | 1.358753 | 1.062979 | 0.287791 | NA |
| | 1.045726 | 0.338916 | 1.94625 | 0.174138 | 0.861757 | NA |
| REPIN1 | 3008.967 | -0.34127 | 0.13443 | -2.53862 | 0.011129 | 0.050549 |
| | 0.162376 | 0.424913 | 4.079006 | 0.104171 | 0.917034 | NA |
| | 0.258578 | 1.233151 | 4.047809 | 0.304646 | 0.760635 | NA |
| RPS11P7 | 0.151697 | -1.546 | 4.075469 | -0.37934 | 0.704433 | NA |
| FBXO16 | 113.6131 | 0.168994 | 0.238539 | 0.708454 | 0.478664 | 0.633256 |
| WEE2 | 0.162376 | 0.424913 | 4.079006 | 0.104171 | 0.917034 | NA |
| PAXIP1-AS2 | 15.65996 | 0.411914 | 0.583137 | 0.706376 | 0.479954 | NA |

| | | | | | | |
|---|---|---|---|---|---|---|
| LDHAP4 | 9.513653 | -0.26056 | 0.688232 | -0.3786 | 0.704986 | NA |
| | 0.46735 | -1.30193 | 3.399671 | -0.38296 | 0.701751 | NA |
| TDPX2 | 0.936915 | 0.819377 | 2.189457 | 0.374237 | 0.708228 | NA |
| TMEM213 | 0.162376 | 0.424913 | 4.079006 | 0.104171 | 0.917034 | NA |
| ST7-OT4 | 0.324752 | 1.53082 | 4.022025 | 0.380609 | 0.703493 | NA |
| | 0.399595 | -1.31595 | 3.579466 | -0.36764 | 0.713143 | NA |
| LINC00998 | 1070.183 | 0.891046 | 0.217767 | 4.091739 | 4.28E-05 | 0.00152 |
| | 1.620196 | 0.149506 | 1.623 | 0.092117 | 0.926605 | NA |
| AZGP1P2 | 0.162376 | 0.424913 | 4.079006 | 0.104171 | 0.917034 | NA |
| FIS1 | 5211.746 | 0.039375 | 0.146154 | 0.26941 | 0.787614 | 0.871252 |
| KRT8P14 | 0.162376 | 0.424913 | 4.079006 | 0.104171 | 0.917034 | NA |
| RSBN1L-AS | 115.9829 | 0.292719 | 0.228588 | 1.28055 | 0.200352 | 0.356417 |
| SPDYE3 | 120.0166 | -0.2633 | 0.21381 | -1.23146 | 0.21815 | 0.377598 |
| MBLAC1 | 71.99407 | -0.01267 | 0.286051 | -0.04431 | 0.964661 | 0.980626 |
| CBX1P2 | 0.163957 | 0.424913 | 4.079006 | 0.104171 | 0.917034 | NA |
| RPS3AP26 | 5.369457 | 0.751802 | 0.972567 | 0.773008 | 0.439518 | NA |
| GNAT3 | 0.162376 | 0.424913 | 4.079006 | 0.104171 | 0.917034 | NA |
| FAM185BP | 7.610177 | 0.158986 | 0.727765 | 0.218458 | 0.827073 | NA |
| GTF2IRD2P | 0.47961 | 0.535796 | 3.39605 | 0.15777 | 0.874638 | NA |
| | 2.991623 | -0.47739 | 1.328363 | -0.35938 | 0.71931 | NA |
| TMEM249 | 2.875743 | 0.71712 | 1.194186 | 0.600509 | 0.548167 | NA |
| | 57.98846 | -0.41183 | 0.290444 | -1.41795 | 0.156206 | 0.302092 |
| | 0.247898 | -0.5901 | 4.074633 | -0.14482 | 0.884851 | NA |
| | 0.488912 | 0.771583 | 2.897678 | 0.266276 | 0.790026 | NA |
| | 11.91114 | 0.731311 | 0.626871 | 1.166605 | 0.24337 | NA |
| | 0.28583 | -2.21358 | 3.97376 | -0.55705 | 0.577494 | NA |
| HNRNPA1P | 0.096202 | 0.408546 | 4.079639 | 0.100143 | 0.920231 | NA |
| SEPT7P2 | 121.4802 | 0.762735 | 0.221969 | 3.436225 | 5.90E-04 | 0.007495 |
| POLR2J4 | 194.4875 | -0.14245 | 0.214993 | -0.6626 | 0.507587 | 0.656489 |
| | 1.269533 | 2.018767 | 2.236927 | 0.902474 | 0.366805 | NA |
| FER1L6 | 0.258578 | 1.233151 | 4.047809 | 0.304646 | 0.760635 | NA |
| MTCP1 | 10.05135 | 0.559032 | 0.679571 | 0.822626 | 0.410721 | NA |
| TPM3P4 | 0.28583 | -2.21358 | 3.97376 | -0.55705 | 0.577494 | NA |
| | 26.25836 | -0.47191 | 0.455465 | -1.0361 | 0.300156 | NA |
| | 0.163957 | 0.424913 | 4.079006 | 0.104171 | 0.917034 | NA |
| LRRC69 | 8.63561 | 0.77943 | 0.74077 | 1.052189 | 0.292713 | NA |
| ISPD | 37.38181 | 0.330544 | 0.381356 | 0.866759 | 0.386074 | NA |
| | 1.66046 | 2.530715 | 1.855025 | 1.364249 | 0.172489 | NA |
| | 0.163957 | 0.424913 | 4.079006 | 0.104171 | 0.917034 | NA |
| DNAJA1P3 | 3.198732 | 1.296258 | 1.275645 | 1.016159 | 0.309554 | NA |
| COL28A1 | 110.081 | 0.992825 | 0.447331 | 2.219441 | 0.026457 | 0.091282 |
| FDPSP5 | 0.151697 | -1.546 | 4.075469 | -0.37934 | 0.704433 | NA |
| GRID2IP | 2.756094 | -0.65166 | 1.292174 | -0.50431 | 0.614044 | NA |
| C9orf38 | 0.937264 | -1.23834 | 2.063944 | -0.59999 | 0.548514 | NA |
| EEF1A1P29 | 3.671006 | 0.67854 | 1.116298 | 0.607848 | 0.543288 | NA |
| CHMP1B2P | 8.02309 | 0.751416 | 0.742603 | 1.011868 | 0.311601 | NA |
| TTC3P1 | 14.23411 | 0.672678 | 0.769391 | 0.8743 | 0.381955 | NA |

| | | | | | | |
|---|---|---|---|---|---|---|
| UBXN2B | 322.348 | 0.111992 | 0.208264 | 0.537739 | 0.590757 | 0.722451 |
| | 5.282941 | 0.563179 | 0.956925 | 0.58853 | 0.556177 | NA |
| MAPK6PS4 | 0.28583 | -2.21358 | 3.97376 | -0.55705 | 0.577494 | NA |
| FAM166B | 1.975362 | -0.16295 | 1.91951 | -0.08489 | 0.932349 | NA |
| YWHAZP6 | 4.020345 | -0.32154 | 0.979386 | -0.3283 | 0.742681 | NA |
| RBMXP2 | 4.883181 | 0.07567 | 0.895923 | 0.08446 | 0.932691 | NA |
| UBA52P6 | 3.170543 | -0.34775 | 1.209171 | -0.28759 | 0.773659 | NA |
| RPL7P33 | 0.324752 | 1.53082 | 4.022025 | 0.380609 | 0.703493 | NA |
| | 0.462046 | 0.589069 | 3.442439 | 0.17112 | 0.86413 | NA |
| KCNU1 | 5.330241 | -0.75246 | 0.862517 | -0.8724 | 0.382988 | NA |
| HMGB3P24 | 0.364467 | -1.20166 | 3.685431 | -0.32606 | 0.744381 | NA |
| | 0.81145 | -0.46945 | 2.476203 | -0.18958 | 0.849635 | NA |
| | 0.258578 | 1.233151 | 4.047809 | 0.304646 | 0.760635 | NA |
| DDX3X | 6349.613 | 0.680412 | 0.180238 | 3.775067 | 1.60E-04 | 0.003412 |
| NPM1P12 | 0.151697 | -1.546 | 4.075469 | -0.37934 | 0.704433 | NA |
| | 2.767216 | -0.79083 | 1.201441 | -0.65824 | 0.510386 | NA |
| | 0.35636 | 1.709252 | 3.651833 | 0.468053 | 0.639746 | NA |
| FAM66B | 5.56359 | -0.51233 | 0.887161 | -0.5775 | 0.563603 | NA |
| PSMA6P1 | 1.663046 | 0.194302 | 1.527979 | 0.127163 | 0.898811 | NA |
| | 12.35173 | 1.792787 | 0.757133 | 2.367864 | 0.017891 | NA |
| TMEM242 | 495.8726 | 0.215311 | 0.171583 | 1.254849 | 0.209534 | 0.367119 |
| MIR450B | 0.448206 | -1.2532 | 3.007315 | -0.41672 | 0.676885 | NA |
| MIR513C | 0.163957 | 0.424913 | 4.079006 | 0.104171 | 0.917034 | NA |
| | 1.003293 | 0.228546 | 2.091683 | 0.109264 | 0.912993 | NA |
| | 0.096202 | 0.408546 | 4.079639 | 0.100143 | 0.920231 | NA |
| GAPDHP72 | 2.946137 | -0.58834 | 1.15719 | -0.50842 | 0.611157 | NA |
| RPS2P55 | 4.684279 | -0.24145 | 0.954108 | -0.25306 | 0.800221 | NA |
| | 54.4764 | 0.246808 | 0.300143 | 0.822302 | 0.410905 | 0.57424 |
| | 0.162376 | 0.424913 | 4.079006 | 0.104171 | 0.917034 | NA |
| | 0.972246 | 0.032713 | 2.31623 | 0.014124 | 0.988731 | NA |
| RPS10P3 | 5.259817 | -0.47796 | 1.021477 | -0.46791 | 0.639851 | NA |
| LDHAL6FP | 0.247898 | -0.5901 | 4.074633 | -0.14482 | 0.884851 | NA |
| | 0.815391 | 0.931626 | 2.360431 | 0.394685 | 0.693076 | NA |
| | 0.163957 | 0.424913 | 4.079006 | 0.104171 | 0.917034 | NA |
| KRT18P48 | 0.096202 | 0.408546 | 4.079639 | 0.100143 | 0.920231 | NA |
| TATDN2P2 | 66.20945 | 0.413769 | 0.357148 | 1.158537 | 0.246645 | 0.41021 |
| | 117.8607 | 0.164668 | 0.231332 | 0.711826 | 0.476572 | 0.631541 |
| | 0.495797 | -0.73607 | 3.324282 | -0.22142 | 0.824763 | NA |
| | 0.870886 | -0.53073 | 2.310896 | -0.22966 | 0.818353 | NA |
| RPS4XP8 | 0.354779 | 1.703954 | 3.657232 | 0.465914 | 0.641277 | NA |
| | 0.326333 | 1.533889 | 3.826878 | 0.40082 | 0.688553 | NA |
| FTH1P19 | 0.489058 | -0.58569 | 3.427543 | -0.17088 | 0.864321 | NA |
| CACYBPP3 | 0.488709 | 2.119056 | 3.307564 | 0.64067 | 0.521737 | NA |
| AMZ2P2 | 0.096202 | 0.408546 | 4.079639 | 0.100143 | 0.920231 | NA |
| HSPA8P15 | 1.05261 | -0.79 | 2.093679 | -0.37733 | 0.705932 | NA |
| RPL23P8 | 6.93998 | 0.936362 | 0.830824 | 1.127028 | 0.259731 | NA |
| TPT1P8 | 0.151697 | -1.546 | 4.075469 | -0.37934 | 0.704433 | NA |

| | | | | | | |
|---|---|---|---|---|---|---|
| FTH1P8 | 169.3963 | -0.02517 | 0.195755 | -0.1286 | 0.897676 | 0.939852 |
| RPA3-AS1 | 404.394 | 0.260893 | 0.153235 | 1.702574 | 0.088648 | 0.205459 |
| | 0.162376 | 0.424913 | 4.079006 | 0.104171 | 0.917034 | NA |
| | 3.559776 | 1.495895 | 1.105153 | 1.353563 | 0.175876 | NA |
| | 28.03707 | -0.01398 | 0.451282 | -0.03098 | 0.975285 | NA |
| | 1.51862 | -1.23018 | 2.037368 | -0.60381 | 0.545971 | NA |
| | 0.163957 | 0.424913 | 4.079006 | 0.104171 | 0.917034 | NA |
| HNRNPH1F | 1.015902 | -0.7221 | 2.105992 | -0.34288 | 0.731689 | NA |
| HSPE1P26 | 1.06294 | 2.033685 | 2.088387 | 0.973807 | 0.330152 | NA |
| IGBP1-AS2 | 0.834389 | 1.638864 | 2.19944 | 0.745128 | 0.456195 | NA |
| HNRNPA1P | 0.134133 | -1.48909 | 4.077584 | -0.36519 | 0.71497 | NA |
| | 0.151697 | -1.546 | 4.075469 | -0.37934 | 0.704433 | NA |
| | 0.594599 | -0.12152 | 2.771051 | -0.04385 | 0.965021 | NA |
| MIR1302-8 | 0.151697 | -1.546 | 4.075469 | -0.37934 | 0.704433 | NA |
| | 1.090541 | -1.58559 | 1.951054 | -0.81268 | 0.4164 | NA |
| | 0.489058 | -0.58569 | 3.427543 | -0.17088 | 0.864321 | NA |
| | 0.258578 | 1.233151 | 4.047809 | 0.304646 | 0.760635 | NA |
| RNU6ATAC | 1.205538 | 0.514777 | 2.000276 | 0.257353 | 0.796906 | NA |
| | 0.151697 | -1.546 | 4.075469 | -0.37934 | 0.704433 | NA |
| SNORA11 | 2.800026 | 0.956112 | 1.233781 | 0.774944 | 0.438372 | NA |
| | 0.151697 | -1.546 | 4.075469 | -0.37934 | 0.704433 | NA |
| SNORD93 | 1.843415 | 2.949948 | 1.749784 | 1.685893 | 0.091816 | NA |
| FANCG | 648.576 | -0.37172 | 0.187251 | -1.98514 | 0.047128 | 0.133662 |
| AP4M1 | 309.5536 | -0.01129 | 0.173666 | -0.06501 | 0.948162 | 0.971147 |
| | 0.70693 | -1.62766 | 2.687154 | -0.60572 | 0.544701 | NA |
| PLXNA4 | 4.785121 | 1.046441 | 0.967633 | 1.081444 | 0.2795 | NA |
| CEBPD | 111.9729 | -0.50455 | 0.324723 | -1.55379 | 0.120235 | 0.25246 |
| TMEM257 | 2.557221 | -0.01842 | 1.231546 | -0.01496 | 0.988065 | NA |
| FAM200A | 175.0403 | 0.711445 | 0.218325 | 3.258651 | 0.001119 | 0.01112 |
| PPP2R2A | 1876.22 | 0.465106 | 0.134615 | 3.455073 | 5.50E-04 | 0.007174 |
| FAM45B | 1.169033 | -0.01411 | 1.776406 | -0.00794 | 0.993662 | NA |
| OR2A25 | 0.163957 | 0.424913 | 4.079006 | 0.104171 | 0.917034 | NA |
| OR2A14 | 0.096202 | 0.408546 | 4.079639 | 0.100143 | 0.920231 | NA |
| XKR9 | 1.081056 | -0.6332 | 2.300921 | -0.2752 | 0.783166 | NA |
| OR2A1 | 0.096202 | 0.408546 | 4.079639 | 0.100143 | 0.920231 | NA |
| ZNF630 | 22.84857 | 0.149297 | 0.486057 | 0.307159 | 0.758722 | NA |
| C7orf71 | 0.247898 | -0.5901 | 4.074633 | -0.14482 | 0.884851 | NA |
| FAM185A | 69.21596 | 0.16966 | 0.301552 | 0.562622 | 0.573692 | 0.709406 |
| | 0.134133 | -1.48909 | 4.077584 | -0.36519 | 0.71497 | NA |
| | 0.35636 | 1.709252 | 3.651833 | 0.468053 | 0.639746 | NA |
| RNU6-892F | 1.007445 | 3.175243 | 2.206758 | 1.438872 | 0.150187 | NA |
| RN7SKP225 | 0.151697 | -1.546 | 4.075469 | -0.37934 | 0.704433 | NA |
| RNU2-5P | 0.096202 | 0.408546 | 4.079639 | 0.100143 | 0.920231 | NA |
| RNU6-285F | 0.096202 | 0.408546 | 4.079639 | 0.100143 | 0.920231 | NA |
| RNU6-628F | 0.314073 | -0.57308 | 3.898934 | -0.14698 | 0.883146 | NA |
| RN7SKP97 | 0.996766 | 1.94108 | 2.070839 | 0.93734 | 0.348584 | NA |
| RN7SKP268 | 0.163957 | 0.424913 | 4.079006 | 0.104171 | 0.917034 | NA |

| | | | | | | |
|---|---|---|---|---|---|---|
| RNU6-722F | 2.325475 | -1.65907 | 1.450969 | -1.14342 | 0.252863 | NA |
| LINC00473 | 1588.331 | 0.045383 | 0.170596 | 0.266027 | 0.790219 | 0.872781 |
| RPL6P21 | 0.151697 | -1.546 | 4.075469 | -0.37934 | 0.704433 | NA |
| | 0.544554 | -2.65036 | 3.220086 | -0.82307 | 0.410468 | NA |
| FAM115B | 8.68415 | 0.469617 | 0.786857 | 0.596827 | 0.550623 | NA |
| CALM1P1 | 0.151697 | -1.546 | 4.075469 | -0.37934 | 0.704433 | NA |
| | 3.011363 | 0.471331 | 1.165742 | 0.404318 | 0.685979 | NA |
| VN1R42P | 0.845418 | 1.095819 | 2.535106 | 0.432258 | 0.665554 | NA |
| | 86.62159 | 0.154083 | 0.247921 | 0.621498 | 0.534272 | 0.677992 |
| | 2.244875 | -0.67674 | 1.551656 | -0.43614 | 0.662733 | NA |
| | 0.382031 | -1.26484 | 3.151989 | -0.40128 | 0.688212 | NA |
| RPL23AP53 | 16.8965 | 0.612143 | 0.534284 | 1.145726 | 0.251909 | NA |
| LINC00630 | 32.59766 | 0.230702 | 0.388133 | 0.59439 | 0.552251 | NA |
| TMSB4XP4 | 1.439667 | 1.722755 | 1.816404 | 0.948443 | 0.342904 | NA |
| | 0.918331 | 0.764804 | 2.118268 | 0.361052 | 0.718061 | NA |
| TNRC18P2 | 0.392857 | -1.546 | 4.075469 | -0.37934 | 0.704433 | NA |
| SUPT20HL2 | 0.134133 | -1.48909 | 4.077584 | -0.36519 | 0.71497 | NA |
| | 0.163957 | 0.424913 | 4.079006 | 0.104171 | 0.917034 | NA |
| IFNWP18 | 0.151697 | -1.546 | 4.075469 | -0.37934 | 0.704433 | NA |
| | 0.422534 | 1.924583 | 3.001758 | 0.641152 | 0.521424 | NA |
| NSUN5P1 | 112.142 | 0.466466 | 0.227045 | 2.054509 | 0.039926 | 0.120287 |
| | 1.21484 | 0.621271 | 1.748506 | 0.355315 | 0.722353 | NA |
| | 0.698908 | -1.96881 | 2.984525 | -0.65967 | 0.509463 | NA |
| | 0.71919 | 0.510978 | 2.728734 | 0.187258 | 0.851458 | NA |
| MIR503HG | 19.98786 | 0.236494 | 0.459118 | 0.515106 | 0.606479 | NA |
| | 0.096202 | 0.408546 | 4.079639 | 0.100143 | 0.920231 | NA |
| | 0.096202 | 0.408546 | 4.079639 | 0.100143 | 0.920231 | NA |
| | 0.247898 | -0.5901 | 4.074633 | -0.14482 | 0.884851 | NA |
| | 0.422737 | 0.518872 | 3.501356 | 0.148192 | 0.882191 | NA |
| | 0.384806 | 1.846216 | 3.999752 | 0.461582 | 0.644381 | NA |
| | 1.809439 | 0.428428 | 1.452082 | 0.295044 | 0.76796 | NA |
| | 0.736607 | 1.395844 | 2.318642 | 0.602009 | 0.547168 | NA |
| | 13.94726 | 0.342654 | 0.577146 | 0.593704 | 0.55271 | NA |
| | 0.162376 | 0.424913 | 4.079006 | 0.104171 | 0.917034 | NA |
| | 0.72077 | 0.527966 | 3.060238 | 0.172524 | 0.863025 | NA |
| MIR181A2F | 36.82095 | 0.791569 | 0.395477 | 2.001556 | 0.045333 | NA |
| | 9.129422 | 0.435586 | 0.673167 | 0.647069 | 0.517587 | NA |
| | 0.410275 | 0.244283 | 3.100889 | 0.078778 | 0.937209 | NA |
| HSPA8P16 | 0.411855 | 0.250456 | 3.097618 | 0.080854 | 0.935558 | NA |
| SRIP2 | 0.163957 | 0.424913 | 4.079006 | 0.104171 | 0.917034 | NA |
| | 0.096202 | 0.408546 | 4.079639 | 0.100143 | 0.920231 | NA |
| | 0.604258 | 1.087325 | 2.707696 | 0.401568 | 0.688002 | NA |
| TTTY22 | 0.622842 | 1.158846 | 2.735323 | 0.42366 | 0.671814 | NA |
| | 0.096202 | 0.408546 | 4.079639 | 0.100143 | 0.920231 | NA |
| CENPVP3 | 0.314073 | -0.57308 | 3.898934 | -0.14698 | 0.883146 | NA |
| | 2.574492 | 0.801658 | 1.270753 | 0.630853 | 0.528137 | NA |
| AMMECR1- | 0.096202 | 0.408546 | 4.079639 | 0.100143 | 0.920231 | NA |

| | | | | | | |
|---|---|---|---|---|---|---|
| SNX18P9 | 0.162376 | 0.424913 | 4.079006 | 0.104171 | 0.917034 | NA |
| PHEX-AS1 | 0.096202 | 0.408546 | 4.079639 | 0.100143 | 0.920231 | NA |
| | 1.626492 | 1.152152 | 1.746252 | 0.659786 | 0.509391 | NA |
| | 0.392857 | -1.546 | 4.075469 | -0.37934 | 0.704433 | NA |
| SLC25A5-A: | 35.91674 | -0.15104 | 0.381243 | -0.39617 | 0.691983 | NA |
| | 1.598655 | -0.27264 | 1.729733 | -0.15762 | 0.874758 | NA |
| | 1.118436 | 1.112087 | 2.263211 | 0.491376 | 0.623161 | NA |
| THAP5P1 | 0.151697 | -1.546 | 4.075469 | -0.37934 | 0.704433 | NA |
| | 1.143158 | 2.223222 | 2.00111 | 1.110994 | 0.266571 | NA |
| | 0.096202 | 0.408546 | 4.079639 | 0.100143 | 0.920231 | NA |
| MTND2P5 | 0.162376 | 0.424913 | 4.079006 | 0.104171 | 0.917034 | NA |
| RPL7P57 | 0.832809 | 1.636407 | 2.200093 | 0.74379 | 0.457004 | NA |
| RPL19P12 | 3.350758 | 0.427096 | 1.068362 | 0.399768 | 0.689328 | NA |
| | 0.296509 | -0.51709 | 3.982155 | -0.12985 | 0.896684 | NA |
| | 0.163957 | 0.424913 | 4.079006 | 0.104171 | 0.917034 | NA |
| TMLHE-AS: | 2.639676 | -0.08936 | 1.24036 | -0.07204 | 0.94257 | NA |
| EIF4BP3 | 23.88482 | 0.186874 | 0.480068 | 0.389266 | 0.69708 | NA |
| | 0.324752 | 1.53082 | 4.022025 | 0.380609 | 0.703493 | NA |
| RPL29P19 | 0.924655 | -0.07191 | 2.416297 | -0.02976 | 0.976257 | NA |
| | 1.045726 | 0.338916 | 1.94625 | 0.174138 | 0.861757 | NA |
| RPL36AP2! | 0.163957 | 0.424913 | 4.079006 | 0.104171 | 0.917034 | NA |
| PRNCR1 | 0.689366 | -1.56115 | 2.700123 | -0.57818 | 0.563145 | NA |
| | 0.315653 | -0.57305 | 3.892639 | -0.14721 | 0.882963 | NA |
| | 0.458885 | 0.577525 | 3.449278 | 0.167433 | 0.867029 | NA |
| HSPA8P7 | 0.574231 | 0.93227 | 2.49998 | 0.372911 | 0.709215 | NA |
| BRAFP1 | 0.392857 | -1.546 | 4.075469 | -0.37934 | 0.704433 | NA |
| EIF4A2P4 | 1.03648 | 0.354657 | 2.004615 | 0.17692 | 0.859571 | NA |
| | 0.134133 | -1.48909 | 4.077584 | -0.36519 | 0.71497 | NA |
| | 0.096202 | 0.408546 | 4.079639 | 0.100143 | 0.920231 | NA |
| | 0.898195 | -1.76199 | 2.351517 | -0.7493 | 0.453676 | NA |
| BTF3P4 | 0.151697 | -1.546 | 4.075469 | -0.37934 | 0.704433 | NA |
| | 0.926438 | -0.78569 | 2.072011 | -0.37919 | 0.704544 | NA |
| | 0.163957 | 0.424913 | 4.079006 | 0.104171 | 0.917034 | NA |
| | 16.803 | 0.179761 | 0.577673 | 0.311181 | 0.755663 | NA |
| C9orf53 | 0.518736 | 2.237423 | 2.7713 | 0.807355 | 0.419462 | NA |
| | 0.296509 | -0.51709 | 3.982155 | -0.12985 | 0.896684 | NA |
| POT1-AS1 | 11.15936 | -0.33971 | 0.638147 | -0.53233 | 0.594497 | NA |
| | 0.134133 | -1.48909 | 4.077584 | -0.36519 | 0.71497 | NA |
| PRRT4 | 18.18703 | -0.65354 | 0.483809 | -1.35082 | 0.176754 | NA |
| | 0.422534 | 1.924583 | 3.001758 | 0.641152 | 0.521424 | NA |
| SRIP3 | 0.315653 | -0.57305 | 3.892639 | -0.14721 | 0.882963 | NA |
| SUMO2P3 | 0.449786 | -1.25325 | 3.00474 | -0.41709 | 0.676613 | NA |
| INE1 | 6.077036 | 0.268476 | 0.864369 | 0.310603 | 0.756102 | NA |
| | 0.151697 | -1.546 | 4.075469 | -0.37934 | 0.704433 | NA |
| | 2.851215 | -0.75396 | 1.280897 | -0.58862 | 0.556115 | NA |
| | 0.096202 | 0.408546 | 4.079639 | 0.100143 | 0.920231 | NA |
| EIF4BP7 | 2.908262 | -0.16928 | 1.202542 | -0.14077 | 0.888055 | NA |

| | | | | | | |
|---|---|---|---|---|---|---|
| | 7.000011 | 0.194282 | 0.742087 | 0.261805 | 0.793472 | NA |
| EIF1AX-AS1 | 0.162376 | 0.424913 | 4.079006 | 0.104171 | 0.917034 | NA |
| | 0.392857 | -1.546 | 4.075469 | -0.37934 | 0.704433 | NA |
| | 0.752028 | 2.752519 | 2.876634 | 0.956854 | 0.338641 | NA |
| RANP4 | 0.754374 | 0.344906 | 2.513743 | 0.137208 | 0.890866 | NA |
| | 0.151697 | -1.546 | 4.075469 | -0.37934 | 0.704433 | NA |
| LINC00092 | 18.54192 | -0.60246 | 0.507478 | -1.18718 | 0.235159 | NA |
| | 0.28583 | -2.21358 | 3.97376 | -0.55705 | 0.577494 | NA |
| DDX26B-AS | 0.327913 | 1.543154 | 4.021063 | 0.383768 | 0.701151 | NA |
| ST13P6 | 4.740465 | 0.084059 | 0.934694 | 0.089932 | 0.928341 | NA |
| ZNRF2P2 | 44.97849 | -0.20123 | 0.327141 | -0.61512 | 0.538478 | NA |
| | 0.260158 | 1.240347 | 4.047124 | 0.306476 | 0.759242 | NA |
| SNX18P3 | 2.746406 | -0.00586 | 1.451304 | -0.00404 | 0.996779 | NA |
| SLC25A5P8 | 0.303394 | -2.27006 | 4.032451 | -0.56295 | 0.57347 | NA |
| | 0.096202 | 0.408546 | 4.079639 | 0.100143 | 0.920231 | NA |
| MIPEPP1 | 0.392857 | -1.546 | 4.075469 | -0.37934 | 0.704433 | NA |
| PPP1R26-A | 2.252458 | 1.370809 | 1.45018 | 0.945268 | 0.344522 | NA |
| CICP8 | 0.46577 | -1.30191 | 3.402879 | -0.38259 | 0.702024 | NA |
| | 3.664812 | -0.82459 | 1.264758 | -0.65197 | 0.514418 | NA |
| | 0.096202 | 0.408546 | 4.079639 | 0.100143 | 0.920231 | NA |
| | 1.366334 | 0.789187 | 1.898748 | 0.415636 | 0.677677 | NA |
| | 1.714964 | -0.37457 | 1.501527 | -0.24946 | 0.803003 | NA |
| | 1.462738 | 0.413906 | 1.637425 | 0.252778 | 0.800439 | NA |
| | 1.202726 | -1.13177 | 2.006914 | -0.56393 | 0.572799 | NA |
| | 0.49187 | 2.136088 | 3.895953 | 0.548284 | 0.583497 | NA |
| | 0.134133 | -1.48909 | 4.077584 | -0.36519 | 0.71497 | NA |
| JPX | 131.8581 | 0.507863 | 0.276769 | 1.834968 | 0.06651 | 0.168462 |
| | 4.492693 | 1.056544 | 1.034645 | 1.021166 | 0.307176 | NA |
| | 0.260158 | 1.240347 | 4.047124 | 0.306476 | 0.759242 | NA |
| | 0.163957 | 0.424913 | 4.079006 | 0.104171 | 0.917034 | NA |
| | 7.868368 | 0.489824 | 0.708176 | 0.69167 | 0.489145 | NA |
| LINC00475 | 6.972103 | 0.554463 | 0.788875 | 0.702852 | 0.482148 | NA |
| | 0.151697 | -1.546 | 4.075469 | -0.37934 | 0.704433 | NA |
| | 0.096202 | 0.408546 | 4.079639 | 0.100143 | 0.920231 | NA |
| | 0.449786 | -1.25325 | 3.00474 | -0.41709 | 0.676613 | NA |
| | 1.04716 | 1.17329 | 2.006462 | 0.584756 | 0.558712 | NA |
| | 0.911832 | -0.02323 | 2.42372 | -0.00958 | 0.992354 | NA |
| | 0.096202 | 0.408546 | 4.079639 | 0.100143 | 0.920231 | NA |
| | 0.162376 | 0.424913 | 4.079006 | 0.104171 | 0.917034 | NA |
| RPL35P5 | 4.613169 | 0.461418 | 1.069121 | 0.431586 | 0.666042 | NA |
| | 0.096202 | 0.408546 | 4.079639 | 0.100143 | 0.920231 | NA |
| | 0.230334 | -0.53318 | 4.076749 | -0.13079 | 0.895945 | NA |
| SBDSP1 | 518.3406 | 0.653398 | 0.188667 | 3.463233 | 5.34E-04 | 0.00703 |
| | 0.162376 | 0.424913 | 4.079006 | 0.104171 | 0.917034 | NA |
| | 0.804712 | -0.7622 | 2.348804 | -0.3245 | 0.745556 | NA |
| | 0.315653 | -0.57305 | 3.892639 | -0.14721 | 0.882963 | NA |
| | 0.247898 | -0.5901 | 4.074633 | -0.14482 | 0.884851 | NA |

| | | | | | | |
|---|---|---|---|---|---|---|
| NPM1P18 | 0.151697 | -1.546 | 4.075469 | -0.37934 | 0.704433 | NA |
| DBH-AS1 | 1.072557 | -2.6908 | 2.651653 | -1.01476 | 0.310218 | NA |
| | 1.10954 | -0.31713 | 1.971981 | -0.16082 | 0.872235 | NA |
| | 0.392857 | -1.546 | 4.075469 | -0.37934 | 0.704433 | NA |
| RBMY2KP | 0.096202 | 0.408546 | 4.079639 | 0.100143 | 0.920231 | NA |
| | 1622.28 | 1.239218 | 0.334169 | 3.708363 | 2.09E-04 | 0.003995 |
| | 0.151697 | -1.546 | 4.075469 | -0.37934 | 0.704433 | NA |
| | 0.3441 | -0.0166 | 3.769682 | -0.0044 | 0.996486 | NA |
| MTND2P9 | 0.163957 | 0.424913 | 4.079006 | 0.104171 | 0.917034 | NA |
| RPS27P16 | 0.555233 | -0.57009 | 3.296947 | -0.17291 | 0.862719 | NA |
| | 0.162376 | 0.424913 | 4.079006 | 0.104171 | 0.917034 | NA |
| RPL7P59 | 0.260158 | 1.240347 | 4.047124 | 0.306476 | 0.759242 | NA |
| PCA3 | 1.159491 | 0.557655 | 1.885778 | 0.295716 | 0.767447 | NA |
| | 0.163957 | 0.424913 | 4.079006 | 0.104171 | 0.917034 | NA |
| ODF2-AS1 | 1.102452 | 1.146114 | 2.069369 | 0.553847 | 0.579683 | NA |
| ELFN1 | 712.1704 | 0.310688 | 0.294972 | 1.053281 | 0.292212 | 0.458987 |
| LINC00035 | 59.26581 | 0.164333 | 0.327733 | 0.501424 | 0.616073 | 0.742059 |
| | 1.550418 | 1.233972 | 1.753317 | 0.703793 | 0.481562 | NA |
| | 2.453115 | 0.249584 | 1.300814 | 0.191867 | 0.847846 | NA |
| | 1.241889 | -0.06002 | 1.948349 | -0.03081 | 0.975424 | NA |
| | 0.392857 | -1.546 | 4.075469 | -0.37934 | 0.704433 | NA |
| | 0.392857 | -1.546 | 4.075469 | -0.37934 | 0.704433 | NA |
| PAICSP1 | 1.781341 | -0.69181 | 1.425442 | -0.48533 | 0.627443 | NA |
| HMGB3P2( | 0.151697 | -1.546 | 4.075469 | -0.37934 | 0.704433 | NA |
| | 0.561971 | -0.49218 | 2.810061 | -0.17515 | 0.860962 | NA |
| SEPT7P3 | 0.35636 | 1.709252 | 3.651833 | 0.468053 | 0.639746 | NA |
| | 0.298089 | -0.51711 | 3.975232 | -0.13008 | 0.896501 | NA |
| | 1.465485 | -0.31895 | 1.726907 | -0.18469 | 0.85347 | NA |
| CICP22 | 0.881915 | -2.2305 | 2.871853 | -0.77668 | 0.437349 | NA |
| | 0.268266 | -2.15396 | 4.038351 | -0.53338 | 0.593773 | NA |
| | 24.25262 | -0.02206 | 0.434253 | -0.05079 | 0.959493 | NA |
| PTP4A1P3 | 0.163957 | 0.424913 | 4.079006 | 0.104171 | 0.917034 | NA |
| PSPHP1 | 92.92984 | 0.25867 | 0.252146 | 1.025875 | 0.304951 | 0.472185 |
| | 0.749217 | 0.7397 | 2.633289 | 0.280903 | 0.778784 | NA |
| | 0.851742 | -0.48655 | 2.547586 | -0.19099 | 0.848537 | NA |
| | 1.747576 | -1.55187 | 1.668065 | -0.93034 | 0.352194 | NA |
| | 1.202377 | 0.510208 | 2.000895 | 0.25499 | 0.798731 | NA |
| | 14.59691 | -0.04901 | 0.588001 | -0.08336 | 0.933568 | NA |
| | 0.230334 | -0.53318 | 4.076749 | -0.13079 | 0.895945 | NA |
| | 0.151697 | -1.546 | 4.075469 | -0.37934 | 0.704433 | NA |
| ST7-AS2 | 0.096202 | 0.408546 | 4.079639 | 0.100143 | 0.920231 | NA |
| MIR29B1 | 65.53028 | 1.039503 | 0.400207 | 2.597415 | 0.009393 | 0.045248 |
| MIR29A | 65.53028 | 1.039503 | 0.400207 | 2.597415 | 0.009393 | 0.045248 |
| IFNA20P | 4.627232 | 0.547759 | 0.934187 | 0.586348 | 0.557642 | NA |
| | 0.192403 | 0.975392 | 4.060716 | 0.240202 | 0.810174 | NA |
| SLC25A5P5 | 0.641986 | 1.117767 | 2.709494 | 0.412537 | 0.679946 | NA |
| | 0.489058 | -0.58569 | 3.427543 | -0.17088 | 0.864321 | NA |

| | | | | | | |
|---|---|---|---|---|---|---|
| | 24.28568 | -0.19041 | 0.435858 | -0.43687 | 0.662207 | NA |
| | 0.096202 | 0.408546 | 4.079639 | 0.100143 | 0.920231 | NA |
| | 1.294855 | 1.463581 | 1.753443 | 0.83469 | 0.403892 | NA |
| | 53.50595 | -0.00886 | 0.308526 | -0.02872 | 0.977087 | 0.987584 |
| | 0.640755 | -1.64826 | 2.754778 | -0.59833 | 0.549621 | NA |
| CYP4F26P | 0.3441 | -0.0166 | 3.769682 | -0.0044 | 0.996486 | NA |
| SPANXA2-( | 0.151697 | -1.546 | 4.075469 | -0.37934 | 0.704433 | NA |
| PAPPA-AS2 | 0.096202 | 0.408546 | 4.079639 | 0.100143 | 0.920231 | NA |
| | 1.527719 | -0.62615 | 1.783612 | -0.35106 | 0.725546 | NA |
| | 0.815391 | 0.931626 | 2.360431 | 0.394685 | 0.693076 | NA |
| | 1.370827 | -0.71211 | 1.970974 | -0.3613 | 0.717877 | NA |
| | 4.06108 | -0.68272 | 1.001859 | -0.68145 | 0.495587 | NA |
| | 3.266231 | -0.57315 | 1.116294 | -0.51344 | 0.607641 | NA |
| | 0.314073 | -0.57308 | 3.898934 | -0.14698 | 0.883146 | NA |
| PSMD5-AS: | 395.245 | 0.423485 | 0.19211 | 2.204387 | 0.027497 | 0.093699 |
| | 0.298089 | -0.51711 | 3.975232 | -0.13008 | 0.896501 | NA |
| TBCAP1 | 2.420459 | -1.39049 | 1.511433 | -0.91998 | 0.357584 | NA |
| TLK1P1 | 0.134133 | -1.48909 | 4.077584 | -0.36519 | 0.71497 | NA |
| PGAM4 | 2.034615 | 0.269551 | 1.473422 | 0.182942 | 0.854843 | NA |
| | 0.326333 | 1.533889 | 3.826878 | 0.40082 | 0.688553 | NA |
| | 3.117612 | -0.10176 | 1.166924 | -0.08721 | 0.930508 | NA |
| | 32.06809 | -0.04082 | 0.37843 | -0.10787 | 0.914102 | NA |
| | 1.968597 | 0.486705 | 1.677224 | 0.290185 | 0.771675 | NA |
| SHROOM2I | 0.151697 | -1.546 | 4.075469 | -0.37934 | 0.704433 | NA |
| LHFPL3-AS: | 1787.656 | 0.244773 | 0.209738 | 1.167042 | 0.243193 | 0.406164 |
| | 0.586491 | 2.39072 | 2.689057 | 0.889055 | 0.373973 | NA |
| H3F3AP5 | 0.151697 | -1.546 | 4.075469 | -0.37934 | 0.704433 | NA |
| | 0.324752 | 1.53082 | 4.022025 | 0.380609 | 0.703493 | NA |
| | 8167.062 | 0.410384 | 0.247622 | 1.657298 | 0.097459 | 0.219362 |
| | 1.235565 | 0.616768 | 2.143708 | 0.287711 | 0.773568 | NA |
| | 0.555087 | 0.973802 | 2.808575 | 0.346725 | 0.728798 | NA |
| MAGI2-AS2 | 0.163957 | 0.424913 | 4.079006 | 0.104171 | 0.917034 | NA |
| | 9.149826 | 0.354133 | 0.667044 | 0.530899 | 0.595489 | NA |
| | 9.140881 | 0.413823 | 0.70106 | 0.590282 | 0.555001 | NA |
| | 0.288605 | 1.443619 | 4.029058 | 0.358302 | 0.720117 | NA |
| | 1.445321 | -0.30112 | 1.586081 | -0.18985 | 0.849425 | NA |
| LINC00629 | 1.743613 | -0.66006 | 1.560343 | -0.42302 | 0.67228 | NA |
| | 0.596179 | -0.09275 | 3.146759 | -0.02948 | 0.976485 | NA |
| SDHDP2 | 0.096202 | 0.408546 | 4.079639 | 0.100143 | 0.920231 | NA |
| | 0.096202 | 0.408546 | 4.079639 | 0.100143 | 0.920231 | NA |
| | 0.260158 | 1.240347 | 4.047124 | 0.306476 | 0.759242 | NA |
| | 1.085266 | 2.142965 | 2.332199 | 0.91886 | 0.358169 | NA |
| | 0.897992 | -1.0518 | 2.16635 | -0.48552 | 0.627309 | NA |
| HDGFP1 | 1.178424 | -1.58299 | 2.140626 | -0.7395 | 0.459604 | NA |
| EPPK1 | 0.653015 | 0.2552 | 2.787103 | 0.091565 | 0.927044 | NA |
| ST7-AS1 | 33.30131 | -0.60788 | 0.405769 | -1.49808 | 0.134111 | NA |
| | 0.987309 | 0.267164 | 2.098144 | 0.127333 | 0.898677 | NA |

| | | | | | | |
|---|---|---|---|---|---|---|
| SUB1P1 | 0.326333 | 1.533889 | 3.826878 | 0.40082 | 0.688553 | NA |
| | 0.162376 | 0.424913 | 4.079006 | 0.104171 | 0.917034 | NA |
| | 3.913603 | 0.617978 | 1.115606 | 0.553939 | 0.57962 | NA |
| | 7.900161 | 0.246268 | 0.719895 | 0.342089 | 0.732284 | NA |
| | 1.336656 | -0.80389 | 1.980439 | -0.40591 | 0.684807 | NA |
| HAUS6P1 | 1.792021 | -0.1506 | 1.434632 | -0.10497 | 0.916398 | NA |
| | 0.163957 | 0.424913 | 4.079006 | 0.104171 | 0.917034 | NA |
| FCF1P1 | 0.830383 | -3.4083 | 2.430438 | -1.40234 | 0.160814 | NA |
| | 0.918331 | 0.764804 | 2.118268 | 0.361052 | 0.718061 | NA |
| | 0.163957 | 0.424913 | 4.079006 | 0.104171 | 0.917034 | NA |
| AKR1B15 | 0.985875 | -1.0033 | 2.40761 | -0.41672 | 0.676883 | NA |
| TSSK5P1 | 0.298089 | -0.51711 | 3.975232 | -0.13008 | 0.896501 | NA |
| | 9.606747 | 0.571481 | 1.065827 | 0.536186 | 0.59183 | NA |
| | 28.16316 | 0.113923 | 0.403181 | 0.28256 | 0.777514 | NA |
| | 1.139152 | -1.36402 | 1.928942 | -0.70714 | 0.479482 | NA |
| SOCS5P4 | 0.3441 | -0.0166 | 3.769682 | -0.0044 | 0.996486 | NA |
| | 0.096202 | 0.408546 | 4.079639 | 0.100143 | 0.920231 | NA |
| | 0.096202 | 0.408546 | 4.079639 | 0.100143 | 0.920231 | NA |
| PGM5P2 | 0.884581 | 1.788183 | 2.221027 | 0.805115 | 0.420753 | NA |
| | 0.880631 | -1.73238 | 2.414456 | -0.7175 | 0.473064 | NA |
| | 1.319092 | -0.78009 | 1.811759 | -0.43057 | 0.666782 | NA |
| | 0.947944 | -0.02704 | 2.20277 | -0.01228 | 0.990206 | NA |
| CYCSP24 | 4.514218 | 1.401912 | 1.043402 | 1.343597 | 0.179079 | NA |
| | 3.353939 | 0.523272 | 1.114215 | 0.469633 | 0.638617 | NA |
| MRPS17P9 | 0.151697 | -1.546 | 4.075469 | -0.37934 | 0.704433 | NA |
| | 0.392857 | -1.546 | 4.075469 | -0.37934 | 0.704433 | NA |
| | 0.192403 | 0.975392 | 4.060716 | 0.240202 | 0.810174 | NA |
| DPY19L2P3 | 84.92435 | -0.28876 | 0.256505 | -1.12575 | 0.260271 | 0.424534 |
| FAM66A | 2.036758 | -0.22167 | 1.464317 | -0.15138 | 0.879675 | NA |
| KRT8P44 | 0.163957 | 0.424913 | 4.079006 | 0.104171 | 0.917034 | NA |
| | 0.86916 | 0.686589 | 2.213986 | 0.310114 | 0.756474 | NA |
| | 3.073275 | 0.02979 | 1.221139 | 0.024395 | 0.980537 | NA |
| | 0.151697 | -1.546 | 4.075469 | -0.37934 | 0.704433 | NA |
| FRMD8P1 | 0.162376 | 0.424913 | 4.079006 | 0.104171 | 0.917034 | NA |
| | 0.163957 | 0.424913 | 4.079006 | 0.104171 | 0.917034 | NA |
| TRIM60P18 | 2.823731 | 0.98814 | 1.278061 | 0.773156 | 0.43943 | NA |
| | 0.411855 | 0.250456 | 3.097618 | 0.080854 | 0.935558 | NA |
| | 0.462046 | 0.589069 | 3.442439 | 0.17112 | 0.86413 | NA |
| CYP1D1P | 0.785714 | -3.2458 | 3.343263 | -0.97085 | 0.331623 | NA |
| VN1R51P | 0.163957 | 0.424913 | 4.079006 | 0.104171 | 0.917034 | NA |
| | 0.260158 | 1.240347 | 4.047124 | 0.306476 | 0.759242 | NA |
| | 0.096202 | 0.408546 | 4.079639 | 0.100143 | 0.920231 | NA |
| | 0.7777 | 0.426209 | 2.574008 | 0.165582 | 0.868486 | NA |
| | 0.163957 | 0.424913 | 4.079006 | 0.104171 | 0.917034 | NA |
| | 0.151697 | -1.546 | 4.075469 | -0.37934 | 0.704433 | NA |
| | 0.392857 | -1.546 | 4.075469 | -0.37934 | 0.704433 | NA |
| | 0.151697 | -1.546 | 4.075469 | -0.37934 | 0.704433 | NA |

| | | | | | | |
|---|---|---|---|---|---|---|
| | 0.392857 | -1.546 | 4.075469 | -0.37934 | 0.704433 | NA |
| TLE1P1 | 12.71137 | -0.27412 | 0.579794 | -0.47278 | 0.636369 | NA |
| | 1.290703 | -0.48664 | 1.714071 | -0.28391 | 0.776479 | NA |
| NACAP1 | 13.85809 | 0.504113 | 0.560346 | 0.899647 | 0.368308 | NA |
| GAPDHP1 | 73.24968 | -0.15788 | 0.310948 | -0.50774 | 0.611639 | 0.738389 |
| | 0.642189 | 0.046766 | 2.378271 | 0.019664 | 0.984311 | NA |
| ORM2 | 0.162376 | 0.424913 | 4.079006 | 0.104171 | 0.917034 | NA |
| | 0.392857 | -1.546 | 4.075469 | -0.37934 | 0.704433 | NA |
| | 0.192403 | 0.975392 | 4.060716 | 0.240202 | 0.810174 | NA |
| | 1.85963 | 0.528221 | 1.500116 | 0.35212 | 0.724748 | NA |
| | 3.405767 | -0.83379 | 1.345831 | -0.61954 | 0.535561 | NA |
| | 90.00887 | -0.02406 | 0.297093 | -0.081 | 0.935441 | 0.963898 |
| | 2.763101 | 0.296533 | 1.27094 | 0.233318 | 0.815515 | NA |
| | 0.192403 | 0.975392 | 4.060716 | 0.240202 | 0.810174 | NA |
| | 0.979202 | 1.972754 | 2.08204 | 0.94751 | 0.343379 | NA |
| | 0.488709 | 2.119056 | 3.307564 | 0.64067 | 0.521737 | NA |
| | 0.096202 | 0.408546 | 4.079639 | 0.100143 | 0.920231 | NA |
| | 0.517155 | 2.232561 | 3.179631 | 0.702145 | 0.482589 | NA |
| LINC01004 | 4.338534 | 0.29845 | 1.038197 | 0.28747 | 0.773753 | NA |
| | 26.73967 | -0.1782 | 0.45509 | -0.39156 | 0.695381 | NA |
| CCT6P1 | 24.39363 | 1.007881 | 0.471456 | 2.137804 | 0.032533 | NA |
| CDY4P | 0.096202 | 0.408546 | 4.079639 | 0.100143 | 0.920231 | NA |
| | 0.151697 | -1.546 | 4.075469 | -0.37934 | 0.704433 | NA |
| | 1.074412 | -0.25414 | 1.986075 | -0.12796 | 0.898179 | NA |
| SDAD1P1 | 19.84906 | 0.578333 | 0.576975 | 1.002353 | 0.316173 | NA |
| CXorf28 | 0.545988 | -0.43913 | 2.541145 | -0.17281 | 0.862803 | NA |
| TUBB4AP1 | 0.096202 | 0.408546 | 4.079639 | 0.100143 | 0.920231 | NA |
| | 0.163957 | 0.424913 | 4.079006 | 0.104171 | 0.917034 | NA |
| RAB11FIP1 | 17.28015 | 0.649882 | 0.511051 | 1.271657 | 0.203495 | NA |
| | 0.163957 | 0.424913 | 4.079006 | 0.104171 | 0.917034 | NA |
| | 0.163957 | 0.424913 | 4.079006 | 0.104171 | 0.917034 | NA |
| | 14.19837 | 0.10584 | 0.561696 | 0.188429 | 0.85054 | NA |
| | 2.346167 | 3.363693 | 1.662039 | 2.023836 | 0.042987 | NA |
| | 0.488709 | 2.119056 | 3.307564 | 0.64067 | 0.521737 | NA |
| CCDC183-A | 10.34289 | -0.43512 | 0.75485 | -0.57643 | 0.564323 | NA |
| | 2.898546 | -0.65887 | 1.146283 | -0.57479 | 0.565435 | NA |
| | 0.096202 | 0.408546 | 4.079639 | 0.100143 | 0.920231 | NA |
| | 0.536503 | 0.87599 | 3.23231 | 0.27101 | 0.786383 | NA |
| HK2P1 | 8.859976 | -0.82336 | 0.690191 | -1.19295 | 0.232888 | NA |
| ZNF883 | 15.89895 | 0.915564 | 0.523494 | 1.74895 | 0.0803 | NA |
| | 88.45364 | 0.480323 | 0.292262 | 1.643469 | 0.100286 | 0.223392 |
| HNRNPCP7 | 2.16679 | 2.373894 | 1.50404 | 1.578345 | 0.114486 | NA |
| | 0.871935 | 2.99121 | 2.649305 | 1.129055 | 0.258875 | NA |
| | 0.298089 | -0.51711 | 3.975232 | -0.13008 | 0.896501 | NA |
| | 0.326333 | 1.533889 | 3.826878 | 0.40082 | 0.688553 | NA |
| | 0.162376 | 0.424913 | 4.079006 | 0.104171 | 0.917034 | NA |
| | 0.096202 | 0.408546 | 4.079639 | 0.100143 | 0.920231 | NA |

| | | | | | | |
|---|---|---|---|---|---|---|
| WEE2-AS1 | 4.165186 | 0.50166 | 0.994086 | 0.504644 | 0.613809 | NA |
| NLGN4Y-AS | 0.966942 | 0.917791 | 1.915974 | 0.47902 | 0.631924 | NA |
| | 28.62882 | -0.11633 | 0.400813 | -0.29023 | 0.771642 | NA |
| ATP5G2P4 | 1.182873 | 0.724533 | 2.160891 | 0.335294 | 0.737403 | NA |
| | 0.392857 | -1.546 | 4.075469 | -0.37934 | 0.704433 | NA |
| | 8.073391 | -0.48023 | 0.768779 | -0.62467 | 0.532186 | NA |
| | 0.163957 | 0.424913 | 4.079006 | 0.104171 | 0.917034 | NA |
| RASA4CP | 16.59955 | -0.03709 | 0.508884 | -0.07288 | 0.9419 | NA |
| | 1.462535 | 0.94103 | 1.806888 | 0.520801 | 0.602505 | NA |
| | 0.399595 | -1.31595 | 3.579466 | -0.36764 | 0.713143 | NA |
| OR2A9P | 3.163862 | -0.27412 | 1.106794 | -0.24767 | 0.80439 | NA |
| | 0.46735 | -1.30193 | 3.399671 | -0.38296 | 0.701751 | NA |
| | 6.807783 | -0.42579 | 0.768516 | -0.55405 | 0.579547 | NA |
| | 71.56834 | -0.26817 | 0.277358 | -0.96688 | 0.333602 | 0.50214 |
| PSMA6P2 | 0.364467 | -1.20166 | 3.685431 | -0.32606 | 0.744381 | NA |
| HSPA8P8 | 1.625261 | -0.02536 | 1.631497 | -0.01554 | 0.987599 | NA |
| | 3.753788 | 0.787275 | 1.097426 | 0.717384 | 0.473137 | NA |
| EIF4A1P10 | 7.003318 | 0.039541 | 0.7505 | 0.052687 | 0.957982 | NA |
| CCDC26 | 4.50375 | 0.240514 | 0.943298 | 0.254971 | 0.798746 | NA |
| | 0.506476 | 0.726418 | 2.864186 | 0.253621 | 0.799788 | NA |
| EPHA1-AS1 | 0.096202 | 0.408546 | 4.079639 | 0.100143 | 0.920231 | NA |
| TCP1P1 | 0.558248 | 0.982076 | 2.804738 | 0.350149 | 0.726227 | NA |
| NAP1L1P2 | 0.096202 | 0.408546 | 4.079639 | 0.100143 | 0.920231 | NA |
| RPL19P20 | 0.422534 | 1.924583 | 3.001758 | 0.641152 | 0.521424 | NA |
| | 0.151697 | -1.546 | 4.075469 | -0.37934 | 0.704433 | NA |
| | 28.37711 | -0.48852 | 0.439744 | -1.11092 | 0.266601 | NA |
| SERPINH1P | 1.713383 | -0.37731 | 1.475815 | -0.25566 | 0.798213 | NA |
| | 0.162376 | 0.424913 | 4.079006 | 0.104171 | 0.917034 | NA |
| LINC00242 | 1.28151 | -2.12613 | 2.235536 | -0.95106 | 0.341574 | NA |
| | 0.722204 | 1.447218 | 2.611582 | 0.554154 | 0.579474 | NA |
| TTTY10 | 0.787002 | 0.596645 | 2.218025 | 0.268998 | 0.787931 | NA |
| HMGN1P37 | 1.007445 | 3.175243 | 2.206758 | 1.438872 | 0.150187 | NA |
| PPP1R12BP | 0.555233 | -0.57009 | 3.296947 | -0.17291 | 0.862719 | NA |
| | 0.392857 | -1.546 | 4.075469 | -0.37934 | 0.704433 | NA |
| HNRNPA1P | 0.613743 | -0.17004 | 2.749758 | -0.06184 | 0.950691 | NA |
| | 0.392857 | -1.546 | 4.075469 | -0.37934 | 0.704433 | NA |
| PES1P2 | 0.303394 | -2.27006 | 4.032451 | -0.56295 | 0.57347 | NA |
| | 39.0412 | -0.14784 | 0.33843 | -0.43683 | 0.662235 | NA |
| | 0.488709 | 2.119056 | 3.307564 | 0.64067 | 0.521737 | NA |
| TUBB8P1 | 1.146882 | 1.181239 | 1.979819 | 0.59664 | 0.550748 | NA |
| | 0.162376 | 0.424913 | 4.079006 | 0.104171 | 0.917034 | NA |
| | 0.134133 | -1.48909 | 4.077584 | -0.36519 | 0.71497 | NA |
| | 0.28583 | -2.21358 | 3.97376 | -0.55705 | 0.577494 | NA |
| GK-IT1 | 0.35636 | 1.709252 | 3.651833 | 0.468053 | 0.639746 | NA |
| | 0.163957 | 0.424913 | 4.079006 | 0.104171 | 0.917034 | NA |
| DPY19L1P1 | 41.18142 | 0.208052 | 0.356108 | 0.58424 | 0.559059 | NA |
| | 0.392857 | -1.546 | 4.075469 | -0.37934 | 0.704433 | NA |

| | | | | | | |
|---|---|---|---|---|---|---|
| HMGB1P16 | 0.163957 | 0.424913 | 4.079006 | 0.104171 | 0.917034 | NA |
| | 3.299161 | -0.78349 | 1.163926 | -0.67315 | 0.500853 | NA |
| RALGAPA1I | 2.486959 | 0.062924 | 1.292567 | 0.048681 | 0.961173 | NA |
| | 0.134133 | -1.48909 | 4.077584 | -0.36519 | 0.71497 | NA |
| PIP5K1P2 | 0.151697 | -1.546 | 4.075469 | -0.37934 | 0.704433 | NA |
| | 7.541203 | 2.707493 | 0.926937 | 2.920904 | 0.00349 | NA |
| ALG13-AS1 | 0.163957 | 0.424913 | 4.079006 | 0.104171 | 0.917034 | NA |
| | 0.162376 | 0.424913 | 4.079006 | 0.104171 | 0.917034 | NA |
| | 0.571659 | -3.00664 | 3.038724 | -0.98944 | 0.322447 | NA |
| | 0.192403 | 0.975392 | 4.060716 | 0.240202 | 0.810174 | NA |
| | 0.411855 | 0.250456 | 3.097618 | 0.080854 | 0.935558 | NA |
| | 0.6882 | 0.151421 | 2.969628 | 0.05099 | 0.959334 | NA |
| | 0.162376 | 0.424913 | 4.079006 | 0.104171 | 0.917034 | NA |
| | 0.913027 | 1.87659 | 2.13519 | 0.878887 | 0.379463 | NA |
| | 1.744641 | 0.799053 | 1.527853 | 0.522991 | 0.600981 | NA |
| | 0.624422 | 1.162588 | 2.733761 | 0.425271 | 0.670639 | NA |
| ISPD-AS1 | 0.78443 | -2.34592 | 2.507874 | -0.93542 | 0.34957 | NA |
| | 0.449786 | -1.25325 | 3.00474 | -0.41709 | 0.676613 | NA |
| | 0.096202 | 0.408546 | 4.079639 | 0.100143 | 0.920231 | NA |
| | 63.34428 | 0.364345 | 0.293954 | 1.239462 | 0.215174 | 0.374251 |
| | 0.163957 | 0.424913 | 4.079006 | 0.104171 | 0.917034 | NA |
| RPL35AP21 | 0.420954 | 1.923376 | 3.462803 | 0.555439 | 0.578594 | NA |
| | 0.476449 | 0.524262 | 3.402674 | 0.154073 | 0.877552 | NA |
| | 0.258578 | 1.233151 | 4.047809 | 0.304646 | 0.760635 | NA |
| KHSRPP1 | 0.908875 | -0.75101 | 2.116968 | -0.35476 | 0.722771 | NA |
| | 0.46577 | -1.30191 | 3.402879 | -0.38259 | 0.702024 | NA |
| SEPT7P7 | 2.479026 | -0.75316 | 1.234838 | -0.60993 | 0.541908 | NA |
| KIF25-AS1 | 0.612365 | -1.10341 | 2.696105 | -0.40926 | 0.682349 | NA |
| | 18.8578 | -0.10888 | 0.515029 | -0.21141 | 0.832566 | NA |
| | 0.162376 | 0.424913 | 4.079006 | 0.104171 | 0.917034 | NA |
| | 0.151697 | -1.546 | 4.075469 | -0.37934 | 0.704433 | NA |
| | 0.364467 | -1.20166 | 3.685431 | -0.32606 | 0.744381 | NA |
| | 0.392857 | -1.546 | 4.075469 | -0.37934 | 0.704433 | NA |
| | 0.47803 | 0.520083 | 2.97881 | 0.174594 | 0.861399 | NA |
| AK3P3 | 4.4957 | 0.686172 | 0.933675 | 0.734915 | 0.462391 | NA |
| | 0.653015 | 0.2552 | 2.787103 | 0.091565 | 0.927044 | NA |
| | 22.75637 | -0.21708 | 0.448113 | -0.48443 | 0.628083 | NA |
| | 4.856307 | -0.49532 | 0.89112 | -0.55584 | 0.578319 | NA |
| | 0.162376 | 0.424913 | 4.079006 | 0.104171 | 0.917034 | NA |
| | 0.690946 | -1.56103 | 2.699004 | -0.57837 | 0.563013 | NA |
| SEPHS1P4 | 0.35636 | 1.709252 | 3.651833 | 0.468053 | 0.639746 | NA |
| | 0.151697 | -1.546 | 4.075469 | -0.37934 | 0.704433 | NA |
| HNRNPA1P | 0.162376 | 0.424913 | 4.079006 | 0.104171 | 0.917034 | NA |
| C9orf147 | 2.250126 | -0.2245 | 1.366657 | -0.16427 | 0.869516 | NA |
| | 0.163957 | 0.424913 | 4.079006 | 0.104171 | 0.917034 | NA |
| | 2.40988 | 1.102715 | 1.472314 | 0.748967 | 0.453877 | NA |
| | 0.096202 | 0.408546 | 4.079639 | 0.100143 | 0.920231 | NA |

| | | | | | | |
|---|---|---|---|---|---|---|
| | 0.247898 | -0.5901 | 4.074633 | -0.14482 | 0.884851 | NA |
| | 0.096202 | 0.408546 | 4.079639 | 0.100143 | 0.920231 | NA |
| RPL4P5 | 7.86189 | -0.33074 | 0.777887 | -0.42517 | 0.670712 | NA |
| HSPB1P2 | 3.282082 | 0.496936 | 1.250864 | 0.397274 | 0.691165 | NA |
| MTND5P14 | 0.163957 | 0.424913 | 4.079006 | 0.104171 | 0.917034 | NA |
| MIRLET7DH | 9.621168 | 1.988758 | 0.785867 | 2.530656 | 0.011385 | NA |
| | 0.402398 | -2.58771 | 4.018369 | -0.64397 | 0.519595 | NA |
| | 1.587715 | -0.76707 | 1.873519 | -0.40943 | 0.682226 | NA |
| | 0.163957 | 0.424913 | 4.079006 | 0.104171 | 0.917034 | NA |
| | 0.526843 | -0.38719 | 2.855892 | -0.13558 | 0.892156 | NA |
| | 1.467029 | -0.42631 | 1.931736 | -0.22069 | 0.825335 | NA |
| FEZF1-AS1 | 0.151697 | -1.546 | 4.075469 | -0.37934 | 0.704433 | NA |
| | 0.460466 | 0.568655 | 3.014642 | 0.188631 | 0.850382 | NA |
| NCAPD2P1 | 0.544407 | -0.44425 | 2.542878 | -0.1747 | 0.861313 | NA |
| | 2.794082 | -0.70759 | 1.295733 | -0.54609 | 0.585002 | NA |
| TPI1P2 | 10.23325 | 0.073946 | 0.65923 | 0.11217 | 0.910688 | NA |
| DDX10P2 | 0.096202 | 0.408546 | 4.079639 | 0.100143 | 0.920231 | NA |
| | 2.128013 | 0.763944 | 1.355313 | 0.563666 | 0.572981 | NA |
| TFAMP1 | 0.151697 | -1.546 | 4.075469 | -0.37934 | 0.704433 | NA |
| ANKRD18B | 11.9382 | 0.481855 | 0.681017 | 0.707553 | 0.479223 | NA |
| PSMG3-AS1 | 394.427 | -0.34012 | 0.164382 | -2.06908 | 0.038538 | 0.117236 |
| | 0.151697 | -1.546 | 4.075469 | -0.37934 | 0.704433 | NA |
| RPL19P21 | 0.3441 | -0.0166 | 3.769682 | -0.0044 | 0.996486 | NA |
| | 0.096202 | 0.408546 | 4.079639 | 0.100143 | 0.920231 | NA |
| | 1.642116 | 1.248619 | 1.784943 | 0.699529 | 0.484222 | NA |
| | 1.534435 | 1.29036 | 1.713176 | 0.753198 | 0.451331 | NA |
| GTF2IRD1P | 4.233107 | -0.97103 | 0.993322 | -0.97755 | 0.328295 | NA |
| FTX | 18.01475 | 0.652848 | 0.517689 | 1.261081 | 0.20728 | NA |
| RPSAP8 | 0.768215 | 1.51173 | 2.494681 | 0.605981 | 0.544527 | NA |
| | 0.392711 | 0.292538 | 3.145221 | 0.09301 | 0.925895 | NA |
| | 9.114956 | -0.18327 | 0.672182 | -0.27265 | 0.785121 | NA |
| RPS23P8 | 119.0942 | 0.300546 | 0.254743 | 1.179802 | 0.238079 | 0.401059 |
| KLHL7-AS1 | 10.34324 | -0.06077 | 0.613747 | -0.09901 | 0.921127 | NA |
| PRKRIRP1 | 0.708364 | 0.219301 | 2.144998 | 0.102238 | 0.918568 | NA |
| | 0.151697 | -1.546 | 4.075469 | -0.37934 | 0.704433 | NA |
| PABPC1P3 | 0.965158 | 1.861755 | 2.440023 | 0.763007 | 0.445459 | NA |
| | 0.449786 | -1.25325 | 3.00474 | -0.41709 | 0.676613 | NA |
| | 0.163957 | 0.424913 | 4.079006 | 0.104171 | 0.917034 | NA |
| | 5.941926 | -0.94213 | 0.851251 | -1.10676 | 0.268396 | NA |
| | 23.92516 | 0.141191 | 0.528709 | 0.267049 | 0.789432 | NA |
| | 368.3144 | -0.75286 | 0.17237 | -4.3677 | 1.26E-05 | 7.08E-04 |
| RPL10P3 | 1.211263 | 0.702208 | 2.02268 | 0.347167 | 0.728466 | NA |
| | 0.748867 | 2.740193 | 2.468928 | 1.109871 | 0.267054 | NA |
| | 0.303394 | -2.27006 | 4.032451 | -0.56295 | 0.57347 | NA |
| YY2 | 29.5446 | 0.572562 | 0.456615 | 1.253927 | 0.209869 | NA |
| | 0.134133 | -1.48909 | 4.077584 | -0.36519 | 0.71497 | NA |
| CBX1P1 | 0.296509 | -0.51709 | 3.982155 | -0.12985 | 0.896684 | NA |

| | | | | | | |
|---|---|---|---|---|---|---|
| | 0.954682 | 0.055803 | 2.069758 | 0.026961 | 0.978491 | NA |
| | 0.163957 | 0.424913 | 4.079006 | 0.104171 | 0.917034 | NA |
| ZNF674-AS | 46.00872 | 0.009657 | 0.317539 | 0.030412 | 0.975738 | NA |
| | 0.392857 | -1.546 | 4.075469 | -0.37934 | 0.704433 | NA |
| | 0.258578 | 1.233151 | 4.047809 | 0.304646 | 0.760635 | NA |
| | 0.460466 | 0.568655 | 3.014642 | 0.188631 | 0.850382 | NA |
| | 0.763859 | -0.6945 | 2.307767 | -0.30094 | 0.76346 | NA |
| | 0.392857 | -1.546 | 4.075469 | -0.37934 | 0.704433 | NA |
| | 0.640755 | -1.64826 | 2.754778 | -0.59833 | 0.549621 | NA |
| | 0.696104 | -0.93034 | 2.354268 | -0.39517 | 0.692715 | NA |
| RPL7P32 | 1.827149 | -0.19241 | 1.433576 | -0.13422 | 0.893232 | NA |
| CDC20P1 | 2.617029 | -1.97566 | 1.296179 | -1.52422 | 0.127454 | NA |
| | 0.162376 | 0.424913 | 4.079006 | 0.104171 | 0.917034 | NA |
| NPM1P9 | 3.149744 | 0.9315 | 1.233268 | 0.75531 | 0.450063 | NA |
| | 0.315653 | -0.57305 | 3.892639 | -0.14721 | 0.882963 | NA |
| MORF4L2-/ | 0.738188 | 1.39874 | 2.317793 | 0.603479 | 0.54619 | NA |
| TRBV26OR! | 0.899333 | -0.26387 | 2.222941 | -0.1187 | 0.905512 | NA |
| YBX1P2 | 8.384506 | 0.294773 | 0.770412 | 0.382618 | 0.702003 | NA |
| | 2.265949 | 3.273852 | 1.662479 | 1.96926 | 0.048923 | NA |
| HNRNPA1P | 1.083657 | -0.53499 | 1.932845 | -0.27679 | 0.781943 | NA |
| | 0.392857 | -1.546 | 4.075469 | -0.37934 | 0.704433 | NA |
| SKP1P1 | 3.05677 | -1.41508 | 1.254334 | -1.12815 | 0.259256 | NA |
| | 0.134133 | -1.48909 | 4.077584 | -0.36519 | 0.71497 | NA |
| | 0.49029 | 2.123092 | 3.304282 | 0.642527 | 0.520531 | NA |
| | 0.096202 | 0.408546 | 4.079639 | 0.100143 | 0.920231 | NA |
| ACTG1P10 | 3.203213 | 1.626988 | 1.197725 | 1.358399 | 0.174337 | NA |
| VDAC1P6 | 0.162376 | 0.424913 | 4.079006 | 0.104171 | 0.917034 | NA |
| | 0.134133 | -1.48909 | 4.077584 | -0.36519 | 0.71497 | NA |
| | 0.192403 | 0.975392 | 4.060716 | 0.240202 | 0.810174 | NA |
| | 0.28583 | -2.21358 | 3.97376 | -0.55705 | 0.577494 | NA |
| LINC00032 | 0.163957 | 0.424913 | 4.079006 | 0.104171 | 0.917034 | NA |
| RPL7L1P2 | 0.134133 | -1.48909 | 4.077584 | -0.36519 | 0.71497 | NA |
| | 0.354779 | 1.703954 | 3.657232 | 0.465914 | 0.641277 | NA |
| FAM58CP | 0.258578 | 1.233151 | 4.047809 | 0.304646 | 0.760635 | NA |
| | 1.023632 | 2.010953 | 2.294312 | 0.876495 | 0.380761 | NA |
| | 0.096202 | 0.408546 | 4.079639 | 0.100143 | 0.920231 | NA |
| | 0.424115 | 1.932531 | 3.45462 | 0.559405 | 0.575885 | NA |
| HINT1P1 | 1.185162 | -1.08022 | 1.836234 | -0.58828 | 0.556345 | NA |
| LINC00278 | 3.004892 | 1.163576 | 1.242345 | 0.936596 | 0.348966 | NA |
| USMG5P1 | 1.476376 | 1.646496 | 1.822583 | 0.903386 | 0.366321 | NA |
| | 0.230334 | -0.53318 | 4.076749 | -0.13079 | 0.895945 | NA |
| | 1.178424 | -1.58299 | 2.140626 | -0.7395 | 0.459604 | NA |
| | 0.134133 | -1.48909 | 4.077584 | -0.36519 | 0.71497 | NA |
| LINC00574 | 0.151697 | -1.546 | 4.075469 | -0.37934 | 0.704433 | NA |
| NANOGP5 | 0.134133 | -1.48909 | 4.077584 | -0.36519 | 0.71497 | NA |
| | 0.517155 | 2.232561 | 3.179631 | 0.702145 | 0.482589 | NA |
| LINC-PINT | 154.2776 | 0.916315 | 0.309875 | 2.957044 | 0.003106 | 0.021663 |

| | | | | | | |
|---|---|---|---|---|---|---|
| | 7.256456 | -0.55103 | 0.743318 | -0.74131 | 0.458505 | NA |
| ARHGEF9-I | 0.162376 | 0.424913 | 4.079006 | 0.104171 | 0.917034 | NA |
| | 0.151697 | -1.546 | 4.075469 | -0.37934 | 0.704433 | NA |
| DLX6-AS1 | 0.096202 | 0.408546 | 4.079639 | 0.100143 | 0.920231 | NA |
| | 0.327913 | 1.543154 | 4.021063 | 0.383768 | 0.701151 | NA |
| | 1.406049 | -0.26601 | 1.627656 | -0.16343 | 0.870178 | NA |
| | 1.751168 | -0.10773 | 1.535374 | -0.07016 | 0.944064 | NA |
| PCAT7 | 2.682015 | -0.33005 | 1.319979 | -0.25005 | 0.802552 | NA |
| | 0.163957 | 0.424913 | 4.079006 | 0.104171 | 0.917034 | NA |
| MTHFD1P1 | 3.033121 | -1.22881 | 1.154648 | -1.06423 | 0.287226 | NA |
| | 0.582339 | -1.69412 | 2.750801 | -0.61586 | 0.537985 | NA |
| MORF4L1P | 0.096202 | 0.408546 | 4.079639 | 0.100143 | 0.920231 | NA |
| | 0.096202 | 0.408546 | 4.079639 | 0.100143 | 0.920231 | NA |
| | 0.163957 | 0.424913 | 4.079006 | 0.104171 | 0.917034 | NA |
| | 0.298089 | -0.51711 | 3.975232 | -0.13008 | 0.896501 | NA |
| HNRNPA1P | 0.260158 | 1.240347 | 4.047124 | 0.306476 | 0.759242 | NA |
| DPY19L1P2 | 1.703752 | 0.798229 | 1.53059 | 0.521517 | 0.602007 | NA |
| | 0.096202 | 0.408546 | 4.079639 | 0.100143 | 0.920231 | NA |
| | 1.22868 | 1.387311 | 1.789047 | 0.775447 | 0.438075 | NA |
| ANXA2P2 | 121.7565 | -0.06141 | 0.26743 | -0.22962 | 0.818388 | 0.891236 |
| | 0.506476 | 0.726418 | 2.864186 | 0.253621 | 0.799788 | NA |
| FAM27D1 | 0.096202 | 0.408546 | 4.079639 | 0.100143 | 0.920231 | NA |
| | 0.151697 | -1.546 | 4.075469 | -0.37934 | 0.704433 | NA |
| | 8.08142 | 0.71832 | 0.889871 | 0.807218 | 0.419541 | NA |
| LSM12P1 | 1.751721 | -1.66434 | 1.711307 | -0.97256 | 0.330773 | NA |
| | 1.161072 | 0.527093 | 2.031776 | 0.259425 | 0.795308 | NA |
| NPM1P34 | 0.327913 | 1.543154 | 4.021063 | 0.383768 | 0.701151 | NA |
| | 0.776119 | 0.401378 | 2.295521 | 0.174853 | 0.861196 | NA |
| | 2.535716 | -0.30891 | 1.198026 | -0.25785 | 0.796522 | NA |
| RPS6KA2-IT | 4.994838 | 0.945059 | 1.101097 | 0.858289 | 0.390733 | NA |
| | 1.0349 | 0.343148 | 1.852366 | 0.185249 | 0.853034 | NA |
| MCTS1 | 725.4422 | 0.155287 | 0.157823 | 0.983931 | 0.32515 | 0.493048 |
| RAP2C-AS1 | 46.3522 | -0.15058 | 0.331989 | -0.45356 | 0.650145 | NA |
| | 0.096202 | 0.408546 | 4.079639 | 0.100143 | 0.920231 | NA |
| | 0.392857 | -1.546 | 4.075469 | -0.37934 | 0.704433 | NA |
| | 2.327972 | 1.366387 | 1.379451 | 0.990529 | 0.321915 | NA |
| MTND4P24 | 0.151697 | -1.546 | 4.075469 | -0.37934 | 0.704433 | NA |
| | 1.1402 | 1.35289 | 2.18259 | 0.619855 | 0.535353 | NA |
| TOMM22P | 0.327913 | 1.543154 | 4.021063 | 0.383768 | 0.701151 | NA |
| SLC25A6P2 | 0.574231 | 0.93227 | 2.49998 | 0.372911 | 0.709215 | NA |
| PHBP15 | 0.192403 | 0.975392 | 4.060716 | 0.240202 | 0.810174 | NA |
| | 0.786799 | 1.578877 | 2.532268 | 0.623503 | 0.532954 | NA |
| ACTR3P2 | 0.394291 | 0.298745 | 3.141711 | 0.09509 | 0.924243 | NA |
| MTND1P32 | 0.096202 | 0.408546 | 4.079639 | 0.100143 | 0.920231 | NA |
| | 0.804769 | -0.21293 | 2.457871 | -0.08663 | 0.930964 | NA |
| | 0.163957 | 0.424913 | 4.079006 | 0.104171 | 0.917034 | NA |
| LINC00279 | 2.984279 | -0.19974 | 1.249842 | -0.15981 | 0.873029 | NA |

| | | | | | | |
|---|---|---|---|---|---|---|
| FTLP2 | 2.262349 | 0.147942 | 1.309881 | 0.112943 | 0.910076 | NA |
| | 0.151697 | -1.546 | 4.075469 | -0.37934 | 0.704433 | NA |
| | 0.70851 | -1.62753 | 2.686091 | -0.60591 | 0.544574 | NA |
| RPS3AP25 | 0.315653 | -0.57305 | 3.892639 | -0.14721 | 0.882963 | NA |
| SKA2P1 | 0.151697 | -1.546 | 4.075469 | -0.37934 | 0.704433 | NA |
| | 0.268266 | -2.15396 | 4.038351 | -0.53338 | 0.593773 | NA |
| | 0.392857 | -1.546 | 4.075469 | -0.37934 | 0.704433 | NA |
| | 0.134133 | -1.48909 | 4.077584 | -0.36519 | 0.71497 | NA |
| TTTY19 | 0.544554 | -2.65036 | 3.220086 | -0.82307 | 0.410468 | NA |
| C9orf172 | 25.18449 | -0.12172 | 0.487388 | -0.24974 | 0.80279 | NA |
| | 53.36584 | 0.285736 | 0.352952 | 0.80956 | 0.418193 | 0.581245 |
| EEF1B2P3 | 29.76573 | 0.3295 | 0.387961 | 0.849313 | 0.395707 | NA |
| | 0.162376 | 0.424913 | 4.079006 | 0.104171 | 0.917034 | NA |
| RPL3P12 | 0.489058 | -0.58569 | 3.427543 | -0.17088 | 0.864321 | NA |
| | 0.151697 | -1.546 | 4.075469 | -0.37934 | 0.704433 | NA |
| | 21.15383 | 0.316378 | 0.482281 | 0.656004 | 0.511822 | NA |
| | 2.421508 | 0.172314 | 1.33497 | 0.129077 | 0.897297 | NA |
| | 0.702989 | -2.12537 | 2.983149 | -0.71246 | 0.476181 | NA |
| | 172.8359 | 0.069793 | 0.216849 | 0.32185 | 0.747567 | 0.841287 |
| GTF2IP1 | 0.601483 | -1.7332 | 2.729494 | -0.63499 | 0.525436 | NA |
| | 0.162376 | 0.424913 | 4.079006 | 0.104171 | 0.917034 | NA |
| LINC01155 | 5.306664 | -0.23683 | 0.885257 | -0.26753 | 0.789062 | NA |
| | 2.617011 | 0.027855 | 1.343955 | 0.020726 | 0.983464 | NA |
| | 4.75071 | 1.198305 | 0.980837 | 1.221717 | 0.221815 | NA |
| RPL31P35 | 0.162376 | 0.424913 | 4.079006 | 0.104171 | 0.917034 | NA |
| PRPS1P2 | 8.420818 | 0.533631 | 0.74607 | 0.715256 | 0.474451 | NA |
| | 8.123472 | -0.27243 | 0.702742 | -0.38766 | 0.698264 | NA |
| PHBP10 | 0.163957 | 0.424913 | 4.079006 | 0.104171 | 0.917034 | NA |
| | 6.920407 | 0.384583 | 0.82259 | 0.467527 | 0.640123 | NA |
| YWHAEP1 | 0.192403 | 0.975392 | 4.060716 | 0.240202 | 0.810174 | NA |
| | 1.209333 | 2.30346 | 2.022464 | 1.138938 | 0.254729 | NA |
| | 1.273139 | -0.4528 | 1.762969 | -0.25684 | 0.797302 | NA |
| | 0.247898 | -0.5901 | 4.074633 | -0.14482 | 0.884851 | NA |
| HNRNPDP1 | 0.544554 | -2.65036 | 3.220086 | -0.82307 | 0.410468 | NA |
| SDCBPP3 | 2.325692 | -0.88281 | 1.45179 | -0.60809 | 0.54313 | NA |
| LINC00537 | 5.179284 | 0.367142 | 0.930139 | 0.394717 | 0.693052 | NA |
| RPS2P32 | 46.62607 | -0.19962 | 0.329637 | -0.60559 | 0.544787 | NA |
| FAM27E3 | 2.68763 | 0.173123 | 1.336216 | 0.129562 | 0.896913 | NA |
| | 0.28583 | -2.21358 | 3.97376 | -0.55705 | 0.577494 | NA |
| | 2.190036 | -0.42804 | 1.29841 | -0.32966 | 0.741654 | NA |
| | 0.134133 | -1.48909 | 4.077584 | -0.36519 | 0.71497 | NA |
| DDX3YP1 | 0.816972 | 0.936457 | 2.589651 | 0.361615 | 0.71764 | NA |
| | 0.162376 | 0.424913 | 4.079006 | 0.104171 | 0.917034 | NA |
| IPO7P1 | 0.399595 | -1.31595 | 3.579466 | -0.36764 | 0.713143 | NA |
| SNHG15 | 684.1616 | 0.053558 | 0.165239 | 0.324127 | 0.745842 | 0.839962 |
| SNHG7 | 688.0776 | 0.107262 | 0.16853 | 0.636459 | 0.524478 | 0.670401 |
| CRYZP1 | 2.381287 | 1.575004 | 1.420285 | 1.108935 | 0.267458 | NA |

| | | | | | | |
|---|---|---|---|---|---|---|
| CASK-AS1 | 0.327913 | 1.543154 | 4.021063 | 0.383768 | 0.701151 | NA |
| | 0.151697 | -1.546 | 4.075469 | -0.37934 | 0.704433 | NA |
| ZFY-AS1 | 1.790798 | 0.241408 | 1.584876 | 0.15232 | 0.878935 | NA |
| | 0.151697 | -1.546 | 4.075469 | -0.37934 | 0.704433 | NA |
| | 0.096202 | 0.408546 | 4.079639 | 0.100143 | 0.920231 | NA |
| LINC00892 | 0.096202 | 0.408546 | 4.079639 | 0.100143 | 0.920231 | NA |
| | 0.230334 | -0.53318 | 4.076749 | -0.13079 | 0.895945 | NA |
| | 0.324752 | 1.53082 | 4.022025 | 0.380609 | 0.703493 | NA |
| RAB1C | 5.073093 | -1.27464 | 0.929345 | -1.37155 | 0.170204 | NA |
| | 586.7355 | 0.62785 | 0.201261 | 3.119587 | 0.001811 | 0.015146 |
| | 14.47898 | 0.638199 | 0.574469 | 1.110937 | 0.266596 | NA |
| | 0.247898 | -0.5901 | 4.074633 | -0.14482 | 0.884851 | NA |
| RNF224 | 2.675427 | -0.02564 | 1.245063 | -0.02059 | 0.98357 | NA |
| | 8.410626 | -0.08045 | 0.72934 | -0.1103 | 0.912172 | NA |
| HNRNPA1P | 1.036424 | 0.067268 | 2.421139 | 0.027784 | 0.977835 | NA |
| | 0.654246 | 2.541085 | 3.009764 | 0.84428 | 0.398513 | NA |
| | 2.765045 | 0.588123 | 1.284676 | 0.457798 | 0.647097 | NA |
| PNPT1P2 | 0.392857 | -1.546 | 4.075469 | -0.37934 | 0.704433 | NA |
| RPS26P2 | 0.326333 | 1.533889 | 3.826878 | 0.40082 | 0.688553 | NA |
| | 1.927145 | 1.097205 | 1.542711 | 0.711219 | 0.476948 | NA |
| RASA4DP | 0.298089 | -0.51711 | 3.975232 | -0.13008 | 0.896501 | NA |
| | 0.096202 | 0.408546 | 4.079639 | 0.100143 | 0.920231 | NA |
| | 40.38454 | 0.119566 | 0.359107 | 0.332954 | 0.739169 | NA |
| NKAPP1 | 13.02204 | 0.085085 | 0.581418 | 0.14634 | 0.883653 | NA |
| | 0.314073 | -0.57308 | 3.898934 | -0.14698 | 0.883146 | NA |
| | 0.163957 | 0.424913 | 4.079006 | 0.104171 | 0.917034 | NA |
| HOTAIRM1 | 29.05322 | 0.548426 | 0.417014 | 1.315127 | 0.188467 | NA |
| | 0.3441 | -0.0166 | 3.769682 | -0.0044 | 0.996486 | NA |
| | 0.163957 | 0.424913 | 4.079006 | 0.104171 | 0.917034 | NA |
| | 0.288605 | 1.443619 | 4.029058 | 0.358302 | 0.720117 | NA |
| EEF1A1P6 | 26.5886 | -0.30731 | 0.42034 | -0.7311 | 0.464716 | NA |
| | 34.08394 | -0.83899 | 0.526283 | -1.59417 | 0.110897 | NA |
| CHTF8P1 | 0.163957 | 0.424913 | 4.079006 | 0.104171 | 0.917034 | NA |
| | 25.67845 | 0.370205 | 0.412458 | 0.897558 | 0.369421 | NA |
| | 7.037985 | 0.877867 | 0.766688 | 1.145012 | 0.252204 | NA |
| | 2.01505 | -0.19267 | 1.448509 | -0.13301 | 0.894184 | NA |
| | 0.151697 | -1.546 | 4.075469 | -0.37934 | 0.704433 | NA |
| | 0.462046 | 0.589069 | 3.442439 | 0.17112 | 0.86413 | NA |
| | 0.296509 | -0.51709 | 3.982155 | -0.12985 | 0.896684 | NA |
| TTTY18 | 0.46577 | -1.30191 | 3.402879 | -0.38259 | 0.702024 | NA |
| SLC26A4-A | 0.096202 | 0.408546 | 4.079639 | 0.100143 | 0.920231 | NA |
| | 0.134133 | -1.48909 | 4.077584 | -0.36519 | 0.71497 | NA |
| | 0.151697 | -1.546 | 4.075469 | -0.37934 | 0.704433 | NA |
| | 0.163957 | 0.424913 | 4.079006 | 0.104171 | 0.917034 | NA |
| | 1.031536 | 1.037604 | 2.133871 | 0.486254 | 0.626787 | NA |
| | 3.814065 | -0.80794 | 1.157875 | -0.69778 | 0.485314 | NA |
| | 0.58526 | 0.001197 | 3.235889 | 3.70E-04 | 0.999705 | NA |

| | | | | | | |
|---|---|---|---|---|---|---|
| | 7.860041 | 0.125654 | 0.825561 | 0.152204 | 0.879026 | NA |
| | 8.015484 | -0.34199 | 0.744764 | -0.4592 | 0.646093 | NA |
| EIF4HP1 | 5.264077 | -0.45647 | 0.985595 | -0.46314 | 0.643261 | NA |
| | 0.134133 | -1.48909 | 4.077584 | -0.36519 | 0.71497 | NA |
| | 2.297302 | -0.69437 | 1.310331 | -0.52992 | 0.596166 | NA |
| POU6F2-AS | 3.988874 | 1.111876 | 1.075088 | 1.034218 | 0.301034 | NA |
| TTTY15 | 100.0109 | 0.406278 | 0.350833 | 1.15804 | 0.246848 | 0.410366 |
| | 0.162376 | 0.424913 | 4.079006 | 0.104171 | 0.917034 | NA |
| GAPDHP68 | 0.46735 | -1.30193 | 3.399671 | -0.38296 | 0.701751 | NA |
| | 0.58333 | 2.389479 | 3.105916 | 0.769331 | 0.441697 | NA |
| EZR-AS1 | 1.073969 | 1.636961 | 2.132951 | 0.767463 | 0.442806 | NA |
| | 1.959701 | -0.41599 | 1.385214 | -0.30031 | 0.763944 | NA |
| RPS15AP4( | 0.296509 | -0.51709 | 3.982155 | -0.12985 | 0.896684 | NA |
| | 1.669156 | -0.9199 | 1.657633 | -0.55495 | 0.578931 | NA |
| | 0.364467 | -1.20166 | 3.685431 | -0.32606 | 0.744381 | NA |
| | 11.10282 | 0.469748 | 0.609132 | 0.771176 | 0.440602 | NA |
| RPSAP14 | 1.33844 | -1.35576 | 1.767672 | -0.76698 | 0.443096 | NA |
| | 3.001778 | -1.40489 | 1.264377 | -1.11113 | 0.266513 | NA |
| | 6.126679 | -0.36439 | 0.840466 | -0.43355 | 0.664613 | NA |
| | 0.163957 | 0.424913 | 4.079006 | 0.104171 | 0.917034 | NA |
| | 11.77442 | -0.30023 | 0.622745 | -0.48211 | 0.629727 | NA |
| | 2.100719 | -1.09489 | 1.364587 | -0.80236 | 0.422343 | NA |
| | 0.757178 | -0.36061 | 2.51554 | -0.14335 | 0.886013 | NA |
| | 24.9096 | 0.013596 | 0.443972 | 0.030623 | 0.97557 | NA |
| PABPC5-AS | 0.430642 | -1.1891 | 3.485782 | -0.34113 | 0.733008 | NA |
| HSPA8P1 | 12.32245 | 0.630182 | 0.634172 | 0.993708 | 0.320365 | NA |
| | 0.382031 | -1.26484 | 3.151989 | -0.40128 | 0.688212 | NA |
| | 0.162376 | 0.424913 | 4.079006 | 0.104171 | 0.917034 | NA |
| | 0.151697 | -1.546 | 4.075469 | -0.37934 | 0.704433 | NA |
| | 0.556813 | -0.57007 | 3.294615 | -0.17303 | 0.862627 | NA |
| | 7.999487 | 1.401825 | 0.828098 | 1.692825 | 0.090489 | NA |
| ZFX-AS1 | 1.266758 | -0.10165 | 1.859165 | -0.05468 | 0.956395 | NA |
| | 0.613743 | -0.17004 | 2.749758 | -0.06184 | 0.950691 | NA |
| | 0.628146 | -0.21495 | 3.102079 | -0.06929 | 0.944756 | NA |
| CDK2AP2P: | 0.151697 | -1.546 | 4.075469 | -0.37934 | 0.704433 | NA |
| | 270.2359 | 0.861161 | 0.193148 | 4.45855 | 8.25E-06 | 5.54E-04 |
| JAZF1-AS1 | 4.514836 | 0.116672 | 0.927886 | 0.12574 | 0.899938 | NA |
| | 2.784685 | 3.595966 | 1.58167 | 2.273525 | 0.022995 | NA |
| | 0.555233 | -0.57009 | 3.296947 | -0.17291 | 0.862719 | NA |
| | 0.192403 | 0.975392 | 4.060716 | 0.240202 | 0.810174 | NA |
| | 0.096202 | 0.408546 | 4.079639 | 0.100143 | 0.920231 | NA |
| | 0.134133 | -1.48909 | 4.077584 | -0.36519 | 0.71497 | NA |
| SNX18P7 | 1.974184 | 3.044169 | 1.775372 | 1.714666 | 0.086406 | NA |
| USP27X-AS | 14.16399 | -0.0821 | 0.602773 | -0.1362 | 0.891664 | NA |
| | 0.460466 | 0.568655 | 3.014642 | 0.188631 | 0.850382 | NA |
| | 1.272119 | -0.46537 | 1.856946 | -0.25061 | 0.802115 | NA |
| | 0.162376 | 0.424913 | 4.079006 | 0.104171 | 0.917034 | NA |

| | | | | | | |
|---|---|---|---|---|---|---|
| | 4.720594 | -0.41233 | 0.895951 | -0.46021 | 0.645365 | NA |
| ZNF736 | 38.60886 | 0.608461 | 0.386819 | 1.572985 | 0.115722 | NA |
| | 40.50167 | 0.050772 | 0.377024 | 0.134665 | 0.892877 | NA |
| | 0.432222 | -1.18916 | 3.482045 | -0.34151 | 0.732718 | NA |
| MAGI2-AS3 | 806.8085 | 0.679272 | 0.168474 | 4.031911 | 5.53E-05 | 0.001749 |
| | 0.462046 | 0.589069 | 3.442439 | 0.17112 | 0.86413 | NA |
| | 0.096202 | 0.408546 | 4.079639 | 0.100143 | 0.920231 | NA |
| | 0.382031 | -1.26484 | 3.151989 | -0.40128 | 0.688212 | NA |
| | 0.163957 | 0.424913 | 4.079006 | 0.104171 | 0.917034 | NA |
| | 0.585114 | 1.12675 | 2.73598 | 0.411827 | 0.680466 | NA |
| | 1.052464 | 0.310783 | 1.847086 | 0.168256 | 0.866382 | NA |
| | 0.162376 | 0.424913 | 4.079006 | 0.104171 | 0.917034 | NA |
| | 3.951066 | -0.23854 | 1.001482 | -0.23818 | 0.811739 | NA |
| FAM133B | 101.122 | 0.983534 | 0.395526 | 2.486651 | 0.012895 | 0.056183 |
| API5P1 | 0.678686 | -3.08037 | 2.621268 | -1.17514 | 0.239937 | NA |
| COX6B1P1 | 0.52699 | -2.57878 | 3.248478 | -0.79384 | 0.427288 | NA |
| CCT6P3 | 12.36322 | 1.436223 | 0.675611 | 2.125813 | 0.033519 | NA |
| JRK | 1100.086 | -0.63666 | 0.1732 | -3.67586 | 2.37E-04 | 0.004317 |
| RPSAP9 | 6.600773 | 0.413321 | 0.777871 | 0.531349 | 0.595177 | NA |
| MED14-AS1 | 16.87979 | 0.58015 | 0.54841 | 1.057877 | 0.290111 | NA |
| | 0.096202 | 0.408546 | 4.079639 | 0.100143 | 0.920231 | NA |
| | 1.078135 | -1.10729 | 1.970896 | -0.56182 | 0.57424 | NA |
| | 0.327913 | 1.543154 | 4.021063 | 0.383768 | 0.701151 | NA |
| HMGA1P4 | 0.432222 | -1.18916 | 3.482045 | -0.34151 | 0.732718 | NA |
| | 0.28583 | -2.21358 | 3.97376 | -0.55705 | 0.577494 | NA |
| UPF3AP3 | 4.192797 | 0.055947 | 1.116726 | 0.050099 | 0.960043 | NA |
| | 0.162376 | 0.424913 | 4.079006 | 0.104171 | 0.917034 | NA |
| | 0.163957 | 0.424913 | 4.079006 | 0.104171 | 0.917034 | NA |
| | 5.531075 | 0.026924 | 0.929752 | 0.028959 | 0.976898 | NA |
| | 8.535569 | 0.22383 | 0.71284 | 0.313998 | 0.753523 | NA |
| | 40.36157 | -0.11282 | 0.361727 | -0.3119 | 0.755114 | NA |
| TPT1P9 | 15.60001 | 0.794179 | 0.546187 | 1.454042 | 0.145935 | NA |
| EEF1GP5 | 0.640609 | 0.042838 | 2.379511 | 0.018003 | 0.985637 | NA |
| | 2.940979 | -0.70255 | 1.209911 | -0.58066 | 0.56147 | NA |
| PCMTD1P3 | 0.096202 | 0.408546 | 4.079639 | 0.100143 | 0.920231 | NA |
| | 0.754374 | 0.344906 | 2.513743 | 0.137208 | 0.890866 | NA |
| XRCC6P2 | 4.606647 | -0.82423 | 0.975921 | -0.84457 | 0.398353 | NA |
| | 0.327913 | 1.543154 | 4.021063 | 0.383768 | 0.701151 | NA |
| | 0.82213 | 0.50433 | 2.197682 | 0.229483 | 0.818494 | NA |
| | 0.096202 | 0.408546 | 4.079639 | 0.100143 | 0.920231 | NA |
| | 0.544407 | -0.44425 | 2.542878 | -0.1747 | 0.861313 | NA |
| EIF4A1P2 | 6.298966 | -0.5764 | 0.816023 | -0.70635 | 0.479972 | NA |
| | 0.588072 | 2.39955 | 3.098804 | 0.774347 | 0.438726 | NA |
| | 0.706784 | 0.225544 | 2.334352 | 0.096619 | 0.923029 | NA |
| | 0.096202 | 0.408546 | 4.079639 | 0.100143 | 0.920231 | NA |
| FNDC1-IT1 | 1.400156 | 2.55627 | 1.88787 | 1.35405 | 0.17572 | NA |
| RPL7L1P3 | 0.462046 | 0.589069 | 3.442439 | 0.17112 | 0.86413 | NA |

| | | | | | | |
|---|---|---|---|---|---|---|
| LINC00094 | 611.0776 | -0.21552 | 0.148064 | -1.45558 | 0.145508 | 0.287692 |
| | 0.684273 | 2.627785 | 2.896325 | 0.907282 | 0.364257 | NA |
| | 1.861919 | -0.61536 | 1.581414 | -0.38912 | 0.697186 | NA |
| FAM203A | 47.38192 | -0.37235 | 0.325987 | -1.14223 | 0.253359 | 0.417475 |
| RPL26P37 | 0.574434 | -0.17781 | 2.756096 | -0.06452 | 0.948559 | NA |
| | 2.529788 | -0.00939 | 1.403175 | -0.0067 | 0.994658 | NA |
| TUBBP6 | 0.163957 | 0.424913 | 4.079006 | 0.104171 | 0.917034 | NA |
| TMSB10P2 | 0.151697 | -1.546 | 4.075469 | -0.37934 | 0.704433 | NA |
| | 2.721031 | 0.567482 | 1.175179 | 0.48289 | 0.629174 | NA |
| TMEM185/ | 22.25215 | -0.33764 | 0.431607 | -0.78229 | 0.434042 | NA |
| KDM5C-IT1 | 0.096202 | 0.408546 | 4.079639 | 0.100143 | 0.920231 | NA |
| FAM103A2 | 21.26373 | 0.157027 | 0.472713 | 0.332183 | 0.739751 | NA |
| | 0.096202 | 0.408546 | 4.079639 | 0.100143 | 0.920231 | NA |
| LINC00957 | 87.15327 | -0.63764 | 0.248088 | -2.57023 | 0.010163 | 0.047698 |
| | 0.3441 | -0.0166 | 3.769682 | -0.0044 | 0.996486 | NA |
| | 0.134133 | -1.48909 | 4.077584 | -0.36519 | 0.71497 | NA |
| | 0.298089 | -0.51711 | 3.975232 | -0.13008 | 0.896501 | NA |
| | 2.449402 | 1.503751 | 1.490012 | 1.009221 | 0.312869 | NA |
| | 10.28122 | 0.085154 | 0.642814 | 0.132471 | 0.894612 | NA |
| | 0.326536 | 0.037056 | 3.842554 | 0.009644 | 0.992306 | NA |
| | 0.151697 | -1.546 | 4.075469 | -0.37934 | 0.704433 | NA |
| | 0.096202 | 0.408546 | 4.079639 | 0.100143 | 0.920231 | NA |
| | 0.162376 | 0.424913 | 4.079006 | 0.104171 | 0.917034 | NA |
| DPY19L2P4 | 2.829667 | 1.857006 | 1.390922 | 1.33509 | 0.181847 | NA |
| | 53.16222 | -0.1454 | 0.373868 | -0.38891 | 0.697342 | 0.805043 |
| | 46.89179 | 1.174317 | 0.36267 | 3.237974 | 0.001204 | NA |
| TOPORS-AS | 46.26207 | 0.043563 | 0.339662 | 0.128255 | 0.897947 | NA |
| | 1.34683 | 0.987709 | 2.09212 | 0.472109 | 0.636849 | NA |
| RPS26P31 | 0.967145 | 0.230519 | 2.075373 | 0.111074 | 0.911558 | NA |
| | 2.057097 | 0.680253 | 1.525887 | 0.445808 | 0.655736 | NA |
| | 288.4057 | 0.220325 | 0.280311 | 0.786003 | 0.431866 | 0.593115 |
| | 0.151697 | -1.546 | 4.075469 | -0.37934 | 0.704433 | NA |
| GAPDHP65 | 5.143522 | 0.00576 | 0.95616 | 0.006024 | 0.995194 | NA |
| RPL36AP33 | 0.096202 | 0.408546 | 4.079639 | 0.100143 | 0.920231 | NA |
| KRT18P66 | 0.192403 | 0.975392 | 4.060716 | 0.240202 | 0.810174 | NA |
| RNF148 | 1.759394 | -0.27571 | 1.458245 | -0.18907 | 0.85004 | NA |
| NUS1P1 | 5.527723 | 1.373485 | 1.022418 | 1.343369 | 0.179153 | NA |
| LINC00484 | 0.586491 | 2.39072 | 2.689057 | 0.889055 | 0.373973 | NA |
| | 3.020913 | 1.099163 | 1.244456 | 0.883248 | 0.377102 | NA |
| | 0.420954 | 1.923376 | 3.462803 | 0.555439 | 0.578594 | NA |
| LINC00894 | 1.790652 | 0.835355 | 1.504443 | 0.555259 | 0.578718 | NA |
| | 0.134133 | -1.48909 | 4.077584 | -0.36519 | 0.71497 | NA |
| MAGI2-IT1 | 0.72077 | 0.527966 | 3.060238 | 0.172524 | 0.863025 | NA |
| | 0.58526 | 0.001197 | 3.235889 | 3.70E-04 | 0.999705 | NA |
| HCFC1-AS1 | 1.282741 | -0.13551 | 1.82613 | -0.07421 | 0.940846 | NA |
| | 4.924109 | 1.037456 | 1.238594 | 0.837608 | 0.402251 | NA |
| | 2.087307 | 0.208675 | 1.354452 | 0.154066 | 0.877558 | NA |

| | | | | | | |
|---|---|---|---|---|---|---|
| CTBP2P1 | 0.163957 | 0.424913 | 4.079006 | 0.104171 | 0.917034 | NA |
| | 29.18887 | 0.611175 | 0.540273 | 1.131233 | 0.257957 | NA |
| GSN-AS1 | 114.3079 | 0.243565 | 0.27902 | 0.872933 | 0.382699 | 0.548751 |
| PKMP2 | 0.163957 | 0.424913 | 4.079006 | 0.104171 | 0.917034 | NA |
| | 0.804712 | -0.7622 | 2.348804 | -0.3245 | 0.745556 | NA |
| | 0.134133 | -1.48909 | 4.077584 | -0.36519 | 0.71497 | NA |
| ZNF815P | 77.94937 | -0.15686 | 0.259401 | -0.6047 | 0.545377 | 0.686936 |
| | 0.364467 | -1.20166 | 3.685431 | -0.32606 | 0.744381 | NA |
| PNMA6C | 10.56548 | -0.18686 | 0.658612 | -0.28372 | 0.776629 | NA |
| | 3.757337 | 1.523724 | 1.118873 | 1.361839 | 0.173249 | NA |
| | 0.709944 | 0.232855 | 2.332364 | 0.099837 | 0.920474 | NA |
| | 0.449786 | -1.25325 | 3.00474 | -0.41709 | 0.676613 | NA |
| HSPB1P1 | 2.773941 | 1.406875 | 1.409514 | 0.998128 | 0.318217 | NA |
| | 1.045523 | 1.534279 | 2.413816 | 0.635624 | 0.525022 | NA |
| OR13E1P | 0.981934 | -1.6022 | 2.093932 | -0.76516 | 0.444173 | NA |
| | 0.151697 | -1.546 | 4.075469 | -0.37934 | 0.704433 | NA |
| | 0.247898 | -0.5901 | 4.074633 | -0.14482 | 0.884851 | NA |
| CHCHD4P3 | 6.314623 | 0.472697 | 0.820917 | 0.575816 | 0.56474 | NA |
| | 0.258578 | 1.233151 | 4.047809 | 0.304646 | 0.760635 | NA |
| TCEA1P4 | 2.937737 | 0.728452 | 1.212779 | 0.600647 | 0.548075 | NA |
| RPL18AP15 | 0.606787 | -3.06078 | 3.567758 | -0.8579 | 0.390947 | NA |
| RPS15AP25 | 0.163957 | 0.424913 | 4.079006 | 0.104171 | 0.917034 | NA |
| | 13.70922 | 0.508265 | 0.568525 | 0.894007 | 0.371318 | NA |
| | 1.799954 | 0.920797 | 1.509688 | 0.609926 | 0.541911 | NA |
| | 0.326333 | 1.533889 | 3.826878 | 0.40082 | 0.688553 | NA |
| | 0.163957 | 0.424913 | 4.079006 | 0.104171 | 0.917034 | NA |
| DIAPH2-AS | 0.260158 | 1.240347 | 4.047124 | 0.306476 | 0.759242 | NA |
| CLEC2L | 2.350051 | -0.47986 | 1.320432 | -0.36341 | 0.716295 | NA |
| | 0.555233 | -0.57009 | 3.296947 | -0.17291 | 0.862719 | NA |
| VN1R31P | 0.096202 | 0.408546 | 4.079639 | 0.100143 | 0.920231 | NA |
| NPM1P8 | 0.47803 | 0.520083 | 2.97881 | 0.174594 | 0.861399 | NA |
| | 5.796326 | -0.35428 | 0.893968 | -0.3963 | 0.691884 | NA |
| | 0.192403 | 0.975392 | 4.060716 | 0.240202 | 0.810174 | NA |
| FMR1-IT1 | 1.079513 | -0.47694 | 2.056893 | -0.23187 | 0.816636 | NA |
| | 1.510358 | -1.07898 | 1.603256 | -0.67299 | 0.500953 | NA |
| | 8.251742 | 0.129386 | 0.726678 | 0.178052 | 0.858682 | NA |
| | 1.426619 | -1.16726 | 1.79493 | -0.65031 | 0.515491 | NA |
| | 0.382031 | -1.26484 | 3.151989 | -0.40128 | 0.688212 | NA |
| | 0.247898 | -0.5901 | 4.074633 | -0.14482 | 0.884851 | NA |
| | 26.68142 | -0.28803 | 0.463866 | -0.62094 | 0.534639 | NA |
| | 0.096202 | 0.408546 | 4.079639 | 0.100143 | 0.920231 | NA |
| | 0.134133 | -1.48909 | 4.077584 | -0.36519 | 0.71497 | NA |
| | 66.26061 | -0.1079 | 0.312731 | -0.34501 | 0.730086 | 0.828802 |
| | 0.151697 | -1.546 | 4.075469 | -0.37934 | 0.704433 | NA |
| | 0.134133 | -1.48909 | 4.077584 | -0.36519 | 0.71497 | NA |
| HNRNPA1P | 0.162376 | 0.424913 | 4.079006 | 0.104171 | 0.917034 | NA |
| | 0.298089 | -0.51711 | 3.975232 | -0.13008 | 0.896501 | NA |

| | | | | | | |
|---|---|---|---|---|---|---|
| ZNF853 | 189.3233 | -0.34243 | 0.212461 | -1.61173 | 0.107021 | 0.233524 |
| | 0.096202 | 0.408546 | 4.079639 | 0.100143 | 0.920231 | NA |
| | 1.332933 | 0.019174 | 1.852079 | 0.010353 | 0.99174 | NA |
| | 0.489058 | -0.58569 | 3.427543 | -0.17088 | 0.864321 | NA |
| RPL31P63 | 1.864188 | 1.113015 | 1.68702 | 0.659752 | 0.509413 | NA |
| | 0.478233 | -0.67301 | 2.937473 | -0.22911 | 0.818782 | NA |
| YRDCP2 | 0.162376 | 0.424913 | 4.079006 | 0.104171 | 0.917034 | NA |
| | 0.544407 | -0.44425 | 2.542878 | -0.1747 | 0.861313 | NA |
| | 0.163957 | 0.424913 | 4.079006 | 0.104171 | 0.917034 | NA |
| MKLN1-AS | 82.93023 | -0.04946 | 0.25579 | -0.19336 | 0.846675 | 0.908806 |
| TRMT112P | 0.163957 | 0.424913 | 4.079006 | 0.104171 | 0.917034 | NA |
| | 0.162376 | 0.424913 | 4.079006 | 0.104171 | 0.917034 | NA |
| RPL24P8 | 0.834389 | 1.638864 | 2.19944 | 0.745128 | 0.456195 | NA |
| BTF3P8 | 0.424115 | 1.932531 | 3.45462 | 0.559405 | 0.575885 | NA |
| | 0.781626 | -1.50265 | 2.506115 | -0.59959 | 0.548777 | NA |
| ANKRD20A | 1.388282 | 0.29126 | 1.854762 | 0.157033 | 0.875219 | NA |
| | 0.296509 | -0.51709 | 3.982155 | -0.12985 | 0.896684 | NA |
| | 8.003714 | 0.09384 | 0.75823 | 0.123762 | 0.901504 | NA |
| | 0.162376 | 0.424913 | 4.079006 | 0.104171 | 0.917034 | NA |
| | 1.393832 | 3.667627 | 1.97519 | 1.856848 | 0.063333 | NA |
| RPS20P24 | 0.134133 | -1.48909 | 4.077584 | -0.36519 | 0.71497 | NA |
| | 0.096202 | 0.408546 | 4.079639 | 0.100143 | 0.920231 | NA |
| MIR600HG | 13.61791 | 0.543931 | 0.608805 | 0.893441 | 0.371621 | NA |
| | 0.392711 | 0.292538 | 3.145221 | 0.09301 | 0.925895 | NA |
| | 0.096202 | 0.408546 | 4.079639 | 0.100143 | 0.920231 | NA |
| | 0.151697 | -1.546 | 4.075469 | -0.37934 | 0.704433 | NA |
| C8orf56 | 2.365029 | 0.495343 | 1.414365 | 0.350223 | 0.726172 | NA |
| | 0.776119 | 0.401378 | 2.295521 | 0.174853 | 0.861196 | NA |
| | 0.192403 | 0.975392 | 4.060716 | 0.240202 | 0.810174 | NA |
| ZNRF2P1 | 84.83647 | 0.151602 | 0.331628 | 0.457145 | 0.647567 | 0.766201 |
| | 0.162376 | 0.424913 | 4.079006 | 0.104171 | 0.917034 | NA |
| | 0.58333 | 2.389479 | 3.105916 | 0.769331 | 0.441697 | NA |
| EIF3IP1 | 0.900767 | 0.808531 | 1.963257 | 0.411831 | 0.680463 | NA |
| | 0.394291 | 0.298745 | 3.141711 | 0.09509 | 0.924243 | NA |
| | 1.572423 | 0.748853 | 1.766392 | 0.423945 | 0.671606 | NA |
| FAM27E1 | 0.833012 | 0.681553 | 2.203157 | 0.309353 | 0.757053 | NA |
| IMPDH1P4 | 0.151697 | -1.546 | 4.075469 | -0.37934 | 0.704433 | NA |
| | 0.151697 | -1.546 | 4.075469 | -0.37934 | 0.704433 | NA |
| BMS1P10 | 37.43125 | 0.736623 | 0.388337 | 1.896864 | 0.057846 | NA |
| | 0.096202 | 0.408546 | 4.079639 | 0.100143 | 0.920231 | NA |
| | 0.151697 | -1.546 | 4.075469 | -0.37934 | 0.704433 | NA |
| FTH1P11 | 7.665121 | -0.93734 | 0.762221 | -1.22975 | 0.218793 | NA |
| SRRM1P3 | 0.392857 | -1.546 | 4.075469 | -0.37934 | 0.704433 | NA |
| | 12.18293 | -0.39359 | 0.582317 | -0.67591 | 0.499098 | NA |
| | 0.162376 | 0.424913 | 4.079006 | 0.104171 | 0.917034 | NA |
| XIAP-AS1 | 0.230334 | -0.53318 | 4.076749 | -0.13079 | 0.895945 | NA |
| | 0.096202 | 0.408546 | 4.079639 | 0.100143 | 0.920231 | NA |

| | | | | | | |
|---|---|---|---|---|---|---|
| | 0.096202 | 0.408546 | 4.079639 | 0.100143 | 0.920231 | NA |
| | 0.134133 | -1.48909 | 4.077584 | -0.36519 | 0.71497 | NA |
| | 0.162376 | 0.424913 | 4.079006 | 0.104171 | 0.917034 | NA |
| | 0.588072 | 2.39955 | 3.098804 | 0.774347 | 0.438726 | NA |
| ASS1P12 | 1.870281 | 2.987889 | 1.726464 | 1.730641 | 0.083516 | NA |
| | 0.585114 | 1.12675 | 2.73598 | 0.411827 | 0.680466 | NA |
| RPSAP15 | 0.818406 | 1.679649 | 2.471195 | 0.679691 | 0.4967 | NA |
| | 0.192403 | 0.975392 | 4.060716 | 0.240202 | 0.810174 | NA |
| TTLL11-IT1 | 0.52699 | -2.57878 | 3.248478 | -0.79384 | 0.427288 | NA |
| | 0.163957 | 0.424913 | 4.079006 | 0.104171 | 0.917034 | NA |
| RAC1P4 | 0.096202 | 0.408546 | 4.079639 | 0.100143 | 0.920231 | NA |
| | 0.326333 | 1.533889 | 3.826878 | 0.40082 | 0.688553 | NA |
| | 2.359782 | 1.022712 | 1.327152 | 0.770607 | 0.44094 | NA |
| | 0.151697 | -1.546 | 4.075469 | -0.37934 | 0.704433 | NA |
| | 0.162376 | 0.424913 | 4.079006 | 0.104171 | 0.917034 | NA |
| S100A11P1 | 0.889885 | 0.644651 | 2.1706 | 0.296992 | 0.766472 | NA |
| | 0.870886 | -0.53073 | 2.310896 | -0.22966 | 0.818353 | NA |
| ERICH1-AS: | 0.452561 | 2.061505 | 3.349456 | 0.615474 | 0.538241 | NA |
| | 0.151697 | -1.546 | 4.075469 | -0.37934 | 0.704433 | NA |
| VDAC1P11 | 0.314073 | -0.57308 | 3.898934 | -0.14698 | 0.883146 | NA |
| | 0.162376 | 0.424913 | 4.079006 | 0.104171 | 0.917034 | NA |
| IFNWP2 | 0.162376 | 0.424913 | 4.079006 | 0.104171 | 0.917034 | NA |
| ATP5JP1 | 0.258578 | 1.233151 | 4.047809 | 0.304646 | 0.760635 | NA |
| | 0.192403 | 0.975392 | 4.060716 | 0.240202 | 0.810174 | NA |
| EEF1A1P28 | 0.162376 | 0.424913 | 4.079006 | 0.104171 | 0.917034 | NA |
| | 1.736765 | -0.06189 | 1.634137 | -0.03787 | 0.96979 | NA |
| | 2.811894 | 0.001672 | 1.279261 | 0.001307 | 0.998957 | NA |
| UQCRBP1 | 0.545988 | -0.43913 | 2.541145 | -0.17281 | 0.862803 | NA |
| EEF1A1P30 | 0.544554 | -2.65036 | 3.220086 | -0.82307 | 0.410468 | NA |
| | 2.483024 | -0.2666 | 1.268993 | -0.21009 | 0.8336 | NA |
| | 0.315653 | -0.57305 | 3.892639 | -0.14721 | 0.882963 | NA |
| RPL18AP16 | 0.28583 | -2.21358 | 3.97376 | -0.55705 | 0.577494 | NA |
| | 28.85631 | 0.349435 | 0.39046 | 0.894932 | 0.370823 | NA |
| | 0.314073 | -0.57308 | 3.898934 | -0.14698 | 0.883146 | NA |
| | 0.382031 | -1.26484 | 3.151989 | -0.40128 | 0.688212 | NA |
| | 0.476449 | 0.524262 | 3.402674 | 0.154073 | 0.877552 | NA |
| RPS3AP29 | 0.192403 | 0.975392 | 4.060716 | 0.240202 | 0.810174 | NA |
| | 1.24776 | -1.38472 | 2.303574 | -0.60112 | 0.547762 | NA |
| | 0.735027 | 1.3952 | 2.594798 | 0.537691 | 0.59079 | NA |
| PA2G4P1 | 1.041581 | 0.164062 | 2.078273 | 0.078942 | 0.937079 | NA |
| PHKA2-AS1 | 1.204161 | -0.07212 | 1.765777 | -0.04084 | 0.967423 | NA |
| | 13.36403 | -0.04315 | 0.569234 | -0.07581 | 0.939573 | NA |
| | 1.02422 | -0.42649 | 1.996186 | -0.21365 | 0.83082 | NA |
| | 9.277318 | 1.036764 | 0.810147 | 1.279724 | 0.200642 | NA |
| | 4.36747 | 0.386628 | 1.086787 | 0.355753 | 0.722025 | NA |
| | 0.95764 | 0.792425 | 2.400829 | 0.330063 | 0.741352 | NA |
| | 2.837326 | -0.36989 | 1.195009 | -0.30953 | 0.756919 | NA |

| | | | | | | |
|---|---|---|---|---|---|---|
| | 0.883203 | 0.854847 | 2.135496 | 0.400304 | 0.688933 | NA |
| | 0.162376 | 0.424913 | 4.079006 | 0.104171 | 0.917034 | NA |
| | 0.681461 | 0.491544 | 3.066278 | 0.160307 | 0.87264 | NA |
| | 0.399595 | -1.31595 | 3.579466 | -0.36764 | 0.713143 | NA |
| TUBG1P | 1.629405 | -0.0684 | 1.843273 | -0.03711 | 0.9704 | NA |
| PTENP1 | 25.72352 | 0.057066 | 0.495654 | 0.115132 | 0.908341 | NA |
| | 0.192403 | 0.975392 | 4.060716 | 0.240202 | 0.810174 | NA |
| | 5.459387 | 0.728424 | 0.90388 | 0.805885 | 0.420309 | NA |
| | 0.382031 | -1.26484 | 3.151989 | -0.40128 | 0.688212 | NA |
| | 1.015699 | 0.142451 | 2.181052 | 0.065313 | 0.947925 | NA |
| RPL9P7 | 1.148462 | 1.182044 | 1.944788 | 0.607801 | 0.54332 | NA |
| | 0.35636 | 1.709252 | 3.651833 | 0.468053 | 0.639746 | NA |
| | 0.52699 | -2.57878 | 3.248478 | -0.79384 | 0.427288 | NA |
| | 14.80412 | 0.225018 | 0.610944 | 0.368311 | 0.712641 | NA |
| MID1IP1-A | 20.27643 | -0.18951 | 0.51783 | -0.36598 | 0.714383 | NA |
| RPS2P35 | 0.949524 | -0.003 | 2.040023 | -0.00147 | 0.998826 | NA |
| | 2.489762 | -0.1774 | 1.284373 | -0.13812 | 0.890145 | NA |
| AHCYP2 | 0.096202 | 0.408546 | 4.079639 | 0.100143 | 0.920231 | NA |
| | 0.096202 | 0.408546 | 4.079639 | 0.100143 | 0.920231 | NA |
| | 0.9197 | -1.22346 | 2.227737 | -0.54919 | 0.582872 | NA |
| MPC1L | 0.162376 | 0.424913 | 4.079006 | 0.104171 | 0.917034 | NA |
| | 0.28583 | -2.21358 | 3.97376 | -0.55705 | 0.577494 | NA |
| MKRN4P | 8.337677 | -0.13329 | 0.699469 | -0.19056 | 0.848867 | NA |
| C9orf69 | 1827.629 | -0.14379 | 0.155023 | -0.92754 | 0.353647 | 0.521648 |
| OR7E7P | 1.885208 | -0.67551 | 1.572884 | -0.42947 | 0.66758 | NA |
| | 0.462046 | 0.589069 | 3.442439 | 0.17112 | 0.86413 | NA |
| | 0.151697 | -1.546 | 4.075469 | -0.37934 | 0.704433 | NA |
| IFNWP19 | 0.35636 | 1.709252 | 3.651833 | 0.468053 | 0.639746 | NA |
| | 0.392857 | -1.546 | 4.075469 | -0.37934 | 0.704433 | NA |
| RNU7-171F | 0.556667 | 0.972496 | 2.522454 | 0.385536 | 0.699841 | NA |
| | 0.151697 | -1.546 | 4.075469 | -0.37934 | 0.704433 | NA |
| | 0.163957 | 0.424913 | 4.079006 | 0.104171 | 0.917034 | NA |
| | 0.096202 | 0.408546 | 4.079639 | 0.100143 | 0.920231 | NA |
| | 1.56052 | 0.572883 | 1.609696 | 0.355896 | 0.721919 | NA |
| | 0.163957 | 0.424913 | 4.079006 | 0.104171 | 0.917034 | NA |
| | 0.162376 | 0.424913 | 4.079006 | 0.104171 | 0.917034 | NA |
| | 0.392711 | 0.292538 | 3.145221 | 0.09301 | 0.925895 | NA |
| | 2.364682 | 0.994261 | 1.976082 | 0.503148 | 0.614861 | NA |
| | 0.758758 | -0.35643 | 2.514365 | -0.14176 | 0.887273 | NA |
| RNA5SP24( | 0.163957 | 0.424913 | 4.079006 | 0.104171 | 0.917034 | NA |
| | 0.260158 | 1.240347 | 4.047124 | 0.306476 | 0.759242 | NA |
| RNU6-1218 | 0.910012 | 1.249754 | 2.467015 | 0.506585 | 0.612446 | NA |
| SNORA84 | 0.652869 | 1.292592 | 2.374873 | 0.544278 | 0.58625 | NA |
| RPL5P22 | 0.192403 | 0.975392 | 4.060716 | 0.240202 | 0.810174 | NA |
| RPS20P22 | 1.28379 | 2.393551 | 1.949457 | 1.227804 | 0.219521 | NA |
| | 0.555087 | 0.973802 | 2.808575 | 0.346725 | 0.728798 | NA |
| | 0.842643 | -2.08432 | 2.306153 | -0.90381 | 0.366097 | NA |

| | | | | | | |
|---|---|---|---|---|---|---|
| | 0.668852 | 1.246964 | 2.610519 | 0.477669 | 0.632886 | NA |
| | 1.247614 | -0.06697 | 1.833256 | -0.03653 | 0.97086 | NA |
| | 0.58526 | 0.001197 | 3.235889 | 3.70E-04 | 0.999705 | NA |
| | 6.920996 | 0.782807 | 0.813912 | 0.961783 | 0.336159 | NA |
| RN7SL535F | 1.2271 | 1.384124 | 1.759418 | 0.786694 | 0.431461 | NA |
| | 0.757324 | -2.08271 | 2.587512 | -0.80491 | 0.420873 | NA |
| | 0.151697 | -1.546 | 4.075469 | -0.37934 | 0.704433 | NA |
| | 68.07618 | -0.14123 | 0.30111 | -0.46904 | 0.639039 | 0.759385 |
| | 4.079721 | -0.52748 | 1.051777 | -0.50151 | 0.616013 | NA |
| KMT2E-AS1 | 60.40096 | 0.182561 | 0.327998 | 0.556592 | 0.577806 | 0.712417 |
| RN7SL388F | 0.096202 | 0.408546 | 4.079639 | 0.100143 | 0.920231 | NA |
| | 5.938079 | -1.05701 | 1.018841 | -1.03746 | 0.299522 | NA |
| MTND1P3 | 0.096202 | 0.408546 | 4.079639 | 0.100143 | 0.920231 | NA |
| | 0.422534 | 1.924583 | 3.001758 | 0.641152 | 0.521424 | NA |
| | 18.30029 | -0.06491 | 0.565446 | -0.11479 | 0.908609 | NA |
| IMP3P2 | 0.163957 | 0.424913 | 4.079006 | 0.104171 | 0.917034 | NA |
| MRPS17 | 91.90325 | -0.27343 | 0.264059 | -1.03551 | 0.300433 | 0.467283 |
| | 8.007241 | 0.036684 | 0.808728 | 0.045361 | 0.96382 | NA |
| GET4 | 90.75258 | -0.09198 | 0.270347 | -0.34024 | 0.733678 | 0.831605 |
| RN7SL75P | 0.765054 | 1.506026 | 2.496832 | 0.603175 | 0.546392 | NA |
| PRKAG2-AS | 33.81938 | 0.506364 | 0.385189 | 1.314586 | 0.188649 | NA |
| RPS10P16 | 1.427204 | 1.633674 | 1.942319 | 0.841095 | 0.400295 | NA |
| SNRPGP3 | 0.641986 | 1.117767 | 2.709494 | 0.412537 | 0.679946 | NA |
| | 0.134133 | -1.48909 | 4.077584 | -0.36519 | 0.71497 | NA |
| | 0.260158 | 1.240347 | 4.047124 | 0.306476 | 0.759242 | NA |
| RPS3AP32 | 0.163957 | 0.424913 | 4.079006 | 0.104171 | 0.917034 | NA |
| | 0.326333 | 1.533889 | 3.826878 | 0.40082 | 0.688553 | NA |
| ETF1P2 | 0.945583 | -0.81197 | 2.098385 | -0.38695 | 0.698792 | NA |
| | 0.151697 | -1.546 | 4.075469 | -0.37934 | 0.704433 | NA |
| SMKR1 | 4.694573 | 0.363682 | 0.90671 | 0.401101 | 0.688346 | NA |
| | 0.134133 | -1.48909 | 4.077584 | -0.36519 | 0.71497 | NA |
| COX19 | 219.8178 | 0.142535 | 0.177238 | 0.804199 | 0.421282 | 0.583992 |
| | 0.151697 | -1.546 | 4.075469 | -0.37934 | 0.704433 | NA |
| RN7SL100F | 0.096202 | 0.408546 | 4.079639 | 0.100143 | 0.920231 | NA |
| RN7SL481F | 0.258578 | 1.233151 | 4.047809 | 0.304646 | 0.760635 | NA |
| RPL5P23 | 3.365646 | 1.666963 | 1.161895 | 1.434694 | 0.151374 | NA |
| | 17.9016 | -0.00874 | 0.511295 | -0.01708 | 0.986369 | NA |
| CDKN2B-AS | 5.195009 | -0.36597 | 0.944858 | -0.38733 | 0.698515 | NA |
| | 0.162376 | 0.424913 | 4.079006 | 0.104171 | 0.917034 | NA |
| RPL34P18 | 2.14898 | -0.00904 | 1.618733 | -0.00559 | 0.995542 | NA |
| | 0.096202 | 0.408546 | 4.079639 | 0.100143 | 0.920231 | NA |
| RPL3P9 | 0.303394 | -2.27006 | 4.032451 | -0.56295 | 0.57347 | NA |
| | 0.45509 | -2.71227 | 3.963744 | -0.68427 | 0.493805 | NA |
| | 0.151697 | -1.546 | 4.075469 | -0.37934 | 0.704433 | NA |
| AQP1 | 3.115672 | -0.00972 | 1.119512 | -0.00868 | 0.993073 | NA |
| SPATA31D5 | 0.326333 | 1.533889 | 3.826878 | 0.40082 | 0.688553 | NA |
| | 3.093624 | -2.10235 | 1.286337 | -1.63437 | 0.102181 | NA |

| | | | | | | |
|---|---|---|---|---|---|---|
| PNMA2 | 534.1055 | 0.13643 | 0.177709 | 0.767717 | 0.442656 | 0.602766 |
| LRRD1 | 0.327913 | 1.543154 | 4.021063 | 0.383768 | 0.701151 | NA |
| | 0.096202 | 0.408546 | 4.079639 | 0.100143 | 0.920231 | NA |
| | 4.240874 | -0.26207 | 0.974287 | -0.26899 | 0.78794 | NA |
| | 0.230334 | -0.53318 | 4.076749 | -0.13079 | 0.895945 | NA |
| | 36.42058 | 0.193168 | 0.366237 | 0.527439 | 0.597889 | NA |
| RN7SL521F | 1.131101 | 0.561912 | 1.807251 | 0.310921 | 0.755861 | NA |
| NDUFB2-AS | 7.740072 | 0.039962 | 0.724643 | 0.055147 | 0.956022 | NA |
| RN7SL798F | 0.623191 | -1.58103 | 2.770593 | -0.57065 | 0.568239 | NA |
| | 0.490493 | 0.776209 | 2.895261 | 0.268096 | 0.788625 | NA |
| | 0.83597 | 1.639551 | 2.453777 | 0.668175 | 0.504022 | NA |
| HOXA11-AS | 2.617771 | 2.690663 | 1.443247 | 1.864313 | 0.062278 | NA |
| SEPT7P6 | 0.448206 | -1.2532 | 3.007315 | -0.41672 | 0.676885 | NA |
| | 0.162376 | 0.424913 | 4.079006 | 0.104171 | 0.917034 | NA |
| GVQW1 | 21.19746 | -0.31842 | 0.544492 | -0.5848 | 0.55868 | NA |
| YAE1D1 | 395.5974 | 0.897309 | 0.24884 | 3.605975 | 3.11E-04 | 0.00505 |
| CRCP | 586.7375 | 0.027514 | 0.173655 | 0.158438 | 0.874111 | 0.925069 |
| | 3.034042 | -1.33034 | 1.202033 | -1.10674 | 0.268407 | NA |
| RPL36A | 45.0659 | 0.802776 | 0.321024 | 2.500672 | 0.012396 | NA |
| | 1.034697 | 1.024735 | 1.902596 | 0.538598 | 0.590164 | NA |
| RN7SL505F | 0.163957 | 0.424913 | 4.079006 | 0.104171 | 0.917034 | NA |
| | 1.398961 | 0.94508 | 1.898234 | 0.497873 | 0.618573 | NA |
| ATP5J2 | 2273.132 | 0.317486 | 0.161498 | 1.965883 | 0.049312 | 0.137358 |
| | 2.288866 | 0.982847 | 1.389703 | 0.707236 | 0.47942 | NA |
| | 0.163957 | 0.424913 | 4.079006 | 0.104171 | 0.917034 | NA |
| RPS15AP24 | 4.205589 | -0.44511 | 1.020526 | -0.43616 | 0.66272 | NA |
| PPP1R2P6 | 0.096202 | 0.408546 | 4.079639 | 0.100143 | 0.920231 | NA |
| ARPC1A | 4480.768 | -0.25199 | 0.126936 | -1.98515 | 0.047128 | 0.133662 |
| TMEFF1 | 4.181712 | 0.199132 | 1.249148 | 0.159414 | 0.873342 | NA |
| FABP5P3 | 0.296509 | -0.51709 | 3.982155 | -0.12985 | 0.896684 | NA |
| | 2.511865 | 0.109044 | 1.47757 | 0.073799 | 0.94117 | NA |
| | 5.222019 | -1.1244 | 0.906722 | -1.24007 | 0.214949 | NA |
| LINC00893 | 2.228812 | 0.894975 | 1.42372 | 0.628617 | 0.5296 | NA |
| | 213.5302 | 0.736789 | 0.337878 | 2.180636 | 0.02921 | 0.097148 |
| C8orf58 | 348.99 | -0.19325 | 0.159217 | -1.21376 | 0.224838 | 0.385906 |
| | 6.59446 | 0.805469 | 0.85057 | 0.946975 | 0.343652 | NA |
| | 0.096202 | 0.408546 | 4.079639 | 0.100143 | 0.920231 | NA |
| RPS20P20 | 0.162376 | 0.424913 | 4.079006 | 0.104171 | 0.917034 | NA |
| RN7SL76P | 0.303394 | -2.27006 | 4.032451 | -0.56295 | 0.57347 | NA |
| AKAP2 | 24.29358 | -0.42599 | 0.431555 | -0.98709 | 0.323596 | NA |
| USP27X | 108.3989 | 0.214106 | 0.247901 | 0.863677 | 0.387765 | 0.553708 |
| | 1.186743 | -1.07729 | 1.835809 | -0.58682 | 0.557324 | NA |
| | 0.134133 | -1.48909 | 4.077584 | -0.36519 | 0.71497 | NA |
| RPS26P35 | 0.163957 | 0.424913 | 4.079006 | 0.104171 | 0.917034 | NA |
| RN7SL306F | 0.163957 | 0.424913 | 4.079006 | 0.104171 | 0.917034 | NA |
| | 72.80187 | 0.126037 | 0.28333 | 0.444841 | 0.656435 | 0.77328 |
| PEG10 | 947.86 | -0.44291 | 0.296851 | -1.49201 | 0.135695 | 0.274564 |

| | | | | | | |
|---|---|---|---|---|---|---|
| STAG3L5P | 9.455454 | -0.06882 | 0.739094 | -0.09312 | 0.925809 | NA |
| UPK3BP1 | 0.096202 | 0.408546 | 4.079639 | 0.100143 | 0.920231 | NA |
| | 11.74804 | 0.251174 | 0.66694 | 0.376607 | 0.706466 | NA |
| DCAF13P1 | 0.508056 | 0.731032 | 2.861912 | 0.255435 | 0.798387 | NA |
| | 26.68914 | -0.13698 | 0.520998 | -0.26292 | 0.792613 | NA |
| RPS3AP34 | 0.845069 | 2.925226 | 2.326646 | 1.257272 | 0.208655 | NA |
| GNG10 | 83.84676 | -0.01295 | 0.272906 | -0.04745 | 0.962154 | 0.97942 |
| | 42.99288 | -1.12356 | 0.337285 | -3.33117 | 8.65E-04 | NA |
| CCT4P1 | 0.134133 | -1.48909 | 4.077584 | -0.36519 | 0.71497 | NA |
| RN7SL806F | 0.096202 | 0.408546 | 4.079639 | 0.100143 | 0.920231 | NA |
| FAM86LP | 0.096202 | 0.408546 | 4.079639 | 0.100143 | 0.920231 | NA |
| RGAG4 | 80.87342 | -0.49726 | 0.374471 | -1.32791 | 0.184207 | 0.337524 |
| | 7.070501 | -0.70808 | 0.893812 | -0.79221 | 0.428241 | NA |
| AP5Z1 | 2513.014 | -0.24482 | 0.140728 | -1.73968 | 0.081915 | 0.194416 |
| EPS15P1 | 0.096202 | 0.408546 | 4.079639 | 0.100143 | 0.920231 | NA |
| ERVW-1 | 0.720827 | 0.448488 | 2.540859 | 0.17651 | 0.859893 | NA |
| | 1.281364 | -0.71591 | 1.774413 | -0.40346 | 0.686607 | NA |
| | 0.296509 | -0.51709 | 3.982155 | -0.12985 | 0.896684 | NA |
| GK-AS1 | 4.855397 | 1.055137 | 0.946518 | 1.114756 | 0.264955 | NA |
| | 0.586491 | 2.39072 | 2.689057 | 0.889055 | 0.373973 | NA |
| RN7SL849F | 0.192403 | 0.975392 | 4.060716 | 0.240202 | 0.810174 | NA |
| | 0.67854 | -0.89371 | 2.366391 | -0.37767 | 0.705677 | NA |
| | 3.442004 | 0.261975 | 1.062754 | 0.246505 | 0.805291 | NA |
| | 0.163957 | 0.424913 | 4.079006 | 0.104171 | 0.917034 | NA |
| | 0.162376 | 0.424913 | 4.079006 | 0.104171 | 0.917034 | NA |
| | 1.080707 | 1.085144 | 1.993395 | 0.54437 | 0.586187 | NA |
| | 1.460595 | 1.079353 | 1.826252 | 0.591021 | 0.554506 | NA |
| PRAF2 | 938.5056 | -0.17424 | 0.151457 | -1.15045 | 0.249959 | 0.41369 |
| | 22.26316 | -0.11206 | 0.489857 | -0.22875 | 0.819061 | NA |
| C7orf73 | 924.0087 | -0.02921 | 0.141102 | -0.20698 | 0.836026 | 0.901643 |
| KCTD7 | 1256.536 | 0.037306 | 0.188683 | 0.197716 | 0.843267 | 0.906757 |
| | 0.096202 | 0.408546 | 4.079639 | 0.100143 | 0.920231 | NA |
| | 1.236585 | 0.682769 | 1.964737 | 0.347512 | 0.728207 | NA |
| PALM2 | 26.78622 | 1.217598 | 0.492536 | 2.472101 | 0.013432 | NA |
| INGX | 0.162376 | 0.424913 | 4.079006 | 0.104171 | 0.917034 | NA |
| UBA52P5 | 2.51272 | -1.38535 | 1.341944 | -1.03234 | 0.301911 | NA |
| RPS23P1 | 0.448206 | -1.2532 | 3.007315 | -0.41672 | 0.676885 | NA |
| | 247.4726 | 0.018231 | 0.182596 | 0.099843 | 0.920469 | 0.953884 |
| UPK3B | 1.331555 | -0.54837 | 1.636667 | -0.33505 | 0.737584 | NA |
| | 0.134133 | -1.48909 | 4.077584 | -0.36519 | 0.71497 | NA |
| | 30.0952 | 0.016759 | 0.478406 | 0.03503 | 0.972056 | NA |
| RN7SL181F | 1.015699 | 0.142451 | 2.181052 | 0.065313 | 0.947925 | NA |
| EEF1A1P10 | 1.387693 | 2.47614 | 1.982457 | 1.249025 | 0.211656 | NA |
| HOTTIP | 0.258578 | 1.233151 | 4.047809 | 0.304646 | 0.760635 | NA |
| OSTCP1 | 0.392857 | -1.546 | 4.075469 | -0.37934 | 0.704433 | NA |
| | 2.683486 | 0.155434 | 1.204205 | 0.129076 | 0.897297 | NA |
| RN7SL30P | 0.096202 | 0.408546 | 4.079639 | 0.100143 | 0.920231 | NA |

| | | | | | | |
|---|---|---|---|---|---|---|
| | 6.094897 | 0.200757 | 0.842303 | 0.238343 | 0.811615 | NA |
| OR2A7 | 0.162376 | 0.424913 | 4.079006 | 0.104171 | 0.917034 | NA |
| RGAG1 | 21.06191 | 0.232138 | 0.547453 | 0.424033 | 0.671542 | NA |
| RPL5P24 | 0.889885 | 0.644651 | 2.1706 | 0.296992 | 0.766472 | NA |
| | 1.912771 | 3.084214 | 1.757567 | 1.75482 | 0.07929 | NA |
| | 4.15329 | 0.478595 | 1.027853 | 0.465626 | 0.641483 | NA |
| | 0.460466 | 0.568655 | 3.014642 | 0.188631 | 0.850382 | NA |
| | 0.70046 | 1.37107 | 2.594868 | 0.528378 | 0.597237 | NA |
| TMEM141 | 1071.45 | -0.13767 | 0.190861 | -0.72134 | 0.470702 | 0.627057 |
| | 3.329602 | -0.33431 | 1.074517 | -0.31113 | 0.755701 | NA |
| RN7SL81P | 3.53782 | 0.909397 | 1.143266 | 0.795438 | 0.426359 | NA |
| | 2.490829 | -0.77677 | 1.447896 | -0.53648 | 0.591628 | NA |
| RN7SL151F | 0.162376 | 0.424913 | 4.079006 | 0.104171 | 0.917034 | NA |
| RPS3AP35 | 0.163957 | 0.424913 | 4.079006 | 0.104171 | 0.917034 | NA |
| C7orf13 | 35.94831 | -0.00195 | 0.387407 | -0.00504 | 0.995978 | NA |
| RN7SL395F | 0.392711 | 0.292538 | 3.145221 | 0.09301 | 0.925895 | NA |
| | 0.134133 | -1.48909 | 4.077584 | -0.36519 | 0.71497 | NA |
| OR2A1-AS1 | 4.834452 | -0.23755 | 0.901134 | -0.26362 | 0.792075 | NA |
| | 6.850858 | 0.231188 | 0.781321 | 0.295893 | 0.767311 | NA |
| RWDD4P1 | 0.096202 | 0.408546 | 4.079639 | 0.100143 | 0.920231 | NA |
| ODCP | 1.563324 | 0.172115 | 1.593555 | 0.108007 | 0.91399 | NA |
| | 13.91402 | 0.72456 | 0.577255 | 1.255182 | 0.209413 | NA |
| | 0.163957 | 0.424913 | 4.079006 | 0.104171 | 0.917034 | NA |
| | 0.303394 | -2.27006 | 4.032451 | -0.56295 | 0.57347 | NA |
| OR2AE1 | 0.151697 | -1.546 | 4.075469 | -0.37934 | 0.704433 | NA |
| RN7SL396F | 0.382031 | -1.26484 | 3.151989 | -0.40128 | 0.688212 | NA |
| | 6.395351 | 1.488206 | 0.910118 | 1.635179 | 0.102012 | NA |
| | 0.424115 | 1.932531 | 3.45462 | 0.559405 | 0.575885 | NA |
| | 6.939943 | -0.43945 | 0.870467 | -0.50484 | 0.613669 | NA |
| MIR3150B | 3.043561 | -0.69013 | 1.134144 | -0.6085 | 0.542856 | NA |
| MIR3150A | 3.043561 | -0.69013 | 1.134144 | -0.6085 | 0.542856 | NA |
| RNF139-AS | 19.29242 | 0.995342 | 0.49844 | 1.996914 | 0.045835 | NA |
| | 13.3943 | -0.11257 | 0.600638 | -0.18741 | 0.851339 | NA |
| | 0.151697 | -1.546 | 4.075469 | -0.37934 | 0.704433 | NA |
| SNHG6 | 2070.207 | 0.393562 | 0.161071 | 2.443402 | 0.01455 | 0.06067 |
| | 15.51702 | 0.07751 | 0.577485 | 0.134221 | 0.893228 | NA |
| | 77.81177 | -0.31943 | 0.25904 | -1.23314 | 0.217522 | 0.376981 |
| | 6.710807 | 0.216946 | 0.7779 | 0.278886 | 0.780332 | NA |
| | 1.736323 | 0.821497 | 1.650262 | 0.497798 | 0.618626 | NA |
| | 4.177685 | 0.5451 | 1.039806 | 0.524233 | 0.600117 | NA |
| EXTL3-AS1 | 1.818895 | 1.382314 | 1.649483 | 0.838028 | 0.402015 | NA |
| | 1.974976 | 0.620222 | 1.421099 | 0.436438 | 0.662519 | NA |
| LINC00968 | 10.01916 | -0.03358 | 1.00816 | -0.03331 | 0.973428 | NA |
| | 3.570456 | 1.887159 | 1.152287 | 1.63775 | 0.101474 | NA |
| | 5.020089 | 0.354929 | 0.890339 | 0.398645 | 0.690155 | NA |
| LINC00535 | 2.148381 | 0.384613 | 1.326055 | 0.290043 | 0.771783 | NA |
| | 2.191616 | -0.42726 | 1.274182 | -0.33532 | 0.737384 | NA |

| | | | | | | |
|---|---|---|---|---|---|---|
| | 0.768215 | 1.51173 | 2.494681 | 0.605981 | 0.544527 | NA |
| | 84.17917 | -0.20598 | 0.258792 | -0.79591 | 0.426082 | 0.587965 |
| | 392.552 | 0.496262 | 0.180888 | 2.743472 | 0.006079 | 0.033748 |
| | 6.950566 | -0.11972 | 0.787174 | -0.15209 | 0.879113 | NA |
| SDCBPP2 | 3.170972 | 0.815033 | 1.30975 | 0.622281 | 0.533757 | NA |
| USP51 | 88.9624 | 0.235869 | 0.400575 | 0.588826 | 0.555978 | 0.695486 |
| HSPA8P11 | 0.489058 | -0.58569 | 3.427543 | -0.17088 | 0.864321 | NA |
| KRT8P4 | 0.134133 | -1.48909 | 4.077584 | -0.36519 | 0.71497 | NA |
| | 0.163957 | 0.424913 | 4.079006 | 0.104171 | 0.917034 | NA |
| | 8.304867 | -0.02623 | 0.70735 | -0.03708 | 0.970421 | NA |
| SBF1P1 | 0.881915 | -2.2305 | 2.871853 | -0.77668 | 0.437349 | NA |
| NDUFA5P1 | 0.151697 | -1.546 | 4.075469 | -0.37934 | 0.704433 | NA |
| | 0.288605 | 1.443619 | 4.029058 | 0.358302 | 0.720117 | NA |
| NPM1P21 | 0.544407 | -0.44425 | 2.542878 | -0.1747 | 0.861313 | NA |
| | 0.802985 | 0.54709 | 2.208262 | 0.247747 | 0.80433 | NA |
| HAS2-AS1 | 10.26225 | -0.13037 | 0.713225 | -0.18279 | 0.854963 | NA |
| | 0.096202 | 0.408546 | 4.079639 | 0.100143 | 0.920231 | NA |
| | 0.096202 | 0.408546 | 4.079639 | 0.100143 | 0.920231 | NA |
| | 1.710434 | 0.70104 | 1.69845 | 0.412753 | 0.679788 | NA |
| | 2.030581 | -0.3493 | 1.438218 | -0.24287 | 0.808106 | NA |
| | 181.7447 | 0.015695 | 0.23121 | 0.067882 | 0.94588 | 0.96985 |
| | 0.096202 | 0.408546 | 4.079639 | 0.100143 | 0.920231 | NA |
| | 0.602678 | 1.083611 | 2.70938 | 0.399948 | 0.689195 | NA |
| | 2.501608 | -0.2508 | 1.431121 | -0.17525 | 0.860886 | NA |
| CASC11 | 0.938845 | -1.23445 | 2.063177 | -0.59832 | 0.549624 | NA |
| | 0.258578 | 1.233151 | 4.047809 | 0.304646 | 0.760635 | NA |
| | 0.82213 | 0.50433 | 2.197682 | 0.229483 | 0.818494 | NA |
| | 0.544554 | -2.65036 | 3.220086 | -0.82307 | 0.410468 | NA |
| PVT1 | 315.4688 | 0.037528 | 0.207717 | 0.180669 | 0.856627 | 0.914811 |
| | 6.037171 | -0.83139 | 0.80761 | -1.02944 | 0.303273 | NA |
| LINC00536 | 0.45509 | -2.71227 | 3.963744 | -0.68427 | 0.493805 | NA |
| TRGV7 | 0.151697 | -1.546 | 4.075469 | -0.37934 | 0.704433 | NA |
| | 0.612162 | -0.19706 | 2.47693 | -0.07956 | 0.936588 | NA |
| | 0.392857 | -1.546 | 4.075469 | -0.37934 | 0.704433 | NA |
| | 0.096202 | 0.408546 | 4.079639 | 0.100143 | 0.920231 | NA |
| | 0.258578 | 1.233151 | 4.047809 | 0.304646 | 0.760635 | NA |
| | 1.388485 | -0.24258 | 1.630363 | -0.14879 | 0.88172 | NA |
| KIAA1456 | 267.393 | 0.015692 | 0.274873 | 0.05709 | 0.954474 | 0.974998 |
| LINC00977 | 0.938698 | 0.081169 | 1.915817 | 0.042368 | 0.966205 | NA |
| KIAA1210 | 2.059495 | 1.207579 | 1.621969 | 0.744515 | 0.456565 | NA |
| WBP1LP2 | 1.323504 | 0.914682 | 1.745605 | 0.523991 | 0.600285 | NA |
| PPM1AP1 | 0.548763 | 2.343695 | 3.135478 | 0.747476 | 0.454776 | NA |
| OR7E59P | 0.162376 | 0.424913 | 4.079006 | 0.104171 | 0.917034 | NA |
| NTAN1P2 | 4.007939 | -0.25814 | 0.975271 | -0.26469 | 0.79125 | NA |
| GLI4 | 88.69015 | -0.10132 | 0.301245 | -0.33635 | 0.736611 | 0.833121 |
| ALG1L12P | 0.151697 | -1.546 | 4.075469 | -0.37934 | 0.704433 | NA |
| | 0.162376 | 0.424913 | 4.079006 | 0.104171 | 0.917034 | NA |

| | | | | | | |
|---|---|---|---|---|---|---|
| | 0.574231 | 0.93227 | 2.49998 | 0.372911 | 0.709215 | NA |
| | 0.096202 | 0.408546 | 4.079639 | 0.100143 | 0.920231 | NA |
| | 34.30879 | 0.849466 | 0.460775 | 1.843558 | 0.065248 | NA |
| GAPDHP62 | 0.163957 | 0.424913 | 4.079006 | 0.104171 | 0.917034 | NA |
| | 32.8263 | 0.17039 | 0.365959 | 0.465597 | 0.641504 | NA |
| | 50.58778 | 0.52343 | 0.352276 | 1.485853 | 0.137318 | 0.276526 |
| LINC00589 | 1181.283 | 0.123596 | 0.136087 | 0.908214 | 0.363765 | 0.531088 |
| ZNF674 | 35.94689 | 0.674747 | 0.40097 | 1.682787 | 0.092416 | NA |
| PSMA2P2 | 0.151697 | -1.546 | 4.075469 | -0.37934 | 0.704433 | NA |
| | 5.472339 | -0.87991 | 0.8549 | -1.02925 | 0.30336 | NA |
| | 0.476449 | 0.524262 | 3.402674 | 0.154073 | 0.877552 | NA |
| | 8.759559 | 0.266613 | 0.702548 | 0.379495 | 0.704321 | NA |
| IMPA1P | 0.163957 | 0.424913 | 4.079006 | 0.104171 | 0.917034 | NA |
| GYG1P1 | 0.096202 | 0.408546 | 4.079639 | 0.100143 | 0.920231 | NA |
| RNA5-8SP6 | 528.6209 | 0.937258 | 0.255588 | 3.667068 | 2.45E-04 | 0.0044 |
| RNU7-20P | 0.162376 | 0.424913 | 4.079006 | 0.104171 | 0.917034 | NA |
| | 0.162376 | 0.424913 | 4.079006 | 0.104171 | 0.917034 | NA |
| SCARNA8 | 0.247898 | -0.5901 | 4.074633 | -0.14482 | 0.884851 | NA |
| RN7SKP242 | 0.151697 | -1.546 | 4.075469 | -0.37934 | 0.704433 | NA |
| RNU6ATAC | 0.151697 | -1.546 | 4.075469 | -0.37934 | 0.704433 | NA |
| | 0.517155 | 2.232561 | 3.179631 | 0.702145 | 0.482589 | NA |
| RNA5SP288 | 0.096202 | 0.408546 | 4.079639 | 0.100143 | 0.920231 | NA |
| RNU4ATAC | 0.260158 | 1.240347 | 4.047124 | 0.306476 | 0.759242 | NA |
| RNU7-154P | 0.096202 | 0.408546 | 4.079639 | 0.100143 | 0.920231 | NA |
| RNU7-109P | 0.163957 | 0.424913 | 4.079006 | 0.104171 | 0.917034 | NA |
| | 0.163957 | 0.424913 | 4.079006 | 0.104171 | 0.917034 | NA |
| | 0.709944 | 0.232855 | 2.332364 | 0.099837 | 0.920474 | NA |
| | 0.163957 | 0.424913 | 4.079006 | 0.104171 | 0.917034 | NA |
| RNU4ATAC | 0.134133 | -1.48909 | 4.077584 | -0.36519 | 0.71497 | NA |
| RNU6-875P | 0.096202 | 0.408546 | 4.079639 | 0.100143 | 0.920231 | NA |
| RNY5 | 1.344617 | -0.99306 | 2.086066 | -0.47604 | 0.634043 | NA |
| | 0.163957 | 0.424913 | 4.079006 | 0.104171 | 0.917034 | NA |
| RNU7-36P | 0.151697 | -1.546 | 4.075469 | -0.37934 | 0.704433 | NA |
| | 0.096202 | 0.408546 | 4.079639 | 0.100143 | 0.920231 | NA |
| SNORD36C | 0.258578 | 1.233151 | 4.047809 | 0.304646 | 0.760635 | NA |
| | 6.721903 | 0.369391 | 0.796754 | 0.46362 | 0.64292 | NA |
| RNU6-1136 | 0.303394 | -2.27006 | 4.032451 | -0.56295 | 0.57347 | NA |
| RNU6-786P | 0.324752 | 1.53082 | 4.022025 | 0.380609 | 0.703493 | NA |
| RNU6-703P | 0.096202 | 0.408546 | 4.079639 | 0.100143 | 0.920231 | NA |
| RNA5SP243 | 0.151697 | -1.546 | 4.075469 | -0.37934 | 0.704433 | NA |
| RNU7-177P | 0.47961 | 0.535796 | 3.39605 | 0.15777 | 0.874638 | NA |
| | 0.162376 | 0.424913 | 4.079006 | 0.104171 | 0.917034 | NA |
| | 21.6142 | 0.185938 | 0.487863 | 0.381127 | 0.703109 | NA |
| | 0.719393 | -0.9644 | 2.652838 | -0.36354 | 0.716204 | NA |
| | 0.163957 | 0.424913 | 4.079006 | 0.104171 | 0.917034 | NA |
| | 0.163957 | 0.424913 | 4.079006 | 0.104171 | 0.917034 | NA |
| | 0.604258 | 1.087325 | 2.707696 | 0.401568 | 0.688002 | NA |

| | | | | | | |
|---|---|---|---|---|---|---|
| | 0.489058 | -0.58569 | 3.427543 | -0.17088 | 0.864321 | NA |
| | 2.011822 | -1.41639 | 1.48563 | -0.95339 | 0.340391 | NA |
| | 10.50725 | -0.00924 | 0.932832 | -0.0099 | 0.992101 | NA |
| | 0.163957 | 0.424913 | 4.079006 | 0.104171 | 0.917034 | NA |
| | 0.52699 | -2.57878 | 3.248478 | -0.79384 | 0.427288 | NA |
| | 0.162376 | 0.424913 | 4.079006 | 0.104171 | 0.917034 | NA |
| KIAA0196-/ | 0.563552 | -0.48645 | 2.808029 | -0.17324 | 0.862465 | NA |
| | 2.179998 | 1.217055 | 1.464584 | 0.83099 | 0.405979 | NA |
| | 0.096202 | 0.408546 | 4.079639 | 0.100143 | 0.920231 | NA |
| | 0.949175 | 1.90243 | 2.405529 | 0.790857 | 0.429027 | NA |
| | 0.163957 | 0.424913 | 4.079006 | 0.104171 | 0.917034 | NA |
| | 117.6898 | -0.08866 | 0.271877 | -0.32611 | 0.744342 | 0.838959 |
| | 0.315653 | -0.57305 | 3.892639 | -0.14721 | 0.882963 | NA |
| | 0.354779 | 1.703954 | 3.657232 | 0.465914 | 0.641277 | NA |
| | 14.74849 | 0.674715 | 0.583523 | 1.156279 | 0.247567 | NA |
| | 6.490346 | -0.32838 | 0.821116 | -0.39992 | 0.689217 | NA |
| | 0.260158 | 1.240347 | 4.047124 | 0.306476 | 0.759242 | NA |
| | 25.38944 | 0.023664 | 0.517128 | 0.045761 | 0.963501 | NA |
| | 0.163957 | 0.424913 | 4.079006 | 0.104171 | 0.917034 | NA |
| C8orf88 | 97.72986 | 0.137903 | 0.246271 | 0.559964 | 0.575504 | 0.710649 |
| | 0.151697 | -1.546 | 4.075469 | -0.37934 | 0.704433 | NA |
| CCDC71L | 477.9722 | -0.02553 | 0.179832 | -0.14196 | 0.887109 | 0.933443 |
| | 0.151697 | -1.546 | 4.075469 | -0.37934 | 0.704433 | NA |
| | 0.422534 | 1.924583 | 3.001758 | 0.641152 | 0.521424 | NA |
| | 5.444057 | -0.23346 | 0.949845 | -0.24579 | 0.805847 | NA |
| HOXA10 | 14.70732 | 0.730178 | 0.561441 | 1.300543 | 0.193415 | NA |
| | 0.392857 | -1.546 | 4.075469 | -0.37934 | 0.704433 | NA |
| STAU2-AS1 | 1.393789 | -0.8924 | 1.663459 | -0.53647 | 0.591633 | NA |
| | 0.35636 | 1.709252 | 3.651833 | 0.468053 | 0.639746 | NA |
| | 0.247898 | -0.5901 | 4.074633 | -0.14482 | 0.884851 | NA |
| | 14.57915 | -0.32081 | 0.538487 | -0.59576 | 0.551338 | NA |
| RAD21-AS1 | 1.010606 | 3.179931 | 2.258787 | 1.407805 | 0.159189 | NA |
| | 2.171826 | 0.668273 | 1.670833 | 0.399964 | 0.689183 | NA |
| | 0.163957 | 0.424913 | 4.079006 | 0.104171 | 0.917034 | NA |
| | 0.324752 | 1.53082 | 4.022025 | 0.380609 | 0.703493 | NA |
| | 0.384806 | 1.846216 | 3.999752 | 0.461582 | 0.644381 | NA |
| | 84.30288 | -0.34112 | 0.257793 | -1.32322 | 0.185763 | 0.339587 |
| | 0.52699 | -2.57878 | 3.248478 | -0.79384 | 0.427288 | NA |
| | 0.315653 | -0.57305 | 3.892639 | -0.14721 | 0.882963 | NA |
| | 5.40212 | 0.209025 | 0.858806 | 0.243391 | 0.807703 | NA |
| | 0.796661 | 1.607053 | 2.50763 | 0.640865 | 0.52161 | NA |
| | 0.096202 | 0.408546 | 4.079639 | 0.100143 | 0.920231 | NA |
| | 0.462046 | 0.589069 | 3.442439 | 0.17112 | 0.86413 | NA |
| | 4.230617 | 0.811005 | 1.194307 | 0.679059 | 0.497101 | NA |
| | 1.577524 | 0.5935 | 1.606623 | 0.369409 | 0.711823 | NA |
| | 0.652869 | 1.292592 | 2.374873 | 0.544278 | 0.58625 | NA |
| | 0.303394 | -2.27006 | 4.032451 | -0.56295 | 0.57347 | NA |

| | | | | | | |
|---|---|---|---|---|---|---|
| | 2.237626 | -0.35039 | 1.302913 | -0.26893 | 0.787986 | NA |
| LINC00534 | 0.096202 | 0.408546 | 4.079639 | 0.100143 | 0.920231 | NA |
| | 2.688782 | -1.24678 | 1.208429 | -1.03173 | 0.302197 | NA |
| | 3.834191 | 1.87694 | 1.122979 | 1.671394 | 0.094644 | NA |
| | 2.027217 | -0.02327 | 1.699442 | -0.01369 | 0.989075 | NA |
| | 0.806146 | 0.553068 | 2.206737 | 0.250627 | 0.802102 | NA |
| | 2.001988 | 0.165454 | 1.339658 | 0.123505 | 0.901707 | NA |
| | 5.783948 | 0.212996 | 0.827572 | 0.257375 | 0.796889 | NA |
| | 0.151697 | -1.546 | 4.075469 | -0.37934 | 0.704433 | NA |
| | 0.151697 | -1.546 | 4.075469 | -0.37934 | 0.704433 | NA |
| | 0.623045 | 0.091811 | 2.666142 | 0.034436 | 0.97253 | NA |
| | 0.70851 | -1.62753 | 2.686091 | -0.60591 | 0.544574 | NA |
| PCAT1 | 1.036424 | 0.067268 | 2.421139 | 0.027784 | 0.977835 | NA |
| | 0.134133 | -1.48909 | 4.077584 | -0.36519 | 0.71497 | NA |
| | 0.162376 | 0.424913 | 4.079006 | 0.104171 | 0.917034 | NA |
| SMIM18 | 0.163957 | 0.424913 | 4.079006 | 0.104171 | 0.917034 | NA |
| HMGB1P19 | 0.419962 | -2.64111 | 3.429364 | -0.77014 | 0.441214 | NA |
| | 0.151697 | -1.546 | 4.075469 | -0.37934 | 0.704433 | NA |
| | 0.134133 | -1.48909 | 4.077584 | -0.36519 | 0.71497 | NA |
| | 1.959984 | 2.251507 | 1.641087 | 1.371961 | 0.170076 | NA |
| | 1.372408 | -0.70922 | 1.970671 | -0.35989 | 0.718932 | NA |
| | 0.327913 | 1.543154 | 4.021063 | 0.383768 | 0.701151 | NA |
| | 2974.621 | 0.522852 | 0.153955 | 3.396141 | 6.83E-04 | 0.008192 |
| | 0.134133 | -1.48909 | 4.077584 | -0.36519 | 0.71497 | NA |
| | 0.324752 | 1.53082 | 4.022025 | 0.380609 | 0.703493 | NA |
| | 0.192403 | 0.975392 | 4.060716 | 0.240202 | 0.810174 | NA |
| HMGB1P41 | 0.670433 | 1.255281 | 2.356401 | 0.532711 | 0.594234 | NA |
| XRCC6P4 | 0.489058 | -0.58569 | 3.427543 | -0.17088 | 0.864321 | NA |
| | 0.096202 | 0.408546 | 4.079639 | 0.100143 | 0.920231 | NA |
| | 0.134133 | -1.48909 | 4.077584 | -0.36519 | 0.71497 | NA |
| | 0.556813 | -0.57007 | 3.294615 | -0.17303 | 0.862627 | NA |
| HOXA-AS2 | 1.878884 | -0.09816 | 1.5128 | -0.06488 | 0.948266 | NA |
| | 0.392857 | -1.546 | 4.075469 | -0.37934 | 0.704433 | NA |
| | 0.392857 | -1.546 | 4.075469 | -0.37934 | 0.704433 | NA |
| | 12.39298 | 0.600784 | 0.697992 | 0.860731 | 0.389386 | NA |
| RNF5P1 | 48.13137 | 0.628053 | 0.339101 | 1.852114 | 0.064009 | 0.16457 |
| | 0.163957 | 0.424913 | 4.079006 | 0.104171 | 0.917034 | NA |
| | 0.533728 | -1.74213 | 2.820432 | -0.61768 | 0.536785 | NA |
| | 0.096202 | 0.408546 | 4.079639 | 0.100143 | 0.920231 | NA |
| | 0.096202 | 0.408546 | 4.079639 | 0.100143 | 0.920231 | NA |
| SLC10A5 | 1.331766 | 0.790302 | 2.090457 | 0.378052 | 0.705392 | NA |
| | 0.354779 | 1.703954 | 3.657232 | 0.465914 | 0.641277 | NA |
| HNRNPA1P | 0.096202 | 0.408546 | 4.079639 | 0.100143 | 0.920231 | NA |
| AFG3L2P1 | 0.163957 | 0.424913 | 4.079006 | 0.104171 | 0.917034 | NA |
| | 0.163957 | 0.424913 | 4.079006 | 0.104171 | 0.917034 | NA |
| | 23.06244 | -0.91595 | 0.624578 | -1.46651 | 0.14251 | NA |
| | 0.563552 | -0.48645 | 2.808029 | -0.17324 | 0.862465 | NA |

| | | | | | | |
|---|---|---|---|---|---|---|
| | 0.163957 | 0.424913 | 4.079006 | 0.104171 | 0.917034 | NA |
| | 1.645831 | -0.89586 | 1.584376 | -0.56543 | 0.571781 | NA |
| | 0.162376 | 0.424913 | 4.079006 | 0.104171 | 0.917034 | NA |
| | 0.725928 | 0.181819 | 2.321433 | 0.078322 | 0.937572 | NA |
| | 0.315653 | -0.57305 | 3.892639 | -0.14721 | 0.882963 | NA |
| PRSS51 | 0.624422 | 1.162588 | 2.733761 | 0.425271 | 0.670639 | NA |
| ZFHX4-AS1 | 0.096202 | 0.408546 | 4.079639 | 0.100143 | 0.920231 | NA |
| | 1.718584 | 3.961632 | 1.844001 | 2.148389 | 0.031683 | NA |
| | 14.03611 | -0.01197 | 0.541985 | -0.02208 | 0.982383 | NA |
| | 0.47961 | 0.535796 | 3.39605 | 0.15777 | 0.874638 | NA |
| TAGLN2P1 | 5.346607 | 0.122649 | 0.924916 | 0.132606 | 0.894505 | NA |
| UBE2HP1 | 1.232404 | 0.600643 | 1.916349 | 0.313431 | 0.753953 | NA |
| | 0.260158 | 1.240347 | 4.047124 | 0.306476 | 0.759242 | NA |
| | 0.260158 | 1.240347 | 4.047124 | 0.306476 | 0.759242 | NA |
| | 0.638825 | 1.117948 | 3.106359 | 0.35989 | 0.718929 | NA |
| | 7.756183 | 0.461994 | 0.723609 | 0.638458 | 0.523176 | NA |
| | 2.46147 | 0.305609 | 1.395903 | 0.218933 | 0.826703 | NA |
| | 0.096202 | 0.408546 | 4.079639 | 0.100143 | 0.920231 | NA |
| | 30.65542 | -0.03338 | 0.465376 | -0.07172 | 0.942826 | NA |
| | 0.247898 | -0.5901 | 4.074633 | -0.14482 | 0.884851 | NA |
| SUMO2P2( | 0.46577 | -1.30191 | 3.402879 | -0.38259 | 0.702024 | NA |
| PRKDC | 5419.84 | -0.11166 | 0.141246 | -0.79052 | 0.429222 | 0.590716 |
| LZTS1-AS1 | 0.247898 | -0.5901 | 4.074633 | -0.14482 | 0.884851 | NA |
| | 0.545988 | -0.43913 | 2.541145 | -0.17281 | 0.862803 | NA |
| | 386.8148 | 0.97592 | 0.281869 | 3.462312 | 5.36E-04 | 0.007047 |
| | 3.85682 | 0.2515 | 1.073495 | 0.234281 | 0.814767 | NA |
| | 0.162376 | 0.424913 | 4.079006 | 0.104171 | 0.917034 | NA |
| | 0.460466 | 0.568655 | 3.014642 | 0.188631 | 0.850382 | NA |
| HMGB1P2: | 0.096202 | 0.408546 | 4.079639 | 0.100143 | 0.920231 | NA |
| | 0.096202 | 0.408546 | 4.079639 | 0.100143 | 0.920231 | NA |
| | 0.49187 | 2.136088 | 3.895953 | 0.548284 | 0.583497 | NA |
| | 0.163957 | 0.424913 | 4.079006 | 0.104171 | 0.917034 | NA |
| LINC01030 | 0.162376 | 0.424913 | 4.079006 | 0.104171 | 0.917034 | NA |
| PSAT1P1 | 0.151697 | -1.546 | 4.075469 | -0.37934 | 0.704433 | NA |
| | 1.952079 | 3.095022 | 1.734086 | 1.784814 | 0.074291 | NA |
| | 0.260158 | 1.240347 | 4.047124 | 0.306476 | 0.759242 | NA |
| | 2.268819 | -0.81876 | 1.305544 | -0.62714 | 0.530565 | NA |
| | 0.788582 | 0.608095 | 2.456267 | 0.247569 | 0.804468 | NA |
| | 0.260158 | 1.240347 | 4.047124 | 0.306476 | 0.759242 | NA |
| | 0.508056 | 0.731032 | 2.861912 | 0.255435 | 0.798387 | NA |
| | 5.802543 | 0.512214 | 0.941362 | 0.54412 | 0.586359 | NA |
| | 0.096202 | 0.408546 | 4.079639 | 0.100143 | 0.920231 | NA |
| | 30.9825 | 0.282868 | 0.38211 | 0.740279 | 0.459131 | NA |
| | 9.182445 | 0.079713 | 0.705827 | 0.112935 | 0.910082 | NA |
| | 0.670636 | 0.219794 | 2.632532 | 0.083491 | 0.933461 | NA |
| FAM85B | 1.683706 | -2.08511 | 1.718893 | -1.21306 | 0.225108 | NA |
| | 0.410275 | 0.244283 | 3.100889 | 0.078778 | 0.937209 | NA |

| | | | | | | |
|---|---|---|---|---|---|---|
| MAPRE1P1 | 0.783638 | 1.586366 | 2.937049 | 0.540122 | 0.589113 | NA |
| PRKRIRP7 | 0.382031 | -1.26484 | 3.151989 | -0.40128 | 0.688212 | NA |
| | 3.082833 | -0.76085 | 1.269309 | -0.59942 | 0.548892 | NA |
| | 0.163957 | 0.424913 | 4.079006 | 0.104171 | 0.917034 | NA |
| | 1.812543 | 0.264744 | 1.695249 | 0.156168 | 0.8759 | NA |
| | 4.357617 | 0.051613 | 0.940348 | 0.054888 | 0.956228 | NA |
| CLDN23 | 10.06901 | -0.82377 | 0.706302 | -1.16632 | 0.243487 | NA |
| | 0.296509 | -0.51709 | 3.982155 | -0.12985 | 0.896684 | NA |
| | 0.096202 | 0.408546 | 4.079639 | 0.100143 | 0.920231 | NA |
| | 0.151697 | -1.546 | 4.075469 | -0.37934 | 0.704433 | NA |
| ALG1L13P | 2.454383 | 1.035727 | 1.450001 | 0.714294 | 0.475046 | NA |
| | 403.2058 | -0.17602 | 0.163138 | -1.07896 | 0.280604 | 0.446698 |
| | 0.760136 | 0.498384 | 2.921408 | 0.170597 | 0.86454 | NA |
| | 1.407389 | 0.233783 | 1.745172 | 0.13396 | 0.893434 | NA |
| | 0.096202 | 0.408546 | 4.079639 | 0.100143 | 0.920231 | NA |
| | 0.879985 | 1.114342 | 2.892066 | 0.38531 | 0.700008 | NA |
| IGHEP2 | 0.324752 | 1.53082 | 4.022025 | 0.380609 | 0.703493 | NA |
| PKMP4 | 0.858627 | -2.14058 | 2.513451 | -0.85165 | 0.394408 | NA |
| | 11.38772 | 0.161733 | 0.631642 | 0.256052 | 0.797911 | NA |
| | 0.411855 | 0.250456 | 3.097618 | 0.080854 | 0.935558 | NA |
| | 0.096202 | 0.408546 | 4.079639 | 0.100143 | 0.920231 | NA |
| | 1.006656 | -0.39276 | 2.040151 | -0.19251 | 0.847339 | NA |
| | 0.151697 | -1.546 | 4.075469 | -0.37934 | 0.704433 | NA |
| | 0.392857 | -1.546 | 4.075469 | -0.37934 | 0.704433 | NA |
| | 0.420954 | 1.923376 | 3.462803 | 0.555439 | 0.578594 | NA |
| | 0.258578 | 1.233151 | 4.047809 | 0.304646 | 0.760635 | NA |
| | 0.096202 | 0.408546 | 4.079639 | 0.100143 | 0.920231 | NA |
| | 15.39839 | -0.53693 | 0.542099 | -0.99047 | 0.321945 | NA |
| | 0.151697 | -1.546 | 4.075469 | -0.37934 | 0.704433 | NA |
| | 0.151697 | -1.546 | 4.075469 | -0.37934 | 0.704433 | NA |
| LYN | 596.0825 | 0.181111 | 0.161617 | 1.120623 | 0.262449 | 0.426852 |
| SLC2A3P4 | 1.033466 | -0.7302 | 1.946441 | -0.37515 | 0.707552 | NA |
| PINX1 | 316.6299 | 0.07424 | 0.226801 | 0.327335 | 0.743414 | 0.838392 |
| | 0.163957 | 0.424913 | 4.079006 | 0.104171 | 0.917034 | NA |
| RBPMS-AS1 | 114.2264 | -0.10327 | 0.269932 | -0.38258 | 0.702031 | 0.808431 |
| | 0.298089 | -0.51711 | 3.975232 | -0.13008 | 0.896501 | NA |
| | 0.163957 | 0.424913 | 4.079006 | 0.104171 | 0.917034 | NA |
| | 0.949727 | -0.95431 | 2.383855 | -0.40032 | 0.688919 | NA |
| | 0.134133 | -1.48909 | 4.077584 | -0.36519 | 0.71497 | NA |
| | 4.192631 | 0.274928 | 1.186868 | 0.231642 | 0.816816 | NA |
| | 2.236621 | 0.831159 | 1.544529 | 0.538131 | 0.590487 | NA |
| | 0.884727 | 1.125991 | 2.88877 | 0.389782 | 0.696698 | NA |
| | 1.075403 | 2.139729 | 2.013922 | 1.062469 | 0.288023 | NA |
| | 0.162376 | 0.424913 | 4.079006 | 0.104171 | 0.917034 | NA |
| TYW1B | 25.88814 | 0.268057 | 0.426415 | 0.628629 | 0.529592 | NA |
| | 0.096202 | 0.408546 | 4.079639 | 0.100143 | 0.920231 | NA |
| | 0.096202 | 0.408546 | 4.079639 | 0.100143 | 0.920231 | NA |

| | | | | | | |
|---|---|---|---|---|---|---|
| | 11.59205 | 0.030437 | 0.650813 | 0.046768 | 0.962698 | NA |
| | 1.353125 | 2.465848 | 1.88624 | 1.307282 | 0.191117 | NA |
| | 0.411855 | 0.250456 | 3.097618 | 0.080854 | 0.935558 | NA |
| | 0.162376 | 0.424913 | 4.079006 | 0.104171 | 0.917034 | NA |
| | 0.162376 | 0.424913 | 4.079006 | 0.104171 | 0.917034 | NA |
| | 1.332896 | -0.16531 | 2.226653 | -0.07424 | 0.94082 | NA |
| | 1.801534 | 0.918548 | 1.480142 | 0.620581 | 0.534875 | NA |
| | 1.65675 | -0.69289 | 1.546651 | -0.44799 | 0.65416 | NA |
| | 0.163957 | 0.424913 | 4.079006 | 0.104171 | 0.917034 | NA |
| | 0.326333 | 1.533889 | 3.826878 | 0.40082 | 0.688553 | NA |
| PAICSP4 | 1.350553 | 0.261011 | 1.647077 | 0.158469 | 0.874087 | NA |
| | 1.556457 | -3.16133 | 2.134773 | -1.48087 | 0.138641 | NA |
| | 0.326333 | 1.533889 | 3.826878 | 0.40082 | 0.688553 | NA |
| | 0.324752 | 1.53082 | 4.022025 | 0.380609 | 0.703493 | NA |
| | 0.258578 | 1.233151 | 4.047809 | 0.304646 | 0.760635 | NA |
| | 0.162376 | 0.424913 | 4.079006 | 0.104171 | 0.917034 | NA |
| ERHP1 | 0.52699 | -2.57878 | 3.248478 | -0.79384 | 0.427288 | NA |
| | 0.096202 | 0.408546 | 4.079639 | 0.100143 | 0.920231 | NA |
| | 0.162376 | 0.424913 | 4.079006 | 0.104171 | 0.917034 | NA |
| | 0.163957 | 0.424913 | 4.079006 | 0.104171 | 0.917034 | NA |
| KRT8P3 | 0.162376 | 0.424913 | 4.079006 | 0.104171 | 0.917034 | NA |
| | 0.134133 | -1.48909 | 4.077584 | -0.36519 | 0.71497 | NA |
| | 3.002343 | 2.630341 | 1.526943 | 1.722619 | 0.084957 | NA |
| | 0.419962 | -2.64111 | 3.429364 | -0.77014 | 0.441214 | NA |
| IMPDH1P6 | 0.326333 | 1.533889 | 3.826878 | 0.40082 | 0.688553 | NA |
| | 9.001122 | 1.531645 | 0.736448 | 2.079775 | 0.037546 | NA |
| | 1.555013 | 1.798836 | 1.661792 | 1.082468 | 0.279045 | NA |
| | 0.162376 | 0.424913 | 4.079006 | 0.104171 | 0.917034 | NA |
| | 0.096202 | 0.408546 | 4.079639 | 0.100143 | 0.920231 | NA |
| CKS1BP7 | 0.47803 | 0.520083 | 2.97881 | 0.174594 | 0.861399 | NA |
| | 7.8512 | -0.67726 | 0.726958 | -0.93164 | 0.351523 | NA |
| | 5.450406 | -0.28419 | 0.941683 | -0.30179 | 0.762813 | NA |
| | 0.82371 | 0.509057 | 2.431911 | 0.209324 | 0.834195 | NA |
| | 1.103996 | 1.740043 | 2.131488 | 0.816351 | 0.414299 | NA |
| | 0.096202 | 0.408546 | 4.079639 | 0.100143 | 0.920231 | NA |
| HOXA-AS3 | 2.580428 | 1.594005 | 1.311578 | 1.215333 | 0.224239 | NA |
| | 0.422534 | 1.924583 | 3.001758 | 0.641152 | 0.521424 | NA |
| LINC00966 | 0.096202 | 0.408546 | 4.079639 | 0.100143 | 0.920231 | NA |
| | 0.163957 | 0.424913 | 4.079006 | 0.104171 | 0.917034 | NA |
| | 0.755955 | 0.348448 | 2.512583 | 0.138681 | 0.889702 | NA |
| RHPN1-AS1 | 4.754123 | 0.502772 | 0.9709 | 0.517841 | 0.604569 | NA |
| | 0.162376 | 0.424913 | 4.079006 | 0.104171 | 0.917034 | NA |
| | 76.19465 | 0.222012 | 0.280404 | 0.791755 | 0.428503 | 0.590275 |
| | 1.542367 | 2.689772 | 1.869697 | 1.438613 | 0.15026 | NA |
| ENPP7P12 | 0.096202 | 0.408546 | 4.079639 | 0.100143 | 0.920231 | NA |
| | 0.151697 | -1.546 | 4.075469 | -0.37934 | 0.704433 | NA |
| | 0.163957 | 0.424913 | 4.079006 | 0.104171 | 0.917034 | NA |

| | | | | | | |
|---|---|---|---|---|---|---|
| | 0.258578 | 1.233151 | 4.047809 | 0.304646 | 0.760635 | NA |
| | 4.520475 | 0.640662 | 1.096531 | 0.584263 | 0.559043 | NA |
| | 144.7405 | 0.408337 | 0.278161 | 1.467989 | 0.142107 | 0.282988 |
| | 0.151697 | -1.546 | 4.075469 | -0.37934 | 0.704433 | NA |
| | 0.096202 | 0.408546 | 4.079639 | 0.100143 | 0.920231 | NA |
| | 8.277172 | 0.356754 | 0.749532 | 0.47597 | 0.634096 | NA |
| | 0.518939 | 0.925178 | 3.271397 | 0.282808 | 0.777324 | NA |
| | 0.684273 | 2.627785 | 2.896325 | 0.907282 | 0.364257 | NA |
| | 29.42327 | -0.45315 | 0.552325 | -0.82045 | 0.411962 | NA |
| | 0.422534 | 1.924583 | 3.001758 | 0.641152 | 0.521424 | NA |
| | 0.424115 | 1.932531 | 3.45462 | 0.559405 | 0.575885 | NA |
| | 0.815391 | 0.931626 | 2.360431 | 0.394685 | 0.693076 | NA |
| | 12.81501 | 0.727128 | 0.614853 | 1.182605 | 0.236966 | NA |
| OR7E15P | 0.268266 | -2.15396 | 4.038351 | -0.53338 | 0.593773 | NA |
| | 0.555233 | -0.57009 | 3.296947 | -0.17291 | 0.862719 | NA |
| OR7E158P | 2.794596 | 1.862285 | 1.508398 | 1.234611 | 0.216975 | NA |
| SLC2A13P1 | 0.163957 | 0.424913 | 4.079006 | 0.104171 | 0.917034 | NA |
| | 2.649213 | -1.26844 | 1.32972 | -0.95391 | 0.340127 | NA |
| | 0.678686 | -3.08037 | 2.621268 | -1.17514 | 0.239937 | NA |
| | 0.134133 | -1.48909 | 4.077584 | -0.36519 | 0.71497 | NA |
| | 0.096202 | 0.408546 | 4.079639 | 0.100143 | 0.920231 | NA |
| | 0.162376 | 0.424913 | 4.079006 | 0.104171 | 0.917034 | NA |
| | 10.51725 | -0.89778 | 0.720338 | -1.24633 | 0.212643 | NA |
| | 0.162376 | 0.424913 | 4.079006 | 0.104171 | 0.917034 | NA |
| FAM66D | 4.631762 | 0.171271 | 0.948978 | 0.180479 | 0.856776 | NA |
| | 1.263946 | -2.05362 | 2.064371 | -0.99479 | 0.319838 | NA |
| | 403.2932 | 0.239388 | 0.276104 | 0.86702 | 0.385931 | 0.551962 |
| | 5.58308 | 0.286377 | 0.824869 | 0.347179 | 0.728457 | NA |
| | 8.263397 | -0.06504 | 0.704531 | -0.09231 | 0.926449 | NA |
| | 0.096202 | 0.408546 | 4.079639 | 0.100143 | 0.920231 | NA |
| | 1.562532 | 2.727447 | 1.849963 | 1.474325 | 0.140394 | NA |
| | 1.160925 | 1.304491 | 1.810948 | 0.720336 | 0.471318 | NA |
| | 0.876676 | 2.997866 | 2.64573 | 1.133096 | 0.257174 | NA |
| | 50.18152 | -0.24305 | 0.318446 | -0.76325 | 0.445317 | 0.605208 |
| | 0.134133 | -1.48909 | 4.077584 | -0.36519 | 0.71497 | NA |
| | 139.1578 | 0.09825 | 0.226508 | 0.43376 | 0.664463 | 0.779567 |
| | 0.096202 | 0.408546 | 4.079639 | 0.100143 | 0.920231 | NA |
| | 0.881915 | -2.2305 | 2.871853 | -0.77668 | 0.437349 | NA |
| | 0.134133 | -1.48909 | 4.077584 | -0.36519 | 0.71497 | NA |
| | 0.314073 | -0.57308 | 3.898934 | -0.14698 | 0.883146 | NA |
| | 0.163957 | 0.424913 | 4.079006 | 0.104171 | 0.917034 | NA |
| LINC00681 | 22.2546 | -0.05307 | 0.506096 | -0.10486 | 0.916487 | NA |
| FAM85A | 4.09878 | -0.04859 | 0.985899 | -0.04928 | 0.960695 | NA |
| ZNF252P-A | 4.085114 | 0.156925 | 1.068784 | 0.146825 | 0.88327 | NA |
| | 0.815391 | 0.931626 | 2.360431 | 0.394685 | 0.693076 | NA |
| | 0.298089 | -0.51711 | 3.975232 | -0.13008 | 0.896501 | NA |
| | 0.162376 | 0.424913 | 4.079006 | 0.104171 | 0.917034 | NA |

| | | | | | | |
|---|---|---|---|---|---|---|
| SDIM1 | 27.33802 | 0.326543 | 0.42658 | 0.765491 | 0.443979 | NA |
| | 1.332547 | 2.108016 | 2.004334 | 1.051729 | 0.292924 | NA |
| TMEM75 | 0.230334 | -0.53318 | 4.076749 | -0.13079 | 0.895945 | NA |
| | 0.162376 | 0.424913 | 4.079006 | 0.104171 | 0.917034 | NA |
| SNRPEP2 | 5.209221 | 0.653395 | 0.923089 | 0.707835 | 0.479047 | NA |
| KIAA1147 | 430.898 | 0.338681 | 0.178926 | 1.892851 | 0.058378 | 0.15417 |
| MGAM | 254.1513 | 0.790971 | 0.692163 | 1.142753 | 0.253141 | 0.417384 |
| | 0.823352 | -0.17011 | 2.472341 | -0.06881 | 0.945144 | NA |
| | 0.096202 | 0.408546 | 4.079639 | 0.100143 | 0.920231 | NA |
| CUX1 | 2419.98 | -0.24252 | 0.194183 | -1.24895 | 0.211683 | 0.370133 |
| | 0.163957 | 0.424913 | 4.079006 | 0.104171 | 0.917034 | NA |
| | 0.096202 | 0.408546 | 4.079639 | 0.100143 | 0.920231 | NA |
| | 0.410275 | 0.244283 | 3.100889 | 0.078778 | 0.937209 | NA |
| | 0.151697 | -1.546 | 4.075469 | -0.37934 | 0.704433 | NA |
| ZHX1-C8OF | 2.357007 | 0.262822 | 1.299782 | 0.202205 | 0.839757 | NA |
| | 41.63668 | -0.07774 | 0.361539 | -0.21502 | 0.829748 | NA |
| | 0.151697 | -1.546 | 4.075469 | -0.37934 | 0.704433 | NA |
| | 0.151697 | -1.546 | 4.075469 | -0.37934 | 0.704433 | NA |
| | 0.47961 | 0.535796 | 3.39605 | 0.15777 | 0.874638 | NA |
| CASC7 | 252.8207 | 0.048709 | 0.215741 | 0.225776 | 0.821376 | 0.892522 |
| | 56.16704 | 0.837506 | 0.355903 | 2.353182 | 0.018614 | 0.071971 |
| | 14.27076 | -0.02455 | 0.55479 | -0.04425 | 0.964705 | NA |
| | 13.26291 | 0.266307 | 0.652878 | 0.407897 | 0.683349 | NA |
| | 10.27132 | -0.10996 | 0.651943 | -0.16866 | 0.866061 | NA |
| | 0.096202 | 0.408546 | 4.079639 | 0.100143 | 0.920231 | NA |
| | 8.149554 | -0.68704 | 0.836532 | -0.8213 | 0.411476 | NA |
| | 493.648 | -0.41748 | 0.173272 | -2.40941 | 0.015978 | 0.064804 |
| | 22.80256 | 0.522229 | 0.461735 | 1.131014 | 0.258049 | NA |
| | 0.327913 | 1.543154 | 4.021063 | 0.383768 | 0.701151 | NA |
| | 0.096202 | 0.408546 | 4.079639 | 0.100143 | 0.920231 | NA |
| | 0.720827 | 0.448488 | 2.540859 | 0.17651 | 0.859893 | NA |
| | 0.478233 | -0.67301 | 2.937473 | -0.22911 | 0.818782 | NA |
| | 8.475365 | 0.121297 | 0.752233 | 0.161249 | 0.871897 | NA |
| | 3.335695 | 0.353987 | 1.268424 | 0.279076 | 0.780186 | NA |
| FRRS1L | 2.844973 | 1.148597 | 1.526542 | 0.752418 | 0.4518 | NA |
| JHDM1D-A | 30.90198 | 0.011703 | 0.391102 | 0.029923 | 0.976129 | NA |
| | 4.507319 | -0.29599 | 0.963202 | -0.3073 | 0.758618 | NA |
| | 2.836591 | 0.954032 | 1.393987 | 0.684391 | 0.493728 | NA |
| | 370.5169 | -0.06714 | 0.249199 | -0.26943 | 0.787597 | 0.871252 |
| | 1.799897 | 0.836302 | 1.571148 | 0.532287 | 0.594527 | NA |
| | 0.708364 | 0.219301 | 2.144998 | 0.102238 | 0.918568 | NA |
| | 35.39698 | 0.870534 | 0.414846 | 2.09845 | 0.035865 | NA |
| | 1.033705 | -1.3133 | 2.272356 | -0.57794 | 0.563302 | NA |
| | 0.230334 | -0.53318 | 4.076749 | -0.13079 | 0.895945 | NA |
| | 0.260158 | 1.240347 | 4.047124 | 0.306476 | 0.759242 | NA |
| | 16.54849 | 0.157438 | 0.548016 | 0.287287 | 0.773892 | NA |
| LINC00890 | 1.044314 | -3.85838 | 2.557513 | -1.50864 | 0.13139 | NA |

| | | | | | | |
|---|---|---|---|---|---|---|
| | 210.2472 | 0.226474 | 0.302297 | 0.749177 | 0.45375 | 0.612629 |
| | 0.163957 | 0.424913 | 4.079006 | 0.104171 | 0.917034 | NA |
| | 15.20701 | -0.63767 | 0.545632 | -1.16867 | 0.242535 | NA |
| | 15.44601 | 0.782459 | 0.52901 | 1.479101 | 0.139113 | NA |
| | 1.467065 | -0.31569 | 1.756543 | -0.17972 | 0.857371 | NA |
| | 0.708364 | 0.219301 | 2.144998 | 0.102238 | 0.918568 | NA |
| | 0.134133 | -1.48909 | 4.077584 | -0.36519 | 0.71497 | NA |
| | 2.419072 | 1.558246 | 1.479862 | 1.052967 | 0.292356 | NA |
| | 0.88142 | 1.783485 | 2.222213 | 0.802571 | 0.422222 | NA |
| | 0.640406 | 1.11402 | 2.710977 | 0.410929 | 0.681124 | NA |
| | 0.151697 | -1.546 | 4.075469 | -0.37934 | 0.704433 | NA |
| | 14.34609 | 0.383753 | 0.586628 | 0.654167 | 0.513004 | NA |
| | 9.807024 | 0.79613 | 0.675859 | 1.177952 | 0.238816 | NA |
| | 6.662849 | 1.140947 | 0.838715 | 1.360351 | 0.173719 | NA |
| TMEM178I | 66.18823 | 0.372363 | 0.427639 | 0.870741 | 0.383895 | 0.550046 |
| | 1.238129 | 0.835091 | 1.848802 | 0.451693 | 0.65149 | NA |
| | 0.151697 | -1.546 | 4.075469 | -0.37934 | 0.704433 | NA |
| | 0.437526 | -2.68242 | 3.384147 | -0.79264 | 0.427986 | NA |
| | 15.87882 | -0.69463 | 0.537909 | -1.29135 | 0.196584 | NA |
| | 4.620079 | 0.316889 | 0.954784 | 0.331896 | 0.739968 | NA |
| | 0.163957 | 0.424913 | 4.079006 | 0.104171 | 0.917034 | NA |
| | 0.913173 | 1.256193 | 2.465498 | 0.509509 | 0.610396 | NA |
| | 0.163957 | 0.424913 | 4.079006 | 0.104171 | 0.917034 | NA |
| | 11.72935 | -0.49862 | 0.719818 | -0.6927 | 0.4885 | NA |
| | 2.264775 | 1.881059 | 1.430818 | 1.314674 | 0.18862 | NA |
| | 1.645977 | -1.91609 | 1.778358 | -1.07745 | 0.281279 | NA |
| | 0.651435 | 0.249629 | 2.788679 | 0.089515 | 0.928672 | NA |
| | 0.886724 | 0.648622 | 2.434305 | 0.26645 | 0.789892 | NA |
| LINC01003 | 111.4379 | -0.11684 | 0.249149 | -0.46895 | 0.639103 | 0.759385 |
| | 43.78054 | -0.40553 | 0.346774 | -1.16944 | 0.242227 | NA |
| | 4.268337 | -0.19233 | 1.054457 | -0.1824 | 0.855273 | NA |
| | 0.151697 | -1.546 | 4.075469 | -0.37934 | 0.704433 | NA |
| | 0.268266 | -2.15396 | 4.038351 | -0.53338 | 0.593773 | NA |
| | 5.492377 | -0.54211 | 0.912008 | -0.59441 | 0.552237 | NA |
| | 0.163957 | 0.424913 | 4.079006 | 0.104171 | 0.917034 | NA |
| | 0.096202 | 0.408546 | 4.079639 | 0.100143 | 0.920231 | NA |
| | 0.392857 | -1.546 | 4.075469 | -0.37934 | 0.704433 | NA |
| MIR548AN | 0.151697 | -1.546 | 4.075469 | -0.37934 | 0.704433 | NA |
| MIR4665 | 0.382031 | -1.26484 | 3.151989 | -0.40128 | 0.688212 | NA |
| | 0.096202 | 0.408546 | 4.079639 | 0.100143 | 0.920231 | NA |
| | 0.260158 | 1.240347 | 4.047124 | 0.306476 | 0.759242 | NA |
| RN7SL777I | 0.326333 | 1.533889 | 3.826878 | 0.40082 | 0.688553 | NA |
| | 0.28583 | -2.21358 | 3.97376 | -0.55705 | 0.577494 | NA |
| MIR4672 | 0.247898 | -0.5901 | 4.074633 | -0.14482 | 0.884851 | NA |
| MIR4666B | 0.460466 | 0.568655 | 3.014642 | 0.188631 | 0.850382 | NA |
| RN7SL474I | 0.151697 | -1.546 | 4.075469 | -0.37934 | 0.704433 | NA |
| RN7SL158I | 1.265177 | -0.09094 | 1.722664 | -0.05279 | 0.957898 | NA |

| | | | | | | |
|---|---|---|---|---|---|---|
| | 2.06401 | -1.06361 | 1.444842 | -0.73614 | 0.461643 | NA |
| RN7SL463F | 0.462046 | 0.589069 | 3.442439 | 0.17112 | 0.86413 | NA |
| MIR4767 | 8.041325 | -0.0805 | 0.705205 | -0.11416 | 0.909114 | NA |
| MIR3911 | 0.930591 | 1.843923 | 2.122799 | 0.868628 | 0.385051 | NA |
| | 1.924728 | 0.392972 | 1.600688 | 0.245502 | 0.806068 | NA |
| MIR4470 | 0.096202 | 0.408546 | 4.079639 | 0.100143 | 0.920231 | NA |
| MIR4653 | 1.342381 | -0.86875 | 1.876521 | -0.46296 | 0.643393 | NA |
| MIR378D2 | 0.151697 | -1.546 | 4.075469 | -0.37934 | 0.704433 | NA |
| | 0.544407 | -0.44425 | 2.542878 | -0.1747 | 0.861313 | NA |
| RN7SL15P | 0.192403 | 0.975392 | 4.060716 | 0.240202 | 0.810174 | NA |
| RN7SL592F | 0.162376 | 0.424913 | 4.079006 | 0.104171 | 0.917034 | NA |
| | 0.096202 | 0.408546 | 4.079639 | 0.100143 | 0.920231 | NA |
| | 0.096202 | 0.408546 | 4.079639 | 0.100143 | 0.920231 | NA |
| | 3.116371 | 1.581272 | 1.24836 | 1.26668 | 0.20527 | NA |
| | 0.260158 | 1.240347 | 4.047124 | 0.306476 | 0.759242 | NA |
| RN7SL262F | 0.682693 | 2.623213 | 2.518338 | 1.041645 | 0.297576 | NA |
| | 0.096202 | 0.408546 | 4.079639 | 0.100143 | 0.920231 | NA |
| MIR4655 | 0.134133 | -1.48909 | 4.077584 | -0.36519 | 0.71497 | NA |
| MIR3153 | 0.738188 | 1.39874 | 2.317793 | 0.603479 | 0.54619 | NA |
| | 5.459683 | 0.602646 | 0.860804 | 0.700097 | 0.483867 | NA |
| RN7SL365F | 0.392857 | -1.546 | 4.075469 | -0.37934 | 0.704433 | NA |
| | 0.547182 | 2.340248 | 3.137689 | 0.745851 | 0.455757 | NA |
| | 0.134133 | -1.48909 | 4.077584 | -0.36519 | 0.71497 | NA |
| MIR4664 | 16.5093 | -0.47602 | 0.552021 | -0.86232 | 0.388511 | NA |
| RN7SL64P | 0.392857 | -1.546 | 4.075469 | -0.37934 | 0.704433 | NA |
| MIR4292 | 7.194047 | 0.061126 | 0.783148 | 0.078052 | 0.937787 | NA |
| FSBP | 7.155257 | 0.48953 | 0.832047 | 0.588345 | 0.556301 | NA |
| MIR3939 | 0.151697 | -1.546 | 4.075469 | -0.37934 | 0.704433 | NA |
| MIR4477B | 0.420954 | 1.923376 | 3.462803 | 0.555439 | 0.578594 | NA |
| MIR4477A | 0.420954 | 1.923376 | 3.462803 | 0.555439 | 0.578594 | NA |
| MIR3609 | 0.315653 | -0.57305 | 3.892639 | -0.14721 | 0.882963 | NA |
| MIR4469 | 0.326333 | 1.533889 | 3.826878 | 0.40082 | 0.688553 | NA |
| RN7SL625F | 0.134133 | -1.48909 | 4.077584 | -0.36519 | 0.71497 | NA |
| | 0.326536 | 0.037056 | 3.842554 | 0.009644 | 0.992306 | NA |
| MIR4668 | 0.163957 | 0.424913 | 4.079006 | 0.104171 | 0.917034 | NA |
| MIR5692A1 | 0.134133 | -1.48909 | 4.077584 | -0.36519 | 0.71497 | NA |
| MIR5692A2 | 0.134133 | -1.48909 | 4.077584 | -0.36519 | 0.71497 | NA |
| | 0.424115 | 1.932531 | 3.45462 | 0.559405 | 0.575885 | NA |
| RN7SL118F | 0.488709 | 2.119056 | 3.307564 | 0.64067 | 0.521737 | NA |
| | 0.660976 | -0.84029 | 2.673548 | -0.3143 | 0.753294 | NA |
| MIR4479 | 1.246236 | -0.68541 | 1.955344 | -0.35053 | 0.72594 | NA |
| RN7SL377F | 0.096202 | 0.408546 | 4.079639 | 0.100143 | 0.920231 | NA |
| MIR5692C2 | 0.151697 | -1.546 | 4.075469 | -0.37934 | 0.704433 | NA |
| RN7SL48P | 0.096202 | 0.408546 | 4.079639 | 0.100143 | 0.920231 | NA |
| | 3.151791 | 0.848041 | 1.266277 | 0.669712 | 0.503041 | NA |
| | 0.364467 | -1.20166 | 3.685431 | -0.32606 | 0.744381 | NA |
| | 0.586694 | 1.130467 | 2.734203 | 0.413454 | 0.679274 | NA |

| | | | | | | |
|---|---|---|---|---|---|---|
| UXT-AS1 | 1.857029 | 0.477319 | 1.439072 | 0.331685 | 0.740127 | NA |
| | 0.785218 | 1.577495 | 2.568228 | 0.614235 | 0.53906 | NA |
| | 0.951104 | -0.01996 | 2.201588 | -0.00907 | 0.992765 | NA |
| | 0.247898 | -0.5901 | 4.074633 | -0.14482 | 0.884851 | NA |
| FMR1-AS1 | 0.162376 | 0.424913 | 4.079006 | 0.104171 | 0.917034 | NA |
| | 0.681112 | 2.620532 | 2.519661 | 1.040033 | 0.298324 | NA |
| AIRN | 0.260158 | 1.240347 | 4.047124 | 0.306476 | 0.759242 | NA |
| SMC5-AS1 | 3.940973 | 0.53923 | 1.043077 | 0.516961 | 0.605183 | NA |
| | 0.3441 | -0.0166 | 3.769682 | -0.0044 | 0.996486 | NA |
| | 0.162376 | 0.424913 | 4.079006 | 0.104171 | 0.917034 | NA |
| | 16.30271 | -0.30075 | 0.547032 | -0.54978 | 0.582473 | NA |
| | 6.027432 | 0.26514 | 0.923464 | 0.287114 | 0.774025 | NA |
| | 0.162376 | 0.424913 | 4.079006 | 0.104171 | 0.917034 | NA |
| | 12.35168 | 0.113156 | 0.659266 | 0.171639 | 0.863721 | NA |
| | 0.296509 | -0.51709 | 3.982155 | -0.12985 | 0.896684 | NA |
| | 0.268266 | -2.15396 | 4.038351 | -0.53338 | 0.593773 | NA |
| | 0.392857 | -1.546 | 4.075469 | -0.37934 | 0.704433 | NA |
| | 0.096202 | 0.408546 | 4.079639 | 0.100143 | 0.920231 | NA |
| | 2.713155 | -0.03478 | 1.21729 | -0.02857 | 0.977205 | NA |
| NA | 0.163957 | 0.424913 | 4.079006 | 0.104171 | 0.917034 | NA |
| NA | 0.096202 | 0.408546 | 4.079639 | 0.100143 | 0.920231 | NA |
| NA | 0.162376 | 0.424913 | 4.079006 | 0.104171 | 0.917034 | NA |
| NA | 0.096202 | 0.408546 | 4.079639 | 0.100143 | 0.920231 | NA |
| NA | 2.705019 | 1.759231 | 1.381449 | 1.273468 | 0.202852 | NA |
| NA | 0.247898 | -0.5901 | 4.074633 | -0.14482 | 0.884851 | NA |
| NA | 0.163957 | 0.424913 | 4.079006 | 0.104171 | 0.917034 | NA |
| NA | 0.432222 | -1.18916 | 3.482045 | -0.34151 | 0.732718 | NA |
| NA | 0.719393 | -0.9644 | 2.652838 | -0.36354 | 0.716204 | NA |
| NA | 1.377216 | 0.878801 | 1.689434 | 0.520175 | 0.602942 | NA |
| NA | 1.066304 | 1.133112 | 2.033868 | 0.557122 | 0.577444 | NA |
| NA | 0.599903 | -1.73313 | 2.730982 | -0.63462 | 0.525678 | NA |
| NA | 3.860386 | -0.1383 | 0.990674 | -0.1396 | 0.888972 | NA |
| NA | 0.872321 | 0.678609 | 2.011629 | 0.337343 | 0.735858 | NA |
| NA | 0.247898 | -0.5901 | 4.074633 | -0.14482 | 0.884851 | NA |
| NA | 0.520316 | 2.239969 | 3.173769 | 0.705776 | 0.480328 | NA |
| NA | 2.826738 | 0.429572 | 1.379874 | 0.311313 | 0.755563 | NA |
| NA | 1.006454 | 0.21027 | 1.900041 | 0.110666 | 0.911881 | NA |
| NA | 5.204194 | 1.168014 | 1.250813 | 0.933803 | 0.350405 | NA |
| NA | 0.410275 | 0.244283 | 3.100889 | 0.078778 | 0.937209 | NA |
| NA | 12.40638 | -0.01832 | 0.634561 | -0.02888 | 0.976963 | NA |
| NA | 0.134133 | -1.48909 | 4.077584 | -0.36519 | 0.71497 | NA |
| NA | 0.096202 | 0.408546 | 4.079639 | 0.100143 | 0.920231 | NA |
| NA | 3.970882 | 0.227571 | 1.064225 | 0.213837 | 0.830674 | NA |
| NA | 0.548763 | 2.343695 | 3.135478 | 0.747476 | 0.454776 | NA |
| NA | 3.977526 | 0.152755 | 1.023836 | 0.149199 | 0.881397 | NA |
| NA | 4.172024 | 0.624185 | 1.077713 | 0.579176 | 0.56247 | NA |
| NA | 9.126216 | -0.04678 | 0.703139 | -0.06653 | 0.946954 | NA |

| | | | | | | |
|---|---|---|---|---|---|---|
| NA | 0.096202 | 0.408546 | 4.079639 | 0.100143 | 0.920231 | NA |
| NA | 2.618709 | 0.525213 | 1.255873 | 0.418205 | 0.675797 | NA |
| NA | 1.169179 | -1.04993 | 1.839034 | -0.57091 | 0.568059 | NA |
| NA | 12.05283 | 0.783165 | 0.624042 | 1.254989 | 0.209483 | NA |
| NA | 0.230334 | -0.53318 | 4.076749 | -0.13079 | 0.895945 | NA |
| NA | 4.033653 | 0.653628 | 0.974224 | 0.670921 | 0.502271 | NA |
| NA | 1.454862 | -0.58515 | 1.616608 | -0.36196 | 0.717381 | NA |
| NA | 0.163957 | 0.424913 | 4.079006 | 0.104171 | 0.917034 | NA |
| NA | 0.288605 | 1.443619 | 4.029058 | 0.358302 | 0.720117 | NA |
| NA | 5.298071 | 0.38444 | 0.928535 | 0.414029 | 0.678853 | NA |
| NA | 0.747636 | 0.735436 | 2.634453 | 0.279161 | 0.780122 | NA |
| NA | 0.844077 | -0.22325 | 2.189502 | -0.10196 | 0.918786 | NA |
| NA | 0.151697 | -1.546 | 4.075469 | -0.37934 | 0.704433 | NA |
| NA | 185.1007 | -0.1563 | 0.245311 | -0.63713 | 0.524038 | 0.670307 |
| NA | 0.192403 | 0.975392 | 4.060716 | 0.240202 | 0.810174 | NA |
| NA | 0.392857 | -1.546 | 4.075469 | -0.37934 | 0.704433 | NA |
| NA | 0.298089 | -0.51711 | 3.975232 | -0.13008 | 0.896501 | NA |
| NA | 0.192403 | 0.975392 | 4.060716 | 0.240202 | 0.810174 | NA |
| NA | 0.258578 | 1.233151 | 4.047809 | 0.304646 | 0.760635 | NA |
| NA | 0.096202 | 0.408546 | 4.079639 | 0.100143 | 0.920231 | NA |
| NA | 0.163957 | 0.424913 | 4.079006 | 0.104171 | 0.917034 | NA |
| NA | 1.801477 | 0.837718 | 1.571131 | 0.533194 | 0.593899 | NA |
| NA | 66.98923 | -0.12418 | 0.283331 | -0.4383 | 0.661172 | 0.776905 |
| NA | 0.163957 | 0.424913 | 4.079006 | 0.104171 | 0.917034 | NA |
| NA | 0.163957 | 0.424913 | 4.079006 | 0.104171 | 0.917034 | NA |
| NA | 0.420954 | 1.923376 | 3.462803 | 0.555439 | 0.578594 | NA |
| NA | 0.970103 | 0.931158 | 2.103793 | 0.442609 | 0.658049 | NA |
| NA | 1.870384 | -1.19022 | 1.504417 | -0.79115 | 0.428855 | NA |
| NA | 0.192403 | 0.975392 | 4.060716 | 0.240202 | 0.810174 | NA |
| NA | 0.163957 | 0.424913 | 4.079006 | 0.104171 | 0.917034 | NA |
| NA | 54.71069 | -0.23842 | 0.399992 | -0.59607 | 0.551128 | 0.69215 |
| NA | 0.192403 | 0.975392 | 4.060716 | 0.240202 | 0.810174 | NA |
| NA | 0.151697 | -1.546 | 4.075469 | -0.37934 | 0.704433 | NA |
| NA | 0.163957 | 0.424913 | 4.079006 | 0.104171 | 0.917034 | NA |
| NA | 1.981474 | 1.060056 | 1.457971 | 0.727076 | 0.467179 | NA |
| NA | 0.392857 | -1.546 | 4.075469 | -0.37934 | 0.704433 | NA |
| NA | 0.949378 | 0.969416 | 2.078125 | 0.466486 | 0.640868 | NA |
| NA | 2.03082 | -0.68062 | 1.438891 | -0.47302 | 0.636199 | NA |
| NA | 0.842643 | -2.08432 | 2.306153 | -0.90381 | 0.366097 | NA |
| NA | 0.58526 | 0.001197 | 3.235889 | 3.70E-04 | 0.999705 | NA |
| NA | 0.881623 | 0.857232 | 2.360567 | 0.363147 | 0.716495 | NA |
| NA | 40.85569 | 0.228931 | 0.348784 | 0.656369 | 0.511587 | NA |
| NA | 7.694195 | -0.5809 | 0.769509 | -0.75489 | 0.450315 | NA |
| NA | 52.01346 | 0.765969 | 0.358387 | 2.137267 | 0.032576 | 0.105021 |
| NA | 0.260158 | 1.240347 | 4.047124 | 0.306476 | 0.759242 | NA |
| NA | 2.887617 | 1.817345 | 1.282066 | 1.417513 | 0.156333 | NA |
| NA | 3.92815 | 0.550405 | 1.016364 | 0.541543 | 0.588133 | NA |

| | | | | | | |
|---|---|---|---|---|---|---|
| NA | 1.717739 | 0.809782 | 1.52915 | 0.529563 | 0.596415 | NA |
| NA | 23.92622 | 0.888599 | 0.504392 | 1.761724 | 0.078116 | NA |
| NA | 0.28583 | -2.21358 | 3.97376 | -0.55705 | 0.577494 | NA |
| NA | 0.644964 | 2.579089 | 2.975624 | 0.866739 | 0.386085 | NA |
| NA | 0.260158 | 1.240347 | 4.047124 | 0.306476 | 0.759242 | NA |
| NA | 1.763575 | -0.25188 | 1.622659 | -0.15523 | 0.876642 | NA |
| NA | 129.827 | 0.458562 | 0.241132 | 1.901705 | 0.05721 | 0.151659 |
| NA | 0.92901 | 1.841773 | 2.123319 | 0.867403 | 0.385721 | NA |
| NA | 0.487129 | 2.123445 | 3.910539 | 0.543006 | 0.587126 | NA |
| NA | 0.490493 | 0.776209 | 2.895261 | 0.268096 | 0.788625 | NA |
| NA | 1.541319 | 0.429663 | 1.664577 | 0.258121 | 0.796313 | NA |
| NA | 0.327913 | 1.543154 | 4.021063 | 0.383768 | 0.701151 | NA |
| NA | 0.260158 | 1.240347 | 4.047124 | 0.306476 | 0.759242 | NA |
| NA | 0.394291 | 0.298745 | 3.141711 | 0.09509 | 0.924243 | NA |
| NA | 5.384528 | -0.3359 | 0.86191 | -0.38972 | 0.696746 | NA |
| NA | 0.163957 | 0.424913 | 4.079006 | 0.104171 | 0.917034 | NA |
| NA | 10.8931 | 0.871916 | 0.645058 | 1.351685 | 0.176476 | NA |
| NA | 0.288605 | 1.443619 | 4.029058 | 0.358302 | 0.720117 | NA |
| NA | 0.651288 | 1.289307 | 2.633717 | 0.489539 | 0.62446 | NA |
| NA | 0.47961 | 0.535796 | 3.39605 | 0.15777 | 0.874638 | NA |
| NA | 11.36492 | -0.65133 | 0.777282 | -0.83796 | 0.40205 | NA |
| NA | 0.392857 | -1.546 | 4.075469 | -0.37934 | 0.704433 | NA |
| NA | 1.108162 | -0.89528 | 2.172518 | -0.41209 | 0.680272 | NA |
| NA | 3.3203 | -0.39736 | 1.14031 | -0.34846 | 0.727492 | NA |
| NA | 6.860052 | -0.18046 | 0.824601 | -0.21884 | 0.826775 | NA |
| NA | 1.425624 | 1.633938 | 1.964017 | 0.831937 | 0.405445 | NA |
| NA | 0.667861 | -2.0782 | 2.593993 | -0.80116 | 0.42304 | NA |
| NA | 12.34881 | 1.020449 | 0.633191 | 1.611598 | 0.10705 | NA |
| NA | 1.197073 | 1.305651 | 1.836533 | 0.710932 | 0.477126 | NA |
| NA | 0.326333 | 1.533889 | 3.826878 | 0.40082 | 0.688553 | NA |
| NA | 0.528424 | -0.38142 | 2.853651 | -0.13366 | 0.893671 | NA |
| NA | 3.254371 | 0.979549 | 1.312002 | 0.746606 | 0.455301 | NA |
| NA | 31.23871 | -0.19563 | 0.385307 | -0.50774 | 0.611639 | NA |
| NA | 0.668852 | 1.246964 | 2.610519 | 0.477669 | 0.632886 | NA |
| NA | 1.312768 | -0.01332 | 1.920498 | -0.00694 | 0.994466 | NA |
| NA | 1.064521 | 2.035642 | 2.088007 | 0.974921 | 0.329599 | NA |
| NA | 0.422534 | 1.924583 | 3.001758 | 0.641152 | 0.521424 | NA |
| NA | 6.074162 | -0.05793 | 0.807125 | -0.07177 | 0.942786 | NA |
| NA | 1.209479 | 1.863531 | 2.324529 | 0.801681 | 0.422737 | NA |
| NA | 0.096202 | 0.408546 | 4.079639 | 0.100143 | 0.920231 | NA |
| NA | 20.57896 | 1.118347 | 0.641882 | 1.742294 | 0.081457 | NA |
| NA | 0.614937 | 2.493639 | 2.617677 | 0.952615 | 0.340785 | NA |
| NA | 2.763261 | 0.946956 | 1.293513 | 0.732081 | 0.464119 | NA |
| NA | 1.896338 | 0.479132 | 1.467658 | 0.32646 | 0.744076 | NA |
| NA | 0.741166 | 2.78101 | 2.851312 | 0.975344 | 0.32939 | NA |
| NA | 4.661431 | 0.05836 | 0.969689 | 0.060184 | 0.952009 | NA |
| NA | 5.977271 | -0.67833 | 0.844564 | -0.80318 | 0.421872 | NA |

| | | | | | | |
|---|---|---|---|---|---|---|
| NA | 6.764013 | 0.405856 | 0.827137 | 0.490675 | 0.623656 | NA |
| NA | 0.815391 | 0.931626 | 2.360431 | 0.394685 | 0.693076 | NA |
| NA | 1.629791 | -1.39133 | 1.699006 | -0.81891 | 0.412837 | NA |
| NA | 0.151697 | -1.546 | 4.075469 | -0.37934 | 0.704433 | NA |
| NA | 0.151697 | -1.546 | 4.075469 | -0.37934 | 0.704433 | NA |
| NA | 2.077473 | 1.630485 | 1.577913 | 1.033317 | 0.301455 | NA |
| NA | 6.480512 | 0.108513 | 0.818263 | 0.132614 | 0.894498 | NA |
| NA | 689.9089 | 0.760902 | 0.204643 | 3.718186 | 2.01E-04 | 0.003868 |
| NA | 0.260158 | 1.240347 | 4.047124 | 0.306476 | 0.759242 | NA |
| NA | 1.344617 | -0.99306 | 2.086066 | -0.47604 | 0.634043 | NA |
| NA | 0.856337 | 0.757304 | 2.453242 | 0.308695 | 0.757553 | NA |
| NA | 1.469637 | 2.274236 | 2.004742 | 1.134429 | 0.256615 | NA |
| NA | 2.828115 | 0.719244 | 1.209705 | 0.594562 | 0.552136 | NA |
| NA | 0.151697 | -1.546 | 4.075469 | -0.37934 | 0.704433 | NA |
| NA | 0.422737 | 0.518872 | 3.501356 | 0.148192 | 0.882191 | NA |
| NA | 0.518736 | 2.237423 | 2.7713 | 0.807355 | 0.419462 | NA |
| NA | 0.096202 | 0.408546 | 4.079639 | 0.100143 | 0.920231 | NA |
| NA | 0.163957 | 0.424913 | 4.079006 | 0.104171 | 0.917034 | NA |
| NA | 0.452561 | 2.061505 | 3.349456 | 0.615474 | 0.538241 | NA |
| NA | 0.55687 | -0.14239 | 3.170029 | -0.04492 | 0.964172 | NA |
| NA | 0.162376 | 0.424913 | 4.079006 | 0.104171 | 0.917034 | NA |
| NA | 1.633368 | -1.08276 | 1.583569 | -0.68375 | 0.494136 | NA |
| NA | 1.466826 | 0.187141 | 1.944883 | 0.096222 | 0.923344 | NA |
| NA | 0.151697 | -1.546 | 4.075469 | -0.37934 | 0.704433 | NA |
| NA | 2.489705 | -0.37598 | 1.208265 | -0.31117 | 0.75567 | NA |
| NA | 0.3441 | -0.0166 | 3.769682 | -0.0044 | 0.996486 | NA |
| NA | 0.506476 | 0.726418 | 2.864186 | 0.253621 | 0.799788 | NA |
| NA | 0.151697 | -1.546 | 4.075469 | -0.37934 | 0.704433 | NA |
| NA | 0.096202 | 0.408546 | 4.079639 | 0.100143 | 0.920231 | NA |
| NA | 94.34982 | 0.494045 | 0.342856 | 1.440971 | 0.149593 | 0.293079 |
| NA | 0.749273 | 0.569822 | 2.889797 | 0.197184 | 0.843683 | NA |
| NA | 0.096202 | 0.408546 | 4.079639 | 0.100143 | 0.920231 | NA |
| NA | 0.134133 | -1.48909 | 4.077584 | -0.36519 | 0.71497 | NA |
| NA | 0.134133 | -1.48909 | 4.077584 | -0.36519 | 0.71497 | NA |
| NA | 0.364467 | -1.20166 | 3.685431 | -0.32606 | 0.744381 | NA |
| NA | 0.247898 | -0.5901 | 4.074633 | -0.14482 | 0.884851 | NA |
| NA | 4.61616 | -0.3491 | 1.012309 | -0.34486 | 0.730203 | NA |
| NA | 2.74626 | 0.50783 | 1.31029 | 0.387571 | 0.698334 | NA |
| NA | 269.4246 | -0.47591 | 0.192236 | -2.47565 | 0.013299 | 0.057374 |
| NA | 0.260158 | 1.240347 | 4.047124 | 0.306476 | 0.759242 | NA |
| NA | 0.394291 | 0.298745 | 3.141711 | 0.09509 | 0.924243 | NA |
| NA | 0.096202 | 0.408546 | 4.079639 | 0.100143 | 0.920231 | NA |
| NA | 6.455308 | 1.764107 | 0.91469 | 1.928639 | 0.053776 | NA |
| NA | 40.77676 | -0.09668 | 0.344558 | -0.28059 | 0.779026 | NA |
| NA | 7.215339 | -0.54205 | 0.761949 | -0.7114 | 0.476835 | NA |
| NA | 2.468353 | 1.148125 | 1.549118 | 0.741147 | 0.458604 | NA |
| NA | 0.392857 | -1.546 | 4.075469 | -0.37934 | 0.704433 | NA |

| | | | | | | | |
|---|---|---|---|---|---|---|---|
| NA | 0.670636 | 0.219794 | 2.632532 | 0.083491 | 0.933461 | NA | |
| NA | 5.435171 | -0.90459 | 1.059064 | -0.85414 | 0.393029 | NA | |
| NA | 0.324752 | 1.53082 | 4.022025 | 0.380609 | 0.703493 | NA | |
| NA | 0.572651 | 0.928649 | 2.780261 | 0.334015 | 0.738368 | NA | |
| NA | 0.163957 | 0.424913 | 4.079006 | 0.104171 | 0.917034 | NA | |
| NA | 0.247898 | -0.5901 | 4.074633 | -0.14482 | 0.884851 | NA | |
| NA | 1.29763 | 3.559825 | 2.014572 | 1.767038 | 0.077222 | NA | |
| NA | 2.746151 | 0.917161 | 1.37454 | 0.667249 | 0.504613 | NA | |
| NA | 0.151697 | -1.546 | 4.075469 | -0.37934 | 0.704433 | NA | |
| NA | 0.670433 | 1.255281 | 2.356401 | 0.532711 | 0.594234 | NA | |
| NA | 9.965292 | -0.03028 | 0.635321 | -0.04766 | 0.96199 | NA | |
| NA | 14.59014 | -0.59489 | 0.542947 | -1.09567 | 0.273223 | NA | |
| NA | 1.763603 | 1.398047 | 1.552819 | 0.900328 | 0.367946 | NA | |
| NA | 2.712778 | 1.662301 | 1.28044 | 1.298226 | 0.19421 | NA | |
| NA | 0.902348 | 0.812045 | 2.125951 | 0.381968 | 0.702485 | NA | |
| NA | 110.0731 | -0.01322 | 0.264613 | -0.04996 | 0.960156 | | 0.9781 |
| NA | 123.0363 | 0.274222 | 0.23431 | 1.17034 | 0.241864 | | 0.404924 |
| NA | 27.20548 | -0.18876 | 0.449203 | -0.42022 | 0.674327 | NA | |
| NA | 11.00476 | 0.255343 | 0.634491 | 0.402438 | 0.687362 | NA | |
| NA | 20.80226 | -0.12054 | 0.488349 | -0.24683 | 0.805043 | NA | |
| NA | 0.162376 | 0.424913 | 4.079006 | 0.104171 | 0.917034 | NA | |
| NA | 25.91129 | 0.581085 | 0.44269 | 1.312623 | 0.18931 | NA | |
| NA | 4.330045 | 0.304836 | 1.061252 | 0.287242 | 0.773927 | NA | |
| NA | 37.79299 | 0.157151 | 0.402879 | 0.39007 | 0.696485 | NA | |
| NA | 551.406 | 0.061909 | 0.214505 | 0.288612 | 0.772879 | | 0.859277 |
| NA | 0.670433 | 1.255281 | 2.356401 | 0.532711 | 0.594234 | NA | |
| NA | 116.3308 | -0.50404 | 0.248651 | -2.02711 | 0.042652 | | 0.125686 |
| NA | 0.163957 | 0.424913 | 4.079006 | 0.104171 | 0.917034 | NA | |
| NA | 0.575812 | 0.936922 | 2.776634 | 0.337431 | 0.735792 | NA | |
| NA | 605.5667 | 0.024185 | 0.18115 | 0.133507 | 0.893792 | | 0.938023 |
| NA | 0.192403 | 0.975392 | 4.060716 | 0.240202 | 0.810174 | NA | |
| NA | 6.009116 | -0.27561 | 0.835548 | -0.32985 | 0.741513 | NA | |
| NA | 3678.611 | -0.36701 | 0.180965 | -2.02805 | 0.042555 | | 0.125496 |
| NA | 0.816825 | 1.673341 | 2.21346 | 0.755984 | 0.449659 | NA | |
| NA | 78.01318 | -0.14291 | 0.278175 | -0.51375 | 0.60743 | | 0.73511 |
| NA | 5.697384 | 0.822691 | 0.939511 | 0.875659 | 0.381216 | NA | |
| NA | 0.162376 | 0.424913 | 4.079006 | 0.104171 | 0.917034 | NA | |
| NA | 2.279296 | -0.03202 | 1.403524 | -0.02282 | 0.981798 | NA | |
| NA | 2.340591 | 1.837885 | 1.729271 | 1.062809 | 0.287869 | NA | |
| NA | 0.151697 | -1.546 | 4.075469 | -0.37934 | 0.704433 | NA | |
| NA | 2.817757 | -1.56943 | 1.297134 | -1.20992 | 0.226309 | NA | |
| NA | 12.55256 | 0.329673 | 0.579379 | 0.569011 | 0.569349 | NA | |
| NA | 1.014118 | 0.121699 | 2.429485 | 0.050092 | 0.960049 | NA | |
| NA | 1.698741 | 0.093534 | 1.670057 | 0.056007 | 0.955336 | NA | |
| NA | 1.1119 | 0.46633 | 1.933926 | 0.241131 | 0.809453 | NA | |
| NA | 0.410275 | 0.244283 | 3.100889 | 0.078778 | 0.937209 | NA | |
| NA | 0.260158 | 1.240347 | 4.047124 | 0.306476 | 0.759242 | NA | |

| | | | | | | | |
|---|---|---|---|---|---|---|---|
| NA | 0.640609 | 0.042838 | 2.379511 | 0.018003 | 0.985637 | NA | |
| NA | 0.623045 | 0.091811 | 2.666142 | 0.034436 | 0.97253 | NA | |
| NA | 1.320527 | 0.156199 | 1.694976 | 0.092154 | 0.926575 | NA | |
| NA | 0.52699 | -2.57878 | 3.248478 | -0.79384 | 0.427288 | NA | |
| NA | 15.44273 | 1.043196 | 0.554299 | 1.882011 | 0.059834 | NA | |
| NA | 0.151697 | -1.546 | 4.075469 | -0.37934 | 0.704433 | NA | |
| NA | 2.266538 | 1.340283 | 1.464905 | 0.914928 | 0.360229 | NA | |
| NA | 2.192957 | -0.09881 | 1.375251 | -0.07185 | 0.942722 | NA | |
| NA | 1.06136 | 2.032712 | 2.134262 | 0.952419 | 0.340884 | NA | |
| NA | 7.157432 | 0.133903 | 0.738937 | 0.18121 | 0.856203 | NA | |
| NA | 4.558091 | -1.4914 | 1.0743 | -1.38825 | 0.16506 | NA | |
| NA | 1.05261 | -0.79 | 2.093679 | -0.37733 | 0.705932 | NA | |
| NA | 1.083803 | -2.45142 | 2.403819 | -1.0198 | 0.307822 | NA | |
| NA | 4.855027 | -0.48401 | 1.018843 | -0.47506 | 0.634747 | NA | |
| NA | 20.40421 | -0.48903 | 0.458709 | -1.06611 | 0.286376 | NA | |
| NA | 0.968319 | 1.86533 | 2.390384 | 0.780348 | 0.435186 | NA | |
| NA | 0.411855 | 0.250456 | 3.097618 | 0.080854 | 0.935558 | NA | |
| NA | 0.966942 | 0.917791 | 1.915974 | 0.47902 | 0.631924 | NA | |
| NA | 51.31068 | -0.23257 | 0.338015 | -0.68803 | 0.491432 | | 0.643291 |
| NA | 9.671725 | 0.398501 | 0.668977 | 0.595687 | 0.551384 | NA | |
| NA | 99.4965 | 0.112892 | 0.247715 | 0.455734 | 0.648581 | | 0.766669 |
| NA | 2.373033 | 3.391224 | 1.631156 | 2.079031 | 0.037614 | NA | |
| NA | 25.09557 | -0.34542 | 0.436329 | -0.79165 | 0.428568 | NA | |
| NA | 0.134133 | -1.48909 | 4.077584 | -0.36519 | 0.71497 | NA | |
| NA | 0.411855 | 0.250456 | 3.097618 | 0.080854 | 0.935558 | NA | |
| NA | 23.9466 | 0.253374 | 0.425809 | 0.595041 | 0.551816 | NA | |
| NA | 0.623191 | -1.58103 | 2.770593 | -0.57065 | 0.568239 | NA | |
| NA | 0.163957 | 0.424913 | 4.079006 | 0.104171 | 0.917034 | NA | |
| NA | 0.096202 | 0.408546 | 4.079639 | 0.100143 | 0.920231 | NA | |
| NA | 632.8943 | 0.082983 | 0.211184 | 0.392942 | 0.694362 | | 0.802993 |
| NA | 0.772958 | 0.415176 | 2.577252 | 0.161092 | 0.872021 | NA | |
| NA | 0.392857 | -1.546 | 4.075469 | -0.37934 | 0.704433 | NA | |
| NA | 0.260158 | 1.240347 | 4.047124 | 0.306476 | 0.759242 | NA | |
| NA | 6.858819 | 0.937486 | 0.790884 | 1.185364 | 0.235874 | NA | |
| NA | 0.162376 | 0.424913 | 4.079006 | 0.104171 | 0.917034 | NA | |
| NA | 1.017482 | -0.71858 | 2.105405 | -0.3413 | 0.732874 | NA | |
| NA | 2.229547 | -0.77402 | 1.281507 | -0.60399 | 0.545851 | NA | |
| NA | 0.134133 | -1.48909 | 4.077584 | -0.36519 | 0.71497 | NA | |
| NA | 6.654176 | -0.01278 | 0.838386 | -0.01525 | 0.987835 | NA | |
| NA | 0.134133 | -1.48909 | 4.077584 | -0.36519 | 0.71497 | NA | |
| NA | 1.621188 | 1.859647 | 1.660407 | 1.119994 | 0.262716 | NA | |
| NA | 1.141927 | 0.609996 | 1.889109 | 0.322901 | 0.74677 | NA | |
| NA | 0.844077 | -0.22325 | 2.189502 | -0.10196 | 0.918786 | NA | |
| NA | 1.078282 | -2.71064 | 2.113412 | -1.28259 | 0.199636 | NA | |
| NA | 0.192403 | 0.975392 | 4.060716 | 0.240202 | 0.810174 | NA | |
| NA | 50.9714 | -0.03072 | 0.396223 | -0.07753 | 0.938205 | | 0.965739 |
| NA | 20.75854 | 0.686502 | 0.583581 | 1.176362 | 0.23945 | NA | |

| | | | | | | |
|---|---|---|---|---|---|---|
| NA | 14.78211 | 0.170224 | 0.534367 | 0.318552 | 0.750066 | NA |
| NA | 0.746295 | -0.65731 | 2.318283 | -0.28353 | 0.776767 | NA |
| NA | 0.70693 | -1.62766 | 2.687154 | -0.60572 | 0.544701 | NA |
| NA | 59.25318 | 0.513192 | 0.321478 | 1.596352 | 0.11041 | 0.238287 |
| NA | 6.30898 | -0.17834 | 0.841406 | -0.21195 | 0.832145 | NA |
| NA | 22.74525 | -0.30077 | 0.426216 | -0.70567 | 0.480393 | NA |
| NA | 0.326333 | 1.533889 | 3.826878 | 0.40082 | 0.688553 | NA |
| NA | 1.129667 | -0.3654 | 1.881429 | -0.19421 | 0.846008 | NA |
| NA | 0.163957 | 0.424913 | 4.079006 | 0.104171 | 0.917034 | NA |
| NA | 0.49029 | 2.123092 | 3.304282 | 0.642527 | 0.520531 | NA |
| NA | 0.813811 | 0.928591 | 2.591569 | 0.358312 | 0.72011 | NA |
| NA | 2.13454 | -0.0628 | 1.316887 | -0.04768 | 0.961968 | NA |
| NA | 0.096202 | 0.408546 | 4.079639 | 0.100143 | 0.920231 | NA |
| NA | 0.163957 | 0.424913 | 4.079006 | 0.104171 | 0.917034 | NA |
| NA | 1.993466 | 0.509146 | 1.640922 | 0.31028 | 0.756348 | NA |
| NA | 0.354779 | 1.703954 | 3.657232 | 0.465914 | 0.641277 | NA |
| NA | 0.792306 | -0.44145 | 2.21354 | -0.19943 | 0.841926 | NA |
| NA | 0.230334 | -0.53318 | 4.076749 | -0.13079 | 0.895945 | NA |
| NA | 19.83694 | 0.603017 | 0.494301 | 1.219941 | 0.222487 | NA |
| NA | 2.850421 | 0.704407 | 1.219022 | 0.577846 | 0.563368 | NA |
| NA | 9.902591 | 0.617972 | 0.662693 | 0.932517 | 0.35107 | NA |
| NA | 0.151697 | -1.546 | 4.075469 | -0.37934 | 0.704433 | NA |
| NA | 0.682896 | 1.412658 | 2.61672 | 0.539858 | 0.589295 | NA |
| NA | 0.430642 | -1.1891 | 3.485782 | -0.34113 | 0.733008 | NA |
| NA | 2.033538 | -0.01636 | 1.699369 | -0.00963 | 0.99232 | NA |
| NA | 1.459015 | 1.061255 | 1.654615 | 0.641391 | 0.521269 | NA |
| NA | 0.162376 | 0.424913 | 4.079006 | 0.104171 | 0.917034 | NA |
| NA | 0.192403 | 0.975392 | 4.060716 | 0.240202 | 0.810174 | NA |
| NA | 39.41562 | -0.39178 | 0.334518 | -1.17119 | 0.241523 | NA |
| NA | 3.424652 | 0.798681 | 1.097644 | 0.727632 | 0.466839 | NA |
| NA | 0.151697 | -1.546 | 4.075469 | -0.37934 | 0.704433 | NA |
| NA | 0.995185 | 1.939099 | 2.104716 | 0.921311 | 0.356888 | NA |
| NA | 6.256826 | 1.244429 | 0.941608 | 1.321599 | 0.186302 | NA |
| NA | 0.86916 | 0.686589 | 2.213986 | 0.310114 | 0.756474 | NA |
| NA | 1.072977 | -1.55799 | 1.954834 | -0.79699 | 0.425455 | NA |
| NA | 3.550623 | 0.80519 | 1.166444 | 0.690295 | 0.490009 | NA |
| NA | 0.670433 | 1.255281 | 2.356401 | 0.532711 | 0.594234 | NA |
| NA | 1.018713 | 1.086497 | 2.092548 | 0.519222 | 0.603606 | NA |
| NA | 49.50061 | 0.220464 | 0.375398 | 0.587282 | 0.557015 | 0.696198 |
| NA | 0.134133 | -1.48909 | 4.077584 | -0.36519 | 0.71497 | NA |
| NA | 0.260158 | 1.240347 | 4.047124 | 0.306476 | 0.759242 | NA |
| NA | 1.389476 | 1.604276 | 1.701484 | 0.942869 | 0.345748 | NA |
| NA | 1.50496 | -0.97023 | 1.870122 | -0.51881 | 0.603895 | NA |
| NA | 4.03916 | 0.225133 | 0.995188 | 0.226221 | 0.821029 | NA |
| NA | 0.096202 | 0.408546 | 4.079639 | 0.100143 | 0.920231 | NA |
| NA | 0.163957 | 0.424913 | 4.079006 | 0.104171 | 0.917034 | NA |
| NA | 0.151697 | -1.546 | 4.075469 | -0.37934 | 0.704433 | NA |

| | | | | | | |
|---|---|---|---|---|---|---|
| NA | 0.298089 | -0.51711 | 3.975232 | -0.13008 | 0.896501 | NA |
| NA | 0.096202 | 0.408546 | 4.079639 | 0.100143 | 0.920231 | NA |
| NA | 1.016048 | -2.46876 | 2.437554 | -1.0128 | 0.311154 | NA |
| NA | 0.556813 | -0.57007 | 3.294615 | -0.17303 | 0.862627 | NA |
| NA | 14.36111 | 0.284085 | 0.57735 | 0.49205 | 0.622684 | NA |
| NA | 0.163957 | 0.424913 | 4.079006 | 0.104171 | 0.917034 | NA |
| NA | 0.163957 | 0.424913 | 4.079006 | 0.104171 | 0.917034 | NA |
| NA | 36.11147 | 0.501859 | 0.389999 | 1.286818 | 0.198158 | NA |
| NA | 1.111957 | 0.613148 | 2.197118 | 0.279069 | 0.780192 | NA |
| NA | 0.966942 | 0.917791 | 1.915974 | 0.47902 | 0.631924 | NA |
| NA | 0.298089 | -0.51711 | 3.975232 | -0.13008 | 0.896501 | NA |
| NA | 1426.602 | -0.20465 | 0.152109 | -1.34542 | 0.17849 | 0.331093 |
| NA | 0.134133 | -1.48909 | 4.077584 | -0.36519 | 0.71497 | NA |
| NA | 0.162376 | 0.424913 | 4.079006 | 0.104171 | 0.917034 | NA |
| NA | 1.574296 | -0.82564 | 1.83021 | -0.45112 | 0.651903 | NA |
| NA | 3.735012 | 1.480977 | 1.136968 | 1.302567 | 0.192723 | NA |
| NA | 0.134133 | -1.48909 | 4.077584 | -0.36519 | 0.71497 | NA |
| NA | 1.015699 | 0.142451 | 2.181052 | 0.065313 | 0.947925 | NA |
| NA | 0.324752 | 1.53082 | 4.022025 | 0.380609 | 0.703493 | NA |
| NA | 10.01836 | 0.659949 | 0.671032 | 0.983484 | 0.325369 | NA |
| NA | 0.162376 | 0.424913 | 4.079006 | 0.104171 | 0.917034 | NA |
| NA | 0.151697 | -1.546 | 4.075469 | -0.37934 | 0.704433 | NA |
| NA | 0.260158 | 1.240347 | 4.047124 | 0.306476 | 0.759242 | NA |
| NA | 2.655068 | 1.193448 | 1.333164 | 0.895199 | 0.37068 | NA |
| NA | 1195.384 | 0.287839 | 0.166458 | 1.729197 | 0.083774 | 0.197146 |
| NA | 0.096202 | 0.408546 | 4.079639 | 0.100143 | 0.920231 | NA |
| NA | 26.76229 | -0.04868 | 0.484541 | -0.10046 | 0.919976 | NA |
| NA | 0.489058 | -0.58569 | 3.427543 | -0.17088 | 0.864321 | NA |
| NA | 9.666108 | -0.18555 | 0.656496 | -0.28264 | 0.777451 | NA |
| NA | 0.192403 | 0.975392 | 4.060716 | 0.240202 | 0.810174 | NA |
| NA | 0.681461 | 0.491544 | 3.066278 | 0.160307 | 0.87264 | NA |
| NA | 0.134133 | -1.48909 | 4.077584 | -0.36519 | 0.71497 | NA |
| NA | 40.81513 | 1.084003 | 0.507967 | 2.134005 | 0.032842 | NA |
| NA | 1.8771 | 0.331367 | 1.625066 | 0.20391 | 0.838424 | NA |
| NA | 3.938498 | 1.65544 | 1.222006 | 1.35469 | 0.175516 | NA |
| NA | 1.581879 | 1.852431 | 1.635532 | 1.132617 | 0.257375 | NA |
| NA | 0.49029 | 2.123092 | 3.304282 | 0.642527 | 0.520531 | NA |
| NA | 0.151697 | -1.546 | 4.075469 | -0.37934 | 0.704433 | NA |
| NA | 0.192403 | 0.975392 | 4.060716 | 0.240202 | 0.810174 | NA |
| NA | 0.508056 | 0.731032 | 2.861912 | 0.255435 | 0.798387 | NA |
| NA | 1.044145 | 0.31036 | 2.099918 | 0.147796 | 0.882504 | NA |
| NA | 92.01408 | -0.06835 | 0.289631 | -0.23599 | 0.81344 | 0.888705 |
| NA | 0.163957 | 0.424913 | 4.079006 | 0.104171 | 0.917034 | NA |
| NA | 1.064724 | 1.125776 | 1.854789 | 0.606957 | 0.54388 | NA |
| NA | 98.45061 | 0.177384 | 0.277017 | 0.640336 | 0.521954 | 0.668506 |
| NA | 0.096202 | 0.408546 | 4.079639 | 0.100143 | 0.920231 | NA |
| NA | 8.898671 | -1.05781 | 0.7523 | -1.4061 | 0.159695 | NA |

| | | | | | | |
|---|---|---|---|---|---|---|
| NA | 1.363805 | 3.621971 | 1.977166 | 1.831901 | 0.066966 | NA |
| NA | 0.192403 | 0.975392 | 4.060716 | 0.240202 | 0.810174 | NA |
| NA | 1.831504 | 0.901042 | 1.589352 | 0.566924 | 0.570766 | NA |
| NA | 0.49029 | 2.123092 | 3.304282 | 0.642527 | 0.520531 | NA |
| NA | 1.178278 | -0.21536 | 1.987735 | -0.10835 | 0.913721 | NA |
| NA | 0.327913 | 1.543154 | 4.021063 | 0.383768 | 0.701151 | NA |
| NA | 57.08682 | 0.1954 | 0.334836 | 0.583571 | 0.559509 | 0.698208 |
| NA | 0.162376 | 0.424913 | 4.079006 | 0.104171 | 0.917034 | NA |
| NA | 21.202 | 0.871565 | 0.496051 | 1.757005 | 0.078917 | NA |
| NA | 0.134133 | -1.48909 | 4.077584 | -0.36519 | 0.71497 | NA |
| NA | 0.096202 | 0.408546 | 4.079639 | 0.100143 | 0.920231 | NA |
| NA | 3.215633 | -0.21847 | 1.078036 | -0.20265 | 0.839406 | NA |
| NA | 98.42462 | -0.27107 | 0.253165 | -1.07071 | 0.284298 | 0.450883 |
| NA | 1.073969 | 1.636961 | 2.132951 | 0.767463 | 0.442806 | NA |
| NA | 2.246965 | -0.2361 | 1.321807 | -0.17862 | 0.858234 | NA |
| NA | 7.866849 | -0.10492 | 0.800035 | -0.13115 | 0.895659 | NA |
| NA | 6.196517 | 1.548583 | 0.855595 | 1.80995 | 0.070304 | NA |
| NA | 26.32478 | 0.088684 | 0.407276 | 0.217748 | 0.827625 | NA |
| NA | 0.163957 | 0.424913 | 4.079006 | 0.104171 | 0.917034 | NA |
| NA | 0.667861 | -2.0782 | 2.593993 | -0.80116 | 0.42304 | NA |
| NA | 5.033554 | 1.303381 | 1.02471 | 1.271951 | 0.20339 | NA |
| NA | 2.208188 | -1.7278 | 1.421078 | -1.21584 | 0.224047 | NA |
| NA | 0.134133 | -1.48909 | 4.077584 | -0.36519 | 0.71497 | NA |
| NA | 0.162376 | 0.424913 | 4.079006 | 0.104171 | 0.917034 | NA |
| NA | 2.914586 | -0.47352 | 1.259235 | -0.37603 | 0.706891 | NA |
| NA | 0.268266 | -2.15396 | 4.038351 | -0.53338 | 0.593773 | NA |
| NA | 0.162376 | 0.424913 | 4.079006 | 0.104171 | 0.917034 | NA |
| NA | 4.016816 | -0.9346 | 1.077665 | -0.86725 | 0.385807 | NA |
| NA | 1.266023 | 3.509235 | 2.049126 | 1.712552 | 0.086795 | NA |
| NA | 5.779178 | -0.30424 | 0.84599 | -0.35962 | 0.719129 | NA |
| NA | 0.162376 | 0.424913 | 4.079006 | 0.104171 | 0.917034 | NA |
| NA | 5.352388 | 0.199463 | 0.879338 | 0.226833 | 0.820553 | NA |
| NA | 3.285603 | 0.415708 | 1.129021 | 0.368202 | 0.712723 | NA |
| NA | 0.298089 | -0.51711 | 3.975232 | -0.13008 | 0.896501 | NA |
| NA | 1.839262 | 0.914215 | 1.464635 | 0.624193 | 0.532501 | NA |
| NA | 0.28583 | -2.21358 | 3.97376 | -0.55705 | 0.577494 | NA |
| NA | 0.151697 | -1.546 | 4.075469 | -0.37934 | 0.704433 | NA |
| NA | 0.652869 | 1.292592 | 2.374873 | 0.544278 | 0.58625 | NA |
| NA | 2.874512 | 0.100918 | 1.13792 | 0.088687 | 0.929331 | NA |
| NA | 1.064724 | 1.125776 | 1.854789 | 0.606957 | 0.54388 | NA |
| NA | 2.581741 | 0.801615 | 1.37506 | 0.582968 | 0.559915 | NA |
| NA | 0.49029 | 2.123092 | 3.304282 | 0.642527 | 0.520531 | NA |
| NA | 1.53895 | -1.90624 | 1.779374 | -1.0713 | 0.284035 | NA |
| NA | 1.295001 | 0.77564 | 1.869139 | 0.414972 | 0.678163 | NA |
| NA | 18.79715 | -0.12846 | 0.518831 | -0.2476 | 0.804445 | NA |
| NA | 0.717463 | 1.437863 | 2.615223 | 0.549805 | 0.582453 | NA |
| NA | 0.260158 | 1.240347 | 4.047124 | 0.306476 | 0.759242 | NA |

| | | | | | | | |
|---|---|---|---|---|---|---|---|
| NA | | 4.432828 | 0.235117 | 1.106731 | 0.212443 | 0.831761 | NA |
| NA | | 2.862252 | -0.21243 | 1.144941 | -0.18554 | 0.852808 | NA |
| NA | | 0.247898 | -0.5901 | 4.074633 | -0.14482 | 0.884851 | NA |
| NA | | 0.437526 | -2.68242 | 3.384147 | -0.79264 | 0.427986 | NA |
| NA | | 1.345424 | 2.489115 | 2.169846 | 1.147139 | 0.251324 | NA |
